# Membrane proteins ClcB, PtsI and YcaM mediate the bactericidal effects of colistin in *Escherichia coli*

**DOI:** 10.1101/2025.07.07.662571

**Authors:** Rhys Donafee, Mohammad Radi, Douglas B. Kell, J. Enrique Salcedo-Sora

## Abstract

Bacteria can be killed very effectively by targeting their first line of protection. The cell membranes (outer and inner membranes) of Gram-negative bacteria are directly targeted by antibiotics, such as polymyxins, via electrostatic interactions with their lipopolysaccharide (LPS) fraction. The downstream effects - preceding the cell death - of the disruptions of the bacterial membranes upon the intercalation of these antibiotics are unknown. By screening a set of *E. coli* membrane protein knockouts, three membrane transporters were shown to mediate the growth-inhibitory effects of colistin. This was corroborated with growth assays on gain-of-function strains, cytotoxic assays and *in vivo* infections in an invertebrate animal model. This is first-time evidence that the disruption caused to membrane proteins, such as the chloride channel ClcB, the sugar transport system component PtsI and the hypothetical Glutamate:GABA antiporter YcaM, is part of the cytotoxic pathway that follows or is concomitant to the electrostatic intercalations of polymyxins with the Gram-negative bacteria membrane.

## INTRODUCTION

Colistin is currently used as a ‘last resort’ antibiotic against infections by multidrug-resistant Gram-negative bacilli, such as carbapenemase-producing *Enterobacteriales, Pseudomonas aeruginosa*, and *Acinetobacter baumanni* (1). The World Health Organization (WHO) has listed colistin as a critically important antimicrobial because of the increasing use of colistin to treat serious infections in humans in many parts of the world, the prevalence of *mcr* (mobile colistin resistance) genes, and the spread of colistin-resistant bacteria via the food chain (2). Polymyxins such as colistin, and other cell membrane-binding antibacterials, are also effective at targeting antibiotic persister bacteria (stochastic phenotypes that survive high concentrations of antibiotics) (3, 4).

Colistin (polymyxin E) is a cyclic lipopolypeptide antibiotic produced using a nonribosomal synthase by *Bacillus polymyxa* var colistinus (previously *B. colistinus*) (5, 6). Colistin is a polycation that binds lipid A in the lipopolysaccharide (LPS) fraction of the cell outer membrane (OM) of Gram-negative bacteria, displaces the divalent cations calcium (Ca^2+^) and magnesium (Mg^2+^), and impairs the structure of LPS while inserting its hydrophobic terminal acyl chain. This causes an expansion of the OM, permeabilization of the OM and a further movement of colistin beyond the OM (7-10). This can lead to an increase in cell permeability and cell barrier disruptions, including destabilization of membranes and loss of cytoplasm (11). This is recognised as the basis for the antimicrobial properties of polymyxins, manifested as an undetectable cell viability after half an hour of exposure to bactericidal concentrations of colistin (12).

Modes of resistance to colistin have substantiated the cell membrane-binding properties of polymyxins. A reduction in the net negative charge of lipid A affects polymyxin’s affinity for the OM. Enzymes that mediate chemical modifications of lipid A and cause a loss of sensitivity to polymyxins include plasmid-encoded *mcr* gene products (13), and two-component regulatory systems and sensor kinase systems such as pmrA/pmrB (14) and phoP/phoQ (11, 15). Inhibiting the production of fatty acids destined for lipid A biosynthesis overcomes resistance to colistin altogether (16).

This said, the mechanisms of the proposed increase in permeability and cell lysis after the interactions of polymyxins with the outer, and the inner membranes (17) are not known. Cells show some visible cell membrane disruptions one (18) and two hours(19) post-exposure. Other morphological post-exposure observations include the formation of bleb-like projections from the cell wall of *E. coli* (8), with cytoplasmic material detectable in the extracellular milieu after half an hour of exposure to polymyxins(20). Further reports show cell aggregation rather than cell membranes disruption of bacteria exposed to bactericidal concentrations of colistin (21). Furthermore, modified versions of polymyxin B that still have cell membrane-binding properties cause leakage of cytoplasmic contents, but have no antibacterial activity (22). Binding the Gram-negative bacteria membranes does not alone make such a molecule bactericidal.

The links between the well-established interactions of polymyxins with the cell membranes of Gram-negative bacteria and the rapid cell death that follows are thus still missing. Part of the answer could be in the direct interaction of colistin with the inner membrane (cell membrane) of Gram-negative bacteria (23) and the proteins within. The often overlooked but essential integral membrane proteins and membrane-associated proteins are the most abundant component by mass of the cell membranes (24), and responsible for the uptake of any number of small molecules (25-27). It thus seems logical to investigate the involvement of membrane proteins in the bactericidal effects of a membrane-binding antibiotic such as colistin. We here describe several membrane-associated proteins mediating cell death caused by colistin. Specifically, the absence of these proteins caused tolerance while their presence sensitised *E. coli* to this antibiotic. The relevance of these findings to *in vivo* infections is illustrated in an invertebrate animal model.

## RESULTS

### The absence of certain membrane transporters in *E. coli* gene knockouts reduced their sensitivity to colistin

A set of 534 strains lacking membrane transporters and other membrane associated proteins from the *E. coli* Keio collection of gene knock outs (KOs) (28) was used to screen for tolerance to colistin. They were exposed to incremental concentrations of colistin up to 5 mg/L in complex media and ranked according to their growth in comparison to that of the reference strain for that collection *E. coli* BW25113 (Supporting table 1). From that initial list, eight KO strains were selected for further growth inhibition assays (IC_50_): *ΔclcB, ΔfocB, ΔhisJ, ΔmalX, ΔptsI, ΔrhtC, ΔyadI* and *ΔycaM*. The IC_50_ data from four or more replicates for each of these strains showed that the KO strains *ΔclcB, ΔptsI, ΔyadI* and *ΔycaM* had IC_50_ median values six to twenty-fold that of the reference strain BW25113 (Figure 1).

**Figure 1.**
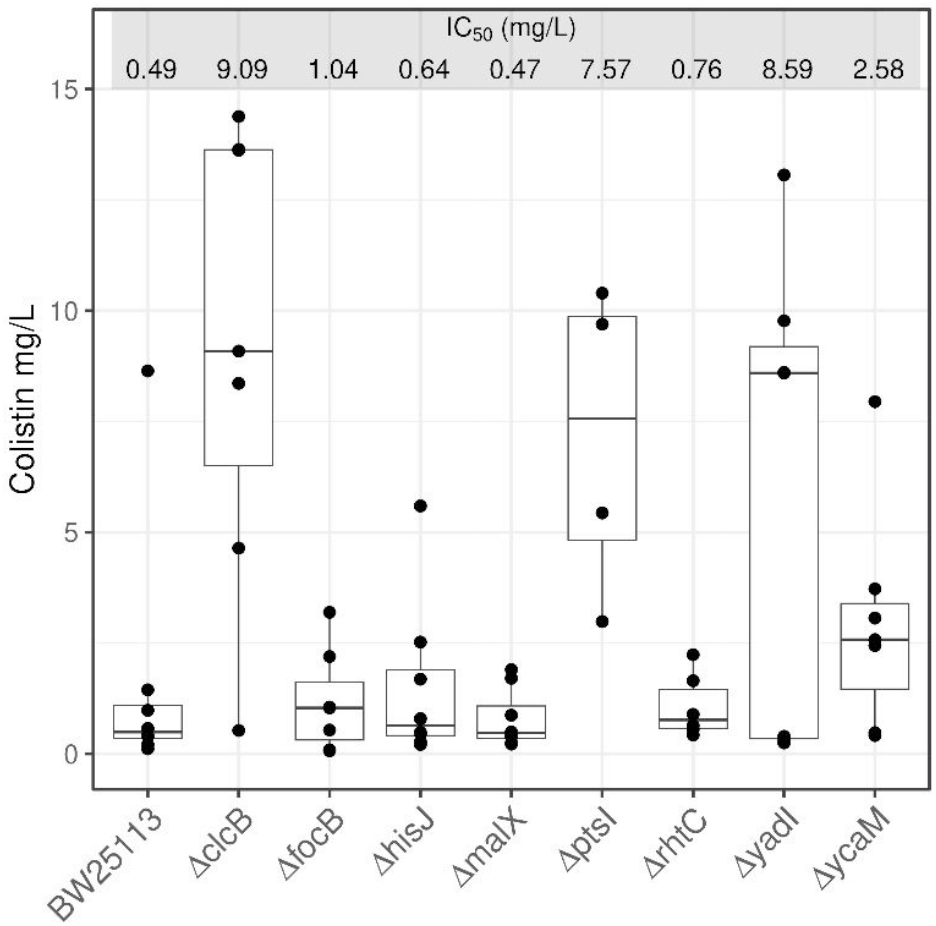
*E. coli* KOs with increased tolerance to colistin. The median for the IC_50_ values of the *E. coli* reference strain BW25113 is presented against those of the eight KO strains initially observed to have different levels of higher tolerance to colistin: *ΔclcB, ΔfocB, ΔhisJ, ΔmalX, ΔptsI, ΔrhtC, ΔyadI* and *ΔycaM*. Data represent at least four biological replicates (black dots). The median values for each IC_50_ range are listed as mg/L. The pairwise comparison between BW25113 against *ΔclcB, ΔptsI, ΔyadI* and *ΔycaM* had *p* values of less than 0.005.

### Expression of membrane transporters *clcB, ptsI* and *ycaM* increased the sensitivity of *E. coli* to colistin

The availability of the ASKA collection of *E. coli* transformed with plasmids expressing most of the protein-encoding genes for these bacteria allowed us to assess if the opposite (increased sensitivity to colistin) was observed in strains expressing the genes *clcB, ptsI, yadI* and *ycaM*. The ASKA collection uses the *E. coli* K-12 derivative strain *AG1* carrying recombinant constructs in an IPTG-inducible *lacO*-T5 promoter expression system in the multicopy plasmid pCA24N (*CmR, lacIq*) (29). The *E. coli* reference strain AG1 as used here was a true control transformed with an empty version of the pCA24N plasmid.

Strains with the recombinant plasmids carrying *clcB, ptsI, yadI* or *ycaM* (denoted as *AG1+pclcB, AG1+pptsI, AG1+ pyadI* and *AG1+pycaM*), respectively) showed IC_50_ values similar to that of the reference AG1 strain (*AG1+pCA24N*) (Figure 2): 0.09 – 0.2 mg/L. However, when 0.1 mM IPTG was added, IC_50_ values decreased to half or less for those strains carrying recombinant plasmids: 0.05 – 0.07 mg/L; except for the strains carrying the empty vector (*AG1+pCA24N*), and the strain expressing *pyadI* (*AG1+pyadI*) (Figure 2).

**Figure 2.**
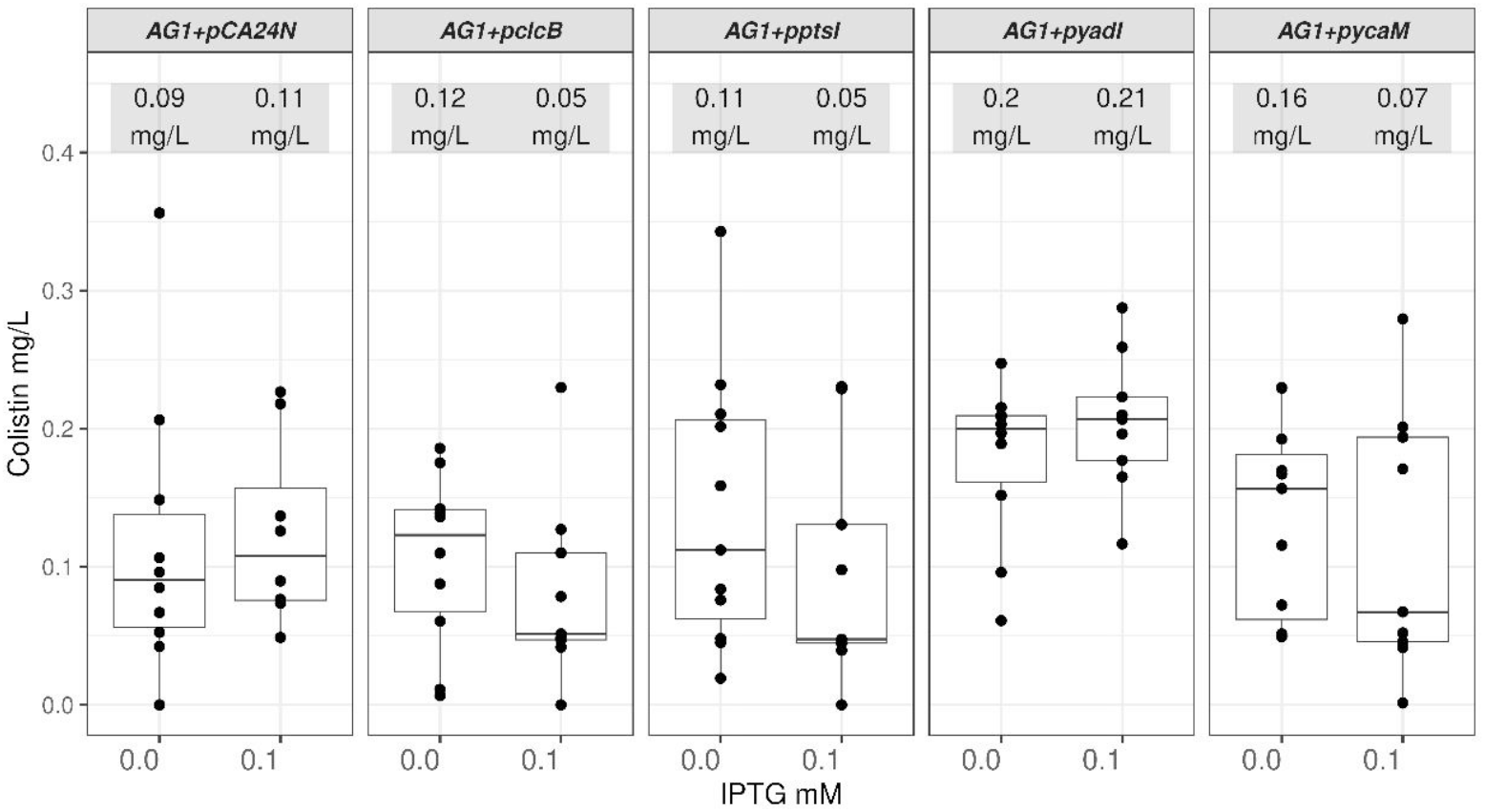
*E. coli* strains expressing *clcB, ptsI* and *ycaM* are more sensitive to colistin. The median values for the IC_50_ of the *E. coli* reference strain AG1 carrying the empty pCA24N (*AG1+pCA24N*) plasmid and those strains containing this plasmid with either of four genes were calculated from samples incubated in the absence or the presence of 0.1 mM IPTG. Three of those strains (*AG1+pclcB, AG1+pptsI* and *AG1+pycaM*) showed a trend of increased sensitivity to colistin when the cognate membrane transporter was induced. Data illustrated originated from at least eight biological repeats (black dots). The median values for each IC_50_ range are listed as mg/L. The pairwise comparison between non-IPTG and 0.1 mM IPTG samples had p values of less than 0.005 for the *AG1+pclcB, AG1+pptsI* and *AG1+pycaM* strains.

The *AG1+pclcB, AG1+pptsI and AG1+pycaM* strains were selected for kinetic growth assays. IPTG (0.05 mM) was added to the relevant samples after four hours of initial growth in complex media (see Methods). Colistin was later added at eight hours. At this time the membrane transporters were expected to have reached detectable protein levels induced by IPTG (30). All three strains had visible lower culture densities (absorbance at OD_600_) than the *AG1+pCA24N* control in the presence of colistin. Those differences widened in the samples upon induction with 0.05 mM IPTG (Figure 3 and Supplementary figure 1), particularly for the *AG1+pclcB* and *AG1+pycaM* samples. The growth of the *AG1+pptsI* samples seemed to have had an intermediate sensitivity to the growth inhibitory effects of colistin (Figure 3).

**Figure 3.**
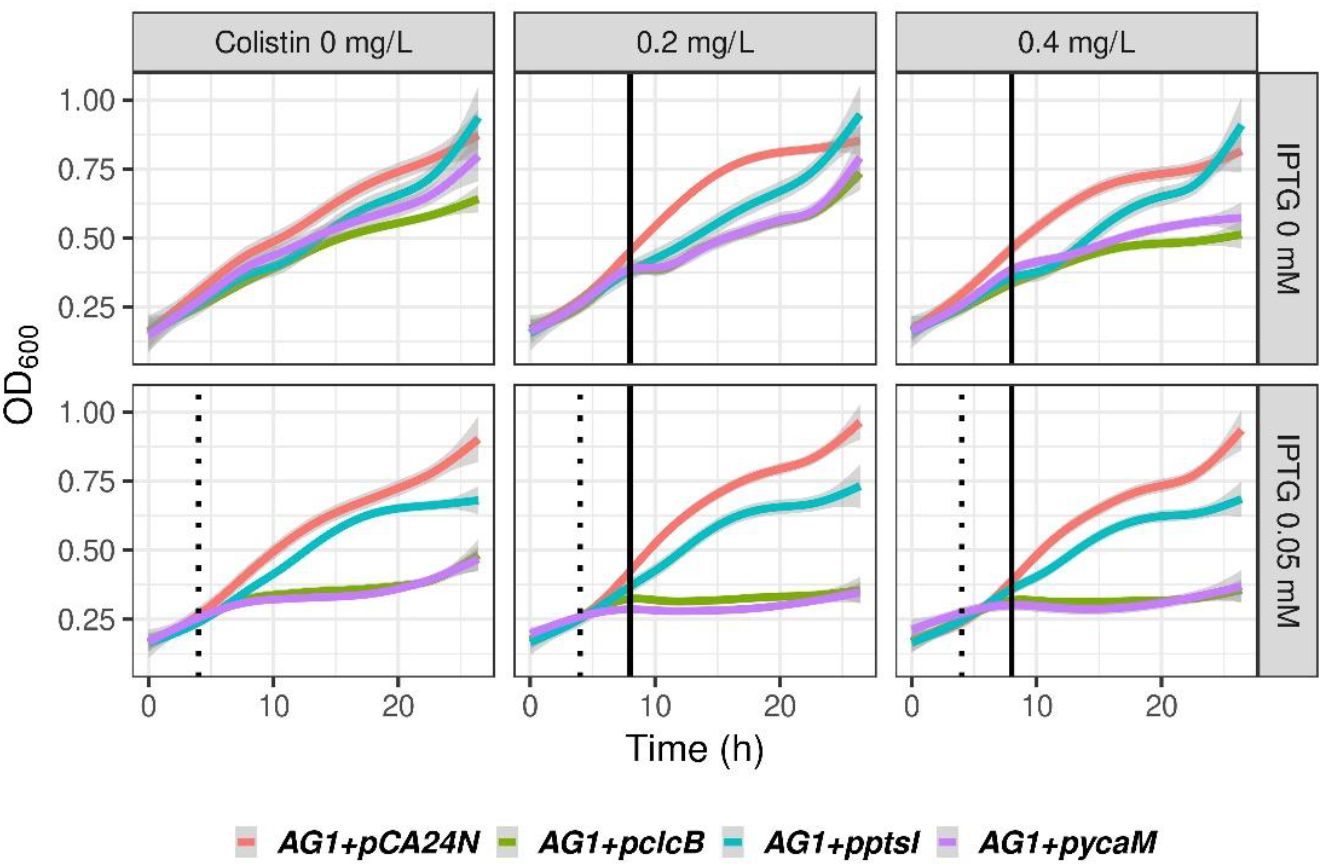
Growth inhibitory effects of colistin in *E. coli* expressing the membrane transporters *clcB, ptsI* and *ycaM*. The abscissa represents time. The ordinate represents the culture media optical densities at 600nm. The AG1 control and the recombinant samples are colour-coded: *AG1+pCA24N* is red, *AG1+pclcB* is green, *AG1+pptsI* is blue and *AG1+pycaM* is magenta. Colistin is represented here in the 0.2 and 0.4 mg/L samples. The full range of colistin concentrations analysed is presented in Supplementary figure 1. The dotted vertical black line represents the addition of 0.05 mM IPTG at 4 hours. The solid vertical black line represents the addition of colistin at 8 hours. The data represented in this figure and the expanded version of it in Supporting figure 1 are tabulated in Supporting table 2.

The overexpression of membrane transporters is often toxic to the bacterial host (31). This was evident in the growth assays where *AG1+pclcB* and *AG1+pycaM* were visibly inhibited in the IPTG-induced samples in the absence of colistin (Figure 3). However, further assays based on growth in solid media substantiated, at two incremental doses, the observed sensitization to colistin caused by the expression of *clcB, ptsI* and *ycaM* additional to the baseline toxicity due to the expression of these proteins upon their induction with IPTG (Figure 4).

**Figure 4.**
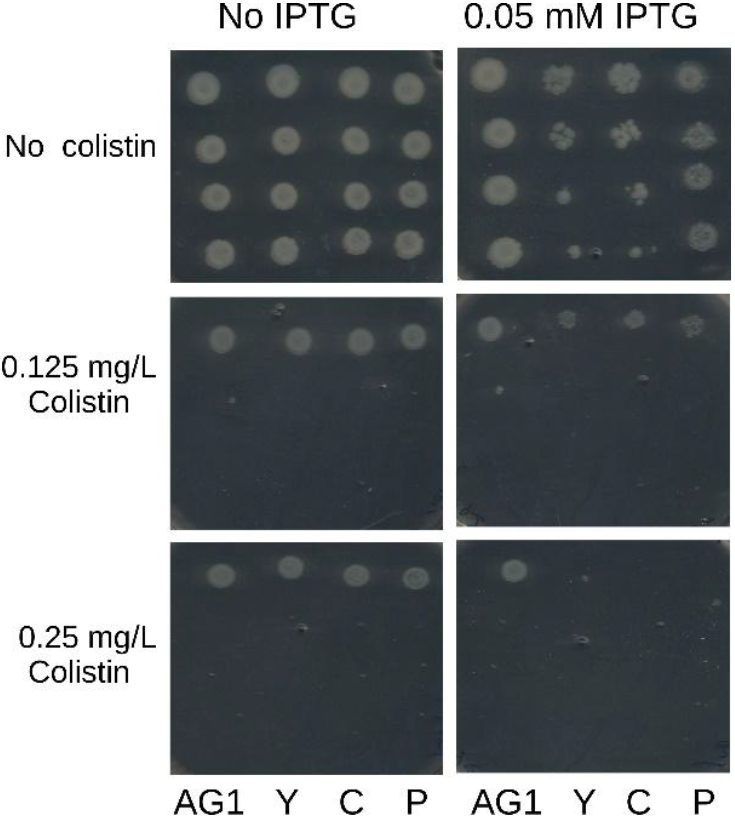
The presence of *clcB, ptsI* and *ycaM* mediated the growth inhibition caused by colistin in *E. coli*. AG1 is the control strain carrying the empty plasmid pCA24N. Y, C, and P represent the AG1 strains carrying the plasmids *pycaM. pclcB* or *pptsI*, respectively. Each strain is represented by four row-wise 1:10 dilutions of culture broth used for spotting. Two different colistin concentrations are shown, in the absence or the presence of IPTG (0.05mM.

### Gene complementation-induced sensitivity to colistin in otherwise tolerant *E. coli* KO strains

Each of the *E. coli* KOs *ΔclcB, ΔptsI* and *ΔycaM* from the Keio collection were transformed with either an empty pCA24N plasmid or with a recombinant plasmid carrying their cognate gene *pclcB, pptsI* or *pycaM*. The identity of the recombinant constructs in pCA24N plasmids as well as the empty plasmid were all confirmed by DNA sequencing (sequence data available upon request). Equally, whole genome sequencing for the *ΔclcB, ΔptsI* and *ΔycaM* strains confirmed their loci to have been replaced by the *S. aureus aadD1* gene encoding for an aminoglycoside O-nucleotidyltransferase that confers the kanamycin resistance marker for the *E. coli* Keio KO collection.

The strains *ΔclcB, ΔptsI* and *ΔycaM* carrying the recombinant plasmids *pclcB, pptsI* and *pycaM*, respectively, had their growth affected in the absence of IPTG (Figure 5B) when compared against the empty plasmid controls (pCA24N, Figure 5A). We interpreted this to be the consequence of a leakier gene expression system in the *E. coli* BW25113 background strain in comparison to the *E. coli* AG1 strains (Figure 3). The BW25113-based strains seemed to have a higher basal lac operon activity, as is sometimes observed when it is assembled with the endogenous bacterial promoter T5 (30), as is the case of the pCA24N plasmid. Growth was further affected by the presence of 0.1 mM IPTG, particularly for the strain carrying *pclcB* which impeded to observe any further growth inhibition upon addition of colistin in this case. Nevertheless, it was still possible to observe the growth inhibitory effect of colistin in relation to the heterologous expression of *ptsI* and *ycaM* (Figure 5D).

**Figure 5.**
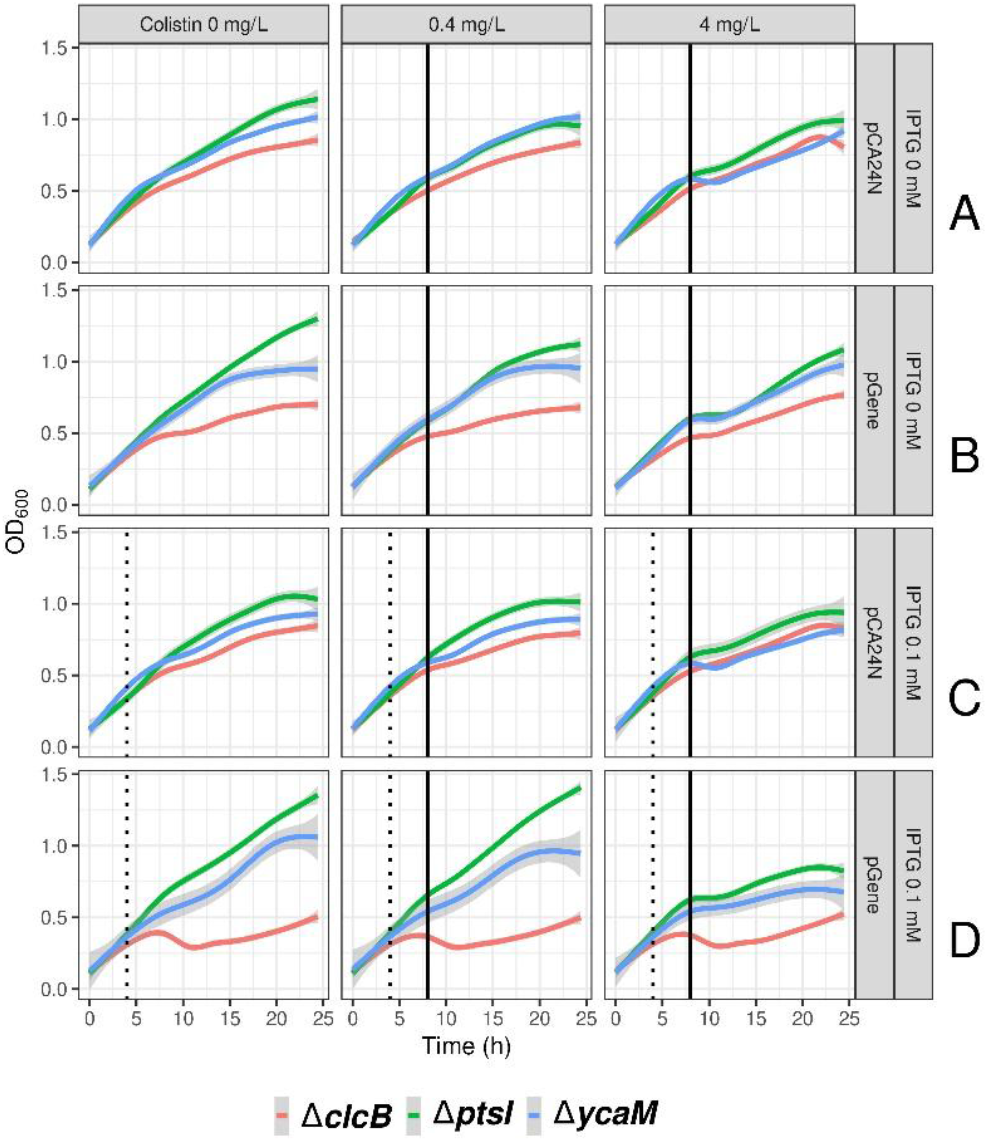
The expression of *ptsI* and *ycaM* re-sensitises *E. coli* KO strains to colistin. No-colistin or colistin samples at 0.4 and 4 mg/L are presented column-wise. Rows A and B represent growth (absorbance at OD_600_) in the absence of IPTG. Rows C and D represent growth in the presence of 0.1 mM IPTG. Strains transformed with the empty pCA24N plasmid are in rows A and C. Strains transformed with the recombinant plasmids expressing their cognate genes (denoted as pGene) are in rows B and D. Each KO strain carrying either the empty plasmid or the recombinant plasmid is colour coded: *ΔclcB* is red, *ΔptsI* is green and *ΔycaM* is blue.

### Short-term exposure to high concentrations of colistin supports the mediation of *clcB, ptsI* and *ycaM* in the action of colistin

Growth-inhibitory effects are usually observed in short-term exposures to bactericidal compounds. We measured the survival of the KOs *ΔclcB, ΔptsI* and *ΔycaM* after half an hour exposure to 5 mg/L of colistin. While the reference strain BW25113 had a 32% survival in relation to the starting population, the KOs had higher and incremental survival levels of up to a median value of 77 % for *ΔptsI*), with *ΔclcB* and *ΔycaM* showing similar survival rates of 72 % and 71 %, respectively (Figure 6).

**Figure 6.**
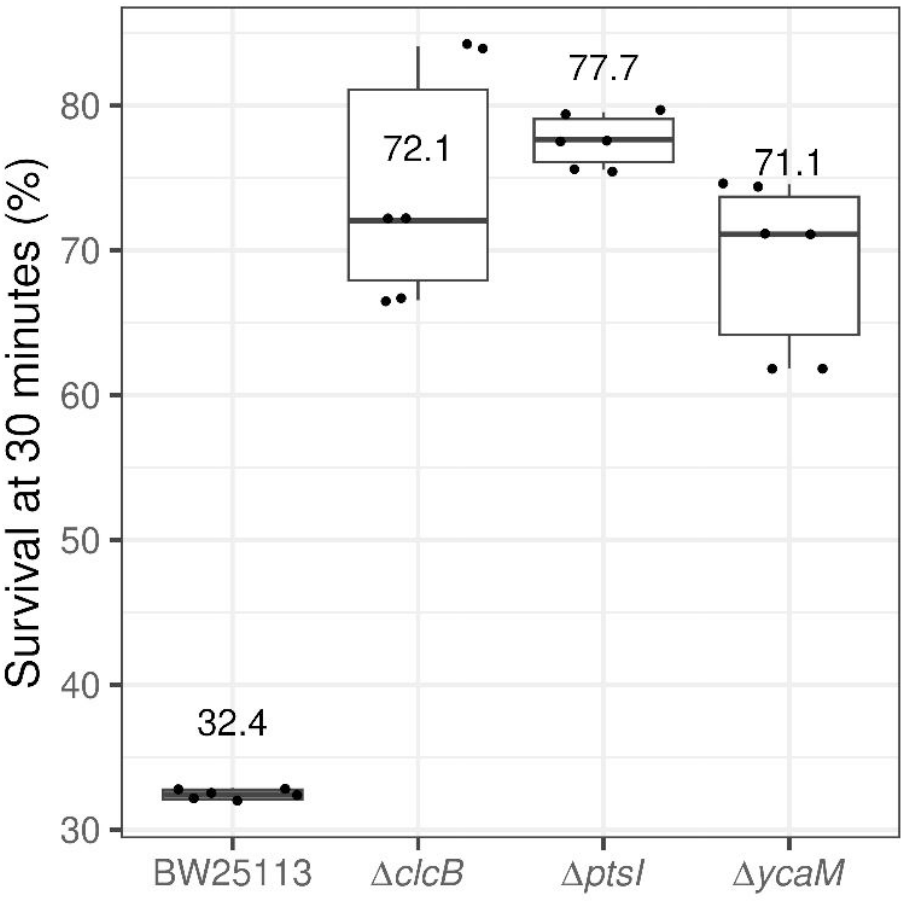
*E. coli* KOs increased survival to short term exposure to high concentrations of colistin. Viable cell counts are presented at time zero and after 30 minutes exposure to 5 mg/L of colistin. Data is presented for the *E. coli* reference strain BW25113 (median 32.4 % survival) and the three KO strains: *ΔclcB* (72.1 %), *ΔptsI* (77.7 %) and *ΔycaM* (71.1 %). Data represent three biological replicates. The pairwise comparison between BW25113 against *ΔclcB, ΔptsI* and *ΔycaM* had *p* values of less than 0.005.

### The absence of *clcB, ptsI*, and *ycaM* reduces the efficacy of colistin *in vivo*

The larvae of the Greater wax moth (*Galleria mellonella*) provide an invertebrate animal model that is often used to study bacterial infections (32-35). A positive correlation between the virulence and immune responses between mammalian models and the innate immune response of *G. mellonella* has been established for several infections (32, 36). We used larvae of *G. mellonella* to determine potential differences, in the absence as well as the presence of colistin, in the survival of the larvae infected with the *E. coli* reference BW25113 strain or the KOs *ΔclcB, ΔptsI* and ΔycaM. The data were represented as time-to-event analysis using Kaplan-Meier curves and using log rank tests to test for significance between strains. Each larva was injected with 50 ng of colistin. At an average weight of 250 mg, each larva received approximately 0.2 mg/Kg of colistin.

In the absence of colistin, at day four the survival probability of larvae infected with *E. coli* BW25113 was 8.2% (Fig. 7A, Supporting table 3). When the infected larvae were also injected with colistin, the survival trend increased to 47 % for BW25113 at day four (Fig. 7A). Thus, the presence of colistin increased the survival of larvae infected with the reference strain by over six-fold.

**Figure 7.**
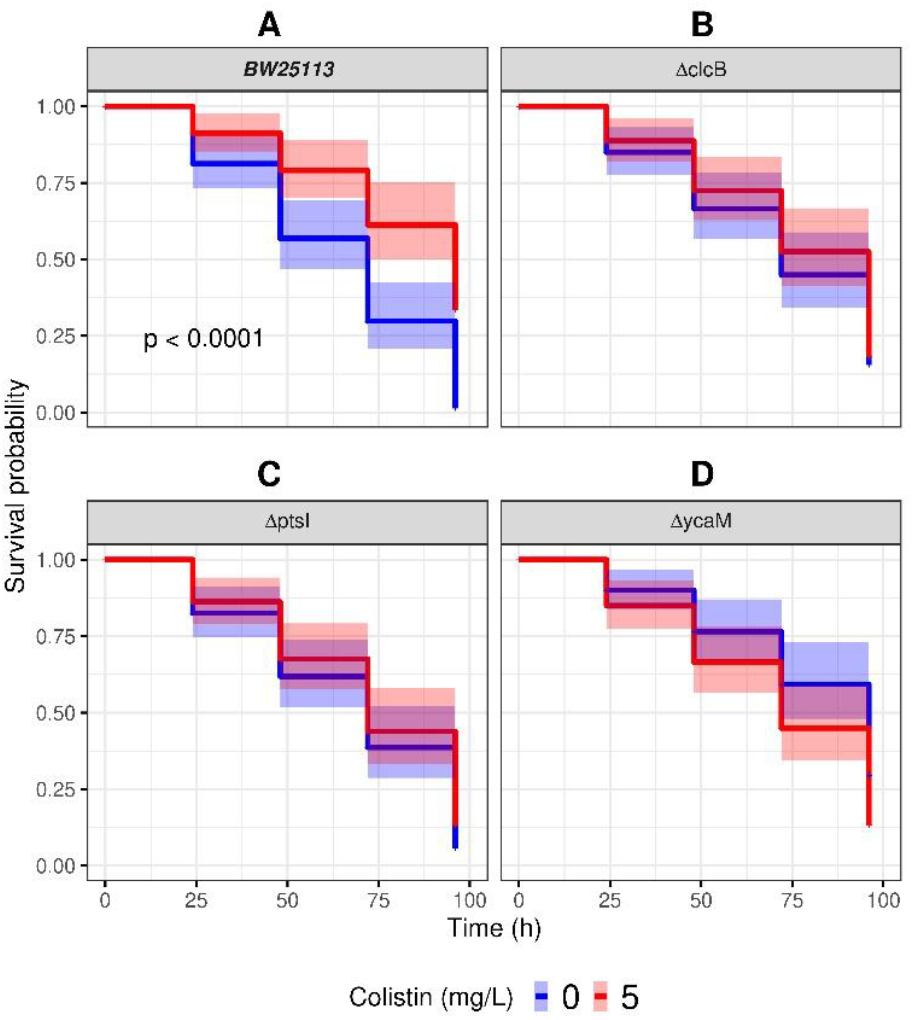
*Galleria mellonella larvae infected E. coli* KOs have reduced survival rates. Kaplan-Meier survival curves are shown for larvae firstly infected with the reference strain BW25113 **(A)** or either of the KOs *ΔclcB* **(B)**, *ΔptsI* **(C)** and *ΔycaM* **(D)**. Colistin control larvae (blue curves) were subsequently injected with PBS. Data of larvae subsequently injected with 5 mg/L of colistin are represented by the red curve. Data represents up to 80 larvae per group, from at least four different biological replicates. Shaded areas cover the 95 % confidence intervals (Supporting table 3).

The probabilities of survival for the larvae injected with the KOs were 14%, 10%, and 18% for *ΔclcB, ΔptsI* and *ΔycaM*, respectively, in the absence of colistin (Figs. 7B - 7D, Supporting table 3). In the presence of colistin, survival rates were 13%, 12%, and 8% for *ΔclcB, ΔptsI* and *ΔycaM*, respectively (Fig. 7B - 7D, Supporting table 3). Therefore, the presence of the antibiotic made no significant difference to those larvae infected with the KOs.

## DISCUSSION

Membrane lipid composition and integrity can modulate the structure and activity of transport proteins (37, 38). The derangement of the bacterial cell membranes after its interactions with polymyxins such as colistin (17-22) are thus expected to alter the integrity and activity of at least some membrane proteins. Colistin also reaches the cytoplasmic side of the inner membrane whereby it can disrupt the initial movement of LPS into the cell membrane. This would be expected to add to the irreversible changes in the integrity of the bacterial membrane (23). This is corroborated by the reports of efflux inhibitors increasing the susceptibility of bacteria to colistin (39, 40). Altogether, disruptions of cellular homeostasis are expected to be a primary cytotoxic effect caused by permeability disruption (19). Our findings here point to specific membrane proteins whose malfunction can affect this disruption of cell homeostasis. Given their location and roles, the dysregulation of PtsI, ClcB and YcaM provides evidence for specific membrane protein dysgenesis as a mediator of the colistin’s lethality.

ClcB is one of the two CLC-type voltage-gated chloride channels found in bacteria, although the function of ClcB is largely unknown (41). It is thought to act as an electrical shunt for an outwardly-directed proton pump that is linked to amino acid decarboxylation, and is of importance in extreme acid resistance responses (42). PtsI is enzyme I of the phosphoenolpyruvate:carbohydrate phosphotransferase system (PTS) (43). This system mediates the transport of sugars and their intracellular phosphorylation in bacteria. PtsI localises to the inner surface of the cytoplasmic membrane where it interacts with sugar-specific inner-membrane permeases. PTS systems regulate carbon utilisation in switching between the hierarchical (diauxic) and co-utilisation (non-specific) use of sugars. YcaM has been classified by sequence similarity as a Glutamate:GABA antiporter, part of the amino acid-polyamine organocation superfamily (44). YcaM function has not been characterised. It seems to be expressed when bacteria are in the presence of abundant nutrients *in vitro* (45).

The Keio collection of *E. coli* gene KO used here has been previously profiled against a set of antibiotics that included colistin (46). However, opposite to our approach, these authors assessed only the sensitivity of KOs to antibiotics (we screened for tolerance to colistin). This could explain why the membrane protein-encoding genes reported here do not appear in the list of genes reported in that work as conferring different levels of sensitivity to colistin (46). A subsequent study, although polymyxins were not included, showed other membrane transporters mediating the growth inhibitory effects of a number of antibiotics in a subset of the Keio collection (47).

The dysregulation of ClcB can affect the pH and ionic homeostasis of bacteria. Alkaline pH has a synergistic effect with colistin across several species of Gram-negative (48).

This observation was extended to intrinsically polymyxin-tolerant Gram-negative bacteria *Burkholderia spp*. (48). PtsI has been shown to be involved in ROS-mediated cellular responses to antimicrobials and other environmental stressors, where a lack of PtsI function allowed tolerance to different types of stressors. Such pan-tolerance in the absence of PtsI seems to be based on a reduced capacity for metabolic shifts between different central carbon metabolism pathways. This offers protection from stress-mediated ATP surges, with a concomitant reduced accumulation of ROS (49). The potential mechanisms by which YcaM would mediate the toxicity of a membrane-acting antibiotic such as colistin will require further investigation, particularly since the function of this membrane protein is itself still to be characterised.

The evidence presented here opens new views regarding the mechanism of action of polymyxins. These antibiotics kill the bacterial cell via a series of discernible steps as opposed to a single event of detergent-like cell lysis. In contrast to detergents, the interactions of polymyxins with the cell membranes, mainly with the LPS fraction, are driven by electrostatic interactions (i.e., not hydrophobic), which cause membrane disorganisation by antibiotic intercalation and self-promoted permeation (50). It is evident that the ensuing cytotoxicity follows then a pathway that includes the disfunction of critical membrane transporters, represented here for *E. coli* by the chloride channel ClcB, the sugar transport system (at least PtsI) and the Glutamate:GABA antiporter YcaM.

## METHODS

### Strains

The *E. coli* Keio collection of gene KO strains was provided by the National Institute of Genetics, Mishima, Shizuoka, Japan (28, 51). A subset of 534 strains from the Keio Collection whose cognate genes are annotated as encoding for membrane associated proteins were selected for this study (Supporting table 1). Individual knockout strains highlighted in this work are as follows: ***clcB*** *(F-, Δ(araD-araB)567, ΔlacZ4787(::rrnB-3), λ-, ΔclcB740::kan, rph-1, Δ(rhaD-rhaB)568, hsdR514);* ***ptsI*** *(F-, Δ(araD-araB)567, ΔlacZ4787(::rrnB-3), λ-, ΔptsI745::kan, rph-1, Δ(rhaD-rhaB)568, hsdR514);* ***ycaM*** *(F-, Δ(araD-araB)567, ΔlacZ4787(::rrnB-3), λ-, rph-1, ΔycaM723::kan, Δ(rhaD-rhaB)568, hsdR514)* ; ***ydaI*** *(F-, Δ(araD-araB)567, ΔlacZ4787(::rrnB-3), λ-, ΔyadI746::kan, rph-1, Δ(rhaD-rhaB)568, hsdR514)*. Whole genome sequencing was used to verify the absence of the cognate genes in the *clcB, ptsI* and *ycaM* KO strains (Plasmidsaurus Inc; Supporting information files). We also used strain overexpressing *clcB, ptsI* and *ycaM* from the ASKA collection: *E. coli* K-12 derivative strain AG1 [*recA1 endA1 gyrA96 thi-1 hsdR17 (r K*− *m K+) supE44 relA1*] carrying recombinant constructs for those genes in the IPTG-inducible and multicopy plasmid pCA24N (*CmR, lacIq*) (29). The composition of those recombinant in pCA24N plasmids was also confirmed by whole plasmid DNA sequencing (Plasmidsaurus Inc; Supporting information files)

### Inhibitory concentrations assays

Bacterial cultures were carried out routinely in complex media (lysogeny broth, Merck LB 110285) (26). Colistin was purchased from MP Biomedicals, cat number 194157, activity ≥15,000 U/mg. The screening of the Keio collection subset of 534 strains (Supporting table 1) to incremental concentrations of colistin was carried out by replicating their glycerol stocks into 384-well plates with 50 µL of LB containing 30 µg/mL of kanamycin. These sealed plates were incubated overnight at 37 °C. The overnight cultures were diluted 1 in 100 in 50 µL of fresh LB without kanamycin in polystyrene 384-well plates with a transparent bottom (Sigma M6936-40EA). End-point growth inhibitory concentrations (IC_50_) were calculated from microtitration assays. Overnight cultures in LB were diluted 1 in 100 in fresh LB. Fifty µL of two-fold dilutions of colistin were prepared in 96-well plates to which 50 µL of fresh bacterial culture were added. Endpoint read outs of the media turbidity at 600 nm (OD_600_) were taken after 24 h at 37 °C. Continuous growth assays to follow the effect of protein expression and subsequent exposure to colistin were carried out with freshly 1 in 100 diluted cultures in 180 µL at 30°C in 96 microwell plates and shaking at 200 rpm.

After four hours protein expression was induced with 50 µM IPTG. At eight hours colistin was added to final concentrations 0.05, 0.1, 0.2, 0.4, 0.8, 1.6, 3.2 mg/L. OD_600_ readouts were collected every 15 minutes in a BMG LabTech CLARIOstar Plus plate reader for 27 hours.

### Short term exposure to colistin

Overnight cultures were diluted 1:500 in 5 mL of complex media and allow to grow for 2h, 37°C and shaking 250 rpm. Number of viable cells were measured from 1:10 dilutions in 0.1 mL of complex media, prepared for each strain row-wise in a 96-well plate. Samples from different and incremental dilutions were plated out in solid complex media and incubated overnight at 37°C. Viable cells were counted manually from plates with colonies clearly visible and separated from each other. These samples were the time 0 data. Subsequently, colistin, at 5 mg/L, was added to an aliquot of each strain and incubated at 37°C, for 30 minutes, shaking 250 rpm. The inoculation size that allowed a measurable number of viable cells in the reference strain BW25113 after half an hour of exposure to colistin was 10_7_ cfu/mL. The number of viable cells were obtained as for the time 0 sample.

### Gene complementation

*E. coli* KO strains *ΔclcB, ΔptsI* and *ΔycaM* were transformed with the recombinant plasmids pCA24N carrying either their cognates genes, or empty plasmids as controls. The presence of those plasmids in their respective strains was verified by enzyme restriction analysis before assessing their antibiotic sensitivity. The IC_50_ assays for colistin were carried out as described above.

### Larval survival assay

Larvae of the Greater wax moth *Galleria mellonella* were purchased from Livefoods UK Ltd. (Somerset, UK). Larvae were injected the same day upon delivery. Healthy large larvae (approximately 250 – 350 mg body weight) were selected. They had a uniform cream colour without spots or marks indicating melanisation. Ten larvae per condition were injected with 10^4^ cells of *E. coli* (for both reference strain and KO strains) in10 µL in either of the penultimate prolegs, using a 10 µL Hamilton 750 syringe (Hamilton Company, UK). After injections, the larvae were placed into Petri dishes and incubated at 37oC. Syringes were cleaned with 70% ethanol between groups of injection. Larvae mortality was followed for 96 hours by counting death individuals (black or dark brown in colour) every 24 hours. Colistin was injected in the opposite penultimate proleg, 30 min after they had been injected with bacteria. The control group was injected with phosphate buffer saline (PBS).

### Data analysis

The inhibitory concentrations of colistin that kills half of the bacteria population (IC_50_) were calculated with the four-parameter logistic model as implemented in the R package *drc* (52, 53) and python package py50 (54), following the guidelines for relative calculations according to the spread of the data (55). Pairwise comparisons were carried out using the built-in pairwise t-test function in R (52).

Visualization and annotation of bacterial genomes and plasmid sequences used Proksee (56). Survival analyses of larvae were carried out with the time to event model Kaplan-Meier which estimates survival differences in survival times between compared groups as implemented in the R packages survival and survminer (57).

## ACKNOWLEDGMENTS

DBK thanks the Novo Nordisk Foundation for financial support (grant NNF20CC0035580). We also thank the University of Liverpool and the Liverpool Shared Research Facilities for their support.

