## Supplementary material for "Membrane proteins ClcB, PtsI and YcaM mediate the bactericidal effects of colistin in *Escherichia coli*": Supp_table_3

| time | n.risk | n.event | n.censor | lower | upper | Strain | Colistin | surv |
|---|---|---|---|---|---|---|---|---|
| 0 | 150 | 0 | 30 | 1 | 1 | BW25113 | No | 1 |
| 24 | 120 | 16 | 14 | 0.807931 | 0.929673 | BW25113 | No | 0.866667 |
| 48 | 90 | 21 | 9 | 0.581205 | 0.759606 | BW25113 | No | 0.664444 |
| 72 | 60 | 23 | 7 | 0.322237 | 0.521007 | BW25113 | No | 0.409741 |
| 96 | 30 | 24 | 6 | 0.038519 | 0.17434 | BW25113 | No | 0.081948 |
| 0 | 150 | 0 | 30 | 1 | 1 | BW25113 | Yes | 1 |
| 24 | 120 | 7 | 23 | 0.900653 | 0.984548 | BW25113 | Yes | 0.941667 |
| 48 | 90 | 9 | 21 | 0.78077 | 0.919933 | BW25113 | Yes | 0.8475 |
| 72 | 60 | 10 | 20 | 0.614137 | 0.812179 | BW25113 | Yes | 0.70625 |
| 96 | 30 | 10 | 20 | 0.352639 | 0.628642 | BW25113 | Yes | 0.470833 |
| 0 | 150 | 0 | 30 | 1 | 1 | clcB | No | 1 |
| 24 | 120 | 15 | 15 | 0.817784 | 0.936219 | clcB | No | 0.875 |
| 48 | 90 | 17 | 13 | 0.629171 | 0.800586 | clcB | No | 0.709722 |
| 72 | 60 | 19 | 11 | 0.393037 | 0.598423 | clcB | No | 0.484977 |
| 96 | 30 | 21 | 9 | 0.081004 | 0.261323 | clcB | No | 0.145493 |
| 0 | 150 | 0 | 30 | 1 | 1 | clcB | Yes | 1 |
| 24 | 120 | 12 | 18 | 0.847893 | 0.955309 | clcB | Yes | 0.9 |
| 48 | 90 | 18 | 12 | 0.639043 | 0.811213 | clcB | Yes | 0.72 |
| 72 | 60 | 18 | 12 | 0.410942 | 0.618131 | clcB | Yes | 0.504 |
| 96 | 30 | 22 | 8 | 0.071757 | 0.25173 | clcB | Yes | 0.1344 |
| 0 | 150 | 0 | 30 | 1 | 1 | ptsI | No | 1 |
| 24 | 120 | 14 | 16 | 0.827724 | 0.942679 | ptsI | No | 0.883333 |
| 48 | 90 | 17 | 13 | 0.636079 | 0.807046 | ptsI | No | 0.716481 |
| 72 | 60 | 19 | 11 | 0.397107 | 0.603626 | ptsI | No | 0.489596 |
| 96 | 30 | 24 | 6 | 0.046454 | 0.2064 | ptsI | No | 0.097919 |
| 0 | 150 | 0 | 30 | 1 | 1 | ptsI | Yes | 1 |
| 24 | 120 | 14 | 16 | 0.827724 | 0.942679 | ptsI | Yes | 0.883333 |
| 48 | 90 | 19 | 11 | 0.614908 | 0.789716 | ptsI | Yes | 0.696852 |
| 72 | 60 | 20 | 10 | 0.373452 | 0.577914 | ptsI | Yes | 0.464568 |
| 96 | 30 | 22 | 8 | 0.065829 | 0.233142 | ptsI | Yes | 0.123885 |
| 0 | 150 | 0 | 30 | 1 | 1 | ycaM | No | 1 |
| 24 | 120 | 13 | 17 | 0.837757 | 0.949046 | ycaM | No | 0.891667 |
| 48 | 90 | 18 | 12 | 0.632249 | 0.804816 | ycaM | No | 0.713333 |
| 72 | 60 | 18 | 12 | 0.406806 | 0.612905 | ycaM | No | 0.499333 |
| 96 | 30 | 19 | 11 | 0.109614 | 0.305814 | ycaM | No | 0.183089 |
| 0 | 150 | 0 | 30 | 1 | 1 | ycaM | Yes | 1 |
| 24 | 120 | 16 | 14 | 0.807931 | 0.929673 | ycaM | Yes | 0.866667 |
| 48 | 90 | 23 | 7 | 0.560943 | 0.742079 | ycaM | Yes | 0.645185 |
| 72 | 60 | 23 | 7 | 0.311824 | 0.507645 | ycaM | Yes | 0.397864 |
| 96 | 30 | 24 | 6 | 0.037362 | 0.169473 | ycaM | Yes | 0.079573 |
