## Supplementary material for "Membrane proteins ClcB, PtsI and YcaM mediate the bactericidal effects of colistin in *Escherichia coli*": Supp_table_2

| Colistin | IPTG | Strain | Repeat | Time | OD600 |
|---|---|---|---|---|---|
| 0 | 0 | ptsI | 2 | 0 | 0.199 |
| 0 | 0 | clcB | 2 | 0 | 0.206 |
| 0 | 0 | ycaM | 2 | 0 | 0.213 |
| 0 | 0 | yadI | 2 | 0 | 0.222 |
| 0 | 0 | AG1 | 2 | 0 | 0.228 |
| 0 | 0 | ptsI | 2 | 0.25 | 0.202 |
| 0 | 0 | clcB | 2 | 0.25 | 0.207 |
| 0 | 0 | ycaM | 2 | 0.25 | 0.206 |
| 0 | 0 | yadI | 2 | 0.25 | 0.213 |
| 0 | 0 | AG1 | 2 | 0.25 | 0.222 |
| 0 | 0 | ptsI | 2 | 0.5 | 0.199 |
| 0 | 0 | clcB | 2 | 0.5 | 0.204 |
| 0 | 0 | ycaM | 2 | 0.5 | 0.205 |
| 0 | 0 | yadI | 2 | 0.5 | 0.214 |
| 0 | 0 | AG1 | 2 | 0.5 | 0.219 |
| 0 | 0 | ptsI | 2 | 0.75 | 0.2 |
| 0 | 0 | clcB | 2 | 0.75 | 0.205 |
| 0 | 0 | ycaM | 2 | 0.75 | 0.206 |
| 0 | 0 | yadI | 2 | 0.75 | 0.215 |
| 0 | 0 | AG1 | 2 | 0.75 | 0.22 |
| 0 | 0 | ptsI | 2 | 1 | 0.209 |
| 0 | 0 | clcB | 2 | 1 | 0.21 |
| 0 | 0 | ycaM | 2 | 1 | 0.212 |
| 0 | 0 | yadI | 2 | 1 | 0.222 |
| 0 | 0 | AG1 | 2 | 1 | 0.225 |
| 0 | 0 | ptsI | 2 | 1.25 | 0.212 |
| 0 | 0 | clcB | 2 | 1.25 | 0.21 |
| 0 | 0 | ycaM | 2 | 1.25 | 0.214 |
| 0 | 0 | yadI | 2 | 1.25 | 0.231 |
| 0 | 0 | AG1 | 2 | 1.25 | 0.23 |
| 0 | 0 | ptsI | 2 | 1.5 | 0.21 |
| 0 | 0 | clcB | 2 | 1.5 | 0.213 |
| 0 | 0 | ycaM | 2 | 1.5 | 0.219 |
| 0 | 0 | yadI | 2 | 1.5 | 0.234 |
| 0 | 0 | AG1 | 2 | 1.5 | 0.238 |
| 0 | 0 | ptsI | 2 | 1.75 | 0.214 |
| 0 | 0 | clcB | 2 | 1.75 | 0.216 |
| 0 | 0 | ycaM | 2 | 1.75 | 0.222 |
| 0 | 0 | yadI | 2 | 1.75 | 0.242 |
| 0 | 0 | AG1 | 2 | 1.75 | 0.246 |
| 0 | 0 | ptsI | 2 | 2 | 0.218 |
| 0 | 0 | clcB | 2 | 2 | 0.219 |
| 0 | 0 | ycaM | 2 | 2 | 0.228 |
| 0 | 0 | yadI | 2 | 2 | 0.252 |
| 0 | 0 | AG1 | 2 | 2 | 0.258 |
| 0 | 0 | ptsI | 2 | 2.25 | 0.224 |
| 0 | 0 | clcB | 2 | 2.25 | 0.224 |

| | | | | | |
|---|---|---|---|---|---|
| 0 | 0 | ycaM | 2 | 2.25 | 0.235 |
| 0 | 0 | yadI | 2 | 2.25 | 0.262 |
| 0 | 0 | AG1 | 2 | 2.25 | 0.274 |
| 0 | 0 | ptsI | 2 | 2.5 | 0.24 |
| 0 | 0 | clcB | 2 | 2.5 | 0.231 |
| 0 | 0 | ycaM | 2 | 2.5 | 0.241 |
| 0 | 0 | yadI | 2 | 2.5 | 0.272 |
| 0 | 0 | AG1 | 2 | 2.5 | 0.287 |
| 0 | 0 | ptsI | 2 | 2.75 | 0.243 |
| 0 | 0 | clcB | 2 | 2.75 | 0.234 |
| 0 | 0 | ycaM | 2 | 2.75 | 0.249 |
| 0 | 0 | yadI | 2 | 2.75 | 0.29 |
| 0 | 0 | AG1 | 2 | 2.75 | 0.303 |
| 0 | 0 | ptsI | 2 | 3 | 0.248 |
| 0 | 0 | clcB | 2 | 3 | 0.238 |
| 0 | 0 | ycaM | 2 | 3 | 0.256 |
| 0 | 0 | yadI | 2 | 3 | 0.3 |
| 0 | 0 | AG1 | 2 | 3 | 0.308 |
| 0 | 0 | ptsI | 2 | 3.25 | 0.26 |
| 0 | 0 | clcB | 2 | 3.25 | 0.246 |
| 0 | 0 | ycaM | 2 | 3.25 | 0.267 |
| 0 | 0 | yadI | 2 | 3.25 | 0.318 |
| 0 | 0 | AG1 | 2 | 3.25 | 0.324 |
| 0 | 0 | ptsI | 2 | 3.5 | 0.258 |
| 0 | 0 | clcB | 2 | 3.5 | 0.254 |
| 0 | 0 | ycaM | 2 | 3.5 | 0.275 |
| 0 | 0 | yadI | 2 | 3.5 | 0.334 |
| 0 | 0 | AG1 | 2 | 3.5 | 0.33 |
| 0 | 0 | ptsI | 2 | 3.75 | 0.271 |
| 0 | 0 | clcB | 2 | 3.75 | 0.26 |
| 0 | 0 | ycaM | 2 | 3.75 | 0.29 |
| 0 | 0 | yadI | 2 | 3.75 | 0.353 |
| 0 | 0 | AG1 | 2 | 3.75 | 0.341 |
| 0 | 0 | ptsI | 2 | 4 | 0.286 |
| 0 | 0 | clcB | 2 | 4 | 0.267 |
| 0 | 0 | ycaM | 2 | 4 | 0.298 |
| 0 | 0 | yadI | 2 | 4 | 0.37 |
| 0 | 0 | AG1 | 2 | 4 | 0.353 |
| 0 | 0 | ptsI | 2 | 4.38333 | 0.291 |
| 0 | 0 | clcB | 2 | 4.38333 | 0.278 |
| 0 | 0 | ycaM | 2 | 4.38333 | 0.317 |
| 0 | 0 | yadI | 2 | 4.38333 | 0.405 |
| 0 | 0 | AG1 | 2 | 4.38333 | 0.378 |
| 0 | 0 | ptsI | 2 | 4.63333 | 0.286 |
| 0 | 0 | clcB | 2 | 4.63333 | 0.27 |
| 0 | 0 | ycaM | 2 | 4.63333 | 0.31 |
| 0 | 0 | yadI | 2 | 4.63333 | 0.411 |
| 0 | 0 | AG1 | 2 | 4.63333 | 0.368 |

| | | | | | |
|---|---|---|---|---|---|
| 0 | 0 | ptsI | 2 | 4.88333 | 0.295 |
| 0 | 0 | clcB | 2 | 4.88333 | 0.275 |
| 0 | 0 | ycaM | 2 | 4.88333 | 0.324 |
| 0 | 0 | yadI | 2 | 4.88333 | 0.43 |
| 0 | 0 | AG1 | 2 | 4.88333 | 0.382 |
| 0 | 0 | ptsI | 2 | 5.13333 | 0.36 |
| 0 | 0 | clcB | 2 | 5.13333 | 0.281 |
| 0 | 0 | ycaM | 2 | 5.13333 | 0.333 |
| 0 | 0 | yadI | 2 | 5.13333 | 0.445 |
| 0 | 0 | AG1 | 2 | 5.13333 | 0.39 |
| 0 | 0 | ptsI | 2 | 5.38333 | 0.308 |
| 0 | 0 | clcB | 2 | 5.38333 | 0.289 |
| 0 | 0 | ycaM | 2 | 5.38333 | 0.342 |
| 0 | 0 | yadI | 2 | 5.38333 | 0.458 |
| 0 | 0 | AG1 | 2 | 5.38333 | 0.401 |
| 0 | 0 | ptsI | 2 | 5.63333 | 0.318 |
| 0 | 0 | clcB | 2 | 5.63333 | 0.302 |
| 0 | 0 | ycaM | 2 | 5.63333 | 0.354 |
| 0 | 0 | yadI | 2 | 5.63333 | 0.473 |
| 0 | 0 | AG1 | 2 | 5.63333 | 0.412 |
| 0 | 0 | ptsI | 2 | 5.88333 | 0.324 |
| 0 | 0 | clcB | 2 | 5.88333 | 0.307 |
| 0 | 0 | ycaM | 2 | 5.88333 | 0.363 |
| 0 | 0 | yadI | 2 | 5.88333 | 0.484 |
| 0 | 0 | AG1 | 2 | 5.88333 | 0.422 |
| 0 | 0 | ptsI | 2 | 6.13333 | 0.333 |
| 0 | 0 | clcB | 2 | 6.13333 | 0.311 |
| 0 | 0 | ycaM | 2 | 6.13333 | 0.373 |
| 0 | 0 | yadI | 2 | 6.13333 | 0.496 |
| 0 | 0 | AG1 | 2 | 6.13333 | 0.435 |
| 0 | 0 | ptsI | 2 | 6.38333 | 0.343 |
| 0 | 0 | clcB | 2 | 6.38333 | 0.321 |
| 0 | 0 | ycaM | 2 | 6.38333 | 0.385 |
| 0 | 0 | yadI | 2 | 6.38333 | 0.51 |
| 0 | 0 | AG1 | 2 | 6.38333 | 0.446 |
| 0 | 0 | ptsI | 2 | 6.63333 | 0.347 |
| 0 | 0 | clcB | 2 | 6.63333 | 0.327 |
| 0 | 0 | ycaM | 2 | 6.63333 | 0.395 |
| 0 | 0 | yadI | 2 | 6.63333 | 0.527 |
| 0 | 0 | AG1 | 2 | 6.63333 | 0.458 |
| 0 | 0 | ptsI | 2 | 6.88333 | 0.359 |
| 0 | 0 | clcB | 2 | 6.88333 | 0.338 |
| 0 | 0 | ycaM | 2 | 6.88333 | 0.414 |
| 0 | 0 | yadI | 2 | 6.88333 | 0.55 |
| 0 | 0 | AG1 | 2 | 6.88333 | 0.474 |
| 0 | 0 | ptsI | 2 | 7.13333 | 0.363 |
| 0 | 0 | clcB | 2 | 7.13333 | 0.338 |
| 0 | 0 | ycaM | 2 | 7.13333 | 0.416 |

| | | | | | |
|---|---|---|---|---|---|
| 0 | 0 | yadI | 2 | 7.13333 | 0.544 |
| 0 | 0 | AG1 | 2 | 7.13333 | 0.476 |
| 0 | 0 | ptsI | 2 | 7.38333 | 0.373 |
| 0 | 0 | clcB | 2 | 7.38333 | 0.35 |
| 0 | 0 | ycaM | 2 | 7.38333 | 0.427 |
| 0 | 0 | yadI | 2 | 7.38333 | 0.558 |
| 0 | 0 | AG1 | 2 | 7.38333 | 0.489 |
| 0 | 0 | ptsI | 2 | 7.63333 | 0.382 |
| 0 | 0 | clcB | 2 | 7.63333 | 0.361 |
| 0 | 0 | ycaM | 2 | 7.63333 | 0.438 |
| 0 | 0 | yadI | 2 | 7.63333 | 0.576 |
| 0 | 0 | AG1 | 2 | 7.63333 | 0.505 |
| 0 | 0 | ptsI | 2 | 7.88333 | 0.404 |
| 0 | 0 | clcB | 2 | 7.88333 | 0.371 |
| 0 | 0 | ycaM | 2 | 7.88333 | 0.446 |
| 0 | 0 | yadI | 2 | 7.88333 | 0.59 |
| 0 | 0 | AG1 | 2 | 7.88333 | 0.515 |
| 0 | 0 | ptsI | 2 | 8.13333 | 0.401 |
| 0 | 0 | clcB | 2 | 8.13333 | 0.372 |
| 0 | 0 | ycaM | 2 | 8.13333 | 0.456 |
| 0 | 0 | yadI | 2 | 8.13333 | 0.598 |
| 0 | 0 | AG1 | 2 | 8.13333 | 0.526 |
| 0 | 0 | ptsI | 2 | 8.38333 | 0.408 |
| 0 | 0 | clcB | 2 | 8.38333 | 0.384 |
| 0 | 0 | ycaM | 2 | 8.38333 | 0.46 |
| 0 | 0 | yadI | 2 | 8.38333 | 0.608 |
| 0 | 0 | AG1 | 2 | 8.38333 | 0.54 |
| 0 | 0 | ptsI | 2 | 8.91667 | 0.423 |
| 0 | 0 | clcB | 2 | 8.91667 | 0.387 |
| 0 | 0 | ycaM | 2 | 8.91667 | 0.489 |
| 0 | 0 | yadI | 2 | 8.91667 | 0.632 |
| 0 | 0 | AG1 | 2 | 8.91667 | 0.361 |
| 0 | 0 | ptsI | 2 | 9.16667 | 0.429 |
| 0 | 0 | clcB | 2 | 9.16667 | 0.394 |
| 0 | 0 | ycaM | 2 | 9.16667 | 0.483 |
| 0 | 0 | yadI | 2 | 9.16667 | 0.613 |
| 0 | 0 | AG1 | 2 | 9.16667 | 0.356 |
| 0 | 0 | ptsI | 2 | 9.41667 | 0.443 |
| 0 | 0 | clcB | 2 | 9.41667 | 0.412 |
| 0 | 0 | ycaM | 2 | 9.41667 | 0.489 |
| 0 | 0 | yadI | 2 | 9.41667 | 0.624 |
| 0 | 0 | AG1 | 2 | 9.41667 | 0.365 |
| 0 | 0 | ptsI | 2 | 9.66667 | 0.454 |
| 0 | 0 | clcB | 2 | 9.66667 | 0.419 |
| 0 | 0 | ycaM | 2 | 9.66667 | 0.498 |
| 0 | 0 | yadI | 2 | 9.66667 | 0.63 |
| 0 | 0 | AG1 | 2 | 9.66667 | 0.371 |
| 0 | 0 | ptsI | 2 | 9.91667 | 0.467 |

| | | | | | |
|---|---|---|---|---|---|
| 0 | 0 | clcB | 2 | 9.91667 | 0.426 |
| 0 | 0 | ycaM | 2 | 9.91667 | 0.508 |
| 0 | 0 | yadI | 2 | 9.91667 | 0.642 |
| 0 | 0 | AG1 | 2 | 9.91667 | 0.379 |
| 0 | 0 | ptsI | 2 | 10.1667 | 0.479 |
| 0 | 0 | clcB | 2 | 10.1667 | 0.434 |
| 0 | 0 | ycaM | 2 | 10.1667 | 0.526 |
| 0 | 0 | yadI | 2 | 10.1667 | 0.658 |
| 0 | 0 | AG1 | 2 | 10.1667 | 0.386 |
| 0 | 0 | ptsI | 2 | 10.4167 | 0.494 |
| 0 | 0 | clcB | 2 | 10.4167 | 0.445 |
| 0 | 0 | ycaM | 2 | 10.4167 | 0.537 |
| 0 | 0 | yadI | 2 | 10.4167 | 0.67 |
| 0 | 0 | AG1 | 2 | 10.4167 | 0.393 |
| 0 | 0 | ptsI | 2 | 10.6667 | 0.494 |
| 0 | 0 | clcB | 2 | 10.6667 | 0.456 |
| 0 | 0 | ycaM | 2 | 10.6667 | 0.548 |
| 0 | 0 | yadI | 2 | 10.6667 | 0.688 |
| 0 | 0 | AG1 | 2 | 10.6667 | 0.399 |
| 0 | 0 | ptsI | 2 | 10.9167 | 0.505 |
| 0 | 0 | clcB | 2 | 10.9167 | 0.459 |
| 0 | 0 | ycaM | 2 | 10.9167 | 0.558 |
| 0 | 0 | yadI | 2 | 10.9167 | 0.695 |
| 0 | 0 | AG1 | 2 | 10.9167 | 0.407 |
| 0 | 0 | ptsI | 2 | 11.1667 | 0.541 |
| 0 | 0 | clcB | 2 | 11.1667 | 0.474 |
| 0 | 0 | ycaM | 2 | 11.1667 | 0.573 |
| 0 | 0 | yadI | 2 | 11.1667 | 0.709 |
| 0 | 0 | AG1 | 2 | 11.1667 | 0.42 |
| 0 | 0 | ptsI | 2 | 11.4167 | 0.517 |
| 0 | 0 | clcB | 2 | 11.4167 | 0.476 |
| 0 | 0 | ycaM | 2 | 11.4167 | 0.587 |
| 0 | 0 | yadI | 2 | 11.4167 | 0.717 |
| 0 | 0 | AG1 | 2 | 11.4167 | 0.417 |
| 0 | 0 | ptsI | 2 | 11.6667 | 0.565 |
| 0 | 0 | clcB | 2 | 11.6667 | 0.486 |
| 0 | 0 | ycaM | 2 | 11.6667 | 0.598 |
| 0 | 0 | yadI | 2 | 11.6667 | 0.731 |
| 0 | 0 | AG1 | 2 | 11.6667 | 0.432 |
| 0 | 0 | ptsI | 2 | 11.9167 | 0.525 |
| 0 | 0 | clcB | 2 | 11.9167 | 0.496 |
| 0 | 0 | ycaM | 2 | 11.9167 | 0.62 |
| 0 | 0 | yadI | 2 | 11.9167 | 0.741 |
| 0 | 0 | AG1 | 2 | 11.9167 | 0.444 |
| 0 | 0 | ptsI | 2 | 12.1667 | 0.554 |
| 0 | 0 | clcB | 2 | 12.1667 | 0.511 |
| 0 | 0 | ycaM | 2 | 12.1667 | 0.642 |
| 0 | 0 | yadI | 2 | 12.1667 | 0.756 |

| | | | | | |
|---|---|---|---|---|---|
| 0 | 0 | AG1 | 2 | 12.1667 | 0.452 |
| 0 | 0 | ptsI | 2 | 12.4167 | 0.576 |
| 0 | 0 | clcB | 2 | 12.4167 | 0.523 |
| 0 | 0 | ycaM | 2 | 12.4167 | 0.654 |
| 0 | 0 | yadI | 2 | 12.4167 | 0.762 |
| 0 | 0 | AG1 | 2 | 12.4167 | 0.471 |
| 0 | 0 | ptsI | 2 | 12.6667 | 0.554 |
| 0 | 0 | clcB | 2 | 12.6667 | 0.521 |
| 0 | 0 | ycaM | 2 | 12.6667 | 0.652 |
| 0 | 0 | yadI | 2 | 12.6667 | 0.772 |
| 0 | 0 | AG1 | 2 | 12.6667 | 0.464 |
| 0 | 0 | ptsI | 2 | 12.9167 | 0.573 |
| 0 | 0 | clcB | 2 | 12.9167 | 0.528 |
| 0 | 0 | ycaM | 2 | 12.9167 | 0.672 |
| 0 | 0 | yadI | 2 | 12.9167 | 0.791 |
| 0 | 0 | AG1 | 2 | 12.9167 | 0.476 |
| 0 | 0 | ptsI | 2 | 13.1667 | 0.601 |
| 0 | 0 | clcB | 2 | 13.1667 | 0.534 |
| 0 | 0 | ycaM | 2 | 13.1667 | 0.687 |
| 0 | 0 | yadI | 2 | 13.1667 | 0.802 |
| 0 | 0 | AG1 | 2 | 13.1667 | 0.502 |
| 0 | 0 | ptsI | 2 | 13.4167 | 0.583 |
| 0 | 0 | clcB | 2 | 13.4167 | 0.527 |
| 0 | 0 | ycaM | 2 | 13.4167 | 0.672 |
| 0 | 0 | yadI | 2 | 13.4167 | 0.803 |
| 0 | 0 | AG1 | 2 | 13.4167 | 0.504 |
| 0 | 0 | ptsI | 2 | 13.6667 | 0.618 |
| 0 | 0 | clcB | 2 | 13.6667 | 0.538 |
| 0 | 0 | ycaM | 2 | 13.6667 | 0.688 |
| 0 | 0 | yadI | 2 | 13.6667 | 0.818 |
| 0 | 0 | AG1 | 2 | 13.6667 | 0.56 |
| 0 | 0 | ptsI | 2 | 13.9167 | 0.633 |
| 0 | 0 | clcB | 2 | 13.9167 | 0.546 |
| 0 | 0 | ycaM | 2 | 13.9167 | 0.675 |
| 0 | 0 | yadI | 2 | 13.9167 | 0.823 |
| 0 | 0 | AG1 | 2 | 13.9167 | 0.553 |
| 0 | 0 | ptsI | 2 | 14.1667 | 0.642 |
| 0 | 0 | clcB | 2 | 14.1667 | 0.542 |
| 0 | 0 | ycaM | 2 | 14.1667 | 0.68 |
| 0 | 0 | yadI | 2 | 14.1667 | 0.836 |
| 0 | 0 | AG1 | 2 | 14.1667 | 0.563 |
| 0 | 0 | ptsI | 2 | 14.4167 | 0.672 |
| 0 | 0 | clcB | 2 | 14.4167 | 0.538 |
| 0 | 0 | ycaM | 2 | 14.4167 | 0.674 |
| 0 | 0 | yadI | 2 | 14.4167 | 0.836 |
| 0 | 0 | AG1 | 2 | 14.4167 | 0.564 |
| 0 | 0 | ptsI | 2 | 14.6667 | 0.686 |
| 0 | 0 | clcB | 2 | 14.6667 | 0.549 |

| | | | | | |
|---|---|---|---|---|---|
| 0 | 0 | ycaM | 2 | 14.6667 | 0.68 |
| 0 | 0 | yadI | 2 | 14.6667 | 0.841 |
| 0 | 0 | AG1 | 2 | 14.6667 | 0.582 |
| 0 | 0 | ptsI | 2 | 14.9167 | 0.701 |
| 0 | 0 | clcB | 2 | 14.9167 | 0.56 |
| 0 | 0 | ycaM | 2 | 14.9167 | 0.662 |
| 0 | 0 | yadI | 2 | 14.9167 | 0.844 |
| 0 | 0 | AG1 | 2 | 14.9167 | 0.587 |
| 0 | 0 | ptsI | 2 | 15.1667 | 0.726 |
| 0 | 0 | clcB | 2 | 15.1667 | 0.543 |
| 0 | 0 | ycaM | 2 | 15.1667 | 0.661 |
| 0 | 0 | yadI | 2 | 15.1667 | 0.862 |
| 0 | 0 | AG1 | 2 | 15.1667 | 0.628 |
| 0 | 0 | ptsI | 2 | 15.4167 | 0.759 |
| 0 | 0 | clcB | 2 | 15.4167 | 0.561 |
| 0 | 0 | ycaM | 2 | 15.4167 | 0.671 |
| 0 | 0 | yadI | 2 | 15.4167 | 0.869 |
| 0 | 0 | AG1 | 2 | 15.4167 | 0.656 |
| 0 | 0 | ptsI | 2 | 15.6667 | 0.742 |
| 0 | 0 | clcB | 2 | 15.6667 | 0.564 |
| 0 | 0 | ycaM | 2 | 15.6667 | 0.662 |
| 0 | 0 | yadI | 2 | 15.6667 | 0.868 |
| 0 | 0 | AG1 | 2 | 15.6667 | 0.672 |
| 0 | 0 | ptsI | 2 | 15.9167 | 0.728 |
| 0 | 0 | clcB | 2 | 15.9167 | 0.571 |
| 0 | 0 | ycaM | 2 | 15.9167 | 0.671 |
| 0 | 0 | yadI | 2 | 15.9167 | 0.878 |
| 0 | 0 | AG1 | 2 | 15.9167 | 0.67 |
| 0 | 0 | ptsI | 2 | 16.1667 | 0.769 |
| 0 | 0 | clcB | 2 | 16.1667 | 0.561 |
| 0 | 0 | ycaM | 2 | 16.1667 | 0.689 |
| 0 | 0 | yadI | 2 | 16.1667 | 0.884 |
| 0 | 0 | AG1 | 2 | 16.1667 | 0.704 |
| 0 | 0 | ptsI | 2 | 16.4167 | 0.754 |
| 0 | 0 | clcB | 2 | 16.4167 | 0.559 |
| 0 | 0 | ycaM | 2 | 16.4167 | 0.697 |
| 0 | 0 | yadI | 2 | 16.4167 | 0.884 |
| 0 | 0 | AG1 | 2 | 16.4167 | 0.706 |
| 0 | 0 | ptsI | 2 | 16.6667 | 0.763 |
| 0 | 0 | clcB | 2 | 16.6667 | 0.587 |
| 0 | 0 | ycaM | 2 | 16.6667 | 0.711 |
| 0 | 0 | yadI | 2 | 16.6667 | 0.889 |
| 0 | 0 | AG1 | 2 | 16.6667 | 0.71 |
| 0 | 0 | ptsI | 2 | 16.9167 | 0.77 |
| 0 | 0 | clcB | 2 | 16.9167 | 0.564 |
| 0 | 0 | ycaM | 2 | 16.9167 | 0.702 |
| 0 | 0 | yadI | 2 | 16.9167 | 0.891 |
| 0 | 0 | AG1 | 2 | 16.9167 | 0.729 |

| | | | | | |
|---|---|---|---|---|---|
| 0 | 0 | ptsI | 2 | 17.1667 | 0.799 |
| 0 | 0 | clcB | 2 | 17.1667 | 0.56 |
| 0 | 0 | ycaM | 2 | 17.1667 | 0.72 |
| 0 | 0 | yadI | 2 | 17.1667 | 0.909 |
| 0 | 0 | AG1 | 2 | 17.1667 | 0.746 |
| 0 | 0 | ptsI | 2 | 17.4167 | 0.786 |
| 0 | 0 | clcB | 2 | 17.4167 | 0.608 |
| 0 | 0 | ycaM | 2 | 17.4167 | 0.727 |
| 0 | 0 | yadI | 2 | 17.4167 | 0.916 |
| 0 | 0 | AG1 | 2 | 17.4167 | 0.761 |
| 0 | 0 | ptsI | 2 | 17.6667 | 0.81 |
| 0 | 0 | clcB | 2 | 17.6667 | 0.588 |
| 0 | 0 | ycaM | 2 | 17.6667 | 0.715 |
| 0 | 0 | yadI | 2 | 17.6667 | 0.916 |
| 0 | 0 | AG1 | 2 | 17.6667 | 0.758 |
| 0 | 0 | ptsI | 2 | 17.9167 | 0.809 |
| 0 | 0 | clcB | 2 | 17.9167 | 0.575 |
| 0 | 0 | ycaM | 2 | 17.9167 | 0.717 |
| 0 | 0 | yadI | 2 | 17.9167 | 0.914 |
| 0 | 0 | AG1 | 2 | 17.9167 | 0.771 |
| 0 | 0 | ptsI | 2 | 18.1667 | 0.835 |
| 0 | 0 | clcB | 2 | 18.1667 | 0.595 |
| 0 | 0 | ycaM | 2 | 18.1667 | 0.724 |
| 0 | 0 | yadI | 2 | 18.1667 | 0.926 |
| 0 | 0 | AG1 | 2 | 18.1667 | 0.768 |
| 0 | 0 | ptsI | 2 | 18.4167 | 0.818 |
| 0 | 0 | clcB | 2 | 18.4167 | 0.586 |
| 0 | 0 | ycaM | 2 | 18.4167 | 0.738 |
| 0 | 0 | yadI | 2 | 18.4167 | 0.945 |
| 0 | 0 | AG1 | 2 | 18.4167 | 0.776 |
| 0 | 0 | ptsI | 2 | 18.6667 | 0.822 |
| 0 | 0 | clcB | 2 | 18.6667 | 0.582 |
| 0 | 0 | ycaM | 2 | 18.6667 | 0.742 |
| 0 | 0 | yadI | 2 | 18.6667 | 0.938 |
| 0 | 0 | AG1 | 2 | 18.6667 | 0.775 |
| 0 | 0 | ptsI | 2 | 18.9167 | 0.837 |
| 0 | 0 | clcB | 2 | 18.9167 | 0.612 |
| 0 | 0 | ycaM | 2 | 18.9167 | 0.748 |
| 0 | 0 | yadI | 2 | 18.9167 | 0.93 |
| 0 | 0 | AG1 | 2 | 18.9167 | 0.781 |
| 0 | 0 | ptsI | 2 | 19.1667 | 0.821 |
| 0 | 0 | clcB | 2 | 19.1667 | 0.6 |
| 0 | 0 | ycaM | 2 | 19.1667 | 0.745 |
| 0 | 0 | yadI | 2 | 19.1667 | 0.939 |
| 0 | 0 | AG1 | 2 | 19.1667 | 0.778 |
| 0 | 0 | ptsI | 2 | 19.4167 | 0.834 |
| 0 | 0 | clcB | 2 | 19.4167 | 0.601 |
| 0 | 0 | ycaM | 2 | 19.4167 | 0.752 |

| | | | | | |
|---|---|---|---|---|---|
| 0 | 0 | yadI | 2 | 19.4167 | 0.948 |
| 0 | 0 | AG1 | 2 | 19.4167 | 0.788 |
| 0 | 0 | ptsI | 2 | 19.6667 | 0.882 |
| 0 | 0 | clcB | 2 | 19.6667 | 0.598 |
| 0 | 0 | ycaM | 2 | 19.6667 | 0.753 |
| 0 | 0 | yadI | 2 | 19.6667 | 0.943 |
| 0 | 0 | AG1 | 2 | 19.6667 | 0.792 |
| 0 | 0 | ptsI | 2 | 19.9167 | 0.862 |
| 0 | 0 | clcB | 2 | 19.9167 | 0.621 |
| 0 | 0 | ycaM | 2 | 19.9167 | 0.77 |
| 0 | 0 | yadI | 2 | 19.9167 | 0.958 |
| 0 | 0 | AG1 | 2 | 19.9167 | 0.815 |
| 0 | 0 | ptsI | 2 | 20.1667 | 0.859 |
| 0 | 0 | clcB | 2 | 20.1667 | 0.598 |
| 0 | 0 | ycaM | 2 | 20.1667 | 0.763 |
| 0 | 0 | yadI | 2 | 20.1667 | 0.954 |
| 0 | 0 | AG1 | 2 | 20.1667 | 0.806 |
| 0 | 0 | ptsI | 2 | 20.4167 | 0.844 |
| 0 | 0 | clcB | 2 | 20.4167 | 0.602 |
| 0 | 0 | ycaM | 2 | 20.4167 | 0.757 |
| 0 | 0 | yadI | 2 | 20.4167 | 0.953 |
| 0 | 0 | AG1 | 2 | 20.4167 | 0.807 |
| 0 | 0 | ptsI | 2 | 20.6667 | 0.854 |
| 0 | 0 | clcB | 2 | 20.6667 | 0.61 |
| 0 | 0 | ycaM | 2 | 20.6667 | 0.768 |
| 0 | 0 | yadI | 2 | 20.6667 | 0.962 |
| 0 | 0 | AG1 | 2 | 20.6667 | 0.811 |
| 0 | 0 | ptsI | 2 | 20.9167 | 0.907 |
| 0 | 0 | clcB | 2 | 20.9167 | 0.604 |
| 0 | 0 | ycaM | 2 | 20.9167 | 0.766 |
| 0 | 0 | yadI | 2 | 20.9167 | 0.96 |
| 0 | 0 | AG1 | 2 | 20.9167 | 0.815 |
| 0 | 0 | ptsI | 2 | 21.1667 | 0.849 |
| 0 | 0 | clcB | 2 | 21.1667 | 0.598 |
| 0 | 0 | ycaM | 2 | 21.1667 | 0.772 |
| 0 | 0 | yadI | 2 | 21.1667 | 0.959 |
| 0 | 0 | AG1 | 2 | 21.1667 | 0.819 |
| 0 | 0 | ptsI | 2 | 21.4167 | 0.856 |
| 0 | 0 | clcB | 2 | 21.4167 | 0.603 |
| 0 | 0 | ycaM | 2 | 21.4167 | 0.78 |
| 0 | 0 | yadI | 2 | 21.4167 | 0.965 |
| 0 | 0 | AG1 | 2 | 21.4167 | 0.834 |
| 0 | 0 | ptsI | 2 | 21.6667 | 0.876 |
| 0 | 0 | clcB | 2 | 21.6667 | 0.614 |
| 0 | 0 | ycaM | 2 | 21.6667 | 0.791 |
| 0 | 0 | yadI | 2 | 21.6667 | 0.979 |
| 0 | 0 | AG1 | 2 | 21.6667 | 0.838 |
| 0 | 0 | ptsI | 2 | 21.9167 | 0.878 |

| | | | | | |
|---|---|---|---|---|---|
| 0 | 0 | clcB | 2 | 21.9167 | 0.622 |
| 0 | 0 | ycaM | 2 | 21.9167 | 0.789 |
| 0 | 0 | yadI | 2 | 21.9167 | 0.98 |
| 0 | 0 | AG1 | 2 | 21.9167 | 0.844 |
| 0 | 0 | ptsI | 2 | 22.1667 | 0.884 |
| 0 | 0 | clcB | 2 | 22.1667 | 0.622 |
| 0 | 0 | ycaM | 2 | 22.1667 | 0.793 |
| 0 | 0 | yadI | 2 | 22.1667 | 0.984 |
| 0 | 0 | AG1 | 2 | 22.1667 | 0.845 |
| 0 | 0 | ptsI | 2 | 22.4167 | 0.884 |
| 0 | 0 | clcB | 2 | 22.4167 | 0.613 |
| 0 | 0 | ycaM | 2 | 22.4167 | 0.793 |
| 0 | 0 | yadI | 2 | 22.4167 | 0.981 |
| 0 | 0 | AG1 | 2 | 22.4167 | 0.852 |
| 0 | 0 | ptsI | 2 | 22.6667 | 0.885 |
| 0 | 0 | clcB | 2 | 22.6667 | 0.626 |
| 0 | 0 | ycaM | 2 | 22.6667 | 0.795 |
| 0 | 0 | yadI | 2 | 22.6667 | 0.979 |
| 0 | 0 | AG1 | 2 | 22.6667 | 0.851 |
| 0 | 0 | ptsI | 2 | 22.9167 | 0.9 |
| 0 | 0 | clcB | 2 | 22.9167 | 0.624 |
| 0 | 0 | ycaM | 2 | 22.9167 | 0.807 |
| 0 | 0 | yadI | 2 | 22.9167 | 0.984 |
| 0 | 0 | AG1 | 2 | 22.9167 | 0.865 |
| 0 | 0 | ptsI | 2 | 23.1667 | 0.895 |
| 0 | 0 | clcB | 2 | 23.1667 | 0.625 |
| 0 | 0 | ycaM | 2 | 23.1667 | 0.797 |
| 0 | 0 | yadI | 2 | 23.1667 | 0.971 |
| 0 | 0 | AG1 | 2 | 23.1667 | 0.844 |
| 0 | 0 | ptsI | 2 | 23.4167 | 0.906 |
| 0 | 0 | clcB | 2 | 23.4167 | 0.632 |
| 0 | 0 | ycaM | 2 | 23.4167 | 0.804 |
| 0 | 0 | yadI | 2 | 23.4167 | 0.988 |
| 0 | 0 | AG1 | 2 | 23.4167 | 0.855 |
| 0 | 0 | ptsI | 2 | 23.6667 | 0.912 |
| 0 | 0 | clcB | 2 | 23.6667 | 0.632 |
| 0 | 0 | ycaM | 2 | 23.6667 | 0.806 |
| 0 | 0 | yadI | 2 | 23.6667 | 0.989 |
| 0 | 0 | AG1 | 2 | 23.6667 | 0.859 |
| 0 | 0 | ptsI | 2 | 23.9167 | 0.955 |
| 0 | 0 | clcB | 2 | 23.9167 | 0.638 |
| 0 | 0 | ycaM | 2 | 23.9167 | 0.812 |
| 0 | 0 | yadI | 2 | 23.9167 | 0.991 |
| 0 | 0 | AG1 | 2 | 23.9167 | 0.863 |
| 0 | 0 | ptsI | 2 | 24.1667 | 0.95 |
| 0 | 0 | clcB | 2 | 24.1667 | 0.635 |
| 0 | 0 | ycaM | 2 | 24.1667 | 0.812 |
| 0 | 0 | yadI | 2 | 24.1667 | 0.991 |

| | | | | | |
|---|---|---|---|---|---|
| 0 | 0 | AG1 | 2 | 24.1667 | 0.864 |
| 0 | 0 | ptsI | 2 | 24.4167 | 0.929 |
| 0 | 0 | clcB | 2 | 24.4167 | 0.639 |
| 0 | 0 | ycaM | 2 | 24.4167 | 0.82 |
| 0 | 0 | yadI | 2 | 24.4167 | 0.996 |
| 0 | 0 | AG1 | 2 | 24.4167 | 0.869 |
| 0 | 0 | ptsI | 2 | 24.6667 | 0.915 |
| 0 | 0 | clcB | 2 | 24.6667 | 0.644 |
| 0 | 0 | ycaM | 2 | 24.6667 | 0.822 |
| 0 | 0 | yadI | 2 | 24.6667 | 0.991 |
| 0 | 0 | AG1 | 2 | 24.6667 | 0.875 |
| 0 | 0 | ptsI | 2 | 24.9167 | 0.964 |
| 0 | 0 | clcB | 2 | 24.9167 | 0.646 |
| 0 | 0 | ycaM | 2 | 24.9167 | 0.82 |
| 0 | 0 | yadI | 2 | 24.9167 | 0.992 |
| 0 | 0 | AG1 | 2 | 24.9167 | 0.88 |
| 0 | 0 | ptsI | 2 | 25.1667 | 0.936 |
| 0 | 0 | clcB | 2 | 25.1667 | 0.646 |
| 0 | 0 | ycaM | 2 | 25.1667 | 0.825 |
| 0 | 0 | yadI | 2 | 25.1667 | 0.999 |
| 0 | 0 | AG1 | 2 | 25.1667 | 0.884 |
| 0 | 0 | ptsI | 2 | 25.4167 | 0.943 |
| 0 | 0 | clcB | 2 | 25.4167 | 0.654 |
| 0 | 0 | ycaM | 2 | 25.4167 | 0.832 |
| 0 | 0 | yadI | 2 | 25.4167 | 0.989 |
| 0 | 0 | AG1 | 2 | 25.4167 | 0.898 |
| 0 | 0 | ptsI | 2 | 25.6667 | 0.953 |
| 0 | 0 | clcB | 2 | 25.6667 | 0.651 |
| 0 | 0 | ycaM | 2 | 25.6667 | 0.831 |
| 0 | 0 | yadI | 2 | 25.6667 | 0.989 |
| 0 | 0 | AG1 | 2 | 25.6667 | 0.897 |
| 0 | 0 | ptsI | 2 | 25.9167 | 0.957 |
| 0 | 0 | clcB | 2 | 25.9167 | 0.648 |
| 0 | 0 | ycaM | 2 | 25.9167 | 0.833 |
| 0 | 0 | yadI | 2 | 25.9167 | 1.008 |
| 0 | 0 | AG1 | 2 | 25.9167 | 0.904 |
| 0 | 0 | ptsI | 2 | 26.1667 | 0.957 |
| 0 | 0 | clcB | 2 | 26.1667 | 0.65 |
| 0 | 0 | ycaM | 2 | 26.1667 | 0.835 |
| 0 | 0 | yadI | 2 | 26.1667 | 0.989 |
| 0 | 0 | AG1 | 2 | 26.1667 | 0.91 |
| 0 | 0 | ptsI | 2 | 26.4167 | 0.964 |
| 0 | 0 | clcB | 2 | 26.4167 | 0.661 |
| 0 | 0 | ycaM | 2 | 26.4167 | 0.844 |
| 0 | 0 | yadI | 2 | 26.4167 | 0.998 |
| 0 | 0 | AG1 | 2 | 26.4167 | 0.925 |
| 0 | 0 | ptsI | 3 | 0 | 0.155 |
| 0 | 0 | clcB | 3 | 0 | 0.162 |

| | | | | | |
|---|---|---|---|---|---|
| 0 | 0 | ycaM | 3 | 0 | 0.151 |
| 0 | 0 | yadI | 3 | 0 | 0.165 |
| 0 | 0 | AG1 | 3 | 0 | 0.165 |
| 0 | 0 | ptsI | 3 | 0.25 | 0.151 |
| 0 | 0 | clcB | 3 | 0.25 | 0.162 |
| 0 | 0 | ycaM | 3 | 0.25 | 0.148 |
| 0 | 0 | yadI | 3 | 0.25 | 0.163 |
| 0 | 0 | AG1 | 3 | 0.25 | 0.165 |
| 0 | 0 | ptsI | 3 | 0.5 | 0.15 |
| 0 | 0 | clcB | 3 | 0.5 | 0.162 |
| 0 | 0 | ycaM | 3 | 0.5 | 0.149 |
| 0 | 0 | yadI | 3 | 0.5 | 0.162 |
| 0 | 0 | AG1 | 3 | 0.5 | 0.166 |
| 0 | 0 | ptsI | 3 | 0.75 | 0.151 |
| 0 | 0 | clcB | 3 | 0.75 | 0.16 |
| 0 | 0 | ycaM | 3 | 0.75 | 0.149 |
| 0 | 0 | yadI | 3 | 0.75 | 0.179 |
| 0 | 0 | AG1 | 3 | 0.75 | 0.166 |
| 0 | 0 | ptsI | 3 | 1 | 0.152 |
| 0 | 0 | clcB | 3 | 1 | 0.162 |
| 0 | 0 | ycaM | 3 | 1 | 0.149 |
| 0 | 0 | yadI | 3 | 1 | 0.163 |
| 0 | 0 | AG1 | 3 | 1 | 0.168 |
| 0 | 0 | ptsI | 3 | 1.25 | 0.152 |
| 0 | 0 | clcB | 3 | 1.25 | 0.162 |
| 0 | 0 | ycaM | 3 | 1.25 | 0.149 |
| 0 | 0 | yadI | 3 | 1.25 | 0.164 |
| 0 | 0 | AG1 | 3 | 1.25 | 0.169 |
| 0 | 0 | ptsI | 3 | 1.5 | 0.154 |
| 0 | 0 | clcB | 3 | 1.5 | 0.163 |
| 0 | 0 | ycaM | 3 | 1.5 | 0.151 |
| 0 | 0 | yadI | 3 | 1.5 | 0.168 |
| 0 | 0 | AG1 | 3 | 1.5 | 0.172 |
| 0 | 0 | ptsI | 3 | 1.75 | 0.155 |
| 0 | 0 | clcB | 3 | 1.75 | 0.164 |
| 0 | 0 | ycaM | 3 | 1.75 | 0.152 |
| 0 | 0 | yadI | 3 | 1.75 | 0.17 |
| 0 | 0 | AG1 | 3 | 1.75 | 0.175 |
| 0 | 0 | ptsI | 3 | 2 | 0.156 |
| 0 | 0 | clcB | 3 | 2 | 0.166 |
| 0 | 0 | ycaM | 3 | 2 | 0.154 |
| 0 | 0 | yadI | 3 | 2 | 0.174 |
| 0 | 0 | AG1 | 3 | 2 | 0.178 |
| 0 | 0 | ptsI | 3 | 2.25 | 0.159 |
| 0 | 0 | clcB | 3 | 2.25 | 0.168 |
| 0 | 0 | ycaM | 3 | 2.25 | 0.156 |
| 0 | 0 | yadI | 3 | 2.25 | 0.178 |
| 0 | 0 | AG1 | 3 | 2.25 | 0.18 |

| | | | | | |
|---|---|---|---|---|---|
| 0 | 0 | ptsI | 3 | 2.5 | 0.16 |
| 0 | 0 | clcB | 3 | 2.5 | 0.168 |
| 0 | 0 | ycaM | 3 | 2.5 | 0.159 |
| 0 | 0 | yadI | 3 | 2.5 | 0.182 |
| 0 | 0 | AG1 | 3 | 2.5 | 0.183 |
| 0 | 0 | ptsI | 3 | 2.75 | 0.164 |
| 0 | 0 | clcB | 3 | 2.75 | 0.17 |
| 0 | 0 | ycaM | 3 | 2.75 | 0.162 |
| 0 | 0 | yadI | 3 | 2.75 | 0.187 |
| 0 | 0 | AG1 | 3 | 2.75 | 0.188 |
| 0 | 0 | ptsI | 3 | 3 | 0.166 |
| 0 | 0 | clcB | 3 | 3 | 0.172 |
| 0 | 0 | ycaM | 3 | 3 | 0.164 |
| 0 | 0 | yadI | 3 | 3 | 0.192 |
| 0 | 0 | AG1 | 3 | 3 | 0.19 |
| 0 | 0 | ptsI | 3 | 3.25 | 0.169 |
| 0 | 0 | clcB | 3 | 3.25 | 0.173 |
| 0 | 0 | ycaM | 3 | 3.25 | 0.165 |
| 0 | 0 | yadI | 3 | 3.25 | 0.197 |
| 0 | 0 | AG1 | 3 | 3.25 | 0.194 |
| 0 | 0 | ptsI | 3 | 3.5 | 0.173 |
| 0 | 0 | clcB | 3 | 3.5 | 0.178 |
| 0 | 0 | ycaM | 3 | 3.5 | 0.169 |
| 0 | 0 | yadI | 3 | 3.5 | 0.205 |
| 0 | 0 | AG1 | 3 | 3.5 | 0.199 |
| 0 | 0 | ptsI | 3 | 3.75 | 0.177 |
| 0 | 0 | clcB | 3 | 3.75 | 0.18 |
| 0 | 0 | ycaM | 3 | 3.75 | 0.173 |
| 0 | 0 | yadI | 3 | 3.75 | 0.212 |
| 0 | 0 | AG1 | 3 | 3.75 | 0.204 |
| 0 | 0 | ptsI | 3 | 4 | 0.181 |
| 0 | 0 | clcB | 3 | 4 | 0.183 |
| 0 | 0 | ycaM | 3 | 4 | 0.176 |
| 0 | 0 | yadI | 3 | 4 | 0.221 |
| 0 | 0 | AG1 | 3 | 4 | 0.213 |
| 0 | 0 | ptsI | 3 | 4.41667 | 0.183 |
| 0 | 0 | clcB | 3 | 4.41667 | 0.196 |
| 0 | 0 | ycaM | 3 | 4.41667 | 0.191 |
| 0 | 0 | yadI | 3 | 4.41667 | 0.232 |
| 0 | 0 | AG1 | 3 | 4.41667 | 0.218 |
| 0 | 0 | ptsI | 3 | 4.66667 | 0.182 |
| 0 | 0 | clcB | 3 | 4.66667 | 0.187 |
| 0 | 0 | ycaM | 3 | 4.66667 | 0.189 |
| 0 | 0 | yadI | 3 | 4.66667 | 0.225 |
| 0 | 0 | AG1 | 3 | 4.66667 | 0.209 |
| 0 | 0 | ptsI | 3 | 4.91667 | 0.187 |
| 0 | 0 | clcB | 3 | 4.91667 | 0.192 |
| 0 | 0 | ycaM | 3 | 4.91667 | 0.19 |

| | | | | | |
|---|---|---|---|---|---|
| 0 | 0 | yadI | 3 | 4.91667 | 0.234 |
| 0 | 0 | AG1 | 3 | 4.91667 | 0.213 |
| 0 | 0 | ptsI | 3 | 5.16667 | 0.191 |
| 0 | 0 | clcB | 3 | 5.16667 | 0.196 |
| 0 | 0 | ycaM | 3 | 5.16667 | 0.195 |
| 0 | 0 | yadI | 3 | 5.16667 | 0.246 |
| 0 | 0 | AG1 | 3 | 5.16667 | 0.218 |
| 0 | 0 | ptsI | 3 | 5.41667 | 0.196 |
| 0 | 0 | clcB | 3 | 5.41667 | 0.202 |
| 0 | 0 | ycaM | 3 | 5.41667 | 0.201 |
| 0 | 0 | yadI | 3 | 5.41667 | 0.262 |
| 0 | 0 | AG1 | 3 | 5.41667 | 0.227 |
| 0 | 0 | ptsI | 3 | 5.66667 | 0.202 |
| 0 | 0 | clcB | 3 | 5.66667 | 0.206 |
| 0 | 0 | ycaM | 3 | 5.66667 | 0.209 |
| 0 | 0 | yadI | 3 | 5.66667 | 0.283 |
| 0 | 0 | AG1 | 3 | 5.66667 | 0.236 |
| 0 | 0 | ptsI | 3 | 5.91667 | 0.209 |
| 0 | 0 | clcB | 3 | 5.91667 | 0.212 |
| 0 | 0 | ycaM | 3 | 5.91667 | 0.218 |
| 0 | 0 | yadI | 3 | 5.91667 | 0.309 |
| 0 | 0 | AG1 | 3 | 5.91667 | 0.252 |
| 0 | 0 | ptsI | 3 | 6.16667 | 0.217 |
| 0 | 0 | clcB | 3 | 6.16667 | 0.22 |
| 0 | 0 | ycaM | 3 | 6.16667 | 0.229 |
| 0 | 0 | yadI | 3 | 6.16667 | 0.33 |
| 0 | 0 | AG1 | 3 | 6.16667 | 0.264 |
| 0 | 0 | ptsI | 3 | 6.41667 | 0.225 |
| 0 | 0 | clcB | 3 | 6.41667 | 0.227 |
| 0 | 0 | ycaM | 3 | 6.41667 | 0.24 |
| 0 | 0 | yadI | 3 | 6.41667 | 0.35 |
| 0 | 0 | AG1 | 3 | 6.41667 | 0.276 |
| 0 | 0 | ptsI | 3 | 6.66667 | 0.238 |
| 0 | 0 | clcB | 3 | 6.66667 | 0.236 |
| 0 | 0 | ycaM | 3 | 6.66667 | 0.256 |
| 0 | 0 | yadI | 3 | 6.66667 | 0.369 |
| 0 | 0 | AG1 | 3 | 6.66667 | 0.29 |
| 0 | 0 | ptsI | 3 | 6.91667 | 0.25 |
| 0 | 0 | clcB | 3 | 6.91667 | 0.245 |
| 0 | 0 | ycaM | 3 | 6.91667 | 0.268 |
| 0 | 0 | yadI | 3 | 6.91667 | 0.383 |
| 0 | 0 | AG1 | 3 | 6.91667 | 0.296 |
| 0 | 0 | ptsI | 3 | 7.16667 | 0.259 |
| 0 | 0 | clcB | 3 | 7.16667 | 0.256 |
| 0 | 0 | ycaM | 3 | 7.16667 | 0.274 |
| 0 | 0 | yadI | 3 | 7.16667 | 0.405 |
| 0 | 0 | AG1 | 3 | 7.16667 | 0.303 |
| 0 | 0 | ptsI | 3 | 7.41667 | 0.268 |

| | | | | | |
|---|---|---|---|---|---|
| 0 | 0 | clcB | 3 | 7.41667 | 0.268 |
| 0 | 0 | ycaM | 3 | 7.41667 | 0.285 |
| 0 | 0 | yadI | 3 | 7.41667 | 0.42 |
| 0 | 0 | AG1 | 3 | 7.41667 | 0.312 |
| 0 | 0 | ptsI | 3 | 7.66667 | 0.28 |
| 0 | 0 | clcB | 3 | 7.66667 | 0.283 |
| 0 | 0 | ycaM | 3 | 7.66667 | 0.295 |
| 0 | 0 | yadI | 3 | 7.66667 | 0.432 |
| 0 | 0 | AG1 | 3 | 7.66667 | 0.324 |
| 0 | 0 | ptsI | 3 | 7.91667 | 0.286 |
| 0 | 0 | clcB | 3 | 7.91667 | 0.294 |
| 0 | 0 | ycaM | 3 | 7.91667 | 0.302 |
| 0 | 0 | yadI | 3 | 7.91667 | 0.45 |
| 0 | 0 | AG1 | 3 | 7.91667 | 0.33 |
| 0 | 0 | ptsI | 3 | 8.16667 | 0.296 |
| 0 | 0 | clcB | 3 | 8.16667 | 0.301 |
| 0 | 0 | ycaM | 3 | 8.16667 | 0.312 |
| 0 | 0 | yadI | 3 | 8.16667 | 0.466 |
| 0 | 0 | AG1 | 3 | 8.16667 | 0.337 |
| 0 | 0 | ptsI | 3 | 8.41667 | 0.31 |
| 0 | 0 | clcB | 3 | 8.41667 | 0.311 |
| 0 | 0 | ycaM | 3 | 8.41667 | 0.324 |
| 0 | 0 | yadI | 3 | 8.41667 | 0.488 |
| 0 | 0 | AG1 | 3 | 8.41667 | 0.349 |
| 0 | 0 | ptsI | 3 | 9.05 | 0.267 |
| 0 | 0 | clcB | 3 | 9.05 | 0.255 |
| 0 | 0 | ycaM | 3 | 9.05 | 0.294 |
| 0 | 0 | yadI | 3 | 9.05 | 0.401 |
| 0 | 0 | AG1 | 3 | 9.05 | 0.339 |
| 0 | 0 | ptsI | 3 | 9.3 | 0.267 |
| 0 | 0 | clcB | 3 | 9.3 | 0.251 |
| 0 | 0 | ycaM | 3 | 9.3 | 0.304 |
| 0 | 0 | yadI | 3 | 9.3 | 0.408 |
| 0 | 0 | AG1 | 3 | 9.3 | 0.34 |
| 0 | 0 | ptsI | 3 | 9.55 | 0.278 |
| 0 | 0 | clcB | 3 | 9.55 | 0.259 |
| 0 | 0 | ycaM | 3 | 9.55 | 0.317 |
| 0 | 0 | yadI | 3 | 9.55 | 0.415 |
| 0 | 0 | AG1 | 3 | 9.55 | 0.35 |
| 0 | 0 | ptsI | 3 | 9.8 | 0.284 |
| 0 | 0 | clcB | 3 | 9.8 | 0.267 |
| 0 | 0 | ycaM | 3 | 9.8 | 0.328 |
| 0 | 0 | yadI | 3 | 9.8 | 0.426 |
| 0 | 0 | AG1 | 3 | 9.8 | 0.357 |
| 0 | 0 | ptsI | 3 | 10.05 | 0.288 |
| 0 | 0 | clcB | 3 | 10.05 | 0.274 |
| 0 | 0 | ycaM | 3 | 10.05 | 0.335 |
| 0 | 0 | yadI | 3 | 10.05 | 0.439 |

| | | | | | |
|---|---|---|---|---|---|
| 0 | 0 | AG1 | 3 | 10.05 | 0.366 |
| 0 | 0 | ptsI | 3 | 10.3 | 0.299 |
| 0 | 0 | clcB | 3 | 10.3 | 0.278 |
| 0 | 0 | ycaM | 3 | 10.3 | 0.343 |
| 0 | 0 | yadI | 3 | 10.3 | 0.451 |
| 0 | 0 | AG1 | 3 | 10.3 | 0.375 |
| 0 | 0 | ptsI | 3 | 10.55 | 0.308 |
| 0 | 0 | clcB | 3 | 10.55 | 0.287 |
| 0 | 0 | ycaM | 3 | 10.55 | 0.356 |
| 0 | 0 | yadI | 3 | 10.55 | 0.47 |
| 0 | 0 | AG1 | 3 | 10.55 | 0.388 |
| 0 | 0 | ptsI | 3 | 10.8 | 0.316 |
| 0 | 0 | clcB | 3 | 10.8 | 0.292 |
| 0 | 0 | ycaM | 3 | 10.8 | 0.359 |
| 0 | 0 | yadI | 3 | 10.8 | 0.475 |
| 0 | 0 | AG1 | 3 | 10.8 | 0.394 |
| 0 | 0 | ptsI | 3 | 11.05 | 0.322 |
| 0 | 0 | clcB | 3 | 11.05 | 0.301 |
| 0 | 0 | ycaM | 3 | 11.05 | 0.366 |
| 0 | 0 | yadI | 3 | 11.05 | 0.487 |
| 0 | 0 | AG1 | 3 | 11.05 | 0.405 |
| 0 | 0 | ptsI | 3 | 11.3 | 0.331 |
| 0 | 0 | clcB | 3 | 11.3 | 0.31 |
| 0 | 0 | ycaM | 3 | 11.3 | 0.37 |
| 0 | 0 | yadI | 3 | 11.3 | 0.514 |
| 0 | 0 | AG1 | 3 | 11.3 | 0.411 |
| 0 | 0 | ptsI | 3 | 11.55 | 0.339 |
| 0 | 0 | clcB | 3 | 11.55 | 0.317 |
| 0 | 0 | ycaM | 3 | 11.55 | 0.372 |
| 0 | 0 | yadI | 3 | 11.55 | 0.517 |
| 0 | 0 | AG1 | 3 | 11.55 | 0.421 |
| 0 | 0 | ptsI | 3 | 11.8 | 0.348 |
| 0 | 0 | clcB | 3 | 11.8 | 0.328 |
| 0 | 0 | ycaM | 3 | 11.8 | 0.374 |
| 0 | 0 | yadI | 3 | 11.8 | 0.519 |
| 0 | 0 | AG1 | 3 | 11.8 | 0.433 |
| 0 | 0 | ptsI | 3 | 12.05 | 0.354 |
| 0 | 0 | clcB | 3 | 12.05 | 0.336 |
| 0 | 0 | ycaM | 3 | 12.05 | 0.378 |
| 0 | 0 | yadI | 3 | 12.05 | 0.537 |
| 0 | 0 | AG1 | 3 | 12.05 | 0.441 |
| 0 | 0 | ptsI | 3 | 12.3 | 0.367 |
| 0 | 0 | clcB | 3 | 12.3 | 0.348 |
| 0 | 0 | ycaM | 3 | 12.3 | 0.39 |
| 0 | 0 | yadI | 3 | 12.3 | 0.551 |
| 0 | 0 | AG1 | 3 | 12.3 | 0.457 |
| 0 | 0 | ptsI | 3 | 12.55 | 0.379 |
| 0 | 0 | clcB | 3 | 12.55 | 0.358 |

| | | | | | |
|---|---|---|---|---|---|
| 0 | 0 | ycaM | 3 | 12.55 | 0.402 |
| 0 | 0 | yadI | 3 | 12.55 | 0.573 |
| 0 | 0 | AG1 | 3 | 12.55 | 0.474 |
| 0 | 0 | ptsI | 3 | 12.8 | 0.385 |
| 0 | 0 | clcB | 3 | 12.8 | 0.364 |
| 0 | 0 | ycaM | 3 | 12.8 | 0.408 |
| 0 | 0 | yadI | 3 | 12.8 | 0.602 |
| 0 | 0 | AG1 | 3 | 12.8 | 0.482 |
| 0 | 0 | ptsI | 3 | 13.05 | 0.392 |
| 0 | 0 | clcB | 3 | 13.05 | 0.362 |
| 0 | 0 | ycaM | 3 | 13.05 | 0.408 |
| 0 | 0 | yadI | 3 | 13.05 | 0.575 |
| 0 | 0 | AG1 | 3 | 13.05 | 0.484 |
| 0 | 0 | ptsI | 3 | 13.3 | 0.396 |
| 0 | 0 | clcB | 3 | 13.3 | 0.365 |
| 0 | 0 | ycaM | 3 | 13.3 | 0.415 |
| 0 | 0 | yadI | 3 | 13.3 | 0.588 |
| 0 | 0 | AG1 | 3 | 13.3 | 0.495 |
| 0 | 0 | ptsI | 3 | 13.55 | 0.41 |
| 0 | 0 | clcB | 3 | 13.55 | 0.377 |
| 0 | 0 | ycaM | 3 | 13.55 | 0.426 |
| 0 | 0 | yadI | 3 | 13.55 | 0.621 |
| 0 | 0 | AG1 | 3 | 13.55 | 0.508 |
| 0 | 0 | ptsI | 3 | 13.8 | 0.423 |
| 0 | 0 | clcB | 3 | 13.8 | 0.39 |
| 0 | 0 | ycaM | 3 | 13.8 | 0.438 |
| 0 | 0 | yadI | 3 | 13.8 | 0.637 |
| 0 | 0 | AG1 | 3 | 13.8 | 0.525 |
| 0 | 0 | ptsI | 3 | 14.05 | 0.426 |
| 0 | 0 | clcB | 3 | 14.05 | 0.39 |
| 0 | 0 | ycaM | 3 | 14.05 | 0.443 |
| 0 | 0 | yadI | 3 | 14.05 | 0.617 |
| 0 | 0 | AG1 | 3 | 14.05 | 0.529 |
| 0 | 0 | ptsI | 3 | 14.3 | 0.437 |
| 0 | 0 | clcB | 3 | 14.3 | 0.403 |
| 0 | 0 | ycaM | 3 | 14.3 | 0.45 |
| 0 | 0 | yadI | 3 | 14.3 | 0.628 |
| 0 | 0 | AG1 | 3 | 14.3 | 0.542 |
| 0 | 0 | ptsI | 3 | 14.55 | 0.441 |
| 0 | 0 | clcB | 3 | 14.55 | 0.411 |
| 0 | 0 | ycaM | 3 | 14.55 | 0.46 |
| 0 | 0 | yadI | 3 | 14.55 | 0.646 |
| 0 | 0 | AG1 | 3 | 14.55 | 0.546 |
| 0 | 0 | ptsI | 3 | 14.8 | 0.466 |
| 0 | 0 | clcB | 3 | 14.8 | 0.431 |
| 0 | 0 | ycaM | 3 | 14.8 | 0.473 |
| 0 | 0 | yadI | 3 | 14.8 | 0.669 |
| 0 | 0 | AG1 | 3 | 14.8 | 0.56 |

| | | | | | |
|---|---|---|---|---|---|
| 0 | 0 | ptsI | 3 | 15.05 | 0.47 |
| 0 | 0 | clcB | 3 | 15.05 | 0.438 |
| 0 | 0 | ycaM | 3 | 15.05 | 0.477 |
| 0 | 0 | yadI | 3 | 15.05 | 0.682 |
| 0 | 0 | AG1 | 3 | 15.05 | 0.568 |
| 0 | 0 | ptsI | 3 | 15.3 | 0.477 |
| 0 | 0 | clcB | 3 | 15.3 | 0.446 |
| 0 | 0 | ycaM | 3 | 15.3 | 0.474 |
| 0 | 0 | yadI | 3 | 15.3 | 0.684 |
| 0 | 0 | AG1 | 3 | 15.3 | 0.573 |
| 0 | 0 | ptsI | 3 | 15.55 | 0.474 |
| 0 | 0 | clcB | 3 | 15.55 | 0.446 |
| 0 | 0 | ycaM | 3 | 15.55 | 0.479 |
| 0 | 0 | yadI | 3 | 15.55 | 0.686 |
| 0 | 0 | AG1 | 3 | 15.55 | 0.574 |
| 0 | 0 | ptsI | 3 | 15.8 | 0.49 |
| 0 | 0 | clcB | 3 | 15.8 | 0.456 |
| 0 | 0 | ycaM | 3 | 15.8 | 0.489 |
| 0 | 0 | yadI | 3 | 15.8 | 0.698 |
| 0 | 0 | AG1 | 3 | 15.8 | 0.599 |
| 0 | 0 | ptsI | 3 | 16.05 | 0.485 |
| 0 | 0 | clcB | 3 | 16.05 | 0.459 |
| 0 | 0 | ycaM | 3 | 16.05 | 0.489 |
| 0 | 0 | yadI | 3 | 16.05 | 0.699 |
| 0 | 0 | AG1 | 3 | 16.05 | 0.586 |
| 0 | 0 | ptsI | 3 | 16.3 | 0.494 |
| 0 | 0 | clcB | 3 | 16.3 | 0.461 |
| 0 | 0 | ycaM | 3 | 16.3 | 0.492 |
| 0 | 0 | yadI | 3 | 16.3 | 0.699 |
| 0 | 0 | AG1 | 3 | 16.3 | 0.589 |
| 0 | 0 | ptsI | 3 | 16.55 | 0.512 |
| 0 | 0 | clcB | 3 | 16.55 | 0.47 |
| 0 | 0 | ycaM | 3 | 16.55 | 0.504 |
| 0 | 0 | yadI | 3 | 16.55 | 0.717 |
| 0 | 0 | AG1 | 3 | 16.55 | 0.609 |
| 0 | 0 | ptsI | 3 | 16.8 | 0.532 |
| 0 | 0 | clcB | 3 | 16.8 | 0.483 |
| 0 | 0 | ycaM | 3 | 16.8 | 0.508 |
| 0 | 0 | yadI | 3 | 16.8 | 0.733 |
| 0 | 0 | AG1 | 3 | 16.8 | 0.627 |
| 0 | 0 | ptsI | 3 | 17.05 | 0.543 |
| 0 | 0 | clcB | 3 | 17.05 | 0.486 |
| 0 | 0 | ycaM | 3 | 17.05 | 0.508 |
| 0 | 0 | yadI | 3 | 17.05 | 0.734 |
| 0 | 0 | AG1 | 3 | 17.05 | 0.642 |
| 0 | 0 | ptsI | 3 | 17.3 | 0.539 |
| 0 | 0 | clcB | 3 | 17.3 | 0.494 |
| 0 | 0 | ycaM | 3 | 17.3 | 0.523 |

| | | | | |
|---|---|---|---|---|
| 0 | 0 | yadI | 3 | 17.3 | 0.741 |
| 0 | 0 | AG1 | 3 | 17.3 | 0.637 |
| 0 | 0 | ptsI | 3 | 17.55 | 0.548 |
| 0 | 0 | clcB | 3 | 17.55 | 0.494 |
| 0 | 0 | ycaM | 3 | 17.55 | 0.516 |
| 0 | 0 | yadI | 3 | 17.55 | 0.747 |
| 0 | 0 | AG1 | 3 | 17.55 | 0.648 |
| 0 | 0 | ptsI | 3 | 17.8 | 0.56 |
| 0 | 0 | clcB | 3 | 17.8 | 0.498 |
| 0 | 0 | ycaM | 3 | 17.8 | 0.522 |
| 0 | 0 | yadI | 3 | 17.8 | 0.755 |
| 0 | 0 | AG1 | 3 | 17.8 | 0.645 |
| 0 | 0 | ptsI | 3 | 18.05 | 0.562 |
| 0 | 0 | clcB | 3 | 18.05 | 0.501 |
| 0 | 0 | ycaM | 3 | 18.05 | 0.527 |
| 0 | 0 | yadI | 3 | 18.05 | 0.781 |
| 0 | 0 | AG1 | 3 | 18.05 | 0.649 |
| 0 | 0 | ptsI | 3 | 18.3 | 0.57 |
| 0 | 0 | clcB | 3 | 18.3 | 0.507 |
| 0 | 0 | ycaM | 3 | 18.3 | 0.535 |
| 0 | 0 | yadI | 3 | 18.3 | 0.778 |
| 0 | 0 | AG1 | 3 | 18.3 | 0.667 |
| 0 | 0 | ptsI | 3 | 18.55 | 0.571 |
| 0 | 0 | clcB | 3 | 18.55 | 0.511 |
| 0 | 0 | ycaM | 3 | 18.55 | 0.542 |
| 0 | 0 | yadI | 3 | 18.55 | 0.772 |
| 0 | 0 | AG1 | 3 | 18.55 | 0.675 |
| 0 | 0 | ptsI | 3 | 18.8 | 0.577 |
| 0 | 0 | clcB | 3 | 18.8 | 0.523 |
| 0 | 0 | ycaM | 3 | 18.8 | 0.547 |
| 0 | 0 | yadI | 3 | 18.8 | 0.799 |
| 0 | 0 | AG1 | 3 | 18.8 | 0.664 |
| 0 | 0 | ptsI | 3 | 19.05 | 0.594 |
| 0 | 0 | clcB | 3 | 19.05 | 0.528 |
| 0 | 0 | ycaM | 3 | 19.05 | 0.556 |
| 0 | 0 | yadI | 3 | 19.05 | 0.79 |
| 0 | 0 | AG1 | 3 | 19.05 | 0.702 |
| 0 | 0 | ptsI | 3 | 19.3 | 0.603 |
| 0 | 0 | clcB | 3 | 19.3 | 0.538 |
| 0 | 0 | ycaM | 3 | 19.3 | 0.578 |
| 0 | 0 | yadI | 3 | 19.3 | 0.804 |
| 0 | 0 | AG1 | 3 | 19.3 | 0.699 |
| 0 | 0 | ptsI | 3 | 19.55 | 0.619 |
| 0 | 0 | clcB | 3 | 19.55 | 0.555 |
| 0 | 0 | ycaM | 3 | 19.55 | 0.595 |
| 0 | 0 | yadI | 3 | 19.55 | 0.843 |
| 0 | 0 | AG1 | 3 | 19.55 | 0.711 |
| 0 | 0 | ptsI | 3 | 19.8 | 0.612 |

| | | | | | |
|---|---|---|---|---|---|
| 0 | 0 | clcB | 3 | 19.8 | 0.551 |
| 0 | 0 | ycaM | 3 | 19.8 | 0.583 |
| 0 | 0 | yadI | 3 | 19.8 | 0.81 |
| 0 | 0 | AG1 | 3 | 19.8 | 0.722 |
| 0 | 0 | ptsI | 3 | 20.05 | 0.63 |
| 0 | 0 | clcB | 3 | 20.05 | 0.562 |
| 0 | 0 | ycaM | 3 | 20.05 | 0.588 |
| 0 | 0 | yadI | 3 | 20.05 | 0.832 |
| 0 | 0 | AG1 | 3 | 20.05 | 0.728 |
| 0 | 0 | ptsI | 3 | 20.3 | 0.621 |
| 0 | 0 | clcB | 3 | 20.3 | 0.554 |
| 0 | 0 | ycaM | 3 | 20.3 | 0.582 |
| 0 | 0 | yadI | 3 | 20.3 | 0.828 |
| 0 | 0 | AG1 | 3 | 20.3 | 0.73 |
| 0 | 0 | ptsI | 3 | 20.55 | 0.63 |
| 0 | 0 | clcB | 3 | 20.55 | 0.567 |
| 0 | 0 | ycaM | 3 | 20.55 | 0.597 |
| 0 | 0 | yadI | 3 | 20.55 | 0.839 |
| 0 | 0 | AG1 | 3 | 20.55 | 0.749 |
| 0 | 0 | ptsI | 3 | 20.8 | 0.638 |
| 0 | 0 | clcB | 3 | 20.8 | 0.573 |
| 0 | 0 | ycaM | 3 | 20.8 | 0.607 |
| 0 | 0 | yadI | 3 | 20.8 | 0.859 |
| 0 | 0 | AG1 | 3 | 20.8 | 0.747 |
| 0 | 0 | ptsI | 3 | 21.05 | 0.648 |
| 0 | 0 | clcB | 3 | 21.05 | 0.572 |
| 0 | 0 | ycaM | 3 | 21.05 | 0.602 |
| 0 | 0 | yadI | 3 | 21.05 | 0.869 |
| 0 | 0 | AG1 | 3 | 21.05 | 0.74 |
| 0 | 0 | ptsI | 3 | 21.3 | 0.65 |
| 0 | 0 | clcB | 3 | 21.3 | 0.574 |
| 0 | 0 | ycaM | 3 | 21.3 | 0.605 |
| 0 | 0 | yadI | 3 | 21.3 | 0.862 |
| 0 | 0 | AG1 | 3 | 21.3 | 0.744 |
| 0 | 0 | ptsI | 3 | 21.55 | 0.674 |
| 0 | 0 | clcB | 3 | 21.55 | 0.584 |
| 0 | 0 | ycaM | 3 | 21.55 | 0.62 |
| 0 | 0 | yadI | 3 | 21.55 | 0.871 |
| 0 | 0 | AG1 | 3 | 21.55 | 0.757 |
| 0 | 0 | ptsI | 3 | 21.8 | 0.683 |
| 0 | 0 | clcB | 3 | 21.8 | 0.587 |
| 0 | 0 | ycaM | 3 | 21.8 | 0.624 |
| 0 | 0 | yadI | 3 | 21.8 | 0.88 |
| 0 | 0 | AG1 | 3 | 21.8 | 0.764 |
| 0 | 0 | ptsI | 3 | 22.05 | 0.704 |
| 0 | 0 | clcB | 3 | 22.05 | 0.586 |
| 0 | 0 | ycaM | 3 | 22.05 | 0.626 |
| 0 | 0 | yadI | 3 | 22.05 | 0.894 |

| | | | | | |
|---|---|---|---|---|---|
| 0 | 0 | AG1 | 3 | 22.05 | 0.773 |
| 0 | 0 | ptsI | 3 | 22.3 | 0.72 |
| 0 | 0 | clcB | 3 | 22.3 | 0.591 |
| 0 | 0 | ycaM | 3 | 22.3 | 0.637 |
| 0 | 0 | yadI | 3 | 22.3 | 0.913 |
| 0 | 0 | AG1 | 3 | 22.3 | 0.783 |
| 0 | 0 | ptsI | 3 | 22.55 | 0.737 |
| 0 | 0 | clcB | 3 | 22.55 | 0.592 |
| 0 | 0 | ycaM | 3 | 22.55 | 0.642 |
| 0 | 0 | yadI | 3 | 22.55 | 0.902 |
| 0 | 0 | AG1 | 3 | 22.55 | 0.793 |
| 0 | 0 | ptsI | 3 | 22.8 | 0.744 |
| 0 | 0 | clcB | 3 | 22.8 | 0.598 |
| 0 | 0 | ycaM | 3 | 22.8 | 0.641 |
| 0 | 0 | yadI | 3 | 22.8 | 0.904 |
| 0 | 0 | AG1 | 3 | 22.8 | 0.801 |
| 0 | 0 | ptsI | 3 | 23.05 | 0.761 |
| 0 | 0 | clcB | 3 | 23.05 | 0.601 |
| 0 | 0 | ycaM | 3 | 23.05 | 0.651 |
| 0 | 0 | yadI | 3 | 23.05 | 0.928 |
| 0 | 0 | AG1 | 3 | 23.05 | 0.804 |
| 0 | 0 | ptsI | 3 | 23.3 | 0.776 |
| 0 | 0 | clcB | 3 | 23.3 | 0.607 |
| 0 | 0 | ycaM | 3 | 23.3 | 0.653 |
| 0 | 0 | yadI | 3 | 23.3 | 0.928 |
| 0 | 0 | AG1 | 3 | 23.3 | 0.814 |
| 0 | 0 | ptsI | 3 | 23.55 | 0.812 |
| 0 | 0 | clcB | 3 | 23.55 | 0.628 |
| 0 | 0 | ycaM | 3 | 23.55 | 0.678 |
| 0 | 0 | yadI | 3 | 23.55 | 0.936 |
| 0 | 0 | AG1 | 3 | 23.55 | 0.827 |
| 0 | 0 | ptsI | 3 | 23.8 | 0.801 |
| 0 | 0 | clcB | 3 | 23.8 | 0.618 |
| 0 | 0 | ycaM | 3 | 23.8 | 0.675 |
| 0 | 0 | yadI | 3 | 23.8 | 0.942 |
| 0 | 0 | AG1 | 3 | 23.8 | 0.826 |
| 0 | 0 | ptsI | 3 | 24.05 | 0.809 |
| 0 | 0 | clcB | 3 | 24.05 | 0.62 |
| 0 | 0 | ycaM | 3 | 24.05 | 0.679 |
| 0 | 0 | yadI | 3 | 24.05 | 0.959 |
| 0 | 0 | AG1 | 3 | 24.05 | 0.83 |
| 0 | 0 | ptsI | 3 | 24.3 | 0.794 |
| 0 | 0 | clcB | 3 | 24.3 | 0.623 |
| 0 | 0 | ycaM | 3 | 24.3 | 0.672 |
| 0 | 0 | yadI | 3 | 24.3 | 0.938 |
| 0 | 0 | AG1 | 3 | 24.3 | 0.815 |
| 0 | 0 | ptsI | 4 | 0 | 0.182 |
| 0 | 0 | clcB | 4 | 0 | 0.196 |

| | | | | | |
|---|---|---|---|---|---|
| 0 | 0 | ycaM | 4 | 0 | 0.195 |
| 0 | 0 | yadI | 4 | 0 | 0.212 |
| 0 | 0 | AG1 | 4 | 0 | 0.202 |
| 0 | 0 | ptsI | 4 | 0.25 | 0.181 |
| 0 | 0 | clcB | 4 | 0.25 | 0.192 |
| 0 | 0 | ycaM | 4 | 0.25 | 0.193 |
| 0 | 0 | yadI | 4 | 0.25 | 0.21 |
| 0 | 0 | AG1 | 4 | 0.25 | 0.204 |
| 0 | 0 | ptsI | 4 | 0.5 | 0.184 |
| 0 | 0 | clcB | 4 | 0.5 | 0.193 |
| 0 | 0 | ycaM | 4 | 0.5 | 0.193 |
| 0 | 0 | yadI | 4 | 0.5 | 0.212 |
| 0 | 0 | AG1 | 4 | 0.5 | 0.207 |
| 0 | 0 | ptsI | 4 | 0.75 | 0.189 |
| 0 | 0 | clcB | 4 | 0.75 | 0.196 |
| 0 | 0 | ycaM | 4 | 0.75 | 0.196 |
| 0 | 0 | yadI | 4 | 0.75 | 0.215 |
| 0 | 0 | AG1 | 4 | 0.75 | 0.211 |
| 0 | 0 | ptsI | 4 | 1 | 0.195 |
| 0 | 0 | clcB | 4 | 1 | 0.201 |
| 0 | 0 | ycaM | 4 | 1 | 0.199 |
| 0 | 0 | yadI | 4 | 1 | 0.219 |
| 0 | 0 | AG1 | 4 | 1 | 0.216 |
| 0 | 0 | ptsI | 4 | 1.25 | 0.206 |
| 0 | 0 | clcB | 4 | 1.25 | 0.207 |
| 0 | 0 | ycaM | 4 | 1.25 | 0.21 |
| 0 | 0 | yadI | 4 | 1.25 | 0.225 |
| 0 | 0 | AG1 | 4 | 1.25 | 0.221 |
| 0 | 0 | ptsI | 4 | 1.5 | 0.205 |
| 0 | 0 | clcB | 4 | 1.5 | 0.212 |
| 0 | 0 | ycaM | 4 | 1.5 | 0.212 |
| 0 | 0 | yadI | 4 | 1.5 | 0.23 |
| 0 | 0 | AG1 | 4 | 1.5 | 0.224 |
| 0 | 0 | ptsI | 4 | 1.75 | 0.22 |
| 0 | 0 | clcB | 4 | 1.75 | 0.224 |
| 0 | 0 | ycaM | 4 | 1.75 | 0.221 |
| 0 | 0 | yadI | 4 | 1.75 | 0.241 |
| 0 | 0 | AG1 | 4 | 1.75 | 0.236 |
| 0 | 0 | ptsI | 4 | 2 | 0.233 |
| 0 | 0 | clcB | 4 | 2 | 0.235 |
| 0 | 0 | ycaM | 4 | 2 | 0.23 |
| 0 | 0 | yadI | 4 | 2 | 0.254 |
| 0 | 0 | AG1 | 4 | 2 | 0.248 |
| 0 | 0 | ptsI | 4 | 2.25 | 0.246 |
| 0 | 0 | clcB | 4 | 2.25 | 0.249 |
| 0 | 0 | ycaM | 4 | 2.25 | 0.239 |
| 0 | 0 | yadI | 4 | 2.25 | 0.266 |
| 0 | 0 | AG1 | 4 | 2.25 | 0.26 |

| | | | | | |
|---|---|---|---|---|---|
| 0 | 0 | ptsI | 4 | 2.5 | 0.254 |
| 0 | 0 | clcB | 4 | 2.5 | 0.259 |
| 0 | 0 | ycaM | 4 | 2.5 | 0.25 |
| 0 | 0 | yadI | 4 | 2.5 | 0.28 |
| 0 | 0 | AG1 | 4 | 2.5 | 0.267 |
| 0 | 0 | ptsI | 4 | 2.75 | 0.272 |
| 0 | 0 | clcB | 4 | 2.75 | 0.269 |
| 0 | 0 | ycaM | 4 | 2.75 | 0.264 |
| 0 | 0 | yadI | 4 | 2.75 | 0.299 |
| 0 | 0 | AG1 | 4 | 2.75 | 0.28 |
| 0 | 0 | ptsI | 4 | 3 | 0.284 |
| 0 | 0 | clcB | 4 | 3 | 0.292 |
| 0 | 0 | ycaM | 4 | 3 | 0.28 |
| 0 | 0 | yadI | 4 | 3 | 0.318 |
| 0 | 0 | AG1 | 4 | 3 | 0.298 |
| 0 | 0 | ptsI | 4 | 3.25 | 0.297 |
| 0 | 0 | clcB | 4 | 3.25 | 0.301 |
| 0 | 0 | ycaM | 4 | 3.25 | 0.297 |
| 0 | 0 | yadI | 4 | 3.25 | 0.332 |
| 0 | 0 | AG1 | 4 | 3.25 | 0.311 |
| 0 | 0 | ptsI | 4 | 3.5 | 0.319 |
| 0 | 0 | clcB | 4 | 3.5 | 0.32 |
| 0 | 0 | ycaM | 4 | 3.5 | 0.312 |
| 0 | 0 | yadI | 4 | 3.5 | 0.355 |
| 0 | 0 | AG1 | 4 | 3.5 | 0.33 |
| 0 | 0 | ptsI | 4 | 3.75 | 0.338 |
| 0 | 0 | clcB | 4 | 3.75 | 0.326 |
| 0 | 0 | ycaM | 4 | 3.75 | 0.333 |
| 0 | 0 | yadI | 4 | 3.75 | 0.372 |
| 0 | 0 | AG1 | 4 | 3.75 | 0.343 |
| 0 | 0 | ptsI | 4 | 4 | 0.351 |
| 0 | 0 | clcB | 4 | 4 | 0.35 |
| 0 | 0 | ycaM | 4 | 4 | 0.35 |
| 0 | 0 | yadI | 4 | 4 | 0.401 |
| 0 | 0 | AG1 | 4 | 4 | 0.364 |
| 0 | 0 | ptsI | 4 | 4.35 | 0.38 |
| 0 | 0 | clcB | 4 | 4.35 | 0.293 |
| 0 | 0 | ycaM | 4 | 4.35 | 0.415 |
| 0 | 0 | yadI | 4 | 4.35 | 0.348 |
| 0 | 0 | AG1 | 4 | 4.35 | 0.406 |
| 0 | 0 | ptsI | 4 | 4.6 | 0.377 |
| 0 | 0 | clcB | 4 | 4.6 | 0.288 |
| 0 | 0 | ycaM | 4 | 4.6 | 0.405 |
| 0 | 0 | yadI | 4 | 4.6 | 0.324 |
| 0 | 0 | AG1 | 4 | 4.6 | 0.407 |
| 0 | 0 | ptsI | 4 | 4.85 | 0.39 |
| 0 | 0 | clcB | 4 | 4.85 | 0.291 |
| 0 | 0 | ycaM | 4 | 4.85 | 0.415 |

| | | | | | |
|---|---|---|---|---|---|
| 0 | 0 | yadI | 4 | 4.85 | 0.332 |
| 0 | 0 | AG1 | 4 | 4.85 | 0.417 |
| 0 | 0 | ptsI | 4 | 5.1 | 0.403 |
| 0 | 0 | clcB | 4 | 5.1 | 0.301 |
| 0 | 0 | ycaM | 4 | 5.1 | 0.428 |
| 0 | 0 | yadI | 4 | 5.1 | 0.344 |
| 0 | 0 | AG1 | 4 | 5.1 | 0.431 |
| 0 | 0 | ptsI | 4 | 5.35 | 0.416 |
| 0 | 0 | clcB | 4 | 5.35 | 0.312 |
| 0 | 0 | ycaM | 4 | 5.35 | 0.44 |
| 0 | 0 | yadI | 4 | 5.35 | 0.359 |
| 0 | 0 | AG1 | 4 | 5.35 | 0.447 |
| 0 | 0 | ptsI | 4 | 5.6 | 0.433 |
| 0 | 0 | clcB | 4 | 5.6 | 0.317 |
| 0 | 0 | ycaM | 4 | 5.6 | 0.455 |
| 0 | 0 | yadI | 4 | 5.6 | 0.369 |
| 0 | 0 | AG1 | 4 | 5.6 | 0.458 |
| 0 | 0 | ptsI | 4 | 5.85 | 0.452 |
| 0 | 0 | clcB | 4 | 5.85 | 0.328 |
| 0 | 0 | ycaM | 4 | 5.85 | 0.466 |
| 0 | 0 | yadI | 4 | 5.85 | 0.383 |
| 0 | 0 | AG1 | 4 | 5.85 | 0.474 |
| 0 | 0 | ptsI | 4 | 6.1 | 0.469 |
| 0 | 0 | clcB | 4 | 6.1 | 0.345 |
| 0 | 0 | ycaM | 4 | 6.1 | 0.485 |
| 0 | 0 | yadI | 4 | 6.1 | 0.399 |
| 0 | 0 | AG1 | 4 | 6.1 | 0.485 |
| 0 | 0 | ptsI | 4 | 6.35 | 0.484 |
| 0 | 0 | clcB | 4 | 6.35 | 0.35 |
| 0 | 0 | ycaM | 4 | 6.35 | 0.496 |
| 0 | 0 | yadI | 4 | 6.35 | 0.414 |
| 0 | 0 | AG1 | 4 | 6.35 | 0.497 |
| 0 | 0 | ptsI | 4 | 6.6 | 0.502 |
| 0 | 0 | clcB | 4 | 6.6 | 0.364 |
| 0 | 0 | ycaM | 4 | 6.6 | 0.509 |
| 0 | 0 | yadI | 4 | 6.6 | 0.433 |
| 0 | 0 | AG1 | 4 | 6.6 | 0.512 |
| 0 | 0 | ptsI | 4 | 6.85 | 0.515 |
| 0 | 0 | clcB | 4 | 6.85 | 0.374 |
| 0 | 0 | ycaM | 4 | 6.85 | 0.519 |
| 0 | 0 | yadI | 4 | 6.85 | 0.441 |
| 0 | 0 | AG1 | 4 | 6.85 | 0.528 |
| 0 | 0 | ptsI | 4 | 7.1 | 0.526 |
| 0 | 0 | clcB | 4 | 7.1 | 0.383 |
| 0 | 0 | ycaM | 4 | 7.1 | 0.531 |
| 0 | 0 | yadI | 4 | 7.1 | 0.456 |
| 0 | 0 | AG1 | 4 | 7.1 | 0.536 |
| 0 | 0 | ptsI | 4 | 7.35 | 0.533 |

| | | | | | |
|---|---|---|---|---|---|
| 0 | 0 | clcB | 4 | 7.35 | 0.388 |
| 0 | 0 | ycaM | 4 | 7.35 | 0.536 |
| 0 | 0 | yadI | 4 | 7.35 | 0.457 |
| 0 | 0 | AG1 | 4 | 7.35 | 0.547 |
| 0 | 0 | ptsI | 4 | 7.6 | 0.541 |
| 0 | 0 | clcB | 4 | 7.6 | 0.398 |
| 0 | 0 | ycaM | 4 | 7.6 | 0.545 |
| 0 | 0 | yadI | 4 | 7.6 | 0.473 |
| 0 | 0 | AG1 | 4 | 7.6 | 0.558 |
| 0 | 0 | ptsI | 4 | 7.85 | 0.546 |
| 0 | 0 | clcB | 4 | 7.85 | 0.404 |
| 0 | 0 | ycaM | 4 | 7.85 | 0.546 |
| 0 | 0 | yadI | 4 | 7.85 | 0.476 |
| 0 | 0 | AG1 | 4 | 7.85 | 0.565 |
| 0 | 0 | ptsI | 4 | 8.1 | 0.56 |
| 0 | 0 | clcB | 4 | 8.1 | 0.422 |
| 0 | 0 | ycaM | 4 | 8.1 | 0.563 |
| 0 | 0 | yadI | 4 | 8.1 | 0.496 |
| 0 | 0 | AG1 | 4 | 8.1 | 0.578 |
| 0 | 0 | ptsI | 4 | 8.35 | 0.572 |
| 0 | 0 | clcB | 4 | 8.35 | 0.421 |
| 0 | 0 | ycaM | 4 | 8.35 | 0.564 |
| 0 | 0 | yadI | 4 | 8.35 | 0.502 |
| 0 | 0 | AG1 | 4 | 8.35 | 0.586 |
| 0 | 0 | ptsI | 4 | 8.83333 | 0.379 |
| 0 | 0 | clcB | 4 | 8.83333 | 0.394 |
| 0 | 0 | ycaM | 4 | 8.83333 | 0.397 |
| 0 | 0 | yadI | 4 | 8.83333 | 0.53 |
| 0 | 0 | AG1 | 4 | 8.83333 | 0.619 |
| 0 | 0 | ptsI | 4 | 9.08333 | 0.37 |
| 0 | 0 | clcB | 4 | 9.08333 | 0.386 |
| 0 | 0 | ycaM | 4 | 9.08333 | 0.389 |
| 0 | 0 | yadI | 4 | 9.08333 | 0.522 |
| 0 | 0 | AG1 | 4 | 9.08333 | 0.62 |
| 0 | 0 | ptsI | 4 | 9.33333 | 0.378 |
| 0 | 0 | clcB | 4 | 9.33333 | 0.392 |
| 0 | 0 | ycaM | 4 | 9.33333 | 0.394 |
| 0 | 0 | yadI | 4 | 9.33333 | 0.533 |
| 0 | 0 | AG1 | 4 | 9.33333 | 0.628 |
| 0 | 0 | ptsI | 4 | 9.58333 | 0.382 |
| 0 | 0 | clcB | 4 | 9.58333 | 0.395 |
| 0 | 0 | ycaM | 4 | 9.58333 | 0.402 |
| 0 | 0 | yadI | 4 | 9.58333 | 0.543 |
| 0 | 0 | AG1 | 4 | 9.58333 | 0.64 |
| 0 | 0 | ptsI | 4 | 9.83333 | 0.389 |
| 0 | 0 | clcB | 4 | 9.83333 | 0.4 |
| 0 | 0 | ycaM | 4 | 9.83333 | 0.4 |
| 0 | 0 | yadI | 4 | 9.83333 | 0.548 |

| | | | | | |
|---|---|---|---|---|---|
| 0 | 0 | AG1 | 4 | 9.83333 | 0.65 |
| 0 | 0 | ptsI | 4 | 10.0833 | 0.397 |
| 0 | 0 | clcB | 4 | 10.0833 | 0.406 |
| 0 | 0 | ycaM | 4 | 10.0833 | 0.407 |
| 0 | 0 | yadI | 4 | 10.0833 | 0.557 |
| 0 | 0 | AG1 | 4 | 10.0833 | 0.66 |
| 0 | 0 | ptsI | 4 | 10.3333 | 0.407 |
| 0 | 0 | clcB | 4 | 10.3333 | 0.414 |
| 0 | 0 | ycaM | 4 | 10.3333 | 0.415 |
| 0 | 0 | yadI | 4 | 10.3333 | 0.566 |
| 0 | 0 | AG1 | 4 | 10.3333 | 0.669 |
| 0 | 0 | ptsI | 4 | 10.5833 | 0.418 |
| 0 | 0 | clcB | 4 | 10.5833 | 0.427 |
| 0 | 0 | ycaM | 4 | 10.5833 | 0.426 |
| 0 | 0 | yadI | 4 | 10.5833 | 0.578 |
| 0 | 0 | AG1 | 4 | 10.5833 | 0.684 |
| 0 | 0 | ptsI | 4 | 10.8333 | 0.417 |
| 0 | 0 | clcB | 4 | 10.8333 | 0.43 |
| 0 | 0 | ycaM | 4 | 10.8333 | 0.425 |
| 0 | 0 | yadI | 4 | 10.8333 | 0.583 |
| 0 | 0 | AG1 | 4 | 10.8333 | 0.686 |
| 0 | 0 | ptsI | 4 | 11.0833 | 0.431 |
| 0 | 0 | clcB | 4 | 11.0833 | 0.425 |
| 0 | 0 | ycaM | 4 | 11.0833 | 0.424 |
| 0 | 0 | yadI | 4 | 11.0833 | 0.584 |
| 0 | 0 | AG1 | 4 | 11.0833 | 0.698 |
| 0 | 0 | ptsI | 4 | 11.3333 | 0.437 |
| 0 | 0 | clcB | 4 | 11.3333 | 0.449 |
| 0 | 0 | ycaM | 4 | 11.3333 | 0.442 |
| 0 | 0 | yadI | 4 | 11.3333 | 0.6 |
| 0 | 0 | AG1 | 4 | 11.3333 | 0.71 |
| 0 | 0 | ptsI | 4 | 11.5833 | 0.436 |
| 0 | 0 | clcB | 4 | 11.5833 | 0.442 |
| 0 | 0 | ycaM | 4 | 11.5833 | 0.439 |
| 0 | 0 | yadI | 4 | 11.5833 | 0.603 |
| 0 | 0 | AG1 | 4 | 11.5833 | 0.713 |
| 0 | 0 | ptsI | 4 | 11.8333 | 0.442 |
| 0 | 0 | clcB | 4 | 11.8333 | 0.443 |
| 0 | 0 | ycaM | 4 | 11.8333 | 0.437 |
| 0 | 0 | yadI | 4 | 11.8333 | 0.602 |
| 0 | 0 | AG1 | 4 | 11.8333 | 0.721 |
| 0 | 0 | ptsI | 4 | 12.0833 | 0.448 |
| 0 | 0 | clcB | 4 | 12.0833 | 0.452 |
| 0 | 0 | ycaM | 4 | 12.0833 | 0.448 |
| 0 | 0 | yadI | 4 | 12.0833 | 0.614 |
| 0 | 0 | AG1 | 4 | 12.0833 | 0.729 |
| 0 | 0 | ptsI | 4 | 12.3333 | 0.456 |
| 0 | 0 | clcB | 4 | 12.3333 | 0.462 |

| | | | | | |
|---|---|---|---|---|---|
| 0 | 0 | ycaM | 4 | 12.3333 | 0.456 |
| 0 | 0 | yadI | 4 | 12.3333 | 0.615 |
| 0 | 0 | AG1 | 4 | 12.3333 | 0.734 |
| 0 | 0 | ptsI | 4 | 12.5833 | 0.468 |
| 0 | 0 | clcB | 4 | 12.5833 | 0.462 |
| 0 | 0 | ycaM | 4 | 12.5833 | 0.458 |
| 0 | 0 | yadI | 4 | 12.5833 | 0.619 |
| 0 | 0 | AG1 | 4 | 12.5833 | 0.738 |
| 0 | 0 | ptsI | 4 | 12.8333 | 0.466 |
| 0 | 0 | clcB | 4 | 12.8333 | 0.463 |
| 0 | 0 | ycaM | 4 | 12.8333 | 0.46 |
| 0 | 0 | yadI | 4 | 12.8333 | 0.623 |
| 0 | 0 | AG1 | 4 | 12.8333 | 0.748 |
| 0 | 0 | ptsI | 4 | 13.0833 | 0.479 |
| 0 | 0 | clcB | 4 | 13.0833 | 0.474 |
| 0 | 0 | ycaM | 4 | 13.0833 | 0.466 |
| 0 | 0 | yadI | 4 | 13.0833 | 0.627 |
| 0 | 0 | AG1 | 4 | 13.0833 | 0.751 |
| 0 | 0 | ptsI | 4 | 13.3333 | 0.491 |
| 0 | 0 | clcB | 4 | 13.3333 | 0.483 |
| 0 | 0 | ycaM | 4 | 13.3333 | 0.474 |
| 0 | 0 | yadI | 4 | 13.3333 | 0.634 |
| 0 | 0 | AG1 | 4 | 13.3333 | 0.758 |
| 0 | 0 | ptsI | 4 | 13.5833 | 0.489 |
| 0 | 0 | clcB | 4 | 13.5833 | 0.481 |
| 0 | 0 | ycaM | 4 | 13.5833 | 0.474 |
| 0 | 0 | yadI | 4 | 13.5833 | 0.634 |
| 0 | 0 | AG1 | 4 | 13.5833 | 0.757 |
| 0 | 0 | ptsI | 4 | 13.8333 | 0.489 |
| 0 | 0 | clcB | 4 | 13.8333 | 0.487 |
| 0 | 0 | ycaM | 4 | 13.8333 | 0.484 |
| 0 | 0 | yadI | 4 | 13.8333 | 0.642 |
| 0 | 0 | AG1 | 4 | 13.8333 | 0.764 |
| 0 | 0 | ptsI | 4 | 14.0833 | 0.49 |
| 0 | 0 | clcB | 4 | 14.0833 | 0.491 |
| 0 | 0 | ycaM | 4 | 14.0833 | 0.487 |
| 0 | 0 | yadI | 4 | 14.0833 | 0.644 |
| 0 | 0 | AG1 | 4 | 14.0833 | 0.769 |
| 0 | 0 | ptsI | 4 | 14.3333 | 0.495 |
| 0 | 0 | clcB | 4 | 14.3333 | 0.496 |
| 0 | 0 | ycaM | 4 | 14.3333 | 0.498 |
| 0 | 0 | yadI | 4 | 14.3333 | 0.654 |
| 0 | 0 | AG1 | 4 | 14.3333 | 0.788 |
| 0 | 0 | ptsI | 4 | 14.5833 | 0.501 |
| 0 | 0 | clcB | 4 | 14.5833 | 0.499 |
| 0 | 0 | ycaM | 4 | 14.5833 | 0.491 |
| 0 | 0 | yadI | 4 | 14.5833 | 0.648 |
| 0 | 0 | AG1 | 4 | 14.5833 | 0.777 |

| | | | | | |
|---|---|---|---|---|---|
| 0 | 0 | ptsI | 4 | 14.8333 | 0.515 |
| 0 | 0 | clcB | 4 | 14.8333 | 0.499 |
| 0 | 0 | ycaM | 4 | 14.8333 | 0.498 |
| 0 | 0 | yadI | 4 | 14.8333 | 0.654 |
| 0 | 0 | AG1 | 4 | 14.8333 | 0.784 |
| 0 | 0 | ptsI | 4 | 15.0833 | 0.513 |
| 0 | 0 | clcB | 4 | 15.0833 | 0.497 |
| 0 | 0 | ycaM | 4 | 15.0833 | 0.497 |
| 0 | 0 | yadI | 4 | 15.0833 | 0.653 |
| 0 | 0 | AG1 | 4 | 15.0833 | 0.783 |
| 0 | 0 | ptsI | 4 | 15.3333 | 0.512 |
| 0 | 0 | clcB | 4 | 15.3333 | 0.496 |
| 0 | 0 | ycaM | 4 | 15.3333 | 0.507 |
| 0 | 0 | yadI | 4 | 15.3333 | 0.663 |
| 0 | 0 | AG1 | 4 | 15.3333 | 0.799 |
| 0 | 0 | ptsI | 4 | 15.5833 | 0.53 |
| 0 | 0 | clcB | 4 | 15.5833 | 0.513 |
| 0 | 0 | ycaM | 4 | 15.5833 | 0.512 |
| 0 | 0 | yadI | 4 | 15.5833 | 0.663 |
| 0 | 0 | AG1 | 4 | 15.5833 | 0.797 |
| 0 | 0 | ptsI | 4 | 15.8333 | 0.514 |
| 0 | 0 | clcB | 4 | 15.8333 | 0.501 |
| 0 | 0 | ycaM | 4 | 15.8333 | 0.512 |
| 0 | 0 | yadI | 4 | 15.8333 | 0.67 |
| 0 | 0 | AG1 | 4 | 15.8333 | 0.796 |
| 0 | 0 | ptsI | 4 | 16.0833 | 0.534 |
| 0 | 0 | clcB | 4 | 16.0833 | 0.517 |
| 0 | 0 | ycaM | 4 | 16.0833 | 0.519 |
| 0 | 0 | yadI | 4 | 16.0833 | 0.668 |
| 0 | 0 | AG1 | 4 | 16.0833 | 0.801 |
| 0 | 0 | ptsI | 4 | 16.3333 | 0.524 |
| 0 | 0 | clcB | 4 | 16.3333 | 0.502 |
| 0 | 0 | ycaM | 4 | 16.3333 | 0.512 |
| 0 | 0 | yadI | 4 | 16.3333 | 0.678 |
| 0 | 0 | AG1 | 4 | 16.3333 | 0.803 |
| 0 | 0 | ptsI | 4 | 16.5833 | 0.524 |
| 0 | 0 | clcB | 4 | 16.5833 | 0.502 |
| 0 | 0 | ycaM | 4 | 16.5833 | 0.517 |
| 0 | 0 | yadI | 4 | 16.5833 | 0.678 |
| 0 | 0 | AG1 | 4 | 16.5833 | 0.81 |
| 0 | 0 | ptsI | 4 | 16.8333 | 0.54 |
| 0 | 0 | clcB | 4 | 16.8333 | 0.515 |
| 0 | 0 | ycaM | 4 | 16.8333 | 0.521 |
| 0 | 0 | yadI | 4 | 16.8333 | 0.674 |
| 0 | 0 | AG1 | 4 | 16.8333 | 0.803 |
| 0 | 0 | ptsI | 4 | 17.0833 | 0.535 |
| 0 | 0 | clcB | 4 | 17.0833 | 0.509 |
| 0 | 0 | ycaM | 4 | 17.0833 | 0.52 |

| | | | | | |
|---|---|---|---|---|---|
| 0 | 0 | yadI | 4 | 17.0833 | 0.676 |
| 0 | 0 | AG1 | 4 | 17.0833 | 0.801 |
| 0 | 0 | ptsI | 4 | 17.3333 | 0.535 |
| 0 | 0 | clcB | 4 | 17.3333 | 0.512 |
| 0 | 0 | ycaM | 4 | 17.3333 | 0.528 |
| 0 | 0 | yadI | 4 | 17.3333 | 0.689 |
| 0 | 0 | AG1 | 4 | 17.3333 | 0.817 |
| 0 | 0 | ptsI | 4 | 17.5833 | 0.542 |
| 0 | 0 | clcB | 4 | 17.5833 | 0.518 |
| 0 | 0 | ycaM | 4 | 17.5833 | 0.525 |
| 0 | 0 | yadI | 4 | 17.5833 | 0.678 |
| 0 | 0 | AG1 | 4 | 17.5833 | 0.807 |
| 0 | 0 | ptsI | 4 | 17.8333 | 0.545 |
| 0 | 0 | clcB | 4 | 17.8333 | 0.515 |
| 0 | 0 | ycaM | 4 | 17.8333 | 0.524 |
| 0 | 0 | yadI | 4 | 17.8333 | 0.678 |
| 0 | 0 | AG1 | 4 | 17.8333 | 0.808 |
| 0 | 0 | ptsI | 4 | 18.0833 | 0.55 |
| 0 | 0 | clcB | 4 | 18.0833 | 0.52 |
| 0 | 0 | ycaM | 4 | 18.0833 | 0.529 |
| 0 | 0 | yadI | 4 | 18.0833 | 0.687 |
| 0 | 0 | AG1 | 4 | 18.0833 | 0.808 |
| 0 | 0 | ptsI | 4 | 18.3333 | 0.545 |
| 0 | 0 | clcB | 4 | 18.3333 | 0.518 |
| 0 | 0 | ycaM | 4 | 18.3333 | 0.537 |
| 0 | 0 | yadI | 4 | 18.3333 | 0.686 |
| 0 | 0 | AG1 | 4 | 18.3333 | 0.814 |
| 0 | 0 | ptsI | 4 | 18.5833 | 0.556 |
| 0 | 0 | clcB | 4 | 18.5833 | 0.523 |
| 0 | 0 | ycaM | 4 | 18.5833 | 0.529 |
| 0 | 0 | yadI | 4 | 18.5833 | 0.689 |
| 0 | 0 | AG1 | 4 | 18.5833 | 0.814 |
| 0 | 0 | ptsI | 4 | 18.8333 | 0.557 |
| 0 | 0 | clcB | 4 | 18.8333 | 0.516 |
| 0 | 0 | ycaM | 4 | 18.8333 | 0.53 |
| 0 | 0 | yadI | 4 | 18.8333 | 0.691 |
| 0 | 0 | AG1 | 4 | 18.8333 | 0.819 |
| 0 | 0 | ptsI | 4 | 19.0833 | 0.549 |
| 0 | 0 | clcB | 4 | 19.0833 | 0.526 |
| 0 | 0 | ycaM | 4 | 19.0833 | 0.532 |
| 0 | 0 | yadI | 4 | 19.0833 | 0.693 |
| 0 | 0 | AG1 | 4 | 19.0833 | 0.818 |
| 0 | 0 | ptsI | 4 | 19.3333 | 0.556 |
| 0 | 0 | clcB | 4 | 19.3333 | 0.519 |
| 0 | 0 | ycaM | 4 | 19.3333 | 0.534 |
| 0 | 0 | yadI | 4 | 19.3333 | 0.707 |
| 0 | 0 | AG1 | 4 | 19.3333 | 0.829 |
| 0 | 0 | ptsI | 4 | 19.5833 | 0.57 |

| | | | | | |
|---|---|---|---|---|---|
| 0 | 0 | clcB | 4 | 19.5833 | 0.536 |
| 0 | 0 | ycaM | 4 | 19.5833 | 0.541 |
| 0 | 0 | yadI | 4 | 19.5833 | 0.704 |
| 0 | 0 | AG1 | 4 | 19.5833 | 0.829 |
| 0 | 0 | ptsI | 4 | 19.8333 | 0.57 |
| 0 | 0 | clcB | 4 | 19.8333 | 0.532 |
| 0 | 0 | ycaM | 4 | 19.8333 | 0.533 |
| 0 | 0 | yadI | 4 | 19.8333 | 0.694 |
| 0 | 0 | AG1 | 4 | 19.8333 | 0.823 |
| 0 | 0 | ptsI | 4 | 20.0833 | 0.566 |
| 0 | 0 | clcB | 4 | 20.0833 | 0.53 |
| 0 | 0 | ycaM | 4 | 20.0833 | 0.544 |
| 0 | 0 | yadI | 4 | 20.0833 | 0.707 |
| 0 | 0 | AG1 | 4 | 20.0833 | 0.824 |
| 0 | 0 | ptsI | 4 | 20.3333 | 0.572 |
| 0 | 0 | clcB | 4 | 20.3333 | 0.534 |
| 0 | 0 | ycaM | 4 | 20.3333 | 0.548 |
| 0 | 0 | yadI | 4 | 20.3333 | 0.706 |
| 0 | 0 | AG1 | 4 | 20.3333 | 0.828 |
| 0 | 0 | ptsI | 4 | 20.5833 | 0.565 |
| 0 | 0 | clcB | 4 | 20.5833 | 0.53 |
| 0 | 0 | ycaM | 4 | 20.5833 | 0.543 |
| 0 | 0 | yadI | 4 | 20.5833 | 0.712 |
| 0 | 0 | AG1 | 4 | 20.5833 | 0.828 |
| 0 | 0 | ptsI | 4 | 20.8333 | 0.568 |
| 0 | 0 | clcB | 4 | 20.8333 | 0.531 |
| 0 | 0 | ycaM | 4 | 20.8333 | 0.551 |
| 0 | 0 | yadI | 4 | 20.8333 | 0.713 |
| 0 | 0 | AG1 | 4 | 20.8333 | 0.839 |
| 0 | 0 | ptsI | 4 | 21.0833 | 0.581 |
| 0 | 0 | clcB | 4 | 21.0833 | 0.536 |
| 0 | 0 | ycaM | 4 | 21.0833 | 0.554 |
| 0 | 0 | yadI | 4 | 21.0833 | 0.711 |
| 0 | 0 | AG1 | 4 | 21.0833 | 0.836 |
| 0 | 0 | ptsI | 4 | 21.3333 | 0.573 |
| 0 | 0 | clcB | 4 | 21.3333 | 0.534 |
| 0 | 0 | ycaM | 4 | 21.3333 | 0.543 |
| 0 | 0 | yadI | 4 | 21.3333 | 0.718 |
| 0 | 0 | AG1 | 4 | 21.3333 | 0.84 |
| 0 | 0 | ptsI | 4 | 21.5833 | 0.582 |
| 0 | 0 | clcB | 4 | 21.5833 | 0.541 |
| 0 | 0 | ycaM | 4 | 21.5833 | 0.558 |
| 0 | 0 | yadI | 4 | 21.5833 | 0.712 |
| 0 | 0 | AG1 | 4 | 21.5833 | 0.842 |
| 0 | 0 | ptsI | 4 | 21.8333 | 0.571 |
| 0 | 0 | clcB | 4 | 21.8333 | 0.529 |
| 0 | 0 | ycaM | 4 | 21.8333 | 0.546 |
| 0 | 0 | yadI | 4 | 21.8333 | 0.708 |

| | | | | | |
|---|---|---|---|---|---|
| 0 | 0 | AG1 | 4 | 21.8333 | 0.837 |
| 0 | 0 | ptsI | 4 | 22.0833 | 0.582 |
| 0 | 0 | clcB | 4 | 22.0833 | 0.539 |
| 0 | 0 | ycaM | 4 | 22.0833 | 0.556 |
| 0 | 0 | yadI | 4 | 22.0833 | 0.717 |
| 0 | 0 | AG1 | 4 | 22.0833 | 0.841 |
| 0 | 0 | ptsI | 4 | 22.3333 | 0.594 |
| 0 | 0 | clcB | 4 | 22.3333 | 0.549 |
| 0 | 0 | ycaM | 4 | 22.3333 | 0.566 |
| 0 | 0 | yadI | 4 | 22.3333 | 0.713 |
| 0 | 0 | AG1 | 4 | 22.3333 | 0.842 |
| 0 | 0 | ptsI | 4 | 22.5833 | 0.579 |
| 0 | 0 | clcB | 4 | 22.5833 | 0.539 |
| 0 | 0 | ycaM | 4 | 22.5833 | 0.561 |
| 0 | 0 | yadI | 4 | 22.5833 | 0.72 |
| 0 | 0 | AG1 | 4 | 22.5833 | 0.838 |
| 0 | 0 | ptsI | 4 | 22.8333 | 0.584 |
| 0 | 0 | clcB | 4 | 22.8333 | 0.543 |
| 0 | 0 | ycaM | 4 | 22.8333 | 0.555 |
| 0 | 0 | yadI | 4 | 22.8333 | 0.719 |
| 0 | 0 | AG1 | 4 | 22.8333 | 0.84 |
| 0 | 0 | ptsI | 4 | 23.0833 | 0.594 |
| 0 | 0 | clcB | 4 | 23.0833 | 0.547 |
| 0 | 0 | ycaM | 4 | 23.0833 | 0.566 |
| 0 | 0 | yadI | 4 | 23.0833 | 0.722 |
| 0 | 0 | AG1 | 4 | 23.0833 | 0.843 |
| 0 | 0 | ptsI | 4 | 23.3333 | 0.583 |
| 0 | 0 | clcB | 4 | 23.3333 | 0.54 |
| 0 | 0 | ycaM | 4 | 23.3333 | 0.558 |
| 0 | 0 | yadI | 4 | 23.3333 | 0.728 |
| 0 | 0 | AG1 | 4 | 23.3333 | 0.848 |
| 0 | 0 | ptsI | 4 | 23.5833 | 0.589 |
| 0 | 0 | clcB | 4 | 23.5833 | 0.541 |
| 0 | 0 | ycaM | 4 | 23.5833 | 0.557 |
| 0 | 0 | yadI | 4 | 23.5833 | 0.73 |
| 0 | 0 | AG1 | 4 | 23.5833 | 0.846 |
| 0 | 0 | ptsI | 4 | 23.8333 | 0.593 |
| 0 | 0 | clcB | 4 | 23.8333 | 0.543 |
| 0 | 0 | ycaM | 4 | 23.8333 | 0.56 |
| 0 | 0 | yadI | 4 | 23.8333 | 0.733 |
| 0 | 0 | AG1 | 4 | 23.8333 | 0.844 |
| 0 | 0 | ptsI | 4 | 24.0833 | 0.596 |
| 0 | 0 | clcB | 4 | 24.0833 | 0.543 |
| 0 | 0 | ycaM | 4 | 24.0833 | 0.561 |
| 0 | 0 | yadI | 4 | 24.0833 | 0.734 |
| 0 | 0 | AG1 | 4 | 24.0833 | 0.848 |
| 0 | 0 | ptsI | 5 | 0 | 0.156 |
| 0 | 0 | clcB | 5 | 0 | 0.163 |

| | | | | | |
|---|---|---|---|---|---|
| 0 | 0 | ycaM | 5 | 0 | 0.158 |
| 0 | 0 | yadI | 5 | 0 | 0.181 |
| 0 | 0 | AG1 | 5 | 0 | 0.171 |
| 0 | 0 | ptsI | 5 | 0.35 | 0.154 |
| 0 | 0 | clcB | 5 | 0.35 | 0.158 |
| 0 | 0 | ycaM | 5 | 0.35 | 0.153 |
| 0 | 0 | yadI | 5 | 0.35 | 0.174 |
| 0 | 0 | AG1 | 5 | 0.35 | 0.168 |
| 0 | 0 | ptsI | 5 | 0.6 | 0.157 |
| 0 | 0 | clcB | 5 | 0.6 | 0.161 |
| 0 | 0 | ycaM | 5 | 0.6 | 0.152 |
| 0 | 0 | yadI | 5 | 0.6 | 0.173 |
| 0 | 0 | AG1 | 5 | 0.6 | 0.17 |
| 0 | 0 | ptsI | 5 | 0.85 | 0.152 |
| 0 | 0 | clcB | 5 | 0.85 | 0.16 |
| 0 | 0 | ycaM | 5 | 0.85 | 0.152 |
| 0 | 0 | yadI | 5 | 0.85 | 0.177 |
| 0 | 0 | AG1 | 5 | 0.85 | 0.17 |
| 0 | 0 | ptsI | 5 | 1.1 | 0.158 |
| 0 | 0 | clcB | 5 | 1.1 | 0.164 |
| 0 | 0 | ycaM | 5 | 1.1 | 0.155 |
| 0 | 0 | yadI | 5 | 1.1 | 0.178 |
| 0 | 0 | AG1 | 5 | 1.1 | 0.172 |
| 0 | 0 | ptsI | 5 | 1.35 | 0.157 |
| 0 | 0 | clcB | 5 | 1.35 | 0.165 |
| 0 | 0 | ycaM | 5 | 1.35 | 0.154 |
| 0 | 0 | yadI | 5 | 1.35 | 0.182 |
| 0 | 0 | AG1 | 5 | 1.35 | 0.175 |
| 0 | 0 | ptsI | 5 | 1.6 | 0.164 |
| 0 | 0 | clcB | 5 | 1.6 | 0.17 |
| 0 | 0 | ycaM | 5 | 1.6 | 0.159 |
| 0 | 0 | yadI | 5 | 1.6 | 0.185 |
| 0 | 0 | AG1 | 5 | 1.6 | 0.18 |
| 0 | 0 | ptsI | 5 | 1.85 | 0.164 |
| 0 | 0 | clcB | 5 | 1.85 | 0.172 |
| 0 | 0 | ycaM | 5 | 1.85 | 0.161 |
| 0 | 0 | yadI | 5 | 1.85 | 0.191 |
| 0 | 0 | AG1 | 5 | 1.85 | 0.185 |
| 0 | 0 | ptsI | 5 | 2.1 | 0.168 |
| 0 | 0 | clcB | 5 | 2.1 | 0.173 |
| 0 | 0 | ycaM | 5 | 2.1 | 0.165 |
| 0 | 0 | yadI | 5 | 2.1 | 0.196 |
| 0 | 0 | AG1 | 5 | 2.1 | 0.188 |
| 0 | 0 | ptsI | 5 | 2.35 | 0.176 |
| 0 | 0 | clcB | 5 | 2.35 | 0.183 |
| 0 | 0 | ycaM | 5 | 2.35 | 0.172 |
| 0 | 0 | yadI | 5 | 2.35 | 0.203 |
| 0 | 0 | AG1 | 5 | 2.35 | 0.201 |

| | | | | | |
|---|---|---|---|---|---|
| 0 | 0 | ptsI | 5 | 2.6 | 0.174 |
| 0 | 0 | clcB | 5 | 2.6 | 0.184 |
| 0 | 0 | ycaM | 5 | 2.6 | 0.176 |
| 0 | 0 | yadI | 5 | 2.6 | 0.212 |
| 0 | 0 | AG1 | 5 | 2.6 | 0.203 |
| 0 | 0 | ptsI | 5 | 2.85 | 0.185 |
| 0 | 0 | clcB | 5 | 2.85 | 0.197 |
| 0 | 0 | ycaM | 5 | 2.85 | 0.184 |
| 0 | 0 | yadI | 5 | 2.85 | 0.219 |
| 0 | 0 | AG1 | 5 | 2.85 | 0.215 |
| 0 | 0 | ptsI | 5 | 3.1 | 0.194 |
| 0 | 0 | clcB | 5 | 3.1 | 0.21 |
| 0 | 0 | ycaM | 5 | 3.1 | 0.191 |
| 0 | 0 | yadI | 5 | 3.1 | 0.228 |
| 0 | 0 | AG1 | 5 | 3.1 | 0.23 |
| 0 | 0 | ptsI | 5 | 3.35 | 0.202 |
| 0 | 0 | clcB | 5 | 3.35 | 0.225 |
| 0 | 0 | ycaM | 5 | 3.35 | 0.201 |
| 0 | 0 | yadI | 5 | 3.35 | 0.241 |
| 0 | 0 | AG1 | 5 | 3.35 | 0.251 |
| 0 | 0 | ptsI | 5 | 3.6 | 0.21 |
| 0 | 0 | clcB | 5 | 3.6 | 0.24 |
| 0 | 0 | ycaM | 5 | 3.6 | 0.209 |
| 0 | 0 | yadI | 5 | 3.6 | 0.259 |
| 0 | 0 | AG1 | 5 | 3.6 | 0.266 |
| 0 | 0 | ptsI | 5 | 3.85 | 0.22 |
| 0 | 0 | clcB | 5 | 3.85 | 0.253 |
| 0 | 0 | ycaM | 5 | 3.85 | 0.223 |
| 0 | 0 | yadI | 5 | 3.85 | 0.276 |
| 0 | 0 | AG1 | 5 | 3.85 | 0.283 |
| 0 | 0 | ptsI | 5 | 4.38333 | 0.214 |
| 0 | 0 | clcB | 5 | 4.38333 | 0.245 |
| 0 | 0 | ycaM | 5 | 4.38333 | 0.242 |
| 0 | 0 | yadI | 5 | 4.38333 | 0.288 |
| 0 | 0 | AG1 | 5 | 4.38333 | 0.289 |
| 0 | 0 | ptsI | 5 | 4.63333 | 0.224 |
| 0 | 0 | clcB | 5 | 4.63333 | 0.249 |
| 0 | 0 | ycaM | 5 | 4.63333 | 0.239 |
| 0 | 0 | yadI | 5 | 4.63333 | 0.281 |
| 0 | 0 | AG1 | 5 | 4.63333 | 0.294 |
| 0 | 0 | ptsI | 5 | 4.88333 | 0.232 |
| 0 | 0 | clcB | 5 | 4.88333 | 0.261 |
| 0 | 0 | ycaM | 5 | 4.88333 | 0.252 |
| 0 | 0 | yadI | 5 | 4.88333 | 0.298 |
| 0 | 0 | AG1 | 5 | 4.88333 | 0.3 |
| 0 | 0 | ptsI | 5 | 5.13333 | 0.24 |
| 0 | 0 | clcB | 5 | 5.13333 | 0.273 |
| 0 | 0 | ycaM | 5 | 5.13333 | 0.264 |

| | | | | | |
|---|---|---|---|---|---|
| 0 | 0 | yadI | 5 | 5.13333 | 0.311 |
| 0 | 0 | AG1 | 5 | 5.13333 | 0.312 |
| 0 | 0 | ptsI | 5 | 5.38333 | 0.248 |
| 0 | 0 | clcB | 5 | 5.38333 | 0.286 |
| 0 | 0 | ycaM | 5 | 5.38333 | 0.279 |
| 0 | 0 | yadI | 5 | 5.38333 | 0.322 |
| 0 | 0 | AG1 | 5 | 5.38333 | 0.324 |
| 0 | 0 | ptsI | 5 | 5.63333 | 0.26 |
| 0 | 0 | clcB | 5 | 5.63333 | 0.301 |
| 0 | 0 | ycaM | 5 | 5.63333 | 0.29 |
| 0 | 0 | yadI | 5 | 5.63333 | 0.338 |
| 0 | 0 | AG1 | 5 | 5.63333 | 0.336 |
| 0 | 0 | ptsI | 5 | 5.88333 | 0.271 |
| 0 | 0 | clcB | 5 | 5.88333 | 0.31 |
| 0 | 0 | ycaM | 5 | 5.88333 | 0.3 |
| 0 | 0 | yadI | 5 | 5.88333 | 0.355 |
| 0 | 0 | AG1 | 5 | 5.88333 | 0.35 |
| 0 | 0 | ptsI | 5 | 6.13333 | 0.29 |
| 0 | 0 | clcB | 5 | 6.13333 | 0.33 |
| 0 | 0 | ycaM | 5 | 6.13333 | 0.317 |
| 0 | 0 | yadI | 5 | 6.13333 | 0.38 |
| 0 | 0 | AG1 | 5 | 6.13333 | 0.375 |
| 0 | 0 | ptsI | 5 | 6.38333 | 0.293 |
| 0 | 0 | clcB | 5 | 6.38333 | 0.336 |
| 0 | 0 | ycaM | 5 | 6.38333 | 0.326 |
| 0 | 0 | yadI | 5 | 6.38333 | 0.401 |
| 0 | 0 | AG1 | 5 | 6.38333 | 0.405 |
| 0 | 0 | ptsI | 5 | 6.63333 | 0.305 |
| 0 | 0 | clcB | 5 | 6.63333 | 0.347 |
| 0 | 0 | ycaM | 5 | 6.63333 | 0.34 |
| 0 | 0 | yadI | 5 | 6.63333 | 0.421 |
| 0 | 0 | AG1 | 5 | 6.63333 | 0.43 |
| 0 | 0 | ptsI | 5 | 6.88333 | 0.316 |
| 0 | 0 | clcB | 5 | 6.88333 | 0.359 |
| 0 | 0 | ycaM | 5 | 6.88333 | 0.356 |
| 0 | 0 | yadI | 5 | 6.88333 | 0.439 |
| 0 | 0 | AG1 | 5 | 6.88333 | 0.454 |
| 0 | 0 | ptsI | 5 | 7.13333 | 0.326 |
| 0 | 0 | clcB | 5 | 7.13333 | 0.37 |
| 0 | 0 | ycaM | 5 | 7.13333 | 0.373 |
| 0 | 0 | yadI | 5 | 7.13333 | 0.46 |
| 0 | 0 | AG1 | 5 | 7.13333 | 0.471 |
| 0 | 0 | ptsI | 5 | 7.38333 | 0.342 |
| 0 | 0 | clcB | 5 | 7.38333 | 0.387 |
| 0 | 0 | ycaM | 5 | 7.38333 | 0.394 |
| 0 | 0 | yadI | 5 | 7.38333 | 0.486 |
| 0 | 0 | AG1 | 5 | 7.38333 | 0.496 |
| 0 | 0 | ptsI | 5 | 7.63333 | 0.353 |

| | | | | | |
|---|---|---|---|---|---|
| 0 | 0 | clcB | 5 | 7.63333 | 0.396 |
| 0 | 0 | ycaM | 5 | 7.63333 | 0.404 |
| 0 | 0 | yadI | 5 | 7.63333 | 0.501 |
| 0 | 0 | AG1 | 5 | 7.63333 | 0.52 |
| 0 | 0 | ptsI | 5 | 7.88333 | 0.364 |
| 0 | 0 | clcB | 5 | 7.88333 | 0.407 |
| 0 | 0 | ycaM | 5 | 7.88333 | 0.416 |
| 0 | 0 | yadI | 5 | 7.88333 | 0.511 |
| 0 | 0 | AG1 | 5 | 7.88333 | 0.531 |
| 0 | 0 | ptsI | 5 | 8.13333 | 0.378 |
| 0 | 0 | clcB | 5 | 8.13333 | 0.414 |
| 0 | 0 | ycaM | 5 | 8.13333 | 0.427 |
| 0 | 0 | yadI | 5 | 8.13333 | 0.528 |
| 0 | 0 | AG1 | 5 | 8.13333 | 0.552 |
| 0 | 0 | ptsI | 5 | 8.38333 | 0.39 |
| 0 | 0 | clcB | 5 | 8.38333 | 0.428 |
| 0 | 0 | ycaM | 5 | 8.38333 | 0.444 |
| 0 | 0 | yadI | 5 | 8.38333 | 0.537 |
| 0 | 0 | AG1 | 5 | 8.38333 | 0.564 |
| 0 | 0 | ptsI | 5 | 8.88333 | 0.318 |
| 0 | 0 | clcB | 5 | 8.88333 | 0.361 |
| 0 | 0 | ycaM | 5 | 8.88333 | 0.382 |
| 0 | 0 | yadI | 5 | 8.88333 | 0.433 |
| 0 | 0 | AG1 | 5 | 8.88333 | 0.402 |
| 0 | 0 | ptsI | 5 | 9.13333 | 0.319 |
| 0 | 0 | clcB | 5 | 9.13333 | 0.364 |
| 0 | 0 | ycaM | 5 | 9.13333 | 0.382 |
| 0 | 0 | yadI | 5 | 9.13333 | 0.436 |
| 0 | 0 | AG1 | 5 | 9.13333 | 0.404 |
| 0 | 0 | ptsI | 5 | 9.38333 | 0.325 |
| 0 | 0 | clcB | 5 | 9.38333 | 0.369 |
| 0 | 0 | ycaM | 5 | 9.38333 | 0.389 |
| 0 | 0 | yadI | 5 | 9.38333 | 0.443 |
| 0 | 0 | AG1 | 5 | 9.38333 | 0.407 |
| 0 | 0 | ptsI | 5 | 9.63333 | 0.339 |
| 0 | 0 | clcB | 5 | 9.63333 | 0.381 |
| 0 | 0 | ycaM | 5 | 9.63333 | 0.4 |
| 0 | 0 | yadI | 5 | 9.63333 | 0.458 |
| 0 | 0 | AG1 | 5 | 9.63333 | 0.424 |
| 0 | 0 | ptsI | 5 | 9.88333 | 0.354 |
| 0 | 0 | clcB | 5 | 9.88333 | 0.394 |
| 0 | 0 | ycaM | 5 | 9.88333 | 0.413 |
| 0 | 0 | yadI | 5 | 9.88333 | 0.474 |
| 0 | 0 | AG1 | 5 | 9.88333 | 0.435 |
| 0 | 0 | ptsI | 5 | 10.1333 | 0.362 |
| 0 | 0 | clcB | 5 | 10.1333 | 0.402 |
| 0 | 0 | ycaM | 5 | 10.1333 | 0.416 |
| 0 | 0 | yadI | 5 | 10.1333 | 0.472 |

| | | | | | |
|---|---|---|---|---|---|
| 0 | 0 | AG1 | 5 | 10.1333 | 0.443 |
| 0 | 0 | ptsI | 5 | 10.3833 | 0.369 |
| 0 | 0 | clcB | 5 | 10.3833 | 0.408 |
| 0 | 0 | ycaM | 5 | 10.3833 | 0.419 |
| 0 | 0 | yadI | 5 | 10.3833 | 0.482 |
| 0 | 0 | AG1 | 5 | 10.3833 | 0.449 |
| 0 | 0 | ptsI | 5 | 10.6333 | 0.384 |
| 0 | 0 | clcB | 5 | 10.6333 | 0.414 |
| 0 | 0 | ycaM | 5 | 10.6333 | 0.423 |
| 0 | 0 | yadI | 5 | 10.6333 | 0.487 |
| 0 | 0 | AG1 | 5 | 10.6333 | 0.457 |
| 0 | 0 | ptsI | 5 | 10.8833 | 0.397 |
| 0 | 0 | clcB | 5 | 10.8833 | 0.427 |
| 0 | 0 | ycaM | 5 | 10.8833 | 0.434 |
| 0 | 0 | yadI | 5 | 10.8833 | 0.503 |
| 0 | 0 | AG1 | 5 | 10.8833 | 0.47 |
| 0 | 0 | ptsI | 5 | 11.1333 | 0.402 |
| 0 | 0 | clcB | 5 | 11.1333 | 0.425 |
| 0 | 0 | ycaM | 5 | 11.1333 | 0.431 |
| 0 | 0 | yadI | 5 | 11.1333 | 0.505 |
| 0 | 0 | AG1 | 5 | 11.1333 | 0.473 |
| 0 | 0 | ptsI | 5 | 11.3833 | 0.411 |
| 0 | 0 | clcB | 5 | 11.3833 | 0.43 |
| 0 | 0 | ycaM | 5 | 11.3833 | 0.435 |
| 0 | 0 | yadI | 5 | 11.3833 | 0.516 |
| 0 | 0 | AG1 | 5 | 11.3833 | 0.486 |
| 0 | 0 | ptsI | 5 | 11.6333 | 0.419 |
| 0 | 0 | clcB | 5 | 11.6333 | 0.438 |
| 0 | 0 | ycaM | 5 | 11.6333 | 0.444 |
| 0 | 0 | yadI | 5 | 11.6333 | 0.52 |
| 0 | 0 | AG1 | 5 | 11.6333 | 0.49 |
| 0 | 0 | ptsI | 5 | 11.8833 | 0.426 |
| 0 | 0 | clcB | 5 | 11.8833 | 0.441 |
| 0 | 0 | ycaM | 5 | 11.8833 | 0.445 |
| 0 | 0 | yadI | 5 | 11.8833 | 0.532 |
| 0 | 0 | AG1 | 5 | 11.8833 | 0.496 |
| 0 | 0 | ptsI | 5 | 12.1333 | 0.444 |
| 0 | 0 | clcB | 5 | 12.1333 | 0.456 |
| 0 | 0 | ycaM | 5 | 12.1333 | 0.458 |
| 0 | 0 | yadI | 5 | 12.1333 | 0.543 |
| 0 | 0 | AG1 | 5 | 12.1333 | 0.511 |
| 0 | 0 | ptsI | 5 | 12.3833 | 0.436 |
| 0 | 0 | clcB | 5 | 12.3833 | 0.449 |
| 0 | 0 | ycaM | 5 | 12.3833 | 0.451 |
| 0 | 0 | yadI | 5 | 12.3833 | 0.538 |
| 0 | 0 | AG1 | 5 | 12.3833 | 0.504 |
| 0 | 0 | ptsI | 5 | 12.6333 | 0.451 |
| 0 | 0 | clcB | 5 | 12.6333 | 0.461 |

| | | | | | |
|---|---|---|---|---|---|
| 0 | 0 | ycaM | 5 | 12.6333 | 0.462 |
| 0 | 0 | yadI | 5 | 12.6333 | 0.545 |
| 0 | 0 | AG1 | 5 | 12.6333 | 0.518 |
| 0 | 0 | ptsI | 5 | 12.8833 | 0.464 |
| 0 | 0 | clcB | 5 | 12.8833 | 0.476 |
| 0 | 0 | ycaM | 5 | 12.8833 | 0.474 |
| 0 | 0 | yadI | 5 | 12.8833 | 0.561 |
| 0 | 0 | AG1 | 5 | 12.8833 | 0.538 |
| 0 | 0 | ptsI | 5 | 13.1333 | 0.463 |
| 0 | 0 | clcB | 5 | 13.1333 | 0.469 |
| 0 | 0 | ycaM | 5 | 13.1333 | 0.467 |
| 0 | 0 | yadI | 5 | 13.1333 | 0.557 |
| 0 | 0 | AG1 | 5 | 13.1333 | 0.531 |
| 0 | 0 | ptsI | 5 | 13.3833 | 0.475 |
| 0 | 0 | clcB | 5 | 13.3833 | 0.478 |
| 0 | 0 | ycaM | 5 | 13.3833 | 0.472 |
| 0 | 0 | yadI | 5 | 13.3833 | 0.561 |
| 0 | 0 | AG1 | 5 | 13.3833 | 0.536 |
| 0 | 0 | ptsI | 5 | 13.6333 | 0.48 |
| 0 | 0 | clcB | 5 | 13.6333 | 0.479 |
| 0 | 0 | ycaM | 5 | 13.6333 | 0.48 |
| 0 | 0 | yadI | 5 | 13.6333 | 0.574 |
| 0 | 0 | AG1 | 5 | 13.6333 | 0.541 |
| 0 | 0 | ptsI | 5 | 13.8833 | 0.484 |
| 0 | 0 | clcB | 5 | 13.8833 | 0.478 |
| 0 | 0 | ycaM | 5 | 13.8833 | 0.474 |
| 0 | 0 | yadI | 5 | 13.8833 | 0.573 |
| 0 | 0 | AG1 | 5 | 13.8833 | 0.543 |
| 0 | 0 | ptsI | 5 | 14.1333 | 0.497 |
| 0 | 0 | clcB | 5 | 14.1333 | 0.485 |
| 0 | 0 | ycaM | 5 | 14.1333 | 0.484 |
| 0 | 0 | yadI | 5 | 14.1333 | 0.582 |
| 0 | 0 | AG1 | 5 | 14.1333 | 0.555 |
| 0 | 0 | ptsI | 5 | 14.3833 | 0.502 |
| 0 | 0 | clcB | 5 | 14.3833 | 0.508 |
| 0 | 0 | ycaM | 5 | 14.3833 | 0.502 |
| 0 | 0 | yadI | 5 | 14.3833 | 0.597 |
| 0 | 0 | AG1 | 5 | 14.3833 | 0.566 |
| 0 | 0 | ptsI | 5 | 14.6333 | 0.508 |
| 0 | 0 | clcB | 5 | 14.6333 | 0.505 |
| 0 | 0 | ycaM | 5 | 14.6333 | 0.501 |
| 0 | 0 | yadI | 5 | 14.6333 | 0.599 |
| 0 | 0 | AG1 | 5 | 14.6333 | 0.568 |
| 0 | 0 | ptsI | 5 | 14.8833 | 0.522 |
| 0 | 0 | clcB | 5 | 14.8833 | 0.508 |
| 0 | 0 | ycaM | 5 | 14.8833 | 0.501 |
| 0 | 0 | yadI | 5 | 14.8833 | 0.6 |
| 0 | 0 | AG1 | 5 | 14.8833 | 0.571 |

| | | | | | |
|---|---|---|---|---|---|
| 0 | 0 | ptsI | 5 | 15.1333 | 0.541 |
| 0 | 0 | clcB | 5 | 15.1333 | 0.514 |
| 0 | 0 | ycaM | 5 | 15.1333 | 0.507 |
| 0 | 0 | yadI | 5 | 15.1333 | 0.609 |
| 0 | 0 | AG1 | 5 | 15.1333 | 0.577 |
| 0 | 0 | ptsI | 5 | 15.3833 | 0.534 |
| 0 | 0 | clcB | 5 | 15.3833 | 0.516 |
| 0 | 0 | ycaM | 5 | 15.3833 | 0.515 |
| 0 | 0 | yadI | 5 | 15.3833 | 0.613 |
| 0 | 0 | AG1 | 5 | 15.3833 | 0.579 |
| 0 | 0 | ptsI | 5 | 15.6333 | 0.543 |
| 0 | 0 | clcB | 5 | 15.6333 | 0.513 |
| 0 | 0 | ycaM | 5 | 15.6333 | 0.513 |
| 0 | 0 | yadI | 5 | 15.6333 | 0.611 |
| 0 | 0 | AG1 | 5 | 15.6333 | 0.577 |
| 0 | 0 | ptsI | 5 | 15.8833 | 0.548 |
| 0 | 0 | clcB | 5 | 15.8833 | 0.513 |
| 0 | 0 | ycaM | 5 | 15.8833 | 0.517 |
| 0 | 0 | yadI | 5 | 15.8833 | 0.615 |
| 0 | 0 | AG1 | 5 | 15.8833 | 0.577 |
| 0 | 0 | ptsI | 5 | 16.1333 | 0.545 |
| 0 | 0 | clcB | 5 | 16.1333 | 0.511 |
| 0 | 0 | ycaM | 5 | 16.1333 | 0.519 |
| 0 | 0 | yadI | 5 | 16.1333 | 0.611 |
| 0 | 0 | AG1 | 5 | 16.1333 | 0.577 |
| 0 | 0 | ptsI | 5 | 16.3833 | 0.542 |
| 0 | 0 | clcB | 5 | 16.3833 | 0.508 |
| 0 | 0 | ycaM | 5 | 16.3833 | 0.517 |
| 0 | 0 | yadI | 5 | 16.3833 | 0.612 |
| 0 | 0 | AG1 | 5 | 16.3833 | 0.581 |
| 0 | 0 | ptsI | 5 | 16.6333 | 0.551 |
| 0 | 0 | clcB | 5 | 16.6333 | 0.524 |
| 0 | 0 | ycaM | 5 | 16.6333 | 0.523 |
| 0 | 0 | yadI | 5 | 16.6333 | 0.62 |
| 0 | 0 | AG1 | 5 | 16.6333 | 0.591 |
| 0 | 0 | ptsI | 5 | 16.8833 | 0.561 |
| 0 | 0 | clcB | 5 | 16.8833 | 0.535 |
| 0 | 0 | ycaM | 5 | 16.8833 | 0.535 |
| 0 | 0 | yadI | 5 | 16.8833 | 0.635 |
| 0 | 0 | AG1 | 5 | 16.8833 | 0.601 |
| 0 | 0 | ptsI | 5 | 17.1333 | 0.565 |
| 0 | 0 | clcB | 5 | 17.1333 | 0.526 |
| 0 | 0 | ycaM | 5 | 17.1333 | 0.537 |
| 0 | 0 | yadI | 5 | 17.1333 | 0.63 |
| 0 | 0 | AG1 | 5 | 17.1333 | 0.595 |
| 0 | 0 | ptsI | 5 | 17.3833 | 0.565 |
| 0 | 0 | clcB | 5 | 17.3833 | 0.525 |
| 0 | 0 | ycaM | 5 | 17.3833 | 0.524 |

| | | | | | |
|---|---|---|---|---|---|
| 0 | 0 | yadI | 5 | 17.3833 | 0.627 |
| 0 | 0 | AG1 | 5 | 17.3833 | 0.594 |
| 0 | 0 | ptsI | 5 | 17.6333 | 0.566 |
| 0 | 0 | clcB | 5 | 17.6333 | 0.525 |
| 0 | 0 | ycaM | 5 | 17.6333 | 0.536 |
| 0 | 0 | yadI | 5 | 17.6333 | 0.632 |
| 0 | 0 | AG1 | 5 | 17.6333 | 0.594 |
| 0 | 0 | ptsI | 5 | 17.8833 | 0.572 |
| 0 | 0 | clcB | 5 | 17.8833 | 0.525 |
| 0 | 0 | ycaM | 5 | 17.8833 | 0.544 |
| 0 | 0 | yadI | 5 | 17.8833 | 0.634 |
| 0 | 0 | AG1 | 5 | 17.8833 | 0.594 |
| 0 | 0 | ptsI | 5 | 18.1333 | 0.576 |
| 0 | 0 | clcB | 5 | 18.1333 | 0.539 |
| 0 | 0 | ycaM | 5 | 18.1333 | 0.543 |
| 0 | 0 | yadI | 5 | 18.1333 | 0.646 |
| 0 | 0 | AG1 | 5 | 18.1333 | 0.608 |
| 0 | 0 | ptsI | 5 | 18.3833 | 0.589 |
| 0 | 0 | clcB | 5 | 18.3833 | 0.527 |
| 0 | 0 | ycaM | 5 | 18.3833 | 0.545 |
| 0 | 0 | yadI | 5 | 18.3833 | 0.637 |
| 0 | 0 | AG1 | 5 | 18.3833 | 0.601 |
| 0 | 0 | ptsI | 5 | 18.6333 | 0.582 |
| 0 | 0 | clcB | 5 | 18.6333 | 0.527 |
| 0 | 0 | ycaM | 5 | 18.6333 | 0.55 |
| 0 | 0 | yadI | 5 | 18.6333 | 0.636 |
| 0 | 0 | AG1 | 5 | 18.6333 | 0.603 |
| 0 | 0 | ptsI | 5 | 18.8833 | 0.583 |
| 0 | 0 | clcB | 5 | 18.8833 | 0.534 |
| 0 | 0 | ycaM | 5 | 18.8833 | 0.546 |
| 0 | 0 | yadI | 5 | 18.8833 | 0.647 |
| 0 | 0 | AG1 | 5 | 18.8833 | 0.609 |
| 0 | 0 | ptsI | 5 | 19.1333 | 0.595 |
| 0 | 0 | clcB | 5 | 19.1333 | 0.532 |
| 0 | 0 | ycaM | 5 | 19.1333 | 0.554 |
| 0 | 0 | yadI | 5 | 19.1333 | 0.645 |
| 0 | 0 | AG1 | 5 | 19.1333 | 0.608 |
| 0 | 0 | ptsI | 5 | 19.3833 | 0.596 |
| 0 | 0 | clcB | 5 | 19.3833 | 0.534 |
| 0 | 0 | ycaM | 5 | 19.3833 | 0.559 |
| 0 | 0 | yadI | 5 | 19.3833 | 0.651 |
| 0 | 0 | AG1 | 5 | 19.3833 | 0.61 |
| 0 | 0 | ptsI | 5 | 19.6333 | 0.595 |
| 0 | 0 | clcB | 5 | 19.6333 | 0.539 |
| 0 | 0 | ycaM | 5 | 19.6333 | 0.554 |
| 0 | 0 | yadI | 5 | 19.6333 | 0.654 |
| 0 | 0 | AG1 | 5 | 19.6333 | 0.616 |
| 0 | 0 | ptsI | 5 | 19.8833 | 0.596 |

| | | | | | |
|---|---|---|---|---|---|
| 0 | 0 | clcB | 5 | 19.8833 | 0.536 |
| 0 | 0 | ycaM | 5 | 19.8833 | 0.557 |
| 0 | 0 | yadI | 5 | 19.8833 | 0.648 |
| 0 | 0 | AG1 | 5 | 19.8833 | 0.61 |
| 0 | 0 | ptsI | 5 | 20.1333 | 0.608 |
| 0 | 0 | clcB | 5 | 20.1333 | 0.536 |
| 0 | 0 | ycaM | 5 | 20.1333 | 0.566 |
| 0 | 0 | yadI | 5 | 20.1333 | 0.65 |
| 0 | 0 | AG1 | 5 | 20.1333 | 0.612 |
| 0 | 0 | ptsI | 5 | 20.3833 | 0.605 |
| 0 | 0 | clcB | 5 | 20.3833 | 0.538 |
| 0 | 0 | ycaM | 5 | 20.3833 | 0.572 |
| 0 | 0 | yadI | 5 | 20.3833 | 0.65 |
| 0 | 0 | AG1 | 5 | 20.3833 | 0.613 |
| 0 | 0 | ptsI | 5 | 20.6333 | 0.604 |
| 0 | 0 | clcB | 5 | 20.6333 | 0.536 |
| 0 | 0 | ycaM | 5 | 20.6333 | 0.565 |
| 0 | 0 | yadI | 5 | 20.6333 | 0.654 |
| 0 | 0 | AG1 | 5 | 20.6333 | 0.618 |
| 0 | 0 | ptsI | 5 | 20.8833 | 0.614 |
| 0 | 0 | clcB | 5 | 20.8833 | 0.538 |
| 0 | 0 | ycaM | 5 | 20.8833 | 0.572 |
| 0 | 0 | yadI | 5 | 20.8833 | 0.656 |
| 0 | 0 | AG1 | 5 | 20.8833 | 0.619 |
| 0 | 0 | ptsI | 5 | 21.1333 | 0.613 |
| 0 | 0 | clcB | 5 | 21.1333 | 0.539 |
| 0 | 0 | ycaM | 5 | 21.1333 | 0.571 |
| 0 | 0 | yadI | 5 | 21.1333 | 0.656 |
| 0 | 0 | AG1 | 5 | 21.1333 | 0.62 |
| 0 | 0 | ptsI | 5 | 21.3833 | 0.623 |
| 0 | 0 | clcB | 5 | 21.3833 | 0.544 |
| 0 | 0 | ycaM | 5 | 21.3833 | 0.575 |
| 0 | 0 | yadI | 5 | 21.3833 | 0.666 |
| 0 | 0 | AG1 | 5 | 21.3833 | 0.626 |
| 0 | 0 | ptsI | 5 | 21.6333 | 0.613 |
| 0 | 0 | clcB | 5 | 21.6333 | 0.553 |
| 0 | 0 | ycaM | 5 | 21.6333 | 0.568 |
| 0 | 0 | yadI | 5 | 21.6333 | 0.673 |
| 0 | 0 | AG1 | 5 | 21.6333 | 0.635 |
| 0 | 0 | ptsI | 5 | 21.8833 | 0.632 |
| 0 | 0 | clcB | 5 | 21.8833 | 0.556 |
| 0 | 0 | ycaM | 5 | 21.8833 | 0.583 |
| 0 | 0 | yadI | 5 | 21.8833 | 0.674 |
| 0 | 0 | AG1 | 5 | 21.8833 | 0.637 |
| 0 | 0 | ptsI | 5 | 22.1333 | 0.623 |
| 0 | 0 | clcB | 5 | 22.1333 | 0.543 |
| 0 | 0 | ycaM | 5 | 22.1333 | 0.579 |
| 0 | 0 | yadI | 5 | 22.1333 | 0.665 |

| | | | | | |
|---|---|---|---|---|---|
| 0 | 0 | AG1 | 5 | 22.1333 | 0.621 |
| 0 | 0 | ptsI | 5 | 22.3833 | 0.628 |
| 0 | 0 | clcB | 5 | 22.3833 | 0.546 |
| 0 | 0 | ycaM | 5 | 22.3833 | 0.58 |
| 0 | 0 | yadI | 5 | 22.3833 | 0.664 |
| 0 | 0 | AG1 | 5 | 22.3833 | 0.623 |
| 0 | 0 | ptsI | 5 | 22.6333 | 0.623 |
| 0 | 0 | clcB | 5 | 22.6333 | 0.555 |
| 0 | 0 | ycaM | 5 | 22.6333 | 0.578 |
| 0 | 0 | yadI | 5 | 22.6333 | 0.681 |
| 0 | 0 | AG1 | 5 | 22.6333 | 0.639 |
| 0 | 0 | ptsI | 5 | 22.8833 | 0.626 |
| 0 | 0 | clcB | 5 | 22.8833 | 0.555 |
| 0 | 0 | ycaM | 5 | 22.8833 | 0.578 |
| 0 | 0 | yadI | 5 | 22.8833 | 0.678 |
| 0 | 0 | AG1 | 5 | 22.8833 | 0.635 |
| 0 | 0 | ptsI | 5 | 23.1333 | 0.626 |
| 0 | 0 | clcB | 5 | 23.1333 | 0.558 |
| 0 | 0 | ycaM | 5 | 23.1333 | 0.582 |
| 0 | 0 | yadI | 5 | 23.1333 | 0.676 |
| 0 | 0 | AG1 | 5 | 23.1333 | 0.636 |
| 0 | 0 | ptsI | 5 | 23.3833 | 0.633 |
| 0 | 0 | clcB | 5 | 23.3833 | 0.548 |
| 0 | 0 | ycaM | 5 | 23.3833 | 0.589 |
| 0 | 0 | yadI | 5 | 23.3833 | 0.685 |
| 0 | 0 | AG1 | 5 | 23.3833 | 0.634 |
| 0 | 0 | ptsI | 5 | 23.6333 | 0.631 |
| 0 | 0 | clcB | 5 | 23.6333 | 0.549 |
| 0 | 0 | ycaM | 5 | 23.6333 | 0.59 |
| 0 | 0 | yadI | 5 | 23.6333 | 0.686 |
| 0 | 0 | AG1 | 5 | 23.6333 | 0.633 |
| 0 | 0 | ptsI | 5 | 23.8833 | 0.632 |
| 0 | 0 | clcB | 5 | 23.8833 | 0.55 |
| 0 | 0 | ycaM | 5 | 23.8833 | 0.589 |
| 0 | 0 | yadI | 5 | 23.8833 | 0.686 |
| 0 | 0 | AG1 | 5 | 23.8833 | 0.633 |
| 0 | 0 | ptsI | 5 | 24.1333 | 0.647 |
| 0 | 0 | clcB | 5 | 24.1333 | 0.554 |
| 0 | 0 | ycaM | 5 | 24.1333 | 0.593 |
| 0 | 0 | yadI | 5 | 24.1333 | 0.689 |
| 0 | 0 | AG1 | 5 | 24.1333 | 0.644 |
| 0 | 0 | ptsI | 5 | 24.3833 | 0.634 |
| 0 | 0 | clcB | 5 | 24.3833 | 0.546 |
| 0 | 0 | ycaM | 5 | 24.3833 | 0.587 |
| 0 | 0 | yadI | 5 | 24.3833 | 0.686 |
| 0 | 0 | AG1 | 5 | 24.3833 | 0.63 |
| 0.05 | 0 | ptsI | 1 | 0 | 0.234 |
| 0.05 | 0 | clcB | 1 | 0 | 0.237 |

| | | | | | |
|---|---|---|---|---|---|
| 0.05 | 0 | ycaM | 1 | 0 | 0.223 |
| 0.05 | 0 | yadI | 1 | 0 | 0.253 |
| 0.05 | 0 | AG1 | 1 | 0 | 0.255 |
| 0.05 | 0 | ptsI | 1 | 0.25 | 0.226 |
| 0.05 | 0 | clcB | 1 | 0.25 | 0.226 |
| 0.05 | 0 | ycaM | 1 | 0.25 | 0.214 |
| 0.05 | 0 | yadI | 1 | 0.25 | 0.245 |
| 0.05 | 0 | AG1 | 1 | 0.25 | 0.251 |
| 0.05 | 0 | ptsI | 1 | 0.5 | 0.223 |
| 0.05 | 0 | clcB | 1 | 0.5 | 0.223 |
| 0.05 | 0 | ycaM | 1 | 0.5 | 0.211 |
| 0.05 | 0 | yadI | 1 | 0.5 | 0.242 |
| 0.05 | 0 | AG1 | 1 | 0.5 | 0.246 |
| 0.05 | 0 | ptsI | 1 | 0.75 | 0.223 |
| 0.05 | 0 | clcB | 1 | 0.75 | 0.221 |
| 0.05 | 0 | ycaM | 1 | 0.75 | 0.21 |
| 0.05 | 0 | yadI | 1 | 0.75 | 0.238 |
| 0.05 | 0 | AG1 | 1 | 0.75 | 0.246 |
| 0.05 | 0 | ptsI | 1 | 1 | 0.23 |
| 0.05 | 0 | clcB | 1 | 1 | 0.222 |
| 0.05 | 0 | ycaM | 1 | 1 | 0.213 |
| 0.05 | 0 | yadI | 1 | 1 | 0.242 |
| 0.05 | 0 | AG1 | 1 | 1 | 0.245 |
| 0.05 | 0 | ptsI | 1 | 1.25 | 0.232 |
| 0.05 | 0 | clcB | 1 | 1.25 | 0.218 |
| 0.05 | 0 | ycaM | 1 | 1.25 | 0.211 |
| 0.05 | 0 | yadI | 1 | 1.25 | 0.245 |
| 0.05 | 0 | AG1 | 1 | 1.25 | 0.247 |
| 0.05 | 0 | ptsI | 1 | 1.5 | 0.238 |
| 0.05 | 0 | clcB | 1 | 1.5 | 0.221 |
| 0.05 | 0 | ycaM | 1 | 1.5 | 0.217 |
| 0.05 | 0 | yadI | 1 | 1.5 | 0.251 |
| 0.05 | 0 | AG1 | 1 | 1.5 | 0.249 |
| 0.05 | 0 | ptsI | 1 | 1.75 | 0.241 |
| 0.05 | 0 | clcB | 1 | 1.75 | 0.218 |
| 0.05 | 0 | ycaM | 1 | 1.75 | 0.22 |
| 0.05 | 0 | yadI | 1 | 1.75 | 0.255 |
| 0.05 | 0 | AG1 | 1 | 1.75 | 0.256 |
| 0.05 | 0 | ptsI | 1 | 2 | 0.246 |
| 0.05 | 0 | clcB | 1 | 2 | 0.22 |
| 0.05 | 0 | ycaM | 1 | 2 | 0.226 |
| 0.05 | 0 | yadI | 1 | 2 | 0.265 |
| 0.05 | 0 | AG1 | 1 | 2 | 0.264 |
| 0.05 | 0 | ptsI | 1 | 2.25 | 0.252 |
| 0.05 | 0 | clcB | 1 | 2.25 | 0.22 |
| 0.05 | 0 | ycaM | 1 | 2.25 | 0.229 |
| 0.05 | 0 | yadI | 1 | 2.25 | 0.272 |
| 0.05 | 0 | AG1 | 1 | 2.25 | 0.272 |

| | | | | | |
|---|---|---|---|---|---|
| 0.05 | 0 | ptsI | 1 | 2.5 | 0.258 |
| 0.05 | 0 | clcB | 1 | 2.5 | 0.223 |
| 0.05 | 0 | ycaM | 1 | 2.5 | 0.236 |
| 0.05 | 0 | yadI | 1 | 2.5 | 0.28 |
| 0.05 | 0 | AG1 | 1 | 2.5 | 0.28 |
| 0.05 | 0 | ptsI | 1 | 2.75 | 0.264 |
| 0.05 | 0 | clcB | 1 | 2.75 | 0.223 |
| 0.05 | 0 | ycaM | 1 | 2.75 | 0.24 |
| 0.05 | 0 | yadI | 1 | 2.75 | 0.294 |
| 0.05 | 0 | AG1 | 1 | 2.75 | 0.291 |
| 0.05 | 0 | ptsI | 1 | 3 | 0.27 |
| 0.05 | 0 | clcB | 1 | 3 | 0.227 |
| 0.05 | 0 | ycaM | 1 | 3 | 0.253 |
| 0.05 | 0 | yadI | 1 | 3 | 0.308 |
| 0.05 | 0 | AG1 | 1 | 3 | 0.301 |
| 0.05 | 0 | ptsI | 1 | 3.25 | 0.278 |
| 0.05 | 0 | clcB | 1 | 3.25 | 0.228 |
| 0.05 | 0 | ycaM | 1 | 3.25 | 0.255 |
| 0.05 | 0 | yadI | 1 | 3.25 | 0.328 |
| 0.05 | 0 | AG1 | 1 | 3.25 | 0.312 |
| 0.05 | 0 | ptsI | 1 | 3.5 | 0.286 |
| 0.05 | 0 | clcB | 1 | 3.5 | 0.23 |
| 0.05 | 0 | ycaM | 1 | 3.5 | 0.262 |
| 0.05 | 0 | yadI | 1 | 3.5 | 0.348 |
| 0.05 | 0 | AG1 | 1 | 3.5 | 0.318 |
| 0.05 | 0 | ptsI | 1 | 3.75 | 0.298 |
| 0.05 | 0 | clcB | 1 | 3.75 | 0.234 |
| 0.05 | 0 | ycaM | 1 | 3.75 | 0.274 |
| 0.05 | 0 | yadI | 1 | 3.75 | 0.374 |
| 0.05 | 0 | AG1 | 1 | 3.75 | 0.333 |
| 0.05 | 0 | ptsI | 1 | 4 | 0.308 |
| 0.05 | 0 | clcB | 1 | 4 | 0.238 |
| 0.05 | 0 | ycaM | 1 | 4 | 0.29 |
| 0.05 | 0 | yadI | 1 | 4 | 0.398 |
| 0.05 | 0 | AG1 | 1 | 4 | 0.331 |
| 0.05 | 0 | ptsI | 1 | 4.25 | 0.321 |
| 0.05 | 0 | clcB | 1 | 4.25 | 0.24 |
| 0.05 | 0 | ycaM | 1 | 4.25 | 0.301 |
| 0.05 | 0 | yadI | 1 | 4.25 | 0.42 |
| 0.05 | 0 | AG1 | 1 | 4.25 | 0.359 |
| 0.05 | 0 | ptsI | 1 | 4.5 | 0.325 |
| 0.05 | 0 | clcB | 1 | 4.5 | 0.243 |
| 0.05 | 0 | ycaM | 1 | 4.5 | 0.311 |
| 0.05 | 0 | yadI | 1 | 4.5 | 0.442 |
| 0.05 | 0 | AG1 | 1 | 4.5 | 0.373 |
| 0.05 | 0 | ptsI | 1 | 4.75 | 0.334 |
| 0.05 | 0 | clcB | 1 | 4.75 | 0.247 |
| 0.05 | 0 | ycaM | 1 | 4.75 | 0.32 |

| | | | | | |
|---|---|---|---|---|---|
| 0.05 | 0 | yadI | 1 | 4.75 | 0.459 |
| 0.05 | 0 | AG1 | 1 | 4.75 | 0.386 |
| 0.05 | 0 | ptsI | 1 | 5 | 0.343 |
| 0.05 | 0 | clcB | 1 | 5 | 0.254 |
| 0.05 | 0 | ycaM | 1 | 5 | 0.333 |
| 0.05 | 0 | yadI | 1 | 5 | 0.479 |
| 0.05 | 0 | AG1 | 1 | 5 | 0.407 |
| 0.05 | 0 | ptsI | 1 | 5.25 | 0.353 |
| 0.05 | 0 | clcB | 1 | 5.25 | 0.258 |
| 0.05 | 0 | ycaM | 1 | 5.25 | 0.344 |
| 0.05 | 0 | yadI | 1 | 5.25 | 0.489 |
| 0.05 | 0 | AG1 | 1 | 5.25 | 0.456 |
| 0.05 | 0 | ptsI | 1 | 5.5 | 0.362 |
| 0.05 | 0 | clcB | 1 | 5.5 | 0.268 |
| 0.05 | 0 | ycaM | 1 | 5.5 | 0.357 |
| 0.05 | 0 | yadI | 1 | 5.5 | 0.493 |
| 0.05 | 0 | AG1 | 1 | 5.5 | 0.416 |
| 0.05 | 0 | ptsI | 1 | 5.75 | 0.37 |
| 0.05 | 0 | clcB | 1 | 5.75 | 0.266 |
| 0.05 | 0 | ycaM | 1 | 5.75 | 0.368 |
| 0.05 | 0 | yadI | 1 | 5.75 | 0.508 |
| 0.05 | 0 | AG1 | 1 | 5.75 | 0.424 |
| 0.05 | 0 | ptsI | 1 | 6 | 0.39 |
| 0.05 | 0 | clcB | 1 | 6 | 0.274 |
| 0.05 | 0 | ycaM | 1 | 6 | 0.377 |
| 0.05 | 0 | yadI | 1 | 6 | 0.517 |
| 0.05 | 0 | AG1 | 1 | 6 | 0.436 |
| 0.05 | 0 | ptsI | 1 | 6.25 | 0.405 |
| 0.05 | 0 | clcB | 1 | 6.25 | 0.285 |
| 0.05 | 0 | ycaM | 1 | 6.25 | 0.404 |
| 0.05 | 0 | yadI | 1 | 6.25 | 0.533 |
| 0.05 | 0 | AG1 | 1 | 6.25 | 0.459 |
| 0.05 | 0 | ptsI | 1 | 6.5 | 0.418 |
| 0.05 | 0 | clcB | 1 | 6.5 | 0.289 |
| 0.05 | 0 | ycaM | 1 | 6.5 | 0.41 |
| 0.05 | 0 | yadI | 1 | 6.5 | 0.538 |
| 0.05 | 0 | AG1 | 1 | 6.5 | 0.462 |
| 0.05 | 0 | ptsI | 1 | 6.75 | 0.419 |
| 0.05 | 0 | clcB | 1 | 6.75 | 0.297 |
| 0.05 | 0 | ycaM | 1 | 6.75 | 0.419 |
| 0.05 | 0 | yadI | 1 | 6.75 | 0.56 |
| 0.05 | 0 | AG1 | 1 | 6.75 | 0.468 |
| 0.05 | 0 | ptsI | 1 | 7 | 0.426 |
| 0.05 | 0 | clcB | 1 | 7 | 0.306 |
| 0.05 | 0 | ycaM | 1 | 7 | 0.429 |
| 0.05 | 0 | yadI | 1 | 7 | 0.573 |
| 0.05 | 0 | AG1 | 1 | 7 | 0.482 |
| 0.05 | 0 | ptsI | 1 | 7.25 | 0.436 |

| | | | | | |
|---|---|---|---|---|---|
| 0.05 | 0 | clcB | 1 | 7.25 | 0.316 |
| 0.05 | 0 | ycaM | 1 | 7.25 | 0.44 |
| 0.05 | 0 | yadI | 1 | 7.25 | 0.579 |
| 0.05 | 0 | AG1 | 1 | 7.25 | 0.492 |
| 0.05 | 0 | ptsI | 1 | 7.5 | 0.443 |
| 0.05 | 0 | clcB | 1 | 7.5 | 0.327 |
| 0.05 | 0 | ycaM | 1 | 7.5 | 0.446 |
| 0.05 | 0 | yadI | 1 | 7.5 | 0.593 |
| 0.05 | 0 | AG1 | 1 | 7.5 | 0.504 |
| 0.05 | 0 | ptsI | 1 | 7.75 | 0.452 |
| 0.05 | 0 | clcB | 1 | 7.75 | 0.339 |
| 0.05 | 0 | ycaM | 1 | 7.75 | 0.455 |
| 0.05 | 0 | yadI | 1 | 7.75 | 0.606 |
| 0.05 | 0 | AG1 | 1 | 7.75 | 0.515 |
| 0.05 | 0 | ptsI | 1 | 8 | 0.478 |
| 0.05 | 0 | clcB | 1 | 8 | 0.348 |
| 0.05 | 0 | ycaM | 1 | 8 | 0.463 |
| 0.05 | 0 | yadI | 1 | 8 | 0.617 |
| 0.05 | 0 | AG1 | 1 | 8 | 0.523 |
| 0.05 | 0 | ptsI | 1 | 8.25 | 0.477 |
| 0.05 | 0 | clcB | 1 | 8.25 | 0.36 |
| 0.05 | 0 | ycaM | 1 | 8.25 | 0.469 |
| 0.05 | 0 | yadI | 1 | 8.25 | 0.629 |
| 0.05 | 0 | AG1 | 1 | 8.25 | 0.534 |
| 0.05 | 0 | ptsI | 1 | 8.5 | 0.493 |
| 0.05 | 0 | clcB | 1 | 8.5 | 0.368 |
| 0.05 | 0 | ycaM | 1 | 8.5 | 0.48 |
| 0.05 | 0 | yadI | 1 | 8.5 | 0.646 |
| 0.05 | 0 | AG1 | 1 | 8.5 | 0.551 |
| 0.05 | 0 | ptsI | 1 | 8.75 | 0.495 |
| 0.05 | 0 | clcB | 1 | 8.75 | 0.375 |
| 0.05 | 0 | ycaM | 1 | 8.75 | 0.482 |
| 0.05 | 0 | yadI | 1 | 8.75 | 0.65 |
| 0.05 | 0 | AG1 | 1 | 8.75 | 0.57 |
| 0.05 | 0 | ptsI | 1 | 9 | 0.507 |
| 0.05 | 0 | clcB | 1 | 9 | 0.382 |
| 0.05 | 0 | ycaM | 1 | 9 | 0.487 |
| 0.05 | 0 | yadI | 1 | 9 | 0.661 |
| 0.05 | 0 | AG1 | 1 | 9 | 0.588 |
| 0.05 | 0 | ptsI | 1 | 9.25 | 0.522 |
| 0.05 | 0 | clcB | 1 | 9.25 | 0.393 |
| 0.05 | 0 | ycaM | 1 | 9.25 | 0.493 |
| 0.05 | 0 | yadI | 1 | 9.25 | 0.678 |
| 0.05 | 0 | AG1 | 1 | 9.25 | 0.618 |
| 0.05 | 0 | ptsI | 1 | 9.5 | 0.551 |
| 0.05 | 0 | clcB | 1 | 9.5 | 0.406 |
| 0.05 | 0 | ycaM | 1 | 9.5 | 0.509 |
| 0.05 | 0 | yadI | 1 | 9.5 | 0.714 |

| | | | | | |
|---|---|---|---|---|---|
| 0.05 | 0 | AG1 | 1 | 9.5 | 0.614 |
| 0.05 | 0 | ptsI | 1 | 9.75 | 0.548 |
| 0.05 | 0 | clcB | 1 | 9.75 | 0.41 |
| 0.05 | 0 | ycaM | 1 | 9.75 | 0.515 |
| 0.05 | 0 | yadI | 1 | 9.75 | 0.714 |
| 0.05 | 0 | AG1 | 1 | 9.75 | 0.623 |
| 0.05 | 0 | ptsI | 1 | 10 | 0.569 |
| 0.05 | 0 | clcB | 1 | 10 | 0.418 |
| 0.05 | 0 | ycaM | 1 | 10 | 0.52 |
| 0.05 | 0 | yadI | 1 | 10 | 0.723 |
| 0.05 | 0 | AG1 | 1 | 10 | 0.625 |
| 0.05 | 0 | ptsI | 1 | 10.25 | 0.575 |
| 0.05 | 0 | clcB | 1 | 10.25 | 0.422 |
| 0.05 | 0 | ycaM | 1 | 10.25 | 0.531 |
| 0.05 | 0 | yadI | 1 | 10.25 | 0.726 |
| 0.05 | 0 | AG1 | 1 | 10.25 | 0.628 |
| 0.05 | 0 | ptsI | 1 | 10.5 | 0.604 |
| 0.05 | 0 | clcB | 1 | 10.5 | 0.425 |
| 0.05 | 0 | ycaM | 1 | 10.5 | 0.539 |
| 0.05 | 0 | yadI | 1 | 10.5 | 0.732 |
| 0.05 | 0 | AG1 | 1 | 10.5 | 0.656 |
| 0.05 | 0 | ptsI | 1 | 10.75 | 0.653 |
| 0.05 | 0 | clcB | 1 | 10.75 | 0.44 |
| 0.05 | 0 | ycaM | 1 | 10.75 | 0.551 |
| 0.05 | 0 | yadI | 1 | 10.75 | 0.749 |
| 0.05 | 0 | AG1 | 1 | 10.75 | 0.625 |
| 0.05 | 0 | ptsI | 1 | 11 | 0.634 |
| 0.05 | 0 | clcB | 1 | 11 | 0.462 |
| 0.05 | 0 | ycaM | 1 | 11 | 0.572 |
| 0.05 | 0 | yadI | 1 | 11 | 0.762 |
| 0.05 | 0 | AG1 | 1 | 11 | 0.632 |
| 0.05 | 0 | ptsI | 1 | 11.25 | 0.659 |
| 0.05 | 0 | clcB | 1 | 11.25 | 0.465 |
| 0.05 | 0 | ycaM | 1 | 11.25 | 0.563 |
| 0.05 | 0 | yadI | 1 | 11.25 | 0.765 |
| 0.05 | 0 | AG1 | 1 | 11.25 | 0.621 |
| 0.05 | 0 | ptsI | 1 | 11.5 | 0.689 |
| 0.05 | 0 | clcB | 1 | 11.5 | 0.481 |
| 0.05 | 0 | ycaM | 1 | 11.5 | 0.56 |
| 0.05 | 0 | yadI | 1 | 11.5 | 0.779 |
| 0.05 | 0 | AG1 | 1 | 11.5 | 0.622 |
| 0.05 | 0 | ptsI | 1 | 11.75 | 0.721 |
| 0.05 | 0 | clcB | 1 | 11.75 | 0.497 |
| 0.05 | 0 | ycaM | 1 | 11.75 | 0.571 |
| 0.05 | 0 | yadI | 1 | 11.75 | 0.809 |
| 0.05 | 0 | AG1 | 1 | 11.75 | 0.636 |
| 0.05 | 0 | ptsI | 1 | 12 | 0.71 |
| 0.05 | 0 | clcB | 1 | 12 | 0.497 |

| | | | | | |
|---|---|---|---|---|---|
| 0.05 | 0 | ycaM | 1 | 12 | 0.58 |
| 0.05 | 0 | yadI | 1 | 12 | 0.796 |
| 0.05 | 0 | AG1 | 1 | 12 | 0.644 |
| 0.05 | 0 | ptsI | 1 | 12.25 | 0.717 |
| 0.05 | 0 | clcB | 1 | 12.25 | 0.517 |
| 0.05 | 0 | ycaM | 1 | 12.25 | 0.58 |
| 0.05 | 0 | yadI | 1 | 12.25 | 0.808 |
| 0.05 | 0 | AG1 | 1 | 12.25 | 0.669 |
| 0.05 | 0 | ptsI | 1 | 12.5 | 0.733 |
| 0.05 | 0 | clcB | 1 | 12.5 | 0.519 |
| 0.05 | 0 | ycaM | 1 | 12.5 | 0.584 |
| 0.05 | 0 | yadI | 1 | 12.5 | 0.823 |
| 0.05 | 0 | AG1 | 1 | 12.5 | 0.668 |
| 0.05 | 0 | ptsI | 1 | 12.75 | 0.745 |
| 0.05 | 0 | clcB | 1 | 12.75 | 0.519 |
| 0.05 | 0 | ycaM | 1 | 12.75 | 0.591 |
| 0.05 | 0 | yadI | 1 | 12.75 | 0.818 |
| 0.05 | 0 | AG1 | 1 | 12.75 | 0.698 |
| 0.05 | 0 | ptsI | 1 | 13 | 0.76 |
| 0.05 | 0 | clcB | 1 | 13 | 0.526 |
| 0.05 | 0 | ycaM | 1 | 13 | 0.598 |
| 0.05 | 0 | yadI | 1 | 13 | 0.825 |
| 0.05 | 0 | AG1 | 1 | 13 | 0.7 |
| 0.05 | 0 | ptsI | 1 | 13.25 | 0.763 |
| 0.05 | 0 | clcB | 1 | 13.25 | 0.543 |
| 0.05 | 0 | ycaM | 1 | 13.25 | 0.606 |
| 0.05 | 0 | yadI | 1 | 13.25 | 0.83 |
| 0.05 | 0 | AG1 | 1 | 13.25 | 0.705 |
| 0.05 | 0 | ptsI | 1 | 13.5 | 0.773 |
| 0.05 | 0 | clcB | 1 | 13.5 | 0.551 |
| 0.05 | 0 | ycaM | 1 | 13.5 | 0.611 |
| 0.05 | 0 | yadI | 1 | 13.5 | 0.847 |
| 0.05 | 0 | AG1 | 1 | 13.5 | 0.726 |
| 0.05 | 0 | ptsI | 1 | 13.75 | 0.783 |
| 0.05 | 0 | clcB | 1 | 13.75 | 0.562 |
| 0.05 | 0 | ycaM | 1 | 13.75 | 0.61 |
| 0.05 | 0 | yadI | 1 | 13.75 | 0.859 |
| 0.05 | 0 | AG1 | 1 | 13.75 | 0.72 |
| 0.05 | 0 | ptsI | 1 | 14 | 0.808 |
| 0.05 | 0 | clcB | 1 | 14 | 0.563 |
| 0.05 | 0 | ycaM | 1 | 14 | 0.613 |
| 0.05 | 0 | yadI | 1 | 14 | 0.871 |
| 0.05 | 0 | AG1 | 1 | 14 | 0.754 |
| 0.05 | 0 | ptsI | 1 | 14.25 | 0.81 |
| 0.05 | 0 | clcB | 1 | 14.25 | 0.566 |
| 0.05 | 0 | ycaM | 1 | 14.25 | 0.622 |
| 0.05 | 0 | yadI | 1 | 14.25 | 0.864 |
| 0.05 | 0 | AG1 | 1 | 14.25 | 0.748 |

| | | | | | |
|---|---|---|---|---|---|
| 0.05 | 0 | ptsI | 1 | 14.5 | 0.816 |
| 0.05 | 0 | clcB | 1 | 14.5 | 0.569 |
| 0.05 | 0 | ycaM | 1 | 14.5 | 0.618 |
| 0.05 | 0 | yadI | 1 | 14.5 | 0.872 |
| 0.05 | 0 | AG1 | 1 | 14.5 | 0.757 |
| 0.05 | 0 | ptsI | 1 | 14.75 | 0.821 |
| 0.05 | 0 | clcB | 1 | 14.75 | 0.579 |
| 0.05 | 0 | ycaM | 1 | 14.75 | 0.619 |
| 0.05 | 0 | yadI | 1 | 14.75 | 0.88 |
| 0.05 | 0 | AG1 | 1 | 14.75 | 0.768 |
| 0.05 | 0 | ptsI | 1 | 15 | 0.811 |
| 0.05 | 0 | clcB | 1 | 15 | 0.59 |
| 0.05 | 0 | ycaM | 1 | 15 | 0.62 |
| 0.05 | 0 | yadI | 1 | 15 | 0.876 |
| 0.05 | 0 | AG1 | 1 | 15 | 0.771 |
| 0.05 | 0 | ptsI | 1 | 15.25 | 0.841 |
| 0.05 | 0 | clcB | 1 | 15.25 | 0.616 |
| 0.05 | 0 | ycaM | 1 | 15.25 | 0.645 |
| 0.05 | 0 | yadI | 1 | 15.25 | 0.873 |
| 0.05 | 0 | AG1 | 1 | 15.25 | 0.782 |
| 0.05 | 0 | ptsI | 1 | 15.5 | 0.838 |
| 0.05 | 0 | clcB | 1 | 15.5 | 0.648 |
| 0.05 | 0 | ycaM | 1 | 15.5 | 0.659 |
| 0.05 | 0 | yadI | 1 | 15.5 | 0.881 |
| 0.05 | 0 | AG1 | 1 | 15.5 | 0.761 |
| 0.05 | 0 | ptsI | 1 | 15.75 | 0.829 |
| 0.05 | 0 | clcB | 1 | 15.75 | 0.636 |
| 0.05 | 0 | ycaM | 1 | 15.75 | 0.644 |
| 0.05 | 0 | yadI | 1 | 15.75 | 0.889 |
| 0.05 | 0 | AG1 | 1 | 15.75 | 0.784 |
| 0.05 | 0 | ptsI | 1 | 16 | 0.831 |
| 0.05 | 0 | clcB | 1 | 16 | 0.676 |
| 0.05 | 0 | ycaM | 1 | 16 | 0.653 |
| 0.05 | 0 | yadI | 1 | 16 | 0.887 |
| 0.05 | 0 | AG1 | 1 | 16 | 0.776 |
| 0.05 | 0 | ptsI | 1 | 16.25 | 0.848 |
| 0.05 | 0 | clcB | 1 | 16.25 | 0.653 |
| 0.05 | 0 | ycaM | 1 | 16.25 | 0.652 |
| 0.05 | 0 | yadI | 1 | 16.25 | 0.88 |
| 0.05 | 0 | AG1 | 1 | 16.25 | 0.81 |
| 0.05 | 0 | ptsI | 1 | 16.5 | 0.856 |
| 0.05 | 0 | clcB | 1 | 16.5 | 0.67 |
| 0.05 | 0 | ycaM | 1 | 16.5 | 0.678 |
| 0.05 | 0 | yadI | 1 | 16.5 | 0.884 |
| 0.05 | 0 | AG1 | 1 | 16.5 | 0.789 |
| 0.05 | 0 | ptsI | 1 | 17.0667 | 0.906 |
| 0.05 | 0 | clcB | 1 | 17.0667 | 0.662 |
| 0.05 | 0 | ycaM | 1 | 17.0667 | 0.744 |

| | | | | | |
|---|---|---|---|---|---|
| 0.05 | 0 | yadI | 1 | 17.0667 | 0.952 |
| 0.05 | 0 | AG1 | 1 | 17.0667 | 0.811 |
| 0.05 | 0 | ptsI | 1 | 17.3167 | 0.81 |
| 0.05 | 0 | clcB | 1 | 17.3167 | 0.642 |
| 0.05 | 0 | ycaM | 1 | 17.3167 | 0.686 |
| 0.05 | 0 | yadI | 1 | 17.3167 | 0.914 |
| 0.05 | 0 | AG1 | 1 | 17.3167 | 0.792 |
| 0.05 | 0 | ptsI | 1 | 17.5667 | 0.823 |
| 0.05 | 0 | clcB | 1 | 17.5667 | 0.658 |
| 0.05 | 0 | ycaM | 1 | 17.5667 | 0.707 |
| 0.05 | 0 | yadI | 1 | 17.5667 | 0.9 |
| 0.05 | 0 | AG1 | 1 | 17.5667 | 0.8 |
| 0.05 | 0 | ptsI | 1 | 17.8167 | 0.853 |
| 0.05 | 0 | clcB | 1 | 17.8167 | 0.736 |
| 0.05 | 0 | ycaM | 1 | 17.8167 | 0.697 |
| 0.05 | 0 | yadI | 1 | 17.8167 | 0.909 |
| 0.05 | 0 | AG1 | 1 | 17.8167 | 0.832 |
| 0.05 | 0 | ptsI | 1 | 18.0667 | 0.85 |
| 0.05 | 0 | clcB | 1 | 18.0667 | 0.644 |
| 0.05 | 0 | ycaM | 1 | 18.0667 | 0.704 |
| 0.05 | 0 | yadI | 1 | 18.0667 | 0.909 |
| 0.05 | 0 | AG1 | 1 | 18.0667 | 0.779 |
| 0.05 | 0 | ptsI | 1 | 18.3167 | 0.853 |
| 0.05 | 0 | clcB | 1 | 18.3167 | 0.668 |
| 0.05 | 0 | ycaM | 1 | 18.3167 | 0.712 |
| 0.05 | 0 | yadI | 1 | 18.3167 | 0.917 |
| 0.05 | 0 | AG1 | 1 | 18.3167 | 0.802 |
| 0.05 | 0 | ptsI | 1 | 18.5667 | 0.866 |
| 0.05 | 0 | clcB | 1 | 18.5667 | 0.697 |
| 0.05 | 0 | ycaM | 1 | 18.5667 | 0.725 |
| 0.05 | 0 | yadI | 1 | 18.5667 | 0.916 |
| 0.05 | 0 | AG1 | 1 | 18.5667 | 0.822 |
| 0.05 | 0 | ptsI | 1 | 18.8167 | 0.872 |
| 0.05 | 0 | clcB | 1 | 18.8167 | 0.711 |
| 0.05 | 0 | ycaM | 1 | 18.8167 | 0.735 |
| 0.05 | 0 | yadI | 1 | 18.8167 | 0.918 |
| 0.05 | 0 | AG1 | 1 | 18.8167 | 0.826 |
| 0.05 | 0 | ptsI | 1 | 19.0667 | 0.88 |
| 0.05 | 0 | clcB | 1 | 19.0667 | 0.708 |
| 0.05 | 0 | ycaM | 1 | 19.0667 | 0.733 |
| 0.05 | 0 | yadI | 1 | 19.0667 | 0.93 |
| 0.05 | 0 | AG1 | 1 | 19.0667 | 0.823 |
| 0.05 | 0 | ptsI | 1 | 19.3167 | 0.887 |
| 0.05 | 0 | clcB | 1 | 19.3167 | 0.709 |
| 0.05 | 0 | ycaM | 1 | 19.3167 | 0.73 |
| 0.05 | 0 | yadI | 1 | 19.3167 | 0.931 |
| 0.05 | 0 | AG1 | 1 | 19.3167 | 0.819 |
| 0.05 | 0 | ptsI | 1 | 19.5667 | 0.894 |

| | | | | | |
|---|---|---|---|---|---|
| 0.05 | 0 | clcB | 1 | 19.5667 | 0.742 |
| 0.05 | 0 | ycaM | 1 | 19.5667 | 0.748 |
| 0.05 | 0 | yadI | 1 | 19.5667 | 0.934 |
| 0.05 | 0 | AG1 | 1 | 19.5667 | 0.838 |
| 0.05 | 0 | ptsI | 1 | 19.8167 | 0.9 |
| 0.05 | 0 | clcB | 1 | 19.8167 | 0.731 |
| 0.05 | 0 | ycaM | 1 | 19.8167 | 0.744 |
| 0.05 | 0 | yadI | 1 | 19.8167 | 0.937 |
| 0.05 | 0 | AG1 | 1 | 19.8167 | 0.835 |
| 0.05 | 0 | ptsI | 1 | 20.0667 | 0.915 |
| 0.05 | 0 | clcB | 1 | 20.0667 | 0.743 |
| 0.05 | 0 | ycaM | 1 | 20.0667 | 0.747 |
| 0.05 | 0 | yadI | 1 | 20.0667 | 0.94 |
| 0.05 | 0 | AG1 | 1 | 20.0667 | 0.84 |
| 0.05 | 0 | ptsI | 1 | 20.3167 | 0.917 |
| 0.05 | 0 | clcB | 1 | 20.3167 | 0.737 |
| 0.05 | 0 | ycaM | 1 | 20.3167 | 0.752 |
| 0.05 | 0 | yadI | 1 | 20.3167 | 0.95 |
| 0.05 | 0 | AG1 | 1 | 20.3167 | 0.831 |
| 0.05 | 0 | ptsI | 1 | 20.5667 | 0.921 |
| 0.05 | 0 | clcB | 1 | 20.5667 | 0.726 |
| 0.05 | 0 | ycaM | 1 | 20.5667 | 0.748 |
| 0.05 | 0 | yadI | 1 | 20.5667 | 0.947 |
| 0.05 | 0 | AG1 | 1 | 20.5667 | 0.826 |
| 0.05 | 0 | ptsI | 1 | 20.8167 | 0.936 |
| 0.05 | 0 | clcB | 1 | 20.8167 | 0.728 |
| 0.05 | 0 | ycaM | 1 | 20.8167 | 0.749 |
| 0.05 | 0 | yadI | 1 | 20.8167 | 0.953 |
| 0.05 | 0 | AG1 | 1 | 20.8167 | 0.817 |
| 0.05 | 0 | ptsI | 1 | 21.0667 | 0.952 |
| 0.05 | 0 | clcB | 1 | 21.0667 | 0.73 |
| 0.05 | 0 | ycaM | 1 | 21.0667 | 0.764 |
| 0.05 | 0 | yadI | 1 | 21.0667 | 0.952 |
| 0.05 | 0 | AG1 | 1 | 21.0667 | 0.811 |
| 0.05 | 0 | ptsI | 1 | 21.3167 | 0.956 |
| 0.05 | 0 | clcB | 1 | 21.3167 | 0.738 |
| 0.05 | 0 | ycaM | 1 | 21.3167 | 0.787 |
| 0.05 | 0 | yadI | 1 | 21.3167 | 0.959 |
| 0.05 | 0 | AG1 | 1 | 21.3167 | 0.82 |
| 0.05 | 0 | ptsI | 1 | 21.5667 | 0.972 |
| 0.05 | 0 | clcB | 1 | 21.5667 | 0.735 |
| 0.05 | 0 | ycaM | 1 | 21.5667 | 0.769 |
| 0.05 | 0 | yadI | 1 | 21.5667 | 0.961 |
| 0.05 | 0 | AG1 | 1 | 21.5667 | 0.823 |
| 0.05 | 0 | ptsI | 1 | 21.8167 | 0.985 |
| 0.05 | 0 | clcB | 1 | 21.8167 | 0.738 |
| 0.05 | 0 | ycaM | 1 | 21.8167 | 0.775 |
| 0.05 | 0 | yadI | 1 | 21.8167 | 0.966 |

| | | | | | |
|---|---|---|---|---|---|
| 0.05 | 0 | AG1 | 1 | 21.8167 | 0.82 |
| 0.05 | 0 | ptsI | 1 | 22.0667 | 1.007 |
| 0.05 | 0 | clcB | 1 | 22.0667 | 0.745 |
| 0.05 | 0 | ycaM | 1 | 22.0667 | 0.792 |
| 0.05 | 0 | yadI | 1 | 22.0667 | 0.97 |
| 0.05 | 0 | AG1 | 1 | 22.0667 | 0.82 |
| 0.05 | 0 | ptsI | 1 | 22.9333 | 1.125 |
| 0.05 | 0 | clcB | 1 | 22.9333 | 0.701 |
| 0.05 | 0 | ycaM | 1 | 22.9333 | 0.744 |
| 0.05 | 0 | yadI | 1 | 22.9333 | 1.088 |
| 0.05 | 0 | AG1 | 1 | 22.9333 | 0.812 |
| 0.05 | 0 | ptsI | 1 | 23.1833 | 1.088 |
| 0.05 | 0 | clcB | 1 | 23.1833 | 0.665 |
| 0.05 | 0 | ycaM | 1 | 23.1833 | 0.716 |
| 0.05 | 0 | yadI | 1 | 23.1833 | 1.039 |
| 0.05 | 0 | AG1 | 1 | 23.1833 | 0.776 |
| 0.05 | 0 | ptsI | 1 | 23.4333 | 1.073 |
| 0.05 | 0 | clcB | 1 | 23.4333 | 0.706 |
| 0.05 | 0 | ycaM | 1 | 23.4333 | 0.745 |
| 0.05 | 0 | yadI | 1 | 23.4333 | 1.035 |
| 0.05 | 0 | AG1 | 1 | 23.4333 | 0.779 |
| 0.05 | 0 | ptsI | 1 | 23.6833 | 1.068 |
| 0.05 | 0 | clcB | 1 | 23.6833 | 0.685 |
| 0.05 | 0 | ycaM | 1 | 23.6833 | 0.739 |
| 0.05 | 0 | yadI | 1 | 23.6833 | 1.043 |
| 0.05 | 0 | AG1 | 1 | 23.6833 | 0.802 |
| 0.05 | 0 | ptsI | 1 | 23.9333 | 1.066 |
| 0.05 | 0 | clcB | 1 | 23.9333 | 0.674 |
| 0.05 | 0 | ycaM | 1 | 23.9333 | 0.732 |
| 0.05 | 0 | yadI | 1 | 23.9333 | 1.042 |
| 0.05 | 0 | AG1 | 1 | 23.9333 | 0.805 |
| 0.05 | 0 | ptsI | 1 | 24.1833 | 1.067 |
| 0.05 | 0 | clcB | 1 | 24.1833 | 0.686 |
| 0.05 | 0 | ycaM | 1 | 24.1833 | 0.748 |
| 0.05 | 0 | yadI | 1 | 24.1833 | 1.052 |
| 0.05 | 0 | AG1 | 1 | 24.1833 | 0.807 |
| 0.05 | 0 | ptsI | 1 | 24.4333 | 1.074 |
| 0.05 | 0 | clcB | 1 | 24.4333 | 0.696 |
| 0.05 | 0 | ycaM | 1 | 24.4333 | 0.751 |
| 0.05 | 0 | yadI | 1 | 24.4333 | 1.051 |
| 0.05 | 0 | AG1 | 1 | 24.4333 | 0.808 |
| 0.05 | 0 | ptsI | 1 | 24.6833 | 1.066 |
| 0.05 | 0 | clcB | 1 | 24.6833 | 0.699 |
| 0.05 | 0 | ycaM | 1 | 24.6833 | 0.75 |
| 0.05 | 0 | yadI | 1 | 24.6833 | 1.053 |
| 0.05 | 0 | AG1 | 1 | 24.6833 | 0.814 |
| 0.05 | 0 | ptsI | 1 | 24.9333 | 1.07 |
| 0.05 | 0 | clcB | 1 | 24.9333 | 0.7 |

| | | | | | |
|---|---|---|---|---|---|
| 0.05 | 0 | ycaM | 1 | 24.9333 | 0.755 |
| 0.05 | 0 | yadI | 1 | 24.9333 | 1.065 |
| 0.05 | 0 | AG1 | 1 | 24.9333 | 0.832 |
| 0.05 | 0 | ptsI | 1 | 25.1833 | 1.072 |
| 0.05 | 0 | clcB | 1 | 25.1833 | 0.706 |
| 0.05 | 0 | ycaM | 1 | 25.1833 | 0.753 |
| 0.05 | 0 | yadI | 1 | 25.1833 | 1.063 |
| 0.05 | 0 | AG1 | 1 | 25.1833 | 0.823 |
| 0.05 | 0 | ptsI | 1 | 25.4333 | 1.079 |
| 0.05 | 0 | clcB | 1 | 25.4333 | 0.715 |
| 0.05 | 0 | ycaM | 1 | 25.4333 | 0.771 |
| 0.05 | 0 | yadI | 1 | 25.4333 | 1.074 |
| 0.05 | 0 | AG1 | 1 | 25.4333 | 0.836 |
| 0.05 | 0 | ptsI | 1 | 25.6833 | 1.08 |
| 0.05 | 0 | clcB | 1 | 25.6833 | 0.719 |
| 0.05 | 0 | ycaM | 1 | 25.6833 | 0.775 |
| 0.05 | 0 | yadI | 1 | 25.6833 | 1.074 |
| 0.05 | 0 | AG1 | 1 | 25.6833 | 0.836 |
| 0.05 | 0 | ptsI | 1 | 25.9333 | 1.078 |
| 0.05 | 0 | clcB | 1 | 25.9333 | 0.731 |
| 0.05 | 0 | ycaM | 1 | 25.9333 | 0.778 |
| 0.05 | 0 | yadI | 1 | 25.9333 | 1.073 |
| 0.05 | 0 | AG1 | 1 | 25.9333 | 0.841 |
| 0.05 | 0 | ptsI | 1 | 26.1833 | 1.088 |
| 0.05 | 0 | clcB | 1 | 26.1833 | 0.746 |
| 0.05 | 0 | ycaM | 1 | 26.1833 | 0.793 |
| 0.05 | 0 | yadI | 1 | 26.1833 | 1.081 |
| 0.05 | 0 | AG1 | 1 | 26.1833 | 0.851 |
| 0.05 | 0 | ptsI | 1 | 26.4333 | 1.087 |
| 0.05 | 0 | clcB | 1 | 26.4333 | 0.752 |
| 0.05 | 0 | ycaM | 1 | 26.4333 | 0.788 |
| 0.05 | 0 | yadI | 1 | 26.4333 | 1.08 |
| 0.05 | 0 | AG1 | 1 | 26.4333 | 0.845 |
| 0.05 | 0 | ptsI | 1 | 26.6833 | 1.091 |
| 0.05 | 0 | clcB | 1 | 26.6833 | 0.754 |
| 0.05 | 0 | ycaM | 1 | 26.6833 | 0.789 |
| 0.05 | 0 | yadI | 1 | 26.6833 | 1.087 |
| 0.05 | 0 | AG1 | 1 | 26.6833 | 0.847 |
| 0.05 | 0 | ptsI | 1 | 26.9333 | 1.092 |
| 0.05 | 0 | clcB | 1 | 26.9333 | 0.762 |
| 0.05 | 0 | ycaM | 1 | 26.9333 | 0.797 |
| 0.05 | 0 | yadI | 1 | 26.9333 | 1.087 |
| 0.05 | 0 | AG1 | 1 | 26.9333 | 0.85 |
| 0.05 | 0 | ptsI | 1 | 27.1833 | 1.095 |
| 0.05 | 0 | clcB | 1 | 27.1833 | 0.768 |
| 0.05 | 0 | ycaM | 1 | 27.1833 | 0.802 |
| 0.05 | 0 | yadI | 1 | 27.1833 | 1.094 |
| 0.05 | 0 | AG1 | 1 | 27.1833 | 0.857 |

| | | | | | |
|---|---|---|---|---|---|
| 0.05 | 0 | ptsI | 1 | 27.4333 | 1.097 |
| 0.05 | 0 | clcB | 1 | 27.4333 | 0.761 |
| 0.05 | 0 | ycaM | 1 | 27.4333 | 0.799 |
| 0.05 | 0 | yadI | 1 | 27.4333 | 1.097 |
| 0.05 | 0 | AG1 | 1 | 27.4333 | 0.85 |
| 0.05 | 0 | ptsI | 1 | 27.6833 | 1.103 |
| 0.05 | 0 | clcB | 1 | 27.6833 | 0.776 |
| 0.05 | 0 | ycaM | 1 | 27.6833 | 0.806 |
| 0.05 | 0 | yadI | 1 | 27.6833 | 1.104 |
| 0.05 | 0 | AG1 | 1 | 27.6833 | 0.861 |
| 0.05 | 0 | ptsI | 1 | 27.9333 | 1.105 |
| 0.05 | 0 | clcB | 1 | 27.9333 | 0.772 |
| 0.05 | 0 | ycaM | 1 | 27.9333 | 0.8 |
| 0.05 | 0 | yadI | 1 | 27.9333 | 1.101 |
| 0.05 | 0 | AG1 | 1 | 27.9333 | 0.853 |
| 0.05 | 0 | ptsI | 1 | 28.1833 | 1.11 |
| 0.05 | 0 | clcB | 1 | 28.1833 | 0.791 |
| 0.05 | 0 | ycaM | 1 | 28.1833 | 0.812 |
| 0.05 | 0 | yadI | 1 | 28.1833 | 1.108 |
| 0.05 | 0 | AG1 | 1 | 28.1833 | 0.865 |
| 0.05 | 0 | ptsI | 1 | 28.4333 | 1.114 |
| 0.05 | 0 | clcB | 1 | 28.4333 | 0.788 |
| 0.05 | 0 | ycaM | 1 | 28.4333 | 0.812 |
| 0.05 | 0 | yadI | 1 | 28.4333 | 1.104 |
| 0.05 | 0 | AG1 | 1 | 28.4333 | 0.868 |
| 0.05 | 0 | ptsI | 1 | 28.6833 | 1.118 |
| 0.05 | 0 | clcB | 1 | 28.6833 | 0.788 |
| 0.05 | 0 | ycaM | 1 | 28.6833 | 0.816 |
| 0.05 | 0 | yadI | 1 | 28.6833 | 1.109 |
| 0.05 | 0 | AG1 | 1 | 28.6833 | 0.868 |
| 0.05 | 0 | ptsI | 1 | 28.9333 | 1.122 |
| 0.05 | 0 | clcB | 1 | 28.9333 | 0.801 |
| 0.05 | 0 | ycaM | 1 | 28.9333 | 0.82 |
| 0.05 | 0 | yadI | 1 | 28.9333 | 1.117 |
| 0.05 | 0 | AG1 | 1 | 28.9333 | 0.875 |
| 0.05 | 0 | ptsI | 1 | 29.1833 | 1.123 |
| 0.05 | 0 | clcB | 1 | 29.1833 | 0.786 |
| 0.05 | 0 | ycaM | 1 | 29.1833 | 0.817 |
| 0.05 | 0 | yadI | 1 | 29.1833 | 1.117 |
| 0.05 | 0 | AG1 | 1 | 29.1833 | 0.873 |
| 0.05 | 0 | ptsI | 1 | 29.4333 | 1.126 |
| 0.05 | 0 | clcB | 1 | 29.4333 | 0.798 |
| 0.05 | 0 | ycaM | 1 | 29.4333 | 0.818 |
| 0.05 | 0 | yadI | 1 | 29.4333 | 1.119 |
| 0.05 | 0 | AG1 | 1 | 29.4333 | 0.876 |
| 0.05 | 0 | ptsI | 1 | 29.6833 | 1.134 |
| 0.05 | 0 | clcB | 1 | 29.6833 | 0.795 |
| 0.05 | 0 | ycaM | 1 | 29.6833 | 0.822 |

| | | | | | |
|---|---|---|---|---|---|
| 0.05 | 0 | yadI | 1 | 29.6833 | 1.124 |
| 0.05 | 0 | AG1 | 1 | 29.6833 | 0.878 |
| 0.05 | 0 | ptsI | 1 | 29.9333 | 1.141 |
| 0.05 | 0 | clcB | 1 | 29.9333 | 0.798 |
| 0.05 | 0 | ycaM | 1 | 29.9333 | 0.825 |
| 0.05 | 0 | yadI | 1 | 29.9333 | 1.131 |
| 0.05 | 0 | AG1 | 1 | 29.9333 | 0.88 |
| 0.05 | 0 | ptsI | 1 | 30.1833 | 1.147 |
| 0.05 | 0 | clcB | 1 | 30.1833 | 0.796 |
| 0.05 | 0 | ycaM | 1 | 30.1833 | 0.824 |
| 0.05 | 0 | yadI | 1 | 30.1833 | 1.133 |
| 0.05 | 0 | AG1 | 1 | 30.1833 | 0.88 |
| 0.05 | 0 | ptsI | 1 | 30.4333 | 1.152 |
| 0.05 | 0 | clcB | 1 | 30.4333 | 0.805 |
| 0.05 | 0 | ycaM | 1 | 30.4333 | 0.829 |
| 0.05 | 0 | yadI | 1 | 30.4333 | 1.137 |
| 0.05 | 0 | AG1 | 1 | 30.4333 | 0.883 |
| 0.05 | 0 | ptsI | 1 | 30.6833 | 1.161 |
| 0.05 | 0 | clcB | 1 | 30.6833 | 0.811 |
| 0.05 | 0 | ycaM | 1 | 30.6833 | 0.826 |
| 0.05 | 0 | yadI | 1 | 30.6833 | 1.146 |
| 0.05 | 0 | AG1 | 1 | 30.6833 | 0.882 |
| 0.05 | 0 | ptsI | 1 | 30.9333 | 1.164 |
| 0.05 | 0 | clcB | 1 | 30.9333 | 0.815 |
| 0.05 | 0 | ycaM | 1 | 30.9333 | 0.833 |
| 0.05 | 0 | yadI | 1 | 30.9333 | 1.151 |
| 0.05 | 0 | AG1 | 1 | 30.9333 | 0.889 |
| 0.05 | 0 | ptsI | 1 | 31.1833 | 1.172 |
| 0.05 | 0 | clcB | 1 | 31.1833 | 0.804 |
| 0.05 | 0 | ycaM | 1 | 31.1833 | 0.832 |
| 0.05 | 0 | yadI | 1 | 31.1833 | 1.152 |
| 0.05 | 0 | AG1 | 1 | 31.1833 | 0.887 |
| 0.05 | 0 | ptsI | 1 | 31.4333 | 1.183 |
| 0.05 | 0 | clcB | 1 | 31.4333 | 0.804 |
| 0.05 | 0 | ycaM | 1 | 31.4333 | 0.843 |
| 0.05 | 0 | yadI | 1 | 31.4333 | 1.158 |
| 0.05 | 0 | AG1 | 1 | 31.4333 | 0.892 |
| 0.05 | 0 | ptsI | 1 | 31.6833 | 1.191 |
| 0.05 | 0 | clcB | 1 | 31.6833 | 0.809 |
| 0.05 | 0 | ycaM | 1 | 31.6833 | 0.844 |
| 0.05 | 0 | yadI | 1 | 31.6833 | 1.16 |
| 0.05 | 0 | AG1 | 1 | 31.6833 | 0.893 |
| 0.05 | 0 | ptsI | 1 | 31.9333 | 1.204 |
| 0.05 | 0 | clcB | 1 | 31.9333 | 0.807 |
| 0.05 | 0 | ycaM | 1 | 31.9333 | 0.847 |
| 0.05 | 0 | yadI | 1 | 31.9333 | 1.163 |
| 0.05 | 0 | AG1 | 1 | 31.9333 | 0.896 |
| 0.05 | 0 | ptsI | 1 | 32.1833 | 1.207 |

| | | | | | |
|---|---|---|---|---|---|
| 0.05 | 0 | clcB | 1 | 32.1833 | 0.808 |
| 0.05 | 0 | ycaM | 1 | 32.1833 | 0.849 |
| 0.05 | 0 | yadI | 1 | 32.1833 | 1.164 |
| 0.05 | 0 | AG1 | 1 | 32.1833 | 0.897 |
| 0.05 | 0 | ptsI | 1 | 32.4333 | 1.227 |
| 0.05 | 0 | clcB | 1 | 32.4333 | 0.805 |
| 0.05 | 0 | ycaM | 1 | 32.4333 | 0.849 |
| 0.05 | 0 | yadI | 1 | 32.4333 | 1.164 |
| 0.05 | 0 | AG1 | 1 | 32.4333 | 0.897 |
| 0.05 | 0 | ptsI | 1 | 32.6833 | 1.223 |
| 0.05 | 0 | clcB | 1 | 32.6833 | 0.816 |
| 0.05 | 0 | ycaM | 1 | 32.6833 | 0.856 |
| 0.05 | 0 | yadI | 1 | 32.6833 | 1.17 |
| 0.05 | 0 | AG1 | 1 | 32.6833 | 0.899 |
| 0.05 | 0 | ptsI | 1 | 32.9333 | 1.236 |
| 0.05 | 0 | clcB | 1 | 32.9333 | 0.824 |
| 0.05 | 0 | ycaM | 1 | 32.9333 | 0.858 |
| 0.05 | 0 | yadI | 1 | 32.9333 | 1.174 |
| 0.05 | 0 | AG1 | 1 | 32.9333 | 0.901 |
| 0.05 | 0 | ptsI | 1 | 33.1833 | 1.245 |
| 0.05 | 0 | clcB | 1 | 33.1833 | 0.826 |
| 0.05 | 0 | ycaM | 1 | 33.1833 | 0.857 |
| 0.05 | 0 | yadI | 1 | 33.1833 | 1.179 |
| 0.05 | 0 | AG1 | 1 | 33.1833 | 0.901 |
| 0.05 | 0 | ptsI | 1 | 33.4333 | 1.254 |
| 0.05 | 0 | clcB | 1 | 33.4333 | 0.838 |
| 0.05 | 0 | ycaM | 1 | 33.4333 | 0.863 |
| 0.05 | 0 | yadI | 1 | 33.4333 | 1.183 |
| 0.05 | 0 | AG1 | 1 | 33.4333 | 0.907 |
| 0.05 | 0 | ptsI | 1 | 33.6833 | 1.263 |
| 0.05 | 0 | clcB | 1 | 33.6833 | 0.832 |
| 0.05 | 0 | ycaM | 1 | 33.6833 | 0.861 |
| 0.05 | 0 | yadI | 1 | 33.6833 | 1.187 |
| 0.05 | 0 | AG1 | 1 | 33.6833 | 0.902 |
| 0.05 | 0 | ptsI | 1 | 33.9333 | 1.273 |
| 0.05 | 0 | clcB | 1 | 33.9333 | 0.846 |
| 0.05 | 0 | ycaM | 1 | 33.9333 | 0.872 |
| 0.05 | 0 | yadI | 1 | 33.9333 | 1.195 |
| 0.05 | 0 | AG1 | 1 | 33.9333 | 0.911 |
| 0.05 | 0 | ptsI | 1 | 34.1833 | 1.282 |
| 0.05 | 0 | clcB | 1 | 34.1833 | 0.842 |
| 0.05 | 0 | ycaM | 1 | 34.1833 | 0.869 |
| 0.05 | 0 | yadI | 1 | 34.1833 | 1.2 |
| 0.05 | 0 | AG1 | 1 | 34.1833 | 0.903 |
| 0.05 | 0 | ptsI | 1 | 34.4333 | 1.298 |
| 0.05 | 0 | clcB | 1 | 34.4333 | 0.842 |
| 0.05 | 0 | ycaM | 1 | 34.4333 | 0.873 |
| 0.05 | 0 | yadI | 1 | 34.4333 | 1.204 |

| | | | | | |
|---|---|---|---|---|---|
| 0.05 | 0 | AG1 | 1 | 34.4333 | 0.912 |
| 0.05 | 0 | ptsI | 1 | 34.6833 | 1.302 |
| 0.05 | 0 | clcB | 1 | 34.6833 | 0.842 |
| 0.05 | 0 | ycaM | 1 | 34.6833 | 0.873 |
| 0.05 | 0 | yadI | 1 | 34.6833 | 1.211 |
| 0.05 | 0 | AG1 | 1 | 34.6833 | 0.918 |
| 0.05 | 0 | ptsI | 1 | 34.9333 | 1.318 |
| 0.05 | 0 | clcB | 1 | 34.9333 | 0.859 |
| 0.05 | 0 | ycaM | 1 | 34.9333 | 0.874 |
| 0.05 | 0 | yadI | 1 | 34.9333 | 1.216 |
| 0.05 | 0 | AG1 | 1 | 34.9333 | 0.918 |
| 0.05 | 0 | ptsI | 1 | 35.1833 | 1.33 |
| 0.05 | 0 | clcB | 1 | 35.1833 | 0.854 |
| 0.05 | 0 | ycaM | 1 | 35.1833 | 0.874 |
| 0.05 | 0 | yadI | 1 | 35.1833 | 1.221 |
| 0.05 | 0 | AG1 | 1 | 35.1833 | 0.917 |
| 0.05 | 0 | ptsI | 1 | 35.4333 | 1.338 |
| 0.05 | 0 | clcB | 1 | 35.4333 | 0.861 |
| 0.05 | 0 | ycaM | 1 | 35.4333 | 0.873 |
| 0.05 | 0 | yadI | 1 | 35.4333 | 1.226 |
| 0.05 | 0 | AG1 | 1 | 35.4333 | 0.924 |
| 0.05 | 0 | ptsI | 1 | 35.6833 | 1.347 |
| 0.05 | 0 | clcB | 1 | 35.6833 | 0.866 |
| 0.05 | 0 | ycaM | 1 | 35.6833 | 0.874 |
| 0.05 | 0 | yadI | 1 | 35.6833 | 1.233 |
| 0.05 | 0 | AG1 | 1 | 35.6833 | 0.93 |
| 0.05 | 0 | ptsI | 1 | 35.9333 | 1.355 |
| 0.05 | 0 | clcB | 1 | 35.9333 | 0.862 |
| 0.05 | 0 | ycaM | 1 | 35.9333 | 0.869 |
| 0.05 | 0 | yadI | 1 | 35.9333 | 1.235 |
| 0.05 | 0 | AG1 | 1 | 35.9333 | 0.924 |
| 0.05 | 0 | ptsI | 1 | 36.1833 | 1.358 |
| 0.05 | 0 | clcB | 1 | 36.1833 | 0.874 |
| 0.05 | 0 | ycaM | 1 | 36.1833 | 0.874 |
| 0.05 | 0 | yadI | 1 | 36.1833 | 1.242 |
| 0.05 | 0 | AG1 | 1 | 36.1833 | 0.928 |
| 0.05 | 0 | ptsI | 1 | 36.4333 | 1.377 |
| 0.05 | 0 | clcB | 1 | 36.4333 | 0.873 |
| 0.05 | 0 | ycaM | 1 | 36.4333 | 0.872 |
| 0.05 | 0 | yadI | 1 | 36.4333 | 1.25 |
| 0.05 | 0 | AG1 | 1 | 36.4333 | 0.932 |
| 0.05 | 0 | ptsI | 1 | 36.6833 | 1.375 |
| 0.05 | 0 | clcB | 1 | 36.6833 | 0.869 |
| 0.05 | 0 | ycaM | 1 | 36.6833 | 0.877 |
| 0.05 | 0 | yadI | 1 | 36.6833 | 1.253 |
| 0.05 | 0 | AG1 | 1 | 36.6833 | 0.927 |
| 0.05 | 0 | ptsI | 2 | 0 | 0.209 |
| 0.05 | 0 | clcB | 2 | 0 | 0.224 |

| | | | | | |
|---|---|---|---|---|---|
| 0.05 | 0 | ycaM | 2 | 0 | 0.225 |
| 0.05 | 0 | yadI | 2 | 0 | 0.222 |
| 0.05 | 0 | AG1 | 2 | 0 | 0.237 |
| 0.05 | 0 | ptsI | 2 | 0.25 | 0.21 |
| 0.05 | 0 | clcB | 2 | 0.25 | 0.218 |
| 0.05 | 0 | ycaM | 2 | 0.25 | 0.217 |
| 0.05 | 0 | yadI | 2 | 0.25 | 0.217 |
| 0.05 | 0 | AG1 | 2 | 0.25 | 0.23 |
| 0.05 | 0 | ptsI | 2 | 0.5 | 0.211 |
| 0.05 | 0 | clcB | 2 | 0.5 | 0.218 |
| 0.05 | 0 | ycaM | 2 | 0.5 | 0.216 |
| 0.05 | 0 | yadI | 2 | 0.5 | 0.214 |
| 0.05 | 0 | AG1 | 2 | 0.5 | 0.226 |
| 0.05 | 0 | ptsI | 2 | 0.75 | 0.212 |
| 0.05 | 0 | clcB | 2 | 0.75 | 0.219 |
| 0.05 | 0 | ycaM | 2 | 0.75 | 0.218 |
| 0.05 | 0 | yadI | 2 | 0.75 | 0.218 |
| 0.05 | 0 | AG1 | 2 | 0.75 | 0.229 |
| 0.05 | 0 | ptsI | 2 | 1 | 0.216 |
| 0.05 | 0 | clcB | 2 | 1 | 0.22 |
| 0.05 | 0 | ycaM | 2 | 1 | 0.22 |
| 0.05 | 0 | yadI | 2 | 1 | 0.223 |
| 0.05 | 0 | AG1 | 2 | 1 | 0.231 |
| 0.05 | 0 | ptsI | 2 | 1.25 | 0.217 |
| 0.05 | 0 | clcB | 2 | 1.25 | 0.223 |
| 0.05 | 0 | ycaM | 2 | 1.25 | 0.222 |
| 0.05 | 0 | yadI | 2 | 1.25 | 0.23 |
| 0.05 | 0 | AG1 | 2 | 1.25 | 0.237 |
| 0.05 | 0 | ptsI | 2 | 1.5 | 0.22 |
| 0.05 | 0 | clcB | 2 | 1.5 | 0.225 |
| 0.05 | 0 | ycaM | 2 | 1.5 | 0.225 |
| 0.05 | 0 | yadI | 2 | 1.5 | 0.236 |
| 0.05 | 0 | AG1 | 2 | 1.5 | 0.244 |
| 0.05 | 0 | ptsI | 2 | 1.75 | 0.224 |
| 0.05 | 0 | clcB | 2 | 1.75 | 0.228 |
| 0.05 | 0 | ycaM | 2 | 1.75 | 0.23 |
| 0.05 | 0 | yadI | 2 | 1.75 | 0.24 |
| 0.05 | 0 | AG1 | 2 | 1.75 | 0.247 |
| 0.05 | 0 | ptsI | 2 | 2 | 0.231 |
| 0.05 | 0 | clcB | 2 | 2 | 0.234 |
| 0.05 | 0 | ycaM | 2 | 2 | 0.236 |
| 0.05 | 0 | yadI | 2 | 2 | 0.248 |
| 0.05 | 0 | AG1 | 2 | 2 | 0.255 |
| 0.05 | 0 | ptsI | 2 | 2.25 | 0.24 |
| 0.05 | 0 | clcB | 2 | 2.25 | 0.241 |
| 0.05 | 0 | ycaM | 2 | 2.25 | 0.243 |
| 0.05 | 0 | yadI | 2 | 2.25 | 0.262 |
| 0.05 | 0 | AG1 | 2 | 2.25 | 0.265 |

| | | | | | |
|---|---|---|---|---|---|
| 0.05 | 0 | ptsI | 2 | 2.5 | 0.248 |
| 0.05 | 0 | clcB | 2 | 2.5 | 0.247 |
| 0.05 | 0 | ycaM | 2 | 2.5 | 0.248 |
| 0.05 | 0 | yadI | 2 | 2.5 | 0.275 |
| 0.05 | 0 | AG1 | 2 | 2.5 | 0.274 |
| 0.05 | 0 | ptsI | 2 | 2.75 | 0.254 |
| 0.05 | 0 | clcB | 2 | 2.75 | 0.252 |
| 0.05 | 0 | ycaM | 2 | 2.75 | 0.257 |
| 0.05 | 0 | yadI | 2 | 2.75 | 0.287 |
| 0.05 | 0 | AG1 | 2 | 2.75 | 0.281 |
| 0.05 | 0 | ptsI | 2 | 3 | 0.265 |
| 0.05 | 0 | clcB | 2 | 3 | 0.258 |
| 0.05 | 0 | ycaM | 2 | 3 | 0.264 |
| 0.05 | 0 | yadI | 2 | 3 | 0.303 |
| 0.05 | 0 | AG1 | 2 | 3 | 0.287 |
| 0.05 | 0 | ptsI | 2 | 3.25 | 0.271 |
| 0.05 | 0 | clcB | 2 | 3.25 | 0.27 |
| 0.05 | 0 | ycaM | 2 | 3.25 | 0.273 |
| 0.05 | 0 | yadI | 2 | 3.25 | 0.32 |
| 0.05 | 0 | AG1 | 2 | 3.25 | 0.3 |
| 0.05 | 0 | ptsI | 2 | 3.5 | 0.292 |
| 0.05 | 0 | clcB | 2 | 3.5 | 0.283 |
| 0.05 | 0 | ycaM | 2 | 3.5 | 0.281 |
| 0.05 | 0 | yadI | 2 | 3.5 | 0.343 |
| 0.05 | 0 | AG1 | 2 | 3.5 | 0.313 |
| 0.05 | 0 | ptsI | 2 | 3.75 | 0.294 |
| 0.05 | 0 | clcB | 2 | 3.75 | 0.282 |
| 0.05 | 0 | ycaM | 2 | 3.75 | 0.291 |
| 0.05 | 0 | yadI | 2 | 3.75 | 0.364 |
| 0.05 | 0 | AG1 | 2 | 3.75 | 0.326 |
| 0.05 | 0 | ptsI | 2 | 4 | 0.302 |
| 0.05 | 0 | clcB | 2 | 4 | 0.291 |
| 0.05 | 0 | ycaM | 2 | 4 | 0.302 |
| 0.05 | 0 | yadI | 2 | 4 | 0.389 |
| 0.05 | 0 | AG1 | 2 | 4 | 0.336 |
| 0.05 | 0 | ptsI | 2 | 4.38333 | 0.325 |
| 0.05 | 0 | clcB | 2 | 4.38333 | 0.307 |
| 0.05 | 0 | ycaM | 2 | 4.38333 | 0.32 |
| 0.05 | 0 | yadI | 2 | 4.38333 | 0.406 |
| 0.05 | 0 | AG1 | 2 | 4.38333 | 0.356 |
| 0.05 | 0 | ptsI | 2 | 4.63333 | 0.315 |
| 0.05 | 0 | clcB | 2 | 4.63333 | 0.3 |
| 0.05 | 0 | ycaM | 2 | 4.63333 | 0.315 |
| 0.05 | 0 | yadI | 2 | 4.63333 | 0.412 |
| 0.05 | 0 | AG1 | 2 | 4.63333 | 0.351 |
| 0.05 | 0 | ptsI | 2 | 4.88333 | 0.322 |
| 0.05 | 0 | clcB | 2 | 4.88333 | 0.305 |
| 0.05 | 0 | ycaM | 2 | 4.88333 | 0.323 |

| | | | | | |
|---|---|---|---|---|---|
| 0.05 | 0 | yadI | 2 | 4.88333 | 0.427 |
| 0.05 | 0 | AG1 | 2 | 4.88333 | 0.361 |
| 0.05 | 0 | ptsI | 2 | 5.13333 | 0.329 |
| 0.05 | 0 | clcB | 2 | 5.13333 | 0.315 |
| 0.05 | 0 | ycaM | 2 | 5.13333 | 0.332 |
| 0.05 | 0 | yadI | 2 | 5.13333 | 0.442 |
| 0.05 | 0 | AG1 | 2 | 5.13333 | 0.373 |
| 0.05 | 0 | ptsI | 2 | 5.38333 | 0.337 |
| 0.05 | 0 | clcB | 2 | 5.38333 | 0.323 |
| 0.05 | 0 | ycaM | 2 | 5.38333 | 0.341 |
| 0.05 | 0 | yadI | 2 | 5.38333 | 0.469 |
| 0.05 | 0 | AG1 | 2 | 5.38333 | 0.383 |
| 0.05 | 0 | ptsI | 2 | 5.63333 | 0.345 |
| 0.05 | 0 | clcB | 2 | 5.63333 | 0.33 |
| 0.05 | 0 | ycaM | 2 | 5.63333 | 0.352 |
| 0.05 | 0 | yadI | 2 | 5.63333 | 0.47 |
| 0.05 | 0 | AG1 | 2 | 5.63333 | 0.392 |
| 0.05 | 0 | ptsI | 2 | 5.88333 | 0.353 |
| 0.05 | 0 | clcB | 2 | 5.88333 | 0.336 |
| 0.05 | 0 | ycaM | 2 | 5.88333 | 0.363 |
| 0.05 | 0 | yadI | 2 | 5.88333 | 0.479 |
| 0.05 | 0 | AG1 | 2 | 5.88333 | 0.401 |
| 0.05 | 0 | ptsI | 2 | 6.13333 | 0.361 |
| 0.05 | 0 | clcB | 2 | 6.13333 | 0.343 |
| 0.05 | 0 | ycaM | 2 | 6.13333 | 0.376 |
| 0.05 | 0 | yadI | 2 | 6.13333 | 0.494 |
| 0.05 | 0 | AG1 | 2 | 6.13333 | 0.417 |
| 0.05 | 0 | ptsI | 2 | 6.38333 | 0.37 |
| 0.05 | 0 | clcB | 2 | 6.38333 | 0.352 |
| 0.05 | 0 | ycaM | 2 | 6.38333 | 0.388 |
| 0.05 | 0 | yadI | 2 | 6.38333 | 0.508 |
| 0.05 | 0 | AG1 | 2 | 6.38333 | 0.43 |
| 0.05 | 0 | ptsI | 2 | 6.63333 | 0.379 |
| 0.05 | 0 | clcB | 2 | 6.63333 | 0.361 |
| 0.05 | 0 | ycaM | 2 | 6.63333 | 0.401 |
| 0.05 | 0 | yadI | 2 | 6.63333 | 0.523 |
| 0.05 | 0 | AG1 | 2 | 6.63333 | 0.439 |
| 0.05 | 0 | ptsI | 2 | 6.88333 | 0.385 |
| 0.05 | 0 | clcB | 2 | 6.88333 | 0.368 |
| 0.05 | 0 | ycaM | 2 | 6.88333 | 0.41 |
| 0.05 | 0 | yadI | 2 | 6.88333 | 0.539 |
| 0.05 | 0 | AG1 | 2 | 6.88333 | 0.454 |
| 0.05 | 0 | ptsI | 2 | 7.13333 | 0.391 |
| 0.05 | 0 | clcB | 2 | 7.13333 | 0.375 |
| 0.05 | 0 | ycaM | 2 | 7.13333 | 0.417 |
| 0.05 | 0 | yadI | 2 | 7.13333 | 0.546 |
| 0.05 | 0 | AG1 | 2 | 7.13333 | 0.462 |
| 0.05 | 0 | ptsI | 2 | 7.38333 | 0.398 |

| | | | | | |
|---|---|---|---|---|---|
| 0.05 | 0 | clcB | 2 | 7.38333 | 0.384 |
| 0.05 | 0 | ycaM | 2 | 7.38333 | 0.429 |
| 0.05 | 0 | yadI | 2 | 7.38333 | 0.565 |
| 0.05 | 0 | AG1 | 2 | 7.38333 | 0.476 |
| 0.05 | 0 | ptsI | 2 | 7.63333 | 0.406 |
| 0.05 | 0 | clcB | 2 | 7.63333 | 0.392 |
| 0.05 | 0 | ycaM | 2 | 7.63333 | 0.437 |
| 0.05 | 0 | yadI | 2 | 7.63333 | 0.576 |
| 0.05 | 0 | AG1 | 2 | 7.63333 | 0.488 |
| 0.05 | 0 | ptsI | 2 | 7.88333 | 0.418 |
| 0.05 | 0 | clcB | 2 | 7.88333 | 0.4 |
| 0.05 | 0 | ycaM | 2 | 7.88333 | 0.446 |
| 0.05 | 0 | yadI | 2 | 7.88333 | 0.594 |
| 0.05 | 0 | AG1 | 2 | 7.88333 | 0.499 |
| 0.05 | 0 | ptsI | 2 | 8.13333 | 0.428 |
| 0.05 | 0 | clcB | 2 | 8.13333 | 0.407 |
| 0.05 | 0 | ycaM | 2 | 8.13333 | 0.452 |
| 0.05 | 0 | yadI | 2 | 8.13333 | 0.606 |
| 0.05 | 0 | AG1 | 2 | 8.13333 | 0.512 |
| 0.05 | 0 | ptsI | 2 | 8.38333 | 0.434 |
| 0.05 | 0 | clcB | 2 | 8.38333 | 0.419 |
| 0.05 | 0 | ycaM | 2 | 8.38333 | 0.456 |
| 0.05 | 0 | yadI | 2 | 8.38333 | 0.624 |
| 0.05 | 0 | AG1 | 2 | 8.38333 | 0.523 |
| 0.05 | 0 | ptsI | 2 | 8.91667 | 0.496 |
| 0.05 | 0 | clcB | 2 | 8.91667 | 0.47 |
| 0.05 | 0 | ycaM | 2 | 8.91667 | 0.492 |
| 0.05 | 0 | yadI | 2 | 8.91667 | 0.669 |
| 0.05 | 0 | AG1 | 2 | 8.91667 | 0.403 |
| 0.05 | 0 | ptsI | 2 | 9.16667 | 0.497 |
| 0.05 | 0 | clcB | 2 | 9.16667 | 0.471 |
| 0.05 | 0 | ycaM | 2 | 9.16667 | 0.489 |
| 0.05 | 0 | yadI | 2 | 9.16667 | 0.66 |
| 0.05 | 0 | AG1 | 2 | 9.16667 | 0.404 |
| 0.05 | 0 | ptsI | 2 | 9.41667 | 0.501 |
| 0.05 | 0 | clcB | 2 | 9.41667 | 0.48 |
| 0.05 | 0 | ycaM | 2 | 9.41667 | 0.494 |
| 0.05 | 0 | yadI | 2 | 9.41667 | 0.667 |
| 0.05 | 0 | AG1 | 2 | 9.41667 | 0.417 |
| 0.05 | 0 | ptsI | 2 | 9.66667 | 0.513 |
| 0.05 | 0 | clcB | 2 | 9.66667 | 0.488 |
| 0.05 | 0 | ycaM | 2 | 9.66667 | 0.503 |
| 0.05 | 0 | yadI | 2 | 9.66667 | 0.675 |
| 0.05 | 0 | AG1 | 2 | 9.66667 | 0.428 |
| 0.05 | 0 | ptsI | 2 | 9.91667 | 0.517 |
| 0.05 | 0 | clcB | 2 | 9.91667 | 0.494 |
| 0.05 | 0 | ycaM | 2 | 9.91667 | 0.511 |
| 0.05 | 0 | yadI | 2 | 9.91667 | 0.685 |

| | | | | | |
|---|---|---|---|---|---|
| 0.05 | 0 | AG1 | 2 | 9.91667 | 0.434 |
| 0.05 | 0 | ptsI | 2 | 10.1667 | 0.525 |
| 0.05 | 0 | clcB | 2 | 10.1667 | 0.503 |
| 0.05 | 0 | ycaM | 2 | 10.1667 | 0.517 |
| 0.05 | 0 | yadI | 2 | 10.1667 | 0.697 |
| 0.05 | 0 | AG1 | 2 | 10.1667 | 0.434 |
| 0.05 | 0 | ptsI | 2 | 10.4167 | 0.528 |
| 0.05 | 0 | clcB | 2 | 10.4167 | 0.513 |
| 0.05 | 0 | ycaM | 2 | 10.4167 | 0.527 |
| 0.05 | 0 | yadI | 2 | 10.4167 | 0.712 |
| 0.05 | 0 | AG1 | 2 | 10.4167 | 0.454 |
| 0.05 | 0 | ptsI | 2 | 10.6667 | 0.533 |
| 0.05 | 0 | clcB | 2 | 10.6667 | 0.52 |
| 0.05 | 0 | ycaM | 2 | 10.6667 | 0.536 |
| 0.05 | 0 | yadI | 2 | 10.6667 | 0.723 |
| 0.05 | 0 | AG1 | 2 | 10.6667 | 0.458 |
| 0.05 | 0 | ptsI | 2 | 10.9167 | 0.561 |
| 0.05 | 0 | clcB | 2 | 10.9167 | 0.528 |
| 0.05 | 0 | ycaM | 2 | 10.9167 | 0.544 |
| 0.05 | 0 | yadI | 2 | 10.9167 | 0.731 |
| 0.05 | 0 | AG1 | 2 | 10.9167 | 0.463 |
| 0.05 | 0 | ptsI | 2 | 11.1667 | 0.581 |
| 0.05 | 0 | clcB | 2 | 11.1667 | 0.541 |
| 0.05 | 0 | ycaM | 2 | 11.1667 | 0.555 |
| 0.05 | 0 | yadI | 2 | 11.1667 | 0.744 |
| 0.05 | 0 | AG1 | 2 | 11.1667 | 0.466 |
| 0.05 | 0 | ptsI | 2 | 11.4167 | 0.575 |
| 0.05 | 0 | clcB | 2 | 11.4167 | 0.545 |
| 0.05 | 0 | ycaM | 2 | 11.4167 | 0.563 |
| 0.05 | 0 | yadI | 2 | 11.4167 | 0.753 |
| 0.05 | 0 | AG1 | 2 | 11.4167 | 0.464 |
| 0.05 | 0 | ptsI | 2 | 11.6667 | 0.569 |
| 0.05 | 0 | clcB | 2 | 11.6667 | 0.56 |
| 0.05 | 0 | ycaM | 2 | 11.6667 | 0.574 |
| 0.05 | 0 | yadI | 2 | 11.6667 | 0.767 |
| 0.05 | 0 | AG1 | 2 | 11.6667 | 0.485 |
| 0.05 | 0 | ptsI | 2 | 11.9167 | 0.548 |
| 0.05 | 0 | clcB | 2 | 11.9167 | 0.564 |
| 0.05 | 0 | ycaM | 2 | 11.9167 | 0.585 |
| 0.05 | 0 | yadI | 2 | 11.9167 | 0.774 |
| 0.05 | 0 | AG1 | 2 | 11.9167 | 0.483 |
| 0.05 | 0 | ptsI | 2 | 12.1667 | 0.587 |
| 0.05 | 0 | clcB | 2 | 12.1667 | 0.58 |
| 0.05 | 0 | ycaM | 2 | 12.1667 | 0.595 |
| 0.05 | 0 | yadI | 2 | 12.1667 | 0.792 |
| 0.05 | 0 | AG1 | 2 | 12.1667 | 0.496 |
| 0.05 | 0 | ptsI | 2 | 12.4167 | 0.568 |
| 0.05 | 0 | clcB | 2 | 12.4167 | 0.596 |

| | | | | | |
|---|---|---|---|---|---|
| 0.05 | 0 | ycaM | 2 | 12.4167 | 0.604 |
| 0.05 | 0 | yadI | 2 | 12.4167 | 0.792 |
| 0.05 | 0 | AG1 | 2 | 12.4167 | 0.496 |
| 0.05 | 0 | ptsI | 2 | 12.6667 | 0.573 |
| 0.05 | 0 | clcB | 2 | 12.6667 | 0.587 |
| 0.05 | 0 | ycaM | 2 | 12.6667 | 0.612 |
| 0.05 | 0 | yadI | 2 | 12.6667 | 0.804 |
| 0.05 | 0 | AG1 | 2 | 12.6667 | 0.5 |
| 0.05 | 0 | ptsI | 2 | 12.9167 | 0.585 |
| 0.05 | 0 | clcB | 2 | 12.9167 | 0.592 |
| 0.05 | 0 | ycaM | 2 | 12.9167 | 0.633 |
| 0.05 | 0 | yadI | 2 | 12.9167 | 0.818 |
| 0.05 | 0 | AG1 | 2 | 12.9167 | 0.504 |
| 0.05 | 0 | ptsI | 2 | 13.1667 | 0.59 |
| 0.05 | 0 | clcB | 2 | 13.1667 | 0.578 |
| 0.05 | 0 | ycaM | 2 | 13.1667 | 0.641 |
| 0.05 | 0 | yadI | 2 | 13.1667 | 0.821 |
| 0.05 | 0 | AG1 | 2 | 13.1667 | 0.509 |
| 0.05 | 0 | ptsI | 2 | 13.4167 | 0.587 |
| 0.05 | 0 | clcB | 2 | 13.4167 | 0.575 |
| 0.05 | 0 | ycaM | 2 | 13.4167 | 0.654 |
| 0.05 | 0 | yadI | 2 | 13.4167 | 0.837 |
| 0.05 | 0 | AG1 | 2 | 13.4167 | 0.507 |
| 0.05 | 0 | ptsI | 2 | 13.6667 | 0.594 |
| 0.05 | 0 | clcB | 2 | 13.6667 | 0.583 |
| 0.05 | 0 | ycaM | 2 | 13.6667 | 0.678 |
| 0.05 | 0 | yadI | 2 | 13.6667 | 0.835 |
| 0.05 | 0 | AG1 | 2 | 13.6667 | 0.539 |
| 0.05 | 0 | ptsI | 2 | 13.9167 | 0.604 |
| 0.05 | 0 | clcB | 2 | 13.9167 | 0.589 |
| 0.05 | 0 | ycaM | 2 | 13.9167 | 0.679 |
| 0.05 | 0 | yadI | 2 | 13.9167 | 0.841 |
| 0.05 | 0 | AG1 | 2 | 13.9167 | 0.543 |
| 0.05 | 0 | ptsI | 2 | 14.1667 | 0.606 |
| 0.05 | 0 | clcB | 2 | 14.1667 | 0.584 |
| 0.05 | 0 | ycaM | 2 | 14.1667 | 0.685 |
| 0.05 | 0 | yadI | 2 | 14.1667 | 0.857 |
| 0.05 | 0 | AG1 | 2 | 14.1667 | 0.534 |
| 0.05 | 0 | ptsI | 2 | 14.4167 | 0.624 |
| 0.05 | 0 | clcB | 2 | 14.4167 | 0.584 |
| 0.05 | 0 | ycaM | 2 | 14.4167 | 0.706 |
| 0.05 | 0 | yadI | 2 | 14.4167 | 0.862 |
| 0.05 | 0 | AG1 | 2 | 14.4167 | 0.564 |
| 0.05 | 0 | ptsI | 2 | 14.6667 | 0.648 |
| 0.05 | 0 | clcB | 2 | 14.6667 | 0.592 |
| 0.05 | 0 | ycaM | 2 | 14.6667 | 0.693 |
| 0.05 | 0 | yadI | 2 | 14.6667 | 0.862 |
| 0.05 | 0 | AG1 | 2 | 14.6667 | 0.586 |

| | | | | | |
|---|---|---|---|---|---|
| 0.05 | 0 | ptsI | 2 | 14.9167 | 0.67 |
| 0.05 | 0 | clcB | 2 | 14.9167 | 0.596 |
| 0.05 | 0 | ycaM | 2 | 14.9167 | 0.696 |
| 0.05 | 0 | yadI | 2 | 14.9167 | 0.865 |
| 0.05 | 0 | AG1 | 2 | 14.9167 | 0.597 |
| 0.05 | 0 | ptsI | 2 | 15.1667 | 0.661 |
| 0.05 | 0 | clcB | 2 | 15.1667 | 0.605 |
| 0.05 | 0 | ycaM | 2 | 15.1667 | 0.69 |
| 0.05 | 0 | yadI | 2 | 15.1667 | 0.873 |
| 0.05 | 0 | AG1 | 2 | 15.1667 | 0.614 |
| 0.05 | 0 | ptsI | 2 | 15.4167 | 0.678 |
| 0.05 | 0 | clcB | 2 | 15.4167 | 0.597 |
| 0.05 | 0 | ycaM | 2 | 15.4167 | 0.681 |
| 0.05 | 0 | yadI | 2 | 15.4167 | 0.878 |
| 0.05 | 0 | AG1 | 2 | 15.4167 | 0.635 |
| 0.05 | 0 | ptsI | 2 | 15.6667 | 0.687 |
| 0.05 | 0 | clcB | 2 | 15.6667 | 0.618 |
| 0.05 | 0 | ycaM | 2 | 15.6667 | 0.691 |
| 0.05 | 0 | yadI | 2 | 15.6667 | 0.879 |
| 0.05 | 0 | AG1 | 2 | 15.6667 | 0.654 |
| 0.05 | 0 | ptsI | 2 | 15.9167 | 0.712 |
| 0.05 | 0 | clcB | 2 | 15.9167 | 0.59 |
| 0.05 | 0 | ycaM | 2 | 15.9167 | 0.679 |
| 0.05 | 0 | yadI | 2 | 15.9167 | 0.889 |
| 0.05 | 0 | AG1 | 2 | 15.9167 | 0.653 |
| 0.05 | 0 | ptsI | 2 | 16.1667 | 0.719 |
| 0.05 | 0 | clcB | 2 | 16.1667 | 0.62 |
| 0.05 | 0 | ycaM | 2 | 16.1667 | 0.676 |
| 0.05 | 0 | yadI | 2 | 16.1667 | 0.894 |
| 0.05 | 0 | AG1 | 2 | 16.1667 | 0.688 |
| 0.05 | 0 | ptsI | 2 | 16.4167 | 0.744 |
| 0.05 | 0 | clcB | 2 | 16.4167 | 0.613 |
| 0.05 | 0 | ycaM | 2 | 16.4167 | 0.685 |
| 0.05 | 0 | yadI | 2 | 16.4167 | 0.896 |
| 0.05 | 0 | AG1 | 2 | 16.4167 | 0.687 |
| 0.05 | 0 | ptsI | 2 | 16.6667 | 0.77 |
| 0.05 | 0 | clcB | 2 | 16.6667 | 0.634 |
| 0.05 | 0 | ycaM | 2 | 16.6667 | 0.679 |
| 0.05 | 0 | yadI | 2 | 16.6667 | 0.895 |
| 0.05 | 0 | AG1 | 2 | 16.6667 | 0.673 |
| 0.05 | 0 | ptsI | 2 | 16.9167 | 0.788 |
| 0.05 | 0 | clcB | 2 | 16.9167 | 0.638 |
| 0.05 | 0 | ycaM | 2 | 16.9167 | 0.685 |
| 0.05 | 0 | yadI | 2 | 16.9167 | 0.902 |
| 0.05 | 0 | AG1 | 2 | 16.9167 | 0.674 |
| 0.05 | 0 | ptsI | 2 | 17.1667 | 0.803 |
| 0.05 | 0 | clcB | 2 | 17.1667 | 0.629 |
| 0.05 | 0 | ycaM | 2 | 17.1667 | 0.681 |

| | | | | | |
|---|---|---|---|---|---|
| 0.05 | 0 | yadI | 2 | 17.1667 | 0.907 |
| 0.05 | 0 | AG1 | 2 | 17.1667 | 0.692 |
| 0.05 | 0 | ptsI | 2 | 17.4167 | 0.811 |
| 0.05 | 0 | clcB | 2 | 17.4167 | 0.639 |
| 0.05 | 0 | ycaM | 2 | 17.4167 | 0.713 |
| 0.05 | 0 | yadI | 2 | 17.4167 | 0.906 |
| 0.05 | 0 | AG1 | 2 | 17.4167 | 0.692 |
| 0.05 | 0 | ptsI | 2 | 17.6667 | 0.819 |
| 0.05 | 0 | clcB | 2 | 17.6667 | 0.62 |
| 0.05 | 0 | ycaM | 2 | 17.6667 | 0.714 |
| 0.05 | 0 | yadI | 2 | 17.6667 | 0.918 |
| 0.05 | 0 | AG1 | 2 | 17.6667 | 0.699 |
| 0.05 | 0 | ptsI | 2 | 17.9167 | 0.825 |
| 0.05 | 0 | clcB | 2 | 17.9167 | 0.614 |
| 0.05 | 0 | ycaM | 2 | 17.9167 | 0.715 |
| 0.05 | 0 | yadI | 2 | 17.9167 | 0.91 |
| 0.05 | 0 | AG1 | 2 | 17.9167 | 0.708 |
| 0.05 | 0 | ptsI | 2 | 18.1667 | 0.83 |
| 0.05 | 0 | clcB | 2 | 18.1667 | 0.62 |
| 0.05 | 0 | ycaM | 2 | 18.1667 | 0.706 |
| 0.05 | 0 | yadI | 2 | 18.1667 | 0.918 |
| 0.05 | 0 | AG1 | 2 | 18.1667 | 0.718 |
| 0.05 | 0 | ptsI | 2 | 18.4167 | 0.842 |
| 0.05 | 0 | clcB | 2 | 18.4167 | 0.596 |
| 0.05 | 0 | ycaM | 2 | 18.4167 | 0.699 |
| 0.05 | 0 | yadI | 2 | 18.4167 | 0.92 |
| 0.05 | 0 | AG1 | 2 | 18.4167 | 0.72 |
| 0.05 | 0 | ptsI | 2 | 18.6667 | 0.839 |
| 0.05 | 0 | clcB | 2 | 18.6667 | 0.6 |
| 0.05 | 0 | ycaM | 2 | 18.6667 | 0.693 |
| 0.05 | 0 | yadI | 2 | 18.6667 | 0.917 |
| 0.05 | 0 | AG1 | 2 | 18.6667 | 0.724 |
| 0.05 | 0 | ptsI | 2 | 18.9167 | 0.84 |
| 0.05 | 0 | clcB | 2 | 18.9167 | 0.604 |
| 0.05 | 0 | ycaM | 2 | 18.9167 | 0.696 |
| 0.05 | 0 | yadI | 2 | 18.9167 | 0.92 |
| 0.05 | 0 | AG1 | 2 | 18.9167 | 0.731 |
| 0.05 | 0 | ptsI | 2 | 19.1667 | 0.856 |
| 0.05 | 0 | clcB | 2 | 19.1667 | 0.611 |
| 0.05 | 0 | ycaM | 2 | 19.1667 | 0.703 |
| 0.05 | 0 | yadI | 2 | 19.1667 | 0.919 |
| 0.05 | 0 | AG1 | 2 | 19.1667 | 0.737 |
| 0.05 | 0 | ptsI | 2 | 19.4167 | 0.857 |
| 0.05 | 0 | clcB | 2 | 19.4167 | 0.611 |
| 0.05 | 0 | ycaM | 2 | 19.4167 | 0.712 |
| 0.05 | 0 | yadI | 2 | 19.4167 | 0.917 |
| 0.05 | 0 | AG1 | 2 | 19.4167 | 0.74 |
| 0.05 | 0 | ptsI | 2 | 19.6667 | 0.865 |

| | | | | | |
|---|---|---|---|---|---|
| 0.05 | 0 | clcB | 2 | 19.6667 | 0.613 |
| 0.05 | 0 | ycaM | 2 | 19.6667 | 0.711 |
| 0.05 | 0 | yadI | 2 | 19.6667 | 0.929 |
| 0.05 | 0 | AG1 | 2 | 19.6667 | 0.748 |
| 0.05 | 0 | ptsI | 2 | 19.9167 | 0.877 |
| 0.05 | 0 | clcB | 2 | 19.9167 | 0.62 |
| 0.05 | 0 | ycaM | 2 | 19.9167 | 0.715 |
| 0.05 | 0 | yadI | 2 | 19.9167 | 0.924 |
| 0.05 | 0 | AG1 | 2 | 19.9167 | 0.76 |
| 0.05 | 0 | ptsI | 2 | 20.1667 | 0.871 |
| 0.05 | 0 | clcB | 2 | 20.1667 | 0.616 |
| 0.05 | 0 | ycaM | 2 | 20.1667 | 0.712 |
| 0.05 | 0 | yadI | 2 | 20.1667 | 0.925 |
| 0.05 | 0 | AG1 | 2 | 20.1667 | 0.753 |
| 0.05 | 0 | ptsI | 2 | 20.4167 | 0.876 |
| 0.05 | 0 | clcB | 2 | 20.4167 | 0.618 |
| 0.05 | 0 | ycaM | 2 | 20.4167 | 0.716 |
| 0.05 | 0 | yadI | 2 | 20.4167 | 0.926 |
| 0.05 | 0 | AG1 | 2 | 20.4167 | 0.751 |
| 0.05 | 0 | ptsI | 2 | 20.6667 | 0.88 |
| 0.05 | 0 | clcB | 2 | 20.6667 | 0.618 |
| 0.05 | 0 | ycaM | 2 | 20.6667 | 0.716 |
| 0.05 | 0 | yadI | 2 | 20.6667 | 0.926 |
| 0.05 | 0 | AG1 | 2 | 20.6667 | 0.754 |
| 0.05 | 0 | ptsI | 2 | 20.9167 | 0.88 |
| 0.05 | 0 | clcB | 2 | 20.9167 | 0.621 |
| 0.05 | 0 | ycaM | 2 | 20.9167 | 0.716 |
| 0.05 | 0 | yadI | 2 | 20.9167 | 0.932 |
| 0.05 | 0 | AG1 | 2 | 20.9167 | 0.756 |
| 0.05 | 0 | ptsI | 2 | 21.1667 | 0.881 |
| 0.05 | 0 | clcB | 2 | 21.1667 | 0.623 |
| 0.05 | 0 | ycaM | 2 | 21.1667 | 0.721 |
| 0.05 | 0 | yadI | 2 | 21.1667 | 0.932 |
| 0.05 | 0 | AG1 | 2 | 21.1667 | 0.755 |
| 0.05 | 0 | ptsI | 2 | 21.4167 | 0.872 |
| 0.05 | 0 | clcB | 2 | 21.4167 | 0.619 |
| 0.05 | 0 | ycaM | 2 | 21.4167 | 0.71 |
| 0.05 | 0 | yadI | 2 | 21.4167 | 0.943 |
| 0.05 | 0 | AG1 | 2 | 21.4167 | 0.753 |
| 0.05 | 0 | ptsI | 2 | 21.6667 | 0.886 |
| 0.05 | 0 | clcB | 2 | 21.6667 | 0.624 |
| 0.05 | 0 | ycaM | 2 | 21.6667 | 0.722 |
| 0.05 | 0 | yadI | 2 | 21.6667 | 0.937 |
| 0.05 | 0 | AG1 | 2 | 21.6667 | 0.757 |
| 0.05 | 0 | ptsI | 2 | 21.9167 | 0.899 |
| 0.05 | 0 | clcB | 2 | 21.9167 | 0.629 |
| 0.05 | 0 | ycaM | 2 | 21.9167 | 0.729 |
| 0.05 | 0 | yadI | 2 | 21.9167 | 0.937 |

| | | | | | |
|---|---|---|---|---|---|
| 0.05 | 0 | AG1 | 2 | 21.9167 | 0.761 |
| 0.05 | 0 | ptsI | 2 | 22.1667 | 0.903 |
| 0.05 | 0 | clcB | 2 | 22.1667 | 0.629 |
| 0.05 | 0 | ycaM | 2 | 22.1667 | 0.726 |
| 0.05 | 0 | yadI | 2 | 22.1667 | 0.936 |
| 0.05 | 0 | AG1 | 2 | 22.1667 | 0.76 |
| 0.05 | 0 | ptsI | 2 | 22.4167 | 0.91 |
| 0.05 | 0 | clcB | 2 | 22.4167 | 0.63 |
| 0.05 | 0 | ycaM | 2 | 22.4167 | 0.723 |
| 0.05 | 0 | yadI | 2 | 22.4167 | 0.941 |
| 0.05 | 0 | AG1 | 2 | 22.4167 | 0.759 |
| 0.05 | 0 | ptsI | 2 | 22.6667 | 0.918 |
| 0.05 | 0 | clcB | 2 | 22.6667 | 0.633 |
| 0.05 | 0 | ycaM | 2 | 22.6667 | 0.725 |
| 0.05 | 0 | yadI | 2 | 22.6667 | 0.939 |
| 0.05 | 0 | AG1 | 2 | 22.6667 | 0.759 |
| 0.05 | 0 | ptsI | 2 | 22.9167 | 0.915 |
| 0.05 | 0 | clcB | 2 | 22.9167 | 0.635 |
| 0.05 | 0 | ycaM | 2 | 22.9167 | 0.728 |
| 0.05 | 0 | yadI | 2 | 22.9167 | 0.943 |
| 0.05 | 0 | AG1 | 2 | 22.9167 | 0.767 |
| 0.05 | 0 | ptsI | 2 | 23.1667 | 0.926 |
| 0.05 | 0 | clcB | 2 | 23.1667 | 0.64 |
| 0.05 | 0 | ycaM | 2 | 23.1667 | 0.725 |
| 0.05 | 0 | yadI | 2 | 23.1667 | 0.936 |
| 0.05 | 0 | AG1 | 2 | 23.1667 | 0.758 |
| 0.05 | 0 | ptsI | 2 | 23.4167 | 0.927 |
| 0.05 | 0 | clcB | 2 | 23.4167 | 0.64 |
| 0.05 | 0 | ycaM | 2 | 23.4167 | 0.729 |
| 0.05 | 0 | yadI | 2 | 23.4167 | 0.942 |
| 0.05 | 0 | AG1 | 2 | 23.4167 | 0.758 |
| 0.05 | 0 | ptsI | 2 | 23.6667 | 0.934 |
| 0.05 | 0 | clcB | 2 | 23.6667 | 0.644 |
| 0.05 | 0 | ycaM | 2 | 23.6667 | 0.729 |
| 0.05 | 0 | yadI | 2 | 23.6667 | 0.94 |
| 0.05 | 0 | AG1 | 2 | 23.6667 | 0.76 |
| 0.05 | 0 | ptsI | 2 | 23.9167 | 0.946 |
| 0.05 | 0 | clcB | 2 | 23.9167 | 0.646 |
| 0.05 | 0 | ycaM | 2 | 23.9167 | 0.731 |
| 0.05 | 0 | yadI | 2 | 23.9167 | 0.943 |
| 0.05 | 0 | AG1 | 2 | 23.9167 | 0.763 |
| 0.05 | 0 | ptsI | 2 | 24.1667 | 0.949 |
| 0.05 | 0 | clcB | 2 | 24.1667 | 0.644 |
| 0.05 | 0 | ycaM | 2 | 24.1667 | 0.73 |
| 0.05 | 0 | yadI | 2 | 24.1667 | 0.942 |
| 0.05 | 0 | AG1 | 2 | 24.1667 | 0.762 |
| 0.05 | 0 | ptsI | 2 | 24.4167 | 0.952 |
| 0.05 | 0 | clcB | 2 | 24.4167 | 0.643 |

| | | | | | |
|---|---|---|---|---|---|
| 0.05 | 0 | ycaM | 2 | 24.4167 | 0.731 |
| 0.05 | 0 | yadI | 2 | 24.4167 | 0.945 |
| 0.05 | 0 | AG1 | 2 | 24.4167 | 0.761 |
| 0.05 | 0 | ptsI | 2 | 24.6667 | 0.965 |
| 0.05 | 0 | clcB | 2 | 24.6667 | 0.649 |
| 0.05 | 0 | ycaM | 2 | 24.6667 | 0.732 |
| 0.05 | 0 | yadI | 2 | 24.6667 | 0.943 |
| 0.05 | 0 | AG1 | 2 | 24.6667 | 0.764 |
| 0.05 | 0 | ptsI | 2 | 24.9167 | 0.977 |
| 0.05 | 0 | clcB | 2 | 24.9167 | 0.651 |
| 0.05 | 0 | ycaM | 2 | 24.9167 | 0.734 |
| 0.05 | 0 | yadI | 2 | 24.9167 | 0.953 |
| 0.05 | 0 | AG1 | 2 | 24.9167 | 0.765 |
| 0.05 | 0 | ptsI | 2 | 25.1667 | 0.983 |
| 0.05 | 0 | clcB | 2 | 25.1667 | 0.648 |
| 0.05 | 0 | ycaM | 2 | 25.1667 | 0.735 |
| 0.05 | 0 | yadI | 2 | 25.1667 | 0.948 |
| 0.05 | 0 | AG1 | 2 | 25.1667 | 0.763 |
| 0.05 | 0 | ptsI | 2 | 25.4167 | 0.996 |
| 0.05 | 0 | clcB | 2 | 25.4167 | 0.655 |
| 0.05 | 0 | ycaM | 2 | 25.4167 | 0.738 |
| 0.05 | 0 | yadI | 2 | 25.4167 | 0.944 |
| 0.05 | 0 | AG1 | 2 | 25.4167 | 0.772 |
| 0.05 | 0 | ptsI | 2 | 25.6667 | 1.002 |
| 0.05 | 0 | clcB | 2 | 25.6667 | 0.652 |
| 0.05 | 0 | ycaM | 2 | 25.6667 | 0.736 |
| 0.05 | 0 | yadI | 2 | 25.6667 | 0.944 |
| 0.05 | 0 | AG1 | 2 | 25.6667 | 0.766 |
| 0.05 | 0 | ptsI | 2 | 25.9167 | 1.015 |
| 0.05 | 0 | clcB | 2 | 25.9167 | 0.651 |
| 0.05 | 0 | ycaM | 2 | 25.9167 | 0.736 |
| 0.05 | 0 | yadI | 2 | 25.9167 | 0.954 |
| 0.05 | 0 | AG1 | 2 | 25.9167 | 0.765 |
| 0.05 | 0 | ptsI | 2 | 26.1667 | 1.02 |
| 0.05 | 0 | clcB | 2 | 26.1667 | 0.656 |
| 0.05 | 0 | ycaM | 2 | 26.1667 | 0.733 |
| 0.05 | 0 | yadI | 2 | 26.1667 | 0.954 |
| 0.05 | 0 | AG1 | 2 | 26.1667 | 0.768 |
| 0.05 | 0 | ptsI | 2 | 26.4167 | 1.038 |
| 0.05 | 0 | clcB | 2 | 26.4167 | 0.661 |
| 0.05 | 0 | ycaM | 2 | 26.4167 | 0.744 |
| 0.05 | 0 | yadI | 2 | 26.4167 | 0.952 |
| 0.05 | 0 | AG1 | 2 | 26.4167 | 0.78 |
| 0.05 | 0 | ptsI | 3 | 0 | 0.254 |
| 0.05 | 0 | clcB | 3 | 0 | 0.153 |
| 0.05 | 0 | ycaM | 3 | 0 | 0.156 |
| 0.05 | 0 | yadI | 3 | 0 | 0.175 |
| 0.05 | 0 | AG1 | 3 | 0 | 0.168 |

| | | | | | |
|---|---|---|---|---|---|
| 0.05 | 0 | ptsI | 3 | 0.25 | 0.149 |
| 0.05 | 0 | clcB | 3 | 0.25 | 0.152 |
| 0.05 | 0 | ycaM | 3 | 0.25 | 0.158 |
| 0.05 | 0 | yadI | 3 | 0.25 | 0.171 |
| 0.05 | 0 | AG1 | 3 | 0.25 | 0.17 |
| 0.05 | 0 | ptsI | 3 | 0.5 | 0.145 |
| 0.05 | 0 | clcB | 3 | 0.5 | 0.152 |
| 0.05 | 0 | ycaM | 3 | 0.5 | 0.159 |
| 0.05 | 0 | yadI | 3 | 0.5 | 0.17 |
| 0.05 | 0 | AG1 | 3 | 0.5 | 0.163 |
| 0.05 | 0 | ptsI | 3 | 0.75 | 0.146 |
| 0.05 | 0 | clcB | 3 | 0.75 | 0.152 |
| 0.05 | 0 | ycaM | 3 | 0.75 | 0.159 |
| 0.05 | 0 | yadI | 3 | 0.75 | 0.17 |
| 0.05 | 0 | AG1 | 3 | 0.75 | 0.165 |
| 0.05 | 0 | ptsI | 3 | 1 | 0.147 |
| 0.05 | 0 | clcB | 3 | 1 | 0.153 |
| 0.05 | 0 | ycaM | 3 | 1 | 0.162 |
| 0.05 | 0 | yadI | 3 | 1 | 0.172 |
| 0.05 | 0 | AG1 | 3 | 1 | 0.167 |
| 0.05 | 0 | ptsI | 3 | 1.25 | 0.149 |
| 0.05 | 0 | clcB | 3 | 1.25 | 0.153 |
| 0.05 | 0 | ycaM | 3 | 1.25 | 0.163 |
| 0.05 | 0 | yadI | 3 | 1.25 | 0.176 |
| 0.05 | 0 | AG1 | 3 | 1.25 | 0.166 |
| 0.05 | 0 | ptsI | 3 | 1.5 | 0.15 |
| 0.05 | 0 | clcB | 3 | 1.5 | 0.156 |
| 0.05 | 0 | ycaM | 3 | 1.5 | 0.165 |
| 0.05 | 0 | yadI | 3 | 1.5 | 0.18 |
| 0.05 | 0 | AG1 | 3 | 1.5 | 0.169 |
| 0.05 | 0 | ptsI | 3 | 1.75 | 0.151 |
| 0.05 | 0 | clcB | 3 | 1.75 | 0.156 |
| 0.05 | 0 | ycaM | 3 | 1.75 | 0.164 |
| 0.05 | 0 | yadI | 3 | 1.75 | 0.183 |
| 0.05 | 0 | AG1 | 3 | 1.75 | 0.17 |
| 0.05 | 0 | ptsI | 3 | 2 | 0.154 |
| 0.05 | 0 | clcB | 3 | 2 | 0.158 |
| 0.05 | 0 | ycaM | 3 | 2 | 0.165 |
| 0.05 | 0 | yadI | 3 | 2 | 0.188 |
| 0.05 | 0 | AG1 | 3 | 2 | 0.172 |
| 0.05 | 0 | ptsI | 3 | 2.25 | 0.157 |
| 0.05 | 0 | clcB | 3 | 2.25 | 0.16 |
| 0.05 | 0 | ycaM | 3 | 2.25 | 0.168 |
| 0.05 | 0 | yadI | 3 | 2.25 | 0.192 |
| 0.05 | 0 | AG1 | 3 | 2.25 | 0.175 |
| 0.05 | 0 | ptsI | 3 | 2.5 | 0.159 |
| 0.05 | 0 | clcB | 3 | 2.5 | 0.162 |
| 0.05 | 0 | ycaM | 3 | 2.5 | 0.17 |

| | | | | | |
|---|---|---|---|---|---|
| 0.05 | 0 | yadI | 3 | 2.5 | 0.197 |
| 0.05 | 0 | AG1 | 3 | 2.5 | 0.179 |
| 0.05 | 0 | ptsI | 3 | 2.75 | 0.164 |
| 0.05 | 0 | clcB | 3 | 2.75 | 0.166 |
| 0.05 | 0 | ycaM | 3 | 2.75 | 0.172 |
| 0.05 | 0 | yadI | 3 | 2.75 | 0.206 |
| 0.05 | 0 | AG1 | 3 | 2.75 | 0.184 |
| 0.05 | 0 | ptsI | 3 | 3 | 0.167 |
| 0.05 | 0 | clcB | 3 | 3 | 0.167 |
| 0.05 | 0 | ycaM | 3 | 3 | 0.174 |
| 0.05 | 0 | yadI | 3 | 3 | 0.208 |
| 0.05 | 0 | AG1 | 3 | 3 | 0.187 |
| 0.05 | 0 | ptsI | 3 | 3.25 | 0.171 |
| 0.05 | 0 | clcB | 3 | 3.25 | 0.172 |
| 0.05 | 0 | ycaM | 3 | 3.25 | 0.177 |
| 0.05 | 0 | yadI | 3 | 3.25 | 0.218 |
| 0.05 | 0 | AG1 | 3 | 3.25 | 0.191 |
| 0.05 | 0 | ptsI | 3 | 3.5 | 0.175 |
| 0.05 | 0 | clcB | 3 | 3.5 | 0.176 |
| 0.05 | 0 | ycaM | 3 | 3.5 | 0.181 |
| 0.05 | 0 | yadI | 3 | 3.5 | 0.228 |
| 0.05 | 0 | AG1 | 3 | 3.5 | 0.194 |
| 0.05 | 0 | ptsI | 3 | 3.75 | 0.18 |
| 0.05 | 0 | clcB | 3 | 3.75 | 0.181 |
| 0.05 | 0 | ycaM | 3 | 3.75 | 0.186 |
| 0.05 | 0 | yadI | 3 | 3.75 | 0.24 |
| 0.05 | 0 | AG1 | 3 | 3.75 | 0.198 |
| 0.05 | 0 | ptsI | 3 | 4 | 0.184 |
| 0.05 | 0 | clcB | 3 | 4 | 0.184 |
| 0.05 | 0 | ycaM | 3 | 4 | 0.19 |
| 0.05 | 0 | yadI | 3 | 4 | 0.253 |
| 0.05 | 0 | AG1 | 3 | 4 | 0.203 |
| 0.05 | 0 | ptsI | 3 | 4.41667 | 0.194 |
| 0.05 | 0 | clcB | 3 | 4.41667 | 0.204 |
| 0.05 | 0 | ycaM | 3 | 4.41667 | 0.201 |
| 0.05 | 0 | yadI | 3 | 4.41667 | 0.27 |
| 0.05 | 0 | AG1 | 3 | 4.41667 | 0.212 |
| 0.05 | 0 | ptsI | 3 | 4.66667 | 0.198 |
| 0.05 | 0 | clcB | 3 | 4.66667 | 0.198 |
| 0.05 | 0 | ycaM | 3 | 4.66667 | 0.197 |
| 0.05 | 0 | yadI | 3 | 4.66667 | 0.266 |
| 0.05 | 0 | AG1 | 3 | 4.66667 | 0.209 |
| 0.05 | 0 | ptsI | 3 | 4.91667 | 0.2 |
| 0.05 | 0 | clcB | 3 | 4.91667 | 0.205 |
| 0.05 | 0 | ycaM | 3 | 4.91667 | 0.199 |
| 0.05 | 0 | yadI | 3 | 4.91667 | 0.277 |
| 0.05 | 0 | AG1 | 3 | 4.91667 | 0.211 |
| 0.05 | 0 | ptsI | 3 | 5.16667 | 0.205 |

| | | | | | |
|---|---|---|---|---|---|
| 0.05 | 0 | clcB | 3 | 5.16667 | 0.21 |
| 0.05 | 0 | ycaM | 3 | 5.16667 | 0.204 |
| 0.05 | 0 | yadI | 3 | 5.16667 | 0.295 |
| 0.05 | 0 | AG1 | 3 | 5.16667 | 0.218 |
| 0.05 | 0 | ptsI | 3 | 5.41667 | 0.211 |
| 0.05 | 0 | clcB | 3 | 5.41667 | 0.22 |
| 0.05 | 0 | ycaM | 3 | 5.41667 | 0.211 |
| 0.05 | 0 | yadI | 3 | 5.41667 | 0.311 |
| 0.05 | 0 | AG1 | 3 | 5.41667 | 0.228 |
| 0.05 | 0 | ptsI | 3 | 5.66667 | 0.218 |
| 0.05 | 0 | clcB | 3 | 5.66667 | 0.229 |
| 0.05 | 0 | ycaM | 3 | 5.66667 | 0.22 |
| 0.05 | 0 | yadI | 3 | 5.66667 | 0.323 |
| 0.05 | 0 | AG1 | 3 | 5.66667 | 0.237 |
| 0.05 | 0 | ptsI | 3 | 5.91667 | 0.225 |
| 0.05 | 0 | clcB | 3 | 5.91667 | 0.243 |
| 0.05 | 0 | ycaM | 3 | 5.91667 | 0.228 |
| 0.05 | 0 | yadI | 3 | 5.91667 | 0.335 |
| 0.05 | 0 | AG1 | 3 | 5.91667 | 0.246 |
| 0.05 | 0 | ptsI | 3 | 6.16667 | 0.236 |
| 0.05 | 0 | clcB | 3 | 6.16667 | 0.258 |
| 0.05 | 0 | ycaM | 3 | 6.16667 | 0.238 |
| 0.05 | 0 | yadI | 3 | 6.16667 | 0.352 |
| 0.05 | 0 | AG1 | 3 | 6.16667 | 0.268 |
| 0.05 | 0 | ptsI | 3 | 6.41667 | 0.243 |
| 0.05 | 0 | clcB | 3 | 6.41667 | 0.272 |
| 0.05 | 0 | ycaM | 3 | 6.41667 | 0.25 |
| 0.05 | 0 | yadI | 3 | 6.41667 | 0.368 |
| 0.05 | 0 | AG1 | 3 | 6.41667 | 0.273 |
| 0.05 | 0 | ptsI | 3 | 6.66667 | 0.255 |
| 0.05 | 0 | clcB | 3 | 6.66667 | 0.289 |
| 0.05 | 0 | ycaM | 3 | 6.66667 | 0.263 |
| 0.05 | 0 | yadI | 3 | 6.66667 | 0.383 |
| 0.05 | 0 | AG1 | 3 | 6.66667 | 0.286 |
| 0.05 | 0 | ptsI | 3 | 6.91667 | 0.264 |
| 0.05 | 0 | clcB | 3 | 6.91667 | 0.291 |
| 0.05 | 0 | ycaM | 3 | 6.91667 | 0.276 |
| 0.05 | 0 | yadI | 3 | 6.91667 | 0.394 |
| 0.05 | 0 | AG1 | 3 | 6.91667 | 0.293 |
| 0.05 | 0 | ptsI | 3 | 7.16667 | 0.274 |
| 0.05 | 0 | clcB | 3 | 7.16667 | 0.3 |
| 0.05 | 0 | ycaM | 3 | 7.16667 | 0.292 |
| 0.05 | 0 | yadI | 3 | 7.16667 | 0.406 |
| 0.05 | 0 | AG1 | 3 | 7.16667 | 0.302 |
| 0.05 | 0 | ptsI | 3 | 7.41667 | 0.284 |
| 0.05 | 0 | clcB | 3 | 7.41667 | 0.309 |
| 0.05 | 0 | ycaM | 3 | 7.41667 | 0.3 |
| 0.05 | 0 | yadI | 3 | 7.41667 | 0.419 |

| | | | | | |
|---|---|---|---|---|---|
| 0.05 | 0 | AG1 | 3 | 7.41667 | 0.313 |
| 0.05 | 0 | ptsI | 3 | 7.66667 | 0.291 |
| 0.05 | 0 | clcB | 3 | 7.66667 | 0.32 |
| 0.05 | 0 | ycaM | 3 | 7.66667 | 0.31 |
| 0.05 | 0 | yadI | 3 | 7.66667 | 0.43 |
| 0.05 | 0 | AG1 | 3 | 7.66667 | 0.323 |
| 0.05 | 0 | ptsI | 3 | 7.91667 | 0.299 |
| 0.05 | 0 | clcB | 3 | 7.91667 | 0.324 |
| 0.05 | 0 | ycaM | 3 | 7.91667 | 0.317 |
| 0.05 | 0 | yadI | 3 | 7.91667 | 0.443 |
| 0.05 | 0 | AG1 | 3 | 7.91667 | 0.331 |
| 0.05 | 0 | ptsI | 3 | 8.16667 | 0.306 |
| 0.05 | 0 | clcB | 3 | 8.16667 | 0.334 |
| 0.05 | 0 | ycaM | 3 | 8.16667 | 0.326 |
| 0.05 | 0 | yadI | 3 | 8.16667 | 0.453 |
| 0.05 | 0 | AG1 | 3 | 8.16667 | 0.341 |
| 0.05 | 0 | ptsI | 3 | 8.41667 | 0.316 |
| 0.05 | 0 | clcB | 3 | 8.41667 | 0.344 |
| 0.05 | 0 | ycaM | 3 | 8.41667 | 0.336 |
| 0.05 | 0 | yadI | 3 | 8.41667 | 0.471 |
| 0.05 | 0 | AG1 | 3 | 8.41667 | 0.354 |
| 0.05 | 0 | ptsI | 3 | 9.05 | 0.278 |
| 0.05 | 0 | clcB | 3 | 9.05 | 0.306 |
| 0.05 | 0 | ycaM | 3 | 9.05 | 0.302 |
| 0.05 | 0 | yadI | 3 | 9.05 | 0.451 |
| 0.05 | 0 | AG1 | 3 | 9.05 | 0.317 |
| 0.05 | 0 | ptsI | 3 | 9.3 | 0.284 |
| 0.05 | 0 | clcB | 3 | 9.3 | 0.31 |
| 0.05 | 0 | ycaM | 3 | 9.3 | 0.306 |
| 0.05 | 0 | yadI | 3 | 9.3 | 0.455 |
| 0.05 | 0 | AG1 | 3 | 9.3 | 0.315 |
| 0.05 | 0 | ptsI | 3 | 9.55 | 0.29 |
| 0.05 | 0 | clcB | 3 | 9.55 | 0.319 |
| 0.05 | 0 | ycaM | 3 | 9.55 | 0.313 |
| 0.05 | 0 | yadI | 3 | 9.55 | 0.464 |
| 0.05 | 0 | AG1 | 3 | 9.55 | 0.323 |
| 0.05 | 0 | ptsI | 3 | 9.8 | 0.298 |
| 0.05 | 0 | clcB | 3 | 9.8 | 0.325 |
| 0.05 | 0 | ycaM | 3 | 9.8 | 0.321 |
| 0.05 | 0 | yadI | 3 | 9.8 | 0.476 |
| 0.05 | 0 | AG1 | 3 | 9.8 | 0.331 |
| 0.05 | 0 | ptsI | 3 | 10.05 | 0.303 |
| 0.05 | 0 | clcB | 3 | 10.05 | 0.333 |
| 0.05 | 0 | ycaM | 3 | 10.05 | 0.328 |
| 0.05 | 0 | yadI | 3 | 10.05 | 0.489 |
| 0.05 | 0 | AG1 | 3 | 10.05 | 0.339 |
| 0.05 | 0 | ptsI | 3 | 10.3 | 0.31 |
| 0.05 | 0 | clcB | 3 | 10.3 | 0.339 |

| | | | | | |
|---|---|---|---|---|---|
| 0.05 | 0 | ycaM | 3 | 10.3 | 0.338 |
| 0.05 | 0 | yadI | 3 | 10.3 | 0.5 |
| 0.05 | 0 | AG1 | 3 | 10.3 | 0.35 |
| 0.05 | 0 | ptsI | 3 | 10.55 | 0.32 |
| 0.05 | 0 | clcB | 3 | 10.55 | 0.344 |
| 0.05 | 0 | ycaM | 3 | 10.55 | 0.348 |
| 0.05 | 0 | yadI | 3 | 10.55 | 0.521 |
| 0.05 | 0 | AG1 | 3 | 10.55 | 0.358 |
| 0.05 | 0 | ptsI | 3 | 10.8 | 0.328 |
| 0.05 | 0 | clcB | 3 | 10.8 | 0.35 |
| 0.05 | 0 | ycaM | 3 | 10.8 | 0.356 |
| 0.05 | 0 | yadI | 3 | 10.8 | 0.534 |
| 0.05 | 0 | AG1 | 3 | 10.8 | 0.366 |
| 0.05 | 0 | ptsI | 3 | 11.05 | 0.336 |
| 0.05 | 0 | clcB | 3 | 11.05 | 0.354 |
| 0.05 | 0 | ycaM | 3 | 11.05 | 0.365 |
| 0.05 | 0 | yadI | 3 | 11.05 | 0.545 |
| 0.05 | 0 | AG1 | 3 | 11.05 | 0.375 |
| 0.05 | 0 | ptsI | 3 | 11.3 | 0.345 |
| 0.05 | 0 | clcB | 3 | 11.3 | 0.36 |
| 0.05 | 0 | ycaM | 3 | 11.3 | 0.373 |
| 0.05 | 0 | yadI | 3 | 11.3 | 0.555 |
| 0.05 | 0 | AG1 | 3 | 11.3 | 0.381 |
| 0.05 | 0 | ptsI | 3 | 11.55 | 0.352 |
| 0.05 | 0 | clcB | 3 | 11.55 | 0.368 |
| 0.05 | 0 | ycaM | 3 | 11.55 | 0.375 |
| 0.05 | 0 | yadI | 3 | 11.55 | 0.572 |
| 0.05 | 0 | AG1 | 3 | 11.55 | 0.388 |
| 0.05 | 0 | ptsI | 3 | 11.8 | 0.36 |
| 0.05 | 0 | clcB | 3 | 11.8 | 0.372 |
| 0.05 | 0 | ycaM | 3 | 11.8 | 0.377 |
| 0.05 | 0 | yadI | 3 | 11.8 | 0.586 |
| 0.05 | 0 | AG1 | 3 | 11.8 | 0.4 |
| 0.05 | 0 | ptsI | 3 | 12.05 | 0.367 |
| 0.05 | 0 | clcB | 3 | 12.05 | 0.381 |
| 0.05 | 0 | ycaM | 3 | 12.05 | 0.381 |
| 0.05 | 0 | yadI | 3 | 12.05 | 0.602 |
| 0.05 | 0 | AG1 | 3 | 12.05 | 0.409 |
| 0.05 | 0 | ptsI | 3 | 12.3 | 0.38 |
| 0.05 | 0 | clcB | 3 | 12.3 | 0.393 |
| 0.05 | 0 | ycaM | 3 | 12.3 | 0.386 |
| 0.05 | 0 | yadI | 3 | 12.3 | 0.617 |
| 0.05 | 0 | AG1 | 3 | 12.3 | 0.421 |
| 0.05 | 0 | ptsI | 3 | 12.55 | 0.387 |
| 0.05 | 0 | clcB | 3 | 12.55 | 0.4 |
| 0.05 | 0 | ycaM | 3 | 12.55 | 0.389 |
| 0.05 | 0 | yadI | 3 | 12.55 | 0.631 |
| 0.05 | 0 | AG1 | 3 | 12.55 | 0.435 |

| | | | | | |
|---|---|---|---|---|---|
| 0.05 | 0 | ptsI | 3 | 12.8 | 0.398 |
| 0.05 | 0 | clcB | 3 | 12.8 | 0.405 |
| 0.05 | 0 | ycaM | 3 | 12.8 | 0.399 |
| 0.05 | 0 | yadI | 3 | 12.8 | 0.648 |
| 0.05 | 0 | AG1 | 3 | 12.8 | 0.448 |
| 0.05 | 0 | ptsI | 3 | 13.05 | 0.409 |
| 0.05 | 0 | clcB | 3 | 13.05 | 0.408 |
| 0.05 | 0 | ycaM | 3 | 13.05 | 0.4 |
| 0.05 | 0 | yadI | 3 | 13.05 | 0.641 |
| 0.05 | 0 | AG1 | 3 | 13.05 | 0.454 |
| 0.05 | 0 | ptsI | 3 | 13.3 | 0.42 |
| 0.05 | 0 | clcB | 3 | 13.3 | 0.414 |
| 0.05 | 0 | ycaM | 3 | 13.3 | 0.408 |
| 0.05 | 0 | yadI | 3 | 13.3 | 0.658 |
| 0.05 | 0 | AG1 | 3 | 13.3 | 0.466 |
| 0.05 | 0 | ptsI | 3 | 13.55 | 0.433 |
| 0.05 | 0 | clcB | 3 | 13.55 | 0.416 |
| 0.05 | 0 | ycaM | 3 | 13.55 | 0.415 |
| 0.05 | 0 | yadI | 3 | 13.55 | 0.674 |
| 0.05 | 0 | AG1 | 3 | 13.55 | 0.478 |
| 0.05 | 0 | ptsI | 3 | 13.8 | 0.444 |
| 0.05 | 0 | clcB | 3 | 13.8 | 0.431 |
| 0.05 | 0 | ycaM | 3 | 13.8 | 0.419 |
| 0.05 | 0 | yadI | 3 | 13.8 | 0.699 |
| 0.05 | 0 | AG1 | 3 | 13.8 | 0.497 |
| 0.05 | 0 | ptsI | 3 | 14.05 | 0.453 |
| 0.05 | 0 | clcB | 3 | 14.05 | 0.438 |
| 0.05 | 0 | ycaM | 3 | 14.05 | 0.431 |
| 0.05 | 0 | yadI | 3 | 14.05 | 0.71 |
| 0.05 | 0 | AG1 | 3 | 14.05 | 0.5 |
| 0.05 | 0 | ptsI | 3 | 14.3 | 0.463 |
| 0.05 | 0 | clcB | 3 | 14.3 | 0.451 |
| 0.05 | 0 | ycaM | 3 | 14.3 | 0.444 |
| 0.05 | 0 | yadI | 3 | 14.3 | 0.717 |
| 0.05 | 0 | AG1 | 3 | 14.3 | 0.514 |
| 0.05 | 0 | ptsI | 3 | 14.55 | 0.475 |
| 0.05 | 0 | clcB | 3 | 14.55 | 0.447 |
| 0.05 | 0 | ycaM | 3 | 14.55 | 0.45 |
| 0.05 | 0 | yadI | 3 | 14.55 | 0.732 |
| 0.05 | 0 | AG1 | 3 | 14.55 | 0.518 |
| 0.05 | 0 | ptsI | 3 | 14.8 | 0.491 |
| 0.05 | 0 | clcB | 3 | 14.8 | 0.443 |
| 0.05 | 0 | ycaM | 3 | 14.8 | 0.459 |
| 0.05 | 0 | yadI | 3 | 14.8 | 0.752 |
| 0.05 | 0 | AG1 | 3 | 14.8 | 0.53 |
| 0.05 | 0 | ptsI | 3 | 15.05 | 0.504 |
| 0.05 | 0 | clcB | 3 | 15.05 | 0.437 |
| 0.05 | 0 | ycaM | 3 | 15.05 | 0.465 |

| | | | | | |
|---|---|---|---|---|---|
| 0.05 | 0 | yadI | 3 | 15.05 | 0.762 |
| 0.05 | 0 | AG1 | 3 | 15.05 | 0.538 |
| 0.05 | 0 | ptsI | 3 | 15.3 | 0.519 |
| 0.05 | 0 | clcB | 3 | 15.3 | 0.456 |
| 0.05 | 0 | ycaM | 3 | 15.3 | 0.474 |
| 0.05 | 0 | yadI | 3 | 15.3 | 0.772 |
| 0.05 | 0 | AG1 | 3 | 15.3 | 0.553 |
| 0.05 | 0 | ptsI | 3 | 15.55 | 0.525 |
| 0.05 | 0 | clcB | 3 | 15.55 | 0.45 |
| 0.05 | 0 | ycaM | 3 | 15.55 | 0.474 |
| 0.05 | 0 | yadI | 3 | 15.55 | 0.78 |
| 0.05 | 0 | AG1 | 3 | 15.55 | 0.555 |
| 0.05 | 0 | ptsI | 3 | 15.8 | 0.53 |
| 0.05 | 0 | clcB | 3 | 15.8 | 0.448 |
| 0.05 | 0 | ycaM | 3 | 15.8 | 0.48 |
| 0.05 | 0 | yadI | 3 | 15.8 | 0.791 |
| 0.05 | 0 | AG1 | 3 | 15.8 | 0.571 |
| 0.05 | 0 | ptsI | 3 | 16.05 | 0.532 |
| 0.05 | 0 | clcB | 3 | 16.05 | 0.456 |
| 0.05 | 0 | ycaM | 3 | 16.05 | 0.49 |
| 0.05 | 0 | yadI | 3 | 16.05 | 0.792 |
| 0.05 | 0 | AG1 | 3 | 16.05 | 0.575 |
| 0.05 | 0 | ptsI | 3 | 16.3 | 0.544 |
| 0.05 | 0 | clcB | 3 | 16.3 | 0.453 |
| 0.05 | 0 | ycaM | 3 | 16.3 | 0.493 |
| 0.05 | 0 | yadI | 3 | 16.3 | 0.801 |
| 0.05 | 0 | AG1 | 3 | 16.3 | 0.584 |
| 0.05 | 0 | ptsI | 3 | 16.55 | 0.55 |
| 0.05 | 0 | clcB | 3 | 16.55 | 0.461 |
| 0.05 | 0 | ycaM | 3 | 16.55 | 0.501 |
| 0.05 | 0 | yadI | 3 | 16.55 | 0.812 |
| 0.05 | 0 | AG1 | 3 | 16.55 | 0.608 |
| 0.05 | 0 | ptsI | 3 | 16.8 | 0.557 |
| 0.05 | 0 | clcB | 3 | 16.8 | 0.468 |
| 0.05 | 0 | ycaM | 3 | 16.8 | 0.506 |
| 0.05 | 0 | yadI | 3 | 16.8 | 0.82 |
| 0.05 | 0 | AG1 | 3 | 16.8 | 0.642 |
| 0.05 | 0 | ptsI | 3 | 17.05 | 0.579 |
| 0.05 | 0 | clcB | 3 | 17.05 | 0.47 |
| 0.05 | 0 | ycaM | 3 | 17.05 | 0.51 |
| 0.05 | 0 | yadI | 3 | 17.05 | 0.825 |
| 0.05 | 0 | AG1 | 3 | 17.05 | 0.653 |
| 0.05 | 0 | ptsI | 3 | 17.3 | 0.589 |
| 0.05 | 0 | clcB | 3 | 17.3 | 0.466 |
| 0.05 | 0 | ycaM | 3 | 17.3 | 0.517 |
| 0.05 | 0 | yadI | 3 | 17.3 | 0.834 |
| 0.05 | 0 | AG1 | 3 | 17.3 | 0.66 |
| 0.05 | 0 | ptsI | 3 | 17.55 | 0.623 |

| | | | | | |
|---|---|---|---|---|---|
| 0.05 | 0 | clcB | 3 | 17.55 | 0.473 |
| 0.05 | 0 | ycaM | 3 | 17.55 | 0.52 |
| 0.05 | 0 | yadI | 3 | 17.55 | 0.842 |
| 0.05 | 0 | AG1 | 3 | 17.55 | 0.686 |
| 0.05 | 0 | ptsI | 3 | 17.8 | 0.647 |
| 0.05 | 0 | clcB | 3 | 17.8 | 0.47 |
| 0.05 | 0 | ycaM | 3 | 17.8 | 0.526 |
| 0.05 | 0 | yadI | 3 | 17.8 | 0.855 |
| 0.05 | 0 | AG1 | 3 | 17.8 | 0.699 |
| 0.05 | 0 | ptsI | 3 | 18.05 | 0.643 |
| 0.05 | 0 | clcB | 3 | 18.05 | 0.474 |
| 0.05 | 0 | ycaM | 3 | 18.05 | 0.533 |
| 0.05 | 0 | yadI | 3 | 18.05 | 0.861 |
| 0.05 | 0 | AG1 | 3 | 18.05 | 0.73 |
| 0.05 | 0 | ptsI | 3 | 18.3 | 0.656 |
| 0.05 | 0 | clcB | 3 | 18.3 | 0.473 |
| 0.05 | 0 | ycaM | 3 | 18.3 | 0.536 |
| 0.05 | 0 | yadI | 3 | 18.3 | 0.868 |
| 0.05 | 0 | AG1 | 3 | 18.3 | 0.738 |
| 0.05 | 0 | ptsI | 3 | 18.55 | 0.628 |
| 0.05 | 0 | clcB | 3 | 18.55 | 0.473 |
| 0.05 | 0 | ycaM | 3 | 18.55 | 0.54 |
| 0.05 | 0 | yadI | 3 | 18.55 | 0.879 |
| 0.05 | 0 | AG1 | 3 | 18.55 | 0.742 |
| 0.05 | 0 | ptsI | 3 | 18.8 | 0.647 |
| 0.05 | 0 | clcB | 3 | 18.8 | 0.473 |
| 0.05 | 0 | ycaM | 3 | 18.8 | 0.549 |
| 0.05 | 0 | yadI | 3 | 18.8 | 0.884 |
| 0.05 | 0 | AG1 | 3 | 18.8 | 0.758 |
| 0.05 | 0 | ptsI | 3 | 19.05 | 0.672 |
| 0.05 | 0 | clcB | 3 | 19.05 | 0.477 |
| 0.05 | 0 | ycaM | 3 | 19.05 | 0.554 |
| 0.05 | 0 | yadI | 3 | 19.05 | 0.891 |
| 0.05 | 0 | AG1 | 3 | 19.05 | 0.778 |
| 0.05 | 0 | ptsI | 3 | 19.3 | 0.688 |
| 0.05 | 0 | clcB | 3 | 19.3 | 0.476 |
| 0.05 | 0 | ycaM | 3 | 19.3 | 0.564 |
| 0.05 | 0 | yadI | 3 | 19.3 | 0.903 |
| 0.05 | 0 | AG1 | 3 | 19.3 | 0.789 |
| 0.05 | 0 | ptsI | 3 | 19.55 | 0.71 |
| 0.05 | 0 | clcB | 3 | 19.55 | 0.485 |
| 0.05 | 0 | ycaM | 3 | 19.55 | 0.57 |
| 0.05 | 0 | yadI | 3 | 19.55 | 0.907 |
| 0.05 | 0 | AG1 | 3 | 19.55 | 0.807 |
| 0.05 | 0 | ptsI | 3 | 19.8 | 0.699 |
| 0.05 | 0 | clcB | 3 | 19.8 | 0.485 |
| 0.05 | 0 | ycaM | 3 | 19.8 | 0.571 |
| 0.05 | 0 | yadI | 3 | 19.8 | 0.912 |

| | | | | | |
|---|---|---|---|---|---|
| 0.05 | 0 | AG1 | 3 | 19.8 | 0.808 |
| 0.05 | 0 | ptsI | 3 | 20.05 | 0.706 |
| 0.05 | 0 | clcB | 3 | 20.05 | 0.481 |
| 0.05 | 0 | ycaM | 3 | 20.05 | 0.576 |
| 0.05 | 0 | yadI | 3 | 20.05 | 0.922 |
| 0.05 | 0 | AG1 | 3 | 20.05 | 0.826 |
| 0.05 | 0 | ptsI | 3 | 20.3 | 0.708 |
| 0.05 | 0 | clcB | 3 | 20.3 | 0.476 |
| 0.05 | 0 | ycaM | 3 | 20.3 | 0.578 |
| 0.05 | 0 | yadI | 3 | 20.3 | 0.925 |
| 0.05 | 0 | AG1 | 3 | 20.3 | 0.825 |
| 0.05 | 0 | ptsI | 3 | 20.55 | 0.73 |
| 0.05 | 0 | clcB | 3 | 20.55 | 0.481 |
| 0.05 | 0 | ycaM | 3 | 20.55 | 0.583 |
| 0.05 | 0 | yadI | 3 | 20.55 | 0.937 |
| 0.05 | 0 | AG1 | 3 | 20.55 | 0.844 |
| 0.05 | 0 | ptsI | 3 | 20.8 | 0.74 |
| 0.05 | 0 | clcB | 3 | 20.8 | 0.482 |
| 0.05 | 0 | ycaM | 3 | 20.8 | 0.589 |
| 0.05 | 0 | yadI | 3 | 20.8 | 0.946 |
| 0.05 | 0 | AG1 | 3 | 20.8 | 0.852 |
| 0.05 | 0 | ptsI | 3 | 21.05 | 0.757 |
| 0.05 | 0 | clcB | 3 | 21.05 | 0.482 |
| 0.05 | 0 | ycaM | 3 | 21.05 | 0.594 |
| 0.05 | 0 | yadI | 3 | 21.05 | 0.954 |
| 0.05 | 0 | AG1 | 3 | 21.05 | 0.859 |
| 0.05 | 0 | ptsI | 3 | 21.3 | 0.766 |
| 0.05 | 0 | clcB | 3 | 21.3 | 0.483 |
| 0.05 | 0 | ycaM | 3 | 21.3 | 0.595 |
| 0.05 | 0 | yadI | 3 | 21.3 | 0.956 |
| 0.05 | 0 | AG1 | 3 | 21.3 | 0.863 |
| 0.05 | 0 | ptsI | 3 | 21.55 | 0.782 |
| 0.05 | 0 | clcB | 3 | 21.55 | 0.486 |
| 0.05 | 0 | ycaM | 3 | 21.55 | 0.603 |
| 0.05 | 0 | yadI | 3 | 21.55 | 0.964 |
| 0.05 | 0 | AG1 | 3 | 21.55 | 0.876 |
| 0.05 | 0 | ptsI | 3 | 21.8 | 0.794 |
| 0.05 | 0 | clcB | 3 | 21.8 | 0.488 |
| 0.05 | 0 | ycaM | 3 | 21.8 | 0.605 |
| 0.05 | 0 | yadI | 3 | 21.8 | 0.968 |
| 0.05 | 0 | AG1 | 3 | 21.8 | 0.89 |
| 0.05 | 0 | ptsI | 3 | 22.05 | 0.802 |
| 0.05 | 0 | clcB | 3 | 22.05 | 0.488 |
| 0.05 | 0 | ycaM | 3 | 22.05 | 0.609 |
| 0.05 | 0 | yadI | 3 | 22.05 | 0.977 |
| 0.05 | 0 | AG1 | 3 | 22.05 | 0.891 |
| 0.05 | 0 | ptsI | 3 | 22.3 | 0.814 |
| 0.05 | 0 | clcB | 3 | 22.3 | 0.491 |

| | | | | | |
|---|---|---|---|---|---|
| 0.05 | 0 | ycaM | 3 | 22.3 | 0.613 |
| 0.05 | 0 | yadI | 3 | 22.3 | 0.978 |
| 0.05 | 0 | AG1 | 3 | 22.3 | 0.899 |
| 0.05 | 0 | ptsI | 3 | 22.55 | 0.824 |
| 0.05 | 0 | clcB | 3 | 22.55 | 0.493 |
| 0.05 | 0 | ycaM | 3 | 22.55 | 0.615 |
| 0.05 | 0 | yadI | 3 | 22.55 | 0.981 |
| 0.05 | 0 | AG1 | 3 | 22.55 | 0.903 |
| 0.05 | 0 | ptsI | 3 | 22.8 | 0.826 |
| 0.05 | 0 | clcB | 3 | 22.8 | 0.493 |
| 0.05 | 0 | ycaM | 3 | 22.8 | 0.614 |
| 0.05 | 0 | yadI | 3 | 22.8 | 0.985 |
| 0.05 | 0 | AG1 | 3 | 22.8 | 0.904 |
| 0.05 | 0 | ptsI | 3 | 23.05 | 0.833 |
| 0.05 | 0 | clcB | 3 | 23.05 | 0.495 |
| 0.05 | 0 | ycaM | 3 | 23.05 | 0.619 |
| 0.05 | 0 | yadI | 3 | 23.05 | 0.988 |
| 0.05 | 0 | AG1 | 3 | 23.05 | 0.909 |
| 0.05 | 0 | ptsI | 3 | 23.3 | 0.836 |
| 0.05 | 0 | clcB | 3 | 23.3 | 0.495 |
| 0.05 | 0 | ycaM | 3 | 23.3 | 0.62 |
| 0.05 | 0 | yadI | 3 | 23.3 | 0.993 |
| 0.05 | 0 | AG1 | 3 | 23.3 | 0.916 |
| 0.05 | 0 | ptsI | 3 | 23.55 | 0.855 |
| 0.05 | 0 | clcB | 3 | 23.55 | 0.501 |
| 0.05 | 0 | ycaM | 3 | 23.55 | 0.625 |
| 0.05 | 0 | yadI | 3 | 23.55 | 1.005 |
| 0.05 | 0 | AG1 | 3 | 23.55 | 0.926 |
| 0.05 | 0 | ptsI | 3 | 23.8 | 0.854 |
| 0.05 | 0 | clcB | 3 | 23.8 | 0.496 |
| 0.05 | 0 | ycaM | 3 | 23.8 | 0.623 |
| 0.05 | 0 | yadI | 3 | 23.8 | 1.004 |
| 0.05 | 0 | AG1 | 3 | 23.8 | 0.925 |
| 0.05 | 0 | ptsI | 3 | 24.05 | 0.862 |
| 0.05 | 0 | clcB | 3 | 24.05 | 0.499 |
| 0.05 | 0 | ycaM | 3 | 24.05 | 0.622 |
| 0.05 | 0 | yadI | 3 | 24.05 | 1.012 |
| 0.05 | 0 | AG1 | 3 | 24.05 | 0.931 |
| 0.05 | 0 | ptsI | 3 | 24.3 | 0.862 |
| 0.05 | 0 | clcB | 3 | 24.3 | 0.492 |
| 0.05 | 0 | ycaM | 3 | 24.3 | 0.62 |
| 0.05 | 0 | yadI | 3 | 24.3 | 1.012 |
| 0.05 | 0 | AG1 | 3 | 24.3 | 0.926 |
| 0.05 | 0 | ptsI | 4 | 0 | 0.202 |
| 0.05 | 0 | clcB | 4 | 0 | 0.207 |
| 0.05 | 0 | ycaM | 4 | 0 | 0.199 |
| 0.05 | 0 | yadI | 4 | 0 | 0.227 |
| 0.05 | 0 | AG1 | 4 | 0 | 0.204 |

| | | | | | |
|---|---|---|---|---|---|
| 0.05 | 0 | ptsI | 4 | 0.25 | 0.201 |
| 0.05 | 0 | clcB | 4 | 0.25 | 0.21 |
| 0.05 | 0 | ycaM | 4 | 0.25 | 0.2 |
| 0.05 | 0 | yadI | 4 | 0.25 | 0.217 |
| 0.05 | 0 | AG1 | 4 | 0.25 | 0.2 |
| 0.05 | 0 | ptsI | 4 | 0.5 | 0.204 |
| 0.05 | 0 | clcB | 4 | 0.5 | 0.214 |
| 0.05 | 0 | ycaM | 4 | 0.5 | 0.2 |
| 0.05 | 0 | yadI | 4 | 0.5 | 0.222 |
| 0.05 | 0 | AG1 | 4 | 0.5 | 0.202 |
| 0.05 | 0 | ptsI | 4 | 0.75 | 0.209 |
| 0.05 | 0 | clcB | 4 | 0.75 | 0.218 |
| 0.05 | 0 | ycaM | 4 | 0.75 | 0.205 |
| 0.05 | 0 | yadI | 4 | 0.75 | 0.222 |
| 0.05 | 0 | AG1 | 4 | 0.75 | 0.205 |
| 0.05 | 0 | ptsI | 4 | 1 | 0.213 |
| 0.05 | 0 | clcB | 4 | 1 | 0.222 |
| 0.05 | 0 | ycaM | 4 | 1 | 0.21 |
| 0.05 | 0 | yadI | 4 | 1 | 0.223 |
| 0.05 | 0 | AG1 | 4 | 1 | 0.208 |
| 0.05 | 0 | ptsI | 4 | 1.25 | 0.22 |
| 0.05 | 0 | clcB | 4 | 1.25 | 0.228 |
| 0.05 | 0 | ycaM | 4 | 1.25 | 0.22 |
| 0.05 | 0 | yadI | 4 | 1.25 | 0.229 |
| 0.05 | 0 | AG1 | 4 | 1.25 | 0.216 |
| 0.05 | 0 | ptsI | 4 | 1.5 | 0.224 |
| 0.05 | 0 | clcB | 4 | 1.5 | 0.232 |
| 0.05 | 0 | ycaM | 4 | 1.5 | 0.224 |
| 0.05 | 0 | yadI | 4 | 1.5 | 0.234 |
| 0.05 | 0 | AG1 | 4 | 1.5 | 0.222 |
| 0.05 | 0 | ptsI | 4 | 1.75 | 0.235 |
| 0.05 | 0 | clcB | 4 | 1.75 | 0.245 |
| 0.05 | 0 | ycaM | 4 | 1.75 | 0.236 |
| 0.05 | 0 | yadI | 4 | 1.75 | 0.243 |
| 0.05 | 0 | AG1 | 4 | 1.75 | 0.231 |
| 0.05 | 0 | ptsI | 4 | 2 | 0.246 |
| 0.05 | 0 | clcB | 4 | 2 | 0.254 |
| 0.05 | 0 | ycaM | 4 | 2 | 0.248 |
| 0.05 | 0 | yadI | 4 | 2 | 0.252 |
| 0.05 | 0 | AG1 | 4 | 2 | 0.245 |
| 0.05 | 0 | ptsI | 4 | 2.25 | 0.26 |
| 0.05 | 0 | clcB | 4 | 2.25 | 0.266 |
| 0.05 | 0 | ycaM | 4 | 2.25 | 0.261 |
| 0.05 | 0 | yadI | 4 | 2.25 | 0.265 |
| 0.05 | 0 | AG1 | 4 | 2.25 | 0.259 |
| 0.05 | 0 | ptsI | 4 | 2.5 | 0.271 |
| 0.05 | 0 | clcB | 4 | 2.5 | 0.278 |
| 0.05 | 0 | ycaM | 4 | 2.5 | 0.27 |

| | | | | | |
|---|---|---|---|---|---|
| 0.05 | 0 | yadI | 4 | 2.5 | 0.276 |
| 0.05 | 0 | AG1 | 4 | 2.5 | 0.272 |
| 0.05 | 0 | ptsI | 4 | 2.75 | 0.283 |
| 0.05 | 0 | clcB | 4 | 2.75 | 0.288 |
| 0.05 | 0 | ycaM | 4 | 2.75 | 0.276 |
| 0.05 | 0 | yadI | 4 | 2.75 | 0.289 |
| 0.05 | 0 | AG1 | 4 | 2.75 | 0.284 |
| 0.05 | 0 | ptsI | 4 | 3 | 0.3 |
| 0.05 | 0 | clcB | 4 | 3 | 0.304 |
| 0.05 | 0 | ycaM | 4 | 3 | 0.296 |
| 0.05 | 0 | yadI | 4 | 3 | 0.31 |
| 0.05 | 0 | AG1 | 4 | 3 | 0.302 |
| 0.05 | 0 | ptsI | 4 | 3.25 | 0.313 |
| 0.05 | 0 | clcB | 4 | 3.25 | 0.312 |
| 0.05 | 0 | ycaM | 4 | 3.25 | 0.306 |
| 0.05 | 0 | yadI | 4 | 3.25 | 0.324 |
| 0.05 | 0 | AG1 | 4 | 3.25 | 0.319 |
| 0.05 | 0 | ptsI | 4 | 3.5 | 0.328 |
| 0.05 | 0 | clcB | 4 | 3.5 | 0.321 |
| 0.05 | 0 | ycaM | 4 | 3.5 | 0.325 |
| 0.05 | 0 | yadI | 4 | 3.5 | 0.345 |
| 0.05 | 0 | AG1 | 4 | 3.5 | 0.337 |
| 0.05 | 0 | ptsI | 4 | 3.75 | 0.345 |
| 0.05 | 0 | clcB | 4 | 3.75 | 0.336 |
| 0.05 | 0 | ycaM | 4 | 3.75 | 0.333 |
| 0.05 | 0 | yadI | 4 | 3.75 | 0.361 |
| 0.05 | 0 | AG1 | 4 | 3.75 | 0.351 |
| 0.05 | 0 | ptsI | 4 | 4 | 0.362 |
| 0.05 | 0 | clcB | 4 | 4 | 0.354 |
| 0.05 | 0 | ycaM | 4 | 4 | 0.356 |
| 0.05 | 0 | yadI | 4 | 4 | 0.39 |
| 0.05 | 0 | AG1 | 4 | 4 | 0.375 |
| 0.05 | 0 | ptsI | 4 | 4.35 | 0.37 |
| 0.05 | 0 | clcB | 4 | 4.35 | 0.29 |
| 0.05 | 0 | ycaM | 4 | 4.35 | 0.397 |
| 0.05 | 0 | yadI | 4 | 4.35 | 0.341 |
| 0.05 | 0 | AG1 | 4 | 4.35 | 0.384 |
| 0.05 | 0 | ptsI | 4 | 4.6 | 0.37 |
| 0.05 | 0 | clcB | 4 | 4.6 | 0.294 |
| 0.05 | 0 | ycaM | 4 | 4.6 | 0.381 |
| 0.05 | 0 | yadI | 4 | 4.6 | 0.315 |
| 0.05 | 0 | AG1 | 4 | 4.6 | 0.365 |
| 0.05 | 0 | ptsI | 4 | 4.85 | 0.385 |
| 0.05 | 0 | clcB | 4 | 4.85 | 0.303 |
| 0.05 | 0 | ycaM | 4 | 4.85 | 0.393 |
| 0.05 | 0 | yadI | 4 | 4.85 | 0.32 |
| 0.05 | 0 | AG1 | 4 | 4.85 | 0.374 |
| 0.05 | 0 | ptsI | 4 | 5.1 | 0.398 |

| | | | | | |
|---|---|---|---|---|---|
| 0.05 | 0 | clcB | 4 | 5.1 | 0.312 |
| 0.05 | 0 | ycaM | 4 | 5.1 | 0.409 |
| 0.05 | 0 | yadI | 4 | 5.1 | 0.332 |
| 0.05 | 0 | AG1 | 4 | 5.1 | 0.385 |
| 0.05 | 0 | ptsI | 4 | 5.35 | 0.417 |
| 0.05 | 0 | clcB | 4 | 5.35 | 0.319 |
| 0.05 | 0 | ycaM | 4 | 5.35 | 0.427 |
| 0.05 | 0 | yadI | 4 | 5.35 | 0.345 |
| 0.05 | 0 | AG1 | 4 | 5.35 | 0.399 |
| 0.05 | 0 | ptsI | 4 | 5.6 | 0.43 |
| 0.05 | 0 | clcB | 4 | 5.6 | 0.327 |
| 0.05 | 0 | ycaM | 4 | 5.6 | 0.445 |
| 0.05 | 0 | yadI | 4 | 5.6 | 0.356 |
| 0.05 | 0 | AG1 | 4 | 5.6 | 0.408 |
| 0.05 | 0 | ptsI | 4 | 5.85 | 0.444 |
| 0.05 | 0 | clcB | 4 | 5.85 | 0.336 |
| 0.05 | 0 | ycaM | 4 | 5.85 | 0.452 |
| 0.05 | 0 | yadI | 4 | 5.85 | 0.369 |
| 0.05 | 0 | AG1 | 4 | 5.85 | 0.422 |
| 0.05 | 0 | ptsI | 4 | 6.1 | 0.459 |
| 0.05 | 0 | clcB | 4 | 6.1 | 0.349 |
| 0.05 | 0 | ycaM | 4 | 6.1 | 0.465 |
| 0.05 | 0 | yadI | 4 | 6.1 | 0.384 |
| 0.05 | 0 | AG1 | 4 | 6.1 | 0.435 |
| 0.05 | 0 | ptsI | 4 | 6.35 | 0.471 |
| 0.05 | 0 | clcB | 4 | 6.35 | 0.356 |
| 0.05 | 0 | ycaM | 4 | 6.35 | 0.473 |
| 0.05 | 0 | yadI | 4 | 6.35 | 0.395 |
| 0.05 | 0 | AG1 | 4 | 6.35 | 0.444 |
| 0.05 | 0 | ptsI | 4 | 6.6 | 0.483 |
| 0.05 | 0 | clcB | 4 | 6.6 | 0.365 |
| 0.05 | 0 | ycaM | 4 | 6.6 | 0.482 |
| 0.05 | 0 | yadI | 4 | 6.6 | 0.409 |
| 0.05 | 0 | AG1 | 4 | 6.6 | 0.455 |
| 0.05 | 0 | ptsI | 4 | 6.85 | 0.495 |
| 0.05 | 0 | clcB | 4 | 6.85 | 0.373 |
| 0.05 | 0 | ycaM | 4 | 6.85 | 0.493 |
| 0.05 | 0 | yadI | 4 | 6.85 | 0.418 |
| 0.05 | 0 | AG1 | 4 | 6.85 | 0.468 |
| 0.05 | 0 | ptsI | 4 | 7.1 | 0.503 |
| 0.05 | 0 | clcB | 4 | 7.1 | 0.38 |
| 0.05 | 0 | ycaM | 4 | 7.1 | 0.506 |
| 0.05 | 0 | yadI | 4 | 7.1 | 0.429 |
| 0.05 | 0 | AG1 | 4 | 7.1 | 0.482 |
| 0.05 | 0 | ptsI | 4 | 7.35 | 0.514 |
| 0.05 | 0 | clcB | 4 | 7.35 | 0.387 |
| 0.05 | 0 | ycaM | 4 | 7.35 | 0.517 |
| 0.05 | 0 | yadI | 4 | 7.35 | 0.435 |

| | | | | | |
|---|---|---|---|---|---|
| 0.05 | 0 | AG1 | 4 | 7.35 | 0.497 |
| 0.05 | 0 | ptsI | 4 | 7.6 | 0.521 |
| 0.05 | 0 | clcB | 4 | 7.6 | 0.397 |
| 0.05 | 0 | ycaM | 4 | 7.6 | 0.524 |
| 0.05 | 0 | yadI | 4 | 7.6 | 0.443 |
| 0.05 | 0 | AG1 | 4 | 7.6 | 0.506 |
| 0.05 | 0 | ptsI | 4 | 7.85 | 0.528 |
| 0.05 | 0 | clcB | 4 | 7.85 | 0.403 |
| 0.05 | 0 | ycaM | 4 | 7.85 | 0.528 |
| 0.05 | 0 | yadI | 4 | 7.85 | 0.448 |
| 0.05 | 0 | AG1 | 4 | 7.85 | 0.516 |
| 0.05 | 0 | ptsI | 4 | 8.1 | 0.537 |
| 0.05 | 0 | clcB | 4 | 8.1 | 0.412 |
| 0.05 | 0 | ycaM | 4 | 8.1 | 0.536 |
| 0.05 | 0 | yadI | 4 | 8.1 | 0.465 |
| 0.05 | 0 | AG1 | 4 | 8.1 | 0.529 |
| 0.05 | 0 | ptsI | 4 | 8.35 | 0.547 |
| 0.05 | 0 | clcB | 4 | 8.35 | 0.419 |
| 0.05 | 0 | ycaM | 4 | 8.35 | 0.54 |
| 0.05 | 0 | yadI | 4 | 8.35 | 0.476 |
| 0.05 | 0 | AG1 | 4 | 8.35 | 0.541 |
| 0.05 | 0 | ptsI | 4 | 8.83333 | 0.39 |
| 0.05 | 0 | clcB | 4 | 8.83333 | 0.412 |
| 0.05 | 0 | ycaM | 4 | 8.83333 | 0.38 |
| 0.05 | 0 | yadI | 4 | 8.83333 | 0.504 |
| 0.05 | 0 | AG1 | 4 | 8.83333 | 0.578 |
| 0.05 | 0 | ptsI | 4 | 9.08333 | 0.392 |
| 0.05 | 0 | clcB | 4 | 9.08333 | 0.409 |
| 0.05 | 0 | ycaM | 4 | 9.08333 | 0.379 |
| 0.05 | 0 | yadI | 4 | 9.08333 | 0.498 |
| 0.05 | 0 | AG1 | 4 | 9.08333 | 0.573 |
| 0.05 | 0 | ptsI | 4 | 9.33333 | 0.402 |
| 0.05 | 0 | clcB | 4 | 9.33333 | 0.414 |
| 0.05 | 0 | ycaM | 4 | 9.33333 | 0.381 |
| 0.05 | 0 | yadI | 4 | 9.33333 | 0.507 |
| 0.05 | 0 | AG1 | 4 | 9.33333 | 0.58 |
| 0.05 | 0 | ptsI | 4 | 9.58333 | 0.407 |
| 0.05 | 0 | clcB | 4 | 9.58333 | 0.42 |
| 0.05 | 0 | ycaM | 4 | 9.58333 | 0.382 |
| 0.05 | 0 | yadI | 4 | 9.58333 | 0.517 |
| 0.05 | 0 | AG1 | 4 | 9.58333 | 0.594 |
| 0.05 | 0 | ptsI | 4 | 9.83333 | 0.417 |
| 0.05 | 0 | clcB | 4 | 9.83333 | 0.425 |
| 0.05 | 0 | ycaM | 4 | 9.83333 | 0.39 |
| 0.05 | 0 | yadI | 4 | 9.83333 | 0.523 |
| 0.05 | 0 | AG1 | 4 | 9.83333 | 0.605 |
| 0.05 | 0 | ptsI | 4 | 10.0833 | 0.423 |
| 0.05 | 0 | clcB | 4 | 10.0833 | 0.43 |

| | | | | | |
|---|---|---|---|---|---|
| 0.05 | 0 | ycaM | 4 | 10.0833 | 0.393 |
| 0.05 | 0 | yadI | 4 | 10.0833 | 0.534 |
| 0.05 | 0 | AG1 | 4 | 10.0833 | 0.614 |
| 0.05 | 0 | ptsI | 4 | 10.3333 | 0.43 |
| 0.05 | 0 | clcB | 4 | 10.3333 | 0.436 |
| 0.05 | 0 | ycaM | 4 | 10.3333 | 0.398 |
| 0.05 | 0 | yadI | 4 | 10.3333 | 0.539 |
| 0.05 | 0 | AG1 | 4 | 10.3333 | 0.624 |
| 0.05 | 0 | ptsI | 4 | 10.5833 | 0.437 |
| 0.05 | 0 | clcB | 4 | 10.5833 | 0.44 |
| 0.05 | 0 | ycaM | 4 | 10.5833 | 0.401 |
| 0.05 | 0 | yadI | 4 | 10.5833 | 0.547 |
| 0.05 | 0 | AG1 | 4 | 10.5833 | 0.63 |
| 0.05 | 0 | ptsI | 4 | 10.8333 | 0.445 |
| 0.05 | 0 | clcB | 4 | 10.8333 | 0.445 |
| 0.05 | 0 | ycaM | 4 | 10.8333 | 0.406 |
| 0.05 | 0 | yadI | 4 | 10.8333 | 0.551 |
| 0.05 | 0 | AG1 | 4 | 10.8333 | 0.642 |
| 0.05 | 0 | ptsI | 4 | 11.0833 | 0.448 |
| 0.05 | 0 | clcB | 4 | 11.0833 | 0.449 |
| 0.05 | 0 | ycaM | 4 | 11.0833 | 0.407 |
| 0.05 | 0 | yadI | 4 | 11.0833 | 0.558 |
| 0.05 | 0 | AG1 | 4 | 11.0833 | 0.649 |
| 0.05 | 0 | ptsI | 4 | 11.3333 | 0.46 |
| 0.05 | 0 | clcB | 4 | 11.3333 | 0.461 |
| 0.05 | 0 | ycaM | 4 | 11.3333 | 0.416 |
| 0.05 | 0 | yadI | 4 | 11.3333 | 0.566 |
| 0.05 | 0 | AG1 | 4 | 11.3333 | 0.66 |
| 0.05 | 0 | ptsI | 4 | 11.5833 | 0.462 |
| 0.05 | 0 | clcB | 4 | 11.5833 | 0.458 |
| 0.05 | 0 | ycaM | 4 | 11.5833 | 0.419 |
| 0.05 | 0 | yadI | 4 | 11.5833 | 0.57 |
| 0.05 | 0 | AG1 | 4 | 11.5833 | 0.663 |
| 0.05 | 0 | ptsI | 4 | 11.8333 | 0.466 |
| 0.05 | 0 | clcB | 4 | 11.8333 | 0.462 |
| 0.05 | 0 | ycaM | 4 | 11.8333 | 0.422 |
| 0.05 | 0 | yadI | 4 | 11.8333 | 0.575 |
| 0.05 | 0 | AG1 | 4 | 11.8333 | 0.67 |
| 0.05 | 0 | ptsI | 4 | 12.0833 | 0.47 |
| 0.05 | 0 | clcB | 4 | 12.0833 | 0.469 |
| 0.05 | 0 | ycaM | 4 | 12.0833 | 0.427 |
| 0.05 | 0 | yadI | 4 | 12.0833 | 0.584 |
| 0.05 | 0 | AG1 | 4 | 12.0833 | 0.679 |
| 0.05 | 0 | ptsI | 4 | 12.3333 | 0.48 |
| 0.05 | 0 | clcB | 4 | 12.3333 | 0.474 |
| 0.05 | 0 | ycaM | 4 | 12.3333 | 0.432 |
| 0.05 | 0 | yadI | 4 | 12.3333 | 0.586 |
| 0.05 | 0 | AG1 | 4 | 12.3333 | 0.69 |

| | | | | | |
|---|---|---|---|---|---|
| 0.05 | 0 | ptsI | 4 | 12.5833 | 0.485 |
| 0.05 | 0 | clcB | 4 | 12.5833 | 0.478 |
| 0.05 | 0 | ycaM | 4 | 12.5833 | 0.437 |
| 0.05 | 0 | yadI | 4 | 12.5833 | 0.589 |
| 0.05 | 0 | AG1 | 4 | 12.5833 | 0.693 |
| 0.05 | 0 | ptsI | 4 | 12.8333 | 0.484 |
| 0.05 | 0 | clcB | 4 | 12.8333 | 0.479 |
| 0.05 | 0 | ycaM | 4 | 12.8333 | 0.443 |
| 0.05 | 0 | yadI | 4 | 12.8333 | 0.595 |
| 0.05 | 0 | AG1 | 4 | 12.8333 | 0.698 |
| 0.05 | 0 | ptsI | 4 | 13.0833 | 0.493 |
| 0.05 | 0 | clcB | 4 | 13.0833 | 0.487 |
| 0.05 | 0 | ycaM | 4 | 13.0833 | 0.446 |
| 0.05 | 0 | yadI | 4 | 13.0833 | 0.598 |
| 0.05 | 0 | AG1 | 4 | 13.0833 | 0.705 |
| 0.05 | 0 | ptsI | 4 | 13.3333 | 0.5 |
| 0.05 | 0 | clcB | 4 | 13.3333 | 0.488 |
| 0.05 | 0 | ycaM | 4 | 13.3333 | 0.452 |
| 0.05 | 0 | yadI | 4 | 13.3333 | 0.603 |
| 0.05 | 0 | AG1 | 4 | 13.3333 | 0.711 |
| 0.05 | 0 | ptsI | 4 | 13.5833 | 0.506 |
| 0.05 | 0 | clcB | 4 | 13.5833 | 0.493 |
| 0.05 | 0 | ycaM | 4 | 13.5833 | 0.455 |
| 0.05 | 0 | yadI | 4 | 13.5833 | 0.608 |
| 0.05 | 0 | AG1 | 4 | 13.5833 | 0.716 |
| 0.05 | 0 | ptsI | 4 | 13.8333 | 0.508 |
| 0.05 | 0 | clcB | 4 | 13.8333 | 0.495 |
| 0.05 | 0 | ycaM | 4 | 13.8333 | 0.458 |
| 0.05 | 0 | yadI | 4 | 13.8333 | 0.612 |
| 0.05 | 0 | AG1 | 4 | 13.8333 | 0.718 |
| 0.05 | 0 | ptsI | 4 | 14.0833 | 0.515 |
| 0.05 | 0 | clcB | 4 | 14.0833 | 0.497 |
| 0.05 | 0 | ycaM | 4 | 14.0833 | 0.462 |
| 0.05 | 0 | yadI | 4 | 14.0833 | 0.614 |
| 0.05 | 0 | AG1 | 4 | 14.0833 | 0.726 |
| 0.05 | 0 | ptsI | 4 | 14.3333 | 0.522 |
| 0.05 | 0 | clcB | 4 | 14.3333 | 0.502 |
| 0.05 | 0 | ycaM | 4 | 14.3333 | 0.468 |
| 0.05 | 0 | yadI | 4 | 14.3333 | 0.621 |
| 0.05 | 0 | AG1 | 4 | 14.3333 | 0.732 |
| 0.05 | 0 | ptsI | 4 | 14.5833 | 0.525 |
| 0.05 | 0 | clcB | 4 | 14.5833 | 0.501 |
| 0.05 | 0 | ycaM | 4 | 14.5833 | 0.469 |
| 0.05 | 0 | yadI | 4 | 14.5833 | 0.619 |
| 0.05 | 0 | AG1 | 4 | 14.5833 | 0.732 |
| 0.05 | 0 | ptsI | 4 | 14.8333 | 0.532 |
| 0.05 | 0 | clcB | 4 | 14.8333 | 0.506 |
| 0.05 | 0 | ycaM | 4 | 14.8333 | 0.473 |

| | | | | | |
|---|---|---|---|---|---|
| 0.05 | 0 | yadI | 4 | 14.8333 | 0.624 |
| 0.05 | 0 | AG1 | 4 | 14.8333 | 0.738 |
| 0.05 | 0 | ptsI | 4 | 15.0833 | 0.534 |
| 0.05 | 0 | clcB | 4 | 15.0833 | 0.506 |
| 0.05 | 0 | ycaM | 4 | 15.0833 | 0.474 |
| 0.05 | 0 | yadI | 4 | 15.0833 | 0.628 |
| 0.05 | 0 | AG1 | 4 | 15.0833 | 0.74 |
| 0.05 | 0 | ptsI | 4 | 15.3333 | 0.541 |
| 0.05 | 0 | clcB | 4 | 15.3333 | 0.508 |
| 0.05 | 0 | ycaM | 4 | 15.3333 | 0.479 |
| 0.05 | 0 | yadI | 4 | 15.3333 | 0.631 |
| 0.05 | 0 | AG1 | 4 | 15.3333 | 0.742 |
| 0.05 | 0 | ptsI | 4 | 15.5833 | 0.548 |
| 0.05 | 0 | clcB | 4 | 15.5833 | 0.512 |
| 0.05 | 0 | ycaM | 4 | 15.5833 | 0.482 |
| 0.05 | 0 | yadI | 4 | 15.5833 | 0.633 |
| 0.05 | 0 | AG1 | 4 | 15.5833 | 0.747 |
| 0.05 | 0 | ptsI | 4 | 15.8333 | 0.545 |
| 0.05 | 0 | clcB | 4 | 15.8333 | 0.511 |
| 0.05 | 0 | ycaM | 4 | 15.8333 | 0.484 |
| 0.05 | 0 | yadI | 4 | 15.8333 | 0.634 |
| 0.05 | 0 | AG1 | 4 | 15.8333 | 0.748 |
| 0.05 | 0 | ptsI | 4 | 16.0833 | 0.558 |
| 0.05 | 0 | clcB | 4 | 16.0833 | 0.516 |
| 0.05 | 0 | ycaM | 4 | 16.0833 | 0.487 |
| 0.05 | 0 | yadI | 4 | 16.0833 | 0.64 |
| 0.05 | 0 | AG1 | 4 | 16.0833 | 0.752 |
| 0.05 | 0 | ptsI | 4 | 16.3333 | 0.552 |
| 0.05 | 0 | clcB | 4 | 16.3333 | 0.512 |
| 0.05 | 0 | ycaM | 4 | 16.3333 | 0.488 |
| 0.05 | 0 | yadI | 4 | 16.3333 | 0.644 |
| 0.05 | 0 | AG1 | 4 | 16.3333 | 0.749 |
| 0.05 | 0 | ptsI | 4 | 16.5833 | 0.559 |
| 0.05 | 0 | clcB | 4 | 16.5833 | 0.516 |
| 0.05 | 0 | ycaM | 4 | 16.5833 | 0.493 |
| 0.05 | 0 | yadI | 4 | 16.5833 | 0.646 |
| 0.05 | 0 | AG1 | 4 | 16.5833 | 0.755 |
| 0.05 | 0 | ptsI | 4 | 16.8333 | 0.566 |
| 0.05 | 0 | clcB | 4 | 16.8333 | 0.517 |
| 0.05 | 0 | ycaM | 4 | 16.8333 | 0.49 |
| 0.05 | 0 | yadI | 4 | 16.8333 | 0.647 |
| 0.05 | 0 | AG1 | 4 | 16.8333 | 0.755 |
| 0.05 | 0 | ptsI | 4 | 17.0833 | 0.568 |
| 0.05 | 0 | clcB | 4 | 17.0833 | 0.519 |
| 0.05 | 0 | ycaM | 4 | 17.0833 | 0.491 |
| 0.05 | 0 | yadI | 4 | 17.0833 | 0.65 |
| 0.05 | 0 | AG1 | 4 | 17.0833 | 0.755 |
| 0.05 | 0 | ptsI | 4 | 17.3333 | 0.574 |

| | | | | | |
|---|---|---|---|---|---|
| 0.05 | 0 | clcB | 4 | 17.3333 | 0.524 |
| 0.05 | 0 | ycaM | 4 | 17.3333 | 0.499 |
| 0.05 | 0 | yadI | 4 | 17.3333 | 0.657 |
| 0.05 | 0 | AG1 | 4 | 17.3333 | 0.759 |
| 0.05 | 0 | ptsI | 4 | 17.5833 | 0.577 |
| 0.05 | 0 | clcB | 4 | 17.5833 | 0.522 |
| 0.05 | 0 | ycaM | 4 | 17.5833 | 0.495 |
| 0.05 | 0 | yadI | 4 | 17.5833 | 0.655 |
| 0.05 | 0 | AG1 | 4 | 17.5833 | 0.757 |
| 0.05 | 0 | ptsI | 4 | 17.8333 | 0.58 |
| 0.05 | 0 | clcB | 4 | 17.8333 | 0.522 |
| 0.05 | 0 | ycaM | 4 | 17.8333 | 0.499 |
| 0.05 | 0 | yadI | 4 | 17.8333 | 0.655 |
| 0.05 | 0 | AG1 | 4 | 17.8333 | 0.762 |
| 0.05 | 0 | ptsI | 4 | 18.0833 | 0.582 |
| 0.05 | 0 | clcB | 4 | 18.0833 | 0.523 |
| 0.05 | 0 | ycaM | 4 | 18.0833 | 0.5 |
| 0.05 | 0 | yadI | 4 | 18.0833 | 0.661 |
| 0.05 | 0 | AG1 | 4 | 18.0833 | 0.762 |
| 0.05 | 0 | ptsI | 4 | 18.3333 | 0.583 |
| 0.05 | 0 | clcB | 4 | 18.3333 | 0.525 |
| 0.05 | 0 | ycaM | 4 | 18.3333 | 0.502 |
| 0.05 | 0 | yadI | 4 | 18.3333 | 0.662 |
| 0.05 | 0 | AG1 | 4 | 18.3333 | 0.764 |
| 0.05 | 0 | ptsI | 4 | 18.5833 | 0.586 |
| 0.05 | 0 | clcB | 4 | 18.5833 | 0.525 |
| 0.05 | 0 | ycaM | 4 | 18.5833 | 0.501 |
| 0.05 | 0 | yadI | 4 | 18.5833 | 0.665 |
| 0.05 | 0 | AG1 | 4 | 18.5833 | 0.766 |
| 0.05 | 0 | ptsI | 4 | 18.8333 | 0.589 |
| 0.05 | 0 | clcB | 4 | 18.8333 | 0.522 |
| 0.05 | 0 | ycaM | 4 | 18.8333 | 0.503 |
| 0.05 | 0 | yadI | 4 | 18.8333 | 0.663 |
| 0.05 | 0 | AG1 | 4 | 18.8333 | 0.766 |
| 0.05 | 0 | ptsI | 4 | 19.0833 | 0.595 |
| 0.05 | 0 | clcB | 4 | 19.0833 | 0.529 |
| 0.05 | 0 | ycaM | 4 | 19.0833 | 0.506 |
| 0.05 | 0 | yadI | 4 | 19.0833 | 0.669 |
| 0.05 | 0 | AG1 | 4 | 19.0833 | 0.769 |
| 0.05 | 0 | ptsI | 4 | 19.3333 | 0.596 |
| 0.05 | 0 | clcB | 4 | 19.3333 | 0.529 |
| 0.05 | 0 | ycaM | 4 | 19.3333 | 0.512 |
| 0.05 | 0 | yadI | 4 | 19.3333 | 0.675 |
| 0.05 | 0 | AG1 | 4 | 19.3333 | 0.77 |
| 0.05 | 0 | ptsI | 4 | 19.5833 | 0.604 |
| 0.05 | 0 | clcB | 4 | 19.5833 | 0.532 |
| 0.05 | 0 | ycaM | 4 | 19.5833 | 0.511 |
| 0.05 | 0 | yadI | 4 | 19.5833 | 0.676 |

| | | | | | |
|---|---|---|---|---|---|
| 0.05 | 0 | AG1 | 4 | 19.5833 | 0.773 |
| 0.05 | 0 | ptsI | 4 | 19.8333 | 0.602 |
| 0.05 | 0 | clcB | 4 | 19.8333 | 0.529 |
| 0.05 | 0 | ycaM | 4 | 19.8333 | 0.509 |
| 0.05 | 0 | yadI | 4 | 19.8333 | 0.672 |
| 0.05 | 0 | AG1 | 4 | 19.8333 | 0.773 |
| 0.05 | 0 | ptsI | 4 | 20.0833 | 0.607 |
| 0.05 | 0 | clcB | 4 | 20.0833 | 0.529 |
| 0.05 | 0 | ycaM | 4 | 20.0833 | 0.513 |
| 0.05 | 0 | yadI | 4 | 20.0833 | 0.678 |
| 0.05 | 0 | AG1 | 4 | 20.0833 | 0.768 |
| 0.05 | 0 | ptsI | 4 | 20.3333 | 0.606 |
| 0.05 | 0 | clcB | 4 | 20.3333 | 0.532 |
| 0.05 | 0 | ycaM | 4 | 20.3333 | 0.516 |
| 0.05 | 0 | yadI | 4 | 20.3333 | 0.676 |
| 0.05 | 0 | AG1 | 4 | 20.3333 | 0.776 |
| 0.05 | 0 | ptsI | 4 | 20.5833 | 0.612 |
| 0.05 | 0 | clcB | 4 | 20.5833 | 0.53 |
| 0.05 | 0 | ycaM | 4 | 20.5833 | 0.516 |
| 0.05 | 0 | yadI | 4 | 20.5833 | 0.682 |
| 0.05 | 0 | AG1 | 4 | 20.5833 | 0.778 |
| 0.05 | 0 | ptsI | 4 | 20.8333 | 0.613 |
| 0.05 | 0 | clcB | 4 | 20.8333 | 0.535 |
| 0.05 | 0 | ycaM | 4 | 20.8333 | 0.523 |
| 0.05 | 0 | yadI | 4 | 20.8333 | 0.681 |
| 0.05 | 0 | AG1 | 4 | 20.8333 | 0.779 |
| 0.05 | 0 | ptsI | 4 | 21.0833 | 0.62 |
| 0.05 | 0 | clcB | 4 | 21.0833 | 0.534 |
| 0.05 | 0 | ycaM | 4 | 21.0833 | 0.522 |
| 0.05 | 0 | yadI | 4 | 21.0833 | 0.681 |
| 0.05 | 0 | AG1 | 4 | 21.0833 | 0.781 |
| 0.05 | 0 | ptsI | 4 | 21.3333 | 0.62 |
| 0.05 | 0 | clcB | 4 | 21.3333 | 0.534 |
| 0.05 | 0 | ycaM | 4 | 21.3333 | 0.52 |
| 0.05 | 0 | yadI | 4 | 21.3333 | 0.681 |
| 0.05 | 0 | AG1 | 4 | 21.3333 | 0.778 |
| 0.05 | 0 | ptsI | 4 | 21.5833 | 0.625 |
| 0.05 | 0 | clcB | 4 | 21.5833 | 0.534 |
| 0.05 | 0 | ycaM | 4 | 21.5833 | 0.525 |
| 0.05 | 0 | yadI | 4 | 21.5833 | 0.684 |
| 0.05 | 0 | AG1 | 4 | 21.5833 | 0.78 |
| 0.05 | 0 | ptsI | 4 | 21.8333 | 0.621 |
| 0.05 | 0 | clcB | 4 | 21.8333 | 0.534 |
| 0.05 | 0 | ycaM | 4 | 21.8333 | 0.525 |
| 0.05 | 0 | yadI | 4 | 21.8333 | 0.687 |
| 0.05 | 0 | AG1 | 4 | 21.8333 | 0.782 |
| 0.05 | 0 | ptsI | 4 | 22.0833 | 0.631 |
| 0.05 | 0 | clcB | 4 | 22.0833 | 0.536 |

| | | | | | |
|---|---|---|---|---|---|
| 0.05 | 0 | ycaM | 4 | 22.0833 | 0.526 |
| 0.05 | 0 | yadI | 4 | 22.0833 | 0.686 |
| 0.05 | 0 | AG1 | 4 | 22.0833 | 0.781 |
| 0.05 | 0 | ptsI | 4 | 22.3333 | 0.637 |
| 0.05 | 0 | clcB | 4 | 22.3333 | 0.534 |
| 0.05 | 0 | ycaM | 4 | 22.3333 | 0.527 |
| 0.05 | 0 | yadI | 4 | 22.3333 | 0.686 |
| 0.05 | 0 | AG1 | 4 | 22.3333 | 0.777 |
| 0.05 | 0 | ptsI | 4 | 22.5833 | 0.635 |
| 0.05 | 0 | clcB | 4 | 22.5833 | 0.533 |
| 0.05 | 0 | ycaM | 4 | 22.5833 | 0.525 |
| 0.05 | 0 | yadI | 4 | 22.5833 | 0.686 |
| 0.05 | 0 | AG1 | 4 | 22.5833 | 0.779 |
| 0.05 | 0 | ptsI | 4 | 22.8333 | 0.638 |
| 0.05 | 0 | clcB | 4 | 22.8333 | 0.534 |
| 0.05 | 0 | ycaM | 4 | 22.8333 | 0.528 |
| 0.05 | 0 | yadI | 4 | 22.8333 | 0.686 |
| 0.05 | 0 | AG1 | 4 | 22.8333 | 0.78 |
| 0.05 | 0 | ptsI | 4 | 23.0833 | 0.644 |
| 0.05 | 0 | clcB | 4 | 23.0833 | 0.536 |
| 0.05 | 0 | ycaM | 4 | 23.0833 | 0.531 |
| 0.05 | 0 | yadI | 4 | 23.0833 | 0.687 |
| 0.05 | 0 | AG1 | 4 | 23.0833 | 0.781 |
| 0.05 | 0 | ptsI | 4 | 23.3333 | 0.643 |
| 0.05 | 0 | clcB | 4 | 23.3333 | 0.535 |
| 0.05 | 0 | ycaM | 4 | 23.3333 | 0.532 |
| 0.05 | 0 | yadI | 4 | 23.3333 | 0.688 |
| 0.05 | 0 | AG1 | 4 | 23.3333 | 0.781 |
| 0.05 | 0 | ptsI | 4 | 23.5833 | 0.649 |
| 0.05 | 0 | clcB | 4 | 23.5833 | 0.535 |
| 0.05 | 0 | ycaM | 4 | 23.5833 | 0.532 |
| 0.05 | 0 | yadI | 4 | 23.5833 | 0.688 |
| 0.05 | 0 | AG1 | 4 | 23.5833 | 0.781 |
| 0.05 | 0 | ptsI | 4 | 23.8333 | 0.655 |
| 0.05 | 0 | clcB | 4 | 23.8333 | 0.535 |
| 0.05 | 0 | ycaM | 4 | 23.8333 | 0.533 |
| 0.05 | 0 | yadI | 4 | 23.8333 | 0.689 |
| 0.05 | 0 | AG1 | 4 | 23.8333 | 0.782 |
| 0.05 | 0 | ptsI | 4 | 24.0833 | 0.657 |
| 0.05 | 0 | clcB | 4 | 24.0833 | 0.535 |
| 0.05 | 0 | ycaM | 4 | 24.0833 | 0.534 |
| 0.05 | 0 | yadI | 4 | 24.0833 | 0.691 |
| 0.05 | 0 | AG1 | 4 | 24.0833 | 0.78 |
| 0.05 | 0 | ptsI | 5 | 0 | 0.186 |
| 0.05 | 0 | clcB | 5 | 0 | 0.163 |
| 0.05 | 0 | ycaM | 5 | 0 | 0.176 |
| 0.05 | 0 | yadI | 5 | 0 | 0.17 |
| 0.05 | 0 | AG1 | 5 | 0 | 0.19 |

| | | | | | |
|---|---|---|---|---|---|
| 0.05 | 0 | ptsI | 5 | 0.35 | 0.175 |
| 0.05 | 0 | clcB | 5 | 0.35 | 0.16 |
| 0.05 | 0 | ycaM | 5 | 0.35 | 0.167 |
| 0.05 | 0 | yadI | 5 | 0.35 | 0.166 |
| 0.05 | 0 | AG1 | 5 | 0.35 | 0.187 |
| 0.05 | 0 | ptsI | 5 | 0.6 | 0.173 |
| 0.05 | 0 | clcB | 5 | 0.6 | 0.161 |
| 0.05 | 0 | ycaM | 5 | 0.6 | 0.168 |
| 0.05 | 0 | yadI | 5 | 0.6 | 0.166 |
| 0.05 | 0 | AG1 | 5 | 0.6 | 0.187 |
| 0.05 | 0 | ptsI | 5 | 0.85 | 0.178 |
| 0.05 | 0 | clcB | 5 | 0.85 | 0.163 |
| 0.05 | 0 | ycaM | 5 | 0.85 | 0.19 |
| 0.05 | 0 | yadI | 5 | 0.85 | 0.167 |
| 0.05 | 0 | AG1 | 5 | 0.85 | 0.188 |
| 0.05 | 0 | ptsI | 5 | 1.1 | 0.18 |
| 0.05 | 0 | clcB | 5 | 1.1 | 0.166 |
| 0.05 | 0 | ycaM | 5 | 1.1 | 0.228 |
| 0.05 | 0 | yadI | 5 | 1.1 | 0.168 |
| 0.05 | 0 | AG1 | 5 | 1.1 | 0.188 |
| 0.05 | 0 | ptsI | 5 | 1.35 | 0.168 |
| 0.05 | 0 | clcB | 5 | 1.35 | 0.168 |
| 0.05 | 0 | ycaM | 5 | 1.35 | 0.172 |
| 0.05 | 0 | yadI | 5 | 1.35 | 0.172 |
| 0.05 | 0 | AG1 | 5 | 1.35 | 0.192 |
| 0.05 | 0 | ptsI | 5 | 1.6 | 0.174 |
| 0.05 | 0 | clcB | 5 | 1.6 | 0.173 |
| 0.05 | 0 | ycaM | 5 | 1.6 | 0.181 |
| 0.05 | 0 | yadI | 5 | 1.6 | 0.174 |
| 0.05 | 0 | AG1 | 5 | 1.6 | 0.196 |
| 0.05 | 0 | ptsI | 5 | 1.85 | 0.192 |
| 0.05 | 0 | clcB | 5 | 1.85 | 0.179 |
| 0.05 | 0 | ycaM | 5 | 1.85 | 0.182 |
| 0.05 | 0 | yadI | 5 | 1.85 | 0.179 |
| 0.05 | 0 | AG1 | 5 | 1.85 | 0.201 |
| 0.05 | 0 | ptsI | 5 | 2.1 | 0.205 |
| 0.05 | 0 | clcB | 5 | 2.1 | 0.186 |
| 0.05 | 0 | ycaM | 5 | 2.1 | 0.184 |
| 0.05 | 0 | yadI | 5 | 2.1 | 0.183 |
| 0.05 | 0 | AG1 | 5 | 2.1 | 0.207 |
| 0.05 | 0 | ptsI | 5 | 2.35 | 0.196 |
| 0.05 | 0 | clcB | 5 | 2.35 | 0.195 |
| 0.05 | 0 | ycaM | 5 | 2.35 | 0.19 |
| 0.05 | 0 | yadI | 5 | 2.35 | 0.19 |
| 0.05 | 0 | AG1 | 5 | 2.35 | 0.216 |
| 0.05 | 0 | ptsI | 5 | 2.6 | 0.209 |
| 0.05 | 0 | clcB | 5 | 2.6 | 0.2 |
| 0.05 | 0 | ycaM | 5 | 2.6 | 0.201 |

| | | | | | |
|---|---|---|---|---|---|
| 0.05 | 0 | yadI | 5 | 2.6 | 0.195 |
| 0.05 | 0 | AG1 | 5 | 2.6 | 0.212 |
| 0.05 | 0 | ptsI | 5 | 2.85 | 0.214 |
| 0.05 | 0 | clcB | 5 | 2.85 | 0.217 |
| 0.05 | 0 | ycaM | 5 | 2.85 | 0.208 |
| 0.05 | 0 | yadI | 5 | 2.85 | 0.205 |
| 0.05 | 0 | AG1 | 5 | 2.85 | 0.228 |
| 0.05 | 0 | ptsI | 5 | 3.1 | 0.22 |
| 0.05 | 0 | clcB | 5 | 3.1 | 0.231 |
| 0.05 | 0 | ycaM | 5 | 3.1 | 0.218 |
| 0.05 | 0 | yadI | 5 | 3.1 | 0.215 |
| 0.05 | 0 | AG1 | 5 | 3.1 | 0.238 |
| 0.05 | 0 | ptsI | 5 | 3.35 | 0.227 |
| 0.05 | 0 | clcB | 5 | 3.35 | 0.244 |
| 0.05 | 0 | ycaM | 5 | 3.35 | 0.245 |
| 0.05 | 0 | yadI | 5 | 3.35 | 0.23 |
| 0.05 | 0 | AG1 | 5 | 3.35 | 0.255 |
| 0.05 | 0 | ptsI | 5 | 3.6 | 0.238 |
| 0.05 | 0 | clcB | 5 | 3.6 | 0.258 |
| 0.05 | 0 | ycaM | 5 | 3.6 | 0.246 |
| 0.05 | 0 | yadI | 5 | 3.6 | 0.246 |
| 0.05 | 0 | AG1 | 5 | 3.6 | 0.277 |
| 0.05 | 0 | ptsI | 5 | 3.85 | 0.253 |
| 0.05 | 0 | clcB | 5 | 3.85 | 0.274 |
| 0.05 | 0 | ycaM | 5 | 3.85 | 0.252 |
| 0.05 | 0 | yadI | 5 | 3.85 | 0.263 |
| 0.05 | 0 | AG1 | 5 | 3.85 | 0.299 |
| 0.05 | 0 | ptsI | 5 | 4.38333 | 0.249 |
| 0.05 | 0 | clcB | 5 | 4.38333 | 0.246 |
| 0.05 | 0 | ycaM | 5 | 4.38333 | 0.259 |
| 0.05 | 0 | yadI | 5 | 4.38333 | 0.273 |
| 0.05 | 0 | AG1 | 5 | 4.38333 | 0.33 |
| 0.05 | 0 | ptsI | 5 | 4.63333 | 0.23 |
| 0.05 | 0 | clcB | 5 | 4.63333 | 0.248 |
| 0.05 | 0 | ycaM | 5 | 4.63333 | 0.261 |
| 0.05 | 0 | yadI | 5 | 4.63333 | 0.276 |
| 0.05 | 0 | AG1 | 5 | 4.63333 | 0.328 |
| 0.05 | 0 | ptsI | 5 | 4.88333 | 0.233 |
| 0.05 | 0 | clcB | 5 | 4.88333 | 0.265 |
| 0.05 | 0 | ycaM | 5 | 4.88333 | 0.271 |
| 0.05 | 0 | yadI | 5 | 4.88333 | 0.287 |
| 0.05 | 0 | AG1 | 5 | 4.88333 | 0.343 |
| 0.05 | 0 | ptsI | 5 | 5.13333 | 0.241 |
| 0.05 | 0 | clcB | 5 | 5.13333 | 0.278 |
| 0.05 | 0 | ycaM | 5 | 5.13333 | 0.28 |
| 0.05 | 0 | yadI | 5 | 5.13333 | 0.299 |
| 0.05 | 0 | AG1 | 5 | 5.13333 | 0.352 |
| 0.05 | 0 | ptsI | 5 | 5.38333 | 0.248 |

| | | | | | |
|---|---|---|---|---|---|
| 0.05 | 0 | clcB | 5 | 5.38333 | 0.293 |
| 0.05 | 0 | ycaM | 5 | 5.38333 | 0.291 |
| 0.05 | 0 | yadI | 5 | 5.38333 | 0.314 |
| 0.05 | 0 | AG1 | 5 | 5.38333 | 0.367 |
| 0.05 | 0 | ptsI | 5 | 5.63333 | 0.257 |
| 0.05 | 0 | clcB | 5 | 5.63333 | 0.304 |
| 0.05 | 0 | ycaM | 5 | 5.63333 | 0.304 |
| 0.05 | 0 | yadI | 5 | 5.63333 | 0.331 |
| 0.05 | 0 | AG1 | 5 | 5.63333 | 0.38 |
| 0.05 | 0 | ptsI | 5 | 5.88333 | 0.267 |
| 0.05 | 0 | clcB | 5 | 5.88333 | 0.314 |
| 0.05 | 0 | ycaM | 5 | 5.88333 | 0.317 |
| 0.05 | 0 | yadI | 5 | 5.88333 | 0.346 |
| 0.05 | 0 | AG1 | 5 | 5.88333 | 0.393 |
| 0.05 | 0 | ptsI | 5 | 6.13333 | 0.277 |
| 0.05 | 0 | clcB | 5 | 6.13333 | 0.326 |
| 0.05 | 0 | ycaM | 5 | 6.13333 | 0.329 |
| 0.05 | 0 | yadI | 5 | 6.13333 | 0.366 |
| 0.05 | 0 | AG1 | 5 | 6.13333 | 0.408 |
| 0.05 | 0 | ptsI | 5 | 6.38333 | 0.288 |
| 0.05 | 0 | clcB | 5 | 6.38333 | 0.336 |
| 0.05 | 0 | ycaM | 5 | 6.38333 | 0.343 |
| 0.05 | 0 | yadI | 5 | 6.38333 | 0.386 |
| 0.05 | 0 | AG1 | 5 | 6.38333 | 0.431 |
| 0.05 | 0 | ptsI | 5 | 6.63333 | 0.304 |
| 0.05 | 0 | clcB | 5 | 6.63333 | 0.348 |
| 0.05 | 0 | ycaM | 5 | 6.63333 | 0.356 |
| 0.05 | 0 | yadI | 5 | 6.63333 | 0.409 |
| 0.05 | 0 | AG1 | 5 | 6.63333 | 0.45 |
| 0.05 | 0 | ptsI | 5 | 6.88333 | 0.317 |
| 0.05 | 0 | clcB | 5 | 6.88333 | 0.359 |
| 0.05 | 0 | ycaM | 5 | 6.88333 | 0.368 |
| 0.05 | 0 | yadI | 5 | 6.88333 | 0.429 |
| 0.05 | 0 | AG1 | 5 | 6.88333 | 0.476 |
| 0.05 | 0 | ptsI | 5 | 7.13333 | 0.329 |
| 0.05 | 0 | clcB | 5 | 7.13333 | 0.371 |
| 0.05 | 0 | ycaM | 5 | 7.13333 | 0.382 |
| 0.05 | 0 | yadI | 5 | 7.13333 | 0.453 |
| 0.05 | 0 | AG1 | 5 | 7.13333 | 0.497 |
| 0.05 | 0 | ptsI | 5 | 7.38333 | 0.348 |
| 0.05 | 0 | clcB | 5 | 7.38333 | 0.382 |
| 0.05 | 0 | ycaM | 5 | 7.38333 | 0.392 |
| 0.05 | 0 | yadI | 5 | 7.38333 | 0.478 |
| 0.05 | 0 | AG1 | 5 | 7.38333 | 0.515 |
| 0.05 | 0 | ptsI | 5 | 7.63333 | 0.364 |
| 0.05 | 0 | clcB | 5 | 7.63333 | 0.395 |
| 0.05 | 0 | ycaM | 5 | 7.63333 | 0.403 |
| 0.05 | 0 | yadI | 5 | 7.63333 | 0.502 |

| | | | | | |
|---|---|---|---|---|---|
| 0.05 | 0 | AG1 | 5 | 7.63333 | 0.537 |
| 0.05 | 0 | ptsI | 5 | 7.88333 | 0.373 |
| 0.05 | 0 | clcB | 5 | 7.88333 | 0.405 |
| 0.05 | 0 | ycaM | 5 | 7.88333 | 0.412 |
| 0.05 | 0 | yadI | 5 | 7.88333 | 0.515 |
| 0.05 | 0 | AG1 | 5 | 7.88333 | 0.549 |
| 0.05 | 0 | ptsI | 5 | 8.13333 | 0.387 |
| 0.05 | 0 | clcB | 5 | 8.13333 | 0.417 |
| 0.05 | 0 | ycaM | 5 | 8.13333 | 0.423 |
| 0.05 | 0 | yadI | 5 | 8.13333 | 0.526 |
| 0.05 | 0 | AG1 | 5 | 8.13333 | 0.568 |
| 0.05 | 0 | ptsI | 5 | 8.38333 | 0.395 |
| 0.05 | 0 | clcB | 5 | 8.38333 | 0.427 |
| 0.05 | 0 | ycaM | 5 | 8.38333 | 0.435 |
| 0.05 | 0 | yadI | 5 | 8.38333 | 0.539 |
| 0.05 | 0 | AG1 | 5 | 8.38333 | 0.581 |
| 0.05 | 0 | ptsI | 5 | 8.88333 | 0.378 |
| 0.05 | 0 | clcB | 5 | 8.88333 | 0.354 |
| 0.05 | 0 | ycaM | 5 | 8.88333 | 0.362 |
| 0.05 | 0 | yadI | 5 | 8.88333 | 0.5 |
| 0.05 | 0 | AG1 | 5 | 8.88333 | 0.609 |
| 0.05 | 0 | ptsI | 5 | 9.13333 | 0.379 |
| 0.05 | 0 | clcB | 5 | 9.13333 | 0.356 |
| 0.05 | 0 | ycaM | 5 | 9.13333 | 0.367 |
| 0.05 | 0 | yadI | 5 | 9.13333 | 0.502 |
| 0.05 | 0 | AG1 | 5 | 9.13333 | 0.608 |
| 0.05 | 0 | ptsI | 5 | 9.38333 | 0.388 |
| 0.05 | 0 | clcB | 5 | 9.38333 | 0.366 |
| 0.05 | 0 | ycaM | 5 | 9.38333 | 0.373 |
| 0.05 | 0 | yadI | 5 | 9.38333 | 0.506 |
| 0.05 | 0 | AG1 | 5 | 9.38333 | 0.614 |
| 0.05 | 0 | ptsI | 5 | 9.63333 | 0.405 |
| 0.05 | 0 | clcB | 5 | 9.63333 | 0.375 |
| 0.05 | 0 | ycaM | 5 | 9.63333 | 0.384 |
| 0.05 | 0 | yadI | 5 | 9.63333 | 0.522 |
| 0.05 | 0 | AG1 | 5 | 9.63333 | 0.637 |
| 0.05 | 0 | ptsI | 5 | 9.88333 | 0.414 |
| 0.05 | 0 | clcB | 5 | 9.88333 | 0.385 |
| 0.05 | 0 | ycaM | 5 | 9.88333 | 0.395 |
| 0.05 | 0 | yadI | 5 | 9.88333 | 0.534 |
| 0.05 | 0 | AG1 | 5 | 9.88333 | 0.652 |
| 0.05 | 0 | ptsI | 5 | 10.1333 | 0.421 |
| 0.05 | 0 | clcB | 5 | 10.1333 | 0.392 |
| 0.05 | 0 | ycaM | 5 | 10.1333 | 0.396 |
| 0.05 | 0 | yadI | 5 | 10.1333 | 0.538 |
| 0.05 | 0 | AG1 | 5 | 10.1333 | 0.662 |
| 0.05 | 0 | ptsI | 5 | 10.3833 | 0.433 |
| 0.05 | 0 | clcB | 5 | 10.3833 | 0.398 |

| | | | | | |
|---|---|---|---|---|---|
| 0.05 | 0 | ycaM | 5 | 10.3833 | 0.403 |
| 0.05 | 0 | yadI | 5 | 10.3833 | 0.547 |
| 0.05 | 0 | AG1 | 5 | 10.3833 | 0.676 |
| 0.05 | 0 | ptsI | 5 | 10.6333 | 0.44 |
| 0.05 | 0 | clcB | 5 | 10.6333 | 0.404 |
| 0.05 | 0 | ycaM | 5 | 10.6333 | 0.403 |
| 0.05 | 0 | yadI | 5 | 10.6333 | 0.556 |
| 0.05 | 0 | AG1 | 5 | 10.6333 | 0.685 |
| 0.05 | 0 | ptsI | 5 | 10.8833 | 0.452 |
| 0.05 | 0 | clcB | 5 | 10.8833 | 0.413 |
| 0.05 | 0 | ycaM | 5 | 10.8833 | 0.408 |
| 0.05 | 0 | yadI | 5 | 10.8833 | 0.569 |
| 0.05 | 0 | AG1 | 5 | 10.8833 | 0.699 |
| 0.05 | 0 | ptsI | 5 | 11.1333 | 0.458 |
| 0.05 | 0 | clcB | 5 | 11.1333 | 0.414 |
| 0.05 | 0 | ycaM | 5 | 11.1333 | 0.417 |
| 0.05 | 0 | yadI | 5 | 11.1333 | 0.576 |
| 0.05 | 0 | AG1 | 5 | 11.1333 | 0.71 |
| 0.05 | 0 | ptsI | 5 | 11.3833 | 0.47 |
| 0.05 | 0 | clcB | 5 | 11.3833 | 0.42 |
| 0.05 | 0 | ycaM | 5 | 11.3833 | 0.415 |
| 0.05 | 0 | yadI | 5 | 11.3833 | 0.583 |
| 0.05 | 0 | AG1 | 5 | 11.3833 | 0.722 |
| 0.05 | 0 | ptsI | 5 | 11.6333 | 0.478 |
| 0.05 | 0 | clcB | 5 | 11.6333 | 0.426 |
| 0.05 | 0 | ycaM | 5 | 11.6333 | 0.42 |
| 0.05 | 0 | yadI | 5 | 11.6333 | 0.593 |
| 0.05 | 0 | AG1 | 5 | 11.6333 | 0.735 |
| 0.05 | 0 | ptsI | 5 | 11.8833 | 0.486 |
| 0.05 | 0 | clcB | 5 | 11.8833 | 0.431 |
| 0.05 | 0 | ycaM | 5 | 11.8833 | 0.422 |
| 0.05 | 0 | yadI | 5 | 11.8833 | 0.601 |
| 0.05 | 0 | AG1 | 5 | 11.8833 | 0.743 |
| 0.05 | 0 | ptsI | 5 | 12.1333 | 0.501 |
| 0.05 | 0 | clcB | 5 | 12.1333 | 0.437 |
| 0.05 | 0 | ycaM | 5 | 12.1333 | 0.429 |
| 0.05 | 0 | yadI | 5 | 12.1333 | 0.611 |
| 0.05 | 0 | AG1 | 5 | 12.1333 | 0.754 |
| 0.05 | 0 | ptsI | 5 | 12.3833 | 0.516 |
| 0.05 | 0 | clcB | 5 | 12.3833 | 0.438 |
| 0.05 | 0 | ycaM | 5 | 12.3833 | 0.429 |
| 0.05 | 0 | yadI | 5 | 12.3833 | 0.603 |
| 0.05 | 0 | AG1 | 5 | 12.3833 | 0.747 |
| 0.05 | 0 | ptsI | 5 | 12.6333 | 0.514 |
| 0.05 | 0 | clcB | 5 | 12.6333 | 0.443 |
| 0.05 | 0 | ycaM | 5 | 12.6333 | 0.438 |
| 0.05 | 0 | yadI | 5 | 12.6333 | 0.621 |
| 0.05 | 0 | AG1 | 5 | 12.6333 | 0.771 |

| | | | | | |
|---|---|---|---|---|---|
| 0.05 | 0 | ptsI | 5 | 12.8833 | 0.529 |
| 0.05 | 0 | clcB | 5 | 12.8833 | 0.451 |
| 0.05 | 0 | ycaM | 5 | 12.8833 | 0.448 |
| 0.05 | 0 | yadI | 5 | 12.8833 | 0.63 |
| 0.05 | 0 | AG1 | 5 | 12.8833 | 0.783 |
| 0.05 | 0 | ptsI | 5 | 13.1333 | 0.533 |
| 0.05 | 0 | clcB | 5 | 13.1333 | 0.455 |
| 0.05 | 0 | ycaM | 5 | 13.1333 | 0.449 |
| 0.05 | 0 | yadI | 5 | 13.1333 | 0.634 |
| 0.05 | 0 | AG1 | 5 | 13.1333 | 0.789 |
| 0.05 | 0 | ptsI | 5 | 13.3833 | 0.543 |
| 0.05 | 0 | clcB | 5 | 13.3833 | 0.457 |
| 0.05 | 0 | ycaM | 5 | 13.3833 | 0.447 |
| 0.05 | 0 | yadI | 5 | 13.3833 | 0.642 |
| 0.05 | 0 | AG1 | 5 | 13.3833 | 0.795 |
| 0.05 | 0 | ptsI | 5 | 13.6333 | 0.554 |
| 0.05 | 0 | clcB | 5 | 13.6333 | 0.463 |
| 0.05 | 0 | ycaM | 5 | 13.6333 | 0.452 |
| 0.05 | 0 | yadI | 5 | 13.6333 | 0.648 |
| 0.05 | 0 | AG1 | 5 | 13.6333 | 0.804 |
| 0.05 | 0 | ptsI | 5 | 13.8833 | 0.555 |
| 0.05 | 0 | clcB | 5 | 13.8833 | 0.466 |
| 0.05 | 0 | ycaM | 5 | 13.8833 | 0.452 |
| 0.05 | 0 | yadI | 5 | 13.8833 | 0.651 |
| 0.05 | 0 | AG1 | 5 | 13.8833 | 0.81 |
| 0.05 | 0 | ptsI | 5 | 14.1333 | 0.567 |
| 0.05 | 0 | clcB | 5 | 14.1333 | 0.472 |
| 0.05 | 0 | ycaM | 5 | 14.1333 | 0.458 |
| 0.05 | 0 | yadI | 5 | 14.1333 | 0.659 |
| 0.05 | 0 | AG1 | 5 | 14.1333 | 0.816 |
| 0.05 | 0 | ptsI | 5 | 14.3833 | 0.58 |
| 0.05 | 0 | clcB | 5 | 14.3833 | 0.479 |
| 0.05 | 0 | ycaM | 5 | 14.3833 | 0.462 |
| 0.05 | 0 | yadI | 5 | 14.3833 | 0.666 |
| 0.05 | 0 | AG1 | 5 | 14.3833 | 0.824 |
| 0.05 | 0 | ptsI | 5 | 14.6333 | 0.584 |
| 0.05 | 0 | clcB | 5 | 14.6333 | 0.482 |
| 0.05 | 0 | ycaM | 5 | 14.6333 | 0.466 |
| 0.05 | 0 | yadI | 5 | 14.6333 | 0.671 |
| 0.05 | 0 | AG1 | 5 | 14.6333 | 0.833 |
| 0.05 | 0 | ptsI | 5 | 14.8833 | 0.589 |
| 0.05 | 0 | clcB | 5 | 14.8833 | 0.485 |
| 0.05 | 0 | ycaM | 5 | 14.8833 | 0.469 |
| 0.05 | 0 | yadI | 5 | 14.8833 | 0.673 |
| 0.05 | 0 | AG1 | 5 | 14.8833 | 0.835 |
| 0.05 | 0 | ptsI | 5 | 15.1333 | 0.602 |
| 0.05 | 0 | clcB | 5 | 15.1333 | 0.487 |
| 0.05 | 0 | ycaM | 5 | 15.1333 | 0.473 |

| | | | | | |
|---|---|---|---|---|---|
| 0.05 | 0 | yadI | 5 | 15.1333 | 0.678 |
| 0.05 | 0 | AG1 | 5 | 15.1333 | 0.841 |
| 0.05 | 0 | ptsI | 5 | 15.3833 | 0.596 |
| 0.05 | 0 | clcB | 5 | 15.3833 | 0.492 |
| 0.05 | 0 | ycaM | 5 | 15.3833 | 0.479 |
| 0.05 | 0 | yadI | 5 | 15.3833 | 0.684 |
| 0.05 | 0 | AG1 | 5 | 15.3833 | 0.849 |
| 0.05 | 0 | ptsI | 5 | 15.6333 | 0.599 |
| 0.05 | 0 | clcB | 5 | 15.6333 | 0.491 |
| 0.05 | 0 | ycaM | 5 | 15.6333 | 0.479 |
| 0.05 | 0 | yadI | 5 | 15.6333 | 0.685 |
| 0.05 | 0 | AG1 | 5 | 15.6333 | 0.848 |
| 0.05 | 0 | ptsI | 5 | 15.8833 | 0.603 |
| 0.05 | 0 | clcB | 5 | 15.8833 | 0.493 |
| 0.05 | 0 | ycaM | 5 | 15.8833 | 0.482 |
| 0.05 | 0 | yadI | 5 | 15.8833 | 0.69 |
| 0.05 | 0 | AG1 | 5 | 15.8833 | 0.854 |
| 0.05 | 0 | ptsI | 5 | 16.1333 | 0.601 |
| 0.05 | 0 | clcB | 5 | 16.1333 | 0.494 |
| 0.05 | 0 | ycaM | 5 | 16.1333 | 0.486 |
| 0.05 | 0 | yadI | 5 | 16.1333 | 0.69 |
| 0.05 | 0 | AG1 | 5 | 16.1333 | 0.848 |
| 0.05 | 0 | ptsI | 5 | 16.3833 | 0.621 |
| 0.05 | 0 | clcB | 5 | 16.3833 | 0.496 |
| 0.05 | 0 | ycaM | 5 | 16.3833 | 0.483 |
| 0.05 | 0 | yadI | 5 | 16.3833 | 0.696 |
| 0.05 | 0 | AG1 | 5 | 16.3833 | 0.861 |
| 0.05 | 0 | ptsI | 5 | 16.6333 | 0.627 |
| 0.05 | 0 | clcB | 5 | 16.6333 | 0.496 |
| 0.05 | 0 | ycaM | 5 | 16.6333 | 0.489 |
| 0.05 | 0 | yadI | 5 | 16.6333 | 0.697 |
| 0.05 | 0 | AG1 | 5 | 16.6333 | 0.864 |
| 0.05 | 0 | ptsI | 5 | 16.8833 | 0.626 |
| 0.05 | 0 | clcB | 5 | 16.8833 | 0.504 |
| 0.05 | 0 | ycaM | 5 | 16.8833 | 0.494 |
| 0.05 | 0 | yadI | 5 | 16.8833 | 0.702 |
| 0.05 | 0 | AG1 | 5 | 16.8833 | 0.872 |
| 0.05 | 0 | ptsI | 5 | 17.1333 | 0.639 |
| 0.05 | 0 | clcB | 5 | 17.1333 | 0.504 |
| 0.05 | 0 | ycaM | 5 | 17.1333 | 0.498 |
| 0.05 | 0 | yadI | 5 | 17.1333 | 0.706 |
| 0.05 | 0 | AG1 | 5 | 17.1333 | 0.875 |
| 0.05 | 0 | ptsI | 5 | 17.3833 | 0.641 |
| 0.05 | 0 | clcB | 5 | 17.3833 | 0.499 |
| 0.05 | 0 | ycaM | 5 | 17.3833 | 0.495 |
| 0.05 | 0 | yadI | 5 | 17.3833 | 0.707 |
| 0.05 | 0 | AG1 | 5 | 17.3833 | 0.877 |
| 0.05 | 0 | ptsI | 5 | 17.6333 | 0.644 |

| | | | | | |
|---|---|---|---|---|---|
| 0.05 | 0 | clcB | 5 | 17.6333 | 0.5 |
| 0.05 | 0 | ycaM | 5 | 17.6333 | 0.501 |
| 0.05 | 0 | yadI | 5 | 17.6333 | 0.708 |
| 0.05 | 0 | AG1 | 5 | 17.6333 | 0.877 |
| 0.05 | 0 | ptsI | 5 | 17.8833 | 0.657 |
| 0.05 | 0 | clcB | 5 | 17.8833 | 0.506 |
| 0.05 | 0 | ycaM | 5 | 17.8833 | 0.507 |
| 0.05 | 0 | yadI | 5 | 17.8833 | 0.701 |
| 0.05 | 0 | AG1 | 5 | 17.8833 | 0.869 |
| 0.05 | 0 | ptsI | 5 | 18.1333 | 0.65 |
| 0.05 | 0 | clcB | 5 | 18.1333 | 0.509 |
| 0.05 | 0 | ycaM | 5 | 18.1333 | 0.51 |
| 0.05 | 0 | yadI | 5 | 18.1333 | 0.714 |
| 0.05 | 0 | AG1 | 5 | 18.1333 | 0.886 |
| 0.05 | 0 | ptsI | 5 | 18.3833 | 0.651 |
| 0.05 | 0 | clcB | 5 | 18.3833 | 0.509 |
| 0.05 | 0 | ycaM | 5 | 18.3833 | 0.51 |
| 0.05 | 0 | yadI | 5 | 18.3833 | 0.718 |
| 0.05 | 0 | AG1 | 5 | 18.3833 | 0.886 |
| 0.05 | 0 | ptsI | 5 | 18.6333 | 0.659 |
| 0.05 | 0 | clcB | 5 | 18.6333 | 0.511 |
| 0.05 | 0 | ycaM | 5 | 18.6333 | 0.511 |
| 0.05 | 0 | yadI | 5 | 18.6333 | 0.717 |
| 0.05 | 0 | AG1 | 5 | 18.6333 | 0.887 |
| 0.05 | 0 | ptsI | 5 | 18.8833 | 0.666 |
| 0.05 | 0 | clcB | 5 | 18.8833 | 0.509 |
| 0.05 | 0 | ycaM | 5 | 18.8833 | 0.518 |
| 0.05 | 0 | yadI | 5 | 18.8833 | 0.717 |
| 0.05 | 0 | AG1 | 5 | 18.8833 | 0.89 |
| 0.05 | 0 | ptsI | 5 | 19.1333 | 0.676 |
| 0.05 | 0 | clcB | 5 | 19.1333 | 0.512 |
| 0.05 | 0 | ycaM | 5 | 19.1333 | 0.516 |
| 0.05 | 0 | yadI | 5 | 19.1333 | 0.722 |
| 0.05 | 0 | AG1 | 5 | 19.1333 | 0.893 |
| 0.05 | 0 | ptsI | 5 | 19.3833 | 0.669 |
| 0.05 | 0 | clcB | 5 | 19.3833 | 0.517 |
| 0.05 | 0 | ycaM | 5 | 19.3833 | 0.524 |
| 0.05 | 0 | yadI | 5 | 19.3833 | 0.724 |
| 0.05 | 0 | AG1 | 5 | 19.3833 | 0.895 |
| 0.05 | 0 | ptsI | 5 | 19.6333 | 0.678 |
| 0.05 | 0 | clcB | 5 | 19.6333 | 0.515 |
| 0.05 | 0 | ycaM | 5 | 19.6333 | 0.53 |
| 0.05 | 0 | yadI | 5 | 19.6333 | 0.726 |
| 0.05 | 0 | AG1 | 5 | 19.6333 | 0.898 |
| 0.05 | 0 | ptsI | 5 | 19.8833 | 0.684 |
| 0.05 | 0 | clcB | 5 | 19.8833 | 0.516 |
| 0.05 | 0 | ycaM | 5 | 19.8833 | 0.518 |
| 0.05 | 0 | yadI | 5 | 19.8833 | 0.729 |

| | | | | | |
|---|---|---|---|---|---|
| 0.05 | 0 | AG1 | 5 | 19.8833 | 0.9 |
| 0.05 | 0 | ptsI | 5 | 20.1333 | 0.686 |
| 0.05 | 0 | clcB | 5 | 20.1333 | 0.517 |
| 0.05 | 0 | ycaM | 5 | 20.1333 | 0.52 |
| 0.05 | 0 | yadI | 5 | 20.1333 | 0.731 |
| 0.05 | 0 | AG1 | 5 | 20.1333 | 0.901 |
| 0.05 | 0 | ptsI | 5 | 20.3833 | 0.693 |
| 0.05 | 0 | clcB | 5 | 20.3833 | 0.517 |
| 0.05 | 0 | ycaM | 5 | 20.3833 | 0.525 |
| 0.05 | 0 | yadI | 5 | 20.3833 | 0.734 |
| 0.05 | 0 | AG1 | 5 | 20.3833 | 0.903 |
| 0.05 | 0 | ptsI | 5 | 20.6333 | 0.699 |
| 0.05 | 0 | clcB | 5 | 20.6333 | 0.52 |
| 0.05 | 0 | ycaM | 5 | 20.6333 | 0.529 |
| 0.05 | 0 | yadI | 5 | 20.6333 | 0.721 |
| 0.05 | 0 | AG1 | 5 | 20.6333 | 0.895 |
| 0.05 | 0 | ptsI | 5 | 20.8833 | 0.704 |
| 0.05 | 0 | clcB | 5 | 20.8833 | 0.518 |
| 0.05 | 0 | ycaM | 5 | 20.8833 | 0.524 |
| 0.05 | 0 | yadI | 5 | 20.8833 | 0.735 |
| 0.05 | 0 | AG1 | 5 | 20.8833 | 0.905 |
| 0.05 | 0 | ptsI | 5 | 21.1333 | 0.709 |
| 0.05 | 0 | clcB | 5 | 21.1333 | 0.521 |
| 0.05 | 0 | ycaM | 5 | 21.1333 | 0.534 |
| 0.05 | 0 | yadI | 5 | 21.1333 | 0.735 |
| 0.05 | 0 | AG1 | 5 | 21.1333 | 0.907 |
| 0.05 | 0 | ptsI | 5 | 21.3833 | 0.71 |
| 0.05 | 0 | clcB | 5 | 21.3833 | 0.521 |
| 0.05 | 0 | ycaM | 5 | 21.3833 | 0.536 |
| 0.05 | 0 | yadI | 5 | 21.3833 | 0.738 |
| 0.05 | 0 | AG1 | 5 | 21.3833 | 0.907 |
| 0.05 | 0 | ptsI | 5 | 21.6333 | 0.714 |
| 0.05 | 0 | clcB | 5 | 21.6333 | 0.523 |
| 0.05 | 0 | ycaM | 5 | 21.6333 | 0.533 |
| 0.05 | 0 | yadI | 5 | 21.6333 | 0.738 |
| 0.05 | 0 | AG1 | 5 | 21.6333 | 0.911 |
| 0.05 | 0 | ptsI | 5 | 21.8833 | 0.71 |
| 0.05 | 0 | clcB | 5 | 21.8833 | 0.524 |
| 0.05 | 0 | ycaM | 5 | 21.8833 | 0.536 |
| 0.05 | 0 | yadI | 5 | 21.8833 | 0.744 |
| 0.05 | 0 | AG1 | 5 | 21.8833 | 0.911 |
| 0.05 | 0 | ptsI | 5 | 22.1333 | 0.73 |
| 0.05 | 0 | clcB | 5 | 22.1333 | 0.523 |
| 0.05 | 0 | ycaM | 5 | 22.1333 | 0.535 |
| 0.05 | 0 | yadI | 5 | 22.1333 | 0.73 |
| 0.05 | 0 | AG1 | 5 | 22.1333 | 0.896 |
| 0.05 | 0 | ptsI | 5 | 22.3833 | 0.727 |
| 0.05 | 0 | clcB | 5 | 22.3833 | 0.522 |

| | | | | | |
|---|---|---|---|---|---|
| 0.05 | 0 | ycaM | 5 | 22.3833 | 0.542 |
| 0.05 | 0 | yadI | 5 | 22.3833 | 0.741 |
| 0.05 | 0 | AG1 | 5 | 22.3833 | 0.91 |
| 0.05 | 0 | ptsI | 5 | 22.6333 | 0.728 |
| 0.05 | 0 | clcB | 5 | 22.6333 | 0.523 |
| 0.05 | 0 | ycaM | 5 | 22.6333 | 0.547 |
| 0.05 | 0 | yadI | 5 | 22.6333 | 0.745 |
| 0.05 | 0 | AG1 | 5 | 22.6333 | 0.913 |
| 0.05 | 0 | ptsI | 5 | 22.8833 | 0.732 |
| 0.05 | 0 | clcB | 5 | 22.8833 | 0.522 |
| 0.05 | 0 | ycaM | 5 | 22.8833 | 0.543 |
| 0.05 | 0 | yadI | 5 | 22.8833 | 0.741 |
| 0.05 | 0 | AG1 | 5 | 22.8833 | 0.913 |
| 0.05 | 0 | ptsI | 5 | 23.1333 | 0.736 |
| 0.05 | 0 | clcB | 5 | 23.1333 | 0.524 |
| 0.05 | 0 | ycaM | 5 | 23.1333 | 0.541 |
| 0.05 | 0 | yadI | 5 | 23.1333 | 0.746 |
| 0.05 | 0 | AG1 | 5 | 23.1333 | 0.915 |
| 0.05 | 0 | ptsI | 5 | 23.3833 | 0.739 |
| 0.05 | 0 | clcB | 5 | 23.3833 | 0.524 |
| 0.05 | 0 | ycaM | 5 | 23.3833 | 0.546 |
| 0.05 | 0 | yadI | 5 | 23.3833 | 0.746 |
| 0.05 | 0 | AG1 | 5 | 23.3833 | 0.917 |
| 0.05 | 0 | ptsI | 5 | 23.6333 | 0.742 |
| 0.05 | 0 | clcB | 5 | 23.6333 | 0.525 |
| 0.05 | 0 | ycaM | 5 | 23.6333 | 0.55 |
| 0.05 | 0 | yadI | 5 | 23.6333 | 0.748 |
| 0.05 | 0 | AG1 | 5 | 23.6333 | 0.916 |
| 0.05 | 0 | ptsI | 5 | 23.8833 | 0.755 |
| 0.05 | 0 | clcB | 5 | 23.8833 | 0.526 |
| 0.05 | 0 | ycaM | 5 | 23.8833 | 0.549 |
| 0.05 | 0 | yadI | 5 | 23.8833 | 0.733 |
| 0.05 | 0 | AG1 | 5 | 23.8833 | 0.902 |
| 0.05 | 0 | ptsI | 5 | 24.1333 | 0.751 |
| 0.05 | 0 | clcB | 5 | 24.1333 | 0.526 |
| 0.05 | 0 | ycaM | 5 | 24.1333 | 0.548 |
| 0.05 | 0 | yadI | 5 | 24.1333 | 0.748 |
| 0.05 | 0 | AG1 | 5 | 24.1333 | 0.918 |
| 0.05 | 0 | ptsI | 5 | 24.3833 | 0.76 |
| 0.05 | 0 | clcB | 5 | 24.3833 | 0.527 |
| 0.05 | 0 | ycaM | 5 | 24.3833 | 0.552 |
| 0.05 | 0 | yadI | 5 | 24.3833 | 0.747 |
| 0.05 | 0 | AG1 | 5 | 24.3833 | 0.915 |
| 0.1 | 0 | ptsI | 1 | 0 | 0.223 |
| 0.1 | 0 | clcB | 1 | 0 | 0.232 |
| 0.1 | 0 | ycaM | 1 | 0 | 0.239 |
| 0.1 | 0 | yadI | 1 | 0 | 0.252 |
| 0.1 | 0 | AG1 | 1 | 0 | 0.236 |

| | | | | | |
|---|---|---|---|---|---|
| 0.1 | 0 | ptsI | 1 | 0.25 | 0.218 |
| 0.1 | 0 | clcB | 1 | 0.25 | 0.224 |
| 0.1 | 0 | ycaM | 1 | 0.25 | 0.231 |
| 0.1 | 0 | yadI | 1 | 0.25 | 0.242 |
| 0.1 | 0 | AG1 | 1 | 0.25 | 0.228 |
| 0.1 | 0 | ptsI | 1 | 0.5 | 0.218 |
| 0.1 | 0 | clcB | 1 | 0.5 | 0.218 |
| 0.1 | 0 | ycaM | 1 | 0.5 | 0.226 |
| 0.1 | 0 | yadI | 1 | 0.5 | 0.24 |
| 0.1 | 0 | AG1 | 1 | 0.5 | 0.224 |
| 0.1 | 0 | ptsI | 1 | 0.75 | 0.22 |
| 0.1 | 0 | clcB | 1 | 0.75 | 0.219 |
| 0.1 | 0 | ycaM | 1 | 0.75 | 0.219 |
| 0.1 | 0 | yadI | 1 | 0.75 | 0.239 |
| 0.1 | 0 | AG1 | 1 | 0.75 | 0.221 |
| 0.1 | 0 | ptsI | 1 | 1 | 0.223 |
| 0.1 | 0 | clcB | 1 | 1 | 0.22 |
| 0.1 | 0 | ycaM | 1 | 1 | 0.221 |
| 0.1 | 0 | yadI | 1 | 1 | 0.242 |
| 0.1 | 0 | AG1 | 1 | 1 | 0.227 |
| 0.1 | 0 | ptsI | 1 | 1.25 | 0.224 |
| 0.1 | 0 | clcB | 1 | 1.25 | 0.217 |
| 0.1 | 0 | ycaM | 1 | 1.25 | 0.223 |
| 0.1 | 0 | yadI | 1 | 1.25 | 0.242 |
| 0.1 | 0 | AG1 | 1 | 1.25 | 0.226 |
| 0.1 | 0 | ptsI | 1 | 1.5 | 0.229 |
| 0.1 | 0 | clcB | 1 | 1.5 | 0.223 |
| 0.1 | 0 | ycaM | 1 | 1.5 | 0.242 |
| 0.1 | 0 | yadI | 1 | 1.5 | 0.256 |
| 0.1 | 0 | AG1 | 1 | 1.5 | 0.229 |
| 0.1 | 0 | ptsI | 1 | 1.75 | 0.237 |
| 0.1 | 0 | clcB | 1 | 1.75 | 0.223 |
| 0.1 | 0 | ycaM | 1 | 1.75 | 0.23 |
| 0.1 | 0 | yadI | 1 | 1.75 | 0.255 |
| 0.1 | 0 | AG1 | 1 | 1.75 | 0.232 |
| 0.1 | 0 | ptsI | 1 | 2 | 0.243 |
| 0.1 | 0 | clcB | 1 | 2 | 0.23 |
| 0.1 | 0 | ycaM | 1 | 2 | 0.238 |
| 0.1 | 0 | yadI | 1 | 2 | 0.262 |
| 0.1 | 0 | AG1 | 1 | 2 | 0.238 |
| 0.1 | 0 | ptsI | 1 | 2.25 | 0.251 |
| 0.1 | 0 | clcB | 1 | 2.25 | 0.228 |
| 0.1 | 0 | ycaM | 1 | 2.25 | 0.243 |
| 0.1 | 0 | yadI | 1 | 2.25 | 0.268 |
| 0.1 | 0 | AG1 | 1 | 2.25 | 0.239 |
| 0.1 | 0 | ptsI | 1 | 2.5 | 0.258 |
| 0.1 | 0 | clcB | 1 | 2.5 | 0.234 |
| 0.1 | 0 | ycaM | 1 | 2.5 | 0.25 |

| | | | | | |
|---|---|---|---|---|---|
| 0.1 | 0 | yadI | 1 | 2.5 | 0.278 |
| 0.1 | 0 | AG1 | 1 | 2.5 | 0.245 |
| 0.1 | 0 | ptsI | 1 | 2.75 | 0.267 |
| 0.1 | 0 | clcB | 1 | 2.75 | 0.235 |
| 0.1 | 0 | ycaM | 1 | 2.75 | 0.256 |
| 0.1 | 0 | yadI | 1 | 2.75 | 0.286 |
| 0.1 | 0 | AG1 | 1 | 2.75 | 0.248 |
| 0.1 | 0 | ptsI | 1 | 3 | 0.274 |
| 0.1 | 0 | clcB | 1 | 3 | 0.245 |
| 0.1 | 0 | ycaM | 1 | 3 | 0.264 |
| 0.1 | 0 | yadI | 1 | 3 | 0.298 |
| 0.1 | 0 | AG1 | 1 | 3 | 0.254 |
| 0.1 | 0 | ptsI | 1 | 3.25 | 0.284 |
| 0.1 | 0 | clcB | 1 | 3.25 | 0.25 |
| 0.1 | 0 | ycaM | 1 | 3.25 | 0.273 |
| 0.1 | 0 | yadI | 1 | 3.25 | 0.314 |
| 0.1 | 0 | AG1 | 1 | 3.25 | 0.26 |
| 0.1 | 0 | ptsI | 1 | 3.5 | 0.307 |
| 0.1 | 0 | clcB | 1 | 3.5 | 0.249 |
| 0.1 | 0 | ycaM | 1 | 3.5 | 0.297 |
| 0.1 | 0 | yadI | 1 | 3.5 | 0.331 |
| 0.1 | 0 | AG1 | 1 | 3.5 | 0.264 |
| 0.1 | 0 | ptsI | 1 | 3.75 | 0.308 |
| 0.1 | 0 | clcB | 1 | 3.75 | 0.259 |
| 0.1 | 0 | ycaM | 1 | 3.75 | 0.3 |
| 0.1 | 0 | yadI | 1 | 3.75 | 0.35 |
| 0.1 | 0 | AG1 | 1 | 3.75 | 0.269 |
| 0.1 | 0 | ptsI | 1 | 4 | 0.323 |
| 0.1 | 0 | clcB | 1 | 4 | 0.264 |
| 0.1 | 0 | ycaM | 1 | 4 | 0.313 |
| 0.1 | 0 | yadI | 1 | 4 | 0.382 |
| 0.1 | 0 | AG1 | 1 | 4 | 0.283 |
| 0.1 | 0 | ptsI | 1 | 4.25 | 0.327 |
| 0.1 | 0 | clcB | 1 | 4.25 | 0.27 |
| 0.1 | 0 | ycaM | 1 | 4.25 | 0.317 |
| 0.1 | 0 | yadI | 1 | 4.25 | 0.406 |
| 0.1 | 0 | AG1 | 1 | 4.25 | 0.287 |
| 0.1 | 0 | ptsI | 1 | 4.5 | 0.337 |
| 0.1 | 0 | clcB | 1 | 4.5 | 0.278 |
| 0.1 | 0 | ycaM | 1 | 4.5 | 0.327 |
| 0.1 | 0 | yadI | 1 | 4.5 | 0.414 |
| 0.1 | 0 | AG1 | 1 | 4.5 | 0.3 |
| 0.1 | 0 | ptsI | 1 | 4.75 | 0.35 |
| 0.1 | 0 | clcB | 1 | 4.75 | 0.284 |
| 0.1 | 0 | ycaM | 1 | 4.75 | 0.338 |
| 0.1 | 0 | yadI | 1 | 4.75 | 0.423 |
| 0.1 | 0 | AG1 | 1 | 4.75 | 0.306 |
| 0.1 | 0 | ptsI | 1 | 5 | 0.359 |

| | | | | | |
|---|---|---|---|---|---|
| 0.1 | 0 | clcB | 1 | 5 | 0.296 |
| 0.1 | 0 | ycaM | 1 | 5 | 0.359 |
| 0.1 | 0 | yadI | 1 | 5 | 0.446 |
| 0.1 | 0 | AG1 | 1 | 5 | 0.314 |
| 0.1 | 0 | ptsI | 1 | 5.25 | 0.371 |
| 0.1 | 0 | clcB | 1 | 5.25 | 0.304 |
| 0.1 | 0 | ycaM | 1 | 5.25 | 0.366 |
| 0.1 | 0 | yadI | 1 | 5.25 | 0.462 |
| 0.1 | 0 | AG1 | 1 | 5.25 | 0.32 |
| 0.1 | 0 | ptsI | 1 | 5.5 | 0.379 |
| 0.1 | 0 | clcB | 1 | 5.5 | 0.316 |
| 0.1 | 0 | ycaM | 1 | 5.5 | 0.368 |
| 0.1 | 0 | yadI | 1 | 5.5 | 0.47 |
| 0.1 | 0 | AG1 | 1 | 5.5 | 0.337 |
| 0.1 | 0 | ptsI | 1 | 5.75 | 0.391 |
| 0.1 | 0 | clcB | 1 | 5.75 | 0.322 |
| 0.1 | 0 | ycaM | 1 | 5.75 | 0.376 |
| 0.1 | 0 | yadI | 1 | 5.75 | 0.483 |
| 0.1 | 0 | AG1 | 1 | 5.75 | 0.348 |
| 0.1 | 0 | ptsI | 1 | 6 | 0.407 |
| 0.1 | 0 | clcB | 1 | 6 | 0.328 |
| 0.1 | 0 | ycaM | 1 | 6 | 0.387 |
| 0.1 | 0 | yadI | 1 | 6 | 0.485 |
| 0.1 | 0 | AG1 | 1 | 6 | 0.353 |
| 0.1 | 0 | ptsI | 1 | 6.25 | 0.422 |
| 0.1 | 0 | clcB | 1 | 6.25 | 0.338 |
| 0.1 | 0 | ycaM | 1 | 6.25 | 0.393 |
| 0.1 | 0 | yadI | 1 | 6.25 | 0.514 |
| 0.1 | 0 | AG1 | 1 | 6.25 | 0.366 |
| 0.1 | 0 | ptsI | 1 | 6.5 | 0.43 |
| 0.1 | 0 | clcB | 1 | 6.5 | 0.348 |
| 0.1 | 0 | ycaM | 1 | 6.5 | 0.401 |
| 0.1 | 0 | yadI | 1 | 6.5 | 0.516 |
| 0.1 | 0 | AG1 | 1 | 6.5 | 0.378 |
| 0.1 | 0 | ptsI | 1 | 6.75 | 0.439 |
| 0.1 | 0 | clcB | 1 | 6.75 | 0.361 |
| 0.1 | 0 | ycaM | 1 | 6.75 | 0.409 |
| 0.1 | 0 | yadI | 1 | 6.75 | 0.534 |
| 0.1 | 0 | AG1 | 1 | 6.75 | 0.395 |
| 0.1 | 0 | ptsI | 1 | 7 | 0.445 |
| 0.1 | 0 | clcB | 1 | 7 | 0.368 |
| 0.1 | 0 | ycaM | 1 | 7 | 0.412 |
| 0.1 | 0 | yadI | 1 | 7 | 0.553 |
| 0.1 | 0 | AG1 | 1 | 7 | 0.406 |
| 0.1 | 0 | ptsI | 1 | 7.25 | 0.451 |
| 0.1 | 0 | clcB | 1 | 7.25 | 0.378 |
| 0.1 | 0 | ycaM | 1 | 7.25 | 0.417 |
| 0.1 | 0 | yadI | 1 | 7.25 | 0.564 |

| | | | | | |
|---|---|---|---|---|---|
| 0.1 | 0 | AG1 | 1 | 7.25 | 0.42 |
| 0.1 | 0 | ptsI | 1 | 7.5 | 0.458 |
| 0.1 | 0 | clcB | 1 | 7.5 | 0.385 |
| 0.1 | 0 | ycaM | 1 | 7.5 | 0.42 |
| 0.1 | 0 | yadI | 1 | 7.5 | 0.57 |
| 0.1 | 0 | AG1 | 1 | 7.5 | 0.429 |
| 0.1 | 0 | ptsI | 1 | 7.75 | 0.474 |
| 0.1 | 0 | clcB | 1 | 7.75 | 0.397 |
| 0.1 | 0 | ycaM | 1 | 7.75 | 0.432 |
| 0.1 | 0 | yadI | 1 | 7.75 | 0.585 |
| 0.1 | 0 | AG1 | 1 | 7.75 | 0.438 |
| 0.1 | 0 | ptsI | 1 | 8 | 0.486 |
| 0.1 | 0 | clcB | 1 | 8 | 0.405 |
| 0.1 | 0 | ycaM | 1 | 8 | 0.442 |
| 0.1 | 0 | yadI | 1 | 8 | 0.6 |
| 0.1 | 0 | AG1 | 1 | 8 | 0.452 |
| 0.1 | 0 | ptsI | 1 | 8.25 | 0.498 |
| 0.1 | 0 | clcB | 1 | 8.25 | 0.415 |
| 0.1 | 0 | ycaM | 1 | 8.25 | 0.449 |
| 0.1 | 0 | yadI | 1 | 8.25 | 0.613 |
| 0.1 | 0 | AG1 | 1 | 8.25 | 0.458 |
| 0.1 | 0 | ptsI | 1 | 8.5 | 0.511 |
| 0.1 | 0 | clcB | 1 | 8.5 | 0.423 |
| 0.1 | 0 | ycaM | 1 | 8.5 | 0.462 |
| 0.1 | 0 | yadI | 1 | 8.5 | 0.624 |
| 0.1 | 0 | AG1 | 1 | 8.5 | 0.474 |
| 0.1 | 0 | ptsI | 1 | 8.75 | 0.519 |
| 0.1 | 0 | clcB | 1 | 8.75 | 0.434 |
| 0.1 | 0 | ycaM | 1 | 8.75 | 0.469 |
| 0.1 | 0 | yadI | 1 | 8.75 | 0.631 |
| 0.1 | 0 | AG1 | 1 | 8.75 | 0.484 |
| 0.1 | 0 | ptsI | 1 | 9 | 0.533 |
| 0.1 | 0 | clcB | 1 | 9 | 0.441 |
| 0.1 | 0 | ycaM | 1 | 9 | 0.481 |
| 0.1 | 0 | yadI | 1 | 9 | 0.647 |
| 0.1 | 0 | AG1 | 1 | 9 | 0.495 |
| 0.1 | 0 | ptsI | 1 | 9.25 | 0.556 |
| 0.1 | 0 | clcB | 1 | 9.25 | 0.454 |
| 0.1 | 0 | ycaM | 1 | 9.25 | 0.492 |
| 0.1 | 0 | yadI | 1 | 9.25 | 0.651 |
| 0.1 | 0 | AG1 | 1 | 9.25 | 0.514 |
| 0.1 | 0 | ptsI | 1 | 9.5 | 0.579 |
| 0.1 | 0 | clcB | 1 | 9.5 | 0.474 |
| 0.1 | 0 | ycaM | 1 | 9.5 | 0.509 |
| 0.1 | 0 | yadI | 1 | 9.5 | 0.659 |
| 0.1 | 0 | AG1 | 1 | 9.5 | 0.531 |
| 0.1 | 0 | ptsI | 1 | 9.75 | 0.567 |
| 0.1 | 0 | clcB | 1 | 9.75 | 0.479 |

| | | | | | |
|---|---|---|---|---|---|
| 0.1 | 0 | ycaM | 1 | 9.75 | 0.536 |
| 0.1 | 0 | yadI | 1 | 9.75 | 0.681 |
| 0.1 | 0 | AG1 | 1 | 9.75 | 0.545 |
| 0.1 | 0 | ptsI | 1 | 10 | 0.591 |
| 0.1 | 0 | clcB | 1 | 10 | 0.486 |
| 0.1 | 0 | ycaM | 1 | 10 | 0.527 |
| 0.1 | 0 | yadI | 1 | 10 | 0.67 |
| 0.1 | 0 | AG1 | 1 | 10 | 0.55 |
| 0.1 | 0 | ptsI | 1 | 10.25 | 0.596 |
| 0.1 | 0 | clcB | 1 | 10.25 | 0.502 |
| 0.1 | 0 | ycaM | 1 | 10.25 | 0.543 |
| 0.1 | 0 | yadI | 1 | 10.25 | 0.692 |
| 0.1 | 0 | AG1 | 1 | 10.25 | 0.565 |
| 0.1 | 0 | ptsI | 1 | 10.5 | 0.606 |
| 0.1 | 0 | clcB | 1 | 10.5 | 0.511 |
| 0.1 | 0 | ycaM | 1 | 10.5 | 0.573 |
| 0.1 | 0 | yadI | 1 | 10.5 | 0.707 |
| 0.1 | 0 | AG1 | 1 | 10.5 | 0.573 |
| 0.1 | 0 | ptsI | 1 | 10.75 | 0.639 |
| 0.1 | 0 | clcB | 1 | 10.75 | 0.543 |
| 0.1 | 0 | ycaM | 1 | 10.75 | 0.558 |
| 0.1 | 0 | yadI | 1 | 10.75 | 0.715 |
| 0.1 | 0 | AG1 | 1 | 10.75 | 0.583 |
| 0.1 | 0 | ptsI | 1 | 11 | 0.652 |
| 0.1 | 0 | clcB | 1 | 11 | 0.558 |
| 0.1 | 0 | ycaM | 1 | 11 | 0.603 |
| 0.1 | 0 | yadI | 1 | 11 | 0.713 |
| 0.1 | 0 | AG1 | 1 | 11 | 0.607 |
| 0.1 | 0 | ptsI | 1 | 11.25 | 0.674 |
| 0.1 | 0 | clcB | 1 | 11.25 | 0.557 |
| 0.1 | 0 | ycaM | 1 | 11.25 | 0.614 |
| 0.1 | 0 | yadI | 1 | 11.25 | 0.729 |
| 0.1 | 0 | AG1 | 1 | 11.25 | 0.626 |
| 0.1 | 0 | ptsI | 1 | 11.5 | 0.694 |
| 0.1 | 0 | clcB | 1 | 11.5 | 0.568 |
| 0.1 | 0 | ycaM | 1 | 11.5 | 0.608 |
| 0.1 | 0 | yadI | 1 | 11.5 | 0.734 |
| 0.1 | 0 | AG1 | 1 | 11.5 | 0.634 |
| 0.1 | 0 | ptsI | 1 | 11.75 | 0.728 |
| 0.1 | 0 | clcB | 1 | 11.75 | 0.589 |
| 0.1 | 0 | ycaM | 1 | 11.75 | 0.628 |
| 0.1 | 0 | yadI | 1 | 11.75 | 0.754 |
| 0.1 | 0 | AG1 | 1 | 11.75 | 0.671 |
| 0.1 | 0 | ptsI | 1 | 12 | 0.713 |
| 0.1 | 0 | clcB | 1 | 12 | 0.575 |
| 0.1 | 0 | ycaM | 1 | 12 | 0.631 |
| 0.1 | 0 | yadI | 1 | 12 | 0.778 |
| 0.1 | 0 | AG1 | 1 | 12 | 0.678 |

| | | | | | |
|---|---|---|---|---|---|
| 0.1 | 0 | ptsI | 1 | 12.25 | 0.726 |
| 0.1 | 0 | clcB | 1 | 12.25 | 0.596 |
| 0.1 | 0 | ycaM | 1 | 12.25 | 0.639 |
| 0.1 | 0 | yadI | 1 | 12.25 | 0.776 |
| 0.1 | 0 | AG1 | 1 | 12.25 | 0.668 |
| 0.1 | 0 | ptsI | 1 | 12.5 | 0.727 |
| 0.1 | 0 | clcB | 1 | 12.5 | 0.608 |
| 0.1 | 0 | ycaM | 1 | 12.5 | 0.648 |
| 0.1 | 0 | yadI | 1 | 12.5 | 0.795 |
| 0.1 | 0 | AG1 | 1 | 12.5 | 0.696 |
| 0.1 | 0 | ptsI | 1 | 12.75 | 0.747 |
| 0.1 | 0 | clcB | 1 | 12.75 | 0.612 |
| 0.1 | 0 | ycaM | 1 | 12.75 | 0.668 |
| 0.1 | 0 | yadI | 1 | 12.75 | 0.812 |
| 0.1 | 0 | AG1 | 1 | 12.75 | 0.715 |
| 0.1 | 0 | ptsI | 1 | 13 | 0.755 |
| 0.1 | 0 | clcB | 1 | 13 | 0.616 |
| 0.1 | 0 | ycaM | 1 | 13 | 0.668 |
| 0.1 | 0 | yadI | 1 | 13 | 0.823 |
| 0.1 | 0 | AG1 | 1 | 13 | 0.732 |
| 0.1 | 0 | ptsI | 1 | 13.25 | 0.758 |
| 0.1 | 0 | clcB | 1 | 13.25 | 0.646 |
| 0.1 | 0 | ycaM | 1 | 13.25 | 0.666 |
| 0.1 | 0 | yadI | 1 | 13.25 | 0.835 |
| 0.1 | 0 | AG1 | 1 | 13.25 | 0.724 |
| 0.1 | 0 | ptsI | 1 | 13.5 | 0.768 |
| 0.1 | 0 | clcB | 1 | 13.5 | 0.651 |
| 0.1 | 0 | ycaM | 1 | 13.5 | 0.698 |
| 0.1 | 0 | yadI | 1 | 13.5 | 0.866 |
| 0.1 | 0 | AG1 | 1 | 13.5 | 0.748 |
| 0.1 | 0 | ptsI | 1 | 13.75 | 0.784 |
| 0.1 | 0 | clcB | 1 | 13.75 | 0.655 |
| 0.1 | 0 | ycaM | 1 | 13.75 | 0.688 |
| 0.1 | 0 | yadI | 1 | 13.75 | 0.856 |
| 0.1 | 0 | AG1 | 1 | 13.75 | 0.744 |
| 0.1 | 0 | ptsI | 1 | 14 | 0.811 |
| 0.1 | 0 | clcB | 1 | 14 | 0.668 |
| 0.1 | 0 | ycaM | 1 | 14 | 0.727 |
| 0.1 | 0 | yadI | 1 | 14 | 0.883 |
| 0.1 | 0 | AG1 | 1 | 14 | 0.779 |
| 0.1 | 0 | ptsI | 1 | 14.25 | 0.812 |
| 0.1 | 0 | clcB | 1 | 14.25 | 0.672 |
| 0.1 | 0 | ycaM | 1 | 14.25 | 0.728 |
| 0.1 | 0 | yadI | 1 | 14.25 | 0.89 |
| 0.1 | 0 | AG1 | 1 | 14.25 | 0.773 |
| 0.1 | 0 | ptsI | 1 | 14.5 | 0.814 |
| 0.1 | 0 | clcB | 1 | 14.5 | 0.682 |
| 0.1 | 0 | ycaM | 1 | 14.5 | 0.69 |

| | | | | | |
|---|---|---|---|---|---|
| 0.1 | 0 | yadI | 1 | 14.5 | 0.898 |
| 0.1 | 0 | AG1 | 1 | 14.5 | 0.778 |
| 0.1 | 0 | ptsI | 1 | 14.75 | 0.808 |
| 0.1 | 0 | clcB | 1 | 14.75 | 0.704 |
| 0.1 | 0 | ycaM | 1 | 14.75 | 0.704 |
| 0.1 | 0 | yadI | 1 | 14.75 | 0.885 |
| 0.1 | 0 | AG1 | 1 | 14.75 | 0.786 |
| 0.1 | 0 | ptsI | 1 | 15 | 0.814 |
| 0.1 | 0 | clcB | 1 | 15 | 0.697 |
| 0.1 | 0 | ycaM | 1 | 15 | 0.716 |
| 0.1 | 0 | yadI | 1 | 15 | 0.896 |
| 0.1 | 0 | AG1 | 1 | 15 | 0.791 |
| 0.1 | 0 | ptsI | 1 | 15.25 | 0.835 |
| 0.1 | 0 | clcB | 1 | 15.25 | 0.718 |
| 0.1 | 0 | ycaM | 1 | 15.25 | 0.713 |
| 0.1 | 0 | yadI | 1 | 15.25 | 0.91 |
| 0.1 | 0 | AG1 | 1 | 15.25 | 0.826 |
| 0.1 | 0 | ptsI | 1 | 15.5 | 0.83 |
| 0.1 | 0 | clcB | 1 | 15.5 | 0.742 |
| 0.1 | 0 | ycaM | 1 | 15.5 | 0.722 |
| 0.1 | 0 | yadI | 1 | 15.5 | 0.921 |
| 0.1 | 0 | AG1 | 1 | 15.5 | 0.799 |
| 0.1 | 0 | ptsI | 1 | 15.75 | 0.832 |
| 0.1 | 0 | clcB | 1 | 15.75 | 0.734 |
| 0.1 | 0 | ycaM | 1 | 15.75 | 0.727 |
| 0.1 | 0 | yadI | 1 | 15.75 | 0.927 |
| 0.1 | 0 | AG1 | 1 | 15.75 | 0.812 |
| 0.1 | 0 | ptsI | 1 | 16 | 0.836 |
| 0.1 | 0 | clcB | 1 | 16 | 0.76 |
| 0.1 | 0 | ycaM | 1 | 16 | 0.735 |
| 0.1 | 0 | yadI | 1 | 16 | 0.931 |
| 0.1 | 0 | AG1 | 1 | 16 | 0.816 |
| 0.1 | 0 | ptsI | 1 | 16.25 | 0.845 |
| 0.1 | 0 | clcB | 1 | 16.25 | 0.779 |
| 0.1 | 0 | ycaM | 1 | 16.25 | 0.743 |
| 0.1 | 0 | yadI | 1 | 16.25 | 0.937 |
| 0.1 | 0 | AG1 | 1 | 16.25 | 0.833 |
| 0.1 | 0 | ptsI | 1 | 16.5 | 0.859 |
| 0.1 | 0 | clcB | 1 | 16.5 | 0.788 |
| 0.1 | 0 | ycaM | 1 | 16.5 | 0.748 |
| 0.1 | 0 | yadI | 1 | 16.5 | 0.933 |
| 0.1 | 0 | AG1 | 1 | 16.5 | 0.838 |
| 0.1 | 0 | ptsI | 1 | 17.0667 | 0.912 |
| 0.1 | 0 | clcB | 1 | 17.0667 | 0.8 |
| 0.1 | 0 | ycaM | 1 | 17.0667 | 0.789 |
| 0.1 | 0 | yadI | 1 | 17.0667 | 1.005 |
| 0.1 | 0 | AG1 | 1 | 17.0667 | 0.829 |
| 0.1 | 0 | ptsI | 1 | 17.3167 | 0.831 |

| | | | | | |
|---|---|---|---|---|---|
| 0.1 | 0 | clcB | 1 | 17.3167 | 0.748 |
| 0.1 | 0 | ycaM | 1 | 17.3167 | 0.736 |
| 0.1 | 0 | yadI | 1 | 17.3167 | 0.968 |
| 0.1 | 0 | AG1 | 1 | 17.3167 | 0.777 |
| 0.1 | 0 | ptsI | 1 | 17.5667 | 0.837 |
| 0.1 | 0 | clcB | 1 | 17.5667 | 0.747 |
| 0.1 | 0 | ycaM | 1 | 17.5667 | 0.732 |
| 0.1 | 0 | yadI | 1 | 17.5667 | 0.959 |
| 0.1 | 0 | AG1 | 1 | 17.5667 | 0.813 |
| 0.1 | 0 | ptsI | 1 | 17.8167 | 0.858 |
| 0.1 | 0 | clcB | 1 | 17.8167 | 0.789 |
| 0.1 | 0 | ycaM | 1 | 17.8167 | 0.75 |
| 0.1 | 0 | yadI | 1 | 17.8167 | 0.964 |
| 0.1 | 0 | AG1 | 1 | 17.8167 | 0.84 |
| 0.1 | 0 | ptsI | 1 | 18.0667 | 0.851 |
| 0.1 | 0 | clcB | 1 | 18.0667 | 0.748 |
| 0.1 | 0 | ycaM | 1 | 18.0667 | 0.748 |
| 0.1 | 0 | yadI | 1 | 18.0667 | 0.968 |
| 0.1 | 0 | AG1 | 1 | 18.0667 | 0.833 |
| 0.1 | 0 | ptsI | 1 | 18.3167 | 0.864 |
| 0.1 | 0 | clcB | 1 | 18.3167 | 0.762 |
| 0.1 | 0 | ycaM | 1 | 18.3167 | 0.751 |
| 0.1 | 0 | yadI | 1 | 18.3167 | 0.965 |
| 0.1 | 0 | AG1 | 1 | 18.3167 | 0.835 |
| 0.1 | 0 | ptsI | 1 | 18.5667 | 0.873 |
| 0.1 | 0 | clcB | 1 | 18.5667 | 0.784 |
| 0.1 | 0 | ycaM | 1 | 18.5667 | 0.76 |
| 0.1 | 0 | yadI | 1 | 18.5667 | 0.97 |
| 0.1 | 0 | AG1 | 1 | 18.5667 | 0.843 |
| 0.1 | 0 | ptsI | 1 | 18.8167 | 0.878 |
| 0.1 | 0 | clcB | 1 | 18.8167 | 0.782 |
| 0.1 | 0 | ycaM | 1 | 18.8167 | 0.759 |
| 0.1 | 0 | yadI | 1 | 18.8167 | 0.974 |
| 0.1 | 0 | AG1 | 1 | 18.8167 | 0.848 |
| 0.1 | 0 | ptsI | 1 | 19.0667 | 0.887 |
| 0.1 | 0 | clcB | 1 | 19.0667 | 0.795 |
| 0.1 | 0 | ycaM | 1 | 19.0667 | 0.764 |
| 0.1 | 0 | yadI | 1 | 19.0667 | 0.981 |
| 0.1 | 0 | AG1 | 1 | 19.0667 | 0.854 |
| 0.1 | 0 | ptsI | 1 | 19.3167 | 0.883 |
| 0.1 | 0 | clcB | 1 | 19.3167 | 0.799 |
| 0.1 | 0 | ycaM | 1 | 19.3167 | 0.766 |
| 0.1 | 0 | yadI | 1 | 19.3167 | 0.985 |
| 0.1 | 0 | AG1 | 1 | 19.3167 | 0.853 |
| 0.1 | 0 | ptsI | 1 | 19.5667 | 0.898 |
| 0.1 | 0 | clcB | 1 | 19.5667 | 0.815 |
| 0.1 | 0 | ycaM | 1 | 19.5667 | 0.779 |
| 0.1 | 0 | yadI | 1 | 19.5667 | 0.985 |

| | | | | | |
|---|---|---|---|---|---|
| 0.1 | 0 | AG1 | 1 | 19.5667 | 0.876 |
| 0.1 | 0 | ptsI | 1 | 19.8167 | 0.907 |
| 0.1 | 0 | clcB | 1 | 19.8167 | 0.811 |
| 0.1 | 0 | ycaM | 1 | 19.8167 | 0.781 |
| 0.1 | 0 | yadI | 1 | 19.8167 | 0.989 |
| 0.1 | 0 | AG1 | 1 | 19.8167 | 0.87 |
| 0.1 | 0 | ptsI | 1 | 20.0667 | 0.922 |
| 0.1 | 0 | clcB | 1 | 20.0667 | 0.818 |
| 0.1 | 0 | ycaM | 1 | 20.0667 | 0.788 |
| 0.1 | 0 | yadI | 1 | 20.0667 | 0.989 |
| 0.1 | 0 | AG1 | 1 | 20.0667 | 0.882 |
| 0.1 | 0 | ptsI | 1 | 20.3167 | 0.925 |
| 0.1 | 0 | clcB | 1 | 20.3167 | 0.819 |
| 0.1 | 0 | ycaM | 1 | 20.3167 | 0.788 |
| 0.1 | 0 | yadI | 1 | 20.3167 | 0.996 |
| 0.1 | 0 | AG1 | 1 | 20.3167 | 0.882 |
| 0.1 | 0 | ptsI | 1 | 20.5667 | 0.924 |
| 0.1 | 0 | clcB | 1 | 20.5667 | 0.813 |
| 0.1 | 0 | ycaM | 1 | 20.5667 | 0.792 |
| 0.1 | 0 | yadI | 1 | 20.5667 | 0.999 |
| 0.1 | 0 | AG1 | 1 | 20.5667 | 0.879 |
| 0.1 | 0 | ptsI | 1 | 20.8167 | 0.935 |
| 0.1 | 0 | clcB | 1 | 20.8167 | 0.821 |
| 0.1 | 0 | ycaM | 1 | 20.8167 | 0.802 |
| 0.1 | 0 | yadI | 1 | 20.8167 | 1.003 |
| 0.1 | 0 | AG1 | 1 | 20.8167 | 0.887 |
| 0.1 | 0 | ptsI | 1 | 21.0667 | 0.951 |
| 0.1 | 0 | clcB | 1 | 21.0667 | 0.827 |
| 0.1 | 0 | ycaM | 1 | 21.0667 | 0.81 |
| 0.1 | 0 | yadI | 1 | 21.0667 | 1.007 |
| 0.1 | 0 | AG1 | 1 | 21.0667 | 0.894 |
| 0.1 | 0 | ptsI | 1 | 21.3167 | 0.953 |
| 0.1 | 0 | clcB | 1 | 21.3167 | 0.836 |
| 0.1 | 0 | ycaM | 1 | 21.3167 | 0.813 |
| 0.1 | 0 | yadI | 1 | 21.3167 | 1.013 |
| 0.1 | 0 | AG1 | 1 | 21.3167 | 0.904 |
| 0.1 | 0 | ptsI | 1 | 21.5667 | 0.957 |
| 0.1 | 0 | clcB | 1 | 21.5667 | 0.827 |
| 0.1 | 0 | ycaM | 1 | 21.5667 | 0.815 |
| 0.1 | 0 | yadI | 1 | 21.5667 | 1.01 |
| 0.1 | 0 | AG1 | 1 | 21.5667 | 0.902 |
| 0.1 | 0 | ptsI | 1 | 21.8167 | 0.968 |
| 0.1 | 0 | clcB | 1 | 21.8167 | 0.835 |
| 0.1 | 0 | ycaM | 1 | 21.8167 | 0.823 |
| 0.1 | 0 | yadI | 1 | 21.8167 | 1.009 |
| 0.1 | 0 | AG1 | 1 | 21.8167 | 0.909 |
| 0.1 | 0 | ptsI | 1 | 22.0667 | 0.99 |
| 0.1 | 0 | clcB | 1 | 22.0667 | 0.839 |

| | | | | | |
|---|---|---|---|---|---|
| 0.1 | 0 | ycaM | 1 | 22.0667 | 0.829 |
| 0.1 | 0 | yadI | 1 | 22.0667 | 1.013 |
| 0.1 | 0 | AG1 | 1 | 22.0667 | 0.913 |
| 0.1 | 0 | ptsI | 1 | 22.9333 | 0.687 |
| 0.1 | 0 | clcB | 1 | 22.9333 | 0.549 |
| 0.1 | 0 | ycaM | 1 | 22.9333 | 0.639 |
| 0.1 | 0 | yadI | 1 | 22.9333 | 0.761 |
| 0.1 | 0 | AG1 | 1 | 22.9333 | 0.84 |
| 0.1 | 0 | ptsI | 1 | 23.1833 | 0.702 |
| 0.1 | 0 | clcB | 1 | 23.1833 | 0.603 |
| 0.1 | 0 | ycaM | 1 | 23.1833 | 0.664 |
| 0.1 | 0 | yadI | 1 | 23.1833 | 0.747 |
| 0.1 | 0 | AG1 | 1 | 23.1833 | 0.776 |
| 0.1 | 0 | ptsI | 1 | 23.4333 | 0.696 |
| 0.1 | 0 | clcB | 1 | 23.4333 | 0.547 |
| 0.1 | 0 | ycaM | 1 | 23.4333 | 0.677 |
| 0.1 | 0 | yadI | 1 | 23.4333 | 0.751 |
| 0.1 | 0 | AG1 | 1 | 23.4333 | 0.787 |
| 0.1 | 0 | ptsI | 1 | 23.6833 | 0.713 |
| 0.1 | 0 | clcB | 1 | 23.6833 | 0.564 |
| 0.1 | 0 | ycaM | 1 | 23.6833 | 0.692 |
| 0.1 | 0 | yadI | 1 | 23.6833 | 0.768 |
| 0.1 | 0 | AG1 | 1 | 23.6833 | 0.804 |
| 0.1 | 0 | ptsI | 1 | 23.9333 | 0.742 |
| 0.1 | 0 | clcB | 1 | 23.9333 | 0.56 |
| 0.1 | 0 | ycaM | 1 | 23.9333 | 0.704 |
| 0.1 | 0 | yadI | 1 | 23.9333 | 0.783 |
| 0.1 | 0 | AG1 | 1 | 23.9333 | 0.803 |
| 0.1 | 0 | ptsI | 1 | 24.1833 | 0.766 |
| 0.1 | 0 | clcB | 1 | 24.1833 | 0.572 |
| 0.1 | 0 | ycaM | 1 | 24.1833 | 0.711 |
| 0.1 | 0 | yadI | 1 | 24.1833 | 0.802 |
| 0.1 | 0 | AG1 | 1 | 24.1833 | 0.814 |
| 0.1 | 0 | ptsI | 1 | 24.4333 | 0.781 |
| 0.1 | 0 | clcB | 1 | 24.4333 | 0.578 |
| 0.1 | 0 | ycaM | 1 | 24.4333 | 0.727 |
| 0.1 | 0 | yadI | 1 | 24.4333 | 0.815 |
| 0.1 | 0 | AG1 | 1 | 24.4333 | 0.823 |
| 0.1 | 0 | ptsI | 1 | 24.6833 | 0.788 |
| 0.1 | 0 | clcB | 1 | 24.6833 | 0.58 |
| 0.1 | 0 | ycaM | 1 | 24.6833 | 0.716 |
| 0.1 | 0 | yadI | 1 | 24.6833 | 0.825 |
| 0.1 | 0 | AG1 | 1 | 24.6833 | 0.823 |
| 0.1 | 0 | ptsI | 1 | 24.9333 | 0.806 |
| 0.1 | 0 | clcB | 1 | 24.9333 | 0.596 |
| 0.1 | 0 | ycaM | 1 | 24.9333 | 0.722 |
| 0.1 | 0 | yadI | 1 | 24.9333 | 0.842 |
| 0.1 | 0 | AG1 | 1 | 24.9333 | 0.829 |

| | | | | | |
|---|---|---|---|---|---|
| 0.1 | 0 | ptsI | 1 | 25.1833 | 0.822 |
| 0.1 | 0 | clcB | 1 | 25.1833 | 0.597 |
| 0.1 | 0 | ycaM | 1 | 25.1833 | 0.731 |
| 0.1 | 0 | yadI | 1 | 25.1833 | 0.845 |
| 0.1 | 0 | AG1 | 1 | 25.1833 | 0.83 |
| 0.1 | 0 | ptsI | 1 | 25.4333 | 0.835 |
| 0.1 | 0 | clcB | 1 | 25.4333 | 0.608 |
| 0.1 | 0 | ycaM | 1 | 25.4333 | 0.738 |
| 0.1 | 0 | yadI | 1 | 25.4333 | 0.854 |
| 0.1 | 0 | AG1 | 1 | 25.4333 | 0.839 |
| 0.1 | 0 | ptsI | 1 | 25.6833 | 0.842 |
| 0.1 | 0 | clcB | 1 | 25.6833 | 0.602 |
| 0.1 | 0 | ycaM | 1 | 25.6833 | 0.744 |
| 0.1 | 0 | yadI | 1 | 25.6833 | 0.862 |
| 0.1 | 0 | AG1 | 1 | 25.6833 | 0.842 |
| 0.1 | 0 | ptsI | 1 | 25.9333 | 0.861 |
| 0.1 | 0 | clcB | 1 | 25.9333 | 0.609 |
| 0.1 | 0 | ycaM | 1 | 25.9333 | 0.75 |
| 0.1 | 0 | yadI | 1 | 25.9333 | 0.871 |
| 0.1 | 0 | AG1 | 1 | 25.9333 | 0.847 |
| 0.1 | 0 | ptsI | 1 | 26.1833 | 0.864 |
| 0.1 | 0 | clcB | 1 | 26.1833 | 0.62 |
| 0.1 | 0 | ycaM | 1 | 26.1833 | 0.759 |
| 0.1 | 0 | yadI | 1 | 26.1833 | 0.882 |
| 0.1 | 0 | AG1 | 1 | 26.1833 | 0.851 |
| 0.1 | 0 | ptsI | 1 | 26.4333 | 0.873 |
| 0.1 | 0 | clcB | 1 | 26.4333 | 0.615 |
| 0.1 | 0 | ycaM | 1 | 26.4333 | 0.761 |
| 0.1 | 0 | yadI | 1 | 26.4333 | 0.885 |
| 0.1 | 0 | AG1 | 1 | 26.4333 | 0.843 |
| 0.1 | 0 | ptsI | 1 | 26.6833 | 0.882 |
| 0.1 | 0 | clcB | 1 | 26.6833 | 0.622 |
| 0.1 | 0 | ycaM | 1 | 26.6833 | 0.774 |
| 0.1 | 0 | yadI | 1 | 26.6833 | 0.892 |
| 0.1 | 0 | AG1 | 1 | 26.6833 | 0.847 |
| 0.1 | 0 | ptsI | 1 | 26.9333 | 0.89 |
| 0.1 | 0 | clcB | 1 | 26.9333 | 0.624 |
| 0.1 | 0 | ycaM | 1 | 26.9333 | 0.782 |
| 0.1 | 0 | yadI | 1 | 26.9333 | 0.899 |
| 0.1 | 0 | AG1 | 1 | 26.9333 | 0.852 |
| 0.1 | 0 | ptsI | 1 | 27.1833 | 0.904 |
| 0.1 | 0 | clcB | 1 | 27.1833 | 0.636 |
| 0.1 | 0 | ycaM | 1 | 27.1833 | 0.79 |
| 0.1 | 0 | yadI | 1 | 27.1833 | 0.906 |
| 0.1 | 0 | AG1 | 1 | 27.1833 | 0.856 |
| 0.1 | 0 | ptsI | 1 | 27.4333 | 0.912 |
| 0.1 | 0 | clcB | 1 | 27.4333 | 0.638 |
| 0.1 | 0 | ycaM | 1 | 27.4333 | 0.798 |

| | | | | | |
|---|---|---|---|---|---|
| 0.1 | 0 | yadI | 1 | 27.4333 | 0.912 |
| 0.1 | 0 | AG1 | 1 | 27.4333 | 0.851 |
| 0.1 | 0 | ptsI | 1 | 27.6833 | 0.922 |
| 0.1 | 0 | clcB | 1 | 27.6833 | 0.641 |
| 0.1 | 0 | ycaM | 1 | 27.6833 | 0.806 |
| 0.1 | 0 | yadI | 1 | 27.6833 | 0.92 |
| 0.1 | 0 | AG1 | 1 | 27.6833 | 0.858 |
| 0.1 | 0 | ptsI | 1 | 27.9333 | 0.931 |
| 0.1 | 0 | clcB | 1 | 27.9333 | 0.642 |
| 0.1 | 0 | ycaM | 1 | 27.9333 | 0.817 |
| 0.1 | 0 | yadI | 1 | 27.9333 | 0.912 |
| 0.1 | 0 | AG1 | 1 | 27.9333 | 0.856 |
| 0.1 | 0 | ptsI | 1 | 28.1833 | 0.938 |
| 0.1 | 0 | clcB | 1 | 28.1833 | 0.666 |
| 0.1 | 0 | ycaM | 1 | 28.1833 | 0.826 |
| 0.1 | 0 | yadI | 1 | 28.1833 | 0.93 |
| 0.1 | 0 | AG1 | 1 | 28.1833 | 0.865 |
| 0.1 | 0 | ptsI | 1 | 28.4333 | 0.947 |
| 0.1 | 0 | clcB | 1 | 28.4333 | 0.659 |
| 0.1 | 0 | ycaM | 1 | 28.4333 | 0.835 |
| 0.1 | 0 | yadI | 1 | 28.4333 | 0.935 |
| 0.1 | 0 | AG1 | 1 | 28.4333 | 0.865 |
| 0.1 | 0 | ptsI | 1 | 28.6833 | 0.955 |
| 0.1 | 0 | clcB | 1 | 28.6833 | 0.67 |
| 0.1 | 0 | ycaM | 1 | 28.6833 | 0.845 |
| 0.1 | 0 | yadI | 1 | 28.6833 | 0.94 |
| 0.1 | 0 | AG1 | 1 | 28.6833 | 0.866 |
| 0.1 | 0 | ptsI | 1 | 28.9333 | 0.966 |
| 0.1 | 0 | clcB | 1 | 28.9333 | 0.685 |
| 0.1 | 0 | ycaM | 1 | 28.9333 | 0.856 |
| 0.1 | 0 | yadI | 1 | 28.9333 | 0.947 |
| 0.1 | 0 | AG1 | 1 | 28.9333 | 0.872 |
| 0.1 | 0 | ptsI | 1 | 29.1833 | 0.969 |
| 0.1 | 0 | clcB | 1 | 29.1833 | 0.679 |
| 0.1 | 0 | ycaM | 1 | 29.1833 | 0.864 |
| 0.1 | 0 | yadI | 1 | 29.1833 | 0.948 |
| 0.1 | 0 | AG1 | 1 | 29.1833 | 0.867 |
| 0.1 | 0 | ptsI | 1 | 29.4333 | 0.984 |
| 0.1 | 0 | clcB | 1 | 29.4333 | 0.695 |
| 0.1 | 0 | ycaM | 1 | 29.4333 | 0.876 |
| 0.1 | 0 | yadI | 1 | 29.4333 | 0.959 |
| 0.1 | 0 | AG1 | 1 | 29.4333 | 0.873 |
| 0.1 | 0 | ptsI | 1 | 29.6833 | 0.996 |
| 0.1 | 0 | clcB | 1 | 29.6833 | 0.707 |
| 0.1 | 0 | ycaM | 1 | 29.6833 | 0.886 |
| 0.1 | 0 | yadI | 1 | 29.6833 | 0.966 |
| 0.1 | 0 | AG1 | 1 | 29.6833 | 0.874 |
| 0.1 | 0 | ptsI | 1 | 29.9333 | 1.01 |

| | | | | | |
|---|---|---|---|---|---|
| 0.1 | 0 | clcB | 1 | 29.9333 | 0.71 |
| 0.1 | 0 | ycaM | 1 | 29.9333 | 0.894 |
| 0.1 | 0 | yadI | 1 | 29.9333 | 0.972 |
| 0.1 | 0 | AG1 | 1 | 29.9333 | 0.876 |
| 0.1 | 0 | ptsI | 1 | 30.1833 | 1.02 |
| 0.1 | 0 | clcB | 1 | 30.1833 | 0.725 |
| 0.1 | 0 | ycaM | 1 | 30.1833 | 0.909 |
| 0.1 | 0 | yadI | 1 | 30.1833 | 0.98 |
| 0.1 | 0 | AG1 | 1 | 30.1833 | 0.878 |
| 0.1 | 0 | ptsI | 1 | 30.4333 | 1.033 |
| 0.1 | 0 | clcB | 1 | 30.4333 | 0.741 |
| 0.1 | 0 | ycaM | 1 | 30.4333 | 0.918 |
| 0.1 | 0 | yadI | 1 | 30.4333 | 0.989 |
| 0.1 | 0 | AG1 | 1 | 30.4333 | 0.879 |
| 0.1 | 0 | ptsI | 1 | 30.6833 | 1.036 |
| 0.1 | 0 | clcB | 1 | 30.6833 | 0.751 |
| 0.1 | 0 | ycaM | 1 | 30.6833 | 0.93 |
| 0.1 | 0 | yadI | 1 | 30.6833 | 0.991 |
| 0.1 | 0 | AG1 | 1 | 30.6833 | 0.876 |
| 0.1 | 0 | ptsI | 1 | 30.9333 | 1.052 |
| 0.1 | 0 | clcB | 1 | 30.9333 | 0.768 |
| 0.1 | 0 | ycaM | 1 | 30.9333 | 0.942 |
| 0.1 | 0 | yadI | 1 | 30.9333 | 1.006 |
| 0.1 | 0 | AG1 | 1 | 30.9333 | 0.879 |
| 0.1 | 0 | ptsI | 1 | 31.1833 | 1.062 |
| 0.1 | 0 | clcB | 1 | 31.1833 | 0.784 |
| 0.1 | 0 | ycaM | 1 | 31.1833 | 0.949 |
| 0.1 | 0 | yadI | 1 | 31.1833 | 1.01 |
| 0.1 | 0 | AG1 | 1 | 31.1833 | 0.877 |
| 0.1 | 0 | ptsI | 1 | 31.4333 | 1.075 |
| 0.1 | 0 | clcB | 1 | 31.4333 | 0.804 |
| 0.1 | 0 | ycaM | 1 | 31.4333 | 0.97 |
| 0.1 | 0 | yadI | 1 | 31.4333 | 1.025 |
| 0.1 | 0 | AG1 | 1 | 31.4333 | 0.884 |
| 0.1 | 0 | ptsI | 1 | 31.6833 | 1.084 |
| 0.1 | 0 | clcB | 1 | 31.6833 | 0.82 |
| 0.1 | 0 | ycaM | 1 | 31.6833 | 0.978 |
| 0.1 | 0 | yadI | 1 | 31.6833 | 1.029 |
| 0.1 | 0 | AG1 | 1 | 31.6833 | 0.882 |
| 0.1 | 0 | ptsI | 1 | 31.9333 | 1.096 |
| 0.1 | 0 | clcB | 1 | 31.9333 | 0.834 |
| 0.1 | 0 | ycaM | 1 | 31.9333 | 0.992 |
| 0.1 | 0 | yadI | 1 | 31.9333 | 1.04 |
| 0.1 | 0 | AG1 | 1 | 31.9333 | 0.885 |
| 0.1 | 0 | ptsI | 1 | 32.1833 | 1.106 |
| 0.1 | 0 | clcB | 1 | 32.1833 | 0.849 |
| 0.1 | 0 | ycaM | 1 | 32.1833 | 1 |
| 0.1 | 0 | yadI | 1 | 32.1833 | 1.05 |

| | | | | | |
|---|---|---|---|---|---|
| 0.1 | 0 | AG1 | 1 | 32.1833 | 0.886 |
| 0.1 | 0 | ptsI | 1 | 32.4333 | 1.12 |
| 0.1 | 0 | clcB | 1 | 32.4333 | 0.859 |
| 0.1 | 0 | ycaM | 1 | 32.4333 | 1.014 |
| 0.1 | 0 | yadI | 1 | 32.4333 | 1.046 |
| 0.1 | 0 | AG1 | 1 | 32.4333 | 0.89 |
| 0.1 | 0 | ptsI | 1 | 32.6833 | 1.123 |
| 0.1 | 0 | clcB | 1 | 32.6833 | 0.871 |
| 0.1 | 0 | ycaM | 1 | 32.6833 | 1.021 |
| 0.1 | 0 | yadI | 1 | 32.6833 | 1.065 |
| 0.1 | 0 | AG1 | 1 | 32.6833 | 0.889 |
| 0.1 | 0 | ptsI | 1 | 32.9333 | 1.133 |
| 0.1 | 0 | clcB | 1 | 32.9333 | 0.886 |
| 0.1 | 0 | ycaM | 1 | 32.9333 | 1.037 |
| 0.1 | 0 | yadI | 1 | 32.9333 | 1.068 |
| 0.1 | 0 | AG1 | 1 | 32.9333 | 0.887 |
| 0.1 | 0 | ptsI | 1 | 33.1833 | 1.148 |
| 0.1 | 0 | clcB | 1 | 33.1833 | 0.894 |
| 0.1 | 0 | ycaM | 1 | 33.1833 | 1.05 |
| 0.1 | 0 | yadI | 1 | 33.1833 | 1.072 |
| 0.1 | 0 | AG1 | 1 | 33.1833 | 0.892 |
| 0.1 | 0 | ptsI | 1 | 33.4333 | 1.16 |
| 0.1 | 0 | clcB | 1 | 33.4333 | 0.906 |
| 0.1 | 0 | ycaM | 1 | 33.4333 | 1.056 |
| 0.1 | 0 | yadI | 1 | 33.4333 | 1.093 |
| 0.1 | 0 | AG1 | 1 | 33.4333 | 0.892 |
| 0.1 | 0 | ptsI | 1 | 33.6833 | 1.163 |
| 0.1 | 0 | clcB | 1 | 33.6833 | 0.922 |
| 0.1 | 0 | ycaM | 1 | 33.6833 | 1.07 |
| 0.1 | 0 | yadI | 1 | 33.6833 | 1.092 |
| 0.1 | 0 | AG1 | 1 | 33.6833 | 0.894 |
| 0.1 | 0 | ptsI | 1 | 33.9333 | 1.172 |
| 0.1 | 0 | clcB | 1 | 33.9333 | 0.932 |
| 0.1 | 0 | ycaM | 1 | 33.9333 | 1.079 |
| 0.1 | 0 | yadI | 1 | 33.9333 | 1.105 |
| 0.1 | 0 | AG1 | 1 | 33.9333 | 0.897 |
| 0.1 | 0 | ptsI | 1 | 34.1833 | 1.182 |
| 0.1 | 0 | clcB | 1 | 34.1833 | 0.944 |
| 0.1 | 0 | ycaM | 1 | 34.1833 | 1.093 |
| 0.1 | 0 | yadI | 1 | 34.1833 | 1.119 |
| 0.1 | 0 | AG1 | 1 | 34.1833 | 0.903 |
| 0.1 | 0 | ptsI | 1 | 34.4333 | 1.194 |
| 0.1 | 0 | clcB | 1 | 34.4333 | 0.947 |
| 0.1 | 0 | ycaM | 1 | 34.4333 | 1.098 |
| 0.1 | 0 | yadI | 1 | 34.4333 | 1.126 |
| 0.1 | 0 | AG1 | 1 | 34.4333 | 0.903 |
| 0.1 | 0 | ptsI | 1 | 34.6833 | 1.205 |
| 0.1 | 0 | clcB | 1 | 34.6833 | 0.964 |

| | | | | | |
|---|---|---|---|---|---|
| 0.1 | 0 | ycaM | 1 | 34.6833 | 1.109 |
| 0.1 | 0 | yadI | 1 | 34.6833 | 1.14 |
| 0.1 | 0 | AG1 | 1 | 34.6833 | 0.909 |
| 0.1 | 0 | ptsI | 1 | 34.9333 | 1.211 |
| 0.1 | 0 | clcB | 1 | 34.9333 | 0.974 |
| 0.1 | 0 | ycaM | 1 | 34.9333 | 1.116 |
| 0.1 | 0 | yadI | 1 | 34.9333 | 1.137 |
| 0.1 | 0 | AG1 | 1 | 34.9333 | 0.905 |
| 0.1 | 0 | ptsI | 1 | 35.1833 | 1.222 |
| 0.1 | 0 | clcB | 1 | 35.1833 | 0.978 |
| 0.1 | 0 | ycaM | 1 | 35.1833 | 1.124 |
| 0.1 | 0 | yadI | 1 | 35.1833 | 1.15 |
| 0.1 | 0 | AG1 | 1 | 35.1833 | 0.906 |
| 0.1 | 0 | ptsI | 1 | 35.4333 | 1.23 |
| 0.1 | 0 | clcB | 1 | 35.4333 | 0.991 |
| 0.1 | 0 | ycaM | 1 | 35.4333 | 1.133 |
| 0.1 | 0 | yadI | 1 | 35.4333 | 1.161 |
| 0.1 | 0 | AG1 | 1 | 35.4333 | 0.908 |
| 0.1 | 0 | ptsI | 1 | 35.6833 | 1.241 |
| 0.1 | 0 | clcB | 1 | 35.6833 | 0.999 |
| 0.1 | 0 | ycaM | 1 | 35.6833 | 1.142 |
| 0.1 | 0 | yadI | 1 | 35.6833 | 1.174 |
| 0.1 | 0 | AG1 | 1 | 35.6833 | 0.918 |
| 0.1 | 0 | ptsI | 1 | 35.9333 | 1.25 |
| 0.1 | 0 | clcB | 1 | 35.9333 | 1.01 |
| 0.1 | 0 | ycaM | 1 | 35.9333 | 1.152 |
| 0.1 | 0 | yadI | 1 | 35.9333 | 1.166 |
| 0.1 | 0 | AG1 | 1 | 35.9333 | 0.917 |
| 0.1 | 0 | ptsI | 1 | 36.1833 | 1.255 |
| 0.1 | 0 | clcB | 1 | 36.1833 | 1.016 |
| 0.1 | 0 | ycaM | 1 | 36.1833 | 1.156 |
| 0.1 | 0 | yadI | 1 | 36.1833 | 1.187 |
| 0.1 | 0 | AG1 | 1 | 36.1833 | 0.922 |
| 0.1 | 0 | ptsI | 1 | 36.4333 | 1.264 |
| 0.1 | 0 | clcB | 1 | 36.4333 | 1.032 |
| 0.1 | 0 | ycaM | 1 | 36.4333 | 1.161 |
| 0.1 | 0 | yadI | 1 | 36.4333 | 1.195 |
| 0.1 | 0 | AG1 | 1 | 36.4333 | 0.921 |
| 0.1 | 0 | ptsI | 1 | 36.6833 | 1.273 |
| 0.1 | 0 | clcB | 1 | 36.6833 | 1.033 |
| 0.1 | 0 | ycaM | 1 | 36.6833 | 1.17 |
| 0.1 | 0 | yadI | 1 | 36.6833 | 1.201 |
| 0.1 | 0 | AG1 | 1 | 36.6833 | 0.916 |
| 0.1 | 0 | ptsI | 2 | 0 | 0.222 |
| 0.1 | 0 | clcB | 2 | 0 | 0.206 |
| 0.1 | 0 | ycaM | 2 | 0 | 0.211 |
| 0.1 | 0 | yadI | 2 | 0 | 0.224 |
| 0.1 | 0 | AG1 | 2 | 0 | 0.226 |

| | | | | | |
|---|---|---|---|---|---|
| 0.1 | 0 | ptsI | 2 | 0.25 | 0.216 |
| 0.1 | 0 | clcB | 2 | 0.25 | 0.199 |
| 0.1 | 0 | ycaM | 2 | 0.25 | 0.21 |
| 0.1 | 0 | yadI | 2 | 0.25 | 0.217 |
| 0.1 | 0 | AG1 | 2 | 0.25 | 0.222 |
| 0.1 | 0 | ptsI | 2 | 0.5 | 0.213 |
| 0.1 | 0 | clcB | 2 | 0.5 | 0.2 |
| 0.1 | 0 | ycaM | 2 | 0.5 | 0.208 |
| 0.1 | 0 | yadI | 2 | 0.5 | 0.214 |
| 0.1 | 0 | AG1 | 2 | 0.5 | 0.223 |
| 0.1 | 0 | ptsI | 2 | 0.75 | 0.217 |
| 0.1 | 0 | clcB | 2 | 0.75 | 0.201 |
| 0.1 | 0 | ycaM | 2 | 0.75 | 0.21 |
| 0.1 | 0 | yadI | 2 | 0.75 | 0.219 |
| 0.1 | 0 | AG1 | 2 | 0.75 | 0.226 |
| 0.1 | 0 | ptsI | 2 | 1 | 0.22 |
| 0.1 | 0 | clcB | 2 | 1 | 0.202 |
| 0.1 | 0 | ycaM | 2 | 1 | 0.215 |
| 0.1 | 0 | yadI | 2 | 1 | 0.224 |
| 0.1 | 0 | AG1 | 2 | 1 | 0.227 |
| 0.1 | 0 | ptsI | 2 | 1.25 | 0.22 |
| 0.1 | 0 | clcB | 2 | 1.25 | 0.204 |
| 0.1 | 0 | ycaM | 2 | 1.25 | 0.217 |
| 0.1 | 0 | yadI | 2 | 1.25 | 0.231 |
| 0.1 | 0 | AG1 | 2 | 1.25 | 0.231 |
| 0.1 | 0 | ptsI | 2 | 1.5 | 0.224 |
| 0.1 | 0 | clcB | 2 | 1.5 | 0.206 |
| 0.1 | 0 | ycaM | 2 | 1.5 | 0.22 |
| 0.1 | 0 | yadI | 2 | 1.5 | 0.239 |
| 0.1 | 0 | AG1 | 2 | 1.5 | 0.236 |
| 0.1 | 0 | ptsI | 2 | 1.75 | 0.228 |
| 0.1 | 0 | clcB | 2 | 1.75 | 0.208 |
| 0.1 | 0 | ycaM | 2 | 1.75 | 0.225 |
| 0.1 | 0 | yadI | 2 | 1.75 | 0.244 |
| 0.1 | 0 | AG1 | 2 | 1.75 | 0.239 |
| 0.1 | 0 | ptsI | 2 | 2 | 0.233 |
| 0.1 | 0 | clcB | 2 | 2 | 0.21 |
| 0.1 | 0 | ycaM | 2 | 2 | 0.23 |
| 0.1 | 0 | yadI | 2 | 2 | 0.251 |
| 0.1 | 0 | AG1 | 2 | 2 | 0.252 |
| 0.1 | 0 | ptsI | 2 | 2.25 | 0.242 |
| 0.1 | 0 | clcB | 2 | 2.25 | 0.212 |
| 0.1 | 0 | ycaM | 2 | 2.25 | 0.235 |
| 0.1 | 0 | yadI | 2 | 2.25 | 0.263 |
| 0.1 | 0 | AG1 | 2 | 2.25 | 0.259 |
| 0.1 | 0 | ptsI | 2 | 2.5 | 0.247 |
| 0.1 | 0 | clcB | 2 | 2.5 | 0.217 |
| 0.1 | 0 | ycaM | 2 | 2.5 | 0.244 |

| | | | | | |
|---|---|---|---|---|---|
| 0.1 | 0 | yadI | 2 | 2.5 | 0.274 |
| 0.1 | 0 | AG1 | 2 | 2.5 | 0.27 |
| 0.1 | 0 | ptsI | 2 | 2.75 | 0.256 |
| 0.1 | 0 | clcB | 2 | 2.75 | 0.222 |
| 0.1 | 0 | ycaM | 2 | 2.75 | 0.25 |
| 0.1 | 0 | yadI | 2 | 2.75 | 0.289 |
| 0.1 | 0 | AG1 | 2 | 2.75 | 0.287 |
| 0.1 | 0 | ptsI | 2 | 3 | 0.267 |
| 0.1 | 0 | clcB | 2 | 3 | 0.23 |
| 0.1 | 0 | ycaM | 2 | 3 | 0.253 |
| 0.1 | 0 | yadI | 2 | 3 | 0.303 |
| 0.1 | 0 | AG1 | 2 | 3 | 0.299 |
| 0.1 | 0 | ptsI | 2 | 3.25 | 0.273 |
| 0.1 | 0 | clcB | 2 | 3.25 | 0.232 |
| 0.1 | 0 | ycaM | 2 | 3.25 | 0.266 |
| 0.1 | 0 | yadI | 2 | 3.25 | 0.325 |
| 0.1 | 0 | AG1 | 2 | 3.25 | 0.31 |
| 0.1 | 0 | ptsI | 2 | 3.5 | 0.282 |
| 0.1 | 0 | clcB | 2 | 3.5 | 0.236 |
| 0.1 | 0 | ycaM | 2 | 3.5 | 0.272 |
| 0.1 | 0 | yadI | 2 | 3.5 | 0.343 |
| 0.1 | 0 | AG1 | 2 | 3.5 | 0.32 |
| 0.1 | 0 | ptsI | 2 | 3.75 | 0.289 |
| 0.1 | 0 | clcB | 2 | 3.75 | 0.241 |
| 0.1 | 0 | ycaM | 2 | 3.75 | 0.284 |
| 0.1 | 0 | yadI | 2 | 3.75 | 0.363 |
| 0.1 | 0 | AG1 | 2 | 3.75 | 0.334 |
| 0.1 | 0 | ptsI | 2 | 4 | 0.293 |
| 0.1 | 0 | clcB | 2 | 4 | 0.246 |
| 0.1 | 0 | ycaM | 2 | 4 | 0.293 |
| 0.1 | 0 | yadI | 2 | 4 | 0.385 |
| 0.1 | 0 | AG1 | 2 | 4 | 0.344 |
| 0.1 | 0 | ptsI | 2 | 4.38333 | 0.303 |
| 0.1 | 0 | clcB | 2 | 4.38333 | 0.259 |
| 0.1 | 0 | ycaM | 2 | 4.38333 | 0.307 |
| 0.1 | 0 | yadI | 2 | 4.38333 | 0.424 |
| 0.1 | 0 | AG1 | 2 | 4.38333 | 0.382 |
| 0.1 | 0 | ptsI | 2 | 4.63333 | 0.296 |
| 0.1 | 0 | clcB | 2 | 4.63333 | 0.256 |
| 0.1 | 0 | ycaM | 2 | 4.63333 | 0.306 |
| 0.1 | 0 | yadI | 2 | 4.63333 | 0.415 |
| 0.1 | 0 | AG1 | 2 | 4.63333 | 0.368 |
| 0.1 | 0 | ptsI | 2 | 4.88333 | 0.302 |
| 0.1 | 0 | clcB | 2 | 4.88333 | 0.26 |
| 0.1 | 0 | ycaM | 2 | 4.88333 | 0.314 |
| 0.1 | 0 | yadI | 2 | 4.88333 | 0.423 |
| 0.1 | 0 | AG1 | 2 | 4.88333 | 0.378 |
| 0.1 | 0 | ptsI | 2 | 5.13333 | 0.309 |

| | | | | | |
|---|---|---|---|---|---|
| 0.1 | 0 | clcB | 2 | 5.13333 | 0.265 |
| 0.1 | 0 | ycaM | 2 | 5.13333 | 0.322 |
| 0.1 | 0 | yadI | 2 | 5.13333 | 0.436 |
| 0.1 | 0 | AG1 | 2 | 5.13333 | 0.387 |
| 0.1 | 0 | ptsI | 2 | 5.38333 | 0.316 |
| 0.1 | 0 | clcB | 2 | 5.38333 | 0.274 |
| 0.1 | 0 | ycaM | 2 | 5.38333 | 0.332 |
| 0.1 | 0 | yadI | 2 | 5.38333 | 0.45 |
| 0.1 | 0 | AG1 | 2 | 5.38333 | 0.398 |
| 0.1 | 0 | ptsI | 2 | 5.63333 | 0.321 |
| 0.1 | 0 | clcB | 2 | 5.63333 | 0.284 |
| 0.1 | 0 | ycaM | 2 | 5.63333 | 0.341 |
| 0.1 | 0 | yadI | 2 | 5.63333 | 0.46 |
| 0.1 | 0 | AG1 | 2 | 5.63333 | 0.407 |
| 0.1 | 0 | ptsI | 2 | 5.88333 | 0.329 |
| 0.1 | 0 | clcB | 2 | 5.88333 | 0.292 |
| 0.1 | 0 | ycaM | 2 | 5.88333 | 0.354 |
| 0.1 | 0 | yadI | 2 | 5.88333 | 0.474 |
| 0.1 | 0 | AG1 | 2 | 5.88333 | 0.419 |
| 0.1 | 0 | ptsI | 2 | 6.13333 | 0.334 |
| 0.1 | 0 | clcB | 2 | 6.13333 | 0.297 |
| 0.1 | 0 | ycaM | 2 | 6.13333 | 0.363 |
| 0.1 | 0 | yadI | 2 | 6.13333 | 0.486 |
| 0.1 | 0 | AG1 | 2 | 6.13333 | 0.431 |
| 0.1 | 0 | ptsI | 2 | 6.38333 | 0.345 |
| 0.1 | 0 | clcB | 2 | 6.38333 | 0.302 |
| 0.1 | 0 | ycaM | 2 | 6.38333 | 0.374 |
| 0.1 | 0 | yadI | 2 | 6.38333 | 0.498 |
| 0.1 | 0 | AG1 | 2 | 6.38333 | 0.445 |
| 0.1 | 0 | ptsI | 2 | 6.63333 | 0.351 |
| 0.1 | 0 | clcB | 2 | 6.63333 | 0.312 |
| 0.1 | 0 | ycaM | 2 | 6.63333 | 0.381 |
| 0.1 | 0 | yadI | 2 | 6.63333 | 0.509 |
| 0.1 | 0 | AG1 | 2 | 6.63333 | 0.456 |
| 0.1 | 0 | ptsI | 2 | 6.88333 | 0.359 |
| 0.1 | 0 | clcB | 2 | 6.88333 | 0.319 |
| 0.1 | 0 | ycaM | 2 | 6.88333 | 0.391 |
| 0.1 | 0 | yadI | 2 | 6.88333 | 0.523 |
| 0.1 | 0 | AG1 | 2 | 6.88333 | 0.47 |
| 0.1 | 0 | ptsI | 2 | 7.13333 | 0.365 |
| 0.1 | 0 | clcB | 2 | 7.13333 | 0.324 |
| 0.1 | 0 | ycaM | 2 | 7.13333 | 0.394 |
| 0.1 | 0 | yadI | 2 | 7.13333 | 0.529 |
| 0.1 | 0 | AG1 | 2 | 7.13333 | 0.48 |
| 0.1 | 0 | ptsI | 2 | 7.38333 | 0.375 |
| 0.1 | 0 | clcB | 2 | 7.38333 | 0.331 |
| 0.1 | 0 | ycaM | 2 | 7.38333 | 0.407 |
| 0.1 | 0 | yadI | 2 | 7.38333 | 0.545 |

| | | | | | |
|---|---|---|---|---|---|
| 0.1 | 0 | AG1 | 2 | 7.38333 | 0.493 |
| 0.1 | 0 | ptsI | 2 | 7.63333 | 0.379 |
| 0.1 | 0 | clcB | 2 | 7.63333 | 0.339 |
| 0.1 | 0 | ycaM | 2 | 7.63333 | 0.417 |
| 0.1 | 0 | yadI | 2 | 7.63333 | 0.561 |
| 0.1 | 0 | AG1 | 2 | 7.63333 | 0.509 |
| 0.1 | 0 | ptsI | 2 | 7.88333 | 0.389 |
| 0.1 | 0 | clcB | 2 | 7.88333 | 0.345 |
| 0.1 | 0 | ycaM | 2 | 7.88333 | 0.426 |
| 0.1 | 0 | yadI | 2 | 7.88333 | 0.576 |
| 0.1 | 0 | AG1 | 2 | 7.88333 | 0.521 |
| 0.1 | 0 | ptsI | 2 | 8.13333 | 0.396 |
| 0.1 | 0 | clcB | 2 | 8.13333 | 0.353 |
| 0.1 | 0 | ycaM | 2 | 8.13333 | 0.433 |
| 0.1 | 0 | yadI | 2 | 8.13333 | 0.582 |
| 0.1 | 0 | AG1 | 2 | 8.13333 | 0.534 |
| 0.1 | 0 | ptsI | 2 | 8.38333 | 0.401 |
| 0.1 | 0 | clcB | 2 | 8.38333 | 0.361 |
| 0.1 | 0 | ycaM | 2 | 8.38333 | 0.439 |
| 0.1 | 0 | yadI | 2 | 8.38333 | 0.599 |
| 0.1 | 0 | AG1 | 2 | 8.38333 | 0.548 |
| 0.1 | 0 | ptsI | 2 | 8.91667 | 0.338 |
| 0.1 | 0 | clcB | 2 | 8.91667 | 0.324 |
| 0.1 | 0 | ycaM | 2 | 8.91667 | 0.361 |
| 0.1 | 0 | yadI | 2 | 8.91667 | 0.568 |
| 0.1 | 0 | AG1 | 2 | 8.91667 | 0.479 |
| 0.1 | 0 | ptsI | 2 | 9.16667 | 0.341 |
| 0.1 | 0 | clcB | 2 | 9.16667 | 0.325 |
| 0.1 | 0 | ycaM | 2 | 9.16667 | 0.364 |
| 0.1 | 0 | yadI | 2 | 9.16667 | 0.555 |
| 0.1 | 0 | AG1 | 2 | 9.16667 | 0.475 |
| 0.1 | 0 | ptsI | 2 | 9.41667 | 0.346 |
| 0.1 | 0 | clcB | 2 | 9.41667 | 0.329 |
| 0.1 | 0 | ycaM | 2 | 9.41667 | 0.364 |
| 0.1 | 0 | yadI | 2 | 9.41667 | 0.564 |
| 0.1 | 0 | AG1 | 2 | 9.41667 | 0.481 |
| 0.1 | 0 | ptsI | 2 | 9.66667 | 0.354 |
| 0.1 | 0 | clcB | 2 | 9.66667 | 0.339 |
| 0.1 | 0 | ycaM | 2 | 9.66667 | 0.37 |
| 0.1 | 0 | yadI | 2 | 9.66667 | 0.574 |
| 0.1 | 0 | AG1 | 2 | 9.66667 | 0.496 |
| 0.1 | 0 | ptsI | 2 | 9.91667 | 0.358 |
| 0.1 | 0 | clcB | 2 | 9.91667 | 0.342 |
| 0.1 | 0 | ycaM | 2 | 9.91667 | 0.377 |
| 0.1 | 0 | yadI | 2 | 9.91667 | 0.582 |
| 0.1 | 0 | AG1 | 2 | 9.91667 | 0.499 |
| 0.1 | 0 | ptsI | 2 | 10.1667 | 0.365 |
| 0.1 | 0 | clcB | 2 | 10.1667 | 0.348 |

| | | | | | |
|---|---|---|---|---|---|
| 0.1 | 0 | ycaM | 2 | 10.1667 | 0.382 |
| 0.1 | 0 | yadI | 2 | 10.1667 | 0.592 |
| 0.1 | 0 | AG1 | 2 | 10.1667 | 0.502 |
| 0.1 | 0 | ptsI | 2 | 10.4167 | 0.368 |
| 0.1 | 0 | clcB | 2 | 10.4167 | 0.352 |
| 0.1 | 0 | ycaM | 2 | 10.4167 | 0.389 |
| 0.1 | 0 | yadI | 2 | 10.4167 | 0.601 |
| 0.1 | 0 | AG1 | 2 | 10.4167 | 0.508 |
| 0.1 | 0 | ptsI | 2 | 10.6667 | 0.377 |
| 0.1 | 0 | clcB | 2 | 10.6667 | 0.358 |
| 0.1 | 0 | ycaM | 2 | 10.6667 | 0.396 |
| 0.1 | 0 | yadI | 2 | 10.6667 | 0.612 |
| 0.1 | 0 | AG1 | 2 | 10.6667 | 0.508 |
| 0.1 | 0 | ptsI | 2 | 10.9167 | 0.384 |
| 0.1 | 0 | clcB | 2 | 10.9167 | 0.362 |
| 0.1 | 0 | ycaM | 2 | 10.9167 | 0.403 |
| 0.1 | 0 | yadI | 2 | 10.9167 | 0.62 |
| 0.1 | 0 | AG1 | 2 | 10.9167 | 0.52 |
| 0.1 | 0 | ptsI | 2 | 11.1667 | 0.392 |
| 0.1 | 0 | clcB | 2 | 11.1667 | 0.367 |
| 0.1 | 0 | ycaM | 2 | 11.1667 | 0.409 |
| 0.1 | 0 | yadI | 2 | 11.1667 | 0.63 |
| 0.1 | 0 | AG1 | 2 | 11.1667 | 0.544 |
| 0.1 | 0 | ptsI | 2 | 11.4167 | 0.399 |
| 0.1 | 0 | clcB | 2 | 11.4167 | 0.372 |
| 0.1 | 0 | ycaM | 2 | 11.4167 | 0.41 |
| 0.1 | 0 | yadI | 2 | 11.4167 | 0.635 |
| 0.1 | 0 | AG1 | 2 | 11.4167 | 0.562 |
| 0.1 | 0 | ptsI | 2 | 11.6667 | 0.409 |
| 0.1 | 0 | clcB | 2 | 11.6667 | 0.381 |
| 0.1 | 0 | ycaM | 2 | 11.6667 | 0.424 |
| 0.1 | 0 | yadI | 2 | 11.6667 | 0.649 |
| 0.1 | 0 | AG1 | 2 | 11.6667 | 0.603 |
| 0.1 | 0 | ptsI | 2 | 11.9167 | 0.411 |
| 0.1 | 0 | clcB | 2 | 11.9167 | 0.385 |
| 0.1 | 0 | ycaM | 2 | 11.9167 | 0.434 |
| 0.1 | 0 | yadI | 2 | 11.9167 | 0.658 |
| 0.1 | 0 | AG1 | 2 | 11.9167 | 0.611 |
| 0.1 | 0 | ptsI | 2 | 12.1667 | 0.424 |
| 0.1 | 0 | clcB | 2 | 12.1667 | 0.392 |
| 0.1 | 0 | ycaM | 2 | 12.1667 | 0.435 |
| 0.1 | 0 | yadI | 2 | 12.1667 | 0.669 |
| 0.1 | 0 | AG1 | 2 | 12.1667 | 0.642 |
| 0.1 | 0 | ptsI | 2 | 12.4167 | 0.458 |
| 0.1 | 0 | clcB | 2 | 12.4167 | 0.397 |
| 0.1 | 0 | ycaM | 2 | 12.4167 | 0.438 |
| 0.1 | 0 | yadI | 2 | 12.4167 | 0.674 |
| 0.1 | 0 | AG1 | 2 | 12.4167 | 0.663 |

| | | | | | |
|---|---|---|---|---|---|
| 0.1 | 0 | ptsI | 2 | 12.6667 | 0.46 |
| 0.1 | 0 | clcB | 2 | 12.6667 | 0.404 |
| 0.1 | 0 | ycaM | 2 | 12.6667 | 0.431 |
| 0.1 | 0 | yadI | 2 | 12.6667 | 0.683 |
| 0.1 | 0 | AG1 | 2 | 12.6667 | 0.681 |
| 0.1 | 0 | ptsI | 2 | 12.9167 | 0.447 |
| 0.1 | 0 | clcB | 2 | 12.9167 | 0.413 |
| 0.1 | 0 | ycaM | 2 | 12.9167 | 0.446 |
| 0.1 | 0 | yadI | 2 | 12.9167 | 0.697 |
| 0.1 | 0 | AG1 | 2 | 12.9167 | 0.698 |
| 0.1 | 0 | ptsI | 2 | 13.1667 | 0.449 |
| 0.1 | 0 | clcB | 2 | 13.1667 | 0.414 |
| 0.1 | 0 | ycaM | 2 | 13.1667 | 0.452 |
| 0.1 | 0 | yadI | 2 | 13.1667 | 0.704 |
| 0.1 | 0 | AG1 | 2 | 13.1667 | 0.698 |
| 0.1 | 0 | ptsI | 2 | 13.4167 | 0.458 |
| 0.1 | 0 | clcB | 2 | 13.4167 | 0.42 |
| 0.1 | 0 | ycaM | 2 | 13.4167 | 0.449 |
| 0.1 | 0 | yadI | 2 | 13.4167 | 0.714 |
| 0.1 | 0 | AG1 | 2 | 13.4167 | 0.717 |
| 0.1 | 0 | ptsI | 2 | 13.6667 | 0.469 |
| 0.1 | 0 | clcB | 2 | 13.6667 | 0.43 |
| 0.1 | 0 | ycaM | 2 | 13.6667 | 0.46 |
| 0.1 | 0 | yadI | 2 | 13.6667 | 0.727 |
| 0.1 | 0 | AG1 | 2 | 13.6667 | 0.758 |
| 0.1 | 0 | ptsI | 2 | 13.9167 | 0.477 |
| 0.1 | 0 | clcB | 2 | 13.9167 | 0.44 |
| 0.1 | 0 | ycaM | 2 | 13.9167 | 0.469 |
| 0.1 | 0 | yadI | 2 | 13.9167 | 0.735 |
| 0.1 | 0 | AG1 | 2 | 13.9167 | 0.764 |
| 0.1 | 0 | ptsI | 2 | 14.1667 | 0.476 |
| 0.1 | 0 | clcB | 2 | 14.1667 | 0.439 |
| 0.1 | 0 | ycaM | 2 | 14.1667 | 0.465 |
| 0.1 | 0 | yadI | 2 | 14.1667 | 0.748 |
| 0.1 | 0 | AG1 | 2 | 14.1667 | 0.764 |
| 0.1 | 0 | ptsI | 2 | 14.4167 | 0.48 |
| 0.1 | 0 | clcB | 2 | 14.4167 | 0.435 |
| 0.1 | 0 | ycaM | 2 | 14.4167 | 0.481 |
| 0.1 | 0 | yadI | 2 | 14.4167 | 0.764 |
| 0.1 | 0 | AG1 | 2 | 14.4167 | 0.777 |
| 0.1 | 0 | ptsI | 2 | 14.6667 | 0.482 |
| 0.1 | 0 | clcB | 2 | 14.6667 | 0.441 |
| 0.1 | 0 | ycaM | 2 | 14.6667 | 0.484 |
| 0.1 | 0 | yadI | 2 | 14.6667 | 0.766 |
| 0.1 | 0 | AG1 | 2 | 14.6667 | 0.78 |
| 0.1 | 0 | ptsI | 2 | 14.9167 | 0.487 |
| 0.1 | 0 | clcB | 2 | 14.9167 | 0.452 |
| 0.1 | 0 | ycaM | 2 | 14.9167 | 0.479 |

| | | | | | |
|---|---|---|---|---|---|
| 0.1 | 0 | yadI | 2 | 14.9167 | 0.779 |
| 0.1 | 0 | AG1 | 2 | 14.9167 | 0.79 |
| 0.1 | 0 | ptsI | 2 | 15.1667 | 0.5 |
| 0.1 | 0 | clcB | 2 | 15.1667 | 0.461 |
| 0.1 | 0 | ycaM | 2 | 15.1667 | 0.499 |
| 0.1 | 0 | yadI | 2 | 15.1667 | 0.783 |
| 0.1 | 0 | AG1 | 2 | 15.1667 | 0.807 |
| 0.1 | 0 | ptsI | 2 | 15.4167 | 0.508 |
| 0.1 | 0 | clcB | 2 | 15.4167 | 0.459 |
| 0.1 | 0 | ycaM | 2 | 15.4167 | 0.515 |
| 0.1 | 0 | yadI | 2 | 15.4167 | 0.803 |
| 0.1 | 0 | AG1 | 2 | 15.4167 | 0.825 |
| 0.1 | 0 | ptsI | 2 | 15.6667 | 0.511 |
| 0.1 | 0 | clcB | 2 | 15.6667 | 0.471 |
| 0.1 | 0 | ycaM | 2 | 15.6667 | 0.514 |
| 0.1 | 0 | yadI | 2 | 15.6667 | 0.804 |
| 0.1 | 0 | AG1 | 2 | 15.6667 | 0.829 |
| 0.1 | 0 | ptsI | 2 | 15.9167 | 0.524 |
| 0.1 | 0 | clcB | 2 | 15.9167 | 0.468 |
| 0.1 | 0 | ycaM | 2 | 15.9167 | 0.527 |
| 0.1 | 0 | yadI | 2 | 15.9167 | 0.813 |
| 0.1 | 0 | AG1 | 2 | 15.9167 | 0.828 |
| 0.1 | 0 | ptsI | 2 | 16.1667 | 0.538 |
| 0.1 | 0 | clcB | 2 | 16.1667 | 0.476 |
| 0.1 | 0 | ycaM | 2 | 16.1667 | 0.551 |
| 0.1 | 0 | yadI | 2 | 16.1667 | 0.824 |
| 0.1 | 0 | AG1 | 2 | 16.1667 | 0.836 |
| 0.1 | 0 | ptsI | 2 | 16.4167 | 0.562 |
| 0.1 | 0 | clcB | 2 | 16.4167 | 0.474 |
| 0.1 | 0 | ycaM | 2 | 16.4167 | 0.536 |
| 0.1 | 0 | yadI | 2 | 16.4167 | 0.829 |
| 0.1 | 0 | AG1 | 2 | 16.4167 | 0.844 |
| 0.1 | 0 | ptsI | 2 | 16.6667 | 0.586 |
| 0.1 | 0 | clcB | 2 | 16.6667 | 0.495 |
| 0.1 | 0 | ycaM | 2 | 16.6667 | 0.552 |
| 0.1 | 0 | yadI | 2 | 16.6667 | 0.836 |
| 0.1 | 0 | AG1 | 2 | 16.6667 | 0.851 |
| 0.1 | 0 | ptsI | 2 | 16.9167 | 0.602 |
| 0.1 | 0 | clcB | 2 | 16.9167 | 0.488 |
| 0.1 | 0 | ycaM | 2 | 16.9167 | 0.562 |
| 0.1 | 0 | yadI | 2 | 16.9167 | 0.843 |
| 0.1 | 0 | AG1 | 2 | 16.9167 | 0.854 |
| 0.1 | 0 | ptsI | 2 | 17.1667 | 0.656 |
| 0.1 | 0 | clcB | 2 | 17.1667 | 0.489 |
| 0.1 | 0 | ycaM | 2 | 17.1667 | 0.549 |
| 0.1 | 0 | yadI | 2 | 17.1667 | 0.852 |
| 0.1 | 0 | AG1 | 2 | 17.1667 | 0.854 |
| 0.1 | 0 | ptsI | 2 | 17.4167 | 0.691 |

| | | | | | |
|---|---|---|---|---|---|
| 0.1 | 0 | clcB | 2 | 17.4167 | 0.502 |
| 0.1 | 0 | ycaM | 2 | 17.4167 | 0.556 |
| 0.1 | 0 | yadI | 2 | 17.4167 | 0.856 |
| 0.1 | 0 | AG1 | 2 | 17.4167 | 0.862 |
| 0.1 | 0 | ptsI | 2 | 17.6667 | 0.713 |
| 0.1 | 0 | clcB | 2 | 17.6667 | 0.5 |
| 0.1 | 0 | ycaM | 2 | 17.6667 | 0.549 |
| 0.1 | 0 | yadI | 2 | 17.6667 | 0.862 |
| 0.1 | 0 | AG1 | 2 | 17.6667 | 0.862 |
| 0.1 | 0 | ptsI | 2 | 17.9167 | 0.713 |
| 0.1 | 0 | clcB | 2 | 17.9167 | 0.495 |
| 0.1 | 0 | ycaM | 2 | 17.9167 | 0.552 |
| 0.1 | 0 | yadI | 2 | 17.9167 | 0.872 |
| 0.1 | 0 | AG1 | 2 | 17.9167 | 0.87 |
| 0.1 | 0 | ptsI | 2 | 18.1667 | 0.726 |
| 0.1 | 0 | clcB | 2 | 18.1667 | 0.504 |
| 0.1 | 0 | ycaM | 2 | 18.1667 | 0.555 |
| 0.1 | 0 | yadI | 2 | 18.1667 | 0.872 |
| 0.1 | 0 | AG1 | 2 | 18.1667 | 0.881 |
| 0.1 | 0 | ptsI | 2 | 18.4167 | 0.742 |
| 0.1 | 0 | clcB | 2 | 18.4167 | 0.508 |
| 0.1 | 0 | ycaM | 2 | 18.4167 | 0.548 |
| 0.1 | 0 | yadI | 2 | 18.4167 | 0.88 |
| 0.1 | 0 | AG1 | 2 | 18.4167 | 0.884 |
| 0.1 | 0 | ptsI | 2 | 18.6667 | 0.748 |
| 0.1 | 0 | clcB | 2 | 18.6667 | 0.495 |
| 0.1 | 0 | ycaM | 2 | 18.6667 | 0.556 |
| 0.1 | 0 | yadI | 2 | 18.6667 | 0.884 |
| 0.1 | 0 | AG1 | 2 | 18.6667 | 0.893 |
| 0.1 | 0 | ptsI | 2 | 18.9167 | 0.773 |
| 0.1 | 0 | clcB | 2 | 18.9167 | 0.498 |
| 0.1 | 0 | ycaM | 2 | 18.9167 | 0.559 |
| 0.1 | 0 | yadI | 2 | 18.9167 | 0.887 |
| 0.1 | 0 | AG1 | 2 | 18.9167 | 0.898 |
| 0.1 | 0 | ptsI | 2 | 19.1667 | 0.775 |
| 0.1 | 0 | clcB | 2 | 19.1667 | 0.499 |
| 0.1 | 0 | ycaM | 2 | 19.1667 | 0.566 |
| 0.1 | 0 | yadI | 2 | 19.1667 | 0.896 |
| 0.1 | 0 | AG1 | 2 | 19.1667 | 0.908 |
| 0.1 | 0 | ptsI | 2 | 19.4167 | 0.776 |
| 0.1 | 0 | clcB | 2 | 19.4167 | 0.506 |
| 0.1 | 0 | ycaM | 2 | 19.4167 | 0.574 |
| 0.1 | 0 | yadI | 2 | 19.4167 | 0.9 |
| 0.1 | 0 | AG1 | 2 | 19.4167 | 0.912 |
| 0.1 | 0 | ptsI | 2 | 19.6667 | 0.784 |
| 0.1 | 0 | clcB | 2 | 19.6667 | 0.502 |
| 0.1 | 0 | ycaM | 2 | 19.6667 | 0.58 |
| 0.1 | 0 | yadI | 2 | 19.6667 | 0.905 |

| | | | | | |
|---|---|---|---|---|---|
| 0.1 | 0 | AG1 | 2 | 19.6667 | 0.911 |
| 0.1 | 0 | ptsI | 2 | 19.9167 | 0.801 |
| 0.1 | 0 | clcB | 2 | 19.9167 | 0.507 |
| 0.1 | 0 | ycaM | 2 | 19.9167 | 0.586 |
| 0.1 | 0 | yadI | 2 | 19.9167 | 0.908 |
| 0.1 | 0 | AG1 | 2 | 19.9167 | 0.922 |
| 0.1 | 0 | ptsI | 2 | 20.1667 | 0.789 |
| 0.1 | 0 | clcB | 2 | 20.1667 | 0.5 |
| 0.1 | 0 | ycaM | 2 | 20.1667 | 0.587 |
| 0.1 | 0 | yadI | 2 | 20.1667 | 0.912 |
| 0.1 | 0 | AG1 | 2 | 20.1667 | 0.914 |
| 0.1 | 0 | ptsI | 2 | 20.4167 | 0.793 |
| 0.1 | 0 | clcB | 2 | 20.4167 | 0.501 |
| 0.1 | 0 | ycaM | 2 | 20.4167 | 0.588 |
| 0.1 | 0 | yadI | 2 | 20.4167 | 0.92 |
| 0.1 | 0 | AG1 | 2 | 20.4167 | 0.916 |
| 0.1 | 0 | ptsI | 2 | 20.6667 | 0.791 |
| 0.1 | 0 | clcB | 2 | 20.6667 | 0.507 |
| 0.1 | 0 | ycaM | 2 | 20.6667 | 0.594 |
| 0.1 | 0 | yadI | 2 | 20.6667 | 0.923 |
| 0.1 | 0 | AG1 | 2 | 20.6667 | 0.922 |
| 0.1 | 0 | ptsI | 2 | 20.9167 | 0.799 |
| 0.1 | 0 | clcB | 2 | 20.9167 | 0.508 |
| 0.1 | 0 | ycaM | 2 | 20.9167 | 0.605 |
| 0.1 | 0 | yadI | 2 | 20.9167 | 0.93 |
| 0.1 | 0 | AG1 | 2 | 20.9167 | 0.924 |
| 0.1 | 0 | ptsI | 2 | 21.1667 | 0.797 |
| 0.1 | 0 | clcB | 2 | 21.1667 | 0.509 |
| 0.1 | 0 | ycaM | 2 | 21.1667 | 0.61 |
| 0.1 | 0 | yadI | 2 | 21.1667 | 0.936 |
| 0.1 | 0 | AG1 | 2 | 21.1667 | 0.925 |
| 0.1 | 0 | ptsI | 2 | 21.4167 | 0.798 |
| 0.1 | 0 | clcB | 2 | 21.4167 | 0.508 |
| 0.1 | 0 | ycaM | 2 | 21.4167 | 0.617 |
| 0.1 | 0 | yadI | 2 | 21.4167 | 0.932 |
| 0.1 | 0 | AG1 | 2 | 21.4167 | 0.929 |
| 0.1 | 0 | ptsI | 2 | 21.6667 | 0.809 |
| 0.1 | 0 | clcB | 2 | 21.6667 | 0.51 |
| 0.1 | 0 | ycaM | 2 | 21.6667 | 0.623 |
| 0.1 | 0 | yadI | 2 | 21.6667 | 0.949 |
| 0.1 | 0 | AG1 | 2 | 21.6667 | 0.941 |
| 0.1 | 0 | ptsI | 2 | 21.9167 | 0.821 |
| 0.1 | 0 | clcB | 2 | 21.9167 | 0.515 |
| 0.1 | 0 | ycaM | 2 | 21.9167 | 0.628 |
| 0.1 | 0 | yadI | 2 | 21.9167 | 0.953 |
| 0.1 | 0 | AG1 | 2 | 21.9167 | 0.948 |
| 0.1 | 0 | ptsI | 2 | 22.1667 | 0.82 |
| 0.1 | 0 | clcB | 2 | 22.1667 | 0.512 |

| | | | | | |
|---|---|---|---|---|---|
| 0.1 | 0 | ycaM | 2 | 22.1667 | 0.633 |
| 0.1 | 0 | yadI | 2 | 22.1667 | 0.956 |
| 0.1 | 0 | AG1 | 2 | 22.1667 | 0.946 |
| 0.1 | 0 | ptsI | 2 | 22.4167 | 0.828 |
| 0.1 | 0 | clcB | 2 | 22.4167 | 0.518 |
| 0.1 | 0 | ycaM | 2 | 22.4167 | 0.635 |
| 0.1 | 0 | yadI | 2 | 22.4167 | 0.962 |
| 0.1 | 0 | AG1 | 2 | 22.4167 | 0.951 |
| 0.1 | 0 | ptsI | 2 | 22.6667 | 0.835 |
| 0.1 | 0 | clcB | 2 | 22.6667 | 0.519 |
| 0.1 | 0 | ycaM | 2 | 22.6667 | 0.643 |
| 0.1 | 0 | yadI | 2 | 22.6667 | 0.965 |
| 0.1 | 0 | AG1 | 2 | 22.6667 | 0.96 |
| 0.1 | 0 | ptsI | 2 | 22.9167 | 0.844 |
| 0.1 | 0 | clcB | 2 | 22.9167 | 0.525 |
| 0.1 | 0 | ycaM | 2 | 22.9167 | 0.648 |
| 0.1 | 0 | yadI | 2 | 22.9167 | 0.972 |
| 0.1 | 0 | AG1 | 2 | 22.9167 | 0.969 |
| 0.1 | 0 | ptsI | 2 | 23.1667 | 0.844 |
| 0.1 | 0 | clcB | 2 | 23.1667 | 0.517 |
| 0.1 | 0 | ycaM | 2 | 23.1667 | 0.645 |
| 0.1 | 0 | yadI | 2 | 23.1667 | 0.976 |
| 0.1 | 0 | AG1 | 2 | 23.1667 | 0.962 |
| 0.1 | 0 | ptsI | 2 | 23.4167 | 0.851 |
| 0.1 | 0 | clcB | 2 | 23.4167 | 0.525 |
| 0.1 | 0 | ycaM | 2 | 23.4167 | 0.654 |
| 0.1 | 0 | yadI | 2 | 23.4167 | 0.978 |
| 0.1 | 0 | AG1 | 2 | 23.4167 | 0.968 |
| 0.1 | 0 | ptsI | 2 | 23.6667 | 0.855 |
| 0.1 | 0 | clcB | 2 | 23.6667 | 0.53 |
| 0.1 | 0 | ycaM | 2 | 23.6667 | 0.657 |
| 0.1 | 0 | yadI | 2 | 23.6667 | 0.986 |
| 0.1 | 0 | AG1 | 2 | 23.6667 | 0.971 |
| 0.1 | 0 | ptsI | 2 | 23.9167 | 0.86 |
| 0.1 | 0 | clcB | 2 | 23.9167 | 0.528 |
| 0.1 | 0 | ycaM | 2 | 23.9167 | 0.665 |
| 0.1 | 0 | yadI | 2 | 23.9167 | 0.99 |
| 0.1 | 0 | AG1 | 2 | 23.9167 | 0.977 |
| 0.1 | 0 | ptsI | 2 | 24.1667 | 0.859 |
| 0.1 | 0 | clcB | 2 | 24.1667 | 0.526 |
| 0.1 | 0 | ycaM | 2 | 24.1667 | 0.667 |
| 0.1 | 0 | yadI | 2 | 24.1667 | 0.993 |
| 0.1 | 0 | AG1 | 2 | 24.1667 | 0.976 |
| 0.1 | 0 | ptsI | 2 | 24.4167 | 0.863 |
| 0.1 | 0 | clcB | 2 | 24.4167 | 0.527 |
| 0.1 | 0 | ycaM | 2 | 24.4167 | 0.673 |
| 0.1 | 0 | yadI | 2 | 24.4167 | 1.003 |
| 0.1 | 0 | AG1 | 2 | 24.4167 | 0.979 |

| | | | | | |
|---|---|---|---|---|---|
| 0.1 | 0 | ptsI | 2 | 24.6667 | 0.871 |
| 0.1 | 0 | clcB | 2 | 24.6667 | 0.535 |
| 0.1 | 0 | ycaM | 2 | 24.6667 | 0.676 |
| 0.1 | 0 | yadI | 2 | 24.6667 | 1.003 |
| 0.1 | 0 | AG1 | 2 | 24.6667 | 0.989 |
| 0.1 | 0 | ptsI | 2 | 24.9167 | 0.884 |
| 0.1 | 0 | clcB | 2 | 24.9167 | 0.54 |
| 0.1 | 0 | ycaM | 2 | 24.9167 | 0.682 |
| 0.1 | 0 | yadI | 2 | 24.9167 | 1.01 |
| 0.1 | 0 | AG1 | 2 | 24.9167 | 0.992 |
| 0.1 | 0 | ptsI | 2 | 25.1667 | 0.878 |
| 0.1 | 0 | clcB | 2 | 25.1667 | 0.531 |
| 0.1 | 0 | ycaM | 2 | 25.1667 | 0.685 |
| 0.1 | 0 | yadI | 2 | 25.1667 | 1.017 |
| 0.1 | 0 | AG1 | 2 | 25.1667 | 0.992 |
| 0.1 | 0 | ptsI | 2 | 25.4167 | 0.895 |
| 0.1 | 0 | clcB | 2 | 25.4167 | 0.54 |
| 0.1 | 0 | ycaM | 2 | 25.4167 | 0.692 |
| 0.1 | 0 | yadI | 2 | 25.4167 | 1.019 |
| 0.1 | 0 | AG1 | 2 | 25.4167 | 1.003 |
| 0.1 | 0 | ptsI | 2 | 25.6667 | 0.894 |
| 0.1 | 0 | clcB | 2 | 25.6667 | 0.542 |
| 0.1 | 0 | ycaM | 2 | 25.6667 | 0.694 |
| 0.1 | 0 | yadI | 2 | 25.6667 | 1.019 |
| 0.1 | 0 | AG1 | 2 | 25.6667 | 1.005 |
| 0.1 | 0 | ptsI | 2 | 25.9167 | 0.899 |
| 0.1 | 0 | clcB | 2 | 25.9167 | 0.537 |
| 0.1 | 0 | ycaM | 2 | 25.9167 | 0.7 |
| 0.1 | 0 | yadI | 2 | 25.9167 | 1.032 |
| 0.1 | 0 | AG1 | 2 | 25.9167 | 1.011 |
| 0.1 | 0 | ptsI | 2 | 26.1667 | 0.907 |
| 0.1 | 0 | clcB | 2 | 26.1667 | 0.542 |
| 0.1 | 0 | ycaM | 2 | 26.1667 | 0.706 |
| 0.1 | 0 | yadI | 2 | 26.1667 | 1.025 |
| 0.1 | 0 | AG1 | 2 | 26.1667 | 1.013 |
| 0.1 | 0 | ptsI | 2 | 26.4167 | 0.922 |
| 0.1 | 0 | clcB | 2 | 26.4167 | 0.546 |
| 0.1 | 0 | ycaM | 2 | 26.4167 | 0.714 |
| 0.1 | 0 | yadI | 2 | 26.4167 | 1.036 |
| 0.1 | 0 | AG1 | 2 | 26.4167 | 1.027 |
| 0.1 | 0 | ptsI | 3 | 0 | 0.147 |
| 0.1 | 0 | clcB | 3 | 0 | 0.161 |
| 0.1 | 0 | ycaM | 3 | 0 | 0.151 |
| 0.1 | 0 | yadI | 3 | 0 | 0.168 |
| 0.1 | 0 | AG1 | 3 | 0 | 0.188 |
| 0.1 | 0 | ptsI | 3 | 0.25 | 0.146 |
| 0.1 | 0 | clcB | 3 | 0.25 | 0.154 |
| 0.1 | 0 | ycaM | 3 | 0.25 | 0.15 |

| | | | | | |
|---|---|---|---|---|---|
| 0.1 | 0 | yadI | 3 | 0.25 | 0.164 |
| 0.1 | 0 | AG1 | 3 | 0.25 | 0.198 |
| 0.1 | 0 | ptsI | 3 | 0.5 | 0.148 |
| 0.1 | 0 | clcB | 3 | 0.5 | 0.152 |
| 0.1 | 0 | ycaM | 3 | 0.5 | 0.148 |
| 0.1 | 0 | yadI | 3 | 0.5 | 0.164 |
| 0.1 | 0 | AG1 | 3 | 0.5 | 0.204 |
| 0.1 | 0 | ptsI | 3 | 0.75 | 0.148 |
| 0.1 | 0 | clcB | 3 | 0.75 | 0.152 |
| 0.1 | 0 | ycaM | 3 | 0.75 | 0.148 |
| 0.1 | 0 | yadI | 3 | 0.75 | 0.165 |
| 0.1 | 0 | AG1 | 3 | 0.75 | 0.21 |
| 0.1 | 0 | ptsI | 3 | 1 | 0.149 |
| 0.1 | 0 | clcB | 3 | 1 | 0.154 |
| 0.1 | 0 | ycaM | 3 | 1 | 0.15 |
| 0.1 | 0 | yadI | 3 | 1 | 0.165 |
| 0.1 | 0 | AG1 | 3 | 1 | 0.206 |
| 0.1 | 0 | ptsI | 3 | 1.25 | 0.148 |
| 0.1 | 0 | clcB | 3 | 1.25 | 0.154 |
| 0.1 | 0 | ycaM | 3 | 1.25 | 0.15 |
| 0.1 | 0 | yadI | 3 | 1.25 | 0.166 |
| 0.1 | 0 | AG1 | 3 | 1.25 | 0.21 |
| 0.1 | 0 | ptsI | 3 | 1.5 | 0.15 |
| 0.1 | 0 | clcB | 3 | 1.5 | 0.156 |
| 0.1 | 0 | ycaM | 3 | 1.5 | 0.153 |
| 0.1 | 0 | yadI | 3 | 1.5 | 0.169 |
| 0.1 | 0 | AG1 | 3 | 1.5 | 0.215 |
| 0.1 | 0 | ptsI | 3 | 1.75 | 0.15 |
| 0.1 | 0 | clcB | 3 | 1.75 | 0.158 |
| 0.1 | 0 | ycaM | 3 | 1.75 | 0.152 |
| 0.1 | 0 | yadI | 3 | 1.75 | 0.169 |
| 0.1 | 0 | AG1 | 3 | 1.75 | 0.221 |
| 0.1 | 0 | ptsI | 3 | 2 | 0.152 |
| 0.1 | 0 | clcB | 3 | 2 | 0.162 |
| 0.1 | 0 | ycaM | 3 | 2 | 0.155 |
| 0.1 | 0 | yadI | 3 | 2 | 0.171 |
| 0.1 | 0 | AG1 | 3 | 2 | 0.218 |
| 0.1 | 0 | ptsI | 3 | 2.25 | 0.154 |
| 0.1 | 0 | clcB | 3 | 2.25 | 0.166 |
| 0.1 | 0 | ycaM | 3 | 2.25 | 0.156 |
| 0.1 | 0 | yadI | 3 | 2.25 | 0.175 |
| 0.1 | 0 | AG1 | 3 | 2.25 | 0.23 |
| 0.1 | 0 | ptsI | 3 | 2.5 | 0.156 |
| 0.1 | 0 | clcB | 3 | 2.5 | 0.167 |
| 0.1 | 0 | ycaM | 3 | 2.5 | 0.159 |
| 0.1 | 0 | yadI | 3 | 2.5 | 0.176 |
| 0.1 | 0 | AG1 | 3 | 2.5 | 0.221 |
| 0.1 | 0 | ptsI | 3 | 2.75 | 0.159 |

| | | | | | |
|---|---|---|---|---|---|
| 0.1 | 0 | clcB | 3 | 2.75 | 0.173 |
| 0.1 | 0 | ycaM | 3 | 2.75 | 0.164 |
| 0.1 | 0 | yadI | 3 | 2.75 | 0.181 |
| 0.1 | 0 | AG1 | 3 | 2.75 | 0.213 |
| 0.1 | 0 | ptsI | 3 | 3 | 0.16 |
| 0.1 | 0 | clcB | 3 | 3 | 0.173 |
| 0.1 | 0 | ycaM | 3 | 3 | 0.162 |
| 0.1 | 0 | yadI | 3 | 3 | 0.184 |
| 0.1 | 0 | AG1 | 3 | 3 | 0.228 |
| 0.1 | 0 | ptsI | 3 | 3.25 | 0.162 |
| 0.1 | 0 | clcB | 3 | 3.25 | 0.175 |
| 0.1 | 0 | ycaM | 3 | 3.25 | 0.165 |
| 0.1 | 0 | yadI | 3 | 3.25 | 0.188 |
| 0.1 | 0 | AG1 | 3 | 3.25 | 0.229 |
| 0.1 | 0 | ptsI | 3 | 3.5 | 0.166 |
| 0.1 | 0 | clcB | 3 | 3.5 | 0.181 |
| 0.1 | 0 | ycaM | 3 | 3.5 | 0.17 |
| 0.1 | 0 | yadI | 3 | 3.5 | 0.196 |
| 0.1 | 0 | AG1 | 3 | 3.5 | 0.204 |
| 0.1 | 0 | ptsI | 3 | 3.75 | 0.17 |
| 0.1 | 0 | clcB | 3 | 3.75 | 0.186 |
| 0.1 | 0 | ycaM | 3 | 3.75 | 0.173 |
| 0.1 | 0 | yadI | 3 | 3.75 | 0.2 |
| 0.1 | 0 | AG1 | 3 | 3.75 | 0.22 |
| 0.1 | 0 | ptsI | 3 | 4 | 0.172 |
| 0.1 | 0 | clcB | 3 | 4 | 0.19 |
| 0.1 | 0 | ycaM | 3 | 4 | 0.177 |
| 0.1 | 0 | yadI | 3 | 4 | 0.208 |
| 0.1 | 0 | AG1 | 3 | 4 | 0.207 |
| 0.1 | 0 | ptsI | 3 | 4.41667 | 0.181 |
| 0.1 | 0 | clcB | 3 | 4.41667 | 0.198 |
| 0.1 | 0 | ycaM | 3 | 4.41667 | 0.194 |
| 0.1 | 0 | yadI | 3 | 4.41667 | 0.218 |
| 0.1 | 0 | AG1 | 3 | 4.41667 | 0.193 |
| 0.1 | 0 | ptsI | 3 | 4.66667 | 0.18 |
| 0.1 | 0 | clcB | 3 | 4.66667 | 0.198 |
| 0.1 | 0 | ycaM | 3 | 4.66667 | 0.187 |
| 0.1 | 0 | yadI | 3 | 4.66667 | 0.205 |
| 0.1 | 0 | AG1 | 3 | 4.66667 | 0.191 |
| 0.1 | 0 | ptsI | 3 | 4.91667 | 0.181 |
| 0.1 | 0 | clcB | 3 | 4.91667 | 0.199 |
| 0.1 | 0 | ycaM | 3 | 4.91667 | 0.188 |
| 0.1 | 0 | yadI | 3 | 4.91667 | 0.206 |
| 0.1 | 0 | AG1 | 3 | 4.91667 | 0.194 |
| 0.1 | 0 | ptsI | 3 | 5.16667 | 0.185 |
| 0.1 | 0 | clcB | 3 | 5.16667 | 0.204 |
| 0.1 | 0 | ycaM | 3 | 5.16667 | 0.193 |
| 0.1 | 0 | yadI | 3 | 5.16667 | 0.213 |

| | | | | | |
|---|---|---|---|---|---|
| 0.1 | 0 | AG1 | 3 | 5.16667 | 0.199 |
| 0.1 | 0 | ptsI | 3 | 5.41667 | 0.191 |
| 0.1 | 0 | clcB | 3 | 5.41667 | 0.21 |
| 0.1 | 0 | ycaM | 3 | 5.41667 | 0.201 |
| 0.1 | 0 | yadI | 3 | 5.41667 | 0.222 |
| 0.1 | 0 | AG1 | 3 | 5.41667 | 0.205 |
| 0.1 | 0 | ptsI | 3 | 5.66667 | 0.196 |
| 0.1 | 0 | clcB | 3 | 5.66667 | 0.216 |
| 0.1 | 0 | ycaM | 3 | 5.66667 | 0.207 |
| 0.1 | 0 | yadI | 3 | 5.66667 | 0.235 |
| 0.1 | 0 | AG1 | 3 | 5.66667 | 0.212 |
| 0.1 | 0 | ptsI | 3 | 5.91667 | 0.202 |
| 0.1 | 0 | clcB | 3 | 5.91667 | 0.223 |
| 0.1 | 0 | ycaM | 3 | 5.91667 | 0.216 |
| 0.1 | 0 | yadI | 3 | 5.91667 | 0.251 |
| 0.1 | 0 | AG1 | 3 | 5.91667 | 0.22 |
| 0.1 | 0 | ptsI | 3 | 6.16667 | 0.209 |
| 0.1 | 0 | clcB | 3 | 6.16667 | 0.232 |
| 0.1 | 0 | ycaM | 3 | 6.16667 | 0.226 |
| 0.1 | 0 | yadI | 3 | 6.16667 | 0.266 |
| 0.1 | 0 | AG1 | 3 | 6.16667 | 0.23 |
| 0.1 | 0 | ptsI | 3 | 6.41667 | 0.215 |
| 0.1 | 0 | clcB | 3 | 6.41667 | 0.243 |
| 0.1 | 0 | ycaM | 3 | 6.41667 | 0.238 |
| 0.1 | 0 | yadI | 3 | 6.41667 | 0.276 |
| 0.1 | 0 | AG1 | 3 | 6.41667 | 0.239 |
| 0.1 | 0 | ptsI | 3 | 6.66667 | 0.225 |
| 0.1 | 0 | clcB | 3 | 6.66667 | 0.254 |
| 0.1 | 0 | ycaM | 3 | 6.66667 | 0.251 |
| 0.1 | 0 | yadI | 3 | 6.66667 | 0.285 |
| 0.1 | 0 | AG1 | 3 | 6.66667 | 0.251 |
| 0.1 | 0 | ptsI | 3 | 6.91667 | 0.236 |
| 0.1 | 0 | clcB | 3 | 6.91667 | 0.266 |
| 0.1 | 0 | ycaM | 3 | 6.91667 | 0.264 |
| 0.1 | 0 | yadI | 3 | 6.91667 | 0.296 |
| 0.1 | 0 | AG1 | 3 | 6.91667 | 0.256 |
| 0.1 | 0 | ptsI | 3 | 7.16667 | 0.246 |
| 0.1 | 0 | clcB | 3 | 7.16667 | 0.278 |
| 0.1 | 0 | ycaM | 3 | 7.16667 | 0.274 |
| 0.1 | 0 | yadI | 3 | 7.16667 | 0.304 |
| 0.1 | 0 | AG1 | 3 | 7.16667 | 0.262 |
| 0.1 | 0 | ptsI | 3 | 7.41667 | 0.259 |
| 0.1 | 0 | clcB | 3 | 7.41667 | 0.291 |
| 0.1 | 0 | ycaM | 3 | 7.41667 | 0.281 |
| 0.1 | 0 | yadI | 3 | 7.41667 | 0.315 |
| 0.1 | 0 | AG1 | 3 | 7.41667 | 0.271 |
| 0.1 | 0 | ptsI | 3 | 7.66667 | 0.268 |
| 0.1 | 0 | clcB | 3 | 7.66667 | 0.301 |

| | | | | | |
|---|---|---|---|---|---|
| 0.1 | 0 | ycaM | 3 | 7.66667 | 0.289 |
| 0.1 | 0 | yadI | 3 | 7.66667 | 0.322 |
| 0.1 | 0 | AG1 | 3 | 7.66667 | 0.277 |
| 0.1 | 0 | ptsI | 3 | 7.91667 | 0.275 |
| 0.1 | 0 | clcB | 3 | 7.91667 | 0.308 |
| 0.1 | 0 | ycaM | 3 | 7.91667 | 0.297 |
| 0.1 | 0 | yadI | 3 | 7.91667 | 0.334 |
| 0.1 | 0 | AG1 | 3 | 7.91667 | 0.283 |
| 0.1 | 0 | ptsI | 3 | 8.16667 | 0.283 |
| 0.1 | 0 | clcB | 3 | 8.16667 | 0.316 |
| 0.1 | 0 | ycaM | 3 | 8.16667 | 0.304 |
| 0.1 | 0 | yadI | 3 | 8.16667 | 0.343 |
| 0.1 | 0 | AG1 | 3 | 8.16667 | 0.291 |
| 0.1 | 0 | ptsI | 3 | 8.41667 | 0.29 |
| 0.1 | 0 | clcB | 3 | 8.41667 | 0.324 |
| 0.1 | 0 | ycaM | 3 | 8.41667 | 0.313 |
| 0.1 | 0 | yadI | 3 | 8.41667 | 0.35 |
| 0.1 | 0 | AG1 | 3 | 8.41667 | 0.297 |
| 0.1 | 0 | ptsI | 3 | 9.05 | 0.28 |
| 0.1 | 0 | clcB | 3 | 9.05 | 0.291 |
| 0.1 | 0 | ycaM | 3 | 9.05 | 0.295 |
| 0.1 | 0 | yadI | 3 | 9.05 | 0.402 |
| 0.1 | 0 | AG1 | 3 | 9.05 | 0.335 |
| 0.1 | 0 | ptsI | 3 | 9.3 | 0.282 |
| 0.1 | 0 | clcB | 3 | 9.3 | 0.293 |
| 0.1 | 0 | ycaM | 3 | 9.3 | 0.299 |
| 0.1 | 0 | yadI | 3 | 9.3 | 0.406 |
| 0.1 | 0 | AG1 | 3 | 9.3 | 0.336 |
| 0.1 | 0 | ptsI | 3 | 9.55 | 0.29 |
| 0.1 | 0 | clcB | 3 | 9.55 | 0.3 |
| 0.1 | 0 | ycaM | 3 | 9.55 | 0.305 |
| 0.1 | 0 | yadI | 3 | 9.55 | 0.415 |
| 0.1 | 0 | AG1 | 3 | 9.55 | 0.345 |
| 0.1 | 0 | ptsI | 3 | 9.8 | 0.298 |
| 0.1 | 0 | clcB | 3 | 9.8 | 0.305 |
| 0.1 | 0 | ycaM | 3 | 9.8 | 0.312 |
| 0.1 | 0 | yadI | 3 | 9.8 | 0.423 |
| 0.1 | 0 | AG1 | 3 | 9.8 | 0.354 |
| 0.1 | 0 | ptsI | 3 | 10.05 | 0.305 |
| 0.1 | 0 | clcB | 3 | 10.05 | 0.313 |
| 0.1 | 0 | ycaM | 3 | 10.05 | 0.32 |
| 0.1 | 0 | yadI | 3 | 10.05 | 0.433 |
| 0.1 | 0 | AG1 | 3 | 10.05 | 0.365 |
| 0.1 | 0 | ptsI | 3 | 10.3 | 0.315 |
| 0.1 | 0 | clcB | 3 | 10.3 | 0.319 |
| 0.1 | 0 | ycaM | 3 | 10.3 | 0.328 |
| 0.1 | 0 | yadI | 3 | 10.3 | 0.444 |
| 0.1 | 0 | AG1 | 3 | 10.3 | 0.376 |

| | | | | | |
|---|---|---|---|---|---|
| 0.1 | 0 | ptsI | 3 | 10.55 | 0.322 |
| 0.1 | 0 | clcB | 3 | 10.55 | 0.328 |
| 0.1 | 0 | ycaM | 3 | 10.55 | 0.336 |
| 0.1 | 0 | yadI | 3 | 10.55 | 0.454 |
| 0.1 | 0 | AG1 | 3 | 10.55 | 0.383 |
| 0.1 | 0 | ptsI | 3 | 10.8 | 0.33 |
| 0.1 | 0 | clcB | 3 | 10.8 | 0.34 |
| 0.1 | 0 | ycaM | 3 | 10.8 | 0.348 |
| 0.1 | 0 | yadI | 3 | 10.8 | 0.462 |
| 0.1 | 0 | AG1 | 3 | 10.8 | 0.391 |
| 0.1 | 0 | ptsI | 3 | 11.05 | 0.339 |
| 0.1 | 0 | clcB | 3 | 11.05 | 0.349 |
| 0.1 | 0 | ycaM | 3 | 11.05 | 0.352 |
| 0.1 | 0 | yadI | 3 | 11.05 | 0.468 |
| 0.1 | 0 | AG1 | 3 | 11.05 | 0.398 |
| 0.1 | 0 | ptsI | 3 | 11.3 | 0.351 |
| 0.1 | 0 | clcB | 3 | 11.3 | 0.356 |
| 0.1 | 0 | ycaM | 3 | 11.3 | 0.359 |
| 0.1 | 0 | yadI | 3 | 11.3 | 0.48 |
| 0.1 | 0 | AG1 | 3 | 11.3 | 0.408 |
| 0.1 | 0 | ptsI | 3 | 11.55 | 0.356 |
| 0.1 | 0 | clcB | 3 | 11.55 | 0.366 |
| 0.1 | 0 | ycaM | 3 | 11.55 | 0.361 |
| 0.1 | 0 | yadI | 3 | 11.55 | 0.488 |
| 0.1 | 0 | AG1 | 3 | 11.55 | 0.415 |
| 0.1 | 0 | ptsI | 3 | 11.8 | 0.363 |
| 0.1 | 0 | clcB | 3 | 11.8 | 0.374 |
| 0.1 | 0 | ycaM | 3 | 11.8 | 0.366 |
| 0.1 | 0 | yadI | 3 | 11.8 | 0.491 |
| 0.1 | 0 | AG1 | 3 | 11.8 | 0.423 |
| 0.1 | 0 | ptsI | 3 | 12.05 | 0.376 |
| 0.1 | 0 | clcB | 3 | 12.05 | 0.376 |
| 0.1 | 0 | ycaM | 3 | 12.05 | 0.368 |
| 0.1 | 0 | yadI | 3 | 12.05 | 0.503 |
| 0.1 | 0 | AG1 | 3 | 12.05 | 0.432 |
| 0.1 | 0 | ptsI | 3 | 12.3 | 0.382 |
| 0.1 | 0 | clcB | 3 | 12.3 | 0.385 |
| 0.1 | 0 | ycaM | 3 | 12.3 | 0.374 |
| 0.1 | 0 | yadI | 3 | 12.3 | 0.506 |
| 0.1 | 0 | AG1 | 3 | 12.3 | 0.444 |
| 0.1 | 0 | ptsI | 3 | 12.55 | 0.391 |
| 0.1 | 0 | clcB | 3 | 12.55 | 0.392 |
| 0.1 | 0 | ycaM | 3 | 12.55 | 0.378 |
| 0.1 | 0 | yadI | 3 | 12.55 | 0.514 |
| 0.1 | 0 | AG1 | 3 | 12.55 | 0.454 |
| 0.1 | 0 | ptsI | 3 | 12.8 | 0.401 |
| 0.1 | 0 | clcB | 3 | 12.8 | 0.399 |
| 0.1 | 0 | ycaM | 3 | 12.8 | 0.383 |

| | | | | | |
|---|---|---|---|---|---|
| 0.1 | 0 | yadI | 3 | 12.8 | 0.522 |
| 0.1 | 0 | AG1 | 3 | 12.8 | 0.472 |
| 0.1 | 0 | ptsI | 3 | 13.05 | 0.415 |
| 0.1 | 0 | clcB | 3 | 13.05 | 0.407 |
| 0.1 | 0 | ycaM | 3 | 13.05 | 0.388 |
| 0.1 | 0 | yadI | 3 | 13.05 | 0.532 |
| 0.1 | 0 | AG1 | 3 | 13.05 | 0.482 |
| 0.1 | 0 | ptsI | 3 | 13.3 | 0.423 |
| 0.1 | 0 | clcB | 3 | 13.3 | 0.424 |
| 0.1 | 0 | ycaM | 3 | 13.3 | 0.397 |
| 0.1 | 0 | yadI | 3 | 13.3 | 0.532 |
| 0.1 | 0 | AG1 | 3 | 13.3 | 0.492 |
| 0.1 | 0 | ptsI | 3 | 13.55 | 0.435 |
| 0.1 | 0 | clcB | 3 | 13.55 | 0.431 |
| 0.1 | 0 | ycaM | 3 | 13.55 | 0.403 |
| 0.1 | 0 | yadI | 3 | 13.55 | 0.536 |
| 0.1 | 0 | AG1 | 3 | 13.55 | 0.508 |
| 0.1 | 0 | ptsI | 3 | 13.8 | 0.442 |
| 0.1 | 0 | clcB | 3 | 13.8 | 0.436 |
| 0.1 | 0 | ycaM | 3 | 13.8 | 0.414 |
| 0.1 | 0 | yadI | 3 | 13.8 | 0.539 |
| 0.1 | 0 | AG1 | 3 | 13.8 | 0.522 |
| 0.1 | 0 | ptsI | 3 | 14.05 | 0.459 |
| 0.1 | 0 | clcB | 3 | 14.05 | 0.447 |
| 0.1 | 0 | ycaM | 3 | 14.05 | 0.417 |
| 0.1 | 0 | yadI | 3 | 14.05 | 0.548 |
| 0.1 | 0 | AG1 | 3 | 14.05 | 0.533 |
| 0.1 | 0 | ptsI | 3 | 14.3 | 0.47 |
| 0.1 | 0 | clcB | 3 | 14.3 | 0.458 |
| 0.1 | 0 | ycaM | 3 | 14.3 | 0.422 |
| 0.1 | 0 | yadI | 3 | 14.3 | 0.552 |
| 0.1 | 0 | AG1 | 3 | 14.3 | 0.542 |
| 0.1 | 0 | ptsI | 3 | 14.55 | 0.48 |
| 0.1 | 0 | clcB | 3 | 14.55 | 0.466 |
| 0.1 | 0 | ycaM | 3 | 14.55 | 0.426 |
| 0.1 | 0 | yadI | 3 | 14.55 | 0.552 |
| 0.1 | 0 | AG1 | 3 | 14.55 | 0.546 |
| 0.1 | 0 | ptsI | 3 | 14.8 | 0.489 |
| 0.1 | 0 | clcB | 3 | 14.8 | 0.478 |
| 0.1 | 0 | ycaM | 3 | 14.8 | 0.435 |
| 0.1 | 0 | yadI | 3 | 14.8 | 0.552 |
| 0.1 | 0 | AG1 | 3 | 14.8 | 0.564 |
| 0.1 | 0 | ptsI | 3 | 15.05 | 0.501 |
| 0.1 | 0 | clcB | 3 | 15.05 | 0.49 |
| 0.1 | 0 | ycaM | 3 | 15.05 | 0.442 |
| 0.1 | 0 | yadI | 3 | 15.05 | 0.555 |
| 0.1 | 0 | AG1 | 3 | 15.05 | 0.576 |
| 0.1 | 0 | ptsI | 3 | 15.3 | 0.514 |

| | | | | | |
|---|---|---|---|---|---|
| 0.1 | 0 | clcB | 3 | 15.3 | 0.494 |
| 0.1 | 0 | ycaM | 3 | 15.3 | 0.444 |
| 0.1 | 0 | yadI | 3 | 15.3 | 0.556 |
| 0.1 | 0 | AG1 | 3 | 15.3 | 0.582 |
| 0.1 | 0 | ptsI | 3 | 15.55 | 0.525 |
| 0.1 | 0 | clcB | 3 | 15.55 | 0.502 |
| 0.1 | 0 | ycaM | 3 | 15.55 | 0.448 |
| 0.1 | 0 | yadI | 3 | 15.55 | 0.555 |
| 0.1 | 0 | AG1 | 3 | 15.55 | 0.591 |
| 0.1 | 0 | ptsI | 3 | 15.8 | 0.532 |
| 0.1 | 0 | clcB | 3 | 15.8 | 0.51 |
| 0.1 | 0 | ycaM | 3 | 15.8 | 0.451 |
| 0.1 | 0 | yadI | 3 | 15.8 | 0.558 |
| 0.1 | 0 | AG1 | 3 | 15.8 | 0.594 |
| 0.1 | 0 | ptsI | 3 | 16.05 | 0.546 |
| 0.1 | 0 | clcB | 3 | 16.05 | 0.518 |
| 0.1 | 0 | ycaM | 3 | 16.05 | 0.456 |
| 0.1 | 0 | yadI | 3 | 16.05 | 0.565 |
| 0.1 | 0 | AG1 | 3 | 16.05 | 0.607 |
| 0.1 | 0 | ptsI | 3 | 16.3 | 0.557 |
| 0.1 | 0 | clcB | 3 | 16.3 | 0.53 |
| 0.1 | 0 | ycaM | 3 | 16.3 | 0.457 |
| 0.1 | 0 | yadI | 3 | 16.3 | 0.563 |
| 0.1 | 0 | AG1 | 3 | 16.3 | 0.643 |
| 0.1 | 0 | ptsI | 3 | 16.55 | 0.564 |
| 0.1 | 0 | clcB | 3 | 16.55 | 0.529 |
| 0.1 | 0 | ycaM | 3 | 16.55 | 0.465 |
| 0.1 | 0 | yadI | 3 | 16.55 | 0.564 |
| 0.1 | 0 | AG1 | 3 | 16.55 | 0.637 |
| 0.1 | 0 | ptsI | 3 | 16.8 | 0.581 |
| 0.1 | 0 | clcB | 3 | 16.8 | 0.539 |
| 0.1 | 0 | ycaM | 3 | 16.8 | 0.47 |
| 0.1 | 0 | yadI | 3 | 16.8 | 0.559 |
| 0.1 | 0 | AG1 | 3 | 16.8 | 0.638 |
| 0.1 | 0 | ptsI | 3 | 17.05 | 0.594 |
| 0.1 | 0 | clcB | 3 | 17.05 | 0.542 |
| 0.1 | 0 | ycaM | 3 | 17.05 | 0.47 |
| 0.1 | 0 | yadI | 3 | 17.05 | 0.567 |
| 0.1 | 0 | AG1 | 3 | 17.05 | 0.629 |
| 0.1 | 0 | ptsI | 3 | 17.3 | 0.607 |
| 0.1 | 0 | clcB | 3 | 17.3 | 0.553 |
| 0.1 | 0 | ycaM | 3 | 17.3 | 0.478 |
| 0.1 | 0 | yadI | 3 | 17.3 | 0.569 |
| 0.1 | 0 | AG1 | 3 | 17.3 | 0.623 |
| 0.1 | 0 | ptsI | 3 | 17.55 | 0.646 |
| 0.1 | 0 | clcB | 3 | 17.55 | 0.56 |
| 0.1 | 0 | ycaM | 3 | 17.55 | 0.483 |
| 0.1 | 0 | yadI | 3 | 17.55 | 0.566 |

| | | | | | |
|---|---|---|---|---|---|
| 0.1 | 0 | AG1 | 3 | 17.55 | 0.625 |
| 0.1 | 0 | ptsI | 3 | 17.8 | 0.668 |
| 0.1 | 0 | clcB | 3 | 17.8 | 0.564 |
| 0.1 | 0 | ycaM | 3 | 17.8 | 0.485 |
| 0.1 | 0 | yadI | 3 | 17.8 | 0.562 |
| 0.1 | 0 | AG1 | 3 | 17.8 | 0.622 |
| 0.1 | 0 | ptsI | 3 | 18.05 | 0.688 |
| 0.1 | 0 | clcB | 3 | 18.05 | 0.579 |
| 0.1 | 0 | ycaM | 3 | 18.05 | 0.488 |
| 0.1 | 0 | yadI | 3 | 18.05 | 0.566 |
| 0.1 | 0 | AG1 | 3 | 18.05 | 0.639 |
| 0.1 | 0 | ptsI | 3 | 18.3 | 0.678 |
| 0.1 | 0 | clcB | 3 | 18.3 | 0.589 |
| 0.1 | 0 | ycaM | 3 | 18.3 | 0.496 |
| 0.1 | 0 | yadI | 3 | 18.3 | 0.564 |
| 0.1 | 0 | AG1 | 3 | 18.3 | 0.647 |
| 0.1 | 0 | ptsI | 3 | 18.55 | 0.674 |
| 0.1 | 0 | clcB | 3 | 18.55 | 0.585 |
| 0.1 | 0 | ycaM | 3 | 18.55 | 0.5 |
| 0.1 | 0 | yadI | 3 | 18.55 | 0.572 |
| 0.1 | 0 | AG1 | 3 | 18.55 | 0.645 |
| 0.1 | 0 | ptsI | 3 | 18.8 | 0.673 |
| 0.1 | 0 | clcB | 3 | 18.8 | 0.586 |
| 0.1 | 0 | ycaM | 3 | 18.8 | 0.505 |
| 0.1 | 0 | yadI | 3 | 18.8 | 0.569 |
| 0.1 | 0 | AG1 | 3 | 18.8 | 0.644 |
| 0.1 | 0 | ptsI | 3 | 19.05 | 0.698 |
| 0.1 | 0 | clcB | 3 | 19.05 | 0.595 |
| 0.1 | 0 | ycaM | 3 | 19.05 | 0.51 |
| 0.1 | 0 | yadI | 3 | 19.05 | 0.566 |
| 0.1 | 0 | AG1 | 3 | 19.05 | 0.642 |
| 0.1 | 0 | ptsI | 3 | 19.3 | 0.714 |
| 0.1 | 0 | clcB | 3 | 19.3 | 0.605 |
| 0.1 | 0 | ycaM | 3 | 19.3 | 0.513 |
| 0.1 | 0 | yadI | 3 | 19.3 | 0.57 |
| 0.1 | 0 | AG1 | 3 | 19.3 | 0.648 |
| 0.1 | 0 | ptsI | 3 | 19.55 | 0.738 |
| 0.1 | 0 | clcB | 3 | 19.55 | 0.628 |
| 0.1 | 0 | ycaM | 3 | 19.55 | 0.518 |
| 0.1 | 0 | yadI | 3 | 19.55 | 0.57 |
| 0.1 | 0 | AG1 | 3 | 19.55 | 0.663 |
| 0.1 | 0 | ptsI | 3 | 19.8 | 0.73 |
| 0.1 | 0 | clcB | 3 | 19.8 | 0.642 |
| 0.1 | 0 | ycaM | 3 | 19.8 | 0.52 |
| 0.1 | 0 | yadI | 3 | 19.8 | 0.569 |
| 0.1 | 0 | AG1 | 3 | 19.8 | 0.649 |
| 0.1 | 0 | ptsI | 3 | 20.05 | 0.729 |
| 0.1 | 0 | clcB | 3 | 20.05 | 0.642 |

| | | | | | |
|---|---|---|---|---|---|
| 0.1 | 0 | ycaM | 3 | 20.05 | 0.525 |
| 0.1 | 0 | yadI | 3 | 20.05 | 0.572 |
| 0.1 | 0 | AG1 | 3 | 20.05 | 0.648 |
| 0.1 | 0 | ptsI | 3 | 20.3 | 0.721 |
| 0.1 | 0 | clcB | 3 | 20.3 | 0.64 |
| 0.1 | 0 | ycaM | 3 | 20.3 | 0.528 |
| 0.1 | 0 | yadI | 3 | 20.3 | 0.572 |
| 0.1 | 0 | AG1 | 3 | 20.3 | 0.639 |
| 0.1 | 0 | ptsI | 3 | 20.55 | 0.73 |
| 0.1 | 0 | clcB | 3 | 20.55 | 0.654 |
| 0.1 | 0 | ycaM | 3 | 20.55 | 0.536 |
| 0.1 | 0 | yadI | 3 | 20.55 | 0.573 |
| 0.1 | 0 | AG1 | 3 | 20.55 | 0.639 |
| 0.1 | 0 | ptsI | 3 | 20.8 | 0.745 |
| 0.1 | 0 | clcB | 3 | 20.8 | 0.66 |
| 0.1 | 0 | ycaM | 3 | 20.8 | 0.541 |
| 0.1 | 0 | yadI | 3 | 20.8 | 0.57 |
| 0.1 | 0 | AG1 | 3 | 20.8 | 0.634 |
| 0.1 | 0 | ptsI | 3 | 21.05 | 0.759 |
| 0.1 | 0 | clcB | 3 | 21.05 | 0.662 |
| 0.1 | 0 | ycaM | 3 | 21.05 | 0.544 |
| 0.1 | 0 | yadI | 3 | 21.05 | 0.578 |
| 0.1 | 0 | AG1 | 3 | 21.05 | 0.637 |
| 0.1 | 0 | ptsI | 3 | 21.3 | 0.769 |
| 0.1 | 0 | clcB | 3 | 21.3 | 0.676 |
| 0.1 | 0 | ycaM | 3 | 21.3 | 0.548 |
| 0.1 | 0 | yadI | 3 | 21.3 | 0.57 |
| 0.1 | 0 | AG1 | 3 | 21.3 | 0.635 |
| 0.1 | 0 | ptsI | 3 | 21.55 | 0.788 |
| 0.1 | 0 | clcB | 3 | 21.55 | 0.684 |
| 0.1 | 0 | ycaM | 3 | 21.55 | 0.555 |
| 0.1 | 0 | yadI | 3 | 21.55 | 0.571 |
| 0.1 | 0 | AG1 | 3 | 21.55 | 0.637 |
| 0.1 | 0 | ptsI | 3 | 21.8 | 0.797 |
| 0.1 | 0 | clcB | 3 | 21.8 | 0.688 |
| 0.1 | 0 | ycaM | 3 | 21.8 | 0.556 |
| 0.1 | 0 | yadI | 3 | 21.8 | 0.567 |
| 0.1 | 0 | AG1 | 3 | 21.8 | 0.64 |
| 0.1 | 0 | ptsI | 3 | 22.05 | 0.808 |
| 0.1 | 0 | clcB | 3 | 22.05 | 0.696 |
| 0.1 | 0 | ycaM | 3 | 22.05 | 0.562 |
| 0.1 | 0 | yadI | 3 | 22.05 | 0.569 |
| 0.1 | 0 | AG1 | 3 | 22.05 | 0.643 |
| 0.1 | 0 | ptsI | 3 | 22.3 | 0.824 |
| 0.1 | 0 | clcB | 3 | 22.3 | 0.701 |
| 0.1 | 0 | ycaM | 3 | 22.3 | 0.563 |
| 0.1 | 0 | yadI | 3 | 22.3 | 0.577 |
| 0.1 | 0 | AG1 | 3 | 22.3 | 0.644 |

| | | | | | |
|---|---|---|---|---|---|
| 0.1 | 0 | ptsI | 3 | 22.55 | 0.833 |
| 0.1 | 0 | clcB | 3 | 22.55 | 0.707 |
| 0.1 | 0 | ycaM | 3 | 22.55 | 0.564 |
| 0.1 | 0 | yadI | 3 | 22.55 | 0.575 |
| 0.1 | 0 | AG1 | 3 | 22.55 | 0.648 |
| 0.1 | 0 | ptsI | 3 | 22.8 | 0.838 |
| 0.1 | 0 | clcB | 3 | 22.8 | 0.702 |
| 0.1 | 0 | ycaM | 3 | 22.8 | 0.565 |
| 0.1 | 0 | yadI | 3 | 22.8 | 0.566 |
| 0.1 | 0 | AG1 | 3 | 22.8 | 0.649 |
| 0.1 | 0 | ptsI | 3 | 23.05 | 0.848 |
| 0.1 | 0 | clcB | 3 | 23.05 | 0.717 |
| 0.1 | 0 | ycaM | 3 | 23.05 | 0.566 |
| 0.1 | 0 | yadI | 3 | 23.05 | 0.567 |
| 0.1 | 0 | AG1 | 3 | 23.05 | 0.65 |
| 0.1 | 0 | ptsI | 3 | 23.3 | 0.856 |
| 0.1 | 0 | clcB | 3 | 23.3 | 0.716 |
| 0.1 | 0 | ycaM | 3 | 23.3 | 0.567 |
| 0.1 | 0 | yadI | 3 | 23.3 | 0.563 |
| 0.1 | 0 | AG1 | 3 | 23.3 | 0.651 |
| 0.1 | 0 | ptsI | 3 | 23.55 | 0.867 |
| 0.1 | 0 | clcB | 3 | 23.55 | 0.735 |
| 0.1 | 0 | ycaM | 3 | 23.55 | 0.573 |
| 0.1 | 0 | yadI | 3 | 23.55 | 0.564 |
| 0.1 | 0 | AG1 | 3 | 23.55 | 0.657 |
| 0.1 | 0 | ptsI | 3 | 23.8 | 0.871 |
| 0.1 | 0 | clcB | 3 | 23.8 | 0.727 |
| 0.1 | 0 | ycaM | 3 | 23.8 | 0.572 |
| 0.1 | 0 | yadI | 3 | 23.8 | 0.562 |
| 0.1 | 0 | AG1 | 3 | 23.8 | 0.652 |
| 0.1 | 0 | ptsI | 3 | 24.05 | 0.878 |
| 0.1 | 0 | clcB | 3 | 24.05 | 0.74 |
| 0.1 | 0 | ycaM | 3 | 24.05 | 0.573 |
| 0.1 | 0 | yadI | 3 | 24.05 | 0.558 |
| 0.1 | 0 | AG1 | 3 | 24.05 | 0.657 |
| 0.1 | 0 | ptsI | 3 | 24.3 | 0.876 |
| 0.1 | 0 | clcB | 3 | 24.3 | 0.755 |
| 0.1 | 0 | ycaM | 3 | 24.3 | 0.57 |
| 0.1 | 0 | yadI | 3 | 24.3 | 0.559 |
| 0.1 | 0 | AG1 | 3 | 24.3 | 0.658 |
| 0.1 | 0 | ptsI | 4 | 0 | 0.192 |
| 0.1 | 0 | clcB | 4 | 0 | 0.196 |
| 0.1 | 0 | ycaM | 4 | 0 | 0.204 |
| 0.1 | 0 | yadI | 4 | 0 | 0.225 |
| 0.1 | 0 | AG1 | 4 | 0 | 0.214 |
| 0.1 | 0 | ptsI | 4 | 0.25 | 0.194 |
| 0.1 | 0 | clcB | 4 | 0.25 | 0.194 |
| 0.1 | 0 | ycaM | 4 | 0.25 | 0.196 |

| | | | | | |
|---|---|---|---|---|---|
| 0.1 | 0 | yadI | 4 | 0.25 | 0.223 |
| 0.1 | 0 | AG1 | 4 | 0.25 | 0.214 |
| 0.1 | 0 | ptsI | 4 | 0.5 | 0.195 |
| 0.1 | 0 | clcB | 4 | 0.5 | 0.194 |
| 0.1 | 0 | ycaM | 4 | 0.5 | 0.199 |
| 0.1 | 0 | yadI | 4 | 0.5 | 0.225 |
| 0.1 | 0 | AG1 | 4 | 0.5 | 0.217 |
| 0.1 | 0 | ptsI | 4 | 0.75 | 0.199 |
| 0.1 | 0 | clcB | 4 | 0.75 | 0.197 |
| 0.1 | 0 | ycaM | 4 | 0.75 | 0.202 |
| 0.1 | 0 | yadI | 4 | 0.75 | 0.23 |
| 0.1 | 0 | AG1 | 4 | 0.75 | 0.22 |
| 0.1 | 0 | ptsI | 4 | 1 | 0.203 |
| 0.1 | 0 | clcB | 4 | 1 | 0.2 |
| 0.1 | 0 | ycaM | 4 | 1 | 0.206 |
| 0.1 | 0 | yadI | 4 | 1 | 0.232 |
| 0.1 | 0 | AG1 | 4 | 1 | 0.223 |
| 0.1 | 0 | ptsI | 4 | 1.25 | 0.21 |
| 0.1 | 0 | clcB | 4 | 1.25 | 0.21 |
| 0.1 | 0 | ycaM | 4 | 1.25 | 0.212 |
| 0.1 | 0 | yadI | 4 | 1.25 | 0.238 |
| 0.1 | 0 | AG1 | 4 | 1.25 | 0.23 |
| 0.1 | 0 | ptsI | 4 | 1.5 | 0.216 |
| 0.1 | 0 | clcB | 4 | 1.5 | 0.213 |
| 0.1 | 0 | ycaM | 4 | 1.5 | 0.212 |
| 0.1 | 0 | yadI | 4 | 1.5 | 0.243 |
| 0.1 | 0 | AG1 | 4 | 1.5 | 0.233 |
| 0.1 | 0 | ptsI | 4 | 1.75 | 0.226 |
| 0.1 | 0 | clcB | 4 | 1.75 | 0.226 |
| 0.1 | 0 | ycaM | 4 | 1.75 | 0.224 |
| 0.1 | 0 | yadI | 4 | 1.75 | 0.252 |
| 0.1 | 0 | AG1 | 4 | 1.75 | 0.242 |
| 0.1 | 0 | ptsI | 4 | 2 | 0.235 |
| 0.1 | 0 | clcB | 4 | 2 | 0.24 |
| 0.1 | 0 | ycaM | 4 | 2 | 0.233 |
| 0.1 | 0 | yadI | 4 | 2 | 0.266 |
| 0.1 | 0 | AG1 | 4 | 2 | 0.252 |
| 0.1 | 0 | ptsI | 4 | 2.25 | 0.248 |
| 0.1 | 0 | clcB | 4 | 2.25 | 0.253 |
| 0.1 | 0 | ycaM | 4 | 2.25 | 0.246 |
| 0.1 | 0 | yadI | 4 | 2.25 | 0.278 |
| 0.1 | 0 | AG1 | 4 | 2.25 | 0.267 |
| 0.1 | 0 | ptsI | 4 | 2.5 | 0.262 |
| 0.1 | 0 | clcB | 4 | 2.5 | 0.262 |
| 0.1 | 0 | ycaM | 4 | 2.5 | 0.253 |
| 0.1 | 0 | yadI | 4 | 2.5 | 0.29 |
| 0.1 | 0 | AG1 | 4 | 2.5 | 0.274 |
| 0.1 | 0 | ptsI | 4 | 2.75 | 0.276 |

| | | | | | |
|---|---|---|---|---|---|
| 0.1 | 0 | clcB | 4 | 2.75 | 0.28 |
| 0.1 | 0 | ycaM | 4 | 2.75 | 0.266 |
| 0.1 | 0 | yadI | 4 | 2.75 | 0.301 |
| 0.1 | 0 | AG1 | 4 | 2.75 | 0.282 |
| 0.1 | 0 | ptsI | 4 | 3 | 0.292 |
| 0.1 | 0 | clcB | 4 | 3 | 0.295 |
| 0.1 | 0 | ycaM | 4 | 3 | 0.284 |
| 0.1 | 0 | yadI | 4 | 3 | 0.321 |
| 0.1 | 0 | AG1 | 4 | 3 | 0.301 |
| 0.1 | 0 | ptsI | 4 | 3.25 | 0.306 |
| 0.1 | 0 | clcB | 4 | 3.25 | 0.311 |
| 0.1 | 0 | ycaM | 4 | 3.25 | 0.298 |
| 0.1 | 0 | yadI | 4 | 3.25 | 0.341 |
| 0.1 | 0 | AG1 | 4 | 3.25 | 0.316 |
| 0.1 | 0 | ptsI | 4 | 3.5 | 0.325 |
| 0.1 | 0 | clcB | 4 | 3.5 | 0.325 |
| 0.1 | 0 | ycaM | 4 | 3.5 | 0.316 |
| 0.1 | 0 | yadI | 4 | 3.5 | 0.362 |
| 0.1 | 0 | AG1 | 4 | 3.5 | 0.334 |
| 0.1 | 0 | ptsI | 4 | 3.75 | 0.341 |
| 0.1 | 0 | clcB | 4 | 3.75 | 0.343 |
| 0.1 | 0 | ycaM | 4 | 3.75 | 0.325 |
| 0.1 | 0 | yadI | 4 | 3.75 | 0.382 |
| 0.1 | 0 | AG1 | 4 | 3.75 | 0.346 |
| 0.1 | 0 | ptsI | 4 | 4 | 0.361 |
| 0.1 | 0 | clcB | 4 | 4 | 0.354 |
| 0.1 | 0 | ycaM | 4 | 4 | 0.343 |
| 0.1 | 0 | yadI | 4 | 4 | 0.403 |
| 0.1 | 0 | AG1 | 4 | 4 | 0.369 |
| 0.1 | 0 | ptsI | 4 | 4.35 | 0.36 |
| 0.1 | 0 | clcB | 4 | 4.35 | 0.318 |
| 0.1 | 0 | ycaM | 4 | 4.35 | 0.373 |
| 0.1 | 0 | yadI | 4 | 4.35 | 0.412 |
| 0.1 | 0 | AG1 | 4 | 4.35 | 0.41 |
| 0.1 | 0 | ptsI | 4 | 4.6 | 0.364 |
| 0.1 | 0 | clcB | 4 | 4.6 | 0.303 |
| 0.1 | 0 | ycaM | 4 | 4.6 | 0.34 |
| 0.1 | 0 | yadI | 4 | 4.6 | 0.386 |
| 0.1 | 0 | AG1 | 4 | 4.6 | 0.392 |
| 0.1 | 0 | ptsI | 4 | 4.85 | 0.377 |
| 0.1 | 0 | clcB | 4 | 4.85 | 0.303 |
| 0.1 | 0 | ycaM | 4 | 4.85 | 0.345 |
| 0.1 | 0 | yadI | 4 | 4.85 | 0.397 |
| 0.1 | 0 | AG1 | 4 | 4.85 | 0.401 |
| 0.1 | 0 | ptsI | 4 | 5.1 | 0.392 |
| 0.1 | 0 | clcB | 4 | 5.1 | 0.312 |
| 0.1 | 0 | ycaM | 4 | 5.1 | 0.356 |
| 0.1 | 0 | yadI | 4 | 5.1 | 0.41 |

| | | | | | |
|---|---|---|---|---|---|
| 0.1 | 0 | AG1 | 4 | 5.1 | 0.417 |
| 0.1 | 0 | ptsI | 4 | 5.35 | 0.414 |
| 0.1 | 0 | clcB | 4 | 5.35 | 0.321 |
| 0.1 | 0 | ycaM | 4 | 5.35 | 0.365 |
| 0.1 | 0 | yadI | 4 | 5.35 | 0.426 |
| 0.1 | 0 | AG1 | 4 | 5.35 | 0.431 |
| 0.1 | 0 | ptsI | 4 | 5.6 | 0.425 |
| 0.1 | 0 | clcB | 4 | 5.6 | 0.331 |
| 0.1 | 0 | ycaM | 4 | 5.6 | 0.376 |
| 0.1 | 0 | yadI | 4 | 5.6 | 0.443 |
| 0.1 | 0 | AG1 | 4 | 5.6 | 0.441 |
| 0.1 | 0 | ptsI | 4 | 5.85 | 0.434 |
| 0.1 | 0 | clcB | 4 | 5.85 | 0.346 |
| 0.1 | 0 | ycaM | 4 | 5.85 | 0.385 |
| 0.1 | 0 | yadI | 4 | 5.85 | 0.456 |
| 0.1 | 0 | AG1 | 4 | 5.85 | 0.451 |
| 0.1 | 0 | ptsI | 4 | 6.1 | 0.449 |
| 0.1 | 0 | clcB | 4 | 6.1 | 0.361 |
| 0.1 | 0 | ycaM | 4 | 6.1 | 0.396 |
| 0.1 | 0 | yadI | 4 | 6.1 | 0.473 |
| 0.1 | 0 | AG1 | 4 | 6.1 | 0.466 |
| 0.1 | 0 | ptsI | 4 | 6.35 | 0.462 |
| 0.1 | 0 | clcB | 4 | 6.35 | 0.371 |
| 0.1 | 0 | ycaM | 4 | 6.35 | 0.405 |
| 0.1 | 0 | yadI | 4 | 6.35 | 0.496 |
| 0.1 | 0 | AG1 | 4 | 6.35 | 0.486 |
| 0.1 | 0 | ptsI | 4 | 6.6 | 0.477 |
| 0.1 | 0 | clcB | 4 | 6.6 | 0.386 |
| 0.1 | 0 | ycaM | 4 | 6.6 | 0.414 |
| 0.1 | 0 | yadI | 4 | 6.6 | 0.512 |
| 0.1 | 0 | AG1 | 4 | 6.6 | 0.492 |
| 0.1 | 0 | ptsI | 4 | 6.85 | 0.486 |
| 0.1 | 0 | clcB | 4 | 6.85 | 0.397 |
| 0.1 | 0 | ycaM | 4 | 6.85 | 0.425 |
| 0.1 | 0 | yadI | 4 | 6.85 | 0.528 |
| 0.1 | 0 | AG1 | 4 | 6.85 | 0.507 |
| 0.1 | 0 | ptsI | 4 | 7.1 | 0.497 |
| 0.1 | 0 | clcB | 4 | 7.1 | 0.408 |
| 0.1 | 0 | ycaM | 4 | 7.1 | 0.432 |
| 0.1 | 0 | yadI | 4 | 7.1 | 0.541 |
| 0.1 | 0 | AG1 | 4 | 7.1 | 0.525 |
| 0.1 | 0 | ptsI | 4 | 7.35 | 0.505 |
| 0.1 | 0 | clcB | 4 | 7.35 | 0.417 |
| 0.1 | 0 | ycaM | 4 | 7.35 | 0.44 |
| 0.1 | 0 | yadI | 4 | 7.35 | 0.549 |
| 0.1 | 0 | AG1 | 4 | 7.35 | 0.538 |
| 0.1 | 0 | ptsI | 4 | 7.6 | 0.513 |
| 0.1 | 0 | clcB | 4 | 7.6 | 0.428 |

| | | | | | |
|---|---|---|---|---|---|
| 0.1 | 0 | ycaM | 4 | 7.6 | 0.451 |
| 0.1 | 0 | yadI | 4 | 7.6 | 0.56 |
| 0.1 | 0 | AG1 | 4 | 7.6 | 0.544 |
| 0.1 | 0 | ptsI | 4 | 7.85 | 0.519 |
| 0.1 | 0 | clcB | 4 | 7.85 | 0.433 |
| 0.1 | 0 | ycaM | 4 | 7.85 | 0.457 |
| 0.1 | 0 | yadI | 4 | 7.85 | 0.57 |
| 0.1 | 0 | AG1 | 4 | 7.85 | 0.558 |
| 0.1 | 0 | ptsI | 4 | 8.1 | 0.529 |
| 0.1 | 0 | clcB | 4 | 8.1 | 0.444 |
| 0.1 | 0 | ycaM | 4 | 8.1 | 0.467 |
| 0.1 | 0 | yadI | 4 | 8.1 | 0.584 |
| 0.1 | 0 | AG1 | 4 | 8.1 | 0.568 |
| 0.1 | 0 | ptsI | 4 | 8.35 | 0.538 |
| 0.1 | 0 | clcB | 4 | 8.35 | 0.447 |
| 0.1 | 0 | ycaM | 4 | 8.35 | 0.472 |
| 0.1 | 0 | yadI | 4 | 8.35 | 0.593 |
| 0.1 | 0 | AG1 | 4 | 8.35 | 0.581 |
| 0.1 | 0 | ptsI | 4 | 8.83333 | 0.378 |
| 0.1 | 0 | clcB | 4 | 8.83333 | 0.414 |
| 0.1 | 0 | ycaM | 4 | 8.83333 | 0.397 |
| 0.1 | 0 | yadI | 4 | 8.83333 | 0.499 |
| 0.1 | 0 | AG1 | 4 | 8.83333 | 0.533 |
| 0.1 | 0 | ptsI | 4 | 9.08333 | 0.382 |
| 0.1 | 0 | clcB | 4 | 9.08333 | 0.41 |
| 0.1 | 0 | ycaM | 4 | 9.08333 | 0.398 |
| 0.1 | 0 | yadI | 4 | 9.08333 | 0.496 |
| 0.1 | 0 | AG1 | 4 | 9.08333 | 0.539 |
| 0.1 | 0 | ptsI | 4 | 9.33333 | 0.389 |
| 0.1 | 0 | clcB | 4 | 9.33333 | 0.413 |
| 0.1 | 0 | ycaM | 4 | 9.33333 | 0.399 |
| 0.1 | 0 | yadI | 4 | 9.33333 | 0.503 |
| 0.1 | 0 | AG1 | 4 | 9.33333 | 0.549 |
| 0.1 | 0 | ptsI | 4 | 9.58333 | 0.399 |
| 0.1 | 0 | clcB | 4 | 9.58333 | 0.422 |
| 0.1 | 0 | ycaM | 4 | 9.58333 | 0.4 |
| 0.1 | 0 | yadI | 4 | 9.58333 | 0.513 |
| 0.1 | 0 | AG1 | 4 | 9.58333 | 0.562 |
| 0.1 | 0 | ptsI | 4 | 9.83333 | 0.407 |
| 0.1 | 0 | clcB | 4 | 9.83333 | 0.428 |
| 0.1 | 0 | ycaM | 4 | 9.83333 | 0.404 |
| 0.1 | 0 | yadI | 4 | 9.83333 | 0.522 |
| 0.1 | 0 | AG1 | 4 | 9.83333 | 0.571 |
| 0.1 | 0 | ptsI | 4 | 10.0833 | 0.417 |
| 0.1 | 0 | clcB | 4 | 10.0833 | 0.435 |
| 0.1 | 0 | ycaM | 4 | 10.0833 | 0.405 |
| 0.1 | 0 | yadI | 4 | 10.0833 | 0.529 |
| 0.1 | 0 | AG1 | 4 | 10.0833 | 0.576 |

| | | | | | |
|---|---|---|---|---|---|
| 0.1 | 0 | ptsI | 4 | 10.3333 | 0.422 |
| 0.1 | 0 | clcB | 4 | 10.3333 | 0.441 |
| 0.1 | 0 | ycaM | 4 | 10.3333 | 0.409 |
| 0.1 | 0 | yadI | 4 | 10.3333 | 0.536 |
| 0.1 | 0 | AG1 | 4 | 10.3333 | 0.584 |
| 0.1 | 0 | ptsI | 4 | 10.5833 | 0.428 |
| 0.1 | 0 | clcB | 4 | 10.5833 | 0.449 |
| 0.1 | 0 | ycaM | 4 | 10.5833 | 0.415 |
| 0.1 | 0 | yadI | 4 | 10.5833 | 0.543 |
| 0.1 | 0 | AG1 | 4 | 10.5833 | 0.592 |
| 0.1 | 0 | ptsI | 4 | 10.8333 | 0.432 |
| 0.1 | 0 | clcB | 4 | 10.8333 | 0.453 |
| 0.1 | 0 | ycaM | 4 | 10.8333 | 0.418 |
| 0.1 | 0 | yadI | 4 | 10.8333 | 0.548 |
| 0.1 | 0 | AG1 | 4 | 10.8333 | 0.603 |
| 0.1 | 0 | ptsI | 4 | 11.0833 | 0.437 |
| 0.1 | 0 | clcB | 4 | 11.0833 | 0.456 |
| 0.1 | 0 | ycaM | 4 | 11.0833 | 0.421 |
| 0.1 | 0 | yadI | 4 | 11.0833 | 0.556 |
| 0.1 | 0 | AG1 | 4 | 11.0833 | 0.61 |
| 0.1 | 0 | ptsI | 4 | 11.3333 | 0.445 |
| 0.1 | 0 | clcB | 4 | 11.3333 | 0.465 |
| 0.1 | 0 | ycaM | 4 | 11.3333 | 0.429 |
| 0.1 | 0 | yadI | 4 | 11.3333 | 0.563 |
| 0.1 | 0 | AG1 | 4 | 11.3333 | 0.62 |
| 0.1 | 0 | ptsI | 4 | 11.5833 | 0.451 |
| 0.1 | 0 | clcB | 4 | 11.5833 | 0.466 |
| 0.1 | 0 | ycaM | 4 | 11.5833 | 0.431 |
| 0.1 | 0 | yadI | 4 | 11.5833 | 0.569 |
| 0.1 | 0 | AG1 | 4 | 11.5833 | 0.624 |
| 0.1 | 0 | ptsI | 4 | 11.8333 | 0.458 |
| 0.1 | 0 | clcB | 4 | 11.8333 | 0.47 |
| 0.1 | 0 | ycaM | 4 | 11.8333 | 0.434 |
| 0.1 | 0 | yadI | 4 | 11.8333 | 0.577 |
| 0.1 | 0 | AG1 | 4 | 11.8333 | 0.632 |
| 0.1 | 0 | ptsI | 4 | 12.0833 | 0.462 |
| 0.1 | 0 | clcB | 4 | 12.0833 | 0.473 |
| 0.1 | 0 | ycaM | 4 | 12.0833 | 0.438 |
| 0.1 | 0 | yadI | 4 | 12.0833 | 0.582 |
| 0.1 | 0 | AG1 | 4 | 12.0833 | 0.636 |
| 0.1 | 0 | ptsI | 4 | 12.3333 | 0.472 |
| 0.1 | 0 | clcB | 4 | 12.3333 | 0.48 |
| 0.1 | 0 | ycaM | 4 | 12.3333 | 0.441 |
| 0.1 | 0 | yadI | 4 | 12.3333 | 0.584 |
| 0.1 | 0 | AG1 | 4 | 12.3333 | 0.644 |
| 0.1 | 0 | ptsI | 4 | 12.5833 | 0.481 |
| 0.1 | 0 | clcB | 4 | 12.5833 | 0.483 |
| 0.1 | 0 | ycaM | 4 | 12.5833 | 0.443 |

| | | | | | |
|---|---|---|---|---|---|
| 0.1 | 0 | yadI | 4 | 12.5833 | 0.591 |
| 0.1 | 0 | AG1 | 4 | 12.5833 | 0.648 |
| 0.1 | 0 | ptsI | 4 | 12.8333 | 0.484 |
| 0.1 | 0 | clcB | 4 | 12.8333 | 0.484 |
| 0.1 | 0 | ycaM | 4 | 12.8333 | 0.449 |
| 0.1 | 0 | yadI | 4 | 12.8333 | 0.598 |
| 0.1 | 0 | AG1 | 4 | 12.8333 | 0.652 |
| 0.1 | 0 | ptsI | 4 | 13.0833 | 0.49 |
| 0.1 | 0 | clcB | 4 | 13.0833 | 0.49 |
| 0.1 | 0 | ycaM | 4 | 13.0833 | 0.452 |
| 0.1 | 0 | yadI | 4 | 13.0833 | 0.6 |
| 0.1 | 0 | AG1 | 4 | 13.0833 | 0.659 |
| 0.1 | 0 | ptsI | 4 | 13.3333 | 0.499 |
| 0.1 | 0 | clcB | 4 | 13.3333 | 0.493 |
| 0.1 | 0 | ycaM | 4 | 13.3333 | 0.455 |
| 0.1 | 0 | yadI | 4 | 13.3333 | 0.604 |
| 0.1 | 0 | AG1 | 4 | 13.3333 | 0.666 |
| 0.1 | 0 | ptsI | 4 | 13.5833 | 0.503 |
| 0.1 | 0 | clcB | 4 | 13.5833 | 0.494 |
| 0.1 | 0 | ycaM | 4 | 13.5833 | 0.459 |
| 0.1 | 0 | yadI | 4 | 13.5833 | 0.61 |
| 0.1 | 0 | AG1 | 4 | 13.5833 | 0.672 |
| 0.1 | 0 | ptsI | 4 | 13.8333 | 0.505 |
| 0.1 | 0 | clcB | 4 | 13.8333 | 0.5 |
| 0.1 | 0 | ycaM | 4 | 13.8333 | 0.464 |
| 0.1 | 0 | yadI | 4 | 13.8333 | 0.615 |
| 0.1 | 0 | AG1 | 4 | 13.8333 | 0.676 |
| 0.1 | 0 | ptsI | 4 | 14.0833 | 0.512 |
| 0.1 | 0 | clcB | 4 | 14.0833 | 0.502 |
| 0.1 | 0 | ycaM | 4 | 14.0833 | 0.467 |
| 0.1 | 0 | yadI | 4 | 14.0833 | 0.618 |
| 0.1 | 0 | AG1 | 4 | 14.0833 | 0.683 |
| 0.1 | 0 | ptsI | 4 | 14.3333 | 0.518 |
| 0.1 | 0 | clcB | 4 | 14.3333 | 0.509 |
| 0.1 | 0 | ycaM | 4 | 14.3333 | 0.471 |
| 0.1 | 0 | yadI | 4 | 14.3333 | 0.626 |
| 0.1 | 0 | AG1 | 4 | 14.3333 | 0.683 |
| 0.1 | 0 | ptsI | 4 | 14.5833 | 0.52 |
| 0.1 | 0 | clcB | 4 | 14.5833 | 0.506 |
| 0.1 | 0 | ycaM | 4 | 14.5833 | 0.471 |
| 0.1 | 0 | yadI | 4 | 14.5833 | 0.626 |
| 0.1 | 0 | AG1 | 4 | 14.5833 | 0.686 |
| 0.1 | 0 | ptsI | 4 | 14.8333 | 0.526 |
| 0.1 | 0 | clcB | 4 | 14.8333 | 0.51 |
| 0.1 | 0 | ycaM | 4 | 14.8333 | 0.478 |
| 0.1 | 0 | yadI | 4 | 14.8333 | 0.629 |
| 0.1 | 0 | AG1 | 4 | 14.8333 | 0.693 |
| 0.1 | 0 | ptsI | 4 | 15.0833 | 0.531 |

| | | | | | |
|---|---|---|---|---|---|
| 0.1 | 0 | clcB | 4 | 15.0833 | 0.51 |
| 0.1 | 0 | ycaM | 4 | 15.0833 | 0.479 |
| 0.1 | 0 | yadI | 4 | 15.0833 | 0.634 |
| 0.1 | 0 | AG1 | 4 | 15.0833 | 0.694 |
| 0.1 | 0 | ptsI | 4 | 15.3333 | 0.534 |
| 0.1 | 0 | clcB | 4 | 15.3333 | 0.514 |
| 0.1 | 0 | ycaM | 4 | 15.3333 | 0.484 |
| 0.1 | 0 | yadI | 4 | 15.3333 | 0.637 |
| 0.1 | 0 | AG1 | 4 | 15.3333 | 0.697 |
| 0.1 | 0 | ptsI | 4 | 15.5833 | 0.543 |
| 0.1 | 0 | clcB | 4 | 15.5833 | 0.516 |
| 0.1 | 0 | ycaM | 4 | 15.5833 | 0.487 |
| 0.1 | 0 | yadI | 4 | 15.5833 | 0.644 |
| 0.1 | 0 | AG1 | 4 | 15.5833 | 0.701 |
| 0.1 | 0 | ptsI | 4 | 15.8333 | 0.54 |
| 0.1 | 0 | clcB | 4 | 15.8333 | 0.516 |
| 0.1 | 0 | ycaM | 4 | 15.8333 | 0.488 |
| 0.1 | 0 | yadI | 4 | 15.8333 | 0.648 |
| 0.1 | 0 | AG1 | 4 | 15.8333 | 0.704 |
| 0.1 | 0 | ptsI | 4 | 16.0833 | 0.548 |
| 0.1 | 0 | clcB | 4 | 16.0833 | 0.521 |
| 0.1 | 0 | ycaM | 4 | 16.0833 | 0.492 |
| 0.1 | 0 | yadI | 4 | 16.0833 | 0.65 |
| 0.1 | 0 | AG1 | 4 | 16.0833 | 0.707 |
| 0.1 | 0 | ptsI | 4 | 16.3333 | 0.546 |
| 0.1 | 0 | clcB | 4 | 16.3333 | 0.517 |
| 0.1 | 0 | ycaM | 4 | 16.3333 | 0.494 |
| 0.1 | 0 | yadI | 4 | 16.3333 | 0.651 |
| 0.1 | 0 | AG1 | 4 | 16.3333 | 0.71 |
| 0.1 | 0 | ptsI | 4 | 16.5833 | 0.551 |
| 0.1 | 0 | clcB | 4 | 16.5833 | 0.516 |
| 0.1 | 0 | ycaM | 4 | 16.5833 | 0.496 |
| 0.1 | 0 | yadI | 4 | 16.5833 | 0.658 |
| 0.1 | 0 | AG1 | 4 | 16.5833 | 0.714 |
| 0.1 | 0 | ptsI | 4 | 16.8333 | 0.559 |
| 0.1 | 0 | clcB | 4 | 16.8333 | 0.52 |
| 0.1 | 0 | ycaM | 4 | 16.8333 | 0.495 |
| 0.1 | 0 | yadI | 4 | 16.8333 | 0.658 |
| 0.1 | 0 | AG1 | 4 | 16.8333 | 0.712 |
| 0.1 | 0 | ptsI | 4 | 17.0833 | 0.562 |
| 0.1 | 0 | clcB | 4 | 17.0833 | 0.519 |
| 0.1 | 0 | ycaM | 4 | 17.0833 | 0.493 |
| 0.1 | 0 | yadI | 4 | 17.0833 | 0.662 |
| 0.1 | 0 | AG1 | 4 | 17.0833 | 0.715 |
| 0.1 | 0 | ptsI | 4 | 17.3333 | 0.559 |
| 0.1 | 0 | clcB | 4 | 17.3333 | 0.523 |
| 0.1 | 0 | ycaM | 4 | 17.3333 | 0.501 |
| 0.1 | 0 | yadI | 4 | 17.3333 | 0.67 |

| | | | | | |
|---|---|---|---|---|---|
| 0.1 | 0 | AG1 | 4 | 17.3333 | 0.717 |
| 0.1 | 0 | ptsI | 4 | 17.5833 | 0.566 |
| 0.1 | 0 | clcB | 4 | 17.5833 | 0.523 |
| 0.1 | 0 | ycaM | 4 | 17.5833 | 0.501 |
| 0.1 | 0 | yadI | 4 | 17.5833 | 0.668 |
| 0.1 | 0 | AG1 | 4 | 17.5833 | 0.722 |
| 0.1 | 0 | ptsI | 4 | 17.8333 | 0.58 |
| 0.1 | 0 | clcB | 4 | 17.8333 | 0.525 |
| 0.1 | 0 | ycaM | 4 | 17.8333 | 0.497 |
| 0.1 | 0 | yadI | 4 | 17.8333 | 0.668 |
| 0.1 | 0 | AG1 | 4 | 17.8333 | 0.729 |
| 0.1 | 0 | ptsI | 4 | 18.0833 | 0.575 |
| 0.1 | 0 | clcB | 4 | 18.0833 | 0.525 |
| 0.1 | 0 | ycaM | 4 | 18.0833 | 0.504 |
| 0.1 | 0 | yadI | 4 | 18.0833 | 0.676 |
| 0.1 | 0 | AG1 | 4 | 18.0833 | 0.726 |
| 0.1 | 0 | ptsI | 4 | 18.3333 | 0.573 |
| 0.1 | 0 | clcB | 4 | 18.3333 | 0.526 |
| 0.1 | 0 | ycaM | 4 | 18.3333 | 0.505 |
| 0.1 | 0 | yadI | 4 | 18.3333 | 0.68 |
| 0.1 | 0 | AG1 | 4 | 18.3333 | 0.728 |
| 0.1 | 0 | ptsI | 4 | 18.5833 | 0.58 |
| 0.1 | 0 | clcB | 4 | 18.5833 | 0.525 |
| 0.1 | 0 | ycaM | 4 | 18.5833 | 0.506 |
| 0.1 | 0 | yadI | 4 | 18.5833 | 0.683 |
| 0.1 | 0 | AG1 | 4 | 18.5833 | 0.732 |
| 0.1 | 0 | ptsI | 4 | 18.8333 | 0.59 |
| 0.1 | 0 | clcB | 4 | 18.8333 | 0.526 |
| 0.1 | 0 | ycaM | 4 | 18.8333 | 0.503 |
| 0.1 | 0 | yadI | 4 | 18.8333 | 0.684 |
| 0.1 | 0 | AG1 | 4 | 18.8333 | 0.739 |
| 0.1 | 0 | ptsI | 4 | 19.0833 | 0.588 |
| 0.1 | 0 | clcB | 4 | 19.0833 | 0.528 |
| 0.1 | 0 | ycaM | 4 | 19.0833 | 0.511 |
| 0.1 | 0 | yadI | 4 | 19.0833 | 0.689 |
| 0.1 | 0 | AG1 | 4 | 19.0833 | 0.738 |
| 0.1 | 0 | ptsI | 4 | 19.3333 | 0.585 |
| 0.1 | 0 | clcB | 4 | 19.3333 | 0.528 |
| 0.1 | 0 | ycaM | 4 | 19.3333 | 0.513 |
| 0.1 | 0 | yadI | 4 | 19.3333 | 0.694 |
| 0.1 | 0 | AG1 | 4 | 19.3333 | 0.738 |
| 0.1 | 0 | ptsI | 4 | 19.5833 | 0.596 |
| 0.1 | 0 | clcB | 4 | 19.5833 | 0.532 |
| 0.1 | 0 | ycaM | 4 | 19.5833 | 0.515 |
| 0.1 | 0 | yadI | 4 | 19.5833 | 0.698 |
| 0.1 | 0 | AG1 | 4 | 19.5833 | 0.741 |
| 0.1 | 0 | ptsI | 4 | 19.8333 | 0.595 |
| 0.1 | 0 | clcB | 4 | 19.8333 | 0.53 |

| | | | | | |
|---|---|---|---|---|---|
| 0.1 | 0 | ycaM | 4 | 19.8333 | 0.513 |
| 0.1 | 0 | yadI | 4 | 19.8333 | 0.697 |
| 0.1 | 0 | AG1 | 4 | 19.8333 | 0.742 |
| 0.1 | 0 | ptsI | 4 | 20.0833 | 0.59 |
| 0.1 | 0 | clcB | 4 | 20.0833 | 0.534 |
| 0.1 | 0 | ycaM | 4 | 20.0833 | 0.521 |
| 0.1 | 0 | yadI | 4 | 20.0833 | 0.703 |
| 0.1 | 0 | AG1 | 4 | 20.0833 | 0.714 |
| 0.1 | 0 | ptsI | 4 | 20.3333 | 0.597 |
| 0.1 | 0 | clcB | 4 | 20.3333 | 0.53 |
| 0.1 | 0 | ycaM | 4 | 20.3333 | 0.519 |
| 0.1 | 0 | yadI | 4 | 20.3333 | 0.703 |
| 0.1 | 0 | AG1 | 4 | 20.3333 | 0.749 |
| 0.1 | 0 | ptsI | 4 | 20.5833 | 0.602 |
| 0.1 | 0 | clcB | 4 | 20.5833 | 0.53 |
| 0.1 | 0 | ycaM | 4 | 20.5833 | 0.515 |
| 0.1 | 0 | yadI | 4 | 20.5833 | 0.709 |
| 0.1 | 0 | AG1 | 4 | 20.5833 | 0.747 |
| 0.1 | 0 | ptsI | 4 | 20.8333 | 0.601 |
| 0.1 | 0 | clcB | 4 | 20.8333 | 0.534 |
| 0.1 | 0 | ycaM | 4 | 20.8333 | 0.52 |
| 0.1 | 0 | yadI | 4 | 20.8333 | 0.71 |
| 0.1 | 0 | AG1 | 4 | 20.8333 | 0.751 |
| 0.1 | 0 | ptsI | 4 | 21.0833 | 0.606 |
| 0.1 | 0 | clcB | 4 | 21.0833 | 0.532 |
| 0.1 | 0 | ycaM | 4 | 21.0833 | 0.52 |
| 0.1 | 0 | yadI | 4 | 21.0833 | 0.712 |
| 0.1 | 0 | AG1 | 4 | 21.0833 | 0.753 |
| 0.1 | 0 | ptsI | 4 | 21.3333 | 0.605 |
| 0.1 | 0 | clcB | 4 | 21.3333 | 0.534 |
| 0.1 | 0 | ycaM | 4 | 21.3333 | 0.521 |
| 0.1 | 0 | yadI | 4 | 21.3333 | 0.714 |
| 0.1 | 0 | AG1 | 4 | 21.3333 | 0.757 |
| 0.1 | 0 | ptsI | 4 | 21.5833 | 0.615 |
| 0.1 | 0 | clcB | 4 | 21.5833 | 0.534 |
| 0.1 | 0 | ycaM | 4 | 21.5833 | 0.523 |
| 0.1 | 0 | yadI | 4 | 21.5833 | 0.719 |
| 0.1 | 0 | AG1 | 4 | 21.5833 | 0.758 |
| 0.1 | 0 | ptsI | 4 | 21.8333 | 0.609 |
| 0.1 | 0 | clcB | 4 | 21.8333 | 0.533 |
| 0.1 | 0 | ycaM | 4 | 21.8333 | 0.526 |
| 0.1 | 0 | yadI | 4 | 21.8333 | 0.722 |
| 0.1 | 0 | AG1 | 4 | 21.8333 | 0.756 |
| 0.1 | 0 | ptsI | 4 | 22.0833 | 0.617 |
| 0.1 | 0 | clcB | 4 | 22.0833 | 0.536 |
| 0.1 | 0 | ycaM | 4 | 22.0833 | 0.525 |
| 0.1 | 0 | yadI | 4 | 22.0833 | 0.721 |
| 0.1 | 0 | AG1 | 4 | 22.0833 | 0.759 |

| | | | | | |
|---|---|---|---|---|---|
| 0.1 | 0 | ptsI | 4 | 22.3333 | 0.628 |
| 0.1 | 0 | clcB | 4 | 22.3333 | 0.536 |
| 0.1 | 0 | ycaM | 4 | 22.3333 | 0.525 |
| 0.1 | 0 | yadI | 4 | 22.3333 | 0.724 |
| 0.1 | 0 | AG1 | 4 | 22.3333 | 0.758 |
| 0.1 | 0 | ptsI | 4 | 22.5833 | 0.628 |
| 0.1 | 0 | clcB | 4 | 22.5833 | 0.532 |
| 0.1 | 0 | ycaM | 4 | 22.5833 | 0.521 |
| 0.1 | 0 | yadI | 4 | 22.5833 | 0.728 |
| 0.1 | 0 | AG1 | 4 | 22.5833 | 0.759 |
| 0.1 | 0 | ptsI | 4 | 22.8333 | 0.63 |
| 0.1 | 0 | clcB | 4 | 22.8333 | 0.536 |
| 0.1 | 0 | ycaM | 4 | 22.8333 | 0.524 |
| 0.1 | 0 | yadI | 4 | 22.8333 | 0.729 |
| 0.1 | 0 | AG1 | 4 | 22.8333 | 0.762 |
| 0.1 | 0 | ptsI | 4 | 23.0833 | 0.638 |
| 0.1 | 0 | clcB | 4 | 23.0833 | 0.538 |
| 0.1 | 0 | ycaM | 4 | 23.0833 | 0.528 |
| 0.1 | 0 | yadI | 4 | 23.0833 | 0.731 |
| 0.1 | 0 | AG1 | 4 | 23.0833 | 0.767 |
| 0.1 | 0 | ptsI | 4 | 23.3333 | 0.632 |
| 0.1 | 0 | clcB | 4 | 23.3333 | 0.538 |
| 0.1 | 0 | ycaM | 4 | 23.3333 | 0.53 |
| 0.1 | 0 | yadI | 4 | 23.3333 | 0.734 |
| 0.1 | 0 | AG1 | 4 | 23.3333 | 0.767 |
| 0.1 | 0 | ptsI | 4 | 23.5833 | 0.639 |
| 0.1 | 0 | clcB | 4 | 23.5833 | 0.536 |
| 0.1 | 0 | ycaM | 4 | 23.5833 | 0.527 |
| 0.1 | 0 | yadI | 4 | 23.5833 | 0.735 |
| 0.1 | 0 | AG1 | 4 | 23.5833 | 0.766 |
| 0.1 | 0 | ptsI | 4 | 23.8333 | 0.645 |
| 0.1 | 0 | clcB | 4 | 23.8333 | 0.537 |
| 0.1 | 0 | ycaM | 4 | 23.8333 | 0.528 |
| 0.1 | 0 | yadI | 4 | 23.8333 | 0.738 |
| 0.1 | 0 | AG1 | 4 | 23.8333 | 0.768 |
| 0.1 | 0 | ptsI | 4 | 24.0833 | 0.645 |
| 0.1 | 0 | clcB | 4 | 24.0833 | 0.538 |
| 0.1 | 0 | ycaM | 4 | 24.0833 | 0.529 |
| 0.1 | 0 | yadI | 4 | 24.0833 | 0.74 |
| 0.1 | 0 | AG1 | 4 | 24.0833 | 0.773 |
| 0.1 | 0 | ptsI | 5 | 0 | 0.157 |
| 0.1 | 0 | clcB | 5 | 0 | 0.166 |
| 0.1 | 0 | ycaM | 5 | 0 | 0.173 |
| 0.1 | 0 | yadI | 5 | 0 | 0.192 |
| 0.1 | 0 | AG1 | 5 | 0 | 0.166 |
| 0.1 | 0 | ptsI | 5 | 0.35 | 0.154 |
| 0.1 | 0 | clcB | 5 | 0.35 | 0.163 |
| 0.1 | 0 | ycaM | 5 | 0.35 | 0.173 |

| | | | | | |
|---|---|---|---|---|---|
| 0.1 | 0 | yadI | 5 | 0.35 | 0.182 |
| 0.1 | 0 | AG1 | 5 | 0.35 | 0.164 |
| 0.1 | 0 | ptsI | 5 | 0.6 | 0.154 |
| 0.1 | 0 | clcB | 5 | 0.6 | 0.165 |
| 0.1 | 0 | ycaM | 5 | 0.6 | 0.168 |
| 0.1 | 0 | yadI | 5 | 0.6 | 0.182 |
| 0.1 | 0 | AG1 | 5 | 0.6 | 0.164 |
| 0.1 | 0 | ptsI | 5 | 0.85 | 0.155 |
| 0.1 | 0 | clcB | 5 | 0.85 | 0.164 |
| 0.1 | 0 | ycaM | 5 | 0.85 | 0.168 |
| 0.1 | 0 | yadI | 5 | 0.85 | 0.184 |
| 0.1 | 0 | AG1 | 5 | 0.85 | 0.166 |
| 0.1 | 0 | ptsI | 5 | 1.1 | 0.157 |
| 0.1 | 0 | clcB | 5 | 1.1 | 0.167 |
| 0.1 | 0 | ycaM | 5 | 1.1 | 0.171 |
| 0.1 | 0 | yadI | 5 | 1.1 | 0.189 |
| 0.1 | 0 | AG1 | 5 | 1.1 | 0.167 |
| 0.1 | 0 | ptsI | 5 | 1.35 | 0.158 |
| 0.1 | 0 | clcB | 5 | 1.35 | 0.167 |
| 0.1 | 0 | ycaM | 5 | 1.35 | 0.175 |
| 0.1 | 0 | yadI | 5 | 1.35 | 0.189 |
| 0.1 | 0 | AG1 | 5 | 1.35 | 0.169 |
| 0.1 | 0 | ptsI | 5 | 1.6 | 0.163 |
| 0.1 | 0 | clcB | 5 | 1.6 | 0.173 |
| 0.1 | 0 | ycaM | 5 | 1.6 | 0.178 |
| 0.1 | 0 | yadI | 5 | 1.6 | 0.196 |
| 0.1 | 0 | AG1 | 5 | 1.6 | 0.175 |
| 0.1 | 0 | ptsI | 5 | 1.85 | 0.164 |
| 0.1 | 0 | clcB | 5 | 1.85 | 0.175 |
| 0.1 | 0 | ycaM | 5 | 1.85 | 0.182 |
| 0.1 | 0 | yadI | 5 | 1.85 | 0.201 |
| 0.1 | 0 | AG1 | 5 | 1.85 | 0.178 |
| 0.1 | 0 | ptsI | 5 | 2.1 | 0.168 |
| 0.1 | 0 | clcB | 5 | 2.1 | 0.181 |
| 0.1 | 0 | ycaM | 5 | 2.1 | 0.186 |
| 0.1 | 0 | yadI | 5 | 2.1 | 0.208 |
| 0.1 | 0 | AG1 | 5 | 2.1 | 0.186 |
| 0.1 | 0 | ptsI | 5 | 2.35 | 0.175 |
| 0.1 | 0 | clcB | 5 | 2.35 | 0.19 |
| 0.1 | 0 | ycaM | 5 | 2.35 | 0.191 |
| 0.1 | 0 | yadI | 5 | 2.35 | 0.219 |
| 0.1 | 0 | AG1 | 5 | 2.35 | 0.194 |
| 0.1 | 0 | ptsI | 5 | 2.6 | 0.178 |
| 0.1 | 0 | clcB | 5 | 2.6 | 0.193 |
| 0.1 | 0 | ycaM | 5 | 2.6 | 0.196 |
| 0.1 | 0 | yadI | 5 | 2.6 | 0.224 |
| 0.1 | 0 | AG1 | 5 | 2.6 | 0.193 |
| 0.1 | 0 | ptsI | 5 | 2.85 | 0.187 |

| | | | | | |
|---|---|---|---|---|---|
| 0.1 | 0 | clcB | 5 | 2.85 | 0.204 |
| 0.1 | 0 | ycaM | 5 | 2.85 | 0.201 |
| 0.1 | 0 | yadI | 5 | 2.85 | 0.238 |
| 0.1 | 0 | AG1 | 5 | 2.85 | 0.209 |
| 0.1 | 0 | ptsI | 5 | 3.1 | 0.196 |
| 0.1 | 0 | clcB | 5 | 3.1 | 0.217 |
| 0.1 | 0 | ycaM | 5 | 3.1 | 0.21 |
| 0.1 | 0 | yadI | 5 | 3.1 | 0.252 |
| 0.1 | 0 | AG1 | 5 | 3.1 | 0.22 |
| 0.1 | 0 | ptsI | 5 | 3.35 | 0.201 |
| 0.1 | 0 | clcB | 5 | 3.35 | 0.227 |
| 0.1 | 0 | ycaM | 5 | 3.35 | 0.218 |
| 0.1 | 0 | yadI | 5 | 3.35 | 0.268 |
| 0.1 | 0 | AG1 | 5 | 3.35 | 0.237 |
| 0.1 | 0 | ptsI | 5 | 3.6 | 0.204 |
| 0.1 | 0 | clcB | 5 | 3.6 | 0.239 |
| 0.1 | 0 | ycaM | 5 | 3.6 | 0.23 |
| 0.1 | 0 | yadI | 5 | 3.6 | 0.287 |
| 0.1 | 0 | AG1 | 5 | 3.6 | 0.256 |
| 0.1 | 0 | ptsI | 5 | 3.85 | 0.211 |
| 0.1 | 0 | clcB | 5 | 3.85 | 0.257 |
| 0.1 | 0 | ycaM | 5 | 3.85 | 0.244 |
| 0.1 | 0 | yadI | 5 | 3.85 | 0.308 |
| 0.1 | 0 | AG1 | 5 | 3.85 | 0.276 |
| 0.1 | 0 | ptsI | 5 | 4.38333 | 0.21 |
| 0.1 | 0 | clcB | 5 | 4.38333 | 0.241 |
| 0.1 | 0 | ycaM | 5 | 4.38333 | 0.27 |
| 0.1 | 0 | yadI | 5 | 4.38333 | 0.323 |
| 0.1 | 0 | AG1 | 5 | 4.38333 | 0.313 |
| 0.1 | 0 | ptsI | 5 | 4.63333 | 0.21 |
| 0.1 | 0 | clcB | 5 | 4.63333 | 0.244 |
| 0.1 | 0 | ycaM | 5 | 4.63333 | 0.268 |
| 0.1 | 0 | yadI | 5 | 4.63333 | 0.316 |
| 0.1 | 0 | AG1 | 5 | 4.63333 | 0.302 |
| 0.1 | 0 | ptsI | 5 | 4.88333 | 0.214 |
| 0.1 | 0 | clcB | 5 | 4.88333 | 0.257 |
| 0.1 | 0 | ycaM | 5 | 4.88333 | 0.28 |
| 0.1 | 0 | yadI | 5 | 4.88333 | 0.328 |
| 0.1 | 0 | AG1 | 5 | 4.88333 | 0.316 |
| 0.1 | 0 | ptsI | 5 | 5.13333 | 0.223 |
| 0.1 | 0 | clcB | 5 | 5.13333 | 0.269 |
| 0.1 | 0 | ycaM | 5 | 5.13333 | 0.291 |
| 0.1 | 0 | yadI | 5 | 5.13333 | 0.342 |
| 0.1 | 0 | AG1 | 5 | 5.13333 | 0.326 |
| 0.1 | 0 | ptsI | 5 | 5.38333 | 0.23 |
| 0.1 | 0 | clcB | 5 | 5.38333 | 0.282 |
| 0.1 | 0 | ycaM | 5 | 5.38333 | 0.306 |
| 0.1 | 0 | yadI | 5 | 5.38333 | 0.354 |

| | | | | | |
|---|---|---|---|---|---|
| 0.1 | 0 | AG1 | 5 | 5.38333 | 0.339 |
| 0.1 | 0 | ptsI | 5 | 5.63333 | 0.238 |
| 0.1 | 0 | clcB | 5 | 5.63333 | 0.295 |
| 0.1 | 0 | ycaM | 5 | 5.63333 | 0.316 |
| 0.1 | 0 | yadI | 5 | 5.63333 | 0.37 |
| 0.1 | 0 | AG1 | 5 | 5.63333 | 0.351 |
| 0.1 | 0 | ptsI | 5 | 5.88333 | 0.248 |
| 0.1 | 0 | clcB | 5 | 5.88333 | 0.307 |
| 0.1 | 0 | ycaM | 5 | 5.88333 | 0.326 |
| 0.1 | 0 | yadI | 5 | 5.88333 | 0.382 |
| 0.1 | 0 | AG1 | 5 | 5.88333 | 0.362 |
| 0.1 | 0 | ptsI | 5 | 6.13333 | 0.258 |
| 0.1 | 0 | clcB | 5 | 6.13333 | 0.322 |
| 0.1 | 0 | ycaM | 5 | 6.13333 | 0.335 |
| 0.1 | 0 | yadI | 5 | 6.13333 | 0.397 |
| 0.1 | 0 | AG1 | 5 | 6.13333 | 0.376 |
| 0.1 | 0 | ptsI | 5 | 6.38333 | 0.268 |
| 0.1 | 0 | clcB | 5 | 6.38333 | 0.332 |
| 0.1 | 0 | ycaM | 5 | 6.38333 | 0.345 |
| 0.1 | 0 | yadI | 5 | 6.38333 | 0.414 |
| 0.1 | 0 | AG1 | 5 | 6.38333 | 0.395 |
| 0.1 | 0 | ptsI | 5 | 6.63333 | 0.281 |
| 0.1 | 0 | clcB | 5 | 6.63333 | 0.345 |
| 0.1 | 0 | ycaM | 5 | 6.63333 | 0.355 |
| 0.1 | 0 | yadI | 5 | 6.63333 | 0.429 |
| 0.1 | 0 | AG1 | 5 | 6.63333 | 0.416 |
| 0.1 | 0 | ptsI | 5 | 6.88333 | 0.292 |
| 0.1 | 0 | clcB | 5 | 6.88333 | 0.357 |
| 0.1 | 0 | ycaM | 5 | 6.88333 | 0.364 |
| 0.1 | 0 | yadI | 5 | 6.88333 | 0.454 |
| 0.1 | 0 | AG1 | 5 | 6.88333 | 0.439 |
| 0.1 | 0 | ptsI | 5 | 7.13333 | 0.303 |
| 0.1 | 0 | clcB | 5 | 7.13333 | 0.367 |
| 0.1 | 0 | ycaM | 5 | 7.13333 | 0.372 |
| 0.1 | 0 | yadI | 5 | 7.13333 | 0.474 |
| 0.1 | 0 | AG1 | 5 | 7.13333 | 0.459 |
| 0.1 | 0 | ptsI | 5 | 7.38333 | 0.316 |
| 0.1 | 0 | clcB | 5 | 7.38333 | 0.381 |
| 0.1 | 0 | ycaM | 5 | 7.38333 | 0.385 |
| 0.1 | 0 | yadI | 5 | 7.38333 | 0.492 |
| 0.1 | 0 | AG1 | 5 | 7.38333 | 0.476 |
| 0.1 | 0 | ptsI | 5 | 7.63333 | 0.326 |
| 0.1 | 0 | clcB | 5 | 7.63333 | 0.392 |
| 0.1 | 0 | ycaM | 5 | 7.63333 | 0.396 |
| 0.1 | 0 | yadI | 5 | 7.63333 | 0.51 |
| 0.1 | 0 | AG1 | 5 | 7.63333 | 0.5 |
| 0.1 | 0 | ptsI | 5 | 7.88333 | 0.336 |
| 0.1 | 0 | clcB | 5 | 7.88333 | 0.404 |

| | | | | | |
|---|---|---|---|---|---|
| 0.1 | 0 | ycaM | 5 | 7.88333 | 0.407 |
| 0.1 | 0 | yadI | 5 | 7.88333 | 0.529 |
| 0.1 | 0 | AG1 | 5 | 7.88333 | 0.516 |
| 0.1 | 0 | ptsI | 5 | 8.13333 | 0.349 |
| 0.1 | 0 | clcB | 5 | 8.13333 | 0.414 |
| 0.1 | 0 | ycaM | 5 | 8.13333 | 0.415 |
| 0.1 | 0 | yadI | 5 | 8.13333 | 0.535 |
| 0.1 | 0 | AG1 | 5 | 8.13333 | 0.532 |
| 0.1 | 0 | ptsI | 5 | 8.38333 | 0.363 |
| 0.1 | 0 | clcB | 5 | 8.38333 | 0.427 |
| 0.1 | 0 | ycaM | 5 | 8.38333 | 0.426 |
| 0.1 | 0 | yadI | 5 | 8.38333 | 0.554 |
| 0.1 | 0 | AG1 | 5 | 8.38333 | 0.546 |
| 0.1 | 0 | ptsI | 5 | 8.88333 | 0.335 |
| 0.1 | 0 | clcB | 5 | 8.88333 | 0.362 |
| 0.1 | 0 | ycaM | 5 | 8.88333 | 0.356 |
| 0.1 | 0 | yadI | 5 | 8.88333 | 0.508 |
| 0.1 | 0 | AG1 | 5 | 8.88333 | 0.599 |
| 0.1 | 0 | ptsI | 5 | 9.13333 | 0.339 |
| 0.1 | 0 | clcB | 5 | 9.13333 | 0.369 |
| 0.1 | 0 | ycaM | 5 | 9.13333 | 0.361 |
| 0.1 | 0 | yadI | 5 | 9.13333 | 0.507 |
| 0.1 | 0 | AG1 | 5 | 9.13333 | 0.597 |
| 0.1 | 0 | ptsI | 5 | 9.38333 | 0.342 |
| 0.1 | 0 | clcB | 5 | 9.38333 | 0.378 |
| 0.1 | 0 | ycaM | 5 | 9.38333 | 0.365 |
| 0.1 | 0 | yadI | 5 | 9.38333 | 0.504 |
| 0.1 | 0 | AG1 | 5 | 9.38333 | 0.616 |
| 0.1 | 0 | ptsI | 5 | 9.63333 | 0.357 |
| 0.1 | 0 | clcB | 5 | 9.63333 | 0.388 |
| 0.1 | 0 | ycaM | 5 | 9.63333 | 0.374 |
| 0.1 | 0 | yadI | 5 | 9.63333 | 0.52 |
| 0.1 | 0 | AG1 | 5 | 9.63333 | 0.624 |
| 0.1 | 0 | ptsI | 5 | 9.88333 | 0.371 |
| 0.1 | 0 | clcB | 5 | 9.88333 | 0.4 |
| 0.1 | 0 | ycaM | 5 | 9.88333 | 0.384 |
| 0.1 | 0 | yadI | 5 | 9.88333 | 0.529 |
| 0.1 | 0 | AG1 | 5 | 9.88333 | 0.638 |
| 0.1 | 0 | ptsI | 5 | 10.1333 | 0.378 |
| 0.1 | 0 | clcB | 5 | 10.1333 | 0.406 |
| 0.1 | 0 | ycaM | 5 | 10.1333 | 0.391 |
| 0.1 | 0 | yadI | 5 | 10.1333 | 0.536 |
| 0.1 | 0 | AG1 | 5 | 10.1333 | 0.646 |
| 0.1 | 0 | ptsI | 5 | 10.3833 | 0.392 |
| 0.1 | 0 | clcB | 5 | 10.3833 | 0.416 |
| 0.1 | 0 | ycaM | 5 | 10.3833 | 0.397 |
| 0.1 | 0 | yadI | 5 | 10.3833 | 0.544 |
| 0.1 | 0 | AG1 | 5 | 10.3833 | 0.66 |

| | | | | | |
|---|---|---|---|---|---|
| 0.1 | 0 | ptsI | 5 | 10.6333 | 0.397 |
| 0.1 | 0 | clcB | 5 | 10.6333 | 0.42 |
| 0.1 | 0 | ycaM | 5 | 10.6333 | 0.399 |
| 0.1 | 0 | yadI | 5 | 10.6333 | 0.551 |
| 0.1 | 0 | AG1 | 5 | 10.6333 | 0.666 |
| 0.1 | 0 | ptsI | 5 | 10.8833 | 0.407 |
| 0.1 | 0 | clcB | 5 | 10.8833 | 0.426 |
| 0.1 | 0 | ycaM | 5 | 10.8833 | 0.405 |
| 0.1 | 0 | yadI | 5 | 10.8833 | 0.562 |
| 0.1 | 0 | AG1 | 5 | 10.8833 | 0.678 |
| 0.1 | 0 | ptsI | 5 | 11.1333 | 0.412 |
| 0.1 | 0 | clcB | 5 | 11.1333 | 0.429 |
| 0.1 | 0 | ycaM | 5 | 11.1333 | 0.403 |
| 0.1 | 0 | yadI | 5 | 11.1333 | 0.564 |
| 0.1 | 0 | AG1 | 5 | 11.1333 | 0.689 |
| 0.1 | 0 | ptsI | 5 | 11.3833 | 0.419 |
| 0.1 | 0 | clcB | 5 | 11.3833 | 0.433 |
| 0.1 | 0 | ycaM | 5 | 11.3833 | 0.406 |
| 0.1 | 0 | yadI | 5 | 11.3833 | 0.573 |
| 0.1 | 0 | AG1 | 5 | 11.3833 | 0.7 |
| 0.1 | 0 | ptsI | 5 | 11.6333 | 0.427 |
| 0.1 | 0 | clcB | 5 | 11.6333 | 0.438 |
| 0.1 | 0 | ycaM | 5 | 11.6333 | 0.411 |
| 0.1 | 0 | yadI | 5 | 11.6333 | 0.58 |
| 0.1 | 0 | AG1 | 5 | 11.6333 | 0.712 |
| 0.1 | 0 | ptsI | 5 | 11.8833 | 0.438 |
| 0.1 | 0 | clcB | 5 | 11.8833 | 0.445 |
| 0.1 | 0 | ycaM | 5 | 11.8833 | 0.416 |
| 0.1 | 0 | yadI | 5 | 11.8833 | 0.589 |
| 0.1 | 0 | AG1 | 5 | 11.8833 | 0.719 |
| 0.1 | 0 | ptsI | 5 | 12.1333 | 0.448 |
| 0.1 | 0 | clcB | 5 | 12.1333 | 0.452 |
| 0.1 | 0 | ycaM | 5 | 12.1333 | 0.42 |
| 0.1 | 0 | yadI | 5 | 12.1333 | 0.598 |
| 0.1 | 0 | AG1 | 5 | 12.1333 | 0.73 |
| 0.1 | 0 | ptsI | 5 | 12.3833 | 0.45 |
| 0.1 | 0 | clcB | 5 | 12.3833 | 0.453 |
| 0.1 | 0 | ycaM | 5 | 12.3833 | 0.422 |
| 0.1 | 0 | yadI | 5 | 12.3833 | 0.599 |
| 0.1 | 0 | AG1 | 5 | 12.3833 | 0.733 |
| 0.1 | 0 | ptsI | 5 | 12.6333 | 0.46 |
| 0.1 | 0 | clcB | 5 | 12.6333 | 0.458 |
| 0.1 | 0 | ycaM | 5 | 12.6333 | 0.426 |
| 0.1 | 0 | yadI | 5 | 12.6333 | 0.607 |
| 0.1 | 0 | AG1 | 5 | 12.6333 | 0.747 |
| 0.1 | 0 | ptsI | 5 | 12.8833 | 0.473 |
| 0.1 | 0 | clcB | 5 | 12.8833 | 0.469 |
| 0.1 | 0 | ycaM | 5 | 12.8833 | 0.431 |

| | | | | | |
|---|---|---|---|---|---|
| 0.1 | 0 | yadI | 5 | 12.8833 | 0.617 |
| 0.1 | 0 | AG1 | 5 | 12.8833 | 0.756 |
| 0.1 | 0 | ptsI | 5 | 13.1333 | 0.478 |
| 0.1 | 0 | clcB | 5 | 13.1333 | 0.47 |
| 0.1 | 0 | ycaM | 5 | 13.1333 | 0.434 |
| 0.1 | 0 | yadI | 5 | 13.1333 | 0.62 |
| 0.1 | 0 | AG1 | 5 | 13.1333 | 0.765 |
| 0.1 | 0 | ptsI | 5 | 13.3833 | 0.482 |
| 0.1 | 0 | clcB | 5 | 13.3833 | 0.473 |
| 0.1 | 0 | ycaM | 5 | 13.3833 | 0.436 |
| 0.1 | 0 | yadI | 5 | 13.3833 | 0.625 |
| 0.1 | 0 | AG1 | 5 | 13.3833 | 0.771 |
| 0.1 | 0 | ptsI | 5 | 13.6333 | 0.491 |
| 0.1 | 0 | clcB | 5 | 13.6333 | 0.481 |
| 0.1 | 0 | ycaM | 5 | 13.6333 | 0.441 |
| 0.1 | 0 | yadI | 5 | 13.6333 | 0.632 |
| 0.1 | 0 | AG1 | 5 | 13.6333 | 0.779 |
| 0.1 | 0 | ptsI | 5 | 13.8833 | 0.497 |
| 0.1 | 0 | clcB | 5 | 13.8833 | 0.481 |
| 0.1 | 0 | ycaM | 5 | 13.8833 | 0.443 |
| 0.1 | 0 | yadI | 5 | 13.8833 | 0.636 |
| 0.1 | 0 | AG1 | 5 | 13.8833 | 0.783 |
| 0.1 | 0 | ptsI | 5 | 14.1333 | 0.51 |
| 0.1 | 0 | clcB | 5 | 14.1333 | 0.487 |
| 0.1 | 0 | ycaM | 5 | 14.1333 | 0.448 |
| 0.1 | 0 | yadI | 5 | 14.1333 | 0.643 |
| 0.1 | 0 | AG1 | 5 | 14.1333 | 0.792 |
| 0.1 | 0 | ptsI | 5 | 14.3833 | 0.517 |
| 0.1 | 0 | clcB | 5 | 14.3833 | 0.492 |
| 0.1 | 0 | ycaM | 5 | 14.3833 | 0.455 |
| 0.1 | 0 | yadI | 5 | 14.3833 | 0.65 |
| 0.1 | 0 | AG1 | 5 | 14.3833 | 0.8 |
| 0.1 | 0 | ptsI | 5 | 14.6333 | 0.521 |
| 0.1 | 0 | clcB | 5 | 14.6333 | 0.499 |
| 0.1 | 0 | ycaM | 5 | 14.6333 | 0.459 |
| 0.1 | 0 | yadI | 5 | 14.6333 | 0.651 |
| 0.1 | 0 | AG1 | 5 | 14.6333 | 0.806 |
| 0.1 | 0 | ptsI | 5 | 14.8833 | 0.529 |
| 0.1 | 0 | clcB | 5 | 14.8833 | 0.498 |
| 0.1 | 0 | ycaM | 5 | 14.8833 | 0.462 |
| 0.1 | 0 | yadI | 5 | 14.8833 | 0.657 |
| 0.1 | 0 | AG1 | 5 | 14.8833 | 0.811 |
| 0.1 | 0 | ptsI | 5 | 15.1333 | 0.537 |
| 0.1 | 0 | clcB | 5 | 15.1333 | 0.505 |
| 0.1 | 0 | ycaM | 5 | 15.1333 | 0.466 |
| 0.1 | 0 | yadI | 5 | 15.1333 | 0.663 |
| 0.1 | 0 | AG1 | 5 | 15.1333 | 0.816 |
| 0.1 | 0 | ptsI | 5 | 15.3833 | 0.539 |

| | | | | | |
|---|---|---|---|---|---|
| 0.1 | 0 | clcB | 5 | 15.3833 | 0.507 |
| 0.1 | 0 | ycaM | 5 | 15.3833 | 0.469 |
| 0.1 | 0 | yadI | 5 | 15.3833 | 0.666 |
| 0.1 | 0 | AG1 | 5 | 15.3833 | 0.819 |
| 0.1 | 0 | ptsI | 5 | 15.6333 | 0.545 |
| 0.1 | 0 | clcB | 5 | 15.6333 | 0.506 |
| 0.1 | 0 | ycaM | 5 | 15.6333 | 0.471 |
| 0.1 | 0 | yadI | 5 | 15.6333 | 0.67 |
| 0.1 | 0 | AG1 | 5 | 15.6333 | 0.821 |
| 0.1 | 0 | ptsI | 5 | 15.8833 | 0.549 |
| 0.1 | 0 | clcB | 5 | 15.8833 | 0.509 |
| 0.1 | 0 | ycaM | 5 | 15.8833 | 0.473 |
| 0.1 | 0 | yadI | 5 | 15.8833 | 0.673 |
| 0.1 | 0 | AG1 | 5 | 15.8833 | 0.826 |
| 0.1 | 0 | ptsI | 5 | 16.1333 | 0.554 |
| 0.1 | 0 | clcB | 5 | 16.1333 | 0.514 |
| 0.1 | 0 | ycaM | 5 | 16.1333 | 0.475 |
| 0.1 | 0 | yadI | 5 | 16.1333 | 0.67 |
| 0.1 | 0 | AG1 | 5 | 16.1333 | 0.844 |
| 0.1 | 0 | ptsI | 5 | 16.3833 | 0.561 |
| 0.1 | 0 | clcB | 5 | 16.3833 | 0.512 |
| 0.1 | 0 | ycaM | 5 | 16.3833 | 0.478 |
| 0.1 | 0 | yadI | 5 | 16.3833 | 0.677 |
| 0.1 | 0 | AG1 | 5 | 16.3833 | 0.834 |
| 0.1 | 0 | ptsI | 5 | 16.6333 | 0.562 |
| 0.1 | 0 | clcB | 5 | 16.6333 | 0.514 |
| 0.1 | 0 | ycaM | 5 | 16.6333 | 0.484 |
| 0.1 | 0 | yadI | 5 | 16.6333 | 0.683 |
| 0.1 | 0 | AG1 | 5 | 16.6333 | 0.84 |
| 0.1 | 0 | ptsI | 5 | 16.8833 | 0.571 |
| 0.1 | 0 | clcB | 5 | 16.8833 | 0.519 |
| 0.1 | 0 | ycaM | 5 | 16.8833 | 0.49 |
| 0.1 | 0 | yadI | 5 | 16.8833 | 0.686 |
| 0.1 | 0 | AG1 | 5 | 16.8833 | 0.843 |
| 0.1 | 0 | ptsI | 5 | 17.1333 | 0.578 |
| 0.1 | 0 | clcB | 5 | 17.1333 | 0.521 |
| 0.1 | 0 | ycaM | 5 | 17.1333 | 0.491 |
| 0.1 | 0 | yadI | 5 | 17.1333 | 0.692 |
| 0.1 | 0 | AG1 | 5 | 17.1333 | 0.847 |
| 0.1 | 0 | ptsI | 5 | 17.3833 | 0.577 |
| 0.1 | 0 | clcB | 5 | 17.3833 | 0.518 |
| 0.1 | 0 | ycaM | 5 | 17.3833 | 0.492 |
| 0.1 | 0 | yadI | 5 | 17.3833 | 0.687 |
| 0.1 | 0 | AG1 | 5 | 17.3833 | 0.848 |
| 0.1 | 0 | ptsI | 5 | 17.6333 | 0.584 |
| 0.1 | 0 | clcB | 5 | 17.6333 | 0.52 |
| 0.1 | 0 | ycaM | 5 | 17.6333 | 0.494 |
| 0.1 | 0 | yadI | 5 | 17.6333 | 0.69 |

| | | | | | |
|---|---|---|---|---|---|
| 0.1 | 0 | AG1 | 5 | 17.6333 | 0.85 |
| 0.1 | 0 | ptsI | 5 | 17.8833 | 0.592 |
| 0.1 | 0 | clcB | 5 | 17.8833 | 0.523 |
| 0.1 | 0 | ycaM | 5 | 17.8833 | 0.499 |
| 0.1 | 0 | yadI | 5 | 17.8833 | 0.694 |
| 0.1 | 0 | AG1 | 5 | 17.8833 | 0.853 |
| 0.1 | 0 | ptsI | 5 | 18.1333 | 0.591 |
| 0.1 | 0 | clcB | 5 | 18.1333 | 0.523 |
| 0.1 | 0 | ycaM | 5 | 18.1333 | 0.503 |
| 0.1 | 0 | yadI | 5 | 18.1333 | 0.698 |
| 0.1 | 0 | AG1 | 5 | 18.1333 | 0.856 |
| 0.1 | 0 | ptsI | 5 | 18.3833 | 0.598 |
| 0.1 | 0 | clcB | 5 | 18.3833 | 0.527 |
| 0.1 | 0 | ycaM | 5 | 18.3833 | 0.504 |
| 0.1 | 0 | yadI | 5 | 18.3833 | 0.702 |
| 0.1 | 0 | AG1 | 5 | 18.3833 | 0.858 |
| 0.1 | 0 | ptsI | 5 | 18.6333 | 0.603 |
| 0.1 | 0 | clcB | 5 | 18.6333 | 0.527 |
| 0.1 | 0 | ycaM | 5 | 18.6333 | 0.504 |
| 0.1 | 0 | yadI | 5 | 18.6333 | 0.702 |
| 0.1 | 0 | AG1 | 5 | 18.6333 | 0.863 |
| 0.1 | 0 | ptsI | 5 | 18.8833 | 0.603 |
| 0.1 | 0 | clcB | 5 | 18.8833 | 0.527 |
| 0.1 | 0 | ycaM | 5 | 18.8833 | 0.507 |
| 0.1 | 0 | yadI | 5 | 18.8833 | 0.705 |
| 0.1 | 0 | AG1 | 5 | 18.8833 | 0.864 |
| 0.1 | 0 | ptsI | 5 | 19.1333 | 0.611 |
| 0.1 | 0 | clcB | 5 | 19.1333 | 0.528 |
| 0.1 | 0 | ycaM | 5 | 19.1333 | 0.508 |
| 0.1 | 0 | yadI | 5 | 19.1333 | 0.706 |
| 0.1 | 0 | AG1 | 5 | 19.1333 | 0.864 |
| 0.1 | 0 | ptsI | 5 | 19.3833 | 0.616 |
| 0.1 | 0 | clcB | 5 | 19.3833 | 0.532 |
| 0.1 | 0 | ycaM | 5 | 19.3833 | 0.51 |
| 0.1 | 0 | yadI | 5 | 19.3833 | 0.713 |
| 0.1 | 0 | AG1 | 5 | 19.3833 | 0.869 |
| 0.1 | 0 | ptsI | 5 | 19.6333 | 0.616 |
| 0.1 | 0 | clcB | 5 | 19.6333 | 0.533 |
| 0.1 | 0 | ycaM | 5 | 19.6333 | 0.516 |
| 0.1 | 0 | yadI | 5 | 19.6333 | 0.713 |
| 0.1 | 0 | AG1 | 5 | 19.6333 | 0.872 |
| 0.1 | 0 | ptsI | 5 | 19.8833 | 0.622 |
| 0.1 | 0 | clcB | 5 | 19.8833 | 0.533 |
| 0.1 | 0 | ycaM | 5 | 19.8833 | 0.515 |
| 0.1 | 0 | yadI | 5 | 19.8833 | 0.714 |
| 0.1 | 0 | AG1 | 5 | 19.8833 | 0.872 |
| 0.1 | 0 | ptsI | 5 | 20.1333 | 0.625 |
| 0.1 | 0 | clcB | 5 | 20.1333 | 0.534 |

| | | | | | |
|---|---|---|---|---|---|
| 0.1 | 0 | ycaM | 5 | 20.1333 | 0.516 |
| 0.1 | 0 | yadI | 5 | 20.1333 | 0.714 |
| 0.1 | 0 | AG1 | 5 | 20.1333 | 0.876 |
| 0.1 | 0 | ptsI | 5 | 20.3833 | 0.629 |
| 0.1 | 0 | clcB | 5 | 20.3833 | 0.535 |
| 0.1 | 0 | ycaM | 5 | 20.3833 | 0.517 |
| 0.1 | 0 | yadI | 5 | 20.3833 | 0.718 |
| 0.1 | 0 | AG1 | 5 | 20.3833 | 0.876 |
| 0.1 | 0 | ptsI | 5 | 20.6333 | 0.628 |
| 0.1 | 0 | clcB | 5 | 20.6333 | 0.531 |
| 0.1 | 0 | ycaM | 5 | 20.6333 | 0.523 |
| 0.1 | 0 | yadI | 5 | 20.6333 | 0.717 |
| 0.1 | 0 | AG1 | 5 | 20.6333 | 0.858 |
| 0.1 | 0 | ptsI | 5 | 20.8833 | 0.633 |
| 0.1 | 0 | clcB | 5 | 20.8833 | 0.538 |
| 0.1 | 0 | ycaM | 5 | 20.8833 | 0.521 |
| 0.1 | 0 | yadI | 5 | 20.8833 | 0.722 |
| 0.1 | 0 | AG1 | 5 | 20.8833 | 0.879 |
| 0.1 | 0 | ptsI | 5 | 21.1333 | 0.634 |
| 0.1 | 0 | clcB | 5 | 21.1333 | 0.535 |
| 0.1 | 0 | ycaM | 5 | 21.1333 | 0.521 |
| 0.1 | 0 | yadI | 5 | 21.1333 | 0.723 |
| 0.1 | 0 | AG1 | 5 | 21.1333 | 0.879 |
| 0.1 | 0 | ptsI | 5 | 21.3833 | 0.636 |
| 0.1 | 0 | clcB | 5 | 21.3833 | 0.539 |
| 0.1 | 0 | ycaM | 5 | 21.3833 | 0.522 |
| 0.1 | 0 | yadI | 5 | 21.3833 | 0.726 |
| 0.1 | 0 | AG1 | 5 | 21.3833 | 0.882 |
| 0.1 | 0 | ptsI | 5 | 21.6333 | 0.639 |
| 0.1 | 0 | clcB | 5 | 21.6333 | 0.539 |
| 0.1 | 0 | ycaM | 5 | 21.6333 | 0.528 |
| 0.1 | 0 | yadI | 5 | 21.6333 | 0.725 |
| 0.1 | 0 | AG1 | 5 | 21.6333 | 0.882 |
| 0.1 | 0 | ptsI | 5 | 21.8833 | 0.646 |
| 0.1 | 0 | clcB | 5 | 21.8833 | 0.54 |
| 0.1 | 0 | ycaM | 5 | 21.8833 | 0.531 |
| 0.1 | 0 | yadI | 5 | 21.8833 | 0.729 |
| 0.1 | 0 | AG1 | 5 | 21.8833 | 0.884 |
| 0.1 | 0 | ptsI | 5 | 22.1333 | 0.64 |
| 0.1 | 0 | clcB | 5 | 22.1333 | 0.538 |
| 0.1 | 0 | ycaM | 5 | 22.1333 | 0.53 |
| 0.1 | 0 | yadI | 5 | 22.1333 | 0.724 |
| 0.1 | 0 | AG1 | 5 | 22.1333 | 0.88 |
| 0.1 | 0 | ptsI | 5 | 22.3833 | 0.65 |
| 0.1 | 0 | clcB | 5 | 22.3833 | 0.539 |
| 0.1 | 0 | ycaM | 5 | 22.3833 | 0.529 |
| 0.1 | 0 | yadI | 5 | 22.3833 | 0.73 |
| 0.1 | 0 | AG1 | 5 | 22.3833 | 0.883 |

| | | | | | |
|---|---|---|---|---|---|
| 0.1 | 0 | ptsI | 5 | 22.6333 | 0.643 |
| 0.1 | 0 | clcB | 5 | 22.6333 | 0.54 |
| 0.1 | 0 | ycaM | 5 | 22.6333 | 0.535 |
| 0.1 | 0 | yadI | 5 | 22.6333 | 0.732 |
| 0.1 | 0 | AG1 | 5 | 22.6333 | 0.886 |
| 0.1 | 0 | ptsI | 5 | 22.8833 | 0.648 |
| 0.1 | 0 | clcB | 5 | 22.8833 | 0.538 |
| 0.1 | 0 | ycaM | 5 | 22.8833 | 0.533 |
| 0.1 | 0 | yadI | 5 | 22.8833 | 0.728 |
| 0.1 | 0 | AG1 | 5 | 22.8833 | 0.884 |
| 0.1 | 0 | ptsI | 5 | 23.1333 | 0.651 |
| 0.1 | 0 | clcB | 5 | 23.1333 | 0.539 |
| 0.1 | 0 | ycaM | 5 | 23.1333 | 0.536 |
| 0.1 | 0 | yadI | 5 | 23.1333 | 0.732 |
| 0.1 | 0 | AG1 | 5 | 23.1333 | 0.888 |
| 0.1 | 0 | ptsI | 5 | 23.3833 | 0.656 |
| 0.1 | 0 | clcB | 5 | 23.3833 | 0.542 |
| 0.1 | 0 | ycaM | 5 | 23.3833 | 0.534 |
| 0.1 | 0 | yadI | 5 | 23.3833 | 0.736 |
| 0.1 | 0 | AG1 | 5 | 23.3833 | 0.887 |
| 0.1 | 0 | ptsI | 5 | 23.6333 | 0.659 |
| 0.1 | 0 | clcB | 5 | 23.6333 | 0.54 |
| 0.1 | 0 | ycaM | 5 | 23.6333 | 0.535 |
| 0.1 | 0 | yadI | 5 | 23.6333 | 0.736 |
| 0.1 | 0 | AG1 | 5 | 23.6333 | 0.89 |
| 0.1 | 0 | ptsI | 5 | 23.8833 | 0.659 |
| 0.1 | 0 | clcB | 5 | 23.8833 | 0.54 |
| 0.1 | 0 | ycaM | 5 | 23.8833 | 0.536 |
| 0.1 | 0 | yadI | 5 | 23.8833 | 0.729 |
| 0.1 | 0 | AG1 | 5 | 23.8833 | 0.884 |
| 0.1 | 0 | ptsI | 5 | 24.1333 | 0.664 |
| 0.1 | 0 | clcB | 5 | 24.1333 | 0.541 |
| 0.1 | 0 | ycaM | 5 | 24.1333 | 0.543 |
| 0.1 | 0 | yadI | 5 | 24.1333 | 0.739 |
| 0.1 | 0 | AG1 | 5 | 24.1333 | 0.889 |
| 0.1 | 0 | ptsI | 5 | 24.3833 | 0.664 |
| 0.1 | 0 | clcB | 5 | 24.3833 | 0.541 |
| 0.1 | 0 | ycaM | 5 | 24.3833 | 0.539 |
| 0.1 | 0 | yadI | 5 | 24.3833 | 0.734 |
| 0.1 | 0 | AG1 | 5 | 24.3833 | 0.889 |
| 0.2 | 0 | ptsI | 1 | 0 | 0.244 |
| 0.2 | 0 | clcB | 1 | 0 | 0.241 |
| 0.2 | 0 | ycaM | 1 | 0 | 0.241 |
| 0.2 | 0 | yadI | 1 | 0 | 0.281 |
| 0.2 | 0 | AG1 | 1 | 0 | 0.283 |
| 0.2 | 0 | ptsI | 1 | 0.25 | 0.233 |
| 0.2 | 0 | clcB | 1 | 0.25 | 0.223 |
| 0.2 | 0 | ycaM | 1 | 0.25 | 0.224 |

| | | | | | |
|---|---|---|---|---|---|
| 0.2 | 0 | yadI | 1 | 0.25 | 0.267 |
| 0.2 | 0 | AG1 | 1 | 0.25 | 0.27 |
| 0.2 | 0 | ptsI | 1 | 0.5 | 0.231 |
| 0.2 | 0 | clcB | 1 | 0.5 | 0.22 |
| 0.2 | 0 | ycaM | 1 | 0.5 | 0.217 |
| 0.2 | 0 | yadI | 1 | 0.5 | 0.263 |
| 0.2 | 0 | AG1 | 1 | 0.5 | 0.265 |
| 0.2 | 0 | ptsI | 1 | 0.75 | 0.232 |
| 0.2 | 0 | clcB | 1 | 0.75 | 0.222 |
| 0.2 | 0 | ycaM | 1 | 0.75 | 0.217 |
| 0.2 | 0 | yadI | 1 | 0.75 | 0.261 |
| 0.2 | 0 | AG1 | 1 | 0.75 | 0.264 |
| 0.2 | 0 | ptsI | 1 | 1 | 0.234 |
| 0.2 | 0 | clcB | 1 | 1 | 0.225 |
| 0.2 | 0 | ycaM | 1 | 1 | 0.217 |
| 0.2 | 0 | yadI | 1 | 1 | 0.262 |
| 0.2 | 0 | AG1 | 1 | 1 | 0.266 |
| 0.2 | 0 | ptsI | 1 | 1.25 | 0.239 |
| 0.2 | 0 | clcB | 1 | 1.25 | 0.228 |
| 0.2 | 0 | ycaM | 1 | 1.25 | 0.218 |
| 0.2 | 0 | yadI | 1 | 1.25 | 0.267 |
| 0.2 | 0 | AG1 | 1 | 1.25 | 0.264 |
| 0.2 | 0 | ptsI | 1 | 1.5 | 0.241 |
| 0.2 | 0 | clcB | 1 | 1.5 | 0.233 |
| 0.2 | 0 | ycaM | 1 | 1.5 | 0.219 |
| 0.2 | 0 | yadI | 1 | 1.5 | 0.27 |
| 0.2 | 0 | AG1 | 1 | 1.5 | 0.267 |
| 0.2 | 0 | ptsI | 1 | 1.75 | 0.25 |
| 0.2 | 0 | clcB | 1 | 1.75 | 0.239 |
| 0.2 | 0 | ycaM | 1 | 1.75 | 0.222 |
| 0.2 | 0 | yadI | 1 | 1.75 | 0.28 |
| 0.2 | 0 | AG1 | 1 | 1.75 | 0.268 |
| 0.2 | 0 | ptsI | 1 | 2 | 0.253 |
| 0.2 | 0 | clcB | 1 | 2 | 0.243 |
| 0.2 | 0 | ycaM | 1 | 2 | 0.226 |
| 0.2 | 0 | yadI | 1 | 2 | 0.284 |
| 0.2 | 0 | AG1 | 1 | 2 | 0.274 |
| 0.2 | 0 | ptsI | 1 | 2.25 | 0.263 |
| 0.2 | 0 | clcB | 1 | 2.25 | 0.248 |
| 0.2 | 0 | ycaM | 1 | 2.25 | 0.234 |
| 0.2 | 0 | yadI | 1 | 2.25 | 0.293 |
| 0.2 | 0 | AG1 | 1 | 2.25 | 0.278 |
| 0.2 | 0 | ptsI | 1 | 2.5 | 0.266 |
| 0.2 | 0 | clcB | 1 | 2.5 | 0.255 |
| 0.2 | 0 | ycaM | 1 | 2.5 | 0.236 |
| 0.2 | 0 | yadI | 1 | 2.5 | 0.299 |
| 0.2 | 0 | AG1 | 1 | 2.5 | 0.284 |
| 0.2 | 0 | ptsI | 1 | 2.75 | 0.278 |

| | | | | | |
|---|---|---|---|---|---|
| 0.2 | 0 | clcB | 1 | 2.75 | 0.26 |
| 0.2 | 0 | ycaM | 1 | 2.75 | 0.246 |
| 0.2 | 0 | yadI | 1 | 2.75 | 0.312 |
| 0.2 | 0 | AG1 | 1 | 2.75 | 0.288 |
| 0.2 | 0 | ptsI | 1 | 3 | 0.284 |
| 0.2 | 0 | clcB | 1 | 3 | 0.268 |
| 0.2 | 0 | ycaM | 1 | 3 | 0.248 |
| 0.2 | 0 | yadI | 1 | 3 | 0.32 |
| 0.2 | 0 | AG1 | 1 | 3 | 0.294 |
| 0.2 | 0 | ptsI | 1 | 3.25 | 0.296 |
| 0.2 | 0 | clcB | 1 | 3.25 | 0.272 |
| 0.2 | 0 | ycaM | 1 | 3.25 | 0.259 |
| 0.2 | 0 | yadI | 1 | 3.25 | 0.334 |
| 0.2 | 0 | AG1 | 1 | 3.25 | 0.304 |
| 0.2 | 0 | ptsI | 1 | 3.5 | 0.307 |
| 0.2 | 0 | clcB | 1 | 3.5 | 0.279 |
| 0.2 | 0 | ycaM | 1 | 3.5 | 0.268 |
| 0.2 | 0 | yadI | 1 | 3.5 | 0.354 |
| 0.2 | 0 | AG1 | 1 | 3.5 | 0.31 |
| 0.2 | 0 | ptsI | 1 | 3.75 | 0.317 |
| 0.2 | 0 | clcB | 1 | 3.75 | 0.286 |
| 0.2 | 0 | ycaM | 1 | 3.75 | 0.275 |
| 0.2 | 0 | yadI | 1 | 3.75 | 0.372 |
| 0.2 | 0 | AG1 | 1 | 3.75 | 0.32 |
| 0.2 | 0 | ptsI | 1 | 4 | 0.331 |
| 0.2 | 0 | clcB | 1 | 4 | 0.291 |
| 0.2 | 0 | ycaM | 1 | 4 | 0.289 |
| 0.2 | 0 | yadI | 1 | 4 | 0.387 |
| 0.2 | 0 | AG1 | 1 | 4 | 0.319 |
| 0.2 | 0 | ptsI | 1 | 4.25 | 0.333 |
| 0.2 | 0 | clcB | 1 | 4.25 | 0.3 |
| 0.2 | 0 | ycaM | 1 | 4.25 | 0.296 |
| 0.2 | 0 | yadI | 1 | 4.25 | 0.385 |
| 0.2 | 0 | AG1 | 1 | 4.25 | 0.338 |
| 0.2 | 0 | ptsI | 1 | 4.5 | 0.342 |
| 0.2 | 0 | clcB | 1 | 4.5 | 0.309 |
| 0.2 | 0 | ycaM | 1 | 4.5 | 0.304 |
| 0.2 | 0 | yadI | 1 | 4.5 | 0.408 |
| 0.2 | 0 | AG1 | 1 | 4.5 | 0.353 |
| 0.2 | 0 | ptsI | 1 | 4.75 | 0.355 |
| 0.2 | 0 | clcB | 1 | 4.75 | 0.314 |
| 0.2 | 0 | ycaM | 1 | 4.75 | 0.318 |
| 0.2 | 0 | yadI | 1 | 4.75 | 0.408 |
| 0.2 | 0 | AG1 | 1 | 4.75 | 0.362 |
| 0.2 | 0 | ptsI | 1 | 5 | 0.365 |
| 0.2 | 0 | clcB | 1 | 5 | 0.326 |
| 0.2 | 0 | ycaM | 1 | 5 | 0.327 |
| 0.2 | 0 | yadI | 1 | 5 | 0.431 |

| | | | | | |
|---|---|---|---|---|---|
| 0.2 | 0 | AG1 | 1 | 5 | 0.37 |
| 0.2 | 0 | ptsI | 1 | 5.25 | 0.372 |
| 0.2 | 0 | clcB | 1 | 5.25 | 0.337 |
| 0.2 | 0 | ycaM | 1 | 5.25 | 0.34 |
| 0.2 | 0 | yadI | 1 | 5.25 | 0.448 |
| 0.2 | 0 | AG1 | 1 | 5.25 | 0.378 |
| 0.2 | 0 | ptsI | 1 | 5.5 | 0.378 |
| 0.2 | 0 | clcB | 1 | 5.5 | 0.341 |
| 0.2 | 0 | ycaM | 1 | 5.5 | 0.348 |
| 0.2 | 0 | yadI | 1 | 5.5 | 0.456 |
| 0.2 | 0 | AG1 | 1 | 5.5 | 0.388 |
| 0.2 | 0 | ptsI | 1 | 5.75 | 0.39 |
| 0.2 | 0 | clcB | 1 | 5.75 | 0.348 |
| 0.2 | 0 | ycaM | 1 | 5.75 | 0.359 |
| 0.2 | 0 | yadI | 1 | 5.75 | 0.47 |
| 0.2 | 0 | AG1 | 1 | 5.75 | 0.397 |
| 0.2 | 0 | ptsI | 1 | 6 | 0.408 |
| 0.2 | 0 | clcB | 1 | 6 | 0.356 |
| 0.2 | 0 | ycaM | 1 | 6 | 0.37 |
| 0.2 | 0 | yadI | 1 | 6 | 0.486 |
| 0.2 | 0 | AG1 | 1 | 6 | 0.414 |
| 0.2 | 0 | ptsI | 1 | 6.25 | 0.417 |
| 0.2 | 0 | clcB | 1 | 6.25 | 0.37 |
| 0.2 | 0 | ycaM | 1 | 6.25 | 0.379 |
| 0.2 | 0 | yadI | 1 | 6.25 | 0.494 |
| 0.2 | 0 | AG1 | 1 | 6.25 | 0.424 |
| 0.2 | 0 | ptsI | 1 | 6.5 | 0.426 |
| 0.2 | 0 | clcB | 1 | 6.5 | 0.382 |
| 0.2 | 0 | ycaM | 1 | 6.5 | 0.389 |
| 0.2 | 0 | yadI | 1 | 6.5 | 0.51 |
| 0.2 | 0 | AG1 | 1 | 6.5 | 0.433 |
| 0.2 | 0 | ptsI | 1 | 6.75 | 0.434 |
| 0.2 | 0 | clcB | 1 | 6.75 | 0.388 |
| 0.2 | 0 | ycaM | 1 | 6.75 | 0.397 |
| 0.2 | 0 | yadI | 1 | 6.75 | 0.519 |
| 0.2 | 0 | AG1 | 1 | 6.75 | 0.443 |
| 0.2 | 0 | ptsI | 1 | 7 | 0.441 |
| 0.2 | 0 | clcB | 1 | 7 | 0.397 |
| 0.2 | 0 | ycaM | 1 | 7 | 0.406 |
| 0.2 | 0 | yadI | 1 | 7 | 0.53 |
| 0.2 | 0 | AG1 | 1 | 7 | 0.454 |
| 0.2 | 0 | ptsI | 1 | 7.25 | 0.448 |
| 0.2 | 0 | clcB | 1 | 7.25 | 0.404 |
| 0.2 | 0 | ycaM | 1 | 7.25 | 0.408 |
| 0.2 | 0 | yadI | 1 | 7.25 | 0.546 |
| 0.2 | 0 | AG1 | 1 | 7.25 | 0.464 |
| 0.2 | 0 | ptsI | 1 | 7.5 | 0.457 |
| 0.2 | 0 | clcB | 1 | 7.5 | 0.416 |

| | | | | | |
|---|---|---|---|---|---|
| 0.2 | 0 | ycaM | 1 | 7.5 | 0.414 |
| 0.2 | 0 | yadI | 1 | 7.5 | 0.554 |
| 0.2 | 0 | AG1 | 1 | 7.5 | 0.468 |
| 0.2 | 0 | ptsI | 1 | 7.75 | 0.488 |
| 0.2 | 0 | clcB | 1 | 7.75 | 0.43 |
| 0.2 | 0 | ycaM | 1 | 7.75 | 0.422 |
| 0.2 | 0 | yadI | 1 | 7.75 | 0.568 |
| 0.2 | 0 | AG1 | 1 | 7.75 | 0.479 |
| 0.2 | 0 | ptsI | 1 | 8 | 0.48 |
| 0.2 | 0 | clcB | 1 | 8 | 0.437 |
| 0.2 | 0 | ycaM | 1 | 8 | 0.427 |
| 0.2 | 0 | yadI | 1 | 8 | 0.574 |
| 0.2 | 0 | AG1 | 1 | 8 | 0.486 |
| 0.2 | 0 | ptsI | 1 | 8.25 | 0.489 |
| 0.2 | 0 | clcB | 1 | 8.25 | 0.447 |
| 0.2 | 0 | ycaM | 1 | 8.25 | 0.436 |
| 0.2 | 0 | yadI | 1 | 8.25 | 0.587 |
| 0.2 | 0 | AG1 | 1 | 8.25 | 0.494 |
| 0.2 | 0 | ptsI | 1 | 8.5 | 0.5 |
| 0.2 | 0 | clcB | 1 | 8.5 | 0.458 |
| 0.2 | 0 | ycaM | 1 | 8.5 | 0.45 |
| 0.2 | 0 | yadI | 1 | 8.5 | 0.596 |
| 0.2 | 0 | AG1 | 1 | 8.5 | 0.503 |
| 0.2 | 0 | ptsI | 1 | 8.75 | 0.509 |
| 0.2 | 0 | clcB | 1 | 8.75 | 0.463 |
| 0.2 | 0 | ycaM | 1 | 8.75 | 0.464 |
| 0.2 | 0 | yadI | 1 | 8.75 | 0.605 |
| 0.2 | 0 | AG1 | 1 | 8.75 | 0.514 |
| 0.2 | 0 | ptsI | 1 | 9 | 0.519 |
| 0.2 | 0 | clcB | 1 | 9 | 0.471 |
| 0.2 | 0 | ycaM | 1 | 9 | 0.483 |
| 0.2 | 0 | yadI | 1 | 9 | 0.619 |
| 0.2 | 0 | AG1 | 1 | 9 | 0.523 |
| 0.2 | 0 | ptsI | 1 | 9.25 | 0.546 |
| 0.2 | 0 | clcB | 1 | 9.25 | 0.482 |
| 0.2 | 0 | ycaM | 1 | 9.25 | 0.488 |
| 0.2 | 0 | yadI | 1 | 9.25 | 0.62 |
| 0.2 | 0 | AG1 | 1 | 9.25 | 0.533 |
| 0.2 | 0 | ptsI | 1 | 9.5 | 0.57 |
| 0.2 | 0 | clcB | 1 | 9.5 | 0.498 |
| 0.2 | 0 | ycaM | 1 | 9.5 | 0.504 |
| 0.2 | 0 | yadI | 1 | 9.5 | 0.634 |
| 0.2 | 0 | AG1 | 1 | 9.5 | 0.548 |
| 0.2 | 0 | ptsI | 1 | 9.75 | 0.557 |
| 0.2 | 0 | clcB | 1 | 9.75 | 0.508 |
| 0.2 | 0 | ycaM | 1 | 9.75 | 0.519 |
| 0.2 | 0 | yadI | 1 | 9.75 | 0.666 |
| 0.2 | 0 | AG1 | 1 | 9.75 | 0.565 |

| | | | | | |
|---|---|---|---|---|---|
| 0.2 | 0 | ptsI | 1 | 10 | 0.588 |
| 0.2 | 0 | clcB | 1 | 10 | 0.517 |
| 0.2 | 0 | ycaM | 1 | 10 | 0.531 |
| 0.2 | 0 | yadI | 1 | 10 | 0.66 |
| 0.2 | 0 | AG1 | 1 | 10 | 0.569 |
| 0.2 | 0 | ptsI | 1 | 10.25 | 0.599 |
| 0.2 | 0 | clcB | 1 | 10.25 | 0.524 |
| 0.2 | 0 | ycaM | 1 | 10.25 | 0.552 |
| 0.2 | 0 | yadI | 1 | 10.25 | 0.671 |
| 0.2 | 0 | AG1 | 1 | 10.25 | 0.58 |
| 0.2 | 0 | ptsI | 1 | 10.5 | 0.616 |
| 0.2 | 0 | clcB | 1 | 10.5 | 0.533 |
| 0.2 | 0 | ycaM | 1 | 10.5 | 0.572 |
| 0.2 | 0 | yadI | 1 | 10.5 | 0.668 |
| 0.2 | 0 | AG1 | 1 | 10.5 | 0.584 |
| 0.2 | 0 | ptsI | 1 | 10.75 | 0.667 |
| 0.2 | 0 | clcB | 1 | 10.75 | 0.545 |
| 0.2 | 0 | ycaM | 1 | 10.75 | 0.567 |
| 0.2 | 0 | yadI | 1 | 10.75 | 0.683 |
| 0.2 | 0 | AG1 | 1 | 10.75 | 0.586 |
| 0.2 | 0 | ptsI | 1 | 11 | 0.68 |
| 0.2 | 0 | clcB | 1 | 11 | 0.566 |
| 0.2 | 0 | ycaM | 1 | 11 | 0.594 |
| 0.2 | 0 | yadI | 1 | 11 | 0.664 |
| 0.2 | 0 | AG1 | 1 | 11 | 0.596 |
| 0.2 | 0 | ptsI | 1 | 11.25 | 0.674 |
| 0.2 | 0 | clcB | 1 | 11.25 | 0.567 |
| 0.2 | 0 | ycaM | 1 | 11.25 | 0.616 |
| 0.2 | 0 | yadI | 1 | 11.25 | 0.685 |
| 0.2 | 0 | AG1 | 1 | 11.25 | 0.602 |
| 0.2 | 0 | ptsI | 1 | 11.5 | 0.675 |
| 0.2 | 0 | clcB | 1 | 11.5 | 0.583 |
| 0.2 | 0 | ycaM | 1 | 11.5 | 0.628 |
| 0.2 | 0 | yadI | 1 | 11.5 | 0.693 |
| 0.2 | 0 | AG1 | 1 | 11.5 | 0.618 |
| 0.2 | 0 | ptsI | 1 | 11.75 | 0.705 |
| 0.2 | 0 | clcB | 1 | 11.75 | 0.594 |
| 0.2 | 0 | ycaM | 1 | 11.75 | 0.643 |
| 0.2 | 0 | yadI | 1 | 11.75 | 0.704 |
| 0.2 | 0 | AG1 | 1 | 11.75 | 0.621 |
| 0.2 | 0 | ptsI | 1 | 12 | 0.721 |
| 0.2 | 0 | clcB | 1 | 12 | 0.595 |
| 0.2 | 0 | ycaM | 1 | 12 | 0.633 |
| 0.2 | 0 | yadI | 1 | 12 | 0.708 |
| 0.2 | 0 | AG1 | 1 | 12 | 0.63 |
| 0.2 | 0 | ptsI | 1 | 12.25 | 0.715 |
| 0.2 | 0 | clcB | 1 | 12.25 | 0.603 |
| 0.2 | 0 | ycaM | 1 | 12.25 | 0.645 |

| | | | | | |
|---|---|---|---|---|---|
| 0.2 | 0 | yadI | 1 | 12.25 | 0.679 |
| 0.2 | 0 | AG1 | 1 | 12.25 | 0.637 |
| 0.2 | 0 | ptsI | 1 | 12.5 | 0.715 |
| 0.2 | 0 | clcB | 1 | 12.5 | 0.609 |
| 0.2 | 0 | ycaM | 1 | 12.5 | 0.629 |
| 0.2 | 0 | yadI | 1 | 12.5 | 0.703 |
| 0.2 | 0 | AG1 | 1 | 12.5 | 0.656 |
| 0.2 | 0 | ptsI | 1 | 12.75 | 0.74 |
| 0.2 | 0 | clcB | 1 | 12.75 | 0.622 |
| 0.2 | 0 | ycaM | 1 | 12.75 | 0.629 |
| 0.2 | 0 | yadI | 1 | 12.75 | 0.708 |
| 0.2 | 0 | AG1 | 1 | 12.75 | 0.67 |
| 0.2 | 0 | ptsI | 1 | 13 | 0.751 |
| 0.2 | 0 | clcB | 1 | 13 | 0.618 |
| 0.2 | 0 | ycaM | 1 | 13 | 0.646 |
| 0.2 | 0 | yadI | 1 | 13 | 0.724 |
| 0.2 | 0 | AG1 | 1 | 13 | 0.683 |
| 0.2 | 0 | ptsI | 1 | 13.25 | 0.756 |
| 0.2 | 0 | clcB | 1 | 13.25 | 0.633 |
| 0.2 | 0 | ycaM | 1 | 13.25 | 0.682 |
| 0.2 | 0 | yadI | 1 | 13.25 | 0.724 |
| 0.2 | 0 | AG1 | 1 | 13.25 | 0.689 |
| 0.2 | 0 | ptsI | 1 | 13.5 | 0.771 |
| 0.2 | 0 | clcB | 1 | 13.5 | 0.637 |
| 0.2 | 0 | ycaM | 1 | 13.5 | 0.69 |
| 0.2 | 0 | yadI | 1 | 13.5 | 0.738 |
| 0.2 | 0 | AG1 | 1 | 13.5 | 0.716 |
| 0.2 | 0 | ptsI | 1 | 13.75 | 0.777 |
| 0.2 | 0 | clcB | 1 | 13.75 | 0.649 |
| 0.2 | 0 | ycaM | 1 | 13.75 | 0.688 |
| 0.2 | 0 | yadI | 1 | 13.75 | 0.745 |
| 0.2 | 0 | AG1 | 1 | 13.75 | 0.725 |
| 0.2 | 0 | ptsI | 1 | 14 | 0.805 |
| 0.2 | 0 | clcB | 1 | 14 | 0.667 |
| 0.2 | 0 | ycaM | 1 | 14 | 0.707 |
| 0.2 | 0 | yadI | 1 | 14 | 0.753 |
| 0.2 | 0 | AG1 | 1 | 14 | 0.753 |
| 0.2 | 0 | ptsI | 1 | 14.25 | 0.806 |
| 0.2 | 0 | clcB | 1 | 14.25 | 0.698 |
| 0.2 | 0 | ycaM | 1 | 14.25 | 0.712 |
| 0.2 | 0 | yadI | 1 | 14.25 | 0.757 |
| 0.2 | 0 | AG1 | 1 | 14.25 | 0.759 |
| 0.2 | 0 | ptsI | 1 | 14.5 | 0.796 |
| 0.2 | 0 | clcB | 1 | 14.5 | 0.724 |
| 0.2 | 0 | ycaM | 1 | 14.5 | 0.713 |
| 0.2 | 0 | yadI | 1 | 14.5 | 0.761 |
| 0.2 | 0 | AG1 | 1 | 14.5 | 0.767 |
| 0.2 | 0 | ptsI | 1 | 14.75 | 0.796 |

| | | | | | |
|---|---|---|---|---|---|
| 0.2 | 0 | clcB | 1 | 14.75 | 0.717 |
| 0.2 | 0 | ycaM | 1 | 14.75 | 0.724 |
| 0.2 | 0 | yadI | 1 | 14.75 | 0.731 |
| 0.2 | 0 | AG1 | 1 | 14.75 | 0.771 |
| 0.2 | 0 | ptsI | 1 | 15 | 0.809 |
| 0.2 | 0 | clcB | 1 | 15 | 0.732 |
| 0.2 | 0 | ycaM | 1 | 15 | 0.73 |
| 0.2 | 0 | yadI | 1 | 15 | 0.739 |
| 0.2 | 0 | AG1 | 1 | 15 | 0.794 |
| 0.2 | 0 | ptsI | 1 | 15.25 | 0.83 |
| 0.2 | 0 | clcB | 1 | 15.25 | 0.736 |
| 0.2 | 0 | ycaM | 1 | 15.25 | 0.741 |
| 0.2 | 0 | yadI | 1 | 15.25 | 0.752 |
| 0.2 | 0 | AG1 | 1 | 15.25 | 0.81 |
| 0.2 | 0 | ptsI | 1 | 15.5 | 0.815 |
| 0.2 | 0 | clcB | 1 | 15.5 | 0.749 |
| 0.2 | 0 | ycaM | 1 | 15.5 | 0.743 |
| 0.2 | 0 | yadI | 1 | 15.5 | 0.737 |
| 0.2 | 0 | AG1 | 1 | 15.5 | 0.807 |
| 0.2 | 0 | ptsI | 1 | 15.75 | 0.808 |
| 0.2 | 0 | clcB | 1 | 15.75 | 0.746 |
| 0.2 | 0 | ycaM | 1 | 15.75 | 0.746 |
| 0.2 | 0 | yadI | 1 | 15.75 | 0.733 |
| 0.2 | 0 | AG1 | 1 | 15.75 | 0.804 |
| 0.2 | 0 | ptsI | 1 | 16 | 0.816 |
| 0.2 | 0 | clcB | 1 | 16 | 0.759 |
| 0.2 | 0 | ycaM | 1 | 16 | 0.752 |
| 0.2 | 0 | yadI | 1 | 16 | 0.729 |
| 0.2 | 0 | AG1 | 1 | 16 | 0.802 |
| 0.2 | 0 | ptsI | 1 | 16.25 | 0.83 |
| 0.2 | 0 | clcB | 1 | 16.25 | 0.774 |
| 0.2 | 0 | ycaM | 1 | 16.25 | 0.763 |
| 0.2 | 0 | yadI | 1 | 16.25 | 0.74 |
| 0.2 | 0 | AG1 | 1 | 16.25 | 0.824 |
| 0.2 | 0 | ptsI | 1 | 16.5 | 0.844 |
| 0.2 | 0 | clcB | 1 | 16.5 | 0.782 |
| 0.2 | 0 | ycaM | 1 | 16.5 | 0.77 |
| 0.2 | 0 | yadI | 1 | 16.5 | 0.747 |
| 0.2 | 0 | AG1 | 1 | 16.5 | 0.843 |
| 0.2 | 0 | ptsI | 1 | 17.0667 | 0.867 |
| 0.2 | 0 | clcB | 1 | 17.0667 | 0.85 |
| 0.2 | 0 | ycaM | 1 | 17.0667 | 0.809 |
| 0.2 | 0 | yadI | 1 | 17.0667 | 0.784 |
| 0.2 | 0 | AG1 | 1 | 17.0667 | 0.854 |
| 0.2 | 0 | ptsI | 1 | 17.3167 | 0.81 |
| 0.2 | 0 | clcB | 1 | 17.3167 | 0.772 |
| 0.2 | 0 | ycaM | 1 | 17.3167 | 0.774 |
| 0.2 | 0 | yadI | 1 | 17.3167 | 0.774 |

| | | | | | |
|---|---|---|---|---|---|
| 0.2 | 0 | AG1 | 1 | 17.3167 | 0.824 |
| 0.2 | 0 | ptsI | 1 | 17.5667 | 0.816 |
| 0.2 | 0 | clcB | 1 | 17.5667 | 0.771 |
| 0.2 | 0 | ycaM | 1 | 17.5667 | 0.762 |
| 0.2 | 0 | yadI | 1 | 17.5667 | 0.772 |
| 0.2 | 0 | AG1 | 1 | 17.5667 | 0.824 |
| 0.2 | 0 | ptsI | 1 | 17.8167 | 0.839 |
| 0.2 | 0 | clcB | 1 | 17.8167 | 0.799 |
| 0.2 | 0 | ycaM | 1 | 17.8167 | 0.766 |
| 0.2 | 0 | yadI | 1 | 17.8167 | 0.779 |
| 0.2 | 0 | AG1 | 1 | 17.8167 | 0.866 |
| 0.2 | 0 | ptsI | 1 | 18.0667 | 0.842 |
| 0.2 | 0 | clcB | 1 | 18.0667 | 0.778 |
| 0.2 | 0 | ycaM | 1 | 18.0667 | 0.774 |
| 0.2 | 0 | yadI | 1 | 18.0667 | 0.78 |
| 0.2 | 0 | AG1 | 1 | 18.0667 | 0.816 |
| 0.2 | 0 | ptsI | 1 | 18.3167 | 0.846 |
| 0.2 | 0 | clcB | 1 | 18.3167 | 0.78 |
| 0.2 | 0 | ycaM | 1 | 18.3167 | 0.773 |
| 0.2 | 0 | yadI | 1 | 18.3167 | 0.781 |
| 0.2 | 0 | AG1 | 1 | 18.3167 | 0.778 |
| 0.2 | 0 | ptsI | 1 | 18.5667 | 0.861 |
| 0.2 | 0 | clcB | 1 | 18.5667 | 0.8 |
| 0.2 | 0 | ycaM | 1 | 18.5667 | 0.78 |
| 0.2 | 0 | yadI | 1 | 18.5667 | 0.802 |
| 0.2 | 0 | AG1 | 1 | 18.5667 | 0.818 |
| 0.2 | 0 | ptsI | 1 | 18.8167 | 0.865 |
| 0.2 | 0 | clcB | 1 | 18.8167 | 0.808 |
| 0.2 | 0 | ycaM | 1 | 18.8167 | 0.785 |
| 0.2 | 0 | yadI | 1 | 18.8167 | 0.79 |
| 0.2 | 0 | AG1 | 1 | 18.8167 | 0.826 |
| 0.2 | 0 | ptsI | 1 | 19.0667 | 0.872 |
| 0.2 | 0 | clcB | 1 | 19.0667 | 0.811 |
| 0.2 | 0 | ycaM | 1 | 19.0667 | 0.79 |
| 0.2 | 0 | yadI | 1 | 19.0667 | 0.798 |
| 0.2 | 0 | AG1 | 1 | 19.0667 | 0.834 |
| 0.2 | 0 | ptsI | 1 | 19.3167 | 0.875 |
| 0.2 | 0 | clcB | 1 | 19.3167 | 0.818 |
| 0.2 | 0 | ycaM | 1 | 19.3167 | 0.789 |
| 0.2 | 0 | yadI | 1 | 19.3167 | 0.81 |
| 0.2 | 0 | AG1 | 1 | 19.3167 | 0.843 |
| 0.2 | 0 | ptsI | 1 | 19.5667 | 0.884 |
| 0.2 | 0 | clcB | 1 | 19.5667 | 0.822 |
| 0.2 | 0 | ycaM | 1 | 19.5667 | 0.789 |
| 0.2 | 0 | yadI | 1 | 19.5667 | 0.824 |
| 0.2 | 0 | AG1 | 1 | 19.5667 | 0.845 |
| 0.2 | 0 | ptsI | 1 | 19.8167 | 0.886 |
| 0.2 | 0 | clcB | 1 | 19.8167 | 0.821 |

| | | | | | |
|---|---|---|---|---|---|
| 0.2 | 0 | ycaM | 1 | 19.8167 | 0.797 |
| 0.2 | 0 | yadI | 1 | 19.8167 | 0.815 |
| 0.2 | 0 | AG1 | 1 | 19.8167 | 0.848 |
| 0.2 | 0 | ptsI | 1 | 20.0667 | 0.897 |
| 0.2 | 0 | clcB | 1 | 20.0667 | 0.83 |
| 0.2 | 0 | ycaM | 1 | 20.0667 | 0.799 |
| 0.2 | 0 | yadI | 1 | 20.0667 | 0.823 |
| 0.2 | 0 | AG1 | 1 | 20.0667 | 0.857 |
| 0.2 | 0 | ptsI | 1 | 20.3167 | 0.902 |
| 0.2 | 0 | clcB | 1 | 20.3167 | 0.83 |
| 0.2 | 0 | ycaM | 1 | 20.3167 | 0.804 |
| 0.2 | 0 | yadI | 1 | 20.3167 | 0.824 |
| 0.2 | 0 | AG1 | 1 | 20.3167 | 0.858 |
| 0.2 | 0 | ptsI | 1 | 20.5667 | 0.898 |
| 0.2 | 0 | clcB | 1 | 20.5667 | 0.822 |
| 0.2 | 0 | ycaM | 1 | 20.5667 | 0.803 |
| 0.2 | 0 | yadI | 1 | 20.5667 | 0.836 |
| 0.2 | 0 | AG1 | 1 | 20.5667 | 0.846 |
| 0.2 | 0 | ptsI | 1 | 20.8167 | 0.91 |
| 0.2 | 0 | clcB | 1 | 20.8167 | 0.828 |
| 0.2 | 0 | ycaM | 1 | 20.8167 | 0.811 |
| 0.2 | 0 | yadI | 1 | 20.8167 | 0.831 |
| 0.2 | 0 | AG1 | 1 | 20.8167 | 0.833 |
| 0.2 | 0 | ptsI | 1 | 21.0667 | 0.926 |
| 0.2 | 0 | clcB | 1 | 21.0667 | 0.835 |
| 0.2 | 0 | ycaM | 1 | 21.0667 | 0.818 |
| 0.2 | 0 | yadI | 1 | 21.0667 | 0.841 |
| 0.2 | 0 | AG1 | 1 | 21.0667 | 0.842 |
| 0.2 | 0 | ptsI | 1 | 21.3167 | 0.935 |
| 0.2 | 0 | clcB | 1 | 21.3167 | 0.84 |
| 0.2 | 0 | ycaM | 1 | 21.3167 | 0.82 |
| 0.2 | 0 | yadI | 1 | 21.3167 | 0.851 |
| 0.2 | 0 | AG1 | 1 | 21.3167 | 0.838 |
| 0.2 | 0 | ptsI | 1 | 21.5667 | 0.933 |
| 0.2 | 0 | clcB | 1 | 21.5667 | 0.836 |
| 0.2 | 0 | ycaM | 1 | 21.5667 | 0.819 |
| 0.2 | 0 | yadI | 1 | 21.5667 | 0.857 |
| 0.2 | 0 | AG1 | 1 | 21.5667 | 0.842 |
| 0.2 | 0 | ptsI | 1 | 21.8167 | 0.943 |
| 0.2 | 0 | clcB | 1 | 21.8167 | 0.835 |
| 0.2 | 0 | ycaM | 1 | 21.8167 | 0.826 |
| 0.2 | 0 | yadI | 1 | 21.8167 | 0.856 |
| 0.2 | 0 | AG1 | 1 | 21.8167 | 0.832 |
| 0.2 | 0 | ptsI | 1 | 22.0667 | 0.963 |
| 0.2 | 0 | clcB | 1 | 22.0667 | 0.842 |
| 0.2 | 0 | ycaM | 1 | 22.0667 | 0.834 |
| 0.2 | 0 | yadI | 1 | 22.0667 | 0.859 |
| 0.2 | 0 | AG1 | 1 | 22.0667 | 0.842 |

| | | | | | |
|---|---|---|---|---|---|
| 0.2 | 0 | ptsI | 1 | 22.9333 | 0.703 |
| 0.2 | 0 | clcB | 1 | 22.9333 | 0.912 |
| 0.2 | 0 | ycaM | 1 | 22.9333 | 0.704 |
| 0.2 | 0 | yadI | 1 | 22.9333 | 0.856 |
| 0.2 | 0 | AG1 | 1 | 22.9333 | 0.844 |
| 0.2 | 0 | ptsI | 1 | 23.1833 | 0.745 |
| 0.2 | 0 | clcB | 1 | 23.1833 | 0.909 |
| 0.2 | 0 | ycaM | 1 | 23.1833 | 0.734 |
| 0.2 | 0 | yadI | 1 | 23.1833 | 0.874 |
| 0.2 | 0 | AG1 | 1 | 23.1833 | 0.833 |
| 0.2 | 0 | ptsI | 1 | 23.4333 | 0.768 |
| 0.2 | 0 | clcB | 1 | 23.4333 | 0.899 |
| 0.2 | 0 | ycaM | 1 | 23.4333 | 0.767 |
| 0.2 | 0 | yadI | 1 | 23.4333 | 0.875 |
| 0.2 | 0 | AG1 | 1 | 23.4333 | 0.845 |
| 0.2 | 0 | ptsI | 1 | 23.6833 | 0.793 |
| 0.2 | 0 | clcB | 1 | 23.6833 | 0.896 |
| 0.2 | 0 | ycaM | 1 | 23.6833 | 0.788 |
| 0.2 | 0 | yadI | 1 | 23.6833 | 0.88 |
| 0.2 | 0 | AG1 | 1 | 23.6833 | 0.83 |
| 0.2 | 0 | ptsI | 1 | 23.9333 | 0.822 |
| 0.2 | 0 | clcB | 1 | 23.9333 | 0.892 |
| 0.2 | 0 | ycaM | 1 | 23.9333 | 0.793 |
| 0.2 | 0 | yadI | 1 | 23.9333 | 0.886 |
| 0.2 | 0 | AG1 | 1 | 23.9333 | 0.851 |
| 0.2 | 0 | ptsI | 1 | 24.1833 | 0.841 |
| 0.2 | 0 | clcB | 1 | 24.1833 | 0.885 |
| 0.2 | 0 | ycaM | 1 | 24.1833 | 0.806 |
| 0.2 | 0 | yadI | 1 | 24.1833 | 0.897 |
| 0.2 | 0 | AG1 | 1 | 24.1833 | 0.865 |
| 0.2 | 0 | ptsI | 1 | 24.4333 | 0.854 |
| 0.2 | 0 | clcB | 1 | 24.4333 | 0.896 |
| 0.2 | 0 | ycaM | 1 | 24.4333 | 0.807 |
| 0.2 | 0 | yadI | 1 | 24.4333 | 0.906 |
| 0.2 | 0 | AG1 | 1 | 24.4333 | 0.859 |
| 0.2 | 0 | ptsI | 1 | 24.6833 | 0.861 |
| 0.2 | 0 | clcB | 1 | 24.6833 | 0.892 |
| 0.2 | 0 | ycaM | 1 | 24.6833 | 0.808 |
| 0.2 | 0 | yadI | 1 | 24.6833 | 0.909 |
| 0.2 | 0 | AG1 | 1 | 24.6833 | 0.864 |
| 0.2 | 0 | ptsI | 1 | 24.9333 | 0.875 |
| 0.2 | 0 | clcB | 1 | 24.9333 | 0.895 |
| 0.2 | 0 | ycaM | 1 | 24.9333 | 0.814 |
| 0.2 | 0 | yadI | 1 | 24.9333 | 0.93 |
| 0.2 | 0 | AG1 | 1 | 24.9333 | 0.861 |
| 0.2 | 0 | ptsI | 1 | 25.1833 | 0.886 |
| 0.2 | 0 | clcB | 1 | 25.1833 | 0.9 |
| 0.2 | 0 | ycaM | 1 | 25.1833 | 0.82 |

| | | | | | |
|---|---|---|---|---|---|
| 0.2 | 0 | yadI | 1 | 25.1833 | 0.926 |
| 0.2 | 0 | AG1 | 1 | 25.1833 | 0.871 |
| 0.2 | 0 | ptsI | 1 | 25.4333 | 0.9 |
| 0.2 | 0 | clcB | 1 | 25.4333 | 0.899 |
| 0.2 | 0 | ycaM | 1 | 25.4333 | 0.832 |
| 0.2 | 0 | yadI | 1 | 25.4333 | 0.935 |
| 0.2 | 0 | AG1 | 1 | 25.4333 | 0.887 |
| 0.2 | 0 | ptsI | 1 | 25.6833 | 0.907 |
| 0.2 | 0 | clcB | 1 | 25.6833 | 0.908 |
| 0.2 | 0 | ycaM | 1 | 25.6833 | 0.845 |
| 0.2 | 0 | yadI | 1 | 25.6833 | 0.939 |
| 0.2 | 0 | AG1 | 1 | 25.6833 | 0.886 |
| 0.2 | 0 | ptsI | 1 | 25.9333 | 0.927 |
| 0.2 | 0 | clcB | 1 | 25.9333 | 0.908 |
| 0.2 | 0 | ycaM | 1 | 25.9333 | 0.854 |
| 0.2 | 0 | yadI | 1 | 25.9333 | 0.96 |
| 0.2 | 0 | AG1 | 1 | 25.9333 | 0.894 |
| 0.2 | 0 | ptsI | 1 | 26.1833 | 0.924 |
| 0.2 | 0 | clcB | 1 | 26.1833 | 0.911 |
| 0.2 | 0 | ycaM | 1 | 26.1833 | 0.866 |
| 0.2 | 0 | yadI | 1 | 26.1833 | 0.955 |
| 0.2 | 0 | AG1 | 1 | 26.1833 | 0.913 |
| 0.2 | 0 | ptsI | 1 | 26.4333 | 0.93 |
| 0.2 | 0 | clcB | 1 | 26.4333 | 0.909 |
| 0.2 | 0 | ycaM | 1 | 26.4333 | 0.871 |
| 0.2 | 0 | yadI | 1 | 26.4333 | 0.961 |
| 0.2 | 0 | AG1 | 1 | 26.4333 | 0.906 |
| 0.2 | 0 | ptsI | 1 | 26.6833 | 0.939 |
| 0.2 | 0 | clcB | 1 | 26.6833 | 0.913 |
| 0.2 | 0 | ycaM | 1 | 26.6833 | 0.882 |
| 0.2 | 0 | yadI | 1 | 26.6833 | 0.962 |
| 0.2 | 0 | AG1 | 1 | 26.6833 | 0.911 |
| 0.2 | 0 | ptsI | 1 | 26.9333 | 0.945 |
| 0.2 | 0 | clcB | 1 | 26.9333 | 0.911 |
| 0.2 | 0 | ycaM | 1 | 26.9333 | 0.889 |
| 0.2 | 0 | yadI | 1 | 26.9333 | 0.968 |
| 0.2 | 0 | AG1 | 1 | 26.9333 | 0.917 |
| 0.2 | 0 | ptsI | 1 | 27.1833 | 0.946 |
| 0.2 | 0 | clcB | 1 | 27.1833 | 0.912 |
| 0.2 | 0 | ycaM | 1 | 27.1833 | 0.9 |
| 0.2 | 0 | yadI | 1 | 27.1833 | 0.979 |
| 0.2 | 0 | AG1 | 1 | 27.1833 | 0.923 |
| 0.2 | 0 | ptsI | 1 | 27.4333 | 0.955 |
| 0.2 | 0 | clcB | 1 | 27.4333 | 0.915 |
| 0.2 | 0 | ycaM | 1 | 27.4333 | 0.906 |
| 0.2 | 0 | yadI | 1 | 27.4333 | 0.974 |
| 0.2 | 0 | AG1 | 1 | 27.4333 | 0.916 |
| 0.2 | 0 | ptsI | 1 | 27.6833 | 0.965 |

| | | | | | |
|---|---|---|---|---|---|
| 0.2 | 0 | clcB | 1 | 27.6833 | 0.919 |
| 0.2 | 0 | ycaM | 1 | 27.6833 | 0.913 |
| 0.2 | 0 | yadI | 1 | 27.6833 | 0.982 |
| 0.2 | 0 | AG1 | 1 | 27.6833 | 0.926 |
| 0.2 | 0 | ptsI | 1 | 27.9333 | 0.967 |
| 0.2 | 0 | clcB | 1 | 27.9333 | 0.909 |
| 0.2 | 0 | ycaM | 1 | 27.9333 | 0.924 |
| 0.2 | 0 | yadI | 1 | 27.9333 | 0.99 |
| 0.2 | 0 | AG1 | 1 | 27.9333 | 0.916 |
| 0.2 | 0 | ptsI | 1 | 28.1833 | 0.972 |
| 0.2 | 0 | clcB | 1 | 28.1833 | 0.916 |
| 0.2 | 0 | ycaM | 1 | 28.1833 | 0.933 |
| 0.2 | 0 | yadI | 1 | 28.1833 | 0.994 |
| 0.2 | 0 | AG1 | 1 | 28.1833 | 0.939 |
| 0.2 | 0 | ptsI | 1 | 28.4333 | 0.985 |
| 0.2 | 0 | clcB | 1 | 28.4333 | 0.926 |
| 0.2 | 0 | ycaM | 1 | 28.4333 | 0.938 |
| 0.2 | 0 | yadI | 1 | 28.4333 | 0.995 |
| 0.2 | 0 | AG1 | 1 | 28.4333 | 0.937 |
| 0.2 | 0 | ptsI | 1 | 28.6833 | 0.992 |
| 0.2 | 0 | clcB | 1 | 28.6833 | 0.928 |
| 0.2 | 0 | ycaM | 1 | 28.6833 | 0.948 |
| 0.2 | 0 | yadI | 1 | 28.6833 | 1.002 |
| 0.2 | 0 | AG1 | 1 | 28.6833 | 0.939 |
| 0.2 | 0 | ptsI | 1 | 28.9333 | 0.997 |
| 0.2 | 0 | clcB | 1 | 28.9333 | 0.933 |
| 0.2 | 0 | ycaM | 1 | 28.9333 | 0.954 |
| 0.2 | 0 | yadI | 1 | 28.9333 | 1.009 |
| 0.2 | 0 | AG1 | 1 | 28.9333 | 0.948 |
| 0.2 | 0 | ptsI | 1 | 29.1833 | 1.005 |
| 0.2 | 0 | clcB | 1 | 29.1833 | 0.936 |
| 0.2 | 0 | ycaM | 1 | 29.1833 | 0.958 |
| 0.2 | 0 | yadI | 1 | 29.1833 | 1.003 |
| 0.2 | 0 | AG1 | 1 | 29.1833 | 0.942 |
| 0.2 | 0 | ptsI | 1 | 29.4333 | 1.012 |
| 0.2 | 0 | clcB | 1 | 29.4333 | 0.94 |
| 0.2 | 0 | ycaM | 1 | 29.4333 | 0.964 |
| 0.2 | 0 | yadI | 1 | 29.4333 | 1.016 |
| 0.2 | 0 | AG1 | 1 | 29.4333 | 0.95 |
| 0.2 | 0 | ptsI | 1 | 29.6833 | 1.019 |
| 0.2 | 0 | clcB | 1 | 29.6833 | 0.944 |
| 0.2 | 0 | ycaM | 1 | 29.6833 | 0.972 |
| 0.2 | 0 | yadI | 1 | 29.6833 | 1.018 |
| 0.2 | 0 | AG1 | 1 | 29.6833 | 0.954 |
| 0.2 | 0 | ptsI | 1 | 29.9333 | 1.025 |
| 0.2 | 0 | clcB | 1 | 29.9333 | 0.945 |
| 0.2 | 0 | ycaM | 1 | 29.9333 | 0.966 |
| 0.2 | 0 | yadI | 1 | 29.9333 | 1.027 |

| | | | | | |
|---|---|---|---|---|---|
| 0.2 | 0 | AG1 | 1 | 29.9333 | 0.961 |
| 0.2 | 0 | ptsI | 1 | 30.1833 | 1.034 |
| 0.2 | 0 | clcB | 1 | 30.1833 | 0.944 |
| 0.2 | 0 | ycaM | 1 | 30.1833 | 0.994 |
| 0.2 | 0 | yadI | 1 | 30.1833 | 1.03 |
| 0.2 | 0 | AG1 | 1 | 30.1833 | 0.964 |
| 0.2 | 0 | ptsI | 1 | 30.4333 | 1.044 |
| 0.2 | 0 | clcB | 1 | 30.4333 | 0.953 |
| 0.2 | 0 | ycaM | 1 | 30.4333 | 0.984 |
| 0.2 | 0 | yadI | 1 | 30.4333 | 1.039 |
| 0.2 | 0 | AG1 | 1 | 30.4333 | 0.968 |
| 0.2 | 0 | ptsI | 1 | 30.6833 | 1.051 |
| 0.2 | 0 | clcB | 1 | 30.6833 | 0.954 |
| 0.2 | 0 | ycaM | 1 | 30.6833 | 0.984 |
| 0.2 | 0 | yadI | 1 | 30.6833 | 1.033 |
| 0.2 | 0 | AG1 | 1 | 30.6833 | 0.968 |
| 0.2 | 0 | ptsI | 1 | 30.9333 | 1.054 |
| 0.2 | 0 | clcB | 1 | 30.9333 | 0.955 |
| 0.2 | 0 | ycaM | 1 | 30.9333 | 1.003 |
| 0.2 | 0 | yadI | 1 | 30.9333 | 1.042 |
| 0.2 | 0 | AG1 | 1 | 30.9333 | 0.972 |
| 0.2 | 0 | ptsI | 1 | 31.1833 | 1.061 |
| 0.2 | 0 | clcB | 1 | 31.1833 | 0.957 |
| 0.2 | 0 | ycaM | 1 | 31.1833 | 0.987 |
| 0.2 | 0 | yadI | 1 | 31.1833 | 1.044 |
| 0.2 | 0 | AG1 | 1 | 31.1833 | 0.976 |
| 0.2 | 0 | ptsI | 1 | 31.4333 | 1.073 |
| 0.2 | 0 | clcB | 1 | 31.4333 | 0.96 |
| 0.2 | 0 | ycaM | 1 | 31.4333 | 1.008 |
| 0.2 | 0 | yadI | 1 | 31.4333 | 1.058 |
| 0.2 | 0 | AG1 | 1 | 31.4333 | 0.989 |
| 0.2 | 0 | ptsI | 1 | 31.6833 | 1.08 |
| 0.2 | 0 | clcB | 1 | 31.6833 | 0.968 |
| 0.2 | 0 | ycaM | 1 | 31.6833 | 1.009 |
| 0.2 | 0 | yadI | 1 | 31.6833 | 1.065 |
| 0.2 | 0 | AG1 | 1 | 31.6833 | 0.99 |
| 0.2 | 0 | ptsI | 1 | 31.9333 | 1.086 |
| 0.2 | 0 | clcB | 1 | 31.9333 | 0.976 |
| 0.2 | 0 | ycaM | 1 | 31.9333 | 1.007 |
| 0.2 | 0 | yadI | 1 | 31.9333 | 1.073 |
| 0.2 | 0 | AG1 | 1 | 31.9333 | 0.998 |
| 0.2 | 0 | ptsI | 1 | 32.1833 | 1.093 |
| 0.2 | 0 | clcB | 1 | 32.1833 | 0.978 |
| 0.2 | 0 | ycaM | 1 | 32.1833 | 1.01 |
| 0.2 | 0 | yadI | 1 | 32.1833 | 1.083 |
| 0.2 | 0 | AG1 | 1 | 32.1833 | 1 |
| 0.2 | 0 | ptsI | 1 | 32.4333 | 1.107 |
| 0.2 | 0 | clcB | 1 | 32.4333 | 0.973 |

| | | | | | |
|---|---|---|---|---|---|
| 0.2 | 0 | ycaM | 1 | 32.4333 | 1.011 |
| 0.2 | 0 | yadI | 1 | 32.4333 | 1.102 |
| 0.2 | 0 | AG1 | 1 | 32.4333 | 0.988 |
| 0.2 | 0 | ptsI | 1 | 32.6833 | 1.109 |
| 0.2 | 0 | clcB | 1 | 32.6833 | 0.99 |
| 0.2 | 0 | ycaM | 1 | 32.6833 | 1.016 |
| 0.2 | 0 | yadI | 1 | 32.6833 | 1.096 |
| 0.2 | 0 | AG1 | 1 | 32.6833 | 1.01 |
| 0.2 | 0 | ptsI | 1 | 32.9333 | 1.117 |
| 0.2 | 0 | clcB | 1 | 32.9333 | 1.005 |
| 0.2 | 0 | ycaM | 1 | 32.9333 | 1.017 |
| 0.2 | 0 | yadI | 1 | 32.9333 | 1.108 |
| 0.2 | 0 | AG1 | 1 | 32.9333 | 1.016 |
| 0.2 | 0 | ptsI | 1 | 33.1833 | 1.129 |
| 0.2 | 0 | clcB | 1 | 33.1833 | 1.005 |
| 0.2 | 0 | ycaM | 1 | 33.1833 | 1.025 |
| 0.2 | 0 | yadI | 1 | 33.1833 | 1.12 |
| 0.2 | 0 | AG1 | 1 | 33.1833 | 1.012 |
| 0.2 | 0 | ptsI | 1 | 33.4333 | 1.135 |
| 0.2 | 0 | clcB | 1 | 33.4333 | 1.022 |
| 0.2 | 0 | ycaM | 1 | 33.4333 | 1.03 |
| 0.2 | 0 | yadI | 1 | 33.4333 | 1.13 |
| 0.2 | 0 | AG1 | 1 | 33.4333 | 1.035 |
| 0.2 | 0 | ptsI | 1 | 33.6833 | 1.148 |
| 0.2 | 0 | clcB | 1 | 33.6833 | 1.018 |
| 0.2 | 0 | ycaM | 1 | 33.6833 | 1.029 |
| 0.2 | 0 | yadI | 1 | 33.6833 | 1.146 |
| 0.2 | 0 | AG1 | 1 | 33.6833 | 1.028 |
| 0.2 | 0 | ptsI | 1 | 33.9333 | 1.149 |
| 0.2 | 0 | clcB | 1 | 33.9333 | 1.038 |
| 0.2 | 0 | ycaM | 1 | 33.9333 | 1.028 |
| 0.2 | 0 | yadI | 1 | 33.9333 | 1.15 |
| 0.2 | 0 | AG1 | 1 | 33.9333 | 1.04 |
| 0.2 | 0 | ptsI | 1 | 34.1833 | 1.156 |
| 0.2 | 0 | clcB | 1 | 34.1833 | 1.031 |
| 0.2 | 0 | ycaM | 1 | 34.1833 | 1.032 |
| 0.2 | 0 | yadI | 1 | 34.1833 | 1.144 |
| 0.2 | 0 | AG1 | 1 | 34.1833 | 1.046 |
| 0.2 | 0 | ptsI | 1 | 34.4333 | 1.17 |
| 0.2 | 0 | clcB | 1 | 34.4333 | 1.045 |
| 0.2 | 0 | ycaM | 1 | 34.4333 | 1.034 |
| 0.2 | 0 | yadI | 1 | 34.4333 | 1.17 |
| 0.2 | 0 | AG1 | 1 | 34.4333 | 1.059 |
| 0.2 | 0 | ptsI | 1 | 34.6833 | 1.185 |
| 0.2 | 0 | clcB | 1 | 34.6833 | 1.043 |
| 0.2 | 0 | ycaM | 1 | 34.6833 | 1.041 |
| 0.2 | 0 | yadI | 1 | 34.6833 | 1.182 |
| 0.2 | 0 | AG1 | 1 | 34.6833 | 1.065 |

| | | | | | |
|---|---|---|---|---|---|
| 0.2 | 0 | ptsI | 1 | 34.9333 | 1.187 |
| 0.2 | 0 | clcB | 1 | 34.9333 | 1.042 |
| 0.2 | 0 | ycaM | 1 | 34.9333 | 1.04 |
| 0.2 | 0 | yadI | 1 | 34.9333 | 1.191 |
| 0.2 | 0 | AG1 | 1 | 34.9333 | 1.071 |
| 0.2 | 0 | ptsI | 1 | 35.1833 | 1.2 |
| 0.2 | 0 | clcB | 1 | 35.1833 | 1.04 |
| 0.2 | 0 | ycaM | 1 | 35.1833 | 1.043 |
| 0.2 | 0 | yadI | 1 | 35.1833 | 1.2 |
| 0.2 | 0 | AG1 | 1 | 35.1833 | 1.077 |
| 0.2 | 0 | ptsI | 1 | 35.4333 | 1.21 |
| 0.2 | 0 | clcB | 1 | 35.4333 | 1.052 |
| 0.2 | 0 | ycaM | 1 | 35.4333 | 1.042 |
| 0.2 | 0 | yadI | 1 | 35.4333 | 1.21 |
| 0.2 | 0 | AG1 | 1 | 35.4333 | 1.091 |
| 0.2 | 0 | ptsI | 1 | 35.6833 | 1.223 |
| 0.2 | 0 | clcB | 1 | 35.6833 | 1.038 |
| 0.2 | 0 | ycaM | 1 | 35.6833 | 1.053 |
| 0.2 | 0 | yadI | 1 | 35.6833 | 1.219 |
| 0.2 | 0 | AG1 | 1 | 35.6833 | 1.096 |
| 0.2 | 0 | ptsI | 1 | 35.9333 | 1.234 |
| 0.2 | 0 | clcB | 1 | 35.9333 | 1.034 |
| 0.2 | 0 | ycaM | 1 | 35.9333 | 1.047 |
| 0.2 | 0 | yadI | 1 | 35.9333 | 1.236 |
| 0.2 | 0 | AG1 | 1 | 35.9333 | 1.098 |
| 0.2 | 0 | ptsI | 1 | 36.1833 | 1.238 |
| 0.2 | 0 | clcB | 1 | 36.1833 | 1.043 |
| 0.2 | 0 | ycaM | 1 | 36.1833 | 1.053 |
| 0.2 | 0 | yadI | 1 | 36.1833 | 1.235 |
| 0.2 | 0 | AG1 | 1 | 36.1833 | 1.113 |
| 0.2 | 0 | ptsI | 1 | 36.4333 | 1.24 |
| 0.2 | 0 | clcB | 1 | 36.4333 | 1.049 |
| 0.2 | 0 | ycaM | 1 | 36.4333 | 1.056 |
| 0.2 | 0 | yadI | 1 | 36.4333 | 1.245 |
| 0.2 | 0 | AG1 | 1 | 36.4333 | 1.122 |
| 0.2 | 0 | ptsI | 1 | 36.6833 | 1.256 |
| 0.2 | 0 | clcB | 1 | 36.6833 | 1.042 |
| 0.2 | 0 | ycaM | 1 | 36.6833 | 1.056 |
| 0.2 | 0 | yadI | 1 | 36.6833 | 1.254 |
| 0.2 | 0 | AG1 | 1 | 36.6833 | 1.129 |
| 0.2 | 0 | ptsI | 2 | 0 | 0.213 |
| 0.2 | 0 | clcB | 2 | 0 | 0.206 |
| 0.2 | 0 | ycaM | 2 | 0 | 0.218 |
| 0.2 | 0 | yadI | 2 | 0 | 0.22 |
| 0.2 | 0 | AG1 | 2 | 0 | 0.226 |
| 0.2 | 0 | ptsI | 2 | 0.25 | 0.213 |
| 0.2 | 0 | clcB | 2 | 0.25 | 0.201 |
| 0.2 | 0 | ycaM | 2 | 0.25 | 0.211 |

| | | | | | |
|---|---|---|---|---|---|
| 0.2 | 0 | yadI | 2 | 0.25 | 0.219 |
| 0.2 | 0 | AG1 | 2 | 0.25 | 0.221 |
| 0.2 | 0 | ptsI | 2 | 0.5 | 0.214 |
| 0.2 | 0 | clcB | 2 | 0.5 | 0.201 |
| 0.2 | 0 | ycaM | 2 | 0.5 | 0.208 |
| 0.2 | 0 | yadI | 2 | 0.5 | 0.219 |
| 0.2 | 0 | AG1 | 2 | 0.5 | 0.222 |
| 0.2 | 0 | ptsI | 2 | 0.75 | 0.217 |
| 0.2 | 0 | clcB | 2 | 0.75 | 0.201 |
| 0.2 | 0 | ycaM | 2 | 0.75 | 0.21 |
| 0.2 | 0 | yadI | 2 | 0.75 | 0.221 |
| 0.2 | 0 | AG1 | 2 | 0.75 | 0.221 |
| 0.2 | 0 | ptsI | 2 | 1 | 0.221 |
| 0.2 | 0 | clcB | 2 | 1 | 0.202 |
| 0.2 | 0 | ycaM | 2 | 1 | 0.213 |
| 0.2 | 0 | yadI | 2 | 1 | 0.226 |
| 0.2 | 0 | AG1 | 2 | 1 | 0.228 |
| 0.2 | 0 | ptsI | 2 | 1.25 | 0.224 |
| 0.2 | 0 | clcB | 2 | 1.25 | 0.204 |
| 0.2 | 0 | ycaM | 2 | 1.25 | 0.216 |
| 0.2 | 0 | yadI | 2 | 1.25 | 0.235 |
| 0.2 | 0 | AG1 | 2 | 1.25 | 0.233 |
| 0.2 | 0 | ptsI | 2 | 1.5 | 0.228 |
| 0.2 | 0 | clcB | 2 | 1.5 | 0.208 |
| 0.2 | 0 | ycaM | 2 | 1.5 | 0.219 |
| 0.2 | 0 | yadI | 2 | 1.5 | 0.239 |
| 0.2 | 0 | AG1 | 2 | 1.5 | 0.236 |
| 0.2 | 0 | ptsI | 2 | 1.75 | 0.235 |
| 0.2 | 0 | clcB | 2 | 1.75 | 0.211 |
| 0.2 | 0 | ycaM | 2 | 1.75 | 0.223 |
| 0.2 | 0 | yadI | 2 | 1.75 | 0.248 |
| 0.2 | 0 | AG1 | 2 | 1.75 | 0.248 |
| 0.2 | 0 | ptsI | 2 | 2 | 0.241 |
| 0.2 | 0 | clcB | 2 | 2 | 0.214 |
| 0.2 | 0 | ycaM | 2 | 2 | 0.229 |
| 0.2 | 0 | yadI | 2 | 2 | 0.255 |
| 0.2 | 0 | AG1 | 2 | 2 | 0.26 |
| 0.2 | 0 | ptsI | 2 | 2.25 | 0.25 |
| 0.2 | 0 | clcB | 2 | 2.25 | 0.216 |
| 0.2 | 0 | ycaM | 2 | 2.25 | 0.234 |
| 0.2 | 0 | yadI | 2 | 2.25 | 0.267 |
| 0.2 | 0 | AG1 | 2 | 2.25 | 0.272 |
| 0.2 | 0 | ptsI | 2 | 2.5 | 0.262 |
| 0.2 | 0 | clcB | 2 | 2.5 | 0.223 |
| 0.2 | 0 | ycaM | 2 | 2.5 | 0.242 |
| 0.2 | 0 | yadI | 2 | 2.5 | 0.282 |
| 0.2 | 0 | AG1 | 2 | 2.5 | 0.285 |
| 0.2 | 0 | ptsI | 2 | 2.75 | 0.272 |

| | | | | | |
|---|---|---|---|---|---|
| 0.2 | 0 | clcB | 2 | 2.75 | 0.227 |
| 0.2 | 0 | ycaM | 2 | 2.75 | 0.248 |
| 0.2 | 0 | yadI | 2 | 2.75 | 0.293 |
| 0.2 | 0 | AG1 | 2 | 2.75 | 0.279 |
| 0.2 | 0 | ptsI | 2 | 3 | 0.28 |
| 0.2 | 0 | clcB | 2 | 3 | 0.234 |
| 0.2 | 0 | ycaM | 2 | 3 | 0.256 |
| 0.2 | 0 | yadI | 2 | 3 | 0.312 |
| 0.2 | 0 | AG1 | 2 | 3 | 0.285 |
| 0.2 | 0 | ptsI | 2 | 3.25 | 0.298 |
| 0.2 | 0 | clcB | 2 | 3.25 | 0.238 |
| 0.2 | 0 | ycaM | 2 | 3.25 | 0.265 |
| 0.2 | 0 | yadI | 2 | 3.25 | 0.33 |
| 0.2 | 0 | AG1 | 2 | 3.25 | 0.296 |
| 0.2 | 0 | ptsI | 2 | 3.5 | 0.305 |
| 0.2 | 0 | clcB | 2 | 3.5 | 0.24 |
| 0.2 | 0 | ycaM | 2 | 3.5 | 0.274 |
| 0.2 | 0 | yadI | 2 | 3.5 | 0.342 |
| 0.2 | 0 | AG1 | 2 | 3.5 | 0.311 |
| 0.2 | 0 | ptsI | 2 | 3.75 | 0.316 |
| 0.2 | 0 | clcB | 2 | 3.75 | 0.247 |
| 0.2 | 0 | ycaM | 2 | 3.75 | 0.285 |
| 0.2 | 0 | yadI | 2 | 3.75 | 0.362 |
| 0.2 | 0 | AG1 | 2 | 3.75 | 0.326 |
| 0.2 | 0 | ptsI | 2 | 4 | 0.329 |
| 0.2 | 0 | clcB | 2 | 4 | 0.254 |
| 0.2 | 0 | ycaM | 2 | 4 | 0.293 |
| 0.2 | 0 | yadI | 2 | 4 | 0.38 |
| 0.2 | 0 | AG1 | 2 | 4 | 0.339 |
| 0.2 | 0 | ptsI | 2 | 4.38333 | 0.346 |
| 0.2 | 0 | clcB | 2 | 4.38333 | 0.312 |
| 0.2 | 0 | ycaM | 2 | 4.38333 | 0.315 |
| 0.2 | 0 | yadI | 2 | 4.38333 | 0.409 |
| 0.2 | 0 | AG1 | 2 | 4.38333 | 0.304 |
| 0.2 | 0 | ptsI | 2 | 4.63333 | 0.341 |
| 0.2 | 0 | clcB | 2 | 4.63333 | 0.306 |
| 0.2 | 0 | ycaM | 2 | 4.63333 | 0.31 |
| 0.2 | 0 | yadI | 2 | 4.63333 | 0.406 |
| 0.2 | 0 | AG1 | 2 | 4.63333 | 0.3 |
| 0.2 | 0 | ptsI | 2 | 4.88333 | 0.345 |
| 0.2 | 0 | clcB | 2 | 4.88333 | 0.308 |
| 0.2 | 0 | ycaM | 2 | 4.88333 | 0.319 |
| 0.2 | 0 | yadI | 2 | 4.88333 | 0.419 |
| 0.2 | 0 | AG1 | 2 | 4.88333 | 0.308 |
| 0.2 | 0 | ptsI | 2 | 5.13333 | 0.355 |
| 0.2 | 0 | clcB | 2 | 5.13333 | 0.313 |
| 0.2 | 0 | ycaM | 2 | 5.13333 | 0.326 |
| 0.2 | 0 | yadI | 2 | 5.13333 | 0.429 |

| | | | | | |
|---|---|---|---|---|---|
| 0.2 | 0 | AG1 | 2 | 5.13333 | 0.316 |
| 0.2 | 0 | ptsI | 2 | 5.38333 | 0.363 |
| 0.2 | 0 | clcB | 2 | 5.38333 | 0.32 |
| 0.2 | 0 | ycaM | 2 | 5.38333 | 0.336 |
| 0.2 | 0 | yadI | 2 | 5.38333 | 0.44 |
| 0.2 | 0 | AG1 | 2 | 5.38333 | 0.322 |
| 0.2 | 0 | ptsI | 2 | 5.63333 | 0.375 |
| 0.2 | 0 | clcB | 2 | 5.63333 | 0.328 |
| 0.2 | 0 | ycaM | 2 | 5.63333 | 0.346 |
| 0.2 | 0 | yadI | 2 | 5.63333 | 0.453 |
| 0.2 | 0 | AG1 | 2 | 5.63333 | 0.332 |
| 0.2 | 0 | ptsI | 2 | 5.88333 | 0.382 |
| 0.2 | 0 | clcB | 2 | 5.88333 | 0.339 |
| 0.2 | 0 | ycaM | 2 | 5.88333 | 0.359 |
| 0.2 | 0 | yadI | 2 | 5.88333 | 0.463 |
| 0.2 | 0 | AG1 | 2 | 5.88333 | 0.34 |
| 0.2 | 0 | ptsI | 2 | 6.13333 | 0.391 |
| 0.2 | 0 | clcB | 2 | 6.13333 | 0.344 |
| 0.2 | 0 | ycaM | 2 | 6.13333 | 0.37 |
| 0.2 | 0 | yadI | 2 | 6.13333 | 0.471 |
| 0.2 | 0 | AG1 | 2 | 6.13333 | 0.353 |
| 0.2 | 0 | ptsI | 2 | 6.38333 | 0.401 |
| 0.2 | 0 | clcB | 2 | 6.38333 | 0.352 |
| 0.2 | 0 | ycaM | 2 | 6.38333 | 0.381 |
| 0.2 | 0 | yadI | 2 | 6.38333 | 0.483 |
| 0.2 | 0 | AG1 | 2 | 6.38333 | 0.364 |
| 0.2 | 0 | ptsI | 2 | 6.63333 | 0.408 |
| 0.2 | 0 | clcB | 2 | 6.63333 | 0.362 |
| 0.2 | 0 | ycaM | 2 | 6.63333 | 0.388 |
| 0.2 | 0 | yadI | 2 | 6.63333 | 0.492 |
| 0.2 | 0 | AG1 | 2 | 6.63333 | 0.373 |
| 0.2 | 0 | ptsI | 2 | 6.88333 | 0.418 |
| 0.2 | 0 | clcB | 2 | 6.88333 | 0.367 |
| 0.2 | 0 | ycaM | 2 | 6.88333 | 0.396 |
| 0.2 | 0 | yadI | 2 | 6.88333 | 0.5 |
| 0.2 | 0 | AG1 | 2 | 6.88333 | 0.383 |
| 0.2 | 0 | ptsI | 2 | 7.13333 | 0.427 |
| 0.2 | 0 | clcB | 2 | 7.13333 | 0.372 |
| 0.2 | 0 | ycaM | 2 | 7.13333 | 0.403 |
| 0.2 | 0 | yadI | 2 | 7.13333 | 0.512 |
| 0.2 | 0 | AG1 | 2 | 7.13333 | 0.394 |
| 0.2 | 0 | ptsI | 2 | 7.38333 | 0.436 |
| 0.2 | 0 | clcB | 2 | 7.38333 | 0.382 |
| 0.2 | 0 | ycaM | 2 | 7.38333 | 0.412 |
| 0.2 | 0 | yadI | 2 | 7.38333 | 0.525 |
| 0.2 | 0 | AG1 | 2 | 7.38333 | 0.4 |
| 0.2 | 0 | ptsI | 2 | 7.63333 | 0.444 |
| 0.2 | 0 | clcB | 2 | 7.63333 | 0.392 |

| | | | | | |
|---|---|---|---|---|---|
| 0.2 | 0 | ycaM | 2 | 7.63333 | 0.422 |
| 0.2 | 0 | yadI | 2 | 7.63333 | 0.535 |
| 0.2 | 0 | AG1 | 2 | 7.63333 | 0.409 |
| 0.2 | 0 | ptsI | 2 | 7.88333 | 0.454 |
| 0.2 | 0 | clcB | 2 | 7.88333 | 0.402 |
| 0.2 | 0 | ycaM | 2 | 7.88333 | 0.428 |
| 0.2 | 0 | yadI | 2 | 7.88333 | 0.548 |
| 0.2 | 0 | AG1 | 2 | 7.88333 | 0.417 |
| 0.2 | 0 | ptsI | 2 | 8.13333 | 0.464 |
| 0.2 | 0 | clcB | 2 | 8.13333 | 0.406 |
| 0.2 | 0 | ycaM | 2 | 8.13333 | 0.433 |
| 0.2 | 0 | yadI | 2 | 8.13333 | 0.557 |
| 0.2 | 0 | AG1 | 2 | 8.13333 | 0.429 |
| 0.2 | 0 | ptsI | 2 | 8.38333 | 0.472 |
| 0.2 | 0 | clcB | 2 | 8.38333 | 0.42 |
| 0.2 | 0 | ycaM | 2 | 8.38333 | 0.438 |
| 0.2 | 0 | yadI | 2 | 8.38333 | 0.57 |
| 0.2 | 0 | AG1 | 2 | 8.38333 | 0.436 |
| 0.2 | 0 | ptsI | 2 | 8.91667 | 0.543 |
| 0.2 | 0 | clcB | 2 | 8.91667 | 0.328 |
| 0.2 | 0 | ycaM | 2 | 8.91667 | 0.384 |
| 0.2 | 0 | yadI | 2 | 8.91667 | 0.464 |
| 0.2 | 0 | AG1 | 2 | 8.91667 | 0.475 |
| 0.2 | 0 | ptsI | 2 | 9.16667 | 0.549 |
| 0.2 | 0 | clcB | 2 | 9.16667 | 0.33 |
| 0.2 | 0 | ycaM | 2 | 9.16667 | 0.389 |
| 0.2 | 0 | yadI | 2 | 9.16667 | 0.462 |
| 0.2 | 0 | AG1 | 2 | 9.16667 | 0.475 |
| 0.2 | 0 | ptsI | 2 | 9.41667 | 0.55 |
| 0.2 | 0 | clcB | 2 | 9.41667 | 0.336 |
| 0.2 | 0 | ycaM | 2 | 9.41667 | 0.392 |
| 0.2 | 0 | yadI | 2 | 9.41667 | 0.469 |
| 0.2 | 0 | AG1 | 2 | 9.41667 | 0.488 |
| 0.2 | 0 | ptsI | 2 | 9.66667 | 0.57 |
| 0.2 | 0 | clcB | 2 | 9.66667 | 0.341 |
| 0.2 | 0 | ycaM | 2 | 9.66667 | 0.398 |
| 0.2 | 0 | yadI | 2 | 9.66667 | 0.481 |
| 0.2 | 0 | AG1 | 2 | 9.66667 | 0.504 |
| 0.2 | 0 | ptsI | 2 | 9.91667 | 0.565 |
| 0.2 | 0 | clcB | 2 | 9.91667 | 0.35 |
| 0.2 | 0 | ycaM | 2 | 9.91667 | 0.404 |
| 0.2 | 0 | yadI | 2 | 9.91667 | 0.49 |
| 0.2 | 0 | AG1 | 2 | 9.91667 | 0.517 |
| 0.2 | 0 | ptsI | 2 | 10.1667 | 0.567 |
| 0.2 | 0 | clcB | 2 | 10.1667 | 0.355 |
| 0.2 | 0 | ycaM | 2 | 10.1667 | 0.406 |
| 0.2 | 0 | yadI | 2 | 10.1667 | 0.496 |
| 0.2 | 0 | AG1 | 2 | 10.1667 | 0.524 |

| | | | | | |
|---|---|---|---|---|---|
| 0.2 | 0 | ptsI | 2 | 10.4167 | 0.603 |
| 0.2 | 0 | clcB | 2 | 10.4167 | 0.358 |
| 0.2 | 0 | ycaM | 2 | 10.4167 | 0.414 |
| 0.2 | 0 | yadI | 2 | 10.4167 | 0.505 |
| 0.2 | 0 | AG1 | 2 | 10.4167 | 0.537 |
| 0.2 | 0 | ptsI | 2 | 10.6667 | 0.573 |
| 0.2 | 0 | clcB | 2 | 10.6667 | 0.369 |
| 0.2 | 0 | ycaM | 2 | 10.6667 | 0.423 |
| 0.2 | 0 | yadI | 2 | 10.6667 | 0.516 |
| 0.2 | 0 | AG1 | 2 | 10.6667 | 0.544 |
| 0.2 | 0 | ptsI | 2 | 10.9167 | 0.58 |
| 0.2 | 0 | clcB | 2 | 10.9167 | 0.372 |
| 0.2 | 0 | ycaM | 2 | 10.9167 | 0.43 |
| 0.2 | 0 | yadI | 2 | 10.9167 | 0.523 |
| 0.2 | 0 | AG1 | 2 | 10.9167 | 0.554 |
| 0.2 | 0 | ptsI | 2 | 11.1667 | 0.586 |
| 0.2 | 0 | clcB | 2 | 11.1667 | 0.378 |
| 0.2 | 0 | ycaM | 2 | 11.1667 | 0.438 |
| 0.2 | 0 | yadI | 2 | 11.1667 | 0.533 |
| 0.2 | 0 | AG1 | 2 | 11.1667 | 0.568 |
| 0.2 | 0 | ptsI | 2 | 11.4167 | 0.584 |
| 0.2 | 0 | clcB | 2 | 11.4167 | 0.379 |
| 0.2 | 0 | ycaM | 2 | 11.4167 | 0.442 |
| 0.2 | 0 | yadI | 2 | 11.4167 | 0.537 |
| 0.2 | 0 | AG1 | 2 | 11.4167 | 0.568 |
| 0.2 | 0 | ptsI | 2 | 11.6667 | 0.605 |
| 0.2 | 0 | clcB | 2 | 11.6667 | 0.386 |
| 0.2 | 0 | ycaM | 2 | 11.6667 | 0.46 |
| 0.2 | 0 | yadI | 2 | 11.6667 | 0.547 |
| 0.2 | 0 | AG1 | 2 | 11.6667 | 0.598 |
| 0.2 | 0 | ptsI | 2 | 11.9167 | 0.581 |
| 0.2 | 0 | clcB | 2 | 11.9167 | 0.394 |
| 0.2 | 0 | ycaM | 2 | 11.9167 | 0.484 |
| 0.2 | 0 | yadI | 2 | 11.9167 | 0.552 |
| 0.2 | 0 | AG1 | 2 | 11.9167 | 0.599 |
| 0.2 | 0 | ptsI | 2 | 12.1667 | 0.6 |
| 0.2 | 0 | clcB | 2 | 12.1667 | 0.4 |
| 0.2 | 0 | ycaM | 2 | 12.1667 | 0.472 |
| 0.2 | 0 | yadI | 2 | 12.1667 | 0.56 |
| 0.2 | 0 | AG1 | 2 | 12.1667 | 0.625 |
| 0.2 | 0 | ptsI | 2 | 12.4167 | 0.585 |
| 0.2 | 0 | clcB | 2 | 12.4167 | 0.409 |
| 0.2 | 0 | ycaM | 2 | 12.4167 | 0.471 |
| 0.2 | 0 | yadI | 2 | 12.4167 | 0.566 |
| 0.2 | 0 | AG1 | 2 | 12.4167 | 0.643 |
| 0.2 | 0 | ptsI | 2 | 12.6667 | 0.592 |
| 0.2 | 0 | clcB | 2 | 12.6667 | 0.419 |
| 0.2 | 0 | ycaM | 2 | 12.6667 | 0.474 |

| | | | | | |
|---|---|---|---|---|---|
| 0.2 | 0 | yadI | 2 | 12.6667 | 0.574 |
| 0.2 | 0 | AG1 | 2 | 12.6667 | 0.639 |
| 0.2 | 0 | ptsI | 2 | 12.9167 | 0.603 |
| 0.2 | 0 | clcB | 2 | 12.9167 | 0.428 |
| 0.2 | 0 | ycaM | 2 | 12.9167 | 0.48 |
| 0.2 | 0 | yadI | 2 | 12.9167 | 0.584 |
| 0.2 | 0 | AG1 | 2 | 12.9167 | 0.653 |
| 0.2 | 0 | ptsI | 2 | 13.1667 | 0.61 |
| 0.2 | 0 | clcB | 2 | 13.1667 | 0.434 |
| 0.2 | 0 | ycaM | 2 | 13.1667 | 0.501 |
| 0.2 | 0 | yadI | 2 | 13.1667 | 0.591 |
| 0.2 | 0 | AG1 | 2 | 13.1667 | 0.664 |
| 0.2 | 0 | ptsI | 2 | 13.4167 | 0.611 |
| 0.2 | 0 | clcB | 2 | 13.4167 | 0.443 |
| 0.2 | 0 | ycaM | 2 | 13.4167 | 0.499 |
| 0.2 | 0 | yadI | 2 | 13.4167 | 0.598 |
| 0.2 | 0 | AG1 | 2 | 13.4167 | 0.678 |
| 0.2 | 0 | ptsI | 2 | 13.6667 | 0.628 |
| 0.2 | 0 | clcB | 2 | 13.6667 | 0.448 |
| 0.2 | 0 | ycaM | 2 | 13.6667 | 0.518 |
| 0.2 | 0 | yadI | 2 | 13.6667 | 0.608 |
| 0.2 | 0 | AG1 | 2 | 13.6667 | 0.726 |
| 0.2 | 0 | ptsI | 2 | 13.9167 | 0.64 |
| 0.2 | 0 | clcB | 2 | 13.9167 | 0.455 |
| 0.2 | 0 | ycaM | 2 | 13.9167 | 0.526 |
| 0.2 | 0 | yadI | 2 | 13.9167 | 0.619 |
| 0.2 | 0 | AG1 | 2 | 13.9167 | 0.704 |
| 0.2 | 0 | ptsI | 2 | 14.1667 | 0.685 |
| 0.2 | 0 | clcB | 2 | 14.1667 | 0.461 |
| 0.2 | 0 | ycaM | 2 | 14.1667 | 0.536 |
| 0.2 | 0 | yadI | 2 | 14.1667 | 0.628 |
| 0.2 | 0 | AG1 | 2 | 14.1667 | 0.707 |
| 0.2 | 0 | ptsI | 2 | 14.4167 | 0.689 |
| 0.2 | 0 | clcB | 2 | 14.4167 | 0.465 |
| 0.2 | 0 | ycaM | 2 | 14.4167 | 0.545 |
| 0.2 | 0 | yadI | 2 | 14.4167 | 0.642 |
| 0.2 | 0 | AG1 | 2 | 14.4167 | 0.698 |
| 0.2 | 0 | ptsI | 2 | 14.6667 | 0.677 |
| 0.2 | 0 | clcB | 2 | 14.6667 | 0.47 |
| 0.2 | 0 | ycaM | 2 | 14.6667 | 0.558 |
| 0.2 | 0 | yadI | 2 | 14.6667 | 0.648 |
| 0.2 | 0 | AG1 | 2 | 14.6667 | 0.711 |
| 0.2 | 0 | ptsI | 2 | 14.9167 | 0.702 |
| 0.2 | 0 | clcB | 2 | 14.9167 | 0.482 |
| 0.2 | 0 | ycaM | 2 | 14.9167 | 0.563 |
| 0.2 | 0 | yadI | 2 | 14.9167 | 0.661 |
| 0.2 | 0 | AG1 | 2 | 14.9167 | 0.733 |
| 0.2 | 0 | ptsI | 2 | 15.1667 | 0.671 |

| | | | | | |
|---|---|---|---|---|---|
| 0.2 | 0 | clcB | 2 | 15.1667 | 0.493 |
| 0.2 | 0 | ycaM | 2 | 15.1667 | 0.566 |
| 0.2 | 0 | yadI | 2 | 15.1667 | 0.667 |
| 0.2 | 0 | AG1 | 2 | 15.1667 | 0.74 |
| 0.2 | 0 | ptsI | 2 | 15.4167 | 0.688 |
| 0.2 | 0 | clcB | 2 | 15.4167 | 0.495 |
| 0.2 | 0 | ycaM | 2 | 15.4167 | 0.589 |
| 0.2 | 0 | yadI | 2 | 15.4167 | 0.688 |
| 0.2 | 0 | AG1 | 2 | 15.4167 | 0.784 |
| 0.2 | 0 | ptsI | 2 | 15.6667 | 0.694 |
| 0.2 | 0 | clcB | 2 | 15.6667 | 0.503 |
| 0.2 | 0 | ycaM | 2 | 15.6667 | 0.589 |
| 0.2 | 0 | yadI | 2 | 15.6667 | 0.69 |
| 0.2 | 0 | AG1 | 2 | 15.6667 | 0.787 |
| 0.2 | 0 | ptsI | 2 | 15.9167 | 0.723 |
| 0.2 | 0 | clcB | 2 | 15.9167 | 0.512 |
| 0.2 | 0 | ycaM | 2 | 15.9167 | 0.596 |
| 0.2 | 0 | yadI | 2 | 15.9167 | 0.7 |
| 0.2 | 0 | AG1 | 2 | 15.9167 | 0.785 |
| 0.2 | 0 | ptsI | 2 | 16.1667 | 0.72 |
| 0.2 | 0 | clcB | 2 | 16.1667 | 0.522 |
| 0.2 | 0 | ycaM | 2 | 16.1667 | 0.581 |
| 0.2 | 0 | yadI | 2 | 16.1667 | 0.717 |
| 0.2 | 0 | AG1 | 2 | 16.1667 | 0.798 |
| 0.2 | 0 | ptsI | 2 | 16.4167 | 0.742 |
| 0.2 | 0 | clcB | 2 | 16.4167 | 0.534 |
| 0.2 | 0 | ycaM | 2 | 16.4167 | 0.592 |
| 0.2 | 0 | yadI | 2 | 16.4167 | 0.722 |
| 0.2 | 0 | AG1 | 2 | 16.4167 | 0.809 |
| 0.2 | 0 | ptsI | 2 | 16.6667 | 0.768 |
| 0.2 | 0 | clcB | 2 | 16.6667 | 0.544 |
| 0.2 | 0 | ycaM | 2 | 16.6667 | 0.603 |
| 0.2 | 0 | yadI | 2 | 16.6667 | 0.729 |
| 0.2 | 0 | AG1 | 2 | 16.6667 | 0.807 |
| 0.2 | 0 | ptsI | 2 | 16.9167 | 0.781 |
| 0.2 | 0 | clcB | 2 | 16.9167 | 0.556 |
| 0.2 | 0 | ycaM | 2 | 16.9167 | 0.608 |
| 0.2 | 0 | yadI | 2 | 16.9167 | 0.735 |
| 0.2 | 0 | AG1 | 2 | 16.9167 | 0.818 |
| 0.2 | 0 | ptsI | 2 | 17.1667 | 0.786 |
| 0.2 | 0 | clcB | 2 | 17.1667 | 0.537 |
| 0.2 | 0 | ycaM | 2 | 17.1667 | 0.609 |
| 0.2 | 0 | yadI | 2 | 17.1667 | 0.746 |
| 0.2 | 0 | AG1 | 2 | 17.1667 | 0.82 |
| 0.2 | 0 | ptsI | 2 | 17.4167 | 0.809 |
| 0.2 | 0 | clcB | 2 | 17.4167 | 0.56 |
| 0.2 | 0 | ycaM | 2 | 17.4167 | 0.62 |
| 0.2 | 0 | yadI | 2 | 17.4167 | 0.748 |

| | | | | | |
|---|---|---|---|---|---|
| 0.2 | 0 | AG1 | 2 | 17.4167 | 0.82 |
| 0.2 | 0 | ptsI | 2 | 17.6667 | 0.824 |
| 0.2 | 0 | clcB | 2 | 17.6667 | 0.559 |
| 0.2 | 0 | ycaM | 2 | 17.6667 | 0.617 |
| 0.2 | 0 | yadI | 2 | 17.6667 | 0.758 |
| 0.2 | 0 | AG1 | 2 | 17.6667 | 0.818 |
| 0.2 | 0 | ptsI | 2 | 17.9167 | 0.827 |
| 0.2 | 0 | clcB | 2 | 17.9167 | 0.56 |
| 0.2 | 0 | ycaM | 2 | 17.9167 | 0.622 |
| 0.2 | 0 | yadI | 2 | 17.9167 | 0.761 |
| 0.2 | 0 | AG1 | 2 | 17.9167 | 0.784 |
| 0.2 | 0 | ptsI | 2 | 18.1667 | 0.842 |
| 0.2 | 0 | clcB | 2 | 18.1667 | 0.571 |
| 0.2 | 0 | ycaM | 2 | 18.1667 | 0.624 |
| 0.2 | 0 | yadI | 2 | 18.1667 | 0.768 |
| 0.2 | 0 | AG1 | 2 | 18.1667 | 0.802 |
| 0.2 | 0 | ptsI | 2 | 18.4167 | 0.844 |
| 0.2 | 0 | clcB | 2 | 18.4167 | 0.566 |
| 0.2 | 0 | ycaM | 2 | 18.4167 | 0.634 |
| 0.2 | 0 | yadI | 2 | 18.4167 | 0.77 |
| 0.2 | 0 | AG1 | 2 | 18.4167 | 0.802 |
| 0.2 | 0 | ptsI | 2 | 18.6667 | 0.84 |
| 0.2 | 0 | clcB | 2 | 18.6667 | 0.569 |
| 0.2 | 0 | ycaM | 2 | 18.6667 | 0.647 |
| 0.2 | 0 | yadI | 2 | 18.6667 | 0.779 |
| 0.2 | 0 | AG1 | 2 | 18.6667 | 0.793 |
| 0.2 | 0 | ptsI | 2 | 18.9167 | 0.837 |
| 0.2 | 0 | clcB | 2 | 18.9167 | 0.569 |
| 0.2 | 0 | ycaM | 2 | 18.9167 | 0.658 |
| 0.2 | 0 | yadI | 2 | 18.9167 | 0.78 |
| 0.2 | 0 | AG1 | 2 | 18.9167 | 0.818 |
| 0.2 | 0 | ptsI | 2 | 19.1667 | 0.856 |
| 0.2 | 0 | clcB | 2 | 19.1667 | 0.578 |
| 0.2 | 0 | ycaM | 2 | 19.1667 | 0.654 |
| 0.2 | 0 | yadI | 2 | 19.1667 | 0.784 |
| 0.2 | 0 | AG1 | 2 | 19.1667 | 0.827 |
| 0.2 | 0 | ptsI | 2 | 19.4167 | 0.86 |
| 0.2 | 0 | clcB | 2 | 19.4167 | 0.582 |
| 0.2 | 0 | ycaM | 2 | 19.4167 | 0.662 |
| 0.2 | 0 | yadI | 2 | 19.4167 | 0.79 |
| 0.2 | 0 | AG1 | 2 | 19.4167 | 0.826 |
| 0.2 | 0 | ptsI | 2 | 19.6667 | 0.864 |
| 0.2 | 0 | clcB | 2 | 19.6667 | 0.585 |
| 0.2 | 0 | ycaM | 2 | 19.6667 | 0.674 |
| 0.2 | 0 | yadI | 2 | 19.6667 | 0.795 |
| 0.2 | 0 | AG1 | 2 | 19.6667 | 0.827 |
| 0.2 | 0 | ptsI | 2 | 19.9167 | 0.872 |
| 0.2 | 0 | clcB | 2 | 19.9167 | 0.599 |

| | | | | | |
|---|---|---|---|---|---|
| 0.2 | 0 | ycaM | 2 | 19.9167 | 0.683 |
| 0.2 | 0 | yadI | 2 | 19.9167 | 0.799 |
| 0.2 | 0 | AG1 | 2 | 19.9167 | 0.844 |
| 0.2 | 0 | ptsI | 2 | 20.1667 | 0.874 |
| 0.2 | 0 | clcB | 2 | 20.1667 | 0.602 |
| 0.2 | 0 | ycaM | 2 | 20.1667 | 0.684 |
| 0.2 | 0 | yadI | 2 | 20.1667 | 0.805 |
| 0.2 | 0 | AG1 | 2 | 20.1667 | 0.837 |
| 0.2 | 0 | ptsI | 2 | 20.4167 | 0.873 |
| 0.2 | 0 | clcB | 2 | 20.4167 | 0.612 |
| 0.2 | 0 | ycaM | 2 | 20.4167 | 0.691 |
| 0.2 | 0 | yadI | 2 | 20.4167 | 0.812 |
| 0.2 | 0 | AG1 | 2 | 20.4167 | 0.84 |
| 0.2 | 0 | ptsI | 2 | 20.6667 | 0.878 |
| 0.2 | 0 | clcB | 2 | 20.6667 | 0.616 |
| 0.2 | 0 | ycaM | 2 | 20.6667 | 0.701 |
| 0.2 | 0 | yadI | 2 | 20.6667 | 0.811 |
| 0.2 | 0 | AG1 | 2 | 20.6667 | 0.835 |
| 0.2 | 0 | ptsI | 2 | 20.9167 | 0.884 |
| 0.2 | 0 | clcB | 2 | 20.9167 | 0.621 |
| 0.2 | 0 | ycaM | 2 | 20.9167 | 0.704 |
| 0.2 | 0 | yadI | 2 | 20.9167 | 0.814 |
| 0.2 | 0 | AG1 | 2 | 20.9167 | 0.843 |
| 0.2 | 0 | ptsI | 2 | 21.1667 | 0.89 |
| 0.2 | 0 | clcB | 2 | 21.1667 | 0.622 |
| 0.2 | 0 | ycaM | 2 | 21.1667 | 0.705 |
| 0.2 | 0 | yadI | 2 | 21.1667 | 0.82 |
| 0.2 | 0 | AG1 | 2 | 21.1667 | 0.84 |
| 0.2 | 0 | ptsI | 2 | 21.4167 | 0.874 |
| 0.2 | 0 | clcB | 2 | 21.4167 | 0.626 |
| 0.2 | 0 | ycaM | 2 | 21.4167 | 0.704 |
| 0.2 | 0 | yadI | 2 | 21.4167 | 0.821 |
| 0.2 | 0 | AG1 | 2 | 21.4167 | 0.84 |
| 0.2 | 0 | ptsI | 2 | 21.6667 | 0.902 |
| 0.2 | 0 | clcB | 2 | 21.6667 | 0.636 |
| 0.2 | 0 | ycaM | 2 | 21.6667 | 0.717 |
| 0.2 | 0 | yadI | 2 | 21.6667 | 0.825 |
| 0.2 | 0 | AG1 | 2 | 21.6667 | 0.839 |
| 0.2 | 0 | ptsI | 2 | 21.9167 | 0.91 |
| 0.2 | 0 | clcB | 2 | 21.9167 | 0.645 |
| 0.2 | 0 | ycaM | 2 | 21.9167 | 0.724 |
| 0.2 | 0 | yadI | 2 | 21.9167 | 0.83 |
| 0.2 | 0 | AG1 | 2 | 21.9167 | 0.844 |
| 0.2 | 0 | ptsI | 2 | 22.1667 | 0.909 |
| 0.2 | 0 | clcB | 2 | 22.1667 | 0.649 |
| 0.2 | 0 | ycaM | 2 | 22.1667 | 0.726 |
| 0.2 | 0 | yadI | 2 | 22.1667 | 0.835 |
| 0.2 | 0 | AG1 | 2 | 22.1667 | 0.844 |

| | | | | | |
|---|---|---|---|---|---|
| 0.2 | 0 | ptsI | 2 | 22.4167 | 0.916 |
| 0.2 | 0 | clcB | 2 | 22.4167 | 0.655 |
| 0.2 | 0 | ycaM | 2 | 22.4167 | 0.731 |
| 0.2 | 0 | yadI | 2 | 22.4167 | 0.836 |
| 0.2 | 0 | AG1 | 2 | 22.4167 | 0.843 |
| 0.2 | 0 | ptsI | 2 | 22.6667 | 0.92 |
| 0.2 | 0 | clcB | 2 | 22.6667 | 0.664 |
| 0.2 | 0 | ycaM | 2 | 22.6667 | 0.732 |
| 0.2 | 0 | yadI | 2 | 22.6667 | 0.84 |
| 0.2 | 0 | AG1 | 2 | 22.6667 | 0.849 |
| 0.2 | 0 | ptsI | 2 | 22.9167 | 0.93 |
| 0.2 | 0 | clcB | 2 | 22.9167 | 0.67 |
| 0.2 | 0 | ycaM | 2 | 22.9167 | 0.735 |
| 0.2 | 0 | yadI | 2 | 22.9167 | 0.843 |
| 0.2 | 0 | AG1 | 2 | 22.9167 | 0.85 |
| 0.2 | 0 | ptsI | 2 | 23.1667 | 0.936 |
| 0.2 | 0 | clcB | 2 | 23.1667 | 0.673 |
| 0.2 | 0 | ycaM | 2 | 23.1667 | 0.739 |
| 0.2 | 0 | yadI | 2 | 23.1667 | 0.845 |
| 0.2 | 0 | AG1 | 2 | 23.1667 | 0.851 |
| 0.2 | 0 | ptsI | 2 | 23.4167 | 0.939 |
| 0.2 | 0 | clcB | 2 | 23.4167 | 0.679 |
| 0.2 | 0 | ycaM | 2 | 23.4167 | 0.741 |
| 0.2 | 0 | yadI | 2 | 23.4167 | 0.85 |
| 0.2 | 0 | AG1 | 2 | 23.4167 | 0.847 |
| 0.2 | 0 | ptsI | 2 | 23.6667 | 0.941 |
| 0.2 | 0 | clcB | 2 | 23.6667 | 0.686 |
| 0.2 | 0 | ycaM | 2 | 23.6667 | 0.755 |
| 0.2 | 0 | yadI | 2 | 23.6667 | 0.852 |
| 0.2 | 0 | AG1 | 2 | 23.6667 | 0.852 |
| 0.2 | 0 | ptsI | 2 | 23.9167 | 0.955 |
| 0.2 | 0 | clcB | 2 | 23.9167 | 0.694 |
| 0.2 | 0 | ycaM | 2 | 23.9167 | 0.755 |
| 0.2 | 0 | yadI | 2 | 23.9167 | 0.851 |
| 0.2 | 0 | AG1 | 2 | 23.9167 | 0.853 |
| 0.2 | 0 | ptsI | 2 | 24.1667 | 0.951 |
| 0.2 | 0 | clcB | 2 | 24.1667 | 0.696 |
| 0.2 | 0 | ycaM | 2 | 24.1667 | 0.758 |
| 0.2 | 0 | yadI | 2 | 24.1667 | 0.854 |
| 0.2 | 0 | AG1 | 2 | 24.1667 | 0.846 |
| 0.2 | 0 | ptsI | 2 | 24.4167 | 0.963 |
| 0.2 | 0 | clcB | 2 | 24.4167 | 0.699 |
| 0.2 | 0 | ycaM | 2 | 24.4167 | 0.762 |
| 0.2 | 0 | yadI | 2 | 24.4167 | 0.858 |
| 0.2 | 0 | AG1 | 2 | 24.4167 | 0.846 |
| 0.2 | 0 | ptsI | 2 | 24.6667 | 0.975 |
| 0.2 | 0 | clcB | 2 | 24.6667 | 0.706 |
| 0.2 | 0 | ycaM | 2 | 24.6667 | 0.765 |

| | | | | | |
|---|---|---|---|---|---|
| 0.2 | 0 | yadI | 2 | 24.6667 | 0.863 |
| 0.2 | 0 | AG1 | 2 | 24.6667 | 0.852 |
| 0.2 | 0 | ptsI | 2 | 24.9167 | 0.977 |
| 0.2 | 0 | clcB | 2 | 24.9167 | 0.71 |
| 0.2 | 0 | ycaM | 2 | 24.9167 | 0.771 |
| 0.2 | 0 | yadI | 2 | 24.9167 | 0.864 |
| 0.2 | 0 | AG1 | 2 | 24.9167 | 0.852 |
| 0.2 | 0 | ptsI | 2 | 25.1667 | 0.972 |
| 0.2 | 0 | clcB | 2 | 25.1667 | 0.716 |
| 0.2 | 0 | ycaM | 2 | 25.1667 | 0.774 |
| 0.2 | 0 | yadI | 2 | 25.1667 | 0.866 |
| 0.2 | 0 | AG1 | 2 | 25.1667 | 0.848 |
| 0.2 | 0 | ptsI | 2 | 25.4167 | 0.983 |
| 0.2 | 0 | clcB | 2 | 25.4167 | 0.723 |
| 0.2 | 0 | ycaM | 2 | 25.4167 | 0.784 |
| 0.2 | 0 | yadI | 2 | 25.4167 | 0.868 |
| 0.2 | 0 | AG1 | 2 | 25.4167 | 0.855 |
| 0.2 | 0 | ptsI | 2 | 25.6667 | 0.982 |
| 0.2 | 0 | clcB | 2 | 25.6667 | 0.724 |
| 0.2 | 0 | ycaM | 2 | 25.6667 | 0.783 |
| 0.2 | 0 | yadI | 2 | 25.6667 | 0.87 |
| 0.2 | 0 | AG1 | 2 | 25.6667 | 0.848 |
| 0.2 | 0 | ptsI | 2 | 25.9167 | 0.988 |
| 0.2 | 0 | clcB | 2 | 25.9167 | 0.729 |
| 0.2 | 0 | ycaM | 2 | 25.9167 | 0.787 |
| 0.2 | 0 | yadI | 2 | 25.9167 | 0.873 |
| 0.2 | 0 | AG1 | 2 | 25.9167 | 0.852 |
| 0.2 | 0 | ptsI | 2 | 26.1667 | 0.977 |
| 0.2 | 0 | clcB | 2 | 26.1667 | 0.727 |
| 0.2 | 0 | ycaM | 2 | 26.1667 | 0.793 |
| 0.2 | 0 | yadI | 2 | 26.1667 | 0.879 |
| 0.2 | 0 | AG1 | 2 | 26.1667 | 0.849 |
| 0.2 | 0 | ptsI | 2 | 26.4167 | 1.003 |
| 0.2 | 0 | clcB | 2 | 26.4167 | 0.741 |
| 0.2 | 0 | ycaM | 2 | 26.4167 | 0.799 |
| 0.2 | 0 | yadI | 2 | 26.4167 | 0.877 |
| 0.2 | 0 | AG1 | 2 | 26.4167 | 0.856 |
| 0.2 | 0 | ptsI | 3 | 0 | 0.154 |
| 0.2 | 0 | clcB | 3 | 0 | 0.163 |
| 0.2 | 0 | ycaM | 3 | 0 | 0.152 |
| 0.2 | 0 | yadI | 3 | 0 | 0.17 |
| 0.2 | 0 | AG1 | 3 | 0 | 0.166 |
| 0.2 | 0 | ptsI | 3 | 0.25 | 0.153 |
| 0.2 | 0 | clcB | 3 | 0.25 | 0.214 |
| 0.2 | 0 | ycaM | 3 | 0.25 | 0.15 |
| 0.2 | 0 | yadI | 3 | 0.25 | 0.167 |
| 0.2 | 0 | AG1 | 3 | 0.25 | 0.165 |
| 0.2 | 0 | ptsI | 3 | 0.5 | 0.151 |

| | | | | | |
|---|---|---|---|---|---|
| 0.2 | 0 | clcB | 3 | 0.5 | 0.154 |
| 0.2 | 0 | ycaM | 3 | 0.5 | 0.147 |
| 0.2 | 0 | yadI | 3 | 0.5 | 0.166 |
| 0.2 | 0 | AG1 | 3 | 0.5 | 0.164 |
| 0.2 | 0 | ptsI | 3 | 0.75 | 0.151 |
| 0.2 | 0 | clcB | 3 | 0.75 | 0.156 |
| 0.2 | 0 | ycaM | 3 | 0.75 | 0.147 |
| 0.2 | 0 | yadI | 3 | 0.75 | 0.168 |
| 0.2 | 0 | AG1 | 3 | 0.75 | 0.164 |
| 0.2 | 0 | ptsI | 3 | 1 | 0.152 |
| 0.2 | 0 | clcB | 3 | 1 | 0.162 |
| 0.2 | 0 | ycaM | 3 | 1 | 0.149 |
| 0.2 | 0 | yadI | 3 | 1 | 0.168 |
| 0.2 | 0 | AG1 | 3 | 1 | 0.165 |
| 0.2 | 0 | ptsI | 3 | 1.25 | 0.152 |
| 0.2 | 0 | clcB | 3 | 1.25 | 0.154 |
| 0.2 | 0 | ycaM | 3 | 1.25 | 0.148 |
| 0.2 | 0 | yadI | 3 | 1.25 | 0.17 |
| 0.2 | 0 | AG1 | 3 | 1.25 | 0.165 |
| 0.2 | 0 | ptsI | 3 | 1.5 | 0.153 |
| 0.2 | 0 | clcB | 3 | 1.5 | 0.154 |
| 0.2 | 0 | ycaM | 3 | 1.5 | 0.15 |
| 0.2 | 0 | yadI | 3 | 1.5 | 0.173 |
| 0.2 | 0 | AG1 | 3 | 1.5 | 0.167 |
| 0.2 | 0 | ptsI | 3 | 1.75 | 0.153 |
| 0.2 | 0 | clcB | 3 | 1.75 | 0.154 |
| 0.2 | 0 | ycaM | 3 | 1.75 | 0.15 |
| 0.2 | 0 | yadI | 3 | 1.75 | 0.176 |
| 0.2 | 0 | AG1 | 3 | 1.75 | 0.167 |
| 0.2 | 0 | ptsI | 3 | 2 | 0.155 |
| 0.2 | 0 | clcB | 3 | 2 | 0.156 |
| 0.2 | 0 | ycaM | 3 | 2 | 0.174 |
| 0.2 | 0 | yadI | 3 | 2 | 0.177 |
| 0.2 | 0 | AG1 | 3 | 2 | 0.169 |
| 0.2 | 0 | ptsI | 3 | 2.25 | 0.156 |
| 0.2 | 0 | clcB | 3 | 2.25 | 0.157 |
| 0.2 | 0 | ycaM | 3 | 2.25 | 0.154 |
| 0.2 | 0 | yadI | 3 | 2.25 | 0.18 |
| 0.2 | 0 | AG1 | 3 | 2.25 | 0.173 |
| 0.2 | 0 | ptsI | 3 | 2.5 | 0.158 |
| 0.2 | 0 | clcB | 3 | 2.5 | 0.159 |
| 0.2 | 0 | ycaM | 3 | 2.5 | 0.154 |
| 0.2 | 0 | yadI | 3 | 2.5 | 0.183 |
| 0.2 | 0 | AG1 | 3 | 2.5 | 0.175 |
| 0.2 | 0 | ptsI | 3 | 2.75 | 0.16 |
| 0.2 | 0 | clcB | 3 | 2.75 | 0.16 |
| 0.2 | 0 | ycaM | 3 | 2.75 | 0.158 |
| 0.2 | 0 | yadI | 3 | 2.75 | 0.185 |

| | | | | | |
|---|---|---|---|---|---|
| 0.2 | 0 | AG1 | 3 | 2.75 | 0.182 |
| 0.2 | 0 | ptsI | 3 | 3 | 0.162 |
| 0.2 | 0 | clcB | 3 | 3 | 0.163 |
| 0.2 | 0 | ycaM | 3 | 3 | 0.157 |
| 0.2 | 0 | yadI | 3 | 3 | 0.192 |
| 0.2 | 0 | AG1 | 3 | 3 | 0.183 |
| 0.2 | 0 | ptsI | 3 | 3.25 | 0.164 |
| 0.2 | 0 | clcB | 3 | 3.25 | 0.162 |
| 0.2 | 0 | ycaM | 3 | 3.25 | 0.16 |
| 0.2 | 0 | yadI | 3 | 3.25 | 0.197 |
| 0.2 | 0 | AG1 | 3 | 3.25 | 0.183 |
| 0.2 | 0 | ptsI | 3 | 3.5 | 0.168 |
| 0.2 | 0 | clcB | 3 | 3.5 | 0.164 |
| 0.2 | 0 | ycaM | 3 | 3.5 | 0.165 |
| 0.2 | 0 | yadI | 3 | 3.5 | 0.202 |
| 0.2 | 0 | AG1 | 3 | 3.5 | 0.188 |
| 0.2 | 0 | ptsI | 3 | 3.75 | 0.172 |
| 0.2 | 0 | clcB | 3 | 3.75 | 0.167 |
| 0.2 | 0 | ycaM | 3 | 3.75 | 0.169 |
| 0.2 | 0 | yadI | 3 | 3.75 | 0.208 |
| 0.2 | 0 | AG1 | 3 | 3.75 | 0.192 |
| 0.2 | 0 | ptsI | 3 | 4 | 0.176 |
| 0.2 | 0 | clcB | 3 | 4 | 0.171 |
| 0.2 | 0 | ycaM | 3 | 4 | 0.171 |
| 0.2 | 0 | yadI | 3 | 4 | 0.22 |
| 0.2 | 0 | AG1 | 3 | 4 | 0.201 |
| 0.2 | 0 | ptsI | 3 | 4.41667 | 0.186 |
| 0.2 | 0 | clcB | 3 | 4.41667 | 0.203 |
| 0.2 | 0 | ycaM | 3 | 4.41667 | 0.192 |
| 0.2 | 0 | yadI | 3 | 4.41667 | 0.218 |
| 0.2 | 0 | AG1 | 3 | 4.41667 | 0.228 |
| 0.2 | 0 | ptsI | 3 | 4.66667 | 0.183 |
| 0.2 | 0 | clcB | 3 | 4.66667 | 0.181 |
| 0.2 | 0 | ycaM | 3 | 4.66667 | 0.189 |
| 0.2 | 0 | yadI | 3 | 4.66667 | 0.212 |
| 0.2 | 0 | AG1 | 3 | 4.66667 | 0.223 |
| 0.2 | 0 | ptsI | 3 | 4.91667 | 0.183 |
| 0.2 | 0 | clcB | 3 | 4.91667 | 0.188 |
| 0.2 | 0 | ycaM | 3 | 4.91667 | 0.189 |
| 0.2 | 0 | yadI | 3 | 4.91667 | 0.222 |
| 0.2 | 0 | AG1 | 3 | 4.91667 | 0.226 |
| 0.2 | 0 | ptsI | 3 | 5.16667 | 0.188 |
| 0.2 | 0 | clcB | 3 | 5.16667 | 0.192 |
| 0.2 | 0 | ycaM | 3 | 5.16667 | 0.192 |
| 0.2 | 0 | yadI | 3 | 5.16667 | 0.234 |
| 0.2 | 0 | AG1 | 3 | 5.16667 | 0.232 |
| 0.2 | 0 | ptsI | 3 | 5.41667 | 0.193 |
| 0.2 | 0 | clcB | 3 | 5.41667 | 0.199 |

| | | | | | |
|---|---|---|---|---|---|
| 0.2 | 0 | ycaM | 3 | 5.41667 | 0.199 |
| 0.2 | 0 | yadI | 3 | 5.41667 | 0.249 |
| 0.2 | 0 | AG1 | 3 | 5.41667 | 0.242 |
| 0.2 | 0 | ptsI | 3 | 5.66667 | 0.199 |
| 0.2 | 0 | clcB | 3 | 5.66667 | 0.208 |
| 0.2 | 0 | ycaM | 3 | 5.66667 | 0.204 |
| 0.2 | 0 | yadI | 3 | 5.66667 | 0.265 |
| 0.2 | 0 | AG1 | 3 | 5.66667 | 0.251 |
| 0.2 | 0 | ptsI | 3 | 5.91667 | 0.206 |
| 0.2 | 0 | clcB | 3 | 5.91667 | 0.216 |
| 0.2 | 0 | ycaM | 3 | 5.91667 | 0.212 |
| 0.2 | 0 | yadI | 3 | 5.91667 | 0.281 |
| 0.2 | 0 | AG1 | 3 | 5.91667 | 0.265 |
| 0.2 | 0 | ptsI | 3 | 6.16667 | 0.214 |
| 0.2 | 0 | clcB | 3 | 6.16667 | 0.23 |
| 0.2 | 0 | ycaM | 3 | 6.16667 | 0.222 |
| 0.2 | 0 | yadI | 3 | 6.16667 | 0.302 |
| 0.2 | 0 | AG1 | 3 | 6.16667 | 0.278 |
| 0.2 | 0 | ptsI | 3 | 6.41667 | 0.222 |
| 0.2 | 0 | clcB | 3 | 6.41667 | 0.247 |
| 0.2 | 0 | ycaM | 3 | 6.41667 | 0.232 |
| 0.2 | 0 | yadI | 3 | 6.41667 | 0.314 |
| 0.2 | 0 | AG1 | 3 | 6.41667 | 0.294 |
| 0.2 | 0 | ptsI | 3 | 6.66667 | 0.232 |
| 0.2 | 0 | clcB | 3 | 6.66667 | 0.265 |
| 0.2 | 0 | ycaM | 3 | 6.66667 | 0.244 |
| 0.2 | 0 | yadI | 3 | 6.66667 | 0.327 |
| 0.2 | 0 | AG1 | 3 | 6.66667 | 0.308 |
| 0.2 | 0 | ptsI | 3 | 6.91667 | 0.24 |
| 0.2 | 0 | clcB | 3 | 6.91667 | 0.276 |
| 0.2 | 0 | ycaM | 3 | 6.91667 | 0.258 |
| 0.2 | 0 | yadI | 3 | 6.91667 | 0.341 |
| 0.2 | 0 | AG1 | 3 | 6.91667 | 0.324 |
| 0.2 | 0 | ptsI | 3 | 7.16667 | 0.252 |
| 0.2 | 0 | clcB | 3 | 7.16667 | 0.291 |
| 0.2 | 0 | ycaM | 3 | 7.16667 | 0.272 |
| 0.2 | 0 | yadI | 3 | 7.16667 | 0.358 |
| 0.2 | 0 | AG1 | 3 | 7.16667 | 0.332 |
| 0.2 | 0 | ptsI | 3 | 7.41667 | 0.265 |
| 0.2 | 0 | clcB | 3 | 7.41667 | 0.299 |
| 0.2 | 0 | ycaM | 3 | 7.41667 | 0.28 |
| 0.2 | 0 | yadI | 3 | 7.41667 | 0.37 |
| 0.2 | 0 | AG1 | 3 | 7.41667 | 0.344 |
| 0.2 | 0 | ptsI | 3 | 7.66667 | 0.274 |
| 0.2 | 0 | clcB | 3 | 7.66667 | 0.309 |
| 0.2 | 0 | ycaM | 3 | 7.66667 | 0.288 |
| 0.2 | 0 | yadI | 3 | 7.66667 | 0.381 |
| 0.2 | 0 | AG1 | 3 | 7.66667 | 0.351 |

| | | | | | |
|---|---|---|---|---|---|
| 0.2 | 0 | ptsI | 3 | 7.91667 | 0.28 |
| 0.2 | 0 | clcB | 3 | 7.91667 | 0.316 |
| 0.2 | 0 | ycaM | 3 | 7.91667 | 0.295 |
| 0.2 | 0 | yadI | 3 | 7.91667 | 0.39 |
| 0.2 | 0 | AG1 | 3 | 7.91667 | 0.364 |
| 0.2 | 0 | ptsI | 3 | 8.16667 | 0.29 |
| 0.2 | 0 | clcB | 3 | 8.16667 | 0.326 |
| 0.2 | 0 | ycaM | 3 | 8.16667 | 0.302 |
| 0.2 | 0 | yadI | 3 | 8.16667 | 0.401 |
| 0.2 | 0 | AG1 | 3 | 8.16667 | 0.371 |
| 0.2 | 0 | ptsI | 3 | 8.41667 | 0.298 |
| 0.2 | 0 | clcB | 3 | 8.41667 | 0.335 |
| 0.2 | 0 | ycaM | 3 | 8.41667 | 0.313 |
| 0.2 | 0 | yadI | 3 | 8.41667 | 0.412 |
| 0.2 | 0 | AG1 | 3 | 8.41667 | 0.381 |
| 0.2 | 0 | ptsI | 3 | 9.05 | 0.282 |
| 0.2 | 0 | clcB | 3 | 9.05 | 0.287 |
| 0.2 | 0 | ycaM | 3 | 9.05 | 0.291 |
| 0.2 | 0 | yadI | 3 | 9.05 | 0.37 |
| 0.2 | 0 | AG1 | 3 | 9.05 | 0.432 |
| 0.2 | 0 | ptsI | 3 | 9.3 | 0.287 |
| 0.2 | 0 | clcB | 3 | 9.3 | 0.292 |
| 0.2 | 0 | ycaM | 3 | 9.3 | 0.297 |
| 0.2 | 0 | yadI | 3 | 9.3 | 0.373 |
| 0.2 | 0 | AG1 | 3 | 9.3 | 0.427 |
| 0.2 | 0 | ptsI | 3 | 9.55 | 0.293 |
| 0.2 | 0 | clcB | 3 | 9.55 | 0.297 |
| 0.2 | 0 | ycaM | 3 | 9.55 | 0.303 |
| 0.2 | 0 | yadI | 3 | 9.55 | 0.382 |
| 0.2 | 0 | AG1 | 3 | 9.55 | 0.438 |
| 0.2 | 0 | ptsI | 3 | 9.8 | 0.301 |
| 0.2 | 0 | clcB | 3 | 9.8 | 0.303 |
| 0.2 | 0 | ycaM | 3 | 9.8 | 0.311 |
| 0.2 | 0 | yadI | 3 | 9.8 | 0.392 |
| 0.2 | 0 | AG1 | 3 | 9.8 | 0.45 |
| 0.2 | 0 | ptsI | 3 | 10.05 | 0.307 |
| 0.2 | 0 | clcB | 3 | 10.05 | 0.308 |
| 0.2 | 0 | ycaM | 3 | 10.05 | 0.316 |
| 0.2 | 0 | yadI | 3 | 10.05 | 0.4 |
| 0.2 | 0 | AG1 | 3 | 10.05 | 0.46 |
| 0.2 | 0 | ptsI | 3 | 10.3 | 0.317 |
| 0.2 | 0 | clcB | 3 | 10.3 | 0.316 |
| 0.2 | 0 | ycaM | 3 | 10.3 | 0.325 |
| 0.2 | 0 | yadI | 3 | 10.3 | 0.41 |
| 0.2 | 0 | AG1 | 3 | 10.3 | 0.472 |
| 0.2 | 0 | ptsI | 3 | 10.55 | 0.325 |
| 0.2 | 0 | clcB | 3 | 10.55 | 0.324 |
| 0.2 | 0 | ycaM | 3 | 10.55 | 0.335 |

| | | | | | |
|---|---|---|---|---|---|
| 0.2 | 0 | yadI | 3 | 10.55 | 0.422 |
| 0.2 | 0 | AG1 | 3 | 10.55 | 0.48 |
| 0.2 | 0 | ptsI | 3 | 10.8 | 0.335 |
| 0.2 | 0 | clcB | 3 | 10.8 | 0.334 |
| 0.2 | 0 | ycaM | 3 | 10.8 | 0.342 |
| 0.2 | 0 | yadI | 3 | 10.8 | 0.434 |
| 0.2 | 0 | AG1 | 3 | 10.8 | 0.492 |
| 0.2 | 0 | ptsI | 3 | 11.05 | 0.343 |
| 0.2 | 0 | clcB | 3 | 11.05 | 0.342 |
| 0.2 | 0 | ycaM | 3 | 11.05 | 0.365 |
| 0.2 | 0 | yadI | 3 | 11.05 | 0.444 |
| 0.2 | 0 | AG1 | 3 | 11.05 | 0.501 |
| 0.2 | 0 | ptsI | 3 | 11.3 | 0.353 |
| 0.2 | 0 | clcB | 3 | 11.3 | 0.349 |
| 0.2 | 0 | ycaM | 3 | 11.3 | 0.357 |
| 0.2 | 0 | yadI | 3 | 11.3 | 0.454 |
| 0.2 | 0 | AG1 | 3 | 11.3 | 0.514 |
| 0.2 | 0 | ptsI | 3 | 11.55 | 0.359 |
| 0.2 | 0 | clcB | 3 | 11.55 | 0.353 |
| 0.2 | 0 | ycaM | 3 | 11.55 | 0.361 |
| 0.2 | 0 | yadI | 3 | 11.55 | 0.465 |
| 0.2 | 0 | AG1 | 3 | 11.55 | 0.525 |
| 0.2 | 0 | ptsI | 3 | 11.8 | 0.368 |
| 0.2 | 0 | clcB | 3 | 11.8 | 0.358 |
| 0.2 | 0 | ycaM | 3 | 11.8 | 0.363 |
| 0.2 | 0 | yadI | 3 | 11.8 | 0.476 |
| 0.2 | 0 | AG1 | 3 | 11.8 | 0.538 |
| 0.2 | 0 | ptsI | 3 | 12.05 | 0.378 |
| 0.2 | 0 | clcB | 3 | 12.05 | 0.361 |
| 0.2 | 0 | ycaM | 3 | 12.05 | 0.366 |
| 0.2 | 0 | yadI | 3 | 12.05 | 0.484 |
| 0.2 | 0 | AG1 | 3 | 12.05 | 0.549 |
| 0.2 | 0 | ptsI | 3 | 12.3 | 0.387 |
| 0.2 | 0 | clcB | 3 | 12.3 | 0.368 |
| 0.2 | 0 | ycaM | 3 | 12.3 | 0.374 |
| 0.2 | 0 | yadI | 3 | 12.3 | 0.5 |
| 0.2 | 0 | AG1 | 3 | 12.3 | 0.57 |
| 0.2 | 0 | ptsI | 3 | 12.55 | 0.394 |
| 0.2 | 0 | clcB | 3 | 12.55 | 0.374 |
| 0.2 | 0 | ycaM | 3 | 12.55 | 0.373 |
| 0.2 | 0 | yadI | 3 | 12.55 | 0.509 |
| 0.2 | 0 | AG1 | 3 | 12.55 | 0.585 |
| 0.2 | 0 | ptsI | 3 | 12.8 | 0.403 |
| 0.2 | 0 | clcB | 3 | 12.8 | 0.381 |
| 0.2 | 0 | ycaM | 3 | 12.8 | 0.381 |
| 0.2 | 0 | yadI | 3 | 12.8 | 0.515 |
| 0.2 | 0 | AG1 | 3 | 12.8 | 0.606 |
| 0.2 | 0 | ptsI | 3 | 13.05 | 0.415 |

| | | | | | |
|---|---|---|---|---|---|
| 0.2 | 0 | clcB | 3 | 13.05 | 0.387 |
| 0.2 | 0 | ycaM | 3 | 13.05 | 0.386 |
| 0.2 | 0 | yadI | 3 | 13.05 | 0.524 |
| 0.2 | 0 | AG1 | 3 | 13.05 | 0.621 |
| 0.2 | 0 | ptsI | 3 | 13.3 | 0.421 |
| 0.2 | 0 | clcB | 3 | 13.3 | 0.396 |
| 0.2 | 0 | ycaM | 3 | 13.3 | 0.391 |
| 0.2 | 0 | yadI | 3 | 13.3 | 0.535 |
| 0.2 | 0 | AG1 | 3 | 13.3 | 0.635 |
| 0.2 | 0 | ptsI | 3 | 13.55 | 0.431 |
| 0.2 | 0 | clcB | 3 | 13.55 | 0.404 |
| 0.2 | 0 | ycaM | 3 | 13.55 | 0.399 |
| 0.2 | 0 | yadI | 3 | 13.55 | 0.548 |
| 0.2 | 0 | AG1 | 3 | 13.55 | 0.65 |
| 0.2 | 0 | ptsI | 3 | 13.8 | 0.438 |
| 0.2 | 0 | clcB | 3 | 13.8 | 0.41 |
| 0.2 | 0 | ycaM | 3 | 13.8 | 0.407 |
| 0.2 | 0 | yadI | 3 | 13.8 | 0.556 |
| 0.2 | 0 | AG1 | 3 | 13.8 | 0.661 |
| 0.2 | 0 | ptsI | 3 | 14.05 | 0.45 |
| 0.2 | 0 | clcB | 3 | 14.05 | 0.421 |
| 0.2 | 0 | ycaM | 3 | 14.05 | 0.413 |
| 0.2 | 0 | yadI | 3 | 14.05 | 0.562 |
| 0.2 | 0 | AG1 | 3 | 14.05 | 0.672 |
| 0.2 | 0 | ptsI | 3 | 14.3 | 0.46 |
| 0.2 | 0 | clcB | 3 | 14.3 | 0.43 |
| 0.2 | 0 | ycaM | 3 | 14.3 | 0.419 |
| 0.2 | 0 | yadI | 3 | 14.3 | 0.57 |
| 0.2 | 0 | AG1 | 3 | 14.3 | 0.681 |
| 0.2 | 0 | ptsI | 3 | 14.55 | 0.473 |
| 0.2 | 0 | clcB | 3 | 14.55 | 0.434 |
| 0.2 | 0 | ycaM | 3 | 14.55 | 0.423 |
| 0.2 | 0 | yadI | 3 | 14.55 | 0.576 |
| 0.2 | 0 | AG1 | 3 | 14.55 | 0.681 |
| 0.2 | 0 | ptsI | 3 | 14.8 | 0.484 |
| 0.2 | 0 | clcB | 3 | 14.8 | 0.439 |
| 0.2 | 0 | ycaM | 3 | 14.8 | 0.433 |
| 0.2 | 0 | yadI | 3 | 14.8 | 0.583 |
| 0.2 | 0 | AG1 | 3 | 14.8 | 0.692 |
| 0.2 | 0 | ptsI | 3 | 15.05 | 0.49 |
| 0.2 | 0 | clcB | 3 | 15.05 | 0.446 |
| 0.2 | 0 | ycaM | 3 | 15.05 | 0.439 |
| 0.2 | 0 | yadI | 3 | 15.05 | 0.589 |
| 0.2 | 0 | AG1 | 3 | 15.05 | 0.699 |
| 0.2 | 0 | ptsI | 3 | 15.3 | 0.504 |
| 0.2 | 0 | clcB | 3 | 15.3 | 0.452 |
| 0.2 | 0 | ycaM | 3 | 15.3 | 0.443 |
| 0.2 | 0 | yadI | 3 | 15.3 | 0.602 |

| | | | | | |
|---|---|---|---|---|---|
| 0.2 | 0 | AG1 | 3 | 15.3 | 0.709 |
| 0.2 | 0 | ptsI | 3 | 15.55 | 0.505 |
| 0.2 | 0 | clcB | 3 | 15.55 | 0.455 |
| 0.2 | 0 | ycaM | 3 | 15.55 | 0.446 |
| 0.2 | 0 | yadI | 3 | 15.55 | 0.604 |
| 0.2 | 0 | AG1 | 3 | 15.55 | 0.711 |
| 0.2 | 0 | ptsI | 3 | 15.8 | 0.515 |
| 0.2 | 0 | clcB | 3 | 15.8 | 0.46 |
| 0.2 | 0 | ycaM | 3 | 15.8 | 0.451 |
| 0.2 | 0 | yadI | 3 | 15.8 | 0.614 |
| 0.2 | 0 | AG1 | 3 | 15.8 | 0.722 |
| 0.2 | 0 | ptsI | 3 | 16.05 | 0.521 |
| 0.2 | 0 | clcB | 3 | 16.05 | 0.467 |
| 0.2 | 0 | ycaM | 3 | 16.05 | 0.458 |
| 0.2 | 0 | yadI | 3 | 16.05 | 0.618 |
| 0.2 | 0 | AG1 | 3 | 16.05 | 0.731 |
| 0.2 | 0 | ptsI | 3 | 16.3 | 0.528 |
| 0.2 | 0 | clcB | 3 | 16.3 | 0.468 |
| 0.2 | 0 | ycaM | 3 | 16.3 | 0.458 |
| 0.2 | 0 | yadI | 3 | 16.3 | 0.623 |
| 0.2 | 0 | AG1 | 3 | 16.3 | 0.744 |
| 0.2 | 0 | ptsI | 3 | 16.55 | 0.535 |
| 0.2 | 0 | clcB | 3 | 16.55 | 0.471 |
| 0.2 | 0 | ycaM | 3 | 16.55 | 0.465 |
| 0.2 | 0 | yadI | 3 | 16.55 | 0.629 |
| 0.2 | 0 | AG1 | 3 | 16.55 | 0.752 |
| 0.2 | 0 | ptsI | 3 | 16.8 | 0.538 |
| 0.2 | 0 | clcB | 3 | 16.8 | 0.478 |
| 0.2 | 0 | ycaM | 3 | 16.8 | 0.474 |
| 0.2 | 0 | yadI | 3 | 16.8 | 0.633 |
| 0.2 | 0 | AG1 | 3 | 16.8 | 0.757 |
| 0.2 | 0 | ptsI | 3 | 17.05 | 0.545 |
| 0.2 | 0 | clcB | 3 | 17.05 | 0.484 |
| 0.2 | 0 | ycaM | 3 | 17.05 | 0.472 |
| 0.2 | 0 | yadI | 3 | 17.05 | 0.64 |
| 0.2 | 0 | AG1 | 3 | 17.05 | 0.776 |
| 0.2 | 0 | ptsI | 3 | 17.3 | 0.551 |
| 0.2 | 0 | clcB | 3 | 17.3 | 0.486 |
| 0.2 | 0 | ycaM | 3 | 17.3 | 0.48 |
| 0.2 | 0 | yadI | 3 | 17.3 | 0.649 |
| 0.2 | 0 | AG1 | 3 | 17.3 | 0.793 |
| 0.2 | 0 | ptsI | 3 | 17.55 | 0.563 |
| 0.2 | 0 | clcB | 3 | 17.55 | 0.491 |
| 0.2 | 0 | ycaM | 3 | 17.55 | 0.48 |
| 0.2 | 0 | yadI | 3 | 17.55 | 0.654 |
| 0.2 | 0 | AG1 | 3 | 17.55 | 0.806 |
| 0.2 | 0 | ptsI | 3 | 17.8 | 0.561 |
| 0.2 | 0 | clcB | 3 | 17.8 | 0.495 |

| | | | | | |
|---|---|---|---|---|---|
| 0.2 | 0 | ycaM | 3 | 17.8 | 0.488 |
| 0.2 | 0 | yadI | 3 | 17.8 | 0.66 |
| 0.2 | 0 | AG1 | 3 | 17.8 | 0.798 |
| 0.2 | 0 | ptsI | 3 | 18.05 | 0.572 |
| 0.2 | 0 | clcB | 3 | 18.05 | 0.5 |
| 0.2 | 0 | ycaM | 3 | 18.05 | 0.492 |
| 0.2 | 0 | yadI | 3 | 18.05 | 0.664 |
| 0.2 | 0 | AG1 | 3 | 18.05 | 0.806 |
| 0.2 | 0 | ptsI | 3 | 18.3 | 0.579 |
| 0.2 | 0 | clcB | 3 | 18.3 | 0.505 |
| 0.2 | 0 | ycaM | 3 | 18.3 | 0.497 |
| 0.2 | 0 | yadI | 3 | 18.3 | 0.674 |
| 0.2 | 0 | AG1 | 3 | 18.3 | 0.806 |
| 0.2 | 0 | ptsI | 3 | 18.55 | 0.584 |
| 0.2 | 0 | clcB | 3 | 18.55 | 0.51 |
| 0.2 | 0 | ycaM | 3 | 18.55 | 0.502 |
| 0.2 | 0 | yadI | 3 | 18.55 | 0.68 |
| 0.2 | 0 | AG1 | 3 | 18.55 | 0.811 |
| 0.2 | 0 | ptsI | 3 | 18.8 | 0.593 |
| 0.2 | 0 | clcB | 3 | 18.8 | 0.51 |
| 0.2 | 0 | ycaM | 3 | 18.8 | 0.504 |
| 0.2 | 0 | yadI | 3 | 18.8 | 0.68 |
| 0.2 | 0 | AG1 | 3 | 18.8 | 0.814 |
| 0.2 | 0 | ptsI | 3 | 19.05 | 0.602 |
| 0.2 | 0 | clcB | 3 | 19.05 | 0.512 |
| 0.2 | 0 | ycaM | 3 | 19.05 | 0.51 |
| 0.2 | 0 | yadI | 3 | 19.05 | 0.686 |
| 0.2 | 0 | AG1 | 3 | 19.05 | 0.826 |
| 0.2 | 0 | ptsI | 3 | 19.3 | 0.611 |
| 0.2 | 0 | clcB | 3 | 19.3 | 0.521 |
| 0.2 | 0 | ycaM | 3 | 19.3 | 0.514 |
| 0.2 | 0 | yadI | 3 | 19.3 | 0.69 |
| 0.2 | 0 | AG1 | 3 | 19.3 | 0.833 |
| 0.2 | 0 | ptsI | 3 | 19.55 | 0.618 |
| 0.2 | 0 | clcB | 3 | 19.55 | 0.529 |
| 0.2 | 0 | ycaM | 3 | 19.55 | 0.518 |
| 0.2 | 0 | yadI | 3 | 19.55 | 0.693 |
| 0.2 | 0 | AG1 | 3 | 19.55 | 0.851 |
| 0.2 | 0 | ptsI | 3 | 19.8 | 0.624 |
| 0.2 | 0 | clcB | 3 | 19.8 | 0.538 |
| 0.2 | 0 | ycaM | 3 | 19.8 | 0.521 |
| 0.2 | 0 | yadI | 3 | 19.8 | 0.698 |
| 0.2 | 0 | AG1 | 3 | 19.8 | 0.856 |
| 0.2 | 0 | ptsI | 3 | 20.05 | 0.622 |
| 0.2 | 0 | clcB | 3 | 20.05 | 0.556 |
| 0.2 | 0 | ycaM | 3 | 20.05 | 0.523 |
| 0.2 | 0 | yadI | 3 | 20.05 | 0.704 |
| 0.2 | 0 | AG1 | 3 | 20.05 | 0.838 |

| | | | | | |
|---|---|---|---|---|---|
| 0.2 | 0 | ptsI | 3 | 20.3 | 0.624 |
| 0.2 | 0 | clcB | 3 | 20.3 | 0.56 |
| 0.2 | 0 | ycaM | 3 | 20.3 | 0.528 |
| 0.2 | 0 | yadI | 3 | 20.3 | 0.707 |
| 0.2 | 0 | AG1 | 3 | 20.3 | 0.819 |
| 0.2 | 0 | ptsI | 3 | 20.55 | 0.634 |
| 0.2 | 0 | clcB | 3 | 20.55 | 0.569 |
| 0.2 | 0 | ycaM | 3 | 20.55 | 0.535 |
| 0.2 | 0 | yadI | 3 | 20.55 | 0.709 |
| 0.2 | 0 | AG1 | 3 | 20.55 | 0.815 |
| 0.2 | 0 | ptsI | 3 | 20.8 | 0.644 |
| 0.2 | 0 | clcB | 3 | 20.8 | 0.571 |
| 0.2 | 0 | ycaM | 3 | 20.8 | 0.535 |
| 0.2 | 0 | yadI | 3 | 20.8 | 0.718 |
| 0.2 | 0 | AG1 | 3 | 20.8 | 0.823 |
| 0.2 | 0 | ptsI | 3 | 21.05 | 0.661 |
| 0.2 | 0 | clcB | 3 | 21.05 | 0.574 |
| 0.2 | 0 | ycaM | 3 | 21.05 | 0.537 |
| 0.2 | 0 | yadI | 3 | 21.05 | 0.72 |
| 0.2 | 0 | AG1 | 3 | 21.05 | 0.824 |
| 0.2 | 0 | ptsI | 3 | 21.3 | 0.678 |
| 0.2 | 0 | clcB | 3 | 21.3 | 0.578 |
| 0.2 | 0 | ycaM | 3 | 21.3 | 0.539 |
| 0.2 | 0 | yadI | 3 | 21.3 | 0.722 |
| 0.2 | 0 | AG1 | 3 | 21.3 | 0.829 |
| 0.2 | 0 | ptsI | 3 | 21.55 | 0.692 |
| 0.2 | 0 | clcB | 3 | 21.55 | 0.58 |
| 0.2 | 0 | ycaM | 3 | 21.55 | 0.547 |
| 0.2 | 0 | yadI | 3 | 21.55 | 0.722 |
| 0.2 | 0 | AG1 | 3 | 21.55 | 0.83 |
| 0.2 | 0 | ptsI | 3 | 21.8 | 0.697 |
| 0.2 | 0 | clcB | 3 | 21.8 | 0.584 |
| 0.2 | 0 | ycaM | 3 | 21.8 | 0.552 |
| 0.2 | 0 | yadI | 3 | 21.8 | 0.727 |
| 0.2 | 0 | AG1 | 3 | 21.8 | 0.832 |
| 0.2 | 0 | ptsI | 3 | 22.05 | 0.698 |
| 0.2 | 0 | clcB | 3 | 22.05 | 0.589 |
| 0.2 | 0 | ycaM | 3 | 22.05 | 0.555 |
| 0.2 | 0 | yadI | 3 | 22.05 | 0.732 |
| 0.2 | 0 | AG1 | 3 | 22.05 | 0.832 |
| 0.2 | 0 | ptsI | 3 | 22.3 | 0.707 |
| 0.2 | 0 | clcB | 3 | 22.3 | 0.594 |
| 0.2 | 0 | ycaM | 3 | 22.3 | 0.561 |
| 0.2 | 0 | yadI | 3 | 22.3 | 0.734 |
| 0.2 | 0 | AG1 | 3 | 22.3 | 0.834 |
| 0.2 | 0 | ptsI | 3 | 22.55 | 0.718 |
| 0.2 | 0 | clcB | 3 | 22.55 | 0.594 |
| 0.2 | 0 | ycaM | 3 | 22.55 | 0.585 |

| | | | | | |
|---|---|---|---|---|---|
| 0.2 | 0 | yadI | 3 | 22.55 | 0.737 |
| 0.2 | 0 | AG1 | 3 | 22.55 | 0.834 |
| 0.2 | 0 | ptsI | 3 | 22.8 | 0.726 |
| 0.2 | 0 | clcB | 3 | 22.8 | 0.594 |
| 0.2 | 0 | ycaM | 3 | 22.8 | 0.565 |
| 0.2 | 0 | yadI | 3 | 22.8 | 0.74 |
| 0.2 | 0 | AG1 | 3 | 22.8 | 0.839 |
| 0.2 | 0 | ptsI | 3 | 23.05 | 0.736 |
| 0.2 | 0 | clcB | 3 | 23.05 | 0.594 |
| 0.2 | 0 | ycaM | 3 | 23.05 | 0.568 |
| 0.2 | 0 | yadI | 3 | 23.05 | 0.743 |
| 0.2 | 0 | AG1 | 3 | 23.05 | 0.835 |
| 0.2 | 0 | ptsI | 3 | 23.3 | 0.75 |
| 0.2 | 0 | clcB | 3 | 23.3 | 0.597 |
| 0.2 | 0 | ycaM | 3 | 23.3 | 0.57 |
| 0.2 | 0 | yadI | 3 | 23.3 | 0.746 |
| 0.2 | 0 | AG1 | 3 | 23.3 | 0.838 |
| 0.2 | 0 | ptsI | 3 | 23.55 | 0.764 |
| 0.2 | 0 | clcB | 3 | 23.55 | 0.598 |
| 0.2 | 0 | ycaM | 3 | 23.55 | 0.573 |
| 0.2 | 0 | yadI | 3 | 23.55 | 0.748 |
| 0.2 | 0 | AG1 | 3 | 23.55 | 0.839 |
| 0.2 | 0 | ptsI | 3 | 23.8 | 0.769 |
| 0.2 | 0 | clcB | 3 | 23.8 | 0.598 |
| 0.2 | 0 | ycaM | 3 | 23.8 | 0.575 |
| 0.2 | 0 | yadI | 3 | 23.8 | 0.748 |
| 0.2 | 0 | AG1 | 3 | 23.8 | 0.841 |
| 0.2 | 0 | ptsI | 3 | 24.05 | 0.782 |
| 0.2 | 0 | clcB | 3 | 24.05 | 0.599 |
| 0.2 | 0 | ycaM | 3 | 24.05 | 0.574 |
| 0.2 | 0 | yadI | 3 | 24.05 | 0.754 |
| 0.2 | 0 | AG1 | 3 | 24.05 | 0.842 |
| 0.2 | 0 | ptsI | 3 | 24.3 | 0.786 |
| 0.2 | 0 | clcB | 3 | 24.3 | 0.605 |
| 0.2 | 0 | ycaM | 3 | 24.3 | 0.576 |
| 0.2 | 0 | yadI | 3 | 24.3 | 0.756 |
| 0.2 | 0 | AG1 | 3 | 24.3 | 0.84 |
| 0.2 | 0 | ptsI | 4 | 0 | 0.183 |
| 0.2 | 0 | clcB | 4 | 0 | 0.202 |
| 0.2 | 0 | ycaM | 4 | 0 | 0.197 |
| 0.2 | 0 | yadI | 4 | 0 | 0.232 |
| 0.2 | 0 | AG1 | 4 | 0 | 0.204 |
| 0.2 | 0 | ptsI | 4 | 0.25 | 0.182 |
| 0.2 | 0 | clcB | 4 | 0.25 | 0.199 |
| 0.2 | 0 | ycaM | 4 | 0.25 | 0.191 |
| 0.2 | 0 | yadI | 4 | 0.25 | 0.221 |
| 0.2 | 0 | AG1 | 4 | 0.25 | 0.202 |
| 0.2 | 0 | ptsI | 4 | 0.5 | 0.183 |

| | | | | | |
|---|---|---|---|---|---|
| 0.2 | 0 | clcB | 4 | 0.5 | 0.199 |
| 0.2 | 0 | ycaM | 4 | 0.5 | 0.191 |
| 0.2 | 0 | yadI | 4 | 0.5 | 0.221 |
| 0.2 | 0 | AG1 | 4 | 0.5 | 0.203 |
| 0.2 | 0 | ptsI | 4 | 0.75 | 0.186 |
| 0.2 | 0 | clcB | 4 | 0.75 | 0.203 |
| 0.2 | 0 | ycaM | 4 | 0.75 | 0.194 |
| 0.2 | 0 | yadI | 4 | 0.75 | 0.224 |
| 0.2 | 0 | AG1 | 4 | 0.75 | 0.206 |
| 0.2 | 0 | ptsI | 4 | 1 | 0.191 |
| 0.2 | 0 | clcB | 4 | 1 | 0.206 |
| 0.2 | 0 | ycaM | 4 | 1 | 0.198 |
| 0.2 | 0 | yadI | 4 | 1 | 0.228 |
| 0.2 | 0 | AG1 | 4 | 1 | 0.212 |
| 0.2 | 0 | ptsI | 4 | 1.25 | 0.198 |
| 0.2 | 0 | clcB | 4 | 1.25 | 0.212 |
| 0.2 | 0 | ycaM | 4 | 1.25 | 0.203 |
| 0.2 | 0 | yadI | 4 | 1.25 | 0.232 |
| 0.2 | 0 | AG1 | 4 | 1.25 | 0.221 |
| 0.2 | 0 | ptsI | 4 | 1.5 | 0.2 |
| 0.2 | 0 | clcB | 4 | 1.5 | 0.212 |
| 0.2 | 0 | ycaM | 4 | 1.5 | 0.207 |
| 0.2 | 0 | yadI | 4 | 1.5 | 0.243 |
| 0.2 | 0 | AG1 | 4 | 1.5 | 0.217 |
| 0.2 | 0 | ptsI | 4 | 1.75 | 0.21 |
| 0.2 | 0 | clcB | 4 | 1.75 | 0.222 |
| 0.2 | 0 | ycaM | 4 | 1.75 | 0.216 |
| 0.2 | 0 | yadI | 4 | 1.75 | 0.255 |
| 0.2 | 0 | AG1 | 4 | 1.75 | 0.23 |
| 0.2 | 0 | ptsI | 4 | 2 | 0.22 |
| 0.2 | 0 | clcB | 4 | 2 | 0.233 |
| 0.2 | 0 | ycaM | 4 | 2 | 0.226 |
| 0.2 | 0 | yadI | 4 | 2 | 0.269 |
| 0.2 | 0 | AG1 | 4 | 2 | 0.247 |
| 0.2 | 0 | ptsI | 4 | 2.25 | 0.232 |
| 0.2 | 0 | clcB | 4 | 2.25 | 0.246 |
| 0.2 | 0 | ycaM | 4 | 2.25 | 0.237 |
| 0.2 | 0 | yadI | 4 | 2.25 | 0.284 |
| 0.2 | 0 | AG1 | 4 | 2.25 | 0.269 |
| 0.2 | 0 | ptsI | 4 | 2.5 | 0.235 |
| 0.2 | 0 | clcB | 4 | 2.5 | 0.249 |
| 0.2 | 0 | ycaM | 4 | 2.5 | 0.249 |
| 0.2 | 0 | yadI | 4 | 2.5 | 0.305 |
| 0.2 | 0 | AG1 | 4 | 2.5 | 0.26 |
| 0.2 | 0 | ptsI | 4 | 2.75 | 0.248 |
| 0.2 | 0 | clcB | 4 | 2.75 | 0.268 |
| 0.2 | 0 | ycaM | 4 | 2.75 | 0.263 |
| 0.2 | 0 | yadI | 4 | 2.75 | 0.322 |

| | | | | | |
|---|---|---|---|---|---|
| 0.2 | 0 | AG1 | 4 | 2.75 | 0.274 |
| 0.2 | 0 | ptsI | 4 | 3 | 0.266 |
| 0.2 | 0 | clcB | 4 | 3 | 0.277 |
| 0.2 | 0 | ycaM | 4 | 3 | 0.273 |
| 0.2 | 0 | yadI | 4 | 3 | 0.342 |
| 0.2 | 0 | AG1 | 4 | 3 | 0.294 |
| 0.2 | 0 | ptsI | 4 | 3.25 | 0.279 |
| 0.2 | 0 | clcB | 4 | 3.25 | 0.288 |
| 0.2 | 0 | ycaM | 4 | 3.25 | 0.289 |
| 0.2 | 0 | yadI | 4 | 3.25 | 0.363 |
| 0.2 | 0 | AG1 | 4 | 3.25 | 0.312 |
| 0.2 | 0 | ptsI | 4 | 3.5 | 0.296 |
| 0.2 | 0 | clcB | 4 | 3.5 | 0.309 |
| 0.2 | 0 | ycaM | 4 | 3.5 | 0.306 |
| 0.2 | 0 | yadI | 4 | 3.5 | 0.385 |
| 0.2 | 0 | AG1 | 4 | 3.5 | 0.338 |
| 0.2 | 0 | ptsI | 4 | 3.75 | 0.313 |
| 0.2 | 0 | clcB | 4 | 3.75 | 0.33 |
| 0.2 | 0 | ycaM | 4 | 3.75 | 0.323 |
| 0.2 | 0 | yadI | 4 | 3.75 | 0.406 |
| 0.2 | 0 | AG1 | 4 | 3.75 | 0.353 |
| 0.2 | 0 | ptsI | 4 | 4 | 0.329 |
| 0.2 | 0 | clcB | 4 | 4 | 0.342 |
| 0.2 | 0 | ycaM | 4 | 4 | 0.335 |
| 0.2 | 0 | yadI | 4 | 4 | 0.439 |
| 0.2 | 0 | AG1 | 4 | 4 | 0.38 |
| 0.2 | 0 | ptsI | 4 | 4.35 | 0.348 |
| 0.2 | 0 | clcB | 4 | 4.35 | 0.324 |
| 0.2 | 0 | ycaM | 4 | 4.35 | 0.388 |
| 0.2 | 0 | yadI | 4 | 4.35 | 0.394 |
| 0.2 | 0 | AG1 | 4 | 4.35 | 0.326 |
| 0.2 | 0 | ptsI | 4 | 4.6 | 0.361 |
| 0.2 | 0 | clcB | 4 | 4.6 | 0.306 |
| 0.2 | 0 | ycaM | 4 | 4.6 | 0.36 |
| 0.2 | 0 | yadI | 4 | 4.6 | 0.375 |
| 0.2 | 0 | AG1 | 4 | 4.6 | 0.347 |
| 0.2 | 0 | ptsI | 4 | 4.85 | 0.373 |
| 0.2 | 0 | clcB | 4 | 4.85 | 0.314 |
| 0.2 | 0 | ycaM | 4 | 4.85 | 0.37 |
| 0.2 | 0 | yadI | 4 | 4.85 | 0.385 |
| 0.2 | 0 | AG1 | 4 | 4.85 | 0.361 |
| 0.2 | 0 | ptsI | 4 | 5.1 | 0.388 |
| 0.2 | 0 | clcB | 4 | 5.1 | 0.326 |
| 0.2 | 0 | ycaM | 4 | 5.1 | 0.386 |
| 0.2 | 0 | yadI | 4 | 5.1 | 0.398 |
| 0.2 | 0 | AG1 | 4 | 5.1 | 0.376 |
| 0.2 | 0 | ptsI | 4 | 5.35 | 0.401 |
| 0.2 | 0 | clcB | 4 | 5.35 | 0.337 |

| | | | | | |
|---|---|---|---|---|---|
| 0.2 | 0 | ycaM | 4 | 5.35 | 0.403 |
| 0.2 | 0 | yadI | 4 | 5.35 | 0.413 |
| 0.2 | 0 | AG1 | 4 | 5.35 | 0.392 |
| 0.2 | 0 | ptsI | 4 | 5.6 | 0.417 |
| 0.2 | 0 | clcB | 4 | 5.6 | 0.342 |
| 0.2 | 0 | ycaM | 4 | 5.6 | 0.417 |
| 0.2 | 0 | yadI | 4 | 5.6 | 0.428 |
| 0.2 | 0 | AG1 | 4 | 5.6 | 0.405 |
| 0.2 | 0 | ptsI | 4 | 5.85 | 0.428 |
| 0.2 | 0 | clcB | 4 | 5.85 | 0.35 |
| 0.2 | 0 | ycaM | 4 | 5.85 | 0.427 |
| 0.2 | 0 | yadI | 4 | 5.85 | 0.441 |
| 0.2 | 0 | AG1 | 4 | 5.85 | 0.424 |
| 0.2 | 0 | ptsI | 4 | 6.1 | 0.437 |
| 0.2 | 0 | clcB | 4 | 6.1 | 0.366 |
| 0.2 | 0 | ycaM | 4 | 6.1 | 0.442 |
| 0.2 | 0 | yadI | 4 | 6.1 | 0.456 |
| 0.2 | 0 | AG1 | 4 | 6.1 | 0.442 |
| 0.2 | 0 | ptsI | 4 | 6.35 | 0.454 |
| 0.2 | 0 | clcB | 4 | 6.35 | 0.374 |
| 0.2 | 0 | ycaM | 4 | 6.35 | 0.458 |
| 0.2 | 0 | yadI | 4 | 6.35 | 0.47 |
| 0.2 | 0 | AG1 | 4 | 6.35 | 0.453 |
| 0.2 | 0 | ptsI | 4 | 6.6 | 0.461 |
| 0.2 | 0 | clcB | 4 | 6.6 | 0.384 |
| 0.2 | 0 | ycaM | 4 | 6.6 | 0.463 |
| 0.2 | 0 | yadI | 4 | 6.6 | 0.484 |
| 0.2 | 0 | AG1 | 4 | 6.6 | 0.468 |
| 0.2 | 0 | ptsI | 4 | 6.85 | 0.475 |
| 0.2 | 0 | clcB | 4 | 6.85 | 0.392 |
| 0.2 | 0 | ycaM | 4 | 6.85 | 0.474 |
| 0.2 | 0 | yadI | 4 | 6.85 | 0.496 |
| 0.2 | 0 | AG1 | 4 | 6.85 | 0.48 |
| 0.2 | 0 | ptsI | 4 | 7.1 | 0.485 |
| 0.2 | 0 | clcB | 4 | 7.1 | 0.403 |
| 0.2 | 0 | ycaM | 4 | 7.1 | 0.483 |
| 0.2 | 0 | yadI | 4 | 7.1 | 0.506 |
| 0.2 | 0 | AG1 | 4 | 7.1 | 0.495 |
| 0.2 | 0 | ptsI | 4 | 7.35 | 0.495 |
| 0.2 | 0 | clcB | 4 | 7.35 | 0.411 |
| 0.2 | 0 | ycaM | 4 | 7.35 | 0.493 |
| 0.2 | 0 | yadI | 4 | 7.35 | 0.514 |
| 0.2 | 0 | AG1 | 4 | 7.35 | 0.503 |
| 0.2 | 0 | ptsI | 4 | 7.6 | 0.506 |
| 0.2 | 0 | clcB | 4 | 7.6 | 0.421 |
| 0.2 | 0 | ycaM | 4 | 7.6 | 0.5 |
| 0.2 | 0 | yadI | 4 | 7.6 | 0.523 |
| 0.2 | 0 | AG1 | 4 | 7.6 | 0.515 |

| | | | | | |
|---|---|---|---|---|---|
| 0.2 | 0 | ptsI | 4 | 7.85 | 0.516 |
| 0.2 | 0 | clcB | 4 | 7.85 | 0.427 |
| 0.2 | 0 | ycaM | 4 | 7.85 | 0.501 |
| 0.2 | 0 | yadI | 4 | 7.85 | 0.531 |
| 0.2 | 0 | AG1 | 4 | 7.85 | 0.52 |
| 0.2 | 0 | ptsI | 4 | 8.1 | 0.525 |
| 0.2 | 0 | clcB | 4 | 8.1 | 0.438 |
| 0.2 | 0 | ycaM | 4 | 8.1 | 0.505 |
| 0.2 | 0 | yadI | 4 | 8.1 | 0.543 |
| 0.2 | 0 | AG1 | 4 | 8.1 | 0.532 |
| 0.2 | 0 | ptsI | 4 | 8.35 | 0.534 |
| 0.2 | 0 | clcB | 4 | 8.35 | 0.444 |
| 0.2 | 0 | ycaM | 4 | 8.35 | 0.511 |
| 0.2 | 0 | yadI | 4 | 8.35 | 0.552 |
| 0.2 | 0 | AG1 | 4 | 8.35 | 0.54 |
| 0.2 | 0 | ptsI | 4 | 8.83333 | 0.383 |
| 0.2 | 0 | clcB | 4 | 8.83333 | 0.426 |
| 0.2 | 0 | ycaM | 4 | 8.83333 | 0.389 |
| 0.2 | 0 | yadI | 4 | 8.83333 | 0.514 |
| 0.2 | 0 | AG1 | 4 | 8.83333 | 0.582 |
| 0.2 | 0 | ptsI | 4 | 9.08333 | 0.386 |
| 0.2 | 0 | clcB | 4 | 9.08333 | 0.422 |
| 0.2 | 0 | ycaM | 4 | 9.08333 | 0.389 |
| 0.2 | 0 | yadI | 4 | 9.08333 | 0.51 |
| 0.2 | 0 | AG1 | 4 | 9.08333 | 0.578 |
| 0.2 | 0 | ptsI | 4 | 9.33333 | 0.393 |
| 0.2 | 0 | clcB | 4 | 9.33333 | 0.426 |
| 0.2 | 0 | ycaM | 4 | 9.33333 | 0.395 |
| 0.2 | 0 | yadI | 4 | 9.33333 | 0.52 |
| 0.2 | 0 | AG1 | 4 | 9.33333 | 0.588 |
| 0.2 | 0 | ptsI | 4 | 9.58333 | 0.406 |
| 0.2 | 0 | clcB | 4 | 9.58333 | 0.434 |
| 0.2 | 0 | ycaM | 4 | 9.58333 | 0.402 |
| 0.2 | 0 | yadI | 4 | 9.58333 | 0.529 |
| 0.2 | 0 | AG1 | 4 | 9.58333 | 0.6 |
| 0.2 | 0 | ptsI | 4 | 9.83333 | 0.412 |
| 0.2 | 0 | clcB | 4 | 9.83333 | 0.438 |
| 0.2 | 0 | ycaM | 4 | 9.83333 | 0.405 |
| 0.2 | 0 | yadI | 4 | 9.83333 | 0.536 |
| 0.2 | 0 | AG1 | 4 | 9.83333 | 0.608 |
| 0.2 | 0 | ptsI | 4 | 10.0833 | 0.42 |
| 0.2 | 0 | clcB | 4 | 10.0833 | 0.446 |
| 0.2 | 0 | ycaM | 4 | 10.0833 | 0.412 |
| 0.2 | 0 | yadI | 4 | 10.0833 | 0.543 |
| 0.2 | 0 | AG1 | 4 | 10.0833 | 0.62 |
| 0.2 | 0 | ptsI | 4 | 10.3333 | 0.426 |
| 0.2 | 0 | clcB | 4 | 10.3333 | 0.45 |
| 0.2 | 0 | ycaM | 4 | 10.3333 | 0.415 |

| | | | | | |
|---|---|---|---|---|---|
| 0.2 | 0 | yadI | 4 | 10.3333 | 0.551 |
| 0.2 | 0 | AG1 | 4 | 10.3333 | 0.629 |
| 0.2 | 0 | ptsI | 4 | 10.5833 | 0.432 |
| 0.2 | 0 | clcB | 4 | 10.5833 | 0.459 |
| 0.2 | 0 | ycaM | 4 | 10.5833 | 0.42 |
| 0.2 | 0 | yadI | 4 | 10.5833 | 0.559 |
| 0.2 | 0 | AG1 | 4 | 10.5833 | 0.638 |
| 0.2 | 0 | ptsI | 4 | 10.8333 | 0.436 |
| 0.2 | 0 | clcB | 4 | 10.8333 | 0.462 |
| 0.2 | 0 | ycaM | 4 | 10.8333 | 0.424 |
| 0.2 | 0 | yadI | 4 | 10.8333 | 0.565 |
| 0.2 | 0 | AG1 | 4 | 10.8333 | 0.645 |
| 0.2 | 0 | ptsI | 4 | 11.0833 | 0.444 |
| 0.2 | 0 | clcB | 4 | 11.0833 | 0.466 |
| 0.2 | 0 | ycaM | 4 | 11.0833 | 0.43 |
| 0.2 | 0 | yadI | 4 | 11.0833 | 0.573 |
| 0.2 | 0 | AG1 | 4 | 11.0833 | 0.657 |
| 0.2 | 0 | ptsI | 4 | 11.3333 | 0.454 |
| 0.2 | 0 | clcB | 4 | 11.3333 | 0.477 |
| 0.2 | 0 | ycaM | 4 | 11.3333 | 0.435 |
| 0.2 | 0 | yadI | 4 | 11.3333 | 0.578 |
| 0.2 | 0 | AG1 | 4 | 11.3333 | 0.665 |
| 0.2 | 0 | ptsI | 4 | 11.5833 | 0.457 |
| 0.2 | 0 | clcB | 4 | 11.5833 | 0.478 |
| 0.2 | 0 | ycaM | 4 | 11.5833 | 0.438 |
| 0.2 | 0 | yadI | 4 | 11.5833 | 0.581 |
| 0.2 | 0 | AG1 | 4 | 11.5833 | 0.672 |
| 0.2 | 0 | ptsI | 4 | 11.8333 | 0.466 |
| 0.2 | 0 | clcB | 4 | 11.8333 | 0.481 |
| 0.2 | 0 | ycaM | 4 | 11.8333 | 0.443 |
| 0.2 | 0 | yadI | 4 | 11.8333 | 0.588 |
| 0.2 | 0 | AG1 | 4 | 11.8333 | 0.68 |
| 0.2 | 0 | ptsI | 4 | 12.0833 | 0.472 |
| 0.2 | 0 | clcB | 4 | 12.0833 | 0.488 |
| 0.2 | 0 | ycaM | 4 | 12.0833 | 0.446 |
| 0.2 | 0 | yadI | 4 | 12.0833 | 0.592 |
| 0.2 | 0 | AG1 | 4 | 12.0833 | 0.686 |
| 0.2 | 0 | ptsI | 4 | 12.3333 | 0.477 |
| 0.2 | 0 | clcB | 4 | 12.3333 | 0.497 |
| 0.2 | 0 | ycaM | 4 | 12.3333 | 0.452 |
| 0.2 | 0 | yadI | 4 | 12.3333 | 0.597 |
| 0.2 | 0 | AG1 | 4 | 12.3333 | 0.694 |
| 0.2 | 0 | ptsI | 4 | 12.5833 | 0.484 |
| 0.2 | 0 | clcB | 4 | 12.5833 | 0.5 |
| 0.2 | 0 | ycaM | 4 | 12.5833 | 0.455 |
| 0.2 | 0 | yadI | 4 | 12.5833 | 0.605 |
| 0.2 | 0 | AG1 | 4 | 12.5833 | 0.701 |
| 0.2 | 0 | ptsI | 4 | 12.8333 | 0.488 |

| | | | | | |
|---|---|---|---|---|---|
| 0.2 | 0 | clcB | 4 | 12.8333 | 0.5 |
| 0.2 | 0 | ycaM | 4 | 12.8333 | 0.458 |
| 0.2 | 0 | yadI | 4 | 12.8333 | 0.609 |
| 0.2 | 0 | AG1 | 4 | 12.8333 | 0.707 |
| 0.2 | 0 | ptsI | 4 | 13.0833 | 0.494 |
| 0.2 | 0 | clcB | 4 | 13.0833 | 0.509 |
| 0.2 | 0 | ycaM | 4 | 13.0833 | 0.464 |
| 0.2 | 0 | yadI | 4 | 13.0833 | 0.614 |
| 0.2 | 0 | AG1 | 4 | 13.0833 | 0.713 |
| 0.2 | 0 | ptsI | 4 | 13.3333 | 0.499 |
| 0.2 | 0 | clcB | 4 | 13.3333 | 0.515 |
| 0.2 | 0 | ycaM | 4 | 13.3333 | 0.47 |
| 0.2 | 0 | yadI | 4 | 13.3333 | 0.616 |
| 0.2 | 0 | AG1 | 4 | 13.3333 | 0.719 |
| 0.2 | 0 | ptsI | 4 | 13.5833 | 0.504 |
| 0.2 | 0 | clcB | 4 | 13.5833 | 0.516 |
| 0.2 | 0 | ycaM | 4 | 13.5833 | 0.473 |
| 0.2 | 0 | yadI | 4 | 13.5833 | 0.626 |
| 0.2 | 0 | AG1 | 4 | 13.5833 | 0.721 |
| 0.2 | 0 | ptsI | 4 | 13.8333 | 0.509 |
| 0.2 | 0 | clcB | 4 | 13.8333 | 0.521 |
| 0.2 | 0 | ycaM | 4 | 13.8333 | 0.483 |
| 0.2 | 0 | yadI | 4 | 13.8333 | 0.623 |
| 0.2 | 0 | AG1 | 4 | 13.8333 | 0.726 |
| 0.2 | 0 | ptsI | 4 | 14.0833 | 0.509 |
| 0.2 | 0 | clcB | 4 | 14.0833 | 0.524 |
| 0.2 | 0 | ycaM | 4 | 14.0833 | 0.485 |
| 0.2 | 0 | yadI | 4 | 14.0833 | 0.629 |
| 0.2 | 0 | AG1 | 4 | 14.0833 | 0.73 |
| 0.2 | 0 | ptsI | 4 | 14.3333 | 0.518 |
| 0.2 | 0 | clcB | 4 | 14.3333 | 0.525 |
| 0.2 | 0 | ycaM | 4 | 14.3333 | 0.488 |
| 0.2 | 0 | yadI | 4 | 14.3333 | 0.632 |
| 0.2 | 0 | AG1 | 4 | 14.3333 | 0.728 |
| 0.2 | 0 | ptsI | 4 | 14.5833 | 0.521 |
| 0.2 | 0 | clcB | 4 | 14.5833 | 0.527 |
| 0.2 | 0 | ycaM | 4 | 14.5833 | 0.491 |
| 0.2 | 0 | yadI | 4 | 14.5833 | 0.633 |
| 0.2 | 0 | AG1 | 4 | 14.5833 | 0.737 |
| 0.2 | 0 | ptsI | 4 | 14.8333 | 0.524 |
| 0.2 | 0 | clcB | 4 | 14.8333 | 0.531 |
| 0.2 | 0 | ycaM | 4 | 14.8333 | 0.499 |
| 0.2 | 0 | yadI | 4 | 14.8333 | 0.638 |
| 0.2 | 0 | AG1 | 4 | 14.8333 | 0.742 |
| 0.2 | 0 | ptsI | 4 | 15.0833 | 0.53 |
| 0.2 | 0 | clcB | 4 | 15.0833 | 0.531 |
| 0.2 | 0 | ycaM | 4 | 15.0833 | 0.5 |
| 0.2 | 0 | yadI | 4 | 15.0833 | 0.644 |

| | | | | | |
|---|---|---|---|---|---|
| 0.2 | 0 | AG1 | 4 | 15.0833 | 0.746 |
| 0.2 | 0 | ptsI | 4 | 15.3333 | 0.536 |
| 0.2 | 0 | clcB | 4 | 15.3333 | 0.536 |
| 0.2 | 0 | ycaM | 4 | 15.3333 | 0.5 |
| 0.2 | 0 | yadI | 4 | 15.3333 | 0.644 |
| 0.2 | 0 | AG1 | 4 | 15.3333 | 0.747 |
| 0.2 | 0 | ptsI | 4 | 15.5833 | 0.54 |
| 0.2 | 0 | clcB | 4 | 15.5833 | 0.541 |
| 0.2 | 0 | ycaM | 4 | 15.5833 | 0.503 |
| 0.2 | 0 | yadI | 4 | 15.5833 | 0.648 |
| 0.2 | 0 | AG1 | 4 | 15.5833 | 0.75 |
| 0.2 | 0 | ptsI | 4 | 15.8333 | 0.542 |
| 0.2 | 0 | clcB | 4 | 15.8333 | 0.537 |
| 0.2 | 0 | ycaM | 4 | 15.8333 | 0.508 |
| 0.2 | 0 | yadI | 4 | 15.8333 | 0.655 |
| 0.2 | 0 | AG1 | 4 | 15.8333 | 0.752 |
| 0.2 | 0 | ptsI | 4 | 16.0833 | 0.542 |
| 0.2 | 0 | clcB | 4 | 16.0833 | 0.546 |
| 0.2 | 0 | ycaM | 4 | 16.0833 | 0.511 |
| 0.2 | 0 | yadI | 4 | 16.0833 | 0.659 |
| 0.2 | 0 | AG1 | 4 | 16.0833 | 0.757 |
| 0.2 | 0 | ptsI | 4 | 16.3333 | 0.546 |
| 0.2 | 0 | clcB | 4 | 16.3333 | 0.546 |
| 0.2 | 0 | ycaM | 4 | 16.3333 | 0.513 |
| 0.2 | 0 | yadI | 4 | 16.3333 | 0.656 |
| 0.2 | 0 | AG1 | 4 | 16.3333 | 0.757 |
| 0.2 | 0 | ptsI | 4 | 16.5833 | 0.55 |
| 0.2 | 0 | clcB | 4 | 16.5833 | 0.545 |
| 0.2 | 0 | ycaM | 4 | 16.5833 | 0.518 |
| 0.2 | 0 | yadI | 4 | 16.5833 | 0.661 |
| 0.2 | 0 | AG1 | 4 | 16.5833 | 0.757 |
| 0.2 | 0 | ptsI | 4 | 16.8333 | 0.552 |
| 0.2 | 0 | clcB | 4 | 16.8333 | 0.546 |
| 0.2 | 0 | ycaM | 4 | 16.8333 | 0.518 |
| 0.2 | 0 | yadI | 4 | 16.8333 | 0.663 |
| 0.2 | 0 | AG1 | 4 | 16.8333 | 0.761 |
| 0.2 | 0 | ptsI | 4 | 17.0833 | 0.559 |
| 0.2 | 0 | clcB | 4 | 17.0833 | 0.551 |
| 0.2 | 0 | ycaM | 4 | 17.0833 | 0.52 |
| 0.2 | 0 | yadI | 4 | 17.0833 | 0.667 |
| 0.2 | 0 | AG1 | 4 | 17.0833 | 0.76 |
| 0.2 | 0 | ptsI | 4 | 17.3333 | 0.563 |
| 0.2 | 0 | clcB | 4 | 17.3333 | 0.557 |
| 0.2 | 0 | ycaM | 4 | 17.3333 | 0.528 |
| 0.2 | 0 | yadI | 4 | 17.3333 | 0.668 |
| 0.2 | 0 | AG1 | 4 | 17.3333 | 0.763 |
| 0.2 | 0 | ptsI | 4 | 17.5833 | 0.56 |
| 0.2 | 0 | clcB | 4 | 17.5833 | 0.551 |

| | | | | | |
|---|---|---|---|---|---|
| 0.2 | 0 | ycaM | 4 | 17.5833 | 0.524 |
| 0.2 | 0 | yadI | 4 | 17.5833 | 0.67 |
| 0.2 | 0 | AG1 | 4 | 17.5833 | 0.762 |
| 0.2 | 0 | ptsI | 4 | 17.8333 | 0.577 |
| 0.2 | 0 | clcB | 4 | 17.8333 | 0.556 |
| 0.2 | 0 | ycaM | 4 | 17.8333 | 0.527 |
| 0.2 | 0 | yadI | 4 | 17.8333 | 0.671 |
| 0.2 | 0 | AG1 | 4 | 17.8333 | 0.77 |
| 0.2 | 0 | ptsI | 4 | 18.0833 | 0.575 |
| 0.2 | 0 | clcB | 4 | 18.0833 | 0.554 |
| 0.2 | 0 | ycaM | 4 | 18.0833 | 0.53 |
| 0.2 | 0 | yadI | 4 | 18.0833 | 0.676 |
| 0.2 | 0 | AG1 | 4 | 18.0833 | 0.764 |
| 0.2 | 0 | ptsI | 4 | 18.3333 | 0.57 |
| 0.2 | 0 | clcB | 4 | 18.3333 | 0.556 |
| 0.2 | 0 | ycaM | 4 | 18.3333 | 0.532 |
| 0.2 | 0 | yadI | 4 | 18.3333 | 0.675 |
| 0.2 | 0 | AG1 | 4 | 18.3333 | 0.769 |
| 0.2 | 0 | ptsI | 4 | 18.5833 | 0.571 |
| 0.2 | 0 | clcB | 4 | 18.5833 | 0.558 |
| 0.2 | 0 | ycaM | 4 | 18.5833 | 0.532 |
| 0.2 | 0 | yadI | 4 | 18.5833 | 0.679 |
| 0.2 | 0 | AG1 | 4 | 18.5833 | 0.77 |
| 0.2 | 0 | ptsI | 4 | 18.8333 | 0.585 |
| 0.2 | 0 | clcB | 4 | 18.8333 | 0.563 |
| 0.2 | 0 | ycaM | 4 | 18.8333 | 0.536 |
| 0.2 | 0 | yadI | 4 | 18.8333 | 0.677 |
| 0.2 | 0 | AG1 | 4 | 18.8333 | 0.775 |
| 0.2 | 0 | ptsI | 4 | 19.0833 | 0.585 |
| 0.2 | 0 | clcB | 4 | 19.0833 | 0.566 |
| 0.2 | 0 | ycaM | 4 | 19.0833 | 0.539 |
| 0.2 | 0 | yadI | 4 | 19.0833 | 0.68 |
| 0.2 | 0 | AG1 | 4 | 19.0833 | 0.777 |
| 0.2 | 0 | ptsI | 4 | 19.3333 | 0.587 |
| 0.2 | 0 | clcB | 4 | 19.3333 | 0.568 |
| 0.2 | 0 | ycaM | 4 | 19.3333 | 0.54 |
| 0.2 | 0 | yadI | 4 | 19.3333 | 0.683 |
| 0.2 | 0 | AG1 | 4 | 19.3333 | 0.776 |
| 0.2 | 0 | ptsI | 4 | 19.5833 | 0.594 |
| 0.2 | 0 | clcB | 4 | 19.5833 | 0.57 |
| 0.2 | 0 | ycaM | 4 | 19.5833 | 0.543 |
| 0.2 | 0 | yadI | 4 | 19.5833 | 0.684 |
| 0.2 | 0 | AG1 | 4 | 19.5833 | 0.776 |
| 0.2 | 0 | ptsI | 4 | 19.8333 | 0.594 |
| 0.2 | 0 | clcB | 4 | 19.8333 | 0.566 |
| 0.2 | 0 | ycaM | 4 | 19.8333 | 0.544 |
| 0.2 | 0 | yadI | 4 | 19.8333 | 0.68 |
| 0.2 | 0 | AG1 | 4 | 19.8333 | 0.777 |

| | | | | | |
|---|---|---|---|---|---|
| 0.2 | 0 | ptsI | 4 | 20.0833 | 0.595 |
| 0.2 | 0 | clcB | 4 | 20.0833 | 0.573 |
| 0.2 | 0 | ycaM | 4 | 20.0833 | 0.548 |
| 0.2 | 0 | yadI | 4 | 20.0833 | 0.686 |
| 0.2 | 0 | AG1 | 4 | 20.0833 | 0.787 |
| 0.2 | 0 | ptsI | 4 | 20.3333 | 0.6 |
| 0.2 | 0 | clcB | 4 | 20.3333 | 0.572 |
| 0.2 | 0 | ycaM | 4 | 20.3333 | 0.549 |
| 0.2 | 0 | yadI | 4 | 20.3333 | 0.686 |
| 0.2 | 0 | AG1 | 4 | 20.3333 | 0.778 |
| 0.2 | 0 | ptsI | 4 | 20.5833 | 0.601 |
| 0.2 | 0 | clcB | 4 | 20.5833 | 0.572 |
| 0.2 | 0 | ycaM | 4 | 20.5833 | 0.552 |
| 0.2 | 0 | yadI | 4 | 20.5833 | 0.686 |
| 0.2 | 0 | AG1 | 4 | 20.5833 | 0.778 |
| 0.2 | 0 | ptsI | 4 | 20.8333 | 0.604 |
| 0.2 | 0 | clcB | 4 | 20.8333 | 0.578 |
| 0.2 | 0 | ycaM | 4 | 20.8333 | 0.557 |
| 0.2 | 0 | yadI | 4 | 20.8333 | 0.687 |
| 0.2 | 0 | AG1 | 4 | 20.8333 | 0.779 |
| 0.2 | 0 | ptsI | 4 | 21.0833 | 0.61 |
| 0.2 | 0 | clcB | 4 | 21.0833 | 0.576 |
| 0.2 | 0 | ycaM | 4 | 21.0833 | 0.557 |
| 0.2 | 0 | yadI | 4 | 21.0833 | 0.69 |
| 0.2 | 0 | AG1 | 4 | 21.0833 | 0.781 |
| 0.2 | 0 | ptsI | 4 | 21.3333 | 0.605 |
| 0.2 | 0 | clcB | 4 | 21.3333 | 0.576 |
| 0.2 | 0 | ycaM | 4 | 21.3333 | 0.558 |
| 0.2 | 0 | yadI | 4 | 21.3333 | 0.687 |
| 0.2 | 0 | AG1 | 4 | 21.3333 | 0.781 |
| 0.2 | 0 | ptsI | 4 | 21.5833 | 0.613 |
| 0.2 | 0 | clcB | 4 | 21.5833 | 0.579 |
| 0.2 | 0 | ycaM | 4 | 21.5833 | 0.564 |
| 0.2 | 0 | yadI | 4 | 21.5833 | 0.689 |
| 0.2 | 0 | AG1 | 4 | 21.5833 | 0.784 |
| 0.2 | 0 | ptsI | 4 | 21.8333 | 0.613 |
| 0.2 | 0 | clcB | 4 | 21.8333 | 0.581 |
| 0.2 | 0 | ycaM | 4 | 21.8333 | 0.565 |
| 0.2 | 0 | yadI | 4 | 21.8333 | 0.691 |
| 0.2 | 0 | AG1 | 4 | 21.8333 | 0.784 |
| 0.2 | 0 | ptsI | 4 | 22.0833 | 0.619 |
| 0.2 | 0 | clcB | 4 | 22.0833 | 0.582 |
| 0.2 | 0 | ycaM | 4 | 22.0833 | 0.565 |
| 0.2 | 0 | yadI | 4 | 22.0833 | 0.69 |
| 0.2 | 0 | AG1 | 4 | 22.0833 | 0.784 |
| 0.2 | 0 | ptsI | 4 | 22.3333 | 0.623 |
| 0.2 | 0 | clcB | 4 | 22.3333 | 0.583 |
| 0.2 | 0 | ycaM | 4 | 22.3333 | 0.567 |

| | | | | | |
|---|---|---|---|---|---|
| 0.2 | 0 | yadI | 4 | 22.3333 | 0.695 |
| 0.2 | 0 | AG1 | 4 | 22.3333 | 0.784 |
| 0.2 | 0 | ptsI | 4 | 22.5833 | 0.623 |
| 0.2 | 0 | clcB | 4 | 22.5833 | 0.581 |
| 0.2 | 0 | ycaM | 4 | 22.5833 | 0.57 |
| 0.2 | 0 | yadI | 4 | 22.5833 | 0.692 |
| 0.2 | 0 | AG1 | 4 | 22.5833 | 0.781 |
| 0.2 | 0 | ptsI | 4 | 22.8333 | 0.628 |
| 0.2 | 0 | clcB | 4 | 22.8333 | 0.586 |
| 0.2 | 0 | ycaM | 4 | 22.8333 | 0.571 |
| 0.2 | 0 | yadI | 4 | 22.8333 | 0.695 |
| 0.2 | 0 | AG1 | 4 | 22.8333 | 0.783 |
| 0.2 | 0 | ptsI | 4 | 23.0833 | 0.639 |
| 0.2 | 0 | clcB | 4 | 23.0833 | 0.588 |
| 0.2 | 0 | ycaM | 4 | 23.0833 | 0.574 |
| 0.2 | 0 | yadI | 4 | 23.0833 | 0.693 |
| 0.2 | 0 | AG1 | 4 | 23.0833 | 0.786 |
| 0.2 | 0 | ptsI | 4 | 23.3333 | 0.638 |
| 0.2 | 0 | clcB | 4 | 23.3333 | 0.592 |
| 0.2 | 0 | ycaM | 4 | 23.3333 | 0.579 |
| 0.2 | 0 | yadI | 4 | 23.3333 | 0.698 |
| 0.2 | 0 | AG1 | 4 | 23.3333 | 0.787 |
| 0.2 | 0 | ptsI | 4 | 23.5833 | 0.64 |
| 0.2 | 0 | clcB | 4 | 23.5833 | 0.59 |
| 0.2 | 0 | ycaM | 4 | 23.5833 | 0.583 |
| 0.2 | 0 | yadI | 4 | 23.5833 | 0.695 |
| 0.2 | 0 | AG1 | 4 | 23.5833 | 0.787 |
| 0.2 | 0 | ptsI | 4 | 23.8333 | 0.645 |
| 0.2 | 0 | clcB | 4 | 23.8333 | 0.591 |
| 0.2 | 0 | ycaM | 4 | 23.8333 | 0.584 |
| 0.2 | 0 | yadI | 4 | 23.8333 | 0.697 |
| 0.2 | 0 | AG1 | 4 | 23.8333 | 0.785 |
| 0.2 | 0 | ptsI | 4 | 24.0833 | 0.651 |
| 0.2 | 0 | clcB | 4 | 24.0833 | 0.594 |
| 0.2 | 0 | ycaM | 4 | 24.0833 | 0.584 |
| 0.2 | 0 | yadI | 4 | 24.0833 | 0.7 |
| 0.2 | 0 | AG1 | 4 | 24.0833 | 0.784 |
| 0.2 | 0 | ptsI | 5 | 0 | 0.155 |
| 0.2 | 0 | clcB | 5 | 0 | 0.161 |
| 0.2 | 0 | ycaM | 5 | 0 | 0.175 |
| 0.2 | 0 | yadI | 5 | 0 | 0.166 |
| 0.2 | 0 | AG1 | 5 | 0 | 0.162 |
| 0.2 | 0 | ptsI | 5 | 0.35 | 0.154 |
| 0.2 | 0 | clcB | 5 | 0.35 | 0.158 |
| 0.2 | 0 | ycaM | 5 | 0.35 | 0.17 |
| 0.2 | 0 | yadI | 5 | 0.35 | 0.161 |
| 0.2 | 0 | AG1 | 5 | 0.35 | 0.157 |
| 0.2 | 0 | ptsI | 5 | 0.6 | 0.155 |

| | | | | | |
|---|---|---|---|---|---|
| 0.2 | 0 | clcB | 5 | 0.6 | 0.158 |
| 0.2 | 0 | ycaM | 5 | 0.6 | 0.169 |
| 0.2 | 0 | yadI | 5 | 0.6 | 0.161 |
| 0.2 | 0 | AG1 | 5 | 0.6 | 0.159 |
| 0.2 | 0 | ptsI | 5 | 0.85 | 0.153 |
| 0.2 | 0 | clcB | 5 | 0.85 | 0.158 |
| 0.2 | 0 | ycaM | 5 | 0.85 | 0.17 |
| 0.2 | 0 | yadI | 5 | 0.85 | 0.161 |
| 0.2 | 0 | AG1 | 5 | 0.85 | 0.16 |
| 0.2 | 0 | ptsI | 5 | 1.1 | 0.157 |
| 0.2 | 0 | clcB | 5 | 1.1 | 0.161 |
| 0.2 | 0 | ycaM | 5 | 1.1 | 0.172 |
| 0.2 | 0 | yadI | 5 | 1.1 | 0.162 |
| 0.2 | 0 | AG1 | 5 | 1.1 | 0.162 |
| 0.2 | 0 | ptsI | 5 | 1.35 | 0.157 |
| 0.2 | 0 | clcB | 5 | 1.35 | 0.162 |
| 0.2 | 0 | ycaM | 5 | 1.35 | 0.174 |
| 0.2 | 0 | yadI | 5 | 1.35 | 0.164 |
| 0.2 | 0 | AG1 | 5 | 1.35 | 0.163 |
| 0.2 | 0 | ptsI | 5 | 1.6 | 0.163 |
| 0.2 | 0 | clcB | 5 | 1.6 | 0.166 |
| 0.2 | 0 | ycaM | 5 | 1.6 | 0.177 |
| 0.2 | 0 | yadI | 5 | 1.6 | 0.166 |
| 0.2 | 0 | AG1 | 5 | 1.6 | 0.169 |
| 0.2 | 0 | ptsI | 5 | 1.85 | 0.165 |
| 0.2 | 0 | clcB | 5 | 1.85 | 0.169 |
| 0.2 | 0 | ycaM | 5 | 1.85 | 0.179 |
| 0.2 | 0 | yadI | 5 | 1.85 | 0.169 |
| 0.2 | 0 | AG1 | 5 | 1.85 | 0.174 |
| 0.2 | 0 | ptsI | 5 | 2.1 | 0.17 |
| 0.2 | 0 | clcB | 5 | 2.1 | 0.175 |
| 0.2 | 0 | ycaM | 5 | 2.1 | 0.184 |
| 0.2 | 0 | yadI | 5 | 2.1 | 0.174 |
| 0.2 | 0 | AG1 | 5 | 2.1 | 0.182 |
| 0.2 | 0 | ptsI | 5 | 2.35 | 0.176 |
| 0.2 | 0 | clcB | 5 | 2.35 | 0.183 |
| 0.2 | 0 | ycaM | 5 | 2.35 | 0.188 |
| 0.2 | 0 | yadI | 5 | 2.35 | 0.179 |
| 0.2 | 0 | AG1 | 5 | 2.35 | 0.191 |
| 0.2 | 0 | ptsI | 5 | 2.6 | 0.18 |
| 0.2 | 0 | clcB | 5 | 2.6 | 0.184 |
| 0.2 | 0 | ycaM | 5 | 2.6 | 0.19 |
| 0.2 | 0 | yadI | 5 | 2.6 | 0.187 |
| 0.2 | 0 | AG1 | 5 | 2.6 | 0.191 |
| 0.2 | 0 | ptsI | 5 | 2.85 | 0.189 |
| 0.2 | 0 | clcB | 5 | 2.85 | 0.198 |
| 0.2 | 0 | ycaM | 5 | 2.85 | 0.204 |
| 0.2 | 0 | yadI | 5 | 2.85 | 0.194 |

| | | | | | |
|---|---|---|---|---|---|
| 0.2 | 0 | AG1 | 5 | 2.85 | 0.21 |
| 0.2 | 0 | ptsI | 5 | 3.1 | 0.196 |
| 0.2 | 0 | clcB | 5 | 3.1 | 0.21 |
| 0.2 | 0 | ycaM | 5 | 3.1 | 0.207 |
| 0.2 | 0 | yadI | 5 | 3.1 | 0.202 |
| 0.2 | 0 | AG1 | 5 | 3.1 | 0.221 |
| 0.2 | 0 | ptsI | 5 | 3.35 | 0.201 |
| 0.2 | 0 | clcB | 5 | 3.35 | 0.225 |
| 0.2 | 0 | ycaM | 5 | 3.35 | 0.217 |
| 0.2 | 0 | yadI | 5 | 3.35 | 0.213 |
| 0.2 | 0 | AG1 | 5 | 3.35 | 0.244 |
| 0.2 | 0 | ptsI | 5 | 3.6 | 0.204 |
| 0.2 | 0 | clcB | 5 | 3.6 | 0.24 |
| 0.2 | 0 | ycaM | 5 | 3.6 | 0.228 |
| 0.2 | 0 | yadI | 5 | 3.6 | 0.224 |
| 0.2 | 0 | AG1 | 5 | 3.6 | 0.263 |
| 0.2 | 0 | ptsI | 5 | 3.85 | 0.212 |
| 0.2 | 0 | clcB | 5 | 3.85 | 0.252 |
| 0.2 | 0 | ycaM | 5 | 3.85 | 0.243 |
| 0.2 | 0 | yadI | 5 | 3.85 | 0.237 |
| 0.2 | 0 | AG1 | 5 | 3.85 | 0.278 |
| 0.2 | 0 | ptsI | 5 | 4.38333 | 0.209 |
| 0.2 | 0 | clcB | 5 | 4.38333 | 0.255 |
| 0.2 | 0 | ycaM | 5 | 4.38333 | 0.256 |
| 0.2 | 0 | yadI | 5 | 4.38333 | 0.263 |
| 0.2 | 0 | AG1 | 5 | 4.38333 | 0.312 |
| 0.2 | 0 | ptsI | 5 | 4.63333 | 0.213 |
| 0.2 | 0 | clcB | 5 | 4.63333 | 0.268 |
| 0.2 | 0 | ycaM | 5 | 4.63333 | 0.258 |
| 0.2 | 0 | yadI | 5 | 4.63333 | 0.266 |
| 0.2 | 0 | AG1 | 5 | 4.63333 | 0.283 |
| 0.2 | 0 | ptsI | 5 | 4.88333 | 0.218 |
| 0.2 | 0 | clcB | 5 | 4.88333 | 0.283 |
| 0.2 | 0 | ycaM | 5 | 4.88333 | 0.269 |
| 0.2 | 0 | yadI | 5 | 4.88333 | 0.277 |
| 0.2 | 0 | AG1 | 5 | 4.88333 | 0.303 |
| 0.2 | 0 | ptsI | 5 | 5.13333 | 0.224 |
| 0.2 | 0 | clcB | 5 | 5.13333 | 0.296 |
| 0.2 | 0 | ycaM | 5 | 5.13333 | 0.281 |
| 0.2 | 0 | yadI | 5 | 5.13333 | 0.29 |
| 0.2 | 0 | AG1 | 5 | 5.13333 | 0.313 |
| 0.2 | 0 | ptsI | 5 | 5.38333 | 0.233 |
| 0.2 | 0 | clcB | 5 | 5.38333 | 0.309 |
| 0.2 | 0 | ycaM | 5 | 5.38333 | 0.296 |
| 0.2 | 0 | yadI | 5 | 5.38333 | 0.304 |
| 0.2 | 0 | AG1 | 5 | 5.38333 | 0.32 |
| 0.2 | 0 | ptsI | 5 | 5.63333 | 0.243 |
| 0.2 | 0 | clcB | 5 | 5.63333 | 0.33 |

| | | | | | |
|---|---|---|---|---|---|
| 0.2 | 0 | ycaM | 5 | 5.63333 | 0.307 |
| 0.2 | 0 | yadI | 5 | 5.63333 | 0.318 |
| 0.2 | 0 | AG1 | 5 | 5.63333 | 0.338 |
| 0.2 | 0 | ptsI | 5 | 5.88333 | 0.254 |
| 0.2 | 0 | clcB | 5 | 5.88333 | 0.346 |
| 0.2 | 0 | ycaM | 5 | 5.88333 | 0.32 |
| 0.2 | 0 | yadI | 5 | 5.88333 | 0.333 |
| 0.2 | 0 | AG1 | 5 | 5.88333 | 0.341 |
| 0.2 | 0 | ptsI | 5 | 6.13333 | 0.266 |
| 0.2 | 0 | clcB | 5 | 6.13333 | 0.36 |
| 0.2 | 0 | ycaM | 5 | 6.13333 | 0.33 |
| 0.2 | 0 | yadI | 5 | 6.13333 | 0.348 |
| 0.2 | 0 | AG1 | 5 | 6.13333 | 0.36 |
| 0.2 | 0 | ptsI | 5 | 6.38333 | 0.276 |
| 0.2 | 0 | clcB | 5 | 6.38333 | 0.371 |
| 0.2 | 0 | ycaM | 5 | 6.38333 | 0.341 |
| 0.2 | 0 | yadI | 5 | 6.38333 | 0.364 |
| 0.2 | 0 | AG1 | 5 | 6.38333 | 0.381 |
| 0.2 | 0 | ptsI | 5 | 6.63333 | 0.288 |
| 0.2 | 0 | clcB | 5 | 6.63333 | 0.385 |
| 0.2 | 0 | ycaM | 5 | 6.63333 | 0.349 |
| 0.2 | 0 | yadI | 5 | 6.63333 | 0.377 |
| 0.2 | 0 | AG1 | 5 | 6.63333 | 0.4 |
| 0.2 | 0 | ptsI | 5 | 6.88333 | 0.299 |
| 0.2 | 0 | clcB | 5 | 6.88333 | 0.398 |
| 0.2 | 0 | ycaM | 5 | 6.88333 | 0.36 |
| 0.2 | 0 | yadI | 5 | 6.88333 | 0.392 |
| 0.2 | 0 | AG1 | 5 | 6.88333 | 0.425 |
| 0.2 | 0 | ptsI | 5 | 7.13333 | 0.312 |
| 0.2 | 0 | clcB | 5 | 7.13333 | 0.413 |
| 0.2 | 0 | ycaM | 5 | 7.13333 | 0.369 |
| 0.2 | 0 | yadI | 5 | 7.13333 | 0.408 |
| 0.2 | 0 | AG1 | 5 | 7.13333 | 0.443 |
| 0.2 | 0 | ptsI | 5 | 7.38333 | 0.324 |
| 0.2 | 0 | clcB | 5 | 7.38333 | 0.426 |
| 0.2 | 0 | ycaM | 5 | 7.38333 | 0.381 |
| 0.2 | 0 | yadI | 5 | 7.38333 | 0.421 |
| 0.2 | 0 | AG1 | 5 | 7.38333 | 0.457 |
| 0.2 | 0 | ptsI | 5 | 7.63333 | 0.335 |
| 0.2 | 0 | clcB | 5 | 7.63333 | 0.438 |
| 0.2 | 0 | ycaM | 5 | 7.63333 | 0.391 |
| 0.2 | 0 | yadI | 5 | 7.63333 | 0.436 |
| 0.2 | 0 | AG1 | 5 | 7.63333 | 0.472 |
| 0.2 | 0 | ptsI | 5 | 7.88333 | 0.346 |
| 0.2 | 0 | clcB | 5 | 7.88333 | 0.454 |
| 0.2 | 0 | ycaM | 5 | 7.88333 | 0.42 |
| 0.2 | 0 | yadI | 5 | 7.88333 | 0.45 |
| 0.2 | 0 | AG1 | 5 | 7.88333 | 0.489 |

| | | | | | |
|---|---|---|---|---|---|
| 0.2 | 0 | ptsI | 5 | 8.13333 | 0.361 |
| 0.2 | 0 | clcB | 5 | 8.13333 | 0.465 |
| 0.2 | 0 | ycaM | 5 | 8.13333 | 0.422 |
| 0.2 | 0 | yadI | 5 | 8.13333 | 0.463 |
| 0.2 | 0 | AG1 | 5 | 8.13333 | 0.502 |
| 0.2 | 0 | ptsI | 5 | 8.38333 | 0.375 |
| 0.2 | 0 | clcB | 5 | 8.38333 | 0.479 |
| 0.2 | 0 | ycaM | 5 | 8.38333 | 0.439 |
| 0.2 | 0 | yadI | 5 | 8.38333 | 0.475 |
| 0.2 | 0 | AG1 | 5 | 8.38333 | 0.515 |
| 0.2 | 0 | ptsI | 5 | 8.88333 | 0.333 |
| 0.2 | 0 | clcB | 5 | 8.88333 | 0.349 |
| 0.2 | 0 | ycaM | 5 | 8.88333 | 0.349 |
| 0.2 | 0 | yadI | 5 | 8.88333 | 0.381 |
| 0.2 | 0 | AG1 | 5 | 8.88333 | 0.536 |
| 0.2 | 0 | ptsI | 5 | 9.13333 | 0.337 |
| 0.2 | 0 | clcB | 5 | 9.13333 | 0.358 |
| 0.2 | 0 | ycaM | 5 | 9.13333 | 0.355 |
| 0.2 | 0 | yadI | 5 | 9.13333 | 0.382 |
| 0.2 | 0 | AG1 | 5 | 9.13333 | 0.536 |
| 0.2 | 0 | ptsI | 5 | 9.38333 | 0.342 |
| 0.2 | 0 | clcB | 5 | 9.38333 | 0.368 |
| 0.2 | 0 | ycaM | 5 | 9.38333 | 0.363 |
| 0.2 | 0 | yadI | 5 | 9.38333 | 0.391 |
| 0.2 | 0 | AG1 | 5 | 9.38333 | 0.546 |
| 0.2 | 0 | ptsI | 5 | 9.63333 | 0.357 |
| 0.2 | 0 | clcB | 5 | 9.63333 | 0.378 |
| 0.2 | 0 | ycaM | 5 | 9.63333 | 0.371 |
| 0.2 | 0 | yadI | 5 | 9.63333 | 0.401 |
| 0.2 | 0 | AG1 | 5 | 9.63333 | 0.56 |
| 0.2 | 0 | ptsI | 5 | 9.88333 | 0.364 |
| 0.2 | 0 | clcB | 5 | 9.88333 | 0.387 |
| 0.2 | 0 | ycaM | 5 | 9.88333 | 0.378 |
| 0.2 | 0 | yadI | 5 | 9.88333 | 0.409 |
| 0.2 | 0 | AG1 | 5 | 9.88333 | 0.575 |
| 0.2 | 0 | ptsI | 5 | 10.1333 | 0.373 |
| 0.2 | 0 | clcB | 5 | 10.1333 | 0.395 |
| 0.2 | 0 | ycaM | 5 | 10.1333 | 0.385 |
| 0.2 | 0 | yadI | 5 | 10.1333 | 0.416 |
| 0.2 | 0 | AG1 | 5 | 10.1333 | 0.583 |
| 0.2 | 0 | ptsI | 5 | 10.3833 | 0.388 |
| 0.2 | 0 | clcB | 5 | 10.3833 | 0.404 |
| 0.2 | 0 | ycaM | 5 | 10.3833 | 0.392 |
| 0.2 | 0 | yadI | 5 | 10.3833 | 0.425 |
| 0.2 | 0 | AG1 | 5 | 10.3833 | 0.595 |
| 0.2 | 0 | ptsI | 5 | 10.6333 | 0.391 |
| 0.2 | 0 | clcB | 5 | 10.6333 | 0.408 |
| 0.2 | 0 | ycaM | 5 | 10.6333 | 0.398 |

| | | | | | |
|---|---|---|---|---|---|
| 0.2 | 0 | yadI | 5 | 10.6333 | 0.429 |
| 0.2 | 0 | AG1 | 5 | 10.6333 | 0.603 |
| 0.2 | 0 | ptsI | 5 | 10.8833 | 0.4 |
| 0.2 | 0 | clcB | 5 | 10.8833 | 0.414 |
| 0.2 | 0 | ycaM | 5 | 10.8833 | 0.401 |
| 0.2 | 0 | yadI | 5 | 10.8833 | 0.438 |
| 0.2 | 0 | AG1 | 5 | 10.8833 | 0.613 |
| 0.2 | 0 | ptsI | 5 | 11.1333 | 0.408 |
| 0.2 | 0 | clcB | 5 | 11.1333 | 0.417 |
| 0.2 | 0 | ycaM | 5 | 11.1333 | 0.404 |
| 0.2 | 0 | yadI | 5 | 11.1333 | 0.44 |
| 0.2 | 0 | AG1 | 5 | 11.1333 | 0.621 |
| 0.2 | 0 | ptsI | 5 | 11.3833 | 0.417 |
| 0.2 | 0 | clcB | 5 | 11.3833 | 0.422 |
| 0.2 | 0 | ycaM | 5 | 11.3833 | 0.405 |
| 0.2 | 0 | yadI | 5 | 11.3833 | 0.447 |
| 0.2 | 0 | AG1 | 5 | 11.3833 | 0.635 |
| 0.2 | 0 | ptsI | 5 | 11.6333 | 0.424 |
| 0.2 | 0 | clcB | 5 | 11.6333 | 0.427 |
| 0.2 | 0 | ycaM | 5 | 11.6333 | 0.408 |
| 0.2 | 0 | yadI | 5 | 11.6333 | 0.454 |
| 0.2 | 0 | AG1 | 5 | 11.6333 | 0.644 |
| 0.2 | 0 | ptsI | 5 | 11.8833 | 0.435 |
| 0.2 | 0 | clcB | 5 | 11.8833 | 0.432 |
| 0.2 | 0 | ycaM | 5 | 11.8833 | 0.411 |
| 0.2 | 0 | yadI | 5 | 11.8833 | 0.462 |
| 0.2 | 0 | AG1 | 5 | 11.8833 | 0.652 |
| 0.2 | 0 | ptsI | 5 | 12.1333 | 0.446 |
| 0.2 | 0 | clcB | 5 | 12.1333 | 0.442 |
| 0.2 | 0 | ycaM | 5 | 12.1333 | 0.416 |
| 0.2 | 0 | yadI | 5 | 12.1333 | 0.469 |
| 0.2 | 0 | AG1 | 5 | 12.1333 | 0.662 |
| 0.2 | 0 | ptsI | 5 | 12.3833 | 0.45 |
| 0.2 | 0 | clcB | 5 | 12.3833 | 0.445 |
| 0.2 | 0 | ycaM | 5 | 12.3833 | 0.414 |
| 0.2 | 0 | yadI | 5 | 12.3833 | 0.476 |
| 0.2 | 0 | AG1 | 5 | 12.3833 | 0.665 |
| 0.2 | 0 | ptsI | 5 | 12.6333 | 0.457 |
| 0.2 | 0 | clcB | 5 | 12.6333 | 0.447 |
| 0.2 | 0 | ycaM | 5 | 12.6333 | 0.419 |
| 0.2 | 0 | yadI | 5 | 12.6333 | 0.479 |
| 0.2 | 0 | AG1 | 5 | 12.6333 | 0.677 |
| 0.2 | 0 | ptsI | 5 | 12.8833 | 0.469 |
| 0.2 | 0 | clcB | 5 | 12.8833 | 0.457 |
| 0.2 | 0 | ycaM | 5 | 12.8833 | 0.427 |
| 0.2 | 0 | yadI | 5 | 12.8833 | 0.487 |
| 0.2 | 0 | AG1 | 5 | 12.8833 | 0.687 |
| 0.2 | 0 | ptsI | 5 | 13.1333 | 0.473 |

| | | | | | |
|---|---|---|---|---|---|
| 0.2 | 0 | clcB | 5 | 13.1333 | 0.459 |
| 0.2 | 0 | ycaM | 5 | 13.1333 | 0.43 |
| 0.2 | 0 | yadI | 5 | 13.1333 | 0.491 |
| 0.2 | 0 | AG1 | 5 | 13.1333 | 0.694 |
| 0.2 | 0 | ptsI | 5 | 13.3833 | 0.477 |
| 0.2 | 0 | clcB | 5 | 13.3833 | 0.463 |
| 0.2 | 0 | ycaM | 5 | 13.3833 | 0.432 |
| 0.2 | 0 | yadI | 5 | 13.3833 | 0.496 |
| 0.2 | 0 | AG1 | 5 | 13.3833 | 0.698 |
| 0.2 | 0 | ptsI | 5 | 13.6333 | 0.488 |
| 0.2 | 0 | clcB | 5 | 13.6333 | 0.468 |
| 0.2 | 0 | ycaM | 5 | 13.6333 | 0.436 |
| 0.2 | 0 | yadI | 5 | 13.6333 | 0.503 |
| 0.2 | 0 | AG1 | 5 | 13.6333 | 0.708 |
| 0.2 | 0 | ptsI | 5 | 13.8833 | 0.494 |
| 0.2 | 0 | clcB | 5 | 13.8833 | 0.471 |
| 0.2 | 0 | ycaM | 5 | 13.8833 | 0.44 |
| 0.2 | 0 | yadI | 5 | 13.8833 | 0.506 |
| 0.2 | 0 | AG1 | 5 | 13.8833 | 0.714 |
| 0.2 | 0 | ptsI | 5 | 14.1333 | 0.503 |
| 0.2 | 0 | clcB | 5 | 14.1333 | 0.478 |
| 0.2 | 0 | ycaM | 5 | 14.1333 | 0.448 |
| 0.2 | 0 | yadI | 5 | 14.1333 | 0.51 |
| 0.2 | 0 | AG1 | 5 | 14.1333 | 0.722 |
| 0.2 | 0 | ptsI | 5 | 14.3833 | 0.509 |
| 0.2 | 0 | clcB | 5 | 14.3833 | 0.486 |
| 0.2 | 0 | ycaM | 5 | 14.3833 | 0.45 |
| 0.2 | 0 | yadI | 5 | 14.3833 | 0.516 |
| 0.2 | 0 | AG1 | 5 | 14.3833 | 0.729 |
| 0.2 | 0 | ptsI | 5 | 14.6333 | 0.517 |
| 0.2 | 0 | clcB | 5 | 14.6333 | 0.487 |
| 0.2 | 0 | ycaM | 5 | 14.6333 | 0.454 |
| 0.2 | 0 | yadI | 5 | 14.6333 | 0.519 |
| 0.2 | 0 | AG1 | 5 | 14.6333 | 0.732 |
| 0.2 | 0 | ptsI | 5 | 14.8833 | 0.519 |
| 0.2 | 0 | clcB | 5 | 14.8833 | 0.488 |
| 0.2 | 0 | ycaM | 5 | 14.8833 | 0.456 |
| 0.2 | 0 | yadI | 5 | 14.8833 | 0.523 |
| 0.2 | 0 | AG1 | 5 | 14.8833 | 0.737 |
| 0.2 | 0 | ptsI | 5 | 15.1333 | 0.524 |
| 0.2 | 0 | clcB | 5 | 15.1333 | 0.492 |
| 0.2 | 0 | ycaM | 5 | 15.1333 | 0.461 |
| 0.2 | 0 | yadI | 5 | 15.1333 | 0.528 |
| 0.2 | 0 | AG1 | 5 | 15.1333 | 0.742 |
| 0.2 | 0 | ptsI | 5 | 15.3833 | 0.53 |
| 0.2 | 0 | clcB | 5 | 15.3833 | 0.495 |
| 0.2 | 0 | ycaM | 5 | 15.3833 | 0.464 |
| 0.2 | 0 | yadI | 5 | 15.3833 | 0.53 |

| | | | | | |
|---|---|---|---|---|---|
| 0.2 | 0 | AG1 | 5 | 15.3833 | 0.745 |
| 0.2 | 0 | ptsI | 5 | 15.6333 | 0.531 |
| 0.2 | 0 | clcB | 5 | 15.6333 | 0.496 |
| 0.2 | 0 | ycaM | 5 | 15.6333 | 0.465 |
| 0.2 | 0 | yadI | 5 | 15.6333 | 0.535 |
| 0.2 | 0 | AG1 | 5 | 15.6333 | 0.75 |
| 0.2 | 0 | ptsI | 5 | 15.8833 | 0.54 |
| 0.2 | 0 | clcB | 5 | 15.8833 | 0.499 |
| 0.2 | 0 | ycaM | 5 | 15.8833 | 0.47 |
| 0.2 | 0 | yadI | 5 | 15.8833 | 0.535 |
| 0.2 | 0 | AG1 | 5 | 15.8833 | 0.754 |
| 0.2 | 0 | ptsI | 5 | 16.1333 | 0.545 |
| 0.2 | 0 | clcB | 5 | 16.1333 | 0.498 |
| 0.2 | 0 | ycaM | 5 | 16.1333 | 0.477 |
| 0.2 | 0 | yadI | 5 | 16.1333 | 0.543 |
| 0.2 | 0 | AG1 | 5 | 16.1333 | 0.759 |
| 0.2 | 0 | ptsI | 5 | 16.3833 | 0.549 |
| 0.2 | 0 | clcB | 5 | 16.3833 | 0.501 |
| 0.2 | 0 | ycaM | 5 | 16.3833 | 0.475 |
| 0.2 | 0 | yadI | 5 | 16.3833 | 0.542 |
| 0.2 | 0 | AG1 | 5 | 16.3833 | 0.761 |
| 0.2 | 0 | ptsI | 5 | 16.6333 | 0.553 |
| 0.2 | 0 | clcB | 5 | 16.6333 | 0.502 |
| 0.2 | 0 | ycaM | 5 | 16.6333 | 0.48 |
| 0.2 | 0 | yadI | 5 | 16.6333 | 0.544 |
| 0.2 | 0 | AG1 | 5 | 16.6333 | 0.764 |
| 0.2 | 0 | ptsI | 5 | 16.8833 | 0.56 |
| 0.2 | 0 | clcB | 5 | 16.8833 | 0.508 |
| 0.2 | 0 | ycaM | 5 | 16.8833 | 0.484 |
| 0.2 | 0 | yadI | 5 | 16.8833 | 0.551 |
| 0.2 | 0 | AG1 | 5 | 16.8833 | 0.771 |
| 0.2 | 0 | ptsI | 5 | 17.1333 | 0.566 |
| 0.2 | 0 | clcB | 5 | 17.1333 | 0.511 |
| 0.2 | 0 | ycaM | 5 | 17.1333 | 0.488 |
| 0.2 | 0 | yadI | 5 | 17.1333 | 0.556 |
| 0.2 | 0 | AG1 | 5 | 17.1333 | 0.775 |
| 0.2 | 0 | ptsI | 5 | 17.3833 | 0.565 |
| 0.2 | 0 | clcB | 5 | 17.3833 | 0.508 |
| 0.2 | 0 | ycaM | 5 | 17.3833 | 0.489 |
| 0.2 | 0 | yadI | 5 | 17.3833 | 0.555 |
| 0.2 | 0 | AG1 | 5 | 17.3833 | 0.776 |
| 0.2 | 0 | ptsI | 5 | 17.6333 | 0.571 |
| 0.2 | 0 | clcB | 5 | 17.6333 | 0.51 |
| 0.2 | 0 | ycaM | 5 | 17.6333 | 0.492 |
| 0.2 | 0 | yadI | 5 | 17.6333 | 0.558 |
| 0.2 | 0 | AG1 | 5 | 17.6333 | 0.778 |
| 0.2 | 0 | ptsI | 5 | 17.8833 | 0.577 |
| 0.2 | 0 | clcB | 5 | 17.8833 | 0.517 |

| | | | | | |
|---|---|---|---|---|---|
| 0.2 | 0 | ycaM | 5 | 17.8833 | 0.492 |
| 0.2 | 0 | yadI | 5 | 17.8833 | 0.558 |
| 0.2 | 0 | AG1 | 5 | 17.8833 | 0.78 |
| 0.2 | 0 | ptsI | 5 | 18.1333 | 0.58 |
| 0.2 | 0 | clcB | 5 | 18.1333 | 0.517 |
| 0.2 | 0 | ycaM | 5 | 18.1333 | 0.499 |
| 0.2 | 0 | yadI | 5 | 18.1333 | 0.564 |
| 0.2 | 0 | AG1 | 5 | 18.1333 | 0.786 |
| 0.2 | 0 | ptsI | 5 | 18.3833 | 0.585 |
| 0.2 | 0 | clcB | 5 | 18.3833 | 0.517 |
| 0.2 | 0 | ycaM | 5 | 18.3833 | 0.499 |
| 0.2 | 0 | yadI | 5 | 18.3833 | 0.566 |
| 0.2 | 0 | AG1 | 5 | 18.3833 | 0.789 |
| 0.2 | 0 | ptsI | 5 | 18.6333 | 0.588 |
| 0.2 | 0 | clcB | 5 | 18.6333 | 0.518 |
| 0.2 | 0 | ycaM | 5 | 18.6333 | 0.503 |
| 0.2 | 0 | yadI | 5 | 18.6333 | 0.568 |
| 0.2 | 0 | AG1 | 5 | 18.6333 | 0.79 |
| 0.2 | 0 | ptsI | 5 | 18.8833 | 0.59 |
| 0.2 | 0 | clcB | 5 | 18.8833 | 0.517 |
| 0.2 | 0 | ycaM | 5 | 18.8833 | 0.507 |
| 0.2 | 0 | yadI | 5 | 18.8833 | 0.569 |
| 0.2 | 0 | AG1 | 5 | 18.8833 | 0.793 |
| 0.2 | 0 | ptsI | 5 | 19.1333 | 0.595 |
| 0.2 | 0 | clcB | 5 | 19.1333 | 0.52 |
| 0.2 | 0 | ycaM | 5 | 19.1333 | 0.508 |
| 0.2 | 0 | yadI | 5 | 19.1333 | 0.572 |
| 0.2 | 0 | AG1 | 5 | 19.1333 | 0.795 |
| 0.2 | 0 | ptsI | 5 | 19.3833 | 0.6 |
| 0.2 | 0 | clcB | 5 | 19.3833 | 0.523 |
| 0.2 | 0 | ycaM | 5 | 19.3833 | 0.509 |
| 0.2 | 0 | yadI | 5 | 19.3833 | 0.577 |
| 0.2 | 0 | AG1 | 5 | 19.3833 | 0.799 |
| 0.2 | 0 | ptsI | 5 | 19.6333 | 0.602 |
| 0.2 | 0 | clcB | 5 | 19.6333 | 0.522 |
| 0.2 | 0 | ycaM | 5 | 19.6333 | 0.515 |
| 0.2 | 0 | yadI | 5 | 19.6333 | 0.582 |
| 0.2 | 0 | AG1 | 5 | 19.6333 | 0.801 |
| 0.2 | 0 | ptsI | 5 | 19.8833 | 0.605 |
| 0.2 | 0 | clcB | 5 | 19.8833 | 0.525 |
| 0.2 | 0 | ycaM | 5 | 19.8833 | 0.513 |
| 0.2 | 0 | yadI | 5 | 19.8833 | 0.582 |
| 0.2 | 0 | AG1 | 5 | 19.8833 | 0.801 |
| 0.2 | 0 | ptsI | 5 | 20.1333 | 0.608 |
| 0.2 | 0 | clcB | 5 | 20.1333 | 0.523 |
| 0.2 | 0 | ycaM | 5 | 20.1333 | 0.514 |
| 0.2 | 0 | yadI | 5 | 20.1333 | 0.584 |
| 0.2 | 0 | AG1 | 5 | 20.1333 | 0.804 |

| | | | | | |
|---|---|---|---|---|---|
| 0.2 | 0 | ptsI | 5 | 20.3833 | 0.613 |
| 0.2 | 0 | clcB | 5 | 20.3833 | 0.525 |
| 0.2 | 0 | ycaM | 5 | 20.3833 | 0.515 |
| 0.2 | 0 | yadI | 5 | 20.3833 | 0.586 |
| 0.2 | 0 | AG1 | 5 | 20.3833 | 0.807 |
| 0.2 | 0 | ptsI | 5 | 20.6333 | 0.614 |
| 0.2 | 0 | clcB | 5 | 20.6333 | 0.53 |
| 0.2 | 0 | ycaM | 5 | 20.6333 | 0.515 |
| 0.2 | 0 | yadI | 5 | 20.6333 | 0.584 |
| 0.2 | 0 | AG1 | 5 | 20.6333 | 0.806 |
| 0.2 | 0 | ptsI | 5 | 20.8833 | 0.617 |
| 0.2 | 0 | clcB | 5 | 20.8833 | 0.526 |
| 0.2 | 0 | ycaM | 5 | 20.8833 | 0.518 |
| 0.2 | 0 | yadI | 5 | 20.8833 | 0.591 |
| 0.2 | 0 | AG1 | 5 | 20.8833 | 0.812 |
| 0.2 | 0 | ptsI | 5 | 21.1333 | 0.619 |
| 0.2 | 0 | clcB | 5 | 21.1333 | 0.527 |
| 0.2 | 0 | ycaM | 5 | 21.1333 | 0.519 |
| 0.2 | 0 | yadI | 5 | 21.1333 | 0.591 |
| 0.2 | 0 | AG1 | 5 | 21.1333 | 0.814 |
| 0.2 | 0 | ptsI | 5 | 21.3833 | 0.619 |
| 0.2 | 0 | clcB | 5 | 21.3833 | 0.527 |
| 0.2 | 0 | ycaM | 5 | 21.3833 | 0.522 |
| 0.2 | 0 | yadI | 5 | 21.3833 | 0.597 |
| 0.2 | 0 | AG1 | 5 | 21.3833 | 0.814 |
| 0.2 | 0 | ptsI | 5 | 21.6333 | 0.622 |
| 0.2 | 0 | clcB | 5 | 21.6333 | 0.529 |
| 0.2 | 0 | ycaM | 5 | 21.6333 | 0.525 |
| 0.2 | 0 | yadI | 5 | 21.6333 | 0.597 |
| 0.2 | 0 | AG1 | 5 | 21.6333 | 0.817 |
| 0.2 | 0 | ptsI | 5 | 21.8833 | 0.629 |
| 0.2 | 0 | clcB | 5 | 21.8833 | 0.53 |
| 0.2 | 0 | ycaM | 5 | 21.8833 | 0.527 |
| 0.2 | 0 | yadI | 5 | 21.8833 | 0.603 |
| 0.2 | 0 | AG1 | 5 | 21.8833 | 0.818 |
| 0.2 | 0 | ptsI | 5 | 22.1333 | 0.627 |
| 0.2 | 0 | clcB | 5 | 22.1333 | 0.531 |
| 0.2 | 0 | ycaM | 5 | 22.1333 | 0.524 |
| 0.2 | 0 | yadI | 5 | 22.1333 | 0.601 |
| 0.2 | 0 | AG1 | 5 | 22.1333 | 0.816 |
| 0.2 | 0 | ptsI | 5 | 22.3833 | 0.63 |
| 0.2 | 0 | clcB | 5 | 22.3833 | 0.528 |
| 0.2 | 0 | ycaM | 5 | 22.3833 | 0.529 |
| 0.2 | 0 | yadI | 5 | 22.3833 | 0.605 |
| 0.2 | 0 | AG1 | 5 | 22.3833 | 0.819 |
| 0.2 | 0 | ptsI | 5 | 22.6333 | 0.629 |
| 0.2 | 0 | clcB | 5 | 22.6333 | 0.53 |
| 0.2 | 0 | ycaM | 5 | 22.6333 | 0.531 |

| | | | | | |
|---|---|---|---|---|---|
| 0.2 | 0 | yadI | 5 | 22.6333 | 0.609 |
| 0.2 | 0 | AG1 | 5 | 22.6333 | 0.821 |
| 0.2 | 0 | ptsI | 5 | 22.8833 | 0.631 |
| 0.2 | 0 | clcB | 5 | 22.8833 | 0.53 |
| 0.2 | 0 | ycaM | 5 | 22.8833 | 0.53 |
| 0.2 | 0 | yadI | 5 | 22.8833 | 0.606 |
| 0.2 | 0 | AG1 | 5 | 22.8833 | 0.821 |
| 0.2 | 0 | ptsI | 5 | 23.1333 | 0.637 |
| 0.2 | 0 | clcB | 5 | 23.1333 | 0.531 |
| 0.2 | 0 | ycaM | 5 | 23.1333 | 0.53 |
| 0.2 | 0 | yadI | 5 | 23.1333 | 0.61 |
| 0.2 | 0 | AG1 | 5 | 23.1333 | 0.822 |
| 0.2 | 0 | ptsI | 5 | 23.3833 | 0.639 |
| 0.2 | 0 | clcB | 5 | 23.3833 | 0.53 |
| 0.2 | 0 | ycaM | 5 | 23.3833 | 0.53 |
| 0.2 | 0 | yadI | 5 | 23.3833 | 0.61 |
| 0.2 | 0 | AG1 | 5 | 23.3833 | 0.821 |
| 0.2 | 0 | ptsI | 5 | 23.6333 | 0.638 |
| 0.2 | 0 | clcB | 5 | 23.6333 | 0.532 |
| 0.2 | 0 | ycaM | 5 | 23.6333 | 0.532 |
| 0.2 | 0 | yadI | 5 | 23.6333 | 0.614 |
| 0.2 | 0 | AG1 | 5 | 23.6333 | 0.824 |
| 0.2 | 0 | ptsI | 5 | 23.8833 | 0.648 |
| 0.2 | 0 | clcB | 5 | 23.8833 | 0.534 |
| 0.2 | 0 | ycaM | 5 | 23.8833 | 0.53 |
| 0.2 | 0 | yadI | 5 | 23.8833 | 0.614 |
| 0.2 | 0 | AG1 | 5 | 23.8833 | 0.822 |
| 0.2 | 0 | ptsI | 5 | 24.1333 | 0.648 |
| 0.2 | 0 | clcB | 5 | 24.1333 | 0.531 |
| 0.2 | 0 | ycaM | 5 | 24.1333 | 0.538 |
| 0.2 | 0 | yadI | 5 | 24.1333 | 0.622 |
| 0.2 | 0 | AG1 | 5 | 24.1333 | 0.829 |
| 0.2 | 0 | ptsI | 5 | 24.3833 | 0.653 |
| 0.2 | 0 | clcB | 5 | 24.3833 | 0.531 |
| 0.2 | 0 | ycaM | 5 | 24.3833 | 0.533 |
| 0.2 | 0 | yadI | 5 | 24.3833 | 0.619 |
| 0.2 | 0 | AG1 | 5 | 24.3833 | 0.825 |
| 0.4 | 0 | ptsI | 1 | 0 | 0.221 |
| 0.4 | 0 | clcB | 1 | 0 | 0.243 |
| 0.4 | 0 | ycaM | 1 | 0 | 0.227 |
| 0.4 | 0 | yadI | 1 | 0 | 0.272 |
| 0.4 | 0 | AG1 | 1 | 0 | 0.237 |
| 0.4 | 0 | ptsI | 1 | 0.25 | 0.218 |
| 0.4 | 0 | clcB | 1 | 0.25 | 0.23 |
| 0.4 | 0 | ycaM | 1 | 0.25 | 0.232 |
| 0.4 | 0 | yadI | 1 | 0.25 | 0.259 |
| 0.4 | 0 | AG1 | 1 | 0.25 | 0.223 |
| 0.4 | 0 | ptsI | 1 | 0.5 | 0.219 |

| | | | | | |
|---|---|---|---|---|---|
| 0.4 | 0 | clcB | 1 | 0.5 | 0.226 |
| 0.4 | 0 | ycaM | 1 | 0.5 | 0.23 |
| 0.4 | 0 | yadI | 1 | 0.5 | 0.255 |
| 0.4 | 0 | AG1 | 1 | 0.5 | 0.222 |
| 0.4 | 0 | ptsI | 1 | 0.75 | 0.22 |
| 0.4 | 0 | clcB | 1 | 0.75 | 0.224 |
| 0.4 | 0 | ycaM | 1 | 0.75 | 0.227 |
| 0.4 | 0 | yadI | 1 | 0.75 | 0.255 |
| 0.4 | 0 | AG1 | 1 | 0.75 | 0.222 |
| 0.4 | 0 | ptsI | 1 | 1 | 0.224 |
| 0.4 | 0 | clcB | 1 | 1 | 0.224 |
| 0.4 | 0 | ycaM | 1 | 1 | 0.229 |
| 0.4 | 0 | yadI | 1 | 1 | 0.257 |
| 0.4 | 0 | AG1 | 1 | 1 | 0.225 |
| 0.4 | 0 | ptsI | 1 | 1.25 | 0.225 |
| 0.4 | 0 | clcB | 1 | 1.25 | 0.226 |
| 0.4 | 0 | ycaM | 1 | 1.25 | 0.23 |
| 0.4 | 0 | yadI | 1 | 1.25 | 0.258 |
| 0.4 | 0 | AG1 | 1 | 1.25 | 0.226 |
| 0.4 | 0 | ptsI | 1 | 1.5 | 0.231 |
| 0.4 | 0 | clcB | 1 | 1.5 | 0.225 |
| 0.4 | 0 | ycaM | 1 | 1.5 | 0.235 |
| 0.4 | 0 | yadI | 1 | 1.5 | 0.261 |
| 0.4 | 0 | AG1 | 1 | 1.5 | 0.228 |
| 0.4 | 0 | ptsI | 1 | 1.75 | 0.235 |
| 0.4 | 0 | clcB | 1 | 1.75 | 0.229 |
| 0.4 | 0 | ycaM | 1 | 1.75 | 0.238 |
| 0.4 | 0 | yadI | 1 | 1.75 | 0.266 |
| 0.4 | 0 | AG1 | 1 | 1.75 | 0.232 |
| 0.4 | 0 | ptsI | 1 | 2 | 0.241 |
| 0.4 | 0 | clcB | 1 | 2 | 0.229 |
| 0.4 | 0 | ycaM | 1 | 2 | 0.247 |
| 0.4 | 0 | yadI | 1 | 2 | 0.272 |
| 0.4 | 0 | AG1 | 1 | 2 | 0.239 |
| 0.4 | 0 | ptsI | 1 | 2.25 | 0.249 |
| 0.4 | 0 | clcB | 1 | 2.25 | 0.236 |
| 0.4 | 0 | ycaM | 1 | 2.25 | 0.253 |
| 0.4 | 0 | yadI | 1 | 2.25 | 0.28 |
| 0.4 | 0 | AG1 | 1 | 2.25 | 0.241 |
| 0.4 | 0 | ptsI | 1 | 2.5 | 0.256 |
| 0.4 | 0 | clcB | 1 | 2.5 | 0.236 |
| 0.4 | 0 | ycaM | 1 | 2.5 | 0.263 |
| 0.4 | 0 | yadI | 1 | 2.5 | 0.286 |
| 0.4 | 0 | AG1 | 1 | 2.5 | 0.253 |
| 0.4 | 0 | ptsI | 1 | 2.75 | 0.266 |
| 0.4 | 0 | clcB | 1 | 2.75 | 0.242 |
| 0.4 | 0 | ycaM | 1 | 2.75 | 0.271 |
| 0.4 | 0 | yadI | 1 | 2.75 | 0.296 |

| | | | | | |
|---|---|---|---|---|---|
| 0.4 | 0 | AG1 | 1 | 2.75 | 0.253 |
| 0.4 | 0 | ptsI | 1 | 3 | 0.27 |
| 0.4 | 0 | clcB | 1 | 3 | 0.246 |
| 0.4 | 0 | ycaM | 1 | 3 | 0.28 |
| 0.4 | 0 | yadI | 1 | 3 | 0.307 |
| 0.4 | 0 | AG1 | 1 | 3 | 0.265 |
| 0.4 | 0 | ptsI | 1 | 3.25 | 0.276 |
| 0.4 | 0 | clcB | 1 | 3.25 | 0.252 |
| 0.4 | 0 | ycaM | 1 | 3.25 | 0.291 |
| 0.4 | 0 | yadI | 1 | 3.25 | 0.327 |
| 0.4 | 0 | AG1 | 1 | 3.25 | 0.279 |
| 0.4 | 0 | ptsI | 1 | 3.5 | 0.289 |
| 0.4 | 0 | clcB | 1 | 3.5 | 0.258 |
| 0.4 | 0 | ycaM | 1 | 3.5 | 0.301 |
| 0.4 | 0 | yadI | 1 | 3.5 | 0.342 |
| 0.4 | 0 | AG1 | 1 | 3.5 | 0.278 |
| 0.4 | 0 | ptsI | 1 | 3.75 | 0.3 |
| 0.4 | 0 | clcB | 1 | 3.75 | 0.26 |
| 0.4 | 0 | ycaM | 1 | 3.75 | 0.317 |
| 0.4 | 0 | yadI | 1 | 3.75 | 0.357 |
| 0.4 | 0 | AG1 | 1 | 3.75 | 0.278 |
| 0.4 | 0 | ptsI | 1 | 4 | 0.315 |
| 0.4 | 0 | clcB | 1 | 4 | 0.262 |
| 0.4 | 0 | ycaM | 1 | 4 | 0.326 |
| 0.4 | 0 | yadI | 1 | 4 | 0.39 |
| 0.4 | 0 | AG1 | 1 | 4 | 0.296 |
| 0.4 | 0 | ptsI | 1 | 4.25 | 0.315 |
| 0.4 | 0 | clcB | 1 | 4.25 | 0.27 |
| 0.4 | 0 | ycaM | 1 | 4.25 | 0.338 |
| 0.4 | 0 | yadI | 1 | 4.25 | 0.394 |
| 0.4 | 0 | AG1 | 1 | 4.25 | 0.302 |
| 0.4 | 0 | ptsI | 1 | 4.5 | 0.322 |
| 0.4 | 0 | clcB | 1 | 4.5 | 0.278 |
| 0.4 | 0 | ycaM | 1 | 4.5 | 0.348 |
| 0.4 | 0 | yadI | 1 | 4.5 | 0.412 |
| 0.4 | 0 | AG1 | 1 | 4.5 | 0.314 |
| 0.4 | 0 | ptsI | 1 | 4.75 | 0.336 |
| 0.4 | 0 | clcB | 1 | 4.75 | 0.286 |
| 0.4 | 0 | ycaM | 1 | 4.75 | 0.376 |
| 0.4 | 0 | yadI | 1 | 4.75 | 0.412 |
| 0.4 | 0 | AG1 | 1 | 4.75 | 0.32 |
| 0.4 | 0 | ptsI | 1 | 5 | 0.344 |
| 0.4 | 0 | clcB | 1 | 5 | 0.294 |
| 0.4 | 0 | ycaM | 1 | 5 | 0.371 |
| 0.4 | 0 | yadI | 1 | 5 | 0.433 |
| 0.4 | 0 | AG1 | 1 | 5 | 0.334 |
| 0.4 | 0 | ptsI | 1 | 5.25 | 0.355 |
| 0.4 | 0 | clcB | 1 | 5.25 | 0.302 |

| | | | | | |
|---|---|---|---|---|---|
| 0.4 | 0 | ycaM | 1 | 5.25 | 0.378 |
| 0.4 | 0 | yadI | 1 | 5.25 | 0.452 |
| 0.4 | 0 | AG1 | 1 | 5.25 | 0.345 |
| 0.4 | 0 | ptsI | 1 | 5.5 | 0.363 |
| 0.4 | 0 | clcB | 1 | 5.5 | 0.309 |
| 0.4 | 0 | ycaM | 1 | 5.5 | 0.386 |
| 0.4 | 0 | yadI | 1 | 5.5 | 0.459 |
| 0.4 | 0 | AG1 | 1 | 5.5 | 0.358 |
| 0.4 | 0 | ptsI | 1 | 5.75 | 0.374 |
| 0.4 | 0 | clcB | 1 | 5.75 | 0.315 |
| 0.4 | 0 | ycaM | 1 | 5.75 | 0.393 |
| 0.4 | 0 | yadI | 1 | 5.75 | 0.474 |
| 0.4 | 0 | AG1 | 1 | 5.75 | 0.369 |
| 0.4 | 0 | ptsI | 1 | 6 | 0.394 |
| 0.4 | 0 | clcB | 1 | 6 | 0.327 |
| 0.4 | 0 | ycaM | 1 | 6 | 0.402 |
| 0.4 | 0 | yadI | 1 | 6 | 0.475 |
| 0.4 | 0 | AG1 | 1 | 6 | 0.378 |
| 0.4 | 0 | ptsI | 1 | 6.25 | 0.405 |
| 0.4 | 0 | clcB | 1 | 6.25 | 0.332 |
| 0.4 | 0 | ycaM | 1 | 6.25 | 0.412 |
| 0.4 | 0 | yadI | 1 | 6.25 | 0.486 |
| 0.4 | 0 | AG1 | 1 | 6.25 | 0.388 |
| 0.4 | 0 | ptsI | 1 | 6.5 | 0.415 |
| 0.4 | 0 | clcB | 1 | 6.5 | 0.349 |
| 0.4 | 0 | ycaM | 1 | 6.5 | 0.418 |
| 0.4 | 0 | yadI | 1 | 6.5 | 0.497 |
| 0.4 | 0 | AG1 | 1 | 6.5 | 0.402 |
| 0.4 | 0 | ptsI | 1 | 6.75 | 0.42 |
| 0.4 | 0 | clcB | 1 | 6.75 | 0.351 |
| 0.4 | 0 | ycaM | 1 | 6.75 | 0.42 |
| 0.4 | 0 | yadI | 1 | 6.75 | 0.514 |
| 0.4 | 0 | AG1 | 1 | 6.75 | 0.419 |
| 0.4 | 0 | ptsI | 1 | 7 | 0.424 |
| 0.4 | 0 | clcB | 1 | 7 | 0.362 |
| 0.4 | 0 | ycaM | 1 | 7 | 0.423 |
| 0.4 | 0 | yadI | 1 | 7 | 0.529 |
| 0.4 | 0 | AG1 | 1 | 7 | 0.431 |
| 0.4 | 0 | ptsI | 1 | 7.25 | 0.434 |
| 0.4 | 0 | clcB | 1 | 7.25 | 0.368 |
| 0.4 | 0 | ycaM | 1 | 7.25 | 0.43 |
| 0.4 | 0 | yadI | 1 | 7.25 | 0.549 |
| 0.4 | 0 | AG1 | 1 | 7.25 | 0.441 |
| 0.4 | 0 | ptsI | 1 | 7.5 | 0.441 |
| 0.4 | 0 | clcB | 1 | 7.5 | 0.377 |
| 0.4 | 0 | ycaM | 1 | 7.5 | 0.437 |
| 0.4 | 0 | yadI | 1 | 7.5 | 0.559 |
| 0.4 | 0 | AG1 | 1 | 7.5 | 0.448 |

| | | | | | |
|---|---|---|---|---|---|
| 0.4 | 0 | ptsI | 1 | 7.75 | 0.456 |
| 0.4 | 0 | clcB | 1 | 7.75 | 0.389 |
| 0.4 | 0 | ycaM | 1 | 7.75 | 0.447 |
| 0.4 | 0 | yadI | 1 | 7.75 | 0.566 |
| 0.4 | 0 | AG1 | 1 | 7.75 | 0.456 |
| 0.4 | 0 | ptsI | 1 | 8 | 0.462 |
| 0.4 | 0 | clcB | 1 | 8 | 0.396 |
| 0.4 | 0 | ycaM | 1 | 8 | 0.457 |
| 0.4 | 0 | yadI | 1 | 8 | 0.583 |
| 0.4 | 0 | AG1 | 1 | 8 | 0.468 |
| 0.4 | 0 | ptsI | 1 | 8.25 | 0.474 |
| 0.4 | 0 | clcB | 1 | 8.25 | 0.403 |
| 0.4 | 0 | ycaM | 1 | 8.25 | 0.466 |
| 0.4 | 0 | yadI | 1 | 8.25 | 0.601 |
| 0.4 | 0 | AG1 | 1 | 8.25 | 0.477 |
| 0.4 | 0 | ptsI | 1 | 8.5 | 0.49 |
| 0.4 | 0 | clcB | 1 | 8.5 | 0.41 |
| 0.4 | 0 | ycaM | 1 | 8.5 | 0.477 |
| 0.4 | 0 | yadI | 1 | 8.5 | 0.604 |
| 0.4 | 0 | AG1 | 1 | 8.5 | 0.492 |
| 0.4 | 0 | ptsI | 1 | 8.75 | 0.496 |
| 0.4 | 0 | clcB | 1 | 8.75 | 0.415 |
| 0.4 | 0 | ycaM | 1 | 8.75 | 0.487 |
| 0.4 | 0 | yadI | 1 | 8.75 | 0.615 |
| 0.4 | 0 | AG1 | 1 | 8.75 | 0.504 |
| 0.4 | 0 | ptsI | 1 | 9 | 0.508 |
| 0.4 | 0 | clcB | 1 | 9 | 0.424 |
| 0.4 | 0 | ycaM | 1 | 9 | 0.498 |
| 0.4 | 0 | yadI | 1 | 9 | 0.621 |
| 0.4 | 0 | AG1 | 1 | 9 | 0.517 |
| 0.4 | 0 | ptsI | 1 | 9.25 | 0.547 |
| 0.4 | 0 | clcB | 1 | 9.25 | 0.435 |
| 0.4 | 0 | ycaM | 1 | 9.25 | 0.509 |
| 0.4 | 0 | yadI | 1 | 9.25 | 0.631 |
| 0.4 | 0 | AG1 | 1 | 9.25 | 0.528 |
| 0.4 | 0 | ptsI | 1 | 9.5 | 0.595 |
| 0.4 | 0 | clcB | 1 | 9.5 | 0.444 |
| 0.4 | 0 | ycaM | 1 | 9.5 | 0.527 |
| 0.4 | 0 | yadI | 1 | 9.5 | 0.638 |
| 0.4 | 0 | AG1 | 1 | 9.5 | 0.554 |
| 0.4 | 0 | ptsI | 1 | 9.75 | 0.58 |
| 0.4 | 0 | clcB | 1 | 9.75 | 0.452 |
| 0.4 | 0 | ycaM | 1 | 9.75 | 0.525 |
| 0.4 | 0 | yadI | 1 | 9.75 | 0.663 |
| 0.4 | 0 | AG1 | 1 | 9.75 | 0.572 |
| 0.4 | 0 | ptsI | 1 | 10 | 0.61 |
| 0.4 | 0 | clcB | 1 | 10 | 0.465 |
| 0.4 | 0 | ycaM | 1 | 10 | 0.54 |

| | | | | | |
|---|---|---|---|---|---|
| 0.4 | 0 | yadI | 1 | 10 | 0.656 |
| 0.4 | 0 | AG1 | 1 | 10 | 0.566 |
| 0.4 | 0 | ptsI | 1 | 10.25 | 0.613 |
| 0.4 | 0 | clcB | 1 | 10.25 | 0.469 |
| 0.4 | 0 | ycaM | 1 | 10.25 | 0.548 |
| 0.4 | 0 | yadI | 1 | 10.25 | 0.673 |
| 0.4 | 0 | AG1 | 1 | 10.25 | 0.577 |
| 0.4 | 0 | ptsI | 1 | 10.5 | 0.608 |
| 0.4 | 0 | clcB | 1 | 10.5 | 0.475 |
| 0.4 | 0 | ycaM | 1 | 10.5 | 0.566 |
| 0.4 | 0 | yadI | 1 | 10.5 | 0.681 |
| 0.4 | 0 | AG1 | 1 | 10.5 | 0.588 |
| 0.4 | 0 | ptsI | 1 | 10.75 | 0.649 |
| 0.4 | 0 | clcB | 1 | 10.75 | 0.493 |
| 0.4 | 0 | ycaM | 1 | 10.75 | 0.563 |
| 0.4 | 0 | yadI | 1 | 10.75 | 0.681 |
| 0.4 | 0 | AG1 | 1 | 10.75 | 0.584 |
| 0.4 | 0 | ptsI | 1 | 11 | 0.687 |
| 0.4 | 0 | clcB | 1 | 11 | 0.502 |
| 0.4 | 0 | ycaM | 1 | 11 | 0.579 |
| 0.4 | 0 | yadI | 1 | 11 | 0.681 |
| 0.4 | 0 | AG1 | 1 | 11 | 0.631 |
| 0.4 | 0 | ptsI | 1 | 11.25 | 0.693 |
| 0.4 | 0 | clcB | 1 | 11.25 | 0.515 |
| 0.4 | 0 | ycaM | 1 | 11.25 | 0.595 |
| 0.4 | 0 | yadI | 1 | 11.25 | 0.7 |
| 0.4 | 0 | AG1 | 1 | 11.25 | 0.636 |
| 0.4 | 0 | ptsI | 1 | 11.5 | 0.703 |
| 0.4 | 0 | clcB | 1 | 11.5 | 0.534 |
| 0.4 | 0 | ycaM | 1 | 11.5 | 0.575 |
| 0.4 | 0 | yadI | 1 | 11.5 | 0.702 |
| 0.4 | 0 | AG1 | 1 | 11.5 | 0.663 |
| 0.4 | 0 | ptsI | 1 | 11.75 | 0.733 |
| 0.4 | 0 | clcB | 1 | 11.75 | 0.548 |
| 0.4 | 0 | ycaM | 1 | 11.75 | 0.601 |
| 0.4 | 0 | yadI | 1 | 11.75 | 0.714 |
| 0.4 | 0 | AG1 | 1 | 11.75 | 0.71 |
| 0.4 | 0 | ptsI | 1 | 12 | 0.715 |
| 0.4 | 0 | clcB | 1 | 12 | 0.547 |
| 0.4 | 0 | ycaM | 1 | 12 | 0.615 |
| 0.4 | 0 | yadI | 1 | 12 | 0.727 |
| 0.4 | 0 | AG1 | 1 | 12 | 0.723 |
| 0.4 | 0 | ptsI | 1 | 12.25 | 0.736 |
| 0.4 | 0 | clcB | 1 | 12.25 | 0.555 |
| 0.4 | 0 | ycaM | 1 | 12.25 | 0.604 |
| 0.4 | 0 | yadI | 1 | 12.25 | 0.721 |
| 0.4 | 0 | AG1 | 1 | 12.25 | 0.733 |
| 0.4 | 0 | ptsI | 1 | 12.5 | 0.725 |

| | | | | | |
|---|---|---|---|---|---|
| 0.4 | 0 | clcB | 1 | 12.5 | 0.561 |
| 0.4 | 0 | ycaM | 1 | 12.5 | 0.629 |
| 0.4 | 0 | yadI | 1 | 12.5 | 0.728 |
| 0.4 | 0 | AG1 | 1 | 12.5 | 0.746 |
| 0.4 | 0 | ptsI | 1 | 12.75 | 0.738 |
| 0.4 | 0 | clcB | 1 | 12.75 | 0.561 |
| 0.4 | 0 | ycaM | 1 | 12.75 | 0.65 |
| 0.4 | 0 | yadI | 1 | 12.75 | 0.732 |
| 0.4 | 0 | AG1 | 1 | 12.75 | 0.745 |
| 0.4 | 0 | ptsI | 1 | 13 | 0.745 |
| 0.4 | 0 | clcB | 1 | 13 | 0.561 |
| 0.4 | 0 | ycaM | 1 | 13 | 0.646 |
| 0.4 | 0 | yadI | 1 | 13 | 0.739 |
| 0.4 | 0 | AG1 | 1 | 13 | 0.754 |
| 0.4 | 0 | ptsI | 1 | 13.25 | 0.759 |
| 0.4 | 0 | clcB | 1 | 13.25 | 0.574 |
| 0.4 | 0 | ycaM | 1 | 13.25 | 0.614 |
| 0.4 | 0 | yadI | 1 | 13.25 | 0.731 |
| 0.4 | 0 | AG1 | 1 | 13.25 | 0.759 |
| 0.4 | 0 | ptsI | 1 | 13.5 | 0.772 |
| 0.4 | 0 | clcB | 1 | 13.5 | 0.58 |
| 0.4 | 0 | ycaM | 1 | 13.5 | 0.617 |
| 0.4 | 0 | yadI | 1 | 13.5 | 0.756 |
| 0.4 | 0 | AG1 | 1 | 13.5 | 0.756 |
| 0.4 | 0 | ptsI | 1 | 13.75 | 0.773 |
| 0.4 | 0 | clcB | 1 | 13.75 | 0.578 |
| 0.4 | 0 | ycaM | 1 | 13.75 | 0.617 |
| 0.4 | 0 | yadI | 1 | 13.75 | 0.76 |
| 0.4 | 0 | AG1 | 1 | 13.75 | 0.762 |
| 0.4 | 0 | ptsI | 1 | 14 | 0.798 |
| 0.4 | 0 | clcB | 1 | 14 | 0.586 |
| 0.4 | 0 | ycaM | 1 | 14 | 0.64 |
| 0.4 | 0 | yadI | 1 | 14 | 0.77 |
| 0.4 | 0 | AG1 | 1 | 14 | 0.747 |
| 0.4 | 0 | ptsI | 1 | 14.25 | 0.805 |
| 0.4 | 0 | clcB | 1 | 14.25 | 0.598 |
| 0.4 | 0 | ycaM | 1 | 14.25 | 0.674 |
| 0.4 | 0 | yadI | 1 | 14.25 | 0.778 |
| 0.4 | 0 | AG1 | 1 | 14.25 | 0.764 |
| 0.4 | 0 | ptsI | 1 | 14.5 | 0.8 |
| 0.4 | 0 | clcB | 1 | 14.5 | 0.601 |
| 0.4 | 0 | ycaM | 1 | 14.5 | 0.681 |
| 0.4 | 0 | yadI | 1 | 14.5 | 0.775 |
| 0.4 | 0 | AG1 | 1 | 14.5 | 0.777 |
| 0.4 | 0 | ptsI | 1 | 14.75 | 0.806 |
| 0.4 | 0 | clcB | 1 | 14.75 | 0.61 |
| 0.4 | 0 | ycaM | 1 | 14.75 | 0.647 |
| 0.4 | 0 | yadI | 1 | 14.75 | 0.785 |

| | | | | | |
|---|---|---|---|---|---|
| 0.4 | 0 | AG1 | 1 | 14.75 | 0.788 |
| 0.4 | 0 | ptsI | 1 | 15 | 0.807 |
| 0.4 | 0 | clcB | 1 | 15 | 0.607 |
| 0.4 | 0 | ycaM | 1 | 15 | 0.637 |
| 0.4 | 0 | yadI | 1 | 15 | 0.79 |
| 0.4 | 0 | AG1 | 1 | 15 | 0.796 |
| 0.4 | 0 | ptsI | 1 | 15.25 | 0.828 |
| 0.4 | 0 | clcB | 1 | 15.25 | 0.618 |
| 0.4 | 0 | ycaM | 1 | 15.25 | 0.681 |
| 0.4 | 0 | yadI | 1 | 15.25 | 0.798 |
| 0.4 | 0 | AG1 | 1 | 15.25 | 0.81 |
| 0.4 | 0 | ptsI | 1 | 15.5 | 0.823 |
| 0.4 | 0 | clcB | 1 | 15.5 | 0.613 |
| 0.4 | 0 | ycaM | 1 | 15.5 | 0.69 |
| 0.4 | 0 | yadI | 1 | 15.5 | 0.805 |
| 0.4 | 0 | AG1 | 1 | 15.5 | 0.799 |
| 0.4 | 0 | ptsI | 1 | 15.75 | 0.813 |
| 0.4 | 0 | clcB | 1 | 15.75 | 0.627 |
| 0.4 | 0 | ycaM | 1 | 15.75 | 0.7 |
| 0.4 | 0 | yadI | 1 | 15.75 | 0.804 |
| 0.4 | 0 | AG1 | 1 | 15.75 | 0.811 |
| 0.4 | 0 | ptsI | 1 | 16 | 0.824 |
| 0.4 | 0 | clcB | 1 | 16 | 0.619 |
| 0.4 | 0 | ycaM | 1 | 16 | 0.697 |
| 0.4 | 0 | yadI | 1 | 16 | 0.805 |
| 0.4 | 0 | AG1 | 1 | 16 | 0.809 |
| 0.4 | 0 | ptsI | 1 | 16.25 | 0.82 |
| 0.4 | 0 | clcB | 1 | 16.25 | 0.628 |
| 0.4 | 0 | ycaM | 1 | 16.25 | 0.722 |
| 0.4 | 0 | yadI | 1 | 16.25 | 0.813 |
| 0.4 | 0 | AG1 | 1 | 16.25 | 0.833 |
| 0.4 | 0 | ptsI | 1 | 16.5 | 0.833 |
| 0.4 | 0 | clcB | 1 | 16.5 | 0.641 |
| 0.4 | 0 | ycaM | 1 | 16.5 | 0.733 |
| 0.4 | 0 | yadI | 1 | 16.5 | 0.808 |
| 0.4 | 0 | AG1 | 1 | 16.5 | 0.833 |
| 0.4 | 0 | ptsI | 1 | 17.0667 | 0.861 |
| 0.4 | 0 | clcB | 1 | 17.0667 | 0.661 |
| 0.4 | 0 | ycaM | 1 | 17.0667 | 0.752 |
| 0.4 | 0 | yadI | 1 | 17.0667 | 0.864 |
| 0.4 | 0 | AG1 | 1 | 17.0667 | 0.866 |
| 0.4 | 0 | ptsI | 1 | 17.3167 | 0.802 |
| 0.4 | 0 | clcB | 1 | 17.3167 | 0.624 |
| 0.4 | 0 | ycaM | 1 | 17.3167 | 0.712 |
| 0.4 | 0 | yadI | 1 | 17.3167 | 0.844 |
| 0.4 | 0 | AG1 | 1 | 17.3167 | 0.837 |
| 0.4 | 0 | ptsI | 1 | 17.5667 | 0.806 |
| 0.4 | 0 | clcB | 1 | 17.5667 | 0.643 |

| | | | | | |
|---|---|---|---|---|---|
| 0.4 | 0 | ycaM | 1 | 17.5667 | 0.702 |
| 0.4 | 0 | yadI | 1 | 17.5667 | 0.841 |
| 0.4 | 0 | AG1 | 1 | 17.5667 | 0.843 |
| 0.4 | 0 | ptsI | 1 | 17.8167 | 0.828 |
| 0.4 | 0 | clcB | 1 | 17.8167 | 0.687 |
| 0.4 | 0 | ycaM | 1 | 17.8167 | 0.716 |
| 0.4 | 0 | yadI | 1 | 17.8167 | 0.851 |
| 0.4 | 0 | AG1 | 1 | 17.8167 | 0.808 |
| 0.4 | 0 | ptsI | 1 | 18.0667 | 0.834 |
| 0.4 | 0 | clcB | 1 | 18.0667 | 0.696 |
| 0.4 | 0 | ycaM | 1 | 18.0667 | 0.716 |
| 0.4 | 0 | yadI | 1 | 18.0667 | 0.851 |
| 0.4 | 0 | AG1 | 1 | 18.0667 | 0.846 |
| 0.4 | 0 | ptsI | 1 | 18.3167 | 0.838 |
| 0.4 | 0 | clcB | 1 | 18.3167 | 0.676 |
| 0.4 | 0 | ycaM | 1 | 18.3167 | 0.713 |
| 0.4 | 0 | yadI | 1 | 18.3167 | 0.834 |
| 0.4 | 0 | AG1 | 1 | 18.3167 | 0.853 |
| 0.4 | 0 | ptsI | 1 | 18.5667 | 0.847 |
| 0.4 | 0 | clcB | 1 | 18.5667 | 0.73 |
| 0.4 | 0 | ycaM | 1 | 18.5667 | 0.718 |
| 0.4 | 0 | yadI | 1 | 18.5667 | 0.844 |
| 0.4 | 0 | AG1 | 1 | 18.5667 | 0.848 |
| 0.4 | 0 | ptsI | 1 | 18.8167 | 0.855 |
| 0.4 | 0 | clcB | 1 | 18.8167 | 0.714 |
| 0.4 | 0 | ycaM | 1 | 18.8167 | 0.716 |
| 0.4 | 0 | yadI | 1 | 18.8167 | 0.849 |
| 0.4 | 0 | AG1 | 1 | 18.8167 | 0.856 |
| 0.4 | 0 | ptsI | 1 | 19.0667 | 0.862 |
| 0.4 | 0 | clcB | 1 | 19.0667 | 0.684 |
| 0.4 | 0 | ycaM | 1 | 19.0667 | 0.721 |
| 0.4 | 0 | yadI | 1 | 19.0667 | 0.857 |
| 0.4 | 0 | AG1 | 1 | 19.0667 | 0.865 |
| 0.4 | 0 | ptsI | 1 | 19.3167 | 0.866 |
| 0.4 | 0 | clcB | 1 | 19.3167 | 0.707 |
| 0.4 | 0 | ycaM | 1 | 19.3167 | 0.723 |
| 0.4 | 0 | yadI | 1 | 19.3167 | 0.853 |
| 0.4 | 0 | AG1 | 1 | 19.3167 | 0.859 |
| 0.4 | 0 | ptsI | 1 | 19.5667 | 0.879 |
| 0.4 | 0 | clcB | 1 | 19.5667 | 0.726 |
| 0.4 | 0 | ycaM | 1 | 19.5667 | 0.735 |
| 0.4 | 0 | yadI | 1 | 19.5667 | 0.868 |
| 0.4 | 0 | AG1 | 1 | 19.5667 | 0.868 |
| 0.4 | 0 | ptsI | 1 | 19.8167 | 0.883 |
| 0.4 | 0 | clcB | 1 | 19.8167 | 0.721 |
| 0.4 | 0 | ycaM | 1 | 19.8167 | 0.734 |
| 0.4 | 0 | yadI | 1 | 19.8167 | 0.868 |
| 0.4 | 0 | AG1 | 1 | 19.8167 | 0.855 |

| | | | | | |
|---|---|---|---|---|---|
| 0.4 | 0 | ptsI | 1 | 20.0667 | 0.895 |
| 0.4 | 0 | clcB | 1 | 20.0667 | 0.713 |
| 0.4 | 0 | ycaM | 1 | 20.0667 | 0.742 |
| 0.4 | 0 | yadI | 1 | 20.0667 | 0.872 |
| 0.4 | 0 | AG1 | 1 | 20.0667 | 0.863 |
| 0.4 | 0 | ptsI | 1 | 20.3167 | 0.9 |
| 0.4 | 0 | clcB | 1 | 20.3167 | 0.719 |
| 0.4 | 0 | ycaM | 1 | 20.3167 | 0.742 |
| 0.4 | 0 | yadI | 1 | 20.3167 | 0.877 |
| 0.4 | 0 | AG1 | 1 | 20.3167 | 0.86 |
| 0.4 | 0 | ptsI | 1 | 20.5667 | 0.902 |
| 0.4 | 0 | clcB | 1 | 20.5667 | 0.718 |
| 0.4 | 0 | ycaM | 1 | 20.5667 | 0.746 |
| 0.4 | 0 | yadI | 1 | 20.5667 | 0.88 |
| 0.4 | 0 | AG1 | 1 | 20.5667 | 0.85 |
| 0.4 | 0 | ptsI | 1 | 20.8167 | 0.912 |
| 0.4 | 0 | clcB | 1 | 20.8167 | 0.725 |
| 0.4 | 0 | ycaM | 1 | 20.8167 | 0.749 |
| 0.4 | 0 | yadI | 1 | 20.8167 | 0.886 |
| 0.4 | 0 | AG1 | 1 | 20.8167 | 0.837 |
| 0.4 | 0 | ptsI | 1 | 21.0667 | 0.928 |
| 0.4 | 0 | clcB | 1 | 21.0667 | 0.726 |
| 0.4 | 0 | ycaM | 1 | 21.0667 | 0.763 |
| 0.4 | 0 | yadI | 1 | 21.0667 | 0.894 |
| 0.4 | 0 | AG1 | 1 | 21.0667 | 0.837 |
| 0.4 | 0 | ptsI | 1 | 21.3167 | 0.933 |
| 0.4 | 0 | clcB | 1 | 21.3167 | 0.725 |
| 0.4 | 0 | ycaM | 1 | 21.3167 | 0.763 |
| 0.4 | 0 | yadI | 1 | 21.3167 | 0.896 |
| 0.4 | 0 | AG1 | 1 | 21.3167 | 0.842 |
| 0.4 | 0 | ptsI | 1 | 21.5667 | 0.943 |
| 0.4 | 0 | clcB | 1 | 21.5667 | 0.722 |
| 0.4 | 0 | ycaM | 1 | 21.5667 | 0.771 |
| 0.4 | 0 | yadI | 1 | 21.5667 | 0.896 |
| 0.4 | 0 | AG1 | 1 | 21.5667 | 0.842 |
| 0.4 | 0 | ptsI | 1 | 21.8167 | 0.945 |
| 0.4 | 0 | clcB | 1 | 21.8167 | 0.723 |
| 0.4 | 0 | ycaM | 1 | 21.8167 | 0.775 |
| 0.4 | 0 | yadI | 1 | 21.8167 | 0.898 |
| 0.4 | 0 | AG1 | 1 | 21.8167 | 0.844 |
| 0.4 | 0 | ptsI | 1 | 22.0667 | 0.963 |
| 0.4 | 0 | clcB | 1 | 22.0667 | 0.728 |
| 0.4 | 0 | ycaM | 1 | 22.0667 | 0.788 |
| 0.4 | 0 | yadI | 1 | 22.0667 | 0.906 |
| 0.4 | 0 | AG1 | 1 | 22.0667 | 0.858 |
| 0.4 | 0 | ptsI | 1 | 22.9333 | 1.107 |
| 0.4 | 0 | clcB | 1 | 22.9333 | 0.601 |
| 0.4 | 0 | ycaM | 1 | 22.9333 | 0.534 |

| | | | | | |
|---|---|---|---|---|---|
| 0.4 | 0 | yadI | 1 | 22.9333 | 0.895 |
| 0.4 | 0 | AG1 | 1 | 22.9333 | 0.798 |
| 0.4 | 0 | ptsI | 1 | 23.1833 | 1.093 |
| 0.4 | 0 | clcB | 1 | 23.1833 | 0.621 |
| 0.4 | 0 | ycaM | 1 | 23.1833 | 0.594 |
| 0.4 | 0 | yadI | 1 | 23.1833 | 0.86 |
| 0.4 | 0 | AG1 | 1 | 23.1833 | 0.777 |
| 0.4 | 0 | ptsI | 1 | 23.4333 | 1.078 |
| 0.4 | 0 | clcB | 1 | 23.4333 | 0.603 |
| 0.4 | 0 | ycaM | 1 | 23.4333 | 0.616 |
| 0.4 | 0 | yadI | 1 | 23.4333 | 0.858 |
| 0.4 | 0 | AG1 | 1 | 23.4333 | 0.815 |
| 0.4 | 0 | ptsI | 1 | 23.6833 | 1.082 |
| 0.4 | 0 | clcB | 1 | 23.6833 | 0.607 |
| 0.4 | 0 | ycaM | 1 | 23.6833 | 0.63 |
| 0.4 | 0 | yadI | 1 | 23.6833 | 0.855 |
| 0.4 | 0 | AG1 | 1 | 23.6833 | 0.83 |
| 0.4 | 0 | ptsI | 1 | 23.9333 | 1.085 |
| 0.4 | 0 | clcB | 1 | 23.9333 | 0.604 |
| 0.4 | 0 | ycaM | 1 | 23.9333 | 0.648 |
| 0.4 | 0 | yadI | 1 | 23.9333 | 0.868 |
| 0.4 | 0 | AG1 | 1 | 23.9333 | 0.832 |
| 0.4 | 0 | ptsI | 1 | 24.1833 | 1.087 |
| 0.4 | 0 | clcB | 1 | 24.1833 | 0.644 |
| 0.4 | 0 | ycaM | 1 | 24.1833 | 0.657 |
| 0.4 | 0 | yadI | 1 | 24.1833 | 0.881 |
| 0.4 | 0 | AG1 | 1 | 24.1833 | 0.838 |
| 0.4 | 0 | ptsI | 1 | 24.4333 | 1.09 |
| 0.4 | 0 | clcB | 1 | 24.4333 | 0.657 |
| 0.4 | 0 | ycaM | 1 | 24.4333 | 0.669 |
| 0.4 | 0 | yadI | 1 | 24.4333 | 0.887 |
| 0.4 | 0 | AG1 | 1 | 24.4333 | 0.829 |
| 0.4 | 0 | ptsI | 1 | 24.6833 | 1.087 |
| 0.4 | 0 | clcB | 1 | 24.6833 | 0.655 |
| 0.4 | 0 | ycaM | 1 | 24.6833 | 0.676 |
| 0.4 | 0 | yadI | 1 | 24.6833 | 0.894 |
| 0.4 | 0 | AG1 | 1 | 24.6833 | 0.836 |
| 0.4 | 0 | ptsI | 1 | 24.9333 | 1.093 |
| 0.4 | 0 | clcB | 1 | 24.9333 | 0.654 |
| 0.4 | 0 | ycaM | 1 | 24.9333 | 0.686 |
| 0.4 | 0 | yadI | 1 | 24.9333 | 0.906 |
| 0.4 | 0 | AG1 | 1 | 24.9333 | 0.845 |
| 0.4 | 0 | ptsI | 1 | 25.1833 | 1.097 |
| 0.4 | 0 | clcB | 1 | 25.1833 | 0.661 |
| 0.4 | 0 | ycaM | 1 | 25.1833 | 0.702 |
| 0.4 | 0 | yadI | 1 | 25.1833 | 0.915 |
| 0.4 | 0 | AG1 | 1 | 25.1833 | 0.843 |
| 0.4 | 0 | ptsI | 1 | 25.4333 | 1.103 |

| | | | | | |
|---|---|---|---|---|---|
| 0.4 | 0 | clcB | 1 | 25.4333 | 0.665 |
| 0.4 | 0 | ycaM | 1 | 25.4333 | 0.708 |
| 0.4 | 0 | yadI | 1 | 25.4333 | 0.928 |
| 0.4 | 0 | AG1 | 1 | 25.4333 | 0.859 |
| 0.4 | 0 | ptsI | 1 | 25.6833 | 1.109 |
| 0.4 | 0 | clcB | 1 | 25.6833 | 0.673 |
| 0.4 | 0 | ycaM | 1 | 25.6833 | 0.712 |
| 0.4 | 0 | yadI | 1 | 25.6833 | 0.933 |
| 0.4 | 0 | AG1 | 1 | 25.6833 | 0.862 |
| 0.4 | 0 | ptsI | 1 | 25.9333 | 1.106 |
| 0.4 | 0 | clcB | 1 | 25.9333 | 0.698 |
| 0.4 | 0 | ycaM | 1 | 25.9333 | 0.722 |
| 0.4 | 0 | yadI | 1 | 25.9333 | 0.927 |
| 0.4 | 0 | AG1 | 1 | 25.9333 | 0.872 |
| 0.4 | 0 | ptsI | 1 | 26.1833 | 1.112 |
| 0.4 | 0 | clcB | 1 | 26.1833 | 0.704 |
| 0.4 | 0 | ycaM | 1 | 26.1833 | 0.727 |
| 0.4 | 0 | yadI | 1 | 26.1833 | 0.949 |
| 0.4 | 0 | AG1 | 1 | 26.1833 | 0.887 |
| 0.4 | 0 | ptsI | 1 | 26.4333 | 1.114 |
| 0.4 | 0 | clcB | 1 | 26.4333 | 0.703 |
| 0.4 | 0 | ycaM | 1 | 26.4333 | 0.725 |
| 0.4 | 0 | yadI | 1 | 26.4333 | 0.951 |
| 0.4 | 0 | AG1 | 1 | 26.4333 | 0.876 |
| 0.4 | 0 | ptsI | 1 | 26.6833 | 1.122 |
| 0.4 | 0 | clcB | 1 | 26.6833 | 0.714 |
| 0.4 | 0 | ycaM | 1 | 26.6833 | 0.73 |
| 0.4 | 0 | yadI | 1 | 26.6833 | 0.954 |
| 0.4 | 0 | AG1 | 1 | 26.6833 | 0.88 |
| 0.4 | 0 | ptsI | 1 | 26.9333 | 1.125 |
| 0.4 | 0 | clcB | 1 | 26.9333 | 0.719 |
| 0.4 | 0 | ycaM | 1 | 26.9333 | 0.735 |
| 0.4 | 0 | yadI | 1 | 26.9333 | 0.959 |
| 0.4 | 0 | AG1 | 1 | 26.9333 | 0.885 |
| 0.4 | 0 | ptsI | 1 | 27.1833 | 1.128 |
| 0.4 | 0 | clcB | 1 | 27.1833 | 0.738 |
| 0.4 | 0 | ycaM | 1 | 27.1833 | 0.738 |
| 0.4 | 0 | yadI | 1 | 27.1833 | 0.96 |
| 0.4 | 0 | AG1 | 1 | 27.1833 | 0.89 |
| 0.4 | 0 | ptsI | 1 | 27.4333 | 1.135 |
| 0.4 | 0 | clcB | 1 | 27.4333 | 0.731 |
| 0.4 | 0 | ycaM | 1 | 27.4333 | 0.73 |
| 0.4 | 0 | yadI | 1 | 27.4333 | 0.965 |
| 0.4 | 0 | AG1 | 1 | 27.4333 | 0.882 |
| 0.4 | 0 | ptsI | 1 | 27.6833 | 1.138 |
| 0.4 | 0 | clcB | 1 | 27.6833 | 0.744 |
| 0.4 | 0 | ycaM | 1 | 27.6833 | 0.74 |
| 0.4 | 0 | yadI | 1 | 27.6833 | 0.971 |

| | | | | | |
|---|---|---|---|---|---|
| 0.4 | 0 | AG1 | 1 | 27.6833 | 0.894 |
| 0.4 | 0 | ptsI | 1 | 27.9333 | 1.145 |
| 0.4 | 0 | clcB | 1 | 27.9333 | 0.741 |
| 0.4 | 0 | ycaM | 1 | 27.9333 | 0.733 |
| 0.4 | 0 | yadI | 1 | 27.9333 | 0.954 |
| 0.4 | 0 | AG1 | 1 | 27.9333 | 0.884 |
| 0.4 | 0 | ptsI | 1 | 28.1833 | 1.151 |
| 0.4 | 0 | clcB | 1 | 28.1833 | 0.762 |
| 0.4 | 0 | ycaM | 1 | 28.1833 | 0.747 |
| 0.4 | 0 | yadI | 1 | 28.1833 | 0.977 |
| 0.4 | 0 | AG1 | 1 | 28.1833 | 0.902 |
| 0.4 | 0 | ptsI | 1 | 28.4333 | 1.16 |
| 0.4 | 0 | clcB | 1 | 28.4333 | 0.754 |
| 0.4 | 0 | ycaM | 1 | 28.4333 | 0.75 |
| 0.4 | 0 | yadI | 1 | 28.4333 | 0.984 |
| 0.4 | 0 | AG1 | 1 | 28.4333 | 0.898 |
| 0.4 | 0 | ptsI | 1 | 28.6833 | 1.164 |
| 0.4 | 0 | clcB | 1 | 28.6833 | 0.76 |
| 0.4 | 0 | ycaM | 1 | 28.6833 | 0.753 |
| 0.4 | 0 | yadI | 1 | 28.6833 | 0.987 |
| 0.4 | 0 | AG1 | 1 | 28.6833 | 0.896 |
| 0.4 | 0 | ptsI | 1 | 28.9333 | 1.171 |
| 0.4 | 0 | clcB | 1 | 28.9333 | 0.768 |
| 0.4 | 0 | ycaM | 1 | 28.9333 | 0.764 |
| 0.4 | 0 | yadI | 1 | 28.9333 | 0.989 |
| 0.4 | 0 | AG1 | 1 | 28.9333 | 0.904 |
| 0.4 | 0 | ptsI | 1 | 29.1833 | 1.174 |
| 0.4 | 0 | clcB | 1 | 29.1833 | 0.755 |
| 0.4 | 0 | ycaM | 1 | 29.1833 | 0.759 |
| 0.4 | 0 | yadI | 1 | 29.1833 | 0.998 |
| 0.4 | 0 | AG1 | 1 | 29.1833 | 0.901 |
| 0.4 | 0 | ptsI | 1 | 29.4333 | 1.18 |
| 0.4 | 0 | clcB | 1 | 29.4333 | 0.768 |
| 0.4 | 0 | ycaM | 1 | 29.4333 | 0.771 |
| 0.4 | 0 | yadI | 1 | 29.4333 | 1 |
| 0.4 | 0 | AG1 | 1 | 29.4333 | 0.908 |
| 0.4 | 0 | ptsI | 1 | 29.6833 | 1.187 |
| 0.4 | 0 | clcB | 1 | 29.6833 | 0.766 |
| 0.4 | 0 | ycaM | 1 | 29.6833 | 0.774 |
| 0.4 | 0 | yadI | 1 | 29.6833 | 1.009 |
| 0.4 | 0 | AG1 | 1 | 29.6833 | 0.907 |
| 0.4 | 0 | ptsI | 1 | 29.9333 | 1.198 |
| 0.4 | 0 | clcB | 1 | 29.9333 | 0.784 |
| 0.4 | 0 | ycaM | 1 | 29.9333 | 0.775 |
| 0.4 | 0 | yadI | 1 | 29.9333 | 1.015 |
| 0.4 | 0 | AG1 | 1 | 29.9333 | 0.911 |
| 0.4 | 0 | ptsI | 1 | 30.1833 | 1.202 |
| 0.4 | 0 | clcB | 1 | 30.1833 | 0.775 |

| | | | | | |
|---|---|---|---|---|---|
| 0.4 | 0 | ycaM | 1 | 30.1833 | 0.784 |
| 0.4 | 0 | yadI | 1 | 30.1833 | 1.018 |
| 0.4 | 0 | AG1 | 1 | 30.1833 | 0.914 |
| 0.4 | 0 | ptsI | 1 | 30.4333 | 1.205 |
| 0.4 | 0 | clcB | 1 | 30.4333 | 0.773 |
| 0.4 | 0 | ycaM | 1 | 30.4333 | 0.794 |
| 0.4 | 0 | yadI | 1 | 30.4333 | 1.019 |
| 0.4 | 0 | AG1 | 1 | 30.4333 | 0.917 |
| 0.4 | 0 | ptsI | 1 | 30.6833 | 1.212 |
| 0.4 | 0 | clcB | 1 | 30.6833 | 0.776 |
| 0.4 | 0 | ycaM | 1 | 30.6833 | 0.783 |
| 0.4 | 0 | yadI | 1 | 30.6833 | 1.035 |
| 0.4 | 0 | AG1 | 1 | 30.6833 | 0.913 |
| 0.4 | 0 | ptsI | 1 | 30.9333 | 1.22 |
| 0.4 | 0 | clcB | 1 | 30.9333 | 0.776 |
| 0.4 | 0 | ycaM | 1 | 30.9333 | 0.79 |
| 0.4 | 0 | yadI | 1 | 30.9333 | 1.04 |
| 0.4 | 0 | AG1 | 1 | 30.9333 | 0.915 |
| 0.4 | 0 | ptsI | 1 | 31.1833 | 1.232 |
| 0.4 | 0 | clcB | 1 | 31.1833 | 0.78 |
| 0.4 | 0 | ycaM | 1 | 31.1833 | 0.794 |
| 0.4 | 0 | yadI | 1 | 31.1833 | 1.048 |
| 0.4 | 0 | AG1 | 1 | 31.1833 | 0.913 |
| 0.4 | 0 | ptsI | 1 | 31.4333 | 1.236 |
| 0.4 | 0 | clcB | 1 | 31.4333 | 0.78 |
| 0.4 | 0 | ycaM | 1 | 31.4333 | 0.81 |
| 0.4 | 0 | yadI | 1 | 31.4333 | 1.051 |
| 0.4 | 0 | AG1 | 1 | 31.4333 | 0.921 |
| 0.4 | 0 | ptsI | 1 | 31.6833 | 1.241 |
| 0.4 | 0 | clcB | 1 | 31.6833 | 0.779 |
| 0.4 | 0 | ycaM | 1 | 31.6833 | 0.806 |
| 0.4 | 0 | yadI | 1 | 31.6833 | 1.053 |
| 0.4 | 0 | AG1 | 1 | 31.6833 | 0.92 |
| 0.4 | 0 | ptsI | 1 | 31.9333 | 1.248 |
| 0.4 | 0 | clcB | 1 | 31.9333 | 0.779 |
| 0.4 | 0 | ycaM | 1 | 31.9333 | 0.808 |
| 0.4 | 0 | yadI | 1 | 31.9333 | 1.058 |
| 0.4 | 0 | AG1 | 1 | 31.9333 | 0.921 |
| 0.4 | 0 | ptsI | 1 | 32.1833 | 1.254 |
| 0.4 | 0 | clcB | 1 | 32.1833 | 0.783 |
| 0.4 | 0 | ycaM | 1 | 32.1833 | 0.81 |
| 0.4 | 0 | yadI | 1 | 32.1833 | 1.062 |
| 0.4 | 0 | AG1 | 1 | 32.1833 | 0.924 |
| 0.4 | 0 | ptsI | 1 | 32.4333 | 1.262 |
| 0.4 | 0 | clcB | 1 | 32.4333 | 0.786 |
| 0.4 | 0 | ycaM | 1 | 32.4333 | 0.804 |
| 0.4 | 0 | yadI | 1 | 32.4333 | 1.041 |
| 0.4 | 0 | AG1 | 1 | 32.4333 | 0.916 |

| | | | | | |
|---|---|---|---|---|---|
| 0.4 | 0 | ptsI | 1 | 32.6833 | 1.266 |
| 0.4 | 0 | clcB | 1 | 32.6833 | 0.781 |
| 0.4 | 0 | ycaM | 1 | 32.6833 | 0.814 |
| 0.4 | 0 | yadI | 1 | 32.6833 | 1.072 |
| 0.4 | 0 | AG1 | 1 | 32.6833 | 0.921 |
| 0.4 | 0 | ptsI | 1 | 32.9333 | 1.286 |
| 0.4 | 0 | clcB | 1 | 32.9333 | 0.787 |
| 0.4 | 0 | ycaM | 1 | 32.9333 | 0.813 |
| 0.4 | 0 | yadI | 1 | 32.9333 | 1.079 |
| 0.4 | 0 | AG1 | 1 | 32.9333 | 0.92 |
| 0.4 | 0 | ptsI | 1 | 33.1833 | 1.288 |
| 0.4 | 0 | clcB | 1 | 33.1833 | 0.781 |
| 0.4 | 0 | ycaM | 1 | 33.1833 | 0.816 |
| 0.4 | 0 | yadI | 1 | 33.1833 | 1.066 |
| 0.4 | 0 | AG1 | 1 | 33.1833 | 0.916 |
| 0.4 | 0 | ptsI | 1 | 33.4333 | 1.301 |
| 0.4 | 0 | clcB | 1 | 33.4333 | 0.786 |
| 0.4 | 0 | ycaM | 1 | 33.4333 | 0.832 |
| 0.4 | 0 | yadI | 1 | 33.4333 | 1.094 |
| 0.4 | 0 | AG1 | 1 | 33.4333 | 0.926 |
| 0.4 | 0 | ptsI | 1 | 33.6833 | 1.302 |
| 0.4 | 0 | clcB | 1 | 33.6833 | 0.782 |
| 0.4 | 0 | ycaM | 1 | 33.6833 | 0.828 |
| 0.4 | 0 | yadI | 1 | 33.6833 | 1.084 |
| 0.4 | 0 | AG1 | 1 | 33.6833 | 0.92 |
| 0.4 | 0 | ptsI | 1 | 33.9333 | 1.303 |
| 0.4 | 0 | clcB | 1 | 33.9333 | 0.787 |
| 0.4 | 0 | ycaM | 1 | 33.9333 | 0.829 |
| 0.4 | 0 | yadI | 1 | 33.9333 | 1.105 |
| 0.4 | 0 | AG1 | 1 | 33.9333 | 0.927 |
| 0.4 | 0 | ptsI | 1 | 34.1833 | 1.313 |
| 0.4 | 0 | clcB | 1 | 34.1833 | 0.779 |
| 0.4 | 0 | ycaM | 1 | 34.1833 | 0.836 |
| 0.4 | 0 | yadI | 1 | 34.1833 | 1.117 |
| 0.4 | 0 | AG1 | 1 | 34.1833 | 0.921 |
| 0.4 | 0 | ptsI | 1 | 34.4333 | 1.32 |
| 0.4 | 0 | clcB | 1 | 34.4333 | 0.793 |
| 0.4 | 0 | ycaM | 1 | 34.4333 | 0.835 |
| 0.4 | 0 | yadI | 1 | 34.4333 | 1.116 |
| 0.4 | 0 | AG1 | 1 | 34.4333 | 0.93 |
| 0.4 | 0 | ptsI | 1 | 34.6833 | 1.332 |
| 0.4 | 0 | clcB | 1 | 34.6833 | 0.796 |
| 0.4 | 0 | ycaM | 1 | 34.6833 | 0.845 |
| 0.4 | 0 | yadI | 1 | 34.6833 | 1.129 |
| 0.4 | 0 | AG1 | 1 | 34.6833 | 0.935 |
| 0.4 | 0 | ptsI | 1 | 34.9333 | 1.341 |
| 0.4 | 0 | clcB | 1 | 34.9333 | 0.805 |
| 0.4 | 0 | ycaM | 1 | 34.9333 | 0.842 |

| | | | | | |
|---|---|---|---|---|---|
| 0.4 | 0 | yadI | 1 | 34.9333 | 1.128 |
| 0.4 | 0 | AG1 | 1 | 34.9333 | 0.935 |
| 0.4 | 0 | ptsI | 1 | 35.1833 | 1.349 |
| 0.4 | 0 | clcB | 1 | 35.1833 | 0.805 |
| 0.4 | 0 | ycaM | 1 | 35.1833 | 0.845 |
| 0.4 | 0 | yadI | 1 | 35.1833 | 1.135 |
| 0.4 | 0 | AG1 | 1 | 35.1833 | 0.936 |
| 0.4 | 0 | ptsI | 1 | 35.4333 | 1.358 |
| 0.4 | 0 | clcB | 1 | 35.4333 | 0.815 |
| 0.4 | 0 | ycaM | 1 | 35.4333 | 0.846 |
| 0.4 | 0 | yadI | 1 | 35.4333 | 1.147 |
| 0.4 | 0 | AG1 | 1 | 35.4333 | 0.936 |
| 0.4 | 0 | ptsI | 1 | 35.6833 | 1.363 |
| 0.4 | 0 | clcB | 1 | 35.6833 | 0.817 |
| 0.4 | 0 | ycaM | 1 | 35.6833 | 0.852 |
| 0.4 | 0 | yadI | 1 | 35.6833 | 1.152 |
| 0.4 | 0 | AG1 | 1 | 35.6833 | 0.942 |
| 0.4 | 0 | ptsI | 1 | 35.9333 | 1.374 |
| 0.4 | 0 | clcB | 1 | 35.9333 | 0.824 |
| 0.4 | 0 | ycaM | 1 | 35.9333 | 0.85 |
| 0.4 | 0 | yadI | 1 | 35.9333 | 1.141 |
| 0.4 | 0 | AG1 | 1 | 35.9333 | 0.939 |
| 0.4 | 0 | ptsI | 1 | 36.1833 | 1.372 |
| 0.4 | 0 | clcB | 1 | 36.1833 | 0.83 |
| 0.4 | 0 | ycaM | 1 | 36.1833 | 0.858 |
| 0.4 | 0 | yadI | 1 | 36.1833 | 1.167 |
| 0.4 | 0 | AG1 | 1 | 36.1833 | 0.947 |
| 0.4 | 0 | ptsI | 1 | 36.4333 | 1.385 |
| 0.4 | 0 | clcB | 1 | 36.4333 | 0.845 |
| 0.4 | 0 | ycaM | 1 | 36.4333 | 0.86 |
| 0.4 | 0 | yadI | 1 | 36.4333 | 1.173 |
| 0.4 | 0 | AG1 | 1 | 36.4333 | 0.95 |
| 0.4 | 0 | ptsI | 1 | 36.6833 | 1.394 |
| 0.4 | 0 | clcB | 1 | 36.6833 | 0.854 |
| 0.4 | 0 | ycaM | 1 | 36.6833 | 0.861 |
| 0.4 | 0 | yadI | 1 | 36.6833 | 1.184 |
| 0.4 | 0 | AG1 | 1 | 36.6833 | 0.956 |
| 0.4 | 0 | ptsI | 2 | 0 | 0.212 |
| 0.4 | 0 | clcB | 2 | 0 | 0.205 |
| 0.4 | 0 | ycaM | 2 | 0 | 0.212 |
| 0.4 | 0 | yadI | 2 | 0 | 0.232 |
| 0.4 | 0 | AG1 | 2 | 0 | 0.244 |
| 0.4 | 0 | ptsI | 2 | 0.25 | 0.214 |
| 0.4 | 0 | clcB | 2 | 0.25 | 0.203 |
| 0.4 | 0 | ycaM | 2 | 0.25 | 0.207 |
| 0.4 | 0 | yadI | 2 | 0.25 | 0.226 |
| 0.4 | 0 | AG1 | 2 | 0.25 | 0.234 |
| 0.4 | 0 | ptsI | 2 | 0.5 | 0.212 |

| | | | | | |
|---|---|---|---|---|---|
| 0.4 | 0 | clcB | 2 | 0.5 | 0.201 |
| 0.4 | 0 | ycaM | 2 | 0.5 | 0.206 |
| 0.4 | 0 | yadI | 2 | 0.5 | 0.224 |
| 0.4 | 0 | AG1 | 2 | 0.5 | 0.232 |
| 0.4 | 0 | ptsI | 2 | 0.75 | 0.215 |
| 0.4 | 0 | clcB | 2 | 0.75 | 0.203 |
| 0.4 | 0 | ycaM | 2 | 0.75 | 0.207 |
| 0.4 | 0 | yadI | 2 | 0.75 | 0.227 |
| 0.4 | 0 | AG1 | 2 | 0.75 | 0.236 |
| 0.4 | 0 | ptsI | 2 | 1 | 0.215 |
| 0.4 | 0 | clcB | 2 | 1 | 0.207 |
| 0.4 | 0 | ycaM | 2 | 1 | 0.213 |
| 0.4 | 0 | yadI | 2 | 1 | 0.234 |
| 0.4 | 0 | AG1 | 2 | 1 | 0.245 |
| 0.4 | 0 | ptsI | 2 | 1.25 | 0.22 |
| 0.4 | 0 | clcB | 2 | 1.25 | 0.205 |
| 0.4 | 0 | ycaM | 2 | 1.25 | 0.215 |
| 0.4 | 0 | yadI | 2 | 1.25 | 0.241 |
| 0.4 | 0 | AG1 | 2 | 1.25 | 0.251 |
| 0.4 | 0 | ptsI | 2 | 1.5 | 0.223 |
| 0.4 | 0 | clcB | 2 | 1.5 | 0.208 |
| 0.4 | 0 | ycaM | 2 | 1.5 | 0.218 |
| 0.4 | 0 | yadI | 2 | 1.5 | 0.247 |
| 0.4 | 0 | AG1 | 2 | 1.5 | 0.256 |
| 0.4 | 0 | ptsI | 2 | 1.75 | 0.226 |
| 0.4 | 0 | clcB | 2 | 1.75 | 0.211 |
| 0.4 | 0 | ycaM | 2 | 1.75 | 0.222 |
| 0.4 | 0 | yadI | 2 | 1.75 | 0.253 |
| 0.4 | 0 | AG1 | 2 | 1.75 | 0.264 |
| 0.4 | 0 | ptsI | 2 | 2 | 0.23 |
| 0.4 | 0 | clcB | 2 | 2 | 0.215 |
| 0.4 | 0 | ycaM | 2 | 2 | 0.229 |
| 0.4 | 0 | yadI | 2 | 2 | 0.261 |
| 0.4 | 0 | AG1 | 2 | 2 | 0.279 |
| 0.4 | 0 | ptsI | 2 | 2.25 | 0.241 |
| 0.4 | 0 | clcB | 2 | 2.25 | 0.219 |
| 0.4 | 0 | ycaM | 2 | 2.25 | 0.232 |
| 0.4 | 0 | yadI | 2 | 2.25 | 0.274 |
| 0.4 | 0 | AG1 | 2 | 2.25 | 0.292 |
| 0.4 | 0 | ptsI | 2 | 2.5 | 0.248 |
| 0.4 | 0 | clcB | 2 | 2.5 | 0.226 |
| 0.4 | 0 | ycaM | 2 | 2.5 | 0.24 |
| 0.4 | 0 | yadI | 2 | 2.5 | 0.289 |
| 0.4 | 0 | AG1 | 2 | 2.5 | 0.307 |
| 0.4 | 0 | ptsI | 2 | 2.75 | 0.252 |
| 0.4 | 0 | clcB | 2 | 2.75 | 0.231 |
| 0.4 | 0 | ycaM | 2 | 2.75 | 0.246 |
| 0.4 | 0 | yadI | 2 | 2.75 | 0.298 |

| | | | | | |
|---|---|---|---|---|---|
| 0.4 | 0 | AG1 | 2 | 2.75 | 0.322 |
| 0.4 | 0 | ptsI | 2 | 3 | 0.258 |
| 0.4 | 0 | clcB | 2 | 3 | 0.238 |
| 0.4 | 0 | ycaM | 2 | 3 | 0.254 |
| 0.4 | 0 | yadI | 2 | 3 | 0.309 |
| 0.4 | 0 | AG1 | 2 | 3 | 0.321 |
| 0.4 | 0 | ptsI | 2 | 3.25 | 0.27 |
| 0.4 | 0 | clcB | 2 | 3.25 | 0.242 |
| 0.4 | 0 | ycaM | 2 | 3.25 | 0.263 |
| 0.4 | 0 | yadI | 2 | 3.25 | 0.332 |
| 0.4 | 0 | AG1 | 2 | 3.25 | 0.345 |
| 0.4 | 0 | ptsI | 2 | 3.5 | 0.277 |
| 0.4 | 0 | clcB | 2 | 3.5 | 0.248 |
| 0.4 | 0 | ycaM | 2 | 3.5 | 0.271 |
| 0.4 | 0 | yadI | 2 | 3.5 | 0.349 |
| 0.4 | 0 | AG1 | 2 | 3.5 | 0.357 |
| 0.4 | 0 | ptsI | 2 | 3.75 | 0.285 |
| 0.4 | 0 | clcB | 2 | 3.75 | 0.253 |
| 0.4 | 0 | ycaM | 2 | 3.75 | 0.285 |
| 0.4 | 0 | yadI | 2 | 3.75 | 0.367 |
| 0.4 | 0 | AG1 | 2 | 3.75 | 0.368 |
| 0.4 | 0 | ptsI | 2 | 4 | 0.292 |
| 0.4 | 0 | clcB | 2 | 4 | 0.259 |
| 0.4 | 0 | ycaM | 2 | 4 | 0.292 |
| 0.4 | 0 | yadI | 2 | 4 | 0.386 |
| 0.4 | 0 | AG1 | 2 | 4 | 0.379 |
| 0.4 | 0 | ptsI | 2 | 4.38333 | 0.309 |
| 0.4 | 0 | clcB | 2 | 4.38333 | 0.269 |
| 0.4 | 0 | ycaM | 2 | 4.38333 | 0.306 |
| 0.4 | 0 | yadI | 2 | 4.38333 | 0.405 |
| 0.4 | 0 | AG1 | 2 | 4.38333 | 0.44 |
| 0.4 | 0 | ptsI | 2 | 4.63333 | 0.303 |
| 0.4 | 0 | clcB | 2 | 4.63333 | 0.265 |
| 0.4 | 0 | ycaM | 2 | 4.63333 | 0.3 |
| 0.4 | 0 | yadI | 2 | 4.63333 | 0.402 |
| 0.4 | 0 | AG1 | 2 | 4.63333 | 0.418 |
| 0.4 | 0 | ptsI | 2 | 4.88333 | 0.307 |
| 0.4 | 0 | clcB | 2 | 4.88333 | 0.269 |
| 0.4 | 0 | ycaM | 2 | 4.88333 | 0.307 |
| 0.4 | 0 | yadI | 2 | 4.88333 | 0.412 |
| 0.4 | 0 | AG1 | 2 | 4.88333 | 0.426 |
| 0.4 | 0 | ptsI | 2 | 5.13333 | 0.315 |
| 0.4 | 0 | clcB | 2 | 5.13333 | 0.275 |
| 0.4 | 0 | ycaM | 2 | 5.13333 | 0.316 |
| 0.4 | 0 | yadI | 2 | 5.13333 | 0.423 |
| 0.4 | 0 | AG1 | 2 | 5.13333 | 0.436 |
| 0.4 | 0 | ptsI | 2 | 5.38333 | 0.323 |
| 0.4 | 0 | clcB | 2 | 5.38333 | 0.283 |

| | | | | | |
|---|---|---|---|---|---|
| 0.4 | 0 | ycaM | 2 | 5.38333 | 0.342 |
| 0.4 | 0 | yadI | 2 | 5.38333 | 0.436 |
| 0.4 | 0 | AG1 | 2 | 5.38333 | 0.443 |
| 0.4 | 0 | ptsI | 2 | 5.63333 | 0.332 |
| 0.4 | 0 | clcB | 2 | 5.63333 | 0.293 |
| 0.4 | 0 | ycaM | 2 | 5.63333 | 0.338 |
| 0.4 | 0 | yadI | 2 | 5.63333 | 0.446 |
| 0.4 | 0 | AG1 | 2 | 5.63333 | 0.454 |
| 0.4 | 0 | ptsI | 2 | 5.88333 | 0.337 |
| 0.4 | 0 | clcB | 2 | 5.88333 | 0.301 |
| 0.4 | 0 | ycaM | 2 | 5.88333 | 0.35 |
| 0.4 | 0 | yadI | 2 | 5.88333 | 0.455 |
| 0.4 | 0 | AG1 | 2 | 5.88333 | 0.465 |
| 0.4 | 0 | ptsI | 2 | 6.13333 | 0.345 |
| 0.4 | 0 | clcB | 2 | 6.13333 | 0.306 |
| 0.4 | 0 | ycaM | 2 | 6.13333 | 0.359 |
| 0.4 | 0 | yadI | 2 | 6.13333 | 0.465 |
| 0.4 | 0 | AG1 | 2 | 6.13333 | 0.478 |
| 0.4 | 0 | ptsI | 2 | 6.38333 | 0.354 |
| 0.4 | 0 | clcB | 2 | 6.38333 | 0.313 |
| 0.4 | 0 | ycaM | 2 | 6.38333 | 0.372 |
| 0.4 | 0 | yadI | 2 | 6.38333 | 0.476 |
| 0.4 | 0 | AG1 | 2 | 6.38333 | 0.49 |
| 0.4 | 0 | ptsI | 2 | 6.63333 | 0.361 |
| 0.4 | 0 | clcB | 2 | 6.63333 | 0.319 |
| 0.4 | 0 | ycaM | 2 | 6.63333 | 0.381 |
| 0.4 | 0 | yadI | 2 | 6.63333 | 0.487 |
| 0.4 | 0 | AG1 | 2 | 6.63333 | 0.5 |
| 0.4 | 0 | ptsI | 2 | 6.88333 | 0.37 |
| 0.4 | 0 | clcB | 2 | 6.88333 | 0.329 |
| 0.4 | 0 | ycaM | 2 | 6.88333 | 0.391 |
| 0.4 | 0 | yadI | 2 | 6.88333 | 0.497 |
| 0.4 | 0 | AG1 | 2 | 6.88333 | 0.51 |
| 0.4 | 0 | ptsI | 2 | 7.13333 | 0.378 |
| 0.4 | 0 | clcB | 2 | 7.13333 | 0.334 |
| 0.4 | 0 | ycaM | 2 | 7.13333 | 0.399 |
| 0.4 | 0 | yadI | 2 | 7.13333 | 0.506 |
| 0.4 | 0 | AG1 | 2 | 7.13333 | 0.524 |
| 0.4 | 0 | ptsI | 2 | 7.38333 | 0.385 |
| 0.4 | 0 | clcB | 2 | 7.38333 | 0.343 |
| 0.4 | 0 | ycaM | 2 | 7.38333 | 0.407 |
| 0.4 | 0 | yadI | 2 | 7.38333 | 0.516 |
| 0.4 | 0 | AG1 | 2 | 7.38333 | 0.532 |
| 0.4 | 0 | ptsI | 2 | 7.63333 | 0.393 |
| 0.4 | 0 | clcB | 2 | 7.63333 | 0.351 |
| 0.4 | 0 | ycaM | 2 | 7.63333 | 0.417 |
| 0.4 | 0 | yadI | 2 | 7.63333 | 0.527 |
| 0.4 | 0 | AG1 | 2 | 7.63333 | 0.546 |

| | | | | | |
|---|---|---|---|---|---|
| 0.4 | 0 | ptsI | 2 | 7.88333 | 0.4 |
| 0.4 | 0 | clcB | 2 | 7.88333 | 0.36 |
| 0.4 | 0 | ycaM | 2 | 7.88333 | 0.423 |
| 0.4 | 0 | yadI | 2 | 7.88333 | 0.542 |
| 0.4 | 0 | AG1 | 2 | 7.88333 | 0.56 |
| 0.4 | 0 | ptsI | 2 | 8.13333 | 0.409 |
| 0.4 | 0 | clcB | 2 | 8.13333 | 0.367 |
| 0.4 | 0 | ycaM | 2 | 8.13333 | 0.431 |
| 0.4 | 0 | yadI | 2 | 8.13333 | 0.55 |
| 0.4 | 0 | AG1 | 2 | 8.13333 | 0.569 |
| 0.4 | 0 | ptsI | 2 | 8.38333 | 0.412 |
| 0.4 | 0 | clcB | 2 | 8.38333 | 0.376 |
| 0.4 | 0 | ycaM | 2 | 8.38333 | 0.436 |
| 0.4 | 0 | yadI | 2 | 8.38333 | 0.562 |
| 0.4 | 0 | AG1 | 2 | 8.38333 | 0.582 |
| 0.4 | 0 | ptsI | 2 | 8.91667 | 0.354 |
| 0.4 | 0 | clcB | 2 | 8.91667 | 0.323 |
| 0.4 | 0 | ycaM | 2 | 8.91667 | 0.356 |
| 0.4 | 0 | yadI | 2 | 8.91667 | 0.652 |
| 0.4 | 0 | AG1 | 2 | 8.91667 | 0.643 |
| 0.4 | 0 | ptsI | 2 | 9.16667 | 0.352 |
| 0.4 | 0 | clcB | 2 | 9.16667 | 0.332 |
| 0.4 | 0 | ycaM | 2 | 9.16667 | 0.356 |
| 0.4 | 0 | yadI | 2 | 9.16667 | 0.652 |
| 0.4 | 0 | AG1 | 2 | 9.16667 | 0.636 |
| 0.4 | 0 | ptsI | 2 | 9.41667 | 0.356 |
| 0.4 | 0 | clcB | 2 | 9.41667 | 0.332 |
| 0.4 | 0 | ycaM | 2 | 9.41667 | 0.36 |
| 0.4 | 0 | yadI | 2 | 9.41667 | 0.659 |
| 0.4 | 0 | AG1 | 2 | 9.41667 | 0.65 |
| 0.4 | 0 | ptsI | 2 | 9.66667 | 0.367 |
| 0.4 | 0 | clcB | 2 | 9.66667 | 0.338 |
| 0.4 | 0 | ycaM | 2 | 9.66667 | 0.365 |
| 0.4 | 0 | yadI | 2 | 9.66667 | 0.672 |
| 0.4 | 0 | AG1 | 2 | 9.66667 | 0.66 |
| 0.4 | 0 | ptsI | 2 | 9.91667 | 0.372 |
| 0.4 | 0 | clcB | 2 | 9.91667 | 0.392 |
| 0.4 | 0 | ycaM | 2 | 9.91667 | 0.37 |
| 0.4 | 0 | yadI | 2 | 9.91667 | 0.685 |
| 0.4 | 0 | AG1 | 2 | 9.91667 | 0.656 |
| 0.4 | 0 | ptsI | 2 | 10.1667 | 0.377 |
| 0.4 | 0 | clcB | 2 | 10.1667 | 0.346 |
| 0.4 | 0 | ycaM | 2 | 10.1667 | 0.374 |
| 0.4 | 0 | yadI | 2 | 10.1667 | 0.696 |
| 0.4 | 0 | AG1 | 2 | 10.1667 | 0.648 |
| 0.4 | 0 | ptsI | 2 | 10.4167 | 0.385 |
| 0.4 | 0 | clcB | 2 | 10.4167 | 0.352 |
| 0.4 | 0 | ycaM | 2 | 10.4167 | 0.38 |

| | | | | | |
|---|---|---|---|---|---|
| 0.4 | 0 | yadI | 2 | 10.4167 | 0.708 |
| 0.4 | 0 | AG1 | 2 | 10.4167 | 0.674 |
| 0.4 | 0 | ptsI | 2 | 10.6667 | 0.392 |
| 0.4 | 0 | clcB | 2 | 10.6667 | 0.358 |
| 0.4 | 0 | ycaM | 2 | 10.6667 | 0.391 |
| 0.4 | 0 | yadI | 2 | 10.6667 | 0.724 |
| 0.4 | 0 | AG1 | 2 | 10.6667 | 0.65 |
| 0.4 | 0 | ptsI | 2 | 10.9167 | 0.399 |
| 0.4 | 0 | clcB | 2 | 10.9167 | 0.36 |
| 0.4 | 0 | ycaM | 2 | 10.9167 | 0.397 |
| 0.4 | 0 | yadI | 2 | 10.9167 | 0.733 |
| 0.4 | 0 | AG1 | 2 | 10.9167 | 0.664 |
| 0.4 | 0 | ptsI | 2 | 11.1667 | 0.405 |
| 0.4 | 0 | clcB | 2 | 11.1667 | 0.366 |
| 0.4 | 0 | ycaM | 2 | 11.1667 | 0.406 |
| 0.4 | 0 | yadI | 2 | 11.1667 | 0.742 |
| 0.4 | 0 | AG1 | 2 | 11.1667 | 0.672 |
| 0.4 | 0 | ptsI | 2 | 11.4167 | 0.425 |
| 0.4 | 0 | clcB | 2 | 11.4167 | 0.368 |
| 0.4 | 0 | ycaM | 2 | 11.4167 | 0.41 |
| 0.4 | 0 | yadI | 2 | 11.4167 | 0.752 |
| 0.4 | 0 | AG1 | 2 | 11.4167 | 0.674 |
| 0.4 | 0 | ptsI | 2 | 11.6667 | 0.431 |
| 0.4 | 0 | clcB | 2 | 11.6667 | 0.378 |
| 0.4 | 0 | ycaM | 2 | 11.6667 | 0.416 |
| 0.4 | 0 | yadI | 2 | 11.6667 | 0.764 |
| 0.4 | 0 | AG1 | 2 | 11.6667 | 0.696 |
| 0.4 | 0 | ptsI | 2 | 11.9167 | 0.46 |
| 0.4 | 0 | clcB | 2 | 11.9167 | 0.386 |
| 0.4 | 0 | ycaM | 2 | 11.9167 | 0.425 |
| 0.4 | 0 | yadI | 2 | 11.9167 | 0.773 |
| 0.4 | 0 | AG1 | 2 | 11.9167 | 0.699 |
| 0.4 | 0 | ptsI | 2 | 12.1667 | 0.45 |
| 0.4 | 0 | clcB | 2 | 12.1667 | 0.397 |
| 0.4 | 0 | ycaM | 2 | 12.1667 | 0.421 |
| 0.4 | 0 | yadI | 2 | 12.1667 | 0.784 |
| 0.4 | 0 | AG1 | 2 | 12.1667 | 0.718 |
| 0.4 | 0 | ptsI | 2 | 12.4167 | 0.452 |
| 0.4 | 0 | clcB | 2 | 12.4167 | 0.401 |
| 0.4 | 0 | ycaM | 2 | 12.4167 | 0.432 |
| 0.4 | 0 | yadI | 2 | 12.4167 | 0.791 |
| 0.4 | 0 | AG1 | 2 | 12.4167 | 0.711 |
| 0.4 | 0 | ptsI | 2 | 12.6667 | 0.464 |
| 0.4 | 0 | clcB | 2 | 12.6667 | 0.407 |
| 0.4 | 0 | ycaM | 2 | 12.6667 | 0.429 |
| 0.4 | 0 | yadI | 2 | 12.6667 | 0.794 |
| 0.4 | 0 | AG1 | 2 | 12.6667 | 0.719 |
| 0.4 | 0 | ptsI | 2 | 12.9167 | 0.477 |

| | | | | | |
|---|---|---|---|---|---|
| 0.4 | 0 | clcB | 2 | 12.9167 | 0.421 |
| 0.4 | 0 | ycaM | 2 | 12.9167 | 0.438 |
| 0.4 | 0 | yadI | 2 | 12.9167 | 0.806 |
| 0.4 | 0 | AG1 | 2 | 12.9167 | 0.724 |
| 0.4 | 0 | ptsI | 2 | 13.1667 | 0.476 |
| 0.4 | 0 | clcB | 2 | 13.1667 | 0.417 |
| 0.4 | 0 | ycaM | 2 | 13.1667 | 0.455 |
| 0.4 | 0 | yadI | 2 | 13.1667 | 0.814 |
| 0.4 | 0 | AG1 | 2 | 13.1667 | 0.733 |
| 0.4 | 0 | ptsI | 2 | 13.4167 | 0.478 |
| 0.4 | 0 | clcB | 2 | 13.4167 | 0.421 |
| 0.4 | 0 | ycaM | 2 | 13.4167 | 0.458 |
| 0.4 | 0 | yadI | 2 | 13.4167 | 0.821 |
| 0.4 | 0 | AG1 | 2 | 13.4167 | 0.746 |
| 0.4 | 0 | ptsI | 2 | 13.6667 | 0.489 |
| 0.4 | 0 | clcB | 2 | 13.6667 | 0.427 |
| 0.4 | 0 | ycaM | 2 | 13.6667 | 0.468 |
| 0.4 | 0 | yadI | 2 | 13.6667 | 0.83 |
| 0.4 | 0 | AG1 | 2 | 13.6667 | 0.756 |
| 0.4 | 0 | ptsI | 2 | 13.9167 | 0.524 |
| 0.4 | 0 | clcB | 2 | 13.9167 | 0.434 |
| 0.4 | 0 | ycaM | 2 | 13.9167 | 0.461 |
| 0.4 | 0 | yadI | 2 | 13.9167 | 0.839 |
| 0.4 | 0 | AG1 | 2 | 13.9167 | 0.779 |
| 0.4 | 0 | ptsI | 2 | 14.1667 | 0.546 |
| 0.4 | 0 | clcB | 2 | 14.1667 | 0.432 |
| 0.4 | 0 | ycaM | 2 | 14.1667 | 0.468 |
| 0.4 | 0 | yadI | 2 | 14.1667 | 0.846 |
| 0.4 | 0 | AG1 | 2 | 14.1667 | 0.792 |
| 0.4 | 0 | ptsI | 2 | 14.4167 | 0.571 |
| 0.4 | 0 | clcB | 2 | 14.4167 | 0.438 |
| 0.4 | 0 | ycaM | 2 | 14.4167 | 0.469 |
| 0.4 | 0 | yadI | 2 | 14.4167 | 0.843 |
| 0.4 | 0 | AG1 | 2 | 14.4167 | 0.801 |
| 0.4 | 0 | ptsI | 2 | 14.6667 | 0.594 |
| 0.4 | 0 | clcB | 2 | 14.6667 | 0.44 |
| 0.4 | 0 | ycaM | 2 | 14.6667 | 0.467 |
| 0.4 | 0 | yadI | 2 | 14.6667 | 0.847 |
| 0.4 | 0 | AG1 | 2 | 14.6667 | 0.797 |
| 0.4 | 0 | ptsI | 2 | 14.9167 | 0.62 |
| 0.4 | 0 | clcB | 2 | 14.9167 | 0.448 |
| 0.4 | 0 | ycaM | 2 | 14.9167 | 0.468 |
| 0.4 | 0 | yadI | 2 | 14.9167 | 0.85 |
| 0.4 | 0 | AG1 | 2 | 14.9167 | 0.79 |
| 0.4 | 0 | ptsI | 2 | 15.1667 | 0.614 |
| 0.4 | 0 | clcB | 2 | 15.1667 | 0.46 |
| 0.4 | 0 | ycaM | 2 | 15.1667 | 0.472 |
| 0.4 | 0 | yadI | 2 | 15.1667 | 0.856 |

| | | | | | |
|---|---|---|---|---|---|
| 0.4 | 0 | AG1 | 2 | 15.1667 | 0.777 |
| 0.4 | 0 | ptsI | 2 | 15.4167 | 0.673 |
| 0.4 | 0 | clcB | 2 | 15.4167 | 0.457 |
| 0.4 | 0 | ycaM | 2 | 15.4167 | 0.485 |
| 0.4 | 0 | yadI | 2 | 15.4167 | 0.865 |
| 0.4 | 0 | AG1 | 2 | 15.4167 | 0.803 |
| 0.4 | 0 | ptsI | 2 | 15.6667 | 0.678 |
| 0.4 | 0 | clcB | 2 | 15.6667 | 0.465 |
| 0.4 | 0 | ycaM | 2 | 15.6667 | 0.482 |
| 0.4 | 0 | yadI | 2 | 15.6667 | 0.862 |
| 0.4 | 0 | AG1 | 2 | 15.6667 | 0.802 |
| 0.4 | 0 | ptsI | 2 | 15.9167 | 0.68 |
| 0.4 | 0 | clcB | 2 | 15.9167 | 0.477 |
| 0.4 | 0 | ycaM | 2 | 15.9167 | 0.49 |
| 0.4 | 0 | yadI | 2 | 15.9167 | 0.868 |
| 0.4 | 0 | AG1 | 2 | 15.9167 | 0.779 |
| 0.4 | 0 | ptsI | 2 | 16.1667 | 0.696 |
| 0.4 | 0 | clcB | 2 | 16.1667 | 0.47 |
| 0.4 | 0 | ycaM | 2 | 16.1667 | 0.499 |
| 0.4 | 0 | yadI | 2 | 16.1667 | 0.873 |
| 0.4 | 0 | AG1 | 2 | 16.1667 | 0.78 |
| 0.4 | 0 | ptsI | 2 | 16.4167 | 0.718 |
| 0.4 | 0 | clcB | 2 | 16.4167 | 0.49 |
| 0.4 | 0 | ycaM | 2 | 16.4167 | 0.495 |
| 0.4 | 0 | yadI | 2 | 16.4167 | 0.877 |
| 0.4 | 0 | AG1 | 2 | 16.4167 | 0.804 |
| 0.4 | 0 | ptsI | 2 | 16.6667 | 0.731 |
| 0.4 | 0 | clcB | 2 | 16.6667 | 0.492 |
| 0.4 | 0 | ycaM | 2 | 16.6667 | 0.508 |
| 0.4 | 0 | yadI | 2 | 16.6667 | 0.882 |
| 0.4 | 0 | AG1 | 2 | 16.6667 | 0.806 |
| 0.4 | 0 | ptsI | 2 | 16.9167 | 0.747 |
| 0.4 | 0 | clcB | 2 | 16.9167 | 0.48 |
| 0.4 | 0 | ycaM | 2 | 16.9167 | 0.525 |
| 0.4 | 0 | yadI | 2 | 16.9167 | 0.882 |
| 0.4 | 0 | AG1 | 2 | 16.9167 | 0.808 |
| 0.4 | 0 | ptsI | 2 | 17.1667 | 0.754 |
| 0.4 | 0 | clcB | 2 | 17.1667 | 0.482 |
| 0.4 | 0 | ycaM | 2 | 17.1667 | 0.526 |
| 0.4 | 0 | yadI | 2 | 17.1667 | 0.885 |
| 0.4 | 0 | AG1 | 2 | 17.1667 | 0.802 |
| 0.4 | 0 | ptsI | 2 | 17.4167 | 0.759 |
| 0.4 | 0 | clcB | 2 | 17.4167 | 0.508 |
| 0.4 | 0 | ycaM | 2 | 17.4167 | 0.535 |
| 0.4 | 0 | yadI | 2 | 17.4167 | 0.887 |
| 0.4 | 0 | AG1 | 2 | 17.4167 | 0.81 |
| 0.4 | 0 | ptsI | 2 | 17.6667 | 0.769 |
| 0.4 | 0 | clcB | 2 | 17.6667 | 0.507 |

| | | | | | |
|---|---|---|---|---|---|
| 0.4 | 0 | ycaM | 2 | 17.6667 | 0.531 |
| 0.4 | 0 | yadI | 2 | 17.6667 | 0.886 |
| 0.4 | 0 | AG1 | 2 | 17.6667 | 0.818 |
| 0.4 | 0 | ptsI | 2 | 17.9167 | 0.773 |
| 0.4 | 0 | clcB | 2 | 17.9167 | 0.502 |
| 0.4 | 0 | ycaM | 2 | 17.9167 | 0.521 |
| 0.4 | 0 | yadI | 2 | 17.9167 | 0.89 |
| 0.4 | 0 | AG1 | 2 | 17.9167 | 0.814 |
| 0.4 | 0 | ptsI | 2 | 18.1667 | 0.786 |
| 0.4 | 0 | clcB | 2 | 18.1667 | 0.501 |
| 0.4 | 0 | ycaM | 2 | 18.1667 | 0.528 |
| 0.4 | 0 | yadI | 2 | 18.1667 | 0.891 |
| 0.4 | 0 | AG1 | 2 | 18.1667 | 0.824 |
| 0.4 | 0 | ptsI | 2 | 18.4167 | 0.792 |
| 0.4 | 0 | clcB | 2 | 18.4167 | 0.495 |
| 0.4 | 0 | ycaM | 2 | 18.4167 | 0.52 |
| 0.4 | 0 | yadI | 2 | 18.4167 | 0.893 |
| 0.4 | 0 | AG1 | 2 | 18.4167 | 0.822 |
| 0.4 | 0 | ptsI | 2 | 18.6667 | 0.787 |
| 0.4 | 0 | clcB | 2 | 18.6667 | 0.498 |
| 0.4 | 0 | ycaM | 2 | 18.6667 | 0.526 |
| 0.4 | 0 | yadI | 2 | 18.6667 | 0.894 |
| 0.4 | 0 | AG1 | 2 | 18.6667 | 0.821 |
| 0.4 | 0 | ptsI | 2 | 18.9167 | 0.813 |
| 0.4 | 0 | clcB | 2 | 18.9167 | 0.488 |
| 0.4 | 0 | ycaM | 2 | 18.9167 | 0.522 |
| 0.4 | 0 | yadI | 2 | 18.9167 | 0.875 |
| 0.4 | 0 | AG1 | 2 | 18.9167 | 0.826 |
| 0.4 | 0 | ptsI | 2 | 19.1667 | 0.805 |
| 0.4 | 0 | clcB | 2 | 19.1667 | 0.49 |
| 0.4 | 0 | ycaM | 2 | 19.1667 | 0.522 |
| 0.4 | 0 | yadI | 2 | 19.1667 | 0.897 |
| 0.4 | 0 | AG1 | 2 | 19.1667 | 0.816 |
| 0.4 | 0 | ptsI | 2 | 19.4167 | 0.81 |
| 0.4 | 0 | clcB | 2 | 19.4167 | 0.49 |
| 0.4 | 0 | ycaM | 2 | 19.4167 | 0.522 |
| 0.4 | 0 | yadI | 2 | 19.4167 | 0.896 |
| 0.4 | 0 | AG1 | 2 | 19.4167 | 0.818 |
| 0.4 | 0 | ptsI | 2 | 19.6667 | 0.813 |
| 0.4 | 0 | clcB | 2 | 19.6667 | 0.488 |
| 0.4 | 0 | ycaM | 2 | 19.6667 | 0.524 |
| 0.4 | 0 | yadI | 2 | 19.6667 | 0.897 |
| 0.4 | 0 | AG1 | 2 | 19.6667 | 0.817 |
| 0.4 | 0 | ptsI | 2 | 19.9167 | 0.826 |
| 0.4 | 0 | clcB | 2 | 19.9167 | 0.5 |
| 0.4 | 0 | ycaM | 2 | 19.9167 | 0.527 |
| 0.4 | 0 | yadI | 2 | 19.9167 | 0.896 |
| 0.4 | 0 | AG1 | 2 | 19.9167 | 0.818 |

| | | | | | |
|---|---|---|---|---|---|
| 0.4 | 0 | ptsI | 2 | 20.1667 | 0.825 |
| 0.4 | 0 | clcB | 2 | 20.1667 | 0.495 |
| 0.4 | 0 | ycaM | 2 | 20.1667 | 0.529 |
| 0.4 | 0 | yadI | 2 | 20.1667 | 0.901 |
| 0.4 | 0 | AG1 | 2 | 20.1667 | 0.815 |
| 0.4 | 0 | ptsI | 2 | 20.4167 | 0.83 |
| 0.4 | 0 | clcB | 2 | 20.4167 | 0.498 |
| 0.4 | 0 | ycaM | 2 | 20.4167 | 0.534 |
| 0.4 | 0 | yadI | 2 | 20.4167 | 0.903 |
| 0.4 | 0 | AG1 | 2 | 20.4167 | 0.824 |
| 0.4 | 0 | ptsI | 2 | 20.6667 | 0.839 |
| 0.4 | 0 | clcB | 2 | 20.6667 | 0.494 |
| 0.4 | 0 | ycaM | 2 | 20.6667 | 0.535 |
| 0.4 | 0 | yadI | 2 | 20.6667 | 0.901 |
| 0.4 | 0 | AG1 | 2 | 20.6667 | 0.82 |
| 0.4 | 0 | ptsI | 2 | 20.9167 | 0.843 |
| 0.4 | 0 | clcB | 2 | 20.9167 | 0.491 |
| 0.4 | 0 | ycaM | 2 | 20.9167 | 0.538 |
| 0.4 | 0 | yadI | 2 | 20.9167 | 0.905 |
| 0.4 | 0 | AG1 | 2 | 20.9167 | 0.824 |
| 0.4 | 0 | ptsI | 2 | 21.1667 | 0.835 |
| 0.4 | 0 | clcB | 2 | 21.1667 | 0.492 |
| 0.4 | 0 | ycaM | 2 | 21.1667 | 0.539 |
| 0.4 | 0 | yadI | 2 | 21.1667 | 0.907 |
| 0.4 | 0 | AG1 | 2 | 21.1667 | 0.823 |
| 0.4 | 0 | ptsI | 2 | 21.4167 | 0.841 |
| 0.4 | 0 | clcB | 2 | 21.4167 | 0.494 |
| 0.4 | 0 | ycaM | 2 | 21.4167 | 0.545 |
| 0.4 | 0 | yadI | 2 | 21.4167 | 0.891 |
| 0.4 | 0 | AG1 | 2 | 21.4167 | 0.828 |
| 0.4 | 0 | ptsI | 2 | 21.6667 | 0.847 |
| 0.4 | 0 | clcB | 2 | 21.6667 | 0.498 |
| 0.4 | 0 | ycaM | 2 | 21.6667 | 0.545 |
| 0.4 | 0 | yadI | 2 | 21.6667 | 0.907 |
| 0.4 | 0 | AG1 | 2 | 21.6667 | 0.83 |
| 0.4 | 0 | ptsI | 2 | 21.9167 | 0.852 |
| 0.4 | 0 | clcB | 2 | 21.9167 | 0.5 |
| 0.4 | 0 | ycaM | 2 | 21.9167 | 0.547 |
| 0.4 | 0 | yadI | 2 | 21.9167 | 0.909 |
| 0.4 | 0 | AG1 | 2 | 21.9167 | 0.834 |
| 0.4 | 0 | ptsI | 2 | 22.1667 | 0.861 |
| 0.4 | 0 | clcB | 2 | 22.1667 | 0.502 |
| 0.4 | 0 | ycaM | 2 | 22.1667 | 0.549 |
| 0.4 | 0 | yadI | 2 | 22.1667 | 0.909 |
| 0.4 | 0 | AG1 | 2 | 22.1667 | 0.835 |
| 0.4 | 0 | ptsI | 2 | 22.4167 | 0.859 |
| 0.4 | 0 | clcB | 2 | 22.4167 | 0.508 |
| 0.4 | 0 | ycaM | 2 | 22.4167 | 0.55 |

| | | | | | |
|---|---|---|---|---|---|
| 0.4 | 0 | yadI | 2 | 22.4167 | 0.912 |
| 0.4 | 0 | AG1 | 2 | 22.4167 | 0.837 |
| 0.4 | 0 | ptsI | 2 | 22.6667 | 0.86 |
| 0.4 | 0 | clcB | 2 | 22.6667 | 0.507 |
| 0.4 | 0 | ycaM | 2 | 22.6667 | 0.554 |
| 0.4 | 0 | yadI | 2 | 22.6667 | 0.912 |
| 0.4 | 0 | AG1 | 2 | 22.6667 | 0.841 |
| 0.4 | 0 | ptsI | 2 | 22.9167 | 0.86 |
| 0.4 | 0 | clcB | 2 | 22.9167 | 0.512 |
| 0.4 | 0 | ycaM | 2 | 22.9167 | 0.553 |
| 0.4 | 0 | yadI | 2 | 22.9167 | 0.917 |
| 0.4 | 0 | AG1 | 2 | 22.9167 | 0.845 |
| 0.4 | 0 | ptsI | 2 | 23.1667 | 0.866 |
| 0.4 | 0 | clcB | 2 | 23.1667 | 0.511 |
| 0.4 | 0 | ycaM | 2 | 23.1667 | 0.554 |
| 0.4 | 0 | yadI | 2 | 23.1667 | 0.916 |
| 0.4 | 0 | AG1 | 2 | 23.1667 | 0.846 |
| 0.4 | 0 | ptsI | 2 | 23.4167 | 0.868 |
| 0.4 | 0 | clcB | 2 | 23.4167 | 0.511 |
| 0.4 | 0 | ycaM | 2 | 23.4167 | 0.554 |
| 0.4 | 0 | yadI | 2 | 23.4167 | 0.916 |
| 0.4 | 0 | AG1 | 2 | 23.4167 | 0.848 |
| 0.4 | 0 | ptsI | 2 | 23.6667 | 0.874 |
| 0.4 | 0 | clcB | 2 | 23.6667 | 0.514 |
| 0.4 | 0 | ycaM | 2 | 23.6667 | 0.561 |
| 0.4 | 0 | yadI | 2 | 23.6667 | 0.916 |
| 0.4 | 0 | AG1 | 2 | 23.6667 | 0.852 |
| 0.4 | 0 | ptsI | 2 | 23.9167 | 0.879 |
| 0.4 | 0 | clcB | 2 | 23.9167 | 0.52 |
| 0.4 | 0 | ycaM | 2 | 23.9167 | 0.561 |
| 0.4 | 0 | yadI | 2 | 23.9167 | 0.919 |
| 0.4 | 0 | AG1 | 2 | 23.9167 | 0.858 |
| 0.4 | 0 | ptsI | 2 | 24.1667 | 0.88 |
| 0.4 | 0 | clcB | 2 | 24.1667 | 0.518 |
| 0.4 | 0 | ycaM | 2 | 24.1667 | 0.56 |
| 0.4 | 0 | yadI | 2 | 24.1667 | 0.919 |
| 0.4 | 0 | AG1 | 2 | 24.1667 | 0.854 |
| 0.4 | 0 | ptsI | 2 | 24.4167 | 0.884 |
| 0.4 | 0 | clcB | 2 | 24.4167 | 0.517 |
| 0.4 | 0 | ycaM | 2 | 24.4167 | 0.563 |
| 0.4 | 0 | yadI | 2 | 24.4167 | 0.92 |
| 0.4 | 0 | AG1 | 2 | 24.4167 | 0.854 |
| 0.4 | 0 | ptsI | 2 | 24.6667 | 0.893 |
| 0.4 | 0 | clcB | 2 | 24.6667 | 0.52 |
| 0.4 | 0 | ycaM | 2 | 24.6667 | 0.564 |
| 0.4 | 0 | yadI | 2 | 24.6667 | 0.924 |
| 0.4 | 0 | AG1 | 2 | 24.6667 | 0.849 |
| 0.4 | 0 | ptsI | 2 | 24.9167 | 0.896 |

| | | | | | |
|---|---|---|---|---|---|
| 0.4 | 0 | clcB | 2 | 24.9167 | 0.523 |
| 0.4 | 0 | ycaM | 2 | 24.9167 | 0.563 |
| 0.4 | 0 | yadI | 2 | 24.9167 | 0.922 |
| 0.4 | 0 | AG1 | 2 | 24.9167 | 0.835 |
| 0.4 | 0 | ptsI | 2 | 25.1667 | 0.896 |
| 0.4 | 0 | clcB | 2 | 25.1667 | 0.521 |
| 0.4 | 0 | ycaM | 2 | 25.1667 | 0.565 |
| 0.4 | 0 | yadI | 2 | 25.1667 | 0.927 |
| 0.4 | 0 | AG1 | 2 | 25.1667 | 0.825 |
| 0.4 | 0 | ptsI | 2 | 25.4167 | 0.906 |
| 0.4 | 0 | clcB | 2 | 25.4167 | 0.524 |
| 0.4 | 0 | ycaM | 2 | 25.4167 | 0.567 |
| 0.4 | 0 | yadI | 2 | 25.4167 | 0.925 |
| 0.4 | 0 | AG1 | 2 | 25.4167 | 0.839 |
| 0.4 | 0 | ptsI | 2 | 25.6667 | 0.909 |
| 0.4 | 0 | clcB | 2 | 25.6667 | 0.527 |
| 0.4 | 0 | ycaM | 2 | 25.6667 | 0.565 |
| 0.4 | 0 | yadI | 2 | 25.6667 | 0.925 |
| 0.4 | 0 | AG1 | 2 | 25.6667 | 0.826 |
| 0.4 | 0 | ptsI | 2 | 25.9167 | 0.914 |
| 0.4 | 0 | clcB | 2 | 25.9167 | 0.529 |
| 0.4 | 0 | ycaM | 2 | 25.9167 | 0.569 |
| 0.4 | 0 | yadI | 2 | 25.9167 | 0.927 |
| 0.4 | 0 | AG1 | 2 | 25.9167 | 0.827 |
| 0.4 | 0 | ptsI | 2 | 26.1667 | 0.92 |
| 0.4 | 0 | clcB | 2 | 26.1667 | 0.527 |
| 0.4 | 0 | ycaM | 2 | 26.1667 | 0.571 |
| 0.4 | 0 | yadI | 2 | 26.1667 | 0.917 |
| 0.4 | 0 | AG1 | 2 | 26.1667 | 0.84 |
| 0.4 | 0 | ptsI | 2 | 26.4167 | 0.933 |
| 0.4 | 0 | clcB | 2 | 26.4167 | 0.533 |
| 0.4 | 0 | ycaM | 2 | 26.4167 | 0.572 |
| 0.4 | 0 | yadI | 2 | 26.4167 | 0.931 |
| 0.4 | 0 | AG1 | 2 | 26.4167 | 0.829 |
| 0.4 | 0 | ptsI | 3 | 0 | 0.149 |
| 0.4 | 0 | clcB | 3 | 0 | 0.157 |
| 0.4 | 0 | ycaM | 3 | 0 | 0.157 |
| 0.4 | 0 | yadI | 3 | 0 | 0.352 |
| 0.4 | 0 | AG1 | 3 | 0 | 0.168 |
| 0.4 | 0 | ptsI | 3 | 0.25 | 0.147 |
| 0.4 | 0 | clcB | 3 | 0.25 | 0.158 |
| 0.4 | 0 | ycaM | 3 | 0.25 | 0.149 |
| 0.4 | 0 | yadI | 3 | 0.25 | 0.341 |
| 0.4 | 0 | AG1 | 3 | 0.25 | 0.163 |
| 0.4 | 0 | ptsI | 3 | 0.5 | 0.147 |
| 0.4 | 0 | clcB | 3 | 0.5 | 0.158 |
| 0.4 | 0 | ycaM | 3 | 0.5 | 0.148 |
| 0.4 | 0 | yadI | 3 | 0.5 | 0.381 |

| | | | | | |
|---|---|---|---|---|---|
| 0.4 | 0 | AG1 | 3 | 0.5 | 0.161 |
| 0.4 | 0 | ptsI | 3 | 0.75 | 0.149 |
| 0.4 | 0 | clcB | 3 | 0.75 | 0.158 |
| 0.4 | 0 | ycaM | 3 | 0.75 | 0.149 |
| 0.4 | 0 | yadI | 3 | 0.75 | 0.359 |
| 0.4 | 0 | AG1 | 3 | 0.75 | 0.161 |
| 0.4 | 0 | ptsI | 3 | 1 | 0.15 |
| 0.4 | 0 | clcB | 3 | 1 | 0.157 |
| 0.4 | 0 | ycaM | 3 | 1 | 0.148 |
| 0.4 | 0 | yadI | 3 | 1 | 0.354 |
| 0.4 | 0 | AG1 | 3 | 1 | 0.163 |
| 0.4 | 0 | ptsI | 3 | 1.25 | 0.15 |
| 0.4 | 0 | clcB | 3 | 1.25 | 0.157 |
| 0.4 | 0 | ycaM | 3 | 1.25 | 0.148 |
| 0.4 | 0 | yadI | 3 | 1.25 | 0.371 |
| 0.4 | 0 | AG1 | 3 | 1.25 | 0.162 |
| 0.4 | 0 | ptsI | 3 | 1.5 | 0.152 |
| 0.4 | 0 | clcB | 3 | 1.5 | 0.158 |
| 0.4 | 0 | ycaM | 3 | 1.5 | 0.149 |
| 0.4 | 0 | yadI | 3 | 1.5 | 0.358 |
| 0.4 | 0 | AG1 | 3 | 1.5 | 0.165 |
| 0.4 | 0 | ptsI | 3 | 1.75 | 0.153 |
| 0.4 | 0 | clcB | 3 | 1.75 | 0.158 |
| 0.4 | 0 | ycaM | 3 | 1.75 | 0.149 |
| 0.4 | 0 | yadI | 3 | 1.75 | 0.377 |
| 0.4 | 0 | AG1 | 3 | 1.75 | 0.164 |
| 0.4 | 0 | ptsI | 3 | 2 | 0.155 |
| 0.4 | 0 | clcB | 3 | 2 | 0.159 |
| 0.4 | 0 | ycaM | 3 | 2 | 0.151 |
| 0.4 | 0 | yadI | 3 | 2 | 0.354 |
| 0.4 | 0 | AG1 | 3 | 2 | 0.167 |
| 0.4 | 0 | ptsI | 3 | 2.25 | 0.159 |
| 0.4 | 0 | clcB | 3 | 2.25 | 0.16 |
| 0.4 | 0 | ycaM | 3 | 2.25 | 0.152 |
| 0.4 | 0 | yadI | 3 | 2.25 | 0.343 |
| 0.4 | 0 | AG1 | 3 | 2.25 | 0.169 |
| 0.4 | 0 | ptsI | 3 | 2.5 | 0.162 |
| 0.4 | 0 | clcB | 3 | 2.5 | 0.161 |
| 0.4 | 0 | ycaM | 3 | 2.5 | 0.153 |
| 0.4 | 0 | yadI | 3 | 2.5 | 0.364 |
| 0.4 | 0 | AG1 | 3 | 2.5 | 0.171 |
| 0.4 | 0 | ptsI | 3 | 2.75 | 0.166 |
| 0.4 | 0 | clcB | 3 | 2.75 | 0.162 |
| 0.4 | 0 | ycaM | 3 | 2.75 | 0.158 |
| 0.4 | 0 | yadI | 3 | 2.75 | 0.363 |
| 0.4 | 0 | AG1 | 3 | 2.75 | 0.179 |
| 0.4 | 0 | ptsI | 3 | 3 | 0.168 |
| 0.4 | 0 | clcB | 3 | 3 | 0.162 |

| | | | | |
|---|---|---|---|---|
| 0.4 | 0 | ycaM | 3 | 3 | 0.157 |
| 0.4 | 0 | yadI | 3 | 3 | 0.364 |
| 0.4 | 0 | AG1 | 3 | 3 | 0.175 |
| 0.4 | 0 | ptsI | 3 | 3.25 | 0.171 |
| 0.4 | 0 | clcB | 3 | 3.25 | 0.164 |
| 0.4 | 0 | ycaM | 3 | 3.25 | 0.158 |
| 0.4 | 0 | yadI | 3 | 3.25 | 0.374 |
| 0.4 | 0 | AG1 | 3 | 3.25 | 0.179 |
| 0.4 | 0 | ptsI | 3 | 3.5 | 0.175 |
| 0.4 | 0 | clcB | 3 | 3.5 | 0.165 |
| 0.4 | 0 | ycaM | 3 | 3.5 | 0.163 |
| 0.4 | 0 | yadI | 3 | 3.5 | 0.365 |
| 0.4 | 0 | AG1 | 3 | 3.5 | 0.186 |
| 0.4 | 0 | ptsI | 3 | 3.75 | 0.18 |
| 0.4 | 0 | clcB | 3 | 3.75 | 0.166 |
| 0.4 | 0 | ycaM | 3 | 3.75 | 0.166 |
| 0.4 | 0 | yadI | 3 | 3.75 | 0.353 |
| 0.4 | 0 | AG1 | 3 | 3.75 | 0.189 |
| 0.4 | 0 | ptsI | 3 | 4 | 0.182 |
| 0.4 | 0 | clcB | 3 | 4 | 0.169 |
| 0.4 | 0 | ycaM | 3 | 4 | 0.169 |
| 0.4 | 0 | yadI | 3 | 4 | 0.365 |
| 0.4 | 0 | AG1 | 3 | 4 | 0.195 |
| 0.4 | 0 | ptsI | 3 | 4.41667 | 0.189 |
| 0.4 | 0 | clcB | 3 | 4.41667 | 0.171 |
| 0.4 | 0 | ycaM | 3 | 4.41667 | 0.188 |
| 0.4 | 0 | yadI | 3 | 4.41667 | 0.606 |
| 0.4 | 0 | AG1 | 3 | 4.41667 | 0.214 |
| 0.4 | 0 | ptsI | 3 | 4.66667 | 0.186 |
| 0.4 | 0 | clcB | 3 | 4.66667 | 0.171 |
| 0.4 | 0 | ycaM | 3 | 4.66667 | 0.184 |
| 0.4 | 0 | yadI | 3 | 4.66667 | 0.526 |
| 0.4 | 0 | AG1 | 3 | 4.66667 | 0.207 |
| 0.4 | 0 | ptsI | 3 | 4.91667 | 0.186 |
| 0.4 | 0 | clcB | 3 | 4.91667 | 0.17 |
| 0.4 | 0 | ycaM | 3 | 4.91667 | 0.184 |
| 0.4 | 0 | yadI | 3 | 4.91667 | 0.399 |
| 0.4 | 0 | AG1 | 3 | 4.91667 | 0.211 |
| 0.4 | 0 | ptsI | 3 | 5.16667 | 0.189 |
| 0.4 | 0 | clcB | 3 | 5.16667 | 0.172 |
| 0.4 | 0 | ycaM | 3 | 5.16667 | 0.188 |
| 0.4 | 0 | yadI | 3 | 5.16667 | 0.422 |
| 0.4 | 0 | AG1 | 3 | 5.16667 | 0.216 |
| 0.4 | 0 | ptsI | 3 | 5.41667 | 0.193 |
| 0.4 | 0 | clcB | 3 | 5.41667 | 0.174 |
| 0.4 | 0 | ycaM | 3 | 5.41667 | 0.193 |
| 0.4 | 0 | yadI | 3 | 5.41667 | 0.433 |
| 0.4 | 0 | AG1 | 3 | 5.41667 | 0.227 |

| | | | | | |
|---|---|---|---|---|---|
| 0.4 | 0 | ptsI | 3 | 5.66667 | 0.198 |
| 0.4 | 0 | clcB | 3 | 5.66667 | 0.178 |
| 0.4 | 0 | ycaM | 3 | 5.66667 | 0.201 |
| 0.4 | 0 | yadI | 3 | 5.66667 | 0.45 |
| 0.4 | 0 | AG1 | 3 | 5.66667 | 0.236 |
| 0.4 | 0 | ptsI | 3 | 5.91667 | 0.203 |
| 0.4 | 0 | clcB | 3 | 5.91667 | 0.18 |
| 0.4 | 0 | ycaM | 3 | 5.91667 | 0.208 |
| 0.4 | 0 | yadI | 3 | 5.91667 | 0.551 |
| 0.4 | 0 | AG1 | 3 | 5.91667 | 0.248 |
| 0.4 | 0 | ptsI | 3 | 6.16667 | 0.209 |
| 0.4 | 0 | clcB | 3 | 6.16667 | 0.186 |
| 0.4 | 0 | ycaM | 3 | 6.16667 | 0.216 |
| 0.4 | 0 | yadI | 3 | 6.16667 | 0.538 |
| 0.4 | 0 | AG1 | 3 | 6.16667 | 0.259 |
| 0.4 | 0 | ptsI | 3 | 6.41667 | 0.216 |
| 0.4 | 0 | clcB | 3 | 6.41667 | 0.191 |
| 0.4 | 0 | ycaM | 3 | 6.41667 | 0.227 |
| 0.4 | 0 | yadI | 3 | 6.41667 | 0.54 |
| 0.4 | 0 | AG1 | 3 | 6.41667 | 0.272 |
| 0.4 | 0 | ptsI | 3 | 6.66667 | 0.223 |
| 0.4 | 0 | clcB | 3 | 6.66667 | 0.195 |
| 0.4 | 0 | ycaM | 3 | 6.66667 | 0.238 |
| 0.4 | 0 | yadI | 3 | 6.66667 | 0.545 |
| 0.4 | 0 | AG1 | 3 | 6.66667 | 0.286 |
| 0.4 | 0 | ptsI | 3 | 6.91667 | 0.232 |
| 0.4 | 0 | clcB | 3 | 6.91667 | 0.202 |
| 0.4 | 0 | ycaM | 3 | 6.91667 | 0.252 |
| 0.4 | 0 | yadI | 3 | 6.91667 | 0.565 |
| 0.4 | 0 | AG1 | 3 | 6.91667 | 0.299 |
| 0.4 | 0 | ptsI | 3 | 7.16667 | 0.241 |
| 0.4 | 0 | clcB | 3 | 7.16667 | 0.211 |
| 0.4 | 0 | ycaM | 3 | 7.16667 | 0.265 |
| 0.4 | 0 | yadI | 3 | 7.16667 | 0.547 |
| 0.4 | 0 | AG1 | 3 | 7.16667 | 0.31 |
| 0.4 | 0 | ptsI | 3 | 7.41667 | 0.253 |
| 0.4 | 0 | clcB | 3 | 7.41667 | 0.218 |
| 0.4 | 0 | ycaM | 3 | 7.41667 | 0.277 |
| 0.4 | 0 | yadI | 3 | 7.41667 | 0.563 |
| 0.4 | 0 | AG1 | 3 | 7.41667 | 0.317 |
| 0.4 | 0 | ptsI | 3 | 7.66667 | 0.263 |
| 0.4 | 0 | clcB | 3 | 7.66667 | 0.228 |
| 0.4 | 0 | ycaM | 3 | 7.66667 | 0.285 |
| 0.4 | 0 | yadI | 3 | 7.66667 | 0.558 |
| 0.4 | 0 | AG1 | 3 | 7.66667 | 0.328 |
| 0.4 | 0 | ptsI | 3 | 7.91667 | 0.27 |
| 0.4 | 0 | clcB | 3 | 7.91667 | 0.236 |
| 0.4 | 0 | ycaM | 3 | 7.91667 | 0.292 |

| | | | | | |
|---|---|---|---|---|---|
| 0.4 | 0 | yadI | 3 | 7.91667 | 0.571 |
| 0.4 | 0 | AG1 | 3 | 7.91667 | 0.334 |
| 0.4 | 0 | ptsI | 3 | 8.16667 | 0.279 |
| 0.4 | 0 | clcB | 3 | 8.16667 | 0.241 |
| 0.4 | 0 | ycaM | 3 | 8.16667 | 0.302 |
| 0.4 | 0 | yadI | 3 | 8.16667 | 0.572 |
| 0.4 | 0 | AG1 | 3 | 8.16667 | 0.343 |
| 0.4 | 0 | ptsI | 3 | 8.41667 | 0.286 |
| 0.4 | 0 | clcB | 3 | 8.41667 | 0.247 |
| 0.4 | 0 | ycaM | 3 | 8.41667 | 0.31 |
| 0.4 | 0 | yadI | 3 | 8.41667 | 0.609 |
| 0.4 | 0 | AG1 | 3 | 8.41667 | 0.353 |
| 0.4 | 0 | ptsI | 3 | 9.05 | 0.26 |
| 0.4 | 0 | clcB | 3 | 9.05 | 0.278 |
| 0.4 | 0 | ycaM | 3 | 9.05 | 0.593 |
| 0.4 | 0 | yadI | 3 | 9.05 | 1.3 |
| 0.4 | 0 | AG1 | 3 | 9.05 | 0.373 |
| 0.4 | 0 | ptsI | 3 | 9.3 | 0.265 |
| 0.4 | 0 | clcB | 3 | 9.3 | 0.285 |
| 0.4 | 0 | ycaM | 3 | 9.3 | 0.527 |
| 0.4 | 0 | yadI | 3 | 9.3 | 1.255 |
| 0.4 | 0 | AG1 | 3 | 9.3 | 0.372 |
| 0.4 | 0 | ptsI | 3 | 9.55 | 0.272 |
| 0.4 | 0 | clcB | 3 | 9.55 | 0.29 |
| 0.4 | 0 | ycaM | 3 | 9.55 | 0.488 |
| 0.4 | 0 | yadI | 3 | 9.55 | 1.237 |
| 0.4 | 0 | AG1 | 3 | 9.55 | 0.381 |
| 0.4 | 0 | ptsI | 3 | 9.8 | 0.278 |
| 0.4 | 0 | clcB | 3 | 9.8 | 0.294 |
| 0.4 | 0 | ycaM | 3 | 9.8 | 0.475 |
| 0.4 | 0 | yadI | 3 | 9.8 | 1.234 |
| 0.4 | 0 | AG1 | 3 | 9.8 | 0.393 |
| 0.4 | 0 | ptsI | 3 | 10.05 | 0.286 |
| 0.4 | 0 | clcB | 3 | 10.05 | 0.298 |
| 0.4 | 0 | ycaM | 3 | 10.05 | 0.472 |
| 0.4 | 0 | yadI | 3 | 10.05 | 1.248 |
| 0.4 | 0 | AG1 | 3 | 10.05 | 0.404 |
| 0.4 | 0 | ptsI | 3 | 10.3 | 0.294 |
| 0.4 | 0 | clcB | 3 | 10.3 | 0.306 |
| 0.4 | 0 | ycaM | 3 | 10.3 | 0.47 |
| 0.4 | 0 | yadI | 3 | 10.3 | 1.249 |
| 0.4 | 0 | AG1 | 3 | 10.3 | 0.413 |
| 0.4 | 0 | ptsI | 3 | 10.55 | 0.298 |
| 0.4 | 0 | clcB | 3 | 10.55 | 0.311 |
| 0.4 | 0 | ycaM | 3 | 10.55 | 0.455 |
| 0.4 | 0 | yadI | 3 | 10.55 | 1.262 |
| 0.4 | 0 | AG1 | 3 | 10.55 | 0.421 |
| 0.4 | 0 | ptsI | 3 | 10.8 | 0.309 |

| | | | | | |
|---|---|---|---|---|---|
| 0.4 | 0 | clcB | 3 | 10.8 | 0.317 |
| 0.4 | 0 | ycaM | 3 | 10.8 | 0.444 |
| 0.4 | 0 | yadI | 3 | 10.8 | 1.26 |
| 0.4 | 0 | AG1 | 3 | 10.8 | 0.429 |
| 0.4 | 0 | ptsI | 3 | 11.05 | 0.317 |
| 0.4 | 0 | clcB | 3 | 11.05 | 0.321 |
| 0.4 | 0 | ycaM | 3 | 11.05 | 0.427 |
| 0.4 | 0 | yadI | 3 | 11.05 | 1.334 |
| 0.4 | 0 | AG1 | 3 | 11.05 | 0.441 |
| 0.4 | 0 | ptsI | 3 | 11.3 | 0.324 |
| 0.4 | 0 | clcB | 3 | 11.3 | 0.328 |
| 0.4 | 0 | ycaM | 3 | 11.3 | 0.424 |
| 0.4 | 0 | yadI | 3 | 11.3 | 1.311 |
| 0.4 | 0 | AG1 | 3 | 11.3 | 0.45 |
| 0.4 | 0 | ptsI | 3 | 11.55 | 0.331 |
| 0.4 | 0 | clcB | 3 | 11.55 | 0.337 |
| 0.4 | 0 | ycaM | 3 | 11.55 | 0.42 |
| 0.4 | 0 | yadI | 3 | 11.55 | 1.323 |
| 0.4 | 0 | AG1 | 3 | 11.55 | 0.46 |
| 0.4 | 0 | ptsI | 3 | 11.8 | 0.339 |
| 0.4 | 0 | clcB | 3 | 11.8 | 0.34 |
| 0.4 | 0 | ycaM | 3 | 11.8 | 0.412 |
| 0.4 | 0 | yadI | 3 | 11.8 | 1.451 |
| 0.4 | 0 | AG1 | 3 | 11.8 | 0.471 |
| 0.4 | 0 | ptsI | 3 | 12.05 | 0.346 |
| 0.4 | 0 | clcB | 3 | 12.05 | 0.35 |
| 0.4 | 0 | ycaM | 3 | 12.05 | 0.403 |
| 0.4 | 0 | yadI | 3 | 12.05 | 1.368 |
| 0.4 | 0 | AG1 | 3 | 12.05 | 0.48 |
| 0.4 | 0 | ptsI | 3 | 12.3 | 0.356 |
| 0.4 | 0 | clcB | 3 | 12.3 | 0.352 |
| 0.4 | 0 | ycaM | 3 | 12.3 | 0.404 |
| 0.4 | 0 | yadI | 3 | 12.3 | 1.4 |
| 0.4 | 0 | AG1 | 3 | 12.3 | 0.496 |
| 0.4 | 0 | ptsI | 3 | 12.55 | 0.363 |
| 0.4 | 0 | clcB | 3 | 12.55 | 0.36 |
| 0.4 | 0 | ycaM | 3 | 12.55 | 0.405 |
| 0.4 | 0 | yadI | 3 | 12.55 | 1.416 |
| 0.4 | 0 | AG1 | 3 | 12.55 | 0.509 |
| 0.4 | 0 | ptsI | 3 | 12.8 | 0.371 |
| 0.4 | 0 | clcB | 3 | 12.8 | 0.364 |
| 0.4 | 0 | ycaM | 3 | 12.8 | 0.41 |
| 0.4 | 0 | yadI | 3 | 12.8 | 1.489 |
| 0.4 | 0 | AG1 | 3 | 12.8 | 0.524 |
| 0.4 | 0 | ptsI | 3 | 13.05 | 0.38 |
| 0.4 | 0 | clcB | 3 | 13.05 | 0.365 |
| 0.4 | 0 | ycaM | 3 | 13.05 | 0.407 |
| 0.4 | 0 | yadI | 3 | 13.05 | 1.456 |

| | | | | | |
|---|---|---|---|---|---|
| 0.4 | 0 | AG1 | 3 | 13.05 | 0.534 |
| 0.4 | 0 | ptsI | 3 | 13.3 | 0.389 |
| 0.4 | 0 | clcB | 3 | 13.3 | 0.363 |
| 0.4 | 0 | ycaM | 3 | 13.3 | 0.411 |
| 0.4 | 0 | yadI | 3 | 13.3 | 1.5 |
| 0.4 | 0 | AG1 | 3 | 13.3 | 0.55 |
| 0.4 | 0 | ptsI | 3 | 13.55 | 0.397 |
| 0.4 | 0 | clcB | 3 | 13.55 | 0.366 |
| 0.4 | 0 | ycaM | 3 | 13.55 | 0.422 |
| 0.4 | 0 | yadI | 3 | 13.55 | 1.532 |
| 0.4 | 0 | AG1 | 3 | 13.55 | 0.567 |
| 0.4 | 0 | ptsI | 3 | 13.8 | 0.404 |
| 0.4 | 0 | clcB | 3 | 13.8 | 0.372 |
| 0.4 | 0 | ycaM | 3 | 13.8 | 0.431 |
| 0.4 | 0 | yadI | 3 | 13.8 | 1.599 |
| 0.4 | 0 | AG1 | 3 | 13.8 | 0.579 |
| 0.4 | 0 | ptsI | 3 | 14.05 | 0.415 |
| 0.4 | 0 | clcB | 3 | 14.05 | 0.376 |
| 0.4 | 0 | ycaM | 3 | 14.05 | 0.435 |
| 0.4 | 0 | yadI | 3 | 14.05 | 1.574 |
| 0.4 | 0 | AG1 | 3 | 14.05 | 0.582 |
| 0.4 | 0 | ptsI | 3 | 14.3 | 0.419 |
| 0.4 | 0 | clcB | 3 | 14.3 | 0.384 |
| 0.4 | 0 | ycaM | 3 | 14.3 | 0.437 |
| 0.4 | 0 | yadI | 3 | 14.3 | 1.563 |
| 0.4 | 0 | AG1 | 3 | 14.3 | 0.589 |
| 0.4 | 0 | ptsI | 3 | 14.55 | 0.431 |
| 0.4 | 0 | clcB | 3 | 14.55 | 0.388 |
| 0.4 | 0 | ycaM | 3 | 14.55 | 0.442 |
| 0.4 | 0 | yadI | 3 | 14.55 | 1.564 |
| 0.4 | 0 | AG1 | 3 | 14.55 | 0.592 |
| 0.4 | 0 | ptsI | 3 | 14.8 | 0.442 |
| 0.4 | 0 | clcB | 3 | 14.8 | 0.392 |
| 0.4 | 0 | ycaM | 3 | 14.8 | 0.448 |
| 0.4 | 0 | yadI | 3 | 14.8 | 1.655 |
| 0.4 | 0 | AG1 | 3 | 14.8 | 0.604 |
| 0.4 | 0 | ptsI | 3 | 15.05 | 0.45 |
| 0.4 | 0 | clcB | 3 | 15.05 | 0.394 |
| 0.4 | 0 | ycaM | 3 | 15.05 | 0.459 |
| 0.4 | 0 | yadI | 3 | 15.05 | 1.698 |
| 0.4 | 0 | AG1 | 3 | 15.05 | 0.612 |
| 0.4 | 0 | ptsI | 3 | 15.3 | 0.461 |
| 0.4 | 0 | clcB | 3 | 15.3 | 0.398 |
| 0.4 | 0 | ycaM | 3 | 15.3 | 0.464 |
| 0.4 | 0 | yadI | 3 | 15.3 | 1.682 |
| 0.4 | 0 | AG1 | 3 | 15.3 | 0.622 |
| 0.4 | 0 | ptsI | 3 | 15.55 | 0.464 |
| 0.4 | 0 | clcB | 3 | 15.55 | 0.398 |

| | | | | | |
|---|---|---|---|---|---|
| 0.4 | 0 | ycaM | 3 | 15.55 | 0.469 |
| 0.4 | 0 | yadI | 3 | 15.55 | 1.716 |
| 0.4 | 0 | AG1 | 3 | 15.55 | 0.628 |
| 0.4 | 0 | ptsI | 3 | 15.8 | 0.476 |
| 0.4 | 0 | clcB | 3 | 15.8 | 0.399 |
| 0.4 | 0 | ycaM | 3 | 15.8 | 0.478 |
| 0.4 | 0 | yadI | 3 | 15.8 | 1.725 |
| 0.4 | 0 | AG1 | 3 | 15.8 | 0.63 |
| 0.4 | 0 | ptsI | 3 | 16.05 | 0.487 |
| 0.4 | 0 | clcB | 3 | 16.05 | 0.403 |
| 0.4 | 0 | ycaM | 3 | 16.05 | 0.483 |
| 0.4 | 0 | yadI | 3 | 16.05 | 1.704 |
| 0.4 | 0 | AG1 | 3 | 16.05 | 0.643 |
| 0.4 | 0 | ptsI | 3 | 16.3 | 0.496 |
| 0.4 | 0 | clcB | 3 | 16.3 | 0.406 |
| 0.4 | 0 | ycaM | 3 | 16.3 | 0.488 |
| 0.4 | 0 | yadI | 3 | 16.3 | 1.645 |
| 0.4 | 0 | AG1 | 3 | 16.3 | 0.644 |
| 0.4 | 0 | ptsI | 3 | 16.55 | 0.498 |
| 0.4 | 0 | clcB | 3 | 16.55 | 0.405 |
| 0.4 | 0 | ycaM | 3 | 16.55 | 0.496 |
| 0.4 | 0 | yadI | 3 | 16.55 | 1.734 |
| 0.4 | 0 | AG1 | 3 | 16.55 | 0.642 |
| 0.4 | 0 | ptsI | 3 | 16.8 | 0.503 |
| 0.4 | 0 | clcB | 3 | 16.8 | 0.406 |
| 0.4 | 0 | ycaM | 3 | 16.8 | 0.502 |
| 0.4 | 0 | yadI | 3 | 16.8 | 1.686 |
| 0.4 | 0 | AG1 | 3 | 16.8 | 0.655 |
| 0.4 | 0 | ptsI | 3 | 17.05 | 0.509 |
| 0.4 | 0 | clcB | 3 | 17.05 | 0.408 |
| 0.4 | 0 | ycaM | 3 | 17.05 | 0.507 |
| 0.4 | 0 | yadI | 3 | 17.05 | 1.694 |
| 0.4 | 0 | AG1 | 3 | 17.05 | 0.662 |
| 0.4 | 0 | ptsI | 3 | 17.3 | 0.519 |
| 0.4 | 0 | clcB | 3 | 17.3 | 0.412 |
| 0.4 | 0 | ycaM | 3 | 17.3 | 0.511 |
| 0.4 | 0 | yadI | 3 | 17.3 | 1.691 |
| 0.4 | 0 | AG1 | 3 | 17.3 | 0.659 |
| 0.4 | 0 | ptsI | 3 | 17.55 | 0.533 |
| 0.4 | 0 | clcB | 3 | 17.55 | 0.411 |
| 0.4 | 0 | ycaM | 3 | 17.55 | 0.513 |
| 0.4 | 0 | yadI | 3 | 17.55 | 1.672 |
| 0.4 | 0 | AG1 | 3 | 17.55 | 0.66 |
| 0.4 | 0 | ptsI | 3 | 17.8 | 0.532 |
| 0.4 | 0 | clcB | 3 | 17.8 | 0.409 |
| 0.4 | 0 | ycaM | 3 | 17.8 | 0.519 |
| 0.4 | 0 | yadI | 3 | 17.8 | 1.672 |
| 0.4 | 0 | AG1 | 3 | 17.8 | 0.666 |

| | | | | | |
|---|---|---|---|---|---|
| 0.4 | 0 | ptsI | 3 | 18.05 | 0.539 |
| 0.4 | 0 | clcB | 3 | 18.05 | 0.41 |
| 0.4 | 0 | ycaM | 3 | 18.05 | 0.524 |
| 0.4 | 0 | yadI | 3 | 18.05 | 1.653 |
| 0.4 | 0 | AG1 | 3 | 18.05 | 0.67 |
| 0.4 | 0 | ptsI | 3 | 18.3 | 0.546 |
| 0.4 | 0 | clcB | 3 | 18.3 | 0.408 |
| 0.4 | 0 | ycaM | 3 | 18.3 | 0.526 |
| 0.4 | 0 | yadI | 3 | 18.3 | 1.627 |
| 0.4 | 0 | AG1 | 3 | 18.3 | 0.664 |
| 0.4 | 0 | ptsI | 3 | 18.55 | 0.552 |
| 0.4 | 0 | clcB | 3 | 18.55 | 0.406 |
| 0.4 | 0 | ycaM | 3 | 18.55 | 0.533 |
| 0.4 | 0 | yadI | 3 | 18.55 | 1.601 |
| 0.4 | 0 | AG1 | 3 | 18.55 | 0.675 |
| 0.4 | 0 | ptsI | 3 | 18.8 | 0.559 |
| 0.4 | 0 | clcB | 3 | 18.8 | 0.41 |
| 0.4 | 0 | ycaM | 3 | 18.8 | 0.538 |
| 0.4 | 0 | yadI | 3 | 18.8 | 1.589 |
| 0.4 | 0 | AG1 | 3 | 18.8 | 0.686 |
| 0.4 | 0 | ptsI | 3 | 19.05 | 0.57 |
| 0.4 | 0 | clcB | 3 | 19.05 | 0.403 |
| 0.4 | 0 | ycaM | 3 | 19.05 | 0.544 |
| 0.4 | 0 | yadI | 3 | 19.05 | 1.426 |
| 0.4 | 0 | AG1 | 3 | 19.05 | 0.691 |
| 0.4 | 0 | ptsI | 3 | 19.3 | 0.585 |
| 0.4 | 0 | clcB | 3 | 19.3 | 0.408 |
| 0.4 | 0 | ycaM | 3 | 19.3 | 0.55 |
| 0.4 | 0 | yadI | 3 | 19.3 | 1.356 |
| 0.4 | 0 | AG1 | 3 | 19.3 | 0.693 |
| 0.4 | 0 | ptsI | 3 | 19.55 | 0.6 |
| 0.4 | 0 | clcB | 3 | 19.55 | 0.406 |
| 0.4 | 0 | ycaM | 3 | 19.55 | 0.551 |
| 0.4 | 0 | yadI | 3 | 19.55 | 1.359 |
| 0.4 | 0 | AG1 | 3 | 19.55 | 0.694 |
| 0.4 | 0 | ptsI | 3 | 19.8 | 0.597 |
| 0.4 | 0 | clcB | 3 | 19.8 | 0.405 |
| 0.4 | 0 | ycaM | 3 | 19.8 | 0.558 |
| 0.4 | 0 | yadI | 3 | 19.8 | 1.346 |
| 0.4 | 0 | AG1 | 3 | 19.8 | 0.694 |
| 0.4 | 0 | ptsI | 3 | 20.05 | 0.597 |
| 0.4 | 0 | clcB | 3 | 20.05 | 0.404 |
| 0.4 | 0 | ycaM | 3 | 20.05 | 0.56 |
| 0.4 | 0 | yadI | 3 | 20.05 | 1.339 |
| 0.4 | 0 | AG1 | 3 | 20.05 | 0.692 |
| 0.4 | 0 | ptsI | 3 | 20.3 | 0.596 |
| 0.4 | 0 | clcB | 3 | 20.3 | 0.406 |
| 0.4 | 0 | ycaM | 3 | 20.3 | 0.566 |

| | | | | | |
|---|---|---|---|---|---|
| 0.4 | 0 | yadI | 3 | 20.3 | 1.215 |
| 0.4 | 0 | AG1 | 3 | 20.3 | 0.694 |
| 0.4 | 0 | ptsI | 3 | 20.55 | 0.609 |
| 0.4 | 0 | clcB | 3 | 20.55 | 0.402 |
| 0.4 | 0 | ycaM | 3 | 20.55 | 0.573 |
| 0.4 | 0 | yadI | 3 | 20.55 | 1.223 |
| 0.4 | 0 | AG1 | 3 | 20.55 | 0.699 |
| 0.4 | 0 | ptsI | 3 | 20.8 | 0.611 |
| 0.4 | 0 | clcB | 3 | 20.8 | 0.404 |
| 0.4 | 0 | ycaM | 3 | 20.8 | 0.574 |
| 0.4 | 0 | yadI | 3 | 20.8 | 1.135 |
| 0.4 | 0 | AG1 | 3 | 20.8 | 0.701 |
| 0.4 | 0 | ptsI | 3 | 21.05 | 0.611 |
| 0.4 | 0 | clcB | 3 | 21.05 | 0.405 |
| 0.4 | 0 | ycaM | 3 | 21.05 | 0.578 |
| 0.4 | 0 | yadI | 3 | 21.05 | 1.101 |
| 0.4 | 0 | AG1 | 3 | 21.05 | 0.687 |
| 0.4 | 0 | ptsI | 3 | 21.3 | 0.607 |
| 0.4 | 0 | clcB | 3 | 21.3 | 0.395 |
| 0.4 | 0 | ycaM | 3 | 21.3 | 0.581 |
| 0.4 | 0 | yadI | 3 | 21.3 | 1.103 |
| 0.4 | 0 | AG1 | 3 | 21.3 | 0.686 |
| 0.4 | 0 | ptsI | 3 | 21.55 | 0.614 |
| 0.4 | 0 | clcB | 3 | 21.55 | 0.394 |
| 0.4 | 0 | ycaM | 3 | 21.55 | 0.586 |
| 0.4 | 0 | yadI | 3 | 21.55 | 1.116 |
| 0.4 | 0 | AG1 | 3 | 21.55 | 0.689 |
| 0.4 | 0 | ptsI | 3 | 21.8 | 0.616 |
| 0.4 | 0 | clcB | 3 | 21.8 | 0.394 |
| 0.4 | 0 | ycaM | 3 | 21.8 | 0.588 |
| 0.4 | 0 | yadI | 3 | 21.8 | 1.102 |
| 0.4 | 0 | AG1 | 3 | 21.8 | 0.693 |
| 0.4 | 0 | ptsI | 3 | 22.05 | 0.618 |
| 0.4 | 0 | clcB | 3 | 22.05 | 0.396 |
| 0.4 | 0 | ycaM | 3 | 22.05 | 0.592 |
| 0.4 | 0 | yadI | 3 | 22.05 | 1.044 |
| 0.4 | 0 | AG1 | 3 | 22.05 | 0.695 |
| 0.4 | 0 | ptsI | 3 | 22.3 | 0.621 |
| 0.4 | 0 | clcB | 3 | 22.3 | 0.395 |
| 0.4 | 0 | ycaM | 3 | 22.3 | 0.596 |
| 0.4 | 0 | yadI | 3 | 22.3 | 1.04 |
| 0.4 | 0 | AG1 | 3 | 22.3 | 0.697 |
| 0.4 | 0 | ptsI | 3 | 22.55 | 0.634 |
| 0.4 | 0 | clcB | 3 | 22.55 | 0.395 |
| 0.4 | 0 | ycaM | 3 | 22.55 | 0.6 |
| 0.4 | 0 | yadI | 3 | 22.55 | 1.002 |
| 0.4 | 0 | AG1 | 3 | 22.55 | 0.698 |
| 0.4 | 0 | ptsI | 3 | 22.8 | 0.643 |

| | | | | | |
|---|---|---|---|---|---|
| 0.4 | 0 | clcB | 3 | 22.8 | 0.394 |
| 0.4 | 0 | ycaM | 3 | 22.8 | 0.6 |
| 0.4 | 0 | yadI | 3 | 22.8 | 1.01 |
| 0.4 | 0 | AG1 | 3 | 22.8 | 0.691 |
| 0.4 | 0 | ptsI | 3 | 23.05 | 0.651 |
| 0.4 | 0 | clcB | 3 | 23.05 | 0.39 |
| 0.4 | 0 | ycaM | 3 | 23.05 | 0.603 |
| 0.4 | 0 | yadI | 3 | 23.05 | 1 |
| 0.4 | 0 | AG1 | 3 | 23.05 | 0.694 |
| 0.4 | 0 | ptsI | 3 | 23.3 | 0.657 |
| 0.4 | 0 | clcB | 3 | 23.3 | 0.39 |
| 0.4 | 0 | ycaM | 3 | 23.3 | 0.608 |
| 0.4 | 0 | yadI | 3 | 23.3 | 0.982 |
| 0.4 | 0 | AG1 | 3 | 23.3 | 0.692 |
| 0.4 | 0 | ptsI | 3 | 23.55 | 0.665 |
| 0.4 | 0 | clcB | 3 | 23.55 | 0.394 |
| 0.4 | 0 | ycaM | 3 | 23.55 | 0.606 |
| 0.4 | 0 | yadI | 3 | 23.55 | 0.91 |
| 0.4 | 0 | AG1 | 3 | 23.55 | 0.695 |
| 0.4 | 0 | ptsI | 3 | 23.8 | 0.665 |
| 0.4 | 0 | clcB | 3 | 23.8 | 0.386 |
| 0.4 | 0 | ycaM | 3 | 23.8 | 0.614 |
| 0.4 | 0 | yadI | 3 | 23.8 | 0.891 |
| 0.4 | 0 | AG1 | 3 | 23.8 | 0.693 |
| 0.4 | 0 | ptsI | 3 | 24.05 | 0.672 |
| 0.4 | 0 | clcB | 3 | 24.05 | 0.381 |
| 0.4 | 0 | ycaM | 3 | 24.05 | 0.615 |
| 0.4 | 0 | yadI | 3 | 24.05 | 0.885 |
| 0.4 | 0 | AG1 | 3 | 24.05 | 0.692 |
| 0.4 | 0 | ptsI | 3 | 24.3 | 0.672 |
| 0.4 | 0 | clcB | 3 | 24.3 | 0.38 |
| 0.4 | 0 | ycaM | 3 | 24.3 | 0.622 |
| 0.4 | 0 | yadI | 3 | 24.3 | 0.763 |
| 0.4 | 0 | AG1 | 3 | 24.3 | 0.704 |
| 0.4 | 0 | ptsI | 4 | 0 | 0.183 |
| 0.4 | 0 | clcB | 4 | 0 | 0.197 |
| 0.4 | 0 | ycaM | 4 | 0 | 0.223 |
| 0.4 | 0 | yadI | 4 | 0 | 0.233 |
| 0.4 | 0 | AG1 | 4 | 0 | 0.22 |
| 0.4 | 0 | ptsI | 4 | 0.25 | 0.184 |
| 0.4 | 0 | clcB | 4 | 0.25 | 0.193 |
| 0.4 | 0 | ycaM | 4 | 0.25 | 0.216 |
| 0.4 | 0 | yadI | 4 | 0.25 | 0.227 |
| 0.4 | 0 | AG1 | 4 | 0.25 | 0.214 |
| 0.4 | 0 | ptsI | 4 | 0.5 | 0.184 |
| 0.4 | 0 | clcB | 4 | 0.5 | 0.193 |
| 0.4 | 0 | ycaM | 4 | 0.5 | 0.216 |
| 0.4 | 0 | yadI | 4 | 0.5 | 0.227 |

| | | | | | |
|---|---|---|---|---|---|
| 0.4 | 0 | AG1 | 4 | 0.5 | 0.217 |
| 0.4 | 0 | ptsI | 4 | 0.75 | 0.188 |
| 0.4 | 0 | clcB | 4 | 0.75 | 0.198 |
| 0.4 | 0 | ycaM | 4 | 0.75 | 0.217 |
| 0.4 | 0 | yadI | 4 | 0.75 | 0.228 |
| 0.4 | 0 | AG1 | 4 | 0.75 | 0.22 |
| 0.4 | 0 | ptsI | 4 | 1 | 0.192 |
| 0.4 | 0 | clcB | 4 | 1 | 0.202 |
| 0.4 | 0 | ycaM | 4 | 1 | 0.219 |
| 0.4 | 0 | yadI | 4 | 1 | 0.231 |
| 0.4 | 0 | AG1 | 4 | 1 | 0.222 |
| 0.4 | 0 | ptsI | 4 | 1.25 | 0.198 |
| 0.4 | 0 | clcB | 4 | 1.25 | 0.208 |
| 0.4 | 0 | ycaM | 4 | 1.25 | 0.225 |
| 0.4 | 0 | yadI | 4 | 1.25 | 0.235 |
| 0.4 | 0 | AG1 | 4 | 1.25 | 0.227 |
| 0.4 | 0 | ptsI | 4 | 1.5 | 0.198 |
| 0.4 | 0 | clcB | 4 | 1.5 | 0.209 |
| 0.4 | 0 | ycaM | 4 | 1.5 | 0.23 |
| 0.4 | 0 | yadI | 4 | 1.5 | 0.24 |
| 0.4 | 0 | AG1 | 4 | 1.5 | 0.231 |
| 0.4 | 0 | ptsI | 4 | 1.75 | 0.21 |
| 0.4 | 0 | clcB | 4 | 1.75 | 0.221 |
| 0.4 | 0 | ycaM | 4 | 1.75 | 0.238 |
| 0.4 | 0 | yadI | 4 | 1.75 | 0.25 |
| 0.4 | 0 | AG1 | 4 | 1.75 | 0.239 |
| 0.4 | 0 | ptsI | 4 | 2 | 0.223 |
| 0.4 | 0 | clcB | 4 | 2 | 0.231 |
| 0.4 | 0 | ycaM | 4 | 2 | 0.247 |
| 0.4 | 0 | yadI | 4 | 2 | 0.26 |
| 0.4 | 0 | AG1 | 4 | 2 | 0.248 |
| 0.4 | 0 | ptsI | 4 | 2.25 | 0.239 |
| 0.4 | 0 | clcB | 4 | 2.25 | 0.244 |
| 0.4 | 0 | ycaM | 4 | 2.25 | 0.259 |
| 0.4 | 0 | yadI | 4 | 2.25 | 0.271 |
| 0.4 | 0 | AG1 | 4 | 2.25 | 0.262 |
| 0.4 | 0 | ptsI | 4 | 2.5 | 0.246 |
| 0.4 | 0 | clcB | 4 | 2.5 | 0.247 |
| 0.4 | 0 | ycaM | 4 | 2.5 | 0.268 |
| 0.4 | 0 | yadI | 4 | 2.5 | 0.282 |
| 0.4 | 0 | AG1 | 4 | 2.5 | 0.267 |
| 0.4 | 0 | ptsI | 4 | 2.75 | 0.26 |
| 0.4 | 0 | clcB | 4 | 2.75 | 0.254 |
| 0.4 | 0 | ycaM | 4 | 2.75 | 0.28 |
| 0.4 | 0 | yadI | 4 | 2.75 | 0.293 |
| 0.4 | 0 | AG1 | 4 | 2.75 | 0.278 |
| 0.4 | 0 | ptsI | 4 | 3 | 0.273 |
| 0.4 | 0 | clcB | 4 | 3 | 0.276 |

| | | | | | |
|---|---|---|---|---|---|
| 0.4 | 0 | ycaM | 4 | 3 | 0.292 |
| 0.4 | 0 | yadI | 4 | 3 | 0.311 |
| 0.4 | 0 | AG1 | 4 | 3 | 0.29 |
| 0.4 | 0 | ptsI | 4 | 3.25 | 0.289 |
| 0.4 | 0 | clcB | 4 | 3.25 | 0.281 |
| 0.4 | 0 | ycaM | 4 | 3.25 | 0.306 |
| 0.4 | 0 | yadI | 4 | 3.25 | 0.329 |
| 0.4 | 0 | AG1 | 4 | 3.25 | 0.301 |
| 0.4 | 0 | ptsI | 4 | 3.5 | 0.305 |
| 0.4 | 0 | clcB | 4 | 3.5 | 0.301 |
| 0.4 | 0 | ycaM | 4 | 3.5 | 0.32 |
| 0.4 | 0 | yadI | 4 | 3.5 | 0.356 |
| 0.4 | 0 | AG1 | 4 | 3.5 | 0.32 |
| 0.4 | 0 | ptsI | 4 | 3.75 | 0.323 |
| 0.4 | 0 | clcB | 4 | 3.75 | 0.307 |
| 0.4 | 0 | ycaM | 4 | 3.75 | 0.335 |
| 0.4 | 0 | yadI | 4 | 3.75 | 0.374 |
| 0.4 | 0 | AG1 | 4 | 3.75 | 0.34 |
| 0.4 | 0 | ptsI | 4 | 4 | 0.338 |
| 0.4 | 0 | clcB | 4 | 4 | 0.329 |
| 0.4 | 0 | ycaM | 4 | 4 | 0.346 |
| 0.4 | 0 | yadI | 4 | 4 | 0.391 |
| 0.4 | 0 | AG1 | 4 | 4 | 0.352 |
| 0.4 | 0 | ptsI | 4 | 4.35 | 0.348 |
| 0.4 | 0 | clcB | 4 | 4.35 | 0.271 |
| 0.4 | 0 | ycaM | 4 | 4.35 | 0.392 |
| 0.4 | 0 | yadI | 4 | 4.35 | 0.402 |
| 0.4 | 0 | AG1 | 4 | 4.35 | 0.346 |
| 0.4 | 0 | ptsI | 4 | 4.6 | 0.355 |
| 0.4 | 0 | clcB | 4 | 4.6 | 0.28 |
| 0.4 | 0 | ycaM | 4 | 4.6 | 0.367 |
| 0.4 | 0 | yadI | 4 | 4.6 | 0.384 |
| 0.4 | 0 | AG1 | 4 | 4.6 | 0.35 |
| 0.4 | 0 | ptsI | 4 | 4.85 | 0.368 |
| 0.4 | 0 | clcB | 4 | 4.85 | 0.291 |
| 0.4 | 0 | ycaM | 4 | 4.85 | 0.378 |
| 0.4 | 0 | yadI | 4 | 4.85 | 0.396 |
| 0.4 | 0 | AG1 | 4 | 4.85 | 0.362 |
| 0.4 | 0 | ptsI | 4 | 5.1 | 0.384 |
| 0.4 | 0 | clcB | 4 | 5.1 | 0.3 |
| 0.4 | 0 | ycaM | 4 | 5.1 | 0.39 |
| 0.4 | 0 | yadI | 4 | 5.1 | 0.411 |
| 0.4 | 0 | AG1 | 4 | 5.1 | 0.374 |
| 0.4 | 0 | ptsI | 4 | 5.35 | 0.403 |
| 0.4 | 0 | clcB | 4 | 5.35 | 0.31 |
| 0.4 | 0 | ycaM | 4 | 5.35 | 0.406 |
| 0.4 | 0 | yadI | 4 | 5.35 | 0.426 |
| 0.4 | 0 | AG1 | 4 | 5.35 | 0.391 |

| | | | | | |
|---|---|---|---|---|---|
| 0.4 | 0 | ptsI | 4 | 5.6 | 0.416 |
| 0.4 | 0 | clcB | 4 | 5.6 | 0.323 |
| 0.4 | 0 | ycaM | 4 | 5.6 | 0.411 |
| 0.4 | 0 | yadI | 4 | 5.6 | 0.448 |
| 0.4 | 0 | AG1 | 4 | 5.6 | 0.403 |
| 0.4 | 0 | ptsI | 4 | 5.85 | 0.428 |
| 0.4 | 0 | clcB | 4 | 5.85 | 0.338 |
| 0.4 | 0 | ycaM | 4 | 5.85 | 0.419 |
| 0.4 | 0 | yadI | 4 | 5.85 | 0.454 |
| 0.4 | 0 | AG1 | 4 | 5.85 | 0.418 |
| 0.4 | 0 | ptsI | 4 | 6.1 | 0.447 |
| 0.4 | 0 | clcB | 4 | 6.1 | 0.353 |
| 0.4 | 0 | ycaM | 4 | 6.1 | 0.428 |
| 0.4 | 0 | yadI | 4 | 6.1 | 0.47 |
| 0.4 | 0 | AG1 | 4 | 6.1 | 0.431 |
| 0.4 | 0 | ptsI | 4 | 6.35 | 0.464 |
| 0.4 | 0 | clcB | 4 | 6.35 | 0.365 |
| 0.4 | 0 | ycaM | 4 | 6.35 | 0.437 |
| 0.4 | 0 | yadI | 4 | 6.35 | 0.485 |
| 0.4 | 0 | AG1 | 4 | 6.35 | 0.439 |
| 0.4 | 0 | ptsI | 4 | 6.6 | 0.477 |
| 0.4 | 0 | clcB | 4 | 6.6 | 0.374 |
| 0.4 | 0 | ycaM | 4 | 6.6 | 0.446 |
| 0.4 | 0 | yadI | 4 | 6.6 | 0.499 |
| 0.4 | 0 | AG1 | 4 | 6.6 | 0.452 |
| 0.4 | 0 | ptsI | 4 | 6.85 | 0.492 |
| 0.4 | 0 | clcB | 4 | 6.85 | 0.385 |
| 0.4 | 0 | ycaM | 4 | 6.85 | 0.451 |
| 0.4 | 0 | yadI | 4 | 6.85 | 0.512 |
| 0.4 | 0 | AG1 | 4 | 6.85 | 0.463 |
| 0.4 | 0 | ptsI | 4 | 7.1 | 0.503 |
| 0.4 | 0 | clcB | 4 | 7.1 | 0.393 |
| 0.4 | 0 | ycaM | 4 | 7.1 | 0.459 |
| 0.4 | 0 | yadI | 4 | 7.1 | 0.522 |
| 0.4 | 0 | AG1 | 4 | 7.1 | 0.477 |
| 0.4 | 0 | ptsI | 4 | 7.35 | 0.511 |
| 0.4 | 0 | clcB | 4 | 7.35 | 0.403 |
| 0.4 | 0 | ycaM | 4 | 7.35 | 0.465 |
| 0.4 | 0 | yadI | 4 | 7.35 | 0.529 |
| 0.4 | 0 | AG1 | 4 | 7.35 | 0.486 |
| 0.4 | 0 | ptsI | 4 | 7.6 | 0.52 |
| 0.4 | 0 | clcB | 4 | 7.6 | 0.412 |
| 0.4 | 0 | ycaM | 4 | 7.6 | 0.469 |
| 0.4 | 0 | yadI | 4 | 7.6 | 0.536 |
| 0.4 | 0 | AG1 | 4 | 7.6 | 0.496 |
| 0.4 | 0 | ptsI | 4 | 7.85 | 0.526 |
| 0.4 | 0 | clcB | 4 | 7.85 | 0.419 |
| 0.4 | 0 | ycaM | 4 | 7.85 | 0.471 |

| | | | | | |
|---|---|---|---|---|---|
| 0.4 | 0 | yadI | 4 | 7.85 | 0.54 |
| 0.4 | 0 | AG1 | 4 | 7.85 | 0.505 |
| 0.4 | 0 | ptsI | 4 | 8.1 | 0.537 |
| 0.4 | 0 | clcB | 4 | 8.1 | 0.43 |
| 0.4 | 0 | ycaM | 4 | 8.1 | 0.477 |
| 0.4 | 0 | yadI | 4 | 8.1 | 0.554 |
| 0.4 | 0 | AG1 | 4 | 8.1 | 0.515 |
| 0.4 | 0 | ptsI | 4 | 8.35 | 0.546 |
| 0.4 | 0 | clcB | 4 | 8.35 | 0.436 |
| 0.4 | 0 | ycaM | 4 | 8.35 | 0.48 |
| 0.4 | 0 | yadI | 4 | 8.35 | 0.564 |
| 0.4 | 0 | AG1 | 4 | 8.35 | 0.52 |
| 0.4 | 0 | ptsI | 4 | 8.83333 | 0.38 |
| 0.4 | 0 | clcB | 4 | 8.83333 | 0.375 |
| 0.4 | 0 | ycaM | 4 | 8.83333 | 0.411 |
| 0.4 | 0 | yadI | 4 | 8.83333 | 0.512 |
| 0.4 | 0 | AG1 | 4 | 8.83333 | 0.537 |
| 0.4 | 0 | ptsI | 4 | 9.08333 | 0.382 |
| 0.4 | 0 | clcB | 4 | 9.08333 | 0.375 |
| 0.4 | 0 | ycaM | 4 | 9.08333 | 0.412 |
| 0.4 | 0 | yadI | 4 | 9.08333 | 0.509 |
| 0.4 | 0 | AG1 | 4 | 9.08333 | 0.531 |
| 0.4 | 0 | ptsI | 4 | 9.33333 | 0.393 |
| 0.4 | 0 | clcB | 4 | 9.33333 | 0.38 |
| 0.4 | 0 | ycaM | 4 | 9.33333 | 0.414 |
| 0.4 | 0 | yadI | 4 | 9.33333 | 0.512 |
| 0.4 | 0 | AG1 | 4 | 9.33333 | 0.543 |
| 0.4 | 0 | ptsI | 4 | 9.58333 | 0.407 |
| 0.4 | 0 | clcB | 4 | 9.58333 | 0.385 |
| 0.4 | 0 | ycaM | 4 | 9.58333 | 0.42 |
| 0.4 | 0 | yadI | 4 | 9.58333 | 0.526 |
| 0.4 | 0 | AG1 | 4 | 9.58333 | 0.557 |
| 0.4 | 0 | ptsI | 4 | 9.83333 | 0.414 |
| 0.4 | 0 | clcB | 4 | 9.83333 | 0.389 |
| 0.4 | 0 | ycaM | 4 | 9.83333 | 0.423 |
| 0.4 | 0 | yadI | 4 | 9.83333 | 0.532 |
| 0.4 | 0 | AG1 | 4 | 9.83333 | 0.566 |
| 0.4 | 0 | ptsI | 4 | 10.0833 | 0.422 |
| 0.4 | 0 | clcB | 4 | 10.0833 | 0.394 |
| 0.4 | 0 | ycaM | 4 | 10.0833 | 0.428 |
| 0.4 | 0 | yadI | 4 | 10.0833 | 0.54 |
| 0.4 | 0 | AG1 | 4 | 10.0833 | 0.572 |
| 0.4 | 0 | ptsI | 4 | 10.3333 | 0.428 |
| 0.4 | 0 | clcB | 4 | 10.3333 | 0.402 |
| 0.4 | 0 | ycaM | 4 | 10.3333 | 0.434 |
| 0.4 | 0 | yadI | 4 | 10.3333 | 0.546 |
| 0.4 | 0 | AG1 | 4 | 10.3333 | 0.581 |
| 0.4 | 0 | ptsI | 4 | 10.5833 | 0.434 |

| | | | | | |
|---|---|---|---|---|---|
| 0.4 | 0 | clcB | 4 | 10.5833 | 0.406 |
| 0.4 | 0 | ycaM | 4 | 10.5833 | 0.439 |
| 0.4 | 0 | yadI | 4 | 10.5833 | 0.554 |
| 0.4 | 0 | AG1 | 4 | 10.5833 | 0.589 |
| 0.4 | 0 | ptsI | 4 | 10.8333 | 0.442 |
| 0.4 | 0 | clcB | 4 | 10.8333 | 0.409 |
| 0.4 | 0 | ycaM | 4 | 10.8333 | 0.44 |
| 0.4 | 0 | yadI | 4 | 10.8333 | 0.558 |
| 0.4 | 0 | AG1 | 4 | 10.8333 | 0.594 |
| 0.4 | 0 | ptsI | 4 | 11.0833 | 0.449 |
| 0.4 | 0 | clcB | 4 | 11.0833 | 0.413 |
| 0.4 | 0 | ycaM | 4 | 11.0833 | 0.446 |
| 0.4 | 0 | yadI | 4 | 11.0833 | 0.563 |
| 0.4 | 0 | AG1 | 4 | 11.0833 | 0.602 |
| 0.4 | 0 | ptsI | 4 | 11.3333 | 0.458 |
| 0.4 | 0 | clcB | 4 | 11.3333 | 0.424 |
| 0.4 | 0 | ycaM | 4 | 11.3333 | 0.453 |
| 0.4 | 0 | yadI | 4 | 11.3333 | 0.57 |
| 0.4 | 0 | AG1 | 4 | 11.3333 | 0.609 |
| 0.4 | 0 | ptsI | 4 | 11.5833 | 0.463 |
| 0.4 | 0 | clcB | 4 | 11.5833 | 0.425 |
| 0.4 | 0 | ycaM | 4 | 11.5833 | 0.458 |
| 0.4 | 0 | yadI | 4 | 11.5833 | 0.576 |
| 0.4 | 0 | AG1 | 4 | 11.5833 | 0.615 |
| 0.4 | 0 | ptsI | 4 | 11.8333 | 0.471 |
| 0.4 | 0 | clcB | 4 | 11.8333 | 0.429 |
| 0.4 | 0 | ycaM | 4 | 11.8333 | 0.461 |
| 0.4 | 0 | yadI | 4 | 11.8333 | 0.58 |
| 0.4 | 0 | AG1 | 4 | 11.8333 | 0.62 |
| 0.4 | 0 | ptsI | 4 | 12.0833 | 0.475 |
| 0.4 | 0 | clcB | 4 | 12.0833 | 0.433 |
| 0.4 | 0 | ycaM | 4 | 12.0833 | 0.467 |
| 0.4 | 0 | yadI | 4 | 12.0833 | 0.586 |
| 0.4 | 0 | AG1 | 4 | 12.0833 | 0.625 |
| 0.4 | 0 | ptsI | 4 | 12.3333 | 0.483 |
| 0.4 | 0 | clcB | 4 | 12.3333 | 0.44 |
| 0.4 | 0 | ycaM | 4 | 12.3333 | 0.469 |
| 0.4 | 0 | yadI | 4 | 12.3333 | 0.59 |
| 0.4 | 0 | AG1 | 4 | 12.3333 | 0.631 |
| 0.4 | 0 | ptsI | 4 | 12.5833 | 0.489 |
| 0.4 | 0 | clcB | 4 | 12.5833 | 0.445 |
| 0.4 | 0 | ycaM | 4 | 12.5833 | 0.475 |
| 0.4 | 0 | yadI | 4 | 12.5833 | 0.595 |
| 0.4 | 0 | AG1 | 4 | 12.5833 | 0.64 |
| 0.4 | 0 | ptsI | 4 | 12.8333 | 0.492 |
| 0.4 | 0 | clcB | 4 | 12.8333 | 0.445 |
| 0.4 | 0 | ycaM | 4 | 12.8333 | 0.479 |
| 0.4 | 0 | yadI | 4 | 12.8333 | 0.599 |

| | | | | | |
|---|---|---|---|---|---|
| 0.4 | 0 | AG1 | 4 | 12.8333 | 0.64 |
| 0.4 | 0 | ptsI | 4 | 13.0833 | 0.498 |
| 0.4 | 0 | clcB | 4 | 13.0833 | 0.449 |
| 0.4 | 0 | ycaM | 4 | 13.0833 | 0.482 |
| 0.4 | 0 | yadI | 4 | 13.0833 | 0.602 |
| 0.4 | 0 | AG1 | 4 | 13.0833 | 0.649 |
| 0.4 | 0 | ptsI | 4 | 13.3333 | 0.507 |
| 0.4 | 0 | clcB | 4 | 13.3333 | 0.452 |
| 0.4 | 0 | ycaM | 4 | 13.3333 | 0.487 |
| 0.4 | 0 | yadI | 4 | 13.3333 | 0.606 |
| 0.4 | 0 | AG1 | 4 | 13.3333 | 0.655 |
| 0.4 | 0 | ptsI | 4 | 13.5833 | 0.509 |
| 0.4 | 0 | clcB | 4 | 13.5833 | 0.454 |
| 0.4 | 0 | ycaM | 4 | 13.5833 | 0.491 |
| 0.4 | 0 | yadI | 4 | 13.5833 | 0.608 |
| 0.4 | 0 | AG1 | 4 | 13.5833 | 0.658 |
| 0.4 | 0 | ptsI | 4 | 13.8333 | 0.515 |
| 0.4 | 0 | clcB | 4 | 13.8333 | 0.457 |
| 0.4 | 0 | ycaM | 4 | 13.8333 | 0.495 |
| 0.4 | 0 | yadI | 4 | 13.8333 | 0.611 |
| 0.4 | 0 | AG1 | 4 | 13.8333 | 0.661 |
| 0.4 | 0 | ptsI | 4 | 14.0833 | 0.519 |
| 0.4 | 0 | clcB | 4 | 14.0833 | 0.456 |
| 0.4 | 0 | ycaM | 4 | 14.0833 | 0.496 |
| 0.4 | 0 | yadI | 4 | 14.0833 | 0.615 |
| 0.4 | 0 | AG1 | 4 | 14.0833 | 0.667 |
| 0.4 | 0 | ptsI | 4 | 14.3333 | 0.527 |
| 0.4 | 0 | clcB | 4 | 14.3333 | 0.461 |
| 0.4 | 0 | ycaM | 4 | 14.3333 | 0.503 |
| 0.4 | 0 | yadI | 4 | 14.3333 | 0.618 |
| 0.4 | 0 | AG1 | 4 | 14.3333 | 0.67 |
| 0.4 | 0 | ptsI | 4 | 14.5833 | 0.527 |
| 0.4 | 0 | clcB | 4 | 14.5833 | 0.462 |
| 0.4 | 0 | ycaM | 4 | 14.5833 | 0.501 |
| 0.4 | 0 | yadI | 4 | 14.5833 | 0.62 |
| 0.4 | 0 | AG1 | 4 | 14.5833 | 0.672 |
| 0.4 | 0 | ptsI | 4 | 14.8333 | 0.532 |
| 0.4 | 0 | clcB | 4 | 14.8333 | 0.464 |
| 0.4 | 0 | ycaM | 4 | 14.8333 | 0.509 |
| 0.4 | 0 | yadI | 4 | 14.8333 | 0.623 |
| 0.4 | 0 | AG1 | 4 | 14.8333 | 0.679 |
| 0.4 | 0 | ptsI | 4 | 15.0833 | 0.532 |
| 0.4 | 0 | clcB | 4 | 15.0833 | 0.465 |
| 0.4 | 0 | ycaM | 4 | 15.0833 | 0.512 |
| 0.4 | 0 | yadI | 4 | 15.0833 | 0.626 |
| 0.4 | 0 | AG1 | 4 | 15.0833 | 0.682 |
| 0.4 | 0 | ptsI | 4 | 15.3333 | 0.541 |
| 0.4 | 0 | clcB | 4 | 15.3333 | 0.471 |

| | | | | | |
|---|---|---|---|---|---|
| 0.4 | 0 | ycaM | 4 | 15.3333 | 0.514 |
| 0.4 | 0 | yadI | 4 | 15.3333 | 0.631 |
| 0.4 | 0 | AG1 | 4 | 15.3333 | 0.68 |
| 0.4 | 0 | ptsI | 4 | 15.5833 | 0.548 |
| 0.4 | 0 | clcB | 4 | 15.5833 | 0.472 |
| 0.4 | 0 | ycaM | 4 | 15.5833 | 0.516 |
| 0.4 | 0 | yadI | 4 | 15.5833 | 0.632 |
| 0.4 | 0 | AG1 | 4 | 15.5833 | 0.685 |
| 0.4 | 0 | ptsI | 4 | 15.8333 | 0.546 |
| 0.4 | 0 | clcB | 4 | 15.8333 | 0.472 |
| 0.4 | 0 | ycaM | 4 | 15.8333 | 0.516 |
| 0.4 | 0 | yadI | 4 | 15.8333 | 0.636 |
| 0.4 | 0 | AG1 | 4 | 15.8333 | 0.683 |
| 0.4 | 0 | ptsI | 4 | 16.0833 | 0.553 |
| 0.4 | 0 | clcB | 4 | 16.0833 | 0.476 |
| 0.4 | 0 | ycaM | 4 | 16.0833 | 0.519 |
| 0.4 | 0 | yadI | 4 | 16.0833 | 0.64 |
| 0.4 | 0 | AG1 | 4 | 16.0833 | 0.69 |
| 0.4 | 0 | ptsI | 4 | 16.3333 | 0.555 |
| 0.4 | 0 | clcB | 4 | 16.3333 | 0.476 |
| 0.4 | 0 | ycaM | 4 | 16.3333 | 0.522 |
| 0.4 | 0 | yadI | 4 | 16.3333 | 0.64 |
| 0.4 | 0 | AG1 | 4 | 16.3333 | 0.686 |
| 0.4 | 0 | ptsI | 4 | 16.5833 | 0.56 |
| 0.4 | 0 | clcB | 4 | 16.5833 | 0.475 |
| 0.4 | 0 | ycaM | 4 | 16.5833 | 0.523 |
| 0.4 | 0 | yadI | 4 | 16.5833 | 0.643 |
| 0.4 | 0 | AG1 | 4 | 16.5833 | 0.69 |
| 0.4 | 0 | ptsI | 4 | 16.8333 | 0.561 |
| 0.4 | 0 | clcB | 4 | 16.8333 | 0.478 |
| 0.4 | 0 | ycaM | 4 | 16.8333 | 0.521 |
| 0.4 | 0 | yadI | 4 | 16.8333 | 0.642 |
| 0.4 | 0 | AG1 | 4 | 16.8333 | 0.694 |
| 0.4 | 0 | ptsI | 4 | 17.0833 | 0.564 |
| 0.4 | 0 | clcB | 4 | 17.0833 | 0.483 |
| 0.4 | 0 | ycaM | 4 | 17.0833 | 0.527 |
| 0.4 | 0 | yadI | 4 | 17.0833 | 0.644 |
| 0.4 | 0 | AG1 | 4 | 17.0833 | 0.696 |
| 0.4 | 0 | ptsI | 4 | 17.3333 | 0.568 |
| 0.4 | 0 | clcB | 4 | 17.3333 | 0.486 |
| 0.4 | 0 | ycaM | 4 | 17.3333 | 0.53 |
| 0.4 | 0 | yadI | 4 | 17.3333 | 0.649 |
| 0.4 | 0 | AG1 | 4 | 17.3333 | 0.696 |
| 0.4 | 0 | ptsI | 4 | 17.5833 | 0.567 |
| 0.4 | 0 | clcB | 4 | 17.5833 | 0.483 |
| 0.4 | 0 | ycaM | 4 | 17.5833 | 0.528 |
| 0.4 | 0 | yadI | 4 | 17.5833 | 0.649 |
| 0.4 | 0 | AG1 | 4 | 17.5833 | 0.701 |

| | | | | | |
|---|---|---|---|---|---|
| 0.4 | 0 | ptsI | 4 | 17.8333 | 0.573 |
| 0.4 | 0 | clcB | 4 | 17.8333 | 0.483 |
| 0.4 | 0 | ycaM | 4 | 17.8333 | 0.533 |
| 0.4 | 0 | yadI | 4 | 17.8333 | 0.647 |
| 0.4 | 0 | AG1 | 4 | 17.8333 | 0.678 |
| 0.4 | 0 | ptsI | 4 | 18.0833 | 0.572 |
| 0.4 | 0 | clcB | 4 | 18.0833 | 0.486 |
| 0.4 | 0 | ycaM | 4 | 18.0833 | 0.535 |
| 0.4 | 0 | yadI | 4 | 18.0833 | 0.654 |
| 0.4 | 0 | AG1 | 4 | 18.0833 | 0.7 |
| 0.4 | 0 | ptsI | 4 | 18.3333 | 0.577 |
| 0.4 | 0 | clcB | 4 | 18.3333 | 0.486 |
| 0.4 | 0 | ycaM | 4 | 18.3333 | 0.536 |
| 0.4 | 0 | yadI | 4 | 18.3333 | 0.659 |
| 0.4 | 0 | AG1 | 4 | 18.3333 | 0.706 |
| 0.4 | 0 | ptsI | 4 | 18.5833 | 0.577 |
| 0.4 | 0 | clcB | 4 | 18.5833 | 0.487 |
| 0.4 | 0 | ycaM | 4 | 18.5833 | 0.536 |
| 0.4 | 0 | yadI | 4 | 18.5833 | 0.656 |
| 0.4 | 0 | AG1 | 4 | 18.5833 | 0.705 |
| 0.4 | 0 | ptsI | 4 | 18.8333 | 0.586 |
| 0.4 | 0 | clcB | 4 | 18.8333 | 0.486 |
| 0.4 | 0 | ycaM | 4 | 18.8333 | 0.54 |
| 0.4 | 0 | yadI | 4 | 18.8333 | 0.654 |
| 0.4 | 0 | AG1 | 4 | 18.8333 | 0.687 |
| 0.4 | 0 | ptsI | 4 | 19.0833 | 0.589 |
| 0.4 | 0 | clcB | 4 | 19.0833 | 0.492 |
| 0.4 | 0 | ycaM | 4 | 19.0833 | 0.54 |
| 0.4 | 0 | yadI | 4 | 19.0833 | 0.658 |
| 0.4 | 0 | AG1 | 4 | 19.0833 | 0.712 |
| 0.4 | 0 | ptsI | 4 | 19.3333 | 0.59 |
| 0.4 | 0 | clcB | 4 | 19.3333 | 0.49 |
| 0.4 | 0 | ycaM | 4 | 19.3333 | 0.543 |
| 0.4 | 0 | yadI | 4 | 19.3333 | 0.66 |
| 0.4 | 0 | AG1 | 4 | 19.3333 | 0.711 |
| 0.4 | 0 | ptsI | 4 | 19.5833 | 0.596 |
| 0.4 | 0 | clcB | 4 | 19.5833 | 0.497 |
| 0.4 | 0 | ycaM | 4 | 19.5833 | 0.545 |
| 0.4 | 0 | yadI | 4 | 19.5833 | 0.664 |
| 0.4 | 0 | AG1 | 4 | 19.5833 | 0.712 |
| 0.4 | 0 | ptsI | 4 | 19.8333 | 0.597 |
| 0.4 | 0 | clcB | 4 | 19.8333 | 0.493 |
| 0.4 | 0 | ycaM | 4 | 19.8333 | 0.541 |
| 0.4 | 0 | yadI | 4 | 19.8333 | 0.663 |
| 0.4 | 0 | AG1 | 4 | 19.8333 | 0.712 |
| 0.4 | 0 | ptsI | 4 | 20.0833 | 0.603 |
| 0.4 | 0 | clcB | 4 | 20.0833 | 0.498 |
| 0.4 | 0 | ycaM | 4 | 20.0833 | 0.549 |

| | | | | | |
|---|---|---|---|---|---|
| 0.4 | 0 | yadI | 4 | 20.0833 | 0.663 |
| 0.4 | 0 | AG1 | 4 | 20.0833 | 0.72 |
| 0.4 | 0 | ptsI | 4 | 20.3333 | 0.602 |
| 0.4 | 0 | clcB | 4 | 20.3333 | 0.492 |
| 0.4 | 0 | ycaM | 4 | 20.3333 | 0.546 |
| 0.4 | 0 | yadI | 4 | 20.3333 | 0.665 |
| 0.4 | 0 | AG1 | 4 | 20.3333 | 0.714 |
| 0.4 | 0 | ptsI | 4 | 20.5833 | 0.605 |
| 0.4 | 0 | clcB | 4 | 20.5833 | 0.493 |
| 0.4 | 0 | ycaM | 4 | 20.5833 | 0.546 |
| 0.4 | 0 | yadI | 4 | 20.5833 | 0.668 |
| 0.4 | 0 | AG1 | 4 | 20.5833 | 0.717 |
| 0.4 | 0 | ptsI | 4 | 20.8333 | 0.608 |
| 0.4 | 0 | clcB | 4 | 20.8333 | 0.5 |
| 0.4 | 0 | ycaM | 4 | 20.8333 | 0.552 |
| 0.4 | 0 | yadI | 4 | 20.8333 | 0.67 |
| 0.4 | 0 | AG1 | 4 | 20.8333 | 0.714 |
| 0.4 | 0 | ptsI | 4 | 21.0833 | 0.611 |
| 0.4 | 0 | clcB | 4 | 21.0833 | 0.498 |
| 0.4 | 0 | ycaM | 4 | 21.0833 | 0.547 |
| 0.4 | 0 | yadI | 4 | 21.0833 | 0.669 |
| 0.4 | 0 | AG1 | 4 | 21.0833 | 0.716 |
| 0.4 | 0 | ptsI | 4 | 21.3333 | 0.61 |
| 0.4 | 0 | clcB | 4 | 21.3333 | 0.498 |
| 0.4 | 0 | ycaM | 4 | 21.3333 | 0.548 |
| 0.4 | 0 | yadI | 4 | 21.3333 | 0.671 |
| 0.4 | 0 | AG1 | 4 | 21.3333 | 0.716 |
| 0.4 | 0 | ptsI | 4 | 21.5833 | 0.613 |
| 0.4 | 0 | clcB | 4 | 21.5833 | 0.497 |
| 0.4 | 0 | ycaM | 4 | 21.5833 | 0.551 |
| 0.4 | 0 | yadI | 4 | 21.5833 | 0.67 |
| 0.4 | 0 | AG1 | 4 | 21.5833 | 0.718 |
| 0.4 | 0 | ptsI | 4 | 21.8333 | 0.618 |
| 0.4 | 0 | clcB | 4 | 21.8333 | 0.498 |
| 0.4 | 0 | ycaM | 4 | 21.8333 | 0.552 |
| 0.4 | 0 | yadI | 4 | 21.8333 | 0.675 |
| 0.4 | 0 | AG1 | 4 | 21.8333 | 0.717 |
| 0.4 | 0 | ptsI | 4 | 22.0833 | 0.624 |
| 0.4 | 0 | clcB | 4 | 22.0833 | 0.5 |
| 0.4 | 0 | ycaM | 4 | 22.0833 | 0.554 |
| 0.4 | 0 | yadI | 4 | 22.0833 | 0.67 |
| 0.4 | 0 | AG1 | 4 | 22.0833 | 0.716 |
| 0.4 | 0 | ptsI | 4 | 22.3333 | 0.626 |
| 0.4 | 0 | clcB | 4 | 22.3333 | 0.504 |
| 0.4 | 0 | ycaM | 4 | 22.3333 | 0.552 |
| 0.4 | 0 | yadI | 4 | 22.3333 | 0.674 |
| 0.4 | 0 | AG1 | 4 | 22.3333 | 0.714 |
| 0.4 | 0 | ptsI | 4 | 22.5833 | 0.627 |

| | | | | | |
|---|---|---|---|---|---|
| 0.4 | 0 | clcB | 4 | 22.5833 | 0.5 |
| 0.4 | 0 | ycaM | 4 | 22.5833 | 0.55 |
| 0.4 | 0 | yadI | 4 | 22.5833 | 0.674 |
| 0.4 | 0 | AG1 | 4 | 22.5833 | 0.718 |
| 0.4 | 0 | ptsI | 4 | 22.8333 | 0.631 |
| 0.4 | 0 | clcB | 4 | 22.8333 | 0.5 |
| 0.4 | 0 | ycaM | 4 | 22.8333 | 0.551 |
| 0.4 | 0 | yadI | 4 | 22.8333 | 0.676 |
| 0.4 | 0 | AG1 | 4 | 22.8333 | 0.72 |
| 0.4 | 0 | ptsI | 4 | 23.0833 | 0.638 |
| 0.4 | 0 | clcB | 4 | 23.0833 | 0.501 |
| 0.4 | 0 | ycaM | 4 | 23.0833 | 0.552 |
| 0.4 | 0 | yadI | 4 | 23.0833 | 0.672 |
| 0.4 | 0 | AG1 | 4 | 23.0833 | 0.72 |
| 0.4 | 0 | ptsI | 4 | 23.3333 | 0.64 |
| 0.4 | 0 | clcB | 4 | 23.3333 | 0.501 |
| 0.4 | 0 | ycaM | 4 | 23.3333 | 0.557 |
| 0.4 | 0 | yadI | 4 | 23.3333 | 0.679 |
| 0.4 | 0 | AG1 | 4 | 23.3333 | 0.72 |
| 0.4 | 0 | ptsI | 4 | 23.5833 | 0.638 |
| 0.4 | 0 | clcB | 4 | 23.5833 | 0.502 |
| 0.4 | 0 | ycaM | 4 | 23.5833 | 0.557 |
| 0.4 | 0 | yadI | 4 | 23.5833 | 0.676 |
| 0.4 | 0 | AG1 | 4 | 23.5833 | 0.719 |
| 0.4 | 0 | ptsI | 4 | 23.8333 | 0.65 |
| 0.4 | 0 | clcB | 4 | 23.8333 | 0.503 |
| 0.4 | 0 | ycaM | 4 | 23.8333 | 0.553 |
| 0.4 | 0 | yadI | 4 | 23.8333 | 0.678 |
| 0.4 | 0 | AG1 | 4 | 23.8333 | 0.722 |
| 0.4 | 0 | ptsI | 4 | 24.0833 | 0.649 |
| 0.4 | 0 | clcB | 4 | 24.0833 | 0.503 |
| 0.4 | 0 | ycaM | 4 | 24.0833 | 0.553 |
| 0.4 | 0 | yadI | 4 | 24.0833 | 0.68 |
| 0.4 | 0 | AG1 | 4 | 24.0833 | 0.718 |
| 0.4 | 0 | ptsI | 5 | 0 | 0.156 |
| 0.4 | 0 | clcB | 5 | 0 | 0.167 |
| 0.4 | 0 | ycaM | 5 | 0 | 0.16 |
| 0.4 | 0 | yadI | 5 | 0 | 0.171 |
| 0.4 | 0 | AG1 | 5 | 0 | 0.166 |
| 0.4 | 0 | ptsI | 5 | 0.35 | 0.158 |
| 0.4 | 0 | clcB | 5 | 0.35 | 0.162 |
| 0.4 | 0 | ycaM | 5 | 0.35 | 0.154 |
| 0.4 | 0 | yadI | 5 | 0.35 | 0.166 |
| 0.4 | 0 | AG1 | 5 | 0.35 | 0.162 |
| 0.4 | 0 | ptsI | 5 | 0.6 | 0.159 |
| 0.4 | 0 | clcB | 5 | 0.6 | 0.162 |
| 0.4 | 0 | ycaM | 5 | 0.6 | 0.153 |
| 0.4 | 0 | yadI | 5 | 0.6 | 0.163 |

| | | | | | |
|---|---|---|---|---|---|
| 0.4 | 0 | AG1 | 5 | 0.6 | 0.163 |
| 0.4 | 0 | ptsI | 5 | 0.85 | 0.159 |
| 0.4 | 0 | clcB | 5 | 0.85 | 0.162 |
| 0.4 | 0 | ycaM | 5 | 0.85 | 0.152 |
| 0.4 | 0 | yadI | 5 | 0.85 | 0.164 |
| 0.4 | 0 | AG1 | 5 | 0.85 | 0.164 |
| 0.4 | 0 | ptsI | 5 | 1.1 | 0.161 |
| 0.4 | 0 | clcB | 5 | 1.1 | 0.164 |
| 0.4 | 0 | ycaM | 5 | 1.1 | 0.154 |
| 0.4 | 0 | yadI | 5 | 1.1 | 0.167 |
| 0.4 | 0 | AG1 | 5 | 1.1 | 0.165 |
| 0.4 | 0 | ptsI | 5 | 1.35 | 0.163 |
| 0.4 | 0 | clcB | 5 | 1.35 | 0.165 |
| 0.4 | 0 | ycaM | 5 | 1.35 | 0.156 |
| 0.4 | 0 | yadI | 5 | 1.35 | 0.167 |
| 0.4 | 0 | AG1 | 5 | 1.35 | 0.168 |
| 0.4 | 0 | ptsI | 5 | 1.6 | 0.164 |
| 0.4 | 0 | clcB | 5 | 1.6 | 0.169 |
| 0.4 | 0 | ycaM | 5 | 1.6 | 0.158 |
| 0.4 | 0 | yadI | 5 | 1.6 | 0.172 |
| 0.4 | 0 | AG1 | 5 | 1.6 | 0.172 |
| 0.4 | 0 | ptsI | 5 | 1.85 | 0.167 |
| 0.4 | 0 | clcB | 5 | 1.85 | 0.171 |
| 0.4 | 0 | ycaM | 5 | 1.85 | 0.161 |
| 0.4 | 0 | yadI | 5 | 1.85 | 0.175 |
| 0.4 | 0 | AG1 | 5 | 1.85 | 0.176 |
| 0.4 | 0 | ptsI | 5 | 2.1 | 0.17 |
| 0.4 | 0 | clcB | 5 | 2.1 | 0.177 |
| 0.4 | 0 | ycaM | 5 | 2.1 | 0.164 |
| 0.4 | 0 | yadI | 5 | 2.1 | 0.183 |
| 0.4 | 0 | AG1 | 5 | 2.1 | 0.184 |
| 0.4 | 0 | ptsI | 5 | 2.35 | 0.177 |
| 0.4 | 0 | clcB | 5 | 2.35 | 0.184 |
| 0.4 | 0 | ycaM | 5 | 2.35 | 0.169 |
| 0.4 | 0 | yadI | 5 | 2.35 | 0.19 |
| 0.4 | 0 | AG1 | 5 | 2.35 | 0.192 |
| 0.4 | 0 | ptsI | 5 | 2.6 | 0.174 |
| 0.4 | 0 | clcB | 5 | 2.6 | 0.186 |
| 0.4 | 0 | ycaM | 5 | 2.6 | 0.176 |
| 0.4 | 0 | yadI | 5 | 2.6 | 0.192 |
| 0.4 | 0 | AG1 | 5 | 2.6 | 0.189 |
| 0.4 | 0 | ptsI | 5 | 2.85 | 0.184 |
| 0.4 | 0 | clcB | 5 | 2.85 | 0.199 |
| 0.4 | 0 | ycaM | 5 | 2.85 | 0.182 |
| 0.4 | 0 | yadI | 5 | 2.85 | 0.205 |
| 0.4 | 0 | AG1 | 5 | 2.85 | 0.205 |
| 0.4 | 0 | ptsI | 5 | 3.1 | 0.192 |
| 0.4 | 0 | clcB | 5 | 3.1 | 0.213 |

| | | | | | |
|---|---|---|---|---|---|
| 0.4 | 0 | ycaM | 5 | 3.1 | 0.19 |
| 0.4 | 0 | yadI | 5 | 3.1 | 0.218 |
| 0.4 | 0 | AG1 | 5 | 3.1 | 0.217 |
| 0.4 | 0 | ptsI | 5 | 3.35 | 0.196 |
| 0.4 | 0 | clcB | 5 | 3.35 | 0.226 |
| 0.4 | 0 | ycaM | 5 | 3.35 | 0.201 |
| 0.4 | 0 | yadI | 5 | 3.35 | 0.234 |
| 0.4 | 0 | AG1 | 5 | 3.35 | 0.236 |
| 0.4 | 0 | ptsI | 5 | 3.6 | 0.201 |
| 0.4 | 0 | clcB | 5 | 3.6 | 0.239 |
| 0.4 | 0 | ycaM | 5 | 3.6 | 0.21 |
| 0.4 | 0 | yadI | 5 | 3.6 | 0.25 |
| 0.4 | 0 | AG1 | 5 | 3.6 | 0.252 |
| 0.4 | 0 | ptsI | 5 | 3.85 | 0.21 |
| 0.4 | 0 | clcB | 5 | 3.85 | 0.257 |
| 0.4 | 0 | ycaM | 5 | 3.85 | 0.223 |
| 0.4 | 0 | yadI | 5 | 3.85 | 0.266 |
| 0.4 | 0 | AG1 | 5 | 3.85 | 0.269 |
| 0.4 | 0 | ptsI | 5 | 4.38333 | 0.209 |
| 0.4 | 0 | clcB | 5 | 4.38333 | 0.264 |
| 0.4 | 0 | ycaM | 5 | 4.38333 | 0.234 |
| 0.4 | 0 | yadI | 5 | 4.38333 | 0.284 |
| 0.4 | 0 | AG1 | 5 | 4.38333 | 0.3 |
| 0.4 | 0 | ptsI | 5 | 4.63333 | 0.21 |
| 0.4 | 0 | clcB | 5 | 4.63333 | 0.258 |
| 0.4 | 0 | ycaM | 5 | 4.63333 | 0.236 |
| 0.4 | 0 | yadI | 5 | 4.63333 | 0.272 |
| 0.4 | 0 | AG1 | 5 | 4.63333 | 0.286 |
| 0.4 | 0 | ptsI | 5 | 4.88333 | 0.216 |
| 0.4 | 0 | clcB | 5 | 4.88333 | 0.262 |
| 0.4 | 0 | ycaM | 5 | 4.88333 | 0.244 |
| 0.4 | 0 | yadI | 5 | 4.88333 | 0.288 |
| 0.4 | 0 | AG1 | 5 | 4.88333 | 0.296 |
| 0.4 | 0 | ptsI | 5 | 5.13333 | 0.219 |
| 0.4 | 0 | clcB | 5 | 5.13333 | 0.27 |
| 0.4 | 0 | ycaM | 5 | 5.13333 | 0.256 |
| 0.4 | 0 | yadI | 5 | 5.13333 | 0.304 |
| 0.4 | 0 | AG1 | 5 | 5.13333 | 0.308 |
| 0.4 | 0 | ptsI | 5 | 5.38333 | 0.226 |
| 0.4 | 0 | clcB | 5 | 5.38333 | 0.28 |
| 0.4 | 0 | ycaM | 5 | 5.38333 | 0.271 |
| 0.4 | 0 | yadI | 5 | 5.38333 | 0.318 |
| 0.4 | 0 | AG1 | 5 | 5.38333 | 0.322 |
| 0.4 | 0 | ptsI | 5 | 5.63333 | 0.236 |
| 0.4 | 0 | clcB | 5 | 5.63333 | 0.294 |
| 0.4 | 0 | ycaM | 5 | 5.63333 | 0.284 |
| 0.4 | 0 | yadI | 5 | 5.63333 | 0.333 |
| 0.4 | 0 | AG1 | 5 | 5.63333 | 0.332 |

| | | | | | |
|---|---|---|---|---|---|
| 0.4 | 0 | ptsI | 5 | 5.88333 | 0.243 |
| 0.4 | 0 | clcB | 5 | 5.88333 | 0.313 |
| 0.4 | 0 | ycaM | 5 | 5.88333 | 0.298 |
| 0.4 | 0 | yadI | 5 | 5.88333 | 0.347 |
| 0.4 | 0 | AG1 | 5 | 5.88333 | 0.34 |
| 0.4 | 0 | ptsI | 5 | 6.13333 | 0.255 |
| 0.4 | 0 | clcB | 5 | 6.13333 | 0.329 |
| 0.4 | 0 | ycaM | 5 | 6.13333 | 0.312 |
| 0.4 | 0 | yadI | 5 | 6.13333 | 0.363 |
| 0.4 | 0 | AG1 | 5 | 6.13333 | 0.353 |
| 0.4 | 0 | ptsI | 5 | 6.38333 | 0.265 |
| 0.4 | 0 | clcB | 5 | 6.38333 | 0.345 |
| 0.4 | 0 | ycaM | 5 | 6.38333 | 0.323 |
| 0.4 | 0 | yadI | 5 | 6.38333 | 0.376 |
| 0.4 | 0 | AG1 | 5 | 6.38333 | 0.37 |
| 0.4 | 0 | ptsI | 5 | 6.63333 | 0.275 |
| 0.4 | 0 | clcB | 5 | 6.63333 | 0.355 |
| 0.4 | 0 | ycaM | 5 | 6.63333 | 0.336 |
| 0.4 | 0 | yadI | 5 | 6.63333 | 0.391 |
| 0.4 | 0 | AG1 | 5 | 6.63333 | 0.388 |
| 0.4 | 0 | ptsI | 5 | 6.88333 | 0.287 |
| 0.4 | 0 | clcB | 5 | 6.88333 | 0.369 |
| 0.4 | 0 | ycaM | 5 | 6.88333 | 0.345 |
| 0.4 | 0 | yadI | 5 | 6.88333 | 0.408 |
| 0.4 | 0 | AG1 | 5 | 6.88333 | 0.404 |
| 0.4 | 0 | ptsI | 5 | 7.13333 | 0.298 |
| 0.4 | 0 | clcB | 5 | 7.13333 | 0.377 |
| 0.4 | 0 | ycaM | 5 | 7.13333 | 0.358 |
| 0.4 | 0 | yadI | 5 | 7.13333 | 0.425 |
| 0.4 | 0 | AG1 | 5 | 7.13333 | 0.417 |
| 0.4 | 0 | ptsI | 5 | 7.38333 | 0.31 |
| 0.4 | 0 | clcB | 5 | 7.38333 | 0.39 |
| 0.4 | 0 | ycaM | 5 | 7.38333 | 0.368 |
| 0.4 | 0 | yadI | 5 | 7.38333 | 0.438 |
| 0.4 | 0 | AG1 | 5 | 7.38333 | 0.433 |
| 0.4 | 0 | ptsI | 5 | 7.63333 | 0.321 |
| 0.4 | 0 | clcB | 5 | 7.63333 | 0.402 |
| 0.4 | 0 | ycaM | 5 | 7.63333 | 0.382 |
| 0.4 | 0 | yadI | 5 | 7.63333 | 0.453 |
| 0.4 | 0 | AG1 | 5 | 7.63333 | 0.45 |
| 0.4 | 0 | ptsI | 5 | 7.88333 | 0.331 |
| 0.4 | 0 | clcB | 5 | 7.88333 | 0.408 |
| 0.4 | 0 | ycaM | 5 | 7.88333 | 0.394 |
| 0.4 | 0 | yadI | 5 | 7.88333 | 0.466 |
| 0.4 | 0 | AG1 | 5 | 7.88333 | 0.462 |
| 0.4 | 0 | ptsI | 5 | 8.13333 | 0.343 |
| 0.4 | 0 | clcB | 5 | 8.13333 | 0.42 |
| 0.4 | 0 | ycaM | 5 | 8.13333 | 0.404 |

| | | | | | |
|---|---|---|---|---|---|
| 0.4 | 0 | yadI | 5 | 8.13333 | 0.48 |
| 0.4 | 0 | AG1 | 5 | 8.13333 | 0.474 |
| 0.4 | 0 | ptsI | 5 | 8.38333 | 0.356 |
| 0.4 | 0 | clcB | 5 | 8.38333 | 0.431 |
| 0.4 | 0 | ycaM | 5 | 8.38333 | 0.415 |
| 0.4 | 0 | yadI | 5 | 8.38333 | 0.493 |
| 0.4 | 0 | AG1 | 5 | 8.38333 | 0.487 |
| 0.4 | 0 | ptsI | 5 | 8.88333 | 0.325 |
| 0.4 | 0 | clcB | 5 | 8.88333 | 0.352 |
| 0.4 | 0 | ycaM | 5 | 8.88333 | 0.337 |
| 0.4 | 0 | yadI | 5 | 8.88333 | 0.39 |
| 0.4 | 0 | AG1 | 5 | 8.88333 | 0.478 |
| 0.4 | 0 | ptsI | 5 | 9.13333 | 0.333 |
| 0.4 | 0 | clcB | 5 | 9.13333 | 0.364 |
| 0.4 | 0 | ycaM | 5 | 9.13333 | 0.344 |
| 0.4 | 0 | yadI | 5 | 9.13333 | 0.39 |
| 0.4 | 0 | AG1 | 5 | 9.13333 | 0.476 |
| 0.4 | 0 | ptsI | 5 | 9.38333 | 0.344 |
| 0.4 | 0 | clcB | 5 | 9.38333 | 0.376 |
| 0.4 | 0 | ycaM | 5 | 9.38333 | 0.355 |
| 0.4 | 0 | yadI | 5 | 9.38333 | 0.399 |
| 0.4 | 0 | AG1 | 5 | 9.38333 | 0.483 |
| 0.4 | 0 | ptsI | 5 | 9.63333 | 0.353 |
| 0.4 | 0 | clcB | 5 | 9.63333 | 0.384 |
| 0.4 | 0 | ycaM | 5 | 9.63333 | 0.361 |
| 0.4 | 0 | yadI | 5 | 9.63333 | 0.412 |
| 0.4 | 0 | AG1 | 5 | 9.63333 | 0.503 |
| 0.4 | 0 | ptsI | 5 | 9.88333 | 0.362 |
| 0.4 | 0 | clcB | 5 | 9.88333 | 0.392 |
| 0.4 | 0 | ycaM | 5 | 9.88333 | 0.37 |
| 0.4 | 0 | yadI | 5 | 9.88333 | 0.42 |
| 0.4 | 0 | AG1 | 5 | 9.88333 | 0.513 |
| 0.4 | 0 | ptsI | 5 | 10.1333 | 0.372 |
| 0.4 | 0 | clcB | 5 | 10.1333 | 0.398 |
| 0.4 | 0 | ycaM | 5 | 10.1333 | 0.379 |
| 0.4 | 0 | yadI | 5 | 10.1333 | 0.428 |
| 0.4 | 0 | AG1 | 5 | 10.1333 | 0.52 |
| 0.4 | 0 | ptsI | 5 | 10.3833 | 0.386 |
| 0.4 | 0 | clcB | 5 | 10.3833 | 0.408 |
| 0.4 | 0 | ycaM | 5 | 10.3833 | 0.386 |
| 0.4 | 0 | yadI | 5 | 10.3833 | 0.438 |
| 0.4 | 0 | AG1 | 5 | 10.3833 | 0.531 |
| 0.4 | 0 | ptsI | 5 | 10.6333 | 0.394 |
| 0.4 | 0 | clcB | 5 | 10.6333 | 0.412 |
| 0.4 | 0 | ycaM | 5 | 10.6333 | 0.392 |
| 0.4 | 0 | yadI | 5 | 10.6333 | 0.441 |
| 0.4 | 0 | AG1 | 5 | 10.6333 | 0.539 |
| 0.4 | 0 | ptsI | 5 | 10.8833 | 0.406 |

| | | | | | |
|---|---|---|---|---|---|
| 0.4 | 0 | clcB | 5 | 10.8833 | 0.422 |
| 0.4 | 0 | ycaM | 5 | 10.8833 | 0.398 |
| 0.4 | 0 | yadI | 5 | 10.8833 | 0.452 |
| 0.4 | 0 | AG1 | 5 | 10.8833 | 0.548 |
| 0.4 | 0 | ptsI | 5 | 11.1333 | 0.409 |
| 0.4 | 0 | clcB | 5 | 11.1333 | 0.421 |
| 0.4 | 0 | ycaM | 5 | 11.1333 | 0.399 |
| 0.4 | 0 | yadI | 5 | 11.1333 | 0.454 |
| 0.4 | 0 | AG1 | 5 | 11.1333 | 0.556 |
| 0.4 | 0 | ptsI | 5 | 11.3833 | 0.418 |
| 0.4 | 0 | clcB | 5 | 11.3833 | 0.426 |
| 0.4 | 0 | ycaM | 5 | 11.3833 | 0.402 |
| 0.4 | 0 | yadI | 5 | 11.3833 | 0.464 |
| 0.4 | 0 | AG1 | 5 | 11.3833 | 0.565 |
| 0.4 | 0 | ptsI | 5 | 11.6333 | 0.428 |
| 0.4 | 0 | clcB | 5 | 11.6333 | 0.429 |
| 0.4 | 0 | ycaM | 5 | 11.6333 | 0.403 |
| 0.4 | 0 | yadI | 5 | 11.6333 | 0.47 |
| 0.4 | 0 | AG1 | 5 | 11.6333 | 0.57 |
| 0.4 | 0 | ptsI | 5 | 11.8833 | 0.435 |
| 0.4 | 0 | clcB | 5 | 11.8833 | 0.437 |
| 0.4 | 0 | ycaM | 5 | 11.8833 | 0.407 |
| 0.4 | 0 | yadI | 5 | 11.8833 | 0.477 |
| 0.4 | 0 | AG1 | 5 | 11.8833 | 0.581 |
| 0.4 | 0 | ptsI | 5 | 12.1333 | 0.447 |
| 0.4 | 0 | clcB | 5 | 12.1333 | 0.444 |
| 0.4 | 0 | ycaM | 5 | 12.1333 | 0.411 |
| 0.4 | 0 | yadI | 5 | 12.1333 | 0.486 |
| 0.4 | 0 | AG1 | 5 | 12.1333 | 0.589 |
| 0.4 | 0 | ptsI | 5 | 12.3833 | 0.45 |
| 0.4 | 0 | clcB | 5 | 12.3833 | 0.443 |
| 0.4 | 0 | ycaM | 5 | 12.3833 | 0.412 |
| 0.4 | 0 | yadI | 5 | 12.3833 | 0.488 |
| 0.4 | 0 | AG1 | 5 | 12.3833 | 0.594 |
| 0.4 | 0 | ptsI | 5 | 12.6333 | 0.458 |
| 0.4 | 0 | clcB | 5 | 12.6333 | 0.45 |
| 0.4 | 0 | ycaM | 5 | 12.6333 | 0.416 |
| 0.4 | 0 | yadI | 5 | 12.6333 | 0.496 |
| 0.4 | 0 | AG1 | 5 | 12.6333 | 0.602 |
| 0.4 | 0 | ptsI | 5 | 12.8833 | 0.47 |
| 0.4 | 0 | clcB | 5 | 12.8833 | 0.459 |
| 0.4 | 0 | ycaM | 5 | 12.8833 | 0.424 |
| 0.4 | 0 | yadI | 5 | 12.8833 | 0.503 |
| 0.4 | 0 | AG1 | 5 | 12.8833 | 0.611 |
| 0.4 | 0 | ptsI | 5 | 13.1333 | 0.478 |
| 0.4 | 0 | clcB | 5 | 13.1333 | 0.461 |
| 0.4 | 0 | ycaM | 5 | 13.1333 | 0.425 |
| 0.4 | 0 | yadI | 5 | 13.1333 | 0.506 |

| | | | | | |
|---|---|---|---|---|---|
| 0.4 | 0 | AG1 | 5 | 13.1333 | 0.617 |
| 0.4 | 0 | ptsI | 5 | 13.3833 | 0.479 |
| 0.4 | 0 | clcB | 5 | 13.3833 | 0.467 |
| 0.4 | 0 | ycaM | 5 | 13.3833 | 0.428 |
| 0.4 | 0 | yadI | 5 | 13.3833 | 0.509 |
| 0.4 | 0 | AG1 | 5 | 13.3833 | 0.622 |
| 0.4 | 0 | ptsI | 5 | 13.6333 | 0.491 |
| 0.4 | 0 | clcB | 5 | 13.6333 | 0.47 |
| 0.4 | 0 | ycaM | 5 | 13.6333 | 0.432 |
| 0.4 | 0 | yadI | 5 | 13.6333 | 0.517 |
| 0.4 | 0 | AG1 | 5 | 13.6333 | 0.631 |
| 0.4 | 0 | ptsI | 5 | 13.8833 | 0.496 |
| 0.4 | 0 | clcB | 5 | 13.8833 | 0.473 |
| 0.4 | 0 | ycaM | 5 | 13.8833 | 0.435 |
| 0.4 | 0 | yadI | 5 | 13.8833 | 0.519 |
| 0.4 | 0 | AG1 | 5 | 13.8833 | 0.636 |
| 0.4 | 0 | ptsI | 5 | 14.1333 | 0.506 |
| 0.4 | 0 | clcB | 5 | 14.1333 | 0.479 |
| 0.4 | 0 | ycaM | 5 | 14.1333 | 0.441 |
| 0.4 | 0 | yadI | 5 | 14.1333 | 0.526 |
| 0.4 | 0 | AG1 | 5 | 14.1333 | 0.64 |
| 0.4 | 0 | ptsI | 5 | 14.3833 | 0.514 |
| 0.4 | 0 | clcB | 5 | 14.3833 | 0.484 |
| 0.4 | 0 | ycaM | 5 | 14.3833 | 0.444 |
| 0.4 | 0 | yadI | 5 | 14.3833 | 0.529 |
| 0.4 | 0 | AG1 | 5 | 14.3833 | 0.646 |
| 0.4 | 0 | ptsI | 5 | 14.6333 | 0.519 |
| 0.4 | 0 | clcB | 5 | 14.6333 | 0.488 |
| 0.4 | 0 | ycaM | 5 | 14.6333 | 0.447 |
| 0.4 | 0 | yadI | 5 | 14.6333 | 0.533 |
| 0.4 | 0 | AG1 | 5 | 14.6333 | 0.65 |
| 0.4 | 0 | ptsI | 5 | 14.8833 | 0.524 |
| 0.4 | 0 | clcB | 5 | 14.8833 | 0.49 |
| 0.4 | 0 | ycaM | 5 | 14.8833 | 0.45 |
| 0.4 | 0 | yadI | 5 | 14.8833 | 0.537 |
| 0.4 | 0 | AG1 | 5 | 14.8833 | 0.655 |
| 0.4 | 0 | ptsI | 5 | 15.1333 | 0.532 |
| 0.4 | 0 | clcB | 5 | 15.1333 | 0.495 |
| 0.4 | 0 | ycaM | 5 | 15.1333 | 0.455 |
| 0.4 | 0 | yadI | 5 | 15.1333 | 0.543 |
| 0.4 | 0 | AG1 | 5 | 15.1333 | 0.661 |
| 0.4 | 0 | ptsI | 5 | 15.3833 | 0.538 |
| 0.4 | 0 | clcB | 5 | 15.3833 | 0.496 |
| 0.4 | 0 | ycaM | 5 | 15.3833 | 0.457 |
| 0.4 | 0 | yadI | 5 | 15.3833 | 0.544 |
| 0.4 | 0 | AG1 | 5 | 15.3833 | 0.664 |
| 0.4 | 0 | ptsI | 5 | 15.6333 | 0.543 |
| 0.4 | 0 | clcB | 5 | 15.6333 | 0.498 |

| | | | | | |
|---|---|---|---|---|---|
| 0.4 | 0 | ycaM | 5 | 15.6333 | 0.461 |
| 0.4 | 0 | yadI | 5 | 15.6333 | 0.547 |
| 0.4 | 0 | AG1 | 5 | 15.6333 | 0.667 |
| 0.4 | 0 | ptsI | 5 | 15.8833 | 0.547 |
| 0.4 | 0 | clcB | 5 | 15.8833 | 0.5 |
| 0.4 | 0 | ycaM | 5 | 15.8833 | 0.464 |
| 0.4 | 0 | yadI | 5 | 15.8833 | 0.551 |
| 0.4 | 0 | AG1 | 5 | 15.8833 | 0.67 |
| 0.4 | 0 | ptsI | 5 | 16.1333 | 0.553 |
| 0.4 | 0 | clcB | 5 | 16.1333 | 0.505 |
| 0.4 | 0 | ycaM | 5 | 16.1333 | 0.472 |
| 0.4 | 0 | yadI | 5 | 16.1333 | 0.559 |
| 0.4 | 0 | AG1 | 5 | 16.1333 | 0.667 |
| 0.4 | 0 | ptsI | 5 | 16.3833 | 0.552 |
| 0.4 | 0 | clcB | 5 | 16.3833 | 0.503 |
| 0.4 | 0 | ycaM | 5 | 16.3833 | 0.469 |
| 0.4 | 0 | yadI | 5 | 16.3833 | 0.557 |
| 0.4 | 0 | AG1 | 5 | 16.3833 | 0.678 |
| 0.4 | 0 | ptsI | 5 | 16.6333 | 0.557 |
| 0.4 | 0 | clcB | 5 | 16.6333 | 0.507 |
| 0.4 | 0 | ycaM | 5 | 16.6333 | 0.476 |
| 0.4 | 0 | yadI | 5 | 16.6333 | 0.562 |
| 0.4 | 0 | AG1 | 5 | 16.6333 | 0.681 |
| 0.4 | 0 | ptsI | 5 | 16.8833 | 0.561 |
| 0.4 | 0 | clcB | 5 | 16.8833 | 0.511 |
| 0.4 | 0 | ycaM | 5 | 16.8833 | 0.478 |
| 0.4 | 0 | yadI | 5 | 16.8833 | 0.562 |
| 0.4 | 0 | AG1 | 5 | 16.8833 | 0.683 |
| 0.4 | 0 | ptsI | 5 | 17.1333 | 0.567 |
| 0.4 | 0 | clcB | 5 | 17.1333 | 0.513 |
| 0.4 | 0 | ycaM | 5 | 17.1333 | 0.481 |
| 0.4 | 0 | yadI | 5 | 17.1333 | 0.565 |
| 0.4 | 0 | AG1 | 5 | 17.1333 | 0.687 |
| 0.4 | 0 | ptsI | 5 | 17.3833 | 0.567 |
| 0.4 | 0 | clcB | 5 | 17.3833 | 0.512 |
| 0.4 | 0 | ycaM | 5 | 17.3833 | 0.484 |
| 0.4 | 0 | yadI | 5 | 17.3833 | 0.567 |
| 0.4 | 0 | AG1 | 5 | 17.3833 | 0.689 |
| 0.4 | 0 | ptsI | 5 | 17.6333 | 0.573 |
| 0.4 | 0 | clcB | 5 | 17.6333 | 0.515 |
| 0.4 | 0 | ycaM | 5 | 17.6333 | 0.487 |
| 0.4 | 0 | yadI | 5 | 17.6333 | 0.568 |
| 0.4 | 0 | AG1 | 5 | 17.6333 | 0.693 |
| 0.4 | 0 | ptsI | 5 | 17.8833 | 0.572 |
| 0.4 | 0 | clcB | 5 | 17.8833 | 0.514 |
| 0.4 | 0 | ycaM | 5 | 17.8833 | 0.489 |
| 0.4 | 0 | yadI | 5 | 17.8833 | 0.567 |
| 0.4 | 0 | AG1 | 5 | 17.8833 | 0.692 |

| | | | | | |
|---|---|---|---|---|---|
| 0.4 | 0 | ptsI | 5 | 18.1333 | 0.58 |
| 0.4 | 0 | clcB | 5 | 18.1333 | 0.52 |
| 0.4 | 0 | ycaM | 5 | 18.1333 | 0.491 |
| 0.4 | 0 | yadI | 5 | 18.1333 | 0.576 |
| 0.4 | 0 | AG1 | 5 | 18.1333 | 0.696 |
| 0.4 | 0 | ptsI | 5 | 18.3833 | 0.582 |
| 0.4 | 0 | clcB | 5 | 18.3833 | 0.52 |
| 0.4 | 0 | ycaM | 5 | 18.3833 | 0.494 |
| 0.4 | 0 | yadI | 5 | 18.3833 | 0.577 |
| 0.4 | 0 | AG1 | 5 | 18.3833 | 0.702 |
| 0.4 | 0 | ptsI | 5 | 18.6333 | 0.586 |
| 0.4 | 0 | clcB | 5 | 18.6333 | 0.522 |
| 0.4 | 0 | ycaM | 5 | 18.6333 | 0.497 |
| 0.4 | 0 | yadI | 5 | 18.6333 | 0.577 |
| 0.4 | 0 | AG1 | 5 | 18.6333 | 0.705 |
| 0.4 | 0 | ptsI | 5 | 18.8833 | 0.584 |
| 0.4 | 0 | clcB | 5 | 18.8833 | 0.521 |
| 0.4 | 0 | ycaM | 5 | 18.8833 | 0.499 |
| 0.4 | 0 | yadI | 5 | 18.8833 | 0.582 |
| 0.4 | 0 | AG1 | 5 | 18.8833 | 0.704 |
| 0.4 | 0 | ptsI | 5 | 19.1333 | 0.588 |
| 0.4 | 0 | clcB | 5 | 19.1333 | 0.524 |
| 0.4 | 0 | ycaM | 5 | 19.1333 | 0.501 |
| 0.4 | 0 | yadI | 5 | 19.1333 | 0.58 |
| 0.4 | 0 | AG1 | 5 | 19.1333 | 0.706 |
| 0.4 | 0 | ptsI | 5 | 19.3833 | 0.596 |
| 0.4 | 0 | clcB | 5 | 19.3833 | 0.526 |
| 0.4 | 0 | ycaM | 5 | 19.3833 | 0.503 |
| 0.4 | 0 | yadI | 5 | 19.3833 | 0.587 |
| 0.4 | 0 | AG1 | 5 | 19.3833 | 0.707 |
| 0.4 | 0 | ptsI | 5 | 19.6333 | 0.595 |
| 0.4 | 0 | clcB | 5 | 19.6333 | 0.529 |
| 0.4 | 0 | ycaM | 5 | 19.6333 | 0.504 |
| 0.4 | 0 | yadI | 5 | 19.6333 | 0.591 |
| 0.4 | 0 | AG1 | 5 | 19.6333 | 0.71 |
| 0.4 | 0 | ptsI | 5 | 19.8833 | 0.595 |
| 0.4 | 0 | clcB | 5 | 19.8833 | 0.527 |
| 0.4 | 0 | ycaM | 5 | 19.8833 | 0.507 |
| 0.4 | 0 | yadI | 5 | 19.8833 | 0.588 |
| 0.4 | 0 | AG1 | 5 | 19.8833 | 0.71 |
| 0.4 | 0 | ptsI | 5 | 20.1333 | 0.597 |
| 0.4 | 0 | clcB | 5 | 20.1333 | 0.528 |
| 0.4 | 0 | ycaM | 5 | 20.1333 | 0.508 |
| 0.4 | 0 | yadI | 5 | 20.1333 | 0.591 |
| 0.4 | 0 | AG1 | 5 | 20.1333 | 0.714 |
| 0.4 | 0 | ptsI | 5 | 20.3833 | 0.603 |
| 0.4 | 0 | clcB | 5 | 20.3833 | 0.53 |
| 0.4 | 0 | ycaM | 5 | 20.3833 | 0.509 |

| | | | | | |
|---|---|---|---|---|---|
| 0.4 | 0 | yadI | 5 | 20.3833 | 0.595 |
| 0.4 | 0 | AG1 | 5 | 20.3833 | 0.714 |
| 0.4 | 0 | ptsI | 5 | 20.6333 | 0.604 |
| 0.4 | 0 | clcB | 5 | 20.6333 | 0.528 |
| 0.4 | 0 | ycaM | 5 | 20.6333 | 0.51 |
| 0.4 | 0 | yadI | 5 | 20.6333 | 0.592 |
| 0.4 | 0 | AG1 | 5 | 20.6333 | 0.723 |
| 0.4 | 0 | ptsI | 5 | 20.8833 | 0.611 |
| 0.4 | 0 | clcB | 5 | 20.8833 | 0.528 |
| 0.4 | 0 | ycaM | 5 | 20.8833 | 0.513 |
| 0.4 | 0 | yadI | 5 | 20.8833 | 0.596 |
| 0.4 | 0 | AG1 | 5 | 20.8833 | 0.716 |
| 0.4 | 0 | ptsI | 5 | 21.1333 | 0.611 |
| 0.4 | 0 | clcB | 5 | 21.1333 | 0.53 |
| 0.4 | 0 | ycaM | 5 | 21.1333 | 0.515 |
| 0.4 | 0 | yadI | 5 | 21.1333 | 0.595 |
| 0.4 | 0 | AG1 | 5 | 21.1333 | 0.718 |
| 0.4 | 0 | ptsI | 5 | 21.3833 | 0.616 |
| 0.4 | 0 | clcB | 5 | 21.3833 | 0.531 |
| 0.4 | 0 | ycaM | 5 | 21.3833 | 0.516 |
| 0.4 | 0 | yadI | 5 | 21.3833 | 0.597 |
| 0.4 | 0 | AG1 | 5 | 21.3833 | 0.719 |
| 0.4 | 0 | ptsI | 5 | 21.6333 | 0.619 |
| 0.4 | 0 | clcB | 5 | 21.6333 | 0.534 |
| 0.4 | 0 | ycaM | 5 | 21.6333 | 0.52 |
| 0.4 | 0 | yadI | 5 | 21.6333 | 0.607 |
| 0.4 | 0 | AG1 | 5 | 21.6333 | 0.722 |
| 0.4 | 0 | ptsI | 5 | 21.8833 | 0.619 |
| 0.4 | 0 | clcB | 5 | 21.8833 | 0.535 |
| 0.4 | 0 | ycaM | 5 | 21.8833 | 0.519 |
| 0.4 | 0 | yadI | 5 | 21.8833 | 0.607 |
| 0.4 | 0 | AG1 | 5 | 21.8833 | 0.721 |
| 0.4 | 0 | ptsI | 5 | 22.1333 | 0.619 |
| 0.4 | 0 | clcB | 5 | 22.1333 | 0.53 |
| 0.4 | 0 | ycaM | 5 | 22.1333 | 0.521 |
| 0.4 | 0 | yadI | 5 | 22.1333 | 0.602 |
| 0.4 | 0 | AG1 | 5 | 22.1333 | 0.72 |
| 0.4 | 0 | ptsI | 5 | 22.3833 | 0.625 |
| 0.4 | 0 | clcB | 5 | 22.3833 | 0.533 |
| 0.4 | 0 | ycaM | 5 | 22.3833 | 0.523 |
| 0.4 | 0 | yadI | 5 | 22.3833 | 0.605 |
| 0.4 | 0 | AG1 | 5 | 22.3833 | 0.722 |
| 0.4 | 0 | ptsI | 5 | 22.6333 | 0.626 |
| 0.4 | 0 | clcB | 5 | 22.6333 | 0.535 |
| 0.4 | 0 | ycaM | 5 | 22.6333 | 0.527 |
| 0.4 | 0 | yadI | 5 | 22.6333 | 0.615 |
| 0.4 | 0 | AG1 | 5 | 22.6333 | 0.727 |
| 0.4 | 0 | ptsI | 5 | 22.8833 | 0.627 |

| | | | | | |
|---|---|---|---|---|---|
| 0.4 | 0 | clcB | 5 | 22.8833 | 0.535 |
| 0.4 | 0 | ycaM | 5 | 22.8833 | 0.526 |
| 0.4 | 0 | yadI | 5 | 22.8833 | 0.613 |
| 0.4 | 0 | AG1 | 5 | 22.8833 | 0.724 |
| 0.4 | 0 | ptsI | 5 | 23.1333 | 0.631 |
| 0.4 | 0 | clcB | 5 | 23.1333 | 0.535 |
| 0.4 | 0 | ycaM | 5 | 23.1333 | 0.528 |
| 0.4 | 0 | yadI | 5 | 23.1333 | 0.611 |
| 0.4 | 0 | AG1 | 5 | 23.1333 | 0.726 |
| 0.4 | 0 | ptsI | 5 | 23.3833 | 0.632 |
| 0.4 | 0 | clcB | 5 | 23.3833 | 0.535 |
| 0.4 | 0 | ycaM | 5 | 23.3833 | 0.525 |
| 0.4 | 0 | yadI | 5 | 23.3833 | 0.61 |
| 0.4 | 0 | AG1 | 5 | 23.3833 | 0.725 |
| 0.4 | 0 | ptsI | 5 | 23.6333 | 0.633 |
| 0.4 | 0 | clcB | 5 | 23.6333 | 0.536 |
| 0.4 | 0 | ycaM | 5 | 23.6333 | 0.527 |
| 0.4 | 0 | yadI | 5 | 23.6333 | 0.607 |
| 0.4 | 0 | AG1 | 5 | 23.6333 | 0.728 |
| 0.4 | 0 | ptsI | 5 | 23.8833 | 0.63 |
| 0.4 | 0 | clcB | 5 | 23.8833 | 0.531 |
| 0.4 | 0 | ycaM | 5 | 23.8833 | 0.528 |
| 0.4 | 0 | yadI | 5 | 23.8833 | 0.609 |
| 0.4 | 0 | AG1 | 5 | 23.8833 | 0.723 |
| 0.4 | 0 | ptsI | 5 | 24.1333 | 0.64 |
| 0.4 | 0 | clcB | 5 | 24.1333 | 0.538 |
| 0.4 | 0 | ycaM | 5 | 24.1333 | 0.531 |
| 0.4 | 0 | yadI | 5 | 24.1333 | 0.618 |
| 0.4 | 0 | AG1 | 5 | 24.1333 | 0.73 |
| 0.4 | 0 | ptsI | 5 | 24.3833 | 0.642 |
| 0.4 | 0 | clcB | 5 | 24.3833 | 0.535 |
| 0.4 | 0 | ycaM | 5 | 24.3833 | 0.53 |
| 0.4 | 0 | yadI | 5 | 24.3833 | 0.615 |
| 0.4 | 0 | AG1 | 5 | 24.3833 | 0.731 |
| 0.8 | 0 | ptsI | 1 | 0 | 0.253 |
| 0.8 | 0 | clcB | 1 | 0 | 0.217 |
| 0.8 | 0 | ycaM | 1 | 0 | 0.233 |
| 0.8 | 0 | yadI | 1 | 0 | 0.247 |
| 0.8 | 0 | AG1 | 1 | 0 | 0.231 |
| 0.8 | 0 | ptsI | 1 | 0.25 | 0.24 |
| 0.8 | 0 | clcB | 1 | 0.25 | 0.21 |
| 0.8 | 0 | ycaM | 1 | 0.25 | 0.219 |
| 0.8 | 0 | yadI | 1 | 0.25 | 0.237 |
| 0.8 | 0 | AG1 | 1 | 0.25 | 0.223 |
| 0.8 | 0 | ptsI | 1 | 0.5 | 0.24 |
| 0.8 | 0 | clcB | 1 | 0.5 | 0.206 |
| 0.8 | 0 | ycaM | 1 | 0.5 | 0.213 |
| 0.8 | 0 | yadI | 1 | 0.5 | 0.233 |

| | | | | | |
|---|---|---|---|---|---|
| 0.8 | 0 | AG1 | 1 | 0.5 | 0.221 |
| 0.8 | 0 | ptsI | 1 | 0.75 | 0.241 |
| 0.8 | 0 | clcB | 1 | 0.75 | 0.209 |
| 0.8 | 0 | ycaM | 1 | 0.75 | 0.213 |
| 0.8 | 0 | yadI | 1 | 0.75 | 0.232 |
| 0.8 | 0 | AG1 | 1 | 0.75 | 0.221 |
| 0.8 | 0 | ptsI | 1 | 1 | 0.244 |
| 0.8 | 0 | clcB | 1 | 1 | 0.209 |
| 0.8 | 0 | ycaM | 1 | 1 | 0.214 |
| 0.8 | 0 | yadI | 1 | 1 | 0.235 |
| 0.8 | 0 | AG1 | 1 | 1 | 0.224 |
| 0.8 | 0 | ptsI | 1 | 1.25 | 0.244 |
| 0.8 | 0 | clcB | 1 | 1.25 | 0.208 |
| 0.8 | 0 | ycaM | 1 | 1.25 | 0.213 |
| 0.8 | 0 | yadI | 1 | 1.25 | 0.239 |
| 0.8 | 0 | AG1 | 1 | 1.25 | 0.226 |
| 0.8 | 0 | ptsI | 1 | 1.5 | 0.254 |
| 0.8 | 0 | clcB | 1 | 1.5 | 0.21 |
| 0.8 | 0 | ycaM | 1 | 1.5 | 0.217 |
| 0.8 | 0 | yadI | 1 | 1.5 | 0.242 |
| 0.8 | 0 | AG1 | 1 | 1.5 | 0.228 |
| 0.8 | 0 | ptsI | 1 | 1.75 | 0.255 |
| 0.8 | 0 | clcB | 1 | 1.75 | 0.214 |
| 0.8 | 0 | ycaM | 1 | 1.75 | 0.221 |
| 0.8 | 0 | yadI | 1 | 1.75 | 0.249 |
| 0.8 | 0 | AG1 | 1 | 1.75 | 0.231 |
| 0.8 | 0 | ptsI | 1 | 2 | 0.264 |
| 0.8 | 0 | clcB | 1 | 2 | 0.218 |
| 0.8 | 0 | ycaM | 1 | 2 | 0.224 |
| 0.8 | 0 | yadI | 1 | 2 | 0.254 |
| 0.8 | 0 | AG1 | 1 | 2 | 0.236 |
| 0.8 | 0 | ptsI | 1 | 2.25 | 0.266 |
| 0.8 | 0 | clcB | 1 | 2.25 | 0.221 |
| 0.8 | 0 | ycaM | 1 | 2.25 | 0.23 |
| 0.8 | 0 | yadI | 1 | 2.25 | 0.266 |
| 0.8 | 0 | AG1 | 1 | 2.25 | 0.242 |
| 0.8 | 0 | ptsI | 1 | 2.5 | 0.276 |
| 0.8 | 0 | clcB | 1 | 2.5 | 0.227 |
| 0.8 | 0 | ycaM | 1 | 2.5 | 0.234 |
| 0.8 | 0 | yadI | 1 | 2.5 | 0.272 |
| 0.8 | 0 | AG1 | 1 | 2.5 | 0.249 |
| 0.8 | 0 | ptsI | 1 | 2.75 | 0.281 |
| 0.8 | 0 | clcB | 1 | 2.75 | 0.23 |
| 0.8 | 0 | ycaM | 1 | 2.75 | 0.243 |
| 0.8 | 0 | yadI | 1 | 2.75 | 0.292 |
| 0.8 | 0 | AG1 | 1 | 2.75 | 0.255 |
| 0.8 | 0 | ptsI | 1 | 3 | 0.294 |
| 0.8 | 0 | clcB | 1 | 3 | 0.238 |

| | | | | | |
|---|---|---|---|---|---|
| 0.8 | 0 | ycaM | 1 | 3 | 0.248 |
| 0.8 | 0 | yadI | 1 | 3 | 0.3 |
| 0.8 | 0 | AG1 | 1 | 3 | 0.26 |
| 0.8 | 0 | ptsI | 1 | 3.25 | 0.304 |
| 0.8 | 0 | clcB | 1 | 3.25 | 0.245 |
| 0.8 | 0 | ycaM | 1 | 3.25 | 0.256 |
| 0.8 | 0 | yadI | 1 | 3.25 | 0.319 |
| 0.8 | 0 | AG1 | 1 | 3.25 | 0.267 |
| 0.8 | 0 | ptsI | 1 | 3.5 | 0.311 |
| 0.8 | 0 | clcB | 1 | 3.5 | 0.251 |
| 0.8 | 0 | ycaM | 1 | 3.5 | 0.264 |
| 0.8 | 0 | yadI | 1 | 3.5 | 0.338 |
| 0.8 | 0 | AG1 | 1 | 3.5 | 0.276 |
| 0.8 | 0 | ptsI | 1 | 3.75 | 0.323 |
| 0.8 | 0 | clcB | 1 | 3.75 | 0.254 |
| 0.8 | 0 | ycaM | 1 | 3.75 | 0.271 |
| 0.8 | 0 | yadI | 1 | 3.75 | 0.366 |
| 0.8 | 0 | AG1 | 1 | 3.75 | 0.274 |
| 0.8 | 0 | ptsI | 1 | 4 | 0.339 |
| 0.8 | 0 | clcB | 1 | 4 | 0.26 |
| 0.8 | 0 | ycaM | 1 | 4 | 0.285 |
| 0.8 | 0 | yadI | 1 | 4 | 0.373 |
| 0.8 | 0 | AG1 | 1 | 4 | 0.283 |
| 0.8 | 0 | ptsI | 1 | 4.25 | 0.346 |
| 0.8 | 0 | clcB | 1 | 4.25 | 0.266 |
| 0.8 | 0 | ycaM | 1 | 4.25 | 0.294 |
| 0.8 | 0 | yadI | 1 | 4.25 | 0.386 |
| 0.8 | 0 | AG1 | 1 | 4.25 | 0.293 |
| 0.8 | 0 | ptsI | 1 | 4.5 | 0.352 |
| 0.8 | 0 | clcB | 1 | 4.5 | 0.275 |
| 0.8 | 0 | ycaM | 1 | 4.5 | 0.301 |
| 0.8 | 0 | yadI | 1 | 4.5 | 0.404 |
| 0.8 | 0 | AG1 | 1 | 4.5 | 0.308 |
| 0.8 | 0 | ptsI | 1 | 4.75 | 0.36 |
| 0.8 | 0 | clcB | 1 | 4.75 | 0.281 |
| 0.8 | 0 | ycaM | 1 | 4.75 | 0.306 |
| 0.8 | 0 | yadI | 1 | 4.75 | 0.4 |
| 0.8 | 0 | AG1 | 1 | 4.75 | 0.313 |
| 0.8 | 0 | ptsI | 1 | 5 | 0.375 |
| 0.8 | 0 | clcB | 1 | 5 | 0.289 |
| 0.8 | 0 | ycaM | 1 | 5 | 0.315 |
| 0.8 | 0 | yadI | 1 | 5 | 0.436 |
| 0.8 | 0 | AG1 | 1 | 5 | 0.325 |
| 0.8 | 0 | ptsI | 1 | 5.25 | 0.386 |
| 0.8 | 0 | clcB | 1 | 5.25 | 0.299 |
| 0.8 | 0 | ycaM | 1 | 5.25 | 0.326 |
| 0.8 | 0 | yadI | 1 | 5.25 | 0.445 |
| 0.8 | 0 | AG1 | 1 | 5.25 | 0.332 |

| | | | | | |
|---|---|---|---|---|---|
| 0.8 | 0 | ptsI | 1 | 5.5 | 0.395 |
| 0.8 | 0 | clcB | 1 | 5.5 | 0.308 |
| 0.8 | 0 | ycaM | 1 | 5.5 | 0.333 |
| 0.8 | 0 | yadI | 1 | 5.5 | 0.456 |
| 0.8 | 0 | AG1 | 1 | 5.5 | 0.345 |
| 0.8 | 0 | ptsI | 1 | 5.75 | 0.404 |
| 0.8 | 0 | clcB | 1 | 5.75 | 0.316 |
| 0.8 | 0 | ycaM | 1 | 5.75 | 0.356 |
| 0.8 | 0 | yadI | 1 | 5.75 | 0.47 |
| 0.8 | 0 | AG1 | 1 | 5.75 | 0.356 |
| 0.8 | 0 | ptsI | 1 | 6 | 0.416 |
| 0.8 | 0 | clcB | 1 | 6 | 0.326 |
| 0.8 | 0 | ycaM | 1 | 6 | 0.346 |
| 0.8 | 0 | yadI | 1 | 6 | 0.475 |
| 0.8 | 0 | AG1 | 1 | 6 | 0.364 |
| 0.8 | 0 | ptsI | 1 | 6.25 | 0.431 |
| 0.8 | 0 | clcB | 1 | 6.25 | 0.338 |
| 0.8 | 0 | ycaM | 1 | 6.25 | 0.359 |
| 0.8 | 0 | yadI | 1 | 6.25 | 0.488 |
| 0.8 | 0 | AG1 | 1 | 6.25 | 0.37 |
| 0.8 | 0 | ptsI | 1 | 6.5 | 0.439 |
| 0.8 | 0 | clcB | 1 | 6.5 | 0.348 |
| 0.8 | 0 | ycaM | 1 | 6.5 | 0.371 |
| 0.8 | 0 | yadI | 1 | 6.5 | 0.498 |
| 0.8 | 0 | AG1 | 1 | 6.5 | 0.384 |
| 0.8 | 0 | ptsI | 1 | 6.75 | 0.454 |
| 0.8 | 0 | clcB | 1 | 6.75 | 0.357 |
| 0.8 | 0 | ycaM | 1 | 6.75 | 0.381 |
| 0.8 | 0 | yadI | 1 | 6.75 | 0.523 |
| 0.8 | 0 | AG1 | 1 | 6.75 | 0.4 |
| 0.8 | 0 | ptsI | 1 | 7 | 0.462 |
| 0.8 | 0 | clcB | 1 | 7 | 0.367 |
| 0.8 | 0 | ycaM | 1 | 7 | 0.391 |
| 0.8 | 0 | yadI | 1 | 7 | 0.539 |
| 0.8 | 0 | AG1 | 1 | 7 | 0.414 |
| 0.8 | 0 | ptsI | 1 | 7.25 | 0.472 |
| 0.8 | 0 | clcB | 1 | 7.25 | 0.374 |
| 0.8 | 0 | ycaM | 1 | 7.25 | 0.4 |
| 0.8 | 0 | yadI | 1 | 7.25 | 0.55 |
| 0.8 | 0 | AG1 | 1 | 7.25 | 0.422 |
| 0.8 | 0 | ptsI | 1 | 7.5 | 0.475 |
| 0.8 | 0 | clcB | 1 | 7.5 | 0.384 |
| 0.8 | 0 | ycaM | 1 | 7.5 | 0.41 |
| 0.8 | 0 | yadI | 1 | 7.5 | 0.571 |
| 0.8 | 0 | AG1 | 1 | 7.5 | 0.431 |
| 0.8 | 0 | ptsI | 1 | 7.75 | 0.487 |
| 0.8 | 0 | clcB | 1 | 7.75 | 0.391 |
| 0.8 | 0 | ycaM | 1 | 7.75 | 0.417 |

| | | | | | |
|---|---|---|---|---|---|
| 0.8 | 0 | yadI | 1 | 7.75 | 0.582 |
| 0.8 | 0 | AG1 | 1 | 7.75 | 0.441 |
| 0.8 | 0 | ptsI | 1 | 8 | 0.502 |
| 0.8 | 0 | clcB | 1 | 8 | 0.399 |
| 0.8 | 0 | ycaM | 1 | 8 | 0.424 |
| 0.8 | 0 | yadI | 1 | 8 | 0.591 |
| 0.8 | 0 | AG1 | 1 | 8 | 0.451 |
| 0.8 | 0 | ptsI | 1 | 8.25 | 0.509 |
| 0.8 | 0 | clcB | 1 | 8.25 | 0.408 |
| 0.8 | 0 | ycaM | 1 | 8.25 | 0.433 |
| 0.8 | 0 | yadI | 1 | 8.25 | 0.607 |
| 0.8 | 0 | AG1 | 1 | 8.25 | 0.461 |
| 0.8 | 0 | ptsI | 1 | 8.5 | 0.525 |
| 0.8 | 0 | clcB | 1 | 8.5 | 0.419 |
| 0.8 | 0 | ycaM | 1 | 8.5 | 0.437 |
| 0.8 | 0 | yadI | 1 | 8.5 | 0.63 |
| 0.8 | 0 | AG1 | 1 | 8.5 | 0.472 |
| 0.8 | 0 | ptsI | 1 | 8.75 | 0.539 |
| 0.8 | 0 | clcB | 1 | 8.75 | 0.421 |
| 0.8 | 0 | ycaM | 1 | 8.75 | 0.442 |
| 0.8 | 0 | yadI | 1 | 8.75 | 0.628 |
| 0.8 | 0 | AG1 | 1 | 8.75 | 0.479 |
| 0.8 | 0 | ptsI | 1 | 9 | 0.557 |
| 0.8 | 0 | clcB | 1 | 9 | 0.428 |
| 0.8 | 0 | ycaM | 1 | 9 | 0.446 |
| 0.8 | 0 | yadI | 1 | 9 | 0.644 |
| 0.8 | 0 | AG1 | 1 | 9 | 0.492 |
| 0.8 | 0 | ptsI | 1 | 9.25 | 0.576 |
| 0.8 | 0 | clcB | 1 | 9.25 | 0.443 |
| 0.8 | 0 | ycaM | 1 | 9.25 | 0.454 |
| 0.8 | 0 | yadI | 1 | 9.25 | 0.635 |
| 0.8 | 0 | AG1 | 1 | 9.25 | 0.501 |
| 0.8 | 0 | ptsI | 1 | 9.5 | 0.611 |
| 0.8 | 0 | clcB | 1 | 9.5 | 0.463 |
| 0.8 | 0 | ycaM | 1 | 9.5 | 0.466 |
| 0.8 | 0 | yadI | 1 | 9.5 | 0.65 |
| 0.8 | 0 | AG1 | 1 | 9.5 | 0.524 |
| 0.8 | 0 | ptsI | 1 | 9.75 | 0.607 |
| 0.8 | 0 | clcB | 1 | 9.75 | 0.465 |
| 0.8 | 0 | ycaM | 1 | 9.75 | 0.477 |
| 0.8 | 0 | yadI | 1 | 9.75 | 0.678 |
| 0.8 | 0 | AG1 | 1 | 9.75 | 0.541 |
| 0.8 | 0 | ptsI | 1 | 10 | 0.611 |
| 0.8 | 0 | clcB | 1 | 10 | 0.473 |
| 0.8 | 0 | ycaM | 1 | 10 | 0.478 |
| 0.8 | 0 | yadI | 1 | 10 | 0.665 |
| 0.8 | 0 | AG1 | 1 | 10 | 0.54 |
| 0.8 | 0 | ptsI | 1 | 10.25 | 0.617 |

| | | | | | |
|---|---|---|---|---|---|
| 0.8 | 0 | clcB | 1 | 10.25 | 0.478 |
| 0.8 | 0 | ycaM | 1 | 10.25 | 0.495 |
| 0.8 | 0 | yadI | 1 | 10.25 | 0.688 |
| 0.8 | 0 | AG1 | 1 | 10.25 | 0.559 |
| 0.8 | 0 | ptsI | 1 | 10.5 | 0.634 |
| 0.8 | 0 | clcB | 1 | 10.5 | 0.481 |
| 0.8 | 0 | ycaM | 1 | 10.5 | 0.505 |
| 0.8 | 0 | yadI | 1 | 10.5 | 0.708 |
| 0.8 | 0 | AG1 | 1 | 10.5 | 0.571 |
| 0.8 | 0 | ptsI | 1 | 10.75 | 0.685 |
| 0.8 | 0 | clcB | 1 | 10.75 | 0.499 |
| 0.8 | 0 | ycaM | 1 | 10.75 | 0.506 |
| 0.8 | 0 | yadI | 1 | 10.75 | 0.694 |
| 0.8 | 0 | AG1 | 1 | 10.75 | 0.568 |
| 0.8 | 0 | ptsI | 1 | 11 | 0.68 |
| 0.8 | 0 | clcB | 1 | 11 | 0.52 |
| 0.8 | 0 | ycaM | 1 | 11 | 0.511 |
| 0.8 | 0 | yadI | 1 | 11 | 0.708 |
| 0.8 | 0 | AG1 | 1 | 11 | 0.611 |
| 0.8 | 0 | ptsI | 1 | 11.25 | 0.677 |
| 0.8 | 0 | clcB | 1 | 11.25 | 0.527 |
| 0.8 | 0 | ycaM | 1 | 11.25 | 0.524 |
| 0.8 | 0 | yadI | 1 | 11.25 | 0.714 |
| 0.8 | 0 | AG1 | 1 | 11.25 | 0.617 |
| 0.8 | 0 | ptsI | 1 | 11.5 | 0.728 |
| 0.8 | 0 | clcB | 1 | 11.5 | 0.54 |
| 0.8 | 0 | ycaM | 1 | 11.5 | 0.526 |
| 0.8 | 0 | yadI | 1 | 11.5 | 0.725 |
| 0.8 | 0 | AG1 | 1 | 11.5 | 0.63 |
| 0.8 | 0 | ptsI | 1 | 11.75 | 0.753 |
| 0.8 | 0 | clcB | 1 | 11.75 | 0.56 |
| 0.8 | 0 | ycaM | 1 | 11.75 | 0.539 |
| 0.8 | 0 | yadI | 1 | 11.75 | 0.753 |
| 0.8 | 0 | AG1 | 1 | 11.75 | 0.667 |
| 0.8 | 0 | ptsI | 1 | 12 | 0.741 |
| 0.8 | 0 | clcB | 1 | 12 | 0.558 |
| 0.8 | 0 | ycaM | 1 | 12 | 0.549 |
| 0.8 | 0 | yadI | 1 | 12 | 0.762 |
| 0.8 | 0 | AG1 | 1 | 12 | 0.675 |
| 0.8 | 0 | ptsI | 1 | 12.25 | 0.774 |
| 0.8 | 0 | clcB | 1 | 12.25 | 0.578 |
| 0.8 | 0 | ycaM | 1 | 12.25 | 0.562 |
| 0.8 | 0 | yadI | 1 | 12.25 | 0.769 |
| 0.8 | 0 | AG1 | 1 | 12.25 | 0.686 |
| 0.8 | 0 | ptsI | 1 | 12.5 | 0.766 |
| 0.8 | 0 | clcB | 1 | 12.5 | 0.573 |
| 0.8 | 0 | ycaM | 1 | 12.5 | 0.572 |
| 0.8 | 0 | yadI | 1 | 12.5 | 0.77 |

| | | | | | |
|---|---|---|---|---|---|
| 0.8 | 0 | AG1 | 1 | 12.5 | 0.684 |
| 0.8 | 0 | ptsI | 1 | 12.75 | 0.766 |
| 0.8 | 0 | clcB | 1 | 12.75 | 0.579 |
| 0.8 | 0 | ycaM | 1 | 12.75 | 0.57 |
| 0.8 | 0 | yadI | 1 | 12.75 | 0.777 |
| 0.8 | 0 | AG1 | 1 | 12.75 | 0.713 |
| 0.8 | 0 | ptsI | 1 | 13 | 0.78 |
| 0.8 | 0 | clcB | 1 | 13 | 0.582 |
| 0.8 | 0 | ycaM | 1 | 13 | 0.566 |
| 0.8 | 0 | yadI | 1 | 13 | 0.779 |
| 0.8 | 0 | AG1 | 1 | 13 | 0.727 |
| 0.8 | 0 | ptsI | 1 | 13.25 | 0.812 |
| 0.8 | 0 | clcB | 1 | 13.25 | 0.587 |
| 0.8 | 0 | ycaM | 1 | 13.25 | 0.584 |
| 0.8 | 0 | yadI | 1 | 13.25 | 0.799 |
| 0.8 | 0 | AG1 | 1 | 13.25 | 0.738 |
| 0.8 | 0 | ptsI | 1 | 13.5 | 0.812 |
| 0.8 | 0 | clcB | 1 | 13.5 | 0.6 |
| 0.8 | 0 | ycaM | 1 | 13.5 | 0.591 |
| 0.8 | 0 | yadI | 1 | 13.5 | 0.803 |
| 0.8 | 0 | AG1 | 1 | 13.5 | 0.737 |
| 0.8 | 0 | ptsI | 1 | 13.75 | 0.821 |
| 0.8 | 0 | clcB | 1 | 13.75 | 0.605 |
| 0.8 | 0 | ycaM | 1 | 13.75 | 0.598 |
| 0.8 | 0 | yadI | 1 | 13.75 | 0.786 |
| 0.8 | 0 | AG1 | 1 | 13.75 | 0.738 |
| 0.8 | 0 | ptsI | 1 | 14 | 0.831 |
| 0.8 | 0 | clcB | 1 | 14 | 0.605 |
| 0.8 | 0 | ycaM | 1 | 14 | 0.597 |
| 0.8 | 0 | yadI | 1 | 14 | 0.804 |
| 0.8 | 0 | AG1 | 1 | 14 | 0.735 |
| 0.8 | 0 | ptsI | 1 | 14.25 | 0.803 |
| 0.8 | 0 | clcB | 1 | 14.25 | 0.612 |
| 0.8 | 0 | ycaM | 1 | 14.25 | 0.61 |
| 0.8 | 0 | yadI | 1 | 14.25 | 0.801 |
| 0.8 | 0 | AG1 | 1 | 14.25 | 0.744 |
| 0.8 | 0 | ptsI | 1 | 14.5 | 0.803 |
| 0.8 | 0 | clcB | 1 | 14.5 | 0.626 |
| 0.8 | 0 | ycaM | 1 | 14.5 | 0.62 |
| 0.8 | 0 | yadI | 1 | 14.5 | 0.802 |
| 0.8 | 0 | AG1 | 1 | 14.5 | 0.772 |
| 0.8 | 0 | ptsI | 1 | 14.75 | 0.836 |
| 0.8 | 0 | clcB | 1 | 14.75 | 0.634 |
| 0.8 | 0 | ycaM | 1 | 14.75 | 0.634 |
| 0.8 | 0 | yadI | 1 | 14.75 | 0.83 |
| 0.8 | 0 | AG1 | 1 | 14.75 | 0.779 |
| 0.8 | 0 | ptsI | 1 | 15 | 0.843 |
| 0.8 | 0 | clcB | 1 | 15 | 0.638 |

| | | | | | |
|---|---|---|---|---|---|
| 0.8 | 0 | ycaM | 1 | 15 | 0.628 |
| 0.8 | 0 | yadI | 1 | 15 | 0.826 |
| 0.8 | 0 | AG1 | 1 | 15 | 0.797 |
| 0.8 | 0 | ptsI | 1 | 15.25 | 0.853 |
| 0.8 | 0 | clcB | 1 | 15.25 | 0.67 |
| 0.8 | 0 | ycaM | 1 | 15.25 | 0.638 |
| 0.8 | 0 | yadI | 1 | 15.25 | 0.827 |
| 0.8 | 0 | AG1 | 1 | 15.25 | 0.812 |
| 0.8 | 0 | ptsI | 1 | 15.5 | 0.852 |
| 0.8 | 0 | clcB | 1 | 15.5 | 0.659 |
| 0.8 | 0 | ycaM | 1 | 15.5 | 0.652 |
| 0.8 | 0 | yadI | 1 | 15.5 | 0.82 |
| 0.8 | 0 | AG1 | 1 | 15.5 | 0.804 |
| 0.8 | 0 | ptsI | 1 | 15.75 | 0.854 |
| 0.8 | 0 | clcB | 1 | 15.75 | 0.665 |
| 0.8 | 0 | ycaM | 1 | 15.75 | 0.658 |
| 0.8 | 0 | yadI | 1 | 15.75 | 0.841 |
| 0.8 | 0 | AG1 | 1 | 15.75 | 0.808 |
| 0.8 | 0 | ptsI | 1 | 16 | 0.864 |
| 0.8 | 0 | clcB | 1 | 16 | 0.672 |
| 0.8 | 0 | ycaM | 1 | 16 | 0.664 |
| 0.8 | 0 | yadI | 1 | 16 | 0.83 |
| 0.8 | 0 | AG1 | 1 | 16 | 0.817 |
| 0.8 | 0 | ptsI | 1 | 16.25 | 0.863 |
| 0.8 | 0 | clcB | 1 | 16.25 | 0.654 |
| 0.8 | 0 | ycaM | 1 | 16.25 | 0.684 |
| 0.8 | 0 | yadI | 1 | 16.25 | 0.826 |
| 0.8 | 0 | AG1 | 1 | 16.25 | 0.829 |
| 0.8 | 0 | ptsI | 1 | 16.5 | 0.876 |
| 0.8 | 0 | clcB | 1 | 16.5 | 0.683 |
| 0.8 | 0 | ycaM | 1 | 16.5 | 0.685 |
| 0.8 | 0 | yadI | 1 | 16.5 | 0.831 |
| 0.8 | 0 | AG1 | 1 | 16.5 | 0.845 |
| 0.8 | 0 | ptsI | 1 | 17.0667 | 0.869 |
| 0.8 | 0 | clcB | 1 | 17.0667 | 0.706 |
| 0.8 | 0 | ycaM | 1 | 17.0667 | 0.705 |
| 0.8 | 0 | yadI | 1 | 17.0667 | 0.882 |
| 0.8 | 0 | AG1 | 1 | 17.0667 | 0.828 |
| 0.8 | 0 | ptsI | 1 | 17.3167 | 0.823 |
| 0.8 | 0 | clcB | 1 | 17.3167 | 0.673 |
| 0.8 | 0 | ycaM | 1 | 17.3167 | 0.684 |
| 0.8 | 0 | yadI | 1 | 17.3167 | 0.843 |
| 0.8 | 0 | AG1 | 1 | 17.3167 | 0.803 |
| 0.8 | 0 | ptsI | 1 | 17.5667 | 0.836 |
| 0.8 | 0 | clcB | 1 | 17.5667 | 0.679 |
| 0.8 | 0 | ycaM | 1 | 17.5667 | 0.684 |
| 0.8 | 0 | yadI | 1 | 17.5667 | 0.83 |
| 0.8 | 0 | AG1 | 1 | 17.5667 | 0.831 |

| | | | | | |
|---|---|---|---|---|---|
| 0.8 | 0 | ptsI | 1 | 17.8167 | 0.865 |
| 0.8 | 0 | clcB | 1 | 17.8167 | 0.744 |
| 0.8 | 0 | ycaM | 1 | 17.8167 | 0.717 |
| 0.8 | 0 | yadI | 1 | 17.8167 | 0.836 |
| 0.8 | 0 | AG1 | 1 | 17.8167 | 0.816 |
| 0.8 | 0 | ptsI | 1 | 18.0667 | 0.863 |
| 0.8 | 0 | clcB | 1 | 18.0667 | 0.713 |
| 0.8 | 0 | ycaM | 1 | 18.0667 | 0.71 |
| 0.8 | 0 | yadI | 1 | 18.0667 | 0.842 |
| 0.8 | 0 | AG1 | 1 | 18.0667 | 0.801 |
| 0.8 | 0 | ptsI | 1 | 18.3167 | 0.866 |
| 0.8 | 0 | clcB | 1 | 18.3167 | 0.72 |
| 0.8 | 0 | ycaM | 1 | 18.3167 | 0.714 |
| 0.8 | 0 | yadI | 1 | 18.3167 | 0.836 |
| 0.8 | 0 | AG1 | 1 | 18.3167 | 0.822 |
| 0.8 | 0 | ptsI | 1 | 18.5667 | 0.879 |
| 0.8 | 0 | clcB | 1 | 18.5667 | 0.726 |
| 0.8 | 0 | ycaM | 1 | 18.5667 | 0.734 |
| 0.8 | 0 | yadI | 1 | 18.5667 | 0.85 |
| 0.8 | 0 | AG1 | 1 | 18.5667 | 0.832 |
| 0.8 | 0 | ptsI | 1 | 18.8167 | 0.885 |
| 0.8 | 0 | clcB | 1 | 18.8167 | 0.73 |
| 0.8 | 0 | ycaM | 1 | 18.8167 | 0.731 |
| 0.8 | 0 | yadI | 1 | 18.8167 | 0.849 |
| 0.8 | 0 | AG1 | 1 | 18.8167 | 0.826 |
| 0.8 | 0 | ptsI | 1 | 19.0667 | 0.895 |
| 0.8 | 0 | clcB | 1 | 19.0667 | 0.712 |
| 0.8 | 0 | ycaM | 1 | 19.0667 | 0.734 |
| 0.8 | 0 | yadI | 1 | 19.0667 | 0.863 |
| 0.8 | 0 | AG1 | 1 | 19.0667 | 0.836 |
| 0.8 | 0 | ptsI | 1 | 19.3167 | 0.903 |
| 0.8 | 0 | clcB | 1 | 19.3167 | 0.707 |
| 0.8 | 0 | ycaM | 1 | 19.3167 | 0.738 |
| 0.8 | 0 | yadI | 1 | 19.3167 | 0.857 |
| 0.8 | 0 | AG1 | 1 | 19.3167 | 0.829 |
| 0.8 | 0 | ptsI | 1 | 19.5667 | 0.91 |
| 0.8 | 0 | clcB | 1 | 19.5667 | 0.72 |
| 0.8 | 0 | ycaM | 1 | 19.5667 | 0.737 |
| 0.8 | 0 | yadI | 1 | 19.5667 | 0.864 |
| 0.8 | 0 | AG1 | 1 | 19.5667 | 0.834 |
| 0.8 | 0 | ptsI | 1 | 19.8167 | 0.919 |
| 0.8 | 0 | clcB | 1 | 19.8167 | 0.713 |
| 0.8 | 0 | ycaM | 1 | 19.8167 | 0.741 |
| 0.8 | 0 | yadI | 1 | 19.8167 | 0.871 |
| 0.8 | 0 | AG1 | 1 | 19.8167 | 0.842 |
| 0.8 | 0 | ptsI | 1 | 20.0667 | 0.931 |
| 0.8 | 0 | clcB | 1 | 20.0667 | 0.734 |
| 0.8 | 0 | ycaM | 1 | 20.0667 | 0.747 |

| | | | | | |
|---|---|---|---|---|---|
| 0.8 | 0 | yadI | 1 | 20.0667 | 0.876 |
| 0.8 | 0 | AG1 | 1 | 20.0667 | 0.837 |
| 0.8 | 0 | ptsI | 1 | 20.3167 | 0.94 |
| 0.8 | 0 | clcB | 1 | 20.3167 | 0.74 |
| 0.8 | 0 | ycaM | 1 | 20.3167 | 0.749 |
| 0.8 | 0 | yadI | 1 | 20.3167 | 0.882 |
| 0.8 | 0 | AG1 | 1 | 20.3167 | 0.838 |
| 0.8 | 0 | ptsI | 1 | 20.5667 | 0.944 |
| 0.8 | 0 | clcB | 1 | 20.5667 | 0.742 |
| 0.8 | 0 | ycaM | 1 | 20.5667 | 0.751 |
| 0.8 | 0 | yadI | 1 | 20.5667 | 0.89 |
| 0.8 | 0 | AG1 | 1 | 20.5667 | 0.823 |
| 0.8 | 0 | ptsI | 1 | 20.8167 | 0.954 |
| 0.8 | 0 | clcB | 1 | 20.8167 | 0.75 |
| 0.8 | 0 | ycaM | 1 | 20.8167 | 0.744 |
| 0.8 | 0 | yadI | 1 | 20.8167 | 0.888 |
| 0.8 | 0 | AG1 | 1 | 20.8167 | 0.814 |
| 0.8 | 0 | ptsI | 1 | 21.0667 | 0.972 |
| 0.8 | 0 | clcB | 1 | 21.0667 | 0.751 |
| 0.8 | 0 | ycaM | 1 | 21.0667 | 0.745 |
| 0.8 | 0 | yadI | 1 | 21.0667 | 0.891 |
| 0.8 | 0 | AG1 | 1 | 21.0667 | 0.819 |
| 0.8 | 0 | ptsI | 1 | 21.3167 | 0.977 |
| 0.8 | 0 | clcB | 1 | 21.3167 | 0.751 |
| 0.8 | 0 | ycaM | 1 | 21.3167 | 0.752 |
| 0.8 | 0 | yadI | 1 | 21.3167 | 0.896 |
| 0.8 | 0 | AG1 | 1 | 21.3167 | 0.822 |
| 0.8 | 0 | ptsI | 1 | 21.5667 | 0.987 |
| 0.8 | 0 | clcB | 1 | 21.5667 | 0.754 |
| 0.8 | 0 | ycaM | 1 | 21.5667 | 0.755 |
| 0.8 | 0 | yadI | 1 | 21.5667 | 0.904 |
| 0.8 | 0 | AG1 | 1 | 21.5667 | 0.823 |
| 0.8 | 0 | ptsI | 1 | 21.8167 | 0.99 |
| 0.8 | 0 | clcB | 1 | 21.8167 | 0.757 |
| 0.8 | 0 | ycaM | 1 | 21.8167 | 0.753 |
| 0.8 | 0 | yadI | 1 | 21.8167 | 0.902 |
| 0.8 | 0 | AG1 | 1 | 21.8167 | 0.821 |
| 0.8 | 0 | ptsI | 1 | 22.0667 | 1.003 |
| 0.8 | 0 | clcB | 1 | 22.0667 | 0.763 |
| 0.8 | 0 | ycaM | 1 | 22.0667 | 0.752 |
| 0.8 | 0 | yadI | 1 | 22.0667 | 0.908 |
| 0.8 | 0 | AG1 | 1 | 22.0667 | 0.83 |
| 0.8 | 0 | ptsI | 1 | 22.9333 | 1.099 |
| 0.8 | 0 | clcB | 1 | 22.9333 | 0.65 |
| 0.8 | 0 | ycaM | 1 | 22.9333 | 0.684 |
| 0.8 | 0 | yadI | 1 | 22.9333 | 1.006 |
| 0.8 | 0 | AG1 | 1 | 22.9333 | 0.726 |
| 0.8 | 0 | ptsI | 1 | 23.1833 | 1.088 |

| | | | | | |
|---|---|---|---|---|---|
| 0.8 | 0 | clcB | 1 | 23.1833 | 0.668 |
| 0.8 | 0 | ycaM | 1 | 23.1833 | 0.686 |
| 0.8 | 0 | yadI | 1 | 23.1833 | 0.983 |
| 0.8 | 0 | AG1 | 1 | 23.1833 | 0.708 |
| 0.8 | 0 | ptsI | 1 | 23.4333 | 1.084 |
| 0.8 | 0 | clcB | 1 | 23.4333 | 0.697 |
| 0.8 | 0 | ycaM | 1 | 23.4333 | 0.624 |
| 0.8 | 0 | yadI | 1 | 23.4333 | 0.984 |
| 0.8 | 0 | AG1 | 1 | 23.4333 | 0.731 |
| 0.8 | 0 | ptsI | 1 | 23.6833 | 1.086 |
| 0.8 | 0 | clcB | 1 | 23.6833 | 0.708 |
| 0.8 | 0 | ycaM | 1 | 23.6833 | 0.666 |
| 0.8 | 0 | yadI | 1 | 23.6833 | 0.98 |
| 0.8 | 0 | AG1 | 1 | 23.6833 | 0.744 |
| 0.8 | 0 | ptsI | 1 | 23.9333 | 1.088 |
| 0.8 | 0 | clcB | 1 | 23.9333 | 0.715 |
| 0.8 | 0 | ycaM | 1 | 23.9333 | 0.676 |
| 0.8 | 0 | yadI | 1 | 23.9333 | 0.985 |
| 0.8 | 0 | AG1 | 1 | 23.9333 | 0.765 |
| 0.8 | 0 | ptsI | 1 | 24.1833 | 1.089 |
| 0.8 | 0 | clcB | 1 | 24.1833 | 0.727 |
| 0.8 | 0 | ycaM | 1 | 24.1833 | 0.701 |
| 0.8 | 0 | yadI | 1 | 24.1833 | 0.993 |
| 0.8 | 0 | AG1 | 1 | 24.1833 | 0.787 |
| 0.8 | 0 | ptsI | 1 | 24.4333 | 1.083 |
| 0.8 | 0 | clcB | 1 | 24.4333 | 0.73 |
| 0.8 | 0 | ycaM | 1 | 24.4333 | 0.728 |
| 0.8 | 0 | yadI | 1 | 24.4333 | 0.996 |
| 0.8 | 0 | AG1 | 1 | 24.4333 | 0.81 |
| 0.8 | 0 | ptsI | 1 | 24.6833 | 1.085 |
| 0.8 | 0 | clcB | 1 | 24.6833 | 0.731 |
| 0.8 | 0 | ycaM | 1 | 24.6833 | 0.731 |
| 0.8 | 0 | yadI | 1 | 24.6833 | 1 |
| 0.8 | 0 | AG1 | 1 | 24.6833 | 0.802 |
| 0.8 | 0 | ptsI | 1 | 24.9333 | 1.095 |
| 0.8 | 0 | clcB | 1 | 24.9333 | 0.741 |
| 0.8 | 0 | ycaM | 1 | 24.9333 | 0.738 |
| 0.8 | 0 | yadI | 1 | 24.9333 | 1.008 |
| 0.8 | 0 | AG1 | 1 | 24.9333 | 0.808 |
| 0.8 | 0 | ptsI | 1 | 25.1833 | 1.094 |
| 0.8 | 0 | clcB | 1 | 25.1833 | 0.744 |
| 0.8 | 0 | ycaM | 1 | 25.1833 | 0.751 |
| 0.8 | 0 | yadI | 1 | 25.1833 | 1.005 |
| 0.8 | 0 | AG1 | 1 | 25.1833 | 0.814 |
| 0.8 | 0 | ptsI | 1 | 25.4333 | 1.104 |
| 0.8 | 0 | clcB | 1 | 25.4333 | 0.748 |
| 0.8 | 0 | ycaM | 1 | 25.4333 | 0.77 |
| 0.8 | 0 | yadI | 1 | 25.4333 | 1.014 |

| | | | | | |
|---|---|---|---|---|---|
| 0.8 | 0 | AG1 | 1 | 25.4333 | 0.823 |
| 0.8 | 0 | ptsI | 1 | 25.6833 | 1.103 |
| 0.8 | 0 | clcB | 1 | 25.6833 | 0.75 |
| 0.8 | 0 | ycaM | 1 | 25.6833 | 0.771 |
| 0.8 | 0 | yadI | 1 | 25.6833 | 1.017 |
| 0.8 | 0 | AG1 | 1 | 25.6833 | 0.821 |
| 0.8 | 0 | ptsI | 1 | 25.9333 | 1.103 |
| 0.8 | 0 | clcB | 1 | 25.9333 | 0.766 |
| 0.8 | 0 | ycaM | 1 | 25.9333 | 0.779 |
| 0.8 | 0 | yadI | 1 | 25.9333 | 1.008 |
| 0.8 | 0 | AG1 | 1 | 25.9333 | 0.83 |
| 0.8 | 0 | ptsI | 1 | 26.1833 | 1.112 |
| 0.8 | 0 | clcB | 1 | 26.1833 | 0.756 |
| 0.8 | 0 | ycaM | 1 | 26.1833 | 0.801 |
| 0.8 | 0 | yadI | 1 | 26.1833 | 1.019 |
| 0.8 | 0 | AG1 | 1 | 26.1833 | 0.835 |
| 0.8 | 0 | ptsI | 1 | 26.4333 | 1.113 |
| 0.8 | 0 | clcB | 1 | 26.4333 | 0.76 |
| 0.8 | 0 | ycaM | 1 | 26.4333 | 0.786 |
| 0.8 | 0 | yadI | 1 | 26.4333 | 1.026 |
| 0.8 | 0 | AG1 | 1 | 26.4333 | 0.83 |
| 0.8 | 0 | ptsI | 1 | 26.6833 | 1.121 |
| 0.8 | 0 | clcB | 1 | 26.6833 | 0.77 |
| 0.8 | 0 | ycaM | 1 | 26.6833 | 0.788 |
| 0.8 | 0 | yadI | 1 | 26.6833 | 1.025 |
| 0.8 | 0 | AG1 | 1 | 26.6833 | 0.833 |
| 0.8 | 0 | ptsI | 1 | 26.9333 | 1.125 |
| 0.8 | 0 | clcB | 1 | 26.9333 | 0.774 |
| 0.8 | 0 | ycaM | 1 | 26.9333 | 0.798 |
| 0.8 | 0 | yadI | 1 | 26.9333 | 1.022 |
| 0.8 | 0 | AG1 | 1 | 26.9333 | 0.839 |
| 0.8 | 0 | ptsI | 1 | 27.1833 | 1.127 |
| 0.8 | 0 | clcB | 1 | 27.1833 | 0.785 |
| 0.8 | 0 | ycaM | 1 | 27.1833 | 0.818 |
| 0.8 | 0 | yadI | 1 | 27.1833 | 1.023 |
| 0.8 | 0 | AG1 | 1 | 27.1833 | 0.844 |
| 0.8 | 0 | ptsI | 1 | 27.4333 | 1.132 |
| 0.8 | 0 | clcB | 1 | 27.4333 | 0.78 |
| 0.8 | 0 | ycaM | 1 | 27.4333 | 0.794 |
| 0.8 | 0 | yadI | 1 | 27.4333 | 1.027 |
| 0.8 | 0 | AG1 | 1 | 27.4333 | 0.836 |
| 0.8 | 0 | ptsI | 1 | 27.6833 | 1.138 |
| 0.8 | 0 | clcB | 1 | 27.6833 | 0.786 |
| 0.8 | 0 | ycaM | 1 | 27.6833 | 0.803 |
| 0.8 | 0 | yadI | 1 | 27.6833 | 1.03 |
| 0.8 | 0 | AG1 | 1 | 27.6833 | 0.848 |
| 0.8 | 0 | ptsI | 1 | 27.9333 | 1.136 |
| 0.8 | 0 | clcB | 1 | 27.9333 | 0.778 |

| | | | | | |
|---|---|---|---|---|---|
| 0.8 | 0 | ycaM | 1 | 27.9333 | 0.801 |
| 0.8 | 0 | yadI | 1 | 27.9333 | 1.016 |
| 0.8 | 0 | AG1 | 1 | 27.9333 | 0.848 |
| 0.8 | 0 | ptsI | 1 | 28.1833 | 1.137 |
| 0.8 | 0 | clcB | 1 | 28.1833 | 0.788 |
| 0.8 | 0 | ycaM | 1 | 28.1833 | 0.816 |
| 0.8 | 0 | yadI | 1 | 28.1833 | 1.028 |
| 0.8 | 0 | AG1 | 1 | 28.1833 | 0.855 |
| 0.8 | 0 | ptsI | 1 | 28.4333 | 1.143 |
| 0.8 | 0 | clcB | 1 | 28.4333 | 0.785 |
| 0.8 | 0 | ycaM | 1 | 28.4333 | 0.814 |
| 0.8 | 0 | yadI | 1 | 28.4333 | 1.03 |
| 0.8 | 0 | AG1 | 1 | 28.4333 | 0.855 |
| 0.8 | 0 | ptsI | 1 | 28.6833 | 1.148 |
| 0.8 | 0 | clcB | 1 | 28.6833 | 0.788 |
| 0.8 | 0 | ycaM | 1 | 28.6833 | 0.821 |
| 0.8 | 0 | yadI | 1 | 28.6833 | 1.03 |
| 0.8 | 0 | AG1 | 1 | 28.6833 | 0.859 |
| 0.8 | 0 | ptsI | 1 | 28.9333 | 1.153 |
| 0.8 | 0 | clcB | 1 | 28.9333 | 0.787 |
| 0.8 | 0 | ycaM | 1 | 28.9333 | 0.831 |
| 0.8 | 0 | yadI | 1 | 28.9333 | 1.028 |
| 0.8 | 0 | AG1 | 1 | 28.9333 | 0.863 |
| 0.8 | 0 | ptsI | 1 | 29.1833 | 1.149 |
| 0.8 | 0 | clcB | 1 | 29.1833 | 0.785 |
| 0.8 | 0 | ycaM | 1 | 29.1833 | 0.828 |
| 0.8 | 0 | yadI | 1 | 29.1833 | 1.03 |
| 0.8 | 0 | AG1 | 1 | 29.1833 | 0.859 |
| 0.8 | 0 | ptsI | 1 | 29.4333 | 1.154 |
| 0.8 | 0 | clcB | 1 | 29.4333 | 0.783 |
| 0.8 | 0 | ycaM | 1 | 29.4333 | 0.84 |
| 0.8 | 0 | yadI | 1 | 29.4333 | 1.03 |
| 0.8 | 0 | AG1 | 1 | 29.4333 | 0.865 |
| 0.8 | 0 | ptsI | 1 | 29.6833 | 1.158 |
| 0.8 | 0 | clcB | 1 | 29.6833 | 0.783 |
| 0.8 | 0 | ycaM | 1 | 29.6833 | 0.842 |
| 0.8 | 0 | yadI | 1 | 29.6833 | 1.035 |
| 0.8 | 0 | AG1 | 1 | 29.6833 | 0.867 |
| 0.8 | 0 | ptsI | 1 | 29.9333 | 1.161 |
| 0.8 | 0 | clcB | 1 | 29.9333 | 0.785 |
| 0.8 | 0 | ycaM | 1 | 29.9333 | 0.856 |
| 0.8 | 0 | yadI | 1 | 29.9333 | 1.036 |
| 0.8 | 0 | AG1 | 1 | 29.9333 | 0.873 |
| 0.8 | 0 | ptsI | 1 | 30.1833 | 1.172 |
| 0.8 | 0 | clcB | 1 | 30.1833 | 0.782 |
| 0.8 | 0 | ycaM | 1 | 30.1833 | 0.86 |
| 0.8 | 0 | yadI | 1 | 30.1833 | 1.038 |
| 0.8 | 0 | AG1 | 1 | 30.1833 | 0.879 |

| | | | | | |
|---|---|---|---|---|---|
| 0.8 | 0 | ptsI | 1 | 30.4333 | 1.169 |
| 0.8 | 0 | clcB | 1 | 30.4333 | 0.783 |
| 0.8 | 0 | ycaM | 1 | 30.4333 | 0.867 |
| 0.8 | 0 | yadI | 1 | 30.4333 | 1.038 |
| 0.8 | 0 | AG1 | 1 | 30.4333 | 0.881 |
| 0.8 | 0 | ptsI | 1 | 30.6833 | 1.174 |
| 0.8 | 0 | clcB | 1 | 30.6833 | 0.787 |
| 0.8 | 0 | ycaM | 1 | 30.6833 | 0.872 |
| 0.8 | 0 | yadI | 1 | 30.6833 | 1.047 |
| 0.8 | 0 | AG1 | 1 | 30.6833 | 0.885 |
| 0.8 | 0 | ptsI | 1 | 30.9333 | 1.176 |
| 0.8 | 0 | clcB | 1 | 30.9333 | 0.784 |
| 0.8 | 0 | ycaM | 1 | 30.9333 | 0.884 |
| 0.8 | 0 | yadI | 1 | 30.9333 | 1.045 |
| 0.8 | 0 | AG1 | 1 | 30.9333 | 0.891 |
| 0.8 | 0 | ptsI | 1 | 31.1833 | 1.19 |
| 0.8 | 0 | clcB | 1 | 31.1833 | 0.783 |
| 0.8 | 0 | ycaM | 1 | 31.1833 | 0.892 |
| 0.8 | 0 | yadI | 1 | 31.1833 | 1.048 |
| 0.8 | 0 | AG1 | 1 | 31.1833 | 0.892 |
| 0.8 | 0 | ptsI | 1 | 31.4333 | 1.193 |
| 0.8 | 0 | clcB | 1 | 31.4333 | 0.787 |
| 0.8 | 0 | ycaM | 1 | 31.4333 | 0.905 |
| 0.8 | 0 | yadI | 1 | 31.4333 | 1.05 |
| 0.8 | 0 | AG1 | 1 | 31.4333 | 0.901 |
| 0.8 | 0 | ptsI | 1 | 31.6833 | 1.2 |
| 0.8 | 0 | clcB | 1 | 31.6833 | 0.785 |
| 0.8 | 0 | ycaM | 1 | 31.6833 | 0.911 |
| 0.8 | 0 | yadI | 1 | 31.6833 | 1.049 |
| 0.8 | 0 | AG1 | 1 | 31.6833 | 0.906 |
| 0.8 | 0 | ptsI | 1 | 31.9333 | 1.206 |
| 0.8 | 0 | clcB | 1 | 31.9333 | 0.786 |
| 0.8 | 0 | ycaM | 1 | 31.9333 | 0.923 |
| 0.8 | 0 | yadI | 1 | 31.9333 | 1.055 |
| 0.8 | 0 | AG1 | 1 | 31.9333 | 0.908 |
| 0.8 | 0 | ptsI | 1 | 32.1833 | 1.205 |
| 0.8 | 0 | clcB | 1 | 32.1833 | 0.788 |
| 0.8 | 0 | ycaM | 1 | 32.1833 | 0.927 |
| 0.8 | 0 | yadI | 1 | 32.1833 | 1.052 |
| 0.8 | 0 | AG1 | 1 | 32.1833 | 0.916 |
| 0.8 | 0 | ptsI | 1 | 32.4333 | 1.222 |
| 0.8 | 0 | clcB | 1 | 32.4333 | 0.8 |
| 0.8 | 0 | ycaM | 1 | 32.4333 | 0.934 |
| 0.8 | 0 | yadI | 1 | 32.4333 | 1.034 |
| 0.8 | 0 | AG1 | 1 | 32.4333 | 0.92 |
| 0.8 | 0 | ptsI | 1 | 32.6833 | 1.226 |
| 0.8 | 0 | clcB | 1 | 32.6833 | 0.794 |
| 0.8 | 0 | ycaM | 1 | 32.6833 | 0.949 |

| | | | | | |
|---|---|---|---|---|---|
| 0.8 | 0 | yadI | 1 | 32.6833 | 1.055 |
| 0.8 | 0 | AG1 | 1 | 32.6833 | 0.922 |
| 0.8 | 0 | ptsI | 1 | 32.9333 | 1.24 |
| 0.8 | 0 | clcB | 1 | 32.9333 | 0.794 |
| 0.8 | 0 | ycaM | 1 | 32.9333 | 0.955 |
| 0.8 | 0 | yadI | 1 | 32.9333 | 1.058 |
| 0.8 | 0 | AG1 | 1 | 32.9333 | 0.926 |
| 0.8 | 0 | ptsI | 1 | 33.1833 | 1.255 |
| 0.8 | 0 | clcB | 1 | 33.1833 | 0.789 |
| 0.8 | 0 | ycaM | 1 | 33.1833 | 0.96 |
| 0.8 | 0 | yadI | 1 | 33.1833 | 1.049 |
| 0.8 | 0 | AG1 | 1 | 33.1833 | 0.934 |
| 0.8 | 0 | ptsI | 1 | 33.4333 | 1.261 |
| 0.8 | 0 | clcB | 1 | 33.4333 | 0.796 |
| 0.8 | 0 | ycaM | 1 | 33.4333 | 0.977 |
| 0.8 | 0 | yadI | 1 | 33.4333 | 1.059 |
| 0.8 | 0 | AG1 | 1 | 33.4333 | 0.943 |
| 0.8 | 0 | ptsI | 1 | 33.6833 | 1.272 |
| 0.8 | 0 | clcB | 1 | 33.6833 | 0.794 |
| 0.8 | 0 | ycaM | 1 | 33.6833 | 0.981 |
| 0.8 | 0 | yadI | 1 | 33.6833 | 1.049 |
| 0.8 | 0 | AG1 | 1 | 33.6833 | 0.952 |
| 0.8 | 0 | ptsI | 1 | 33.9333 | 1.276 |
| 0.8 | 0 | clcB | 1 | 33.9333 | 0.805 |
| 0.8 | 0 | ycaM | 1 | 33.9333 | 0.991 |
| 0.8 | 0 | yadI | 1 | 33.9333 | 1.059 |
| 0.8 | 0 | AG1 | 1 | 33.9333 | 0.952 |
| 0.8 | 0 | ptsI | 1 | 34.1833 | 1.287 |
| 0.8 | 0 | clcB | 1 | 34.1833 | 0.798 |
| 0.8 | 0 | ycaM | 1 | 34.1833 | 0.997 |
| 0.8 | 0 | yadI | 1 | 34.1833 | 1.072 |
| 0.8 | 0 | AG1 | 1 | 34.1833 | 0.954 |
| 0.8 | 0 | ptsI | 1 | 34.4333 | 1.295 |
| 0.8 | 0 | clcB | 1 | 34.4333 | 0.817 |
| 0.8 | 0 | ycaM | 1 | 34.4333 | 1.009 |
| 0.8 | 0 | yadI | 1 | 34.4333 | 1.066 |
| 0.8 | 0 | AG1 | 1 | 34.4333 | 0.967 |
| 0.8 | 0 | ptsI | 1 | 34.6833 | 1.298 |
| 0.8 | 0 | clcB | 1 | 34.6833 | 0.812 |
| 0.8 | 0 | ycaM | 1 | 34.6833 | 1.022 |
| 0.8 | 0 | yadI | 1 | 34.6833 | 1.07 |
| 0.8 | 0 | AG1 | 1 | 34.6833 | 0.976 |
| 0.8 | 0 | ptsI | 1 | 34.9333 | 1.308 |
| 0.8 | 0 | clcB | 1 | 34.9333 | 0.826 |
| 0.8 | 0 | ycaM | 1 | 34.9333 | 1.026 |
| 0.8 | 0 | yadI | 1 | 34.9333 | 1.069 |
| 0.8 | 0 | AG1 | 1 | 34.9333 | 0.976 |
| 0.8 | 0 | ptsI | 1 | 35.1833 | 1.313 |

| | | | | | |
|---|---|---|---|---|---|
| 0.8 | 0 | clcB | 1 | 35.1833 | 0.829 |
| 0.8 | 0 | ycaM | 1 | 35.1833 | 1.035 |
| 0.8 | 0 | yadI | 1 | 35.1833 | 1.072 |
| 0.8 | 0 | AG1 | 1 | 35.1833 | 0.981 |
| 0.8 | 0 | ptsI | 1 | 35.4333 | 1.326 |
| 0.8 | 0 | clcB | 1 | 35.4333 | 0.84 |
| 0.8 | 0 | ycaM | 1 | 35.4333 | 1.042 |
| 0.8 | 0 | yadI | 1 | 35.4333 | 1.077 |
| 0.8 | 0 | AG1 | 1 | 35.4333 | 0.99 |
| 0.8 | 0 | ptsI | 1 | 35.6833 | 1.329 |
| 0.8 | 0 | clcB | 1 | 35.6833 | 0.834 |
| 0.8 | 0 | ycaM | 1 | 35.6833 | 1.053 |
| 0.8 | 0 | yadI | 1 | 35.6833 | 1.08 |
| 0.8 | 0 | AG1 | 1 | 35.6833 | 0.997 |
| 0.8 | 0 | ptsI | 1 | 35.9333 | 1.342 |
| 0.8 | 0 | clcB | 1 | 35.9333 | 0.831 |
| 0.8 | 0 | ycaM | 1 | 35.9333 | 1.057 |
| 0.8 | 0 | yadI | 1 | 35.9333 | 1.074 |
| 0.8 | 0 | AG1 | 1 | 35.9333 | 1.008 |
| 0.8 | 0 | ptsI | 1 | 36.1833 | 1.339 |
| 0.8 | 0 | clcB | 1 | 36.1833 | 0.834 |
| 0.8 | 0 | ycaM | 1 | 36.1833 | 1.071 |
| 0.8 | 0 | yadI | 1 | 36.1833 | 1.085 |
| 0.8 | 0 | AG1 | 1 | 36.1833 | 1.014 |
| 0.8 | 0 | ptsI | 1 | 36.4333 | 1.356 |
| 0.8 | 0 | clcB | 1 | 36.4333 | 0.835 |
| 0.8 | 0 | ycaM | 1 | 36.4333 | 1.083 |
| 0.8 | 0 | yadI | 1 | 36.4333 | 1.089 |
| 0.8 | 0 | AG1 | 1 | 36.4333 | 1.022 |
| 0.8 | 0 | ptsI | 1 | 36.6833 | 1.366 |
| 0.8 | 0 | clcB | 1 | 36.6833 | 0.836 |
| 0.8 | 0 | ycaM | 1 | 36.6833 | 1.098 |
| 0.8 | 0 | yadI | 1 | 36.6833 | 1.091 |
| 0.8 | 0 | AG1 | 1 | 36.6833 | 1.03 |
| 0.8 | 0 | ptsI | 2 | 0 | 0.206 |
| 0.8 | 0 | clcB | 2 | 0 | 0.202 |
| 0.8 | 0 | ycaM | 2 | 0 | 0.218 |
| 0.8 | 0 | yadI | 2 | 0 | 0.221 |
| 0.8 | 0 | AG1 | 2 | 0 | 0.245 |
| 0.8 | 0 | ptsI | 2 | 0.25 | 0.208 |
| 0.8 | 0 | clcB | 2 | 0.25 | 0.2 |
| 0.8 | 0 | ycaM | 2 | 0.25 | 0.209 |
| 0.8 | 0 | yadI | 2 | 0.25 | 0.217 |
| 0.8 | 0 | AG1 | 2 | 0.25 | 0.241 |
| 0.8 | 0 | ptsI | 2 | 0.5 | 0.213 |
| 0.8 | 0 | clcB | 2 | 0.5 | 0.2 |
| 0.8 | 0 | ycaM | 2 | 0.5 | 0.21 |
| 0.8 | 0 | yadI | 2 | 0.5 | 0.214 |

| | | | | | |
|---|---|---|---|---|---|
| 0.8 | 0 | AG1 | 2 | 0.5 | 0.24 |
| 0.8 | 0 | ptsI | 2 | 0.75 | 0.214 |
| 0.8 | 0 | clcB | 2 | 0.75 | 0.201 |
| 0.8 | 0 | ycaM | 2 | 0.75 | 0.213 |
| 0.8 | 0 | yadI | 2 | 0.75 | 0.216 |
| 0.8 | 0 | AG1 | 2 | 0.75 | 0.243 |
| 0.8 | 0 | ptsI | 2 | 1 | 0.218 |
| 0.8 | 0 | clcB | 2 | 1 | 0.203 |
| 0.8 | 0 | ycaM | 2 | 1 | 0.22 |
| 0.8 | 0 | yadI | 2 | 1 | 0.223 |
| 0.8 | 0 | AG1 | 2 | 1 | 0.247 |
| 0.8 | 0 | ptsI | 2 | 1.25 | 0.222 |
| 0.8 | 0 | clcB | 2 | 1.25 | 0.206 |
| 0.8 | 0 | ycaM | 2 | 1.25 | 0.253 |
| 0.8 | 0 | yadI | 2 | 1.25 | 0.229 |
| 0.8 | 0 | AG1 | 2 | 1.25 | 0.254 |
| 0.8 | 0 | ptsI | 2 | 1.5 | 0.225 |
| 0.8 | 0 | clcB | 2 | 1.5 | 0.206 |
| 0.8 | 0 | ycaM | 2 | 1.5 | 0.254 |
| 0.8 | 0 | yadI | 2 | 1.5 | 0.237 |
| 0.8 | 0 | AG1 | 2 | 1.5 | 0.255 |
| 0.8 | 0 | ptsI | 2 | 1.75 | 0.232 |
| 0.8 | 0 | clcB | 2 | 1.75 | 0.208 |
| 0.8 | 0 | ycaM | 2 | 1.75 | 0.228 |
| 0.8 | 0 | yadI | 2 | 1.75 | 0.243 |
| 0.8 | 0 | AG1 | 2 | 1.75 | 0.261 |
| 0.8 | 0 | ptsI | 2 | 2 | 0.238 |
| 0.8 | 0 | clcB | 2 | 2 | 0.212 |
| 0.8 | 0 | ycaM | 2 | 2 | 0.28 |
| 0.8 | 0 | yadI | 2 | 2 | 0.251 |
| 0.8 | 0 | AG1 | 2 | 2 | 0.275 |
| 0.8 | 0 | ptsI | 2 | 2.25 | 0.249 |
| 0.8 | 0 | clcB | 2 | 2.25 | 0.215 |
| 0.8 | 0 | ycaM | 2 | 2.25 | 0.241 |
| 0.8 | 0 | yadI | 2 | 2.25 | 0.263 |
| 0.8 | 0 | AG1 | 2 | 2.25 | 0.285 |
| 0.8 | 0 | ptsI | 2 | 2.5 | 0.26 |
| 0.8 | 0 | clcB | 2 | 2.5 | 0.22 |
| 0.8 | 0 | ycaM | 2 | 2.5 | 0.253 |
| 0.8 | 0 | yadI | 2 | 2.5 | 0.278 |
| 0.8 | 0 | AG1 | 2 | 2.5 | 0.294 |
| 0.8 | 0 | ptsI | 2 | 2.75 | 0.266 |
| 0.8 | 0 | clcB | 2 | 2.75 | 0.226 |
| 0.8 | 0 | ycaM | 2 | 2.75 | 0.256 |
| 0.8 | 0 | yadI | 2 | 2.75 | 0.291 |
| 0.8 | 0 | AG1 | 2 | 2.75 | 0.298 |
| 0.8 | 0 | ptsI | 2 | 3 | 0.272 |
| 0.8 | 0 | clcB | 2 | 3 | 0.231 |

| | | | | | |
|---|---|---|---|---|---|
| 0.8 | 0 | ycaM | 2 | 3 | 0.258 |
| 0.8 | 0 | yadI | 2 | 3 | 0.304 |
| 0.8 | 0 | AG1 | 2 | 3 | 0.301 |
| 0.8 | 0 | ptsI | 2 | 3.25 | 0.281 |
| 0.8 | 0 | clcB | 2 | 3.25 | 0.236 |
| 0.8 | 0 | ycaM | 2 | 3.25 | 0.27 |
| 0.8 | 0 | yadI | 2 | 3.25 | 0.324 |
| 0.8 | 0 | AG1 | 2 | 3.25 | 0.321 |
| 0.8 | 0 | ptsI | 2 | 3.5 | 0.285 |
| 0.8 | 0 | clcB | 2 | 3.5 | 0.242 |
| 0.8 | 0 | ycaM | 2 | 3.5 | 0.276 |
| 0.8 | 0 | yadI | 2 | 3.5 | 0.344 |
| 0.8 | 0 | AG1 | 2 | 3.5 | 0.324 |
| 0.8 | 0 | ptsI | 2 | 3.75 | 0.294 |
| 0.8 | 0 | clcB | 2 | 3.75 | 0.246 |
| 0.8 | 0 | ycaM | 2 | 3.75 | 0.287 |
| 0.8 | 0 | yadI | 2 | 3.75 | 0.362 |
| 0.8 | 0 | AG1 | 2 | 3.75 | 0.338 |
| 0.8 | 0 | ptsI | 2 | 4 | 0.306 |
| 0.8 | 0 | clcB | 2 | 4 | 0.255 |
| 0.8 | 0 | ycaM | 2 | 4 | 0.299 |
| 0.8 | 0 | yadI | 2 | 4 | 0.38 |
| 0.8 | 0 | AG1 | 2 | 4 | 0.349 |
| 0.8 | 0 | ptsI | 2 | 4.38333 | 0.321 |
| 0.8 | 0 | clcB | 2 | 4.38333 | 0.261 |
| 0.8 | 0 | ycaM | 2 | 4.38333 | 0.32 |
| 0.8 | 0 | yadI | 2 | 4.38333 | 0.39 |
| 0.8 | 0 | AG1 | 2 | 4.38333 | 0.302 |
| 0.8 | 0 | ptsI | 2 | 4.63333 | 0.315 |
| 0.8 | 0 | clcB | 2 | 4.63333 | 0.255 |
| 0.8 | 0 | ycaM | 2 | 4.63333 | 0.362 |
| 0.8 | 0 | yadI | 2 | 4.63333 | 0.386 |
| 0.8 | 0 | AG1 | 2 | 4.63333 | 0.303 |
| 0.8 | 0 | ptsI | 2 | 4.88333 | 0.322 |
| 0.8 | 0 | clcB | 2 | 4.88333 | 0.261 |
| 0.8 | 0 | ycaM | 2 | 4.88333 | 0.317 |
| 0.8 | 0 | yadI | 2 | 4.88333 | 0.395 |
| 0.8 | 0 | AG1 | 2 | 4.88333 | 0.314 |
| 0.8 | 0 | ptsI | 2 | 5.13333 | 0.327 |
| 0.8 | 0 | clcB | 2 | 5.13333 | 0.266 |
| 0.8 | 0 | ycaM | 2 | 5.13333 | 0.32 |
| 0.8 | 0 | yadI | 2 | 5.13333 | 0.407 |
| 0.8 | 0 | AG1 | 2 | 5.13333 | 0.322 |
| 0.8 | 0 | ptsI | 2 | 5.38333 | 0.336 |
| 0.8 | 0 | clcB | 2 | 5.38333 | 0.274 |
| 0.8 | 0 | ycaM | 2 | 5.38333 | 0.33 |
| 0.8 | 0 | yadI | 2 | 5.38333 | 0.419 |
| 0.8 | 0 | AG1 | 2 | 5.38333 | 0.329 |

| | | | | | |
|---|---|---|---|---|---|
| 0.8 | 0 | ptsI | 2 | 5.63333 | 0.345 |
| 0.8 | 0 | clcB | 2 | 5.63333 | 0.285 |
| 0.8 | 0 | ycaM | 2 | 5.63333 | 0.339 |
| 0.8 | 0 | yadI | 2 | 5.63333 | 0.428 |
| 0.8 | 0 | AG1 | 2 | 5.63333 | 0.337 |
| 0.8 | 0 | ptsI | 2 | 5.88333 | 0.353 |
| 0.8 | 0 | clcB | 2 | 5.88333 | 0.291 |
| 0.8 | 0 | ycaM | 2 | 5.88333 | 0.349 |
| 0.8 | 0 | yadI | 2 | 5.88333 | 0.435 |
| 0.8 | 0 | AG1 | 2 | 5.88333 | 0.348 |
| 0.8 | 0 | ptsI | 2 | 6.13333 | 0.362 |
| 0.8 | 0 | clcB | 2 | 6.13333 | 0.298 |
| 0.8 | 0 | ycaM | 2 | 6.13333 | 0.358 |
| 0.8 | 0 | yadI | 2 | 6.13333 | 0.448 |
| 0.8 | 0 | AG1 | 2 | 6.13333 | 0.357 |
| 0.8 | 0 | ptsI | 2 | 6.38333 | 0.369 |
| 0.8 | 0 | clcB | 2 | 6.38333 | 0.306 |
| 0.8 | 0 | ycaM | 2 | 6.38333 | 0.377 |
| 0.8 | 0 | yadI | 2 | 6.38333 | 0.457 |
| 0.8 | 0 | AG1 | 2 | 6.38333 | 0.364 |
| 0.8 | 0 | ptsI | 2 | 6.63333 | 0.379 |
| 0.8 | 0 | clcB | 2 | 6.63333 | 0.312 |
| 0.8 | 0 | ycaM | 2 | 6.63333 | 0.376 |
| 0.8 | 0 | yadI | 2 | 6.63333 | 0.466 |
| 0.8 | 0 | AG1 | 2 | 6.63333 | 0.372 |
| 0.8 | 0 | ptsI | 2 | 6.88333 | 0.387 |
| 0.8 | 0 | clcB | 2 | 6.88333 | 0.32 |
| 0.8 | 0 | ycaM | 2 | 6.88333 | 0.392 |
| 0.8 | 0 | yadI | 2 | 6.88333 | 0.477 |
| 0.8 | 0 | AG1 | 2 | 6.88333 | 0.379 |
| 0.8 | 0 | ptsI | 2 | 7.13333 | 0.394 |
| 0.8 | 0 | clcB | 2 | 7.13333 | 0.326 |
| 0.8 | 0 | ycaM | 2 | 7.13333 | 0.401 |
| 0.8 | 0 | yadI | 2 | 7.13333 | 0.485 |
| 0.8 | 0 | AG1 | 2 | 7.13333 | 0.383 |
| 0.8 | 0 | ptsI | 2 | 7.38333 | 0.403 |
| 0.8 | 0 | clcB | 2 | 7.38333 | 0.332 |
| 0.8 | 0 | ycaM | 2 | 7.38333 | 0.402 |
| 0.8 | 0 | yadI | 2 | 7.38333 | 0.496 |
| 0.8 | 0 | AG1 | 2 | 7.38333 | 0.391 |
| 0.8 | 0 | ptsI | 2 | 7.63333 | 0.412 |
| 0.8 | 0 | clcB | 2 | 7.63333 | 0.342 |
| 0.8 | 0 | ycaM | 2 | 7.63333 | 0.408 |
| 0.8 | 0 | yadI | 2 | 7.63333 | 0.508 |
| 0.8 | 0 | AG1 | 2 | 7.63333 | 0.395 |
| 0.8 | 0 | ptsI | 2 | 7.88333 | 0.418 |
| 0.8 | 0 | clcB | 2 | 7.88333 | 0.35 |
| 0.8 | 0 | ycaM | 2 | 7.88333 | 0.415 |

| | | | | | |
|---|---|---|---|---|---|
| 0.8 | 0 | yadI | 2 | 7.88333 | 0.519 |
| 0.8 | 0 | AG1 | 2 | 7.88333 | 0.405 |
| 0.8 | 0 | ptsI | 2 | 8.13333 | 0.428 |
| 0.8 | 0 | clcB | 2 | 8.13333 | 0.356 |
| 0.8 | 0 | ycaM | 2 | 8.13333 | 0.419 |
| 0.8 | 0 | yadI | 2 | 8.13333 | 0.526 |
| 0.8 | 0 | AG1 | 2 | 8.13333 | 0.413 |
| 0.8 | 0 | ptsI | 2 | 8.38333 | 0.438 |
| 0.8 | 0 | clcB | 2 | 8.38333 | 0.366 |
| 0.8 | 0 | ycaM | 2 | 8.38333 | 0.422 |
| 0.8 | 0 | yadI | 2 | 8.38333 | 0.54 |
| 0.8 | 0 | AG1 | 2 | 8.38333 | 0.422 |
| 0.8 | 0 | ptsI | 2 | 8.91667 | 0.353 |
| 0.8 | 0 | clcB | 2 | 8.91667 | 0.314 |
| 0.8 | 0 | ycaM | 2 | 8.91667 | 0.354 |
| 0.8 | 0 | yadI | 2 | 8.91667 | 0.61 |
| 0.8 | 0 | AG1 | 2 | 8.91667 | 0.481 |
| 0.8 | 0 | ptsI | 2 | 9.16667 | 0.356 |
| 0.8 | 0 | clcB | 2 | 9.16667 | 0.319 |
| 0.8 | 0 | ycaM | 2 | 9.16667 | 0.358 |
| 0.8 | 0 | yadI | 2 | 9.16667 | 0.612 |
| 0.8 | 0 | AG1 | 2 | 9.16667 | 0.484 |
| 0.8 | 0 | ptsI | 2 | 9.41667 | 0.364 |
| 0.8 | 0 | clcB | 2 | 9.41667 | 0.325 |
| 0.8 | 0 | ycaM | 2 | 9.41667 | 0.363 |
| 0.8 | 0 | yadI | 2 | 9.41667 | 0.625 |
| 0.8 | 0 | AG1 | 2 | 9.41667 | 0.496 |
| 0.8 | 0 | ptsI | 2 | 9.66667 | 0.372 |
| 0.8 | 0 | clcB | 2 | 9.66667 | 0.333 |
| 0.8 | 0 | ycaM | 2 | 9.66667 | 0.37 |
| 0.8 | 0 | yadI | 2 | 9.66667 | 0.636 |
| 0.8 | 0 | AG1 | 2 | 9.66667 | 0.508 |
| 0.8 | 0 | ptsI | 2 | 9.91667 | 0.38 |
| 0.8 | 0 | clcB | 2 | 9.91667 | 0.338 |
| 0.8 | 0 | ycaM | 2 | 9.91667 | 0.372 |
| 0.8 | 0 | yadI | 2 | 9.91667 | 0.646 |
| 0.8 | 0 | AG1 | 2 | 9.91667 | 0.516 |
| 0.8 | 0 | ptsI | 2 | 10.1667 | 0.39 |
| 0.8 | 0 | clcB | 2 | 10.1667 | 0.342 |
| 0.8 | 0 | ycaM | 2 | 10.1667 | 0.379 |
| 0.8 | 0 | yadI | 2 | 10.1667 | 0.657 |
| 0.8 | 0 | AG1 | 2 | 10.1667 | 0.524 |
| 0.8 | 0 | ptsI | 2 | 10.4167 | 0.4 |
| 0.8 | 0 | clcB | 2 | 10.4167 | 0.349 |
| 0.8 | 0 | ycaM | 2 | 10.4167 | 0.385 |
| 0.8 | 0 | yadI | 2 | 10.4167 | 0.67 |
| 0.8 | 0 | AG1 | 2 | 10.4167 | 0.548 |
| 0.8 | 0 | ptsI | 2 | 10.6667 | 0.424 |

| | | | | | |
|---|---|---|---|---|---|
| 0.8 | 0 | clcB | 2 | 10.6667 | 0.354 |
| 0.8 | 0 | ycaM | 2 | 10.6667 | 0.392 |
| 0.8 | 0 | yadI | 2 | 10.6667 | 0.68 |
| 0.8 | 0 | AG1 | 2 | 10.6667 | 0.541 |
| 0.8 | 0 | ptsI | 2 | 10.9167 | 0.454 |
| 0.8 | 0 | clcB | 2 | 10.9167 | 0.355 |
| 0.8 | 0 | ycaM | 2 | 10.9167 | 0.398 |
| 0.8 | 0 | yadI | 2 | 10.9167 | 0.69 |
| 0.8 | 0 | AG1 | 2 | 10.9167 | 0.546 |
| 0.8 | 0 | ptsI | 2 | 11.1667 | 0.447 |
| 0.8 | 0 | clcB | 2 | 11.1667 | 0.36 |
| 0.8 | 0 | ycaM | 2 | 11.1667 | 0.408 |
| 0.8 | 0 | yadI | 2 | 11.1667 | 0.7 |
| 0.8 | 0 | AG1 | 2 | 11.1667 | 0.556 |
| 0.8 | 0 | ptsI | 2 | 11.4167 | 0.468 |
| 0.8 | 0 | clcB | 2 | 11.4167 | 0.364 |
| 0.8 | 0 | ycaM | 2 | 11.4167 | 0.413 |
| 0.8 | 0 | yadI | 2 | 11.4167 | 0.708 |
| 0.8 | 0 | AG1 | 2 | 11.4167 | 0.561 |
| 0.8 | 0 | ptsI | 2 | 11.6667 | 0.501 |
| 0.8 | 0 | clcB | 2 | 11.6667 | 0.372 |
| 0.8 | 0 | ycaM | 2 | 11.6667 | 0.417 |
| 0.8 | 0 | yadI | 2 | 11.6667 | 0.717 |
| 0.8 | 0 | AG1 | 2 | 11.6667 | 0.593 |
| 0.8 | 0 | ptsI | 2 | 11.9167 | 0.505 |
| 0.8 | 0 | clcB | 2 | 11.9167 | 0.378 |
| 0.8 | 0 | ycaM | 2 | 11.9167 | 0.422 |
| 0.8 | 0 | yadI | 2 | 11.9167 | 0.723 |
| 0.8 | 0 | AG1 | 2 | 11.9167 | 0.583 |
| 0.8 | 0 | ptsI | 2 | 12.1667 | 0.527 |
| 0.8 | 0 | clcB | 2 | 12.1667 | 0.384 |
| 0.8 | 0 | ycaM | 2 | 12.1667 | 0.44 |
| 0.8 | 0 | yadI | 2 | 12.1667 | 0.726 |
| 0.8 | 0 | AG1 | 2 | 12.1667 | 0.6 |
| 0.8 | 0 | ptsI | 2 | 12.4167 | 0.52 |
| 0.8 | 0 | clcB | 2 | 12.4167 | 0.395 |
| 0.8 | 0 | ycaM | 2 | 12.4167 | 0.434 |
| 0.8 | 0 | yadI | 2 | 12.4167 | 0.731 |
| 0.8 | 0 | AG1 | 2 | 12.4167 | 0.604 |
| 0.8 | 0 | ptsI | 2 | 12.6667 | 0.512 |
| 0.8 | 0 | clcB | 2 | 12.6667 | 0.399 |
| 0.8 | 0 | ycaM | 2 | 12.6667 | 0.463 |
| 0.8 | 0 | yadI | 2 | 12.6667 | 0.742 |
| 0.8 | 0 | AG1 | 2 | 12.6667 | 0.595 |
| 0.8 | 0 | ptsI | 2 | 12.9167 | 0.52 |
| 0.8 | 0 | clcB | 2 | 12.9167 | 0.403 |
| 0.8 | 0 | ycaM | 2 | 12.9167 | 0.465 |
| 0.8 | 0 | yadI | 2 | 12.9167 | 0.755 |

| | | | | | |
|---|---|---|---|---|---|
| 0.8 | 0 | AG1 | 2 | 12.9167 | 0.613 |
| 0.8 | 0 | ptsI | 2 | 13.1667 | 0.518 |
| 0.8 | 0 | clcB | 2 | 13.1667 | 0.404 |
| 0.8 | 0 | ycaM | 2 | 13.1667 | 0.47 |
| 0.8 | 0 | yadI | 2 | 13.1667 | 0.758 |
| 0.8 | 0 | AG1 | 2 | 13.1667 | 0.613 |
| 0.8 | 0 | ptsI | 2 | 13.4167 | 0.537 |
| 0.8 | 0 | clcB | 2 | 13.4167 | 0.41 |
| 0.8 | 0 | ycaM | 2 | 13.4167 | 0.47 |
| 0.8 | 0 | yadI | 2 | 13.4167 | 0.767 |
| 0.8 | 0 | AG1 | 2 | 13.4167 | 0.61 |
| 0.8 | 0 | ptsI | 2 | 13.6667 | 0.566 |
| 0.8 | 0 | clcB | 2 | 13.6667 | 0.42 |
| 0.8 | 0 | ycaM | 2 | 13.6667 | 0.504 |
| 0.8 | 0 | yadI | 2 | 13.6667 | 0.769 |
| 0.8 | 0 | AG1 | 2 | 13.6667 | 0.611 |
| 0.8 | 0 | ptsI | 2 | 13.9167 | 0.597 |
| 0.8 | 0 | clcB | 2 | 13.9167 | 0.421 |
| 0.8 | 0 | ycaM | 2 | 13.9167 | 0.487 |
| 0.8 | 0 | yadI | 2 | 13.9167 | 0.772 |
| 0.8 | 0 | AG1 | 2 | 13.9167 | 0.606 |
| 0.8 | 0 | ptsI | 2 | 14.1667 | 0.609 |
| 0.8 | 0 | clcB | 2 | 14.1667 | 0.422 |
| 0.8 | 0 | ycaM | 2 | 14.1667 | 0.483 |
| 0.8 | 0 | yadI | 2 | 14.1667 | 0.779 |
| 0.8 | 0 | AG1 | 2 | 14.1667 | 0.624 |
| 0.8 | 0 | ptsI | 2 | 14.4167 | 0.625 |
| 0.8 | 0 | clcB | 2 | 14.4167 | 0.431 |
| 0.8 | 0 | ycaM | 2 | 14.4167 | 0.486 |
| 0.8 | 0 | yadI | 2 | 14.4167 | 0.783 |
| 0.8 | 0 | AG1 | 2 | 14.4167 | 0.642 |
| 0.8 | 0 | ptsI | 2 | 14.6667 | 0.655 |
| 0.8 | 0 | clcB | 2 | 14.6667 | 0.432 |
| 0.8 | 0 | ycaM | 2 | 14.6667 | 0.502 |
| 0.8 | 0 | yadI | 2 | 14.6667 | 0.789 |
| 0.8 | 0 | AG1 | 2 | 14.6667 | 0.654 |
| 0.8 | 0 | ptsI | 2 | 14.9167 | 0.648 |
| 0.8 | 0 | clcB | 2 | 14.9167 | 0.442 |
| 0.8 | 0 | ycaM | 2 | 14.9167 | 0.494 |
| 0.8 | 0 | yadI | 2 | 14.9167 | 0.797 |
| 0.8 | 0 | AG1 | 2 | 14.9167 | 0.67 |
| 0.8 | 0 | ptsI | 2 | 15.1667 | 0.665 |
| 0.8 | 0 | clcB | 2 | 15.1667 | 0.44 |
| 0.8 | 0 | ycaM | 2 | 15.1667 | 0.512 |
| 0.8 | 0 | yadI | 2 | 15.1667 | 0.808 |
| 0.8 | 0 | AG1 | 2 | 15.1667 | 0.656 |
| 0.8 | 0 | ptsI | 2 | 15.4167 | 0.669 |
| 0.8 | 0 | clcB | 2 | 15.4167 | 0.437 |

| | | | | | |
|---|---|---|---|---|---|
| 0.8 | 0 | ycaM | 2 | 15.4167 | 0.519 |
| 0.8 | 0 | yadI | 2 | 15.4167 | 0.807 |
| 0.8 | 0 | AG1 | 2 | 15.4167 | 0.677 |
| 0.8 | 0 | ptsI | 2 | 15.6667 | 0.678 |
| 0.8 | 0 | clcB | 2 | 15.6667 | 0.471 |
| 0.8 | 0 | ycaM | 2 | 15.6667 | 0.525 |
| 0.8 | 0 | yadI | 2 | 15.6667 | 0.813 |
| 0.8 | 0 | AG1 | 2 | 15.6667 | 0.7 |
| 0.8 | 0 | ptsI | 2 | 15.9167 | 0.676 |
| 0.8 | 0 | clcB | 2 | 15.9167 | 0.465 |
| 0.8 | 0 | ycaM | 2 | 15.9167 | 0.51 |
| 0.8 | 0 | yadI | 2 | 15.9167 | 0.815 |
| 0.8 | 0 | AG1 | 2 | 15.9167 | 0.702 |
| 0.8 | 0 | ptsI | 2 | 16.1667 | 0.719 |
| 0.8 | 0 | clcB | 2 | 16.1667 | 0.453 |
| 0.8 | 0 | ycaM | 2 | 16.1667 | 0.528 |
| 0.8 | 0 | yadI | 2 | 16.1667 | 0.816 |
| 0.8 | 0 | AG1 | 2 | 16.1667 | 0.702 |
| 0.8 | 0 | ptsI | 2 | 16.4167 | 0.738 |
| 0.8 | 0 | clcB | 2 | 16.4167 | 0.466 |
| 0.8 | 0 | ycaM | 2 | 16.4167 | 0.53 |
| 0.8 | 0 | yadI | 2 | 16.4167 | 0.821 |
| 0.8 | 0 | AG1 | 2 | 16.4167 | 0.73 |
| 0.8 | 0 | ptsI | 2 | 16.6667 | 0.752 |
| 0.8 | 0 | clcB | 2 | 16.6667 | 0.478 |
| 0.8 | 0 | ycaM | 2 | 16.6667 | 0.538 |
| 0.8 | 0 | yadI | 2 | 16.6667 | 0.825 |
| 0.8 | 0 | AG1 | 2 | 16.6667 | 0.739 |
| 0.8 | 0 | ptsI | 2 | 16.9167 | 0.742 |
| 0.8 | 0 | clcB | 2 | 16.9167 | 0.466 |
| 0.8 | 0 | ycaM | 2 | 16.9167 | 0.531 |
| 0.8 | 0 | yadI | 2 | 16.9167 | 0.826 |
| 0.8 | 0 | AG1 | 2 | 16.9167 | 0.746 |
| 0.8 | 0 | ptsI | 2 | 17.1667 | 0.74 |
| 0.8 | 0 | clcB | 2 | 17.1667 | 0.467 |
| 0.8 | 0 | ycaM | 2 | 17.1667 | 0.545 |
| 0.8 | 0 | yadI | 2 | 17.1667 | 0.825 |
| 0.8 | 0 | AG1 | 2 | 17.1667 | 0.747 |
| 0.8 | 0 | ptsI | 2 | 17.4167 | 0.754 |
| 0.8 | 0 | clcB | 2 | 17.4167 | 0.479 |
| 0.8 | 0 | ycaM | 2 | 17.4167 | 0.544 |
| 0.8 | 0 | yadI | 2 | 17.4167 | 0.83 |
| 0.8 | 0 | AG1 | 2 | 17.4167 | 0.749 |
| 0.8 | 0 | ptsI | 2 | 17.6667 | 0.746 |
| 0.8 | 0 | clcB | 2 | 17.6667 | 0.473 |
| 0.8 | 0 | ycaM | 2 | 17.6667 | 0.544 |
| 0.8 | 0 | yadI | 2 | 17.6667 | 0.832 |
| 0.8 | 0 | AG1 | 2 | 17.6667 | 0.756 |

| | | | | | |
|---|---|---|---|---|---|
| 0.8 | 0 | ptsI | 2 | 17.9167 | 0.767 |
| 0.8 | 0 | clcB | 2 | 17.9167 | 0.473 |
| 0.8 | 0 | ycaM | 2 | 17.9167 | 0.539 |
| 0.8 | 0 | yadI | 2 | 17.9167 | 0.834 |
| 0.8 | 0 | AG1 | 2 | 17.9167 | 0.757 |
| 0.8 | 0 | ptsI | 2 | 18.1667 | 0.767 |
| 0.8 | 0 | clcB | 2 | 18.1667 | 0.474 |
| 0.8 | 0 | ycaM | 2 | 18.1667 | 0.532 |
| 0.8 | 0 | yadI | 2 | 18.1667 | 0.836 |
| 0.8 | 0 | AG1 | 2 | 18.1667 | 0.767 |
| 0.8 | 0 | ptsI | 2 | 18.4167 | 0.778 |
| 0.8 | 0 | clcB | 2 | 18.4167 | 0.477 |
| 0.8 | 0 | ycaM | 2 | 18.4167 | 0.534 |
| 0.8 | 0 | yadI | 2 | 18.4167 | 0.838 |
| 0.8 | 0 | AG1 | 2 | 18.4167 | 0.767 |
| 0.8 | 0 | ptsI | 2 | 18.6667 | 0.785 |
| 0.8 | 0 | clcB | 2 | 18.6667 | 0.476 |
| 0.8 | 0 | ycaM | 2 | 18.6667 | 0.532 |
| 0.8 | 0 | yadI | 2 | 18.6667 | 0.839 |
| 0.8 | 0 | AG1 | 2 | 18.6667 | 0.764 |
| 0.8 | 0 | ptsI | 2 | 18.9167 | 0.786 |
| 0.8 | 0 | clcB | 2 | 18.9167 | 0.466 |
| 0.8 | 0 | ycaM | 2 | 18.9167 | 0.534 |
| 0.8 | 0 | yadI | 2 | 18.9167 | 0.839 |
| 0.8 | 0 | AG1 | 2 | 18.9167 | 0.78 |
| 0.8 | 0 | ptsI | 2 | 19.1667 | 0.799 |
| 0.8 | 0 | clcB | 2 | 19.1667 | 0.473 |
| 0.8 | 0 | ycaM | 2 | 19.1667 | 0.541 |
| 0.8 | 0 | yadI | 2 | 19.1667 | 0.843 |
| 0.8 | 0 | AG1 | 2 | 19.1667 | 0.766 |
| 0.8 | 0 | ptsI | 2 | 19.4167 | 0.798 |
| 0.8 | 0 | clcB | 2 | 19.4167 | 0.47 |
| 0.8 | 0 | ycaM | 2 | 19.4167 | 0.543 |
| 0.8 | 0 | yadI | 2 | 19.4167 | 0.844 |
| 0.8 | 0 | AG1 | 2 | 19.4167 | 0.762 |
| 0.8 | 0 | ptsI | 2 | 19.6667 | 0.801 |
| 0.8 | 0 | clcB | 2 | 19.6667 | 0.463 |
| 0.8 | 0 | ycaM | 2 | 19.6667 | 0.55 |
| 0.8 | 0 | yadI | 2 | 19.6667 | 0.847 |
| 0.8 | 0 | AG1 | 2 | 19.6667 | 0.767 |
| 0.8 | 0 | ptsI | 2 | 19.9167 | 0.804 |
| 0.8 | 0 | clcB | 2 | 19.9167 | 0.468 |
| 0.8 | 0 | ycaM | 2 | 19.9167 | 0.547 |
| 0.8 | 0 | yadI | 2 | 19.9167 | 0.848 |
| 0.8 | 0 | AG1 | 2 | 19.9167 | 0.77 |
| 0.8 | 0 | ptsI | 2 | 20.1667 | 0.801 |
| 0.8 | 0 | clcB | 2 | 20.1667 | 0.469 |
| 0.8 | 0 | ycaM | 2 | 20.1667 | 0.549 |

| | | | | | |
|---|---|---|---|---|---|
| 0.8 | 0 | yadI | 2 | 20.1667 | 0.847 |
| 0.8 | 0 | AG1 | 2 | 20.1667 | 0.757 |
| 0.8 | 0 | ptsI | 2 | 20.4167 | 0.809 |
| 0.8 | 0 | clcB | 2 | 20.4167 | 0.476 |
| 0.8 | 0 | ycaM | 2 | 20.4167 | 0.555 |
| 0.8 | 0 | yadI | 2 | 20.4167 | 0.85 |
| 0.8 | 0 | AG1 | 2 | 20.4167 | 0.755 |
| 0.8 | 0 | ptsI | 2 | 20.6667 | 0.809 |
| 0.8 | 0 | clcB | 2 | 20.6667 | 0.468 |
| 0.8 | 0 | ycaM | 2 | 20.6667 | 0.56 |
| 0.8 | 0 | yadI | 2 | 20.6667 | 0.849 |
| 0.8 | 0 | AG1 | 2 | 20.6667 | 0.747 |
| 0.8 | 0 | ptsI | 2 | 20.9167 | 0.815 |
| 0.8 | 0 | clcB | 2 | 20.9167 | 0.469 |
| 0.8 | 0 | ycaM | 2 | 20.9167 | 0.566 |
| 0.8 | 0 | yadI | 2 | 20.9167 | 0.851 |
| 0.8 | 0 | AG1 | 2 | 20.9167 | 0.746 |
| 0.8 | 0 | ptsI | 2 | 21.1667 | 0.808 |
| 0.8 | 0 | clcB | 2 | 21.1667 | 0.472 |
| 0.8 | 0 | ycaM | 2 | 21.1667 | 0.572 |
| 0.8 | 0 | yadI | 2 | 21.1667 | 0.85 |
| 0.8 | 0 | AG1 | 2 | 21.1667 | 0.749 |
| 0.8 | 0 | ptsI | 2 | 21.4167 | 0.813 |
| 0.8 | 0 | clcB | 2 | 21.4167 | 0.472 |
| 0.8 | 0 | ycaM | 2 | 21.4167 | 0.574 |
| 0.8 | 0 | yadI | 2 | 21.4167 | 0.858 |
| 0.8 | 0 | AG1 | 2 | 21.4167 | 0.764 |
| 0.8 | 0 | ptsI | 2 | 21.6667 | 0.816 |
| 0.8 | 0 | clcB | 2 | 21.6667 | 0.475 |
| 0.8 | 0 | ycaM | 2 | 21.6667 | 0.578 |
| 0.8 | 0 | yadI | 2 | 21.6667 | 0.853 |
| 0.8 | 0 | AG1 | 2 | 21.6667 | 0.753 |
| 0.8 | 0 | ptsI | 2 | 21.9167 | 0.821 |
| 0.8 | 0 | clcB | 2 | 21.9167 | 0.481 |
| 0.8 | 0 | ycaM | 2 | 21.9167 | 0.583 |
| 0.8 | 0 | yadI | 2 | 21.9167 | 0.855 |
| 0.8 | 0 | AG1 | 2 | 21.9167 | 0.755 |
| 0.8 | 0 | ptsI | 2 | 22.1667 | 0.826 |
| 0.8 | 0 | clcB | 2 | 22.1667 | 0.48 |
| 0.8 | 0 | ycaM | 2 | 22.1667 | 0.586 |
| 0.8 | 0 | yadI | 2 | 22.1667 | 0.854 |
| 0.8 | 0 | AG1 | 2 | 22.1667 | 0.758 |
| 0.8 | 0 | ptsI | 2 | 22.4167 | 0.826 |
| 0.8 | 0 | clcB | 2 | 22.4167 | 0.482 |
| 0.8 | 0 | ycaM | 2 | 22.4167 | 0.587 |
| 0.8 | 0 | yadI | 2 | 22.4167 | 0.854 |
| 0.8 | 0 | AG1 | 2 | 22.4167 | 0.76 |
| 0.8 | 0 | ptsI | 2 | 22.6667 | 0.835 |

| | | | | | |
|---|---|---|---|---|---|
| 0.8 | 0 | clcB | 2 | 22.6667 | 0.485 |
| 0.8 | 0 | ycaM | 2 | 22.6667 | 0.59 |
| 0.8 | 0 | yadI | 2 | 22.6667 | 0.856 |
| 0.8 | 0 | AG1 | 2 | 22.6667 | 0.759 |
| 0.8 | 0 | ptsI | 2 | 22.9167 | 0.844 |
| 0.8 | 0 | clcB | 2 | 22.9167 | 0.487 |
| 0.8 | 0 | ycaM | 2 | 22.9167 | 0.593 |
| 0.8 | 0 | yadI | 2 | 22.9167 | 0.862 |
| 0.8 | 0 | AG1 | 2 | 22.9167 | 0.761 |
| 0.8 | 0 | ptsI | 2 | 23.1667 | 0.85 |
| 0.8 | 0 | clcB | 2 | 23.1667 | 0.489 |
| 0.8 | 0 | ycaM | 2 | 23.1667 | 0.595 |
| 0.8 | 0 | yadI | 2 | 23.1667 | 0.863 |
| 0.8 | 0 | AG1 | 2 | 23.1667 | 0.764 |
| 0.8 | 0 | ptsI | 2 | 23.4167 | 0.852 |
| 0.8 | 0 | clcB | 2 | 23.4167 | 0.491 |
| 0.8 | 0 | ycaM | 2 | 23.4167 | 0.597 |
| 0.8 | 0 | yadI | 2 | 23.4167 | 0.863 |
| 0.8 | 0 | AG1 | 2 | 23.4167 | 0.762 |
| 0.8 | 0 | ptsI | 2 | 23.6667 | 0.864 |
| 0.8 | 0 | clcB | 2 | 23.6667 | 0.494 |
| 0.8 | 0 | ycaM | 2 | 23.6667 | 0.6 |
| 0.8 | 0 | yadI | 2 | 23.6667 | 0.864 |
| 0.8 | 0 | AG1 | 2 | 23.6667 | 0.768 |
| 0.8 | 0 | ptsI | 2 | 23.9167 | 0.869 |
| 0.8 | 0 | clcB | 2 | 23.9167 | 0.496 |
| 0.8 | 0 | ycaM | 2 | 23.9167 | 0.6 |
| 0.8 | 0 | yadI | 2 | 23.9167 | 0.866 |
| 0.8 | 0 | AG1 | 2 | 23.9167 | 0.764 |
| 0.8 | 0 | ptsI | 2 | 24.1667 | 0.869 |
| 0.8 | 0 | clcB | 2 | 24.1667 | 0.497 |
| 0.8 | 0 | ycaM | 2 | 24.1667 | 0.599 |
| 0.8 | 0 | yadI | 2 | 24.1667 | 0.866 |
| 0.8 | 0 | AG1 | 2 | 24.1667 | 0.764 |
| 0.8 | 0 | ptsI | 2 | 24.4167 | 0.872 |
| 0.8 | 0 | clcB | 2 | 24.4167 | 0.499 |
| 0.8 | 0 | ycaM | 2 | 24.4167 | 0.605 |
| 0.8 | 0 | yadI | 2 | 24.4167 | 0.868 |
| 0.8 | 0 | AG1 | 2 | 24.4167 | 0.766 |
| 0.8 | 0 | ptsI | 2 | 24.6667 | 0.877 |
| 0.8 | 0 | clcB | 2 | 24.6667 | 0.502 |
| 0.8 | 0 | ycaM | 2 | 24.6667 | 0.607 |
| 0.8 | 0 | yadI | 2 | 24.6667 | 0.867 |
| 0.8 | 0 | AG1 | 2 | 24.6667 | 0.77 |
| 0.8 | 0 | ptsI | 2 | 24.9167 | 0.882 |
| 0.8 | 0 | clcB | 2 | 24.9167 | 0.505 |
| 0.8 | 0 | ycaM | 2 | 24.9167 | 0.609 |
| 0.8 | 0 | yadI | 2 | 24.9167 | 0.87 |

| | | | | | |
|---|---|---|---|---|---|
| 0.8 | 0 | AG1 | 2 | 24.9167 | 0.768 |
| 0.8 | 0 | ptsI | 2 | 25.1667 | 0.886 |
| 0.8 | 0 | clcB | 2 | 25.1667 | 0.506 |
| 0.8 | 0 | ycaM | 2 | 25.1667 | 0.61 |
| 0.8 | 0 | yadI | 2 | 25.1667 | 0.872 |
| 0.8 | 0 | AG1 | 2 | 25.1667 | 0.768 |
| 0.8 | 0 | ptsI | 2 | 25.4167 | 0.892 |
| 0.8 | 0 | clcB | 2 | 25.4167 | 0.507 |
| 0.8 | 0 | ycaM | 2 | 25.4167 | 0.611 |
| 0.8 | 0 | yadI | 2 | 25.4167 | 0.873 |
| 0.8 | 0 | AG1 | 2 | 25.4167 | 0.772 |
| 0.8 | 0 | ptsI | 2 | 25.6667 | 0.895 |
| 0.8 | 0 | clcB | 2 | 25.6667 | 0.509 |
| 0.8 | 0 | ycaM | 2 | 25.6667 | 0.614 |
| 0.8 | 0 | yadI | 2 | 25.6667 | 0.87 |
| 0.8 | 0 | AG1 | 2 | 25.6667 | 0.771 |
| 0.8 | 0 | ptsI | 2 | 25.9167 | 0.896 |
| 0.8 | 0 | clcB | 2 | 25.9167 | 0.511 |
| 0.8 | 0 | ycaM | 2 | 25.9167 | 0.616 |
| 0.8 | 0 | yadI | 2 | 25.9167 | 0.876 |
| 0.8 | 0 | AG1 | 2 | 25.9167 | 0.775 |
| 0.8 | 0 | ptsI | 2 | 26.1667 | 0.904 |
| 0.8 | 0 | clcB | 2 | 26.1667 | 0.516 |
| 0.8 | 0 | ycaM | 2 | 26.1667 | 0.617 |
| 0.8 | 0 | yadI | 2 | 26.1667 | 0.88 |
| 0.8 | 0 | AG1 | 2 | 26.1667 | 0.784 |
| 0.8 | 0 | ptsI | 2 | 26.4167 | 0.911 |
| 0.8 | 0 | clcB | 2 | 26.4167 | 0.516 |
| 0.8 | 0 | ycaM | 2 | 26.4167 | 0.62 |
| 0.8 | 0 | yadI | 2 | 26.4167 | 0.878 |
| 0.8 | 0 | AG1 | 2 | 26.4167 | 0.772 |
| 0.8 | 0 | ptsI | 3 | 0 | 0.153 |
| 0.8 | 0 | clcB | 3 | 0 | 0.155 |
| 0.8 | 0 | ycaM | 3 | 0 | 0.153 |
| 0.8 | 0 | yadI | 3 | 0 | 0.167 |
| 0.8 | 0 | AG1 | 3 | 0 | 0.163 |
| 0.8 | 0 | ptsI | 3 | 0.25 | 0.152 |
| 0.8 | 0 | clcB | 3 | 0.25 | 0.152 |
| 0.8 | 0 | ycaM | 3 | 0.25 | 0.149 |
| 0.8 | 0 | yadI | 3 | 0.25 | 0.163 |
| 0.8 | 0 | AG1 | 3 | 0.25 | 0.159 |
| 0.8 | 0 | ptsI | 3 | 0.5 | 0.151 |
| 0.8 | 0 | clcB | 3 | 0.5 | 0.152 |
| 0.8 | 0 | ycaM | 3 | 0.5 | 0.148 |
| 0.8 | 0 | yadI | 3 | 0.5 | 0.163 |
| 0.8 | 0 | AG1 | 3 | 0.5 | 0.158 |
| 0.8 | 0 | ptsI | 3 | 0.75 | 0.152 |
| 0.8 | 0 | clcB | 3 | 0.75 | 0.153 |

| | | | | | |
|---|---|---|---|---|---|
| 0.8 | 0 | ycaM | 3 | 0.75 | 0.15 |
| 0.8 | 0 | yadI | 3 | 0.75 | 0.168 |
| 0.8 | 0 | AG1 | 3 | 0.75 | 0.16 |
| 0.8 | 0 | ptsI | 3 | 1 | 0.153 |
| 0.8 | 0 | clcB | 3 | 1 | 0.154 |
| 0.8 | 0 | ycaM | 3 | 1 | 0.15 |
| 0.8 | 0 | yadI | 3 | 1 | 0.162 |
| 0.8 | 0 | AG1 | 3 | 1 | 0.16 |
| 0.8 | 0 | ptsI | 3 | 1.25 | 0.154 |
| 0.8 | 0 | clcB | 3 | 1.25 | 0.154 |
| 0.8 | 0 | ycaM | 3 | 1.25 | 0.148 |
| 0.8 | 0 | yadI | 3 | 1.25 | 0.169 |
| 0.8 | 0 | AG1 | 3 | 1.25 | 0.159 |
| 0.8 | 0 | ptsI | 3 | 1.5 | 0.154 |
| 0.8 | 0 | clcB | 3 | 1.5 | 0.156 |
| 0.8 | 0 | ycaM | 3 | 1.5 | 0.15 |
| 0.8 | 0 | yadI | 3 | 1.5 | 0.164 |
| 0.8 | 0 | AG1 | 3 | 1.5 | 0.16 |
| 0.8 | 0 | ptsI | 3 | 1.75 | 0.155 |
| 0.8 | 0 | clcB | 3 | 1.75 | 0.156 |
| 0.8 | 0 | ycaM | 3 | 1.75 | 0.151 |
| 0.8 | 0 | yadI | 3 | 1.75 | 0.164 |
| 0.8 | 0 | AG1 | 3 | 1.75 | 0.16 |
| 0.8 | 0 | ptsI | 3 | 2 | 0.157 |
| 0.8 | 0 | clcB | 3 | 2 | 0.157 |
| 0.8 | 0 | ycaM | 3 | 2 | 0.152 |
| 0.8 | 0 | yadI | 3 | 2 | 0.171 |
| 0.8 | 0 | AG1 | 3 | 2 | 0.162 |
| 0.8 | 0 | ptsI | 3 | 2.25 | 0.159 |
| 0.8 | 0 | clcB | 3 | 2.25 | 0.16 |
| 0.8 | 0 | ycaM | 3 | 2.25 | 0.154 |
| 0.8 | 0 | yadI | 3 | 2.25 | 0.17 |
| 0.8 | 0 | AG1 | 3 | 2.25 | 0.164 |
| 0.8 | 0 | ptsI | 3 | 2.5 | 0.161 |
| 0.8 | 0 | clcB | 3 | 2.5 | 0.162 |
| 0.8 | 0 | ycaM | 3 | 2.5 | 0.159 |
| 0.8 | 0 | yadI | 3 | 2.5 | 0.168 |
| 0.8 | 0 | AG1 | 3 | 2.5 | 0.166 |
| 0.8 | 0 | ptsI | 3 | 2.75 | 0.163 |
| 0.8 | 0 | clcB | 3 | 2.75 | 0.164 |
| 0.8 | 0 | ycaM | 3 | 2.75 | 0.158 |
| 0.8 | 0 | yadI | 3 | 2.75 | 0.172 |
| 0.8 | 0 | AG1 | 3 | 2.75 | 0.171 |
| 0.8 | 0 | ptsI | 3 | 3 | 0.164 |
| 0.8 | 0 | clcB | 3 | 3 | 0.165 |
| 0.8 | 0 | ycaM | 3 | 3 | 0.158 |
| 0.8 | 0 | yadI | 3 | 3 | 0.173 |
| 0.8 | 0 | AG1 | 3 | 3 | 0.171 |

| | | | | | |
|---|---|---|---|---|---|
| 0.8 | 0 | ptsI | 3 | 3.25 | 0.167 |
| 0.8 | 0 | clcB | 3 | 3.25 | 0.169 |
| 0.8 | 0 | ycaM | 3 | 3.25 | 0.16 |
| 0.8 | 0 | yadI | 3 | 3.25 | 0.177 |
| 0.8 | 0 | AG1 | 3 | 3.25 | 0.172 |
| 0.8 | 0 | ptsI | 3 | 3.5 | 0.17 |
| 0.8 | 0 | clcB | 3 | 3.5 | 0.171 |
| 0.8 | 0 | ycaM | 3 | 3.5 | 0.166 |
| 0.8 | 0 | yadI | 3 | 3.5 | 0.181 |
| 0.8 | 0 | AG1 | 3 | 3.5 | 0.178 |
| 0.8 | 0 | ptsI | 3 | 3.75 | 0.174 |
| 0.8 | 0 | clcB | 3 | 3.75 | 0.175 |
| 0.8 | 0 | ycaM | 3 | 3.75 | 0.17 |
| 0.8 | 0 | yadI | 3 | 3.75 | 0.185 |
| 0.8 | 0 | AG1 | 3 | 3.75 | 0.181 |
| 0.8 | 0 | ptsI | 3 | 4 | 0.182 |
| 0.8 | 0 | clcB | 3 | 4 | 0.178 |
| 0.8 | 0 | ycaM | 3 | 4 | 0.174 |
| 0.8 | 0 | yadI | 3 | 4 | 0.191 |
| 0.8 | 0 | AG1 | 3 | 4 | 0.186 |
| 0.8 | 0 | ptsI | 3 | 4.41667 | 0.183 |
| 0.8 | 0 | clcB | 3 | 4.41667 | 0.194 |
| 0.8 | 0 | ycaM | 3 | 4.41667 | 0.194 |
| 0.8 | 0 | yadI | 3 | 4.41667 | 0.194 |
| 0.8 | 0 | AG1 | 3 | 4.41667 | 0.201 |
| 0.8 | 0 | ptsI | 3 | 4.66667 | 0.183 |
| 0.8 | 0 | clcB | 3 | 4.66667 | 0.184 |
| 0.8 | 0 | ycaM | 3 | 4.66667 | 0.197 |
| 0.8 | 0 | yadI | 3 | 4.66667 | 0.186 |
| 0.8 | 0 | AG1 | 3 | 4.66667 | 0.2 |
| 0.8 | 0 | ptsI | 3 | 4.91667 | 0.184 |
| 0.8 | 0 | clcB | 3 | 4.91667 | 0.18 |
| 0.8 | 0 | ycaM | 3 | 4.91667 | 0.193 |
| 0.8 | 0 | yadI | 3 | 4.91667 | 0.19 |
| 0.8 | 0 | AG1 | 3 | 4.91667 | 0.199 |
| 0.8 | 0 | ptsI | 3 | 5.16667 | 0.188 |
| 0.8 | 0 | clcB | 3 | 5.16667 | 0.186 |
| 0.8 | 0 | ycaM | 3 | 5.16667 | 0.198 |
| 0.8 | 0 | yadI | 3 | 5.16667 | 0.2 |
| 0.8 | 0 | AG1 | 3 | 5.16667 | 0.204 |
| 0.8 | 0 | ptsI | 3 | 5.41667 | 0.191 |
| 0.8 | 0 | clcB | 3 | 5.41667 | 0.19 |
| 0.8 | 0 | ycaM | 3 | 5.41667 | 0.205 |
| 0.8 | 0 | yadI | 3 | 5.41667 | 0.212 |
| 0.8 | 0 | AG1 | 3 | 5.41667 | 0.212 |
| 0.8 | 0 | ptsI | 3 | 5.66667 | 0.197 |
| 0.8 | 0 | clcB | 3 | 5.66667 | 0.196 |
| 0.8 | 0 | ycaM | 3 | 5.66667 | 0.216 |

| | | | | | |
|---|---|---|---|---|---|
| 0.8 | 0 | yadI | 3 | 5.66667 | 0.226 |
| 0.8 | 0 | AG1 | 3 | 5.66667 | 0.221 |
| 0.8 | 0 | ptsI | 3 | 5.91667 | 0.201 |
| 0.8 | 0 | clcB | 3 | 5.91667 | 0.222 |
| 0.8 | 0 | ycaM | 3 | 5.91667 | 0.227 |
| 0.8 | 0 | yadI | 3 | 5.91667 | 0.24 |
| 0.8 | 0 | AG1 | 3 | 5.91667 | 0.231 |
| 0.8 | 0 | ptsI | 3 | 6.16667 | 0.208 |
| 0.8 | 0 | clcB | 3 | 6.16667 | 0.212 |
| 0.8 | 0 | ycaM | 3 | 6.16667 | 0.242 |
| 0.8 | 0 | yadI | 3 | 6.16667 | 0.258 |
| 0.8 | 0 | AG1 | 3 | 6.16667 | 0.242 |
| 0.8 | 0 | ptsI | 3 | 6.41667 | 0.214 |
| 0.8 | 0 | clcB | 3 | 6.41667 | 0.22 |
| 0.8 | 0 | ycaM | 3 | 6.41667 | 0.257 |
| 0.8 | 0 | yadI | 3 | 6.41667 | 0.27 |
| 0.8 | 0 | AG1 | 3 | 6.41667 | 0.252 |
| 0.8 | 0 | ptsI | 3 | 6.66667 | 0.222 |
| 0.8 | 0 | clcB | 3 | 6.66667 | 0.23 |
| 0.8 | 0 | ycaM | 3 | 6.66667 | 0.272 |
| 0.8 | 0 | yadI | 3 | 6.66667 | 0.282 |
| 0.8 | 0 | AG1 | 3 | 6.66667 | 0.266 |
| 0.8 | 0 | ptsI | 3 | 6.91667 | 0.231 |
| 0.8 | 0 | clcB | 3 | 6.91667 | 0.254 |
| 0.8 | 0 | ycaM | 3 | 6.91667 | 0.281 |
| 0.8 | 0 | yadI | 3 | 6.91667 | 0.296 |
| 0.8 | 0 | AG1 | 3 | 6.91667 | 0.277 |
| 0.8 | 0 | ptsI | 3 | 7.16667 | 0.242 |
| 0.8 | 0 | clcB | 3 | 7.16667 | 0.257 |
| 0.8 | 0 | ycaM | 3 | 7.16667 | 0.291 |
| 0.8 | 0 | yadI | 3 | 7.16667 | 0.318 |
| 0.8 | 0 | AG1 | 3 | 7.16667 | 0.288 |
| 0.8 | 0 | ptsI | 3 | 7.41667 | 0.254 |
| 0.8 | 0 | clcB | 3 | 7.41667 | 0.265 |
| 0.8 | 0 | ycaM | 3 | 7.41667 | 0.301 |
| 0.8 | 0 | yadI | 3 | 7.41667 | 0.328 |
| 0.8 | 0 | AG1 | 3 | 7.41667 | 0.294 |
| 0.8 | 0 | ptsI | 3 | 7.66667 | 0.263 |
| 0.8 | 0 | clcB | 3 | 7.66667 | 0.273 |
| 0.8 | 0 | ycaM | 3 | 7.66667 | 0.311 |
| 0.8 | 0 | yadI | 3 | 7.66667 | 0.339 |
| 0.8 | 0 | AG1 | 3 | 7.66667 | 0.303 |
| 0.8 | 0 | ptsI | 3 | 7.91667 | 0.27 |
| 0.8 | 0 | clcB | 3 | 7.91667 | 0.28 |
| 0.8 | 0 | ycaM | 3 | 7.91667 | 0.322 |
| 0.8 | 0 | yadI | 3 | 7.91667 | 0.355 |
| 0.8 | 0 | AG1 | 3 | 7.91667 | 0.312 |
| 0.8 | 0 | ptsI | 3 | 8.16667 | 0.28 |

| | | | | | |
|---|---|---|---|---|---|
| 0.8 | 0 | clcB | 3 | 8.16667 | 0.289 |
| 0.8 | 0 | ycaM | 3 | 8.16667 | 0.333 |
| 0.8 | 0 | yadI | 3 | 8.16667 | 0.382 |
| 0.8 | 0 | AG1 | 3 | 8.16667 | 0.32 |
| 0.8 | 0 | ptsI | 3 | 8.41667 | 0.286 |
| 0.8 | 0 | clcB | 3 | 8.41667 | 0.298 |
| 0.8 | 0 | ycaM | 3 | 8.41667 | 0.354 |
| 0.8 | 0 | yadI | 3 | 8.41667 | 0.385 |
| 0.8 | 0 | AG1 | 3 | 8.41667 | 0.328 |
| 0.8 | 0 | ptsI | 3 | 9.05 | 0.263 |
| 0.8 | 0 | clcB | 3 | 9.05 | 0.281 |
| 0.8 | 0 | ycaM | 3 | 9.05 | 0.307 |
| 0.8 | 0 | yadI | 3 | 9.05 | 0.452 |
| 0.8 | 0 | AG1 | 3 | 9.05 | 0.358 |
| 0.8 | 0 | ptsI | 3 | 9.3 | 0.27 |
| 0.8 | 0 | clcB | 3 | 9.3 | 0.289 |
| 0.8 | 0 | ycaM | 3 | 9.3 | 0.31 |
| 0.8 | 0 | yadI | 3 | 9.3 | 0.448 |
| 0.8 | 0 | AG1 | 3 | 9.3 | 0.356 |
| 0.8 | 0 | ptsI | 3 | 9.55 | 0.28 |
| 0.8 | 0 | clcB | 3 | 9.55 | 0.298 |
| 0.8 | 0 | ycaM | 3 | 9.55 | 0.321 |
| 0.8 | 0 | yadI | 3 | 9.55 | 0.462 |
| 0.8 | 0 | AG1 | 3 | 9.55 | 0.368 |
| 0.8 | 0 | ptsI | 3 | 9.8 | 0.287 |
| 0.8 | 0 | clcB | 3 | 9.8 | 0.308 |
| 0.8 | 0 | ycaM | 3 | 9.8 | 0.33 |
| 0.8 | 0 | yadI | 3 | 9.8 | 0.47 |
| 0.8 | 0 | AG1 | 3 | 9.8 | 0.38 |
| 0.8 | 0 | ptsI | 3 | 10.05 | 0.296 |
| 0.8 | 0 | clcB | 3 | 10.05 | 0.316 |
| 0.8 | 0 | ycaM | 3 | 10.05 | 0.342 |
| 0.8 | 0 | yadI | 3 | 10.05 | 0.485 |
| 0.8 | 0 | AG1 | 3 | 10.05 | 0.392 |
| 0.8 | 0 | ptsI | 3 | 10.3 | 0.304 |
| 0.8 | 0 | clcB | 3 | 10.3 | 0.327 |
| 0.8 | 0 | ycaM | 3 | 10.3 | 0.356 |
| 0.8 | 0 | yadI | 3 | 10.3 | 0.497 |
| 0.8 | 0 | AG1 | 3 | 10.3 | 0.398 |
| 0.8 | 0 | ptsI | 3 | 10.55 | 0.31 |
| 0.8 | 0 | clcB | 3 | 10.55 | 0.335 |
| 0.8 | 0 | ycaM | 3 | 10.55 | 0.363 |
| 0.8 | 0 | yadI | 3 | 10.55 | 0.503 |
| 0.8 | 0 | AG1 | 3 | 10.55 | 0.406 |
| 0.8 | 0 | ptsI | 3 | 10.8 | 0.319 |
| 0.8 | 0 | clcB | 3 | 10.8 | 0.344 |
| 0.8 | 0 | ycaM | 3 | 10.8 | 0.371 |
| 0.8 | 0 | yadI | 3 | 10.8 | 0.516 |

| | | | | | |
|---|---|---|---|---|---|
| 0.8 | 0 | AG1 | 3 | 10.8 | 0.414 |
| 0.8 | 0 | ptsI | 3 | 11.05 | 0.326 |
| 0.8 | 0 | clcB | 3 | 11.05 | 0.354 |
| 0.8 | 0 | ycaM | 3 | 11.05 | 0.38 |
| 0.8 | 0 | yadI | 3 | 11.05 | 0.534 |
| 0.8 | 0 | AG1 | 3 | 11.05 | 0.427 |
| 0.8 | 0 | ptsI | 3 | 11.3 | 0.336 |
| 0.8 | 0 | clcB | 3 | 11.3 | 0.361 |
| 0.8 | 0 | ycaM | 3 | 11.3 | 0.38 |
| 0.8 | 0 | yadI | 3 | 11.3 | 0.542 |
| 0.8 | 0 | AG1 | 3 | 11.3 | 0.435 |
| 0.8 | 0 | ptsI | 3 | 11.55 | 0.344 |
| 0.8 | 0 | clcB | 3 | 11.55 | 0.363 |
| 0.8 | 0 | ycaM | 3 | 11.55 | 0.39 |
| 0.8 | 0 | yadI | 3 | 11.55 | 0.556 |
| 0.8 | 0 | AG1 | 3 | 11.55 | 0.445 |
| 0.8 | 0 | ptsI | 3 | 11.8 | 0.354 |
| 0.8 | 0 | clcB | 3 | 11.8 | 0.368 |
| 0.8 | 0 | ycaM | 3 | 11.8 | 0.402 |
| 0.8 | 0 | yadI | 3 | 11.8 | 0.564 |
| 0.8 | 0 | AG1 | 3 | 11.8 | 0.453 |
| 0.8 | 0 | ptsI | 3 | 12.05 | 0.361 |
| 0.8 | 0 | clcB | 3 | 12.05 | 0.37 |
| 0.8 | 0 | ycaM | 3 | 12.05 | 0.407 |
| 0.8 | 0 | yadI | 3 | 12.05 | 0.569 |
| 0.8 | 0 | AG1 | 3 | 12.05 | 0.463 |
| 0.8 | 0 | ptsI | 3 | 12.3 | 0.373 |
| 0.8 | 0 | clcB | 3 | 12.3 | 0.377 |
| 0.8 | 0 | ycaM | 3 | 12.3 | 0.415 |
| 0.8 | 0 | yadI | 3 | 12.3 | 0.589 |
| 0.8 | 0 | AG1 | 3 | 12.3 | 0.478 |
| 0.8 | 0 | ptsI | 3 | 12.55 | 0.378 |
| 0.8 | 0 | clcB | 3 | 12.55 | 0.384 |
| 0.8 | 0 | ycaM | 3 | 12.55 | 0.423 |
| 0.8 | 0 | yadI | 3 | 12.55 | 0.597 |
| 0.8 | 0 | AG1 | 3 | 12.55 | 0.491 |
| 0.8 | 0 | ptsI | 3 | 12.8 | 0.388 |
| 0.8 | 0 | clcB | 3 | 12.8 | 0.39 |
| 0.8 | 0 | ycaM | 3 | 12.8 | 0.432 |
| 0.8 | 0 | yadI | 3 | 12.8 | 0.609 |
| 0.8 | 0 | AG1 | 3 | 12.8 | 0.502 |
| 0.8 | 0 | ptsI | 3 | 13.05 | 0.392 |
| 0.8 | 0 | clcB | 3 | 13.05 | 0.394 |
| 0.8 | 0 | ycaM | 3 | 13.05 | 0.43 |
| 0.8 | 0 | yadI | 3 | 13.05 | 0.612 |
| 0.8 | 0 | AG1 | 3 | 13.05 | 0.512 |
| 0.8 | 0 | ptsI | 3 | 13.3 | 0.405 |
| 0.8 | 0 | clcB | 3 | 13.3 | 0.402 |

| | | | | | |
|---|---|---|---|---|---|
| 0.8 | 0 | ycaM | 3 | 13.3 | 0.438 |
| 0.8 | 0 | yadI | 3 | 13.3 | 0.633 |
| 0.8 | 0 | AG1 | 3 | 13.3 | 0.529 |
| 0.8 | 0 | ptsI | 3 | 13.55 | 0.412 |
| 0.8 | 0 | clcB | 3 | 13.55 | 0.414 |
| 0.8 | 0 | ycaM | 3 | 13.55 | 0.445 |
| 0.8 | 0 | yadI | 3 | 13.55 | 0.627 |
| 0.8 | 0 | AG1 | 3 | 13.55 | 0.544 |
| 0.8 | 0 | ptsI | 3 | 13.8 | 0.421 |
| 0.8 | 0 | clcB | 3 | 13.8 | 0.414 |
| 0.8 | 0 | ycaM | 3 | 13.8 | 0.452 |
| 0.8 | 0 | yadI | 3 | 13.8 | 0.647 |
| 0.8 | 0 | AG1 | 3 | 13.8 | 0.555 |
| 0.8 | 0 | ptsI | 3 | 14.05 | 0.429 |
| 0.8 | 0 | clcB | 3 | 14.05 | 0.42 |
| 0.8 | 0 | ycaM | 3 | 14.05 | 0.463 |
| 0.8 | 0 | yadI | 3 | 14.05 | 0.644 |
| 0.8 | 0 | AG1 | 3 | 14.05 | 0.568 |
| 0.8 | 0 | ptsI | 3 | 14.3 | 0.432 |
| 0.8 | 0 | clcB | 3 | 14.3 | 0.427 |
| 0.8 | 0 | ycaM | 3 | 14.3 | 0.47 |
| 0.8 | 0 | yadI | 3 | 14.3 | 0.651 |
| 0.8 | 0 | AG1 | 3 | 14.3 | 0.575 |
| 0.8 | 0 | ptsI | 3 | 14.55 | 0.44 |
| 0.8 | 0 | clcB | 3 | 14.55 | 0.428 |
| 0.8 | 0 | ycaM | 3 | 14.55 | 0.474 |
| 0.8 | 0 | yadI | 3 | 14.55 | 0.665 |
| 0.8 | 0 | AG1 | 3 | 14.55 | 0.579 |
| 0.8 | 0 | ptsI | 3 | 14.8 | 0.45 |
| 0.8 | 0 | clcB | 3 | 14.8 | 0.436 |
| 0.8 | 0 | ycaM | 3 | 14.8 | 0.486 |
| 0.8 | 0 | yadI | 3 | 14.8 | 0.672 |
| 0.8 | 0 | AG1 | 3 | 14.8 | 0.583 |
| 0.8 | 0 | ptsI | 3 | 15.05 | 0.458 |
| 0.8 | 0 | clcB | 3 | 15.05 | 0.44 |
| 0.8 | 0 | ycaM | 3 | 15.05 | 0.488 |
| 0.8 | 0 | yadI | 3 | 15.05 | 0.679 |
| 0.8 | 0 | AG1 | 3 | 15.05 | 0.615 |
| 0.8 | 0 | ptsI | 3 | 15.3 | 0.466 |
| 0.8 | 0 | clcB | 3 | 15.3 | 0.445 |
| 0.8 | 0 | ycaM | 3 | 15.3 | 0.499 |
| 0.8 | 0 | yadI | 3 | 15.3 | 0.698 |
| 0.8 | 0 | AG1 | 3 | 15.3 | 0.632 |
| 0.8 | 0 | ptsI | 3 | 15.55 | 0.48 |
| 0.8 | 0 | clcB | 3 | 15.55 | 0.448 |
| 0.8 | 0 | ycaM | 3 | 15.55 | 0.507 |
| 0.8 | 0 | yadI | 3 | 15.55 | 0.694 |
| 0.8 | 0 | AG1 | 3 | 15.55 | 0.639 |

| | | | | | |
|---|---|---|---|---|---|
| 0.8 | 0 | ptsI | 3 | 15.8 | 0.488 |
| 0.8 | 0 | clcB | 3 | 15.8 | 0.45 |
| 0.8 | 0 | ycaM | 3 | 15.8 | 0.524 |
| 0.8 | 0 | yadI | 3 | 15.8 | 0.703 |
| 0.8 | 0 | AG1 | 3 | 15.8 | 0.646 |
| 0.8 | 0 | ptsI | 3 | 16.05 | 0.5 |
| 0.8 | 0 | clcB | 3 | 16.05 | 0.451 |
| 0.8 | 0 | ycaM | 3 | 16.05 | 0.528 |
| 0.8 | 0 | yadI | 3 | 16.05 | 0.71 |
| 0.8 | 0 | AG1 | 3 | 16.05 | 0.646 |
| 0.8 | 0 | ptsI | 3 | 16.3 | 0.506 |
| 0.8 | 0 | clcB | 3 | 16.3 | 0.458 |
| 0.8 | 0 | ycaM | 3 | 16.3 | 0.531 |
| 0.8 | 0 | yadI | 3 | 16.3 | 0.692 |
| 0.8 | 0 | AG1 | 3 | 16.3 | 0.638 |
| 0.8 | 0 | ptsI | 3 | 16.55 | 0.513 |
| 0.8 | 0 | clcB | 3 | 16.55 | 0.456 |
| 0.8 | 0 | ycaM | 3 | 16.55 | 0.542 |
| 0.8 | 0 | yadI | 3 | 16.55 | 0.72 |
| 0.8 | 0 | AG1 | 3 | 16.55 | 0.661 |
| 0.8 | 0 | ptsI | 3 | 16.8 | 0.522 |
| 0.8 | 0 | clcB | 3 | 16.8 | 0.464 |
| 0.8 | 0 | ycaM | 3 | 16.8 | 0.547 |
| 0.8 | 0 | yadI | 3 | 16.8 | 0.75 |
| 0.8 | 0 | AG1 | 3 | 16.8 | 0.678 |
| 0.8 | 0 | ptsI | 3 | 17.05 | 0.52 |
| 0.8 | 0 | clcB | 3 | 17.05 | 0.466 |
| 0.8 | 0 | ycaM | 3 | 17.05 | 0.549 |
| 0.8 | 0 | yadI | 3 | 17.05 | 0.738 |
| 0.8 | 0 | AG1 | 3 | 17.05 | 0.675 |
| 0.8 | 0 | ptsI | 3 | 17.3 | 0.531 |
| 0.8 | 0 | clcB | 3 | 17.3 | 0.467 |
| 0.8 | 0 | ycaM | 3 | 17.3 | 0.552 |
| 0.8 | 0 | yadI | 3 | 17.3 | 0.742 |
| 0.8 | 0 | AG1 | 3 | 17.3 | 0.682 |
| 0.8 | 0 | ptsI | 3 | 17.55 | 0.537 |
| 0.8 | 0 | clcB | 3 | 17.55 | 0.47 |
| 0.8 | 0 | ycaM | 3 | 17.55 | 0.568 |
| 0.8 | 0 | yadI | 3 | 17.55 | 0.749 |
| 0.8 | 0 | AG1 | 3 | 17.55 | 0.678 |
| 0.8 | 0 | ptsI | 3 | 17.8 | 0.541 |
| 0.8 | 0 | clcB | 3 | 17.8 | 0.474 |
| 0.8 | 0 | ycaM | 3 | 17.8 | 0.577 |
| 0.8 | 0 | yadI | 3 | 17.8 | 0.755 |
| 0.8 | 0 | AG1 | 3 | 17.8 | 0.701 |
| 0.8 | 0 | ptsI | 3 | 18.05 | 0.552 |
| 0.8 | 0 | clcB | 3 | 18.05 | 0.475 |
| 0.8 | 0 | ycaM | 3 | 18.05 | 0.574 |

| | | | | | |
|---|---|---|---|---|---|
| 0.8 | 0 | yadI | 3 | 18.05 | 0.763 |
| 0.8 | 0 | AG1 | 3 | 18.05 | 0.693 |
| 0.8 | 0 | ptsI | 3 | 18.3 | 0.554 |
| 0.8 | 0 | clcB | 3 | 18.3 | 0.477 |
| 0.8 | 0 | ycaM | 3 | 18.3 | 0.592 |
| 0.8 | 0 | yadI | 3 | 18.3 | 0.765 |
| 0.8 | 0 | AG1 | 3 | 18.3 | 0.708 |
| 0.8 | 0 | ptsI | 3 | 18.55 | 0.561 |
| 0.8 | 0 | clcB | 3 | 18.55 | 0.48 |
| 0.8 | 0 | ycaM | 3 | 18.55 | 0.584 |
| 0.8 | 0 | yadI | 3 | 18.55 | 0.772 |
| 0.8 | 0 | AG1 | 3 | 18.55 | 0.721 |
| 0.8 | 0 | ptsI | 3 | 18.8 | 0.567 |
| 0.8 | 0 | clcB | 3 | 18.8 | 0.48 |
| 0.8 | 0 | ycaM | 3 | 18.8 | 0.579 |
| 0.8 | 0 | yadI | 3 | 18.8 | 0.781 |
| 0.8 | 0 | AG1 | 3 | 18.8 | 0.71 |
| 0.8 | 0 | ptsI | 3 | 19.05 | 0.576 |
| 0.8 | 0 | clcB | 3 | 19.05 | 0.487 |
| 0.8 | 0 | ycaM | 3 | 19.05 | 0.579 |
| 0.8 | 0 | yadI | 3 | 19.05 | 0.785 |
| 0.8 | 0 | AG1 | 3 | 19.05 | 0.715 |
| 0.8 | 0 | ptsI | 3 | 19.3 | 0.588 |
| 0.8 | 0 | clcB | 3 | 19.3 | 0.489 |
| 0.8 | 0 | ycaM | 3 | 19.3 | 0.574 |
| 0.8 | 0 | yadI | 3 | 19.3 | 0.796 |
| 0.8 | 0 | AG1 | 3 | 19.3 | 0.716 |
| 0.8 | 0 | ptsI | 3 | 19.55 | 0.607 |
| 0.8 | 0 | clcB | 3 | 19.55 | 0.496 |
| 0.8 | 0 | ycaM | 3 | 19.55 | 0.572 |
| 0.8 | 0 | yadI | 3 | 19.55 | 0.804 |
| 0.8 | 0 | AG1 | 3 | 19.55 | 0.715 |
| 0.8 | 0 | ptsI | 3 | 19.8 | 0.597 |
| 0.8 | 0 | clcB | 3 | 19.8 | 0.503 |
| 0.8 | 0 | ycaM | 3 | 19.8 | 0.575 |
| 0.8 | 0 | yadI | 3 | 19.8 | 0.805 |
| 0.8 | 0 | AG1 | 3 | 19.8 | 0.734 |
| 0.8 | 0 | ptsI | 3 | 20.05 | 0.604 |
| 0.8 | 0 | clcB | 3 | 20.05 | 0.507 |
| 0.8 | 0 | ycaM | 3 | 20.05 | 0.58 |
| 0.8 | 0 | yadI | 3 | 20.05 | 0.818 |
| 0.8 | 0 | AG1 | 3 | 20.05 | 0.747 |
| 0.8 | 0 | ptsI | 3 | 20.3 | 0.593 |
| 0.8 | 0 | clcB | 3 | 20.3 | 0.503 |
| 0.8 | 0 | ycaM | 3 | 20.3 | 0.571 |
| 0.8 | 0 | yadI | 3 | 20.3 | 0.857 |
| 0.8 | 0 | AG1 | 3 | 20.3 | 0.75 |
| 0.8 | 0 | ptsI | 3 | 20.55 | 0.601 |

| | | | | | |
|---|---|---|---|---|---|
| 0.8 | 0 | clcB | 3 | 20.55 | 0.509 |
| 0.8 | 0 | ycaM | 3 | 20.55 | 0.584 |
| 0.8 | 0 | yadI | 3 | 20.55 | 0.828 |
| 0.8 | 0 | AG1 | 3 | 20.55 | 0.757 |
| 0.8 | 0 | ptsI | 3 | 20.8 | 0.601 |
| 0.8 | 0 | clcB | 3 | 20.8 | 0.517 |
| 0.8 | 0 | ycaM | 3 | 20.8 | 0.594 |
| 0.8 | 0 | yadI | 3 | 20.8 | 0.834 |
| 0.8 | 0 | AG1 | 3 | 20.8 | 0.76 |
| 0.8 | 0 | ptsI | 3 | 21.05 | 0.598 |
| 0.8 | 0 | clcB | 3 | 21.05 | 0.529 |
| 0.8 | 0 | ycaM | 3 | 21.05 | 0.601 |
| 0.8 | 0 | yadI | 3 | 21.05 | 0.843 |
| 0.8 | 0 | AG1 | 3 | 21.05 | 0.769 |
| 0.8 | 0 | ptsI | 3 | 21.3 | 0.598 |
| 0.8 | 0 | clcB | 3 | 21.3 | 0.536 |
| 0.8 | 0 | ycaM | 3 | 21.3 | 0.604 |
| 0.8 | 0 | yadI | 3 | 21.3 | 0.848 |
| 0.8 | 0 | AG1 | 3 | 21.3 | 0.774 |
| 0.8 | 0 | ptsI | 3 | 21.55 | 0.598 |
| 0.8 | 0 | clcB | 3 | 21.55 | 0.537 |
| 0.8 | 0 | ycaM | 3 | 21.55 | 0.606 |
| 0.8 | 0 | yadI | 3 | 21.55 | 0.852 |
| 0.8 | 0 | AG1 | 3 | 21.55 | 0.772 |
| 0.8 | 0 | ptsI | 3 | 21.8 | 0.594 |
| 0.8 | 0 | clcB | 3 | 21.8 | 0.542 |
| 0.8 | 0 | ycaM | 3 | 21.8 | 0.611 |
| 0.8 | 0 | yadI | 3 | 21.8 | 0.854 |
| 0.8 | 0 | AG1 | 3 | 21.8 | 0.774 |
| 0.8 | 0 | ptsI | 3 | 22.05 | 0.606 |
| 0.8 | 0 | clcB | 3 | 22.05 | 0.537 |
| 0.8 | 0 | ycaM | 3 | 22.05 | 0.611 |
| 0.8 | 0 | yadI | 3 | 22.05 | 0.863 |
| 0.8 | 0 | AG1 | 3 | 22.05 | 0.784 |
| 0.8 | 0 | ptsI | 3 | 22.3 | 0.602 |
| 0.8 | 0 | clcB | 3 | 22.3 | 0.537 |
| 0.8 | 0 | ycaM | 3 | 22.3 | 0.618 |
| 0.8 | 0 | yadI | 3 | 22.3 | 0.87 |
| 0.8 | 0 | AG1 | 3 | 22.3 | 0.786 |
| 0.8 | 0 | ptsI | 3 | 22.55 | 0.604 |
| 0.8 | 0 | clcB | 3 | 22.55 | 0.537 |
| 0.8 | 0 | ycaM | 3 | 22.55 | 0.625 |
| 0.8 | 0 | yadI | 3 | 22.55 | 0.876 |
| 0.8 | 0 | AG1 | 3 | 22.55 | 0.79 |
| 0.8 | 0 | ptsI | 3 | 22.8 | 0.608 |
| 0.8 | 0 | clcB | 3 | 22.8 | 0.55 |
| 0.8 | 0 | ycaM | 3 | 22.8 | 0.636 |
| 0.8 | 0 | yadI | 3 | 22.8 | 0.881 |

| | | | | | |
|---|---|---|---|---|---|
| 0.8 | 0 | AG1 | 3 | 22.8 | 0.79 |
| 0.8 | 0 | ptsI | 3 | 23.05 | 0.606 |
| 0.8 | 0 | clcB | 3 | 23.05 | 0.541 |
| 0.8 | 0 | ycaM | 3 | 23.05 | 0.636 |
| 0.8 | 0 | yadI | 3 | 23.05 | 0.883 |
| 0.8 | 0 | AG1 | 3 | 23.05 | 0.802 |
| 0.8 | 0 | ptsI | 3 | 23.3 | 0.609 |
| 0.8 | 0 | clcB | 3 | 23.3 | 0.542 |
| 0.8 | 0 | ycaM | 3 | 23.3 | 0.642 |
| 0.8 | 0 | yadI | 3 | 23.3 | 0.925 |
| 0.8 | 0 | AG1 | 3 | 23.3 | 0.807 |
| 0.8 | 0 | ptsI | 3 | 23.55 | 0.615 |
| 0.8 | 0 | clcB | 3 | 23.55 | 0.544 |
| 0.8 | 0 | ycaM | 3 | 23.55 | 0.646 |
| 0.8 | 0 | yadI | 3 | 23.55 | 0.92 |
| 0.8 | 0 | AG1 | 3 | 23.55 | 0.81 |
| 0.8 | 0 | ptsI | 3 | 23.8 | 0.616 |
| 0.8 | 0 | clcB | 3 | 23.8 | 0.546 |
| 0.8 | 0 | ycaM | 3 | 23.8 | 0.649 |
| 0.8 | 0 | yadI | 3 | 23.8 | 0.948 |
| 0.8 | 0 | AG1 | 3 | 23.8 | 0.82 |
| 0.8 | 0 | ptsI | 3 | 24.05 | 0.617 |
| 0.8 | 0 | clcB | 3 | 24.05 | 0.547 |
| 0.8 | 0 | ycaM | 3 | 24.05 | 0.656 |
| 0.8 | 0 | yadI | 3 | 24.05 | 0.915 |
| 0.8 | 0 | AG1 | 3 | 24.05 | 0.825 |
| 0.8 | 0 | ptsI | 3 | 24.3 | 0.63 |
| 0.8 | 0 | clcB | 3 | 24.3 | 0.555 |
| 0.8 | 0 | ycaM | 3 | 24.3 | 0.666 |
| 0.8 | 0 | yadI | 3 | 24.3 | 0.946 |
| 0.8 | 0 | AG1 | 3 | 24.3 | 0.832 |
| 0.8 | 0 | ptsI | 4 | 0 | 0.197 |
| 0.8 | 0 | clcB | 4 | 0 | 0.211 |
| 0.8 | 0 | ycaM | 4 | 0 | 0.218 |
| 0.8 | 0 | yadI | 4 | 0 | 0.235 |
| 0.8 | 0 | AG1 | 4 | 0 | 0.203 |
| 0.8 | 0 | ptsI | 4 | 0.25 | 0.198 |
| 0.8 | 0 | clcB | 4 | 0.25 | 0.207 |
| 0.8 | 0 | ycaM | 4 | 0.25 | 0.212 |
| 0.8 | 0 | yadI | 4 | 0.25 | 0.236 |
| 0.8 | 0 | AG1 | 4 | 0.25 | 0.201 |
| 0.8 | 0 | ptsI | 4 | 0.5 | 0.196 |
| 0.8 | 0 | clcB | 4 | 0.5 | 0.207 |
| 0.8 | 0 | ycaM | 4 | 0.5 | 0.213 |
| 0.8 | 0 | yadI | 4 | 0.5 | 0.236 |
| 0.8 | 0 | AG1 | 4 | 0.5 | 0.202 |
| 0.8 | 0 | ptsI | 4 | 0.75 | 0.203 |
| 0.8 | 0 | clcB | 4 | 0.75 | 0.212 |

| | | | | | |
|---|---|---|---|---|---|
| 0.8 | 0 | ycaM | 4 | 0.75 | 0.216 |
| 0.8 | 0 | yadI | 4 | 0.75 | 0.239 |
| 0.8 | 0 | AG1 | 4 | 0.75 | 0.207 |
| 0.8 | 0 | ptsI | 4 | 1 | 0.206 |
| 0.8 | 0 | clcB | 4 | 1 | 0.215 |
| 0.8 | 0 | ycaM | 4 | 1 | 0.219 |
| 0.8 | 0 | yadI | 4 | 1 | 0.24 |
| 0.8 | 0 | AG1 | 4 | 1 | 0.207 |
| 0.8 | 0 | ptsI | 4 | 1.25 | 0.214 |
| 0.8 | 0 | clcB | 4 | 1.25 | 0.222 |
| 0.8 | 0 | ycaM | 4 | 1.25 | 0.226 |
| 0.8 | 0 | yadI | 4 | 1.25 | 0.243 |
| 0.8 | 0 | AG1 | 4 | 1.25 | 0.213 |
| 0.8 | 0 | ptsI | 4 | 1.5 | 0.212 |
| 0.8 | 0 | clcB | 4 | 1.5 | 0.224 |
| 0.8 | 0 | ycaM | 4 | 1.5 | 0.228 |
| 0.8 | 0 | yadI | 4 | 1.5 | 0.251 |
| 0.8 | 0 | AG1 | 4 | 1.5 | 0.214 |
| 0.8 | 0 | ptsI | 4 | 1.75 | 0.224 |
| 0.8 | 0 | clcB | 4 | 1.75 | 0.237 |
| 0.8 | 0 | ycaM | 4 | 1.75 | 0.234 |
| 0.8 | 0 | yadI | 4 | 1.75 | 0.259 |
| 0.8 | 0 | AG1 | 4 | 1.75 | 0.223 |
| 0.8 | 0 | ptsI | 4 | 2 | 0.235 |
| 0.8 | 0 | clcB | 4 | 2 | 0.251 |
| 0.8 | 0 | ycaM | 4 | 2 | 0.249 |
| 0.8 | 0 | yadI | 4 | 2 | 0.269 |
| 0.8 | 0 | AG1 | 4 | 2 | 0.234 |
| 0.8 | 0 | ptsI | 4 | 2.25 | 0.25 |
| 0.8 | 0 | clcB | 4 | 2.25 | 0.262 |
| 0.8 | 0 | ycaM | 4 | 2.25 | 0.259 |
| 0.8 | 0 | yadI | 4 | 2.25 | 0.281 |
| 0.8 | 0 | AG1 | 4 | 2.25 | 0.244 |
| 0.8 | 0 | ptsI | 4 | 2.5 | 0.248 |
| 0.8 | 0 | clcB | 4 | 2.5 | 0.273 |
| 0.8 | 0 | ycaM | 4 | 2.5 | 0.266 |
| 0.8 | 0 | yadI | 4 | 2.5 | 0.292 |
| 0.8 | 0 | AG1 | 4 | 2.5 | 0.252 |
| 0.8 | 0 | ptsI | 4 | 2.75 | 0.258 |
| 0.8 | 0 | clcB | 4 | 2.75 | 0.289 |
| 0.8 | 0 | ycaM | 4 | 2.75 | 0.28 |
| 0.8 | 0 | yadI | 4 | 2.75 | 0.311 |
| 0.8 | 0 | AG1 | 4 | 2.75 | 0.261 |
| 0.8 | 0 | ptsI | 4 | 3 | 0.28 |
| 0.8 | 0 | clcB | 4 | 3 | 0.309 |
| 0.8 | 0 | ycaM | 4 | 3 | 0.285 |
| 0.8 | 0 | yadI | 4 | 3 | 0.325 |
| 0.8 | 0 | AG1 | 4 | 3 | 0.274 |

| | | | | | |
|---|---|---|---|---|---|
| 0.8 | 0 | ptsI | 4 | 3.25 | 0.3 |
| 0.8 | 0 | clcB | 4 | 3.25 | 0.326 |
| 0.8 | 0 | ycaM | 4 | 3.25 | 0.299 |
| 0.8 | 0 | yadI | 4 | 3.25 | 0.341 |
| 0.8 | 0 | AG1 | 4 | 3.25 | 0.287 |
| 0.8 | 0 | ptsI | 4 | 3.5 | 0.319 |
| 0.8 | 0 | clcB | 4 | 3.5 | 0.341 |
| 0.8 | 0 | ycaM | 4 | 3.5 | 0.314 |
| 0.8 | 0 | yadI | 4 | 3.5 | 0.361 |
| 0.8 | 0 | AG1 | 4 | 3.5 | 0.306 |
| 0.8 | 0 | ptsI | 4 | 3.75 | 0.327 |
| 0.8 | 0 | clcB | 4 | 3.75 | 0.352 |
| 0.8 | 0 | ycaM | 4 | 3.75 | 0.333 |
| 0.8 | 0 | yadI | 4 | 3.75 | 0.381 |
| 0.8 | 0 | AG1 | 4 | 3.75 | 0.322 |
| 0.8 | 0 | ptsI | 4 | 4 | 0.347 |
| 0.8 | 0 | clcB | 4 | 4 | 0.365 |
| 0.8 | 0 | ycaM | 4 | 4 | 0.345 |
| 0.8 | 0 | yadI | 4 | 4 | 0.394 |
| 0.8 | 0 | AG1 | 4 | 4 | 0.338 |
| 0.8 | 0 | ptsI | 4 | 4.35 | 0.339 |
| 0.8 | 0 | clcB | 4 | 4.35 | 0.281 |
| 0.8 | 0 | ycaM | 4 | 4.35 | 0.387 |
| 0.8 | 0 | yadI | 4 | 4.35 | 0.417 |
| 0.8 | 0 | AG1 | 4 | 4.35 | 0.305 |
| 0.8 | 0 | ptsI | 4 | 4.6 | 0.352 |
| 0.8 | 0 | clcB | 4 | 4.6 | 0.278 |
| 0.8 | 0 | ycaM | 4 | 4.6 | 0.372 |
| 0.8 | 0 | yadI | 4 | 4.6 | 0.4 |
| 0.8 | 0 | AG1 | 4 | 4.6 | 0.287 |
| 0.8 | 0 | ptsI | 4 | 4.85 | 0.366 |
| 0.8 | 0 | clcB | 4 | 4.85 | 0.284 |
| 0.8 | 0 | ycaM | 4 | 4.85 | 0.384 |
| 0.8 | 0 | yadI | 4 | 4.85 | 0.412 |
| 0.8 | 0 | AG1 | 4 | 4.85 | 0.294 |
| 0.8 | 0 | ptsI | 4 | 5.1 | 0.379 |
| 0.8 | 0 | clcB | 4 | 5.1 | 0.295 |
| 0.8 | 0 | ycaM | 4 | 5.1 | 0.397 |
| 0.8 | 0 | yadI | 4 | 5.1 | 0.423 |
| 0.8 | 0 | AG1 | 4 | 5.1 | 0.304 |
| 0.8 | 0 | ptsI | 4 | 5.35 | 0.391 |
| 0.8 | 0 | clcB | 4 | 5.35 | 0.303 |
| 0.8 | 0 | ycaM | 4 | 5.35 | 0.411 |
| 0.8 | 0 | yadI | 4 | 5.35 | 0.438 |
| 0.8 | 0 | AG1 | 4 | 5.35 | 0.314 |
| 0.8 | 0 | ptsI | 4 | 5.6 | 0.406 |
| 0.8 | 0 | clcB | 4 | 5.6 | 0.309 |
| 0.8 | 0 | ycaM | 4 | 5.6 | 0.419 |

| | | | | | |
|---|---|---|---|---|---|
| 0.8 | 0 | yadI | 4 | 5.6 | 0.45 |
| 0.8 | 0 | AG1 | 4 | 5.6 | 0.323 |
| 0.8 | 0 | ptsI | 4 | 5.85 | 0.411 |
| 0.8 | 0 | clcB | 4 | 5.85 | 0.317 |
| 0.8 | 0 | ycaM | 4 | 5.85 | 0.427 |
| 0.8 | 0 | yadI | 4 | 5.85 | 0.468 |
| 0.8 | 0 | AG1 | 4 | 5.85 | 0.334 |
| 0.8 | 0 | ptsI | 4 | 6.1 | 0.425 |
| 0.8 | 0 | clcB | 4 | 6.1 | 0.329 |
| 0.8 | 0 | ycaM | 4 | 6.1 | 0.434 |
| 0.8 | 0 | yadI | 4 | 6.1 | 0.484 |
| 0.8 | 0 | AG1 | 4 | 6.1 | 0.344 |
| 0.8 | 0 | ptsI | 4 | 6.35 | 0.437 |
| 0.8 | 0 | clcB | 4 | 6.35 | 0.336 |
| 0.8 | 0 | ycaM | 4 | 6.35 | 0.447 |
| 0.8 | 0 | yadI | 4 | 6.35 | 0.498 |
| 0.8 | 0 | AG1 | 4 | 6.35 | 0.354 |
| 0.8 | 0 | ptsI | 4 | 6.6 | 0.449 |
| 0.8 | 0 | clcB | 4 | 6.6 | 0.346 |
| 0.8 | 0 | ycaM | 4 | 6.6 | 0.456 |
| 0.8 | 0 | yadI | 4 | 6.6 | 0.512 |
| 0.8 | 0 | AG1 | 4 | 6.6 | 0.363 |
| 0.8 | 0 | ptsI | 4 | 6.85 | 0.462 |
| 0.8 | 0 | clcB | 4 | 6.85 | 0.353 |
| 0.8 | 0 | ycaM | 4 | 6.85 | 0.467 |
| 0.8 | 0 | yadI | 4 | 6.85 | 0.524 |
| 0.8 | 0 | AG1 | 4 | 6.85 | 0.372 |
| 0.8 | 0 | ptsI | 4 | 7.1 | 0.472 |
| 0.8 | 0 | clcB | 4 | 7.1 | 0.364 |
| 0.8 | 0 | ycaM | 4 | 7.1 | 0.476 |
| 0.8 | 0 | yadI | 4 | 7.1 | 0.531 |
| 0.8 | 0 | AG1 | 4 | 7.1 | 0.381 |
| 0.8 | 0 | ptsI | 4 | 7.35 | 0.484 |
| 0.8 | 0 | clcB | 4 | 7.35 | 0.369 |
| 0.8 | 0 | ycaM | 4 | 7.35 | 0.478 |
| 0.8 | 0 | yadI | 4 | 7.35 | 0.542 |
| 0.8 | 0 | AG1 | 4 | 7.35 | 0.39 |
| 0.8 | 0 | ptsI | 4 | 7.6 | 0.492 |
| 0.8 | 0 | clcB | 4 | 7.6 | 0.376 |
| 0.8 | 0 | ycaM | 4 | 7.6 | 0.484 |
| 0.8 | 0 | yadI | 4 | 7.6 | 0.549 |
| 0.8 | 0 | AG1 | 4 | 7.6 | 0.401 |
| 0.8 | 0 | ptsI | 4 | 7.85 | 0.5 |
| 0.8 | 0 | clcB | 4 | 7.85 | 0.387 |
| 0.8 | 0 | ycaM | 4 | 7.85 | 0.487 |
| 0.8 | 0 | yadI | 4 | 7.85 | 0.559 |
| 0.8 | 0 | AG1 | 4 | 7.85 | 0.406 |
| 0.8 | 0 | ptsI | 4 | 8.1 | 0.51 |

| | | | | | |
|---|---|---|---|---|---|
| 0.8 | 0 | clcB | 4 | 8.1 | 0.396 |
| 0.8 | 0 | ycaM | 4 | 8.1 | 0.49 |
| 0.8 | 0 | yadI | 4 | 8.1 | 0.566 |
| 0.8 | 0 | AG1 | 4 | 8.1 | 0.415 |
| 0.8 | 0 | ptsI | 4 | 8.35 | 0.52 |
| 0.8 | 0 | clcB | 4 | 8.35 | 0.402 |
| 0.8 | 0 | ycaM | 4 | 8.35 | 0.492 |
| 0.8 | 0 | yadI | 4 | 8.35 | 0.574 |
| 0.8 | 0 | AG1 | 4 | 8.35 | 0.421 |
| 0.8 | 0 | ptsI | 4 | 8.83333 | 0.416 |
| 0.8 | 0 | clcB | 4 | 8.83333 | 0.4 |
| 0.8 | 0 | ycaM | 4 | 8.83333 | 0.438 |
| 0.8 | 0 | yadI | 4 | 8.83333 | 0.553 |
| 0.8 | 0 | AG1 | 4 | 8.83333 | 0.465 |
| 0.8 | 0 | ptsI | 4 | 9.08333 | 0.416 |
| 0.8 | 0 | clcB | 4 | 9.08333 | 0.397 |
| 0.8 | 0 | ycaM | 4 | 9.08333 | 0.436 |
| 0.8 | 0 | yadI | 4 | 9.08333 | 0.552 |
| 0.8 | 0 | AG1 | 4 | 9.08333 | 0.462 |
| 0.8 | 0 | ptsI | 4 | 9.33333 | 0.427 |
| 0.8 | 0 | clcB | 4 | 9.33333 | 0.4 |
| 0.8 | 0 | ycaM | 4 | 9.33333 | 0.44 |
| 0.8 | 0 | yadI | 4 | 9.33333 | 0.555 |
| 0.8 | 0 | AG1 | 4 | 9.33333 | 0.471 |
| 0.8 | 0 | ptsI | 4 | 9.58333 | 0.439 |
| 0.8 | 0 | clcB | 4 | 9.58333 | 0.401 |
| 0.8 | 0 | ycaM | 4 | 9.58333 | 0.45 |
| 0.8 | 0 | yadI | 4 | 9.58333 | 0.566 |
| 0.8 | 0 | AG1 | 4 | 9.58333 | 0.483 |
| 0.8 | 0 | ptsI | 4 | 9.83333 | 0.447 |
| 0.8 | 0 | clcB | 4 | 9.83333 | 0.406 |
| 0.8 | 0 | ycaM | 4 | 9.83333 | 0.456 |
| 0.8 | 0 | yadI | 4 | 9.83333 | 0.577 |
| 0.8 | 0 | AG1 | 4 | 9.83333 | 0.495 |
| 0.8 | 0 | ptsI | 4 | 10.0833 | 0.453 |
| 0.8 | 0 | clcB | 4 | 10.0833 | 0.412 |
| 0.8 | 0 | ycaM | 4 | 10.0833 | 0.462 |
| 0.8 | 0 | yadI | 4 | 10.0833 | 0.583 |
| 0.8 | 0 | AG1 | 4 | 10.0833 | 0.502 |
| 0.8 | 0 | ptsI | 4 | 10.3333 | 0.459 |
| 0.8 | 0 | clcB | 4 | 10.3333 | 0.412 |
| 0.8 | 0 | ycaM | 4 | 10.3333 | 0.466 |
| 0.8 | 0 | yadI | 4 | 10.3333 | 0.59 |
| 0.8 | 0 | AG1 | 4 | 10.3333 | 0.51 |
| 0.8 | 0 | ptsI | 4 | 10.5833 | 0.466 |
| 0.8 | 0 | clcB | 4 | 10.5833 | 0.418 |
| 0.8 | 0 | ycaM | 4 | 10.5833 | 0.473 |
| 0.8 | 0 | yadI | 4 | 10.5833 | 0.596 |

| | | | | | |
|---|---|---|---|---|---|
| 0.8 | 0 | AG1 | 4 | 10.5833 | 0.519 |
| 0.8 | 0 | ptsI | 4 | 10.8333 | 0.473 |
| 0.8 | 0 | clcB | 4 | 10.8333 | 0.42 |
| 0.8 | 0 | ycaM | 4 | 10.8333 | 0.477 |
| 0.8 | 0 | yadI | 4 | 10.8333 | 0.602 |
| 0.8 | 0 | AG1 | 4 | 10.8333 | 0.525 |
| 0.8 | 0 | ptsI | 4 | 11.0833 | 0.482 |
| 0.8 | 0 | clcB | 4 | 11.0833 | 0.423 |
| 0.8 | 0 | ycaM | 4 | 11.0833 | 0.481 |
| 0.8 | 0 | yadI | 4 | 11.0833 | 0.608 |
| 0.8 | 0 | AG1 | 4 | 11.0833 | 0.526 |
| 0.8 | 0 | ptsI | 4 | 11.3333 | 0.489 |
| 0.8 | 0 | clcB | 4 | 11.3333 | 0.432 |
| 0.8 | 0 | ycaM | 4 | 11.3333 | 0.486 |
| 0.8 | 0 | yadI | 4 | 11.3333 | 0.615 |
| 0.8 | 0 | AG1 | 4 | 11.3333 | 0.536 |
| 0.8 | 0 | ptsI | 4 | 11.5833 | 0.497 |
| 0.8 | 0 | clcB | 4 | 11.5833 | 0.433 |
| 0.8 | 0 | ycaM | 4 | 11.5833 | 0.489 |
| 0.8 | 0 | yadI | 4 | 11.5833 | 0.622 |
| 0.8 | 0 | AG1 | 4 | 11.5833 | 0.539 |
| 0.8 | 0 | ptsI | 4 | 11.8333 | 0.505 |
| 0.8 | 0 | clcB | 4 | 11.8333 | 0.433 |
| 0.8 | 0 | ycaM | 4 | 11.8333 | 0.492 |
| 0.8 | 0 | yadI | 4 | 11.8333 | 0.625 |
| 0.8 | 0 | AG1 | 4 | 11.8333 | 0.542 |
| 0.8 | 0 | ptsI | 4 | 12.0833 | 0.511 |
| 0.8 | 0 | clcB | 4 | 12.0833 | 0.438 |
| 0.8 | 0 | ycaM | 4 | 12.0833 | 0.498 |
| 0.8 | 0 | yadI | 4 | 12.0833 | 0.631 |
| 0.8 | 0 | AG1 | 4 | 12.0833 | 0.549 |
| 0.8 | 0 | ptsI | 4 | 12.3333 | 0.516 |
| 0.8 | 0 | clcB | 4 | 12.3333 | 0.443 |
| 0.8 | 0 | ycaM | 4 | 12.3333 | 0.504 |
| 0.8 | 0 | yadI | 4 | 12.3333 | 0.639 |
| 0.8 | 0 | AG1 | 4 | 12.3333 | 0.557 |
| 0.8 | 0 | ptsI | 4 | 12.5833 | 0.521 |
| 0.8 | 0 | clcB | 4 | 12.5833 | 0.444 |
| 0.8 | 0 | ycaM | 4 | 12.5833 | 0.508 |
| 0.8 | 0 | yadI | 4 | 12.5833 | 0.64 |
| 0.8 | 0 | AG1 | 4 | 12.5833 | 0.561 |
| 0.8 | 0 | ptsI | 4 | 12.8333 | 0.528 |
| 0.8 | 0 | clcB | 4 | 12.8333 | 0.445 |
| 0.8 | 0 | ycaM | 4 | 12.8333 | 0.512 |
| 0.8 | 0 | yadI | 4 | 12.8333 | 0.643 |
| 0.8 | 0 | AG1 | 4 | 12.8333 | 0.567 |
| 0.8 | 0 | ptsI | 4 | 13.0833 | 0.536 |
| 0.8 | 0 | clcB | 4 | 13.0833 | 0.448 |

| | | | | | |
|---|---|---|---|---|---|
| 0.8 | 0 | ycaM | 4 | 13.0833 | 0.516 |
| 0.8 | 0 | yadI | 4 | 13.0833 | 0.65 |
| 0.8 | 0 | AG1 | 4 | 13.0833 | 0.572 |
| 0.8 | 0 | ptsI | 4 | 13.3333 | 0.541 |
| 0.8 | 0 | clcB | 4 | 13.3333 | 0.45 |
| 0.8 | 0 | ycaM | 4 | 13.3333 | 0.523 |
| 0.8 | 0 | yadI | 4 | 13.3333 | 0.652 |
| 0.8 | 0 | AG1 | 4 | 13.3333 | 0.578 |
| 0.8 | 0 | ptsI | 4 | 13.5833 | 0.544 |
| 0.8 | 0 | clcB | 4 | 13.5833 | 0.455 |
| 0.8 | 0 | ycaM | 4 | 13.5833 | 0.524 |
| 0.8 | 0 | yadI | 4 | 13.5833 | 0.657 |
| 0.8 | 0 | AG1 | 4 | 13.5833 | 0.582 |
| 0.8 | 0 | ptsI | 4 | 13.8333 | 0.552 |
| 0.8 | 0 | clcB | 4 | 13.8333 | 0.456 |
| 0.8 | 0 | ycaM | 4 | 13.8333 | 0.53 |
| 0.8 | 0 | yadI | 4 | 13.8333 | 0.657 |
| 0.8 | 0 | AG1 | 4 | 13.8333 | 0.582 |
| 0.8 | 0 | ptsI | 4 | 14.0833 | 0.555 |
| 0.8 | 0 | clcB | 4 | 14.0833 | 0.457 |
| 0.8 | 0 | ycaM | 4 | 14.0833 | 0.533 |
| 0.8 | 0 | yadI | 4 | 14.0833 | 0.663 |
| 0.8 | 0 | AG1 | 4 | 14.0833 | 0.587 |
| 0.8 | 0 | ptsI | 4 | 14.3333 | 0.56 |
| 0.8 | 0 | clcB | 4 | 14.3333 | 0.462 |
| 0.8 | 0 | ycaM | 4 | 14.3333 | 0.533 |
| 0.8 | 0 | yadI | 4 | 14.3333 | 0.663 |
| 0.8 | 0 | AG1 | 4 | 14.3333 | 0.589 |
| 0.8 | 0 | ptsI | 4 | 14.5833 | 0.562 |
| 0.8 | 0 | clcB | 4 | 14.5833 | 0.46 |
| 0.8 | 0 | ycaM | 4 | 14.5833 | 0.535 |
| 0.8 | 0 | yadI | 4 | 14.5833 | 0.667 |
| 0.8 | 0 | AG1 | 4 | 14.5833 | 0.592 |
| 0.8 | 0 | ptsI | 4 | 14.8333 | 0.567 |
| 0.8 | 0 | clcB | 4 | 14.8333 | 0.462 |
| 0.8 | 0 | ycaM | 4 | 14.8333 | 0.539 |
| 0.8 | 0 | yadI | 4 | 14.8333 | 0.667 |
| 0.8 | 0 | AG1 | 4 | 14.8333 | 0.596 |
| 0.8 | 0 | ptsI | 4 | 15.0833 | 0.572 |
| 0.8 | 0 | clcB | 4 | 15.0833 | 0.464 |
| 0.8 | 0 | ycaM | 4 | 15.0833 | 0.539 |
| 0.8 | 0 | yadI | 4 | 15.0833 | 0.669 |
| 0.8 | 0 | AG1 | 4 | 15.0833 | 0.597 |
| 0.8 | 0 | ptsI | 4 | 15.3333 | 0.575 |
| 0.8 | 0 | clcB | 4 | 15.3333 | 0.467 |
| 0.8 | 0 | ycaM | 4 | 15.3333 | 0.545 |
| 0.8 | 0 | yadI | 4 | 15.3333 | 0.67 |
| 0.8 | 0 | AG1 | 4 | 15.3333 | 0.597 |

| | | | | | |
|---|---|---|---|---|---|
| 0.8 | 0 | ptsI | 4 | 15.5833 | 0.576 |
| 0.8 | 0 | clcB | 4 | 15.5833 | 0.47 |
| 0.8 | 0 | ycaM | 4 | 15.5833 | 0.544 |
| 0.8 | 0 | yadI | 4 | 15.5833 | 0.675 |
| 0.8 | 0 | AG1 | 4 | 15.5833 | 0.6 |
| 0.8 | 0 | ptsI | 4 | 15.8333 | 0.583 |
| 0.8 | 0 | clcB | 4 | 15.8333 | 0.472 |
| 0.8 | 0 | ycaM | 4 | 15.8333 | 0.547 |
| 0.8 | 0 | yadI | 4 | 15.8333 | 0.678 |
| 0.8 | 0 | AG1 | 4 | 15.8333 | 0.602 |
| 0.8 | 0 | ptsI | 4 | 16.0833 | 0.588 |
| 0.8 | 0 | clcB | 4 | 16.0833 | 0.475 |
| 0.8 | 0 | ycaM | 4 | 16.0833 | 0.55 |
| 0.8 | 0 | yadI | 4 | 16.0833 | 0.677 |
| 0.8 | 0 | AG1 | 4 | 16.0833 | 0.605 |
| 0.8 | 0 | ptsI | 4 | 16.3333 | 0.589 |
| 0.8 | 0 | clcB | 4 | 16.3333 | 0.473 |
| 0.8 | 0 | ycaM | 4 | 16.3333 | 0.551 |
| 0.8 | 0 | yadI | 4 | 16.3333 | 0.682 |
| 0.8 | 0 | AG1 | 4 | 16.3333 | 0.604 |
| 0.8 | 0 | ptsI | 4 | 16.5833 | 0.594 |
| 0.8 | 0 | clcB | 4 | 16.5833 | 0.473 |
| 0.8 | 0 | ycaM | 4 | 16.5833 | 0.553 |
| 0.8 | 0 | yadI | 4 | 16.5833 | 0.685 |
| 0.8 | 0 | AG1 | 4 | 16.5833 | 0.608 |
| 0.8 | 0 | ptsI | 4 | 16.8333 | 0.596 |
| 0.8 | 0 | clcB | 4 | 16.8333 | 0.476 |
| 0.8 | 0 | ycaM | 4 | 16.8333 | 0.555 |
| 0.8 | 0 | yadI | 4 | 16.8333 | 0.686 |
| 0.8 | 0 | AG1 | 4 | 16.8333 | 0.607 |
| 0.8 | 0 | ptsI | 4 | 17.0833 | 0.601 |
| 0.8 | 0 | clcB | 4 | 17.0833 | 0.478 |
| 0.8 | 0 | ycaM | 4 | 17.0833 | 0.558 |
| 0.8 | 0 | yadI | 4 | 17.0833 | 0.688 |
| 0.8 | 0 | AG1 | 4 | 17.0833 | 0.606 |
| 0.8 | 0 | ptsI | 4 | 17.3333 | 0.603 |
| 0.8 | 0 | clcB | 4 | 17.3333 | 0.477 |
| 0.8 | 0 | ycaM | 4 | 17.3333 | 0.563 |
| 0.8 | 0 | yadI | 4 | 17.3333 | 0.694 |
| 0.8 | 0 | AG1 | 4 | 17.3333 | 0.608 |
| 0.8 | 0 | ptsI | 4 | 17.5833 | 0.607 |
| 0.8 | 0 | clcB | 4 | 17.5833 | 0.481 |
| 0.8 | 0 | ycaM | 4 | 17.5833 | 0.559 |
| 0.8 | 0 | yadI | 4 | 17.5833 | 0.694 |
| 0.8 | 0 | AG1 | 4 | 17.5833 | 0.611 |
| 0.8 | 0 | ptsI | 4 | 17.8333 | 0.609 |
| 0.8 | 0 | clcB | 4 | 17.8333 | 0.481 |
| 0.8 | 0 | ycaM | 4 | 17.8333 | 0.558 |

| | | | | | |
|---|---|---|---|---|---|
| 0.8 | 0 | yadI | 4 | 17.8333 | 0.693 |
| 0.8 | 0 | AG1 | 4 | 17.8333 | 0.61 |
| 0.8 | 0 | ptsI | 4 | 18.0833 | 0.617 |
| 0.8 | 0 | clcB | 4 | 18.0833 | 0.481 |
| 0.8 | 0 | ycaM | 4 | 18.0833 | 0.563 |
| 0.8 | 0 | yadI | 4 | 18.0833 | 0.698 |
| 0.8 | 0 | AG1 | 4 | 18.0833 | 0.612 |
| 0.8 | 0 | ptsI | 4 | 18.3333 | 0.62 |
| 0.8 | 0 | clcB | 4 | 18.3333 | 0.482 |
| 0.8 | 0 | ycaM | 4 | 18.3333 | 0.565 |
| 0.8 | 0 | yadI | 4 | 18.3333 | 0.701 |
| 0.8 | 0 | AG1 | 4 | 18.3333 | 0.613 |
| 0.8 | 0 | ptsI | 4 | 18.5833 | 0.62 |
| 0.8 | 0 | clcB | 4 | 18.5833 | 0.482 |
| 0.8 | 0 | ycaM | 4 | 18.5833 | 0.566 |
| 0.8 | 0 | yadI | 4 | 18.5833 | 0.704 |
| 0.8 | 0 | AG1 | 4 | 18.5833 | 0.613 |
| 0.8 | 0 | ptsI | 4 | 18.8333 | 0.624 |
| 0.8 | 0 | clcB | 4 | 18.8333 | 0.483 |
| 0.8 | 0 | ycaM | 4 | 18.8333 | 0.563 |
| 0.8 | 0 | yadI | 4 | 18.8333 | 0.703 |
| 0.8 | 0 | AG1 | 4 | 18.8333 | 0.614 |
| 0.8 | 0 | ptsI | 4 | 19.0833 | 0.628 |
| 0.8 | 0 | clcB | 4 | 19.0833 | 0.486 |
| 0.8 | 0 | ycaM | 4 | 19.0833 | 0.568 |
| 0.8 | 0 | yadI | 4 | 19.0833 | 0.705 |
| 0.8 | 0 | AG1 | 4 | 19.0833 | 0.619 |
| 0.8 | 0 | ptsI | 4 | 19.3333 | 0.628 |
| 0.8 | 0 | clcB | 4 | 19.3333 | 0.484 |
| 0.8 | 0 | ycaM | 4 | 19.3333 | 0.572 |
| 0.8 | 0 | yadI | 4 | 19.3333 | 0.709 |
| 0.8 | 0 | AG1 | 4 | 19.3333 | 0.618 |
| 0.8 | 0 | ptsI | 4 | 19.5833 | 0.628 |
| 0.8 | 0 | clcB | 4 | 19.5833 | 0.489 |
| 0.8 | 0 | ycaM | 4 | 19.5833 | 0.57 |
| 0.8 | 0 | yadI | 4 | 19.5833 | 0.708 |
| 0.8 | 0 | AG1 | 4 | 19.5833 | 0.616 |
| 0.8 | 0 | ptsI | 4 | 19.8333 | 0.632 |
| 0.8 | 0 | clcB | 4 | 19.8333 | 0.487 |
| 0.8 | 0 | ycaM | 4 | 19.8333 | 0.57 |
| 0.8 | 0 | yadI | 4 | 19.8333 | 0.706 |
| 0.8 | 0 | AG1 | 4 | 19.8333 | 0.617 |
| 0.8 | 0 | ptsI | 4 | 20.0833 | 0.626 |
| 0.8 | 0 | clcB | 4 | 20.0833 | 0.489 |
| 0.8 | 0 | ycaM | 4 | 20.0833 | 0.572 |
| 0.8 | 0 | yadI | 4 | 20.0833 | 0.703 |
| 0.8 | 0 | AG1 | 4 | 20.0833 | 0.626 |
| 0.8 | 0 | ptsI | 4 | 20.3333 | 0.633 |

| | | | | | |
|---|---|---|---|---|---|
| 0.8 | 0 | clcB | 4 | 20.3333 | 0.487 |
| 0.8 | 0 | ycaM | 4 | 20.3333 | 0.575 |
| 0.8 | 0 | yadI | 4 | 20.3333 | 0.71 |
| 0.8 | 0 | AG1 | 4 | 20.3333 | 0.621 |
| 0.8 | 0 | ptsI | 4 | 20.5833 | 0.637 |
| 0.8 | 0 | clcB | 4 | 20.5833 | 0.49 |
| 0.8 | 0 | ycaM | 4 | 20.5833 | 0.574 |
| 0.8 | 0 | yadI | 4 | 20.5833 | 0.71 |
| 0.8 | 0 | AG1 | 4 | 20.5833 | 0.624 |
| 0.8 | 0 | ptsI | 4 | 20.8333 | 0.64 |
| 0.8 | 0 | clcB | 4 | 20.8333 | 0.486 |
| 0.8 | 0 | ycaM | 4 | 20.8333 | 0.577 |
| 0.8 | 0 | yadI | 4 | 20.8333 | 0.711 |
| 0.8 | 0 | AG1 | 4 | 20.8333 | 0.621 |
| 0.8 | 0 | ptsI | 4 | 21.0833 | 0.638 |
| 0.8 | 0 | clcB | 4 | 21.0833 | 0.492 |
| 0.8 | 0 | ycaM | 4 | 21.0833 | 0.578 |
| 0.8 | 0 | yadI | 4 | 21.0833 | 0.709 |
| 0.8 | 0 | AG1 | 4 | 21.0833 | 0.621 |
| 0.8 | 0 | ptsI | 4 | 21.3333 | 0.64 |
| 0.8 | 0 | clcB | 4 | 21.3333 | 0.493 |
| 0.8 | 0 | ycaM | 4 | 21.3333 | 0.58 |
| 0.8 | 0 | yadI | 4 | 21.3333 | 0.708 |
| 0.8 | 0 | AG1 | 4 | 21.3333 | 0.623 |
| 0.8 | 0 | ptsI | 4 | 21.5833 | 0.644 |
| 0.8 | 0 | clcB | 4 | 21.5833 | 0.496 |
| 0.8 | 0 | ycaM | 4 | 21.5833 | 0.577 |
| 0.8 | 0 | yadI | 4 | 21.5833 | 0.71 |
| 0.8 | 0 | AG1 | 4 | 21.5833 | 0.623 |
| 0.8 | 0 | ptsI | 4 | 21.8333 | 0.644 |
| 0.8 | 0 | clcB | 4 | 21.8333 | 0.497 |
| 0.8 | 0 | ycaM | 4 | 21.8333 | 0.582 |
| 0.8 | 0 | yadI | 4 | 21.8333 | 0.713 |
| 0.8 | 0 | AG1 | 4 | 21.8333 | 0.624 |
| 0.8 | 0 | ptsI | 4 | 22.0833 | 0.646 |
| 0.8 | 0 | clcB | 4 | 22.0833 | 0.498 |
| 0.8 | 0 | ycaM | 4 | 22.0833 | 0.581 |
| 0.8 | 0 | yadI | 4 | 22.0833 | 0.713 |
| 0.8 | 0 | AG1 | 4 | 22.0833 | 0.622 |
| 0.8 | 0 | ptsI | 4 | 22.3333 | 0.65 |
| 0.8 | 0 | clcB | 4 | 22.3333 | 0.501 |
| 0.8 | 0 | ycaM | 4 | 22.3333 | 0.581 |
| 0.8 | 0 | yadI | 4 | 22.3333 | 0.711 |
| 0.8 | 0 | AG1 | 4 | 22.3333 | 0.625 |
| 0.8 | 0 | ptsI | 4 | 22.5833 | 0.651 |
| 0.8 | 0 | clcB | 4 | 22.5833 | 0.501 |
| 0.8 | 0 | ycaM | 4 | 22.5833 | 0.582 |
| 0.8 | 0 | yadI | 4 | 22.5833 | 0.713 |

| | | | | | |
|---|---|---|---|---|---|
| 0.8 | 0 | AG1 | 4 | 22.5833 | 0.624 |
| 0.8 | 0 | ptsI | 4 | 22.8333 | 0.65 |
| 0.8 | 0 | clcB | 4 | 22.8333 | 0.504 |
| 0.8 | 0 | ycaM | 4 | 22.8333 | 0.584 |
| 0.8 | 0 | yadI | 4 | 22.8333 | 0.709 |
| 0.8 | 0 | AG1 | 4 | 22.8333 | 0.625 |
| 0.8 | 0 | ptsI | 4 | 23.0833 | 0.657 |
| 0.8 | 0 | clcB | 4 | 23.0833 | 0.508 |
| 0.8 | 0 | ycaM | 4 | 23.0833 | 0.581 |
| 0.8 | 0 | yadI | 4 | 23.0833 | 0.713 |
| 0.8 | 0 | AG1 | 4 | 23.0833 | 0.624 |
| 0.8 | 0 | ptsI | 4 | 23.3333 | 0.655 |
| 0.8 | 0 | clcB | 4 | 23.3333 | 0.511 |
| 0.8 | 0 | ycaM | 4 | 23.3333 | 0.583 |
| 0.8 | 0 | yadI | 4 | 23.3333 | 0.712 |
| 0.8 | 0 | AG1 | 4 | 23.3333 | 0.629 |
| 0.8 | 0 | ptsI | 4 | 23.5833 | 0.66 |
| 0.8 | 0 | clcB | 4 | 23.5833 | 0.509 |
| 0.8 | 0 | ycaM | 4 | 23.5833 | 0.583 |
| 0.8 | 0 | yadI | 4 | 23.5833 | 0.712 |
| 0.8 | 0 | AG1 | 4 | 23.5833 | 0.626 |
| 0.8 | 0 | ptsI | 4 | 23.8333 | 0.668 |
| 0.8 | 0 | clcB | 4 | 23.8333 | 0.512 |
| 0.8 | 0 | ycaM | 4 | 23.8333 | 0.584 |
| 0.8 | 0 | yadI | 4 | 23.8333 | 0.712 |
| 0.8 | 0 | AG1 | 4 | 23.8333 | 0.627 |
| 0.8 | 0 | ptsI | 4 | 24.0833 | 0.668 |
| 0.8 | 0 | clcB | 4 | 24.0833 | 0.514 |
| 0.8 | 0 | ycaM | 4 | 24.0833 | 0.582 |
| 0.8 | 0 | yadI | 4 | 24.0833 | 0.713 |
| 0.8 | 0 | AG1 | 4 | 24.0833 | 0.629 |
| 0.8 | 0 | ptsI | 5 | 0 | 0.156 |
| 0.8 | 0 | clcB | 5 | 0 | 0.188 |
| 0.8 | 0 | ycaM | 5 | 0 | 0.163 |
| 0.8 | 0 | yadI | 5 | 0 | 0.166 |
| 0.8 | 0 | AG1 | 5 | 0 | 0.167 |
| 0.8 | 0 | ptsI | 5 | 0.35 | 0.156 |
| 0.8 | 0 | clcB | 5 | 0.35 | 0.187 |
| 0.8 | 0 | ycaM | 5 | 0.35 | 0.156 |
| 0.8 | 0 | yadI | 5 | 0.35 | 0.164 |
| 0.8 | 0 | AG1 | 5 | 0.35 | 0.163 |
| 0.8 | 0 | ptsI | 5 | 0.6 | 0.156 |
| 0.8 | 0 | clcB | 5 | 0.6 | 0.188 |
| 0.8 | 0 | ycaM | 5 | 0.6 | 0.156 |
| 0.8 | 0 | yadI | 5 | 0.6 | 0.164 |
| 0.8 | 0 | AG1 | 5 | 0.6 | 0.163 |
| 0.8 | 0 | ptsI | 5 | 0.85 | 0.155 |
| 0.8 | 0 | clcB | 5 | 0.85 | 0.187 |

| | | | | | |
|---|---|---|---|---|---|
| 0.8 | 0 | ycaM | 5 | 0.85 | 0.156 |
| 0.8 | 0 | yadI | 5 | 0.85 | 0.165 |
| 0.8 | 0 | AG1 | 5 | 0.85 | 0.163 |
| 0.8 | 0 | ptsI | 5 | 1.1 | 0.158 |
| 0.8 | 0 | clcB | 5 | 1.1 | 0.188 |
| 0.8 | 0 | ycaM | 5 | 1.1 | 0.158 |
| 0.8 | 0 | yadI | 5 | 1.1 | 0.167 |
| 0.8 | 0 | AG1 | 5 | 1.1 | 0.166 |
| 0.8 | 0 | ptsI | 5 | 1.35 | 0.158 |
| 0.8 | 0 | clcB | 5 | 1.35 | 0.189 |
| 0.8 | 0 | ycaM | 5 | 1.35 | 0.16 |
| 0.8 | 0 | yadI | 5 | 1.35 | 0.166 |
| 0.8 | 0 | AG1 | 5 | 1.35 | 0.167 |
| 0.8 | 0 | ptsI | 5 | 1.6 | 0.162 |
| 0.8 | 0 | clcB | 5 | 1.6 | 0.192 |
| 0.8 | 0 | ycaM | 5 | 1.6 | 0.163 |
| 0.8 | 0 | yadI | 5 | 1.6 | 0.172 |
| 0.8 | 0 | AG1 | 5 | 1.6 | 0.17 |
| 0.8 | 0 | ptsI | 5 | 1.85 | 0.164 |
| 0.8 | 0 | clcB | 5 | 1.85 | 0.195 |
| 0.8 | 0 | ycaM | 5 | 1.85 | 0.167 |
| 0.8 | 0 | yadI | 5 | 1.85 | 0.175 |
| 0.8 | 0 | AG1 | 5 | 1.85 | 0.174 |
| 0.8 | 0 | ptsI | 5 | 2.1 | 0.167 |
| 0.8 | 0 | clcB | 5 | 2.1 | 0.198 |
| 0.8 | 0 | ycaM | 5 | 2.1 | 0.172 |
| 0.8 | 0 | yadI | 5 | 2.1 | 0.179 |
| 0.8 | 0 | AG1 | 5 | 2.1 | 0.181 |
| 0.8 | 0 | ptsI | 5 | 2.35 | 0.172 |
| 0.8 | 0 | clcB | 5 | 2.35 | 0.203 |
| 0.8 | 0 | ycaM | 5 | 2.35 | 0.178 |
| 0.8 | 0 | yadI | 5 | 2.35 | 0.184 |
| 0.8 | 0 | AG1 | 5 | 2.35 | 0.188 |
| 0.8 | 0 | ptsI | 5 | 2.6 | 0.173 |
| 0.8 | 0 | clcB | 5 | 2.6 | 0.207 |
| 0.8 | 0 | ycaM | 5 | 2.6 | 0.182 |
| 0.8 | 0 | yadI | 5 | 2.6 | 0.186 |
| 0.8 | 0 | AG1 | 5 | 2.6 | 0.187 |
| 0.8 | 0 | ptsI | 5 | 2.85 | 0.175 |
| 0.8 | 0 | clcB | 5 | 2.85 | 0.214 |
| 0.8 | 0 | ycaM | 5 | 2.85 | 0.191 |
| 0.8 | 0 | yadI | 5 | 2.85 | 0.196 |
| 0.8 | 0 | AG1 | 5 | 2.85 | 0.2 |
| 0.8 | 0 | ptsI | 5 | 3.1 | 0.179 |
| 0.8 | 0 | clcB | 5 | 3.1 | 0.223 |
| 0.8 | 0 | ycaM | 5 | 3.1 | 0.202 |
| 0.8 | 0 | yadI | 5 | 3.1 | 0.206 |
| 0.8 | 0 | AG1 | 5 | 3.1 | 0.212 |

| | | | | | |
|---|---|---|---|---|---|
| 0.8 | 0 | ptsI | 5 | 3.35 | 0.185 |
| 0.8 | 0 | clcB | 5 | 3.35 | 0.234 |
| 0.8 | 0 | ycaM | 5 | 3.35 | 0.214 |
| 0.8 | 0 | yadI | 5 | 3.35 | 0.218 |
| 0.8 | 0 | AG1 | 5 | 3.35 | 0.229 |
| 0.8 | 0 | ptsI | 5 | 3.6 | 0.192 |
| 0.8 | 0 | clcB | 5 | 3.6 | 0.248 |
| 0.8 | 0 | ycaM | 5 | 3.6 | 0.226 |
| 0.8 | 0 | yadI | 5 | 3.6 | 0.23 |
| 0.8 | 0 | AG1 | 5 | 3.6 | 0.246 |
| 0.8 | 0 | ptsI | 5 | 3.85 | 0.201 |
| 0.8 | 0 | clcB | 5 | 3.85 | 0.26 |
| 0.8 | 0 | ycaM | 5 | 3.85 | 0.237 |
| 0.8 | 0 | yadI | 5 | 3.85 | 0.244 |
| 0.8 | 0 | AG1 | 5 | 3.85 | 0.263 |
| 0.8 | 0 | ptsI | 5 | 4.38333 | 0.212 |
| 0.8 | 0 | clcB | 5 | 4.38333 | 0.25 |
| 0.8 | 0 | ycaM | 5 | 4.38333 | 0.244 |
| 0.8 | 0 | yadI | 5 | 4.38333 | 0.258 |
| 0.8 | 0 | AG1 | 5 | 4.38333 | 0.28 |
| 0.8 | 0 | ptsI | 5 | 4.63333 | 0.212 |
| 0.8 | 0 | clcB | 5 | 4.63333 | 0.24 |
| 0.8 | 0 | ycaM | 5 | 4.63333 | 0.247 |
| 0.8 | 0 | yadI | 5 | 4.63333 | 0.251 |
| 0.8 | 0 | AG1 | 5 | 4.63333 | 0.28 |
| 0.8 | 0 | ptsI | 5 | 4.88333 | 0.217 |
| 0.8 | 0 | clcB | 5 | 4.88333 | 0.241 |
| 0.8 | 0 | ycaM | 5 | 4.88333 | 0.254 |
| 0.8 | 0 | yadI | 5 | 4.88333 | 0.266 |
| 0.8 | 0 | AG1 | 5 | 4.88333 | 0.29 |
| 0.8 | 0 | ptsI | 5 | 5.13333 | 0.225 |
| 0.8 | 0 | clcB | 5 | 5.13333 | 0.248 |
| 0.8 | 0 | ycaM | 5 | 5.13333 | 0.266 |
| 0.8 | 0 | yadI | 5 | 5.13333 | 0.28 |
| 0.8 | 0 | AG1 | 5 | 5.13333 | 0.3 |
| 0.8 | 0 | ptsI | 5 | 5.38333 | 0.233 |
| 0.8 | 0 | clcB | 5 | 5.38333 | 0.256 |
| 0.8 | 0 | ycaM | 5 | 5.38333 | 0.279 |
| 0.8 | 0 | yadI | 5 | 5.38333 | 0.295 |
| 0.8 | 0 | AG1 | 5 | 5.38333 | 0.314 |
| 0.8 | 0 | ptsI | 5 | 5.63333 | 0.244 |
| 0.8 | 0 | clcB | 5 | 5.63333 | 0.264 |
| 0.8 | 0 | ycaM | 5 | 5.63333 | 0.29 |
| 0.8 | 0 | yadI | 5 | 5.63333 | 0.31 |
| 0.8 | 0 | AG1 | 5 | 5.63333 | 0.326 |
| 0.8 | 0 | ptsI | 5 | 5.88333 | 0.25 |
| 0.8 | 0 | clcB | 5 | 5.88333 | 0.276 |
| 0.8 | 0 | ycaM | 5 | 5.88333 | 0.304 |

| | | | | | |
|---|---|---|---|---|---|
| 0.8 | 0 | yadI | 5 | 5.88333 | 0.324 |
| 0.8 | 0 | AG1 | 5 | 5.88333 | 0.336 |
| 0.8 | 0 | ptsI | 5 | 6.13333 | 0.26 |
| 0.8 | 0 | clcB | 5 | 6.13333 | 0.287 |
| 0.8 | 0 | ycaM | 5 | 6.13333 | 0.315 |
| 0.8 | 0 | yadI | 5 | 6.13333 | 0.34 |
| 0.8 | 0 | AG1 | 5 | 6.13333 | 0.346 |
| 0.8 | 0 | ptsI | 5 | 6.38333 | 0.268 |
| 0.8 | 0 | clcB | 5 | 6.38333 | 0.304 |
| 0.8 | 0 | ycaM | 5 | 6.38333 | 0.327 |
| 0.8 | 0 | yadI | 5 | 6.38333 | 0.355 |
| 0.8 | 0 | AG1 | 5 | 6.38333 | 0.36 |
| 0.8 | 0 | ptsI | 5 | 6.63333 | 0.278 |
| 0.8 | 0 | clcB | 5 | 6.63333 | 0.32 |
| 0.8 | 0 | ycaM | 5 | 6.63333 | 0.338 |
| 0.8 | 0 | yadI | 5 | 6.63333 | 0.367 |
| 0.8 | 0 | AG1 | 5 | 6.63333 | 0.374 |
| 0.8 | 0 | ptsI | 5 | 6.88333 | 0.288 |
| 0.8 | 0 | clcB | 5 | 6.88333 | 0.334 |
| 0.8 | 0 | ycaM | 5 | 6.88333 | 0.351 |
| 0.8 | 0 | yadI | 5 | 6.88333 | 0.38 |
| 0.8 | 0 | AG1 | 5 | 6.88333 | 0.389 |
| 0.8 | 0 | ptsI | 5 | 7.13333 | 0.295 |
| 0.8 | 0 | clcB | 5 | 7.13333 | 0.349 |
| 0.8 | 0 | ycaM | 5 | 7.13333 | 0.364 |
| 0.8 | 0 | yadI | 5 | 7.13333 | 0.396 |
| 0.8 | 0 | AG1 | 5 | 7.13333 | 0.405 |
| 0.8 | 0 | ptsI | 5 | 7.38333 | 0.304 |
| 0.8 | 0 | clcB | 5 | 7.38333 | 0.365 |
| 0.8 | 0 | ycaM | 5 | 7.38333 | 0.373 |
| 0.8 | 0 | yadI | 5 | 7.38333 | 0.415 |
| 0.8 | 0 | AG1 | 5 | 7.38333 | 0.42 |
| 0.8 | 0 | ptsI | 5 | 7.63333 | 0.314 |
| 0.8 | 0 | clcB | 5 | 7.63333 | 0.382 |
| 0.8 | 0 | ycaM | 5 | 7.63333 | 0.385 |
| 0.8 | 0 | yadI | 5 | 7.63333 | 0.446 |
| 0.8 | 0 | AG1 | 5 | 7.63333 | 0.43 |
| 0.8 | 0 | ptsI | 5 | 7.88333 | 0.325 |
| 0.8 | 0 | clcB | 5 | 7.88333 | 0.39 |
| 0.8 | 0 | ycaM | 5 | 7.88333 | 0.392 |
| 0.8 | 0 | yadI | 5 | 7.88333 | 0.461 |
| 0.8 | 0 | AG1 | 5 | 7.88333 | 0.443 |
| 0.8 | 0 | ptsI | 5 | 8.13333 | 0.338 |
| 0.8 | 0 | clcB | 5 | 8.13333 | 0.4 |
| 0.8 | 0 | ycaM | 5 | 8.13333 | 0.404 |
| 0.8 | 0 | yadI | 5 | 8.13333 | 0.468 |
| 0.8 | 0 | AG1 | 5 | 8.13333 | 0.455 |
| 0.8 | 0 | ptsI | 5 | 8.38333 | 0.347 |

| | | | | | |
|---|---|---|---|---|---|
| 0.8 | 0 | clcB | 5 | 8.38333 | 0.41 |
| 0.8 | 0 | ycaM | 5 | 8.38333 | 0.415 |
| 0.8 | 0 | yadI | 5 | 8.38333 | 0.478 |
| 0.8 | 0 | AG1 | 5 | 8.38333 | 0.47 |
| 0.8 | 0 | ptsI | 5 | 8.88333 | 0.342 |
| 0.8 | 0 | clcB | 5 | 8.88333 | 0.362 |
| 0.8 | 0 | ycaM | 5 | 8.88333 | 0.34 |
| 0.8 | 0 | yadI | 5 | 8.88333 | 0.395 |
| 0.8 | 0 | AG1 | 5 | 8.88333 | 0.525 |
| 0.8 | 0 | ptsI | 5 | 9.13333 | 0.345 |
| 0.8 | 0 | clcB | 5 | 9.13333 | 0.371 |
| 0.8 | 0 | ycaM | 5 | 9.13333 | 0.346 |
| 0.8 | 0 | yadI | 5 | 9.13333 | 0.394 |
| 0.8 | 0 | AG1 | 5 | 9.13333 | 0.519 |
| 0.8 | 0 | ptsI | 5 | 9.38333 | 0.362 |
| 0.8 | 0 | clcB | 5 | 9.38333 | 0.38 |
| 0.8 | 0 | ycaM | 5 | 9.38333 | 0.359 |
| 0.8 | 0 | yadI | 5 | 9.38333 | 0.41 |
| 0.8 | 0 | AG1 | 5 | 9.38333 | 0.534 |
| 0.8 | 0 | ptsI | 5 | 9.63333 | 0.366 |
| 0.8 | 0 | clcB | 5 | 9.63333 | 0.396 |
| 0.8 | 0 | ycaM | 5 | 9.63333 | 0.367 |
| 0.8 | 0 | yadI | 5 | 9.63333 | 0.416 |
| 0.8 | 0 | AG1 | 5 | 9.63333 | 0.551 |
| 0.8 | 0 | ptsI | 5 | 9.88333 | 0.38 |
| 0.8 | 0 | clcB | 5 | 9.88333 | 0.407 |
| 0.8 | 0 | ycaM | 5 | 9.88333 | 0.376 |
| 0.8 | 0 | yadI | 5 | 9.88333 | 0.426 |
| 0.8 | 0 | AG1 | 5 | 9.88333 | 0.564 |
| 0.8 | 0 | ptsI | 5 | 10.1333 | 0.39 |
| 0.8 | 0 | clcB | 5 | 10.1333 | 0.416 |
| 0.8 | 0 | ycaM | 5 | 10.1333 | 0.387 |
| 0.8 | 0 | yadI | 5 | 10.1333 | 0.432 |
| 0.8 | 0 | AG1 | 5 | 10.1333 | 0.574 |
| 0.8 | 0 | ptsI | 5 | 10.3833 | 0.402 |
| 0.8 | 0 | clcB | 5 | 10.3833 | 0.425 |
| 0.8 | 0 | ycaM | 5 | 10.3833 | 0.394 |
| 0.8 | 0 | yadI | 5 | 10.3833 | 0.442 |
| 0.8 | 0 | AG1 | 5 | 10.3833 | 0.585 |
| 0.8 | 0 | ptsI | 5 | 10.6333 | 0.412 |
| 0.8 | 0 | clcB | 5 | 10.6333 | 0.43 |
| 0.8 | 0 | ycaM | 5 | 10.6333 | 0.399 |
| 0.8 | 0 | yadI | 5 | 10.6333 | 0.446 |
| 0.8 | 0 | AG1 | 5 | 10.6333 | 0.591 |
| 0.8 | 0 | ptsI | 5 | 10.8833 | 0.422 |
| 0.8 | 0 | clcB | 5 | 10.8833 | 0.433 |
| 0.8 | 0 | ycaM | 5 | 10.8833 | 0.404 |
| 0.8 | 0 | yadI | 5 | 10.8833 | 0.456 |

| | | | | | |
|---|---|---|---|---|---|
| 0.8 | 0 | AG1 | 5 | 10.8833 | 0.602 |
| 0.8 | 0 | ptsI | 5 | 11.1333 | 0.429 |
| 0.8 | 0 | clcB | 5 | 11.1333 | 0.438 |
| 0.8 | 0 | ycaM | 5 | 11.1333 | 0.404 |
| 0.8 | 0 | yadI | 5 | 11.1333 | 0.458 |
| 0.8 | 0 | AG1 | 5 | 11.1333 | 0.608 |
| 0.8 | 0 | ptsI | 5 | 11.3833 | 0.439 |
| 0.8 | 0 | clcB | 5 | 11.3833 | 0.441 |
| 0.8 | 0 | ycaM | 5 | 11.3833 | 0.408 |
| 0.8 | 0 | yadI | 5 | 11.3833 | 0.469 |
| 0.8 | 0 | AG1 | 5 | 11.3833 | 0.615 |
| 0.8 | 0 | ptsI | 5 | 11.6333 | 0.447 |
| 0.8 | 0 | clcB | 5 | 11.6333 | 0.445 |
| 0.8 | 0 | ycaM | 5 | 11.6333 | 0.411 |
| 0.8 | 0 | yadI | 5 | 11.6333 | 0.474 |
| 0.8 | 0 | AG1 | 5 | 11.6333 | 0.624 |
| 0.8 | 0 | ptsI | 5 | 11.8833 | 0.455 |
| 0.8 | 0 | clcB | 5 | 11.8833 | 0.45 |
| 0.8 | 0 | ycaM | 5 | 11.8833 | 0.414 |
| 0.8 | 0 | yadI | 5 | 11.8833 | 0.479 |
| 0.8 | 0 | AG1 | 5 | 11.8833 | 0.634 |
| 0.8 | 0 | ptsI | 5 | 12.1333 | 0.465 |
| 0.8 | 0 | clcB | 5 | 12.1333 | 0.455 |
| 0.8 | 0 | ycaM | 5 | 12.1333 | 0.42 |
| 0.8 | 0 | yadI | 5 | 12.1333 | 0.486 |
| 0.8 | 0 | AG1 | 5 | 12.1333 | 0.64 |
| 0.8 | 0 | ptsI | 5 | 12.3833 | 0.477 |
| 0.8 | 0 | clcB | 5 | 12.3833 | 0.457 |
| 0.8 | 0 | ycaM | 5 | 12.3833 | 0.419 |
| 0.8 | 0 | yadI | 5 | 12.3833 | 0.49 |
| 0.8 | 0 | AG1 | 5 | 12.3833 | 0.65 |
| 0.8 | 0 | ptsI | 5 | 12.6333 | 0.478 |
| 0.8 | 0 | clcB | 5 | 12.6333 | 0.463 |
| 0.8 | 0 | ycaM | 5 | 12.6333 | 0.427 |
| 0.8 | 0 | yadI | 5 | 12.6333 | 0.496 |
| 0.8 | 0 | AG1 | 5 | 12.6333 | 0.655 |
| 0.8 | 0 | ptsI | 5 | 12.8833 | 0.483 |
| 0.8 | 0 | clcB | 5 | 12.8833 | 0.468 |
| 0.8 | 0 | ycaM | 5 | 12.8833 | 0.429 |
| 0.8 | 0 | yadI | 5 | 12.8833 | 0.502 |
| 0.8 | 0 | AG1 | 5 | 12.8833 | 0.663 |
| 0.8 | 0 | ptsI | 5 | 13.1333 | 0.491 |
| 0.8 | 0 | clcB | 5 | 13.1333 | 0.472 |
| 0.8 | 0 | ycaM | 5 | 13.1333 | 0.434 |
| 0.8 | 0 | yadI | 5 | 13.1333 | 0.507 |
| 0.8 | 0 | AG1 | 5 | 13.1333 | 0.668 |
| 0.8 | 0 | ptsI | 5 | 13.3833 | 0.495 |
| 0.8 | 0 | clcB | 5 | 13.3833 | 0.476 |

| | | | | | |
|---|---|---|---|---|---|
| 0.8 | 0 | ycaM | 5 | 13.3833 | 0.436 |
| 0.8 | 0 | yadI | 5 | 13.3833 | 0.511 |
| 0.8 | 0 | AG1 | 5 | 13.3833 | 0.674 |
| 0.8 | 0 | ptsI | 5 | 13.6333 | 0.5 |
| 0.8 | 0 | clcB | 5 | 13.6333 | 0.48 |
| 0.8 | 0 | ycaM | 5 | 13.6333 | 0.44 |
| 0.8 | 0 | yadI | 5 | 13.6333 | 0.516 |
| 0.8 | 0 | AG1 | 5 | 13.6333 | 0.679 |
| 0.8 | 0 | ptsI | 5 | 13.8833 | 0.502 |
| 0.8 | 0 | clcB | 5 | 13.8833 | 0.484 |
| 0.8 | 0 | ycaM | 5 | 13.8833 | 0.441 |
| 0.8 | 0 | yadI | 5 | 13.8833 | 0.518 |
| 0.8 | 0 | AG1 | 5 | 13.8833 | 0.684 |
| 0.8 | 0 | ptsI | 5 | 14.1333 | 0.507 |
| 0.8 | 0 | clcB | 5 | 14.1333 | 0.485 |
| 0.8 | 0 | ycaM | 5 | 14.1333 | 0.447 |
| 0.8 | 0 | yadI | 5 | 14.1333 | 0.527 |
| 0.8 | 0 | AG1 | 5 | 14.1333 | 0.69 |
| 0.8 | 0 | ptsI | 5 | 14.3833 | 0.504 |
| 0.8 | 0 | clcB | 5 | 14.3833 | 0.486 |
| 0.8 | 0 | ycaM | 5 | 14.3833 | 0.446 |
| 0.8 | 0 | yadI | 5 | 14.3833 | 0.532 |
| 0.8 | 0 | AG1 | 5 | 14.3833 | 0.691 |
| 0.8 | 0 | ptsI | 5 | 14.6333 | 0.505 |
| 0.8 | 0 | clcB | 5 | 14.6333 | 0.491 |
| 0.8 | 0 | ycaM | 5 | 14.6333 | 0.448 |
| 0.8 | 0 | yadI | 5 | 14.6333 | 0.528 |
| 0.8 | 0 | AG1 | 5 | 14.6333 | 0.696 |
| 0.8 | 0 | ptsI | 5 | 14.8833 | 0.513 |
| 0.8 | 0 | clcB | 5 | 14.8833 | 0.495 |
| 0.8 | 0 | ycaM | 5 | 14.8833 | 0.452 |
| 0.8 | 0 | yadI | 5 | 14.8833 | 0.533 |
| 0.8 | 0 | AG1 | 5 | 14.8833 | 0.705 |
| 0.8 | 0 | ptsI | 5 | 15.1333 | 0.515 |
| 0.8 | 0 | clcB | 5 | 15.1333 | 0.496 |
| 0.8 | 0 | ycaM | 5 | 15.1333 | 0.453 |
| 0.8 | 0 | yadI | 5 | 15.1333 | 0.539 |
| 0.8 | 0 | AG1 | 5 | 15.1333 | 0.709 |
| 0.8 | 0 | ptsI | 5 | 15.3833 | 0.521 |
| 0.8 | 0 | clcB | 5 | 15.3833 | 0.499 |
| 0.8 | 0 | ycaM | 5 | 15.3833 | 0.459 |
| 0.8 | 0 | yadI | 5 | 15.3833 | 0.54 |
| 0.8 | 0 | AG1 | 5 | 15.3833 | 0.711 |
| 0.8 | 0 | ptsI | 5 | 15.6333 | 0.521 |
| 0.8 | 0 | clcB | 5 | 15.6333 | 0.502 |
| 0.8 | 0 | ycaM | 5 | 15.6333 | 0.459 |
| 0.8 | 0 | yadI | 5 | 15.6333 | 0.541 |
| 0.8 | 0 | AG1 | 5 | 15.6333 | 0.718 |

| | | | | | |
|---|---|---|---|---|---|
| 0.8 | 0 | ptsI | 5 | 15.8833 | 0.526 |
| 0.8 | 0 | clcB | 5 | 15.8833 | 0.503 |
| 0.8 | 0 | ycaM | 5 | 15.8833 | 0.461 |
| 0.8 | 0 | yadI | 5 | 15.8833 | 0.544 |
| 0.8 | 0 | AG1 | 5 | 15.8833 | 0.72 |
| 0.8 | 0 | ptsI | 5 | 16.1333 | 0.536 |
| 0.8 | 0 | clcB | 5 | 16.1333 | 0.5 |
| 0.8 | 0 | ycaM | 5 | 16.1333 | 0.466 |
| 0.8 | 0 | yadI | 5 | 16.1333 | 0.553 |
| 0.8 | 0 | AG1 | 5 | 16.1333 | 0.725 |
| 0.8 | 0 | ptsI | 5 | 16.3833 | 0.534 |
| 0.8 | 0 | clcB | 5 | 16.3833 | 0.508 |
| 0.8 | 0 | ycaM | 5 | 16.3833 | 0.466 |
| 0.8 | 0 | yadI | 5 | 16.3833 | 0.549 |
| 0.8 | 0 | AG1 | 5 | 16.3833 | 0.725 |
| 0.8 | 0 | ptsI | 5 | 16.6333 | 0.533 |
| 0.8 | 0 | clcB | 5 | 16.6333 | 0.509 |
| 0.8 | 0 | ycaM | 5 | 16.6333 | 0.47 |
| 0.8 | 0 | yadI | 5 | 16.6333 | 0.551 |
| 0.8 | 0 | AG1 | 5 | 16.6333 | 0.732 |
| 0.8 | 0 | ptsI | 5 | 16.8833 | 0.538 |
| 0.8 | 0 | clcB | 5 | 16.8833 | 0.51 |
| 0.8 | 0 | ycaM | 5 | 16.8833 | 0.47 |
| 0.8 | 0 | yadI | 5 | 16.8833 | 0.555 |
| 0.8 | 0 | AG1 | 5 | 16.8833 | 0.733 |
| 0.8 | 0 | ptsI | 5 | 17.1333 | 0.541 |
| 0.8 | 0 | clcB | 5 | 17.1333 | 0.511 |
| 0.8 | 0 | ycaM | 5 | 17.1333 | 0.47 |
| 0.8 | 0 | yadI | 5 | 17.1333 | 0.551 |
| 0.8 | 0 | AG1 | 5 | 17.1333 | 0.741 |
| 0.8 | 0 | ptsI | 5 | 17.3833 | 0.541 |
| 0.8 | 0 | clcB | 5 | 17.3833 | 0.513 |
| 0.8 | 0 | ycaM | 5 | 17.3833 | 0.475 |
| 0.8 | 0 | yadI | 5 | 17.3833 | 0.556 |
| 0.8 | 0 | AG1 | 5 | 17.3833 | 0.742 |
| 0.8 | 0 | ptsI | 5 | 17.6333 | 0.543 |
| 0.8 | 0 | clcB | 5 | 17.6333 | 0.512 |
| 0.8 | 0 | ycaM | 5 | 17.6333 | 0.475 |
| 0.8 | 0 | yadI | 5 | 17.6333 | 0.556 |
| 0.8 | 0 | AG1 | 5 | 17.6333 | 0.748 |
| 0.8 | 0 | ptsI | 5 | 17.8833 | 0.556 |
| 0.8 | 0 | clcB | 5 | 17.8833 | 0.514 |
| 0.8 | 0 | ycaM | 5 | 17.8833 | 0.477 |
| 0.8 | 0 | yadI | 5 | 17.8833 | 0.558 |
| 0.8 | 0 | AG1 | 5 | 17.8833 | 0.75 |
| 0.8 | 0 | ptsI | 5 | 18.1333 | 0.55 |
| 0.8 | 0 | clcB | 5 | 18.1333 | 0.518 |
| 0.8 | 0 | ycaM | 5 | 18.1333 | 0.482 |

| | | | | | |
|---|---|---|---|---|---|
| 0.8 | 0 | yadI | 5 | 18.1333 | 0.564 |
| 0.8 | 0 | AG1 | 5 | 18.1333 | 0.752 |
| 0.8 | 0 | ptsI | 5 | 18.3833 | 0.551 |
| 0.8 | 0 | clcB | 5 | 18.3833 | 0.518 |
| 0.8 | 0 | ycaM | 5 | 18.3833 | 0.482 |
| 0.8 | 0 | yadI | 5 | 18.3833 | 0.561 |
| 0.8 | 0 | AG1 | 5 | 18.3833 | 0.752 |
| 0.8 | 0 | ptsI | 5 | 18.6333 | 0.549 |
| 0.8 | 0 | clcB | 5 | 18.6333 | 0.52 |
| 0.8 | 0 | ycaM | 5 | 18.6333 | 0.484 |
| 0.8 | 0 | yadI | 5 | 18.6333 | 0.564 |
| 0.8 | 0 | AG1 | 5 | 18.6333 | 0.756 |
| 0.8 | 0 | ptsI | 5 | 18.8833 | 0.546 |
| 0.8 | 0 | clcB | 5 | 18.8833 | 0.521 |
| 0.8 | 0 | ycaM | 5 | 18.8833 | 0.487 |
| 0.8 | 0 | yadI | 5 | 18.8833 | 0.562 |
| 0.8 | 0 | AG1 | 5 | 18.8833 | 0.759 |
| 0.8 | 0 | ptsI | 5 | 19.1333 | 0.554 |
| 0.8 | 0 | clcB | 5 | 19.1333 | 0.52 |
| 0.8 | 0 | ycaM | 5 | 19.1333 | 0.488 |
| 0.8 | 0 | yadI | 5 | 19.1333 | 0.561 |
| 0.8 | 0 | AG1 | 5 | 19.1333 | 0.76 |
| 0.8 | 0 | ptsI | 5 | 19.3833 | 0.556 |
| 0.8 | 0 | clcB | 5 | 19.3833 | 0.526 |
| 0.8 | 0 | ycaM | 5 | 19.3833 | 0.49 |
| 0.8 | 0 | yadI | 5 | 19.3833 | 0.567 |
| 0.8 | 0 | AG1 | 5 | 19.3833 | 0.763 |
| 0.8 | 0 | ptsI | 5 | 19.6333 | 0.547 |
| 0.8 | 0 | clcB | 5 | 19.6333 | 0.527 |
| 0.8 | 0 | ycaM | 5 | 19.6333 | 0.491 |
| 0.8 | 0 | yadI | 5 | 19.6333 | 0.568 |
| 0.8 | 0 | AG1 | 5 | 19.6333 | 0.765 |
| 0.8 | 0 | ptsI | 5 | 19.8833 | 0.555 |
| 0.8 | 0 | clcB | 5 | 19.8833 | 0.524 |
| 0.8 | 0 | ycaM | 5 | 19.8833 | 0.494 |
| 0.8 | 0 | yadI | 5 | 19.8833 | 0.566 |
| 0.8 | 0 | AG1 | 5 | 19.8833 | 0.765 |
| 0.8 | 0 | ptsI | 5 | 20.1333 | 0.555 |
| 0.8 | 0 | clcB | 5 | 20.1333 | 0.524 |
| 0.8 | 0 | ycaM | 5 | 20.1333 | 0.495 |
| 0.8 | 0 | yadI | 5 | 20.1333 | 0.567 |
| 0.8 | 0 | AG1 | 5 | 20.1333 | 0.77 |
| 0.8 | 0 | ptsI | 5 | 20.3833 | 0.553 |
| 0.8 | 0 | clcB | 5 | 20.3833 | 0.525 |
| 0.8 | 0 | ycaM | 5 | 20.3833 | 0.495 |
| 0.8 | 0 | yadI | 5 | 20.3833 | 0.566 |
| 0.8 | 0 | AG1 | 5 | 20.3833 | 0.771 |
| 0.8 | 0 | ptsI | 5 | 20.6333 | 0.556 |

| | | | | | |
|---|---|---|---|---|---|
| 0.8 | 0 | clcB | 5 | 20.6333 | 0.53 |
| 0.8 | 0 | ycaM | 5 | 20.6333 | 0.488 |
| 0.8 | 0 | yadI | 5 | 20.6333 | 0.569 |
| 0.8 | 0 | AG1 | 5 | 20.6333 | 0.767 |
| 0.8 | 0 | ptsI | 5 | 20.8833 | 0.554 |
| 0.8 | 0 | clcB | 5 | 20.8833 | 0.532 |
| 0.8 | 0 | ycaM | 5 | 20.8833 | 0.499 |
| 0.8 | 0 | yadI | 5 | 20.8833 | 0.572 |
| 0.8 | 0 | AG1 | 5 | 20.8833 | 0.775 |
| 0.8 | 0 | ptsI | 5 | 21.1333 | 0.556 |
| 0.8 | 0 | clcB | 5 | 21.1333 | 0.531 |
| 0.8 | 0 | ycaM | 5 | 21.1333 | 0.5 |
| 0.8 | 0 | yadI | 5 | 21.1333 | 0.571 |
| 0.8 | 0 | AG1 | 5 | 21.1333 | 0.775 |
| 0.8 | 0 | ptsI | 5 | 21.3833 | 0.553 |
| 0.8 | 0 | clcB | 5 | 21.3833 | 0.531 |
| 0.8 | 0 | ycaM | 5 | 21.3833 | 0.5 |
| 0.8 | 0 | yadI | 5 | 21.3833 | 0.572 |
| 0.8 | 0 | AG1 | 5 | 21.3833 | 0.775 |
| 0.8 | 0 | ptsI | 5 | 21.6333 | 0.552 |
| 0.8 | 0 | clcB | 5 | 21.6333 | 0.532 |
| 0.8 | 0 | ycaM | 5 | 21.6333 | 0.501 |
| 0.8 | 0 | yadI | 5 | 21.6333 | 0.573 |
| 0.8 | 0 | AG1 | 5 | 21.6333 | 0.78 |
| 0.8 | 0 | ptsI | 5 | 21.8833 | 0.556 |
| 0.8 | 0 | clcB | 5 | 21.8833 | 0.532 |
| 0.8 | 0 | ycaM | 5 | 21.8833 | 0.503 |
| 0.8 | 0 | yadI | 5 | 21.8833 | 0.573 |
| 0.8 | 0 | AG1 | 5 | 21.8833 | 0.78 |
| 0.8 | 0 | ptsI | 5 | 22.1333 | 0.563 |
| 0.8 | 0 | clcB | 5 | 22.1333 | 0.534 |
| 0.8 | 0 | ycaM | 5 | 22.1333 | 0.502 |
| 0.8 | 0 | yadI | 5 | 22.1333 | 0.577 |
| 0.8 | 0 | AG1 | 5 | 22.1333 | 0.784 |
| 0.8 | 0 | ptsI | 5 | 22.3833 | 0.555 |
| 0.8 | 0 | clcB | 5 | 22.3833 | 0.535 |
| 0.8 | 0 | ycaM | 5 | 22.3833 | 0.503 |
| 0.8 | 0 | yadI | 5 | 22.3833 | 0.577 |
| 0.8 | 0 | AG1 | 5 | 22.3833 | 0.782 |
| 0.8 | 0 | ptsI | 5 | 22.6333 | 0.555 |
| 0.8 | 0 | clcB | 5 | 22.6333 | 0.536 |
| 0.8 | 0 | ycaM | 5 | 22.6333 | 0.504 |
| 0.8 | 0 | yadI | 5 | 22.6333 | 0.576 |
| 0.8 | 0 | AG1 | 5 | 22.6333 | 0.781 |
| 0.8 | 0 | ptsI | 5 | 22.8833 | 0.557 |
| 0.8 | 0 | clcB | 5 | 22.8833 | 0.534 |
| 0.8 | 0 | ycaM | 5 | 22.8833 | 0.503 |
| 0.8 | 0 | yadI | 5 | 22.8833 | 0.58 |

| | | | | | |
|---|---|---|---|---|---|
| 0.8 | 0 | AG1 | 5 | 22.8833 | 0.783 |
| 0.8 | 0 | ptsI | 5 | 23.1333 | 0.558 |
| 0.8 | 0 | clcB | 5 | 23.1333 | 0.534 |
| 0.8 | 0 | ycaM | 5 | 23.1333 | 0.504 |
| 0.8 | 0 | yadI | 5 | 23.1333 | 0.583 |
| 0.8 | 0 | AG1 | 5 | 23.1333 | 0.788 |
| 0.8 | 0 | ptsI | 5 | 23.3833 | 0.559 |
| 0.8 | 0 | clcB | 5 | 23.3833 | 0.535 |
| 0.8 | 0 | ycaM | 5 | 23.3833 | 0.504 |
| 0.8 | 0 | yadI | 5 | 23.3833 | 0.585 |
| 0.8 | 0 | AG1 | 5 | 23.3833 | 0.784 |
| 0.8 | 0 | ptsI | 5 | 23.6333 | 0.563 |
| 0.8 | 0 | clcB | 5 | 23.6333 | 0.536 |
| 0.8 | 0 | ycaM | 5 | 23.6333 | 0.504 |
| 0.8 | 0 | yadI | 5 | 23.6333 | 0.582 |
| 0.8 | 0 | AG1 | 5 | 23.6333 | 0.789 |
| 0.8 | 0 | ptsI | 5 | 23.8833 | 0.576 |
| 0.8 | 0 | clcB | 5 | 23.8833 | 0.533 |
| 0.8 | 0 | ycaM | 5 | 23.8833 | 0.502 |
| 0.8 | 0 | yadI | 5 | 23.8833 | 0.585 |
| 0.8 | 0 | AG1 | 5 | 23.8833 | 0.787 |
| 0.8 | 0 | ptsI | 5 | 24.1333 | 0.567 |
| 0.8 | 0 | clcB | 5 | 24.1333 | 0.537 |
| 0.8 | 0 | ycaM | 5 | 24.1333 | 0.506 |
| 0.8 | 0 | yadI | 5 | 24.1333 | 0.587 |
| 0.8 | 0 | AG1 | 5 | 24.1333 | 0.787 |
| 0.8 | 0 | ptsI | 5 | 24.3833 | 0.575 |
| 0.8 | 0 | clcB | 5 | 24.3833 | 0.536 |
| 0.8 | 0 | ycaM | 5 | 24.3833 | 0.506 |
| 0.8 | 0 | yadI | 5 | 24.3833 | 0.589 |
| 0.8 | 0 | AG1 | 5 | 24.3833 | 0.79 |
| 1.6 | 0 | ptsI | 1 | 0 | 0.219 |
| 1.6 | 0 | clcB | 1 | 0 | 0.229 |
| 1.6 | 0 | ycaM | 1 | 0 | 0.234 |
| 1.6 | 0 | yadI | 1 | 0 | 0.269 |
| 1.6 | 0 | AG1 | 1 | 0 | 0.243 |
| 1.6 | 0 | ptsI | 1 | 0.25 | 0.217 |
| 1.6 | 0 | clcB | 1 | 0.25 | 0.221 |
| 1.6 | 0 | ycaM | 1 | 0.25 | 0.227 |
| 1.6 | 0 | yadI | 1 | 0.25 | 0.26 |
| 1.6 | 0 | AG1 | 1 | 0.25 | 0.228 |
| 1.6 | 0 | ptsI | 1 | 0.5 | 0.216 |
| 1.6 | 0 | clcB | 1 | 0.5 | 0.22 |
| 1.6 | 0 | ycaM | 1 | 0.5 | 0.222 |
| 1.6 | 0 | yadI | 1 | 0.5 | 0.257 |
| 1.6 | 0 | AG1 | 1 | 0.5 | 0.228 |
| 1.6 | 0 | ptsI | 1 | 0.75 | 0.218 |
| 1.6 | 0 | clcB | 1 | 0.75 | 0.219 |

| | | | | | |
|---|---|---|---|---|---|
| 1.6 | 0 | ycaM | 1 | 0.75 | 0.22 |
| 1.6 | 0 | yadI | 1 | 0.75 | 0.256 |
| 1.6 | 0 | AG1 | 1 | 0.75 | 0.228 |
| 1.6 | 0 | ptsI | 1 | 1 | 0.224 |
| 1.6 | 0 | clcB | 1 | 1 | 0.221 |
| 1.6 | 0 | ycaM | 1 | 1 | 0.22 |
| 1.6 | 0 | yadI | 1 | 1 | 0.26 |
| 1.6 | 0 | AG1 | 1 | 1 | 0.229 |
| 1.6 | 0 | ptsI | 1 | 1.25 | 0.226 |
| 1.6 | 0 | clcB | 1 | 1.25 | 0.222 |
| 1.6 | 0 | ycaM | 1 | 1.25 | 0.221 |
| 1.6 | 0 | yadI | 1 | 1.25 | 0.263 |
| 1.6 | 0 | AG1 | 1 | 1.25 | 0.231 |
| 1.6 | 0 | ptsI | 1 | 1.5 | 0.23 |
| 1.6 | 0 | clcB | 1 | 1.5 | 0.222 |
| 1.6 | 0 | ycaM | 1 | 1.5 | 0.222 |
| 1.6 | 0 | yadI | 1 | 1.5 | 0.264 |
| 1.6 | 0 | AG1 | 1 | 1.5 | 0.232 |
| 1.6 | 0 | ptsI | 1 | 1.75 | 0.236 |
| 1.6 | 0 | clcB | 1 | 1.75 | 0.226 |
| 1.6 | 0 | ycaM | 1 | 1.75 | 0.231 |
| 1.6 | 0 | yadI | 1 | 1.75 | 0.275 |
| 1.6 | 0 | AG1 | 1 | 1.75 | 0.236 |
| 1.6 | 0 | ptsI | 1 | 2 | 0.242 |
| 1.6 | 0 | clcB | 1 | 2 | 0.226 |
| 1.6 | 0 | ycaM | 1 | 2 | 0.234 |
| 1.6 | 0 | yadI | 1 | 2 | 0.274 |
| 1.6 | 0 | AG1 | 1 | 2 | 0.242 |
| 1.6 | 0 | ptsI | 1 | 2.25 | 0.248 |
| 1.6 | 0 | clcB | 1 | 2.25 | 0.229 |
| 1.6 | 0 | ycaM | 1 | 2.25 | 0.242 |
| 1.6 | 0 | yadI | 1 | 2.25 | 0.289 |
| 1.6 | 0 | AG1 | 1 | 2.25 | 0.25 |
| 1.6 | 0 | ptsI | 1 | 2.5 | 0.253 |
| 1.6 | 0 | clcB | 1 | 2.5 | 0.236 |
| 1.6 | 0 | ycaM | 1 | 2.5 | 0.245 |
| 1.6 | 0 | yadI | 1 | 2.5 | 0.294 |
| 1.6 | 0 | AG1 | 1 | 2.5 | 0.254 |
| 1.6 | 0 | ptsI | 1 | 2.75 | 0.259 |
| 1.6 | 0 | clcB | 1 | 2.75 | 0.242 |
| 1.6 | 0 | ycaM | 1 | 2.75 | 0.255 |
| 1.6 | 0 | yadI | 1 | 2.75 | 0.309 |
| 1.6 | 0 | AG1 | 1 | 2.75 | 0.26 |
| 1.6 | 0 | ptsI | 1 | 3 | 0.268 |
| 1.6 | 0 | clcB | 1 | 3 | 0.248 |
| 1.6 | 0 | ycaM | 1 | 3 | 0.26 |
| 1.6 | 0 | yadI | 1 | 3 | 0.317 |
| 1.6 | 0 | AG1 | 1 | 3 | 0.268 |

| | | | | | |
|---|---|---|---|---|---|
| 1.6 | 0 | ptsI | 1 | 3.25 | 0.274 |
| 1.6 | 0 | clcB | 1 | 3.25 | 0.253 |
| 1.6 | 0 | ycaM | 1 | 3.25 | 0.266 |
| 1.6 | 0 | yadI | 1 | 3.25 | 0.329 |
| 1.6 | 0 | AG1 | 1 | 3.25 | 0.279 |
| 1.6 | 0 | ptsI | 1 | 3.5 | 0.28 |
| 1.6 | 0 | clcB | 1 | 3.5 | 0.261 |
| 1.6 | 0 | ycaM | 1 | 3.5 | 0.273 |
| 1.6 | 0 | yadI | 1 | 3.5 | 0.348 |
| 1.6 | 0 | AG1 | 1 | 3.5 | 0.291 |
| 1.6 | 0 | ptsI | 1 | 3.75 | 0.292 |
| 1.6 | 0 | clcB | 1 | 3.75 | 0.264 |
| 1.6 | 0 | ycaM | 1 | 3.75 | 0.281 |
| 1.6 | 0 | yadI | 1 | 3.75 | 0.362 |
| 1.6 | 0 | AG1 | 1 | 3.75 | 0.292 |
| 1.6 | 0 | ptsI | 1 | 4 | 0.285 |
| 1.6 | 0 | clcB | 1 | 4 | 0.276 |
| 1.6 | 0 | ycaM | 1 | 4 | 0.287 |
| 1.6 | 0 | yadI | 1 | 4 | 0.391 |
| 1.6 | 0 | AG1 | 1 | 4 | 0.314 |
| 1.6 | 0 | ptsI | 1 | 4.25 | 0.304 |
| 1.6 | 0 | clcB | 1 | 4.25 | 0.278 |
| 1.6 | 0 | ycaM | 1 | 4.25 | 0.298 |
| 1.6 | 0 | yadI | 1 | 4.25 | 0.397 |
| 1.6 | 0 | AG1 | 1 | 4.25 | 0.314 |
| 1.6 | 0 | ptsI | 1 | 4.5 | 0.308 |
| 1.6 | 0 | clcB | 1 | 4.5 | 0.284 |
| 1.6 | 0 | ycaM | 1 | 4.5 | 0.304 |
| 1.6 | 0 | yadI | 1 | 4.5 | 0.413 |
| 1.6 | 0 | AG1 | 1 | 4.5 | 0.322 |
| 1.6 | 0 | ptsI | 1 | 4.75 | 0.327 |
| 1.6 | 0 | clcB | 1 | 4.75 | 0.29 |
| 1.6 | 0 | ycaM | 1 | 4.75 | 0.31 |
| 1.6 | 0 | yadI | 1 | 4.75 | 0.417 |
| 1.6 | 0 | AG1 | 1 | 4.75 | 0.331 |
| 1.6 | 0 | ptsI | 1 | 5 | 0.339 |
| 1.6 | 0 | clcB | 1 | 5 | 0.299 |
| 1.6 | 0 | ycaM | 1 | 5 | 0.32 |
| 1.6 | 0 | yadI | 1 | 5 | 0.432 |
| 1.6 | 0 | AG1 | 1 | 5 | 0.338 |
| 1.6 | 0 | ptsI | 1 | 5.25 | 0.346 |
| 1.6 | 0 | clcB | 1 | 5.25 | 0.305 |
| 1.6 | 0 | ycaM | 1 | 5.25 | 0.329 |
| 1.6 | 0 | yadI | 1 | 5.25 | 0.445 |
| 1.6 | 0 | AG1 | 1 | 5.25 | 0.348 |
| 1.6 | 0 | ptsI | 1 | 5.5 | 0.358 |
| 1.6 | 0 | clcB | 1 | 5.5 | 0.312 |
| 1.6 | 0 | ycaM | 1 | 5.5 | 0.34 |

| | | | | | |
|---|---|---|---|---|---|
| 1.6 | 0 | yadI | 1 | 5.5 | 0.457 |
| 1.6 | 0 | AG1 | 1 | 5.5 | 0.357 |
| 1.6 | 0 | ptsI | 1 | 5.75 | 0.366 |
| 1.6 | 0 | clcB | 1 | 5.75 | 0.318 |
| 1.6 | 0 | ycaM | 1 | 5.75 | 0.35 |
| 1.6 | 0 | yadI | 1 | 5.75 | 0.465 |
| 1.6 | 0 | AG1 | 1 | 5.75 | 0.366 |
| 1.6 | 0 | ptsI | 1 | 6 | 0.386 |
| 1.6 | 0 | clcB | 1 | 6 | 0.333 |
| 1.6 | 0 | ycaM | 1 | 6 | 0.357 |
| 1.6 | 0 | yadI | 1 | 6 | 0.469 |
| 1.6 | 0 | AG1 | 1 | 6 | 0.382 |
| 1.6 | 0 | ptsI | 1 | 6.25 | 0.402 |
| 1.6 | 0 | clcB | 1 | 6.25 | 0.342 |
| 1.6 | 0 | ycaM | 1 | 6.25 | 0.365 |
| 1.6 | 0 | yadI | 1 | 6.25 | 0.48 |
| 1.6 | 0 | AG1 | 1 | 6.25 | 0.397 |
| 1.6 | 0 | ptsI | 1 | 6.5 | 0.408 |
| 1.6 | 0 | clcB | 1 | 6.5 | 0.353 |
| 1.6 | 0 | ycaM | 1 | 6.5 | 0.375 |
| 1.6 | 0 | yadI | 1 | 6.5 | 0.488 |
| 1.6 | 0 | AG1 | 1 | 6.5 | 0.408 |
| 1.6 | 0 | ptsI | 1 | 6.75 | 0.42 |
| 1.6 | 0 | clcB | 1 | 6.75 | 0.356 |
| 1.6 | 0 | ycaM | 1 | 6.75 | 0.386 |
| 1.6 | 0 | yadI | 1 | 6.75 | 0.505 |
| 1.6 | 0 | AG1 | 1 | 6.75 | 0.419 |
| 1.6 | 0 | ptsI | 1 | 7 | 0.428 |
| 1.6 | 0 | clcB | 1 | 7 | 0.362 |
| 1.6 | 0 | ycaM | 1 | 7 | 0.399 |
| 1.6 | 0 | yadI | 1 | 7 | 0.518 |
| 1.6 | 0 | AG1 | 1 | 7 | 0.427 |
| 1.6 | 0 | ptsI | 1 | 7.25 | 0.434 |
| 1.6 | 0 | clcB | 1 | 7.25 | 0.366 |
| 1.6 | 0 | ycaM | 1 | 7.25 | 0.4 |
| 1.6 | 0 | yadI | 1 | 7.25 | 0.532 |
| 1.6 | 0 | AG1 | 1 | 7.25 | 0.434 |
| 1.6 | 0 | ptsI | 1 | 7.5 | 0.443 |
| 1.6 | 0 | clcB | 1 | 7.5 | 0.374 |
| 1.6 | 0 | ycaM | 1 | 7.5 | 0.407 |
| 1.6 | 0 | yadI | 1 | 7.5 | 0.541 |
| 1.6 | 0 | AG1 | 1 | 7.5 | 0.442 |
| 1.6 | 0 | ptsI | 1 | 7.75 | 0.445 |
| 1.6 | 0 | clcB | 1 | 7.75 | 0.381 |
| 1.6 | 0 | ycaM | 1 | 7.75 | 0.41 |
| 1.6 | 0 | yadI | 1 | 7.75 | 0.557 |
| 1.6 | 0 | AG1 | 1 | 7.75 | 0.452 |
| 1.6 | 0 | ptsI | 1 | 8 | 0.464 |

| | | | | | |
|---|---|---|---|---|---|
| 1.6 | 0 | clcB | 1 | 8 | 0.39 |
| 1.6 | 0 | ycaM | 1 | 8 | 0.412 |
| 1.6 | 0 | yadI | 1 | 8 | 0.56 |
| 1.6 | 0 | AG1 | 1 | 8 | 0.46 |
| 1.6 | 0 | ptsI | 1 | 8.25 | 0.47 |
| 1.6 | 0 | clcB | 1 | 8.25 | 0.403 |
| 1.6 | 0 | ycaM | 1 | 8.25 | 0.416 |
| 1.6 | 0 | yadI | 1 | 8.25 | 0.568 |
| 1.6 | 0 | AG1 | 1 | 8.25 | 0.474 |
| 1.6 | 0 | ptsI | 1 | 8.5 | 0.486 |
| 1.6 | 0 | clcB | 1 | 8.5 | 0.405 |
| 1.6 | 0 | ycaM | 1 | 8.5 | 0.424 |
| 1.6 | 0 | yadI | 1 | 8.5 | 0.578 |
| 1.6 | 0 | AG1 | 1 | 8.5 | 0.484 |
| 1.6 | 0 | ptsI | 1 | 8.75 | 0.499 |
| 1.6 | 0 | clcB | 1 | 8.75 | 0.407 |
| 1.6 | 0 | ycaM | 1 | 8.75 | 0.428 |
| 1.6 | 0 | yadI | 1 | 8.75 | 0.588 |
| 1.6 | 0 | AG1 | 1 | 8.75 | 0.495 |
| 1.6 | 0 | ptsI | 1 | 9 | 0.516 |
| 1.6 | 0 | clcB | 1 | 9 | 0.413 |
| 1.6 | 0 | ycaM | 1 | 9 | 0.432 |
| 1.6 | 0 | yadI | 1 | 9 | 0.591 |
| 1.6 | 0 | AG1 | 1 | 9 | 0.507 |
| 1.6 | 0 | ptsI | 1 | 9.25 | 0.542 |
| 1.6 | 0 | clcB | 1 | 9.25 | 0.428 |
| 1.6 | 0 | ycaM | 1 | 9.25 | 0.444 |
| 1.6 | 0 | yadI | 1 | 9.25 | 0.597 |
| 1.6 | 0 | AG1 | 1 | 9.25 | 0.524 |
| 1.6 | 0 | ptsI | 1 | 9.5 | 0.571 |
| 1.6 | 0 | clcB | 1 | 9.5 | 0.449 |
| 1.6 | 0 | ycaM | 1 | 9.5 | 0.455 |
| 1.6 | 0 | yadI | 1 | 9.5 | 0.605 |
| 1.6 | 0 | AG1 | 1 | 9.5 | 0.547 |
| 1.6 | 0 | ptsI | 1 | 9.75 | 0.562 |
| 1.6 | 0 | clcB | 1 | 9.75 | 0.442 |
| 1.6 | 0 | ycaM | 1 | 9.75 | 0.455 |
| 1.6 | 0 | yadI | 1 | 9.75 | 0.62 |
| 1.6 | 0 | AG1 | 1 | 9.75 | 0.556 |
| 1.6 | 0 | ptsI | 1 | 10 | 0.535 |
| 1.6 | 0 | clcB | 1 | 10 | 0.461 |
| 1.6 | 0 | ycaM | 1 | 10 | 0.467 |
| 1.6 | 0 | yadI | 1 | 10 | 0.617 |
| 1.6 | 0 | AG1 | 1 | 10 | 0.561 |
| 1.6 | 0 | ptsI | 1 | 10.25 | 0.552 |
| 1.6 | 0 | clcB | 1 | 10.25 | 0.463 |
| 1.6 | 0 | ycaM | 1 | 10.25 | 0.471 |
| 1.6 | 0 | yadI | 1 | 10.25 | 0.632 |

| | | | | | |
|---|---|---|---|---|---|
| 1.6 | 0 | AG1 | 1 | 10.25 | 0.572 |
| 1.6 | 0 | ptsI | 1 | 10.5 | 0.554 |
| 1.6 | 0 | clcB | 1 | 10.5 | 0.469 |
| 1.6 | 0 | ycaM | 1 | 10.5 | 0.487 |
| 1.6 | 0 | yadI | 1 | 10.5 | 0.645 |
| 1.6 | 0 | AG1 | 1 | 10.5 | 0.57 |
| 1.6 | 0 | ptsI | 1 | 10.75 | 0.555 |
| 1.6 | 0 | clcB | 1 | 10.75 | 0.49 |
| 1.6 | 0 | ycaM | 1 | 10.75 | 0.491 |
| 1.6 | 0 | yadI | 1 | 10.75 | 0.644 |
| 1.6 | 0 | AG1 | 1 | 10.75 | 0.592 |
| 1.6 | 0 | ptsI | 1 | 11 | 0.567 |
| 1.6 | 0 | clcB | 1 | 11 | 0.518 |
| 1.6 | 0 | ycaM | 1 | 11 | 0.497 |
| 1.6 | 0 | yadI | 1 | 11 | 0.648 |
| 1.6 | 0 | AG1 | 1 | 11 | 0.61 |
| 1.6 | 0 | ptsI | 1 | 11.25 | 0.564 |
| 1.6 | 0 | clcB | 1 | 11.25 | 0.516 |
| 1.6 | 0 | ycaM | 1 | 11.25 | 0.515 |
| 1.6 | 0 | yadI | 1 | 11.25 | 0.673 |
| 1.6 | 0 | AG1 | 1 | 11.25 | 0.601 |
| 1.6 | 0 | ptsI | 1 | 11.5 | 0.603 |
| 1.6 | 0 | clcB | 1 | 11.5 | 0.547 |
| 1.6 | 0 | ycaM | 1 | 11.5 | 0.514 |
| 1.6 | 0 | yadI | 1 | 11.5 | 0.69 |
| 1.6 | 0 | AG1 | 1 | 11.5 | 0.624 |
| 1.6 | 0 | ptsI | 1 | 11.75 | 0.649 |
| 1.6 | 0 | clcB | 1 | 11.75 | 0.56 |
| 1.6 | 0 | ycaM | 1 | 11.75 | 0.52 |
| 1.6 | 0 | yadI | 1 | 11.75 | 0.688 |
| 1.6 | 0 | AG1 | 1 | 11.75 | 0.65 |
| 1.6 | 0 | ptsI | 1 | 12 | 0.64 |
| 1.6 | 0 | clcB | 1 | 12 | 0.548 |
| 1.6 | 0 | ycaM | 1 | 12 | 0.526 |
| 1.6 | 0 | yadI | 1 | 12 | 0.7 |
| 1.6 | 0 | AG1 | 1 | 12 | 0.651 |
| 1.6 | 0 | ptsI | 1 | 12.25 | 0.676 |
| 1.6 | 0 | clcB | 1 | 12.25 | 0.564 |
| 1.6 | 0 | ycaM | 1 | 12.25 | 0.533 |
| 1.6 | 0 | yadI | 1 | 12.25 | 0.706 |
| 1.6 | 0 | AG1 | 1 | 12.25 | 0.683 |
| 1.6 | 0 | ptsI | 1 | 12.5 | 0.674 |
| 1.6 | 0 | clcB | 1 | 12.5 | 0.561 |
| 1.6 | 0 | ycaM | 1 | 12.5 | 0.532 |
| 1.6 | 0 | yadI | 1 | 12.5 | 0.699 |
| 1.6 | 0 | AG1 | 1 | 12.5 | 0.673 |
| 1.6 | 0 | ptsI | 1 | 12.75 | 0.678 |
| 1.6 | 0 | clcB | 1 | 12.75 | 0.559 |

| | | | | | |
|---|---|---|---|---|---|
| 1.6 | 0 | ycaM | 1 | 12.75 | 0.528 |
| 1.6 | 0 | yadI | 1 | 12.75 | 0.694 |
| 1.6 | 0 | AG1 | 1 | 12.75 | 0.666 |
| 1.6 | 0 | ptsI | 1 | 13 | 0.697 |
| 1.6 | 0 | clcB | 1 | 13 | 0.564 |
| 1.6 | 0 | ycaM | 1 | 13 | 0.538 |
| 1.6 | 0 | yadI | 1 | 13 | 0.701 |
| 1.6 | 0 | AG1 | 1 | 13 | 0.66 |
| 1.6 | 0 | ptsI | 1 | 13.25 | 0.725 |
| 1.6 | 0 | clcB | 1 | 13.25 | 0.573 |
| 1.6 | 0 | ycaM | 1 | 13.25 | 0.554 |
| 1.6 | 0 | yadI | 1 | 13.25 | 0.65 |
| 1.6 | 0 | AG1 | 1 | 13.25 | 0.677 |
| 1.6 | 0 | ptsI | 1 | 13.5 | 0.732 |
| 1.6 | 0 | clcB | 1 | 13.5 | 0.564 |
| 1.6 | 0 | ycaM | 1 | 13.5 | 0.553 |
| 1.6 | 0 | yadI | 1 | 13.5 | 0.662 |
| 1.6 | 0 | AG1 | 1 | 13.5 | 0.698 |
| 1.6 | 0 | ptsI | 1 | 13.75 | 0.752 |
| 1.6 | 0 | clcB | 1 | 13.75 | 0.566 |
| 1.6 | 0 | ycaM | 1 | 13.75 | 0.553 |
| 1.6 | 0 | yadI | 1 | 13.75 | 0.658 |
| 1.6 | 0 | AG1 | 1 | 13.75 | 0.687 |
| 1.6 | 0 | ptsI | 1 | 14 | 0.778 |
| 1.6 | 0 | clcB | 1 | 14 | 0.566 |
| 1.6 | 0 | ycaM | 1 | 14 | 0.561 |
| 1.6 | 0 | yadI | 1 | 14 | 0.659 |
| 1.6 | 0 | AG1 | 1 | 14 | 0.684 |
| 1.6 | 0 | ptsI | 1 | 14.25 | 0.793 |
| 1.6 | 0 | clcB | 1 | 14.25 | 0.564 |
| 1.6 | 0 | ycaM | 1 | 14.25 | 0.569 |
| 1.6 | 0 | yadI | 1 | 14.25 | 0.659 |
| 1.6 | 0 | AG1 | 1 | 14.25 | 0.679 |
| 1.6 | 0 | ptsI | 1 | 14.5 | 0.794 |
| 1.6 | 0 | clcB | 1 | 14.5 | 0.577 |
| 1.6 | 0 | ycaM | 1 | 14.5 | 0.572 |
| 1.6 | 0 | yadI | 1 | 14.5 | 0.666 |
| 1.6 | 0 | AG1 | 1 | 14.5 | 0.669 |
| 1.6 | 0 | ptsI | 1 | 14.75 | 0.788 |
| 1.6 | 0 | clcB | 1 | 14.75 | 0.612 |
| 1.6 | 0 | ycaM | 1 | 14.75 | 0.592 |
| 1.6 | 0 | yadI | 1 | 14.75 | 0.669 |
| 1.6 | 0 | AG1 | 1 | 14.75 | 0.685 |
| 1.6 | 0 | ptsI | 1 | 15 | 0.78 |
| 1.6 | 0 | clcB | 1 | 15 | 0.614 |
| 1.6 | 0 | ycaM | 1 | 15 | 0.586 |
| 1.6 | 0 | yadI | 1 | 15 | 0.662 |
| 1.6 | 0 | AG1 | 1 | 15 | 0.712 |

| | | | | | |
|---|---|---|---|---|---|
| 1.6 | 0 | ptsI | 1 | 15.25 | 0.788 |
| 1.6 | 0 | clcB | 1 | 15.25 | 0.651 |
| 1.6 | 0 | ycaM | 1 | 15.25 | 0.616 |
| 1.6 | 0 | yadI | 1 | 15.25 | 0.672 |
| 1.6 | 0 | AG1 | 1 | 15.25 | 0.713 |
| 1.6 | 0 | ptsI | 1 | 15.5 | 0.776 |
| 1.6 | 0 | clcB | 1 | 15.5 | 0.648 |
| 1.6 | 0 | ycaM | 1 | 15.5 | 0.624 |
| 1.6 | 0 | yadI | 1 | 15.5 | 0.678 |
| 1.6 | 0 | AG1 | 1 | 15.5 | 0.694 |
| 1.6 | 0 | ptsI | 1 | 15.75 | 0.737 |
| 1.6 | 0 | clcB | 1 | 15.75 | 0.658 |
| 1.6 | 0 | ycaM | 1 | 15.75 | 0.638 |
| 1.6 | 0 | yadI | 1 | 15.75 | 0.691 |
| 1.6 | 0 | AG1 | 1 | 15.75 | 0.716 |
| 1.6 | 0 | ptsI | 1 | 16 | 0.723 |
| 1.6 | 0 | clcB | 1 | 16 | 0.671 |
| 1.6 | 0 | ycaM | 1 | 16 | 0.654 |
| 1.6 | 0 | yadI | 1 | 16 | 0.689 |
| 1.6 | 0 | AG1 | 1 | 16 | 0.714 |
| 1.6 | 0 | ptsI | 1 | 16.25 | 0.757 |
| 1.6 | 0 | clcB | 1 | 16.25 | 0.669 |
| 1.6 | 0 | ycaM | 1 | 16.25 | 0.666 |
| 1.6 | 0 | yadI | 1 | 16.25 | 0.689 |
| 1.6 | 0 | AG1 | 1 | 16.25 | 0.721 |
| 1.6 | 0 | ptsI | 1 | 16.5 | 0.778 |
| 1.6 | 0 | clcB | 1 | 16.5 | 0.694 |
| 1.6 | 0 | ycaM | 1 | 16.5 | 0.671 |
| 1.6 | 0 | yadI | 1 | 16.5 | 0.68 |
| 1.6 | 0 | AG1 | 1 | 16.5 | 0.736 |
| 1.6 | 0 | ptsI | 1 | 17.0667 | 0.768 |
| 1.6 | 0 | clcB | 1 | 17.0667 | 0.684 |
| 1.6 | 0 | ycaM | 1 | 17.0667 | 0.697 |
| 1.6 | 0 | yadI | 1 | 17.0667 | 0.759 |
| 1.6 | 0 | AG1 | 1 | 17.0667 | 0.706 |
| 1.6 | 0 | ptsI | 1 | 17.3167 | 0.771 |
| 1.6 | 0 | clcB | 1 | 17.3167 | 0.669 |
| 1.6 | 0 | ycaM | 1 | 17.3167 | 0.678 |
| 1.6 | 0 | yadI | 1 | 17.3167 | 0.714 |
| 1.6 | 0 | AG1 | 1 | 17.3167 | 0.696 |
| 1.6 | 0 | ptsI | 1 | 17.5667 | 0.796 |
| 1.6 | 0 | clcB | 1 | 17.5667 | 0.664 |
| 1.6 | 0 | ycaM | 1 | 17.5667 | 0.678 |
| 1.6 | 0 | yadI | 1 | 17.5667 | 0.711 |
| 1.6 | 0 | AG1 | 1 | 17.5667 | 0.696 |
| 1.6 | 0 | ptsI | 1 | 17.8167 | 0.814 |
| 1.6 | 0 | clcB | 1 | 17.8167 | 0.721 |
| 1.6 | 0 | ycaM | 1 | 17.8167 | 0.69 |

| | | | | | |
|---|---|---|---|---|---|
| 1.6 | 0 | yadI | 1 | 17.8167 | 0.72 |
| 1.6 | 0 | AG1 | 1 | 17.8167 | 0.698 |
| 1.6 | 0 | ptsI | 1 | 18.0667 | 0.827 |
| 1.6 | 0 | clcB | 1 | 18.0667 | 0.702 |
| 1.6 | 0 | ycaM | 1 | 18.0667 | 0.686 |
| 1.6 | 0 | yadI | 1 | 18.0667 | 0.721 |
| 1.6 | 0 | AG1 | 1 | 18.0667 | 0.695 |
| 1.6 | 0 | ptsI | 1 | 18.3167 | 0.834 |
| 1.6 | 0 | clcB | 1 | 18.3167 | 0.705 |
| 1.6 | 0 | ycaM | 1 | 18.3167 | 0.69 |
| 1.6 | 0 | yadI | 1 | 18.3167 | 0.727 |
| 1.6 | 0 | AG1 | 1 | 18.3167 | 0.732 |
| 1.6 | 0 | ptsI | 1 | 18.5667 | 0.847 |
| 1.6 | 0 | clcB | 1 | 18.5667 | 0.712 |
| 1.6 | 0 | ycaM | 1 | 18.5667 | 0.702 |
| 1.6 | 0 | yadI | 1 | 18.5667 | 0.73 |
| 1.6 | 0 | AG1 | 1 | 18.5667 | 0.737 |
| 1.6 | 0 | ptsI | 1 | 18.8167 | 0.856 |
| 1.6 | 0 | clcB | 1 | 18.8167 | 0.712 |
| 1.6 | 0 | ycaM | 1 | 18.8167 | 0.703 |
| 1.6 | 0 | yadI | 1 | 18.8167 | 0.728 |
| 1.6 | 0 | AG1 | 1 | 18.8167 | 0.733 |
| 1.6 | 0 | ptsI | 1 | 19.0667 | 0.862 |
| 1.6 | 0 | clcB | 1 | 19.0667 | 0.727 |
| 1.6 | 0 | ycaM | 1 | 19.0667 | 0.703 |
| 1.6 | 0 | yadI | 1 | 19.0667 | 0.742 |
| 1.6 | 0 | AG1 | 1 | 19.0667 | 0.761 |
| 1.6 | 0 | ptsI | 1 | 19.3167 | 0.862 |
| 1.6 | 0 | clcB | 1 | 19.3167 | 0.734 |
| 1.6 | 0 | ycaM | 1 | 19.3167 | 0.706 |
| 1.6 | 0 | yadI | 1 | 19.3167 | 0.753 |
| 1.6 | 0 | AG1 | 1 | 19.3167 | 0.76 |
| 1.6 | 0 | ptsI | 1 | 19.5667 | 0.874 |
| 1.6 | 0 | clcB | 1 | 19.5667 | 0.742 |
| 1.6 | 0 | ycaM | 1 | 19.5667 | 0.714 |
| 1.6 | 0 | yadI | 1 | 19.5667 | 0.748 |
| 1.6 | 0 | AG1 | 1 | 19.5667 | 0.763 |
| 1.6 | 0 | ptsI | 1 | 19.8167 | 0.88 |
| 1.6 | 0 | clcB | 1 | 19.8167 | 0.74 |
| 1.6 | 0 | ycaM | 1 | 19.8167 | 0.713 |
| 1.6 | 0 | yadI | 1 | 19.8167 | 0.752 |
| 1.6 | 0 | AG1 | 1 | 19.8167 | 0.773 |
| 1.6 | 0 | ptsI | 1 | 20.0667 | 0.893 |
| 1.6 | 0 | clcB | 1 | 20.0667 | 0.748 |
| 1.6 | 0 | ycaM | 1 | 20.0667 | 0.723 |
| 1.6 | 0 | yadI | 1 | 20.0667 | 0.758 |
| 1.6 | 0 | AG1 | 1 | 20.0667 | 0.773 |
| 1.6 | 0 | ptsI | 1 | 20.3167 | 0.902 |

| | | | | | |
|---|---|---|---|---|---|
| 1.6 | 0 | clcB | 1 | 20.3167 | 0.754 |
| 1.6 | 0 | ycaM | 1 | 20.3167 | 0.723 |
| 1.6 | 0 | yadI | 1 | 20.3167 | 0.765 |
| 1.6 | 0 | AG1 | 1 | 20.3167 | 0.769 |
| 1.6 | 0 | ptsI | 1 | 20.5667 | 0.905 |
| 1.6 | 0 | clcB | 1 | 20.5667 | 0.751 |
| 1.6 | 0 | ycaM | 1 | 20.5667 | 0.714 |
| 1.6 | 0 | yadI | 1 | 20.5667 | 0.782 |
| 1.6 | 0 | AG1 | 1 | 20.5667 | 0.763 |
| 1.6 | 0 | ptsI | 1 | 20.8167 | 0.909 |
| 1.6 | 0 | clcB | 1 | 20.8167 | 0.752 |
| 1.6 | 0 | ycaM | 1 | 20.8167 | 0.716 |
| 1.6 | 0 | yadI | 1 | 20.8167 | 0.785 |
| 1.6 | 0 | AG1 | 1 | 20.8167 | 0.759 |
| 1.6 | 0 | ptsI | 1 | 21.0667 | 0.928 |
| 1.6 | 0 | clcB | 1 | 21.0667 | 0.757 |
| 1.6 | 0 | ycaM | 1 | 21.0667 | 0.72 |
| 1.6 | 0 | yadI | 1 | 21.0667 | 0.784 |
| 1.6 | 0 | AG1 | 1 | 21.0667 | 0.764 |
| 1.6 | 0 | ptsI | 1 | 21.3167 | 0.925 |
| 1.6 | 0 | clcB | 1 | 21.3167 | 0.76 |
| 1.6 | 0 | ycaM | 1 | 21.3167 | 0.726 |
| 1.6 | 0 | yadI | 1 | 21.3167 | 0.784 |
| 1.6 | 0 | AG1 | 1 | 21.3167 | 0.761 |
| 1.6 | 0 | ptsI | 1 | 21.5667 | 0.936 |
| 1.6 | 0 | clcB | 1 | 21.5667 | 0.762 |
| 1.6 | 0 | ycaM | 1 | 21.5667 | 0.72 |
| 1.6 | 0 | yadI | 1 | 21.5667 | 0.806 |
| 1.6 | 0 | AG1 | 1 | 21.5667 | 0.761 |
| 1.6 | 0 | ptsI | 1 | 21.8167 | 0.942 |
| 1.6 | 0 | clcB | 1 | 21.8167 | 0.76 |
| 1.6 | 0 | ycaM | 1 | 21.8167 | 0.721 |
| 1.6 | 0 | yadI | 1 | 21.8167 | 0.803 |
| 1.6 | 0 | AG1 | 1 | 21.8167 | 0.749 |
| 1.6 | 0 | ptsI | 1 | 22.0667 | 0.952 |
| 1.6 | 0 | clcB | 1 | 22.0667 | 0.766 |
| 1.6 | 0 | ycaM | 1 | 22.0667 | 0.72 |
| 1.6 | 0 | yadI | 1 | 22.0667 | 0.806 |
| 1.6 | 0 | AG1 | 1 | 22.0667 | 0.758 |
| 1.6 | 0 | ptsI | 1 | 22.9333 | 1.05 |
| 1.6 | 0 | clcB | 1 | 22.9333 | 0.736 |
| 1.6 | 0 | ycaM | 1 | 22.9333 | 0.675 |
| 1.6 | 0 | yadI | 1 | 22.9333 | 0.878 |
| 1.6 | 0 | AG1 | 1 | 22.9333 | 0.71 |
| 1.6 | 0 | ptsI | 1 | 23.1833 | 1.054 |
| 1.6 | 0 | clcB | 1 | 23.1833 | 0.721 |
| 1.6 | 0 | ycaM | 1 | 23.1833 | 0.689 |
| 1.6 | 0 | yadI | 1 | 23.1833 | 0.851 |

| | | | | | |
|---|---|---|---|---|---|
| 1.6 | 0 | AG1 | 1 | 23.1833 | 0.692 |
| 1.6 | 0 | ptsI | 1 | 23.4333 | 1.056 |
| 1.6 | 0 | clcB | 1 | 23.4333 | 0.737 |
| 1.6 | 0 | ycaM | 1 | 23.4333 | 0.714 |
| 1.6 | 0 | yadI | 1 | 23.4333 | 0.853 |
| 1.6 | 0 | AG1 | 1 | 23.4333 | 0.701 |
| 1.6 | 0 | ptsI | 1 | 23.6833 | 1.065 |
| 1.6 | 0 | clcB | 1 | 23.6833 | 0.746 |
| 1.6 | 0 | ycaM | 1 | 23.6833 | 0.722 |
| 1.6 | 0 | yadI | 1 | 23.6833 | 0.857 |
| 1.6 | 0 | AG1 | 1 | 23.6833 | 0.706 |
| 1.6 | 0 | ptsI | 1 | 23.9333 | 1.073 |
| 1.6 | 0 | clcB | 1 | 23.9333 | 0.746 |
| 1.6 | 0 | ycaM | 1 | 23.9333 | 0.731 |
| 1.6 | 0 | yadI | 1 | 23.9333 | 0.866 |
| 1.6 | 0 | AG1 | 1 | 23.9333 | 0.717 |
| 1.6 | 0 | ptsI | 1 | 24.1833 | 1.076 |
| 1.6 | 0 | clcB | 1 | 24.1833 | 0.745 |
| 1.6 | 0 | ycaM | 1 | 24.1833 | 0.729 |
| 1.6 | 0 | yadI | 1 | 24.1833 | 0.879 |
| 1.6 | 0 | AG1 | 1 | 24.1833 | 0.716 |
| 1.6 | 0 | ptsI | 1 | 24.4333 | 1.077 |
| 1.6 | 0 | clcB | 1 | 24.4333 | 0.744 |
| 1.6 | 0 | ycaM | 1 | 24.4333 | 0.741 |
| 1.6 | 0 | yadI | 1 | 24.4333 | 0.88 |
| 1.6 | 0 | AG1 | 1 | 24.4333 | 0.725 |
| 1.6 | 0 | ptsI | 1 | 24.6833 | 1.082 |
| 1.6 | 0 | clcB | 1 | 24.6833 | 0.739 |
| 1.6 | 0 | ycaM | 1 | 24.6833 | 0.736 |
| 1.6 | 0 | yadI | 1 | 24.6833 | 0.878 |
| 1.6 | 0 | AG1 | 1 | 24.6833 | 0.72 |
| 1.6 | 0 | ptsI | 1 | 24.9333 | 1.091 |
| 1.6 | 0 | clcB | 1 | 24.9333 | 0.744 |
| 1.6 | 0 | ycaM | 1 | 24.9333 | 0.733 |
| 1.6 | 0 | yadI | 1 | 24.9333 | 0.897 |
| 1.6 | 0 | AG1 | 1 | 24.9333 | 0.717 |
| 1.6 | 0 | ptsI | 1 | 25.1833 | 1.099 |
| 1.6 | 0 | clcB | 1 | 25.1833 | 0.744 |
| 1.6 | 0 | ycaM | 1 | 25.1833 | 0.743 |
| 1.6 | 0 | yadI | 1 | 25.1833 | 0.896 |
| 1.6 | 0 | AG1 | 1 | 25.1833 | 0.736 |
| 1.6 | 0 | ptsI | 1 | 25.4333 | 1.104 |
| 1.6 | 0 | clcB | 1 | 25.4333 | 0.744 |
| 1.6 | 0 | ycaM | 1 | 25.4333 | 0.757 |
| 1.6 | 0 | yadI | 1 | 25.4333 | 0.896 |
| 1.6 | 0 | AG1 | 1 | 25.4333 | 0.751 |
| 1.6 | 0 | ptsI | 1 | 25.6833 | 1.111 |
| 1.6 | 0 | clcB | 1 | 25.6833 | 0.746 |

| | | | | | |
|---|---|---|---|---|---|
| 1.6 | 0 | ycaM | 1 | 25.6833 | 0.751 |
| 1.6 | 0 | yadI | 1 | 25.6833 | 0.904 |
| 1.6 | 0 | AG1 | 1 | 25.6833 | 0.746 |
| 1.6 | 0 | ptsI | 1 | 25.9333 | 1.111 |
| 1.6 | 0 | clcB | 1 | 25.9333 | 0.755 |
| 1.6 | 0 | ycaM | 1 | 25.9333 | 0.762 |
| 1.6 | 0 | yadI | 1 | 25.9333 | 0.888 |
| 1.6 | 0 | AG1 | 1 | 25.9333 | 0.762 |
| 1.6 | 0 | ptsI | 1 | 26.1833 | 1.115 |
| 1.6 | 0 | clcB | 1 | 26.1833 | 0.751 |
| 1.6 | 0 | ycaM | 1 | 26.1833 | 0.774 |
| 1.6 | 0 | yadI | 1 | 26.1833 | 0.906 |
| 1.6 | 0 | AG1 | 1 | 26.1833 | 0.768 |
| 1.6 | 0 | ptsI | 1 | 26.4333 | 1.119 |
| 1.6 | 0 | clcB | 1 | 26.4333 | 0.75 |
| 1.6 | 0 | ycaM | 1 | 26.4333 | 0.774 |
| 1.6 | 0 | yadI | 1 | 26.4333 | 0.91 |
| 1.6 | 0 | AG1 | 1 | 26.4333 | 0.762 |
| 1.6 | 0 | ptsI | 1 | 26.6833 | 1.126 |
| 1.6 | 0 | clcB | 1 | 26.6833 | 0.757 |
| 1.6 | 0 | ycaM | 1 | 26.6833 | 0.776 |
| 1.6 | 0 | yadI | 1 | 26.6833 | 0.915 |
| 1.6 | 0 | AG1 | 1 | 26.6833 | 0.764 |
| 1.6 | 0 | ptsI | 1 | 26.9333 | 1.13 |
| 1.6 | 0 | clcB | 1 | 26.9333 | 0.759 |
| 1.6 | 0 | ycaM | 1 | 26.9333 | 0.78 |
| 1.6 | 0 | yadI | 1 | 26.9333 | 0.915 |
| 1.6 | 0 | AG1 | 1 | 26.9333 | 0.772 |
| 1.6 | 0 | ptsI | 1 | 27.1833 | 1.13 |
| 1.6 | 0 | clcB | 1 | 27.1833 | 0.761 |
| 1.6 | 0 | ycaM | 1 | 27.1833 | 0.783 |
| 1.6 | 0 | yadI | 1 | 27.1833 | 0.916 |
| 1.6 | 0 | AG1 | 1 | 27.1833 | 0.775 |
| 1.6 | 0 | ptsI | 1 | 27.4333 | 1.136 |
| 1.6 | 0 | clcB | 1 | 27.4333 | 0.768 |
| 1.6 | 0 | ycaM | 1 | 27.4333 | 0.775 |
| 1.6 | 0 | yadI | 1 | 27.4333 | 0.917 |
| 1.6 | 0 | AG1 | 1 | 27.4333 | 0.769 |
| 1.6 | 0 | ptsI | 1 | 27.6833 | 1.141 |
| 1.6 | 0 | clcB | 1 | 27.6833 | 0.763 |
| 1.6 | 0 | ycaM | 1 | 27.6833 | 0.781 |
| 1.6 | 0 | yadI | 1 | 27.6833 | 0.92 |
| 1.6 | 0 | AG1 | 1 | 27.6833 | 0.78 |
| 1.6 | 0 | ptsI | 1 | 27.9333 | 1.145 |
| 1.6 | 0 | clcB | 1 | 27.9333 | 0.756 |
| 1.6 | 0 | ycaM | 1 | 27.9333 | 0.782 |
| 1.6 | 0 | yadI | 1 | 27.9333 | 0.922 |
| 1.6 | 0 | AG1 | 1 | 27.9333 | 0.778 |

| | | | | | |
|---|---|---|---|---|---|
| 1.6 | 0 | ptsI | 1 | 28.1833 | 1.142 |
| 1.6 | 0 | clcB | 1 | 28.1833 | 0.762 |
| 1.6 | 0 | ycaM | 1 | 28.1833 | 0.792 |
| 1.6 | 0 | yadI | 1 | 28.1833 | 0.927 |
| 1.6 | 0 | AG1 | 1 | 28.1833 | 0.79 |
| 1.6 | 0 | ptsI | 1 | 28.4333 | 1.148 |
| 1.6 | 0 | clcB | 1 | 28.4333 | 0.762 |
| 1.6 | 0 | ycaM | 1 | 28.4333 | 0.793 |
| 1.6 | 0 | yadI | 1 | 28.4333 | 0.926 |
| 1.6 | 0 | AG1 | 1 | 28.4333 | 0.79 |
| 1.6 | 0 | ptsI | 1 | 28.6833 | 1.156 |
| 1.6 | 0 | clcB | 1 | 28.6833 | 0.763 |
| 1.6 | 0 | ycaM | 1 | 28.6833 | 0.795 |
| 1.6 | 0 | yadI | 1 | 28.6833 | 0.932 |
| 1.6 | 0 | AG1 | 1 | 28.6833 | 0.792 |
| 1.6 | 0 | ptsI | 1 | 28.9333 | 1.154 |
| 1.6 | 0 | clcB | 1 | 28.9333 | 0.76 |
| 1.6 | 0 | ycaM | 1 | 28.9333 | 0.8 |
| 1.6 | 0 | yadI | 1 | 28.9333 | 0.929 |
| 1.6 | 0 | AG1 | 1 | 28.9333 | 0.796 |
| 1.6 | 0 | ptsI | 1 | 29.1833 | 1.153 |
| 1.6 | 0 | clcB | 1 | 29.1833 | 0.761 |
| 1.6 | 0 | ycaM | 1 | 29.1833 | 0.793 |
| 1.6 | 0 | yadI | 1 | 29.1833 | 0.936 |
| 1.6 | 0 | AG1 | 1 | 29.1833 | 0.792 |
| 1.6 | 0 | ptsI | 1 | 29.4333 | 1.154 |
| 1.6 | 0 | clcB | 1 | 29.4333 | 0.762 |
| 1.6 | 0 | ycaM | 1 | 29.4333 | 0.803 |
| 1.6 | 0 | yadI | 1 | 29.4333 | 0.943 |
| 1.6 | 0 | AG1 | 1 | 29.4333 | 0.797 |
| 1.6 | 0 | ptsI | 1 | 29.6833 | 1.156 |
| 1.6 | 0 | clcB | 1 | 29.6833 | 0.762 |
| 1.6 | 0 | ycaM | 1 | 29.6833 | 0.805 |
| 1.6 | 0 | yadI | 1 | 29.6833 | 0.949 |
| 1.6 | 0 | AG1 | 1 | 29.6833 | 0.796 |
| 1.6 | 0 | ptsI | 1 | 29.9333 | 1.162 |
| 1.6 | 0 | clcB | 1 | 29.9333 | 0.762 |
| 1.6 | 0 | ycaM | 1 | 29.9333 | 0.81 |
| 1.6 | 0 | yadI | 1 | 29.9333 | 0.952 |
| 1.6 | 0 | AG1 | 1 | 29.9333 | 0.798 |
| 1.6 | 0 | ptsI | 1 | 30.1833 | 1.173 |
| 1.6 | 0 | clcB | 1 | 30.1833 | 0.759 |
| 1.6 | 0 | ycaM | 1 | 30.1833 | 0.808 |
| 1.6 | 0 | yadI | 1 | 30.1833 | 0.957 |
| 1.6 | 0 | AG1 | 1 | 30.1833 | 0.802 |
| 1.6 | 0 | ptsI | 1 | 30.4333 | 1.173 |
| 1.6 | 0 | clcB | 1 | 30.4333 | 0.764 |
| 1.6 | 0 | ycaM | 1 | 30.4333 | 0.816 |

| | | | | | |
|---|---|---|---|---|---|
| 1.6 | 0 | yadI | 1 | 30.4333 | 0.96 |
| 1.6 | 0 | AG1 | 1 | 30.4333 | 0.805 |
| 1.6 | 0 | ptsI | 1 | 30.6833 | 1.185 |
| 1.6 | 0 | clcB | 1 | 30.6833 | 0.764 |
| 1.6 | 0 | ycaM | 1 | 30.6833 | 0.818 |
| 1.6 | 0 | yadI | 1 | 30.6833 | 0.974 |
| 1.6 | 0 | AG1 | 1 | 30.6833 | 0.802 |
| 1.6 | 0 | ptsI | 1 | 30.9333 | 1.19 |
| 1.6 | 0 | clcB | 1 | 30.9333 | 0.766 |
| 1.6 | 0 | ycaM | 1 | 30.9333 | 0.816 |
| 1.6 | 0 | yadI | 1 | 30.9333 | 0.977 |
| 1.6 | 0 | AG1 | 1 | 30.9333 | 0.809 |
| 1.6 | 0 | ptsI | 1 | 31.1833 | 1.198 |
| 1.6 | 0 | clcB | 1 | 31.1833 | 0.77 |
| 1.6 | 0 | ycaM | 1 | 31.1833 | 0.821 |
| 1.6 | 0 | yadI | 1 | 31.1833 | 0.984 |
| 1.6 | 0 | AG1 | 1 | 31.1833 | 0.806 |
| 1.6 | 0 | ptsI | 1 | 31.4333 | 1.2 |
| 1.6 | 0 | clcB | 1 | 31.4333 | 0.774 |
| 1.6 | 0 | ycaM | 1 | 31.4333 | 0.824 |
| 1.6 | 0 | yadI | 1 | 31.4333 | 0.98 |
| 1.6 | 0 | AG1 | 1 | 31.4333 | 0.812 |
| 1.6 | 0 | ptsI | 1 | 31.6833 | 1.209 |
| 1.6 | 0 | clcB | 1 | 31.6833 | 0.776 |
| 1.6 | 0 | ycaM | 1 | 31.6833 | 0.828 |
| 1.6 | 0 | yadI | 1 | 31.6833 | 0.986 |
| 1.6 | 0 | AG1 | 1 | 31.6833 | 0.815 |
| 1.6 | 0 | ptsI | 1 | 31.9333 | 1.214 |
| 1.6 | 0 | clcB | 1 | 31.9333 | 0.776 |
| 1.6 | 0 | ycaM | 1 | 31.9333 | 0.828 |
| 1.6 | 0 | yadI | 1 | 31.9333 | 0.987 |
| 1.6 | 0 | AG1 | 1 | 31.9333 | 0.812 |
| 1.6 | 0 | ptsI | 1 | 32.1833 | 1.22 |
| 1.6 | 0 | clcB | 1 | 32.1833 | 0.778 |
| 1.6 | 0 | ycaM | 1 | 32.1833 | 0.836 |
| 1.6 | 0 | yadI | 1 | 32.1833 | 0.994 |
| 1.6 | 0 | AG1 | 1 | 32.1833 | 0.82 |
| 1.6 | 0 | ptsI | 1 | 32.4333 | 1.236 |
| 1.6 | 0 | clcB | 1 | 32.4333 | 0.786 |
| 1.6 | 0 | ycaM | 1 | 32.4333 | 0.836 |
| 1.6 | 0 | yadI | 1 | 32.4333 | 0.986 |
| 1.6 | 0 | AG1 | 1 | 32.4333 | 0.821 |
| 1.6 | 0 | ptsI | 1 | 32.6833 | 1.235 |
| 1.6 | 0 | clcB | 1 | 32.6833 | 0.788 |
| 1.6 | 0 | ycaM | 1 | 32.6833 | 0.842 |
| 1.6 | 0 | yadI | 1 | 32.6833 | 1.012 |
| 1.6 | 0 | AG1 | 1 | 32.6833 | 0.824 |
| 1.6 | 0 | ptsI | 1 | 32.9333 | 1.246 |

| | | | | | |
|---|---|---|---|---|---|
| 1.6 | 0 | clcB | 1 | 32.9333 | 0.791 |
| 1.6 | 0 | ycaM | 1 | 32.9333 | 0.847 |
| 1.6 | 0 | yadI | 1 | 32.9333 | 1.022 |
| 1.6 | 0 | AG1 | 1 | 32.9333 | 0.823 |
| 1.6 | 0 | ptsI | 1 | 33.1833 | 1.263 |
| 1.6 | 0 | clcB | 1 | 33.1833 | 0.784 |
| 1.6 | 0 | ycaM | 1 | 33.1833 | 0.85 |
| 1.6 | 0 | yadI | 1 | 33.1833 | 1.019 |
| 1.6 | 0 | AG1 | 1 | 33.1833 | 0.821 |
| 1.6 | 0 | ptsI | 1 | 33.4333 | 1.268 |
| 1.6 | 0 | clcB | 1 | 33.4333 | 0.794 |
| 1.6 | 0 | ycaM | 1 | 33.4333 | 0.85 |
| 1.6 | 0 | yadI | 1 | 33.4333 | 1.03 |
| 1.6 | 0 | AG1 | 1 | 33.4333 | 0.828 |
| 1.6 | 0 | ptsI | 1 | 33.6833 | 1.28 |
| 1.6 | 0 | clcB | 1 | 33.6833 | 0.79 |
| 1.6 | 0 | ycaM | 1 | 33.6833 | 0.857 |
| 1.6 | 0 | yadI | 1 | 33.6833 | 1.03 |
| 1.6 | 0 | AG1 | 1 | 33.6833 | 0.823 |
| 1.6 | 0 | ptsI | 1 | 33.9333 | 1.285 |
| 1.6 | 0 | clcB | 1 | 33.9333 | 0.806 |
| 1.6 | 0 | ycaM | 1 | 33.9333 | 0.859 |
| 1.6 | 0 | yadI | 1 | 33.9333 | 1.042 |
| 1.6 | 0 | AG1 | 1 | 33.9333 | 0.827 |
| 1.6 | 0 | ptsI | 1 | 34.1833 | 1.292 |
| 1.6 | 0 | clcB | 1 | 34.1833 | 0.796 |
| 1.6 | 0 | ycaM | 1 | 34.1833 | 0.861 |
| 1.6 | 0 | yadI | 1 | 34.1833 | 1.069 |
| 1.6 | 0 | AG1 | 1 | 34.1833 | 0.83 |
| 1.6 | 0 | ptsI | 1 | 34.4333 | 1.306 |
| 1.6 | 0 | clcB | 1 | 34.4333 | 0.813 |
| 1.6 | 0 | ycaM | 1 | 34.4333 | 0.864 |
| 1.6 | 0 | yadI | 1 | 34.4333 | 1.061 |
| 1.6 | 0 | AG1 | 1 | 34.4333 | 0.831 |
| 1.6 | 0 | ptsI | 1 | 34.6833 | 1.312 |
| 1.6 | 0 | clcB | 1 | 34.6833 | 0.817 |
| 1.6 | 0 | ycaM | 1 | 34.6833 | 0.864 |
| 1.6 | 0 | yadI | 1 | 34.6833 | 1.068 |
| 1.6 | 0 | AG1 | 1 | 34.6833 | 0.834 |
| 1.6 | 0 | ptsI | 1 | 34.9333 | 1.328 |
| 1.6 | 0 | clcB | 1 | 34.9333 | 0.822 |
| 1.6 | 0 | ycaM | 1 | 34.9333 | 0.862 |
| 1.6 | 0 | yadI | 1 | 34.9333 | 1.075 |
| 1.6 | 0 | AG1 | 1 | 34.9333 | 0.838 |
| 1.6 | 0 | ptsI | 1 | 35.1833 | 1.334 |
| 1.6 | 0 | clcB | 1 | 35.1833 | 0.832 |
| 1.6 | 0 | ycaM | 1 | 35.1833 | 0.859 |
| 1.6 | 0 | yadI | 1 | 35.1833 | 1.082 |

| | | | | | |
|---|---|---|---|---|---|
| 1.6 | 0 | AG1 | 1 | 35.1833 | 0.838 |
| 1.6 | 0 | ptsI | 1 | 35.4333 | 1.342 |
| 1.6 | 0 | clcB | 1 | 35.4333 | 0.836 |
| 1.6 | 0 | ycaM | 1 | 35.4333 | 0.86 |
| 1.6 | 0 | yadI | 1 | 35.4333 | 1.097 |
| 1.6 | 0 | AG1 | 1 | 35.4333 | 0.84 |
| 1.6 | 0 | ptsI | 1 | 35.6833 | 1.355 |
| 1.6 | 0 | clcB | 1 | 35.6833 | 0.834 |
| 1.6 | 0 | ycaM | 1 | 35.6833 | 0.858 |
| 1.6 | 0 | yadI | 1 | 35.6833 | 1.102 |
| 1.6 | 0 | AG1 | 1 | 35.6833 | 0.841 |
| 1.6 | 0 | ptsI | 1 | 35.9333 | 1.37 |
| 1.6 | 0 | clcB | 1 | 35.9333 | 0.83 |
| 1.6 | 0 | ycaM | 1 | 35.9333 | 0.853 |
| 1.6 | 0 | yadI | 1 | 35.9333 | 1.109 |
| 1.6 | 0 | AG1 | 1 | 35.9333 | 0.843 |
| 1.6 | 0 | ptsI | 1 | 36.1833 | 1.357 |
| 1.6 | 0 | clcB | 1 | 36.1833 | 0.835 |
| 1.6 | 0 | ycaM | 1 | 36.1833 | 0.855 |
| 1.6 | 0 | yadI | 1 | 36.1833 | 1.118 |
| 1.6 | 0 | AG1 | 1 | 36.1833 | 0.846 |
| 1.6 | 0 | ptsI | 1 | 36.4333 | 1.382 |
| 1.6 | 0 | clcB | 1 | 36.4333 | 0.838 |
| 1.6 | 0 | ycaM | 1 | 36.4333 | 0.854 |
| 1.6 | 0 | yadI | 1 | 36.4333 | 1.132 |
| 1.6 | 0 | AG1 | 1 | 36.4333 | 0.851 |
| 1.6 | 0 | ptsI | 1 | 36.6833 | 1.39 |
| 1.6 | 0 | clcB | 1 | 36.6833 | 0.839 |
| 1.6 | 0 | ycaM | 1 | 36.6833 | 0.853 |
| 1.6 | 0 | yadI | 1 | 36.6833 | 1.136 |
| 1.6 | 0 | AG1 | 1 | 36.6833 | 0.852 |
| 1.6 | 0 | ptsI | 2 | 0 | 0.22 |
| 1.6 | 0 | clcB | 2 | 0 | 0.204 |
| 1.6 | 0 | ycaM | 2 | 0 | 0.227 |
| 1.6 | 0 | yadI | 2 | 0 | 0.216 |
| 1.6 | 0 | AG1 | 2 | 0 | 0.218 |
| 1.6 | 0 | ptsI | 2 | 0.25 | 0.217 |
| 1.6 | 0 | clcB | 2 | 0.25 | 0.2 |
| 1.6 | 0 | ycaM | 2 | 0.25 | 0.218 |
| 1.6 | 0 | yadI | 2 | 0.25 | 0.212 |
| 1.6 | 0 | AG1 | 2 | 0.25 | 0.211 |
| 1.6 | 0 | ptsI | 2 | 0.5 | 0.222 |
| 1.6 | 0 | clcB | 2 | 0.5 | 0.205 |
| 1.6 | 0 | ycaM | 2 | 0.5 | 0.218 |
| 1.6 | 0 | yadI | 2 | 0.5 | 0.213 |
| 1.6 | 0 | AG1 | 2 | 0.5 | 0.213 |
| 1.6 | 0 | ptsI | 2 | 0.75 | 0.224 |
| 1.6 | 0 | clcB | 2 | 0.75 | 0.201 |

| | | | | | |
|---|---|---|---|---|---|
| 1.6 | 0 | ycaM | 2 | 0.75 | 0.221 |
| 1.6 | 0 | yadI | 2 | 0.75 | 0.216 |
| 1.6 | 0 | AG1 | 2 | 0.75 | 0.216 |
| 1.6 | 0 | ptsI | 2 | 1 | 0.228 |
| 1.6 | 0 | clcB | 2 | 1 | 0.205 |
| 1.6 | 0 | ycaM | 2 | 1 | 0.226 |
| 1.6 | 0 | yadI | 2 | 1 | 0.223 |
| 1.6 | 0 | AG1 | 2 | 1 | 0.221 |
| 1.6 | 0 | ptsI | 2 | 1.25 | 0.229 |
| 1.6 | 0 | clcB | 2 | 1.25 | 0.204 |
| 1.6 | 0 | ycaM | 2 | 1.25 | 0.228 |
| 1.6 | 0 | yadI | 2 | 1.25 | 0.232 |
| 1.6 | 0 | AG1 | 2 | 1.25 | 0.224 |
| 1.6 | 0 | ptsI | 2 | 1.5 | 0.231 |
| 1.6 | 0 | clcB | 2 | 1.5 | 0.209 |
| 1.6 | 0 | ycaM | 2 | 1.5 | 0.231 |
| 1.6 | 0 | yadI | 2 | 1.5 | 0.238 |
| 1.6 | 0 | AG1 | 2 | 1.5 | 0.228 |
| 1.6 | 0 | ptsI | 2 | 1.75 | 0.238 |
| 1.6 | 0 | clcB | 2 | 1.75 | 0.21 |
| 1.6 | 0 | ycaM | 2 | 1.75 | 0.236 |
| 1.6 | 0 | yadI | 2 | 1.75 | 0.243 |
| 1.6 | 0 | AG1 | 2 | 1.75 | 0.234 |
| 1.6 | 0 | ptsI | 2 | 2 | 0.244 |
| 1.6 | 0 | clcB | 2 | 2 | 0.212 |
| 1.6 | 0 | ycaM | 2 | 2 | 0.243 |
| 1.6 | 0 | yadI | 2 | 2 | 0.251 |
| 1.6 | 0 | AG1 | 2 | 2 | 0.245 |
| 1.6 | 0 | ptsI | 2 | 2.25 | 0.252 |
| 1.6 | 0 | clcB | 2 | 2.25 | 0.221 |
| 1.6 | 0 | ycaM | 2 | 2.25 | 0.249 |
| 1.6 | 0 | yadI | 2 | 2.25 | 0.262 |
| 1.6 | 0 | AG1 | 2 | 2.25 | 0.255 |
| 1.6 | 0 | ptsI | 2 | 2.5 | 0.263 |
| 1.6 | 0 | clcB | 2 | 2.5 | 0.222 |
| 1.6 | 0 | ycaM | 2 | 2.5 | 0.258 |
| 1.6 | 0 | yadI | 2 | 2.5 | 0.277 |
| 1.6 | 0 | AG1 | 2 | 2.5 | 0.268 |
| 1.6 | 0 | ptsI | 2 | 2.75 | 0.266 |
| 1.6 | 0 | clcB | 2 | 2.75 | 0.23 |
| 1.6 | 0 | ycaM | 2 | 2.75 | 0.274 |
| 1.6 | 0 | yadI | 2 | 2.75 | 0.306 |
| 1.6 | 0 | AG1 | 2 | 2.75 | 0.279 |
| 1.6 | 0 | ptsI | 2 | 3 | 0.274 |
| 1.6 | 0 | clcB | 2 | 3 | 0.237 |
| 1.6 | 0 | ycaM | 2 | 3 | 0.267 |
| 1.6 | 0 | yadI | 2 | 3 | 0.302 |
| 1.6 | 0 | AG1 | 2 | 3 | 0.29 |

| | | | | | |
|---|---|---|---|---|---|
| 1.6 | 0 | ptsI | 2 | 3.25 | 0.286 |
| 1.6 | 0 | clcB | 2 | 3.25 | 0.243 |
| 1.6 | 0 | ycaM | 2 | 3.25 | 0.279 |
| 1.6 | 0 | yadI | 2 | 3.25 | 0.326 |
| 1.6 | 0 | AG1 | 2 | 3.25 | 0.302 |
| 1.6 | 0 | ptsI | 2 | 3.5 | 0.294 |
| 1.6 | 0 | clcB | 2 | 3.5 | 0.251 |
| 1.6 | 0 | ycaM | 2 | 3.5 | 0.288 |
| 1.6 | 0 | yadI | 2 | 3.5 | 0.344 |
| 1.6 | 0 | AG1 | 2 | 3.5 | 0.314 |
| 1.6 | 0 | ptsI | 2 | 3.75 | 0.306 |
| 1.6 | 0 | clcB | 2 | 3.75 | 0.257 |
| 1.6 | 0 | ycaM | 2 | 3.75 | 0.298 |
| 1.6 | 0 | yadI | 2 | 3.75 | 0.361 |
| 1.6 | 0 | AG1 | 2 | 3.75 | 0.327 |
| 1.6 | 0 | ptsI | 2 | 4 | 0.316 |
| 1.6 | 0 | clcB | 2 | 4 | 0.261 |
| 1.6 | 0 | ycaM | 2 | 4 | 0.306 |
| 1.6 | 0 | yadI | 2 | 4 | 0.383 |
| 1.6 | 0 | AG1 | 2 | 4 | 0.335 |
| 1.6 | 0 | ptsI | 2 | 4.38333 | 0.318 |
| 1.6 | 0 | clcB | 2 | 4.38333 | 0.267 |
| 1.6 | 0 | ycaM | 2 | 4.38333 | 0.342 |
| 1.6 | 0 | yadI | 2 | 4.38333 | 0.396 |
| 1.6 | 0 | AG1 | 2 | 4.38333 | 0.298 |
| 1.6 | 0 | ptsI | 2 | 4.63333 | 0.316 |
| 1.6 | 0 | clcB | 2 | 4.63333 | 0.264 |
| 1.6 | 0 | ycaM | 2 | 4.63333 | 0.334 |
| 1.6 | 0 | yadI | 2 | 4.63333 | 0.394 |
| 1.6 | 0 | AG1 | 2 | 4.63333 | 0.291 |
| 1.6 | 0 | ptsI | 2 | 4.88333 | 0.323 |
| 1.6 | 0 | clcB | 2 | 4.88333 | 0.266 |
| 1.6 | 0 | ycaM | 2 | 4.88333 | 0.343 |
| 1.6 | 0 | yadI | 2 | 4.88333 | 0.407 |
| 1.6 | 0 | AG1 | 2 | 4.88333 | 0.297 |
| 1.6 | 0 | ptsI | 2 | 5.13333 | 0.334 |
| 1.6 | 0 | clcB | 2 | 5.13333 | 0.274 |
| 1.6 | 0 | ycaM | 2 | 5.13333 | 0.354 |
| 1.6 | 0 | yadI | 2 | 5.13333 | 0.418 |
| 1.6 | 0 | AG1 | 2 | 5.13333 | 0.304 |
| 1.6 | 0 | ptsI | 2 | 5.38333 | 0.341 |
| 1.6 | 0 | clcB | 2 | 5.38333 | 0.281 |
| 1.6 | 0 | ycaM | 2 | 5.38333 | 0.36 |
| 1.6 | 0 | yadI | 2 | 5.38333 | 0.428 |
| 1.6 | 0 | AG1 | 2 | 5.38333 | 0.309 |
| 1.6 | 0 | ptsI | 2 | 5.63333 | 0.348 |
| 1.6 | 0 | clcB | 2 | 5.63333 | 0.29 |
| 1.6 | 0 | ycaM | 2 | 5.63333 | 0.372 |

| | | | | | |
|---|---|---|---|---|---|
| 1.6 | 0 | yadI | 2 | 5.63333 | 0.438 |
| 1.6 | 0 | AG1 | 2 | 5.63333 | 0.318 |
| 1.6 | 0 | ptsI | 2 | 5.88333 | 0.358 |
| 1.6 | 0 | clcB | 2 | 5.88333 | 0.297 |
| 1.6 | 0 | ycaM | 2 | 5.88333 | 0.381 |
| 1.6 | 0 | yadI | 2 | 5.88333 | 0.447 |
| 1.6 | 0 | AG1 | 2 | 5.88333 | 0.329 |
| 1.6 | 0 | ptsI | 2 | 6.13333 | 0.364 |
| 1.6 | 0 | clcB | 2 | 6.13333 | 0.305 |
| 1.6 | 0 | ycaM | 2 | 6.13333 | 0.389 |
| 1.6 | 0 | yadI | 2 | 6.13333 | 0.458 |
| 1.6 | 0 | AG1 | 2 | 6.13333 | 0.339 |
| 1.6 | 0 | ptsI | 2 | 6.38333 | 0.371 |
| 1.6 | 0 | clcB | 2 | 6.38333 | 0.31 |
| 1.6 | 0 | ycaM | 2 | 6.38333 | 0.401 |
| 1.6 | 0 | yadI | 2 | 6.38333 | 0.468 |
| 1.6 | 0 | AG1 | 2 | 6.38333 | 0.349 |
| 1.6 | 0 | ptsI | 2 | 6.63333 | 0.378 |
| 1.6 | 0 | clcB | 2 | 6.63333 | 0.317 |
| 1.6 | 0 | ycaM | 2 | 6.63333 | 0.412 |
| 1.6 | 0 | yadI | 2 | 6.63333 | 0.478 |
| 1.6 | 0 | AG1 | 2 | 6.63333 | 0.356 |
| 1.6 | 0 | ptsI | 2 | 6.88333 | 0.388 |
| 1.6 | 0 | clcB | 2 | 6.88333 | 0.324 |
| 1.6 | 0 | ycaM | 2 | 6.88333 | 0.418 |
| 1.6 | 0 | yadI | 2 | 6.88333 | 0.487 |
| 1.6 | 0 | AG1 | 2 | 6.88333 | 0.364 |
| 1.6 | 0 | ptsI | 2 | 7.13333 | 0.397 |
| 1.6 | 0 | clcB | 2 | 7.13333 | 0.332 |
| 1.6 | 0 | ycaM | 2 | 7.13333 | 0.426 |
| 1.6 | 0 | yadI | 2 | 7.13333 | 0.498 |
| 1.6 | 0 | AG1 | 2 | 7.13333 | 0.372 |
| 1.6 | 0 | ptsI | 2 | 7.38333 | 0.404 |
| 1.6 | 0 | clcB | 2 | 7.38333 | 0.34 |
| 1.6 | 0 | ycaM | 2 | 7.38333 | 0.435 |
| 1.6 | 0 | yadI | 2 | 7.38333 | 0.507 |
| 1.6 | 0 | AG1 | 2 | 7.38333 | 0.382 |
| 1.6 | 0 | ptsI | 2 | 7.63333 | 0.412 |
| 1.6 | 0 | clcB | 2 | 7.63333 | 0.347 |
| 1.6 | 0 | ycaM | 2 | 7.63333 | 0.443 |
| 1.6 | 0 | yadI | 2 | 7.63333 | 0.519 |
| 1.6 | 0 | AG1 | 2 | 7.63333 | 0.388 |
| 1.6 | 0 | ptsI | 2 | 7.88333 | 0.423 |
| 1.6 | 0 | clcB | 2 | 7.88333 | 0.354 |
| 1.6 | 0 | ycaM | 2 | 7.88333 | 0.451 |
| 1.6 | 0 | yadI | 2 | 7.88333 | 0.532 |
| 1.6 | 0 | AG1 | 2 | 7.88333 | 0.395 |
| 1.6 | 0 | ptsI | 2 | 8.13333 | 0.429 |

| | | | | | |
|---|---|---|---|---|---|
| 1.6 | 0 | clcB | 2 | 8.13333 | 0.361 |
| 1.6 | 0 | ycaM | 2 | 8.13333 | 0.455 |
| 1.6 | 0 | yadI | 2 | 8.13333 | 0.541 |
| 1.6 | 0 | AG1 | 2 | 8.13333 | 0.403 |
| 1.6 | 0 | ptsI | 2 | 8.38333 | 0.437 |
| 1.6 | 0 | clcB | 2 | 8.38333 | 0.372 |
| 1.6 | 0 | ycaM | 2 | 8.38333 | 0.463 |
| 1.6 | 0 | yadI | 2 | 8.38333 | 0.552 |
| 1.6 | 0 | AG1 | 2 | 8.38333 | 0.409 |
| 1.6 | 0 | ptsI | 2 | 8.91667 | 0.358 |
| 1.6 | 0 | clcB | 2 | 8.91667 | 0.315 |
| 1.6 | 0 | ycaM | 2 | 8.91667 | 0.344 |
| 1.6 | 0 | yadI | 2 | 8.91667 | 0.617 |
| 1.6 | 0 | AG1 | 2 | 8.91667 | 0.456 |
| 1.6 | 0 | ptsI | 2 | 9.16667 | 0.36 |
| 1.6 | 0 | clcB | 2 | 9.16667 | 0.322 |
| 1.6 | 0 | ycaM | 2 | 9.16667 | 0.36 |
| 1.6 | 0 | yadI | 2 | 9.16667 | 0.629 |
| 1.6 | 0 | AG1 | 2 | 9.16667 | 0.46 |
| 1.6 | 0 | ptsI | 2 | 9.41667 | 0.368 |
| 1.6 | 0 | clcB | 2 | 9.41667 | 0.33 |
| 1.6 | 0 | ycaM | 2 | 9.41667 | 0.352 |
| 1.6 | 0 | yadI | 2 | 9.41667 | 0.64 |
| 1.6 | 0 | AG1 | 2 | 9.41667 | 0.479 |
| 1.6 | 0 | ptsI | 2 | 9.66667 | 0.376 |
| 1.6 | 0 | clcB | 2 | 9.66667 | 0.337 |
| 1.6 | 0 | ycaM | 2 | 9.66667 | 0.352 |
| 1.6 | 0 | yadI | 2 | 9.66667 | 0.657 |
| 1.6 | 0 | AG1 | 2 | 9.66667 | 0.494 |
| 1.6 | 0 | ptsI | 2 | 9.91667 | 0.38 |
| 1.6 | 0 | clcB | 2 | 9.91667 | 0.343 |
| 1.6 | 0 | ycaM | 2 | 9.91667 | 0.355 |
| 1.6 | 0 | yadI | 2 | 9.91667 | 0.666 |
| 1.6 | 0 | AG1 | 2 | 9.91667 | 0.503 |
| 1.6 | 0 | ptsI | 2 | 10.1667 | 0.388 |
| 1.6 | 0 | clcB | 2 | 10.1667 | 0.351 |
| 1.6 | 0 | ycaM | 2 | 10.1667 | 0.36 |
| 1.6 | 0 | yadI | 2 | 10.1667 | 0.673 |
| 1.6 | 0 | AG1 | 2 | 10.1667 | 0.506 |
| 1.6 | 0 | ptsI | 2 | 10.4167 | 0.396 |
| 1.6 | 0 | clcB | 2 | 10.4167 | 0.358 |
| 1.6 | 0 | ycaM | 2 | 10.4167 | 0.368 |
| 1.6 | 0 | yadI | 2 | 10.4167 | 0.687 |
| 1.6 | 0 | AG1 | 2 | 10.4167 | 0.518 |
| 1.6 | 0 | ptsI | 2 | 10.6667 | 0.407 |
| 1.6 | 0 | clcB | 2 | 10.6667 | 0.365 |
| 1.6 | 0 | ycaM | 2 | 10.6667 | 0.372 |
| 1.6 | 0 | yadI | 2 | 10.6667 | 0.695 |

| | | | | | |
|---|---|---|---|---|---|
| 1.6 | 0 | AG1 | 2 | 10.6667 | 0.523 |
| 1.6 | 0 | ptsI | 2 | 10.9167 | 0.41 |
| 1.6 | 0 | clcB | 2 | 10.9167 | 0.367 |
| 1.6 | 0 | ycaM | 2 | 10.9167 | 0.376 |
| 1.6 | 0 | yadI | 2 | 10.9167 | 0.694 |
| 1.6 | 0 | AG1 | 2 | 10.9167 | 0.53 |
| 1.6 | 0 | ptsI | 2 | 11.1667 | 0.421 |
| 1.6 | 0 | clcB | 2 | 11.1667 | 0.372 |
| 1.6 | 0 | ycaM | 2 | 11.1667 | 0.378 |
| 1.6 | 0 | yadI | 2 | 11.1667 | 0.703 |
| 1.6 | 0 | AG1 | 2 | 11.1667 | 0.538 |
| 1.6 | 0 | ptsI | 2 | 11.4167 | 0.439 |
| 1.6 | 0 | clcB | 2 | 11.4167 | 0.37 |
| 1.6 | 0 | ycaM | 2 | 11.4167 | 0.382 |
| 1.6 | 0 | yadI | 2 | 11.4167 | 0.711 |
| 1.6 | 0 | AG1 | 2 | 11.4167 | 0.551 |
| 1.6 | 0 | ptsI | 2 | 11.6667 | 0.434 |
| 1.6 | 0 | clcB | 2 | 11.6667 | 0.37 |
| 1.6 | 0 | ycaM | 2 | 11.6667 | 0.382 |
| 1.6 | 0 | yadI | 2 | 11.6667 | 0.722 |
| 1.6 | 0 | AG1 | 2 | 11.6667 | 0.54 |
| 1.6 | 0 | ptsI | 2 | 11.9167 | 0.444 |
| 1.6 | 0 | clcB | 2 | 11.9167 | 0.375 |
| 1.6 | 0 | ycaM | 2 | 11.9167 | 0.389 |
| 1.6 | 0 | yadI | 2 | 11.9167 | 0.73 |
| 1.6 | 0 | AG1 | 2 | 11.9167 | 0.555 |
| 1.6 | 0 | ptsI | 2 | 12.1667 | 0.439 |
| 1.6 | 0 | clcB | 2 | 12.1667 | 0.378 |
| 1.6 | 0 | ycaM | 2 | 12.1667 | 0.39 |
| 1.6 | 0 | yadI | 2 | 12.1667 | 0.736 |
| 1.6 | 0 | AG1 | 2 | 12.1667 | 0.546 |
| 1.6 | 0 | ptsI | 2 | 12.4167 | 0.446 |
| 1.6 | 0 | clcB | 2 | 12.4167 | 0.377 |
| 1.6 | 0 | ycaM | 2 | 12.4167 | 0.393 |
| 1.6 | 0 | yadI | 2 | 12.4167 | 0.742 |
| 1.6 | 0 | AG1 | 2 | 12.4167 | 0.544 |
| 1.6 | 0 | ptsI | 2 | 12.6667 | 0.457 |
| 1.6 | 0 | clcB | 2 | 12.6667 | 0.381 |
| 1.6 | 0 | ycaM | 2 | 12.6667 | 0.394 |
| 1.6 | 0 | yadI | 2 | 12.6667 | 0.746 |
| 1.6 | 0 | AG1 | 2 | 12.6667 | 0.549 |
| 1.6 | 0 | ptsI | 2 | 12.9167 | 0.455 |
| 1.6 | 0 | clcB | 2 | 12.9167 | 0.38 |
| 1.6 | 0 | ycaM | 2 | 12.9167 | 0.399 |
| 1.6 | 0 | yadI | 2 | 12.9167 | 0.756 |
| 1.6 | 0 | AG1 | 2 | 12.9167 | 0.549 |
| 1.6 | 0 | ptsI | 2 | 13.1667 | 0.453 |
| 1.6 | 0 | clcB | 2 | 13.1667 | 0.383 |

| | | | | | |
|---|---|---|---|---|---|
| 1.6 | 0 | ycaM | 2 | 13.1667 | 0.399 |
| 1.6 | 0 | yadI | 2 | 13.1667 | 0.761 |
| 1.6 | 0 | AG1 | 2 | 13.1667 | 0.543 |
| 1.6 | 0 | ptsI | 2 | 13.4167 | 0.458 |
| 1.6 | 0 | clcB | 2 | 13.4167 | 0.387 |
| 1.6 | 0 | ycaM | 2 | 13.4167 | 0.404 |
| 1.6 | 0 | yadI | 2 | 13.4167 | 0.765 |
| 1.6 | 0 | AG1 | 2 | 13.4167 | 0.542 |
| 1.6 | 0 | ptsI | 2 | 13.6667 | 0.468 |
| 1.6 | 0 | clcB | 2 | 13.6667 | 0.39 |
| 1.6 | 0 | ycaM | 2 | 13.6667 | 0.402 |
| 1.6 | 0 | yadI | 2 | 13.6667 | 0.773 |
| 1.6 | 0 | AG1 | 2 | 13.6667 | 0.546 |
| 1.6 | 0 | ptsI | 2 | 13.9167 | 0.469 |
| 1.6 | 0 | clcB | 2 | 13.9167 | 0.399 |
| 1.6 | 0 | ycaM | 2 | 13.9167 | 0.404 |
| 1.6 | 0 | yadI | 2 | 13.9167 | 0.775 |
| 1.6 | 0 | AG1 | 2 | 13.9167 | 0.548 |
| 1.6 | 0 | ptsI | 2 | 14.1667 | 0.48 |
| 1.6 | 0 | clcB | 2 | 14.1667 | 0.4 |
| 1.6 | 0 | ycaM | 2 | 14.1667 | 0.408 |
| 1.6 | 0 | yadI | 2 | 14.1667 | 0.781 |
| 1.6 | 0 | AG1 | 2 | 14.1667 | 0.55 |
| 1.6 | 0 | ptsI | 2 | 14.4167 | 0.481 |
| 1.6 | 0 | clcB | 2 | 14.4167 | 0.401 |
| 1.6 | 0 | ycaM | 2 | 14.4167 | 0.401 |
| 1.6 | 0 | yadI | 2 | 14.4167 | 0.786 |
| 1.6 | 0 | AG1 | 2 | 14.4167 | 0.555 |
| 1.6 | 0 | ptsI | 2 | 14.6667 | 0.483 |
| 1.6 | 0 | clcB | 2 | 14.6667 | 0.408 |
| 1.6 | 0 | ycaM | 2 | 14.6667 | 0.413 |
| 1.6 | 0 | yadI | 2 | 14.6667 | 0.789 |
| 1.6 | 0 | AG1 | 2 | 14.6667 | 0.556 |
| 1.6 | 0 | ptsI | 2 | 14.9167 | 0.496 |
| 1.6 | 0 | clcB | 2 | 14.9167 | 0.409 |
| 1.6 | 0 | ycaM | 2 | 14.9167 | 0.416 |
| 1.6 | 0 | yadI | 2 | 14.9167 | 0.794 |
| 1.6 | 0 | AG1 | 2 | 14.9167 | 0.561 |
| 1.6 | 0 | ptsI | 2 | 15.1667 | 0.506 |
| 1.6 | 0 | clcB | 2 | 15.1667 | 0.416 |
| 1.6 | 0 | ycaM | 2 | 15.1667 | 0.419 |
| 1.6 | 0 | yadI | 2 | 15.1667 | 0.806 |
| 1.6 | 0 | AG1 | 2 | 15.1667 | 0.578 |
| 1.6 | 0 | ptsI | 2 | 15.4167 | 0.517 |
| 1.6 | 0 | clcB | 2 | 15.4167 | 0.418 |
| 1.6 | 0 | ycaM | 2 | 15.4167 | 0.411 |
| 1.6 | 0 | yadI | 2 | 15.4167 | 0.806 |
| 1.6 | 0 | AG1 | 2 | 15.4167 | 0.564 |

| | | | | | |
|---|---|---|---|---|---|
| 1.6 | 0 | ptsI | 2 | 15.6667 | 0.53 |
| 1.6 | 0 | clcB | 2 | 15.6667 | 0.421 |
| 1.6 | 0 | ycaM | 2 | 15.6667 | 0.421 |
| 1.6 | 0 | yadI | 2 | 15.6667 | 0.804 |
| 1.6 | 0 | AG1 | 2 | 15.6667 | 0.572 |
| 1.6 | 0 | ptsI | 2 | 15.9167 | 0.537 |
| 1.6 | 0 | clcB | 2 | 15.9167 | 0.427 |
| 1.6 | 0 | ycaM | 2 | 15.9167 | 0.435 |
| 1.6 | 0 | yadI | 2 | 15.9167 | 0.808 |
| 1.6 | 0 | AG1 | 2 | 15.9167 | 0.568 |
| 1.6 | 0 | ptsI | 2 | 16.1667 | 0.533 |
| 1.6 | 0 | clcB | 2 | 16.1667 | 0.432 |
| 1.6 | 0 | ycaM | 2 | 16.1667 | 0.428 |
| 1.6 | 0 | yadI | 2 | 16.1667 | 0.81 |
| 1.6 | 0 | AG1 | 2 | 16.1667 | 0.568 |
| 1.6 | 0 | ptsI | 2 | 16.4167 | 0.541 |
| 1.6 | 0 | clcB | 2 | 16.4167 | 0.438 |
| 1.6 | 0 | ycaM | 2 | 16.4167 | 0.431 |
| 1.6 | 0 | yadI | 2 | 16.4167 | 0.815 |
| 1.6 | 0 | AG1 | 2 | 16.4167 | 0.566 |
| 1.6 | 0 | ptsI | 2 | 16.6667 | 0.542 |
| 1.6 | 0 | clcB | 2 | 16.6667 | 0.449 |
| 1.6 | 0 | ycaM | 2 | 16.6667 | 0.436 |
| 1.6 | 0 | yadI | 2 | 16.6667 | 0.818 |
| 1.6 | 0 | AG1 | 2 | 16.6667 | 0.553 |
| 1.6 | 0 | ptsI | 2 | 16.9167 | 0.566 |
| 1.6 | 0 | clcB | 2 | 16.9167 | 0.451 |
| 1.6 | 0 | ycaM | 2 | 16.9167 | 0.429 |
| 1.6 | 0 | yadI | 2 | 16.9167 | 0.821 |
| 1.6 | 0 | AG1 | 2 | 16.9167 | 0.559 |
| 1.6 | 0 | ptsI | 2 | 17.1667 | 0.573 |
| 1.6 | 0 | clcB | 2 | 17.1667 | 0.467 |
| 1.6 | 0 | ycaM | 2 | 17.1667 | 0.436 |
| 1.6 | 0 | yadI | 2 | 17.1667 | 0.824 |
| 1.6 | 0 | AG1 | 2 | 17.1667 | 0.591 |
| 1.6 | 0 | ptsI | 2 | 17.4167 | 0.56 |
| 1.6 | 0 | clcB | 2 | 17.4167 | 0.466 |
| 1.6 | 0 | ycaM | 2 | 17.4167 | 0.433 |
| 1.6 | 0 | yadI | 2 | 17.4167 | 0.823 |
| 1.6 | 0 | AG1 | 2 | 17.4167 | 0.586 |
| 1.6 | 0 | ptsI | 2 | 17.6667 | 0.563 |
| 1.6 | 0 | clcB | 2 | 17.6667 | 0.474 |
| 1.6 | 0 | ycaM | 2 | 17.6667 | 0.443 |
| 1.6 | 0 | yadI | 2 | 17.6667 | 0.822 |
| 1.6 | 0 | AG1 | 2 | 17.6667 | 0.572 |
| 1.6 | 0 | ptsI | 2 | 17.9167 | 0.554 |
| 1.6 | 0 | clcB | 2 | 17.9167 | 0.473 |
| 1.6 | 0 | ycaM | 2 | 17.9167 | 0.439 |

| | | | | | |
|---|---|---|---|---|---|
| 1.6 | 0 | yadI | 2 | 17.9167 | 0.824 |
| 1.6 | 0 | AG1 | 2 | 17.9167 | 0.579 |
| 1.6 | 0 | ptsI | 2 | 18.1667 | 0.571 |
| 1.6 | 0 | clcB | 2 | 18.1667 | 0.47 |
| 1.6 | 0 | ycaM | 2 | 18.1667 | 0.443 |
| 1.6 | 0 | yadI | 2 | 18.1667 | 0.822 |
| 1.6 | 0 | AG1 | 2 | 18.1667 | 0.582 |
| 1.6 | 0 | ptsI | 2 | 18.4167 | 0.567 |
| 1.6 | 0 | clcB | 2 | 18.4167 | 0.471 |
| 1.6 | 0 | ycaM | 2 | 18.4167 | 0.442 |
| 1.6 | 0 | yadI | 2 | 18.4167 | 0.824 |
| 1.6 | 0 | AG1 | 2 | 18.4167 | 0.57 |
| 1.6 | 0 | ptsI | 2 | 18.6667 | 0.568 |
| 1.6 | 0 | clcB | 2 | 18.6667 | 0.484 |
| 1.6 | 0 | ycaM | 2 | 18.6667 | 0.453 |
| 1.6 | 0 | yadI | 2 | 18.6667 | 0.825 |
| 1.6 | 0 | AG1 | 2 | 18.6667 | 0.566 |
| 1.6 | 0 | ptsI | 2 | 18.9167 | 0.555 |
| 1.6 | 0 | clcB | 2 | 18.9167 | 0.472 |
| 1.6 | 0 | ycaM | 2 | 18.9167 | 0.449 |
| 1.6 | 0 | yadI | 2 | 18.9167 | 0.811 |
| 1.6 | 0 | AG1 | 2 | 18.9167 | 0.561 |
| 1.6 | 0 | ptsI | 2 | 19.1667 | 0.567 |
| 1.6 | 0 | clcB | 2 | 19.1667 | 0.477 |
| 1.6 | 0 | ycaM | 2 | 19.1667 | 0.449 |
| 1.6 | 0 | yadI | 2 | 19.1667 | 0.822 |
| 1.6 | 0 | AG1 | 2 | 19.1667 | 0.561 |
| 1.6 | 0 | ptsI | 2 | 19.4167 | 0.572 |
| 1.6 | 0 | clcB | 2 | 19.4167 | 0.468 |
| 1.6 | 0 | ycaM | 2 | 19.4167 | 0.458 |
| 1.6 | 0 | yadI | 2 | 19.4167 | 0.825 |
| 1.6 | 0 | AG1 | 2 | 19.4167 | 0.566 |
| 1.6 | 0 | ptsI | 2 | 19.6667 | 0.57 |
| 1.6 | 0 | clcB | 2 | 19.6667 | 0.476 |
| 1.6 | 0 | ycaM | 2 | 19.6667 | 0.45 |
| 1.6 | 0 | yadI | 2 | 19.6667 | 0.83 |
| 1.6 | 0 | AG1 | 2 | 19.6667 | 0.562 |
| 1.6 | 0 | ptsI | 2 | 19.9167 | 0.572 |
| 1.6 | 0 | clcB | 2 | 19.9167 | 0.455 |
| 1.6 | 0 | ycaM | 2 | 19.9167 | 0.461 |
| 1.6 | 0 | yadI | 2 | 19.9167 | 0.827 |
| 1.6 | 0 | AG1 | 2 | 19.9167 | 0.577 |
| 1.6 | 0 | ptsI | 2 | 20.1667 | 0.581 |
| 1.6 | 0 | clcB | 2 | 20.1667 | 0.468 |
| 1.6 | 0 | ycaM | 2 | 20.1667 | 0.461 |
| 1.6 | 0 | yadI | 2 | 20.1667 | 0.827 |
| 1.6 | 0 | AG1 | 2 | 20.1667 | 0.575 |
| 1.6 | 0 | ptsI | 2 | 20.4167 | 0.583 |

| | | | | | |
|---|---|---|---|---|---|
| 1.6 | 0 | clcB | 2 | 20.4167 | 0.466 |
| 1.6 | 0 | ycaM | 2 | 20.4167 | 0.456 |
| 1.6 | 0 | yadI | 2 | 20.4167 | 0.83 |
| 1.6 | 0 | AG1 | 2 | 20.4167 | 0.577 |
| 1.6 | 0 | ptsI | 2 | 20.6667 | 0.584 |
| 1.6 | 0 | clcB | 2 | 20.6667 | 0.468 |
| 1.6 | 0 | ycaM | 2 | 20.6667 | 0.462 |
| 1.6 | 0 | yadI | 2 | 20.6667 | 0.832 |
| 1.6 | 0 | AG1 | 2 | 20.6667 | 0.571 |
| 1.6 | 0 | ptsI | 2 | 20.9167 | 0.595 |
| 1.6 | 0 | clcB | 2 | 20.9167 | 0.482 |
| 1.6 | 0 | ycaM | 2 | 20.9167 | 0.458 |
| 1.6 | 0 | yadI | 2 | 20.9167 | 0.833 |
| 1.6 | 0 | AG1 | 2 | 20.9167 | 0.571 |
| 1.6 | 0 | ptsI | 2 | 21.1667 | 0.597 |
| 1.6 | 0 | clcB | 2 | 21.1667 | 0.464 |
| 1.6 | 0 | ycaM | 2 | 21.1667 | 0.453 |
| 1.6 | 0 | yadI | 2 | 21.1667 | 0.829 |
| 1.6 | 0 | AG1 | 2 | 21.1667 | 0.57 |
| 1.6 | 0 | ptsI | 2 | 21.4167 | 0.602 |
| 1.6 | 0 | clcB | 2 | 21.4167 | 0.463 |
| 1.6 | 0 | ycaM | 2 | 21.4167 | 0.454 |
| 1.6 | 0 | yadI | 2 | 21.4167 | 0.826 |
| 1.6 | 0 | AG1 | 2 | 21.4167 | 0.574 |
| 1.6 | 0 | ptsI | 2 | 21.6667 | 0.608 |
| 1.6 | 0 | clcB | 2 | 21.6667 | 0.465 |
| 1.6 | 0 | ycaM | 2 | 21.6667 | 0.453 |
| 1.6 | 0 | yadI | 2 | 21.6667 | 0.828 |
| 1.6 | 0 | AG1 | 2 | 21.6667 | 0.565 |
| 1.6 | 0 | ptsI | 2 | 21.9167 | 0.619 |
| 1.6 | 0 | clcB | 2 | 21.9167 | 0.466 |
| 1.6 | 0 | ycaM | 2 | 21.9167 | 0.453 |
| 1.6 | 0 | yadI | 2 | 21.9167 | 0.827 |
| 1.6 | 0 | AG1 | 2 | 21.9167 | 0.572 |
| 1.6 | 0 | ptsI | 2 | 22.1667 | 0.618 |
| 1.6 | 0 | clcB | 2 | 22.1667 | 0.46 |
| 1.6 | 0 | ycaM | 2 | 22.1667 | 0.453 |
| 1.6 | 0 | yadI | 2 | 22.1667 | 0.829 |
| 1.6 | 0 | AG1 | 2 | 22.1667 | 0.573 |
| 1.6 | 0 | ptsI | 2 | 22.4167 | 0.622 |
| 1.6 | 0 | clcB | 2 | 22.4167 | 0.443 |
| 1.6 | 0 | ycaM | 2 | 22.4167 | 0.452 |
| 1.6 | 0 | yadI | 2 | 22.4167 | 0.829 |
| 1.6 | 0 | AG1 | 2 | 22.4167 | 0.564 |
| 1.6 | 0 | ptsI | 2 | 22.6667 | 0.626 |
| 1.6 | 0 | clcB | 2 | 22.6667 | 0.452 |
| 1.6 | 0 | ycaM | 2 | 22.6667 | 0.45 |
| 1.6 | 0 | yadI | 2 | 22.6667 | 0.829 |

| | | | | | |
|---|---|---|---|---|---|
| 1.6 | 0 | AG1 | 2 | 22.6667 | 0.565 |
| 1.6 | 0 | ptsI | 2 | 22.9167 | 0.634 |
| 1.6 | 0 | clcB | 2 | 22.9167 | 0.439 |
| 1.6 | 0 | ycaM | 2 | 22.9167 | 0.452 |
| 1.6 | 0 | yadI | 2 | 22.9167 | 0.833 |
| 1.6 | 0 | AG1 | 2 | 22.9167 | 0.565 |
| 1.6 | 0 | ptsI | 2 | 23.1667 | 0.639 |
| 1.6 | 0 | clcB | 2 | 23.1667 | 0.436 |
| 1.6 | 0 | ycaM | 2 | 23.1667 | 0.452 |
| 1.6 | 0 | yadI | 2 | 23.1667 | 0.831 |
| 1.6 | 0 | AG1 | 2 | 23.1667 | 0.567 |
| 1.6 | 0 | ptsI | 2 | 23.4167 | 0.644 |
| 1.6 | 0 | clcB | 2 | 23.4167 | 0.439 |
| 1.6 | 0 | ycaM | 2 | 23.4167 | 0.451 |
| 1.6 | 0 | yadI | 2 | 23.4167 | 0.832 |
| 1.6 | 0 | AG1 | 2 | 23.4167 | 0.566 |
| 1.6 | 0 | ptsI | 2 | 23.6667 | 0.649 |
| 1.6 | 0 | clcB | 2 | 23.6667 | 0.444 |
| 1.6 | 0 | ycaM | 2 | 23.6667 | 0.454 |
| 1.6 | 0 | yadI | 2 | 23.6667 | 0.832 |
| 1.6 | 0 | AG1 | 2 | 23.6667 | 0.566 |
| 1.6 | 0 | ptsI | 2 | 23.9167 | 0.654 |
| 1.6 | 0 | clcB | 2 | 23.9167 | 0.443 |
| 1.6 | 0 | ycaM | 2 | 23.9167 | 0.451 |
| 1.6 | 0 | yadI | 2 | 23.9167 | 0.833 |
| 1.6 | 0 | AG1 | 2 | 23.9167 | 0.563 |
| 1.6 | 0 | ptsI | 2 | 24.1667 | 0.663 |
| 1.6 | 0 | clcB | 2 | 24.1667 | 0.448 |
| 1.6 | 0 | ycaM | 2 | 24.1667 | 0.452 |
| 1.6 | 0 | yadI | 2 | 24.1667 | 0.83 |
| 1.6 | 0 | AG1 | 2 | 24.1667 | 0.561 |
| 1.6 | 0 | ptsI | 2 | 24.4167 | 0.675 |
| 1.6 | 0 | clcB | 2 | 24.4167 | 0.451 |
| 1.6 | 0 | ycaM | 2 | 24.4167 | 0.455 |
| 1.6 | 0 | yadI | 2 | 24.4167 | 0.837 |
| 1.6 | 0 | AG1 | 2 | 24.4167 | 0.563 |
| 1.6 | 0 | ptsI | 2 | 24.6667 | 0.68 |
| 1.6 | 0 | clcB | 2 | 24.6667 | 0.455 |
| 1.6 | 0 | ycaM | 2 | 24.6667 | 0.453 |
| 1.6 | 0 | yadI | 2 | 24.6667 | 0.833 |
| 1.6 | 0 | AG1 | 2 | 24.6667 | 0.565 |
| 1.6 | 0 | ptsI | 2 | 24.9167 | 0.69 |
| 1.6 | 0 | clcB | 2 | 24.9167 | 0.453 |
| 1.6 | 0 | ycaM | 2 | 24.9167 | 0.457 |
| 1.6 | 0 | yadI | 2 | 24.9167 | 0.837 |
| 1.6 | 0 | AG1 | 2 | 24.9167 | 0.568 |
| 1.6 | 0 | ptsI | 2 | 25.1667 | 0.697 |
| 1.6 | 0 | clcB | 2 | 25.1667 | 0.454 |

| | | | | | |
|---|---|---|---|---|---|
| 1.6 | 0 | ycaM | 2 | 25.1667 | 0.456 |
| 1.6 | 0 | yadI | 2 | 25.1667 | 0.834 |
| 1.6 | 0 | AG1 | 2 | 25.1667 | 0.569 |
| 1.6 | 0 | ptsI | 2 | 25.4167 | 0.701 |
| 1.6 | 0 | clcB | 2 | 25.4167 | 0.459 |
| 1.6 | 0 | ycaM | 2 | 25.4167 | 0.458 |
| 1.6 | 0 | yadI | 2 | 25.4167 | 0.833 |
| 1.6 | 0 | AG1 | 2 | 25.4167 | 0.574 |
| 1.6 | 0 | ptsI | 2 | 25.6667 | 0.708 |
| 1.6 | 0 | clcB | 2 | 25.6667 | 0.456 |
| 1.6 | 0 | ycaM | 2 | 25.6667 | 0.457 |
| 1.6 | 0 | yadI | 2 | 25.6667 | 0.831 |
| 1.6 | 0 | AG1 | 2 | 25.6667 | 0.577 |
| 1.6 | 0 | ptsI | 2 | 25.9167 | 0.713 |
| 1.6 | 0 | clcB | 2 | 25.9167 | 0.459 |
| 1.6 | 0 | ycaM | 2 | 25.9167 | 0.458 |
| 1.6 | 0 | yadI | 2 | 25.9167 | 0.833 |
| 1.6 | 0 | AG1 | 2 | 25.9167 | 0.578 |
| 1.6 | 0 | ptsI | 2 | 26.1667 | 0.724 |
| 1.6 | 0 | clcB | 2 | 26.1667 | 0.462 |
| 1.6 | 0 | ycaM | 2 | 26.1667 | 0.462 |
| 1.6 | 0 | yadI | 2 | 26.1667 | 0.83 |
| 1.6 | 0 | AG1 | 2 | 26.1667 | 0.589 |
| 1.6 | 0 | ptsI | 2 | 26.4167 | 0.732 |
| 1.6 | 0 | clcB | 2 | 26.4167 | 0.462 |
| 1.6 | 0 | ycaM | 2 | 26.4167 | 0.463 |
| 1.6 | 0 | yadI | 2 | 26.4167 | 0.835 |
| 1.6 | 0 | AG1 | 2 | 26.4167 | 0.588 |
| 1.6 | 0 | ptsI | 3 | 0 | 0.15 |
| 1.6 | 0 | clcB | 3 | 0 | 0.155 |
| 1.6 | 0 | ycaM | 3 | 0 | 0.154 |
| 1.6 | 0 | yadI | 3 | 0 | 0.166 |
| 1.6 | 0 | AG1 | 3 | 0 | 0.167 |
| 1.6 | 0 | ptsI | 3 | 0.25 | 0.147 |
| 1.6 | 0 | clcB | 3 | 0.25 | 0.152 |
| 1.6 | 0 | ycaM | 3 | 0.25 | 0.15 |
| 1.6 | 0 | yadI | 3 | 0.25 | 0.163 |
| 1.6 | 0 | AG1 | 3 | 0.25 | 0.167 |
| 1.6 | 0 | ptsI | 3 | 0.5 | 0.148 |
| 1.6 | 0 | clcB | 3 | 0.5 | 0.151 |
| 1.6 | 0 | ycaM | 3 | 0.5 | 0.148 |
| 1.6 | 0 | yadI | 3 | 0.5 | 0.162 |
| 1.6 | 0 | AG1 | 3 | 0.5 | 0.161 |
| 1.6 | 0 | ptsI | 3 | 0.75 | 0.149 |
| 1.6 | 0 | clcB | 3 | 0.75 | 0.151 |
| 1.6 | 0 | ycaM | 3 | 0.75 | 0.148 |
| 1.6 | 0 | yadI | 3 | 0.75 | 0.163 |
| 1.6 | 0 | AG1 | 3 | 0.75 | 0.16 |

| | | | | | |
|---|---|---|---|---|---|
| 1.6 | 0 | ptsI | 3 | 1 | 0.162 |
| 1.6 | 0 | clcB | 3 | 1 | 0.152 |
| 1.6 | 0 | ycaM | 3 | 1 | 0.15 |
| 1.6 | 0 | yadI | 3 | 1 | 0.163 |
| 1.6 | 0 | AG1 | 3 | 1 | 0.162 |
| 1.6 | 0 | ptsI | 3 | 1.25 | 0.15 |
| 1.6 | 0 | clcB | 3 | 1.25 | 0.152 |
| 1.6 | 0 | ycaM | 3 | 1.25 | 0.15 |
| 1.6 | 0 | yadI | 3 | 1.25 | 0.163 |
| 1.6 | 0 | AG1 | 3 | 1.25 | 0.164 |
| 1.6 | 0 | ptsI | 3 | 1.5 | 0.151 |
| 1.6 | 0 | clcB | 3 | 1.5 | 0.153 |
| 1.6 | 0 | ycaM | 3 | 1.5 | 0.151 |
| 1.6 | 0 | yadI | 3 | 1.5 | 0.164 |
| 1.6 | 0 | AG1 | 3 | 1.5 | 0.166 |
| 1.6 | 0 | ptsI | 3 | 1.75 | 0.152 |
| 1.6 | 0 | clcB | 3 | 1.75 | 0.153 |
| 1.6 | 0 | ycaM | 3 | 1.75 | 0.151 |
| 1.6 | 0 | yadI | 3 | 1.75 | 0.164 |
| 1.6 | 0 | AG1 | 3 | 1.75 | 0.165 |
| 1.6 | 0 | ptsI | 3 | 2 | 0.153 |
| 1.6 | 0 | clcB | 3 | 2 | 0.156 |
| 1.6 | 0 | ycaM | 3 | 2 | 0.153 |
| 1.6 | 0 | yadI | 3 | 2 | 0.165 |
| 1.6 | 0 | AG1 | 3 | 2 | 0.167 |
| 1.6 | 0 | ptsI | 3 | 2.25 | 0.155 |
| 1.6 | 0 | clcB | 3 | 2.25 | 0.157 |
| 1.6 | 0 | ycaM | 3 | 2.25 | 0.154 |
| 1.6 | 0 | yadI | 3 | 2.25 | 0.169 |
| 1.6 | 0 | AG1 | 3 | 2.25 | 0.169 |
| 1.6 | 0 | ptsI | 3 | 2.5 | 0.156 |
| 1.6 | 0 | clcB | 3 | 2.5 | 0.16 |
| 1.6 | 0 | ycaM | 3 | 2.5 | 0.157 |
| 1.6 | 0 | yadI | 3 | 2.5 | 0.169 |
| 1.6 | 0 | AG1 | 3 | 2.5 | 0.17 |
| 1.6 | 0 | ptsI | 3 | 2.75 | 0.159 |
| 1.6 | 0 | clcB | 3 | 2.75 | 0.163 |
| 1.6 | 0 | ycaM | 3 | 2.75 | 0.161 |
| 1.6 | 0 | yadI | 3 | 2.75 | 0.177 |
| 1.6 | 0 | AG1 | 3 | 2.75 | 0.176 |
| 1.6 | 0 | ptsI | 3 | 3 | 0.16 |
| 1.6 | 0 | clcB | 3 | 3 | 0.164 |
| 1.6 | 0 | ycaM | 3 | 3 | 0.159 |
| 1.6 | 0 | yadI | 3 | 3 | 0.178 |
| 1.6 | 0 | AG1 | 3 | 3 | 0.176 |
| 1.6 | 0 | ptsI | 3 | 3.25 | 0.162 |
| 1.6 | 0 | clcB | 3 | 3.25 | 0.165 |
| 1.6 | 0 | ycaM | 3 | 3.25 | 0.161 |

| | | | | | |
|---|---|---|---|---|---|
| 1.6 | 0 | yadI | 3 | 3.25 | 0.18 |
| 1.6 | 0 | AG1 | 3 | 3.25 | 0.178 |
| 1.6 | 0 | ptsI | 3 | 3.5 | 0.166 |
| 1.6 | 0 | clcB | 3 | 3.5 | 0.168 |
| 1.6 | 0 | ycaM | 3 | 3.5 | 0.166 |
| 1.6 | 0 | yadI | 3 | 3.5 | 0.187 |
| 1.6 | 0 | AG1 | 3 | 3.5 | 0.182 |
| 1.6 | 0 | ptsI | 3 | 3.75 | 0.167 |
| 1.6 | 0 | clcB | 3 | 3.75 | 0.172 |
| 1.6 | 0 | ycaM | 3 | 3.75 | 0.17 |
| 1.6 | 0 | yadI | 3 | 3.75 | 0.195 |
| 1.6 | 0 | AG1 | 3 | 3.75 | 0.186 |
| 1.6 | 0 | ptsI | 3 | 4 | 0.171 |
| 1.6 | 0 | clcB | 3 | 4 | 0.174 |
| 1.6 | 0 | ycaM | 3 | 4 | 0.173 |
| 1.6 | 0 | yadI | 3 | 4 | 0.205 |
| 1.6 | 0 | AG1 | 3 | 4 | 0.19 |
| 1.6 | 0 | ptsI | 3 | 4.41667 | 0.192 |
| 1.6 | 0 | clcB | 3 | 4.41667 | 0.164 |
| 1.6 | 0 | ycaM | 3 | 4.41667 | 0.184 |
| 1.6 | 0 | yadI | 3 | 4.41667 | 0.201 |
| 1.6 | 0 | AG1 | 3 | 4.41667 | 0.204 |
| 1.6 | 0 | ptsI | 3 | 4.66667 | 0.2 |
| 1.6 | 0 | clcB | 3 | 4.66667 | 0.164 |
| 1.6 | 0 | ycaM | 3 | 4.66667 | 0.18 |
| 1.6 | 0 | yadI | 3 | 4.66667 | 0.201 |
| 1.6 | 0 | AG1 | 3 | 4.66667 | 0.193 |
| 1.6 | 0 | ptsI | 3 | 4.91667 | 0.201 |
| 1.6 | 0 | clcB | 3 | 4.91667 | 0.163 |
| 1.6 | 0 | ycaM | 3 | 4.91667 | 0.184 |
| 1.6 | 0 | yadI | 3 | 4.91667 | 0.206 |
| 1.6 | 0 | AG1 | 3 | 4.91667 | 0.196 |
| 1.6 | 0 | ptsI | 3 | 5.16667 | 0.204 |
| 1.6 | 0 | clcB | 3 | 5.16667 | 0.164 |
| 1.6 | 0 | ycaM | 3 | 5.16667 | 0.189 |
| 1.6 | 0 | yadI | 3 | 5.16667 | 0.217 |
| 1.6 | 0 | AG1 | 3 | 5.16667 | 0.2 |
| 1.6 | 0 | ptsI | 3 | 5.41667 | 0.206 |
| 1.6 | 0 | clcB | 3 | 5.41667 | 0.168 |
| 1.6 | 0 | ycaM | 3 | 5.41667 | 0.194 |
| 1.6 | 0 | yadI | 3 | 5.41667 | 0.227 |
| 1.6 | 0 | AG1 | 3 | 5.41667 | 0.208 |
| 1.6 | 0 | ptsI | 3 | 5.66667 | 0.211 |
| 1.6 | 0 | clcB | 3 | 5.66667 | 0.171 |
| 1.6 | 0 | ycaM | 3 | 5.66667 | 0.202 |
| 1.6 | 0 | yadI | 3 | 5.66667 | 0.24 |
| 1.6 | 0 | AG1 | 3 | 5.66667 | 0.214 |
| 1.6 | 0 | ptsI | 3 | 5.91667 | 0.215 |

| | | | | | |
|---|---|---|---|---|---|
| 1.6 | 0 | clcB | 3 | 5.91667 | 0.175 |
| 1.6 | 0 | ycaM | 3 | 5.91667 | 0.209 |
| 1.6 | 0 | yadI | 3 | 5.91667 | 0.254 |
| 1.6 | 0 | AG1 | 3 | 5.91667 | 0.223 |
| 1.6 | 0 | ptsI | 3 | 6.16667 | 0.219 |
| 1.6 | 0 | clcB | 3 | 6.16667 | 0.181 |
| 1.6 | 0 | ycaM | 3 | 6.16667 | 0.219 |
| 1.6 | 0 | yadI | 3 | 6.16667 | 0.267 |
| 1.6 | 0 | AG1 | 3 | 6.16667 | 0.232 |
| 1.6 | 0 | ptsI | 3 | 6.41667 | 0.227 |
| 1.6 | 0 | clcB | 3 | 6.41667 | 0.188 |
| 1.6 | 0 | ycaM | 3 | 6.41667 | 0.231 |
| 1.6 | 0 | yadI | 3 | 6.41667 | 0.277 |
| 1.6 | 0 | AG1 | 3 | 6.41667 | 0.242 |
| 1.6 | 0 | ptsI | 3 | 6.66667 | 0.237 |
| 1.6 | 0 | clcB | 3 | 6.66667 | 0.192 |
| 1.6 | 0 | ycaM | 3 | 6.66667 | 0.243 |
| 1.6 | 0 | yadI | 3 | 6.66667 | 0.289 |
| 1.6 | 0 | AG1 | 3 | 6.66667 | 0.255 |
| 1.6 | 0 | ptsI | 3 | 6.91667 | 0.246 |
| 1.6 | 0 | clcB | 3 | 6.91667 | 0.198 |
| 1.6 | 0 | ycaM | 3 | 6.91667 | 0.257 |
| 1.6 | 0 | yadI | 3 | 6.91667 | 0.3 |
| 1.6 | 0 | AG1 | 3 | 6.91667 | 0.265 |
| 1.6 | 0 | ptsI | 3 | 7.16667 | 0.256 |
| 1.6 | 0 | clcB | 3 | 7.16667 | 0.206 |
| 1.6 | 0 | ycaM | 3 | 7.16667 | 0.27 |
| 1.6 | 0 | yadI | 3 | 7.16667 | 0.313 |
| 1.6 | 0 | AG1 | 3 | 7.16667 | 0.274 |
| 1.6 | 0 | ptsI | 3 | 7.41667 | 0.268 |
| 1.6 | 0 | clcB | 3 | 7.41667 | 0.214 |
| 1.6 | 0 | ycaM | 3 | 7.41667 | 0.278 |
| 1.6 | 0 | yadI | 3 | 7.41667 | 0.324 |
| 1.6 | 0 | AG1 | 3 | 7.41667 | 0.279 |
| 1.6 | 0 | ptsI | 3 | 7.66667 | 0.275 |
| 1.6 | 0 | clcB | 3 | 7.66667 | 0.224 |
| 1.6 | 0 | ycaM | 3 | 7.66667 | 0.286 |
| 1.6 | 0 | yadI | 3 | 7.66667 | 0.337 |
| 1.6 | 0 | AG1 | 3 | 7.66667 | 0.288 |
| 1.6 | 0 | ptsI | 3 | 7.91667 | 0.285 |
| 1.6 | 0 | clcB | 3 | 7.91667 | 0.235 |
| 1.6 | 0 | ycaM | 3 | 7.91667 | 0.294 |
| 1.6 | 0 | yadI | 3 | 7.91667 | 0.348 |
| 1.6 | 0 | AG1 | 3 | 7.91667 | 0.295 |
| 1.6 | 0 | ptsI | 3 | 8.16667 | 0.291 |
| 1.6 | 0 | clcB | 3 | 8.16667 | 0.245 |
| 1.6 | 0 | ycaM | 3 | 8.16667 | 0.303 |
| 1.6 | 0 | yadI | 3 | 8.16667 | 0.358 |

| | | | | | |
|---|---|---|---|---|---|
| 1.6 | 0 | AG1 | 3 | 8.16667 | 0.303 |
| 1.6 | 0 | ptsI | 3 | 8.41667 | 0.3 |
| 1.6 | 0 | clcB | 3 | 8.41667 | 0.252 |
| 1.6 | 0 | ycaM | 3 | 8.41667 | 0.311 |
| 1.6 | 0 | yadI | 3 | 8.41667 | 0.367 |
| 1.6 | 0 | AG1 | 3 | 8.41667 | 0.31 |
| 1.6 | 0 | ptsI | 3 | 9.05 | 0.261 |
| 1.6 | 0 | clcB | 3 | 9.05 | 0.247 |
| 1.6 | 0 | ycaM | 3 | 9.05 | 0.303 |
| 1.6 | 0 | yadI | 3 | 9.05 | 0.401 |
| 1.6 | 0 | AG1 | 3 | 9.05 | 0.342 |
| 1.6 | 0 | ptsI | 3 | 9.3 | 0.272 |
| 1.6 | 0 | clcB | 3 | 9.3 | 0.255 |
| 1.6 | 0 | ycaM | 3 | 9.3 | 0.312 |
| 1.6 | 0 | yadI | 3 | 9.3 | 0.414 |
| 1.6 | 0 | AG1 | 3 | 9.3 | 0.341 |
| 1.6 | 0 | ptsI | 3 | 9.55 | 0.284 |
| 1.6 | 0 | clcB | 3 | 9.55 | 0.261 |
| 1.6 | 0 | ycaM | 3 | 9.55 | 0.322 |
| 1.6 | 0 | yadI | 3 | 9.55 | 0.423 |
| 1.6 | 0 | AG1 | 3 | 9.55 | 0.355 |
| 1.6 | 0 | ptsI | 3 | 9.8 | 0.291 |
| 1.6 | 0 | clcB | 3 | 9.8 | 0.268 |
| 1.6 | 0 | ycaM | 3 | 9.8 | 0.329 |
| 1.6 | 0 | yadI | 3 | 9.8 | 0.434 |
| 1.6 | 0 | AG1 | 3 | 9.8 | 0.367 |
| 1.6 | 0 | ptsI | 3 | 10.05 | 0.302 |
| 1.6 | 0 | clcB | 3 | 10.05 | 0.273 |
| 1.6 | 0 | ycaM | 3 | 10.05 | 0.338 |
| 1.6 | 0 | yadI | 3 | 10.05 | 0.442 |
| 1.6 | 0 | AG1 | 3 | 10.05 | 0.376 |
| 1.6 | 0 | ptsI | 3 | 10.3 | 0.311 |
| 1.6 | 0 | clcB | 3 | 10.3 | 0.28 |
| 1.6 | 0 | ycaM | 3 | 10.3 | 0.345 |
| 1.6 | 0 | yadI | 3 | 10.3 | 0.451 |
| 1.6 | 0 | AG1 | 3 | 10.3 | 0.387 |
| 1.6 | 0 | ptsI | 3 | 10.55 | 0.321 |
| 1.6 | 0 | clcB | 3 | 10.55 | 0.288 |
| 1.6 | 0 | ycaM | 3 | 10.55 | 0.353 |
| 1.6 | 0 | yadI | 3 | 10.55 | 0.458 |
| 1.6 | 0 | AG1 | 3 | 10.55 | 0.397 |
| 1.6 | 0 | ptsI | 3 | 10.8 | 0.328 |
| 1.6 | 0 | clcB | 3 | 10.8 | 0.294 |
| 1.6 | 0 | ycaM | 3 | 10.8 | 0.36 |
| 1.6 | 0 | yadI | 3 | 10.8 | 0.468 |
| 1.6 | 0 | AG1 | 3 | 10.8 | 0.404 |
| 1.6 | 0 | ptsI | 3 | 11.05 | 0.339 |
| 1.6 | 0 | clcB | 3 | 11.05 | 0.301 |

| | | | | | |
|---|---|---|---|---|---|
| 1.6 | 0 | ycaM | 3 | 11.05 | 0.371 |
| 1.6 | 0 | yadI | 3 | 11.05 | 0.474 |
| 1.6 | 0 | AG1 | 3 | 11.05 | 0.414 |
| 1.6 | 0 | ptsI | 3 | 11.3 | 0.346 |
| 1.6 | 0 | clcB | 3 | 11.3 | 0.292 |
| 1.6 | 0 | ycaM | 3 | 11.3 | 0.369 |
| 1.6 | 0 | yadI | 3 | 11.3 | 0.478 |
| 1.6 | 0 | AG1 | 3 | 11.3 | 0.414 |
| 1.6 | 0 | ptsI | 3 | 11.55 | 0.356 |
| 1.6 | 0 | clcB | 3 | 11.55 | 0.3 |
| 1.6 | 0 | ycaM | 3 | 11.55 | 0.369 |
| 1.6 | 0 | yadI | 3 | 11.55 | 0.487 |
| 1.6 | 0 | AG1 | 3 | 11.55 | 0.419 |
| 1.6 | 0 | ptsI | 3 | 11.8 | 0.356 |
| 1.6 | 0 | clcB | 3 | 11.8 | 0.288 |
| 1.6 | 0 | ycaM | 3 | 11.8 | 0.378 |
| 1.6 | 0 | yadI | 3 | 11.8 | 0.495 |
| 1.6 | 0 | AG1 | 3 | 11.8 | 0.429 |
| 1.6 | 0 | ptsI | 3 | 12.05 | 0.349 |
| 1.6 | 0 | clcB | 3 | 12.05 | 0.304 |
| 1.6 | 0 | ycaM | 3 | 12.05 | 0.375 |
| 1.6 | 0 | yadI | 3 | 12.05 | 0.501 |
| 1.6 | 0 | AG1 | 3 | 12.05 | 0.429 |
| 1.6 | 0 | ptsI | 3 | 12.3 | 0.358 |
| 1.6 | 0 | clcB | 3 | 12.3 | 0.308 |
| 1.6 | 0 | ycaM | 3 | 12.3 | 0.376 |
| 1.6 | 0 | yadI | 3 | 12.3 | 0.506 |
| 1.6 | 0 | AG1 | 3 | 12.3 | 0.434 |
| 1.6 | 0 | ptsI | 3 | 12.55 | 0.361 |
| 1.6 | 0 | clcB | 3 | 12.55 | 0.292 |
| 1.6 | 0 | ycaM | 3 | 12.55 | 0.377 |
| 1.6 | 0 | yadI | 3 | 12.55 | 0.51 |
| 1.6 | 0 | AG1 | 3 | 12.55 | 0.448 |
| 1.6 | 0 | ptsI | 3 | 12.8 | 0.366 |
| 1.6 | 0 | clcB | 3 | 12.8 | 0.33 |
| 1.6 | 0 | ycaM | 3 | 12.8 | 0.376 |
| 1.6 | 0 | yadI | 3 | 12.8 | 0.515 |
| 1.6 | 0 | AG1 | 3 | 12.8 | 0.449 |
| 1.6 | 0 | ptsI | 3 | 13.05 | 0.389 |
| 1.6 | 0 | clcB | 3 | 13.05 | 0.317 |
| 1.6 | 0 | ycaM | 3 | 13.05 | 0.388 |
| 1.6 | 0 | yadI | 3 | 13.05 | 0.514 |
| 1.6 | 0 | AG1 | 3 | 13.05 | 0.452 |
| 1.6 | 0 | ptsI | 3 | 13.3 | 0.382 |
| 1.6 | 0 | clcB | 3 | 13.3 | 0.323 |
| 1.6 | 0 | ycaM | 3 | 13.3 | 0.392 |
| 1.6 | 0 | yadI | 3 | 13.3 | 0.516 |
| 1.6 | 0 | AG1 | 3 | 13.3 | 0.457 |

| | | | | | |
|---|---|---|---|---|---|
| 1.6 | 0 | ptsI | 3 | 13.55 | 0.382 |
| 1.6 | 0 | clcB | 3 | 13.55 | 0.321 |
| 1.6 | 0 | ycaM | 3 | 13.55 | 0.393 |
| 1.6 | 0 | yadI | 3 | 13.55 | 0.521 |
| 1.6 | 0 | AG1 | 3 | 13.55 | 0.463 |
| 1.6 | 0 | ptsI | 3 | 13.8 | 0.404 |
| 1.6 | 0 | clcB | 3 | 13.8 | 0.312 |
| 1.6 | 0 | ycaM | 3 | 13.8 | 0.398 |
| 1.6 | 0 | yadI | 3 | 13.8 | 0.523 |
| 1.6 | 0 | AG1 | 3 | 13.8 | 0.47 |
| 1.6 | 0 | ptsI | 3 | 14.05 | 0.405 |
| 1.6 | 0 | clcB | 3 | 14.05 | 0.315 |
| 1.6 | 0 | ycaM | 3 | 14.05 | 0.4 |
| 1.6 | 0 | yadI | 3 | 14.05 | 0.53 |
| 1.6 | 0 | AG1 | 3 | 14.05 | 0.482 |
| 1.6 | 0 | ptsI | 3 | 14.3 | 0.418 |
| 1.6 | 0 | clcB | 3 | 14.3 | 0.326 |
| 1.6 | 0 | ycaM | 3 | 14.3 | 0.41 |
| 1.6 | 0 | yadI | 3 | 14.3 | 0.534 |
| 1.6 | 0 | AG1 | 3 | 14.3 | 0.485 |
| 1.6 | 0 | ptsI | 3 | 14.55 | 0.419 |
| 1.6 | 0 | clcB | 3 | 14.55 | 0.33 |
| 1.6 | 0 | ycaM | 3 | 14.55 | 0.416 |
| 1.6 | 0 | yadI | 3 | 14.55 | 0.539 |
| 1.6 | 0 | AG1 | 3 | 14.55 | 0.496 |
| 1.6 | 0 | ptsI | 3 | 14.8 | 0.424 |
| 1.6 | 0 | clcB | 3 | 14.8 | 0.502 |
| 1.6 | 0 | ycaM | 3 | 14.8 | 0.422 |
| 1.6 | 0 | yadI | 3 | 14.8 | 0.543 |
| 1.6 | 0 | AG1 | 3 | 14.8 | 0.5 |
| 1.6 | 0 | ptsI | 3 | 15.05 | 0.428 |
| 1.6 | 0 | clcB | 3 | 15.05 | 0.337 |
| 1.6 | 0 | ycaM | 3 | 15.05 | 0.427 |
| 1.6 | 0 | yadI | 3 | 15.05 | 0.548 |
| 1.6 | 0 | AG1 | 3 | 15.05 | 0.504 |
| 1.6 | 0 | ptsI | 3 | 15.3 | 0.443 |
| 1.6 | 0 | clcB | 3 | 15.3 | 0.353 |
| 1.6 | 0 | ycaM | 3 | 15.3 | 0.434 |
| 1.6 | 0 | yadI | 3 | 15.3 | 0.551 |
| 1.6 | 0 | AG1 | 3 | 15.3 | 0.513 |
| 1.6 | 0 | ptsI | 3 | 15.55 | 0.446 |
| 1.6 | 0 | clcB | 3 | 15.55 | 0.357 |
| 1.6 | 0 | ycaM | 3 | 15.55 | 0.439 |
| 1.6 | 0 | yadI | 3 | 15.55 | 0.554 |
| 1.6 | 0 | AG1 | 3 | 15.55 | 0.519 |
| 1.6 | 0 | ptsI | 3 | 15.8 | 0.454 |
| 1.6 | 0 | clcB | 3 | 15.8 | 0.359 |
| 1.6 | 0 | ycaM | 3 | 15.8 | 0.446 |

| | | | | | |
|---|---|---|---|---|---|
| 1.6 | 0 | yadI | 3 | 15.8 | 0.557 |
| 1.6 | 0 | AG1 | 3 | 15.8 | 0.529 |
| 1.6 | 0 | ptsI | 3 | 16.05 | 0.466 |
| 1.6 | 0 | clcB | 3 | 16.05 | 0.353 |
| 1.6 | 0 | ycaM | 3 | 16.05 | 0.452 |
| 1.6 | 0 | yadI | 3 | 16.05 | 0.563 |
| 1.6 | 0 | AG1 | 3 | 16.05 | 0.539 |
| 1.6 | 0 | ptsI | 3 | 16.3 | 0.471 |
| 1.6 | 0 | clcB | 3 | 16.3 | 0.356 |
| 1.6 | 0 | ycaM | 3 | 16.3 | 0.456 |
| 1.6 | 0 | yadI | 3 | 16.3 | 0.557 |
| 1.6 | 0 | AG1 | 3 | 16.3 | 0.553 |
| 1.6 | 0 | ptsI | 3 | 16.55 | 0.475 |
| 1.6 | 0 | clcB | 3 | 16.55 | 0.365 |
| 1.6 | 0 | ycaM | 3 | 16.55 | 0.466 |
| 1.6 | 0 | yadI | 3 | 16.55 | 0.568 |
| 1.6 | 0 | AG1 | 3 | 16.55 | 0.556 |
| 1.6 | 0 | ptsI | 3 | 16.8 | 0.477 |
| 1.6 | 0 | clcB | 3 | 16.8 | 0.374 |
| 1.6 | 0 | ycaM | 3 | 16.8 | 0.472 |
| 1.6 | 0 | yadI | 3 | 16.8 | 0.569 |
| 1.6 | 0 | AG1 | 3 | 16.8 | 0.571 |
| 1.6 | 0 | ptsI | 3 | 17.05 | 0.486 |
| 1.6 | 0 | clcB | 3 | 17.05 | 0.378 |
| 1.6 | 0 | ycaM | 3 | 17.05 | 0.476 |
| 1.6 | 0 | yadI | 3 | 17.05 | 0.571 |
| 1.6 | 0 | AG1 | 3 | 17.05 | 0.581 |
| 1.6 | 0 | ptsI | 3 | 17.3 | 0.487 |
| 1.6 | 0 | clcB | 3 | 17.3 | 0.384 |
| 1.6 | 0 | ycaM | 3 | 17.3 | 0.482 |
| 1.6 | 0 | yadI | 3 | 17.3 | 0.571 |
| 1.6 | 0 | AG1 | 3 | 17.3 | 0.593 |
| 1.6 | 0 | ptsI | 3 | 17.55 | 0.49 |
| 1.6 | 0 | clcB | 3 | 17.55 | 0.388 |
| 1.6 | 0 | ycaM | 3 | 17.55 | 0.484 |
| 1.6 | 0 | yadI | 3 | 17.55 | 0.578 |
| 1.6 | 0 | AG1 | 3 | 17.55 | 0.589 |
| 1.6 | 0 | ptsI | 3 | 17.8 | 0.492 |
| 1.6 | 0 | clcB | 3 | 17.8 | 0.391 |
| 1.6 | 0 | ycaM | 3 | 17.8 | 0.495 |
| 1.6 | 0 | yadI | 3 | 17.8 | 0.58 |
| 1.6 | 0 | AG1 | 3 | 17.8 | 0.609 |
| 1.6 | 0 | ptsI | 3 | 18.05 | 0.496 |
| 1.6 | 0 | clcB | 3 | 18.05 | 0.403 |
| 1.6 | 0 | ycaM | 3 | 18.05 | 0.499 |
| 1.6 | 0 | yadI | 3 | 18.05 | 0.579 |
| 1.6 | 0 | AG1 | 3 | 18.05 | 0.608 |
| 1.6 | 0 | ptsI | 3 | 18.3 | 0.5 |

| | | | | | |
|---|---|---|---|---|---|
| 1.6 | 0 | clcB | 3 | 18.3 | 0.404 |
| 1.6 | 0 | ycaM | 3 | 18.3 | 0.502 |
| 1.6 | 0 | yadI | 3 | 18.3 | 0.581 |
| 1.6 | 0 | AG1 | 3 | 18.3 | 0.597 |
| 1.6 | 0 | ptsI | 3 | 18.55 | 0.505 |
| 1.6 | 0 | clcB | 3 | 18.55 | 0.409 |
| 1.6 | 0 | ycaM | 3 | 18.55 | 0.504 |
| 1.6 | 0 | yadI | 3 | 18.55 | 0.587 |
| 1.6 | 0 | AG1 | 3 | 18.55 | 0.598 |
| 1.6 | 0 | ptsI | 3 | 18.8 | 0.517 |
| 1.6 | 0 | clcB | 3 | 18.8 | 0.41 |
| 1.6 | 0 | ycaM | 3 | 18.8 | 0.51 |
| 1.6 | 0 | yadI | 3 | 18.8 | 0.589 |
| 1.6 | 0 | AG1 | 3 | 18.8 | 0.611 |
| 1.6 | 0 | ptsI | 3 | 19.05 | 0.519 |
| 1.6 | 0 | clcB | 3 | 19.05 | 0.414 |
| 1.6 | 0 | ycaM | 3 | 19.05 | 0.514 |
| 1.6 | 0 | yadI | 3 | 19.05 | 0.588 |
| 1.6 | 0 | AG1 | 3 | 19.05 | 0.604 |
| 1.6 | 0 | ptsI | 3 | 19.3 | 0.531 |
| 1.6 | 0 | clcB | 3 | 19.3 | 0.411 |
| 1.6 | 0 | ycaM | 3 | 19.3 | 0.518 |
| 1.6 | 0 | yadI | 3 | 19.3 | 0.59 |
| 1.6 | 0 | AG1 | 3 | 19.3 | 0.59 |
| 1.6 | 0 | ptsI | 3 | 19.55 | 0.542 |
| 1.6 | 0 | clcB | 3 | 19.55 | 0.421 |
| 1.6 | 0 | ycaM | 3 | 19.55 | 0.516 |
| 1.6 | 0 | yadI | 3 | 19.55 | 0.592 |
| 1.6 | 0 | AG1 | 3 | 19.55 | 0.609 |
| 1.6 | 0 | ptsI | 3 | 19.8 | 0.557 |
| 1.6 | 0 | clcB | 3 | 19.8 | 0.425 |
| 1.6 | 0 | ycaM | 3 | 19.8 | 0.516 |
| 1.6 | 0 | yadI | 3 | 19.8 | 0.593 |
| 1.6 | 0 | AG1 | 3 | 19.8 | 0.602 |
| 1.6 | 0 | ptsI | 3 | 20.05 | 0.565 |
| 1.6 | 0 | clcB | 3 | 20.05 | 0.414 |
| 1.6 | 0 | ycaM | 3 | 20.05 | 0.52 |
| 1.6 | 0 | yadI | 3 | 20.05 | 0.591 |
| 1.6 | 0 | AG1 | 3 | 20.05 | 0.598 |
| 1.6 | 0 | ptsI | 3 | 20.3 | 0.563 |
| 1.6 | 0 | clcB | 3 | 20.3 | 0.424 |
| 1.6 | 0 | ycaM | 3 | 20.3 | 0.526 |
| 1.6 | 0 | yadI | 3 | 20.3 | 0.591 |
| 1.6 | 0 | AG1 | 3 | 20.3 | 0.58 |
| 1.6 | 0 | ptsI | 3 | 20.55 | 0.564 |
| 1.6 | 0 | clcB | 3 | 20.55 | 0.427 |
| 1.6 | 0 | ycaM | 3 | 20.55 | 0.532 |
| 1.6 | 0 | yadI | 3 | 20.55 | 0.594 |

| | | | | | |
|---|---|---|---|---|---|
| 1.6 | 0 | AG1 | 3 | 20.55 | 0.581 |
| 1.6 | 0 | ptsI | 3 | 20.8 | 0.571 |
| 1.6 | 0 | clcB | 3 | 20.8 | 0.424 |
| 1.6 | 0 | ycaM | 3 | 20.8 | 0.533 |
| 1.6 | 0 | yadI | 3 | 20.8 | 0.594 |
| 1.6 | 0 | AG1 | 3 | 20.8 | 0.569 |
| 1.6 | 0 | ptsI | 3 | 21.05 | 0.57 |
| 1.6 | 0 | clcB | 3 | 21.05 | 0.434 |
| 1.6 | 0 | ycaM | 3 | 21.05 | 0.543 |
| 1.6 | 0 | yadI | 3 | 21.05 | 0.596 |
| 1.6 | 0 | AG1 | 3 | 21.05 | 0.561 |
| 1.6 | 0 | ptsI | 3 | 21.3 | 0.584 |
| 1.6 | 0 | clcB | 3 | 21.3 | 0.433 |
| 1.6 | 0 | ycaM | 3 | 21.3 | 0.538 |
| 1.6 | 0 | yadI | 3 | 21.3 | 0.594 |
| 1.6 | 0 | AG1 | 3 | 21.3 | 0.555 |
| 1.6 | 0 | ptsI | 3 | 21.55 | 0.576 |
| 1.6 | 0 | clcB | 3 | 21.55 | 0.44 |
| 1.6 | 0 | ycaM | 3 | 21.55 | 0.546 |
| 1.6 | 0 | yadI | 3 | 21.55 | 0.6 |
| 1.6 | 0 | AG1 | 3 | 21.55 | 0.565 |
| 1.6 | 0 | ptsI | 3 | 21.8 | 0.58 |
| 1.6 | 0 | clcB | 3 | 21.8 | 0.444 |
| 1.6 | 0 | ycaM | 3 | 21.8 | 0.551 |
| 1.6 | 0 | yadI | 3 | 21.8 | 0.6 |
| 1.6 | 0 | AG1 | 3 | 21.8 | 0.565 |
| 1.6 | 0 | ptsI | 3 | 22.05 | 0.574 |
| 1.6 | 0 | clcB | 3 | 22.05 | 0.45 |
| 1.6 | 0 | ycaM | 3 | 22.05 | 0.541 |
| 1.6 | 0 | yadI | 3 | 22.05 | 0.6 |
| 1.6 | 0 | AG1 | 3 | 22.05 | 0.564 |
| 1.6 | 0 | ptsI | 3 | 22.3 | 0.576 |
| 1.6 | 0 | clcB | 3 | 22.3 | 0.448 |
| 1.6 | 0 | ycaM | 3 | 22.3 | 0.555 |
| 1.6 | 0 | yadI | 3 | 22.3 | 0.601 |
| 1.6 | 0 | AG1 | 3 | 22.3 | 0.552 |
| 1.6 | 0 | ptsI | 3 | 22.55 | 0.58 |
| 1.6 | 0 | clcB | 3 | 22.55 | 0.461 |
| 1.6 | 0 | ycaM | 3 | 22.55 | 0.556 |
| 1.6 | 0 | yadI | 3 | 22.55 | 0.603 |
| 1.6 | 0 | AG1 | 3 | 22.55 | 0.56 |
| 1.6 | 0 | ptsI | 3 | 22.8 | 0.572 |
| 1.6 | 0 | clcB | 3 | 22.8 | 0.471 |
| 1.6 | 0 | ycaM | 3 | 22.8 | 0.555 |
| 1.6 | 0 | yadI | 3 | 22.8 | 0.602 |
| 1.6 | 0 | AG1 | 3 | 22.8 | 0.56 |
| 1.6 | 0 | ptsI | 3 | 23.05 | 0.57 |
| 1.6 | 0 | clcB | 3 | 23.05 | 0.464 |

| | | | | | |
|---|---|---|---|---|---|
| 1.6 | 0 | ycaM | 3 | 23.05 | 0.564 |
| 1.6 | 0 | yadI | 3 | 23.05 | 0.6 |
| 1.6 | 0 | AG1 | 3 | 23.05 | 0.576 |
| 1.6 | 0 | ptsI | 3 | 23.3 | 0.574 |
| 1.6 | 0 | clcB | 3 | 23.3 | 0.461 |
| 1.6 | 0 | ycaM | 3 | 23.3 | 0.57 |
| 1.6 | 0 | yadI | 3 | 23.3 | 0.602 |
| 1.6 | 0 | AG1 | 3 | 23.3 | 0.576 |
| 1.6 | 0 | ptsI | 3 | 23.55 | 0.564 |
| 1.6 | 0 | clcB | 3 | 23.55 | 0.455 |
| 1.6 | 0 | ycaM | 3 | 23.55 | 0.58 |
| 1.6 | 0 | yadI | 3 | 23.55 | 0.601 |
| 1.6 | 0 | AG1 | 3 | 23.55 | 0.559 |
| 1.6 | 0 | ptsI | 3 | 23.8 | 0.575 |
| 1.6 | 0 | clcB | 3 | 23.8 | 0.457 |
| 1.6 | 0 | ycaM | 3 | 23.8 | 0.591 |
| 1.6 | 0 | yadI | 3 | 23.8 | 0.602 |
| 1.6 | 0 | AG1 | 3 | 23.8 | 0.569 |
| 1.6 | 0 | ptsI | 3 | 24.05 | 0.585 |
| 1.6 | 0 | clcB | 3 | 24.05 | 0.461 |
| 1.6 | 0 | ycaM | 3 | 24.05 | 0.588 |
| 1.6 | 0 | yadI | 3 | 24.05 | 0.601 |
| 1.6 | 0 | AG1 | 3 | 24.05 | 0.592 |
| 1.6 | 0 | ptsI | 3 | 24.3 | 0.587 |
| 1.6 | 0 | clcB | 3 | 24.3 | 0.467 |
| 1.6 | 0 | ycaM | 3 | 24.3 | 0.59 |
| 1.6 | 0 | yadI | 3 | 24.3 | 0.598 |
| 1.6 | 0 | AG1 | 3 | 24.3 | 0.584 |
| 1.6 | 0 | ptsI | 4 | 0 | 0.192 |
| 1.6 | 0 | clcB | 4 | 0 | 0.2 |
| 1.6 | 0 | ycaM | 4 | 0 | 0.278 |
| 1.6 | 0 | yadI | 4 | 0 | 0.248 |
| 1.6 | 0 | AG1 | 4 | 0 | 0.233 |
| 1.6 | 0 | ptsI | 4 | 0.25 | 0.19 |
| 1.6 | 0 | clcB | 4 | 0.25 | 0.197 |
| 1.6 | 0 | ycaM | 4 | 0.25 | 0.271 |
| 1.6 | 0 | yadI | 4 | 0.25 | 0.247 |
| 1.6 | 0 | AG1 | 4 | 0.25 | 0.229 |
| 1.6 | 0 | ptsI | 4 | 0.5 | 0.192 |
| 1.6 | 0 | clcB | 4 | 0.5 | 0.197 |
| 1.6 | 0 | ycaM | 4 | 0.5 | 0.271 |
| 1.6 | 0 | yadI | 4 | 0.5 | 0.257 |
| 1.6 | 0 | AG1 | 4 | 0.5 | 0.232 |
| 1.6 | 0 | ptsI | 4 | 0.75 | 0.195 |
| 1.6 | 0 | clcB | 4 | 0.75 | 0.2 |
| 1.6 | 0 | ycaM | 4 | 0.75 | 0.274 |
| 1.6 | 0 | yadI | 4 | 0.75 | 0.255 |
| 1.6 | 0 | AG1 | 4 | 0.75 | 0.236 |

| | | | | | |
|---|---|---|---|---|---|
| 1.6 | 0 | ptsI | 4 | 1 | 0.201 |
| 1.6 | 0 | clcB | 4 | 1 | 0.201 |
| 1.6 | 0 | ycaM | 4 | 1 | 0.276 |
| 1.6 | 0 | yadI | 4 | 1 | 0.252 |
| 1.6 | 0 | AG1 | 4 | 1 | 0.241 |
| 1.6 | 0 | ptsI | 4 | 1.25 | 0.206 |
| 1.6 | 0 | clcB | 4 | 1.25 | 0.209 |
| 1.6 | 0 | ycaM | 4 | 1.25 | 0.28 |
| 1.6 | 0 | yadI | 4 | 1.25 | 0.258 |
| 1.6 | 0 | AG1 | 4 | 1.25 | 0.248 |
| 1.6 | 0 | ptsI | 4 | 1.5 | 0.205 |
| 1.6 | 0 | clcB | 4 | 1.5 | 0.208 |
| 1.6 | 0 | ycaM | 4 | 1.5 | 0.285 |
| 1.6 | 0 | yadI | 4 | 1.5 | 0.264 |
| 1.6 | 0 | AG1 | 4 | 1.5 | 0.243 |
| 1.6 | 0 | ptsI | 4 | 1.75 | 0.215 |
| 1.6 | 0 | clcB | 4 | 1.75 | 0.219 |
| 1.6 | 0 | ycaM | 4 | 1.75 | 0.292 |
| 1.6 | 0 | yadI | 4 | 1.75 | 0.272 |
| 1.6 | 0 | AG1 | 4 | 1.75 | 0.259 |
| 1.6 | 0 | ptsI | 4 | 2 | 0.227 |
| 1.6 | 0 | clcB | 4 | 2 | 0.232 |
| 1.6 | 0 | ycaM | 4 | 2 | 0.301 |
| 1.6 | 0 | yadI | 4 | 2 | 0.283 |
| 1.6 | 0 | AG1 | 4 | 2 | 0.272 |
| 1.6 | 0 | ptsI | 4 | 2.25 | 0.243 |
| 1.6 | 0 | clcB | 4 | 2.25 | 0.242 |
| 1.6 | 0 | ycaM | 4 | 2.25 | 0.312 |
| 1.6 | 0 | yadI | 4 | 2.25 | 0.292 |
| 1.6 | 0 | AG1 | 4 | 2.25 | 0.289 |
| 1.6 | 0 | ptsI | 4 | 2.5 | 0.24 |
| 1.6 | 0 | clcB | 4 | 2.5 | 0.247 |
| 1.6 | 0 | ycaM | 4 | 2.5 | 0.321 |
| 1.6 | 0 | yadI | 4 | 2.5 | 0.301 |
| 1.6 | 0 | AG1 | 4 | 2.5 | 0.292 |
| 1.6 | 0 | ptsI | 4 | 2.75 | 0.25 |
| 1.6 | 0 | clcB | 4 | 2.75 | 0.26 |
| 1.6 | 0 | ycaM | 4 | 2.75 | 0.327 |
| 1.6 | 0 | yadI | 4 | 2.75 | 0.312 |
| 1.6 | 0 | AG1 | 4 | 2.75 | 0.304 |
| 1.6 | 0 | ptsI | 4 | 3 | 0.264 |
| 1.6 | 0 | clcB | 4 | 3 | 0.275 |
| 1.6 | 0 | ycaM | 4 | 3 | 0.342 |
| 1.6 | 0 | yadI | 4 | 3 | 0.329 |
| 1.6 | 0 | AG1 | 4 | 3 | 0.32 |
| 1.6 | 0 | ptsI | 4 | 3.25 | 0.277 |
| 1.6 | 0 | clcB | 4 | 3.25 | 0.284 |
| 1.6 | 0 | ycaM | 4 | 3.25 | 0.354 |

| | | | | | |
|---|---|---|---|---|---|
| 1.6 | 0 | yadI | 4 | 3.25 | 0.349 |
| 1.6 | 0 | AG1 | 4 | 3.25 | 0.337 |
| 1.6 | 0 | ptsI | 4 | 3.5 | 0.297 |
| 1.6 | 0 | clcB | 4 | 3.5 | 0.3 |
| 1.6 | 0 | ycaM | 4 | 3.5 | 0.362 |
| 1.6 | 0 | yadI | 4 | 3.5 | 0.366 |
| 1.6 | 0 | AG1 | 4 | 3.5 | 0.356 |
| 1.6 | 0 | ptsI | 4 | 3.75 | 0.314 |
| 1.6 | 0 | clcB | 4 | 3.75 | 0.314 |
| 1.6 | 0 | ycaM | 4 | 3.75 | 0.368 |
| 1.6 | 0 | yadI | 4 | 3.75 | 0.385 |
| 1.6 | 0 | AG1 | 4 | 3.75 | 0.368 |
| 1.6 | 0 | ptsI | 4 | 4 | 0.331 |
| 1.6 | 0 | clcB | 4 | 4 | 0.33 |
| 1.6 | 0 | ycaM | 4 | 4 | 0.388 |
| 1.6 | 0 | yadI | 4 | 4 | 0.405 |
| 1.6 | 0 | AG1 | 4 | 4 | 0.39 |
| 1.6 | 0 | ptsI | 4 | 4.35 | 0.334 |
| 1.6 | 0 | clcB | 4 | 4.35 | 0.27 |
| 1.6 | 0 | ycaM | 4 | 4.35 | 0.392 |
| 1.6 | 0 | yadI | 4 | 4.35 | 0.39 |
| 1.6 | 0 | AG1 | 4 | 4.35 | 0.298 |
| 1.6 | 0 | ptsI | 4 | 4.6 | 0.343 |
| 1.6 | 0 | clcB | 4 | 4.6 | 0.273 |
| 1.6 | 0 | ycaM | 4 | 4.6 | 0.386 |
| 1.6 | 0 | yadI | 4 | 4.6 | 0.394 |
| 1.6 | 0 | AG1 | 4 | 4.6 | 0.307 |
| 1.6 | 0 | ptsI | 4 | 4.85 | 0.347 |
| 1.6 | 0 | clcB | 4 | 4.85 | 0.28 |
| 1.6 | 0 | ycaM | 4 | 4.85 | 0.398 |
| 1.6 | 0 | yadI | 4 | 4.85 | 0.408 |
| 1.6 | 0 | AG1 | 4 | 4.85 | 0.316 |
| 1.6 | 0 | ptsI | 4 | 5.1 | 0.353 |
| 1.6 | 0 | clcB | 4 | 5.1 | 0.288 |
| 1.6 | 0 | ycaM | 4 | 5.1 | 0.41 |
| 1.6 | 0 | yadI | 4 | 5.1 | 0.422 |
| 1.6 | 0 | AG1 | 4 | 5.1 | 0.325 |
| 1.6 | 0 | ptsI | 4 | 5.35 | 0.37 |
| 1.6 | 0 | clcB | 4 | 5.35 | 0.298 |
| 1.6 | 0 | ycaM | 4 | 5.35 | 0.42 |
| 1.6 | 0 | yadI | 4 | 5.35 | 0.435 |
| 1.6 | 0 | AG1 | 4 | 5.35 | 0.337 |
| 1.6 | 0 | ptsI | 4 | 5.6 | 0.377 |
| 1.6 | 0 | clcB | 4 | 5.6 | 0.307 |
| 1.6 | 0 | ycaM | 4 | 5.6 | 0.424 |
| 1.6 | 0 | yadI | 4 | 5.6 | 0.448 |
| 1.6 | 0 | AG1 | 4 | 5.6 | 0.346 |
| 1.6 | 0 | ptsI | 4 | 5.85 | 0.382 |

| | | | | | |
|---|---|---|---|---|---|
| 1.6 | 0 | clcB | 4 | 5.85 | 0.317 |
| 1.6 | 0 | ycaM | 4 | 5.85 | 0.431 |
| 1.6 | 0 | yadI | 4 | 5.85 | 0.464 |
| 1.6 | 0 | AG1 | 4 | 5.85 | 0.352 |
| 1.6 | 0 | ptsI | 4 | 6.1 | 0.396 |
| 1.6 | 0 | clcB | 4 | 6.1 | 0.327 |
| 1.6 | 0 | ycaM | 4 | 6.1 | 0.443 |
| 1.6 | 0 | yadI | 4 | 6.1 | 0.477 |
| 1.6 | 0 | AG1 | 4 | 6.1 | 0.361 |
| 1.6 | 0 | ptsI | 4 | 6.35 | 0.407 |
| 1.6 | 0 | clcB | 4 | 6.35 | 0.338 |
| 1.6 | 0 | ycaM | 4 | 6.35 | 0.452 |
| 1.6 | 0 | yadI | 4 | 6.35 | 0.491 |
| 1.6 | 0 | AG1 | 4 | 6.35 | 0.369 |
| 1.6 | 0 | ptsI | 4 | 6.6 | 0.419 |
| 1.6 | 0 | clcB | 4 | 6.6 | 0.347 |
| 1.6 | 0 | ycaM | 4 | 6.6 | 0.461 |
| 1.6 | 0 | yadI | 4 | 6.6 | 0.501 |
| 1.6 | 0 | AG1 | 4 | 6.6 | 0.378 |
| 1.6 | 0 | ptsI | 4 | 6.85 | 0.43 |
| 1.6 | 0 | clcB | 4 | 6.85 | 0.352 |
| 1.6 | 0 | ycaM | 4 | 6.85 | 0.468 |
| 1.6 | 0 | yadI | 4 | 6.85 | 0.512 |
| 1.6 | 0 | AG1 | 4 | 6.85 | 0.386 |
| 1.6 | 0 | ptsI | 4 | 7.1 | 0.438 |
| 1.6 | 0 | clcB | 4 | 7.1 | 0.361 |
| 1.6 | 0 | ycaM | 4 | 7.1 | 0.478 |
| 1.6 | 0 | yadI | 4 | 7.1 | 0.522 |
| 1.6 | 0 | AG1 | 4 | 7.1 | 0.396 |
| 1.6 | 0 | ptsI | 4 | 7.35 | 0.446 |
| 1.6 | 0 | clcB | 4 | 7.35 | 0.369 |
| 1.6 | 0 | ycaM | 4 | 7.35 | 0.482 |
| 1.6 | 0 | yadI | 4 | 7.35 | 0.532 |
| 1.6 | 0 | AG1 | 4 | 7.35 | 0.402 |
| 1.6 | 0 | ptsI | 4 | 7.6 | 0.455 |
| 1.6 | 0 | clcB | 4 | 7.6 | 0.377 |
| 1.6 | 0 | ycaM | 4 | 7.6 | 0.485 |
| 1.6 | 0 | yadI | 4 | 7.6 | 0.541 |
| 1.6 | 0 | AG1 | 4 | 7.6 | 0.406 |
| 1.6 | 0 | ptsI | 4 | 7.85 | 0.462 |
| 1.6 | 0 | clcB | 4 | 7.85 | 0.383 |
| 1.6 | 0 | ycaM | 4 | 7.85 | 0.486 |
| 1.6 | 0 | yadI | 4 | 7.85 | 0.545 |
| 1.6 | 0 | AG1 | 4 | 7.85 | 0.417 |
| 1.6 | 0 | ptsI | 4 | 8.1 | 0.473 |
| 1.6 | 0 | clcB | 4 | 8.1 | 0.391 |
| 1.6 | 0 | ycaM | 4 | 8.1 | 0.493 |
| 1.6 | 0 | yadI | 4 | 8.1 | 0.559 |

| | | | | | |
|---|---|---|---|---|---|
| 1.6 | 0 | AG1 | 4 | 8.1 | 0.426 |
| 1.6 | 0 | ptsI | 4 | 8.35 | 0.483 |
| 1.6 | 0 | clcB | 4 | 8.35 | 0.396 |
| 1.6 | 0 | ycaM | 4 | 8.35 | 0.495 |
| 1.6 | 0 | yadI | 4 | 8.35 | 0.566 |
| 1.6 | 0 | AG1 | 4 | 8.35 | 0.433 |
| 1.6 | 0 | ptsI | 4 | 8.83333 | 0.403 |
| 1.6 | 0 | clcB | 4 | 8.83333 | 0.388 |
| 1.6 | 0 | ycaM | 4 | 8.83333 | 0.434 |
| 1.6 | 0 | yadI | 4 | 8.83333 | 0.543 |
| 1.6 | 0 | AG1 | 4 | 8.83333 | 0.425 |
| 1.6 | 0 | ptsI | 4 | 9.08333 | 0.406 |
| 1.6 | 0 | clcB | 4 | 9.08333 | 0.389 |
| 1.6 | 0 | ycaM | 4 | 9.08333 | 0.429 |
| 1.6 | 0 | yadI | 4 | 9.08333 | 0.533 |
| 1.6 | 0 | AG1 | 4 | 9.08333 | 0.427 |
| 1.6 | 0 | ptsI | 4 | 9.33333 | 0.416 |
| 1.6 | 0 | clcB | 4 | 9.33333 | 0.399 |
| 1.6 | 0 | ycaM | 4 | 9.33333 | 0.431 |
| 1.6 | 0 | yadI | 4 | 9.33333 | 0.534 |
| 1.6 | 0 | AG1 | 4 | 9.33333 | 0.437 |
| 1.6 | 0 | ptsI | 4 | 9.58333 | 0.428 |
| 1.6 | 0 | clcB | 4 | 9.58333 | 0.404 |
| 1.6 | 0 | ycaM | 4 | 9.58333 | 0.434 |
| 1.6 | 0 | yadI | 4 | 9.58333 | 0.546 |
| 1.6 | 0 | AG1 | 4 | 9.58333 | 0.446 |
| 1.6 | 0 | ptsI | 4 | 9.83333 | 0.434 |
| 1.6 | 0 | clcB | 4 | 9.83333 | 0.409 |
| 1.6 | 0 | ycaM | 4 | 9.83333 | 0.442 |
| 1.6 | 0 | yadI | 4 | 9.83333 | 0.553 |
| 1.6 | 0 | AG1 | 4 | 9.83333 | 0.455 |
| 1.6 | 0 | ptsI | 4 | 10.0833 | 0.439 |
| 1.6 | 0 | clcB | 4 | 10.0833 | 0.412 |
| 1.6 | 0 | ycaM | 4 | 10.0833 | 0.447 |
| 1.6 | 0 | yadI | 4 | 10.0833 | 0.559 |
| 1.6 | 0 | AG1 | 4 | 10.0833 | 0.461 |
| 1.6 | 0 | ptsI | 4 | 10.3333 | 0.445 |
| 1.6 | 0 | clcB | 4 | 10.3333 | 0.414 |
| 1.6 | 0 | ycaM | 4 | 10.3333 | 0.45 |
| 1.6 | 0 | yadI | 4 | 10.3333 | 0.563 |
| 1.6 | 0 | AG1 | 4 | 10.3333 | 0.465 |
| 1.6 | 0 | ptsI | 4 | 10.5833 | 0.449 |
| 1.6 | 0 | clcB | 4 | 10.5833 | 0.415 |
| 1.6 | 0 | ycaM | 4 | 10.5833 | 0.452 |
| 1.6 | 0 | yadI | 4 | 10.5833 | 0.57 |
| 1.6 | 0 | AG1 | 4 | 10.5833 | 0.472 |
| 1.6 | 0 | ptsI | 4 | 10.8333 | 0.454 |
| 1.6 | 0 | clcB | 4 | 10.8333 | 0.422 |

| | | | | | |
|---|---|---|---|---|---|
| 1.6 | 0 | ycaM | 4 | 10.8333 | 0.456 |
| 1.6 | 0 | yadI | 4 | 10.8333 | 0.572 |
| 1.6 | 0 | AG1 | 4 | 10.8333 | 0.477 |
| 1.6 | 0 | ptsI | 4 | 11.0833 | 0.457 |
| 1.6 | 0 | clcB | 4 | 11.0833 | 0.424 |
| 1.6 | 0 | ycaM | 4 | 11.0833 | 0.459 |
| 1.6 | 0 | yadI | 4 | 11.0833 | 0.578 |
| 1.6 | 0 | AG1 | 4 | 11.0833 | 0.477 |
| 1.6 | 0 | ptsI | 4 | 11.3333 | 0.462 |
| 1.6 | 0 | clcB | 4 | 11.3333 | 0.426 |
| 1.6 | 0 | ycaM | 4 | 11.3333 | 0.464 |
| 1.6 | 0 | yadI | 4 | 11.3333 | 0.584 |
| 1.6 | 0 | AG1 | 4 | 11.3333 | 0.482 |
| 1.6 | 0 | ptsI | 4 | 11.5833 | 0.467 |
| 1.6 | 0 | clcB | 4 | 11.5833 | 0.428 |
| 1.6 | 0 | ycaM | 4 | 11.5833 | 0.466 |
| 1.6 | 0 | yadI | 4 | 11.5833 | 0.589 |
| 1.6 | 0 | AG1 | 4 | 11.5833 | 0.481 |
| 1.6 | 0 | ptsI | 4 | 11.8333 | 0.47 |
| 1.6 | 0 | clcB | 4 | 11.8333 | 0.43 |
| 1.6 | 0 | ycaM | 4 | 11.8333 | 0.468 |
| 1.6 | 0 | yadI | 4 | 11.8333 | 0.591 |
| 1.6 | 0 | AG1 | 4 | 11.8333 | 0.483 |
| 1.6 | 0 | ptsI | 4 | 12.0833 | 0.473 |
| 1.6 | 0 | clcB | 4 | 12.0833 | 0.432 |
| 1.6 | 0 | ycaM | 4 | 12.0833 | 0.473 |
| 1.6 | 0 | yadI | 4 | 12.0833 | 0.597 |
| 1.6 | 0 | AG1 | 4 | 12.0833 | 0.489 |
| 1.6 | 0 | ptsI | 4 | 12.3333 | 0.474 |
| 1.6 | 0 | clcB | 4 | 12.3333 | 0.436 |
| 1.6 | 0 | ycaM | 4 | 12.3333 | 0.474 |
| 1.6 | 0 | yadI | 4 | 12.3333 | 0.603 |
| 1.6 | 0 | AG1 | 4 | 12.3333 | 0.492 |
| 1.6 | 0 | ptsI | 4 | 12.5833 | 0.478 |
| 1.6 | 0 | clcB | 4 | 12.5833 | 0.432 |
| 1.6 | 0 | ycaM | 4 | 12.5833 | 0.475 |
| 1.6 | 0 | yadI | 4 | 12.5833 | 0.604 |
| 1.6 | 0 | AG1 | 4 | 12.5833 | 0.487 |
| 1.6 | 0 | ptsI | 4 | 12.8333 | 0.482 |
| 1.6 | 0 | clcB | 4 | 12.8333 | 0.439 |
| 1.6 | 0 | ycaM | 4 | 12.8333 | 0.48 |
| 1.6 | 0 | yadI | 4 | 12.8333 | 0.609 |
| 1.6 | 0 | AG1 | 4 | 12.8333 | 0.494 |
| 1.6 | 0 | ptsI | 4 | 13.0833 | 0.483 |
| 1.6 | 0 | clcB | 4 | 13.0833 | 0.437 |
| 1.6 | 0 | ycaM | 4 | 13.0833 | 0.485 |
| 1.6 | 0 | yadI | 4 | 13.0833 | 0.611 |
| 1.6 | 0 | AG1 | 4 | 13.0833 | 0.499 |

| | | | | | |
|---|---|---|---|---|---|
| 1.6 | 0 | ptsI | 4 | 13.3333 | 0.487 |
| 1.6 | 0 | clcB | 4 | 13.3333 | 0.44 |
| 1.6 | 0 | ycaM | 4 | 13.3333 | 0.486 |
| 1.6 | 0 | yadI | 4 | 13.3333 | 0.613 |
| 1.6 | 0 | AG1 | 4 | 13.3333 | 0.499 |
| 1.6 | 0 | ptsI | 4 | 13.5833 | 0.485 |
| 1.6 | 0 | clcB | 4 | 13.5833 | 0.44 |
| 1.6 | 0 | ycaM | 4 | 13.5833 | 0.487 |
| 1.6 | 0 | yadI | 4 | 13.5833 | 0.615 |
| 1.6 | 0 | AG1 | 4 | 13.5833 | 0.5 |
| 1.6 | 0 | ptsI | 4 | 13.8333 | 0.487 |
| 1.6 | 0 | clcB | 4 | 13.8333 | 0.442 |
| 1.6 | 0 | ycaM | 4 | 13.8333 | 0.492 |
| 1.6 | 0 | yadI | 4 | 13.8333 | 0.618 |
| 1.6 | 0 | AG1 | 4 | 13.8333 | 0.505 |
| 1.6 | 0 | ptsI | 4 | 14.0833 | 0.49 |
| 1.6 | 0 | clcB | 4 | 14.0833 | 0.444 |
| 1.6 | 0 | ycaM | 4 | 14.0833 | 0.49 |
| 1.6 | 0 | yadI | 4 | 14.0833 | 0.619 |
| 1.6 | 0 | AG1 | 4 | 14.0833 | 0.503 |
| 1.6 | 0 | ptsI | 4 | 14.3333 | 0.489 |
| 1.6 | 0 | clcB | 4 | 14.3333 | 0.447 |
| 1.6 | 0 | ycaM | 4 | 14.3333 | 0.492 |
| 1.6 | 0 | yadI | 4 | 14.3333 | 0.624 |
| 1.6 | 0 | AG1 | 4 | 14.3333 | 0.506 |
| 1.6 | 0 | ptsI | 4 | 14.5833 | 0.493 |
| 1.6 | 0 | clcB | 4 | 14.5833 | 0.45 |
| 1.6 | 0 | ycaM | 4 | 14.5833 | 0.495 |
| 1.6 | 0 | yadI | 4 | 14.5833 | 0.626 |
| 1.6 | 0 | AG1 | 4 | 14.5833 | 0.505 |
| 1.6 | 0 | ptsI | 4 | 14.8333 | 0.499 |
| 1.6 | 0 | clcB | 4 | 14.8333 | 0.451 |
| 1.6 | 0 | ycaM | 4 | 14.8333 | 0.5 |
| 1.6 | 0 | yadI | 4 | 14.8333 | 0.624 |
| 1.6 | 0 | AG1 | 4 | 14.8333 | 0.508 |
| 1.6 | 0 | ptsI | 4 | 15.0833 | 0.5 |
| 1.6 | 0 | clcB | 4 | 15.0833 | 0.452 |
| 1.6 | 0 | ycaM | 4 | 15.0833 | 0.499 |
| 1.6 | 0 | yadI | 4 | 15.0833 | 0.624 |
| 1.6 | 0 | AG1 | 4 | 15.0833 | 0.508 |
| 1.6 | 0 | ptsI | 4 | 15.3333 | 0.499 |
| 1.6 | 0 | clcB | 4 | 15.3333 | 0.451 |
| 1.6 | 0 | ycaM | 4 | 15.3333 | 0.498 |
| 1.6 | 0 | yadI | 4 | 15.3333 | 0.625 |
| 1.6 | 0 | AG1 | 4 | 15.3333 | 0.509 |
| 1.6 | 0 | ptsI | 4 | 15.5833 | 0.503 |
| 1.6 | 0 | clcB | 4 | 15.5833 | 0.454 |
| 1.6 | 0 | ycaM | 4 | 15.5833 | 0.5 |

| | | | | | |
|---|---|---|---|---|---|
| 1.6 | 0 | yadI | 4 | 15.5833 | 0.629 |
| 1.6 | 0 | AG1 | 4 | 15.5833 | 0.509 |
| 1.6 | 0 | ptsI | 4 | 15.8333 | 0.509 |
| 1.6 | 0 | clcB | 4 | 15.8333 | 0.459 |
| 1.6 | 0 | ycaM | 4 | 15.8333 | 0.503 |
| 1.6 | 0 | yadI | 4 | 15.8333 | 0.632 |
| 1.6 | 0 | AG1 | 4 | 15.8333 | 0.514 |
| 1.6 | 0 | ptsI | 4 | 16.0833 | 0.511 |
| 1.6 | 0 | clcB | 4 | 16.0833 | 0.454 |
| 1.6 | 0 | ycaM | 4 | 16.0833 | 0.504 |
| 1.6 | 0 | yadI | 4 | 16.0833 | 0.63 |
| 1.6 | 0 | AG1 | 4 | 16.0833 | 0.513 |
| 1.6 | 0 | ptsI | 4 | 16.3333 | 0.514 |
| 1.6 | 0 | clcB | 4 | 16.3333 | 0.46 |
| 1.6 | 0 | ycaM | 4 | 16.3333 | 0.508 |
| 1.6 | 0 | yadI | 4 | 16.3333 | 0.634 |
| 1.6 | 0 | AG1 | 4 | 16.3333 | 0.516 |
| 1.6 | 0 | ptsI | 4 | 16.5833 | 0.513 |
| 1.6 | 0 | clcB | 4 | 16.5833 | 0.458 |
| 1.6 | 0 | ycaM | 4 | 16.5833 | 0.507 |
| 1.6 | 0 | yadI | 4 | 16.5833 | 0.634 |
| 1.6 | 0 | AG1 | 4 | 16.5833 | 0.515 |
| 1.6 | 0 | ptsI | 4 | 16.8333 | 0.517 |
| 1.6 | 0 | clcB | 4 | 16.8333 | 0.463 |
| 1.6 | 0 | ycaM | 4 | 16.8333 | 0.511 |
| 1.6 | 0 | yadI | 4 | 16.8333 | 0.635 |
| 1.6 | 0 | AG1 | 4 | 16.8333 | 0.516 |
| 1.6 | 0 | ptsI | 4 | 17.0833 | 0.518 |
| 1.6 | 0 | clcB | 4 | 17.0833 | 0.465 |
| 1.6 | 0 | ycaM | 4 | 17.0833 | 0.513 |
| 1.6 | 0 | yadI | 4 | 17.0833 | 0.639 |
| 1.6 | 0 | AG1 | 4 | 17.0833 | 0.517 |
| 1.6 | 0 | ptsI | 4 | 17.3333 | 0.517 |
| 1.6 | 0 | clcB | 4 | 17.3333 | 0.464 |
| 1.6 | 0 | ycaM | 4 | 17.3333 | 0.516 |
| 1.6 | 0 | yadI | 4 | 17.3333 | 0.641 |
| 1.6 | 0 | AG1 | 4 | 17.3333 | 0.52 |
| 1.6 | 0 | ptsI | 4 | 17.5833 | 0.523 |
| 1.6 | 0 | clcB | 4 | 17.5833 | 0.468 |
| 1.6 | 0 | ycaM | 4 | 17.5833 | 0.516 |
| 1.6 | 0 | yadI | 4 | 17.5833 | 0.645 |
| 1.6 | 0 | AG1 | 4 | 17.5833 | 0.522 |
| 1.6 | 0 | ptsI | 4 | 17.8333 | 0.521 |
| 1.6 | 0 | clcB | 4 | 17.8333 | 0.468 |
| 1.6 | 0 | ycaM | 4 | 17.8333 | 0.513 |
| 1.6 | 0 | yadI | 4 | 17.8333 | 0.642 |
| 1.6 | 0 | AG1 | 4 | 17.8333 | 0.522 |
| 1.6 | 0 | ptsI | 4 | 18.0833 | 0.522 |

| | | | | | |
|---|---|---|---|---|---|
| 1.6 | 0 | clcB | 4 | 18.0833 | 0.47 |
| 1.6 | 0 | ycaM | 4 | 18.0833 | 0.518 |
| 1.6 | 0 | yadI | 4 | 18.0833 | 0.646 |
| 1.6 | 0 | AG1 | 4 | 18.0833 | 0.524 |
| 1.6 | 0 | ptsI | 4 | 18.3333 | 0.519 |
| 1.6 | 0 | clcB | 4 | 18.3333 | 0.472 |
| 1.6 | 0 | ycaM | 4 | 18.3333 | 0.519 |
| 1.6 | 0 | yadI | 4 | 18.3333 | 0.644 |
| 1.6 | 0 | AG1 | 4 | 18.3333 | 0.523 |
| 1.6 | 0 | ptsI | 4 | 18.5833 | 0.518 |
| 1.6 | 0 | clcB | 4 | 18.5833 | 0.472 |
| 1.6 | 0 | ycaM | 4 | 18.5833 | 0.52 |
| 1.6 | 0 | yadI | 4 | 18.5833 | 0.644 |
| 1.6 | 0 | AG1 | 4 | 18.5833 | 0.526 |
| 1.6 | 0 | ptsI | 4 | 18.8333 | 0.518 |
| 1.6 | 0 | clcB | 4 | 18.8333 | 0.474 |
| 1.6 | 0 | ycaM | 4 | 18.8333 | 0.517 |
| 1.6 | 0 | yadI | 4 | 18.8333 | 0.644 |
| 1.6 | 0 | AG1 | 4 | 18.8333 | 0.525 |
| 1.6 | 0 | ptsI | 4 | 19.0833 | 0.523 |
| 1.6 | 0 | clcB | 4 | 19.0833 | 0.474 |
| 1.6 | 0 | ycaM | 4 | 19.0833 | 0.522 |
| 1.6 | 0 | yadI | 4 | 19.0833 | 0.643 |
| 1.6 | 0 | AG1 | 4 | 19.0833 | 0.526 |
| 1.6 | 0 | ptsI | 4 | 19.3333 | 0.523 |
| 1.6 | 0 | clcB | 4 | 19.3333 | 0.477 |
| 1.6 | 0 | ycaM | 4 | 19.3333 | 0.523 |
| 1.6 | 0 | yadI | 4 | 19.3333 | 0.647 |
| 1.6 | 0 | AG1 | 4 | 19.3333 | 0.528 |
| 1.6 | 0 | ptsI | 4 | 19.5833 | 0.524 |
| 1.6 | 0 | clcB | 4 | 19.5833 | 0.479 |
| 1.6 | 0 | ycaM | 4 | 19.5833 | 0.526 |
| 1.6 | 0 | yadI | 4 | 19.5833 | 0.65 |
| 1.6 | 0 | AG1 | 4 | 19.5833 | 0.532 |
| 1.6 | 0 | ptsI | 4 | 19.8333 | 0.527 |
| 1.6 | 0 | clcB | 4 | 19.8333 | 0.481 |
| 1.6 | 0 | ycaM | 4 | 19.8333 | 0.523 |
| 1.6 | 0 | yadI | 4 | 19.8333 | 0.649 |
| 1.6 | 0 | AG1 | 4 | 19.8333 | 0.532 |
| 1.6 | 0 | ptsI | 4 | 20.0833 | 0.53 |
| 1.6 | 0 | clcB | 4 | 20.0833 | 0.478 |
| 1.6 | 0 | ycaM | 4 | 20.0833 | 0.524 |
| 1.6 | 0 | yadI | 4 | 20.0833 | 0.65 |
| 1.6 | 0 | AG1 | 4 | 20.0833 | 0.532 |
| 1.6 | 0 | ptsI | 4 | 20.3333 | 0.532 |
| 1.6 | 0 | clcB | 4 | 20.3333 | 0.481 |
| 1.6 | 0 | ycaM | 4 | 20.3333 | 0.528 |
| 1.6 | 0 | yadI | 4 | 20.3333 | 0.647 |

| | | | | | |
|---|---|---|---|---|---|
| 1.6 | 0 | AG1 | 4 | 20.3333 | 0.531 |
| 1.6 | 0 | ptsI | 4 | 20.5833 | 0.536 |
| 1.6 | 0 | clcB | 4 | 20.5833 | 0.485 |
| 1.6 | 0 | ycaM | 4 | 20.5833 | 0.53 |
| 1.6 | 0 | yadI | 4 | 20.5833 | 0.654 |
| 1.6 | 0 | AG1 | 4 | 20.5833 | 0.538 |
| 1.6 | 0 | ptsI | 4 | 20.8333 | 0.539 |
| 1.6 | 0 | clcB | 4 | 20.8333 | 0.486 |
| 1.6 | 0 | ycaM | 4 | 20.8333 | 0.531 |
| 1.6 | 0 | yadI | 4 | 20.8333 | 0.649 |
| 1.6 | 0 | AG1 | 4 | 20.8333 | 0.537 |
| 1.6 | 0 | ptsI | 4 | 21.0833 | 0.541 |
| 1.6 | 0 | clcB | 4 | 21.0833 | 0.485 |
| 1.6 | 0 | ycaM | 4 | 21.0833 | 0.53 |
| 1.6 | 0 | yadI | 4 | 21.0833 | 0.651 |
| 1.6 | 0 | AG1 | 4 | 21.0833 | 0.533 |
| 1.6 | 0 | ptsI | 4 | 21.3333 | 0.542 |
| 1.6 | 0 | clcB | 4 | 21.3333 | 0.488 |
| 1.6 | 0 | ycaM | 4 | 21.3333 | 0.532 |
| 1.6 | 0 | yadI | 4 | 21.3333 | 0.656 |
| 1.6 | 0 | AG1 | 4 | 21.3333 | 0.535 |
| 1.6 | 0 | ptsI | 4 | 21.5833 | 0.546 |
| 1.6 | 0 | clcB | 4 | 21.5833 | 0.49 |
| 1.6 | 0 | ycaM | 4 | 21.5833 | 0.534 |
| 1.6 | 0 | yadI | 4 | 21.5833 | 0.652 |
| 1.6 | 0 | AG1 | 4 | 21.5833 | 0.539 |
| 1.6 | 0 | ptsI | 4 | 21.8333 | 0.55 |
| 1.6 | 0 | clcB | 4 | 21.8333 | 0.49 |
| 1.6 | 0 | ycaM | 4 | 21.8333 | 0.534 |
| 1.6 | 0 | yadI | 4 | 21.8333 | 0.655 |
| 1.6 | 0 | AG1 | 4 | 21.8333 | 0.542 |
| 1.6 | 0 | ptsI | 4 | 22.0833 | 0.551 |
| 1.6 | 0 | clcB | 4 | 22.0833 | 0.49 |
| 1.6 | 0 | ycaM | 4 | 22.0833 | 0.532 |
| 1.6 | 0 | yadI | 4 | 22.0833 | 0.652 |
| 1.6 | 0 | AG1 | 4 | 22.0833 | 0.54 |
| 1.6 | 0 | ptsI | 4 | 22.3333 | 0.554 |
| 1.6 | 0 | clcB | 4 | 22.3333 | 0.493 |
| 1.6 | 0 | ycaM | 4 | 22.3333 | 0.535 |
| 1.6 | 0 | yadI | 4 | 22.3333 | 0.656 |
| 1.6 | 0 | AG1 | 4 | 22.3333 | 0.545 |
| 1.6 | 0 | ptsI | 4 | 22.5833 | 0.559 |
| 1.6 | 0 | clcB | 4 | 22.5833 | 0.493 |
| 1.6 | 0 | ycaM | 4 | 22.5833 | 0.536 |
| 1.6 | 0 | yadI | 4 | 22.5833 | 0.658 |
| 1.6 | 0 | AG1 | 4 | 22.5833 | 0.543 |
| 1.6 | 0 | ptsI | 4 | 22.8333 | 0.559 |
| 1.6 | 0 | clcB | 4 | 22.8333 | 0.494 |

| | | | | | |
|---|---|---|---|---|---|
| 1.6 | 0 | ycaM | 4 | 22.8333 | 0.538 |
| 1.6 | 0 | yadI | 4 | 22.8333 | 0.655 |
| 1.6 | 0 | AG1 | 4 | 22.8333 | 0.545 |
| 1.6 | 0 | ptsI | 4 | 23.0833 | 0.563 |
| 1.6 | 0 | clcB | 4 | 23.0833 | 0.495 |
| 1.6 | 0 | ycaM | 4 | 23.0833 | 0.538 |
| 1.6 | 0 | yadI | 4 | 23.0833 | 0.658 |
| 1.6 | 0 | AG1 | 4 | 23.0833 | 0.545 |
| 1.6 | 0 | ptsI | 4 | 23.3333 | 0.57 |
| 1.6 | 0 | clcB | 4 | 23.3333 | 0.495 |
| 1.6 | 0 | ycaM | 4 | 23.3333 | 0.538 |
| 1.6 | 0 | yadI | 4 | 23.3333 | 0.657 |
| 1.6 | 0 | AG1 | 4 | 23.3333 | 0.546 |
| 1.6 | 0 | ptsI | 4 | 23.5833 | 0.572 |
| 1.6 | 0 | clcB | 4 | 23.5833 | 0.499 |
| 1.6 | 0 | ycaM | 4 | 23.5833 | 0.539 |
| 1.6 | 0 | yadI | 4 | 23.5833 | 0.658 |
| 1.6 | 0 | AG1 | 4 | 23.5833 | 0.545 |
| 1.6 | 0 | ptsI | 4 | 23.8333 | 0.574 |
| 1.6 | 0 | clcB | 4 | 23.8333 | 0.499 |
| 1.6 | 0 | ycaM | 4 | 23.8333 | 0.538 |
| 1.6 | 0 | yadI | 4 | 23.8333 | 0.657 |
| 1.6 | 0 | AG1 | 4 | 23.8333 | 0.546 |
| 1.6 | 0 | ptsI | 4 | 24.0833 | 0.577 |
| 1.6 | 0 | clcB | 4 | 24.0833 | 0.5 |
| 1.6 | 0 | ycaM | 4 | 24.0833 | 0.538 |
| 1.6 | 0 | yadI | 4 | 24.0833 | 0.659 |
| 1.6 | 0 | AG1 | 4 | 24.0833 | 0.546 |
| 1.6 | 0 | ptsI | 5 | 0 | 0.159 |
| 1.6 | 0 | clcB | 5 | 0 | 0.163 |
| 1.6 | 0 | ycaM | 5 | 0 | 0.168 |
| 1.6 | 0 | yadI | 5 | 0 | 0.161 |
| 1.6 | 0 | AG1 | 5 | 0 | 0.161 |
| 1.6 | 0 | ptsI | 5 | 0.35 | 0.157 |
| 1.6 | 0 | clcB | 5 | 0.35 | 0.159 |
| 1.6 | 0 | ycaM | 5 | 0.35 | 0.162 |
| 1.6 | 0 | yadI | 5 | 0.35 | 0.158 |
| 1.6 | 0 | AG1 | 5 | 0.35 | 0.16 |
| 1.6 | 0 | ptsI | 5 | 0.6 | 0.158 |
| 1.6 | 0 | clcB | 5 | 0.6 | 0.161 |
| 1.6 | 0 | ycaM | 5 | 0.6 | 0.161 |
| 1.6 | 0 | yadI | 5 | 0.6 | 0.16 |
| 1.6 | 0 | AG1 | 5 | 0.6 | 0.161 |
| 1.6 | 0 | ptsI | 5 | 0.85 | 0.158 |
| 1.6 | 0 | clcB | 5 | 0.85 | 0.16 |
| 1.6 | 0 | ycaM | 5 | 0.85 | 0.161 |
| 1.6 | 0 | yadI | 5 | 0.85 | 0.16 |
| 1.6 | 0 | AG1 | 5 | 0.85 | 0.16 |

| | | | | | |
|---|---|---|---|---|---|
| 1.6 | 0 | ptsI | 5 | 1.1 | 0.16 |
| 1.6 | 0 | clcB | 5 | 1.1 | 0.16 |
| 1.6 | 0 | ycaM | 5 | 1.1 | 0.163 |
| 1.6 | 0 | yadI | 5 | 1.1 | 0.161 |
| 1.6 | 0 | AG1 | 5 | 1.1 | 0.161 |
| 1.6 | 0 | ptsI | 5 | 1.35 | 0.162 |
| 1.6 | 0 | clcB | 5 | 1.35 | 0.162 |
| 1.6 | 0 | ycaM | 5 | 1.35 | 0.164 |
| 1.6 | 0 | yadI | 5 | 1.35 | 0.162 |
| 1.6 | 0 | AG1 | 5 | 1.35 | 0.162 |
| 1.6 | 0 | ptsI | 5 | 1.6 | 0.164 |
| 1.6 | 0 | clcB | 5 | 1.6 | 0.164 |
| 1.6 | 0 | ycaM | 5 | 1.6 | 0.167 |
| 1.6 | 0 | yadI | 5 | 1.6 | 0.165 |
| 1.6 | 0 | AG1 | 5 | 1.6 | 0.167 |
| 1.6 | 0 | ptsI | 5 | 1.85 | 0.174 |
| 1.6 | 0 | clcB | 5 | 1.85 | 0.169 |
| 1.6 | 0 | ycaM | 5 | 1.85 | 0.171 |
| 1.6 | 0 | yadI | 5 | 1.85 | 0.168 |
| 1.6 | 0 | AG1 | 5 | 1.85 | 0.171 |
| 1.6 | 0 | ptsI | 5 | 2.1 | 0.168 |
| 1.6 | 0 | clcB | 5 | 2.1 | 0.172 |
| 1.6 | 0 | ycaM | 5 | 2.1 | 0.176 |
| 1.6 | 0 | yadI | 5 | 2.1 | 0.173 |
| 1.6 | 0 | AG1 | 5 | 2.1 | 0.17 |
| 1.6 | 0 | ptsI | 5 | 2.35 | 0.17 |
| 1.6 | 0 | clcB | 5 | 2.35 | 0.176 |
| 1.6 | 0 | ycaM | 5 | 2.35 | 0.18 |
| 1.6 | 0 | yadI | 5 | 2.35 | 0.178 |
| 1.6 | 0 | AG1 | 5 | 2.35 | 0.174 |
| 1.6 | 0 | ptsI | 5 | 2.6 | 0.172 |
| 1.6 | 0 | clcB | 5 | 2.6 | 0.178 |
| 1.6 | 0 | ycaM | 5 | 2.6 | 0.183 |
| 1.6 | 0 | yadI | 5 | 2.6 | 0.185 |
| 1.6 | 0 | AG1 | 5 | 2.6 | 0.184 |
| 1.6 | 0 | ptsI | 5 | 2.85 | 0.176 |
| 1.6 | 0 | clcB | 5 | 2.85 | 0.186 |
| 1.6 | 0 | ycaM | 5 | 2.85 | 0.189 |
| 1.6 | 0 | yadI | 5 | 2.85 | 0.19 |
| 1.6 | 0 | AG1 | 5 | 2.85 | 0.192 |
| 1.6 | 0 | ptsI | 5 | 3.1 | 0.184 |
| 1.6 | 0 | clcB | 5 | 3.1 | 0.198 |
| 1.6 | 0 | ycaM | 5 | 3.1 | 0.198 |
| 1.6 | 0 | yadI | 5 | 3.1 | 0.197 |
| 1.6 | 0 | AG1 | 5 | 3.1 | 0.206 |
| 1.6 | 0 | ptsI | 5 | 3.35 | 0.186 |
| 1.6 | 0 | clcB | 5 | 3.35 | 0.207 |
| 1.6 | 0 | ycaM | 5 | 3.35 | 0.207 |

| | | | | | |
|---|---|---|---|---|---|
| 1.6 | 0 | yadI | 5 | 3.35 | 0.209 |
| 1.6 | 0 | AG1 | 5 | 3.35 | 0.204 |
| 1.6 | 0 | ptsI | 5 | 3.6 | 0.193 |
| 1.6 | 0 | clcB | 5 | 3.6 | 0.214 |
| 1.6 | 0 | ycaM | 5 | 3.6 | 0.219 |
| 1.6 | 0 | yadI | 5 | 3.6 | 0.223 |
| 1.6 | 0 | AG1 | 5 | 3.6 | 0.211 |
| 1.6 | 0 | ptsI | 5 | 3.85 | 0.206 |
| 1.6 | 0 | clcB | 5 | 3.85 | 0.225 |
| 1.6 | 0 | ycaM | 5 | 3.85 | 0.228 |
| 1.6 | 0 | yadI | 5 | 3.85 | 0.236 |
| 1.6 | 0 | AG1 | 5 | 3.85 | 0.217 |
| 1.6 | 0 | ptsI | 5 | 4.38333 | 0.202 |
| 1.6 | 0 | clcB | 5 | 4.38333 | 0.254 |
| 1.6 | 0 | ycaM | 5 | 4.38333 | 0.236 |
| 1.6 | 0 | yadI | 5 | 4.38333 | 0.24 |
| 1.6 | 0 | AG1 | 5 | 4.38333 | 0.232 |
| 1.6 | 0 | ptsI | 5 | 4.63333 | 0.198 |
| 1.6 | 0 | clcB | 5 | 4.63333 | 0.27 |
| 1.6 | 0 | ycaM | 5 | 4.63333 | 0.24 |
| 1.6 | 0 | yadI | 5 | 4.63333 | 0.231 |
| 1.6 | 0 | AG1 | 5 | 4.63333 | 0.226 |
| 1.6 | 0 | ptsI | 5 | 4.88333 | 0.204 |
| 1.6 | 0 | clcB | 5 | 4.88333 | 0.282 |
| 1.6 | 0 | ycaM | 5 | 4.88333 | 0.251 |
| 1.6 | 0 | yadI | 5 | 4.88333 | 0.245 |
| 1.6 | 0 | AG1 | 5 | 4.88333 | 0.229 |
| 1.6 | 0 | ptsI | 5 | 5.13333 | 0.211 |
| 1.6 | 0 | clcB | 5 | 5.13333 | 0.295 |
| 1.6 | 0 | ycaM | 5 | 5.13333 | 0.257 |
| 1.6 | 0 | yadI | 5 | 5.13333 | 0.258 |
| 1.6 | 0 | AG1 | 5 | 5.13333 | 0.237 |
| 1.6 | 0 | ptsI | 5 | 5.38333 | 0.218 |
| 1.6 | 0 | clcB | 5 | 5.38333 | 0.312 |
| 1.6 | 0 | ycaM | 5 | 5.38333 | 0.267 |
| 1.6 | 0 | yadI | 5 | 5.38333 | 0.273 |
| 1.6 | 0 | AG1 | 5 | 5.38333 | 0.242 |
| 1.6 | 0 | ptsI | 5 | 5.63333 | 0.226 |
| 1.6 | 0 | clcB | 5 | 5.63333 | 0.324 |
| 1.6 | 0 | ycaM | 5 | 5.63333 | 0.277 |
| 1.6 | 0 | yadI | 5 | 5.63333 | 0.288 |
| 1.6 | 0 | AG1 | 5 | 5.63333 | 0.249 |
| 1.6 | 0 | ptsI | 5 | 5.88333 | 0.232 |
| 1.6 | 0 | clcB | 5 | 5.88333 | 0.339 |
| 1.6 | 0 | ycaM | 5 | 5.88333 | 0.288 |
| 1.6 | 0 | yadI | 5 | 5.88333 | 0.301 |
| 1.6 | 0 | AG1 | 5 | 5.88333 | 0.259 |
| 1.6 | 0 | ptsI | 5 | 6.13333 | 0.241 |

| | | | | | |
|---|---|---|---|---|---|
| 1.6 | 0 | clcB | 5 | 6.13333 | 0.348 |
| 1.6 | 0 | ycaM | 5 | 6.13333 | 0.3 |
| 1.6 | 0 | yadI | 5 | 6.13333 | 0.316 |
| 1.6 | 0 | AG1 | 5 | 6.13333 | 0.273 |
| 1.6 | 0 | ptsI | 5 | 6.38333 | 0.252 |
| 1.6 | 0 | clcB | 5 | 6.38333 | 0.36 |
| 1.6 | 0 | ycaM | 5 | 6.38333 | 0.31 |
| 1.6 | 0 | yadI | 5 | 6.38333 | 0.326 |
| 1.6 | 0 | AG1 | 5 | 6.38333 | 0.278 |
| 1.6 | 0 | ptsI | 5 | 6.63333 | 0.261 |
| 1.6 | 0 | clcB | 5 | 6.63333 | 0.369 |
| 1.6 | 0 | ycaM | 5 | 6.63333 | 0.322 |
| 1.6 | 0 | yadI | 5 | 6.63333 | 0.335 |
| 1.6 | 0 | AG1 | 5 | 6.63333 | 0.282 |
| 1.6 | 0 | ptsI | 5 | 6.88333 | 0.271 |
| 1.6 | 0 | clcB | 5 | 6.88333 | 0.379 |
| 1.6 | 0 | ycaM | 5 | 6.88333 | 0.332 |
| 1.6 | 0 | yadI | 5 | 6.88333 | 0.348 |
| 1.6 | 0 | AG1 | 5 | 6.88333 | 0.288 |
| 1.6 | 0 | ptsI | 5 | 7.13333 | 0.281 |
| 1.6 | 0 | clcB | 5 | 7.13333 | 0.391 |
| 1.6 | 0 | ycaM | 5 | 7.13333 | 0.343 |
| 1.6 | 0 | yadI | 5 | 7.13333 | 0.364 |
| 1.6 | 0 | AG1 | 5 | 7.13333 | 0.299 |
| 1.6 | 0 | ptsI | 5 | 7.38333 | 0.295 |
| 1.6 | 0 | clcB | 5 | 7.38333 | 0.402 |
| 1.6 | 0 | ycaM | 5 | 7.38333 | 0.356 |
| 1.6 | 0 | yadI | 5 | 7.38333 | 0.378 |
| 1.6 | 0 | AG1 | 5 | 7.38333 | 0.312 |
| 1.6 | 0 | ptsI | 5 | 7.63333 | 0.307 |
| 1.6 | 0 | clcB | 5 | 7.63333 | 0.418 |
| 1.6 | 0 | ycaM | 5 | 7.63333 | 0.366 |
| 1.6 | 0 | yadI | 5 | 7.63333 | 0.39 |
| 1.6 | 0 | AG1 | 5 | 7.63333 | 0.322 |
| 1.6 | 0 | ptsI | 5 | 7.88333 | 0.318 |
| 1.6 | 0 | clcB | 5 | 7.88333 | 0.43 |
| 1.6 | 0 | ycaM | 5 | 7.88333 | 0.375 |
| 1.6 | 0 | yadI | 5 | 7.88333 | 0.401 |
| 1.6 | 0 | AG1 | 5 | 7.88333 | 0.326 |
| 1.6 | 0 | ptsI | 5 | 8.13333 | 0.329 |
| 1.6 | 0 | clcB | 5 | 8.13333 | 0.443 |
| 1.6 | 0 | ycaM | 5 | 8.13333 | 0.385 |
| 1.6 | 0 | yadI | 5 | 8.13333 | 0.414 |
| 1.6 | 0 | AG1 | 5 | 8.13333 | 0.34 |
| 1.6 | 0 | ptsI | 5 | 8.38333 | 0.342 |
| 1.6 | 0 | clcB | 5 | 8.38333 | 0.458 |
| 1.6 | 0 | ycaM | 5 | 8.38333 | 0.395 |
| 1.6 | 0 | yadI | 5 | 8.38333 | 0.424 |

| | | | | | |
|---|---|---|---|---|---|
| 1.6 | 0 | AG1 | 5 | 8.38333 | 0.35 |
| 1.6 | 0 | ptsI | 5 | 8.88333 | 0.32 |
| 1.6 | 0 | clcB | 5 | 8.88333 | 0.367 |
| 1.6 | 0 | ycaM | 5 | 8.88333 | 0.353 |
| 1.6 | 0 | yadI | 5 | 8.88333 | 0.427 |
| 1.6 | 0 | AG1 | 5 | 8.88333 | 0.384 |
| 1.6 | 0 | ptsI | 5 | 9.13333 | 0.332 |
| 1.6 | 0 | clcB | 5 | 9.13333 | 0.373 |
| 1.6 | 0 | ycaM | 5 | 9.13333 | 0.358 |
| 1.6 | 0 | yadI | 5 | 9.13333 | 0.42 |
| 1.6 | 0 | AG1 | 5 | 9.13333 | 0.389 |
| 1.6 | 0 | ptsI | 5 | 9.38333 | 0.345 |
| 1.6 | 0 | clcB | 5 | 9.38333 | 0.383 |
| 1.6 | 0 | ycaM | 5 | 9.38333 | 0.368 |
| 1.6 | 0 | yadI | 5 | 9.38333 | 0.428 |
| 1.6 | 0 | AG1 | 5 | 9.38333 | 0.399 |
| 1.6 | 0 | ptsI | 5 | 9.63333 | 0.36 |
| 1.6 | 0 | clcB | 5 | 9.63333 | 0.398 |
| 1.6 | 0 | ycaM | 5 | 9.63333 | 0.379 |
| 1.6 | 0 | yadI | 5 | 9.63333 | 0.437 |
| 1.6 | 0 | AG1 | 5 | 9.63333 | 0.418 |
| 1.6 | 0 | ptsI | 5 | 9.88333 | 0.369 |
| 1.6 | 0 | clcB | 5 | 9.88333 | 0.408 |
| 1.6 | 0 | ycaM | 5 | 9.88333 | 0.389 |
| 1.6 | 0 | yadI | 5 | 9.88333 | 0.443 |
| 1.6 | 0 | AG1 | 5 | 9.88333 | 0.43 |
| 1.6 | 0 | ptsI | 5 | 10.1333 | 0.379 |
| 1.6 | 0 | clcB | 5 | 10.1333 | 0.414 |
| 1.6 | 0 | ycaM | 5 | 10.1333 | 0.394 |
| 1.6 | 0 | yadI | 5 | 10.1333 | 0.448 |
| 1.6 | 0 | AG1 | 5 | 10.1333 | 0.435 |
| 1.6 | 0 | ptsI | 5 | 10.3833 | 0.39 |
| 1.6 | 0 | clcB | 5 | 10.3833 | 0.42 |
| 1.6 | 0 | ycaM | 5 | 10.3833 | 0.401 |
| 1.6 | 0 | yadI | 5 | 10.3833 | 0.458 |
| 1.6 | 0 | AG1 | 5 | 10.3833 | 0.446 |
| 1.6 | 0 | ptsI | 5 | 10.6333 | 0.395 |
| 1.6 | 0 | clcB | 5 | 10.6333 | 0.424 |
| 1.6 | 0 | ycaM | 5 | 10.6333 | 0.401 |
| 1.6 | 0 | yadI | 5 | 10.6333 | 0.459 |
| 1.6 | 0 | AG1 | 5 | 10.6333 | 0.452 |
| 1.6 | 0 | ptsI | 5 | 10.8833 | 0.402 |
| 1.6 | 0 | clcB | 5 | 10.8833 | 0.428 |
| 1.6 | 0 | ycaM | 5 | 10.8833 | 0.407 |
| 1.6 | 0 | yadI | 5 | 10.8833 | 0.466 |
| 1.6 | 0 | AG1 | 5 | 10.8833 | 0.458 |
| 1.6 | 0 | ptsI | 5 | 11.1333 | 0.407 |
| 1.6 | 0 | clcB | 5 | 11.1333 | 0.431 |

| | | | | | |
|---|---|---|---|---|---|
| 1.6 | 0 | ycaM | 5 | 11.1333 | 0.406 |
| 1.6 | 0 | yadI | 5 | 11.1333 | 0.471 |
| 1.6 | 0 | AG1 | 5 | 11.1333 | 0.459 |
| 1.6 | 0 | ptsI | 5 | 11.3833 | 0.413 |
| 1.6 | 0 | clcB | 5 | 11.3833 | 0.431 |
| 1.6 | 0 | ycaM | 5 | 11.3833 | 0.409 |
| 1.6 | 0 | yadI | 5 | 11.3833 | 0.475 |
| 1.6 | 0 | AG1 | 5 | 11.3833 | 0.463 |
| 1.6 | 0 | ptsI | 5 | 11.6333 | 0.417 |
| 1.6 | 0 | clcB | 5 | 11.6333 | 0.433 |
| 1.6 | 0 | ycaM | 5 | 11.6333 | 0.412 |
| 1.6 | 0 | yadI | 5 | 11.6333 | 0.48 |
| 1.6 | 0 | AG1 | 5 | 11.6333 | 0.468 |
| 1.6 | 0 | ptsI | 5 | 11.8833 | 0.424 |
| 1.6 | 0 | clcB | 5 | 11.8833 | 0.44 |
| 1.6 | 0 | ycaM | 5 | 11.8833 | 0.412 |
| 1.6 | 0 | yadI | 5 | 11.8833 | 0.484 |
| 1.6 | 0 | AG1 | 5 | 11.8833 | 0.471 |
| 1.6 | 0 | ptsI | 5 | 12.1333 | 0.43 |
| 1.6 | 0 | clcB | 5 | 12.1333 | 0.442 |
| 1.6 | 0 | ycaM | 5 | 12.1333 | 0.416 |
| 1.6 | 0 | yadI | 5 | 12.1333 | 0.489 |
| 1.6 | 0 | AG1 | 5 | 12.1333 | 0.474 |
| 1.6 | 0 | ptsI | 5 | 12.3833 | 0.432 |
| 1.6 | 0 | clcB | 5 | 12.3833 | 0.445 |
| 1.6 | 0 | ycaM | 5 | 12.3833 | 0.416 |
| 1.6 | 0 | yadI | 5 | 12.3833 | 0.492 |
| 1.6 | 0 | AG1 | 5 | 12.3833 | 0.478 |
| 1.6 | 0 | ptsI | 5 | 12.6333 | 0.44 |
| 1.6 | 0 | clcB | 5 | 12.6333 | 0.448 |
| 1.6 | 0 | ycaM | 5 | 12.6333 | 0.42 |
| 1.6 | 0 | yadI | 5 | 12.6333 | 0.5 |
| 1.6 | 0 | AG1 | 5 | 12.6333 | 0.483 |
| 1.6 | 0 | ptsI | 5 | 12.8833 | 0.441 |
| 1.6 | 0 | clcB | 5 | 12.8833 | 0.452 |
| 1.6 | 0 | ycaM | 5 | 12.8833 | 0.42 |
| 1.6 | 0 | yadI | 5 | 12.8833 | 0.504 |
| 1.6 | 0 | AG1 | 5 | 12.8833 | 0.487 |
| 1.6 | 0 | ptsI | 5 | 13.1333 | 0.444 |
| 1.6 | 0 | clcB | 5 | 13.1333 | 0.454 |
| 1.6 | 0 | ycaM | 5 | 13.1333 | 0.424 |
| 1.6 | 0 | yadI | 5 | 13.1333 | 0.506 |
| 1.6 | 0 | AG1 | 5 | 13.1333 | 0.491 |
| 1.6 | 0 | ptsI | 5 | 13.3833 | 0.447 |
| 1.6 | 0 | clcB | 5 | 13.3833 | 0.457 |
| 1.6 | 0 | ycaM | 5 | 13.3833 | 0.427 |
| 1.6 | 0 | yadI | 5 | 13.3833 | 0.514 |
| 1.6 | 0 | AG1 | 5 | 13.3833 | 0.497 |

| | | | | | |
|---|---|---|---|---|---|
| 1.6 | 0 | ptsI | 5 | 13.6333 | 0.451 |
| 1.6 | 0 | clcB | 5 | 13.6333 | 0.462 |
| 1.6 | 0 | ycaM | 5 | 13.6333 | 0.43 |
| 1.6 | 0 | yadI | 5 | 13.6333 | 0.517 |
| 1.6 | 0 | AG1 | 5 | 13.6333 | 0.5 |
| 1.6 | 0 | ptsI | 5 | 13.8833 | 0.456 |
| 1.6 | 0 | clcB | 5 | 13.8833 | 0.461 |
| 1.6 | 0 | ycaM | 5 | 13.8833 | 0.433 |
| 1.6 | 0 | yadI | 5 | 13.8833 | 0.518 |
| 1.6 | 0 | AG1 | 5 | 13.8833 | 0.506 |
| 1.6 | 0 | ptsI | 5 | 14.1333 | 0.46 |
| 1.6 | 0 | clcB | 5 | 14.1333 | 0.467 |
| 1.6 | 0 | ycaM | 5 | 14.1333 | 0.436 |
| 1.6 | 0 | yadI | 5 | 14.1333 | 0.525 |
| 1.6 | 0 | AG1 | 5 | 14.1333 | 0.509 |
| 1.6 | 0 | ptsI | 5 | 14.3833 | 0.464 |
| 1.6 | 0 | clcB | 5 | 14.3833 | 0.464 |
| 1.6 | 0 | ycaM | 5 | 14.3833 | 0.435 |
| 1.6 | 0 | yadI | 5 | 14.3833 | 0.527 |
| 1.6 | 0 | AG1 | 5 | 14.3833 | 0.509 |
| 1.6 | 0 | ptsI | 5 | 14.6333 | 0.468 |
| 1.6 | 0 | clcB | 5 | 14.6333 | 0.468 |
| 1.6 | 0 | ycaM | 5 | 14.6333 | 0.439 |
| 1.6 | 0 | yadI | 5 | 14.6333 | 0.528 |
| 1.6 | 0 | AG1 | 5 | 14.6333 | 0.517 |
| 1.6 | 0 | ptsI | 5 | 14.8833 | 0.468 |
| 1.6 | 0 | clcB | 5 | 14.8833 | 0.472 |
| 1.6 | 0 | ycaM | 5 | 14.8833 | 0.443 |
| 1.6 | 0 | yadI | 5 | 14.8833 | 0.534 |
| 1.6 | 0 | AG1 | 5 | 14.8833 | 0.518 |
| 1.6 | 0 | ptsI | 5 | 15.1333 | 0.475 |
| 1.6 | 0 | clcB | 5 | 15.1333 | 0.473 |
| 1.6 | 0 | ycaM | 5 | 15.1333 | 0.444 |
| 1.6 | 0 | yadI | 5 | 15.1333 | 0.536 |
| 1.6 | 0 | AG1 | 5 | 15.1333 | 0.524 |
| 1.6 | 0 | ptsI | 5 | 15.3833 | 0.482 |
| 1.6 | 0 | clcB | 5 | 15.3833 | 0.476 |
| 1.6 | 0 | ycaM | 5 | 15.3833 | 0.447 |
| 1.6 | 0 | yadI | 5 | 15.3833 | 0.537 |
| 1.6 | 0 | AG1 | 5 | 15.3833 | 0.531 |
| 1.6 | 0 | ptsI | 5 | 15.6333 | 0.479 |
| 1.6 | 0 | clcB | 5 | 15.6333 | 0.479 |
| 1.6 | 0 | ycaM | 5 | 15.6333 | 0.45 |
| 1.6 | 0 | yadI | 5 | 15.6333 | 0.538 |
| 1.6 | 0 | AG1 | 5 | 15.6333 | 0.532 |
| 1.6 | 0 | ptsI | 5 | 15.8833 | 0.486 |
| 1.6 | 0 | clcB | 5 | 15.8833 | 0.479 |
| 1.6 | 0 | ycaM | 5 | 15.8833 | 0.452 |

| | | | | | |
|---|---|---|---|---|---|
| 1.6 | 0 | yadI | 5 | 15.8833 | 0.545 |
| 1.6 | 0 | AG1 | 5 | 15.8833 | 0.531 |
| 1.6 | 0 | ptsI | 5 | 16.1333 | 0.49 |
| 1.6 | 0 | clcB | 5 | 16.1333 | 0.48 |
| 1.6 | 0 | ycaM | 5 | 16.1333 | 0.453 |
| 1.6 | 0 | yadI | 5 | 16.1333 | 0.554 |
| 1.6 | 0 | AG1 | 5 | 16.1333 | 0.521 |
| 1.6 | 0 | ptsI | 5 | 16.3833 | 0.495 |
| 1.6 | 0 | clcB | 5 | 16.3833 | 0.484 |
| 1.6 | 0 | ycaM | 5 | 16.3833 | 0.46 |
| 1.6 | 0 | yadI | 5 | 16.3833 | 0.558 |
| 1.6 | 0 | AG1 | 5 | 16.3833 | 0.537 |
| 1.6 | 0 | ptsI | 5 | 16.6333 | 0.501 |
| 1.6 | 0 | clcB | 5 | 16.6333 | 0.485 |
| 1.6 | 0 | ycaM | 5 | 16.6333 | 0.462 |
| 1.6 | 0 | yadI | 5 | 16.6333 | 0.561 |
| 1.6 | 0 | AG1 | 5 | 16.6333 | 0.539 |
| 1.6 | 0 | ptsI | 5 | 16.8833 | 0.501 |
| 1.6 | 0 | clcB | 5 | 16.8833 | 0.487 |
| 1.6 | 0 | ycaM | 5 | 16.8833 | 0.462 |
| 1.6 | 0 | yadI | 5 | 16.8833 | 0.562 |
| 1.6 | 0 | AG1 | 5 | 16.8833 | 0.541 |
| 1.6 | 0 | ptsI | 5 | 17.1333 | 0.5 |
| 1.6 | 0 | clcB | 5 | 17.1333 | 0.489 |
| 1.6 | 0 | ycaM | 5 | 17.1333 | 0.461 |
| 1.6 | 0 | yadI | 5 | 17.1333 | 0.562 |
| 1.6 | 0 | AG1 | 5 | 17.1333 | 0.544 |
| 1.6 | 0 | ptsI | 5 | 17.3833 | 0.507 |
| 1.6 | 0 | clcB | 5 | 17.3833 | 0.49 |
| 1.6 | 0 | ycaM | 5 | 17.3833 | 0.462 |
| 1.6 | 0 | yadI | 5 | 17.3833 | 0.57 |
| 1.6 | 0 | AG1 | 5 | 17.3833 | 0.547 |
| 1.6 | 0 | ptsI | 5 | 17.6333 | 0.511 |
| 1.6 | 0 | clcB | 5 | 17.6333 | 0.493 |
| 1.6 | 0 | ycaM | 5 | 17.6333 | 0.466 |
| 1.6 | 0 | yadI | 5 | 17.6333 | 0.573 |
| 1.6 | 0 | AG1 | 5 | 17.6333 | 0.551 |
| 1.6 | 0 | ptsI | 5 | 17.8833 | 0.512 |
| 1.6 | 0 | clcB | 5 | 17.8833 | 0.494 |
| 1.6 | 0 | ycaM | 5 | 17.8833 | 0.47 |
| 1.6 | 0 | yadI | 5 | 17.8833 | 0.577 |
| 1.6 | 0 | AG1 | 5 | 17.8833 | 0.561 |
| 1.6 | 0 | ptsI | 5 | 18.1333 | 0.514 |
| 1.6 | 0 | clcB | 5 | 18.1333 | 0.492 |
| 1.6 | 0 | ycaM | 5 | 18.1333 | 0.467 |
| 1.6 | 0 | yadI | 5 | 18.1333 | 0.58 |
| 1.6 | 0 | AG1 | 5 | 18.1333 | 0.573 |
| 1.6 | 0 | ptsI | 5 | 18.3833 | 0.517 |

| | | | | | |
|---|---|---|---|---|---|
| 1.6 | 0 | clcB | 5 | 18.3833 | 0.495 |
| 1.6 | 0 | ycaM | 5 | 18.3833 | 0.469 |
| 1.6 | 0 | yadI | 5 | 18.3833 | 0.584 |
| 1.6 | 0 | AG1 | 5 | 18.3833 | 0.583 |
| 1.6 | 0 | ptsI | 5 | 18.6333 | 0.516 |
| 1.6 | 0 | clcB | 5 | 18.6333 | 0.493 |
| 1.6 | 0 | ycaM | 5 | 18.6333 | 0.47 |
| 1.6 | 0 | yadI | 5 | 18.6333 | 0.588 |
| 1.6 | 0 | AG1 | 5 | 18.6333 | 0.589 |
| 1.6 | 0 | ptsI | 5 | 18.8833 | 0.521 |
| 1.6 | 0 | clcB | 5 | 18.8833 | 0.498 |
| 1.6 | 0 | ycaM | 5 | 18.8833 | 0.474 |
| 1.6 | 0 | yadI | 5 | 18.8833 | 0.592 |
| 1.6 | 0 | AG1 | 5 | 18.8833 | 0.596 |
| 1.6 | 0 | ptsI | 5 | 19.1333 | 0.521 |
| 1.6 | 0 | clcB | 5 | 19.1333 | 0.495 |
| 1.6 | 0 | ycaM | 5 | 19.1333 | 0.472 |
| 1.6 | 0 | yadI | 5 | 19.1333 | 0.588 |
| 1.6 | 0 | AG1 | 5 | 19.1333 | 0.603 |
| 1.6 | 0 | ptsI | 5 | 19.3833 | 0.523 |
| 1.6 | 0 | clcB | 5 | 19.3833 | 0.499 |
| 1.6 | 0 | ycaM | 5 | 19.3833 | 0.472 |
| 1.6 | 0 | yadI | 5 | 19.3833 | 0.59 |
| 1.6 | 0 | AG1 | 5 | 19.3833 | 0.603 |
| 1.6 | 0 | ptsI | 5 | 19.6333 | 0.524 |
| 1.6 | 0 | clcB | 5 | 19.6333 | 0.498 |
| 1.6 | 0 | ycaM | 5 | 19.6333 | 0.474 |
| 1.6 | 0 | yadI | 5 | 19.6333 | 0.596 |
| 1.6 | 0 | AG1 | 5 | 19.6333 | 0.604 |
| 1.6 | 0 | ptsI | 5 | 19.8833 | 0.529 |
| 1.6 | 0 | clcB | 5 | 19.8833 | 0.496 |
| 1.6 | 0 | ycaM | 5 | 19.8833 | 0.474 |
| 1.6 | 0 | yadI | 5 | 19.8833 | 0.591 |
| 1.6 | 0 | AG1 | 5 | 19.8833 | 0.604 |
| 1.6 | 0 | ptsI | 5 | 20.1333 | 0.532 |
| 1.6 | 0 | clcB | 5 | 20.1333 | 0.501 |
| 1.6 | 0 | ycaM | 5 | 20.1333 | 0.474 |
| 1.6 | 0 | yadI | 5 | 20.1333 | 0.594 |
| 1.6 | 0 | AG1 | 5 | 20.1333 | 0.607 |
| 1.6 | 0 | ptsI | 5 | 20.3833 | 0.533 |
| 1.6 | 0 | clcB | 5 | 20.3833 | 0.497 |
| 1.6 | 0 | ycaM | 5 | 20.3833 | 0.476 |
| 1.6 | 0 | yadI | 5 | 20.3833 | 0.594 |
| 1.6 | 0 | AG1 | 5 | 20.3833 | 0.608 |
| 1.6 | 0 | ptsI | 5 | 20.6333 | 0.536 |
| 1.6 | 0 | clcB | 5 | 20.6333 | 0.502 |
| 1.6 | 0 | ycaM | 5 | 20.6333 | 0.477 |
| 1.6 | 0 | yadI | 5 | 20.6333 | 0.594 |

| | | | | | |
|---|---|---|---|---|---|
| 1.6 | 0 | AG1 | 5 | 20.6333 | 0.619 |
| 1.6 | 0 | ptsI | 5 | 20.8833 | 0.536 |
| 1.6 | 0 | clcB | 5 | 20.8833 | 0.503 |
| 1.6 | 0 | ycaM | 5 | 20.8833 | 0.479 |
| 1.6 | 0 | yadI | 5 | 20.8833 | 0.597 |
| 1.6 | 0 | AG1 | 5 | 20.8833 | 0.606 |
| 1.6 | 0 | ptsI | 5 | 21.1333 | 0.542 |
| 1.6 | 0 | clcB | 5 | 21.1333 | 0.499 |
| 1.6 | 0 | ycaM | 5 | 21.1333 | 0.481 |
| 1.6 | 0 | yadI | 5 | 21.1333 | 0.597 |
| 1.6 | 0 | AG1 | 5 | 21.1333 | 0.606 |
| 1.6 | 0 | ptsI | 5 | 21.3833 | 0.539 |
| 1.6 | 0 | clcB | 5 | 21.3833 | 0.502 |
| 1.6 | 0 | ycaM | 5 | 21.3833 | 0.479 |
| 1.6 | 0 | yadI | 5 | 21.3833 | 0.592 |
| 1.6 | 0 | AG1 | 5 | 21.3833 | 0.605 |
| 1.6 | 0 | ptsI | 5 | 21.6333 | 0.543 |
| 1.6 | 0 | clcB | 5 | 21.6333 | 0.502 |
| 1.6 | 0 | ycaM | 5 | 21.6333 | 0.481 |
| 1.6 | 0 | yadI | 5 | 21.6333 | 0.597 |
| 1.6 | 0 | AG1 | 5 | 21.6333 | 0.607 |
| 1.6 | 0 | ptsI | 5 | 21.8833 | 0.542 |
| 1.6 | 0 | clcB | 5 | 21.8833 | 0.501 |
| 1.6 | 0 | ycaM | 5 | 21.8833 | 0.482 |
| 1.6 | 0 | yadI | 5 | 21.8833 | 0.594 |
| 1.6 | 0 | AG1 | 5 | 21.8833 | 0.61 |
| 1.6 | 0 | ptsI | 5 | 22.1333 | 0.55 |
| 1.6 | 0 | clcB | 5 | 22.1333 | 0.503 |
| 1.6 | 0 | ycaM | 5 | 22.1333 | 0.483 |
| 1.6 | 0 | yadI | 5 | 22.1333 | 0.594 |
| 1.6 | 0 | AG1 | 5 | 22.1333 | 0.608 |
| 1.6 | 0 | ptsI | 5 | 22.3833 | 0.55 |
| 1.6 | 0 | clcB | 5 | 22.3833 | 0.501 |
| 1.6 | 0 | ycaM | 5 | 22.3833 | 0.481 |
| 1.6 | 0 | yadI | 5 | 22.3833 | 0.596 |
| 1.6 | 0 | AG1 | 5 | 22.3833 | 0.61 |
| 1.6 | 0 | ptsI | 5 | 22.6333 | 0.556 |
| 1.6 | 0 | clcB | 5 | 22.6333 | 0.504 |
| 1.6 | 0 | ycaM | 5 | 22.6333 | 0.484 |
| 1.6 | 0 | yadI | 5 | 22.6333 | 0.594 |
| 1.6 | 0 | AG1 | 5 | 22.6333 | 0.613 |
| 1.6 | 0 | ptsI | 5 | 22.8833 | 0.556 |
| 1.6 | 0 | clcB | 5 | 22.8833 | 0.504 |
| 1.6 | 0 | ycaM | 5 | 22.8833 | 0.486 |
| 1.6 | 0 | yadI | 5 | 22.8833 | 0.597 |
| 1.6 | 0 | AG1 | 5 | 22.8833 | 0.615 |
| 1.6 | 0 | ptsI | 5 | 23.1333 | 0.558 |
| 1.6 | 0 | clcB | 5 | 23.1333 | 0.504 |

| | | | | | |
|---|---|---|---|---|---|
| 1.6 | 0 | ycaM | 5 | 23.1333 | 0.488 |
| 1.6 | 0 | yadI | 5 | 23.1333 | 0.596 |
| 1.6 | 0 | AG1 | 5 | 23.1333 | 0.614 |
| 1.6 | 0 | ptsI | 5 | 23.3833 | 0.559 |
| 1.6 | 0 | clcB | 5 | 23.3833 | 0.505 |
| 1.6 | 0 | ycaM | 5 | 23.3833 | 0.487 |
| 1.6 | 0 | yadI | 5 | 23.3833 | 0.598 |
| 1.6 | 0 | AG1 | 5 | 23.3833 | 0.612 |
| 1.6 | 0 | ptsI | 5 | 23.6333 | 0.559 |
| 1.6 | 0 | clcB | 5 | 23.6333 | 0.506 |
| 1.6 | 0 | ycaM | 5 | 23.6333 | 0.487 |
| 1.6 | 0 | yadI | 5 | 23.6333 | 0.594 |
| 1.6 | 0 | AG1 | 5 | 23.6333 | 0.612 |
| 1.6 | 0 | ptsI | 5 | 23.8833 | 0.563 |
| 1.6 | 0 | clcB | 5 | 23.8833 | 0.506 |
| 1.6 | 0 | ycaM | 5 | 23.8833 | 0.486 |
| 1.6 | 0 | yadI | 5 | 23.8833 | 0.595 |
| 1.6 | 0 | AG1 | 5 | 23.8833 | 0.609 |
| 1.6 | 0 | ptsI | 5 | 24.1333 | 0.565 |
| 1.6 | 0 | clcB | 5 | 24.1333 | 0.505 |
| 1.6 | 0 | ycaM | 5 | 24.1333 | 0.488 |
| 1.6 | 0 | yadI | 5 | 24.1333 | 0.594 |
| 1.6 | 0 | AG1 | 5 | 24.1333 | 0.614 |
| 1.6 | 0 | ptsI | 5 | 24.3833 | 0.566 |
| 1.6 | 0 | clcB | 5 | 24.3833 | 0.506 |
| 1.6 | 0 | ycaM | 5 | 24.3833 | 0.492 |
| 1.6 | 0 | yadI | 5 | 24.3833 | 0.596 |
| 1.6 | 0 | AG1 | 5 | 24.3833 | 0.613 |
| 3.2 | 0 | ptsI | 1 | 0 | 0.219 |
| 3.2 | 0 | clcB | 1 | 0 | 0.22 |
| 3.2 | 0 | ycaM | 1 | 0 | 0.223 |
| 3.2 | 0 | yadI | 1 | 0 | 0.264 |
| 3.2 | 0 | AG1 | 1 | 0 | 0.246 |
| 3.2 | 0 | ptsI | 1 | 0.25 | 0.221 |
| 3.2 | 0 | clcB | 1 | 0.25 | 0.215 |
| 3.2 | 0 | ycaM | 1 | 0.25 | 0.215 |
| 3.2 | 0 | yadI | 1 | 0.25 | 0.26 |
| 3.2 | 0 | AG1 | 1 | 0.25 | 0.233 |
| 3.2 | 0 | ptsI | 1 | 0.5 | 0.224 |
| 3.2 | 0 | clcB | 1 | 0.5 | 0.214 |
| 3.2 | 0 | ycaM | 1 | 0.5 | 0.214 |
| 3.2 | 0 | yadI | 1 | 0.5 | 0.257 |
| 3.2 | 0 | AG1 | 1 | 0.5 | 0.233 |
| 3.2 | 0 | ptsI | 1 | 0.75 | 0.222 |
| 3.2 | 0 | clcB | 1 | 0.75 | 0.214 |
| 3.2 | 0 | ycaM | 1 | 0.75 | 0.21 |
| 3.2 | 0 | yadI | 1 | 0.75 | 0.255 |
| 3.2 | 0 | AG1 | 1 | 0.75 | 0.232 |

| | | | | | |
|---|---|---|---|---|---|
| 3.2 | 0 | ptsI | 1 | 1 | 0.225 |
| 3.2 | 0 | clcB | 1 | 1 | 0.216 |
| 3.2 | 0 | ycaM | 1 | 1 | 0.212 |
| 3.2 | 0 | yadI | 1 | 1 | 0.259 |
| 3.2 | 0 | AG1 | 1 | 1 | 0.234 |
| 3.2 | 0 | ptsI | 1 | 1.25 | 0.229 |
| 3.2 | 0 | clcB | 1 | 1.25 | 0.216 |
| 3.2 | 0 | ycaM | 1 | 1.25 | 0.212 |
| 3.2 | 0 | yadI | 1 | 1.25 | 0.256 |
| 3.2 | 0 | AG1 | 1 | 1.25 | 0.231 |
| 3.2 | 0 | ptsI | 1 | 1.5 | 0.235 |
| 3.2 | 0 | clcB | 1 | 1.5 | 0.217 |
| 3.2 | 0 | ycaM | 1 | 1.5 | 0.214 |
| 3.2 | 0 | yadI | 1 | 1.5 | 0.262 |
| 3.2 | 0 | AG1 | 1 | 1.5 | 0.235 |
| 3.2 | 0 | ptsI | 1 | 1.75 | 0.242 |
| 3.2 | 0 | clcB | 1 | 1.75 | 0.217 |
| 3.2 | 0 | ycaM | 1 | 1.75 | 0.22 |
| 3.2 | 0 | yadI | 1 | 1.75 | 0.266 |
| 3.2 | 0 | AG1 | 1 | 1.75 | 0.236 |
| 3.2 | 0 | ptsI | 1 | 2 | 0.245 |
| 3.2 | 0 | clcB | 1 | 2 | 0.22 |
| 3.2 | 0 | ycaM | 1 | 2 | 0.223 |
| 3.2 | 0 | yadI | 1 | 2 | 0.274 |
| 3.2 | 0 | AG1 | 1 | 2 | 0.238 |
| 3.2 | 0 | ptsI | 1 | 2.25 | 0.255 |
| 3.2 | 0 | clcB | 1 | 2.25 | 0.224 |
| 3.2 | 0 | ycaM | 1 | 2.25 | 0.231 |
| 3.2 | 0 | yadI | 1 | 2.25 | 0.285 |
| 3.2 | 0 | AG1 | 1 | 2.25 | 0.245 |
| 3.2 | 0 | ptsI | 1 | 2.5 | 0.259 |
| 3.2 | 0 | clcB | 1 | 2.5 | 0.228 |
| 3.2 | 0 | ycaM | 1 | 2.5 | 0.236 |
| 3.2 | 0 | yadI | 1 | 2.5 | 0.292 |
| 3.2 | 0 | AG1 | 1 | 2.5 | 0.25 |
| 3.2 | 0 | ptsI | 1 | 2.75 | 0.269 |
| 3.2 | 0 | clcB | 1 | 2.75 | 0.235 |
| 3.2 | 0 | ycaM | 1 | 2.75 | 0.244 |
| 3.2 | 0 | yadI | 1 | 2.75 | 0.301 |
| 3.2 | 0 | AG1 | 1 | 2.75 | 0.256 |
| 3.2 | 0 | ptsI | 1 | 3 | 0.28 |
| 3.2 | 0 | clcB | 1 | 3 | 0.241 |
| 3.2 | 0 | ycaM | 1 | 3 | 0.252 |
| 3.2 | 0 | yadI | 1 | 3 | 0.313 |
| 3.2 | 0 | AG1 | 1 | 3 | 0.262 |
| 3.2 | 0 | ptsI | 1 | 3.25 | 0.29 |
| 3.2 | 0 | clcB | 1 | 3.25 | 0.247 |
| 3.2 | 0 | ycaM | 1 | 3.25 | 0.258 |

| | | | | | |
|---|---|---|---|---|---|
| 3.2 | 0 | yadI | 1 | 3.25 | 0.33 |
| 3.2 | 0 | AG1 | 1 | 3.25 | 0.273 |
| 3.2 | 0 | ptsI | 1 | 3.5 | 0.296 |
| 3.2 | 0 | clcB | 1 | 3.5 | 0.251 |
| 3.2 | 0 | ycaM | 1 | 3.5 | 0.266 |
| 3.2 | 0 | yadI | 1 | 3.5 | 0.343 |
| 3.2 | 0 | AG1 | 1 | 3.5 | 0.276 |
| 3.2 | 0 | ptsI | 1 | 3.75 | 0.307 |
| 3.2 | 0 | clcB | 1 | 3.75 | 0.253 |
| 3.2 | 0 | ycaM | 1 | 3.75 | 0.276 |
| 3.2 | 0 | yadI | 1 | 3.75 | 0.353 |
| 3.2 | 0 | AG1 | 1 | 3.75 | 0.274 |
| 3.2 | 0 | ptsI | 1 | 4 | 0.313 |
| 3.2 | 0 | clcB | 1 | 4 | 0.267 |
| 3.2 | 0 | ycaM | 1 | 4 | 0.288 |
| 3.2 | 0 | yadI | 1 | 4 | 0.386 |
| 3.2 | 0 | AG1 | 1 | 4 | 0.286 |
| 3.2 | 0 | ptsI | 1 | 4.25 | 0.318 |
| 3.2 | 0 | clcB | 1 | 4.25 | 0.266 |
| 3.2 | 0 | ycaM | 1 | 4.25 | 0.298 |
| 3.2 | 0 | yadI | 1 | 4.25 | 0.389 |
| 3.2 | 0 | AG1 | 1 | 4.25 | 0.295 |
| 3.2 | 0 | ptsI | 1 | 4.5 | 0.323 |
| 3.2 | 0 | clcB | 1 | 4.5 | 0.271 |
| 3.2 | 0 | ycaM | 1 | 4.5 | 0.304 |
| 3.2 | 0 | yadI | 1 | 4.5 | 0.412 |
| 3.2 | 0 | AG1 | 1 | 4.5 | 0.308 |
| 3.2 | 0 | ptsI | 1 | 4.75 | 0.332 |
| 3.2 | 0 | clcB | 1 | 4.75 | 0.273 |
| 3.2 | 0 | ycaM | 1 | 4.75 | 0.305 |
| 3.2 | 0 | yadI | 1 | 4.75 | 0.396 |
| 3.2 | 0 | AG1 | 1 | 4.75 | 0.308 |
| 3.2 | 0 | ptsI | 1 | 5 | 0.34 |
| 3.2 | 0 | clcB | 1 | 5 | 0.286 |
| 3.2 | 0 | ycaM | 1 | 5 | 0.318 |
| 3.2 | 0 | yadI | 1 | 5 | 0.43 |
| 3.2 | 0 | AG1 | 1 | 5 | 0.317 |
| 3.2 | 0 | ptsI | 1 | 5.25 | 0.351 |
| 3.2 | 0 | clcB | 1 | 5.25 | 0.292 |
| 3.2 | 0 | ycaM | 1 | 5.25 | 0.329 |
| 3.2 | 0 | yadI | 1 | 5.25 | 0.448 |
| 3.2 | 0 | AG1 | 1 | 5.25 | 0.329 |
| 3.2 | 0 | ptsI | 1 | 5.5 | 0.356 |
| 3.2 | 0 | clcB | 1 | 5.5 | 0.301 |
| 3.2 | 0 | ycaM | 1 | 5.5 | 0.34 |
| 3.2 | 0 | yadI | 1 | 5.5 | 0.464 |
| 3.2 | 0 | AG1 | 1 | 5.5 | 0.34 |
| 3.2 | 0 | ptsI | 1 | 5.75 | 0.362 |

| | | | | | |
|---|---|---|---|---|---|
| 3.2 | 0 | clcB | 1 | 5.75 | 0.307 |
| 3.2 | 0 | ycaM | 1 | 5.75 | 0.351 |
| 3.2 | 0 | yadI | 1 | 5.75 | 0.481 |
| 3.2 | 0 | AG1 | 1 | 5.75 | 0.354 |
| 3.2 | 0 | ptsI | 1 | 6 | 0.378 |
| 3.2 | 0 | clcB | 1 | 6 | 0.313 |
| 3.2 | 0 | ycaM | 1 | 6 | 0.351 |
| 3.2 | 0 | yadI | 1 | 6 | 0.476 |
| 3.2 | 0 | AG1 | 1 | 6 | 0.357 |
| 3.2 | 0 | ptsI | 1 | 6.25 | 0.394 |
| 3.2 | 0 | clcB | 1 | 6.25 | 0.322 |
| 3.2 | 0 | ycaM | 1 | 6.25 | 0.362 |
| 3.2 | 0 | yadI | 1 | 6.25 | 0.495 |
| 3.2 | 0 | AG1 | 1 | 6.25 | 0.364 |
| 3.2 | 0 | ptsI | 1 | 6.5 | 0.402 |
| 3.2 | 0 | clcB | 1 | 6.5 | 0.327 |
| 3.2 | 0 | ycaM | 1 | 6.5 | 0.372 |
| 3.2 | 0 | yadI | 1 | 6.5 | 0.493 |
| 3.2 | 0 | AG1 | 1 | 6.5 | 0.368 |
| 3.2 | 0 | ptsI | 1 | 6.75 | 0.414 |
| 3.2 | 0 | clcB | 1 | 6.75 | 0.344 |
| 3.2 | 0 | ycaM | 1 | 6.75 | 0.393 |
| 3.2 | 0 | yadI | 1 | 6.75 | 0.525 |
| 3.2 | 0 | AG1 | 1 | 6.75 | 0.391 |
| 3.2 | 0 | ptsI | 1 | 7 | 0.417 |
| 3.2 | 0 | clcB | 1 | 7 | 0.35 |
| 3.2 | 0 | ycaM | 1 | 7 | 0.403 |
| 3.2 | 0 | yadI | 1 | 7 | 0.544 |
| 3.2 | 0 | AG1 | 1 | 7 | 0.411 |
| 3.2 | 0 | ptsI | 1 | 7.25 | 0.422 |
| 3.2 | 0 | clcB | 1 | 7.25 | 0.356 |
| 3.2 | 0 | ycaM | 1 | 7.25 | 0.411 |
| 3.2 | 0 | yadI | 1 | 7.25 | 0.558 |
| 3.2 | 0 | AG1 | 1 | 7.25 | 0.419 |
| 3.2 | 0 | ptsI | 1 | 7.5 | 0.429 |
| 3.2 | 0 | clcB | 1 | 7.5 | 0.366 |
| 3.2 | 0 | ycaM | 1 | 7.5 | 0.417 |
| 3.2 | 0 | yadI | 1 | 7.5 | 0.573 |
| 3.2 | 0 | AG1 | 1 | 7.5 | 0.43 |
| 3.2 | 0 | ptsI | 1 | 7.75 | 0.439 |
| 3.2 | 0 | clcB | 1 | 7.75 | 0.374 |
| 3.2 | 0 | ycaM | 1 | 7.75 | 0.425 |
| 3.2 | 0 | yadI | 1 | 7.75 | 0.582 |
| 3.2 | 0 | AG1 | 1 | 7.75 | 0.44 |
| 3.2 | 0 | ptsI | 1 | 8 | 0.451 |
| 3.2 | 0 | clcB | 1 | 8 | 0.382 |
| 3.2 | 0 | ycaM | 1 | 8 | 0.434 |
| 3.2 | 0 | yadI | 1 | 8 | 0.599 |

| | | | | | |
|---|---|---|---|---|---|
| 3.2 | 0 | AG1 | 1 | 8 | 0.454 |
| 3.2 | 0 | ptsI | 1 | 8.25 | 0.459 |
| 3.2 | 0 | clcB | 1 | 8.25 | 0.391 |
| 3.2 | 0 | ycaM | 1 | 8.25 | 0.438 |
| 3.2 | 0 | yadI | 1 | 8.25 | 0.611 |
| 3.2 | 0 | AG1 | 1 | 8.25 | 0.465 |
| 3.2 | 0 | ptsI | 1 | 8.5 | 0.469 |
| 3.2 | 0 | clcB | 1 | 8.5 | 0.396 |
| 3.2 | 0 | ycaM | 1 | 8.5 | 0.44 |
| 3.2 | 0 | yadI | 1 | 8.5 | 0.627 |
| 3.2 | 0 | AG1 | 1 | 8.5 | 0.479 |
| 3.2 | 0 | ptsI | 1 | 8.75 | 0.481 |
| 3.2 | 0 | clcB | 1 | 8.75 | 0.403 |
| 3.2 | 0 | ycaM | 1 | 8.75 | 0.446 |
| 3.2 | 0 | yadI | 1 | 8.75 | 0.632 |
| 3.2 | 0 | AG1 | 1 | 8.75 | 0.494 |
| 3.2 | 0 | ptsI | 1 | 9 | 0.493 |
| 3.2 | 0 | clcB | 1 | 9 | 0.416 |
| 3.2 | 0 | ycaM | 1 | 9 | 0.452 |
| 3.2 | 0 | yadI | 1 | 9 | 0.648 |
| 3.2 | 0 | AG1 | 1 | 9 | 0.505 |
| 3.2 | 0 | ptsI | 1 | 9.25 | 0.516 |
| 3.2 | 0 | clcB | 1 | 9.25 | 0.424 |
| 3.2 | 0 | ycaM | 1 | 9.25 | 0.466 |
| 3.2 | 0 | yadI | 1 | 9.25 | 0.648 |
| 3.2 | 0 | AG1 | 1 | 9.25 | 0.521 |
| 3.2 | 0 | ptsI | 1 | 9.5 | 0.552 |
| 3.2 | 0 | clcB | 1 | 9.5 | 0.448 |
| 3.2 | 0 | ycaM | 1 | 9.5 | 0.485 |
| 3.2 | 0 | yadI | 1 | 9.5 | 0.667 |
| 3.2 | 0 | AG1 | 1 | 9.5 | 0.547 |
| 3.2 | 0 | ptsI | 1 | 9.75 | 0.545 |
| 3.2 | 0 | clcB | 1 | 9.75 | 0.45 |
| 3.2 | 0 | ycaM | 1 | 9.75 | 0.494 |
| 3.2 | 0 | yadI | 1 | 9.75 | 0.694 |
| 3.2 | 0 | AG1 | 1 | 9.75 | 0.567 |
| 3.2 | 0 | ptsI | 1 | 10 | 0.504 |
| 3.2 | 0 | clcB | 1 | 10 | 0.46 |
| 3.2 | 0 | ycaM | 1 | 10 | 0.504 |
| 3.2 | 0 | yadI | 1 | 10 | 0.678 |
| 3.2 | 0 | AG1 | 1 | 10 | 0.562 |
| 3.2 | 0 | ptsI | 1 | 10.25 | 0.526 |
| 3.2 | 0 | clcB | 1 | 10.25 | 0.466 |
| 3.2 | 0 | ycaM | 1 | 10.25 | 0.515 |
| 3.2 | 0 | yadI | 1 | 10.25 | 0.696 |
| 3.2 | 0 | AG1 | 1 | 10.25 | 0.576 |
| 3.2 | 0 | ptsI | 1 | 10.5 | 0.526 |
| 3.2 | 0 | clcB | 1 | 10.5 | 0.465 |

| | | | | | |
|---|---|---|---|---|---|
| 3.2 | 0 | ycaM | 1 | 10.5 | 0.523 |
| 3.2 | 0 | yadI | 1 | 10.5 | 0.718 |
| 3.2 | 0 | AG1 | 1 | 10.5 | 0.576 |
| 3.2 | 0 | ptsI | 1 | 10.75 | 0.515 |
| 3.2 | 0 | clcB | 1 | 10.75 | 0.488 |
| 3.2 | 0 | ycaM | 1 | 10.75 | 0.538 |
| 3.2 | 0 | yadI | 1 | 10.75 | 0.704 |
| 3.2 | 0 | AG1 | 1 | 10.75 | 0.585 |
| 3.2 | 0 | ptsI | 1 | 11 | 0.519 |
| 3.2 | 0 | clcB | 1 | 11 | 0.505 |
| 3.2 | 0 | ycaM | 1 | 11 | 0.557 |
| 3.2 | 0 | yadI | 1 | 11 | 0.728 |
| 3.2 | 0 | AG1 | 1 | 11 | 0.643 |
| 3.2 | 0 | ptsI | 1 | 11.25 | 0.55 |
| 3.2 | 0 | clcB | 1 | 11.25 | 0.517 |
| 3.2 | 0 | ycaM | 1 | 11.25 | 0.568 |
| 3.2 | 0 | yadI | 1 | 11.25 | 0.749 |
| 3.2 | 0 | AG1 | 1 | 11.25 | 0.645 |
| 3.2 | 0 | ptsI | 1 | 11.5 | 0.564 |
| 3.2 | 0 | clcB | 1 | 11.5 | 0.533 |
| 3.2 | 0 | ycaM | 1 | 11.5 | 0.577 |
| 3.2 | 0 | yadI | 1 | 11.5 | 0.73 |
| 3.2 | 0 | AG1 | 1 | 11.5 | 0.675 |
| 3.2 | 0 | ptsI | 1 | 11.75 | 0.585 |
| 3.2 | 0 | clcB | 1 | 11.75 | 0.555 |
| 3.2 | 0 | ycaM | 1 | 11.75 | 0.586 |
| 3.2 | 0 | yadI | 1 | 11.75 | 0.657 |
| 3.2 | 0 | AG1 | 1 | 11.75 | 0.692 |
| 3.2 | 0 | ptsI | 1 | 12 | 0.609 |
| 3.2 | 0 | clcB | 1 | 12 | 0.554 |
| 3.2 | 0 | ycaM | 1 | 12 | 0.58 |
| 3.2 | 0 | yadI | 1 | 12 | 0.677 |
| 3.2 | 0 | AG1 | 1 | 12 | 0.682 |
| 3.2 | 0 | ptsI | 1 | 12.25 | 0.639 |
| 3.2 | 0 | clcB | 1 | 12.25 | 0.565 |
| 3.2 | 0 | ycaM | 1 | 12.25 | 0.601 |
| 3.2 | 0 | yadI | 1 | 12.25 | 0.609 |
| 3.2 | 0 | AG1 | 1 | 12.25 | 0.696 |
| 3.2 | 0 | ptsI | 1 | 12.5 | 0.633 |
| 3.2 | 0 | clcB | 1 | 12.5 | 0.573 |
| 3.2 | 0 | ycaM | 1 | 12.5 | 0.6 |
| 3.2 | 0 | yadI | 1 | 12.5 | 0.643 |
| 3.2 | 0 | AG1 | 1 | 12.5 | 0.706 |
| 3.2 | 0 | ptsI | 1 | 12.75 | 0.654 |
| 3.2 | 0 | clcB | 1 | 12.75 | 0.564 |
| 3.2 | 0 | ycaM | 1 | 12.75 | 0.594 |
| 3.2 | 0 | yadI | 1 | 12.75 | 0.655 |
| 3.2 | 0 | AG1 | 1 | 12.75 | 0.704 |

| | | | | | |
|---|---|---|---|---|---|
| 3.2 | 0 | ptsI | 1 | 13 | 0.665 |
| 3.2 | 0 | clcB | 1 | 13 | 0.568 |
| 3.2 | 0 | ycaM | 1 | 13 | 0.602 |
| 3.2 | 0 | yadI | 1 | 13 | 0.655 |
| 3.2 | 0 | AG1 | 1 | 13 | 0.715 |
| 3.2 | 0 | ptsI | 1 | 13.25 | 0.721 |
| 3.2 | 0 | clcB | 1 | 13.25 | 0.576 |
| 3.2 | 0 | ycaM | 1 | 13.25 | 0.62 |
| 3.2 | 0 | yadI | 1 | 13.25 | 0.639 |
| 3.2 | 0 | AG1 | 1 | 13.25 | 0.748 |
| 3.2 | 0 | ptsI | 1 | 13.5 | 0.724 |
| 3.2 | 0 | clcB | 1 | 13.5 | 0.574 |
| 3.2 | 0 | ycaM | 1 | 13.5 | 0.617 |
| 3.2 | 0 | yadI | 1 | 13.5 | 0.65 |
| 3.2 | 0 | AG1 | 1 | 13.5 | 0.744 |
| 3.2 | 0 | ptsI | 1 | 13.75 | 0.757 |
| 3.2 | 0 | clcB | 1 | 13.75 | 0.578 |
| 3.2 | 0 | ycaM | 1 | 13.75 | 0.626 |
| 3.2 | 0 | yadI | 1 | 13.75 | 0.66 |
| 3.2 | 0 | AG1 | 1 | 13.75 | 0.763 |
| 3.2 | 0 | ptsI | 1 | 14 | 0.782 |
| 3.2 | 0 | clcB | 1 | 14 | 0.579 |
| 3.2 | 0 | ycaM | 1 | 14 | 0.618 |
| 3.2 | 0 | yadI | 1 | 14 | 0.665 |
| 3.2 | 0 | AG1 | 1 | 14 | 0.776 |
| 3.2 | 0 | ptsI | 1 | 14.25 | 0.798 |
| 3.2 | 0 | clcB | 1 | 14.25 | 0.583 |
| 3.2 | 0 | ycaM | 1 | 14.25 | 0.615 |
| 3.2 | 0 | yadI | 1 | 14.25 | 0.668 |
| 3.2 | 0 | AG1 | 1 | 14.25 | 0.779 |
| 3.2 | 0 | ptsI | 1 | 14.5 | 0.807 |
| 3.2 | 0 | clcB | 1 | 14.5 | 0.581 |
| 3.2 | 0 | ycaM | 1 | 14.5 | 0.641 |
| 3.2 | 0 | yadI | 1 | 14.5 | 0.67 |
| 3.2 | 0 | AG1 | 1 | 14.5 | 0.792 |
| 3.2 | 0 | ptsI | 1 | 14.75 | 0.844 |
| 3.2 | 0 | clcB | 1 | 14.75 | 0.596 |
| 3.2 | 0 | ycaM | 1 | 14.75 | 0.683 |
| 3.2 | 0 | yadI | 1 | 14.75 | 0.667 |
| 3.2 | 0 | AG1 | 1 | 14.75 | 0.818 |
| 3.2 | 0 | ptsI | 1 | 15 | 0.824 |
| 3.2 | 0 | clcB | 1 | 15 | 0.599 |
| 3.2 | 0 | ycaM | 1 | 15 | 0.663 |
| 3.2 | 0 | yadI | 1 | 15 | 0.695 |
| 3.2 | 0 | AG1 | 1 | 15 | 0.814 |
| 3.2 | 0 | ptsI | 1 | 15.25 | 0.826 |
| 3.2 | 0 | clcB | 1 | 15.25 | 0.606 |
| 3.2 | 0 | ycaM | 1 | 15.25 | 0.677 |

| | | | | | |
|---|---|---|---|---|---|
| 3.2 | 0 | yadI | 1 | 15.25 | 0.706 |
| 3.2 | 0 | AG1 | 1 | 15.25 | 0.828 |
| 3.2 | 0 | ptsI | 1 | 15.5 | 0.814 |
| 3.2 | 0 | clcB | 1 | 15.5 | 0.629 |
| 3.2 | 0 | ycaM | 1 | 15.5 | 0.674 |
| 3.2 | 0 | yadI | 1 | 15.5 | 0.701 |
| 3.2 | 0 | AG1 | 1 | 15.5 | 0.825 |
| 3.2 | 0 | ptsI | 1 | 15.75 | 0.74 |
| 3.2 | 0 | clcB | 1 | 15.75 | 0.654 |
| 3.2 | 0 | ycaM | 1 | 15.75 | 0.678 |
| 3.2 | 0 | yadI | 1 | 15.75 | 0.691 |
| 3.2 | 0 | AG1 | 1 | 15.75 | 0.813 |
| 3.2 | 0 | ptsI | 1 | 16 | 0.707 |
| 3.2 | 0 | clcB | 1 | 16 | 0.678 |
| 3.2 | 0 | ycaM | 1 | 16 | 0.695 |
| 3.2 | 0 | yadI | 1 | 16 | 0.683 |
| 3.2 | 0 | AG1 | 1 | 16 | 0.812 |
| 3.2 | 0 | ptsI | 1 | 16.25 | 0.722 |
| 3.2 | 0 | clcB | 1 | 16.25 | 0.672 |
| 3.2 | 0 | ycaM | 1 | 16.25 | 0.688 |
| 3.2 | 0 | yadI | 1 | 16.25 | 0.709 |
| 3.2 | 0 | AG1 | 1 | 16.25 | 0.807 |
| 3.2 | 0 | ptsI | 1 | 16.5 | 0.751 |
| 3.2 | 0 | clcB | 1 | 16.5 | 0.675 |
| 3.2 | 0 | ycaM | 1 | 16.5 | 0.695 |
| 3.2 | 0 | yadI | 1 | 16.5 | 0.716 |
| 3.2 | 0 | AG1 | 1 | 16.5 | 0.802 |
| 3.2 | 0 | ptsI | 1 | 17.0667 | 0.684 |
| 3.2 | 0 | clcB | 1 | 17.0667 | 0.677 |
| 3.2 | 0 | ycaM | 1 | 17.0667 | 0.734 |
| 3.2 | 0 | yadI | 1 | 17.0667 | 0.725 |
| 3.2 | 0 | AG1 | 1 | 17.0667 | 0.84 |
| 3.2 | 0 | ptsI | 1 | 17.3167 | 0.722 |
| 3.2 | 0 | clcB | 1 | 17.3167 | 0.675 |
| 3.2 | 0 | ycaM | 1 | 17.3167 | 0.715 |
| 3.2 | 0 | yadI | 1 | 17.3167 | 0.716 |
| 3.2 | 0 | AG1 | 1 | 17.3167 | 0.812 |
| 3.2 | 0 | ptsI | 1 | 17.5667 | 0.763 |
| 3.2 | 0 | clcB | 1 | 17.5667 | 0.678 |
| 3.2 | 0 | ycaM | 1 | 17.5667 | 0.711 |
| 3.2 | 0 | yadI | 1 | 17.5667 | 0.724 |
| 3.2 | 0 | AG1 | 1 | 17.5667 | 0.796 |
| 3.2 | 0 | ptsI | 1 | 17.8167 | 0.778 |
| 3.2 | 0 | clcB | 1 | 17.8167 | 0.677 |
| 3.2 | 0 | ycaM | 1 | 17.8167 | 0.717 |
| 3.2 | 0 | yadI | 1 | 17.8167 | 0.726 |
| 3.2 | 0 | AG1 | 1 | 17.8167 | 0.802 |
| 3.2 | 0 | ptsI | 1 | 18.0667 | 0.801 |

| | | | | | |
|---|---|---|---|---|---|
| 3.2 | 0 | clcB | 1 | 18.0667 | 0.678 |
| 3.2 | 0 | ycaM | 1 | 18.0667 | 0.715 |
| 3.2 | 0 | yadI | 1 | 18.0667 | 0.729 |
| 3.2 | 0 | AG1 | 1 | 18.0667 | 0.802 |
| 3.2 | 0 | ptsI | 1 | 18.3167 | 0.795 |
| 3.2 | 0 | clcB | 1 | 18.3167 | 0.679 |
| 3.2 | 0 | ycaM | 1 | 18.3167 | 0.721 |
| 3.2 | 0 | yadI | 1 | 18.3167 | 0.721 |
| 3.2 | 0 | AG1 | 1 | 18.3167 | 0.812 |
| 3.2 | 0 | ptsI | 1 | 18.5667 | 0.813 |
| 3.2 | 0 | clcB | 1 | 18.5667 | 0.69 |
| 3.2 | 0 | ycaM | 1 | 18.5667 | 0.729 |
| 3.2 | 0 | yadI | 1 | 18.5667 | 0.741 |
| 3.2 | 0 | AG1 | 1 | 18.5667 | 0.811 |
| 3.2 | 0 | ptsI | 1 | 18.8167 | 0.825 |
| 3.2 | 0 | clcB | 1 | 18.8167 | 0.692 |
| 3.2 | 0 | ycaM | 1 | 18.8167 | 0.734 |
| 3.2 | 0 | yadI | 1 | 18.8167 | 0.737 |
| 3.2 | 0 | AG1 | 1 | 18.8167 | 0.822 |
| 3.2 | 0 | ptsI | 1 | 19.0667 | 0.836 |
| 3.2 | 0 | clcB | 1 | 19.0667 | 0.695 |
| 3.2 | 0 | ycaM | 1 | 19.0667 | 0.747 |
| 3.2 | 0 | yadI | 1 | 19.0667 | 0.744 |
| 3.2 | 0 | AG1 | 1 | 19.0667 | 0.831 |
| 3.2 | 0 | ptsI | 1 | 19.3167 | 0.84 |
| 3.2 | 0 | clcB | 1 | 19.3167 | 0.706 |
| 3.2 | 0 | ycaM | 1 | 19.3167 | 0.751 |
| 3.2 | 0 | yadI | 1 | 19.3167 | 0.747 |
| 3.2 | 0 | AG1 | 1 | 19.3167 | 0.836 |
| 3.2 | 0 | ptsI | 1 | 19.5667 | 0.845 |
| 3.2 | 0 | clcB | 1 | 19.5667 | 0.71 |
| 3.2 | 0 | ycaM | 1 | 19.5667 | 0.756 |
| 3.2 | 0 | yadI | 1 | 19.5667 | 0.747 |
| 3.2 | 0 | AG1 | 1 | 19.5667 | 0.842 |
| 3.2 | 0 | ptsI | 1 | 19.8167 | 0.846 |
| 3.2 | 0 | clcB | 1 | 19.8167 | 0.715 |
| 3.2 | 0 | ycaM | 1 | 19.8167 | 0.759 |
| 3.2 | 0 | yadI | 1 | 19.8167 | 0.744 |
| 3.2 | 0 | AG1 | 1 | 19.8167 | 0.849 |
| 3.2 | 0 | ptsI | 1 | 20.0667 | 0.871 |
| 3.2 | 0 | clcB | 1 | 20.0667 | 0.723 |
| 3.2 | 0 | ycaM | 1 | 20.0667 | 0.768 |
| 3.2 | 0 | yadI | 1 | 20.0667 | 0.745 |
| 3.2 | 0 | AG1 | 1 | 20.0667 | 0.856 |
| 3.2 | 0 | ptsI | 1 | 20.3167 | 0.881 |
| 3.2 | 0 | clcB | 1 | 20.3167 | 0.725 |
| 3.2 | 0 | ycaM | 1 | 20.3167 | 0.768 |
| 3.2 | 0 | yadI | 1 | 20.3167 | 0.744 |

| | | | | | |
|---|---|---|---|---|---|
| 3.2 | 0 | AG1 | 1 | 20.3167 | 0.855 |
| 3.2 | 0 | ptsI | 1 | 20.5667 | 0.894 |
| 3.2 | 0 | clcB | 1 | 20.5667 | 0.734 |
| 3.2 | 0 | ycaM | 1 | 20.5667 | 0.78 |
| 3.2 | 0 | yadI | 1 | 20.5667 | 0.765 |
| 3.2 | 0 | AG1 | 1 | 20.5667 | 0.861 |
| 3.2 | 0 | ptsI | 1 | 20.8167 | 0.906 |
| 3.2 | 0 | clcB | 1 | 20.8167 | 0.739 |
| 3.2 | 0 | ycaM | 1 | 20.8167 | 0.783 |
| 3.2 | 0 | yadI | 1 | 20.8167 | 0.773 |
| 3.2 | 0 | AG1 | 1 | 20.8167 | 0.865 |
| 3.2 | 0 | ptsI | 1 | 21.0667 | 0.911 |
| 3.2 | 0 | clcB | 1 | 21.0667 | 0.748 |
| 3.2 | 0 | ycaM | 1 | 21.0667 | 0.79 |
| 3.2 | 0 | yadI | 1 | 21.0667 | 0.771 |
| 3.2 | 0 | AG1 | 1 | 21.0667 | 0.879 |
| 3.2 | 0 | ptsI | 1 | 21.3167 | 0.918 |
| 3.2 | 0 | clcB | 1 | 21.3167 | 0.751 |
| 3.2 | 0 | ycaM | 1 | 21.3167 | 0.794 |
| 3.2 | 0 | yadI | 1 | 21.3167 | 0.769 |
| 3.2 | 0 | AG1 | 1 | 21.3167 | 0.883 |
| 3.2 | 0 | ptsI | 1 | 21.5667 | 0.921 |
| 3.2 | 0 | clcB | 1 | 21.5667 | 0.754 |
| 3.2 | 0 | ycaM | 1 | 21.5667 | 0.797 |
| 3.2 | 0 | yadI | 1 | 21.5667 | 0.782 |
| 3.2 | 0 | AG1 | 1 | 21.5667 | 0.887 |
| 3.2 | 0 | ptsI | 1 | 21.8167 | 0.926 |
| 3.2 | 0 | clcB | 1 | 21.8167 | 0.76 |
| 3.2 | 0 | ycaM | 1 | 21.8167 | 0.796 |
| 3.2 | 0 | yadI | 1 | 21.8167 | 0.773 |
| 3.2 | 0 | AG1 | 1 | 21.8167 | 0.887 |
| 3.2 | 0 | ptsI | 1 | 22.0667 | 0.95 |
| 3.2 | 0 | clcB | 1 | 22.0667 | 0.766 |
| 3.2 | 0 | ycaM | 1 | 22.0667 | 0.796 |
| 3.2 | 0 | yadI | 1 | 22.0667 | 0.77 |
| 3.2 | 0 | AG1 | 1 | 22.0667 | 0.894 |
| 3.2 | 0 | ptsI | 1 | 22.9333 | 0.946 |
| 3.2 | 0 | clcB | 1 | 22.9333 | 0.738 |
| 3.2 | 0 | ycaM | 1 | 22.9333 | 0.895 |
| 3.2 | 0 | yadI | 1 | 22.9333 | 0.879 |
| 3.2 | 0 | AG1 | 1 | 22.9333 | 0.931 |
| 3.2 | 0 | ptsI | 1 | 23.1833 | 0.96 |
| 3.2 | 0 | clcB | 1 | 23.1833 | 0.716 |
| 3.2 | 0 | ycaM | 1 | 23.1833 | 0.863 |
| 3.2 | 0 | yadI | 1 | 23.1833 | 0.866 |
| 3.2 | 0 | AG1 | 1 | 23.1833 | 0.867 |
| 3.2 | 0 | ptsI | 1 | 23.4333 | 0.952 |
| 3.2 | 0 | clcB | 1 | 23.4333 | 0.715 |

| | | | | | |
|---|---|---|---|---|---|
| 3.2 | 0 | ycaM | 1 | 23.4333 | 0.839 |
| 3.2 | 0 | yadI | 1 | 23.4333 | 0.878 |
| 3.2 | 0 | AG1 | 1 | 23.4333 | 0.854 |
| 3.2 | 0 | ptsI | 1 | 23.6833 | 0.977 |
| 3.2 | 0 | clcB | 1 | 23.6833 | 0.718 |
| 3.2 | 0 | ycaM | 1 | 23.6833 | 0.824 |
| 3.2 | 0 | yadI | 1 | 23.6833 | 0.895 |
| 3.2 | 0 | AG1 | 1 | 23.6833 | 0.86 |
| 3.2 | 0 | ptsI | 1 | 23.9333 | 0.986 |
| 3.2 | 0 | clcB | 1 | 23.9333 | 0.719 |
| 3.2 | 0 | ycaM | 1 | 23.9333 | 0.827 |
| 3.2 | 0 | yadI | 1 | 23.9333 | 0.899 |
| 3.2 | 0 | AG1 | 1 | 23.9333 | 0.866 |
| 3.2 | 0 | ptsI | 1 | 24.1833 | 0.99 |
| 3.2 | 0 | clcB | 1 | 24.1833 | 0.715 |
| 3.2 | 0 | ycaM | 1 | 24.1833 | 0.828 |
| 3.2 | 0 | yadI | 1 | 24.1833 | 0.897 |
| 3.2 | 0 | AG1 | 1 | 24.1833 | 0.876 |
| 3.2 | 0 | ptsI | 1 | 24.4333 | 0.958 |
| 3.2 | 0 | clcB | 1 | 24.4333 | 0.714 |
| 3.2 | 0 | ycaM | 1 | 24.4333 | 0.831 |
| 3.2 | 0 | yadI | 1 | 24.4333 | 0.894 |
| 3.2 | 0 | AG1 | 1 | 24.4333 | 0.886 |
| 3.2 | 0 | ptsI | 1 | 24.6833 | 0.999 |
| 3.2 | 0 | clcB | 1 | 24.6833 | 0.715 |
| 3.2 | 0 | ycaM | 1 | 24.6833 | 0.827 |
| 3.2 | 0 | yadI | 1 | 24.6833 | 0.89 |
| 3.2 | 0 | AG1 | 1 | 24.6833 | 0.886 |
| 3.2 | 0 | ptsI | 1 | 24.9333 | 0.968 |
| 3.2 | 0 | clcB | 1 | 24.9333 | 0.72 |
| 3.2 | 0 | ycaM | 1 | 24.9333 | 0.831 |
| 3.2 | 0 | yadI | 1 | 24.9333 | 0.896 |
| 3.2 | 0 | AG1 | 1 | 24.9333 | 0.887 |
| 3.2 | 0 | ptsI | 1 | 25.1833 | 0.997 |
| 3.2 | 0 | clcB | 1 | 25.1833 | 0.723 |
| 3.2 | 0 | ycaM | 1 | 25.1833 | 0.836 |
| 3.2 | 0 | yadI | 1 | 25.1833 | 0.901 |
| 3.2 | 0 | AG1 | 1 | 25.1833 | 0.889 |
| 3.2 | 0 | ptsI | 1 | 25.4333 | 0.995 |
| 3.2 | 0 | clcB | 1 | 25.4333 | 0.735 |
| 3.2 | 0 | ycaM | 1 | 25.4333 | 0.835 |
| 3.2 | 0 | yadI | 1 | 25.4333 | 0.895 |
| 3.2 | 0 | AG1 | 1 | 25.4333 | 0.888 |
| 3.2 | 0 | ptsI | 1 | 25.6833 | 1.003 |
| 3.2 | 0 | clcB | 1 | 25.6833 | 0.73 |
| 3.2 | 0 | ycaM | 1 | 25.6833 | 0.836 |
| 3.2 | 0 | yadI | 1 | 25.6833 | 0.899 |
| 3.2 | 0 | AG1 | 1 | 25.6833 | 0.895 |

| | | | | | |
|---|---|---|---|---|---|
| 3.2 | 0 | ptsI | 1 | 25.9333 | 1.013 |
| 3.2 | 0 | clcB | 1 | 25.9333 | 0.72 |
| 3.2 | 0 | ycaM | 1 | 25.9333 | 0.838 |
| 3.2 | 0 | yadI | 1 | 25.9333 | 0.893 |
| 3.2 | 0 | AG1 | 1 | 25.9333 | 0.889 |
| 3.2 | 0 | ptsI | 1 | 26.1833 | 0.999 |
| 3.2 | 0 | clcB | 1 | 26.1833 | 0.737 |
| 3.2 | 0 | ycaM | 1 | 26.1833 | 0.85 |
| 3.2 | 0 | yadI | 1 | 26.1833 | 0.898 |
| 3.2 | 0 | AG1 | 1 | 26.1833 | 0.897 |
| 3.2 | 0 | ptsI | 1 | 26.4333 | 1.02 |
| 3.2 | 0 | clcB | 1 | 26.4333 | 0.747 |
| 3.2 | 0 | ycaM | 1 | 26.4333 | 0.851 |
| 3.2 | 0 | yadI | 1 | 26.4333 | 0.905 |
| 3.2 | 0 | AG1 | 1 | 26.4333 | 0.899 |
| 3.2 | 0 | ptsI | 1 | 26.6833 | 1.016 |
| 3.2 | 0 | clcB | 1 | 26.6833 | 0.741 |
| 3.2 | 0 | ycaM | 1 | 26.6833 | 0.857 |
| 3.2 | 0 | yadI | 1 | 26.6833 | 0.908 |
| 3.2 | 0 | AG1 | 1 | 26.6833 | 0.898 |
| 3.2 | 0 | ptsI | 1 | 26.9333 | 1.041 |
| 3.2 | 0 | clcB | 1 | 26.9333 | 0.737 |
| 3.2 | 0 | ycaM | 1 | 26.9333 | 0.862 |
| 3.2 | 0 | yadI | 1 | 26.9333 | 0.908 |
| 3.2 | 0 | AG1 | 1 | 26.9333 | 0.897 |
| 3.2 | 0 | ptsI | 1 | 27.1833 | 1.046 |
| 3.2 | 0 | clcB | 1 | 27.1833 | 0.741 |
| 3.2 | 0 | ycaM | 1 | 27.1833 | 0.868 |
| 3.2 | 0 | yadI | 1 | 27.1833 | 0.91 |
| 3.2 | 0 | AG1 | 1 | 27.1833 | 0.898 |
| 3.2 | 0 | ptsI | 1 | 27.4333 | 1.054 |
| 3.2 | 0 | clcB | 1 | 27.4333 | 0.744 |
| 3.2 | 0 | ycaM | 1 | 27.4333 | 0.876 |
| 3.2 | 0 | yadI | 1 | 27.4333 | 0.929 |
| 3.2 | 0 | AG1 | 1 | 27.4333 | 0.901 |
| 3.2 | 0 | ptsI | 1 | 27.6833 | 1.058 |
| 3.2 | 0 | clcB | 1 | 27.6833 | 0.747 |
| 3.2 | 0 | ycaM | 1 | 27.6833 | 0.881 |
| 3.2 | 0 | yadI | 1 | 27.6833 | 0.932 |
| 3.2 | 0 | AG1 | 1 | 27.6833 | 0.901 |
| 3.2 | 0 | ptsI | 1 | 27.9333 | 1.058 |
| 3.2 | 0 | clcB | 1 | 27.9333 | 0.76 |
| 3.2 | 0 | ycaM | 1 | 27.9333 | 0.886 |
| 3.2 | 0 | yadI | 1 | 27.9333 | 0.94 |
| 3.2 | 0 | AG1 | 1 | 27.9333 | 0.905 |
| 3.2 | 0 | ptsI | 1 | 28.1833 | 1.077 |
| 3.2 | 0 | clcB | 1 | 28.1833 | 0.749 |
| 3.2 | 0 | ycaM | 1 | 28.1833 | 0.893 |

| | | | | | |
|---|---|---|---|---|---|
| 3.2 | 0 | yadI | 1 | 28.1833 | 0.937 |
| 3.2 | 0 | AG1 | 1 | 28.1833 | 0.899 |
| 3.2 | 0 | ptsI | 1 | 28.4333 | 1.089 |
| 3.2 | 0 | clcB | 1 | 28.4333 | 0.755 |
| 3.2 | 0 | ycaM | 1 | 28.4333 | 0.9 |
| 3.2 | 0 | yadI | 1 | 28.4333 | 0.946 |
| 3.2 | 0 | AG1 | 1 | 28.4333 | 0.907 |
| 3.2 | 0 | ptsI | 1 | 28.6833 | 1.099 |
| 3.2 | 0 | clcB | 1 | 28.6833 | 0.757 |
| 3.2 | 0 | ycaM | 1 | 28.6833 | 0.909 |
| 3.2 | 0 | yadI | 1 | 28.6833 | 0.954 |
| 3.2 | 0 | AG1 | 1 | 28.6833 | 0.906 |
| 3.2 | 0 | ptsI | 1 | 28.9333 | 1.104 |
| 3.2 | 0 | clcB | 1 | 28.9333 | 0.765 |
| 3.2 | 0 | ycaM | 1 | 28.9333 | 0.917 |
| 3.2 | 0 | yadI | 1 | 28.9333 | 0.956 |
| 3.2 | 0 | AG1 | 1 | 28.9333 | 0.911 |
| 3.2 | 0 | ptsI | 1 | 29.1833 | 1.111 |
| 3.2 | 0 | clcB | 1 | 29.1833 | 0.768 |
| 3.2 | 0 | ycaM | 1 | 29.1833 | 0.928 |
| 3.2 | 0 | yadI | 1 | 29.1833 | 0.965 |
| 3.2 | 0 | AG1 | 1 | 29.1833 | 0.911 |
| 3.2 | 0 | ptsI | 1 | 29.4333 | 1.111 |
| 3.2 | 0 | clcB | 1 | 29.4333 | 0.774 |
| 3.2 | 0 | ycaM | 1 | 29.4333 | 0.936 |
| 3.2 | 0 | yadI | 1 | 29.4333 | 0.968 |
| 3.2 | 0 | AG1 | 1 | 29.4333 | 0.913 |
| 3.2 | 0 | ptsI | 1 | 29.6833 | 1.123 |
| 3.2 | 0 | clcB | 1 | 29.6833 | 0.779 |
| 3.2 | 0 | ycaM | 1 | 29.6833 | 0.944 |
| 3.2 | 0 | yadI | 1 | 29.6833 | 0.971 |
| 3.2 | 0 | AG1 | 1 | 29.6833 | 0.917 |
| 3.2 | 0 | ptsI | 1 | 29.9333 | 1.13 |
| 3.2 | 0 | clcB | 1 | 29.9333 | 0.784 |
| 3.2 | 0 | ycaM | 1 | 29.9333 | 0.952 |
| 3.2 | 0 | yadI | 1 | 29.9333 | 0.984 |
| 3.2 | 0 | AG1 | 1 | 29.9333 | 0.92 |
| 3.2 | 0 | ptsI | 1 | 30.1833 | 1.148 |
| 3.2 | 0 | clcB | 1 | 30.1833 | 0.79 |
| 3.2 | 0 | ycaM | 1 | 30.1833 | 0.963 |
| 3.2 | 0 | yadI | 1 | 30.1833 | 0.992 |
| 3.2 | 0 | AG1 | 1 | 30.1833 | 0.927 |
| 3.2 | 0 | ptsI | 1 | 30.4333 | 1.164 |
| 3.2 | 0 | clcB | 1 | 30.4333 | 0.798 |
| 3.2 | 0 | ycaM | 1 | 30.4333 | 0.97 |
| 3.2 | 0 | yadI | 1 | 30.4333 | 0.995 |
| 3.2 | 0 | AG1 | 1 | 30.4333 | 0.929 |
| 3.2 | 0 | ptsI | 1 | 30.6833 | 1.19 |

| | | | | | |
|---|---|---|---|---|---|
| 3.2 | 0 | clcB | 1 | 30.6833 | 0.803 |
| 3.2 | 0 | ycaM | 1 | 30.6833 | 0.979 |
| 3.2 | 0 | yadI | 1 | 30.6833 | 1.01 |
| 3.2 | 0 | AG1 | 1 | 30.6833 | 0.936 |
| 3.2 | 0 | ptsI | 1 | 30.9333 | 1.178 |
| 3.2 | 0 | clcB | 1 | 30.9333 | 0.808 |
| 3.2 | 0 | ycaM | 1 | 30.9333 | 0.988 |
| 3.2 | 0 | yadI | 1 | 30.9333 | 1.012 |
| 3.2 | 0 | AG1 | 1 | 30.9333 | 0.938 |
| 3.2 | 0 | ptsI | 1 | 31.1833 | 1.216 |
| 3.2 | 0 | clcB | 1 | 31.1833 | 0.813 |
| 3.2 | 0 | ycaM | 1 | 31.1833 | 0.997 |
| 3.2 | 0 | yadI | 1 | 31.1833 | 1.017 |
| 3.2 | 0 | AG1 | 1 | 31.1833 | 0.942 |
| 3.2 | 0 | ptsI | 1 | 31.4333 | 1.211 |
| 3.2 | 0 | clcB | 1 | 31.4333 | 0.821 |
| 3.2 | 0 | ycaM | 1 | 31.4333 | 1.004 |
| 3.2 | 0 | yadI | 1 | 31.4333 | 1.022 |
| 3.2 | 0 | AG1 | 1 | 31.4333 | 0.951 |
| 3.2 | 0 | ptsI | 1 | 31.6833 | 1.224 |
| 3.2 | 0 | clcB | 1 | 31.6833 | 0.828 |
| 3.2 | 0 | ycaM | 1 | 31.6833 | 1.012 |
| 3.2 | 0 | yadI | 1 | 31.6833 | 1.028 |
| 3.2 | 0 | AG1 | 1 | 31.6833 | 0.952 |
| 3.2 | 0 | ptsI | 1 | 31.9333 | 1.217 |
| 3.2 | 0 | clcB | 1 | 31.9333 | 0.834 |
| 3.2 | 0 | ycaM | 1 | 31.9333 | 1.022 |
| 3.2 | 0 | yadI | 1 | 31.9333 | 1.033 |
| 3.2 | 0 | AG1 | 1 | 31.9333 | 0.961 |
| 3.2 | 0 | ptsI | 1 | 32.1833 | 1.244 |
| 3.2 | 0 | clcB | 1 | 32.1833 | 0.839 |
| 3.2 | 0 | ycaM | 1 | 32.1833 | 1.026 |
| 3.2 | 0 | yadI | 1 | 32.1833 | 1.039 |
| 3.2 | 0 | AG1 | 1 | 32.1833 | 0.968 |
| 3.2 | 0 | ptsI | 1 | 32.4333 | 1.273 |
| 3.2 | 0 | clcB | 1 | 32.4333 | 0.855 |
| 3.2 | 0 | ycaM | 1 | 32.4333 | 1.033 |
| 3.2 | 0 | yadI | 1 | 32.4333 | 1.051 |
| 3.2 | 0 | AG1 | 1 | 32.4333 | 0.97 |
| 3.2 | 0 | ptsI | 1 | 32.6833 | 1.263 |
| 3.2 | 0 | clcB | 1 | 32.6833 | 0.848 |
| 3.2 | 0 | ycaM | 1 | 32.6833 | 1.044 |
| 3.2 | 0 | yadI | 1 | 32.6833 | 1.053 |
| 3.2 | 0 | AG1 | 1 | 32.6833 | 0.976 |
| 3.2 | 0 | ptsI | 1 | 32.9333 | 1.289 |
| 3.2 | 0 | clcB | 1 | 32.9333 | 0.857 |
| 3.2 | 0 | ycaM | 1 | 32.9333 | 1.052 |
| 3.2 | 0 | yadI | 1 | 32.9333 | 1.064 |

| | | | | | |
|---|---|---|---|---|---|
| 3.2 | 0 | AG1 | 1 | 32.9333 | 0.984 |
| 3.2 | 0 | ptsI | 1 | 33.1833 | 1.277 |
| 3.2 | 0 | clcB | 1 | 33.1833 | 0.871 |
| 3.2 | 0 | ycaM | 1 | 33.1833 | 1.06 |
| 3.2 | 0 | yadI | 1 | 33.1833 | 1.072 |
| 3.2 | 0 | AG1 | 1 | 33.1833 | 0.993 |
| 3.2 | 0 | ptsI | 1 | 33.4333 | 1.282 |
| 3.2 | 0 | clcB | 1 | 33.4333 | 0.867 |
| 3.2 | 0 | ycaM | 1 | 33.4333 | 1.067 |
| 3.2 | 0 | yadI | 1 | 33.4333 | 1.074 |
| 3.2 | 0 | AG1 | 1 | 33.4333 | 0.998 |
| 3.2 | 0 | ptsI | 1 | 33.6833 | 1.297 |
| 3.2 | 0 | clcB | 1 | 33.6833 | 0.882 |
| 3.2 | 0 | ycaM | 1 | 33.6833 | 1.08 |
| 3.2 | 0 | yadI | 1 | 33.6833 | 1.086 |
| 3.2 | 0 | AG1 | 1 | 33.6833 | 1.007 |
| 3.2 | 0 | ptsI | 1 | 33.9333 | 1.313 |
| 3.2 | 0 | clcB | 1 | 33.9333 | 0.879 |
| 3.2 | 0 | ycaM | 1 | 33.9333 | 1.086 |
| 3.2 | 0 | yadI | 1 | 33.9333 | 1.091 |
| 3.2 | 0 | AG1 | 1 | 33.9333 | 1.01 |
| 3.2 | 0 | ptsI | 1 | 34.1833 | 1.3 |
| 3.2 | 0 | clcB | 1 | 34.1833 | 0.892 |
| 3.2 | 0 | ycaM | 1 | 34.1833 | 1.098 |
| 3.2 | 0 | yadI | 1 | 34.1833 | 1.105 |
| 3.2 | 0 | AG1 | 1 | 34.1833 | 1.016 |
| 3.2 | 0 | ptsI | 1 | 34.4333 | 1.33 |
| 3.2 | 0 | clcB | 1 | 34.4333 | 0.889 |
| 3.2 | 0 | ycaM | 1 | 34.4333 | 1.109 |
| 3.2 | 0 | yadI | 1 | 34.4333 | 1.117 |
| 3.2 | 0 | AG1 | 1 | 34.4333 | 1.022 |
| 3.2 | 0 | ptsI | 1 | 34.6833 | 1.346 |
| 3.2 | 0 | clcB | 1 | 34.6833 | 0.893 |
| 3.2 | 0 | ycaM | 1 | 34.6833 | 1.116 |
| 3.2 | 0 | yadI | 1 | 34.6833 | 1.121 |
| 3.2 | 0 | AG1 | 1 | 34.6833 | 1.031 |
| 3.2 | 0 | ptsI | 1 | 34.9333 | 1.343 |
| 3.2 | 0 | clcB | 1 | 34.9333 | 0.9 |
| 3.2 | 0 | ycaM | 1 | 34.9333 | 1.126 |
| 3.2 | 0 | yadI | 1 | 34.9333 | 1.131 |
| 3.2 | 0 | AG1 | 1 | 34.9333 | 1.038 |
| 3.2 | 0 | ptsI | 1 | 35.1833 | 1.346 |
| 3.2 | 0 | clcB | 1 | 35.1833 | 0.903 |
| 3.2 | 0 | ycaM | 1 | 35.1833 | 1.132 |
| 3.2 | 0 | yadI | 1 | 35.1833 | 1.138 |
| 3.2 | 0 | AG1 | 1 | 35.1833 | 1.043 |
| 3.2 | 0 | ptsI | 1 | 35.4333 | 1.352 |
| 3.2 | 0 | clcB | 1 | 35.4333 | 0.91 |

| | | | | | |
|---|---|---|---|---|---|
| 3.2 | 0 | ycaM | 1 | 35.4333 | 1.144 |
| 3.2 | 0 | yadI | 1 | 35.4333 | 1.147 |
| 3.2 | 0 | AG1 | 1 | 35.4333 | 1.05 |
| 3.2 | 0 | ptsI | 1 | 35.6833 | 1.368 |
| 3.2 | 0 | clcB | 1 | 35.6833 | 0.914 |
| 3.2 | 0 | ycaM | 1 | 35.6833 | 1.149 |
| 3.2 | 0 | yadI | 1 | 35.6833 | 1.155 |
| 3.2 | 0 | AG1 | 1 | 35.6833 | 1.055 |
| 3.2 | 0 | ptsI | 1 | 35.9333 | 1.368 |
| 3.2 | 0 | clcB | 1 | 35.9333 | 0.929 |
| 3.2 | 0 | ycaM | 1 | 35.9333 | 1.151 |
| 3.2 | 0 | yadI | 1 | 35.9333 | 1.169 |
| 3.2 | 0 | AG1 | 1 | 35.9333 | 1.066 |
| 3.2 | 0 | ptsI | 1 | 36.1833 | 1.398 |
| 3.2 | 0 | clcB | 1 | 36.1833 | 0.921 |
| 3.2 | 0 | ycaM | 1 | 36.1833 | 1.152 |
| 3.2 | 0 | yadI | 1 | 36.1833 | 1.172 |
| 3.2 | 0 | AG1 | 1 | 36.1833 | 1.075 |
| 3.2 | 0 | ptsI | 1 | 36.4333 | 1.388 |
| 3.2 | 0 | clcB | 1 | 36.4333 | 0.928 |
| 3.2 | 0 | ycaM | 1 | 36.4333 | 1.157 |
| 3.2 | 0 | yadI | 1 | 36.4333 | 1.185 |
| 3.2 | 0 | AG1 | 1 | 36.4333 | 1.084 |
| 3.2 | 0 | ptsI | 1 | 36.6833 | 1.396 |
| 3.2 | 0 | clcB | 1 | 36.6833 | 0.934 |
| 3.2 | 0 | ycaM | 1 | 36.6833 | 1.163 |
| 3.2 | 0 | yadI | 1 | 36.6833 | 1.191 |
| 3.2 | 0 | AG1 | 1 | 36.6833 | 1.088 |
| 3.2 | 0 | ptsI | 2 | 0 | 0.211 |
| 3.2 | 0 | clcB | 2 | 0 | 0.214 |
| 3.2 | 0 | ycaM | 2 | 0 | 0.216 |
| 3.2 | 0 | yadI | 2 | 0 | 0.215 |
| 3.2 | 0 | AG1 | 2 | 0 | 0.238 |
| 3.2 | 0 | ptsI | 2 | 0.25 | 0.216 |
| 3.2 | 0 | clcB | 2 | 0.25 | 0.204 |
| 3.2 | 0 | ycaM | 2 | 0.25 | 0.213 |
| 3.2 | 0 | yadI | 2 | 0.25 | 0.212 |
| 3.2 | 0 | AG1 | 2 | 0.25 | 0.235 |
| 3.2 | 0 | ptsI | 2 | 0.5 | 0.22 |
| 3.2 | 0 | clcB | 2 | 0.5 | 0.205 |
| 3.2 | 0 | ycaM | 2 | 0.5 | 0.215 |
| 3.2 | 0 | yadI | 2 | 0.5 | 0.212 |
| 3.2 | 0 | AG1 | 2 | 0.5 | 0.238 |
| 3.2 | 0 | ptsI | 2 | 0.75 | 0.226 |
| 3.2 | 0 | clcB | 2 | 0.75 | 0.211 |
| 3.2 | 0 | ycaM | 2 | 0.75 | 0.215 |
| 3.2 | 0 | yadI | 2 | 0.75 | 0.216 |
| 3.2 | 0 | AG1 | 2 | 0.75 | 0.24 |

| | | | | | |
|---|---|---|---|---|---|
| 3.2 | 0 | ptsI | 2 | 1 | 0.228 |
| 3.2 | 0 | clcB | 2 | 1 | 0.21 |
| 3.2 | 0 | ycaM | 2 | 1 | 0.22 |
| 3.2 | 0 | yadI | 2 | 1 | 0.224 |
| 3.2 | 0 | AG1 | 2 | 1 | 0.247 |
| 3.2 | 0 | ptsI | 2 | 1.25 | 0.23 |
| 3.2 | 0 | clcB | 2 | 1.25 | 0.21 |
| 3.2 | 0 | ycaM | 2 | 1.25 | 0.222 |
| 3.2 | 0 | yadI | 2 | 1.25 | 0.231 |
| 3.2 | 0 | AG1 | 2 | 1.25 | 0.25 |
| 3.2 | 0 | ptsI | 2 | 1.5 | 0.236 |
| 3.2 | 0 | clcB | 2 | 1.5 | 0.209 |
| 3.2 | 0 | ycaM | 2 | 1.5 | 0.227 |
| 3.2 | 0 | yadI | 2 | 1.5 | 0.236 |
| 3.2 | 0 | AG1 | 2 | 1.5 | 0.255 |
| 3.2 | 0 | ptsI | 2 | 1.75 | 0.24 |
| 3.2 | 0 | clcB | 2 | 1.75 | 0.212 |
| 3.2 | 0 | ycaM | 2 | 1.75 | 0.233 |
| 3.2 | 0 | yadI | 2 | 1.75 | 0.243 |
| 3.2 | 0 | AG1 | 2 | 1.75 | 0.264 |
| 3.2 | 0 | ptsI | 2 | 2 | 0.247 |
| 3.2 | 0 | clcB | 2 | 2 | 0.215 |
| 3.2 | 0 | ycaM | 2 | 2 | 0.24 |
| 3.2 | 0 | yadI | 2 | 2 | 0.252 |
| 3.2 | 0 | AG1 | 2 | 2 | 0.277 |
| 3.2 | 0 | ptsI | 2 | 2.25 | 0.256 |
| 3.2 | 0 | clcB | 2 | 2.25 | 0.219 |
| 3.2 | 0 | ycaM | 2 | 2.25 | 0.244 |
| 3.2 | 0 | yadI | 2 | 2.25 | 0.263 |
| 3.2 | 0 | AG1 | 2 | 2.25 | 0.289 |
| 3.2 | 0 | ptsI | 2 | 2.5 | 0.268 |
| 3.2 | 0 | clcB | 2 | 2.5 | 0.223 |
| 3.2 | 0 | ycaM | 2 | 2.5 | 0.252 |
| 3.2 | 0 | yadI | 2 | 2.5 | 0.278 |
| 3.2 | 0 | AG1 | 2 | 2.5 | 0.302 |
| 3.2 | 0 | ptsI | 2 | 2.75 | 0.272 |
| 3.2 | 0 | clcB | 2 | 2.75 | 0.228 |
| 3.2 | 0 | ycaM | 2 | 2.75 | 0.257 |
| 3.2 | 0 | yadI | 2 | 2.75 | 0.289 |
| 3.2 | 0 | AG1 | 2 | 2.75 | 0.294 |
| 3.2 | 0 | ptsI | 2 | 3 | 0.27 |
| 3.2 | 0 | clcB | 2 | 3 | 0.238 |
| 3.2 | 0 | ycaM | 2 | 3 | 0.261 |
| 3.2 | 0 | yadI | 2 | 3 | 0.304 |
| 3.2 | 0 | AG1 | 2 | 3 | 0.294 |
| 3.2 | 0 | ptsI | 2 | 3.25 | 0.286 |
| 3.2 | 0 | clcB | 2 | 3.25 | 0.24 |
| 3.2 | 0 | ycaM | 2 | 3.25 | 0.272 |

| | | | | | |
|---|---|---|---|---|---|
| 3.2 | 0 | yadI | 2 | 3.25 | 0.326 |
| 3.2 | 0 | AG1 | 2 | 3.25 | 0.311 |
| 3.2 | 0 | ptsI | 2 | 3.5 | 0.294 |
| 3.2 | 0 | clcB | 2 | 3.5 | 0.245 |
| 3.2 | 0 | ycaM | 2 | 3.5 | 0.277 |
| 3.2 | 0 | yadI | 2 | 3.5 | 0.342 |
| 3.2 | 0 | AG1 | 2 | 3.5 | 0.318 |
| 3.2 | 0 | ptsI | 2 | 3.75 | 0.303 |
| 3.2 | 0 | clcB | 2 | 3.75 | 0.252 |
| 3.2 | 0 | ycaM | 2 | 3.75 | 0.287 |
| 3.2 | 0 | yadI | 2 | 3.75 | 0.363 |
| 3.2 | 0 | AG1 | 2 | 3.75 | 0.329 |
| 3.2 | 0 | ptsI | 2 | 4 | 0.314 |
| 3.2 | 0 | clcB | 2 | 4 | 0.259 |
| 3.2 | 0 | ycaM | 2 | 4 | 0.292 |
| 3.2 | 0 | yadI | 2 | 4 | 0.381 |
| 3.2 | 0 | AG1 | 2 | 4 | 0.339 |
| 3.2 | 0 | ptsI | 2 | 4.38333 | 0.325 |
| 3.2 | 0 | clcB | 2 | 4.38333 | 0.263 |
| 3.2 | 0 | ycaM | 2 | 4.38333 | 0.305 |
| 3.2 | 0 | yadI | 2 | 4.38333 | 0.383 |
| 3.2 | 0 | AG1 | 2 | 4.38333 | 0.275 |
| 3.2 | 0 | ptsI | 2 | 4.63333 | 0.317 |
| 3.2 | 0 | clcB | 2 | 4.63333 | 0.255 |
| 3.2 | 0 | ycaM | 2 | 4.63333 | 0.289 |
| 3.2 | 0 | yadI | 2 | 4.63333 | 0.388 |
| 3.2 | 0 | AG1 | 2 | 4.63333 | 0.283 |
| 3.2 | 0 | ptsI | 2 | 4.88333 | 0.324 |
| 3.2 | 0 | clcB | 2 | 4.88333 | 0.259 |
| 3.2 | 0 | ycaM | 2 | 4.88333 | 0.292 |
| 3.2 | 0 | yadI | 2 | 4.88333 | 0.402 |
| 3.2 | 0 | AG1 | 2 | 4.88333 | 0.295 |
| 3.2 | 0 | ptsI | 2 | 5.13333 | 0.331 |
| 3.2 | 0 | clcB | 2 | 5.13333 | 0.266 |
| 3.2 | 0 | ycaM | 2 | 5.13333 | 0.298 |
| 3.2 | 0 | yadI | 2 | 5.13333 | 0.414 |
| 3.2 | 0 | AG1 | 2 | 5.13333 | 0.303 |
| 3.2 | 0 | ptsI | 2 | 5.38333 | 0.339 |
| 3.2 | 0 | clcB | 2 | 5.38333 | 0.274 |
| 3.2 | 0 | ycaM | 2 | 5.38333 | 0.303 |
| 3.2 | 0 | yadI | 2 | 5.38333 | 0.426 |
| 3.2 | 0 | AG1 | 2 | 5.38333 | 0.313 |
| 3.2 | 0 | ptsI | 2 | 5.63333 | 0.347 |
| 3.2 | 0 | clcB | 2 | 5.63333 | 0.28 |
| 3.2 | 0 | ycaM | 2 | 5.63333 | 0.314 |
| 3.2 | 0 | yadI | 2 | 5.63333 | 0.436 |
| 3.2 | 0 | AG1 | 2 | 5.63333 | 0.321 |
| 3.2 | 0 | ptsI | 2 | 5.88333 | 0.354 |

| | | | | | |
|---|---|---|---|---|---|
| 3.2 | 0 | clcB | 2 | 5.88333 | 0.286 |
| 3.2 | 0 | ycaM | 2 | 5.88333 | 0.32 |
| 3.2 | 0 | yadI | 2 | 5.88333 | 0.443 |
| 3.2 | 0 | AG1 | 2 | 5.88333 | 0.331 |
| 3.2 | 0 | ptsI | 2 | 6.13333 | 0.361 |
| 3.2 | 0 | clcB | 2 | 6.13333 | 0.294 |
| 3.2 | 0 | ycaM | 2 | 6.13333 | 0.332 |
| 3.2 | 0 | yadI | 2 | 6.13333 | 0.453 |
| 3.2 | 0 | AG1 | 2 | 6.13333 | 0.339 |
| 3.2 | 0 | ptsI | 2 | 6.38333 | 0.372 |
| 3.2 | 0 | clcB | 2 | 6.38333 | 0.301 |
| 3.2 | 0 | ycaM | 2 | 6.38333 | 0.34 |
| 3.2 | 0 | yadI | 2 | 6.38333 | 0.467 |
| 3.2 | 0 | AG1 | 2 | 6.38333 | 0.349 |
| 3.2 | 0 | ptsI | 2 | 6.63333 | 0.38 |
| 3.2 | 0 | clcB | 2 | 6.63333 | 0.307 |
| 3.2 | 0 | ycaM | 2 | 6.63333 | 0.348 |
| 3.2 | 0 | yadI | 2 | 6.63333 | 0.477 |
| 3.2 | 0 | AG1 | 2 | 6.63333 | 0.357 |
| 3.2 | 0 | ptsI | 2 | 6.88333 | 0.388 |
| 3.2 | 0 | clcB | 2 | 6.88333 | 0.314 |
| 3.2 | 0 | ycaM | 2 | 6.88333 | 0.356 |
| 3.2 | 0 | yadI | 2 | 6.88333 | 0.487 |
| 3.2 | 0 | AG1 | 2 | 6.88333 | 0.365 |
| 3.2 | 0 | ptsI | 2 | 7.13333 | 0.393 |
| 3.2 | 0 | clcB | 2 | 7.13333 | 0.322 |
| 3.2 | 0 | ycaM | 2 | 7.13333 | 0.364 |
| 3.2 | 0 | yadI | 2 | 7.13333 | 0.498 |
| 3.2 | 0 | AG1 | 2 | 7.13333 | 0.375 |
| 3.2 | 0 | ptsI | 2 | 7.38333 | 0.402 |
| 3.2 | 0 | clcB | 2 | 7.38333 | 0.327 |
| 3.2 | 0 | ycaM | 2 | 7.38333 | 0.372 |
| 3.2 | 0 | yadI | 2 | 7.38333 | 0.512 |
| 3.2 | 0 | AG1 | 2 | 7.38333 | 0.382 |
| 3.2 | 0 | ptsI | 2 | 7.63333 | 0.408 |
| 3.2 | 0 | clcB | 2 | 7.63333 | 0.336 |
| 3.2 | 0 | ycaM | 2 | 7.63333 | 0.379 |
| 3.2 | 0 | yadI | 2 | 7.63333 | 0.523 |
| 3.2 | 0 | AG1 | 2 | 7.63333 | 0.388 |
| 3.2 | 0 | ptsI | 2 | 7.88333 | 0.419 |
| 3.2 | 0 | clcB | 2 | 7.88333 | 0.343 |
| 3.2 | 0 | ycaM | 2 | 7.88333 | 0.384 |
| 3.2 | 0 | yadI | 2 | 7.88333 | 0.534 |
| 3.2 | 0 | AG1 | 2 | 7.88333 | 0.397 |
| 3.2 | 0 | ptsI | 2 | 8.13333 | 0.43 |
| 3.2 | 0 | clcB | 2 | 8.13333 | 0.351 |
| 3.2 | 0 | ycaM | 2 | 8.13333 | 0.388 |
| 3.2 | 0 | yadI | 2 | 8.13333 | 0.55 |

| | | | | | |
|---|---|---|---|---|---|
| 3.2 | 0 | AG1 | 2 | 8.13333 | 0.404 |
| 3.2 | 0 | ptsI | 2 | 8.38333 | 0.437 |
| 3.2 | 0 | clcB | 2 | 8.38333 | 0.36 |
| 3.2 | 0 | ycaM | 2 | 8.38333 | 0.391 |
| 3.2 | 0 | yadI | 2 | 8.38333 | 0.561 |
| 3.2 | 0 | AG1 | 2 | 8.38333 | 0.414 |
| 3.2 | 0 | ptsI | 2 | 8.91667 | 0.368 |
| 3.2 | 0 | clcB | 2 | 8.91667 | 0.319 |
| 3.2 | 0 | ycaM | 2 | 8.91667 | 0.388 |
| 3.2 | 0 | yadI | 2 | 8.91667 | 0.554 |
| 3.2 | 0 | AG1 | 2 | 8.91667 | 0.475 |
| 3.2 | 0 | ptsI | 2 | 9.16667 | 0.378 |
| 3.2 | 0 | clcB | 2 | 9.16667 | 0.333 |
| 3.2 | 0 | ycaM | 2 | 9.16667 | 0.391 |
| 3.2 | 0 | yadI | 2 | 9.16667 | 0.574 |
| 3.2 | 0 | AG1 | 2 | 9.16667 | 0.484 |
| 3.2 | 0 | ptsI | 2 | 9.41667 | 0.381 |
| 3.2 | 0 | clcB | 2 | 9.41667 | 0.34 |
| 3.2 | 0 | ycaM | 2 | 9.41667 | 0.393 |
| 3.2 | 0 | yadI | 2 | 9.41667 | 0.58 |
| 3.2 | 0 | AG1 | 2 | 9.41667 | 0.484 |
| 3.2 | 0 | ptsI | 2 | 9.66667 | 0.389 |
| 3.2 | 0 | clcB | 2 | 9.66667 | 0.345 |
| 3.2 | 0 | ycaM | 2 | 9.66667 | 0.399 |
| 3.2 | 0 | yadI | 2 | 9.66667 | 0.586 |
| 3.2 | 0 | AG1 | 2 | 9.66667 | 0.491 |
| 3.2 | 0 | ptsI | 2 | 9.91667 | 0.393 |
| 3.2 | 0 | clcB | 2 | 9.91667 | 0.352 |
| 3.2 | 0 | ycaM | 2 | 9.91667 | 0.397 |
| 3.2 | 0 | yadI | 2 | 9.91667 | 0.588 |
| 3.2 | 0 | AG1 | 2 | 9.91667 | 0.498 |
| 3.2 | 0 | ptsI | 2 | 10.1667 | 0.402 |
| 3.2 | 0 | clcB | 2 | 10.1667 | 0.356 |
| 3.2 | 0 | ycaM | 2 | 10.1667 | 0.402 |
| 3.2 | 0 | yadI | 2 | 10.1667 | 0.592 |
| 3.2 | 0 | AG1 | 2 | 10.1667 | 0.52 |
| 3.2 | 0 | ptsI | 2 | 10.4167 | 0.402 |
| 3.2 | 0 | clcB | 2 | 10.4167 | 0.36 |
| 3.2 | 0 | ycaM | 2 | 10.4167 | 0.418 |
| 3.2 | 0 | yadI | 2 | 10.4167 | 0.605 |
| 3.2 | 0 | AG1 | 2 | 10.4167 | 0.526 |
| 3.2 | 0 | ptsI | 2 | 10.6667 | 0.409 |
| 3.2 | 0 | clcB | 2 | 10.6667 | 0.366 |
| 3.2 | 0 | ycaM | 2 | 10.6667 | 0.419 |
| 3.2 | 0 | yadI | 2 | 10.6667 | 0.61 |
| 3.2 | 0 | AG1 | 2 | 10.6667 | 0.534 |
| 3.2 | 0 | ptsI | 2 | 10.9167 | 0.41 |
| 3.2 | 0 | clcB | 2 | 10.9167 | 0.37 |

| | | | | | |
|---|---|---|---|---|---|
| 3.2 | 0 | ycaM | 2 | 10.9167 | 0.426 |
| 3.2 | 0 | yadI | 2 | 10.9167 | 0.612 |
| 3.2 | 0 | AG1 | 2 | 10.9167 | 0.54 |
| 3.2 | 0 | ptsI | 2 | 11.1667 | 0.409 |
| 3.2 | 0 | clcB | 2 | 11.1667 | 0.372 |
| 3.2 | 0 | ycaM | 2 | 11.1667 | 0.431 |
| 3.2 | 0 | yadI | 2 | 11.1667 | 0.619 |
| 3.2 | 0 | AG1 | 2 | 11.1667 | 0.537 |
| 3.2 | 0 | ptsI | 2 | 11.4167 | 0.419 |
| 3.2 | 0 | clcB | 2 | 11.4167 | 0.377 |
| 3.2 | 0 | ycaM | 2 | 11.4167 | 0.429 |
| 3.2 | 0 | yadI | 2 | 11.4167 | 0.627 |
| 3.2 | 0 | AG1 | 2 | 11.4167 | 0.542 |
| 3.2 | 0 | ptsI | 2 | 11.6667 | 0.431 |
| 3.2 | 0 | clcB | 2 | 11.6667 | 0.374 |
| 3.2 | 0 | ycaM | 2 | 11.6667 | 0.439 |
| 3.2 | 0 | yadI | 2 | 11.6667 | 0.634 |
| 3.2 | 0 | AG1 | 2 | 11.6667 | 0.543 |
| 3.2 | 0 | ptsI | 2 | 11.9167 | 0.424 |
| 3.2 | 0 | clcB | 2 | 11.9167 | 0.38 |
| 3.2 | 0 | ycaM | 2 | 11.9167 | 0.447 |
| 3.2 | 0 | yadI | 2 | 11.9167 | 0.635 |
| 3.2 | 0 | AG1 | 2 | 11.9167 | 0.543 |
| 3.2 | 0 | ptsI | 2 | 12.1667 | 0.434 |
| 3.2 | 0 | clcB | 2 | 12.1667 | 0.385 |
| 3.2 | 0 | ycaM | 2 | 12.1667 | 0.448 |
| 3.2 | 0 | yadI | 2 | 12.1667 | 0.645 |
| 3.2 | 0 | AG1 | 2 | 12.1667 | 0.548 |
| 3.2 | 0 | ptsI | 2 | 12.4167 | 0.429 |
| 3.2 | 0 | clcB | 2 | 12.4167 | 0.385 |
| 3.2 | 0 | ycaM | 2 | 12.4167 | 0.445 |
| 3.2 | 0 | yadI | 2 | 12.4167 | 0.652 |
| 3.2 | 0 | AG1 | 2 | 12.4167 | 0.553 |
| 3.2 | 0 | ptsI | 2 | 12.6667 | 0.432 |
| 3.2 | 0 | clcB | 2 | 12.6667 | 0.396 |
| 3.2 | 0 | ycaM | 2 | 12.6667 | 0.456 |
| 3.2 | 0 | yadI | 2 | 12.6667 | 0.656 |
| 3.2 | 0 | AG1 | 2 | 12.6667 | 0.556 |
| 3.2 | 0 | ptsI | 2 | 12.9167 | 0.43 |
| 3.2 | 0 | clcB | 2 | 12.9167 | 0.396 |
| 3.2 | 0 | ycaM | 2 | 12.9167 | 0.458 |
| 3.2 | 0 | yadI | 2 | 12.9167 | 0.66 |
| 3.2 | 0 | AG1 | 2 | 12.9167 | 0.562 |
| 3.2 | 0 | ptsI | 2 | 13.1667 | 0.429 |
| 3.2 | 0 | clcB | 2 | 13.1667 | 0.397 |
| 3.2 | 0 | ycaM | 2 | 13.1667 | 0.467 |
| 3.2 | 0 | yadI | 2 | 13.1667 | 0.667 |
| 3.2 | 0 | AG1 | 2 | 13.1667 | 0.563 |

| | | | | | |
|---|---|---|---|---|---|
| 3.2 | 0 | ptsI | 2 | 13.4167 | 0.432 |
| 3.2 | 0 | clcB | 2 | 13.4167 | 0.401 |
| 3.2 | 0 | ycaM | 2 | 13.4167 | 0.469 |
| 3.2 | 0 | yadI | 2 | 13.4167 | 0.672 |
| 3.2 | 0 | AG1 | 2 | 13.4167 | 0.564 |
| 3.2 | 0 | ptsI | 2 | 13.6667 | 0.436 |
| 3.2 | 0 | clcB | 2 | 13.6667 | 0.402 |
| 3.2 | 0 | ycaM | 2 | 13.6667 | 0.471 |
| 3.2 | 0 | yadI | 2 | 13.6667 | 0.68 |
| 3.2 | 0 | AG1 | 2 | 13.6667 | 0.565 |
| 3.2 | 0 | ptsI | 2 | 13.9167 | 0.439 |
| 3.2 | 0 | clcB | 2 | 13.9167 | 0.408 |
| 3.2 | 0 | ycaM | 2 | 13.9167 | 0.476 |
| 3.2 | 0 | yadI | 2 | 13.9167 | 0.683 |
| 3.2 | 0 | AG1 | 2 | 13.9167 | 0.57 |
| 3.2 | 0 | ptsI | 2 | 14.1667 | 0.437 |
| 3.2 | 0 | clcB | 2 | 14.1667 | 0.413 |
| 3.2 | 0 | ycaM | 2 | 14.1667 | 0.48 |
| 3.2 | 0 | yadI | 2 | 14.1667 | 0.691 |
| 3.2 | 0 | AG1 | 2 | 14.1667 | 0.573 |
| 3.2 | 0 | ptsI | 2 | 14.4167 | 0.443 |
| 3.2 | 0 | clcB | 2 | 14.4167 | 0.416 |
| 3.2 | 0 | ycaM | 2 | 14.4167 | 0.478 |
| 3.2 | 0 | yadI | 2 | 14.4167 | 0.694 |
| 3.2 | 0 | AG1 | 2 | 14.4167 | 0.582 |
| 3.2 | 0 | ptsI | 2 | 14.6667 | 0.446 |
| 3.2 | 0 | clcB | 2 | 14.6667 | 0.423 |
| 3.2 | 0 | ycaM | 2 | 14.6667 | 0.485 |
| 3.2 | 0 | yadI | 2 | 14.6667 | 0.704 |
| 3.2 | 0 | AG1 | 2 | 14.6667 | 0.58 |
| 3.2 | 0 | ptsI | 2 | 14.9167 | 0.446 |
| 3.2 | 0 | clcB | 2 | 14.9167 | 0.426 |
| 3.2 | 0 | ycaM | 2 | 14.9167 | 0.484 |
| 3.2 | 0 | yadI | 2 | 14.9167 | 0.709 |
| 3.2 | 0 | AG1 | 2 | 14.9167 | 0.582 |
| 3.2 | 0 | ptsI | 2 | 15.1667 | 0.45 |
| 3.2 | 0 | clcB | 2 | 15.1667 | 0.431 |
| 3.2 | 0 | ycaM | 2 | 15.1667 | 0.483 |
| 3.2 | 0 | yadI | 2 | 15.1667 | 0.701 |
| 3.2 | 0 | AG1 | 2 | 15.1667 | 0.595 |
| 3.2 | 0 | ptsI | 2 | 15.4167 | 0.454 |
| 3.2 | 0 | clcB | 2 | 15.4167 | 0.427 |
| 3.2 | 0 | ycaM | 2 | 15.4167 | 0.488 |
| 3.2 | 0 | yadI | 2 | 15.4167 | 0.721 |
| 3.2 | 0 | AG1 | 2 | 15.4167 | 0.591 |
| 3.2 | 0 | ptsI | 2 | 15.6667 | 0.454 |
| 3.2 | 0 | clcB | 2 | 15.6667 | 0.441 |
| 3.2 | 0 | ycaM | 2 | 15.6667 | 0.492 |

| | | | | | |
|---|---|---|---|---|---|
| 3.2 | 0 | yadI | 2 | 15.6667 | 0.724 |
| 3.2 | 0 | AG1 | 2 | 15.6667 | 0.594 |
| 3.2 | 0 | ptsI | 2 | 15.9167 | 0.461 |
| 3.2 | 0 | clcB | 2 | 15.9167 | 0.444 |
| 3.2 | 0 | ycaM | 2 | 15.9167 | 0.493 |
| 3.2 | 0 | yadI | 2 | 15.9167 | 0.73 |
| 3.2 | 0 | AG1 | 2 | 15.9167 | 0.59 |
| 3.2 | 0 | ptsI | 2 | 16.1667 | 0.465 |
| 3.2 | 0 | clcB | 2 | 16.1667 | 0.444 |
| 3.2 | 0 | ycaM | 2 | 16.1667 | 0.494 |
| 3.2 | 0 | yadI | 2 | 16.1667 | 0.734 |
| 3.2 | 0 | AG1 | 2 | 16.1667 | 0.597 |
| 3.2 | 0 | ptsI | 2 | 16.4167 | 0.462 |
| 3.2 | 0 | clcB | 2 | 16.4167 | 0.452 |
| 3.2 | 0 | ycaM | 2 | 16.4167 | 0.497 |
| 3.2 | 0 | yadI | 2 | 16.4167 | 0.737 |
| 3.2 | 0 | AG1 | 2 | 16.4167 | 0.604 |
| 3.2 | 0 | ptsI | 2 | 16.6667 | 0.467 |
| 3.2 | 0 | clcB | 2 | 16.6667 | 0.453 |
| 3.2 | 0 | ycaM | 2 | 16.6667 | 0.499 |
| 3.2 | 0 | yadI | 2 | 16.6667 | 0.746 |
| 3.2 | 0 | AG1 | 2 | 16.6667 | 0.6 |
| 3.2 | 0 | ptsI | 2 | 16.9167 | 0.47 |
| 3.2 | 0 | clcB | 2 | 16.9167 | 0.455 |
| 3.2 | 0 | ycaM | 2 | 16.9167 | 0.502 |
| 3.2 | 0 | yadI | 2 | 16.9167 | 0.746 |
| 3.2 | 0 | AG1 | 2 | 16.9167 | 0.603 |
| 3.2 | 0 | ptsI | 2 | 17.1667 | 0.478 |
| 3.2 | 0 | clcB | 2 | 17.1667 | 0.457 |
| 3.2 | 0 | ycaM | 2 | 17.1667 | 0.505 |
| 3.2 | 0 | yadI | 2 | 17.1667 | 0.749 |
| 3.2 | 0 | AG1 | 2 | 17.1667 | 0.605 |
| 3.2 | 0 | ptsI | 2 | 17.4167 | 0.48 |
| 3.2 | 0 | clcB | 2 | 17.4167 | 0.464 |
| 3.2 | 0 | ycaM | 2 | 17.4167 | 0.507 |
| 3.2 | 0 | yadI | 2 | 17.4167 | 0.751 |
| 3.2 | 0 | AG1 | 2 | 17.4167 | 0.609 |
| 3.2 | 0 | ptsI | 2 | 17.6667 | 0.484 |
| 3.2 | 0 | clcB | 2 | 17.6667 | 0.466 |
| 3.2 | 0 | ycaM | 2 | 17.6667 | 0.51 |
| 3.2 | 0 | yadI | 2 | 17.6667 | 0.757 |
| 3.2 | 0 | AG1 | 2 | 17.6667 | 0.612 |
| 3.2 | 0 | ptsI | 2 | 17.9167 | 0.49 |
| 3.2 | 0 | clcB | 2 | 17.9167 | 0.475 |
| 3.2 | 0 | ycaM | 2 | 17.9167 | 0.51 |
| 3.2 | 0 | yadI | 2 | 17.9167 | 0.762 |
| 3.2 | 0 | AG1 | 2 | 17.9167 | 0.618 |
| 3.2 | 0 | ptsI | 2 | 18.1667 | 0.494 |

| | | | | | |
|---|---|---|---|---|---|
| 3.2 | 0 | clcB | 2 | 18.1667 | 0.479 |
| 3.2 | 0 | ycaM | 2 | 18.1667 | 0.514 |
| 3.2 | 0 | yadI | 2 | 18.1667 | 0.767 |
| 3.2 | 0 | AG1 | 2 | 18.1667 | 0.618 |
| 3.2 | 0 | ptsI | 2 | 18.4167 | 0.496 |
| 3.2 | 0 | clcB | 2 | 18.4167 | 0.474 |
| 3.2 | 0 | ycaM | 2 | 18.4167 | 0.514 |
| 3.2 | 0 | yadI | 2 | 18.4167 | 0.766 |
| 3.2 | 0 | AG1 | 2 | 18.4167 | 0.619 |
| 3.2 | 0 | ptsI | 2 | 18.6667 | 0.501 |
| 3.2 | 0 | clcB | 2 | 18.6667 | 0.487 |
| 3.2 | 0 | ycaM | 2 | 18.6667 | 0.517 |
| 3.2 | 0 | yadI | 2 | 18.6667 | 0.769 |
| 3.2 | 0 | AG1 | 2 | 18.6667 | 0.622 |
| 3.2 | 0 | ptsI | 2 | 18.9167 | 0.504 |
| 3.2 | 0 | clcB | 2 | 18.9167 | 0.487 |
| 3.2 | 0 | ycaM | 2 | 18.9167 | 0.517 |
| 3.2 | 0 | yadI | 2 | 18.9167 | 0.76 |
| 3.2 | 0 | AG1 | 2 | 18.9167 | 0.608 |
| 3.2 | 0 | ptsI | 2 | 19.1667 | 0.509 |
| 3.2 | 0 | clcB | 2 | 19.1667 | 0.493 |
| 3.2 | 0 | ycaM | 2 | 19.1667 | 0.524 |
| 3.2 | 0 | yadI | 2 | 19.1667 | 0.772 |
| 3.2 | 0 | AG1 | 2 | 19.1667 | 0.626 |
| 3.2 | 0 | ptsI | 2 | 19.4167 | 0.512 |
| 3.2 | 0 | clcB | 2 | 19.4167 | 0.496 |
| 3.2 | 0 | ycaM | 2 | 19.4167 | 0.526 |
| 3.2 | 0 | yadI | 2 | 19.4167 | 0.778 |
| 3.2 | 0 | AG1 | 2 | 19.4167 | 0.628 |
| 3.2 | 0 | ptsI | 2 | 19.6667 | 0.518 |
| 3.2 | 0 | clcB | 2 | 19.6667 | 0.502 |
| 3.2 | 0 | ycaM | 2 | 19.6667 | 0.529 |
| 3.2 | 0 | yadI | 2 | 19.6667 | 0.776 |
| 3.2 | 0 | AG1 | 2 | 19.6667 | 0.63 |
| 3.2 | 0 | ptsI | 2 | 19.9167 | 0.519 |
| 3.2 | 0 | clcB | 2 | 19.9167 | 0.509 |
| 3.2 | 0 | ycaM | 2 | 19.9167 | 0.531 |
| 3.2 | 0 | yadI | 2 | 19.9167 | 0.779 |
| 3.2 | 0 | AG1 | 2 | 19.9167 | 0.63 |
| 3.2 | 0 | ptsI | 2 | 20.1667 | 0.526 |
| 3.2 | 0 | clcB | 2 | 20.1667 | 0.504 |
| 3.2 | 0 | ycaM | 2 | 20.1667 | 0.534 |
| 3.2 | 0 | yadI | 2 | 20.1667 | 0.784 |
| 3.2 | 0 | AG1 | 2 | 20.1667 | 0.634 |
| 3.2 | 0 | ptsI | 2 | 20.4167 | 0.532 |
| 3.2 | 0 | clcB | 2 | 20.4167 | 0.517 |
| 3.2 | 0 | ycaM | 2 | 20.4167 | 0.533 |
| 3.2 | 0 | yadI | 2 | 20.4167 | 0.784 |

| | | | | | |
|---|---|---|---|---|---|
| 3.2 | 0 | AG1 | 2 | 20.4167 | 0.634 |
| 3.2 | 0 | ptsI | 2 | 20.6667 | 0.535 |
| 3.2 | 0 | clcB | 2 | 20.6667 | 0.523 |
| 3.2 | 0 | ycaM | 2 | 20.6667 | 0.538 |
| 3.2 | 0 | yadI | 2 | 20.6667 | 0.785 |
| 3.2 | 0 | AG1 | 2 | 20.6667 | 0.637 |
| 3.2 | 0 | ptsI | 2 | 20.9167 | 0.539 |
| 3.2 | 0 | clcB | 2 | 20.9167 | 0.525 |
| 3.2 | 0 | ycaM | 2 | 20.9167 | 0.539 |
| 3.2 | 0 | yadI | 2 | 20.9167 | 0.785 |
| 3.2 | 0 | AG1 | 2 | 20.9167 | 0.641 |
| 3.2 | 0 | ptsI | 2 | 21.1667 | 0.542 |
| 3.2 | 0 | clcB | 2 | 21.1667 | 0.532 |
| 3.2 | 0 | ycaM | 2 | 21.1667 | 0.54 |
| 3.2 | 0 | yadI | 2 | 21.1667 | 0.785 |
| 3.2 | 0 | AG1 | 2 | 21.1667 | 0.641 |
| 3.2 | 0 | ptsI | 2 | 21.4167 | 0.549 |
| 3.2 | 0 | clcB | 2 | 21.4167 | 0.536 |
| 3.2 | 0 | ycaM | 2 | 21.4167 | 0.538 |
| 3.2 | 0 | yadI | 2 | 21.4167 | 0.763 |
| 3.2 | 0 | AG1 | 2 | 21.4167 | 0.64 |
| 3.2 | 0 | ptsI | 2 | 21.6667 | 0.553 |
| 3.2 | 0 | clcB | 2 | 21.6667 | 0.542 |
| 3.2 | 0 | ycaM | 2 | 21.6667 | 0.543 |
| 3.2 | 0 | yadI | 2 | 21.6667 | 0.79 |
| 3.2 | 0 | AG1 | 2 | 21.6667 | 0.646 |
| 3.2 | 0 | ptsI | 2 | 21.9167 | 0.56 |
| 3.2 | 0 | clcB | 2 | 21.9167 | 0.549 |
| 3.2 | 0 | ycaM | 2 | 21.9167 | 0.546 |
| 3.2 | 0 | yadI | 2 | 21.9167 | 0.79 |
| 3.2 | 0 | AG1 | 2 | 21.9167 | 0.651 |
| 3.2 | 0 | ptsI | 2 | 22.1667 | 0.563 |
| 3.2 | 0 | clcB | 2 | 22.1667 | 0.552 |
| 3.2 | 0 | ycaM | 2 | 22.1667 | 0.546 |
| 3.2 | 0 | yadI | 2 | 22.1667 | 0.791 |
| 3.2 | 0 | AG1 | 2 | 22.1667 | 0.652 |
| 3.2 | 0 | ptsI | 2 | 22.4167 | 0.571 |
| 3.2 | 0 | clcB | 2 | 22.4167 | 0.556 |
| 3.2 | 0 | ycaM | 2 | 22.4167 | 0.548 |
| 3.2 | 0 | yadI | 2 | 22.4167 | 0.791 |
| 3.2 | 0 | AG1 | 2 | 22.4167 | 0.65 |
| 3.2 | 0 | ptsI | 2 | 22.6667 | 0.574 |
| 3.2 | 0 | clcB | 2 | 22.6667 | 0.56 |
| 3.2 | 0 | ycaM | 2 | 22.6667 | 0.549 |
| 3.2 | 0 | yadI | 2 | 22.6667 | 0.794 |
| 3.2 | 0 | AG1 | 2 | 22.6667 | 0.65 |
| 3.2 | 0 | ptsI | 2 | 22.9167 | 0.577 |
| 3.2 | 0 | clcB | 2 | 22.9167 | 0.559 |

| | | | | | |
|---|---|---|---|---|---|
| 3.2 | 0 | ycaM | 2 | 22.9167 | 0.548 |
| 3.2 | 0 | yadI | 2 | 22.9167 | 0.793 |
| 3.2 | 0 | AG1 | 2 | 22.9167 | 0.652 |
| 3.2 | 0 | ptsI | 2 | 23.1667 | 0.58 |
| 3.2 | 0 | clcB | 2 | 23.1667 | 0.568 |
| 3.2 | 0 | ycaM | 2 | 23.1667 | 0.549 |
| 3.2 | 0 | yadI | 2 | 23.1667 | 0.792 |
| 3.2 | 0 | AG1 | 2 | 23.1667 | 0.655 |
| 3.2 | 0 | ptsI | 2 | 23.4167 | 0.587 |
| 3.2 | 0 | clcB | 2 | 23.4167 | 0.575 |
| 3.2 | 0 | ycaM | 2 | 23.4167 | 0.548 |
| 3.2 | 0 | yadI | 2 | 23.4167 | 0.792 |
| 3.2 | 0 | AG1 | 2 | 23.4167 | 0.656 |
| 3.2 | 0 | ptsI | 2 | 23.6667 | 0.587 |
| 3.2 | 0 | clcB | 2 | 23.6667 | 0.575 |
| 3.2 | 0 | ycaM | 2 | 23.6667 | 0.545 |
| 3.2 | 0 | yadI | 2 | 23.6667 | 0.791 |
| 3.2 | 0 | AG1 | 2 | 23.6667 | 0.66 |
| 3.2 | 0 | ptsI | 2 | 23.9167 | 0.594 |
| 3.2 | 0 | clcB | 2 | 23.9167 | 0.586 |
| 3.2 | 0 | ycaM | 2 | 23.9167 | 0.548 |
| 3.2 | 0 | yadI | 2 | 23.9167 | 0.794 |
| 3.2 | 0 | AG1 | 2 | 23.9167 | 0.659 |
| 3.2 | 0 | ptsI | 2 | 24.1667 | 0.592 |
| 3.2 | 0 | clcB | 2 | 24.1667 | 0.587 |
| 3.2 | 0 | ycaM | 2 | 24.1667 | 0.547 |
| 3.2 | 0 | yadI | 2 | 24.1667 | 0.794 |
| 3.2 | 0 | AG1 | 2 | 24.1667 | 0.662 |
| 3.2 | 0 | ptsI | 2 | 24.4167 | 0.602 |
| 3.2 | 0 | clcB | 2 | 24.4167 | 0.59 |
| 3.2 | 0 | ycaM | 2 | 24.4167 | 0.544 |
| 3.2 | 0 | yadI | 2 | 24.4167 | 0.794 |
| 3.2 | 0 | AG1 | 2 | 24.4167 | 0.666 |
| 3.2 | 0 | ptsI | 2 | 24.6667 | 0.605 |
| 3.2 | 0 | clcB | 2 | 24.6667 | 0.597 |
| 3.2 | 0 | ycaM | 2 | 24.6667 | 0.545 |
| 3.2 | 0 | yadI | 2 | 24.6667 | 0.795 |
| 3.2 | 0 | AG1 | 2 | 24.6667 | 0.668 |
| 3.2 | 0 | ptsI | 2 | 24.9167 | 0.608 |
| 3.2 | 0 | clcB | 2 | 24.9167 | 0.599 |
| 3.2 | 0 | ycaM | 2 | 24.9167 | 0.547 |
| 3.2 | 0 | yadI | 2 | 24.9167 | 0.794 |
| 3.2 | 0 | AG1 | 2 | 24.9167 | 0.667 |
| 3.2 | 0 | ptsI | 2 | 25.1667 | 0.611 |
| 3.2 | 0 | clcB | 2 | 25.1667 | 0.607 |
| 3.2 | 0 | ycaM | 2 | 25.1667 | 0.547 |
| 3.2 | 0 | yadI | 2 | 25.1667 | 0.792 |
| 3.2 | 0 | AG1 | 2 | 25.1667 | 0.67 |

| | | | | | |
|---|---|---|---|---|---|
| 3.2 | 0 | ptsI | 2 | 25.4167 | 0.616 |
| 3.2 | 0 | clcB | 2 | 25.4167 | 0.614 |
| 3.2 | 0 | ycaM | 2 | 25.4167 | 0.546 |
| 3.2 | 0 | yadI | 2 | 25.4167 | 0.793 |
| 3.2 | 0 | AG1 | 2 | 25.4167 | 0.676 |
| 3.2 | 0 | ptsI | 2 | 25.6667 | 0.619 |
| 3.2 | 0 | clcB | 2 | 25.6667 | 0.618 |
| 3.2 | 0 | ycaM | 2 | 25.6667 | 0.547 |
| 3.2 | 0 | yadI | 2 | 25.6667 | 0.79 |
| 3.2 | 0 | AG1 | 2 | 25.6667 | 0.675 |
| 3.2 | 0 | ptsI | 2 | 25.9167 | 0.626 |
| 3.2 | 0 | clcB | 2 | 25.9167 | 0.63 |
| 3.2 | 0 | ycaM | 2 | 25.9167 | 0.546 |
| 3.2 | 0 | yadI | 2 | 25.9167 | 0.79 |
| 3.2 | 0 | AG1 | 2 | 25.9167 | 0.678 |
| 3.2 | 0 | ptsI | 2 | 26.1667 | 0.626 |
| 3.2 | 0 | clcB | 2 | 26.1667 | 0.637 |
| 3.2 | 0 | ycaM | 2 | 26.1667 | 0.544 |
| 3.2 | 0 | yadI | 2 | 26.1667 | 0.768 |
| 3.2 | 0 | AG1 | 2 | 26.1667 | 0.674 |
| 3.2 | 0 | ptsI | 2 | 26.4167 | 0.631 |
| 3.2 | 0 | clcB | 2 | 26.4167 | 0.642 |
| 3.2 | 0 | ycaM | 2 | 26.4167 | 0.546 |
| 3.2 | 0 | yadI | 2 | 26.4167 | 0.792 |
| 3.2 | 0 | AG1 | 2 | 26.4167 | 0.684 |
| 3.2 | 0 | ptsI | 3 | 0 | 0.164 |
| 3.2 | 0 | clcB | 3 | 0 | 0.156 |
| 3.2 | 0 | ycaM | 3 | 0 | 0.161 |
| 3.2 | 0 | yadI | 3 | 0 | 0.16 |
| 3.2 | 0 | AG1 | 3 | 0 | 0.18 |
| 3.2 | 0 | ptsI | 3 | 0.25 | 0.162 |
| 3.2 | 0 | clcB | 3 | 0.25 | 0.153 |
| 3.2 | 0 | ycaM | 3 | 0.25 | 0.16 |
| 3.2 | 0 | yadI | 3 | 0.25 | 0.158 |
| 3.2 | 0 | AG1 | 3 | 0.25 | 0.178 |
| 3.2 | 0 | ptsI | 3 | 0.5 | 0.163 |
| 3.2 | 0 | clcB | 3 | 0.5 | 0.152 |
| 3.2 | 0 | ycaM | 3 | 0.5 | 0.16 |
| 3.2 | 0 | yadI | 3 | 0.5 | 0.158 |
| 3.2 | 0 | AG1 | 3 | 0.5 | 0.178 |
| 3.2 | 0 | ptsI | 3 | 0.75 | 0.164 |
| 3.2 | 0 | clcB | 3 | 0.75 | 0.154 |
| 3.2 | 0 | ycaM | 3 | 0.75 | 0.159 |
| 3.2 | 0 | yadI | 3 | 0.75 | 0.159 |
| 3.2 | 0 | AG1 | 3 | 0.75 | 0.177 |
| 3.2 | 0 | ptsI | 3 | 1 | 0.164 |
| 3.2 | 0 | clcB | 3 | 1 | 0.154 |
| 3.2 | 0 | ycaM | 3 | 1 | 0.16 |

| | | | | | |
|---|---|---|---|---|---|
| 3.2 | 0 | yadI | 3 | 1 | 0.162 |
| 3.2 | 0 | AG1 | 3 | 1 | 0.18 |
| 3.2 | 0 | ptsI | 3 | 1.25 | 0.164 |
| 3.2 | 0 | clcB | 3 | 1.25 | 0.156 |
| 3.2 | 0 | ycaM | 3 | 1.25 | 0.162 |
| 3.2 | 0 | yadI | 3 | 1.25 | 0.163 |
| 3.2 | 0 | AG1 | 3 | 1.25 | 0.181 |
| 3.2 | 0 | ptsI | 3 | 1.5 | 0.166 |
| 3.2 | 0 | clcB | 3 | 1.5 | 0.157 |
| 3.2 | 0 | ycaM | 3 | 1.5 | 0.163 |
| 3.2 | 0 | yadI | 3 | 1.5 | 0.164 |
| 3.2 | 0 | AG1 | 3 | 1.5 | 0.181 |
| 3.2 | 0 | ptsI | 3 | 1.75 | 0.165 |
| 3.2 | 0 | clcB | 3 | 1.75 | 0.158 |
| 3.2 | 0 | ycaM | 3 | 1.75 | 0.162 |
| 3.2 | 0 | yadI | 3 | 1.75 | 0.163 |
| 3.2 | 0 | AG1 | 3 | 1.75 | 0.179 |
| 3.2 | 0 | ptsI | 3 | 2 | 0.168 |
| 3.2 | 0 | clcB | 3 | 2 | 0.16 |
| 3.2 | 0 | ycaM | 3 | 2 | 0.165 |
| 3.2 | 0 | yadI | 3 | 2 | 0.167 |
| 3.2 | 0 | AG1 | 3 | 2 | 0.183 |
| 3.2 | 0 | ptsI | 3 | 2.25 | 0.172 |
| 3.2 | 0 | clcB | 3 | 2.25 | 0.161 |
| 3.2 | 0 | ycaM | 3 | 2.25 | 0.167 |
| 3.2 | 0 | yadI | 3 | 2.25 | 0.169 |
| 3.2 | 0 | AG1 | 3 | 2.25 | 0.187 |
| 3.2 | 0 | ptsI | 3 | 2.5 | 0.174 |
| 3.2 | 0 | clcB | 3 | 2.5 | 0.163 |
| 3.2 | 0 | ycaM | 3 | 2.5 | 0.168 |
| 3.2 | 0 | yadI | 3 | 2.5 | 0.172 |
| 3.2 | 0 | AG1 | 3 | 2.5 | 0.189 |
| 3.2 | 0 | ptsI | 3 | 2.75 | 0.176 |
| 3.2 | 0 | clcB | 3 | 2.75 | 0.162 |
| 3.2 | 0 | ycaM | 3 | 2.75 | 0.175 |
| 3.2 | 0 | yadI | 3 | 2.75 | 0.178 |
| 3.2 | 0 | AG1 | 3 | 2.75 | 0.197 |
| 3.2 | 0 | ptsI | 3 | 3 | 0.177 |
| 3.2 | 0 | clcB | 3 | 3 | 0.164 |
| 3.2 | 0 | ycaM | 3 | 3 | 0.174 |
| 3.2 | 0 | yadI | 3 | 3 | 0.178 |
| 3.2 | 0 | AG1 | 3 | 3 | 0.194 |
| 3.2 | 0 | ptsI | 3 | 3.25 | 0.178 |
| 3.2 | 0 | clcB | 3 | 3.25 | 0.168 |
| 3.2 | 0 | ycaM | 3 | 3.25 | 0.177 |
| 3.2 | 0 | yadI | 3 | 3.25 | 0.182 |
| 3.2 | 0 | AG1 | 3 | 3.25 | 0.195 |
| 3.2 | 0 | ptsI | 3 | 3.5 | 0.182 |

| | | | | | |
|---|---|---|---|---|---|
| 3.2 | 0 | clcB | 3 | 3.5 | 0.17 |
| 3.2 | 0 | ycaM | 3 | 3.5 | 0.18 |
| 3.2 | 0 | yadI | 3 | 3.5 | 0.192 |
| 3.2 | 0 | AG1 | 3 | 3.5 | 0.201 |
| 3.2 | 0 | ptsI | 3 | 3.75 | 0.186 |
| 3.2 | 0 | clcB | 3 | 3.75 | 0.171 |
| 3.2 | 0 | ycaM | 3 | 3.75 | 0.185 |
| 3.2 | 0 | yadI | 3 | 3.75 | 0.202 |
| 3.2 | 0 | AG1 | 3 | 3.75 | 0.208 |
| 3.2 | 0 | ptsI | 3 | 4 | 0.188 |
| 3.2 | 0 | clcB | 3 | 4 | 0.174 |
| 3.2 | 0 | ycaM | 3 | 4 | 0.19 |
| 3.2 | 0 | yadI | 3 | 4 | 0.214 |
| 3.2 | 0 | AG1 | 3 | 4 | 0.215 |
| 3.2 | 0 | ptsI | 3 | 4.41667 | 0.195 |
| 3.2 | 0 | clcB | 3 | 4.41667 | 0.17 |
| 3.2 | 0 | ycaM | 3 | 4.41667 | 0.195 |
| 3.2 | 0 | yadI | 3 | 4.41667 | 0.193 |
| 3.2 | 0 | AG1 | 3 | 4.41667 | 0.222 |
| 3.2 | 0 | ptsI | 3 | 4.66667 | 0.189 |
| 3.2 | 0 | clcB | 3 | 4.66667 | 0.17 |
| 3.2 | 0 | ycaM | 3 | 4.66667 | 0.193 |
| 3.2 | 0 | yadI | 3 | 4.66667 | 0.193 |
| 3.2 | 0 | AG1 | 3 | 4.66667 | 0.217 |
| 3.2 | 0 | ptsI | 3 | 4.91667 | 0.189 |
| 3.2 | 0 | clcB | 3 | 4.91667 | 0.17 |
| 3.2 | 0 | ycaM | 3 | 4.91667 | 0.195 |
| 3.2 | 0 | yadI | 3 | 4.91667 | 0.203 |
| 3.2 | 0 | AG1 | 3 | 4.91667 | 0.228 |
| 3.2 | 0 | ptsI | 3 | 5.16667 | 0.193 |
| 3.2 | 0 | clcB | 3 | 5.16667 | 0.174 |
| 3.2 | 0 | ycaM | 3 | 5.16667 | 0.2 |
| 3.2 | 0 | yadI | 3 | 5.16667 | 0.213 |
| 3.2 | 0 | AG1 | 3 | 5.16667 | 0.236 |
| 3.2 | 0 | ptsI | 3 | 5.41667 | 0.198 |
| 3.2 | 0 | clcB | 3 | 5.41667 | 0.176 |
| 3.2 | 0 | ycaM | 3 | 5.41667 | 0.207 |
| 3.2 | 0 | yadI | 3 | 5.41667 | 0.227 |
| 3.2 | 0 | AG1 | 3 | 5.41667 | 0.246 |
| 3.2 | 0 | ptsI | 3 | 5.66667 | 0.204 |
| 3.2 | 0 | clcB | 3 | 5.66667 | 0.18 |
| 3.2 | 0 | ycaM | 3 | 5.66667 | 0.215 |
| 3.2 | 0 | yadI | 3 | 5.66667 | 0.24 |
| 3.2 | 0 | AG1 | 3 | 5.66667 | 0.259 |
| 3.2 | 0 | ptsI | 3 | 5.91667 | 0.21 |
| 3.2 | 0 | clcB | 3 | 5.91667 | 0.185 |
| 3.2 | 0 | ycaM | 3 | 5.91667 | 0.226 |
| 3.2 | 0 | yadI | 3 | 5.91667 | 0.255 |

| | | | | | |
|---|---|---|---|---|---|
| 3.2 | 0 | AG1 | 3 | 5.91667 | 0.27 |
| 3.2 | 0 | ptsI | 3 | 6.16667 | 0.217 |
| 3.2 | 0 | clcB | 3 | 6.16667 | 0.19 |
| 3.2 | 0 | ycaM | 3 | 6.16667 | 0.236 |
| 3.2 | 0 | yadI | 3 | 6.16667 | 0.266 |
| 3.2 | 0 | AG1 | 3 | 6.16667 | 0.281 |
| 3.2 | 0 | ptsI | 3 | 6.41667 | 0.224 |
| 3.2 | 0 | clcB | 3 | 6.41667 | 0.195 |
| 3.2 | 0 | ycaM | 3 | 6.41667 | 0.25 |
| 3.2 | 0 | yadI | 3 | 6.41667 | 0.278 |
| 3.2 | 0 | AG1 | 3 | 6.41667 | 0.288 |
| 3.2 | 0 | ptsI | 3 | 6.66667 | 0.234 |
| 3.2 | 0 | clcB | 3 | 6.66667 | 0.202 |
| 3.2 | 0 | ycaM | 3 | 6.66667 | 0.262 |
| 3.2 | 0 | yadI | 3 | 6.66667 | 0.292 |
| 3.2 | 0 | AG1 | 3 | 6.66667 | 0.298 |
| 3.2 | 0 | ptsI | 3 | 6.91667 | 0.243 |
| 3.2 | 0 | clcB | 3 | 6.91667 | 0.208 |
| 3.2 | 0 | ycaM | 3 | 6.91667 | 0.272 |
| 3.2 | 0 | yadI | 3 | 6.91667 | 0.302 |
| 3.2 | 0 | AG1 | 3 | 6.91667 | 0.307 |
| 3.2 | 0 | ptsI | 3 | 7.16667 | 0.253 |
| 3.2 | 0 | clcB | 3 | 7.16667 | 0.217 |
| 3.2 | 0 | ycaM | 3 | 7.16667 | 0.281 |
| 3.2 | 0 | yadI | 3 | 7.16667 | 0.316 |
| 3.2 | 0 | AG1 | 3 | 7.16667 | 0.315 |
| 3.2 | 0 | ptsI | 3 | 7.41667 | 0.262 |
| 3.2 | 0 | clcB | 3 | 7.41667 | 0.226 |
| 3.2 | 0 | ycaM | 3 | 7.41667 | 0.288 |
| 3.2 | 0 | yadI | 3 | 7.41667 | 0.329 |
| 3.2 | 0 | AG1 | 3 | 7.41667 | 0.325 |
| 3.2 | 0 | ptsI | 3 | 7.66667 | 0.271 |
| 3.2 | 0 | clcB | 3 | 7.66667 | 0.235 |
| 3.2 | 0 | ycaM | 3 | 7.66667 | 0.297 |
| 3.2 | 0 | yadI | 3 | 7.66667 | 0.341 |
| 3.2 | 0 | AG1 | 3 | 7.66667 | 0.332 |
| 3.2 | 0 | ptsI | 3 | 7.91667 | 0.278 |
| 3.2 | 0 | clcB | 3 | 7.91667 | 0.244 |
| 3.2 | 0 | ycaM | 3 | 7.91667 | 0.305 |
| 3.2 | 0 | yadI | 3 | 7.91667 | 0.352 |
| 3.2 | 0 | AG1 | 3 | 7.91667 | 0.34 |
| 3.2 | 0 | ptsI | 3 | 8.16667 | 0.286 |
| 3.2 | 0 | clcB | 3 | 8.16667 | 0.251 |
| 3.2 | 0 | ycaM | 3 | 8.16667 | 0.314 |
| 3.2 | 0 | yadI | 3 | 8.16667 | 0.361 |
| 3.2 | 0 | AG1 | 3 | 8.16667 | 0.35 |
| 3.2 | 0 | ptsI | 3 | 8.41667 | 0.293 |
| 3.2 | 0 | clcB | 3 | 8.41667 | 0.259 |

| | | | | | |
|---|---|---|---|---|---|
| 3.2 | 0 | ycaM | 3 | 8.41667 | 0.325 |
| 3.2 | 0 | yadI | 3 | 8.41667 | 0.372 |
| 3.2 | 0 | AG1 | 3 | 8.41667 | 0.36 |
| 3.2 | 0 | ptsI | 3 | 9.05 | 0.283 |
| 3.2 | 0 | clcB | 3 | 9.05 | 0.266 |
| 3.2 | 0 | ycaM | 3 | 9.05 | 0.313 |
| 3.2 | 0 | yadI | 3 | 9.05 | 0.396 |
| 3.2 | 0 | AG1 | 3 | 9.05 | 0.374 |
| 3.2 | 0 | ptsI | 3 | 9.3 | 0.286 |
| 3.2 | 0 | clcB | 3 | 9.3 | 0.265 |
| 3.2 | 0 | ycaM | 3 | 9.3 | 0.315 |
| 3.2 | 0 | yadI | 3 | 9.3 | 0.415 |
| 3.2 | 0 | AG1 | 3 | 9.3 | 0.383 |
| 3.2 | 0 | ptsI | 3 | 9.55 | 0.286 |
| 3.2 | 0 | clcB | 3 | 9.55 | 0.263 |
| 3.2 | 0 | ycaM | 3 | 9.55 | 0.311 |
| 3.2 | 0 | yadI | 3 | 9.55 | 0.423 |
| 3.2 | 0 | AG1 | 3 | 9.55 | 0.389 |
| 3.2 | 0 | ptsI | 3 | 9.8 | 0.285 |
| 3.2 | 0 | clcB | 3 | 9.8 | 0.26 |
| 3.2 | 0 | ycaM | 3 | 9.8 | 0.309 |
| 3.2 | 0 | yadI | 3 | 9.8 | 0.43 |
| 3.2 | 0 | AG1 | 3 | 9.8 | 0.389 |
| 3.2 | 0 | ptsI | 3 | 10.05 | 0.279 |
| 3.2 | 0 | clcB | 3 | 10.05 | 0.253 |
| 3.2 | 0 | ycaM | 3 | 10.05 | 0.304 |
| 3.2 | 0 | yadI | 3 | 10.05 | 0.434 |
| 3.2 | 0 | AG1 | 3 | 10.05 | 0.391 |
| 3.2 | 0 | ptsI | 3 | 10.3 | 0.278 |
| 3.2 | 0 | clcB | 3 | 10.3 | 0.251 |
| 3.2 | 0 | ycaM | 3 | 10.3 | 0.304 |
| 3.2 | 0 | yadI | 3 | 10.3 | 0.438 |
| 3.2 | 0 | AG1 | 3 | 10.3 | 0.393 |
| 3.2 | 0 | ptsI | 3 | 10.55 | 0.269 |
| 3.2 | 0 | clcB | 3 | 10.55 | 0.248 |
| 3.2 | 0 | ycaM | 3 | 10.55 | 0.306 |
| 3.2 | 0 | yadI | 3 | 10.55 | 0.447 |
| 3.2 | 0 | AG1 | 3 | 10.55 | 0.393 |
| 3.2 | 0 | ptsI | 3 | 10.8 | 0.264 |
| 3.2 | 0 | clcB | 3 | 10.8 | 0.242 |
| 3.2 | 0 | ycaM | 3 | 10.8 | 0.303 |
| 3.2 | 0 | yadI | 3 | 10.8 | 0.449 |
| 3.2 | 0 | AG1 | 3 | 10.8 | 0.4 |
| 3.2 | 0 | ptsI | 3 | 11.05 | 0.261 |
| 3.2 | 0 | clcB | 3 | 11.05 | 0.242 |
| 3.2 | 0 | ycaM | 3 | 11.05 | 0.3 |
| 3.2 | 0 | yadI | 3 | 11.05 | 0.456 |
| 3.2 | 0 | AG1 | 3 | 11.05 | 0.402 |

| | | | | | |
|---|---|---|---|---|---|
| 3.2 | 0 | ptsI | 3 | 11.3 | 0.259 |
| 3.2 | 0 | clcB | 3 | 11.3 | 0.237 |
| 3.2 | 0 | ycaM | 3 | 11.3 | 0.298 |
| 3.2 | 0 | yadI | 3 | 11.3 | 0.457 |
| 3.2 | 0 | AG1 | 3 | 11.3 | 0.402 |
| 3.2 | 0 | ptsI | 3 | 11.55 | 0.254 |
| 3.2 | 0 | clcB | 3 | 11.55 | 0.237 |
| 3.2 | 0 | ycaM | 3 | 11.55 | 0.298 |
| 3.2 | 0 | yadI | 3 | 11.55 | 0.465 |
| 3.2 | 0 | AG1 | 3 | 11.55 | 0.398 |
| 3.2 | 0 | ptsI | 3 | 11.8 | 0.253 |
| 3.2 | 0 | clcB | 3 | 11.8 | 0.234 |
| 3.2 | 0 | ycaM | 3 | 11.8 | 0.297 |
| 3.2 | 0 | yadI | 3 | 11.8 | 0.474 |
| 3.2 | 0 | AG1 | 3 | 11.8 | 0.403 |
| 3.2 | 0 | ptsI | 3 | 12.05 | 0.249 |
| 3.2 | 0 | clcB | 3 | 12.05 | 0.234 |
| 3.2 | 0 | ycaM | 3 | 12.05 | 0.298 |
| 3.2 | 0 | yadI | 3 | 12.05 | 0.476 |
| 3.2 | 0 | AG1 | 3 | 12.05 | 0.4 |
| 3.2 | 0 | ptsI | 3 | 12.3 | 0.247 |
| 3.2 | 0 | clcB | 3 | 12.3 | 0.23 |
| 3.2 | 0 | ycaM | 3 | 12.3 | 0.294 |
| 3.2 | 0 | yadI | 3 | 12.3 | 0.48 |
| 3.2 | 0 | AG1 | 3 | 12.3 | 0.401 |
| 3.2 | 0 | ptsI | 3 | 12.55 | 0.243 |
| 3.2 | 0 | clcB | 3 | 12.55 | 0.229 |
| 3.2 | 0 | ycaM | 3 | 12.55 | 0.294 |
| 3.2 | 0 | yadI | 3 | 12.55 | 0.481 |
| 3.2 | 0 | AG1 | 3 | 12.55 | 0.408 |
| 3.2 | 0 | ptsI | 3 | 12.8 | 0.243 |
| 3.2 | 0 | clcB | 3 | 12.8 | 0.226 |
| 3.2 | 0 | ycaM | 3 | 12.8 | 0.29 |
| 3.2 | 0 | yadI | 3 | 12.8 | 0.486 |
| 3.2 | 0 | AG1 | 3 | 12.8 | 0.409 |
| 3.2 | 0 | ptsI | 3 | 13.05 | 0.242 |
| 3.2 | 0 | clcB | 3 | 13.05 | 0.226 |
| 3.2 | 0 | ycaM | 3 | 13.05 | 0.291 |
| 3.2 | 0 | yadI | 3 | 13.05 | 0.496 |
| 3.2 | 0 | AG1 | 3 | 13.05 | 0.412 |
| 3.2 | 0 | ptsI | 3 | 13.3 | 0.241 |
| 3.2 | 0 | clcB | 3 | 13.3 | 0.224 |
| 3.2 | 0 | ycaM | 3 | 13.3 | 0.289 |
| 3.2 | 0 | yadI | 3 | 13.3 | 0.5 |
| 3.2 | 0 | AG1 | 3 | 13.3 | 0.414 |
| 3.2 | 0 | ptsI | 3 | 13.55 | 0.238 |
| 3.2 | 0 | clcB | 3 | 13.55 | 0.223 |
| 3.2 | 0 | ycaM | 3 | 13.55 | 0.292 |

| | | | | | |
|---|---|---|---|---|---|
| 3.2 | 0 | yadI | 3 | 13.55 | 0.503 |
| 3.2 | 0 | AG1 | 3 | 13.55 | 0.412 |
| 3.2 | 0 | ptsI | 3 | 13.8 | 0.239 |
| 3.2 | 0 | clcB | 3 | 13.8 | 0.221 |
| 3.2 | 0 | ycaM | 3 | 13.8 | 0.287 |
| 3.2 | 0 | yadI | 3 | 13.8 | 0.509 |
| 3.2 | 0 | AG1 | 3 | 13.8 | 0.414 |
| 3.2 | 0 | ptsI | 3 | 14.05 | 0.237 |
| 3.2 | 0 | clcB | 3 | 14.05 | 0.221 |
| 3.2 | 0 | ycaM | 3 | 14.05 | 0.287 |
| 3.2 | 0 | yadI | 3 | 14.05 | 0.514 |
| 3.2 | 0 | AG1 | 3 | 14.05 | 0.412 |
| 3.2 | 0 | ptsI | 3 | 14.3 | 0.237 |
| 3.2 | 0 | clcB | 3 | 14.3 | 0.222 |
| 3.2 | 0 | ycaM | 3 | 14.3 | 0.288 |
| 3.2 | 0 | yadI | 3 | 14.3 | 0.52 |
| 3.2 | 0 | AG1 | 3 | 14.3 | 0.421 |
| 3.2 | 0 | ptsI | 3 | 14.55 | 0.235 |
| 3.2 | 0 | clcB | 3 | 14.55 | 0.221 |
| 3.2 | 0 | ycaM | 3 | 14.55 | 0.287 |
| 3.2 | 0 | yadI | 3 | 14.55 | 0.525 |
| 3.2 | 0 | AG1 | 3 | 14.55 | 0.427 |
| 3.2 | 0 | ptsI | 3 | 14.8 | 0.235 |
| 3.2 | 0 | clcB | 3 | 14.8 | 0.22 |
| 3.2 | 0 | ycaM | 3 | 14.8 | 0.286 |
| 3.2 | 0 | yadI | 3 | 14.8 | 0.536 |
| 3.2 | 0 | AG1 | 3 | 14.8 | 0.431 |
| 3.2 | 0 | ptsI | 3 | 15.05 | 0.234 |
| 3.2 | 0 | clcB | 3 | 15.05 | 0.22 |
| 3.2 | 0 | ycaM | 3 | 15.05 | 0.284 |
| 3.2 | 0 | yadI | 3 | 15.05 | 0.538 |
| 3.2 | 0 | AG1 | 3 | 15.05 | 0.434 |
| 3.2 | 0 | ptsI | 3 | 15.3 | 0.234 |
| 3.2 | 0 | clcB | 3 | 15.3 | 0.22 |
| 3.2 | 0 | ycaM | 3 | 15.3 | 0.285 |
| 3.2 | 0 | yadI | 3 | 15.3 | 0.545 |
| 3.2 | 0 | AG1 | 3 | 15.3 | 0.437 |
| 3.2 | 0 | ptsI | 3 | 15.55 | 0.232 |
| 3.2 | 0 | clcB | 3 | 15.55 | 0.221 |
| 3.2 | 0 | ycaM | 3 | 15.55 | 0.286 |
| 3.2 | 0 | yadI | 3 | 15.55 | 0.54 |
| 3.2 | 0 | AG1 | 3 | 15.55 | 0.44 |
| 3.2 | 0 | ptsI | 3 | 15.8 | 0.232 |
| 3.2 | 0 | clcB | 3 | 15.8 | 0.235 |
| 3.2 | 0 | ycaM | 3 | 15.8 | 0.284 |
| 3.2 | 0 | yadI | 3 | 15.8 | 0.545 |
| 3.2 | 0 | AG1 | 3 | 15.8 | 0.441 |
| 3.2 | 0 | ptsI | 3 | 16.05 | 0.232 |

| | | | | | |
|---|---|---|---|---|---|
| 3.2 | 0 | clcB | 3 | 16.05 | 0.218 |
| 3.2 | 0 | ycaM | 3 | 16.05 | 0.282 |
| 3.2 | 0 | yadI | 3 | 16.05 | 0.555 |
| 3.2 | 0 | AG1 | 3 | 16.05 | 0.446 |
| 3.2 | 0 | ptsI | 3 | 16.3 | 0.229 |
| 3.2 | 0 | clcB | 3 | 16.3 | 0.215 |
| 3.2 | 0 | ycaM | 3 | 16.3 | 0.282 |
| 3.2 | 0 | yadI | 3 | 16.3 | 0.552 |
| 3.2 | 0 | AG1 | 3 | 16.3 | 0.446 |
| 3.2 | 0 | ptsI | 3 | 16.55 | 0.23 |
| 3.2 | 0 | clcB | 3 | 16.55 | 0.216 |
| 3.2 | 0 | ycaM | 3 | 16.55 | 0.281 |
| 3.2 | 0 | yadI | 3 | 16.55 | 0.557 |
| 3.2 | 0 | AG1 | 3 | 16.55 | 0.452 |
| 3.2 | 0 | ptsI | 3 | 16.8 | 0.229 |
| 3.2 | 0 | clcB | 3 | 16.8 | 0.215 |
| 3.2 | 0 | ycaM | 3 | 16.8 | 0.281 |
| 3.2 | 0 | yadI | 3 | 16.8 | 0.556 |
| 3.2 | 0 | AG1 | 3 | 16.8 | 0.456 |
| 3.2 | 0 | ptsI | 3 | 17.05 | 0.229 |
| 3.2 | 0 | clcB | 3 | 17.05 | 0.217 |
| 3.2 | 0 | ycaM | 3 | 17.05 | 0.281 |
| 3.2 | 0 | yadI | 3 | 17.05 | 0.564 |
| 3.2 | 0 | AG1 | 3 | 17.05 | 0.462 |
| 3.2 | 0 | ptsI | 3 | 17.3 | 0.23 |
| 3.2 | 0 | clcB | 3 | 17.3 | 0.219 |
| 3.2 | 0 | ycaM | 3 | 17.3 | 0.281 |
| 3.2 | 0 | yadI | 3 | 17.3 | 0.568 |
| 3.2 | 0 | AG1 | 3 | 17.3 | 0.466 |
| 3.2 | 0 | ptsI | 3 | 17.55 | 0.229 |
| 3.2 | 0 | clcB | 3 | 17.55 | 0.218 |
| 3.2 | 0 | ycaM | 3 | 17.55 | 0.282 |
| 3.2 | 0 | yadI | 3 | 17.55 | 0.572 |
| 3.2 | 0 | AG1 | 3 | 17.55 | 0.466 |
| 3.2 | 0 | ptsI | 3 | 17.8 | 0.229 |
| 3.2 | 0 | clcB | 3 | 17.8 | 0.218 |
| 3.2 | 0 | ycaM | 3 | 17.8 | 0.282 |
| 3.2 | 0 | yadI | 3 | 17.8 | 0.578 |
| 3.2 | 0 | AG1 | 3 | 17.8 | 0.474 |
| 3.2 | 0 | ptsI | 3 | 18.05 | 0.229 |
| 3.2 | 0 | clcB | 3 | 18.05 | 0.219 |
| 3.2 | 0 | ycaM | 3 | 18.05 | 0.284 |
| 3.2 | 0 | yadI | 3 | 18.05 | 0.588 |
| 3.2 | 0 | AG1 | 3 | 18.05 | 0.48 |
| 3.2 | 0 | ptsI | 3 | 18.3 | 0.23 |
| 3.2 | 0 | clcB | 3 | 18.3 | 0.219 |
| 3.2 | 0 | ycaM | 3 | 18.3 | 0.283 |
| 3.2 | 0 | yadI | 3 | 18.3 | 0.593 |

| | | | | | |
|---|---|---|---|---|---|
| 3.2 | 0 | AG1 | 3 | 18.3 | 0.485 |
| 3.2 | 0 | ptsI | 3 | 18.55 | 0.229 |
| 3.2 | 0 | clcB | 3 | 18.55 | 0.22 |
| 3.2 | 0 | ycaM | 3 | 18.55 | 0.286 |
| 3.2 | 0 | yadI | 3 | 18.55 | 0.598 |
| 3.2 | 0 | AG1 | 3 | 18.55 | 0.487 |
| 3.2 | 0 | ptsI | 3 | 18.8 | 0.23 |
| 3.2 | 0 | clcB | 3 | 18.8 | 0.221 |
| 3.2 | 0 | ycaM | 3 | 18.8 | 0.285 |
| 3.2 | 0 | yadI | 3 | 18.8 | 0.601 |
| 3.2 | 0 | AG1 | 3 | 18.8 | 0.488 |
| 3.2 | 0 | ptsI | 3 | 19.05 | 0.231 |
| 3.2 | 0 | clcB | 3 | 19.05 | 0.22 |
| 3.2 | 0 | ycaM | 3 | 19.05 | 0.286 |
| 3.2 | 0 | yadI | 3 | 19.05 | 0.612 |
| 3.2 | 0 | AG1 | 3 | 19.05 | 0.494 |
| 3.2 | 0 | ptsI | 3 | 19.3 | 0.232 |
| 3.2 | 0 | clcB | 3 | 19.3 | 0.223 |
| 3.2 | 0 | ycaM | 3 | 19.3 | 0.287 |
| 3.2 | 0 | yadI | 3 | 19.3 | 0.615 |
| 3.2 | 0 | AG1 | 3 | 19.3 | 0.496 |
| 3.2 | 0 | ptsI | 3 | 19.55 | 0.231 |
| 3.2 | 0 | clcB | 3 | 19.55 | 0.225 |
| 3.2 | 0 | ycaM | 3 | 19.55 | 0.29 |
| 3.2 | 0 | yadI | 3 | 19.55 | 0.617 |
| 3.2 | 0 | AG1 | 3 | 19.55 | 0.499 |
| 3.2 | 0 | ptsI | 3 | 19.8 | 0.232 |
| 3.2 | 0 | clcB | 3 | 19.8 | 0.228 |
| 3.2 | 0 | ycaM | 3 | 19.8 | 0.29 |
| 3.2 | 0 | yadI | 3 | 19.8 | 0.622 |
| 3.2 | 0 | AG1 | 3 | 19.8 | 0.503 |
| 3.2 | 0 | ptsI | 3 | 20.05 | 0.234 |
| 3.2 | 0 | clcB | 3 | 20.05 | 0.232 |
| 3.2 | 0 | ycaM | 3 | 20.05 | 0.293 |
| 3.2 | 0 | yadI | 3 | 20.05 | 0.628 |
| 3.2 | 0 | AG1 | 3 | 20.05 | 0.51 |
| 3.2 | 0 | ptsI | 3 | 20.3 | 0.235 |
| 3.2 | 0 | clcB | 3 | 20.3 | 0.239 |
| 3.2 | 0 | ycaM | 3 | 20.3 | 0.294 |
| 3.2 | 0 | yadI | 3 | 20.3 | 0.634 |
| 3.2 | 0 | AG1 | 3 | 20.3 | 0.511 |
| 3.2 | 0 | ptsI | 3 | 20.55 | 0.236 |
| 3.2 | 0 | clcB | 3 | 20.55 | 0.243 |
| 3.2 | 0 | ycaM | 3 | 20.55 | 0.296 |
| 3.2 | 0 | yadI | 3 | 20.55 | 0.639 |
| 3.2 | 0 | AG1 | 3 | 20.55 | 0.514 |
| 3.2 | 0 | ptsI | 3 | 20.8 | 0.237 |
| 3.2 | 0 | clcB | 3 | 20.8 | 0.249 |

| | | | | | |
|---|---|---|---|---|---|
| 3.2 | 0 | ycaM | 3 | 20.8 | 0.299 |
| 3.2 | 0 | yadI | 3 | 20.8 | 0.642 |
| 3.2 | 0 | AG1 | 3 | 20.8 | 0.52 |
| 3.2 | 0 | ptsI | 3 | 21.05 | 0.238 |
| 3.2 | 0 | clcB | 3 | 21.05 | 0.253 |
| 3.2 | 0 | ycaM | 3 | 21.05 | 0.302 |
| 3.2 | 0 | yadI | 3 | 21.05 | 0.643 |
| 3.2 | 0 | AG1 | 3 | 21.05 | 0.52 |
| 3.2 | 0 | ptsI | 3 | 21.3 | 0.239 |
| 3.2 | 0 | clcB | 3 | 21.3 | 0.259 |
| 3.2 | 0 | ycaM | 3 | 21.3 | 0.303 |
| 3.2 | 0 | yadI | 3 | 21.3 | 0.646 |
| 3.2 | 0 | AG1 | 3 | 21.3 | 0.525 |
| 3.2 | 0 | ptsI | 3 | 21.55 | 0.24 |
| 3.2 | 0 | clcB | 3 | 21.55 | 0.268 |
| 3.2 | 0 | ycaM | 3 | 21.55 | 0.326 |
| 3.2 | 0 | yadI | 3 | 21.55 | 0.652 |
| 3.2 | 0 | AG1 | 3 | 21.55 | 0.529 |
| 3.2 | 0 | ptsI | 3 | 21.8 | 0.242 |
| 3.2 | 0 | clcB | 3 | 21.8 | 0.275 |
| 3.2 | 0 | ycaM | 3 | 21.8 | 0.319 |
| 3.2 | 0 | yadI | 3 | 21.8 | 0.651 |
| 3.2 | 0 | AG1 | 3 | 21.8 | 0.533 |
| 3.2 | 0 | ptsI | 3 | 22.05 | 0.242 |
| 3.2 | 0 | clcB | 3 | 22.05 | 0.282 |
| 3.2 | 0 | ycaM | 3 | 22.05 | 0.32 |
| 3.2 | 0 | yadI | 3 | 22.05 | 0.652 |
| 3.2 | 0 | AG1 | 3 | 22.05 | 0.534 |
| 3.2 | 0 | ptsI | 3 | 22.3 | 0.244 |
| 3.2 | 0 | clcB | 3 | 22.3 | 0.289 |
| 3.2 | 0 | ycaM | 3 | 22.3 | 0.322 |
| 3.2 | 0 | yadI | 3 | 22.3 | 0.659 |
| 3.2 | 0 | AG1 | 3 | 22.3 | 0.537 |
| 3.2 | 0 | ptsI | 3 | 22.55 | 0.244 |
| 3.2 | 0 | clcB | 3 | 22.55 | 0.294 |
| 3.2 | 0 | ycaM | 3 | 22.55 | 0.324 |
| 3.2 | 0 | yadI | 3 | 22.55 | 0.666 |
| 3.2 | 0 | AG1 | 3 | 22.55 | 0.543 |
| 3.2 | 0 | ptsI | 3 | 22.8 | 0.247 |
| 3.2 | 0 | clcB | 3 | 22.8 | 0.301 |
| 3.2 | 0 | ycaM | 3 | 22.8 | 0.326 |
| 3.2 | 0 | yadI | 3 | 22.8 | 0.671 |
| 3.2 | 0 | AG1 | 3 | 22.8 | 0.546 |
| 3.2 | 0 | ptsI | 3 | 23.05 | 0.249 |
| 3.2 | 0 | clcB | 3 | 23.05 | 0.308 |
| 3.2 | 0 | ycaM | 3 | 23.05 | 0.326 |
| 3.2 | 0 | yadI | 3 | 23.05 | 0.666 |
| 3.2 | 0 | AG1 | 3 | 23.05 | 0.551 |

| | | | | | |
|---|---|---|---|---|---|
| 3.2 | 0 | ptsI | 3 | 23.3 | 0.249 |
| 3.2 | 0 | clcB | 3 | 23.3 | 0.312 |
| 3.2 | 0 | ycaM | 3 | 23.3 | 0.328 |
| 3.2 | 0 | yadI | 3 | 23.3 | 0.672 |
| 3.2 | 0 | AG1 | 3 | 23.3 | 0.554 |
| 3.2 | 0 | ptsI | 3 | 23.55 | 0.251 |
| 3.2 | 0 | clcB | 3 | 23.55 | 0.316 |
| 3.2 | 0 | ycaM | 3 | 23.55 | 0.328 |
| 3.2 | 0 | yadI | 3 | 23.55 | 0.682 |
| 3.2 | 0 | AG1 | 3 | 23.55 | 0.558 |
| 3.2 | 0 | ptsI | 3 | 23.8 | 0.253 |
| 3.2 | 0 | clcB | 3 | 23.8 | 0.32 |
| 3.2 | 0 | ycaM | 3 | 23.8 | 0.331 |
| 3.2 | 0 | yadI | 3 | 23.8 | 0.687 |
| 3.2 | 0 | AG1 | 3 | 23.8 | 0.562 |
| 3.2 | 0 | ptsI | 3 | 24.05 | 0.255 |
| 3.2 | 0 | clcB | 3 | 24.05 | 0.322 |
| 3.2 | 0 | ycaM | 3 | 24.05 | 0.333 |
| 3.2 | 0 | yadI | 3 | 24.05 | 0.687 |
| 3.2 | 0 | AG1 | 3 | 24.05 | 0.57 |
| 3.2 | 0 | ptsI | 3 | 24.3 | 0.254 |
| 3.2 | 0 | clcB | 3 | 24.3 | 0.32 |
| 3.2 | 0 | ycaM | 3 | 24.3 | 0.344 |
| 3.2 | 0 | yadI | 3 | 24.3 | 0.695 |
| 3.2 | 0 | AG1 | 3 | 24.3 | 0.569 |
| 3.2 | 0 | ptsI | 4 | 0 | 0.176 |
| 3.2 | 0 | clcB | 4 | 0 | 0.193 |
| 3.2 | 0 | ycaM | 4 | 0 | 0.197 |
| 3.2 | 0 | yadI | 4 | 0 | 0.215 |
| 3.2 | 0 | AG1 | 4 | 0 | 0.238 |
| 3.2 | 0 | ptsI | 4 | 0.25 | 0.181 |
| 3.2 | 0 | clcB | 4 | 0.25 | 0.192 |
| 3.2 | 0 | ycaM | 4 | 0.25 | 0.197 |
| 3.2 | 0 | yadI | 4 | 0.25 | 0.216 |
| 3.2 | 0 | AG1 | 4 | 0.25 | 0.239 |
| 3.2 | 0 | ptsI | 4 | 0.5 | 0.182 |
| 3.2 | 0 | clcB | 4 | 0.5 | 0.195 |
| 3.2 | 0 | ycaM | 4 | 0.5 | 0.199 |
| 3.2 | 0 | yadI | 4 | 0.5 | 0.219 |
| 3.2 | 0 | AG1 | 4 | 0.5 | 0.24 |
| 3.2 | 0 | ptsI | 4 | 0.75 | 0.188 |
| 3.2 | 0 | clcB | 4 | 0.75 | 0.199 |
| 3.2 | 0 | ycaM | 4 | 0.75 | 0.201 |
| 3.2 | 0 | yadI | 4 | 0.75 | 0.224 |
| 3.2 | 0 | AG1 | 4 | 0.75 | 0.244 |
| 3.2 | 0 | ptsI | 4 | 1 | 0.19 |
| 3.2 | 0 | clcB | 4 | 1 | 0.203 |
| 3.2 | 0 | ycaM | 4 | 1 | 0.203 |

| | | | | | |
|---|---|---|---|---|---|
| 3.2 | 0 | yadI | 4 | 1 | 0.224 |
| 3.2 | 0 | AG1 | 4 | 1 | 0.246 |
| 3.2 | 0 | ptsI | 4 | 1.25 | 0.2 |
| 3.2 | 0 | clcB | 4 | 1.25 | 0.209 |
| 3.2 | 0 | ycaM | 4 | 1.25 | 0.207 |
| 3.2 | 0 | yadI | 4 | 1.25 | 0.23 |
| 3.2 | 0 | AG1 | 4 | 1.25 | 0.255 |
| 3.2 | 0 | ptsI | 4 | 1.5 | 0.201 |
| 3.2 | 0 | clcB | 4 | 1.5 | 0.205 |
| 3.2 | 0 | ycaM | 4 | 1.5 | 0.209 |
| 3.2 | 0 | yadI | 4 | 1.5 | 0.229 |
| 3.2 | 0 | AG1 | 4 | 1.5 | 0.254 |
| 3.2 | 0 | ptsI | 4 | 1.75 | 0.213 |
| 3.2 | 0 | clcB | 4 | 1.75 | 0.215 |
| 3.2 | 0 | ycaM | 4 | 1.75 | 0.217 |
| 3.2 | 0 | yadI | 4 | 1.75 | 0.239 |
| 3.2 | 0 | AG1 | 4 | 1.75 | 0.263 |
| 3.2 | 0 | ptsI | 4 | 2 | 0.223 |
| 3.2 | 0 | clcB | 4 | 2 | 0.229 |
| 3.2 | 0 | ycaM | 4 | 2 | 0.226 |
| 3.2 | 0 | yadI | 4 | 2 | 0.247 |
| 3.2 | 0 | AG1 | 4 | 2 | 0.272 |
| 3.2 | 0 | ptsI | 4 | 2.25 | 0.234 |
| 3.2 | 0 | clcB | 4 | 2.25 | 0.244 |
| 3.2 | 0 | ycaM | 4 | 2.25 | 0.236 |
| 3.2 | 0 | yadI | 4 | 2.25 | 0.263 |
| 3.2 | 0 | AG1 | 4 | 2.25 | 0.288 |
| 3.2 | 0 | ptsI | 4 | 2.5 | 0.247 |
| 3.2 | 0 | clcB | 4 | 2.5 | 0.24 |
| 3.2 | 0 | ycaM | 4 | 2.5 | 0.24 |
| 3.2 | 0 | yadI | 4 | 2.5 | 0.269 |
| 3.2 | 0 | AG1 | 4 | 2.5 | 0.292 |
| 3.2 | 0 | ptsI | 4 | 2.75 | 0.262 |
| 3.2 | 0 | clcB | 4 | 2.75 | 0.251 |
| 3.2 | 0 | ycaM | 4 | 2.75 | 0.25 |
| 3.2 | 0 | yadI | 4 | 2.75 | 0.282 |
| 3.2 | 0 | AG1 | 4 | 2.75 | 0.302 |
| 3.2 | 0 | ptsI | 4 | 3 | 0.28 |
| 3.2 | 0 | clcB | 4 | 3 | 0.272 |
| 3.2 | 0 | ycaM | 4 | 3 | 0.264 |
| 3.2 | 0 | yadI | 4 | 3 | 0.301 |
| 3.2 | 0 | AG1 | 4 | 3 | 0.321 |
| 3.2 | 0 | ptsI | 4 | 3.25 | 0.297 |
| 3.2 | 0 | clcB | 4 | 3.25 | 0.287 |
| 3.2 | 0 | ycaM | 4 | 3.25 | 0.273 |
| 3.2 | 0 | yadI | 4 | 3.25 | 0.319 |
| 3.2 | 0 | AG1 | 4 | 3.25 | 0.337 |
| 3.2 | 0 | ptsI | 4 | 3.5 | 0.306 |

| | | | | | |
|---|---|---|---|---|---|
| 3.2 | 0 | clcB | 4 | 3.5 | 0.309 |
| 3.2 | 0 | ycaM | 4 | 3.5 | 0.289 |
| 3.2 | 0 | yadI | 4 | 3.5 | 0.342 |
| 3.2 | 0 | AG1 | 4 | 3.5 | 0.352 |
| 3.2 | 0 | ptsI | 4 | 3.75 | 0.322 |
| 3.2 | 0 | clcB | 4 | 3.75 | 0.32 |
| 3.2 | 0 | ycaM | 4 | 3.75 | 0.299 |
| 3.2 | 0 | yadI | 4 | 3.75 | 0.358 |
| 3.2 | 0 | AG1 | 4 | 3.75 | 0.369 |
| 3.2 | 0 | ptsI | 4 | 4 | 0.338 |
| 3.2 | 0 | clcB | 4 | 4 | 0.341 |
| 3.2 | 0 | ycaM | 4 | 4 | 0.317 |
| 3.2 | 0 | yadI | 4 | 4 | 0.379 |
| 3.2 | 0 | AG1 | 4 | 4 | 0.392 |
| 3.2 | 0 | ptsI | 4 | 4.35 | 0.324 |
| 3.2 | 0 | clcB | 4 | 4.35 | 0.262 |
| 3.2 | 0 | ycaM | 4 | 4.35 | 0.345 |
| 3.2 | 0 | yadI | 4 | 4.35 | 0.362 |
| 3.2 | 0 | AG1 | 4 | 4.35 | 0.28 |
| 3.2 | 0 | ptsI | 4 | 4.6 | 0.362 |
| 3.2 | 0 | clcB | 4 | 4.6 | 0.268 |
| 3.2 | 0 | ycaM | 4 | 4.6 | 0.348 |
| 3.2 | 0 | yadI | 4 | 4.6 | 0.375 |
| 3.2 | 0 | AG1 | 4 | 4.6 | 0.289 |
| 3.2 | 0 | ptsI | 4 | 4.85 | 0.379 |
| 3.2 | 0 | clcB | 4 | 4.85 | 0.276 |
| 3.2 | 0 | ycaM | 4 | 4.85 | 0.358 |
| 3.2 | 0 | yadI | 4 | 4.85 | 0.39 |
| 3.2 | 0 | AG1 | 4 | 4.85 | 0.3 |
| 3.2 | 0 | ptsI | 4 | 5.1 | 0.393 |
| 3.2 | 0 | clcB | 4 | 5.1 | 0.282 |
| 3.2 | 0 | ycaM | 4 | 5.1 | 0.373 |
| 3.2 | 0 | yadI | 4 | 5.1 | 0.405 |
| 3.2 | 0 | AG1 | 4 | 5.1 | 0.311 |
| 3.2 | 0 | ptsI | 4 | 5.35 | 0.407 |
| 3.2 | 0 | clcB | 4 | 5.35 | 0.29 |
| 3.2 | 0 | ycaM | 4 | 5.35 | 0.389 |
| 3.2 | 0 | yadI | 4 | 5.35 | 0.42 |
| 3.2 | 0 | AG1 | 4 | 5.35 | 0.322 |
| 3.2 | 0 | ptsI | 4 | 5.6 | 0.429 |
| 3.2 | 0 | clcB | 4 | 5.6 | 0.298 |
| 3.2 | 0 | ycaM | 4 | 5.6 | 0.401 |
| 3.2 | 0 | yadI | 4 | 5.6 | 0.433 |
| 3.2 | 0 | AG1 | 4 | 5.6 | 0.329 |
| 3.2 | 0 | ptsI | 4 | 5.85 | 0.438 |
| 3.2 | 0 | clcB | 4 | 5.85 | 0.307 |
| 3.2 | 0 | ycaM | 4 | 5.85 | 0.411 |
| 3.2 | 0 | yadI | 4 | 5.85 | 0.449 |

| | | | | | |
|---|---|---|---|---|---|
| 3.2 | 0 | AG1 | 4 | 5.85 | 0.335 |
| 3.2 | 0 | ptsI | 4 | 6.1 | 0.453 |
| 3.2 | 0 | clcB | 4 | 6.1 | 0.317 |
| 3.2 | 0 | ycaM | 4 | 6.1 | 0.422 |
| 3.2 | 0 | yadI | 4 | 6.1 | 0.466 |
| 3.2 | 0 | AG1 | 4 | 6.1 | 0.345 |
| 3.2 | 0 | ptsI | 4 | 6.35 | 0.466 |
| 3.2 | 0 | clcB | 4 | 6.35 | 0.326 |
| 3.2 | 0 | ycaM | 4 | 6.35 | 0.435 |
| 3.2 | 0 | yadI | 4 | 6.35 | 0.48 |
| 3.2 | 0 | AG1 | 4 | 6.35 | 0.356 |
| 3.2 | 0 | ptsI | 4 | 6.6 | 0.475 |
| 3.2 | 0 | clcB | 4 | 6.6 | 0.337 |
| 3.2 | 0 | ycaM | 4 | 6.6 | 0.445 |
| 3.2 | 0 | yadI | 4 | 6.6 | 0.492 |
| 3.2 | 0 | AG1 | 4 | 6.6 | 0.364 |
| 3.2 | 0 | ptsI | 4 | 6.85 | 0.487 |
| 3.2 | 0 | clcB | 4 | 6.85 | 0.347 |
| 3.2 | 0 | ycaM | 4 | 6.85 | 0.455 |
| 3.2 | 0 | yadI | 4 | 6.85 | 0.506 |
| 3.2 | 0 | AG1 | 4 | 6.85 | 0.375 |
| 3.2 | 0 | ptsI | 4 | 7.1 | 0.497 |
| 3.2 | 0 | clcB | 4 | 7.1 | 0.354 |
| 3.2 | 0 | ycaM | 4 | 7.1 | 0.465 |
| 3.2 | 0 | yadI | 4 | 7.1 | 0.517 |
| 3.2 | 0 | AG1 | 4 | 7.1 | 0.384 |
| 3.2 | 0 | ptsI | 4 | 7.35 | 0.507 |
| 3.2 | 0 | clcB | 4 | 7.35 | 0.363 |
| 3.2 | 0 | ycaM | 4 | 7.35 | 0.473 |
| 3.2 | 0 | yadI | 4 | 7.35 | 0.525 |
| 3.2 | 0 | AG1 | 4 | 7.35 | 0.393 |
| 3.2 | 0 | ptsI | 4 | 7.6 | 0.514 |
| 3.2 | 0 | clcB | 4 | 7.6 | 0.372 |
| 3.2 | 0 | ycaM | 4 | 7.6 | 0.48 |
| 3.2 | 0 | yadI | 4 | 7.6 | 0.536 |
| 3.2 | 0 | AG1 | 4 | 7.6 | 0.4 |
| 3.2 | 0 | ptsI | 4 | 7.85 | 0.524 |
| 3.2 | 0 | clcB | 4 | 7.85 | 0.383 |
| 3.2 | 0 | ycaM | 4 | 7.85 | 0.486 |
| 3.2 | 0 | yadI | 4 | 7.85 | 0.544 |
| 3.2 | 0 | AG1 | 4 | 7.85 | 0.409 |
| 3.2 | 0 | ptsI | 4 | 8.1 | 0.532 |
| 3.2 | 0 | clcB | 4 | 8.1 | 0.394 |
| 3.2 | 0 | ycaM | 4 | 8.1 | 0.492 |
| 3.2 | 0 | yadI | 4 | 8.1 | 0.556 |
| 3.2 | 0 | AG1 | 4 | 8.1 | 0.416 |
| 3.2 | 0 | ptsI | 4 | 8.35 | 0.541 |
| 3.2 | 0 | clcB | 4 | 8.35 | 0.401 |

| | | | | | |
|---|---|---|---|---|---|
| 3.2 | 0 | ycaM | 4 | 8.35 | 0.494 |
| 3.2 | 0 | yadI | 4 | 8.35 | 0.564 |
| 3.2 | 0 | AG1 | 4 | 8.35 | 0.425 |
| 3.2 | 0 | ptsI | 4 | 8.83333 | 0.391 |
| 3.2 | 0 | clcB | 4 | 8.83333 | 0.399 |
| 3.2 | 0 | ycaM | 4 | 8.83333 | 0.398 |
| 3.2 | 0 | yadI | 4 | 8.83333 | 0.564 |
| 3.2 | 0 | AG1 | 4 | 8.83333 | 0.467 |
| 3.2 | 0 | ptsI | 4 | 9.08333 | 0.414 |
| 3.2 | 0 | clcB | 4 | 9.08333 | 0.417 |
| 3.2 | 0 | ycaM | 4 | 9.08333 | 0.407 |
| 3.2 | 0 | yadI | 4 | 9.08333 | 0.568 |
| 3.2 | 0 | AG1 | 4 | 9.08333 | 0.49 |
| 3.2 | 0 | ptsI | 4 | 9.33333 | 0.434 |
| 3.2 | 0 | clcB | 4 | 9.33333 | 0.437 |
| 3.2 | 0 | ycaM | 4 | 9.33333 | 0.42 |
| 3.2 | 0 | yadI | 4 | 9.33333 | 0.582 |
| 3.2 | 0 | AG1 | 4 | 9.33333 | 0.509 |
| 3.2 | 0 | ptsI | 4 | 9.58333 | 0.437 |
| 3.2 | 0 | clcB | 4 | 9.58333 | 0.439 |
| 3.2 | 0 | ycaM | 4 | 9.58333 | 0.427 |
| 3.2 | 0 | yadI | 4 | 9.58333 | 0.597 |
| 3.2 | 0 | AG1 | 4 | 9.58333 | 0.514 |
| 3.2 | 0 | ptsI | 4 | 9.83333 | 0.443 |
| 3.2 | 0 | clcB | 4 | 9.83333 | 0.434 |
| 3.2 | 0 | ycaM | 4 | 9.83333 | 0.426 |
| 3.2 | 0 | yadI | 4 | 9.83333 | 0.601 |
| 3.2 | 0 | AG1 | 4 | 9.83333 | 0.517 |
| 3.2 | 0 | ptsI | 4 | 10.0833 | 0.443 |
| 3.2 | 0 | clcB | 4 | 10.0833 | 0.44 |
| 3.2 | 0 | ycaM | 4 | 10.0833 | 0.43 |
| 3.2 | 0 | yadI | 4 | 10.0833 | 0.605 |
| 3.2 | 0 | AG1 | 4 | 10.0833 | 0.521 |
| 3.2 | 0 | ptsI | 4 | 10.3333 | 0.441 |
| 3.2 | 0 | clcB | 4 | 10.3333 | 0.437 |
| 3.2 | 0 | ycaM | 4 | 10.3333 | 0.433 |
| 3.2 | 0 | yadI | 4 | 10.3333 | 0.613 |
| 3.2 | 0 | AG1 | 4 | 10.3333 | 0.529 |
| 3.2 | 0 | ptsI | 4 | 10.5833 | 0.447 |
| 3.2 | 0 | clcB | 4 | 10.5833 | 0.443 |
| 3.2 | 0 | ycaM | 4 | 10.5833 | 0.438 |
| 3.2 | 0 | yadI | 4 | 10.5833 | 0.615 |
| 3.2 | 0 | AG1 | 4 | 10.5833 | 0.534 |
| 3.2 | 0 | ptsI | 4 | 10.8333 | 0.445 |
| 3.2 | 0 | clcB | 4 | 10.8333 | 0.442 |
| 3.2 | 0 | ycaM | 4 | 10.8333 | 0.438 |
| 3.2 | 0 | yadI | 4 | 10.8333 | 0.622 |
| 3.2 | 0 | AG1 | 4 | 10.8333 | 0.532 |

| | | | | | |
|---|---|---|---|---|---|
| 3.2 | 0 | ptsI | 4 | 11.0833 | 0.456 |
| 3.2 | 0 | clcB | 4 | 11.0833 | 0.444 |
| 3.2 | 0 | ycaM | 4 | 11.0833 | 0.442 |
| 3.2 | 0 | yadI | 4 | 11.0833 | 0.626 |
| 3.2 | 0 | AG1 | 4 | 11.0833 | 0.535 |
| 3.2 | 0 | ptsI | 4 | 11.3333 | 0.461 |
| 3.2 | 0 | clcB | 4 | 11.3333 | 0.443 |
| 3.2 | 0 | ycaM | 4 | 11.3333 | 0.449 |
| 3.2 | 0 | yadI | 4 | 11.3333 | 0.63 |
| 3.2 | 0 | AG1 | 4 | 11.3333 | 0.538 |
| 3.2 | 0 | ptsI | 4 | 11.5833 | 0.464 |
| 3.2 | 0 | clcB | 4 | 11.5833 | 0.45 |
| 3.2 | 0 | ycaM | 4 | 11.5833 | 0.451 |
| 3.2 | 0 | yadI | 4 | 11.5833 | 0.636 |
| 3.2 | 0 | AG1 | 4 | 11.5833 | 0.543 |
| 3.2 | 0 | ptsI | 4 | 11.8333 | 0.471 |
| 3.2 | 0 | clcB | 4 | 11.8333 | 0.454 |
| 3.2 | 0 | ycaM | 4 | 11.8333 | 0.459 |
| 3.2 | 0 | yadI | 4 | 11.8333 | 0.645 |
| 3.2 | 0 | AG1 | 4 | 11.8333 | 0.544 |
| 3.2 | 0 | ptsI | 4 | 12.0833 | 0.474 |
| 3.2 | 0 | clcB | 4 | 12.0833 | 0.458 |
| 3.2 | 0 | ycaM | 4 | 12.0833 | 0.462 |
| 3.2 | 0 | yadI | 4 | 12.0833 | 0.646 |
| 3.2 | 0 | AG1 | 4 | 12.0833 | 0.553 |
| 3.2 | 0 | ptsI | 4 | 12.3333 | 0.481 |
| 3.2 | 0 | clcB | 4 | 12.3333 | 0.456 |
| 3.2 | 0 | ycaM | 4 | 12.3333 | 0.47 |
| 3.2 | 0 | yadI | 4 | 12.3333 | 0.656 |
| 3.2 | 0 | AG1 | 4 | 12.3333 | 0.555 |
| 3.2 | 0 | ptsI | 4 | 12.5833 | 0.48 |
| 3.2 | 0 | clcB | 4 | 12.5833 | 0.462 |
| 3.2 | 0 | ycaM | 4 | 12.5833 | 0.467 |
| 3.2 | 0 | yadI | 4 | 12.5833 | 0.658 |
| 3.2 | 0 | AG1 | 4 | 12.5833 | 0.561 |
| 3.2 | 0 | ptsI | 4 | 12.8333 | 0.488 |
| 3.2 | 0 | clcB | 4 | 12.8333 | 0.463 |
| 3.2 | 0 | ycaM | 4 | 12.8333 | 0.475 |
| 3.2 | 0 | yadI | 4 | 12.8333 | 0.663 |
| 3.2 | 0 | AG1 | 4 | 12.8333 | 0.564 |
| 3.2 | 0 | ptsI | 4 | 13.0833 | 0.499 |
| 3.2 | 0 | clcB | 4 | 13.0833 | 0.465 |
| 3.2 | 0 | ycaM | 4 | 13.0833 | 0.48 |
| 3.2 | 0 | yadI | 4 | 13.0833 | 0.668 |
| 3.2 | 0 | AG1 | 4 | 13.0833 | 0.568 |
| 3.2 | 0 | ptsI | 4 | 13.3333 | 0.504 |
| 3.2 | 0 | clcB | 4 | 13.3333 | 0.47 |
| 3.2 | 0 | ycaM | 4 | 13.3333 | 0.484 |

| | | | | | |
|---|---|---|---|---|---|
| 3.2 | 0 | yadI | 4 | 13.3333 | 0.672 |
| 3.2 | 0 | AG1 | 4 | 13.3333 | 0.575 |
| 3.2 | 0 | ptsI | 4 | 13.5833 | 0.512 |
| 3.2 | 0 | clcB | 4 | 13.5833 | 0.473 |
| 3.2 | 0 | ycaM | 4 | 13.5833 | 0.487 |
| 3.2 | 0 | yadI | 4 | 13.5833 | 0.675 |
| 3.2 | 0 | AG1 | 4 | 13.5833 | 0.573 |
| 3.2 | 0 | ptsI | 4 | 13.8333 | 0.518 |
| 3.2 | 0 | clcB | 4 | 13.8333 | 0.475 |
| 3.2 | 0 | ycaM | 4 | 13.8333 | 0.493 |
| 3.2 | 0 | yadI | 4 | 13.8333 | 0.682 |
| 3.2 | 0 | AG1 | 4 | 13.8333 | 0.584 |
| 3.2 | 0 | ptsI | 4 | 14.0833 | 0.519 |
| 3.2 | 0 | clcB | 4 | 14.0833 | 0.483 |
| 3.2 | 0 | ycaM | 4 | 14.0833 | 0.495 |
| 3.2 | 0 | yadI | 4 | 14.0833 | 0.687 |
| 3.2 | 0 | AG1 | 4 | 14.0833 | 0.586 |
| 3.2 | 0 | ptsI | 4 | 14.3333 | 0.52 |
| 3.2 | 0 | clcB | 4 | 14.3333 | 0.486 |
| 3.2 | 0 | ycaM | 4 | 14.3333 | 0.498 |
| 3.2 | 0 | yadI | 4 | 14.3333 | 0.689 |
| 3.2 | 0 | AG1 | 4 | 14.3333 | 0.589 |
| 3.2 | 0 | ptsI | 4 | 14.5833 | 0.528 |
| 3.2 | 0 | clcB | 4 | 14.5833 | 0.488 |
| 3.2 | 0 | ycaM | 4 | 14.5833 | 0.508 |
| 3.2 | 0 | yadI | 4 | 14.5833 | 0.698 |
| 3.2 | 0 | AG1 | 4 | 14.5833 | 0.592 |
| 3.2 | 0 | ptsI | 4 | 14.8333 | 0.531 |
| 3.2 | 0 | clcB | 4 | 14.8333 | 0.487 |
| 3.2 | 0 | ycaM | 4 | 14.8333 | 0.511 |
| 3.2 | 0 | yadI | 4 | 14.8333 | 0.696 |
| 3.2 | 0 | AG1 | 4 | 14.8333 | 0.597 |
| 3.2 | 0 | ptsI | 4 | 15.0833 | 0.537 |
| 3.2 | 0 | clcB | 4 | 15.0833 | 0.491 |
| 3.2 | 0 | ycaM | 4 | 15.0833 | 0.507 |
| 3.2 | 0 | yadI | 4 | 15.0833 | 0.698 |
| 3.2 | 0 | AG1 | 4 | 15.0833 | 0.599 |
| 3.2 | 0 | ptsI | 4 | 15.3333 | 0.546 |
| 3.2 | 0 | clcB | 4 | 15.3333 | 0.494 |
| 3.2 | 0 | ycaM | 4 | 15.3333 | 0.514 |
| 3.2 | 0 | yadI | 4 | 15.3333 | 0.706 |
| 3.2 | 0 | AG1 | 4 | 15.3333 | 0.606 |
| 3.2 | 0 | ptsI | 4 | 15.5833 | 0.545 |
| 3.2 | 0 | clcB | 4 | 15.5833 | 0.496 |
| 3.2 | 0 | ycaM | 4 | 15.5833 | 0.521 |
| 3.2 | 0 | yadI | 4 | 15.5833 | 0.707 |
| 3.2 | 0 | AG1 | 4 | 15.5833 | 0.608 |
| 3.2 | 0 | ptsI | 4 | 15.8333 | 0.552 |

| | | | | | |
|---|---|---|---|---|---|
| 3.2 | 0 | clcB | 4 | 15.8333 | 0.499 |
| 3.2 | 0 | ycaM | 4 | 15.8333 | 0.521 |
| 3.2 | 0 | yadI | 4 | 15.8333 | 0.714 |
| 3.2 | 0 | AG1 | 4 | 15.8333 | 0.614 |
| 3.2 | 0 | ptsI | 4 | 16.0833 | 0.559 |
| 3.2 | 0 | clcB | 4 | 16.0833 | 0.505 |
| 3.2 | 0 | ycaM | 4 | 16.0833 | 0.524 |
| 3.2 | 0 | yadI | 4 | 16.0833 | 0.716 |
| 3.2 | 0 | AG1 | 4 | 16.0833 | 0.614 |
| 3.2 | 0 | ptsI | 4 | 16.3333 | 0.561 |
| 3.2 | 0 | clcB | 4 | 16.3333 | 0.506 |
| 3.2 | 0 | ycaM | 4 | 16.3333 | 0.529 |
| 3.2 | 0 | yadI | 4 | 16.3333 | 0.722 |
| 3.2 | 0 | AG1 | 4 | 16.3333 | 0.621 |
| 3.2 | 0 | ptsI | 4 | 16.5833 | 0.56 |
| 3.2 | 0 | clcB | 4 | 16.5833 | 0.506 |
| 3.2 | 0 | ycaM | 4 | 16.5833 | 0.53 |
| 3.2 | 0 | yadI | 4 | 16.5833 | 0.725 |
| 3.2 | 0 | AG1 | 4 | 16.5833 | 0.623 |
| 3.2 | 0 | ptsI | 4 | 16.8333 | 0.573 |
| 3.2 | 0 | clcB | 4 | 16.8333 | 0.512 |
| 3.2 | 0 | ycaM | 4 | 16.8333 | 0.534 |
| 3.2 | 0 | yadI | 4 | 16.8333 | 0.727 |
| 3.2 | 0 | AG1 | 4 | 16.8333 | 0.627 |
| 3.2 | 0 | ptsI | 4 | 17.0833 | 0.578 |
| 3.2 | 0 | clcB | 4 | 17.0833 | 0.517 |
| 3.2 | 0 | ycaM | 4 | 17.0833 | 0.537 |
| 3.2 | 0 | yadI | 4 | 17.0833 | 0.728 |
| 3.2 | 0 | AG1 | 4 | 17.0833 | 0.629 |
| 3.2 | 0 | ptsI | 4 | 17.3333 | 0.583 |
| 3.2 | 0 | clcB | 4 | 17.3333 | 0.52 |
| 3.2 | 0 | ycaM | 4 | 17.3333 | 0.54 |
| 3.2 | 0 | yadI | 4 | 17.3333 | 0.732 |
| 3.2 | 0 | AG1 | 4 | 17.3333 | 0.63 |
| 3.2 | 0 | ptsI | 4 | 17.5833 | 0.592 |
| 3.2 | 0 | clcB | 4 | 17.5833 | 0.519 |
| 3.2 | 0 | ycaM | 4 | 17.5833 | 0.544 |
| 3.2 | 0 | yadI | 4 | 17.5833 | 0.733 |
| 3.2 | 0 | AG1 | 4 | 17.5833 | 0.636 |
| 3.2 | 0 | ptsI | 4 | 17.8333 | 0.598 |
| 3.2 | 0 | clcB | 4 | 17.8333 | 0.522 |
| 3.2 | 0 | ycaM | 4 | 17.8333 | 0.548 |
| 3.2 | 0 | yadI | 4 | 17.8333 | 0.719 |
| 3.2 | 0 | AG1 | 4 | 17.8333 | 0.64 |
| 3.2 | 0 | ptsI | 4 | 18.0833 | 0.599 |
| 3.2 | 0 | clcB | 4 | 18.0833 | 0.525 |
| 3.2 | 0 | ycaM | 4 | 18.0833 | 0.551 |
| 3.2 | 0 | yadI | 4 | 18.0833 | 0.738 |

| | | | | | |
|---|---|---|---|---|---|
| 3.2 | 0 | AG1 | 4 | 18.0833 | 0.64 |
| 3.2 | 0 | ptsI | 4 | 18.3333 | 0.608 |
| 3.2 | 0 | clcB | 4 | 18.3333 | 0.527 |
| 3.2 | 0 | ycaM | 4 | 18.3333 | 0.554 |
| 3.2 | 0 | yadI | 4 | 18.3333 | 0.743 |
| 3.2 | 0 | AG1 | 4 | 18.3333 | 0.643 |
| 3.2 | 0 | ptsI | 4 | 18.5833 | 0.609 |
| 3.2 | 0 | clcB | 4 | 18.5833 | 0.527 |
| 3.2 | 0 | ycaM | 4 | 18.5833 | 0.555 |
| 3.2 | 0 | yadI | 4 | 18.5833 | 0.744 |
| 3.2 | 0 | AG1 | 4 | 18.5833 | 0.645 |
| 3.2 | 0 | ptsI | 4 | 18.8333 | 0.613 |
| 3.2 | 0 | clcB | 4 | 18.8333 | 0.528 |
| 3.2 | 0 | ycaM | 4 | 18.8333 | 0.56 |
| 3.2 | 0 | yadI | 4 | 18.8333 | 0.73 |
| 3.2 | 0 | AG1 | 4 | 18.8333 | 0.648 |
| 3.2 | 0 | ptsI | 4 | 19.0833 | 0.62 |
| 3.2 | 0 | clcB | 4 | 19.0833 | 0.533 |
| 3.2 | 0 | ycaM | 4 | 19.0833 | 0.56 |
| 3.2 | 0 | yadI | 4 | 19.0833 | 0.75 |
| 3.2 | 0 | AG1 | 4 | 19.0833 | 0.65 |
| 3.2 | 0 | ptsI | 4 | 19.3333 | 0.621 |
| 3.2 | 0 | clcB | 4 | 19.3333 | 0.534 |
| 3.2 | 0 | ycaM | 4 | 19.3333 | 0.564 |
| 3.2 | 0 | yadI | 4 | 19.3333 | 0.754 |
| 3.2 | 0 | AG1 | 4 | 19.3333 | 0.652 |
| 3.2 | 0 | ptsI | 4 | 19.5833 | 0.624 |
| 3.2 | 0 | clcB | 4 | 19.5833 | 0.538 |
| 3.2 | 0 | ycaM | 4 | 19.5833 | 0.568 |
| 3.2 | 0 | yadI | 4 | 19.5833 | 0.756 |
| 3.2 | 0 | AG1 | 4 | 19.5833 | 0.656 |
| 3.2 | 0 | ptsI | 4 | 19.8333 | 0.629 |
| 3.2 | 0 | clcB | 4 | 19.8333 | 0.535 |
| 3.2 | 0 | ycaM | 4 | 19.8333 | 0.569 |
| 3.2 | 0 | yadI | 4 | 19.8333 | 0.758 |
| 3.2 | 0 | AG1 | 4 | 19.8333 | 0.662 |
| 3.2 | 0 | ptsI | 4 | 20.0833 | 0.644 |
| 3.2 | 0 | clcB | 4 | 20.0833 | 0.541 |
| 3.2 | 0 | ycaM | 4 | 20.0833 | 0.573 |
| 3.2 | 0 | yadI | 4 | 20.0833 | 0.75 |
| 3.2 | 0 | AG1 | 4 | 20.0833 | 0.663 |
| 3.2 | 0 | ptsI | 4 | 20.3333 | 0.638 |
| 3.2 | 0 | clcB | 4 | 20.3333 | 0.54 |
| 3.2 | 0 | ycaM | 4 | 20.3333 | 0.574 |
| 3.2 | 0 | yadI | 4 | 20.3333 | 0.767 |
| 3.2 | 0 | AG1 | 4 | 20.3333 | 0.663 |
| 3.2 | 0 | ptsI | 4 | 20.5833 | 0.642 |
| 3.2 | 0 | clcB | 4 | 20.5833 | 0.546 |

| | | | | | |
|---|---|---|---|---|---|
| 3.2 | 0 | ycaM | 4 | 20.5833 | 0.576 |
| 3.2 | 0 | yadI | 4 | 20.5833 | 0.769 |
| 3.2 | 0 | AG1 | 4 | 20.5833 | 0.669 |
| 3.2 | 0 | ptsI | 4 | 20.8333 | 0.648 |
| 3.2 | 0 | clcB | 4 | 20.8333 | 0.548 |
| 3.2 | 0 | ycaM | 4 | 20.8333 | 0.58 |
| 3.2 | 0 | yadI | 4 | 20.8333 | 0.771 |
| 3.2 | 0 | AG1 | 4 | 20.8333 | 0.669 |
| 3.2 | 0 | ptsI | 4 | 21.0833 | 0.653 |
| 3.2 | 0 | clcB | 4 | 21.0833 | 0.547 |
| 3.2 | 0 | ycaM | 4 | 21.0833 | 0.578 |
| 3.2 | 0 | yadI | 4 | 21.0833 | 0.774 |
| 3.2 | 0 | AG1 | 4 | 21.0833 | 0.668 |
| 3.2 | 0 | ptsI | 4 | 21.3333 | 0.654 |
| 3.2 | 0 | clcB | 4 | 21.3333 | 0.548 |
| 3.2 | 0 | ycaM | 4 | 21.3333 | 0.584 |
| 3.2 | 0 | yadI | 4 | 21.3333 | 0.777 |
| 3.2 | 0 | AG1 | 4 | 21.3333 | 0.672 |
| 3.2 | 0 | ptsI | 4 | 21.5833 | 0.658 |
| 3.2 | 0 | clcB | 4 | 21.5833 | 0.552 |
| 3.2 | 0 | ycaM | 4 | 21.5833 | 0.582 |
| 3.2 | 0 | yadI | 4 | 21.5833 | 0.775 |
| 3.2 | 0 | AG1 | 4 | 21.5833 | 0.671 |
| 3.2 | 0 | ptsI | 4 | 21.8333 | 0.658 |
| 3.2 | 0 | clcB | 4 | 21.8333 | 0.556 |
| 3.2 | 0 | ycaM | 4 | 21.8333 | 0.586 |
| 3.2 | 0 | yadI | 4 | 21.8333 | 0.782 |
| 3.2 | 0 | AG1 | 4 | 21.8333 | 0.676 |
| 3.2 | 0 | ptsI | 4 | 22.0833 | 0.666 |
| 3.2 | 0 | clcB | 4 | 22.0833 | 0.557 |
| 3.2 | 0 | ycaM | 4 | 22.0833 | 0.587 |
| 3.2 | 0 | yadI | 4 | 22.0833 | 0.784 |
| 3.2 | 0 | AG1 | 4 | 22.0833 | 0.679 |
| 3.2 | 0 | ptsI | 4 | 22.3333 | 0.666 |
| 3.2 | 0 | clcB | 4 | 22.3333 | 0.56 |
| 3.2 | 0 | ycaM | 4 | 22.3333 | 0.59 |
| 3.2 | 0 | yadI | 4 | 22.3333 | 0.786 |
| 3.2 | 0 | AG1 | 4 | 22.3333 | 0.683 |
| 3.2 | 0 | ptsI | 4 | 22.5833 | 0.665 |
| 3.2 | 0 | clcB | 4 | 22.5833 | 0.561 |
| 3.2 | 0 | ycaM | 4 | 22.5833 | 0.592 |
| 3.2 | 0 | yadI | 4 | 22.5833 | 0.786 |
| 3.2 | 0 | AG1 | 4 | 22.5833 | 0.684 |
| 3.2 | 0 | ptsI | 4 | 22.8333 | 0.672 |
| 3.2 | 0 | clcB | 4 | 22.8333 | 0.561 |
| 3.2 | 0 | ycaM | 4 | 22.8333 | 0.593 |
| 3.2 | 0 | yadI | 4 | 22.8333 | 0.789 |
| 3.2 | 0 | AG1 | 4 | 22.8333 | 0.686 |

| | | | | | |
|---|---|---|---|---|---|
| 3.2 | 0 | ptsI | 4 | 23.0833 | 0.675 |
| 3.2 | 0 | clcB | 4 | 23.0833 | 0.561 |
| 3.2 | 0 | ycaM | 4 | 23.0833 | 0.593 |
| 3.2 | 0 | yadI | 4 | 23.0833 | 0.789 |
| 3.2 | 0 | AG1 | 4 | 23.0833 | 0.683 |
| 3.2 | 0 | ptsI | 4 | 23.3333 | 0.679 |
| 3.2 | 0 | clcB | 4 | 23.3333 | 0.565 |
| 3.2 | 0 | ycaM | 4 | 23.3333 | 0.596 |
| 3.2 | 0 | yadI | 4 | 23.3333 | 0.803 |
| 3.2 | 0 | AG1 | 4 | 23.3333 | 0.688 |
| 3.2 | 0 | ptsI | 4 | 23.5833 | 0.683 |
| 3.2 | 0 | clcB | 4 | 23.5833 | 0.566 |
| 3.2 | 0 | ycaM | 4 | 23.5833 | 0.6 |
| 3.2 | 0 | yadI | 4 | 23.5833 | 0.796 |
| 3.2 | 0 | AG1 | 4 | 23.5833 | 0.692 |
| 3.2 | 0 | ptsI | 4 | 23.8333 | 0.684 |
| 3.2 | 0 | clcB | 4 | 23.8333 | 0.571 |
| 3.2 | 0 | ycaM | 4 | 23.8333 | 0.596 |
| 3.2 | 0 | yadI | 4 | 23.8333 | 0.793 |
| 3.2 | 0 | AG1 | 4 | 23.8333 | 0.693 |
| 3.2 | 0 | ptsI | 4 | 24.0833 | 0.689 |
| 3.2 | 0 | clcB | 4 | 24.0833 | 0.57 |
| 3.2 | 0 | ycaM | 4 | 24.0833 | 0.601 |
| 3.2 | 0 | yadI | 4 | 24.0833 | 0.799 |
| 3.2 | 0 | AG1 | 4 | 24.0833 | 0.695 |
| 3.2 | 0 | ptsI | 5 | 0 | 0.163 |
| 3.2 | 0 | clcB | 5 | 0 | 0.163 |
| 3.2 | 0 | ycaM | 5 | 0 | 0.165 |
| 3.2 | 0 | yadI | 5 | 0 | 0.154 |
| 3.2 | 0 | AG1 | 5 | 0 | 0.156 |
| 3.2 | 0 | ptsI | 5 | 0.35 | 0.161 |
| 3.2 | 0 | clcB | 5 | 0.35 | 0.162 |
| 3.2 | 0 | ycaM | 5 | 0.35 | 0.163 |
| 3.2 | 0 | yadI | 5 | 0.35 | 0.157 |
| 3.2 | 0 | AG1 | 5 | 0.35 | 0.156 |
| 3.2 | 0 | ptsI | 5 | 0.6 | 0.165 |
| 3.2 | 0 | clcB | 5 | 0.6 | 0.163 |
| 3.2 | 0 | ycaM | 5 | 0.6 | 0.164 |
| 3.2 | 0 | yadI | 5 | 0.6 | 0.158 |
| 3.2 | 0 | AG1 | 5 | 0.6 | 0.156 |
| 3.2 | 0 | ptsI | 5 | 0.85 | 0.164 |
| 3.2 | 0 | clcB | 5 | 0.85 | 0.163 |
| 3.2 | 0 | ycaM | 5 | 0.85 | 0.162 |
| 3.2 | 0 | yadI | 5 | 0.85 | 0.157 |
| 3.2 | 0 | AG1 | 5 | 0.85 | 0.156 |
| 3.2 | 0 | ptsI | 5 | 1.1 | 0.164 |
| 3.2 | 0 | clcB | 5 | 1.1 | 0.163 |
| 3.2 | 0 | ycaM | 5 | 1.1 | 0.165 |

| | | | | | |
|---|---|---|---|---|---|
| 3.2 | 0 | yadI | 5 | 1.1 | 0.158 |
| 3.2 | 0 | AG1 | 5 | 1.1 | 0.157 |
| 3.2 | 0 | ptsI | 5 | 1.35 | 0.168 |
| 3.2 | 0 | clcB | 5 | 1.35 | 0.165 |
| 3.2 | 0 | ycaM | 5 | 1.35 | 0.166 |
| 3.2 | 0 | yadI | 5 | 1.35 | 0.161 |
| 3.2 | 0 | AG1 | 5 | 1.35 | 0.158 |
| 3.2 | 0 | ptsI | 5 | 1.6 | 0.166 |
| 3.2 | 0 | clcB | 5 | 1.6 | 0.169 |
| 3.2 | 0 | ycaM | 5 | 1.6 | 0.17 |
| 3.2 | 0 | yadI | 5 | 1.6 | 0.163 |
| 3.2 | 0 | AG1 | 5 | 1.6 | 0.161 |
| 3.2 | 0 | ptsI | 5 | 1.85 | 0.171 |
| 3.2 | 0 | clcB | 5 | 1.85 | 0.172 |
| 3.2 | 0 | ycaM | 5 | 1.85 | 0.173 |
| 3.2 | 0 | yadI | 5 | 1.85 | 0.167 |
| 3.2 | 0 | AG1 | 5 | 1.85 | 0.165 |
| 3.2 | 0 | ptsI | 5 | 2.1 | 0.172 |
| 3.2 | 0 | clcB | 5 | 2.1 | 0.177 |
| 3.2 | 0 | ycaM | 5 | 2.1 | 0.175 |
| 3.2 | 0 | yadI | 5 | 2.1 | 0.173 |
| 3.2 | 0 | AG1 | 5 | 2.1 | 0.172 |
| 3.2 | 0 | ptsI | 5 | 2.35 | 0.173 |
| 3.2 | 0 | clcB | 5 | 2.35 | 0.183 |
| 3.2 | 0 | ycaM | 5 | 2.35 | 0.18 |
| 3.2 | 0 | yadI | 5 | 2.35 | 0.178 |
| 3.2 | 0 | AG1 | 5 | 2.35 | 0.177 |
| 3.2 | 0 | ptsI | 5 | 2.6 | 0.179 |
| 3.2 | 0 | clcB | 5 | 2.6 | 0.189 |
| 3.2 | 0 | ycaM | 5 | 2.6 | 0.183 |
| 3.2 | 0 | yadI | 5 | 2.6 | 0.179 |
| 3.2 | 0 | AG1 | 5 | 2.6 | 0.174 |
| 3.2 | 0 | ptsI | 5 | 2.85 | 0.185 |
| 3.2 | 0 | clcB | 5 | 2.85 | 0.199 |
| 3.2 | 0 | ycaM | 5 | 2.85 | 0.19 |
| 3.2 | 0 | yadI | 5 | 2.85 | 0.189 |
| 3.2 | 0 | AG1 | 5 | 2.85 | 0.189 |
| 3.2 | 0 | ptsI | 5 | 3.1 | 0.188 |
| 3.2 | 0 | clcB | 5 | 3.1 | 0.21 |
| 3.2 | 0 | ycaM | 5 | 3.1 | 0.199 |
| 3.2 | 0 | yadI | 5 | 3.1 | 0.199 |
| 3.2 | 0 | AG1 | 5 | 3.1 | 0.198 |
| 3.2 | 0 | ptsI | 5 | 3.35 | 0.193 |
| 3.2 | 0 | clcB | 5 | 3.35 | 0.222 |
| 3.2 | 0 | ycaM | 5 | 3.35 | 0.209 |
| 3.2 | 0 | yadI | 5 | 3.35 | 0.211 |
| 3.2 | 0 | AG1 | 5 | 3.35 | 0.213 |
| 3.2 | 0 | ptsI | 5 | 3.6 | 0.197 |

| | | | | | |
|---|---|---|---|---|---|
| 3.2 | 0 | clcB | 5 | 3.6 | 0.229 |
| 3.2 | 0 | ycaM | 5 | 3.6 | 0.216 |
| 3.2 | 0 | yadI | 5 | 3.6 | 0.224 |
| 3.2 | 0 | AG1 | 5 | 3.6 | 0.227 |
| 3.2 | 0 | ptsI | 5 | 3.85 | 0.205 |
| 3.2 | 0 | clcB | 5 | 3.85 | 0.24 |
| 3.2 | 0 | ycaM | 5 | 3.85 | 0.226 |
| 3.2 | 0 | yadI | 5 | 3.85 | 0.238 |
| 3.2 | 0 | AG1 | 5 | 3.85 | 0.244 |
| 3.2 | 0 | ptsI | 5 | 4.38333 | 0.202 |
| 3.2 | 0 | clcB | 5 | 4.38333 | 0.22 |
| 3.2 | 0 | ycaM | 5 | 4.38333 | 0.235 |
| 3.2 | 0 | yadI | 5 | 4.38333 | 0.198 |
| 3.2 | 0 | AG1 | 5 | 4.38333 | 0.201 |
| 3.2 | 0 | ptsI | 5 | 4.63333 | 0.2 |
| 3.2 | 0 | clcB | 5 | 4.63333 | 0.213 |
| 3.2 | 0 | ycaM | 5 | 4.63333 | 0.237 |
| 3.2 | 0 | yadI | 5 | 4.63333 | 0.201 |
| 3.2 | 0 | AG1 | 5 | 4.63333 | 0.208 |
| 3.2 | 0 | ptsI | 5 | 4.88333 | 0.207 |
| 3.2 | 0 | clcB | 5 | 4.88333 | 0.22 |
| 3.2 | 0 | ycaM | 5 | 4.88333 | 0.249 |
| 3.2 | 0 | yadI | 5 | 4.88333 | 0.209 |
| 3.2 | 0 | AG1 | 5 | 4.88333 | 0.223 |
| 3.2 | 0 | ptsI | 5 | 5.13333 | 0.214 |
| 3.2 | 0 | clcB | 5 | 5.13333 | 0.227 |
| 3.2 | 0 | ycaM | 5 | 5.13333 | 0.26 |
| 3.2 | 0 | yadI | 5 | 5.13333 | 0.219 |
| 3.2 | 0 | AG1 | 5 | 5.13333 | 0.236 |
| 3.2 | 0 | ptsI | 5 | 5.38333 | 0.221 |
| 3.2 | 0 | clcB | 5 | 5.38333 | 0.236 |
| 3.2 | 0 | ycaM | 5 | 5.38333 | 0.268 |
| 3.2 | 0 | yadI | 5 | 5.38333 | 0.229 |
| 3.2 | 0 | AG1 | 5 | 5.38333 | 0.249 |
| 3.2 | 0 | ptsI | 5 | 5.63333 | 0.226 |
| 3.2 | 0 | clcB | 5 | 5.63333 | 0.246 |
| 3.2 | 0 | ycaM | 5 | 5.63333 | 0.277 |
| 3.2 | 0 | yadI | 5 | 5.63333 | 0.241 |
| 3.2 | 0 | AG1 | 5 | 5.63333 | 0.264 |
| 3.2 | 0 | ptsI | 5 | 5.88333 | 0.233 |
| 3.2 | 0 | clcB | 5 | 5.88333 | 0.255 |
| 3.2 | 0 | ycaM | 5 | 5.88333 | 0.29 |
| 3.2 | 0 | yadI | 5 | 5.88333 | 0.249 |
| 3.2 | 0 | AG1 | 5 | 5.88333 | 0.276 |
| 3.2 | 0 | ptsI | 5 | 6.13333 | 0.244 |
| 3.2 | 0 | clcB | 5 | 6.13333 | 0.271 |
| 3.2 | 0 | ycaM | 5 | 6.13333 | 0.3 |
| 3.2 | 0 | yadI | 5 | 6.13333 | 0.261 |

| | | | | | |
|---|---|---|---|---|---|
| 3.2 | 0 | AG1 | 5 | 6.13333 | 0.294 |
| 3.2 | 0 | ptsI | 5 | 6.38333 | 0.252 |
| 3.2 | 0 | clcB | 5 | 6.38333 | 0.287 |
| 3.2 | 0 | ycaM | 5 | 6.38333 | 0.308 |
| 3.2 | 0 | yadI | 5 | 6.38333 | 0.276 |
| 3.2 | 0 | AG1 | 5 | 6.38333 | 0.314 |
| 3.2 | 0 | ptsI | 5 | 6.63333 | 0.258 |
| 3.2 | 0 | clcB | 5 | 6.63333 | 0.306 |
| 3.2 | 0 | ycaM | 5 | 6.63333 | 0.322 |
| 3.2 | 0 | yadI | 5 | 6.63333 | 0.292 |
| 3.2 | 0 | AG1 | 5 | 6.63333 | 0.337 |
| 3.2 | 0 | ptsI | 5 | 6.88333 | 0.268 |
| 3.2 | 0 | clcB | 5 | 6.88333 | 0.322 |
| 3.2 | 0 | ycaM | 5 | 6.88333 | 0.332 |
| 3.2 | 0 | yadI | 5 | 6.88333 | 0.31 |
| 3.2 | 0 | AG1 | 5 | 6.88333 | 0.353 |
| 3.2 | 0 | ptsI | 5 | 7.13333 | 0.276 |
| 3.2 | 0 | clcB | 5 | 7.13333 | 0.341 |
| 3.2 | 0 | ycaM | 5 | 7.13333 | 0.344 |
| 3.2 | 0 | yadI | 5 | 7.13333 | 0.325 |
| 3.2 | 0 | AG1 | 5 | 7.13333 | 0.376 |
| 3.2 | 0 | ptsI | 5 | 7.38333 | 0.284 |
| 3.2 | 0 | clcB | 5 | 7.38333 | 0.36 |
| 3.2 | 0 | ycaM | 5 | 7.38333 | 0.357 |
| 3.2 | 0 | yadI | 5 | 7.38333 | 0.343 |
| 3.2 | 0 | AG1 | 5 | 7.38333 | 0.396 |
| 3.2 | 0 | ptsI | 5 | 7.63333 | 0.293 |
| 3.2 | 0 | clcB | 5 | 7.63333 | 0.383 |
| 3.2 | 0 | ycaM | 5 | 7.63333 | 0.371 |
| 3.2 | 0 | yadI | 5 | 7.63333 | 0.367 |
| 3.2 | 0 | AG1 | 5 | 7.63333 | 0.413 |
| 3.2 | 0 | ptsI | 5 | 7.88333 | 0.302 |
| 3.2 | 0 | clcB | 5 | 7.88333 | 0.396 |
| 3.2 | 0 | ycaM | 5 | 7.88333 | 0.38 |
| 3.2 | 0 | yadI | 5 | 7.88333 | 0.389 |
| 3.2 | 0 | AG1 | 5 | 7.88333 | 0.435 |
| 3.2 | 0 | ptsI | 5 | 8.13333 | 0.312 |
| 3.2 | 0 | clcB | 5 | 8.13333 | 0.407 |
| 3.2 | 0 | ycaM | 5 | 8.13333 | 0.388 |
| 3.2 | 0 | yadI | 5 | 8.13333 | 0.402 |
| 3.2 | 0 | AG1 | 5 | 8.13333 | 0.445 |
| 3.2 | 0 | ptsI | 5 | 8.38333 | 0.318 |
| 3.2 | 0 | clcB | 5 | 8.38333 | 0.425 |
| 3.2 | 0 | ycaM | 5 | 8.38333 | 0.401 |
| 3.2 | 0 | yadI | 5 | 8.38333 | 0.42 |
| 3.2 | 0 | AG1 | 5 | 8.38333 | 0.458 |
| 3.2 | 0 | ptsI | 5 | 8.88333 | 0.324 |
| 3.2 | 0 | clcB | 5 | 8.88333 | 0.346 |

| | | | | | |
|---|---|---|---|---|---|
| 3.2 | 0 | ycaM | 5 | 8.88333 | 0.338 |
| 3.2 | 0 | yadI | 5 | 8.88333 | 0.39 |
| 3.2 | 0 | AG1 | 5 | 8.88333 | 0.455 |
| 3.2 | 0 | ptsI | 5 | 9.13333 | 0.343 |
| 3.2 | 0 | clcB | 5 | 9.13333 | 0.372 |
| 3.2 | 0 | ycaM | 5 | 9.13333 | 0.361 |
| 3.2 | 0 | yadI | 5 | 9.13333 | 0.408 |
| 3.2 | 0 | AG1 | 5 | 9.13333 | 0.463 |
| 3.2 | 0 | ptsI | 5 | 9.38333 | 0.35 |
| 3.2 | 0 | clcB | 5 | 9.38333 | 0.389 |
| 3.2 | 0 | ycaM | 5 | 9.38333 | 0.373 |
| 3.2 | 0 | yadI | 5 | 9.38333 | 0.424 |
| 3.2 | 0 | AG1 | 5 | 9.38333 | 0.482 |
| 3.2 | 0 | ptsI | 5 | 9.63333 | 0.357 |
| 3.2 | 0 | clcB | 5 | 9.63333 | 0.388 |
| 3.2 | 0 | ycaM | 5 | 9.63333 | 0.379 |
| 3.2 | 0 | yadI | 5 | 9.63333 | 0.431 |
| 3.2 | 0 | AG1 | 5 | 9.63333 | 0.487 |
| 3.2 | 0 | ptsI | 5 | 9.88333 | 0.361 |
| 3.2 | 0 | clcB | 5 | 9.88333 | 0.389 |
| 3.2 | 0 | ycaM | 5 | 9.88333 | 0.386 |
| 3.2 | 0 | yadI | 5 | 9.88333 | 0.438 |
| 3.2 | 0 | AG1 | 5 | 9.88333 | 0.494 |
| 3.2 | 0 | ptsI | 5 | 10.1333 | 0.361 |
| 3.2 | 0 | clcB | 5 | 10.1333 | 0.393 |
| 3.2 | 0 | ycaM | 5 | 10.1333 | 0.381 |
| 3.2 | 0 | yadI | 5 | 10.1333 | 0.434 |
| 3.2 | 0 | AG1 | 5 | 10.1333 | 0.503 |
| 3.2 | 0 | ptsI | 5 | 10.3833 | 0.372 |
| 3.2 | 0 | clcB | 5 | 10.3833 | 0.4 |
| 3.2 | 0 | ycaM | 5 | 10.3833 | 0.389 |
| 3.2 | 0 | yadI | 5 | 10.3833 | 0.444 |
| 3.2 | 0 | AG1 | 5 | 10.3833 | 0.51 |
| 3.2 | 0 | ptsI | 5 | 10.6333 | 0.372 |
| 3.2 | 0 | clcB | 5 | 10.6333 | 0.399 |
| 3.2 | 0 | ycaM | 5 | 10.6333 | 0.388 |
| 3.2 | 0 | yadI | 5 | 10.6333 | 0.444 |
| 3.2 | 0 | AG1 | 5 | 10.6333 | 0.512 |
| 3.2 | 0 | ptsI | 5 | 10.8833 | 0.378 |
| 3.2 | 0 | clcB | 5 | 10.8833 | 0.407 |
| 3.2 | 0 | ycaM | 5 | 10.8833 | 0.39 |
| 3.2 | 0 | yadI | 5 | 10.8833 | 0.451 |
| 3.2 | 0 | AG1 | 5 | 10.8833 | 0.525 |
| 3.2 | 0 | ptsI | 5 | 11.1333 | 0.379 |
| 3.2 | 0 | clcB | 5 | 11.1333 | 0.408 |
| 3.2 | 0 | ycaM | 5 | 11.1333 | 0.391 |
| 3.2 | 0 | yadI | 5 | 11.1333 | 0.451 |
| 3.2 | 0 | AG1 | 5 | 11.1333 | 0.53 |

| | | | | | |
|---|---|---|---|---|---|
| 3.2 | 0 | ptsI | 5 | 11.3833 | 0.386 |
| 3.2 | 0 | clcB | 5 | 11.3833 | 0.414 |
| 3.2 | 0 | ycaM | 5 | 11.3833 | 0.396 |
| 3.2 | 0 | yadI | 5 | 11.3833 | 0.461 |
| 3.2 | 0 | AG1 | 5 | 11.3833 | 0.537 |
| 3.2 | 0 | ptsI | 5 | 11.6333 | 0.388 |
| 3.2 | 0 | clcB | 5 | 11.6333 | 0.418 |
| 3.2 | 0 | ycaM | 5 | 11.6333 | 0.396 |
| 3.2 | 0 | yadI | 5 | 11.6333 | 0.461 |
| 3.2 | 0 | AG1 | 5 | 11.6333 | 0.543 |
| 3.2 | 0 | ptsI | 5 | 11.8833 | 0.395 |
| 3.2 | 0 | clcB | 5 | 11.8833 | 0.418 |
| 3.2 | 0 | ycaM | 5 | 11.8833 | 0.396 |
| 3.2 | 0 | yadI | 5 | 11.8833 | 0.466 |
| 3.2 | 0 | AG1 | 5 | 11.8833 | 0.549 |
| 3.2 | 0 | ptsI | 5 | 12.1333 | 0.398 |
| 3.2 | 0 | clcB | 5 | 12.1333 | 0.423 |
| 3.2 | 0 | ycaM | 5 | 12.1333 | 0.402 |
| 3.2 | 0 | yadI | 5 | 12.1333 | 0.472 |
| 3.2 | 0 | AG1 | 5 | 12.1333 | 0.56 |
| 3.2 | 0 | ptsI | 5 | 12.3833 | 0.401 |
| 3.2 | 0 | clcB | 5 | 12.3833 | 0.426 |
| 3.2 | 0 | ycaM | 5 | 12.3833 | 0.405 |
| 3.2 | 0 | yadI | 5 | 12.3833 | 0.472 |
| 3.2 | 0 | AG1 | 5 | 12.3833 | 0.539 |
| 3.2 | 0 | ptsI | 5 | 12.6333 | 0.408 |
| 3.2 | 0 | clcB | 5 | 12.6333 | 0.43 |
| 3.2 | 0 | ycaM | 5 | 12.6333 | 0.41 |
| 3.2 | 0 | yadI | 5 | 12.6333 | 0.479 |
| 3.2 | 0 | AG1 | 5 | 12.6333 | 0.57 |
| 3.2 | 0 | ptsI | 5 | 12.8833 | 0.412 |
| 3.2 | 0 | clcB | 5 | 12.8833 | 0.435 |
| 3.2 | 0 | ycaM | 5 | 12.8833 | 0.412 |
| 3.2 | 0 | yadI | 5 | 12.8833 | 0.486 |
| 3.2 | 0 | AG1 | 5 | 12.8833 | 0.578 |
| 3.2 | 0 | ptsI | 5 | 13.1333 | 0.41 |
| 3.2 | 0 | clcB | 5 | 13.1333 | 0.44 |
| 3.2 | 0 | ycaM | 5 | 13.1333 | 0.416 |
| 3.2 | 0 | yadI | 5 | 13.1333 | 0.484 |
| 3.2 | 0 | AG1 | 5 | 13.1333 | 0.585 |
| 3.2 | 0 | ptsI | 5 | 13.3833 | 0.414 |
| 3.2 | 0 | clcB | 5 | 13.3833 | 0.443 |
| 3.2 | 0 | ycaM | 5 | 13.3833 | 0.422 |
| 3.2 | 0 | yadI | 5 | 13.3833 | 0.493 |
| 3.2 | 0 | AG1 | 5 | 13.3833 | 0.591 |
| 3.2 | 0 | ptsI | 5 | 13.6333 | 0.421 |
| 3.2 | 0 | clcB | 5 | 13.6333 | 0.443 |
| 3.2 | 0 | ycaM | 5 | 13.6333 | 0.427 |

| | | | | | |
|---|---|---|---|---|---|
| 3.2 | 0 | yadI | 5 | 13.6333 | 0.498 |
| 3.2 | 0 | AG1 | 5 | 13.6333 | 0.596 |
| 3.2 | 0 | ptsI | 5 | 13.8833 | 0.422 |
| 3.2 | 0 | clcB | 5 | 13.8833 | 0.446 |
| 3.2 | 0 | ycaM | 5 | 13.8833 | 0.43 |
| 3.2 | 0 | yadI | 5 | 13.8833 | 0.503 |
| 3.2 | 0 | AG1 | 5 | 13.8833 | 0.599 |
| 3.2 | 0 | ptsI | 5 | 14.1333 | 0.424 |
| 3.2 | 0 | clcB | 5 | 14.1333 | 0.454 |
| 3.2 | 0 | ycaM | 5 | 14.1333 | 0.431 |
| 3.2 | 0 | yadI | 5 | 14.1333 | 0.507 |
| 3.2 | 0 | AG1 | 5 | 14.1333 | 0.607 |
| 3.2 | 0 | ptsI | 5 | 14.3833 | 0.437 |
| 3.2 | 0 | clcB | 5 | 14.3833 | 0.454 |
| 3.2 | 0 | ycaM | 5 | 14.3833 | 0.438 |
| 3.2 | 0 | yadI | 5 | 14.3833 | 0.511 |
| 3.2 | 0 | AG1 | 5 | 14.3833 | 0.615 |
| 3.2 | 0 | ptsI | 5 | 14.6333 | 0.438 |
| 3.2 | 0 | clcB | 5 | 14.6333 | 0.463 |
| 3.2 | 0 | ycaM | 5 | 14.6333 | 0.445 |
| 3.2 | 0 | yadI | 5 | 14.6333 | 0.512 |
| 3.2 | 0 | AG1 | 5 | 14.6333 | 0.618 |
| 3.2 | 0 | ptsI | 5 | 14.8833 | 0.437 |
| 3.2 | 0 | clcB | 5 | 14.8833 | 0.463 |
| 3.2 | 0 | ycaM | 5 | 14.8833 | 0.442 |
| 3.2 | 0 | yadI | 5 | 14.8833 | 0.52 |
| 3.2 | 0 | AG1 | 5 | 14.8833 | 0.622 |
| 3.2 | 0 | ptsI | 5 | 15.1333 | 0.444 |
| 3.2 | 0 | clcB | 5 | 15.1333 | 0.47 |
| 3.2 | 0 | ycaM | 5 | 15.1333 | 0.452 |
| 3.2 | 0 | yadI | 5 | 15.1333 | 0.523 |
| 3.2 | 0 | AG1 | 5 | 15.1333 | 0.632 |
| 3.2 | 0 | ptsI | 5 | 15.3833 | 0.449 |
| 3.2 | 0 | clcB | 5 | 15.3833 | 0.472 |
| 3.2 | 0 | ycaM | 5 | 15.3833 | 0.453 |
| 3.2 | 0 | yadI | 5 | 15.3833 | 0.531 |
| 3.2 | 0 | AG1 | 5 | 15.3833 | 0.638 |
| 3.2 | 0 | ptsI | 5 | 15.6333 | 0.45 |
| 3.2 | 0 | clcB | 5 | 15.6333 | 0.476 |
| 3.2 | 0 | ycaM | 5 | 15.6333 | 0.457 |
| 3.2 | 0 | yadI | 5 | 15.6333 | 0.53 |
| 3.2 | 0 | AG1 | 5 | 15.6333 | 0.638 |
| 3.2 | 0 | ptsI | 5 | 15.8833 | 0.448 |
| 3.2 | 0 | clcB | 5 | 15.8833 | 0.479 |
| 3.2 | 0 | ycaM | 5 | 15.8833 | 0.458 |
| 3.2 | 0 | yadI | 5 | 15.8833 | 0.539 |
| 3.2 | 0 | AG1 | 5 | 15.8833 | 0.646 |
| 3.2 | 0 | ptsI | 5 | 16.1333 | 0.458 |

| | | | | | |
|---|---|---|---|---|---|
| 3.2 | 0 | clcB | 5 | 16.1333 | 0.483 |
| 3.2 | 0 | ycaM | 5 | 16.1333 | 0.463 |
| 3.2 | 0 | yadI | 5 | 16.1333 | 0.541 |
| 3.2 | 0 | AG1 | 5 | 16.1333 | 0.652 |
| 3.2 | 0 | ptsI | 5 | 16.3833 | 0.459 |
| 3.2 | 0 | clcB | 5 | 16.3833 | 0.485 |
| 3.2 | 0 | ycaM | 5 | 16.3833 | 0.464 |
| 3.2 | 0 | yadI | 5 | 16.3833 | 0.547 |
| 3.2 | 0 | AG1 | 5 | 16.3833 | 0.658 |
| 3.2 | 0 | ptsI | 5 | 16.6333 | 0.464 |
| 3.2 | 0 | clcB | 5 | 16.6333 | 0.488 |
| 3.2 | 0 | ycaM | 5 | 16.6333 | 0.472 |
| 3.2 | 0 | yadI | 5 | 16.6333 | 0.555 |
| 3.2 | 0 | AG1 | 5 | 16.6333 | 0.665 |
| 3.2 | 0 | ptsI | 5 | 16.8833 | 0.47 |
| 3.2 | 0 | clcB | 5 | 16.8833 | 0.491 |
| 3.2 | 0 | ycaM | 5 | 16.8833 | 0.478 |
| 3.2 | 0 | yadI | 5 | 16.8833 | 0.558 |
| 3.2 | 0 | AG1 | 5 | 16.8833 | 0.667 |
| 3.2 | 0 | ptsI | 5 | 17.1333 | 0.468 |
| 3.2 | 0 | clcB | 5 | 17.1333 | 0.493 |
| 3.2 | 0 | ycaM | 5 | 17.1333 | 0.476 |
| 3.2 | 0 | yadI | 5 | 17.1333 | 0.558 |
| 3.2 | 0 | AG1 | 5 | 17.1333 | 0.67 |
| 3.2 | 0 | ptsI | 5 | 17.3833 | 0.48 |
| 3.2 | 0 | clcB | 5 | 17.3833 | 0.497 |
| 3.2 | 0 | ycaM | 5 | 17.3833 | 0.483 |
| 3.2 | 0 | yadI | 5 | 17.3833 | 0.56 |
| 3.2 | 0 | AG1 | 5 | 17.3833 | 0.68 |
| 3.2 | 0 | ptsI | 5 | 17.6333 | 0.479 |
| 3.2 | 0 | clcB | 5 | 17.6333 | 0.5 |
| 3.2 | 0 | ycaM | 5 | 17.6333 | 0.483 |
| 3.2 | 0 | yadI | 5 | 17.6333 | 0.567 |
| 3.2 | 0 | AG1 | 5 | 17.6333 | 0.682 |
| 3.2 | 0 | ptsI | 5 | 17.8833 | 0.478 |
| 3.2 | 0 | clcB | 5 | 17.8833 | 0.505 |
| 3.2 | 0 | ycaM | 5 | 17.8833 | 0.487 |
| 3.2 | 0 | yadI | 5 | 17.8833 | 0.566 |
| 3.2 | 0 | AG1 | 5 | 17.8833 | 0.648 |
| 3.2 | 0 | ptsI | 5 | 18.1333 | 0.485 |
| 3.2 | 0 | clcB | 5 | 18.1333 | 0.503 |
| 3.2 | 0 | ycaM | 5 | 18.1333 | 0.491 |
| 3.2 | 0 | yadI | 5 | 18.1333 | 0.572 |
| 3.2 | 0 | AG1 | 5 | 18.1333 | 0.691 |
| 3.2 | 0 | ptsI | 5 | 18.3833 | 0.487 |
| 3.2 | 0 | clcB | 5 | 18.3833 | 0.509 |
| 3.2 | 0 | ycaM | 5 | 18.3833 | 0.497 |
| 3.2 | 0 | yadI | 5 | 18.3833 | 0.577 |

| | | | | | |
|---|---|---|---|---|---|
| 3.2 | 0 | AG1 | 5 | 18.3833 | 0.69 |
| 3.2 | 0 | ptsI | 5 | 18.6333 | 0.494 |
| 3.2 | 0 | clcB | 5 | 18.6333 | 0.506 |
| 3.2 | 0 | ycaM | 5 | 18.6333 | 0.5 |
| 3.2 | 0 | yadI | 5 | 18.6333 | 0.583 |
| 3.2 | 0 | AG1 | 5 | 18.6333 | 0.698 |
| 3.2 | 0 | ptsI | 5 | 18.8833 | 0.496 |
| 3.2 | 0 | clcB | 5 | 18.8833 | 0.512 |
| 3.2 | 0 | ycaM | 5 | 18.8833 | 0.506 |
| 3.2 | 0 | yadI | 5 | 18.8833 | 0.59 |
| 3.2 | 0 | AG1 | 5 | 18.8833 | 0.704 |
| 3.2 | 0 | ptsI | 5 | 19.1333 | 0.494 |
| 3.2 | 0 | clcB | 5 | 19.1333 | 0.513 |
| 3.2 | 0 | ycaM | 5 | 19.1333 | 0.505 |
| 3.2 | 0 | yadI | 5 | 19.1333 | 0.591 |
| 3.2 | 0 | AG1 | 5 | 19.1333 | 0.705 |
| 3.2 | 0 | ptsI | 5 | 19.3833 | 0.508 |
| 3.2 | 0 | clcB | 5 | 19.3833 | 0.512 |
| 3.2 | 0 | ycaM | 5 | 19.3833 | 0.502 |
| 3.2 | 0 | yadI | 5 | 19.3833 | 0.601 |
| 3.2 | 0 | AG1 | 5 | 19.3833 | 0.712 |
| 3.2 | 0 | ptsI | 5 | 19.6333 | 0.507 |
| 3.2 | 0 | clcB | 5 | 19.6333 | 0.519 |
| 3.2 | 0 | ycaM | 5 | 19.6333 | 0.513 |
| 3.2 | 0 | yadI | 5 | 19.6333 | 0.603 |
| 3.2 | 0 | AG1 | 5 | 19.6333 | 0.717 |
| 3.2 | 0 | ptsI | 5 | 19.8833 | 0.511 |
| 3.2 | 0 | clcB | 5 | 19.8833 | 0.519 |
| 3.2 | 0 | ycaM | 5 | 19.8833 | 0.516 |
| 3.2 | 0 | yadI | 5 | 19.8833 | 0.603 |
| 3.2 | 0 | AG1 | 5 | 19.8833 | 0.718 |
| 3.2 | 0 | ptsI | 5 | 20.1333 | 0.517 |
| 3.2 | 0 | clcB | 5 | 20.1333 | 0.521 |
| 3.2 | 0 | ycaM | 5 | 20.1333 | 0.517 |
| 3.2 | 0 | yadI | 5 | 20.1333 | 0.611 |
| 3.2 | 0 | AG1 | 5 | 20.1333 | 0.723 |
| 3.2 | 0 | ptsI | 5 | 20.3833 | 0.52 |
| 3.2 | 0 | clcB | 5 | 20.3833 | 0.521 |
| 3.2 | 0 | ycaM | 5 | 20.3833 | 0.52 |
| 3.2 | 0 | yadI | 5 | 20.3833 | 0.617 |
| 3.2 | 0 | AG1 | 5 | 20.3833 | 0.726 |
| 3.2 | 0 | ptsI | 5 | 20.6333 | 0.521 |
| 3.2 | 0 | clcB | 5 | 20.6333 | 0.522 |
| 3.2 | 0 | ycaM | 5 | 20.6333 | 0.524 |
| 3.2 | 0 | yadI | 5 | 20.6333 | 0.625 |
| 3.2 | 0 | AG1 | 5 | 20.6333 | 0.69 |
| 3.2 | 0 | ptsI | 5 | 20.8833 | 0.524 |
| 3.2 | 0 | clcB | 5 | 20.8833 | 0.532 |

| | | | | | |
|---|---|---|---|---|---|
| 3.2 | 0 | ycaM | 5 | 20.8833 | 0.529 |
| 3.2 | 0 | yadI | 5 | 20.8833 | 0.624 |
| 3.2 | 0 | AG1 | 5 | 20.8833 | 0.733 |
| 3.2 | 0 | ptsI | 5 | 21.1333 | 0.531 |
| 3.2 | 0 | clcB | 5 | 21.1333 | 0.531 |
| 3.2 | 0 | ycaM | 5 | 21.1333 | 0.529 |
| 3.2 | 0 | yadI | 5 | 21.1333 | 0.629 |
| 3.2 | 0 | AG1 | 5 | 21.1333 | 0.738 |
| 3.2 | 0 | ptsI | 5 | 21.3833 | 0.529 |
| 3.2 | 0 | clcB | 5 | 21.3833 | 0.533 |
| 3.2 | 0 | ycaM | 5 | 21.3833 | 0.533 |
| 3.2 | 0 | yadI | 5 | 21.3833 | 0.629 |
| 3.2 | 0 | AG1 | 5 | 21.3833 | 0.743 |
| 3.2 | 0 | ptsI | 5 | 21.6333 | 0.534 |
| 3.2 | 0 | clcB | 5 | 21.6333 | 0.537 |
| 3.2 | 0 | ycaM | 5 | 21.6333 | 0.538 |
| 3.2 | 0 | yadI | 5 | 21.6333 | 0.634 |
| 3.2 | 0 | AG1 | 5 | 21.6333 | 0.746 |
| 3.2 | 0 | ptsI | 5 | 21.8833 | 0.535 |
| 3.2 | 0 | clcB | 5 | 21.8833 | 0.539 |
| 3.2 | 0 | ycaM | 5 | 21.8833 | 0.539 |
| 3.2 | 0 | yadI | 5 | 21.8833 | 0.639 |
| 3.2 | 0 | AG1 | 5 | 21.8833 | 0.746 |
| 3.2 | 0 | ptsI | 5 | 22.1333 | 0.537 |
| 3.2 | 0 | clcB | 5 | 22.1333 | 0.545 |
| 3.2 | 0 | ycaM | 5 | 22.1333 | 0.539 |
| 3.2 | 0 | yadI | 5 | 22.1333 | 0.643 |
| 3.2 | 0 | AG1 | 5 | 22.1333 | 0.718 |
| 3.2 | 0 | ptsI | 5 | 22.3833 | 0.539 |
| 3.2 | 0 | clcB | 5 | 22.3833 | 0.543 |
| 3.2 | 0 | ycaM | 5 | 22.3833 | 0.543 |
| 3.2 | 0 | yadI | 5 | 22.3833 | 0.647 |
| 3.2 | 0 | AG1 | 5 | 22.3833 | 0.764 |
| 3.2 | 0 | ptsI | 5 | 22.6333 | 0.548 |
| 3.2 | 0 | clcB | 5 | 22.6333 | 0.546 |
| 3.2 | 0 | ycaM | 5 | 22.6333 | 0.546 |
| 3.2 | 0 | yadI | 5 | 22.6333 | 0.655 |
| 3.2 | 0 | AG1 | 5 | 22.6333 | 0.765 |
| 3.2 | 0 | ptsI | 5 | 22.8833 | 0.553 |
| 3.2 | 0 | clcB | 5 | 22.8833 | 0.55 |
| 3.2 | 0 | ycaM | 5 | 22.8833 | 0.55 |
| 3.2 | 0 | yadI | 5 | 22.8833 | 0.658 |
| 3.2 | 0 | AG1 | 5 | 22.8833 | 0.768 |
| 3.2 | 0 | ptsI | 5 | 23.1333 | 0.557 |
| 3.2 | 0 | clcB | 5 | 23.1333 | 0.556 |
| 3.2 | 0 | ycaM | 5 | 23.1333 | 0.549 |
| 3.2 | 0 | yadI | 5 | 23.1333 | 0.666 |
| 3.2 | 0 | AG1 | 5 | 23.1333 | 0.775 |

| | | | | | |
|---|---|---|---|---|---|
| 3.2 | 0 | ptsI | 5 | 23.3833 | 0.558 |
| 3.2 | 0 | clcB | 5 | 23.3833 | 0.552 |
| 3.2 | 0 | ycaM | 5 | 23.3833 | 0.554 |
| 3.2 | 0 | yadI | 5 | 23.3833 | 0.666 |
| 3.2 | 0 | AG1 | 5 | 23.3833 | 0.773 |
| 3.2 | 0 | ptsI | 5 | 23.6333 | 0.568 |
| 3.2 | 0 | clcB | 5 | 23.6333 | 0.556 |
| 3.2 | 0 | ycaM | 5 | 23.6333 | 0.554 |
| 3.2 | 0 | yadI | 5 | 23.6333 | 0.669 |
| 3.2 | 0 | AG1 | 5 | 23.6333 | 0.772 |
| 3.2 | 0 | ptsI | 5 | 23.8833 | 0.575 |
| 3.2 | 0 | clcB | 5 | 23.8833 | 0.558 |
| 3.2 | 0 | ycaM | 5 | 23.8833 | 0.554 |
| 3.2 | 0 | yadI | 5 | 23.8833 | 0.668 |
| 3.2 | 0 | AG1 | 5 | 23.8833 | 0.734 |
| 3.2 | 0 | ptsI | 5 | 24.1333 | 0.582 |
| 3.2 | 0 | clcB | 5 | 24.1333 | 0.561 |
| 3.2 | 0 | ycaM | 5 | 24.1333 | 0.56 |
| 3.2 | 0 | yadI | 5 | 24.1333 | 0.677 |
| 3.2 | 0 | AG1 | 5 | 24.1333 | 0.777 |
| 3.2 | 0 | ptsI | 5 | 24.3833 | 0.586 |
| 3.2 | 0 | clcB | 5 | 24.3833 | 0.566 |
| 3.2 | 0 | ycaM | 5 | 24.3833 | 0.563 |
| 3.2 | 0 | yadI | 5 | 24.3833 | 0.68 |
| 3.2 | 0 | AG1 | 5 | 24.3833 | 0.778 |
| 0 | 0.05 | ptsI | 2 | 0 | 0.206 |
| 0 | 0.05 | clcB | 2 | 0 | 0.204 |
| 0 | 0.05 | ycaM | 2 | 0 | 0.214 |
| 0 | 0.05 | yadI | 2 | 0 | 0.216 |
| 0 | 0.05 | AG1 | 2 | 0 | 0.233 |
| 0 | 0.05 | ptsI | 2 | 0.25 | 0.204 |
| 0 | 0.05 | clcB | 2 | 0.25 | 0.204 |
| 0 | 0.05 | ycaM | 2 | 0.25 | 0.211 |
| 0 | 0.05 | yadI | 2 | 0.25 | 0.21 |
| 0 | 0.05 | AG1 | 2 | 0.25 | 0.223 |
| 0 | 0.05 | ptsI | 2 | 0.5 | 0.203 |
| 0 | 0.05 | clcB | 2 | 0.5 | 0.203 |
| 0 | 0.05 | ycaM | 2 | 0.5 | 0.209 |
| 0 | 0.05 | yadI | 2 | 0.5 | 0.209 |
| 0 | 0.05 | AG1 | 2 | 0.5 | 0.221 |
| 0 | 0.05 | ptsI | 2 | 0.75 | 0.207 |
| 0 | 0.05 | clcB | 2 | 0.75 | 0.205 |
| 0 | 0.05 | ycaM | 2 | 0.75 | 0.21 |
| 0 | 0.05 | yadI | 2 | 0.75 | 0.211 |
| 0 | 0.05 | AG1 | 2 | 0.75 | 0.221 |
| 0 | 0.05 | ptsI | 2 | 1 | 0.206 |
| 0 | 0.05 | clcB | 2 | 1 | 0.207 |
| 0 | 0.05 | ycaM | 2 | 1 | 0.214 |

| | | | | | |
|---|---|---|---|---|---|
| 0 | 0.05 | yadI | 2 | 1 | 0.216 |
| 0 | 0.05 | AG1 | 2 | 1 | 0.227 |
| 0 | 0.05 | ptsI | 2 | 1.25 | 0.209 |
| 0 | 0.05 | clcB | 2 | 1.25 | 0.208 |
| 0 | 0.05 | ycaM | 2 | 1.25 | 0.218 |
| 0 | 0.05 | yadI | 2 | 1.25 | 0.224 |
| 0 | 0.05 | AG1 | 2 | 1.25 | 0.231 |
| 0 | 0.05 | ptsI | 2 | 1.5 | 0.21 |
| 0 | 0.05 | clcB | 2 | 1.5 | 0.21 |
| 0 | 0.05 | ycaM | 2 | 1.5 | 0.22 |
| 0 | 0.05 | yadI | 2 | 1.5 | 0.228 |
| 0 | 0.05 | AG1 | 2 | 1.5 | 0.235 |
| 0 | 0.05 | ptsI | 2 | 1.75 | 0.216 |
| 0 | 0.05 | clcB | 2 | 1.75 | 0.212 |
| 0 | 0.05 | ycaM | 2 | 1.75 | 0.224 |
| 0 | 0.05 | yadI | 2 | 1.75 | 0.236 |
| 0 | 0.05 | AG1 | 2 | 1.75 | 0.239 |
| 0 | 0.05 | ptsI | 2 | 2 | 0.217 |
| 0 | 0.05 | clcB | 2 | 2 | 0.214 |
| 0 | 0.05 | ycaM | 2 | 2 | 0.232 |
| 0 | 0.05 | yadI | 2 | 2 | 0.243 |
| 0 | 0.05 | AG1 | 2 | 2 | 0.246 |
| 0 | 0.05 | ptsI | 2 | 2.25 | 0.223 |
| 0 | 0.05 | clcB | 2 | 2.25 | 0.217 |
| 0 | 0.05 | ycaM | 2 | 2.25 | 0.239 |
| 0 | 0.05 | yadI | 2 | 2.25 | 0.251 |
| 0 | 0.05 | AG1 | 2 | 2.25 | 0.251 |
| 0 | 0.05 | ptsI | 2 | 2.5 | 0.231 |
| 0 | 0.05 | clcB | 2 | 2.5 | 0.223 |
| 0 | 0.05 | ycaM | 2 | 2.5 | 0.247 |
| 0 | 0.05 | yadI | 2 | 2.5 | 0.266 |
| 0 | 0.05 | AG1 | 2 | 2.5 | 0.259 |
| 0 | 0.05 | ptsI | 2 | 2.75 | 0.235 |
| 0 | 0.05 | clcB | 2 | 2.75 | 0.232 |
| 0 | 0.05 | ycaM | 2 | 2.75 | 0.259 |
| 0 | 0.05 | yadI | 2 | 2.75 | 0.277 |
| 0 | 0.05 | AG1 | 2 | 2.75 | 0.268 |
| 0 | 0.05 | ptsI | 2 | 3 | 0.241 |
| 0 | 0.05 | clcB | 2 | 3 | 0.233 |
| 0 | 0.05 | ycaM | 2 | 3 | 0.26 |
| 0 | 0.05 | yadI | 2 | 3 | 0.286 |
| 0 | 0.05 | AG1 | 2 | 3 | 0.265 |
| 0 | 0.05 | ptsI | 2 | 3.25 | 0.247 |
| 0 | 0.05 | clcB | 2 | 3.25 | 0.241 |
| 0 | 0.05 | ycaM | 2 | 3.25 | 0.276 |
| 0 | 0.05 | yadI | 2 | 3.25 | 0.302 |
| 0 | 0.05 | AG1 | 2 | 3.25 | 0.281 |
| 0 | 0.05 | ptsI | 2 | 3.5 | 0.257 |

| | | | | | |
|---|---|---|---|---|---|
| 0 | 0.05 | clcB | 2 | 3.5 | 0.243 |
| 0 | 0.05 | ycaM | 2 | 3.5 | 0.284 |
| 0 | 0.05 | yadI | 2 | 3.5 | 0.312 |
| 0 | 0.05 | AG1 | 2 | 3.5 | 0.289 |
| 0 | 0.05 | ptsI | 2 | 3.75 | 0.263 |
| 0 | 0.05 | clcB | 2 | 3.75 | 0.247 |
| 0 | 0.05 | ycaM | 2 | 3.75 | 0.294 |
| 0 | 0.05 | yadI | 2 | 3.75 | 0.326 |
| 0 | 0.05 | AG1 | 2 | 3.75 | 0.3 |
| 0 | 0.05 | ptsI | 2 | 4 | 0.271 |
| 0 | 0.05 | clcB | 2 | 4 | 0.253 |
| 0 | 0.05 | ycaM | 2 | 4 | 0.306 |
| 0 | 0.05 | yadI | 2 | 4 | 0.343 |
| 0 | 0.05 | AG1 | 2 | 4 | 0.311 |
| 0 | 0.05 | ptsI | 2 | 4.38333 | 0.282 |
| 0 | 0.05 | clcB | 2 | 4.38333 | 0.27 |
| 0 | 0.05 | ycaM | 2 | 4.38333 | 0.297 |
| 0 | 0.05 | yadI | 2 | 4.38333 | 0.377 |
| 0 | 0.05 | AG1 | 2 | 4.38333 | 0.34 |
| 0 | 0.05 | ptsI | 2 | 4.63333 | 0.28 |
| 0 | 0.05 | clcB | 2 | 4.63333 | 0.265 |
| 0 | 0.05 | ycaM | 2 | 4.63333 | 0.299 |
| 0 | 0.05 | yadI | 2 | 4.63333 | 0.386 |
| 0 | 0.05 | AG1 | 2 | 4.63333 | 0.339 |
| 0 | 0.05 | ptsI | 2 | 4.88333 | 0.288 |
| 0 | 0.05 | clcB | 2 | 4.88333 | 0.27 |
| 0 | 0.05 | ycaM | 2 | 4.88333 | 0.313 |
| 0 | 0.05 | yadI | 2 | 4.88333 | 0.401 |
| 0 | 0.05 | AG1 | 2 | 4.88333 | 0.349 |
| 0 | 0.05 | ptsI | 2 | 5.13333 | 0.293 |
| 0 | 0.05 | clcB | 2 | 5.13333 | 0.276 |
| 0 | 0.05 | ycaM | 2 | 5.13333 | 0.316 |
| 0 | 0.05 | yadI | 2 | 5.13333 | 0.417 |
| 0 | 0.05 | AG1 | 2 | 5.13333 | 0.357 |
| 0 | 0.05 | ptsI | 2 | 5.38333 | 0.299 |
| 0 | 0.05 | clcB | 2 | 5.38333 | 0.284 |
| 0 | 0.05 | ycaM | 2 | 5.38333 | 0.322 |
| 0 | 0.05 | yadI | 2 | 5.38333 | 0.434 |
| 0 | 0.05 | AG1 | 2 | 5.38333 | 0.368 |
| 0 | 0.05 | ptsI | 2 | 5.63333 | 0.307 |
| 0 | 0.05 | clcB | 2 | 5.63333 | 0.29 |
| 0 | 0.05 | ycaM | 2 | 5.63333 | 0.324 |
| 0 | 0.05 | yadI | 2 | 5.63333 | 0.45 |
| 0 | 0.05 | AG1 | 2 | 5.63333 | 0.378 |
| 0 | 0.05 | ptsI | 2 | 5.88333 | 0.314 |
| 0 | 0.05 | clcB | 2 | 5.88333 | 0.296 |
| 0 | 0.05 | ycaM | 2 | 5.88333 | 0.326 |
| 0 | 0.05 | yadI | 2 | 5.88333 | 0.466 |

| | | | | | |
|---|---|---|---|---|---|
| 0 | 0.05 | AG1 | 2 | 5.88333 | 0.39 |
| 0 | 0.05 | ptsI | 2 | 6.13333 | 0.317 |
| 0 | 0.05 | clcB | 2 | 6.13333 | 0.305 |
| 0 | 0.05 | ycaM | 2 | 6.13333 | 0.37 |
| 0 | 0.05 | yadI | 2 | 6.13333 | 0.459 |
| 0 | 0.05 | AG1 | 2 | 6.13333 | 0.4 |
| 0 | 0.05 | ptsI | 2 | 6.38333 | 0.326 |
| 0 | 0.05 | clcB | 2 | 6.38333 | 0.309 |
| 0 | 0.05 | ycaM | 2 | 6.38333 | 0.327 |
| 0 | 0.05 | yadI | 2 | 6.38333 | 0.47 |
| 0 | 0.05 | AG1 | 2 | 6.38333 | 0.41 |
| 0 | 0.05 | ptsI | 2 | 6.63333 | 0.334 |
| 0 | 0.05 | clcB | 2 | 6.63333 | 0.315 |
| 0 | 0.05 | ycaM | 2 | 6.63333 | 0.331 |
| 0 | 0.05 | yadI | 2 | 6.63333 | 0.482 |
| 0 | 0.05 | AG1 | 2 | 6.63333 | 0.424 |
| 0 | 0.05 | ptsI | 2 | 6.88333 | 0.344 |
| 0 | 0.05 | clcB | 2 | 6.88333 | 0.324 |
| 0 | 0.05 | ycaM | 2 | 6.88333 | 0.37 |
| 0 | 0.05 | yadI | 2 | 6.88333 | 0.492 |
| 0 | 0.05 | AG1 | 2 | 6.88333 | 0.438 |
| 0 | 0.05 | ptsI | 2 | 7.13333 | 0.348 |
| 0 | 0.05 | clcB | 2 | 7.13333 | 0.322 |
| 0 | 0.05 | ycaM | 2 | 7.13333 | 0.335 |
| 0 | 0.05 | yadI | 2 | 7.13333 | 0.514 |
| 0 | 0.05 | AG1 | 2 | 7.13333 | 0.438 |
| 0 | 0.05 | ptsI | 2 | 7.38333 | 0.357 |
| 0 | 0.05 | clcB | 2 | 7.38333 | 0.331 |
| 0 | 0.05 | ycaM | 2 | 7.38333 | 0.339 |
| 0 | 0.05 | yadI | 2 | 7.38333 | 0.506 |
| 0 | 0.05 | AG1 | 2 | 7.38333 | 0.45 |
| 0 | 0.05 | ptsI | 2 | 7.63333 | 0.362 |
| 0 | 0.05 | clcB | 2 | 7.63333 | 0.34 |
| 0 | 0.05 | ycaM | 2 | 7.63333 | 0.347 |
| 0 | 0.05 | yadI | 2 | 7.63333 | 0.518 |
| 0 | 0.05 | AG1 | 2 | 7.63333 | 0.464 |
| 0 | 0.05 | ptsI | 2 | 7.88333 | 0.374 |
| 0 | 0.05 | clcB | 2 | 7.88333 | 0.346 |
| 0 | 0.05 | ycaM | 2 | 7.88333 | 0.344 |
| 0 | 0.05 | yadI | 2 | 7.88333 | 0.528 |
| 0 | 0.05 | AG1 | 2 | 7.88333 | 0.47 |
| 0 | 0.05 | ptsI | 2 | 8.13333 | 0.383 |
| 0 | 0.05 | clcB | 2 | 8.13333 | 0.349 |
| 0 | 0.05 | ycaM | 2 | 8.13333 | 0.344 |
| 0 | 0.05 | yadI | 2 | 8.13333 | 0.536 |
| 0 | 0.05 | AG1 | 2 | 8.13333 | 0.476 |
| 0 | 0.05 | ptsI | 2 | 8.38333 | 0.39 |
| 0 | 0.05 | clcB | 2 | 8.38333 | 0.355 |

| | | | | | |
|---|---|---|---|---|---|
| 0 | 0.05 | ycaM | 2 | 8.38333 | 0.348 |
| 0 | 0.05 | yadI | 2 | 8.38333 | 0.55 |
| 0 | 0.05 | AG1 | 2 | 8.38333 | 0.491 |
| 0 | 0.05 | ptsI | 2 | 8.91667 | 0.424 |
| 0 | 0.05 | clcB | 2 | 8.91667 | 0.358 |
| 0 | 0.05 | ycaM | 2 | 8.91667 | 0.374 |
| 0 | 0.05 | yadI | 2 | 8.91667 | 0.578 |
| 0 | 0.05 | AG1 | 2 | 8.91667 | 0.558 |
| 0 | 0.05 | ptsI | 2 | 9.16667 | 0.425 |
| 0 | 0.05 | clcB | 2 | 9.16667 | 0.362 |
| 0 | 0.05 | ycaM | 2 | 9.16667 | 0.351 |
| 0 | 0.05 | yadI | 2 | 9.16667 | 0.59 |
| 0 | 0.05 | AG1 | 2 | 9.16667 | 0.573 |
| 0 | 0.05 | ptsI | 2 | 9.41667 | 0.432 |
| 0 | 0.05 | clcB | 2 | 9.41667 | 0.371 |
| 0 | 0.05 | ycaM | 2 | 9.41667 | 0.387 |
| 0 | 0.05 | yadI | 2 | 9.41667 | 0.606 |
| 0 | 0.05 | AG1 | 2 | 9.41667 | 0.59 |
| 0 | 0.05 | ptsI | 2 | 9.66667 | 0.445 |
| 0 | 0.05 | clcB | 2 | 9.66667 | 0.373 |
| 0 | 0.05 | ycaM | 2 | 9.66667 | 0.355 |
| 0 | 0.05 | yadI | 2 | 9.66667 | 0.614 |
| 0 | 0.05 | AG1 | 2 | 9.66667 | 0.599 |
| 0 | 0.05 | ptsI | 2 | 9.91667 | 0.454 |
| 0 | 0.05 | clcB | 2 | 9.91667 | 0.382 |
| 0 | 0.05 | ycaM | 2 | 9.91667 | 0.357 |
| 0 | 0.05 | yadI | 2 | 9.91667 | 0.641 |
| 0 | 0.05 | AG1 | 2 | 9.91667 | 0.613 |
| 0 | 0.05 | ptsI | 2 | 10.1667 | 0.463 |
| 0 | 0.05 | clcB | 2 | 10.1667 | 0.384 |
| 0 | 0.05 | ycaM | 2 | 10.1667 | 0.409 |
| 0 | 0.05 | yadI | 2 | 10.1667 | 0.645 |
| 0 | 0.05 | AG1 | 2 | 10.1667 | 0.621 |
| 0 | 0.05 | ptsI | 2 | 10.4167 | 0.476 |
| 0 | 0.05 | clcB | 2 | 10.4167 | 0.388 |
| 0 | 0.05 | ycaM | 2 | 10.4167 | 0.358 |
| 0 | 0.05 | yadI | 2 | 10.4167 | 0.654 |
| 0 | 0.05 | AG1 | 2 | 10.4167 | 0.626 |
| 0 | 0.05 | ptsI | 2 | 10.6667 | 0.484 |
| 0 | 0.05 | clcB | 2 | 10.6667 | 0.399 |
| 0 | 0.05 | ycaM | 2 | 10.6667 | 0.362 |
| 0 | 0.05 | yadI | 2 | 10.6667 | 0.673 |
| 0 | 0.05 | AG1 | 2 | 10.6667 | 0.638 |
| 0 | 0.05 | ptsI | 2 | 10.9167 | 0.494 |
| 0 | 0.05 | clcB | 2 | 10.9167 | 0.405 |
| 0 | 0.05 | ycaM | 2 | 10.9167 | 0.364 |
| 0 | 0.05 | yadI | 2 | 10.9167 | 0.678 |
| 0 | 0.05 | AG1 | 2 | 10.9167 | 0.649 |

| | | | | | |
|---|---|---|---|---|---|
| 0 | 0.05 | ptsI | 2 | 11.1667 | 0.502 |
| 0 | 0.05 | clcB | 2 | 11.1667 | 0.401 |
| 0 | 0.05 | ycaM | 2 | 11.1667 | 0.361 |
| 0 | 0.05 | yadI | 2 | 11.1667 | 0.693 |
| 0 | 0.05 | AG1 | 2 | 11.1667 | 0.659 |
| 0 | 0.05 | ptsI | 2 | 11.4167 | 0.512 |
| 0 | 0.05 | clcB | 2 | 11.4167 | 0.406 |
| 0 | 0.05 | ycaM | 2 | 11.4167 | 0.363 |
| 0 | 0.05 | yadI | 2 | 11.4167 | 0.7 |
| 0 | 0.05 | AG1 | 2 | 11.4167 | 0.657 |
| 0 | 0.05 | ptsI | 2 | 11.6667 | 0.532 |
| 0 | 0.05 | clcB | 2 | 11.6667 | 0.406 |
| 0 | 0.05 | ycaM | 2 | 11.6667 | 0.363 |
| 0 | 0.05 | yadI | 2 | 11.6667 | 0.706 |
| 0 | 0.05 | AG1 | 2 | 11.6667 | 0.673 |
| 0 | 0.05 | ptsI | 2 | 11.9167 | 0.53 |
| 0 | 0.05 | clcB | 2 | 11.9167 | 0.417 |
| 0 | 0.05 | ycaM | 2 | 11.9167 | 0.367 |
| 0 | 0.05 | yadI | 2 | 11.9167 | 0.723 |
| 0 | 0.05 | AG1 | 2 | 11.9167 | 0.671 |
| 0 | 0.05 | ptsI | 2 | 12.1667 | 0.548 |
| 0 | 0.05 | clcB | 2 | 12.1667 | 0.417 |
| 0 | 0.05 | ycaM | 2 | 12.1667 | 0.366 |
| 0 | 0.05 | yadI | 2 | 12.1667 | 0.723 |
| 0 | 0.05 | AG1 | 2 | 12.1667 | 0.688 |
| 0 | 0.05 | ptsI | 2 | 12.4167 | 0.548 |
| 0 | 0.05 | clcB | 2 | 12.4167 | 0.414 |
| 0 | 0.05 | ycaM | 2 | 12.4167 | 0.368 |
| 0 | 0.05 | yadI | 2 | 12.4167 | 0.732 |
| 0 | 0.05 | AG1 | 2 | 12.4167 | 0.688 |
| 0 | 0.05 | ptsI | 2 | 12.6667 | 0.54 |
| 0 | 0.05 | clcB | 2 | 12.6667 | 0.411 |
| 0 | 0.05 | ycaM | 2 | 12.6667 | 0.362 |
| 0 | 0.05 | yadI | 2 | 12.6667 | 0.74 |
| 0 | 0.05 | AG1 | 2 | 12.6667 | 0.689 |
| 0 | 0.05 | ptsI | 2 | 12.9167 | 0.548 |
| 0 | 0.05 | clcB | 2 | 12.9167 | 0.412 |
| 0 | 0.05 | ycaM | 2 | 12.9167 | 0.367 |
| 0 | 0.05 | yadI | 2 | 12.9167 | 0.754 |
| 0 | 0.05 | AG1 | 2 | 12.9167 | 0.692 |
| 0 | 0.05 | ptsI | 2 | 13.1667 | 0.558 |
| 0 | 0.05 | clcB | 2 | 13.1667 | 0.417 |
| 0 | 0.05 | ycaM | 2 | 13.1667 | 0.367 |
| 0 | 0.05 | yadI | 2 | 13.1667 | 0.772 |
| 0 | 0.05 | AG1 | 2 | 13.1667 | 0.692 |
| 0 | 0.05 | ptsI | 2 | 13.4167 | 0.557 |
| 0 | 0.05 | clcB | 2 | 13.4167 | 0.412 |
| 0 | 0.05 | ycaM | 2 | 13.4167 | 0.367 |

| | | | | | |
|---|---|---|---|---|---|
| 0 | 0.05 | yadI | 2 | 13.4167 | 0.784 |
| 0 | 0.05 | AG1 | 2 | 13.4167 | 0.695 |
| 0 | 0.05 | ptsI | 2 | 13.6667 | 0.576 |
| 0 | 0.05 | clcB | 2 | 13.6667 | 0.412 |
| 0 | 0.05 | ycaM | 2 | 13.6667 | 0.369 |
| 0 | 0.05 | yadI | 2 | 13.6667 | 0.791 |
| 0 | 0.05 | AG1 | 2 | 13.6667 | 0.735 |
| 0 | 0.05 | ptsI | 2 | 13.9167 | 0.577 |
| 0 | 0.05 | clcB | 2 | 13.9167 | 0.414 |
| 0 | 0.05 | ycaM | 2 | 13.9167 | 0.37 |
| 0 | 0.05 | yadI | 2 | 13.9167 | 0.797 |
| 0 | 0.05 | AG1 | 2 | 13.9167 | 0.709 |
| 0 | 0.05 | ptsI | 2 | 14.1667 | 0.587 |
| 0 | 0.05 | clcB | 2 | 14.1667 | 0.416 |
| 0 | 0.05 | ycaM | 2 | 14.1667 | 0.375 |
| 0 | 0.05 | yadI | 2 | 14.1667 | 0.82 |
| 0 | 0.05 | AG1 | 2 | 14.1667 | 0.727 |
| 0 | 0.05 | ptsI | 2 | 14.4167 | 0.583 |
| 0 | 0.05 | clcB | 2 | 14.4167 | 0.414 |
| 0 | 0.05 | ycaM | 2 | 14.4167 | 0.368 |
| 0 | 0.05 | yadI | 2 | 14.4167 | 0.826 |
| 0 | 0.05 | AG1 | 2 | 14.4167 | 0.71 |
| 0 | 0.05 | ptsI | 2 | 14.6667 | 0.587 |
| 0 | 0.05 | clcB | 2 | 14.6667 | 0.418 |
| 0 | 0.05 | ycaM | 2 | 14.6667 | 0.365 |
| 0 | 0.05 | yadI | 2 | 14.6667 | 0.821 |
| 0 | 0.05 | AG1 | 2 | 14.6667 | 0.701 |
| 0 | 0.05 | ptsI | 2 | 14.9167 | 0.589 |
| 0 | 0.05 | clcB | 2 | 14.9167 | 0.419 |
| 0 | 0.05 | ycaM | 2 | 14.9167 | 0.363 |
| 0 | 0.05 | yadI | 2 | 14.9167 | 0.839 |
| 0 | 0.05 | AG1 | 2 | 14.9167 | 0.702 |
| 0 | 0.05 | ptsI | 2 | 15.1667 | 0.611 |
| 0 | 0.05 | clcB | 2 | 15.1667 | 0.425 |
| 0 | 0.05 | ycaM | 2 | 15.1667 | 0.374 |
| 0 | 0.05 | yadI | 2 | 15.1667 | 0.853 |
| 0 | 0.05 | AG1 | 2 | 15.1667 | 0.705 |
| 0 | 0.05 | ptsI | 2 | 15.4167 | 0.614 |
| 0 | 0.05 | clcB | 2 | 15.4167 | 0.425 |
| 0 | 0.05 | ycaM | 2 | 15.4167 | 0.368 |
| 0 | 0.05 | yadI | 2 | 15.4167 | 0.854 |
| 0 | 0.05 | AG1 | 2 | 15.4167 | 0.697 |
| 0 | 0.05 | ptsI | 2 | 15.6667 | 0.603 |
| 0 | 0.05 | clcB | 2 | 15.6667 | 0.427 |
| 0 | 0.05 | ycaM | 2 | 15.6667 | 0.363 |
| 0 | 0.05 | yadI | 2 | 15.6667 | 0.856 |
| 0 | 0.05 | AG1 | 2 | 15.6667 | 0.7 |
| 0 | 0.05 | ptsI | 2 | 15.9167 | 0.612 |

| | | | | | |
|---|---|---|---|---|---|
| 0 | 0.05 | clcB | 2 | 15.9167 | 0.432 |
| 0 | 0.05 | ycaM | 2 | 15.9167 | 0.366 |
| 0 | 0.05 | yadI | 2 | 15.9167 | 0.865 |
| 0 | 0.05 | AG1 | 2 | 15.9167 | 0.717 |
| 0 | 0.05 | ptsI | 2 | 16.1667 | 0.632 |
| 0 | 0.05 | clcB | 2 | 16.1667 | 0.433 |
| 0 | 0.05 | ycaM | 2 | 16.1667 | 0.372 |
| 0 | 0.05 | yadI | 2 | 16.1667 | 0.885 |
| 0 | 0.05 | AG1 | 2 | 16.1667 | 0.735 |
| 0 | 0.05 | ptsI | 2 | 16.4167 | 0.625 |
| 0 | 0.05 | clcB | 2 | 16.4167 | 0.437 |
| 0 | 0.05 | ycaM | 2 | 16.4167 | 0.369 |
| 0 | 0.05 | yadI | 2 | 16.4167 | 0.889 |
| 0 | 0.05 | AG1 | 2 | 16.4167 | 0.73 |
| 0 | 0.05 | ptsI | 2 | 16.6667 | 0.625 |
| 0 | 0.05 | clcB | 2 | 16.6667 | 0.436 |
| 0 | 0.05 | ycaM | 2 | 16.6667 | 0.371 |
| 0 | 0.05 | yadI | 2 | 16.6667 | 0.893 |
| 0 | 0.05 | AG1 | 2 | 16.6667 | 0.703 |
| 0 | 0.05 | ptsI | 2 | 16.9167 | 0.64 |
| 0 | 0.05 | clcB | 2 | 16.9167 | 0.441 |
| 0 | 0.05 | ycaM | 2 | 16.9167 | 0.374 |
| 0 | 0.05 | yadI | 2 | 16.9167 | 0.901 |
| 0 | 0.05 | AG1 | 2 | 16.9167 | 0.712 |
| 0 | 0.05 | ptsI | 2 | 17.1667 | 0.649 |
| 0 | 0.05 | clcB | 2 | 17.1667 | 0.445 |
| 0 | 0.05 | ycaM | 2 | 17.1667 | 0.38 |
| 0 | 0.05 | yadI | 2 | 17.1667 | 0.925 |
| 0 | 0.05 | AG1 | 2 | 17.1667 | 0.726 |
| 0 | 0.05 | ptsI | 2 | 17.4167 | 0.657 |
| 0 | 0.05 | clcB | 2 | 17.4167 | 0.446 |
| 0 | 0.05 | ycaM | 2 | 17.4167 | 0.386 |
| 0 | 0.05 | yadI | 2 | 17.4167 | 0.933 |
| 0 | 0.05 | AG1 | 2 | 17.4167 | 0.733 |
| 0 | 0.05 | ptsI | 2 | 17.6667 | 0.656 |
| 0 | 0.05 | clcB | 2 | 17.6667 | 0.445 |
| 0 | 0.05 | ycaM | 2 | 17.6667 | 0.385 |
| 0 | 0.05 | yadI | 2 | 17.6667 | 0.933 |
| 0 | 0.05 | AG1 | 2 | 17.6667 | 0.73 |
| 0 | 0.05 | ptsI | 2 | 17.9167 | 0.65 |
| 0 | 0.05 | clcB | 2 | 17.9167 | 0.45 |
| 0 | 0.05 | ycaM | 2 | 17.9167 | 0.393 |
| 0 | 0.05 | yadI | 2 | 17.9167 | 0.947 |
| 0 | 0.05 | AG1 | 2 | 17.9167 | 0.734 |
| 0 | 0.05 | ptsI | 2 | 18.1667 | 0.669 |
| 0 | 0.05 | clcB | 2 | 18.1667 | 0.447 |
| 0 | 0.05 | ycaM | 2 | 18.1667 | 0.388 |
| 0 | 0.05 | yadI | 2 | 18.1667 | 0.942 |

| | | | | | |
|---|---|---|---|---|---|
| 0 | 0.05 | AG1 | 2 | 18.1667 | 0.757 |
| 0 | 0.05 | ptsI | 2 | 18.4167 | 0.682 |
| 0 | 0.05 | clcB | 2 | 18.4167 | 0.452 |
| 0 | 0.05 | ycaM | 2 | 18.4167 | 0.398 |
| 0 | 0.05 | yadI | 2 | 18.4167 | 0.959 |
| 0 | 0.05 | AG1 | 2 | 18.4167 | 0.763 |
| 0 | 0.05 | ptsI | 2 | 18.6667 | 0.665 |
| 0 | 0.05 | clcB | 2 | 18.6667 | 0.458 |
| 0 | 0.05 | ycaM | 2 | 18.6667 | 0.396 |
| 0 | 0.05 | yadI | 2 | 18.6667 | 0.962 |
| 0 | 0.05 | AG1 | 2 | 18.6667 | 0.765 |
| 0 | 0.05 | ptsI | 2 | 18.9167 | 0.633 |
| 0 | 0.05 | clcB | 2 | 18.9167 | 0.45 |
| 0 | 0.05 | ycaM | 2 | 18.9167 | 0.392 |
| 0 | 0.05 | yadI | 2 | 18.9167 | 0.956 |
| 0 | 0.05 | AG1 | 2 | 18.9167 | 0.773 |
| 0 | 0.05 | ptsI | 2 | 19.1667 | 0.68 |
| 0 | 0.05 | clcB | 2 | 19.1667 | 0.459 |
| 0 | 0.05 | ycaM | 2 | 19.1667 | 0.402 |
| 0 | 0.05 | yadI | 2 | 19.1667 | 0.969 |
| 0 | 0.05 | AG1 | 2 | 19.1667 | 0.792 |
| 0 | 0.05 | ptsI | 2 | 19.4167 | 0.663 |
| 0 | 0.05 | clcB | 2 | 19.4167 | 0.459 |
| 0 | 0.05 | ycaM | 2 | 19.4167 | 0.4 |
| 0 | 0.05 | yadI | 2 | 19.4167 | 0.972 |
| 0 | 0.05 | AG1 | 2 | 19.4167 | 0.787 |
| 0 | 0.05 | ptsI | 2 | 19.6667 | 0.668 |
| 0 | 0.05 | clcB | 2 | 19.6667 | 0.465 |
| 0 | 0.05 | ycaM | 2 | 19.6667 | 0.406 |
| 0 | 0.05 | yadI | 2 | 19.6667 | 0.983 |
| 0 | 0.05 | AG1 | 2 | 19.6667 | 0.795 |
| 0 | 0.05 | ptsI | 2 | 19.9167 | 0.696 |
| 0 | 0.05 | clcB | 2 | 19.9167 | 0.466 |
| 0 | 0.05 | ycaM | 2 | 19.9167 | 0.404 |
| 0 | 0.05 | yadI | 2 | 19.9167 | 0.992 |
| 0 | 0.05 | AG1 | 2 | 19.9167 | 0.822 |
| 0 | 0.05 | ptsI | 2 | 20.1667 | 0.665 |
| 0 | 0.05 | clcB | 2 | 20.1667 | 0.469 |
| 0 | 0.05 | ycaM | 2 | 20.1667 | 0.412 |
| 0 | 0.05 | yadI | 2 | 20.1667 | 0.994 |
| 0 | 0.05 | AG1 | 2 | 20.1667 | 0.81 |
| 0 | 0.05 | ptsI | 2 | 20.4167 | 0.679 |
| 0 | 0.05 | clcB | 2 | 20.4167 | 0.469 |
| 0 | 0.05 | ycaM | 2 | 20.4167 | 0.406 |
| 0 | 0.05 | yadI | 2 | 20.4167 | 0.996 |
| 0 | 0.05 | AG1 | 2 | 20.4167 | 0.819 |
| 0 | 0.05 | ptsI | 2 | 20.6667 | 0.666 |
| 0 | 0.05 | clcB | 2 | 20.6667 | 0.473 |

| | | | | | |
|---|---|---|---|---|---|
| 0 | 0.05 | ycaM | 2 | 20.6667 | 0.416 |
| 0 | 0.05 | yadI | 2 | 20.6667 | 0.995 |
| 0 | 0.05 | AG1 | 2 | 20.6667 | 0.816 |
| 0 | 0.05 | ptsI | 2 | 20.9167 | 0.675 |
| 0 | 0.05 | clcB | 2 | 20.9167 | 0.477 |
| 0 | 0.05 | ycaM | 2 | 20.9167 | 0.416 |
| 0 | 0.05 | yadI | 2 | 20.9167 | 1.004 |
| 0 | 0.05 | AG1 | 2 | 20.9167 | 0.822 |
| 0 | 0.05 | ptsI | 2 | 21.1667 | 0.676 |
| 0 | 0.05 | clcB | 2 | 21.1667 | 0.478 |
| 0 | 0.05 | ycaM | 2 | 21.1667 | 0.425 |
| 0 | 0.05 | yadI | 2 | 21.1667 | 1.018 |
| 0 | 0.05 | AG1 | 2 | 21.1667 | 0.829 |
| 0 | 0.05 | ptsI | 2 | 21.4167 | 0.662 |
| 0 | 0.05 | clcB | 2 | 21.4167 | 0.472 |
| 0 | 0.05 | ycaM | 2 | 21.4167 | 0.429 |
| 0 | 0.05 | yadI | 2 | 21.4167 | 1.021 |
| 0 | 0.05 | AG1 | 2 | 21.4167 | 0.834 |
| 0 | 0.05 | ptsI | 2 | 21.6667 | 0.687 |
| 0 | 0.05 | clcB | 2 | 21.6667 | 0.482 |
| 0 | 0.05 | ycaM | 2 | 21.6667 | 0.433 |
| 0 | 0.05 | yadI | 2 | 21.6667 | 1.033 |
| 0 | 0.05 | AG1 | 2 | 21.6667 | 0.842 |
| 0 | 0.05 | ptsI | 2 | 21.9167 | 0.698 |
| 0 | 0.05 | clcB | 2 | 21.9167 | 0.481 |
| 0 | 0.05 | ycaM | 2 | 21.9167 | 0.436 |
| 0 | 0.05 | yadI | 2 | 21.9167 | 1.038 |
| 0 | 0.05 | AG1 | 2 | 21.9167 | 0.845 |
| 0 | 0.05 | ptsI | 2 | 22.1667 | 0.692 |
| 0 | 0.05 | clcB | 2 | 22.1667 | 0.485 |
| 0 | 0.05 | ycaM | 2 | 22.1667 | 0.442 |
| 0 | 0.05 | yadI | 2 | 22.1667 | 1.043 |
| 0 | 0.05 | AG1 | 2 | 22.1667 | 0.845 |
| 0 | 0.05 | ptsI | 2 | 22.4167 | 0.684 |
| 0 | 0.05 | clcB | 2 | 22.4167 | 0.484 |
| 0 | 0.05 | ycaM | 2 | 22.4167 | 0.445 |
| 0 | 0.05 | yadI | 2 | 22.4167 | 1.046 |
| 0 | 0.05 | AG1 | 2 | 22.4167 | 0.852 |
| 0 | 0.05 | ptsI | 2 | 22.6667 | 0.683 |
| 0 | 0.05 | clcB | 2 | 22.6667 | 0.485 |
| 0 | 0.05 | ycaM | 2 | 22.6667 | 0.446 |
| 0 | 0.05 | yadI | 2 | 22.6667 | 1.052 |
| 0 | 0.05 | AG1 | 2 | 22.6667 | 0.857 |
| 0 | 0.05 | ptsI | 2 | 22.9167 | 0.689 |
| 0 | 0.05 | clcB | 2 | 22.9167 | 0.486 |
| 0 | 0.05 | ycaM | 2 | 22.9167 | 0.45 |
| 0 | 0.05 | yadI | 2 | 22.9167 | 1.059 |
| 0 | 0.05 | AG1 | 2 | 22.9167 | 0.867 |

| | | | | | |
|---|---|---|---|---|---|
| 0 | 0.05 | ptsI | 2 | 23.1667 | 0.68 |
| 0 | 0.05 | clcB | 2 | 23.1667 | 0.492 |
| 0 | 0.05 | ycaM | 2 | 23.1667 | 0.44 |
| 0 | 0.05 | yadI | 2 | 23.1667 | 1.054 |
| 0 | 0.05 | AG1 | 2 | 23.1667 | 0.868 |
| 0 | 0.05 | ptsI | 2 | 23.4167 | 0.685 |
| 0 | 0.05 | clcB | 2 | 23.4167 | 0.485 |
| 0 | 0.05 | ycaM | 2 | 23.4167 | 0.45 |
| 0 | 0.05 | yadI | 2 | 23.4167 | 1.066 |
| 0 | 0.05 | AG1 | 2 | 23.4167 | 0.868 |
| 0 | 0.05 | ptsI | 2 | 23.6667 | 0.691 |
| 0 | 0.05 | clcB | 2 | 23.6667 | 0.491 |
| 0 | 0.05 | ycaM | 2 | 23.6667 | 0.45 |
| 0 | 0.05 | yadI | 2 | 23.6667 | 1.069 |
| 0 | 0.05 | AG1 | 2 | 23.6667 | 0.874 |
| 0 | 0.05 | ptsI | 2 | 23.9167 | 0.691 |
| 0 | 0.05 | clcB | 2 | 23.9167 | 0.49 |
| 0 | 0.05 | ycaM | 2 | 23.9167 | 0.457 |
| 0 | 0.05 | yadI | 2 | 23.9167 | 1.078 |
| 0 | 0.05 | AG1 | 2 | 23.9167 | 0.883 |
| 0 | 0.05 | ptsI | 2 | 24.1667 | 0.685 |
| 0 | 0.05 | clcB | 2 | 24.1667 | 0.486 |
| 0 | 0.05 | ycaM | 2 | 24.1667 | 0.458 |
| 0 | 0.05 | yadI | 2 | 24.1667 | 1.079 |
| 0 | 0.05 | AG1 | 2 | 24.1667 | 0.881 |
| 0 | 0.05 | ptsI | 2 | 24.4167 | 0.688 |
| 0 | 0.05 | clcB | 2 | 24.4167 | 0.488 |
| 0 | 0.05 | ycaM | 2 | 24.4167 | 0.462 |
| 0 | 0.05 | yadI | 2 | 24.4167 | 1.084 |
| 0 | 0.05 | AG1 | 2 | 24.4167 | 0.888 |
| 0 | 0.05 | ptsI | 2 | 24.6667 | 0.685 |
| 0 | 0.05 | clcB | 2 | 24.6667 | 0.493 |
| 0 | 0.05 | ycaM | 2 | 24.6667 | 0.46 |
| 0 | 0.05 | yadI | 2 | 24.6667 | 1.084 |
| 0 | 0.05 | AG1 | 2 | 24.6667 | 0.897 |
| 0 | 0.05 | ptsI | 2 | 24.9167 | 0.685 |
| 0 | 0.05 | clcB | 2 | 24.9167 | 0.493 |
| 0 | 0.05 | ycaM | 2 | 24.9167 | 0.464 |
| 0 | 0.05 | yadI | 2 | 24.9167 | 1.09 |
| 0 | 0.05 | AG1 | 2 | 24.9167 | 0.906 |
| 0 | 0.05 | ptsI | 2 | 25.1667 | 0.688 |
| 0 | 0.05 | clcB | 2 | 25.1667 | 0.497 |
| 0 | 0.05 | ycaM | 2 | 25.1667 | 0.468 |
| 0 | 0.05 | yadI | 2 | 25.1667 | 1.103 |
| 0 | 0.05 | AG1 | 2 | 25.1667 | 0.908 |
| 0 | 0.05 | ptsI | 2 | 25.4167 | 0.684 |
| 0 | 0.05 | clcB | 2 | 25.4167 | 0.493 |
| 0 | 0.05 | ycaM | 2 | 25.4167 | 0.469 |

| | | | | | |
|---|---|---|---|---|---|
| 0 | 0.05 | yadI | 2 | 25.4167 | 1.102 |
| 0 | 0.05 | AG1 | 2 | 25.4167 | 0.92 |
| 0 | 0.05 | ptsI | 2 | 25.6667 | 0.682 |
| 0 | 0.05 | clcB | 2 | 25.6667 | 0.495 |
| 0 | 0.05 | ycaM | 2 | 25.6667 | 0.474 |
| 0 | 0.05 | yadI | 2 | 25.6667 | 1.104 |
| 0 | 0.05 | AG1 | 2 | 25.6667 | 0.922 |
| 0 | 0.05 | ptsI | 2 | 25.9167 | 0.69 |
| 0 | 0.05 | clcB | 2 | 25.9167 | 0.499 |
| 0 | 0.05 | ycaM | 2 | 25.9167 | 0.481 |
| 0 | 0.05 | yadI | 2 | 25.9167 | 1.118 |
| 0 | 0.05 | AG1 | 2 | 25.9167 | 0.932 |
| 0 | 0.05 | ptsI | 2 | 26.1667 | 0.665 |
| 0 | 0.05 | clcB | 2 | 26.1667 | 0.494 |
| 0 | 0.05 | ycaM | 2 | 26.1667 | 0.474 |
| 0 | 0.05 | yadI | 2 | 26.1667 | 1.116 |
| 0 | 0.05 | AG1 | 2 | 26.1667 | 0.939 |
| 0 | 0.05 | ptsI | 2 | 26.4167 | 0.685 |
| 0 | 0.05 | clcB | 2 | 26.4167 | 0.499 |
| 0 | 0.05 | ycaM | 2 | 26.4167 | 0.483 |
| 0 | 0.05 | yadI | 2 | 26.4167 | 1.116 |
| 0 | 0.05 | AG1 | 2 | 26.4167 | 0.953 |
| 0 | 0.05 | ptsI | 3 | 0 | 0.16 |
| 0 | 0.05 | clcB | 3 | 0 | 0.162 |
| 0 | 0.05 | ycaM | 3 | 0 | 0.156 |
| 0 | 0.05 | yadI | 3 | 0 | 0.159 |
| 0 | 0.05 | AG1 | 3 | 0 | 0.159 |
| 0 | 0.05 | ptsI | 3 | 0.25 | 0.156 |
| 0 | 0.05 | clcB | 3 | 0.25 | 0.161 |
| 0 | 0.05 | ycaM | 3 | 0.25 | 0.153 |
| 0 | 0.05 | yadI | 3 | 0.25 | 0.167 |
| 0 | 0.05 | AG1 | 3 | 0.25 | 0.158 |
| 0 | 0.05 | ptsI | 3 | 0.5 | 0.156 |
| 0 | 0.05 | clcB | 3 | 0.5 | 0.16 |
| 0 | 0.05 | ycaM | 3 | 0.5 | 0.153 |
| 0 | 0.05 | yadI | 3 | 0.5 | 0.158 |
| 0 | 0.05 | AG1 | 3 | 0.5 | 0.157 |
| 0 | 0.05 | ptsI | 3 | 0.75 | 0.156 |
| 0 | 0.05 | clcB | 3 | 0.75 | 0.161 |
| 0 | 0.05 | ycaM | 3 | 0.75 | 0.152 |
| 0 | 0.05 | yadI | 3 | 0.75 | 0.159 |
| 0 | 0.05 | AG1 | 3 | 0.75 | 0.157 |
| 0 | 0.05 | ptsI | 3 | 1 | 0.156 |
| 0 | 0.05 | clcB | 3 | 1 | 0.161 |
| 0 | 0.05 | ycaM | 3 | 1 | 0.154 |
| 0 | 0.05 | yadI | 3 | 1 | 0.158 |
| 0 | 0.05 | AG1 | 3 | 1 | 0.158 |
| 0 | 0.05 | ptsI | 3 | 1.25 | 0.157 |

| | | | | | |
|---|---|---|---|---|---|
| 0 | 0.05 | clcB | 3 | 1.25 | 0.162 |
| 0 | 0.05 | ycaM | 3 | 1.25 | 0.153 |
| 0 | 0.05 | yadI | 3 | 1.25 | 0.16 |
| 0 | 0.05 | AG1 | 3 | 1.25 | 0.159 |
| 0 | 0.05 | ptsI | 3 | 1.5 | 0.158 |
| 0 | 0.05 | clcB | 3 | 1.5 | 0.166 |
| 0 | 0.05 | ycaM | 3 | 1.5 | 0.155 |
| 0 | 0.05 | yadI | 3 | 1.5 | 0.163 |
| 0 | 0.05 | AG1 | 3 | 1.5 | 0.161 |
| 0 | 0.05 | ptsI | 3 | 1.75 | 0.159 |
| 0 | 0.05 | clcB | 3 | 1.75 | 0.163 |
| 0 | 0.05 | ycaM | 3 | 1.75 | 0.156 |
| 0 | 0.05 | yadI | 3 | 1.75 | 0.163 |
| 0 | 0.05 | AG1 | 3 | 1.75 | 0.162 |
| 0 | 0.05 | ptsI | 3 | 2 | 0.161 |
| 0 | 0.05 | clcB | 3 | 2 | 0.162 |
| 0 | 0.05 | ycaM | 3 | 2 | 0.157 |
| 0 | 0.05 | yadI | 3 | 2 | 0.164 |
| 0 | 0.05 | AG1 | 3 | 2 | 0.164 |
| 0 | 0.05 | ptsI | 3 | 2.25 | 0.162 |
| 0 | 0.05 | clcB | 3 | 2.25 | 0.164 |
| 0 | 0.05 | ycaM | 3 | 2.25 | 0.158 |
| 0 | 0.05 | yadI | 3 | 2.25 | 0.166 |
| 0 | 0.05 | AG1 | 3 | 2.25 | 0.165 |
| 0 | 0.05 | ptsI | 3 | 2.5 | 0.162 |
| 0 | 0.05 | clcB | 3 | 2.5 | 0.166 |
| 0 | 0.05 | ycaM | 3 | 2.5 | 0.159 |
| 0 | 0.05 | yadI | 3 | 2.5 | 0.169 |
| 0 | 0.05 | AG1 | 3 | 2.5 | 0.167 |
| 0 | 0.05 | ptsI | 3 | 2.75 | 0.165 |
| 0 | 0.05 | clcB | 3 | 2.75 | 0.168 |
| 0 | 0.05 | ycaM | 3 | 2.75 | 0.161 |
| 0 | 0.05 | yadI | 3 | 2.75 | 0.171 |
| 0 | 0.05 | AG1 | 3 | 2.75 | 0.168 |
| 0 | 0.05 | ptsI | 3 | 3 | 0.165 |
| 0 | 0.05 | clcB | 3 | 3 | 0.168 |
| 0 | 0.05 | ycaM | 3 | 3 | 0.163 |
| 0 | 0.05 | yadI | 3 | 3 | 0.175 |
| 0 | 0.05 | AG1 | 3 | 3 | 0.17 |
| 0 | 0.05 | ptsI | 3 | 3.25 | 0.168 |
| 0 | 0.05 | clcB | 3 | 3.25 | 0.171 |
| 0 | 0.05 | ycaM | 3 | 3.25 | 0.165 |
| 0 | 0.05 | yadI | 3 | 3.25 | 0.178 |
| 0 | 0.05 | AG1 | 3 | 3.25 | 0.172 |
| 0 | 0.05 | ptsI | 3 | 3.5 | 0.171 |
| 0 | 0.05 | clcB | 3 | 3.5 | 0.171 |
| 0 | 0.05 | ycaM | 3 | 3.5 | 0.168 |
| 0 | 0.05 | yadI | 3 | 3.5 | 0.182 |

| | | | | | |
|---|---|---|---|---|---|
| 0 | 0.05 | AG1 | 3 | 3.5 | 0.174 |
| 0 | 0.05 | ptsI | 3 | 3.75 | 0.173 |
| 0 | 0.05 | clcB | 3 | 3.75 | 0.175 |
| 0 | 0.05 | ycaM | 3 | 3.75 | 0.171 |
| 0 | 0.05 | yadI | 3 | 3.75 | 0.186 |
| 0 | 0.05 | AG1 | 3 | 3.75 | 0.182 |
| 0 | 0.05 | ptsI | 3 | 4 | 0.177 |
| 0 | 0.05 | clcB | 3 | 4 | 0.178 |
| 0 | 0.05 | ycaM | 3 | 4 | 0.174 |
| 0 | 0.05 | yadI | 3 | 4 | 0.194 |
| 0 | 0.05 | AG1 | 3 | 4 | 0.18 |
| 0 | 0.05 | ptsI | 3 | 4.41667 | 0.197 |
| 0 | 0.05 | clcB | 3 | 4.41667 | 0.196 |
| 0 | 0.05 | ycaM | 3 | 4.41667 | 0.183 |
| 0 | 0.05 | yadI | 3 | 4.41667 | 0.218 |
| 0 | 0.05 | AG1 | 3 | 4.41667 | 0.208 |
| 0 | 0.05 | ptsI | 3 | 4.66667 | 0.197 |
| 0 | 0.05 | clcB | 3 | 4.66667 | 0.193 |
| 0 | 0.05 | ycaM | 3 | 4.66667 | 0.184 |
| 0 | 0.05 | yadI | 3 | 4.66667 | 0.226 |
| 0 | 0.05 | AG1 | 3 | 4.66667 | 0.201 |
| 0 | 0.05 | ptsI | 3 | 4.91667 | 0.198 |
| 0 | 0.05 | clcB | 3 | 4.91667 | 0.197 |
| 0 | 0.05 | ycaM | 3 | 4.91667 | 0.19 |
| 0 | 0.05 | yadI | 3 | 4.91667 | 0.237 |
| 0 | 0.05 | AG1 | 3 | 4.91667 | 0.208 |
| 0 | 0.05 | ptsI | 3 | 5.16667 | 0.203 |
| 0 | 0.05 | clcB | 3 | 5.16667 | 0.2 |
| 0 | 0.05 | ycaM | 3 | 5.16667 | 0.195 |
| 0 | 0.05 | yadI | 3 | 5.16667 | 0.252 |
| 0 | 0.05 | AG1 | 3 | 5.16667 | 0.214 |
| 0 | 0.05 | ptsI | 3 | 5.41667 | 0.209 |
| 0 | 0.05 | clcB | 3 | 5.41667 | 0.207 |
| 0 | 0.05 | ycaM | 3 | 5.41667 | 0.201 |
| 0 | 0.05 | yadI | 3 | 5.41667 | 0.268 |
| 0 | 0.05 | AG1 | 3 | 5.41667 | 0.221 |
| 0 | 0.05 | ptsI | 3 | 5.66667 | 0.212 |
| 0 | 0.05 | clcB | 3 | 5.66667 | 0.232 |
| 0 | 0.05 | ycaM | 3 | 5.66667 | 0.21 |
| 0 | 0.05 | yadI | 3 | 5.66667 | 0.287 |
| 0 | 0.05 | AG1 | 3 | 5.66667 | 0.23 |
| 0 | 0.05 | ptsI | 3 | 5.91667 | 0.221 |
| 0 | 0.05 | clcB | 3 | 5.91667 | 0.22 |
| 0 | 0.05 | ycaM | 3 | 5.91667 | 0.218 |
| 0 | 0.05 | yadI | 3 | 5.91667 | 0.304 |
| 0 | 0.05 | AG1 | 3 | 5.91667 | 0.241 |
| 0 | 0.05 | ptsI | 3 | 6.16667 | 0.228 |
| 0 | 0.05 | clcB | 3 | 6.16667 | 0.23 |

| | | | | | |
|---|---|---|---|---|---|
| 0 | 0.05 | ycaM | 3 | 6.16667 | 0.226 |
| 0 | 0.05 | yadI | 3 | 6.16667 | 0.318 |
| 0 | 0.05 | AG1 | 3 | 6.16667 | 0.25 |
| 0 | 0.05 | ptsI | 3 | 6.41667 | 0.235 |
| 0 | 0.05 | clcB | 3 | 6.41667 | 0.236 |
| 0 | 0.05 | ycaM | 3 | 6.41667 | 0.233 |
| 0 | 0.05 | yadI | 3 | 6.41667 | 0.325 |
| 0 | 0.05 | AG1 | 3 | 6.41667 | 0.264 |
| 0 | 0.05 | ptsI | 3 | 6.66667 | 0.246 |
| 0 | 0.05 | clcB | 3 | 6.66667 | 0.247 |
| 0 | 0.05 | ycaM | 3 | 6.66667 | 0.243 |
| 0 | 0.05 | yadI | 3 | 6.66667 | 0.338 |
| 0 | 0.05 | AG1 | 3 | 6.66667 | 0.276 |
| 0 | 0.05 | ptsI | 3 | 6.91667 | 0.258 |
| 0 | 0.05 | clcB | 3 | 6.91667 | 0.261 |
| 0 | 0.05 | ycaM | 3 | 6.91667 | 0.252 |
| 0 | 0.05 | yadI | 3 | 6.91667 | 0.347 |
| 0 | 0.05 | AG1 | 3 | 6.91667 | 0.29 |
| 0 | 0.05 | ptsI | 3 | 7.16667 | 0.263 |
| 0 | 0.05 | clcB | 3 | 7.16667 | 0.268 |
| 0 | 0.05 | ycaM | 3 | 7.16667 | 0.253 |
| 0 | 0.05 | yadI | 3 | 7.16667 | 0.357 |
| 0 | 0.05 | AG1 | 3 | 7.16667 | 0.298 |
| 0 | 0.05 | ptsI | 3 | 7.41667 | 0.275 |
| 0 | 0.05 | clcB | 3 | 7.41667 | 0.278 |
| 0 | 0.05 | ycaM | 3 | 7.41667 | 0.259 |
| 0 | 0.05 | yadI | 3 | 7.41667 | 0.372 |
| 0 | 0.05 | AG1 | 3 | 7.41667 | 0.308 |
| 0 | 0.05 | ptsI | 3 | 7.66667 | 0.28 |
| 0 | 0.05 | clcB | 3 | 7.66667 | 0.292 |
| 0 | 0.05 | ycaM | 3 | 7.66667 | 0.261 |
| 0 | 0.05 | yadI | 3 | 7.66667 | 0.385 |
| 0 | 0.05 | AG1 | 3 | 7.66667 | 0.317 |
| 0 | 0.05 | ptsI | 3 | 7.91667 | 0.287 |
| 0 | 0.05 | clcB | 3 | 7.91667 | 0.29 |
| 0 | 0.05 | ycaM | 3 | 7.91667 | 0.264 |
| 0 | 0.05 | yadI | 3 | 7.91667 | 0.393 |
| 0 | 0.05 | AG1 | 3 | 7.91667 | 0.323 |
| 0 | 0.05 | ptsI | 3 | 8.16667 | 0.296 |
| 0 | 0.05 | clcB | 3 | 8.16667 | 0.296 |
| 0 | 0.05 | ycaM | 3 | 8.16667 | 0.266 |
| 0 | 0.05 | yadI | 3 | 8.16667 | 0.403 |
| 0 | 0.05 | AG1 | 3 | 8.16667 | 0.332 |
| 0 | 0.05 | ptsI | 3 | 8.41667 | 0.306 |
| 0 | 0.05 | clcB | 3 | 8.41667 | 0.305 |
| 0 | 0.05 | ycaM | 3 | 8.41667 | 0.269 |
| 0 | 0.05 | yadI | 3 | 8.41667 | 0.418 |
| 0 | 0.05 | AG1 | 3 | 8.41667 | 0.343 |

| | | | | | |
|---|---|---|---|---|---|
| 0 | 0.05 | ptsI | 3 | 9.05 | 0.29 |
| 0 | 0.05 | clcB | 3 | 9.05 | 0.39 |
| 0 | 0.05 | ycaM | 3 | 9.05 | 0.295 |
| 0 | 0.05 | yadI | 3 | 9.05 | 0.433 |
| 0 | 0.05 | AG1 | 3 | 9.05 | 0.374 |
| 0 | 0.05 | ptsI | 3 | 9.3 | 0.292 |
| 0 | 0.05 | clcB | 3 | 9.3 | 0.283 |
| 0 | 0.05 | ycaM | 3 | 9.3 | 0.29 |
| 0 | 0.05 | yadI | 3 | 9.3 | 0.442 |
| 0 | 0.05 | AG1 | 3 | 9.3 | 0.378 |
| 0 | 0.05 | ptsI | 3 | 9.55 | 0.3 |
| 0 | 0.05 | clcB | 3 | 9.55 | 0.294 |
| 0 | 0.05 | ycaM | 3 | 9.55 | 0.296 |
| 0 | 0.05 | yadI | 3 | 9.55 | 0.456 |
| 0 | 0.05 | AG1 | 3 | 9.55 | 0.388 |
| 0 | 0.05 | ptsI | 3 | 9.8 | 0.309 |
| 0 | 0.05 | clcB | 3 | 9.8 | 0.299 |
| 0 | 0.05 | ycaM | 3 | 9.8 | 0.298 |
| 0 | 0.05 | yadI | 3 | 9.8 | 0.472 |
| 0 | 0.05 | AG1 | 3 | 9.8 | 0.396 |
| 0 | 0.05 | ptsI | 3 | 10.05 | 0.318 |
| 0 | 0.05 | clcB | 3 | 10.05 | 0.307 |
| 0 | 0.05 | ycaM | 3 | 10.05 | 0.304 |
| 0 | 0.05 | yadI | 3 | 10.05 | 0.486 |
| 0 | 0.05 | AG1 | 3 | 10.05 | 0.409 |
| 0 | 0.05 | ptsI | 3 | 10.3 | 0.326 |
| 0 | 0.05 | clcB | 3 | 10.3 | 0.314 |
| 0 | 0.05 | ycaM | 3 | 10.3 | 0.306 |
| 0 | 0.05 | yadI | 3 | 10.3 | 0.498 |
| 0 | 0.05 | AG1 | 3 | 10.3 | 0.422 |
| 0 | 0.05 | ptsI | 3 | 10.55 | 0.343 |
| 0 | 0.05 | clcB | 3 | 10.55 | 0.32 |
| 0 | 0.05 | ycaM | 3 | 10.55 | 0.311 |
| 0 | 0.05 | yadI | 3 | 10.55 | 0.516 |
| 0 | 0.05 | AG1 | 3 | 10.55 | 0.435 |
| 0 | 0.05 | ptsI | 3 | 10.8 | 0.345 |
| 0 | 0.05 | clcB | 3 | 10.8 | 0.316 |
| 0 | 0.05 | ycaM | 3 | 10.8 | 0.309 |
| 0 | 0.05 | yadI | 3 | 10.8 | 0.522 |
| 0 | 0.05 | AG1 | 3 | 10.8 | 0.442 |
| 0 | 0.05 | ptsI | 3 | 11.05 | 0.358 |
| 0 | 0.05 | clcB | 3 | 11.05 | 0.328 |
| 0 | 0.05 | ycaM | 3 | 11.05 | 0.315 |
| 0 | 0.05 | yadI | 3 | 11.05 | 0.538 |
| 0 | 0.05 | AG1 | 3 | 11.05 | 0.458 |
| 0 | 0.05 | ptsI | 3 | 11.3 | 0.362 |
| 0 | 0.05 | clcB | 3 | 11.3 | 0.321 |
| 0 | 0.05 | ycaM | 3 | 11.3 | 0.31 |

| | | | | | |
|---|---|---|---|---|---|
| 0 | 0.05 | yadI | 3 | 11.3 | 0.54 |
| 0 | 0.05 | AG1 | 3 | 11.3 | 0.462 |
| 0 | 0.05 | ptsI | 3 | 11.55 | 0.373 |
| 0 | 0.05 | clcB | 3 | 11.55 | 0.328 |
| 0 | 0.05 | ycaM | 3 | 11.55 | 0.312 |
| 0 | 0.05 | yadI | 3 | 11.55 | 0.557 |
| 0 | 0.05 | AG1 | 3 | 11.55 | 0.476 |
| 0 | 0.05 | ptsI | 3 | 11.8 | 0.391 |
| 0 | 0.05 | clcB | 3 | 11.8 | 0.343 |
| 0 | 0.05 | ycaM | 3 | 11.8 | 0.322 |
| 0 | 0.05 | yadI | 3 | 11.8 | 0.579 |
| 0 | 0.05 | AG1 | 3 | 11.8 | 0.494 |
| 0 | 0.05 | ptsI | 3 | 12.05 | 0.392 |
| 0 | 0.05 | clcB | 3 | 12.05 | 0.333 |
| 0 | 0.05 | ycaM | 3 | 12.05 | 0.315 |
| 0 | 0.05 | yadI | 3 | 12.05 | 0.578 |
| 0 | 0.05 | AG1 | 3 | 12.05 | 0.495 |
| 0 | 0.05 | ptsI | 3 | 12.3 | 0.406 |
| 0 | 0.05 | clcB | 3 | 12.3 | 0.392 |
| 0 | 0.05 | ycaM | 3 | 12.3 | 0.325 |
| 0 | 0.05 | yadI | 3 | 12.3 | 0.596 |
| 0 | 0.05 | AG1 | 3 | 12.3 | 0.512 |
| 0 | 0.05 | ptsI | 3 | 12.55 | 0.424 |
| 0 | 0.05 | clcB | 3 | 12.55 | 0.342 |
| 0 | 0.05 | ycaM | 3 | 12.55 | 0.326 |
| 0 | 0.05 | yadI | 3 | 12.55 | 0.62 |
| 0 | 0.05 | AG1 | 3 | 12.55 | 0.532 |
| 0 | 0.05 | ptsI | 3 | 12.8 | 0.437 |
| 0 | 0.05 | clcB | 3 | 12.8 | 0.337 |
| 0 | 0.05 | ycaM | 3 | 12.8 | 0.328 |
| 0 | 0.05 | yadI | 3 | 12.8 | 0.635 |
| 0 | 0.05 | AG1 | 3 | 12.8 | 0.548 |
| 0 | 0.05 | ptsI | 3 | 13.05 | 0.428 |
| 0 | 0.05 | clcB | 3 | 13.05 | 0.331 |
| 0 | 0.05 | ycaM | 3 | 13.05 | 0.316 |
| 0 | 0.05 | yadI | 3 | 13.05 | 0.627 |
| 0 | 0.05 | AG1 | 3 | 13.05 | 0.549 |
| 0 | 0.05 | ptsI | 3 | 13.3 | 0.443 |
| 0 | 0.05 | clcB | 3 | 13.3 | 0.341 |
| 0 | 0.05 | ycaM | 3 | 13.3 | 0.326 |
| 0 | 0.05 | yadI | 3 | 13.3 | 0.645 |
| 0 | 0.05 | AG1 | 3 | 13.3 | 0.574 |
| 0 | 0.05 | ptsI | 3 | 13.55 | 0.455 |
| 0 | 0.05 | clcB | 3 | 13.55 | 0.336 |
| 0 | 0.05 | ycaM | 3 | 13.55 | 0.322 |
| 0 | 0.05 | yadI | 3 | 13.55 | 0.657 |
| 0 | 0.05 | AG1 | 3 | 13.55 | 0.586 |
| 0 | 0.05 | ptsI | 3 | 13.8 | 0.472 |

| | | | | | |
|---|---|---|---|---|---|
| 0 | 0.05 | clcB | 3 | 13.8 | 0.34 |
| 0 | 0.05 | ycaM | 3 | 13.8 | 0.325 |
| 0 | 0.05 | yadI | 3 | 13.8 | 0.67 |
| 0 | 0.05 | AG1 | 3 | 13.8 | 0.604 |
| 0 | 0.05 | ptsI | 3 | 14.05 | 0.473 |
| 0 | 0.05 | clcB | 3 | 14.05 | 0.334 |
| 0 | 0.05 | ycaM | 3 | 14.05 | 0.326 |
| 0 | 0.05 | yadI | 3 | 14.05 | 0.676 |
| 0 | 0.05 | AG1 | 3 | 14.05 | 0.616 |
| 0 | 0.05 | ptsI | 3 | 14.3 | 0.481 |
| 0 | 0.05 | clcB | 3 | 14.3 | 0.336 |
| 0 | 0.05 | ycaM | 3 | 14.3 | 0.319 |
| 0 | 0.05 | yadI | 3 | 14.3 | 0.682 |
| 0 | 0.05 | AG1 | 3 | 14.3 | 0.626 |
| 0 | 0.05 | ptsI | 3 | 14.55 | 0.491 |
| 0 | 0.05 | clcB | 3 | 14.55 | 0.338 |
| 0 | 0.05 | ycaM | 3 | 14.55 | 0.317 |
| 0 | 0.05 | yadI | 3 | 14.55 | 0.693 |
| 0 | 0.05 | AG1 | 3 | 14.55 | 0.637 |
| 0 | 0.05 | ptsI | 3 | 14.8 | 0.512 |
| 0 | 0.05 | clcB | 3 | 14.8 | 0.34 |
| 0 | 0.05 | ycaM | 3 | 14.8 | 0.333 |
| 0 | 0.05 | yadI | 3 | 14.8 | 0.721 |
| 0 | 0.05 | AG1 | 3 | 14.8 | 0.656 |
| 0 | 0.05 | ptsI | 3 | 15.05 | 0.53 |
| 0 | 0.05 | clcB | 3 | 15.05 | 0.336 |
| 0 | 0.05 | ycaM | 3 | 15.05 | 0.334 |
| 0 | 0.05 | yadI | 3 | 15.05 | 0.734 |
| 0 | 0.05 | AG1 | 3 | 15.05 | 0.657 |
| 0 | 0.05 | ptsI | 3 | 15.3 | 0.527 |
| 0 | 0.05 | clcB | 3 | 15.3 | 0.332 |
| 0 | 0.05 | ycaM | 3 | 15.3 | 0.325 |
| 0 | 0.05 | yadI | 3 | 15.3 | 0.737 |
| 0 | 0.05 | AG1 | 3 | 15.3 | 0.668 |
| 0 | 0.05 | ptsI | 3 | 15.55 | 0.532 |
| 0 | 0.05 | clcB | 3 | 15.55 | 0.336 |
| 0 | 0.05 | ycaM | 3 | 15.55 | 0.328 |
| 0 | 0.05 | yadI | 3 | 15.55 | 0.745 |
| 0 | 0.05 | AG1 | 3 | 15.55 | 0.684 |
| 0 | 0.05 | ptsI | 3 | 15.8 | 0.549 |
| 0 | 0.05 | clcB | 3 | 15.8 | 0.34 |
| 0 | 0.05 | ycaM | 3 | 15.8 | 0.327 |
| 0 | 0.05 | yadI | 3 | 15.8 | 0.766 |
| 0 | 0.05 | AG1 | 3 | 15.8 | 0.693 |
| 0 | 0.05 | ptsI | 3 | 16.05 | 0.54 |
| 0 | 0.05 | clcB | 3 | 16.05 | 0.328 |
| 0 | 0.05 | ycaM | 3 | 16.05 | 0.322 |
| 0 | 0.05 | yadI | 3 | 16.05 | 0.756 |

| | | | | | |
|---|---|---|---|---|---|
| 0 | 0.05 | AG1 | 3 | 16.05 | 0.686 |
| 0 | 0.05 | ptsI | 3 | 16.3 | 0.553 |
| 0 | 0.05 | clcB | 3 | 16.3 | 0.329 |
| 0 | 0.05 | ycaM | 3 | 16.3 | 0.318 |
| 0 | 0.05 | yadI | 3 | 16.3 | 0.766 |
| 0 | 0.05 | AG1 | 3 | 16.3 | 0.706 |
| 0 | 0.05 | ptsI | 3 | 16.55 | 0.569 |
| 0 | 0.05 | clcB | 3 | 16.55 | 0.332 |
| 0 | 0.05 | ycaM | 3 | 16.55 | 0.322 |
| 0 | 0.05 | yadI | 3 | 16.55 | 0.786 |
| 0 | 0.05 | AG1 | 3 | 16.55 | 0.707 |
| 0 | 0.05 | ptsI | 3 | 16.8 | 0.576 |
| 0 | 0.05 | clcB | 3 | 16.8 | 0.338 |
| 0 | 0.05 | ycaM | 3 | 16.8 | 0.32 |
| 0 | 0.05 | yadI | 3 | 16.8 | 0.807 |
| 0 | 0.05 | AG1 | 3 | 16.8 | 0.734 |
| 0 | 0.05 | ptsI | 3 | 17.05 | 0.574 |
| 0 | 0.05 | clcB | 3 | 17.05 | 0.329 |
| 0 | 0.05 | ycaM | 3 | 17.05 | 0.316 |
| 0 | 0.05 | yadI | 3 | 17.05 | 0.809 |
| 0 | 0.05 | AG1 | 3 | 17.05 | 0.72 |
| 0 | 0.05 | ptsI | 3 | 17.3 | 0.581 |
| 0 | 0.05 | clcB | 3 | 17.3 | 0.327 |
| 0 | 0.05 | ycaM | 3 | 17.3 | 0.323 |
| 0 | 0.05 | yadI | 3 | 17.3 | 0.822 |
| 0 | 0.05 | AG1 | 3 | 17.3 | 0.724 |
| 0 | 0.05 | ptsI | 3 | 17.55 | 0.587 |
| 0 | 0.05 | clcB | 3 | 17.55 | 0.333 |
| 0 | 0.05 | ycaM | 3 | 17.55 | 0.318 |
| 0 | 0.05 | yadI | 3 | 17.55 | 0.828 |
| 0 | 0.05 | AG1 | 3 | 17.55 | 0.734 |
| 0 | 0.05 | ptsI | 3 | 17.8 | 0.599 |
| 0 | 0.05 | clcB | 3 | 17.8 | 0.332 |
| 0 | 0.05 | ycaM | 3 | 17.8 | 0.321 |
| 0 | 0.05 | yadI | 3 | 17.8 | 0.838 |
| 0 | 0.05 | AG1 | 3 | 17.8 | 0.76 |
| 0 | 0.05 | ptsI | 3 | 18.05 | 0.599 |
| 0 | 0.05 | clcB | 3 | 18.05 | 0.334 |
| 0 | 0.05 | ycaM | 3 | 18.05 | 0.316 |
| 0 | 0.05 | yadI | 3 | 18.05 | 0.837 |
| 0 | 0.05 | AG1 | 3 | 18.05 | 0.748 |
| 0 | 0.05 | ptsI | 3 | 18.3 | 0.595 |
| 0 | 0.05 | clcB | 3 | 18.3 | 0.34 |
| 0 | 0.05 | ycaM | 3 | 18.3 | 0.318 |
| 0 | 0.05 | yadI | 3 | 18.3 | 0.856 |
| 0 | 0.05 | AG1 | 3 | 18.3 | 0.755 |
| 0 | 0.05 | ptsI | 3 | 18.55 | 0.598 |
| 0 | 0.05 | clcB | 3 | 18.55 | 0.336 |

| | | | | | |
|---|---|---|---|---|---|
| 0 | 0.05 | ycaM | 3 | 18.55 | 0.318 |
| 0 | 0.05 | yadI | 3 | 18.55 | 0.86 |
| 0 | 0.05 | AG1 | 3 | 18.55 | 0.755 |
| 0 | 0.05 | ptsI | 3 | 18.8 | 0.602 |
| 0 | 0.05 | clcB | 3 | 18.8 | 0.33 |
| 0 | 0.05 | ycaM | 3 | 18.8 | 0.319 |
| 0 | 0.05 | yadI | 3 | 18.8 | 0.866 |
| 0 | 0.05 | AG1 | 3 | 18.8 | 0.774 |
| 0 | 0.05 | ptsI | 3 | 19.05 | 0.615 |
| 0 | 0.05 | clcB | 3 | 19.05 | 0.333 |
| 0 | 0.05 | ycaM | 3 | 19.05 | 0.32 |
| 0 | 0.05 | yadI | 3 | 19.05 | 0.881 |
| 0 | 0.05 | AG1 | 3 | 19.05 | 0.805 |
| 0 | 0.05 | ptsI | 3 | 19.3 | 0.613 |
| 0 | 0.05 | clcB | 3 | 19.3 | 0.334 |
| 0 | 0.05 | ycaM | 3 | 19.3 | 0.323 |
| 0 | 0.05 | yadI | 3 | 19.3 | 0.885 |
| 0 | 0.05 | AG1 | 3 | 19.3 | 0.802 |
| 0 | 0.05 | ptsI | 3 | 19.55 | 0.625 |
| 0 | 0.05 | clcB | 3 | 19.55 | 0.335 |
| 0 | 0.05 | ycaM | 3 | 19.55 | 0.321 |
| 0 | 0.05 | yadI | 3 | 19.55 | 0.903 |
| 0 | 0.05 | AG1 | 3 | 19.55 | 0.822 |
| 0 | 0.05 | ptsI | 3 | 19.8 | 0.607 |
| 0 | 0.05 | clcB | 3 | 19.8 | 0.333 |
| 0 | 0.05 | ycaM | 3 | 19.8 | 0.32 |
| 0 | 0.05 | yadI | 3 | 19.8 | 0.89 |
| 0 | 0.05 | AG1 | 3 | 19.8 | 0.83 |
| 0 | 0.05 | ptsI | 3 | 20.05 | 0.611 |
| 0 | 0.05 | clcB | 3 | 20.05 | 0.332 |
| 0 | 0.05 | ycaM | 3 | 20.05 | 0.325 |
| 0 | 0.05 | yadI | 3 | 20.05 | 0.9 |
| 0 | 0.05 | AG1 | 3 | 20.05 | 0.839 |
| 0 | 0.05 | ptsI | 3 | 20.3 | 0.611 |
| 0 | 0.05 | clcB | 3 | 20.3 | 0.336 |
| 0 | 0.05 | ycaM | 3 | 20.3 | 0.323 |
| 0 | 0.05 | yadI | 3 | 20.3 | 0.917 |
| 0 | 0.05 | AG1 | 3 | 20.3 | 0.839 |
| 0 | 0.05 | ptsI | 3 | 20.55 | 0.615 |
| 0 | 0.05 | clcB | 3 | 20.55 | 0.334 |
| 0 | 0.05 | ycaM | 3 | 20.55 | 0.327 |
| 0 | 0.05 | yadI | 3 | 20.55 | 0.915 |
| 0 | 0.05 | AG1 | 3 | 20.55 | 0.849 |
| 0 | 0.05 | ptsI | 3 | 20.8 | 0.629 |
| 0 | 0.05 | clcB | 3 | 20.8 | 0.335 |
| 0 | 0.05 | ycaM | 3 | 20.8 | 0.329 |
| 0 | 0.05 | yadI | 3 | 20.8 | 0.947 |
| 0 | 0.05 | AG1 | 3 | 20.8 | 0.857 |

| | | | | | |
|---|---|---|---|---|---|
| 0 | 0.05 | ptsI | 3 | 21.05 | 0.62 |
| 0 | 0.05 | clcB | 3 | 21.05 | 0.333 |
| 0 | 0.05 | ycaM | 3 | 21.05 | 0.332 |
| 0 | 0.05 | yadI | 3 | 21.05 | 0.933 |
| 0 | 0.05 | AG1 | 3 | 21.05 | 0.865 |
| 0 | 0.05 | ptsI | 3 | 21.3 | 0.617 |
| 0 | 0.05 | clcB | 3 | 21.3 | 0.334 |
| 0 | 0.05 | ycaM | 3 | 21.3 | 0.334 |
| 0 | 0.05 | yadI | 3 | 21.3 | 0.942 |
| 0 | 0.05 | AG1 | 3 | 21.3 | 0.879 |
| 0 | 0.05 | ptsI | 3 | 21.55 | 0.63 |
| 0 | 0.05 | clcB | 3 | 21.55 | 0.338 |
| 0 | 0.05 | ycaM | 3 | 21.55 | 0.334 |
| 0 | 0.05 | yadI | 3 | 21.55 | 0.947 |
| 0 | 0.05 | AG1 | 3 | 21.55 | 0.888 |
| 0 | 0.05 | ptsI | 3 | 21.8 | 0.628 |
| 0 | 0.05 | clcB | 3 | 21.8 | 0.339 |
| 0 | 0.05 | ycaM | 3 | 21.8 | 0.333 |
| 0 | 0.05 | yadI | 3 | 21.8 | 0.959 |
| 0 | 0.05 | AG1 | 3 | 21.8 | 0.896 |
| 0 | 0.05 | ptsI | 3 | 22.05 | 0.622 |
| 0 | 0.05 | clcB | 3 | 22.05 | 0.336 |
| 0 | 0.05 | ycaM | 3 | 22.05 | 0.336 |
| 0 | 0.05 | yadI | 3 | 22.05 | 0.964 |
| 0 | 0.05 | AG1 | 3 | 22.05 | 0.894 |
| 0 | 0.05 | ptsI | 3 | 22.3 | 0.625 |
| 0 | 0.05 | clcB | 3 | 22.3 | 0.341 |
| 0 | 0.05 | ycaM | 3 | 22.3 | 0.341 |
| 0 | 0.05 | yadI | 3 | 22.3 | 0.971 |
| 0 | 0.05 | AG1 | 3 | 22.3 | 0.902 |
| 0 | 0.05 | ptsI | 3 | 22.55 | 0.621 |
| 0 | 0.05 | clcB | 3 | 22.55 | 0.339 |
| 0 | 0.05 | ycaM | 3 | 22.55 | 0.342 |
| 0 | 0.05 | yadI | 3 | 22.55 | 0.976 |
| 0 | 0.05 | AG1 | 3 | 22.55 | 0.913 |
| 0 | 0.05 | ptsI | 3 | 22.8 | 0.618 |
| 0 | 0.05 | clcB | 3 | 22.8 | 0.339 |
| 0 | 0.05 | ycaM | 3 | 22.8 | 0.344 |
| 0 | 0.05 | yadI | 3 | 22.8 | 0.986 |
| 0 | 0.05 | AG1 | 3 | 22.8 | 0.916 |
| 0 | 0.05 | ptsI | 3 | 23.05 | 0.625 |
| 0 | 0.05 | clcB | 3 | 23.05 | 0.339 |
| 0 | 0.05 | ycaM | 3 | 23.05 | 0.349 |
| 0 | 0.05 | yadI | 3 | 23.05 | 0.997 |
| 0 | 0.05 | AG1 | 3 | 23.05 | 0.919 |
| 0 | 0.05 | ptsI | 3 | 23.3 | 0.62 |
| 0 | 0.05 | clcB | 3 | 23.3 | 0.338 |
| 0 | 0.05 | ycaM | 3 | 23.3 | 0.354 |

| | | | | | |
|---|---|---|---|---|---|
| 0 | 0.05 | yadI | 3 | 23.3 | 1.001 |
| 0 | 0.05 | AG1 | 3 | 23.3 | 0.924 |
| 0 | 0.05 | ptsI | 3 | 23.55 | 0.632 |
| 0 | 0.05 | clcB | 3 | 23.55 | 0.34 |
| 0 | 0.05 | ycaM | 3 | 23.55 | 0.369 |
| 0 | 0.05 | yadI | 3 | 23.55 | 1.018 |
| 0 | 0.05 | AG1 | 3 | 23.55 | 0.937 |
| 0 | 0.05 | ptsI | 3 | 23.8 | 0.625 |
| 0 | 0.05 | clcB | 3 | 23.8 | 0.345 |
| 0 | 0.05 | ycaM | 3 | 23.8 | 0.366 |
| 0 | 0.05 | yadI | 3 | 23.8 | 1.024 |
| 0 | 0.05 | AG1 | 3 | 23.8 | 0.937 |
| 0 | 0.05 | ptsI | 3 | 24.05 | 0.623 |
| 0 | 0.05 | clcB | 3 | 24.05 | 0.343 |
| 0 | 0.05 | ycaM | 3 | 24.05 | 0.369 |
| 0 | 0.05 | yadI | 3 | 24.05 | 1.034 |
| 0 | 0.05 | AG1 | 3 | 24.05 | 0.943 |
| 0 | 0.05 | ptsI | 3 | 24.3 | 0.616 |
| 0 | 0.05 | clcB | 3 | 24.3 | 0.341 |
| 0 | 0.05 | ycaM | 3 | 24.3 | 0.373 |
| 0 | 0.05 | yadI | 3 | 24.3 | 1.028 |
| 0 | 0.05 | AG1 | 3 | 24.3 | 0.935 |
| 0 | 0.05 | ptsI | 4 | 0 | 0.194 |
| 0 | 0.05 | clcB | 4 | 0 | 0.2 |
| 0 | 0.05 | ycaM | 4 | 0 | 0.224 |
| 0 | 0.05 | yadI | 4 | 0 | 0.21 |
| 0 | 0.05 | AG1 | 4 | 0 | 0.197 |
| 0 | 0.05 | ptsI | 4 | 0.25 | 0.194 |
| 0 | 0.05 | clcB | 4 | 0.25 | 0.205 |
| 0 | 0.05 | ycaM | 4 | 0.25 | 0.214 |
| 0 | 0.05 | yadI | 4 | 0.25 | 0.205 |
| 0 | 0.05 | AG1 | 4 | 0.25 | 0.196 |
| 0 | 0.05 | ptsI | 4 | 0.5 | 0.194 |
| 0 | 0.05 | clcB | 4 | 0.5 | 0.209 |
| 0 | 0.05 | ycaM | 4 | 0.5 | 0.212 |
| 0 | 0.05 | yadI | 4 | 0.5 | 0.206 |
| 0 | 0.05 | AG1 | 4 | 0.5 | 0.198 |
| 0 | 0.05 | ptsI | 4 | 0.75 | 0.197 |
| 0 | 0.05 | clcB | 4 | 0.75 | 0.213 |
| 0 | 0.05 | ycaM | 4 | 0.75 | 0.219 |
| 0 | 0.05 | yadI | 4 | 0.75 | 0.209 |
| 0 | 0.05 | AG1 | 4 | 0.75 | 0.202 |
| 0 | 0.05 | ptsI | 4 | 1 | 0.201 |
| 0 | 0.05 | clcB | 4 | 1 | 0.219 |
| 0 | 0.05 | ycaM | 4 | 1 | 0.223 |
| 0 | 0.05 | yadI | 4 | 1 | 0.212 |
| 0 | 0.05 | AG1 | 4 | 1 | 0.207 |
| 0 | 0.05 | ptsI | 4 | 1.25 | 0.205 |

| | | | | | |
|---|---|---|---|---|---|
| 0 | 0.05 | clcB | 4 | 1.25 | 0.223 |
| 0 | 0.05 | ycaM | 4 | 1.25 | 0.227 |
| 0 | 0.05 | yadI | 4 | 1.25 | 0.218 |
| 0 | 0.05 | AG1 | 4 | 1.25 | 0.211 |
| 0 | 0.05 | ptsI | 4 | 1.5 | 0.21 |
| 0 | 0.05 | clcB | 4 | 1.5 | 0.227 |
| 0 | 0.05 | ycaM | 4 | 1.5 | 0.235 |
| 0 | 0.05 | yadI | 4 | 1.5 | 0.224 |
| 0 | 0.05 | AG1 | 4 | 1.5 | 0.215 |
| 0 | 0.05 | ptsI | 4 | 1.75 | 0.218 |
| 0 | 0.05 | clcB | 4 | 1.75 | 0.236 |
| 0 | 0.05 | ycaM | 4 | 1.75 | 0.242 |
| 0 | 0.05 | yadI | 4 | 1.75 | 0.232 |
| 0 | 0.05 | AG1 | 4 | 1.75 | 0.222 |
| 0 | 0.05 | ptsI | 4 | 2 | 0.223 |
| 0 | 0.05 | clcB | 4 | 2 | 0.246 |
| 0 | 0.05 | ycaM | 4 | 2 | 0.252 |
| 0 | 0.05 | yadI | 4 | 2 | 0.239 |
| 0 | 0.05 | AG1 | 4 | 2 | 0.234 |
| 0 | 0.05 | ptsI | 4 | 2.25 | 0.23 |
| 0 | 0.05 | clcB | 4 | 2.25 | 0.255 |
| 0 | 0.05 | ycaM | 4 | 2.25 | 0.26 |
| 0 | 0.05 | yadI | 4 | 2.25 | 0.248 |
| 0 | 0.05 | AG1 | 4 | 2.25 | 0.238 |
| 0 | 0.05 | ptsI | 4 | 2.5 | 0.24 |
| 0 | 0.05 | clcB | 4 | 2.5 | 0.258 |
| 0 | 0.05 | ycaM | 4 | 2.5 | 0.269 |
| 0 | 0.05 | yadI | 4 | 2.5 | 0.257 |
| 0 | 0.05 | AG1 | 4 | 2.5 | 0.248 |
| 0 | 0.05 | ptsI | 4 | 2.75 | 0.248 |
| 0 | 0.05 | clcB | 4 | 2.75 | 0.264 |
| 0 | 0.05 | ycaM | 4 | 2.75 | 0.28 |
| 0 | 0.05 | yadI | 4 | 2.75 | 0.264 |
| 0 | 0.05 | AG1 | 4 | 2.75 | 0.259 |
| 0 | 0.05 | ptsI | 4 | 3 | 0.256 |
| 0 | 0.05 | clcB | 4 | 3 | 0.276 |
| 0 | 0.05 | ycaM | 4 | 3 | 0.29 |
| 0 | 0.05 | yadI | 4 | 3 | 0.274 |
| 0 | 0.05 | AG1 | 4 | 3 | 0.272 |
| 0 | 0.05 | ptsI | 4 | 3.25 | 0.265 |
| 0 | 0.05 | clcB | 4 | 3.25 | 0.286 |
| 0 | 0.05 | ycaM | 4 | 3.25 | 0.301 |
| 0 | 0.05 | yadI | 4 | 3.25 | 0.285 |
| 0 | 0.05 | AG1 | 4 | 3.25 | 0.281 |
| 0 | 0.05 | ptsI | 4 | 3.5 | 0.274 |
| 0 | 0.05 | clcB | 4 | 3.5 | 0.298 |
| 0 | 0.05 | ycaM | 4 | 3.5 | 0.313 |
| 0 | 0.05 | yadI | 4 | 3.5 | 0.298 |

| | | | | | |
|---|---|---|---|---|---|
| 0 | 0.05 | AG1 | 4 | 3.5 | 0.292 |
| 0 | 0.05 | ptsI | 4 | 3.75 | 0.282 |
| 0 | 0.05 | clcB | 4 | 3.75 | 0.305 |
| 0 | 0.05 | ycaM | 4 | 3.75 | 0.321 |
| 0 | 0.05 | yadI | 4 | 3.75 | 0.305 |
| 0 | 0.05 | AG1 | 4 | 3.75 | 0.299 |
| 0 | 0.05 | ptsI | 4 | 4 | 0.291 |
| 0 | 0.05 | clcB | 4 | 4 | 0.32 |
| 0 | 0.05 | ycaM | 4 | 4 | 0.34 |
| 0 | 0.05 | yadI | 4 | 4 | 0.321 |
| 0 | 0.05 | AG1 | 4 | 4 | 0.314 |
| 0 | 0.05 | ptsI | 4 | 4.35 | 0.287 |
| 0 | 0.05 | clcB | 4 | 4.35 | 0.311 |
| 0 | 0.05 | ycaM | 4 | 4.35 | 0.343 |
| 0 | 0.05 | yadI | 4 | 4.35 | 0.316 |
| 0 | 0.05 | AG1 | 4 | 4.35 | 0.293 |
| 0 | 0.05 | ptsI | 4 | 4.6 | 0.291 |
| 0 | 0.05 | clcB | 4 | 4.6 | 0.315 |
| 0 | 0.05 | ycaM | 4 | 4.6 | 0.336 |
| 0 | 0.05 | yadI | 4 | 4.6 | 0.323 |
| 0 | 0.05 | AG1 | 4 | 4.6 | 0.307 |
| 0 | 0.05 | ptsI | 4 | 4.85 | 0.3 |
| 0 | 0.05 | clcB | 4 | 4.85 | 0.32 |
| 0 | 0.05 | ycaM | 4 | 4.85 | 0.332 |
| 0 | 0.05 | yadI | 4 | 4.85 | 0.338 |
| 0 | 0.05 | AG1 | 4 | 4.85 | 0.316 |
| 0 | 0.05 | ptsI | 4 | 5.1 | 0.308 |
| 0 | 0.05 | clcB | 4 | 5.1 | 0.322 |
| 0 | 0.05 | ycaM | 4 | 5.1 | 0.333 |
| 0 | 0.05 | yadI | 4 | 5.1 | 0.35 |
| 0 | 0.05 | AG1 | 4 | 5.1 | 0.328 |
| 0 | 0.05 | ptsI | 4 | 5.35 | 0.319 |
| 0 | 0.05 | clcB | 4 | 5.35 | 0.324 |
| 0 | 0.05 | ycaM | 4 | 5.35 | 0.334 |
| 0 | 0.05 | yadI | 4 | 5.35 | 0.363 |
| 0 | 0.05 | AG1 | 4 | 5.35 | 0.338 |
| 0 | 0.05 | ptsI | 4 | 5.6 | 0.331 |
| 0 | 0.05 | clcB | 4 | 5.6 | 0.324 |
| 0 | 0.05 | ycaM | 4 | 5.6 | 0.33 |
| 0 | 0.05 | yadI | 4 | 5.6 | 0.374 |
| 0 | 0.05 | AG1 | 4 | 5.6 | 0.349 |
| 0 | 0.05 | ptsI | 4 | 5.85 | 0.348 |
| 0 | 0.05 | clcB | 4 | 5.85 | 0.325 |
| 0 | 0.05 | ycaM | 4 | 5.85 | 0.33 |
| 0 | 0.05 | yadI | 4 | 5.85 | 0.387 |
| 0 | 0.05 | AG1 | 4 | 5.85 | 0.359 |
| 0 | 0.05 | ptsI | 4 | 6.1 | 0.363 |
| 0 | 0.05 | clcB | 4 | 6.1 | 0.328 |

| | | | | | |
|---|---|---|---|---|---|
| 0 | 0.05 | ycaM | 4 | 6.1 | 0.332 |
| 0 | 0.05 | yadI | 4 | 6.1 | 0.398 |
| 0 | 0.05 | AG1 | 4 | 6.1 | 0.368 |
| 0 | 0.05 | ptsI | 4 | 6.35 | 0.372 |
| 0 | 0.05 | clcB | 4 | 6.35 | 0.329 |
| 0 | 0.05 | ycaM | 4 | 6.35 | 0.331 |
| 0 | 0.05 | yadI | 4 | 6.35 | 0.411 |
| 0 | 0.05 | AG1 | 4 | 6.35 | 0.377 |
| 0 | 0.05 | ptsI | 4 | 6.6 | 0.385 |
| 0 | 0.05 | clcB | 4 | 6.6 | 0.329 |
| 0 | 0.05 | ycaM | 4 | 6.6 | 0.328 |
| 0 | 0.05 | yadI | 4 | 6.6 | 0.417 |
| 0 | 0.05 | AG1 | 4 | 6.6 | 0.384 |
| 0 | 0.05 | ptsI | 4 | 6.85 | 0.398 |
| 0 | 0.05 | clcB | 4 | 6.85 | 0.33 |
| 0 | 0.05 | ycaM | 4 | 6.85 | 0.328 |
| 0 | 0.05 | yadI | 4 | 6.85 | 0.424 |
| 0 | 0.05 | AG1 | 4 | 6.85 | 0.397 |
| 0 | 0.05 | ptsI | 4 | 7.1 | 0.405 |
| 0 | 0.05 | clcB | 4 | 7.1 | 0.329 |
| 0 | 0.05 | ycaM | 4 | 7.1 | 0.328 |
| 0 | 0.05 | yadI | 4 | 7.1 | 0.431 |
| 0 | 0.05 | AG1 | 4 | 7.1 | 0.406 |
| 0 | 0.05 | ptsI | 4 | 7.35 | 0.415 |
| 0 | 0.05 | clcB | 4 | 7.35 | 0.333 |
| 0 | 0.05 | ycaM | 4 | 7.35 | 0.33 |
| 0 | 0.05 | yadI | 4 | 7.35 | 0.439 |
| 0 | 0.05 | AG1 | 4 | 7.35 | 0.416 |
| 0 | 0.05 | ptsI | 4 | 7.6 | 0.421 |
| 0 | 0.05 | clcB | 4 | 7.6 | 0.337 |
| 0 | 0.05 | ycaM | 4 | 7.6 | 0.334 |
| 0 | 0.05 | yadI | 4 | 7.6 | 0.446 |
| 0 | 0.05 | AG1 | 4 | 7.6 | 0.423 |
| 0 | 0.05 | ptsI | 4 | 7.85 | 0.427 |
| 0 | 0.05 | clcB | 4 | 7.85 | 0.338 |
| 0 | 0.05 | ycaM | 4 | 7.85 | 0.339 |
| 0 | 0.05 | yadI | 4 | 7.85 | 0.452 |
| 0 | 0.05 | AG1 | 4 | 7.85 | 0.43 |
| 0 | 0.05 | ptsI | 4 | 8.1 | 0.44 |
| 0 | 0.05 | clcB | 4 | 8.1 | 0.34 |
| 0 | 0.05 | ycaM | 4 | 8.1 | 0.342 |
| 0 | 0.05 | yadI | 4 | 8.1 | 0.461 |
| 0 | 0.05 | AG1 | 4 | 8.1 | 0.44 |
| 0 | 0.05 | ptsI | 4 | 8.35 | 0.449 |
| 0 | 0.05 | clcB | 4 | 8.35 | 0.343 |
| 0 | 0.05 | ycaM | 4 | 8.35 | 0.345 |
| 0 | 0.05 | yadI | 4 | 8.35 | 0.466 |
| 0 | 0.05 | AG1 | 4 | 8.35 | 0.448 |

| | | | | | |
|---|---|---|---|---|---|
| 0 | 0.05 | ptsI | 4 | 8.83333 | 0.427 |
| 0 | 0.05 | clcB | 4 | 8.83333 | 0.316 |
| 0 | 0.05 | ycaM | 4 | 8.83333 | 0.311 |
| 0 | 0.05 | yadI | 4 | 8.83333 | 0.51 |
| 0 | 0.05 | AG1 | 4 | 8.83333 | 0.488 |
| 0 | 0.05 | ptsI | 4 | 9.08333 | 0.43 |
| 0 | 0.05 | clcB | 4 | 9.08333 | 0.314 |
| 0 | 0.05 | ycaM | 4 | 9.08333 | 0.316 |
| 0 | 0.05 | yadI | 4 | 9.08333 | 0.513 |
| 0 | 0.05 | AG1 | 4 | 9.08333 | 0.49 |
| 0 | 0.05 | ptsI | 4 | 9.33333 | 0.438 |
| 0 | 0.05 | clcB | 4 | 9.33333 | 0.316 |
| 0 | 0.05 | ycaM | 4 | 9.33333 | 0.316 |
| 0 | 0.05 | yadI | 4 | 9.33333 | 0.518 |
| 0 | 0.05 | AG1 | 4 | 9.33333 | 0.503 |
| 0 | 0.05 | ptsI | 4 | 9.58333 | 0.45 |
| 0 | 0.05 | clcB | 4 | 9.58333 | 0.327 |
| 0 | 0.05 | ycaM | 4 | 9.58333 | 0.321 |
| 0 | 0.05 | yadI | 4 | 9.58333 | 0.529 |
| 0 | 0.05 | AG1 | 4 | 9.58333 | 0.513 |
| 0 | 0.05 | ptsI | 4 | 9.83333 | 0.459 |
| 0 | 0.05 | clcB | 4 | 9.83333 | 0.318 |
| 0 | 0.05 | ycaM | 4 | 9.83333 | 0.33 |
| 0 | 0.05 | yadI | 4 | 9.83333 | 0.544 |
| 0 | 0.05 | AG1 | 4 | 9.83333 | 0.52 |
| 0 | 0.05 | ptsI | 4 | 10.0833 | 0.47 |
| 0 | 0.05 | clcB | 4 | 10.0833 | 0.32 |
| 0 | 0.05 | ycaM | 4 | 10.0833 | 0.328 |
| 0 | 0.05 | yadI | 4 | 10.0833 | 0.551 |
| 0 | 0.05 | AG1 | 4 | 10.0833 | 0.526 |
| 0 | 0.05 | ptsI | 4 | 10.3333 | 0.481 |
| 0 | 0.05 | clcB | 4 | 10.3333 | 0.326 |
| 0 | 0.05 | ycaM | 4 | 10.3333 | 0.328 |
| 0 | 0.05 | yadI | 4 | 10.3333 | 0.554 |
| 0 | 0.05 | AG1 | 4 | 10.3333 | 0.536 |
| 0 | 0.05 | ptsI | 4 | 10.5833 | 0.491 |
| 0 | 0.05 | clcB | 4 | 10.5833 | 0.324 |
| 0 | 0.05 | ycaM | 4 | 10.5833 | 0.331 |
| 0 | 0.05 | yadI | 4 | 10.5833 | 0.566 |
| 0 | 0.05 | AG1 | 4 | 10.5833 | 0.544 |
| 0 | 0.05 | ptsI | 4 | 10.8333 | 0.493 |
| 0 | 0.05 | clcB | 4 | 10.8333 | 0.341 |
| 0 | 0.05 | ycaM | 4 | 10.8333 | 0.331 |
| 0 | 0.05 | yadI | 4 | 10.8333 | 0.57 |
| 0 | 0.05 | AG1 | 4 | 10.8333 | 0.552 |
| 0 | 0.05 | ptsI | 4 | 11.0833 | 0.503 |
| 0 | 0.05 | clcB | 4 | 11.0833 | 0.329 |
| 0 | 0.05 | ycaM | 4 | 11.0833 | 0.328 |

| | | | | | |
|---|---|---|---|---|---|
| 0 | 0.05 | yadI | 4 | 11.0833 | 0.579 |
| 0 | 0.05 | AG1 | 4 | 11.0833 | 0.559 |
| 0 | 0.05 | ptsI | 4 | 11.3333 | 0.513 |
| 0 | 0.05 | clcB | 4 | 11.3333 | 0.336 |
| 0 | 0.05 | ycaM | 4 | 11.3333 | 0.335 |
| 0 | 0.05 | yadI | 4 | 11.3333 | 0.586 |
| 0 | 0.05 | AG1 | 4 | 11.3333 | 0.567 |
| 0 | 0.05 | ptsI | 4 | 11.5833 | 0.52 |
| 0 | 0.05 | clcB | 4 | 11.5833 | 0.327 |
| 0 | 0.05 | ycaM | 4 | 11.5833 | 0.335 |
| 0 | 0.05 | yadI | 4 | 11.5833 | 0.594 |
| 0 | 0.05 | AG1 | 4 | 11.5833 | 0.575 |
| 0 | 0.05 | ptsI | 4 | 11.8333 | 0.528 |
| 0 | 0.05 | clcB | 4 | 11.8333 | 0.336 |
| 0 | 0.05 | ycaM | 4 | 11.8333 | 0.329 |
| 0 | 0.05 | yadI | 4 | 11.8333 | 0.598 |
| 0 | 0.05 | AG1 | 4 | 11.8333 | 0.575 |
| 0 | 0.05 | ptsI | 4 | 12.0833 | 0.538 |
| 0 | 0.05 | clcB | 4 | 12.0833 | 0.322 |
| 0 | 0.05 | ycaM | 4 | 12.0833 | 0.329 |
| 0 | 0.05 | yadI | 4 | 12.0833 | 0.606 |
| 0 | 0.05 | AG1 | 4 | 12.0833 | 0.582 |
| 0 | 0.05 | ptsI | 4 | 12.3333 | 0.544 |
| 0 | 0.05 | clcB | 4 | 12.3333 | 0.341 |
| 0 | 0.05 | ycaM | 4 | 12.3333 | 0.328 |
| 0 | 0.05 | yadI | 4 | 12.3333 | 0.608 |
| 0 | 0.05 | AG1 | 4 | 12.3333 | 0.586 |
| 0 | 0.05 | ptsI | 4 | 12.5833 | 0.55 |
| 0 | 0.05 | clcB | 4 | 12.5833 | 0.338 |
| 0 | 0.05 | ycaM | 4 | 12.5833 | 0.336 |
| 0 | 0.05 | yadI | 4 | 12.5833 | 0.616 |
| 0 | 0.05 | AG1 | 4 | 12.5833 | 0.593 |
| 0 | 0.05 | ptsI | 4 | 12.8333 | 0.557 |
| 0 | 0.05 | clcB | 4 | 12.8333 | 0.336 |
| 0 | 0.05 | ycaM | 4 | 12.8333 | 0.333 |
| 0 | 0.05 | yadI | 4 | 12.8333 | 0.621 |
| 0 | 0.05 | AG1 | 4 | 12.8333 | 0.597 |
| 0 | 0.05 | ptsI | 4 | 13.0833 | 0.559 |
| 0 | 0.05 | clcB | 4 | 13.0833 | 0.342 |
| 0 | 0.05 | ycaM | 4 | 13.0833 | 0.334 |
| 0 | 0.05 | yadI | 4 | 13.0833 | 0.623 |
| 0 | 0.05 | AG1 | 4 | 13.0833 | 0.6 |
| 0 | 0.05 | ptsI | 4 | 13.3333 | 0.571 |
| 0 | 0.05 | clcB | 4 | 13.3333 | 0.341 |
| 0 | 0.05 | ycaM | 4 | 13.3333 | 0.354 |
| 0 | 0.05 | yadI | 4 | 13.3333 | 0.629 |
| 0 | 0.05 | AG1 | 4 | 13.3333 | 0.607 |
| 0 | 0.05 | ptsI | 4 | 13.5833 | 0.574 |

| | | | | | |
|---|---|---|---|---|---|
| 0 | 0.05 | clcB | 4 | 13.5833 | 0.34 |
| 0 | 0.05 | ycaM | 4 | 13.5833 | 0.354 |
| 0 | 0.05 | yadI | 4 | 13.5833 | 0.635 |
| 0 | 0.05 | AG1 | 4 | 13.5833 | 0.61 |
| 0 | 0.05 | ptsI | 4 | 13.8333 | 0.583 |
| 0 | 0.05 | clcB | 4 | 13.8333 | 0.34 |
| 0 | 0.05 | ycaM | 4 | 13.8333 | 0.334 |
| 0 | 0.05 | yadI | 4 | 13.8333 | 0.636 |
| 0 | 0.05 | AG1 | 4 | 13.8333 | 0.614 |
| 0 | 0.05 | ptsI | 4 | 14.0833 | 0.589 |
| 0 | 0.05 | clcB | 4 | 14.0833 | 0.338 |
| 0 | 0.05 | ycaM | 4 | 14.0833 | 0.33 |
| 0 | 0.05 | yadI | 4 | 14.0833 | 0.643 |
| 0 | 0.05 | AG1 | 4 | 14.0833 | 0.616 |
| 0 | 0.05 | ptsI | 4 | 14.3333 | 0.597 |
| 0 | 0.05 | clcB | 4 | 14.3333 | 0.334 |
| 0 | 0.05 | ycaM | 4 | 14.3333 | 0.33 |
| 0 | 0.05 | yadI | 4 | 14.3333 | 0.651 |
| 0 | 0.05 | AG1 | 4 | 14.3333 | 0.626 |
| 0 | 0.05 | ptsI | 4 | 14.5833 | 0.595 |
| 0 | 0.05 | clcB | 4 | 14.5833 | 0.342 |
| 0 | 0.05 | ycaM | 4 | 14.5833 | 0.332 |
| 0 | 0.05 | yadI | 4 | 14.5833 | 0.649 |
| 0 | 0.05 | AG1 | 4 | 14.5833 | 0.62 |
| 0 | 0.05 | ptsI | 4 | 14.8333 | 0.604 |
| 0 | 0.05 | clcB | 4 | 14.8333 | 0.338 |
| 0 | 0.05 | ycaM | 4 | 14.8333 | 0.331 |
| 0 | 0.05 | yadI | 4 | 14.8333 | 0.657 |
| 0 | 0.05 | AG1 | 4 | 14.8333 | 0.629 |
| 0 | 0.05 | ptsI | 4 | 15.0833 | 0.606 |
| 0 | 0.05 | clcB | 4 | 15.0833 | 0.34 |
| 0 | 0.05 | ycaM | 4 | 15.0833 | 0.33 |
| 0 | 0.05 | yadI | 4 | 15.0833 | 0.66 |
| 0 | 0.05 | AG1 | 4 | 15.0833 | 0.628 |
| 0 | 0.05 | ptsI | 4 | 15.3333 | 0.622 |
| 0 | 0.05 | clcB | 4 | 15.3333 | 0.345 |
| 0 | 0.05 | ycaM | 4 | 15.3333 | 0.332 |
| 0 | 0.05 | yadI | 4 | 15.3333 | 0.667 |
| 0 | 0.05 | AG1 | 4 | 15.3333 | 0.635 |
| 0 | 0.05 | ptsI | 4 | 15.5833 | 0.622 |
| 0 | 0.05 | clcB | 4 | 15.5833 | 0.341 |
| 0 | 0.05 | ycaM | 4 | 15.5833 | 0.335 |
| 0 | 0.05 | yadI | 4 | 15.5833 | 0.671 |
| 0 | 0.05 | AG1 | 4 | 15.5833 | 0.636 |
| 0 | 0.05 | ptsI | 4 | 15.8333 | 0.629 |
| 0 | 0.05 | clcB | 4 | 15.8333 | 0.342 |
| 0 | 0.05 | ycaM | 4 | 15.8333 | 0.345 |
| 0 | 0.05 | yadI | 4 | 15.8333 | 0.678 |

| | | | | | |
|---|---|---|---|---|---|
| 0 | 0.05 | AG1 | 4 | 15.8333 | 0.639 |
| 0 | 0.05 | ptsI | 4 | 16.0833 | 0.632 |
| 0 | 0.05 | clcB | 4 | 16.0833 | 0.342 |
| 0 | 0.05 | ycaM | 4 | 16.0833 | 0.35 |
| 0 | 0.05 | yadI | 4 | 16.0833 | 0.679 |
| 0 | 0.05 | AG1 | 4 | 16.0833 | 0.641 |
| 0 | 0.05 | ptsI | 4 | 16.3333 | 0.635 |
| 0 | 0.05 | clcB | 4 | 16.3333 | 0.345 |
| 0 | 0.05 | ycaM | 4 | 16.3333 | 0.339 |
| 0 | 0.05 | yadI | 4 | 16.3333 | 0.682 |
| 0 | 0.05 | AG1 | 4 | 16.3333 | 0.642 |
| 0 | 0.05 | ptsI | 4 | 16.5833 | 0.642 |
| 0 | 0.05 | clcB | 4 | 16.5833 | 0.349 |
| 0 | 0.05 | ycaM | 4 | 16.5833 | 0.341 |
| 0 | 0.05 | yadI | 4 | 16.5833 | 0.686 |
| 0 | 0.05 | AG1 | 4 | 16.5833 | 0.646 |
| 0 | 0.05 | ptsI | 4 | 16.8333 | 0.637 |
| 0 | 0.05 | clcB | 4 | 16.8333 | 0.347 |
| 0 | 0.05 | ycaM | 4 | 16.8333 | 0.339 |
| 0 | 0.05 | yadI | 4 | 16.8333 | 0.687 |
| 0 | 0.05 | AG1 | 4 | 16.8333 | 0.648 |
| 0 | 0.05 | ptsI | 4 | 17.0833 | 0.639 |
| 0 | 0.05 | clcB | 4 | 17.0833 | 0.349 |
| 0 | 0.05 | ycaM | 4 | 17.0833 | 0.344 |
| 0 | 0.05 | yadI | 4 | 17.0833 | 0.692 |
| 0 | 0.05 | AG1 | 4 | 17.0833 | 0.65 |
| 0 | 0.05 | ptsI | 4 | 17.3333 | 0.65 |
| 0 | 0.05 | clcB | 4 | 17.3333 | 0.378 |
| 0 | 0.05 | ycaM | 4 | 17.3333 | 0.348 |
| 0 | 0.05 | yadI | 4 | 17.3333 | 0.698 |
| 0 | 0.05 | AG1 | 4 | 17.3333 | 0.657 |
| 0 | 0.05 | ptsI | 4 | 17.5833 | 0.642 |
| 0 | 0.05 | clcB | 4 | 17.5833 | 0.362 |
| 0 | 0.05 | ycaM | 4 | 17.5833 | 0.348 |
| 0 | 0.05 | yadI | 4 | 17.5833 | 0.694 |
| 0 | 0.05 | AG1 | 4 | 17.5833 | 0.651 |
| 0 | 0.05 | ptsI | 4 | 17.8333 | 0.64 |
| 0 | 0.05 | clcB | 4 | 17.8333 | 0.347 |
| 0 | 0.05 | ycaM | 4 | 17.8333 | 0.349 |
| 0 | 0.05 | yadI | 4 | 17.8333 | 0.697 |
| 0 | 0.05 | AG1 | 4 | 17.8333 | 0.645 |
| 0 | 0.05 | ptsI | 4 | 18.0833 | 0.644 |
| 0 | 0.05 | clcB | 4 | 18.0833 | 0.351 |
| 0 | 0.05 | ycaM | 4 | 18.0833 | 0.36 |
| 0 | 0.05 | yadI | 4 | 18.0833 | 0.707 |
| 0 | 0.05 | AG1 | 4 | 18.0833 | 0.659 |
| 0 | 0.05 | ptsI | 4 | 18.3333 | 0.649 |
| 0 | 0.05 | clcB | 4 | 18.3333 | 0.363 |

| | | | | | |
|---|---|---|---|---|---|
| 0 | 0.05 | ycaM | 4 | 18.3333 | 0.362 |
| 0 | 0.05 | yadI | 4 | 18.3333 | 0.707 |
| 0 | 0.05 | AG1 | 4 | 18.3333 | 0.657 |
| 0 | 0.05 | ptsI | 4 | 18.5833 | 0.645 |
| 0 | 0.05 | clcB | 4 | 18.5833 | 0.352 |
| 0 | 0.05 | ycaM | 4 | 18.5833 | 0.365 |
| 0 | 0.05 | yadI | 4 | 18.5833 | 0.71 |
| 0 | 0.05 | AG1 | 4 | 18.5833 | 0.663 |
| 0 | 0.05 | ptsI | 4 | 18.8333 | 0.645 |
| 0 | 0.05 | clcB | 4 | 18.8333 | 0.351 |
| 0 | 0.05 | ycaM | 4 | 18.8333 | 0.362 |
| 0 | 0.05 | yadI | 4 | 18.8333 | 0.714 |
| 0 | 0.05 | AG1 | 4 | 18.8333 | 0.659 |
| 0 | 0.05 | ptsI | 4 | 19.0833 | 0.644 |
| 0 | 0.05 | clcB | 4 | 19.0833 | 0.354 |
| 0 | 0.05 | ycaM | 4 | 19.0833 | 0.366 |
| 0 | 0.05 | yadI | 4 | 19.0833 | 0.717 |
| 0 | 0.05 | AG1 | 4 | 19.0833 | 0.667 |
| 0 | 0.05 | ptsI | 4 | 19.3333 | 0.658 |
| 0 | 0.05 | clcB | 4 | 19.3333 | 0.383 |
| 0 | 0.05 | ycaM | 4 | 19.3333 | 0.382 |
| 0 | 0.05 | yadI | 4 | 19.3333 | 0.724 |
| 0 | 0.05 | AG1 | 4 | 19.3333 | 0.674 |
| 0 | 0.05 | ptsI | 4 | 19.5833 | 0.651 |
| 0 | 0.05 | clcB | 4 | 19.5833 | 0.351 |
| 0 | 0.05 | ycaM | 4 | 19.5833 | 0.375 |
| 0 | 0.05 | yadI | 4 | 19.5833 | 0.723 |
| 0 | 0.05 | AG1 | 4 | 19.5833 | 0.67 |
| 0 | 0.05 | ptsI | 4 | 19.8333 | 0.652 |
| 0 | 0.05 | clcB | 4 | 19.8333 | 0.344 |
| 0 | 0.05 | ycaM | 4 | 19.8333 | 0.38 |
| 0 | 0.05 | yadI | 4 | 19.8333 | 0.717 |
| 0 | 0.05 | AG1 | 4 | 19.8333 | 0.668 |
| 0 | 0.05 | ptsI | 4 | 20.0833 | 0.657 |
| 0 | 0.05 | clcB | 4 | 20.0833 | 0.378 |
| 0 | 0.05 | ycaM | 4 | 20.0833 | 0.394 |
| 0 | 0.05 | yadI | 4 | 20.0833 | 0.727 |
| 0 | 0.05 | AG1 | 4 | 20.0833 | 0.678 |
| 0 | 0.05 | ptsI | 4 | 20.3333 | 0.654 |
| 0 | 0.05 | clcB | 4 | 20.3333 | 0.4 |
| 0 | 0.05 | ycaM | 4 | 20.3333 | 0.401 |
| 0 | 0.05 | yadI | 4 | 20.3333 | 0.722 |
| 0 | 0.05 | AG1 | 4 | 20.3333 | 0.675 |
| 0 | 0.05 | ptsI | 4 | 20.5833 | 0.653 |
| 0 | 0.05 | clcB | 4 | 20.5833 | 0.378 |
| 0 | 0.05 | ycaM | 4 | 20.5833 | 0.395 |
| 0 | 0.05 | yadI | 4 | 20.5833 | 0.74 |
| 0 | 0.05 | AG1 | 4 | 20.5833 | 0.681 |

| | | | | | |
|---|---|---|---|---|---|
| 0 | 0.05 | ptsI | 4 | 20.8333 | 0.657 |
| 0 | 0.05 | clcB | 4 | 20.8333 | 0.381 |
| 0 | 0.05 | ycaM | 4 | 20.8333 | 0.404 |
| 0 | 0.05 | yadI | 4 | 20.8333 | 0.73 |
| 0 | 0.05 | AG1 | 4 | 20.8333 | 0.681 |
| 0 | 0.05 | ptsI | 4 | 21.0833 | 0.66 |
| 0 | 0.05 | clcB | 4 | 21.0833 | 0.396 |
| 0 | 0.05 | ycaM | 4 | 21.0833 | 0.405 |
| 0 | 0.05 | yadI | 4 | 21.0833 | 0.73 |
| 0 | 0.05 | AG1 | 4 | 21.0833 | 0.68 |
| 0 | 0.05 | ptsI | 4 | 21.3333 | 0.656 |
| 0 | 0.05 | clcB | 4 | 21.3333 | 0.371 |
| 0 | 0.05 | ycaM | 4 | 21.3333 | 0.424 |
| 0 | 0.05 | yadI | 4 | 21.3333 | 0.732 |
| 0 | 0.05 | AG1 | 4 | 21.3333 | 0.679 |
| 0 | 0.05 | ptsI | 4 | 21.5833 | 0.662 |
| 0 | 0.05 | clcB | 4 | 21.5833 | 0.365 |
| 0 | 0.05 | ycaM | 4 | 21.5833 | 0.42 |
| 0 | 0.05 | yadI | 4 | 21.5833 | 0.734 |
| 0 | 0.05 | AG1 | 4 | 21.5833 | 0.68 |
| 0 | 0.05 | ptsI | 4 | 21.8333 | 0.659 |
| 0 | 0.05 | clcB | 4 | 21.8333 | 0.388 |
| 0 | 0.05 | ycaM | 4 | 21.8333 | 0.433 |
| 0 | 0.05 | yadI | 4 | 21.8333 | 0.744 |
| 0 | 0.05 | AG1 | 4 | 21.8333 | 0.688 |
| 0 | 0.05 | ptsI | 4 | 22.0833 | 0.664 |
| 0 | 0.05 | clcB | 4 | 22.0833 | 0.392 |
| 0 | 0.05 | ycaM | 4 | 22.0833 | 0.43 |
| 0 | 0.05 | yadI | 4 | 22.0833 | 0.738 |
| 0 | 0.05 | AG1 | 4 | 22.0833 | 0.686 |
| 0 | 0.05 | ptsI | 4 | 22.3333 | 0.663 |
| 0 | 0.05 | clcB | 4 | 22.3333 | 0.358 |
| 0 | 0.05 | ycaM | 4 | 22.3333 | 0.426 |
| 0 | 0.05 | yadI | 4 | 22.3333 | 0.74 |
| 0 | 0.05 | AG1 | 4 | 22.3333 | 0.682 |
| 0 | 0.05 | ptsI | 4 | 22.5833 | 0.666 |
| 0 | 0.05 | clcB | 4 | 22.5833 | 0.354 |
| 0 | 0.05 | ycaM | 4 | 22.5833 | 0.43 |
| 0 | 0.05 | yadI | 4 | 22.5833 | 0.742 |
| 0 | 0.05 | AG1 | 4 | 22.5833 | 0.688 |
| 0 | 0.05 | ptsI | 4 | 22.8333 | 0.655 |
| 0 | 0.05 | clcB | 4 | 22.8333 | 0.376 |
| 0 | 0.05 | ycaM | 4 | 22.8333 | 0.434 |
| 0 | 0.05 | yadI | 4 | 22.8333 | 0.743 |
| 0 | 0.05 | AG1 | 4 | 22.8333 | 0.686 |
| 0 | 0.05 | ptsI | 4 | 23.0833 | 0.668 |
| 0 | 0.05 | clcB | 4 | 23.0833 | 0.37 |
| 0 | 0.05 | ycaM | 4 | 23.0833 | 0.43 |

| | | | | | |
|---|---|---|---|---|---|
| 0 | 0.05 | yadI | 4 | 23.0833 | 0.744 |
| 0 | 0.05 | AG1 | 4 | 23.0833 | 0.688 |
| 0 | 0.05 | ptsI | 4 | 23.3333 | 0.673 |
| 0 | 0.05 | clcB | 4 | 23.3333 | 0.379 |
| 0 | 0.05 | ycaM | 4 | 23.3333 | 0.432 |
| 0 | 0.05 | yadI | 4 | 23.3333 | 0.75 |
| 0 | 0.05 | AG1 | 4 | 23.3333 | 0.689 |
| 0 | 0.05 | ptsI | 4 | 23.5833 | 0.66 |
| 0 | 0.05 | clcB | 4 | 23.5833 | 0.389 |
| 0 | 0.05 | ycaM | 4 | 23.5833 | 0.433 |
| 0 | 0.05 | yadI | 4 | 23.5833 | 0.75 |
| 0 | 0.05 | AG1 | 4 | 23.5833 | 0.69 |
| 0 | 0.05 | ptsI | 4 | 23.8333 | 0.665 |
| 0 | 0.05 | clcB | 4 | 23.8333 | 0.389 |
| 0 | 0.05 | ycaM | 4 | 23.8333 | 0.429 |
| 0 | 0.05 | yadI | 4 | 23.8333 | 0.752 |
| 0 | 0.05 | AG1 | 4 | 23.8333 | 0.693 |
| 0 | 0.05 | ptsI | 4 | 24.0833 | 0.665 |
| 0 | 0.05 | clcB | 4 | 24.0833 | 0.394 |
| 0 | 0.05 | ycaM | 4 | 24.0833 | 0.424 |
| 0 | 0.05 | yadI | 4 | 24.0833 | 0.75 |
| 0 | 0.05 | AG1 | 4 | 24.0833 | 0.691 |
| 0 | 0.05 | ptsI | 5 | 0 | 0.159 |
| 0 | 0.05 | clcB | 5 | 0 | 0.161 |
| 0 | 0.05 | ycaM | 5 | 0 | 0.16 |
| 0 | 0.05 | yadI | 5 | 0 | 0.188 |
| 0 | 0.05 | AG1 | 5 | 0 | 0.16 |
| 0 | 0.05 | ptsI | 5 | 0.35 | 0.153 |
| 0 | 0.05 | clcB | 5 | 0.35 | 0.158 |
| 0 | 0.05 | ycaM | 5 | 0.35 | 0.157 |
| 0 | 0.05 | yadI | 5 | 0.35 | 0.16 |
| 0 | 0.05 | AG1 | 5 | 0.35 | 0.16 |
| 0 | 0.05 | ptsI | 5 | 0.6 | 0.153 |
| 0 | 0.05 | clcB | 5 | 0.6 | 0.159 |
| 0 | 0.05 | ycaM | 5 | 0.6 | 0.157 |
| 0 | 0.05 | yadI | 5 | 0.6 | 0.163 |
| 0 | 0.05 | AG1 | 5 | 0.6 | 0.159 |
| 0 | 0.05 | ptsI | 5 | 0.85 | 0.153 |
| 0 | 0.05 | clcB | 5 | 0.85 | 0.158 |
| 0 | 0.05 | ycaM | 5 | 0.85 | 0.157 |
| 0 | 0.05 | yadI | 5 | 0.85 | 0.168 |
| 0 | 0.05 | AG1 | 5 | 0.85 | 0.159 |
| 0 | 0.05 | ptsI | 5 | 1.1 | 0.154 |
| 0 | 0.05 | clcB | 5 | 1.1 | 0.16 |
| 0 | 0.05 | ycaM | 5 | 1.1 | 0.158 |
| 0 | 0.05 | yadI | 5 | 1.1 | 0.166 |
| 0 | 0.05 | AG1 | 5 | 1.1 | 0.161 |
| 0 | 0.05 | ptsI | 5 | 1.35 | 0.157 |

| | | | | | |
|---|---|---|---|---|---|
| 0 | 0.05 | clcB | 5 | 1.35 | 0.162 |
| 0 | 0.05 | ycaM | 5 | 1.35 | 0.163 |
| 0 | 0.05 | yadI | 5 | 1.35 | 0.187 |
| 0 | 0.05 | AG1 | 5 | 1.35 | 0.164 |
| 0 | 0.05 | ptsI | 5 | 1.6 | 0.159 |
| 0 | 0.05 | clcB | 5 | 1.6 | 0.166 |
| 0 | 0.05 | ycaM | 5 | 1.6 | 0.165 |
| 0 | 0.05 | yadI | 5 | 1.6 | 0.174 |
| 0 | 0.05 | AG1 | 5 | 1.6 | 0.166 |
| 0 | 0.05 | ptsI | 5 | 1.85 | 0.164 |
| 0 | 0.05 | clcB | 5 | 1.85 | 0.169 |
| 0 | 0.05 | ycaM | 5 | 1.85 | 0.168 |
| 0 | 0.05 | yadI | 5 | 1.85 | 0.178 |
| 0 | 0.05 | AG1 | 5 | 1.85 | 0.17 |
| 0 | 0.05 | ptsI | 5 | 2.1 | 0.164 |
| 0 | 0.05 | clcB | 5 | 2.1 | 0.172 |
| 0 | 0.05 | ycaM | 5 | 2.1 | 0.171 |
| 0 | 0.05 | yadI | 5 | 2.1 | 0.183 |
| 0 | 0.05 | AG1 | 5 | 2.1 | 0.172 |
| 0 | 0.05 | ptsI | 5 | 2.35 | 0.168 |
| 0 | 0.05 | clcB | 5 | 2.35 | 0.179 |
| 0 | 0.05 | ycaM | 5 | 2.35 | 0.175 |
| 0 | 0.05 | yadI | 5 | 2.35 | 0.188 |
| 0 | 0.05 | AG1 | 5 | 2.35 | 0.176 |
| 0 | 0.05 | ptsI | 5 | 2.6 | 0.173 |
| 0 | 0.05 | clcB | 5 | 2.6 | 0.178 |
| 0 | 0.05 | ycaM | 5 | 2.6 | 0.181 |
| 0 | 0.05 | yadI | 5 | 2.6 | 0.198 |
| 0 | 0.05 | AG1 | 5 | 2.6 | 0.181 |
| 0 | 0.05 | ptsI | 5 | 2.85 | 0.177 |
| 0 | 0.05 | clcB | 5 | 2.85 | 0.186 |
| 0 | 0.05 | ycaM | 5 | 2.85 | 0.187 |
| 0 | 0.05 | yadI | 5 | 2.85 | 0.202 |
| 0 | 0.05 | AG1 | 5 | 2.85 | 0.186 |
| 0 | 0.05 | ptsI | 5 | 3.1 | 0.181 |
| 0 | 0.05 | clcB | 5 | 3.1 | 0.195 |
| 0 | 0.05 | ycaM | 5 | 3.1 | 0.192 |
| 0 | 0.05 | yadI | 5 | 3.1 | 0.211 |
| 0 | 0.05 | AG1 | 5 | 3.1 | 0.192 |
| 0 | 0.05 | ptsI | 5 | 3.35 | 0.188 |
| 0 | 0.05 | clcB | 5 | 3.35 | 0.207 |
| 0 | 0.05 | ycaM | 5 | 3.35 | 0.203 |
| 0 | 0.05 | yadI | 5 | 3.35 | 0.223 |
| 0 | 0.05 | AG1 | 5 | 3.35 | 0.198 |
| 0 | 0.05 | ptsI | 5 | 3.6 | 0.2 |
| 0 | 0.05 | clcB | 5 | 3.6 | 0.22 |
| 0 | 0.05 | ycaM | 5 | 3.6 | 0.212 |
| 0 | 0.05 | yadI | 5 | 3.6 | 0.229 |

| | | | | | |
|---|---|---|---|---|---|
| 0 | 0.05 | AG1 | 5 | 3.6 | 0.207 |
| 0 | 0.05 | ptsI | 5 | 3.85 | 0.209 |
| 0 | 0.05 | clcB | 5 | 3.85 | 0.228 |
| 0 | 0.05 | ycaM | 5 | 3.85 | 0.221 |
| 0 | 0.05 | yadI | 5 | 3.85 | 0.239 |
| 0 | 0.05 | AG1 | 5 | 3.85 | 0.215 |
| 0 | 0.05 | ptsI | 5 | 4.38333 | 0.254 |
| 0 | 0.05 | clcB | 5 | 4.38333 | 0.255 |
| 0 | 0.05 | ycaM | 5 | 4.38333 | 0.262 |
| 0 | 0.05 | yadI | 5 | 4.38333 | 0.319 |
| 0 | 0.05 | AG1 | 5 | 4.38333 | 0.249 |
| 0 | 0.05 | ptsI | 5 | 4.63333 | 0.233 |
| 0 | 0.05 | clcB | 5 | 4.63333 | 0.252 |
| 0 | 0.05 | ycaM | 5 | 4.63333 | 0.266 |
| 0 | 0.05 | yadI | 5 | 4.63333 | 0.314 |
| 0 | 0.05 | AG1 | 5 | 4.63333 | 0.266 |
| 0 | 0.05 | ptsI | 5 | 4.88333 | 0.234 |
| 0 | 0.05 | clcB | 5 | 4.88333 | 0.27 |
| 0 | 0.05 | ycaM | 5 | 4.88333 | 0.274 |
| 0 | 0.05 | yadI | 5 | 4.88333 | 0.332 |
| 0 | 0.05 | AG1 | 5 | 4.88333 | 0.287 |
| 0 | 0.05 | ptsI | 5 | 5.13333 | 0.244 |
| 0 | 0.05 | clcB | 5 | 5.13333 | 0.281 |
| 0 | 0.05 | ycaM | 5 | 5.13333 | 0.278 |
| 0 | 0.05 | yadI | 5 | 5.13333 | 0.353 |
| 0 | 0.05 | AG1 | 5 | 5.13333 | 0.302 |
| 0 | 0.05 | ptsI | 5 | 5.38333 | 0.254 |
| 0 | 0.05 | clcB | 5 | 5.38333 | 0.292 |
| 0 | 0.05 | ycaM | 5 | 5.38333 | 0.281 |
| 0 | 0.05 | yadI | 5 | 5.38333 | 0.377 |
| 0 | 0.05 | AG1 | 5 | 5.38333 | 0.321 |
| 0 | 0.05 | ptsI | 5 | 5.63333 | 0.262 |
| 0 | 0.05 | clcB | 5 | 5.63333 | 0.303 |
| 0 | 0.05 | ycaM | 5 | 5.63333 | 0.286 |
| 0 | 0.05 | yadI | 5 | 5.63333 | 0.391 |
| 0 | 0.05 | AG1 | 5 | 5.63333 | 0.335 |
| 0 | 0.05 | ptsI | 5 | 5.88333 | 0.271 |
| 0 | 0.05 | clcB | 5 | 5.88333 | 0.313 |
| 0 | 0.05 | ycaM | 5 | 5.88333 | 0.286 |
| 0 | 0.05 | yadI | 5 | 5.88333 | 0.407 |
| 0 | 0.05 | AG1 | 5 | 5.88333 | 0.341 |
| 0 | 0.05 | ptsI | 5 | 6.13333 | 0.282 |
| 0 | 0.05 | clcB | 5 | 6.13333 | 0.319 |
| 0 | 0.05 | ycaM | 5 | 6.13333 | 0.289 |
| 0 | 0.05 | yadI | 5 | 6.13333 | 0.418 |
| 0 | 0.05 | AG1 | 5 | 6.13333 | 0.356 |
| 0 | 0.05 | ptsI | 5 | 6.38333 | 0.295 |
| 0 | 0.05 | clcB | 5 | 6.38333 | 0.328 |

| | | | | | |
|---|---|---|---|---|---|
| 0 | 0.05 | ycaM | 5 | 6.38333 | 0.293 |
| 0 | 0.05 | yadI | 5 | 6.38333 | 0.436 |
| 0 | 0.05 | AG1 | 5 | 6.38333 | 0.371 |
| 0 | 0.05 | ptsI | 5 | 6.63333 | 0.309 |
| 0 | 0.05 | clcB | 5 | 6.63333 | 0.332 |
| 0 | 0.05 | ycaM | 5 | 6.63333 | 0.297 |
| 0 | 0.05 | yadI | 5 | 6.63333 | 0.45 |
| 0 | 0.05 | AG1 | 5 | 6.63333 | 0.384 |
| 0 | 0.05 | ptsI | 5 | 6.88333 | 0.323 |
| 0 | 0.05 | clcB | 5 | 6.88333 | 0.339 |
| 0 | 0.05 | ycaM | 5 | 6.88333 | 0.299 |
| 0 | 0.05 | yadI | 5 | 6.88333 | 0.466 |
| 0 | 0.05 | AG1 | 5 | 6.88333 | 0.398 |
| 0 | 0.05 | ptsI | 5 | 7.13333 | 0.331 |
| 0 | 0.05 | clcB | 5 | 7.13333 | 0.345 |
| 0 | 0.05 | ycaM | 5 | 7.13333 | 0.3 |
| 0 | 0.05 | yadI | 5 | 7.13333 | 0.476 |
| 0 | 0.05 | AG1 | 5 | 7.13333 | 0.412 |
| 0 | 0.05 | ptsI | 5 | 7.38333 | 0.343 |
| 0 | 0.05 | clcB | 5 | 7.38333 | 0.353 |
| 0 | 0.05 | ycaM | 5 | 7.38333 | 0.303 |
| 0 | 0.05 | yadI | 5 | 7.38333 | 0.49 |
| 0 | 0.05 | AG1 | 5 | 7.38333 | 0.423 |
| 0 | 0.05 | ptsI | 5 | 7.63333 | 0.352 |
| 0 | 0.05 | clcB | 5 | 7.63333 | 0.359 |
| 0 | 0.05 | ycaM | 5 | 7.63333 | 0.306 |
| 0 | 0.05 | yadI | 5 | 7.63333 | 0.499 |
| 0 | 0.05 | AG1 | 5 | 7.63333 | 0.437 |
| 0 | 0.05 | ptsI | 5 | 7.88333 | 0.366 |
| 0 | 0.05 | clcB | 5 | 7.88333 | 0.366 |
| 0 | 0.05 | ycaM | 5 | 7.88333 | 0.312 |
| 0 | 0.05 | yadI | 5 | 7.88333 | 0.512 |
| 0 | 0.05 | AG1 | 5 | 7.88333 | 0.451 |
| 0 | 0.05 | ptsI | 5 | 8.13333 | 0.378 |
| 0 | 0.05 | clcB | 5 | 8.13333 | 0.369 |
| 0 | 0.05 | ycaM | 5 | 8.13333 | 0.312 |
| 0 | 0.05 | yadI | 5 | 8.13333 | 0.517 |
| 0 | 0.05 | AG1 | 5 | 8.13333 | 0.466 |
| 0 | 0.05 | ptsI | 5 | 8.38333 | 0.396 |
| 0 | 0.05 | clcB | 5 | 8.38333 | 0.375 |
| 0 | 0.05 | ycaM | 5 | 8.38333 | 0.313 |
| 0 | 0.05 | yadI | 5 | 8.38333 | 0.525 |
| 0 | 0.05 | AG1 | 5 | 8.38333 | 0.48 |
| 0 | 0.05 | ptsI | 5 | 8.88333 | 0.337 |
| 0 | 0.05 | clcB | 5 | 8.88333 | 0.294 |
| 0 | 0.05 | ycaM | 5 | 8.88333 | 0.27 |
| 0 | 0.05 | yadI | 5 | 8.88333 | 0.531 |
| 0 | 0.05 | AG1 | 5 | 8.88333 | 0.386 |

| | | | | | |
|---|---|---|---|---|---|
| 0 | 0.05 | ptsI | 5 | 9.13333 | 0.34 |
| 0 | 0.05 | clcB | 5 | 9.13333 | 0.288 |
| 0 | 0.05 | ycaM | 5 | 9.13333 | 0.28 |
| 0 | 0.05 | yadI | 5 | 9.13333 | 0.533 |
| 0 | 0.05 | AG1 | 5 | 9.13333 | 0.386 |
| 0 | 0.05 | ptsI | 5 | 9.38333 | 0.35 |
| 0 | 0.05 | clcB | 5 | 9.38333 | 0.294 |
| 0 | 0.05 | ycaM | 5 | 9.38333 | 0.283 |
| 0 | 0.05 | yadI | 5 | 9.38333 | 0.556 |
| 0 | 0.05 | AG1 | 5 | 9.38333 | 0.402 |
| 0 | 0.05 | ptsI | 5 | 9.63333 | 0.362 |
| 0 | 0.05 | clcB | 5 | 9.63333 | 0.295 |
| 0 | 0.05 | ycaM | 5 | 9.63333 | 0.282 |
| 0 | 0.05 | yadI | 5 | 9.63333 | 0.564 |
| 0 | 0.05 | AG1 | 5 | 9.63333 | 0.406 |
| 0 | 0.05 | ptsI | 5 | 9.88333 | 0.374 |
| 0 | 0.05 | clcB | 5 | 9.88333 | 0.296 |
| 0 | 0.05 | ycaM | 5 | 9.88333 | 0.283 |
| 0 | 0.05 | yadI | 5 | 9.88333 | 0.576 |
| 0 | 0.05 | AG1 | 5 | 9.88333 | 0.42 |
| 0 | 0.05 | ptsI | 5 | 10.1333 | 0.386 |
| 0 | 0.05 | clcB | 5 | 10.1333 | 0.304 |
| 0 | 0.05 | ycaM | 5 | 10.1333 | 0.286 |
| 0 | 0.05 | yadI | 5 | 10.1333 | 0.581 |
| 0 | 0.05 | AG1 | 5 | 10.1333 | 0.431 |
| 0 | 0.05 | ptsI | 5 | 10.3833 | 0.395 |
| 0 | 0.05 | clcB | 5 | 10.3833 | 0.305 |
| 0 | 0.05 | ycaM | 5 | 10.3833 | 0.282 |
| 0 | 0.05 | yadI | 5 | 10.3833 | 0.59 |
| 0 | 0.05 | AG1 | 5 | 10.3833 | 0.441 |
| 0 | 0.05 | ptsI | 5 | 10.6333 | 0.403 |
| 0 | 0.05 | clcB | 5 | 10.6333 | 0.304 |
| 0 | 0.05 | ycaM | 5 | 10.6333 | 0.281 |
| 0 | 0.05 | yadI | 5 | 10.6333 | 0.598 |
| 0 | 0.05 | AG1 | 5 | 10.6333 | 0.449 |
| 0 | 0.05 | ptsI | 5 | 10.8833 | 0.419 |
| 0 | 0.05 | clcB | 5 | 10.8833 | 0.307 |
| 0 | 0.05 | ycaM | 5 | 10.8833 | 0.281 |
| 0 | 0.05 | yadI | 5 | 10.8833 | 0.602 |
| 0 | 0.05 | AG1 | 5 | 10.8833 | 0.458 |
| 0 | 0.05 | ptsI | 5 | 11.1333 | 0.424 |
| 0 | 0.05 | clcB | 5 | 11.1333 | 0.302 |
| 0 | 0.05 | ycaM | 5 | 11.1333 | 0.281 |
| 0 | 0.05 | yadI | 5 | 11.1333 | 0.611 |
| 0 | 0.05 | AG1 | 5 | 11.1333 | 0.466 |
| 0 | 0.05 | ptsI | 5 | 11.3833 | 0.438 |
| 0 | 0.05 | clcB | 5 | 11.3833 | 0.31 |
| 0 | 0.05 | ycaM | 5 | 11.3833 | 0.283 |

| | | | | | |
|---|---|---|---|---|---|
| 0 | 0.05 | yadI | 5 | 11.3833 | 0.618 |
| 0 | 0.05 | AG1 | 5 | 11.3833 | 0.473 |
| 0 | 0.05 | ptsI | 5 | 11.6333 | 0.448 |
| 0 | 0.05 | clcB | 5 | 11.6333 | 0.313 |
| 0 | 0.05 | ycaM | 5 | 11.6333 | 0.285 |
| 0 | 0.05 | yadI | 5 | 11.6333 | 0.625 |
| 0 | 0.05 | AG1 | 5 | 11.6333 | 0.483 |
| 0 | 0.05 | ptsI | 5 | 11.8833 | 0.455 |
| 0 | 0.05 | clcB | 5 | 11.8833 | 0.311 |
| 0 | 0.05 | ycaM | 5 | 11.8833 | 0.285 |
| 0 | 0.05 | yadI | 5 | 11.8833 | 0.631 |
| 0 | 0.05 | AG1 | 5 | 11.8833 | 0.489 |
| 0 | 0.05 | ptsI | 5 | 12.1333 | 0.47 |
| 0 | 0.05 | clcB | 5 | 12.1333 | 0.308 |
| 0 | 0.05 | ycaM | 5 | 12.1333 | 0.287 |
| 0 | 0.05 | yadI | 5 | 12.1333 | 0.638 |
| 0 | 0.05 | AG1 | 5 | 12.1333 | 0.497 |
| 0 | 0.05 | ptsI | 5 | 12.3833 | 0.471 |
| 0 | 0.05 | clcB | 5 | 12.3833 | 0.299 |
| 0 | 0.05 | ycaM | 5 | 12.3833 | 0.288 |
| 0 | 0.05 | yadI | 5 | 12.3833 | 0.631 |
| 0 | 0.05 | AG1 | 5 | 12.3833 | 0.5 |
| 0 | 0.05 | ptsI | 5 | 12.6333 | 0.486 |
| 0 | 0.05 | clcB | 5 | 12.6333 | 0.312 |
| 0 | 0.05 | ycaM | 5 | 12.6333 | 0.292 |
| 0 | 0.05 | yadI | 5 | 12.6333 | 0.648 |
| 0 | 0.05 | AG1 | 5 | 12.6333 | 0.512 |
| 0 | 0.05 | ptsI | 5 | 12.8833 | 0.503 |
| 0 | 0.05 | clcB | 5 | 12.8833 | 0.305 |
| 0 | 0.05 | ycaM | 5 | 12.8833 | 0.294 |
| 0 | 0.05 | yadI | 5 | 12.8833 | 0.66 |
| 0 | 0.05 | AG1 | 5 | 12.8833 | 0.517 |
| 0 | 0.05 | ptsI | 5 | 13.1333 | 0.506 |
| 0 | 0.05 | clcB | 5 | 13.1333 | 0.316 |
| 0 | 0.05 | ycaM | 5 | 13.1333 | 0.295 |
| 0 | 0.05 | yadI | 5 | 13.1333 | 0.662 |
| 0 | 0.05 | AG1 | 5 | 13.1333 | 0.524 |
| 0 | 0.05 | ptsI | 5 | 13.3833 | 0.513 |
| 0 | 0.05 | clcB | 5 | 13.3833 | 0.312 |
| 0 | 0.05 | ycaM | 5 | 13.3833 | 0.291 |
| 0 | 0.05 | yadI | 5 | 13.3833 | 0.663 |
| 0 | 0.05 | AG1 | 5 | 13.3833 | 0.528 |
| 0 | 0.05 | ptsI | 5 | 13.6333 | 0.523 |
| 0 | 0.05 | clcB | 5 | 13.6333 | 0.305 |
| 0 | 0.05 | ycaM | 5 | 13.6333 | 0.292 |
| 0 | 0.05 | yadI | 5 | 13.6333 | 0.671 |
| 0 | 0.05 | AG1 | 5 | 13.6333 | 0.534 |
| 0 | 0.05 | ptsI | 5 | 13.8833 | 0.528 |

| | | | | | |
|---|---|---|---|---|---|
| 0 | 0.05 | clcB | 5 | 13.8833 | 0.303 |
| 0 | 0.05 | ycaM | 5 | 13.8833 | 0.295 |
| 0 | 0.05 | yadI | 5 | 13.8833 | 0.679 |
| 0 | 0.05 | AG1 | 5 | 13.8833 | 0.54 |
| 0 | 0.05 | ptsI | 5 | 14.1333 | 0.537 |
| 0 | 0.05 | clcB | 5 | 14.1333 | 0.305 |
| 0 | 0.05 | ycaM | 5 | 14.1333 | 0.298 |
| 0 | 0.05 | yadI | 5 | 14.1333 | 0.682 |
| 0 | 0.05 | AG1 | 5 | 14.1333 | 0.541 |
| 0 | 0.05 | ptsI | 5 | 14.3833 | 0.55 |
| 0 | 0.05 | clcB | 5 | 14.3833 | 0.304 |
| 0 | 0.05 | ycaM | 5 | 14.3833 | 0.3 |
| 0 | 0.05 | yadI | 5 | 14.3833 | 0.691 |
| 0 | 0.05 | AG1 | 5 | 14.3833 | 0.548 |
| 0 | 0.05 | ptsI | 5 | 14.6333 | 0.563 |
| 0 | 0.05 | clcB | 5 | 14.6333 | 0.312 |
| 0 | 0.05 | ycaM | 5 | 14.6333 | 0.304 |
| 0 | 0.05 | yadI | 5 | 14.6333 | 0.693 |
| 0 | 0.05 | AG1 | 5 | 14.6333 | 0.554 |
| 0 | 0.05 | ptsI | 5 | 14.8833 | 0.567 |
| 0 | 0.05 | clcB | 5 | 14.8833 | 0.312 |
| 0 | 0.05 | ycaM | 5 | 14.8833 | 0.3 |
| 0 | 0.05 | yadI | 5 | 14.8833 | 0.691 |
| 0 | 0.05 | AG1 | 5 | 14.8833 | 0.556 |
| 0 | 0.05 | ptsI | 5 | 15.1333 | 0.578 |
| 0 | 0.05 | clcB | 5 | 15.1333 | 0.307 |
| 0 | 0.05 | ycaM | 5 | 15.1333 | 0.301 |
| 0 | 0.05 | yadI | 5 | 15.1333 | 0.696 |
| 0 | 0.05 | AG1 | 5 | 15.1333 | 0.56 |
| 0 | 0.05 | ptsI | 5 | 15.3833 | 0.585 |
| 0 | 0.05 | clcB | 5 | 15.3833 | 0.31 |
| 0 | 0.05 | ycaM | 5 | 15.3833 | 0.303 |
| 0 | 0.05 | yadI | 5 | 15.3833 | 0.698 |
| 0 | 0.05 | AG1 | 5 | 15.3833 | 0.563 |
| 0 | 0.05 | ptsI | 5 | 15.6333 | 0.592 |
| 0 | 0.05 | clcB | 5 | 15.6333 | 0.308 |
| 0 | 0.05 | ycaM | 5 | 15.6333 | 0.305 |
| 0 | 0.05 | yadI | 5 | 15.6333 | 0.702 |
| 0 | 0.05 | AG1 | 5 | 15.6333 | 0.568 |
| 0 | 0.05 | ptsI | 5 | 15.8833 | 0.598 |
| 0 | 0.05 | clcB | 5 | 15.8833 | 0.307 |
| 0 | 0.05 | ycaM | 5 | 15.8833 | 0.306 |
| 0 | 0.05 | yadI | 5 | 15.8833 | 0.706 |
| 0 | 0.05 | AG1 | 5 | 15.8833 | 0.57 |
| 0 | 0.05 | ptsI | 5 | 16.1333 | 0.597 |
| 0 | 0.05 | clcB | 5 | 16.1333 | 0.317 |
| 0 | 0.05 | ycaM | 5 | 16.1333 | 0.303 |
| 0 | 0.05 | yadI | 5 | 16.1333 | 0.713 |

| | | | | | |
|---|---|---|---|---|---|
| 0 | 0.05 | AG1 | 5 | 16.1333 | 0.584 |
| 0 | 0.05 | ptsI | 5 | 16.3833 | 0.6 |
| 0 | 0.05 | clcB | 5 | 16.3833 | 0.309 |
| 0 | 0.05 | ycaM | 5 | 16.3833 | 0.3 |
| 0 | 0.05 | yadI | 5 | 16.3833 | 0.71 |
| 0 | 0.05 | AG1 | 5 | 16.3833 | 0.574 |
| 0 | 0.05 | ptsI | 5 | 16.6333 | 0.607 |
| 0 | 0.05 | clcB | 5 | 16.6333 | 0.308 |
| 0 | 0.05 | ycaM | 5 | 16.6333 | 0.305 |
| 0 | 0.05 | yadI | 5 | 16.6333 | 0.715 |
| 0 | 0.05 | AG1 | 5 | 16.6333 | 0.576 |
| 0 | 0.05 | ptsI | 5 | 16.8833 | 0.625 |
| 0 | 0.05 | clcB | 5 | 16.8833 | 0.312 |
| 0 | 0.05 | ycaM | 5 | 16.8833 | 0.304 |
| 0 | 0.05 | yadI | 5 | 16.8833 | 0.722 |
| 0 | 0.05 | AG1 | 5 | 16.8833 | 0.583 |
| 0 | 0.05 | ptsI | 5 | 17.1333 | 0.624 |
| 0 | 0.05 | clcB | 5 | 17.1333 | 0.312 |
| 0 | 0.05 | ycaM | 5 | 17.1333 | 0.306 |
| 0 | 0.05 | yadI | 5 | 17.1333 | 0.72 |
| 0 | 0.05 | AG1 | 5 | 17.1333 | 0.583 |
| 0 | 0.05 | ptsI | 5 | 17.3833 | 0.625 |
| 0 | 0.05 | clcB | 5 | 17.3833 | 0.308 |
| 0 | 0.05 | ycaM | 5 | 17.3833 | 0.306 |
| 0 | 0.05 | yadI | 5 | 17.3833 | 0.718 |
| 0 | 0.05 | AG1 | 5 | 17.3833 | 0.583 |
| 0 | 0.05 | ptsI | 5 | 17.6333 | 0.629 |
| 0 | 0.05 | clcB | 5 | 17.6333 | 0.309 |
| 0 | 0.05 | ycaM | 5 | 17.6333 | 0.308 |
| 0 | 0.05 | yadI | 5 | 17.6333 | 0.719 |
| 0 | 0.05 | AG1 | 5 | 17.6333 | 0.586 |
| 0 | 0.05 | ptsI | 5 | 17.8833 | 0.635 |
| 0 | 0.05 | clcB | 5 | 17.8833 | 0.31 |
| 0 | 0.05 | ycaM | 5 | 17.8833 | 0.302 |
| 0 | 0.05 | yadI | 5 | 17.8833 | 0.709 |
| 0 | 0.05 | AG1 | 5 | 17.8833 | 0.586 |
| 0 | 0.05 | ptsI | 5 | 18.1333 | 0.65 |
| 0 | 0.05 | clcB | 5 | 18.1333 | 0.31 |
| 0 | 0.05 | ycaM | 5 | 18.1333 | 0.31 |
| 0 | 0.05 | yadI | 5 | 18.1333 | 0.727 |
| 0 | 0.05 | AG1 | 5 | 18.1333 | 0.596 |
| 0 | 0.05 | ptsI | 5 | 18.3833 | 0.641 |
| 0 | 0.05 | clcB | 5 | 18.3833 | 0.313 |
| 0 | 0.05 | ycaM | 5 | 18.3833 | 0.309 |
| 0 | 0.05 | yadI | 5 | 18.3833 | 0.726 |
| 0 | 0.05 | AG1 | 5 | 18.3833 | 0.594 |
| 0 | 0.05 | ptsI | 5 | 18.6333 | 0.644 |
| 0 | 0.05 | clcB | 5 | 18.6333 | 0.315 |

| | | | | | |
|---|---|---|---|---|---|
| 0 | 0.05 | ycaM | 5 | 18.6333 | 0.307 |
| 0 | 0.05 | yadI | 5 | 18.6333 | 0.733 |
| 0 | 0.05 | AG1 | 5 | 18.6333 | 0.598 |
| 0 | 0.05 | ptsI | 5 | 18.8833 | 0.653 |
| 0 | 0.05 | clcB | 5 | 18.8833 | 0.307 |
| 0 | 0.05 | ycaM | 5 | 18.8833 | 0.313 |
| 0 | 0.05 | yadI | 5 | 18.8833 | 0.732 |
| 0 | 0.05 | AG1 | 5 | 18.8833 | 0.597 |
| 0 | 0.05 | ptsI | 5 | 19.1333 | 0.651 |
| 0 | 0.05 | clcB | 5 | 19.1333 | 0.317 |
| 0 | 0.05 | ycaM | 5 | 19.1333 | 0.31 |
| 0 | 0.05 | yadI | 5 | 19.1333 | 0.743 |
| 0 | 0.05 | AG1 | 5 | 19.1333 | 0.604 |
| 0 | 0.05 | ptsI | 5 | 19.3833 | 0.669 |
| 0 | 0.05 | clcB | 5 | 19.3833 | 0.316 |
| 0 | 0.05 | ycaM | 5 | 19.3833 | 0.318 |
| 0 | 0.05 | yadI | 5 | 19.3833 | 0.737 |
| 0 | 0.05 | AG1 | 5 | 19.3833 | 0.607 |
| 0 | 0.05 | ptsI | 5 | 19.6333 | 0.668 |
| 0 | 0.05 | clcB | 5 | 19.6333 | 0.309 |
| 0 | 0.05 | ycaM | 5 | 19.6333 | 0.319 |
| 0 | 0.05 | yadI | 5 | 19.6333 | 0.736 |
| 0 | 0.05 | AG1 | 5 | 19.6333 | 0.606 |
| 0 | 0.05 | ptsI | 5 | 19.8833 | 0.661 |
| 0 | 0.05 | clcB | 5 | 19.8833 | 0.323 |
| 0 | 0.05 | ycaM | 5 | 19.8833 | 0.319 |
| 0 | 0.05 | yadI | 5 | 19.8833 | 0.74 |
| 0 | 0.05 | AG1 | 5 | 19.8833 | 0.608 |
| 0 | 0.05 | ptsI | 5 | 20.1333 | 0.661 |
| 0 | 0.05 | clcB | 5 | 20.1333 | 0.326 |
| 0 | 0.05 | ycaM | 5 | 20.1333 | 0.324 |
| 0 | 0.05 | yadI | 5 | 20.1333 | 0.744 |
| 0 | 0.05 | AG1 | 5 | 20.1333 | 0.615 |
| 0 | 0.05 | ptsI | 5 | 20.3833 | 0.66 |
| 0 | 0.05 | clcB | 5 | 20.3833 | 0.327 |
| 0 | 0.05 | ycaM | 5 | 20.3833 | 0.327 |
| 0 | 0.05 | yadI | 5 | 20.3833 | 0.741 |
| 0 | 0.05 | AG1 | 5 | 20.3833 | 0.616 |
| 0 | 0.05 | ptsI | 5 | 20.6333 | 0.665 |
| 0 | 0.05 | clcB | 5 | 20.6333 | 0.33 |
| 0 | 0.05 | ycaM | 5 | 20.6333 | 0.327 |
| 0 | 0.05 | yadI | 5 | 20.6333 | 0.727 |
| 0 | 0.05 | AG1 | 5 | 20.6333 | 0.613 |
| 0 | 0.05 | ptsI | 5 | 20.8833 | 0.675 |
| 0 | 0.05 | clcB | 5 | 20.8833 | 0.328 |
| 0 | 0.05 | ycaM | 5 | 20.8833 | 0.338 |
| 0 | 0.05 | yadI | 5 | 20.8833 | 0.744 |
| 0 | 0.05 | AG1 | 5 | 20.8833 | 0.618 |

| | | | | | |
|---|---|---|---|---|---|
| 0 | 0.05 | ptsI | 5 | 21.1333 | 0.668 |
| 0 | 0.05 | clcB | 5 | 21.1333 | 0.332 |
| 0 | 0.05 | ycaM | 5 | 21.1333 | 0.336 |
| 0 | 0.05 | yadI | 5 | 21.1333 | 0.751 |
| 0 | 0.05 | AG1 | 5 | 21.1333 | 0.621 |
| 0 | 0.05 | ptsI | 5 | 21.3833 | 0.677 |
| 0 | 0.05 | clcB | 5 | 21.3833 | 0.335 |
| 0 | 0.05 | ycaM | 5 | 21.3833 | 0.343 |
| 0 | 0.05 | yadI | 5 | 21.3833 | 0.748 |
| 0 | 0.05 | AG1 | 5 | 21.3833 | 0.618 |
| 0 | 0.05 | ptsI | 5 | 21.6333 | 0.671 |
| 0 | 0.05 | clcB | 5 | 21.6333 | 0.325 |
| 0 | 0.05 | ycaM | 5 | 21.6333 | 0.348 |
| 0 | 0.05 | yadI | 5 | 21.6333 | 0.749 |
| 0 | 0.05 | AG1 | 5 | 21.6333 | 0.624 |
| 0 | 0.05 | ptsI | 5 | 21.8833 | 0.68 |
| 0 | 0.05 | clcB | 5 | 21.8833 | 0.341 |
| 0 | 0.05 | ycaM | 5 | 21.8833 | 0.347 |
| 0 | 0.05 | yadI | 5 | 21.8833 | 0.75 |
| 0 | 0.05 | AG1 | 5 | 21.8833 | 0.629 |
| 0 | 0.05 | ptsI | 5 | 22.1333 | 0.668 |
| 0 | 0.05 | clcB | 5 | 22.1333 | 0.337 |
| 0 | 0.05 | ycaM | 5 | 22.1333 | 0.34 |
| 0 | 0.05 | yadI | 5 | 22.1333 | 0.731 |
| 0 | 0.05 | AG1 | 5 | 22.1333 | 0.615 |
| 0 | 0.05 | ptsI | 5 | 22.3833 | 0.674 |
| 0 | 0.05 | clcB | 5 | 22.3833 | 0.34 |
| 0 | 0.05 | ycaM | 5 | 22.3833 | 0.353 |
| 0 | 0.05 | yadI | 5 | 22.3833 | 0.75 |
| 0 | 0.05 | AG1 | 5 | 22.3833 | 0.622 |
| 0 | 0.05 | ptsI | 5 | 22.6333 | 0.664 |
| 0 | 0.05 | clcB | 5 | 22.6333 | 0.332 |
| 0 | 0.05 | ycaM | 5 | 22.6333 | 0.361 |
| 0 | 0.05 | yadI | 5 | 22.6333 | 0.753 |
| 0 | 0.05 | AG1 | 5 | 22.6333 | 0.63 |
| 0 | 0.05 | ptsI | 5 | 22.8833 | 0.671 |
| 0 | 0.05 | clcB | 5 | 22.8833 | 0.344 |
| 0 | 0.05 | ycaM | 5 | 22.8833 | 0.366 |
| 0 | 0.05 | yadI | 5 | 22.8833 | 0.746 |
| 0 | 0.05 | AG1 | 5 | 22.8833 | 0.631 |
| 0 | 0.05 | ptsI | 5 | 23.1333 | 0.659 |
| 0 | 0.05 | clcB | 5 | 23.1333 | 0.336 |
| 0 | 0.05 | ycaM | 5 | 23.1333 | 0.367 |
| 0 | 0.05 | yadI | 5 | 23.1333 | 0.754 |
| 0 | 0.05 | AG1 | 5 | 23.1333 | 0.635 |
| 0 | 0.05 | ptsI | 5 | 23.3833 | 0.682 |
| 0 | 0.05 | clcB | 5 | 23.3833 | 0.361 |
| 0 | 0.05 | ycaM | 5 | 23.3833 | 0.364 |

| | | | | | |
|---|---|---|---|---|---|
| 0 | 0.05 | yadI | 5 | 23.3833 | 0.751 |
| 0 | 0.05 | AG1 | 5 | 23.3833 | 0.63 |
| 0 | 0.05 | ptsI | 5 | 23.6333 | 0.677 |
| 0 | 0.05 | clcB | 5 | 23.6333 | 0.367 |
| 0 | 0.05 | ycaM | 5 | 23.6333 | 0.37 |
| 0 | 0.05 | yadI | 5 | 23.6333 | 0.751 |
| 0 | 0.05 | AG1 | 5 | 23.6333 | 0.632 |
| 0 | 0.05 | ptsI | 5 | 23.8833 | 0.684 |
| 0 | 0.05 | clcB | 5 | 23.8833 | 0.373 |
| 0 | 0.05 | ycaM | 5 | 23.8833 | 0.363 |
| 0 | 0.05 | yadI | 5 | 23.8833 | 0.732 |
| 0 | 0.05 | AG1 | 5 | 23.8833 | 0.627 |
| 0 | 0.05 | ptsI | 5 | 24.1333 | 0.667 |
| 0 | 0.05 | clcB | 5 | 24.1333 | 0.364 |
| 0 | 0.05 | ycaM | 5 | 24.1333 | 0.391 |
| 0 | 0.05 | yadI | 5 | 24.1333 | 0.752 |
| 0 | 0.05 | AG1 | 5 | 24.1333 | 0.637 |
| 0 | 0.05 | ptsI | 5 | 24.3833 | 0.676 |
| 0 | 0.05 | clcB | 5 | 24.3833 | 0.373 |
| 0 | 0.05 | ycaM | 5 | 24.3833 | 0.372 |
| 0 | 0.05 | yadI | 5 | 24.3833 | 0.75 |
| 0 | 0.05 | AG1 | 5 | 24.3833 | 0.63 |
| 0.05 | 0.05 | ptsI | 2 | 0 | 0.222 |
| 0.05 | 0.05 | clcB | 2 | 0 | 0.207 |
| 0.05 | 0.05 | ycaM | 2 | 0 | 0.214 |
| 0.05 | 0.05 | yadI | 2 | 0 | 0.213 |
| 0.05 | 0.05 | AG1 | 2 | 0 | 0.225 |
| 0.05 | 0.05 | ptsI | 2 | 0.25 | 0.219 |
| 0.05 | 0.05 | clcB | 2 | 0.25 | 0.201 |
| 0.05 | 0.05 | ycaM | 2 | 0.25 | 0.209 |
| 0.05 | 0.05 | yadI | 2 | 0.25 | 0.206 |
| 0.05 | 0.05 | AG1 | 2 | 0.25 | 0.216 |
| 0.05 | 0.05 | ptsI | 2 | 0.5 | 0.219 |
| 0.05 | 0.05 | clcB | 2 | 0.5 | 0.2 |
| 0.05 | 0.05 | ycaM | 2 | 0.5 | 0.208 |
| 0.05 | 0.05 | yadI | 2 | 0.5 | 0.205 |
| 0.05 | 0.05 | AG1 | 2 | 0.5 | 0.214 |
| 0.05 | 0.05 | ptsI | 2 | 0.75 | 0.222 |
| 0.05 | 0.05 | clcB | 2 | 0.75 | 0.202 |
| 0.05 | 0.05 | ycaM | 2 | 0.75 | 0.213 |
| 0.05 | 0.05 | yadI | 2 | 0.75 | 0.209 |
| 0.05 | 0.05 | AG1 | 2 | 0.75 | 0.216 |
| 0.05 | 0.05 | ptsI | 2 | 1 | 0.222 |
| 0.05 | 0.05 | clcB | 2 | 1 | 0.199 |
| 0.05 | 0.05 | ycaM | 2 | 1 | 0.216 |
| 0.05 | 0.05 | yadI | 2 | 1 | 0.213 |
| 0.05 | 0.05 | AG1 | 2 | 1 | 0.223 |
| 0.05 | 0.05 | ptsI | 2 | 1.25 | 0.225 |

| | | | | | |
|---|---|---|---|---|---|
| 0.05 | 0.05 | clcB | 2 | 1.25 | 0.202 |
| 0.05 | 0.05 | ycaM | 2 | 1.25 | 0.22 |
| 0.05 | 0.05 | yadI | 2 | 1.25 | 0.221 |
| 0.05 | 0.05 | AG1 | 2 | 1.25 | 0.226 |
| 0.05 | 0.05 | ptsI | 2 | 1.5 | 0.225 |
| 0.05 | 0.05 | clcB | 2 | 1.5 | 0.204 |
| 0.05 | 0.05 | ycaM | 2 | 1.5 | 0.223 |
| 0.05 | 0.05 | yadI | 2 | 1.5 | 0.225 |
| 0.05 | 0.05 | AG1 | 2 | 1.5 | 0.227 |
| 0.05 | 0.05 | ptsI | 2 | 1.75 | 0.23 |
| 0.05 | 0.05 | clcB | 2 | 1.75 | 0.204 |
| 0.05 | 0.05 | ycaM | 2 | 1.75 | 0.228 |
| 0.05 | 0.05 | yadI | 2 | 1.75 | 0.232 |
| 0.05 | 0.05 | AG1 | 2 | 1.75 | 0.232 |
| 0.05 | 0.05 | ptsI | 2 | 2 | 0.234 |
| 0.05 | 0.05 | clcB | 2 | 2 | 0.206 |
| 0.05 | 0.05 | ycaM | 2 | 2 | 0.234 |
| 0.05 | 0.05 | yadI | 2 | 2 | 0.237 |
| 0.05 | 0.05 | AG1 | 2 | 2 | 0.242 |
| 0.05 | 0.05 | ptsI | 2 | 2.25 | 0.24 |
| 0.05 | 0.05 | clcB | 2 | 2.25 | 0.209 |
| 0.05 | 0.05 | ycaM | 2 | 2.25 | 0.238 |
| 0.05 | 0.05 | yadI | 2 | 2.25 | 0.245 |
| 0.05 | 0.05 | AG1 | 2 | 2.25 | 0.249 |
| 0.05 | 0.05 | ptsI | 2 | 2.5 | 0.247 |
| 0.05 | 0.05 | clcB | 2 | 2.5 | 0.214 |
| 0.05 | 0.05 | ycaM | 2 | 2.5 | 0.248 |
| 0.05 | 0.05 | yadI | 2 | 2.5 | 0.263 |
| 0.05 | 0.05 | AG1 | 2 | 2.5 | 0.261 |
| 0.05 | 0.05 | ptsI | 2 | 2.75 | 0.251 |
| 0.05 | 0.05 | clcB | 2 | 2.75 | 0.215 |
| 0.05 | 0.05 | ycaM | 2 | 2.75 | 0.253 |
| 0.05 | 0.05 | yadI | 2 | 2.75 | 0.269 |
| 0.05 | 0.05 | AG1 | 2 | 2.75 | 0.267 |
| 0.05 | 0.05 | ptsI | 2 | 3 | 0.257 |
| 0.05 | 0.05 | clcB | 2 | 3 | 0.217 |
| 0.05 | 0.05 | ycaM | 2 | 3 | 0.256 |
| 0.05 | 0.05 | yadI | 2 | 3 | 0.278 |
| 0.05 | 0.05 | AG1 | 2 | 3 | 0.265 |
| 0.05 | 0.05 | ptsI | 2 | 3.25 | 0.267 |
| 0.05 | 0.05 | clcB | 2 | 3.25 | 0.225 |
| 0.05 | 0.05 | ycaM | 2 | 3.25 | 0.266 |
| 0.05 | 0.05 | yadI | 2 | 3.25 | 0.292 |
| 0.05 | 0.05 | AG1 | 2 | 3.25 | 0.282 |
| 0.05 | 0.05 | ptsI | 2 | 3.5 | 0.271 |
| 0.05 | 0.05 | clcB | 2 | 3.5 | 0.23 |
| 0.05 | 0.05 | ycaM | 2 | 3.5 | 0.273 |
| 0.05 | 0.05 | yadI | 2 | 3.5 | 0.304 |

| | | | | | |
|---|---|---|---|---|---|
| 0.05 | 0.05 | AG1 | 2 | 3.5 | 0.287 |
| 0.05 | 0.05 | ptsI | 2 | 3.75 | 0.279 |
| 0.05 | 0.05 | clcB | 2 | 3.75 | 0.231 |
| 0.05 | 0.05 | ycaM | 2 | 3.75 | 0.282 |
| 0.05 | 0.05 | yadI | 2 | 3.75 | 0.315 |
| 0.05 | 0.05 | AG1 | 2 | 3.75 | 0.3 |
| 0.05 | 0.05 | ptsI | 2 | 4 | 0.286 |
| 0.05 | 0.05 | clcB | 2 | 4 | 0.252 |
| 0.05 | 0.05 | ycaM | 2 | 4 | 0.293 |
| 0.05 | 0.05 | yadI | 2 | 4 | 0.328 |
| 0.05 | 0.05 | AG1 | 2 | 4 | 0.315 |
| 0.05 | 0.05 | ptsI | 2 | 4.38333 | 0.292 |
| 0.05 | 0.05 | clcB | 2 | 4.38333 | 0.254 |
| 0.05 | 0.05 | ycaM | 2 | 4.38333 | 0.294 |
| 0.05 | 0.05 | yadI | 2 | 4.38333 | 0.38 |
| 0.05 | 0.05 | AG1 | 2 | 4.38333 | 0.287 |
| 0.05 | 0.05 | ptsI | 2 | 4.63333 | 0.287 |
| 0.05 | 0.05 | clcB | 2 | 4.63333 | 0.248 |
| 0.05 | 0.05 | ycaM | 2 | 4.63333 | 0.29 |
| 0.05 | 0.05 | yadI | 2 | 4.63333 | 0.389 |
| 0.05 | 0.05 | AG1 | 2 | 4.63333 | 0.289 |
| 0.05 | 0.05 | ptsI | 2 | 4.88333 | 0.285 |
| 0.05 | 0.05 | clcB | 2 | 4.88333 | 0.251 |
| 0.05 | 0.05 | ycaM | 2 | 4.88333 | 0.299 |
| 0.05 | 0.05 | yadI | 2 | 4.88333 | 0.404 |
| 0.05 | 0.05 | AG1 | 2 | 4.88333 | 0.299 |
| 0.05 | 0.05 | ptsI | 2 | 5.13333 | 0.287 |
| 0.05 | 0.05 | clcB | 2 | 5.13333 | 0.255 |
| 0.05 | 0.05 | ycaM | 2 | 5.13333 | 0.298 |
| 0.05 | 0.05 | yadI | 2 | 5.13333 | 0.417 |
| 0.05 | 0.05 | AG1 | 2 | 5.13333 | 0.306 |
| 0.05 | 0.05 | ptsI | 2 | 5.38333 | 0.29 |
| 0.05 | 0.05 | clcB | 2 | 5.38333 | 0.261 |
| 0.05 | 0.05 | ycaM | 2 | 5.38333 | 0.3 |
| 0.05 | 0.05 | yadI | 2 | 5.38333 | 0.434 |
| 0.05 | 0.05 | AG1 | 2 | 5.38333 | 0.318 |
| 0.05 | 0.05 | ptsI | 2 | 5.63333 | 0.296 |
| 0.05 | 0.05 | clcB | 2 | 5.63333 | 0.269 |
| 0.05 | 0.05 | ycaM | 2 | 5.63333 | 0.301 |
| 0.05 | 0.05 | yadI | 2 | 5.63333 | 0.444 |
| 0.05 | 0.05 | AG1 | 2 | 5.63333 | 0.324 |
| 0.05 | 0.05 | ptsI | 2 | 5.88333 | 0.301 |
| 0.05 | 0.05 | clcB | 2 | 5.88333 | 0.276 |
| 0.05 | 0.05 | ycaM | 2 | 5.88333 | 0.303 |
| 0.05 | 0.05 | yadI | 2 | 5.88333 | 0.455 |
| 0.05 | 0.05 | AG1 | 2 | 5.88333 | 0.333 |
| 0.05 | 0.05 | ptsI | 2 | 6.13333 | 0.305 |
| 0.05 | 0.05 | clcB | 2 | 6.13333 | 0.283 |

| | | | | | |
|---|---|---|---|---|---|
| 0.05 | 0.05 | ycaM | 2 | 6.13333 | 0.302 |
| 0.05 | 0.05 | yadI | 2 | 6.13333 | 0.465 |
| 0.05 | 0.05 | AG1 | 2 | 6.13333 | 0.341 |
| 0.05 | 0.05 | ptsI | 2 | 6.38333 | 0.316 |
| 0.05 | 0.05 | clcB | 2 | 6.38333 | 0.286 |
| 0.05 | 0.05 | ycaM | 2 | 6.38333 | 0.305 |
| 0.05 | 0.05 | yadI | 2 | 6.38333 | 0.47 |
| 0.05 | 0.05 | AG1 | 2 | 6.38333 | 0.348 |
| 0.05 | 0.05 | ptsI | 2 | 6.63333 | 0.319 |
| 0.05 | 0.05 | clcB | 2 | 6.63333 | 0.294 |
| 0.05 | 0.05 | ycaM | 2 | 6.63333 | 0.308 |
| 0.05 | 0.05 | yadI | 2 | 6.63333 | 0.48 |
| 0.05 | 0.05 | AG1 | 2 | 6.63333 | 0.358 |
| 0.05 | 0.05 | ptsI | 2 | 6.88333 | 0.328 |
| 0.05 | 0.05 | clcB | 2 | 6.88333 | 0.299 |
| 0.05 | 0.05 | ycaM | 2 | 6.88333 | 0.311 |
| 0.05 | 0.05 | yadI | 2 | 6.88333 | 0.492 |
| 0.05 | 0.05 | AG1 | 2 | 6.88333 | 0.369 |
| 0.05 | 0.05 | ptsI | 2 | 7.13333 | 0.332 |
| 0.05 | 0.05 | clcB | 2 | 7.13333 | 0.301 |
| 0.05 | 0.05 | ycaM | 2 | 7.13333 | 0.307 |
| 0.05 | 0.05 | yadI | 2 | 7.13333 | 0.494 |
| 0.05 | 0.05 | AG1 | 2 | 7.13333 | 0.374 |
| 0.05 | 0.05 | ptsI | 2 | 7.38333 | 0.342 |
| 0.05 | 0.05 | clcB | 2 | 7.38333 | 0.308 |
| 0.05 | 0.05 | ycaM | 2 | 7.38333 | 0.308 |
| 0.05 | 0.05 | yadI | 2 | 7.38333 | 0.503 |
| 0.05 | 0.05 | AG1 | 2 | 7.38333 | 0.382 |
| 0.05 | 0.05 | ptsI | 2 | 7.63333 | 0.347 |
| 0.05 | 0.05 | clcB | 2 | 7.63333 | 0.311 |
| 0.05 | 0.05 | ycaM | 2 | 7.63333 | 0.31 |
| 0.05 | 0.05 | yadI | 2 | 7.63333 | 0.513 |
| 0.05 | 0.05 | AG1 | 2 | 7.63333 | 0.393 |
| 0.05 | 0.05 | ptsI | 2 | 7.88333 | 0.354 |
| 0.05 | 0.05 | clcB | 2 | 7.88333 | 0.315 |
| 0.05 | 0.05 | ycaM | 2 | 7.88333 | 0.311 |
| 0.05 | 0.05 | yadI | 2 | 7.88333 | 0.529 |
| 0.05 | 0.05 | AG1 | 2 | 7.88333 | 0.399 |
| 0.05 | 0.05 | ptsI | 2 | 8.13333 | 0.362 |
| 0.05 | 0.05 | clcB | 2 | 8.13333 | 0.317 |
| 0.05 | 0.05 | ycaM | 2 | 8.13333 | 0.311 |
| 0.05 | 0.05 | yadI | 2 | 8.13333 | 0.528 |
| 0.05 | 0.05 | AG1 | 2 | 8.13333 | 0.41 |
| 0.05 | 0.05 | ptsI | 2 | 8.38333 | 0.372 |
| 0.05 | 0.05 | clcB | 2 | 8.38333 | 0.322 |
| 0.05 | 0.05 | ycaM | 2 | 8.38333 | 0.311 |
| 0.05 | 0.05 | yadI | 2 | 8.38333 | 0.538 |
| 0.05 | 0.05 | AG1 | 2 | 8.38333 | 0.417 |

| | | | | | |
|---|---|---|---|---|---|
| 0.05 | 0.05 | ptsI | 2 | 8.91667 | 0.369 |
| 0.05 | 0.05 | clcB | 2 | 8.91667 | 0.293 |
| 0.05 | 0.05 | ycaM | 2 | 8.91667 | 0.276 |
| 0.05 | 0.05 | yadI | 2 | 8.91667 | 0.563 |
| 0.05 | 0.05 | AG1 | 2 | 8.91667 | 0.46 |
| 0.05 | 0.05 | ptsI | 2 | 9.16667 | 0.374 |
| 0.05 | 0.05 | clcB | 2 | 9.16667 | 0.286 |
| 0.05 | 0.05 | ycaM | 2 | 9.16667 | 0.275 |
| 0.05 | 0.05 | yadI | 2 | 9.16667 | 0.574 |
| 0.05 | 0.05 | AG1 | 2 | 9.16667 | 0.466 |
| 0.05 | 0.05 | ptsI | 2 | 9.41667 | 0.381 |
| 0.05 | 0.05 | clcB | 2 | 9.41667 | 0.298 |
| 0.05 | 0.05 | ycaM | 2 | 9.41667 | 0.279 |
| 0.05 | 0.05 | yadI | 2 | 9.41667 | 0.59 |
| 0.05 | 0.05 | AG1 | 2 | 9.41667 | 0.48 |
| 0.05 | 0.05 | ptsI | 2 | 9.66667 | 0.389 |
| 0.05 | 0.05 | clcB | 2 | 9.66667 | 0.303 |
| 0.05 | 0.05 | ycaM | 2 | 9.66667 | 0.283 |
| 0.05 | 0.05 | yadI | 2 | 9.66667 | 0.605 |
| 0.05 | 0.05 | AG1 | 2 | 9.66667 | 0.494 |
| 0.05 | 0.05 | ptsI | 2 | 9.91667 | 0.394 |
| 0.05 | 0.05 | clcB | 2 | 9.91667 | 0.309 |
| 0.05 | 0.05 | ycaM | 2 | 9.91667 | 0.282 |
| 0.05 | 0.05 | yadI | 2 | 9.91667 | 0.619 |
| 0.05 | 0.05 | AG1 | 2 | 9.91667 | 0.51 |
| 0.05 | 0.05 | ptsI | 2 | 10.1667 | 0.401 |
| 0.05 | 0.05 | clcB | 2 | 10.1667 | 0.314 |
| 0.05 | 0.05 | ycaM | 2 | 10.1667 | 0.285 |
| 0.05 | 0.05 | yadI | 2 | 10.1667 | 0.631 |
| 0.05 | 0.05 | AG1 | 2 | 10.1667 | 0.517 |
| 0.05 | 0.05 | ptsI | 2 | 10.4167 | 0.411 |
| 0.05 | 0.05 | clcB | 2 | 10.4167 | 0.315 |
| 0.05 | 0.05 | ycaM | 2 | 10.4167 | 0.286 |
| 0.05 | 0.05 | yadI | 2 | 10.4167 | 0.644 |
| 0.05 | 0.05 | AG1 | 2 | 10.4167 | 0.529 |
| 0.05 | 0.05 | ptsI | 2 | 10.6667 | 0.414 |
| 0.05 | 0.05 | clcB | 2 | 10.6667 | 0.324 |
| 0.05 | 0.05 | ycaM | 2 | 10.6667 | 0.288 |
| 0.05 | 0.05 | yadI | 2 | 10.6667 | 0.656 |
| 0.05 | 0.05 | AG1 | 2 | 10.6667 | 0.54 |
| 0.05 | 0.05 | ptsI | 2 | 10.9167 | 0.423 |
| 0.05 | 0.05 | clcB | 2 | 10.9167 | 0.325 |
| 0.05 | 0.05 | ycaM | 2 | 10.9167 | 0.286 |
| 0.05 | 0.05 | yadI | 2 | 10.9167 | 0.664 |
| 0.05 | 0.05 | AG1 | 2 | 10.9167 | 0.546 |
| 0.05 | 0.05 | ptsI | 2 | 11.1667 | 0.433 |
| 0.05 | 0.05 | clcB | 2 | 11.1667 | 0.346 |
| 0.05 | 0.05 | ycaM | 2 | 11.1667 | 0.29 |

| | | | | | |
|---|---|---|---|---|---|
| 0.05 | 0.05 | yadI | 2 | 11.1667 | 0.673 |
| 0.05 | 0.05 | AG1 | 2 | 11.1667 | 0.564 |
| 0.05 | 0.05 | ptsI | 2 | 11.4167 | 0.439 |
| 0.05 | 0.05 | clcB | 2 | 11.4167 | 0.334 |
| 0.05 | 0.05 | ycaM | 2 | 11.4167 | 0.286 |
| 0.05 | 0.05 | yadI | 2 | 11.4167 | 0.682 |
| 0.05 | 0.05 | AG1 | 2 | 11.4167 | 0.588 |
| 0.05 | 0.05 | ptsI | 2 | 11.6667 | 0.446 |
| 0.05 | 0.05 | clcB | 2 | 11.6667 | 0.327 |
| 0.05 | 0.05 | ycaM | 2 | 11.6667 | 0.287 |
| 0.05 | 0.05 | yadI | 2 | 11.6667 | 0.693 |
| 0.05 | 0.05 | AG1 | 2 | 11.6667 | 0.622 |
| 0.05 | 0.05 | ptsI | 2 | 11.9167 | 0.452 |
| 0.05 | 0.05 | clcB | 2 | 11.9167 | 0.336 |
| 0.05 | 0.05 | ycaM | 2 | 11.9167 | 0.29 |
| 0.05 | 0.05 | yadI | 2 | 11.9167 | 0.699 |
| 0.05 | 0.05 | AG1 | 2 | 11.9167 | 0.617 |
| 0.05 | 0.05 | ptsI | 2 | 12.1667 | 0.459 |
| 0.05 | 0.05 | clcB | 2 | 12.1667 | 0.33 |
| 0.05 | 0.05 | ycaM | 2 | 12.1667 | 0.289 |
| 0.05 | 0.05 | yadI | 2 | 12.1667 | 0.71 |
| 0.05 | 0.05 | AG1 | 2 | 12.1667 | 0.656 |
| 0.05 | 0.05 | ptsI | 2 | 12.4167 | 0.47 |
| 0.05 | 0.05 | clcB | 2 | 12.4167 | 0.33 |
| 0.05 | 0.05 | ycaM | 2 | 12.4167 | 0.286 |
| 0.05 | 0.05 | yadI | 2 | 12.4167 | 0.72 |
| 0.05 | 0.05 | AG1 | 2 | 12.4167 | 0.664 |
| 0.05 | 0.05 | ptsI | 2 | 12.6667 | 0.471 |
| 0.05 | 0.05 | clcB | 2 | 12.6667 | 0.329 |
| 0.05 | 0.05 | ycaM | 2 | 12.6667 | 0.286 |
| 0.05 | 0.05 | yadI | 2 | 12.6667 | 0.723 |
| 0.05 | 0.05 | AG1 | 2 | 12.6667 | 0.672 |
| 0.05 | 0.05 | ptsI | 2 | 12.9167 | 0.482 |
| 0.05 | 0.05 | clcB | 2 | 12.9167 | 0.333 |
| 0.05 | 0.05 | ycaM | 2 | 12.9167 | 0.29 |
| 0.05 | 0.05 | yadI | 2 | 12.9167 | 0.738 |
| 0.05 | 0.05 | AG1 | 2 | 12.9167 | 0.662 |
| 0.05 | 0.05 | ptsI | 2 | 13.1667 | 0.481 |
| 0.05 | 0.05 | clcB | 2 | 13.1667 | 0.334 |
| 0.05 | 0.05 | ycaM | 2 | 13.1667 | 0.291 |
| 0.05 | 0.05 | yadI | 2 | 13.1667 | 0.749 |
| 0.05 | 0.05 | AG1 | 2 | 13.1667 | 0.662 |
| 0.05 | 0.05 | ptsI | 2 | 13.4167 | 0.485 |
| 0.05 | 0.05 | clcB | 2 | 13.4167 | 0.332 |
| 0.05 | 0.05 | ycaM | 2 | 13.4167 | 0.289 |
| 0.05 | 0.05 | yadI | 2 | 13.4167 | 0.752 |
| 0.05 | 0.05 | AG1 | 2 | 13.4167 | 0.659 |
| 0.05 | 0.05 | ptsI | 2 | 13.6667 | 0.491 |

| | | | | | |
|---|---|---|---|---|---|
| 0.05 | 0.05 | clcB | 2 | 13.6667 | 0.338 |
| 0.05 | 0.05 | ycaM | 2 | 13.6667 | 0.289 |
| 0.05 | 0.05 | yadI | 2 | 13.6667 | 0.762 |
| 0.05 | 0.05 | AG1 | 2 | 13.6667 | 0.684 |
| 0.05 | 0.05 | ptsI | 2 | 13.9167 | 0.493 |
| 0.05 | 0.05 | clcB | 2 | 13.9167 | 0.331 |
| 0.05 | 0.05 | ycaM | 2 | 13.9167 | 0.289 |
| 0.05 | 0.05 | yadI | 2 | 13.9167 | 0.77 |
| 0.05 | 0.05 | AG1 | 2 | 13.9167 | 0.663 |
| 0.05 | 0.05 | ptsI | 2 | 14.1667 | 0.502 |
| 0.05 | 0.05 | clcB | 2 | 14.1667 | 0.339 |
| 0.05 | 0.05 | ycaM | 2 | 14.1667 | 0.29 |
| 0.05 | 0.05 | yadI | 2 | 14.1667 | 0.776 |
| 0.05 | 0.05 | AG1 | 2 | 14.1667 | 0.705 |
| 0.05 | 0.05 | ptsI | 2 | 14.4167 | 0.504 |
| 0.05 | 0.05 | clcB | 2 | 14.4167 | 0.332 |
| 0.05 | 0.05 | ycaM | 2 | 14.4167 | 0.286 |
| 0.05 | 0.05 | yadI | 2 | 14.4167 | 0.78 |
| 0.05 | 0.05 | AG1 | 2 | 14.4167 | 0.674 |
| 0.05 | 0.05 | ptsI | 2 | 14.6667 | 0.502 |
| 0.05 | 0.05 | clcB | 2 | 14.6667 | 0.334 |
| 0.05 | 0.05 | ycaM | 2 | 14.6667 | 0.284 |
| 0.05 | 0.05 | yadI | 2 | 14.6667 | 0.782 |
| 0.05 | 0.05 | AG1 | 2 | 14.6667 | 0.684 |
| 0.05 | 0.05 | ptsI | 2 | 14.9167 | 0.511 |
| 0.05 | 0.05 | clcB | 2 | 14.9167 | 0.338 |
| 0.05 | 0.05 | ycaM | 2 | 14.9167 | 0.285 |
| 0.05 | 0.05 | yadI | 2 | 14.9167 | 0.788 |
| 0.05 | 0.05 | AG1 | 2 | 14.9167 | 0.692 |
| 0.05 | 0.05 | ptsI | 2 | 15.1667 | 0.515 |
| 0.05 | 0.05 | clcB | 2 | 15.1667 | 0.333 |
| 0.05 | 0.05 | ycaM | 2 | 15.1667 | 0.286 |
| 0.05 | 0.05 | yadI | 2 | 15.1667 | 0.801 |
| 0.05 | 0.05 | AG1 | 2 | 15.1667 | 0.709 |
| 0.05 | 0.05 | ptsI | 2 | 15.4167 | 0.525 |
| 0.05 | 0.05 | clcB | 2 | 15.4167 | 0.332 |
| 0.05 | 0.05 | ycaM | 2 | 15.4167 | 0.282 |
| 0.05 | 0.05 | yadI | 2 | 15.4167 | 0.803 |
| 0.05 | 0.05 | AG1 | 2 | 15.4167 | 0.701 |
| 0.05 | 0.05 | ptsI | 2 | 15.6667 | 0.521 |
| 0.05 | 0.05 | clcB | 2 | 15.6667 | 0.334 |
| 0.05 | 0.05 | ycaM | 2 | 15.6667 | 0.282 |
| 0.05 | 0.05 | yadI | 2 | 15.6667 | 0.799 |
| 0.05 | 0.05 | AG1 | 2 | 15.6667 | 0.702 |
| 0.05 | 0.05 | ptsI | 2 | 15.9167 | 0.519 |
| 0.05 | 0.05 | clcB | 2 | 15.9167 | 0.334 |
| 0.05 | 0.05 | ycaM | 2 | 15.9167 | 0.281 |
| 0.05 | 0.05 | yadI | 2 | 15.9167 | 0.801 |

| | | | | | |
|---|---|---|---|---|---|
| 0.05 | 0.05 | AG1 | 2 | 15.9167 | 0.708 |
| 0.05 | 0.05 | ptsI | 2 | 16.1667 | 0.529 |
| 0.05 | 0.05 | clcB | 2 | 16.1667 | 0.346 |
| 0.05 | 0.05 | ycaM | 2 | 16.1667 | 0.281 |
| 0.05 | 0.05 | yadI | 2 | 16.1667 | 0.821 |
| 0.05 | 0.05 | AG1 | 2 | 16.1667 | 0.727 |
| 0.05 | 0.05 | ptsI | 2 | 16.4167 | 0.526 |
| 0.05 | 0.05 | clcB | 2 | 16.4167 | 0.332 |
| 0.05 | 0.05 | ycaM | 2 | 16.4167 | 0.284 |
| 0.05 | 0.05 | yadI | 2 | 16.4167 | 0.818 |
| 0.05 | 0.05 | AG1 | 2 | 16.4167 | 0.719 |
| 0.05 | 0.05 | ptsI | 2 | 16.6667 | 0.523 |
| 0.05 | 0.05 | clcB | 2 | 16.6667 | 0.332 |
| 0.05 | 0.05 | ycaM | 2 | 16.6667 | 0.288 |
| 0.05 | 0.05 | yadI | 2 | 16.6667 | 0.822 |
| 0.05 | 0.05 | AG1 | 2 | 16.6667 | 0.717 |
| 0.05 | 0.05 | ptsI | 2 | 16.9167 | 0.521 |
| 0.05 | 0.05 | clcB | 2 | 16.9167 | 0.338 |
| 0.05 | 0.05 | ycaM | 2 | 16.9167 | 0.287 |
| 0.05 | 0.05 | yadI | 2 | 16.9167 | 0.826 |
| 0.05 | 0.05 | AG1 | 2 | 16.9167 | 0.732 |
| 0.05 | 0.05 | ptsI | 2 | 17.1667 | 0.53 |
| 0.05 | 0.05 | clcB | 2 | 17.1667 | 0.338 |
| 0.05 | 0.05 | ycaM | 2 | 17.1667 | 0.287 |
| 0.05 | 0.05 | yadI | 2 | 17.1667 | 0.839 |
| 0.05 | 0.05 | AG1 | 2 | 17.1667 | 0.754 |
| 0.05 | 0.05 | ptsI | 2 | 17.4167 | 0.533 |
| 0.05 | 0.05 | clcB | 2 | 17.4167 | 0.337 |
| 0.05 | 0.05 | ycaM | 2 | 17.4167 | 0.289 |
| 0.05 | 0.05 | yadI | 2 | 17.4167 | 0.843 |
| 0.05 | 0.05 | AG1 | 2 | 17.4167 | 0.764 |
| 0.05 | 0.05 | ptsI | 2 | 17.6667 | 0.537 |
| 0.05 | 0.05 | clcB | 2 | 17.6667 | 0.335 |
| 0.05 | 0.05 | ycaM | 2 | 17.6667 | 0.289 |
| 0.05 | 0.05 | yadI | 2 | 17.6667 | 0.845 |
| 0.05 | 0.05 | AG1 | 2 | 17.6667 | 0.774 |
| 0.05 | 0.05 | ptsI | 2 | 17.9167 | 0.53 |
| 0.05 | 0.05 | clcB | 2 | 17.9167 | 0.346 |
| 0.05 | 0.05 | ycaM | 2 | 17.9167 | 0.293 |
| 0.05 | 0.05 | yadI | 2 | 17.9167 | 0.844 |
| 0.05 | 0.05 | AG1 | 2 | 17.9167 | 0.763 |
| 0.05 | 0.05 | ptsI | 2 | 18.1667 | 0.54 |
| 0.05 | 0.05 | clcB | 2 | 18.1667 | 0.329 |
| 0.05 | 0.05 | ycaM | 2 | 18.1667 | 0.29 |
| 0.05 | 0.05 | yadI | 2 | 18.1667 | 0.848 |
| 0.05 | 0.05 | AG1 | 2 | 18.1667 | 0.775 |
| 0.05 | 0.05 | ptsI | 2 | 18.4167 | 0.542 |
| 0.05 | 0.05 | clcB | 2 | 18.4167 | 0.333 |

| | | | | | |
|---|---|---|---|---|---|
| 0.05 | 0.05 | ycaM | 2 | 18.4167 | 0.294 |
| 0.05 | 0.05 | yadI | 2 | 18.4167 | 0.856 |
| 0.05 | 0.05 | AG1 | 2 | 18.4167 | 0.775 |
| 0.05 | 0.05 | ptsI | 2 | 18.6667 | 0.535 |
| 0.05 | 0.05 | clcB | 2 | 18.6667 | 0.336 |
| 0.05 | 0.05 | ycaM | 2 | 18.6667 | 0.291 |
| 0.05 | 0.05 | yadI | 2 | 18.6667 | 0.853 |
| 0.05 | 0.05 | AG1 | 2 | 18.6667 | 0.776 |
| 0.05 | 0.05 | ptsI | 2 | 18.9167 | 0.521 |
| 0.05 | 0.05 | clcB | 2 | 18.9167 | 0.33 |
| 0.05 | 0.05 | ycaM | 2 | 18.9167 | 0.294 |
| 0.05 | 0.05 | yadI | 2 | 18.9167 | 0.844 |
| 0.05 | 0.05 | AG1 | 2 | 18.9167 | 0.778 |
| 0.05 | 0.05 | ptsI | 2 | 19.1667 | 0.542 |
| 0.05 | 0.05 | clcB | 2 | 19.1667 | 0.327 |
| 0.05 | 0.05 | ycaM | 2 | 19.1667 | 0.292 |
| 0.05 | 0.05 | yadI | 2 | 19.1667 | 0.857 |
| 0.05 | 0.05 | AG1 | 2 | 19.1667 | 0.794 |
| 0.05 | 0.05 | ptsI | 2 | 19.4167 | 0.534 |
| 0.05 | 0.05 | clcB | 2 | 19.4167 | 0.332 |
| 0.05 | 0.05 | ycaM | 2 | 19.4167 | 0.298 |
| 0.05 | 0.05 | yadI | 2 | 19.4167 | 0.848 |
| 0.05 | 0.05 | AG1 | 2 | 19.4167 | 0.793 |
| 0.05 | 0.05 | ptsI | 2 | 19.6667 | 0.538 |
| 0.05 | 0.05 | clcB | 2 | 19.6667 | 0.33 |
| 0.05 | 0.05 | ycaM | 2 | 19.6667 | 0.298 |
| 0.05 | 0.05 | yadI | 2 | 19.6667 | 0.863 |
| 0.05 | 0.05 | AG1 | 2 | 19.6667 | 0.796 |
| 0.05 | 0.05 | ptsI | 2 | 19.9167 | 0.539 |
| 0.05 | 0.05 | clcB | 2 | 19.9167 | 0.328 |
| 0.05 | 0.05 | ycaM | 2 | 19.9167 | 0.3 |
| 0.05 | 0.05 | yadI | 2 | 19.9167 | 0.864 |
| 0.05 | 0.05 | AG1 | 2 | 19.9167 | 0.819 |
| 0.05 | 0.05 | ptsI | 2 | 20.1667 | 0.539 |
| 0.05 | 0.05 | clcB | 2 | 20.1667 | 0.329 |
| 0.05 | 0.05 | ycaM | 2 | 20.1667 | 0.304 |
| 0.05 | 0.05 | yadI | 2 | 20.1667 | 0.865 |
| 0.05 | 0.05 | AG1 | 2 | 20.1667 | 0.806 |
| 0.05 | 0.05 | ptsI | 2 | 20.4167 | 0.541 |
| 0.05 | 0.05 | clcB | 2 | 20.4167 | 0.329 |
| 0.05 | 0.05 | ycaM | 2 | 20.4167 | 0.304 |
| 0.05 | 0.05 | yadI | 2 | 20.4167 | 0.864 |
| 0.05 | 0.05 | AG1 | 2 | 20.4167 | 0.811 |
| 0.05 | 0.05 | ptsI | 2 | 20.6667 | 0.536 |
| 0.05 | 0.05 | clcB | 2 | 20.6667 | 0.328 |
| 0.05 | 0.05 | ycaM | 2 | 20.6667 | 0.307 |
| 0.05 | 0.05 | yadI | 2 | 20.6667 | 0.861 |
| 0.05 | 0.05 | AG1 | 2 | 20.6667 | 0.811 |

| | | | | | |
|---|---|---|---|---|---|
| 0.05 | 0.05 | ptsI | 2 | 20.9167 | 0.536 |
| 0.05 | 0.05 | clcB | 2 | 20.9167 | 0.327 |
| 0.05 | 0.05 | ycaM | 2 | 20.9167 | 0.309 |
| 0.05 | 0.05 | yadI | 2 | 20.9167 | 0.864 |
| 0.05 | 0.05 | AG1 | 2 | 20.9167 | 0.813 |
| 0.05 | 0.05 | ptsI | 2 | 21.1667 | 0.534 |
| 0.05 | 0.05 | clcB | 2 | 21.1667 | 0.327 |
| 0.05 | 0.05 | ycaM | 2 | 21.1667 | 0.312 |
| 0.05 | 0.05 | yadI | 2 | 21.1667 | 0.877 |
| 0.05 | 0.05 | AG1 | 2 | 21.1667 | 0.819 |
| 0.05 | 0.05 | ptsI | 2 | 21.4167 | 0.527 |
| 0.05 | 0.05 | clcB | 2 | 21.4167 | 0.325 |
| 0.05 | 0.05 | ycaM | 2 | 21.4167 | 0.316 |
| 0.05 | 0.05 | yadI | 2 | 21.4167 | 0.87 |
| 0.05 | 0.05 | AG1 | 2 | 21.4167 | 0.832 |
| 0.05 | 0.05 | ptsI | 2 | 21.6667 | 0.538 |
| 0.05 | 0.05 | clcB | 2 | 21.6667 | 0.329 |
| 0.05 | 0.05 | ycaM | 2 | 21.6667 | 0.32 |
| 0.05 | 0.05 | yadI | 2 | 21.6667 | 0.874 |
| 0.05 | 0.05 | AG1 | 2 | 21.6667 | 0.835 |
| 0.05 | 0.05 | ptsI | 2 | 21.9167 | 0.538 |
| 0.05 | 0.05 | clcB | 2 | 21.9167 | 0.329 |
| 0.05 | 0.05 | ycaM | 2 | 21.9167 | 0.32 |
| 0.05 | 0.05 | yadI | 2 | 21.9167 | 0.882 |
| 0.05 | 0.05 | AG1 | 2 | 21.9167 | 0.838 |
| 0.05 | 0.05 | ptsI | 2 | 22.1667 | 0.539 |
| 0.05 | 0.05 | clcB | 2 | 22.1667 | 0.327 |
| 0.05 | 0.05 | ycaM | 2 | 22.1667 | 0.326 |
| 0.05 | 0.05 | yadI | 2 | 22.1667 | 0.882 |
| 0.05 | 0.05 | AG1 | 2 | 22.1667 | 0.84 |
| 0.05 | 0.05 | ptsI | 2 | 22.4167 | 0.535 |
| 0.05 | 0.05 | clcB | 2 | 22.4167 | 0.329 |
| 0.05 | 0.05 | ycaM | 2 | 22.4167 | 0.325 |
| 0.05 | 0.05 | yadI | 2 | 22.4167 | 0.884 |
| 0.05 | 0.05 | AG1 | 2 | 22.4167 | 0.847 |
| 0.05 | 0.05 | ptsI | 2 | 22.6667 | 0.534 |
| 0.05 | 0.05 | clcB | 2 | 22.6667 | 0.327 |
| 0.05 | 0.05 | ycaM | 2 | 22.6667 | 0.33 |
| 0.05 | 0.05 | yadI | 2 | 22.6667 | 0.887 |
| 0.05 | 0.05 | AG1 | 2 | 22.6667 | 0.85 |
| 0.05 | 0.05 | ptsI | 2 | 22.9167 | 0.543 |
| 0.05 | 0.05 | clcB | 2 | 22.9167 | 0.33 |
| 0.05 | 0.05 | ycaM | 2 | 22.9167 | 0.333 |
| 0.05 | 0.05 | yadI | 2 | 22.9167 | 0.885 |
| 0.05 | 0.05 | AG1 | 2 | 22.9167 | 0.859 |
| 0.05 | 0.05 | ptsI | 2 | 23.1667 | 0.537 |
| 0.05 | 0.05 | clcB | 2 | 23.1667 | 0.329 |
| 0.05 | 0.05 | ycaM | 2 | 23.1667 | 0.332 |

| | | | | | |
|---|---|---|---|---|---|
| 0.05 | 0.05 | yadI | 2 | 23.1667 | 0.882 |
| 0.05 | 0.05 | AG1 | 2 | 23.1667 | 0.861 |
| 0.05 | 0.05 | ptsI | 2 | 23.4167 | 0.537 |
| 0.05 | 0.05 | clcB | 2 | 23.4167 | 0.329 |
| 0.05 | 0.05 | ycaM | 2 | 23.4167 | 0.336 |
| 0.05 | 0.05 | yadI | 2 | 23.4167 | 0.891 |
| 0.05 | 0.05 | AG1 | 2 | 23.4167 | 0.865 |
| 0.05 | 0.05 | ptsI | 2 | 23.6667 | 0.537 |
| 0.05 | 0.05 | clcB | 2 | 23.6667 | 0.33 |
| 0.05 | 0.05 | ycaM | 2 | 23.6667 | 0.34 |
| 0.05 | 0.05 | yadI | 2 | 23.6667 | 0.89 |
| 0.05 | 0.05 | AG1 | 2 | 23.6667 | 0.87 |
| 0.05 | 0.05 | ptsI | 2 | 23.9167 | 0.539 |
| 0.05 | 0.05 | clcB | 2 | 23.9167 | 0.332 |
| 0.05 | 0.05 | ycaM | 2 | 23.9167 | 0.337 |
| 0.05 | 0.05 | yadI | 2 | 23.9167 | 0.897 |
| 0.05 | 0.05 | AG1 | 2 | 23.9167 | 0.878 |
| 0.05 | 0.05 | ptsI | 2 | 24.1667 | 0.538 |
| 0.05 | 0.05 | clcB | 2 | 24.1667 | 0.333 |
| 0.05 | 0.05 | ycaM | 2 | 24.1667 | 0.337 |
| 0.05 | 0.05 | yadI | 2 | 24.1667 | 0.89 |
| 0.05 | 0.05 | AG1 | 2 | 24.1667 | 0.882 |
| 0.05 | 0.05 | ptsI | 2 | 24.4167 | 0.54 |
| 0.05 | 0.05 | clcB | 2 | 24.4167 | 0.333 |
| 0.05 | 0.05 | ycaM | 2 | 24.4167 | 0.344 |
| 0.05 | 0.05 | yadI | 2 | 24.4167 | 0.89 |
| 0.05 | 0.05 | AG1 | 2 | 24.4167 | 0.89 |
| 0.05 | 0.05 | ptsI | 2 | 24.6667 | 0.543 |
| 0.05 | 0.05 | clcB | 2 | 24.6667 | 0.332 |
| 0.05 | 0.05 | ycaM | 2 | 24.6667 | 0.343 |
| 0.05 | 0.05 | yadI | 2 | 24.6667 | 0.889 |
| 0.05 | 0.05 | AG1 | 2 | 24.6667 | 0.898 |
| 0.05 | 0.05 | ptsI | 2 | 24.9167 | 0.541 |
| 0.05 | 0.05 | clcB | 2 | 24.9167 | 0.334 |
| 0.05 | 0.05 | ycaM | 2 | 24.9167 | 0.344 |
| 0.05 | 0.05 | yadI | 2 | 24.9167 | 0.89 |
| 0.05 | 0.05 | AG1 | 2 | 24.9167 | 0.91 |
| 0.05 | 0.05 | ptsI | 2 | 25.1667 | 0.542 |
| 0.05 | 0.05 | clcB | 2 | 25.1667 | 0.335 |
| 0.05 | 0.05 | ycaM | 2 | 25.1667 | 0.347 |
| 0.05 | 0.05 | yadI | 2 | 25.1667 | 0.9 |
| 0.05 | 0.05 | AG1 | 2 | 25.1667 | 0.913 |
| 0.05 | 0.05 | ptsI | 2 | 25.4167 | 0.546 |
| 0.05 | 0.05 | clcB | 2 | 25.4167 | 0.338 |
| 0.05 | 0.05 | ycaM | 2 | 25.4167 | 0.348 |
| 0.05 | 0.05 | yadI | 2 | 25.4167 | 0.896 |
| 0.05 | 0.05 | AG1 | 2 | 25.4167 | 0.924 |
| 0.05 | 0.05 | ptsI | 2 | 25.6667 | 0.54 |

| | | | | | |
|---|---|---|---|---|---|
| 0.05 | 0.05 | clcB | 2 | 25.6667 | 0.335 |
| 0.05 | 0.05 | ycaM | 2 | 25.6667 | 0.348 |
| 0.05 | 0.05 | yadI | 2 | 25.6667 | 0.898 |
| 0.05 | 0.05 | AG1 | 2 | 25.6667 | 0.93 |
| 0.05 | 0.05 | ptsI | 2 | 25.9167 | 0.542 |
| 0.05 | 0.05 | clcB | 2 | 25.9167 | 0.339 |
| 0.05 | 0.05 | ycaM | 2 | 25.9167 | 0.355 |
| 0.05 | 0.05 | yadI | 2 | 25.9167 | 0.905 |
| 0.05 | 0.05 | AG1 | 2 | 25.9167 | 0.94 |
| 0.05 | 0.05 | ptsI | 2 | 26.1667 | 0.538 |
| 0.05 | 0.05 | clcB | 2 | 26.1667 | 0.335 |
| 0.05 | 0.05 | ycaM | 2 | 26.1667 | 0.356 |
| 0.05 | 0.05 | yadI | 2 | 26.1667 | 0.891 |
| 0.05 | 0.05 | AG1 | 2 | 26.1667 | 0.954 |
| 0.05 | 0.05 | ptsI | 2 | 26.4167 | 0.54 |
| 0.05 | 0.05 | clcB | 2 | 26.4167 | 0.336 |
| 0.05 | 0.05 | ycaM | 2 | 26.4167 | 0.357 |
| 0.05 | 0.05 | yadI | 2 | 26.4167 | 0.9 |
| 0.05 | 0.05 | AG1 | 2 | 26.4167 | 0.963 |
| 0.05 | 0.05 | ptsI | 3 | 0 | 0.155 |
| 0.05 | 0.05 | clcB | 3 | 0 | 0.164 |
| 0.05 | 0.05 | ycaM | 3 | 0 | 0.162 |
| 0.05 | 0.05 | yadI | 3 | 0 | 0.162 |
| 0.05 | 0.05 | AG1 | 3 | 0 | 0.163 |
| 0.05 | 0.05 | ptsI | 3 | 0.25 | 0.152 |
| 0.05 | 0.05 | clcB | 3 | 0.25 | 0.161 |
| 0.05 | 0.05 | ycaM | 3 | 0.25 | 0.158 |
| 0.05 | 0.05 | yadI | 3 | 0.25 | 0.159 |
| 0.05 | 0.05 | AG1 | 3 | 0.25 | 0.159 |
| 0.05 | 0.05 | ptsI | 3 | 0.5 | 0.151 |
| 0.05 | 0.05 | clcB | 3 | 0.5 | 0.16 |
| 0.05 | 0.05 | ycaM | 3 | 0.5 | 0.156 |
| 0.05 | 0.05 | yadI | 3 | 0.5 | 0.158 |
| 0.05 | 0.05 | AG1 | 3 | 0.5 | 0.158 |
| 0.05 | 0.05 | ptsI | 3 | 0.75 | 0.152 |
| 0.05 | 0.05 | clcB | 3 | 0.75 | 0.16 |
| 0.05 | 0.05 | ycaM | 3 | 0.75 | 0.156 |
| 0.05 | 0.05 | yadI | 3 | 0.75 | 0.16 |
| 0.05 | 0.05 | AG1 | 3 | 0.75 | 0.159 |
| 0.05 | 0.05 | ptsI | 3 | 1 | 0.152 |
| 0.05 | 0.05 | clcB | 3 | 1 | 0.16 |
| 0.05 | 0.05 | ycaM | 3 | 1 | 0.157 |
| 0.05 | 0.05 | yadI | 3 | 1 | 0.161 |
| 0.05 | 0.05 | AG1 | 3 | 1 | 0.159 |
| 0.05 | 0.05 | ptsI | 3 | 1.25 | 0.153 |
| 0.05 | 0.05 | clcB | 3 | 1.25 | 0.161 |
| 0.05 | 0.05 | ycaM | 3 | 1.25 | 0.157 |
| 0.05 | 0.05 | yadI | 3 | 1.25 | 0.161 |

| | | | | | |
|---|---|---|---|---|---|
| 0.05 | 0.05 | AG1 | 3 | 1.25 | 0.161 |
| 0.05 | 0.05 | ptsI | 3 | 1.5 | 0.166 |
| 0.05 | 0.05 | clcB | 3 | 1.5 | 0.16 |
| 0.05 | 0.05 | ycaM | 3 | 1.5 | 0.158 |
| 0.05 | 0.05 | yadI | 3 | 1.5 | 0.163 |
| 0.05 | 0.05 | AG1 | 3 | 1.5 | 0.162 |
| 0.05 | 0.05 | ptsI | 3 | 1.75 | 0.155 |
| 0.05 | 0.05 | clcB | 3 | 1.75 | 0.161 |
| 0.05 | 0.05 | ycaM | 3 | 1.75 | 0.16 |
| 0.05 | 0.05 | yadI | 3 | 1.75 | 0.164 |
| 0.05 | 0.05 | AG1 | 3 | 1.75 | 0.165 |
| 0.05 | 0.05 | ptsI | 3 | 2 | 0.158 |
| 0.05 | 0.05 | clcB | 3 | 2 | 0.161 |
| 0.05 | 0.05 | ycaM | 3 | 2 | 0.161 |
| 0.05 | 0.05 | yadI | 3 | 2 | 0.166 |
| 0.05 | 0.05 | AG1 | 3 | 2 | 0.164 |
| 0.05 | 0.05 | ptsI | 3 | 2.25 | 0.159 |
| 0.05 | 0.05 | clcB | 3 | 2.25 | 0.162 |
| 0.05 | 0.05 | ycaM | 3 | 2.25 | 0.163 |
| 0.05 | 0.05 | yadI | 3 | 2.25 | 0.168 |
| 0.05 | 0.05 | AG1 | 3 | 2.25 | 0.168 |
| 0.05 | 0.05 | ptsI | 3 | 2.5 | 0.16 |
| 0.05 | 0.05 | clcB | 3 | 2.5 | 0.164 |
| 0.05 | 0.05 | ycaM | 3 | 2.5 | 0.163 |
| 0.05 | 0.05 | yadI | 3 | 2.5 | 0.172 |
| 0.05 | 0.05 | AG1 | 3 | 2.5 | 0.17 |
| 0.05 | 0.05 | ptsI | 3 | 2.75 | 0.164 |
| 0.05 | 0.05 | clcB | 3 | 2.75 | 0.167 |
| 0.05 | 0.05 | ycaM | 3 | 2.75 | 0.168 |
| 0.05 | 0.05 | yadI | 3 | 2.75 | 0.174 |
| 0.05 | 0.05 | AG1 | 3 | 2.75 | 0.173 |
| 0.05 | 0.05 | ptsI | 3 | 3 | 0.165 |
| 0.05 | 0.05 | clcB | 3 | 3 | 0.167 |
| 0.05 | 0.05 | ycaM | 3 | 3 | 0.167 |
| 0.05 | 0.05 | yadI | 3 | 3 | 0.18 |
| 0.05 | 0.05 | AG1 | 3 | 3 | 0.176 |
| 0.05 | 0.05 | ptsI | 3 | 3.25 | 0.167 |
| 0.05 | 0.05 | clcB | 3 | 3.25 | 0.169 |
| 0.05 | 0.05 | ycaM | 3 | 3.25 | 0.17 |
| 0.05 | 0.05 | yadI | 3 | 3.25 | 0.186 |
| 0.05 | 0.05 | AG1 | 3 | 3.25 | 0.179 |
| 0.05 | 0.05 | ptsI | 3 | 3.5 | 0.17 |
| 0.05 | 0.05 | clcB | 3 | 3.5 | 0.172 |
| 0.05 | 0.05 | ycaM | 3 | 3.5 | 0.174 |
| 0.05 | 0.05 | yadI | 3 | 3.5 | 0.19 |
| 0.05 | 0.05 | AG1 | 3 | 3.5 | 0.186 |
| 0.05 | 0.05 | ptsI | 3 | 3.75 | 0.174 |
| 0.05 | 0.05 | clcB | 3 | 3.75 | 0.174 |

| | | | | | |
|---|---|---|---|---|---|
| 0.05 | 0.05 | ycaM | 3 | 3.75 | 0.175 |
| 0.05 | 0.05 | yadI | 3 | 3.75 | 0.197 |
| 0.05 | 0.05 | AG1 | 3 | 3.75 | 0.185 |
| 0.05 | 0.05 | ptsI | 3 | 4 | 0.176 |
| 0.05 | 0.05 | clcB | 3 | 4 | 0.177 |
| 0.05 | 0.05 | ycaM | 3 | 4 | 0.178 |
| 0.05 | 0.05 | yadI | 3 | 4 | 0.204 |
| 0.05 | 0.05 | AG1 | 3 | 4 | 0.191 |
| 0.05 | 0.05 | ptsI | 3 | 4.41667 | 0.195 |
| 0.05 | 0.05 | clcB | 3 | 4.41667 | 0.202 |
| 0.05 | 0.05 | ycaM | 3 | 4.41667 | 0.206 |
| 0.05 | 0.05 | yadI | 3 | 4.41667 | 0.221 |
| 0.05 | 0.05 | AG1 | 3 | 4.41667 | 0.197 |
| 0.05 | 0.05 | ptsI | 3 | 4.66667 | 0.186 |
| 0.05 | 0.05 | clcB | 3 | 4.66667 | 0.2 |
| 0.05 | 0.05 | ycaM | 3 | 4.66667 | 0.194 |
| 0.05 | 0.05 | yadI | 3 | 4.66667 | 0.218 |
| 0.05 | 0.05 | AG1 | 3 | 4.66667 | 0.194 |
| 0.05 | 0.05 | ptsI | 3 | 4.91667 | 0.189 |
| 0.05 | 0.05 | clcB | 3 | 4.91667 | 0.198 |
| 0.05 | 0.05 | ycaM | 3 | 4.91667 | 0.202 |
| 0.05 | 0.05 | yadI | 3 | 4.91667 | 0.232 |
| 0.05 | 0.05 | AG1 | 3 | 4.91667 | 0.194 |
| 0.05 | 0.05 | ptsI | 3 | 5.16667 | 0.192 |
| 0.05 | 0.05 | clcB | 3 | 5.16667 | 0.203 |
| 0.05 | 0.05 | ycaM | 3 | 5.16667 | 0.207 |
| 0.05 | 0.05 | yadI | 3 | 5.16667 | 0.244 |
| 0.05 | 0.05 | AG1 | 3 | 5.16667 | 0.2 |
| 0.05 | 0.05 | ptsI | 3 | 5.41667 | 0.197 |
| 0.05 | 0.05 | clcB | 3 | 5.41667 | 0.212 |
| 0.05 | 0.05 | ycaM | 3 | 5.41667 | 0.215 |
| 0.05 | 0.05 | yadI | 3 | 5.41667 | 0.26 |
| 0.05 | 0.05 | AG1 | 3 | 5.41667 | 0.206 |
| 0.05 | 0.05 | ptsI | 3 | 5.66667 | 0.201 |
| 0.05 | 0.05 | clcB | 3 | 5.66667 | 0.219 |
| 0.05 | 0.05 | ycaM | 3 | 5.66667 | 0.221 |
| 0.05 | 0.05 | yadI | 3 | 5.66667 | 0.277 |
| 0.05 | 0.05 | AG1 | 3 | 5.66667 | 0.214 |
| 0.05 | 0.05 | ptsI | 3 | 5.91667 | 0.206 |
| 0.05 | 0.05 | clcB | 3 | 5.91667 | 0.227 |
| 0.05 | 0.05 | ycaM | 3 | 5.91667 | 0.23 |
| 0.05 | 0.05 | yadI | 3 | 5.91667 | 0.297 |
| 0.05 | 0.05 | AG1 | 3 | 5.91667 | 0.229 |
| 0.05 | 0.05 | ptsI | 3 | 6.16667 | 0.212 |
| 0.05 | 0.05 | clcB | 3 | 6.16667 | 0.237 |
| 0.05 | 0.05 | ycaM | 3 | 6.16667 | 0.238 |
| 0.05 | 0.05 | yadI | 3 | 6.16667 | 0.316 |
| 0.05 | 0.05 | AG1 | 3 | 6.16667 | 0.24 |

| | | | | | |
|---|---|---|---|---|---|
| 0.05 | 0.05 | ptsI | 3 | 6.41667 | 0.218 |
| 0.05 | 0.05 | clcB | 3 | 6.41667 | 0.245 |
| 0.05 | 0.05 | ycaM | 3 | 6.41667 | 0.245 |
| 0.05 | 0.05 | yadI | 3 | 6.41667 | 0.326 |
| 0.05 | 0.05 | AG1 | 3 | 6.41667 | 0.251 |
| 0.05 | 0.05 | ptsI | 3 | 6.66667 | 0.226 |
| 0.05 | 0.05 | clcB | 3 | 6.66667 | 0.256 |
| 0.05 | 0.05 | ycaM | 3 | 6.66667 | 0.255 |
| 0.05 | 0.05 | yadI | 3 | 6.66667 | 0.336 |
| 0.05 | 0.05 | AG1 | 3 | 6.66667 | 0.266 |
| 0.05 | 0.05 | ptsI | 3 | 6.91667 | 0.234 |
| 0.05 | 0.05 | clcB | 3 | 6.91667 | 0.266 |
| 0.05 | 0.05 | ycaM | 3 | 6.91667 | 0.262 |
| 0.05 | 0.05 | yadI | 3 | 6.91667 | 0.348 |
| 0.05 | 0.05 | AG1 | 3 | 6.91667 | 0.281 |
| 0.05 | 0.05 | ptsI | 3 | 7.16667 | 0.242 |
| 0.05 | 0.05 | clcB | 3 | 7.16667 | 0.276 |
| 0.05 | 0.05 | ycaM | 3 | 7.16667 | 0.264 |
| 0.05 | 0.05 | yadI | 3 | 7.16667 | 0.36 |
| 0.05 | 0.05 | AG1 | 3 | 7.16667 | 0.29 |
| 0.05 | 0.05 | ptsI | 3 | 7.41667 | 0.254 |
| 0.05 | 0.05 | clcB | 3 | 7.41667 | 0.283 |
| 0.05 | 0.05 | ycaM | 3 | 7.41667 | 0.266 |
| 0.05 | 0.05 | yadI | 3 | 7.41667 | 0.374 |
| 0.05 | 0.05 | AG1 | 3 | 7.41667 | 0.298 |
| 0.05 | 0.05 | ptsI | 3 | 7.66667 | 0.264 |
| 0.05 | 0.05 | clcB | 3 | 7.66667 | 0.289 |
| 0.05 | 0.05 | ycaM | 3 | 7.66667 | 0.27 |
| 0.05 | 0.05 | yadI | 3 | 7.66667 | 0.389 |
| 0.05 | 0.05 | AG1 | 3 | 7.66667 | 0.306 |
| 0.05 | 0.05 | ptsI | 3 | 7.91667 | 0.272 |
| 0.05 | 0.05 | clcB | 3 | 7.91667 | 0.293 |
| 0.05 | 0.05 | ycaM | 3 | 7.91667 | 0.271 |
| 0.05 | 0.05 | yadI | 3 | 7.91667 | 0.398 |
| 0.05 | 0.05 | AG1 | 3 | 7.91667 | 0.316 |
| 0.05 | 0.05 | ptsI | 3 | 8.16667 | 0.278 |
| 0.05 | 0.05 | clcB | 3 | 8.16667 | 0.299 |
| 0.05 | 0.05 | ycaM | 3 | 8.16667 | 0.272 |
| 0.05 | 0.05 | yadI | 3 | 8.16667 | 0.41 |
| 0.05 | 0.05 | AG1 | 3 | 8.16667 | 0.325 |
| 0.05 | 0.05 | ptsI | 3 | 8.41667 | 0.287 |
| 0.05 | 0.05 | clcB | 3 | 8.41667 | 0.306 |
| 0.05 | 0.05 | ycaM | 3 | 8.41667 | 0.275 |
| 0.05 | 0.05 | yadI | 3 | 8.41667 | 0.418 |
| 0.05 | 0.05 | AG1 | 3 | 8.41667 | 0.334 |
| 0.05 | 0.05 | ptsI | 3 | 9.05 | 0.279 |
| 0.05 | 0.05 | clcB | 3 | 9.05 | 0.27 |
| 0.05 | 0.05 | ycaM | 3 | 9.05 | 0.252 |

| | | | | | |
|---|---|---|---|---|---|
| 0.05 | 0.05 | yadI | 3 | 9.05 | 0.453 |
| 0.05 | 0.05 | AG1 | 3 | 9.05 | 0.363 |
| 0.05 | 0.05 | ptsI | 3 | 9.3 | 0.279 |
| 0.05 | 0.05 | clcB | 3 | 9.3 | 0.269 |
| 0.05 | 0.05 | ycaM | 3 | 9.3 | 0.251 |
| 0.05 | 0.05 | yadI | 3 | 9.3 | 0.463 |
| 0.05 | 0.05 | AG1 | 3 | 9.3 | 0.368 |
| 0.05 | 0.05 | ptsI | 3 | 9.55 | 0.286 |
| 0.05 | 0.05 | clcB | 3 | 9.55 | 0.278 |
| 0.05 | 0.05 | ycaM | 3 | 9.55 | 0.255 |
| 0.05 | 0.05 | yadI | 3 | 9.55 | 0.476 |
| 0.05 | 0.05 | AG1 | 3 | 9.55 | 0.377 |
| 0.05 | 0.05 | ptsI | 3 | 9.8 | 0.293 |
| 0.05 | 0.05 | clcB | 3 | 9.8 | 0.282 |
| 0.05 | 0.05 | ycaM | 3 | 9.8 | 0.256 |
| 0.05 | 0.05 | yadI | 3 | 9.8 | 0.492 |
| 0.05 | 0.05 | AG1 | 3 | 9.8 | 0.387 |
| 0.05 | 0.05 | ptsI | 3 | 10.05 | 0.302 |
| 0.05 | 0.05 | clcB | 3 | 10.05 | 0.288 |
| 0.05 | 0.05 | ycaM | 3 | 10.05 | 0.262 |
| 0.05 | 0.05 | yadI | 3 | 10.05 | 0.511 |
| 0.05 | 0.05 | AG1 | 3 | 10.05 | 0.4 |
| 0.05 | 0.05 | ptsI | 3 | 10.3 | 0.31 |
| 0.05 | 0.05 | clcB | 3 | 10.3 | 0.295 |
| 0.05 | 0.05 | ycaM | 3 | 10.3 | 0.263 |
| 0.05 | 0.05 | yadI | 3 | 10.3 | 0.524 |
| 0.05 | 0.05 | AG1 | 3 | 10.3 | 0.411 |
| 0.05 | 0.05 | ptsI | 3 | 10.55 | 0.319 |
| 0.05 | 0.05 | clcB | 3 | 10.55 | 0.296 |
| 0.05 | 0.05 | ycaM | 3 | 10.55 | 0.263 |
| 0.05 | 0.05 | yadI | 3 | 10.55 | 0.536 |
| 0.05 | 0.05 | AG1 | 3 | 10.55 | 0.424 |
| 0.05 | 0.05 | ptsI | 3 | 10.8 | 0.327 |
| 0.05 | 0.05 | clcB | 3 | 10.8 | 0.297 |
| 0.05 | 0.05 | ycaM | 3 | 10.8 | 0.265 |
| 0.05 | 0.05 | yadI | 3 | 10.8 | 0.548 |
| 0.05 | 0.05 | AG1 | 3 | 10.8 | 0.431 |
| 0.05 | 0.05 | ptsI | 3 | 11.05 | 0.338 |
| 0.05 | 0.05 | clcB | 3 | 11.05 | 0.308 |
| 0.05 | 0.05 | ycaM | 3 | 11.05 | 0.267 |
| 0.05 | 0.05 | yadI | 3 | 11.05 | 0.562 |
| 0.05 | 0.05 | AG1 | 3 | 11.05 | 0.445 |
| 0.05 | 0.05 | ptsI | 3 | 11.3 | 0.345 |
| 0.05 | 0.05 | clcB | 3 | 11.3 | 0.3 |
| 0.05 | 0.05 | ycaM | 3 | 11.3 | 0.264 |
| 0.05 | 0.05 | yadI | 3 | 11.3 | 0.571 |
| 0.05 | 0.05 | AG1 | 3 | 11.3 | 0.451 |
| 0.05 | 0.05 | ptsI | 3 | 11.55 | 0.356 |

| | | | | | |
|---|---|---|---|---|---|
| 0.05 | 0.05 | clcB | 3 | 11.55 | 0.301 |
| 0.05 | 0.05 | ycaM | 3 | 11.55 | 0.265 |
| 0.05 | 0.05 | yadI | 3 | 11.55 | 0.582 |
| 0.05 | 0.05 | AG1 | 3 | 11.55 | 0.463 |
| 0.05 | 0.05 | ptsI | 3 | 11.8 | 0.366 |
| 0.05 | 0.05 | clcB | 3 | 11.8 | 0.311 |
| 0.05 | 0.05 | ycaM | 3 | 11.8 | 0.269 |
| 0.05 | 0.05 | yadI | 3 | 11.8 | 0.599 |
| 0.05 | 0.05 | AG1 | 3 | 11.8 | 0.479 |
| 0.05 | 0.05 | ptsI | 3 | 12.05 | 0.371 |
| 0.05 | 0.05 | clcB | 3 | 12.05 | 0.299 |
| 0.05 | 0.05 | ycaM | 3 | 12.05 | 0.264 |
| 0.05 | 0.05 | yadI | 3 | 12.05 | 0.605 |
| 0.05 | 0.05 | AG1 | 3 | 12.05 | 0.487 |
| 0.05 | 0.05 | ptsI | 3 | 12.3 | 0.386 |
| 0.05 | 0.05 | clcB | 3 | 12.3 | 0.31 |
| 0.05 | 0.05 | ycaM | 3 | 12.3 | 0.268 |
| 0.05 | 0.05 | yadI | 3 | 12.3 | 0.621 |
| 0.05 | 0.05 | AG1 | 3 | 12.3 | 0.5 |
| 0.05 | 0.05 | ptsI | 3 | 12.55 | 0.396 |
| 0.05 | 0.05 | clcB | 3 | 12.55 | 0.304 |
| 0.05 | 0.05 | ycaM | 3 | 12.55 | 0.269 |
| 0.05 | 0.05 | yadI | 3 | 12.55 | 0.638 |
| 0.05 | 0.05 | AG1 | 3 | 12.55 | 0.519 |
| 0.05 | 0.05 | ptsI | 3 | 12.8 | 0.408 |
| 0.05 | 0.05 | clcB | 3 | 12.8 | 0.298 |
| 0.05 | 0.05 | ycaM | 3 | 12.8 | 0.268 |
| 0.05 | 0.05 | yadI | 3 | 12.8 | 0.655 |
| 0.05 | 0.05 | AG1 | 3 | 12.8 | 0.539 |
| 0.05 | 0.05 | ptsI | 3 | 13.05 | 0.41 |
| 0.05 | 0.05 | clcB | 3 | 13.05 | 0.296 |
| 0.05 | 0.05 | ycaM | 3 | 13.05 | 0.264 |
| 0.05 | 0.05 | yadI | 3 | 13.05 | 0.654 |
| 0.05 | 0.05 | AG1 | 3 | 13.05 | 0.546 |
| 0.05 | 0.05 | ptsI | 3 | 13.3 | 0.417 |
| 0.05 | 0.05 | clcB | 3 | 13.3 | 0.305 |
| 0.05 | 0.05 | ycaM | 3 | 13.3 | 0.267 |
| 0.05 | 0.05 | yadI | 3 | 13.3 | 0.671 |
| 0.05 | 0.05 | AG1 | 3 | 13.3 | 0.562 |
| 0.05 | 0.05 | ptsI | 3 | 13.55 | 0.43 |
| 0.05 | 0.05 | clcB | 3 | 13.55 | 0.301 |
| 0.05 | 0.05 | ycaM | 3 | 13.55 | 0.266 |
| 0.05 | 0.05 | yadI | 3 | 13.55 | 0.682 |
| 0.05 | 0.05 | AG1 | 3 | 13.55 | 0.574 |
| 0.05 | 0.05 | ptsI | 3 | 13.8 | 0.438 |
| 0.05 | 0.05 | clcB | 3 | 13.8 | 0.296 |
| 0.05 | 0.05 | ycaM | 3 | 13.8 | 0.266 |
| 0.05 | 0.05 | yadI | 3 | 13.8 | 0.695 |

| | | | | | |
|---|---|---|---|---|---|
| 0.05 | 0.05 | AG1 | 3 | 13.8 | 0.592 |
| 0.05 | 0.05 | ptsI | 3 | 14.05 | 0.446 |
| 0.05 | 0.05 | clcB | 3 | 14.05 | 0.304 |
| 0.05 | 0.05 | ycaM | 3 | 14.05 | 0.267 |
| 0.05 | 0.05 | yadI | 3 | 14.05 | 0.695 |
| 0.05 | 0.05 | AG1 | 3 | 14.05 | 0.598 |
| 0.05 | 0.05 | ptsI | 3 | 14.3 | 0.45 |
| 0.05 | 0.05 | clcB | 3 | 14.3 | 0.303 |
| 0.05 | 0.05 | ycaM | 3 | 14.3 | 0.266 |
| 0.05 | 0.05 | yadI | 3 | 14.3 | 0.704 |
| 0.05 | 0.05 | AG1 | 3 | 14.3 | 0.607 |
| 0.05 | 0.05 | ptsI | 3 | 14.55 | 0.459 |
| 0.05 | 0.05 | clcB | 3 | 14.55 | 0.296 |
| 0.05 | 0.05 | ycaM | 3 | 14.55 | 0.261 |
| 0.05 | 0.05 | yadI | 3 | 14.55 | 0.711 |
| 0.05 | 0.05 | AG1 | 3 | 14.55 | 0.618 |
| 0.05 | 0.05 | ptsI | 3 | 14.8 | 0.471 |
| 0.05 | 0.05 | clcB | 3 | 14.8 | 0.296 |
| 0.05 | 0.05 | ycaM | 3 | 14.8 | 0.264 |
| 0.05 | 0.05 | yadI | 3 | 14.8 | 0.733 |
| 0.05 | 0.05 | AG1 | 3 | 14.8 | 0.632 |
| 0.05 | 0.05 | ptsI | 3 | 15.05 | 0.486 |
| 0.05 | 0.05 | clcB | 3 | 15.05 | 0.297 |
| 0.05 | 0.05 | ycaM | 3 | 15.05 | 0.265 |
| 0.05 | 0.05 | yadI | 3 | 15.05 | 0.752 |
| 0.05 | 0.05 | AG1 | 3 | 15.05 | 0.65 |
| 0.05 | 0.05 | ptsI | 3 | 15.3 | 0.486 |
| 0.05 | 0.05 | clcB | 3 | 15.3 | 0.297 |
| 0.05 | 0.05 | ycaM | 3 | 15.3 | 0.261 |
| 0.05 | 0.05 | yadI | 3 | 15.3 | 0.745 |
| 0.05 | 0.05 | AG1 | 3 | 15.3 | 0.653 |
| 0.05 | 0.05 | ptsI | 3 | 15.55 | 0.49 |
| 0.05 | 0.05 | clcB | 3 | 15.55 | 0.294 |
| 0.05 | 0.05 | ycaM | 3 | 15.55 | 0.261 |
| 0.05 | 0.05 | yadI | 3 | 15.55 | 0.757 |
| 0.05 | 0.05 | AG1 | 3 | 15.55 | 0.665 |
| 0.05 | 0.05 | ptsI | 3 | 15.8 | 0.499 |
| 0.05 | 0.05 | clcB | 3 | 15.8 | 0.295 |
| 0.05 | 0.05 | ycaM | 3 | 15.8 | 0.261 |
| 0.05 | 0.05 | yadI | 3 | 15.8 | 0.761 |
| 0.05 | 0.05 | AG1 | 3 | 15.8 | 0.685 |
| 0.05 | 0.05 | ptsI | 3 | 16.05 | 0.501 |
| 0.05 | 0.05 | clcB | 3 | 16.05 | 0.292 |
| 0.05 | 0.05 | ycaM | 3 | 16.05 | 0.258 |
| 0.05 | 0.05 | yadI | 3 | 16.05 | 0.764 |
| 0.05 | 0.05 | AG1 | 3 | 16.05 | 0.696 |
| 0.05 | 0.05 | ptsI | 3 | 16.3 | 0.51 |
| 0.05 | 0.05 | clcB | 3 | 16.3 | 0.295 |

| | | | | | |
|---|---|---|---|---|---|
| 0.05 | 0.05 | ycaM | 3 | 16.3 | 0.258 |
| 0.05 | 0.05 | yadI | 3 | 16.3 | 0.758 |
| 0.05 | 0.05 | AG1 | 3 | 16.3 | 0.725 |
| 0.05 | 0.05 | ptsI | 3 | 16.55 | 0.517 |
| 0.05 | 0.05 | clcB | 3 | 16.55 | 0.296 |
| 0.05 | 0.05 | ycaM | 3 | 16.55 | 0.259 |
| 0.05 | 0.05 | yadI | 3 | 16.55 | 0.783 |
| 0.05 | 0.05 | AG1 | 3 | 16.55 | 0.718 |
| 0.05 | 0.05 | ptsI | 3 | 16.8 | 0.526 |
| 0.05 | 0.05 | clcB | 3 | 16.8 | 0.294 |
| 0.05 | 0.05 | ycaM | 3 | 16.8 | 0.257 |
| 0.05 | 0.05 | yadI | 3 | 16.8 | 0.789 |
| 0.05 | 0.05 | AG1 | 3 | 16.8 | 0.749 |
| 0.05 | 0.05 | ptsI | 3 | 17.05 | 0.531 |
| 0.05 | 0.05 | clcB | 3 | 17.05 | 0.294 |
| 0.05 | 0.05 | ycaM | 3 | 17.05 | 0.257 |
| 0.05 | 0.05 | yadI | 3 | 17.05 | 0.789 |
| 0.05 | 0.05 | AG1 | 3 | 17.05 | 0.752 |
| 0.05 | 0.05 | ptsI | 3 | 17.3 | 0.53 |
| 0.05 | 0.05 | clcB | 3 | 17.3 | 0.293 |
| 0.05 | 0.05 | ycaM | 3 | 17.3 | 0.258 |
| 0.05 | 0.05 | yadI | 3 | 17.3 | 0.795 |
| 0.05 | 0.05 | AG1 | 3 | 17.3 | 0.748 |
| 0.05 | 0.05 | ptsI | 3 | 17.55 | 0.538 |
| 0.05 | 0.05 | clcB | 3 | 17.55 | 0.296 |
| 0.05 | 0.05 | ycaM | 3 | 17.55 | 0.259 |
| 0.05 | 0.05 | yadI | 3 | 17.55 | 0.796 |
| 0.05 | 0.05 | AG1 | 3 | 17.55 | 0.756 |
| 0.05 | 0.05 | ptsI | 3 | 17.8 | 0.545 |
| 0.05 | 0.05 | clcB | 3 | 17.8 | 0.293 |
| 0.05 | 0.05 | ycaM | 3 | 17.8 | 0.259 |
| 0.05 | 0.05 | yadI | 3 | 17.8 | 0.806 |
| 0.05 | 0.05 | AG1 | 3 | 17.8 | 0.771 |
| 0.05 | 0.05 | ptsI | 3 | 18.05 | 0.545 |
| 0.05 | 0.05 | clcB | 3 | 18.05 | 0.295 |
| 0.05 | 0.05 | ycaM | 3 | 18.05 | 0.259 |
| 0.05 | 0.05 | yadI | 3 | 18.05 | 0.803 |
| 0.05 | 0.05 | AG1 | 3 | 18.05 | 0.778 |
| 0.05 | 0.05 | ptsI | 3 | 18.3 | 0.545 |
| 0.05 | 0.05 | clcB | 3 | 18.3 | 0.294 |
| 0.05 | 0.05 | ycaM | 3 | 18.3 | 0.259 |
| 0.05 | 0.05 | yadI | 3 | 18.3 | 0.813 |
| 0.05 | 0.05 | AG1 | 3 | 18.3 | 0.777 |
| 0.05 | 0.05 | ptsI | 3 | 18.55 | 0.544 |
| 0.05 | 0.05 | clcB | 3 | 18.55 | 0.296 |
| 0.05 | 0.05 | ycaM | 3 | 18.55 | 0.26 |
| 0.05 | 0.05 | yadI | 3 | 18.55 | 0.818 |
| 0.05 | 0.05 | AG1 | 3 | 18.55 | 0.757 |

| | | | | | |
|---|---|---|---|---|---|
| 0.05 | 0.05 | ptsI | 3 | 18.8 | 0.546 |
| 0.05 | 0.05 | clcB | 3 | 18.8 | 0.296 |
| 0.05 | 0.05 | ycaM | 3 | 18.8 | 0.264 |
| 0.05 | 0.05 | yadI | 3 | 18.8 | 0.817 |
| 0.05 | 0.05 | AG1 | 3 | 18.8 | 0.747 |
| 0.05 | 0.05 | ptsI | 3 | 19.05 | 0.549 |
| 0.05 | 0.05 | clcB | 3 | 19.05 | 0.298 |
| 0.05 | 0.05 | ycaM | 3 | 19.05 | 0.263 |
| 0.05 | 0.05 | yadI | 3 | 19.05 | 0.829 |
| 0.05 | 0.05 | AG1 | 3 | 19.05 | 0.759 |
| 0.05 | 0.05 | ptsI | 3 | 19.3 | 0.552 |
| 0.05 | 0.05 | clcB | 3 | 19.3 | 0.297 |
| 0.05 | 0.05 | ycaM | 3 | 19.3 | 0.266 |
| 0.05 | 0.05 | yadI | 3 | 19.3 | 0.832 |
| 0.05 | 0.05 | AG1 | 3 | 19.3 | 0.774 |
| 0.05 | 0.05 | ptsI | 3 | 19.55 | 0.56 |
| 0.05 | 0.05 | clcB | 3 | 19.55 | 0.298 |
| 0.05 | 0.05 | ycaM | 3 | 19.55 | 0.264 |
| 0.05 | 0.05 | yadI | 3 | 19.55 | 0.828 |
| 0.05 | 0.05 | AG1 | 3 | 19.55 | 0.772 |
| 0.05 | 0.05 | ptsI | 3 | 19.8 | 0.55 |
| 0.05 | 0.05 | clcB | 3 | 19.8 | 0.296 |
| 0.05 | 0.05 | ycaM | 3 | 19.8 | 0.266 |
| 0.05 | 0.05 | yadI | 3 | 19.8 | 0.824 |
| 0.05 | 0.05 | AG1 | 3 | 19.8 | 0.769 |
| 0.05 | 0.05 | ptsI | 3 | 20.05 | 0.554 |
| 0.05 | 0.05 | clcB | 3 | 20.05 | 0.296 |
| 0.05 | 0.05 | ycaM | 3 | 20.05 | 0.267 |
| 0.05 | 0.05 | yadI | 3 | 20.05 | 0.833 |
| 0.05 | 0.05 | AG1 | 3 | 20.05 | 0.757 |
| 0.05 | 0.05 | ptsI | 3 | 20.3 | 0.556 |
| 0.05 | 0.05 | clcB | 3 | 20.3 | 0.295 |
| 0.05 | 0.05 | ycaM | 3 | 20.3 | 0.269 |
| 0.05 | 0.05 | yadI | 3 | 20.3 | 0.832 |
| 0.05 | 0.05 | AG1 | 3 | 20.3 | 0.758 |
| 0.05 | 0.05 | ptsI | 3 | 20.55 | 0.553 |
| 0.05 | 0.05 | clcB | 3 | 20.55 | 0.296 |
| 0.05 | 0.05 | ycaM | 3 | 20.55 | 0.271 |
| 0.05 | 0.05 | yadI | 3 | 20.55 | 0.831 |
| 0.05 | 0.05 | AG1 | 3 | 20.55 | 0.767 |
| 0.05 | 0.05 | ptsI | 3 | 20.8 | 0.557 |
| 0.05 | 0.05 | clcB | 3 | 20.8 | 0.296 |
| 0.05 | 0.05 | ycaM | 3 | 20.8 | 0.272 |
| 0.05 | 0.05 | yadI | 3 | 20.8 | 0.839 |
| 0.05 | 0.05 | AG1 | 3 | 20.8 | 0.781 |
| 0.05 | 0.05 | ptsI | 3 | 21.05 | 0.554 |
| 0.05 | 0.05 | clcB | 3 | 21.05 | 0.296 |
| 0.05 | 0.05 | ycaM | 3 | 21.05 | 0.275 |

| | | | | | |
|---|---|---|---|---|---|
| 0.05 | 0.05 | yadI | 3 | 21.05 | 0.837 |
| 0.05 | 0.05 | AG1 | 3 | 21.05 | 0.782 |
| 0.05 | 0.05 | ptsI | 3 | 21.3 | 0.558 |
| 0.05 | 0.05 | clcB | 3 | 21.3 | 0.296 |
| 0.05 | 0.05 | ycaM | 3 | 21.3 | 0.277 |
| 0.05 | 0.05 | yadI | 3 | 21.3 | 0.84 |
| 0.05 | 0.05 | AG1 | 3 | 21.3 | 0.793 |
| 0.05 | 0.05 | ptsI | 3 | 21.55 | 0.557 |
| 0.05 | 0.05 | clcB | 3 | 21.55 | 0.297 |
| 0.05 | 0.05 | ycaM | 3 | 21.55 | 0.278 |
| 0.05 | 0.05 | yadI | 3 | 21.55 | 0.842 |
| 0.05 | 0.05 | AG1 | 3 | 21.55 | 0.802 |
| 0.05 | 0.05 | ptsI | 3 | 21.8 | 0.559 |
| 0.05 | 0.05 | clcB | 3 | 21.8 | 0.299 |
| 0.05 | 0.05 | ycaM | 3 | 21.8 | 0.279 |
| 0.05 | 0.05 | yadI | 3 | 21.8 | 0.844 |
| 0.05 | 0.05 | AG1 | 3 | 21.8 | 0.809 |
| 0.05 | 0.05 | ptsI | 3 | 22.05 | 0.557 |
| 0.05 | 0.05 | clcB | 3 | 22.05 | 0.297 |
| 0.05 | 0.05 | ycaM | 3 | 22.05 | 0.28 |
| 0.05 | 0.05 | yadI | 3 | 22.05 | 0.846 |
| 0.05 | 0.05 | AG1 | 3 | 22.05 | 0.811 |
| 0.05 | 0.05 | ptsI | 3 | 22.3 | 0.557 |
| 0.05 | 0.05 | clcB | 3 | 22.3 | 0.299 |
| 0.05 | 0.05 | ycaM | 3 | 22.3 | 0.285 |
| 0.05 | 0.05 | yadI | 3 | 22.3 | 0.843 |
| 0.05 | 0.05 | AG1 | 3 | 22.3 | 0.822 |
| 0.05 | 0.05 | ptsI | 3 | 22.55 | 0.559 |
| 0.05 | 0.05 | clcB | 3 | 22.55 | 0.301 |
| 0.05 | 0.05 | ycaM | 3 | 22.55 | 0.288 |
| 0.05 | 0.05 | yadI | 3 | 22.55 | 0.851 |
| 0.05 | 0.05 | AG1 | 3 | 22.55 | 0.828 |
| 0.05 | 0.05 | ptsI | 3 | 22.8 | 0.558 |
| 0.05 | 0.05 | clcB | 3 | 22.8 | 0.3 |
| 0.05 | 0.05 | ycaM | 3 | 22.8 | 0.29 |
| 0.05 | 0.05 | yadI | 3 | 22.8 | 0.848 |
| 0.05 | 0.05 | AG1 | 3 | 22.8 | 0.836 |
| 0.05 | 0.05 | ptsI | 3 | 23.05 | 0.558 |
| 0.05 | 0.05 | clcB | 3 | 23.05 | 0.301 |
| 0.05 | 0.05 | ycaM | 3 | 23.05 | 0.292 |
| 0.05 | 0.05 | yadI | 3 | 23.05 | 0.855 |
| 0.05 | 0.05 | AG1 | 3 | 23.05 | 0.841 |
| 0.05 | 0.05 | ptsI | 3 | 23.3 | 0.562 |
| 0.05 | 0.05 | clcB | 3 | 23.3 | 0.302 |
| 0.05 | 0.05 | ycaM | 3 | 23.3 | 0.294 |
| 0.05 | 0.05 | yadI | 3 | 23.3 | 0.854 |
| 0.05 | 0.05 | AG1 | 3 | 23.3 | 0.848 |
| 0.05 | 0.05 | ptsI | 3 | 23.55 | 0.567 |

| | | | | | |
|---|---|---|---|---|---|
| 0.05 | 0.05 | clcB | 3 | 23.55 | 0.306 |
| 0.05 | 0.05 | ycaM | 3 | 23.55 | 0.304 |
| 0.05 | 0.05 | yadI | 3 | 23.55 | 0.862 |
| 0.05 | 0.05 | AG1 | 3 | 23.55 | 0.865 |
| 0.05 | 0.05 | ptsI | 3 | 23.8 | 0.565 |
| 0.05 | 0.05 | clcB | 3 | 23.8 | 0.31 |
| 0.05 | 0.05 | ycaM | 3 | 23.8 | 0.306 |
| 0.05 | 0.05 | yadI | 3 | 23.8 | 0.86 |
| 0.05 | 0.05 | AG1 | 3 | 23.8 | 0.866 |
| 0.05 | 0.05 | ptsI | 3 | 24.05 | 0.569 |
| 0.05 | 0.05 | clcB | 3 | 24.05 | 0.308 |
| 0.05 | 0.05 | ycaM | 3 | 24.05 | 0.306 |
| 0.05 | 0.05 | yadI | 3 | 24.05 | 0.861 |
| 0.05 | 0.05 | AG1 | 3 | 24.05 | 0.87 |
| 0.05 | 0.05 | ptsI | 3 | 24.3 | 0.562 |
| 0.05 | 0.05 | clcB | 3 | 24.3 | 0.305 |
| 0.05 | 0.05 | ycaM | 3 | 24.3 | 0.305 |
| 0.05 | 0.05 | yadI | 3 | 24.3 | 0.851 |
| 0.05 | 0.05 | AG1 | 3 | 24.3 | 0.881 |
| 0.05 | 0.05 | ptsI | 4 | 0 | 0.207 |
| 0.05 | 0.05 | clcB | 4 | 0 | 0.198 |
| 0.05 | 0.05 | ycaM | 4 | 0 | 0.194 |
| 0.05 | 0.05 | yadI | 4 | 0 | 0.254 |
| 0.05 | 0.05 | AG1 | 4 | 0 | 0.196 |
| 0.05 | 0.05 | ptsI | 4 | 0.25 | 0.204 |
| 0.05 | 0.05 | clcB | 4 | 0.25 | 0.196 |
| 0.05 | 0.05 | ycaM | 4 | 0.25 | 0.19 |
| 0.05 | 0.05 | yadI | 4 | 0.25 | 0.257 |
| 0.05 | 0.05 | AG1 | 4 | 0.25 | 0.194 |
| 0.05 | 0.05 | ptsI | 4 | 0.5 | 0.204 |
| 0.05 | 0.05 | clcB | 4 | 0.5 | 0.197 |
| 0.05 | 0.05 | ycaM | 4 | 0.5 | 0.19 |
| 0.05 | 0.05 | yadI | 4 | 0.5 | 0.255 |
| 0.05 | 0.05 | AG1 | 4 | 0.5 | 0.195 |
| 0.05 | 0.05 | ptsI | 4 | 0.75 | 0.202 |
| 0.05 | 0.05 | clcB | 4 | 0.75 | 0.199 |
| 0.05 | 0.05 | ycaM | 4 | 0.75 | 0.193 |
| 0.05 | 0.05 | yadI | 4 | 0.75 | 0.259 |
| 0.05 | 0.05 | AG1 | 4 | 0.75 | 0.197 |
| 0.05 | 0.05 | ptsI | 4 | 1 | 0.205 |
| 0.05 | 0.05 | clcB | 4 | 1 | 0.202 |
| 0.05 | 0.05 | ycaM | 4 | 1 | 0.194 |
| 0.05 | 0.05 | yadI | 4 | 1 | 0.262 |
| 0.05 | 0.05 | AG1 | 4 | 1 | 0.2 |
| 0.05 | 0.05 | ptsI | 4 | 1.25 | 0.209 |
| 0.05 | 0.05 | clcB | 4 | 1.25 | 0.206 |
| 0.05 | 0.05 | ycaM | 4 | 1.25 | 0.201 |
| 0.05 | 0.05 | yadI | 4 | 1.25 | 0.265 |

| | | | | | |
|---|---|---|---|---|---|
| 0.05 | 0.05 | AG1 | 4 | 1.25 | 0.205 |
| 0.05 | 0.05 | ptsI | 4 | 1.5 | 0.213 |
| 0.05 | 0.05 | clcB | 4 | 1.5 | 0.212 |
| 0.05 | 0.05 | ycaM | 4 | 1.5 | 0.204 |
| 0.05 | 0.05 | yadI | 4 | 1.5 | 0.272 |
| 0.05 | 0.05 | AG1 | 4 | 1.5 | 0.213 |
| 0.05 | 0.05 | ptsI | 4 | 1.75 | 0.221 |
| 0.05 | 0.05 | clcB | 4 | 1.75 | 0.218 |
| 0.05 | 0.05 | ycaM | 4 | 1.75 | 0.211 |
| 0.05 | 0.05 | yadI | 4 | 1.75 | 0.277 |
| 0.05 | 0.05 | AG1 | 4 | 1.75 | 0.218 |
| 0.05 | 0.05 | ptsI | 4 | 2 | 0.226 |
| 0.05 | 0.05 | clcB | 4 | 2 | 0.226 |
| 0.05 | 0.05 | ycaM | 4 | 2 | 0.219 |
| 0.05 | 0.05 | yadI | 4 | 2 | 0.287 |
| 0.05 | 0.05 | AG1 | 4 | 2 | 0.226 |
| 0.05 | 0.05 | ptsI | 4 | 2.25 | 0.236 |
| 0.05 | 0.05 | clcB | 4 | 2.25 | 0.234 |
| 0.05 | 0.05 | ycaM | 4 | 2.25 | 0.228 |
| 0.05 | 0.05 | yadI | 4 | 2.25 | 0.298 |
| 0.05 | 0.05 | AG1 | 4 | 2.25 | 0.235 |
| 0.05 | 0.05 | ptsI | 4 | 2.5 | 0.243 |
| 0.05 | 0.05 | clcB | 4 | 2.5 | 0.241 |
| 0.05 | 0.05 | ycaM | 4 | 2.5 | 0.236 |
| 0.05 | 0.05 | yadI | 4 | 2.5 | 0.311 |
| 0.05 | 0.05 | AG1 | 4 | 2.5 | 0.244 |
| 0.05 | 0.05 | ptsI | 4 | 2.75 | 0.249 |
| 0.05 | 0.05 | clcB | 4 | 2.75 | 0.247 |
| 0.05 | 0.05 | ycaM | 4 | 2.75 | 0.245 |
| 0.05 | 0.05 | yadI | 4 | 2.75 | 0.33 |
| 0.05 | 0.05 | AG1 | 4 | 2.75 | 0.252 |
| 0.05 | 0.05 | ptsI | 4 | 3 | 0.26 |
| 0.05 | 0.05 | clcB | 4 | 3 | 0.259 |
| 0.05 | 0.05 | ycaM | 4 | 3 | 0.254 |
| 0.05 | 0.05 | yadI | 4 | 3 | 0.324 |
| 0.05 | 0.05 | AG1 | 4 | 3 | 0.261 |
| 0.05 | 0.05 | ptsI | 4 | 3.25 | 0.265 |
| 0.05 | 0.05 | clcB | 4 | 3.25 | 0.27 |
| 0.05 | 0.05 | ycaM | 4 | 3.25 | 0.263 |
| 0.05 | 0.05 | yadI | 4 | 3.25 | 0.334 |
| 0.05 | 0.05 | AG1 | 4 | 3.25 | 0.272 |
| 0.05 | 0.05 | ptsI | 4 | 3.5 | 0.277 |
| 0.05 | 0.05 | clcB | 4 | 3.5 | 0.278 |
| 0.05 | 0.05 | ycaM | 4 | 3.5 | 0.276 |
| 0.05 | 0.05 | yadI | 4 | 3.5 | 0.347 |
| 0.05 | 0.05 | AG1 | 4 | 3.5 | 0.281 |
| 0.05 | 0.05 | ptsI | 4 | 3.75 | 0.285 |
| 0.05 | 0.05 | clcB | 4 | 3.75 | 0.288 |

| | | | | | |
|---|---|---|---|---|---|
| 0.05 | 0.05 | ycaM | 4 | 3.75 | 0.283 |
| 0.05 | 0.05 | yadI | 4 | 3.75 | 0.372 |
| 0.05 | 0.05 | AG1 | 4 | 3.75 | 0.29 |
| 0.05 | 0.05 | ptsI | 4 | 4 | 0.296 |
| 0.05 | 0.05 | clcB | 4 | 4 | 0.301 |
| 0.05 | 0.05 | ycaM | 4 | 4 | 0.295 |
| 0.05 | 0.05 | yadI | 4 | 4 | 0.37 |
| 0.05 | 0.05 | AG1 | 4 | 4 | 0.301 |
| 0.05 | 0.05 | ptsI | 4 | 4.35 | 0.307 |
| 0.05 | 0.05 | clcB | 4 | 4.35 | 0.312 |
| 0.05 | 0.05 | ycaM | 4 | 4.35 | 0.39 |
| 0.05 | 0.05 | yadI | 4 | 4.35 | 0.338 |
| 0.05 | 0.05 | AG1 | 4 | 4.35 | 0.329 |
| 0.05 | 0.05 | ptsI | 4 | 4.6 | 0.3 |
| 0.05 | 0.05 | clcB | 4 | 4.6 | 0.302 |
| 0.05 | 0.05 | ycaM | 4 | 4.6 | 0.344 |
| 0.05 | 0.05 | yadI | 4 | 4.6 | 0.342 |
| 0.05 | 0.05 | AG1 | 4 | 4.6 | 0.342 |
| 0.05 | 0.05 | ptsI | 4 | 4.85 | 0.31 |
| 0.05 | 0.05 | clcB | 4 | 4.85 | 0.307 |
| 0.05 | 0.05 | ycaM | 4 | 4.85 | 0.344 |
| 0.05 | 0.05 | yadI | 4 | 4.85 | 0.355 |
| 0.05 | 0.05 | AG1 | 4 | 4.85 | 0.356 |
| 0.05 | 0.05 | ptsI | 4 | 5.1 | 0.321 |
| 0.05 | 0.05 | clcB | 4 | 5.1 | 0.309 |
| 0.05 | 0.05 | ycaM | 4 | 5.1 | 0.345 |
| 0.05 | 0.05 | yadI | 4 | 5.1 | 0.366 |
| 0.05 | 0.05 | AG1 | 4 | 5.1 | 0.369 |
| 0.05 | 0.05 | ptsI | 4 | 5.35 | 0.333 |
| 0.05 | 0.05 | clcB | 4 | 5.35 | 0.312 |
| 0.05 | 0.05 | ycaM | 4 | 5.35 | 0.347 |
| 0.05 | 0.05 | yadI | 4 | 5.35 | 0.378 |
| 0.05 | 0.05 | AG1 | 4 | 5.35 | 0.385 |
| 0.05 | 0.05 | ptsI | 4 | 5.6 | 0.347 |
| 0.05 | 0.05 | clcB | 4 | 5.6 | 0.311 |
| 0.05 | 0.05 | ycaM | 4 | 5.6 | 0.346 |
| 0.05 | 0.05 | yadI | 4 | 5.6 | 0.396 |
| 0.05 | 0.05 | AG1 | 4 | 5.6 | 0.394 |
| 0.05 | 0.05 | ptsI | 4 | 5.85 | 0.36 |
| 0.05 | 0.05 | clcB | 4 | 5.85 | 0.315 |
| 0.05 | 0.05 | ycaM | 4 | 5.85 | 0.426 |
| 0.05 | 0.05 | yadI | 4 | 5.85 | 0.406 |
| 0.05 | 0.05 | AG1 | 4 | 5.85 | 0.405 |
| 0.05 | 0.05 | ptsI | 4 | 6.1 | 0.375 |
| 0.05 | 0.05 | clcB | 4 | 6.1 | 0.322 |
| 0.05 | 0.05 | ycaM | 4 | 6.1 | 0.362 |
| 0.05 | 0.05 | yadI | 4 | 6.1 | 0.416 |
| 0.05 | 0.05 | AG1 | 4 | 6.1 | 0.415 |

| | | | | | |
|---|---|---|---|---|---|
| 0.05 | 0.05 | ptsI | 4 | 6.35 | 0.384 |
| 0.05 | 0.05 | clcB | 4 | 6.35 | 0.325 |
| 0.05 | 0.05 | ycaM | 4 | 6.35 | 0.365 |
| 0.05 | 0.05 | yadI | 4 | 6.35 | 0.428 |
| 0.05 | 0.05 | AG1 | 4 | 6.35 | 0.425 |
| 0.05 | 0.05 | ptsI | 4 | 6.6 | 0.398 |
| 0.05 | 0.05 | clcB | 4 | 6.6 | 0.326 |
| 0.05 | 0.05 | ycaM | 4 | 6.6 | 0.373 |
| 0.05 | 0.05 | yadI | 4 | 6.6 | 0.428 |
| 0.05 | 0.05 | AG1 | 4 | 6.6 | 0.436 |
| 0.05 | 0.05 | ptsI | 4 | 6.85 | 0.405 |
| 0.05 | 0.05 | clcB | 4 | 6.85 | 0.329 |
| 0.05 | 0.05 | ycaM | 4 | 6.85 | 0.453 |
| 0.05 | 0.05 | yadI | 4 | 6.85 | 0.432 |
| 0.05 | 0.05 | AG1 | 4 | 6.85 | 0.447 |
| 0.05 | 0.05 | ptsI | 4 | 7.1 | 0.413 |
| 0.05 | 0.05 | clcB | 4 | 7.1 | 0.331 |
| 0.05 | 0.05 | ycaM | 4 | 7.1 | 0.447 |
| 0.05 | 0.05 | yadI | 4 | 7.1 | 0.438 |
| 0.05 | 0.05 | AG1 | 4 | 7.1 | 0.461 |
| 0.05 | 0.05 | ptsI | 4 | 7.35 | 0.42 |
| 0.05 | 0.05 | clcB | 4 | 7.35 | 0.332 |
| 0.05 | 0.05 | ycaM | 4 | 7.35 | 0.508 |
| 0.05 | 0.05 | yadI | 4 | 7.35 | 0.445 |
| 0.05 | 0.05 | AG1 | 4 | 7.35 | 0.472 |
| 0.05 | 0.05 | ptsI | 4 | 7.6 | 0.426 |
| 0.05 | 0.05 | clcB | 4 | 7.6 | 0.335 |
| 0.05 | 0.05 | ycaM | 4 | 7.6 | 0.521 |
| 0.05 | 0.05 | yadI | 4 | 7.6 | 0.451 |
| 0.05 | 0.05 | AG1 | 4 | 7.6 | 0.478 |
| 0.05 | 0.05 | ptsI | 4 | 7.85 | 0.435 |
| 0.05 | 0.05 | clcB | 4 | 7.85 | 0.336 |
| 0.05 | 0.05 | ycaM | 4 | 7.85 | 0.363 |
| 0.05 | 0.05 | yadI | 4 | 7.85 | 0.459 |
| 0.05 | 0.05 | AG1 | 4 | 7.85 | 0.487 |
| 0.05 | 0.05 | ptsI | 4 | 8.1 | 0.443 |
| 0.05 | 0.05 | clcB | 4 | 8.1 | 0.337 |
| 0.05 | 0.05 | ycaM | 4 | 8.1 | 0.366 |
| 0.05 | 0.05 | yadI | 4 | 8.1 | 0.46 |
| 0.05 | 0.05 | AG1 | 4 | 8.1 | 0.497 |
| 0.05 | 0.05 | ptsI | 4 | 8.35 | 0.452 |
| 0.05 | 0.05 | clcB | 4 | 8.35 | 0.342 |
| 0.05 | 0.05 | ycaM | 4 | 8.35 | 0.371 |
| 0.05 | 0.05 | yadI | 4 | 8.35 | 0.467 |
| 0.05 | 0.05 | AG1 | 4 | 8.35 | 0.507 |
| 0.05 | 0.05 | ptsI | 4 | 8.83333 | 0.438 |
| 0.05 | 0.05 | clcB | 4 | 8.83333 | 0.306 |
| 0.05 | 0.05 | ycaM | 4 | 8.83333 | 0.284 |

| | | | | | |
|---|---|---|---|---|---|
| 0.05 | 0.05 | yadI | 4 | 8.83333 | 0.521 |
| 0.05 | 0.05 | AG1 | 4 | 8.83333 | 0.536 |
| 0.05 | 0.05 | ptsI | 4 | 9.08333 | 0.437 |
| 0.05 | 0.05 | clcB | 4 | 9.08333 | 0.299 |
| 0.05 | 0.05 | ycaM | 4 | 9.08333 | 0.279 |
| 0.05 | 0.05 | yadI | 4 | 9.08333 | 0.522 |
| 0.05 | 0.05 | AG1 | 4 | 9.08333 | 0.541 |
| 0.05 | 0.05 | ptsI | 4 | 9.33333 | 0.445 |
| 0.05 | 0.05 | clcB | 4 | 9.33333 | 0.297 |
| 0.05 | 0.05 | ycaM | 4 | 9.33333 | 0.276 |
| 0.05 | 0.05 | yadI | 4 | 9.33333 | 0.529 |
| 0.05 | 0.05 | AG1 | 4 | 9.33333 | 0.554 |
| 0.05 | 0.05 | ptsI | 4 | 9.58333 | 0.457 |
| 0.05 | 0.05 | clcB | 4 | 9.58333 | 0.303 |
| 0.05 | 0.05 | ycaM | 4 | 9.58333 | 0.279 |
| 0.05 | 0.05 | yadI | 4 | 9.58333 | 0.534 |
| 0.05 | 0.05 | AG1 | 4 | 9.58333 | 0.565 |
| 0.05 | 0.05 | ptsI | 4 | 9.83333 | 0.471 |
| 0.05 | 0.05 | clcB | 4 | 9.83333 | 0.3 |
| 0.05 | 0.05 | ycaM | 4 | 9.83333 | 0.279 |
| 0.05 | 0.05 | yadI | 4 | 9.83333 | 0.536 |
| 0.05 | 0.05 | AG1 | 4 | 9.83333 | 0.577 |
| 0.05 | 0.05 | ptsI | 4 | 10.0833 | 0.478 |
| 0.05 | 0.05 | clcB | 4 | 10.0833 | 0.301 |
| 0.05 | 0.05 | ycaM | 4 | 10.0833 | 0.279 |
| 0.05 | 0.05 | yadI | 4 | 10.0833 | 0.541 |
| 0.05 | 0.05 | AG1 | 4 | 10.0833 | 0.585 |
| 0.05 | 0.05 | ptsI | 4 | 10.3333 | 0.488 |
| 0.05 | 0.05 | clcB | 4 | 10.3333 | 0.3 |
| 0.05 | 0.05 | ycaM | 4 | 10.3333 | 0.278 |
| 0.05 | 0.05 | yadI | 4 | 10.3333 | 0.548 |
| 0.05 | 0.05 | AG1 | 4 | 10.3333 | 0.596 |
| 0.05 | 0.05 | ptsI | 4 | 10.5833 | 0.494 |
| 0.05 | 0.05 | clcB | 4 | 10.5833 | 0.3 |
| 0.05 | 0.05 | ycaM | 4 | 10.5833 | 0.281 |
| 0.05 | 0.05 | yadI | 4 | 10.5833 | 0.55 |
| 0.05 | 0.05 | AG1 | 4 | 10.5833 | 0.604 |
| 0.05 | 0.05 | ptsI | 4 | 10.8333 | 0.503 |
| 0.05 | 0.05 | clcB | 4 | 10.8333 | 0.3 |
| 0.05 | 0.05 | ycaM | 4 | 10.8333 | 0.283 |
| 0.05 | 0.05 | yadI | 4 | 10.8333 | 0.561 |
| 0.05 | 0.05 | AG1 | 4 | 10.8333 | 0.613 |
| 0.05 | 0.05 | ptsI | 4 | 11.0833 | 0.51 |
| 0.05 | 0.05 | clcB | 4 | 11.0833 | 0.301 |
| 0.05 | 0.05 | ycaM | 4 | 11.0833 | 0.275 |
| 0.05 | 0.05 | yadI | 4 | 11.0833 | 0.561 |
| 0.05 | 0.05 | AG1 | 4 | 11.0833 | 0.62 |
| 0.05 | 0.05 | ptsI | 4 | 11.3333 | 0.52 |

| | | | | | |
|---|---|---|---|---|---|
| 0.05 | 0.05 | clcB | 4 | 11.3333 | 0.302 |
| 0.05 | 0.05 | ycaM | 4 | 11.3333 | 0.283 |
| 0.05 | 0.05 | yadI | 4 | 11.3333 | 0.559 |
| 0.05 | 0.05 | AG1 | 4 | 11.3333 | 0.627 |
| 0.05 | 0.05 | ptsI | 4 | 11.5833 | 0.529 |
| 0.05 | 0.05 | clcB | 4 | 11.5833 | 0.304 |
| 0.05 | 0.05 | ycaM | 4 | 11.5833 | 0.28 |
| 0.05 | 0.05 | yadI | 4 | 11.5833 | 0.565 |
| 0.05 | 0.05 | AG1 | 4 | 11.5833 | 0.632 |
| 0.05 | 0.05 | ptsI | 4 | 11.8333 | 0.534 |
| 0.05 | 0.05 | clcB | 4 | 11.8333 | 0.31 |
| 0.05 | 0.05 | ycaM | 4 | 11.8333 | 0.279 |
| 0.05 | 0.05 | yadI | 4 | 11.8333 | 0.567 |
| 0.05 | 0.05 | AG1 | 4 | 11.8333 | 0.639 |
| 0.05 | 0.05 | ptsI | 4 | 12.0833 | 0.545 |
| 0.05 | 0.05 | clcB | 4 | 12.0833 | 0.31 |
| 0.05 | 0.05 | ycaM | 4 | 12.0833 | 0.278 |
| 0.05 | 0.05 | yadI | 4 | 12.0833 | 0.566 |
| 0.05 | 0.05 | AG1 | 4 | 12.0833 | 0.642 |
| 0.05 | 0.05 | ptsI | 4 | 12.3333 | 0.553 |
| 0.05 | 0.05 | clcB | 4 | 12.3333 | 0.307 |
| 0.05 | 0.05 | ycaM | 4 | 12.3333 | 0.279 |
| 0.05 | 0.05 | yadI | 4 | 12.3333 | 0.572 |
| 0.05 | 0.05 | AG1 | 4 | 12.3333 | 0.647 |
| 0.05 | 0.05 | ptsI | 4 | 12.5833 | 0.56 |
| 0.05 | 0.05 | clcB | 4 | 12.5833 | 0.315 |
| 0.05 | 0.05 | ycaM | 4 | 12.5833 | 0.285 |
| 0.05 | 0.05 | yadI | 4 | 12.5833 | 0.572 |
| 0.05 | 0.05 | AG1 | 4 | 12.5833 | 0.655 |
| 0.05 | 0.05 | ptsI | 4 | 12.8333 | 0.566 |
| 0.05 | 0.05 | clcB | 4 | 12.8333 | 0.304 |
| 0.05 | 0.05 | ycaM | 4 | 12.8333 | 0.284 |
| 0.05 | 0.05 | yadI | 4 | 12.8333 | 0.574 |
| 0.05 | 0.05 | AG1 | 4 | 12.8333 | 0.66 |
| 0.05 | 0.05 | ptsI | 4 | 13.0833 | 0.571 |
| 0.05 | 0.05 | clcB | 4 | 13.0833 | 0.31 |
| 0.05 | 0.05 | ycaM | 4 | 13.0833 | 0.283 |
| 0.05 | 0.05 | yadI | 4 | 13.0833 | 0.571 |
| 0.05 | 0.05 | AG1 | 4 | 13.0833 | 0.664 |
| 0.05 | 0.05 | ptsI | 4 | 13.3333 | 0.579 |
| 0.05 | 0.05 | clcB | 4 | 13.3333 | 0.315 |
| 0.05 | 0.05 | ycaM | 4 | 13.3333 | 0.286 |
| 0.05 | 0.05 | yadI | 4 | 13.3333 | 0.572 |
| 0.05 | 0.05 | AG1 | 4 | 13.3333 | 0.674 |
| 0.05 | 0.05 | ptsI | 4 | 13.5833 | 0.586 |
| 0.05 | 0.05 | clcB | 4 | 13.5833 | 0.31 |
| 0.05 | 0.05 | ycaM | 4 | 13.5833 | 0.284 |
| 0.05 | 0.05 | yadI | 4 | 13.5833 | 0.573 |

| | | | | | |
|---|---|---|---|---|---|
| 0.05 | 0.05 | AG1 | 4 | 13.5833 | 0.676 |
| 0.05 | 0.05 | ptsI | 4 | 13.8333 | 0.592 |
| 0.05 | 0.05 | clcB | 4 | 13.8333 | 0.327 |
| 0.05 | 0.05 | ycaM | 4 | 13.8333 | 0.285 |
| 0.05 | 0.05 | yadI | 4 | 13.8333 | 0.571 |
| 0.05 | 0.05 | AG1 | 4 | 13.8333 | 0.681 |
| 0.05 | 0.05 | ptsI | 4 | 14.0833 | 0.599 |
| 0.05 | 0.05 | clcB | 4 | 14.0833 | 0.317 |
| 0.05 | 0.05 | ycaM | 4 | 14.0833 | 0.286 |
| 0.05 | 0.05 | yadI | 4 | 14.0833 | 0.577 |
| 0.05 | 0.05 | AG1 | 4 | 14.0833 | 0.684 |
| 0.05 | 0.05 | ptsI | 4 | 14.3333 | 0.604 |
| 0.05 | 0.05 | clcB | 4 | 14.3333 | 0.315 |
| 0.05 | 0.05 | ycaM | 4 | 14.3333 | 0.288 |
| 0.05 | 0.05 | yadI | 4 | 14.3333 | 0.573 |
| 0.05 | 0.05 | AG1 | 4 | 14.3333 | 0.692 |
| 0.05 | 0.05 | ptsI | 4 | 14.5833 | 0.61 |
| 0.05 | 0.05 | clcB | 4 | 14.5833 | 0.315 |
| 0.05 | 0.05 | ycaM | 4 | 14.5833 | 0.288 |
| 0.05 | 0.05 | yadI | 4 | 14.5833 | 0.579 |
| 0.05 | 0.05 | AG1 | 4 | 14.5833 | 0.693 |
| 0.05 | 0.05 | ptsI | 4 | 14.8333 | 0.618 |
| 0.05 | 0.05 | clcB | 4 | 14.8333 | 0.313 |
| 0.05 | 0.05 | ycaM | 4 | 14.8333 | 0.286 |
| 0.05 | 0.05 | yadI | 4 | 14.8333 | 0.578 |
| 0.05 | 0.05 | AG1 | 4 | 14.8333 | 0.699 |
| 0.05 | 0.05 | ptsI | 4 | 15.0833 | 0.619 |
| 0.05 | 0.05 | clcB | 4 | 15.0833 | 0.315 |
| 0.05 | 0.05 | ycaM | 4 | 15.0833 | 0.287 |
| 0.05 | 0.05 | yadI | 4 | 15.0833 | 0.58 |
| 0.05 | 0.05 | AG1 | 4 | 15.0833 | 0.7 |
| 0.05 | 0.05 | ptsI | 4 | 15.3333 | 0.625 |
| 0.05 | 0.05 | clcB | 4 | 15.3333 | 0.316 |
| 0.05 | 0.05 | ycaM | 4 | 15.3333 | 0.292 |
| 0.05 | 0.05 | yadI | 4 | 15.3333 | 0.568 |
| 0.05 | 0.05 | AG1 | 4 | 15.3333 | 0.705 |
| 0.05 | 0.05 | ptsI | 4 | 15.5833 | 0.629 |
| 0.05 | 0.05 | clcB | 4 | 15.5833 | 0.316 |
| 0.05 | 0.05 | ycaM | 4 | 15.5833 | 0.292 |
| 0.05 | 0.05 | yadI | 4 | 15.5833 | 0.572 |
| 0.05 | 0.05 | AG1 | 4 | 15.5833 | 0.707 |
| 0.05 | 0.05 | ptsI | 4 | 15.8333 | 0.63 |
| 0.05 | 0.05 | clcB | 4 | 15.8333 | 0.318 |
| 0.05 | 0.05 | ycaM | 4 | 15.8333 | 0.292 |
| 0.05 | 0.05 | yadI | 4 | 15.8333 | 0.573 |
| 0.05 | 0.05 | AG1 | 4 | 15.8333 | 0.71 |
| 0.05 | 0.05 | ptsI | 4 | 16.0833 | 0.635 |
| 0.05 | 0.05 | clcB | 4 | 16.0833 | 0.324 |

| | | | | | |
|---|---|---|---|---|---|
| 0.05 | 0.05 | ycaM | 4 | 16.0833 | 0.291 |
| 0.05 | 0.05 | yadI | 4 | 16.0833 | 0.576 |
| 0.05 | 0.05 | AG1 | 4 | 16.0833 | 0.712 |
| 0.05 | 0.05 | ptsI | 4 | 16.3333 | 0.634 |
| 0.05 | 0.05 | clcB | 4 | 16.3333 | 0.322 |
| 0.05 | 0.05 | ycaM | 4 | 16.3333 | 0.295 |
| 0.05 | 0.05 | yadI | 4 | 16.3333 | 0.57 |
| 0.05 | 0.05 | AG1 | 4 | 16.3333 | 0.716 |
| 0.05 | 0.05 | ptsI | 4 | 16.5833 | 0.637 |
| 0.05 | 0.05 | clcB | 4 | 16.5833 | 0.327 |
| 0.05 | 0.05 | ycaM | 4 | 16.5833 | 0.299 |
| 0.05 | 0.05 | yadI | 4 | 16.5833 | 0.575 |
| 0.05 | 0.05 | AG1 | 4 | 16.5833 | 0.719 |
| 0.05 | 0.05 | ptsI | 4 | 16.8333 | 0.637 |
| 0.05 | 0.05 | clcB | 4 | 16.8333 | 0.328 |
| 0.05 | 0.05 | ycaM | 4 | 16.8333 | 0.296 |
| 0.05 | 0.05 | yadI | 4 | 16.8333 | 0.576 |
| 0.05 | 0.05 | AG1 | 4 | 16.8333 | 0.718 |
| 0.05 | 0.05 | ptsI | 4 | 17.0833 | 0.637 |
| 0.05 | 0.05 | clcB | 4 | 17.0833 | 0.327 |
| 0.05 | 0.05 | ycaM | 4 | 17.0833 | 0.299 |
| 0.05 | 0.05 | yadI | 4 | 17.0833 | 0.568 |
| 0.05 | 0.05 | AG1 | 4 | 17.0833 | 0.724 |
| 0.05 | 0.05 | ptsI | 4 | 17.3333 | 0.643 |
| 0.05 | 0.05 | clcB | 4 | 17.3333 | 0.332 |
| 0.05 | 0.05 | ycaM | 4 | 17.3333 | 0.302 |
| 0.05 | 0.05 | yadI | 4 | 17.3333 | 0.566 |
| 0.05 | 0.05 | AG1 | 4 | 17.3333 | 0.727 |
| 0.05 | 0.05 | ptsI | 4 | 17.5833 | 0.642 |
| 0.05 | 0.05 | clcB | 4 | 17.5833 | 0.337 |
| 0.05 | 0.05 | ycaM | 4 | 17.5833 | 0.305 |
| 0.05 | 0.05 | yadI | 4 | 17.5833 | 0.572 |
| 0.05 | 0.05 | AG1 | 4 | 17.5833 | 0.727 |
| 0.05 | 0.05 | ptsI | 4 | 17.8333 | 0.638 |
| 0.05 | 0.05 | clcB | 4 | 17.8333 | 0.339 |
| 0.05 | 0.05 | ycaM | 4 | 17.8333 | 0.299 |
| 0.05 | 0.05 | yadI | 4 | 17.8333 | 0.577 |
| 0.05 | 0.05 | AG1 | 4 | 17.8333 | 0.734 |
| 0.05 | 0.05 | ptsI | 4 | 18.0833 | 0.642 |
| 0.05 | 0.05 | clcB | 4 | 18.0833 | 0.328 |
| 0.05 | 0.05 | ycaM | 4 | 18.0833 | 0.306 |
| 0.05 | 0.05 | yadI | 4 | 18.0833 | 0.571 |
| 0.05 | 0.05 | AG1 | 4 | 18.0833 | 0.738 |
| 0.05 | 0.05 | ptsI | 4 | 18.3333 | 0.644 |
| 0.05 | 0.05 | clcB | 4 | 18.3333 | 0.328 |
| 0.05 | 0.05 | ycaM | 4 | 18.3333 | 0.312 |
| 0.05 | 0.05 | yadI | 4 | 18.3333 | 0.572 |
| 0.05 | 0.05 | AG1 | 4 | 18.3333 | 0.734 |

| | | | | | |
|---|---|---|---|---|---|
| 0.05 | 0.05 | ptsI | 4 | 18.5833 | 0.645 |
| 0.05 | 0.05 | clcB | 4 | 18.5833 | 0.329 |
| 0.05 | 0.05 | ycaM | 4 | 18.5833 | 0.313 |
| 0.05 | 0.05 | yadI | 4 | 18.5833 | 0.569 |
| 0.05 | 0.05 | AG1 | 4 | 18.5833 | 0.74 |
| 0.05 | 0.05 | ptsI | 4 | 18.8333 | 0.644 |
| 0.05 | 0.05 | clcB | 4 | 18.8333 | 0.329 |
| 0.05 | 0.05 | ycaM | 4 | 18.8333 | 0.311 |
| 0.05 | 0.05 | yadI | 4 | 18.8333 | 0.57 |
| 0.05 | 0.05 | AG1 | 4 | 18.8333 | 0.753 |
| 0.05 | 0.05 | ptsI | 4 | 19.0833 | 0.649 |
| 0.05 | 0.05 | clcB | 4 | 19.0833 | 0.338 |
| 0.05 | 0.05 | ycaM | 4 | 19.0833 | 0.31 |
| 0.05 | 0.05 | yadI | 4 | 19.0833 | 0.567 |
| 0.05 | 0.05 | AG1 | 4 | 19.0833 | 0.749 |
| 0.05 | 0.05 | ptsI | 4 | 19.3333 | 0.649 |
| 0.05 | 0.05 | clcB | 4 | 19.3333 | 0.319 |
| 0.05 | 0.05 | ycaM | 4 | 19.3333 | 0.329 |
| 0.05 | 0.05 | yadI | 4 | 19.3333 | 0.557 |
| 0.05 | 0.05 | AG1 | 4 | 19.3333 | 0.756 |
| 0.05 | 0.05 | ptsI | 4 | 19.5833 | 0.65 |
| 0.05 | 0.05 | clcB | 4 | 19.5833 | 0.337 |
| 0.05 | 0.05 | ycaM | 4 | 19.5833 | 0.313 |
| 0.05 | 0.05 | yadI | 4 | 19.5833 | 0.555 |
| 0.05 | 0.05 | AG1 | 4 | 19.5833 | 0.755 |
| 0.05 | 0.05 | ptsI | 4 | 19.8333 | 0.65 |
| 0.05 | 0.05 | clcB | 4 | 19.8333 | 0.33 |
| 0.05 | 0.05 | ycaM | 4 | 19.8333 | 0.315 |
| 0.05 | 0.05 | yadI | 4 | 19.8333 | 0.564 |
| 0.05 | 0.05 | AG1 | 4 | 19.8333 | 0.754 |
| 0.05 | 0.05 | ptsI | 4 | 20.0833 | 0.65 |
| 0.05 | 0.05 | clcB | 4 | 20.0833 | 0.328 |
| 0.05 | 0.05 | ycaM | 4 | 20.0833 | 0.324 |
| 0.05 | 0.05 | yadI | 4 | 20.0833 | 0.565 |
| 0.05 | 0.05 | AG1 | 4 | 20.0833 | 0.765 |
| 0.05 | 0.05 | ptsI | 4 | 20.3333 | 0.649 |
| 0.05 | 0.05 | clcB | 4 | 20.3333 | 0.317 |
| 0.05 | 0.05 | ycaM | 4 | 20.3333 | 0.341 |
| 0.05 | 0.05 | yadI | 4 | 20.3333 | 0.555 |
| 0.05 | 0.05 | AG1 | 4 | 20.3333 | 0.757 |
| 0.05 | 0.05 | ptsI | 4 | 20.5833 | 0.648 |
| 0.05 | 0.05 | clcB | 4 | 20.5833 | 0.317 |
| 0.05 | 0.05 | ycaM | 4 | 20.5833 | 0.328 |
| 0.05 | 0.05 | yadI | 4 | 20.5833 | 0.56 |
| 0.05 | 0.05 | AG1 | 4 | 20.5833 | 0.767 |
| 0.05 | 0.05 | ptsI | 4 | 20.8333 | 0.652 |
| 0.05 | 0.05 | clcB | 4 | 20.8333 | 0.322 |
| 0.05 | 0.05 | ycaM | 4 | 20.8333 | 0.36 |

| | | | | | |
|---|---|---|---|---|---|
| 0.05 | 0.05 | yadI | 4 | 20.8333 | 0.552 |
| 0.05 | 0.05 | AG1 | 4 | 20.8333 | 0.765 |
| 0.05 | 0.05 | ptsI | 4 | 21.0833 | 0.648 |
| 0.05 | 0.05 | clcB | 4 | 21.0833 | 0.323 |
| 0.05 | 0.05 | ycaM | 4 | 21.0833 | 0.336 |
| 0.05 | 0.05 | yadI | 4 | 21.0833 | 0.55 |
| 0.05 | 0.05 | AG1 | 4 | 21.0833 | 0.767 |
| 0.05 | 0.05 | ptsI | 4 | 21.3333 | 0.653 |
| 0.05 | 0.05 | clcB | 4 | 21.3333 | 0.324 |
| 0.05 | 0.05 | ycaM | 4 | 21.3333 | 0.34 |
| 0.05 | 0.05 | yadI | 4 | 21.3333 | 0.548 |
| 0.05 | 0.05 | AG1 | 4 | 21.3333 | 0.769 |
| 0.05 | 0.05 | ptsI | 4 | 21.5833 | 0.655 |
| 0.05 | 0.05 | clcB | 4 | 21.5833 | 0.335 |
| 0.05 | 0.05 | ycaM | 4 | 21.5833 | 0.33 |
| 0.05 | 0.05 | yadI | 4 | 21.5833 | 0.549 |
| 0.05 | 0.05 | AG1 | 4 | 21.5833 | 0.774 |
| 0.05 | 0.05 | ptsI | 4 | 21.8333 | 0.648 |
| 0.05 | 0.05 | clcB | 4 | 21.8333 | 0.331 |
| 0.05 | 0.05 | ycaM | 4 | 21.8333 | 0.344 |
| 0.05 | 0.05 | yadI | 4 | 21.8333 | 0.539 |
| 0.05 | 0.05 | AG1 | 4 | 21.8333 | 0.777 |
| 0.05 | 0.05 | ptsI | 4 | 22.0833 | 0.652 |
| 0.05 | 0.05 | clcB | 4 | 22.0833 | 0.33 |
| 0.05 | 0.05 | ycaM | 4 | 22.0833 | 0.335 |
| 0.05 | 0.05 | yadI | 4 | 22.0833 | 0.547 |
| 0.05 | 0.05 | AG1 | 4 | 22.0833 | 0.776 |
| 0.05 | 0.05 | ptsI | 4 | 22.3333 | 0.653 |
| 0.05 | 0.05 | clcB | 4 | 22.3333 | 0.331 |
| 0.05 | 0.05 | ycaM | 4 | 22.3333 | 0.324 |
| 0.05 | 0.05 | yadI | 4 | 22.3333 | 0.543 |
| 0.05 | 0.05 | AG1 | 4 | 22.3333 | 0.78 |
| 0.05 | 0.05 | ptsI | 4 | 22.5833 | 0.647 |
| 0.05 | 0.05 | clcB | 4 | 22.5833 | 0.334 |
| 0.05 | 0.05 | ycaM | 4 | 22.5833 | 0.324 |
| 0.05 | 0.05 | yadI | 4 | 22.5833 | 0.54 |
| 0.05 | 0.05 | AG1 | 4 | 22.5833 | 0.783 |
| 0.05 | 0.05 | ptsI | 4 | 22.8333 | 0.64 |
| 0.05 | 0.05 | clcB | 4 | 22.8333 | 0.335 |
| 0.05 | 0.05 | ycaM | 4 | 22.8333 | 0.328 |
| 0.05 | 0.05 | yadI | 4 | 22.8333 | 0.539 |
| 0.05 | 0.05 | AG1 | 4 | 22.8333 | 0.784 |
| 0.05 | 0.05 | ptsI | 4 | 23.0833 | 0.652 |
| 0.05 | 0.05 | clcB | 4 | 23.0833 | 0.344 |
| 0.05 | 0.05 | ycaM | 4 | 23.0833 | 0.326 |
| 0.05 | 0.05 | yadI | 4 | 23.0833 | 0.542 |
| 0.05 | 0.05 | AG1 | 4 | 23.0833 | 0.789 |
| 0.05 | 0.05 | ptsI | 4 | 23.3333 | 0.65 |

| | | | | | |
|---|---|---|---|---|---|
| 0.05 | 0.05 | clcB | 4 | 23.3333 | 0.343 |
| 0.05 | 0.05 | ycaM | 4 | 23.3333 | 0.331 |
| 0.05 | 0.05 | yadI | 4 | 23.3333 | 0.539 |
| 0.05 | 0.05 | AG1 | 4 | 23.3333 | 0.789 |
| 0.05 | 0.05 | ptsI | 4 | 23.5833 | 0.64 |
| 0.05 | 0.05 | clcB | 4 | 23.5833 | 0.336 |
| 0.05 | 0.05 | ycaM | 4 | 23.5833 | 0.332 |
| 0.05 | 0.05 | yadI | 4 | 23.5833 | 0.542 |
| 0.05 | 0.05 | AG1 | 4 | 23.5833 | 0.79 |
| 0.05 | 0.05 | ptsI | 4 | 23.8333 | 0.638 |
| 0.05 | 0.05 | clcB | 4 | 23.8333 | 0.345 |
| 0.05 | 0.05 | ycaM | 4 | 23.8333 | 0.333 |
| 0.05 | 0.05 | yadI | 4 | 23.8333 | 0.543 |
| 0.05 | 0.05 | AG1 | 4 | 23.8333 | 0.794 |
| 0.05 | 0.05 | ptsI | 4 | 24.0833 | 0.642 |
| 0.05 | 0.05 | clcB | 4 | 24.0833 | 0.348 |
| 0.05 | 0.05 | ycaM | 4 | 24.0833 | 0.334 |
| 0.05 | 0.05 | yadI | 4 | 24.0833 | 0.541 |
| 0.05 | 0.05 | AG1 | 4 | 24.0833 | 0.794 |
| 0.05 | 0.05 | ptsI | 5 | 0 | 0.161 |
| 0.05 | 0.05 | clcB | 5 | 0 | 0.157 |
| 0.05 | 0.05 | ycaM | 5 | 0 | 0.165 |
| 0.05 | 0.05 | yadI | 5 | 0 | 0.17 |
| 0.05 | 0.05 | AG1 | 5 | 0 | 0.163 |
| 0.05 | 0.05 | ptsI | 5 | 0.35 | 0.158 |
| 0.05 | 0.05 | clcB | 5 | 0.35 | 0.154 |
| 0.05 | 0.05 | ycaM | 5 | 0.35 | 0.159 |
| 0.05 | 0.05 | yadI | 5 | 0.35 | 0.167 |
| 0.05 | 0.05 | AG1 | 5 | 0.35 | 0.158 |
| 0.05 | 0.05 | ptsI | 5 | 0.6 | 0.158 |
| 0.05 | 0.05 | clcB | 5 | 0.6 | 0.156 |
| 0.05 | 0.05 | ycaM | 5 | 0.6 | 0.159 |
| 0.05 | 0.05 | yadI | 5 | 0.6 | 0.167 |
| 0.05 | 0.05 | AG1 | 5 | 0.6 | 0.157 |
| 0.05 | 0.05 | ptsI | 5 | 0.85 | 0.157 |
| 0.05 | 0.05 | clcB | 5 | 0.85 | 0.156 |
| 0.05 | 0.05 | ycaM | 5 | 0.85 | 0.159 |
| 0.05 | 0.05 | yadI | 5 | 0.85 | 0.167 |
| 0.05 | 0.05 | AG1 | 5 | 0.85 | 0.159 |
| 0.05 | 0.05 | ptsI | 5 | 1.1 | 0.159 |
| 0.05 | 0.05 | clcB | 5 | 1.1 | 0.158 |
| 0.05 | 0.05 | ycaM | 5 | 1.1 | 0.162 |
| 0.05 | 0.05 | yadI | 5 | 1.1 | 0.168 |
| 0.05 | 0.05 | AG1 | 5 | 1.1 | 0.162 |
| 0.05 | 0.05 | ptsI | 5 | 1.35 | 0.161 |
| 0.05 | 0.05 | clcB | 5 | 1.35 | 0.16 |
| 0.05 | 0.05 | ycaM | 5 | 1.35 | 0.163 |
| 0.05 | 0.05 | yadI | 5 | 1.35 | 0.17 |

| | | | | | |
|---|---|---|---|---|---|
| 0.05 | 0.05 | AG1 | 5 | 1.35 | 0.163 |
| 0.05 | 0.05 | ptsI | 5 | 1.6 | 0.163 |
| 0.05 | 0.05 | clcB | 5 | 1.6 | 0.16 |
| 0.05 | 0.05 | ycaM | 5 | 1.6 | 0.165 |
| 0.05 | 0.05 | yadI | 5 | 1.6 | 0.173 |
| 0.05 | 0.05 | AG1 | 5 | 1.6 | 0.167 |
| 0.05 | 0.05 | ptsI | 5 | 1.85 | 0.166 |
| 0.05 | 0.05 | clcB | 5 | 1.85 | 0.164 |
| 0.05 | 0.05 | ycaM | 5 | 1.85 | 0.17 |
| 0.05 | 0.05 | yadI | 5 | 1.85 | 0.176 |
| 0.05 | 0.05 | AG1 | 5 | 1.85 | 0.169 |
| 0.05 | 0.05 | ptsI | 5 | 2.1 | 0.17 |
| 0.05 | 0.05 | clcB | 5 | 2.1 | 0.167 |
| 0.05 | 0.05 | ycaM | 5 | 2.1 | 0.175 |
| 0.05 | 0.05 | yadI | 5 | 2.1 | 0.181 |
| 0.05 | 0.05 | AG1 | 5 | 2.1 | 0.174 |
| 0.05 | 0.05 | ptsI | 5 | 2.35 | 0.175 |
| 0.05 | 0.05 | clcB | 5 | 2.35 | 0.172 |
| 0.05 | 0.05 | ycaM | 5 | 2.35 | 0.18 |
| 0.05 | 0.05 | yadI | 5 | 2.35 | 0.185 |
| 0.05 | 0.05 | AG1 | 5 | 2.35 | 0.176 |
| 0.05 | 0.05 | ptsI | 5 | 2.6 | 0.175 |
| 0.05 | 0.05 | clcB | 5 | 2.6 | 0.174 |
| 0.05 | 0.05 | ycaM | 5 | 2.6 | 0.184 |
| 0.05 | 0.05 | yadI | 5 | 2.6 | 0.187 |
| 0.05 | 0.05 | AG1 | 5 | 2.6 | 0.182 |
| 0.05 | 0.05 | ptsI | 5 | 2.85 | 0.183 |
| 0.05 | 0.05 | clcB | 5 | 2.85 | 0.181 |
| 0.05 | 0.05 | ycaM | 5 | 2.85 | 0.192 |
| 0.05 | 0.05 | yadI | 5 | 2.85 | 0.196 |
| 0.05 | 0.05 | AG1 | 5 | 2.85 | 0.186 |
| 0.05 | 0.05 | ptsI | 5 | 3.1 | 0.187 |
| 0.05 | 0.05 | clcB | 5 | 3.1 | 0.189 |
| 0.05 | 0.05 | ycaM | 5 | 3.1 | 0.198 |
| 0.05 | 0.05 | yadI | 5 | 3.1 | 0.203 |
| 0.05 | 0.05 | AG1 | 5 | 3.1 | 0.192 |
| 0.05 | 0.05 | ptsI | 5 | 3.35 | 0.196 |
| 0.05 | 0.05 | clcB | 5 | 3.35 | 0.198 |
| 0.05 | 0.05 | ycaM | 5 | 3.35 | 0.209 |
| 0.05 | 0.05 | yadI | 5 | 3.35 | 0.212 |
| 0.05 | 0.05 | AG1 | 5 | 3.35 | 0.199 |
| 0.05 | 0.05 | ptsI | 5 | 3.6 | 0.205 |
| 0.05 | 0.05 | clcB | 5 | 3.6 | 0.209 |
| 0.05 | 0.05 | ycaM | 5 | 3.6 | 0.22 |
| 0.05 | 0.05 | yadI | 5 | 3.6 | 0.224 |
| 0.05 | 0.05 | AG1 | 5 | 3.6 | 0.206 |
| 0.05 | 0.05 | ptsI | 5 | 3.85 | 0.221 |
| 0.05 | 0.05 | clcB | 5 | 3.85 | 0.219 |

| | | | | | |
|---|---|---|---|---|---|
| 0.05 | 0.05 | ycaM | 5 | 3.85 | 0.232 |
| 0.05 | 0.05 | yadI | 5 | 3.85 | 0.233 |
| 0.05 | 0.05 | AG1 | 5 | 3.85 | 0.214 |
| 0.05 | 0.05 | ptsI | 5 | 4.38333 | 0.233 |
| 0.05 | 0.05 | clcB | 5 | 4.38333 | 0.267 |
| 0.05 | 0.05 | ycaM | 5 | 4.38333 | 0.262 |
| 0.05 | 0.05 | yadI | 5 | 4.38333 | 0.278 |
| 0.05 | 0.05 | AG1 | 5 | 4.38333 | 0.252 |
| 0.05 | 0.05 | ptsI | 5 | 4.63333 | 0.239 |
| 0.05 | 0.05 | clcB | 5 | 4.63333 | 0.261 |
| 0.05 | 0.05 | ycaM | 5 | 4.63333 | 0.254 |
| 0.05 | 0.05 | yadI | 5 | 4.63333 | 0.279 |
| 0.05 | 0.05 | AG1 | 5 | 4.63333 | 0.263 |
| 0.05 | 0.05 | ptsI | 5 | 4.88333 | 0.249 |
| 0.05 | 0.05 | clcB | 5 | 4.88333 | 0.284 |
| 0.05 | 0.05 | ycaM | 5 | 4.88333 | 0.268 |
| 0.05 | 0.05 | yadI | 5 | 4.88333 | 0.296 |
| 0.05 | 0.05 | AG1 | 5 | 4.88333 | 0.284 |
| 0.05 | 0.05 | ptsI | 5 | 5.13333 | 0.256 |
| 0.05 | 0.05 | clcB | 5 | 5.13333 | 0.297 |
| 0.05 | 0.05 | ycaM | 5 | 5.13333 | 0.273 |
| 0.05 | 0.05 | yadI | 5 | 5.13333 | 0.312 |
| 0.05 | 0.05 | AG1 | 5 | 5.13333 | 0.299 |
| 0.05 | 0.05 | ptsI | 5 | 5.38333 | 0.269 |
| 0.05 | 0.05 | clcB | 5 | 5.38333 | 0.306 |
| 0.05 | 0.05 | ycaM | 5 | 5.38333 | 0.277 |
| 0.05 | 0.05 | yadI | 5 | 5.38333 | 0.329 |
| 0.05 | 0.05 | AG1 | 5 | 5.38333 | 0.315 |
| 0.05 | 0.05 | ptsI | 5 | 5.63333 | 0.28 |
| 0.05 | 0.05 | clcB | 5 | 5.63333 | 0.313 |
| 0.05 | 0.05 | ycaM | 5 | 5.63333 | 0.284 |
| 0.05 | 0.05 | yadI | 5 | 5.63333 | 0.342 |
| 0.05 | 0.05 | AG1 | 5 | 5.63333 | 0.329 |
| 0.05 | 0.05 | ptsI | 5 | 5.88333 | 0.291 |
| 0.05 | 0.05 | clcB | 5 | 5.88333 | 0.321 |
| 0.05 | 0.05 | ycaM | 5 | 5.88333 | 0.288 |
| 0.05 | 0.05 | yadI | 5 | 5.88333 | 0.357 |
| 0.05 | 0.05 | AG1 | 5 | 5.88333 | 0.34 |
| 0.05 | 0.05 | ptsI | 5 | 6.13333 | 0.303 |
| 0.05 | 0.05 | clcB | 5 | 6.13333 | 0.33 |
| 0.05 | 0.05 | ycaM | 5 | 6.13333 | 0.292 |
| 0.05 | 0.05 | yadI | 5 | 6.13333 | 0.37 |
| 0.05 | 0.05 | AG1 | 5 | 6.13333 | 0.35 |
| 0.05 | 0.05 | ptsI | 5 | 6.38333 | 0.316 |
| 0.05 | 0.05 | clcB | 5 | 6.38333 | 0.334 |
| 0.05 | 0.05 | ycaM | 5 | 6.38333 | 0.298 |
| 0.05 | 0.05 | yadI | 5 | 6.38333 | 0.384 |
| 0.05 | 0.05 | AG1 | 5 | 6.38333 | 0.366 |

| | | | | | |
|---|---|---|---|---|---|
| 0.05 | 0.05 | ptsI | 5 | 6.63333 | 0.326 |
| 0.05 | 0.05 | clcB | 5 | 6.63333 | 0.34 |
| 0.05 | 0.05 | ycaM | 5 | 6.63333 | 0.3 |
| 0.05 | 0.05 | yadI | 5 | 6.63333 | 0.398 |
| 0.05 | 0.05 | AG1 | 5 | 6.63333 | 0.381 |
| 0.05 | 0.05 | ptsI | 5 | 6.88333 | 0.339 |
| 0.05 | 0.05 | clcB | 5 | 6.88333 | 0.343 |
| 0.05 | 0.05 | ycaM | 5 | 6.88333 | 0.302 |
| 0.05 | 0.05 | yadI | 5 | 6.88333 | 0.413 |
| 0.05 | 0.05 | AG1 | 5 | 6.88333 | 0.394 |
| 0.05 | 0.05 | ptsI | 5 | 7.13333 | 0.352 |
| 0.05 | 0.05 | clcB | 5 | 7.13333 | 0.348 |
| 0.05 | 0.05 | ycaM | 5 | 7.13333 | 0.306 |
| 0.05 | 0.05 | yadI | 5 | 7.13333 | 0.428 |
| 0.05 | 0.05 | AG1 | 5 | 7.13333 | 0.409 |
| 0.05 | 0.05 | ptsI | 5 | 7.38333 | 0.362 |
| 0.05 | 0.05 | clcB | 5 | 7.38333 | 0.351 |
| 0.05 | 0.05 | ycaM | 5 | 7.38333 | 0.309 |
| 0.05 | 0.05 | yadI | 5 | 7.38333 | 0.443 |
| 0.05 | 0.05 | AG1 | 5 | 7.38333 | 0.425 |
| 0.05 | 0.05 | ptsI | 5 | 7.63333 | 0.374 |
| 0.05 | 0.05 | clcB | 5 | 7.63333 | 0.354 |
| 0.05 | 0.05 | ycaM | 5 | 7.63333 | 0.31 |
| 0.05 | 0.05 | yadI | 5 | 7.63333 | 0.456 |
| 0.05 | 0.05 | AG1 | 5 | 7.63333 | 0.438 |
| 0.05 | 0.05 | ptsI | 5 | 7.88333 | 0.384 |
| 0.05 | 0.05 | clcB | 5 | 7.88333 | 0.358 |
| 0.05 | 0.05 | ycaM | 5 | 7.88333 | 0.313 |
| 0.05 | 0.05 | yadI | 5 | 7.88333 | 0.468 |
| 0.05 | 0.05 | AG1 | 5 | 7.88333 | 0.454 |
| 0.05 | 0.05 | ptsI | 5 | 8.13333 | 0.398 |
| 0.05 | 0.05 | clcB | 5 | 8.13333 | 0.361 |
| 0.05 | 0.05 | ycaM | 5 | 8.13333 | 0.317 |
| 0.05 | 0.05 | yadI | 5 | 8.13333 | 0.477 |
| 0.05 | 0.05 | AG1 | 5 | 8.13333 | 0.468 |
| 0.05 | 0.05 | ptsI | 5 | 8.38333 | 0.409 |
| 0.05 | 0.05 | clcB | 5 | 8.38333 | 0.364 |
| 0.05 | 0.05 | ycaM | 5 | 8.38333 | 0.316 |
| 0.05 | 0.05 | yadI | 5 | 8.38333 | 0.486 |
| 0.05 | 0.05 | AG1 | 5 | 8.38333 | 0.484 |
| 0.05 | 0.05 | ptsI | 5 | 8.88333 | 0.34 |
| 0.05 | 0.05 | clcB | 5 | 8.88333 | 0.299 |
| 0.05 | 0.05 | ycaM | 5 | 8.88333 | 0.312 |
| 0.05 | 0.05 | yadI | 5 | 8.88333 | 0.442 |
| 0.05 | 0.05 | AG1 | 5 | 8.88333 | 0.468 |
| 0.05 | 0.05 | ptsI | 5 | 9.13333 | 0.349 |
| 0.05 | 0.05 | clcB | 5 | 9.13333 | 0.294 |
| 0.05 | 0.05 | ycaM | 5 | 9.13333 | 0.308 |

| | | | | | |
|---|---|---|---|---|---|
| 0.05 | 0.05 | yadI | 5 | 9.13333 | 0.438 |
| 0.05 | 0.05 | AG1 | 5 | 9.13333 | 0.475 |
| 0.05 | 0.05 | ptsI | 5 | 9.38333 | 0.361 |
| 0.05 | 0.05 | clcB | 5 | 9.38333 | 0.298 |
| 0.05 | 0.05 | ycaM | 5 | 9.38333 | 0.309 |
| 0.05 | 0.05 | yadI | 5 | 9.38333 | 0.446 |
| 0.05 | 0.05 | AG1 | 5 | 9.38333 | 0.463 |
| 0.05 | 0.05 | ptsI | 5 | 9.63333 | 0.372 |
| 0.05 | 0.05 | clcB | 5 | 9.63333 | 0.307 |
| 0.05 | 0.05 | ycaM | 5 | 9.63333 | 0.311 |
| 0.05 | 0.05 | yadI | 5 | 9.63333 | 0.46 |
| 0.05 | 0.05 | AG1 | 5 | 9.63333 | 0.501 |
| 0.05 | 0.05 | ptsI | 5 | 9.88333 | 0.378 |
| 0.05 | 0.05 | clcB | 5 | 9.88333 | 0.305 |
| 0.05 | 0.05 | ycaM | 5 | 9.88333 | 0.311 |
| 0.05 | 0.05 | yadI | 5 | 9.88333 | 0.47 |
| 0.05 | 0.05 | AG1 | 5 | 9.88333 | 0.511 |
| 0.05 | 0.05 | ptsI | 5 | 10.1333 | 0.389 |
| 0.05 | 0.05 | clcB | 5 | 10.1333 | 0.308 |
| 0.05 | 0.05 | ycaM | 5 | 10.1333 | 0.309 |
| 0.05 | 0.05 | yadI | 5 | 10.1333 | 0.478 |
| 0.05 | 0.05 | AG1 | 5 | 10.1333 | 0.518 |
| 0.05 | 0.05 | ptsI | 5 | 10.3833 | 0.398 |
| 0.05 | 0.05 | clcB | 5 | 10.3833 | 0.313 |
| 0.05 | 0.05 | ycaM | 5 | 10.3833 | 0.31 |
| 0.05 | 0.05 | yadI | 5 | 10.3833 | 0.487 |
| 0.05 | 0.05 | AG1 | 5 | 10.3833 | 0.53 |
| 0.05 | 0.05 | ptsI | 5 | 10.6333 | 0.406 |
| 0.05 | 0.05 | clcB | 5 | 10.6333 | 0.311 |
| 0.05 | 0.05 | ycaM | 5 | 10.6333 | 0.312 |
| 0.05 | 0.05 | yadI | 5 | 10.6333 | 0.497 |
| 0.05 | 0.05 | AG1 | 5 | 10.6333 | 0.545 |
| 0.05 | 0.05 | ptsI | 5 | 10.8833 | 0.418 |
| 0.05 | 0.05 | clcB | 5 | 10.8833 | 0.316 |
| 0.05 | 0.05 | ycaM | 5 | 10.8833 | 0.312 |
| 0.05 | 0.05 | yadI | 5 | 10.8833 | 0.504 |
| 0.05 | 0.05 | AG1 | 5 | 10.8833 | 0.556 |
| 0.05 | 0.05 | ptsI | 5 | 11.1333 | 0.425 |
| 0.05 | 0.05 | clcB | 5 | 11.1333 | 0.31 |
| 0.05 | 0.05 | ycaM | 5 | 11.1333 | 0.31 |
| 0.05 | 0.05 | yadI | 5 | 11.1333 | 0.51 |
| 0.05 | 0.05 | AG1 | 5 | 11.1333 | 0.566 |
| 0.05 | 0.05 | ptsI | 5 | 11.3833 | 0.435 |
| 0.05 | 0.05 | clcB | 5 | 11.3833 | 0.31 |
| 0.05 | 0.05 | ycaM | 5 | 11.3833 | 0.309 |
| 0.05 | 0.05 | yadI | 5 | 11.3833 | 0.518 |
| 0.05 | 0.05 | AG1 | 5 | 11.3833 | 0.577 |
| 0.05 | 0.05 | ptsI | 5 | 11.6333 | 0.443 |

| | | | | | |
|---|---|---|---|---|---|
| 0.05 | 0.05 | clcB | 5 | 11.6333 | 0.309 |
| 0.05 | 0.05 | ycaM | 5 | 11.6333 | 0.31 |
| 0.05 | 0.05 | yadI | 5 | 11.6333 | 0.526 |
| 0.05 | 0.05 | AG1 | 5 | 11.6333 | 0.589 |
| 0.05 | 0.05 | ptsI | 5 | 11.8833 | 0.451 |
| 0.05 | 0.05 | clcB | 5 | 11.8833 | 0.315 |
| 0.05 | 0.05 | ycaM | 5 | 11.8833 | 0.312 |
| 0.05 | 0.05 | yadI | 5 | 11.8833 | 0.532 |
| 0.05 | 0.05 | AG1 | 5 | 11.8833 | 0.601 |
| 0.05 | 0.05 | ptsI | 5 | 12.1333 | 0.458 |
| 0.05 | 0.05 | clcB | 5 | 12.1333 | 0.312 |
| 0.05 | 0.05 | ycaM | 5 | 12.1333 | 0.313 |
| 0.05 | 0.05 | yadI | 5 | 12.1333 | 0.54 |
| 0.05 | 0.05 | AG1 | 5 | 12.1333 | 0.612 |
| 0.05 | 0.05 | ptsI | 5 | 12.3833 | 0.465 |
| 0.05 | 0.05 | clcB | 5 | 12.3833 | 0.311 |
| 0.05 | 0.05 | ycaM | 5 | 12.3833 | 0.311 |
| 0.05 | 0.05 | yadI | 5 | 12.3833 | 0.539 |
| 0.05 | 0.05 | AG1 | 5 | 12.3833 | 0.612 |
| 0.05 | 0.05 | ptsI | 5 | 12.6333 | 0.473 |
| 0.05 | 0.05 | clcB | 5 | 12.6333 | 0.312 |
| 0.05 | 0.05 | ycaM | 5 | 12.6333 | 0.314 |
| 0.05 | 0.05 | yadI | 5 | 12.6333 | 0.558 |
| 0.05 | 0.05 | AG1 | 5 | 12.6333 | 0.631 |
| 0.05 | 0.05 | ptsI | 5 | 12.8833 | 0.484 |
| 0.05 | 0.05 | clcB | 5 | 12.8833 | 0.311 |
| 0.05 | 0.05 | ycaM | 5 | 12.8833 | 0.316 |
| 0.05 | 0.05 | yadI | 5 | 12.8833 | 0.561 |
| 0.05 | 0.05 | AG1 | 5 | 12.8833 | 0.642 |
| 0.05 | 0.05 | ptsI | 5 | 13.1333 | 0.493 |
| 0.05 | 0.05 | clcB | 5 | 13.1333 | 0.314 |
| 0.05 | 0.05 | ycaM | 5 | 13.1333 | 0.315 |
| 0.05 | 0.05 | yadI | 5 | 13.1333 | 0.566 |
| 0.05 | 0.05 | AG1 | 5 | 13.1333 | 0.648 |
| 0.05 | 0.05 | ptsI | 5 | 13.3833 | 0.496 |
| 0.05 | 0.05 | clcB | 5 | 13.3833 | 0.31 |
| 0.05 | 0.05 | ycaM | 5 | 13.3833 | 0.314 |
| 0.05 | 0.05 | yadI | 5 | 13.3833 | 0.572 |
| 0.05 | 0.05 | AG1 | 5 | 13.3833 | 0.655 |
| 0.05 | 0.05 | ptsI | 5 | 13.6333 | 0.508 |
| 0.05 | 0.05 | clcB | 5 | 13.6333 | 0.316 |
| 0.05 | 0.05 | ycaM | 5 | 13.6333 | 0.317 |
| 0.05 | 0.05 | yadI | 5 | 13.6333 | 0.577 |
| 0.05 | 0.05 | AG1 | 5 | 13.6333 | 0.66 |
| 0.05 | 0.05 | ptsI | 5 | 13.8833 | 0.512 |
| 0.05 | 0.05 | clcB | 5 | 13.8833 | 0.318 |
| 0.05 | 0.05 | ycaM | 5 | 13.8833 | 0.318 |
| 0.05 | 0.05 | yadI | 5 | 13.8833 | 0.585 |

| | | | | | |
|---|---|---|---|---|---|
| 0.05 | 0.05 | AG1 | 5 | 13.8833 | 0.672 |
| 0.05 | 0.05 | ptsI | 5 | 14.1333 | 0.522 |
| 0.05 | 0.05 | clcB | 5 | 14.1333 | 0.314 |
| 0.05 | 0.05 | ycaM | 5 | 14.1333 | 0.317 |
| 0.05 | 0.05 | yadI | 5 | 14.1333 | 0.589 |
| 0.05 | 0.05 | AG1 | 5 | 14.1333 | 0.676 |
| 0.05 | 0.05 | ptsI | 5 | 14.3833 | 0.53 |
| 0.05 | 0.05 | clcB | 5 | 14.3833 | 0.309 |
| 0.05 | 0.05 | ycaM | 5 | 14.3833 | 0.32 |
| 0.05 | 0.05 | yadI | 5 | 14.3833 | 0.596 |
| 0.05 | 0.05 | AG1 | 5 | 14.3833 | 0.682 |
| 0.05 | 0.05 | ptsI | 5 | 14.6333 | 0.542 |
| 0.05 | 0.05 | clcB | 5 | 14.6333 | 0.312 |
| 0.05 | 0.05 | ycaM | 5 | 14.6333 | 0.32 |
| 0.05 | 0.05 | yadI | 5 | 14.6333 | 0.602 |
| 0.05 | 0.05 | AG1 | 5 | 14.6333 | 0.687 |
| 0.05 | 0.05 | ptsI | 5 | 14.8833 | 0.543 |
| 0.05 | 0.05 | clcB | 5 | 14.8833 | 0.314 |
| 0.05 | 0.05 | ycaM | 5 | 14.8833 | 0.32 |
| 0.05 | 0.05 | yadI | 5 | 14.8833 | 0.603 |
| 0.05 | 0.05 | AG1 | 5 | 14.8833 | 0.692 |
| 0.05 | 0.05 | ptsI | 5 | 15.1333 | 0.55 |
| 0.05 | 0.05 | clcB | 5 | 15.1333 | 0.313 |
| 0.05 | 0.05 | ycaM | 5 | 15.1333 | 0.322 |
| 0.05 | 0.05 | yadI | 5 | 15.1333 | 0.607 |
| 0.05 | 0.05 | AG1 | 5 | 15.1333 | 0.697 |
| 0.05 | 0.05 | ptsI | 5 | 15.3833 | 0.557 |
| 0.05 | 0.05 | clcB | 5 | 15.3833 | 0.312 |
| 0.05 | 0.05 | ycaM | 5 | 15.3833 | 0.322 |
| 0.05 | 0.05 | yadI | 5 | 15.3833 | 0.61 |
| 0.05 | 0.05 | AG1 | 5 | 15.3833 | 0.703 |
| 0.05 | 0.05 | ptsI | 5 | 15.6333 | 0.561 |
| 0.05 | 0.05 | clcB | 5 | 15.6333 | 0.306 |
| 0.05 | 0.05 | ycaM | 5 | 15.6333 | 0.322 |
| 0.05 | 0.05 | yadI | 5 | 15.6333 | 0.614 |
| 0.05 | 0.05 | AG1 | 5 | 15.6333 | 0.709 |
| 0.05 | 0.05 | ptsI | 5 | 15.8833 | 0.567 |
| 0.05 | 0.05 | clcB | 5 | 15.8833 | 0.31 |
| 0.05 | 0.05 | ycaM | 5 | 15.8833 | 0.323 |
| 0.05 | 0.05 | yadI | 5 | 15.8833 | 0.618 |
| 0.05 | 0.05 | AG1 | 5 | 15.8833 | 0.713 |
| 0.05 | 0.05 | ptsI | 5 | 16.1333 | 0.575 |
| 0.05 | 0.05 | clcB | 5 | 16.1333 | 0.315 |
| 0.05 | 0.05 | ycaM | 5 | 16.1333 | 0.322 |
| 0.05 | 0.05 | yadI | 5 | 16.1333 | 0.621 |
| 0.05 | 0.05 | AG1 | 5 | 16.1333 | 0.685 |
| 0.05 | 0.05 | ptsI | 5 | 16.3833 | 0.578 |
| 0.05 | 0.05 | clcB | 5 | 16.3833 | 0.312 |

| | | | | | |
|---|---|---|---|---|---|
| 0.05 | 0.05 | ycaM | 5 | 16.3833 | 0.323 |
| 0.05 | 0.05 | yadI | 5 | 16.3833 | 0.628 |
| 0.05 | 0.05 | AG1 | 5 | 16.3833 | 0.723 |
| 0.05 | 0.05 | ptsI | 5 | 16.6333 | 0.583 |
| 0.05 | 0.05 | clcB | 5 | 16.6333 | 0.313 |
| 0.05 | 0.05 | ycaM | 5 | 16.6333 | 0.325 |
| 0.05 | 0.05 | yadI | 5 | 16.6333 | 0.629 |
| 0.05 | 0.05 | AG1 | 5 | 16.6333 | 0.725 |
| 0.05 | 0.05 | ptsI | 5 | 16.8833 | 0.591 |
| 0.05 | 0.05 | clcB | 5 | 16.8833 | 0.319 |
| 0.05 | 0.05 | ycaM | 5 | 16.8833 | 0.326 |
| 0.05 | 0.05 | yadI | 5 | 16.8833 | 0.634 |
| 0.05 | 0.05 | AG1 | 5 | 16.8833 | 0.729 |
| 0.05 | 0.05 | ptsI | 5 | 17.1333 | 0.594 |
| 0.05 | 0.05 | clcB | 5 | 17.1333 | 0.317 |
| 0.05 | 0.05 | ycaM | 5 | 17.1333 | 0.325 |
| 0.05 | 0.05 | yadI | 5 | 17.1333 | 0.637 |
| 0.05 | 0.05 | AG1 | 5 | 17.1333 | 0.738 |
| 0.05 | 0.05 | ptsI | 5 | 17.3833 | 0.595 |
| 0.05 | 0.05 | clcB | 5 | 17.3833 | 0.318 |
| 0.05 | 0.05 | ycaM | 5 | 17.3833 | 0.327 |
| 0.05 | 0.05 | yadI | 5 | 17.3833 | 0.64 |
| 0.05 | 0.05 | AG1 | 5 | 17.3833 | 0.736 |
| 0.05 | 0.05 | ptsI | 5 | 17.6333 | 0.601 |
| 0.05 | 0.05 | clcB | 5 | 17.6333 | 0.324 |
| 0.05 | 0.05 | ycaM | 5 | 17.6333 | 0.325 |
| 0.05 | 0.05 | yadI | 5 | 17.6333 | 0.642 |
| 0.05 | 0.05 | AG1 | 5 | 17.6333 | 0.742 |
| 0.05 | 0.05 | ptsI | 5 | 17.8833 | 0.606 |
| 0.05 | 0.05 | clcB | 5 | 17.8833 | 0.325 |
| 0.05 | 0.05 | ycaM | 5 | 17.8833 | 0.328 |
| 0.05 | 0.05 | yadI | 5 | 17.8833 | 0.634 |
| 0.05 | 0.05 | AG1 | 5 | 17.8833 | 0.734 |
| 0.05 | 0.05 | ptsI | 5 | 18.1333 | 0.612 |
| 0.05 | 0.05 | clcB | 5 | 18.1333 | 0.332 |
| 0.05 | 0.05 | ycaM | 5 | 18.1333 | 0.335 |
| 0.05 | 0.05 | yadI | 5 | 18.1333 | 0.646 |
| 0.05 | 0.05 | AG1 | 5 | 18.1333 | 0.751 |
| 0.05 | 0.05 | ptsI | 5 | 18.3833 | 0.618 |
| 0.05 | 0.05 | clcB | 5 | 18.3833 | 0.333 |
| 0.05 | 0.05 | ycaM | 5 | 18.3833 | 0.332 |
| 0.05 | 0.05 | yadI | 5 | 18.3833 | 0.653 |
| 0.05 | 0.05 | AG1 | 5 | 18.3833 | 0.753 |
| 0.05 | 0.05 | ptsI | 5 | 18.6333 | 0.622 |
| 0.05 | 0.05 | clcB | 5 | 18.6333 | 0.333 |
| 0.05 | 0.05 | ycaM | 5 | 18.6333 | 0.34 |
| 0.05 | 0.05 | yadI | 5 | 18.6333 | 0.653 |
| 0.05 | 0.05 | AG1 | 5 | 18.6333 | 0.756 |

| | | | | | |
|---|---|---|---|---|---|
| 0.05 | 0.05 | ptsI | 5 | 18.8833 | 0.625 |
| 0.05 | 0.05 | clcB | 5 | 18.8833 | 0.334 |
| 0.05 | 0.05 | ycaM | 5 | 18.8833 | 0.343 |
| 0.05 | 0.05 | yadI | 5 | 18.8833 | 0.655 |
| 0.05 | 0.05 | AG1 | 5 | 18.8833 | 0.76 |
| 0.05 | 0.05 | ptsI | 5 | 19.1333 | 0.63 |
| 0.05 | 0.05 | clcB | 5 | 19.1333 | 0.338 |
| 0.05 | 0.05 | ycaM | 5 | 19.1333 | 0.343 |
| 0.05 | 0.05 | yadI | 5 | 19.1333 | 0.664 |
| 0.05 | 0.05 | AG1 | 5 | 19.1333 | 0.768 |
| 0.05 | 0.05 | ptsI | 5 | 19.3833 | 0.626 |
| 0.05 | 0.05 | clcB | 5 | 19.3833 | 0.34 |
| 0.05 | 0.05 | ycaM | 5 | 19.3833 | 0.34 |
| 0.05 | 0.05 | yadI | 5 | 19.3833 | 0.661 |
| 0.05 | 0.05 | AG1 | 5 | 19.3833 | 0.77 |
| 0.05 | 0.05 | ptsI | 5 | 19.6333 | 0.632 |
| 0.05 | 0.05 | clcB | 5 | 19.6333 | 0.341 |
| 0.05 | 0.05 | ycaM | 5 | 19.6333 | 0.351 |
| 0.05 | 0.05 | yadI | 5 | 19.6333 | 0.661 |
| 0.05 | 0.05 | AG1 | 5 | 19.6333 | 0.768 |
| 0.05 | 0.05 | ptsI | 5 | 19.8833 | 0.633 |
| 0.05 | 0.05 | clcB | 5 | 19.8833 | 0.342 |
| 0.05 | 0.05 | ycaM | 5 | 19.8833 | 0.352 |
| 0.05 | 0.05 | yadI | 5 | 19.8833 | 0.666 |
| 0.05 | 0.05 | AG1 | 5 | 19.8833 | 0.774 |
| 0.05 | 0.05 | ptsI | 5 | 20.1333 | 0.638 |
| 0.05 | 0.05 | clcB | 5 | 20.1333 | 0.345 |
| 0.05 | 0.05 | ycaM | 5 | 20.1333 | 0.358 |
| 0.05 | 0.05 | yadI | 5 | 20.1333 | 0.663 |
| 0.05 | 0.05 | AG1 | 5 | 20.1333 | 0.774 |
| 0.05 | 0.05 | ptsI | 5 | 20.3833 | 0.639 |
| 0.05 | 0.05 | clcB | 5 | 20.3833 | 0.345 |
| 0.05 | 0.05 | ycaM | 5 | 20.3833 | 0.358 |
| 0.05 | 0.05 | yadI | 5 | 20.3833 | 0.67 |
| 0.05 | 0.05 | AG1 | 5 | 20.3833 | 0.776 |
| 0.05 | 0.05 | ptsI | 5 | 20.6333 | 0.642 |
| 0.05 | 0.05 | clcB | 5 | 20.6333 | 0.345 |
| 0.05 | 0.05 | ycaM | 5 | 20.6333 | 0.356 |
| 0.05 | 0.05 | yadI | 5 | 20.6333 | 0.662 |
| 0.05 | 0.05 | AG1 | 5 | 20.6333 | 0.812 |
| 0.05 | 0.05 | ptsI | 5 | 20.8833 | 0.643 |
| 0.05 | 0.05 | clcB | 5 | 20.8833 | 0.348 |
| 0.05 | 0.05 | ycaM | 5 | 20.8833 | 0.36 |
| 0.05 | 0.05 | yadI | 5 | 20.8833 | 0.67 |
| 0.05 | 0.05 | AG1 | 5 | 20.8833 | 0.778 |
| 0.05 | 0.05 | ptsI | 5 | 21.1333 | 0.644 |
| 0.05 | 0.05 | clcB | 5 | 21.1333 | 0.358 |
| 0.05 | 0.05 | ycaM | 5 | 21.1333 | 0.366 |

| | | | | | |
|---|---|---|---|---|---|
| 0.05 | 0.05 | yadI | 5 | 21.1333 | 0.676 |
| 0.05 | 0.05 | AG1 | 5 | 21.1333 | 0.784 |
| 0.05 | 0.05 | ptsI | 5 | 21.3833 | 0.64 |
| 0.05 | 0.05 | clcB | 5 | 21.3833 | 0.362 |
| 0.05 | 0.05 | ycaM | 5 | 21.3833 | 0.366 |
| 0.05 | 0.05 | yadI | 5 | 21.3833 | 0.671 |
| 0.05 | 0.05 | AG1 | 5 | 21.3833 | 0.783 |
| 0.05 | 0.05 | ptsI | 5 | 21.6333 | 0.636 |
| 0.05 | 0.05 | clcB | 5 | 21.6333 | 0.367 |
| 0.05 | 0.05 | ycaM | 5 | 21.6333 | 0.379 |
| 0.05 | 0.05 | yadI | 5 | 21.6333 | 0.672 |
| 0.05 | 0.05 | AG1 | 5 | 21.6333 | 0.785 |
| 0.05 | 0.05 | ptsI | 5 | 21.8833 | 0.638 |
| 0.05 | 0.05 | clcB | 5 | 21.8833 | 0.386 |
| 0.05 | 0.05 | ycaM | 5 | 21.8833 | 0.382 |
| 0.05 | 0.05 | yadI | 5 | 21.8833 | 0.674 |
| 0.05 | 0.05 | AG1 | 5 | 21.8833 | 0.789 |
| 0.05 | 0.05 | ptsI | 5 | 22.1333 | 0.634 |
| 0.05 | 0.05 | clcB | 5 | 22.1333 | 0.379 |
| 0.05 | 0.05 | ycaM | 5 | 22.1333 | 0.381 |
| 0.05 | 0.05 | yadI | 5 | 22.1333 | 0.665 |
| 0.05 | 0.05 | AG1 | 5 | 22.1333 | 0.778 |
| 0.05 | 0.05 | ptsI | 5 | 22.3833 | 0.635 |
| 0.05 | 0.05 | clcB | 5 | 22.3833 | 0.383 |
| 0.05 | 0.05 | ycaM | 5 | 22.3833 | 0.388 |
| 0.05 | 0.05 | yadI | 5 | 22.3833 | 0.674 |
| 0.05 | 0.05 | AG1 | 5 | 22.3833 | 0.792 |
| 0.05 | 0.05 | ptsI | 5 | 22.6333 | 0.638 |
| 0.05 | 0.05 | clcB | 5 | 22.6333 | 0.376 |
| 0.05 | 0.05 | ycaM | 5 | 22.6333 | 0.393 |
| 0.05 | 0.05 | yadI | 5 | 22.6333 | 0.676 |
| 0.05 | 0.05 | AG1 | 5 | 22.6333 | 0.798 |
| 0.05 | 0.05 | ptsI | 5 | 22.8833 | 0.641 |
| 0.05 | 0.05 | clcB | 5 | 22.8833 | 0.377 |
| 0.05 | 0.05 | ycaM | 5 | 22.8833 | 0.399 |
| 0.05 | 0.05 | yadI | 5 | 22.8833 | 0.676 |
| 0.05 | 0.05 | AG1 | 5 | 22.8833 | 0.795 |
| 0.05 | 0.05 | ptsI | 5 | 23.1333 | 0.646 |
| 0.05 | 0.05 | clcB | 5 | 23.1333 | 0.381 |
| 0.05 | 0.05 | ycaM | 5 | 23.1333 | 0.404 |
| 0.05 | 0.05 | yadI | 5 | 23.1333 | 0.679 |
| 0.05 | 0.05 | AG1 | 5 | 23.1333 | 0.802 |
| 0.05 | 0.05 | ptsI | 5 | 23.3833 | 0.643 |
| 0.05 | 0.05 | clcB | 5 | 23.3833 | 0.379 |
| 0.05 | 0.05 | ycaM | 5 | 23.3833 | 0.396 |
| 0.05 | 0.05 | yadI | 5 | 23.3833 | 0.679 |
| 0.05 | 0.05 | AG1 | 5 | 23.3833 | 0.8 |
| 0.05 | 0.05 | ptsI | 5 | 23.6333 | 0.647 |

| | | | | | |
|---|---|---|---|---|---|
| 0.05 | 0.05 | clcB | 5 | 23.6333 | 0.379 |
| 0.05 | 0.05 | ycaM | 5 | 23.6333 | 0.404 |
| 0.05 | 0.05 | yadI | 5 | 23.6333 | 0.679 |
| 0.05 | 0.05 | AG1 | 5 | 23.6333 | 0.804 |
| 0.05 | 0.05 | ptsI | 5 | 23.8833 | 0.643 |
| 0.05 | 0.05 | clcB | 5 | 23.8833 | 0.388 |
| 0.05 | 0.05 | ycaM | 5 | 23.8833 | 0.406 |
| 0.05 | 0.05 | yadI | 5 | 23.8833 | 0.672 |
| 0.05 | 0.05 | AG1 | 5 | 23.8833 | 0.79 |
| 0.05 | 0.05 | ptsI | 5 | 24.1333 | 0.642 |
| 0.05 | 0.05 | clcB | 5 | 24.1333 | 0.39 |
| 0.05 | 0.05 | ycaM | 5 | 24.1333 | 0.424 |
| 0.05 | 0.05 | yadI | 5 | 24.1333 | 0.68 |
| 0.05 | 0.05 | AG1 | 5 | 24.1333 | 0.806 |
| 0.05 | 0.05 | ptsI | 5 | 24.3833 | 0.647 |
| 0.05 | 0.05 | clcB | 5 | 24.3833 | 0.398 |
| 0.05 | 0.05 | ycaM | 5 | 24.3833 | 0.421 |
| 0.05 | 0.05 | yadI | 5 | 24.3833 | 0.685 |
| 0.05 | 0.05 | AG1 | 5 | 24.3833 | 0.811 |
| 0.1 | 0.05 | ptsI | 2 | 0 | 0.216 |
| 0.1 | 0.05 | clcB | 2 | 0 | 0.226 |
| 0.1 | 0.05 | ycaM | 2 | 0 | 0.218 |
| 0.1 | 0.05 | yadI | 2 | 0 | 0.222 |
| 0.1 | 0.05 | AG1 | 2 | 0 | 0.224 |
| 0.1 | 0.05 | ptsI | 2 | 0.25 | 0.212 |
| 0.1 | 0.05 | clcB | 2 | 0.25 | 0.221 |
| 0.1 | 0.05 | ycaM | 2 | 0.25 | 0.213 |
| 0.1 | 0.05 | yadI | 2 | 0.25 | 0.215 |
| 0.1 | 0.05 | AG1 | 2 | 0.25 | 0.22 |
| 0.1 | 0.05 | ptsI | 2 | 0.5 | 0.209 |
| 0.1 | 0.05 | clcB | 2 | 0.5 | 0.215 |
| 0.1 | 0.05 | ycaM | 2 | 0.5 | 0.213 |
| 0.1 | 0.05 | yadI | 2 | 0.5 | 0.214 |
| 0.1 | 0.05 | AG1 | 2 | 0.5 | 0.217 |
| 0.1 | 0.05 | ptsI | 2 | 0.75 | 0.211 |
| 0.1 | 0.05 | clcB | 2 | 0.75 | 0.217 |
| 0.1 | 0.05 | ycaM | 2 | 0.75 | 0.216 |
| 0.1 | 0.05 | yadI | 2 | 0.75 | 0.218 |
| 0.1 | 0.05 | AG1 | 2 | 0.75 | 0.218 |
| 0.1 | 0.05 | ptsI | 2 | 1 | 0.213 |
| 0.1 | 0.05 | clcB | 2 | 1 | 0.219 |
| 0.1 | 0.05 | ycaM | 2 | 1 | 0.215 |
| 0.1 | 0.05 | yadI | 2 | 1 | 0.225 |
| 0.1 | 0.05 | AG1 | 2 | 1 | 0.225 |
| 0.1 | 0.05 | ptsI | 2 | 1.25 | 0.214 |
| 0.1 | 0.05 | clcB | 2 | 1.25 | 0.217 |
| 0.1 | 0.05 | ycaM | 2 | 1.25 | 0.218 |
| 0.1 | 0.05 | yadI | 2 | 1.25 | 0.232 |

| | | | | | |
|---|---|---|---|---|---|
| 0.1 | 0.05 | AG1 | 2 | 1.25 | 0.227 |
| 0.1 | 0.05 | ptsI | 2 | 1.5 | 0.215 |
| 0.1 | 0.05 | clcB | 2 | 1.5 | 0.219 |
| 0.1 | 0.05 | ycaM | 2 | 1.5 | 0.221 |
| 0.1 | 0.05 | yadI | 2 | 1.5 | 0.24 |
| 0.1 | 0.05 | AG1 | 2 | 1.5 | 0.233 |
| 0.1 | 0.05 | ptsI | 2 | 1.75 | 0.219 |
| 0.1 | 0.05 | clcB | 2 | 1.75 | 0.22 |
| 0.1 | 0.05 | ycaM | 2 | 1.75 | 0.226 |
| 0.1 | 0.05 | yadI | 2 | 1.75 | 0.246 |
| 0.1 | 0.05 | AG1 | 2 | 1.75 | 0.236 |
| 0.1 | 0.05 | ptsI | 2 | 2 | 0.221 |
| 0.1 | 0.05 | clcB | 2 | 2 | 0.222 |
| 0.1 | 0.05 | ycaM | 2 | 2 | 0.232 |
| 0.1 | 0.05 | yadI | 2 | 2 | 0.253 |
| 0.1 | 0.05 | AG1 | 2 | 2 | 0.246 |
| 0.1 | 0.05 | ptsI | 2 | 2.25 | 0.226 |
| 0.1 | 0.05 | clcB | 2 | 2.25 | 0.227 |
| 0.1 | 0.05 | ycaM | 2 | 2.25 | 0.24 |
| 0.1 | 0.05 | yadI | 2 | 2.25 | 0.264 |
| 0.1 | 0.05 | AG1 | 2 | 2.25 | 0.256 |
| 0.1 | 0.05 | ptsI | 2 | 2.5 | 0.231 |
| 0.1 | 0.05 | clcB | 2 | 2.5 | 0.232 |
| 0.1 | 0.05 | ycaM | 2 | 2.5 | 0.246 |
| 0.1 | 0.05 | yadI | 2 | 2.5 | 0.278 |
| 0.1 | 0.05 | AG1 | 2 | 2.5 | 0.268 |
| 0.1 | 0.05 | ptsI | 2 | 2.75 | 0.237 |
| 0.1 | 0.05 | clcB | 2 | 2.75 | 0.236 |
| 0.1 | 0.05 | ycaM | 2 | 2.75 | 0.247 |
| 0.1 | 0.05 | yadI | 2 | 2.75 | 0.287 |
| 0.1 | 0.05 | AG1 | 2 | 2.75 | 0.277 |
| 0.1 | 0.05 | ptsI | 2 | 3 | 0.243 |
| 0.1 | 0.05 | clcB | 2 | 3 | 0.236 |
| 0.1 | 0.05 | ycaM | 2 | 3 | 0.252 |
| 0.1 | 0.05 | yadI | 2 | 3 | 0.294 |
| 0.1 | 0.05 | AG1 | 2 | 3 | 0.28 |
| 0.1 | 0.05 | ptsI | 2 | 3.25 | 0.248 |
| 0.1 | 0.05 | clcB | 2 | 3.25 | 0.243 |
| 0.1 | 0.05 | ycaM | 2 | 3.25 | 0.259 |
| 0.1 | 0.05 | yadI | 2 | 3.25 | 0.311 |
| 0.1 | 0.05 | AG1 | 2 | 3.25 | 0.297 |
| 0.1 | 0.05 | ptsI | 2 | 3.5 | 0.254 |
| 0.1 | 0.05 | clcB | 2 | 3.5 | 0.245 |
| 0.1 | 0.05 | ycaM | 2 | 3.5 | 0.264 |
| 0.1 | 0.05 | yadI | 2 | 3.5 | 0.32 |
| 0.1 | 0.05 | AG1 | 2 | 3.5 | 0.303 |
| 0.1 | 0.05 | ptsI | 2 | 3.75 | 0.26 |
| 0.1 | 0.05 | clcB | 2 | 3.75 | 0.25 |

| | | | | | |
|---|---|---|---|---|---|
| 0.1 | 0.05 | ycaM | 2 | 3.75 | 0.274 |
| 0.1 | 0.05 | yadI | 2 | 3.75 | 0.332 |
| 0.1 | 0.05 | AG1 | 2 | 3.75 | 0.317 |
| 0.1 | 0.05 | ptsI | 2 | 4 | 0.265 |
| 0.1 | 0.05 | clcB | 2 | 4 | 0.254 |
| 0.1 | 0.05 | ycaM | 2 | 4 | 0.286 |
| 0.1 | 0.05 | yadI | 2 | 4 | 0.347 |
| 0.1 | 0.05 | AG1 | 2 | 4 | 0.329 |
| 0.1 | 0.05 | ptsI | 2 | 4.38333 | 0.306 |
| 0.1 | 0.05 | clcB | 2 | 4.38333 | 0.244 |
| 0.1 | 0.05 | ycaM | 2 | 4.38333 | 0.299 |
| 0.1 | 0.05 | yadI | 2 | 4.38333 | 0.372 |
| 0.1 | 0.05 | AG1 | 2 | 4.38333 | 0.306 |
| 0.1 | 0.05 | ptsI | 2 | 4.63333 | 0.298 |
| 0.1 | 0.05 | clcB | 2 | 4.63333 | 0.241 |
| 0.1 | 0.05 | ycaM | 2 | 4.63333 | 0.296 |
| 0.1 | 0.05 | yadI | 2 | 4.63333 | 0.378 |
| 0.1 | 0.05 | AG1 | 2 | 4.63333 | 0.303 |
| 0.1 | 0.05 | ptsI | 2 | 4.88333 | 0.304 |
| 0.1 | 0.05 | clcB | 2 | 4.88333 | 0.244 |
| 0.1 | 0.05 | ycaM | 2 | 4.88333 | 0.302 |
| 0.1 | 0.05 | yadI | 2 | 4.88333 | 0.39 |
| 0.1 | 0.05 | AG1 | 2 | 4.88333 | 0.312 |
| 0.1 | 0.05 | ptsI | 2 | 5.13333 | 0.311 |
| 0.1 | 0.05 | clcB | 2 | 5.13333 | 0.247 |
| 0.1 | 0.05 | ycaM | 2 | 5.13333 | 0.304 |
| 0.1 | 0.05 | yadI | 2 | 5.13333 | 0.406 |
| 0.1 | 0.05 | AG1 | 2 | 5.13333 | 0.32 |
| 0.1 | 0.05 | ptsI | 2 | 5.38333 | 0.319 |
| 0.1 | 0.05 | clcB | 2 | 5.38333 | 0.259 |
| 0.1 | 0.05 | ycaM | 2 | 5.38333 | 0.306 |
| 0.1 | 0.05 | yadI | 2 | 5.38333 | 0.417 |
| 0.1 | 0.05 | AG1 | 2 | 5.38333 | 0.329 |
| 0.1 | 0.05 | ptsI | 2 | 5.63333 | 0.325 |
| 0.1 | 0.05 | clcB | 2 | 5.63333 | 0.266 |
| 0.1 | 0.05 | ycaM | 2 | 5.63333 | 0.309 |
| 0.1 | 0.05 | yadI | 2 | 5.63333 | 0.428 |
| 0.1 | 0.05 | AG1 | 2 | 5.63333 | 0.339 |
| 0.1 | 0.05 | ptsI | 2 | 5.88333 | 0.331 |
| 0.1 | 0.05 | clcB | 2 | 5.88333 | 0.274 |
| 0.1 | 0.05 | ycaM | 2 | 5.88333 | 0.309 |
| 0.1 | 0.05 | yadI | 2 | 5.88333 | 0.437 |
| 0.1 | 0.05 | AG1 | 2 | 5.88333 | 0.348 |
| 0.1 | 0.05 | ptsI | 2 | 6.13333 | 0.338 |
| 0.1 | 0.05 | clcB | 2 | 6.13333 | 0.281 |
| 0.1 | 0.05 | ycaM | 2 | 6.13333 | 0.31 |
| 0.1 | 0.05 | yadI | 2 | 6.13333 | 0.445 |
| 0.1 | 0.05 | AG1 | 2 | 6.13333 | 0.358 |

| | | | | | |
|---|---|---|---|---|---|
| 0.1 | 0.05 | ptsI | 2 | 6.38333 | 0.345 |
| 0.1 | 0.05 | clcB | 2 | 6.38333 | 0.287 |
| 0.1 | 0.05 | ycaM | 2 | 6.38333 | 0.311 |
| 0.1 | 0.05 | yadI | 2 | 6.38333 | 0.458 |
| 0.1 | 0.05 | AG1 | 2 | 6.38333 | 0.366 |
| 0.1 | 0.05 | ptsI | 2 | 6.63333 | 0.354 |
| 0.1 | 0.05 | clcB | 2 | 6.63333 | 0.293 |
| 0.1 | 0.05 | ycaM | 2 | 6.63333 | 0.313 |
| 0.1 | 0.05 | yadI | 2 | 6.63333 | 0.466 |
| 0.1 | 0.05 | AG1 | 2 | 6.63333 | 0.378 |
| 0.1 | 0.05 | ptsI | 2 | 6.88333 | 0.362 |
| 0.1 | 0.05 | clcB | 2 | 6.88333 | 0.302 |
| 0.1 | 0.05 | ycaM | 2 | 6.88333 | 0.315 |
| 0.1 | 0.05 | yadI | 2 | 6.88333 | 0.484 |
| 0.1 | 0.05 | AG1 | 2 | 6.88333 | 0.387 |
| 0.1 | 0.05 | ptsI | 2 | 7.13333 | 0.369 |
| 0.1 | 0.05 | clcB | 2 | 7.13333 | 0.303 |
| 0.1 | 0.05 | ycaM | 2 | 7.13333 | 0.313 |
| 0.1 | 0.05 | yadI | 2 | 7.13333 | 0.487 |
| 0.1 | 0.05 | AG1 | 2 | 7.13333 | 0.398 |
| 0.1 | 0.05 | ptsI | 2 | 7.38333 | 0.377 |
| 0.1 | 0.05 | clcB | 2 | 7.38333 | 0.309 |
| 0.1 | 0.05 | ycaM | 2 | 7.38333 | 0.313 |
| 0.1 | 0.05 | yadI | 2 | 7.38333 | 0.498 |
| 0.1 | 0.05 | AG1 | 2 | 7.38333 | 0.409 |
| 0.1 | 0.05 | ptsI | 2 | 7.63333 | 0.386 |
| 0.1 | 0.05 | clcB | 2 | 7.63333 | 0.317 |
| 0.1 | 0.05 | ycaM | 2 | 7.63333 | 0.314 |
| 0.1 | 0.05 | yadI | 2 | 7.63333 | 0.507 |
| 0.1 | 0.05 | AG1 | 2 | 7.63333 | 0.417 |
| 0.1 | 0.05 | ptsI | 2 | 7.88333 | 0.393 |
| 0.1 | 0.05 | clcB | 2 | 7.88333 | 0.322 |
| 0.1 | 0.05 | ycaM | 2 | 7.88333 | 0.314 |
| 0.1 | 0.05 | yadI | 2 | 7.88333 | 0.524 |
| 0.1 | 0.05 | AG1 | 2 | 7.88333 | 0.427 |
| 0.1 | 0.05 | ptsI | 2 | 8.13333 | 0.401 |
| 0.1 | 0.05 | clcB | 2 | 8.13333 | 0.321 |
| 0.1 | 0.05 | ycaM | 2 | 8.13333 | 0.315 |
| 0.1 | 0.05 | yadI | 2 | 8.13333 | 0.53 |
| 0.1 | 0.05 | AG1 | 2 | 8.13333 | 0.434 |
| 0.1 | 0.05 | ptsI | 2 | 8.38333 | 0.409 |
| 0.1 | 0.05 | clcB | 2 | 8.38333 | 0.329 |
| 0.1 | 0.05 | ycaM | 2 | 8.38333 | 0.315 |
| 0.1 | 0.05 | yadI | 2 | 8.38333 | 0.546 |
| 0.1 | 0.05 | AG1 | 2 | 8.38333 | 0.446 |
| 0.1 | 0.05 | ptsI | 2 | 8.91667 | 0.357 |
| 0.1 | 0.05 | clcB | 2 | 8.91667 | 0.302 |
| 0.1 | 0.05 | ycaM | 2 | 8.91667 | 0.275 |

| | | | | | |
|---|---|---|---|---|---|
| 0.1 | 0.05 | yadI | 2 | 8.91667 | 0.481 |
| 0.1 | 0.05 | AG1 | 2 | 8.91667 | 0.472 |
| 0.1 | 0.05 | ptsI | 2 | 9.16667 | 0.356 |
| 0.1 | 0.05 | clcB | 2 | 9.16667 | 0.298 |
| 0.1 | 0.05 | ycaM | 2 | 9.16667 | 0.274 |
| 0.1 | 0.05 | yadI | 2 | 9.16667 | 0.483 |
| 0.1 | 0.05 | AG1 | 2 | 9.16667 | 0.484 |
| 0.1 | 0.05 | ptsI | 2 | 9.41667 | 0.365 |
| 0.1 | 0.05 | clcB | 2 | 9.41667 | 0.304 |
| 0.1 | 0.05 | ycaM | 2 | 9.41667 | 0.275 |
| 0.1 | 0.05 | yadI | 2 | 9.41667 | 0.495 |
| 0.1 | 0.05 | AG1 | 2 | 9.41667 | 0.505 |
| 0.1 | 0.05 | ptsI | 2 | 9.66667 | 0.369 |
| 0.1 | 0.05 | clcB | 2 | 9.66667 | 0.31 |
| 0.1 | 0.05 | ycaM | 2 | 9.66667 | 0.274 |
| 0.1 | 0.05 | yadI | 2 | 9.66667 | 0.505 |
| 0.1 | 0.05 | AG1 | 2 | 9.66667 | 0.516 |
| 0.1 | 0.05 | ptsI | 2 | 9.91667 | 0.378 |
| 0.1 | 0.05 | clcB | 2 | 9.91667 | 0.315 |
| 0.1 | 0.05 | ycaM | 2 | 9.91667 | 0.279 |
| 0.1 | 0.05 | yadI | 2 | 9.91667 | 0.516 |
| 0.1 | 0.05 | AG1 | 2 | 9.91667 | 0.528 |
| 0.1 | 0.05 | ptsI | 2 | 10.1667 | 0.385 |
| 0.1 | 0.05 | clcB | 2 | 10.1667 | 0.32 |
| 0.1 | 0.05 | ycaM | 2 | 10.1667 | 0.28 |
| 0.1 | 0.05 | yadI | 2 | 10.1667 | 0.523 |
| 0.1 | 0.05 | AG1 | 2 | 10.1667 | 0.53 |
| 0.1 | 0.05 | ptsI | 2 | 10.4167 | 0.395 |
| 0.1 | 0.05 | clcB | 2 | 10.4167 | 0.321 |
| 0.1 | 0.05 | ycaM | 2 | 10.4167 | 0.28 |
| 0.1 | 0.05 | yadI | 2 | 10.4167 | 0.534 |
| 0.1 | 0.05 | AG1 | 2 | 10.4167 | 0.543 |
| 0.1 | 0.05 | ptsI | 2 | 10.6667 | 0.402 |
| 0.1 | 0.05 | clcB | 2 | 10.6667 | 0.327 |
| 0.1 | 0.05 | ycaM | 2 | 10.6667 | 0.281 |
| 0.1 | 0.05 | yadI | 2 | 10.6667 | 0.543 |
| 0.1 | 0.05 | AG1 | 2 | 10.6667 | 0.556 |
| 0.1 | 0.05 | ptsI | 2 | 10.9167 | 0.413 |
| 0.1 | 0.05 | clcB | 2 | 10.9167 | 0.328 |
| 0.1 | 0.05 | ycaM | 2 | 10.9167 | 0.282 |
| 0.1 | 0.05 | yadI | 2 | 10.9167 | 0.55 |
| 0.1 | 0.05 | AG1 | 2 | 10.9167 | 0.574 |
| 0.1 | 0.05 | ptsI | 2 | 11.1667 | 0.423 |
| 0.1 | 0.05 | clcB | 2 | 11.1667 | 0.334 |
| 0.1 | 0.05 | ycaM | 2 | 11.1667 | 0.281 |
| 0.1 | 0.05 | yadI | 2 | 11.1667 | 0.564 |
| 0.1 | 0.05 | AG1 | 2 | 11.1667 | 0.59 |
| 0.1 | 0.05 | ptsI | 2 | 11.4167 | 0.432 |

| | | | | | |
|---|---|---|---|---|---|
| 0.1 | 0.05 | clcB | 2 | 11.4167 | 0.335 |
| 0.1 | 0.05 | ycaM | 2 | 11.4167 | 0.283 |
| 0.1 | 0.05 | yadI | 2 | 11.4167 | 0.565 |
| 0.1 | 0.05 | AG1 | 2 | 11.4167 | 0.597 |
| 0.1 | 0.05 | ptsI | 2 | 11.6667 | 0.439 |
| 0.1 | 0.05 | clcB | 2 | 11.6667 | 0.337 |
| 0.1 | 0.05 | ycaM | 2 | 11.6667 | 0.282 |
| 0.1 | 0.05 | yadI | 2 | 11.6667 | 0.572 |
| 0.1 | 0.05 | AG1 | 2 | 11.6667 | 0.626 |
| 0.1 | 0.05 | ptsI | 2 | 11.9167 | 0.446 |
| 0.1 | 0.05 | clcB | 2 | 11.9167 | 0.337 |
| 0.1 | 0.05 | ycaM | 2 | 11.9167 | 0.282 |
| 0.1 | 0.05 | yadI | 2 | 11.9167 | 0.579 |
| 0.1 | 0.05 | AG1 | 2 | 11.9167 | 0.626 |
| 0.1 | 0.05 | ptsI | 2 | 12.1667 | 0.458 |
| 0.1 | 0.05 | clcB | 2 | 12.1667 | 0.331 |
| 0.1 | 0.05 | ycaM | 2 | 12.1667 | 0.278 |
| 0.1 | 0.05 | yadI | 2 | 12.1667 | 0.588 |
| 0.1 | 0.05 | AG1 | 2 | 12.1667 | 0.648 |
| 0.1 | 0.05 | ptsI | 2 | 12.4167 | 0.466 |
| 0.1 | 0.05 | clcB | 2 | 12.4167 | 0.33 |
| 0.1 | 0.05 | ycaM | 2 | 12.4167 | 0.276 |
| 0.1 | 0.05 | yadI | 2 | 12.4167 | 0.59 |
| 0.1 | 0.05 | AG1 | 2 | 12.4167 | 0.662 |
| 0.1 | 0.05 | ptsI | 2 | 12.6667 | 0.465 |
| 0.1 | 0.05 | clcB | 2 | 12.6667 | 0.33 |
| 0.1 | 0.05 | ycaM | 2 | 12.6667 | 0.274 |
| 0.1 | 0.05 | yadI | 2 | 12.6667 | 0.599 |
| 0.1 | 0.05 | AG1 | 2 | 12.6667 | 0.671 |
| 0.1 | 0.05 | ptsI | 2 | 12.9167 | 0.477 |
| 0.1 | 0.05 | clcB | 2 | 12.9167 | 0.325 |
| 0.1 | 0.05 | ycaM | 2 | 12.9167 | 0.277 |
| 0.1 | 0.05 | yadI | 2 | 12.9167 | 0.609 |
| 0.1 | 0.05 | AG1 | 2 | 12.9167 | 0.677 |
| 0.1 | 0.05 | ptsI | 2 | 13.1667 | 0.485 |
| 0.1 | 0.05 | clcB | 2 | 13.1667 | 0.327 |
| 0.1 | 0.05 | ycaM | 2 | 13.1667 | 0.279 |
| 0.1 | 0.05 | yadI | 2 | 13.1667 | 0.616 |
| 0.1 | 0.05 | AG1 | 2 | 13.1667 | 0.684 |
| 0.1 | 0.05 | ptsI | 2 | 13.4167 | 0.486 |
| 0.1 | 0.05 | clcB | 2 | 13.4167 | 0.327 |
| 0.1 | 0.05 | ycaM | 2 | 13.4167 | 0.278 |
| 0.1 | 0.05 | yadI | 2 | 13.4167 | 0.618 |
| 0.1 | 0.05 | AG1 | 2 | 13.4167 | 0.682 |
| 0.1 | 0.05 | ptsI | 2 | 13.6667 | 0.494 |
| 0.1 | 0.05 | clcB | 2 | 13.6667 | 0.328 |
| 0.1 | 0.05 | ycaM | 2 | 13.6667 | 0.278 |
| 0.1 | 0.05 | yadI | 2 | 13.6667 | 0.628 |

| | | | | | |
|---|---|---|---|---|---|
| 0.1 | 0.05 | AG1 | 2 | 13.6667 | 0.707 |
| 0.1 | 0.05 | ptsI | 2 | 13.9167 | 0.5 |
| 0.1 | 0.05 | clcB | 2 | 13.9167 | 0.325 |
| 0.1 | 0.05 | ycaM | 2 | 13.9167 | 0.282 |
| 0.1 | 0.05 | yadI | 2 | 13.9167 | 0.637 |
| 0.1 | 0.05 | AG1 | 2 | 13.9167 | 0.704 |
| 0.1 | 0.05 | ptsI | 2 | 14.1667 | 0.503 |
| 0.1 | 0.05 | clcB | 2 | 14.1667 | 0.321 |
| 0.1 | 0.05 | ycaM | 2 | 14.1667 | 0.281 |
| 0.1 | 0.05 | yadI | 2 | 14.1667 | 0.636 |
| 0.1 | 0.05 | AG1 | 2 | 14.1667 | 0.714 |
| 0.1 | 0.05 | ptsI | 2 | 14.4167 | 0.506 |
| 0.1 | 0.05 | clcB | 2 | 14.4167 | 0.325 |
| 0.1 | 0.05 | ycaM | 2 | 14.4167 | 0.279 |
| 0.1 | 0.05 | yadI | 2 | 14.4167 | 0.645 |
| 0.1 | 0.05 | AG1 | 2 | 14.4167 | 0.711 |
| 0.1 | 0.05 | ptsI | 2 | 14.6667 | 0.504 |
| 0.1 | 0.05 | clcB | 2 | 14.6667 | 0.327 |
| 0.1 | 0.05 | ycaM | 2 | 14.6667 | 0.278 |
| 0.1 | 0.05 | yadI | 2 | 14.6667 | 0.648 |
| 0.1 | 0.05 | AG1 | 2 | 14.6667 | 0.7 |
| 0.1 | 0.05 | ptsI | 2 | 14.9167 | 0.512 |
| 0.1 | 0.05 | clcB | 2 | 14.9167 | 0.328 |
| 0.1 | 0.05 | ycaM | 2 | 14.9167 | 0.275 |
| 0.1 | 0.05 | yadI | 2 | 14.9167 | 0.66 |
| 0.1 | 0.05 | AG1 | 2 | 14.9167 | 0.704 |
| 0.1 | 0.05 | ptsI | 2 | 15.1667 | 0.529 |
| 0.1 | 0.05 | clcB | 2 | 15.1667 | 0.328 |
| 0.1 | 0.05 | ycaM | 2 | 15.1667 | 0.279 |
| 0.1 | 0.05 | yadI | 2 | 15.1667 | 0.663 |
| 0.1 | 0.05 | AG1 | 2 | 15.1667 | 0.703 |
| 0.1 | 0.05 | ptsI | 2 | 15.4167 | 0.529 |
| 0.1 | 0.05 | clcB | 2 | 15.4167 | 0.327 |
| 0.1 | 0.05 | ycaM | 2 | 15.4167 | 0.277 |
| 0.1 | 0.05 | yadI | 2 | 15.4167 | 0.67 |
| 0.1 | 0.05 | AG1 | 2 | 15.4167 | 0.704 |
| 0.1 | 0.05 | ptsI | 2 | 15.6667 | 0.517 |
| 0.1 | 0.05 | clcB | 2 | 15.6667 | 0.329 |
| 0.1 | 0.05 | ycaM | 2 | 15.6667 | 0.272 |
| 0.1 | 0.05 | yadI | 2 | 15.6667 | 0.672 |
| 0.1 | 0.05 | AG1 | 2 | 15.6667 | 0.706 |
| 0.1 | 0.05 | ptsI | 2 | 15.9167 | 0.516 |
| 0.1 | 0.05 | clcB | 2 | 15.9167 | 0.33 |
| 0.1 | 0.05 | ycaM | 2 | 15.9167 | 0.274 |
| 0.1 | 0.05 | yadI | 2 | 15.9167 | 0.678 |
| 0.1 | 0.05 | AG1 | 2 | 15.9167 | 0.716 |
| 0.1 | 0.05 | ptsI | 2 | 16.1667 | 0.528 |
| 0.1 | 0.05 | clcB | 2 | 16.1667 | 0.33 |

| | | | | | |
|---|---|---|---|---|---|
| 0.1 | 0.05 | ycaM | 2 | 16.1667 | 0.273 |
| 0.1 | 0.05 | yadI | 2 | 16.1667 | 0.682 |
| 0.1 | 0.05 | AG1 | 2 | 16.1667 | 0.727 |
| 0.1 | 0.05 | ptsI | 2 | 16.4167 | 0.527 |
| 0.1 | 0.05 | clcB | 2 | 16.4167 | 0.33 |
| 0.1 | 0.05 | ycaM | 2 | 16.4167 | 0.275 |
| 0.1 | 0.05 | yadI | 2 | 16.4167 | 0.69 |
| 0.1 | 0.05 | AG1 | 2 | 16.4167 | 0.72 |
| 0.1 | 0.05 | ptsI | 2 | 16.6667 | 0.528 |
| 0.1 | 0.05 | clcB | 2 | 16.6667 | 0.328 |
| 0.1 | 0.05 | ycaM | 2 | 16.6667 | 0.274 |
| 0.1 | 0.05 | yadI | 2 | 16.6667 | 0.694 |
| 0.1 | 0.05 | AG1 | 2 | 16.6667 | 0.729 |
| 0.1 | 0.05 | ptsI | 2 | 16.9167 | 0.53 |
| 0.1 | 0.05 | clcB | 2 | 16.9167 | 0.33 |
| 0.1 | 0.05 | ycaM | 2 | 16.9167 | 0.275 |
| 0.1 | 0.05 | yadI | 2 | 16.9167 | 0.699 |
| 0.1 | 0.05 | AG1 | 2 | 16.9167 | 0.741 |
| 0.1 | 0.05 | ptsI | 2 | 17.1667 | 0.539 |
| 0.1 | 0.05 | clcB | 2 | 17.1667 | 0.331 |
| 0.1 | 0.05 | ycaM | 2 | 17.1667 | 0.274 |
| 0.1 | 0.05 | yadI | 2 | 17.1667 | 0.708 |
| 0.1 | 0.05 | AG1 | 2 | 17.1667 | 0.731 |
| 0.1 | 0.05 | ptsI | 2 | 17.4167 | 0.542 |
| 0.1 | 0.05 | clcB | 2 | 17.4167 | 0.328 |
| 0.1 | 0.05 | ycaM | 2 | 17.4167 | 0.275 |
| 0.1 | 0.05 | yadI | 2 | 17.4167 | 0.711 |
| 0.1 | 0.05 | AG1 | 2 | 17.4167 | 0.739 |
| 0.1 | 0.05 | ptsI | 2 | 17.6667 | 0.552 |
| 0.1 | 0.05 | clcB | 2 | 17.6667 | 0.332 |
| 0.1 | 0.05 | ycaM | 2 | 17.6667 | 0.276 |
| 0.1 | 0.05 | yadI | 2 | 17.6667 | 0.717 |
| 0.1 | 0.05 | AG1 | 2 | 17.6667 | 0.726 |
| 0.1 | 0.05 | ptsI | 2 | 17.9167 | 0.545 |
| 0.1 | 0.05 | clcB | 2 | 17.9167 | 0.331 |
| 0.1 | 0.05 | ycaM | 2 | 17.9167 | 0.273 |
| 0.1 | 0.05 | yadI | 2 | 17.9167 | 0.722 |
| 0.1 | 0.05 | AG1 | 2 | 17.9167 | 0.727 |
| 0.1 | 0.05 | ptsI | 2 | 18.1667 | 0.55 |
| 0.1 | 0.05 | clcB | 2 | 18.1667 | 0.33 |
| 0.1 | 0.05 | ycaM | 2 | 18.1667 | 0.277 |
| 0.1 | 0.05 | yadI | 2 | 18.1667 | 0.718 |
| 0.1 | 0.05 | AG1 | 2 | 18.1667 | 0.74 |
| 0.1 | 0.05 | ptsI | 2 | 18.4167 | 0.556 |
| 0.1 | 0.05 | clcB | 2 | 18.4167 | 0.333 |
| 0.1 | 0.05 | ycaM | 2 | 18.4167 | 0.278 |
| 0.1 | 0.05 | yadI | 2 | 18.4167 | 0.728 |
| 0.1 | 0.05 | AG1 | 2 | 18.4167 | 0.737 |

| | | | | | |
|---|---|---|---|---|---|
| 0.1 | 0.05 | ptsI | 2 | 18.6667 | 0.547 |
| 0.1 | 0.05 | clcB | 2 | 18.6667 | 0.333 |
| 0.1 | 0.05 | ycaM | 2 | 18.6667 | 0.278 |
| 0.1 | 0.05 | yadI | 2 | 18.6667 | 0.729 |
| 0.1 | 0.05 | AG1 | 2 | 18.6667 | 0.743 |
| 0.1 | 0.05 | ptsI | 2 | 18.9167 | 0.536 |
| 0.1 | 0.05 | clcB | 2 | 18.9167 | 0.33 |
| 0.1 | 0.05 | ycaM | 2 | 18.9167 | 0.278 |
| 0.1 | 0.05 | yadI | 2 | 18.9167 | 0.727 |
| 0.1 | 0.05 | AG1 | 2 | 18.9167 | 0.744 |
| 0.1 | 0.05 | ptsI | 2 | 19.1667 | 0.556 |
| 0.1 | 0.05 | clcB | 2 | 19.1667 | 0.331 |
| 0.1 | 0.05 | ycaM | 2 | 19.1667 | 0.28 |
| 0.1 | 0.05 | yadI | 2 | 19.1667 | 0.727 |
| 0.1 | 0.05 | AG1 | 2 | 19.1667 | 0.76 |
| 0.1 | 0.05 | ptsI | 2 | 19.4167 | 0.553 |
| 0.1 | 0.05 | clcB | 2 | 19.4167 | 0.333 |
| 0.1 | 0.05 | ycaM | 2 | 19.4167 | 0.28 |
| 0.1 | 0.05 | yadI | 2 | 19.4167 | 0.729 |
| 0.1 | 0.05 | AG1 | 2 | 19.4167 | 0.76 |
| 0.1 | 0.05 | ptsI | 2 | 19.6667 | 0.556 |
| 0.1 | 0.05 | clcB | 2 | 19.6667 | 0.332 |
| 0.1 | 0.05 | ycaM | 2 | 19.6667 | 0.281 |
| 0.1 | 0.05 | yadI | 2 | 19.6667 | 0.74 |
| 0.1 | 0.05 | AG1 | 2 | 19.6667 | 0.764 |
| 0.1 | 0.05 | ptsI | 2 | 19.9167 | 0.559 |
| 0.1 | 0.05 | clcB | 2 | 19.9167 | 0.329 |
| 0.1 | 0.05 | ycaM | 2 | 19.9167 | 0.283 |
| 0.1 | 0.05 | yadI | 2 | 19.9167 | 0.736 |
| 0.1 | 0.05 | AG1 | 2 | 19.9167 | 0.784 |
| 0.1 | 0.05 | ptsI | 2 | 20.1667 | 0.557 |
| 0.1 | 0.05 | clcB | 2 | 20.1667 | 0.334 |
| 0.1 | 0.05 | ycaM | 2 | 20.1667 | 0.286 |
| 0.1 | 0.05 | yadI | 2 | 20.1667 | 0.749 |
| 0.1 | 0.05 | AG1 | 2 | 20.1667 | 0.777 |
| 0.1 | 0.05 | ptsI | 2 | 20.4167 | 0.56 |
| 0.1 | 0.05 | clcB | 2 | 20.4167 | 0.333 |
| 0.1 | 0.05 | ycaM | 2 | 20.4167 | 0.287 |
| 0.1 | 0.05 | yadI | 2 | 20.4167 | 0.75 |
| 0.1 | 0.05 | AG1 | 2 | 20.4167 | 0.788 |
| 0.1 | 0.05 | ptsI | 2 | 20.6667 | 0.556 |
| 0.1 | 0.05 | clcB | 2 | 20.6667 | 0.334 |
| 0.1 | 0.05 | ycaM | 2 | 20.6667 | 0.291 |
| 0.1 | 0.05 | yadI | 2 | 20.6667 | 0.749 |
| 0.1 | 0.05 | AG1 | 2 | 20.6667 | 0.786 |
| 0.1 | 0.05 | ptsI | 2 | 20.9167 | 0.564 |
| 0.1 | 0.05 | clcB | 2 | 20.9167 | 0.332 |
| 0.1 | 0.05 | ycaM | 2 | 20.9167 | 0.294 |

| | | | | | |
|---|---|---|---|---|---|
| 0.1 | 0.05 | yadI | 2 | 20.9167 | 0.752 |
| 0.1 | 0.05 | AG1 | 2 | 20.9167 | 0.792 |
| 0.1 | 0.05 | ptsI | 2 | 21.1667 | 0.572 |
| 0.1 | 0.05 | clcB | 2 | 21.1667 | 0.332 |
| 0.1 | 0.05 | ycaM | 2 | 21.1667 | 0.297 |
| 0.1 | 0.05 | yadI | 2 | 21.1667 | 0.762 |
| 0.1 | 0.05 | AG1 | 2 | 21.1667 | 0.801 |
| 0.1 | 0.05 | ptsI | 2 | 21.4167 | 0.571 |
| 0.1 | 0.05 | clcB | 2 | 21.4167 | 0.33 |
| 0.1 | 0.05 | ycaM | 2 | 21.4167 | 0.297 |
| 0.1 | 0.05 | yadI | 2 | 21.4167 | 0.761 |
| 0.1 | 0.05 | AG1 | 2 | 21.4167 | 0.809 |
| 0.1 | 0.05 | ptsI | 2 | 21.6667 | 0.581 |
| 0.1 | 0.05 | clcB | 2 | 21.6667 | 0.333 |
| 0.1 | 0.05 | ycaM | 2 | 21.6667 | 0.299 |
| 0.1 | 0.05 | yadI | 2 | 21.6667 | 0.768 |
| 0.1 | 0.05 | AG1 | 2 | 21.6667 | 0.81 |
| 0.1 | 0.05 | ptsI | 2 | 21.9167 | 0.589 |
| 0.1 | 0.05 | clcB | 2 | 21.9167 | 0.333 |
| 0.1 | 0.05 | ycaM | 2 | 21.9167 | 0.299 |
| 0.1 | 0.05 | yadI | 2 | 21.9167 | 0.771 |
| 0.1 | 0.05 | AG1 | 2 | 21.9167 | 0.818 |
| 0.1 | 0.05 | ptsI | 2 | 22.1667 | 0.593 |
| 0.1 | 0.05 | clcB | 2 | 22.1667 | 0.333 |
| 0.1 | 0.05 | ycaM | 2 | 22.1667 | 0.302 |
| 0.1 | 0.05 | yadI | 2 | 22.1667 | 0.768 |
| 0.1 | 0.05 | AG1 | 2 | 22.1667 | 0.823 |
| 0.1 | 0.05 | ptsI | 2 | 22.4167 | 0.588 |
| 0.1 | 0.05 | clcB | 2 | 22.4167 | 0.332 |
| 0.1 | 0.05 | ycaM | 2 | 22.4167 | 0.306 |
| 0.1 | 0.05 | yadI | 2 | 22.4167 | 0.773 |
| 0.1 | 0.05 | AG1 | 2 | 22.4167 | 0.828 |
| 0.1 | 0.05 | ptsI | 2 | 22.6667 | 0.59 |
| 0.1 | 0.05 | clcB | 2 | 22.6667 | 0.332 |
| 0.1 | 0.05 | ycaM | 2 | 22.6667 | 0.306 |
| 0.1 | 0.05 | yadI | 2 | 22.6667 | 0.776 |
| 0.1 | 0.05 | AG1 | 2 | 22.6667 | 0.834 |
| 0.1 | 0.05 | ptsI | 2 | 22.9167 | 0.596 |
| 0.1 | 0.05 | clcB | 2 | 22.9167 | 0.333 |
| 0.1 | 0.05 | ycaM | 2 | 22.9167 | 0.31 |
| 0.1 | 0.05 | yadI | 2 | 22.9167 | 0.779 |
| 0.1 | 0.05 | AG1 | 2 | 22.9167 | 0.84 |
| 0.1 | 0.05 | ptsI | 2 | 23.1667 | 0.596 |
| 0.1 | 0.05 | clcB | 2 | 23.1667 | 0.333 |
| 0.1 | 0.05 | ycaM | 2 | 23.1667 | 0.308 |
| 0.1 | 0.05 | yadI | 2 | 23.1667 | 0.777 |
| 0.1 | 0.05 | AG1 | 2 | 23.1667 | 0.843 |
| 0.1 | 0.05 | ptsI | 2 | 23.4167 | 0.593 |

| | | | | | |
|---|---|---|---|---|---|
| 0.1 | 0.05 | clcB | 2 | 23.4167 | 0.334 |
| 0.1 | 0.05 | ycaM | 2 | 23.4167 | 0.31 |
| 0.1 | 0.05 | yadI | 2 | 23.4167 | 0.784 |
| 0.1 | 0.05 | AG1 | 2 | 23.4167 | 0.848 |
| 0.1 | 0.05 | ptsI | 2 | 23.6667 | 0.598 |
| 0.1 | 0.05 | clcB | 2 | 23.6667 | 0.335 |
| 0.1 | 0.05 | ycaM | 2 | 23.6667 | 0.312 |
| 0.1 | 0.05 | yadI | 2 | 23.6667 | 0.785 |
| 0.1 | 0.05 | AG1 | 2 | 23.6667 | 0.857 |
| 0.1 | 0.05 | ptsI | 2 | 23.9167 | 0.6 |
| 0.1 | 0.05 | clcB | 2 | 23.9167 | 0.335 |
| 0.1 | 0.05 | ycaM | 2 | 23.9167 | 0.314 |
| 0.1 | 0.05 | yadI | 2 | 23.9167 | 0.79 |
| 0.1 | 0.05 | AG1 | 2 | 23.9167 | 0.865 |
| 0.1 | 0.05 | ptsI | 2 | 24.1667 | 0.596 |
| 0.1 | 0.05 | clcB | 2 | 24.1667 | 0.338 |
| 0.1 | 0.05 | ycaM | 2 | 24.1667 | 0.317 |
| 0.1 | 0.05 | yadI | 2 | 24.1667 | 0.794 |
| 0.1 | 0.05 | AG1 | 2 | 24.1667 | 0.866 |
| 0.1 | 0.05 | ptsI | 2 | 24.4167 | 0.597 |
| 0.1 | 0.05 | clcB | 2 | 24.4167 | 0.34 |
| 0.1 | 0.05 | ycaM | 2 | 24.4167 | 0.321 |
| 0.1 | 0.05 | yadI | 2 | 24.4167 | 0.794 |
| 0.1 | 0.05 | AG1 | 2 | 24.4167 | 0.873 |
| 0.1 | 0.05 | ptsI | 2 | 24.6667 | 0.597 |
| 0.1 | 0.05 | clcB | 2 | 24.6667 | 0.341 |
| 0.1 | 0.05 | ycaM | 2 | 24.6667 | 0.324 |
| 0.1 | 0.05 | yadI | 2 | 24.6667 | 0.797 |
| 0.1 | 0.05 | AG1 | 2 | 24.6667 | 0.883 |
| 0.1 | 0.05 | ptsI | 2 | 24.9167 | 0.599 |
| 0.1 | 0.05 | clcB | 2 | 24.9167 | 0.342 |
| 0.1 | 0.05 | ycaM | 2 | 24.9167 | 0.323 |
| 0.1 | 0.05 | yadI | 2 | 24.9167 | 0.802 |
| 0.1 | 0.05 | AG1 | 2 | 24.9167 | 0.893 |
| 0.1 | 0.05 | ptsI | 2 | 25.1667 | 0.599 |
| 0.1 | 0.05 | clcB | 2 | 25.1667 | 0.344 |
| 0.1 | 0.05 | ycaM | 2 | 25.1667 | 0.331 |
| 0.1 | 0.05 | yadI | 2 | 25.1667 | 0.809 |
| 0.1 | 0.05 | AG1 | 2 | 25.1667 | 0.897 |
| 0.1 | 0.05 | ptsI | 2 | 25.4167 | 0.6 |
| 0.1 | 0.05 | clcB | 2 | 25.4167 | 0.347 |
| 0.1 | 0.05 | ycaM | 2 | 25.4167 | 0.336 |
| 0.1 | 0.05 | yadI | 2 | 25.4167 | 0.811 |
| 0.1 | 0.05 | AG1 | 2 | 25.4167 | 0.906 |
| 0.1 | 0.05 | ptsI | 2 | 25.6667 | 0.597 |
| 0.1 | 0.05 | clcB | 2 | 25.6667 | 0.345 |
| 0.1 | 0.05 | ycaM | 2 | 25.6667 | 0.336 |
| 0.1 | 0.05 | yadI | 2 | 25.6667 | 0.811 |

| | | | | | |
|---|---|---|---|---|---|
| 0.1 | 0.05 | AG1 | 2 | 25.6667 | 0.912 |
| 0.1 | 0.05 | ptsI | 2 | 25.9167 | 0.598 |
| 0.1 | 0.05 | clcB | 2 | 25.9167 | 0.346 |
| 0.1 | 0.05 | ycaM | 2 | 25.9167 | 0.345 |
| 0.1 | 0.05 | yadI | 2 | 25.9167 | 0.817 |
| 0.1 | 0.05 | AG1 | 2 | 25.9167 | 0.923 |
| 0.1 | 0.05 | ptsI | 2 | 26.1667 | 0.594 |
| 0.1 | 0.05 | clcB | 2 | 26.1667 | 0.345 |
| 0.1 | 0.05 | ycaM | 2 | 26.1667 | 0.344 |
| 0.1 | 0.05 | yadI | 2 | 26.1667 | 0.819 |
| 0.1 | 0.05 | AG1 | 2 | 26.1667 | 0.934 |
| 0.1 | 0.05 | ptsI | 2 | 26.4167 | 0.598 |
| 0.1 | 0.05 | clcB | 2 | 26.4167 | 0.347 |
| 0.1 | 0.05 | ycaM | 2 | 26.4167 | 0.353 |
| 0.1 | 0.05 | yadI | 2 | 26.4167 | 0.822 |
| 0.1 | 0.05 | AG1 | 2 | 26.4167 | 0.946 |
| 0.1 | 0.05 | ptsI | 3 | 0 | 0.176 |
| 0.1 | 0.05 | clcB | 3 | 0 | 0.161 |
| 0.1 | 0.05 | ycaM | 3 | 0 | 0.162 |
| 0.1 | 0.05 | yadI | 3 | 0 | 0.176 |
| 0.1 | 0.05 | AG1 | 3 | 0 | 0.351 |
| 0.1 | 0.05 | ptsI | 3 | 0.25 | 0.172 |
| 0.1 | 0.05 | clcB | 3 | 0.25 | 0.156 |
| 0.1 | 0.05 | ycaM | 3 | 0.25 | 0.159 |
| 0.1 | 0.05 | yadI | 3 | 0.25 | 0.174 |
| 0.1 | 0.05 | AG1 | 3 | 0.25 | 0.359 |
| 0.1 | 0.05 | ptsI | 3 | 0.5 | 0.169 |
| 0.1 | 0.05 | clcB | 3 | 0.5 | 0.154 |
| 0.1 | 0.05 | ycaM | 3 | 0.5 | 0.158 |
| 0.1 | 0.05 | yadI | 3 | 0.5 | 0.174 |
| 0.1 | 0.05 | AG1 | 3 | 0.5 | 0.358 |
| 0.1 | 0.05 | ptsI | 3 | 0.75 | 0.17 |
| 0.1 | 0.05 | clcB | 3 | 0.75 | 0.154 |
| 0.1 | 0.05 | ycaM | 3 | 0.75 | 0.158 |
| 0.1 | 0.05 | yadI | 3 | 0.75 | 0.173 |
| 0.1 | 0.05 | AG1 | 3 | 0.75 | 0.361 |
| 0.1 | 0.05 | ptsI | 3 | 1 | 0.17 |
| 0.1 | 0.05 | clcB | 3 | 1 | 0.153 |
| 0.1 | 0.05 | ycaM | 3 | 1 | 0.157 |
| 0.1 | 0.05 | yadI | 3 | 1 | 0.174 |
| 0.1 | 0.05 | AG1 | 3 | 1 | 0.368 |
| 0.1 | 0.05 | ptsI | 3 | 1.25 | 0.17 |
| 0.1 | 0.05 | clcB | 3 | 1.25 | 0.154 |
| 0.1 | 0.05 | ycaM | 3 | 1.25 | 0.159 |
| 0.1 | 0.05 | yadI | 3 | 1.25 | 0.174 |
| 0.1 | 0.05 | AG1 | 3 | 1.25 | 0.366 |
| 0.1 | 0.05 | ptsI | 3 | 1.5 | 0.17 |
| 0.1 | 0.05 | clcB | 3 | 1.5 | 0.153 |

| | | | | | |
|---|---|---|---|---|---|
| 0.1 | 0.05 | ycaM | 3 | 1.5 | 0.158 |
| 0.1 | 0.05 | yadI | 3 | 1.5 | 0.176 |
| 0.1 | 0.05 | AG1 | 3 | 1.5 | 0.359 |
| 0.1 | 0.05 | ptsI | 3 | 1.75 | 0.172 |
| 0.1 | 0.05 | clcB | 3 | 1.75 | 0.154 |
| 0.1 | 0.05 | ycaM | 3 | 1.75 | 0.158 |
| 0.1 | 0.05 | yadI | 3 | 1.75 | 0.19 |
| 0.1 | 0.05 | AG1 | 3 | 1.75 | 0.364 |
| 0.1 | 0.05 | ptsI | 3 | 2 | 0.174 |
| 0.1 | 0.05 | clcB | 3 | 2 | 0.16 |
| 0.1 | 0.05 | ycaM | 3 | 2 | 0.161 |
| 0.1 | 0.05 | yadI | 3 | 2 | 0.176 |
| 0.1 | 0.05 | AG1 | 3 | 2 | 0.356 |
| 0.1 | 0.05 | ptsI | 3 | 2.25 | 0.174 |
| 0.1 | 0.05 | clcB | 3 | 2.25 | 0.155 |
| 0.1 | 0.05 | ycaM | 3 | 2.25 | 0.161 |
| 0.1 | 0.05 | yadI | 3 | 2.25 | 0.178 |
| 0.1 | 0.05 | AG1 | 3 | 2.25 | 0.357 |
| 0.1 | 0.05 | ptsI | 3 | 2.5 | 0.177 |
| 0.1 | 0.05 | clcB | 3 | 2.5 | 0.156 |
| 0.1 | 0.05 | ycaM | 3 | 2.5 | 0.162 |
| 0.1 | 0.05 | yadI | 3 | 2.5 | 0.178 |
| 0.1 | 0.05 | AG1 | 3 | 2.5 | 0.347 |
| 0.1 | 0.05 | ptsI | 3 | 2.75 | 0.178 |
| 0.1 | 0.05 | clcB | 3 | 2.75 | 0.158 |
| 0.1 | 0.05 | ycaM | 3 | 2.75 | 0.165 |
| 0.1 | 0.05 | yadI | 3 | 2.75 | 0.179 |
| 0.1 | 0.05 | AG1 | 3 | 2.75 | 0.329 |
| 0.1 | 0.05 | ptsI | 3 | 3 | 0.18 |
| 0.1 | 0.05 | clcB | 3 | 3 | 0.159 |
| 0.1 | 0.05 | ycaM | 3 | 3 | 0.166 |
| 0.1 | 0.05 | yadI | 3 | 3 | 0.183 |
| 0.1 | 0.05 | AG1 | 3 | 3 | 0.297 |
| 0.1 | 0.05 | ptsI | 3 | 3.25 | 0.182 |
| 0.1 | 0.05 | clcB | 3 | 3.25 | 0.16 |
| 0.1 | 0.05 | ycaM | 3 | 3.25 | 0.167 |
| 0.1 | 0.05 | yadI | 3 | 3.25 | 0.186 |
| 0.1 | 0.05 | AG1 | 3 | 3.25 | 0.269 |
| 0.1 | 0.05 | ptsI | 3 | 3.5 | 0.184 |
| 0.1 | 0.05 | clcB | 3 | 3.5 | 0.161 |
| 0.1 | 0.05 | ycaM | 3 | 3.5 | 0.172 |
| 0.1 | 0.05 | yadI | 3 | 3.5 | 0.189 |
| 0.1 | 0.05 | AG1 | 3 | 3.5 | 0.249 |
| 0.1 | 0.05 | ptsI | 3 | 3.75 | 0.186 |
| 0.1 | 0.05 | clcB | 3 | 3.75 | 0.164 |
| 0.1 | 0.05 | ycaM | 3 | 3.75 | 0.174 |
| 0.1 | 0.05 | yadI | 3 | 3.75 | 0.191 |
| 0.1 | 0.05 | AG1 | 3 | 3.75 | 0.236 |

| | | | | | |
|---|---|---|---|---|---|
| 0.1 | 0.05 | ptsI | 3 | 4 | 0.189 |
| 0.1 | 0.05 | clcB | 3 | 4 | 0.166 |
| 0.1 | 0.05 | ycaM | 3 | 4 | 0.176 |
| 0.1 | 0.05 | yadI | 3 | 4 | 0.197 |
| 0.1 | 0.05 | AG1 | 3 | 4 | 0.227 |
| 0.1 | 0.05 | ptsI | 3 | 4.41667 | 0.195 |
| 0.1 | 0.05 | clcB | 3 | 4.41667 | 0.192 |
| 0.1 | 0.05 | ycaM | 3 | 4.41667 | 0.193 |
| 0.1 | 0.05 | yadI | 3 | 4.41667 | 0.216 |
| 0.1 | 0.05 | AG1 | 3 | 4.41667 | 0.204 |
| 0.1 | 0.05 | ptsI | 3 | 4.66667 | 0.185 |
| 0.1 | 0.05 | clcB | 3 | 4.66667 | 0.186 |
| 0.1 | 0.05 | ycaM | 3 | 4.66667 | 0.193 |
| 0.1 | 0.05 | yadI | 3 | 4.66667 | 0.22 |
| 0.1 | 0.05 | AG1 | 3 | 4.66667 | 0.191 |
| 0.1 | 0.05 | ptsI | 3 | 4.91667 | 0.188 |
| 0.1 | 0.05 | clcB | 3 | 4.91667 | 0.188 |
| 0.1 | 0.05 | ycaM | 3 | 4.91667 | 0.193 |
| 0.1 | 0.05 | yadI | 3 | 4.91667 | 0.232 |
| 0.1 | 0.05 | AG1 | 3 | 4.91667 | 0.199 |
| 0.1 | 0.05 | ptsI | 3 | 5.16667 | 0.191 |
| 0.1 | 0.05 | clcB | 3 | 5.16667 | 0.193 |
| 0.1 | 0.05 | ycaM | 3 | 5.16667 | 0.198 |
| 0.1 | 0.05 | yadI | 3 | 5.16667 | 0.246 |
| 0.1 | 0.05 | AG1 | 3 | 5.16667 | 0.198 |
| 0.1 | 0.05 | ptsI | 3 | 5.41667 | 0.195 |
| 0.1 | 0.05 | clcB | 3 | 5.41667 | 0.198 |
| 0.1 | 0.05 | ycaM | 3 | 5.41667 | 0.204 |
| 0.1 | 0.05 | yadI | 3 | 5.41667 | 0.264 |
| 0.1 | 0.05 | AG1 | 3 | 5.41667 | 0.198 |
| 0.1 | 0.05 | ptsI | 3 | 5.66667 | 0.2 |
| 0.1 | 0.05 | clcB | 3 | 5.66667 | 0.205 |
| 0.1 | 0.05 | ycaM | 3 | 5.66667 | 0.212 |
| 0.1 | 0.05 | yadI | 3 | 5.66667 | 0.284 |
| 0.1 | 0.05 | AG1 | 3 | 5.66667 | 0.199 |
| 0.1 | 0.05 | ptsI | 3 | 5.91667 | 0.205 |
| 0.1 | 0.05 | clcB | 3 | 5.91667 | 0.213 |
| 0.1 | 0.05 | ycaM | 3 | 5.91667 | 0.218 |
| 0.1 | 0.05 | yadI | 3 | 5.91667 | 0.286 |
| 0.1 | 0.05 | AG1 | 3 | 5.91667 | 0.204 |
| 0.1 | 0.05 | ptsI | 3 | 6.16667 | 0.212 |
| 0.1 | 0.05 | clcB | 3 | 6.16667 | 0.22 |
| 0.1 | 0.05 | ycaM | 3 | 6.16667 | 0.228 |
| 0.1 | 0.05 | yadI | 3 | 6.16667 | 0.316 |
| 0.1 | 0.05 | AG1 | 3 | 6.16667 | 0.211 |
| 0.1 | 0.05 | ptsI | 3 | 6.41667 | 0.217 |
| 0.1 | 0.05 | clcB | 3 | 6.41667 | 0.229 |
| 0.1 | 0.05 | ycaM | 3 | 6.41667 | 0.24 |

| | | | | | |
|---|---|---|---|---|---|
| 0.1 | 0.05 | yadI | 3 | 6.41667 | 0.307 |
| 0.1 | 0.05 | AG1 | 3 | 6.41667 | 0.219 |
| 0.1 | 0.05 | ptsI | 3 | 6.66667 | 0.226 |
| 0.1 | 0.05 | clcB | 3 | 6.66667 | 0.239 |
| 0.1 | 0.05 | ycaM | 3 | 6.66667 | 0.249 |
| 0.1 | 0.05 | yadI | 3 | 6.66667 | 0.317 |
| 0.1 | 0.05 | AG1 | 3 | 6.66667 | 0.23 |
| 0.1 | 0.05 | ptsI | 3 | 6.91667 | 0.237 |
| 0.1 | 0.05 | clcB | 3 | 6.91667 | 0.25 |
| 0.1 | 0.05 | ycaM | 3 | 6.91667 | 0.259 |
| 0.1 | 0.05 | yadI | 3 | 6.91667 | 0.33 |
| 0.1 | 0.05 | AG1 | 3 | 6.91667 | 0.239 |
| 0.1 | 0.05 | ptsI | 3 | 7.16667 | 0.246 |
| 0.1 | 0.05 | clcB | 3 | 7.16667 | 0.26 |
| 0.1 | 0.05 | ycaM | 3 | 7.16667 | 0.265 |
| 0.1 | 0.05 | yadI | 3 | 7.16667 | 0.342 |
| 0.1 | 0.05 | AG1 | 3 | 7.16667 | 0.251 |
| 0.1 | 0.05 | ptsI | 3 | 7.41667 | 0.259 |
| 0.1 | 0.05 | clcB | 3 | 7.41667 | 0.267 |
| 0.1 | 0.05 | ycaM | 3 | 7.41667 | 0.269 |
| 0.1 | 0.05 | yadI | 3 | 7.41667 | 0.358 |
| 0.1 | 0.05 | AG1 | 3 | 7.41667 | 0.258 |
| 0.1 | 0.05 | ptsI | 3 | 7.66667 | 0.269 |
| 0.1 | 0.05 | clcB | 3 | 7.66667 | 0.276 |
| 0.1 | 0.05 | ycaM | 3 | 7.66667 | 0.274 |
| 0.1 | 0.05 | yadI | 3 | 7.66667 | 0.358 |
| 0.1 | 0.05 | AG1 | 3 | 7.66667 | 0.266 |
| 0.1 | 0.05 | ptsI | 3 | 7.91667 | 0.276 |
| 0.1 | 0.05 | clcB | 3 | 7.91667 | 0.281 |
| 0.1 | 0.05 | ycaM | 3 | 7.91667 | 0.275 |
| 0.1 | 0.05 | yadI | 3 | 7.91667 | 0.366 |
| 0.1 | 0.05 | AG1 | 3 | 7.91667 | 0.273 |
| 0.1 | 0.05 | ptsI | 3 | 8.16667 | 0.284 |
| 0.1 | 0.05 | clcB | 3 | 8.16667 | 0.286 |
| 0.1 | 0.05 | ycaM | 3 | 8.16667 | 0.274 |
| 0.1 | 0.05 | yadI | 3 | 8.16667 | 0.379 |
| 0.1 | 0.05 | AG1 | 3 | 8.16667 | 0.28 |
| 0.1 | 0.05 | ptsI | 3 | 8.41667 | 0.294 |
| 0.1 | 0.05 | clcB | 3 | 8.41667 | 0.294 |
| 0.1 | 0.05 | ycaM | 3 | 8.41667 | 0.275 |
| 0.1 | 0.05 | yadI | 3 | 8.41667 | 0.388 |
| 0.1 | 0.05 | AG1 | 3 | 8.41667 | 0.293 |
| 0.1 | 0.05 | ptsI | 3 | 9.05 | 0.283 |
| 0.1 | 0.05 | clcB | 3 | 9.05 | 0.28 |
| 0.1 | 0.05 | ycaM | 3 | 9.05 | 0.292 |
| 0.1 | 0.05 | yadI | 3 | 9.05 | 0.412 |
| 0.1 | 0.05 | AG1 | 3 | 9.05 | 0.299 |
| 0.1 | 0.05 | ptsI | 3 | 9.3 | 0.288 |

| | | | | | |
|---|---|---|---|---|---|
| 0.1 | 0.05 | clcB | 3 | 9.3 | 0.282 |
| 0.1 | 0.05 | ycaM | 3 | 9.3 | 0.287 |
| 0.1 | 0.05 | yadI | 3 | 9.3 | 0.421 |
| 0.1 | 0.05 | AG1 | 3 | 9.3 | 0.313 |
| 0.1 | 0.05 | ptsI | 3 | 9.55 | 0.294 |
| 0.1 | 0.05 | clcB | 3 | 9.55 | 0.286 |
| 0.1 | 0.05 | ycaM | 3 | 9.55 | 0.286 |
| 0.1 | 0.05 | yadI | 3 | 9.55 | 0.435 |
| 0.1 | 0.05 | AG1 | 3 | 9.55 | 0.325 |
| 0.1 | 0.05 | ptsI | 3 | 9.8 | 0.302 |
| 0.1 | 0.05 | clcB | 3 | 9.8 | 0.295 |
| 0.1 | 0.05 | ycaM | 3 | 9.8 | 0.289 |
| 0.1 | 0.05 | yadI | 3 | 9.8 | 0.448 |
| 0.1 | 0.05 | AG1 | 3 | 9.8 | 0.331 |
| 0.1 | 0.05 | ptsI | 3 | 10.05 | 0.309 |
| 0.1 | 0.05 | clcB | 3 | 10.05 | 0.298 |
| 0.1 | 0.05 | ycaM | 3 | 10.05 | 0.29 |
| 0.1 | 0.05 | yadI | 3 | 10.05 | 0.461 |
| 0.1 | 0.05 | AG1 | 3 | 10.05 | 0.342 |
| 0.1 | 0.05 | ptsI | 3 | 10.3 | 0.32 |
| 0.1 | 0.05 | clcB | 3 | 10.3 | 0.3 |
| 0.1 | 0.05 | ycaM | 3 | 10.3 | 0.293 |
| 0.1 | 0.05 | yadI | 3 | 10.3 | 0.472 |
| 0.1 | 0.05 | AG1 | 3 | 10.3 | 0.349 |
| 0.1 | 0.05 | ptsI | 3 | 10.55 | 0.328 |
| 0.1 | 0.05 | clcB | 3 | 10.55 | 0.305 |
| 0.1 | 0.05 | ycaM | 3 | 10.55 | 0.292 |
| 0.1 | 0.05 | yadI | 3 | 10.55 | 0.486 |
| 0.1 | 0.05 | AG1 | 3 | 10.55 | 0.356 |
| 0.1 | 0.05 | ptsI | 3 | 10.8 | 0.332 |
| 0.1 | 0.05 | clcB | 3 | 10.8 | 0.306 |
| 0.1 | 0.05 | ycaM | 3 | 10.8 | 0.292 |
| 0.1 | 0.05 | yadI | 3 | 10.8 | 0.495 |
| 0.1 | 0.05 | AG1 | 3 | 10.8 | 0.361 |
| 0.1 | 0.05 | ptsI | 3 | 11.05 | 0.347 |
| 0.1 | 0.05 | clcB | 3 | 11.05 | 0.311 |
| 0.1 | 0.05 | ycaM | 3 | 11.05 | 0.295 |
| 0.1 | 0.05 | yadI | 3 | 11.05 | 0.506 |
| 0.1 | 0.05 | AG1 | 3 | 11.05 | 0.37 |
| 0.1 | 0.05 | ptsI | 3 | 11.3 | 0.353 |
| 0.1 | 0.05 | clcB | 3 | 11.3 | 0.316 |
| 0.1 | 0.05 | ycaM | 3 | 11.3 | 0.294 |
| 0.1 | 0.05 | yadI | 3 | 11.3 | 0.513 |
| 0.1 | 0.05 | AG1 | 3 | 11.3 | 0.376 |
| 0.1 | 0.05 | ptsI | 3 | 11.55 | 0.363 |
| 0.1 | 0.05 | clcB | 3 | 11.55 | 0.312 |
| 0.1 | 0.05 | ycaM | 3 | 11.55 | 0.292 |
| 0.1 | 0.05 | yadI | 3 | 11.55 | 0.525 |

| | | | | | |
|---|---|---|---|---|---|
| 0.1 | 0.05 | AG1 | 3 | 11.55 | 0.383 |
| 0.1 | 0.05 | ptsI | 3 | 11.8 | 0.373 |
| 0.1 | 0.05 | clcB | 3 | 11.8 | 0.31 |
| 0.1 | 0.05 | ycaM | 3 | 11.8 | 0.293 |
| 0.1 | 0.05 | yadI | 3 | 11.8 | 0.54 |
| 0.1 | 0.05 | AG1 | 3 | 11.8 | 0.396 |
| 0.1 | 0.05 | ptsI | 3 | 12.05 | 0.38 |
| 0.1 | 0.05 | clcB | 3 | 12.05 | 0.317 |
| 0.1 | 0.05 | ycaM | 3 | 12.05 | 0.294 |
| 0.1 | 0.05 | yadI | 3 | 12.05 | 0.547 |
| 0.1 | 0.05 | AG1 | 3 | 12.05 | 0.397 |
| 0.1 | 0.05 | ptsI | 3 | 12.3 | 0.394 |
| 0.1 | 0.05 | clcB | 3 | 12.3 | 0.314 |
| 0.1 | 0.05 | ycaM | 3 | 12.3 | 0.291 |
| 0.1 | 0.05 | yadI | 3 | 12.3 | 0.558 |
| 0.1 | 0.05 | AG1 | 3 | 12.3 | 0.406 |
| 0.1 | 0.05 | ptsI | 3 | 12.55 | 0.403 |
| 0.1 | 0.05 | clcB | 3 | 12.55 | 0.307 |
| 0.1 | 0.05 | ycaM | 3 | 12.55 | 0.294 |
| 0.1 | 0.05 | yadI | 3 | 12.55 | 0.569 |
| 0.1 | 0.05 | AG1 | 3 | 12.55 | 0.42 |
| 0.1 | 0.05 | ptsI | 3 | 12.8 | 0.414 |
| 0.1 | 0.05 | clcB | 3 | 12.8 | 0.305 |
| 0.1 | 0.05 | ycaM | 3 | 12.8 | 0.29 |
| 0.1 | 0.05 | yadI | 3 | 12.8 | 0.582 |
| 0.1 | 0.05 | AG1 | 3 | 12.8 | 0.43 |
| 0.1 | 0.05 | ptsI | 3 | 13.05 | 0.418 |
| 0.1 | 0.05 | clcB | 3 | 13.05 | 0.302 |
| 0.1 | 0.05 | ycaM | 3 | 13.05 | 0.289 |
| 0.1 | 0.05 | yadI | 3 | 13.05 | 0.586 |
| 0.1 | 0.05 | AG1 | 3 | 13.05 | 0.434 |
| 0.1 | 0.05 | ptsI | 3 | 13.3 | 0.427 |
| 0.1 | 0.05 | clcB | 3 | 13.3 | 0.317 |
| 0.1 | 0.05 | ycaM | 3 | 13.3 | 0.287 |
| 0.1 | 0.05 | yadI | 3 | 13.3 | 0.597 |
| 0.1 | 0.05 | AG1 | 3 | 13.3 | 0.447 |
| 0.1 | 0.05 | ptsI | 3 | 13.55 | 0.44 |
| 0.1 | 0.05 | clcB | 3 | 13.55 | 0.321 |
| 0.1 | 0.05 | ycaM | 3 | 13.55 | 0.288 |
| 0.1 | 0.05 | yadI | 3 | 13.55 | 0.606 |
| 0.1 | 0.05 | AG1 | 3 | 13.55 | 0.46 |
| 0.1 | 0.05 | ptsI | 3 | 13.8 | 0.451 |
| 0.1 | 0.05 | clcB | 3 | 13.8 | 0.304 |
| 0.1 | 0.05 | ycaM | 3 | 13.8 | 0.284 |
| 0.1 | 0.05 | yadI | 3 | 13.8 | 0.617 |
| 0.1 | 0.05 | AG1 | 3 | 13.8 | 0.474 |
| 0.1 | 0.05 | ptsI | 3 | 14.05 | 0.459 |
| 0.1 | 0.05 | clcB | 3 | 14.05 | 0.31 |

| | | | | | |
|---|---|---|---|---|---|
| 0.1 | 0.05 | ycaM | 3 | 14.05 | 0.286 |
| 0.1 | 0.05 | yadI | 3 | 14.05 | 0.625 |
| 0.1 | 0.05 | AG1 | 3 | 14.05 | 0.484 |
| 0.1 | 0.05 | ptsI | 3 | 14.3 | 0.464 |
| 0.1 | 0.05 | clcB | 3 | 14.3 | 0.318 |
| 0.1 | 0.05 | ycaM | 3 | 14.3 | 0.284 |
| 0.1 | 0.05 | yadI | 3 | 14.3 | 0.63 |
| 0.1 | 0.05 | AG1 | 3 | 14.3 | 0.489 |
| 0.1 | 0.05 | ptsI | 3 | 14.55 | 0.472 |
| 0.1 | 0.05 | clcB | 3 | 14.55 | 0.31 |
| 0.1 | 0.05 | ycaM | 3 | 14.55 | 0.286 |
| 0.1 | 0.05 | yadI | 3 | 14.55 | 0.639 |
| 0.1 | 0.05 | AG1 | 3 | 14.55 | 0.501 |
| 0.1 | 0.05 | ptsI | 3 | 14.8 | 0.486 |
| 0.1 | 0.05 | clcB | 3 | 14.8 | 0.304 |
| 0.1 | 0.05 | ycaM | 3 | 14.8 | 0.285 |
| 0.1 | 0.05 | yadI | 3 | 14.8 | 0.652 |
| 0.1 | 0.05 | AG1 | 3 | 14.8 | 0.522 |
| 0.1 | 0.05 | ptsI | 3 | 15.05 | 0.497 |
| 0.1 | 0.05 | clcB | 3 | 15.05 | 0.308 |
| 0.1 | 0.05 | ycaM | 3 | 15.05 | 0.286 |
| 0.1 | 0.05 | yadI | 3 | 15.05 | 0.67 |
| 0.1 | 0.05 | AG1 | 3 | 15.05 | 0.542 |
| 0.1 | 0.05 | ptsI | 3 | 15.3 | 0.501 |
| 0.1 | 0.05 | clcB | 3 | 15.3 | 0.309 |
| 0.1 | 0.05 | ycaM | 3 | 15.3 | 0.28 |
| 0.1 | 0.05 | yadI | 3 | 15.3 | 0.67 |
| 0.1 | 0.05 | AG1 | 3 | 15.3 | 0.542 |
| 0.1 | 0.05 | ptsI | 3 | 15.55 | 0.505 |
| 0.1 | 0.05 | clcB | 3 | 15.55 | 0.307 |
| 0.1 | 0.05 | ycaM | 3 | 15.55 | 0.284 |
| 0.1 | 0.05 | yadI | 3 | 15.55 | 0.675 |
| 0.1 | 0.05 | AG1 | 3 | 15.55 | 0.551 |
| 0.1 | 0.05 | ptsI | 3 | 15.8 | 0.511 |
| 0.1 | 0.05 | clcB | 3 | 15.8 | 0.305 |
| 0.1 | 0.05 | ycaM | 3 | 15.8 | 0.279 |
| 0.1 | 0.05 | yadI | 3 | 15.8 | 0.685 |
| 0.1 | 0.05 | AG1 | 3 | 15.8 | 0.564 |
| 0.1 | 0.05 | ptsI | 3 | 16.05 | 0.514 |
| 0.1 | 0.05 | clcB | 3 | 16.05 | 0.305 |
| 0.1 | 0.05 | ycaM | 3 | 16.05 | 0.278 |
| 0.1 | 0.05 | yadI | 3 | 16.05 | 0.681 |
| 0.1 | 0.05 | AG1 | 3 | 16.05 | 0.567 |
| 0.1 | 0.05 | ptsI | 3 | 16.3 | 0.521 |
| 0.1 | 0.05 | clcB | 3 | 16.3 | 0.308 |
| 0.1 | 0.05 | ycaM | 3 | 16.3 | 0.275 |
| 0.1 | 0.05 | yadI | 3 | 16.3 | 0.689 |
| 0.1 | 0.05 | AG1 | 3 | 16.3 | 0.591 |

| | | | | | |
|---|---|---|---|---|---|
| 0.1 | 0.05 | ptsI | 3 | 16.55 | 0.527 |
| 0.1 | 0.05 | clcB | 3 | 16.55 | 0.305 |
| 0.1 | 0.05 | ycaM | 3 | 16.55 | 0.278 |
| 0.1 | 0.05 | yadI | 3 | 16.55 | 0.703 |
| 0.1 | 0.05 | AG1 | 3 | 16.55 | 0.598 |
| 0.1 | 0.05 | ptsI | 3 | 16.8 | 0.536 |
| 0.1 | 0.05 | clcB | 3 | 16.8 | 0.306 |
| 0.1 | 0.05 | ycaM | 3 | 16.8 | 0.278 |
| 0.1 | 0.05 | yadI | 3 | 16.8 | 0.71 |
| 0.1 | 0.05 | AG1 | 3 | 16.8 | 0.62 |
| 0.1 | 0.05 | ptsI | 3 | 17.05 | 0.536 |
| 0.1 | 0.05 | clcB | 3 | 17.05 | 0.305 |
| 0.1 | 0.05 | ycaM | 3 | 17.05 | 0.278 |
| 0.1 | 0.05 | yadI | 3 | 17.05 | 0.714 |
| 0.1 | 0.05 | AG1 | 3 | 17.05 | 0.633 |
| 0.1 | 0.05 | ptsI | 3 | 17.3 | 0.542 |
| 0.1 | 0.05 | clcB | 3 | 17.3 | 0.307 |
| 0.1 | 0.05 | ycaM | 3 | 17.3 | 0.277 |
| 0.1 | 0.05 | yadI | 3 | 17.3 | 0.715 |
| 0.1 | 0.05 | AG1 | 3 | 17.3 | 0.64 |
| 0.1 | 0.05 | ptsI | 3 | 17.55 | 0.547 |
| 0.1 | 0.05 | clcB | 3 | 17.55 | 0.305 |
| 0.1 | 0.05 | ycaM | 3 | 17.55 | 0.276 |
| 0.1 | 0.05 | yadI | 3 | 17.55 | 0.718 |
| 0.1 | 0.05 | AG1 | 3 | 17.55 | 0.653 |
| 0.1 | 0.05 | ptsI | 3 | 17.8 | 0.552 |
| 0.1 | 0.05 | clcB | 3 | 17.8 | 0.306 |
| 0.1 | 0.05 | ycaM | 3 | 17.8 | 0.276 |
| 0.1 | 0.05 | yadI | 3 | 17.8 | 0.729 |
| 0.1 | 0.05 | AG1 | 3 | 17.8 | 0.678 |
| 0.1 | 0.05 | ptsI | 3 | 18.05 | 0.55 |
| 0.1 | 0.05 | clcB | 3 | 18.05 | 0.305 |
| 0.1 | 0.05 | ycaM | 3 | 18.05 | 0.278 |
| 0.1 | 0.05 | yadI | 3 | 18.05 | 0.736 |
| 0.1 | 0.05 | AG1 | 3 | 18.05 | 0.684 |
| 0.1 | 0.05 | ptsI | 3 | 18.3 | 0.554 |
| 0.1 | 0.05 | clcB | 3 | 18.3 | 0.305 |
| 0.1 | 0.05 | ycaM | 3 | 18.3 | 0.278 |
| 0.1 | 0.05 | yadI | 3 | 18.3 | 0.742 |
| 0.1 | 0.05 | AG1 | 3 | 18.3 | 0.689 |
| 0.1 | 0.05 | ptsI | 3 | 18.55 | 0.558 |
| 0.1 | 0.05 | clcB | 3 | 18.55 | 0.305 |
| 0.1 | 0.05 | ycaM | 3 | 18.55 | 0.281 |
| 0.1 | 0.05 | yadI | 3 | 18.55 | 0.748 |
| 0.1 | 0.05 | AG1 | 3 | 18.55 | 0.678 |
| 0.1 | 0.05 | ptsI | 3 | 18.8 | 0.561 |
| 0.1 | 0.05 | clcB | 3 | 18.8 | 0.302 |
| 0.1 | 0.05 | ycaM | 3 | 18.8 | 0.281 |

| | | | | | |
|---|---|---|---|---|---|
| 0.1 | 0.05 | yadI | 3 | 18.8 | 0.754 |
| 0.1 | 0.05 | AG1 | 3 | 18.8 | 0.673 |
| 0.1 | 0.05 | ptsI | 3 | 19.05 | 0.569 |
| 0.1 | 0.05 | clcB | 3 | 19.05 | 0.302 |
| 0.1 | 0.05 | ycaM | 3 | 19.05 | 0.28 |
| 0.1 | 0.05 | yadI | 3 | 19.05 | 0.76 |
| 0.1 | 0.05 | AG1 | 3 | 19.05 | 0.677 |
| 0.1 | 0.05 | ptsI | 3 | 19.3 | 0.57 |
| 0.1 | 0.05 | clcB | 3 | 19.3 | 0.304 |
| 0.1 | 0.05 | ycaM | 3 | 19.3 | 0.281 |
| 0.1 | 0.05 | yadI | 3 | 19.3 | 0.762 |
| 0.1 | 0.05 | AG1 | 3 | 19.3 | 0.683 |
| 0.1 | 0.05 | ptsI | 3 | 19.55 | 0.565 |
| 0.1 | 0.05 | clcB | 3 | 19.55 | 0.3 |
| 0.1 | 0.05 | ycaM | 3 | 19.55 | 0.281 |
| 0.1 | 0.05 | yadI | 3 | 19.55 | 0.764 |
| 0.1 | 0.05 | AG1 | 3 | 19.55 | 0.677 |
| 0.1 | 0.05 | ptsI | 3 | 19.8 | 0.568 |
| 0.1 | 0.05 | clcB | 3 | 19.8 | 0.302 |
| 0.1 | 0.05 | ycaM | 3 | 19.8 | 0.283 |
| 0.1 | 0.05 | yadI | 3 | 19.8 | 0.765 |
| 0.1 | 0.05 | AG1 | 3 | 19.8 | 0.68 |
| 0.1 | 0.05 | ptsI | 3 | 20.05 | 0.572 |
| 0.1 | 0.05 | clcB | 3 | 20.05 | 0.301 |
| 0.1 | 0.05 | ycaM | 3 | 20.05 | 0.284 |
| 0.1 | 0.05 | yadI | 3 | 20.05 | 0.774 |
| 0.1 | 0.05 | AG1 | 3 | 20.05 | 0.675 |
| 0.1 | 0.05 | ptsI | 3 | 20.3 | 0.573 |
| 0.1 | 0.05 | clcB | 3 | 20.3 | 0.302 |
| 0.1 | 0.05 | ycaM | 3 | 20.3 | 0.284 |
| 0.1 | 0.05 | yadI | 3 | 20.3 | 0.77 |
| 0.1 | 0.05 | AG1 | 3 | 20.3 | 0.667 |
| 0.1 | 0.05 | ptsI | 3 | 20.55 | 0.573 |
| 0.1 | 0.05 | clcB | 3 | 20.55 | 0.304 |
| 0.1 | 0.05 | ycaM | 3 | 20.55 | 0.285 |
| 0.1 | 0.05 | yadI | 3 | 20.55 | 0.774 |
| 0.1 | 0.05 | AG1 | 3 | 20.55 | 0.68 |
| 0.1 | 0.05 | ptsI | 3 | 20.8 | 0.582 |
| 0.1 | 0.05 | clcB | 3 | 20.8 | 0.303 |
| 0.1 | 0.05 | ycaM | 3 | 20.8 | 0.286 |
| 0.1 | 0.05 | yadI | 3 | 20.8 | 0.783 |
| 0.1 | 0.05 | AG1 | 3 | 20.8 | 0.686 |
| 0.1 | 0.05 | ptsI | 3 | 21.05 | 0.578 |
| 0.1 | 0.05 | clcB | 3 | 21.05 | 0.302 |
| 0.1 | 0.05 | ycaM | 3 | 21.05 | 0.29 |
| 0.1 | 0.05 | yadI | 3 | 21.05 | 0.779 |
| 0.1 | 0.05 | AG1 | 3 | 21.05 | 0.69 |
| 0.1 | 0.05 | ptsI | 3 | 21.3 | 0.581 |

| | | | | | |
|---|---|---|---|---|---|
| 0.1 | 0.05 | clcB | 3 | 21.3 | 0.302 |
| 0.1 | 0.05 | ycaM | 3 | 21.3 | 0.291 |
| 0.1 | 0.05 | yadI | 3 | 21.3 | 0.778 |
| 0.1 | 0.05 | AG1 | 3 | 21.3 | 0.696 |
| 0.1 | 0.05 | ptsI | 3 | 21.55 | 0.582 |
| 0.1 | 0.05 | clcB | 3 | 21.55 | 0.304 |
| 0.1 | 0.05 | ycaM | 3 | 21.55 | 0.295 |
| 0.1 | 0.05 | yadI | 3 | 21.55 | 0.787 |
| 0.1 | 0.05 | AG1 | 3 | 21.55 | 0.709 |
| 0.1 | 0.05 | ptsI | 3 | 21.8 | 0.585 |
| 0.1 | 0.05 | clcB | 3 | 21.8 | 0.305 |
| 0.1 | 0.05 | ycaM | 3 | 21.8 | 0.299 |
| 0.1 | 0.05 | yadI | 3 | 21.8 | 0.787 |
| 0.1 | 0.05 | AG1 | 3 | 21.8 | 0.714 |
| 0.1 | 0.05 | ptsI | 3 | 22.05 | 0.588 |
| 0.1 | 0.05 | clcB | 3 | 22.05 | 0.304 |
| 0.1 | 0.05 | ycaM | 3 | 22.05 | 0.3 |
| 0.1 | 0.05 | yadI | 3 | 22.05 | 0.792 |
| 0.1 | 0.05 | AG1 | 3 | 22.05 | 0.722 |
| 0.1 | 0.05 | ptsI | 3 | 22.3 | 0.591 |
| 0.1 | 0.05 | clcB | 3 | 22.3 | 0.307 |
| 0.1 | 0.05 | ycaM | 3 | 22.3 | 0.306 |
| 0.1 | 0.05 | yadI | 3 | 22.3 | 0.792 |
| 0.1 | 0.05 | AG1 | 3 | 22.3 | 0.729 |
| 0.1 | 0.05 | ptsI | 3 | 22.55 | 0.59 |
| 0.1 | 0.05 | clcB | 3 | 22.55 | 0.309 |
| 0.1 | 0.05 | ycaM | 3 | 22.55 | 0.311 |
| 0.1 | 0.05 | yadI | 3 | 22.55 | 0.797 |
| 0.1 | 0.05 | AG1 | 3 | 22.55 | 0.738 |
| 0.1 | 0.05 | ptsI | 3 | 22.8 | 0.59 |
| 0.1 | 0.05 | clcB | 3 | 22.8 | 0.308 |
| 0.1 | 0.05 | ycaM | 3 | 22.8 | 0.315 |
| 0.1 | 0.05 | yadI | 3 | 22.8 | 0.796 |
| 0.1 | 0.05 | AG1 | 3 | 22.8 | 0.741 |
| 0.1 | 0.05 | ptsI | 3 | 23.05 | 0.59 |
| 0.1 | 0.05 | clcB | 3 | 23.05 | 0.31 |
| 0.1 | 0.05 | ycaM | 3 | 23.05 | 0.321 |
| 0.1 | 0.05 | yadI | 3 | 23.05 | 0.801 |
| 0.1 | 0.05 | AG1 | 3 | 23.05 | 0.752 |
| 0.1 | 0.05 | ptsI | 3 | 23.3 | 0.594 |
| 0.1 | 0.05 | clcB | 3 | 23.3 | 0.31 |
| 0.1 | 0.05 | ycaM | 3 | 23.3 | 0.327 |
| 0.1 | 0.05 | yadI | 3 | 23.3 | 0.802 |
| 0.1 | 0.05 | AG1 | 3 | 23.3 | 0.757 |
| 0.1 | 0.05 | ptsI | 3 | 23.55 | 0.594 |
| 0.1 | 0.05 | clcB | 3 | 23.55 | 0.309 |
| 0.1 | 0.05 | ycaM | 3 | 23.55 | 0.33 |
| 0.1 | 0.05 | yadI | 3 | 23.55 | 0.81 |

| | | | | | |
|---|---|---|---|---|---|
| 0.1 | 0.05 | AG1 | 3 | 23.55 | 0.768 |
| 0.1 | 0.05 | ptsI | 3 | 23.8 | 0.593 |
| 0.1 | 0.05 | clcB | 3 | 23.8 | 0.311 |
| 0.1 | 0.05 | ycaM | 3 | 23.8 | 0.336 |
| 0.1 | 0.05 | yadI | 3 | 23.8 | 0.807 |
| 0.1 | 0.05 | AG1 | 3 | 23.8 | 0.773 |
| 0.1 | 0.05 | ptsI | 3 | 24.05 | 0.593 |
| 0.1 | 0.05 | clcB | 3 | 24.05 | 0.311 |
| 0.1 | 0.05 | ycaM | 3 | 24.05 | 0.34 |
| 0.1 | 0.05 | yadI | 3 | 24.05 | 0.814 |
| 0.1 | 0.05 | AG1 | 3 | 24.05 | 0.779 |
| 0.1 | 0.05 | ptsI | 3 | 24.3 | 0.593 |
| 0.1 | 0.05 | clcB | 3 | 24.3 | 0.312 |
| 0.1 | 0.05 | ycaM | 3 | 24.3 | 0.345 |
| 0.1 | 0.05 | yadI | 3 | 24.3 | 0.813 |
| 0.1 | 0.05 | AG1 | 3 | 24.3 | 0.791 |
| 0.1 | 0.05 | ptsI | 4 | 0 | 0.192 |
| 0.1 | 0.05 | clcB | 4 | 0 | 0.212 |
| 0.1 | 0.05 | ycaM | 4 | 0 | 0.224 |
| 0.1 | 0.05 | yadI | 4 | 0 | 0.232 |
| 0.1 | 0.05 | AG1 | 4 | 0 | 0.198 |
| 0.1 | 0.05 | ptsI | 4 | 0.25 | 0.191 |
| 0.1 | 0.05 | clcB | 4 | 0.25 | 0.208 |
| 0.1 | 0.05 | ycaM | 4 | 0.25 | 0.218 |
| 0.1 | 0.05 | yadI | 4 | 0.25 | 0.222 |
| 0.1 | 0.05 | AG1 | 4 | 0.25 | 0.196 |
| 0.1 | 0.05 | ptsI | 4 | 0.5 | 0.187 |
| 0.1 | 0.05 | clcB | 4 | 0.5 | 0.208 |
| 0.1 | 0.05 | ycaM | 4 | 0.5 | 0.218 |
| 0.1 | 0.05 | yadI | 4 | 0.5 | 0.225 |
| 0.1 | 0.05 | AG1 | 4 | 0.5 | 0.198 |
| 0.1 | 0.05 | ptsI | 4 | 0.75 | 0.19 |
| 0.1 | 0.05 | clcB | 4 | 0.75 | 0.214 |
| 0.1 | 0.05 | ycaM | 4 | 0.75 | 0.22 |
| 0.1 | 0.05 | yadI | 4 | 0.75 | 0.228 |
| 0.1 | 0.05 | AG1 | 4 | 0.75 | 0.199 |
| 0.1 | 0.05 | ptsI | 4 | 1 | 0.193 |
| 0.1 | 0.05 | clcB | 4 | 1 | 0.217 |
| 0.1 | 0.05 | ycaM | 4 | 1 | 0.222 |
| 0.1 | 0.05 | yadI | 4 | 1 | 0.228 |
| 0.1 | 0.05 | AG1 | 4 | 1 | 0.201 |
| 0.1 | 0.05 | ptsI | 4 | 1.25 | 0.197 |
| 0.1 | 0.05 | clcB | 4 | 1.25 | 0.223 |
| 0.1 | 0.05 | ycaM | 4 | 1.25 | 0.228 |
| 0.1 | 0.05 | yadI | 4 | 1.25 | 0.231 |
| 0.1 | 0.05 | AG1 | 4 | 1.25 | 0.205 |
| 0.1 | 0.05 | ptsI | 4 | 1.5 | 0.201 |
| 0.1 | 0.05 | clcB | 4 | 1.5 | 0.224 |

| | | | | | |
|---|---|---|---|---|---|
| 0.1 | 0.05 | ycaM | 4 | 1.5 | 0.233 |
| 0.1 | 0.05 | yadI | 4 | 1.5 | 0.235 |
| 0.1 | 0.05 | AG1 | 4 | 1.5 | 0.212 |
| 0.1 | 0.05 | ptsI | 4 | 1.75 | 0.208 |
| 0.1 | 0.05 | clcB | 4 | 1.75 | 0.229 |
| 0.1 | 0.05 | ycaM | 4 | 1.75 | 0.237 |
| 0.1 | 0.05 | yadI | 4 | 1.75 | 0.242 |
| 0.1 | 0.05 | AG1 | 4 | 1.75 | 0.22 |
| 0.1 | 0.05 | ptsI | 4 | 2 | 0.215 |
| 0.1 | 0.05 | clcB | 4 | 2 | 0.24 |
| 0.1 | 0.05 | ycaM | 4 | 2 | 0.248 |
| 0.1 | 0.05 | yadI | 4 | 2 | 0.249 |
| 0.1 | 0.05 | AG1 | 4 | 2 | 0.228 |
| 0.1 | 0.05 | ptsI | 4 | 2.25 | 0.224 |
| 0.1 | 0.05 | clcB | 4 | 2.25 | 0.249 |
| 0.1 | 0.05 | ycaM | 4 | 2.25 | 0.253 |
| 0.1 | 0.05 | yadI | 4 | 2.25 | 0.258 |
| 0.1 | 0.05 | AG1 | 4 | 2.25 | 0.239 |
| 0.1 | 0.05 | ptsI | 4 | 2.5 | 0.229 |
| 0.1 | 0.05 | clcB | 4 | 2.5 | 0.254 |
| 0.1 | 0.05 | ycaM | 4 | 2.5 | 0.263 |
| 0.1 | 0.05 | yadI | 4 | 2.5 | 0.265 |
| 0.1 | 0.05 | AG1 | 4 | 2.5 | 0.247 |
| 0.1 | 0.05 | ptsI | 4 | 2.75 | 0.24 |
| 0.1 | 0.05 | clcB | 4 | 2.75 | 0.26 |
| 0.1 | 0.05 | ycaM | 4 | 2.75 | 0.272 |
| 0.1 | 0.05 | yadI | 4 | 2.75 | 0.269 |
| 0.1 | 0.05 | AG1 | 4 | 2.75 | 0.256 |
| 0.1 | 0.05 | ptsI | 4 | 3 | 0.247 |
| 0.1 | 0.05 | clcB | 4 | 3 | 0.268 |
| 0.1 | 0.05 | ycaM | 4 | 3 | 0.276 |
| 0.1 | 0.05 | yadI | 4 | 3 | 0.282 |
| 0.1 | 0.05 | AG1 | 4 | 3 | 0.27 |
| 0.1 | 0.05 | ptsI | 4 | 3.25 | 0.258 |
| 0.1 | 0.05 | clcB | 4 | 3.25 | 0.278 |
| 0.1 | 0.05 | ycaM | 4 | 3.25 | 0.288 |
| 0.1 | 0.05 | yadI | 4 | 3.25 | 0.293 |
| 0.1 | 0.05 | AG1 | 4 | 3.25 | 0.28 |
| 0.1 | 0.05 | ptsI | 4 | 3.5 | 0.266 |
| 0.1 | 0.05 | clcB | 4 | 3.5 | 0.293 |
| 0.1 | 0.05 | ycaM | 4 | 3.5 | 0.298 |
| 0.1 | 0.05 | yadI | 4 | 3.5 | 0.302 |
| 0.1 | 0.05 | AG1 | 4 | 3.5 | 0.296 |
| 0.1 | 0.05 | ptsI | 4 | 3.75 | 0.275 |
| 0.1 | 0.05 | clcB | 4 | 3.75 | 0.298 |
| 0.1 | 0.05 | ycaM | 4 | 3.75 | 0.31 |
| 0.1 | 0.05 | yadI | 4 | 3.75 | 0.308 |
| 0.1 | 0.05 | AG1 | 4 | 3.75 | 0.304 |

| | | | | | |
|---|---|---|---|---|---|
| 0.1 | 0.05 | ptsI | 4 | 4 | 0.292 |
| 0.1 | 0.05 | clcB | 4 | 4 | 0.312 |
| 0.1 | 0.05 | ycaM | 4 | 4 | 0.322 |
| 0.1 | 0.05 | yadI | 4 | 4 | 0.327 |
| 0.1 | 0.05 | AG1 | 4 | 4 | 0.318 |
| 0.1 | 0.05 | ptsI | 4 | 4.35 | 0.307 |
| 0.1 | 0.05 | clcB | 4 | 4.35 | 0.312 |
| 0.1 | 0.05 | ycaM | 4 | 4.35 | 0.408 |
| 0.1 | 0.05 | yadI | 4 | 4.35 | 0.308 |
| 0.1 | 0.05 | AG1 | 4 | 4.35 | 0.318 |
| 0.1 | 0.05 | ptsI | 4 | 4.6 | 0.289 |
| 0.1 | 0.05 | clcB | 4 | 4.6 | 0.297 |
| 0.1 | 0.05 | ycaM | 4 | 4.6 | 0.371 |
| 0.1 | 0.05 | yadI | 4 | 4.6 | 0.32 |
| 0.1 | 0.05 | AG1 | 4 | 4.6 | 0.334 |
| 0.1 | 0.05 | ptsI | 4 | 4.85 | 0.297 |
| 0.1 | 0.05 | clcB | 4 | 4.85 | 0.304 |
| 0.1 | 0.05 | ycaM | 4 | 4.85 | 0.37 |
| 0.1 | 0.05 | yadI | 4 | 4.85 | 0.332 |
| 0.1 | 0.05 | AG1 | 4 | 4.85 | 0.344 |
| 0.1 | 0.05 | ptsI | 4 | 5.1 | 0.305 |
| 0.1 | 0.05 | clcB | 4 | 5.1 | 0.308 |
| 0.1 | 0.05 | ycaM | 4 | 5.1 | 0.37 |
| 0.1 | 0.05 | yadI | 4 | 5.1 | 0.346 |
| 0.1 | 0.05 | AG1 | 4 | 5.1 | 0.355 |
| 0.1 | 0.05 | ptsI | 4 | 5.35 | 0.315 |
| 0.1 | 0.05 | clcB | 4 | 5.35 | 0.309 |
| 0.1 | 0.05 | ycaM | 4 | 5.35 | 0.375 |
| 0.1 | 0.05 | yadI | 4 | 5.35 | 0.359 |
| 0.1 | 0.05 | AG1 | 4 | 5.35 | 0.372 |
| 0.1 | 0.05 | ptsI | 4 | 5.6 | 0.325 |
| 0.1 | 0.05 | clcB | 4 | 5.6 | 0.308 |
| 0.1 | 0.05 | ycaM | 4 | 5.6 | 0.376 |
| 0.1 | 0.05 | yadI | 4 | 5.6 | 0.37 |
| 0.1 | 0.05 | AG1 | 4 | 5.6 | 0.383 |
| 0.1 | 0.05 | ptsI | 4 | 5.85 | 0.335 |
| 0.1 | 0.05 | clcB | 4 | 5.85 | 0.31 |
| 0.1 | 0.05 | ycaM | 4 | 5.85 | 0.376 |
| 0.1 | 0.05 | yadI | 4 | 5.85 | 0.381 |
| 0.1 | 0.05 | AG1 | 4 | 5.85 | 0.392 |
| 0.1 | 0.05 | ptsI | 4 | 6.1 | 0.349 |
| 0.1 | 0.05 | clcB | 4 | 6.1 | 0.315 |
| 0.1 | 0.05 | ycaM | 4 | 6.1 | 0.377 |
| 0.1 | 0.05 | yadI | 4 | 6.1 | 0.394 |
| 0.1 | 0.05 | AG1 | 4 | 6.1 | 0.408 |
| 0.1 | 0.05 | ptsI | 4 | 6.35 | 0.365 |
| 0.1 | 0.05 | clcB | 4 | 6.35 | 0.32 |
| 0.1 | 0.05 | ycaM | 4 | 6.35 | 0.38 |

| | | | | | |
|---|---|---|---|---|---|
| 0.1 | 0.05 | yadI | 4 | 6.35 | 0.406 |
| 0.1 | 0.05 | AG1 | 4 | 6.35 | 0.417 |
| 0.1 | 0.05 | ptsI | 4 | 6.6 | 0.376 |
| 0.1 | 0.05 | clcB | 4 | 6.6 | 0.319 |
| 0.1 | 0.05 | ycaM | 4 | 6.6 | 0.376 |
| 0.1 | 0.05 | yadI | 4 | 6.6 | 0.41 |
| 0.1 | 0.05 | AG1 | 4 | 6.6 | 0.428 |
| 0.1 | 0.05 | ptsI | 4 | 6.85 | 0.388 |
| 0.1 | 0.05 | clcB | 4 | 6.85 | 0.322 |
| 0.1 | 0.05 | ycaM | 4 | 6.85 | 0.38 |
| 0.1 | 0.05 | yadI | 4 | 6.85 | 0.418 |
| 0.1 | 0.05 | AG1 | 4 | 6.85 | 0.44 |
| 0.1 | 0.05 | ptsI | 4 | 7.1 | 0.397 |
| 0.1 | 0.05 | clcB | 4 | 7.1 | 0.322 |
| 0.1 | 0.05 | ycaM | 4 | 7.1 | 0.375 |
| 0.1 | 0.05 | yadI | 4 | 7.1 | 0.423 |
| 0.1 | 0.05 | AG1 | 4 | 7.1 | 0.452 |
| 0.1 | 0.05 | ptsI | 4 | 7.35 | 0.406 |
| 0.1 | 0.05 | clcB | 4 | 7.35 | 0.326 |
| 0.1 | 0.05 | ycaM | 4 | 7.35 | 0.392 |
| 0.1 | 0.05 | yadI | 4 | 7.35 | 0.43 |
| 0.1 | 0.05 | AG1 | 4 | 7.35 | 0.464 |
| 0.1 | 0.05 | ptsI | 4 | 7.6 | 0.413 |
| 0.1 | 0.05 | clcB | 4 | 7.6 | 0.326 |
| 0.1 | 0.05 | ycaM | 4 | 7.6 | 0.392 |
| 0.1 | 0.05 | yadI | 4 | 7.6 | 0.436 |
| 0.1 | 0.05 | AG1 | 4 | 7.6 | 0.472 |
| 0.1 | 0.05 | ptsI | 4 | 7.85 | 0.418 |
| 0.1 | 0.05 | clcB | 4 | 7.85 | 0.327 |
| 0.1 | 0.05 | ycaM | 4 | 7.85 | 0.369 |
| 0.1 | 0.05 | yadI | 4 | 7.85 | 0.441 |
| 0.1 | 0.05 | AG1 | 4 | 7.85 | 0.478 |
| 0.1 | 0.05 | ptsI | 4 | 8.1 | 0.43 |
| 0.1 | 0.05 | clcB | 4 | 8.1 | 0.326 |
| 0.1 | 0.05 | ycaM | 4 | 8.1 | 0.378 |
| 0.1 | 0.05 | yadI | 4 | 8.1 | 0.45 |
| 0.1 | 0.05 | AG1 | 4 | 8.1 | 0.49 |
| 0.1 | 0.05 | ptsI | 4 | 8.35 | 0.439 |
| 0.1 | 0.05 | clcB | 4 | 8.35 | 0.332 |
| 0.1 | 0.05 | ycaM | 4 | 8.35 | 0.389 |
| 0.1 | 0.05 | yadI | 4 | 8.35 | 0.455 |
| 0.1 | 0.05 | AG1 | 4 | 8.35 | 0.5 |
| 0.1 | 0.05 | ptsI | 4 | 8.83333 | 0.416 |
| 0.1 | 0.05 | clcB | 4 | 8.83333 | 0.293 |
| 0.1 | 0.05 | ycaM | 4 | 8.83333 | 0.286 |
| 0.1 | 0.05 | yadI | 4 | 8.83333 | 0.484 |
| 0.1 | 0.05 | AG1 | 4 | 8.83333 | 0.529 |
| 0.1 | 0.05 | ptsI | 4 | 9.08333 | 0.415 |

| | | | | | |
|---|---|---|---|---|---|
| 0.1 | 0.05 | clcB | 4 | 9.08333 | 0.288 |
| 0.1 | 0.05 | ycaM | 4 | 9.08333 | 0.283 |
| 0.1 | 0.05 | yadI | 4 | 9.08333 | 0.482 |
| 0.1 | 0.05 | AG1 | 4 | 9.08333 | 0.534 |
| 0.1 | 0.05 | ptsI | 4 | 9.33333 | 0.422 |
| 0.1 | 0.05 | clcB | 4 | 9.33333 | 0.285 |
| 0.1 | 0.05 | ycaM | 4 | 9.33333 | 0.279 |
| 0.1 | 0.05 | yadI | 4 | 9.33333 | 0.489 |
| 0.1 | 0.05 | AG1 | 4 | 9.33333 | 0.549 |
| 0.1 | 0.05 | ptsI | 4 | 9.58333 | 0.434 |
| 0.1 | 0.05 | clcB | 4 | 9.58333 | 0.287 |
| 0.1 | 0.05 | ycaM | 4 | 9.58333 | 0.28 |
| 0.1 | 0.05 | yadI | 4 | 9.58333 | 0.501 |
| 0.1 | 0.05 | AG1 | 4 | 9.58333 | 0.562 |
| 0.1 | 0.05 | ptsI | 4 | 9.83333 | 0.444 |
| 0.1 | 0.05 | clcB | 4 | 9.83333 | 0.286 |
| 0.1 | 0.05 | ycaM | 4 | 9.83333 | 0.28 |
| 0.1 | 0.05 | yadI | 4 | 9.83333 | 0.508 |
| 0.1 | 0.05 | AG1 | 4 | 9.83333 | 0.571 |
| 0.1 | 0.05 | ptsI | 4 | 10.0833 | 0.451 |
| 0.1 | 0.05 | clcB | 4 | 10.0833 | 0.286 |
| 0.1 | 0.05 | ycaM | 4 | 10.0833 | 0.282 |
| 0.1 | 0.05 | yadI | 4 | 10.0833 | 0.514 |
| 0.1 | 0.05 | AG1 | 4 | 10.0833 | 0.581 |
| 0.1 | 0.05 | ptsI | 4 | 10.3333 | 0.461 |
| 0.1 | 0.05 | clcB | 4 | 10.3333 | 0.285 |
| 0.1 | 0.05 | ycaM | 4 | 10.3333 | 0.28 |
| 0.1 | 0.05 | yadI | 4 | 10.3333 | 0.522 |
| 0.1 | 0.05 | AG1 | 4 | 10.3333 | 0.592 |
| 0.1 | 0.05 | ptsI | 4 | 10.5833 | 0.471 |
| 0.1 | 0.05 | clcB | 4 | 10.5833 | 0.286 |
| 0.1 | 0.05 | ycaM | 4 | 10.5833 | 0.284 |
| 0.1 | 0.05 | yadI | 4 | 10.5833 | 0.532 |
| 0.1 | 0.05 | AG1 | 4 | 10.5833 | 0.598 |
| 0.1 | 0.05 | ptsI | 4 | 10.8333 | 0.48 |
| 0.1 | 0.05 | clcB | 4 | 10.8333 | 0.288 |
| 0.1 | 0.05 | ycaM | 4 | 10.8333 | 0.282 |
| 0.1 | 0.05 | yadI | 4 | 10.8333 | 0.535 |
| 0.1 | 0.05 | AG1 | 4 | 10.8333 | 0.607 |
| 0.1 | 0.05 | ptsI | 4 | 11.0833 | 0.488 |
| 0.1 | 0.05 | clcB | 4 | 11.0833 | 0.279 |
| 0.1 | 0.05 | ycaM | 4 | 11.0833 | 0.282 |
| 0.1 | 0.05 | yadI | 4 | 11.0833 | 0.541 |
| 0.1 | 0.05 | AG1 | 4 | 11.0833 | 0.613 |
| 0.1 | 0.05 | ptsI | 4 | 11.3333 | 0.499 |
| 0.1 | 0.05 | clcB | 4 | 11.3333 | 0.292 |
| 0.1 | 0.05 | ycaM | 4 | 11.3333 | 0.287 |
| 0.1 | 0.05 | yadI | 4 | 11.3333 | 0.547 |

| | | | | | |
|---|---|---|---|---|---|
| 0.1 | 0.05 | AG1 | 4 | 11.3333 | 0.619 |
| 0.1 | 0.05 | ptsI | 4 | 11.5833 | 0.504 |
| 0.1 | 0.05 | clcB | 4 | 11.5833 | 0.285 |
| 0.1 | 0.05 | ycaM | 4 | 11.5833 | 0.286 |
| 0.1 | 0.05 | yadI | 4 | 11.5833 | 0.555 |
| 0.1 | 0.05 | AG1 | 4 | 11.5833 | 0.628 |
| 0.1 | 0.05 | ptsI | 4 | 11.8333 | 0.51 |
| 0.1 | 0.05 | clcB | 4 | 11.8333 | 0.29 |
| 0.1 | 0.05 | ycaM | 4 | 11.8333 | 0.29 |
| 0.1 | 0.05 | yadI | 4 | 11.8333 | 0.558 |
| 0.1 | 0.05 | AG1 | 4 | 11.8333 | 0.632 |
| 0.1 | 0.05 | ptsI | 4 | 12.0833 | 0.519 |
| 0.1 | 0.05 | clcB | 4 | 12.0833 | 0.291 |
| 0.1 | 0.05 | ycaM | 4 | 12.0833 | 0.29 |
| 0.1 | 0.05 | yadI | 4 | 12.0833 | 0.564 |
| 0.1 | 0.05 | AG1 | 4 | 12.0833 | 0.637 |
| 0.1 | 0.05 | ptsI | 4 | 12.3333 | 0.528 |
| 0.1 | 0.05 | clcB | 4 | 12.3333 | 0.292 |
| 0.1 | 0.05 | ycaM | 4 | 12.3333 | 0.289 |
| 0.1 | 0.05 | yadI | 4 | 12.3333 | 0.569 |
| 0.1 | 0.05 | AG1 | 4 | 12.3333 | 0.646 |
| 0.1 | 0.05 | ptsI | 4 | 12.5833 | 0.535 |
| 0.1 | 0.05 | clcB | 4 | 12.5833 | 0.3 |
| 0.1 | 0.05 | ycaM | 4 | 12.5833 | 0.289 |
| 0.1 | 0.05 | yadI | 4 | 12.5833 | 0.574 |
| 0.1 | 0.05 | AG1 | 4 | 12.5833 | 0.653 |
| 0.1 | 0.05 | ptsI | 4 | 12.8333 | 0.54 |
| 0.1 | 0.05 | clcB | 4 | 12.8333 | 0.299 |
| 0.1 | 0.05 | ycaM | 4 | 12.8333 | 0.289 |
| 0.1 | 0.05 | yadI | 4 | 12.8333 | 0.58 |
| 0.1 | 0.05 | AG1 | 4 | 12.8333 | 0.656 |
| 0.1 | 0.05 | ptsI | 4 | 13.0833 | 0.545 |
| 0.1 | 0.05 | clcB | 4 | 13.0833 | 0.299 |
| 0.1 | 0.05 | ycaM | 4 | 13.0833 | 0.291 |
| 0.1 | 0.05 | yadI | 4 | 13.0833 | 0.586 |
| 0.1 | 0.05 | AG1 | 4 | 13.0833 | 0.663 |
| 0.1 | 0.05 | ptsI | 4 | 13.3333 | 0.553 |
| 0.1 | 0.05 | clcB | 4 | 13.3333 | 0.304 |
| 0.1 | 0.05 | ycaM | 4 | 13.3333 | 0.291 |
| 0.1 | 0.05 | yadI | 4 | 13.3333 | 0.588 |
| 0.1 | 0.05 | AG1 | 4 | 13.3333 | 0.667 |
| 0.1 | 0.05 | ptsI | 4 | 13.5833 | 0.558 |
| 0.1 | 0.05 | clcB | 4 | 13.5833 | 0.301 |
| 0.1 | 0.05 | ycaM | 4 | 13.5833 | 0.302 |
| 0.1 | 0.05 | yadI | 4 | 13.5833 | 0.592 |
| 0.1 | 0.05 | AG1 | 4 | 13.5833 | 0.672 |
| 0.1 | 0.05 | ptsI | 4 | 13.8333 | 0.565 |
| 0.1 | 0.05 | clcB | 4 | 13.8333 | 0.304 |

| | | | | | |
|---|---|---|---|---|---|
| 0.1 | 0.05 | ycaM | 4 | 13.8333 | 0.292 |
| 0.1 | 0.05 | yadI | 4 | 13.8333 | 0.595 |
| 0.1 | 0.05 | AG1 | 4 | 13.8333 | 0.675 |
| 0.1 | 0.05 | ptsI | 4 | 14.0833 | 0.571 |
| 0.1 | 0.05 | clcB | 4 | 14.0833 | 0.303 |
| 0.1 | 0.05 | ycaM | 4 | 14.0833 | 0.304 |
| 0.1 | 0.05 | yadI | 4 | 14.0833 | 0.602 |
| 0.1 | 0.05 | AG1 | 4 | 14.0833 | 0.679 |
| 0.1 | 0.05 | ptsI | 4 | 14.3333 | 0.577 |
| 0.1 | 0.05 | clcB | 4 | 14.3333 | 0.305 |
| 0.1 | 0.05 | ycaM | 4 | 14.3333 | 0.304 |
| 0.1 | 0.05 | yadI | 4 | 14.3333 | 0.603 |
| 0.1 | 0.05 | AG1 | 4 | 14.3333 | 0.68 |
| 0.1 | 0.05 | ptsI | 4 | 14.5833 | 0.582 |
| 0.1 | 0.05 | clcB | 4 | 14.5833 | 0.307 |
| 0.1 | 0.05 | ycaM | 4 | 14.5833 | 0.288 |
| 0.1 | 0.05 | yadI | 4 | 14.5833 | 0.609 |
| 0.1 | 0.05 | AG1 | 4 | 14.5833 | 0.686 |
| 0.1 | 0.05 | ptsI | 4 | 14.8333 | 0.588 |
| 0.1 | 0.05 | clcB | 4 | 14.8333 | 0.31 |
| 0.1 | 0.05 | ycaM | 4 | 14.8333 | 0.322 |
| 0.1 | 0.05 | yadI | 4 | 14.8333 | 0.612 |
| 0.1 | 0.05 | AG1 | 4 | 14.8333 | 0.693 |
| 0.1 | 0.05 | ptsI | 4 | 15.0833 | 0.59 |
| 0.1 | 0.05 | clcB | 4 | 15.0833 | 0.308 |
| 0.1 | 0.05 | ycaM | 4 | 15.0833 | 0.319 |
| 0.1 | 0.05 | yadI | 4 | 15.0833 | 0.617 |
| 0.1 | 0.05 | AG1 | 4 | 15.0833 | 0.691 |
| 0.1 | 0.05 | ptsI | 4 | 15.3333 | 0.596 |
| 0.1 | 0.05 | clcB | 4 | 15.3333 | 0.31 |
| 0.1 | 0.05 | ycaM | 4 | 15.3333 | 0.324 |
| 0.1 | 0.05 | yadI | 4 | 15.3333 | 0.616 |
| 0.1 | 0.05 | AG1 | 4 | 15.3333 | 0.696 |
| 0.1 | 0.05 | ptsI | 4 | 15.5833 | 0.602 |
| 0.1 | 0.05 | clcB | 4 | 15.5833 | 0.314 |
| 0.1 | 0.05 | ycaM | 4 | 15.5833 | 0.326 |
| 0.1 | 0.05 | yadI | 4 | 15.5833 | 0.623 |
| 0.1 | 0.05 | AG1 | 4 | 15.5833 | 0.698 |
| 0.1 | 0.05 | ptsI | 4 | 15.8333 | 0.606 |
| 0.1 | 0.05 | clcB | 4 | 15.8333 | 0.313 |
| 0.1 | 0.05 | ycaM | 4 | 15.8333 | 0.326 |
| 0.1 | 0.05 | yadI | 4 | 15.8333 | 0.628 |
| 0.1 | 0.05 | AG1 | 4 | 15.8333 | 0.702 |
| 0.1 | 0.05 | ptsI | 4 | 16.0833 | 0.611 |
| 0.1 | 0.05 | clcB | 4 | 16.0833 | 0.316 |
| 0.1 | 0.05 | ycaM | 4 | 16.0833 | 0.328 |
| 0.1 | 0.05 | yadI | 4 | 16.0833 | 0.629 |
| 0.1 | 0.05 | AG1 | 4 | 16.0833 | 0.705 |

| | | | | | |
|---|---|---|---|---|---|
| 0.1 | 0.05 | ptsI | 4 | 16.3333 | 0.616 |
| 0.1 | 0.05 | clcB | 4 | 16.3333 | 0.316 |
| 0.1 | 0.05 | ycaM | 4 | 16.3333 | 0.331 |
| 0.1 | 0.05 | yadI | 4 | 16.3333 | 0.631 |
| 0.1 | 0.05 | AG1 | 4 | 16.3333 | 0.708 |
| 0.1 | 0.05 | ptsI | 4 | 16.5833 | 0.619 |
| 0.1 | 0.05 | clcB | 4 | 16.5833 | 0.319 |
| 0.1 | 0.05 | ycaM | 4 | 16.5833 | 0.331 |
| 0.1 | 0.05 | yadI | 4 | 16.5833 | 0.635 |
| 0.1 | 0.05 | AG1 | 4 | 16.5833 | 0.711 |
| 0.1 | 0.05 | ptsI | 4 | 16.8333 | 0.623 |
| 0.1 | 0.05 | clcB | 4 | 16.8333 | 0.323 |
| 0.1 | 0.05 | ycaM | 4 | 16.8333 | 0.335 |
| 0.1 | 0.05 | yadI | 4 | 16.8333 | 0.641 |
| 0.1 | 0.05 | AG1 | 4 | 16.8333 | 0.717 |
| 0.1 | 0.05 | ptsI | 4 | 17.0833 | 0.619 |
| 0.1 | 0.05 | clcB | 4 | 17.0833 | 0.325 |
| 0.1 | 0.05 | ycaM | 4 | 17.0833 | 0.339 |
| 0.1 | 0.05 | yadI | 4 | 17.0833 | 0.64 |
| 0.1 | 0.05 | AG1 | 4 | 17.0833 | 0.719 |
| 0.1 | 0.05 | ptsI | 4 | 17.3333 | 0.625 |
| 0.1 | 0.05 | clcB | 4 | 17.3333 | 0.338 |
| 0.1 | 0.05 | ycaM | 4 | 17.3333 | 0.341 |
| 0.1 | 0.05 | yadI | 4 | 17.3333 | 0.642 |
| 0.1 | 0.05 | AG1 | 4 | 17.3333 | 0.719 |
| 0.1 | 0.05 | ptsI | 4 | 17.5833 | 0.625 |
| 0.1 | 0.05 | clcB | 4 | 17.5833 | 0.339 |
| 0.1 | 0.05 | ycaM | 4 | 17.5833 | 0.344 |
| 0.1 | 0.05 | yadI | 4 | 17.5833 | 0.647 |
| 0.1 | 0.05 | AG1 | 4 | 17.5833 | 0.722 |
| 0.1 | 0.05 | ptsI | 4 | 17.8333 | 0.628 |
| 0.1 | 0.05 | clcB | 4 | 17.8333 | 0.339 |
| 0.1 | 0.05 | ycaM | 4 | 17.8333 | 0.352 |
| 0.1 | 0.05 | yadI | 4 | 17.8333 | 0.644 |
| 0.1 | 0.05 | AG1 | 4 | 17.8333 | 0.716 |
| 0.1 | 0.05 | ptsI | 4 | 18.0833 | 0.629 |
| 0.1 | 0.05 | clcB | 4 | 18.0833 | 0.339 |
| 0.1 | 0.05 | ycaM | 4 | 18.0833 | 0.351 |
| 0.1 | 0.05 | yadI | 4 | 18.0833 | 0.649 |
| 0.1 | 0.05 | AG1 | 4 | 18.0833 | 0.727 |
| 0.1 | 0.05 | ptsI | 4 | 18.3333 | 0.63 |
| 0.1 | 0.05 | clcB | 4 | 18.3333 | 0.331 |
| 0.1 | 0.05 | ycaM | 4 | 18.3333 | 0.354 |
| 0.1 | 0.05 | yadI | 4 | 18.3333 | 0.654 |
| 0.1 | 0.05 | AG1 | 4 | 18.3333 | 0.729 |
| 0.1 | 0.05 | ptsI | 4 | 18.5833 | 0.63 |
| 0.1 | 0.05 | clcB | 4 | 18.5833 | 0.33 |
| 0.1 | 0.05 | ycaM | 4 | 18.5833 | 0.361 |

| | | | | | |
|---|---|---|---|---|---|
| 0.1 | 0.05 | yadI | 4 | 18.5833 | 0.656 |
| 0.1 | 0.05 | AG1 | 4 | 18.5833 | 0.735 |
| 0.1 | 0.05 | ptsI | 4 | 18.8333 | 0.634 |
| 0.1 | 0.05 | clcB | 4 | 18.8333 | 0.322 |
| 0.1 | 0.05 | ycaM | 4 | 18.8333 | 0.37 |
| 0.1 | 0.05 | yadI | 4 | 18.8333 | 0.657 |
| 0.1 | 0.05 | AG1 | 4 | 18.8333 | 0.731 |
| 0.1 | 0.05 | ptsI | 4 | 19.0833 | 0.633 |
| 0.1 | 0.05 | clcB | 4 | 19.0833 | 0.331 |
| 0.1 | 0.05 | ycaM | 4 | 19.0833 | 0.366 |
| 0.1 | 0.05 | yadI | 4 | 19.0833 | 0.664 |
| 0.1 | 0.05 | AG1 | 4 | 19.0833 | 0.743 |
| 0.1 | 0.05 | ptsI | 4 | 19.3333 | 0.629 |
| 0.1 | 0.05 | clcB | 4 | 19.3333 | 0.318 |
| 0.1 | 0.05 | ycaM | 4 | 19.3333 | 0.371 |
| 0.1 | 0.05 | yadI | 4 | 19.3333 | 0.66 |
| 0.1 | 0.05 | AG1 | 4 | 19.3333 | 0.752 |
| 0.1 | 0.05 | ptsI | 4 | 19.5833 | 0.628 |
| 0.1 | 0.05 | clcB | 4 | 19.5833 | 0.334 |
| 0.1 | 0.05 | ycaM | 4 | 19.5833 | 0.373 |
| 0.1 | 0.05 | yadI | 4 | 19.5833 | 0.66 |
| 0.1 | 0.05 | AG1 | 4 | 19.5833 | 0.76 |
| 0.1 | 0.05 | ptsI | 4 | 19.8333 | 0.628 |
| 0.1 | 0.05 | clcB | 4 | 19.8333 | 0.323 |
| 0.1 | 0.05 | ycaM | 4 | 19.8333 | 0.378 |
| 0.1 | 0.05 | yadI | 4 | 19.8333 | 0.662 |
| 0.1 | 0.05 | AG1 | 4 | 19.8333 | 0.765 |
| 0.1 | 0.05 | ptsI | 4 | 20.0833 | 0.631 |
| 0.1 | 0.05 | clcB | 4 | 20.0833 | 0.327 |
| 0.1 | 0.05 | ycaM | 4 | 20.0833 | 0.387 |
| 0.1 | 0.05 | yadI | 4 | 20.0833 | 0.665 |
| 0.1 | 0.05 | AG1 | 4 | 20.0833 | 0.765 |
| 0.1 | 0.05 | ptsI | 4 | 20.3333 | 0.633 |
| 0.1 | 0.05 | clcB | 4 | 20.3333 | 0.32 |
| 0.1 | 0.05 | ycaM | 4 | 20.3333 | 0.387 |
| 0.1 | 0.05 | yadI | 4 | 20.3333 | 0.667 |
| 0.1 | 0.05 | AG1 | 4 | 20.3333 | 0.769 |
| 0.1 | 0.05 | ptsI | 4 | 20.5833 | 0.627 |
| 0.1 | 0.05 | clcB | 4 | 20.5833 | 0.326 |
| 0.1 | 0.05 | ycaM | 4 | 20.5833 | 0.393 |
| 0.1 | 0.05 | yadI | 4 | 20.5833 | 0.668 |
| 0.1 | 0.05 | AG1 | 4 | 20.5833 | 0.775 |
| 0.1 | 0.05 | ptsI | 4 | 20.8333 | 0.637 |
| 0.1 | 0.05 | clcB | 4 | 20.8333 | 0.327 |
| 0.1 | 0.05 | ycaM | 4 | 20.8333 | 0.397 |
| 0.1 | 0.05 | yadI | 4 | 20.8333 | 0.669 |
| 0.1 | 0.05 | AG1 | 4 | 20.8333 | 0.774 |
| 0.1 | 0.05 | ptsI | 4 | 21.0833 | 0.634 |

| | | | | | |
|---|---|---|---|---|---|
| 0.1 | 0.05 | clcB | 4 | 21.0833 | 0.327 |
| 0.1 | 0.05 | ycaM | 4 | 21.0833 | 0.396 |
| 0.1 | 0.05 | yadI | 4 | 21.0833 | 0.669 |
| 0.1 | 0.05 | AG1 | 4 | 21.0833 | 0.778 |
| 0.1 | 0.05 | ptsI | 4 | 21.3333 | 0.631 |
| 0.1 | 0.05 | clcB | 4 | 21.3333 | 0.326 |
| 0.1 | 0.05 | ycaM | 4 | 21.3333 | 0.399 |
| 0.1 | 0.05 | yadI | 4 | 21.3333 | 0.67 |
| 0.1 | 0.05 | AG1 | 4 | 21.3333 | 0.778 |
| 0.1 | 0.05 | ptsI | 4 | 21.5833 | 0.636 |
| 0.1 | 0.05 | clcB | 4 | 21.5833 | 0.33 |
| 0.1 | 0.05 | ycaM | 4 | 21.5833 | 0.401 |
| 0.1 | 0.05 | yadI | 4 | 21.5833 | 0.671 |
| 0.1 | 0.05 | AG1 | 4 | 21.5833 | 0.781 |
| 0.1 | 0.05 | ptsI | 4 | 21.8333 | 0.638 |
| 0.1 | 0.05 | clcB | 4 | 21.8333 | 0.325 |
| 0.1 | 0.05 | ycaM | 4 | 21.8333 | 0.401 |
| 0.1 | 0.05 | yadI | 4 | 21.8333 | 0.671 |
| 0.1 | 0.05 | AG1 | 4 | 21.8333 | 0.781 |
| 0.1 | 0.05 | ptsI | 4 | 22.0833 | 0.634 |
| 0.1 | 0.05 | clcB | 4 | 22.0833 | 0.325 |
| 0.1 | 0.05 | ycaM | 4 | 22.0833 | 0.403 |
| 0.1 | 0.05 | yadI | 4 | 22.0833 | 0.671 |
| 0.1 | 0.05 | AG1 | 4 | 22.0833 | 0.784 |
| 0.1 | 0.05 | ptsI | 4 | 22.3333 | 0.637 |
| 0.1 | 0.05 | clcB | 4 | 22.3333 | 0.331 |
| 0.1 | 0.05 | ycaM | 4 | 22.3333 | 0.397 |
| 0.1 | 0.05 | yadI | 4 | 22.3333 | 0.672 |
| 0.1 | 0.05 | AG1 | 4 | 22.3333 | 0.786 |
| 0.1 | 0.05 | ptsI | 4 | 22.5833 | 0.637 |
| 0.1 | 0.05 | clcB | 4 | 22.5833 | 0.327 |
| 0.1 | 0.05 | ycaM | 4 | 22.5833 | 0.399 |
| 0.1 | 0.05 | yadI | 4 | 22.5833 | 0.674 |
| 0.1 | 0.05 | AG1 | 4 | 22.5833 | 0.788 |
| 0.1 | 0.05 | ptsI | 4 | 22.8333 | 0.634 |
| 0.1 | 0.05 | clcB | 4 | 22.8333 | 0.326 |
| 0.1 | 0.05 | ycaM | 4 | 22.8333 | 0.4 |
| 0.1 | 0.05 | yadI | 4 | 22.8333 | 0.674 |
| 0.1 | 0.05 | AG1 | 4 | 22.8333 | 0.792 |
| 0.1 | 0.05 | ptsI | 4 | 23.0833 | 0.633 |
| 0.1 | 0.05 | clcB | 4 | 23.0833 | 0.334 |
| 0.1 | 0.05 | ycaM | 4 | 23.0833 | 0.401 |
| 0.1 | 0.05 | yadI | 4 | 23.0833 | 0.674 |
| 0.1 | 0.05 | AG1 | 4 | 23.0833 | 0.793 |
| 0.1 | 0.05 | ptsI | 4 | 23.3333 | 0.636 |
| 0.1 | 0.05 | clcB | 4 | 23.3333 | 0.33 |
| 0.1 | 0.05 | ycaM | 4 | 23.3333 | 0.406 |
| 0.1 | 0.05 | yadI | 4 | 23.3333 | 0.677 |

| | | | | | |
|---|---|---|---|---|---|
| 0.1 | 0.05 | AG1 | 4 | 23.3333 | 0.796 |
| 0.1 | 0.05 | ptsI | 4 | 23.5833 | 0.634 |
| 0.1 | 0.05 | clcB | 4 | 23.5833 | 0.328 |
| 0.1 | 0.05 | ycaM | 4 | 23.5833 | 0.407 |
| 0.1 | 0.05 | yadI | 4 | 23.5833 | 0.678 |
| 0.1 | 0.05 | AG1 | 4 | 23.5833 | 0.798 |
| 0.1 | 0.05 | ptsI | 4 | 23.8333 | 0.63 |
| 0.1 | 0.05 | clcB | 4 | 23.8333 | 0.336 |
| 0.1 | 0.05 | ycaM | 4 | 23.8333 | 0.408 |
| 0.1 | 0.05 | yadI | 4 | 23.8333 | 0.678 |
| 0.1 | 0.05 | AG1 | 4 | 23.8333 | 0.801 |
| 0.1 | 0.05 | ptsI | 4 | 24.0833 | 0.635 |
| 0.1 | 0.05 | clcB | 4 | 24.0833 | 0.338 |
| 0.1 | 0.05 | ycaM | 4 | 24.0833 | 0.41 |
| 0.1 | 0.05 | yadI | 4 | 24.0833 | 0.679 |
| 0.1 | 0.05 | AG1 | 4 | 24.0833 | 0.8 |
| 0.1 | 0.05 | ptsI | 5 | 0 | 0.18 |
| 0.1 | 0.05 | clcB | 5 | 0 | 0.172 |
| 0.1 | 0.05 | ycaM | 5 | 0 | 0.176 |
| 0.1 | 0.05 | yadI | 5 | 0 | 0.18 |
| 0.1 | 0.05 | AG1 | 5 | 0 | 0.174 |
| 0.1 | 0.05 | ptsI | 5 | 0.35 | 0.176 |
| 0.1 | 0.05 | clcB | 5 | 0.35 | 0.166 |
| 0.1 | 0.05 | ycaM | 5 | 0.35 | 0.167 |
| 0.1 | 0.05 | yadI | 5 | 0.35 | 0.173 |
| 0.1 | 0.05 | AG1 | 5 | 0.35 | 0.17 |
| 0.1 | 0.05 | ptsI | 5 | 0.6 | 0.176 |
| 0.1 | 0.05 | clcB | 5 | 0.6 | 0.166 |
| 0.1 | 0.05 | ycaM | 5 | 0.6 | 0.168 |
| 0.1 | 0.05 | yadI | 5 | 0.6 | 0.173 |
| 0.1 | 0.05 | AG1 | 5 | 0.6 | 0.171 |
| 0.1 | 0.05 | ptsI | 5 | 0.85 | 0.175 |
| 0.1 | 0.05 | clcB | 5 | 0.85 | 0.166 |
| 0.1 | 0.05 | ycaM | 5 | 0.85 | 0.168 |
| 0.1 | 0.05 | yadI | 5 | 0.85 | 0.176 |
| 0.1 | 0.05 | AG1 | 5 | 0.85 | 0.171 |
| 0.1 | 0.05 | ptsI | 5 | 1.1 | 0.178 |
| 0.1 | 0.05 | clcB | 5 | 1.1 | 0.166 |
| 0.1 | 0.05 | ycaM | 5 | 1.1 | 0.17 |
| 0.1 | 0.05 | yadI | 5 | 1.1 | 0.176 |
| 0.1 | 0.05 | AG1 | 5 | 1.1 | 0.172 |
| 0.1 | 0.05 | ptsI | 5 | 1.35 | 0.179 |
| 0.1 | 0.05 | clcB | 5 | 1.35 | 0.167 |
| 0.1 | 0.05 | ycaM | 5 | 1.35 | 0.172 |
| 0.1 | 0.05 | yadI | 5 | 1.35 | 0.18 |
| 0.1 | 0.05 | AG1 | 5 | 1.35 | 0.175 |
| 0.1 | 0.05 | ptsI | 5 | 1.6 | 0.182 |
| 0.1 | 0.05 | clcB | 5 | 1.6 | 0.168 |

| | | | | | |
|---|---|---|---|---|---|
| 0.1 | 0.05 | ycaM | 5 | 1.6 | 0.174 |
| 0.1 | 0.05 | yadI | 5 | 1.6 | 0.18 |
| 0.1 | 0.05 | AG1 | 5 | 1.6 | 0.177 |
| 0.1 | 0.05 | ptsI | 5 | 1.85 | 0.185 |
| 0.1 | 0.05 | clcB | 5 | 1.85 | 0.171 |
| 0.1 | 0.05 | ycaM | 5 | 1.85 | 0.178 |
| 0.1 | 0.05 | yadI | 5 | 1.85 | 0.184 |
| 0.1 | 0.05 | AG1 | 5 | 1.85 | 0.18 |
| 0.1 | 0.05 | ptsI | 5 | 2.1 | 0.191 |
| 0.1 | 0.05 | clcB | 5 | 2.1 | 0.181 |
| 0.1 | 0.05 | ycaM | 5 | 2.1 | 0.182 |
| 0.1 | 0.05 | yadI | 5 | 2.1 | 0.187 |
| 0.1 | 0.05 | AG1 | 5 | 2.1 | 0.185 |
| 0.1 | 0.05 | ptsI | 5 | 2.35 | 0.198 |
| 0.1 | 0.05 | clcB | 5 | 2.35 | 0.188 |
| 0.1 | 0.05 | ycaM | 5 | 2.35 | 0.187 |
| 0.1 | 0.05 | yadI | 5 | 2.35 | 0.192 |
| 0.1 | 0.05 | AG1 | 5 | 2.35 | 0.186 |
| 0.1 | 0.05 | ptsI | 5 | 2.6 | 0.194 |
| 0.1 | 0.05 | clcB | 5 | 2.6 | 0.183 |
| 0.1 | 0.05 | ycaM | 5 | 2.6 | 0.19 |
| 0.1 | 0.05 | yadI | 5 | 2.6 | 0.192 |
| 0.1 | 0.05 | AG1 | 5 | 2.6 | 0.183 |
| 0.1 | 0.05 | ptsI | 5 | 2.85 | 0.203 |
| 0.1 | 0.05 | clcB | 5 | 2.85 | 0.193 |
| 0.1 | 0.05 | ycaM | 5 | 2.85 | 0.196 |
| 0.1 | 0.05 | yadI | 5 | 2.85 | 0.203 |
| 0.1 | 0.05 | AG1 | 5 | 2.85 | 0.194 |
| 0.1 | 0.05 | ptsI | 5 | 3.1 | 0.209 |
| 0.1 | 0.05 | clcB | 5 | 3.1 | 0.2 |
| 0.1 | 0.05 | ycaM | 5 | 3.1 | 0.2 |
| 0.1 | 0.05 | yadI | 5 | 3.1 | 0.211 |
| 0.1 | 0.05 | AG1 | 5 | 3.1 | 0.2 |
| 0.1 | 0.05 | ptsI | 5 | 3.35 | 0.22 |
| 0.1 | 0.05 | clcB | 5 | 3.35 | 0.216 |
| 0.1 | 0.05 | ycaM | 5 | 3.35 | 0.209 |
| 0.1 | 0.05 | yadI | 5 | 3.35 | 0.22 |
| 0.1 | 0.05 | AG1 | 5 | 3.35 | 0.205 |
| 0.1 | 0.05 | ptsI | 5 | 3.6 | 0.233 |
| 0.1 | 0.05 | clcB | 5 | 3.6 | 0.234 |
| 0.1 | 0.05 | ycaM | 5 | 3.6 | 0.219 |
| 0.1 | 0.05 | yadI | 5 | 3.6 | 0.228 |
| 0.1 | 0.05 | AG1 | 5 | 3.6 | 0.214 |
| 0.1 | 0.05 | ptsI | 5 | 3.85 | 0.241 |
| 0.1 | 0.05 | clcB | 5 | 3.85 | 0.248 |
| 0.1 | 0.05 | ycaM | 5 | 3.85 | 0.227 |
| 0.1 | 0.05 | yadI | 5 | 3.85 | 0.239 |
| 0.1 | 0.05 | AG1 | 5 | 3.85 | 0.225 |

| | | | | | |
|---|---|---|---|---|---|
| 0.1 | 0.05 | ptsI | 5 | 4.38333 | 0.263 |
| 0.1 | 0.05 | clcB | 5 | 4.38333 | 0.286 |
| 0.1 | 0.05 | ycaM | 5 | 4.38333 | 0.246 |
| 0.1 | 0.05 | yadI | 5 | 4.38333 | 0.267 |
| 0.1 | 0.05 | AG1 | 5 | 4.38333 | 0.257 |
| 0.1 | 0.05 | ptsI | 5 | 4.63333 | 0.264 |
| 0.1 | 0.05 | clcB | 5 | 4.63333 | 0.292 |
| 0.1 | 0.05 | ycaM | 5 | 4.63333 | 0.257 |
| 0.1 | 0.05 | yadI | 5 | 4.63333 | 0.269 |
| 0.1 | 0.05 | AG1 | 5 | 4.63333 | 0.257 |
| 0.1 | 0.05 | ptsI | 5 | 4.88333 | 0.271 |
| 0.1 | 0.05 | clcB | 5 | 4.88333 | 0.311 |
| 0.1 | 0.05 | ycaM | 5 | 4.88333 | 0.27 |
| 0.1 | 0.05 | yadI | 5 | 4.88333 | 0.288 |
| 0.1 | 0.05 | AG1 | 5 | 4.88333 | 0.268 |
| 0.1 | 0.05 | ptsI | 5 | 5.13333 | 0.279 |
| 0.1 | 0.05 | clcB | 5 | 5.13333 | 0.324 |
| 0.1 | 0.05 | ycaM | 5 | 5.13333 | 0.273 |
| 0.1 | 0.05 | yadI | 5 | 5.13333 | 0.3 |
| 0.1 | 0.05 | AG1 | 5 | 5.13333 | 0.281 |
| 0.1 | 0.05 | ptsI | 5 | 5.38333 | 0.289 |
| 0.1 | 0.05 | clcB | 5 | 5.38333 | 0.336 |
| 0.1 | 0.05 | ycaM | 5 | 5.38333 | 0.275 |
| 0.1 | 0.05 | yadI | 5 | 5.38333 | 0.313 |
| 0.1 | 0.05 | AG1 | 5 | 5.38333 | 0.297 |
| 0.1 | 0.05 | ptsI | 5 | 5.63333 | 0.302 |
| 0.1 | 0.05 | clcB | 5 | 5.63333 | 0.348 |
| 0.1 | 0.05 | ycaM | 5 | 5.63333 | 0.282 |
| 0.1 | 0.05 | yadI | 5 | 5.63333 | 0.326 |
| 0.1 | 0.05 | AG1 | 5 | 5.63333 | 0.312 |
| 0.1 | 0.05 | ptsI | 5 | 5.88333 | 0.314 |
| 0.1 | 0.05 | clcB | 5 | 5.88333 | 0.355 |
| 0.1 | 0.05 | ycaM | 5 | 5.88333 | 0.286 |
| 0.1 | 0.05 | yadI | 5 | 5.88333 | 0.336 |
| 0.1 | 0.05 | AG1 | 5 | 5.88333 | 0.324 |
| 0.1 | 0.05 | ptsI | 5 | 6.13333 | 0.328 |
| 0.1 | 0.05 | clcB | 5 | 6.13333 | 0.364 |
| 0.1 | 0.05 | ycaM | 5 | 6.13333 | 0.286 |
| 0.1 | 0.05 | yadI | 5 | 6.13333 | 0.351 |
| 0.1 | 0.05 | AG1 | 5 | 6.13333 | 0.338 |
| 0.1 | 0.05 | ptsI | 5 | 6.38333 | 0.342 |
| 0.1 | 0.05 | clcB | 5 | 6.38333 | 0.368 |
| 0.1 | 0.05 | ycaM | 5 | 6.38333 | 0.289 |
| 0.1 | 0.05 | yadI | 5 | 6.38333 | 0.364 |
| 0.1 | 0.05 | AG1 | 5 | 6.38333 | 0.351 |
| 0.1 | 0.05 | ptsI | 5 | 6.63333 | 0.361 |
| 0.1 | 0.05 | clcB | 5 | 6.63333 | 0.373 |
| 0.1 | 0.05 | ycaM | 5 | 6.63333 | 0.29 |

| | | | | | |
|---|---|---|---|---|---|
| 0.1 | 0.05 | yadI | 5 | 6.63333 | 0.377 |
| 0.1 | 0.05 | AG1 | 5 | 6.63333 | 0.365 |
| 0.1 | 0.05 | ptsI | 5 | 6.88333 | 0.376 |
| 0.1 | 0.05 | clcB | 5 | 6.88333 | 0.381 |
| 0.1 | 0.05 | ycaM | 5 | 6.88333 | 0.291 |
| 0.1 | 0.05 | yadI | 5 | 6.88333 | 0.39 |
| 0.1 | 0.05 | AG1 | 5 | 6.88333 | 0.378 |
| 0.1 | 0.05 | ptsI | 5 | 7.13333 | 0.392 |
| 0.1 | 0.05 | clcB | 5 | 7.13333 | 0.384 |
| 0.1 | 0.05 | ycaM | 5 | 7.13333 | 0.294 |
| 0.1 | 0.05 | yadI | 5 | 7.13333 | 0.406 |
| 0.1 | 0.05 | AG1 | 5 | 7.13333 | 0.392 |
| 0.1 | 0.05 | ptsI | 5 | 7.38333 | 0.405 |
| 0.1 | 0.05 | clcB | 5 | 7.38333 | 0.385 |
| 0.1 | 0.05 | ycaM | 5 | 7.38333 | 0.297 |
| 0.1 | 0.05 | yadI | 5 | 7.38333 | 0.42 |
| 0.1 | 0.05 | AG1 | 5 | 7.38333 | 0.404 |
| 0.1 | 0.05 | ptsI | 5 | 7.63333 | 0.414 |
| 0.1 | 0.05 | clcB | 5 | 7.63333 | 0.388 |
| 0.1 | 0.05 | ycaM | 5 | 7.63333 | 0.298 |
| 0.1 | 0.05 | yadI | 5 | 7.63333 | 0.429 |
| 0.1 | 0.05 | AG1 | 5 | 7.63333 | 0.415 |
| 0.1 | 0.05 | ptsI | 5 | 7.88333 | 0.432 |
| 0.1 | 0.05 | clcB | 5 | 7.88333 | 0.393 |
| 0.1 | 0.05 | ycaM | 5 | 7.88333 | 0.297 |
| 0.1 | 0.05 | yadI | 5 | 7.88333 | 0.443 |
| 0.1 | 0.05 | AG1 | 5 | 7.88333 | 0.428 |
| 0.1 | 0.05 | ptsI | 5 | 8.13333 | 0.452 |
| 0.1 | 0.05 | clcB | 5 | 8.13333 | 0.392 |
| 0.1 | 0.05 | ycaM | 5 | 8.13333 | 0.301 |
| 0.1 | 0.05 | yadI | 5 | 8.13333 | 0.452 |
| 0.1 | 0.05 | AG1 | 5 | 8.13333 | 0.444 |
| 0.1 | 0.05 | ptsI | 5 | 8.38333 | 0.465 |
| 0.1 | 0.05 | clcB | 5 | 8.38333 | 0.397 |
| 0.1 | 0.05 | ycaM | 5 | 8.38333 | 0.299 |
| 0.1 | 0.05 | yadI | 5 | 8.38333 | 0.46 |
| 0.1 | 0.05 | AG1 | 5 | 8.38333 | 0.455 |
| 0.1 | 0.05 | ptsI | 5 | 8.88333 | 0.355 |
| 0.1 | 0.05 | clcB | 5 | 8.88333 | 0.296 |
| 0.1 | 0.05 | ycaM | 5 | 8.88333 | 0.266 |
| 0.1 | 0.05 | yadI | 5 | 8.88333 | 0.459 |
| 0.1 | 0.05 | AG1 | 5 | 8.88333 | 0.503 |
| 0.1 | 0.05 | ptsI | 5 | 9.13333 | 0.365 |
| 0.1 | 0.05 | clcB | 5 | 9.13333 | 0.295 |
| 0.1 | 0.05 | ycaM | 5 | 9.13333 | 0.265 |
| 0.1 | 0.05 | yadI | 5 | 9.13333 | 0.457 |
| 0.1 | 0.05 | AG1 | 5 | 9.13333 | 0.506 |
| 0.1 | 0.05 | ptsI | 5 | 9.38333 | 0.372 |

| | | | | | |
|---|---|---|---|---|---|
| 0.1 | 0.05 | clcB | 5 | 9.38333 | 0.298 |
| 0.1 | 0.05 | ycaM | 5 | 9.38333 | 0.263 |
| 0.1 | 0.05 | yadI | 5 | 9.38333 | 0.472 |
| 0.1 | 0.05 | AG1 | 5 | 9.38333 | 0.526 |
| 0.1 | 0.05 | ptsI | 5 | 9.63333 | 0.384 |
| 0.1 | 0.05 | clcB | 5 | 9.63333 | 0.301 |
| 0.1 | 0.05 | ycaM | 5 | 9.63333 | 0.266 |
| 0.1 | 0.05 | yadI | 5 | 9.63333 | 0.482 |
| 0.1 | 0.05 | AG1 | 5 | 9.63333 | 0.54 |
| 0.1 | 0.05 | ptsI | 5 | 9.88333 | 0.392 |
| 0.1 | 0.05 | clcB | 5 | 9.88333 | 0.301 |
| 0.1 | 0.05 | ycaM | 5 | 9.88333 | 0.266 |
| 0.1 | 0.05 | yadI | 5 | 9.88333 | 0.491 |
| 0.1 | 0.05 | AG1 | 5 | 9.88333 | 0.554 |
| 0.1 | 0.05 | ptsI | 5 | 10.1333 | 0.402 |
| 0.1 | 0.05 | clcB | 5 | 10.1333 | 0.304 |
| 0.1 | 0.05 | ycaM | 5 | 10.1333 | 0.264 |
| 0.1 | 0.05 | yadI | 5 | 10.1333 | 0.499 |
| 0.1 | 0.05 | AG1 | 5 | 10.1333 | 0.563 |
| 0.1 | 0.05 | ptsI | 5 | 10.3833 | 0.414 |
| 0.1 | 0.05 | clcB | 5 | 10.3833 | 0.305 |
| 0.1 | 0.05 | ycaM | 5 | 10.3833 | 0.27 |
| 0.1 | 0.05 | yadI | 5 | 10.3833 | 0.509 |
| 0.1 | 0.05 | AG1 | 5 | 10.3833 | 0.578 |
| 0.1 | 0.05 | ptsI | 5 | 10.6333 | 0.42 |
| 0.1 | 0.05 | clcB | 5 | 10.6333 | 0.304 |
| 0.1 | 0.05 | ycaM | 5 | 10.6333 | 0.268 |
| 0.1 | 0.05 | yadI | 5 | 10.6333 | 0.515 |
| 0.1 | 0.05 | AG1 | 5 | 10.6333 | 0.588 |
| 0.1 | 0.05 | ptsI | 5 | 10.8833 | 0.433 |
| 0.1 | 0.05 | clcB | 5 | 10.8833 | 0.309 |
| 0.1 | 0.05 | ycaM | 5 | 10.8833 | 0.269 |
| 0.1 | 0.05 | yadI | 5 | 10.8833 | 0.523 |
| 0.1 | 0.05 | AG1 | 5 | 10.8833 | 0.599 |
| 0.1 | 0.05 | ptsI | 5 | 11.1333 | 0.441 |
| 0.1 | 0.05 | clcB | 5 | 11.1333 | 0.306 |
| 0.1 | 0.05 | ycaM | 5 | 11.1333 | 0.267 |
| 0.1 | 0.05 | yadI | 5 | 11.1333 | 0.526 |
| 0.1 | 0.05 | AG1 | 5 | 11.1333 | 0.609 |
| 0.1 | 0.05 | ptsI | 5 | 11.3833 | 0.45 |
| 0.1 | 0.05 | clcB | 5 | 11.3833 | 0.306 |
| 0.1 | 0.05 | ycaM | 5 | 11.3833 | 0.268 |
| 0.1 | 0.05 | yadI | 5 | 11.3833 | 0.534 |
| 0.1 | 0.05 | AG1 | 5 | 11.3833 | 0.618 |
| 0.1 | 0.05 | ptsI | 5 | 11.6333 | 0.461 |
| 0.1 | 0.05 | clcB | 5 | 11.6333 | 0.305 |
| 0.1 | 0.05 | ycaM | 5 | 11.6333 | 0.268 |
| 0.1 | 0.05 | yadI | 5 | 11.6333 | 0.544 |

| | | | | | |
|---|---|---|---|---|---|
| 0.1 | 0.05 | AG1 | 5 | 11.6333 | 0.632 |
| 0.1 | 0.05 | ptsI | 5 | 11.8833 | 0.466 |
| 0.1 | 0.05 | clcB | 5 | 11.8833 | 0.304 |
| 0.1 | 0.05 | ycaM | 5 | 11.8833 | 0.268 |
| 0.1 | 0.05 | yadI | 5 | 11.8833 | 0.549 |
| 0.1 | 0.05 | AG1 | 5 | 11.8833 | 0.64 |
| 0.1 | 0.05 | ptsI | 5 | 12.1333 | 0.478 |
| 0.1 | 0.05 | clcB | 5 | 12.1333 | 0.303 |
| 0.1 | 0.05 | ycaM | 5 | 12.1333 | 0.271 |
| 0.1 | 0.05 | yadI | 5 | 12.1333 | 0.556 |
| 0.1 | 0.05 | AG1 | 5 | 12.1333 | 0.652 |
| 0.1 | 0.05 | ptsI | 5 | 12.3833 | 0.481 |
| 0.1 | 0.05 | clcB | 5 | 12.3833 | 0.298 |
| 0.1 | 0.05 | ycaM | 5 | 12.3833 | 0.268 |
| 0.1 | 0.05 | yadI | 5 | 12.3833 | 0.558 |
| 0.1 | 0.05 | AG1 | 5 | 12.3833 | 0.658 |
| 0.1 | 0.05 | ptsI | 5 | 12.6333 | 0.496 |
| 0.1 | 0.05 | clcB | 5 | 12.6333 | 0.309 |
| 0.1 | 0.05 | ycaM | 5 | 12.6333 | 0.271 |
| 0.1 | 0.05 | yadI | 5 | 12.6333 | 0.568 |
| 0.1 | 0.05 | AG1 | 5 | 12.6333 | 0.671 |
| 0.1 | 0.05 | ptsI | 5 | 12.8833 | 0.503 |
| 0.1 | 0.05 | clcB | 5 | 12.8833 | 0.316 |
| 0.1 | 0.05 | ycaM | 5 | 12.8833 | 0.273 |
| 0.1 | 0.05 | yadI | 5 | 12.8833 | 0.575 |
| 0.1 | 0.05 | AG1 | 5 | 12.8833 | 0.681 |
| 0.1 | 0.05 | ptsI | 5 | 13.1333 | 0.512 |
| 0.1 | 0.05 | clcB | 5 | 13.1333 | 0.317 |
| 0.1 | 0.05 | ycaM | 5 | 13.1333 | 0.275 |
| 0.1 | 0.05 | yadI | 5 | 13.1333 | 0.581 |
| 0.1 | 0.05 | AG1 | 5 | 13.1333 | 0.69 |
| 0.1 | 0.05 | ptsI | 5 | 13.3833 | 0.521 |
| 0.1 | 0.05 | clcB | 5 | 13.3833 | 0.312 |
| 0.1 | 0.05 | ycaM | 5 | 13.3833 | 0.273 |
| 0.1 | 0.05 | yadI | 5 | 13.3833 | 0.584 |
| 0.1 | 0.05 | AG1 | 5 | 13.3833 | 0.696 |
| 0.1 | 0.05 | ptsI | 5 | 13.6333 | 0.527 |
| 0.1 | 0.05 | clcB | 5 | 13.6333 | 0.316 |
| 0.1 | 0.05 | ycaM | 5 | 13.6333 | 0.275 |
| 0.1 | 0.05 | yadI | 5 | 13.6333 | 0.592 |
| 0.1 | 0.05 | AG1 | 5 | 13.6333 | 0.707 |
| 0.1 | 0.05 | ptsI | 5 | 13.8833 | 0.533 |
| 0.1 | 0.05 | clcB | 5 | 13.8833 | 0.318 |
| 0.1 | 0.05 | ycaM | 5 | 13.8833 | 0.278 |
| 0.1 | 0.05 | yadI | 5 | 13.8833 | 0.595 |
| 0.1 | 0.05 | AG1 | 5 | 13.8833 | 0.715 |
| 0.1 | 0.05 | ptsI | 5 | 14.1333 | 0.543 |
| 0.1 | 0.05 | clcB | 5 | 14.1333 | 0.319 |

| | | | | | |
|---|---|---|---|---|---|
| 0.1 | 0.05 | ycaM | 5 | 14.1333 | 0.28 |
| 0.1 | 0.05 | yadI | 5 | 14.1333 | 0.601 |
| 0.1 | 0.05 | AG1 | 5 | 14.1333 | 0.717 |
| 0.1 | 0.05 | ptsI | 5 | 14.3833 | 0.552 |
| 0.1 | 0.05 | clcB | 5 | 14.3833 | 0.317 |
| 0.1 | 0.05 | ycaM | 5 | 14.3833 | 0.278 |
| 0.1 | 0.05 | yadI | 5 | 14.3833 | 0.605 |
| 0.1 | 0.05 | AG1 | 5 | 14.3833 | 0.724 |
| 0.1 | 0.05 | ptsI | 5 | 14.6333 | 0.558 |
| 0.1 | 0.05 | clcB | 5 | 14.6333 | 0.32 |
| 0.1 | 0.05 | ycaM | 5 | 14.6333 | 0.279 |
| 0.1 | 0.05 | yadI | 5 | 14.6333 | 0.609 |
| 0.1 | 0.05 | AG1 | 5 | 14.6333 | 0.73 |
| 0.1 | 0.05 | ptsI | 5 | 14.8833 | 0.567 |
| 0.1 | 0.05 | clcB | 5 | 14.8833 | 0.316 |
| 0.1 | 0.05 | ycaM | 5 | 14.8833 | 0.28 |
| 0.1 | 0.05 | yadI | 5 | 14.8833 | 0.612 |
| 0.1 | 0.05 | AG1 | 5 | 14.8833 | 0.738 |
| 0.1 | 0.05 | ptsI | 5 | 15.1333 | 0.574 |
| 0.1 | 0.05 | clcB | 5 | 15.1333 | 0.317 |
| 0.1 | 0.05 | ycaM | 5 | 15.1333 | 0.282 |
| 0.1 | 0.05 | yadI | 5 | 15.1333 | 0.616 |
| 0.1 | 0.05 | AG1 | 5 | 15.1333 | 0.742 |
| 0.1 | 0.05 | ptsI | 5 | 15.3833 | 0.581 |
| 0.1 | 0.05 | clcB | 5 | 15.3833 | 0.318 |
| 0.1 | 0.05 | ycaM | 5 | 15.3833 | 0.281 |
| 0.1 | 0.05 | yadI | 5 | 15.3833 | 0.62 |
| 0.1 | 0.05 | AG1 | 5 | 15.3833 | 0.745 |
| 0.1 | 0.05 | ptsI | 5 | 15.6333 | 0.584 |
| 0.1 | 0.05 | clcB | 5 | 15.6333 | 0.313 |
| 0.1 | 0.05 | ycaM | 5 | 15.6333 | 0.282 |
| 0.1 | 0.05 | yadI | 5 | 15.6333 | 0.626 |
| 0.1 | 0.05 | AG1 | 5 | 15.6333 | 0.753 |
| 0.1 | 0.05 | ptsI | 5 | 15.8833 | 0.591 |
| 0.1 | 0.05 | clcB | 5 | 15.8833 | 0.314 |
| 0.1 | 0.05 | ycaM | 5 | 15.8833 | 0.283 |
| 0.1 | 0.05 | yadI | 5 | 15.8833 | 0.629 |
| 0.1 | 0.05 | AG1 | 5 | 15.8833 | 0.755 |
| 0.1 | 0.05 | ptsI | 5 | 16.1333 | 0.6 |
| 0.1 | 0.05 | clcB | 5 | 16.1333 | 0.317 |
| 0.1 | 0.05 | ycaM | 5 | 16.1333 | 0.287 |
| 0.1 | 0.05 | yadI | 5 | 16.1333 | 0.639 |
| 0.1 | 0.05 | AG1 | 5 | 16.1333 | 0.765 |
| 0.1 | 0.05 | ptsI | 5 | 16.3833 | 0.602 |
| 0.1 | 0.05 | clcB | 5 | 16.3833 | 0.324 |
| 0.1 | 0.05 | ycaM | 5 | 16.3833 | 0.284 |
| 0.1 | 0.05 | yadI | 5 | 16.3833 | 0.634 |
| 0.1 | 0.05 | AG1 | 5 | 16.3833 | 0.766 |

| | | | | | |
|---|---|---|---|---|---|
| 0.1 | 0.05 | ptsI | 5 | 16.6333 | 0.609 |
| 0.1 | 0.05 | clcB | 5 | 16.6333 | 0.324 |
| 0.1 | 0.05 | ycaM | 5 | 16.6333 | 0.288 |
| 0.1 | 0.05 | yadI | 5 | 16.6333 | 0.637 |
| 0.1 | 0.05 | AG1 | 5 | 16.6333 | 0.768 |
| 0.1 | 0.05 | ptsI | 5 | 16.8833 | 0.614 |
| 0.1 | 0.05 | clcB | 5 | 16.8833 | 0.331 |
| 0.1 | 0.05 | ycaM | 5 | 16.8833 | 0.289 |
| 0.1 | 0.05 | yadI | 5 | 16.8833 | 0.64 |
| 0.1 | 0.05 | AG1 | 5 | 16.8833 | 0.775 |
| 0.1 | 0.05 | ptsI | 5 | 17.1333 | 0.624 |
| 0.1 | 0.05 | clcB | 5 | 17.1333 | 0.327 |
| 0.1 | 0.05 | ycaM | 5 | 17.1333 | 0.291 |
| 0.1 | 0.05 | yadI | 5 | 17.1333 | 0.647 |
| 0.1 | 0.05 | AG1 | 5 | 17.1333 | 0.784 |
| 0.1 | 0.05 | ptsI | 5 | 17.3833 | 0.627 |
| 0.1 | 0.05 | clcB | 5 | 17.3833 | 0.331 |
| 0.1 | 0.05 | ycaM | 5 | 17.3833 | 0.294 |
| 0.1 | 0.05 | yadI | 5 | 17.3833 | 0.645 |
| 0.1 | 0.05 | AG1 | 5 | 17.3833 | 0.786 |
| 0.1 | 0.05 | ptsI | 5 | 17.6333 | 0.629 |
| 0.1 | 0.05 | clcB | 5 | 17.6333 | 0.331 |
| 0.1 | 0.05 | ycaM | 5 | 17.6333 | 0.297 |
| 0.1 | 0.05 | yadI | 5 | 17.6333 | 0.647 |
| 0.1 | 0.05 | AG1 | 5 | 17.6333 | 0.79 |
| 0.1 | 0.05 | ptsI | 5 | 17.8833 | 0.634 |
| 0.1 | 0.05 | clcB | 5 | 17.8833 | 0.334 |
| 0.1 | 0.05 | ycaM | 5 | 17.8833 | 0.292 |
| 0.1 | 0.05 | yadI | 5 | 17.8833 | 0.648 |
| 0.1 | 0.05 | AG1 | 5 | 17.8833 | 0.789 |
| 0.1 | 0.05 | ptsI | 5 | 18.1333 | 0.643 |
| 0.1 | 0.05 | clcB | 5 | 18.1333 | 0.344 |
| 0.1 | 0.05 | ycaM | 5 | 18.1333 | 0.298 |
| 0.1 | 0.05 | yadI | 5 | 18.1333 | 0.653 |
| 0.1 | 0.05 | AG1 | 5 | 18.1333 | 0.794 |
| 0.1 | 0.05 | ptsI | 5 | 18.3833 | 0.648 |
| 0.1 | 0.05 | clcB | 5 | 18.3833 | 0.348 |
| 0.1 | 0.05 | ycaM | 5 | 18.3833 | 0.301 |
| 0.1 | 0.05 | yadI | 5 | 18.3833 | 0.658 |
| 0.1 | 0.05 | AG1 | 5 | 18.3833 | 0.802 |
| 0.1 | 0.05 | ptsI | 5 | 18.6333 | 0.65 |
| 0.1 | 0.05 | clcB | 5 | 18.6333 | 0.351 |
| 0.1 | 0.05 | ycaM | 5 | 18.6333 | 0.303 |
| 0.1 | 0.05 | yadI | 5 | 18.6333 | 0.659 |
| 0.1 | 0.05 | AG1 | 5 | 18.6333 | 0.805 |
| 0.1 | 0.05 | ptsI | 5 | 18.8833 | 0.656 |
| 0.1 | 0.05 | clcB | 5 | 18.8833 | 0.355 |
| 0.1 | 0.05 | ycaM | 5 | 18.8833 | 0.307 |

| | | | | | |
|---|---|---|---|---|---|
| 0.1 | 0.05 | yadI | 5 | 18.8833 | 0.662 |
| 0.1 | 0.05 | AG1 | 5 | 18.8833 | 0.807 |
| 0.1 | 0.05 | ptsI | 5 | 19.1333 | 0.658 |
| 0.1 | 0.05 | clcB | 5 | 19.1333 | 0.362 |
| 0.1 | 0.05 | ycaM | 5 | 19.1333 | 0.31 |
| 0.1 | 0.05 | yadI | 5 | 19.1333 | 0.67 |
| 0.1 | 0.05 | AG1 | 5 | 19.1333 | 0.813 |
| 0.1 | 0.05 | ptsI | 5 | 19.3833 | 0.66 |
| 0.1 | 0.05 | clcB | 5 | 19.3833 | 0.366 |
| 0.1 | 0.05 | ycaM | 5 | 19.3833 | 0.313 |
| 0.1 | 0.05 | yadI | 5 | 19.3833 | 0.67 |
| 0.1 | 0.05 | AG1 | 5 | 19.3833 | 0.817 |
| 0.1 | 0.05 | ptsI | 5 | 19.6333 | 0.667 |
| 0.1 | 0.05 | clcB | 5 | 19.6333 | 0.366 |
| 0.1 | 0.05 | ycaM | 5 | 19.6333 | 0.318 |
| 0.1 | 0.05 | yadI | 5 | 19.6333 | 0.667 |
| 0.1 | 0.05 | AG1 | 5 | 19.6333 | 0.817 |
| 0.1 | 0.05 | ptsI | 5 | 19.8833 | 0.669 |
| 0.1 | 0.05 | clcB | 5 | 19.8833 | 0.371 |
| 0.1 | 0.05 | ycaM | 5 | 19.8833 | 0.326 |
| 0.1 | 0.05 | yadI | 5 | 19.8833 | 0.673 |
| 0.1 | 0.05 | AG1 | 5 | 19.8833 | 0.82 |
| 0.1 | 0.05 | ptsI | 5 | 20.1333 | 0.668 |
| 0.1 | 0.05 | clcB | 5 | 20.1333 | 0.377 |
| 0.1 | 0.05 | ycaM | 5 | 20.1333 | 0.329 |
| 0.1 | 0.05 | yadI | 5 | 20.1333 | 0.674 |
| 0.1 | 0.05 | AG1 | 5 | 20.1333 | 0.827 |
| 0.1 | 0.05 | ptsI | 5 | 20.3833 | 0.667 |
| 0.1 | 0.05 | clcB | 5 | 20.3833 | 0.38 |
| 0.1 | 0.05 | ycaM | 5 | 20.3833 | 0.336 |
| 0.1 | 0.05 | yadI | 5 | 20.3833 | 0.677 |
| 0.1 | 0.05 | AG1 | 5 | 20.3833 | 0.828 |
| 0.1 | 0.05 | ptsI | 5 | 20.6333 | 0.663 |
| 0.1 | 0.05 | clcB | 5 | 20.6333 | 0.376 |
| 0.1 | 0.05 | ycaM | 5 | 20.6333 | 0.333 |
| 0.1 | 0.05 | yadI | 5 | 20.6333 | 0.67 |
| 0.1 | 0.05 | AG1 | 5 | 20.6333 | 0.816 |
| 0.1 | 0.05 | ptsI | 5 | 20.8833 | 0.669 |
| 0.1 | 0.05 | clcB | 5 | 20.8833 | 0.387 |
| 0.1 | 0.05 | ycaM | 5 | 20.8833 | 0.344 |
| 0.1 | 0.05 | yadI | 5 | 20.8833 | 0.682 |
| 0.1 | 0.05 | AG1 | 5 | 20.8833 | 0.835 |
| 0.1 | 0.05 | ptsI | 5 | 21.1333 | 0.667 |
| 0.1 | 0.05 | clcB | 5 | 21.1333 | 0.383 |
| 0.1 | 0.05 | ycaM | 5 | 21.1333 | 0.355 |
| 0.1 | 0.05 | yadI | 5 | 21.1333 | 0.684 |
| 0.1 | 0.05 | AG1 | 5 | 21.1333 | 0.837 |
| 0.1 | 0.05 | ptsI | 5 | 21.3833 | 0.665 |

| | | | | | |
|---|---|---|---|---|---|
| 0.1 | 0.05 | clcB | 5 | 21.3833 | 0.383 |
| 0.1 | 0.05 | ycaM | 5 | 21.3833 | 0.358 |
| 0.1 | 0.05 | yadI | 5 | 21.3833 | 0.687 |
| 0.1 | 0.05 | AG1 | 5 | 21.3833 | 0.834 |
| 0.1 | 0.05 | ptsI | 5 | 21.6333 | 0.668 |
| 0.1 | 0.05 | clcB | 5 | 21.6333 | 0.382 |
| 0.1 | 0.05 | ycaM | 5 | 21.6333 | 0.366 |
| 0.1 | 0.05 | yadI | 5 | 21.6333 | 0.684 |
| 0.1 | 0.05 | AG1 | 5 | 21.6333 | 0.833 |
| 0.1 | 0.05 | ptsI | 5 | 21.8833 | 0.667 |
| 0.1 | 0.05 | clcB | 5 | 21.8833 | 0.387 |
| 0.1 | 0.05 | ycaM | 5 | 21.8833 | 0.373 |
| 0.1 | 0.05 | yadI | 5 | 21.8833 | 0.688 |
| 0.1 | 0.05 | AG1 | 5 | 21.8833 | 0.836 |
| 0.1 | 0.05 | ptsI | 5 | 22.1333 | 0.665 |
| 0.1 | 0.05 | clcB | 5 | 22.1333 | 0.384 |
| 0.1 | 0.05 | ycaM | 5 | 22.1333 | 0.378 |
| 0.1 | 0.05 | yadI | 5 | 22.1333 | 0.684 |
| 0.1 | 0.05 | AG1 | 5 | 22.1333 | 0.836 |
| 0.1 | 0.05 | ptsI | 5 | 22.3833 | 0.662 |
| 0.1 | 0.05 | clcB | 5 | 22.3833 | 0.387 |
| 0.1 | 0.05 | ycaM | 5 | 22.3833 | 0.396 |
| 0.1 | 0.05 | yadI | 5 | 22.3833 | 0.691 |
| 0.1 | 0.05 | AG1 | 5 | 22.3833 | 0.848 |
| 0.1 | 0.05 | ptsI | 5 | 22.6333 | 0.664 |
| 0.1 | 0.05 | clcB | 5 | 22.6333 | 0.381 |
| 0.1 | 0.05 | ycaM | 5 | 22.6333 | 0.395 |
| 0.1 | 0.05 | yadI | 5 | 22.6333 | 0.69 |
| 0.1 | 0.05 | AG1 | 5 | 22.6333 | 0.842 |
| 0.1 | 0.05 | ptsI | 5 | 22.8833 | 0.661 |
| 0.1 | 0.05 | clcB | 5 | 22.8833 | 0.379 |
| 0.1 | 0.05 | ycaM | 5 | 22.8833 | 0.405 |
| 0.1 | 0.05 | yadI | 5 | 22.8833 | 0.69 |
| 0.1 | 0.05 | AG1 | 5 | 22.8833 | 0.847 |
| 0.1 | 0.05 | ptsI | 5 | 23.1333 | 0.664 |
| 0.1 | 0.05 | clcB | 5 | 23.1333 | 0.376 |
| 0.1 | 0.05 | ycaM | 5 | 23.1333 | 0.41 |
| 0.1 | 0.05 | yadI | 5 | 23.1333 | 0.688 |
| 0.1 | 0.05 | AG1 | 5 | 23.1333 | 0.845 |
| 0.1 | 0.05 | ptsI | 5 | 23.3833 | 0.666 |
| 0.1 | 0.05 | clcB | 5 | 23.3833 | 0.377 |
| 0.1 | 0.05 | ycaM | 5 | 23.3833 | 0.417 |
| 0.1 | 0.05 | yadI | 5 | 23.3833 | 0.694 |
| 0.1 | 0.05 | AG1 | 5 | 23.3833 | 0.852 |
| 0.1 | 0.05 | ptsI | 5 | 23.6333 | 0.664 |
| 0.1 | 0.05 | clcB | 5 | 23.6333 | 0.37 |
| 0.1 | 0.05 | ycaM | 5 | 23.6333 | 0.418 |
| 0.1 | 0.05 | yadI | 5 | 23.6333 | 0.698 |

| | | | | | |
|---|---|---|---|---|---|
| 0.1 | 0.05 | AG1 | 5 | 23.6333 | 0.858 |
| 0.1 | 0.05 | ptsI | 5 | 23.8833 | 0.657 |
| 0.1 | 0.05 | clcB | 5 | 23.8833 | 0.369 |
| 0.1 | 0.05 | ycaM | 5 | 23.8833 | 0.416 |
| 0.1 | 0.05 | yadI | 5 | 23.8833 | 0.693 |
| 0.1 | 0.05 | AG1 | 5 | 23.8833 | 0.851 |
| 0.1 | 0.05 | ptsI | 5 | 24.1333 | 0.658 |
| 0.1 | 0.05 | clcB | 5 | 24.1333 | 0.362 |
| 0.1 | 0.05 | ycaM | 5 | 24.1333 | 0.422 |
| 0.1 | 0.05 | yadI | 5 | 24.1333 | 0.698 |
| 0.1 | 0.05 | AG1 | 5 | 24.1333 | 0.859 |
| 0.1 | 0.05 | ptsI | 5 | 24.3833 | 0.654 |
| 0.1 | 0.05 | clcB | 5 | 24.3833 | 0.362 |
| 0.1 | 0.05 | ycaM | 5 | 24.3833 | 0.43 |
| 0.1 | 0.05 | yadI | 5 | 24.3833 | 0.698 |
| 0.1 | 0.05 | AG1 | 5 | 24.3833 | 0.862 |
| 0.2 | 0.05 | ptsI | 2 | 0 | 0.216 |
| 0.2 | 0.05 | clcB | 2 | 0 | 0.229 |
| 0.2 | 0.05 | ycaM | 2 | 0 | 0.218 |
| 0.2 | 0.05 | yadI | 2 | 0 | 0.222 |
| 0.2 | 0.05 | AG1 | 2 | 0 | 0.234 |
| 0.2 | 0.05 | ptsI | 2 | 0.25 | 0.218 |
| 0.2 | 0.05 | clcB | 2 | 0.25 | 0.226 |
| 0.2 | 0.05 | ycaM | 2 | 0.25 | 0.213 |
| 0.2 | 0.05 | yadI | 2 | 0.25 | 0.218 |
| 0.2 | 0.05 | AG1 | 2 | 0.25 | 0.226 |
| 0.2 | 0.05 | ptsI | 2 | 0.5 | 0.213 |
| 0.2 | 0.05 | clcB | 2 | 0.5 | 0.214 |
| 0.2 | 0.05 | ycaM | 2 | 0.5 | 0.212 |
| 0.2 | 0.05 | yadI | 2 | 0.5 | 0.216 |
| 0.2 | 0.05 | AG1 | 2 | 0.5 | 0.221 |
| 0.2 | 0.05 | ptsI | 2 | 0.75 | 0.214 |
| 0.2 | 0.05 | clcB | 2 | 0.75 | 0.214 |
| 0.2 | 0.05 | ycaM | 2 | 0.75 | 0.213 |
| 0.2 | 0.05 | yadI | 2 | 0.75 | 0.218 |
| 0.2 | 0.05 | AG1 | 2 | 0.75 | 0.225 |
| 0.2 | 0.05 | ptsI | 2 | 1 | 0.218 |
| 0.2 | 0.05 | clcB | 2 | 1 | 0.218 |
| 0.2 | 0.05 | ycaM | 2 | 1 | 0.216 |
| 0.2 | 0.05 | yadI | 2 | 1 | 0.224 |
| 0.2 | 0.05 | AG1 | 2 | 1 | 0.227 |
| 0.2 | 0.05 | ptsI | 2 | 1.25 | 0.22 |
| 0.2 | 0.05 | clcB | 2 | 1.25 | 0.219 |
| 0.2 | 0.05 | ycaM | 2 | 1.25 | 0.217 |
| 0.2 | 0.05 | yadI | 2 | 1.25 | 0.229 |
| 0.2 | 0.05 | AG1 | 2 | 1.25 | 0.229 |
| 0.2 | 0.05 | ptsI | 2 | 1.5 | 0.22 |
| 0.2 | 0.05 | clcB | 2 | 1.5 | 0.222 |

| | | | | | |
|---|---|---|---|---|---|
| 0.2 | 0.05 | ycaM | 2 | 1.5 | 0.219 |
| 0.2 | 0.05 | yadI | 2 | 1.5 | 0.236 |
| 0.2 | 0.05 | AG1 | 2 | 1.5 | 0.23 |
| 0.2 | 0.05 | ptsI | 2 | 1.75 | 0.224 |
| 0.2 | 0.05 | clcB | 2 | 1.75 | 0.226 |
| 0.2 | 0.05 | ycaM | 2 | 1.75 | 0.223 |
| 0.2 | 0.05 | yadI | 2 | 1.75 | 0.241 |
| 0.2 | 0.05 | AG1 | 2 | 1.75 | 0.239 |
| 0.2 | 0.05 | ptsI | 2 | 2 | 0.226 |
| 0.2 | 0.05 | clcB | 2 | 2 | 0.229 |
| 0.2 | 0.05 | ycaM | 2 | 2 | 0.23 |
| 0.2 | 0.05 | yadI | 2 | 2 | 0.249 |
| 0.2 | 0.05 | AG1 | 2 | 2 | 0.244 |
| 0.2 | 0.05 | ptsI | 2 | 2.25 | 0.235 |
| 0.2 | 0.05 | clcB | 2 | 2.25 | 0.236 |
| 0.2 | 0.05 | ycaM | 2 | 2.25 | 0.234 |
| 0.2 | 0.05 | yadI | 2 | 2.25 | 0.259 |
| 0.2 | 0.05 | AG1 | 2 | 2.25 | 0.252 |
| 0.2 | 0.05 | ptsI | 2 | 2.5 | 0.242 |
| 0.2 | 0.05 | clcB | 2 | 2.5 | 0.243 |
| 0.2 | 0.05 | ycaM | 2 | 2.5 | 0.241 |
| 0.2 | 0.05 | yadI | 2 | 2.5 | 0.27 |
| 0.2 | 0.05 | AG1 | 2 | 2.5 | 0.26 |
| 0.2 | 0.05 | ptsI | 2 | 2.75 | 0.243 |
| 0.2 | 0.05 | clcB | 2 | 2.75 | 0.251 |
| 0.2 | 0.05 | ycaM | 2 | 2.75 | 0.244 |
| 0.2 | 0.05 | yadI | 2 | 2.75 | 0.281 |
| 0.2 | 0.05 | AG1 | 2 | 2.75 | 0.258 |
| 0.2 | 0.05 | ptsI | 2 | 3 | 0.248 |
| 0.2 | 0.05 | clcB | 2 | 3 | 0.249 |
| 0.2 | 0.05 | ycaM | 2 | 3 | 0.246 |
| 0.2 | 0.05 | yadI | 2 | 3 | 0.287 |
| 0.2 | 0.05 | AG1 | 2 | 3 | 0.258 |
| 0.2 | 0.05 | ptsI | 2 | 3.25 | 0.255 |
| 0.2 | 0.05 | clcB | 2 | 3.25 | 0.265 |
| 0.2 | 0.05 | ycaM | 2 | 3.25 | 0.254 |
| 0.2 | 0.05 | yadI | 2 | 3.25 | 0.305 |
| 0.2 | 0.05 | AG1 | 2 | 3.25 | 0.269 |
| 0.2 | 0.05 | ptsI | 2 | 3.5 | 0.264 |
| 0.2 | 0.05 | clcB | 2 | 3.5 | 0.271 |
| 0.2 | 0.05 | ycaM | 2 | 3.5 | 0.258 |
| 0.2 | 0.05 | yadI | 2 | 3.5 | 0.318 |
| 0.2 | 0.05 | AG1 | 2 | 3.5 | 0.281 |
| 0.2 | 0.05 | ptsI | 2 | 3.75 | 0.27 |
| 0.2 | 0.05 | clcB | 2 | 3.75 | 0.274 |
| 0.2 | 0.05 | ycaM | 2 | 3.75 | 0.268 |
| 0.2 | 0.05 | yadI | 2 | 3.75 | 0.332 |
| 0.2 | 0.05 | AG1 | 2 | 3.75 | 0.289 |

| | | | | | |
|---|---|---|---|---|---|
| 0.2 | 0.05 | ptsI | 2 | 4 | 0.28 |
| 0.2 | 0.05 | clcB | 2 | 4 | 0.282 |
| 0.2 | 0.05 | ycaM | 2 | 4 | 0.279 |
| 0.2 | 0.05 | yadI | 2 | 4 | 0.344 |
| 0.2 | 0.05 | AG1 | 2 | 4 | 0.3 |
| 0.2 | 0.05 | ptsI | 2 | 4.38333 | 0.318 |
| 0.2 | 0.05 | clcB | 2 | 4.38333 | 0.262 |
| 0.2 | 0.05 | ycaM | 2 | 4.38333 | 0.288 |
| 0.2 | 0.05 | yadI | 2 | 4.38333 | 0.32 |
| 0.2 | 0.05 | AG1 | 2 | 4.38333 | 0.294 |
| 0.2 | 0.05 | ptsI | 2 | 4.63333 | 0.311 |
| 0.2 | 0.05 | clcB | 2 | 4.63333 | 0.252 |
| 0.2 | 0.05 | ycaM | 2 | 4.63333 | 0.285 |
| 0.2 | 0.05 | yadI | 2 | 4.63333 | 0.33 |
| 0.2 | 0.05 | AG1 | 2 | 4.63333 | 0.293 |
| 0.2 | 0.05 | ptsI | 2 | 4.88333 | 0.316 |
| 0.2 | 0.05 | clcB | 2 | 4.88333 | 0.252 |
| 0.2 | 0.05 | ycaM | 2 | 4.88333 | 0.289 |
| 0.2 | 0.05 | yadI | 2 | 4.88333 | 0.34 |
| 0.2 | 0.05 | AG1 | 2 | 4.88333 | 0.304 |
| 0.2 | 0.05 | ptsI | 2 | 5.13333 | 0.324 |
| 0.2 | 0.05 | clcB | 2 | 5.13333 | 0.255 |
| 0.2 | 0.05 | ycaM | 2 | 5.13333 | 0.293 |
| 0.2 | 0.05 | yadI | 2 | 5.13333 | 0.357 |
| 0.2 | 0.05 | AG1 | 2 | 5.13333 | 0.311 |
| 0.2 | 0.05 | ptsI | 2 | 5.38333 | 0.334 |
| 0.2 | 0.05 | clcB | 2 | 5.38333 | 0.259 |
| 0.2 | 0.05 | ycaM | 2 | 5.38333 | 0.295 |
| 0.2 | 0.05 | yadI | 2 | 5.38333 | 0.368 |
| 0.2 | 0.05 | AG1 | 2 | 5.38333 | 0.323 |
| 0.2 | 0.05 | ptsI | 2 | 5.63333 | 0.342 |
| 0.2 | 0.05 | clcB | 2 | 5.63333 | 0.269 |
| 0.2 | 0.05 | ycaM | 2 | 5.63333 | 0.297 |
| 0.2 | 0.05 | yadI | 2 | 5.63333 | 0.382 |
| 0.2 | 0.05 | AG1 | 2 | 5.63333 | 0.332 |
| 0.2 | 0.05 | ptsI | 2 | 5.88333 | 0.35 |
| 0.2 | 0.05 | clcB | 2 | 5.88333 | 0.274 |
| 0.2 | 0.05 | ycaM | 2 | 5.88333 | 0.298 |
| 0.2 | 0.05 | yadI | 2 | 5.88333 | 0.387 |
| 0.2 | 0.05 | AG1 | 2 | 5.88333 | 0.34 |
| 0.2 | 0.05 | ptsI | 2 | 6.13333 | 0.362 |
| 0.2 | 0.05 | clcB | 2 | 6.13333 | 0.277 |
| 0.2 | 0.05 | ycaM | 2 | 6.13333 | 0.299 |
| 0.2 | 0.05 | yadI | 2 | 6.13333 | 0.396 |
| 0.2 | 0.05 | AG1 | 2 | 6.13333 | 0.35 |
| 0.2 | 0.05 | ptsI | 2 | 6.38333 | 0.371 |
| 0.2 | 0.05 | clcB | 2 | 6.38333 | 0.282 |
| 0.2 | 0.05 | ycaM | 2 | 6.38333 | 0.302 |

| | | | | | |
|---|---|---|---|---|---|
| 0.2 | 0.05 | yadI | 2 | 6.38333 | 0.404 |
| 0.2 | 0.05 | AG1 | 2 | 6.38333 | 0.356 |
| 0.2 | 0.05 | ptsI | 2 | 6.63333 | 0.381 |
| 0.2 | 0.05 | clcB | 2 | 6.63333 | 0.287 |
| 0.2 | 0.05 | ycaM | 2 | 6.63333 | 0.301 |
| 0.2 | 0.05 | yadI | 2 | 6.63333 | 0.414 |
| 0.2 | 0.05 | AG1 | 2 | 6.63333 | 0.366 |
| 0.2 | 0.05 | ptsI | 2 | 6.88333 | 0.394 |
| 0.2 | 0.05 | clcB | 2 | 6.88333 | 0.293 |
| 0.2 | 0.05 | ycaM | 2 | 6.88333 | 0.303 |
| 0.2 | 0.05 | yadI | 2 | 6.88333 | 0.424 |
| 0.2 | 0.05 | AG1 | 2 | 6.88333 | 0.376 |
| 0.2 | 0.05 | ptsI | 2 | 7.13333 | 0.406 |
| 0.2 | 0.05 | clcB | 2 | 7.13333 | 0.294 |
| 0.2 | 0.05 | ycaM | 2 | 7.13333 | 0.304 |
| 0.2 | 0.05 | yadI | 2 | 7.13333 | 0.431 |
| 0.2 | 0.05 | AG1 | 2 | 7.13333 | 0.385 |
| 0.2 | 0.05 | ptsI | 2 | 7.38333 | 0.415 |
| 0.2 | 0.05 | clcB | 2 | 7.38333 | 0.303 |
| 0.2 | 0.05 | ycaM | 2 | 7.38333 | 0.305 |
| 0.2 | 0.05 | yadI | 2 | 7.38333 | 0.441 |
| 0.2 | 0.05 | AG1 | 2 | 7.38333 | 0.393 |
| 0.2 | 0.05 | ptsI | 2 | 7.63333 | 0.422 |
| 0.2 | 0.05 | clcB | 2 | 7.63333 | 0.304 |
| 0.2 | 0.05 | ycaM | 2 | 7.63333 | 0.303 |
| 0.2 | 0.05 | yadI | 2 | 7.63333 | 0.443 |
| 0.2 | 0.05 | AG1 | 2 | 7.63333 | 0.399 |
| 0.2 | 0.05 | ptsI | 2 | 7.88333 | 0.435 |
| 0.2 | 0.05 | clcB | 2 | 7.88333 | 0.306 |
| 0.2 | 0.05 | ycaM | 2 | 7.88333 | 0.305 |
| 0.2 | 0.05 | yadI | 2 | 7.88333 | 0.456 |
| 0.2 | 0.05 | AG1 | 2 | 7.88333 | 0.408 |
| 0.2 | 0.05 | ptsI | 2 | 8.13333 | 0.448 |
| 0.2 | 0.05 | clcB | 2 | 8.13333 | 0.309 |
| 0.2 | 0.05 | ycaM | 2 | 8.13333 | 0.306 |
| 0.2 | 0.05 | yadI | 2 | 8.13333 | 0.462 |
| 0.2 | 0.05 | AG1 | 2 | 8.13333 | 0.418 |
| 0.2 | 0.05 | ptsI | 2 | 8.38333 | 0.455 |
| 0.2 | 0.05 | clcB | 2 | 8.38333 | 0.318 |
| 0.2 | 0.05 | ycaM | 2 | 8.38333 | 0.306 |
| 0.2 | 0.05 | yadI | 2 | 8.38333 | 0.472 |
| 0.2 | 0.05 | AG1 | 2 | 8.38333 | 0.426 |
| 0.2 | 0.05 | ptsI | 2 | 8.91667 | 0.425 |
| 0.2 | 0.05 | clcB | 2 | 8.91667 | 0.328 |
| 0.2 | 0.05 | ycaM | 2 | 8.91667 | 0.286 |
| 0.2 | 0.05 | yadI | 2 | 8.91667 | 0.503 |
| 0.2 | 0.05 | AG1 | 2 | 8.91667 | 0.463 |
| 0.2 | 0.05 | ptsI | 2 | 9.16667 | 0.424 |

| | | | | | |
|---|---|---|---|---|---|
| 0.2 | 0.05 | clcB | 2 | 9.16667 | 0.327 |
| 0.2 | 0.05 | ycaM | 2 | 9.16667 | 0.287 |
| 0.2 | 0.05 | yadI | 2 | 9.16667 | 0.512 |
| 0.2 | 0.05 | AG1 | 2 | 9.16667 | 0.476 |
| 0.2 | 0.05 | ptsI | 2 | 9.41667 | 0.432 |
| 0.2 | 0.05 | clcB | 2 | 9.41667 | 0.332 |
| 0.2 | 0.05 | ycaM | 2 | 9.41667 | 0.286 |
| 0.2 | 0.05 | yadI | 2 | 9.41667 | 0.524 |
| 0.2 | 0.05 | AG1 | 2 | 9.41667 | 0.492 |
| 0.2 | 0.05 | ptsI | 2 | 9.66667 | 0.445 |
| 0.2 | 0.05 | clcB | 2 | 9.66667 | 0.338 |
| 0.2 | 0.05 | ycaM | 2 | 9.66667 | 0.289 |
| 0.2 | 0.05 | yadI | 2 | 9.66667 | 0.538 |
| 0.2 | 0.05 | AG1 | 2 | 9.66667 | 0.51 |
| 0.2 | 0.05 | ptsI | 2 | 9.91667 | 0.456 |
| 0.2 | 0.05 | clcB | 2 | 9.91667 | 0.342 |
| 0.2 | 0.05 | ycaM | 2 | 9.91667 | 0.29 |
| 0.2 | 0.05 | yadI | 2 | 9.91667 | 0.549 |
| 0.2 | 0.05 | AG1 | 2 | 9.91667 | 0.528 |
| 0.2 | 0.05 | ptsI | 2 | 10.1667 | 0.47 |
| 0.2 | 0.05 | clcB | 2 | 10.1667 | 0.342 |
| 0.2 | 0.05 | ycaM | 2 | 10.1667 | 0.29 |
| 0.2 | 0.05 | yadI | 2 | 10.1667 | 0.555 |
| 0.2 | 0.05 | AG1 | 2 | 10.1667 | 0.536 |
| 0.2 | 0.05 | ptsI | 2 | 10.4167 | 0.487 |
| 0.2 | 0.05 | clcB | 2 | 10.4167 | 0.349 |
| 0.2 | 0.05 | ycaM | 2 | 10.4167 | 0.294 |
| 0.2 | 0.05 | yadI | 2 | 10.4167 | 0.567 |
| 0.2 | 0.05 | AG1 | 2 | 10.4167 | 0.559 |
| 0.2 | 0.05 | ptsI | 2 | 10.6667 | 0.502 |
| 0.2 | 0.05 | clcB | 2 | 10.6667 | 0.353 |
| 0.2 | 0.05 | ycaM | 2 | 10.6667 | 0.295 |
| 0.2 | 0.05 | yadI | 2 | 10.6667 | 0.574 |
| 0.2 | 0.05 | AG1 | 2 | 10.6667 | 0.571 |
| 0.2 | 0.05 | ptsI | 2 | 10.9167 | 0.518 |
| 0.2 | 0.05 | clcB | 2 | 10.9167 | 0.361 |
| 0.2 | 0.05 | ycaM | 2 | 10.9167 | 0.294 |
| 0.2 | 0.05 | yadI | 2 | 10.9167 | 0.585 |
| 0.2 | 0.05 | AG1 | 2 | 10.9167 | 0.585 |
| 0.2 | 0.05 | ptsI | 2 | 11.1667 | 0.539 |
| 0.2 | 0.05 | clcB | 2 | 11.1667 | 0.352 |
| 0.2 | 0.05 | ycaM | 2 | 11.1667 | 0.296 |
| 0.2 | 0.05 | yadI | 2 | 11.1667 | 0.592 |
| 0.2 | 0.05 | AG1 | 2 | 11.1667 | 0.597 |
| 0.2 | 0.05 | ptsI | 2 | 11.4167 | 0.551 |
| 0.2 | 0.05 | clcB | 2 | 11.4167 | 0.352 |
| 0.2 | 0.05 | ycaM | 2 | 11.4167 | 0.296 |
| 0.2 | 0.05 | yadI | 2 | 11.4167 | 0.6 |

| | | | | | |
|---|---|---|---|---|---|
| 0.2 | 0.05 | AG1 | 2 | 11.4167 | 0.596 |
| 0.2 | 0.05 | ptsI | 2 | 11.6667 | 0.564 |
| 0.2 | 0.05 | clcB | 2 | 11.6667 | 0.353 |
| 0.2 | 0.05 | ycaM | 2 | 11.6667 | 0.296 |
| 0.2 | 0.05 | yadI | 2 | 11.6667 | 0.613 |
| 0.2 | 0.05 | AG1 | 2 | 11.6667 | 0.628 |
| 0.2 | 0.05 | ptsI | 2 | 11.9167 | 0.578 |
| 0.2 | 0.05 | clcB | 2 | 11.9167 | 0.351 |
| 0.2 | 0.05 | ycaM | 2 | 11.9167 | 0.296 |
| 0.2 | 0.05 | yadI | 2 | 11.9167 | 0.614 |
| 0.2 | 0.05 | AG1 | 2 | 11.9167 | 0.615 |
| 0.2 | 0.05 | ptsI | 2 | 12.1667 | 0.598 |
| 0.2 | 0.05 | clcB | 2 | 12.1667 | 0.352 |
| 0.2 | 0.05 | ycaM | 2 | 12.1667 | 0.296 |
| 0.2 | 0.05 | yadI | 2 | 12.1667 | 0.625 |
| 0.2 | 0.05 | AG1 | 2 | 12.1667 | 0.647 |
| 0.2 | 0.05 | ptsI | 2 | 12.4167 | 0.6 |
| 0.2 | 0.05 | clcB | 2 | 12.4167 | 0.347 |
| 0.2 | 0.05 | ycaM | 2 | 12.4167 | 0.296 |
| 0.2 | 0.05 | yadI | 2 | 12.4167 | 0.631 |
| 0.2 | 0.05 | AG1 | 2 | 12.4167 | 0.637 |
| 0.2 | 0.05 | ptsI | 2 | 12.6667 | 0.609 |
| 0.2 | 0.05 | clcB | 2 | 12.6667 | 0.347 |
| 0.2 | 0.05 | ycaM | 2 | 12.6667 | 0.298 |
| 0.2 | 0.05 | yadI | 2 | 12.6667 | 0.635 |
| 0.2 | 0.05 | AG1 | 2 | 12.6667 | 0.644 |
| 0.2 | 0.05 | ptsI | 2 | 12.9167 | 0.629 |
| 0.2 | 0.05 | clcB | 2 | 12.9167 | 0.338 |
| 0.2 | 0.05 | ycaM | 2 | 12.9167 | 0.297 |
| 0.2 | 0.05 | yadI | 2 | 12.9167 | 0.644 |
| 0.2 | 0.05 | AG1 | 2 | 12.9167 | 0.638 |
| 0.2 | 0.05 | ptsI | 2 | 13.1667 | 0.637 |
| 0.2 | 0.05 | clcB | 2 | 13.1667 | 0.343 |
| 0.2 | 0.05 | ycaM | 2 | 13.1667 | 0.297 |
| 0.2 | 0.05 | yadI | 2 | 13.1667 | 0.65 |
| 0.2 | 0.05 | AG1 | 2 | 13.1667 | 0.645 |
| 0.2 | 0.05 | ptsI | 2 | 13.4167 | 0.646 |
| 0.2 | 0.05 | clcB | 2 | 13.4167 | 0.336 |
| 0.2 | 0.05 | ycaM | 2 | 13.4167 | 0.293 |
| 0.2 | 0.05 | yadI | 2 | 13.4167 | 0.656 |
| 0.2 | 0.05 | AG1 | 2 | 13.4167 | 0.638 |
| 0.2 | 0.05 | ptsI | 2 | 13.6667 | 0.668 |
| 0.2 | 0.05 | clcB | 2 | 13.6667 | 0.341 |
| 0.2 | 0.05 | ycaM | 2 | 13.6667 | 0.293 |
| 0.2 | 0.05 | yadI | 2 | 13.6667 | 0.669 |
| 0.2 | 0.05 | AG1 | 2 | 13.6667 | 0.663 |
| 0.2 | 0.05 | ptsI | 2 | 13.9167 | 0.674 |
| 0.2 | 0.05 | clcB | 2 | 13.9167 | 0.35 |

| | | | | | |
|---|---|---|---|---|---|
| 0.2 | 0.05 | ycaM | 2 | 13.9167 | 0.294 |
| 0.2 | 0.05 | yadI | 2 | 13.9167 | 0.672 |
| 0.2 | 0.05 | AG1 | 2 | 13.9167 | 0.666 |
| 0.2 | 0.05 | ptsI | 2 | 14.1667 | 0.676 |
| 0.2 | 0.05 | clcB | 2 | 14.1667 | 0.34 |
| 0.2 | 0.05 | ycaM | 2 | 14.1667 | 0.296 |
| 0.2 | 0.05 | yadI | 2 | 14.1667 | 0.677 |
| 0.2 | 0.05 | AG1 | 2 | 14.1667 | 0.669 |
| 0.2 | 0.05 | ptsI | 2 | 14.4167 | 0.69 |
| 0.2 | 0.05 | clcB | 2 | 14.4167 | 0.343 |
| 0.2 | 0.05 | ycaM | 2 | 14.4167 | 0.295 |
| 0.2 | 0.05 | yadI | 2 | 14.4167 | 0.681 |
| 0.2 | 0.05 | AG1 | 2 | 14.4167 | 0.655 |
| 0.2 | 0.05 | ptsI | 2 | 14.6667 | 0.69 |
| 0.2 | 0.05 | clcB | 2 | 14.6667 | 0.342 |
| 0.2 | 0.05 | ycaM | 2 | 14.6667 | 0.294 |
| 0.2 | 0.05 | yadI | 2 | 14.6667 | 0.69 |
| 0.2 | 0.05 | AG1 | 2 | 14.6667 | 0.659 |
| 0.2 | 0.05 | ptsI | 2 | 14.9167 | 0.708 |
| 0.2 | 0.05 | clcB | 2 | 14.9167 | 0.345 |
| 0.2 | 0.05 | ycaM | 2 | 14.9167 | 0.294 |
| 0.2 | 0.05 | yadI | 2 | 14.9167 | 0.703 |
| 0.2 | 0.05 | AG1 | 2 | 14.9167 | 0.68 |
| 0.2 | 0.05 | ptsI | 2 | 15.1667 | 0.709 |
| 0.2 | 0.05 | clcB | 2 | 15.1667 | 0.353 |
| 0.2 | 0.05 | ycaM | 2 | 15.1667 | 0.296 |
| 0.2 | 0.05 | yadI | 2 | 15.1667 | 0.712 |
| 0.2 | 0.05 | AG1 | 2 | 15.1667 | 0.703 |
| 0.2 | 0.05 | ptsI | 2 | 15.4167 | 0.722 |
| 0.2 | 0.05 | clcB | 2 | 15.4167 | 0.358 |
| 0.2 | 0.05 | ycaM | 2 | 15.4167 | 0.292 |
| 0.2 | 0.05 | yadI | 2 | 15.4167 | 0.713 |
| 0.2 | 0.05 | AG1 | 2 | 15.4167 | 0.696 |
| 0.2 | 0.05 | ptsI | 2 | 15.6667 | 0.719 |
| 0.2 | 0.05 | clcB | 2 | 15.6667 | 0.35 |
| 0.2 | 0.05 | ycaM | 2 | 15.6667 | 0.29 |
| 0.2 | 0.05 | yadI | 2 | 15.6667 | 0.716 |
| 0.2 | 0.05 | AG1 | 2 | 15.6667 | 0.704 |
| 0.2 | 0.05 | ptsI | 2 | 15.9167 | 0.719 |
| 0.2 | 0.05 | clcB | 2 | 15.9167 | 0.355 |
| 0.2 | 0.05 | ycaM | 2 | 15.9167 | 0.29 |
| 0.2 | 0.05 | yadI | 2 | 15.9167 | 0.719 |
| 0.2 | 0.05 | AG1 | 2 | 15.9167 | 0.727 |
| 0.2 | 0.05 | ptsI | 2 | 16.1667 | 0.739 |
| 0.2 | 0.05 | clcB | 2 | 16.1667 | 0.352 |
| 0.2 | 0.05 | ycaM | 2 | 16.1667 | 0.29 |
| 0.2 | 0.05 | yadI | 2 | 16.1667 | 0.731 |
| 0.2 | 0.05 | AG1 | 2 | 16.1667 | 0.733 |

| | | | | | |
|---|---|---|---|---|---|
| 0.2 | 0.05 | ptsI | 2 | 16.4167 | 0.731 |
| 0.2 | 0.05 | clcB | 2 | 16.4167 | 0.353 |
| 0.2 | 0.05 | ycaM | 2 | 16.4167 | 0.29 |
| 0.2 | 0.05 | yadI | 2 | 16.4167 | 0.733 |
| 0.2 | 0.05 | AG1 | 2 | 16.4167 | 0.752 |
| 0.2 | 0.05 | ptsI | 2 | 16.6667 | 0.732 |
| 0.2 | 0.05 | clcB | 2 | 16.6667 | 0.353 |
| 0.2 | 0.05 | ycaM | 2 | 16.6667 | 0.291 |
| 0.2 | 0.05 | yadI | 2 | 16.6667 | 0.738 |
| 0.2 | 0.05 | AG1 | 2 | 16.6667 | 0.748 |
| 0.2 | 0.05 | ptsI | 2 | 16.9167 | 0.742 |
| 0.2 | 0.05 | clcB | 2 | 16.9167 | 0.357 |
| 0.2 | 0.05 | ycaM | 2 | 16.9167 | 0.29 |
| 0.2 | 0.05 | yadI | 2 | 16.9167 | 0.739 |
| 0.2 | 0.05 | AG1 | 2 | 16.9167 | 0.747 |
| 0.2 | 0.05 | ptsI | 2 | 17.1667 | 0.748 |
| 0.2 | 0.05 | clcB | 2 | 17.1667 | 0.354 |
| 0.2 | 0.05 | ycaM | 2 | 17.1667 | 0.291 |
| 0.2 | 0.05 | yadI | 2 | 17.1667 | 0.756 |
| 0.2 | 0.05 | AG1 | 2 | 17.1667 | 0.734 |
| 0.2 | 0.05 | ptsI | 2 | 17.4167 | 0.755 |
| 0.2 | 0.05 | clcB | 2 | 17.4167 | 0.354 |
| 0.2 | 0.05 | ycaM | 2 | 17.4167 | 0.291 |
| 0.2 | 0.05 | yadI | 2 | 17.4167 | 0.757 |
| 0.2 | 0.05 | AG1 | 2 | 17.4167 | 0.753 |
| 0.2 | 0.05 | ptsI | 2 | 17.6667 | 0.758 |
| 0.2 | 0.05 | clcB | 2 | 17.6667 | 0.356 |
| 0.2 | 0.05 | ycaM | 2 | 17.6667 | 0.291 |
| 0.2 | 0.05 | yadI | 2 | 17.6667 | 0.763 |
| 0.2 | 0.05 | AG1 | 2 | 17.6667 | 0.768 |
| 0.2 | 0.05 | ptsI | 2 | 17.9167 | 0.755 |
| 0.2 | 0.05 | clcB | 2 | 17.9167 | 0.354 |
| 0.2 | 0.05 | ycaM | 2 | 17.9167 | 0.292 |
| 0.2 | 0.05 | yadI | 2 | 17.9167 | 0.764 |
| 0.2 | 0.05 | AG1 | 2 | 17.9167 | 0.753 |
| 0.2 | 0.05 | ptsI | 2 | 18.1667 | 0.765 |
| 0.2 | 0.05 | clcB | 2 | 18.1667 | 0.355 |
| 0.2 | 0.05 | ycaM | 2 | 18.1667 | 0.291 |
| 0.2 | 0.05 | yadI | 2 | 18.1667 | 0.761 |
| 0.2 | 0.05 | AG1 | 2 | 18.1667 | 0.781 |
| 0.2 | 0.05 | ptsI | 2 | 18.4167 | 0.772 |
| 0.2 | 0.05 | clcB | 2 | 18.4167 | 0.354 |
| 0.2 | 0.05 | ycaM | 2 | 18.4167 | 0.293 |
| 0.2 | 0.05 | yadI | 2 | 18.4167 | 0.766 |
| 0.2 | 0.05 | AG1 | 2 | 18.4167 | 0.776 |
| 0.2 | 0.05 | ptsI | 2 | 18.6667 | 0.758 |
| 0.2 | 0.05 | clcB | 2 | 18.6667 | 0.354 |
| 0.2 | 0.05 | ycaM | 2 | 18.6667 | 0.293 |

| | | | | | |
|---|---|---|---|---|---|
| 0.2 | 0.05 | yadI | 2 | 18.6667 | 0.767 |
| 0.2 | 0.05 | AG1 | 2 | 18.6667 | 0.782 |
| 0.2 | 0.05 | ptsI | 2 | 18.9167 | 0.744 |
| 0.2 | 0.05 | clcB | 2 | 18.9167 | 0.355 |
| 0.2 | 0.05 | ycaM | 2 | 18.9167 | 0.289 |
| 0.2 | 0.05 | yadI | 2 | 18.9167 | 0.769 |
| 0.2 | 0.05 | AG1 | 2 | 18.9167 | 0.791 |
| 0.2 | 0.05 | ptsI | 2 | 19.1667 | 0.77 |
| 0.2 | 0.05 | clcB | 2 | 19.1667 | 0.355 |
| 0.2 | 0.05 | ycaM | 2 | 19.1667 | 0.294 |
| 0.2 | 0.05 | yadI | 2 | 19.1667 | 0.767 |
| 0.2 | 0.05 | AG1 | 2 | 19.1667 | 0.802 |
| 0.2 | 0.05 | ptsI | 2 | 19.4167 | 0.772 |
| 0.2 | 0.05 | clcB | 2 | 19.4167 | 0.356 |
| 0.2 | 0.05 | ycaM | 2 | 19.4167 | 0.294 |
| 0.2 | 0.05 | yadI | 2 | 19.4167 | 0.773 |
| 0.2 | 0.05 | AG1 | 2 | 19.4167 | 0.801 |
| 0.2 | 0.05 | ptsI | 2 | 19.6667 | 0.776 |
| 0.2 | 0.05 | clcB | 2 | 19.6667 | 0.356 |
| 0.2 | 0.05 | ycaM | 2 | 19.6667 | 0.295 |
| 0.2 | 0.05 | yadI | 2 | 19.6667 | 0.785 |
| 0.2 | 0.05 | AG1 | 2 | 19.6667 | 0.804 |
| 0.2 | 0.05 | ptsI | 2 | 19.9167 | 0.782 |
| 0.2 | 0.05 | clcB | 2 | 19.9167 | 0.353 |
| 0.2 | 0.05 | ycaM | 2 | 19.9167 | 0.297 |
| 0.2 | 0.05 | yadI | 2 | 19.9167 | 0.778 |
| 0.2 | 0.05 | AG1 | 2 | 19.9167 | 0.819 |
| 0.2 | 0.05 | ptsI | 2 | 20.1667 | 0.775 |
| 0.2 | 0.05 | clcB | 2 | 20.1667 | 0.356 |
| 0.2 | 0.05 | ycaM | 2 | 20.1667 | 0.297 |
| 0.2 | 0.05 | yadI | 2 | 20.1667 | 0.79 |
| 0.2 | 0.05 | AG1 | 2 | 20.1667 | 0.816 |
| 0.2 | 0.05 | ptsI | 2 | 20.4167 | 0.782 |
| 0.2 | 0.05 | clcB | 2 | 20.4167 | 0.353 |
| 0.2 | 0.05 | ycaM | 2 | 20.4167 | 0.297 |
| 0.2 | 0.05 | yadI | 2 | 20.4167 | 0.787 |
| 0.2 | 0.05 | AG1 | 2 | 20.4167 | 0.825 |
| 0.2 | 0.05 | ptsI | 2 | 20.6667 | 0.774 |
| 0.2 | 0.05 | clcB | 2 | 20.6667 | 0.354 |
| 0.2 | 0.05 | ycaM | 2 | 20.6667 | 0.3 |
| 0.2 | 0.05 | yadI | 2 | 20.6667 | 0.786 |
| 0.2 | 0.05 | AG1 | 2 | 20.6667 | 0.824 |
| 0.2 | 0.05 | ptsI | 2 | 20.9167 | 0.782 |
| 0.2 | 0.05 | clcB | 2 | 20.9167 | 0.354 |
| 0.2 | 0.05 | ycaM | 2 | 20.9167 | 0.302 |
| 0.2 | 0.05 | yadI | 2 | 20.9167 | 0.797 |
| 0.2 | 0.05 | AG1 | 2 | 20.9167 | 0.832 |
| 0.2 | 0.05 | ptsI | 2 | 21.1667 | 0.784 |

| | | | | | |
|---|---|---|---|---|---|
| 0.2 | 0.05 | clcB | 2 | 21.1667 | 0.351 |
| 0.2 | 0.05 | ycaM | 2 | 21.1667 | 0.304 |
| 0.2 | 0.05 | yadI | 2 | 21.1667 | 0.799 |
| 0.2 | 0.05 | AG1 | 2 | 21.1667 | 0.837 |
| 0.2 | 0.05 | ptsI | 2 | 21.4167 | 0.768 |
| 0.2 | 0.05 | clcB | 2 | 21.4167 | 0.353 |
| 0.2 | 0.05 | ycaM | 2 | 21.4167 | 0.304 |
| 0.2 | 0.05 | yadI | 2 | 21.4167 | 0.8 |
| 0.2 | 0.05 | AG1 | 2 | 21.4167 | 0.84 |
| 0.2 | 0.05 | ptsI | 2 | 21.6667 | 0.773 |
| 0.2 | 0.05 | clcB | 2 | 21.6667 | 0.352 |
| 0.2 | 0.05 | ycaM | 2 | 21.6667 | 0.307 |
| 0.2 | 0.05 | yadI | 2 | 21.6667 | 0.798 |
| 0.2 | 0.05 | AG1 | 2 | 21.6667 | 0.849 |
| 0.2 | 0.05 | ptsI | 2 | 21.9167 | 0.766 |
| 0.2 | 0.05 | clcB | 2 | 21.9167 | 0.352 |
| 0.2 | 0.05 | ycaM | 2 | 21.9167 | 0.308 |
| 0.2 | 0.05 | yadI | 2 | 21.9167 | 0.806 |
| 0.2 | 0.05 | AG1 | 2 | 21.9167 | 0.851 |
| 0.2 | 0.05 | ptsI | 2 | 22.1667 | 0.769 |
| 0.2 | 0.05 | clcB | 2 | 22.1667 | 0.355 |
| 0.2 | 0.05 | ycaM | 2 | 22.1667 | 0.31 |
| 0.2 | 0.05 | yadI | 2 | 22.1667 | 0.801 |
| 0.2 | 0.05 | AG1 | 2 | 22.1667 | 0.858 |
| 0.2 | 0.05 | ptsI | 2 | 22.4167 | 0.776 |
| 0.2 | 0.05 | clcB | 2 | 22.4167 | 0.355 |
| 0.2 | 0.05 | ycaM | 2 | 22.4167 | 0.313 |
| 0.2 | 0.05 | yadI | 2 | 22.4167 | 0.803 |
| 0.2 | 0.05 | AG1 | 2 | 22.4167 | 0.862 |
| 0.2 | 0.05 | ptsI | 2 | 22.6667 | 0.766 |
| 0.2 | 0.05 | clcB | 2 | 22.6667 | 0.354 |
| 0.2 | 0.05 | ycaM | 2 | 22.6667 | 0.314 |
| 0.2 | 0.05 | yadI | 2 | 22.6667 | 0.81 |
| 0.2 | 0.05 | AG1 | 2 | 22.6667 | 0.868 |
| 0.2 | 0.05 | ptsI | 2 | 22.9167 | 0.778 |
| 0.2 | 0.05 | clcB | 2 | 22.9167 | 0.352 |
| 0.2 | 0.05 | ycaM | 2 | 22.9167 | 0.316 |
| 0.2 | 0.05 | yadI | 2 | 22.9167 | 0.809 |
| 0.2 | 0.05 | AG1 | 2 | 22.9167 | 0.878 |
| 0.2 | 0.05 | ptsI | 2 | 23.1667 | 0.767 |
| 0.2 | 0.05 | clcB | 2 | 23.1667 | 0.352 |
| 0.2 | 0.05 | ycaM | 2 | 23.1667 | 0.318 |
| 0.2 | 0.05 | yadI | 2 | 23.1667 | 0.809 |
| 0.2 | 0.05 | AG1 | 2 | 23.1667 | 0.882 |
| 0.2 | 0.05 | ptsI | 2 | 23.4167 | 0.773 |
| 0.2 | 0.05 | clcB | 2 | 23.4167 | 0.352 |
| 0.2 | 0.05 | ycaM | 2 | 23.4167 | 0.321 |
| 0.2 | 0.05 | yadI | 2 | 23.4167 | 0.813 |

| | | | | | |
|---|---|---|---|---|---|
| 0.2 | 0.05 | AG1 | 2 | 23.4167 | 0.883 |
| 0.2 | 0.05 | ptsI | 2 | 23.6667 | 0.775 |
| 0.2 | 0.05 | clcB | 2 | 23.6667 | 0.351 |
| 0.2 | 0.05 | ycaM | 2 | 23.6667 | 0.322 |
| 0.2 | 0.05 | yadI | 2 | 23.6667 | 0.821 |
| 0.2 | 0.05 | AG1 | 2 | 23.6667 | 0.889 |
| 0.2 | 0.05 | ptsI | 2 | 23.9167 | 0.773 |
| 0.2 | 0.05 | clcB | 2 | 23.9167 | 0.352 |
| 0.2 | 0.05 | ycaM | 2 | 23.9167 | 0.327 |
| 0.2 | 0.05 | yadI | 2 | 23.9167 | 0.822 |
| 0.2 | 0.05 | AG1 | 2 | 23.9167 | 0.899 |
| 0.2 | 0.05 | ptsI | 2 | 24.1667 | 0.77 |
| 0.2 | 0.05 | clcB | 2 | 24.1667 | 0.352 |
| 0.2 | 0.05 | ycaM | 2 | 24.1667 | 0.327 |
| 0.2 | 0.05 | yadI | 2 | 24.1667 | 0.824 |
| 0.2 | 0.05 | AG1 | 2 | 24.1667 | 0.907 |
| 0.2 | 0.05 | ptsI | 2 | 24.4167 | 0.764 |
| 0.2 | 0.05 | clcB | 2 | 24.4167 | 0.353 |
| 0.2 | 0.05 | ycaM | 2 | 24.4167 | 0.329 |
| 0.2 | 0.05 | yadI | 2 | 24.4167 | 0.825 |
| 0.2 | 0.05 | AG1 | 2 | 24.4167 | 0.912 |
| 0.2 | 0.05 | ptsI | 2 | 24.6667 | 0.763 |
| 0.2 | 0.05 | clcB | 2 | 24.6667 | 0.355 |
| 0.2 | 0.05 | ycaM | 2 | 24.6667 | 0.332 |
| 0.2 | 0.05 | yadI | 2 | 24.6667 | 0.826 |
| 0.2 | 0.05 | AG1 | 2 | 24.6667 | 0.917 |
| 0.2 | 0.05 | ptsI | 2 | 24.9167 | 0.764 |
| 0.2 | 0.05 | clcB | 2 | 24.9167 | 0.354 |
| 0.2 | 0.05 | ycaM | 2 | 24.9167 | 0.333 |
| 0.2 | 0.05 | yadI | 2 | 24.9167 | 0.829 |
| 0.2 | 0.05 | AG1 | 2 | 24.9167 | 0.925 |
| 0.2 | 0.05 | ptsI | 2 | 25.1667 | 0.764 |
| 0.2 | 0.05 | clcB | 2 | 25.1667 | 0.355 |
| 0.2 | 0.05 | ycaM | 2 | 25.1667 | 0.339 |
| 0.2 | 0.05 | yadI | 2 | 25.1667 | 0.837 |
| 0.2 | 0.05 | AG1 | 2 | 25.1667 | 0.932 |
| 0.2 | 0.05 | ptsI | 2 | 25.4167 | 0.762 |
| 0.2 | 0.05 | clcB | 2 | 25.4167 | 0.36 |
| 0.2 | 0.05 | ycaM | 2 | 25.4167 | 0.343 |
| 0.2 | 0.05 | yadI | 2 | 25.4167 | 0.833 |
| 0.2 | 0.05 | AG1 | 2 | 25.4167 | 0.941 |
| 0.2 | 0.05 | ptsI | 2 | 25.6667 | 0.756 |
| 0.2 | 0.05 | clcB | 2 | 25.6667 | 0.359 |
| 0.2 | 0.05 | ycaM | 2 | 25.6667 | 0.343 |
| 0.2 | 0.05 | yadI | 2 | 25.6667 | 0.84 |
| 0.2 | 0.05 | AG1 | 2 | 25.6667 | 0.95 |
| 0.2 | 0.05 | ptsI | 2 | 25.9167 | 0.759 |
| 0.2 | 0.05 | clcB | 2 | 25.9167 | 0.361 |

| | | | | | |
|---|---|---|---|---|---|
| 0.2 | 0.05 | ycaM | 2 | 25.9167 | 0.348 |
| 0.2 | 0.05 | yadI | 2 | 25.9167 | 0.845 |
| 0.2 | 0.05 | AG1 | 2 | 25.9167 | 0.959 |
| 0.2 | 0.05 | ptsI | 2 | 26.1667 | 0.734 |
| 0.2 | 0.05 | clcB | 2 | 26.1667 | 0.357 |
| 0.2 | 0.05 | ycaM | 2 | 26.1667 | 0.347 |
| 0.2 | 0.05 | yadI | 2 | 26.1667 | 0.846 |
| 0.2 | 0.05 | AG1 | 2 | 26.1667 | 0.972 |
| 0.2 | 0.05 | ptsI | 2 | 26.4167 | 0.752 |
| 0.2 | 0.05 | clcB | 2 | 26.4167 | 0.362 |
| 0.2 | 0.05 | ycaM | 2 | 26.4167 | 0.35 |
| 0.2 | 0.05 | yadI | 2 | 26.4167 | 0.844 |
| 0.2 | 0.05 | AG1 | 2 | 26.4167 | 0.984 |
| 0.2 | 0.05 | ptsI | 3 | 0 | 0.161 |
| 0.2 | 0.05 | clcB | 3 | 0 | 0.163 |
| 0.2 | 0.05 | ycaM | 3 | 0 | 0.155 |
| 0.2 | 0.05 | yadI | 3 | 0 | 0.168 |
| 0.2 | 0.05 | AG1 | 3 | 0 | 0.163 |
| 0.2 | 0.05 | ptsI | 3 | 0.25 | 0.153 |
| 0.2 | 0.05 | clcB | 3 | 0.25 | 0.161 |
| 0.2 | 0.05 | ycaM | 3 | 0.25 | 0.152 |
| 0.2 | 0.05 | yadI | 3 | 0.25 | 0.164 |
| 0.2 | 0.05 | AG1 | 3 | 0.25 | 0.159 |
| 0.2 | 0.05 | ptsI | 3 | 0.5 | 0.152 |
| 0.2 | 0.05 | clcB | 3 | 0.5 | 0.159 |
| 0.2 | 0.05 | ycaM | 3 | 0.5 | 0.151 |
| 0.2 | 0.05 | yadI | 3 | 0.5 | 0.164 |
| 0.2 | 0.05 | AG1 | 3 | 0.5 | 0.156 |
| 0.2 | 0.05 | ptsI | 3 | 0.75 | 0.152 |
| 0.2 | 0.05 | clcB | 3 | 0.75 | 0.159 |
| 0.2 | 0.05 | ycaM | 3 | 0.75 | 0.151 |
| 0.2 | 0.05 | yadI | 3 | 0.75 | 0.164 |
| 0.2 | 0.05 | AG1 | 3 | 0.75 | 0.156 |
| 0.2 | 0.05 | ptsI | 3 | 1 | 0.152 |
| 0.2 | 0.05 | clcB | 3 | 1 | 0.16 |
| 0.2 | 0.05 | ycaM | 3 | 1 | 0.152 |
| 0.2 | 0.05 | yadI | 3 | 1 | 0.164 |
| 0.2 | 0.05 | AG1 | 3 | 1 | 0.158 |
| 0.2 | 0.05 | ptsI | 3 | 1.25 | 0.152 |
| 0.2 | 0.05 | clcB | 3 | 1.25 | 0.16 |
| 0.2 | 0.05 | ycaM | 3 | 1.25 | 0.152 |
| 0.2 | 0.05 | yadI | 3 | 1.25 | 0.164 |
| 0.2 | 0.05 | AG1 | 3 | 1.25 | 0.157 |
| 0.2 | 0.05 | ptsI | 3 | 1.5 | 0.154 |
| 0.2 | 0.05 | clcB | 3 | 1.5 | 0.161 |
| 0.2 | 0.05 | ycaM | 3 | 1.5 | 0.153 |
| 0.2 | 0.05 | yadI | 3 | 1.5 | 0.167 |
| 0.2 | 0.05 | AG1 | 3 | 1.5 | 0.158 |

| | | | | | |
|---|---|---|---|---|---|
| 0.2 | 0.05 | ptsI | 3 | 1.75 | 0.154 |
| 0.2 | 0.05 | clcB | 3 | 1.75 | 0.161 |
| 0.2 | 0.05 | ycaM | 3 | 1.75 | 0.153 |
| 0.2 | 0.05 | yadI | 3 | 1.75 | 0.167 |
| 0.2 | 0.05 | AG1 | 3 | 1.75 | 0.158 |
| 0.2 | 0.05 | ptsI | 3 | 2 | 0.156 |
| 0.2 | 0.05 | clcB | 3 | 2 | 0.161 |
| 0.2 | 0.05 | ycaM | 3 | 2 | 0.154 |
| 0.2 | 0.05 | yadI | 3 | 2 | 0.168 |
| 0.2 | 0.05 | AG1 | 3 | 2 | 0.159 |
| 0.2 | 0.05 | ptsI | 3 | 2.25 | 0.158 |
| 0.2 | 0.05 | clcB | 3 | 2.25 | 0.164 |
| 0.2 | 0.05 | ycaM | 3 | 2.25 | 0.156 |
| 0.2 | 0.05 | yadI | 3 | 2.25 | 0.17 |
| 0.2 | 0.05 | AG1 | 3 | 2.25 | 0.159 |
| 0.2 | 0.05 | ptsI | 3 | 2.5 | 0.16 |
| 0.2 | 0.05 | clcB | 3 | 2.5 | 0.164 |
| 0.2 | 0.05 | ycaM | 3 | 2.5 | 0.156 |
| 0.2 | 0.05 | yadI | 3 | 2.5 | 0.172 |
| 0.2 | 0.05 | AG1 | 3 | 2.5 | 0.162 |
| 0.2 | 0.05 | ptsI | 3 | 2.75 | 0.164 |
| 0.2 | 0.05 | clcB | 3 | 2.75 | 0.167 |
| 0.2 | 0.05 | ycaM | 3 | 2.75 | 0.159 |
| 0.2 | 0.05 | yadI | 3 | 2.75 | 0.177 |
| 0.2 | 0.05 | AG1 | 3 | 2.75 | 0.165 |
| 0.2 | 0.05 | ptsI | 3 | 3 | 0.164 |
| 0.2 | 0.05 | clcB | 3 | 3 | 0.167 |
| 0.2 | 0.05 | ycaM | 3 | 3 | 0.16 |
| 0.2 | 0.05 | yadI | 3 | 3 | 0.181 |
| 0.2 | 0.05 | AG1 | 3 | 3 | 0.164 |
| 0.2 | 0.05 | ptsI | 3 | 3.25 | 0.166 |
| 0.2 | 0.05 | clcB | 3 | 3.25 | 0.168 |
| 0.2 | 0.05 | ycaM | 3 | 3.25 | 0.162 |
| 0.2 | 0.05 | yadI | 3 | 3.25 | 0.185 |
| 0.2 | 0.05 | AG1 | 3 | 3.25 | 0.167 |
| 0.2 | 0.05 | ptsI | 3 | 3.5 | 0.17 |
| 0.2 | 0.05 | clcB | 3 | 3.5 | 0.172 |
| 0.2 | 0.05 | ycaM | 3 | 3.5 | 0.164 |
| 0.2 | 0.05 | yadI | 3 | 3.5 | 0.19 |
| 0.2 | 0.05 | AG1 | 3 | 3.5 | 0.17 |
| 0.2 | 0.05 | ptsI | 3 | 3.75 | 0.172 |
| 0.2 | 0.05 | clcB | 3 | 3.75 | 0.173 |
| 0.2 | 0.05 | ycaM | 3 | 3.75 | 0.167 |
| 0.2 | 0.05 | yadI | 3 | 3.75 | 0.194 |
| 0.2 | 0.05 | AG1 | 3 | 3.75 | 0.172 |
| 0.2 | 0.05 | ptsI | 3 | 4 | 0.175 |
| 0.2 | 0.05 | clcB | 3 | 4 | 0.176 |
| 0.2 | 0.05 | ycaM | 3 | 4 | 0.169 |

| | | | | | |
|---|---|---|---|---|---|
| 0.2 | 0.05 | yadI | 3 | 4 | 0.199 |
| 0.2 | 0.05 | AG1 | 3 | 4 | 0.173 |
| 0.2 | 0.05 | ptsI | 3 | 4.41667 | 0.208 |
| 0.2 | 0.05 | clcB | 3 | 4.41667 | 0.223 |
| 0.2 | 0.05 | ycaM | 3 | 4.41667 | 0.183 |
| 0.2 | 0.05 | yadI | 3 | 4.41667 | 0.235 |
| 0.2 | 0.05 | AG1 | 3 | 4.41667 | 0.187 |
| 0.2 | 0.05 | ptsI | 3 | 4.66667 | 0.193 |
| 0.2 | 0.05 | clcB | 3 | 4.66667 | 0.182 |
| 0.2 | 0.05 | ycaM | 3 | 4.66667 | 0.177 |
| 0.2 | 0.05 | yadI | 3 | 4.66667 | 0.236 |
| 0.2 | 0.05 | AG1 | 3 | 4.66667 | 0.196 |
| 0.2 | 0.05 | ptsI | 3 | 4.91667 | 0.191 |
| 0.2 | 0.05 | clcB | 3 | 4.91667 | 0.183 |
| 0.2 | 0.05 | ycaM | 3 | 4.91667 | 0.18 |
| 0.2 | 0.05 | yadI | 3 | 4.91667 | 0.249 |
| 0.2 | 0.05 | AG1 | 3 | 4.91667 | 0.206 |
| 0.2 | 0.05 | ptsI | 3 | 5.16667 | 0.196 |
| 0.2 | 0.05 | clcB | 3 | 5.16667 | 0.186 |
| 0.2 | 0.05 | ycaM | 3 | 5.16667 | 0.184 |
| 0.2 | 0.05 | yadI | 3 | 5.16667 | 0.264 |
| 0.2 | 0.05 | AG1 | 3 | 5.16667 | 0.212 |
| 0.2 | 0.05 | ptsI | 3 | 5.41667 | 0.2 |
| 0.2 | 0.05 | clcB | 3 | 5.41667 | 0.189 |
| 0.2 | 0.05 | ycaM | 3 | 5.41667 | 0.19 |
| 0.2 | 0.05 | yadI | 3 | 5.41667 | 0.282 |
| 0.2 | 0.05 | AG1 | 3 | 5.41667 | 0.214 |
| 0.2 | 0.05 | ptsI | 3 | 5.66667 | 0.205 |
| 0.2 | 0.05 | clcB | 3 | 5.66667 | 0.195 |
| 0.2 | 0.05 | ycaM | 3 | 5.66667 | 0.196 |
| 0.2 | 0.05 | yadI | 3 | 5.66667 | 0.301 |
| 0.2 | 0.05 | AG1 | 3 | 5.66667 | 0.221 |
| 0.2 | 0.05 | ptsI | 3 | 5.91667 | 0.212 |
| 0.2 | 0.05 | clcB | 3 | 5.91667 | 0.201 |
| 0.2 | 0.05 | ycaM | 3 | 5.91667 | 0.203 |
| 0.2 | 0.05 | yadI | 3 | 5.91667 | 0.321 |
| 0.2 | 0.05 | AG1 | 3 | 5.91667 | 0.222 |
| 0.2 | 0.05 | ptsI | 3 | 6.16667 | 0.218 |
| 0.2 | 0.05 | clcB | 3 | 6.16667 | 0.21 |
| 0.2 | 0.05 | ycaM | 3 | 6.16667 | 0.212 |
| 0.2 | 0.05 | yadI | 3 | 6.16667 | 0.336 |
| 0.2 | 0.05 | AG1 | 3 | 6.16667 | 0.23 |
| 0.2 | 0.05 | ptsI | 3 | 6.41667 | 0.228 |
| 0.2 | 0.05 | clcB | 3 | 6.41667 | 0.217 |
| 0.2 | 0.05 | ycaM | 3 | 6.41667 | 0.219 |
| 0.2 | 0.05 | yadI | 3 | 6.41667 | 0.345 |
| 0.2 | 0.05 | AG1 | 3 | 6.41667 | 0.239 |
| 0.2 | 0.05 | ptsI | 3 | 6.66667 | 0.238 |

| | | | | | |
|---|---|---|---|---|---|
| 0.2 | 0.05 | clcB | 3 | 6.66667 | 0.227 |
| 0.2 | 0.05 | ycaM | 3 | 6.66667 | 0.227 |
| 0.2 | 0.05 | yadI | 3 | 6.66667 | 0.357 |
| 0.2 | 0.05 | AG1 | 3 | 6.66667 | 0.251 |
| 0.2 | 0.05 | ptsI | 3 | 6.91667 | 0.249 |
| 0.2 | 0.05 | clcB | 3 | 6.91667 | 0.236 |
| 0.2 | 0.05 | ycaM | 3 | 6.91667 | 0.235 |
| 0.2 | 0.05 | yadI | 3 | 6.91667 | 0.368 |
| 0.2 | 0.05 | AG1 | 3 | 6.91667 | 0.262 |
| 0.2 | 0.05 | ptsI | 3 | 7.16667 | 0.258 |
| 0.2 | 0.05 | clcB | 3 | 7.16667 | 0.247 |
| 0.2 | 0.05 | ycaM | 3 | 7.16667 | 0.241 |
| 0.2 | 0.05 | yadI | 3 | 7.16667 | 0.381 |
| 0.2 | 0.05 | AG1 | 3 | 7.16667 | 0.271 |
| 0.2 | 0.05 | ptsI | 3 | 7.41667 | 0.268 |
| 0.2 | 0.05 | clcB | 3 | 7.41667 | 0.258 |
| 0.2 | 0.05 | ycaM | 3 | 7.41667 | 0.245 |
| 0.2 | 0.05 | yadI | 3 | 7.41667 | 0.393 |
| 0.2 | 0.05 | AG1 | 3 | 7.41667 | 0.279 |
| 0.2 | 0.05 | ptsI | 3 | 7.66667 | 0.278 |
| 0.2 | 0.05 | clcB | 3 | 7.66667 | 0.267 |
| 0.2 | 0.05 | ycaM | 3 | 7.66667 | 0.249 |
| 0.2 | 0.05 | yadI | 3 | 7.66667 | 0.405 |
| 0.2 | 0.05 | AG1 | 3 | 7.66667 | 0.287 |
| 0.2 | 0.05 | ptsI | 3 | 7.91667 | 0.287 |
| 0.2 | 0.05 | clcB | 3 | 7.91667 | 0.272 |
| 0.2 | 0.05 | ycaM | 3 | 7.91667 | 0.249 |
| 0.2 | 0.05 | yadI | 3 | 7.91667 | 0.416 |
| 0.2 | 0.05 | AG1 | 3 | 7.91667 | 0.295 |
| 0.2 | 0.05 | ptsI | 3 | 8.16667 | 0.3 |
| 0.2 | 0.05 | clcB | 3 | 8.16667 | 0.28 |
| 0.2 | 0.05 | ycaM | 3 | 8.16667 | 0.252 |
| 0.2 | 0.05 | yadI | 3 | 8.16667 | 0.426 |
| 0.2 | 0.05 | AG1 | 3 | 8.16667 | 0.302 |
| 0.2 | 0.05 | ptsI | 3 | 8.41667 | 0.31 |
| 0.2 | 0.05 | clcB | 3 | 8.41667 | 0.285 |
| 0.2 | 0.05 | ycaM | 3 | 8.41667 | 0.254 |
| 0.2 | 0.05 | yadI | 3 | 8.41667 | 0.437 |
| 0.2 | 0.05 | AG1 | 3 | 8.41667 | 0.31 |
| 0.2 | 0.05 | ptsI | 3 | 9.05 | 0.29 |
| 0.2 | 0.05 | clcB | 3 | 9.05 | 0.273 |
| 0.2 | 0.05 | ycaM | 3 | 9.05 | 0.252 |
| 0.2 | 0.05 | yadI | 3 | 9.05 | 0.443 |
| 0.2 | 0.05 | AG1 | 3 | 9.05 | 0.372 |
| 0.2 | 0.05 | ptsI | 3 | 9.3 | 0.294 |
| 0.2 | 0.05 | clcB | 3 | 9.3 | 0.275 |
| 0.2 | 0.05 | ycaM | 3 | 9.3 | 0.249 |
| 0.2 | 0.05 | yadI | 3 | 9.3 | 0.45 |

| | | | | | |
|---|---|---|---|---|---|
| 0.2 | 0.05 | AG1 | 3 | 9.3 | 0.382 |
| 0.2 | 0.05 | ptsI | 3 | 9.55 | 0.302 |
| 0.2 | 0.05 | clcB | 3 | 9.55 | 0.282 |
| 0.2 | 0.05 | ycaM | 3 | 9.55 | 0.252 |
| 0.2 | 0.05 | yadI | 3 | 9.55 | 0.467 |
| 0.2 | 0.05 | AG1 | 3 | 9.55 | 0.394 |
| 0.2 | 0.05 | ptsI | 3 | 9.8 | 0.31 |
| 0.2 | 0.05 | clcB | 3 | 9.8 | 0.289 |
| 0.2 | 0.05 | ycaM | 3 | 9.8 | 0.254 |
| 0.2 | 0.05 | yadI | 3 | 9.8 | 0.48 |
| 0.2 | 0.05 | AG1 | 3 | 9.8 | 0.406 |
| 0.2 | 0.05 | ptsI | 3 | 10.05 | 0.32 |
| 0.2 | 0.05 | clcB | 3 | 10.05 | 0.296 |
| 0.2 | 0.05 | ycaM | 3 | 10.05 | 0.256 |
| 0.2 | 0.05 | yadI | 3 | 10.05 | 0.494 |
| 0.2 | 0.05 | AG1 | 3 | 10.05 | 0.418 |
| 0.2 | 0.05 | ptsI | 3 | 10.3 | 0.325 |
| 0.2 | 0.05 | clcB | 3 | 10.3 | 0.299 |
| 0.2 | 0.05 | ycaM | 3 | 10.3 | 0.256 |
| 0.2 | 0.05 | yadI | 3 | 10.3 | 0.507 |
| 0.2 | 0.05 | AG1 | 3 | 10.3 | 0.429 |
| 0.2 | 0.05 | ptsI | 3 | 10.55 | 0.336 |
| 0.2 | 0.05 | clcB | 3 | 10.55 | 0.304 |
| 0.2 | 0.05 | ycaM | 3 | 10.55 | 0.257 |
| 0.2 | 0.05 | yadI | 3 | 10.55 | 0.518 |
| 0.2 | 0.05 | AG1 | 3 | 10.55 | 0.439 |
| 0.2 | 0.05 | ptsI | 3 | 10.8 | 0.343 |
| 0.2 | 0.05 | clcB | 3 | 10.8 | 0.301 |
| 0.2 | 0.05 | ycaM | 3 | 10.8 | 0.259 |
| 0.2 | 0.05 | yadI | 3 | 10.8 | 0.53 |
| 0.2 | 0.05 | AG1 | 3 | 10.8 | 0.446 |
| 0.2 | 0.05 | ptsI | 3 | 11.05 | 0.355 |
| 0.2 | 0.05 | clcB | 3 | 11.05 | 0.309 |
| 0.2 | 0.05 | ycaM | 3 | 11.05 | 0.262 |
| 0.2 | 0.05 | yadI | 3 | 11.05 | 0.544 |
| 0.2 | 0.05 | AG1 | 3 | 11.05 | 0.458 |
| 0.2 | 0.05 | ptsI | 3 | 11.3 | 0.362 |
| 0.2 | 0.05 | clcB | 3 | 11.3 | 0.307 |
| 0.2 | 0.05 | ycaM | 3 | 11.3 | 0.263 |
| 0.2 | 0.05 | yadI | 3 | 11.3 | 0.553 |
| 0.2 | 0.05 | AG1 | 3 | 11.3 | 0.468 |
| 0.2 | 0.05 | ptsI | 3 | 11.55 | 0.373 |
| 0.2 | 0.05 | clcB | 3 | 11.55 | 0.308 |
| 0.2 | 0.05 | ycaM | 3 | 11.55 | 0.262 |
| 0.2 | 0.05 | yadI | 3 | 11.55 | 0.566 |
| 0.2 | 0.05 | AG1 | 3 | 11.55 | 0.479 |
| 0.2 | 0.05 | ptsI | 3 | 11.8 | 0.383 |
| 0.2 | 0.05 | clcB | 3 | 11.8 | 0.317 |

| | | | | | |
|---|---|---|---|---|---|
| 0.2 | 0.05 | ycaM | 3 | 11.8 | 0.266 |
| 0.2 | 0.05 | yadI | 3 | 11.8 | 0.578 |
| 0.2 | 0.05 | AG1 | 3 | 11.8 | 0.493 |
| 0.2 | 0.05 | ptsI | 3 | 12.05 | 0.394 |
| 0.2 | 0.05 | clcB | 3 | 12.05 | 0.31 |
| 0.2 | 0.05 | ycaM | 3 | 12.05 | 0.261 |
| 0.2 | 0.05 | yadI | 3 | 12.05 | 0.589 |
| 0.2 | 0.05 | AG1 | 3 | 12.05 | 0.504 |
| 0.2 | 0.05 | ptsI | 3 | 12.3 | 0.405 |
| 0.2 | 0.05 | clcB | 3 | 12.3 | 0.315 |
| 0.2 | 0.05 | ycaM | 3 | 12.3 | 0.27 |
| 0.2 | 0.05 | yadI | 3 | 12.3 | 0.602 |
| 0.2 | 0.05 | AG1 | 3 | 12.3 | 0.519 |
| 0.2 | 0.05 | ptsI | 3 | 12.55 | 0.413 |
| 0.2 | 0.05 | clcB | 3 | 12.55 | 0.313 |
| 0.2 | 0.05 | ycaM | 3 | 12.55 | 0.268 |
| 0.2 | 0.05 | yadI | 3 | 12.55 | 0.615 |
| 0.2 | 0.05 | AG1 | 3 | 12.55 | 0.538 |
| 0.2 | 0.05 | ptsI | 3 | 12.8 | 0.423 |
| 0.2 | 0.05 | clcB | 3 | 12.8 | 0.316 |
| 0.2 | 0.05 | ycaM | 3 | 12.8 | 0.269 |
| 0.2 | 0.05 | yadI | 3 | 12.8 | 0.626 |
| 0.2 | 0.05 | AG1 | 3 | 12.8 | 0.557 |
| 0.2 | 0.05 | ptsI | 3 | 13.05 | 0.43 |
| 0.2 | 0.05 | clcB | 3 | 13.05 | 0.302 |
| 0.2 | 0.05 | ycaM | 3 | 13.05 | 0.264 |
| 0.2 | 0.05 | yadI | 3 | 13.05 | 0.633 |
| 0.2 | 0.05 | AG1 | 3 | 13.05 | 0.565 |
| 0.2 | 0.05 | ptsI | 3 | 13.3 | 0.438 |
| 0.2 | 0.05 | clcB | 3 | 13.3 | 0.301 |
| 0.2 | 0.05 | ycaM | 3 | 13.3 | 0.266 |
| 0.2 | 0.05 | yadI | 3 | 13.3 | 0.644 |
| 0.2 | 0.05 | AG1 | 3 | 13.3 | 0.583 |
| 0.2 | 0.05 | ptsI | 3 | 13.55 | 0.45 |
| 0.2 | 0.05 | clcB | 3 | 13.55 | 0.309 |
| 0.2 | 0.05 | ycaM | 3 | 13.55 | 0.268 |
| 0.2 | 0.05 | yadI | 3 | 13.55 | 0.659 |
| 0.2 | 0.05 | AG1 | 3 | 13.55 | 0.601 |
| 0.2 | 0.05 | ptsI | 3 | 13.8 | 0.461 |
| 0.2 | 0.05 | clcB | 3 | 13.8 | 0.304 |
| 0.2 | 0.05 | ycaM | 3 | 13.8 | 0.269 |
| 0.2 | 0.05 | yadI | 3 | 13.8 | 0.665 |
| 0.2 | 0.05 | AG1 | 3 | 13.8 | 0.611 |
| 0.2 | 0.05 | ptsI | 3 | 14.05 | 0.465 |
| 0.2 | 0.05 | clcB | 3 | 14.05 | 0.316 |
| 0.2 | 0.05 | ycaM | 3 | 14.05 | 0.27 |
| 0.2 | 0.05 | yadI | 3 | 14.05 | 0.672 |
| 0.2 | 0.05 | AG1 | 3 | 14.05 | 0.62 |

| | | | | | |
|---|---|---|---|---|---|
| 0.2 | 0.05 | ptsI | 3 | 14.3 | 0.469 |
| 0.2 | 0.05 | clcB | 3 | 14.3 | 0.299 |
| 0.2 | 0.05 | ycaM | 3 | 14.3 | 0.265 |
| 0.2 | 0.05 | yadI | 3 | 14.3 | 0.679 |
| 0.2 | 0.05 | AG1 | 3 | 14.3 | 0.63 |
| 0.2 | 0.05 | ptsI | 3 | 14.55 | 0.473 |
| 0.2 | 0.05 | clcB | 3 | 14.55 | 0.298 |
| 0.2 | 0.05 | ycaM | 3 | 14.55 | 0.262 |
| 0.2 | 0.05 | yadI | 3 | 14.55 | 0.688 |
| 0.2 | 0.05 | AG1 | 3 | 14.55 | 0.642 |
| 0.2 | 0.05 | ptsI | 3 | 14.8 | 0.489 |
| 0.2 | 0.05 | clcB | 3 | 14.8 | 0.295 |
| 0.2 | 0.05 | ycaM | 3 | 14.8 | 0.265 |
| 0.2 | 0.05 | yadI | 3 | 14.8 | 0.703 |
| 0.2 | 0.05 | AG1 | 3 | 14.8 | 0.656 |
| 0.2 | 0.05 | ptsI | 3 | 15.05 | 0.503 |
| 0.2 | 0.05 | clcB | 3 | 15.05 | 0.295 |
| 0.2 | 0.05 | ycaM | 3 | 15.05 | 0.266 |
| 0.2 | 0.05 | yadI | 3 | 15.05 | 0.717 |
| 0.2 | 0.05 | AG1 | 3 | 15.05 | 0.663 |
| 0.2 | 0.05 | ptsI | 3 | 15.3 | 0.501 |
| 0.2 | 0.05 | clcB | 3 | 15.3 | 0.296 |
| 0.2 | 0.05 | ycaM | 3 | 15.3 | 0.262 |
| 0.2 | 0.05 | yadI | 3 | 15.3 | 0.717 |
| 0.2 | 0.05 | AG1 | 3 | 15.3 | 0.682 |
| 0.2 | 0.05 | ptsI | 3 | 15.55 | 0.506 |
| 0.2 | 0.05 | clcB | 3 | 15.55 | 0.298 |
| 0.2 | 0.05 | ycaM | 3 | 15.55 | 0.261 |
| 0.2 | 0.05 | yadI | 3 | 15.55 | 0.72 |
| 0.2 | 0.05 | AG1 | 3 | 15.55 | 0.686 |
| 0.2 | 0.05 | ptsI | 3 | 15.8 | 0.512 |
| 0.2 | 0.05 | clcB | 3 | 15.8 | 0.298 |
| 0.2 | 0.05 | ycaM | 3 | 15.8 | 0.258 |
| 0.2 | 0.05 | yadI | 3 | 15.8 | 0.73 |
| 0.2 | 0.05 | AG1 | 3 | 15.8 | 0.697 |
| 0.2 | 0.05 | ptsI | 3 | 16.05 | 0.519 |
| 0.2 | 0.05 | clcB | 3 | 16.05 | 0.298 |
| 0.2 | 0.05 | ycaM | 3 | 16.05 | 0.257 |
| 0.2 | 0.05 | yadI | 3 | 16.05 | 0.733 |
| 0.2 | 0.05 | AG1 | 3 | 16.05 | 0.693 |
| 0.2 | 0.05 | ptsI | 3 | 16.3 | 0.522 |
| 0.2 | 0.05 | clcB | 3 | 16.3 | 0.302 |
| 0.2 | 0.05 | ycaM | 3 | 16.3 | 0.255 |
| 0.2 | 0.05 | yadI | 3 | 16.3 | 0.735 |
| 0.2 | 0.05 | AG1 | 3 | 16.3 | 0.696 |
| 0.2 | 0.05 | ptsI | 3 | 16.55 | 0.534 |
| 0.2 | 0.05 | clcB | 3 | 16.55 | 0.306 |
| 0.2 | 0.05 | ycaM | 3 | 16.55 | 0.259 |

| | | | | | |
|---|---|---|---|---|---|
| 0.2 | 0.05 | yadI | 3 | 16.55 | 0.748 |
| 0.2 | 0.05 | AG1 | 3 | 16.55 | 0.711 |
| 0.2 | 0.05 | ptsI | 3 | 16.8 | 0.542 |
| 0.2 | 0.05 | clcB | 3 | 16.8 | 0.303 |
| 0.2 | 0.05 | ycaM | 3 | 16.8 | 0.256 |
| 0.2 | 0.05 | yadI | 3 | 16.8 | 0.759 |
| 0.2 | 0.05 | AG1 | 3 | 16.8 | 0.734 |
| 0.2 | 0.05 | ptsI | 3 | 17.05 | 0.544 |
| 0.2 | 0.05 | clcB | 3 | 17.05 | 0.305 |
| 0.2 | 0.05 | ycaM | 3 | 17.05 | 0.256 |
| 0.2 | 0.05 | yadI | 3 | 17.05 | 0.762 |
| 0.2 | 0.05 | AG1 | 3 | 17.05 | 0.727 |
| 0.2 | 0.05 | ptsI | 3 | 17.3 | 0.544 |
| 0.2 | 0.05 | clcB | 3 | 17.3 | 0.304 |
| 0.2 | 0.05 | ycaM | 3 | 17.3 | 0.257 |
| 0.2 | 0.05 | yadI | 3 | 17.3 | 0.77 |
| 0.2 | 0.05 | AG1 | 3 | 17.3 | 0.733 |
| 0.2 | 0.05 | ptsI | 3 | 17.55 | 0.554 |
| 0.2 | 0.05 | clcB | 3 | 17.55 | 0.303 |
| 0.2 | 0.05 | ycaM | 3 | 17.55 | 0.256 |
| 0.2 | 0.05 | yadI | 3 | 17.55 | 0.785 |
| 0.2 | 0.05 | AG1 | 3 | 17.55 | 0.743 |
| 0.2 | 0.05 | ptsI | 3 | 17.8 | 0.559 |
| 0.2 | 0.05 | clcB | 3 | 17.8 | 0.302 |
| 0.2 | 0.05 | ycaM | 3 | 17.8 | 0.259 |
| 0.2 | 0.05 | yadI | 3 | 17.8 | 0.79 |
| 0.2 | 0.05 | AG1 | 3 | 17.8 | 0.753 |
| 0.2 | 0.05 | ptsI | 3 | 18.05 | 0.56 |
| 0.2 | 0.05 | clcB | 3 | 18.05 | 0.3 |
| 0.2 | 0.05 | ycaM | 3 | 18.05 | 0.26 |
| 0.2 | 0.05 | yadI | 3 | 18.05 | 0.791 |
| 0.2 | 0.05 | AG1 | 3 | 18.05 | 0.77 |
| 0.2 | 0.05 | ptsI | 3 | 18.3 | 0.562 |
| 0.2 | 0.05 | clcB | 3 | 18.3 | 0.3 |
| 0.2 | 0.05 | ycaM | 3 | 18.3 | 0.26 |
| 0.2 | 0.05 | yadI | 3 | 18.3 | 0.796 |
| 0.2 | 0.05 | AG1 | 3 | 18.3 | 0.766 |
| 0.2 | 0.05 | ptsI | 3 | 18.55 | 0.563 |
| 0.2 | 0.05 | clcB | 3 | 18.55 | 0.3 |
| 0.2 | 0.05 | ycaM | 3 | 18.55 | 0.262 |
| 0.2 | 0.05 | yadI | 3 | 18.55 | 0.798 |
| 0.2 | 0.05 | AG1 | 3 | 18.55 | 0.757 |
| 0.2 | 0.05 | ptsI | 3 | 18.8 | 0.565 |
| 0.2 | 0.05 | clcB | 3 | 18.8 | 0.298 |
| 0.2 | 0.05 | ycaM | 3 | 18.8 | 0.263 |
| 0.2 | 0.05 | yadI | 3 | 18.8 | 0.795 |
| 0.2 | 0.05 | AG1 | 3 | 18.8 | 0.76 |
| 0.2 | 0.05 | ptsI | 3 | 19.05 | 0.566 |

| | | | | | |
|---|---|---|---|---|---|
| 0.2 | 0.05 | clcB | 3 | 19.05 | 0.297 |
| 0.2 | 0.05 | ycaM | 3 | 19.05 | 0.264 |
| 0.2 | 0.05 | yadI | 3 | 19.05 | 0.802 |
| 0.2 | 0.05 | AG1 | 3 | 19.05 | 0.774 |
| 0.2 | 0.05 | ptsI | 3 | 19.3 | 0.567 |
| 0.2 | 0.05 | clcB | 3 | 19.3 | 0.298 |
| 0.2 | 0.05 | ycaM | 3 | 19.3 | 0.264 |
| 0.2 | 0.05 | yadI | 3 | 19.3 | 0.803 |
| 0.2 | 0.05 | AG1 | 3 | 19.3 | 0.77 |
| 0.2 | 0.05 | ptsI | 3 | 19.55 | 0.566 |
| 0.2 | 0.05 | clcB | 3 | 19.55 | 0.299 |
| 0.2 | 0.05 | ycaM | 3 | 19.55 | 0.265 |
| 0.2 | 0.05 | yadI | 3 | 19.55 | 0.81 |
| 0.2 | 0.05 | AG1 | 3 | 19.55 | 0.778 |
| 0.2 | 0.05 | ptsI | 3 | 19.8 | 0.568 |
| 0.2 | 0.05 | clcB | 3 | 19.8 | 0.299 |
| 0.2 | 0.05 | ycaM | 3 | 19.8 | 0.266 |
| 0.2 | 0.05 | yadI | 3 | 19.8 | 0.806 |
| 0.2 | 0.05 | AG1 | 3 | 19.8 | 0.768 |
| 0.2 | 0.05 | ptsI | 3 | 20.05 | 0.569 |
| 0.2 | 0.05 | clcB | 3 | 20.05 | 0.299 |
| 0.2 | 0.05 | ycaM | 3 | 20.05 | 0.268 |
| 0.2 | 0.05 | yadI | 3 | 20.05 | 0.814 |
| 0.2 | 0.05 | AG1 | 3 | 20.05 | 0.76 |
| 0.2 | 0.05 | ptsI | 3 | 20.3 | 0.568 |
| 0.2 | 0.05 | clcB | 3 | 20.3 | 0.298 |
| 0.2 | 0.05 | ycaM | 3 | 20.3 | 0.268 |
| 0.2 | 0.05 | yadI | 3 | 20.3 | 0.812 |
| 0.2 | 0.05 | AG1 | 3 | 20.3 | 0.754 |
| 0.2 | 0.05 | ptsI | 3 | 20.55 | 0.563 |
| 0.2 | 0.05 | clcB | 3 | 20.55 | 0.299 |
| 0.2 | 0.05 | ycaM | 3 | 20.55 | 0.269 |
| 0.2 | 0.05 | yadI | 3 | 20.55 | 0.81 |
| 0.2 | 0.05 | AG1 | 3 | 20.55 | 0.768 |
| 0.2 | 0.05 | ptsI | 3 | 20.8 | 0.566 |
| 0.2 | 0.05 | clcB | 3 | 20.8 | 0.298 |
| 0.2 | 0.05 | ycaM | 3 | 20.8 | 0.27 |
| 0.2 | 0.05 | yadI | 3 | 20.8 | 0.817 |
| 0.2 | 0.05 | AG1 | 3 | 20.8 | 0.777 |
| 0.2 | 0.05 | ptsI | 3 | 21.05 | 0.563 |
| 0.2 | 0.05 | clcB | 3 | 21.05 | 0.298 |
| 0.2 | 0.05 | ycaM | 3 | 21.05 | 0.271 |
| 0.2 | 0.05 | yadI | 3 | 21.05 | 0.813 |
| 0.2 | 0.05 | AG1 | 3 | 21.05 | 0.779 |
| 0.2 | 0.05 | ptsI | 3 | 21.3 | 0.561 |
| 0.2 | 0.05 | clcB | 3 | 21.3 | 0.298 |
| 0.2 | 0.05 | ycaM | 3 | 21.3 | 0.274 |
| 0.2 | 0.05 | yadI | 3 | 21.3 | 0.818 |

| | | | | | |
|---|---|---|---|---|---|
| 0.2 | 0.05 | AG1 | 3 | 21.3 | 0.784 |
| 0.2 | 0.05 | ptsI | 3 | 21.55 | 0.561 |
| 0.2 | 0.05 | clcB | 3 | 21.55 | 0.3 |
| 0.2 | 0.05 | ycaM | 3 | 21.55 | 0.276 |
| 0.2 | 0.05 | yadI | 3 | 21.55 | 0.817 |
| 0.2 | 0.05 | AG1 | 3 | 21.55 | 0.796 |
| 0.2 | 0.05 | ptsI | 3 | 21.8 | 0.561 |
| 0.2 | 0.05 | clcB | 3 | 21.8 | 0.299 |
| 0.2 | 0.05 | ycaM | 3 | 21.8 | 0.278 |
| 0.2 | 0.05 | yadI | 3 | 21.8 | 0.819 |
| 0.2 | 0.05 | AG1 | 3 | 21.8 | 0.804 |
| 0.2 | 0.05 | ptsI | 3 | 22.05 | 0.562 |
| 0.2 | 0.05 | clcB | 3 | 22.05 | 0.298 |
| 0.2 | 0.05 | ycaM | 3 | 22.05 | 0.278 |
| 0.2 | 0.05 | yadI | 3 | 22.05 | 0.822 |
| 0.2 | 0.05 | AG1 | 3 | 22.05 | 0.806 |
| 0.2 | 0.05 | ptsI | 3 | 22.3 | 0.558 |
| 0.2 | 0.05 | clcB | 3 | 22.3 | 0.298 |
| 0.2 | 0.05 | ycaM | 3 | 22.3 | 0.283 |
| 0.2 | 0.05 | yadI | 3 | 22.3 | 0.822 |
| 0.2 | 0.05 | AG1 | 3 | 22.3 | 0.814 |
| 0.2 | 0.05 | ptsI | 3 | 22.55 | 0.562 |
| 0.2 | 0.05 | clcB | 3 | 22.55 | 0.302 |
| 0.2 | 0.05 | ycaM | 3 | 22.55 | 0.284 |
| 0.2 | 0.05 | yadI | 3 | 22.55 | 0.828 |
| 0.2 | 0.05 | AG1 | 3 | 22.55 | 0.819 |
| 0.2 | 0.05 | ptsI | 3 | 22.8 | 0.564 |
| 0.2 | 0.05 | clcB | 3 | 22.8 | 0.302 |
| 0.2 | 0.05 | ycaM | 3 | 22.8 | 0.286 |
| 0.2 | 0.05 | yadI | 3 | 22.8 | 0.831 |
| 0.2 | 0.05 | AG1 | 3 | 22.8 | 0.829 |
| 0.2 | 0.05 | ptsI | 3 | 23.05 | 0.566 |
| 0.2 | 0.05 | clcB | 3 | 23.05 | 0.302 |
| 0.2 | 0.05 | ycaM | 3 | 23.05 | 0.291 |
| 0.2 | 0.05 | yadI | 3 | 23.05 | 0.833 |
| 0.2 | 0.05 | AG1 | 3 | 23.05 | 0.834 |
| 0.2 | 0.05 | ptsI | 3 | 23.3 | 0.565 |
| 0.2 | 0.05 | clcB | 3 | 23.3 | 0.301 |
| 0.2 | 0.05 | ycaM | 3 | 23.3 | 0.293 |
| 0.2 | 0.05 | yadI | 3 | 23.3 | 0.829 |
| 0.2 | 0.05 | AG1 | 3 | 23.3 | 0.839 |
| 0.2 | 0.05 | ptsI | 3 | 23.55 | 0.569 |
| 0.2 | 0.05 | clcB | 3 | 23.55 | 0.306 |
| 0.2 | 0.05 | ycaM | 3 | 23.55 | 0.297 |
| 0.2 | 0.05 | yadI | 3 | 23.55 | 0.834 |
| 0.2 | 0.05 | AG1 | 3 | 23.55 | 0.851 |
| 0.2 | 0.05 | ptsI | 3 | 23.8 | 0.564 |
| 0.2 | 0.05 | clcB | 3 | 23.8 | 0.307 |

| | | | | | |
|---|---|---|---|---|---|
| 0.2 | 0.05 | ycaM | 3 | 23.8 | 0.3 |
| 0.2 | 0.05 | yadI | 3 | 23.8 | 0.829 |
| 0.2 | 0.05 | AG1 | 3 | 23.8 | 0.859 |
| 0.2 | 0.05 | ptsI | 3 | 24.05 | 0.566 |
| 0.2 | 0.05 | clcB | 3 | 24.05 | 0.309 |
| 0.2 | 0.05 | ycaM | 3 | 24.05 | 0.305 |
| 0.2 | 0.05 | yadI | 3 | 24.05 | 0.829 |
| 0.2 | 0.05 | AG1 | 3 | 24.05 | 0.87 |
| 0.2 | 0.05 | ptsI | 3 | 24.3 | 0.566 |
| 0.2 | 0.05 | clcB | 3 | 24.3 | 0.308 |
| 0.2 | 0.05 | ycaM | 3 | 24.3 | 0.307 |
| 0.2 | 0.05 | yadI | 3 | 24.3 | 0.824 |
| 0.2 | 0.05 | AG1 | 3 | 24.3 | 0.874 |
| 0.2 | 0.05 | ptsI | 4 | 0 | 0.212 |
| 0.2 | 0.05 | clcB | 4 | 0 | 0.205 |
| 0.2 | 0.05 | ycaM | 4 | 0 | 0.204 |
| 0.2 | 0.05 | yadI | 4 | 0 | 0.215 |
| 0.2 | 0.05 | AG1 | 4 | 0 | 0.205 |
| 0.2 | 0.05 | ptsI | 4 | 0.25 | 0.205 |
| 0.2 | 0.05 | clcB | 4 | 0.25 | 0.198 |
| 0.2 | 0.05 | ycaM | 4 | 0.25 | 0.199 |
| 0.2 | 0.05 | yadI | 4 | 0.25 | 0.217 |
| 0.2 | 0.05 | AG1 | 4 | 0.25 | 0.204 |
| 0.2 | 0.05 | ptsI | 4 | 0.5 | 0.203 |
| 0.2 | 0.05 | clcB | 4 | 0.5 | 0.197 |
| 0.2 | 0.05 | ycaM | 4 | 0.5 | 0.2 |
| 0.2 | 0.05 | yadI | 4 | 0.5 | 0.217 |
| 0.2 | 0.05 | AG1 | 4 | 0.5 | 0.204 |
| 0.2 | 0.05 | ptsI | 4 | 0.75 | 0.205 |
| 0.2 | 0.05 | clcB | 4 | 0.75 | 0.2 |
| 0.2 | 0.05 | ycaM | 4 | 0.75 | 0.2 |
| 0.2 | 0.05 | yadI | 4 | 0.75 | 0.217 |
| 0.2 | 0.05 | AG1 | 4 | 0.75 | 0.206 |
| 0.2 | 0.05 | ptsI | 4 | 1 | 0.207 |
| 0.2 | 0.05 | clcB | 4 | 1 | 0.204 |
| 0.2 | 0.05 | ycaM | 4 | 1 | 0.204 |
| 0.2 | 0.05 | yadI | 4 | 1 | 0.218 |
| 0.2 | 0.05 | AG1 | 4 | 1 | 0.21 |
| 0.2 | 0.05 | ptsI | 4 | 1.25 | 0.212 |
| 0.2 | 0.05 | clcB | 4 | 1.25 | 0.209 |
| 0.2 | 0.05 | ycaM | 4 | 1.25 | 0.208 |
| 0.2 | 0.05 | yadI | 4 | 1.25 | 0.224 |
| 0.2 | 0.05 | AG1 | 4 | 1.25 | 0.214 |
| 0.2 | 0.05 | ptsI | 4 | 1.5 | 0.221 |
| 0.2 | 0.05 | clcB | 4 | 1.5 | 0.212 |
| 0.2 | 0.05 | ycaM | 4 | 1.5 | 0.214 |
| 0.2 | 0.05 | yadI | 4 | 1.5 | 0.226 |
| 0.2 | 0.05 | AG1 | 4 | 1.5 | 0.213 |

| | | | | | |
|---|---|---|---|---|---|
| 0.2 | 0.05 | ptsI | 4 | 1.75 | 0.225 |
| 0.2 | 0.05 | clcB | 4 | 1.75 | 0.218 |
| 0.2 | 0.05 | ycaM | 4 | 1.75 | 0.221 |
| 0.2 | 0.05 | yadI | 4 | 1.75 | 0.233 |
| 0.2 | 0.05 | AG1 | 4 | 1.75 | 0.221 |
| 0.2 | 0.05 | ptsI | 4 | 2 | 0.236 |
| 0.2 | 0.05 | clcB | 4 | 2 | 0.229 |
| 0.2 | 0.05 | ycaM | 4 | 2 | 0.23 |
| 0.2 | 0.05 | yadI | 4 | 2 | 0.242 |
| 0.2 | 0.05 | AG1 | 4 | 2 | 0.228 |
| 0.2 | 0.05 | ptsI | 4 | 2.25 | 0.245 |
| 0.2 | 0.05 | clcB | 4 | 2.25 | 0.236 |
| 0.2 | 0.05 | ycaM | 4 | 2.25 | 0.238 |
| 0.2 | 0.05 | yadI | 4 | 2.25 | 0.254 |
| 0.2 | 0.05 | AG1 | 4 | 2.25 | 0.238 |
| 0.2 | 0.05 | ptsI | 4 | 2.5 | 0.254 |
| 0.2 | 0.05 | clcB | 4 | 2.5 | 0.24 |
| 0.2 | 0.05 | ycaM | 4 | 2.5 | 0.244 |
| 0.2 | 0.05 | yadI | 4 | 2.5 | 0.257 |
| 0.2 | 0.05 | AG1 | 4 | 2.5 | 0.241 |
| 0.2 | 0.05 | ptsI | 4 | 2.75 | 0.264 |
| 0.2 | 0.05 | clcB | 4 | 2.75 | 0.252 |
| 0.2 | 0.05 | ycaM | 4 | 2.75 | 0.253 |
| 0.2 | 0.05 | yadI | 4 | 2.75 | 0.267 |
| 0.2 | 0.05 | AG1 | 4 | 2.75 | 0.25 |
| 0.2 | 0.05 | ptsI | 4 | 3 | 0.272 |
| 0.2 | 0.05 | clcB | 4 | 3 | 0.257 |
| 0.2 | 0.05 | ycaM | 4 | 3 | 0.26 |
| 0.2 | 0.05 | yadI | 4 | 3 | 0.276 |
| 0.2 | 0.05 | AG1 | 4 | 3 | 0.26 |
| 0.2 | 0.05 | ptsI | 4 | 3.25 | 0.282 |
| 0.2 | 0.05 | clcB | 4 | 3.25 | 0.268 |
| 0.2 | 0.05 | ycaM | 4 | 3.25 | 0.271 |
| 0.2 | 0.05 | yadI | 4 | 3.25 | 0.285 |
| 0.2 | 0.05 | AG1 | 4 | 3.25 | 0.268 |
| 0.2 | 0.05 | ptsI | 4 | 3.5 | 0.298 |
| 0.2 | 0.05 | clcB | 4 | 3.5 | 0.283 |
| 0.2 | 0.05 | ycaM | 4 | 3.5 | 0.279 |
| 0.2 | 0.05 | yadI | 4 | 3.5 | 0.296 |
| 0.2 | 0.05 | AG1 | 4 | 3.5 | 0.282 |
| 0.2 | 0.05 | ptsI | 4 | 3.75 | 0.31 |
| 0.2 | 0.05 | clcB | 4 | 3.75 | 0.296 |
| 0.2 | 0.05 | ycaM | 4 | 3.75 | 0.289 |
| 0.2 | 0.05 | yadI | 4 | 3.75 | 0.304 |
| 0.2 | 0.05 | AG1 | 4 | 3.75 | 0.289 |
| 0.2 | 0.05 | ptsI | 4 | 4 | 0.328 |
| 0.2 | 0.05 | clcB | 4 | 4 | 0.309 |
| 0.2 | 0.05 | ycaM | 4 | 4 | 0.297 |

| | | | | | |
|---|---|---|---|---|---|
| 0.2 | 0.05 | yadI | 4 | 4 | 0.318 |
| 0.2 | 0.05 | AG1 | 4 | 4 | 0.307 |
| 0.2 | 0.05 | ptsI | 4 | 4.35 | 0.331 |
| 0.2 | 0.05 | clcB | 4 | 4.35 | 0.406 |
| 0.2 | 0.05 | ycaM | 4 | 4.35 | 0.378 |
| 0.2 | 0.05 | yadI | 4 | 4.35 | 0.323 |
| 0.2 | 0.05 | AG1 | 4 | 4.35 | 0.339 |
| 0.2 | 0.05 | ptsI | 4 | 4.6 | 0.31 |
| 0.2 | 0.05 | clcB | 4 | 4.6 | 0.388 |
| 0.2 | 0.05 | ycaM | 4 | 4.6 | 0.358 |
| 0.2 | 0.05 | yadI | 4 | 4.6 | 0.33 |
| 0.2 | 0.05 | AG1 | 4 | 4.6 | 0.35 |
| 0.2 | 0.05 | ptsI | 4 | 4.85 | 0.314 |
| 0.2 | 0.05 | clcB | 4 | 4.85 | 0.39 |
| 0.2 | 0.05 | ycaM | 4 | 4.85 | 0.354 |
| 0.2 | 0.05 | yadI | 4 | 4.85 | 0.346 |
| 0.2 | 0.05 | AG1 | 4 | 4.85 | 0.363 |
| 0.2 | 0.05 | ptsI | 4 | 5.1 | 0.323 |
| 0.2 | 0.05 | clcB | 4 | 5.1 | 0.392 |
| 0.2 | 0.05 | ycaM | 4 | 5.1 | 0.357 |
| 0.2 | 0.05 | yadI | 4 | 5.1 | 0.359 |
| 0.2 | 0.05 | AG1 | 4 | 5.1 | 0.373 |
| 0.2 | 0.05 | ptsI | 4 | 5.35 | 0.338 |
| 0.2 | 0.05 | clcB | 4 | 5.35 | 0.396 |
| 0.2 | 0.05 | ycaM | 4 | 5.35 | 0.36 |
| 0.2 | 0.05 | yadI | 4 | 5.35 | 0.374 |
| 0.2 | 0.05 | AG1 | 4 | 5.35 | 0.39 |
| 0.2 | 0.05 | ptsI | 4 | 5.6 | 0.347 |
| 0.2 | 0.05 | clcB | 4 | 5.6 | 0.392 |
| 0.2 | 0.05 | ycaM | 4 | 5.6 | 0.36 |
| 0.2 | 0.05 | yadI | 4 | 5.6 | 0.384 |
| 0.2 | 0.05 | AG1 | 4 | 5.6 | 0.408 |
| 0.2 | 0.05 | ptsI | 4 | 5.85 | 0.358 |
| 0.2 | 0.05 | clcB | 4 | 5.85 | 0.394 |
| 0.2 | 0.05 | ycaM | 4 | 5.85 | 0.359 |
| 0.2 | 0.05 | yadI | 4 | 5.85 | 0.395 |
| 0.2 | 0.05 | AG1 | 4 | 5.85 | 0.413 |
| 0.2 | 0.05 | ptsI | 4 | 6.1 | 0.376 |
| 0.2 | 0.05 | clcB | 4 | 6.1 | 0.399 |
| 0.2 | 0.05 | ycaM | 4 | 6.1 | 0.359 |
| 0.2 | 0.05 | yadI | 4 | 6.1 | 0.408 |
| 0.2 | 0.05 | AG1 | 4 | 6.1 | 0.423 |
| 0.2 | 0.05 | ptsI | 4 | 6.35 | 0.39 |
| 0.2 | 0.05 | clcB | 4 | 6.35 | 0.402 |
| 0.2 | 0.05 | ycaM | 4 | 6.35 | 0.361 |
| 0.2 | 0.05 | yadI | 4 | 6.35 | 0.421 |
| 0.2 | 0.05 | AG1 | 4 | 6.35 | 0.435 |
| 0.2 | 0.05 | ptsI | 4 | 6.6 | 0.404 |

| | | | | | |
|---|---|---|---|---|---|
| 0.2 | 0.05 | clcB | 4 | 6.6 | 0.406 |
| 0.2 | 0.05 | ycaM | 4 | 6.6 | 0.36 |
| 0.2 | 0.05 | yadI | 4 | 6.6 | 0.426 |
| 0.2 | 0.05 | AG1 | 4 | 6.6 | 0.447 |
| 0.2 | 0.05 | ptsI | 4 | 6.85 | 0.414 |
| 0.2 | 0.05 | clcB | 4 | 6.85 | 0.407 |
| 0.2 | 0.05 | ycaM | 4 | 6.85 | 0.364 |
| 0.2 | 0.05 | yadI | 4 | 6.85 | 0.435 |
| 0.2 | 0.05 | AG1 | 4 | 6.85 | 0.46 |
| 0.2 | 0.05 | ptsI | 4 | 7.1 | 0.42 |
| 0.2 | 0.05 | clcB | 4 | 7.1 | 0.408 |
| 0.2 | 0.05 | ycaM | 4 | 7.1 | 0.364 |
| 0.2 | 0.05 | yadI | 4 | 7.1 | 0.444 |
| 0.2 | 0.05 | AG1 | 4 | 7.1 | 0.468 |
| 0.2 | 0.05 | ptsI | 4 | 7.35 | 0.431 |
| 0.2 | 0.05 | clcB | 4 | 7.35 | 0.412 |
| 0.2 | 0.05 | ycaM | 4 | 7.35 | 0.373 |
| 0.2 | 0.05 | yadI | 4 | 7.35 | 0.453 |
| 0.2 | 0.05 | AG1 | 4 | 7.35 | 0.481 |
| 0.2 | 0.05 | ptsI | 4 | 7.6 | 0.441 |
| 0.2 | 0.05 | clcB | 4 | 7.6 | 0.414 |
| 0.2 | 0.05 | ycaM | 4 | 7.6 | 0.38 |
| 0.2 | 0.05 | yadI | 4 | 7.6 | 0.463 |
| 0.2 | 0.05 | AG1 | 4 | 7.6 | 0.49 |
| 0.2 | 0.05 | ptsI | 4 | 7.85 | 0.446 |
| 0.2 | 0.05 | clcB | 4 | 7.85 | 0.407 |
| 0.2 | 0.05 | ycaM | 4 | 7.85 | 0.356 |
| 0.2 | 0.05 | yadI | 4 | 7.85 | 0.47 |
| 0.2 | 0.05 | AG1 | 4 | 7.85 | 0.497 |
| 0.2 | 0.05 | ptsI | 4 | 8.1 | 0.458 |
| 0.2 | 0.05 | clcB | 4 | 8.1 | 0.412 |
| 0.2 | 0.05 | ycaM | 4 | 8.1 | 0.364 |
| 0.2 | 0.05 | yadI | 4 | 8.1 | 0.479 |
| 0.2 | 0.05 | AG1 | 4 | 8.1 | 0.503 |
| 0.2 | 0.05 | ptsI | 4 | 8.35 | 0.468 |
| 0.2 | 0.05 | clcB | 4 | 8.35 | 0.416 |
| 0.2 | 0.05 | ycaM | 4 | 8.35 | 0.376 |
| 0.2 | 0.05 | yadI | 4 | 8.35 | 0.488 |
| 0.2 | 0.05 | AG1 | 4 | 8.35 | 0.516 |
| 0.2 | 0.05 | ptsI | 4 | 8.83333 | 0.431 |
| 0.2 | 0.05 | clcB | 4 | 8.83333 | 0.306 |
| 0.2 | 0.05 | ycaM | 4 | 8.83333 | 0.282 |
| 0.2 | 0.05 | yadI | 4 | 8.83333 | 0.512 |
| 0.2 | 0.05 | AG1 | 4 | 8.83333 | 0.56 |
| 0.2 | 0.05 | ptsI | 4 | 9.08333 | 0.436 |
| 0.2 | 0.05 | clcB | 4 | 9.08333 | 0.307 |
| 0.2 | 0.05 | ycaM | 4 | 9.08333 | 0.286 |
| 0.2 | 0.05 | yadI | 4 | 9.08333 | 0.508 |

| | | | | | |
|---|---|---|---|---|---|
| 0.2 | 0.05 | AG1 | 4 | 9.08333 | 0.563 |
| 0.2 | 0.05 | ptsI | 4 | 9.33333 | 0.443 |
| 0.2 | 0.05 | clcB | 4 | 9.33333 | 0.303 |
| 0.2 | 0.05 | ycaM | 4 | 9.33333 | 0.28 |
| 0.2 | 0.05 | yadI | 4 | 9.33333 | 0.518 |
| 0.2 | 0.05 | AG1 | 4 | 9.33333 | 0.579 |
| 0.2 | 0.05 | ptsI | 4 | 9.58333 | 0.451 |
| 0.2 | 0.05 | clcB | 4 | 9.58333 | 0.302 |
| 0.2 | 0.05 | ycaM | 4 | 9.58333 | 0.284 |
| 0.2 | 0.05 | yadI | 4 | 9.58333 | 0.529 |
| 0.2 | 0.05 | AG1 | 4 | 9.58333 | 0.592 |
| 0.2 | 0.05 | ptsI | 4 | 9.83333 | 0.463 |
| 0.2 | 0.05 | clcB | 4 | 9.83333 | 0.302 |
| 0.2 | 0.05 | ycaM | 4 | 9.83333 | 0.282 |
| 0.2 | 0.05 | yadI | 4 | 9.83333 | 0.539 |
| 0.2 | 0.05 | AG1 | 4 | 9.83333 | 0.601 |
| 0.2 | 0.05 | ptsI | 4 | 10.0833 | 0.471 |
| 0.2 | 0.05 | clcB | 4 | 10.0833 | 0.301 |
| 0.2 | 0.05 | ycaM | 4 | 10.0833 | 0.282 |
| 0.2 | 0.05 | yadI | 4 | 10.0833 | 0.543 |
| 0.2 | 0.05 | AG1 | 4 | 10.0833 | 0.607 |
| 0.2 | 0.05 | ptsI | 4 | 10.3333 | 0.481 |
| 0.2 | 0.05 | clcB | 4 | 10.3333 | 0.304 |
| 0.2 | 0.05 | ycaM | 4 | 10.3333 | 0.282 |
| 0.2 | 0.05 | yadI | 4 | 10.3333 | 0.551 |
| 0.2 | 0.05 | AG1 | 4 | 10.3333 | 0.619 |
| 0.2 | 0.05 | ptsI | 4 | 10.5833 | 0.492 |
| 0.2 | 0.05 | clcB | 4 | 10.5833 | 0.302 |
| 0.2 | 0.05 | ycaM | 4 | 10.5833 | 0.284 |
| 0.2 | 0.05 | yadI | 4 | 10.5833 | 0.56 |
| 0.2 | 0.05 | AG1 | 4 | 10.5833 | 0.626 |
| 0.2 | 0.05 | ptsI | 4 | 10.8333 | 0.501 |
| 0.2 | 0.05 | clcB | 4 | 10.8333 | 0.297 |
| 0.2 | 0.05 | ycaM | 4 | 10.8333 | 0.285 |
| 0.2 | 0.05 | yadI | 4 | 10.8333 | 0.566 |
| 0.2 | 0.05 | AG1 | 4 | 10.8333 | 0.636 |
| 0.2 | 0.05 | ptsI | 4 | 11.0833 | 0.508 |
| 0.2 | 0.05 | clcB | 4 | 11.0833 | 0.299 |
| 0.2 | 0.05 | ycaM | 4 | 11.0833 | 0.281 |
| 0.2 | 0.05 | yadI | 4 | 11.0833 | 0.57 |
| 0.2 | 0.05 | AG1 | 4 | 11.0833 | 0.641 |
| 0.2 | 0.05 | ptsI | 4 | 11.3333 | 0.518 |
| 0.2 | 0.05 | clcB | 4 | 11.3333 | 0.303 |
| 0.2 | 0.05 | ycaM | 4 | 11.3333 | 0.288 |
| 0.2 | 0.05 | yadI | 4 | 11.3333 | 0.58 |
| 0.2 | 0.05 | AG1 | 4 | 11.3333 | 0.651 |
| 0.2 | 0.05 | ptsI | 4 | 11.5833 | 0.524 |
| 0.2 | 0.05 | clcB | 4 | 11.5833 | 0.298 |

| | | | | | |
|---|---|---|---|---|---|
| 0.2 | 0.05 | ycaM | 4 | 11.5833 | 0.281 |
| 0.2 | 0.05 | yadI | 4 | 11.5833 | 0.587 |
| 0.2 | 0.05 | AG1 | 4 | 11.5833 | 0.656 |
| 0.2 | 0.05 | ptsI | 4 | 11.8333 | 0.534 |
| 0.2 | 0.05 | clcB | 4 | 11.8333 | 0.292 |
| 0.2 | 0.05 | ycaM | 4 | 11.8333 | 0.29 |
| 0.2 | 0.05 | yadI | 4 | 11.8333 | 0.589 |
| 0.2 | 0.05 | AG1 | 4 | 11.8333 | 0.658 |
| 0.2 | 0.05 | ptsI | 4 | 12.0833 | 0.539 |
| 0.2 | 0.05 | clcB | 4 | 12.0833 | 0.299 |
| 0.2 | 0.05 | ycaM | 4 | 12.0833 | 0.288 |
| 0.2 | 0.05 | yadI | 4 | 12.0833 | 0.595 |
| 0.2 | 0.05 | AG1 | 4 | 12.0833 | 0.665 |
| 0.2 | 0.05 | ptsI | 4 | 12.3333 | 0.549 |
| 0.2 | 0.05 | clcB | 4 | 12.3333 | 0.304 |
| 0.2 | 0.05 | ycaM | 4 | 12.3333 | 0.286 |
| 0.2 | 0.05 | yadI | 4 | 12.3333 | 0.601 |
| 0.2 | 0.05 | AG1 | 4 | 12.3333 | 0.672 |
| 0.2 | 0.05 | ptsI | 4 | 12.5833 | 0.555 |
| 0.2 | 0.05 | clcB | 4 | 12.5833 | 0.307 |
| 0.2 | 0.05 | ycaM | 4 | 12.5833 | 0.287 |
| 0.2 | 0.05 | yadI | 4 | 12.5833 | 0.605 |
| 0.2 | 0.05 | AG1 | 4 | 12.5833 | 0.678 |
| 0.2 | 0.05 | ptsI | 4 | 12.8333 | 0.56 |
| 0.2 | 0.05 | clcB | 4 | 12.8333 | 0.299 |
| 0.2 | 0.05 | ycaM | 4 | 12.8333 | 0.286 |
| 0.2 | 0.05 | yadI | 4 | 12.8333 | 0.612 |
| 0.2 | 0.05 | AG1 | 4 | 12.8333 | 0.683 |
| 0.2 | 0.05 | ptsI | 4 | 13.0833 | 0.569 |
| 0.2 | 0.05 | clcB | 4 | 13.0833 | 0.303 |
| 0.2 | 0.05 | ycaM | 4 | 13.0833 | 0.291 |
| 0.2 | 0.05 | yadI | 4 | 13.0833 | 0.616 |
| 0.2 | 0.05 | AG1 | 4 | 13.0833 | 0.69 |
| 0.2 | 0.05 | ptsI | 4 | 13.3333 | 0.579 |
| 0.2 | 0.05 | clcB | 4 | 13.3333 | 0.302 |
| 0.2 | 0.05 | ycaM | 4 | 13.3333 | 0.29 |
| 0.2 | 0.05 | yadI | 4 | 13.3333 | 0.619 |
| 0.2 | 0.05 | AG1 | 4 | 13.3333 | 0.696 |
| 0.2 | 0.05 | ptsI | 4 | 13.5833 | 0.585 |
| 0.2 | 0.05 | clcB | 4 | 13.5833 | 0.306 |
| 0.2 | 0.05 | ycaM | 4 | 13.5833 | 0.284 |
| 0.2 | 0.05 | yadI | 4 | 13.5833 | 0.624 |
| 0.2 | 0.05 | AG1 | 4 | 13.5833 | 0.699 |
| 0.2 | 0.05 | ptsI | 4 | 13.8333 | 0.592 |
| 0.2 | 0.05 | clcB | 4 | 13.8333 | 0.311 |
| 0.2 | 0.05 | ycaM | 4 | 13.8333 | 0.29 |
| 0.2 | 0.05 | yadI | 4 | 13.8333 | 0.627 |
| 0.2 | 0.05 | AG1 | 4 | 13.8333 | 0.7 |

| | | | | | |
|---|---|---|---|---|---|
| 0.2 | 0.05 | ptsI | 4 | 14.0833 | 0.6 |
| 0.2 | 0.05 | clcB | 4 | 14.0833 | 0.311 |
| 0.2 | 0.05 | ycaM | 4 | 14.0833 | 0.29 |
| 0.2 | 0.05 | yadI | 4 | 14.0833 | 0.634 |
| 0.2 | 0.05 | AG1 | 4 | 14.0833 | 0.704 |
| 0.2 | 0.05 | ptsI | 4 | 14.3333 | 0.606 |
| 0.2 | 0.05 | clcB | 4 | 14.3333 | 0.309 |
| 0.2 | 0.05 | ycaM | 4 | 14.3333 | 0.292 |
| 0.2 | 0.05 | yadI | 4 | 14.3333 | 0.632 |
| 0.2 | 0.05 | AG1 | 4 | 14.3333 | 0.708 |
| 0.2 | 0.05 | ptsI | 4 | 14.5833 | 0.609 |
| 0.2 | 0.05 | clcB | 4 | 14.5833 | 0.311 |
| 0.2 | 0.05 | ycaM | 4 | 14.5833 | 0.302 |
| 0.2 | 0.05 | yadI | 4 | 14.5833 | 0.638 |
| 0.2 | 0.05 | AG1 | 4 | 14.5833 | 0.711 |
| 0.2 | 0.05 | ptsI | 4 | 14.8333 | 0.614 |
| 0.2 | 0.05 | clcB | 4 | 14.8333 | 0.311 |
| 0.2 | 0.05 | ycaM | 4 | 14.8333 | 0.292 |
| 0.2 | 0.05 | yadI | 4 | 14.8333 | 0.644 |
| 0.2 | 0.05 | AG1 | 4 | 14.8333 | 0.718 |
| 0.2 | 0.05 | ptsI | 4 | 15.0833 | 0.622 |
| 0.2 | 0.05 | clcB | 4 | 15.0833 | 0.308 |
| 0.2 | 0.05 | ycaM | 4 | 15.0833 | 0.292 |
| 0.2 | 0.05 | yadI | 4 | 15.0833 | 0.645 |
| 0.2 | 0.05 | AG1 | 4 | 15.0833 | 0.72 |
| 0.2 | 0.05 | ptsI | 4 | 15.3333 | 0.626 |
| 0.2 | 0.05 | clcB | 4 | 15.3333 | 0.312 |
| 0.2 | 0.05 | ycaM | 4 | 15.3333 | 0.295 |
| 0.2 | 0.05 | yadI | 4 | 15.3333 | 0.647 |
| 0.2 | 0.05 | AG1 | 4 | 15.3333 | 0.722 |
| 0.2 | 0.05 | ptsI | 4 | 15.5833 | 0.632 |
| 0.2 | 0.05 | clcB | 4 | 15.5833 | 0.313 |
| 0.2 | 0.05 | ycaM | 4 | 15.5833 | 0.298 |
| 0.2 | 0.05 | yadI | 4 | 15.5833 | 0.65 |
| 0.2 | 0.05 | AG1 | 4 | 15.5833 | 0.727 |
| 0.2 | 0.05 | ptsI | 4 | 15.8333 | 0.632 |
| 0.2 | 0.05 | clcB | 4 | 15.8333 | 0.316 |
| 0.2 | 0.05 | ycaM | 4 | 15.8333 | 0.299 |
| 0.2 | 0.05 | yadI | 4 | 15.8333 | 0.658 |
| 0.2 | 0.05 | AG1 | 4 | 15.8333 | 0.73 |
| 0.2 | 0.05 | ptsI | 4 | 16.0833 | 0.638 |
| 0.2 | 0.05 | clcB | 4 | 16.0833 | 0.314 |
| 0.2 | 0.05 | ycaM | 4 | 16.0833 | 0.299 |
| 0.2 | 0.05 | yadI | 4 | 16.0833 | 0.658 |
| 0.2 | 0.05 | AG1 | 4 | 16.0833 | 0.734 |
| 0.2 | 0.05 | ptsI | 4 | 16.3333 | 0.636 |
| 0.2 | 0.05 | clcB | 4 | 16.3333 | 0.315 |
| 0.2 | 0.05 | ycaM | 4 | 16.3333 | 0.301 |

| | | | | | |
|---|---|---|---|---|---|
| 0.2 | 0.05 | yadI | 4 | 16.3333 | 0.66 |
| 0.2 | 0.05 | AG1 | 4 | 16.3333 | 0.737 |
| 0.2 | 0.05 | ptsI | 4 | 16.5833 | 0.637 |
| 0.2 | 0.05 | clcB | 4 | 16.5833 | 0.314 |
| 0.2 | 0.05 | ycaM | 4 | 16.5833 | 0.301 |
| 0.2 | 0.05 | yadI | 4 | 16.5833 | 0.665 |
| 0.2 | 0.05 | AG1 | 4 | 16.5833 | 0.744 |
| 0.2 | 0.05 | ptsI | 4 | 16.8333 | 0.645 |
| 0.2 | 0.05 | clcB | 4 | 16.8333 | 0.316 |
| 0.2 | 0.05 | ycaM | 4 | 16.8333 | 0.298 |
| 0.2 | 0.05 | yadI | 4 | 16.8333 | 0.67 |
| 0.2 | 0.05 | AG1 | 4 | 16.8333 | 0.746 |
| 0.2 | 0.05 | ptsI | 4 | 17.0833 | 0.644 |
| 0.2 | 0.05 | clcB | 4 | 17.0833 | 0.317 |
| 0.2 | 0.05 | ycaM | 4 | 17.0833 | 0.303 |
| 0.2 | 0.05 | yadI | 4 | 17.0833 | 0.67 |
| 0.2 | 0.05 | AG1 | 4 | 17.0833 | 0.746 |
| 0.2 | 0.05 | ptsI | 4 | 17.3333 | 0.644 |
| 0.2 | 0.05 | clcB | 4 | 17.3333 | 0.322 |
| 0.2 | 0.05 | ycaM | 4 | 17.3333 | 0.3 |
| 0.2 | 0.05 | yadI | 4 | 17.3333 | 0.672 |
| 0.2 | 0.05 | AG1 | 4 | 17.3333 | 0.746 |
| 0.2 | 0.05 | ptsI | 4 | 17.5833 | 0.649 |
| 0.2 | 0.05 | clcB | 4 | 17.5833 | 0.319 |
| 0.2 | 0.05 | ycaM | 4 | 17.5833 | 0.305 |
| 0.2 | 0.05 | yadI | 4 | 17.5833 | 0.677 |
| 0.2 | 0.05 | AG1 | 4 | 17.5833 | 0.752 |
| 0.2 | 0.05 | ptsI | 4 | 17.8333 | 0.645 |
| 0.2 | 0.05 | clcB | 4 | 17.8333 | 0.313 |
| 0.2 | 0.05 | ycaM | 4 | 17.8333 | 0.307 |
| 0.2 | 0.05 | yadI | 4 | 17.8333 | 0.679 |
| 0.2 | 0.05 | AG1 | 4 | 17.8333 | 0.759 |
| 0.2 | 0.05 | ptsI | 4 | 18.0833 | 0.648 |
| 0.2 | 0.05 | clcB | 4 | 18.0833 | 0.314 |
| 0.2 | 0.05 | ycaM | 4 | 18.0833 | 0.305 |
| 0.2 | 0.05 | yadI | 4 | 18.0833 | 0.679 |
| 0.2 | 0.05 | AG1 | 4 | 18.0833 | 0.758 |
| 0.2 | 0.05 | ptsI | 4 | 18.3333 | 0.653 |
| 0.2 | 0.05 | clcB | 4 | 18.3333 | 0.31 |
| 0.2 | 0.05 | ycaM | 4 | 18.3333 | 0.311 |
| 0.2 | 0.05 | yadI | 4 | 18.3333 | 0.683 |
| 0.2 | 0.05 | AG1 | 4 | 18.3333 | 0.759 |
| 0.2 | 0.05 | ptsI | 4 | 18.5833 | 0.652 |
| 0.2 | 0.05 | clcB | 4 | 18.5833 | 0.309 |
| 0.2 | 0.05 | ycaM | 4 | 18.5833 | 0.308 |
| 0.2 | 0.05 | yadI | 4 | 18.5833 | 0.682 |
| 0.2 | 0.05 | AG1 | 4 | 18.5833 | 0.76 |
| 0.2 | 0.05 | ptsI | 4 | 18.8333 | 0.652 |

| | | | | | |
|---|---|---|---|---|---|
| 0.2 | 0.05 | clcB | 4 | 18.8333 | 0.303 |
| 0.2 | 0.05 | ycaM | 4 | 18.8333 | 0.317 |
| 0.2 | 0.05 | yadI | 4 | 18.8333 | 0.686 |
| 0.2 | 0.05 | AG1 | 4 | 18.8333 | 0.767 |
| 0.2 | 0.05 | ptsI | 4 | 19.0833 | 0.653 |
| 0.2 | 0.05 | clcB | 4 | 19.0833 | 0.304 |
| 0.2 | 0.05 | ycaM | 4 | 19.0833 | 0.315 |
| 0.2 | 0.05 | yadI | 4 | 19.0833 | 0.69 |
| 0.2 | 0.05 | AG1 | 4 | 19.0833 | 0.765 |
| 0.2 | 0.05 | ptsI | 4 | 19.3333 | 0.65 |
| 0.2 | 0.05 | clcB | 4 | 19.3333 | 0.306 |
| 0.2 | 0.05 | ycaM | 4 | 19.3333 | 0.32 |
| 0.2 | 0.05 | yadI | 4 | 19.3333 | 0.685 |
| 0.2 | 0.05 | AG1 | 4 | 19.3333 | 0.763 |
| 0.2 | 0.05 | ptsI | 4 | 19.5833 | 0.654 |
| 0.2 | 0.05 | clcB | 4 | 19.5833 | 0.307 |
| 0.2 | 0.05 | ycaM | 4 | 19.5833 | 0.322 |
| 0.2 | 0.05 | yadI | 4 | 19.5833 | 0.687 |
| 0.2 | 0.05 | AG1 | 4 | 19.5833 | 0.764 |
| 0.2 | 0.05 | ptsI | 4 | 19.8333 | 0.654 |
| 0.2 | 0.05 | clcB | 4 | 19.8333 | 0.305 |
| 0.2 | 0.05 | ycaM | 4 | 19.8333 | 0.323 |
| 0.2 | 0.05 | yadI | 4 | 19.8333 | 0.686 |
| 0.2 | 0.05 | AG1 | 4 | 19.8333 | 0.767 |
| 0.2 | 0.05 | ptsI | 4 | 20.0833 | 0.66 |
| 0.2 | 0.05 | clcB | 4 | 20.0833 | 0.304 |
| 0.2 | 0.05 | ycaM | 4 | 20.0833 | 0.326 |
| 0.2 | 0.05 | yadI | 4 | 20.0833 | 0.688 |
| 0.2 | 0.05 | AG1 | 4 | 20.0833 | 0.772 |
| 0.2 | 0.05 | ptsI | 4 | 20.3333 | 0.654 |
| 0.2 | 0.05 | clcB | 4 | 20.3333 | 0.302 |
| 0.2 | 0.05 | ycaM | 4 | 20.3333 | 0.325 |
| 0.2 | 0.05 | yadI | 4 | 20.3333 | 0.69 |
| 0.2 | 0.05 | AG1 | 4 | 20.3333 | 0.769 |
| 0.2 | 0.05 | ptsI | 4 | 20.5833 | 0.653 |
| 0.2 | 0.05 | clcB | 4 | 20.5833 | 0.3 |
| 0.2 | 0.05 | ycaM | 4 | 20.5833 | 0.328 |
| 0.2 | 0.05 | yadI | 4 | 20.5833 | 0.693 |
| 0.2 | 0.05 | AG1 | 4 | 20.5833 | 0.778 |
| 0.2 | 0.05 | ptsI | 4 | 20.8333 | 0.662 |
| 0.2 | 0.05 | clcB | 4 | 20.8333 | 0.303 |
| 0.2 | 0.05 | ycaM | 4 | 20.8333 | 0.333 |
| 0.2 | 0.05 | yadI | 4 | 20.8333 | 0.692 |
| 0.2 | 0.05 | AG1 | 4 | 20.8333 | 0.774 |
| 0.2 | 0.05 | ptsI | 4 | 21.0833 | 0.659 |
| 0.2 | 0.05 | clcB | 4 | 21.0833 | 0.301 |
| 0.2 | 0.05 | ycaM | 4 | 21.0833 | 0.332 |
| 0.2 | 0.05 | yadI | 4 | 21.0833 | 0.694 |

| | | | | | |
|---|---|---|---|---|---|
| 0.2 | 0.05 | AG1 | 4 | 21.0833 | 0.777 |
| 0.2 | 0.05 | ptsI | 4 | 21.3333 | 0.664 |
| 0.2 | 0.05 | clcB | 4 | 21.3333 | 0.303 |
| 0.2 | 0.05 | ycaM | 4 | 21.3333 | 0.33 |
| 0.2 | 0.05 | yadI | 4 | 21.3333 | 0.692 |
| 0.2 | 0.05 | AG1 | 4 | 21.3333 | 0.779 |
| 0.2 | 0.05 | ptsI | 4 | 21.5833 | 0.659 |
| 0.2 | 0.05 | clcB | 4 | 21.5833 | 0.304 |
| 0.2 | 0.05 | ycaM | 4 | 21.5833 | 0.328 |
| 0.2 | 0.05 | yadI | 4 | 21.5833 | 0.693 |
| 0.2 | 0.05 | AG1 | 4 | 21.5833 | 0.782 |
| 0.2 | 0.05 | ptsI | 4 | 21.8333 | 0.66 |
| 0.2 | 0.05 | clcB | 4 | 21.8333 | 0.307 |
| 0.2 | 0.05 | ycaM | 4 | 21.8333 | 0.333 |
| 0.2 | 0.05 | yadI | 4 | 21.8333 | 0.694 |
| 0.2 | 0.05 | AG1 | 4 | 21.8333 | 0.782 |
| 0.2 | 0.05 | ptsI | 4 | 22.0833 | 0.662 |
| 0.2 | 0.05 | clcB | 4 | 22.0833 | 0.308 |
| 0.2 | 0.05 | ycaM | 4 | 22.0833 | 0.33 |
| 0.2 | 0.05 | yadI | 4 | 22.0833 | 0.694 |
| 0.2 | 0.05 | AG1 | 4 | 22.0833 | 0.786 |
| 0.2 | 0.05 | ptsI | 4 | 22.3333 | 0.657 |
| 0.2 | 0.05 | clcB | 4 | 22.3333 | 0.307 |
| 0.2 | 0.05 | ycaM | 4 | 22.3333 | 0.328 |
| 0.2 | 0.05 | yadI | 4 | 22.3333 | 0.695 |
| 0.2 | 0.05 | AG1 | 4 | 22.3333 | 0.787 |
| 0.2 | 0.05 | ptsI | 4 | 22.5833 | 0.659 |
| 0.2 | 0.05 | clcB | 4 | 22.5833 | 0.304 |
| 0.2 | 0.05 | ycaM | 4 | 22.5833 | 0.328 |
| 0.2 | 0.05 | yadI | 4 | 22.5833 | 0.695 |
| 0.2 | 0.05 | AG1 | 4 | 22.5833 | 0.789 |
| 0.2 | 0.05 | ptsI | 4 | 22.8333 | 0.663 |
| 0.2 | 0.05 | clcB | 4 | 22.8333 | 0.308 |
| 0.2 | 0.05 | ycaM | 4 | 22.8333 | 0.332 |
| 0.2 | 0.05 | yadI | 4 | 22.8333 | 0.699 |
| 0.2 | 0.05 | AG1 | 4 | 22.8333 | 0.789 |
| 0.2 | 0.05 | ptsI | 4 | 23.0833 | 0.658 |
| 0.2 | 0.05 | clcB | 4 | 23.0833 | 0.312 |
| 0.2 | 0.05 | ycaM | 4 | 23.0833 | 0.329 |
| 0.2 | 0.05 | yadI | 4 | 23.0833 | 0.696 |
| 0.2 | 0.05 | AG1 | 4 | 23.0833 | 0.792 |
| 0.2 | 0.05 | ptsI | 4 | 23.3333 | 0.663 |
| 0.2 | 0.05 | clcB | 4 | 23.3333 | 0.316 |
| 0.2 | 0.05 | ycaM | 4 | 23.3333 | 0.335 |
| 0.2 | 0.05 | yadI | 4 | 23.3333 | 0.7 |
| 0.2 | 0.05 | AG1 | 4 | 23.3333 | 0.794 |
| 0.2 | 0.05 | ptsI | 4 | 23.5833 | 0.662 |
| 0.2 | 0.05 | clcB | 4 | 23.5833 | 0.315 |

| | | | | | |
|---|---|---|---|---|---|
| 0.2 | 0.05 | ycaM | 4 | 23.5833 | 0.334 |
| 0.2 | 0.05 | yadI | 4 | 23.5833 | 0.702 |
| 0.2 | 0.05 | AG1 | 4 | 23.5833 | 0.795 |
| 0.2 | 0.05 | ptsI | 4 | 23.8333 | 0.659 |
| 0.2 | 0.05 | clcB | 4 | 23.8333 | 0.316 |
| 0.2 | 0.05 | ycaM | 4 | 23.8333 | 0.334 |
| 0.2 | 0.05 | yadI | 4 | 23.8333 | 0.701 |
| 0.2 | 0.05 | AG1 | 4 | 23.8333 | 0.801 |
| 0.2 | 0.05 | ptsI | 4 | 24.0833 | 0.661 |
| 0.2 | 0.05 | clcB | 4 | 24.0833 | 0.318 |
| 0.2 | 0.05 | ycaM | 4 | 24.0833 | 0.34 |
| 0.2 | 0.05 | yadI | 4 | 24.0833 | 0.702 |
| 0.2 | 0.05 | AG1 | 4 | 24.0833 | 0.801 |
| 0.2 | 0.05 | ptsI | 5 | 0 | 0.167 |
| 0.2 | 0.05 | clcB | 5 | 0 | 0.21 |
| 0.2 | 0.05 | ycaM | 5 | 0 | 0.256 |
| 0.2 | 0.05 | yadI | 5 | 0 | 0.186 |
| 0.2 | 0.05 | AG1 | 5 | 0 | 0.169 |
| 0.2 | 0.05 | ptsI | 5 | 0.35 | 0.159 |
| 0.2 | 0.05 | clcB | 5 | 0.35 | 0.206 |
| 0.2 | 0.05 | ycaM | 5 | 0.35 | 0.257 |
| 0.2 | 0.05 | yadI | 5 | 0.35 | 0.182 |
| 0.2 | 0.05 | AG1 | 5 | 0.35 | 0.167 |
| 0.2 | 0.05 | ptsI | 5 | 0.6 | 0.161 |
| 0.2 | 0.05 | clcB | 5 | 0.6 | 0.205 |
| 0.2 | 0.05 | ycaM | 5 | 0.6 | 0.26 |
| 0.2 | 0.05 | yadI | 5 | 0.6 | 0.183 |
| 0.2 | 0.05 | AG1 | 5 | 0.6 | 0.167 |
| 0.2 | 0.05 | ptsI | 5 | 0.85 | 0.161 |
| 0.2 | 0.05 | clcB | 5 | 0.85 | 0.207 |
| 0.2 | 0.05 | ycaM | 5 | 0.85 | 0.265 |
| 0.2 | 0.05 | yadI | 5 | 0.85 | 0.183 |
| 0.2 | 0.05 | AG1 | 5 | 0.85 | 0.168 |
| 0.2 | 0.05 | ptsI | 5 | 1.1 | 0.161 |
| 0.2 | 0.05 | clcB | 5 | 1.1 | 0.205 |
| 0.2 | 0.05 | ycaM | 5 | 1.1 | 0.267 |
| 0.2 | 0.05 | yadI | 5 | 1.1 | 0.186 |
| 0.2 | 0.05 | AG1 | 5 | 1.1 | 0.169 |
| 0.2 | 0.05 | ptsI | 5 | 1.35 | 0.162 |
| 0.2 | 0.05 | clcB | 5 | 1.35 | 0.207 |
| 0.2 | 0.05 | ycaM | 5 | 1.35 | 0.267 |
| 0.2 | 0.05 | yadI | 5 | 1.35 | 0.186 |
| 0.2 | 0.05 | AG1 | 5 | 1.35 | 0.171 |
| 0.2 | 0.05 | ptsI | 5 | 1.6 | 0.164 |
| 0.2 | 0.05 | clcB | 5 | 1.6 | 0.209 |
| 0.2 | 0.05 | ycaM | 5 | 1.6 | 0.275 |
| 0.2 | 0.05 | yadI | 5 | 1.6 | 0.189 |
| 0.2 | 0.05 | AG1 | 5 | 1.6 | 0.172 |

| | | | | | |
|---|---|---|---|---|---|
| 0.2 | 0.05 | ptsI | 5 | 1.85 | 0.168 |
| 0.2 | 0.05 | clcB | 5 | 1.85 | 0.211 |
| 0.2 | 0.05 | ycaM | 5 | 1.85 | 0.284 |
| 0.2 | 0.05 | yadI | 5 | 1.85 | 0.191 |
| 0.2 | 0.05 | AG1 | 5 | 1.85 | 0.175 |
| 0.2 | 0.05 | ptsI | 5 | 2.1 | 0.174 |
| 0.2 | 0.05 | clcB | 5 | 2.1 | 0.215 |
| 0.2 | 0.05 | ycaM | 5 | 2.1 | 0.29 |
| 0.2 | 0.05 | yadI | 5 | 2.1 | 0.195 |
| 0.2 | 0.05 | AG1 | 5 | 2.1 | 0.179 |
| 0.2 | 0.05 | ptsI | 5 | 2.35 | 0.177 |
| 0.2 | 0.05 | clcB | 5 | 2.35 | 0.221 |
| 0.2 | 0.05 | ycaM | 5 | 2.35 | 0.299 |
| 0.2 | 0.05 | yadI | 5 | 2.35 | 0.2 |
| 0.2 | 0.05 | AG1 | 5 | 2.35 | 0.182 |
| 0.2 | 0.05 | ptsI | 5 | 2.6 | 0.174 |
| 0.2 | 0.05 | clcB | 5 | 2.6 | 0.225 |
| 0.2 | 0.05 | ycaM | 5 | 2.6 | 0.263 |
| 0.2 | 0.05 | yadI | 5 | 2.6 | 0.207 |
| 0.2 | 0.05 | AG1 | 5 | 2.6 | 0.187 |
| 0.2 | 0.05 | ptsI | 5 | 2.85 | 0.185 |
| 0.2 | 0.05 | clcB | 5 | 2.85 | 0.23 |
| 0.2 | 0.05 | ycaM | 5 | 2.85 | 0.316 |
| 0.2 | 0.05 | yadI | 5 | 2.85 | 0.208 |
| 0.2 | 0.05 | AG1 | 5 | 2.85 | 0.19 |
| 0.2 | 0.05 | ptsI | 5 | 3.1 | 0.19 |
| 0.2 | 0.05 | clcB | 5 | 3.1 | 0.235 |
| 0.2 | 0.05 | ycaM | 5 | 3.1 | 0.321 |
| 0.2 | 0.05 | yadI | 5 | 3.1 | 0.215 |
| 0.2 | 0.05 | AG1 | 5 | 3.1 | 0.193 |
| 0.2 | 0.05 | ptsI | 5 | 3.35 | 0.204 |
| 0.2 | 0.05 | clcB | 5 | 3.35 | 0.244 |
| 0.2 | 0.05 | ycaM | 5 | 3.35 | 0.335 |
| 0.2 | 0.05 | yadI | 5 | 3.35 | 0.223 |
| 0.2 | 0.05 | AG1 | 5 | 3.35 | 0.202 |
| 0.2 | 0.05 | ptsI | 5 | 3.6 | 0.211 |
| 0.2 | 0.05 | clcB | 5 | 3.6 | 0.256 |
| 0.2 | 0.05 | ycaM | 5 | 3.6 | 0.347 |
| 0.2 | 0.05 | yadI | 5 | 3.6 | 0.23 |
| 0.2 | 0.05 | AG1 | 5 | 3.6 | 0.21 |
| 0.2 | 0.05 | ptsI | 5 | 3.85 | 0.22 |
| 0.2 | 0.05 | clcB | 5 | 3.85 | 0.268 |
| 0.2 | 0.05 | ycaM | 5 | 3.85 | 0.353 |
| 0.2 | 0.05 | yadI | 5 | 3.85 | 0.241 |
| 0.2 | 0.05 | AG1 | 5 | 3.85 | 0.221 |
| 0.2 | 0.05 | ptsI | 5 | 4.38333 | 0.232 |
| 0.2 | 0.05 | clcB | 5 | 4.38333 | 0.283 |
| 0.2 | 0.05 | ycaM | 5 | 4.38333 | 0.227 |

| | | | | | |
|---|---|---|---|---|---|
| 0.2 | 0.05 | yadI | 5 | 4.38333 | 0.282 |
| 0.2 | 0.05 | AG1 | 5 | 4.38333 | 0.251 |
| 0.2 | 0.05 | ptsI | 5 | 4.63333 | 0.23 |
| 0.2 | 0.05 | clcB | 5 | 4.63333 | 0.273 |
| 0.2 | 0.05 | ycaM | 5 | 4.63333 | 0.23 |
| 0.2 | 0.05 | yadI | 5 | 4.63333 | 0.288 |
| 0.2 | 0.05 | AG1 | 5 | 4.63333 | 0.261 |
| 0.2 | 0.05 | ptsI | 5 | 4.88333 | 0.239 |
| 0.2 | 0.05 | clcB | 5 | 4.88333 | 0.285 |
| 0.2 | 0.05 | ycaM | 5 | 4.88333 | 0.24 |
| 0.2 | 0.05 | yadI | 5 | 4.88333 | 0.305 |
| 0.2 | 0.05 | AG1 | 5 | 4.88333 | 0.272 |
| 0.2 | 0.05 | ptsI | 5 | 5.13333 | 0.246 |
| 0.2 | 0.05 | clcB | 5 | 5.13333 | 0.296 |
| 0.2 | 0.05 | ycaM | 5 | 5.13333 | 0.245 |
| 0.2 | 0.05 | yadI | 5 | 5.13333 | 0.319 |
| 0.2 | 0.05 | AG1 | 5 | 5.13333 | 0.287 |
| 0.2 | 0.05 | ptsI | 5 | 5.38333 | 0.256 |
| 0.2 | 0.05 | clcB | 5 | 5.38333 | 0.307 |
| 0.2 | 0.05 | ycaM | 5 | 5.38333 | 0.249 |
| 0.2 | 0.05 | yadI | 5 | 5.38333 | 0.334 |
| 0.2 | 0.05 | AG1 | 5 | 5.38333 | 0.304 |
| 0.2 | 0.05 | ptsI | 5 | 5.63333 | 0.266 |
| 0.2 | 0.05 | clcB | 5 | 5.63333 | 0.317 |
| 0.2 | 0.05 | ycaM | 5 | 5.63333 | 0.249 |
| 0.2 | 0.05 | yadI | 5 | 5.63333 | 0.348 |
| 0.2 | 0.05 | AG1 | 5 | 5.63333 | 0.317 |
| 0.2 | 0.05 | ptsI | 5 | 5.88333 | 0.276 |
| 0.2 | 0.05 | clcB | 5 | 5.88333 | 0.328 |
| 0.2 | 0.05 | ycaM | 5 | 5.88333 | 0.252 |
| 0.2 | 0.05 | yadI | 5 | 5.88333 | 0.363 |
| 0.2 | 0.05 | AG1 | 5 | 5.88333 | 0.328 |
| 0.2 | 0.05 | ptsI | 5 | 6.13333 | 0.287 |
| 0.2 | 0.05 | clcB | 5 | 6.13333 | 0.338 |
| 0.2 | 0.05 | ycaM | 5 | 6.13333 | 0.256 |
| 0.2 | 0.05 | yadI | 5 | 6.13333 | 0.375 |
| 0.2 | 0.05 | AG1 | 5 | 6.13333 | 0.344 |
| 0.2 | 0.05 | ptsI | 5 | 6.38333 | 0.299 |
| 0.2 | 0.05 | clcB | 5 | 6.38333 | 0.344 |
| 0.2 | 0.05 | ycaM | 5 | 6.38333 | 0.256 |
| 0.2 | 0.05 | yadI | 5 | 6.38333 | 0.386 |
| 0.2 | 0.05 | AG1 | 5 | 6.38333 | 0.357 |
| 0.2 | 0.05 | ptsI | 5 | 6.63333 | 0.308 |
| 0.2 | 0.05 | clcB | 5 | 6.63333 | 0.351 |
| 0.2 | 0.05 | ycaM | 5 | 6.63333 | 0.258 |
| 0.2 | 0.05 | yadI | 5 | 6.63333 | 0.4 |
| 0.2 | 0.05 | AG1 | 5 | 6.63333 | 0.371 |
| 0.2 | 0.05 | ptsI | 5 | 6.88333 | 0.32 |

| | | | | | |
|---|---|---|---|---|---|
| 0.2 | 0.05 | clcB | 5 | 6.88333 | 0.356 |
| 0.2 | 0.05 | ycaM | 5 | 6.88333 | 0.261 |
| 0.2 | 0.05 | yadI | 5 | 6.88333 | 0.414 |
| 0.2 | 0.05 | AG1 | 5 | 6.88333 | 0.385 |
| 0.2 | 0.05 | ptsI | 5 | 7.13333 | 0.336 |
| 0.2 | 0.05 | clcB | 5 | 7.13333 | 0.363 |
| 0.2 | 0.05 | ycaM | 5 | 7.13333 | 0.262 |
| 0.2 | 0.05 | yadI | 5 | 7.13333 | 0.427 |
| 0.2 | 0.05 | AG1 | 5 | 7.13333 | 0.399 |
| 0.2 | 0.05 | ptsI | 5 | 7.38333 | 0.348 |
| 0.2 | 0.05 | clcB | 5 | 7.38333 | 0.366 |
| 0.2 | 0.05 | ycaM | 5 | 7.38333 | 0.265 |
| 0.2 | 0.05 | yadI | 5 | 7.38333 | 0.44 |
| 0.2 | 0.05 | AG1 | 5 | 7.38333 | 0.411 |
| 0.2 | 0.05 | ptsI | 5 | 7.63333 | 0.362 |
| 0.2 | 0.05 | clcB | 5 | 7.63333 | 0.369 |
| 0.2 | 0.05 | ycaM | 5 | 7.63333 | 0.268 |
| 0.2 | 0.05 | yadI | 5 | 7.63333 | 0.45 |
| 0.2 | 0.05 | AG1 | 5 | 7.63333 | 0.422 |
| 0.2 | 0.05 | ptsI | 5 | 7.88333 | 0.372 |
| 0.2 | 0.05 | clcB | 5 | 7.88333 | 0.372 |
| 0.2 | 0.05 | ycaM | 5 | 7.88333 | 0.269 |
| 0.2 | 0.05 | yadI | 5 | 7.88333 | 0.462 |
| 0.2 | 0.05 | AG1 | 5 | 7.88333 | 0.432 |
| 0.2 | 0.05 | ptsI | 5 | 8.13333 | 0.388 |
| 0.2 | 0.05 | clcB | 5 | 8.13333 | 0.38 |
| 0.2 | 0.05 | ycaM | 5 | 8.13333 | 0.273 |
| 0.2 | 0.05 | yadI | 5 | 8.13333 | 0.471 |
| 0.2 | 0.05 | AG1 | 5 | 8.13333 | 0.448 |
| 0.2 | 0.05 | ptsI | 5 | 8.38333 | 0.405 |
| 0.2 | 0.05 | clcB | 5 | 8.38333 | 0.384 |
| 0.2 | 0.05 | ycaM | 5 | 8.38333 | 0.271 |
| 0.2 | 0.05 | yadI | 5 | 8.38333 | 0.476 |
| 0.2 | 0.05 | AG1 | 5 | 8.38333 | 0.46 |
| 0.2 | 0.05 | ptsI | 5 | 8.88333 | 0.35 |
| 0.2 | 0.05 | clcB | 5 | 8.88333 | 0.296 |
| 0.2 | 0.05 | ycaM | 5 | 8.88333 | 0.272 |
| 0.2 | 0.05 | yadI | 5 | 8.88333 | 0.48 |
| 0.2 | 0.05 | AG1 | 5 | 8.88333 | 0.499 |
| 0.2 | 0.05 | ptsI | 5 | 9.13333 | 0.356 |
| 0.2 | 0.05 | clcB | 5 | 9.13333 | 0.294 |
| 0.2 | 0.05 | ycaM | 5 | 9.13333 | 0.268 |
| 0.2 | 0.05 | yadI | 5 | 9.13333 | 0.482 |
| 0.2 | 0.05 | AG1 | 5 | 9.13333 | 0.503 |
| 0.2 | 0.05 | ptsI | 5 | 9.38333 | 0.37 |
| 0.2 | 0.05 | clcB | 5 | 9.38333 | 0.296 |
| 0.2 | 0.05 | ycaM | 5 | 9.38333 | 0.267 |
| 0.2 | 0.05 | yadI | 5 | 9.38333 | 0.493 |

| | | | | | |
|---|---|---|---|---|---|
| 0.2 | 0.05 | AG1 | 5 | 9.38333 | 0.525 |
| 0.2 | 0.05 | ptsI | 5 | 9.63333 | 0.375 |
| 0.2 | 0.05 | clcB | 5 | 9.63333 | 0.298 |
| 0.2 | 0.05 | ycaM | 5 | 9.63333 | 0.266 |
| 0.2 | 0.05 | yadI | 5 | 9.63333 | 0.504 |
| 0.2 | 0.05 | AG1 | 5 | 9.63333 | 0.536 |
| 0.2 | 0.05 | ptsI | 5 | 9.88333 | 0.385 |
| 0.2 | 0.05 | clcB | 5 | 9.88333 | 0.3 |
| 0.2 | 0.05 | ycaM | 5 | 9.88333 | 0.266 |
| 0.2 | 0.05 | yadI | 5 | 9.88333 | 0.513 |
| 0.2 | 0.05 | AG1 | 5 | 9.88333 | 0.55 |
| 0.2 | 0.05 | ptsI | 5 | 10.1333 | 0.393 |
| 0.2 | 0.05 | clcB | 5 | 10.1333 | 0.303 |
| 0.2 | 0.05 | ycaM | 5 | 10.1333 | 0.267 |
| 0.2 | 0.05 | yadI | 5 | 10.1333 | 0.521 |
| 0.2 | 0.05 | AG1 | 5 | 10.1333 | 0.562 |
| 0.2 | 0.05 | ptsI | 5 | 10.3833 | 0.404 |
| 0.2 | 0.05 | clcB | 5 | 10.3833 | 0.306 |
| 0.2 | 0.05 | ycaM | 5 | 10.3833 | 0.268 |
| 0.2 | 0.05 | yadI | 5 | 10.3833 | 0.534 |
| 0.2 | 0.05 | AG1 | 5 | 10.3833 | 0.575 |
| 0.2 | 0.05 | ptsI | 5 | 10.6333 | 0.411 |
| 0.2 | 0.05 | clcB | 5 | 10.6333 | 0.304 |
| 0.2 | 0.05 | ycaM | 5 | 10.6333 | 0.266 |
| 0.2 | 0.05 | yadI | 5 | 10.6333 | 0.536 |
| 0.2 | 0.05 | AG1 | 5 | 10.6333 | 0.585 |
| 0.2 | 0.05 | ptsI | 5 | 10.8833 | 0.421 |
| 0.2 | 0.05 | clcB | 5 | 10.8833 | 0.307 |
| 0.2 | 0.05 | ycaM | 5 | 10.8833 | 0.27 |
| 0.2 | 0.05 | yadI | 5 | 10.8833 | 0.546 |
| 0.2 | 0.05 | AG1 | 5 | 10.8833 | 0.597 |
| 0.2 | 0.05 | ptsI | 5 | 11.1333 | 0.428 |
| 0.2 | 0.05 | clcB | 5 | 11.1333 | 0.304 |
| 0.2 | 0.05 | ycaM | 5 | 11.1333 | 0.268 |
| 0.2 | 0.05 | yadI | 5 | 11.1333 | 0.552 |
| 0.2 | 0.05 | AG1 | 5 | 11.1333 | 0.609 |
| 0.2 | 0.05 | ptsI | 5 | 11.3833 | 0.436 |
| 0.2 | 0.05 | clcB | 5 | 11.3833 | 0.302 |
| 0.2 | 0.05 | ycaM | 5 | 11.3833 | 0.269 |
| 0.2 | 0.05 | yadI | 5 | 11.3833 | 0.559 |
| 0.2 | 0.05 | AG1 | 5 | 11.3833 | 0.62 |
| 0.2 | 0.05 | ptsI | 5 | 11.6333 | 0.447 |
| 0.2 | 0.05 | clcB | 5 | 11.6333 | 0.306 |
| 0.2 | 0.05 | ycaM | 5 | 11.6333 | 0.269 |
| 0.2 | 0.05 | yadI | 5 | 11.6333 | 0.568 |
| 0.2 | 0.05 | AG1 | 5 | 11.6333 | 0.63 |
| 0.2 | 0.05 | ptsI | 5 | 11.8833 | 0.454 |
| 0.2 | 0.05 | clcB | 5 | 11.8833 | 0.306 |

| | | | | | |
|---|---|---|---|---|---|
| 0.2 | 0.05 | ycaM | 5 | 11.8833 | 0.269 |
| 0.2 | 0.05 | yadI | 5 | 11.8833 | 0.576 |
| 0.2 | 0.05 | AG1 | 5 | 11.8833 | 0.64 |
| 0.2 | 0.05 | ptsI | 5 | 12.1333 | 0.466 |
| 0.2 | 0.05 | clcB | 5 | 12.1333 | 0.303 |
| 0.2 | 0.05 | ycaM | 5 | 12.1333 | 0.271 |
| 0.2 | 0.05 | yadI | 5 | 12.1333 | 0.583 |
| 0.2 | 0.05 | AG1 | 5 | 12.1333 | 0.653 |
| 0.2 | 0.05 | ptsI | 5 | 12.3833 | 0.467 |
| 0.2 | 0.05 | clcB | 5 | 12.3833 | 0.295 |
| 0.2 | 0.05 | ycaM | 5 | 12.3833 | 0.275 |
| 0.2 | 0.05 | yadI | 5 | 12.3833 | 0.596 |
| 0.2 | 0.05 | AG1 | 5 | 12.3833 | 0.661 |
| 0.2 | 0.05 | ptsI | 5 | 12.6333 | 0.479 |
| 0.2 | 0.05 | clcB | 5 | 12.6333 | 0.301 |
| 0.2 | 0.05 | ycaM | 5 | 12.6333 | 0.272 |
| 0.2 | 0.05 | yadI | 5 | 12.6333 | 0.594 |
| 0.2 | 0.05 | AG1 | 5 | 12.6333 | 0.672 |
| 0.2 | 0.05 | ptsI | 5 | 12.8833 | 0.491 |
| 0.2 | 0.05 | clcB | 5 | 12.8833 | 0.309 |
| 0.2 | 0.05 | ycaM | 5 | 12.8833 | 0.276 |
| 0.2 | 0.05 | yadI | 5 | 12.8833 | 0.601 |
| 0.2 | 0.05 | AG1 | 5 | 12.8833 | 0.679 |
| 0.2 | 0.05 | ptsI | 5 | 13.1333 | 0.497 |
| 0.2 | 0.05 | clcB | 5 | 13.1333 | 0.311 |
| 0.2 | 0.05 | ycaM | 5 | 13.1333 | 0.276 |
| 0.2 | 0.05 | yadI | 5 | 13.1333 | 0.607 |
| 0.2 | 0.05 | AG1 | 5 | 13.1333 | 0.692 |
| 0.2 | 0.05 | ptsI | 5 | 13.3833 | 0.505 |
| 0.2 | 0.05 | clcB | 5 | 13.3833 | 0.307 |
| 0.2 | 0.05 | ycaM | 5 | 13.3833 | 0.274 |
| 0.2 | 0.05 | yadI | 5 | 13.3833 | 0.612 |
| 0.2 | 0.05 | AG1 | 5 | 13.3833 | 0.699 |
| 0.2 | 0.05 | ptsI | 5 | 13.6333 | 0.513 |
| 0.2 | 0.05 | clcB | 5 | 13.6333 | 0.31 |
| 0.2 | 0.05 | ycaM | 5 | 13.6333 | 0.277 |
| 0.2 | 0.05 | yadI | 5 | 13.6333 | 0.618 |
| 0.2 | 0.05 | AG1 | 5 | 13.6333 | 0.707 |
| 0.2 | 0.05 | ptsI | 5 | 13.8833 | 0.518 |
| 0.2 | 0.05 | clcB | 5 | 13.8833 | 0.307 |
| 0.2 | 0.05 | ycaM | 5 | 13.8833 | 0.277 |
| 0.2 | 0.05 | yadI | 5 | 13.8833 | 0.618 |
| 0.2 | 0.05 | AG1 | 5 | 13.8833 | 0.712 |
| 0.2 | 0.05 | ptsI | 5 | 14.1333 | 0.525 |
| 0.2 | 0.05 | clcB | 5 | 14.1333 | 0.319 |
| 0.2 | 0.05 | ycaM | 5 | 14.1333 | 0.279 |
| 0.2 | 0.05 | yadI | 5 | 14.1333 | 0.625 |
| 0.2 | 0.05 | AG1 | 5 | 14.1333 | 0.72 |

| | | | | | |
|---|---|---|---|---|---|
| 0.2 | 0.05 | ptsI | 5 | 14.3833 | 0.536 |
| 0.2 | 0.05 | clcB | 5 | 14.3833 | 0.316 |
| 0.2 | 0.05 | ycaM | 5 | 14.3833 | 0.282 |
| 0.2 | 0.05 | yadI | 5 | 14.3833 | 0.63 |
| 0.2 | 0.05 | AG1 | 5 | 14.3833 | 0.725 |
| 0.2 | 0.05 | ptsI | 5 | 14.6333 | 0.545 |
| 0.2 | 0.05 | clcB | 5 | 14.6333 | 0.313 |
| 0.2 | 0.05 | ycaM | 5 | 14.6333 | 0.283 |
| 0.2 | 0.05 | yadI | 5 | 14.6333 | 0.635 |
| 0.2 | 0.05 | AG1 | 5 | 14.6333 | 0.729 |
| 0.2 | 0.05 | ptsI | 5 | 14.8833 | 0.549 |
| 0.2 | 0.05 | clcB | 5 | 14.8833 | 0.316 |
| 0.2 | 0.05 | ycaM | 5 | 14.8833 | 0.283 |
| 0.2 | 0.05 | yadI | 5 | 14.8833 | 0.641 |
| 0.2 | 0.05 | AG1 | 5 | 14.8833 | 0.739 |
| 0.2 | 0.05 | ptsI | 5 | 15.1333 | 0.555 |
| 0.2 | 0.05 | clcB | 5 | 15.1333 | 0.317 |
| 0.2 | 0.05 | ycaM | 5 | 15.1333 | 0.284 |
| 0.2 | 0.05 | yadI | 5 | 15.1333 | 0.646 |
| 0.2 | 0.05 | AG1 | 5 | 15.1333 | 0.742 |
| 0.2 | 0.05 | ptsI | 5 | 15.3833 | 0.56 |
| 0.2 | 0.05 | clcB | 5 | 15.3833 | 0.315 |
| 0.2 | 0.05 | ycaM | 5 | 15.3833 | 0.286 |
| 0.2 | 0.05 | yadI | 5 | 15.3833 | 0.647 |
| 0.2 | 0.05 | AG1 | 5 | 15.3833 | 0.749 |
| 0.2 | 0.05 | ptsI | 5 | 15.6333 | 0.568 |
| 0.2 | 0.05 | clcB | 5 | 15.6333 | 0.317 |
| 0.2 | 0.05 | ycaM | 5 | 15.6333 | 0.286 |
| 0.2 | 0.05 | yadI | 5 | 15.6333 | 0.654 |
| 0.2 | 0.05 | AG1 | 5 | 15.6333 | 0.755 |
| 0.2 | 0.05 | ptsI | 5 | 15.8833 | 0.574 |
| 0.2 | 0.05 | clcB | 5 | 15.8833 | 0.312 |
| 0.2 | 0.05 | ycaM | 5 | 15.8833 | 0.285 |
| 0.2 | 0.05 | yadI | 5 | 15.8833 | 0.658 |
| 0.2 | 0.05 | AG1 | 5 | 15.8833 | 0.76 |
| 0.2 | 0.05 | ptsI | 5 | 16.1333 | 0.585 |
| 0.2 | 0.05 | clcB | 5 | 16.1333 | 0.321 |
| 0.2 | 0.05 | ycaM | 5 | 16.1333 | 0.289 |
| 0.2 | 0.05 | yadI | 5 | 16.1333 | 0.662 |
| 0.2 | 0.05 | AG1 | 5 | 16.1333 | 0.772 |
| 0.2 | 0.05 | ptsI | 5 | 16.3833 | 0.586 |
| 0.2 | 0.05 | clcB | 5 | 16.3833 | 0.325 |
| 0.2 | 0.05 | ycaM | 5 | 16.3833 | 0.289 |
| 0.2 | 0.05 | yadI | 5 | 16.3833 | 0.659 |
| 0.2 | 0.05 | AG1 | 5 | 16.3833 | 0.77 |
| 0.2 | 0.05 | ptsI | 5 | 16.6333 | 0.592 |
| 0.2 | 0.05 | clcB | 5 | 16.6333 | 0.326 |
| 0.2 | 0.05 | ycaM | 5 | 16.6333 | 0.291 |

| | | | | | |
|---|---|---|---|---|---|
| 0.2 | 0.05 | yadI | 5 | 16.6333 | 0.665 |
| 0.2 | 0.05 | AG1 | 5 | 16.6333 | 0.776 |
| 0.2 | 0.05 | ptsI | 5 | 16.8833 | 0.595 |
| 0.2 | 0.05 | clcB | 5 | 16.8833 | 0.329 |
| 0.2 | 0.05 | ycaM | 5 | 16.8833 | 0.292 |
| 0.2 | 0.05 | yadI | 5 | 16.8833 | 0.668 |
| 0.2 | 0.05 | AG1 | 5 | 16.8833 | 0.777 |
| 0.2 | 0.05 | ptsI | 5 | 17.1333 | 0.602 |
| 0.2 | 0.05 | clcB | 5 | 17.1333 | 0.326 |
| 0.2 | 0.05 | ycaM | 5 | 17.1333 | 0.293 |
| 0.2 | 0.05 | yadI | 5 | 17.1333 | 0.674 |
| 0.2 | 0.05 | AG1 | 5 | 17.1333 | 0.785 |
| 0.2 | 0.05 | ptsI | 5 | 17.3833 | 0.604 |
| 0.2 | 0.05 | clcB | 5 | 17.3833 | 0.331 |
| 0.2 | 0.05 | ycaM | 5 | 17.3833 | 0.294 |
| 0.2 | 0.05 | yadI | 5 | 17.3833 | 0.675 |
| 0.2 | 0.05 | AG1 | 5 | 17.3833 | 0.791 |
| 0.2 | 0.05 | ptsI | 5 | 17.6333 | 0.61 |
| 0.2 | 0.05 | clcB | 5 | 17.6333 | 0.341 |
| 0.2 | 0.05 | ycaM | 5 | 17.6333 | 0.294 |
| 0.2 | 0.05 | yadI | 5 | 17.6333 | 0.678 |
| 0.2 | 0.05 | AG1 | 5 | 17.6333 | 0.793 |
| 0.2 | 0.05 | ptsI | 5 | 17.8833 | 0.612 |
| 0.2 | 0.05 | clcB | 5 | 17.8833 | 0.34 |
| 0.2 | 0.05 | ycaM | 5 | 17.8833 | 0.299 |
| 0.2 | 0.05 | yadI | 5 | 17.8833 | 0.689 |
| 0.2 | 0.05 | AG1 | 5 | 17.8833 | 0.796 |
| 0.2 | 0.05 | ptsI | 5 | 18.1333 | 0.621 |
| 0.2 | 0.05 | clcB | 5 | 18.1333 | 0.349 |
| 0.2 | 0.05 | ycaM | 5 | 18.1333 | 0.299 |
| 0.2 | 0.05 | yadI | 5 | 18.1333 | 0.68 |
| 0.2 | 0.05 | AG1 | 5 | 18.1333 | 0.798 |
| 0.2 | 0.05 | ptsI | 5 | 18.3833 | 0.625 |
| 0.2 | 0.05 | clcB | 5 | 18.3833 | 0.351 |
| 0.2 | 0.05 | ycaM | 5 | 18.3833 | 0.298 |
| 0.2 | 0.05 | yadI | 5 | 18.3833 | 0.688 |
| 0.2 | 0.05 | AG1 | 5 | 18.3833 | 0.804 |
| 0.2 | 0.05 | ptsI | 5 | 18.6333 | 0.626 |
| 0.2 | 0.05 | clcB | 5 | 18.6333 | 0.354 |
| 0.2 | 0.05 | ycaM | 5 | 18.6333 | 0.302 |
| 0.2 | 0.05 | yadI | 5 | 18.6333 | 0.689 |
| 0.2 | 0.05 | AG1 | 5 | 18.6333 | 0.806 |
| 0.2 | 0.05 | ptsI | 5 | 18.8833 | 0.634 |
| 0.2 | 0.05 | clcB | 5 | 18.8833 | 0.359 |
| 0.2 | 0.05 | ycaM | 5 | 18.8833 | 0.304 |
| 0.2 | 0.05 | yadI | 5 | 18.8833 | 0.69 |
| 0.2 | 0.05 | AG1 | 5 | 18.8833 | 0.814 |
| 0.2 | 0.05 | ptsI | 5 | 19.1333 | 0.637 |

| | | | | | |
|---|---|---|---|---|---|
| 0.2 | 0.05 | clcB | 5 | 19.1333 | 0.362 |
| 0.2 | 0.05 | ycaM | 5 | 19.1333 | 0.306 |
| 0.2 | 0.05 | yadI | 5 | 19.1333 | 0.694 |
| 0.2 | 0.05 | AG1 | 5 | 19.1333 | 0.816 |
| 0.2 | 0.05 | ptsI | 5 | 19.3833 | 0.637 |
| 0.2 | 0.05 | clcB | 5 | 19.3833 | 0.366 |
| 0.2 | 0.05 | ycaM | 5 | 19.3833 | 0.303 |
| 0.2 | 0.05 | yadI | 5 | 19.3833 | 0.699 |
| 0.2 | 0.05 | AG1 | 5 | 19.3833 | 0.818 |
| 0.2 | 0.05 | ptsI | 5 | 19.6333 | 0.639 |
| 0.2 | 0.05 | clcB | 5 | 19.6333 | 0.37 |
| 0.2 | 0.05 | ycaM | 5 | 19.6333 | 0.309 |
| 0.2 | 0.05 | yadI | 5 | 19.6333 | 0.702 |
| 0.2 | 0.05 | AG1 | 5 | 19.6333 | 0.821 |
| 0.2 | 0.05 | ptsI | 5 | 19.8833 | 0.643 |
| 0.2 | 0.05 | clcB | 5 | 19.8833 | 0.375 |
| 0.2 | 0.05 | ycaM | 5 | 19.8833 | 0.31 |
| 0.2 | 0.05 | yadI | 5 | 19.8833 | 0.701 |
| 0.2 | 0.05 | AG1 | 5 | 19.8833 | 0.826 |
| 0.2 | 0.05 | ptsI | 5 | 20.1333 | 0.642 |
| 0.2 | 0.05 | clcB | 5 | 20.1333 | 0.378 |
| 0.2 | 0.05 | ycaM | 5 | 20.1333 | 0.315 |
| 0.2 | 0.05 | yadI | 5 | 20.1333 | 0.704 |
| 0.2 | 0.05 | AG1 | 5 | 20.1333 | 0.829 |
| 0.2 | 0.05 | ptsI | 5 | 20.3833 | 0.645 |
| 0.2 | 0.05 | clcB | 5 | 20.3833 | 0.376 |
| 0.2 | 0.05 | ycaM | 5 | 20.3833 | 0.314 |
| 0.2 | 0.05 | yadI | 5 | 20.3833 | 0.706 |
| 0.2 | 0.05 | AG1 | 5 | 20.3833 | 0.832 |
| 0.2 | 0.05 | ptsI | 5 | 20.6333 | 0.638 |
| 0.2 | 0.05 | clcB | 5 | 20.6333 | 0.372 |
| 0.2 | 0.05 | ycaM | 5 | 20.6333 | 0.315 |
| 0.2 | 0.05 | yadI | 5 | 20.6333 | 0.714 |
| 0.2 | 0.05 | AG1 | 5 | 20.6333 | 0.828 |
| 0.2 | 0.05 | ptsI | 5 | 20.8833 | 0.648 |
| 0.2 | 0.05 | clcB | 5 | 20.8833 | 0.387 |
| 0.2 | 0.05 | ycaM | 5 | 20.8833 | 0.317 |
| 0.2 | 0.05 | yadI | 5 | 20.8833 | 0.712 |
| 0.2 | 0.05 | AG1 | 5 | 20.8833 | 0.836 |
| 0.2 | 0.05 | ptsI | 5 | 21.1333 | 0.65 |
| 0.2 | 0.05 | clcB | 5 | 21.1333 | 0.385 |
| 0.2 | 0.05 | ycaM | 5 | 21.1333 | 0.32 |
| 0.2 | 0.05 | yadI | 5 | 21.1333 | 0.712 |
| 0.2 | 0.05 | AG1 | 5 | 21.1333 | 0.838 |
| 0.2 | 0.05 | ptsI | 5 | 21.3833 | 0.65 |
| 0.2 | 0.05 | clcB | 5 | 21.3833 | 0.394 |
| 0.2 | 0.05 | ycaM | 5 | 21.3833 | 0.322 |
| 0.2 | 0.05 | yadI | 5 | 21.3833 | 0.713 |

| | | | | | |
|---|---|---|---|---|---|
| 0.2 | 0.05 | AG1 | 5 | 21.3833 | 0.839 |
| 0.2 | 0.05 | ptsI | 5 | 21.6333 | 0.652 |
| 0.2 | 0.05 | clcB | 5 | 21.6333 | 0.379 |
| 0.2 | 0.05 | ycaM | 5 | 21.6333 | 0.325 |
| 0.2 | 0.05 | yadI | 5 | 21.6333 | 0.716 |
| 0.2 | 0.05 | AG1 | 5 | 21.6333 | 0.839 |
| 0.2 | 0.05 | ptsI | 5 | 21.8833 | 0.651 |
| 0.2 | 0.05 | clcB | 5 | 21.8833 | 0.398 |
| 0.2 | 0.05 | ycaM | 5 | 21.8833 | 0.328 |
| 0.2 | 0.05 | yadI | 5 | 21.8833 | 0.715 |
| 0.2 | 0.05 | AG1 | 5 | 21.8833 | 0.839 |
| 0.2 | 0.05 | ptsI | 5 | 22.1333 | 0.648 |
| 0.2 | 0.05 | clcB | 5 | 22.1333 | 0.375 |
| 0.2 | 0.05 | ycaM | 5 | 22.1333 | 0.33 |
| 0.2 | 0.05 | yadI | 5 | 22.1333 | 0.726 |
| 0.2 | 0.05 | AG1 | 5 | 22.1333 | 0.84 |
| 0.2 | 0.05 | ptsI | 5 | 22.3833 | 0.638 |
| 0.2 | 0.05 | clcB | 5 | 22.3833 | 0.383 |
| 0.2 | 0.05 | ycaM | 5 | 22.3833 | 0.329 |
| 0.2 | 0.05 | yadI | 5 | 22.3833 | 0.724 |
| 0.2 | 0.05 | AG1 | 5 | 22.3833 | 0.845 |
| 0.2 | 0.05 | ptsI | 5 | 22.6333 | 0.638 |
| 0.2 | 0.05 | clcB | 5 | 22.6333 | 0.377 |
| 0.2 | 0.05 | ycaM | 5 | 22.6333 | 0.334 |
| 0.2 | 0.05 | yadI | 5 | 22.6333 | 0.718 |
| 0.2 | 0.05 | AG1 | 5 | 22.6333 | 0.846 |
| 0.2 | 0.05 | ptsI | 5 | 22.8833 | 0.642 |
| 0.2 | 0.05 | clcB | 5 | 22.8833 | 0.379 |
| 0.2 | 0.05 | ycaM | 5 | 22.8833 | 0.337 |
| 0.2 | 0.05 | yadI | 5 | 22.8833 | 0.721 |
| 0.2 | 0.05 | AG1 | 5 | 22.8833 | 0.848 |
| 0.2 | 0.05 | ptsI | 5 | 23.1333 | 0.647 |
| 0.2 | 0.05 | clcB | 5 | 23.1333 | 0.366 |
| 0.2 | 0.05 | ycaM | 5 | 23.1333 | 0.338 |
| 0.2 | 0.05 | yadI | 5 | 23.1333 | 0.719 |
| 0.2 | 0.05 | AG1 | 5 | 23.1333 | 0.85 |
| 0.2 | 0.05 | ptsI | 5 | 23.3833 | 0.653 |
| 0.2 | 0.05 | clcB | 5 | 23.3833 | 0.373 |
| 0.2 | 0.05 | ycaM | 5 | 23.3833 | 0.341 |
| 0.2 | 0.05 | yadI | 5 | 23.3833 | 0.724 |
| 0.2 | 0.05 | AG1 | 5 | 23.3833 | 0.852 |
| 0.2 | 0.05 | ptsI | 5 | 23.6333 | 0.651 |
| 0.2 | 0.05 | clcB | 5 | 23.6333 | 0.369 |
| 0.2 | 0.05 | ycaM | 5 | 23.6333 | 0.342 |
| 0.2 | 0.05 | yadI | 5 | 23.6333 | 0.727 |
| 0.2 | 0.05 | AG1 | 5 | 23.6333 | 0.855 |
| 0.2 | 0.05 | ptsI | 5 | 23.8833 | 0.651 |
| 0.2 | 0.05 | clcB | 5 | 23.8833 | 0.356 |

| | | | | | |
|---|---|---|---|---|---|
| 0.2 | 0.05 | ycaM | 5 | 23.8833 | 0.34 |
| 0.2 | 0.05 | yadI | 5 | 23.8833 | 0.732 |
| 0.2 | 0.05 | AG1 | 5 | 23.8833 | 0.851 |
| 0.2 | 0.05 | ptsI | 5 | 24.1333 | 0.65 |
| 0.2 | 0.05 | clcB | 5 | 24.1333 | 0.371 |
| 0.2 | 0.05 | ycaM | 5 | 24.1333 | 0.345 |
| 0.2 | 0.05 | yadI | 5 | 24.1333 | 0.723 |
| 0.2 | 0.05 | AG1 | 5 | 24.1333 | 0.858 |
| 0.2 | 0.05 | ptsI | 5 | 24.3833 | 0.656 |
| 0.2 | 0.05 | clcB | 5 | 24.3833 | 0.365 |
| 0.2 | 0.05 | ycaM | 5 | 24.3833 | 0.345 |
| 0.2 | 0.05 | yadI | 5 | 24.3833 | 0.728 |
| 0.2 | 0.05 | AG1 | 5 | 24.3833 | 0.856 |
| 0.4 | 0.05 | ptsI | 2 | 0 | 0.211 |
| 0.4 | 0.05 | clcB | 2 | 0 | 0.228 |
| 0.4 | 0.05 | ycaM | 2 | 0 | 0.217 |
| 0.4 | 0.05 | yadI | 2 | 0 | 0.221 |
| 0.4 | 0.05 | AG1 | 2 | 0 | 0.227 |
| 0.4 | 0.05 | ptsI | 2 | 0.25 | 0.211 |
| 0.4 | 0.05 | clcB | 2 | 0.25 | 0.22 |
| 0.4 | 0.05 | ycaM | 2 | 0.25 | 0.212 |
| 0.4 | 0.05 | yadI | 2 | 0.25 | 0.215 |
| 0.4 | 0.05 | AG1 | 2 | 0.25 | 0.22 |
| 0.4 | 0.05 | ptsI | 2 | 0.5 | 0.21 |
| 0.4 | 0.05 | clcB | 2 | 0.5 | 0.218 |
| 0.4 | 0.05 | ycaM | 2 | 0.5 | 0.211 |
| 0.4 | 0.05 | yadI | 2 | 0.5 | 0.214 |
| 0.4 | 0.05 | AG1 | 2 | 0.5 | 0.221 |
| 0.4 | 0.05 | ptsI | 2 | 0.75 | 0.214 |
| 0.4 | 0.05 | clcB | 2 | 0.75 | 0.218 |
| 0.4 | 0.05 | ycaM | 2 | 0.75 | 0.212 |
| 0.4 | 0.05 | yadI | 2 | 0.75 | 0.216 |
| 0.4 | 0.05 | AG1 | 2 | 0.75 | 0.221 |
| 0.4 | 0.05 | ptsI | 2 | 1 | 0.217 |
| 0.4 | 0.05 | clcB | 2 | 1 | 0.22 |
| 0.4 | 0.05 | ycaM | 2 | 1 | 0.214 |
| 0.4 | 0.05 | yadI | 2 | 1 | 0.223 |
| 0.4 | 0.05 | AG1 | 2 | 1 | 0.223 |
| 0.4 | 0.05 | ptsI | 2 | 1.25 | 0.219 |
| 0.4 | 0.05 | clcB | 2 | 1.25 | 0.222 |
| 0.4 | 0.05 | ycaM | 2 | 1.25 | 0.214 |
| 0.4 | 0.05 | yadI | 2 | 1.25 | 0.227 |
| 0.4 | 0.05 | AG1 | 2 | 1.25 | 0.227 |
| 0.4 | 0.05 | ptsI | 2 | 1.5 | 0.224 |
| 0.4 | 0.05 | clcB | 2 | 1.5 | 0.229 |
| 0.4 | 0.05 | ycaM | 2 | 1.5 | 0.214 |
| 0.4 | 0.05 | yadI | 2 | 1.5 | 0.233 |
| 0.4 | 0.05 | AG1 | 2 | 1.5 | 0.229 |

| | | | | | |
|---|---|---|---|---|---|
| 0.4 | 0.05 | ptsI | 2 | 1.75 | 0.23 |
| 0.4 | 0.05 | clcB | 2 | 1.75 | 0.23 |
| 0.4 | 0.05 | ycaM | 2 | 1.75 | 0.217 |
| 0.4 | 0.05 | yadI | 2 | 1.75 | 0.239 |
| 0.4 | 0.05 | AG1 | 2 | 1.75 | 0.234 |
| 0.4 | 0.05 | ptsI | 2 | 2 | 0.24 |
| 0.4 | 0.05 | clcB | 2 | 2 | 0.232 |
| 0.4 | 0.05 | ycaM | 2 | 2 | 0.225 |
| 0.4 | 0.05 | yadI | 2 | 2 | 0.247 |
| 0.4 | 0.05 | AG1 | 2 | 2 | 0.24 |
| 0.4 | 0.05 | ptsI | 2 | 2.25 | 0.249 |
| 0.4 | 0.05 | clcB | 2 | 2.25 | 0.235 |
| 0.4 | 0.05 | ycaM | 2 | 2.25 | 0.227 |
| 0.4 | 0.05 | yadI | 2 | 2.25 | 0.256 |
| 0.4 | 0.05 | AG1 | 2 | 2.25 | 0.248 |
| 0.4 | 0.05 | ptsI | 2 | 2.5 | 0.26 |
| 0.4 | 0.05 | clcB | 2 | 2.5 | 0.24 |
| 0.4 | 0.05 | ycaM | 2 | 2.5 | 0.235 |
| 0.4 | 0.05 | yadI | 2 | 2.5 | 0.266 |
| 0.4 | 0.05 | AG1 | 2 | 2.5 | 0.259 |
| 0.4 | 0.05 | ptsI | 2 | 2.75 | 0.256 |
| 0.4 | 0.05 | clcB | 2 | 2.75 | 0.244 |
| 0.4 | 0.05 | ycaM | 2 | 2.75 | 0.236 |
| 0.4 | 0.05 | yadI | 2 | 2.75 | 0.278 |
| 0.4 | 0.05 | AG1 | 2 | 2.75 | 0.253 |
| 0.4 | 0.05 | ptsI | 2 | 3 | 0.262 |
| 0.4 | 0.05 | clcB | 2 | 3 | 0.25 |
| 0.4 | 0.05 | ycaM | 2 | 3 | 0.24 |
| 0.4 | 0.05 | yadI | 2 | 3 | 0.284 |
| 0.4 | 0.05 | AG1 | 2 | 3 | 0.256 |
| 0.4 | 0.05 | ptsI | 2 | 3.25 | 0.286 |
| 0.4 | 0.05 | clcB | 2 | 3.25 | 0.256 |
| 0.4 | 0.05 | ycaM | 2 | 3.25 | 0.249 |
| 0.4 | 0.05 | yadI | 2 | 3.25 | 0.3 |
| 0.4 | 0.05 | AG1 | 2 | 3.25 | 0.269 |
| 0.4 | 0.05 | ptsI | 2 | 3.5 | 0.297 |
| 0.4 | 0.05 | clcB | 2 | 3.5 | 0.261 |
| 0.4 | 0.05 | ycaM | 2 | 3.5 | 0.255 |
| 0.4 | 0.05 | yadI | 2 | 3.5 | 0.311 |
| 0.4 | 0.05 | AG1 | 2 | 3.5 | 0.278 |
| 0.4 | 0.05 | ptsI | 2 | 3.75 | 0.311 |
| 0.4 | 0.05 | clcB | 2 | 3.75 | 0.269 |
| 0.4 | 0.05 | ycaM | 2 | 3.75 | 0.264 |
| 0.4 | 0.05 | yadI | 2 | 3.75 | 0.325 |
| 0.4 | 0.05 | AG1 | 2 | 3.75 | 0.287 |
| 0.4 | 0.05 | ptsI | 2 | 4 | 0.326 |
| 0.4 | 0.05 | clcB | 2 | 4 | 0.274 |
| 0.4 | 0.05 | ycaM | 2 | 4 | 0.275 |

| | | | | | |
|---|---|---|---|---|---|
| 0.4 | 0.05 | yadI | 2 | 4 | 0.339 |
| 0.4 | 0.05 | AG1 | 2 | 4 | 0.298 |
| 0.4 | 0.05 | ptsI | 2 | 4.38333 | 0.299 |
| 0.4 | 0.05 | clcB | 2 | 4.38333 | 0.247 |
| 0.4 | 0.05 | ycaM | 2 | 4.38333 | 0.292 |
| 0.4 | 0.05 | yadI | 2 | 4.38333 | 0.331 |
| 0.4 | 0.05 | AG1 | 2 | 4.38333 | 0.288 |
| 0.4 | 0.05 | ptsI | 2 | 4.63333 | 0.299 |
| 0.4 | 0.05 | clcB | 2 | 4.63333 | 0.245 |
| 0.4 | 0.05 | ycaM | 2 | 4.63333 | 0.29 |
| 0.4 | 0.05 | yadI | 2 | 4.63333 | 0.336 |
| 0.4 | 0.05 | AG1 | 2 | 4.63333 | 0.286 |
| 0.4 | 0.05 | ptsI | 2 | 4.88333 | 0.307 |
| 0.4 | 0.05 | clcB | 2 | 4.88333 | 0.247 |
| 0.4 | 0.05 | ycaM | 2 | 4.88333 | 0.29 |
| 0.4 | 0.05 | yadI | 2 | 4.88333 | 0.35 |
| 0.4 | 0.05 | AG1 | 2 | 4.88333 | 0.298 |
| 0.4 | 0.05 | ptsI | 2 | 5.13333 | 0.317 |
| 0.4 | 0.05 | clcB | 2 | 5.13333 | 0.252 |
| 0.4 | 0.05 | ycaM | 2 | 5.13333 | 0.294 |
| 0.4 | 0.05 | yadI | 2 | 5.13333 | 0.365 |
| 0.4 | 0.05 | AG1 | 2 | 5.13333 | 0.304 |
| 0.4 | 0.05 | ptsI | 2 | 5.38333 | 0.326 |
| 0.4 | 0.05 | clcB | 2 | 5.38333 | 0.258 |
| 0.4 | 0.05 | ycaM | 2 | 5.38333 | 0.293 |
| 0.4 | 0.05 | yadI | 2 | 5.38333 | 0.375 |
| 0.4 | 0.05 | AG1 | 2 | 5.38333 | 0.313 |
| 0.4 | 0.05 | ptsI | 2 | 5.63333 | 0.336 |
| 0.4 | 0.05 | clcB | 2 | 5.63333 | 0.268 |
| 0.4 | 0.05 | ycaM | 2 | 5.63333 | 0.295 |
| 0.4 | 0.05 | yadI | 2 | 5.63333 | 0.387 |
| 0.4 | 0.05 | AG1 | 2 | 5.63333 | 0.324 |
| 0.4 | 0.05 | ptsI | 2 | 5.88333 | 0.349 |
| 0.4 | 0.05 | clcB | 2 | 5.88333 | 0.275 |
| 0.4 | 0.05 | ycaM | 2 | 5.88333 | 0.294 |
| 0.4 | 0.05 | yadI | 2 | 5.88333 | 0.393 |
| 0.4 | 0.05 | AG1 | 2 | 5.88333 | 0.334 |
| 0.4 | 0.05 | ptsI | 2 | 6.13333 | 0.362 |
| 0.4 | 0.05 | clcB | 2 | 6.13333 | 0.28 |
| 0.4 | 0.05 | ycaM | 2 | 6.13333 | 0.297 |
| 0.4 | 0.05 | yadI | 2 | 6.13333 | 0.402 |
| 0.4 | 0.05 | AG1 | 2 | 6.13333 | 0.34 |
| 0.4 | 0.05 | ptsI | 2 | 6.38333 | 0.382 |
| 0.4 | 0.05 | clcB | 2 | 6.38333 | 0.286 |
| 0.4 | 0.05 | ycaM | 2 | 6.38333 | 0.299 |
| 0.4 | 0.05 | yadI | 2 | 6.38333 | 0.411 |
| 0.4 | 0.05 | AG1 | 2 | 6.38333 | 0.351 |
| 0.4 | 0.05 | ptsI | 2 | 6.63333 | 0.396 |

| | | | | | |
|---|---|---|---|---|---|
| 0.4 | 0.05 | clcB | 2 | 6.63333 | 0.291 |
| 0.4 | 0.05 | ycaM | 2 | 6.63333 | 0.298 |
| 0.4 | 0.05 | yadI | 2 | 6.63333 | 0.421 |
| 0.4 | 0.05 | AG1 | 2 | 6.63333 | 0.358 |
| 0.4 | 0.05 | ptsI | 2 | 6.88333 | 0.411 |
| 0.4 | 0.05 | clcB | 2 | 6.88333 | 0.298 |
| 0.4 | 0.05 | ycaM | 2 | 6.88333 | 0.299 |
| 0.4 | 0.05 | yadI | 2 | 6.88333 | 0.428 |
| 0.4 | 0.05 | AG1 | 2 | 6.88333 | 0.371 |
| 0.4 | 0.05 | ptsI | 2 | 7.13333 | 0.425 |
| 0.4 | 0.05 | clcB | 2 | 7.13333 | 0.302 |
| 0.4 | 0.05 | ycaM | 2 | 7.13333 | 0.299 |
| 0.4 | 0.05 | yadI | 2 | 7.13333 | 0.434 |
| 0.4 | 0.05 | AG1 | 2 | 7.13333 | 0.397 |
| 0.4 | 0.05 | ptsI | 2 | 7.38333 | 0.436 |
| 0.4 | 0.05 | clcB | 2 | 7.38333 | 0.308 |
| 0.4 | 0.05 | ycaM | 2 | 7.38333 | 0.301 |
| 0.4 | 0.05 | yadI | 2 | 7.38333 | 0.443 |
| 0.4 | 0.05 | AG1 | 2 | 7.38333 | 0.389 |
| 0.4 | 0.05 | ptsI | 2 | 7.63333 | 0.434 |
| 0.4 | 0.05 | clcB | 2 | 7.63333 | 0.312 |
| 0.4 | 0.05 | ycaM | 2 | 7.63333 | 0.298 |
| 0.4 | 0.05 | yadI | 2 | 7.63333 | 0.448 |
| 0.4 | 0.05 | AG1 | 2 | 7.63333 | 0.392 |
| 0.4 | 0.05 | ptsI | 2 | 7.88333 | 0.452 |
| 0.4 | 0.05 | clcB | 2 | 7.88333 | 0.317 |
| 0.4 | 0.05 | ycaM | 2 | 7.88333 | 0.299 |
| 0.4 | 0.05 | yadI | 2 | 7.88333 | 0.468 |
| 0.4 | 0.05 | AG1 | 2 | 7.88333 | 0.402 |
| 0.4 | 0.05 | ptsI | 2 | 8.13333 | 0.466 |
| 0.4 | 0.05 | clcB | 2 | 8.13333 | 0.318 |
| 0.4 | 0.05 | ycaM | 2 | 8.13333 | 0.3 |
| 0.4 | 0.05 | yadI | 2 | 8.13333 | 0.462 |
| 0.4 | 0.05 | AG1 | 2 | 8.13333 | 0.412 |
| 0.4 | 0.05 | ptsI | 2 | 8.38333 | 0.473 |
| 0.4 | 0.05 | clcB | 2 | 8.38333 | 0.33 |
| 0.4 | 0.05 | ycaM | 2 | 8.38333 | 0.3 |
| 0.4 | 0.05 | yadI | 2 | 8.38333 | 0.471 |
| 0.4 | 0.05 | AG1 | 2 | 8.38333 | 0.42 |
| 0.4 | 0.05 | ptsI | 2 | 8.91667 | 0.376 |
| 0.4 | 0.05 | clcB | 2 | 8.91667 | 0.342 |
| 0.4 | 0.05 | ycaM | 2 | 8.91667 | 0.324 |
| 0.4 | 0.05 | yadI | 2 | 8.91667 | 0.52 |
| 0.4 | 0.05 | AG1 | 2 | 8.91667 | 0.466 |
| 0.4 | 0.05 | ptsI | 2 | 9.16667 | 0.376 |
| 0.4 | 0.05 | clcB | 2 | 9.16667 | 0.341 |
| 0.4 | 0.05 | ycaM | 2 | 9.16667 | 0.322 |
| 0.4 | 0.05 | yadI | 2 | 9.16667 | 0.524 |

| | | | | | |
|---|---|---|---|---|---|
| 0.4 | 0.05 | AG1 | 2 | 9.16667 | 0.474 |
| 0.4 | 0.05 | ptsI | 2 | 9.41667 | 0.384 |
| 0.4 | 0.05 | clcB | 2 | 9.41667 | 0.342 |
| 0.4 | 0.05 | ycaM | 2 | 9.41667 | 0.321 |
| 0.4 | 0.05 | yadI | 2 | 9.41667 | 0.543 |
| 0.4 | 0.05 | AG1 | 2 | 9.41667 | 0.494 |
| 0.4 | 0.05 | ptsI | 2 | 9.66667 | 0.394 |
| 0.4 | 0.05 | clcB | 2 | 9.66667 | 0.345 |
| 0.4 | 0.05 | ycaM | 2 | 9.66667 | 0.322 |
| 0.4 | 0.05 | yadI | 2 | 9.66667 | 0.546 |
| 0.4 | 0.05 | AG1 | 2 | 9.66667 | 0.511 |
| 0.4 | 0.05 | ptsI | 2 | 9.91667 | 0.403 |
| 0.4 | 0.05 | clcB | 2 | 9.91667 | 0.35 |
| 0.4 | 0.05 | ycaM | 2 | 9.91667 | 0.322 |
| 0.4 | 0.05 | yadI | 2 | 9.91667 | 0.548 |
| 0.4 | 0.05 | AG1 | 2 | 9.91667 | 0.523 |
| 0.4 | 0.05 | ptsI | 2 | 10.1667 | 0.407 |
| 0.4 | 0.05 | clcB | 2 | 10.1667 | 0.353 |
| 0.4 | 0.05 | ycaM | 2 | 10.1667 | 0.321 |
| 0.4 | 0.05 | yadI | 2 | 10.1667 | 0.568 |
| 0.4 | 0.05 | AG1 | 2 | 10.1667 | 0.533 |
| 0.4 | 0.05 | ptsI | 2 | 10.4167 | 0.418 |
| 0.4 | 0.05 | clcB | 2 | 10.4167 | 0.36 |
| 0.4 | 0.05 | ycaM | 2 | 10.4167 | 0.326 |
| 0.4 | 0.05 | yadI | 2 | 10.4167 | 0.578 |
| 0.4 | 0.05 | AG1 | 2 | 10.4167 | 0.545 |
| 0.4 | 0.05 | ptsI | 2 | 10.6667 | 0.43 |
| 0.4 | 0.05 | clcB | 2 | 10.6667 | 0.362 |
| 0.4 | 0.05 | ycaM | 2 | 10.6667 | 0.325 |
| 0.4 | 0.05 | yadI | 2 | 10.6667 | 0.578 |
| 0.4 | 0.05 | AG1 | 2 | 10.6667 | 0.572 |
| 0.4 | 0.05 | ptsI | 2 | 10.9167 | 0.441 |
| 0.4 | 0.05 | clcB | 2 | 10.9167 | 0.361 |
| 0.4 | 0.05 | ycaM | 2 | 10.9167 | 0.324 |
| 0.4 | 0.05 | yadI | 2 | 10.9167 | 0.584 |
| 0.4 | 0.05 | AG1 | 2 | 10.9167 | 0.587 |
| 0.4 | 0.05 | ptsI | 2 | 11.1667 | 0.448 |
| 0.4 | 0.05 | clcB | 2 | 11.1667 | 0.36 |
| 0.4 | 0.05 | ycaM | 2 | 11.1667 | 0.324 |
| 0.4 | 0.05 | yadI | 2 | 11.1667 | 0.594 |
| 0.4 | 0.05 | AG1 | 2 | 11.1667 | 0.598 |
| 0.4 | 0.05 | ptsI | 2 | 11.4167 | 0.46 |
| 0.4 | 0.05 | clcB | 2 | 11.4167 | 0.363 |
| 0.4 | 0.05 | ycaM | 2 | 11.4167 | 0.324 |
| 0.4 | 0.05 | yadI | 2 | 11.4167 | 0.595 |
| 0.4 | 0.05 | AG1 | 2 | 11.4167 | 0.606 |
| 0.4 | 0.05 | ptsI | 2 | 11.6667 | 0.473 |
| 0.4 | 0.05 | clcB | 2 | 11.6667 | 0.363 |

| | | | | | |
|---|---|---|---|---|---|
| 0.4 | 0.05 | ycaM | 2 | 11.6667 | 0.325 |
| 0.4 | 0.05 | yadI | 2 | 11.6667 | 0.606 |
| 0.4 | 0.05 | AG1 | 2 | 11.6667 | 0.632 |
| 0.4 | 0.05 | ptsI | 2 | 11.9167 | 0.479 |
| 0.4 | 0.05 | clcB | 2 | 11.9167 | 0.356 |
| 0.4 | 0.05 | ycaM | 2 | 11.9167 | 0.324 |
| 0.4 | 0.05 | yadI | 2 | 11.9167 | 0.61 |
| 0.4 | 0.05 | AG1 | 2 | 11.9167 | 0.632 |
| 0.4 | 0.05 | ptsI | 2 | 12.1667 | 0.492 |
| 0.4 | 0.05 | clcB | 2 | 12.1667 | 0.359 |
| 0.4 | 0.05 | ycaM | 2 | 12.1667 | 0.323 |
| 0.4 | 0.05 | yadI | 2 | 12.1667 | 0.619 |
| 0.4 | 0.05 | AG1 | 2 | 12.1667 | 0.656 |
| 0.4 | 0.05 | ptsI | 2 | 12.4167 | 0.5 |
| 0.4 | 0.05 | clcB | 2 | 12.4167 | 0.356 |
| 0.4 | 0.05 | ycaM | 2 | 12.4167 | 0.325 |
| 0.4 | 0.05 | yadI | 2 | 12.4167 | 0.626 |
| 0.4 | 0.05 | AG1 | 2 | 12.4167 | 0.674 |
| 0.4 | 0.05 | ptsI | 2 | 12.6667 | 0.502 |
| 0.4 | 0.05 | clcB | 2 | 12.6667 | 0.345 |
| 0.4 | 0.05 | ycaM | 2 | 12.6667 | 0.322 |
| 0.4 | 0.05 | yadI | 2 | 12.6667 | 0.63 |
| 0.4 | 0.05 | AG1 | 2 | 12.6667 | 0.669 |
| 0.4 | 0.05 | ptsI | 2 | 12.9167 | 0.523 |
| 0.4 | 0.05 | clcB | 2 | 12.9167 | 0.344 |
| 0.4 | 0.05 | ycaM | 2 | 12.9167 | 0.325 |
| 0.4 | 0.05 | yadI | 2 | 12.9167 | 0.639 |
| 0.4 | 0.05 | AG1 | 2 | 12.9167 | 0.669 |
| 0.4 | 0.05 | ptsI | 2 | 13.1667 | 0.541 |
| 0.4 | 0.05 | clcB | 2 | 13.1667 | 0.342 |
| 0.4 | 0.05 | ycaM | 2 | 13.1667 | 0.322 |
| 0.4 | 0.05 | yadI | 2 | 13.1667 | 0.648 |
| 0.4 | 0.05 | AG1 | 2 | 13.1667 | 0.672 |
| 0.4 | 0.05 | ptsI | 2 | 13.4167 | 0.555 |
| 0.4 | 0.05 | clcB | 2 | 13.4167 | 0.343 |
| 0.4 | 0.05 | ycaM | 2 | 13.4167 | 0.319 |
| 0.4 | 0.05 | yadI | 2 | 13.4167 | 0.663 |
| 0.4 | 0.05 | AG1 | 2 | 13.4167 | 0.649 |
| 0.4 | 0.05 | ptsI | 2 | 13.6667 | 0.57 |
| 0.4 | 0.05 | clcB | 2 | 13.6667 | 0.346 |
| 0.4 | 0.05 | ycaM | 2 | 13.6667 | 0.319 |
| 0.4 | 0.05 | yadI | 2 | 13.6667 | 0.662 |
| 0.4 | 0.05 | AG1 | 2 | 13.6667 | 0.68 |
| 0.4 | 0.05 | ptsI | 2 | 13.9167 | 0.575 |
| 0.4 | 0.05 | clcB | 2 | 13.9167 | 0.348 |
| 0.4 | 0.05 | ycaM | 2 | 13.9167 | 0.32 |
| 0.4 | 0.05 | yadI | 2 | 13.9167 | 0.67 |
| 0.4 | 0.05 | AG1 | 2 | 13.9167 | 0.706 |

| | | | | | |
|---|---|---|---|---|---|
| 0.4 | 0.05 | ptsI | 2 | 14.1667 | 0.577 |
| 0.4 | 0.05 | clcB | 2 | 14.1667 | 0.349 |
| 0.4 | 0.05 | ycaM | 2 | 14.1667 | 0.319 |
| 0.4 | 0.05 | yadI | 2 | 14.1667 | 0.664 |
| 0.4 | 0.05 | AG1 | 2 | 14.1667 | 0.695 |
| 0.4 | 0.05 | ptsI | 2 | 14.4167 | 0.59 |
| 0.4 | 0.05 | clcB | 2 | 14.4167 | 0.345 |
| 0.4 | 0.05 | ycaM | 2 | 14.4167 | 0.318 |
| 0.4 | 0.05 | yadI | 2 | 14.4167 | 0.68 |
| 0.4 | 0.05 | AG1 | 2 | 14.4167 | 0.671 |
| 0.4 | 0.05 | ptsI | 2 | 14.6667 | 0.596 |
| 0.4 | 0.05 | clcB | 2 | 14.6667 | 0.34 |
| 0.4 | 0.05 | ycaM | 2 | 14.6667 | 0.317 |
| 0.4 | 0.05 | yadI | 2 | 14.6667 | 0.697 |
| 0.4 | 0.05 | AG1 | 2 | 14.6667 | 0.679 |
| 0.4 | 0.05 | ptsI | 2 | 14.9167 | 0.608 |
| 0.4 | 0.05 | clcB | 2 | 14.9167 | 0.346 |
| 0.4 | 0.05 | ycaM | 2 | 14.9167 | 0.319 |
| 0.4 | 0.05 | yadI | 2 | 14.9167 | 0.711 |
| 0.4 | 0.05 | AG1 | 2 | 14.9167 | 0.687 |
| 0.4 | 0.05 | ptsI | 2 | 15.1667 | 0.617 |
| 0.4 | 0.05 | clcB | 2 | 15.1667 | 0.349 |
| 0.4 | 0.05 | ycaM | 2 | 15.1667 | 0.319 |
| 0.4 | 0.05 | yadI | 2 | 15.1667 | 0.694 |
| 0.4 | 0.05 | AG1 | 2 | 15.1667 | 0.682 |
| 0.4 | 0.05 | ptsI | 2 | 15.4167 | 0.635 |
| 0.4 | 0.05 | clcB | 2 | 15.4167 | 0.346 |
| 0.4 | 0.05 | ycaM | 2 | 15.4167 | 0.315 |
| 0.4 | 0.05 | yadI | 2 | 15.4167 | 0.709 |
| 0.4 | 0.05 | AG1 | 2 | 15.4167 | 0.712 |
| 0.4 | 0.05 | ptsI | 2 | 15.6667 | 0.617 |
| 0.4 | 0.05 | clcB | 2 | 15.6667 | 0.344 |
| 0.4 | 0.05 | ycaM | 2 | 15.6667 | 0.312 |
| 0.4 | 0.05 | yadI | 2 | 15.6667 | 0.702 |
| 0.4 | 0.05 | AG1 | 2 | 15.6667 | 0.693 |
| 0.4 | 0.05 | ptsI | 2 | 15.9167 | 0.63 |
| 0.4 | 0.05 | clcB | 2 | 15.9167 | 0.344 |
| 0.4 | 0.05 | ycaM | 2 | 15.9167 | 0.315 |
| 0.4 | 0.05 | yadI | 2 | 15.9167 | 0.706 |
| 0.4 | 0.05 | AG1 | 2 | 15.9167 | 0.717 |
| 0.4 | 0.05 | ptsI | 2 | 16.1667 | 0.641 |
| 0.4 | 0.05 | clcB | 2 | 16.1667 | 0.341 |
| 0.4 | 0.05 | ycaM | 2 | 16.1667 | 0.315 |
| 0.4 | 0.05 | yadI | 2 | 16.1667 | 0.738 |
| 0.4 | 0.05 | AG1 | 2 | 16.1667 | 0.729 |
| 0.4 | 0.05 | ptsI | 2 | 16.4167 | 0.643 |
| 0.4 | 0.05 | clcB | 2 | 16.4167 | 0.351 |
| 0.4 | 0.05 | ycaM | 2 | 16.4167 | 0.316 |

| | | | | | |
|---|---|---|---|---|---|
| 0.4 | 0.05 | yadI | 2 | 16.4167 | 0.714 |
| 0.4 | 0.05 | AG1 | 2 | 16.4167 | 0.743 |
| 0.4 | 0.05 | ptsI | 2 | 16.6667 | 0.645 |
| 0.4 | 0.05 | clcB | 2 | 16.6667 | 0.351 |
| 0.4 | 0.05 | ycaM | 2 | 16.6667 | 0.313 |
| 0.4 | 0.05 | yadI | 2 | 16.6667 | 0.718 |
| 0.4 | 0.05 | AG1 | 2 | 16.6667 | 0.751 |
| 0.4 | 0.05 | ptsI | 2 | 16.9167 | 0.65 |
| 0.4 | 0.05 | clcB | 2 | 16.9167 | 0.35 |
| 0.4 | 0.05 | ycaM | 2 | 16.9167 | 0.314 |
| 0.4 | 0.05 | yadI | 2 | 16.9167 | 0.735 |
| 0.4 | 0.05 | AG1 | 2 | 16.9167 | 0.766 |
| 0.4 | 0.05 | ptsI | 2 | 17.1667 | 0.664 |
| 0.4 | 0.05 | clcB | 2 | 17.1667 | 0.35 |
| 0.4 | 0.05 | ycaM | 2 | 17.1667 | 0.315 |
| 0.4 | 0.05 | yadI | 2 | 17.1667 | 0.731 |
| 0.4 | 0.05 | AG1 | 2 | 17.1667 | 0.767 |
| 0.4 | 0.05 | ptsI | 2 | 17.4167 | 0.669 |
| 0.4 | 0.05 | clcB | 2 | 17.4167 | 0.342 |
| 0.4 | 0.05 | ycaM | 2 | 17.4167 | 0.314 |
| 0.4 | 0.05 | yadI | 2 | 17.4167 | 0.748 |
| 0.4 | 0.05 | AG1 | 2 | 17.4167 | 0.775 |
| 0.4 | 0.05 | ptsI | 2 | 17.6667 | 0.676 |
| 0.4 | 0.05 | clcB | 2 | 17.6667 | 0.344 |
| 0.4 | 0.05 | ycaM | 2 | 17.6667 | 0.314 |
| 0.4 | 0.05 | yadI | 2 | 17.6667 | 0.75 |
| 0.4 | 0.05 | AG1 | 2 | 17.6667 | 0.771 |
| 0.4 | 0.05 | ptsI | 2 | 17.9167 | 0.673 |
| 0.4 | 0.05 | clcB | 2 | 17.9167 | 0.343 |
| 0.4 | 0.05 | ycaM | 2 | 17.9167 | 0.314 |
| 0.4 | 0.05 | yadI | 2 | 17.9167 | 0.746 |
| 0.4 | 0.05 | AG1 | 2 | 17.9167 | 0.764 |
| 0.4 | 0.05 | ptsI | 2 | 18.1667 | 0.679 |
| 0.4 | 0.05 | clcB | 2 | 18.1667 | 0.345 |
| 0.4 | 0.05 | ycaM | 2 | 18.1667 | 0.314 |
| 0.4 | 0.05 | yadI | 2 | 18.1667 | 0.753 |
| 0.4 | 0.05 | AG1 | 2 | 18.1667 | 0.766 |
| 0.4 | 0.05 | ptsI | 2 | 18.4167 | 0.685 |
| 0.4 | 0.05 | clcB | 2 | 18.4167 | 0.344 |
| 0.4 | 0.05 | ycaM | 2 | 18.4167 | 0.317 |
| 0.4 | 0.05 | yadI | 2 | 18.4167 | 0.744 |
| 0.4 | 0.05 | AG1 | 2 | 18.4167 | 0.774 |
| 0.4 | 0.05 | ptsI | 2 | 18.6667 | 0.687 |
| 0.4 | 0.05 | clcB | 2 | 18.6667 | 0.343 |
| 0.4 | 0.05 | ycaM | 2 | 18.6667 | 0.315 |
| 0.4 | 0.05 | yadI | 2 | 18.6667 | 0.744 |
| 0.4 | 0.05 | AG1 | 2 | 18.6667 | 0.774 |
| 0.4 | 0.05 | ptsI | 2 | 18.9167 | 0.688 |

| | | | | | |
|---|---|---|---|---|---|
| 0.4 | 0.05 | clcB | 2 | 18.9167 | 0.341 |
| 0.4 | 0.05 | ycaM | 2 | 18.9167 | 0.309 |
| 0.4 | 0.05 | yadI | 2 | 18.9167 | 0.73 |
| 0.4 | 0.05 | AG1 | 2 | 18.9167 | 0.772 |
| 0.4 | 0.05 | ptsI | 2 | 19.1667 | 0.684 |
| 0.4 | 0.05 | clcB | 2 | 19.1667 | 0.344 |
| 0.4 | 0.05 | ycaM | 2 | 19.1667 | 0.314 |
| 0.4 | 0.05 | yadI | 2 | 19.1667 | 0.748 |
| 0.4 | 0.05 | AG1 | 2 | 19.1667 | 0.778 |
| 0.4 | 0.05 | ptsI | 2 | 19.4167 | 0.69 |
| 0.4 | 0.05 | clcB | 2 | 19.4167 | 0.345 |
| 0.4 | 0.05 | ycaM | 2 | 19.4167 | 0.315 |
| 0.4 | 0.05 | yadI | 2 | 19.4167 | 0.753 |
| 0.4 | 0.05 | AG1 | 2 | 19.4167 | 0.783 |
| 0.4 | 0.05 | ptsI | 2 | 19.6667 | 0.693 |
| 0.4 | 0.05 | clcB | 2 | 19.6667 | 0.342 |
| 0.4 | 0.05 | ycaM | 2 | 19.6667 | 0.318 |
| 0.4 | 0.05 | yadI | 2 | 19.6667 | 0.756 |
| 0.4 | 0.05 | AG1 | 2 | 19.6667 | 0.783 |
| 0.4 | 0.05 | ptsI | 2 | 19.9167 | 0.693 |
| 0.4 | 0.05 | clcB | 2 | 19.9167 | 0.343 |
| 0.4 | 0.05 | ycaM | 2 | 19.9167 | 0.32 |
| 0.4 | 0.05 | yadI | 2 | 19.9167 | 0.756 |
| 0.4 | 0.05 | AG1 | 2 | 19.9167 | 0.79 |
| 0.4 | 0.05 | ptsI | 2 | 20.1667 | 0.691 |
| 0.4 | 0.05 | clcB | 2 | 20.1667 | 0.344 |
| 0.4 | 0.05 | ycaM | 2 | 20.1667 | 0.321 |
| 0.4 | 0.05 | yadI | 2 | 20.1667 | 0.766 |
| 0.4 | 0.05 | AG1 | 2 | 20.1667 | 0.793 |
| 0.4 | 0.05 | ptsI | 2 | 20.4167 | 0.693 |
| 0.4 | 0.05 | clcB | 2 | 20.4167 | 0.343 |
| 0.4 | 0.05 | ycaM | 2 | 20.4167 | 0.322 |
| 0.4 | 0.05 | yadI | 2 | 20.4167 | 0.762 |
| 0.4 | 0.05 | AG1 | 2 | 20.4167 | 0.8 |
| 0.4 | 0.05 | ptsI | 2 | 20.6667 | 0.693 |
| 0.4 | 0.05 | clcB | 2 | 20.6667 | 0.344 |
| 0.4 | 0.05 | ycaM | 2 | 20.6667 | 0.323 |
| 0.4 | 0.05 | yadI | 2 | 20.6667 | 0.776 |
| 0.4 | 0.05 | AG1 | 2 | 20.6667 | 0.799 |
| 0.4 | 0.05 | ptsI | 2 | 20.9167 | 0.698 |
| 0.4 | 0.05 | clcB | 2 | 20.9167 | 0.344 |
| 0.4 | 0.05 | ycaM | 2 | 20.9167 | 0.324 |
| 0.4 | 0.05 | yadI | 2 | 20.9167 | 0.768 |
| 0.4 | 0.05 | AG1 | 2 | 20.9167 | 0.806 |
| 0.4 | 0.05 | ptsI | 2 | 21.1667 | 0.702 |
| 0.4 | 0.05 | clcB | 2 | 21.1667 | 0.346 |
| 0.4 | 0.05 | ycaM | 2 | 21.1667 | 0.331 |
| 0.4 | 0.05 | yadI | 2 | 21.1667 | 0.788 |

| | | | | | |
|---|---|---|---|---|---|
| 0.4 | 0.05 | AG1 | 2 | 21.1667 | 0.818 |
| 0.4 | 0.05 | ptsI | 2 | 21.4167 | 0.698 |
| 0.4 | 0.05 | clcB | 2 | 21.4167 | 0.344 |
| 0.4 | 0.05 | ycaM | 2 | 21.4167 | 0.326 |
| 0.4 | 0.05 | yadI | 2 | 21.4167 | 0.783 |
| 0.4 | 0.05 | AG1 | 2 | 21.4167 | 0.817 |
| 0.4 | 0.05 | ptsI | 2 | 21.6667 | 0.693 |
| 0.4 | 0.05 | clcB | 2 | 21.6667 | 0.347 |
| 0.4 | 0.05 | ycaM | 2 | 21.6667 | 0.333 |
| 0.4 | 0.05 | yadI | 2 | 21.6667 | 0.789 |
| 0.4 | 0.05 | AG1 | 2 | 21.6667 | 0.833 |
| 0.4 | 0.05 | ptsI | 2 | 21.9167 | 0.69 |
| 0.4 | 0.05 | clcB | 2 | 21.9167 | 0.347 |
| 0.4 | 0.05 | ycaM | 2 | 21.9167 | 0.335 |
| 0.4 | 0.05 | yadI | 2 | 21.9167 | 0.791 |
| 0.4 | 0.05 | AG1 | 2 | 21.9167 | 0.836 |
| 0.4 | 0.05 | ptsI | 2 | 22.1667 | 0.701 |
| 0.4 | 0.05 | clcB | 2 | 22.1667 | 0.347 |
| 0.4 | 0.05 | ycaM | 2 | 22.1667 | 0.339 |
| 0.4 | 0.05 | yadI | 2 | 22.1667 | 0.797 |
| 0.4 | 0.05 | AG1 | 2 | 22.1667 | 0.834 |
| 0.4 | 0.05 | ptsI | 2 | 22.4167 | 0.706 |
| 0.4 | 0.05 | clcB | 2 | 22.4167 | 0.346 |
| 0.4 | 0.05 | ycaM | 2 | 22.4167 | 0.344 |
| 0.4 | 0.05 | yadI | 2 | 22.4167 | 0.8 |
| 0.4 | 0.05 | AG1 | 2 | 22.4167 | 0.843 |
| 0.4 | 0.05 | ptsI | 2 | 22.6667 | 0.7 |
| 0.4 | 0.05 | clcB | 2 | 22.6667 | 0.345 |
| 0.4 | 0.05 | ycaM | 2 | 22.6667 | 0.347 |
| 0.4 | 0.05 | yadI | 2 | 22.6667 | 0.798 |
| 0.4 | 0.05 | AG1 | 2 | 22.6667 | 0.839 |
| 0.4 | 0.05 | ptsI | 2 | 22.9167 | 0.707 |
| 0.4 | 0.05 | clcB | 2 | 22.9167 | 0.349 |
| 0.4 | 0.05 | ycaM | 2 | 22.9167 | 0.347 |
| 0.4 | 0.05 | yadI | 2 | 22.9167 | 0.803 |
| 0.4 | 0.05 | AG1 | 2 | 22.9167 | 0.862 |
| 0.4 | 0.05 | ptsI | 2 | 23.1667 | 0.699 |
| 0.4 | 0.05 | clcB | 2 | 23.1667 | 0.348 |
| 0.4 | 0.05 | ycaM | 2 | 23.1667 | 0.35 |
| 0.4 | 0.05 | yadI | 2 | 23.1667 | 0.806 |
| 0.4 | 0.05 | AG1 | 2 | 23.1667 | 0.851 |
| 0.4 | 0.05 | ptsI | 2 | 23.4167 | 0.705 |
| 0.4 | 0.05 | clcB | 2 | 23.4167 | 0.352 |
| 0.4 | 0.05 | ycaM | 2 | 23.4167 | 0.352 |
| 0.4 | 0.05 | yadI | 2 | 23.4167 | 0.812 |
| 0.4 | 0.05 | AG1 | 2 | 23.4167 | 0.857 |
| 0.4 | 0.05 | ptsI | 2 | 23.6667 | 0.704 |
| 0.4 | 0.05 | clcB | 2 | 23.6667 | 0.348 |

| | | | | | |
|---|---|---|---|---|---|
| 0.4 | 0.05 | ycaM | 2 | 23.6667 | 0.353 |
| 0.4 | 0.05 | yadI | 2 | 23.6667 | 0.811 |
| 0.4 | 0.05 | AG1 | 2 | 23.6667 | 0.862 |
| 0.4 | 0.05 | ptsI | 2 | 23.9167 | 0.702 |
| 0.4 | 0.05 | clcB | 2 | 23.9167 | 0.347 |
| 0.4 | 0.05 | ycaM | 2 | 23.9167 | 0.357 |
| 0.4 | 0.05 | yadI | 2 | 23.9167 | 0.81 |
| 0.4 | 0.05 | AG1 | 2 | 23.9167 | 0.868 |
| 0.4 | 0.05 | ptsI | 2 | 24.1667 | 0.702 |
| 0.4 | 0.05 | clcB | 2 | 24.1667 | 0.352 |
| 0.4 | 0.05 | ycaM | 2 | 24.1667 | 0.36 |
| 0.4 | 0.05 | yadI | 2 | 24.1667 | 0.814 |
| 0.4 | 0.05 | AG1 | 2 | 24.1667 | 0.877 |
| 0.4 | 0.05 | ptsI | 2 | 24.4167 | 0.7 |
| 0.4 | 0.05 | clcB | 2 | 24.4167 | 0.353 |
| 0.4 | 0.05 | ycaM | 2 | 24.4167 | 0.361 |
| 0.4 | 0.05 | yadI | 2 | 24.4167 | 0.821 |
| 0.4 | 0.05 | AG1 | 2 | 24.4167 | 0.884 |
| 0.4 | 0.05 | ptsI | 2 | 24.6667 | 0.701 |
| 0.4 | 0.05 | clcB | 2 | 24.6667 | 0.352 |
| 0.4 | 0.05 | ycaM | 2 | 24.6667 | 0.36 |
| 0.4 | 0.05 | yadI | 2 | 24.6667 | 0.818 |
| 0.4 | 0.05 | AG1 | 2 | 24.6667 | 0.892 |
| 0.4 | 0.05 | ptsI | 2 | 24.9167 | 0.706 |
| 0.4 | 0.05 | clcB | 2 | 24.9167 | 0.352 |
| 0.4 | 0.05 | ycaM | 2 | 24.9167 | 0.36 |
| 0.4 | 0.05 | yadI | 2 | 24.9167 | 0.82 |
| 0.4 | 0.05 | AG1 | 2 | 24.9167 | 0.9 |
| 0.4 | 0.05 | ptsI | 2 | 25.1667 | 0.703 |
| 0.4 | 0.05 | clcB | 2 | 25.1667 | 0.354 |
| 0.4 | 0.05 | ycaM | 2 | 25.1667 | 0.364 |
| 0.4 | 0.05 | yadI | 2 | 25.1667 | 0.825 |
| 0.4 | 0.05 | AG1 | 2 | 25.1667 | 0.908 |
| 0.4 | 0.05 | ptsI | 2 | 25.4167 | 0.698 |
| 0.4 | 0.05 | clcB | 2 | 25.4167 | 0.357 |
| 0.4 | 0.05 | ycaM | 2 | 25.4167 | 0.366 |
| 0.4 | 0.05 | yadI | 2 | 25.4167 | 0.828 |
| 0.4 | 0.05 | AG1 | 2 | 25.4167 | 0.914 |
| 0.4 | 0.05 | ptsI | 2 | 25.6667 | 0.698 |
| 0.4 | 0.05 | clcB | 2 | 25.6667 | 0.358 |
| 0.4 | 0.05 | ycaM | 2 | 25.6667 | 0.367 |
| 0.4 | 0.05 | yadI | 2 | 25.6667 | 0.835 |
| 0.4 | 0.05 | AG1 | 2 | 25.6667 | 0.925 |
| 0.4 | 0.05 | ptsI | 2 | 25.9167 | 0.698 |
| 0.4 | 0.05 | clcB | 2 | 25.9167 | 0.364 |
| 0.4 | 0.05 | ycaM | 2 | 25.9167 | 0.374 |
| 0.4 | 0.05 | yadI | 2 | 25.9167 | 0.83 |
| 0.4 | 0.05 | AG1 | 2 | 25.9167 | 0.935 |

| | | | | | |
|---|---|---|---|---|---|
| 0.4 | 0.05 | ptsI | 2 | 26.1667 | 0.688 |
| 0.4 | 0.05 | clcB | 2 | 26.1667 | 0.362 |
| 0.4 | 0.05 | ycaM | 2 | 26.1667 | 0.372 |
| 0.4 | 0.05 | yadI | 2 | 26.1667 | 0.827 |
| 0.4 | 0.05 | AG1 | 2 | 26.1667 | 0.936 |
| 0.4 | 0.05 | ptsI | 2 | 26.4167 | 0.697 |
| 0.4 | 0.05 | clcB | 2 | 26.4167 | 0.362 |
| 0.4 | 0.05 | ycaM | 2 | 26.4167 | 0.379 |
| 0.4 | 0.05 | yadI | 2 | 26.4167 | 0.831 |
| 0.4 | 0.05 | AG1 | 2 | 26.4167 | 0.957 |
| 0.4 | 0.05 | ptsI | 3 | 0 | 0.166 |
| 0.4 | 0.05 | clcB | 3 | 0 | 0.16 |
| 0.4 | 0.05 | ycaM | 3 | 0 | 0.158 |
| 0.4 | 0.05 | yadI | 3 | 0 | 0.158 |
| 0.4 | 0.05 | AG1 | 3 | 0 | 0.159 |
| 0.4 | 0.05 | ptsI | 3 | 0.25 | 0.161 |
| 0.4 | 0.05 | clcB | 3 | 0.25 | 0.159 |
| 0.4 | 0.05 | ycaM | 3 | 0.25 | 0.154 |
| 0.4 | 0.05 | yadI | 3 | 0.25 | 0.155 |
| 0.4 | 0.05 | AG1 | 3 | 0.25 | 0.155 |
| 0.4 | 0.05 | ptsI | 3 | 0.5 | 0.16 |
| 0.4 | 0.05 | clcB | 3 | 0.5 | 0.158 |
| 0.4 | 0.05 | ycaM | 3 | 0.5 | 0.153 |
| 0.4 | 0.05 | yadI | 3 | 0.5 | 0.153 |
| 0.4 | 0.05 | AG1 | 3 | 0.5 | 0.155 |
| 0.4 | 0.05 | ptsI | 3 | 0.75 | 0.16 |
| 0.4 | 0.05 | clcB | 3 | 0.75 | 0.159 |
| 0.4 | 0.05 | ycaM | 3 | 0.75 | 0.152 |
| 0.4 | 0.05 | yadI | 3 | 0.75 | 0.169 |
| 0.4 | 0.05 | AG1 | 3 | 0.75 | 0.155 |
| 0.4 | 0.05 | ptsI | 3 | 1 | 0.162 |
| 0.4 | 0.05 | clcB | 3 | 1 | 0.16 |
| 0.4 | 0.05 | ycaM | 3 | 1 | 0.152 |
| 0.4 | 0.05 | yadI | 3 | 1 | 0.153 |
| 0.4 | 0.05 | AG1 | 3 | 1 | 0.155 |
| 0.4 | 0.05 | ptsI | 3 | 1.25 | 0.162 |
| 0.4 | 0.05 | clcB | 3 | 1.25 | 0.16 |
| 0.4 | 0.05 | ycaM | 3 | 1.25 | 0.152 |
| 0.4 | 0.05 | yadI | 3 | 1.25 | 0.153 |
| 0.4 | 0.05 | AG1 | 3 | 1.25 | 0.154 |
| 0.4 | 0.05 | ptsI | 3 | 1.5 | 0.163 |
| 0.4 | 0.05 | clcB | 3 | 1.5 | 0.16 |
| 0.4 | 0.05 | ycaM | 3 | 1.5 | 0.153 |
| 0.4 | 0.05 | yadI | 3 | 1.5 | 0.154 |
| 0.4 | 0.05 | AG1 | 3 | 1.5 | 0.155 |
| 0.4 | 0.05 | ptsI | 3 | 1.75 | 0.166 |
| 0.4 | 0.05 | clcB | 3 | 1.75 | 0.162 |
| 0.4 | 0.05 | ycaM | 3 | 1.75 | 0.152 |

| | | | | | |
|---|---|---|---|---|---|
| 0.4 | 0.05 | yadI | 3 | 1.75 | 0.155 |
| 0.4 | 0.05 | AG1 | 3 | 1.75 | 0.155 |
| 0.4 | 0.05 | ptsI | 3 | 2 | 0.167 |
| 0.4 | 0.05 | clcB | 3 | 2 | 0.165 |
| 0.4 | 0.05 | ycaM | 3 | 2 | 0.154 |
| 0.4 | 0.05 | yadI | 3 | 2 | 0.157 |
| 0.4 | 0.05 | AG1 | 3 | 2 | 0.157 |
| 0.4 | 0.05 | ptsI | 3 | 2.25 | 0.168 |
| 0.4 | 0.05 | clcB | 3 | 2.25 | 0.165 |
| 0.4 | 0.05 | ycaM | 3 | 2.25 | 0.155 |
| 0.4 | 0.05 | yadI | 3 | 2.25 | 0.158 |
| 0.4 | 0.05 | AG1 | 3 | 2.25 | 0.158 |
| 0.4 | 0.05 | ptsI | 3 | 2.5 | 0.17 |
| 0.4 | 0.05 | clcB | 3 | 2.5 | 0.17 |
| 0.4 | 0.05 | ycaM | 3 | 2.5 | 0.157 |
| 0.4 | 0.05 | yadI | 3 | 2.5 | 0.16 |
| 0.4 | 0.05 | AG1 | 3 | 2.5 | 0.16 |
| 0.4 | 0.05 | ptsI | 3 | 2.75 | 0.17 |
| 0.4 | 0.05 | clcB | 3 | 2.75 | 0.174 |
| 0.4 | 0.05 | ycaM | 3 | 2.75 | 0.159 |
| 0.4 | 0.05 | yadI | 3 | 2.75 | 0.163 |
| 0.4 | 0.05 | AG1 | 3 | 2.75 | 0.161 |
| 0.4 | 0.05 | ptsI | 3 | 3 | 0.171 |
| 0.4 | 0.05 | clcB | 3 | 3 | 0.175 |
| 0.4 | 0.05 | ycaM | 3 | 3 | 0.159 |
| 0.4 | 0.05 | yadI | 3 | 3 | 0.166 |
| 0.4 | 0.05 | AG1 | 3 | 3 | 0.163 |
| 0.4 | 0.05 | ptsI | 3 | 3.25 | 0.176 |
| 0.4 | 0.05 | clcB | 3 | 3.25 | 0.175 |
| 0.4 | 0.05 | ycaM | 3 | 3.25 | 0.161 |
| 0.4 | 0.05 | yadI | 3 | 3.25 | 0.168 |
| 0.4 | 0.05 | AG1 | 3 | 3.25 | 0.164 |
| 0.4 | 0.05 | ptsI | 3 | 3.5 | 0.18 |
| 0.4 | 0.05 | clcB | 3 | 3.5 | 0.18 |
| 0.4 | 0.05 | ycaM | 3 | 3.5 | 0.164 |
| 0.4 | 0.05 | yadI | 3 | 3.5 | 0.173 |
| 0.4 | 0.05 | AG1 | 3 | 3.5 | 0.166 |
| 0.4 | 0.05 | ptsI | 3 | 3.75 | 0.182 |
| 0.4 | 0.05 | clcB | 3 | 3.75 | 0.183 |
| 0.4 | 0.05 | ycaM | 3 | 3.75 | 0.166 |
| 0.4 | 0.05 | yadI | 3 | 3.75 | 0.178 |
| 0.4 | 0.05 | AG1 | 3 | 3.75 | 0.168 |
| 0.4 | 0.05 | ptsI | 3 | 4 | 0.184 |
| 0.4 | 0.05 | clcB | 3 | 4 | 0.186 |
| 0.4 | 0.05 | ycaM | 3 | 4 | 0.168 |
| 0.4 | 0.05 | yadI | 3 | 4 | 0.183 |
| 0.4 | 0.05 | AG1 | 3 | 4 | 0.17 |
| 0.4 | 0.05 | ptsI | 3 | 4.41667 | 0.206 |

| | | | | | |
|---|---|---|---|---|---|
| 0.4 | 0.05 | clcB | 3 | 4.41667 | 0.21 |
| 0.4 | 0.05 | ycaM | 3 | 4.41667 | 0.172 |
| 0.4 | 0.05 | yadI | 3 | 4.41667 | 0.214 |
| 0.4 | 0.05 | AG1 | 3 | 4.41667 | 0.192 |
| 0.4 | 0.05 | ptsI | 3 | 4.66667 | 0.195 |
| 0.4 | 0.05 | clcB | 3 | 4.66667 | 0.186 |
| 0.4 | 0.05 | ycaM | 3 | 4.66667 | 0.17 |
| 0.4 | 0.05 | yadI | 3 | 4.66667 | 0.22 |
| 0.4 | 0.05 | AG1 | 3 | 4.66667 | 0.192 |
| 0.4 | 0.05 | ptsI | 3 | 4.91667 | 0.192 |
| 0.4 | 0.05 | clcB | 3 | 4.91667 | 0.191 |
| 0.4 | 0.05 | ycaM | 3 | 4.91667 | 0.171 |
| 0.4 | 0.05 | yadI | 3 | 4.91667 | 0.24 |
| 0.4 | 0.05 | AG1 | 3 | 4.91667 | 0.197 |
| 0.4 | 0.05 | ptsI | 3 | 5.16667 | 0.195 |
| 0.4 | 0.05 | clcB | 3 | 5.16667 | 0.196 |
| 0.4 | 0.05 | ycaM | 3 | 5.16667 | 0.175 |
| 0.4 | 0.05 | yadI | 3 | 5.16667 | 0.254 |
| 0.4 | 0.05 | AG1 | 3 | 5.16667 | 0.202 |
| 0.4 | 0.05 | ptsI | 3 | 5.41667 | 0.2 |
| 0.4 | 0.05 | clcB | 3 | 5.41667 | 0.2 |
| 0.4 | 0.05 | ycaM | 3 | 5.41667 | 0.181 |
| 0.4 | 0.05 | yadI | 3 | 5.41667 | 0.262 |
| 0.4 | 0.05 | AG1 | 3 | 5.41667 | 0.206 |
| 0.4 | 0.05 | ptsI | 3 | 5.66667 | 0.205 |
| 0.4 | 0.05 | clcB | 3 | 5.66667 | 0.208 |
| 0.4 | 0.05 | ycaM | 3 | 5.66667 | 0.187 |
| 0.4 | 0.05 | yadI | 3 | 5.66667 | 0.276 |
| 0.4 | 0.05 | AG1 | 3 | 5.66667 | 0.212 |
| 0.4 | 0.05 | ptsI | 3 | 5.91667 | 0.208 |
| 0.4 | 0.05 | clcB | 3 | 5.91667 | 0.213 |
| 0.4 | 0.05 | ycaM | 3 | 5.91667 | 0.193 |
| 0.4 | 0.05 | yadI | 3 | 5.91667 | 0.282 |
| 0.4 | 0.05 | AG1 | 3 | 5.91667 | 0.214 |
| 0.4 | 0.05 | ptsI | 3 | 6.16667 | 0.216 |
| 0.4 | 0.05 | clcB | 3 | 6.16667 | 0.219 |
| 0.4 | 0.05 | ycaM | 3 | 6.16667 | 0.201 |
| 0.4 | 0.05 | yadI | 3 | 6.16667 | 0.29 |
| 0.4 | 0.05 | AG1 | 3 | 6.16667 | 0.221 |
| 0.4 | 0.05 | ptsI | 3 | 6.41667 | 0.223 |
| 0.4 | 0.05 | clcB | 3 | 6.41667 | 0.227 |
| 0.4 | 0.05 | ycaM | 3 | 6.41667 | 0.21 |
| 0.4 | 0.05 | yadI | 3 | 6.41667 | 0.299 |
| 0.4 | 0.05 | AG1 | 3 | 6.41667 | 0.23 |
| 0.4 | 0.05 | ptsI | 3 | 6.66667 | 0.233 |
| 0.4 | 0.05 | clcB | 3 | 6.66667 | 0.235 |
| 0.4 | 0.05 | ycaM | 3 | 6.66667 | 0.219 |
| 0.4 | 0.05 | yadI | 3 | 6.66667 | 0.307 |

| | | | | | |
|---|---|---|---|---|---|
| 0.4 | 0.05 | AG1 | 3 | 6.66667 | 0.24 |
| 0.4 | 0.05 | ptsI | 3 | 6.91667 | 0.243 |
| 0.4 | 0.05 | clcB | 3 | 6.91667 | 0.242 |
| 0.4 | 0.05 | ycaM | 3 | 6.91667 | 0.226 |
| 0.4 | 0.05 | yadI | 3 | 6.91667 | 0.315 |
| 0.4 | 0.05 | AG1 | 3 | 6.91667 | 0.253 |
| 0.4 | 0.05 | ptsI | 3 | 7.16667 | 0.254 |
| 0.4 | 0.05 | clcB | 3 | 7.16667 | 0.254 |
| 0.4 | 0.05 | ycaM | 3 | 7.16667 | 0.233 |
| 0.4 | 0.05 | yadI | 3 | 7.16667 | 0.325 |
| 0.4 | 0.05 | AG1 | 3 | 7.16667 | 0.26 |
| 0.4 | 0.05 | ptsI | 3 | 7.41667 | 0.263 |
| 0.4 | 0.05 | clcB | 3 | 7.41667 | 0.266 |
| 0.4 | 0.05 | ycaM | 3 | 7.41667 | 0.236 |
| 0.4 | 0.05 | yadI | 3 | 7.41667 | 0.336 |
| 0.4 | 0.05 | AG1 | 3 | 7.41667 | 0.269 |
| 0.4 | 0.05 | ptsI | 3 | 7.66667 | 0.272 |
| 0.4 | 0.05 | clcB | 3 | 7.66667 | 0.274 |
| 0.4 | 0.05 | ycaM | 3 | 7.66667 | 0.24 |
| 0.4 | 0.05 | yadI | 3 | 7.66667 | 0.347 |
| 0.4 | 0.05 | AG1 | 3 | 7.66667 | 0.274 |
| 0.4 | 0.05 | ptsI | 3 | 7.91667 | 0.278 |
| 0.4 | 0.05 | clcB | 3 | 7.91667 | 0.28 |
| 0.4 | 0.05 | ycaM | 3 | 7.91667 | 0.241 |
| 0.4 | 0.05 | yadI | 3 | 7.91667 | 0.357 |
| 0.4 | 0.05 | AG1 | 3 | 7.91667 | 0.28 |
| 0.4 | 0.05 | ptsI | 3 | 8.16667 | 0.289 |
| 0.4 | 0.05 | clcB | 3 | 8.16667 | 0.286 |
| 0.4 | 0.05 | ycaM | 3 | 8.16667 | 0.244 |
| 0.4 | 0.05 | yadI | 3 | 8.16667 | 0.364 |
| 0.4 | 0.05 | AG1 | 3 | 8.16667 | 0.288 |
| 0.4 | 0.05 | ptsI | 3 | 8.41667 | 0.298 |
| 0.4 | 0.05 | clcB | 3 | 8.41667 | 0.292 |
| 0.4 | 0.05 | ycaM | 3 | 8.41667 | 0.246 |
| 0.4 | 0.05 | yadI | 3 | 8.41667 | 0.373 |
| 0.4 | 0.05 | AG1 | 3 | 8.41667 | 0.296 |
| 0.4 | 0.05 | ptsI | 3 | 9.05 | 0.29 |
| 0.4 | 0.05 | clcB | 3 | 9.05 | 0.281 |
| 0.4 | 0.05 | ycaM | 3 | 9.05 | 0.257 |
| 0.4 | 0.05 | yadI | 3 | 9.05 | 0.466 |
| 0.4 | 0.05 | AG1 | 3 | 9.05 | 0.358 |
| 0.4 | 0.05 | ptsI | 3 | 9.3 | 0.296 |
| 0.4 | 0.05 | clcB | 3 | 9.3 | 0.281 |
| 0.4 | 0.05 | ycaM | 3 | 9.3 | 0.252 |
| 0.4 | 0.05 | yadI | 3 | 9.3 | 0.473 |
| 0.4 | 0.05 | AG1 | 3 | 9.3 | 0.372 |
| 0.4 | 0.05 | ptsI | 3 | 9.55 | 0.305 |
| 0.4 | 0.05 | clcB | 3 | 9.55 | 0.289 |

| | | | | | |
|---|---|---|---|---|---|
| 0.4 | 0.05 | ycaM | 3 | 9.55 | 0.257 |
| 0.4 | 0.05 | yadI | 3 | 9.55 | 0.49 |
| 0.4 | 0.05 | AG1 | 3 | 9.55 | 0.386 |
| 0.4 | 0.05 | ptsI | 3 | 9.8 | 0.31 |
| 0.4 | 0.05 | clcB | 3 | 9.8 | 0.29 |
| 0.4 | 0.05 | ycaM | 3 | 9.8 | 0.255 |
| 0.4 | 0.05 | yadI | 3 | 9.8 | 0.505 |
| 0.4 | 0.05 | AG1 | 3 | 9.8 | 0.396 |
| 0.4 | 0.05 | ptsI | 3 | 10.05 | 0.321 |
| 0.4 | 0.05 | clcB | 3 | 10.05 | 0.295 |
| 0.4 | 0.05 | ycaM | 3 | 10.05 | 0.256 |
| 0.4 | 0.05 | yadI | 3 | 10.05 | 0.519 |
| 0.4 | 0.05 | AG1 | 3 | 10.05 | 0.406 |
| 0.4 | 0.05 | ptsI | 3 | 10.3 | 0.326 |
| 0.4 | 0.05 | clcB | 3 | 10.3 | 0.297 |
| 0.4 | 0.05 | ycaM | 3 | 10.3 | 0.26 |
| 0.4 | 0.05 | yadI | 3 | 10.3 | 0.531 |
| 0.4 | 0.05 | AG1 | 3 | 10.3 | 0.415 |
| 0.4 | 0.05 | ptsI | 3 | 10.55 | 0.337 |
| 0.4 | 0.05 | clcB | 3 | 10.55 | 0.303 |
| 0.4 | 0.05 | ycaM | 3 | 10.55 | 0.258 |
| 0.4 | 0.05 | yadI | 3 | 10.55 | 0.539 |
| 0.4 | 0.05 | AG1 | 3 | 10.55 | 0.422 |
| 0.4 | 0.05 | ptsI | 3 | 10.8 | 0.343 |
| 0.4 | 0.05 | clcB | 3 | 10.8 | 0.301 |
| 0.4 | 0.05 | ycaM | 3 | 10.8 | 0.26 |
| 0.4 | 0.05 | yadI | 3 | 10.8 | 0.553 |
| 0.4 | 0.05 | AG1 | 3 | 10.8 | 0.431 |
| 0.4 | 0.05 | ptsI | 3 | 11.05 | 0.356 |
| 0.4 | 0.05 | clcB | 3 | 11.05 | 0.306 |
| 0.4 | 0.05 | ycaM | 3 | 11.05 | 0.262 |
| 0.4 | 0.05 | yadI | 3 | 11.05 | 0.566 |
| 0.4 | 0.05 | AG1 | 3 | 11.05 | 0.444 |
| 0.4 | 0.05 | ptsI | 3 | 11.3 | 0.364 |
| 0.4 | 0.05 | clcB | 3 | 11.3 | 0.308 |
| 0.4 | 0.05 | ycaM | 3 | 11.3 | 0.262 |
| 0.4 | 0.05 | yadI | 3 | 11.3 | 0.572 |
| 0.4 | 0.05 | AG1 | 3 | 11.3 | 0.45 |
| 0.4 | 0.05 | ptsI | 3 | 11.55 | 0.374 |
| 0.4 | 0.05 | clcB | 3 | 11.55 | 0.313 |
| 0.4 | 0.05 | ycaM | 3 | 11.55 | 0.264 |
| 0.4 | 0.05 | yadI | 3 | 11.55 | 0.583 |
| 0.4 | 0.05 | AG1 | 3 | 11.55 | 0.461 |
| 0.4 | 0.05 | ptsI | 3 | 11.8 | 0.384 |
| 0.4 | 0.05 | clcB | 3 | 11.8 | 0.305 |
| 0.4 | 0.05 | ycaM | 3 | 11.8 | 0.264 |
| 0.4 | 0.05 | yadI | 3 | 11.8 | 0.599 |
| 0.4 | 0.05 | AG1 | 3 | 11.8 | 0.472 |

| | | | | | |
|---|---|---|---|---|---|
| 0.4 | 0.05 | ptsI | 3 | 12.05 | 0.391 |
| 0.4 | 0.05 | clcB | 3 | 12.05 | 0.299 |
| 0.4 | 0.05 | ycaM | 3 | 12.05 | 0.265 |
| 0.4 | 0.05 | yadI | 3 | 12.05 | 0.604 |
| 0.4 | 0.05 | AG1 | 3 | 12.05 | 0.481 |
| 0.4 | 0.05 | ptsI | 3 | 12.3 | 0.4 |
| 0.4 | 0.05 | clcB | 3 | 12.3 | 0.305 |
| 0.4 | 0.05 | ycaM | 3 | 12.3 | 0.263 |
| 0.4 | 0.05 | yadI | 3 | 12.3 | 0.621 |
| 0.4 | 0.05 | AG1 | 3 | 12.3 | 0.496 |
| 0.4 | 0.05 | ptsI | 3 | 12.55 | 0.406 |
| 0.4 | 0.05 | clcB | 3 | 12.55 | 0.302 |
| 0.4 | 0.05 | ycaM | 3 | 12.55 | 0.262 |
| 0.4 | 0.05 | yadI | 3 | 12.55 | 0.634 |
| 0.4 | 0.05 | AG1 | 3 | 12.55 | 0.509 |
| 0.4 | 0.05 | ptsI | 3 | 12.8 | 0.414 |
| 0.4 | 0.05 | clcB | 3 | 12.8 | 0.305 |
| 0.4 | 0.05 | ycaM | 3 | 12.8 | 0.262 |
| 0.4 | 0.05 | yadI | 3 | 12.8 | 0.643 |
| 0.4 | 0.05 | AG1 | 3 | 12.8 | 0.52 |
| 0.4 | 0.05 | ptsI | 3 | 13.05 | 0.42 |
| 0.4 | 0.05 | clcB | 3 | 13.05 | 0.296 |
| 0.4 | 0.05 | ycaM | 3 | 13.05 | 0.259 |
| 0.4 | 0.05 | yadI | 3 | 13.05 | 0.641 |
| 0.4 | 0.05 | AG1 | 3 | 13.05 | 0.53 |
| 0.4 | 0.05 | ptsI | 3 | 13.3 | 0.43 |
| 0.4 | 0.05 | clcB | 3 | 13.3 | 0.298 |
| 0.4 | 0.05 | ycaM | 3 | 13.3 | 0.26 |
| 0.4 | 0.05 | yadI | 3 | 13.3 | 0.649 |
| 0.4 | 0.05 | AG1 | 3 | 13.3 | 0.547 |
| 0.4 | 0.05 | ptsI | 3 | 13.55 | 0.439 |
| 0.4 | 0.05 | clcB | 3 | 13.55 | 0.292 |
| 0.4 | 0.05 | ycaM | 3 | 13.55 | 0.258 |
| 0.4 | 0.05 | yadI | 3 | 13.55 | 0.668 |
| 0.4 | 0.05 | AG1 | 3 | 13.55 | 0.562 |
| 0.4 | 0.05 | ptsI | 3 | 13.8 | 0.443 |
| 0.4 | 0.05 | clcB | 3 | 13.8 | 0.29 |
| 0.4 | 0.05 | ycaM | 3 | 13.8 | 0.264 |
| 0.4 | 0.05 | yadI | 3 | 13.8 | 0.664 |
| 0.4 | 0.05 | AG1 | 3 | 13.8 | 0.573 |
| 0.4 | 0.05 | ptsI | 3 | 14.05 | 0.45 |
| 0.4 | 0.05 | clcB | 3 | 14.05 | 0.29 |
| 0.4 | 0.05 | ycaM | 3 | 14.05 | 0.259 |
| 0.4 | 0.05 | yadI | 3 | 14.05 | 0.678 |
| 0.4 | 0.05 | AG1 | 3 | 14.05 | 0.588 |
| 0.4 | 0.05 | ptsI | 3 | 14.3 | 0.453 |
| 0.4 | 0.05 | clcB | 3 | 14.3 | 0.286 |
| 0.4 | 0.05 | ycaM | 3 | 14.3 | 0.255 |

| | | | | | |
|---|---|---|---|---|---|
| 0.4 | 0.05 | yadI | 3 | 14.3 | 0.686 |
| 0.4 | 0.05 | AG1 | 3 | 14.3 | 0.59 |
| 0.4 | 0.05 | ptsI | 3 | 14.55 | 0.462 |
| 0.4 | 0.05 | clcB | 3 | 14.55 | 0.285 |
| 0.4 | 0.05 | ycaM | 3 | 14.55 | 0.254 |
| 0.4 | 0.05 | yadI | 3 | 14.55 | 0.682 |
| 0.4 | 0.05 | AG1 | 3 | 14.55 | 0.604 |
| 0.4 | 0.05 | ptsI | 3 | 14.8 | 0.471 |
| 0.4 | 0.05 | clcB | 3 | 14.8 | 0.286 |
| 0.4 | 0.05 | ycaM | 3 | 14.8 | 0.254 |
| 0.4 | 0.05 | yadI | 3 | 14.8 | 0.691 |
| 0.4 | 0.05 | AG1 | 3 | 14.8 | 0.621 |
| 0.4 | 0.05 | ptsI | 3 | 15.05 | 0.476 |
| 0.4 | 0.05 | clcB | 3 | 15.05 | 0.288 |
| 0.4 | 0.05 | ycaM | 3 | 15.05 | 0.256 |
| 0.4 | 0.05 | yadI | 3 | 15.05 | 0.712 |
| 0.4 | 0.05 | AG1 | 3 | 15.05 | 0.632 |
| 0.4 | 0.05 | ptsI | 3 | 15.3 | 0.479 |
| 0.4 | 0.05 | clcB | 3 | 15.3 | 0.288 |
| 0.4 | 0.05 | ycaM | 3 | 15.3 | 0.257 |
| 0.4 | 0.05 | yadI | 3 | 15.3 | 0.703 |
| 0.4 | 0.05 | AG1 | 3 | 15.3 | 0.632 |
| 0.4 | 0.05 | ptsI | 3 | 15.55 | 0.484 |
| 0.4 | 0.05 | clcB | 3 | 15.55 | 0.285 |
| 0.4 | 0.05 | ycaM | 3 | 15.55 | 0.257 |
| 0.4 | 0.05 | yadI | 3 | 15.55 | 0.72 |
| 0.4 | 0.05 | AG1 | 3 | 15.55 | 0.648 |
| 0.4 | 0.05 | ptsI | 3 | 15.8 | 0.49 |
| 0.4 | 0.05 | clcB | 3 | 15.8 | 0.284 |
| 0.4 | 0.05 | ycaM | 3 | 15.8 | 0.252 |
| 0.4 | 0.05 | yadI | 3 | 15.8 | 0.722 |
| 0.4 | 0.05 | AG1 | 3 | 15.8 | 0.648 |
| 0.4 | 0.05 | ptsI | 3 | 16.05 | 0.49 |
| 0.4 | 0.05 | clcB | 3 | 16.05 | 0.284 |
| 0.4 | 0.05 | ycaM | 3 | 16.05 | 0.256 |
| 0.4 | 0.05 | yadI | 3 | 16.05 | 0.718 |
| 0.4 | 0.05 | AG1 | 3 | 16.05 | 0.655 |
| 0.4 | 0.05 | ptsI | 3 | 16.3 | 0.494 |
| 0.4 | 0.05 | clcB | 3 | 16.3 | 0.284 |
| 0.4 | 0.05 | ycaM | 3 | 16.3 | 0.256 |
| 0.4 | 0.05 | yadI | 3 | 16.3 | 0.722 |
| 0.4 | 0.05 | AG1 | 3 | 16.3 | 0.662 |
| 0.4 | 0.05 | ptsI | 3 | 16.55 | 0.501 |
| 0.4 | 0.05 | clcB | 3 | 16.55 | 0.286 |
| 0.4 | 0.05 | ycaM | 3 | 16.55 | 0.26 |
| 0.4 | 0.05 | yadI | 3 | 16.55 | 0.741 |
| 0.4 | 0.05 | AG1 | 3 | 16.55 | 0.68 |
| 0.4 | 0.05 | ptsI | 3 | 16.8 | 0.503 |

| | | | | | |
|---|---|---|---|---|---|
| 0.4 | 0.05 | clcB | 3 | 16.8 | 0.286 |
| 0.4 | 0.05 | ycaM | 3 | 16.8 | 0.261 |
| 0.4 | 0.05 | yadI | 3 | 16.8 | 0.753 |
| 0.4 | 0.05 | AG1 | 3 | 16.8 | 0.703 |
| 0.4 | 0.05 | ptsI | 3 | 17.05 | 0.509 |
| 0.4 | 0.05 | clcB | 3 | 17.05 | 0.284 |
| 0.4 | 0.05 | ycaM | 3 | 17.05 | 0.262 |
| 0.4 | 0.05 | yadI | 3 | 17.05 | 0.748 |
| 0.4 | 0.05 | AG1 | 3 | 17.05 | 0.694 |
| 0.4 | 0.05 | ptsI | 3 | 17.3 | 0.517 |
| 0.4 | 0.05 | clcB | 3 | 17.3 | 0.282 |
| 0.4 | 0.05 | ycaM | 3 | 17.3 | 0.26 |
| 0.4 | 0.05 | yadI | 3 | 17.3 | 0.759 |
| 0.4 | 0.05 | AG1 | 3 | 17.3 | 0.708 |
| 0.4 | 0.05 | ptsI | 3 | 17.55 | 0.52 |
| 0.4 | 0.05 | clcB | 3 | 17.55 | 0.281 |
| 0.4 | 0.05 | ycaM | 3 | 17.55 | 0.258 |
| 0.4 | 0.05 | yadI | 3 | 17.55 | 0.75 |
| 0.4 | 0.05 | AG1 | 3 | 17.55 | 0.716 |
| 0.4 | 0.05 | ptsI | 3 | 17.8 | 0.524 |
| 0.4 | 0.05 | clcB | 3 | 17.8 | 0.28 |
| 0.4 | 0.05 | ycaM | 3 | 17.8 | 0.258 |
| 0.4 | 0.05 | yadI | 3 | 17.8 | 0.759 |
| 0.4 | 0.05 | AG1 | 3 | 17.8 | 0.733 |
| 0.4 | 0.05 | ptsI | 3 | 18.05 | 0.525 |
| 0.4 | 0.05 | clcB | 3 | 18.05 | 0.283 |
| 0.4 | 0.05 | ycaM | 3 | 18.05 | 0.261 |
| 0.4 | 0.05 | yadI | 3 | 18.05 | 0.761 |
| 0.4 | 0.05 | AG1 | 3 | 18.05 | 0.739 |
| 0.4 | 0.05 | ptsI | 3 | 18.3 | 0.531 |
| 0.4 | 0.05 | clcB | 3 | 18.3 | 0.283 |
| 0.4 | 0.05 | ycaM | 3 | 18.3 | 0.261 |
| 0.4 | 0.05 | yadI | 3 | 18.3 | 0.766 |
| 0.4 | 0.05 | AG1 | 3 | 18.3 | 0.745 |
| 0.4 | 0.05 | ptsI | 3 | 18.55 | 0.538 |
| 0.4 | 0.05 | clcB | 3 | 18.55 | 0.283 |
| 0.4 | 0.05 | ycaM | 3 | 18.55 | 0.264 |
| 0.4 | 0.05 | yadI | 3 | 18.55 | 0.779 |
| 0.4 | 0.05 | AG1 | 3 | 18.55 | 0.728 |
| 0.4 | 0.05 | ptsI | 3 | 18.8 | 0.541 |
| 0.4 | 0.05 | clcB | 3 | 18.8 | 0.282 |
| 0.4 | 0.05 | ycaM | 3 | 18.8 | 0.263 |
| 0.4 | 0.05 | yadI | 3 | 18.8 | 0.776 |
| 0.4 | 0.05 | AG1 | 3 | 18.8 | 0.725 |
| 0.4 | 0.05 | ptsI | 3 | 19.05 | 0.543 |
| 0.4 | 0.05 | clcB | 3 | 19.05 | 0.284 |
| 0.4 | 0.05 | ycaM | 3 | 19.05 | 0.265 |
| 0.4 | 0.05 | yadI | 3 | 19.05 | 0.783 |

| | | | | | |
|---|---|---|---|---|---|
| 0.4 | 0.05 | AG1 | 3 | 19.05 | 0.739 |
| 0.4 | 0.05 | ptsI | 3 | 19.3 | 0.543 |
| 0.4 | 0.05 | clcB | 3 | 19.3 | 0.284 |
| 0.4 | 0.05 | ycaM | 3 | 19.3 | 0.266 |
| 0.4 | 0.05 | yadI | 3 | 19.3 | 0.782 |
| 0.4 | 0.05 | AG1 | 3 | 19.3 | 0.752 |
| 0.4 | 0.05 | ptsI | 3 | 19.55 | 0.549 |
| 0.4 | 0.05 | clcB | 3 | 19.55 | 0.283 |
| 0.4 | 0.05 | ycaM | 3 | 19.55 | 0.267 |
| 0.4 | 0.05 | yadI | 3 | 19.55 | 0.791 |
| 0.4 | 0.05 | AG1 | 3 | 19.55 | 0.756 |
| 0.4 | 0.05 | ptsI | 3 | 19.8 | 0.548 |
| 0.4 | 0.05 | clcB | 3 | 19.8 | 0.284 |
| 0.4 | 0.05 | ycaM | 3 | 19.8 | 0.268 |
| 0.4 | 0.05 | yadI | 3 | 19.8 | 0.792 |
| 0.4 | 0.05 | AG1 | 3 | 19.8 | 0.747 |
| 0.4 | 0.05 | ptsI | 3 | 20.05 | 0.549 |
| 0.4 | 0.05 | clcB | 3 | 20.05 | 0.285 |
| 0.4 | 0.05 | ycaM | 3 | 20.05 | 0.269 |
| 0.4 | 0.05 | yadI | 3 | 20.05 | 0.796 |
| 0.4 | 0.05 | AG1 | 3 | 20.05 | 0.754 |
| 0.4 | 0.05 | ptsI | 3 | 20.3 | 0.553 |
| 0.4 | 0.05 | clcB | 3 | 20.3 | 0.287 |
| 0.4 | 0.05 | ycaM | 3 | 20.3 | 0.271 |
| 0.4 | 0.05 | yadI | 3 | 20.3 | 0.801 |
| 0.4 | 0.05 | AG1 | 3 | 20.3 | 0.747 |
| 0.4 | 0.05 | ptsI | 3 | 20.55 | 0.55 |
| 0.4 | 0.05 | clcB | 3 | 20.55 | 0.287 |
| 0.4 | 0.05 | ycaM | 3 | 20.55 | 0.276 |
| 0.4 | 0.05 | yadI | 3 | 20.55 | 0.795 |
| 0.4 | 0.05 | AG1 | 3 | 20.55 | 0.75 |
| 0.4 | 0.05 | ptsI | 3 | 20.8 | 0.552 |
| 0.4 | 0.05 | clcB | 3 | 20.8 | 0.291 |
| 0.4 | 0.05 | ycaM | 3 | 20.8 | 0.277 |
| 0.4 | 0.05 | yadI | 3 | 20.8 | 0.805 |
| 0.4 | 0.05 | AG1 | 3 | 20.8 | 0.758 |
| 0.4 | 0.05 | ptsI | 3 | 21.05 | 0.555 |
| 0.4 | 0.05 | clcB | 3 | 21.05 | 0.291 |
| 0.4 | 0.05 | ycaM | 3 | 21.05 | 0.279 |
| 0.4 | 0.05 | yadI | 3 | 21.05 | 0.803 |
| 0.4 | 0.05 | AG1 | 3 | 21.05 | 0.757 |
| 0.4 | 0.05 | ptsI | 3 | 21.3 | 0.552 |
| 0.4 | 0.05 | clcB | 3 | 21.3 | 0.291 |
| 0.4 | 0.05 | ycaM | 3 | 21.3 | 0.281 |
| 0.4 | 0.05 | yadI | 3 | 21.3 | 0.805 |
| 0.4 | 0.05 | AG1 | 3 | 21.3 | 0.77 |
| 0.4 | 0.05 | ptsI | 3 | 21.55 | 0.55 |
| 0.4 | 0.05 | clcB | 3 | 21.55 | 0.292 |

| | | | | | |
|---|---|---|---|---|---|
| 0.4 | 0.05 | ycaM | 3 | 21.55 | 0.283 |
| 0.4 | 0.05 | yadI | 3 | 21.55 | 0.815 |
| 0.4 | 0.05 | AG1 | 3 | 21.55 | 0.77 |
| 0.4 | 0.05 | ptsI | 3 | 21.8 | 0.549 |
| 0.4 | 0.05 | clcB | 3 | 21.8 | 0.295 |
| 0.4 | 0.05 | ycaM | 3 | 21.8 | 0.285 |
| 0.4 | 0.05 | yadI | 3 | 21.8 | 0.813 |
| 0.4 | 0.05 | AG1 | 3 | 21.8 | 0.78 |
| 0.4 | 0.05 | ptsI | 3 | 22.05 | 0.545 |
| 0.4 | 0.05 | clcB | 3 | 22.05 | 0.296 |
| 0.4 | 0.05 | ycaM | 3 | 22.05 | 0.286 |
| 0.4 | 0.05 | yadI | 3 | 22.05 | 0.813 |
| 0.4 | 0.05 | AG1 | 3 | 22.05 | 0.79 |
| 0.4 | 0.05 | ptsI | 3 | 22.3 | 0.543 |
| 0.4 | 0.05 | clcB | 3 | 22.3 | 0.296 |
| 0.4 | 0.05 | ycaM | 3 | 22.3 | 0.288 |
| 0.4 | 0.05 | yadI | 3 | 22.3 | 0.817 |
| 0.4 | 0.05 | AG1 | 3 | 22.3 | 0.791 |
| 0.4 | 0.05 | ptsI | 3 | 22.55 | 0.539 |
| 0.4 | 0.05 | clcB | 3 | 22.55 | 0.301 |
| 0.4 | 0.05 | ycaM | 3 | 22.55 | 0.294 |
| 0.4 | 0.05 | yadI | 3 | 22.55 | 0.816 |
| 0.4 | 0.05 | AG1 | 3 | 22.55 | 0.797 |
| 0.4 | 0.05 | ptsI | 3 | 22.8 | 0.537 |
| 0.4 | 0.05 | clcB | 3 | 22.8 | 0.299 |
| 0.4 | 0.05 | ycaM | 3 | 22.8 | 0.296 |
| 0.4 | 0.05 | yadI | 3 | 22.8 | 0.817 |
| 0.4 | 0.05 | AG1 | 3 | 22.8 | 0.809 |
| 0.4 | 0.05 | ptsI | 3 | 23.05 | 0.539 |
| 0.4 | 0.05 | clcB | 3 | 23.05 | 0.301 |
| 0.4 | 0.05 | ycaM | 3 | 23.05 | 0.299 |
| 0.4 | 0.05 | yadI | 3 | 23.05 | 0.818 |
| 0.4 | 0.05 | AG1 | 3 | 23.05 | 0.812 |
| 0.4 | 0.05 | ptsI | 3 | 23.3 | 0.535 |
| 0.4 | 0.05 | clcB | 3 | 23.3 | 0.302 |
| 0.4 | 0.05 | ycaM | 3 | 23.3 | 0.3 |
| 0.4 | 0.05 | yadI | 3 | 23.3 | 0.821 |
| 0.4 | 0.05 | AG1 | 3 | 23.3 | 0.821 |
| 0.4 | 0.05 | ptsI | 3 | 23.55 | 0.535 |
| 0.4 | 0.05 | clcB | 3 | 23.55 | 0.303 |
| 0.4 | 0.05 | ycaM | 3 | 23.55 | 0.305 |
| 0.4 | 0.05 | yadI | 3 | 23.55 | 0.835 |
| 0.4 | 0.05 | AG1 | 3 | 23.55 | 0.83 |
| 0.4 | 0.05 | ptsI | 3 | 23.8 | 0.538 |
| 0.4 | 0.05 | clcB | 3 | 23.8 | 0.303 |
| 0.4 | 0.05 | ycaM | 3 | 23.8 | 0.307 |
| 0.4 | 0.05 | yadI | 3 | 23.8 | 0.838 |
| 0.4 | 0.05 | AG1 | 3 | 23.8 | 0.834 |

| | | | | | |
|---|---|---|---|---|---|
| 0.4 | 0.05 | ptsI | 3 | 24.05 | 0.537 |
| 0.4 | 0.05 | clcB | 3 | 24.05 | 0.306 |
| 0.4 | 0.05 | ycaM | 3 | 24.05 | 0.313 |
| 0.4 | 0.05 | yadI | 3 | 24.05 | 0.836 |
| 0.4 | 0.05 | AG1 | 3 | 24.05 | 0.843 |
| 0.4 | 0.05 | ptsI | 3 | 24.3 | 0.535 |
| 0.4 | 0.05 | clcB | 3 | 24.3 | 0.308 |
| 0.4 | 0.05 | ycaM | 3 | 24.3 | 0.316 |
| 0.4 | 0.05 | yadI | 3 | 24.3 | 0.837 |
| 0.4 | 0.05 | AG1 | 3 | 24.3 | 0.842 |
| 0.4 | 0.05 | ptsI | 4 | 0 | 0.186 |
| 0.4 | 0.05 | clcB | 4 | 0 | 0.203 |
| 0.4 | 0.05 | ycaM | 4 | 0 | 0.353 |
| 0.4 | 0.05 | yadI | 4 | 0 | 0.208 |
| 0.4 | 0.05 | AG1 | 4 | 0 | 0.2 |
| 0.4 | 0.05 | ptsI | 4 | 0.25 | 0.186 |
| 0.4 | 0.05 | clcB | 4 | 0.25 | 0.2 |
| 0.4 | 0.05 | ycaM | 4 | 0.25 | 0.346 |
| 0.4 | 0.05 | yadI | 4 | 0.25 | 0.206 |
| 0.4 | 0.05 | AG1 | 4 | 0.25 | 0.196 |
| 0.4 | 0.05 | ptsI | 4 | 0.5 | 0.186 |
| 0.4 | 0.05 | clcB | 4 | 0.5 | 0.201 |
| 0.4 | 0.05 | ycaM | 4 | 0.5 | 0.338 |
| 0.4 | 0.05 | yadI | 4 | 0.5 | 0.207 |
| 0.4 | 0.05 | AG1 | 4 | 0.5 | 0.196 |
| 0.4 | 0.05 | ptsI | 4 | 0.75 | 0.184 |
| 0.4 | 0.05 | clcB | 4 | 0.75 | 0.204 |
| 0.4 | 0.05 | ycaM | 4 | 0.75 | 0.338 |
| 0.4 | 0.05 | yadI | 4 | 0.75 | 0.212 |
| 0.4 | 0.05 | AG1 | 4 | 0.75 | 0.2 |
| 0.4 | 0.05 | ptsI | 4 | 1 | 0.188 |
| 0.4 | 0.05 | clcB | 4 | 1 | 0.206 |
| 0.4 | 0.05 | ycaM | 4 | 1 | 0.336 |
| 0.4 | 0.05 | yadI | 4 | 1 | 0.213 |
| 0.4 | 0.05 | AG1 | 4 | 1 | 0.202 |
| 0.4 | 0.05 | ptsI | 4 | 1.25 | 0.194 |
| 0.4 | 0.05 | clcB | 4 | 1.25 | 0.21 |
| 0.4 | 0.05 | ycaM | 4 | 1.25 | 0.344 |
| 0.4 | 0.05 | yadI | 4 | 1.25 | 0.218 |
| 0.4 | 0.05 | AG1 | 4 | 1.25 | 0.205 |
| 0.4 | 0.05 | ptsI | 4 | 1.5 | 0.193 |
| 0.4 | 0.05 | clcB | 4 | 1.5 | 0.208 |
| 0.4 | 0.05 | ycaM | 4 | 1.5 | 0.358 |
| 0.4 | 0.05 | yadI | 4 | 1.5 | 0.219 |
| 0.4 | 0.05 | AG1 | 4 | 1.5 | 0.205 |
| 0.4 | 0.05 | ptsI | 4 | 1.75 | 0.203 |
| 0.4 | 0.05 | clcB | 4 | 1.75 | 0.218 |
| 0.4 | 0.05 | ycaM | 4 | 1.75 | 0.368 |

| | | | | | |
|---|---|---|---|---|---|
| 0.4 | 0.05 | yadI | 4 | 1.75 | 0.225 |
| 0.4 | 0.05 | AG1 | 4 | 1.75 | 0.213 |
| 0.4 | 0.05 | ptsI | 4 | 2 | 0.211 |
| 0.4 | 0.05 | clcB | 4 | 2 | 0.226 |
| 0.4 | 0.05 | ycaM | 4 | 2 | 0.376 |
| 0.4 | 0.05 | yadI | 4 | 2 | 0.231 |
| 0.4 | 0.05 | AG1 | 4 | 2 | 0.222 |
| 0.4 | 0.05 | ptsI | 4 | 2.25 | 0.22 |
| 0.4 | 0.05 | clcB | 4 | 2.25 | 0.233 |
| 0.4 | 0.05 | ycaM | 4 | 2.25 | 0.378 |
| 0.4 | 0.05 | yadI | 4 | 2.25 | 0.241 |
| 0.4 | 0.05 | AG1 | 4 | 2.25 | 0.23 |
| 0.4 | 0.05 | ptsI | 4 | 2.5 | 0.224 |
| 0.4 | 0.05 | clcB | 4 | 2.5 | 0.238 |
| 0.4 | 0.05 | ycaM | 4 | 2.5 | 0.386 |
| 0.4 | 0.05 | yadI | 4 | 2.5 | 0.246 |
| 0.4 | 0.05 | AG1 | 4 | 2.5 | 0.233 |
| 0.4 | 0.05 | ptsI | 4 | 2.75 | 0.229 |
| 0.4 | 0.05 | clcB | 4 | 2.75 | 0.248 |
| 0.4 | 0.05 | ycaM | 4 | 2.75 | 0.462 |
| 0.4 | 0.05 | yadI | 4 | 2.75 | 0.252 |
| 0.4 | 0.05 | AG1 | 4 | 2.75 | 0.241 |
| 0.4 | 0.05 | ptsI | 4 | 3 | 0.242 |
| 0.4 | 0.05 | clcB | 4 | 3 | 0.256 |
| 0.4 | 0.05 | ycaM | 4 | 3 | 0.403 |
| 0.4 | 0.05 | yadI | 4 | 3 | 0.262 |
| 0.4 | 0.05 | AG1 | 4 | 3 | 0.249 |
| 0.4 | 0.05 | ptsI | 4 | 3.25 | 0.251 |
| 0.4 | 0.05 | clcB | 4 | 3.25 | 0.268 |
| 0.4 | 0.05 | ycaM | 4 | 3.25 | 0.418 |
| 0.4 | 0.05 | yadI | 4 | 3.25 | 0.269 |
| 0.4 | 0.05 | AG1 | 4 | 3.25 | 0.256 |
| 0.4 | 0.05 | ptsI | 4 | 3.5 | 0.264 |
| 0.4 | 0.05 | clcB | 4 | 3.5 | 0.281 |
| 0.4 | 0.05 | ycaM | 4 | 3.5 | 0.426 |
| 0.4 | 0.05 | yadI | 4 | 3.5 | 0.282 |
| 0.4 | 0.05 | AG1 | 4 | 3.5 | 0.268 |
| 0.4 | 0.05 | ptsI | 4 | 3.75 | 0.273 |
| 0.4 | 0.05 | clcB | 4 | 3.75 | 0.29 |
| 0.4 | 0.05 | ycaM | 4 | 3.75 | 0.507 |
| 0.4 | 0.05 | yadI | 4 | 3.75 | 0.29 |
| 0.4 | 0.05 | AG1 | 4 | 3.75 | 0.276 |
| 0.4 | 0.05 | ptsI | 4 | 4 | 0.292 |
| 0.4 | 0.05 | clcB | 4 | 4 | 0.301 |
| 0.4 | 0.05 | ycaM | 4 | 4 | 0.444 |
| 0.4 | 0.05 | yadI | 4 | 4 | 0.301 |
| 0.4 | 0.05 | AG1 | 4 | 4 | 0.288 |
| 0.4 | 0.05 | ptsI | 4 | 4.35 | 0.34 |

| | | | | | |
|---|---|---|---|---|---|
| 0.4 | 0.05 | clcB | 4 | 4.35 | 0.366 |
| 0.4 | 0.05 | ycaM | 4 | 4.35 | 0.474 |
| 0.4 | 0.05 | yadI | 4 | 4.35 | 0.362 |
| 0.4 | 0.05 | AG1 | 4 | 4.35 | 0.277 |
| 0.4 | 0.05 | ptsI | 4 | 4.6 | 0.307 |
| 0.4 | 0.05 | clcB | 4 | 4.6 | 0.344 |
| 0.4 | 0.05 | ycaM | 4 | 4.6 | 0.43 |
| 0.4 | 0.05 | yadI | 4 | 4.6 | 0.359 |
| 0.4 | 0.05 | AG1 | 4 | 4.6 | 0.287 |
| 0.4 | 0.05 | ptsI | 4 | 4.85 | 0.304 |
| 0.4 | 0.05 | clcB | 4 | 4.85 | 0.35 |
| 0.4 | 0.05 | ycaM | 4 | 4.85 | 0.424 |
| 0.4 | 0.05 | yadI | 4 | 4.85 | 0.376 |
| 0.4 | 0.05 | AG1 | 4 | 4.85 | 0.301 |
| 0.4 | 0.05 | ptsI | 4 | 5.1 | 0.309 |
| 0.4 | 0.05 | clcB | 4 | 5.1 | 0.352 |
| 0.4 | 0.05 | ycaM | 4 | 5.1 | 0.428 |
| 0.4 | 0.05 | yadI | 4 | 5.1 | 0.389 |
| 0.4 | 0.05 | AG1 | 4 | 5.1 | 0.312 |
| 0.4 | 0.05 | ptsI | 4 | 5.35 | 0.319 |
| 0.4 | 0.05 | clcB | 4 | 5.35 | 0.356 |
| 0.4 | 0.05 | ycaM | 4 | 5.35 | 0.431 |
| 0.4 | 0.05 | yadI | 4 | 5.35 | 0.406 |
| 0.4 | 0.05 | AG1 | 4 | 5.35 | 0.322 |
| 0.4 | 0.05 | ptsI | 4 | 5.6 | 0.321 |
| 0.4 | 0.05 | clcB | 4 | 5.6 | 0.353 |
| 0.4 | 0.05 | ycaM | 4 | 5.6 | 0.429 |
| 0.4 | 0.05 | yadI | 4 | 5.6 | 0.414 |
| 0.4 | 0.05 | AG1 | 4 | 5.6 | 0.331 |
| 0.4 | 0.05 | ptsI | 4 | 5.85 | 0.328 |
| 0.4 | 0.05 | clcB | 4 | 5.85 | 0.354 |
| 0.4 | 0.05 | ycaM | 4 | 5.85 | 0.437 |
| 0.4 | 0.05 | yadI | 4 | 5.85 | 0.427 |
| 0.4 | 0.05 | AG1 | 4 | 5.85 | 0.335 |
| 0.4 | 0.05 | ptsI | 4 | 6.1 | 0.34 |
| 0.4 | 0.05 | clcB | 4 | 6.1 | 0.361 |
| 0.4 | 0.05 | ycaM | 4 | 6.1 | 0.441 |
| 0.4 | 0.05 | yadI | 4 | 6.1 | 0.44 |
| 0.4 | 0.05 | AG1 | 4 | 6.1 | 0.346 |
| 0.4 | 0.05 | ptsI | 4 | 6.35 | 0.352 |
| 0.4 | 0.05 | clcB | 4 | 6.35 | 0.364 |
| 0.4 | 0.05 | ycaM | 4 | 6.35 | 0.444 |
| 0.4 | 0.05 | yadI | 4 | 6.35 | 0.45 |
| 0.4 | 0.05 | AG1 | 4 | 6.35 | 0.354 |
| 0.4 | 0.05 | ptsI | 4 | 6.6 | 0.363 |
| 0.4 | 0.05 | clcB | 4 | 6.6 | 0.366 |
| 0.4 | 0.05 | ycaM | 4 | 6.6 | 0.447 |
| 0.4 | 0.05 | yadI | 4 | 6.6 | 0.457 |

| | | | | | |
|---|---|---|---|---|---|
| 0.4 | 0.05 | AG1 | 4 | 6.6 | 0.364 |
| 0.4 | 0.05 | ptsI | 4 | 6.85 | 0.376 |
| 0.4 | 0.05 | clcB | 4 | 6.85 | 0.367 |
| 0.4 | 0.05 | ycaM | 4 | 6.85 | 0.448 |
| 0.4 | 0.05 | yadI | 4 | 6.85 | 0.465 |
| 0.4 | 0.05 | AG1 | 4 | 6.85 | 0.374 |
| 0.4 | 0.05 | ptsI | 4 | 7.1 | 0.384 |
| 0.4 | 0.05 | clcB | 4 | 7.1 | 0.369 |
| 0.4 | 0.05 | ycaM | 4 | 7.1 | 0.448 |
| 0.4 | 0.05 | yadI | 4 | 7.1 | 0.472 |
| 0.4 | 0.05 | AG1 | 4 | 7.1 | 0.383 |
| 0.4 | 0.05 | ptsI | 4 | 7.35 | 0.393 |
| 0.4 | 0.05 | clcB | 4 | 7.35 | 0.37 |
| 0.4 | 0.05 | ycaM | 4 | 7.35 | 0.45 |
| 0.4 | 0.05 | yadI | 4 | 7.35 | 0.48 |
| 0.4 | 0.05 | AG1 | 4 | 7.35 | 0.393 |
| 0.4 | 0.05 | ptsI | 4 | 7.6 | 0.401 |
| 0.4 | 0.05 | clcB | 4 | 7.6 | 0.371 |
| 0.4 | 0.05 | ycaM | 4 | 7.6 | 0.448 |
| 0.4 | 0.05 | yadI | 4 | 7.6 | 0.488 |
| 0.4 | 0.05 | AG1 | 4 | 7.6 | 0.401 |
| 0.4 | 0.05 | ptsI | 4 | 7.85 | 0.403 |
| 0.4 | 0.05 | clcB | 4 | 7.85 | 0.371 |
| 0.4 | 0.05 | ycaM | 4 | 7.85 | 0.446 |
| 0.4 | 0.05 | yadI | 4 | 7.85 | 0.49 |
| 0.4 | 0.05 | AG1 | 4 | 7.85 | 0.408 |
| 0.4 | 0.05 | ptsI | 4 | 8.1 | 0.413 |
| 0.4 | 0.05 | clcB | 4 | 8.1 | 0.374 |
| 0.4 | 0.05 | ycaM | 4 | 8.1 | 0.453 |
| 0.4 | 0.05 | yadI | 4 | 8.1 | 0.496 |
| 0.4 | 0.05 | AG1 | 4 | 8.1 | 0.416 |
| 0.4 | 0.05 | ptsI | 4 | 8.35 | 0.424 |
| 0.4 | 0.05 | clcB | 4 | 8.35 | 0.379 |
| 0.4 | 0.05 | ycaM | 4 | 8.35 | 0.455 |
| 0.4 | 0.05 | yadI | 4 | 8.35 | 0.503 |
| 0.4 | 0.05 | AG1 | 4 | 8.35 | 0.427 |
| 0.4 | 0.05 | ptsI | 4 | 8.83333 | 0.422 |
| 0.4 | 0.05 | clcB | 4 | 8.83333 | 0.312 |
| 0.4 | 0.05 | ycaM | 4 | 8.83333 | 0.417 |
| 0.4 | 0.05 | yadI | 4 | 8.83333 | 0.498 |
| 0.4 | 0.05 | AG1 | 4 | 8.83333 | 0.452 |
| 0.4 | 0.05 | ptsI | 4 | 9.08333 | 0.422 |
| 0.4 | 0.05 | clcB | 4 | 9.08333 | 0.303 |
| 0.4 | 0.05 | ycaM | 4 | 9.08333 | 0.383 |
| 0.4 | 0.05 | yadI | 4 | 9.08333 | 0.496 |
| 0.4 | 0.05 | AG1 | 4 | 9.08333 | 0.454 |
| 0.4 | 0.05 | ptsI | 4 | 9.33333 | 0.431 |
| 0.4 | 0.05 | clcB | 4 | 9.33333 | 0.3 |

| | | | | | |
|---|---|---|---|---|---|
| 0.4 | 0.05 | ycaM | 4 | 9.33333 | 0.364 |
| 0.4 | 0.05 | yadI | 4 | 9.33333 | 0.503 |
| 0.4 | 0.05 | AG1 | 4 | 9.33333 | 0.47 |
| 0.4 | 0.05 | ptsI | 4 | 9.58333 | 0.441 |
| 0.4 | 0.05 | clcB | 4 | 9.58333 | 0.302 |
| 0.4 | 0.05 | ycaM | 4 | 9.58333 | 0.351 |
| 0.4 | 0.05 | yadI | 4 | 9.58333 | 0.514 |
| 0.4 | 0.05 | AG1 | 4 | 9.58333 | 0.48 |
| 0.4 | 0.05 | ptsI | 4 | 9.83333 | 0.45 |
| 0.4 | 0.05 | clcB | 4 | 9.83333 | 0.302 |
| 0.4 | 0.05 | ycaM | 4 | 9.83333 | 0.338 |
| 0.4 | 0.05 | yadI | 4 | 9.83333 | 0.525 |
| 0.4 | 0.05 | AG1 | 4 | 9.83333 | 0.487 |
| 0.4 | 0.05 | ptsI | 4 | 10.0833 | 0.457 |
| 0.4 | 0.05 | clcB | 4 | 10.0833 | 0.301 |
| 0.4 | 0.05 | ycaM | 4 | 10.0833 | 0.325 |
| 0.4 | 0.05 | yadI | 4 | 10.0833 | 0.53 |
| 0.4 | 0.05 | AG1 | 4 | 10.0833 | 0.493 |
| 0.4 | 0.05 | ptsI | 4 | 10.3333 | 0.468 |
| 0.4 | 0.05 | clcB | 4 | 10.3333 | 0.304 |
| 0.4 | 0.05 | ycaM | 4 | 10.3333 | 0.314 |
| 0.4 | 0.05 | yadI | 4 | 10.3333 | 0.537 |
| 0.4 | 0.05 | AG1 | 4 | 10.3333 | 0.502 |
| 0.4 | 0.05 | ptsI | 4 | 10.5833 | 0.475 |
| 0.4 | 0.05 | clcB | 4 | 10.5833 | 0.302 |
| 0.4 | 0.05 | ycaM | 4 | 10.5833 | 0.3 |
| 0.4 | 0.05 | yadI | 4 | 10.5833 | 0.545 |
| 0.4 | 0.05 | AG1 | 4 | 10.5833 | 0.508 |
| 0.4 | 0.05 | ptsI | 4 | 10.8333 | 0.482 |
| 0.4 | 0.05 | clcB | 4 | 10.8333 | 0.302 |
| 0.4 | 0.05 | ycaM | 4 | 10.8333 | 0.297 |
| 0.4 | 0.05 | yadI | 4 | 10.8333 | 0.55 |
| 0.4 | 0.05 | AG1 | 4 | 10.8333 | 0.513 |
| 0.4 | 0.05 | ptsI | 4 | 11.0833 | 0.487 |
| 0.4 | 0.05 | clcB | 4 | 11.0833 | 0.302 |
| 0.4 | 0.05 | ycaM | 4 | 11.0833 | 0.293 |
| 0.4 | 0.05 | yadI | 4 | 11.0833 | 0.558 |
| 0.4 | 0.05 | AG1 | 4 | 11.0833 | 0.518 |
| 0.4 | 0.05 | ptsI | 4 | 11.3333 | 0.497 |
| 0.4 | 0.05 | clcB | 4 | 11.3333 | 0.305 |
| 0.4 | 0.05 | ycaM | 4 | 11.3333 | 0.298 |
| 0.4 | 0.05 | yadI | 4 | 11.3333 | 0.562 |
| 0.4 | 0.05 | AG1 | 4 | 11.3333 | 0.524 |
| 0.4 | 0.05 | ptsI | 4 | 11.5833 | 0.504 |
| 0.4 | 0.05 | clcB | 4 | 11.5833 | 0.3 |
| 0.4 | 0.05 | ycaM | 4 | 11.5833 | 0.293 |
| 0.4 | 0.05 | yadI | 4 | 11.5833 | 0.569 |
| 0.4 | 0.05 | AG1 | 4 | 11.5833 | 0.528 |

| | | | | | |
|---|---|---|---|---|---|
| 0.4 | 0.05 | ptsI | 4 | 11.8333 | 0.507 |
| 0.4 | 0.05 | clcB | 4 | 11.8333 | 0.304 |
| 0.4 | 0.05 | ycaM | 4 | 11.8333 | 0.29 |
| 0.4 | 0.05 | yadI | 4 | 11.8333 | 0.572 |
| 0.4 | 0.05 | AG1 | 4 | 11.8333 | 0.53 |
| 0.4 | 0.05 | ptsI | 4 | 12.0833 | 0.517 |
| 0.4 | 0.05 | clcB | 4 | 12.0833 | 0.313 |
| 0.4 | 0.05 | ycaM | 4 | 12.0833 | 0.29 |
| 0.4 | 0.05 | yadI | 4 | 12.0833 | 0.575 |
| 0.4 | 0.05 | AG1 | 4 | 12.0833 | 0.536 |
| 0.4 | 0.05 | ptsI | 4 | 12.3333 | 0.523 |
| 0.4 | 0.05 | clcB | 4 | 12.3333 | 0.312 |
| 0.4 | 0.05 | ycaM | 4 | 12.3333 | 0.299 |
| 0.4 | 0.05 | yadI | 4 | 12.3333 | 0.581 |
| 0.4 | 0.05 | AG1 | 4 | 12.3333 | 0.54 |
| 0.4 | 0.05 | ptsI | 4 | 12.5833 | 0.533 |
| 0.4 | 0.05 | clcB | 4 | 12.5833 | 0.319 |
| 0.4 | 0.05 | ycaM | 4 | 12.5833 | 0.295 |
| 0.4 | 0.05 | yadI | 4 | 12.5833 | 0.586 |
| 0.4 | 0.05 | AG1 | 4 | 12.5833 | 0.544 |
| 0.4 | 0.05 | ptsI | 4 | 12.8333 | 0.54 |
| 0.4 | 0.05 | clcB | 4 | 12.8333 | 0.316 |
| 0.4 | 0.05 | ycaM | 4 | 12.8333 | 0.296 |
| 0.4 | 0.05 | yadI | 4 | 12.8333 | 0.592 |
| 0.4 | 0.05 | AG1 | 4 | 12.8333 | 0.547 |
| 0.4 | 0.05 | ptsI | 4 | 13.0833 | 0.549 |
| 0.4 | 0.05 | clcB | 4 | 13.0833 | 0.315 |
| 0.4 | 0.05 | ycaM | 4 | 13.0833 | 0.298 |
| 0.4 | 0.05 | yadI | 4 | 13.0833 | 0.596 |
| 0.4 | 0.05 | AG1 | 4 | 13.0833 | 0.555 |
| 0.4 | 0.05 | ptsI | 4 | 13.3333 | 0.556 |
| 0.4 | 0.05 | clcB | 4 | 13.3333 | 0.326 |
| 0.4 | 0.05 | ycaM | 4 | 13.3333 | 0.3 |
| 0.4 | 0.05 | yadI | 4 | 13.3333 | 0.598 |
| 0.4 | 0.05 | AG1 | 4 | 13.3333 | 0.561 |
| 0.4 | 0.05 | ptsI | 4 | 13.5833 | 0.562 |
| 0.4 | 0.05 | clcB | 4 | 13.5833 | 0.326 |
| 0.4 | 0.05 | ycaM | 4 | 13.5833 | 0.3 |
| 0.4 | 0.05 | yadI | 4 | 13.5833 | 0.604 |
| 0.4 | 0.05 | AG1 | 4 | 13.5833 | 0.564 |
| 0.4 | 0.05 | ptsI | 4 | 13.8333 | 0.566 |
| 0.4 | 0.05 | clcB | 4 | 13.8333 | 0.328 |
| 0.4 | 0.05 | ycaM | 4 | 13.8333 | 0.303 |
| 0.4 | 0.05 | yadI | 4 | 13.8333 | 0.607 |
| 0.4 | 0.05 | AG1 | 4 | 13.8333 | 0.569 |
| 0.4 | 0.05 | ptsI | 4 | 14.0833 | 0.569 |
| 0.4 | 0.05 | clcB | 4 | 14.0833 | 0.326 |
| 0.4 | 0.05 | ycaM | 4 | 14.0833 | 0.303 |

| | | | | | |
|---|---|---|---|---|---|
| 0.4 | 0.05 | yadI | 4 | 14.0833 | 0.614 |
| 0.4 | 0.05 | AG1 | 4 | 14.0833 | 0.573 |
| 0.4 | 0.05 | ptsI | 4 | 14.3333 | 0.577 |
| 0.4 | 0.05 | clcB | 4 | 14.3333 | 0.32 |
| 0.4 | 0.05 | ycaM | 4 | 14.3333 | 0.302 |
| 0.4 | 0.05 | yadI | 4 | 14.3333 | 0.611 |
| 0.4 | 0.05 | AG1 | 4 | 14.3333 | 0.577 |
| 0.4 | 0.05 | ptsI | 4 | 14.5833 | 0.58 |
| 0.4 | 0.05 | clcB | 4 | 14.5833 | 0.325 |
| 0.4 | 0.05 | ycaM | 4 | 14.5833 | 0.307 |
| 0.4 | 0.05 | yadI | 4 | 14.5833 | 0.617 |
| 0.4 | 0.05 | AG1 | 4 | 14.5833 | 0.579 |
| 0.4 | 0.05 | ptsI | 4 | 14.8333 | 0.587 |
| 0.4 | 0.05 | clcB | 4 | 14.8333 | 0.324 |
| 0.4 | 0.05 | ycaM | 4 | 14.8333 | 0.304 |
| 0.4 | 0.05 | yadI | 4 | 14.8333 | 0.62 |
| 0.4 | 0.05 | AG1 | 4 | 14.8333 | 0.582 |
| 0.4 | 0.05 | ptsI | 4 | 15.0833 | 0.59 |
| 0.4 | 0.05 | clcB | 4 | 15.0833 | 0.323 |
| 0.4 | 0.05 | ycaM | 4 | 15.0833 | 0.302 |
| 0.4 | 0.05 | yadI | 4 | 15.0833 | 0.624 |
| 0.4 | 0.05 | AG1 | 4 | 15.0833 | 0.587 |
| 0.4 | 0.05 | ptsI | 4 | 15.3333 | 0.595 |
| 0.4 | 0.05 | clcB | 4 | 15.3333 | 0.327 |
| 0.4 | 0.05 | ycaM | 4 | 15.3333 | 0.305 |
| 0.4 | 0.05 | yadI | 4 | 15.3333 | 0.624 |
| 0.4 | 0.05 | AG1 | 4 | 15.3333 | 0.588 |
| 0.4 | 0.05 | ptsI | 4 | 15.5833 | 0.602 |
| 0.4 | 0.05 | clcB | 4 | 15.5833 | 0.327 |
| 0.4 | 0.05 | ycaM | 4 | 15.5833 | 0.308 |
| 0.4 | 0.05 | yadI | 4 | 15.5833 | 0.627 |
| 0.4 | 0.05 | AG1 | 4 | 15.5833 | 0.592 |
| 0.4 | 0.05 | ptsI | 4 | 15.8333 | 0.604 |
| 0.4 | 0.05 | clcB | 4 | 15.8333 | 0.327 |
| 0.4 | 0.05 | ycaM | 4 | 15.8333 | 0.306 |
| 0.4 | 0.05 | yadI | 4 | 15.8333 | 0.633 |
| 0.4 | 0.05 | AG1 | 4 | 15.8333 | 0.594 |
| 0.4 | 0.05 | ptsI | 4 | 16.0833 | 0.609 |
| 0.4 | 0.05 | clcB | 4 | 16.0833 | 0.328 |
| 0.4 | 0.05 | ycaM | 4 | 16.0833 | 0.324 |
| 0.4 | 0.05 | yadI | 4 | 16.0833 | 0.635 |
| 0.4 | 0.05 | AG1 | 4 | 16.0833 | 0.6 |
| 0.4 | 0.05 | ptsI | 4 | 16.3333 | 0.61 |
| 0.4 | 0.05 | clcB | 4 | 16.3333 | 0.332 |
| 0.4 | 0.05 | ycaM | 4 | 16.3333 | 0.33 |
| 0.4 | 0.05 | yadI | 4 | 16.3333 | 0.636 |
| 0.4 | 0.05 | AG1 | 4 | 16.3333 | 0.602 |
| 0.4 | 0.05 | ptsI | 4 | 16.5833 | 0.612 |

| | | | | | |
|---|---|---|---|---|---|
| 0.4 | 0.05 | clcB | 4 | 16.5833 | 0.331 |
| 0.4 | 0.05 | ycaM | 4 | 16.5833 | 0.333 |
| 0.4 | 0.05 | yadI | 4 | 16.5833 | 0.64 |
| 0.4 | 0.05 | AG1 | 4 | 16.5833 | 0.606 |
| 0.4 | 0.05 | ptsI | 4 | 16.8333 | 0.615 |
| 0.4 | 0.05 | clcB | 4 | 16.8333 | 0.33 |
| 0.4 | 0.05 | ycaM | 4 | 16.8333 | 0.323 |
| 0.4 | 0.05 | yadI | 4 | 16.8333 | 0.645 |
| 0.4 | 0.05 | AG1 | 4 | 16.8333 | 0.608 |
| 0.4 | 0.05 | ptsI | 4 | 17.0833 | 0.614 |
| 0.4 | 0.05 | clcB | 4 | 17.0833 | 0.333 |
| 0.4 | 0.05 | ycaM | 4 | 17.0833 | 0.323 |
| 0.4 | 0.05 | yadI | 4 | 17.0833 | 0.644 |
| 0.4 | 0.05 | AG1 | 4 | 17.0833 | 0.607 |
| 0.4 | 0.05 | ptsI | 4 | 17.3333 | 0.617 |
| 0.4 | 0.05 | clcB | 4 | 17.3333 | 0.34 |
| 0.4 | 0.05 | ycaM | 4 | 17.3333 | 0.327 |
| 0.4 | 0.05 | yadI | 4 | 17.3333 | 0.648 |
| 0.4 | 0.05 | AG1 | 4 | 17.3333 | 0.612 |
| 0.4 | 0.05 | ptsI | 4 | 17.5833 | 0.62 |
| 0.4 | 0.05 | clcB | 4 | 17.5833 | 0.342 |
| 0.4 | 0.05 | ycaM | 4 | 17.5833 | 0.327 |
| 0.4 | 0.05 | yadI | 4 | 17.5833 | 0.649 |
| 0.4 | 0.05 | AG1 | 4 | 17.5833 | 0.617 |
| 0.4 | 0.05 | ptsI | 4 | 17.8333 | 0.619 |
| 0.4 | 0.05 | clcB | 4 | 17.8333 | 0.335 |
| 0.4 | 0.05 | ycaM | 4 | 17.8333 | 0.325 |
| 0.4 | 0.05 | yadI | 4 | 17.8333 | 0.648 |
| 0.4 | 0.05 | AG1 | 4 | 17.8333 | 0.62 |
| 0.4 | 0.05 | ptsI | 4 | 18.0833 | 0.619 |
| 0.4 | 0.05 | clcB | 4 | 18.0833 | 0.342 |
| 0.4 | 0.05 | ycaM | 4 | 18.0833 | 0.333 |
| 0.4 | 0.05 | yadI | 4 | 18.0833 | 0.654 |
| 0.4 | 0.05 | AG1 | 4 | 18.0833 | 0.62 |
| 0.4 | 0.05 | ptsI | 4 | 18.3333 | 0.622 |
| 0.4 | 0.05 | clcB | 4 | 18.3333 | 0.337 |
| 0.4 | 0.05 | ycaM | 4 | 18.3333 | 0.338 |
| 0.4 | 0.05 | yadI | 4 | 18.3333 | 0.66 |
| 0.4 | 0.05 | AG1 | 4 | 18.3333 | 0.623 |
| 0.4 | 0.05 | ptsI | 4 | 18.5833 | 0.62 |
| 0.4 | 0.05 | clcB | 4 | 18.5833 | 0.331 |
| 0.4 | 0.05 | ycaM | 4 | 18.5833 | 0.341 |
| 0.4 | 0.05 | yadI | 4 | 18.5833 | 0.66 |
| 0.4 | 0.05 | AG1 | 4 | 18.5833 | 0.624 |
| 0.4 | 0.05 | ptsI | 4 | 18.8333 | 0.62 |
| 0.4 | 0.05 | clcB | 4 | 18.8333 | 0.323 |
| 0.4 | 0.05 | ycaM | 4 | 18.8333 | 0.336 |
| 0.4 | 0.05 | yadI | 4 | 18.8333 | 0.66 |

| | | | | | |
|---|---|---|---|---|---|
| 0.4 | 0.05 | AG1 | 4 | 18.8333 | 0.628 |
| 0.4 | 0.05 | ptsI | 4 | 19.0833 | 0.624 |
| 0.4 | 0.05 | clcB | 4 | 19.0833 | 0.326 |
| 0.4 | 0.05 | ycaM | 4 | 19.0833 | 0.348 |
| 0.4 | 0.05 | yadI | 4 | 19.0833 | 0.664 |
| 0.4 | 0.05 | AG1 | 4 | 19.0833 | 0.629 |
| 0.4 | 0.05 | ptsI | 4 | 19.3333 | 0.622 |
| 0.4 | 0.05 | clcB | 4 | 19.3333 | 0.324 |
| 0.4 | 0.05 | ycaM | 4 | 19.3333 | 0.345 |
| 0.4 | 0.05 | yadI | 4 | 19.3333 | 0.66 |
| 0.4 | 0.05 | AG1 | 4 | 19.3333 | 0.628 |
| 0.4 | 0.05 | ptsI | 4 | 19.5833 | 0.624 |
| 0.4 | 0.05 | clcB | 4 | 19.5833 | 0.321 |
| 0.4 | 0.05 | ycaM | 4 | 19.5833 | 0.347 |
| 0.4 | 0.05 | yadI | 4 | 19.5833 | 0.662 |
| 0.4 | 0.05 | AG1 | 4 | 19.5833 | 0.628 |
| 0.4 | 0.05 | ptsI | 4 | 19.8333 | 0.624 |
| 0.4 | 0.05 | clcB | 4 | 19.8333 | 0.318 |
| 0.4 | 0.05 | ycaM | 4 | 19.8333 | 0.349 |
| 0.4 | 0.05 | yadI | 4 | 19.8333 | 0.665 |
| 0.4 | 0.05 | AG1 | 4 | 19.8333 | 0.634 |
| 0.4 | 0.05 | ptsI | 4 | 20.0833 | 0.635 |
| 0.4 | 0.05 | clcB | 4 | 20.0833 | 0.32 |
| 0.4 | 0.05 | ycaM | 4 | 20.0833 | 0.363 |
| 0.4 | 0.05 | yadI | 4 | 20.0833 | 0.664 |
| 0.4 | 0.05 | AG1 | 4 | 20.0833 | 0.636 |
| 0.4 | 0.05 | ptsI | 4 | 20.3333 | 0.63 |
| 0.4 | 0.05 | clcB | 4 | 20.3333 | 0.316 |
| 0.4 | 0.05 | ycaM | 4 | 20.3333 | 0.349 |
| 0.4 | 0.05 | yadI | 4 | 20.3333 | 0.664 |
| 0.4 | 0.05 | AG1 | 4 | 20.3333 | 0.633 |
| 0.4 | 0.05 | ptsI | 4 | 20.5833 | 0.628 |
| 0.4 | 0.05 | clcB | 4 | 20.5833 | 0.314 |
| 0.4 | 0.05 | ycaM | 4 | 20.5833 | 0.352 |
| 0.4 | 0.05 | yadI | 4 | 20.5833 | 0.671 |
| 0.4 | 0.05 | AG1 | 4 | 20.5833 | 0.641 |
| 0.4 | 0.05 | ptsI | 4 | 20.8333 | 0.628 |
| 0.4 | 0.05 | clcB | 4 | 20.8333 | 0.318 |
| 0.4 | 0.05 | ycaM | 4 | 20.8333 | 0.356 |
| 0.4 | 0.05 | yadI | 4 | 20.8333 | 0.67 |
| 0.4 | 0.05 | AG1 | 4 | 20.8333 | 0.638 |
| 0.4 | 0.05 | ptsI | 4 | 21.0833 | 0.627 |
| 0.4 | 0.05 | clcB | 4 | 21.0833 | 0.316 |
| 0.4 | 0.05 | ycaM | 4 | 21.0833 | 0.357 |
| 0.4 | 0.05 | yadI | 4 | 21.0833 | 0.67 |
| 0.4 | 0.05 | AG1 | 4 | 21.0833 | 0.642 |
| 0.4 | 0.05 | ptsI | 4 | 21.3333 | 0.629 |
| 0.4 | 0.05 | clcB | 4 | 21.3333 | 0.32 |

| | | | | | |
|---|---|---|---|---|---|
| 0.4 | 0.05 | ycaM | 4 | 21.3333 | 0.357 |
| 0.4 | 0.05 | yadI | 4 | 21.3333 | 0.67 |
| 0.4 | 0.05 | AG1 | 4 | 21.3333 | 0.643 |
| 0.4 | 0.05 | ptsI | 4 | 21.5833 | 0.628 |
| 0.4 | 0.05 | clcB | 4 | 21.5833 | 0.318 |
| 0.4 | 0.05 | ycaM | 4 | 21.5833 | 0.357 |
| 0.4 | 0.05 | yadI | 4 | 21.5833 | 0.673 |
| 0.4 | 0.05 | AG1 | 4 | 21.5833 | 0.645 |
| 0.4 | 0.05 | ptsI | 4 | 21.8333 | 0.63 |
| 0.4 | 0.05 | clcB | 4 | 21.8333 | 0.319 |
| 0.4 | 0.05 | ycaM | 4 | 21.8333 | 0.363 |
| 0.4 | 0.05 | yadI | 4 | 21.8333 | 0.672 |
| 0.4 | 0.05 | AG1 | 4 | 21.8333 | 0.644 |
| 0.4 | 0.05 | ptsI | 4 | 22.0833 | 0.624 |
| 0.4 | 0.05 | clcB | 4 | 22.0833 | 0.319 |
| 0.4 | 0.05 | ycaM | 4 | 22.0833 | 0.36 |
| 0.4 | 0.05 | yadI | 4 | 22.0833 | 0.674 |
| 0.4 | 0.05 | AG1 | 4 | 22.0833 | 0.648 |
| 0.4 | 0.05 | ptsI | 4 | 22.3333 | 0.627 |
| 0.4 | 0.05 | clcB | 4 | 22.3333 | 0.32 |
| 0.4 | 0.05 | ycaM | 4 | 22.3333 | 0.359 |
| 0.4 | 0.05 | yadI | 4 | 22.3333 | 0.674 |
| 0.4 | 0.05 | AG1 | 4 | 22.3333 | 0.648 |
| 0.4 | 0.05 | ptsI | 4 | 22.5833 | 0.628 |
| 0.4 | 0.05 | clcB | 4 | 22.5833 | 0.319 |
| 0.4 | 0.05 | ycaM | 4 | 22.5833 | 0.361 |
| 0.4 | 0.05 | yadI | 4 | 22.5833 | 0.68 |
| 0.4 | 0.05 | AG1 | 4 | 22.5833 | 0.651 |
| 0.4 | 0.05 | ptsI | 4 | 22.8333 | 0.625 |
| 0.4 | 0.05 | clcB | 4 | 22.8333 | 0.321 |
| 0.4 | 0.05 | ycaM | 4 | 22.8333 | 0.36 |
| 0.4 | 0.05 | yadI | 4 | 22.8333 | 0.68 |
| 0.4 | 0.05 | AG1 | 4 | 22.8333 | 0.655 |
| 0.4 | 0.05 | ptsI | 4 | 23.0833 | 0.625 |
| 0.4 | 0.05 | clcB | 4 | 23.0833 | 0.324 |
| 0.4 | 0.05 | ycaM | 4 | 23.0833 | 0.363 |
| 0.4 | 0.05 | yadI | 4 | 23.0833 | 0.68 |
| 0.4 | 0.05 | AG1 | 4 | 23.0833 | 0.653 |
| 0.4 | 0.05 | ptsI | 4 | 23.3333 | 0.627 |
| 0.4 | 0.05 | clcB | 4 | 23.3333 | 0.327 |
| 0.4 | 0.05 | ycaM | 4 | 23.3333 | 0.366 |
| 0.4 | 0.05 | yadI | 4 | 23.3333 | 0.683 |
| 0.4 | 0.05 | AG1 | 4 | 23.3333 | 0.657 |
| 0.4 | 0.05 | ptsI | 4 | 23.5833 | 0.63 |
| 0.4 | 0.05 | clcB | 4 | 23.5833 | 0.327 |
| 0.4 | 0.05 | ycaM | 4 | 23.5833 | 0.367 |
| 0.4 | 0.05 | yadI | 4 | 23.5833 | 0.684 |
| 0.4 | 0.05 | AG1 | 4 | 23.5833 | 0.658 |

| | | | | | |
|---|---|---|---|---|---|
| 0.4 | 0.05 | ptsI | 4 | 23.8333 | 0.626 |
| 0.4 | 0.05 | clcB | 4 | 23.8333 | 0.33 |
| 0.4 | 0.05 | ycaM | 4 | 23.8333 | 0.37 |
| 0.4 | 0.05 | yadI | 4 | 23.8333 | 0.682 |
| 0.4 | 0.05 | AG1 | 4 | 23.8333 | 0.658 |
| 0.4 | 0.05 | ptsI | 4 | 24.0833 | 0.628 |
| 0.4 | 0.05 | clcB | 4 | 24.0833 | 0.332 |
| 0.4 | 0.05 | ycaM | 4 | 24.0833 | 0.371 |
| 0.4 | 0.05 | yadI | 4 | 24.0833 | 0.682 |
| 0.4 | 0.05 | AG1 | 4 | 24.0833 | 0.661 |
| 0.4 | 0.05 | ptsI | 5 | 0 | 0.165 |
| 0.4 | 0.05 | clcB | 5 | 0 | 0.168 |
| 0.4 | 0.05 | ycaM | 5 | 0 | 0.177 |
| 0.4 | 0.05 | yadI | 5 | 0 | 0.17 |
| 0.4 | 0.05 | AG1 | 5 | 0 | 0.161 |
| 0.4 | 0.05 | ptsI | 5 | 0.35 | 0.162 |
| 0.4 | 0.05 | clcB | 5 | 0.35 | 0.164 |
| 0.4 | 0.05 | ycaM | 5 | 0.35 | 0.171 |
| 0.4 | 0.05 | yadI | 5 | 0.35 | 0.163 |
| 0.4 | 0.05 | AG1 | 5 | 0.35 | 0.158 |
| 0.4 | 0.05 | ptsI | 5 | 0.6 | 0.161 |
| 0.4 | 0.05 | clcB | 5 | 0.6 | 0.164 |
| 0.4 | 0.05 | ycaM | 5 | 0.6 | 0.17 |
| 0.4 | 0.05 | yadI | 5 | 0.6 | 0.164 |
| 0.4 | 0.05 | AG1 | 5 | 0.6 | 0.159 |
| 0.4 | 0.05 | ptsI | 5 | 0.85 | 0.16 |
| 0.4 | 0.05 | clcB | 5 | 0.85 | 0.163 |
| 0.4 | 0.05 | ycaM | 5 | 0.85 | 0.169 |
| 0.4 | 0.05 | yadI | 5 | 0.85 | 0.163 |
| 0.4 | 0.05 | AG1 | 5 | 0.85 | 0.156 |
| 0.4 | 0.05 | ptsI | 5 | 1.1 | 0.162 |
| 0.4 | 0.05 | clcB | 5 | 1.1 | 0.165 |
| 0.4 | 0.05 | ycaM | 5 | 1.1 | 0.171 |
| 0.4 | 0.05 | yadI | 5 | 1.1 | 0.165 |
| 0.4 | 0.05 | AG1 | 5 | 1.1 | 0.159 |
| 0.4 | 0.05 | ptsI | 5 | 1.35 | 0.163 |
| 0.4 | 0.05 | clcB | 5 | 1.35 | 0.166 |
| 0.4 | 0.05 | ycaM | 5 | 1.35 | 0.172 |
| 0.4 | 0.05 | yadI | 5 | 1.35 | 0.167 |
| 0.4 | 0.05 | AG1 | 5 | 1.35 | 0.16 |
| 0.4 | 0.05 | ptsI | 5 | 1.6 | 0.164 |
| 0.4 | 0.05 | clcB | 5 | 1.6 | 0.168 |
| 0.4 | 0.05 | ycaM | 5 | 1.6 | 0.174 |
| 0.4 | 0.05 | yadI | 5 | 1.6 | 0.169 |
| 0.4 | 0.05 | AG1 | 5 | 1.6 | 0.162 |
| 0.4 | 0.05 | ptsI | 5 | 1.85 | 0.167 |
| 0.4 | 0.05 | clcB | 5 | 1.85 | 0.171 |
| 0.4 | 0.05 | ycaM | 5 | 1.85 | 0.176 |

| | | | | | |
|---|---|---|---|---|---|
| 0.4 | 0.05 | yadI | 5 | 1.85 | 0.171 |
| 0.4 | 0.05 | AG1 | 5 | 1.85 | 0.164 |
| 0.4 | 0.05 | ptsI | 5 | 2.1 | 0.17 |
| 0.4 | 0.05 | clcB | 5 | 2.1 | 0.176 |
| 0.4 | 0.05 | ycaM | 5 | 2.1 | 0.179 |
| 0.4 | 0.05 | yadI | 5 | 2.1 | 0.175 |
| 0.4 | 0.05 | AG1 | 5 | 2.1 | 0.169 |
| 0.4 | 0.05 | ptsI | 5 | 2.35 | 0.173 |
| 0.4 | 0.05 | clcB | 5 | 2.35 | 0.18 |
| 0.4 | 0.05 | ycaM | 5 | 2.35 | 0.184 |
| 0.4 | 0.05 | yadI | 5 | 2.35 | 0.18 |
| 0.4 | 0.05 | AG1 | 5 | 2.35 | 0.172 |
| 0.4 | 0.05 | ptsI | 5 | 2.6 | 0.175 |
| 0.4 | 0.05 | clcB | 5 | 2.6 | 0.179 |
| 0.4 | 0.05 | ycaM | 5 | 2.6 | 0.183 |
| 0.4 | 0.05 | yadI | 5 | 2.6 | 0.18 |
| 0.4 | 0.05 | AG1 | 5 | 2.6 | 0.173 |
| 0.4 | 0.05 | ptsI | 5 | 2.85 | 0.181 |
| 0.4 | 0.05 | clcB | 5 | 2.85 | 0.189 |
| 0.4 | 0.05 | ycaM | 5 | 2.85 | 0.191 |
| 0.4 | 0.05 | yadI | 5 | 2.85 | 0.188 |
| 0.4 | 0.05 | AG1 | 5 | 2.85 | 0.178 |
| 0.4 | 0.05 | ptsI | 5 | 3.1 | 0.186 |
| 0.4 | 0.05 | clcB | 5 | 3.1 | 0.195 |
| 0.4 | 0.05 | ycaM | 5 | 3.1 | 0.198 |
| 0.4 | 0.05 | yadI | 5 | 3.1 | 0.194 |
| 0.4 | 0.05 | AG1 | 5 | 3.1 | 0.183 |
| 0.4 | 0.05 | ptsI | 5 | 3.35 | 0.195 |
| 0.4 | 0.05 | clcB | 5 | 3.35 | 0.205 |
| 0.4 | 0.05 | ycaM | 5 | 3.35 | 0.217 |
| 0.4 | 0.05 | yadI | 5 | 3.35 | 0.203 |
| 0.4 | 0.05 | AG1 | 5 | 3.35 | 0.192 |
| 0.4 | 0.05 | ptsI | 5 | 3.6 | 0.206 |
| 0.4 | 0.05 | clcB | 5 | 3.6 | 0.218 |
| 0.4 | 0.05 | ycaM | 5 | 3.6 | 0.225 |
| 0.4 | 0.05 | yadI | 5 | 3.6 | 0.212 |
| 0.4 | 0.05 | AG1 | 5 | 3.6 | 0.197 |
| 0.4 | 0.05 | ptsI | 5 | 3.85 | 0.22 |
| 0.4 | 0.05 | clcB | 5 | 3.85 | 0.229 |
| 0.4 | 0.05 | ycaM | 5 | 3.85 | 0.234 |
| 0.4 | 0.05 | yadI | 5 | 3.85 | 0.222 |
| 0.4 | 0.05 | AG1 | 5 | 3.85 | 0.207 |
| 0.4 | 0.05 | ptsI | 5 | 4.38333 | 0.231 |
| 0.4 | 0.05 | clcB | 5 | 4.38333 | 0.311 |
| 0.4 | 0.05 | ycaM | 5 | 4.38333 | 0.236 |
| 0.4 | 0.05 | yadI | 5 | 4.38333 | 0.262 |
| 0.4 | 0.05 | AG1 | 5 | 4.38333 | 0.246 |
| 0.4 | 0.05 | ptsI | 5 | 4.63333 | 0.22 |

| | | | | | |
|---|---|---|---|---|---|
| 0.4 | 0.05 | clcB | 5 | 4.63333 | 0.262 |
| 0.4 | 0.05 | ycaM | 5 | 4.63333 | 0.225 |
| 0.4 | 0.05 | yadI | 5 | 4.63333 | 0.26 |
| 0.4 | 0.05 | AG1 | 5 | 4.63333 | 0.242 |
| 0.4 | 0.05 | ptsI | 5 | 4.88333 | 0.23 |
| 0.4 | 0.05 | clcB | 5 | 4.88333 | 0.274 |
| 0.4 | 0.05 | ycaM | 5 | 4.88333 | 0.235 |
| 0.4 | 0.05 | yadI | 5 | 4.88333 | 0.276 |
| 0.4 | 0.05 | AG1 | 5 | 4.88333 | 0.257 |
| 0.4 | 0.05 | ptsI | 5 | 5.13333 | 0.238 |
| 0.4 | 0.05 | clcB | 5 | 5.13333 | 0.286 |
| 0.4 | 0.05 | ycaM | 5 | 5.13333 | 0.239 |
| 0.4 | 0.05 | yadI | 5 | 5.13333 | 0.29 |
| 0.4 | 0.05 | AG1 | 5 | 5.13333 | 0.268 |
| 0.4 | 0.05 | ptsI | 5 | 5.38333 | 0.249 |
| 0.4 | 0.05 | clcB | 5 | 5.38333 | 0.295 |
| 0.4 | 0.05 | ycaM | 5 | 5.38333 | 0.242 |
| 0.4 | 0.05 | yadI | 5 | 5.38333 | 0.307 |
| 0.4 | 0.05 | AG1 | 5 | 5.38333 | 0.282 |
| 0.4 | 0.05 | ptsI | 5 | 5.63333 | 0.26 |
| 0.4 | 0.05 | clcB | 5 | 5.63333 | 0.304 |
| 0.4 | 0.05 | ycaM | 5 | 5.63333 | 0.243 |
| 0.4 | 0.05 | yadI | 5 | 5.63333 | 0.319 |
| 0.4 | 0.05 | AG1 | 5 | 5.63333 | 0.296 |
| 0.4 | 0.05 | ptsI | 5 | 5.88333 | 0.27 |
| 0.4 | 0.05 | clcB | 5 | 5.88333 | 0.312 |
| 0.4 | 0.05 | ycaM | 5 | 5.88333 | 0.245 |
| 0.4 | 0.05 | yadI | 5 | 5.88333 | 0.331 |
| 0.4 | 0.05 | AG1 | 5 | 5.88333 | 0.307 |
| 0.4 | 0.05 | ptsI | 5 | 6.13333 | 0.284 |
| 0.4 | 0.05 | clcB | 5 | 6.13333 | 0.322 |
| 0.4 | 0.05 | ycaM | 5 | 6.13333 | 0.247 |
| 0.4 | 0.05 | yadI | 5 | 6.13333 | 0.343 |
| 0.4 | 0.05 | AG1 | 5 | 6.13333 | 0.32 |
| 0.4 | 0.05 | ptsI | 5 | 6.38333 | 0.297 |
| 0.4 | 0.05 | clcB | 5 | 6.38333 | 0.328 |
| 0.4 | 0.05 | ycaM | 5 | 6.38333 | 0.248 |
| 0.4 | 0.05 | yadI | 5 | 6.38333 | 0.356 |
| 0.4 | 0.05 | AG1 | 5 | 6.38333 | 0.333 |
| 0.4 | 0.05 | ptsI | 5 | 6.63333 | 0.31 |
| 0.4 | 0.05 | clcB | 5 | 6.63333 | 0.332 |
| 0.4 | 0.05 | ycaM | 5 | 6.63333 | 0.25 |
| 0.4 | 0.05 | yadI | 5 | 6.63333 | 0.368 |
| 0.4 | 0.05 | AG1 | 5 | 6.63333 | 0.346 |
| 0.4 | 0.05 | ptsI | 5 | 6.88333 | 0.324 |
| 0.4 | 0.05 | clcB | 5 | 6.88333 | 0.34 |
| 0.4 | 0.05 | ycaM | 5 | 6.88333 | 0.251 |
| 0.4 | 0.05 | yadI | 5 | 6.88333 | 0.383 |

| | | | | | |
|---|---|---|---|---|---|
| 0.4 | 0.05 | AG1 | 5 | 6.88333 | 0.357 |
| 0.4 | 0.05 | ptsI | 5 | 7.13333 | 0.34 |
| 0.4 | 0.05 | clcB | 5 | 7.13333 | 0.346 |
| 0.4 | 0.05 | ycaM | 5 | 7.13333 | 0.251 |
| 0.4 | 0.05 | yadI | 5 | 7.13333 | 0.397 |
| 0.4 | 0.05 | AG1 | 5 | 7.13333 | 0.369 |
| 0.4 | 0.05 | ptsI | 5 | 7.38333 | 0.356 |
| 0.4 | 0.05 | clcB | 5 | 7.38333 | 0.351 |
| 0.4 | 0.05 | ycaM | 5 | 7.38333 | 0.255 |
| 0.4 | 0.05 | yadI | 5 | 7.38333 | 0.411 |
| 0.4 | 0.05 | AG1 | 5 | 7.38333 | 0.382 |
| 0.4 | 0.05 | ptsI | 5 | 7.63333 | 0.372 |
| 0.4 | 0.05 | clcB | 5 | 7.63333 | 0.354 |
| 0.4 | 0.05 | ycaM | 5 | 7.63333 | 0.256 |
| 0.4 | 0.05 | yadI | 5 | 7.63333 | 0.422 |
| 0.4 | 0.05 | AG1 | 5 | 7.63333 | 0.392 |
| 0.4 | 0.05 | ptsI | 5 | 7.88333 | 0.387 |
| 0.4 | 0.05 | clcB | 5 | 7.88333 | 0.358 |
| 0.4 | 0.05 | ycaM | 5 | 7.88333 | 0.261 |
| 0.4 | 0.05 | yadI | 5 | 7.88333 | 0.434 |
| 0.4 | 0.05 | AG1 | 5 | 7.88333 | 0.4 |
| 0.4 | 0.05 | ptsI | 5 | 8.13333 | 0.403 |
| 0.4 | 0.05 | clcB | 5 | 8.13333 | 0.362 |
| 0.4 | 0.05 | ycaM | 5 | 8.13333 | 0.265 |
| 0.4 | 0.05 | yadI | 5 | 8.13333 | 0.443 |
| 0.4 | 0.05 | AG1 | 5 | 8.13333 | 0.413 |
| 0.4 | 0.05 | ptsI | 5 | 8.38333 | 0.421 |
| 0.4 | 0.05 | clcB | 5 | 8.38333 | 0.366 |
| 0.4 | 0.05 | ycaM | 5 | 8.38333 | 0.267 |
| 0.4 | 0.05 | yadI | 5 | 8.38333 | 0.452 |
| 0.4 | 0.05 | AG1 | 5 | 8.38333 | 0.425 |
| 0.4 | 0.05 | ptsI | 5 | 8.88333 | 0.339 |
| 0.4 | 0.05 | clcB | 5 | 8.88333 | 0.29 |
| 0.4 | 0.05 | ycaM | 5 | 8.88333 | 0.262 |
| 0.4 | 0.05 | yadI | 5 | 8.88333 | 0.437 |
| 0.4 | 0.05 | AG1 | 5 | 8.88333 | 0.467 |
| 0.4 | 0.05 | ptsI | 5 | 9.13333 | 0.343 |
| 0.4 | 0.05 | clcB | 5 | 9.13333 | 0.288 |
| 0.4 | 0.05 | ycaM | 5 | 9.13333 | 0.26 |
| 0.4 | 0.05 | yadI | 5 | 9.13333 | 0.436 |
| 0.4 | 0.05 | AG1 | 5 | 9.13333 | 0.472 |
| 0.4 | 0.05 | ptsI | 5 | 9.38333 | 0.356 |
| 0.4 | 0.05 | clcB | 5 | 9.38333 | 0.288 |
| 0.4 | 0.05 | ycaM | 5 | 9.38333 | 0.257 |
| 0.4 | 0.05 | yadI | 5 | 9.38333 | 0.444 |
| 0.4 | 0.05 | AG1 | 5 | 9.38333 | 0.488 |
| 0.4 | 0.05 | ptsI | 5 | 9.63333 | 0.366 |
| 0.4 | 0.05 | clcB | 5 | 9.63333 | 0.29 |

| | | | | | |
|---|---|---|---|---|---|
| 0.4 | 0.05 | ycaM | 5 | 9.63333 | 0.257 |
| 0.4 | 0.05 | yadI | 5 | 9.63333 | 0.455 |
| 0.4 | 0.05 | AG1 | 5 | 9.63333 | 0.506 |
| 0.4 | 0.05 | ptsI | 5 | 9.88333 | 0.375 |
| 0.4 | 0.05 | clcB | 5 | 9.88333 | 0.29 |
| 0.4 | 0.05 | ycaM | 5 | 9.88333 | 0.257 |
| 0.4 | 0.05 | yadI | 5 | 9.88333 | 0.463 |
| 0.4 | 0.05 | AG1 | 5 | 9.88333 | 0.516 |
| 0.4 | 0.05 | ptsI | 5 | 10.1333 | 0.386 |
| 0.4 | 0.05 | clcB | 5 | 10.1333 | 0.295 |
| 0.4 | 0.05 | ycaM | 5 | 10.1333 | 0.259 |
| 0.4 | 0.05 | yadI | 5 | 10.1333 | 0.472 |
| 0.4 | 0.05 | AG1 | 5 | 10.1333 | 0.522 |
| 0.4 | 0.05 | ptsI | 5 | 10.3833 | 0.396 |
| 0.4 | 0.05 | clcB | 5 | 10.3833 | 0.294 |
| 0.4 | 0.05 | ycaM | 5 | 10.3833 | 0.259 |
| 0.4 | 0.05 | yadI | 5 | 10.3833 | 0.48 |
| 0.4 | 0.05 | AG1 | 5 | 10.3833 | 0.536 |
| 0.4 | 0.05 | ptsI | 5 | 10.6333 | 0.406 |
| 0.4 | 0.05 | clcB | 5 | 10.6333 | 0.294 |
| 0.4 | 0.05 | ycaM | 5 | 10.6333 | 0.259 |
| 0.4 | 0.05 | yadI | 5 | 10.6333 | 0.485 |
| 0.4 | 0.05 | AG1 | 5 | 10.6333 | 0.543 |
| 0.4 | 0.05 | ptsI | 5 | 10.8833 | 0.416 |
| 0.4 | 0.05 | clcB | 5 | 10.8833 | 0.296 |
| 0.4 | 0.05 | ycaM | 5 | 10.8833 | 0.258 |
| 0.4 | 0.05 | yadI | 5 | 10.8833 | 0.492 |
| 0.4 | 0.05 | AG1 | 5 | 10.8833 | 0.557 |
| 0.4 | 0.05 | ptsI | 5 | 11.1333 | 0.427 |
| 0.4 | 0.05 | clcB | 5 | 11.1333 | 0.295 |
| 0.4 | 0.05 | ycaM | 5 | 11.1333 | 0.258 |
| 0.4 | 0.05 | yadI | 5 | 11.1333 | 0.498 |
| 0.4 | 0.05 | AG1 | 5 | 11.1333 | 0.566 |
| 0.4 | 0.05 | ptsI | 5 | 11.3833 | 0.434 |
| 0.4 | 0.05 | clcB | 5 | 11.3833 | 0.296 |
| 0.4 | 0.05 | ycaM | 5 | 11.3833 | 0.259 |
| 0.4 | 0.05 | yadI | 5 | 11.3833 | 0.507 |
| 0.4 | 0.05 | AG1 | 5 | 11.3833 | 0.578 |
| 0.4 | 0.05 | ptsI | 5 | 11.6333 | 0.441 |
| 0.4 | 0.05 | clcB | 5 | 11.6333 | 0.296 |
| 0.4 | 0.05 | ycaM | 5 | 11.6333 | 0.259 |
| 0.4 | 0.05 | yadI | 5 | 11.6333 | 0.514 |
| 0.4 | 0.05 | AG1 | 5 | 11.6333 | 0.586 |
| 0.4 | 0.05 | ptsI | 5 | 11.8833 | 0.449 |
| 0.4 | 0.05 | clcB | 5 | 11.8833 | 0.295 |
| 0.4 | 0.05 | ycaM | 5 | 11.8833 | 0.26 |
| 0.4 | 0.05 | yadI | 5 | 11.8833 | 0.521 |
| 0.4 | 0.05 | AG1 | 5 | 11.8833 | 0.596 |

| | | | | | |
|---|---|---|---|---|---|
| 0.4 | 0.05 | ptsI | 5 | 12.1333 | 0.46 |
| 0.4 | 0.05 | clcB | 5 | 12.1333 | 0.294 |
| 0.4 | 0.05 | ycaM | 5 | 12.1333 | 0.26 |
| 0.4 | 0.05 | yadI | 5 | 12.1333 | 0.527 |
| 0.4 | 0.05 | AG1 | 5 | 12.1333 | 0.605 |
| 0.4 | 0.05 | ptsI | 5 | 12.3833 | 0.463 |
| 0.4 | 0.05 | clcB | 5 | 12.3833 | 0.296 |
| 0.4 | 0.05 | ycaM | 5 | 12.3833 | 0.256 |
| 0.4 | 0.05 | yadI | 5 | 12.3833 | 0.523 |
| 0.4 | 0.05 | AG1 | 5 | 12.3833 | 0.602 |
| 0.4 | 0.05 | ptsI | 5 | 12.6333 | 0.474 |
| 0.4 | 0.05 | clcB | 5 | 12.6333 | 0.294 |
| 0.4 | 0.05 | ycaM | 5 | 12.6333 | 0.257 |
| 0.4 | 0.05 | yadI | 5 | 12.6333 | 0.54 |
| 0.4 | 0.05 | AG1 | 5 | 12.6333 | 0.623 |
| 0.4 | 0.05 | ptsI | 5 | 12.8833 | 0.483 |
| 0.4 | 0.05 | clcB | 5 | 12.8833 | 0.296 |
| 0.4 | 0.05 | ycaM | 5 | 12.8833 | 0.262 |
| 0.4 | 0.05 | yadI | 5 | 12.8833 | 0.546 |
| 0.4 | 0.05 | AG1 | 5 | 12.8833 | 0.632 |
| 0.4 | 0.05 | ptsI | 5 | 13.1333 | 0.491 |
| 0.4 | 0.05 | clcB | 5 | 13.1333 | 0.293 |
| 0.4 | 0.05 | ycaM | 5 | 13.1333 | 0.263 |
| 0.4 | 0.05 | yadI | 5 | 13.1333 | 0.549 |
| 0.4 | 0.05 | AG1 | 5 | 13.1333 | 0.64 |
| 0.4 | 0.05 | ptsI | 5 | 13.3833 | 0.499 |
| 0.4 | 0.05 | clcB | 5 | 13.3833 | 0.292 |
| 0.4 | 0.05 | ycaM | 5 | 13.3833 | 0.26 |
| 0.4 | 0.05 | yadI | 5 | 13.3833 | 0.552 |
| 0.4 | 0.05 | AG1 | 5 | 13.3833 | 0.645 |
| 0.4 | 0.05 | ptsI | 5 | 13.6333 | 0.506 |
| 0.4 | 0.05 | clcB | 5 | 13.6333 | 0.292 |
| 0.4 | 0.05 | ycaM | 5 | 13.6333 | 0.262 |
| 0.4 | 0.05 | yadI | 5 | 13.6333 | 0.557 |
| 0.4 | 0.05 | AG1 | 5 | 13.6333 | 0.652 |
| 0.4 | 0.05 | ptsI | 5 | 13.8833 | 0.511 |
| 0.4 | 0.05 | clcB | 5 | 13.8833 | 0.294 |
| 0.4 | 0.05 | ycaM | 5 | 13.8833 | 0.262 |
| 0.4 | 0.05 | yadI | 5 | 13.8833 | 0.563 |
| 0.4 | 0.05 | AG1 | 5 | 13.8833 | 0.656 |
| 0.4 | 0.05 | ptsI | 5 | 14.1333 | 0.52 |
| 0.4 | 0.05 | clcB | 5 | 14.1333 | 0.296 |
| 0.4 | 0.05 | ycaM | 5 | 14.1333 | 0.261 |
| 0.4 | 0.05 | yadI | 5 | 14.1333 | 0.568 |
| 0.4 | 0.05 | AG1 | 5 | 14.1333 | 0.663 |
| 0.4 | 0.05 | ptsI | 5 | 14.3833 | 0.523 |
| 0.4 | 0.05 | clcB | 5 | 14.3833 | 0.296 |
| 0.4 | 0.05 | ycaM | 5 | 14.3833 | 0.264 |

| | | | | | |
|---|---|---|---|---|---|
| 0.4 | 0.05 | yadI | 5 | 14.3833 | 0.57 |
| 0.4 | 0.05 | AG1 | 5 | 14.3833 | 0.669 |
| 0.4 | 0.05 | ptsI | 5 | 14.6333 | 0.53 |
| 0.4 | 0.05 | clcB | 5 | 14.6333 | 0.297 |
| 0.4 | 0.05 | ycaM | 5 | 14.6333 | 0.267 |
| 0.4 | 0.05 | yadI | 5 | 14.6333 | 0.575 |
| 0.4 | 0.05 | AG1 | 5 | 14.6333 | 0.674 |
| 0.4 | 0.05 | ptsI | 5 | 14.8833 | 0.544 |
| 0.4 | 0.05 | clcB | 5 | 14.8833 | 0.297 |
| 0.4 | 0.05 | ycaM | 5 | 14.8833 | 0.265 |
| 0.4 | 0.05 | yadI | 5 | 14.8833 | 0.579 |
| 0.4 | 0.05 | AG1 | 5 | 14.8833 | 0.681 |
| 0.4 | 0.05 | ptsI | 5 | 15.1333 | 0.547 |
| 0.4 | 0.05 | clcB | 5 | 15.1333 | 0.297 |
| 0.4 | 0.05 | ycaM | 5 | 15.1333 | 0.268 |
| 0.4 | 0.05 | yadI | 5 | 15.1333 | 0.581 |
| 0.4 | 0.05 | AG1 | 5 | 15.1333 | 0.684 |
| 0.4 | 0.05 | ptsI | 5 | 15.3833 | 0.551 |
| 0.4 | 0.05 | clcB | 5 | 15.3833 | 0.299 |
| 0.4 | 0.05 | ycaM | 5 | 15.3833 | 0.267 |
| 0.4 | 0.05 | yadI | 5 | 15.3833 | 0.586 |
| 0.4 | 0.05 | AG1 | 5 | 15.3833 | 0.69 |
| 0.4 | 0.05 | ptsI | 5 | 15.6333 | 0.562 |
| 0.4 | 0.05 | clcB | 5 | 15.6333 | 0.298 |
| 0.4 | 0.05 | ycaM | 5 | 15.6333 | 0.268 |
| 0.4 | 0.05 | yadI | 5 | 15.6333 | 0.592 |
| 0.4 | 0.05 | AG1 | 5 | 15.6333 | 0.697 |
| 0.4 | 0.05 | ptsI | 5 | 15.8833 | 0.567 |
| 0.4 | 0.05 | clcB | 5 | 15.8833 | 0.298 |
| 0.4 | 0.05 | ycaM | 5 | 15.8833 | 0.268 |
| 0.4 | 0.05 | yadI | 5 | 15.8833 | 0.596 |
| 0.4 | 0.05 | AG1 | 5 | 15.8833 | 0.702 |
| 0.4 | 0.05 | ptsI | 5 | 16.1333 | 0.576 |
| 0.4 | 0.05 | clcB | 5 | 16.1333 | 0.303 |
| 0.4 | 0.05 | ycaM | 5 | 16.1333 | 0.272 |
| 0.4 | 0.05 | yadI | 5 | 16.1333 | 0.593 |
| 0.4 | 0.05 | AG1 | 5 | 16.1333 | 0.709 |
| 0.4 | 0.05 | ptsI | 5 | 16.3833 | 0.577 |
| 0.4 | 0.05 | clcB | 5 | 16.3833 | 0.299 |
| 0.4 | 0.05 | ycaM | 5 | 16.3833 | 0.269 |
| 0.4 | 0.05 | yadI | 5 | 16.3833 | 0.595 |
| 0.4 | 0.05 | AG1 | 5 | 16.3833 | 0.708 |
| 0.4 | 0.05 | ptsI | 5 | 16.6333 | 0.582 |
| 0.4 | 0.05 | clcB | 5 | 16.6333 | 0.298 |
| 0.4 | 0.05 | ycaM | 5 | 16.6333 | 0.273 |
| 0.4 | 0.05 | yadI | 5 | 16.6333 | 0.605 |
| 0.4 | 0.05 | AG1 | 5 | 16.6333 | 0.713 |
| 0.4 | 0.05 | ptsI | 5 | 16.8833 | 0.589 |

| | | | | | |
|---|---|---|---|---|---|
| 0.4 | 0.05 | clcB | 5 | 16.8833 | 0.303 |
| 0.4 | 0.05 | ycaM | 5 | 16.8833 | 0.274 |
| 0.4 | 0.05 | yadI | 5 | 16.8833 | 0.602 |
| 0.4 | 0.05 | AG1 | 5 | 16.8833 | 0.72 |
| 0.4 | 0.05 | ptsI | 5 | 17.1333 | 0.597 |
| 0.4 | 0.05 | clcB | 5 | 17.1333 | 0.301 |
| 0.4 | 0.05 | ycaM | 5 | 17.1333 | 0.276 |
| 0.4 | 0.05 | yadI | 5 | 17.1333 | 0.604 |
| 0.4 | 0.05 | AG1 | 5 | 17.1333 | 0.728 |
| 0.4 | 0.05 | ptsI | 5 | 17.3833 | 0.599 |
| 0.4 | 0.05 | clcB | 5 | 17.3833 | 0.303 |
| 0.4 | 0.05 | ycaM | 5 | 17.3833 | 0.275 |
| 0.4 | 0.05 | yadI | 5 | 17.3833 | 0.608 |
| 0.4 | 0.05 | AG1 | 5 | 17.3833 | 0.729 |
| 0.4 | 0.05 | ptsI | 5 | 17.6333 | 0.603 |
| 0.4 | 0.05 | clcB | 5 | 17.6333 | 0.304 |
| 0.4 | 0.05 | ycaM | 5 | 17.6333 | 0.276 |
| 0.4 | 0.05 | yadI | 5 | 17.6333 | 0.61 |
| 0.4 | 0.05 | AG1 | 5 | 17.6333 | 0.738 |
| 0.4 | 0.05 | ptsI | 5 | 17.8833 | 0.607 |
| 0.4 | 0.05 | clcB | 5 | 17.8833 | 0.304 |
| 0.4 | 0.05 | ycaM | 5 | 17.8833 | 0.278 |
| 0.4 | 0.05 | yadI | 5 | 17.8833 | 0.605 |
| 0.4 | 0.05 | AG1 | 5 | 17.8833 | 0.723 |
| 0.4 | 0.05 | ptsI | 5 | 18.1333 | 0.613 |
| 0.4 | 0.05 | clcB | 5 | 18.1333 | 0.309 |
| 0.4 | 0.05 | ycaM | 5 | 18.1333 | 0.282 |
| 0.4 | 0.05 | yadI | 5 | 18.1333 | 0.613 |
| 0.4 | 0.05 | AG1 | 5 | 18.1333 | 0.739 |
| 0.4 | 0.05 | ptsI | 5 | 18.3833 | 0.618 |
| 0.4 | 0.05 | clcB | 5 | 18.3833 | 0.312 |
| 0.4 | 0.05 | ycaM | 5 | 18.3833 | 0.282 |
| 0.4 | 0.05 | yadI | 5 | 18.3833 | 0.615 |
| 0.4 | 0.05 | AG1 | 5 | 18.3833 | 0.748 |
| 0.4 | 0.05 | ptsI | 5 | 18.6333 | 0.621 |
| 0.4 | 0.05 | clcB | 5 | 18.6333 | 0.314 |
| 0.4 | 0.05 | ycaM | 5 | 18.6333 | 0.283 |
| 0.4 | 0.05 | yadI | 5 | 18.6333 | 0.618 |
| 0.4 | 0.05 | AG1 | 5 | 18.6333 | 0.746 |
| 0.4 | 0.05 | ptsI | 5 | 18.8833 | 0.623 |
| 0.4 | 0.05 | clcB | 5 | 18.8833 | 0.317 |
| 0.4 | 0.05 | ycaM | 5 | 18.8833 | 0.287 |
| 0.4 | 0.05 | yadI | 5 | 18.8833 | 0.621 |
| 0.4 | 0.05 | AG1 | 5 | 18.8833 | 0.748 |
| 0.4 | 0.05 | ptsI | 5 | 19.1333 | 0.626 |
| 0.4 | 0.05 | clcB | 5 | 19.1333 | 0.316 |
| 0.4 | 0.05 | ycaM | 5 | 19.1333 | 0.285 |
| 0.4 | 0.05 | yadI | 5 | 19.1333 | 0.622 |

| | | | | | |
|---|---|---|---|---|---|
| 0.4 | 0.05 | AG1 | 5 | 19.1333 | 0.754 |
| 0.4 | 0.05 | ptsI | 5 | 19.3833 | 0.634 |
| 0.4 | 0.05 | clcB | 5 | 19.3833 | 0.314 |
| 0.4 | 0.05 | ycaM | 5 | 19.3833 | 0.291 |
| 0.4 | 0.05 | yadI | 5 | 19.3833 | 0.625 |
| 0.4 | 0.05 | AG1 | 5 | 19.3833 | 0.755 |
| 0.4 | 0.05 | ptsI | 5 | 19.6333 | 0.636 |
| 0.4 | 0.05 | clcB | 5 | 19.6333 | 0.318 |
| 0.4 | 0.05 | ycaM | 5 | 19.6333 | 0.294 |
| 0.4 | 0.05 | yadI | 5 | 19.6333 | 0.627 |
| 0.4 | 0.05 | AG1 | 5 | 19.6333 | 0.758 |
| 0.4 | 0.05 | ptsI | 5 | 19.8833 | 0.634 |
| 0.4 | 0.05 | clcB | 5 | 19.8833 | 0.319 |
| 0.4 | 0.05 | ycaM | 5 | 19.8833 | 0.292 |
| 0.4 | 0.05 | yadI | 5 | 19.8833 | 0.627 |
| 0.4 | 0.05 | AG1 | 5 | 19.8833 | 0.764 |
| 0.4 | 0.05 | ptsI | 5 | 20.1333 | 0.635 |
| 0.4 | 0.05 | clcB | 5 | 20.1333 | 0.323 |
| 0.4 | 0.05 | ycaM | 5 | 20.1333 | 0.293 |
| 0.4 | 0.05 | yadI | 5 | 20.1333 | 0.627 |
| 0.4 | 0.05 | AG1 | 5 | 20.1333 | 0.768 |
| 0.4 | 0.05 | ptsI | 5 | 20.3833 | 0.636 |
| 0.4 | 0.05 | clcB | 5 | 20.3833 | 0.33 |
| 0.4 | 0.05 | ycaM | 5 | 20.3833 | 0.295 |
| 0.4 | 0.05 | yadI | 5 | 20.3833 | 0.629 |
| 0.4 | 0.05 | AG1 | 5 | 20.3833 | 0.769 |
| 0.4 | 0.05 | ptsI | 5 | 20.6333 | 0.633 |
| 0.4 | 0.05 | clcB | 5 | 20.6333 | 0.33 |
| 0.4 | 0.05 | ycaM | 5 | 20.6333 | 0.292 |
| 0.4 | 0.05 | yadI | 5 | 20.6333 | 0.621 |
| 0.4 | 0.05 | AG1 | 5 | 20.6333 | 0.757 |
| 0.4 | 0.05 | ptsI | 5 | 20.8833 | 0.637 |
| 0.4 | 0.05 | clcB | 5 | 20.8833 | 0.334 |
| 0.4 | 0.05 | ycaM | 5 | 20.8833 | 0.3 |
| 0.4 | 0.05 | yadI | 5 | 20.8833 | 0.637 |
| 0.4 | 0.05 | AG1 | 5 | 20.8833 | 0.775 |
| 0.4 | 0.05 | ptsI | 5 | 21.1333 | 0.639 |
| 0.4 | 0.05 | clcB | 5 | 21.1333 | 0.335 |
| 0.4 | 0.05 | ycaM | 5 | 21.1333 | 0.3 |
| 0.4 | 0.05 | yadI | 5 | 21.1333 | 0.638 |
| 0.4 | 0.05 | AG1 | 5 | 21.1333 | 0.774 |
| 0.4 | 0.05 | ptsI | 5 | 21.3833 | 0.639 |
| 0.4 | 0.05 | clcB | 5 | 21.3833 | 0.338 |
| 0.4 | 0.05 | ycaM | 5 | 21.3833 | 0.306 |
| 0.4 | 0.05 | yadI | 5 | 21.3833 | 0.64 |
| 0.4 | 0.05 | AG1 | 5 | 21.3833 | 0.774 |
| 0.4 | 0.05 | ptsI | 5 | 21.6333 | 0.64 |
| 0.4 | 0.05 | clcB | 5 | 21.6333 | 0.344 |

| | | | | | |
|---|---|---|---|---|---|
| 0.4 | 0.05 | ycaM | 5 | 21.6333 | 0.31 |
| 0.4 | 0.05 | yadI | 5 | 21.6333 | 0.64 |
| 0.4 | 0.05 | AG1 | 5 | 21.6333 | 0.773 |
| 0.4 | 0.05 | ptsI | 5 | 21.8833 | 0.642 |
| 0.4 | 0.05 | clcB | 5 | 21.8833 | 0.343 |
| 0.4 | 0.05 | ycaM | 5 | 21.8833 | 0.312 |
| 0.4 | 0.05 | yadI | 5 | 21.8833 | 0.639 |
| 0.4 | 0.05 | AG1 | 5 | 21.8833 | 0.775 |
| 0.4 | 0.05 | ptsI | 5 | 22.1333 | 0.636 |
| 0.4 | 0.05 | clcB | 5 | 22.1333 | 0.34 |
| 0.4 | 0.05 | ycaM | 5 | 22.1333 | 0.313 |
| 0.4 | 0.05 | yadI | 5 | 22.1333 | 0.628 |
| 0.4 | 0.05 | AG1 | 5 | 22.1333 | 0.768 |
| 0.4 | 0.05 | ptsI | 5 | 22.3833 | 0.642 |
| 0.4 | 0.05 | clcB | 5 | 22.3833 | 0.337 |
| 0.4 | 0.05 | ycaM | 5 | 22.3833 | 0.319 |
| 0.4 | 0.05 | yadI | 5 | 22.3833 | 0.646 |
| 0.4 | 0.05 | AG1 | 5 | 22.3833 | 0.79 |
| 0.4 | 0.05 | ptsI | 5 | 22.6333 | 0.644 |
| 0.4 | 0.05 | clcB | 5 | 22.6333 | 0.334 |
| 0.4 | 0.05 | ycaM | 5 | 22.6333 | 0.324 |
| 0.4 | 0.05 | yadI | 5 | 22.6333 | 0.648 |
| 0.4 | 0.05 | AG1 | 5 | 22.6333 | 0.788 |
| 0.4 | 0.05 | ptsI | 5 | 22.8833 | 0.641 |
| 0.4 | 0.05 | clcB | 5 | 22.8833 | 0.329 |
| 0.4 | 0.05 | ycaM | 5 | 22.8833 | 0.326 |
| 0.4 | 0.05 | yadI | 5 | 22.8833 | 0.647 |
| 0.4 | 0.05 | AG1 | 5 | 22.8833 | 0.784 |
| 0.4 | 0.05 | ptsI | 5 | 23.1333 | 0.639 |
| 0.4 | 0.05 | clcB | 5 | 23.1333 | 0.334 |
| 0.4 | 0.05 | ycaM | 5 | 23.1333 | 0.328 |
| 0.4 | 0.05 | yadI | 5 | 23.1333 | 0.649 |
| 0.4 | 0.05 | AG1 | 5 | 23.1333 | 0.788 |
| 0.4 | 0.05 | ptsI | 5 | 23.3833 | 0.641 |
| 0.4 | 0.05 | clcB | 5 | 23.3833 | 0.325 |
| 0.4 | 0.05 | ycaM | 5 | 23.3833 | 0.331 |
| 0.4 | 0.05 | yadI | 5 | 23.3833 | 0.649 |
| 0.4 | 0.05 | AG1 | 5 | 23.3833 | 0.791 |
| 0.4 | 0.05 | ptsI | 5 | 23.6333 | 0.641 |
| 0.4 | 0.05 | clcB | 5 | 23.6333 | 0.323 |
| 0.4 | 0.05 | ycaM | 5 | 23.6333 | 0.333 |
| 0.4 | 0.05 | yadI | 5 | 23.6333 | 0.646 |
| 0.4 | 0.05 | AG1 | 5 | 23.6333 | 0.794 |
| 0.4 | 0.05 | ptsI | 5 | 23.8833 | 0.638 |
| 0.4 | 0.05 | clcB | 5 | 23.8833 | 0.316 |
| 0.4 | 0.05 | ycaM | 5 | 23.8833 | 0.335 |
| 0.4 | 0.05 | yadI | 5 | 23.8833 | 0.641 |
| 0.4 | 0.05 | AG1 | 5 | 23.8833 | 0.782 |

| | | | | | |
|---|---|---|---|---|---|
| 0.4 | 0.05 | ptsI | 5 | 24.1333 | 0.647 |
| 0.4 | 0.05 | clcB | 5 | 24.1333 | 0.318 |
| 0.4 | 0.05 | ycaM | 5 | 24.1333 | 0.341 |
| 0.4 | 0.05 | yadI | 5 | 24.1333 | 0.648 |
| 0.4 | 0.05 | AG1 | 5 | 24.1333 | 0.796 |
| 0.4 | 0.05 | ptsI | 5 | 24.3833 | 0.647 |
| 0.4 | 0.05 | clcB | 5 | 24.3833 | 0.314 |
| 0.4 | 0.05 | ycaM | 5 | 24.3833 | 0.341 |
| 0.4 | 0.05 | yadI | 5 | 24.3833 | 0.653 |
| 0.4 | 0.05 | AG1 | 5 | 24.3833 | 0.799 |
| 0.8 | 0.05 | ptsI | 2 | 0 | 0.221 |
| 0.8 | 0.05 | clcB | 2 | 0 | 0.224 |
| 0.8 | 0.05 | ycaM | 2 | 0 | 0.217 |
| 0.8 | 0.05 | yadI | 2 | 0 | 0.223 |
| 0.8 | 0.05 | AG1 | 2 | 0 | 0.232 |
| 0.8 | 0.05 | ptsI | 2 | 0.25 | 0.215 |
| 0.8 | 0.05 | clcB | 2 | 0.25 | 0.218 |
| 0.8 | 0.05 | ycaM | 2 | 0.25 | 0.213 |
| 0.8 | 0.05 | yadI | 2 | 0.25 | 0.217 |
| 0.8 | 0.05 | AG1 | 2 | 0.25 | 0.224 |
| 0.8 | 0.05 | ptsI | 2 | 0.5 | 0.215 |
| 0.8 | 0.05 | clcB | 2 | 0.5 | 0.219 |
| 0.8 | 0.05 | ycaM | 2 | 0.5 | 0.212 |
| 0.8 | 0.05 | yadI | 2 | 0.5 | 0.215 |
| 0.8 | 0.05 | AG1 | 2 | 0.5 | 0.223 |
| 0.8 | 0.05 | ptsI | 2 | 0.75 | 0.216 |
| 0.8 | 0.05 | clcB | 2 | 0.75 | 0.22 |
| 0.8 | 0.05 | ycaM | 2 | 0.75 | 0.212 |
| 0.8 | 0.05 | yadI | 2 | 0.75 | 0.218 |
| 0.8 | 0.05 | AG1 | 2 | 0.75 | 0.224 |
| 0.8 | 0.05 | ptsI | 2 | 1 | 0.219 |
| 0.8 | 0.05 | clcB | 2 | 1 | 0.22 |
| 0.8 | 0.05 | ycaM | 2 | 1 | 0.214 |
| 0.8 | 0.05 | yadI | 2 | 1 | 0.222 |
| 0.8 | 0.05 | AG1 | 2 | 1 | 0.229 |
| 0.8 | 0.05 | ptsI | 2 | 1.25 | 0.22 |
| 0.8 | 0.05 | clcB | 2 | 1.25 | 0.222 |
| 0.8 | 0.05 | ycaM | 2 | 1.25 | 0.212 |
| 0.8 | 0.05 | yadI | 2 | 1.25 | 0.229 |
| 0.8 | 0.05 | AG1 | 2 | 1.25 | 0.23 |
| 0.8 | 0.05 | ptsI | 2 | 1.5 | 0.222 |
| 0.8 | 0.05 | clcB | 2 | 1.5 | 0.221 |
| 0.8 | 0.05 | ycaM | 2 | 1.5 | 0.214 |
| 0.8 | 0.05 | yadI | 2 | 1.5 | 0.234 |
| 0.8 | 0.05 | AG1 | 2 | 1.5 | 0.234 |
| 0.8 | 0.05 | ptsI | 2 | 1.75 | 0.227 |
| 0.8 | 0.05 | clcB | 2 | 1.75 | 0.224 |
| 0.8 | 0.05 | ycaM | 2 | 1.75 | 0.217 |

| | | | | | |
|---|---|---|---|---|---|
| 0.8 | 0.05 | yadI | 2 | 1.75 | 0.239 |
| 0.8 | 0.05 | AG1 | 2 | 1.75 | 0.237 |
| 0.8 | 0.05 | ptsI | 2 | 2 | 0.233 |
| 0.8 | 0.05 | clcB | 2 | 2 | 0.228 |
| 0.8 | 0.05 | ycaM | 2 | 2 | 0.223 |
| 0.8 | 0.05 | yadI | 2 | 2 | 0.244 |
| 0.8 | 0.05 | AG1 | 2 | 2 | 0.243 |
| 0.8 | 0.05 | ptsI | 2 | 2.25 | 0.239 |
| 0.8 | 0.05 | clcB | 2 | 2.25 | 0.234 |
| 0.8 | 0.05 | ycaM | 2 | 2.25 | 0.228 |
| 0.8 | 0.05 | yadI | 2 | 2.25 | 0.256 |
| 0.8 | 0.05 | AG1 | 2 | 2.25 | 0.254 |
| 0.8 | 0.05 | ptsI | 2 | 2.5 | 0.247 |
| 0.8 | 0.05 | clcB | 2 | 2.5 | 0.24 |
| 0.8 | 0.05 | ycaM | 2 | 2.5 | 0.234 |
| 0.8 | 0.05 | yadI | 2 | 2.5 | 0.265 |
| 0.8 | 0.05 | AG1 | 2 | 2.5 | 0.259 |
| 0.8 | 0.05 | ptsI | 2 | 2.75 | 0.246 |
| 0.8 | 0.05 | clcB | 2 | 2.75 | 0.246 |
| 0.8 | 0.05 | ycaM | 2 | 2.75 | 0.233 |
| 0.8 | 0.05 | yadI | 2 | 2.75 | 0.274 |
| 0.8 | 0.05 | AG1 | 2 | 2.75 | 0.256 |
| 0.8 | 0.05 | ptsI | 2 | 3 | 0.251 |
| 0.8 | 0.05 | clcB | 2 | 3 | 0.246 |
| 0.8 | 0.05 | ycaM | 2 | 3 | 0.235 |
| 0.8 | 0.05 | yadI | 2 | 3 | 0.281 |
| 0.8 | 0.05 | AG1 | 2 | 3 | 0.259 |
| 0.8 | 0.05 | ptsI | 2 | 3.25 | 0.257 |
| 0.8 | 0.05 | clcB | 2 | 3.25 | 0.248 |
| 0.8 | 0.05 | ycaM | 2 | 3.25 | 0.244 |
| 0.8 | 0.05 | yadI | 2 | 3.25 | 0.292 |
| 0.8 | 0.05 | AG1 | 2 | 3.25 | 0.271 |
| 0.8 | 0.05 | ptsI | 2 | 3.5 | 0.263 |
| 0.8 | 0.05 | clcB | 2 | 3.5 | 0.257 |
| 0.8 | 0.05 | ycaM | 2 | 3.5 | 0.249 |
| 0.8 | 0.05 | yadI | 2 | 3.5 | 0.3 |
| 0.8 | 0.05 | AG1 | 2 | 3.5 | 0.28 |
| 0.8 | 0.05 | ptsI | 2 | 3.75 | 0.273 |
| 0.8 | 0.05 | clcB | 2 | 3.75 | 0.265 |
| 0.8 | 0.05 | ycaM | 2 | 3.75 | 0.259 |
| 0.8 | 0.05 | yadI | 2 | 3.75 | 0.312 |
| 0.8 | 0.05 | AG1 | 2 | 3.75 | 0.289 |
| 0.8 | 0.05 | ptsI | 2 | 4 | 0.282 |
| 0.8 | 0.05 | clcB | 2 | 4 | 0.275 |
| 0.8 | 0.05 | ycaM | 2 | 4 | 0.267 |
| 0.8 | 0.05 | yadI | 2 | 4 | 0.323 |
| 0.8 | 0.05 | AG1 | 2 | 4 | 0.299 |
| 0.8 | 0.05 | ptsI | 2 | 4.38333 | 0.291 |

| | | | | | |
|---|---|---|---|---|---|
| 0.8 | 0.05 | clcB | 2 | 4.38333 | 0.32 |
| 0.8 | 0.05 | ycaM | 2 | 4.38333 | 0.284 |
| 0.8 | 0.05 | yadI | 2 | 4.38333 | 0.333 |
| 0.8 | 0.05 | AG1 | 2 | 4.38333 | 0.314 |
| 0.8 | 0.05 | ptsI | 2 | 4.63333 | 0.287 |
| 0.8 | 0.05 | clcB | 2 | 4.63333 | 0.316 |
| 0.8 | 0.05 | ycaM | 2 | 4.63333 | 0.284 |
| 0.8 | 0.05 | yadI | 2 | 4.63333 | 0.34 |
| 0.8 | 0.05 | AG1 | 2 | 4.63333 | 0.307 |
| 0.8 | 0.05 | ptsI | 2 | 4.88333 | 0.292 |
| 0.8 | 0.05 | clcB | 2 | 4.88333 | 0.324 |
| 0.8 | 0.05 | ycaM | 2 | 4.88333 | 0.288 |
| 0.8 | 0.05 | yadI | 2 | 4.88333 | 0.356 |
| 0.8 | 0.05 | AG1 | 2 | 4.88333 | 0.317 |
| 0.8 | 0.05 | ptsI | 2 | 5.13333 | 0.299 |
| 0.8 | 0.05 | clcB | 2 | 5.13333 | 0.334 |
| 0.8 | 0.05 | ycaM | 2 | 5.13333 | 0.291 |
| 0.8 | 0.05 | yadI | 2 | 5.13333 | 0.371 |
| 0.8 | 0.05 | AG1 | 2 | 5.13333 | 0.325 |
| 0.8 | 0.05 | ptsI | 2 | 5.38333 | 0.305 |
| 0.8 | 0.05 | clcB | 2 | 5.38333 | 0.34 |
| 0.8 | 0.05 | ycaM | 2 | 5.38333 | 0.293 |
| 0.8 | 0.05 | yadI | 2 | 5.38333 | 0.38 |
| 0.8 | 0.05 | AG1 | 2 | 5.38333 | 0.335 |
| 0.8 | 0.05 | ptsI | 2 | 5.63333 | 0.312 |
| 0.8 | 0.05 | clcB | 2 | 5.63333 | 0.346 |
| 0.8 | 0.05 | ycaM | 2 | 5.63333 | 0.294 |
| 0.8 | 0.05 | yadI | 2 | 5.63333 | 0.39 |
| 0.8 | 0.05 | AG1 | 2 | 5.63333 | 0.343 |
| 0.8 | 0.05 | ptsI | 2 | 5.88333 | 0.319 |
| 0.8 | 0.05 | clcB | 2 | 5.88333 | 0.353 |
| 0.8 | 0.05 | ycaM | 2 | 5.88333 | 0.296 |
| 0.8 | 0.05 | yadI | 2 | 5.88333 | 0.398 |
| 0.8 | 0.05 | AG1 | 2 | 5.88333 | 0.352 |
| 0.8 | 0.05 | ptsI | 2 | 6.13333 | 0.326 |
| 0.8 | 0.05 | clcB | 2 | 6.13333 | 0.359 |
| 0.8 | 0.05 | ycaM | 2 | 6.13333 | 0.299 |
| 0.8 | 0.05 | yadI | 2 | 6.13333 | 0.404 |
| 0.8 | 0.05 | AG1 | 2 | 6.13333 | 0.359 |
| 0.8 | 0.05 | ptsI | 2 | 6.38333 | 0.334 |
| 0.8 | 0.05 | clcB | 2 | 6.38333 | 0.366 |
| 0.8 | 0.05 | ycaM | 2 | 6.38333 | 0.299 |
| 0.8 | 0.05 | yadI | 2 | 6.38333 | 0.412 |
| 0.8 | 0.05 | AG1 | 2 | 6.38333 | 0.368 |
| 0.8 | 0.05 | ptsI | 2 | 6.63333 | 0.343 |
| 0.8 | 0.05 | clcB | 2 | 6.63333 | 0.37 |
| 0.8 | 0.05 | ycaM | 2 | 6.63333 | 0.3 |
| 0.8 | 0.05 | yadI | 2 | 6.63333 | 0.42 |

| | | | | | |
|---|---|---|---|---|---|
| 0.8 | 0.05 | AG1 | 2 | 6.63333 | 0.377 |
| 0.8 | 0.05 | ptsI | 2 | 6.88333 | 0.35 |
| 0.8 | 0.05 | clcB | 2 | 6.88333 | 0.375 |
| 0.8 | 0.05 | ycaM | 2 | 6.88333 | 0.302 |
| 0.8 | 0.05 | yadI | 2 | 6.88333 | 0.427 |
| 0.8 | 0.05 | AG1 | 2 | 6.88333 | 0.386 |
| 0.8 | 0.05 | ptsI | 2 | 7.13333 | 0.36 |
| 0.8 | 0.05 | clcB | 2 | 7.13333 | 0.381 |
| 0.8 | 0.05 | ycaM | 2 | 7.13333 | 0.302 |
| 0.8 | 0.05 | yadI | 2 | 7.13333 | 0.435 |
| 0.8 | 0.05 | AG1 | 2 | 7.13333 | 0.394 |
| 0.8 | 0.05 | ptsI | 2 | 7.38333 | 0.368 |
| 0.8 | 0.05 | clcB | 2 | 7.38333 | 0.386 |
| 0.8 | 0.05 | ycaM | 2 | 7.38333 | 0.302 |
| 0.8 | 0.05 | yadI | 2 | 7.38333 | 0.441 |
| 0.8 | 0.05 | AG1 | 2 | 7.38333 | 0.399 |
| 0.8 | 0.05 | ptsI | 2 | 7.63333 | 0.375 |
| 0.8 | 0.05 | clcB | 2 | 7.63333 | 0.392 |
| 0.8 | 0.05 | ycaM | 2 | 7.63333 | 0.303 |
| 0.8 | 0.05 | yadI | 2 | 7.63333 | 0.445 |
| 0.8 | 0.05 | AG1 | 2 | 7.63333 | 0.405 |
| 0.8 | 0.05 | ptsI | 2 | 7.88333 | 0.382 |
| 0.8 | 0.05 | clcB | 2 | 7.88333 | 0.395 |
| 0.8 | 0.05 | ycaM | 2 | 7.88333 | 0.305 |
| 0.8 | 0.05 | yadI | 2 | 7.88333 | 0.453 |
| 0.8 | 0.05 | AG1 | 2 | 7.88333 | 0.411 |
| 0.8 | 0.05 | ptsI | 2 | 8.13333 | 0.389 |
| 0.8 | 0.05 | clcB | 2 | 8.13333 | 0.396 |
| 0.8 | 0.05 | ycaM | 2 | 8.13333 | 0.306 |
| 0.8 | 0.05 | yadI | 2 | 8.13333 | 0.463 |
| 0.8 | 0.05 | AG1 | 2 | 8.13333 | 0.419 |
| 0.8 | 0.05 | ptsI | 2 | 8.38333 | 0.397 |
| 0.8 | 0.05 | clcB | 2 | 8.38333 | 0.405 |
| 0.8 | 0.05 | ycaM | 2 | 8.38333 | 0.304 |
| 0.8 | 0.05 | yadI | 2 | 8.38333 | 0.471 |
| 0.8 | 0.05 | AG1 | 2 | 8.38333 | 0.427 |
| 0.8 | 0.05 | ptsI | 2 | 8.91667 | 0.36 |
| 0.8 | 0.05 | clcB | 2 | 8.91667 | 0.309 |
| 0.8 | 0.05 | ycaM | 2 | 8.91667 | 0.274 |
| 0.8 | 0.05 | yadI | 2 | 8.91667 | 0.505 |
| 0.8 | 0.05 | AG1 | 2 | 8.91667 | 0.438 |
| 0.8 | 0.05 | ptsI | 2 | 9.16667 | 0.361 |
| 0.8 | 0.05 | clcB | 2 | 9.16667 | 0.304 |
| 0.8 | 0.05 | ycaM | 2 | 9.16667 | 0.272 |
| 0.8 | 0.05 | yadI | 2 | 9.16667 | 0.511 |
| 0.8 | 0.05 | AG1 | 2 | 9.16667 | 0.443 |
| 0.8 | 0.05 | ptsI | 2 | 9.41667 | 0.367 |
| 0.8 | 0.05 | clcB | 2 | 9.41667 | 0.309 |

| | | | | | |
|---|---|---|---|---|---|
| 0.8 | 0.05 | ycaM | 2 | 9.41667 | 0.273 |
| 0.8 | 0.05 | yadI | 2 | 9.41667 | 0.523 |
| 0.8 | 0.05 | AG1 | 2 | 9.41667 | 0.46 |
| 0.8 | 0.05 | ptsI | 2 | 9.66667 | 0.374 |
| 0.8 | 0.05 | clcB | 2 | 9.66667 | 0.314 |
| 0.8 | 0.05 | ycaM | 2 | 9.66667 | 0.276 |
| 0.8 | 0.05 | yadI | 2 | 9.66667 | 0.535 |
| 0.8 | 0.05 | AG1 | 2 | 9.66667 | 0.474 |
| 0.8 | 0.05 | ptsI | 2 | 9.91667 | 0.38 |
| 0.8 | 0.05 | clcB | 2 | 9.91667 | 0.311 |
| 0.8 | 0.05 | ycaM | 2 | 9.91667 | 0.277 |
| 0.8 | 0.05 | yadI | 2 | 9.91667 | 0.543 |
| 0.8 | 0.05 | AG1 | 2 | 9.91667 | 0.476 |
| 0.8 | 0.05 | ptsI | 2 | 10.1667 | 0.385 |
| 0.8 | 0.05 | clcB | 2 | 10.1667 | 0.319 |
| 0.8 | 0.05 | ycaM | 2 | 10.1667 | 0.278 |
| 0.8 | 0.05 | yadI | 2 | 10.1667 | 0.552 |
| 0.8 | 0.05 | AG1 | 2 | 10.1667 | 0.48 |
| 0.8 | 0.05 | ptsI | 2 | 10.4167 | 0.392 |
| 0.8 | 0.05 | clcB | 2 | 10.4167 | 0.319 |
| 0.8 | 0.05 | ycaM | 2 | 10.4167 | 0.28 |
| 0.8 | 0.05 | yadI | 2 | 10.4167 | 0.562 |
| 0.8 | 0.05 | AG1 | 2 | 10.4167 | 0.488 |
| 0.8 | 0.05 | ptsI | 2 | 10.6667 | 0.401 |
| 0.8 | 0.05 | clcB | 2 | 10.6667 | 0.328 |
| 0.8 | 0.05 | ycaM | 2 | 10.6667 | 0.28 |
| 0.8 | 0.05 | yadI | 2 | 10.6667 | 0.57 |
| 0.8 | 0.05 | AG1 | 2 | 10.6667 | 0.5 |
| 0.8 | 0.05 | ptsI | 2 | 10.9167 | 0.406 |
| 0.8 | 0.05 | clcB | 2 | 10.9167 | 0.328 |
| 0.8 | 0.05 | ycaM | 2 | 10.9167 | 0.282 |
| 0.8 | 0.05 | yadI | 2 | 10.9167 | 0.579 |
| 0.8 | 0.05 | AG1 | 2 | 10.9167 | 0.512 |
| 0.8 | 0.05 | ptsI | 2 | 11.1667 | 0.416 |
| 0.8 | 0.05 | clcB | 2 | 11.1667 | 0.32 |
| 0.8 | 0.05 | ycaM | 2 | 11.1667 | 0.284 |
| 0.8 | 0.05 | yadI | 2 | 11.1667 | 0.587 |
| 0.8 | 0.05 | AG1 | 2 | 11.1667 | 0.52 |
| 0.8 | 0.05 | ptsI | 2 | 11.4167 | 0.422 |
| 0.8 | 0.05 | clcB | 2 | 11.4167 | 0.324 |
| 0.8 | 0.05 | ycaM | 2 | 11.4167 | 0.283 |
| 0.8 | 0.05 | yadI | 2 | 11.4167 | 0.593 |
| 0.8 | 0.05 | AG1 | 2 | 11.4167 | 0.52 |
| 0.8 | 0.05 | ptsI | 2 | 11.6667 | 0.432 |
| 0.8 | 0.05 | clcB | 2 | 11.6667 | 0.315 |
| 0.8 | 0.05 | ycaM | 2 | 11.6667 | 0.285 |
| 0.8 | 0.05 | yadI | 2 | 11.6667 | 0.598 |
| 0.8 | 0.05 | AG1 | 2 | 11.6667 | 0.537 |

| | | | | | |
|---|---|---|---|---|---|
| 0.8 | 0.05 | ptsI | 2 | 11.9167 | 0.437 |
| 0.8 | 0.05 | clcB | 2 | 11.9167 | 0.317 |
| 0.8 | 0.05 | ycaM | 2 | 11.9167 | 0.285 |
| 0.8 | 0.05 | yadI | 2 | 11.9167 | 0.6 |
| 0.8 | 0.05 | AG1 | 2 | 11.9167 | 0.533 |
| 0.8 | 0.05 | ptsI | 2 | 12.1667 | 0.447 |
| 0.8 | 0.05 | clcB | 2 | 12.1667 | 0.314 |
| 0.8 | 0.05 | ycaM | 2 | 12.1667 | 0.285 |
| 0.8 | 0.05 | yadI | 2 | 12.1667 | 0.61 |
| 0.8 | 0.05 | AG1 | 2 | 12.1667 | 0.552 |
| 0.8 | 0.05 | ptsI | 2 | 12.4167 | 0.454 |
| 0.8 | 0.05 | clcB | 2 | 12.4167 | 0.316 |
| 0.8 | 0.05 | ycaM | 2 | 12.4167 | 0.284 |
| 0.8 | 0.05 | yadI | 2 | 12.4167 | 0.618 |
| 0.8 | 0.05 | AG1 | 2 | 12.4167 | 0.563 |
| 0.8 | 0.05 | ptsI | 2 | 12.6667 | 0.46 |
| 0.8 | 0.05 | clcB | 2 | 12.6667 | 0.319 |
| 0.8 | 0.05 | ycaM | 2 | 12.6667 | 0.281 |
| 0.8 | 0.05 | yadI | 2 | 12.6667 | 0.624 |
| 0.8 | 0.05 | AG1 | 2 | 12.6667 | 0.564 |
| 0.8 | 0.05 | ptsI | 2 | 12.9167 | 0.469 |
| 0.8 | 0.05 | clcB | 2 | 12.9167 | 0.313 |
| 0.8 | 0.05 | ycaM | 2 | 12.9167 | 0.287 |
| 0.8 | 0.05 | yadI | 2 | 12.9167 | 0.635 |
| 0.8 | 0.05 | AG1 | 2 | 12.9167 | 0.574 |
| 0.8 | 0.05 | ptsI | 2 | 13.1667 | 0.478 |
| 0.8 | 0.05 | clcB | 2 | 13.1667 | 0.316 |
| 0.8 | 0.05 | ycaM | 2 | 13.1667 | 0.282 |
| 0.8 | 0.05 | yadI | 2 | 13.1667 | 0.636 |
| 0.8 | 0.05 | AG1 | 2 | 13.1667 | 0.575 |
| 0.8 | 0.05 | ptsI | 2 | 13.4167 | 0.482 |
| 0.8 | 0.05 | clcB | 2 | 13.4167 | 0.317 |
| 0.8 | 0.05 | ycaM | 2 | 13.4167 | 0.281 |
| 0.8 | 0.05 | yadI | 2 | 13.4167 | 0.645 |
| 0.8 | 0.05 | AG1 | 2 | 13.4167 | 0.572 |
| 0.8 | 0.05 | ptsI | 2 | 13.6667 | 0.493 |
| 0.8 | 0.05 | clcB | 2 | 13.6667 | 0.311 |
| 0.8 | 0.05 | ycaM | 2 | 13.6667 | 0.281 |
| 0.8 | 0.05 | yadI | 2 | 13.6667 | 0.645 |
| 0.8 | 0.05 | AG1 | 2 | 13.6667 | 0.602 |
| 0.8 | 0.05 | ptsI | 2 | 13.9167 | 0.496 |
| 0.8 | 0.05 | clcB | 2 | 13.9167 | 0.312 |
| 0.8 | 0.05 | ycaM | 2 | 13.9167 | 0.279 |
| 0.8 | 0.05 | yadI | 2 | 13.9167 | 0.644 |
| 0.8 | 0.05 | AG1 | 2 | 13.9167 | 0.596 |
| 0.8 | 0.05 | ptsI | 2 | 14.1667 | 0.502 |
| 0.8 | 0.05 | clcB | 2 | 14.1667 | 0.313 |
| 0.8 | 0.05 | ycaM | 2 | 14.1667 | 0.282 |

| | | | | | |
|---|---|---|---|---|---|
| 0.8 | 0.05 | yadI | 2 | 14.1667 | 0.648 |
| 0.8 | 0.05 | AG1 | 2 | 14.1667 | 0.587 |
| 0.8 | 0.05 | ptsI | 2 | 14.4167 | 0.51 |
| 0.8 | 0.05 | clcB | 2 | 14.4167 | 0.32 |
| 0.8 | 0.05 | ycaM | 2 | 14.4167 | 0.283 |
| 0.8 | 0.05 | yadI | 2 | 14.4167 | 0.652 |
| 0.8 | 0.05 | AG1 | 2 | 14.4167 | 0.592 |
| 0.8 | 0.05 | ptsI | 2 | 14.6667 | 0.517 |
| 0.8 | 0.05 | clcB | 2 | 14.6667 | 0.323 |
| 0.8 | 0.05 | ycaM | 2 | 14.6667 | 0.281 |
| 0.8 | 0.05 | yadI | 2 | 14.6667 | 0.659 |
| 0.8 | 0.05 | AG1 | 2 | 14.6667 | 0.604 |
| 0.8 | 0.05 | ptsI | 2 | 14.9167 | 0.518 |
| 0.8 | 0.05 | clcB | 2 | 14.9167 | 0.325 |
| 0.8 | 0.05 | ycaM | 2 | 14.9167 | 0.282 |
| 0.8 | 0.05 | yadI | 2 | 14.9167 | 0.663 |
| 0.8 | 0.05 | AG1 | 2 | 14.9167 | 0.61 |
| 0.8 | 0.05 | ptsI | 2 | 15.1667 | 0.53 |
| 0.8 | 0.05 | clcB | 2 | 15.1667 | 0.33 |
| 0.8 | 0.05 | ycaM | 2 | 15.1667 | 0.281 |
| 0.8 | 0.05 | yadI | 2 | 15.1667 | 0.666 |
| 0.8 | 0.05 | AG1 | 2 | 15.1667 | 0.608 |
| 0.8 | 0.05 | ptsI | 2 | 15.4167 | 0.534 |
| 0.8 | 0.05 | clcB | 2 | 15.4167 | 0.323 |
| 0.8 | 0.05 | ycaM | 2 | 15.4167 | 0.278 |
| 0.8 | 0.05 | yadI | 2 | 15.4167 | 0.667 |
| 0.8 | 0.05 | AG1 | 2 | 15.4167 | 0.634 |
| 0.8 | 0.05 | ptsI | 2 | 15.6667 | 0.525 |
| 0.8 | 0.05 | clcB | 2 | 15.6667 | 0.33 |
| 0.8 | 0.05 | ycaM | 2 | 15.6667 | 0.276 |
| 0.8 | 0.05 | yadI | 2 | 15.6667 | 0.671 |
| 0.8 | 0.05 | AG1 | 2 | 15.6667 | 0.626 |
| 0.8 | 0.05 | ptsI | 2 | 15.9167 | 0.53 |
| 0.8 | 0.05 | clcB | 2 | 15.9167 | 0.326 |
| 0.8 | 0.05 | ycaM | 2 | 15.9167 | 0.278 |
| 0.8 | 0.05 | yadI | 2 | 15.9167 | 0.676 |
| 0.8 | 0.05 | AG1 | 2 | 15.9167 | 0.638 |
| 0.8 | 0.05 | ptsI | 2 | 16.1667 | 0.536 |
| 0.8 | 0.05 | clcB | 2 | 16.1667 | 0.334 |
| 0.8 | 0.05 | ycaM | 2 | 16.1667 | 0.277 |
| 0.8 | 0.05 | yadI | 2 | 16.1667 | 0.68 |
| 0.8 | 0.05 | AG1 | 2 | 16.1667 | 0.653 |
| 0.8 | 0.05 | ptsI | 2 | 16.4167 | 0.54 |
| 0.8 | 0.05 | clcB | 2 | 16.4167 | 0.33 |
| 0.8 | 0.05 | ycaM | 2 | 16.4167 | 0.277 |
| 0.8 | 0.05 | yadI | 2 | 16.4167 | 0.682 |
| 0.8 | 0.05 | AG1 | 2 | 16.4167 | 0.655 |
| 0.8 | 0.05 | ptsI | 2 | 16.6667 | 0.541 |

| | | | | | |
|---|---|---|---|---|---|
| 0.8 | 0.05 | clcB | 2 | 16.6667 | 0.327 |
| 0.8 | 0.05 | ycaM | 2 | 16.6667 | 0.278 |
| 0.8 | 0.05 | yadI | 2 | 16.6667 | 0.683 |
| 0.8 | 0.05 | AG1 | 2 | 16.6667 | 0.646 |
| 0.8 | 0.05 | ptsI | 2 | 16.9167 | 0.551 |
| 0.8 | 0.05 | clcB | 2 | 16.9167 | 0.324 |
| 0.8 | 0.05 | ycaM | 2 | 16.9167 | 0.28 |
| 0.8 | 0.05 | yadI | 2 | 16.9167 | 0.689 |
| 0.8 | 0.05 | AG1 | 2 | 16.9167 | 0.658 |
| 0.8 | 0.05 | ptsI | 2 | 17.1667 | 0.558 |
| 0.8 | 0.05 | clcB | 2 | 17.1667 | 0.326 |
| 0.8 | 0.05 | ycaM | 2 | 17.1667 | 0.279 |
| 0.8 | 0.05 | yadI | 2 | 17.1667 | 0.69 |
| 0.8 | 0.05 | AG1 | 2 | 17.1667 | 0.66 |
| 0.8 | 0.05 | ptsI | 2 | 17.4167 | 0.563 |
| 0.8 | 0.05 | clcB | 2 | 17.4167 | 0.331 |
| 0.8 | 0.05 | ycaM | 2 | 17.4167 | 0.278 |
| 0.8 | 0.05 | yadI | 2 | 17.4167 | 0.696 |
| 0.8 | 0.05 | AG1 | 2 | 17.4167 | 0.662 |
| 0.8 | 0.05 | ptsI | 2 | 17.6667 | 0.561 |
| 0.8 | 0.05 | clcB | 2 | 17.6667 | 0.328 |
| 0.8 | 0.05 | ycaM | 2 | 17.6667 | 0.28 |
| 0.8 | 0.05 | yadI | 2 | 17.6667 | 0.698 |
| 0.8 | 0.05 | AG1 | 2 | 17.6667 | 0.664 |
| 0.8 | 0.05 | ptsI | 2 | 17.9167 | 0.564 |
| 0.8 | 0.05 | clcB | 2 | 17.9167 | 0.326 |
| 0.8 | 0.05 | ycaM | 2 | 17.9167 | 0.28 |
| 0.8 | 0.05 | yadI | 2 | 17.9167 | 0.7 |
| 0.8 | 0.05 | AG1 | 2 | 17.9167 | 0.687 |
| 0.8 | 0.05 | ptsI | 2 | 18.1667 | 0.564 |
| 0.8 | 0.05 | clcB | 2 | 18.1667 | 0.325 |
| 0.8 | 0.05 | ycaM | 2 | 18.1667 | 0.282 |
| 0.8 | 0.05 | yadI | 2 | 18.1667 | 0.708 |
| 0.8 | 0.05 | AG1 | 2 | 18.1667 | 0.701 |
| 0.8 | 0.05 | ptsI | 2 | 18.4167 | 0.567 |
| 0.8 | 0.05 | clcB | 2 | 18.4167 | 0.326 |
| 0.8 | 0.05 | ycaM | 2 | 18.4167 | 0.282 |
| 0.8 | 0.05 | yadI | 2 | 18.4167 | 0.707 |
| 0.8 | 0.05 | AG1 | 2 | 18.4167 | 0.72 |
| 0.8 | 0.05 | ptsI | 2 | 18.6667 | 0.57 |
| 0.8 | 0.05 | clcB | 2 | 18.6667 | 0.328 |
| 0.8 | 0.05 | ycaM | 2 | 18.6667 | 0.285 |
| 0.8 | 0.05 | yadI | 2 | 18.6667 | 0.71 |
| 0.8 | 0.05 | AG1 | 2 | 18.6667 | 0.716 |
| 0.8 | 0.05 | ptsI | 2 | 18.9167 | 0.579 |
| 0.8 | 0.05 | clcB | 2 | 18.9167 | 0.326 |
| 0.8 | 0.05 | ycaM | 2 | 18.9167 | 0.284 |
| 0.8 | 0.05 | yadI | 2 | 18.9167 | 0.708 |

| | | | | | |
|---|---|---|---|---|---|
| 0.8 | 0.05 | AG1 | 2 | 18.9167 | 0.715 |
| 0.8 | 0.05 | ptsI | 2 | 19.1667 | 0.571 |
| 0.8 | 0.05 | clcB | 2 | 19.1667 | 0.328 |
| 0.8 | 0.05 | ycaM | 2 | 19.1667 | 0.29 |
| 0.8 | 0.05 | yadI | 2 | 19.1667 | 0.717 |
| 0.8 | 0.05 | AG1 | 2 | 19.1667 | 0.735 |
| 0.8 | 0.05 | ptsI | 2 | 19.4167 | 0.577 |
| 0.8 | 0.05 | clcB | 2 | 19.4167 | 0.329 |
| 0.8 | 0.05 | ycaM | 2 | 19.4167 | 0.29 |
| 0.8 | 0.05 | yadI | 2 | 19.4167 | 0.721 |
| 0.8 | 0.05 | AG1 | 2 | 19.4167 | 0.734 |
| 0.8 | 0.05 | ptsI | 2 | 19.6667 | 0.582 |
| 0.8 | 0.05 | clcB | 2 | 19.6667 | 0.331 |
| 0.8 | 0.05 | ycaM | 2 | 19.6667 | 0.292 |
| 0.8 | 0.05 | yadI | 2 | 19.6667 | 0.723 |
| 0.8 | 0.05 | AG1 | 2 | 19.6667 | 0.728 |
| 0.8 | 0.05 | ptsI | 2 | 19.9167 | 0.588 |
| 0.8 | 0.05 | clcB | 2 | 19.9167 | 0.329 |
| 0.8 | 0.05 | ycaM | 2 | 19.9167 | 0.293 |
| 0.8 | 0.05 | yadI | 2 | 19.9167 | 0.725 |
| 0.8 | 0.05 | AG1 | 2 | 19.9167 | 0.727 |
| 0.8 | 0.05 | ptsI | 2 | 20.1667 | 0.587 |
| 0.8 | 0.05 | clcB | 2 | 20.1667 | 0.328 |
| 0.8 | 0.05 | ycaM | 2 | 20.1667 | 0.293 |
| 0.8 | 0.05 | yadI | 2 | 20.1667 | 0.727 |
| 0.8 | 0.05 | AG1 | 2 | 20.1667 | 0.726 |
| 0.8 | 0.05 | ptsI | 2 | 20.4167 | 0.589 |
| 0.8 | 0.05 | clcB | 2 | 20.4167 | 0.333 |
| 0.8 | 0.05 | ycaM | 2 | 20.4167 | 0.295 |
| 0.8 | 0.05 | yadI | 2 | 20.4167 | 0.73 |
| 0.8 | 0.05 | AG1 | 2 | 20.4167 | 0.732 |
| 0.8 | 0.05 | ptsI | 2 | 20.6667 | 0.592 |
| 0.8 | 0.05 | clcB | 2 | 20.6667 | 0.326 |
| 0.8 | 0.05 | ycaM | 2 | 20.6667 | 0.297 |
| 0.8 | 0.05 | yadI | 2 | 20.6667 | 0.729 |
| 0.8 | 0.05 | AG1 | 2 | 20.6667 | 0.732 |
| 0.8 | 0.05 | ptsI | 2 | 20.9167 | 0.593 |
| 0.8 | 0.05 | clcB | 2 | 20.9167 | 0.326 |
| 0.8 | 0.05 | ycaM | 2 | 20.9167 | 0.3 |
| 0.8 | 0.05 | yadI | 2 | 20.9167 | 0.735 |
| 0.8 | 0.05 | AG1 | 2 | 20.9167 | 0.736 |
| 0.8 | 0.05 | ptsI | 2 | 21.1667 | 0.593 |
| 0.8 | 0.05 | clcB | 2 | 21.1667 | 0.326 |
| 0.8 | 0.05 | ycaM | 2 | 21.1667 | 0.3 |
| 0.8 | 0.05 | yadI | 2 | 21.1667 | 0.735 |
| 0.8 | 0.05 | AG1 | 2 | 21.1667 | 0.736 |
| 0.8 | 0.05 | ptsI | 2 | 21.4167 | 0.598 |
| 0.8 | 0.05 | clcB | 2 | 21.4167 | 0.329 |

| | | | | | |
|---|---|---|---|---|---|
| 0.8 | 0.05 | ycaM | 2 | 21.4167 | 0.299 |
| 0.8 | 0.05 | yadI | 2 | 21.4167 | 0.735 |
| 0.8 | 0.05 | AG1 | 2 | 21.4167 | 0.748 |
| 0.8 | 0.05 | ptsI | 2 | 21.6667 | 0.599 |
| 0.8 | 0.05 | clcB | 2 | 21.6667 | 0.328 |
| 0.8 | 0.05 | ycaM | 2 | 21.6667 | 0.299 |
| 0.8 | 0.05 | yadI | 2 | 21.6667 | 0.737 |
| 0.8 | 0.05 | AG1 | 2 | 21.6667 | 0.758 |
| 0.8 | 0.05 | ptsI | 2 | 21.9167 | 0.599 |
| 0.8 | 0.05 | clcB | 2 | 21.9167 | 0.329 |
| 0.8 | 0.05 | ycaM | 2 | 21.9167 | 0.301 |
| 0.8 | 0.05 | yadI | 2 | 21.9167 | 0.74 |
| 0.8 | 0.05 | AG1 | 2 | 21.9167 | 0.758 |
| 0.8 | 0.05 | ptsI | 2 | 22.1667 | 0.602 |
| 0.8 | 0.05 | clcB | 2 | 22.1667 | 0.328 |
| 0.8 | 0.05 | ycaM | 2 | 22.1667 | 0.304 |
| 0.8 | 0.05 | yadI | 2 | 22.1667 | 0.745 |
| 0.8 | 0.05 | AG1 | 2 | 22.1667 | 0.761 |
| 0.8 | 0.05 | ptsI | 2 | 22.4167 | 0.596 |
| 0.8 | 0.05 | clcB | 2 | 22.4167 | 0.331 |
| 0.8 | 0.05 | ycaM | 2 | 22.4167 | 0.305 |
| 0.8 | 0.05 | yadI | 2 | 22.4167 | 0.747 |
| 0.8 | 0.05 | AG1 | 2 | 22.4167 | 0.77 |
| 0.8 | 0.05 | ptsI | 2 | 22.6667 | 0.598 |
| 0.8 | 0.05 | clcB | 2 | 22.6667 | 0.332 |
| 0.8 | 0.05 | ycaM | 2 | 22.6667 | 0.307 |
| 0.8 | 0.05 | yadI | 2 | 22.6667 | 0.746 |
| 0.8 | 0.05 | AG1 | 2 | 22.6667 | 0.768 |
| 0.8 | 0.05 | ptsI | 2 | 22.9167 | 0.604 |
| 0.8 | 0.05 | clcB | 2 | 22.9167 | 0.331 |
| 0.8 | 0.05 | ycaM | 2 | 22.9167 | 0.307 |
| 0.8 | 0.05 | yadI | 2 | 22.9167 | 0.746 |
| 0.8 | 0.05 | AG1 | 2 | 22.9167 | 0.792 |
| 0.8 | 0.05 | ptsI | 2 | 23.1667 | 0.603 |
| 0.8 | 0.05 | clcB | 2 | 23.1667 | 0.331 |
| 0.8 | 0.05 | ycaM | 2 | 23.1667 | 0.309 |
| 0.8 | 0.05 | yadI | 2 | 23.1667 | 0.752 |
| 0.8 | 0.05 | AG1 | 2 | 23.1667 | 0.779 |
| 0.8 | 0.05 | ptsI | 2 | 23.4167 | 0.599 |
| 0.8 | 0.05 | clcB | 2 | 23.4167 | 0.331 |
| 0.8 | 0.05 | ycaM | 2 | 23.4167 | 0.312 |
| 0.8 | 0.05 | yadI | 2 | 23.4167 | 0.757 |
| 0.8 | 0.05 | AG1 | 2 | 23.4167 | 0.783 |
| 0.8 | 0.05 | ptsI | 2 | 23.6667 | 0.601 |
| 0.8 | 0.05 | clcB | 2 | 23.6667 | 0.333 |
| 0.8 | 0.05 | ycaM | 2 | 23.6667 | 0.314 |
| 0.8 | 0.05 | yadI | 2 | 23.6667 | 0.76 |
| 0.8 | 0.05 | AG1 | 2 | 23.6667 | 0.787 |

| | | | | | |
|---|---|---|---|---|---|
| 0.8 | 0.05 | ptsI | 2 | 23.9167 | 0.605 |
| 0.8 | 0.05 | clcB | 2 | 23.9167 | 0.331 |
| 0.8 | 0.05 | ycaM | 2 | 23.9167 | 0.313 |
| 0.8 | 0.05 | yadI | 2 | 23.9167 | 0.76 |
| 0.8 | 0.05 | AG1 | 2 | 23.9167 | 0.793 |
| 0.8 | 0.05 | ptsI | 2 | 24.1667 | 0.603 |
| 0.8 | 0.05 | clcB | 2 | 24.1667 | 0.328 |
| 0.8 | 0.05 | ycaM | 2 | 24.1667 | 0.314 |
| 0.8 | 0.05 | yadI | 2 | 24.1667 | 0.759 |
| 0.8 | 0.05 | AG1 | 2 | 24.1667 | 0.799 |
| 0.8 | 0.05 | ptsI | 2 | 24.4167 | 0.6 |
| 0.8 | 0.05 | clcB | 2 | 24.4167 | 0.327 |
| 0.8 | 0.05 | ycaM | 2 | 24.4167 | 0.314 |
| 0.8 | 0.05 | yadI | 2 | 24.4167 | 0.763 |
| 0.8 | 0.05 | AG1 | 2 | 24.4167 | 0.805 |
| 0.8 | 0.05 | ptsI | 2 | 24.6667 | 0.599 |
| 0.8 | 0.05 | clcB | 2 | 24.6667 | 0.327 |
| 0.8 | 0.05 | ycaM | 2 | 24.6667 | 0.315 |
| 0.8 | 0.05 | yadI | 2 | 24.6667 | 0.763 |
| 0.8 | 0.05 | AG1 | 2 | 24.6667 | 0.811 |
| 0.8 | 0.05 | ptsI | 2 | 24.9167 | 0.598 |
| 0.8 | 0.05 | clcB | 2 | 24.9167 | 0.324 |
| 0.8 | 0.05 | ycaM | 2 | 24.9167 | 0.317 |
| 0.8 | 0.05 | yadI | 2 | 24.9167 | 0.77 |
| 0.8 | 0.05 | AG1 | 2 | 24.9167 | 0.82 |
| 0.8 | 0.05 | ptsI | 2 | 25.1667 | 0.6 |
| 0.8 | 0.05 | clcB | 2 | 25.1667 | 0.325 |
| 0.8 | 0.05 | ycaM | 2 | 25.1667 | 0.318 |
| 0.8 | 0.05 | yadI | 2 | 25.1667 | 0.769 |
| 0.8 | 0.05 | AG1 | 2 | 25.1667 | 0.829 |
| 0.8 | 0.05 | ptsI | 2 | 25.4167 | 0.595 |
| 0.8 | 0.05 | clcB | 2 | 25.4167 | 0.326 |
| 0.8 | 0.05 | ycaM | 2 | 25.4167 | 0.32 |
| 0.8 | 0.05 | yadI | 2 | 25.4167 | 0.772 |
| 0.8 | 0.05 | AG1 | 2 | 25.4167 | 0.832 |
| 0.8 | 0.05 | ptsI | 2 | 25.6667 | 0.594 |
| 0.8 | 0.05 | clcB | 2 | 25.6667 | 0.326 |
| 0.8 | 0.05 | ycaM | 2 | 25.6667 | 0.323 |
| 0.8 | 0.05 | yadI | 2 | 25.6667 | 0.773 |
| 0.8 | 0.05 | AG1 | 2 | 25.6667 | 0.839 |
| 0.8 | 0.05 | ptsI | 2 | 25.9167 | 0.593 |
| 0.8 | 0.05 | clcB | 2 | 25.9167 | 0.327 |
| 0.8 | 0.05 | ycaM | 2 | 25.9167 | 0.326 |
| 0.8 | 0.05 | yadI | 2 | 25.9167 | 0.776 |
| 0.8 | 0.05 | AG1 | 2 | 25.9167 | 0.856 |
| 0.8 | 0.05 | ptsI | 2 | 26.1667 | 0.591 |
| 0.8 | 0.05 | clcB | 2 | 26.1667 | 0.328 |
| 0.8 | 0.05 | ycaM | 2 | 26.1667 | 0.328 |

| | | | | | |
|---|---|---|---|---|---|
| 0.8 | 0.05 | yadI | 2 | 26.1667 | 0.778 |
| 0.8 | 0.05 | AG1 | 2 | 26.1667 | 0.865 |
| 0.8 | 0.05 | ptsI | 2 | 26.4167 | 0.59 |
| 0.8 | 0.05 | clcB | 2 | 26.4167 | 0.326 |
| 0.8 | 0.05 | ycaM | 2 | 26.4167 | 0.329 |
| 0.8 | 0.05 | yadI | 2 | 26.4167 | 0.779 |
| 0.8 | 0.05 | AG1 | 2 | 26.4167 | 0.88 |
| 0.8 | 0.05 | ptsI | 3 | 0 | 0.158 |
| 0.8 | 0.05 | clcB | 3 | 0 | 0.155 |
| 0.8 | 0.05 | ycaM | 3 | 0 | 0.154 |
| 0.8 | 0.05 | yadI | 3 | 0 | 0.167 |
| 0.8 | 0.05 | AG1 | 3 | 0 | 0.163 |
| 0.8 | 0.05 | ptsI | 3 | 0.25 | 0.157 |
| 0.8 | 0.05 | clcB | 3 | 0.25 | 0.153 |
| 0.8 | 0.05 | ycaM | 3 | 0.25 | 0.153 |
| 0.8 | 0.05 | yadI | 3 | 0.25 | 0.158 |
| 0.8 | 0.05 | AG1 | 3 | 0.25 | 0.162 |
| 0.8 | 0.05 | ptsI | 3 | 0.5 | 0.156 |
| 0.8 | 0.05 | clcB | 3 | 0.5 | 0.153 |
| 0.8 | 0.05 | ycaM | 3 | 0.5 | 0.152 |
| 0.8 | 0.05 | yadI | 3 | 0.5 | 0.156 |
| 0.8 | 0.05 | AG1 | 3 | 0.5 | 0.159 |
| 0.8 | 0.05 | ptsI | 3 | 0.75 | 0.156 |
| 0.8 | 0.05 | clcB | 3 | 0.75 | 0.152 |
| 0.8 | 0.05 | ycaM | 3 | 0.75 | 0.152 |
| 0.8 | 0.05 | yadI | 3 | 0.75 | 0.156 |
| 0.8 | 0.05 | AG1 | 3 | 0.75 | 0.16 |
| 0.8 | 0.05 | ptsI | 3 | 1 | 0.157 |
| 0.8 | 0.05 | clcB | 3 | 1 | 0.166 |
| 0.8 | 0.05 | ycaM | 3 | 1 | 0.153 |
| 0.8 | 0.05 | yadI | 3 | 1 | 0.156 |
| 0.8 | 0.05 | AG1 | 3 | 1 | 0.16 |
| 0.8 | 0.05 | ptsI | 3 | 1.25 | 0.157 |
| 0.8 | 0.05 | clcB | 3 | 1.25 | 0.154 |
| 0.8 | 0.05 | ycaM | 3 | 1.25 | 0.153 |
| 0.8 | 0.05 | yadI | 3 | 1.25 | 0.156 |
| 0.8 | 0.05 | AG1 | 3 | 1.25 | 0.16 |
| 0.8 | 0.05 | ptsI | 3 | 1.5 | 0.158 |
| 0.8 | 0.05 | clcB | 3 | 1.5 | 0.154 |
| 0.8 | 0.05 | ycaM | 3 | 1.5 | 0.154 |
| 0.8 | 0.05 | yadI | 3 | 1.5 | 0.156 |
| 0.8 | 0.05 | AG1 | 3 | 1.5 | 0.16 |
| 0.8 | 0.05 | ptsI | 3 | 1.75 | 0.158 |
| 0.8 | 0.05 | clcB | 3 | 1.75 | 0.154 |
| 0.8 | 0.05 | ycaM | 3 | 1.75 | 0.154 |
| 0.8 | 0.05 | yadI | 3 | 1.75 | 0.157 |
| 0.8 | 0.05 | AG1 | 3 | 1.75 | 0.161 |
| 0.8 | 0.05 | ptsI | 3 | 2 | 0.159 |

| | | | | | |
|---|---|---|---|---|---|
| 0.8 | 0.05 | clcB | 3 | 2 | 0.158 |
| 0.8 | 0.05 | ycaM | 3 | 2 | 0.155 |
| 0.8 | 0.05 | yadI | 3 | 2 | 0.158 |
| 0.8 | 0.05 | AG1 | 3 | 2 | 0.161 |
| 0.8 | 0.05 | ptsI | 3 | 2.25 | 0.16 |
| 0.8 | 0.05 | clcB | 3 | 2.25 | 0.157 |
| 0.8 | 0.05 | ycaM | 3 | 2.25 | 0.157 |
| 0.8 | 0.05 | yadI | 3 | 2.25 | 0.161 |
| 0.8 | 0.05 | AG1 | 3 | 2.25 | 0.162 |
| 0.8 | 0.05 | ptsI | 3 | 2.5 | 0.162 |
| 0.8 | 0.05 | clcB | 3 | 2.5 | 0.16 |
| 0.8 | 0.05 | ycaM | 3 | 2.5 | 0.157 |
| 0.8 | 0.05 | yadI | 3 | 2.5 | 0.162 |
| 0.8 | 0.05 | AG1 | 3 | 2.5 | 0.164 |
| 0.8 | 0.05 | ptsI | 3 | 2.75 | 0.165 |
| 0.8 | 0.05 | clcB | 3 | 2.75 | 0.161 |
| 0.8 | 0.05 | ycaM | 3 | 2.75 | 0.16 |
| 0.8 | 0.05 | yadI | 3 | 2.75 | 0.165 |
| 0.8 | 0.05 | AG1 | 3 | 2.75 | 0.166 |
| 0.8 | 0.05 | ptsI | 3 | 3 | 0.166 |
| 0.8 | 0.05 | clcB | 3 | 3 | 0.162 |
| 0.8 | 0.05 | ycaM | 3 | 3 | 0.16 |
| 0.8 | 0.05 | yadI | 3 | 3 | 0.166 |
| 0.8 | 0.05 | AG1 | 3 | 3 | 0.167 |
| 0.8 | 0.05 | ptsI | 3 | 3.25 | 0.168 |
| 0.8 | 0.05 | clcB | 3 | 3.25 | 0.164 |
| 0.8 | 0.05 | ycaM | 3 | 3.25 | 0.162 |
| 0.8 | 0.05 | yadI | 3 | 3.25 | 0.169 |
| 0.8 | 0.05 | AG1 | 3 | 3.25 | 0.168 |
| 0.8 | 0.05 | ptsI | 3 | 3.5 | 0.172 |
| 0.8 | 0.05 | clcB | 3 | 3.5 | 0.165 |
| 0.8 | 0.05 | ycaM | 3 | 3.5 | 0.165 |
| 0.8 | 0.05 | yadI | 3 | 3.5 | 0.173 |
| 0.8 | 0.05 | AG1 | 3 | 3.5 | 0.17 |
| 0.8 | 0.05 | ptsI | 3 | 3.75 | 0.174 |
| 0.8 | 0.05 | clcB | 3 | 3.75 | 0.169 |
| 0.8 | 0.05 | ycaM | 3 | 3.75 | 0.167 |
| 0.8 | 0.05 | yadI | 3 | 3.75 | 0.177 |
| 0.8 | 0.05 | AG1 | 3 | 3.75 | 0.171 |
| 0.8 | 0.05 | ptsI | 3 | 4 | 0.178 |
| 0.8 | 0.05 | clcB | 3 | 4 | 0.171 |
| 0.8 | 0.05 | ycaM | 3 | 4 | 0.17 |
| 0.8 | 0.05 | yadI | 3 | 4 | 0.182 |
| 0.8 | 0.05 | AG1 | 3 | 4 | 0.174 |
| 0.8 | 0.05 | ptsI | 3 | 4.41667 | 0.192 |
| 0.8 | 0.05 | clcB | 3 | 4.41667 | 0.219 |
| 0.8 | 0.05 | ycaM | 3 | 4.41667 | 0.172 |
| 0.8 | 0.05 | yadI | 3 | 4.41667 | 0.22 |

| | | | | | |
|---|---|---|---|---|---|
| 0.8 | 0.05 | AG1 | 3 | 4.41667 | 0.188 |
| 0.8 | 0.05 | ptsI | 3 | 4.66667 | 0.171 |
| 0.8 | 0.05 | clcB | 3 | 4.66667 | 0.182 |
| 0.8 | 0.05 | ycaM | 3 | 4.66667 | 0.17 |
| 0.8 | 0.05 | yadI | 3 | 4.66667 | 0.221 |
| 0.8 | 0.05 | AG1 | 3 | 4.66667 | 0.188 |
| 0.8 | 0.05 | ptsI | 3 | 4.91667 | 0.172 |
| 0.8 | 0.05 | clcB | 3 | 4.91667 | 0.183 |
| 0.8 | 0.05 | ycaM | 3 | 4.91667 | 0.171 |
| 0.8 | 0.05 | yadI | 3 | 4.91667 | 0.228 |
| 0.8 | 0.05 | AG1 | 3 | 4.91667 | 0.192 |
| 0.8 | 0.05 | ptsI | 3 | 5.16667 | 0.175 |
| 0.8 | 0.05 | clcB | 3 | 5.16667 | 0.186 |
| 0.8 | 0.05 | ycaM | 3 | 5.16667 | 0.175 |
| 0.8 | 0.05 | yadI | 3 | 5.16667 | 0.24 |
| 0.8 | 0.05 | AG1 | 3 | 5.16667 | 0.197 |
| 0.8 | 0.05 | ptsI | 3 | 5.41667 | 0.178 |
| 0.8 | 0.05 | clcB | 3 | 5.41667 | 0.189 |
| 0.8 | 0.05 | ycaM | 3 | 5.41667 | 0.181 |
| 0.8 | 0.05 | yadI | 3 | 5.41667 | 0.254 |
| 0.8 | 0.05 | AG1 | 3 | 5.41667 | 0.204 |
| 0.8 | 0.05 | ptsI | 3 | 5.66667 | 0.182 |
| 0.8 | 0.05 | clcB | 3 | 5.66667 | 0.195 |
| 0.8 | 0.05 | ycaM | 3 | 5.66667 | 0.187 |
| 0.8 | 0.05 | yadI | 3 | 5.66667 | 0.272 |
| 0.8 | 0.05 | AG1 | 3 | 5.66667 | 0.212 |
| 0.8 | 0.05 | ptsI | 3 | 5.91667 | 0.186 |
| 0.8 | 0.05 | clcB | 3 | 5.91667 | 0.197 |
| 0.8 | 0.05 | ycaM | 3 | 5.91667 | 0.194 |
| 0.8 | 0.05 | yadI | 3 | 5.91667 | 0.274 |
| 0.8 | 0.05 | AG1 | 3 | 5.91667 | 0.214 |
| 0.8 | 0.05 | ptsI | 3 | 6.16667 | 0.192 |
| 0.8 | 0.05 | clcB | 3 | 6.16667 | 0.204 |
| 0.8 | 0.05 | ycaM | 3 | 6.16667 | 0.201 |
| 0.8 | 0.05 | yadI | 3 | 6.16667 | 0.292 |
| 0.8 | 0.05 | AG1 | 3 | 6.16667 | 0.222 |
| 0.8 | 0.05 | ptsI | 3 | 6.41667 | 0.198 |
| 0.8 | 0.05 | clcB | 3 | 6.41667 | 0.212 |
| 0.8 | 0.05 | ycaM | 3 | 6.41667 | 0.209 |
| 0.8 | 0.05 | yadI | 3 | 6.41667 | 0.301 |
| 0.8 | 0.05 | AG1 | 3 | 6.41667 | 0.231 |
| 0.8 | 0.05 | ptsI | 3 | 6.66667 | 0.206 |
| 0.8 | 0.05 | clcB | 3 | 6.66667 | 0.22 |
| 0.8 | 0.05 | ycaM | 3 | 6.66667 | 0.219 |
| 0.8 | 0.05 | yadI | 3 | 6.66667 | 0.309 |
| 0.8 | 0.05 | AG1 | 3 | 6.66667 | 0.242 |
| 0.8 | 0.05 | ptsI | 3 | 6.91667 | 0.214 |
| 0.8 | 0.05 | clcB | 3 | 6.91667 | 0.229 |

| | | | | | |
|---|---|---|---|---|---|
| 0.8 | 0.05 | ycaM | 3 | 6.91667 | 0.227 |
| 0.8 | 0.05 | yadI | 3 | 6.91667 | 0.317 |
| 0.8 | 0.05 | AG1 | 3 | 6.91667 | 0.25 |
| 0.8 | 0.05 | ptsI | 3 | 7.16667 | 0.224 |
| 0.8 | 0.05 | clcB | 3 | 7.16667 | 0.238 |
| 0.8 | 0.05 | ycaM | 3 | 7.16667 | 0.233 |
| 0.8 | 0.05 | yadI | 3 | 7.16667 | 0.328 |
| 0.8 | 0.05 | AG1 | 3 | 7.16667 | 0.259 |
| 0.8 | 0.05 | ptsI | 3 | 7.41667 | 0.233 |
| 0.8 | 0.05 | clcB | 3 | 7.41667 | 0.25 |
| 0.8 | 0.05 | ycaM | 3 | 7.41667 | 0.236 |
| 0.8 | 0.05 | yadI | 3 | 7.41667 | 0.339 |
| 0.8 | 0.05 | AG1 | 3 | 7.41667 | 0.267 |
| 0.8 | 0.05 | ptsI | 3 | 7.66667 | 0.242 |
| 0.8 | 0.05 | clcB | 3 | 7.66667 | 0.26 |
| 0.8 | 0.05 | ycaM | 3 | 7.66667 | 0.238 |
| 0.8 | 0.05 | yadI | 3 | 7.66667 | 0.352 |
| 0.8 | 0.05 | AG1 | 3 | 7.66667 | 0.272 |
| 0.8 | 0.05 | ptsI | 3 | 7.91667 | 0.25 |
| 0.8 | 0.05 | clcB | 3 | 7.91667 | 0.262 |
| 0.8 | 0.05 | ycaM | 3 | 7.91667 | 0.24 |
| 0.8 | 0.05 | yadI | 3 | 7.91667 | 0.362 |
| 0.8 | 0.05 | AG1 | 3 | 7.91667 | 0.281 |
| 0.8 | 0.05 | ptsI | 3 | 8.16667 | 0.258 |
| 0.8 | 0.05 | clcB | 3 | 8.16667 | 0.266 |
| 0.8 | 0.05 | ycaM | 3 | 8.16667 | 0.245 |
| 0.8 | 0.05 | yadI | 3 | 8.16667 | 0.371 |
| 0.8 | 0.05 | AG1 | 3 | 8.16667 | 0.289 |
| 0.8 | 0.05 | ptsI | 3 | 8.41667 | 0.267 |
| 0.8 | 0.05 | clcB | 3 | 8.41667 | 0.27 |
| 0.8 | 0.05 | ycaM | 3 | 8.41667 | 0.245 |
| 0.8 | 0.05 | yadI | 3 | 8.41667 | 0.38 |
| 0.8 | 0.05 | AG1 | 3 | 8.41667 | 0.296 |
| 0.8 | 0.05 | ptsI | 3 | 9.05 | 0.285 |
| 0.8 | 0.05 | clcB | 3 | 9.05 | 0.28 |
| 0.8 | 0.05 | ycaM | 3 | 9.05 | 0.233 |
| 0.8 | 0.05 | yadI | 3 | 9.05 | 0.368 |
| 0.8 | 0.05 | AG1 | 3 | 9.05 | 0.353 |
| 0.8 | 0.05 | ptsI | 3 | 9.3 | 0.288 |
| 0.8 | 0.05 | clcB | 3 | 9.3 | 0.282 |
| 0.8 | 0.05 | ycaM | 3 | 9.3 | 0.229 |
| 0.8 | 0.05 | yadI | 3 | 9.3 | 0.374 |
| 0.8 | 0.05 | AG1 | 3 | 9.3 | 0.362 |
| 0.8 | 0.05 | ptsI | 3 | 9.55 | 0.298 |
| 0.8 | 0.05 | clcB | 3 | 9.55 | 0.292 |
| 0.8 | 0.05 | ycaM | 3 | 9.55 | 0.232 |
| 0.8 | 0.05 | yadI | 3 | 9.55 | 0.387 |
| 0.8 | 0.05 | AG1 | 3 | 9.55 | 0.377 |

| | | | | | |
|---|---|---|---|---|---|
| 0.8 | 0.05 | ptsI | 3 | 9.8 | 0.306 |
| 0.8 | 0.05 | clcB | 3 | 9.8 | 0.296 |
| 0.8 | 0.05 | ycaM | 3 | 9.8 | 0.232 |
| 0.8 | 0.05 | yadI | 3 | 9.8 | 0.401 |
| 0.8 | 0.05 | AG1 | 3 | 9.8 | 0.387 |
| 0.8 | 0.05 | ptsI | 3 | 10.05 | 0.313 |
| 0.8 | 0.05 | clcB | 3 | 10.05 | 0.302 |
| 0.8 | 0.05 | ycaM | 3 | 10.05 | 0.234 |
| 0.8 | 0.05 | yadI | 3 | 10.05 | 0.413 |
| 0.8 | 0.05 | AG1 | 3 | 10.05 | 0.396 |
| 0.8 | 0.05 | ptsI | 3 | 10.3 | 0.321 |
| 0.8 | 0.05 | clcB | 3 | 10.3 | 0.307 |
| 0.8 | 0.05 | ycaM | 3 | 10.3 | 0.236 |
| 0.8 | 0.05 | yadI | 3 | 10.3 | 0.422 |
| 0.8 | 0.05 | AG1 | 3 | 10.3 | 0.404 |
| 0.8 | 0.05 | ptsI | 3 | 10.55 | 0.33 |
| 0.8 | 0.05 | clcB | 3 | 10.55 | 0.312 |
| 0.8 | 0.05 | ycaM | 3 | 10.55 | 0.235 |
| 0.8 | 0.05 | yadI | 3 | 10.55 | 0.431 |
| 0.8 | 0.05 | AG1 | 3 | 10.55 | 0.415 |
| 0.8 | 0.05 | ptsI | 3 | 10.8 | 0.34 |
| 0.8 | 0.05 | clcB | 3 | 10.8 | 0.309 |
| 0.8 | 0.05 | ycaM | 3 | 10.8 | 0.238 |
| 0.8 | 0.05 | yadI | 3 | 10.8 | 0.438 |
| 0.8 | 0.05 | AG1 | 3 | 10.8 | 0.423 |
| 0.8 | 0.05 | ptsI | 3 | 11.05 | 0.352 |
| 0.8 | 0.05 | clcB | 3 | 11.05 | 0.316 |
| 0.8 | 0.05 | ycaM | 3 | 11.05 | 0.24 |
| 0.8 | 0.05 | yadI | 3 | 11.05 | 0.45 |
| 0.8 | 0.05 | AG1 | 3 | 11.05 | 0.435 |
| 0.8 | 0.05 | ptsI | 3 | 11.3 | 0.359 |
| 0.8 | 0.05 | clcB | 3 | 11.3 | 0.318 |
| 0.8 | 0.05 | ycaM | 3 | 11.3 | 0.24 |
| 0.8 | 0.05 | yadI | 3 | 11.3 | 0.456 |
| 0.8 | 0.05 | AG1 | 3 | 11.3 | 0.444 |
| 0.8 | 0.05 | ptsI | 3 | 11.55 | 0.373 |
| 0.8 | 0.05 | clcB | 3 | 11.55 | 0.318 |
| 0.8 | 0.05 | ycaM | 3 | 11.55 | 0.242 |
| 0.8 | 0.05 | yadI | 3 | 11.55 | 0.47 |
| 0.8 | 0.05 | AG1 | 3 | 11.55 | 0.45 |
| 0.8 | 0.05 | ptsI | 3 | 11.8 | 0.38 |
| 0.8 | 0.05 | clcB | 3 | 11.8 | 0.314 |
| 0.8 | 0.05 | ycaM | 3 | 11.8 | 0.241 |
| 0.8 | 0.05 | yadI | 3 | 11.8 | 0.479 |
| 0.8 | 0.05 | AG1 | 3 | 11.8 | 0.46 |
| 0.8 | 0.05 | ptsI | 3 | 12.05 | 0.386 |
| 0.8 | 0.05 | clcB | 3 | 12.05 | 0.311 |
| 0.8 | 0.05 | ycaM | 3 | 12.05 | 0.236 |

| | | | | | |
|---|---|---|---|---|---|
| 0.8 | 0.05 | yadI | 3 | 12.05 | 0.479 |
| 0.8 | 0.05 | AG1 | 3 | 12.05 | 0.467 |
| 0.8 | 0.05 | ptsI | 3 | 12.3 | 0.398 |
| 0.8 | 0.05 | clcB | 3 | 12.3 | 0.305 |
| 0.8 | 0.05 | ycaM | 3 | 12.3 | 0.24 |
| 0.8 | 0.05 | yadI | 3 | 12.3 | 0.494 |
| 0.8 | 0.05 | AG1 | 3 | 12.3 | 0.476 |
| 0.8 | 0.05 | ptsI | 3 | 12.55 | 0.408 |
| 0.8 | 0.05 | clcB | 3 | 12.55 | 0.302 |
| 0.8 | 0.05 | ycaM | 3 | 12.55 | 0.238 |
| 0.8 | 0.05 | yadI | 3 | 12.55 | 0.501 |
| 0.8 | 0.05 | AG1 | 3 | 12.55 | 0.489 |
| 0.8 | 0.05 | ptsI | 3 | 12.8 | 0.417 |
| 0.8 | 0.05 | clcB | 3 | 12.8 | 0.301 |
| 0.8 | 0.05 | ycaM | 3 | 12.8 | 0.239 |
| 0.8 | 0.05 | yadI | 3 | 12.8 | 0.504 |
| 0.8 | 0.05 | AG1 | 3 | 12.8 | 0.499 |
| 0.8 | 0.05 | ptsI | 3 | 13.05 | 0.417 |
| 0.8 | 0.05 | clcB | 3 | 13.05 | 0.296 |
| 0.8 | 0.05 | ycaM | 3 | 13.05 | 0.237 |
| 0.8 | 0.05 | yadI | 3 | 13.05 | 0.505 |
| 0.8 | 0.05 | AG1 | 3 | 13.05 | 0.504 |
| 0.8 | 0.05 | ptsI | 3 | 13.3 | 0.429 |
| 0.8 | 0.05 | clcB | 3 | 13.3 | 0.292 |
| 0.8 | 0.05 | ycaM | 3 | 13.3 | 0.236 |
| 0.8 | 0.05 | yadI | 3 | 13.3 | 0.513 |
| 0.8 | 0.05 | AG1 | 3 | 13.3 | 0.524 |
| 0.8 | 0.05 | ptsI | 3 | 13.55 | 0.439 |
| 0.8 | 0.05 | clcB | 3 | 13.55 | 0.293 |
| 0.8 | 0.05 | ycaM | 3 | 13.55 | 0.237 |
| 0.8 | 0.05 | yadI | 3 | 13.55 | 0.52 |
| 0.8 | 0.05 | AG1 | 3 | 13.55 | 0.534 |
| 0.8 | 0.05 | ptsI | 3 | 13.8 | 0.446 |
| 0.8 | 0.05 | clcB | 3 | 13.8 | 0.284 |
| 0.8 | 0.05 | ycaM | 3 | 13.8 | 0.236 |
| 0.8 | 0.05 | yadI | 3 | 13.8 | 0.526 |
| 0.8 | 0.05 | AG1 | 3 | 13.8 | 0.542 |
| 0.8 | 0.05 | ptsI | 3 | 14.05 | 0.454 |
| 0.8 | 0.05 | clcB | 3 | 14.05 | 0.284 |
| 0.8 | 0.05 | ycaM | 3 | 14.05 | 0.236 |
| 0.8 | 0.05 | yadI | 3 | 14.05 | 0.532 |
| 0.8 | 0.05 | AG1 | 3 | 14.05 | 0.549 |
| 0.8 | 0.05 | ptsI | 3 | 14.3 | 0.454 |
| 0.8 | 0.05 | clcB | 3 | 14.3 | 0.284 |
| 0.8 | 0.05 | ycaM | 3 | 14.3 | 0.238 |
| 0.8 | 0.05 | yadI | 3 | 14.3 | 0.535 |
| 0.8 | 0.05 | AG1 | 3 | 14.3 | 0.552 |
| 0.8 | 0.05 | ptsI | 3 | 14.55 | 0.456 |

| | | | | | |
|---|---|---|---|---|---|
| 0.8 | 0.05 | clcB | 3 | 14.55 | 0.286 |
| 0.8 | 0.05 | ycaM | 3 | 14.55 | 0.235 |
| 0.8 | 0.05 | yadI | 3 | 14.55 | 0.538 |
| 0.8 | 0.05 | AG1 | 3 | 14.55 | 0.558 |
| 0.8 | 0.05 | ptsI | 3 | 14.8 | 0.465 |
| 0.8 | 0.05 | clcB | 3 | 14.8 | 0.289 |
| 0.8 | 0.05 | ycaM | 3 | 14.8 | 0.235 |
| 0.8 | 0.05 | yadI | 3 | 14.8 | 0.541 |
| 0.8 | 0.05 | AG1 | 3 | 14.8 | 0.573 |
| 0.8 | 0.05 | ptsI | 3 | 15.05 | 0.473 |
| 0.8 | 0.05 | clcB | 3 | 15.05 | 0.288 |
| 0.8 | 0.05 | ycaM | 3 | 15.05 | 0.233 |
| 0.8 | 0.05 | yadI | 3 | 15.05 | 0.546 |
| 0.8 | 0.05 | AG1 | 3 | 15.05 | 0.586 |
| 0.8 | 0.05 | ptsI | 3 | 15.3 | 0.48 |
| 0.8 | 0.05 | clcB | 3 | 15.3 | 0.293 |
| 0.8 | 0.05 | ycaM | 3 | 15.3 | 0.235 |
| 0.8 | 0.05 | yadI | 3 | 15.3 | 0.55 |
| 0.8 | 0.05 | AG1 | 3 | 15.3 | 0.582 |
| 0.8 | 0.05 | ptsI | 3 | 15.55 | 0.489 |
| 0.8 | 0.05 | clcB | 3 | 15.55 | 0.284 |
| 0.8 | 0.05 | ycaM | 3 | 15.55 | 0.234 |
| 0.8 | 0.05 | yadI | 3 | 15.55 | 0.555 |
| 0.8 | 0.05 | AG1 | 3 | 15.55 | 0.596 |
| 0.8 | 0.05 | ptsI | 3 | 15.8 | 0.498 |
| 0.8 | 0.05 | clcB | 3 | 15.8 | 0.284 |
| 0.8 | 0.05 | ycaM | 3 | 15.8 | 0.234 |
| 0.8 | 0.05 | yadI | 3 | 15.8 | 0.561 |
| 0.8 | 0.05 | AG1 | 3 | 15.8 | 0.61 |
| 0.8 | 0.05 | ptsI | 3 | 16.05 | 0.498 |
| 0.8 | 0.05 | clcB | 3 | 16.05 | 0.286 |
| 0.8 | 0.05 | ycaM | 3 | 16.05 | 0.234 |
| 0.8 | 0.05 | yadI | 3 | 16.05 | 0.562 |
| 0.8 | 0.05 | AG1 | 3 | 16.05 | 0.618 |
| 0.8 | 0.05 | ptsI | 3 | 16.3 | 0.501 |
| 0.8 | 0.05 | clcB | 3 | 16.3 | 0.284 |
| 0.8 | 0.05 | ycaM | 3 | 16.3 | 0.233 |
| 0.8 | 0.05 | yadI | 3 | 16.3 | 0.564 |
| 0.8 | 0.05 | AG1 | 3 | 16.3 | 0.615 |
| 0.8 | 0.05 | ptsI | 3 | 16.55 | 0.508 |
| 0.8 | 0.05 | clcB | 3 | 16.55 | 0.281 |
| 0.8 | 0.05 | ycaM | 3 | 16.55 | 0.233 |
| 0.8 | 0.05 | yadI | 3 | 16.55 | 0.569 |
| 0.8 | 0.05 | AG1 | 3 | 16.55 | 0.631 |
| 0.8 | 0.05 | ptsI | 3 | 16.8 | 0.515 |
| 0.8 | 0.05 | clcB | 3 | 16.8 | 0.28 |
| 0.8 | 0.05 | ycaM | 3 | 16.8 | 0.232 |
| 0.8 | 0.05 | yadI | 3 | 16.8 | 0.577 |

| | | | | | |
|---|---|---|---|---|---|
| 0.8 | 0.05 | AG1 | 3 | 16.8 | 0.642 |
| 0.8 | 0.05 | ptsI | 3 | 17.05 | 0.511 |
| 0.8 | 0.05 | clcB | 3 | 17.05 | 0.281 |
| 0.8 | 0.05 | ycaM | 3 | 17.05 | 0.232 |
| 0.8 | 0.05 | yadI | 3 | 17.05 | 0.583 |
| 0.8 | 0.05 | AG1 | 3 | 17.05 | 0.658 |
| 0.8 | 0.05 | ptsI | 3 | 17.3 | 0.513 |
| 0.8 | 0.05 | clcB | 3 | 17.3 | 0.283 |
| 0.8 | 0.05 | ycaM | 3 | 17.3 | 0.234 |
| 0.8 | 0.05 | yadI | 3 | 17.3 | 0.585 |
| 0.8 | 0.05 | AG1 | 3 | 17.3 | 0.663 |
| 0.8 | 0.05 | ptsI | 3 | 17.55 | 0.519 |
| 0.8 | 0.05 | clcB | 3 | 17.55 | 0.281 |
| 0.8 | 0.05 | ycaM | 3 | 17.55 | 0.234 |
| 0.8 | 0.05 | yadI | 3 | 17.55 | 0.591 |
| 0.8 | 0.05 | AG1 | 3 | 17.55 | 0.66 |
| 0.8 | 0.05 | ptsI | 3 | 17.8 | 0.518 |
| 0.8 | 0.05 | clcB | 3 | 17.8 | 0.282 |
| 0.8 | 0.05 | ycaM | 3 | 17.8 | 0.234 |
| 0.8 | 0.05 | yadI | 3 | 17.8 | 0.595 |
| 0.8 | 0.05 | AG1 | 3 | 17.8 | 0.667 |
| 0.8 | 0.05 | ptsI | 3 | 18.05 | 0.526 |
| 0.8 | 0.05 | clcB | 3 | 18.05 | 0.283 |
| 0.8 | 0.05 | ycaM | 3 | 18.05 | 0.236 |
| 0.8 | 0.05 | yadI | 3 | 18.05 | 0.598 |
| 0.8 | 0.05 | AG1 | 3 | 18.05 | 0.686 |
| 0.8 | 0.05 | ptsI | 3 | 18.3 | 0.524 |
| 0.8 | 0.05 | clcB | 3 | 18.3 | 0.282 |
| 0.8 | 0.05 | ycaM | 3 | 18.3 | 0.235 |
| 0.8 | 0.05 | yadI | 3 | 18.3 | 0.601 |
| 0.8 | 0.05 | AG1 | 3 | 18.3 | 0.682 |
| 0.8 | 0.05 | ptsI | 3 | 18.55 | 0.528 |
| 0.8 | 0.05 | clcB | 3 | 18.55 | 0.283 |
| 0.8 | 0.05 | ycaM | 3 | 18.55 | 0.237 |
| 0.8 | 0.05 | yadI | 3 | 18.55 | 0.603 |
| 0.8 | 0.05 | AG1 | 3 | 18.55 | 0.673 |
| 0.8 | 0.05 | ptsI | 3 | 18.8 | 0.533 |
| 0.8 | 0.05 | clcB | 3 | 18.8 | 0.281 |
| 0.8 | 0.05 | ycaM | 3 | 18.8 | 0.237 |
| 0.8 | 0.05 | yadI | 3 | 18.8 | 0.606 |
| 0.8 | 0.05 | AG1 | 3 | 18.8 | 0.675 |
| 0.8 | 0.05 | ptsI | 3 | 19.05 | 0.539 |
| 0.8 | 0.05 | clcB | 3 | 19.05 | 0.282 |
| 0.8 | 0.05 | ycaM | 3 | 19.05 | 0.237 |
| 0.8 | 0.05 | yadI | 3 | 19.05 | 0.611 |
| 0.8 | 0.05 | AG1 | 3 | 19.05 | 0.661 |
| 0.8 | 0.05 | ptsI | 3 | 19.3 | 0.54 |
| 0.8 | 0.05 | clcB | 3 | 19.3 | 0.283 |

| | | | | | |
|---|---|---|---|---|---|
| 0.8 | 0.05 | ycaM | 3 | 19.3 | 0.239 |
| 0.8 | 0.05 | yadI | 3 | 19.3 | 0.614 |
| 0.8 | 0.05 | AG1 | 3 | 19.3 | 0.665 |
| 0.8 | 0.05 | ptsI | 3 | 19.55 | 0.546 |
| 0.8 | 0.05 | clcB | 3 | 19.55 | 0.281 |
| 0.8 | 0.05 | ycaM | 3 | 19.55 | 0.24 |
| 0.8 | 0.05 | yadI | 3 | 19.55 | 0.615 |
| 0.8 | 0.05 | AG1 | 3 | 19.55 | 0.665 |
| 0.8 | 0.05 | ptsI | 3 | 19.8 | 0.546 |
| 0.8 | 0.05 | clcB | 3 | 19.8 | 0.282 |
| 0.8 | 0.05 | ycaM | 3 | 19.8 | 0.24 |
| 0.8 | 0.05 | yadI | 3 | 19.8 | 0.619 |
| 0.8 | 0.05 | AG1 | 3 | 19.8 | 0.681 |
| 0.8 | 0.05 | ptsI | 3 | 20.05 | 0.546 |
| 0.8 | 0.05 | clcB | 3 | 20.05 | 0.28 |
| 0.8 | 0.05 | ycaM | 3 | 20.05 | 0.243 |
| 0.8 | 0.05 | yadI | 3 | 20.05 | 0.622 |
| 0.8 | 0.05 | AG1 | 3 | 20.05 | 0.693 |
| 0.8 | 0.05 | ptsI | 3 | 20.3 | 0.55 |
| 0.8 | 0.05 | clcB | 3 | 20.3 | 0.283 |
| 0.8 | 0.05 | ycaM | 3 | 20.3 | 0.246 |
| 0.8 | 0.05 | yadI | 3 | 20.3 | 0.624 |
| 0.8 | 0.05 | AG1 | 3 | 20.3 | 0.701 |
| 0.8 | 0.05 | ptsI | 3 | 20.55 | 0.548 |
| 0.8 | 0.05 | clcB | 3 | 20.55 | 0.283 |
| 0.8 | 0.05 | ycaM | 3 | 20.55 | 0.246 |
| 0.8 | 0.05 | yadI | 3 | 20.55 | 0.628 |
| 0.8 | 0.05 | AG1 | 3 | 20.55 | 0.709 |
| 0.8 | 0.05 | ptsI | 3 | 20.8 | 0.545 |
| 0.8 | 0.05 | clcB | 3 | 20.8 | 0.285 |
| 0.8 | 0.05 | ycaM | 3 | 20.8 | 0.248 |
| 0.8 | 0.05 | yadI | 3 | 20.8 | 0.63 |
| 0.8 | 0.05 | AG1 | 3 | 20.8 | 0.713 |
| 0.8 | 0.05 | ptsI | 3 | 21.05 | 0.546 |
| 0.8 | 0.05 | clcB | 3 | 21.05 | 0.285 |
| 0.8 | 0.05 | ycaM | 3 | 21.05 | 0.248 |
| 0.8 | 0.05 | yadI | 3 | 21.05 | 0.632 |
| 0.8 | 0.05 | AG1 | 3 | 21.05 | 0.714 |
| 0.8 | 0.05 | ptsI | 3 | 21.3 | 0.545 |
| 0.8 | 0.05 | clcB | 3 | 21.3 | 0.284 |
| 0.8 | 0.05 | ycaM | 3 | 21.3 | 0.249 |
| 0.8 | 0.05 | yadI | 3 | 21.3 | 0.633 |
| 0.8 | 0.05 | AG1 | 3 | 21.3 | 0.717 |
| 0.8 | 0.05 | ptsI | 3 | 21.55 | 0.545 |
| 0.8 | 0.05 | clcB | 3 | 21.55 | 0.284 |
| 0.8 | 0.05 | ycaM | 3 | 21.55 | 0.251 |
| 0.8 | 0.05 | yadI | 3 | 21.55 | 0.634 |
| 0.8 | 0.05 | AG1 | 3 | 21.55 | 0.725 |

| | | | | | |
|---|---|---|---|---|---|
| 0.8 | 0.05 | ptsI | 3 | 21.8 | 0.542 |
| 0.8 | 0.05 | clcB | 3 | 21.8 | 0.286 |
| 0.8 | 0.05 | ycaM | 3 | 21.8 | 0.253 |
| 0.8 | 0.05 | yadI | 3 | 21.8 | 0.64 |
| 0.8 | 0.05 | AG1 | 3 | 21.8 | 0.735 |
| 0.8 | 0.05 | ptsI | 3 | 22.05 | 0.544 |
| 0.8 | 0.05 | clcB | 3 | 22.05 | 0.286 |
| 0.8 | 0.05 | ycaM | 3 | 22.05 | 0.254 |
| 0.8 | 0.05 | yadI | 3 | 22.05 | 0.639 |
| 0.8 | 0.05 | AG1 | 3 | 22.05 | 0.744 |
| 0.8 | 0.05 | ptsI | 3 | 22.3 | 0.548 |
| 0.8 | 0.05 | clcB | 3 | 22.3 | 0.286 |
| 0.8 | 0.05 | ycaM | 3 | 22.3 | 0.257 |
| 0.8 | 0.05 | yadI | 3 | 22.3 | 0.639 |
| 0.8 | 0.05 | AG1 | 3 | 22.3 | 0.751 |
| 0.8 | 0.05 | ptsI | 3 | 22.55 | 0.55 |
| 0.8 | 0.05 | clcB | 3 | 22.55 | 0.286 |
| 0.8 | 0.05 | ycaM | 3 | 22.55 | 0.259 |
| 0.8 | 0.05 | yadI | 3 | 22.55 | 0.647 |
| 0.8 | 0.05 | AG1 | 3 | 22.55 | 0.755 |
| 0.8 | 0.05 | ptsI | 3 | 22.8 | 0.544 |
| 0.8 | 0.05 | clcB | 3 | 22.8 | 0.288 |
| 0.8 | 0.05 | ycaM | 3 | 22.8 | 0.259 |
| 0.8 | 0.05 | yadI | 3 | 22.8 | 0.65 |
| 0.8 | 0.05 | AG1 | 3 | 22.8 | 0.766 |
| 0.8 | 0.05 | ptsI | 3 | 23.05 | 0.541 |
| 0.8 | 0.05 | clcB | 3 | 23.05 | 0.291 |
| 0.8 | 0.05 | ycaM | 3 | 23.05 | 0.263 |
| 0.8 | 0.05 | yadI | 3 | 23.05 | 0.651 |
| 0.8 | 0.05 | AG1 | 3 | 23.05 | 0.774 |
| 0.8 | 0.05 | ptsI | 3 | 23.3 | 0.541 |
| 0.8 | 0.05 | clcB | 3 | 23.3 | 0.292 |
| 0.8 | 0.05 | ycaM | 3 | 23.3 | 0.266 |
| 0.8 | 0.05 | yadI | 3 | 23.3 | 0.655 |
| 0.8 | 0.05 | AG1 | 3 | 23.3 | 0.78 |
| 0.8 | 0.05 | ptsI | 3 | 23.55 | 0.548 |
| 0.8 | 0.05 | clcB | 3 | 23.55 | 0.292 |
| 0.8 | 0.05 | ycaM | 3 | 23.55 | 0.268 |
| 0.8 | 0.05 | yadI | 3 | 23.55 | 0.654 |
| 0.8 | 0.05 | AG1 | 3 | 23.55 | 0.791 |
| 0.8 | 0.05 | ptsI | 3 | 23.8 | 0.543 |
| 0.8 | 0.05 | clcB | 3 | 23.8 | 0.295 |
| 0.8 | 0.05 | ycaM | 3 | 23.8 | 0.272 |
| 0.8 | 0.05 | yadI | 3 | 23.8 | 0.656 |
| 0.8 | 0.05 | AG1 | 3 | 23.8 | 0.797 |
| 0.8 | 0.05 | ptsI | 3 | 24.05 | 0.546 |
| 0.8 | 0.05 | clcB | 3 | 24.05 | 0.296 |
| 0.8 | 0.05 | ycaM | 3 | 24.05 | 0.273 |

| | | | | | |
|---|---|---|---|---|---|
| 0.8 | 0.05 | yadI | 3 | 24.05 | 0.658 |
| 0.8 | 0.05 | AG1 | 3 | 24.05 | 0.802 |
| 0.8 | 0.05 | ptsI | 3 | 24.3 | 0.548 |
| 0.8 | 0.05 | clcB | 3 | 24.3 | 0.293 |
| 0.8 | 0.05 | ycaM | 3 | 24.3 | 0.276 |
| 0.8 | 0.05 | yadI | 3 | 24.3 | 0.658 |
| 0.8 | 0.05 | AG1 | 3 | 24.3 | 0.813 |
| 0.8 | 0.05 | ptsI | 4 | 0 | 0.203 |
| 0.8 | 0.05 | clcB | 4 | 0 | 0.196 |
| 0.8 | 0.05 | ycaM | 4 | 0 | 0.212 |
| 0.8 | 0.05 | yadI | 4 | 0 | 0.217 |
| 0.8 | 0.05 | AG1 | 4 | 0 | 0.208 |
| 0.8 | 0.05 | ptsI | 4 | 0.25 | 0.2 |
| 0.8 | 0.05 | clcB | 4 | 0.25 | 0.192 |
| 0.8 | 0.05 | ycaM | 4 | 0.25 | 0.203 |
| 0.8 | 0.05 | yadI | 4 | 0.25 | 0.211 |
| 0.8 | 0.05 | AG1 | 4 | 0.25 | 0.204 |
| 0.8 | 0.05 | ptsI | 4 | 0.5 | 0.202 |
| 0.8 | 0.05 | clcB | 4 | 0.5 | 0.193 |
| 0.8 | 0.05 | ycaM | 4 | 0.5 | 0.204 |
| 0.8 | 0.05 | yadI | 4 | 0.5 | 0.213 |
| 0.8 | 0.05 | AG1 | 4 | 0.5 | 0.206 |
| 0.8 | 0.05 | ptsI | 4 | 0.75 | 0.203 |
| 0.8 | 0.05 | clcB | 4 | 0.75 | 0.194 |
| 0.8 | 0.05 | ycaM | 4 | 0.75 | 0.207 |
| 0.8 | 0.05 | yadI | 4 | 0.75 | 0.214 |
| 0.8 | 0.05 | AG1 | 4 | 0.75 | 0.208 |
| 0.8 | 0.05 | ptsI | 4 | 1 | 0.205 |
| 0.8 | 0.05 | clcB | 4 | 1 | 0.199 |
| 0.8 | 0.05 | ycaM | 4 | 1 | 0.211 |
| 0.8 | 0.05 | yadI | 4 | 1 | 0.211 |
| 0.8 | 0.05 | AG1 | 4 | 1 | 0.212 |
| 0.8 | 0.05 | ptsI | 4 | 1.25 | 0.21 |
| 0.8 | 0.05 | clcB | 4 | 1.25 | 0.205 |
| 0.8 | 0.05 | ycaM | 4 | 1.25 | 0.218 |
| 0.8 | 0.05 | yadI | 4 | 1.25 | 0.218 |
| 0.8 | 0.05 | AG1 | 4 | 1.25 | 0.216 |
| 0.8 | 0.05 | ptsI | 4 | 1.5 | 0.211 |
| 0.8 | 0.05 | clcB | 4 | 1.5 | 0.202 |
| 0.8 | 0.05 | ycaM | 4 | 1.5 | 0.213 |
| 0.8 | 0.05 | yadI | 4 | 1.5 | 0.221 |
| 0.8 | 0.05 | AG1 | 4 | 1.5 | 0.214 |
| 0.8 | 0.05 | ptsI | 4 | 1.75 | 0.22 |
| 0.8 | 0.05 | clcB | 4 | 1.75 | 0.207 |
| 0.8 | 0.05 | ycaM | 4 | 1.75 | 0.222 |
| 0.8 | 0.05 | yadI | 4 | 1.75 | 0.227 |
| 0.8 | 0.05 | AG1 | 4 | 1.75 | 0.223 |
| 0.8 | 0.05 | ptsI | 4 | 2 | 0.23 |

| | | | | | |
|---|---|---|---|---|---|
| 0.8 | 0.05 | clcB | 4 | 2 | 0.216 |
| 0.8 | 0.05 | ycaM | 4 | 2 | 0.229 |
| 0.8 | 0.05 | yadI | 4 | 2 | 0.234 |
| 0.8 | 0.05 | AG1 | 4 | 2 | 0.23 |
| 0.8 | 0.05 | ptsI | 4 | 2.25 | 0.238 |
| 0.8 | 0.05 | clcB | 4 | 2.25 | 0.224 |
| 0.8 | 0.05 | ycaM | 4 | 2.25 | 0.241 |
| 0.8 | 0.05 | yadI | 4 | 2.25 | 0.243 |
| 0.8 | 0.05 | AG1 | 4 | 2.25 | 0.238 |
| 0.8 | 0.05 | ptsI | 4 | 2.5 | 0.242 |
| 0.8 | 0.05 | clcB | 4 | 2.5 | 0.229 |
| 0.8 | 0.05 | ycaM | 4 | 2.5 | 0.24 |
| 0.8 | 0.05 | yadI | 4 | 2.5 | 0.25 |
| 0.8 | 0.05 | AG1 | 4 | 2.5 | 0.24 |
| 0.8 | 0.05 | ptsI | 4 | 2.75 | 0.244 |
| 0.8 | 0.05 | clcB | 4 | 2.75 | 0.235 |
| 0.8 | 0.05 | ycaM | 4 | 2.75 | 0.248 |
| 0.8 | 0.05 | yadI | 4 | 2.75 | 0.256 |
| 0.8 | 0.05 | AG1 | 4 | 2.75 | 0.249 |
| 0.8 | 0.05 | ptsI | 4 | 3 | 0.261 |
| 0.8 | 0.05 | clcB | 4 | 3 | 0.246 |
| 0.8 | 0.05 | ycaM | 4 | 3 | 0.257 |
| 0.8 | 0.05 | yadI | 4 | 3 | 0.264 |
| 0.8 | 0.05 | AG1 | 4 | 3 | 0.257 |
| 0.8 | 0.05 | ptsI | 4 | 3.25 | 0.27 |
| 0.8 | 0.05 | clcB | 4 | 3.25 | 0.255 |
| 0.8 | 0.05 | ycaM | 4 | 3.25 | 0.267 |
| 0.8 | 0.05 | yadI | 4 | 3.25 | 0.274 |
| 0.8 | 0.05 | AG1 | 4 | 3.25 | 0.267 |
| 0.8 | 0.05 | ptsI | 4 | 3.5 | 0.286 |
| 0.8 | 0.05 | clcB | 4 | 3.5 | 0.264 |
| 0.8 | 0.05 | ycaM | 4 | 3.5 | 0.277 |
| 0.8 | 0.05 | yadI | 4 | 3.5 | 0.284 |
| 0.8 | 0.05 | AG1 | 4 | 3.5 | 0.279 |
| 0.8 | 0.05 | ptsI | 4 | 3.75 | 0.292 |
| 0.8 | 0.05 | clcB | 4 | 3.75 | 0.275 |
| 0.8 | 0.05 | ycaM | 4 | 3.75 | 0.284 |
| 0.8 | 0.05 | yadI | 4 | 3.75 | 0.296 |
| 0.8 | 0.05 | AG1 | 4 | 3.75 | 0.287 |
| 0.8 | 0.05 | ptsI | 4 | 4 | 0.321 |
| 0.8 | 0.05 | clcB | 4 | 4 | 0.291 |
| 0.8 | 0.05 | ycaM | 4 | 4 | 0.3 |
| 0.8 | 0.05 | yadI | 4 | 4 | 0.309 |
| 0.8 | 0.05 | AG1 | 4 | 4 | 0.298 |
| 0.8 | 0.05 | ptsI | 4 | 4.35 | 0.313 |
| 0.8 | 0.05 | clcB | 4 | 4.35 | 0.359 |
| 0.8 | 0.05 | ycaM | 4 | 4.35 | 0.383 |
| 0.8 | 0.05 | yadI | 4 | 4.35 | 0.36 |

| | | | | | |
|---|---|---|---|---|---|
| 0.8 | 0.05 | AG1 | 4 | 4.35 | 0.295 |
| 0.8 | 0.05 | ptsI | 4 | 4.6 | 0.312 |
| 0.8 | 0.05 | clcB | 4 | 4.6 | 0.341 |
| 0.8 | 0.05 | ycaM | 4 | 4.6 | 0.351 |
| 0.8 | 0.05 | yadI | 4 | 4.6 | 0.365 |
| 0.8 | 0.05 | AG1 | 4 | 4.6 | 0.303 |
| 0.8 | 0.05 | ptsI | 4 | 4.85 | 0.322 |
| 0.8 | 0.05 | clcB | 4 | 4.85 | 0.346 |
| 0.8 | 0.05 | ycaM | 4 | 4.85 | 0.343 |
| 0.8 | 0.05 | yadI | 4 | 4.85 | 0.385 |
| 0.8 | 0.05 | AG1 | 4 | 4.85 | 0.316 |
| 0.8 | 0.05 | ptsI | 4 | 5.1 | 0.332 |
| 0.8 | 0.05 | clcB | 4 | 5.1 | 0.349 |
| 0.8 | 0.05 | ycaM | 4 | 5.1 | 0.344 |
| 0.8 | 0.05 | yadI | 4 | 5.1 | 0.4 |
| 0.8 | 0.05 | AG1 | 4 | 5.1 | 0.327 |
| 0.8 | 0.05 | ptsI | 4 | 5.35 | 0.347 |
| 0.8 | 0.05 | clcB | 4 | 5.35 | 0.351 |
| 0.8 | 0.05 | ycaM | 4 | 5.35 | 0.347 |
| 0.8 | 0.05 | yadI | 4 | 5.35 | 0.418 |
| 0.8 | 0.05 | AG1 | 4 | 5.35 | 0.338 |
| 0.8 | 0.05 | ptsI | 4 | 5.6 | 0.359 |
| 0.8 | 0.05 | clcB | 4 | 5.6 | 0.346 |
| 0.8 | 0.05 | ycaM | 4 | 5.6 | 0.345 |
| 0.8 | 0.05 | yadI | 4 | 5.6 | 0.427 |
| 0.8 | 0.05 | AG1 | 4 | 5.6 | 0.344 |
| 0.8 | 0.05 | ptsI | 4 | 5.85 | 0.368 |
| 0.8 | 0.05 | clcB | 4 | 5.85 | 0.344 |
| 0.8 | 0.05 | ycaM | 4 | 5.85 | 0.341 |
| 0.8 | 0.05 | yadI | 4 | 5.85 | 0.441 |
| 0.8 | 0.05 | AG1 | 4 | 5.85 | 0.349 |
| 0.8 | 0.05 | ptsI | 4 | 6.1 | 0.383 |
| 0.8 | 0.05 | clcB | 4 | 6.1 | 0.35 |
| 0.8 | 0.05 | ycaM | 4 | 6.1 | 0.342 |
| 0.8 | 0.05 | yadI | 4 | 6.1 | 0.457 |
| 0.8 | 0.05 | AG1 | 4 | 6.1 | 0.36 |
| 0.8 | 0.05 | ptsI | 4 | 6.35 | 0.396 |
| 0.8 | 0.05 | clcB | 4 | 6.35 | 0.354 |
| 0.8 | 0.05 | ycaM | 4 | 6.35 | 0.344 |
| 0.8 | 0.05 | yadI | 4 | 6.35 | 0.466 |
| 0.8 | 0.05 | AG1 | 4 | 6.35 | 0.37 |
| 0.8 | 0.05 | ptsI | 4 | 6.6 | 0.408 |
| 0.8 | 0.05 | clcB | 4 | 6.6 | 0.358 |
| 0.8 | 0.05 | ycaM | 4 | 6.6 | 0.346 |
| 0.8 | 0.05 | yadI | 4 | 6.6 | 0.472 |
| 0.8 | 0.05 | AG1 | 4 | 6.6 | 0.38 |
| 0.8 | 0.05 | ptsI | 4 | 6.85 | 0.417 |
| 0.8 | 0.05 | clcB | 4 | 6.85 | 0.361 |

| | | | | | |
|---|---|---|---|---|---|
| 0.8 | 0.05 | ycaM | 4 | 6.85 | 0.348 |
| 0.8 | 0.05 | yadI | 4 | 6.85 | 0.482 |
| 0.8 | 0.05 | AG1 | 4 | 6.85 | 0.388 |
| 0.8 | 0.05 | ptsI | 4 | 7.1 | 0.427 |
| 0.8 | 0.05 | clcB | 4 | 7.1 | 0.362 |
| 0.8 | 0.05 | ycaM | 4 | 7.1 | 0.349 |
| 0.8 | 0.05 | yadI | 4 | 7.1 | 0.489 |
| 0.8 | 0.05 | AG1 | 4 | 7.1 | 0.401 |
| 0.8 | 0.05 | ptsI | 4 | 7.35 | 0.435 |
| 0.8 | 0.05 | clcB | 4 | 7.35 | 0.365 |
| 0.8 | 0.05 | ycaM | 4 | 7.35 | 0.356 |
| 0.8 | 0.05 | yadI | 4 | 7.35 | 0.497 |
| 0.8 | 0.05 | AG1 | 4 | 7.35 | 0.408 |
| 0.8 | 0.05 | ptsI | 4 | 7.6 | 0.443 |
| 0.8 | 0.05 | clcB | 4 | 7.6 | 0.367 |
| 0.8 | 0.05 | ycaM | 4 | 7.6 | 0.369 |
| 0.8 | 0.05 | yadI | 4 | 7.6 | 0.507 |
| 0.8 | 0.05 | AG1 | 4 | 7.6 | 0.415 |
| 0.8 | 0.05 | ptsI | 4 | 7.85 | 0.446 |
| 0.8 | 0.05 | clcB | 4 | 7.85 | 0.364 |
| 0.8 | 0.05 | ycaM | 4 | 7.85 | 0.347 |
| 0.8 | 0.05 | yadI | 4 | 7.85 | 0.508 |
| 0.8 | 0.05 | AG1 | 4 | 7.85 | 0.419 |
| 0.8 | 0.05 | ptsI | 4 | 8.1 | 0.454 |
| 0.8 | 0.05 | clcB | 4 | 8.1 | 0.37 |
| 0.8 | 0.05 | ycaM | 4 | 8.1 | 0.35 |
| 0.8 | 0.05 | yadI | 4 | 8.1 | 0.514 |
| 0.8 | 0.05 | AG1 | 4 | 8.1 | 0.428 |
| 0.8 | 0.05 | ptsI | 4 | 8.35 | 0.463 |
| 0.8 | 0.05 | clcB | 4 | 8.35 | 0.372 |
| 0.8 | 0.05 | ycaM | 4 | 8.35 | 0.352 |
| 0.8 | 0.05 | yadI | 4 | 8.35 | 0.522 |
| 0.8 | 0.05 | AG1 | 4 | 8.35 | 0.439 |
| 0.8 | 0.05 | ptsI | 4 | 8.83333 | 0.436 |
| 0.8 | 0.05 | clcB | 4 | 8.83333 | 0.32 |
| 0.8 | 0.05 | ycaM | 4 | 8.83333 | 0.31 |
| 0.8 | 0.05 | yadI | 4 | 8.83333 | 0.513 |
| 0.8 | 0.05 | AG1 | 4 | 8.83333 | 0.457 |
| 0.8 | 0.05 | ptsI | 4 | 9.08333 | 0.438 |
| 0.8 | 0.05 | clcB | 4 | 9.08333 | 0.314 |
| 0.8 | 0.05 | ycaM | 4 | 9.08333 | 0.306 |
| 0.8 | 0.05 | yadI | 4 | 9.08333 | 0.507 |
| 0.8 | 0.05 | AG1 | 4 | 9.08333 | 0.456 |
| 0.8 | 0.05 | ptsI | 4 | 9.33333 | 0.446 |
| 0.8 | 0.05 | clcB | 4 | 9.33333 | 0.311 |
| 0.8 | 0.05 | ycaM | 4 | 9.33333 | 0.301 |
| 0.8 | 0.05 | yadI | 4 | 9.33333 | 0.515 |
| 0.8 | 0.05 | AG1 | 4 | 9.33333 | 0.468 |

| | | | | | |
|---|---|---|---|---|---|
| 0.8 | 0.05 | ptsI | 4 | 9.58333 | 0.456 |
| 0.8 | 0.05 | clcB | 4 | 9.58333 | 0.313 |
| 0.8 | 0.05 | ycaM | 4 | 9.58333 | 0.3 |
| 0.8 | 0.05 | yadI | 4 | 9.58333 | 0.528 |
| 0.8 | 0.05 | AG1 | 4 | 9.58333 | 0.479 |
| 0.8 | 0.05 | ptsI | 4 | 9.83333 | 0.465 |
| 0.8 | 0.05 | clcB | 4 | 9.83333 | 0.312 |
| 0.8 | 0.05 | ycaM | 4 | 9.83333 | 0.297 |
| 0.8 | 0.05 | yadI | 4 | 9.83333 | 0.537 |
| 0.8 | 0.05 | AG1 | 4 | 9.83333 | 0.488 |
| 0.8 | 0.05 | ptsI | 4 | 10.0833 | 0.474 |
| 0.8 | 0.05 | clcB | 4 | 10.0833 | 0.31 |
| 0.8 | 0.05 | ycaM | 4 | 10.0833 | 0.301 |
| 0.8 | 0.05 | yadI | 4 | 10.0833 | 0.545 |
| 0.8 | 0.05 | AG1 | 4 | 10.0833 | 0.495 |
| 0.8 | 0.05 | ptsI | 4 | 10.3333 | 0.481 |
| 0.8 | 0.05 | clcB | 4 | 10.3333 | 0.311 |
| 0.8 | 0.05 | ycaM | 4 | 10.3333 | 0.301 |
| 0.8 | 0.05 | yadI | 4 | 10.3333 | 0.55 |
| 0.8 | 0.05 | AG1 | 4 | 10.3333 | 0.499 |
| 0.8 | 0.05 | ptsI | 4 | 10.5833 | 0.491 |
| 0.8 | 0.05 | clcB | 4 | 10.5833 | 0.313 |
| 0.8 | 0.05 | ycaM | 4 | 10.5833 | 0.302 |
| 0.8 | 0.05 | yadI | 4 | 10.5833 | 0.555 |
| 0.8 | 0.05 | AG1 | 4 | 10.5833 | 0.506 |
| 0.8 | 0.05 | ptsI | 4 | 10.8333 | 0.498 |
| 0.8 | 0.05 | clcB | 4 | 10.8333 | 0.312 |
| 0.8 | 0.05 | ycaM | 4 | 10.8333 | 0.301 |
| 0.8 | 0.05 | yadI | 4 | 10.8333 | 0.563 |
| 0.8 | 0.05 | AG1 | 4 | 10.8333 | 0.513 |
| 0.8 | 0.05 | ptsI | 4 | 11.0833 | 0.504 |
| 0.8 | 0.05 | clcB | 4 | 11.0833 | 0.314 |
| 0.8 | 0.05 | ycaM | 4 | 11.0833 | 0.295 |
| 0.8 | 0.05 | yadI | 4 | 11.0833 | 0.565 |
| 0.8 | 0.05 | AG1 | 4 | 11.0833 | 0.519 |
| 0.8 | 0.05 | ptsI | 4 | 11.3333 | 0.511 |
| 0.8 | 0.05 | clcB | 4 | 11.3333 | 0.311 |
| 0.8 | 0.05 | ycaM | 4 | 11.3333 | 0.297 |
| 0.8 | 0.05 | yadI | 4 | 11.3333 | 0.571 |
| 0.8 | 0.05 | AG1 | 4 | 11.3333 | 0.525 |
| 0.8 | 0.05 | ptsI | 4 | 11.5833 | 0.52 |
| 0.8 | 0.05 | clcB | 4 | 11.5833 | 0.314 |
| 0.8 | 0.05 | ycaM | 4 | 11.5833 | 0.297 |
| 0.8 | 0.05 | yadI | 4 | 11.5833 | 0.579 |
| 0.8 | 0.05 | AG1 | 4 | 11.5833 | 0.528 |
| 0.8 | 0.05 | ptsI | 4 | 11.8333 | 0.526 |
| 0.8 | 0.05 | clcB | 4 | 11.8333 | 0.308 |
| 0.8 | 0.05 | ycaM | 4 | 11.8333 | 0.296 |

| | | | | | |
|---|---|---|---|---|---|
| 0.8 | 0.05 | yadI | 4 | 11.8333 | 0.583 |
| 0.8 | 0.05 | AG1 | 4 | 11.8333 | 0.532 |
| 0.8 | 0.05 | ptsI | 4 | 12.0833 | 0.531 |
| 0.8 | 0.05 | clcB | 4 | 12.0833 | 0.315 |
| 0.8 | 0.05 | ycaM | 4 | 12.0833 | 0.295 |
| 0.8 | 0.05 | yadI | 4 | 12.0833 | 0.589 |
| 0.8 | 0.05 | AG1 | 4 | 12.0833 | 0.539 |
| 0.8 | 0.05 | ptsI | 4 | 12.3333 | 0.54 |
| 0.8 | 0.05 | clcB | 4 | 12.3333 | 0.316 |
| 0.8 | 0.05 | ycaM | 4 | 12.3333 | 0.302 |
| 0.8 | 0.05 | yadI | 4 | 12.3333 | 0.594 |
| 0.8 | 0.05 | AG1 | 4 | 12.3333 | 0.543 |
| 0.8 | 0.05 | ptsI | 4 | 12.5833 | 0.548 |
| 0.8 | 0.05 | clcB | 4 | 12.5833 | 0.318 |
| 0.8 | 0.05 | ycaM | 4 | 12.5833 | 0.304 |
| 0.8 | 0.05 | yadI | 4 | 12.5833 | 0.592 |
| 0.8 | 0.05 | AG1 | 4 | 12.5833 | 0.548 |
| 0.8 | 0.05 | ptsI | 4 | 12.8333 | 0.551 |
| 0.8 | 0.05 | clcB | 4 | 12.8333 | 0.318 |
| 0.8 | 0.05 | ycaM | 4 | 12.8333 | 0.308 |
| 0.8 | 0.05 | yadI | 4 | 12.8333 | 0.601 |
| 0.8 | 0.05 | AG1 | 4 | 12.8333 | 0.553 |
| 0.8 | 0.05 | ptsI | 4 | 13.0833 | 0.557 |
| 0.8 | 0.05 | clcB | 4 | 13.0833 | 0.321 |
| 0.8 | 0.05 | ycaM | 4 | 13.0833 | 0.304 |
| 0.8 | 0.05 | yadI | 4 | 13.0833 | 0.605 |
| 0.8 | 0.05 | AG1 | 4 | 13.0833 | 0.559 |
| 0.8 | 0.05 | ptsI | 4 | 13.3333 | 0.562 |
| 0.8 | 0.05 | clcB | 4 | 13.3333 | 0.324 |
| 0.8 | 0.05 | ycaM | 4 | 13.3333 | 0.304 |
| 0.8 | 0.05 | yadI | 4 | 13.3333 | 0.608 |
| 0.8 | 0.05 | AG1 | 4 | 13.3333 | 0.564 |
| 0.8 | 0.05 | ptsI | 4 | 13.5833 | 0.567 |
| 0.8 | 0.05 | clcB | 4 | 13.5833 | 0.323 |
| 0.8 | 0.05 | ycaM | 4 | 13.5833 | 0.314 |
| 0.8 | 0.05 | yadI | 4 | 13.5833 | 0.61 |
| 0.8 | 0.05 | AG1 | 4 | 13.5833 | 0.568 |
| 0.8 | 0.05 | ptsI | 4 | 13.8333 | 0.568 |
| 0.8 | 0.05 | clcB | 4 | 13.8333 | 0.329 |
| 0.8 | 0.05 | ycaM | 4 | 13.8333 | 0.312 |
| 0.8 | 0.05 | yadI | 4 | 13.8333 | 0.612 |
| 0.8 | 0.05 | AG1 | 4 | 13.8333 | 0.573 |
| 0.8 | 0.05 | ptsI | 4 | 14.0833 | 0.575 |
| 0.8 | 0.05 | clcB | 4 | 14.0833 | 0.329 |
| 0.8 | 0.05 | ycaM | 4 | 14.0833 | 0.312 |
| 0.8 | 0.05 | yadI | 4 | 14.0833 | 0.62 |
| 0.8 | 0.05 | AG1 | 4 | 14.0833 | 0.579 |
| 0.8 | 0.05 | ptsI | 4 | 14.3333 | 0.579 |

| | | | | | |
|---|---|---|---|---|---|
| 0.8 | 0.05 | clcB | 4 | 14.3333 | 0.331 |
| 0.8 | 0.05 | ycaM | 4 | 14.3333 | 0.316 |
| 0.8 | 0.05 | yadI | 4 | 14.3333 | 0.618 |
| 0.8 | 0.05 | AG1 | 4 | 14.3333 | 0.58 |
| 0.8 | 0.05 | ptsI | 4 | 14.5833 | 0.583 |
| 0.8 | 0.05 | clcB | 4 | 14.5833 | 0.332 |
| 0.8 | 0.05 | ycaM | 4 | 14.5833 | 0.315 |
| 0.8 | 0.05 | yadI | 4 | 14.5833 | 0.622 |
| 0.8 | 0.05 | AG1 | 4 | 14.5833 | 0.588 |
| 0.8 | 0.05 | ptsI | 4 | 14.8333 | 0.586 |
| 0.8 | 0.05 | clcB | 4 | 14.8333 | 0.332 |
| 0.8 | 0.05 | ycaM | 4 | 14.8333 | 0.317 |
| 0.8 | 0.05 | yadI | 4 | 14.8333 | 0.621 |
| 0.8 | 0.05 | AG1 | 4 | 14.8333 | 0.591 |
| 0.8 | 0.05 | ptsI | 4 | 15.0833 | 0.588 |
| 0.8 | 0.05 | clcB | 4 | 15.0833 | 0.332 |
| 0.8 | 0.05 | ycaM | 4 | 15.0833 | 0.32 |
| 0.8 | 0.05 | yadI | 4 | 15.0833 | 0.622 |
| 0.8 | 0.05 | AG1 | 4 | 15.0833 | 0.594 |
| 0.8 | 0.05 | ptsI | 4 | 15.3333 | 0.592 |
| 0.8 | 0.05 | clcB | 4 | 15.3333 | 0.338 |
| 0.8 | 0.05 | ycaM | 4 | 15.3333 | 0.32 |
| 0.8 | 0.05 | yadI | 4 | 15.3333 | 0.624 |
| 0.8 | 0.05 | AG1 | 4 | 15.3333 | 0.595 |
| 0.8 | 0.05 | ptsI | 4 | 15.5833 | 0.595 |
| 0.8 | 0.05 | clcB | 4 | 15.5833 | 0.339 |
| 0.8 | 0.05 | ycaM | 4 | 15.5833 | 0.322 |
| 0.8 | 0.05 | yadI | 4 | 15.5833 | 0.627 |
| 0.8 | 0.05 | AG1 | 4 | 15.5833 | 0.599 |
| 0.8 | 0.05 | ptsI | 4 | 15.8333 | 0.599 |
| 0.8 | 0.05 | clcB | 4 | 15.8333 | 0.339 |
| 0.8 | 0.05 | ycaM | 4 | 15.8333 | 0.324 |
| 0.8 | 0.05 | yadI | 4 | 15.8333 | 0.629 |
| 0.8 | 0.05 | AG1 | 4 | 15.8333 | 0.604 |
| 0.8 | 0.05 | ptsI | 4 | 16.0833 | 0.599 |
| 0.8 | 0.05 | clcB | 4 | 16.0833 | 0.339 |
| 0.8 | 0.05 | ycaM | 4 | 16.0833 | 0.32 |
| 0.8 | 0.05 | yadI | 4 | 16.0833 | 0.631 |
| 0.8 | 0.05 | AG1 | 4 | 16.0833 | 0.603 |
| 0.8 | 0.05 | ptsI | 4 | 16.3333 | 0.6 |
| 0.8 | 0.05 | clcB | 4 | 16.3333 | 0.34 |
| 0.8 | 0.05 | ycaM | 4 | 16.3333 | 0.325 |
| 0.8 | 0.05 | yadI | 4 | 16.3333 | 0.634 |
| 0.8 | 0.05 | AG1 | 4 | 16.3333 | 0.608 |
| 0.8 | 0.05 | ptsI | 4 | 16.5833 | 0.603 |
| 0.8 | 0.05 | clcB | 4 | 16.5833 | 0.34 |
| 0.8 | 0.05 | ycaM | 4 | 16.5833 | 0.328 |
| 0.8 | 0.05 | yadI | 4 | 16.5833 | 0.635 |

| | | | | | |
|---|---|---|---|---|---|
| 0.8 | 0.05 | AG1 | 4 | 16.5833 | 0.613 |
| 0.8 | 0.05 | ptsI | 4 | 16.8333 | 0.601 |
| 0.8 | 0.05 | clcB | 4 | 16.8333 | 0.337 |
| 0.8 | 0.05 | ycaM | 4 | 16.8333 | 0.326 |
| 0.8 | 0.05 | yadI | 4 | 16.8333 | 0.639 |
| 0.8 | 0.05 | AG1 | 4 | 16.8333 | 0.61 |
| 0.8 | 0.05 | ptsI | 4 | 17.0833 | 0.602 |
| 0.8 | 0.05 | clcB | 4 | 17.0833 | 0.337 |
| 0.8 | 0.05 | ycaM | 4 | 17.0833 | 0.325 |
| 0.8 | 0.05 | yadI | 4 | 17.0833 | 0.641 |
| 0.8 | 0.05 | AG1 | 4 | 17.0833 | 0.612 |
| 0.8 | 0.05 | ptsI | 4 | 17.3333 | 0.606 |
| 0.8 | 0.05 | clcB | 4 | 17.3333 | 0.343 |
| 0.8 | 0.05 | ycaM | 4 | 17.3333 | 0.327 |
| 0.8 | 0.05 | yadI | 4 | 17.3333 | 0.644 |
| 0.8 | 0.05 | AG1 | 4 | 17.3333 | 0.616 |
| 0.8 | 0.05 | ptsI | 4 | 17.5833 | 0.604 |
| 0.8 | 0.05 | clcB | 4 | 17.5833 | 0.34 |
| 0.8 | 0.05 | ycaM | 4 | 17.5833 | 0.325 |
| 0.8 | 0.05 | yadI | 4 | 17.5833 | 0.647 |
| 0.8 | 0.05 | AG1 | 4 | 17.5833 | 0.62 |
| 0.8 | 0.05 | ptsI | 4 | 17.8333 | 0.599 |
| 0.8 | 0.05 | clcB | 4 | 17.8333 | 0.34 |
| 0.8 | 0.05 | ycaM | 4 | 17.8333 | 0.326 |
| 0.8 | 0.05 | yadI | 4 | 17.8333 | 0.647 |
| 0.8 | 0.05 | AG1 | 4 | 17.8333 | 0.616 |
| 0.8 | 0.05 | ptsI | 4 | 18.0833 | 0.605 |
| 0.8 | 0.05 | clcB | 4 | 18.0833 | 0.34 |
| 0.8 | 0.05 | ycaM | 4 | 18.0833 | 0.325 |
| 0.8 | 0.05 | yadI | 4 | 18.0833 | 0.648 |
| 0.8 | 0.05 | AG1 | 4 | 18.0833 | 0.622 |
| 0.8 | 0.05 | ptsI | 4 | 18.3333 | 0.605 |
| 0.8 | 0.05 | clcB | 4 | 18.3333 | 0.338 |
| 0.8 | 0.05 | ycaM | 4 | 18.3333 | 0.328 |
| 0.8 | 0.05 | yadI | 4 | 18.3333 | 0.65 |
| 0.8 | 0.05 | AG1 | 4 | 18.3333 | 0.625 |
| 0.8 | 0.05 | ptsI | 4 | 18.5833 | 0.605 |
| 0.8 | 0.05 | clcB | 4 | 18.5833 | 0.342 |
| 0.8 | 0.05 | ycaM | 4 | 18.5833 | 0.328 |
| 0.8 | 0.05 | yadI | 4 | 18.5833 | 0.652 |
| 0.8 | 0.05 | AG1 | 4 | 18.5833 | 0.628 |
| 0.8 | 0.05 | ptsI | 4 | 18.8333 | 0.599 |
| 0.8 | 0.05 | clcB | 4 | 18.8333 | 0.339 |
| 0.8 | 0.05 | ycaM | 4 | 18.8333 | 0.325 |
| 0.8 | 0.05 | yadI | 4 | 18.8333 | 0.651 |
| 0.8 | 0.05 | AG1 | 4 | 18.8333 | 0.625 |
| 0.8 | 0.05 | ptsI | 4 | 19.0833 | 0.603 |
| 0.8 | 0.05 | clcB | 4 | 19.0833 | 0.343 |

| | | | | | |
|---|---|---|---|---|---|
| 0.8 | 0.05 | ycaM | 4 | 19.0833 | 0.324 |
| 0.8 | 0.05 | yadI | 4 | 19.0833 | 0.654 |
| 0.8 | 0.05 | AG1 | 4 | 19.0833 | 0.632 |
| 0.8 | 0.05 | ptsI | 4 | 19.3333 | 0.601 |
| 0.8 | 0.05 | clcB | 4 | 19.3333 | 0.342 |
| 0.8 | 0.05 | ycaM | 4 | 19.3333 | 0.329 |
| 0.8 | 0.05 | yadI | 4 | 19.3333 | 0.654 |
| 0.8 | 0.05 | AG1 | 4 | 19.3333 | 0.631 |
| 0.8 | 0.05 | ptsI | 4 | 19.5833 | 0.603 |
| 0.8 | 0.05 | clcB | 4 | 19.5833 | 0.342 |
| 0.8 | 0.05 | ycaM | 4 | 19.5833 | 0.331 |
| 0.8 | 0.05 | yadI | 4 | 19.5833 | 0.658 |
| 0.8 | 0.05 | AG1 | 4 | 19.5833 | 0.635 |
| 0.8 | 0.05 | ptsI | 4 | 19.8333 | 0.604 |
| 0.8 | 0.05 | clcB | 4 | 19.8333 | 0.335 |
| 0.8 | 0.05 | ycaM | 4 | 19.8333 | 0.332 |
| 0.8 | 0.05 | yadI | 4 | 19.8333 | 0.654 |
| 0.8 | 0.05 | AG1 | 4 | 19.8333 | 0.641 |
| 0.8 | 0.05 | ptsI | 4 | 20.0833 | 0.605 |
| 0.8 | 0.05 | clcB | 4 | 20.0833 | 0.342 |
| 0.8 | 0.05 | ycaM | 4 | 20.0833 | 0.337 |
| 0.8 | 0.05 | yadI | 4 | 20.0833 | 0.657 |
| 0.8 | 0.05 | AG1 | 4 | 20.0833 | 0.642 |
| 0.8 | 0.05 | ptsI | 4 | 20.3333 | 0.6 |
| 0.8 | 0.05 | clcB | 4 | 20.3333 | 0.333 |
| 0.8 | 0.05 | ycaM | 4 | 20.3333 | 0.338 |
| 0.8 | 0.05 | yadI | 4 | 20.3333 | 0.654 |
| 0.8 | 0.05 | AG1 | 4 | 20.3333 | 0.641 |
| 0.8 | 0.05 | ptsI | 4 | 20.5833 | 0.603 |
| 0.8 | 0.05 | clcB | 4 | 20.5833 | 0.335 |
| 0.8 | 0.05 | ycaM | 4 | 20.5833 | 0.339 |
| 0.8 | 0.05 | yadI | 4 | 20.5833 | 0.659 |
| 0.8 | 0.05 | AG1 | 4 | 20.5833 | 0.649 |
| 0.8 | 0.05 | ptsI | 4 | 20.8333 | 0.6 |
| 0.8 | 0.05 | clcB | 4 | 20.8333 | 0.336 |
| 0.8 | 0.05 | ycaM | 4 | 20.8333 | 0.343 |
| 0.8 | 0.05 | yadI | 4 | 20.8333 | 0.655 |
| 0.8 | 0.05 | AG1 | 4 | 20.8333 | 0.647 |
| 0.8 | 0.05 | ptsI | 4 | 21.0833 | 0.6 |
| 0.8 | 0.05 | clcB | 4 | 21.0833 | 0.336 |
| 0.8 | 0.05 | ycaM | 4 | 21.0833 | 0.34 |
| 0.8 | 0.05 | yadI | 4 | 21.0833 | 0.659 |
| 0.8 | 0.05 | AG1 | 4 | 21.0833 | 0.654 |
| 0.8 | 0.05 | ptsI | 4 | 21.3333 | 0.596 |
| 0.8 | 0.05 | clcB | 4 | 21.3333 | 0.34 |
| 0.8 | 0.05 | ycaM | 4 | 21.3333 | 0.342 |
| 0.8 | 0.05 | yadI | 4 | 21.3333 | 0.656 |
| 0.8 | 0.05 | AG1 | 4 | 21.3333 | 0.652 |

| | | | | | |
|---|---|---|---|---|---|
| 0.8 | 0.05 | ptsI | 4 | 21.5833 | 0.595 |
| 0.8 | 0.05 | clcB | 4 | 21.5833 | 0.338 |
| 0.8 | 0.05 | ycaM | 4 | 21.5833 | 0.344 |
| 0.8 | 0.05 | yadI | 4 | 21.5833 | 0.66 |
| 0.8 | 0.05 | AG1 | 4 | 21.5833 | 0.656 |
| 0.8 | 0.05 | ptsI | 4 | 21.8333 | 0.602 |
| 0.8 | 0.05 | clcB | 4 | 21.8333 | 0.335 |
| 0.8 | 0.05 | ycaM | 4 | 21.8333 | 0.345 |
| 0.8 | 0.05 | yadI | 4 | 21.8333 | 0.662 |
| 0.8 | 0.05 | AG1 | 4 | 21.8333 | 0.655 |
| 0.8 | 0.05 | ptsI | 4 | 22.0833 | 0.592 |
| 0.8 | 0.05 | clcB | 4 | 22.0833 | 0.333 |
| 0.8 | 0.05 | ycaM | 4 | 22.0833 | 0.346 |
| 0.8 | 0.05 | yadI | 4 | 22.0833 | 0.66 |
| 0.8 | 0.05 | AG1 | 4 | 22.0833 | 0.654 |
| 0.8 | 0.05 | ptsI | 4 | 22.3333 | 0.597 |
| 0.8 | 0.05 | clcB | 4 | 22.3333 | 0.331 |
| 0.8 | 0.05 | ycaM | 4 | 22.3333 | 0.345 |
| 0.8 | 0.05 | yadI | 4 | 22.3333 | 0.662 |
| 0.8 | 0.05 | AG1 | 4 | 22.3333 | 0.654 |
| 0.8 | 0.05 | ptsI | 4 | 22.5833 | 0.597 |
| 0.8 | 0.05 | clcB | 4 | 22.5833 | 0.33 |
| 0.8 | 0.05 | ycaM | 4 | 22.5833 | 0.348 |
| 0.8 | 0.05 | yadI | 4 | 22.5833 | 0.667 |
| 0.8 | 0.05 | AG1 | 4 | 22.5833 | 0.658 |
| 0.8 | 0.05 | ptsI | 4 | 22.8333 | 0.595 |
| 0.8 | 0.05 | clcB | 4 | 22.8333 | 0.331 |
| 0.8 | 0.05 | ycaM | 4 | 22.8333 | 0.351 |
| 0.8 | 0.05 | yadI | 4 | 22.8333 | 0.666 |
| 0.8 | 0.05 | AG1 | 4 | 22.8333 | 0.662 |
| 0.8 | 0.05 | ptsI | 4 | 23.0833 | 0.593 |
| 0.8 | 0.05 | clcB | 4 | 23.0833 | 0.335 |
| 0.8 | 0.05 | ycaM | 4 | 23.0833 | 0.35 |
| 0.8 | 0.05 | yadI | 4 | 23.0833 | 0.668 |
| 0.8 | 0.05 | AG1 | 4 | 23.0833 | 0.661 |
| 0.8 | 0.05 | ptsI | 4 | 23.3333 | 0.602 |
| 0.8 | 0.05 | clcB | 4 | 23.3333 | 0.337 |
| 0.8 | 0.05 | ycaM | 4 | 23.3333 | 0.355 |
| 0.8 | 0.05 | yadI | 4 | 23.3333 | 0.668 |
| 0.8 | 0.05 | AG1 | 4 | 23.3333 | 0.664 |
| 0.8 | 0.05 | ptsI | 4 | 23.5833 | 0.603 |
| 0.8 | 0.05 | clcB | 4 | 23.5833 | 0.338 |
| 0.8 | 0.05 | ycaM | 4 | 23.5833 | 0.354 |
| 0.8 | 0.05 | yadI | 4 | 23.5833 | 0.669 |
| 0.8 | 0.05 | AG1 | 4 | 23.5833 | 0.667 |
| 0.8 | 0.05 | ptsI | 4 | 23.8333 | 0.6 |
| 0.8 | 0.05 | clcB | 4 | 23.8333 | 0.335 |
| 0.8 | 0.05 | ycaM | 4 | 23.8333 | 0.354 |

| | | | | | |
|---|---|---|---|---|---|
| 0.8 | 0.05 | yadI | 4 | 23.8333 | 0.668 |
| 0.8 | 0.05 | AG1 | 4 | 23.8333 | 0.668 |
| 0.8 | 0.05 | ptsI | 4 | 24.0833 | 0.594 |
| 0.8 | 0.05 | clcB | 4 | 24.0833 | 0.339 |
| 0.8 | 0.05 | ycaM | 4 | 24.0833 | 0.354 |
| 0.8 | 0.05 | yadI | 4 | 24.0833 | 0.665 |
| 0.8 | 0.05 | AG1 | 4 | 24.0833 | 0.669 |
| 0.8 | 0.05 | ptsI | 5 | 0 | 0.164 |
| 0.8 | 0.05 | clcB | 5 | 0 | 0.171 |
| 0.8 | 0.05 | ycaM | 5 | 0 | 0.163 |
| 0.8 | 0.05 | yadI | 5 | 0 | 0.17 |
| 0.8 | 0.05 | AG1 | 5 | 0 | 0.162 |
| 0.8 | 0.05 | ptsI | 5 | 0.35 | 0.157 |
| 0.8 | 0.05 | clcB | 5 | 0.35 | 0.167 |
| 0.8 | 0.05 | ycaM | 5 | 0.35 | 0.156 |
| 0.8 | 0.05 | yadI | 5 | 0.35 | 0.166 |
| 0.8 | 0.05 | AG1 | 5 | 0.35 | 0.161 |
| 0.8 | 0.05 | ptsI | 5 | 0.6 | 0.157 |
| 0.8 | 0.05 | clcB | 5 | 0.6 | 0.168 |
| 0.8 | 0.05 | ycaM | 5 | 0.6 | 0.156 |
| 0.8 | 0.05 | yadI | 5 | 0.6 | 0.168 |
| 0.8 | 0.05 | AG1 | 5 | 0.6 | 0.161 |
| 0.8 | 0.05 | ptsI | 5 | 0.85 | 0.155 |
| 0.8 | 0.05 | clcB | 5 | 0.85 | 0.165 |
| 0.8 | 0.05 | ycaM | 5 | 0.85 | 0.154 |
| 0.8 | 0.05 | yadI | 5 | 0.85 | 0.167 |
| 0.8 | 0.05 | AG1 | 5 | 0.85 | 0.162 |
| 0.8 | 0.05 | ptsI | 5 | 1.1 | 0.156 |
| 0.8 | 0.05 | clcB | 5 | 1.1 | 0.17 |
| 0.8 | 0.05 | ycaM | 5 | 1.1 | 0.157 |
| 0.8 | 0.05 | yadI | 5 | 1.1 | 0.168 |
| 0.8 | 0.05 | AG1 | 5 | 1.1 | 0.165 |
| 0.8 | 0.05 | ptsI | 5 | 1.35 | 0.157 |
| 0.8 | 0.05 | clcB | 5 | 1.35 | 0.17 |
| 0.8 | 0.05 | ycaM | 5 | 1.35 | 0.156 |
| 0.8 | 0.05 | yadI | 5 | 1.35 | 0.17 |
| 0.8 | 0.05 | AG1 | 5 | 1.35 | 0.164 |
| 0.8 | 0.05 | ptsI | 5 | 1.6 | 0.16 |
| 0.8 | 0.05 | clcB | 5 | 1.6 | 0.172 |
| 0.8 | 0.05 | ycaM | 5 | 1.6 | 0.158 |
| 0.8 | 0.05 | yadI | 5 | 1.6 | 0.173 |
| 0.8 | 0.05 | AG1 | 5 | 1.6 | 0.166 |
| 0.8 | 0.05 | ptsI | 5 | 1.85 | 0.162 |
| 0.8 | 0.05 | clcB | 5 | 1.85 | 0.176 |
| 0.8 | 0.05 | ycaM | 5 | 1.85 | 0.162 |
| 0.8 | 0.05 | yadI | 5 | 1.85 | 0.178 |
| 0.8 | 0.05 | AG1 | 5 | 1.85 | 0.168 |
| 0.8 | 0.05 | ptsI | 5 | 2.1 | 0.166 |

| | | | | | |
|---|---|---|---|---|---|
| 0.8 | 0.05 | clcB | 5 | 2.1 | 0.182 |
| 0.8 | 0.05 | ycaM | 5 | 2.1 | 0.167 |
| 0.8 | 0.05 | yadI | 5 | 2.1 | 0.18 |
| 0.8 | 0.05 | AG1 | 5 | 2.1 | 0.173 |
| 0.8 | 0.05 | ptsI | 5 | 2.35 | 0.169 |
| 0.8 | 0.05 | clcB | 5 | 2.35 | 0.187 |
| 0.8 | 0.05 | ycaM | 5 | 2.35 | 0.171 |
| 0.8 | 0.05 | yadI | 5 | 2.35 | 0.187 |
| 0.8 | 0.05 | AG1 | 5 | 2.35 | 0.177 |
| 0.8 | 0.05 | ptsI | 5 | 2.6 | 0.165 |
| 0.8 | 0.05 | clcB | 5 | 2.6 | 0.186 |
| 0.8 | 0.05 | ycaM | 5 | 2.6 | 0.176 |
| 0.8 | 0.05 | yadI | 5 | 2.6 | 0.194 |
| 0.8 | 0.05 | AG1 | 5 | 2.6 | 0.175 |
| 0.8 | 0.05 | ptsI | 5 | 2.85 | 0.176 |
| 0.8 | 0.05 | clcB | 5 | 2.85 | 0.196 |
| 0.8 | 0.05 | ycaM | 5 | 2.85 | 0.18 |
| 0.8 | 0.05 | yadI | 5 | 2.85 | 0.199 |
| 0.8 | 0.05 | AG1 | 5 | 2.85 | 0.182 |
| 0.8 | 0.05 | ptsI | 5 | 3.1 | 0.181 |
| 0.8 | 0.05 | clcB | 5 | 3.1 | 0.206 |
| 0.8 | 0.05 | ycaM | 5 | 3.1 | 0.186 |
| 0.8 | 0.05 | yadI | 5 | 3.1 | 0.204 |
| 0.8 | 0.05 | AG1 | 5 | 3.1 | 0.189 |
| 0.8 | 0.05 | ptsI | 5 | 3.35 | 0.191 |
| 0.8 | 0.05 | clcB | 5 | 3.35 | 0.217 |
| 0.8 | 0.05 | ycaM | 5 | 3.35 | 0.196 |
| 0.8 | 0.05 | yadI | 5 | 3.35 | 0.213 |
| 0.8 | 0.05 | AG1 | 5 | 3.35 | 0.195 |
| 0.8 | 0.05 | ptsI | 5 | 3.6 | 0.199 |
| 0.8 | 0.05 | clcB | 5 | 3.6 | 0.231 |
| 0.8 | 0.05 | ycaM | 5 | 3.6 | 0.209 |
| 0.8 | 0.05 | yadI | 5 | 3.6 | 0.221 |
| 0.8 | 0.05 | AG1 | 5 | 3.6 | 0.203 |
| 0.8 | 0.05 | ptsI | 5 | 3.85 | 0.209 |
| 0.8 | 0.05 | clcB | 5 | 3.85 | 0.246 |
| 0.8 | 0.05 | ycaM | 5 | 3.85 | 0.222 |
| 0.8 | 0.05 | yadI | 5 | 3.85 | 0.229 |
| 0.8 | 0.05 | AG1 | 5 | 3.85 | 0.21 |
| 0.8 | 0.05 | ptsI | 5 | 4.38333 | 0.214 |
| 0.8 | 0.05 | clcB | 5 | 4.38333 | 0.317 |
| 0.8 | 0.05 | ycaM | 5 | 4.38333 | 0.224 |
| 0.8 | 0.05 | yadI | 5 | 4.38333 | 0.283 |
| 0.8 | 0.05 | AG1 | 5 | 4.38333 | 0.248 |
| 0.8 | 0.05 | ptsI | 5 | 4.63333 | 0.212 |
| 0.8 | 0.05 | clcB | 5 | 4.63333 | 0.271 |
| 0.8 | 0.05 | ycaM | 5 | 4.63333 | 0.216 |
| 0.8 | 0.05 | yadI | 5 | 4.63333 | 0.281 |

| | | | | | |
|---|---|---|---|---|---|
| 0.8 | 0.05 | AG1 | 5 | 4.63333 | 0.24 |
| 0.8 | 0.05 | ptsI | 5 | 4.88333 | 0.22 |
| 0.8 | 0.05 | clcB | 5 | 4.88333 | 0.286 |
| 0.8 | 0.05 | ycaM | 5 | 4.88333 | 0.226 |
| 0.8 | 0.05 | yadI | 5 | 4.88333 | 0.299 |
| 0.8 | 0.05 | AG1 | 5 | 4.88333 | 0.255 |
| 0.8 | 0.05 | ptsI | 5 | 5.13333 | 0.229 |
| 0.8 | 0.05 | clcB | 5 | 5.13333 | 0.295 |
| 0.8 | 0.05 | ycaM | 5 | 5.13333 | 0.231 |
| 0.8 | 0.05 | yadI | 5 | 5.13333 | 0.313 |
| 0.8 | 0.05 | AG1 | 5 | 5.13333 | 0.264 |
| 0.8 | 0.05 | ptsI | 5 | 5.38333 | 0.239 |
| 0.8 | 0.05 | clcB | 5 | 5.38333 | 0.302 |
| 0.8 | 0.05 | ycaM | 5 | 5.38333 | 0.232 |
| 0.8 | 0.05 | yadI | 5 | 5.38333 | 0.333 |
| 0.8 | 0.05 | AG1 | 5 | 5.38333 | 0.28 |
| 0.8 | 0.05 | ptsI | 5 | 5.63333 | 0.248 |
| 0.8 | 0.05 | clcB | 5 | 5.63333 | 0.311 |
| 0.8 | 0.05 | ycaM | 5 | 5.63333 | 0.236 |
| 0.8 | 0.05 | yadI | 5 | 5.63333 | 0.348 |
| 0.8 | 0.05 | AG1 | 5 | 5.63333 | 0.295 |
| 0.8 | 0.05 | ptsI | 5 | 5.88333 | 0.258 |
| 0.8 | 0.05 | clcB | 5 | 5.88333 | 0.315 |
| 0.8 | 0.05 | ycaM | 5 | 5.88333 | 0.236 |
| 0.8 | 0.05 | yadI | 5 | 5.88333 | 0.361 |
| 0.8 | 0.05 | AG1 | 5 | 5.88333 | 0.304 |
| 0.8 | 0.05 | ptsI | 5 | 6.13333 | 0.271 |
| 0.8 | 0.05 | clcB | 5 | 6.13333 | 0.33 |
| 0.8 | 0.05 | ycaM | 5 | 6.13333 | 0.241 |
| 0.8 | 0.05 | yadI | 5 | 6.13333 | 0.374 |
| 0.8 | 0.05 | AG1 | 5 | 6.13333 | 0.317 |
| 0.8 | 0.05 | ptsI | 5 | 6.38333 | 0.286 |
| 0.8 | 0.05 | clcB | 5 | 6.38333 | 0.342 |
| 0.8 | 0.05 | ycaM | 5 | 6.38333 | 0.242 |
| 0.8 | 0.05 | yadI | 5 | 6.38333 | 0.388 |
| 0.8 | 0.05 | AG1 | 5 | 6.38333 | 0.329 |
| 0.8 | 0.05 | ptsI | 5 | 6.63333 | 0.297 |
| 0.8 | 0.05 | clcB | 5 | 6.63333 | 0.351 |
| 0.8 | 0.05 | ycaM | 5 | 6.63333 | 0.245 |
| 0.8 | 0.05 | yadI | 5 | 6.63333 | 0.4 |
| 0.8 | 0.05 | AG1 | 5 | 6.63333 | 0.342 |
| 0.8 | 0.05 | ptsI | 5 | 6.88333 | 0.31 |
| 0.8 | 0.05 | clcB | 5 | 6.88333 | 0.359 |
| 0.8 | 0.05 | ycaM | 5 | 6.88333 | 0.248 |
| 0.8 | 0.05 | yadI | 5 | 6.88333 | 0.413 |
| 0.8 | 0.05 | AG1 | 5 | 6.88333 | 0.354 |
| 0.8 | 0.05 | ptsI | 5 | 7.13333 | 0.324 |
| 0.8 | 0.05 | clcB | 5 | 7.13333 | 0.368 |

| | | | | | |
|---|---|---|---|---|---|
| 0.8 | 0.05 | ycaM | 5 | 7.13333 | 0.25 |
| 0.8 | 0.05 | yadI | 5 | 7.13333 | 0.425 |
| 0.8 | 0.05 | AG1 | 5 | 7.13333 | 0.363 |
| 0.8 | 0.05 | ptsI | 5 | 7.38333 | 0.337 |
| 0.8 | 0.05 | clcB | 5 | 7.38333 | 0.373 |
| 0.8 | 0.05 | ycaM | 5 | 7.38333 | 0.251 |
| 0.8 | 0.05 | yadI | 5 | 7.38333 | 0.438 |
| 0.8 | 0.05 | AG1 | 5 | 7.38333 | 0.377 |
| 0.8 | 0.05 | ptsI | 5 | 7.63333 | 0.352 |
| 0.8 | 0.05 | clcB | 5 | 7.63333 | 0.38 |
| 0.8 | 0.05 | ycaM | 5 | 7.63333 | 0.253 |
| 0.8 | 0.05 | yadI | 5 | 7.63333 | 0.448 |
| 0.8 | 0.05 | AG1 | 5 | 7.63333 | 0.388 |
| 0.8 | 0.05 | ptsI | 5 | 7.88333 | 0.369 |
| 0.8 | 0.05 | clcB | 5 | 7.88333 | 0.391 |
| 0.8 | 0.05 | ycaM | 5 | 7.88333 | 0.257 |
| 0.8 | 0.05 | yadI | 5 | 7.88333 | 0.457 |
| 0.8 | 0.05 | AG1 | 5 | 7.88333 | 0.399 |
| 0.8 | 0.05 | ptsI | 5 | 8.13333 | 0.382 |
| 0.8 | 0.05 | clcB | 5 | 8.13333 | 0.392 |
| 0.8 | 0.05 | ycaM | 5 | 8.13333 | 0.261 |
| 0.8 | 0.05 | yadI | 5 | 8.13333 | 0.468 |
| 0.8 | 0.05 | AG1 | 5 | 8.13333 | 0.412 |
| 0.8 | 0.05 | ptsI | 5 | 8.38333 | 0.4 |
| 0.8 | 0.05 | clcB | 5 | 8.38333 | 0.398 |
| 0.8 | 0.05 | ycaM | 5 | 8.38333 | 0.262 |
| 0.8 | 0.05 | yadI | 5 | 8.38333 | 0.473 |
| 0.8 | 0.05 | AG1 | 5 | 8.38333 | 0.423 |
| 0.8 | 0.05 | ptsI | 5 | 8.88333 | 0.353 |
| 0.8 | 0.05 | clcB | 5 | 8.88333 | 0.291 |
| 0.8 | 0.05 | ycaM | 5 | 8.88333 | 0.25 |
| 0.8 | 0.05 | yadI | 5 | 8.88333 | 0.495 |
| 0.8 | 0.05 | AG1 | 5 | 8.88333 | 0.468 |
| 0.8 | 0.05 | ptsI | 5 | 9.13333 | 0.357 |
| 0.8 | 0.05 | clcB | 5 | 9.13333 | 0.288 |
| 0.8 | 0.05 | ycaM | 5 | 9.13333 | 0.245 |
| 0.8 | 0.05 | yadI | 5 | 9.13333 | 0.486 |
| 0.8 | 0.05 | AG1 | 5 | 9.13333 | 0.475 |
| 0.8 | 0.05 | ptsI | 5 | 9.38333 | 0.372 |
| 0.8 | 0.05 | clcB | 5 | 9.38333 | 0.29 |
| 0.8 | 0.05 | ycaM | 5 | 9.38333 | 0.243 |
| 0.8 | 0.05 | yadI | 5 | 9.38333 | 0.495 |
| 0.8 | 0.05 | AG1 | 5 | 9.38333 | 0.498 |
| 0.8 | 0.05 | ptsI | 5 | 9.63333 | 0.383 |
| 0.8 | 0.05 | clcB | 5 | 9.63333 | 0.29 |
| 0.8 | 0.05 | ycaM | 5 | 9.63333 | 0.246 |
| 0.8 | 0.05 | yadI | 5 | 9.63333 | 0.507 |
| 0.8 | 0.05 | AG1 | 5 | 9.63333 | 0.511 |

| | | | | | |
|---|---|---|---|---|---|
| 0.8 | 0.05 | ptsI | 5 | 9.88333 | 0.392 |
| 0.8 | 0.05 | clcB | 5 | 9.88333 | 0.292 |
| 0.8 | 0.05 | ycaM | 5 | 9.88333 | 0.247 |
| 0.8 | 0.05 | yadI | 5 | 9.88333 | 0.52 |
| 0.8 | 0.05 | AG1 | 5 | 9.88333 | 0.523 |
| 0.8 | 0.05 | ptsI | 5 | 10.1333 | 0.403 |
| 0.8 | 0.05 | clcB | 5 | 10.1333 | 0.292 |
| 0.8 | 0.05 | ycaM | 5 | 10.1333 | 0.247 |
| 0.8 | 0.05 | yadI | 5 | 10.1333 | 0.53 |
| 0.8 | 0.05 | AG1 | 5 | 10.1333 | 0.533 |
| 0.8 | 0.05 | ptsI | 5 | 10.3833 | 0.412 |
| 0.8 | 0.05 | clcB | 5 | 10.3833 | 0.294 |
| 0.8 | 0.05 | ycaM | 5 | 10.3833 | 0.248 |
| 0.8 | 0.05 | yadI | 5 | 10.3833 | 0.537 |
| 0.8 | 0.05 | AG1 | 5 | 10.3833 | 0.545 |
| 0.8 | 0.05 | ptsI | 5 | 10.6333 | 0.42 |
| 0.8 | 0.05 | clcB | 5 | 10.6333 | 0.294 |
| 0.8 | 0.05 | ycaM | 5 | 10.6333 | 0.248 |
| 0.8 | 0.05 | yadI | 5 | 10.6333 | 0.544 |
| 0.8 | 0.05 | AG1 | 5 | 10.6333 | 0.552 |
| 0.8 | 0.05 | ptsI | 5 | 10.8833 | 0.429 |
| 0.8 | 0.05 | clcB | 5 | 10.8833 | 0.296 |
| 0.8 | 0.05 | ycaM | 5 | 10.8833 | 0.249 |
| 0.8 | 0.05 | yadI | 5 | 10.8833 | 0.552 |
| 0.8 | 0.05 | AG1 | 5 | 10.8833 | 0.562 |
| 0.8 | 0.05 | ptsI | 5 | 11.1333 | 0.436 |
| 0.8 | 0.05 | clcB | 5 | 11.1333 | 0.294 |
| 0.8 | 0.05 | ycaM | 5 | 11.1333 | 0.247 |
| 0.8 | 0.05 | yadI | 5 | 11.1333 | 0.557 |
| 0.8 | 0.05 | AG1 | 5 | 11.1333 | 0.569 |
| 0.8 | 0.05 | ptsI | 5 | 11.3833 | 0.445 |
| 0.8 | 0.05 | clcB | 5 | 11.3833 | 0.298 |
| 0.8 | 0.05 | ycaM | 5 | 11.3833 | 0.251 |
| 0.8 | 0.05 | yadI | 5 | 11.3833 | 0.564 |
| 0.8 | 0.05 | AG1 | 5 | 11.3833 | 0.577 |
| 0.8 | 0.05 | ptsI | 5 | 11.6333 | 0.453 |
| 0.8 | 0.05 | clcB | 5 | 11.6333 | 0.297 |
| 0.8 | 0.05 | ycaM | 5 | 11.6333 | 0.252 |
| 0.8 | 0.05 | yadI | 5 | 11.6333 | 0.571 |
| 0.8 | 0.05 | AG1 | 5 | 11.6333 | 0.585 |
| 0.8 | 0.05 | ptsI | 5 | 11.8833 | 0.46 |
| 0.8 | 0.05 | clcB | 5 | 11.8833 | 0.296 |
| 0.8 | 0.05 | ycaM | 5 | 11.8833 | 0.25 |
| 0.8 | 0.05 | yadI | 5 | 11.8833 | 0.578 |
| 0.8 | 0.05 | AG1 | 5 | 11.8833 | 0.595 |
| 0.8 | 0.05 | ptsI | 5 | 12.1333 | 0.468 |
| 0.8 | 0.05 | clcB | 5 | 12.1333 | 0.299 |
| 0.8 | 0.05 | ycaM | 5 | 12.1333 | 0.252 |

| | | | | | |
|---|---|---|---|---|---|
| 0.8 | 0.05 | yadI | 5 | 12.1333 | 0.58 |
| 0.8 | 0.05 | AG1 | 5 | 12.1333 | 0.605 |
| 0.8 | 0.05 | ptsI | 5 | 12.3833 | 0.475 |
| 0.8 | 0.05 | clcB | 5 | 12.3833 | 0.295 |
| 0.8 | 0.05 | ycaM | 5 | 12.3833 | 0.25 |
| 0.8 | 0.05 | yadI | 5 | 12.3833 | 0.57 |
| 0.8 | 0.05 | AG1 | 5 | 12.3833 | 0.613 |
| 0.8 | 0.05 | ptsI | 5 | 12.6333 | 0.481 |
| 0.8 | 0.05 | clcB | 5 | 12.6333 | 0.299 |
| 0.8 | 0.05 | ycaM | 5 | 12.6333 | 0.251 |
| 0.8 | 0.05 | yadI | 5 | 12.6333 | 0.594 |
| 0.8 | 0.05 | AG1 | 5 | 12.6333 | 0.624 |
| 0.8 | 0.05 | ptsI | 5 | 12.8833 | 0.487 |
| 0.8 | 0.05 | clcB | 5 | 12.8833 | 0.297 |
| 0.8 | 0.05 | ycaM | 5 | 12.8833 | 0.251 |
| 0.8 | 0.05 | yadI | 5 | 12.8833 | 0.597 |
| 0.8 | 0.05 | AG1 | 5 | 12.8833 | 0.627 |
| 0.8 | 0.05 | ptsI | 5 | 13.1333 | 0.492 |
| 0.8 | 0.05 | clcB | 5 | 13.1333 | 0.294 |
| 0.8 | 0.05 | ycaM | 5 | 13.1333 | 0.255 |
| 0.8 | 0.05 | yadI | 5 | 13.1333 | 0.603 |
| 0.8 | 0.05 | AG1 | 5 | 13.1333 | 0.637 |
| 0.8 | 0.05 | ptsI | 5 | 13.3833 | 0.5 |
| 0.8 | 0.05 | clcB | 5 | 13.3833 | 0.296 |
| 0.8 | 0.05 | ycaM | 5 | 13.3833 | 0.255 |
| 0.8 | 0.05 | yadI | 5 | 13.3833 | 0.606 |
| 0.8 | 0.05 | AG1 | 5 | 13.3833 | 0.638 |
| 0.8 | 0.05 | ptsI | 5 | 13.6333 | 0.509 |
| 0.8 | 0.05 | clcB | 5 | 13.6333 | 0.293 |
| 0.8 | 0.05 | ycaM | 5 | 13.6333 | 0.256 |
| 0.8 | 0.05 | yadI | 5 | 13.6333 | 0.612 |
| 0.8 | 0.05 | AG1 | 5 | 13.6333 | 0.647 |
| 0.8 | 0.05 | ptsI | 5 | 13.8833 | 0.511 |
| 0.8 | 0.05 | clcB | 5 | 13.8833 | 0.294 |
| 0.8 | 0.05 | ycaM | 5 | 13.8833 | 0.256 |
| 0.8 | 0.05 | yadI | 5 | 13.8833 | 0.617 |
| 0.8 | 0.05 | AG1 | 5 | 13.8833 | 0.655 |
| 0.8 | 0.05 | ptsI | 5 | 14.1333 | 0.517 |
| 0.8 | 0.05 | clcB | 5 | 14.1333 | 0.293 |
| 0.8 | 0.05 | ycaM | 5 | 14.1333 | 0.257 |
| 0.8 | 0.05 | yadI | 5 | 14.1333 | 0.623 |
| 0.8 | 0.05 | AG1 | 5 | 14.1333 | 0.664 |
| 0.8 | 0.05 | ptsI | 5 | 14.3833 | 0.523 |
| 0.8 | 0.05 | clcB | 5 | 14.3833 | 0.296 |
| 0.8 | 0.05 | ycaM | 5 | 14.3833 | 0.259 |
| 0.8 | 0.05 | yadI | 5 | 14.3833 | 0.621 |
| 0.8 | 0.05 | AG1 | 5 | 14.3833 | 0.666 |
| 0.8 | 0.05 | ptsI | 5 | 14.6333 | 0.526 |

| | | | | | |
|---|---|---|---|---|---|
| 0.8 | 0.05 | clcB | 5 | 14.6333 | 0.292 |
| 0.8 | 0.05 | ycaM | 5 | 14.6333 | 0.257 |
| 0.8 | 0.05 | yadI | 5 | 14.6333 | 0.622 |
| 0.8 | 0.05 | AG1 | 5 | 14.6333 | 0.675 |
| 0.8 | 0.05 | ptsI | 5 | 14.8833 | 0.535 |
| 0.8 | 0.05 | clcB | 5 | 14.8833 | 0.292 |
| 0.8 | 0.05 | ycaM | 5 | 14.8833 | 0.259 |
| 0.8 | 0.05 | yadI | 5 | 14.8833 | 0.628 |
| 0.8 | 0.05 | AG1 | 5 | 14.8833 | 0.682 |
| 0.8 | 0.05 | ptsI | 5 | 15.1333 | 0.535 |
| 0.8 | 0.05 | clcB | 5 | 15.1333 | 0.294 |
| 0.8 | 0.05 | ycaM | 5 | 15.1333 | 0.258 |
| 0.8 | 0.05 | yadI | 5 | 15.1333 | 0.627 |
| 0.8 | 0.05 | AG1 | 5 | 15.1333 | 0.686 |
| 0.8 | 0.05 | ptsI | 5 | 15.3833 | 0.542 |
| 0.8 | 0.05 | clcB | 5 | 15.3833 | 0.293 |
| 0.8 | 0.05 | ycaM | 5 | 15.3833 | 0.259 |
| 0.8 | 0.05 | yadI | 5 | 15.3833 | 0.629 |
| 0.8 | 0.05 | AG1 | 5 | 15.3833 | 0.689 |
| 0.8 | 0.05 | ptsI | 5 | 15.6333 | 0.546 |
| 0.8 | 0.05 | clcB | 5 | 15.6333 | 0.295 |
| 0.8 | 0.05 | ycaM | 5 | 15.6333 | 0.259 |
| 0.8 | 0.05 | yadI | 5 | 15.6333 | 0.635 |
| 0.8 | 0.05 | AG1 | 5 | 15.6333 | 0.698 |
| 0.8 | 0.05 | ptsI | 5 | 15.8833 | 0.551 |
| 0.8 | 0.05 | clcB | 5 | 15.8833 | 0.296 |
| 0.8 | 0.05 | ycaM | 5 | 15.8833 | 0.26 |
| 0.8 | 0.05 | yadI | 5 | 15.8833 | 0.635 |
| 0.8 | 0.05 | AG1 | 5 | 15.8833 | 0.696 |
| 0.8 | 0.05 | ptsI | 5 | 16.1333 | 0.56 |
| 0.8 | 0.05 | clcB | 5 | 16.1333 | 0.299 |
| 0.8 | 0.05 | ycaM | 5 | 16.1333 | 0.261 |
| 0.8 | 0.05 | yadI | 5 | 16.1333 | 0.638 |
| 0.8 | 0.05 | AG1 | 5 | 16.1333 | 0.71 |
| 0.8 | 0.05 | ptsI | 5 | 16.3833 | 0.559 |
| 0.8 | 0.05 | clcB | 5 | 16.3833 | 0.296 |
| 0.8 | 0.05 | ycaM | 5 | 16.3833 | 0.261 |
| 0.8 | 0.05 | yadI | 5 | 16.3833 | 0.642 |
| 0.8 | 0.05 | AG1 | 5 | 16.3833 | 0.704 |
| 0.8 | 0.05 | ptsI | 5 | 16.6333 | 0.564 |
| 0.8 | 0.05 | clcB | 5 | 16.6333 | 0.3 |
| 0.8 | 0.05 | ycaM | 5 | 16.6333 | 0.262 |
| 0.8 | 0.05 | yadI | 5 | 16.6333 | 0.643 |
| 0.8 | 0.05 | AG1 | 5 | 16.6333 | 0.71 |
| 0.8 | 0.05 | ptsI | 5 | 16.8833 | 0.567 |
| 0.8 | 0.05 | clcB | 5 | 16.8833 | 0.297 |
| 0.8 | 0.05 | ycaM | 5 | 16.8833 | 0.262 |
| 0.8 | 0.05 | yadI | 5 | 16.8833 | 0.647 |

| | | | | | |
|---|---|---|---|---|---|
| 0.8 | 0.05 | AG1 | 5 | 16.8833 | 0.714 |
| 0.8 | 0.05 | ptsI | 5 | 17.1333 | 0.572 |
| 0.8 | 0.05 | clcB | 5 | 17.1333 | 0.298 |
| 0.8 | 0.05 | ycaM | 5 | 17.1333 | 0.261 |
| 0.8 | 0.05 | yadI | 5 | 17.1333 | 0.643 |
| 0.8 | 0.05 | AG1 | 5 | 17.1333 | 0.72 |
| 0.8 | 0.05 | ptsI | 5 | 17.3833 | 0.577 |
| 0.8 | 0.05 | clcB | 5 | 17.3833 | 0.301 |
| 0.8 | 0.05 | ycaM | 5 | 17.3833 | 0.265 |
| 0.8 | 0.05 | yadI | 5 | 17.3833 | 0.647 |
| 0.8 | 0.05 | AG1 | 5 | 17.3833 | 0.723 |
| 0.8 | 0.05 | ptsI | 5 | 17.6333 | 0.58 |
| 0.8 | 0.05 | clcB | 5 | 17.6333 | 0.302 |
| 0.8 | 0.05 | ycaM | 5 | 17.6333 | 0.266 |
| 0.8 | 0.05 | yadI | 5 | 17.6333 | 0.648 |
| 0.8 | 0.05 | AG1 | 5 | 17.6333 | 0.729 |
| 0.8 | 0.05 | ptsI | 5 | 17.8833 | 0.58 |
| 0.8 | 0.05 | clcB | 5 | 17.8833 | 0.3 |
| 0.8 | 0.05 | ycaM | 5 | 17.8833 | 0.266 |
| 0.8 | 0.05 | yadI | 5 | 17.8833 | 0.631 |
| 0.8 | 0.05 | AG1 | 5 | 17.8833 | 0.727 |
| 0.8 | 0.05 | ptsI | 5 | 18.1333 | 0.587 |
| 0.8 | 0.05 | clcB | 5 | 18.1333 | 0.305 |
| 0.8 | 0.05 | ycaM | 5 | 18.1333 | 0.268 |
| 0.8 | 0.05 | yadI | 5 | 18.1333 | 0.654 |
| 0.8 | 0.05 | AG1 | 5 | 18.1333 | 0.733 |
| 0.8 | 0.05 | ptsI | 5 | 18.3833 | 0.588 |
| 0.8 | 0.05 | clcB | 5 | 18.3833 | 0.306 |
| 0.8 | 0.05 | ycaM | 5 | 18.3833 | 0.268 |
| 0.8 | 0.05 | yadI | 5 | 18.3833 | 0.654 |
| 0.8 | 0.05 | AG1 | 5 | 18.3833 | 0.735 |
| 0.8 | 0.05 | ptsI | 5 | 18.6333 | 0.588 |
| 0.8 | 0.05 | clcB | 5 | 18.6333 | 0.304 |
| 0.8 | 0.05 | ycaM | 5 | 18.6333 | 0.27 |
| 0.8 | 0.05 | yadI | 5 | 18.6333 | 0.658 |
| 0.8 | 0.05 | AG1 | 5 | 18.6333 | 0.736 |
| 0.8 | 0.05 | ptsI | 5 | 18.8833 | 0.593 |
| 0.8 | 0.05 | clcB | 5 | 18.8833 | 0.306 |
| 0.8 | 0.05 | ycaM | 5 | 18.8833 | 0.272 |
| 0.8 | 0.05 | yadI | 5 | 18.8833 | 0.66 |
| 0.8 | 0.05 | AG1 | 5 | 18.8833 | 0.741 |
| 0.8 | 0.05 | ptsI | 5 | 19.1333 | 0.598 |
| 0.8 | 0.05 | clcB | 5 | 19.1333 | 0.309 |
| 0.8 | 0.05 | ycaM | 5 | 19.1333 | 0.272 |
| 0.8 | 0.05 | yadI | 5 | 19.1333 | 0.66 |
| 0.8 | 0.05 | AG1 | 5 | 19.1333 | 0.745 |
| 0.8 | 0.05 | ptsI | 5 | 19.3833 | 0.601 |
| 0.8 | 0.05 | clcB | 5 | 19.3833 | 0.312 |

| | | | | | |
|---|---|---|---|---|---|
| 0.8 | 0.05 | ycaM | 5 | 19.3833 | 0.274 |
| 0.8 | 0.05 | yadI | 5 | 19.3833 | 0.665 |
| 0.8 | 0.05 | AG1 | 5 | 19.3833 | 0.751 |
| 0.8 | 0.05 | ptsI | 5 | 19.6333 | 0.604 |
| 0.8 | 0.05 | clcB | 5 | 19.6333 | 0.316 |
| 0.8 | 0.05 | ycaM | 5 | 19.6333 | 0.276 |
| 0.8 | 0.05 | yadI | 5 | 19.6333 | 0.662 |
| 0.8 | 0.05 | AG1 | 5 | 19.6333 | 0.752 |
| 0.8 | 0.05 | ptsI | 5 | 19.8833 | 0.601 |
| 0.8 | 0.05 | clcB | 5 | 19.8833 | 0.317 |
| 0.8 | 0.05 | ycaM | 5 | 19.8833 | 0.277 |
| 0.8 | 0.05 | yadI | 5 | 19.8833 | 0.66 |
| 0.8 | 0.05 | AG1 | 5 | 19.8833 | 0.756 |
| 0.8 | 0.05 | ptsI | 5 | 20.1333 | 0.603 |
| 0.8 | 0.05 | clcB | 5 | 20.1333 | 0.32 |
| 0.8 | 0.05 | ycaM | 5 | 20.1333 | 0.278 |
| 0.8 | 0.05 | yadI | 5 | 20.1333 | 0.664 |
| 0.8 | 0.05 | AG1 | 5 | 20.1333 | 0.761 |
| 0.8 | 0.05 | ptsI | 5 | 20.3833 | 0.604 |
| 0.8 | 0.05 | clcB | 5 | 20.3833 | 0.324 |
| 0.8 | 0.05 | ycaM | 5 | 20.3833 | 0.28 |
| 0.8 | 0.05 | yadI | 5 | 20.3833 | 0.666 |
| 0.8 | 0.05 | AG1 | 5 | 20.3833 | 0.761 |
| 0.8 | 0.05 | ptsI | 5 | 20.6333 | 0.607 |
| 0.8 | 0.05 | clcB | 5 | 20.6333 | 0.321 |
| 0.8 | 0.05 | ycaM | 5 | 20.6333 | 0.281 |
| 0.8 | 0.05 | yadI | 5 | 20.6333 | 0.644 |
| 0.8 | 0.05 | AG1 | 5 | 20.6333 | 0.756 |
| 0.8 | 0.05 | ptsI | 5 | 20.8833 | 0.606 |
| 0.8 | 0.05 | clcB | 5 | 20.8833 | 0.323 |
| 0.8 | 0.05 | ycaM | 5 | 20.8833 | 0.284 |
| 0.8 | 0.05 | yadI | 5 | 20.8833 | 0.67 |
| 0.8 | 0.05 | AG1 | 5 | 20.8833 | 0.77 |
| 0.8 | 0.05 | ptsI | 5 | 21.1333 | 0.607 |
| 0.8 | 0.05 | clcB | 5 | 21.1333 | 0.324 |
| 0.8 | 0.05 | ycaM | 5 | 21.1333 | 0.286 |
| 0.8 | 0.05 | yadI | 5 | 21.1333 | 0.672 |
| 0.8 | 0.05 | AG1 | 5 | 21.1333 | 0.772 |
| 0.8 | 0.05 | ptsI | 5 | 21.3833 | 0.609 |
| 0.8 | 0.05 | clcB | 5 | 21.3833 | 0.319 |
| 0.8 | 0.05 | ycaM | 5 | 21.3833 | 0.29 |
| 0.8 | 0.05 | yadI | 5 | 21.3833 | 0.671 |
| 0.8 | 0.05 | AG1 | 5 | 21.3833 | 0.772 |
| 0.8 | 0.05 | ptsI | 5 | 21.6333 | 0.611 |
| 0.8 | 0.05 | clcB | 5 | 21.6333 | 0.324 |
| 0.8 | 0.05 | ycaM | 5 | 21.6333 | 0.292 |
| 0.8 | 0.05 | yadI | 5 | 21.6333 | 0.67 |
| 0.8 | 0.05 | AG1 | 5 | 21.6333 | 0.775 |

| | | | | | |
|---|---|---|---|---|---|
| 0.8 | 0.05 | ptsI | 5 | 21.8833 | 0.607 |
| 0.8 | 0.05 | clcB | 5 | 21.8833 | 0.324 |
| 0.8 | 0.05 | ycaM | 5 | 21.8833 | 0.297 |
| 0.8 | 0.05 | yadI | 5 | 21.8833 | 0.668 |
| 0.8 | 0.05 | AG1 | 5 | 21.8833 | 0.772 |
| 0.8 | 0.05 | ptsI | 5 | 22.1333 | 0.61 |
| 0.8 | 0.05 | clcB | 5 | 22.1333 | 0.323 |
| 0.8 | 0.05 | ycaM | 5 | 22.1333 | 0.3 |
| 0.8 | 0.05 | yadI | 5 | 22.1333 | 0.648 |
| 0.8 | 0.05 | AG1 | 5 | 22.1333 | 0.771 |
| 0.8 | 0.05 | ptsI | 5 | 22.3833 | 0.609 |
| 0.8 | 0.05 | clcB | 5 | 22.3833 | 0.324 |
| 0.8 | 0.05 | ycaM | 5 | 22.3833 | 0.301 |
| 0.8 | 0.05 | yadI | 5 | 22.3833 | 0.672 |
| 0.8 | 0.05 | AG1 | 5 | 22.3833 | 0.778 |
| 0.8 | 0.05 | ptsI | 5 | 22.6333 | 0.609 |
| 0.8 | 0.05 | clcB | 5 | 22.6333 | 0.321 |
| 0.8 | 0.05 | ycaM | 5 | 22.6333 | 0.306 |
| 0.8 | 0.05 | yadI | 5 | 22.6333 | 0.674 |
| 0.8 | 0.05 | AG1 | 5 | 22.6333 | 0.781 |
| 0.8 | 0.05 | ptsI | 5 | 22.8833 | 0.61 |
| 0.8 | 0.05 | clcB | 5 | 22.8833 | 0.319 |
| 0.8 | 0.05 | ycaM | 5 | 22.8833 | 0.308 |
| 0.8 | 0.05 | yadI | 5 | 22.8833 | 0.672 |
| 0.8 | 0.05 | AG1 | 5 | 22.8833 | 0.777 |
| 0.8 | 0.05 | ptsI | 5 | 23.1333 | 0.611 |
| 0.8 | 0.05 | clcB | 5 | 23.1333 | 0.32 |
| 0.8 | 0.05 | ycaM | 5 | 23.1333 | 0.311 |
| 0.8 | 0.05 | yadI | 5 | 23.1333 | 0.676 |
| 0.8 | 0.05 | AG1 | 5 | 23.1333 | 0.784 |
| 0.8 | 0.05 | ptsI | 5 | 23.3833 | 0.612 |
| 0.8 | 0.05 | clcB | 5 | 23.3833 | 0.323 |
| 0.8 | 0.05 | ycaM | 5 | 23.3833 | 0.314 |
| 0.8 | 0.05 | yadI | 5 | 23.3833 | 0.674 |
| 0.8 | 0.05 | AG1 | 5 | 23.3833 | 0.784 |
| 0.8 | 0.05 | ptsI | 5 | 23.6333 | 0.612 |
| 0.8 | 0.05 | clcB | 5 | 23.6333 | 0.324 |
| 0.8 | 0.05 | ycaM | 5 | 23.6333 | 0.317 |
| 0.8 | 0.05 | yadI | 5 | 23.6333 | 0.676 |
| 0.8 | 0.05 | AG1 | 5 | 23.6333 | 0.785 |
| 0.8 | 0.05 | ptsI | 5 | 23.8833 | 0.617 |
| 0.8 | 0.05 | clcB | 5 | 23.8833 | 0.321 |
| 0.8 | 0.05 | ycaM | 5 | 23.8833 | 0.323 |
| 0.8 | 0.05 | yadI | 5 | 23.8833 | 0.654 |
| 0.8 | 0.05 | AG1 | 5 | 23.8833 | 0.781 |
| 0.8 | 0.05 | ptsI | 5 | 24.1333 | 0.612 |
| 0.8 | 0.05 | clcB | 5 | 24.1333 | 0.324 |
| 0.8 | 0.05 | ycaM | 5 | 24.1333 | 0.325 |

| | | | | | |
|---|---|---|---|---|---|
| 0.8 | 0.05 | yadI | 5 | 24.1333 | 0.675 |
| 0.8 | 0.05 | AG1 | 5 | 24.1333 | 0.79 |
| 0.8 | 0.05 | ptsI | 5 | 24.3833 | 0.611 |
| 0.8 | 0.05 | clcB | 5 | 24.3833 | 0.319 |
| 0.8 | 0.05 | ycaM | 5 | 24.3833 | 0.328 |
| 0.8 | 0.05 | yadI | 5 | 24.3833 | 0.674 |
| 0.8 | 0.05 | AG1 | 5 | 24.3833 | 0.79 |
| 1.6 | 0.05 | ptsI | 2 | 0 | 0.22 |
| 1.6 | 0.05 | clcB | 2 | 0 | 0.201 |
| 1.6 | 0.05 | ycaM | 2 | 0 | 0.21 |
| 1.6 | 0.05 | yadI | 2 | 0 | 0.222 |
| 1.6 | 0.05 | AG1 | 2 | 0 | 0.233 |
| 1.6 | 0.05 | ptsI | 2 | 0.25 | 0.218 |
| 1.6 | 0.05 | clcB | 2 | 0.25 | 0.2 |
| 1.6 | 0.05 | ycaM | 2 | 0.25 | 0.204 |
| 1.6 | 0.05 | yadI | 2 | 0.25 | 0.218 |
| 1.6 | 0.05 | AG1 | 2 | 0.25 | 0.225 |
| 1.6 | 0.05 | ptsI | 2 | 0.5 | 0.216 |
| 1.6 | 0.05 | clcB | 2 | 0.5 | 0.199 |
| 1.6 | 0.05 | ycaM | 2 | 0.5 | 0.206 |
| 1.6 | 0.05 | yadI | 2 | 0.5 | 0.219 |
| 1.6 | 0.05 | AG1 | 2 | 0.5 | 0.228 |
| 1.6 | 0.05 | ptsI | 2 | 0.75 | 0.218 |
| 1.6 | 0.05 | clcB | 2 | 0.75 | 0.2 |
| 1.6 | 0.05 | ycaM | 2 | 0.75 | 0.207 |
| 1.6 | 0.05 | yadI | 2 | 0.75 | 0.223 |
| 1.6 | 0.05 | AG1 | 2 | 0.75 | 0.229 |
| 1.6 | 0.05 | ptsI | 2 | 1 | 0.219 |
| 1.6 | 0.05 | clcB | 2 | 1 | 0.201 |
| 1.6 | 0.05 | ycaM | 2 | 1 | 0.21 |
| 1.6 | 0.05 | yadI | 2 | 1 | 0.23 |
| 1.6 | 0.05 | AG1 | 2 | 1 | 0.232 |
| 1.6 | 0.05 | ptsI | 2 | 1.25 | 0.222 |
| 1.6 | 0.05 | clcB | 2 | 1.25 | 0.202 |
| 1.6 | 0.05 | ycaM | 2 | 1.25 | 0.21 |
| 1.6 | 0.05 | yadI | 2 | 1.25 | 0.234 |
| 1.6 | 0.05 | AG1 | 2 | 1.25 | 0.233 |
| 1.6 | 0.05 | ptsI | 2 | 1.5 | 0.222 |
| 1.6 | 0.05 | clcB | 2 | 1.5 | 0.207 |
| 1.6 | 0.05 | ycaM | 2 | 1.5 | 0.212 |
| 1.6 | 0.05 | yadI | 2 | 1.5 | 0.24 |
| 1.6 | 0.05 | AG1 | 2 | 1.5 | 0.235 |
| 1.6 | 0.05 | ptsI | 2 | 1.75 | 0.227 |
| 1.6 | 0.05 | clcB | 2 | 1.75 | 0.206 |
| 1.6 | 0.05 | ycaM | 2 | 1.75 | 0.216 |
| 1.6 | 0.05 | yadI | 2 | 1.75 | 0.246 |
| 1.6 | 0.05 | AG1 | 2 | 1.75 | 0.24 |
| 1.6 | 0.05 | ptsI | 2 | 2 | 0.231 |

| | | | | | |
|---|---|---|---|---|---|
| 1.6 | 0.05 | clcB | 2 | 2 | 0.21 |
| 1.6 | 0.05 | ycaM | 2 | 2 | 0.224 |
| 1.6 | 0.05 | yadI | 2 | 2 | 0.253 |
| 1.6 | 0.05 | AG1 | 2 | 2 | 0.247 |
| 1.6 | 0.05 | ptsI | 2 | 2.25 | 0.238 |
| 1.6 | 0.05 | clcB | 2 | 2.25 | 0.214 |
| 1.6 | 0.05 | ycaM | 2 | 2.25 | 0.23 |
| 1.6 | 0.05 | yadI | 2 | 2.25 | 0.263 |
| 1.6 | 0.05 | AG1 | 2 | 2.25 | 0.254 |
| 1.6 | 0.05 | ptsI | 2 | 2.5 | 0.245 |
| 1.6 | 0.05 | clcB | 2 | 2.5 | 0.218 |
| 1.6 | 0.05 | ycaM | 2 | 2.5 | 0.239 |
| 1.6 | 0.05 | yadI | 2 | 2.5 | 0.277 |
| 1.6 | 0.05 | AG1 | 2 | 2.5 | 0.262 |
| 1.6 | 0.05 | ptsI | 2 | 2.75 | 0.244 |
| 1.6 | 0.05 | clcB | 2 | 2.75 | 0.218 |
| 1.6 | 0.05 | ycaM | 2 | 2.75 | 0.238 |
| 1.6 | 0.05 | yadI | 2 | 2.75 | 0.283 |
| 1.6 | 0.05 | AG1 | 2 | 2.75 | 0.259 |
| 1.6 | 0.05 | ptsI | 2 | 3 | 0.249 |
| 1.6 | 0.05 | clcB | 2 | 3 | 0.22 |
| 1.6 | 0.05 | ycaM | 2 | 3 | 0.239 |
| 1.6 | 0.05 | yadI | 2 | 3 | 0.29 |
| 1.6 | 0.05 | AG1 | 2 | 3 | 0.259 |
| 1.6 | 0.05 | ptsI | 2 | 3.25 | 0.255 |
| 1.6 | 0.05 | clcB | 2 | 3.25 | 0.229 |
| 1.6 | 0.05 | ycaM | 2 | 3.25 | 0.249 |
| 1.6 | 0.05 | yadI | 2 | 3.25 | 0.308 |
| 1.6 | 0.05 | AG1 | 2 | 3.25 | 0.268 |
| 1.6 | 0.05 | ptsI | 2 | 3.5 | 0.26 |
| 1.6 | 0.05 | clcB | 2 | 3.5 | 0.232 |
| 1.6 | 0.05 | ycaM | 2 | 3.5 | 0.256 |
| 1.6 | 0.05 | yadI | 2 | 3.5 | 0.319 |
| 1.6 | 0.05 | AG1 | 2 | 3.5 | 0.273 |
| 1.6 | 0.05 | ptsI | 2 | 3.75 | 0.269 |
| 1.6 | 0.05 | clcB | 2 | 3.75 | 0.238 |
| 1.6 | 0.05 | ycaM | 2 | 3.75 | 0.265 |
| 1.6 | 0.05 | yadI | 2 | 3.75 | 0.332 |
| 1.6 | 0.05 | AG1 | 2 | 3.75 | 0.285 |
| 1.6 | 0.05 | ptsI | 2 | 4 | 0.277 |
| 1.6 | 0.05 | clcB | 2 | 4 | 0.252 |
| 1.6 | 0.05 | ycaM | 2 | 4 | 0.279 |
| 1.6 | 0.05 | yadI | 2 | 4 | 0.348 |
| 1.6 | 0.05 | AG1 | 2 | 4 | 0.295 |
| 1.6 | 0.05 | ptsI | 2 | 4.38333 | 0.301 |
| 1.6 | 0.05 | clcB | 2 | 4.38333 | 0.274 |
| 1.6 | 0.05 | ycaM | 2 | 4.38333 | 0.288 |
| 1.6 | 0.05 | yadI | 2 | 4.38333 | 0.333 |

| | | | | | |
|---|---|---|---|---|---|
| 1.6 | 0.05 | AG1 | 2 | 4.38333 | 0.312 |
| 1.6 | 0.05 | ptsI | 2 | 4.63333 | 0.301 |
| 1.6 | 0.05 | clcB | 2 | 4.63333 | 0.267 |
| 1.6 | 0.05 | ycaM | 2 | 4.63333 | 0.287 |
| 1.6 | 0.05 | yadI | 2 | 4.63333 | 0.336 |
| 1.6 | 0.05 | AG1 | 2 | 4.63333 | 0.307 |
| 1.6 | 0.05 | ptsI | 2 | 4.88333 | 0.305 |
| 1.6 | 0.05 | clcB | 2 | 4.88333 | 0.272 |
| 1.6 | 0.05 | ycaM | 2 | 4.88333 | 0.292 |
| 1.6 | 0.05 | yadI | 2 | 4.88333 | 0.35 |
| 1.6 | 0.05 | AG1 | 2 | 4.88333 | 0.317 |
| 1.6 | 0.05 | ptsI | 2 | 5.13333 | 0.313 |
| 1.6 | 0.05 | clcB | 2 | 5.13333 | 0.278 |
| 1.6 | 0.05 | ycaM | 2 | 5.13333 | 0.296 |
| 1.6 | 0.05 | yadI | 2 | 5.13333 | 0.365 |
| 1.6 | 0.05 | AG1 | 2 | 5.13333 | 0.327 |
| 1.6 | 0.05 | ptsI | 2 | 5.38333 | 0.32 |
| 1.6 | 0.05 | clcB | 2 | 5.38333 | 0.284 |
| 1.6 | 0.05 | ycaM | 2 | 5.38333 | 0.296 |
| 1.6 | 0.05 | yadI | 2 | 5.38333 | 0.375 |
| 1.6 | 0.05 | AG1 | 2 | 5.38333 | 0.338 |
| 1.6 | 0.05 | ptsI | 2 | 5.63333 | 0.327 |
| 1.6 | 0.05 | clcB | 2 | 5.63333 | 0.292 |
| 1.6 | 0.05 | ycaM | 2 | 5.63333 | 0.298 |
| 1.6 | 0.05 | yadI | 2 | 5.63333 | 0.386 |
| 1.6 | 0.05 | AG1 | 2 | 5.63333 | 0.346 |
| 1.6 | 0.05 | ptsI | 2 | 5.88333 | 0.335 |
| 1.6 | 0.05 | clcB | 2 | 5.88333 | 0.299 |
| 1.6 | 0.05 | ycaM | 2 | 5.88333 | 0.3 |
| 1.6 | 0.05 | yadI | 2 | 5.88333 | 0.393 |
| 1.6 | 0.05 | AG1 | 2 | 5.88333 | 0.355 |
| 1.6 | 0.05 | ptsI | 2 | 6.13333 | 0.344 |
| 1.6 | 0.05 | clcB | 2 | 6.13333 | 0.304 |
| 1.6 | 0.05 | ycaM | 2 | 6.13333 | 0.302 |
| 1.6 | 0.05 | yadI | 2 | 6.13333 | 0.401 |
| 1.6 | 0.05 | AG1 | 2 | 6.13333 | 0.365 |
| 1.6 | 0.05 | ptsI | 2 | 6.38333 | 0.353 |
| 1.6 | 0.05 | clcB | 2 | 6.38333 | 0.312 |
| 1.6 | 0.05 | ycaM | 2 | 6.38333 | 0.302 |
| 1.6 | 0.05 | yadI | 2 | 6.38333 | 0.41 |
| 1.6 | 0.05 | AG1 | 2 | 6.38333 | 0.374 |
| 1.6 | 0.05 | ptsI | 2 | 6.63333 | 0.361 |
| 1.6 | 0.05 | clcB | 2 | 6.63333 | 0.317 |
| 1.6 | 0.05 | ycaM | 2 | 6.63333 | 0.304 |
| 1.6 | 0.05 | yadI | 2 | 6.63333 | 0.413 |
| 1.6 | 0.05 | AG1 | 2 | 6.63333 | 0.382 |
| 1.6 | 0.05 | ptsI | 2 | 6.88333 | 0.37 |
| 1.6 | 0.05 | clcB | 2 | 6.88333 | 0.323 |

| | | | | | |
|---|---|---|---|---|---|
| 1.6 | 0.05 | ycaM | 2 | 6.88333 | 0.304 |
| 1.6 | 0.05 | yadI | 2 | 6.88333 | 0.423 |
| 1.6 | 0.05 | AG1 | 2 | 6.88333 | 0.391 |
| 1.6 | 0.05 | ptsI | 2 | 7.13333 | 0.38 |
| 1.6 | 0.05 | clcB | 2 | 7.13333 | 0.329 |
| 1.6 | 0.05 | ycaM | 2 | 7.13333 | 0.307 |
| 1.6 | 0.05 | yadI | 2 | 7.13333 | 0.434 |
| 1.6 | 0.05 | AG1 | 2 | 7.13333 | 0.401 |
| 1.6 | 0.05 | ptsI | 2 | 7.38333 | 0.392 |
| 1.6 | 0.05 | clcB | 2 | 7.38333 | 0.336 |
| 1.6 | 0.05 | ycaM | 2 | 7.38333 | 0.309 |
| 1.6 | 0.05 | yadI | 2 | 7.38333 | 0.441 |
| 1.6 | 0.05 | AG1 | 2 | 7.38333 | 0.411 |
| 1.6 | 0.05 | ptsI | 2 | 7.63333 | 0.396 |
| 1.6 | 0.05 | clcB | 2 | 7.63333 | 0.341 |
| 1.6 | 0.05 | ycaM | 2 | 7.63333 | 0.308 |
| 1.6 | 0.05 | yadI | 2 | 7.63333 | 0.439 |
| 1.6 | 0.05 | AG1 | 2 | 7.63333 | 0.418 |
| 1.6 | 0.05 | ptsI | 2 | 7.88333 | 0.405 |
| 1.6 | 0.05 | clcB | 2 | 7.88333 | 0.347 |
| 1.6 | 0.05 | ycaM | 2 | 7.88333 | 0.309 |
| 1.6 | 0.05 | yadI | 2 | 7.88333 | 0.448 |
| 1.6 | 0.05 | AG1 | 2 | 7.88333 | 0.424 |
| 1.6 | 0.05 | ptsI | 2 | 8.13333 | 0.417 |
| 1.6 | 0.05 | clcB | 2 | 8.13333 | 0.349 |
| 1.6 | 0.05 | ycaM | 2 | 8.13333 | 0.312 |
| 1.6 | 0.05 | yadI | 2 | 8.13333 | 0.456 |
| 1.6 | 0.05 | AG1 | 2 | 8.13333 | 0.433 |
| 1.6 | 0.05 | ptsI | 2 | 8.38333 | 0.424 |
| 1.6 | 0.05 | clcB | 2 | 8.38333 | 0.355 |
| 1.6 | 0.05 | ycaM | 2 | 8.38333 | 0.31 |
| 1.6 | 0.05 | yadI | 2 | 8.38333 | 0.463 |
| 1.6 | 0.05 | AG1 | 2 | 8.38333 | 0.441 |
| 1.6 | 0.05 | ptsI | 2 | 8.91667 | 0.348 |
| 1.6 | 0.05 | clcB | 2 | 8.91667 | 0.295 |
| 1.6 | 0.05 | ycaM | 2 | 8.91667 | 0.282 |
| 1.6 | 0.05 | yadI | 2 | 8.91667 | 0.503 |
| 1.6 | 0.05 | AG1 | 2 | 8.91667 | 0.482 |
| 1.6 | 0.05 | ptsI | 2 | 9.16667 | 0.349 |
| 1.6 | 0.05 | clcB | 2 | 9.16667 | 0.297 |
| 1.6 | 0.05 | ycaM | 2 | 9.16667 | 0.279 |
| 1.6 | 0.05 | yadI | 2 | 9.16667 | 0.512 |
| 1.6 | 0.05 | AG1 | 2 | 9.16667 | 0.498 |
| 1.6 | 0.05 | ptsI | 2 | 9.41667 | 0.356 |
| 1.6 | 0.05 | clcB | 2 | 9.41667 | 0.304 |
| 1.6 | 0.05 | ycaM | 2 | 9.41667 | 0.281 |
| 1.6 | 0.05 | yadI | 2 | 9.41667 | 0.525 |
| 1.6 | 0.05 | AG1 | 2 | 9.41667 | 0.515 |

| | | | | | |
|---|---|---|---|---|---|
| 1.6 | 0.05 | ptsI | 2 | 9.66667 | 0.364 |
| 1.6 | 0.05 | clcB | 2 | 9.66667 | 0.309 |
| 1.6 | 0.05 | ycaM | 2 | 9.66667 | 0.281 |
| 1.6 | 0.05 | yadI | 2 | 9.66667 | 0.537 |
| 1.6 | 0.05 | AG1 | 2 | 9.66667 | 0.532 |
| 1.6 | 0.05 | ptsI | 2 | 9.91667 | 0.372 |
| 1.6 | 0.05 | clcB | 2 | 9.91667 | 0.312 |
| 1.6 | 0.05 | ycaM | 2 | 9.91667 | 0.281 |
| 1.6 | 0.05 | yadI | 2 | 9.91667 | 0.546 |
| 1.6 | 0.05 | AG1 | 2 | 9.91667 | 0.541 |
| 1.6 | 0.05 | ptsI | 2 | 10.1667 | 0.379 |
| 1.6 | 0.05 | clcB | 2 | 10.1667 | 0.312 |
| 1.6 | 0.05 | ycaM | 2 | 10.1667 | 0.283 |
| 1.6 | 0.05 | yadI | 2 | 10.1667 | 0.548 |
| 1.6 | 0.05 | AG1 | 2 | 10.1667 | 0.548 |
| 1.6 | 0.05 | ptsI | 2 | 10.4167 | 0.387 |
| 1.6 | 0.05 | clcB | 2 | 10.4167 | 0.317 |
| 1.6 | 0.05 | ycaM | 2 | 10.4167 | 0.281 |
| 1.6 | 0.05 | yadI | 2 | 10.4167 | 0.563 |
| 1.6 | 0.05 | AG1 | 2 | 10.4167 | 0.563 |
| 1.6 | 0.05 | ptsI | 2 | 10.6667 | 0.392 |
| 1.6 | 0.05 | clcB | 2 | 10.6667 | 0.315 |
| 1.6 | 0.05 | ycaM | 2 | 10.6667 | 0.285 |
| 1.6 | 0.05 | yadI | 2 | 10.6667 | 0.563 |
| 1.6 | 0.05 | AG1 | 2 | 10.6667 | 0.57 |
| 1.6 | 0.05 | ptsI | 2 | 10.9167 | 0.4 |
| 1.6 | 0.05 | clcB | 2 | 10.9167 | 0.319 |
| 1.6 | 0.05 | ycaM | 2 | 10.9167 | 0.285 |
| 1.6 | 0.05 | yadI | 2 | 10.9167 | 0.559 |
| 1.6 | 0.05 | AG1 | 2 | 10.9167 | 0.57 |
| 1.6 | 0.05 | ptsI | 2 | 11.1667 | 0.404 |
| 1.6 | 0.05 | clcB | 2 | 11.1667 | 0.317 |
| 1.6 | 0.05 | ycaM | 2 | 11.1667 | 0.282 |
| 1.6 | 0.05 | yadI | 2 | 11.1667 | 0.564 |
| 1.6 | 0.05 | AG1 | 2 | 11.1667 | 0.573 |
| 1.6 | 0.05 | ptsI | 2 | 11.4167 | 0.409 |
| 1.6 | 0.05 | clcB | 2 | 11.4167 | 0.316 |
| 1.6 | 0.05 | ycaM | 2 | 11.4167 | 0.283 |
| 1.6 | 0.05 | yadI | 2 | 11.4167 | 0.566 |
| 1.6 | 0.05 | AG1 | 2 | 11.4167 | 0.574 |
| 1.6 | 0.05 | ptsI | 2 | 11.6667 | 0.413 |
| 1.6 | 0.05 | clcB | 2 | 11.6667 | 0.309 |
| 1.6 | 0.05 | ycaM | 2 | 11.6667 | 0.282 |
| 1.6 | 0.05 | yadI | 2 | 11.6667 | 0.571 |
| 1.6 | 0.05 | AG1 | 2 | 11.6667 | 0.586 |
| 1.6 | 0.05 | ptsI | 2 | 11.9167 | 0.417 |
| 1.6 | 0.05 | clcB | 2 | 11.9167 | 0.309 |
| 1.6 | 0.05 | ycaM | 2 | 11.9167 | 0.28 |

| | | | | | |
|---|---|---|---|---|---|
| 1.6 | 0.05 | yadI | 2 | 11.9167 | 0.573 |
| 1.6 | 0.05 | AG1 | 2 | 11.9167 | 0.592 |
| 1.6 | 0.05 | ptsI | 2 | 12.1667 | 0.421 |
| 1.6 | 0.05 | clcB | 2 | 12.1667 | 0.31 |
| 1.6 | 0.05 | ycaM | 2 | 12.1667 | 0.279 |
| 1.6 | 0.05 | yadI | 2 | 12.1667 | 0.579 |
| 1.6 | 0.05 | AG1 | 2 | 12.1667 | 0.588 |
| 1.6 | 0.05 | ptsI | 2 | 12.4167 | 0.431 |
| 1.6 | 0.05 | clcB | 2 | 12.4167 | 0.309 |
| 1.6 | 0.05 | ycaM | 2 | 12.4167 | 0.278 |
| 1.6 | 0.05 | yadI | 2 | 12.4167 | 0.582 |
| 1.6 | 0.05 | AG1 | 2 | 12.4167 | 0.6 |
| 1.6 | 0.05 | ptsI | 2 | 12.6667 | 0.436 |
| 1.6 | 0.05 | clcB | 2 | 12.6667 | 0.304 |
| 1.6 | 0.05 | ycaM | 2 | 12.6667 | 0.274 |
| 1.6 | 0.05 | yadI | 2 | 12.6667 | 0.576 |
| 1.6 | 0.05 | AG1 | 2 | 12.6667 | 0.601 |
| 1.6 | 0.05 | ptsI | 2 | 12.9167 | 0.446 |
| 1.6 | 0.05 | clcB | 2 | 12.9167 | 0.3 |
| 1.6 | 0.05 | ycaM | 2 | 12.9167 | 0.274 |
| 1.6 | 0.05 | yadI | 2 | 12.9167 | 0.577 |
| 1.6 | 0.05 | AG1 | 2 | 12.9167 | 0.595 |
| 1.6 | 0.05 | ptsI | 2 | 13.1667 | 0.45 |
| 1.6 | 0.05 | clcB | 2 | 13.1667 | 0.302 |
| 1.6 | 0.05 | ycaM | 2 | 13.1667 | 0.273 |
| 1.6 | 0.05 | yadI | 2 | 13.1667 | 0.584 |
| 1.6 | 0.05 | AG1 | 2 | 13.1667 | 0.606 |
| 1.6 | 0.05 | ptsI | 2 | 13.4167 | 0.451 |
| 1.6 | 0.05 | clcB | 2 | 13.4167 | 0.3 |
| 1.6 | 0.05 | ycaM | 2 | 13.4167 | 0.274 |
| 1.6 | 0.05 | yadI | 2 | 13.4167 | 0.585 |
| 1.6 | 0.05 | AG1 | 2 | 13.4167 | 0.629 |
| 1.6 | 0.05 | ptsI | 2 | 13.6667 | 0.453 |
| 1.6 | 0.05 | clcB | 2 | 13.6667 | 0.301 |
| 1.6 | 0.05 | ycaM | 2 | 13.6667 | 0.272 |
| 1.6 | 0.05 | yadI | 2 | 13.6667 | 0.587 |
| 1.6 | 0.05 | AG1 | 2 | 13.6667 | 0.626 |
| 1.6 | 0.05 | ptsI | 2 | 13.9167 | 0.454 |
| 1.6 | 0.05 | clcB | 2 | 13.9167 | 0.299 |
| 1.6 | 0.05 | ycaM | 2 | 13.9167 | 0.27 |
| 1.6 | 0.05 | yadI | 2 | 13.9167 | 0.59 |
| 1.6 | 0.05 | AG1 | 2 | 13.9167 | 0.662 |
| 1.6 | 0.05 | ptsI | 2 | 14.1667 | 0.459 |
| 1.6 | 0.05 | clcB | 2 | 14.1667 | 0.298 |
| 1.6 | 0.05 | ycaM | 2 | 14.1667 | 0.27 |
| 1.6 | 0.05 | yadI | 2 | 14.1667 | 0.592 |
| 1.6 | 0.05 | AG1 | 2 | 14.1667 | 0.649 |
| 1.6 | 0.05 | ptsI | 2 | 14.4167 | 0.46 |

| | | | | | |
|---|---|---|---|---|---|
| 1.6 | 0.05 | clcB | 2 | 14.4167 | 0.295 |
| 1.6 | 0.05 | ycaM | 2 | 14.4167 | 0.272 |
| 1.6 | 0.05 | yadI | 2 | 14.4167 | 0.597 |
| 1.6 | 0.05 | AG1 | 2 | 14.4167 | 0.661 |
| 1.6 | 0.05 | ptsI | 2 | 14.6667 | 0.459 |
| 1.6 | 0.05 | clcB | 2 | 14.6667 | 0.294 |
| 1.6 | 0.05 | ycaM | 2 | 14.6667 | 0.273 |
| 1.6 | 0.05 | yadI | 2 | 14.6667 | 0.598 |
| 1.6 | 0.05 | AG1 | 2 | 14.6667 | 0.687 |
| 1.6 | 0.05 | ptsI | 2 | 14.9167 | 0.463 |
| 1.6 | 0.05 | clcB | 2 | 14.9167 | 0.292 |
| 1.6 | 0.05 | ycaM | 2 | 14.9167 | 0.271 |
| 1.6 | 0.05 | yadI | 2 | 14.9167 | 0.598 |
| 1.6 | 0.05 | AG1 | 2 | 14.9167 | 0.688 |
| 1.6 | 0.05 | ptsI | 2 | 15.1667 | 0.469 |
| 1.6 | 0.05 | clcB | 2 | 15.1667 | 0.289 |
| 1.6 | 0.05 | ycaM | 2 | 15.1667 | 0.272 |
| 1.6 | 0.05 | yadI | 2 | 15.1667 | 0.607 |
| 1.6 | 0.05 | AG1 | 2 | 15.1667 | 0.73 |
| 1.6 | 0.05 | ptsI | 2 | 15.4167 | 0.47 |
| 1.6 | 0.05 | clcB | 2 | 15.4167 | 0.29 |
| 1.6 | 0.05 | ycaM | 2 | 15.4167 | 0.271 |
| 1.6 | 0.05 | yadI | 2 | 15.4167 | 0.604 |
| 1.6 | 0.05 | AG1 | 2 | 15.4167 | 0.702 |
| 1.6 | 0.05 | ptsI | 2 | 15.6667 | 0.476 |
| 1.6 | 0.05 | clcB | 2 | 15.6667 | 0.289 |
| 1.6 | 0.05 | ycaM | 2 | 15.6667 | 0.27 |
| 1.6 | 0.05 | yadI | 2 | 15.6667 | 0.606 |
| 1.6 | 0.05 | AG1 | 2 | 15.6667 | 0.725 |
| 1.6 | 0.05 | ptsI | 2 | 15.9167 | 0.478 |
| 1.6 | 0.05 | clcB | 2 | 15.9167 | 0.288 |
| 1.6 | 0.05 | ycaM | 2 | 15.9167 | 0.272 |
| 1.6 | 0.05 | yadI | 2 | 15.9167 | 0.607 |
| 1.6 | 0.05 | AG1 | 2 | 15.9167 | 0.73 |
| 1.6 | 0.05 | ptsI | 2 | 16.1667 | 0.476 |
| 1.6 | 0.05 | clcB | 2 | 16.1667 | 0.288 |
| 1.6 | 0.05 | ycaM | 2 | 16.1667 | 0.273 |
| 1.6 | 0.05 | yadI | 2 | 16.1667 | 0.613 |
| 1.6 | 0.05 | AG1 | 2 | 16.1667 | 0.703 |
| 1.6 | 0.05 | ptsI | 2 | 16.4167 | 0.48 |
| 1.6 | 0.05 | clcB | 2 | 16.4167 | 0.287 |
| 1.6 | 0.05 | ycaM | 2 | 16.4167 | 0.269 |
| 1.6 | 0.05 | yadI | 2 | 16.4167 | 0.614 |
| 1.6 | 0.05 | AG1 | 2 | 16.4167 | 0.713 |
| 1.6 | 0.05 | ptsI | 2 | 16.6667 | 0.48 |
| 1.6 | 0.05 | clcB | 2 | 16.6667 | 0.287 |
| 1.6 | 0.05 | ycaM | 2 | 16.6667 | 0.271 |
| 1.6 | 0.05 | yadI | 2 | 16.6667 | 0.616 |

| | | | | | |
|---|---|---|---|---|---|
| 1.6 | 0.05 | AG1 | 2 | 16.6667 | 0.726 |
| 1.6 | 0.05 | ptsI | 2 | 16.9167 | 0.482 |
| 1.6 | 0.05 | clcB | 2 | 16.9167 | 0.286 |
| 1.6 | 0.05 | ycaM | 2 | 16.9167 | 0.272 |
| 1.6 | 0.05 | yadI | 2 | 16.9167 | 0.621 |
| 1.6 | 0.05 | AG1 | 2 | 16.9167 | 0.744 |
| 1.6 | 0.05 | ptsI | 2 | 17.1667 | 0.487 |
| 1.6 | 0.05 | clcB | 2 | 17.1667 | 0.287 |
| 1.6 | 0.05 | ycaM | 2 | 17.1667 | 0.27 |
| 1.6 | 0.05 | yadI | 2 | 17.1667 | 0.622 |
| 1.6 | 0.05 | AG1 | 2 | 17.1667 | 0.753 |
| 1.6 | 0.05 | ptsI | 2 | 17.4167 | 0.487 |
| 1.6 | 0.05 | clcB | 2 | 17.4167 | 0.288 |
| 1.6 | 0.05 | ycaM | 2 | 17.4167 | 0.271 |
| 1.6 | 0.05 | yadI | 2 | 17.4167 | 0.625 |
| 1.6 | 0.05 | AG1 | 2 | 17.4167 | 0.739 |
| 1.6 | 0.05 | ptsI | 2 | 17.6667 | 0.489 |
| 1.6 | 0.05 | clcB | 2 | 17.6667 | 0.287 |
| 1.6 | 0.05 | ycaM | 2 | 17.6667 | 0.271 |
| 1.6 | 0.05 | yadI | 2 | 17.6667 | 0.625 |
| 1.6 | 0.05 | AG1 | 2 | 17.6667 | 0.731 |
| 1.6 | 0.05 | ptsI | 2 | 17.9167 | 0.487 |
| 1.6 | 0.05 | clcB | 2 | 17.9167 | 0.287 |
| 1.6 | 0.05 | ycaM | 2 | 17.9167 | 0.272 |
| 1.6 | 0.05 | yadI | 2 | 17.9167 | 0.627 |
| 1.6 | 0.05 | AG1 | 2 | 17.9167 | 0.728 |
| 1.6 | 0.05 | ptsI | 2 | 18.1667 | 0.493 |
| 1.6 | 0.05 | clcB | 2 | 18.1667 | 0.286 |
| 1.6 | 0.05 | ycaM | 2 | 18.1667 | 0.272 |
| 1.6 | 0.05 | yadI | 2 | 18.1667 | 0.63 |
| 1.6 | 0.05 | AG1 | 2 | 18.1667 | 0.739 |
| 1.6 | 0.05 | ptsI | 2 | 18.4167 | 0.496 |
| 1.6 | 0.05 | clcB | 2 | 18.4167 | 0.286 |
| 1.6 | 0.05 | ycaM | 2 | 18.4167 | 0.274 |
| 1.6 | 0.05 | yadI | 2 | 18.4167 | 0.63 |
| 1.6 | 0.05 | AG1 | 2 | 18.4167 | 0.756 |
| 1.6 | 0.05 | ptsI | 2 | 18.6667 | 0.496 |
| 1.6 | 0.05 | clcB | 2 | 18.6667 | 0.285 |
| 1.6 | 0.05 | ycaM | 2 | 18.6667 | 0.275 |
| 1.6 | 0.05 | yadI | 2 | 18.6667 | 0.633 |
| 1.6 | 0.05 | AG1 | 2 | 18.6667 | 0.744 |
| 1.6 | 0.05 | ptsI | 2 | 18.9167 | 0.494 |
| 1.6 | 0.05 | clcB | 2 | 18.9167 | 0.286 |
| 1.6 | 0.05 | ycaM | 2 | 18.9167 | 0.277 |
| 1.6 | 0.05 | yadI | 2 | 18.9167 | 0.626 |
| 1.6 | 0.05 | AG1 | 2 | 18.9167 | 0.753 |
| 1.6 | 0.05 | ptsI | 2 | 19.1667 | 0.498 |
| 1.6 | 0.05 | clcB | 2 | 19.1667 | 0.285 |

| | | | | | |
|---|---|---|---|---|---|
| 1.6 | 0.05 | ycaM | 2 | 19.1667 | 0.278 |
| 1.6 | 0.05 | yadI | 2 | 19.1667 | 0.637 |
| 1.6 | 0.05 | AG1 | 2 | 19.1667 | 0.759 |
| 1.6 | 0.05 | ptsI | 2 | 19.4167 | 0.501 |
| 1.6 | 0.05 | clcB | 2 | 19.4167 | 0.286 |
| 1.6 | 0.05 | ycaM | 2 | 19.4167 | 0.279 |
| 1.6 | 0.05 | yadI | 2 | 19.4167 | 0.639 |
| 1.6 | 0.05 | AG1 | 2 | 19.4167 | 0.751 |
| 1.6 | 0.05 | ptsI | 2 | 19.6667 | 0.503 |
| 1.6 | 0.05 | clcB | 2 | 19.6667 | 0.286 |
| 1.6 | 0.05 | ycaM | 2 | 19.6667 | 0.281 |
| 1.6 | 0.05 | yadI | 2 | 19.6667 | 0.641 |
| 1.6 | 0.05 | AG1 | 2 | 19.6667 | 0.754 |
| 1.6 | 0.05 | ptsI | 2 | 19.9167 | 0.503 |
| 1.6 | 0.05 | clcB | 2 | 19.9167 | 0.286 |
| 1.6 | 0.05 | ycaM | 2 | 19.9167 | 0.282 |
| 1.6 | 0.05 | yadI | 2 | 19.9167 | 0.642 |
| 1.6 | 0.05 | AG1 | 2 | 19.9167 | 0.749 |
| 1.6 | 0.05 | ptsI | 2 | 20.1667 | 0.504 |
| 1.6 | 0.05 | clcB | 2 | 20.1667 | 0.287 |
| 1.6 | 0.05 | ycaM | 2 | 20.1667 | 0.284 |
| 1.6 | 0.05 | yadI | 2 | 20.1667 | 0.645 |
| 1.6 | 0.05 | AG1 | 2 | 20.1667 | 0.75 |
| 1.6 | 0.05 | ptsI | 2 | 20.4167 | 0.504 |
| 1.6 | 0.05 | clcB | 2 | 20.4167 | 0.287 |
| 1.6 | 0.05 | ycaM | 2 | 20.4167 | 0.286 |
| 1.6 | 0.05 | yadI | 2 | 20.4167 | 0.648 |
| 1.6 | 0.05 | AG1 | 2 | 20.4167 | 0.754 |
| 1.6 | 0.05 | ptsI | 2 | 20.6667 | 0.507 |
| 1.6 | 0.05 | clcB | 2 | 20.6667 | 0.287 |
| 1.6 | 0.05 | ycaM | 2 | 20.6667 | 0.288 |
| 1.6 | 0.05 | yadI | 2 | 20.6667 | 0.649 |
| 1.6 | 0.05 | AG1 | 2 | 20.6667 | 0.753 |
| 1.6 | 0.05 | ptsI | 2 | 20.9167 | 0.509 |
| 1.6 | 0.05 | clcB | 2 | 20.9167 | 0.289 |
| 1.6 | 0.05 | ycaM | 2 | 20.9167 | 0.29 |
| 1.6 | 0.05 | yadI | 2 | 20.9167 | 0.651 |
| 1.6 | 0.05 | AG1 | 2 | 20.9167 | 0.759 |
| 1.6 | 0.05 | ptsI | 2 | 21.1667 | 0.508 |
| 1.6 | 0.05 | clcB | 2 | 21.1667 | 0.29 |
| 1.6 | 0.05 | ycaM | 2 | 21.1667 | 0.291 |
| 1.6 | 0.05 | yadI | 2 | 21.1667 | 0.65 |
| 1.6 | 0.05 | AG1 | 2 | 21.1667 | 0.762 |
| 1.6 | 0.05 | ptsI | 2 | 21.4167 | 0.509 |
| 1.6 | 0.05 | clcB | 2 | 21.4167 | 0.29 |
| 1.6 | 0.05 | ycaM | 2 | 21.4167 | 0.293 |
| 1.6 | 0.05 | yadI | 2 | 21.4167 | 0.647 |
| 1.6 | 0.05 | AG1 | 2 | 21.4167 | 0.782 |

| | | | | | |
|---|---|---|---|---|---|
| 1.6 | 0.05 | ptsI | 2 | 21.6667 | 0.508 |
| 1.6 | 0.05 | clcB | 2 | 21.6667 | 0.291 |
| 1.6 | 0.05 | ycaM | 2 | 21.6667 | 0.295 |
| 1.6 | 0.05 | yadI | 2 | 21.6667 | 0.656 |
| 1.6 | 0.05 | AG1 | 2 | 21.6667 | 0.774 |
| 1.6 | 0.05 | ptsI | 2 | 21.9167 | 0.511 |
| 1.6 | 0.05 | clcB | 2 | 21.9167 | 0.292 |
| 1.6 | 0.05 | ycaM | 2 | 21.9167 | 0.297 |
| 1.6 | 0.05 | yadI | 2 | 21.9167 | 0.655 |
| 1.6 | 0.05 | AG1 | 2 | 21.9167 | 0.78 |
| 1.6 | 0.05 | ptsI | 2 | 22.1667 | 0.51 |
| 1.6 | 0.05 | clcB | 2 | 22.1667 | 0.293 |
| 1.6 | 0.05 | ycaM | 2 | 22.1667 | 0.298 |
| 1.6 | 0.05 | yadI | 2 | 22.1667 | 0.655 |
| 1.6 | 0.05 | AG1 | 2 | 22.1667 | 0.782 |
| 1.6 | 0.05 | ptsI | 2 | 22.4167 | 0.509 |
| 1.6 | 0.05 | clcB | 2 | 22.4167 | 0.296 |
| 1.6 | 0.05 | ycaM | 2 | 22.4167 | 0.3 |
| 1.6 | 0.05 | yadI | 2 | 22.4167 | 0.653 |
| 1.6 | 0.05 | AG1 | 2 | 22.4167 | 0.785 |
| 1.6 | 0.05 | ptsI | 2 | 22.6667 | 0.507 |
| 1.6 | 0.05 | clcB | 2 | 22.6667 | 0.296 |
| 1.6 | 0.05 | ycaM | 2 | 22.6667 | 0.302 |
| 1.6 | 0.05 | yadI | 2 | 22.6667 | 0.656 |
| 1.6 | 0.05 | AG1 | 2 | 22.6667 | 0.792 |
| 1.6 | 0.05 | ptsI | 2 | 22.9167 | 0.513 |
| 1.6 | 0.05 | clcB | 2 | 22.9167 | 0.295 |
| 1.6 | 0.05 | ycaM | 2 | 22.9167 | 0.302 |
| 1.6 | 0.05 | yadI | 2 | 22.9167 | 0.656 |
| 1.6 | 0.05 | AG1 | 2 | 22.9167 | 0.798 |
| 1.6 | 0.05 | ptsI | 2 | 23.1667 | 0.51 |
| 1.6 | 0.05 | clcB | 2 | 23.1667 | 0.295 |
| 1.6 | 0.05 | ycaM | 2 | 23.1667 | 0.305 |
| 1.6 | 0.05 | yadI | 2 | 23.1667 | 0.659 |
| 1.6 | 0.05 | AG1 | 2 | 23.1667 | 0.802 |
| 1.6 | 0.05 | ptsI | 2 | 23.4167 | 0.511 |
| 1.6 | 0.05 | clcB | 2 | 23.4167 | 0.297 |
| 1.6 | 0.05 | ycaM | 2 | 23.4167 | 0.307 |
| 1.6 | 0.05 | yadI | 2 | 23.4167 | 0.66 |
| 1.6 | 0.05 | AG1 | 2 | 23.4167 | 0.811 |
| 1.6 | 0.05 | ptsI | 2 | 23.6667 | 0.511 |
| 1.6 | 0.05 | clcB | 2 | 23.6667 | 0.298 |
| 1.6 | 0.05 | ycaM | 2 | 23.6667 | 0.311 |
| 1.6 | 0.05 | yadI | 2 | 23.6667 | 0.659 |
| 1.6 | 0.05 | AG1 | 2 | 23.6667 | 0.816 |
| 1.6 | 0.05 | ptsI | 2 | 23.9167 | 0.509 |
| 1.6 | 0.05 | clcB | 2 | 23.9167 | 0.3 |
| 1.6 | 0.05 | ycaM | 2 | 23.9167 | 0.313 |

| | | | | | |
|---|---|---|---|---|---|
| 1.6 | 0.05 | yadI | 2 | 23.9167 | 0.662 |
| 1.6 | 0.05 | AG1 | 2 | 23.9167 | 0.821 |
| 1.6 | 0.05 | ptsI | 2 | 24.1667 | 0.511 |
| 1.6 | 0.05 | clcB | 2 | 24.1667 | 0.302 |
| 1.6 | 0.05 | ycaM | 2 | 24.1667 | 0.316 |
| 1.6 | 0.05 | yadI | 2 | 24.1667 | 0.662 |
| 1.6 | 0.05 | AG1 | 2 | 24.1667 | 0.83 |
| 1.6 | 0.05 | ptsI | 2 | 24.4167 | 0.511 |
| 1.6 | 0.05 | clcB | 2 | 24.4167 | 0.302 |
| 1.6 | 0.05 | ycaM | 2 | 24.4167 | 0.32 |
| 1.6 | 0.05 | yadI | 2 | 24.4167 | 0.667 |
| 1.6 | 0.05 | AG1 | 2 | 24.4167 | 0.839 |
| 1.6 | 0.05 | ptsI | 2 | 24.6667 | 0.511 |
| 1.6 | 0.05 | clcB | 2 | 24.6667 | 0.305 |
| 1.6 | 0.05 | ycaM | 2 | 24.6667 | 0.323 |
| 1.6 | 0.05 | yadI | 2 | 24.6667 | 0.665 |
| 1.6 | 0.05 | AG1 | 2 | 24.6667 | 0.842 |
| 1.6 | 0.05 | ptsI | 2 | 24.9167 | 0.509 |
| 1.6 | 0.05 | clcB | 2 | 24.9167 | 0.307 |
| 1.6 | 0.05 | ycaM | 2 | 24.9167 | 0.327 |
| 1.6 | 0.05 | yadI | 2 | 24.9167 | 0.666 |
| 1.6 | 0.05 | AG1 | 2 | 24.9167 | 0.853 |
| 1.6 | 0.05 | ptsI | 2 | 25.1667 | 0.512 |
| 1.6 | 0.05 | clcB | 2 | 25.1667 | 0.309 |
| 1.6 | 0.05 | ycaM | 2 | 25.1667 | 0.326 |
| 1.6 | 0.05 | yadI | 2 | 25.1667 | 0.667 |
| 1.6 | 0.05 | AG1 | 2 | 25.1667 | 0.86 |
| 1.6 | 0.05 | ptsI | 2 | 25.4167 | 0.511 |
| 1.6 | 0.05 | clcB | 2 | 25.4167 | 0.311 |
| 1.6 | 0.05 | ycaM | 2 | 25.4167 | 0.331 |
| 1.6 | 0.05 | yadI | 2 | 25.4167 | 0.67 |
| 1.6 | 0.05 | AG1 | 2 | 25.4167 | 0.871 |
| 1.6 | 0.05 | ptsI | 2 | 25.6667 | 0.512 |
| 1.6 | 0.05 | clcB | 2 | 25.6667 | 0.313 |
| 1.6 | 0.05 | ycaM | 2 | 25.6667 | 0.333 |
| 1.6 | 0.05 | yadI | 2 | 25.6667 | 0.67 |
| 1.6 | 0.05 | AG1 | 2 | 25.6667 | 0.877 |
| 1.6 | 0.05 | ptsI | 2 | 25.9167 | 0.514 |
| 1.6 | 0.05 | clcB | 2 | 25.9167 | 0.315 |
| 1.6 | 0.05 | ycaM | 2 | 25.9167 | 0.337 |
| 1.6 | 0.05 | yadI | 2 | 25.9167 | 0.669 |
| 1.6 | 0.05 | AG1 | 2 | 25.9167 | 0.885 |
| 1.6 | 0.05 | ptsI | 2 | 26.1667 | 0.516 |
| 1.6 | 0.05 | clcB | 2 | 26.1667 | 0.318 |
| 1.6 | 0.05 | ycaM | 2 | 26.1667 | 0.339 |
| 1.6 | 0.05 | yadI | 2 | 26.1667 | 0.672 |
| 1.6 | 0.05 | AG1 | 2 | 26.1667 | 0.904 |
| 1.6 | 0.05 | ptsI | 2 | 26.4167 | 0.514 |

| | | | | | |
|---|---|---|---|---|---|
| 1.6 | 0.05 | clcB | 2 | 26.4167 | 0.318 |
| 1.6 | 0.05 | ycaM | 2 | 26.4167 | 0.34 |
| 1.6 | 0.05 | yadI | 2 | 26.4167 | 0.674 |
| 1.6 | 0.05 | AG1 | 2 | 26.4167 | 0.906 |
| 1.6 | 0.05 | ptsI | 3 | 0 | 0.161 |
| 1.6 | 0.05 | clcB | 3 | 0 | 0.162 |
| 1.6 | 0.05 | ycaM | 3 | 0 | 0.157 |
| 1.6 | 0.05 | yadI | 3 | 0 | 0.158 |
| 1.6 | 0.05 | AG1 | 3 | 0 | 0.16 |
| 1.6 | 0.05 | ptsI | 3 | 0.25 | 0.157 |
| 1.6 | 0.05 | clcB | 3 | 0.25 | 0.158 |
| 1.6 | 0.05 | ycaM | 3 | 0.25 | 0.154 |
| 1.6 | 0.05 | yadI | 3 | 0.25 | 0.156 |
| 1.6 | 0.05 | AG1 | 3 | 0.25 | 0.157 |
| 1.6 | 0.05 | ptsI | 3 | 0.5 | 0.158 |
| 1.6 | 0.05 | clcB | 3 | 0.5 | 0.158 |
| 1.6 | 0.05 | ycaM | 3 | 0.5 | 0.151 |
| 1.6 | 0.05 | yadI | 3 | 0.5 | 0.154 |
| 1.6 | 0.05 | AG1 | 3 | 0.5 | 0.155 |
| 1.6 | 0.05 | ptsI | 3 | 0.75 | 0.157 |
| 1.6 | 0.05 | clcB | 3 | 0.75 | 0.157 |
| 1.6 | 0.05 | ycaM | 3 | 0.75 | 0.152 |
| 1.6 | 0.05 | yadI | 3 | 0.75 | 0.153 |
| 1.6 | 0.05 | AG1 | 3 | 0.75 | 0.156 |
| 1.6 | 0.05 | ptsI | 3 | 1 | 0.159 |
| 1.6 | 0.05 | clcB | 3 | 1 | 0.172 |
| 1.6 | 0.05 | ycaM | 3 | 1 | 0.158 |
| 1.6 | 0.05 | yadI | 3 | 1 | 0.155 |
| 1.6 | 0.05 | AG1 | 3 | 1 | 0.156 |
| 1.6 | 0.05 | ptsI | 3 | 1.25 | 0.158 |
| 1.6 | 0.05 | clcB | 3 | 1.25 | 0.157 |
| 1.6 | 0.05 | ycaM | 3 | 1.25 | 0.151 |
| 1.6 | 0.05 | yadI | 3 | 1.25 | 0.155 |
| 1.6 | 0.05 | AG1 | 3 | 1.25 | 0.156 |
| 1.6 | 0.05 | ptsI | 3 | 1.5 | 0.159 |
| 1.6 | 0.05 | clcB | 3 | 1.5 | 0.157 |
| 1.6 | 0.05 | ycaM | 3 | 1.5 | 0.152 |
| 1.6 | 0.05 | yadI | 3 | 1.5 | 0.154 |
| 1.6 | 0.05 | AG1 | 3 | 1.5 | 0.156 |
| 1.6 | 0.05 | ptsI | 3 | 1.75 | 0.161 |
| 1.6 | 0.05 | clcB | 3 | 1.75 | 0.156 |
| 1.6 | 0.05 | ycaM | 3 | 1.75 | 0.152 |
| 1.6 | 0.05 | yadI | 3 | 1.75 | 0.155 |
| 1.6 | 0.05 | AG1 | 3 | 1.75 | 0.157 |
| 1.6 | 0.05 | ptsI | 3 | 2 | 0.161 |
| 1.6 | 0.05 | clcB | 3 | 2 | 0.189 |
| 1.6 | 0.05 | ycaM | 3 | 2 | 0.154 |
| 1.6 | 0.05 | yadI | 3 | 2 | 0.157 |

| | | | | | |
|---|---|---|---|---|---|
| 1.6 | 0.05 | AG1 | 3 | 2 | 0.157 |
| 1.6 | 0.05 | ptsI | 3 | 2.25 | 0.162 |
| 1.6 | 0.05 | clcB | 3 | 2.25 | 0.176 |
| 1.6 | 0.05 | ycaM | 3 | 2.25 | 0.156 |
| 1.6 | 0.05 | yadI | 3 | 2.25 | 0.157 |
| 1.6 | 0.05 | AG1 | 3 | 2.25 | 0.159 |
| 1.6 | 0.05 | ptsI | 3 | 2.5 | 0.163 |
| 1.6 | 0.05 | clcB | 3 | 2.5 | 0.172 |
| 1.6 | 0.05 | ycaM | 3 | 2.5 | 0.157 |
| 1.6 | 0.05 | yadI | 3 | 2.5 | 0.159 |
| 1.6 | 0.05 | AG1 | 3 | 2.5 | 0.16 |
| 1.6 | 0.05 | ptsI | 3 | 2.75 | 0.166 |
| 1.6 | 0.05 | clcB | 3 | 2.75 | 0.191 |
| 1.6 | 0.05 | ycaM | 3 | 2.75 | 0.16 |
| 1.6 | 0.05 | yadI | 3 | 2.75 | 0.162 |
| 1.6 | 0.05 | AG1 | 3 | 2.75 | 0.164 |
| 1.6 | 0.05 | ptsI | 3 | 3 | 0.165 |
| 1.6 | 0.05 | clcB | 3 | 3 | 0.163 |
| 1.6 | 0.05 | ycaM | 3 | 3 | 0.16 |
| 1.6 | 0.05 | yadI | 3 | 3 | 0.163 |
| 1.6 | 0.05 | AG1 | 3 | 3 | 0.163 |
| 1.6 | 0.05 | ptsI | 3 | 3.25 | 0.167 |
| 1.6 | 0.05 | clcB | 3 | 3.25 | 0.165 |
| 1.6 | 0.05 | ycaM | 3 | 3.25 | 0.162 |
| 1.6 | 0.05 | yadI | 3 | 3.25 | 0.165 |
| 1.6 | 0.05 | AG1 | 3 | 3.25 | 0.163 |
| 1.6 | 0.05 | ptsI | 3 | 3.5 | 0.17 |
| 1.6 | 0.05 | clcB | 3 | 3.5 | 0.168 |
| 1.6 | 0.05 | ycaM | 3 | 3.5 | 0.166 |
| 1.6 | 0.05 | yadI | 3 | 3.5 | 0.17 |
| 1.6 | 0.05 | AG1 | 3 | 3.5 | 0.165 |
| 1.6 | 0.05 | ptsI | 3 | 3.75 | 0.172 |
| 1.6 | 0.05 | clcB | 3 | 3.75 | 0.169 |
| 1.6 | 0.05 | ycaM | 3 | 3.75 | 0.17 |
| 1.6 | 0.05 | yadI | 3 | 3.75 | 0.173 |
| 1.6 | 0.05 | AG1 | 3 | 3.75 | 0.168 |
| 1.6 | 0.05 | ptsI | 3 | 4 | 0.174 |
| 1.6 | 0.05 | clcB | 3 | 4 | 0.171 |
| 1.6 | 0.05 | ycaM | 3 | 4 | 0.172 |
| 1.6 | 0.05 | yadI | 3 | 4 | 0.178 |
| 1.6 | 0.05 | AG1 | 3 | 4 | 0.171 |
| 1.6 | 0.05 | ptsI | 3 | 4.41667 | 0.174 |
| 1.6 | 0.05 | clcB | 3 | 4.41667 | 0.214 |
| 1.6 | 0.05 | ycaM | 3 | 4.41667 | 0.179 |
| 1.6 | 0.05 | yadI | 3 | 4.41667 | 0.193 |
| 1.6 | 0.05 | AG1 | 3 | 4.41667 | 0.189 |
| 1.6 | 0.05 | ptsI | 3 | 4.66667 | 0.167 |
| 1.6 | 0.05 | clcB | 3 | 4.66667 | 0.187 |

| | | | | | |
|---|---|---|---|---|---|
| 1.6 | 0.05 | ycaM | 3 | 4.66667 | 0.171 |
| 1.6 | 0.05 | yadI | 3 | 4.66667 | 0.193 |
| 1.6 | 0.05 | AG1 | 3 | 4.66667 | 0.19 |
| 1.6 | 0.05 | ptsI | 3 | 4.91667 | 0.168 |
| 1.6 | 0.05 | clcB | 3 | 4.91667 | 0.192 |
| 1.6 | 0.05 | ycaM | 3 | 4.91667 | 0.179 |
| 1.6 | 0.05 | yadI | 3 | 4.91667 | 0.2 |
| 1.6 | 0.05 | AG1 | 3 | 4.91667 | 0.196 |
| 1.6 | 0.05 | ptsI | 3 | 5.16667 | 0.169 |
| 1.6 | 0.05 | clcB | 3 | 5.16667 | 0.197 |
| 1.6 | 0.05 | ycaM | 3 | 5.16667 | 0.184 |
| 1.6 | 0.05 | yadI | 3 | 5.16667 | 0.209 |
| 1.6 | 0.05 | AG1 | 3 | 5.16667 | 0.202 |
| 1.6 | 0.05 | ptsI | 3 | 5.41667 | 0.173 |
| 1.6 | 0.05 | clcB | 3 | 5.41667 | 0.202 |
| 1.6 | 0.05 | ycaM | 3 | 5.41667 | 0.188 |
| 1.6 | 0.05 | yadI | 3 | 5.41667 | 0.219 |
| 1.6 | 0.05 | AG1 | 3 | 5.41667 | 0.207 |
| 1.6 | 0.05 | ptsI | 3 | 5.66667 | 0.177 |
| 1.6 | 0.05 | clcB | 3 | 5.66667 | 0.208 |
| 1.6 | 0.05 | ycaM | 3 | 5.66667 | 0.196 |
| 1.6 | 0.05 | yadI | 3 | 5.66667 | 0.232 |
| 1.6 | 0.05 | AG1 | 3 | 5.66667 | 0.214 |
| 1.6 | 0.05 | ptsI | 3 | 5.91667 | 0.181 |
| 1.6 | 0.05 | clcB | 3 | 5.91667 | 0.209 |
| 1.6 | 0.05 | ycaM | 3 | 5.91667 | 0.202 |
| 1.6 | 0.05 | yadI | 3 | 5.91667 | 0.238 |
| 1.6 | 0.05 | AG1 | 3 | 5.91667 | 0.214 |
| 1.6 | 0.05 | ptsI | 3 | 6.16667 | 0.187 |
| 1.6 | 0.05 | clcB | 3 | 6.16667 | 0.216 |
| 1.6 | 0.05 | ycaM | 3 | 6.16667 | 0.212 |
| 1.6 | 0.05 | yadI | 3 | 6.16667 | 0.253 |
| 1.6 | 0.05 | AG1 | 3 | 6.16667 | 0.222 |
| 1.6 | 0.05 | ptsI | 3 | 6.41667 | 0.193 |
| 1.6 | 0.05 | clcB | 3 | 6.41667 | 0.225 |
| 1.6 | 0.05 | ycaM | 3 | 6.41667 | 0.219 |
| 1.6 | 0.05 | yadI | 3 | 6.41667 | 0.264 |
| 1.6 | 0.05 | AG1 | 3 | 6.41667 | 0.23 |
| 1.6 | 0.05 | ptsI | 3 | 6.66667 | 0.2 |
| 1.6 | 0.05 | clcB | 3 | 6.66667 | 0.233 |
| 1.6 | 0.05 | ycaM | 3 | 6.66667 | 0.228 |
| 1.6 | 0.05 | yadI | 3 | 6.66667 | 0.272 |
| 1.6 | 0.05 | AG1 | 3 | 6.66667 | 0.241 |
| 1.6 | 0.05 | ptsI | 3 | 6.91667 | 0.207 |
| 1.6 | 0.05 | clcB | 3 | 6.91667 | 0.244 |
| 1.6 | 0.05 | ycaM | 3 | 6.91667 | 0.235 |
| 1.6 | 0.05 | yadI | 3 | 6.91667 | 0.282 |
| 1.6 | 0.05 | AG1 | 3 | 6.91667 | 0.25 |

| | | | | | |
|---|---|---|---|---|---|
| 1.6 | 0.05 | ptsI | 3 | 7.16667 | 0.217 |
| 1.6 | 0.05 | clcB | 3 | 7.16667 | 0.255 |
| 1.6 | 0.05 | ycaM | 3 | 7.16667 | 0.239 |
| 1.6 | 0.05 | yadI | 3 | 7.16667 | 0.292 |
| 1.6 | 0.05 | AG1 | 3 | 7.16667 | 0.258 |
| 1.6 | 0.05 | ptsI | 3 | 7.41667 | 0.225 |
| 1.6 | 0.05 | clcB | 3 | 7.41667 | 0.267 |
| 1.6 | 0.05 | ycaM | 3 | 7.41667 | 0.243 |
| 1.6 | 0.05 | yadI | 3 | 7.41667 | 0.301 |
| 1.6 | 0.05 | AG1 | 3 | 7.41667 | 0.265 |
| 1.6 | 0.05 | ptsI | 3 | 7.66667 | 0.234 |
| 1.6 | 0.05 | clcB | 3 | 7.66667 | 0.275 |
| 1.6 | 0.05 | ycaM | 3 | 7.66667 | 0.244 |
| 1.6 | 0.05 | yadI | 3 | 7.66667 | 0.312 |
| 1.6 | 0.05 | AG1 | 3 | 7.66667 | 0.272 |
| 1.6 | 0.05 | ptsI | 3 | 7.91667 | 0.242 |
| 1.6 | 0.05 | clcB | 3 | 7.91667 | 0.281 |
| 1.6 | 0.05 | ycaM | 3 | 7.91667 | 0.246 |
| 1.6 | 0.05 | yadI | 3 | 7.91667 | 0.321 |
| 1.6 | 0.05 | AG1 | 3 | 7.91667 | 0.281 |
| 1.6 | 0.05 | ptsI | 3 | 8.16667 | 0.252 |
| 1.6 | 0.05 | clcB | 3 | 8.16667 | 0.288 |
| 1.6 | 0.05 | ycaM | 3 | 8.16667 | 0.248 |
| 1.6 | 0.05 | yadI | 3 | 8.16667 | 0.33 |
| 1.6 | 0.05 | AG1 | 3 | 8.16667 | 0.286 |
| 1.6 | 0.05 | ptsI | 3 | 8.41667 | 0.259 |
| 1.6 | 0.05 | clcB | 3 | 8.41667 | 0.294 |
| 1.6 | 0.05 | ycaM | 3 | 8.41667 | 0.25 |
| 1.6 | 0.05 | yadI | 3 | 8.41667 | 0.339 |
| 1.6 | 0.05 | AG1 | 3 | 8.41667 | 0.295 |
| 1.6 | 0.05 | ptsI | 3 | 9.05 | 0.262 |
| 1.6 | 0.05 | clcB | 3 | 9.05 | 0.268 |
| 1.6 | 0.05 | ycaM | 3 | 9.05 | 0.241 |
| 1.6 | 0.05 | yadI | 3 | 9.05 | 0.376 |
| 1.6 | 0.05 | AG1 | 3 | 9.05 | 0.314 |
| 1.6 | 0.05 | ptsI | 3 | 9.3 | 0.27 |
| 1.6 | 0.05 | clcB | 3 | 9.3 | 0.271 |
| 1.6 | 0.05 | ycaM | 3 | 9.3 | 0.236 |
| 1.6 | 0.05 | yadI | 3 | 9.3 | 0.384 |
| 1.6 | 0.05 | AG1 | 3 | 9.3 | 0.324 |
| 1.6 | 0.05 | ptsI | 3 | 9.55 | 0.281 |
| 1.6 | 0.05 | clcB | 3 | 9.55 | 0.275 |
| 1.6 | 0.05 | ycaM | 3 | 9.55 | 0.236 |
| 1.6 | 0.05 | yadI | 3 | 9.55 | 0.399 |
| 1.6 | 0.05 | AG1 | 3 | 9.55 | 0.342 |
| 1.6 | 0.05 | ptsI | 3 | 9.8 | 0.292 |
| 1.6 | 0.05 | clcB | 3 | 9.8 | 0.279 |
| 1.6 | 0.05 | ycaM | 3 | 9.8 | 0.233 |

| | | | | | |
|---|---|---|---|---|---|
| 1.6 | 0.05 | yadI | 3 | 9.8 | 0.411 |
| 1.6 | 0.05 | AG1 | 3 | 9.8 | 0.35 |
| 1.6 | 0.05 | ptsI | 3 | 10.05 | 0.299 |
| 1.6 | 0.05 | clcB | 3 | 10.05 | 0.282 |
| 1.6 | 0.05 | ycaM | 3 | 10.05 | 0.233 |
| 1.6 | 0.05 | yadI | 3 | 10.05 | 0.42 |
| 1.6 | 0.05 | AG1 | 3 | 10.05 | 0.359 |
| 1.6 | 0.05 | ptsI | 3 | 10.3 | 0.307 |
| 1.6 | 0.05 | clcB | 3 | 10.3 | 0.286 |
| 1.6 | 0.05 | ycaM | 3 | 10.3 | 0.233 |
| 1.6 | 0.05 | yadI | 3 | 10.3 | 0.425 |
| 1.6 | 0.05 | AG1 | 3 | 10.3 | 0.366 |
| 1.6 | 0.05 | ptsI | 3 | 10.55 | 0.317 |
| 1.6 | 0.05 | clcB | 3 | 10.55 | 0.283 |
| 1.6 | 0.05 | ycaM | 3 | 10.55 | 0.235 |
| 1.6 | 0.05 | yadI | 3 | 10.55 | 0.43 |
| 1.6 | 0.05 | AG1 | 3 | 10.55 | 0.373 |
| 1.6 | 0.05 | ptsI | 3 | 10.8 | 0.323 |
| 1.6 | 0.05 | clcB | 3 | 10.8 | 0.282 |
| 1.6 | 0.05 | ycaM | 3 | 10.8 | 0.234 |
| 1.6 | 0.05 | yadI | 3 | 10.8 | 0.437 |
| 1.6 | 0.05 | AG1 | 3 | 10.8 | 0.382 |
| 1.6 | 0.05 | ptsI | 3 | 11.05 | 0.331 |
| 1.6 | 0.05 | clcB | 3 | 11.05 | 0.288 |
| 1.6 | 0.05 | ycaM | 3 | 11.05 | 0.234 |
| 1.6 | 0.05 | yadI | 3 | 11.05 | 0.444 |
| 1.6 | 0.05 | AG1 | 3 | 11.05 | 0.387 |
| 1.6 | 0.05 | ptsI | 3 | 11.3 | 0.325 |
| 1.6 | 0.05 | clcB | 3 | 11.3 | 0.281 |
| 1.6 | 0.05 | ycaM | 3 | 11.3 | 0.232 |
| 1.6 | 0.05 | yadI | 3 | 11.3 | 0.452 |
| 1.6 | 0.05 | AG1 | 3 | 11.3 | 0.396 |
| 1.6 | 0.05 | ptsI | 3 | 11.55 | 0.33 |
| 1.6 | 0.05 | clcB | 3 | 11.55 | 0.282 |
| 1.6 | 0.05 | ycaM | 3 | 11.55 | 0.232 |
| 1.6 | 0.05 | yadI | 3 | 11.55 | 0.457 |
| 1.6 | 0.05 | AG1 | 3 | 11.55 | 0.398 |
| 1.6 | 0.05 | ptsI | 3 | 11.8 | 0.335 |
| 1.6 | 0.05 | clcB | 3 | 11.8 | 0.286 |
| 1.6 | 0.05 | ycaM | 3 | 11.8 | 0.23 |
| 1.6 | 0.05 | yadI | 3 | 11.8 | 0.46 |
| 1.6 | 0.05 | AG1 | 3 | 11.8 | 0.399 |
| 1.6 | 0.05 | ptsI | 3 | 12.05 | 0.336 |
| 1.6 | 0.05 | clcB | 3 | 12.05 | 0.278 |
| 1.6 | 0.05 | ycaM | 3 | 12.05 | 0.226 |
| 1.6 | 0.05 | yadI | 3 | 12.05 | 0.46 |
| 1.6 | 0.05 | AG1 | 3 | 12.05 | 0.401 |
| 1.6 | 0.05 | ptsI | 3 | 12.3 | 0.343 |

| | | | | | |
|---|---|---|---|---|---|
| 1.6 | 0.05 | clcB | 3 | 12.3 | 0.279 |
| 1.6 | 0.05 | ycaM | 3 | 12.3 | 0.225 |
| 1.6 | 0.05 | yadI | 3 | 12.3 | 0.465 |
| 1.6 | 0.05 | AG1 | 3 | 12.3 | 0.409 |
| 1.6 | 0.05 | ptsI | 3 | 12.55 | 0.351 |
| 1.6 | 0.05 | clcB | 3 | 12.55 | 0.28 |
| 1.6 | 0.05 | ycaM | 3 | 12.55 | 0.227 |
| 1.6 | 0.05 | yadI | 3 | 12.55 | 0.472 |
| 1.6 | 0.05 | AG1 | 3 | 12.55 | 0.408 |
| 1.6 | 0.05 | ptsI | 3 | 12.8 | 0.356 |
| 1.6 | 0.05 | clcB | 3 | 12.8 | 0.281 |
| 1.6 | 0.05 | ycaM | 3 | 12.8 | 0.226 |
| 1.6 | 0.05 | yadI | 3 | 12.8 | 0.474 |
| 1.6 | 0.05 | AG1 | 3 | 12.8 | 0.411 |
| 1.6 | 0.05 | ptsI | 3 | 13.05 | 0.36 |
| 1.6 | 0.05 | clcB | 3 | 13.05 | 0.278 |
| 1.6 | 0.05 | ycaM | 3 | 13.05 | 0.226 |
| 1.6 | 0.05 | yadI | 3 | 13.05 | 0.472 |
| 1.6 | 0.05 | AG1 | 3 | 13.05 | 0.411 |
| 1.6 | 0.05 | ptsI | 3 | 13.3 | 0.368 |
| 1.6 | 0.05 | clcB | 3 | 13.3 | 0.276 |
| 1.6 | 0.05 | ycaM | 3 | 13.3 | 0.223 |
| 1.6 | 0.05 | yadI | 3 | 13.3 | 0.476 |
| 1.6 | 0.05 | AG1 | 3 | 13.3 | 0.42 |
| 1.6 | 0.05 | ptsI | 3 | 13.55 | 0.367 |
| 1.6 | 0.05 | clcB | 3 | 13.55 | 0.276 |
| 1.6 | 0.05 | ycaM | 3 | 13.55 | 0.221 |
| 1.6 | 0.05 | yadI | 3 | 13.55 | 0.48 |
| 1.6 | 0.05 | AG1 | 3 | 13.55 | 0.425 |
| 1.6 | 0.05 | ptsI | 3 | 13.8 | 0.378 |
| 1.6 | 0.05 | clcB | 3 | 13.8 | 0.274 |
| 1.6 | 0.05 | ycaM | 3 | 13.8 | 0.224 |
| 1.6 | 0.05 | yadI | 3 | 13.8 | 0.484 |
| 1.6 | 0.05 | AG1 | 3 | 13.8 | 0.427 |
| 1.6 | 0.05 | ptsI | 3 | 14.05 | 0.382 |
| 1.6 | 0.05 | clcB | 3 | 14.05 | 0.276 |
| 1.6 | 0.05 | ycaM | 3 | 14.05 | 0.222 |
| 1.6 | 0.05 | yadI | 3 | 14.05 | 0.487 |
| 1.6 | 0.05 | AG1 | 3 | 14.05 | 0.436 |
| 1.6 | 0.05 | ptsI | 3 | 14.3 | 0.388 |
| 1.6 | 0.05 | clcB | 3 | 14.3 | 0.275 |
| 1.6 | 0.05 | ycaM | 3 | 14.3 | 0.221 |
| 1.6 | 0.05 | yadI | 3 | 14.3 | 0.492 |
| 1.6 | 0.05 | AG1 | 3 | 14.3 | 0.44 |
| 1.6 | 0.05 | ptsI | 3 | 14.55 | 0.39 |
| 1.6 | 0.05 | clcB | 3 | 14.55 | 0.274 |
| 1.6 | 0.05 | ycaM | 3 | 14.55 | 0.219 |
| 1.6 | 0.05 | yadI | 3 | 14.55 | 0.49 |

| | | | | | |
|---|---|---|---|---|---|
| 1.6 | 0.05 | AG1 | 3 | 14.55 | 0.445 |
| 1.6 | 0.05 | ptsI | 3 | 14.8 | 0.394 |
| 1.6 | 0.05 | clcB | 3 | 14.8 | 0.272 |
| 1.6 | 0.05 | ycaM | 3 | 14.8 | 0.22 |
| 1.6 | 0.05 | yadI | 3 | 14.8 | 0.494 |
| 1.6 | 0.05 | AG1 | 3 | 14.8 | 0.45 |
| 1.6 | 0.05 | ptsI | 3 | 15.05 | 0.402 |
| 1.6 | 0.05 | clcB | 3 | 15.05 | 0.272 |
| 1.6 | 0.05 | ycaM | 3 | 15.05 | 0.219 |
| 1.6 | 0.05 | yadI | 3 | 15.05 | 0.497 |
| 1.6 | 0.05 | AG1 | 3 | 15.05 | 0.454 |
| 1.6 | 0.05 | ptsI | 3 | 15.3 | 0.405 |
| 1.6 | 0.05 | clcB | 3 | 15.3 | 0.272 |
| 1.6 | 0.05 | ycaM | 3 | 15.3 | 0.218 |
| 1.6 | 0.05 | yadI | 3 | 15.3 | 0.497 |
| 1.6 | 0.05 | AG1 | 3 | 15.3 | 0.46 |
| 1.6 | 0.05 | ptsI | 3 | 15.55 | 0.414 |
| 1.6 | 0.05 | clcB | 3 | 15.55 | 0.272 |
| 1.6 | 0.05 | ycaM | 3 | 15.55 | 0.217 |
| 1.6 | 0.05 | yadI | 3 | 15.55 | 0.501 |
| 1.6 | 0.05 | AG1 | 3 | 15.55 | 0.467 |
| 1.6 | 0.05 | ptsI | 3 | 15.8 | 0.418 |
| 1.6 | 0.05 | clcB | 3 | 15.8 | 0.271 |
| 1.6 | 0.05 | ycaM | 3 | 15.8 | 0.217 |
| 1.6 | 0.05 | yadI | 3 | 15.8 | 0.498 |
| 1.6 | 0.05 | AG1 | 3 | 15.8 | 0.471 |
| 1.6 | 0.05 | ptsI | 3 | 16.05 | 0.427 |
| 1.6 | 0.05 | clcB | 3 | 16.05 | 0.271 |
| 1.6 | 0.05 | ycaM | 3 | 16.05 | 0.215 |
| 1.6 | 0.05 | yadI | 3 | 16.05 | 0.5 |
| 1.6 | 0.05 | AG1 | 3 | 16.05 | 0.475 |
| 1.6 | 0.05 | ptsI | 3 | 16.3 | 0.43 |
| 1.6 | 0.05 | clcB | 3 | 16.3 | 0.269 |
| 1.6 | 0.05 | ycaM | 3 | 16.3 | 0.214 |
| 1.6 | 0.05 | yadI | 3 | 16.3 | 0.498 |
| 1.6 | 0.05 | AG1 | 3 | 16.3 | 0.48 |
| 1.6 | 0.05 | ptsI | 3 | 16.55 | 0.432 |
| 1.6 | 0.05 | clcB | 3 | 16.55 | 0.27 |
| 1.6 | 0.05 | ycaM | 3 | 16.55 | 0.216 |
| 1.6 | 0.05 | yadI | 3 | 16.55 | 0.497 |
| 1.6 | 0.05 | AG1 | 3 | 16.55 | 0.485 |
| 1.6 | 0.05 | ptsI | 3 | 16.8 | 0.438 |
| 1.6 | 0.05 | clcB | 3 | 16.8 | 0.269 |
| 1.6 | 0.05 | ycaM | 3 | 16.8 | 0.214 |
| 1.6 | 0.05 | yadI | 3 | 16.8 | 0.496 |
| 1.6 | 0.05 | AG1 | 3 | 16.8 | 0.491 |
| 1.6 | 0.05 | ptsI | 3 | 17.05 | 0.438 |
| 1.6 | 0.05 | clcB | 3 | 17.05 | 0.268 |

| | | | | | |
|---|---|---|---|---|---|
| 1.6 | 0.05 | ycaM | 3 | 17.05 | 0.214 |
| 1.6 | 0.05 | yadI | 3 | 17.05 | 0.501 |
| 1.6 | 0.05 | AG1 | 3 | 17.05 | 0.495 |
| 1.6 | 0.05 | ptsI | 3 | 17.3 | 0.444 |
| 1.6 | 0.05 | clcB | 3 | 17.3 | 0.265 |
| 1.6 | 0.05 | ycaM | 3 | 17.3 | 0.214 |
| 1.6 | 0.05 | yadI | 3 | 17.3 | 0.502 |
| 1.6 | 0.05 | AG1 | 3 | 17.3 | 0.502 |
| 1.6 | 0.05 | ptsI | 3 | 17.55 | 0.446 |
| 1.6 | 0.05 | clcB | 3 | 17.55 | 0.266 |
| 1.6 | 0.05 | ycaM | 3 | 17.55 | 0.214 |
| 1.6 | 0.05 | yadI | 3 | 17.55 | 0.504 |
| 1.6 | 0.05 | AG1 | 3 | 17.55 | 0.512 |
| 1.6 | 0.05 | ptsI | 3 | 17.8 | 0.45 |
| 1.6 | 0.05 | clcB | 3 | 17.8 | 0.266 |
| 1.6 | 0.05 | ycaM | 3 | 17.8 | 0.215 |
| 1.6 | 0.05 | yadI | 3 | 17.8 | 0.505 |
| 1.6 | 0.05 | AG1 | 3 | 17.8 | 0.522 |
| 1.6 | 0.05 | ptsI | 3 | 18.05 | 0.45 |
| 1.6 | 0.05 | clcB | 3 | 18.05 | 0.266 |
| 1.6 | 0.05 | ycaM | 3 | 18.05 | 0.214 |
| 1.6 | 0.05 | yadI | 3 | 18.05 | 0.506 |
| 1.6 | 0.05 | AG1 | 3 | 18.05 | 0.534 |
| 1.6 | 0.05 | ptsI | 3 | 18.3 | 0.454 |
| 1.6 | 0.05 | clcB | 3 | 18.3 | 0.267 |
| 1.6 | 0.05 | ycaM | 3 | 18.3 | 0.213 |
| 1.6 | 0.05 | yadI | 3 | 18.3 | 0.508 |
| 1.6 | 0.05 | AG1 | 3 | 18.3 | 0.554 |
| 1.6 | 0.05 | ptsI | 3 | 18.55 | 0.458 |
| 1.6 | 0.05 | clcB | 3 | 18.55 | 0.267 |
| 1.6 | 0.05 | ycaM | 3 | 18.55 | 0.214 |
| 1.6 | 0.05 | yadI | 3 | 18.55 | 0.511 |
| 1.6 | 0.05 | AG1 | 3 | 18.55 | 0.551 |
| 1.6 | 0.05 | ptsI | 3 | 18.8 | 0.461 |
| 1.6 | 0.05 | clcB | 3 | 18.8 | 0.266 |
| 1.6 | 0.05 | ycaM | 3 | 18.8 | 0.212 |
| 1.6 | 0.05 | yadI | 3 | 18.8 | 0.512 |
| 1.6 | 0.05 | AG1 | 3 | 18.8 | 0.578 |
| 1.6 | 0.05 | ptsI | 3 | 19.05 | 0.462 |
| 1.6 | 0.05 | clcB | 3 | 19.05 | 0.266 |
| 1.6 | 0.05 | ycaM | 3 | 19.05 | 0.213 |
| 1.6 | 0.05 | yadI | 3 | 19.05 | 0.514 |
| 1.6 | 0.05 | AG1 | 3 | 19.05 | 0.574 |
| 1.6 | 0.05 | ptsI | 3 | 19.3 | 0.459 |
| 1.6 | 0.05 | clcB | 3 | 19.3 | 0.265 |
| 1.6 | 0.05 | ycaM | 3 | 19.3 | 0.213 |
| 1.6 | 0.05 | yadI | 3 | 19.3 | 0.516 |
| 1.6 | 0.05 | AG1 | 3 | 19.3 | 0.61 |

| | | | | | |
|---|---|---|---|---|---|
| 1.6 | 0.05 | ptsI | 3 | 19.55 | 0.467 |
| 1.6 | 0.05 | clcB | 3 | 19.55 | 0.265 |
| 1.6 | 0.05 | ycaM | 3 | 19.55 | 0.212 |
| 1.6 | 0.05 | yadI | 3 | 19.55 | 0.518 |
| 1.6 | 0.05 | AG1 | 3 | 19.55 | 0.644 |
| 1.6 | 0.05 | ptsI | 3 | 19.8 | 0.469 |
| 1.6 | 0.05 | clcB | 3 | 19.8 | 0.265 |
| 1.6 | 0.05 | ycaM | 3 | 19.8 | 0.212 |
| 1.6 | 0.05 | yadI | 3 | 19.8 | 0.517 |
| 1.6 | 0.05 | AG1 | 3 | 19.8 | 0.675 |
| 1.6 | 0.05 | ptsI | 3 | 20.05 | 0.472 |
| 1.6 | 0.05 | clcB | 3 | 20.05 | 0.265 |
| 1.6 | 0.05 | ycaM | 3 | 20.05 | 0.213 |
| 1.6 | 0.05 | yadI | 3 | 20.05 | 0.523 |
| 1.6 | 0.05 | AG1 | 3 | 20.05 | 0.662 |
| 1.6 | 0.05 | ptsI | 3 | 20.3 | 0.474 |
| 1.6 | 0.05 | clcB | 3 | 20.3 | 0.264 |
| 1.6 | 0.05 | ycaM | 3 | 20.3 | 0.212 |
| 1.6 | 0.05 | yadI | 3 | 20.3 | 0.524 |
| 1.6 | 0.05 | AG1 | 3 | 20.3 | 0.676 |
| 1.6 | 0.05 | ptsI | 3 | 20.55 | 0.477 |
| 1.6 | 0.05 | clcB | 3 | 20.55 | 0.264 |
| 1.6 | 0.05 | ycaM | 3 | 20.55 | 0.212 |
| 1.6 | 0.05 | yadI | 3 | 20.55 | 0.527 |
| 1.6 | 0.05 | AG1 | 3 | 20.55 | 0.656 |
| 1.6 | 0.05 | ptsI | 3 | 20.8 | 0.48 |
| 1.6 | 0.05 | clcB | 3 | 20.8 | 0.264 |
| 1.6 | 0.05 | ycaM | 3 | 20.8 | 0.212 |
| 1.6 | 0.05 | yadI | 3 | 20.8 | 0.526 |
| 1.6 | 0.05 | AG1 | 3 | 20.8 | 0.654 |
| 1.6 | 0.05 | ptsI | 3 | 21.05 | 0.485 |
| 1.6 | 0.05 | clcB | 3 | 21.05 | 0.263 |
| 1.6 | 0.05 | ycaM | 3 | 21.05 | 0.212 |
| 1.6 | 0.05 | yadI | 3 | 21.05 | 0.528 |
| 1.6 | 0.05 | AG1 | 3 | 21.05 | 0.712 |
| 1.6 | 0.05 | ptsI | 3 | 21.3 | 0.488 |
| 1.6 | 0.05 | clcB | 3 | 21.3 | 0.263 |
| 1.6 | 0.05 | ycaM | 3 | 21.3 | 0.211 |
| 1.6 | 0.05 | yadI | 3 | 21.3 | 0.531 |
| 1.6 | 0.05 | AG1 | 3 | 21.3 | 0.718 |
| 1.6 | 0.05 | ptsI | 3 | 21.55 | 0.49 |
| 1.6 | 0.05 | clcB | 3 | 21.55 | 0.264 |
| 1.6 | 0.05 | ycaM | 3 | 21.55 | 0.212 |
| 1.6 | 0.05 | yadI | 3 | 21.55 | 0.531 |
| 1.6 | 0.05 | AG1 | 3 | 21.55 | 0.749 |
| 1.6 | 0.05 | ptsI | 3 | 21.8 | 0.493 |
| 1.6 | 0.05 | clcB | 3 | 21.8 | 0.263 |
| 1.6 | 0.05 | ycaM | 3 | 21.8 | 0.212 |

| | | | | | |
|---|---|---|---|---|---|
| 1.6 | 0.05 | yadI | 3 | 21.8 | 0.536 |
| 1.6 | 0.05 | AG1 | 3 | 21.8 | 0.759 |
| 1.6 | 0.05 | ptsI | 3 | 22.05 | 0.495 |
| 1.6 | 0.05 | clcB | 3 | 22.05 | 0.264 |
| 1.6 | 0.05 | ycaM | 3 | 22.05 | 0.213 |
| 1.6 | 0.05 | yadI | 3 | 22.05 | 0.537 |
| 1.6 | 0.05 | AG1 | 3 | 22.05 | 0.76 |
| 1.6 | 0.05 | ptsI | 3 | 22.3 | 0.5 |
| 1.6 | 0.05 | clcB | 3 | 22.3 | 0.264 |
| 1.6 | 0.05 | ycaM | 3 | 22.3 | 0.212 |
| 1.6 | 0.05 | yadI | 3 | 22.3 | 0.539 |
| 1.6 | 0.05 | AG1 | 3 | 22.3 | 0.762 |
| 1.6 | 0.05 | ptsI | 3 | 22.55 | 0.502 |
| 1.6 | 0.05 | clcB | 3 | 22.55 | 0.265 |
| 1.6 | 0.05 | ycaM | 3 | 22.55 | 0.212 |
| 1.6 | 0.05 | yadI | 3 | 22.55 | 0.542 |
| 1.6 | 0.05 | AG1 | 3 | 22.55 | 0.771 |
| 1.6 | 0.05 | ptsI | 3 | 22.8 | 0.502 |
| 1.6 | 0.05 | clcB | 3 | 22.8 | 0.265 |
| 1.6 | 0.05 | ycaM | 3 | 22.8 | 0.212 |
| 1.6 | 0.05 | yadI | 3 | 22.8 | 0.543 |
| 1.6 | 0.05 | AG1 | 3 | 22.8 | 0.777 |
| 1.6 | 0.05 | ptsI | 3 | 23.05 | 0.502 |
| 1.6 | 0.05 | clcB | 3 | 23.05 | 0.264 |
| 1.6 | 0.05 | ycaM | 3 | 23.05 | 0.212 |
| 1.6 | 0.05 | yadI | 3 | 23.05 | 0.545 |
| 1.6 | 0.05 | AG1 | 3 | 23.05 | 0.776 |
| 1.6 | 0.05 | ptsI | 3 | 23.3 | 0.502 |
| 1.6 | 0.05 | clcB | 3 | 23.3 | 0.265 |
| 1.6 | 0.05 | ycaM | 3 | 23.3 | 0.213 |
| 1.6 | 0.05 | yadI | 3 | 23.3 | 0.547 |
| 1.6 | 0.05 | AG1 | 3 | 23.3 | 0.779 |
| 1.6 | 0.05 | ptsI | 3 | 23.55 | 0.503 |
| 1.6 | 0.05 | clcB | 3 | 23.55 | 0.264 |
| 1.6 | 0.05 | ycaM | 3 | 23.55 | 0.212 |
| 1.6 | 0.05 | yadI | 3 | 23.55 | 0.545 |
| 1.6 | 0.05 | AG1 | 3 | 23.55 | 0.762 |
| 1.6 | 0.05 | ptsI | 3 | 23.8 | 0.505 |
| 1.6 | 0.05 | clcB | 3 | 23.8 | 0.266 |
| 1.6 | 0.05 | ycaM | 3 | 23.8 | 0.212 |
| 1.6 | 0.05 | yadI | 3 | 23.8 | 0.548 |
| 1.6 | 0.05 | AG1 | 3 | 23.8 | 0.776 |
| 1.6 | 0.05 | ptsI | 3 | 24.05 | 0.512 |
| 1.6 | 0.05 | clcB | 3 | 24.05 | 0.267 |
| 1.6 | 0.05 | ycaM | 3 | 24.05 | 0.212 |
| 1.6 | 0.05 | yadI | 3 | 24.05 | 0.552 |
| 1.6 | 0.05 | AG1 | 3 | 24.05 | 0.774 |
| 1.6 | 0.05 | ptsI | 3 | 24.3 | 0.513 |

| | | | | | |
|---|---|---|---|---|---|
| 1.6 | 0.05 | clcB | 3 | 24.3 | 0.266 |
| 1.6 | 0.05 | ycaM | 3 | 24.3 | 0.211 |
| 1.6 | 0.05 | yadI | 3 | 24.3 | 0.554 |
| 1.6 | 0.05 | AG1 | 3 | 24.3 | 0.795 |
| 1.6 | 0.05 | ptsI | 4 | 0 | 0.211 |
| 1.6 | 0.05 | clcB | 4 | 0 | 0.191 |
| 1.6 | 0.05 | ycaM | 4 | 0 | 0.214 |
| 1.6 | 0.05 | yadI | 4 | 0 | 0.2 |
| 1.6 | 0.05 | AG1 | 4 | 0 | 0.211 |
| 1.6 | 0.05 | ptsI | 4 | 0.25 | 0.205 |
| 1.6 | 0.05 | clcB | 4 | 0.25 | 0.188 |
| 1.6 | 0.05 | ycaM | 4 | 0.25 | 0.205 |
| 1.6 | 0.05 | yadI | 4 | 0.25 | 0.205 |
| 1.6 | 0.05 | AG1 | 4 | 0.25 | 0.205 |
| 1.6 | 0.05 | ptsI | 4 | 0.5 | 0.207 |
| 1.6 | 0.05 | clcB | 4 | 0.5 | 0.19 |
| 1.6 | 0.05 | ycaM | 4 | 0.5 | 0.206 |
| 1.6 | 0.05 | yadI | 4 | 0.5 | 0.206 |
| 1.6 | 0.05 | AG1 | 4 | 0.5 | 0.205 |
| 1.6 | 0.05 | ptsI | 4 | 0.75 | 0.208 |
| 1.6 | 0.05 | clcB | 4 | 0.75 | 0.191 |
| 1.6 | 0.05 | ycaM | 4 | 0.75 | 0.207 |
| 1.6 | 0.05 | yadI | 4 | 0.75 | 0.209 |
| 1.6 | 0.05 | AG1 | 4 | 0.75 | 0.21 |
| 1.6 | 0.05 | ptsI | 4 | 1 | 0.214 |
| 1.6 | 0.05 | clcB | 4 | 1 | 0.195 |
| 1.6 | 0.05 | ycaM | 4 | 1 | 0.211 |
| 1.6 | 0.05 | yadI | 4 | 1 | 0.21 |
| 1.6 | 0.05 | AG1 | 4 | 1 | 0.216 |
| 1.6 | 0.05 | ptsI | 4 | 1.25 | 0.221 |
| 1.6 | 0.05 | clcB | 4 | 1.25 | 0.201 |
| 1.6 | 0.05 | ycaM | 4 | 1.25 | 0.217 |
| 1.6 | 0.05 | yadI | 4 | 1.25 | 0.214 |
| 1.6 | 0.05 | AG1 | 4 | 1.25 | 0.225 |
| 1.6 | 0.05 | ptsI | 4 | 1.5 | 0.222 |
| 1.6 | 0.05 | clcB | 4 | 1.5 | 0.197 |
| 1.6 | 0.05 | ycaM | 4 | 1.5 | 0.216 |
| 1.6 | 0.05 | yadI | 4 | 1.5 | 0.213 |
| 1.6 | 0.05 | AG1 | 4 | 1.5 | 0.22 |
| 1.6 | 0.05 | ptsI | 4 | 1.75 | 0.229 |
| 1.6 | 0.05 | clcB | 4 | 1.75 | 0.205 |
| 1.6 | 0.05 | ycaM | 4 | 1.75 | 0.225 |
| 1.6 | 0.05 | yadI | 4 | 1.75 | 0.217 |
| 1.6 | 0.05 | AG1 | 4 | 1.75 | 0.226 |
| 1.6 | 0.05 | ptsI | 4 | 2 | 0.24 |
| 1.6 | 0.05 | clcB | 4 | 2 | 0.212 |
| 1.6 | 0.05 | ycaM | 4 | 2 | 0.232 |
| 1.6 | 0.05 | yadI | 4 | 2 | 0.226 |

| | | | | | |
|---|---|---|---|---|---|
| 1.6 | 0.05 | AG1 | 4 | 2 | 0.236 |
| 1.6 | 0.05 | ptsI | 4 | 2.25 | 0.25 |
| 1.6 | 0.05 | clcB | 4 | 2.25 | 0.222 |
| 1.6 | 0.05 | ycaM | 4 | 2.25 | 0.244 |
| 1.6 | 0.05 | yadI | 4 | 2.25 | 0.234 |
| 1.6 | 0.05 | AG1 | 4 | 2.25 | 0.247 |
| 1.6 | 0.05 | ptsI | 4 | 2.5 | 0.252 |
| 1.6 | 0.05 | clcB | 4 | 2.5 | 0.221 |
| 1.6 | 0.05 | ycaM | 4 | 2.5 | 0.245 |
| 1.6 | 0.05 | yadI | 4 | 2.5 | 0.236 |
| 1.6 | 0.05 | AG1 | 4 | 2.5 | 0.242 |
| 1.6 | 0.05 | ptsI | 4 | 2.75 | 0.257 |
| 1.6 | 0.05 | clcB | 4 | 2.75 | 0.228 |
| 1.6 | 0.05 | ycaM | 4 | 2.75 | 0.252 |
| 1.6 | 0.05 | yadI | 4 | 2.75 | 0.247 |
| 1.6 | 0.05 | AG1 | 4 | 2.75 | 0.249 |
| 1.6 | 0.05 | ptsI | 4 | 3 | 0.27 |
| 1.6 | 0.05 | clcB | 4 | 3 | 0.243 |
| 1.6 | 0.05 | ycaM | 4 | 3 | 0.264 |
| 1.6 | 0.05 | yadI | 4 | 3 | 0.255 |
| 1.6 | 0.05 | AG1 | 4 | 3 | 0.26 |
| 1.6 | 0.05 | ptsI | 4 | 3.25 | 0.283 |
| 1.6 | 0.05 | clcB | 4 | 3.25 | 0.25 |
| 1.6 | 0.05 | ycaM | 4 | 3.25 | 0.272 |
| 1.6 | 0.05 | yadI | 4 | 3.25 | 0.263 |
| 1.6 | 0.05 | AG1 | 4 | 3.25 | 0.268 |
| 1.6 | 0.05 | ptsI | 4 | 3.5 | 0.299 |
| 1.6 | 0.05 | clcB | 4 | 3.5 | 0.268 |
| 1.6 | 0.05 | ycaM | 4 | 3.5 | 0.285 |
| 1.6 | 0.05 | yadI | 4 | 3.5 | 0.273 |
| 1.6 | 0.05 | AG1 | 4 | 3.5 | 0.283 |
| 1.6 | 0.05 | ptsI | 4 | 3.75 | 0.311 |
| 1.6 | 0.05 | clcB | 4 | 3.75 | 0.272 |
| 1.6 | 0.05 | ycaM | 4 | 3.75 | 0.296 |
| 1.6 | 0.05 | yadI | 4 | 3.75 | 0.284 |
| 1.6 | 0.05 | AG1 | 4 | 3.75 | 0.289 |
| 1.6 | 0.05 | ptsI | 4 | 4 | 0.329 |
| 1.6 | 0.05 | clcB | 4 | 4 | 0.301 |
| 1.6 | 0.05 | ycaM | 4 | 4 | 0.315 |
| 1.6 | 0.05 | yadI | 4 | 4 | 0.293 |
| 1.6 | 0.05 | AG1 | 4 | 4 | 0.305 |
| 1.6 | 0.05 | ptsI | 4 | 4.35 | 0.346 |
| 1.6 | 0.05 | clcB | 4 | 4.35 | 0.355 |
| 1.6 | 0.05 | ycaM | 4 | 4.35 | 0.346 |
| 1.6 | 0.05 | yadI | 4 | 4.35 | 0.34 |
| 1.6 | 0.05 | AG1 | 4 | 4.35 | 0.28 |
| 1.6 | 0.05 | ptsI | 4 | 4.6 | 0.339 |
| 1.6 | 0.05 | clcB | 4 | 4.6 | 0.352 |

| | | | | | |
|---|---|---|---|---|---|
| 1.6 | 0.05 | ycaM | 4 | 4.6 | 0.34 |
| 1.6 | 0.05 | yadI | 4 | 4.6 | 0.349 |
| 1.6 | 0.05 | AG1 | 4 | 4.6 | 0.292 |
| 1.6 | 0.05 | ptsI | 4 | 4.85 | 0.35 |
| 1.6 | 0.05 | clcB | 4 | 4.85 | 0.358 |
| 1.6 | 0.05 | ycaM | 4 | 4.85 | 0.336 |
| 1.6 | 0.05 | yadI | 4 | 4.85 | 0.358 |
| 1.6 | 0.05 | AG1 | 4 | 4.85 | 0.304 |
| 1.6 | 0.05 | ptsI | 4 | 5.1 | 0.36 |
| 1.6 | 0.05 | clcB | 4 | 5.1 | 0.358 |
| 1.6 | 0.05 | ycaM | 4 | 5.1 | 0.337 |
| 1.6 | 0.05 | yadI | 4 | 5.1 | 0.37 |
| 1.6 | 0.05 | AG1 | 4 | 5.1 | 0.313 |
| 1.6 | 0.05 | ptsI | 4 | 5.35 | 0.373 |
| 1.6 | 0.05 | clcB | 4 | 5.35 | 0.364 |
| 1.6 | 0.05 | ycaM | 4 | 5.35 | 0.34 |
| 1.6 | 0.05 | yadI | 4 | 5.35 | 0.389 |
| 1.6 | 0.05 | AG1 | 4 | 5.35 | 0.324 |
| 1.6 | 0.05 | ptsI | 4 | 5.6 | 0.38 |
| 1.6 | 0.05 | clcB | 4 | 5.6 | 0.355 |
| 1.6 | 0.05 | ycaM | 4 | 5.6 | 0.335 |
| 1.6 | 0.05 | yadI | 4 | 5.6 | 0.393 |
| 1.6 | 0.05 | AG1 | 4 | 5.6 | 0.329 |
| 1.6 | 0.05 | ptsI | 4 | 5.85 | 0.387 |
| 1.6 | 0.05 | clcB | 4 | 5.85 | 0.336 |
| 1.6 | 0.05 | ycaM | 4 | 5.85 | 0.331 |
| 1.6 | 0.05 | yadI | 4 | 5.85 | 0.391 |
| 1.6 | 0.05 | AG1 | 4 | 5.85 | 0.333 |
| 1.6 | 0.05 | ptsI | 4 | 6.1 | 0.401 |
| 1.6 | 0.05 | clcB | 4 | 6.1 | 0.341 |
| 1.6 | 0.05 | ycaM | 4 | 6.1 | 0.337 |
| 1.6 | 0.05 | yadI | 4 | 6.1 | 0.407 |
| 1.6 | 0.05 | AG1 | 4 | 6.1 | 0.342 |
| 1.6 | 0.05 | ptsI | 4 | 6.35 | 0.41 |
| 1.6 | 0.05 | clcB | 4 | 6.35 | 0.35 |
| 1.6 | 0.05 | ycaM | 4 | 6.35 | 0.338 |
| 1.6 | 0.05 | yadI | 4 | 6.35 | 0.418 |
| 1.6 | 0.05 | AG1 | 4 | 6.35 | 0.353 |
| 1.6 | 0.05 | ptsI | 4 | 6.6 | 0.419 |
| 1.6 | 0.05 | clcB | 4 | 6.6 | 0.355 |
| 1.6 | 0.05 | ycaM | 4 | 6.6 | 0.336 |
| 1.6 | 0.05 | yadI | 4 | 6.6 | 0.425 |
| 1.6 | 0.05 | AG1 | 4 | 6.6 | 0.361 |
| 1.6 | 0.05 | ptsI | 4 | 6.85 | 0.428 |
| 1.6 | 0.05 | clcB | 4 | 6.85 | 0.361 |
| 1.6 | 0.05 | ycaM | 4 | 6.85 | 0.34 |
| 1.6 | 0.05 | yadI | 4 | 6.85 | 0.434 |
| 1.6 | 0.05 | AG1 | 4 | 6.85 | 0.371 |

| | | | | | |
|---|---|---|---|---|---|
| 1.6 | 0.05 | ptsI | 4 | 7.1 | 0.436 |
| 1.6 | 0.05 | clcB | 4 | 7.1 | 0.365 |
| 1.6 | 0.05 | ycaM | 4 | 7.1 | 0.341 |
| 1.6 | 0.05 | yadI | 4 | 7.1 | 0.439 |
| 1.6 | 0.05 | AG1 | 4 | 7.1 | 0.38 |
| 1.6 | 0.05 | ptsI | 4 | 7.35 | 0.447 |
| 1.6 | 0.05 | clcB | 4 | 7.35 | 0.37 |
| 1.6 | 0.05 | ycaM | 4 | 7.35 | 0.344 |
| 1.6 | 0.05 | yadI | 4 | 7.35 | 0.447 |
| 1.6 | 0.05 | AG1 | 4 | 7.35 | 0.386 |
| 1.6 | 0.05 | ptsI | 4 | 7.6 | 0.456 |
| 1.6 | 0.05 | clcB | 4 | 7.6 | 0.375 |
| 1.6 | 0.05 | ycaM | 4 | 7.6 | 0.352 |
| 1.6 | 0.05 | yadI | 4 | 7.6 | 0.454 |
| 1.6 | 0.05 | AG1 | 4 | 7.6 | 0.395 |
| 1.6 | 0.05 | ptsI | 4 | 7.85 | 0.456 |
| 1.6 | 0.05 | clcB | 4 | 7.85 | 0.361 |
| 1.6 | 0.05 | ycaM | 4 | 7.85 | 0.336 |
| 1.6 | 0.05 | yadI | 4 | 7.85 | 0.456 |
| 1.6 | 0.05 | AG1 | 4 | 7.85 | 0.394 |
| 1.6 | 0.05 | ptsI | 4 | 8.1 | 0.469 |
| 1.6 | 0.05 | clcB | 4 | 8.1 | 0.366 |
| 1.6 | 0.05 | ycaM | 4 | 8.1 | 0.34 |
| 1.6 | 0.05 | yadI | 4 | 8.1 | 0.463 |
| 1.6 | 0.05 | AG1 | 4 | 8.1 | 0.403 |
| 1.6 | 0.05 | ptsI | 4 | 8.35 | 0.476 |
| 1.6 | 0.05 | clcB | 4 | 8.35 | 0.371 |
| 1.6 | 0.05 | ycaM | 4 | 8.35 | 0.343 |
| 1.6 | 0.05 | yadI | 4 | 8.35 | 0.47 |
| 1.6 | 0.05 | AG1 | 4 | 8.35 | 0.412 |
| 1.6 | 0.05 | ptsI | 4 | 8.83333 | 0.428 |
| 1.6 | 0.05 | clcB | 4 | 8.83333 | 0.295 |
| 1.6 | 0.05 | ycaM | 4 | 8.83333 | 0.28 |
| 1.6 | 0.05 | yadI | 4 | 8.83333 | 0.46 |
| 1.6 | 0.05 | AG1 | 4 | 8.83333 | 0.45 |
| 1.6 | 0.05 | ptsI | 4 | 9.08333 | 0.427 |
| 1.6 | 0.05 | clcB | 4 | 9.08333 | 0.292 |
| 1.6 | 0.05 | ycaM | 4 | 9.08333 | 0.277 |
| 1.6 | 0.05 | yadI | 4 | 9.08333 | 0.459 |
| 1.6 | 0.05 | AG1 | 4 | 9.08333 | 0.456 |
| 1.6 | 0.05 | ptsI | 4 | 9.33333 | 0.436 |
| 1.6 | 0.05 | clcB | 4 | 9.33333 | 0.29 |
| 1.6 | 0.05 | ycaM | 4 | 9.33333 | 0.278 |
| 1.6 | 0.05 | yadI | 4 | 9.33333 | 0.465 |
| 1.6 | 0.05 | AG1 | 4 | 9.33333 | 0.467 |
| 1.6 | 0.05 | ptsI | 4 | 9.58333 | 0.444 |
| 1.6 | 0.05 | clcB | 4 | 9.58333 | 0.292 |
| 1.6 | 0.05 | ycaM | 4 | 9.58333 | 0.281 |

| | | | | | |
|---|---|---|---|---|---|
| 1.6 | 0.05 | yadI | 4 | 9.58333 | 0.475 |
| 1.6 | 0.05 | AG1 | 4 | 9.58333 | 0.478 |
| 1.6 | 0.05 | ptsI | 4 | 9.83333 | 0.449 |
| 1.6 | 0.05 | clcB | 4 | 9.83333 | 0.294 |
| 1.6 | 0.05 | ycaM | 4 | 9.83333 | 0.28 |
| 1.6 | 0.05 | yadI | 4 | 9.83333 | 0.485 |
| 1.6 | 0.05 | AG1 | 4 | 9.83333 | 0.488 |
| 1.6 | 0.05 | ptsI | 4 | 10.0833 | 0.454 |
| 1.6 | 0.05 | clcB | 4 | 10.0833 | 0.293 |
| 1.6 | 0.05 | ycaM | 4 | 10.0833 | 0.283 |
| 1.6 | 0.05 | yadI | 4 | 10.0833 | 0.491 |
| 1.6 | 0.05 | AG1 | 4 | 10.0833 | 0.498 |
| 1.6 | 0.05 | ptsI | 4 | 10.3333 | 0.458 |
| 1.6 | 0.05 | clcB | 4 | 10.3333 | 0.294 |
| 1.6 | 0.05 | ycaM | 4 | 10.3333 | 0.285 |
| 1.6 | 0.05 | yadI | 4 | 10.3333 | 0.496 |
| 1.6 | 0.05 | AG1 | 4 | 10.3333 | 0.503 |
| 1.6 | 0.05 | ptsI | 4 | 10.5833 | 0.464 |
| 1.6 | 0.05 | clcB | 4 | 10.5833 | 0.295 |
| 1.6 | 0.05 | ycaM | 4 | 10.5833 | 0.286 |
| 1.6 | 0.05 | yadI | 4 | 10.5833 | 0.5 |
| 1.6 | 0.05 | AG1 | 4 | 10.5833 | 0.509 |
| 1.6 | 0.05 | ptsI | 4 | 10.8333 | 0.471 |
| 1.6 | 0.05 | clcB | 4 | 10.8333 | 0.294 |
| 1.6 | 0.05 | ycaM | 4 | 10.8333 | 0.285 |
| 1.6 | 0.05 | yadI | 4 | 10.8333 | 0.505 |
| 1.6 | 0.05 | AG1 | 4 | 10.8333 | 0.512 |
| 1.6 | 0.05 | ptsI | 4 | 11.0833 | 0.475 |
| 1.6 | 0.05 | clcB | 4 | 11.0833 | 0.295 |
| 1.6 | 0.05 | ycaM | 4 | 11.0833 | 0.286 |
| 1.6 | 0.05 | yadI | 4 | 11.0833 | 0.508 |
| 1.6 | 0.05 | AG1 | 4 | 11.0833 | 0.514 |
| 1.6 | 0.05 | ptsI | 4 | 11.3333 | 0.479 |
| 1.6 | 0.05 | clcB | 4 | 11.3333 | 0.294 |
| 1.6 | 0.05 | ycaM | 4 | 11.3333 | 0.285 |
| 1.6 | 0.05 | yadI | 4 | 11.3333 | 0.512 |
| 1.6 | 0.05 | AG1 | 4 | 11.3333 | 0.52 |
| 1.6 | 0.05 | ptsI | 4 | 11.5833 | 0.481 |
| 1.6 | 0.05 | clcB | 4 | 11.5833 | 0.293 |
| 1.6 | 0.05 | ycaM | 4 | 11.5833 | 0.285 |
| 1.6 | 0.05 | yadI | 4 | 11.5833 | 0.514 |
| 1.6 | 0.05 | AG1 | 4 | 11.5833 | 0.525 |
| 1.6 | 0.05 | ptsI | 4 | 11.8333 | 0.482 |
| 1.6 | 0.05 | clcB | 4 | 11.8333 | 0.292 |
| 1.6 | 0.05 | ycaM | 4 | 11.8333 | 0.283 |
| 1.6 | 0.05 | yadI | 4 | 11.8333 | 0.516 |
| 1.6 | 0.05 | AG1 | 4 | 11.8333 | 0.522 |
| 1.6 | 0.05 | ptsI | 4 | 12.0833 | 0.483 |

| | | | | | |
|---|---|---|---|---|---|
| 1.6 | 0.05 | clcB | 4 | 12.0833 | 0.292 |
| 1.6 | 0.05 | ycaM | 4 | 12.0833 | 0.281 |
| 1.6 | 0.05 | yadI | 4 | 12.0833 | 0.519 |
| 1.6 | 0.05 | AG1 | 4 | 12.0833 | 0.532 |
| 1.6 | 0.05 | ptsI | 4 | 12.3333 | 0.482 |
| 1.6 | 0.05 | clcB | 4 | 12.3333 | 0.291 |
| 1.6 | 0.05 | ycaM | 4 | 12.3333 | 0.281 |
| 1.6 | 0.05 | yadI | 4 | 12.3333 | 0.521 |
| 1.6 | 0.05 | AG1 | 4 | 12.3333 | 0.532 |
| 1.6 | 0.05 | ptsI | 4 | 12.5833 | 0.483 |
| 1.6 | 0.05 | clcB | 4 | 12.5833 | 0.291 |
| 1.6 | 0.05 | ycaM | 4 | 12.5833 | 0.279 |
| 1.6 | 0.05 | yadI | 4 | 12.5833 | 0.527 |
| 1.6 | 0.05 | AG1 | 4 | 12.5833 | 0.532 |
| 1.6 | 0.05 | ptsI | 4 | 12.8333 | 0.486 |
| 1.6 | 0.05 | clcB | 4 | 12.8333 | 0.29 |
| 1.6 | 0.05 | ycaM | 4 | 12.8333 | 0.282 |
| 1.6 | 0.05 | yadI | 4 | 12.8333 | 0.528 |
| 1.6 | 0.05 | AG1 | 4 | 12.8333 | 0.541 |
| 1.6 | 0.05 | ptsI | 4 | 13.0833 | 0.488 |
| 1.6 | 0.05 | clcB | 4 | 13.0833 | 0.29 |
| 1.6 | 0.05 | ycaM | 4 | 13.0833 | 0.277 |
| 1.6 | 0.05 | yadI | 4 | 13.0833 | 0.532 |
| 1.6 | 0.05 | AG1 | 4 | 13.0833 | 0.544 |
| 1.6 | 0.05 | ptsI | 4 | 13.3333 | 0.491 |
| 1.6 | 0.05 | clcB | 4 | 13.3333 | 0.288 |
| 1.6 | 0.05 | ycaM | 4 | 13.3333 | 0.276 |
| 1.6 | 0.05 | yadI | 4 | 13.3333 | 0.534 |
| 1.6 | 0.05 | AG1 | 4 | 13.3333 | 0.541 |
| 1.6 | 0.05 | ptsI | 4 | 13.5833 | 0.494 |
| 1.6 | 0.05 | clcB | 4 | 13.5833 | 0.286 |
| 1.6 | 0.05 | ycaM | 4 | 13.5833 | 0.276 |
| 1.6 | 0.05 | yadI | 4 | 13.5833 | 0.532 |
| 1.6 | 0.05 | AG1 | 4 | 13.5833 | 0.545 |
| 1.6 | 0.05 | ptsI | 4 | 13.8333 | 0.494 |
| 1.6 | 0.05 | clcB | 4 | 13.8333 | 0.288 |
| 1.6 | 0.05 | ycaM | 4 | 13.8333 | 0.276 |
| 1.6 | 0.05 | yadI | 4 | 13.8333 | 0.537 |
| 1.6 | 0.05 | AG1 | 4 | 13.8333 | 0.55 |
| 1.6 | 0.05 | ptsI | 4 | 14.0833 | 0.495 |
| 1.6 | 0.05 | clcB | 4 | 14.0833 | 0.286 |
| 1.6 | 0.05 | ycaM | 4 | 14.0833 | 0.274 |
| 1.6 | 0.05 | yadI | 4 | 14.0833 | 0.538 |
| 1.6 | 0.05 | AG1 | 4 | 14.0833 | 0.554 |
| 1.6 | 0.05 | ptsI | 4 | 14.3333 | 0.5 |
| 1.6 | 0.05 | clcB | 4 | 14.3333 | 0.285 |
| 1.6 | 0.05 | ycaM | 4 | 14.3333 | 0.274 |
| 1.6 | 0.05 | yadI | 4 | 14.3333 | 0.536 |

| | | | | | |
|---|---|---|---|---|---|
| 1.6 | 0.05 | AG1 | 4 | 14.3333 | 0.552 |
| 1.6 | 0.05 | ptsI | 4 | 14.5833 | 0.504 |
| 1.6 | 0.05 | clcB | 4 | 14.5833 | 0.284 |
| 1.6 | 0.05 | ycaM | 4 | 14.5833 | 0.273 |
| 1.6 | 0.05 | yadI | 4 | 14.5833 | 0.541 |
| 1.6 | 0.05 | AG1 | 4 | 14.5833 | 0.557 |
| 1.6 | 0.05 | ptsI | 4 | 14.8333 | 0.507 |
| 1.6 | 0.05 | clcB | 4 | 14.8333 | 0.286 |
| 1.6 | 0.05 | ycaM | 4 | 14.8333 | 0.274 |
| 1.6 | 0.05 | yadI | 4 | 14.8333 | 0.541 |
| 1.6 | 0.05 | AG1 | 4 | 14.8333 | 0.561 |
| 1.6 | 0.05 | ptsI | 4 | 15.0833 | 0.507 |
| 1.6 | 0.05 | clcB | 4 | 15.0833 | 0.285 |
| 1.6 | 0.05 | ycaM | 4 | 15.0833 | 0.274 |
| 1.6 | 0.05 | yadI | 4 | 15.0833 | 0.544 |
| 1.6 | 0.05 | AG1 | 4 | 15.0833 | 0.564 |
| 1.6 | 0.05 | ptsI | 4 | 15.3333 | 0.512 |
| 1.6 | 0.05 | clcB | 4 | 15.3333 | 0.284 |
| 1.6 | 0.05 | ycaM | 4 | 15.3333 | 0.272 |
| 1.6 | 0.05 | yadI | 4 | 15.3333 | 0.542 |
| 1.6 | 0.05 | AG1 | 4 | 15.3333 | 0.561 |
| 1.6 | 0.05 | ptsI | 4 | 15.5833 | 0.512 |
| 1.6 | 0.05 | clcB | 4 | 15.5833 | 0.285 |
| 1.6 | 0.05 | ycaM | 4 | 15.5833 | 0.273 |
| 1.6 | 0.05 | yadI | 4 | 15.5833 | 0.542 |
| 1.6 | 0.05 | AG1 | 4 | 15.5833 | 0.568 |
| 1.6 | 0.05 | ptsI | 4 | 15.8333 | 0.512 |
| 1.6 | 0.05 | clcB | 4 | 15.8333 | 0.283 |
| 1.6 | 0.05 | ycaM | 4 | 15.8333 | 0.272 |
| 1.6 | 0.05 | yadI | 4 | 15.8333 | 0.545 |
| 1.6 | 0.05 | AG1 | 4 | 15.8333 | 0.567 |
| 1.6 | 0.05 | ptsI | 4 | 16.0833 | 0.514 |
| 1.6 | 0.05 | clcB | 4 | 16.0833 | 0.282 |
| 1.6 | 0.05 | ycaM | 4 | 16.0833 | 0.272 |
| 1.6 | 0.05 | yadI | 4 | 16.0833 | 0.544 |
| 1.6 | 0.05 | AG1 | 4 | 16.0833 | 0.57 |
| 1.6 | 0.05 | ptsI | 4 | 16.3333 | 0.518 |
| 1.6 | 0.05 | clcB | 4 | 16.3333 | 0.281 |
| 1.6 | 0.05 | ycaM | 4 | 16.3333 | 0.272 |
| 1.6 | 0.05 | yadI | 4 | 16.3333 | 0.543 |
| 1.6 | 0.05 | AG1 | 4 | 16.3333 | 0.568 |
| 1.6 | 0.05 | ptsI | 4 | 16.5833 | 0.515 |
| 1.6 | 0.05 | clcB | 4 | 16.5833 | 0.281 |
| 1.6 | 0.05 | ycaM | 4 | 16.5833 | 0.271 |
| 1.6 | 0.05 | yadI | 4 | 16.5833 | 0.542 |
| 1.6 | 0.05 | AG1 | 4 | 16.5833 | 0.57 |
| 1.6 | 0.05 | ptsI | 4 | 16.8333 | 0.515 |
| 1.6 | 0.05 | clcB | 4 | 16.8333 | 0.281 |

| | | | | | |
|---|---|---|---|---|---|
| 1.6 | 0.05 | ycaM | 4 | 16.8333 | 0.271 |
| 1.6 | 0.05 | yadI | 4 | 16.8333 | 0.543 |
| 1.6 | 0.05 | AG1 | 4 | 16.8333 | 0.573 |
| 1.6 | 0.05 | ptsI | 4 | 17.0833 | 0.52 |
| 1.6 | 0.05 | clcB | 4 | 17.0833 | 0.281 |
| 1.6 | 0.05 | ycaM | 4 | 17.0833 | 0.271 |
| 1.6 | 0.05 | yadI | 4 | 17.0833 | 0.545 |
| 1.6 | 0.05 | AG1 | 4 | 17.0833 | 0.572 |
| 1.6 | 0.05 | ptsI | 4 | 17.3333 | 0.523 |
| 1.6 | 0.05 | clcB | 4 | 17.3333 | 0.28 |
| 1.6 | 0.05 | ycaM | 4 | 17.3333 | 0.27 |
| 1.6 | 0.05 | yadI | 4 | 17.3333 | 0.543 |
| 1.6 | 0.05 | AG1 | 4 | 17.3333 | 0.573 |
| 1.6 | 0.05 | ptsI | 4 | 17.5833 | 0.517 |
| 1.6 | 0.05 | clcB | 4 | 17.5833 | 0.28 |
| 1.6 | 0.05 | ycaM | 4 | 17.5833 | 0.271 |
| 1.6 | 0.05 | yadI | 4 | 17.5833 | 0.544 |
| 1.6 | 0.05 | AG1 | 4 | 17.5833 | 0.573 |
| 1.6 | 0.05 | ptsI | 4 | 17.8333 | 0.52 |
| 1.6 | 0.05 | clcB | 4 | 17.8333 | 0.28 |
| 1.6 | 0.05 | ycaM | 4 | 17.8333 | 0.271 |
| 1.6 | 0.05 | yadI | 4 | 17.8333 | 0.544 |
| 1.6 | 0.05 | AG1 | 4 | 17.8333 | 0.573 |
| 1.6 | 0.05 | ptsI | 4 | 18.0833 | 0.523 |
| 1.6 | 0.05 | clcB | 4 | 18.0833 | 0.28 |
| 1.6 | 0.05 | ycaM | 4 | 18.0833 | 0.27 |
| 1.6 | 0.05 | yadI | 4 | 18.0833 | 0.544 |
| 1.6 | 0.05 | AG1 | 4 | 18.0833 | 0.576 |
| 1.6 | 0.05 | ptsI | 4 | 18.3333 | 0.522 |
| 1.6 | 0.05 | clcB | 4 | 18.3333 | 0.28 |
| 1.6 | 0.05 | ycaM | 4 | 18.3333 | 0.27 |
| 1.6 | 0.05 | yadI | 4 | 18.3333 | 0.544 |
| 1.6 | 0.05 | AG1 | 4 | 18.3333 | 0.578 |
| 1.6 | 0.05 | ptsI | 4 | 18.5833 | 0.523 |
| 1.6 | 0.05 | clcB | 4 | 18.5833 | 0.28 |
| 1.6 | 0.05 | ycaM | 4 | 18.5833 | 0.269 |
| 1.6 | 0.05 | yadI | 4 | 18.5833 | 0.542 |
| 1.6 | 0.05 | AG1 | 4 | 18.5833 | 0.577 |
| 1.6 | 0.05 | ptsI | 4 | 18.8333 | 0.525 |
| 1.6 | 0.05 | clcB | 4 | 18.8333 | 0.28 |
| 1.6 | 0.05 | ycaM | 4 | 18.8333 | 0.268 |
| 1.6 | 0.05 | yadI | 4 | 18.8333 | 0.54 |
| 1.6 | 0.05 | AG1 | 4 | 18.8333 | 0.574 |
| 1.6 | 0.05 | ptsI | 4 | 19.0833 | 0.526 |
| 1.6 | 0.05 | clcB | 4 | 19.0833 | 0.279 |
| 1.6 | 0.05 | ycaM | 4 | 19.0833 | 0.267 |
| 1.6 | 0.05 | yadI | 4 | 19.0833 | 0.54 |
| 1.6 | 0.05 | AG1 | 4 | 19.0833 | 0.582 |

| | | | | | |
|---|---|---|---|---|---|
| 1.6 | 0.05 | ptsI | 4 | 19.3333 | 0.529 |
| 1.6 | 0.05 | clcB | 4 | 19.3333 | 0.279 |
| 1.6 | 0.05 | ycaM | 4 | 19.3333 | 0.266 |
| 1.6 | 0.05 | yadI | 4 | 19.3333 | 0.541 |
| 1.6 | 0.05 | AG1 | 4 | 19.3333 | 0.581 |
| 1.6 | 0.05 | ptsI | 4 | 19.5833 | 0.529 |
| 1.6 | 0.05 | clcB | 4 | 19.5833 | 0.278 |
| 1.6 | 0.05 | ycaM | 4 | 19.5833 | 0.267 |
| 1.6 | 0.05 | yadI | 4 | 19.5833 | 0.541 |
| 1.6 | 0.05 | AG1 | 4 | 19.5833 | 0.587 |
| 1.6 | 0.05 | ptsI | 4 | 19.8333 | 0.529 |
| 1.6 | 0.05 | clcB | 4 | 19.8333 | 0.278 |
| 1.6 | 0.05 | ycaM | 4 | 19.8333 | 0.266 |
| 1.6 | 0.05 | yadI | 4 | 19.8333 | 0.541 |
| 1.6 | 0.05 | AG1 | 4 | 19.8333 | 0.584 |
| 1.6 | 0.05 | ptsI | 4 | 20.0833 | 0.531 |
| 1.6 | 0.05 | clcB | 4 | 20.0833 | 0.279 |
| 1.6 | 0.05 | ycaM | 4 | 20.0833 | 0.267 |
| 1.6 | 0.05 | yadI | 4 | 20.0833 | 0.544 |
| 1.6 | 0.05 | AG1 | 4 | 20.0833 | 0.588 |
| 1.6 | 0.05 | ptsI | 4 | 20.3333 | 0.527 |
| 1.6 | 0.05 | clcB | 4 | 20.3333 | 0.278 |
| 1.6 | 0.05 | ycaM | 4 | 20.3333 | 0.265 |
| 1.6 | 0.05 | yadI | 4 | 20.3333 | 0.539 |
| 1.6 | 0.05 | AG1 | 4 | 20.3333 | 0.588 |
| 1.6 | 0.05 | ptsI | 4 | 20.5833 | 0.535 |
| 1.6 | 0.05 | clcB | 4 | 20.5833 | 0.277 |
| 1.6 | 0.05 | ycaM | 4 | 20.5833 | 0.267 |
| 1.6 | 0.05 | yadI | 4 | 20.5833 | 0.545 |
| 1.6 | 0.05 | AG1 | 4 | 20.5833 | 0.591 |
| 1.6 | 0.05 | ptsI | 4 | 20.8333 | 0.536 |
| 1.6 | 0.05 | clcB | 4 | 20.8333 | 0.277 |
| 1.6 | 0.05 | ycaM | 4 | 20.8333 | 0.266 |
| 1.6 | 0.05 | yadI | 4 | 20.8333 | 0.547 |
| 1.6 | 0.05 | AG1 | 4 | 20.8333 | 0.592 |
| 1.6 | 0.05 | ptsI | 4 | 21.0833 | 0.534 |
| 1.6 | 0.05 | clcB | 4 | 21.0833 | 0.277 |
| 1.6 | 0.05 | ycaM | 4 | 21.0833 | 0.266 |
| 1.6 | 0.05 | yadI | 4 | 21.0833 | 0.543 |
| 1.6 | 0.05 | AG1 | 4 | 21.0833 | 0.593 |
| 1.6 | 0.05 | ptsI | 4 | 21.3333 | 0.537 |
| 1.6 | 0.05 | clcB | 4 | 21.3333 | 0.276 |
| 1.6 | 0.05 | ycaM | 4 | 21.3333 | 0.265 |
| 1.6 | 0.05 | yadI | 4 | 21.3333 | 0.546 |
| 1.6 | 0.05 | AG1 | 4 | 21.3333 | 0.592 |
| 1.6 | 0.05 | ptsI | 4 | 21.5833 | 0.537 |
| 1.6 | 0.05 | clcB | 4 | 21.5833 | 0.276 |
| 1.6 | 0.05 | ycaM | 4 | 21.5833 | 0.266 |

| | | | | | |
|---|---|---|---|---|---|
| 1.6 | 0.05 | yadI | 4 | 21.5833 | 0.546 |
| 1.6 | 0.05 | AG1 | 4 | 21.5833 | 0.594 |
| 1.6 | 0.05 | ptsI | 4 | 21.8333 | 0.541 |
| 1.6 | 0.05 | clcB | 4 | 21.8333 | 0.277 |
| 1.6 | 0.05 | ycaM | 4 | 21.8333 | 0.265 |
| 1.6 | 0.05 | yadI | 4 | 21.8333 | 0.547 |
| 1.6 | 0.05 | AG1 | 4 | 21.8333 | 0.597 |
| 1.6 | 0.05 | ptsI | 4 | 22.0833 | 0.539 |
| 1.6 | 0.05 | clcB | 4 | 22.0833 | 0.276 |
| 1.6 | 0.05 | ycaM | 4 | 22.0833 | 0.266 |
| 1.6 | 0.05 | yadI | 4 | 22.0833 | 0.546 |
| 1.6 | 0.05 | AG1 | 4 | 22.0833 | 0.596 |
| 1.6 | 0.05 | ptsI | 4 | 22.3333 | 0.543 |
| 1.6 | 0.05 | clcB | 4 | 22.3333 | 0.276 |
| 1.6 | 0.05 | ycaM | 4 | 22.3333 | 0.265 |
| 1.6 | 0.05 | yadI | 4 | 22.3333 | 0.547 |
| 1.6 | 0.05 | AG1 | 4 | 22.3333 | 0.601 |
| 1.6 | 0.05 | ptsI | 4 | 22.5833 | 0.542 |
| 1.6 | 0.05 | clcB | 4 | 22.5833 | 0.276 |
| 1.6 | 0.05 | ycaM | 4 | 22.5833 | 0.265 |
| 1.6 | 0.05 | yadI | 4 | 22.5833 | 0.549 |
| 1.6 | 0.05 | AG1 | 4 | 22.5833 | 0.603 |
| 1.6 | 0.05 | ptsI | 4 | 22.8333 | 0.542 |
| 1.6 | 0.05 | clcB | 4 | 22.8333 | 0.276 |
| 1.6 | 0.05 | ycaM | 4 | 22.8333 | 0.264 |
| 1.6 | 0.05 | yadI | 4 | 22.8333 | 0.551 |
| 1.6 | 0.05 | AG1 | 4 | 22.8333 | 0.605 |
| 1.6 | 0.05 | ptsI | 4 | 23.0833 | 0.546 |
| 1.6 | 0.05 | clcB | 4 | 23.0833 | 0.275 |
| 1.6 | 0.05 | ycaM | 4 | 23.0833 | 0.264 |
| 1.6 | 0.05 | yadI | 4 | 23.0833 | 0.553 |
| 1.6 | 0.05 | AG1 | 4 | 23.0833 | 0.606 |
| 1.6 | 0.05 | ptsI | 4 | 23.3333 | 0.544 |
| 1.6 | 0.05 | clcB | 4 | 23.3333 | 0.277 |
| 1.6 | 0.05 | ycaM | 4 | 23.3333 | 0.264 |
| 1.6 | 0.05 | yadI | 4 | 23.3333 | 0.551 |
| 1.6 | 0.05 | AG1 | 4 | 23.3333 | 0.606 |
| 1.6 | 0.05 | ptsI | 4 | 23.5833 | 0.542 |
| 1.6 | 0.05 | clcB | 4 | 23.5833 | 0.276 |
| 1.6 | 0.05 | ycaM | 4 | 23.5833 | 0.264 |
| 1.6 | 0.05 | yadI | 4 | 23.5833 | 0.552 |
| 1.6 | 0.05 | AG1 | 4 | 23.5833 | 0.607 |
| 1.6 | 0.05 | ptsI | 4 | 23.8333 | 0.543 |
| 1.6 | 0.05 | clcB | 4 | 23.8333 | 0.276 |
| 1.6 | 0.05 | ycaM | 4 | 23.8333 | 0.263 |
| 1.6 | 0.05 | yadI | 4 | 23.8333 | 0.549 |
| 1.6 | 0.05 | AG1 | 4 | 23.8333 | 0.608 |
| 1.6 | 0.05 | ptsI | 4 | 24.0833 | 0.542 |

| | | | | | |
|---|---|---|---|---|---|
| 1.6 | 0.05 | clcB | 4 | 24.0833 | 0.275 |
| 1.6 | 0.05 | ycaM | 4 | 24.0833 | 0.262 |
| 1.6 | 0.05 | yadI | 4 | 24.0833 | 0.551 |
| 1.6 | 0.05 | AG1 | 4 | 24.0833 | 0.61 |
| 1.6 | 0.05 | ptsI | 5 | 0 | 0.168 |
| 1.6 | 0.05 | clcB | 5 | 0 | 0.167 |
| 1.6 | 0.05 | ycaM | 5 | 0 | 0.167 |
| 1.6 | 0.05 | yadI | 5 | 0 | 0.176 |
| 1.6 | 0.05 | AG1 | 5 | 0 | 0.167 |
| 1.6 | 0.05 | ptsI | 5 | 0.35 | 0.166 |
| 1.6 | 0.05 | clcB | 5 | 0.35 | 0.164 |
| 1.6 | 0.05 | ycaM | 5 | 0.35 | 0.173 |
| 1.6 | 0.05 | yadI | 5 | 0.35 | 0.173 |
| 1.6 | 0.05 | AG1 | 5 | 0.35 | 0.164 |
| 1.6 | 0.05 | ptsI | 5 | 0.6 | 0.166 |
| 1.6 | 0.05 | clcB | 5 | 0.6 | 0.16 |
| 1.6 | 0.05 | ycaM | 5 | 0.6 | 0.158 |
| 1.6 | 0.05 | yadI | 5 | 0.6 | 0.171 |
| 1.6 | 0.05 | AG1 | 5 | 0.6 | 0.162 |
| 1.6 | 0.05 | ptsI | 5 | 0.85 | 0.164 |
| 1.6 | 0.05 | clcB | 5 | 0.85 | 0.16 |
| 1.6 | 0.05 | ycaM | 5 | 0.85 | 0.158 |
| 1.6 | 0.05 | yadI | 5 | 0.85 | 0.172 |
| 1.6 | 0.05 | AG1 | 5 | 0.85 | 0.163 |
| 1.6 | 0.05 | ptsI | 5 | 1.1 | 0.168 |
| 1.6 | 0.05 | clcB | 5 | 1.1 | 0.165 |
| 1.6 | 0.05 | ycaM | 5 | 1.1 | 0.163 |
| 1.6 | 0.05 | yadI | 5 | 1.1 | 0.174 |
| 1.6 | 0.05 | AG1 | 5 | 1.1 | 0.164 |
| 1.6 | 0.05 | ptsI | 5 | 1.35 | 0.167 |
| 1.6 | 0.05 | clcB | 5 | 1.35 | 0.163 |
| 1.6 | 0.05 | ycaM | 5 | 1.35 | 0.161 |
| 1.6 | 0.05 | yadI | 5 | 1.35 | 0.177 |
| 1.6 | 0.05 | AG1 | 5 | 1.35 | 0.165 |
| 1.6 | 0.05 | ptsI | 5 | 1.6 | 0.169 |
| 1.6 | 0.05 | clcB | 5 | 1.6 | 0.167 |
| 1.6 | 0.05 | ycaM | 5 | 1.6 | 0.166 |
| 1.6 | 0.05 | yadI | 5 | 1.6 | 0.179 |
| 1.6 | 0.05 | AG1 | 5 | 1.6 | 0.168 |
| 1.6 | 0.05 | ptsI | 5 | 1.85 | 0.17 |
| 1.6 | 0.05 | clcB | 5 | 1.85 | 0.166 |
| 1.6 | 0.05 | ycaM | 5 | 1.85 | 0.171 |
| 1.6 | 0.05 | yadI | 5 | 1.85 | 0.183 |
| 1.6 | 0.05 | AG1 | 5 | 1.85 | 0.171 |
| 1.6 | 0.05 | ptsI | 5 | 2.1 | 0.176 |
| 1.6 | 0.05 | clcB | 5 | 2.1 | 0.175 |
| 1.6 | 0.05 | ycaM | 5 | 2.1 | 0.175 |
| 1.6 | 0.05 | yadI | 5 | 2.1 | 0.193 |

| | | | | | |
|---|---|---|---|---|---|
| 1.6 | 0.05 | AG1 | 5 | 2.1 | 0.176 |
| 1.6 | 0.05 | ptsI | 5 | 2.35 | 0.18 |
| 1.6 | 0.05 | clcB | 5 | 2.35 | 0.182 |
| 1.6 | 0.05 | ycaM | 5 | 2.35 | 0.184 |
| 1.6 | 0.05 | yadI | 5 | 2.35 | 0.199 |
| 1.6 | 0.05 | AG1 | 5 | 2.35 | 0.182 |
| 1.6 | 0.05 | ptsI | 5 | 2.6 | 0.177 |
| 1.6 | 0.05 | clcB | 5 | 2.6 | 0.175 |
| 1.6 | 0.05 | ycaM | 5 | 2.6 | 0.177 |
| 1.6 | 0.05 | yadI | 5 | 2.6 | 0.2 |
| 1.6 | 0.05 | AG1 | 5 | 2.6 | 0.18 |
| 1.6 | 0.05 | ptsI | 5 | 2.85 | 0.183 |
| 1.6 | 0.05 | clcB | 5 | 2.85 | 0.187 |
| 1.6 | 0.05 | ycaM | 5 | 2.85 | 0.19 |
| 1.6 | 0.05 | yadI | 5 | 2.85 | 0.207 |
| 1.6 | 0.05 | AG1 | 5 | 2.85 | 0.188 |
| 1.6 | 0.05 | ptsI | 5 | 3.1 | 0.189 |
| 1.6 | 0.05 | clcB | 5 | 3.1 | 0.197 |
| 1.6 | 0.05 | ycaM | 5 | 3.1 | 0.202 |
| 1.6 | 0.05 | yadI | 5 | 3.1 | 0.214 |
| 1.6 | 0.05 | AG1 | 5 | 3.1 | 0.197 |
| 1.6 | 0.05 | ptsI | 5 | 3.35 | 0.198 |
| 1.6 | 0.05 | clcB | 5 | 3.35 | 0.205 |
| 1.6 | 0.05 | ycaM | 5 | 3.35 | 0.216 |
| 1.6 | 0.05 | yadI | 5 | 3.35 | 0.224 |
| 1.6 | 0.05 | AG1 | 5 | 3.35 | 0.205 |
| 1.6 | 0.05 | ptsI | 5 | 3.6 | 0.211 |
| 1.6 | 0.05 | clcB | 5 | 3.6 | 0.219 |
| 1.6 | 0.05 | ycaM | 5 | 3.6 | 0.232 |
| 1.6 | 0.05 | yadI | 5 | 3.6 | 0.236 |
| 1.6 | 0.05 | AG1 | 5 | 3.6 | 0.215 |
| 1.6 | 0.05 | ptsI | 5 | 3.85 | 0.222 |
| 1.6 | 0.05 | clcB | 5 | 3.85 | 0.234 |
| 1.6 | 0.05 | ycaM | 5 | 3.85 | 0.247 |
| 1.6 | 0.05 | yadI | 5 | 3.85 | 0.248 |
| 1.6 | 0.05 | AG1 | 5 | 3.85 | 0.222 |
| 1.6 | 0.05 | ptsI | 5 | 4.38333 | 0.223 |
| 1.6 | 0.05 | clcB | 5 | 4.38333 | 0.263 |
| 1.6 | 0.05 | ycaM | 5 | 4.38333 | 0.216 |
| 1.6 | 0.05 | yadI | 5 | 4.38333 | 0.287 |
| 1.6 | 0.05 | AG1 | 5 | 4.38333 | 0.257 |
| 1.6 | 0.05 | ptsI | 5 | 4.63333 | 0.221 |
| 1.6 | 0.05 | clcB | 5 | 4.63333 | 0.24 |
| 1.6 | 0.05 | ycaM | 5 | 4.63333 | 0.22 |
| 1.6 | 0.05 | yadI | 5 | 4.63333 | 0.29 |
| 1.6 | 0.05 | AG1 | 5 | 4.63333 | 0.26 |
| 1.6 | 0.05 | ptsI | 5 | 4.88333 | 0.23 |
| 1.6 | 0.05 | clcB | 5 | 4.88333 | 0.253 |

| | | | | | |
|---|---|---|---|---|---|
| 1.6 | 0.05 | ycaM | 5 | 4.88333 | 0.23 |
| 1.6 | 0.05 | yadI | 5 | 4.88333 | 0.312 |
| 1.6 | 0.05 | AG1 | 5 | 4.88333 | 0.28 |
| 1.6 | 0.05 | ptsI | 5 | 5.13333 | 0.238 |
| 1.6 | 0.05 | clcB | 5 | 5.13333 | 0.262 |
| 1.6 | 0.05 | ycaM | 5 | 5.13333 | 0.235 |
| 1.6 | 0.05 | yadI | 5 | 5.13333 | 0.324 |
| 1.6 | 0.05 | AG1 | 5 | 5.13333 | 0.291 |
| 1.6 | 0.05 | ptsI | 5 | 5.38333 | 0.25 |
| 1.6 | 0.05 | clcB | 5 | 5.38333 | 0.27 |
| 1.6 | 0.05 | ycaM | 5 | 5.38333 | 0.239 |
| 1.6 | 0.05 | yadI | 5 | 5.38333 | 0.336 |
| 1.6 | 0.05 | AG1 | 5 | 5.38333 | 0.307 |
| 1.6 | 0.05 | ptsI | 5 | 5.63333 | 0.257 |
| 1.6 | 0.05 | clcB | 5 | 5.63333 | 0.281 |
| 1.6 | 0.05 | ycaM | 5 | 5.63333 | 0.241 |
| 1.6 | 0.05 | yadI | 5 | 5.63333 | 0.347 |
| 1.6 | 0.05 | AG1 | 5 | 5.63333 | 0.32 |
| 1.6 | 0.05 | ptsI | 5 | 5.88333 | 0.27 |
| 1.6 | 0.05 | clcB | 5 | 5.88333 | 0.288 |
| 1.6 | 0.05 | ycaM | 5 | 5.88333 | 0.243 |
| 1.6 | 0.05 | yadI | 5 | 5.88333 | 0.351 |
| 1.6 | 0.05 | AG1 | 5 | 5.88333 | 0.328 |
| 1.6 | 0.05 | ptsI | 5 | 6.13333 | 0.282 |
| 1.6 | 0.05 | clcB | 5 | 6.13333 | 0.3 |
| 1.6 | 0.05 | ycaM | 5 | 6.13333 | 0.246 |
| 1.6 | 0.05 | yadI | 5 | 6.13333 | 0.363 |
| 1.6 | 0.05 | AG1 | 5 | 6.13333 | 0.341 |
| 1.6 | 0.05 | ptsI | 5 | 6.38333 | 0.297 |
| 1.6 | 0.05 | clcB | 5 | 6.38333 | 0.311 |
| 1.6 | 0.05 | ycaM | 5 | 6.38333 | 0.249 |
| 1.6 | 0.05 | yadI | 5 | 6.38333 | 0.374 |
| 1.6 | 0.05 | AG1 | 5 | 6.38333 | 0.352 |
| 1.6 | 0.05 | ptsI | 5 | 6.63333 | 0.314 |
| 1.6 | 0.05 | clcB | 5 | 6.63333 | 0.321 |
| 1.6 | 0.05 | ycaM | 5 | 6.63333 | 0.251 |
| 1.6 | 0.05 | yadI | 5 | 6.63333 | 0.385 |
| 1.6 | 0.05 | AG1 | 5 | 6.63333 | 0.364 |
| 1.6 | 0.05 | ptsI | 5 | 6.88333 | 0.33 |
| 1.6 | 0.05 | clcB | 5 | 6.88333 | 0.33 |
| 1.6 | 0.05 | ycaM | 5 | 6.88333 | 0.253 |
| 1.6 | 0.05 | yadI | 5 | 6.88333 | 0.396 |
| 1.6 | 0.05 | AG1 | 5 | 6.88333 | 0.376 |
| 1.6 | 0.05 | ptsI | 5 | 7.13333 | 0.346 |
| 1.6 | 0.05 | clcB | 5 | 7.13333 | 0.337 |
| 1.6 | 0.05 | ycaM | 5 | 7.13333 | 0.253 |
| 1.6 | 0.05 | yadI | 5 | 7.13333 | 0.409 |
| 1.6 | 0.05 | AG1 | 5 | 7.13333 | 0.388 |

| | | | | | |
|---|---|---|---|---|---|
| 1.6 | 0.05 | ptsI | 5 | 7.38333 | 0.362 |
| 1.6 | 0.05 | clcB | 5 | 7.38333 | 0.341 |
| 1.6 | 0.05 | ycaM | 5 | 7.38333 | 0.256 |
| 1.6 | 0.05 | yadI | 5 | 7.38333 | 0.415 |
| 1.6 | 0.05 | AG1 | 5 | 7.38333 | 0.398 |
| 1.6 | 0.05 | ptsI | 5 | 7.63333 | 0.374 |
| 1.6 | 0.05 | clcB | 5 | 7.63333 | 0.346 |
| 1.6 | 0.05 | ycaM | 5 | 7.63333 | 0.256 |
| 1.6 | 0.05 | yadI | 5 | 7.63333 | 0.422 |
| 1.6 | 0.05 | AG1 | 5 | 7.63333 | 0.408 |
| 1.6 | 0.05 | ptsI | 5 | 7.88333 | 0.391 |
| 1.6 | 0.05 | clcB | 5 | 7.88333 | 0.353 |
| 1.6 | 0.05 | ycaM | 5 | 7.88333 | 0.257 |
| 1.6 | 0.05 | yadI | 5 | 7.88333 | 0.427 |
| 1.6 | 0.05 | AG1 | 5 | 7.88333 | 0.418 |
| 1.6 | 0.05 | ptsI | 5 | 8.13333 | 0.407 |
| 1.6 | 0.05 | clcB | 5 | 8.13333 | 0.355 |
| 1.6 | 0.05 | ycaM | 5 | 8.13333 | 0.259 |
| 1.6 | 0.05 | yadI | 5 | 8.13333 | 0.431 |
| 1.6 | 0.05 | AG1 | 5 | 8.13333 | 0.429 |
| 1.6 | 0.05 | ptsI | 5 | 8.38333 | 0.424 |
| 1.6 | 0.05 | clcB | 5 | 8.38333 | 0.36 |
| 1.6 | 0.05 | ycaM | 5 | 8.38333 | 0.261 |
| 1.6 | 0.05 | yadI | 5 | 8.38333 | 0.44 |
| 1.6 | 0.05 | AG1 | 5 | 8.38333 | 0.44 |
| 1.6 | 0.05 | ptsI | 5 | 8.88333 | 0.353 |
| 1.6 | 0.05 | clcB | 5 | 8.88333 | 0.282 |
| 1.6 | 0.05 | ycaM | 5 | 8.88333 | 0.262 |
| 1.6 | 0.05 | yadI | 5 | 8.88333 | 0.431 |
| 1.6 | 0.05 | AG1 | 5 | 8.88333 | 0.435 |
| 1.6 | 0.05 | ptsI | 5 | 9.13333 | 0.359 |
| 1.6 | 0.05 | clcB | 5 | 9.13333 | 0.28 |
| 1.6 | 0.05 | ycaM | 5 | 9.13333 | 0.259 |
| 1.6 | 0.05 | yadI | 5 | 9.13333 | 0.424 |
| 1.6 | 0.05 | AG1 | 5 | 9.13333 | 0.441 |
| 1.6 | 0.05 | ptsI | 5 | 9.38333 | 0.375 |
| 1.6 | 0.05 | clcB | 5 | 9.38333 | 0.282 |
| 1.6 | 0.05 | ycaM | 5 | 9.38333 | 0.258 |
| 1.6 | 0.05 | yadI | 5 | 9.38333 | 0.433 |
| 1.6 | 0.05 | AG1 | 5 | 9.38333 | 0.459 |
| 1.6 | 0.05 | ptsI | 5 | 9.63333 | 0.38 |
| 1.6 | 0.05 | clcB | 5 | 9.63333 | 0.279 |
| 1.6 | 0.05 | ycaM | 5 | 9.63333 | 0.258 |
| 1.6 | 0.05 | yadI | 5 | 9.63333 | 0.441 |
| 1.6 | 0.05 | AG1 | 5 | 9.63333 | 0.471 |
| 1.6 | 0.05 | ptsI | 5 | 9.88333 | 0.39 |
| 1.6 | 0.05 | clcB | 5 | 9.88333 | 0.28 |
| 1.6 | 0.05 | ycaM | 5 | 9.88333 | 0.259 |

| | | | | | |
|---|---|---|---|---|---|
| 1.6 | 0.05 | yadI | 5 | 9.88333 | 0.448 |
| 1.6 | 0.05 | AG1 | 5 | 9.88333 | 0.479 |
| 1.6 | 0.05 | ptsI | 5 | 10.1333 | 0.396 |
| 1.6 | 0.05 | clcB | 5 | 10.1333 | 0.279 |
| 1.6 | 0.05 | ycaM | 5 | 10.1333 | 0.258 |
| 1.6 | 0.05 | yadI | 5 | 10.1333 | 0.455 |
| 1.6 | 0.05 | AG1 | 5 | 10.1333 | 0.484 |
| 1.6 | 0.05 | ptsI | 5 | 10.3833 | 0.406 |
| 1.6 | 0.05 | clcB | 5 | 10.3833 | 0.278 |
| 1.6 | 0.05 | ycaM | 5 | 10.3833 | 0.259 |
| 1.6 | 0.05 | yadI | 5 | 10.3833 | 0.462 |
| 1.6 | 0.05 | AG1 | 5 | 10.3833 | 0.492 |
| 1.6 | 0.05 | ptsI | 5 | 10.6333 | 0.41 |
| 1.6 | 0.05 | clcB | 5 | 10.6333 | 0.277 |
| 1.6 | 0.05 | ycaM | 5 | 10.6333 | 0.258 |
| 1.6 | 0.05 | yadI | 5 | 10.6333 | 0.466 |
| 1.6 | 0.05 | AG1 | 5 | 10.6333 | 0.499 |
| 1.6 | 0.05 | ptsI | 5 | 10.8833 | 0.416 |
| 1.6 | 0.05 | clcB | 5 | 10.8833 | 0.279 |
| 1.6 | 0.05 | ycaM | 5 | 10.8833 | 0.258 |
| 1.6 | 0.05 | yadI | 5 | 10.8833 | 0.473 |
| 1.6 | 0.05 | AG1 | 5 | 10.8833 | 0.507 |
| 1.6 | 0.05 | ptsI | 5 | 11.1333 | 0.421 |
| 1.6 | 0.05 | clcB | 5 | 11.1333 | 0.277 |
| 1.6 | 0.05 | ycaM | 5 | 11.1333 | 0.256 |
| 1.6 | 0.05 | yadI | 5 | 11.1333 | 0.476 |
| 1.6 | 0.05 | AG1 | 5 | 11.1333 | 0.511 |
| 1.6 | 0.05 | ptsI | 5 | 11.3833 | 0.423 |
| 1.6 | 0.05 | clcB | 5 | 11.3833 | 0.278 |
| 1.6 | 0.05 | ycaM | 5 | 11.3833 | 0.26 |
| 1.6 | 0.05 | yadI | 5 | 11.3833 | 0.482 |
| 1.6 | 0.05 | AG1 | 5 | 11.3833 | 0.518 |
| 1.6 | 0.05 | ptsI | 5 | 11.6333 | 0.428 |
| 1.6 | 0.05 | clcB | 5 | 11.6333 | 0.275 |
| 1.6 | 0.05 | ycaM | 5 | 11.6333 | 0.257 |
| 1.6 | 0.05 | yadI | 5 | 11.6333 | 0.488 |
| 1.6 | 0.05 | AG1 | 5 | 11.6333 | 0.525 |
| 1.6 | 0.05 | ptsI | 5 | 11.8833 | 0.434 |
| 1.6 | 0.05 | clcB | 5 | 11.8833 | 0.277 |
| 1.6 | 0.05 | ycaM | 5 | 11.8833 | 0.257 |
| 1.6 | 0.05 | yadI | 5 | 11.8833 | 0.49 |
| 1.6 | 0.05 | AG1 | 5 | 11.8833 | 0.531 |
| 1.6 | 0.05 | ptsI | 5 | 12.1333 | 0.439 |
| 1.6 | 0.05 | clcB | 5 | 12.1333 | 0.279 |
| 1.6 | 0.05 | ycaM | 5 | 12.1333 | 0.256 |
| 1.6 | 0.05 | yadI | 5 | 12.1333 | 0.494 |
| 1.6 | 0.05 | AG1 | 5 | 12.1333 | 0.54 |
| 1.6 | 0.05 | ptsI | 5 | 12.3833 | 0.44 |

| | | | | | |
|---|---|---|---|---|---|
| 1.6 | 0.05 | clcB | 5 | 12.3833 | 0.278 |
| 1.6 | 0.05 | ycaM | 5 | 12.3833 | 0.252 |
| 1.6 | 0.05 | yadI | 5 | 12.3833 | 0.494 |
| 1.6 | 0.05 | AG1 | 5 | 12.3833 | 0.536 |
| 1.6 | 0.05 | ptsI | 5 | 12.6333 | 0.446 |
| 1.6 | 0.05 | clcB | 5 | 12.6333 | 0.275 |
| 1.6 | 0.05 | ycaM | 5 | 12.6333 | 0.255 |
| 1.6 | 0.05 | yadI | 5 | 12.6333 | 0.503 |
| 1.6 | 0.05 | AG1 | 5 | 12.6333 | 0.555 |
| 1.6 | 0.05 | ptsI | 5 | 12.8833 | 0.448 |
| 1.6 | 0.05 | clcB | 5 | 12.8833 | 0.276 |
| 1.6 | 0.05 | ycaM | 5 | 12.8833 | 0.252 |
| 1.6 | 0.05 | yadI | 5 | 12.8833 | 0.506 |
| 1.6 | 0.05 | AG1 | 5 | 12.8833 | 0.556 |
| 1.6 | 0.05 | ptsI | 5 | 13.1333 | 0.453 |
| 1.6 | 0.05 | clcB | 5 | 13.1333 | 0.277 |
| 1.6 | 0.05 | ycaM | 5 | 13.1333 | 0.251 |
| 1.6 | 0.05 | yadI | 5 | 13.1333 | 0.51 |
| 1.6 | 0.05 | AG1 | 5 | 13.1333 | 0.57 |
| 1.6 | 0.05 | ptsI | 5 | 13.3833 | 0.455 |
| 1.6 | 0.05 | clcB | 5 | 13.3833 | 0.275 |
| 1.6 | 0.05 | ycaM | 5 | 13.3833 | 0.251 |
| 1.6 | 0.05 | yadI | 5 | 13.3833 | 0.51 |
| 1.6 | 0.05 | AG1 | 5 | 13.3833 | 0.573 |
| 1.6 | 0.05 | ptsI | 5 | 13.6333 | 0.455 |
| 1.6 | 0.05 | clcB | 5 | 13.6333 | 0.277 |
| 1.6 | 0.05 | ycaM | 5 | 13.6333 | 0.25 |
| 1.6 | 0.05 | yadI | 5 | 13.6333 | 0.514 |
| 1.6 | 0.05 | AG1 | 5 | 13.6333 | 0.578 |
| 1.6 | 0.05 | ptsI | 5 | 13.8833 | 0.464 |
| 1.6 | 0.05 | clcB | 5 | 13.8833 | 0.274 |
| 1.6 | 0.05 | ycaM | 5 | 13.8833 | 0.248 |
| 1.6 | 0.05 | yadI | 5 | 13.8833 | 0.519 |
| 1.6 | 0.05 | AG1 | 5 | 13.8833 | 0.583 |
| 1.6 | 0.05 | ptsI | 5 | 14.1333 | 0.467 |
| 1.6 | 0.05 | clcB | 5 | 14.1333 | 0.273 |
| 1.6 | 0.05 | ycaM | 5 | 14.1333 | 0.248 |
| 1.6 | 0.05 | yadI | 5 | 14.1333 | 0.522 |
| 1.6 | 0.05 | AG1 | 5 | 14.1333 | 0.585 |
| 1.6 | 0.05 | ptsI | 5 | 14.3833 | 0.468 |
| 1.6 | 0.05 | clcB | 5 | 14.3833 | 0.273 |
| 1.6 | 0.05 | ycaM | 5 | 14.3833 | 0.248 |
| 1.6 | 0.05 | yadI | 5 | 14.3833 | 0.522 |
| 1.6 | 0.05 | AG1 | 5 | 14.3833 | 0.594 |
| 1.6 | 0.05 | ptsI | 5 | 14.6333 | 0.471 |
| 1.6 | 0.05 | clcB | 5 | 14.6333 | 0.273 |
| 1.6 | 0.05 | ycaM | 5 | 14.6333 | 0.247 |
| 1.6 | 0.05 | yadI | 5 | 14.6333 | 0.524 |

| | | | | | |
|---|---|---|---|---|---|
| 1.6 | 0.05 | AG1 | 5 | 14.6333 | 0.599 |
| 1.6 | 0.05 | ptsI | 5 | 14.8833 | 0.474 |
| 1.6 | 0.05 | clcB | 5 | 14.8833 | 0.271 |
| 1.6 | 0.05 | ycaM | 5 | 14.8833 | 0.246 |
| 1.6 | 0.05 | yadI | 5 | 14.8833 | 0.523 |
| 1.6 | 0.05 | AG1 | 5 | 14.8833 | 0.608 |
| 1.6 | 0.05 | ptsI | 5 | 15.1333 | 0.477 |
| 1.6 | 0.05 | clcB | 5 | 15.1333 | 0.272 |
| 1.6 | 0.05 | ycaM | 5 | 15.1333 | 0.246 |
| 1.6 | 0.05 | yadI | 5 | 15.1333 | 0.527 |
| 1.6 | 0.05 | AG1 | 5 | 15.1333 | 0.608 |
| 1.6 | 0.05 | ptsI | 5 | 15.3833 | 0.481 |
| 1.6 | 0.05 | clcB | 5 | 15.3833 | 0.271 |
| 1.6 | 0.05 | ycaM | 5 | 15.3833 | 0.246 |
| 1.6 | 0.05 | yadI | 5 | 15.3833 | 0.528 |
| 1.6 | 0.05 | AG1 | 5 | 15.3833 | 0.616 |
| 1.6 | 0.05 | ptsI | 5 | 15.6333 | 0.479 |
| 1.6 | 0.05 | clcB | 5 | 15.6333 | 0.27 |
| 1.6 | 0.05 | ycaM | 5 | 15.6333 | 0.247 |
| 1.6 | 0.05 | yadI | 5 | 15.6333 | 0.528 |
| 1.6 | 0.05 | AG1 | 5 | 15.6333 | 0.616 |
| 1.6 | 0.05 | ptsI | 5 | 15.8833 | 0.486 |
| 1.6 | 0.05 | clcB | 5 | 15.8833 | 0.27 |
| 1.6 | 0.05 | ycaM | 5 | 15.8833 | 0.247 |
| 1.6 | 0.05 | yadI | 5 | 15.8833 | 0.532 |
| 1.6 | 0.05 | AG1 | 5 | 15.8833 | 0.622 |
| 1.6 | 0.05 | ptsI | 5 | 16.1333 | 0.49 |
| 1.6 | 0.05 | clcB | 5 | 16.1333 | 0.272 |
| 1.6 | 0.05 | ycaM | 5 | 16.1333 | 0.246 |
| 1.6 | 0.05 | yadI | 5 | 16.1333 | 0.533 |
| 1.6 | 0.05 | AG1 | 5 | 16.1333 | 0.631 |
| 1.6 | 0.05 | ptsI | 5 | 16.3833 | 0.495 |
| 1.6 | 0.05 | clcB | 5 | 16.3833 | 0.271 |
| 1.6 | 0.05 | ycaM | 5 | 16.3833 | 0.247 |
| 1.6 | 0.05 | yadI | 5 | 16.3833 | 0.536 |
| 1.6 | 0.05 | AG1 | 5 | 16.3833 | 0.629 |
| 1.6 | 0.05 | ptsI | 5 | 16.6333 | 0.499 |
| 1.6 | 0.05 | clcB | 5 | 16.6333 | 0.27 |
| 1.6 | 0.05 | ycaM | 5 | 16.6333 | 0.247 |
| 1.6 | 0.05 | yadI | 5 | 16.6333 | 0.536 |
| 1.6 | 0.05 | AG1 | 5 | 16.6333 | 0.634 |
| 1.6 | 0.05 | ptsI | 5 | 16.8833 | 0.499 |
| 1.6 | 0.05 | clcB | 5 | 16.8833 | 0.27 |
| 1.6 | 0.05 | ycaM | 5 | 16.8833 | 0.247 |
| 1.6 | 0.05 | yadI | 5 | 16.8833 | 0.533 |
| 1.6 | 0.05 | AG1 | 5 | 16.8833 | 0.635 |
| 1.6 | 0.05 | ptsI | 5 | 17.1333 | 0.49 |
| 1.6 | 0.05 | clcB | 5 | 17.1333 | 0.268 |

| | | | | | |
|---|---|---|---|---|---|
| 1.6 | 0.05 | ycaM | 5 | 17.1333 | 0.248 |
| 1.6 | 0.05 | yadI | 5 | 17.1333 | 0.533 |
| 1.6 | 0.05 | AG1 | 5 | 17.1333 | 0.637 |
| 1.6 | 0.05 | ptsI | 5 | 17.3833 | 0.502 |
| 1.6 | 0.05 | clcB | 5 | 17.3833 | 0.27 |
| 1.6 | 0.05 | ycaM | 5 | 17.3833 | 0.248 |
| 1.6 | 0.05 | yadI | 5 | 17.3833 | 0.534 |
| 1.6 | 0.05 | AG1 | 5 | 17.3833 | 0.636 |
| 1.6 | 0.05 | ptsI | 5 | 17.6333 | 0.504 |
| 1.6 | 0.05 | clcB | 5 | 17.6333 | 0.27 |
| 1.6 | 0.05 | ycaM | 5 | 17.6333 | 0.248 |
| 1.6 | 0.05 | yadI | 5 | 17.6333 | 0.534 |
| 1.6 | 0.05 | AG1 | 5 | 17.6333 | 0.64 |
| 1.6 | 0.05 | ptsI | 5 | 17.8833 | 0.5 |
| 1.6 | 0.05 | clcB | 5 | 17.8833 | 0.269 |
| 1.6 | 0.05 | ycaM | 5 | 17.8833 | 0.247 |
| 1.6 | 0.05 | yadI | 5 | 17.8833 | 0.526 |
| 1.6 | 0.05 | AG1 | 5 | 17.8833 | 0.634 |
| 1.6 | 0.05 | ptsI | 5 | 18.1333 | 0.511 |
| 1.6 | 0.05 | clcB | 5 | 18.1333 | 0.27 |
| 1.6 | 0.05 | ycaM | 5 | 18.1333 | 0.25 |
| 1.6 | 0.05 | yadI | 5 | 18.1333 | 0.53 |
| 1.6 | 0.05 | AG1 | 5 | 18.1333 | 0.643 |
| 1.6 | 0.05 | ptsI | 5 | 18.3833 | 0.51 |
| 1.6 | 0.05 | clcB | 5 | 18.3833 | 0.269 |
| 1.6 | 0.05 | ycaM | 5 | 18.3833 | 0.25 |
| 1.6 | 0.05 | yadI | 5 | 18.3833 | 0.531 |
| 1.6 | 0.05 | AG1 | 5 | 18.3833 | 0.643 |
| 1.6 | 0.05 | ptsI | 5 | 18.6333 | 0.511 |
| 1.6 | 0.05 | clcB | 5 | 18.6333 | 0.27 |
| 1.6 | 0.05 | ycaM | 5 | 18.6333 | 0.251 |
| 1.6 | 0.05 | yadI | 5 | 18.6333 | 0.527 |
| 1.6 | 0.05 | AG1 | 5 | 18.6333 | 0.644 |
| 1.6 | 0.05 | ptsI | 5 | 18.8833 | 0.516 |
| 1.6 | 0.05 | clcB | 5 | 18.8833 | 0.27 |
| 1.6 | 0.05 | ycaM | 5 | 18.8833 | 0.251 |
| 1.6 | 0.05 | yadI | 5 | 18.8833 | 0.527 |
| 1.6 | 0.05 | AG1 | 5 | 18.8833 | 0.644 |
| 1.6 | 0.05 | ptsI | 5 | 19.1333 | 0.514 |
| 1.6 | 0.05 | clcB | 5 | 19.1333 | 0.269 |
| 1.6 | 0.05 | ycaM | 5 | 19.1333 | 0.251 |
| 1.6 | 0.05 | yadI | 5 | 19.1333 | 0.526 |
| 1.6 | 0.05 | AG1 | 5 | 19.1333 | 0.647 |
| 1.6 | 0.05 | ptsI | 5 | 19.3833 | 0.518 |
| 1.6 | 0.05 | clcB | 5 | 19.3833 | 0.27 |
| 1.6 | 0.05 | ycaM | 5 | 19.3833 | 0.252 |
| 1.6 | 0.05 | yadI | 5 | 19.3833 | 0.528 |
| 1.6 | 0.05 | AG1 | 5 | 19.3833 | 0.652 |

| | | | | | |
|---|---|---|---|---|---|
| 1.6 | 0.05 | ptsI | 5 | 19.6333 | 0.521 |
| 1.6 | 0.05 | clcB | 5 | 19.6333 | 0.27 |
| 1.6 | 0.05 | ycaM | 5 | 19.6333 | 0.253 |
| 1.6 | 0.05 | yadI | 5 | 19.6333 | 0.525 |
| 1.6 | 0.05 | AG1 | 5 | 19.6333 | 0.65 |
| 1.6 | 0.05 | ptsI | 5 | 19.8833 | 0.523 |
| 1.6 | 0.05 | clcB | 5 | 19.8833 | 0.269 |
| 1.6 | 0.05 | ycaM | 5 | 19.8833 | 0.252 |
| 1.6 | 0.05 | yadI | 5 | 19.8833 | 0.525 |
| 1.6 | 0.05 | AG1 | 5 | 19.8833 | 0.65 |
| 1.6 | 0.05 | ptsI | 5 | 20.1333 | 0.521 |
| 1.6 | 0.05 | clcB | 5 | 20.1333 | 0.268 |
| 1.6 | 0.05 | ycaM | 5 | 20.1333 | 0.253 |
| 1.6 | 0.05 | yadI | 5 | 20.1333 | 0.528 |
| 1.6 | 0.05 | AG1 | 5 | 20.1333 | 0.652 |
| 1.6 | 0.05 | ptsI | 5 | 20.3833 | 0.525 |
| 1.6 | 0.05 | clcB | 5 | 20.3833 | 0.268 |
| 1.6 | 0.05 | ycaM | 5 | 20.3833 | 0.254 |
| 1.6 | 0.05 | yadI | 5 | 20.3833 | 0.526 |
| 1.6 | 0.05 | AG1 | 5 | 20.3833 | 0.652 |
| 1.6 | 0.05 | ptsI | 5 | 20.6333 | 0.52 |
| 1.6 | 0.05 | clcB | 5 | 20.6333 | 0.27 |
| 1.6 | 0.05 | ycaM | 5 | 20.6333 | 0.249 |
| 1.6 | 0.05 | yadI | 5 | 20.6333 | 0.527 |
| 1.6 | 0.05 | AG1 | 5 | 20.6333 | 0.645 |
| 1.6 | 0.05 | ptsI | 5 | 20.8833 | 0.533 |
| 1.6 | 0.05 | clcB | 5 | 20.8833 | 0.267 |
| 1.6 | 0.05 | ycaM | 5 | 20.8833 | 0.253 |
| 1.6 | 0.05 | yadI | 5 | 20.8833 | 0.529 |
| 1.6 | 0.05 | AG1 | 5 | 20.8833 | 0.654 |
| 1.6 | 0.05 | ptsI | 5 | 21.1333 | 0.532 |
| 1.6 | 0.05 | clcB | 5 | 21.1333 | 0.268 |
| 1.6 | 0.05 | ycaM | 5 | 21.1333 | 0.252 |
| 1.6 | 0.05 | yadI | 5 | 21.1333 | 0.53 |
| 1.6 | 0.05 | AG1 | 5 | 21.1333 | 0.659 |
| 1.6 | 0.05 | ptsI | 5 | 21.3833 | 0.536 |
| 1.6 | 0.05 | clcB | 5 | 21.3833 | 0.267 |
| 1.6 | 0.05 | ycaM | 5 | 21.3833 | 0.253 |
| 1.6 | 0.05 | yadI | 5 | 21.3833 | 0.531 |
| 1.6 | 0.05 | AG1 | 5 | 21.3833 | 0.659 |
| 1.6 | 0.05 | ptsI | 5 | 21.6333 | 0.538 |
| 1.6 | 0.05 | clcB | 5 | 21.6333 | 0.268 |
| 1.6 | 0.05 | ycaM | 5 | 21.6333 | 0.252 |
| 1.6 | 0.05 | yadI | 5 | 21.6333 | 0.53 |
| 1.6 | 0.05 | AG1 | 5 | 21.6333 | 0.662 |
| 1.6 | 0.05 | ptsI | 5 | 21.8833 | 0.54 |
| 1.6 | 0.05 | clcB | 5 | 21.8833 | 0.268 |
| 1.6 | 0.05 | ycaM | 5 | 21.8833 | 0.252 |

| | | | | | |
|---|---|---|---|---|---|
| 1.6 | 0.05 | yadI | 5 | 21.8833 | 0.53 |
| 1.6 | 0.05 | AG1 | 5 | 21.8833 | 0.666 |
| 1.6 | 0.05 | ptsI | 5 | 22.1333 | 0.539 |
| 1.6 | 0.05 | clcB | 5 | 22.1333 | 0.266 |
| 1.6 | 0.05 | ycaM | 5 | 22.1333 | 0.251 |
| 1.6 | 0.05 | yadI | 5 | 22.1333 | 0.526 |
| 1.6 | 0.05 | AG1 | 5 | 22.1333 | 0.656 |
| 1.6 | 0.05 | ptsI | 5 | 22.3833 | 0.544 |
| 1.6 | 0.05 | clcB | 5 | 22.3833 | 0.267 |
| 1.6 | 0.05 | ycaM | 5 | 22.3833 | 0.252 |
| 1.6 | 0.05 | yadI | 5 | 22.3833 | 0.532 |
| 1.6 | 0.05 | AG1 | 5 | 22.3833 | 0.671 |
| 1.6 | 0.05 | ptsI | 5 | 22.6333 | 0.55 |
| 1.6 | 0.05 | clcB | 5 | 22.6333 | 0.268 |
| 1.6 | 0.05 | ycaM | 5 | 22.6333 | 0.253 |
| 1.6 | 0.05 | yadI | 5 | 22.6333 | 0.534 |
| 1.6 | 0.05 | AG1 | 5 | 22.6333 | 0.675 |
| 1.6 | 0.05 | ptsI | 5 | 22.8833 | 0.548 |
| 1.6 | 0.05 | clcB | 5 | 22.8833 | 0.267 |
| 1.6 | 0.05 | ycaM | 5 | 22.8833 | 0.252 |
| 1.6 | 0.05 | yadI | 5 | 22.8833 | 0.534 |
| 1.6 | 0.05 | AG1 | 5 | 22.8833 | 0.677 |
| 1.6 | 0.05 | ptsI | 5 | 23.1333 | 0.552 |
| 1.6 | 0.05 | clcB | 5 | 23.1333 | 0.267 |
| 1.6 | 0.05 | ycaM | 5 | 23.1333 | 0.252 |
| 1.6 | 0.05 | yadI | 5 | 23.1333 | 0.536 |
| 1.6 | 0.05 | AG1 | 5 | 23.1333 | 0.68 |
| 1.6 | 0.05 | ptsI | 5 | 23.3833 | 0.551 |
| 1.6 | 0.05 | clcB | 5 | 23.3833 | 0.267 |
| 1.6 | 0.05 | ycaM | 5 | 23.3833 | 0.251 |
| 1.6 | 0.05 | yadI | 5 | 23.3833 | 0.536 |
| 1.6 | 0.05 | AG1 | 5 | 23.3833 | 0.682 |
| 1.6 | 0.05 | ptsI | 5 | 23.6333 | 0.551 |
| 1.6 | 0.05 | clcB | 5 | 23.6333 | 0.266 |
| 1.6 | 0.05 | ycaM | 5 | 23.6333 | 0.251 |
| 1.6 | 0.05 | yadI | 5 | 23.6333 | 0.534 |
| 1.6 | 0.05 | AG1 | 5 | 23.6333 | 0.682 |
| 1.6 | 0.05 | ptsI | 5 | 23.8833 | 0.55 |
| 1.6 | 0.05 | clcB | 5 | 23.8833 | 0.266 |
| 1.6 | 0.05 | ycaM | 5 | 23.8833 | 0.249 |
| 1.6 | 0.05 | yadI | 5 | 23.8833 | 0.533 |
| 1.6 | 0.05 | AG1 | 5 | 23.8833 | 0.672 |
| 1.6 | 0.05 | ptsI | 5 | 24.1333 | 0.555 |
| 1.6 | 0.05 | clcB | 5 | 24.1333 | 0.266 |
| 1.6 | 0.05 | ycaM | 5 | 24.1333 | 0.251 |
| 1.6 | 0.05 | yadI | 5 | 24.1333 | 0.537 |
| 1.6 | 0.05 | AG1 | 5 | 24.1333 | 0.69 |
| 1.6 | 0.05 | ptsI | 5 | 24.3833 | 0.553 |

| | | | | | |
|---|---|---|---|---|---|
| 1.6 | 0.05 | clcB | 5 | 24.3833 | 0.267 |
| 1.6 | 0.05 | ycaM | 5 | 24.3833 | 0.249 |
| 1.6 | 0.05 | yadI | 5 | 24.3833 | 0.538 |
| 1.6 | 0.05 | AG1 | 5 | 24.3833 | 0.688 |
| 3.2 | 0.05 | ptsI | 2 | 0 | 0.212 |
| 3.2 | 0.05 | clcB | 2 | 0 | 0.202 |
| 3.2 | 0.05 | ycaM | 2 | 0 | 0.212 |
| 3.2 | 0.05 | yadI | 2 | 0 | 1.942 |
| 3.2 | 0.05 | AG1 | 2 | 0 | 0.233 |
| 3.2 | 0.05 | ptsI | 2 | 0.25 | 0.21 |
| 3.2 | 0.05 | clcB | 2 | 0.25 | 0.198 |
| 3.2 | 0.05 | ycaM | 2 | 0.25 | 0.209 |
| 3.2 | 0.05 | yadI | 2 | 0.25 | 1.93 |
| 3.2 | 0.05 | AG1 | 2 | 0.25 | 0.229 |
| 3.2 | 0.05 | ptsI | 2 | 0.5 | 0.209 |
| 3.2 | 0.05 | clcB | 2 | 0.5 | 0.2 |
| 3.2 | 0.05 | ycaM | 2 | 0.5 | 0.21 |
| 3.2 | 0.05 | yadI | 2 | 0.5 | 1.9 |
| 3.2 | 0.05 | AG1 | 2 | 0.5 | 0.232 |
| 3.2 | 0.05 | ptsI | 2 | 0.75 | 0.215 |
| 3.2 | 0.05 | clcB | 2 | 0.75 | 0.199 |
| 3.2 | 0.05 | ycaM | 2 | 0.75 | 0.215 |
| 3.2 | 0.05 | yadI | 2 | 0.75 | 1.896 |
| 3.2 | 0.05 | AG1 | 2 | 0.75 | 0.235 |
| 3.2 | 0.05 | ptsI | 2 | 1 | 0.214 |
| 3.2 | 0.05 | clcB | 2 | 1 | 0.201 |
| 3.2 | 0.05 | ycaM | 2 | 1 | 0.214 |
| 3.2 | 0.05 | yadI | 2 | 1 | 1.851 |
| 3.2 | 0.05 | AG1 | 2 | 1 | 0.242 |
| 3.2 | 0.05 | ptsI | 2 | 1.25 | 0.214 |
| 3.2 | 0.05 | clcB | 2 | 1.25 | 0.199 |
| 3.2 | 0.05 | ycaM | 2 | 1.25 | 0.216 |
| 3.2 | 0.05 | yadI | 2 | 1.25 | 1.782 |
| 3.2 | 0.05 | AG1 | 2 | 1.25 | 0.236 |
| 3.2 | 0.05 | ptsI | 2 | 1.5 | 0.218 |
| 3.2 | 0.05 | clcB | 2 | 1.5 | 0.202 |
| 3.2 | 0.05 | ycaM | 2 | 1.5 | 0.217 |
| 3.2 | 0.05 | yadI | 2 | 1.5 | 1.692 |
| 3.2 | 0.05 | AG1 | 2 | 1.5 | 0.242 |
| 3.2 | 0.05 | ptsI | 2 | 1.75 | 0.22 |
| 3.2 | 0.05 | clcB | 2 | 1.75 | 0.203 |
| 3.2 | 0.05 | ycaM | 2 | 1.75 | 0.222 |
| 3.2 | 0.05 | yadI | 2 | 1.75 | 1.495 |
| 3.2 | 0.05 | AG1 | 2 | 1.75 | 0.248 |
| 3.2 | 0.05 | ptsI | 2 | 2 | 0.227 |
| 3.2 | 0.05 | clcB | 2 | 2 | 0.207 |
| 3.2 | 0.05 | ycaM | 2 | 2 | 0.229 |
| 3.2 | 0.05 | yadI | 2 | 2 | 1.418 |

| | | | | | |
|---|---|---|---|---|---|
| 3.2 | 0.05 | AG1 | 2 | 2 | 0.257 |
| 3.2 | 0.05 | ptsI | 2 | 2.25 | 0.233 |
| 3.2 | 0.05 | clcB | 2 | 2.25 | 0.209 |
| 3.2 | 0.05 | ycaM | 2 | 2.25 | 0.233 |
| 3.2 | 0.05 | yadI | 2 | 2.25 | 1.288 |
| 3.2 | 0.05 | AG1 | 2 | 2.25 | 0.264 |
| 3.2 | 0.05 | ptsI | 2 | 2.5 | 0.242 |
| 3.2 | 0.05 | clcB | 2 | 2.5 | 0.212 |
| 3.2 | 0.05 | ycaM | 2 | 2.5 | 0.242 |
| 3.2 | 0.05 | yadI | 2 | 2.5 | 1.254 |
| 3.2 | 0.05 | AG1 | 2 | 2.5 | 0.274 |
| 3.2 | 0.05 | ptsI | 2 | 2.75 | 0.238 |
| 3.2 | 0.05 | clcB | 2 | 2.75 | 0.212 |
| 3.2 | 0.05 | ycaM | 2 | 2.75 | 0.244 |
| 3.2 | 0.05 | yadI | 2 | 2.75 | 1.162 |
| 3.2 | 0.05 | AG1 | 2 | 2.75 | 0.261 |
| 3.2 | 0.05 | ptsI | 2 | 3 | 0.239 |
| 3.2 | 0.05 | clcB | 2 | 3 | 0.217 |
| 3.2 | 0.05 | ycaM | 2 | 3 | 0.247 |
| 3.2 | 0.05 | yadI | 2 | 3 | 0.907 |
| 3.2 | 0.05 | AG1 | 2 | 3 | 0.267 |
| 3.2 | 0.05 | ptsI | 2 | 3.25 | 0.25 |
| 3.2 | 0.05 | clcB | 2 | 3.25 | 0.222 |
| 3.2 | 0.05 | ycaM | 2 | 3.25 | 0.254 |
| 3.2 | 0.05 | yadI | 2 | 3.25 | 1.06 |
| 3.2 | 0.05 | AG1 | 2 | 3.25 | 0.277 |
| 3.2 | 0.05 | ptsI | 2 | 3.5 | 0.258 |
| 3.2 | 0.05 | clcB | 2 | 3.5 | 0.227 |
| 3.2 | 0.05 | ycaM | 2 | 3.5 | 0.262 |
| 3.2 | 0.05 | yadI | 2 | 3.5 | 0.936 |
| 3.2 | 0.05 | AG1 | 2 | 3.5 | 0.286 |
| 3.2 | 0.05 | ptsI | 2 | 3.75 | 0.266 |
| 3.2 | 0.05 | clcB | 2 | 3.75 | 0.234 |
| 3.2 | 0.05 | ycaM | 2 | 3.75 | 0.268 |
| 3.2 | 0.05 | yadI | 2 | 3.75 | 0.838 |
| 3.2 | 0.05 | AG1 | 2 | 3.75 | 0.298 |
| 3.2 | 0.05 | ptsI | 2 | 4 | 0.274 |
| 3.2 | 0.05 | clcB | 2 | 4 | 0.239 |
| 3.2 | 0.05 | ycaM | 2 | 4 | 0.277 |
| 3.2 | 0.05 | yadI | 2 | 4 | 0.814 |
| 3.2 | 0.05 | AG1 | 2 | 4 | 0.307 |
| 3.2 | 0.05 | ptsI | 2 | 4.38333 | 0.302 |
| 3.2 | 0.05 | clcB | 2 | 4.38333 | 0.231 |
| 3.2 | 0.05 | ycaM | 2 | 4.38333 | 0.3 |
| 3.2 | 0.05 | yadI | 2 | 4.38333 | 0.4 |
| 3.2 | 0.05 | AG1 | 2 | 4.38333 | 0.279 |
| 3.2 | 0.05 | ptsI | 2 | 4.63333 | 0.298 |
| 3.2 | 0.05 | clcB | 2 | 4.63333 | 0.238 |

| | | | | | |
|---|---|---|---|---|---|
| 3.2 | 0.05 | ycaM | 2 | 4.63333 | 0.297 |
| 3.2 | 0.05 | yadI | 2 | 4.63333 | 0.4 |
| 3.2 | 0.05 | AG1 | 2 | 4.63333 | 0.286 |
| 3.2 | 0.05 | ptsI | 2 | 4.88333 | 0.302 |
| 3.2 | 0.05 | clcB | 2 | 4.88333 | 0.227 |
| 3.2 | 0.05 | ycaM | 2 | 4.88333 | 0.304 |
| 3.2 | 0.05 | yadI | 2 | 4.88333 | 0.418 |
| 3.2 | 0.05 | AG1 | 2 | 4.88333 | 0.3 |
| 3.2 | 0.05 | ptsI | 2 | 5.13333 | 0.309 |
| 3.2 | 0.05 | clcB | 2 | 5.13333 | 0.232 |
| 3.2 | 0.05 | ycaM | 2 | 5.13333 | 0.309 |
| 3.2 | 0.05 | yadI | 2 | 5.13333 | 0.43 |
| 3.2 | 0.05 | AG1 | 2 | 5.13333 | 0.308 |
| 3.2 | 0.05 | ptsI | 2 | 5.38333 | 0.316 |
| 3.2 | 0.05 | clcB | 2 | 5.38333 | 0.238 |
| 3.2 | 0.05 | ycaM | 2 | 5.38333 | 0.31 |
| 3.2 | 0.05 | yadI | 2 | 5.38333 | 0.445 |
| 3.2 | 0.05 | AG1 | 2 | 5.38333 | 0.317 |
| 3.2 | 0.05 | ptsI | 2 | 5.63333 | 0.322 |
| 3.2 | 0.05 | clcB | 2 | 5.63333 | 0.244 |
| 3.2 | 0.05 | ycaM | 2 | 5.63333 | 0.312 |
| 3.2 | 0.05 | yadI | 2 | 5.63333 | 0.444 |
| 3.2 | 0.05 | AG1 | 2 | 5.63333 | 0.325 |
| 3.2 | 0.05 | ptsI | 2 | 5.88333 | 0.328 |
| 3.2 | 0.05 | clcB | 2 | 5.88333 | 0.252 |
| 3.2 | 0.05 | ycaM | 2 | 5.88333 | 0.314 |
| 3.2 | 0.05 | yadI | 2 | 5.88333 | 0.451 |
| 3.2 | 0.05 | AG1 | 2 | 5.88333 | 0.334 |
| 3.2 | 0.05 | ptsI | 2 | 6.13333 | 0.338 |
| 3.2 | 0.05 | clcB | 2 | 6.13333 | 0.258 |
| 3.2 | 0.05 | ycaM | 2 | 6.13333 | 0.317 |
| 3.2 | 0.05 | yadI | 2 | 6.13333 | 0.459 |
| 3.2 | 0.05 | AG1 | 2 | 6.13333 | 0.342 |
| 3.2 | 0.05 | ptsI | 2 | 6.38333 | 0.345 |
| 3.2 | 0.05 | clcB | 2 | 6.38333 | 0.266 |
| 3.2 | 0.05 | ycaM | 2 | 6.38333 | 0.318 |
| 3.2 | 0.05 | yadI | 2 | 6.38333 | 0.47 |
| 3.2 | 0.05 | AG1 | 2 | 6.38333 | 0.351 |
| 3.2 | 0.05 | ptsI | 2 | 6.63333 | 0.353 |
| 3.2 | 0.05 | clcB | 2 | 6.63333 | 0.271 |
| 3.2 | 0.05 | ycaM | 2 | 6.63333 | 0.321 |
| 3.2 | 0.05 | yadI | 2 | 6.63333 | 0.485 |
| 3.2 | 0.05 | AG1 | 2 | 6.63333 | 0.362 |
| 3.2 | 0.05 | ptsI | 2 | 6.88333 | 0.363 |
| 3.2 | 0.05 | clcB | 2 | 6.88333 | 0.278 |
| 3.2 | 0.05 | ycaM | 2 | 6.88333 | 0.32 |
| 3.2 | 0.05 | yadI | 2 | 6.88333 | 0.5 |
| 3.2 | 0.05 | AG1 | 2 | 6.88333 | 0.37 |

| | | | | | |
|---|---|---|---|---|---|
| 3.2 | 0.05 | ptsI | 2 | 7.13333 | 0.371 |
| 3.2 | 0.05 | clcB | 2 | 7.13333 | 0.282 |
| 3.2 | 0.05 | ycaM | 2 | 7.13333 | 0.327 |
| 3.2 | 0.05 | yadI | 2 | 7.13333 | 0.51 |
| 3.2 | 0.05 | AG1 | 2 | 7.13333 | 0.379 |
| 3.2 | 0.05 | ptsI | 2 | 7.38333 | 0.378 |
| 3.2 | 0.05 | clcB | 2 | 7.38333 | 0.29 |
| 3.2 | 0.05 | ycaM | 2 | 7.38333 | 0.328 |
| 3.2 | 0.05 | yadI | 2 | 7.38333 | 0.521 |
| 3.2 | 0.05 | AG1 | 2 | 7.38333 | 0.387 |
| 3.2 | 0.05 | ptsI | 2 | 7.63333 | 0.39 |
| 3.2 | 0.05 | clcB | 2 | 7.63333 | 0.298 |
| 3.2 | 0.05 | ycaM | 2 | 7.63333 | 0.327 |
| 3.2 | 0.05 | yadI | 2 | 7.63333 | 0.556 |
| 3.2 | 0.05 | AG1 | 2 | 7.63333 | 0.393 |
| 3.2 | 0.05 | ptsI | 2 | 7.88333 | 0.398 |
| 3.2 | 0.05 | clcB | 2 | 7.88333 | 0.303 |
| 3.2 | 0.05 | ycaM | 2 | 7.88333 | 0.33 |
| 3.2 | 0.05 | yadI | 2 | 7.88333 | 0.58 |
| 3.2 | 0.05 | AG1 | 2 | 7.88333 | 0.402 |
| 3.2 | 0.05 | ptsI | 2 | 8.13333 | 0.409 |
| 3.2 | 0.05 | clcB | 2 | 8.13333 | 0.311 |
| 3.2 | 0.05 | ycaM | 2 | 8.13333 | 0.333 |
| 3.2 | 0.05 | yadI | 2 | 8.13333 | 0.579 |
| 3.2 | 0.05 | AG1 | 2 | 8.13333 | 0.411 |
| 3.2 | 0.05 | ptsI | 2 | 8.38333 | 0.418 |
| 3.2 | 0.05 | clcB | 2 | 8.38333 | 0.317 |
| 3.2 | 0.05 | ycaM | 2 | 8.38333 | 0.335 |
| 3.2 | 0.05 | yadI | 2 | 8.38333 | 0.608 |
| 3.2 | 0.05 | AG1 | 2 | 8.38333 | 0.42 |
| 3.2 | 0.05 | ptsI | 2 | 8.91667 | 0.371 |
| 3.2 | 0.05 | clcB | 2 | 8.91667 | 0.3 |
| 3.2 | 0.05 | ycaM | 2 | 8.91667 | 0.326 |
| 3.2 | 0.05 | yadI | 2 | 8.91667 | 0.541 |
| 3.2 | 0.05 | AG1 | 2 | 8.91667 | 0.469 |
| 3.2 | 0.05 | ptsI | 2 | 9.16667 | 0.381 |
| 3.2 | 0.05 | clcB | 2 | 9.16667 | 0.296 |
| 3.2 | 0.05 | ycaM | 2 | 9.16667 | 0.321 |
| 3.2 | 0.05 | yadI | 2 | 9.16667 | 0.57 |
| 3.2 | 0.05 | AG1 | 2 | 9.16667 | 0.486 |
| 3.2 | 0.05 | ptsI | 2 | 9.41667 | 0.38 |
| 3.2 | 0.05 | clcB | 2 | 9.41667 | 0.295 |
| 3.2 | 0.05 | ycaM | 2 | 9.41667 | 0.317 |
| 3.2 | 0.05 | yadI | 2 | 9.41667 | 0.581 |
| 3.2 | 0.05 | AG1 | 2 | 9.41667 | 0.486 |
| 3.2 | 0.05 | ptsI | 2 | 9.66667 | 0.385 |
| 3.2 | 0.05 | clcB | 2 | 9.66667 | 0.291 |
| 3.2 | 0.05 | ycaM | 2 | 9.66667 | 0.322 |

| | | | | | |
|---|---|---|---|---|---|
| 3.2 | 0.05 | yadI | 2 | 9.66667 | 0.593 |
| 3.2 | 0.05 | AG1 | 2 | 9.66667 | 0.504 |
| 3.2 | 0.05 | ptsI | 2 | 9.91667 | 0.383 |
| 3.2 | 0.05 | clcB | 2 | 9.91667 | 0.289 |
| 3.2 | 0.05 | ycaM | 2 | 9.91667 | 0.316 |
| 3.2 | 0.05 | yadI | 2 | 9.91667 | 0.593 |
| 3.2 | 0.05 | AG1 | 2 | 9.91667 | 0.499 |
| 3.2 | 0.05 | ptsI | 2 | 10.1667 | 0.385 |
| 3.2 | 0.05 | clcB | 2 | 10.1667 | 0.283 |
| 3.2 | 0.05 | ycaM | 2 | 10.1667 | 0.319 |
| 3.2 | 0.05 | yadI | 2 | 10.1667 | 0.596 |
| 3.2 | 0.05 | AG1 | 2 | 10.1667 | 0.501 |
| 3.2 | 0.05 | ptsI | 2 | 10.4167 | 0.392 |
| 3.2 | 0.05 | clcB | 2 | 10.4167 | 0.287 |
| 3.2 | 0.05 | ycaM | 2 | 10.4167 | 0.32 |
| 3.2 | 0.05 | yadI | 2 | 10.4167 | 0.614 |
| 3.2 | 0.05 | AG1 | 2 | 10.4167 | 0.515 |
| 3.2 | 0.05 | ptsI | 2 | 10.6667 | 0.395 |
| 3.2 | 0.05 | clcB | 2 | 10.6667 | 0.281 |
| 3.2 | 0.05 | ycaM | 2 | 10.6667 | 0.321 |
| 3.2 | 0.05 | yadI | 2 | 10.6667 | 0.614 |
| 3.2 | 0.05 | AG1 | 2 | 10.6667 | 0.519 |
| 3.2 | 0.05 | ptsI | 2 | 10.9167 | 0.399 |
| 3.2 | 0.05 | clcB | 2 | 10.9167 | 0.282 |
| 3.2 | 0.05 | ycaM | 2 | 10.9167 | 0.322 |
| 3.2 | 0.05 | yadI | 2 | 10.9167 | 0.614 |
| 3.2 | 0.05 | AG1 | 2 | 10.9167 | 0.524 |
| 3.2 | 0.05 | ptsI | 2 | 11.1667 | 0.405 |
| 3.2 | 0.05 | clcB | 2 | 11.1667 | 0.284 |
| 3.2 | 0.05 | ycaM | 2 | 11.1667 | 0.324 |
| 3.2 | 0.05 | yadI | 2 | 11.1667 | 0.63 |
| 3.2 | 0.05 | AG1 | 2 | 11.1667 | 0.521 |
| 3.2 | 0.05 | ptsI | 2 | 11.4167 | 0.415 |
| 3.2 | 0.05 | clcB | 2 | 11.4167 | 0.28 |
| 3.2 | 0.05 | ycaM | 2 | 11.4167 | 0.328 |
| 3.2 | 0.05 | yadI | 2 | 11.4167 | 0.633 |
| 3.2 | 0.05 | AG1 | 2 | 11.4167 | 0.519 |
| 3.2 | 0.05 | ptsI | 2 | 11.6667 | 0.414 |
| 3.2 | 0.05 | clcB | 2 | 11.6667 | 0.28 |
| 3.2 | 0.05 | ycaM | 2 | 11.6667 | 0.316 |
| 3.2 | 0.05 | yadI | 2 | 11.6667 | 0.645 |
| 3.2 | 0.05 | AG1 | 2 | 11.6667 | 0.525 |
| 3.2 | 0.05 | ptsI | 2 | 11.9167 | 0.415 |
| 3.2 | 0.05 | clcB | 2 | 11.9167 | 0.28 |
| 3.2 | 0.05 | ycaM | 2 | 11.9167 | 0.321 |
| 3.2 | 0.05 | yadI | 2 | 11.9167 | 0.651 |
| 3.2 | 0.05 | AG1 | 2 | 11.9167 | 0.529 |
| 3.2 | 0.05 | ptsI | 2 | 12.1667 | 0.424 |

| | | | | | |
|---|---|---|---|---|---|
| 3.2 | 0.05 | clcB | 2 | 12.1667 | 0.278 |
| 3.2 | 0.05 | ycaM | 2 | 12.1667 | 0.315 |
| 3.2 | 0.05 | yadI | 2 | 12.1667 | 0.658 |
| 3.2 | 0.05 | AG1 | 2 | 12.1667 | 0.537 |
| 3.2 | 0.05 | ptsI | 2 | 12.4167 | 0.425 |
| 3.2 | 0.05 | clcB | 2 | 12.4167 | 0.279 |
| 3.2 | 0.05 | ycaM | 2 | 12.4167 | 0.305 |
| 3.2 | 0.05 | yadI | 2 | 12.4167 | 0.662 |
| 3.2 | 0.05 | AG1 | 2 | 12.4167 | 0.538 |
| 3.2 | 0.05 | ptsI | 2 | 12.6667 | 0.424 |
| 3.2 | 0.05 | clcB | 2 | 12.6667 | 0.272 |
| 3.2 | 0.05 | ycaM | 2 | 12.6667 | 0.307 |
| 3.2 | 0.05 | yadI | 2 | 12.6667 | 0.664 |
| 3.2 | 0.05 | AG1 | 2 | 12.6667 | 0.539 |
| 3.2 | 0.05 | ptsI | 2 | 12.9167 | 0.426 |
| 3.2 | 0.05 | clcB | 2 | 12.9167 | 0.272 |
| 3.2 | 0.05 | ycaM | 2 | 12.9167 | 0.303 |
| 3.2 | 0.05 | yadI | 2 | 12.9167 | 0.669 |
| 3.2 | 0.05 | AG1 | 2 | 12.9167 | 0.542 |
| 3.2 | 0.05 | ptsI | 2 | 13.1667 | 0.427 |
| 3.2 | 0.05 | clcB | 2 | 13.1667 | 0.268 |
| 3.2 | 0.05 | ycaM | 2 | 13.1667 | 0.502 |
| 3.2 | 0.05 | yadI | 2 | 13.1667 | 0.675 |
| 3.2 | 0.05 | AG1 | 2 | 13.1667 | 0.547 |
| 3.2 | 0.05 | ptsI | 2 | 13.4167 | 0.428 |
| 3.2 | 0.05 | clcB | 2 | 13.4167 | 0.267 |
| 3.2 | 0.05 | ycaM | 2 | 13.4167 | 0.487 |
| 3.2 | 0.05 | yadI | 2 | 13.4167 | 0.686 |
| 3.2 | 0.05 | AG1 | 2 | 13.4167 | 0.549 |
| 3.2 | 0.05 | ptsI | 2 | 13.6667 | 0.43 |
| 3.2 | 0.05 | clcB | 2 | 13.6667 | 0.264 |
| 3.2 | 0.05 | ycaM | 2 | 13.6667 | 0.466 |
| 3.2 | 0.05 | yadI | 2 | 13.6667 | 0.69 |
| 3.2 | 0.05 | AG1 | 2 | 13.6667 | 0.555 |
| 3.2 | 0.05 | ptsI | 2 | 13.9167 | 0.429 |
| 3.2 | 0.05 | clcB | 2 | 13.9167 | 0.264 |
| 3.2 | 0.05 | ycaM | 2 | 13.9167 | 0.456 |
| 3.2 | 0.05 | yadI | 2 | 13.9167 | 0.696 |
| 3.2 | 0.05 | AG1 | 2 | 13.9167 | 0.559 |
| 3.2 | 0.05 | ptsI | 2 | 14.1667 | 0.431 |
| 3.2 | 0.05 | clcB | 2 | 14.1667 | 0.262 |
| 3.2 | 0.05 | ycaM | 2 | 14.1667 | 0.418 |
| 3.2 | 0.05 | yadI | 2 | 14.1667 | 0.699 |
| 3.2 | 0.05 | AG1 | 2 | 14.1667 | 0.562 |
| 3.2 | 0.05 | ptsI | 2 | 14.4167 | 0.432 |
| 3.2 | 0.05 | clcB | 2 | 14.4167 | 0.26 |
| 3.2 | 0.05 | ycaM | 2 | 14.4167 | 0.384 |
| 3.2 | 0.05 | yadI | 2 | 14.4167 | 0.712 |

| | | | | | |
|---|---|---|---|---|---|
| 3.2 | 0.05 | AG1 | 2 | 14.4167 | 0.568 |
| 3.2 | 0.05 | ptsI | 2 | 14.6667 | 0.432 |
| 3.2 | 0.05 | clcB | 2 | 14.6667 | 0.26 |
| 3.2 | 0.05 | ycaM | 2 | 14.6667 | 0.39 |
| 3.2 | 0.05 | yadI | 2 | 14.6667 | 0.717 |
| 3.2 | 0.05 | AG1 | 2 | 14.6667 | 0.576 |
| 3.2 | 0.05 | ptsI | 2 | 14.9167 | 0.433 |
| 3.2 | 0.05 | clcB | 2 | 14.9167 | 0.259 |
| 3.2 | 0.05 | ycaM | 2 | 14.9167 | 0.39 |
| 3.2 | 0.05 | yadI | 2 | 14.9167 | 0.724 |
| 3.2 | 0.05 | AG1 | 2 | 14.9167 | 0.575 |
| 3.2 | 0.05 | ptsI | 2 | 15.1667 | 0.436 |
| 3.2 | 0.05 | clcB | 2 | 15.1667 | 0.257 |
| 3.2 | 0.05 | ycaM | 2 | 15.1667 | 0.436 |
| 3.2 | 0.05 | yadI | 2 | 15.1667 | 0.693 |
| 3.2 | 0.05 | AG1 | 2 | 15.1667 | 0.592 |
| 3.2 | 0.05 | ptsI | 2 | 15.4167 | 0.437 |
| 3.2 | 0.05 | clcB | 2 | 15.4167 | 0.257 |
| 3.2 | 0.05 | ycaM | 2 | 15.4167 | 0.376 |
| 3.2 | 0.05 | yadI | 2 | 15.4167 | 0.741 |
| 3.2 | 0.05 | AG1 | 2 | 15.4167 | 0.582 |
| 3.2 | 0.05 | ptsI | 2 | 15.6667 | 0.437 |
| 3.2 | 0.05 | clcB | 2 | 15.6667 | 0.257 |
| 3.2 | 0.05 | ycaM | 2 | 15.6667 | 0.381 |
| 3.2 | 0.05 | yadI | 2 | 15.6667 | 0.743 |
| 3.2 | 0.05 | AG1 | 2 | 15.6667 | 0.586 |
| 3.2 | 0.05 | ptsI | 2 | 15.9167 | 0.442 |
| 3.2 | 0.05 | clcB | 2 | 15.9167 | 0.257 |
| 3.2 | 0.05 | ycaM | 2 | 15.9167 | 0.381 |
| 3.2 | 0.05 | yadI | 2 | 15.9167 | 0.751 |
| 3.2 | 0.05 | AG1 | 2 | 15.9167 | 0.587 |
| 3.2 | 0.05 | ptsI | 2 | 16.1667 | 0.444 |
| 3.2 | 0.05 | clcB | 2 | 16.1667 | 0.255 |
| 3.2 | 0.05 | ycaM | 2 | 16.1667 | 0.375 |
| 3.2 | 0.05 | yadI | 2 | 16.1667 | 0.758 |
| 3.2 | 0.05 | AG1 | 2 | 16.1667 | 0.594 |
| 3.2 | 0.05 | ptsI | 2 | 16.4167 | 0.443 |
| 3.2 | 0.05 | clcB | 2 | 16.4167 | 0.254 |
| 3.2 | 0.05 | ycaM | 2 | 16.4167 | 0.375 |
| 3.2 | 0.05 | yadI | 2 | 16.4167 | 0.756 |
| 3.2 | 0.05 | AG1 | 2 | 16.4167 | 0.596 |
| 3.2 | 0.05 | ptsI | 2 | 16.6667 | 0.445 |
| 3.2 | 0.05 | clcB | 2 | 16.6667 | 0.253 |
| 3.2 | 0.05 | ycaM | 2 | 16.6667 | 0.376 |
| 3.2 | 0.05 | yadI | 2 | 16.6667 | 0.759 |
| 3.2 | 0.05 | AG1 | 2 | 16.6667 | 0.601 |
| 3.2 | 0.05 | ptsI | 2 | 16.9167 | 0.445 |
| 3.2 | 0.05 | clcB | 2 | 16.9167 | 0.252 |

| | | | | | |
|---|---|---|---|---|---|
| 3.2 | 0.05 | ycaM | 2 | 16.9167 | 0.377 |
| 3.2 | 0.05 | yadI | 2 | 16.9167 | 0.774 |
| 3.2 | 0.05 | AG1 | 2 | 16.9167 | 0.602 |
| 3.2 | 0.05 | ptsI | 2 | 17.1667 | 0.451 |
| 3.2 | 0.05 | clcB | 2 | 17.1667 | 0.253 |
| 3.2 | 0.05 | ycaM | 2 | 17.1667 | 0.372 |
| 3.2 | 0.05 | yadI | 2 | 17.1667 | 0.773 |
| 3.2 | 0.05 | AG1 | 2 | 17.1667 | 0.608 |
| 3.2 | 0.05 | ptsI | 2 | 17.4167 | 0.451 |
| 3.2 | 0.05 | clcB | 2 | 17.4167 | 0.251 |
| 3.2 | 0.05 | ycaM | 2 | 17.4167 | 0.368 |
| 3.2 | 0.05 | yadI | 2 | 17.4167 | 0.783 |
| 3.2 | 0.05 | AG1 | 2 | 17.4167 | 0.612 |
| 3.2 | 0.05 | ptsI | 2 | 17.6667 | 0.45 |
| 3.2 | 0.05 | clcB | 2 | 17.6667 | 0.249 |
| 3.2 | 0.05 | ycaM | 2 | 17.6667 | 0.357 |
| 3.2 | 0.05 | yadI | 2 | 17.6667 | 0.791 |
| 3.2 | 0.05 | AG1 | 2 | 17.6667 | 0.615 |
| 3.2 | 0.05 | ptsI | 2 | 17.9167 | 0.451 |
| 3.2 | 0.05 | clcB | 2 | 17.9167 | 0.248 |
| 3.2 | 0.05 | ycaM | 2 | 17.9167 | 0.358 |
| 3.2 | 0.05 | yadI | 2 | 17.9167 | 0.787 |
| 3.2 | 0.05 | AG1 | 2 | 17.9167 | 0.618 |
| 3.2 | 0.05 | ptsI | 2 | 18.1667 | 0.451 |
| 3.2 | 0.05 | clcB | 2 | 18.1667 | 0.247 |
| 3.2 | 0.05 | ycaM | 2 | 18.1667 | 0.362 |
| 3.2 | 0.05 | yadI | 2 | 18.1667 | 0.797 |
| 3.2 | 0.05 | AG1 | 2 | 18.1667 | 0.623 |
| 3.2 | 0.05 | ptsI | 2 | 18.4167 | 0.452 |
| 3.2 | 0.05 | clcB | 2 | 18.4167 | 0.247 |
| 3.2 | 0.05 | ycaM | 2 | 18.4167 | 0.353 |
| 3.2 | 0.05 | yadI | 2 | 18.4167 | 0.81 |
| 3.2 | 0.05 | AG1 | 2 | 18.4167 | 0.623 |
| 3.2 | 0.05 | ptsI | 2 | 18.6667 | 0.455 |
| 3.2 | 0.05 | clcB | 2 | 18.6667 | 0.247 |
| 3.2 | 0.05 | ycaM | 2 | 18.6667 | 0.348 |
| 3.2 | 0.05 | yadI | 2 | 18.6667 | 0.809 |
| 3.2 | 0.05 | AG1 | 2 | 18.6667 | 0.629 |
| 3.2 | 0.05 | ptsI | 2 | 18.9167 | 0.453 |
| 3.2 | 0.05 | clcB | 2 | 18.9167 | 0.246 |
| 3.2 | 0.05 | ycaM | 2 | 18.9167 | 0.319 |
| 3.2 | 0.05 | yadI | 2 | 18.9167 | 0.88 |
| 3.2 | 0.05 | AG1 | 2 | 18.9167 | 0.619 |
| 3.2 | 0.05 | ptsI | 2 | 19.1667 | 0.458 |
| 3.2 | 0.05 | clcB | 2 | 19.1667 | 0.243 |
| 3.2 | 0.05 | ycaM | 2 | 19.1667 | 0.345 |
| 3.2 | 0.05 | yadI | 2 | 19.1667 | 0.808 |
| 3.2 | 0.05 | AG1 | 2 | 19.1667 | 0.634 |

| | | | | | |
|---|---|---|---|---|---|
| 3.2 | 0.05 | ptsI | 2 | 19.4167 | 0.457 |
| 3.2 | 0.05 | clcB | 2 | 19.4167 | 0.244 |
| 3.2 | 0.05 | ycaM | 2 | 19.4167 | 0.349 |
| 3.2 | 0.05 | yadI | 2 | 19.4167 | 0.813 |
| 3.2 | 0.05 | AG1 | 2 | 19.4167 | 0.637 |
| 3.2 | 0.05 | ptsI | 2 | 19.6667 | 0.459 |
| 3.2 | 0.05 | clcB | 2 | 19.6667 | 0.242 |
| 3.2 | 0.05 | ycaM | 2 | 19.6667 | 0.348 |
| 3.2 | 0.05 | yadI | 2 | 19.6667 | 0.815 |
| 3.2 | 0.05 | AG1 | 2 | 19.6667 | 0.641 |
| 3.2 | 0.05 | ptsI | 2 | 19.9167 | 0.462 |
| 3.2 | 0.05 | clcB | 2 | 19.9167 | 0.242 |
| 3.2 | 0.05 | ycaM | 2 | 19.9167 | 0.347 |
| 3.2 | 0.05 | yadI | 2 | 19.9167 | 0.819 |
| 3.2 | 0.05 | AG1 | 2 | 19.9167 | 0.646 |
| 3.2 | 0.05 | ptsI | 2 | 20.1667 | 0.46 |
| 3.2 | 0.05 | clcB | 2 | 20.1667 | 0.242 |
| 3.2 | 0.05 | ycaM | 2 | 20.1667 | 0.344 |
| 3.2 | 0.05 | yadI | 2 | 20.1667 | 0.827 |
| 3.2 | 0.05 | AG1 | 2 | 20.1667 | 0.65 |
| 3.2 | 0.05 | ptsI | 2 | 20.4167 | 0.462 |
| 3.2 | 0.05 | clcB | 2 | 20.4167 | 0.24 |
| 3.2 | 0.05 | ycaM | 2 | 20.4167 | 0.341 |
| 3.2 | 0.05 | yadI | 2 | 20.4167 | 0.826 |
| 3.2 | 0.05 | AG1 | 2 | 20.4167 | 0.651 |
| 3.2 | 0.05 | ptsI | 2 | 20.6667 | 0.464 |
| 3.2 | 0.05 | clcB | 2 | 20.6667 | 0.24 |
| 3.2 | 0.05 | ycaM | 2 | 20.6667 | 0.34 |
| 3.2 | 0.05 | yadI | 2 | 20.6667 | 0.827 |
| 3.2 | 0.05 | AG1 | 2 | 20.6667 | 0.658 |
| 3.2 | 0.05 | ptsI | 2 | 20.9167 | 0.467 |
| 3.2 | 0.05 | clcB | 2 | 20.9167 | 0.239 |
| 3.2 | 0.05 | ycaM | 2 | 20.9167 | 0.339 |
| 3.2 | 0.05 | yadI | 2 | 20.9167 | 0.829 |
| 3.2 | 0.05 | AG1 | 2 | 20.9167 | 0.657 |
| 3.2 | 0.05 | ptsI | 2 | 21.1667 | 0.466 |
| 3.2 | 0.05 | clcB | 2 | 21.1667 | 0.238 |
| 3.2 | 0.05 | ycaM | 2 | 21.1667 | 0.336 |
| 3.2 | 0.05 | yadI | 2 | 21.1667 | 0.835 |
| 3.2 | 0.05 | AG1 | 2 | 21.1667 | 0.662 |
| 3.2 | 0.05 | ptsI | 2 | 21.4167 | 0.466 |
| 3.2 | 0.05 | clcB | 2 | 21.4167 | 0.238 |
| 3.2 | 0.05 | ycaM | 2 | 21.4167 | 0.332 |
| 3.2 | 0.05 | yadI | 2 | 21.4167 | 0.872 |
| 3.2 | 0.05 | AG1 | 2 | 21.4167 | 0.667 |
| 3.2 | 0.05 | ptsI | 2 | 21.6667 | 0.469 |
| 3.2 | 0.05 | clcB | 2 | 21.6667 | 0.238 |
| 3.2 | 0.05 | ycaM | 2 | 21.6667 | 0.325 |

| | | | | | |
|---|---|---|---|---|---|
| 3.2 | 0.05 | yadI | 2 | 21.6667 | 0.845 |
| 3.2 | 0.05 | AG1 | 2 | 21.6667 | 0.675 |
| 3.2 | 0.05 | ptsI | 2 | 21.9167 | 0.468 |
| 3.2 | 0.05 | clcB | 2 | 21.9167 | 0.237 |
| 3.2 | 0.05 | ycaM | 2 | 21.9167 | 0.333 |
| 3.2 | 0.05 | yadI | 2 | 21.9167 | 0.845 |
| 3.2 | 0.05 | AG1 | 2 | 21.9167 | 0.676 |
| 3.2 | 0.05 | ptsI | 2 | 22.1667 | 0.472 |
| 3.2 | 0.05 | clcB | 2 | 22.1667 | 0.235 |
| 3.2 | 0.05 | ycaM | 2 | 22.1667 | 0.335 |
| 3.2 | 0.05 | yadI | 2 | 22.1667 | 0.849 |
| 3.2 | 0.05 | AG1 | 2 | 22.1667 | 0.683 |
| 3.2 | 0.05 | ptsI | 2 | 22.4167 | 0.47 |
| 3.2 | 0.05 | clcB | 2 | 22.4167 | 0.235 |
| 3.2 | 0.05 | ycaM | 2 | 22.4167 | 0.329 |
| 3.2 | 0.05 | yadI | 2 | 22.4167 | 0.853 |
| 3.2 | 0.05 | AG1 | 2 | 22.4167 | 0.687 |
| 3.2 | 0.05 | ptsI | 2 | 22.6667 | 0.472 |
| 3.2 | 0.05 | clcB | 2 | 22.6667 | 0.233 |
| 3.2 | 0.05 | ycaM | 2 | 22.6667 | 0.331 |
| 3.2 | 0.05 | yadI | 2 | 22.6667 | 0.856 |
| 3.2 | 0.05 | AG1 | 2 | 22.6667 | 0.689 |
| 3.2 | 0.05 | ptsI | 2 | 22.9167 | 0.474 |
| 3.2 | 0.05 | clcB | 2 | 22.9167 | 0.234 |
| 3.2 | 0.05 | ycaM | 2 | 22.9167 | 0.32 |
| 3.2 | 0.05 | yadI | 2 | 22.9167 | 0.853 |
| 3.2 | 0.05 | AG1 | 2 | 22.9167 | 0.694 |
| 3.2 | 0.05 | ptsI | 2 | 23.1667 | 0.476 |
| 3.2 | 0.05 | clcB | 2 | 23.1667 | 0.233 |
| 3.2 | 0.05 | ycaM | 2 | 23.1667 | 0.323 |
| 3.2 | 0.05 | yadI | 2 | 23.1667 | 0.859 |
| 3.2 | 0.05 | AG1 | 2 | 23.1667 | 0.698 |
| 3.2 | 0.05 | ptsI | 2 | 23.4167 | 0.479 |
| 3.2 | 0.05 | clcB | 2 | 23.4167 | 0.231 |
| 3.2 | 0.05 | ycaM | 2 | 23.4167 | 0.32 |
| 3.2 | 0.05 | yadI | 2 | 23.4167 | 0.857 |
| 3.2 | 0.05 | AG1 | 2 | 23.4167 | 0.705 |
| 3.2 | 0.05 | ptsI | 2 | 23.6667 | 0.48 |
| 3.2 | 0.05 | clcB | 2 | 23.6667 | 0.23 |
| 3.2 | 0.05 | ycaM | 2 | 23.6667 | 0.325 |
| 3.2 | 0.05 | yadI | 2 | 23.6667 | 0.858 |
| 3.2 | 0.05 | AG1 | 2 | 23.6667 | 0.71 |
| 3.2 | 0.05 | ptsI | 2 | 23.9167 | 0.479 |
| 3.2 | 0.05 | clcB | 2 | 23.9167 | 0.229 |
| 3.2 | 0.05 | ycaM | 2 | 23.9167 | 0.323 |
| 3.2 | 0.05 | yadI | 2 | 23.9167 | 0.865 |
| 3.2 | 0.05 | AG1 | 2 | 23.9167 | 0.716 |
| 3.2 | 0.05 | ptsI | 2 | 24.1667 | 0.481 |

| | | | | | |
|---|---|---|---|---|---|
| 3.2 | 0.05 | clcB | 2 | 24.1667 | 0.229 |
| 3.2 | 0.05 | ycaM | 2 | 24.1667 | 0.322 |
| 3.2 | 0.05 | yadI | 2 | 24.1667 | 0.869 |
| 3.2 | 0.05 | AG1 | 2 | 24.1667 | 0.722 |
| 3.2 | 0.05 | ptsI | 2 | 24.4167 | 0.485 |
| 3.2 | 0.05 | clcB | 2 | 24.4167 | 0.229 |
| 3.2 | 0.05 | ycaM | 2 | 24.4167 | 0.32 |
| 3.2 | 0.05 | yadI | 2 | 24.4167 | 0.874 |
| 3.2 | 0.05 | AG1 | 2 | 24.4167 | 0.732 |
| 3.2 | 0.05 | ptsI | 2 | 24.6667 | 0.483 |
| 3.2 | 0.05 | clcB | 2 | 24.6667 | 0.229 |
| 3.2 | 0.05 | ycaM | 2 | 24.6667 | 0.32 |
| 3.2 | 0.05 | yadI | 2 | 24.6667 | 0.878 |
| 3.2 | 0.05 | AG1 | 2 | 24.6667 | 0.739 |
| 3.2 | 0.05 | ptsI | 2 | 24.9167 | 0.483 |
| 3.2 | 0.05 | clcB | 2 | 24.9167 | 0.226 |
| 3.2 | 0.05 | ycaM | 2 | 24.9167 | 0.319 |
| 3.2 | 0.05 | yadI | 2 | 24.9167 | 0.876 |
| 3.2 | 0.05 | AG1 | 2 | 24.9167 | 0.744 |
| 3.2 | 0.05 | ptsI | 2 | 25.1667 | 0.489 |
| 3.2 | 0.05 | clcB | 2 | 25.1667 | 0.226 |
| 3.2 | 0.05 | ycaM | 2 | 25.1667 | 0.316 |
| 3.2 | 0.05 | yadI | 2 | 25.1667 | 0.875 |
| 3.2 | 0.05 | AG1 | 2 | 25.1667 | 0.752 |
| 3.2 | 0.05 | ptsI | 2 | 25.4167 | 0.488 |
| 3.2 | 0.05 | clcB | 2 | 25.4167 | 0.226 |
| 3.2 | 0.05 | ycaM | 2 | 25.4167 | 0.317 |
| 3.2 | 0.05 | yadI | 2 | 25.4167 | 0.875 |
| 3.2 | 0.05 | AG1 | 2 | 25.4167 | 0.761 |
| 3.2 | 0.05 | ptsI | 2 | 25.6667 | 0.491 |
| 3.2 | 0.05 | clcB | 2 | 25.6667 | 0.225 |
| 3.2 | 0.05 | ycaM | 2 | 25.6667 | 0.314 |
| 3.2 | 0.05 | yadI | 2 | 25.6667 | 0.877 |
| 3.2 | 0.05 | AG1 | 2 | 25.6667 | 0.764 |
| 3.2 | 0.05 | ptsI | 2 | 25.9167 | 0.493 |
| 3.2 | 0.05 | clcB | 2 | 25.9167 | 0.225 |
| 3.2 | 0.05 | ycaM | 2 | 25.9167 | 0.312 |
| 3.2 | 0.05 | yadI | 2 | 25.9167 | 0.885 |
| 3.2 | 0.05 | AG1 | 2 | 25.9167 | 0.771 |
| 3.2 | 0.05 | ptsI | 2 | 26.1667 | 0.494 |
| 3.2 | 0.05 | clcB | 2 | 26.1667 | 0.224 |
| 3.2 | 0.05 | ycaM | 2 | 26.1667 | 0.305 |
| 3.2 | 0.05 | yadI | 2 | 26.1667 | 0.916 |
| 3.2 | 0.05 | AG1 | 2 | 26.1667 | 0.777 |
| 3.2 | 0.05 | ptsI | 2 | 26.4167 | 0.495 |
| 3.2 | 0.05 | clcB | 2 | 26.4167 | 0.223 |
| 3.2 | 0.05 | ycaM | 2 | 26.4167 | 0.31 |
| 3.2 | 0.05 | yadI | 2 | 26.4167 | 0.89 |

| | | | | | |
|---|---|---|---|---|---|
| 3.2 | 0.05 | AG1 | 2 | 26.4167 | 0.789 |
| 3.2 | 0.05 | ptsI | 3 | 0 | 0.168 |
| 3.2 | 0.05 | clcB | 3 | 0 | 0.161 |
| 3.2 | 0.05 | ycaM | 3 | 0 | 0.158 |
| 3.2 | 0.05 | yadI | 3 | 0 | 0.164 |
| 3.2 | 0.05 | AG1 | 3 | 0 | 0.165 |
| 3.2 | 0.05 | ptsI | 3 | 0.25 | 0.156 |
| 3.2 | 0.05 | clcB | 3 | 0.25 | 0.159 |
| 3.2 | 0.05 | ycaM | 3 | 0.25 | 0.154 |
| 3.2 | 0.05 | yadI | 3 | 0.25 | 0.163 |
| 3.2 | 0.05 | AG1 | 3 | 0.25 | 0.165 |
| 3.2 | 0.05 | ptsI | 3 | 0.5 | 0.155 |
| 3.2 | 0.05 | clcB | 3 | 0.5 | 0.158 |
| 3.2 | 0.05 | ycaM | 3 | 0.5 | 0.153 |
| 3.2 | 0.05 | yadI | 3 | 0.5 | 0.161 |
| 3.2 | 0.05 | AG1 | 3 | 0.5 | 0.163 |
| 3.2 | 0.05 | ptsI | 3 | 0.75 | 0.155 |
| 3.2 | 0.05 | clcB | 3 | 0.75 | 0.158 |
| 3.2 | 0.05 | ycaM | 3 | 0.75 | 0.152 |
| 3.2 | 0.05 | yadI | 3 | 0.75 | 0.159 |
| 3.2 | 0.05 | AG1 | 3 | 0.75 | 0.161 |
| 3.2 | 0.05 | ptsI | 3 | 1 | 0.157 |
| 3.2 | 0.05 | clcB | 3 | 1 | 0.158 |
| 3.2 | 0.05 | ycaM | 3 | 1 | 0.154 |
| 3.2 | 0.05 | yadI | 3 | 1 | 0.16 |
| 3.2 | 0.05 | AG1 | 3 | 1 | 0.163 |
| 3.2 | 0.05 | ptsI | 3 | 1.25 | 0.157 |
| 3.2 | 0.05 | clcB | 3 | 1.25 | 0.158 |
| 3.2 | 0.05 | ycaM | 3 | 1.25 | 0.154 |
| 3.2 | 0.05 | yadI | 3 | 1.25 | 0.16 |
| 3.2 | 0.05 | AG1 | 3 | 1.25 | 0.162 |
| 3.2 | 0.05 | ptsI | 3 | 1.5 | 0.159 |
| 3.2 | 0.05 | clcB | 3 | 1.5 | 0.157 |
| 3.2 | 0.05 | ycaM | 3 | 1.5 | 0.154 |
| 3.2 | 0.05 | yadI | 3 | 1.5 | 0.16 |
| 3.2 | 0.05 | AG1 | 3 | 1.5 | 0.162 |
| 3.2 | 0.05 | ptsI | 3 | 1.75 | 0.16 |
| 3.2 | 0.05 | clcB | 3 | 1.75 | 0.158 |
| 3.2 | 0.05 | ycaM | 3 | 1.75 | 0.154 |
| 3.2 | 0.05 | yadI | 3 | 1.75 | 0.161 |
| 3.2 | 0.05 | AG1 | 3 | 1.75 | 0.163 |
| 3.2 | 0.05 | ptsI | 3 | 2 | 0.161 |
| 3.2 | 0.05 | clcB | 3 | 2 | 0.159 |
| 3.2 | 0.05 | ycaM | 3 | 2 | 0.156 |
| 3.2 | 0.05 | yadI | 3 | 2 | 0.163 |
| 3.2 | 0.05 | AG1 | 3 | 2 | 0.164 |
| 3.2 | 0.05 | ptsI | 3 | 2.25 | 0.163 |
| 3.2 | 0.05 | clcB | 3 | 2.25 | 0.16 |

| | | | | | |
|---|---|---|---|---|---|
| 3.2 | 0.05 | ycaM | 3 | 2.25 | 0.158 |
| 3.2 | 0.05 | yadI | 3 | 2.25 | 0.165 |
| 3.2 | 0.05 | AG1 | 3 | 2.25 | 0.166 |
| 3.2 | 0.05 | ptsI | 3 | 2.5 | 0.165 |
| 3.2 | 0.05 | clcB | 3 | 2.5 | 0.163 |
| 3.2 | 0.05 | ycaM | 3 | 2.5 | 0.159 |
| 3.2 | 0.05 | yadI | 3 | 2.5 | 0.166 |
| 3.2 | 0.05 | AG1 | 3 | 2.5 | 0.168 |
| 3.2 | 0.05 | ptsI | 3 | 2.75 | 0.172 |
| 3.2 | 0.05 | clcB | 3 | 2.75 | 0.165 |
| 3.2 | 0.05 | ycaM | 3 | 2.75 | 0.163 |
| 3.2 | 0.05 | yadI | 3 | 2.75 | 0.17 |
| 3.2 | 0.05 | AG1 | 3 | 2.75 | 0.173 |
| 3.2 | 0.05 | ptsI | 3 | 3 | 0.167 |
| 3.2 | 0.05 | clcB | 3 | 3 | 0.165 |
| 3.2 | 0.05 | ycaM | 3 | 3 | 0.162 |
| 3.2 | 0.05 | yadI | 3 | 3 | 0.172 |
| 3.2 | 0.05 | AG1 | 3 | 3 | 0.173 |
| 3.2 | 0.05 | ptsI | 3 | 3.25 | 0.17 |
| 3.2 | 0.05 | clcB | 3 | 3.25 | 0.166 |
| 3.2 | 0.05 | ycaM | 3 | 3.25 | 0.164 |
| 3.2 | 0.05 | yadI | 3 | 3.25 | 0.175 |
| 3.2 | 0.05 | AG1 | 3 | 3.25 | 0.174 |
| 3.2 | 0.05 | ptsI | 3 | 3.5 | 0.176 |
| 3.2 | 0.05 | clcB | 3 | 3.5 | 0.169 |
| 3.2 | 0.05 | ycaM | 3 | 3.5 | 0.168 |
| 3.2 | 0.05 | yadI | 3 | 3.5 | 0.18 |
| 3.2 | 0.05 | AG1 | 3 | 3.5 | 0.177 |
| 3.2 | 0.05 | ptsI | 3 | 3.75 | 0.18 |
| 3.2 | 0.05 | clcB | 3 | 3.75 | 0.171 |
| 3.2 | 0.05 | ycaM | 3 | 3.75 | 0.171 |
| 3.2 | 0.05 | yadI | 3 | 3.75 | 0.184 |
| 3.2 | 0.05 | AG1 | 3 | 3.75 | 0.18 |
| 3.2 | 0.05 | ptsI | 3 | 4 | 0.184 |
| 3.2 | 0.05 | clcB | 3 | 4 | 0.174 |
| 3.2 | 0.05 | ycaM | 3 | 4 | 0.174 |
| 3.2 | 0.05 | yadI | 3 | 4 | 0.191 |
| 3.2 | 0.05 | AG1 | 3 | 4 | 0.183 |
| 3.2 | 0.05 | ptsI | 3 | 4.41667 | 0.198 |
| 3.2 | 0.05 | clcB | 3 | 4.41667 | 0.2 |
| 3.2 | 0.05 | ycaM | 3 | 4.41667 | 0.174 |
| 3.2 | 0.05 | yadI | 3 | 4.41667 | 0.268 |
| 3.2 | 0.05 | AG1 | 3 | 4.41667 | 0.199 |
| 3.2 | 0.05 | ptsI | 3 | 4.66667 | 0.188 |
| 3.2 | 0.05 | clcB | 3 | 4.66667 | 0.187 |
| 3.2 | 0.05 | ycaM | 3 | 4.66667 | 0.17 |
| 3.2 | 0.05 | yadI | 3 | 4.66667 | 0.27 |
| 3.2 | 0.05 | AG1 | 3 | 4.66667 | 0.199 |

| | | | | | |
|---|---|---|---|---|---|
| 3.2 | 0.05 | ptsI | 3 | 4.91667 | 0.193 |
| 3.2 | 0.05 | clcB | 3 | 4.91667 | 0.189 |
| 3.2 | 0.05 | ycaM | 3 | 4.91667 | 0.174 |
| 3.2 | 0.05 | yadI | 3 | 4.91667 | 0.282 |
| 3.2 | 0.05 | AG1 | 3 | 4.91667 | 0.207 |
| 3.2 | 0.05 | ptsI | 3 | 5.16667 | 0.195 |
| 3.2 | 0.05 | clcB | 3 | 5.16667 | 0.194 |
| 3.2 | 0.05 | ycaM | 3 | 5.16667 | 0.178 |
| 3.2 | 0.05 | yadI | 3 | 5.16667 | 0.294 |
| 3.2 | 0.05 | AG1 | 3 | 5.16667 | 0.214 |
| 3.2 | 0.05 | ptsI | 3 | 5.41667 | 0.198 |
| 3.2 | 0.05 | clcB | 3 | 5.41667 | 0.2 |
| 3.2 | 0.05 | ycaM | 3 | 5.41667 | 0.184 |
| 3.2 | 0.05 | yadI | 3 | 5.41667 | 0.307 |
| 3.2 | 0.05 | AG1 | 3 | 5.41667 | 0.221 |
| 3.2 | 0.05 | ptsI | 3 | 5.66667 | 0.204 |
| 3.2 | 0.05 | clcB | 3 | 5.66667 | 0.206 |
| 3.2 | 0.05 | ycaM | 3 | 5.66667 | 0.191 |
| 3.2 | 0.05 | yadI | 3 | 5.66667 | 0.321 |
| 3.2 | 0.05 | AG1 | 3 | 5.66667 | 0.23 |
| 3.2 | 0.05 | ptsI | 3 | 5.91667 | 0.208 |
| 3.2 | 0.05 | clcB | 3 | 5.91667 | 0.205 |
| 3.2 | 0.05 | ycaM | 3 | 5.91667 | 0.197 |
| 3.2 | 0.05 | yadI | 3 | 5.91667 | 0.33 |
| 3.2 | 0.05 | AG1 | 3 | 5.91667 | 0.235 |
| 3.2 | 0.05 | ptsI | 3 | 6.16667 | 0.215 |
| 3.2 | 0.05 | clcB | 3 | 6.16667 | 0.214 |
| 3.2 | 0.05 | ycaM | 3 | 6.16667 | 0.204 |
| 3.2 | 0.05 | yadI | 3 | 6.16667 | 0.336 |
| 3.2 | 0.05 | AG1 | 3 | 6.16667 | 0.247 |
| 3.2 | 0.05 | ptsI | 3 | 6.41667 | 0.223 |
| 3.2 | 0.05 | clcB | 3 | 6.41667 | 0.223 |
| 3.2 | 0.05 | ycaM | 3 | 6.41667 | 0.215 |
| 3.2 | 0.05 | yadI | 3 | 6.41667 | 0.347 |
| 3.2 | 0.05 | AG1 | 3 | 6.41667 | 0.254 |
| 3.2 | 0.05 | ptsI | 3 | 6.66667 | 0.234 |
| 3.2 | 0.05 | clcB | 3 | 6.66667 | 0.235 |
| 3.2 | 0.05 | ycaM | 3 | 6.66667 | 0.223 |
| 3.2 | 0.05 | yadI | 3 | 6.66667 | 0.357 |
| 3.2 | 0.05 | AG1 | 3 | 6.66667 | 0.262 |
| 3.2 | 0.05 | ptsI | 3 | 6.91667 | 0.24 |
| 3.2 | 0.05 | clcB | 3 | 6.91667 | 0.245 |
| 3.2 | 0.05 | ycaM | 3 | 6.91667 | 0.23 |
| 3.2 | 0.05 | yadI | 3 | 6.91667 | 0.368 |
| 3.2 | 0.05 | AG1 | 3 | 6.91667 | 0.267 |
| 3.2 | 0.05 | ptsI | 3 | 7.16667 | 0.25 |
| 3.2 | 0.05 | clcB | 3 | 7.16667 | 0.26 |
| 3.2 | 0.05 | ycaM | 3 | 7.16667 | 0.235 |

| | | | | | |
|---|---|---|---|---|---|
| 3.2 | 0.05 | yadI | 3 | 7.16667 | 0.379 |
| 3.2 | 0.05 | AG1 | 3 | 7.16667 | 0.274 |
| 3.2 | 0.05 | ptsI | 3 | 7.41667 | 0.258 |
| 3.2 | 0.05 | clcB | 3 | 7.41667 | 0.273 |
| 3.2 | 0.05 | ycaM | 3 | 7.41667 | 0.239 |
| 3.2 | 0.05 | yadI | 3 | 7.41667 | 0.391 |
| 3.2 | 0.05 | AG1 | 3 | 7.41667 | 0.282 |
| 3.2 | 0.05 | ptsI | 3 | 7.66667 | 0.266 |
| 3.2 | 0.05 | clcB | 3 | 7.66667 | 0.284 |
| 3.2 | 0.05 | ycaM | 3 | 7.66667 | 0.243 |
| 3.2 | 0.05 | yadI | 3 | 7.66667 | 0.403 |
| 3.2 | 0.05 | AG1 | 3 | 7.66667 | 0.289 |
| 3.2 | 0.05 | ptsI | 3 | 7.91667 | 0.276 |
| 3.2 | 0.05 | clcB | 3 | 7.91667 | 0.291 |
| 3.2 | 0.05 | ycaM | 3 | 7.91667 | 0.246 |
| 3.2 | 0.05 | yadI | 3 | 7.91667 | 0.417 |
| 3.2 | 0.05 | AG1 | 3 | 7.91667 | 0.297 |
| 3.2 | 0.05 | ptsI | 3 | 8.16667 | 0.288 |
| 3.2 | 0.05 | clcB | 3 | 8.16667 | 0.299 |
| 3.2 | 0.05 | ycaM | 3 | 8.16667 | 0.249 |
| 3.2 | 0.05 | yadI | 3 | 8.16667 | 0.425 |
| 3.2 | 0.05 | AG1 | 3 | 8.16667 | 0.304 |
| 3.2 | 0.05 | ptsI | 3 | 8.41667 | 0.295 |
| 3.2 | 0.05 | clcB | 3 | 8.41667 | 0.308 |
| 3.2 | 0.05 | ycaM | 3 | 8.41667 | 0.252 |
| 3.2 | 0.05 | yadI | 3 | 8.41667 | 0.441 |
| 3.2 | 0.05 | AG1 | 3 | 8.41667 | 0.31 |
| 3.2 | 0.05 | ptsI | 3 | 9.05 | 0.315 |
| 3.2 | 0.05 | clcB | 3 | 9.05 | 0.29 |
| 3.2 | 0.05 | ycaM | 3 | 9.05 | 0.257 |
| 3.2 | 0.05 | yadI | 3 | 9.05 | 0.428 |
| 3.2 | 0.05 | AG1 | 3 | 9.05 | 0.354 |
| 3.2 | 0.05 | ptsI | 3 | 9.3 | 0.315 |
| 3.2 | 0.05 | clcB | 3 | 9.3 | 0.283 |
| 3.2 | 0.05 | ycaM | 3 | 9.3 | 0.252 |
| 3.2 | 0.05 | yadI | 3 | 9.3 | 0.446 |
| 3.2 | 0.05 | AG1 | 3 | 9.3 | 0.363 |
| 3.2 | 0.05 | ptsI | 3 | 9.55 | 0.309 |
| 3.2 | 0.05 | clcB | 3 | 9.55 | 0.279 |
| 3.2 | 0.05 | ycaM | 3 | 9.55 | 0.25 |
| 3.2 | 0.05 | yadI | 3 | 9.55 | 0.461 |
| 3.2 | 0.05 | AG1 | 3 | 9.55 | 0.371 |
| 3.2 | 0.05 | ptsI | 3 | 9.8 | 0.308 |
| 3.2 | 0.05 | clcB | 3 | 9.8 | 0.276 |
| 3.2 | 0.05 | ycaM | 3 | 9.8 | 0.25 |
| 3.2 | 0.05 | yadI | 3 | 9.8 | 0.466 |
| 3.2 | 0.05 | AG1 | 3 | 9.8 | 0.367 |
| 3.2 | 0.05 | ptsI | 3 | 10.05 | 0.303 |

| | | | | | |
|---|---|---|---|---|---|
| 3.2 | 0.05 | clcB | 3 | 10.05 | 0.278 |
| 3.2 | 0.05 | ycaM | 3 | 10.05 | 0.249 |
| 3.2 | 0.05 | yadI | 3 | 10.05 | 0.47 |
| 3.2 | 0.05 | AG1 | 3 | 10.05 | 0.369 |
| 3.2 | 0.05 | ptsI | 3 | 10.3 | 0.302 |
| 3.2 | 0.05 | clcB | 3 | 10.3 | 0.276 |
| 3.2 | 0.05 | ycaM | 3 | 10.3 | 0.246 |
| 3.2 | 0.05 | yadI | 3 | 10.3 | 0.477 |
| 3.2 | 0.05 | AG1 | 3 | 10.3 | 0.368 |
| 3.2 | 0.05 | ptsI | 3 | 10.55 | 0.293 |
| 3.2 | 0.05 | clcB | 3 | 10.55 | 0.273 |
| 3.2 | 0.05 | ycaM | 3 | 10.55 | 0.241 |
| 3.2 | 0.05 | yadI | 3 | 10.55 | 0.484 |
| 3.2 | 0.05 | AG1 | 3 | 10.55 | 0.371 |
| 3.2 | 0.05 | ptsI | 3 | 10.8 | 0.286 |
| 3.2 | 0.05 | clcB | 3 | 10.8 | 0.275 |
| 3.2 | 0.05 | ycaM | 3 | 10.8 | 0.242 |
| 3.2 | 0.05 | yadI | 3 | 10.8 | 0.493 |
| 3.2 | 0.05 | AG1 | 3 | 10.8 | 0.375 |
| 3.2 | 0.05 | ptsI | 3 | 11.05 | 0.284 |
| 3.2 | 0.05 | clcB | 3 | 11.05 | 0.276 |
| 3.2 | 0.05 | ycaM | 3 | 11.05 | 0.242 |
| 3.2 | 0.05 | yadI | 3 | 11.05 | 0.492 |
| 3.2 | 0.05 | AG1 | 3 | 11.05 | 0.372 |
| 3.2 | 0.05 | ptsI | 3 | 11.3 | 0.281 |
| 3.2 | 0.05 | clcB | 3 | 11.3 | 0.274 |
| 3.2 | 0.05 | ycaM | 3 | 11.3 | 0.245 |
| 3.2 | 0.05 | yadI | 3 | 11.3 | 0.496 |
| 3.2 | 0.05 | AG1 | 3 | 11.3 | 0.376 |
| 3.2 | 0.05 | ptsI | 3 | 11.55 | 0.28 |
| 3.2 | 0.05 | clcB | 3 | 11.55 | 0.27 |
| 3.2 | 0.05 | ycaM | 3 | 11.55 | 0.247 |
| 3.2 | 0.05 | yadI | 3 | 11.55 | 0.498 |
| 3.2 | 0.05 | AG1 | 3 | 11.55 | 0.375 |
| 3.2 | 0.05 | ptsI | 3 | 11.8 | 0.273 |
| 3.2 | 0.05 | clcB | 3 | 11.8 | 0.27 |
| 3.2 | 0.05 | ycaM | 3 | 11.8 | 0.245 |
| 3.2 | 0.05 | yadI | 3 | 11.8 | 0.505 |
| 3.2 | 0.05 | AG1 | 3 | 11.8 | 0.376 |
| 3.2 | 0.05 | ptsI | 3 | 12.05 | 0.27 |
| 3.2 | 0.05 | clcB | 3 | 12.05 | 0.27 |
| 3.2 | 0.05 | ycaM | 3 | 12.05 | 0.241 |
| 3.2 | 0.05 | yadI | 3 | 12.05 | 0.505 |
| 3.2 | 0.05 | AG1 | 3 | 12.05 | 0.377 |
| 3.2 | 0.05 | ptsI | 3 | 12.3 | 0.263 |
| 3.2 | 0.05 | clcB | 3 | 12.3 | 0.267 |
| 3.2 | 0.05 | ycaM | 3 | 12.3 | 0.244 |
| 3.2 | 0.05 | yadI | 3 | 12.3 | 0.503 |

| | | | | | |
|---|---|---|---|---|---|
| 3.2 | 0.05 | AG1 | 3 | 12.3 | 0.379 |
| 3.2 | 0.05 | ptsI | 3 | 12.55 | 0.26 |
| 3.2 | 0.05 | clcB | 3 | 12.55 | 0.271 |
| 3.2 | 0.05 | ycaM | 3 | 12.55 | 0.242 |
| 3.2 | 0.05 | yadI | 3 | 12.55 | 0.51 |
| 3.2 | 0.05 | AG1 | 3 | 12.55 | 0.378 |
| 3.2 | 0.05 | ptsI | 3 | 12.8 | 0.26 |
| 3.2 | 0.05 | clcB | 3 | 12.8 | 0.268 |
| 3.2 | 0.05 | ycaM | 3 | 12.8 | 0.241 |
| 3.2 | 0.05 | yadI | 3 | 12.8 | 0.513 |
| 3.2 | 0.05 | AG1 | 3 | 12.8 | 0.375 |
| 3.2 | 0.05 | ptsI | 3 | 13.05 | 0.257 |
| 3.2 | 0.05 | clcB | 3 | 13.05 | 0.266 |
| 3.2 | 0.05 | ycaM | 3 | 13.05 | 0.242 |
| 3.2 | 0.05 | yadI | 3 | 13.05 | 0.516 |
| 3.2 | 0.05 | AG1 | 3 | 13.05 | 0.375 |
| 3.2 | 0.05 | ptsI | 3 | 13.3 | 0.256 |
| 3.2 | 0.05 | clcB | 3 | 13.3 | 0.266 |
| 3.2 | 0.05 | ycaM | 3 | 13.3 | 0.241 |
| 3.2 | 0.05 | yadI | 3 | 13.3 | 0.52 |
| 3.2 | 0.05 | AG1 | 3 | 13.3 | 0.382 |
| 3.2 | 0.05 | ptsI | 3 | 13.55 | 0.256 |
| 3.2 | 0.05 | clcB | 3 | 13.55 | 0.264 |
| 3.2 | 0.05 | ycaM | 3 | 13.55 | 0.24 |
| 3.2 | 0.05 | yadI | 3 | 13.55 | 0.524 |
| 3.2 | 0.05 | AG1 | 3 | 13.55 | 0.379 |
| 3.2 | 0.05 | ptsI | 3 | 13.8 | 0.254 |
| 3.2 | 0.05 | clcB | 3 | 13.8 | 0.262 |
| 3.2 | 0.05 | ycaM | 3 | 13.8 | 0.239 |
| 3.2 | 0.05 | yadI | 3 | 13.8 | 0.524 |
| 3.2 | 0.05 | AG1 | 3 | 13.8 | 0.382 |
| 3.2 | 0.05 | ptsI | 3 | 14.05 | 0.253 |
| 3.2 | 0.05 | clcB | 3 | 14.05 | 0.262 |
| 3.2 | 0.05 | ycaM | 3 | 14.05 | 0.238 |
| 3.2 | 0.05 | yadI | 3 | 14.05 | 0.531 |
| 3.2 | 0.05 | AG1 | 3 | 14.05 | 0.382 |
| 3.2 | 0.05 | ptsI | 3 | 14.3 | 0.252 |
| 3.2 | 0.05 | clcB | 3 | 14.3 | 0.262 |
| 3.2 | 0.05 | ycaM | 3 | 14.3 | 0.236 |
| 3.2 | 0.05 | yadI | 3 | 14.3 | 0.534 |
| 3.2 | 0.05 | AG1 | 3 | 14.3 | 0.384 |
| 3.2 | 0.05 | ptsI | 3 | 14.55 | 0.252 |
| 3.2 | 0.05 | clcB | 3 | 14.55 | 0.259 |
| 3.2 | 0.05 | ycaM | 3 | 14.55 | 0.238 |
| 3.2 | 0.05 | yadI | 3 | 14.55 | 0.541 |
| 3.2 | 0.05 | AG1 | 3 | 14.55 | 0.384 |
| 3.2 | 0.05 | ptsI | 3 | 14.8 | 0.25 |
| 3.2 | 0.05 | clcB | 3 | 14.8 | 0.259 |

| | | | | | |
|---|---|---|---|---|---|
| 3.2 | 0.05 | ycaM | 3 | 14.8 | 0.236 |
| 3.2 | 0.05 | yadI | 3 | 14.8 | 0.547 |
| 3.2 | 0.05 | AG1 | 3 | 14.8 | 0.387 |
| 3.2 | 0.05 | ptsI | 3 | 15.05 | 0.249 |
| 3.2 | 0.05 | clcB | 3 | 15.05 | 0.26 |
| 3.2 | 0.05 | ycaM | 3 | 15.05 | 0.235 |
| 3.2 | 0.05 | yadI | 3 | 15.05 | 0.551 |
| 3.2 | 0.05 | AG1 | 3 | 15.05 | 0.389 |
| 3.2 | 0.05 | ptsI | 3 | 15.3 | 0.25 |
| 3.2 | 0.05 | clcB | 3 | 15.3 | 0.259 |
| 3.2 | 0.05 | ycaM | 3 | 15.3 | 0.234 |
| 3.2 | 0.05 | yadI | 3 | 15.3 | 0.556 |
| 3.2 | 0.05 | AG1 | 3 | 15.3 | 0.389 |
| 3.2 | 0.05 | ptsI | 3 | 15.55 | 0.247 |
| 3.2 | 0.05 | clcB | 3 | 15.55 | 0.258 |
| 3.2 | 0.05 | ycaM | 3 | 15.55 | 0.233 |
| 3.2 | 0.05 | yadI | 3 | 15.55 | 0.556 |
| 3.2 | 0.05 | AG1 | 3 | 15.55 | 0.389 |
| 3.2 | 0.05 | ptsI | 3 | 15.8 | 0.247 |
| 3.2 | 0.05 | clcB | 3 | 15.8 | 0.257 |
| 3.2 | 0.05 | ycaM | 3 | 15.8 | 0.232 |
| 3.2 | 0.05 | yadI | 3 | 15.8 | 0.561 |
| 3.2 | 0.05 | AG1 | 3 | 15.8 | 0.389 |
| 3.2 | 0.05 | ptsI | 3 | 16.05 | 0.247 |
| 3.2 | 0.05 | clcB | 3 | 16.05 | 0.256 |
| 3.2 | 0.05 | ycaM | 3 | 16.05 | 0.234 |
| 3.2 | 0.05 | yadI | 3 | 16.05 | 0.564 |
| 3.2 | 0.05 | AG1 | 3 | 16.05 | 0.39 |
| 3.2 | 0.05 | ptsI | 3 | 16.3 | 0.242 |
| 3.2 | 0.05 | clcB | 3 | 16.3 | 0.256 |
| 3.2 | 0.05 | ycaM | 3 | 16.3 | 0.229 |
| 3.2 | 0.05 | yadI | 3 | 16.3 | 0.567 |
| 3.2 | 0.05 | AG1 | 3 | 16.3 | 0.389 |
| 3.2 | 0.05 | ptsI | 3 | 16.55 | 0.245 |
| 3.2 | 0.05 | clcB | 3 | 16.55 | 0.255 |
| 3.2 | 0.05 | ycaM | 3 | 16.55 | 0.228 |
| 3.2 | 0.05 | yadI | 3 | 16.55 | 0.571 |
| 3.2 | 0.05 | AG1 | 3 | 16.55 | 0.391 |
| 3.2 | 0.05 | ptsI | 3 | 16.8 | 0.243 |
| 3.2 | 0.05 | clcB | 3 | 16.8 | 0.255 |
| 3.2 | 0.05 | ycaM | 3 | 16.8 | 0.228 |
| 3.2 | 0.05 | yadI | 3 | 16.8 | 0.573 |
| 3.2 | 0.05 | AG1 | 3 | 16.8 | 0.392 |
| 3.2 | 0.05 | ptsI | 3 | 17.05 | 0.242 |
| 3.2 | 0.05 | clcB | 3 | 17.05 | 0.254 |
| 3.2 | 0.05 | ycaM | 3 | 17.05 | 0.227 |
| 3.2 | 0.05 | yadI | 3 | 17.05 | 0.578 |
| 3.2 | 0.05 | AG1 | 3 | 17.05 | 0.394 |

| | | | | | |
|---|---|---|---|---|---|
| 3.2 | 0.05 | ptsI | 3 | 17.3 | 0.242 |
| 3.2 | 0.05 | clcB | 3 | 17.3 | 0.253 |
| 3.2 | 0.05 | ycaM | 3 | 17.3 | 0.228 |
| 3.2 | 0.05 | yadI | 3 | 17.3 | 0.584 |
| 3.2 | 0.05 | AG1 | 3 | 17.3 | 0.398 |
| 3.2 | 0.05 | ptsI | 3 | 17.55 | 0.24 |
| 3.2 | 0.05 | clcB | 3 | 17.55 | 0.253 |
| 3.2 | 0.05 | ycaM | 3 | 17.55 | 0.226 |
| 3.2 | 0.05 | yadI | 3 | 17.55 | 0.586 |
| 3.2 | 0.05 | AG1 | 3 | 17.55 | 0.398 |
| 3.2 | 0.05 | ptsI | 3 | 17.8 | 0.241 |
| 3.2 | 0.05 | clcB | 3 | 17.8 | 0.25 |
| 3.2 | 0.05 | ycaM | 3 | 17.8 | 0.225 |
| 3.2 | 0.05 | yadI | 3 | 17.8 | 0.591 |
| 3.2 | 0.05 | AG1 | 3 | 17.8 | 0.403 |
| 3.2 | 0.05 | ptsI | 3 | 18.05 | 0.241 |
| 3.2 | 0.05 | clcB | 3 | 18.05 | 0.249 |
| 3.2 | 0.05 | ycaM | 3 | 18.05 | 0.224 |
| 3.2 | 0.05 | yadI | 3 | 18.05 | 0.595 |
| 3.2 | 0.05 | AG1 | 3 | 18.05 | 0.403 |
| 3.2 | 0.05 | ptsI | 3 | 18.3 | 0.24 |
| 3.2 | 0.05 | clcB | 3 | 18.3 | 0.249 |
| 3.2 | 0.05 | ycaM | 3 | 18.3 | 0.223 |
| 3.2 | 0.05 | yadI | 3 | 18.3 | 0.603 |
| 3.2 | 0.05 | AG1 | 3 | 18.3 | 0.406 |
| 3.2 | 0.05 | ptsI | 3 | 18.55 | 0.24 |
| 3.2 | 0.05 | clcB | 3 | 18.55 | 0.248 |
| 3.2 | 0.05 | ycaM | 3 | 18.55 | 0.223 |
| 3.2 | 0.05 | yadI | 3 | 18.55 | 0.603 |
| 3.2 | 0.05 | AG1 | 3 | 18.55 | 0.41 |
| 3.2 | 0.05 | ptsI | 3 | 18.8 | 0.24 |
| 3.2 | 0.05 | clcB | 3 | 18.8 | 0.246 |
| 3.2 | 0.05 | ycaM | 3 | 18.8 | 0.222 |
| 3.2 | 0.05 | yadI | 3 | 18.8 | 0.611 |
| 3.2 | 0.05 | AG1 | 3 | 18.8 | 0.406 |
| 3.2 | 0.05 | ptsI | 3 | 19.05 | 0.238 |
| 3.2 | 0.05 | clcB | 3 | 19.05 | 0.247 |
| 3.2 | 0.05 | ycaM | 3 | 19.05 | 0.22 |
| 3.2 | 0.05 | yadI | 3 | 19.05 | 0.61 |
| 3.2 | 0.05 | AG1 | 3 | 19.05 | 0.408 |
| 3.2 | 0.05 | ptsI | 3 | 19.3 | 0.238 |
| 3.2 | 0.05 | clcB | 3 | 19.3 | 0.245 |
| 3.2 | 0.05 | ycaM | 3 | 19.3 | 0.22 |
| 3.2 | 0.05 | yadI | 3 | 19.3 | 0.616 |
| 3.2 | 0.05 | AG1 | 3 | 19.3 | 0.413 |
| 3.2 | 0.05 | ptsI | 3 | 19.55 | 0.239 |
| 3.2 | 0.05 | clcB | 3 | 19.55 | 0.243 |
| 3.2 | 0.05 | ycaM | 3 | 19.55 | 0.219 |

| | | | | | |
|---|---|---|---|---|---|
| 3.2 | 0.05 | yadI | 3 | 19.55 | 0.619 |
| 3.2 | 0.05 | AG1 | 3 | 19.55 | 0.413 |
| 3.2 | 0.05 | ptsI | 3 | 19.8 | 0.237 |
| 3.2 | 0.05 | clcB | 3 | 19.8 | 0.243 |
| 3.2 | 0.05 | ycaM | 3 | 19.8 | 0.218 |
| 3.2 | 0.05 | yadI | 3 | 19.8 | 0.625 |
| 3.2 | 0.05 | AG1 | 3 | 19.8 | 0.414 |
| 3.2 | 0.05 | ptsI | 3 | 20.05 | 0.239 |
| 3.2 | 0.05 | clcB | 3 | 20.05 | 0.242 |
| 3.2 | 0.05 | ycaM | 3 | 20.05 | 0.217 |
| 3.2 | 0.05 | yadI | 3 | 20.05 | 0.632 |
| 3.2 | 0.05 | AG1 | 3 | 20.05 | 0.417 |
| 3.2 | 0.05 | ptsI | 3 | 20.3 | 0.237 |
| 3.2 | 0.05 | clcB | 3 | 20.3 | 0.242 |
| 3.2 | 0.05 | ycaM | 3 | 20.3 | 0.216 |
| 3.2 | 0.05 | yadI | 3 | 20.3 | 0.634 |
| 3.2 | 0.05 | AG1 | 3 | 20.3 | 0.418 |
| 3.2 | 0.05 | ptsI | 3 | 20.55 | 0.238 |
| 3.2 | 0.05 | clcB | 3 | 20.55 | 0.242 |
| 3.2 | 0.05 | ycaM | 3 | 20.55 | 0.216 |
| 3.2 | 0.05 | yadI | 3 | 20.55 | 0.639 |
| 3.2 | 0.05 | AG1 | 3 | 20.55 | 0.422 |
| 3.2 | 0.05 | ptsI | 3 | 20.8 | 0.238 |
| 3.2 | 0.05 | clcB | 3 | 20.8 | 0.24 |
| 3.2 | 0.05 | ycaM | 3 | 20.8 | 0.215 |
| 3.2 | 0.05 | yadI | 3 | 20.8 | 0.644 |
| 3.2 | 0.05 | AG1 | 3 | 20.8 | 0.422 |
| 3.2 | 0.05 | ptsI | 3 | 21.05 | 0.237 |
| 3.2 | 0.05 | clcB | 3 | 21.05 | 0.239 |
| 3.2 | 0.05 | ycaM | 3 | 21.05 | 0.213 |
| 3.2 | 0.05 | yadI | 3 | 21.05 | 0.645 |
| 3.2 | 0.05 | AG1 | 3 | 21.05 | 0.423 |
| 3.2 | 0.05 | ptsI | 3 | 21.3 | 0.236 |
| 3.2 | 0.05 | clcB | 3 | 21.3 | 0.238 |
| 3.2 | 0.05 | ycaM | 3 | 21.3 | 0.212 |
| 3.2 | 0.05 | yadI | 3 | 21.3 | 0.65 |
| 3.2 | 0.05 | AG1 | 3 | 21.3 | 0.425 |
| 3.2 | 0.05 | ptsI | 3 | 21.55 | 0.238 |
| 3.2 | 0.05 | clcB | 3 | 21.55 | 0.237 |
| 3.2 | 0.05 | ycaM | 3 | 21.55 | 0.211 |
| 3.2 | 0.05 | yadI | 3 | 21.55 | 0.652 |
| 3.2 | 0.05 | AG1 | 3 | 21.55 | 0.426 |
| 3.2 | 0.05 | ptsI | 3 | 21.8 | 0.237 |
| 3.2 | 0.05 | clcB | 3 | 21.8 | 0.237 |
| 3.2 | 0.05 | ycaM | 3 | 21.8 | 0.21 |
| 3.2 | 0.05 | yadI | 3 | 21.8 | 0.654 |
| 3.2 | 0.05 | AG1 | 3 | 21.8 | 0.432 |
| 3.2 | 0.05 | ptsI | 3 | 22.05 | 0.237 |

| | | | | | |
|---|---|---|---|---|---|
| 3.2 | 0.05 | clcB | 3 | 22.05 | 0.236 |
| 3.2 | 0.05 | ycaM | 3 | 22.05 | 0.209 |
| 3.2 | 0.05 | yadI | 3 | 22.05 | 0.659 |
| 3.2 | 0.05 | AG1 | 3 | 22.05 | 0.43 |
| 3.2 | 0.05 | ptsI | 3 | 22.3 | 0.236 |
| 3.2 | 0.05 | clcB | 3 | 22.3 | 0.235 |
| 3.2 | 0.05 | ycaM | 3 | 22.3 | 0.208 |
| 3.2 | 0.05 | yadI | 3 | 22.3 | 0.661 |
| 3.2 | 0.05 | AG1 | 3 | 22.3 | 0.434 |
| 3.2 | 0.05 | ptsI | 3 | 22.55 | 0.236 |
| 3.2 | 0.05 | clcB | 3 | 22.55 | 0.234 |
| 3.2 | 0.05 | ycaM | 3 | 22.55 | 0.207 |
| 3.2 | 0.05 | yadI | 3 | 22.55 | 0.67 |
| 3.2 | 0.05 | AG1 | 3 | 22.55 | 0.436 |
| 3.2 | 0.05 | ptsI | 3 | 22.8 | 0.236 |
| 3.2 | 0.05 | clcB | 3 | 22.8 | 0.232 |
| 3.2 | 0.05 | ycaM | 3 | 22.8 | 0.206 |
| 3.2 | 0.05 | yadI | 3 | 22.8 | 0.67 |
| 3.2 | 0.05 | AG1 | 3 | 22.8 | 0.436 |
| 3.2 | 0.05 | ptsI | 3 | 23.05 | 0.235 |
| 3.2 | 0.05 | clcB | 3 | 23.05 | 0.232 |
| 3.2 | 0.05 | ycaM | 3 | 23.05 | 0.206 |
| 3.2 | 0.05 | yadI | 3 | 23.05 | 0.677 |
| 3.2 | 0.05 | AG1 | 3 | 23.05 | 0.438 |
| 3.2 | 0.05 | ptsI | 3 | 23.3 | 0.235 |
| 3.2 | 0.05 | clcB | 3 | 23.3 | 0.231 |
| 3.2 | 0.05 | ycaM | 3 | 23.3 | 0.204 |
| 3.2 | 0.05 | yadI | 3 | 23.3 | 0.685 |
| 3.2 | 0.05 | AG1 | 3 | 23.3 | 0.441 |
| 3.2 | 0.05 | ptsI | 3 | 23.55 | 0.235 |
| 3.2 | 0.05 | clcB | 3 | 23.55 | 0.229 |
| 3.2 | 0.05 | ycaM | 3 | 23.55 | 0.202 |
| 3.2 | 0.05 | yadI | 3 | 23.55 | 0.687 |
| 3.2 | 0.05 | AG1 | 3 | 23.55 | 0.442 |
| 3.2 | 0.05 | ptsI | 3 | 23.8 | 0.234 |
| 3.2 | 0.05 | clcB | 3 | 23.8 | 0.229 |
| 3.2 | 0.05 | ycaM | 3 | 23.8 | 0.202 |
| 3.2 | 0.05 | yadI | 3 | 23.8 | 0.688 |
| 3.2 | 0.05 | AG1 | 3 | 23.8 | 0.444 |
| 3.2 | 0.05 | ptsI | 3 | 24.05 | 0.235 |
| 3.2 | 0.05 | clcB | 3 | 24.05 | 0.227 |
| 3.2 | 0.05 | ycaM | 3 | 24.05 | 0.201 |
| 3.2 | 0.05 | yadI | 3 | 24.05 | 0.697 |
| 3.2 | 0.05 | AG1 | 3 | 24.05 | 0.445 |
| 3.2 | 0.05 | ptsI | 3 | 24.3 | 0.231 |
| 3.2 | 0.05 | clcB | 3 | 24.3 | 0.229 |
| 3.2 | 0.05 | ycaM | 3 | 24.3 | 0.199 |
| 3.2 | 0.05 | yadI | 3 | 24.3 | 0.699 |

| | | | | | |
|---|---|---|---|---|---|
| 3.2 | 0.05 | AG1 | 3 | 24.3 | 0.443 |
| 3.2 | 0.05 | ptsI | 4 | 0 | 0.208 |
| 3.2 | 0.05 | clcB | 4 | 0 | 0.19 |
| 3.2 | 0.05 | ycaM | 4 | 0 | 0.223 |
| 3.2 | 0.05 | yadI | 4 | 0 | 0.221 |
| 3.2 | 0.05 | AG1 | 4 | 0 | 0.203 |
| 3.2 | 0.05 | ptsI | 4 | 0.25 | 0.207 |
| 3.2 | 0.05 | clcB | 4 | 0.25 | 0.19 |
| 3.2 | 0.05 | ycaM | 4 | 0.25 | 0.209 |
| 3.2 | 0.05 | yadI | 4 | 0.25 | 0.222 |
| 3.2 | 0.05 | AG1 | 4 | 0.25 | 0.199 |
| 3.2 | 0.05 | ptsI | 4 | 0.5 | 0.208 |
| 3.2 | 0.05 | clcB | 4 | 0.5 | 0.192 |
| 3.2 | 0.05 | ycaM | 4 | 0.5 | 0.215 |
| 3.2 | 0.05 | yadI | 4 | 0.5 | 0.224 |
| 3.2 | 0.05 | AG1 | 4 | 0.5 | 0.202 |
| 3.2 | 0.05 | ptsI | 4 | 0.75 | 0.212 |
| 3.2 | 0.05 | clcB | 4 | 0.75 | 0.193 |
| 3.2 | 0.05 | ycaM | 4 | 0.75 | 0.215 |
| 3.2 | 0.05 | yadI | 4 | 0.75 | 0.228 |
| 3.2 | 0.05 | AG1 | 4 | 0.75 | 0.204 |
| 3.2 | 0.05 | ptsI | 4 | 1 | 0.216 |
| 3.2 | 0.05 | clcB | 4 | 1 | 0.195 |
| 3.2 | 0.05 | ycaM | 4 | 1 | 0.22 |
| 3.2 | 0.05 | yadI | 4 | 1 | 0.23 |
| 3.2 | 0.05 | AG1 | 4 | 1 | 0.21 |
| 3.2 | 0.05 | ptsI | 4 | 1.25 | 0.218 |
| 3.2 | 0.05 | clcB | 4 | 1.25 | 0.201 |
| 3.2 | 0.05 | ycaM | 4 | 1.25 | 0.225 |
| 3.2 | 0.05 | yadI | 4 | 1.25 | 0.234 |
| 3.2 | 0.05 | AG1 | 4 | 1.25 | 0.215 |
| 3.2 | 0.05 | ptsI | 4 | 1.5 | 0.218 |
| 3.2 | 0.05 | clcB | 4 | 1.5 | 0.199 |
| 3.2 | 0.05 | ycaM | 4 | 1.5 | 0.221 |
| 3.2 | 0.05 | yadI | 4 | 1.5 | 0.233 |
| 3.2 | 0.05 | AG1 | 4 | 1.5 | 0.208 |
| 3.2 | 0.05 | ptsI | 4 | 1.75 | 0.225 |
| 3.2 | 0.05 | clcB | 4 | 1.75 | 0.207 |
| 3.2 | 0.05 | ycaM | 4 | 1.75 | 0.23 |
| 3.2 | 0.05 | yadI | 4 | 1.75 | 0.242 |
| 3.2 | 0.05 | AG1 | 4 | 1.75 | 0.215 |
| 3.2 | 0.05 | ptsI | 4 | 2 | 0.234 |
| 3.2 | 0.05 | clcB | 4 | 2 | 0.213 |
| 3.2 | 0.05 | ycaM | 4 | 2 | 0.239 |
| 3.2 | 0.05 | yadI | 4 | 2 | 0.25 |
| 3.2 | 0.05 | AG1 | 4 | 2 | 0.223 |
| 3.2 | 0.05 | ptsI | 4 | 2.25 | 0.242 |
| 3.2 | 0.05 | clcB | 4 | 2.25 | 0.225 |

| | | | | | |
|---|---|---|---|---|---|
| 3.2 | 0.05 | ycaM | 4 | 2.25 | 0.252 |
| 3.2 | 0.05 | yadI | 4 | 2.25 | 0.261 |
| 3.2 | 0.05 | AG1 | 4 | 2.25 | 0.233 |
| 3.2 | 0.05 | ptsI | 4 | 2.5 | 0.246 |
| 3.2 | 0.05 | clcB | 4 | 2.5 | 0.225 |
| 3.2 | 0.05 | ycaM | 4 | 2.5 | 0.25 |
| 3.2 | 0.05 | yadI | 4 | 2.5 | 0.258 |
| 3.2 | 0.05 | AG1 | 4 | 2.5 | 0.23 |
| 3.2 | 0.05 | ptsI | 4 | 2.75 | 0.254 |
| 3.2 | 0.05 | clcB | 4 | 2.75 | 0.234 |
| 3.2 | 0.05 | ycaM | 4 | 2.75 | 0.255 |
| 3.2 | 0.05 | yadI | 4 | 2.75 | 0.265 |
| 3.2 | 0.05 | AG1 | 4 | 2.75 | 0.237 |
| 3.2 | 0.05 | ptsI | 4 | 3 | 0.263 |
| 3.2 | 0.05 | clcB | 4 | 3 | 0.247 |
| 3.2 | 0.05 | ycaM | 4 | 3 | 0.274 |
| 3.2 | 0.05 | yadI | 4 | 3 | 0.276 |
| 3.2 | 0.05 | AG1 | 4 | 3 | 0.244 |
| 3.2 | 0.05 | ptsI | 4 | 3.25 | 0.274 |
| 3.2 | 0.05 | clcB | 4 | 3.25 | 0.261 |
| 3.2 | 0.05 | ycaM | 4 | 3.25 | 0.284 |
| 3.2 | 0.05 | yadI | 4 | 3.25 | 0.286 |
| 3.2 | 0.05 | AG1 | 4 | 3.25 | 0.254 |
| 3.2 | 0.05 | ptsI | 4 | 3.5 | 0.288 |
| 3.2 | 0.05 | clcB | 4 | 3.5 | 0.277 |
| 3.2 | 0.05 | ycaM | 4 | 3.5 | 0.301 |
| 3.2 | 0.05 | yadI | 4 | 3.5 | 0.301 |
| 3.2 | 0.05 | AG1 | 4 | 3.5 | 0.266 |
| 3.2 | 0.05 | ptsI | 4 | 3.75 | 0.296 |
| 3.2 | 0.05 | clcB | 4 | 3.75 | 0.291 |
| 3.2 | 0.05 | ycaM | 4 | 3.75 | 0.311 |
| 3.2 | 0.05 | yadI | 4 | 3.75 | 0.308 |
| 3.2 | 0.05 | AG1 | 4 | 3.75 | 0.276 |
| 3.2 | 0.05 | ptsI | 4 | 4 | 0.314 |
| 3.2 | 0.05 | clcB | 4 | 4 | 0.316 |
| 3.2 | 0.05 | ycaM | 4 | 4 | 0.341 |
| 3.2 | 0.05 | yadI | 4 | 4 | 0.32 |
| 3.2 | 0.05 | AG1 | 4 | 4 | 0.287 |
| 3.2 | 0.05 | ptsI | 4 | 4.35 | 0.327 |
| 3.2 | 0.05 | clcB | 4 | 4.35 | 0.322 |
| 3.2 | 0.05 | ycaM | 4 | 4.35 | 0.338 |
| 3.2 | 0.05 | yadI | 4 | 4.35 | 0.366 |
| 3.2 | 0.05 | AG1 | 4 | 4.35 | 0.262 |
| 3.2 | 0.05 | ptsI | 4 | 4.6 | 0.331 |
| 3.2 | 0.05 | clcB | 4 | 4.6 | 0.328 |
| 3.2 | 0.05 | ycaM | 4 | 4.6 | 0.341 |
| 3.2 | 0.05 | yadI | 4 | 4.6 | 0.383 |
| 3.2 | 0.05 | AG1 | 4 | 4.6 | 0.277 |

| | | | | | |
|---|---|---|---|---|---|
| 3.2 | 0.05 | ptsI | 4 | 4.85 | 0.341 |
| 3.2 | 0.05 | clcB | 4 | 4.85 | 0.329 |
| 3.2 | 0.05 | ycaM | 4 | 4.85 | 0.339 |
| 3.2 | 0.05 | yadI | 4 | 4.85 | 0.402 |
| 3.2 | 0.05 | AG1 | 4 | 4.85 | 0.29 |
| 3.2 | 0.05 | ptsI | 4 | 5.1 | 0.351 |
| 3.2 | 0.05 | clcB | 4 | 5.1 | 0.33 |
| 3.2 | 0.05 | ycaM | 4 | 5.1 | 0.338 |
| 3.2 | 0.05 | yadI | 4 | 5.1 | 0.411 |
| 3.2 | 0.05 | AG1 | 4 | 5.1 | 0.297 |
| 3.2 | 0.05 | ptsI | 4 | 5.35 | 0.363 |
| 3.2 | 0.05 | clcB | 4 | 5.35 | 0.336 |
| 3.2 | 0.05 | ycaM | 4 | 5.35 | 0.344 |
| 3.2 | 0.05 | yadI | 4 | 5.35 | 0.427 |
| 3.2 | 0.05 | AG1 | 4 | 5.35 | 0.307 |
| 3.2 | 0.05 | ptsI | 4 | 5.6 | 0.368 |
| 3.2 | 0.05 | clcB | 4 | 5.6 | 0.33 |
| 3.2 | 0.05 | ycaM | 4 | 5.6 | 0.338 |
| 3.2 | 0.05 | yadI | 4 | 5.6 | 0.457 |
| 3.2 | 0.05 | AG1 | 4 | 5.6 | 0.312 |
| 3.2 | 0.05 | ptsI | 4 | 5.85 | 0.371 |
| 3.2 | 0.05 | clcB | 4 | 5.85 | 0.32 |
| 3.2 | 0.05 | ycaM | 4 | 5.85 | 0.331 |
| 3.2 | 0.05 | yadI | 4 | 5.85 | 0.446 |
| 3.2 | 0.05 | AG1 | 4 | 5.85 | 0.317 |
| 3.2 | 0.05 | ptsI | 4 | 6.1 | 0.388 |
| 3.2 | 0.05 | clcB | 4 | 6.1 | 0.324 |
| 3.2 | 0.05 | ycaM | 4 | 6.1 | 0.336 |
| 3.2 | 0.05 | yadI | 4 | 6.1 | 0.46 |
| 3.2 | 0.05 | AG1 | 4 | 6.1 | 0.328 |
| 3.2 | 0.05 | ptsI | 4 | 6.35 | 0.404 |
| 3.2 | 0.05 | clcB | 4 | 6.35 | 0.33 |
| 3.2 | 0.05 | ycaM | 4 | 6.35 | 0.337 |
| 3.2 | 0.05 | yadI | 4 | 6.35 | 0.482 |
| 3.2 | 0.05 | AG1 | 4 | 6.35 | 0.338 |
| 3.2 | 0.05 | ptsI | 4 | 6.6 | 0.417 |
| 3.2 | 0.05 | clcB | 4 | 6.6 | 0.336 |
| 3.2 | 0.05 | ycaM | 4 | 6.6 | 0.342 |
| 3.2 | 0.05 | yadI | 4 | 6.6 | 0.474 |
| 3.2 | 0.05 | AG1 | 4 | 6.6 | 0.347 |
| 3.2 | 0.05 | ptsI | 4 | 6.85 | 0.429 |
| 3.2 | 0.05 | clcB | 4 | 6.85 | 0.339 |
| 3.2 | 0.05 | ycaM | 4 | 6.85 | 0.345 |
| 3.2 | 0.05 | yadI | 4 | 6.85 | 0.482 |
| 3.2 | 0.05 | AG1 | 4 | 6.85 | 0.355 |
| 3.2 | 0.05 | ptsI | 4 | 7.1 | 0.441 |
| 3.2 | 0.05 | clcB | 4 | 7.1 | 0.343 |
| 3.2 | 0.05 | ycaM | 4 | 7.1 | 0.346 |

| | | | | | |
|---|---|---|---|---|---|
| 3.2 | 0.05 | yadI | 4 | 7.1 | 0.489 |
| 3.2 | 0.05 | AG1 | 4 | 7.1 | 0.363 |
| 3.2 | 0.05 | ptsI | 4 | 7.35 | 0.45 |
| 3.2 | 0.05 | clcB | 4 | 7.35 | 0.348 |
| 3.2 | 0.05 | ycaM | 4 | 7.35 | 0.352 |
| 3.2 | 0.05 | yadI | 4 | 7.35 | 0.499 |
| 3.2 | 0.05 | AG1 | 4 | 7.35 | 0.372 |
| 3.2 | 0.05 | ptsI | 4 | 7.6 | 0.46 |
| 3.2 | 0.05 | clcB | 4 | 7.6 | 0.35 |
| 3.2 | 0.05 | ycaM | 4 | 7.6 | 0.37 |
| 3.2 | 0.05 | yadI | 4 | 7.6 | 0.506 |
| 3.2 | 0.05 | AG1 | 4 | 7.6 | 0.379 |
| 3.2 | 0.05 | ptsI | 4 | 7.85 | 0.459 |
| 3.2 | 0.05 | clcB | 4 | 7.85 | 0.347 |
| 3.2 | 0.05 | ycaM | 4 | 7.85 | 0.338 |
| 3.2 | 0.05 | yadI | 4 | 7.85 | 0.507 |
| 3.2 | 0.05 | AG1 | 4 | 7.85 | 0.381 |
| 3.2 | 0.05 | ptsI | 4 | 8.1 | 0.474 |
| 3.2 | 0.05 | clcB | 4 | 8.1 | 0.354 |
| 3.2 | 0.05 | ycaM | 4 | 8.1 | 0.346 |
| 3.2 | 0.05 | yadI | 4 | 8.1 | 0.524 |
| 3.2 | 0.05 | AG1 | 4 | 8.1 | 0.39 |
| 3.2 | 0.05 | ptsI | 4 | 8.35 | 0.487 |
| 3.2 | 0.05 | clcB | 4 | 8.35 | 0.358 |
| 3.2 | 0.05 | ycaM | 4 | 8.35 | 0.357 |
| 3.2 | 0.05 | yadI | 4 | 8.35 | 0.526 |
| 3.2 | 0.05 | AG1 | 4 | 8.35 | 0.4 |
| 3.2 | 0.05 | ptsI | 4 | 8.83333 | 0.424 |
| 3.2 | 0.05 | clcB | 4 | 8.83333 | 0.304 |
| 3.2 | 0.05 | ycaM | 4 | 8.83333 | 0.306 |
| 3.2 | 0.05 | yadI | 4 | 8.83333 | 0.511 |
| 3.2 | 0.05 | AG1 | 4 | 8.83333 | 0.439 |
| 3.2 | 0.05 | ptsI | 4 | 9.08333 | 0.434 |
| 3.2 | 0.05 | clcB | 4 | 9.08333 | 0.303 |
| 3.2 | 0.05 | ycaM | 4 | 9.08333 | 0.305 |
| 3.2 | 0.05 | yadI | 4 | 9.08333 | 0.522 |
| 3.2 | 0.05 | AG1 | 4 | 9.08333 | 0.468 |
| 3.2 | 0.05 | ptsI | 4 | 9.33333 | 0.446 |
| 3.2 | 0.05 | clcB | 4 | 9.33333 | 0.3 |
| 3.2 | 0.05 | ycaM | 4 | 9.33333 | 0.304 |
| 3.2 | 0.05 | yadI | 4 | 9.33333 | 0.539 |
| 3.2 | 0.05 | AG1 | 4 | 9.33333 | 0.488 |
| 3.2 | 0.05 | ptsI | 4 | 9.58333 | 0.452 |
| 3.2 | 0.05 | clcB | 4 | 9.58333 | 0.301 |
| 3.2 | 0.05 | ycaM | 4 | 9.58333 | 0.307 |
| 3.2 | 0.05 | yadI | 4 | 9.58333 | 0.561 |
| 3.2 | 0.05 | AG1 | 4 | 9.58333 | 0.492 |
| 3.2 | 0.05 | ptsI | 4 | 9.83333 | 0.452 |

| | | | | | |
|---|---|---|---|---|---|
| 3.2 | 0.05 | clcB | 4 | 9.83333 | 0.299 |
| 3.2 | 0.05 | ycaM | 4 | 9.83333 | 0.306 |
| 3.2 | 0.05 | yadI | 4 | 9.83333 | 0.572 |
| 3.2 | 0.05 | AG1 | 4 | 9.83333 | 0.494 |
| 3.2 | 0.05 | ptsI | 4 | 10.0833 | 0.451 |
| 3.2 | 0.05 | clcB | 4 | 10.0833 | 0.298 |
| 3.2 | 0.05 | ycaM | 4 | 10.0833 | 0.307 |
| 3.2 | 0.05 | yadI | 4 | 10.0833 | 0.587 |
| 3.2 | 0.05 | AG1 | 4 | 10.0833 | 0.5 |
| 3.2 | 0.05 | ptsI | 4 | 10.3333 | 0.455 |
| 3.2 | 0.05 | clcB | 4 | 10.3333 | 0.296 |
| 3.2 | 0.05 | ycaM | 4 | 10.3333 | 0.306 |
| 3.2 | 0.05 | yadI | 4 | 10.3333 | 0.555 |
| 3.2 | 0.05 | AG1 | 4 | 10.3333 | 0.504 |
| 3.2 | 0.05 | ptsI | 4 | 10.5833 | 0.464 |
| 3.2 | 0.05 | clcB | 4 | 10.5833 | 0.297 |
| 3.2 | 0.05 | ycaM | 4 | 10.5833 | 0.304 |
| 3.2 | 0.05 | yadI | 4 | 10.5833 | 0.558 |
| 3.2 | 0.05 | AG1 | 4 | 10.5833 | 0.505 |
| 3.2 | 0.05 | ptsI | 4 | 10.8333 | 0.467 |
| 3.2 | 0.05 | clcB | 4 | 10.8333 | 0.299 |
| 3.2 | 0.05 | ycaM | 4 | 10.8333 | 0.306 |
| 3.2 | 0.05 | yadI | 4 | 10.8333 | 0.578 |
| 3.2 | 0.05 | AG1 | 4 | 10.8333 | 0.512 |
| 3.2 | 0.05 | ptsI | 4 | 11.0833 | 0.469 |
| 3.2 | 0.05 | clcB | 4 | 11.0833 | 0.298 |
| 3.2 | 0.05 | ycaM | 4 | 11.0833 | 0.307 |
| 3.2 | 0.05 | yadI | 4 | 11.0833 | 0.583 |
| 3.2 | 0.05 | AG1 | 4 | 11.0833 | 0.518 |
| 3.2 | 0.05 | ptsI | 4 | 11.3333 | 0.475 |
| 3.2 | 0.05 | clcB | 4 | 11.3333 | 0.3 |
| 3.2 | 0.05 | ycaM | 4 | 11.3333 | 0.307 |
| 3.2 | 0.05 | yadI | 4 | 11.3333 | 0.592 |
| 3.2 | 0.05 | AG1 | 4 | 11.3333 | 0.521 |
| 3.2 | 0.05 | ptsI | 4 | 11.5833 | 0.48 |
| 3.2 | 0.05 | clcB | 4 | 11.5833 | 0.3 |
| 3.2 | 0.05 | ycaM | 4 | 11.5833 | 0.308 |
| 3.2 | 0.05 | yadI | 4 | 11.5833 | 0.577 |
| 3.2 | 0.05 | AG1 | 4 | 11.5833 | 0.527 |
| 3.2 | 0.05 | ptsI | 4 | 11.8333 | 0.486 |
| 3.2 | 0.05 | clcB | 4 | 11.8333 | 0.299 |
| 3.2 | 0.05 | ycaM | 4 | 11.8333 | 0.31 |
| 3.2 | 0.05 | yadI | 4 | 11.8333 | 0.583 |
| 3.2 | 0.05 | AG1 | 4 | 11.8333 | 0.528 |
| 3.2 | 0.05 | ptsI | 4 | 12.0833 | 0.494 |
| 3.2 | 0.05 | clcB | 4 | 12.0833 | 0.3 |
| 3.2 | 0.05 | ycaM | 4 | 12.0833 | 0.31 |
| 3.2 | 0.05 | yadI | 4 | 12.0833 | 0.582 |

| | | | | | |
|---|---|---|---|---|---|
| 3.2 | 0.05 | AG1 | 4 | 12.0833 | 0.532 |
| 3.2 | 0.05 | ptsI | 4 | 12.3333 | 0.496 |
| 3.2 | 0.05 | clcB | 4 | 12.3333 | 0.3 |
| 3.2 | 0.05 | ycaM | 4 | 12.3333 | 0.308 |
| 3.2 | 0.05 | yadI | 4 | 12.3333 | 0.595 |
| 3.2 | 0.05 | AG1 | 4 | 12.3333 | 0.537 |
| 3.2 | 0.05 | ptsI | 4 | 12.5833 | 0.502 |
| 3.2 | 0.05 | clcB | 4 | 12.5833 | 0.302 |
| 3.2 | 0.05 | ycaM | 4 | 12.5833 | 0.308 |
| 3.2 | 0.05 | yadI | 4 | 12.5833 | 0.592 |
| 3.2 | 0.05 | AG1 | 4 | 12.5833 | 0.539 |
| 3.2 | 0.05 | ptsI | 4 | 12.8333 | 0.509 |
| 3.2 | 0.05 | clcB | 4 | 12.8333 | 0.3 |
| 3.2 | 0.05 | ycaM | 4 | 12.8333 | 0.307 |
| 3.2 | 0.05 | yadI | 4 | 12.8333 | 0.618 |
| 3.2 | 0.05 | AG1 | 4 | 12.8333 | 0.544 |
| 3.2 | 0.05 | ptsI | 4 | 13.0833 | 0.517 |
| 3.2 | 0.05 | clcB | 4 | 13.0833 | 0.301 |
| 3.2 | 0.05 | ycaM | 4 | 13.0833 | 0.307 |
| 3.2 | 0.05 | yadI | 4 | 13.0833 | 0.602 |
| 3.2 | 0.05 | AG1 | 4 | 13.0833 | 0.546 |
| 3.2 | 0.05 | ptsI | 4 | 13.3333 | 0.519 |
| 3.2 | 0.05 | clcB | 4 | 13.3333 | 0.3 |
| 3.2 | 0.05 | ycaM | 4 | 13.3333 | 0.306 |
| 3.2 | 0.05 | yadI | 4 | 13.3333 | 0.636 |
| 3.2 | 0.05 | AG1 | 4 | 13.3333 | 0.552 |
| 3.2 | 0.05 | ptsI | 4 | 13.5833 | 0.523 |
| 3.2 | 0.05 | clcB | 4 | 13.5833 | 0.301 |
| 3.2 | 0.05 | ycaM | 4 | 13.5833 | 0.306 |
| 3.2 | 0.05 | yadI | 4 | 13.5833 | 0.603 |
| 3.2 | 0.05 | AG1 | 4 | 13.5833 | 0.555 |
| 3.2 | 0.05 | ptsI | 4 | 13.8333 | 0.532 |
| 3.2 | 0.05 | clcB | 4 | 13.8333 | 0.3 |
| 3.2 | 0.05 | ycaM | 4 | 13.8333 | 0.306 |
| 3.2 | 0.05 | yadI | 4 | 13.8333 | 0.61 |
| 3.2 | 0.05 | AG1 | 4 | 13.8333 | 0.557 |
| 3.2 | 0.05 | ptsI | 4 | 14.0833 | 0.53 |
| 3.2 | 0.05 | clcB | 4 | 14.0833 | 0.3 |
| 3.2 | 0.05 | ycaM | 4 | 14.0833 | 0.306 |
| 3.2 | 0.05 | yadI | 4 | 14.0833 | 0.611 |
| 3.2 | 0.05 | AG1 | 4 | 14.0833 | 0.561 |
| 3.2 | 0.05 | ptsI | 4 | 14.3333 | 0.539 |
| 3.2 | 0.05 | clcB | 4 | 14.3333 | 0.302 |
| 3.2 | 0.05 | ycaM | 4 | 14.3333 | 0.306 |
| 3.2 | 0.05 | yadI | 4 | 14.3333 | 0.615 |
| 3.2 | 0.05 | AG1 | 4 | 14.3333 | 0.563 |
| 3.2 | 0.05 | ptsI | 4 | 14.5833 | 0.54 |
| 3.2 | 0.05 | clcB | 4 | 14.5833 | 0.301 |

| | | | | | |
|---|---|---|---|---|---|
| 3.2 | 0.05 | ycaM | 4 | 14.5833 | 0.305 |
| 3.2 | 0.05 | yadI | 4 | 14.5833 | 0.618 |
| 3.2 | 0.05 | AG1 | 4 | 14.5833 | 0.566 |
| 3.2 | 0.05 | ptsI | 4 | 14.8333 | 0.542 |
| 3.2 | 0.05 | clcB | 4 | 14.8333 | 0.301 |
| 3.2 | 0.05 | ycaM | 4 | 14.8333 | 0.306 |
| 3.2 | 0.05 | yadI | 4 | 14.8333 | 0.625 |
| 3.2 | 0.05 | AG1 | 4 | 14.8333 | 0.566 |
| 3.2 | 0.05 | ptsI | 4 | 15.0833 | 0.552 |
| 3.2 | 0.05 | clcB | 4 | 15.0833 | 0.3 |
| 3.2 | 0.05 | ycaM | 4 | 15.0833 | 0.307 |
| 3.2 | 0.05 | yadI | 4 | 15.0833 | 0.638 |
| 3.2 | 0.05 | AG1 | 4 | 15.0833 | 0.573 |
| 3.2 | 0.05 | ptsI | 4 | 15.3333 | 0.553 |
| 3.2 | 0.05 | clcB | 4 | 15.3333 | 0.303 |
| 3.2 | 0.05 | ycaM | 4 | 15.3333 | 0.306 |
| 3.2 | 0.05 | yadI | 4 | 15.3333 | 0.626 |
| 3.2 | 0.05 | AG1 | 4 | 15.3333 | 0.576 |
| 3.2 | 0.05 | ptsI | 4 | 15.5833 | 0.559 |
| 3.2 | 0.05 | clcB | 4 | 15.5833 | 0.302 |
| 3.2 | 0.05 | ycaM | 4 | 15.5833 | 0.305 |
| 3.2 | 0.05 | yadI | 4 | 15.5833 | 0.631 |
| 3.2 | 0.05 | AG1 | 4 | 15.5833 | 0.581 |
| 3.2 | 0.05 | ptsI | 4 | 15.8333 | 0.562 |
| 3.2 | 0.05 | clcB | 4 | 15.8333 | 0.302 |
| 3.2 | 0.05 | ycaM | 4 | 15.8333 | 0.307 |
| 3.2 | 0.05 | yadI | 4 | 15.8333 | 0.634 |
| 3.2 | 0.05 | AG1 | 4 | 15.8333 | 0.581 |
| 3.2 | 0.05 | ptsI | 4 | 16.0833 | 0.567 |
| 3.2 | 0.05 | clcB | 4 | 16.0833 | 0.303 |
| 3.2 | 0.05 | ycaM | 4 | 16.0833 | 0.306 |
| 3.2 | 0.05 | yadI | 4 | 16.0833 | 0.633 |
| 3.2 | 0.05 | AG1 | 4 | 16.0833 | 0.585 |
| 3.2 | 0.05 | ptsI | 4 | 16.3333 | 0.572 |
| 3.2 | 0.05 | clcB | 4 | 16.3333 | 0.302 |
| 3.2 | 0.05 | ycaM | 4 | 16.3333 | 0.306 |
| 3.2 | 0.05 | yadI | 4 | 16.3333 | 0.64 |
| 3.2 | 0.05 | AG1 | 4 | 16.3333 | 0.588 |
| 3.2 | 0.05 | ptsI | 4 | 16.5833 | 0.571 |
| 3.2 | 0.05 | clcB | 4 | 16.5833 | 0.302 |
| 3.2 | 0.05 | ycaM | 4 | 16.5833 | 0.306 |
| 3.2 | 0.05 | yadI | 4 | 16.5833 | 0.642 |
| 3.2 | 0.05 | AG1 | 4 | 16.5833 | 0.589 |
| 3.2 | 0.05 | ptsI | 4 | 16.8333 | 0.577 |
| 3.2 | 0.05 | clcB | 4 | 16.8333 | 0.301 |
| 3.2 | 0.05 | ycaM | 4 | 16.8333 | 0.307 |
| 3.2 | 0.05 | yadI | 4 | 16.8333 | 0.644 |
| 3.2 | 0.05 | AG1 | 4 | 16.8333 | 0.594 |

| | | | | | |
|---|---|---|---|---|---|
| 3.2 | 0.05 | ptsI | 4 | 17.0833 | 0.58 |
| 3.2 | 0.05 | clcB | 4 | 17.0833 | 0.302 |
| 3.2 | 0.05 | ycaM | 4 | 17.0833 | 0.307 |
| 3.2 | 0.05 | yadI | 4 | 17.0833 | 0.647 |
| 3.2 | 0.05 | AG1 | 4 | 17.0833 | 0.595 |
| 3.2 | 0.05 | ptsI | 4 | 17.3333 | 0.586 |
| 3.2 | 0.05 | clcB | 4 | 17.3333 | 0.3 |
| 3.2 | 0.05 | ycaM | 4 | 17.3333 | 0.307 |
| 3.2 | 0.05 | yadI | 4 | 17.3333 | 0.649 |
| 3.2 | 0.05 | AG1 | 4 | 17.3333 | 0.597 |
| 3.2 | 0.05 | ptsI | 4 | 17.5833 | 0.588 |
| 3.2 | 0.05 | clcB | 4 | 17.5833 | 0.301 |
| 3.2 | 0.05 | ycaM | 4 | 17.5833 | 0.305 |
| 3.2 | 0.05 | yadI | 4 | 17.5833 | 0.649 |
| 3.2 | 0.05 | AG1 | 4 | 17.5833 | 0.601 |
| 3.2 | 0.05 | ptsI | 4 | 17.8333 | 0.593 |
| 3.2 | 0.05 | clcB | 4 | 17.8333 | 0.302 |
| 3.2 | 0.05 | ycaM | 4 | 17.8333 | 0.306 |
| 3.2 | 0.05 | yadI | 4 | 17.8333 | 0.65 |
| 3.2 | 0.05 | AG1 | 4 | 17.8333 | 0.602 |
| 3.2 | 0.05 | ptsI | 4 | 18.0833 | 0.597 |
| 3.2 | 0.05 | clcB | 4 | 18.0833 | 0.302 |
| 3.2 | 0.05 | ycaM | 4 | 18.0833 | 0.306 |
| 3.2 | 0.05 | yadI | 4 | 18.0833 | 0.656 |
| 3.2 | 0.05 | AG1 | 4 | 18.0833 | 0.604 |
| 3.2 | 0.05 | ptsI | 4 | 18.3333 | 0.599 |
| 3.2 | 0.05 | clcB | 4 | 18.3333 | 0.302 |
| 3.2 | 0.05 | ycaM | 4 | 18.3333 | 0.305 |
| 3.2 | 0.05 | yadI | 4 | 18.3333 | 0.656 |
| 3.2 | 0.05 | AG1 | 4 | 18.3333 | 0.605 |
| 3.2 | 0.05 | ptsI | 4 | 18.5833 | 0.602 |
| 3.2 | 0.05 | clcB | 4 | 18.5833 | 0.301 |
| 3.2 | 0.05 | ycaM | 4 | 18.5833 | 0.305 |
| 3.2 | 0.05 | yadI | 4 | 18.5833 | 0.658 |
| 3.2 | 0.05 | AG1 | 4 | 18.5833 | 0.608 |
| 3.2 | 0.05 | ptsI | 4 | 18.8333 | 0.609 |
| 3.2 | 0.05 | clcB | 4 | 18.8333 | 0.301 |
| 3.2 | 0.05 | ycaM | 4 | 18.8333 | 0.305 |
| 3.2 | 0.05 | yadI | 4 | 18.8333 | 0.66 |
| 3.2 | 0.05 | AG1 | 4 | 18.8333 | 0.609 |
| 3.2 | 0.05 | ptsI | 4 | 19.0833 | 0.613 |
| 3.2 | 0.05 | clcB | 4 | 19.0833 | 0.301 |
| 3.2 | 0.05 | ycaM | 4 | 19.0833 | 0.304 |
| 3.2 | 0.05 | yadI | 4 | 19.0833 | 0.662 |
| 3.2 | 0.05 | AG1 | 4 | 19.0833 | 0.613 |
| 3.2 | 0.05 | ptsI | 4 | 19.3333 | 0.618 |
| 3.2 | 0.05 | clcB | 4 | 19.3333 | 0.301 |
| 3.2 | 0.05 | ycaM | 4 | 19.3333 | 0.305 |

| | | | | | |
|---|---|---|---|---|---|
| 3.2 | 0.05 | yadI | 4 | 19.3333 | 0.666 |
| 3.2 | 0.05 | AG1 | 4 | 19.3333 | 0.617 |
| 3.2 | 0.05 | ptsI | 4 | 19.5833 | 0.62 |
| 3.2 | 0.05 | clcB | 4 | 19.5833 | 0.302 |
| 3.2 | 0.05 | ycaM | 4 | 19.5833 | 0.305 |
| 3.2 | 0.05 | yadI | 4 | 19.5833 | 0.67 |
| 3.2 | 0.05 | AG1 | 4 | 19.5833 | 0.619 |
| 3.2 | 0.05 | ptsI | 4 | 19.8333 | 0.625 |
| 3.2 | 0.05 | clcB | 4 | 19.8333 | 0.301 |
| 3.2 | 0.05 | ycaM | 4 | 19.8333 | 0.304 |
| 3.2 | 0.05 | yadI | 4 | 19.8333 | 0.672 |
| 3.2 | 0.05 | AG1 | 4 | 19.8333 | 0.622 |
| 3.2 | 0.05 | ptsI | 4 | 20.0833 | 0.63 |
| 3.2 | 0.05 | clcB | 4 | 20.0833 | 0.3 |
| 3.2 | 0.05 | ycaM | 4 | 20.0833 | 0.303 |
| 3.2 | 0.05 | yadI | 4 | 20.0833 | 0.671 |
| 3.2 | 0.05 | AG1 | 4 | 20.0833 | 0.62 |
| 3.2 | 0.05 | ptsI | 4 | 20.3333 | 0.631 |
| 3.2 | 0.05 | clcB | 4 | 20.3333 | 0.301 |
| 3.2 | 0.05 | ycaM | 4 | 20.3333 | 0.304 |
| 3.2 | 0.05 | yadI | 4 | 20.3333 | 0.675 |
| 3.2 | 0.05 | AG1 | 4 | 20.3333 | 0.628 |
| 3.2 | 0.05 | ptsI | 4 | 20.5833 | 0.635 |
| 3.2 | 0.05 | clcB | 4 | 20.5833 | 0.301 |
| 3.2 | 0.05 | ycaM | 4 | 20.5833 | 0.304 |
| 3.2 | 0.05 | yadI | 4 | 20.5833 | 0.679 |
| 3.2 | 0.05 | AG1 | 4 | 20.5833 | 0.633 |
| 3.2 | 0.05 | ptsI | 4 | 20.8333 | 0.638 |
| 3.2 | 0.05 | clcB | 4 | 20.8333 | 0.3 |
| 3.2 | 0.05 | ycaM | 4 | 20.8333 | 0.304 |
| 3.2 | 0.05 | yadI | 4 | 20.8333 | 0.682 |
| 3.2 | 0.05 | AG1 | 4 | 20.8333 | 0.636 |
| 3.2 | 0.05 | ptsI | 4 | 21.0833 | 0.639 |
| 3.2 | 0.05 | clcB | 4 | 21.0833 | 0.3 |
| 3.2 | 0.05 | ycaM | 4 | 21.0833 | 0.303 |
| 3.2 | 0.05 | yadI | 4 | 21.0833 | 0.678 |
| 3.2 | 0.05 | AG1 | 4 | 21.0833 | 0.638 |
| 3.2 | 0.05 | ptsI | 4 | 21.3333 | 0.643 |
| 3.2 | 0.05 | clcB | 4 | 21.3333 | 0.301 |
| 3.2 | 0.05 | ycaM | 4 | 21.3333 | 0.303 |
| 3.2 | 0.05 | yadI | 4 | 21.3333 | 0.683 |
| 3.2 | 0.05 | AG1 | 4 | 21.3333 | 0.642 |
| 3.2 | 0.05 | ptsI | 4 | 21.5833 | 0.649 |
| 3.2 | 0.05 | clcB | 4 | 21.5833 | 0.299 |
| 3.2 | 0.05 | ycaM | 4 | 21.5833 | 0.302 |
| 3.2 | 0.05 | yadI | 4 | 21.5833 | 0.686 |
| 3.2 | 0.05 | AG1 | 4 | 21.5833 | 0.644 |
| 3.2 | 0.05 | ptsI | 4 | 21.8333 | 0.651 |

| | | | | | |
|---|---|---|---|---|---|
| 3.2 | 0.05 | clcB | 4 | 21.8333 | 0.299 |
| 3.2 | 0.05 | ycaM | 4 | 21.8333 | 0.302 |
| 3.2 | 0.05 | yadI | 4 | 21.8333 | 0.689 |
| 3.2 | 0.05 | AG1 | 4 | 21.8333 | 0.647 |
| 3.2 | 0.05 | ptsI | 4 | 22.0833 | 0.656 |
| 3.2 | 0.05 | clcB | 4 | 22.0833 | 0.3 |
| 3.2 | 0.05 | ycaM | 4 | 22.0833 | 0.301 |
| 3.2 | 0.05 | yadI | 4 | 22.0833 | 0.689 |
| 3.2 | 0.05 | AG1 | 4 | 22.0833 | 0.648 |
| 3.2 | 0.05 | ptsI | 4 | 22.3333 | 0.661 |
| 3.2 | 0.05 | clcB | 4 | 22.3333 | 0.299 |
| 3.2 | 0.05 | ycaM | 4 | 22.3333 | 0.301 |
| 3.2 | 0.05 | yadI | 4 | 22.3333 | 0.692 |
| 3.2 | 0.05 | AG1 | 4 | 22.3333 | 0.65 |
| 3.2 | 0.05 | ptsI | 4 | 22.5833 | 0.662 |
| 3.2 | 0.05 | clcB | 4 | 22.5833 | 0.298 |
| 3.2 | 0.05 | ycaM | 4 | 22.5833 | 0.3 |
| 3.2 | 0.05 | yadI | 4 | 22.5833 | 0.694 |
| 3.2 | 0.05 | AG1 | 4 | 22.5833 | 0.65 |
| 3.2 | 0.05 | ptsI | 4 | 22.8333 | 0.668 |
| 3.2 | 0.05 | clcB | 4 | 22.8333 | 0.297 |
| 3.2 | 0.05 | ycaM | 4 | 22.8333 | 0.3 |
| 3.2 | 0.05 | yadI | 4 | 22.8333 | 0.694 |
| 3.2 | 0.05 | AG1 | 4 | 22.8333 | 0.658 |
| 3.2 | 0.05 | ptsI | 4 | 23.0833 | 0.668 |
| 3.2 | 0.05 | clcB | 4 | 23.0833 | 0.296 |
| 3.2 | 0.05 | ycaM | 4 | 23.0833 | 0.3 |
| 3.2 | 0.05 | yadI | 4 | 23.0833 | 0.697 |
| 3.2 | 0.05 | AG1 | 4 | 23.0833 | 0.656 |
| 3.2 | 0.05 | ptsI | 4 | 23.3333 | 0.674 |
| 3.2 | 0.05 | clcB | 4 | 23.3333 | 0.297 |
| 3.2 | 0.05 | ycaM | 4 | 23.3333 | 0.3 |
| 3.2 | 0.05 | yadI | 4 | 23.3333 | 0.701 |
| 3.2 | 0.05 | AG1 | 4 | 23.3333 | 0.661 |
| 3.2 | 0.05 | ptsI | 4 | 23.5833 | 0.677 |
| 3.2 | 0.05 | clcB | 4 | 23.5833 | 0.296 |
| 3.2 | 0.05 | ycaM | 4 | 23.5833 | 0.3 |
| 3.2 | 0.05 | yadI | 4 | 23.5833 | 0.702 |
| 3.2 | 0.05 | AG1 | 4 | 23.5833 | 0.663 |
| 3.2 | 0.05 | ptsI | 4 | 23.8333 | 0.681 |
| 3.2 | 0.05 | clcB | 4 | 23.8333 | 0.296 |
| 3.2 | 0.05 | ycaM | 4 | 23.8333 | 0.299 |
| 3.2 | 0.05 | yadI | 4 | 23.8333 | 0.703 |
| 3.2 | 0.05 | AG1 | 4 | 23.8333 | 0.665 |
| 3.2 | 0.05 | ptsI | 4 | 24.0833 | 0.682 |
| 3.2 | 0.05 | clcB | 4 | 24.0833 | 0.296 |
| 3.2 | 0.05 | ycaM | 4 | 24.0833 | 0.297 |
| 3.2 | 0.05 | yadI | 4 | 24.0833 | 0.705 |

| | | | | | |
|---|---|---|---|---|---|
| 3.2 | 0.05 | AG1 | 4 | 24.0833 | 0.667 |
| 3.2 | 0.05 | ptsI | 5 | 0 | 0.182 |
| 3.2 | 0.05 | clcB | 5 | 0 | 0.158 |
| 3.2 | 0.05 | ycaM | 5 | 0 | 0.154 |
| 3.2 | 0.05 | yadI | 5 | 0 | 0.164 |
| 3.2 | 0.05 | AG1 | 5 | 0 | 0.155 |
| 3.2 | 0.05 | ptsI | 5 | 0.35 | 0.181 |
| 3.2 | 0.05 | clcB | 5 | 0.35 | 0.158 |
| 3.2 | 0.05 | ycaM | 5 | 0.35 | 0.152 |
| 3.2 | 0.05 | yadI | 5 | 0.35 | 0.163 |
| 3.2 | 0.05 | AG1 | 5 | 0.35 | 0.157 |
| 3.2 | 0.05 | ptsI | 5 | 0.6 | 0.175 |
| 3.2 | 0.05 | clcB | 5 | 0.6 | 0.157 |
| 3.2 | 0.05 | ycaM | 5 | 0.6 | 0.152 |
| 3.2 | 0.05 | yadI | 5 | 0.6 | 0.165 |
| 3.2 | 0.05 | AG1 | 5 | 0.6 | 0.156 |
| 3.2 | 0.05 | ptsI | 5 | 0.85 | 0.178 |
| 3.2 | 0.05 | clcB | 5 | 0.85 | 0.157 |
| 3.2 | 0.05 | ycaM | 5 | 0.85 | 0.152 |
| 3.2 | 0.05 | yadI | 5 | 0.85 | 0.165 |
| 3.2 | 0.05 | AG1 | 5 | 0.85 | 0.155 |
| 3.2 | 0.05 | ptsI | 5 | 1.1 | 0.18 |
| 3.2 | 0.05 | clcB | 5 | 1.1 | 0.159 |
| 3.2 | 0.05 | ycaM | 5 | 1.1 | 0.154 |
| 3.2 | 0.05 | yadI | 5 | 1.1 | 0.168 |
| 3.2 | 0.05 | AG1 | 5 | 1.1 | 0.158 |
| 3.2 | 0.05 | ptsI | 5 | 1.35 | 0.185 |
| 3.2 | 0.05 | clcB | 5 | 1.35 | 0.16 |
| 3.2 | 0.05 | ycaM | 5 | 1.35 | 0.155 |
| 3.2 | 0.05 | yadI | 5 | 1.35 | 0.169 |
| 3.2 | 0.05 | AG1 | 5 | 1.35 | 0.157 |
| 3.2 | 0.05 | ptsI | 5 | 1.6 | 0.185 |
| 3.2 | 0.05 | clcB | 5 | 1.6 | 0.161 |
| 3.2 | 0.05 | ycaM | 5 | 1.6 | 0.158 |
| 3.2 | 0.05 | yadI | 5 | 1.6 | 0.171 |
| 3.2 | 0.05 | AG1 | 5 | 1.6 | 0.16 |
| 3.2 | 0.05 | ptsI | 5 | 1.85 | 0.186 |
| 3.2 | 0.05 | clcB | 5 | 1.85 | 0.165 |
| 3.2 | 0.05 | ycaM | 5 | 1.85 | 0.16 |
| 3.2 | 0.05 | yadI | 5 | 1.85 | 0.174 |
| 3.2 | 0.05 | AG1 | 5 | 1.85 | 0.161 |
| 3.2 | 0.05 | ptsI | 5 | 2.1 | 0.194 |
| 3.2 | 0.05 | clcB | 5 | 2.1 | 0.17 |
| 3.2 | 0.05 | ycaM | 5 | 2.1 | 0.167 |
| 3.2 | 0.05 | yadI | 5 | 2.1 | 0.18 |
| 3.2 | 0.05 | AG1 | 5 | 2.1 | 0.166 |
| 3.2 | 0.05 | ptsI | 5 | 2.35 | 0.196 |
| 3.2 | 0.05 | clcB | 5 | 2.35 | 0.174 |

| | | | | | |
|---|---|---|---|---|---|
| 3.2 | 0.05 | ycaM | 5 | 2.35 | 0.17 |
| 3.2 | 0.05 | yadI | 5 | 2.35 | 0.186 |
| 3.2 | 0.05 | AG1 | 5 | 2.35 | 0.169 |
| 3.2 | 0.05 | ptsI | 5 | 2.6 | 0.19 |
| 3.2 | 0.05 | clcB | 5 | 2.6 | 0.176 |
| 3.2 | 0.05 | ycaM | 5 | 2.6 | 0.17 |
| 3.2 | 0.05 | yadI | 5 | 2.6 | 0.185 |
| 3.2 | 0.05 | AG1 | 5 | 2.6 | 0.168 |
| 3.2 | 0.05 | ptsI | 5 | 2.85 | 0.201 |
| 3.2 | 0.05 | clcB | 5 | 2.85 | 0.183 |
| 3.2 | 0.05 | ycaM | 5 | 2.85 | 0.179 |
| 3.2 | 0.05 | yadI | 5 | 2.85 | 0.194 |
| 3.2 | 0.05 | AG1 | 5 | 2.85 | 0.175 |
| 3.2 | 0.05 | ptsI | 5 | 3.1 | 0.208 |
| 3.2 | 0.05 | clcB | 5 | 3.1 | 0.192 |
| 3.2 | 0.05 | ycaM | 5 | 3.1 | 0.188 |
| 3.2 | 0.05 | yadI | 5 | 3.1 | 0.202 |
| 3.2 | 0.05 | AG1 | 5 | 3.1 | 0.181 |
| 3.2 | 0.05 | ptsI | 5 | 3.35 | 0.22 |
| 3.2 | 0.05 | clcB | 5 | 3.35 | 0.2 |
| 3.2 | 0.05 | ycaM | 5 | 3.35 | 0.2 |
| 3.2 | 0.05 | yadI | 5 | 3.35 | 0.217 |
| 3.2 | 0.05 | AG1 | 5 | 3.35 | 0.187 |
| 3.2 | 0.05 | ptsI | 5 | 3.6 | 0.233 |
| 3.2 | 0.05 | clcB | 5 | 3.6 | 0.215 |
| 3.2 | 0.05 | ycaM | 5 | 3.6 | 0.217 |
| 3.2 | 0.05 | yadI | 5 | 3.6 | 0.239 |
| 3.2 | 0.05 | AG1 | 5 | 3.6 | 0.197 |
| 3.2 | 0.05 | ptsI | 5 | 3.85 | 0.243 |
| 3.2 | 0.05 | clcB | 5 | 3.85 | 0.23 |
| 3.2 | 0.05 | ycaM | 5 | 3.85 | 0.23 |
| 3.2 | 0.05 | yadI | 5 | 3.85 | 0.269 |
| 3.2 | 0.05 | AG1 | 5 | 3.85 | 0.208 |
| 3.2 | 0.05 | ptsI | 5 | 4.38333 | 0.246 |
| 3.2 | 0.05 | clcB | 5 | 4.38333 | 0.247 |
| 3.2 | 0.05 | ycaM | 5 | 4.38333 | 0.209 |
| 3.2 | 0.05 | yadI | 5 | 4.38333 | 0.294 |
| 3.2 | 0.05 | AG1 | 5 | 4.38333 | 0.228 |
| 3.2 | 0.05 | ptsI | 5 | 4.63333 | 0.233 |
| 3.2 | 0.05 | clcB | 5 | 4.63333 | 0.245 |
| 3.2 | 0.05 | ycaM | 5 | 4.63333 | 0.211 |
| 3.2 | 0.05 | yadI | 5 | 4.63333 | 0.277 |
| 3.2 | 0.05 | AG1 | 5 | 4.63333 | 0.216 |
| 3.2 | 0.05 | ptsI | 5 | 4.88333 | 0.239 |
| 3.2 | 0.05 | clcB | 5 | 4.88333 | 0.259 |
| 3.2 | 0.05 | ycaM | 5 | 4.88333 | 0.225 |
| 3.2 | 0.05 | yadI | 5 | 4.88333 | 0.3 |
| 3.2 | 0.05 | AG1 | 5 | 4.88333 | 0.232 |

| | | | | | |
|---|---|---|---|---|---|
| 3.2 | 0.05 | ptsI | 5 | 5.13333 | 0.247 |
| 3.2 | 0.05 | clcB | 5 | 5.13333 | 0.269 |
| 3.2 | 0.05 | ycaM | 5 | 5.13333 | 0.234 |
| 3.2 | 0.05 | yadI | 5 | 5.13333 | 0.312 |
| 3.2 | 0.05 | AG1 | 5 | 5.13333 | 0.244 |
| 3.2 | 0.05 | ptsI | 5 | 5.38333 | 0.256 |
| 3.2 | 0.05 | clcB | 5 | 5.38333 | 0.275 |
| 3.2 | 0.05 | ycaM | 5 | 5.38333 | 0.236 |
| 3.2 | 0.05 | yadI | 5 | 5.38333 | 0.327 |
| 3.2 | 0.05 | AG1 | 5 | 5.38333 | 0.253 |
| 3.2 | 0.05 | ptsI | 5 | 5.63333 | 0.269 |
| 3.2 | 0.05 | clcB | 5 | 5.63333 | 0.285 |
| 3.2 | 0.05 | ycaM | 5 | 5.63333 | 0.239 |
| 3.2 | 0.05 | yadI | 5 | 5.63333 | 0.346 |
| 3.2 | 0.05 | AG1 | 5 | 5.63333 | 0.263 |
| 3.2 | 0.05 | ptsI | 5 | 5.88333 | 0.283 |
| 3.2 | 0.05 | clcB | 5 | 5.88333 | 0.29 |
| 3.2 | 0.05 | ycaM | 5 | 5.88333 | 0.239 |
| 3.2 | 0.05 | yadI | 5 | 5.88333 | 0.364 |
| 3.2 | 0.05 | AG1 | 5 | 5.88333 | 0.262 |
| 3.2 | 0.05 | ptsI | 5 | 6.13333 | 0.295 |
| 3.2 | 0.05 | clcB | 5 | 6.13333 | 0.301 |
| 3.2 | 0.05 | ycaM | 5 | 6.13333 | 0.244 |
| 3.2 | 0.05 | yadI | 5 | 6.13333 | 0.381 |
| 3.2 | 0.05 | AG1 | 5 | 6.13333 | 0.27 |
| 3.2 | 0.05 | ptsI | 5 | 6.38333 | 0.308 |
| 3.2 | 0.05 | clcB | 5 | 6.38333 | 0.313 |
| 3.2 | 0.05 | ycaM | 5 | 6.38333 | 0.247 |
| 3.2 | 0.05 | yadI | 5 | 6.38333 | 0.397 |
| 3.2 | 0.05 | AG1 | 5 | 6.38333 | 0.28 |
| 3.2 | 0.05 | ptsI | 5 | 6.63333 | 0.322 |
| 3.2 | 0.05 | clcB | 5 | 6.63333 | 0.322 |
| 3.2 | 0.05 | ycaM | 5 | 6.63333 | 0.252 |
| 3.2 | 0.05 | yadI | 5 | 6.63333 | 0.411 |
| 3.2 | 0.05 | AG1 | 5 | 6.63333 | 0.29 |
| 3.2 | 0.05 | ptsI | 5 | 6.88333 | 0.331 |
| 3.2 | 0.05 | clcB | 5 | 6.88333 | 0.334 |
| 3.2 | 0.05 | ycaM | 5 | 6.88333 | 0.255 |
| 3.2 | 0.05 | yadI | 5 | 6.88333 | 0.434 |
| 3.2 | 0.05 | AG1 | 5 | 6.88333 | 0.296 |
| 3.2 | 0.05 | ptsI | 5 | 7.13333 | 0.343 |
| 3.2 | 0.05 | clcB | 5 | 7.13333 | 0.337 |
| 3.2 | 0.05 | ycaM | 5 | 7.13333 | 0.257 |
| 3.2 | 0.05 | yadI | 5 | 7.13333 | 0.453 |
| 3.2 | 0.05 | AG1 | 5 | 7.13333 | 0.307 |
| 3.2 | 0.05 | ptsI | 5 | 7.38333 | 0.356 |
| 3.2 | 0.05 | clcB | 5 | 7.38333 | 0.339 |
| 3.2 | 0.05 | ycaM | 5 | 7.38333 | 0.263 |

| | | | | | |
|---|---|---|---|---|---|
| 3.2 | 0.05 | yadI | 5 | 7.38333 | 0.471 |
| 3.2 | 0.05 | AG1 | 5 | 7.38333 | 0.314 |
| 3.2 | 0.05 | ptsI | 5 | 7.63333 | 0.371 |
| 3.2 | 0.05 | clcB | 5 | 7.63333 | 0.343 |
| 3.2 | 0.05 | ycaM | 5 | 7.63333 | 0.265 |
| 3.2 | 0.05 | yadI | 5 | 7.63333 | 0.482 |
| 3.2 | 0.05 | AG1 | 5 | 7.63333 | 0.324 |
| 3.2 | 0.05 | ptsI | 5 | 7.88333 | 0.382 |
| 3.2 | 0.05 | clcB | 5 | 7.88333 | 0.354 |
| 3.2 | 0.05 | ycaM | 5 | 7.88333 | 0.271 |
| 3.2 | 0.05 | yadI | 5 | 7.88333 | 0.492 |
| 3.2 | 0.05 | AG1 | 5 | 7.88333 | 0.335 |
| 3.2 | 0.05 | ptsI | 5 | 8.13333 | 0.402 |
| 3.2 | 0.05 | clcB | 5 | 8.13333 | 0.355 |
| 3.2 | 0.05 | ycaM | 5 | 8.13333 | 0.272 |
| 3.2 | 0.05 | yadI | 5 | 8.13333 | 0.502 |
| 3.2 | 0.05 | AG1 | 5 | 8.13333 | 0.343 |
| 3.2 | 0.05 | ptsI | 5 | 8.38333 | 0.415 |
| 3.2 | 0.05 | clcB | 5 | 8.38333 | 0.358 |
| 3.2 | 0.05 | ycaM | 5 | 8.38333 | 0.274 |
| 3.2 | 0.05 | yadI | 5 | 8.38333 | 0.512 |
| 3.2 | 0.05 | AG1 | 5 | 8.38333 | 0.353 |
| 3.2 | 0.05 | ptsI | 5 | 8.88333 | 0.369 |
| 3.2 | 0.05 | clcB | 5 | 8.88333 | 0.311 |
| 3.2 | 0.05 | ycaM | 5 | 8.88333 | 0.264 |
| 3.2 | 0.05 | yadI | 5 | 8.88333 | 0.444 |
| 3.2 | 0.05 | AG1 | 5 | 8.88333 | 0.367 |
| 3.2 | 0.05 | ptsI | 5 | 9.13333 | 0.39 |
| 3.2 | 0.05 | clcB | 5 | 9.13333 | 0.309 |
| 3.2 | 0.05 | ycaM | 5 | 9.13333 | 0.263 |
| 3.2 | 0.05 | yadI | 5 | 9.13333 | 0.448 |
| 3.2 | 0.05 | AG1 | 5 | 9.13333 | 0.394 |
| 3.2 | 0.05 | ptsI | 5 | 9.38333 | 0.394 |
| 3.2 | 0.05 | clcB | 5 | 9.38333 | 0.306 |
| 3.2 | 0.05 | ycaM | 5 | 9.38333 | 0.261 |
| 3.2 | 0.05 | yadI | 5 | 9.38333 | 0.468 |
| 3.2 | 0.05 | AG1 | 5 | 9.38333 | 0.422 |
| 3.2 | 0.05 | ptsI | 5 | 9.63333 | 0.398 |
| 3.2 | 0.05 | clcB | 5 | 9.63333 | 0.306 |
| 3.2 | 0.05 | ycaM | 5 | 9.63333 | 0.258 |
| 3.2 | 0.05 | yadI | 5 | 9.63333 | 0.476 |
| 3.2 | 0.05 | AG1 | 5 | 9.63333 | 0.413 |
| 3.2 | 0.05 | ptsI | 5 | 9.88333 | 0.405 |
| 3.2 | 0.05 | clcB | 5 | 9.88333 | 0.306 |
| 3.2 | 0.05 | ycaM | 5 | 9.88333 | 0.258 |
| 3.2 | 0.05 | yadI | 5 | 9.88333 | 0.482 |
| 3.2 | 0.05 | AG1 | 5 | 9.88333 | 0.419 |
| 3.2 | 0.05 | ptsI | 5 | 10.1333 | 0.407 |

| | | | | | |
|---|---|---|---|---|---|
| 3.2 | 0.05 | clcB | 5 | 10.1333 | 0.302 |
| 3.2 | 0.05 | ycaM | 5 | 10.1333 | 0.256 |
| 3.2 | 0.05 | yadI | 5 | 10.1333 | 0.485 |
| 3.2 | 0.05 | AG1 | 5 | 10.1333 | 0.42 |
| 3.2 | 0.05 | ptsI | 5 | 10.3833 | 0.412 |
| 3.2 | 0.05 | clcB | 5 | 10.3833 | 0.304 |
| 3.2 | 0.05 | ycaM | 5 | 10.3833 | 0.255 |
| 3.2 | 0.05 | yadI | 5 | 10.3833 | 0.492 |
| 3.2 | 0.05 | AG1 | 5 | 10.3833 | 0.428 |
| 3.2 | 0.05 | ptsI | 5 | 10.6333 | 0.414 |
| 3.2 | 0.05 | clcB | 5 | 10.6333 | 0.301 |
| 3.2 | 0.05 | ycaM | 5 | 10.6333 | 0.255 |
| 3.2 | 0.05 | yadI | 5 | 10.6333 | 0.49 |
| 3.2 | 0.05 | AG1 | 5 | 10.6333 | 0.433 |
| 3.2 | 0.05 | ptsI | 5 | 10.8833 | 0.424 |
| 3.2 | 0.05 | clcB | 5 | 10.8833 | 0.304 |
| 3.2 | 0.05 | ycaM | 5 | 10.8833 | 0.256 |
| 3.2 | 0.05 | yadI | 5 | 10.8833 | 0.498 |
| 3.2 | 0.05 | AG1 | 5 | 10.8833 | 0.435 |
| 3.2 | 0.05 | ptsI | 5 | 11.1333 | 0.425 |
| 3.2 | 0.05 | clcB | 5 | 11.1333 | 0.303 |
| 3.2 | 0.05 | ycaM | 5 | 11.1333 | 0.255 |
| 3.2 | 0.05 | yadI | 5 | 11.1333 | 0.502 |
| 3.2 | 0.05 | AG1 | 5 | 11.1333 | 0.466 |
| 3.2 | 0.05 | ptsI | 5 | 11.3833 | 0.428 |
| 3.2 | 0.05 | clcB | 5 | 11.3833 | 0.301 |
| 3.2 | 0.05 | ycaM | 5 | 11.3833 | 0.255 |
| 3.2 | 0.05 | yadI | 5 | 11.3833 | 0.506 |
| 3.2 | 0.05 | AG1 | 5 | 11.3833 | 0.454 |
| 3.2 | 0.05 | ptsI | 5 | 11.6333 | 0.433 |
| 3.2 | 0.05 | clcB | 5 | 11.6333 | 0.299 |
| 3.2 | 0.05 | ycaM | 5 | 11.6333 | 0.26 |
| 3.2 | 0.05 | yadI | 5 | 11.6333 | 0.513 |
| 3.2 | 0.05 | AG1 | 5 | 11.6333 | 0.459 |
| 3.2 | 0.05 | ptsI | 5 | 11.8833 | 0.436 |
| 3.2 | 0.05 | clcB | 5 | 11.8833 | 0.302 |
| 3.2 | 0.05 | ycaM | 5 | 11.8833 | 0.26 |
| 3.2 | 0.05 | yadI | 5 | 11.8833 | 0.517 |
| 3.2 | 0.05 | AG1 | 5 | 11.8833 | 0.451 |
| 3.2 | 0.05 | ptsI | 5 | 12.1333 | 0.44 |
| 3.2 | 0.05 | clcB | 5 | 12.1333 | 0.302 |
| 3.2 | 0.05 | ycaM | 5 | 12.1333 | 0.257 |
| 3.2 | 0.05 | yadI | 5 | 12.1333 | 0.519 |
| 3.2 | 0.05 | AG1 | 5 | 12.1333 | 0.458 |
| 3.2 | 0.05 | ptsI | 5 | 12.3833 | 0.444 |
| 3.2 | 0.05 | clcB | 5 | 12.3833 | 0.304 |
| 3.2 | 0.05 | ycaM | 5 | 12.3833 | 0.255 |
| 3.2 | 0.05 | yadI | 5 | 12.3833 | 0.523 |

| | | | | | |
|---|---|---|---|---|---|
| 3.2 | 0.05 | AG1 | 5 | 12.3833 | 0.456 |
| 3.2 | 0.05 | ptsI | 5 | 12.6333 | 0.452 |
| 3.2 | 0.05 | clcB | 5 | 12.6333 | 0.304 |
| 3.2 | 0.05 | ycaM | 5 | 12.6333 | 0.256 |
| 3.2 | 0.05 | yadI | 5 | 12.6333 | 0.531 |
| 3.2 | 0.05 | AG1 | 5 | 12.6333 | 0.468 |
| 3.2 | 0.05 | ptsI | 5 | 12.8833 | 0.455 |
| 3.2 | 0.05 | clcB | 5 | 12.8833 | 0.301 |
| 3.2 | 0.05 | ycaM | 5 | 12.8833 | 0.259 |
| 3.2 | 0.05 | yadI | 5 | 12.8833 | 0.532 |
| 3.2 | 0.05 | AG1 | 5 | 12.8833 | 0.472 |
| 3.2 | 0.05 | ptsI | 5 | 13.1333 | 0.46 |
| 3.2 | 0.05 | clcB | 5 | 13.1333 | 0.302 |
| 3.2 | 0.05 | ycaM | 5 | 13.1333 | 0.257 |
| 3.2 | 0.05 | yadI | 5 | 13.1333 | 0.542 |
| 3.2 | 0.05 | AG1 | 5 | 13.1333 | 0.478 |
| 3.2 | 0.05 | ptsI | 5 | 13.3833 | 0.462 |
| 3.2 | 0.05 | clcB | 5 | 13.3833 | 0.302 |
| 3.2 | 0.05 | ycaM | 5 | 13.3833 | 0.258 |
| 3.2 | 0.05 | yadI | 5 | 13.3833 | 0.544 |
| 3.2 | 0.05 | AG1 | 5 | 13.3833 | 0.483 |
| 3.2 | 0.05 | ptsI | 5 | 13.6333 | 0.466 |
| 3.2 | 0.05 | clcB | 5 | 13.6333 | 0.3 |
| 3.2 | 0.05 | ycaM | 5 | 13.6333 | 0.257 |
| 3.2 | 0.05 | yadI | 5 | 13.6333 | 0.548 |
| 3.2 | 0.05 | AG1 | 5 | 13.6333 | 0.486 |
| 3.2 | 0.05 | ptsI | 5 | 13.8833 | 0.472 |
| 3.2 | 0.05 | clcB | 5 | 13.8833 | 0.301 |
| 3.2 | 0.05 | ycaM | 5 | 13.8833 | 0.257 |
| 3.2 | 0.05 | yadI | 5 | 13.8833 | 0.553 |
| 3.2 | 0.05 | AG1 | 5 | 13.8833 | 0.489 |
| 3.2 | 0.05 | ptsI | 5 | 14.1333 | 0.466 |
| 3.2 | 0.05 | clcB | 5 | 14.1333 | 0.298 |
| 3.2 | 0.05 | ycaM | 5 | 14.1333 | 0.256 |
| 3.2 | 0.05 | yadI | 5 | 14.1333 | 0.556 |
| 3.2 | 0.05 | AG1 | 5 | 14.1333 | 0.499 |
| 3.2 | 0.05 | ptsI | 5 | 14.3833 | 0.479 |
| 3.2 | 0.05 | clcB | 5 | 14.3833 | 0.3 |
| 3.2 | 0.05 | ycaM | 5 | 14.3833 | 0.255 |
| 3.2 | 0.05 | yadI | 5 | 14.3833 | 0.558 |
| 3.2 | 0.05 | AG1 | 5 | 14.3833 | 0.5 |
| 3.2 | 0.05 | ptsI | 5 | 14.6333 | 0.485 |
| 3.2 | 0.05 | clcB | 5 | 14.6333 | 0.301 |
| 3.2 | 0.05 | ycaM | 5 | 14.6333 | 0.254 |
| 3.2 | 0.05 | yadI | 5 | 14.6333 | 0.561 |
| 3.2 | 0.05 | AG1 | 5 | 14.6333 | 0.506 |
| 3.2 | 0.05 | ptsI | 5 | 14.8833 | 0.486 |
| 3.2 | 0.05 | clcB | 5 | 14.8833 | 0.3 |

| | | | | | |
|---|---|---|---|---|---|
| 3.2 | 0.05 | ycaM | 5 | 14.8833 | 0.255 |
| 3.2 | 0.05 | yadI | 5 | 14.8833 | 0.568 |
| 3.2 | 0.05 | AG1 | 5 | 14.8833 | 0.511 |
| 3.2 | 0.05 | ptsI | 5 | 15.1333 | 0.496 |
| 3.2 | 0.05 | clcB | 5 | 15.1333 | 0.301 |
| 3.2 | 0.05 | ycaM | 5 | 15.1333 | 0.256 |
| 3.2 | 0.05 | yadI | 5 | 15.1333 | 0.572 |
| 3.2 | 0.05 | AG1 | 5 | 15.1333 | 0.512 |
| 3.2 | 0.05 | ptsI | 5 | 15.3833 | 0.501 |
| 3.2 | 0.05 | clcB | 5 | 15.3833 | 0.3 |
| 3.2 | 0.05 | ycaM | 5 | 15.3833 | 0.256 |
| 3.2 | 0.05 | yadI | 5 | 15.3833 | 0.575 |
| 3.2 | 0.05 | AG1 | 5 | 15.3833 | 0.518 |
| 3.2 | 0.05 | ptsI | 5 | 15.6333 | 0.503 |
| 3.2 | 0.05 | clcB | 5 | 15.6333 | 0.299 |
| 3.2 | 0.05 | ycaM | 5 | 15.6333 | 0.253 |
| 3.2 | 0.05 | yadI | 5 | 15.6333 | 0.575 |
| 3.2 | 0.05 | AG1 | 5 | 15.6333 | 0.519 |
| 3.2 | 0.05 | ptsI | 5 | 15.8833 | 0.502 |
| 3.2 | 0.05 | clcB | 5 | 15.8833 | 0.3 |
| 3.2 | 0.05 | ycaM | 5 | 15.8833 | 0.253 |
| 3.2 | 0.05 | yadI | 5 | 15.8833 | 0.579 |
| 3.2 | 0.05 | AG1 | 5 | 15.8833 | 0.523 |
| 3.2 | 0.05 | ptsI | 5 | 16.1333 | 0.51 |
| 3.2 | 0.05 | clcB | 5 | 16.1333 | 0.296 |
| 3.2 | 0.05 | ycaM | 5 | 16.1333 | 0.253 |
| 3.2 | 0.05 | yadI | 5 | 16.1333 | 0.575 |
| 3.2 | 0.05 | AG1 | 5 | 16.1333 | 0.529 |
| 3.2 | 0.05 | ptsI | 5 | 16.3833 | 0.513 |
| 3.2 | 0.05 | clcB | 5 | 16.3833 | 0.299 |
| 3.2 | 0.05 | ycaM | 5 | 16.3833 | 0.255 |
| 3.2 | 0.05 | yadI | 5 | 16.3833 | 0.588 |
| 3.2 | 0.05 | AG1 | 5 | 16.3833 | 0.533 |
| 3.2 | 0.05 | ptsI | 5 | 16.6333 | 0.525 |
| 3.2 | 0.05 | clcB | 5 | 16.6333 | 0.301 |
| 3.2 | 0.05 | ycaM | 5 | 16.6333 | 0.254 |
| 3.2 | 0.05 | yadI | 5 | 16.6333 | 0.591 |
| 3.2 | 0.05 | AG1 | 5 | 16.6333 | 0.538 |
| 3.2 | 0.05 | ptsI | 5 | 16.8833 | 0.528 |
| 3.2 | 0.05 | clcB | 5 | 16.8833 | 0.301 |
| 3.2 | 0.05 | ycaM | 5 | 16.8833 | 0.254 |
| 3.2 | 0.05 | yadI | 5 | 16.8833 | 0.592 |
| 3.2 | 0.05 | AG1 | 5 | 16.8833 | 0.538 |
| 3.2 | 0.05 | ptsI | 5 | 17.1333 | 0.532 |
| 3.2 | 0.05 | clcB | 5 | 17.1333 | 0.299 |
| 3.2 | 0.05 | ycaM | 5 | 17.1333 | 0.254 |
| 3.2 | 0.05 | yadI | 5 | 17.1333 | 0.591 |
| 3.2 | 0.05 | AG1 | 5 | 17.1333 | 0.546 |

| | | | | | |
|---|---|---|---|---|---|
| 3.2 | 0.05 | ptsI | 5 | 17.3833 | 0.539 |
| 3.2 | 0.05 | clcB | 5 | 17.3833 | 0.3 |
| 3.2 | 0.05 | ycaM | 5 | 17.3833 | 0.254 |
| 3.2 | 0.05 | yadI | 5 | 17.3833 | 0.594 |
| 3.2 | 0.05 | AG1 | 5 | 17.3833 | 0.546 |
| 3.2 | 0.05 | ptsI | 5 | 17.6333 | 0.542 |
| 3.2 | 0.05 | clcB | 5 | 17.6333 | 0.299 |
| 3.2 | 0.05 | ycaM | 5 | 17.6333 | 0.254 |
| 3.2 | 0.05 | yadI | 5 | 17.6333 | 0.598 |
| 3.2 | 0.05 | AG1 | 5 | 17.6333 | 0.559 |
| 3.2 | 0.05 | ptsI | 5 | 17.8833 | 0.545 |
| 3.2 | 0.05 | clcB | 5 | 17.8833 | 0.3 |
| 3.2 | 0.05 | ycaM | 5 | 17.8833 | 0.253 |
| 3.2 | 0.05 | yadI | 5 | 17.8833 | 0.597 |
| 3.2 | 0.05 | AG1 | 5 | 17.8833 | 0.554 |
| 3.2 | 0.05 | ptsI | 5 | 18.1333 | 0.553 |
| 3.2 | 0.05 | clcB | 5 | 18.1333 | 0.298 |
| 3.2 | 0.05 | ycaM | 5 | 18.1333 | 0.253 |
| 3.2 | 0.05 | yadI | 5 | 18.1333 | 0.603 |
| 3.2 | 0.05 | AG1 | 5 | 18.1333 | 0.572 |
| 3.2 | 0.05 | ptsI | 5 | 18.3833 | 0.556 |
| 3.2 | 0.05 | clcB | 5 | 18.3833 | 0.298 |
| 3.2 | 0.05 | ycaM | 5 | 18.3833 | 0.253 |
| 3.2 | 0.05 | yadI | 5 | 18.3833 | 0.605 |
| 3.2 | 0.05 | AG1 | 5 | 18.3833 | 0.576 |
| 3.2 | 0.05 | ptsI | 5 | 18.6333 | 0.559 |
| 3.2 | 0.05 | clcB | 5 | 18.6333 | 0.297 |
| 3.2 | 0.05 | ycaM | 5 | 18.6333 | 0.253 |
| 3.2 | 0.05 | yadI | 5 | 18.6333 | 0.605 |
| 3.2 | 0.05 | AG1 | 5 | 18.6333 | 0.591 |
| 3.2 | 0.05 | ptsI | 5 | 18.8833 | 0.567 |
| 3.2 | 0.05 | clcB | 5 | 18.8833 | 0.297 |
| 3.2 | 0.05 | ycaM | 5 | 18.8833 | 0.253 |
| 3.2 | 0.05 | yadI | 5 | 18.8833 | 0.607 |
| 3.2 | 0.05 | AG1 | 5 | 18.8833 | 0.597 |
| 3.2 | 0.05 | ptsI | 5 | 19.1333 | 0.569 |
| 3.2 | 0.05 | clcB | 5 | 19.1333 | 0.297 |
| 3.2 | 0.05 | ycaM | 5 | 19.1333 | 0.252 |
| 3.2 | 0.05 | yadI | 5 | 19.1333 | 0.608 |
| 3.2 | 0.05 | AG1 | 5 | 19.1333 | 0.591 |
| 3.2 | 0.05 | ptsI | 5 | 19.3833 | 0.576 |
| 3.2 | 0.05 | clcB | 5 | 19.3833 | 0.297 |
| 3.2 | 0.05 | ycaM | 5 | 19.3833 | 0.252 |
| 3.2 | 0.05 | yadI | 5 | 19.3833 | 0.614 |
| 3.2 | 0.05 | AG1 | 5 | 19.3833 | 0.614 |
| 3.2 | 0.05 | ptsI | 5 | 19.6333 | 0.584 |
| 3.2 | 0.05 | clcB | 5 | 19.6333 | 0.298 |
| 3.2 | 0.05 | ycaM | 5 | 19.6333 | 0.252 |

| | | | | | |
|---|---|---|---|---|---|
| 3.2 | 0.05 | yadI | 5 | 19.6333 | 0.608 |
| 3.2 | 0.05 | AG1 | 5 | 19.6333 | 0.618 |
| 3.2 | 0.05 | ptsI | 5 | 19.8833 | 0.586 |
| 3.2 | 0.05 | clcB | 5 | 19.8833 | 0.297 |
| 3.2 | 0.05 | ycaM | 5 | 19.8833 | 0.252 |
| 3.2 | 0.05 | yadI | 5 | 19.8833 | 0.613 |
| 3.2 | 0.05 | AG1 | 5 | 19.8833 | 0.589 |
| 3.2 | 0.05 | ptsI | 5 | 20.1333 | 0.589 |
| 3.2 | 0.05 | clcB | 5 | 20.1333 | 0.296 |
| 3.2 | 0.05 | ycaM | 5 | 20.1333 | 0.251 |
| 3.2 | 0.05 | yadI | 5 | 20.1333 | 0.613 |
| 3.2 | 0.05 | AG1 | 5 | 20.1333 | 0.581 |
| 3.2 | 0.05 | ptsI | 5 | 20.3833 | 0.593 |
| 3.2 | 0.05 | clcB | 5 | 20.3833 | 0.296 |
| 3.2 | 0.05 | ycaM | 5 | 20.3833 | 0.251 |
| 3.2 | 0.05 | yadI | 5 | 20.3833 | 0.615 |
| 3.2 | 0.05 | AG1 | 5 | 20.3833 | 0.579 |
| 3.2 | 0.05 | ptsI | 5 | 20.6333 | 0.596 |
| 3.2 | 0.05 | clcB | 5 | 20.6333 | 0.3 |
| 3.2 | 0.05 | ycaM | 5 | 20.6333 | 0.251 |
| 3.2 | 0.05 | yadI | 5 | 20.6333 | 0.614 |
| 3.2 | 0.05 | AG1 | 5 | 20.6333 | 0.583 |
| 3.2 | 0.05 | ptsI | 5 | 20.8833 | 0.609 |
| 3.2 | 0.05 | clcB | 5 | 20.8833 | 0.296 |
| 3.2 | 0.05 | ycaM | 5 | 20.8833 | 0.252 |
| 3.2 | 0.05 | yadI | 5 | 20.8833 | 0.616 |
| 3.2 | 0.05 | AG1 | 5 | 20.8833 | 0.583 |
| 3.2 | 0.05 | ptsI | 5 | 21.1333 | 0.608 |
| 3.2 | 0.05 | clcB | 5 | 21.1333 | 0.296 |
| 3.2 | 0.05 | ycaM | 5 | 21.1333 | 0.251 |
| 3.2 | 0.05 | yadI | 5 | 21.1333 | 0.616 |
| 3.2 | 0.05 | AG1 | 5 | 21.1333 | 0.586 |
| 3.2 | 0.05 | ptsI | 5 | 21.3833 | 0.616 |
| 3.2 | 0.05 | clcB | 5 | 21.3833 | 0.296 |
| 3.2 | 0.05 | ycaM | 5 | 21.3833 | 0.25 |
| 3.2 | 0.05 | yadI | 5 | 21.3833 | 0.62 |
| 3.2 | 0.05 | AG1 | 5 | 21.3833 | 0.589 |
| 3.2 | 0.05 | ptsI | 5 | 21.6333 | 0.619 |
| 3.2 | 0.05 | clcB | 5 | 21.6333 | 0.296 |
| 3.2 | 0.05 | ycaM | 5 | 21.6333 | 0.25 |
| 3.2 | 0.05 | yadI | 5 | 21.6333 | 0.625 |
| 3.2 | 0.05 | AG1 | 5 | 21.6333 | 0.595 |
| 3.2 | 0.05 | ptsI | 5 | 21.8833 | 0.62 |
| 3.2 | 0.05 | clcB | 5 | 21.8833 | 0.295 |
| 3.2 | 0.05 | ycaM | 5 | 21.8833 | 0.25 |
| 3.2 | 0.05 | yadI | 5 | 21.8833 | 0.625 |
| 3.2 | 0.05 | AG1 | 5 | 21.8833 | 0.596 |
| 3.2 | 0.05 | ptsI | 5 | 22.1333 | 0.626 |

| | | | | | |
|---|---|---|---|---|---|
| 3.2 | 0.05 | clcB | 5 | 22.1333 | 0.297 |
| 3.2 | 0.05 | ycaM | 5 | 22.1333 | 0.249 |
| 3.2 | 0.05 | yadI | 5 | 22.1333 | 0.625 |
| 3.2 | 0.05 | AG1 | 5 | 22.1333 | 0.598 |
| 3.2 | 0.05 | ptsI | 5 | 22.3833 | 0.631 |
| 3.2 | 0.05 | clcB | 5 | 22.3833 | 0.294 |
| 3.2 | 0.05 | ycaM | 5 | 22.3833 | 0.249 |
| 3.2 | 0.05 | yadI | 5 | 22.3833 | 0.63 |
| 3.2 | 0.05 | AG1 | 5 | 22.3833 | 0.602 |
| 3.2 | 0.05 | ptsI | 5 | 22.6333 | 0.635 |
| 3.2 | 0.05 | clcB | 5 | 22.6333 | 0.296 |
| 3.2 | 0.05 | ycaM | 5 | 22.6333 | 0.25 |
| 3.2 | 0.05 | yadI | 5 | 22.6333 | 0.632 |
| 3.2 | 0.05 | AG1 | 5 | 22.6333 | 0.607 |
| 3.2 | 0.05 | ptsI | 5 | 22.8833 | 0.643 |
| 3.2 | 0.05 | clcB | 5 | 22.8833 | 0.295 |
| 3.2 | 0.05 | ycaM | 5 | 22.8833 | 0.249 |
| 3.2 | 0.05 | yadI | 5 | 22.8833 | 0.633 |
| 3.2 | 0.05 | AG1 | 5 | 22.8833 | 0.608 |
| 3.2 | 0.05 | ptsI | 5 | 23.1333 | 0.644 |
| 3.2 | 0.05 | clcB | 5 | 23.1333 | 0.295 |
| 3.2 | 0.05 | ycaM | 5 | 23.1333 | 0.249 |
| 3.2 | 0.05 | yadI | 5 | 23.1333 | 0.632 |
| 3.2 | 0.05 | AG1 | 5 | 23.1333 | 0.61 |
| 3.2 | 0.05 | ptsI | 5 | 23.3833 | 0.647 |
| 3.2 | 0.05 | clcB | 5 | 23.3833 | 0.295 |
| 3.2 | 0.05 | ycaM | 5 | 23.3833 | 0.248 |
| 3.2 | 0.05 | yadI | 5 | 23.3833 | 0.634 |
| 3.2 | 0.05 | AG1 | 5 | 23.3833 | 0.615 |
| 3.2 | 0.05 | ptsI | 5 | 23.6333 | 0.649 |
| 3.2 | 0.05 | clcB | 5 | 23.6333 | 0.294 |
| 3.2 | 0.05 | ycaM | 5 | 23.6333 | 0.249 |
| 3.2 | 0.05 | yadI | 5 | 23.6333 | 0.633 |
| 3.2 | 0.05 | AG1 | 5 | 23.6333 | 0.616 |
| 3.2 | 0.05 | ptsI | 5 | 23.8833 | 0.658 |
| 3.2 | 0.05 | clcB | 5 | 23.8833 | 0.296 |
| 3.2 | 0.05 | ycaM | 5 | 23.8833 | 0.248 |
| 3.2 | 0.05 | yadI | 5 | 23.8833 | 0.633 |
| 3.2 | 0.05 | AG1 | 5 | 23.8833 | 0.617 |
| 3.2 | 0.05 | ptsI | 5 | 24.1333 | 0.661 |
| 3.2 | 0.05 | clcB | 5 | 24.1333 | 0.295 |
| 3.2 | 0.05 | ycaM | 5 | 24.1333 | 0.248 |
| 3.2 | 0.05 | yadI | 5 | 24.1333 | 0.637 |
| 3.2 | 0.05 | AG1 | 5 | 24.1333 | 0.623 |
| 3.2 | 0.05 | ptsI | 5 | 24.3833 | 0.666 |
| 3.2 | 0.05 | clcB | 5 | 24.3833 | 0.294 |
| 3.2 | 0.05 | ycaM | 5 | 24.3833 | 0.248 |
| 3.2 | 0.05 | yadI | 5 | 24.3833 | 0.636 |

| | | | | | |
|---|---|---|---|---|---|
| 3.2 | 0.05 | AG1 | 5 | 24.3833 | 0.621 |
| 0 | 0.1 | ptsI | 1 | 0 | 0.242 |
| 0 | 0.1 | clcB | 1 | 0 | 0.224 |
| 0 | 0.1 | ycaM | 1 | 0 | 0.232 |
| 0 | 0.1 | yadI | 1 | 0 | 0.25 |
| 0 | 0.1 | AG1 | 1 | 0 | 0.241 |
| 0 | 0.1 | ptsI | 1 | 0.25 | 0.226 |
| 0 | 0.1 | clcB | 1 | 0.25 | 0.215 |
| 0 | 0.1 | ycaM | 1 | 0.25 | 0.221 |
| 0 | 0.1 | yadI | 1 | 0.25 | 0.236 |
| 0 | 0.1 | AG1 | 1 | 0.25 | 0.229 |
| 0 | 0.1 | ptsI | 1 | 0.5 | 0.225 |
| 0 | 0.1 | clcB | 1 | 0.5 | 0.214 |
| 0 | 0.1 | ycaM | 1 | 0.5 | 0.217 |
| 0 | 0.1 | yadI | 1 | 0.5 | 0.236 |
| 0 | 0.1 | AG1 | 1 | 0.5 | 0.227 |
| 0 | 0.1 | ptsI | 1 | 0.75 | 0.224 |
| 0 | 0.1 | clcB | 1 | 0.75 | 0.211 |
| 0 | 0.1 | ycaM | 1 | 0.75 | 0.216 |
| 0 | 0.1 | yadI | 1 | 0.75 | 0.231 |
| 0 | 0.1 | AG1 | 1 | 0.75 | 0.223 |
| 0 | 0.1 | ptsI | 1 | 1 | 0.235 |
| 0 | 0.1 | clcB | 1 | 1 | 0.215 |
| 0 | 0.1 | ycaM | 1 | 1 | 0.216 |
| 0 | 0.1 | yadI | 1 | 1 | 0.237 |
| 0 | 0.1 | AG1 | 1 | 1 | 0.227 |
| 0 | 0.1 | ptsI | 1 | 1.25 | 0.24 |
| 0 | 0.1 | clcB | 1 | 1.25 | 0.218 |
| 0 | 0.1 | ycaM | 1 | 1.25 | 0.22 |
| 0 | 0.1 | yadI | 1 | 1.25 | 0.24 |
| 0 | 0.1 | AG1 | 1 | 1.25 | 0.228 |
| 0 | 0.1 | ptsI | 1 | 1.5 | 0.246 |
| 0 | 0.1 | clcB | 1 | 1.5 | 0.218 |
| 0 | 0.1 | ycaM | 1 | 1.5 | 0.22 |
| 0 | 0.1 | yadI | 1 | 1.5 | 0.243 |
| 0 | 0.1 | AG1 | 1 | 1.5 | 0.234 |
| 0 | 0.1 | ptsI | 1 | 1.75 | 0.251 |
| 0 | 0.1 | clcB | 1 | 1.75 | 0.222 |
| 0 | 0.1 | ycaM | 1 | 1.75 | 0.224 |
| 0 | 0.1 | yadI | 1 | 1.75 | 0.25 |
| 0 | 0.1 | AG1 | 1 | 1.75 | 0.234 |
| 0 | 0.1 | ptsI | 1 | 2 | 0.26 |
| 0 | 0.1 | clcB | 1 | 2 | 0.224 |
| 0 | 0.1 | ycaM | 1 | 2 | 0.224 |
| 0 | 0.1 | yadI | 1 | 2 | 0.253 |
| 0 | 0.1 | AG1 | 1 | 2 | 0.239 |
| 0 | 0.1 | ptsI | 1 | 2.25 | 0.276 |
| 0 | 0.1 | clcB | 1 | 2.25 | 0.232 |

| | | | | | |
|---|---|---|---|---|---|
| 0 | 0.1 | ycaM | 1 | 2.25 | 0.233 |
| 0 | 0.1 | yadI | 1 | 2.25 | 0.262 |
| 0 | 0.1 | AG1 | 1 | 2.25 | 0.243 |
| 0 | 0.1 | ptsI | 1 | 2.5 | 0.288 |
| 0 | 0.1 | clcB | 1 | 2.5 | 0.233 |
| 0 | 0.1 | ycaM | 1 | 2.5 | 0.232 |
| 0 | 0.1 | yadI | 1 | 2.5 | 0.266 |
| 0 | 0.1 | AG1 | 1 | 2.5 | 0.244 |
| 0 | 0.1 | ptsI | 1 | 2.75 | 0.309 |
| 0 | 0.1 | clcB | 1 | 2.75 | 0.243 |
| 0 | 0.1 | ycaM | 1 | 2.75 | 0.243 |
| 0 | 0.1 | yadI | 1 | 2.75 | 0.278 |
| 0 | 0.1 | AG1 | 1 | 2.75 | 0.253 |
| 0 | 0.1 | ptsI | 1 | 3 | 0.324 |
| 0 | 0.1 | clcB | 1 | 3 | 0.244 |
| 0 | 0.1 | ycaM | 1 | 3 | 0.247 |
| 0 | 0.1 | yadI | 1 | 3 | 0.285 |
| 0 | 0.1 | AG1 | 1 | 3 | 0.261 |
| 0 | 0.1 | ptsI | 1 | 3.25 | 0.341 |
| 0 | 0.1 | clcB | 1 | 3.25 | 0.249 |
| 0 | 0.1 | ycaM | 1 | 3.25 | 0.253 |
| 0 | 0.1 | yadI | 1 | 3.25 | 0.296 |
| 0 | 0.1 | AG1 | 1 | 3.25 | 0.268 |
| 0 | 0.1 | ptsI | 1 | 3.5 | 0.351 |
| 0 | 0.1 | clcB | 1 | 3.5 | 0.259 |
| 0 | 0.1 | ycaM | 1 | 3.5 | 0.263 |
| 0 | 0.1 | yadI | 1 | 3.5 | 0.311 |
| 0 | 0.1 | AG1 | 1 | 3.5 | 0.277 |
| 0 | 0.1 | ptsI | 1 | 3.75 | 0.37 |
| 0 | 0.1 | clcB | 1 | 3.75 | 0.266 |
| 0 | 0.1 | ycaM | 1 | 3.75 | 0.272 |
| 0 | 0.1 | yadI | 1 | 3.75 | 0.326 |
| 0 | 0.1 | AG1 | 1 | 3.75 | 0.29 |
| 0 | 0.1 | ptsI | 1 | 4 | 0.383 |
| 0 | 0.1 | clcB | 1 | 4 | 0.275 |
| 0 | 0.1 | ycaM | 1 | 4 | 0.278 |
| 0 | 0.1 | yadI | 1 | 4 | 0.333 |
| 0 | 0.1 | AG1 | 1 | 4 | 0.301 |
| 0 | 0.1 | ptsI | 1 | 4.25 | 0.391 |
| 0 | 0.1 | clcB | 1 | 4.25 | 0.282 |
| 0 | 0.1 | ycaM | 1 | 4.25 | 0.29 |
| 0 | 0.1 | yadI | 1 | 4.25 | 0.348 |
| 0 | 0.1 | AG1 | 1 | 4.25 | 0.311 |
| 0 | 0.1 | ptsI | 1 | 4.5 | 0.393 |
| 0 | 0.1 | clcB | 1 | 4.5 | 0.295 |
| 0 | 0.1 | ycaM | 1 | 4.5 | 0.298 |
| 0 | 0.1 | yadI | 1 | 4.5 | 0.362 |
| 0 | 0.1 | AG1 | 1 | 4.5 | 0.319 |

| | | | | | |
|---|---|---|---|---|---|
| 0 | 0.1 | ptsI | 1 | 4.75 | 0.402 |
| 0 | 0.1 | clcB | 1 | 4.75 | 0.302 |
| 0 | 0.1 | ycaM | 1 | 4.75 | 0.306 |
| 0 | 0.1 | yadI | 1 | 4.75 | 0.375 |
| 0 | 0.1 | AG1 | 1 | 4.75 | 0.334 |
| 0 | 0.1 | ptsI | 1 | 5 | 0.41 |
| 0 | 0.1 | clcB | 1 | 5 | 0.306 |
| 0 | 0.1 | ycaM | 1 | 5 | 0.311 |
| 0 | 0.1 | yadI | 1 | 5 | 0.386 |
| 0 | 0.1 | AG1 | 1 | 5 | 0.342 |
| 0 | 0.1 | ptsI | 1 | 5.25 | 0.414 |
| 0 | 0.1 | clcB | 1 | 5.25 | 0.314 |
| 0 | 0.1 | ycaM | 1 | 5.25 | 0.32 |
| 0 | 0.1 | yadI | 1 | 5.25 | 0.401 |
| 0 | 0.1 | AG1 | 1 | 5.25 | 0.352 |
| 0 | 0.1 | ptsI | 1 | 5.5 | 0.42 |
| 0 | 0.1 | clcB | 1 | 5.5 | 0.319 |
| 0 | 0.1 | ycaM | 1 | 5.5 | 0.32 |
| 0 | 0.1 | yadI | 1 | 5.5 | 0.41 |
| 0 | 0.1 | AG1 | 1 | 5.5 | 0.36 |
| 0 | 0.1 | ptsI | 1 | 5.75 | 0.425 |
| 0 | 0.1 | clcB | 1 | 5.75 | 0.324 |
| 0 | 0.1 | ycaM | 1 | 5.75 | 0.328 |
| 0 | 0.1 | yadI | 1 | 5.75 | 0.418 |
| 0 | 0.1 | AG1 | 1 | 5.75 | 0.37 |
| 0 | 0.1 | ptsI | 1 | 6 | 0.434 |
| 0 | 0.1 | clcB | 1 | 6 | 0.34 |
| 0 | 0.1 | ycaM | 1 | 6 | 0.347 |
| 0 | 0.1 | yadI | 1 | 6 | 0.434 |
| 0 | 0.1 | AG1 | 1 | 6 | 0.387 |
| 0 | 0.1 | ptsI | 1 | 6.25 | 0.444 |
| 0 | 0.1 | clcB | 1 | 6.25 | 0.344 |
| 0 | 0.1 | ycaM | 1 | 6.25 | 0.356 |
| 0 | 0.1 | yadI | 1 | 6.25 | 0.444 |
| 0 | 0.1 | AG1 | 1 | 6.25 | 0.398 |
| 0 | 0.1 | ptsI | 1 | 6.5 | 0.451 |
| 0 | 0.1 | clcB | 1 | 6.5 | 0.354 |
| 0 | 0.1 | ycaM | 1 | 6.5 | 0.369 |
| 0 | 0.1 | yadI | 1 | 6.5 | 0.467 |
| 0 | 0.1 | AG1 | 1 | 6.5 | 0.416 |
| 0 | 0.1 | ptsI | 1 | 6.75 | 0.462 |
| 0 | 0.1 | clcB | 1 | 6.75 | 0.363 |
| 0 | 0.1 | ycaM | 1 | 6.75 | 0.378 |
| 0 | 0.1 | yadI | 1 | 6.75 | 0.483 |
| 0 | 0.1 | AG1 | 1 | 6.75 | 0.423 |
| 0 | 0.1 | ptsI | 1 | 7 | 0.466 |
| 0 | 0.1 | clcB | 1 | 7 | 0.371 |
| 0 | 0.1 | ycaM | 1 | 7 | 0.381 |

| | | | | | |
|---|---|---|---|---|---|
| 0 | 0.1 | yadI | 1 | 7 | 0.493 |
| 0 | 0.1 | AG1 | 1 | 7 | 0.432 |
| 0 | 0.1 | ptsI | 1 | 7.25 | 0.472 |
| 0 | 0.1 | clcB | 1 | 7.25 | 0.376 |
| 0 | 0.1 | ycaM | 1 | 7.25 | 0.386 |
| 0 | 0.1 | yadI | 1 | 7.25 | 0.504 |
| 0 | 0.1 | AG1 | 1 | 7.25 | 0.439 |
| 0 | 0.1 | ptsI | 1 | 7.5 | 0.472 |
| 0 | 0.1 | clcB | 1 | 7.5 | 0.386 |
| 0 | 0.1 | ycaM | 1 | 7.5 | 0.395 |
| 0 | 0.1 | yadI | 1 | 7.5 | 0.514 |
| 0 | 0.1 | AG1 | 1 | 7.5 | 0.449 |
| 0 | 0.1 | ptsI | 1 | 7.75 | 0.482 |
| 0 | 0.1 | clcB | 1 | 7.75 | 0.394 |
| 0 | 0.1 | ycaM | 1 | 7.75 | 0.408 |
| 0 | 0.1 | yadI | 1 | 7.75 | 0.532 |
| 0 | 0.1 | AG1 | 1 | 7.75 | 0.461 |
| 0 | 0.1 | ptsI | 1 | 8 | 0.486 |
| 0 | 0.1 | clcB | 1 | 8 | 0.398 |
| 0 | 0.1 | ycaM | 1 | 8 | 0.41 |
| 0 | 0.1 | yadI | 1 | 8 | 0.54 |
| 0 | 0.1 | AG1 | 1 | 8 | 0.47 |
| 0 | 0.1 | ptsI | 1 | 8.25 | 0.485 |
| 0 | 0.1 | clcB | 1 | 8.25 | 0.409 |
| 0 | 0.1 | ycaM | 1 | 8.25 | 0.416 |
| 0 | 0.1 | yadI | 1 | 8.25 | 0.55 |
| 0 | 0.1 | AG1 | 1 | 8.25 | 0.483 |
| 0 | 0.1 | ptsI | 1 | 8.5 | 0.504 |
| 0 | 0.1 | clcB | 1 | 8.5 | 0.419 |
| 0 | 0.1 | ycaM | 1 | 8.5 | 0.422 |
| 0 | 0.1 | yadI | 1 | 8.5 | 0.563 |
| 0 | 0.1 | AG1 | 1 | 8.5 | 0.495 |
| 0 | 0.1 | ptsI | 1 | 8.75 | 0.51 |
| 0 | 0.1 | clcB | 1 | 8.75 | 0.422 |
| 0 | 0.1 | ycaM | 1 | 8.75 | 0.425 |
| 0 | 0.1 | yadI | 1 | 8.75 | 0.572 |
| 0 | 0.1 | AG1 | 1 | 8.75 | 0.503 |
| 0 | 0.1 | ptsI | 1 | 9 | 0.534 |
| 0 | 0.1 | clcB | 1 | 9 | 0.432 |
| 0 | 0.1 | ycaM | 1 | 9 | 0.433 |
| 0 | 0.1 | yadI | 1 | 9 | 0.586 |
| 0 | 0.1 | AG1 | 1 | 9 | 0.519 |
| 0 | 0.1 | ptsI | 1 | 9.25 | 0.541 |
| 0 | 0.1 | clcB | 1 | 9.25 | 0.445 |
| 0 | 0.1 | ycaM | 1 | 9.25 | 0.447 |
| 0 | 0.1 | yadI | 1 | 9.25 | 0.589 |
| 0 | 0.1 | AG1 | 1 | 9.25 | 0.54 |
| 0 | 0.1 | ptsI | 1 | 9.5 | 0.586 |

| | | | | | |
|---|---|---|---|---|---|
| 0 | 0.1 | clcB | 1 | 9.5 | 0.463 |
| 0 | 0.1 | ycaM | 1 | 9.5 | 0.462 |
| 0 | 0.1 | yadI | 1 | 9.5 | 0.634 |
| 0 | 0.1 | AG1 | 1 | 9.5 | 0.574 |
| 0 | 0.1 | ptsI | 1 | 9.75 | 0.571 |
| 0 | 0.1 | clcB | 1 | 9.75 | 0.477 |
| 0 | 0.1 | ycaM | 1 | 9.75 | 0.474 |
| 0 | 0.1 | yadI | 1 | 9.75 | 0.647 |
| 0 | 0.1 | AG1 | 1 | 9.75 | 0.577 |
| 0 | 0.1 | ptsI | 1 | 10 | 0.569 |
| 0 | 0.1 | clcB | 1 | 10 | 0.487 |
| 0 | 0.1 | ycaM | 1 | 10 | 0.484 |
| 0 | 0.1 | yadI | 1 | 10 | 0.636 |
| 0 | 0.1 | AG1 | 1 | 10 | 0.575 |
| 0 | 0.1 | ptsI | 1 | 10.25 | 0.559 |
| 0 | 0.1 | clcB | 1 | 10.25 | 0.495 |
| 0 | 0.1 | ycaM | 1 | 10.25 | 0.493 |
| 0 | 0.1 | yadI | 1 | 10.25 | 0.649 |
| 0 | 0.1 | AG1 | 1 | 10.25 | 0.582 |
| 0 | 0.1 | ptsI | 1 | 10.5 | 0.575 |
| 0 | 0.1 | clcB | 1 | 10.5 | 0.517 |
| 0 | 0.1 | ycaM | 1 | 10.5 | 0.503 |
| 0 | 0.1 | yadI | 1 | 10.5 | 0.657 |
| 0 | 0.1 | AG1 | 1 | 10.5 | 0.595 |
| 0 | 0.1 | ptsI | 1 | 10.75 | 0.614 |
| 0 | 0.1 | clcB | 1 | 10.75 | 0.519 |
| 0 | 0.1 | ycaM | 1 | 10.75 | 0.513 |
| 0 | 0.1 | yadI | 1 | 10.75 | 0.669 |
| 0 | 0.1 | AG1 | 1 | 10.75 | 0.598 |
| 0 | 0.1 | ptsI | 1 | 11 | 0.659 |
| 0 | 0.1 | clcB | 1 | 11 | 0.548 |
| 0 | 0.1 | ycaM | 1 | 11 | 0.531 |
| 0 | 0.1 | yadI | 1 | 11 | 0.72 |
| 0 | 0.1 | AG1 | 1 | 11 | 0.643 |
| 0 | 0.1 | ptsI | 1 | 11.25 | 0.707 |
| 0 | 0.1 | clcB | 1 | 11.25 | 0.553 |
| 0 | 0.1 | ycaM | 1 | 11.25 | 0.525 |
| 0 | 0.1 | yadI | 1 | 11.25 | 0.725 |
| 0 | 0.1 | AG1 | 1 | 11.25 | 0.642 |
| 0 | 0.1 | ptsI | 1 | 11.5 | 0.693 |
| 0 | 0.1 | clcB | 1 | 11.5 | 0.55 |
| 0 | 0.1 | ycaM | 1 | 11.5 | 0.544 |
| 0 | 0.1 | yadI | 1 | 11.5 | 0.772 |
| 0 | 0.1 | AG1 | 1 | 11.5 | 0.674 |
| 0 | 0.1 | ptsI | 1 | 11.75 | 0.711 |
| 0 | 0.1 | clcB | 1 | 11.75 | 0.563 |
| 0 | 0.1 | ycaM | 1 | 11.75 | 0.552 |
| 0 | 0.1 | yadI | 1 | 11.75 | 0.828 |

| | | | | | |
|---|---|---|---|---|---|
| 0 | 0.1 | AG1 | 1 | 11.75 | 0.723 |
| 0 | 0.1 | ptsI | 1 | 12 | 0.728 |
| 0 | 0.1 | clcB | 1 | 12 | 0.552 |
| 0 | 0.1 | ycaM | 1 | 12 | 0.553 |
| 0 | 0.1 | yadI | 1 | 12 | 0.834 |
| 0 | 0.1 | AG1 | 1 | 12 | 0.722 |
| 0 | 0.1 | ptsI | 1 | 12.25 | 0.711 |
| 0 | 0.1 | clcB | 1 | 12.25 | 0.58 |
| 0 | 0.1 | ycaM | 1 | 12.25 | 0.564 |
| 0 | 0.1 | yadI | 1 | 12.25 | 0.888 |
| 0 | 0.1 | AG1 | 1 | 12.25 | 0.741 |
| 0 | 0.1 | ptsI | 1 | 12.5 | 0.744 |
| 0 | 0.1 | clcB | 1 | 12.5 | 0.595 |
| 0 | 0.1 | ycaM | 1 | 12.5 | 0.562 |
| 0 | 0.1 | yadI | 1 | 12.5 | 0.906 |
| 0 | 0.1 | AG1 | 1 | 12.5 | 0.748 |
| 0 | 0.1 | ptsI | 1 | 12.75 | 0.759 |
| 0 | 0.1 | clcB | 1 | 12.75 | 0.609 |
| 0 | 0.1 | ycaM | 1 | 12.75 | 0.564 |
| 0 | 0.1 | yadI | 1 | 12.75 | 0.912 |
| 0 | 0.1 | AG1 | 1 | 12.75 | 0.766 |
| 0 | 0.1 | ptsI | 1 | 13 | 0.752 |
| 0 | 0.1 | clcB | 1 | 13 | 0.624 |
| 0 | 0.1 | ycaM | 1 | 13 | 0.573 |
| 0 | 0.1 | yadI | 1 | 13 | 0.917 |
| 0 | 0.1 | AG1 | 1 | 13 | 0.788 |
| 0 | 0.1 | ptsI | 1 | 13.25 | 0.75 |
| 0 | 0.1 | clcB | 1 | 13.25 | 0.639 |
| 0 | 0.1 | ycaM | 1 | 13.25 | 0.594 |
| 0 | 0.1 | yadI | 1 | 13.25 | 0.946 |
| 0 | 0.1 | AG1 | 1 | 13.25 | 0.797 |
| 0 | 0.1 | ptsI | 1 | 13.5 | 0.796 |
| 0 | 0.1 | clcB | 1 | 13.5 | 0.675 |
| 0 | 0.1 | ycaM | 1 | 13.5 | 0.622 |
| 0 | 0.1 | yadI | 1 | 13.5 | 0.952 |
| 0 | 0.1 | AG1 | 1 | 13.5 | 0.811 |
| 0 | 0.1 | ptsI | 1 | 13.75 | 0.783 |
| 0 | 0.1 | clcB | 1 | 13.75 | 0.677 |
| 0 | 0.1 | ycaM | 1 | 13.75 | 0.631 |
| 0 | 0.1 | yadI | 1 | 13.75 | 0.943 |
| 0 | 0.1 | AG1 | 1 | 13.75 | 0.822 |
| 0 | 0.1 | ptsI | 1 | 14 | 0.789 |
| 0 | 0.1 | clcB | 1 | 14 | 0.688 |
| 0 | 0.1 | ycaM | 1 | 14 | 0.666 |
| 0 | 0.1 | yadI | 1 | 14 | 0.947 |
| 0 | 0.1 | AG1 | 1 | 14 | 0.847 |
| 0 | 0.1 | ptsI | 1 | 14.25 | 0.805 |
| 0 | 0.1 | clcB | 1 | 14.25 | 0.656 |

| | | | | | |
|---|---|---|---|---|---|
| 0 | 0.1 | ycaM | 1 | 14.25 | 0.653 |
| 0 | 0.1 | yadI | 1 | 14.25 | 0.952 |
| 0 | 0.1 | AG1 | 1 | 14.25 | 0.842 |
| 0 | 0.1 | ptsI | 1 | 14.5 | 0.796 |
| 0 | 0.1 | clcB | 1 | 14.5 | 0.678 |
| 0 | 0.1 | ycaM | 1 | 14.5 | 0.693 |
| 0 | 0.1 | yadI | 1 | 14.5 | 0.956 |
| 0 | 0.1 | AG1 | 1 | 14.5 | 0.844 |
| 0 | 0.1 | ptsI | 1 | 14.75 | 0.773 |
| 0 | 0.1 | clcB | 1 | 14.75 | 0.704 |
| 0 | 0.1 | ycaM | 1 | 14.75 | 0.72 |
| 0 | 0.1 | yadI | 1 | 14.75 | 0.973 |
| 0 | 0.1 | AG1 | 1 | 14.75 | 0.847 |
| 0 | 0.1 | ptsI | 1 | 15 | 0.806 |
| 0 | 0.1 | clcB | 1 | 15 | 0.714 |
| 0 | 0.1 | ycaM | 1 | 15 | 0.718 |
| 0 | 0.1 | yadI | 1 | 15 | 0.964 |
| 0 | 0.1 | AG1 | 1 | 15 | 0.851 |
| 0 | 0.1 | ptsI | 1 | 15.25 | 0.821 |
| 0 | 0.1 | clcB | 1 | 15.25 | 0.736 |
| 0 | 0.1 | ycaM | 1 | 15.25 | 0.736 |
| 0 | 0.1 | yadI | 1 | 15.25 | 0.958 |
| 0 | 0.1 | AG1 | 1 | 15.25 | 0.877 |
| 0 | 0.1 | ptsI | 1 | 15.5 | 0.806 |
| 0 | 0.1 | clcB | 1 | 15.5 | 0.748 |
| 0 | 0.1 | ycaM | 1 | 15.5 | 0.69 |
| 0 | 0.1 | yadI | 1 | 15.5 | 0.954 |
| 0 | 0.1 | AG1 | 1 | 15.5 | 0.855 |
| 0 | 0.1 | ptsI | 1 | 15.75 | 0.817 |
| 0 | 0.1 | clcB | 1 | 15.75 | 0.722 |
| 0 | 0.1 | ycaM | 1 | 15.75 | 0.705 |
| 0 | 0.1 | yadI | 1 | 15.75 | 0.947 |
| 0 | 0.1 | AG1 | 1 | 15.75 | 0.85 |
| 0 | 0.1 | ptsI | 1 | 16 | 0.8 |
| 0 | 0.1 | clcB | 1 | 16 | 0.708 |
| 0 | 0.1 | ycaM | 1 | 16 | 0.725 |
| 0 | 0.1 | yadI | 1 | 16 | 0.948 |
| 0 | 0.1 | AG1 | 1 | 16 | 0.876 |
| 0 | 0.1 | ptsI | 1 | 16.25 | 0.83 |
| 0 | 0.1 | clcB | 1 | 16.25 | 0.733 |
| 0 | 0.1 | ycaM | 1 | 16.25 | 0.738 |
| 0 | 0.1 | yadI | 1 | 16.25 | 0.943 |
| 0 | 0.1 | AG1 | 1 | 16.25 | 0.896 |
| 0 | 0.1 | ptsI | 1 | 16.5 | 0.827 |
| 0 | 0.1 | clcB | 1 | 16.5 | 0.736 |
| 0 | 0.1 | ycaM | 1 | 16.5 | 0.739 |
| 0 | 0.1 | yadI | 1 | 16.5 | 0.94 |
| 0 | 0.1 | AG1 | 1 | 16.5 | 0.866 |

| | | | | | |
|---|---|---|---|---|---|
| 0 | 0.1 | ptsI | 1 | 17.0667 | 0.808 |
| 0 | 0.1 | clcB | 1 | 17.0667 | 0.569 |
| 0 | 0.1 | ycaM | 1 | 17.0667 | 0.592 |
| 0 | 0.1 | yadI | 1 | 17.0667 | 0.669 |
| 0 | 0.1 | AG1 | 1 | 17.0667 | 0.706 |
| 0 | 0.1 | ptsI | 1 | 17.3167 | 0.784 |
| 0 | 0.1 | clcB | 1 | 17.3167 | 0.522 |
| 0 | 0.1 | ycaM | 1 | 17.3167 | 0.542 |
| 0 | 0.1 | yadI | 1 | 17.3167 | 0.651 |
| 0 | 0.1 | AG1 | 1 | 17.3167 | 0.714 |
| 0 | 0.1 | ptsI | 1 | 17.5667 | 0.793 |
| 0 | 0.1 | clcB | 1 | 17.5667 | 0.512 |
| 0 | 0.1 | ycaM | 1 | 17.5667 | 0.543 |
| 0 | 0.1 | yadI | 1 | 17.5667 | 0.66 |
| 0 | 0.1 | AG1 | 1 | 17.5667 | 0.738 |
| 0 | 0.1 | ptsI | 1 | 17.8167 | 0.834 |
| 0 | 0.1 | clcB | 1 | 17.8167 | 0.515 |
| 0 | 0.1 | ycaM | 1 | 17.8167 | 0.545 |
| 0 | 0.1 | yadI | 1 | 17.8167 | 0.678 |
| 0 | 0.1 | AG1 | 1 | 17.8167 | 0.797 |
| 0 | 0.1 | ptsI | 1 | 18.0667 | 0.819 |
| 0 | 0.1 | clcB | 1 | 18.0667 | 0.517 |
| 0 | 0.1 | ycaM | 1 | 18.0667 | 0.555 |
| 0 | 0.1 | yadI | 1 | 18.0667 | 0.692 |
| 0 | 0.1 | AG1 | 1 | 18.0667 | 0.784 |
| 0 | 0.1 | ptsI | 1 | 18.3167 | 0.829 |
| 0 | 0.1 | clcB | 1 | 18.3167 | 0.526 |
| 0 | 0.1 | ycaM | 1 | 18.3167 | 0.569 |
| 0 | 0.1 | yadI | 1 | 18.3167 | 0.695 |
| 0 | 0.1 | AG1 | 1 | 18.3167 | 0.818 |
| 0 | 0.1 | ptsI | 1 | 18.5667 | 0.837 |
| 0 | 0.1 | clcB | 1 | 18.5667 | 0.534 |
| 0 | 0.1 | ycaM | 1 | 18.5667 | 0.575 |
| 0 | 0.1 | yadI | 1 | 18.5667 | 0.706 |
| 0 | 0.1 | AG1 | 1 | 18.5667 | 0.832 |
| 0 | 0.1 | ptsI | 1 | 18.8167 | 0.844 |
| 0 | 0.1 | clcB | 1 | 18.8167 | 0.538 |
| 0 | 0.1 | ycaM | 1 | 18.8167 | 0.578 |
| 0 | 0.1 | yadI | 1 | 18.8167 | 0.721 |
| 0 | 0.1 | AG1 | 1 | 18.8167 | 0.845 |
| 0 | 0.1 | ptsI | 1 | 19.0667 | 0.847 |
| 0 | 0.1 | clcB | 1 | 19.0667 | 0.542 |
| 0 | 0.1 | ycaM | 1 | 19.0667 | 0.597 |
| 0 | 0.1 | yadI | 1 | 19.0667 | 0.743 |
| 0 | 0.1 | AG1 | 1 | 19.0667 | 0.876 |
| 0 | 0.1 | ptsI | 1 | 19.3167 | 0.849 |
| 0 | 0.1 | clcB | 1 | 19.3167 | 0.55 |
| 0 | 0.1 | ycaM | 1 | 19.3167 | 0.578 |

| | | | | | |
|---|---|---|---|---|---|
| 0 | 0.1 | yadI | 1 | 19.3167 | 0.739 |
| 0 | 0.1 | AG1 | 1 | 19.3167 | 0.853 |
| 0 | 0.1 | ptsI | 1 | 19.5667 | 0.856 |
| 0 | 0.1 | clcB | 1 | 19.5667 | 0.552 |
| 0 | 0.1 | ycaM | 1 | 19.5667 | 0.576 |
| 0 | 0.1 | yadI | 1 | 19.5667 | 0.741 |
| 0 | 0.1 | AG1 | 1 | 19.5667 | 0.858 |
| 0 | 0.1 | ptsI | 1 | 19.8167 | 0.864 |
| 0 | 0.1 | clcB | 1 | 19.8167 | 0.551 |
| 0 | 0.1 | ycaM | 1 | 19.8167 | 0.576 |
| 0 | 0.1 | yadI | 1 | 19.8167 | 0.751 |
| 0 | 0.1 | AG1 | 1 | 19.8167 | 0.863 |
| 0 | 0.1 | ptsI | 1 | 20.0667 | 0.863 |
| 0 | 0.1 | clcB | 1 | 20.0667 | 0.548 |
| 0 | 0.1 | ycaM | 1 | 20.0667 | 0.571 |
| 0 | 0.1 | yadI | 1 | 20.0667 | 0.759 |
| 0 | 0.1 | AG1 | 1 | 20.0667 | 0.86 |
| 0 | 0.1 | ptsI | 1 | 20.3167 | 0.867 |
| 0 | 0.1 | clcB | 1 | 20.3167 | 0.54 |
| 0 | 0.1 | ycaM | 1 | 20.3167 | 0.574 |
| 0 | 0.1 | yadI | 1 | 20.3167 | 0.761 |
| 0 | 0.1 | AG1 | 1 | 20.3167 | 0.864 |
| 0 | 0.1 | ptsI | 1 | 20.5667 | 0.851 |
| 0 | 0.1 | clcB | 1 | 20.5667 | 0.551 |
| 0 | 0.1 | ycaM | 1 | 20.5667 | 0.574 |
| 0 | 0.1 | yadI | 1 | 20.5667 | 0.771 |
| 0 | 0.1 | AG1 | 1 | 20.5667 | 0.852 |
| 0 | 0.1 | ptsI | 1 | 20.8167 | 0.864 |
| 0 | 0.1 | clcB | 1 | 20.8167 | 0.56 |
| 0 | 0.1 | ycaM | 1 | 20.8167 | 0.567 |
| 0 | 0.1 | yadI | 1 | 20.8167 | 0.772 |
| 0 | 0.1 | AG1 | 1 | 20.8167 | 0.86 |
| 0 | 0.1 | ptsI | 1 | 21.0667 | 0.877 |
| 0 | 0.1 | clcB | 1 | 21.0667 | 0.552 |
| 0 | 0.1 | ycaM | 1 | 21.0667 | 0.565 |
| 0 | 0.1 | yadI | 1 | 21.0667 | 0.774 |
| 0 | 0.1 | AG1 | 1 | 21.0667 | 0.877 |
| 0 | 0.1 | ptsI | 1 | 21.3167 | 0.878 |
| 0 | 0.1 | clcB | 1 | 21.3167 | 0.562 |
| 0 | 0.1 | ycaM | 1 | 21.3167 | 0.563 |
| 0 | 0.1 | yadI | 1 | 21.3167 | 0.786 |
| 0 | 0.1 | AG1 | 1 | 21.3167 | 0.884 |
| 0 | 0.1 | ptsI | 1 | 21.5667 | 0.882 |
| 0 | 0.1 | clcB | 1 | 21.5667 | 0.571 |
| 0 | 0.1 | ycaM | 1 | 21.5667 | 0.566 |
| 0 | 0.1 | yadI | 1 | 21.5667 | 0.794 |
| 0 | 0.1 | AG1 | 1 | 21.5667 | 0.885 |
| 0 | 0.1 | ptsI | 1 | 21.8167 | 0.883 |

| | | | | | |
|---|---|---|---|---|---|
| 0 | 0.1 | clcB | 1 | 21.8167 | 0.562 |
| 0 | 0.1 | ycaM | 1 | 21.8167 | 0.558 |
| 0 | 0.1 | yadI | 1 | 21.8167 | 0.801 |
| 0 | 0.1 | AG1 | 1 | 21.8167 | 0.897 |
| 0 | 0.1 | ptsI | 1 | 22.0667 | 0.894 |
| 0 | 0.1 | clcB | 1 | 22.0667 | 0.558 |
| 0 | 0.1 | ycaM | 1 | 22.0667 | 0.557 |
| 0 | 0.1 | yadI | 1 | 22.0667 | 0.796 |
| 0 | 0.1 | AG1 | 1 | 22.0667 | 0.907 |
| 0 | 0.1 | ptsI | 1 | 22.9333 | 0.96 |
| 0 | 0.1 | clcB | 1 | 22.9333 | 0.527 |
| 0 | 0.1 | ycaM | 1 | 22.9333 | 0.639 |
| 0 | 0.1 | yadI | 1 | 22.9333 | 0.86 |
| 0 | 0.1 | AG1 | 1 | 22.9333 | 1.037 |
| 0 | 0.1 | ptsI | 1 | 23.1833 | 0.954 |
| 0 | 0.1 | clcB | 1 | 23.1833 | 0.525 |
| 0 | 0.1 | ycaM | 1 | 23.1833 | 0.602 |
| 0 | 0.1 | yadI | 1 | 23.1833 | 0.86 |
| 0 | 0.1 | AG1 | 1 | 23.1833 | 0.998 |
| 0 | 0.1 | ptsI | 1 | 23.4333 | 0.91 |
| 0 | 0.1 | clcB | 1 | 23.4333 | 0.528 |
| 0 | 0.1 | ycaM | 1 | 23.4333 | 0.605 |
| 0 | 0.1 | yadI | 1 | 23.4333 | 0.884 |
| 0 | 0.1 | AG1 | 1 | 23.4333 | 0.975 |
| 0 | 0.1 | ptsI | 1 | 23.6833 | 0.904 |
| 0 | 0.1 | clcB | 1 | 23.6833 | 0.525 |
| 0 | 0.1 | ycaM | 1 | 23.6833 | 0.603 |
| 0 | 0.1 | yadI | 1 | 23.6833 | 0.898 |
| 0 | 0.1 | AG1 | 1 | 23.6833 | 0.972 |
| 0 | 0.1 | ptsI | 1 | 23.9333 | 0.907 |
| 0 | 0.1 | clcB | 1 | 23.9333 | 0.527 |
| 0 | 0.1 | ycaM | 1 | 23.9333 | 0.603 |
| 0 | 0.1 | yadI | 1 | 23.9333 | 0.891 |
| 0 | 0.1 | AG1 | 1 | 23.9333 | 0.958 |
| 0 | 0.1 | ptsI | 1 | 24.1833 | 0.905 |
| 0 | 0.1 | clcB | 1 | 24.1833 | 0.526 |
| 0 | 0.1 | ycaM | 1 | 24.1833 | 0.608 |
| 0 | 0.1 | yadI | 1 | 24.1833 | 0.898 |
| 0 | 0.1 | AG1 | 1 | 24.1833 | 0.958 |
| 0 | 0.1 | ptsI | 1 | 24.4333 | 0.917 |
| 0 | 0.1 | clcB | 1 | 24.4333 | 0.524 |
| 0 | 0.1 | ycaM | 1 | 24.4333 | 0.6 |
| 0 | 0.1 | yadI | 1 | 24.4333 | 0.898 |
| 0 | 0.1 | AG1 | 1 | 24.4333 | 0.96 |
| 0 | 0.1 | ptsI | 1 | 24.6833 | 0.906 |
| 0 | 0.1 | clcB | 1 | 24.6833 | 0.527 |
| 0 | 0.1 | ycaM | 1 | 24.6833 | 0.604 |
| 0 | 0.1 | yadI | 1 | 24.6833 | 0.906 |

| | | | | | |
|---|---|---|---|---|---|
| 0 | 0.1 | AG1 | 1 | 24.6833 | 0.962 |
| 0 | 0.1 | ptsI | 1 | 24.9333 | 0.912 |
| 0 | 0.1 | clcB | 1 | 24.9333 | 0.521 |
| 0 | 0.1 | ycaM | 1 | 24.9333 | 0.608 |
| 0 | 0.1 | yadI | 1 | 24.9333 | 0.92 |
| 0 | 0.1 | AG1 | 1 | 24.9333 | 0.969 |
| 0 | 0.1 | ptsI | 1 | 25.1833 | 0.91 |
| 0 | 0.1 | clcB | 1 | 25.1833 | 0.516 |
| 0 | 0.1 | ycaM | 1 | 25.1833 | 0.606 |
| 0 | 0.1 | yadI | 1 | 25.1833 | 0.91 |
| 0 | 0.1 | AG1 | 1 | 25.1833 | 0.961 |
| 0 | 0.1 | ptsI | 1 | 25.4333 | 0.91 |
| 0 | 0.1 | clcB | 1 | 25.4333 | 0.523 |
| 0 | 0.1 | ycaM | 1 | 25.4333 | 0.609 |
| 0 | 0.1 | yadI | 1 | 25.4333 | 0.918 |
| 0 | 0.1 | AG1 | 1 | 25.4333 | 0.962 |
| 0 | 0.1 | ptsI | 1 | 25.6833 | 0.907 |
| 0 | 0.1 | clcB | 1 | 25.6833 | 0.515 |
| 0 | 0.1 | ycaM | 1 | 25.6833 | 0.608 |
| 0 | 0.1 | yadI | 1 | 25.6833 | 0.917 |
| 0 | 0.1 | AG1 | 1 | 25.6833 | 0.965 |
| 0 | 0.1 | ptsI | 1 | 25.9333 | 0.908 |
| 0 | 0.1 | clcB | 1 | 25.9333 | 0.514 |
| 0 | 0.1 | ycaM | 1 | 25.9333 | 0.605 |
| 0 | 0.1 | yadI | 1 | 25.9333 | 0.909 |
| 0 | 0.1 | AG1 | 1 | 25.9333 | 0.966 |
| 0 | 0.1 | ptsI | 1 | 26.1833 | 0.915 |
| 0 | 0.1 | clcB | 1 | 26.1833 | 0.513 |
| 0 | 0.1 | ycaM | 1 | 26.1833 | 0.605 |
| 0 | 0.1 | yadI | 1 | 26.1833 | 0.922 |
| 0 | 0.1 | AG1 | 1 | 26.1833 | 0.967 |
| 0 | 0.1 | ptsI | 1 | 26.4333 | 0.908 |
| 0 | 0.1 | clcB | 1 | 26.4333 | 0.518 |
| 0 | 0.1 | ycaM | 1 | 26.4333 | 0.609 |
| 0 | 0.1 | yadI | 1 | 26.4333 | 0.919 |
| 0 | 0.1 | AG1 | 1 | 26.4333 | 0.966 |
| 0 | 0.1 | ptsI | 1 | 26.6833 | 0.904 |
| 0 | 0.1 | clcB | 1 | 26.6833 | 0.516 |
| 0 | 0.1 | ycaM | 1 | 26.6833 | 0.608 |
| 0 | 0.1 | yadI | 1 | 26.6833 | 0.93 |
| 0 | 0.1 | AG1 | 1 | 26.6833 | 0.969 |
| 0 | 0.1 | ptsI | 1 | 26.9333 | 0.909 |
| 0 | 0.1 | clcB | 1 | 26.9333 | 0.511 |
| 0 | 0.1 | ycaM | 1 | 26.9333 | 0.604 |
| 0 | 0.1 | yadI | 1 | 26.9333 | 0.922 |
| 0 | 0.1 | AG1 | 1 | 26.9333 | 0.969 |
| 0 | 0.1 | ptsI | 1 | 27.1833 | 0.914 |
| 0 | 0.1 | clcB | 1 | 27.1833 | 0.513 |

| | | | | | |
|---|---|---|---|---|---|
| 0 | 0.1 | ycaM | 1 | 27.1833 | 0.605 |
| 0 | 0.1 | yadI | 1 | 27.1833 | 0.926 |
| 0 | 0.1 | AG1 | 1 | 27.1833 | 0.973 |
| 0 | 0.1 | ptsI | 1 | 27.4333 | 0.909 |
| 0 | 0.1 | clcB | 1 | 27.4333 | 0.511 |
| 0 | 0.1 | ycaM | 1 | 27.4333 | 0.606 |
| 0 | 0.1 | yadI | 1 | 27.4333 | 0.936 |
| 0 | 0.1 | AG1 | 1 | 27.4333 | 0.968 |
| 0 | 0.1 | ptsI | 1 | 27.6833 | 0.91 |
| 0 | 0.1 | clcB | 1 | 27.6833 | 0.518 |
| 0 | 0.1 | ycaM | 1 | 27.6833 | 0.604 |
| 0 | 0.1 | yadI | 1 | 27.6833 | 0.94 |
| 0 | 0.1 | AG1 | 1 | 27.6833 | 0.972 |
| 0 | 0.1 | ptsI | 1 | 27.9333 | 0.907 |
| 0 | 0.1 | clcB | 1 | 27.9333 | 0.508 |
| 0 | 0.1 | ycaM | 1 | 27.9333 | 0.595 |
| 0 | 0.1 | yadI | 1 | 27.9333 | 0.931 |
| 0 | 0.1 | AG1 | 1 | 27.9333 | 0.968 |
| 0 | 0.1 | ptsI | 1 | 28.1833 | 0.91 |
| 0 | 0.1 | clcB | 1 | 28.1833 | 0.516 |
| 0 | 0.1 | ycaM | 1 | 28.1833 | 0.602 |
| 0 | 0.1 | yadI | 1 | 28.1833 | 0.942 |
| 0 | 0.1 | AG1 | 1 | 28.1833 | 0.979 |
| 0 | 0.1 | ptsI | 1 | 28.4333 | 0.916 |
| 0 | 0.1 | clcB | 1 | 28.4333 | 0.515 |
| 0 | 0.1 | ycaM | 1 | 28.4333 | 0.6 |
| 0 | 0.1 | yadI | 1 | 28.4333 | 0.94 |
| 0 | 0.1 | AG1 | 1 | 28.4333 | 0.977 |
| 0 | 0.1 | ptsI | 1 | 28.6833 | 0.917 |
| 0 | 0.1 | clcB | 1 | 28.6833 | 0.515 |
| 0 | 0.1 | ycaM | 1 | 28.6833 | 0.6 |
| 0 | 0.1 | yadI | 1 | 28.6833 | 0.943 |
| 0 | 0.1 | AG1 | 1 | 28.6833 | 0.977 |
| 0 | 0.1 | ptsI | 1 | 28.9333 | 0.921 |
| 0 | 0.1 | clcB | 1 | 28.9333 | 0.52 |
| 0 | 0.1 | ycaM | 1 | 28.9333 | 0.603 |
| 0 | 0.1 | yadI | 1 | 28.9333 | 0.954 |
| 0 | 0.1 | AG1 | 1 | 28.9333 | 0.983 |
| 0 | 0.1 | ptsI | 1 | 29.1833 | 0.917 |
| 0 | 0.1 | clcB | 1 | 29.1833 | 0.516 |
| 0 | 0.1 | ycaM | 1 | 29.1833 | 0.6 |
| 0 | 0.1 | yadI | 1 | 29.1833 | 0.956 |
| 0 | 0.1 | AG1 | 1 | 29.1833 | 0.98 |
| 0 | 0.1 | ptsI | 1 | 29.4333 | 0.915 |
| 0 | 0.1 | clcB | 1 | 29.4333 | 0.518 |
| 0 | 0.1 | ycaM | 1 | 29.4333 | 0.597 |
| 0 | 0.1 | yadI | 1 | 29.4333 | 0.956 |
| 0 | 0.1 | AG1 | 1 | 29.4333 | 0.981 |

| | | | | | |
|---|---|---|---|---|---|
| 0 | 0.1 | ptsI | 1 | 29.6833 | 0.924 |
| 0 | 0.1 | clcB | 1 | 29.6833 | 0.517 |
| 0 | 0.1 | ycaM | 1 | 29.6833 | 0.602 |
| 0 | 0.1 | yadI | 1 | 29.6833 | 0.958 |
| 0 | 0.1 | AG1 | 1 | 29.6833 | 0.98 |
| 0 | 0.1 | ptsI | 1 | 29.9333 | 0.92 |
| 0 | 0.1 | clcB | 1 | 29.9333 | 0.518 |
| 0 | 0.1 | ycaM | 1 | 29.9333 | 0.6 |
| 0 | 0.1 | yadI | 1 | 29.9333 | 0.966 |
| 0 | 0.1 | AG1 | 1 | 29.9333 | 0.98 |
| 0 | 0.1 | ptsI | 1 | 30.1833 | 0.926 |
| 0 | 0.1 | clcB | 1 | 30.1833 | 0.517 |
| 0 | 0.1 | ycaM | 1 | 30.1833 | 0.602 |
| 0 | 0.1 | yadI | 1 | 30.1833 | 0.966 |
| 0 | 0.1 | AG1 | 1 | 30.1833 | 0.983 |
| 0 | 0.1 | ptsI | 1 | 30.4333 | 0.934 |
| 0 | 0.1 | clcB | 1 | 30.4333 | 0.521 |
| 0 | 0.1 | ycaM | 1 | 30.4333 | 0.6 |
| 0 | 0.1 | yadI | 1 | 30.4333 | 0.972 |
| 0 | 0.1 | AG1 | 1 | 30.4333 | 0.984 |
| 0 | 0.1 | ptsI | 1 | 30.6833 | 0.93 |
| 0 | 0.1 | clcB | 1 | 30.6833 | 0.517 |
| 0 | 0.1 | ycaM | 1 | 30.6833 | 0.602 |
| 0 | 0.1 | yadI | 1 | 30.6833 | 0.978 |
| 0 | 0.1 | AG1 | 1 | 30.6833 | 0.989 |
| 0 | 0.1 | ptsI | 1 | 30.9333 | 0.932 |
| 0 | 0.1 | clcB | 1 | 30.9333 | 0.522 |
| 0 | 0.1 | ycaM | 1 | 30.9333 | 0.602 |
| 0 | 0.1 | yadI | 1 | 30.9333 | 0.985 |
| 0 | 0.1 | AG1 | 1 | 30.9333 | 0.986 |
| 0 | 0.1 | ptsI | 1 | 31.1833 | 0.935 |
| 0 | 0.1 | clcB | 1 | 31.1833 | 0.514 |
| 0 | 0.1 | ycaM | 1 | 31.1833 | 0.602 |
| 0 | 0.1 | yadI | 1 | 31.1833 | 0.986 |
| 0 | 0.1 | AG1 | 1 | 31.1833 | 0.985 |
| 0 | 0.1 | ptsI | 1 | 31.4333 | 0.94 |
| 0 | 0.1 | clcB | 1 | 31.4333 | 0.521 |
| 0 | 0.1 | ycaM | 1 | 31.4333 | 0.602 |
| 0 | 0.1 | yadI | 1 | 31.4333 | 0.994 |
| 0 | 0.1 | AG1 | 1 | 31.4333 | 0.988 |
| 0 | 0.1 | ptsI | 1 | 31.6833 | 0.94 |
| 0 | 0.1 | clcB | 1 | 31.6833 | 0.518 |
| 0 | 0.1 | ycaM | 1 | 31.6833 | 0.602 |
| 0 | 0.1 | yadI | 1 | 31.6833 | 1 |
| 0 | 0.1 | AG1 | 1 | 31.6833 | 0.99 |
| 0 | 0.1 | ptsI | 1 | 31.9333 | 0.942 |
| 0 | 0.1 | clcB | 1 | 31.9333 | 0.521 |
| 0 | 0.1 | ycaM | 1 | 31.9333 | 0.603 |

| | | | | | |
|---|---|---|---|---|---|
| 0 | 0.1 | yadI | 1 | 31.9333 | 0.994 |
| 0 | 0.1 | AG1 | 1 | 31.9333 | 0.99 |
| 0 | 0.1 | ptsI | 1 | 32.1833 | 0.941 |
| 0 | 0.1 | clcB | 1 | 32.1833 | 0.52 |
| 0 | 0.1 | ycaM | 1 | 32.1833 | 0.605 |
| 0 | 0.1 | yadI | 1 | 32.1833 | 1.003 |
| 0 | 0.1 | AG1 | 1 | 32.1833 | 0.993 |
| 0 | 0.1 | ptsI | 1 | 32.4333 | 0.929 |
| 0 | 0.1 | clcB | 1 | 32.4333 | 0.503 |
| 0 | 0.1 | ycaM | 1 | 32.4333 | 0.596 |
| 0 | 0.1 | yadI | 1 | 32.4333 | 0.991 |
| 0 | 0.1 | AG1 | 1 | 32.4333 | 0.994 |
| 0 | 0.1 | ptsI | 1 | 32.6833 | 0.938 |
| 0 | 0.1 | clcB | 1 | 32.6833 | 0.522 |
| 0 | 0.1 | ycaM | 1 | 32.6833 | 0.604 |
| 0 | 0.1 | yadI | 1 | 32.6833 | 1.013 |
| 0 | 0.1 | AG1 | 1 | 32.6833 | 0.994 |
| 0 | 0.1 | ptsI | 1 | 32.9333 | 0.942 |
| 0 | 0.1 | clcB | 1 | 32.9333 | 0.516 |
| 0 | 0.1 | ycaM | 1 | 32.9333 | 0.604 |
| 0 | 0.1 | yadI | 1 | 32.9333 | 1.01 |
| 0 | 0.1 | AG1 | 1 | 32.9333 | 0.994 |
| 0 | 0.1 | ptsI | 1 | 33.1833 | 0.932 |
| 0 | 0.1 | clcB | 1 | 33.1833 | 0.514 |
| 0 | 0.1 | ycaM | 1 | 33.1833 | 0.601 |
| 0 | 0.1 | yadI | 1 | 33.1833 | 1.013 |
| 0 | 0.1 | AG1 | 1 | 33.1833 | 0.995 |
| 0 | 0.1 | ptsI | 1 | 33.4333 | 0.942 |
| 0 | 0.1 | clcB | 1 | 33.4333 | 0.525 |
| 0 | 0.1 | ycaM | 1 | 33.4333 | 0.608 |
| 0 | 0.1 | yadI | 1 | 33.4333 | 1.022 |
| 0 | 0.1 | AG1 | 1 | 33.4333 | 0.999 |
| 0 | 0.1 | ptsI | 1 | 33.6833 | 0.941 |
| 0 | 0.1 | clcB | 1 | 33.6833 | 0.515 |
| 0 | 0.1 | ycaM | 1 | 33.6833 | 0.607 |
| 0 | 0.1 | yadI | 1 | 33.6833 | 1.013 |
| 0 | 0.1 | AG1 | 1 | 33.6833 | 0.995 |
| 0 | 0.1 | ptsI | 1 | 33.9333 | 0.942 |
| 0 | 0.1 | clcB | 1 | 33.9333 | 0.523 |
| 0 | 0.1 | ycaM | 1 | 33.9333 | 0.607 |
| 0 | 0.1 | yadI | 1 | 33.9333 | 1.023 |
| 0 | 0.1 | AG1 | 1 | 33.9333 | 1.001 |
| 0 | 0.1 | ptsI | 1 | 34.1833 | 0.946 |
| 0 | 0.1 | clcB | 1 | 34.1833 | 0.536 |
| 0 | 0.1 | ycaM | 1 | 34.1833 | 0.61 |
| 0 | 0.1 | yadI | 1 | 34.1833 | 1.033 |
| 0 | 0.1 | AG1 | 1 | 34.1833 | 0.992 |
| 0 | 0.1 | ptsI | 1 | 34.4333 | 0.939 |

| | | | | | |
|---|---|---|---|---|---|
| 0 | 0.1 | clcB | 1 | 34.4333 | 0.527 |
| 0 | 0.1 | ycaM | 1 | 34.4333 | 0.611 |
| 0 | 0.1 | yadI | 1 | 34.4333 | 1.027 |
| 0 | 0.1 | AG1 | 1 | 34.4333 | 1.004 |
| 0 | 0.1 | ptsI | 1 | 34.6833 | 0.944 |
| 0 | 0.1 | clcB | 1 | 34.6833 | 0.528 |
| 0 | 0.1 | ycaM | 1 | 34.6833 | 0.615 |
| 0 | 0.1 | yadI | 1 | 34.6833 | 1.037 |
| 0 | 0.1 | AG1 | 1 | 34.6833 | 1.01 |
| 0 | 0.1 | ptsI | 1 | 34.9333 | 0.939 |
| 0 | 0.1 | clcB | 1 | 34.9333 | 0.528 |
| 0 | 0.1 | ycaM | 1 | 34.9333 | 0.611 |
| 0 | 0.1 | yadI | 1 | 34.9333 | 1.029 |
| 0 | 0.1 | AG1 | 1 | 34.9333 | 1.011 |
| 0 | 0.1 | ptsI | 1 | 35.1833 | 0.939 |
| 0 | 0.1 | clcB | 1 | 35.1833 | 0.524 |
| 0 | 0.1 | ycaM | 1 | 35.1833 | 0.612 |
| 0 | 0.1 | yadI | 1 | 35.1833 | 1.033 |
| 0 | 0.1 | AG1 | 1 | 35.1833 | 1.012 |
| 0 | 0.1 | ptsI | 1 | 35.4333 | 0.939 |
| 0 | 0.1 | clcB | 1 | 35.4333 | 0.528 |
| 0 | 0.1 | ycaM | 1 | 35.4333 | 0.613 |
| 0 | 0.1 | yadI | 1 | 35.4333 | 1.035 |
| 0 | 0.1 | AG1 | 1 | 35.4333 | 1.016 |
| 0 | 0.1 | ptsI | 1 | 35.6833 | 0.942 |
| 0 | 0.1 | clcB | 1 | 35.6833 | 0.529 |
| 0 | 0.1 | ycaM | 1 | 35.6833 | 0.617 |
| 0 | 0.1 | yadI | 1 | 35.6833 | 1.036 |
| 0 | 0.1 | AG1 | 1 | 35.6833 | 1.022 |
| 0 | 0.1 | ptsI | 1 | 35.9333 | 0.932 |
| 0 | 0.1 | clcB | 1 | 35.9333 | 0.522 |
| 0 | 0.1 | ycaM | 1 | 35.9333 | 0.611 |
| 0 | 0.1 | yadI | 1 | 35.9333 | 1.026 |
| 0 | 0.1 | AG1 | 1 | 35.9333 | 1.02 |
| 0 | 0.1 | ptsI | 1 | 36.1833 | 0.943 |
| 0 | 0.1 | clcB | 1 | 36.1833 | 0.536 |
| 0 | 0.1 | ycaM | 1 | 36.1833 | 0.622 |
| 0 | 0.1 | yadI | 1 | 36.1833 | 1.04 |
| 0 | 0.1 | AG1 | 1 | 36.1833 | 1.023 |
| 0 | 0.1 | ptsI | 1 | 36.4333 | 0.938 |
| 0 | 0.1 | clcB | 1 | 36.4333 | 0.525 |
| 0 | 0.1 | ycaM | 1 | 36.4333 | 0.617 |
| 0 | 0.1 | yadI | 1 | 36.4333 | 1.037 |
| 0 | 0.1 | AG1 | 1 | 36.4333 | 1.027 |
| 0 | 0.1 | ptsI | 1 | 36.6833 | 0.937 |
| 0 | 0.1 | clcB | 1 | 36.6833 | 0.522 |
| 0 | 0.1 | ycaM | 1 | 36.6833 | 0.618 |
| 0 | 0.1 | yadI | 1 | 36.6833 | 1.039 |

| | | | | | |
|---|---|---|---|---|---|
| 0 | 0.1 | AG1 | 1 | 36.6833 | 1.03 |
| 0.05 | 0.1 | ptsI | 1 | 0 | 0.241 |
| 0.05 | 0.1 | clcB | 1 | 0 | 0.242 |
| 0.05 | 0.1 | ycaM | 1 | 0 | 0.231 |
| 0.05 | 0.1 | yadI | 1 | 0 | 0.254 |
| 0.05 | 0.1 | AG1 | 1 | 0 | 0.232 |
| 0.05 | 0.1 | ptsI | 1 | 0.25 | 0.236 |
| 0.05 | 0.1 | clcB | 1 | 0.25 | 0.238 |
| 0.05 | 0.1 | ycaM | 1 | 0.25 | 0.221 |
| 0.05 | 0.1 | yadI | 1 | 0.25 | 0.24 |
| 0.05 | 0.1 | AG1 | 1 | 0.25 | 0.221 |
| 0.05 | 0.1 | ptsI | 1 | 0.5 | 0.234 |
| 0.05 | 0.1 | clcB | 1 | 0.5 | 0.236 |
| 0.05 | 0.1 | ycaM | 1 | 0.5 | 0.218 |
| 0.05 | 0.1 | yadI | 1 | 0.5 | 0.236 |
| 0.05 | 0.1 | AG1 | 1 | 0.5 | 0.219 |
| 0.05 | 0.1 | ptsI | 1 | 0.75 | 0.232 |
| 0.05 | 0.1 | clcB | 1 | 0.75 | 0.232 |
| 0.05 | 0.1 | ycaM | 1 | 0.75 | 0.216 |
| 0.05 | 0.1 | yadI | 1 | 0.75 | 0.237 |
| 0.05 | 0.1 | AG1 | 1 | 0.75 | 0.227 |
| 0.05 | 0.1 | ptsI | 1 | 1 | 0.234 |
| 0.05 | 0.1 | clcB | 1 | 1 | 0.232 |
| 0.05 | 0.1 | ycaM | 1 | 1 | 0.215 |
| 0.05 | 0.1 | yadI | 1 | 1 | 0.239 |
| 0.05 | 0.1 | AG1 | 1 | 1 | 0.218 |
| 0.05 | 0.1 | ptsI | 1 | 1.25 | 0.23 |
| 0.05 | 0.1 | clcB | 1 | 1.25 | 0.23 |
| 0.05 | 0.1 | ycaM | 1 | 1.25 | 0.218 |
| 0.05 | 0.1 | yadI | 1 | 1.25 | 0.243 |
| 0.05 | 0.1 | AG1 | 1 | 1.25 | 0.221 |
| 0.05 | 0.1 | ptsI | 1 | 1.5 | 0.237 |
| 0.05 | 0.1 | clcB | 1 | 1.5 | 0.234 |
| 0.05 | 0.1 | ycaM | 1 | 1.5 | 0.221 |
| 0.05 | 0.1 | yadI | 1 | 1.5 | 0.244 |
| 0.05 | 0.1 | AG1 | 1 | 1.5 | 0.225 |
| 0.05 | 0.1 | ptsI | 1 | 1.75 | 0.239 |
| 0.05 | 0.1 | clcB | 1 | 1.75 | 0.234 |
| 0.05 | 0.1 | ycaM | 1 | 1.75 | 0.224 |
| 0.05 | 0.1 | yadI | 1 | 1.75 | 0.248 |
| 0.05 | 0.1 | AG1 | 1 | 1.75 | 0.23 |
| 0.05 | 0.1 | ptsI | 1 | 2 | 0.246 |
| 0.05 | 0.1 | clcB | 1 | 2 | 0.24 |
| 0.05 | 0.1 | ycaM | 1 | 2 | 0.226 |
| 0.05 | 0.1 | yadI | 1 | 2 | 0.253 |
| 0.05 | 0.1 | AG1 | 1 | 2 | 0.232 |
| 0.05 | 0.1 | ptsI | 1 | 2.25 | 0.251 |
| 0.05 | 0.1 | clcB | 1 | 2.25 | 0.239 |

| | | | | | |
|---|---|---|---|---|---|
| 0.05 | 0.1 | ycaM | 1 | 2.25 | 0.232 |
| 0.05 | 0.1 | yadI | 1 | 2.25 | 0.26 |
| 0.05 | 0.1 | AG1 | 1 | 2.25 | 0.238 |
| 0.05 | 0.1 | ptsI | 1 | 2.5 | 0.256 |
| 0.05 | 0.1 | clcB | 1 | 2.5 | 0.243 |
| 0.05 | 0.1 | ycaM | 1 | 2.5 | 0.238 |
| 0.05 | 0.1 | yadI | 1 | 2.5 | 0.265 |
| 0.05 | 0.1 | AG1 | 1 | 2.5 | 0.242 |
| 0.05 | 0.1 | ptsI | 1 | 2.75 | 0.26 |
| 0.05 | 0.1 | clcB | 1 | 2.75 | 0.248 |
| 0.05 | 0.1 | ycaM | 1 | 2.75 | 0.243 |
| 0.05 | 0.1 | yadI | 1 | 2.75 | 0.275 |
| 0.05 | 0.1 | AG1 | 1 | 2.75 | 0.247 |
| 0.05 | 0.1 | ptsI | 1 | 3 | 0.272 |
| 0.05 | 0.1 | clcB | 1 | 3 | 0.254 |
| 0.05 | 0.1 | ycaM | 1 | 3 | 0.248 |
| 0.05 | 0.1 | yadI | 1 | 3 | 0.285 |
| 0.05 | 0.1 | AG1 | 1 | 3 | 0.255 |
| 0.05 | 0.1 | ptsI | 1 | 3.25 | 0.277 |
| 0.05 | 0.1 | clcB | 1 | 3.25 | 0.256 |
| 0.05 | 0.1 | ycaM | 1 | 3.25 | 0.253 |
| 0.05 | 0.1 | yadI | 1 | 3.25 | 0.295 |
| 0.05 | 0.1 | AG1 | 1 | 3.25 | 0.263 |
| 0.05 | 0.1 | ptsI | 1 | 3.5 | 0.285 |
| 0.05 | 0.1 | clcB | 1 | 3.5 | 0.26 |
| 0.05 | 0.1 | ycaM | 1 | 3.5 | 0.26 |
| 0.05 | 0.1 | yadI | 1 | 3.5 | 0.307 |
| 0.05 | 0.1 | AG1 | 1 | 3.5 | 0.272 |
| 0.05 | 0.1 | ptsI | 1 | 3.75 | 0.297 |
| 0.05 | 0.1 | clcB | 1 | 3.75 | 0.267 |
| 0.05 | 0.1 | ycaM | 1 | 3.75 | 0.269 |
| 0.05 | 0.1 | yadI | 1 | 3.75 | 0.327 |
| 0.05 | 0.1 | AG1 | 1 | 3.75 | 0.28 |
| 0.05 | 0.1 | ptsI | 1 | 4 | 0.303 |
| 0.05 | 0.1 | clcB | 1 | 4 | 0.272 |
| 0.05 | 0.1 | ycaM | 1 | 4 | 0.275 |
| 0.05 | 0.1 | yadI | 1 | 4 | 0.338 |
| 0.05 | 0.1 | AG1 | 1 | 4 | 0.291 |
| 0.05 | 0.1 | ptsI | 1 | 4.25 | 0.316 |
| 0.05 | 0.1 | clcB | 1 | 4.25 | 0.281 |
| 0.05 | 0.1 | ycaM | 1 | 4.25 | 0.286 |
| 0.05 | 0.1 | yadI | 1 | 4.25 | 0.346 |
| 0.05 | 0.1 | AG1 | 1 | 4.25 | 0.3 |
| 0.05 | 0.1 | ptsI | 1 | 4.5 | 0.327 |
| 0.05 | 0.1 | clcB | 1 | 4.5 | 0.29 |
| 0.05 | 0.1 | ycaM | 1 | 4.5 | 0.294 |
| 0.05 | 0.1 | yadI | 1 | 4.5 | 0.358 |
| 0.05 | 0.1 | AG1 | 1 | 4.5 | 0.308 |

| | | | | | |
|---|---|---|---|---|---|
| 0.05 | 0.1 | ptsI | 1 | 4.75 | 0.338 |
| 0.05 | 0.1 | clcB | 1 | 4.75 | 0.297 |
| 0.05 | 0.1 | ycaM | 1 | 4.75 | 0.303 |
| 0.05 | 0.1 | yadI | 1 | 4.75 | 0.372 |
| 0.05 | 0.1 | AG1 | 1 | 4.75 | 0.323 |
| 0.05 | 0.1 | ptsI | 1 | 5 | 0.355 |
| 0.05 | 0.1 | clcB | 1 | 5 | 0.305 |
| 0.05 | 0.1 | ycaM | 1 | 5 | 0.307 |
| 0.05 | 0.1 | yadI | 1 | 5 | 0.383 |
| 0.05 | 0.1 | AG1 | 1 | 5 | 0.33 |
| 0.05 | 0.1 | ptsI | 1 | 5.25 | 0.358 |
| 0.05 | 0.1 | clcB | 1 | 5.25 | 0.306 |
| 0.05 | 0.1 | ycaM | 1 | 5.25 | 0.316 |
| 0.05 | 0.1 | yadI | 1 | 5.25 | 0.399 |
| 0.05 | 0.1 | AG1 | 1 | 5.25 | 0.338 |
| 0.05 | 0.1 | ptsI | 1 | 5.5 | 0.375 |
| 0.05 | 0.1 | clcB | 1 | 5.5 | 0.317 |
| 0.05 | 0.1 | ycaM | 1 | 5.5 | 0.322 |
| 0.05 | 0.1 | yadI | 1 | 5.5 | 0.408 |
| 0.05 | 0.1 | AG1 | 1 | 5.5 | 0.35 |
| 0.05 | 0.1 | ptsI | 1 | 5.75 | 0.377 |
| 0.05 | 0.1 | clcB | 1 | 5.75 | 0.323 |
| 0.05 | 0.1 | ycaM | 1 | 5.75 | 0.328 |
| 0.05 | 0.1 | yadI | 1 | 5.75 | 0.418 |
| 0.05 | 0.1 | AG1 | 1 | 5.75 | 0.356 |
| 0.05 | 0.1 | ptsI | 1 | 6 | 0.379 |
| 0.05 | 0.1 | clcB | 1 | 6 | 0.332 |
| 0.05 | 0.1 | ycaM | 1 | 6 | 0.339 |
| 0.05 | 0.1 | yadI | 1 | 6 | 0.429 |
| 0.05 | 0.1 | AG1 | 1 | 6 | 0.374 |
| 0.05 | 0.1 | ptsI | 1 | 6.25 | 0.384 |
| 0.05 | 0.1 | clcB | 1 | 6.25 | 0.335 |
| 0.05 | 0.1 | ycaM | 1 | 6.25 | 0.343 |
| 0.05 | 0.1 | yadI | 1 | 6.25 | 0.441 |
| 0.05 | 0.1 | AG1 | 1 | 6.25 | 0.38 |
| 0.05 | 0.1 | ptsI | 1 | 6.5 | 0.397 |
| 0.05 | 0.1 | clcB | 1 | 6.5 | 0.348 |
| 0.05 | 0.1 | ycaM | 1 | 6.5 | 0.358 |
| 0.05 | 0.1 | yadI | 1 | 6.5 | 0.46 |
| 0.05 | 0.1 | AG1 | 1 | 6.5 | 0.399 |
| 0.05 | 0.1 | ptsI | 1 | 6.75 | 0.414 |
| 0.05 | 0.1 | clcB | 1 | 6.75 | 0.352 |
| 0.05 | 0.1 | ycaM | 1 | 6.75 | 0.366 |
| 0.05 | 0.1 | yadI | 1 | 6.75 | 0.476 |
| 0.05 | 0.1 | AG1 | 1 | 6.75 | 0.403 |
| 0.05 | 0.1 | ptsI | 1 | 7 | 0.424 |
| 0.05 | 0.1 | clcB | 1 | 7 | 0.36 |
| 0.05 | 0.1 | ycaM | 1 | 7 | 0.372 |

| | | | | | |
|---|---|---|---|---|---|
| 0.05 | 0.1 | yadI | 1 | 7 | 0.483 |
| 0.05 | 0.1 | AG1 | 1 | 7 | 0.413 |
| 0.05 | 0.1 | ptsI | 1 | 7.25 | 0.431 |
| 0.05 | 0.1 | clcB | 1 | 7.25 | 0.368 |
| 0.05 | 0.1 | ycaM | 1 | 7.25 | 0.379 |
| 0.05 | 0.1 | yadI | 1 | 7.25 | 0.494 |
| 0.05 | 0.1 | AG1 | 1 | 7.25 | 0.421 |
| 0.05 | 0.1 | ptsI | 1 | 7.5 | 0.44 |
| 0.05 | 0.1 | clcB | 1 | 7.5 | 0.371 |
| 0.05 | 0.1 | ycaM | 1 | 7.5 | 0.387 |
| 0.05 | 0.1 | yadI | 1 | 7.5 | 0.506 |
| 0.05 | 0.1 | AG1 | 1 | 7.5 | 0.429 |
| 0.05 | 0.1 | ptsI | 1 | 7.75 | 0.446 |
| 0.05 | 0.1 | clcB | 1 | 7.75 | 0.378 |
| 0.05 | 0.1 | ycaM | 1 | 7.75 | 0.392 |
| 0.05 | 0.1 | yadI | 1 | 7.75 | 0.524 |
| 0.05 | 0.1 | AG1 | 1 | 7.75 | 0.444 |
| 0.05 | 0.1 | ptsI | 1 | 8 | 0.458 |
| 0.05 | 0.1 | clcB | 1 | 8 | 0.387 |
| 0.05 | 0.1 | ycaM | 1 | 8 | 0.396 |
| 0.05 | 0.1 | yadI | 1 | 8 | 0.531 |
| 0.05 | 0.1 | AG1 | 1 | 8 | 0.449 |
| 0.05 | 0.1 | ptsI | 1 | 8.25 | 0.466 |
| 0.05 | 0.1 | clcB | 1 | 8.25 | 0.389 |
| 0.05 | 0.1 | ycaM | 1 | 8.25 | 0.401 |
| 0.05 | 0.1 | yadI | 1 | 8.25 | 0.539 |
| 0.05 | 0.1 | AG1 | 1 | 8.25 | 0.46 |
| 0.05 | 0.1 | ptsI | 1 | 8.5 | 0.475 |
| 0.05 | 0.1 | clcB | 1 | 8.5 | 0.397 |
| 0.05 | 0.1 | ycaM | 1 | 8.5 | 0.404 |
| 0.05 | 0.1 | yadI | 1 | 8.5 | 0.549 |
| 0.05 | 0.1 | AG1 | 1 | 8.5 | 0.471 |
| 0.05 | 0.1 | ptsI | 1 | 8.75 | 0.488 |
| 0.05 | 0.1 | clcB | 1 | 8.75 | 0.406 |
| 0.05 | 0.1 | ycaM | 1 | 8.75 | 0.404 |
| 0.05 | 0.1 | yadI | 1 | 8.75 | 0.557 |
| 0.05 | 0.1 | AG1 | 1 | 8.75 | 0.481 |
| 0.05 | 0.1 | ptsI | 1 | 9 | 0.499 |
| 0.05 | 0.1 | clcB | 1 | 9 | 0.411 |
| 0.05 | 0.1 | ycaM | 1 | 9 | 0.408 |
| 0.05 | 0.1 | yadI | 1 | 9 | 0.566 |
| 0.05 | 0.1 | AG1 | 1 | 9 | 0.498 |
| 0.05 | 0.1 | ptsI | 1 | 9.25 | 0.52 |
| 0.05 | 0.1 | clcB | 1 | 9.25 | 0.415 |
| 0.05 | 0.1 | ycaM | 1 | 9.25 | 0.414 |
| 0.05 | 0.1 | yadI | 1 | 9.25 | 0.584 |
| 0.05 | 0.1 | AG1 | 1 | 9.25 | 0.513 |
| 0.05 | 0.1 | ptsI | 1 | 9.5 | 0.569 |

| | | | | | |
|---|---|---|---|---|---|
| 0.05 | 0.1 | clcB | 1 | 9.5 | 0.424 |
| 0.05 | 0.1 | ycaM | 1 | 9.5 | 0.43 |
| 0.05 | 0.1 | yadI | 1 | 9.5 | 0.641 |
| 0.05 | 0.1 | AG1 | 1 | 9.5 | 0.548 |
| 0.05 | 0.1 | ptsI | 1 | 9.75 | 0.556 |
| 0.05 | 0.1 | clcB | 1 | 9.75 | 0.424 |
| 0.05 | 0.1 | ycaM | 1 | 9.75 | 0.436 |
| 0.05 | 0.1 | yadI | 1 | 9.75 | 0.636 |
| 0.05 | 0.1 | AG1 | 1 | 9.75 | 0.555 |
| 0.05 | 0.1 | ptsI | 1 | 10 | 0.546 |
| 0.05 | 0.1 | clcB | 1 | 10 | 0.429 |
| 0.05 | 0.1 | ycaM | 1 | 10 | 0.439 |
| 0.05 | 0.1 | yadI | 1 | 10 | 0.621 |
| 0.05 | 0.1 | AG1 | 1 | 10 | 0.552 |
| 0.05 | 0.1 | ptsI | 1 | 10.25 | 0.558 |
| 0.05 | 0.1 | clcB | 1 | 10.25 | 0.437 |
| 0.05 | 0.1 | ycaM | 1 | 10.25 | 0.444 |
| 0.05 | 0.1 | yadI | 1 | 10.25 | 0.626 |
| 0.05 | 0.1 | AG1 | 1 | 10.25 | 0.566 |
| 0.05 | 0.1 | ptsI | 1 | 10.5 | 0.597 |
| 0.05 | 0.1 | clcB | 1 | 10.5 | 0.45 |
| 0.05 | 0.1 | ycaM | 1 | 10.5 | 0.45 |
| 0.05 | 0.1 | yadI | 1 | 10.5 | 0.622 |
| 0.05 | 0.1 | AG1 | 1 | 10.5 | 0.572 |
| 0.05 | 0.1 | ptsI | 1 | 10.75 | 0.597 |
| 0.05 | 0.1 | clcB | 1 | 10.75 | 0.456 |
| 0.05 | 0.1 | ycaM | 1 | 10.75 | 0.457 |
| 0.05 | 0.1 | yadI | 1 | 10.75 | 0.622 |
| 0.05 | 0.1 | AG1 | 1 | 10.75 | 0.577 |
| 0.05 | 0.1 | ptsI | 1 | 11 | 0.636 |
| 0.05 | 0.1 | clcB | 1 | 11 | 0.473 |
| 0.05 | 0.1 | ycaM | 1 | 11 | 0.472 |
| 0.05 | 0.1 | yadI | 1 | 11 | 0.682 |
| 0.05 | 0.1 | AG1 | 1 | 11 | 0.634 |
| 0.05 | 0.1 | ptsI | 1 | 11.25 | 0.666 |
| 0.05 | 0.1 | clcB | 1 | 11.25 | 0.488 |
| 0.05 | 0.1 | ycaM | 1 | 11.25 | 0.485 |
| 0.05 | 0.1 | yadI | 1 | 11.25 | 0.673 |
| 0.05 | 0.1 | AG1 | 1 | 11.25 | 0.642 |
| 0.05 | 0.1 | ptsI | 1 | 11.5 | 0.623 |
| 0.05 | 0.1 | clcB | 1 | 11.5 | 0.49 |
| 0.05 | 0.1 | ycaM | 1 | 11.5 | 0.498 |
| 0.05 | 0.1 | yadI | 1 | 11.5 | 0.684 |
| 0.05 | 0.1 | AG1 | 1 | 11.5 | 0.688 |
| 0.05 | 0.1 | ptsI | 1 | 11.75 | 0.638 |
| 0.05 | 0.1 | clcB | 1 | 11.75 | 0.5 |
| 0.05 | 0.1 | ycaM | 1 | 11.75 | 0.516 |
| 0.05 | 0.1 | yadI | 1 | 11.75 | 0.735 |

| | | | | | |
|---|---|---|---|---|---|
| 0.05 | 0.1 | AG1 | 1 | 11.75 | 0.749 |
| 0.05 | 0.1 | ptsI | 1 | 12 | 0.626 |
| 0.05 | 0.1 | clcB | 1 | 12 | 0.505 |
| 0.05 | 0.1 | ycaM | 1 | 12 | 0.524 |
| 0.05 | 0.1 | yadI | 1 | 12 | 0.74 |
| 0.05 | 0.1 | AG1 | 1 | 12 | 0.745 |
| 0.05 | 0.1 | ptsI | 1 | 12.25 | 0.604 |
| 0.05 | 0.1 | clcB | 1 | 12.25 | 0.516 |
| 0.05 | 0.1 | ycaM | 1 | 12.25 | 0.518 |
| 0.05 | 0.1 | yadI | 1 | 12.25 | 0.761 |
| 0.05 | 0.1 | AG1 | 1 | 12.25 | 0.768 |
| 0.05 | 0.1 | ptsI | 1 | 12.5 | 0.626 |
| 0.05 | 0.1 | clcB | 1 | 12.5 | 0.519 |
| 0.05 | 0.1 | ycaM | 1 | 12.5 | 0.524 |
| 0.05 | 0.1 | yadI | 1 | 12.5 | 0.789 |
| 0.05 | 0.1 | AG1 | 1 | 12.5 | 0.761 |
| 0.05 | 0.1 | ptsI | 1 | 12.75 | 0.617 |
| 0.05 | 0.1 | clcB | 1 | 12.75 | 0.526 |
| 0.05 | 0.1 | ycaM | 1 | 12.75 | 0.529 |
| 0.05 | 0.1 | yadI | 1 | 12.75 | 0.807 |
| 0.05 | 0.1 | AG1 | 1 | 12.75 | 0.776 |
| 0.05 | 0.1 | ptsI | 1 | 13 | 0.631 |
| 0.05 | 0.1 | clcB | 1 | 13 | 0.539 |
| 0.05 | 0.1 | ycaM | 1 | 13 | 0.537 |
| 0.05 | 0.1 | yadI | 1 | 13 | 0.812 |
| 0.05 | 0.1 | AG1 | 1 | 13 | 0.798 |
| 0.05 | 0.1 | ptsI | 1 | 13.25 | 0.594 |
| 0.05 | 0.1 | clcB | 1 | 13.25 | 0.528 |
| 0.05 | 0.1 | ycaM | 1 | 13.25 | 0.54 |
| 0.05 | 0.1 | yadI | 1 | 13.25 | 0.83 |
| 0.05 | 0.1 | AG1 | 1 | 13.25 | 0.808 |
| 0.05 | 0.1 | ptsI | 1 | 13.5 | 0.617 |
| 0.05 | 0.1 | clcB | 1 | 13.5 | 0.524 |
| 0.05 | 0.1 | ycaM | 1 | 13.5 | 0.541 |
| 0.05 | 0.1 | yadI | 1 | 13.5 | 0.845 |
| 0.05 | 0.1 | AG1 | 1 | 13.5 | 0.815 |
| 0.05 | 0.1 | ptsI | 1 | 13.75 | 0.608 |
| 0.05 | 0.1 | clcB | 1 | 13.75 | 0.534 |
| 0.05 | 0.1 | ycaM | 1 | 13.75 | 0.544 |
| 0.05 | 0.1 | yadI | 1 | 13.75 | 0.841 |
| 0.05 | 0.1 | AG1 | 1 | 13.75 | 0.81 |
| 0.05 | 0.1 | ptsI | 1 | 14 | 0.643 |
| 0.05 | 0.1 | clcB | 1 | 14 | 0.559 |
| 0.05 | 0.1 | ycaM | 1 | 14 | 0.547 |
| 0.05 | 0.1 | yadI | 1 | 14 | 0.846 |
| 0.05 | 0.1 | AG1 | 1 | 14 | 0.847 |
| 0.05 | 0.1 | ptsI | 1 | 14.25 | 0.687 |
| 0.05 | 0.1 | clcB | 1 | 14.25 | 0.576 |

| | | | | | |
|---|---|---|---|---|---|
| 0.05 | 0.1 | ycaM | 1 | 14.25 | 0.551 |
| 0.05 | 0.1 | yadI | 1 | 14.25 | 0.846 |
| 0.05 | 0.1 | AG1 | 1 | 14.25 | 0.85 |
| 0.05 | 0.1 | ptsI | 1 | 14.5 | 0.676 |
| 0.05 | 0.1 | clcB | 1 | 14.5 | 0.578 |
| 0.05 | 0.1 | ycaM | 1 | 14.5 | 0.564 |
| 0.05 | 0.1 | yadI | 1 | 14.5 | 0.85 |
| 0.05 | 0.1 | AG1 | 1 | 14.5 | 0.847 |
| 0.05 | 0.1 | ptsI | 1 | 14.75 | 0.706 |
| 0.05 | 0.1 | clcB | 1 | 14.75 | 0.585 |
| 0.05 | 0.1 | ycaM | 1 | 14.75 | 0.575 |
| 0.05 | 0.1 | yadI | 1 | 14.75 | 0.85 |
| 0.05 | 0.1 | AG1 | 1 | 14.75 | 0.841 |
| 0.05 | 0.1 | ptsI | 1 | 15 | 0.749 |
| 0.05 | 0.1 | clcB | 1 | 15 | 0.619 |
| 0.05 | 0.1 | ycaM | 1 | 15 | 0.598 |
| 0.05 | 0.1 | yadI | 1 | 15 | 0.852 |
| 0.05 | 0.1 | AG1 | 1 | 15 | 0.861 |
| 0.05 | 0.1 | ptsI | 1 | 15.25 | 0.721 |
| 0.05 | 0.1 | clcB | 1 | 15.25 | 0.656 |
| 0.05 | 0.1 | ycaM | 1 | 15.25 | 0.632 |
| 0.05 | 0.1 | yadI | 1 | 15.25 | 0.852 |
| 0.05 | 0.1 | AG1 | 1 | 15.25 | 0.884 |
| 0.05 | 0.1 | ptsI | 1 | 15.5 | 0.705 |
| 0.05 | 0.1 | clcB | 1 | 15.5 | 0.661 |
| 0.05 | 0.1 | ycaM | 1 | 15.5 | 0.624 |
| 0.05 | 0.1 | yadI | 1 | 15.5 | 0.841 |
| 0.05 | 0.1 | AG1 | 1 | 15.5 | 0.858 |
| 0.05 | 0.1 | ptsI | 1 | 15.75 | 0.741 |
| 0.05 | 0.1 | clcB | 1 | 15.75 | 0.642 |
| 0.05 | 0.1 | ycaM | 1 | 15.75 | 0.647 |
| 0.05 | 0.1 | yadI | 1 | 15.75 | 0.844 |
| 0.05 | 0.1 | AG1 | 1 | 15.75 | 0.864 |
| 0.05 | 0.1 | ptsI | 1 | 16 | 0.746 |
| 0.05 | 0.1 | clcB | 1 | 16 | 0.646 |
| 0.05 | 0.1 | ycaM | 1 | 16 | 0.654 |
| 0.05 | 0.1 | yadI | 1 | 16 | 0.839 |
| 0.05 | 0.1 | AG1 | 1 | 16 | 0.867 |
| 0.05 | 0.1 | ptsI | 1 | 16.25 | 0.777 |
| 0.05 | 0.1 | clcB | 1 | 16.25 | 0.615 |
| 0.05 | 0.1 | ycaM | 1 | 16.25 | 0.662 |
| 0.05 | 0.1 | yadI | 1 | 16.25 | 0.846 |
| 0.05 | 0.1 | AG1 | 1 | 16.25 | 0.889 |
| 0.05 | 0.1 | ptsI | 1 | 16.5 | 0.752 |
| 0.05 | 0.1 | clcB | 1 | 16.5 | 0.596 |
| 0.05 | 0.1 | ycaM | 1 | 16.5 | 0.621 |
| 0.05 | 0.1 | yadI | 1 | 16.5 | 0.845 |
| 0.05 | 0.1 | AG1 | 1 | 16.5 | 0.885 |

| | | | | | |
|---|---|---|---|---|---|
| 0.05 | 0.1 | ptsI | 1 | 17.0667 | 0.609 |
| 0.05 | 0.1 | clcB | 1 | 17.0667 | 0.496 |
| 0.05 | 0.1 | ycaM | 1 | 17.0667 | 0.481 |
| 0.05 | 0.1 | yadI | 1 | 17.0667 | 0.586 |
| 0.05 | 0.1 | AG1 | 1 | 17.0667 | 0.872 |
| 0.05 | 0.1 | ptsI | 1 | 17.3167 | 0.575 |
| 0.05 | 0.1 | clcB | 1 | 17.3167 | 0.435 |
| 0.05 | 0.1 | ycaM | 1 | 17.3167 | 0.444 |
| 0.05 | 0.1 | yadI | 1 | 17.3167 | 0.562 |
| 0.05 | 0.1 | AG1 | 1 | 17.3167 | 0.766 |
| 0.05 | 0.1 | ptsI | 1 | 17.5667 | 0.604 |
| 0.05 | 0.1 | clcB | 1 | 17.5667 | 0.433 |
| 0.05 | 0.1 | ycaM | 1 | 17.5667 | 0.443 |
| 0.05 | 0.1 | yadI | 1 | 17.5667 | 0.576 |
| 0.05 | 0.1 | AG1 | 1 | 17.5667 | 0.762 |
| 0.05 | 0.1 | ptsI | 1 | 17.8167 | 0.607 |
| 0.05 | 0.1 | clcB | 1 | 17.8167 | 0.436 |
| 0.05 | 0.1 | ycaM | 1 | 17.8167 | 0.448 |
| 0.05 | 0.1 | yadI | 1 | 17.8167 | 0.587 |
| 0.05 | 0.1 | AG1 | 1 | 17.8167 | 0.814 |
| 0.05 | 0.1 | ptsI | 1 | 18.0667 | 0.63 |
| 0.05 | 0.1 | clcB | 1 | 18.0667 | 0.435 |
| 0.05 | 0.1 | ycaM | 1 | 18.0667 | 0.449 |
| 0.05 | 0.1 | yadI | 1 | 18.0667 | 0.605 |
| 0.05 | 0.1 | AG1 | 1 | 18.0667 | 0.791 |
| 0.05 | 0.1 | ptsI | 1 | 18.3167 | 0.648 |
| 0.05 | 0.1 | clcB | 1 | 18.3167 | 0.438 |
| 0.05 | 0.1 | ycaM | 1 | 18.3167 | 0.46 |
| 0.05 | 0.1 | yadI | 1 | 18.3167 | 0.611 |
| 0.05 | 0.1 | AG1 | 1 | 18.3167 | 0.81 |
| 0.05 | 0.1 | ptsI | 1 | 18.5667 | 0.707 |
| 0.05 | 0.1 | clcB | 1 | 18.5667 | 0.446 |
| 0.05 | 0.1 | ycaM | 1 | 18.5667 | 0.452 |
| 0.05 | 0.1 | yadI | 1 | 18.5667 | 0.627 |
| 0.05 | 0.1 | AG1 | 1 | 18.5667 | 0.825 |
| 0.05 | 0.1 | ptsI | 1 | 18.8167 | 0.725 |
| 0.05 | 0.1 | clcB | 1 | 18.8167 | 0.45 |
| 0.05 | 0.1 | ycaM | 1 | 18.8167 | 0.459 |
| 0.05 | 0.1 | yadI | 1 | 18.8167 | 0.636 |
| 0.05 | 0.1 | AG1 | 1 | 18.8167 | 0.829 |
| 0.05 | 0.1 | ptsI | 1 | 19.0667 | 0.75 |
| 0.05 | 0.1 | clcB | 1 | 19.0667 | 0.452 |
| 0.05 | 0.1 | ycaM | 1 | 19.0667 | 0.463 |
| 0.05 | 0.1 | yadI | 1 | 19.0667 | 0.655 |
| 0.05 | 0.1 | AG1 | 1 | 19.0667 | 0.857 |
| 0.05 | 0.1 | ptsI | 1 | 19.3167 | 0.743 |
| 0.05 | 0.1 | clcB | 1 | 19.3167 | 0.462 |
| 0.05 | 0.1 | ycaM | 1 | 19.3167 | 0.466 |

| | | | | | |
|---|---|---|---|---|---|
| 0.05 | 0.1 | yadI | 1 | 19.3167 | 0.653 |
| 0.05 | 0.1 | AG1 | 1 | 19.3167 | 0.843 |
| 0.05 | 0.1 | ptsI | 1 | 19.5667 | 0.752 |
| 0.05 | 0.1 | clcB | 1 | 19.5667 | 0.465 |
| 0.05 | 0.1 | ycaM | 1 | 19.5667 | 0.469 |
| 0.05 | 0.1 | yadI | 1 | 19.5667 | 0.664 |
| 0.05 | 0.1 | AG1 | 1 | 19.5667 | 0.848 |
| 0.05 | 0.1 | ptsI | 1 | 19.8167 | 0.748 |
| 0.05 | 0.1 | clcB | 1 | 19.8167 | 0.447 |
| 0.05 | 0.1 | ycaM | 1 | 19.8167 | 0.47 |
| 0.05 | 0.1 | yadI | 1 | 19.8167 | 0.668 |
| 0.05 | 0.1 | AG1 | 1 | 19.8167 | 0.856 |
| 0.05 | 0.1 | ptsI | 1 | 20.0667 | 0.746 |
| 0.05 | 0.1 | clcB | 1 | 20.0667 | 0.443 |
| 0.05 | 0.1 | ycaM | 1 | 20.0667 | 0.473 |
| 0.05 | 0.1 | yadI | 1 | 20.0667 | 0.672 |
| 0.05 | 0.1 | AG1 | 1 | 20.0667 | 0.866 |
| 0.05 | 0.1 | ptsI | 1 | 20.3167 | 0.75 |
| 0.05 | 0.1 | clcB | 1 | 20.3167 | 0.434 |
| 0.05 | 0.1 | ycaM | 1 | 20.3167 | 0.475 |
| 0.05 | 0.1 | yadI | 1 | 20.3167 | 0.676 |
| 0.05 | 0.1 | AG1 | 1 | 20.3167 | 0.869 |
| 0.05 | 0.1 | ptsI | 1 | 20.5667 | 0.744 |
| 0.05 | 0.1 | clcB | 1 | 20.5667 | 0.428 |
| 0.05 | 0.1 | ycaM | 1 | 20.5667 | 0.474 |
| 0.05 | 0.1 | yadI | 1 | 20.5667 | 0.687 |
| 0.05 | 0.1 | AG1 | 1 | 20.5667 | 0.868 |
| 0.05 | 0.1 | ptsI | 1 | 20.8167 | 0.75 |
| 0.05 | 0.1 | clcB | 1 | 20.8167 | 0.422 |
| 0.05 | 0.1 | ycaM | 1 | 20.8167 | 0.482 |
| 0.05 | 0.1 | yadI | 1 | 20.8167 | 0.688 |
| 0.05 | 0.1 | AG1 | 1 | 20.8167 | 0.871 |
| 0.05 | 0.1 | ptsI | 1 | 21.0667 | 0.762 |
| 0.05 | 0.1 | clcB | 1 | 21.0667 | 0.421 |
| 0.05 | 0.1 | ycaM | 1 | 21.0667 | 0.482 |
| 0.05 | 0.1 | yadI | 1 | 21.0667 | 0.696 |
| 0.05 | 0.1 | AG1 | 1 | 21.0667 | 0.888 |
| 0.05 | 0.1 | ptsI | 1 | 21.3167 | 0.767 |
| 0.05 | 0.1 | clcB | 1 | 21.3167 | 0.418 |
| 0.05 | 0.1 | ycaM | 1 | 21.3167 | 0.483 |
| 0.05 | 0.1 | yadI | 1 | 21.3167 | 0.71 |
| 0.05 | 0.1 | AG1 | 1 | 21.3167 | 0.893 |
| 0.05 | 0.1 | ptsI | 1 | 21.5667 | 0.772 |
| 0.05 | 0.1 | clcB | 1 | 21.5667 | 0.427 |
| 0.05 | 0.1 | ycaM | 1 | 21.5667 | 0.489 |
| 0.05 | 0.1 | yadI | 1 | 21.5667 | 0.722 |
| 0.05 | 0.1 | AG1 | 1 | 21.5667 | 0.89 |
| 0.05 | 0.1 | ptsI | 1 | 21.8167 | 0.78 |

| | | | | | |
|---|---|---|---|---|---|
| 0.05 | 0.1 | clcB | 1 | 21.8167 | 0.42 |
| 0.05 | 0.1 | ycaM | 1 | 21.8167 | 0.491 |
| 0.05 | 0.1 | yadI | 1 | 21.8167 | 0.729 |
| 0.05 | 0.1 | AG1 | 1 | 21.8167 | 0.905 |
| 0.05 | 0.1 | ptsI | 1 | 22.0667 | 0.787 |
| 0.05 | 0.1 | clcB | 1 | 22.0667 | 0.42 |
| 0.05 | 0.1 | ycaM | 1 | 22.0667 | 0.491 |
| 0.05 | 0.1 | yadI | 1 | 22.0667 | 0.726 |
| 0.05 | 0.1 | AG1 | 1 | 22.0667 | 0.908 |
| 0.05 | 0.1 | ptsI | 1 | 22.9333 | 0.78 |
| 0.05 | 0.1 | clcB | 1 | 22.9333 | 0.46 |
| 0.05 | 0.1 | ycaM | 1 | 22.9333 | 0.411 |
| 0.05 | 0.1 | yadI | 1 | 22.9333 | 0.745 |
| 0.05 | 0.1 | AG1 | 1 | 22.9333 | 0.779 |
| 0.05 | 0.1 | ptsI | 1 | 23.1833 | 0.768 |
| 0.05 | 0.1 | clcB | 1 | 23.1833 | 0.441 |
| 0.05 | 0.1 | ycaM | 1 | 23.1833 | 0.402 |
| 0.05 | 0.1 | yadI | 1 | 23.1833 | 0.729 |
| 0.05 | 0.1 | AG1 | 1 | 23.1833 | 0.762 |
| 0.05 | 0.1 | ptsI | 1 | 23.4333 | 0.746 |
| 0.05 | 0.1 | clcB | 1 | 23.4333 | 0.44 |
| 0.05 | 0.1 | ycaM | 1 | 23.4333 | 0.404 |
| 0.05 | 0.1 | yadI | 1 | 23.4333 | 0.738 |
| 0.05 | 0.1 | AG1 | 1 | 23.4333 | 0.783 |
| 0.05 | 0.1 | ptsI | 1 | 23.6833 | 0.756 |
| 0.05 | 0.1 | clcB | 1 | 23.6833 | 0.441 |
| 0.05 | 0.1 | ycaM | 1 | 23.6833 | 0.406 |
| 0.05 | 0.1 | yadI | 1 | 23.6833 | 0.756 |
| 0.05 | 0.1 | AG1 | 1 | 23.6833 | 0.815 |
| 0.05 | 0.1 | ptsI | 1 | 23.9333 | 0.735 |
| 0.05 | 0.1 | clcB | 1 | 23.9333 | 0.447 |
| 0.05 | 0.1 | ycaM | 1 | 23.9333 | 0.407 |
| 0.05 | 0.1 | yadI | 1 | 23.9333 | 0.757 |
| 0.05 | 0.1 | AG1 | 1 | 23.9333 | 0.807 |
| 0.05 | 0.1 | ptsI | 1 | 24.1833 | 0.744 |
| 0.05 | 0.1 | clcB | 1 | 24.1833 | 0.436 |
| 0.05 | 0.1 | ycaM | 1 | 24.1833 | 0.406 |
| 0.05 | 0.1 | yadI | 1 | 24.1833 | 0.761 |
| 0.05 | 0.1 | AG1 | 1 | 24.1833 | 0.816 |
| 0.05 | 0.1 | ptsI | 1 | 24.4333 | 0.773 |
| 0.05 | 0.1 | clcB | 1 | 24.4333 | 0.427 |
| 0.05 | 0.1 | ycaM | 1 | 24.4333 | 0.404 |
| 0.05 | 0.1 | yadI | 1 | 24.4333 | 0.767 |
| 0.05 | 0.1 | AG1 | 1 | 24.4333 | 0.816 |
| 0.05 | 0.1 | ptsI | 1 | 24.6833 | 0.772 |
| 0.05 | 0.1 | clcB | 1 | 24.6833 | 0.43 |
| 0.05 | 0.1 | ycaM | 1 | 24.6833 | 0.405 |
| 0.05 | 0.1 | yadI | 1 | 24.6833 | 0.775 |

| | | | | | |
|---|---|---|---|---|---|
| 0.05 | 0.1 | AG1 | 1 | 24.6833 | 0.818 |
| 0.05 | 0.1 | ptsI | 1 | 24.9333 | 0.783 |
| 0.05 | 0.1 | clcB | 1 | 24.9333 | 0.425 |
| 0.05 | 0.1 | ycaM | 1 | 24.9333 | 0.406 |
| 0.05 | 0.1 | yadI | 1 | 24.9333 | 0.782 |
| 0.05 | 0.1 | AG1 | 1 | 24.9333 | 0.838 |
| 0.05 | 0.1 | ptsI | 1 | 25.1833 | 0.788 |
| 0.05 | 0.1 | clcB | 1 | 25.1833 | 0.426 |
| 0.05 | 0.1 | ycaM | 1 | 25.1833 | 0.408 |
| 0.05 | 0.1 | yadI | 1 | 25.1833 | 0.779 |
| 0.05 | 0.1 | AG1 | 1 | 25.1833 | 0.826 |
| 0.05 | 0.1 | ptsI | 1 | 25.4333 | 0.802 |
| 0.05 | 0.1 | clcB | 1 | 25.4333 | 0.428 |
| 0.05 | 0.1 | ycaM | 1 | 25.4333 | 0.406 |
| 0.05 | 0.1 | yadI | 1 | 25.4333 | 0.787 |
| 0.05 | 0.1 | AG1 | 1 | 25.4333 | 0.833 |
| 0.05 | 0.1 | ptsI | 1 | 25.6833 | 0.804 |
| 0.05 | 0.1 | clcB | 1 | 25.6833 | 0.429 |
| 0.05 | 0.1 | ycaM | 1 | 25.6833 | 0.409 |
| 0.05 | 0.1 | yadI | 1 | 25.6833 | 0.793 |
| 0.05 | 0.1 | AG1 | 1 | 25.6833 | 0.83 |
| 0.05 | 0.1 | ptsI | 1 | 25.9333 | 0.813 |
| 0.05 | 0.1 | clcB | 1 | 25.9333 | 0.429 |
| 0.05 | 0.1 | ycaM | 1 | 25.9333 | 0.403 |
| 0.05 | 0.1 | yadI | 1 | 25.9333 | 0.794 |
| 0.05 | 0.1 | AG1 | 1 | 25.9333 | 0.835 |
| 0.05 | 0.1 | ptsI | 1 | 26.1833 | 0.823 |
| 0.05 | 0.1 | clcB | 1 | 26.1833 | 0.43 |
| 0.05 | 0.1 | ycaM | 1 | 26.1833 | 0.406 |
| 0.05 | 0.1 | yadI | 1 | 26.1833 | 0.804 |
| 0.05 | 0.1 | AG1 | 1 | 26.1833 | 0.844 |
| 0.05 | 0.1 | ptsI | 1 | 26.4333 | 0.818 |
| 0.05 | 0.1 | clcB | 1 | 26.4333 | 0.428 |
| 0.05 | 0.1 | ycaM | 1 | 26.4333 | 0.409 |
| 0.05 | 0.1 | yadI | 1 | 26.4333 | 0.804 |
| 0.05 | 0.1 | AG1 | 1 | 26.4333 | 0.84 |
| 0.05 | 0.1 | ptsI | 1 | 26.6833 | 0.821 |
| 0.05 | 0.1 | clcB | 1 | 26.6833 | 0.426 |
| 0.05 | 0.1 | ycaM | 1 | 26.6833 | 0.411 |
| 0.05 | 0.1 | yadI | 1 | 26.6833 | 0.81 |
| 0.05 | 0.1 | AG1 | 1 | 26.6833 | 0.846 |
| 0.05 | 0.1 | ptsI | 1 | 26.9333 | 0.826 |
| 0.05 | 0.1 | clcB | 1 | 26.9333 | 0.427 |
| 0.05 | 0.1 | ycaM | 1 | 26.9333 | 0.408 |
| 0.05 | 0.1 | yadI | 1 | 26.9333 | 0.81 |
| 0.05 | 0.1 | AG1 | 1 | 26.9333 | 0.849 |
| 0.05 | 0.1 | ptsI | 1 | 27.1833 | 0.836 |
| 0.05 | 0.1 | clcB | 1 | 27.1833 | 0.434 |

| | | | | | |
|---|---|---|---|---|---|
| 0.05 | 0.1 | ycaM | 1 | 27.1833 | 0.409 |
| 0.05 | 0.1 | yadI | 1 | 27.1833 | 0.818 |
| 0.05 | 0.1 | AG1 | 1 | 27.1833 | 0.854 |
| 0.05 | 0.1 | ptsI | 1 | 27.4333 | 0.826 |
| 0.05 | 0.1 | clcB | 1 | 27.4333 | 0.433 |
| 0.05 | 0.1 | ycaM | 1 | 27.4333 | 0.41 |
| 0.05 | 0.1 | yadI | 1 | 27.4333 | 0.821 |
| 0.05 | 0.1 | AG1 | 1 | 27.4333 | 0.855 |
| 0.05 | 0.1 | ptsI | 1 | 27.6833 | 0.837 |
| 0.05 | 0.1 | clcB | 1 | 27.6833 | 0.435 |
| 0.05 | 0.1 | ycaM | 1 | 27.6833 | 0.41 |
| 0.05 | 0.1 | yadI | 1 | 27.6833 | 0.822 |
| 0.05 | 0.1 | AG1 | 1 | 27.6833 | 0.862 |
| 0.05 | 0.1 | ptsI | 1 | 27.9333 | 0.832 |
| 0.05 | 0.1 | clcB | 1 | 27.9333 | 0.428 |
| 0.05 | 0.1 | ycaM | 1 | 27.9333 | 0.405 |
| 0.05 | 0.1 | yadI | 1 | 27.9333 | 0.818 |
| 0.05 | 0.1 | AG1 | 1 | 27.9333 | 0.854 |
| 0.05 | 0.1 | ptsI | 1 | 28.1833 | 0.847 |
| 0.05 | 0.1 | clcB | 1 | 28.1833 | 0.434 |
| 0.05 | 0.1 | ycaM | 1 | 28.1833 | 0.411 |
| 0.05 | 0.1 | yadI | 1 | 28.1833 | 0.827 |
| 0.05 | 0.1 | AG1 | 1 | 28.1833 | 0.871 |
| 0.05 | 0.1 | ptsI | 1 | 28.4333 | 0.844 |
| 0.05 | 0.1 | clcB | 1 | 28.4333 | 0.434 |
| 0.05 | 0.1 | ycaM | 1 | 28.4333 | 0.411 |
| 0.05 | 0.1 | yadI | 1 | 28.4333 | 0.832 |
| 0.05 | 0.1 | AG1 | 1 | 28.4333 | 0.872 |
| 0.05 | 0.1 | ptsI | 1 | 28.6833 | 0.84 |
| 0.05 | 0.1 | clcB | 1 | 28.6833 | 0.44 |
| 0.05 | 0.1 | ycaM | 1 | 28.6833 | 0.41 |
| 0.05 | 0.1 | yadI | 1 | 28.6833 | 0.838 |
| 0.05 | 0.1 | AG1 | 1 | 28.6833 | 0.871 |
| 0.05 | 0.1 | ptsI | 1 | 28.9333 | 0.851 |
| 0.05 | 0.1 | clcB | 1 | 28.9333 | 0.439 |
| 0.05 | 0.1 | ycaM | 1 | 28.9333 | 0.411 |
| 0.05 | 0.1 | yadI | 1 | 28.9333 | 0.841 |
| 0.05 | 0.1 | AG1 | 1 | 28.9333 | 0.881 |
| 0.05 | 0.1 | ptsI | 1 | 29.1833 | 0.843 |
| 0.05 | 0.1 | clcB | 1 | 29.1833 | 0.438 |
| 0.05 | 0.1 | ycaM | 1 | 29.1833 | 0.412 |
| 0.05 | 0.1 | yadI | 1 | 29.1833 | 0.849 |
| 0.05 | 0.1 | AG1 | 1 | 29.1833 | 0.879 |
| 0.05 | 0.1 | ptsI | 1 | 29.4333 | 0.849 |
| 0.05 | 0.1 | clcB | 1 | 29.4333 | 0.442 |
| 0.05 | 0.1 | ycaM | 1 | 29.4333 | 0.414 |
| 0.05 | 0.1 | yadI | 1 | 29.4333 | 0.846 |
| 0.05 | 0.1 | AG1 | 1 | 29.4333 | 0.884 |

| | | | | | |
|---|---|---|---|---|---|
| 0.05 | 0.1 | ptsI | 1 | 29.6833 | 0.852 |
| 0.05 | 0.1 | clcB | 1 | 29.6833 | 0.443 |
| 0.05 | 0.1 | ycaM | 1 | 29.6833 | 0.415 |
| 0.05 | 0.1 | yadI | 1 | 29.6833 | 0.848 |
| 0.05 | 0.1 | AG1 | 1 | 29.6833 | 0.888 |
| 0.05 | 0.1 | ptsI | 1 | 29.9333 | 0.858 |
| 0.05 | 0.1 | clcB | 1 | 29.9333 | 0.444 |
| 0.05 | 0.1 | ycaM | 1 | 29.9333 | 0.412 |
| 0.05 | 0.1 | yadI | 1 | 29.9333 | 0.852 |
| 0.05 | 0.1 | AG1 | 1 | 29.9333 | 0.894 |
| 0.05 | 0.1 | ptsI | 1 | 30.1833 | 0.852 |
| 0.05 | 0.1 | clcB | 1 | 30.1833 | 0.446 |
| 0.05 | 0.1 | ycaM | 1 | 30.1833 | 0.415 |
| 0.05 | 0.1 | yadI | 1 | 30.1833 | 0.856 |
| 0.05 | 0.1 | AG1 | 1 | 30.1833 | 0.896 |
| 0.05 | 0.1 | ptsI | 1 | 30.4333 | 0.857 |
| 0.05 | 0.1 | clcB | 1 | 30.4333 | 0.444 |
| 0.05 | 0.1 | ycaM | 1 | 30.4333 | 0.414 |
| 0.05 | 0.1 | yadI | 1 | 30.4333 | 0.858 |
| 0.05 | 0.1 | AG1 | 1 | 30.4333 | 0.903 |
| 0.05 | 0.1 | ptsI | 1 | 30.6833 | 0.852 |
| 0.05 | 0.1 | clcB | 1 | 30.6833 | 0.446 |
| 0.05 | 0.1 | ycaM | 1 | 30.6833 | 0.418 |
| 0.05 | 0.1 | yadI | 1 | 30.6833 | 0.873 |
| 0.05 | 0.1 | AG1 | 1 | 30.6833 | 0.904 |
| 0.05 | 0.1 | ptsI | 1 | 30.9333 | 0.858 |
| 0.05 | 0.1 | clcB | 1 | 30.9333 | 0.448 |
| 0.05 | 0.1 | ycaM | 1 | 30.9333 | 0.417 |
| 0.05 | 0.1 | yadI | 1 | 30.9333 | 0.874 |
| 0.05 | 0.1 | AG1 | 1 | 30.9333 | 0.907 |
| 0.05 | 0.1 | ptsI | 1 | 31.1833 | 0.857 |
| 0.05 | 0.1 | clcB | 1 | 31.1833 | 0.445 |
| 0.05 | 0.1 | ycaM | 1 | 31.1833 | 0.42 |
| 0.05 | 0.1 | yadI | 1 | 31.1833 | 0.876 |
| 0.05 | 0.1 | AG1 | 1 | 31.1833 | 0.911 |
| 0.05 | 0.1 | ptsI | 1 | 31.4333 | 0.866 |
| 0.05 | 0.1 | clcB | 1 | 31.4333 | 0.456 |
| 0.05 | 0.1 | ycaM | 1 | 31.4333 | 0.418 |
| 0.05 | 0.1 | yadI | 1 | 31.4333 | 0.871 |
| 0.05 | 0.1 | AG1 | 1 | 31.4333 | 0.918 |
| 0.05 | 0.1 | ptsI | 1 | 31.6833 | 0.863 |
| 0.05 | 0.1 | clcB | 1 | 31.6833 | 0.461 |
| 0.05 | 0.1 | ycaM | 1 | 31.6833 | 0.42 |
| 0.05 | 0.1 | yadI | 1 | 31.6833 | 0.876 |
| 0.05 | 0.1 | AG1 | 1 | 31.6833 | 0.926 |
| 0.05 | 0.1 | ptsI | 1 | 31.9333 | 0.867 |
| 0.05 | 0.1 | clcB | 1 | 31.9333 | 0.451 |
| 0.05 | 0.1 | ycaM | 1 | 31.9333 | 0.42 |

| | | | | | |
|---|---|---|---|---|---|
| 0.05 | 0.1 | yadI | 1 | 31.9333 | 0.875 |
| 0.05 | 0.1 | AG1 | 1 | 31.9333 | 0.928 |
| 0.05 | 0.1 | ptsI | 1 | 32.1833 | 0.869 |
| 0.05 | 0.1 | clcB | 1 | 32.1833 | 0.457 |
| 0.05 | 0.1 | ycaM | 1 | 32.1833 | 0.422 |
| 0.05 | 0.1 | yadI | 1 | 32.1833 | 0.881 |
| 0.05 | 0.1 | AG1 | 1 | 32.1833 | 0.933 |
| 0.05 | 0.1 | ptsI | 1 | 32.4333 | 0.864 |
| 0.05 | 0.1 | clcB | 1 | 32.4333 | 0.448 |
| 0.05 | 0.1 | ycaM | 1 | 32.4333 | 0.418 |
| 0.05 | 0.1 | yadI | 1 | 32.4333 | 0.879 |
| 0.05 | 0.1 | AG1 | 1 | 32.4333 | 0.928 |
| 0.05 | 0.1 | ptsI | 1 | 32.6833 | 0.864 |
| 0.05 | 0.1 | clcB | 1 | 32.6833 | 0.461 |
| 0.05 | 0.1 | ycaM | 1 | 32.6833 | 0.432 |
| 0.05 | 0.1 | yadI | 1 | 32.6833 | 0.89 |
| 0.05 | 0.1 | AG1 | 1 | 32.6833 | 0.944 |
| 0.05 | 0.1 | ptsI | 1 | 32.9333 | 0.866 |
| 0.05 | 0.1 | clcB | 1 | 32.9333 | 0.446 |
| 0.05 | 0.1 | ycaM | 1 | 32.9333 | 0.425 |
| 0.05 | 0.1 | yadI | 1 | 32.9333 | 0.89 |
| 0.05 | 0.1 | AG1 | 1 | 32.9333 | 0.952 |
| 0.05 | 0.1 | ptsI | 1 | 33.1833 | 0.869 |
| 0.05 | 0.1 | clcB | 1 | 33.1833 | 0.448 |
| 0.05 | 0.1 | ycaM | 1 | 33.1833 | 0.423 |
| 0.05 | 0.1 | yadI | 1 | 33.1833 | 0.89 |
| 0.05 | 0.1 | AG1 | 1 | 33.1833 | 0.95 |
| 0.05 | 0.1 | ptsI | 1 | 33.4333 | 0.872 |
| 0.05 | 0.1 | clcB | 1 | 33.4333 | 0.454 |
| 0.05 | 0.1 | ycaM | 1 | 33.4333 | 0.428 |
| 0.05 | 0.1 | yadI | 1 | 33.4333 | 0.888 |
| 0.05 | 0.1 | AG1 | 1 | 33.4333 | 0.966 |
| 0.05 | 0.1 | ptsI | 1 | 33.6833 | 0.868 |
| 0.05 | 0.1 | clcB | 1 | 33.6833 | 0.458 |
| 0.05 | 0.1 | ycaM | 1 | 33.6833 | 0.425 |
| 0.05 | 0.1 | yadI | 1 | 33.6833 | 0.886 |
| 0.05 | 0.1 | AG1 | 1 | 33.6833 | 0.965 |
| 0.05 | 0.1 | ptsI | 1 | 33.9333 | 0.872 |
| 0.05 | 0.1 | clcB | 1 | 33.9333 | 0.453 |
| 0.05 | 0.1 | ycaM | 1 | 33.9333 | 0.428 |
| 0.05 | 0.1 | yadI | 1 | 33.9333 | 0.896 |
| 0.05 | 0.1 | AG1 | 1 | 33.9333 | 0.978 |
| 0.05 | 0.1 | ptsI | 1 | 34.1833 | 0.876 |
| 0.05 | 0.1 | clcB | 1 | 34.1833 | 0.451 |
| 0.05 | 0.1 | ycaM | 1 | 34.1833 | 0.434 |
| 0.05 | 0.1 | yadI | 1 | 34.1833 | 0.902 |
| 0.05 | 0.1 | AG1 | 1 | 34.1833 | 0.986 |
| 0.05 | 0.1 | ptsI | 1 | 34.4333 | 0.871 |

| | | | | | |
|---|---|---|---|---|---|
| 0.05 | 0.1 | clcB | 1 | 34.4333 | 0.457 |
| 0.05 | 0.1 | ycaM | 1 | 34.4333 | 0.43 |
| 0.05 | 0.1 | yadI | 1 | 34.4333 | 0.903 |
| 0.05 | 0.1 | AG1 | 1 | 34.4333 | 0.994 |
| 0.05 | 0.1 | ptsI | 1 | 34.6833 | 0.879 |
| 0.05 | 0.1 | clcB | 1 | 34.6833 | 0.461 |
| 0.05 | 0.1 | ycaM | 1 | 34.6833 | 0.433 |
| 0.05 | 0.1 | yadI | 1 | 34.6833 | 0.903 |
| 0.05 | 0.1 | AG1 | 1 | 34.6833 | 1.008 |
| 0.05 | 0.1 | ptsI | 1 | 34.9333 | 0.881 |
| 0.05 | 0.1 | clcB | 1 | 34.9333 | 0.453 |
| 0.05 | 0.1 | ycaM | 1 | 34.9333 | 0.431 |
| 0.05 | 0.1 | yadI | 1 | 34.9333 | 0.908 |
| 0.05 | 0.1 | AG1 | 1 | 34.9333 | 1.01 |
| 0.05 | 0.1 | ptsI | 1 | 35.1833 | 0.872 |
| 0.05 | 0.1 | clcB | 1 | 35.1833 | 0.455 |
| 0.05 | 0.1 | ycaM | 1 | 35.1833 | 0.432 |
| 0.05 | 0.1 | yadI | 1 | 35.1833 | 0.911 |
| 0.05 | 0.1 | AG1 | 1 | 35.1833 | 1.015 |
| 0.05 | 0.1 | ptsI | 1 | 35.4333 | 0.876 |
| 0.05 | 0.1 | clcB | 1 | 35.4333 | 0.453 |
| 0.05 | 0.1 | ycaM | 1 | 35.4333 | 0.433 |
| 0.05 | 0.1 | yadI | 1 | 35.4333 | 0.913 |
| 0.05 | 0.1 | AG1 | 1 | 35.4333 | 1.023 |
| 0.05 | 0.1 | ptsI | 1 | 35.6833 | 0.879 |
| 0.05 | 0.1 | clcB | 1 | 35.6833 | 0.463 |
| 0.05 | 0.1 | ycaM | 1 | 35.6833 | 0.438 |
| 0.05 | 0.1 | yadI | 1 | 35.6833 | 0.911 |
| 0.05 | 0.1 | AG1 | 1 | 35.6833 | 1.035 |
| 0.05 | 0.1 | ptsI | 1 | 35.9333 | 0.881 |
| 0.05 | 0.1 | clcB | 1 | 35.9333 | 0.457 |
| 0.05 | 0.1 | ycaM | 1 | 35.9333 | 0.435 |
| 0.05 | 0.1 | yadI | 1 | 35.9333 | 0.912 |
| 0.05 | 0.1 | AG1 | 1 | 35.9333 | 1.032 |
| 0.05 | 0.1 | ptsI | 1 | 36.1833 | 0.877 |
| 0.05 | 0.1 | clcB | 1 | 36.1833 | 0.46 |
| 0.05 | 0.1 | ycaM | 1 | 36.1833 | 0.446 |
| 0.05 | 0.1 | yadI | 1 | 36.1833 | 0.915 |
| 0.05 | 0.1 | AG1 | 1 | 36.1833 | 1.047 |
| 0.05 | 0.1 | ptsI | 1 | 36.4333 | 0.885 |
| 0.05 | 0.1 | clcB | 1 | 36.4333 | 0.456 |
| 0.05 | 0.1 | ycaM | 1 | 36.4333 | 0.437 |
| 0.05 | 0.1 | yadI | 1 | 36.4333 | 0.918 |
| 0.05 | 0.1 | AG1 | 1 | 36.4333 | 1.05 |
| 0.05 | 0.1 | ptsI | 1 | 36.6833 | 0.883 |
| 0.05 | 0.1 | clcB | 1 | 36.6833 | 0.463 |
| 0.05 | 0.1 | ycaM | 1 | 36.6833 | 0.435 |
| 0.05 | 0.1 | yadI | 1 | 36.6833 | 0.921 |

| | | | | | |
|---|---|---|---|---|---|
| 0.05 | 0.1 | AG1 | 1 | 36.6833 | 1.06 |
| 0.1 | 0.1 | ptsI | 1 | 0 | 0.235 |
| 0.1 | 0.1 | clcB | 1 | 0 | 0.249 |
| 0.1 | 0.1 | ycaM | 1 | 0 | 0.228 |
| 0.1 | 0.1 | yadI | 1 | 0 | 0.242 |
| 0.1 | 0.1 | AG1 | 1 | 0 | 0.235 |
| 0.1 | 0.1 | ptsI | 1 | 0.25 | 0.222 |
| 0.1 | 0.1 | clcB | 1 | 0.25 | 0.24 |
| 0.1 | 0.1 | ycaM | 1 | 0.25 | 0.212 |
| 0.1 | 0.1 | yadI | 1 | 0.25 | 0.234 |
| 0.1 | 0.1 | AG1 | 1 | 0.25 | 0.224 |
| 0.1 | 0.1 | ptsI | 1 | 0.5 | 0.219 |
| 0.1 | 0.1 | clcB | 1 | 0.5 | 0.236 |
| 0.1 | 0.1 | ycaM | 1 | 0.5 | 0.209 |
| 0.1 | 0.1 | yadI | 1 | 0.5 | 0.231 |
| 0.1 | 0.1 | AG1 | 1 | 0.5 | 0.22 |
| 0.1 | 0.1 | ptsI | 1 | 0.75 | 0.218 |
| 0.1 | 0.1 | clcB | 1 | 0.75 | 0.232 |
| 0.1 | 0.1 | ycaM | 1 | 0.75 | 0.207 |
| 0.1 | 0.1 | yadI | 1 | 0.75 | 0.236 |
| 0.1 | 0.1 | AG1 | 1 | 0.75 | 0.221 |
| 0.1 | 0.1 | ptsI | 1 | 1 | 0.22 |
| 0.1 | 0.1 | clcB | 1 | 1 | 0.234 |
| 0.1 | 0.1 | ycaM | 1 | 1 | 0.207 |
| 0.1 | 0.1 | yadI | 1 | 1 | 0.235 |
| 0.1 | 0.1 | AG1 | 1 | 1 | 0.221 |
| 0.1 | 0.1 | ptsI | 1 | 1.25 | 0.222 |
| 0.1 | 0.1 | clcB | 1 | 1.25 | 0.232 |
| 0.1 | 0.1 | ycaM | 1 | 1.25 | 0.206 |
| 0.1 | 0.1 | yadI | 1 | 1.25 | 0.238 |
| 0.1 | 0.1 | AG1 | 1 | 1.25 | 0.227 |
| 0.1 | 0.1 | ptsI | 1 | 1.5 | 0.227 |
| 0.1 | 0.1 | clcB | 1 | 1.5 | 0.232 |
| 0.1 | 0.1 | ycaM | 1 | 1.5 | 0.207 |
| 0.1 | 0.1 | yadI | 1 | 1.5 | 0.241 |
| 0.1 | 0.1 | AG1 | 1 | 1.5 | 0.227 |
| 0.1 | 0.1 | ptsI | 1 | 1.75 | 0.23 |
| 0.1 | 0.1 | clcB | 1 | 1.75 | 0.235 |
| 0.1 | 0.1 | ycaM | 1 | 1.75 | 0.211 |
| 0.1 | 0.1 | yadI | 1 | 1.75 | 0.247 |
| 0.1 | 0.1 | AG1 | 1 | 1.75 | 0.232 |
| 0.1 | 0.1 | ptsI | 1 | 2 | 0.236 |
| 0.1 | 0.1 | clcB | 1 | 2 | 0.239 |
| 0.1 | 0.1 | ycaM | 1 | 2 | 0.215 |
| 0.1 | 0.1 | yadI | 1 | 2 | 0.251 |
| 0.1 | 0.1 | AG1 | 1 | 2 | 0.235 |
| 0.1 | 0.1 | ptsI | 1 | 2.25 | 0.242 |
| 0.1 | 0.1 | clcB | 1 | 2.25 | 0.244 |

| | | | | | |
|---|---|---|---|---|---|
| 0.1 | 0.1 | ycaM | 1 | 2.25 | 0.218 |
| 0.1 | 0.1 | yadI | 1 | 2.25 | 0.259 |
| 0.1 | 0.1 | AG1 | 1 | 2.25 | 0.239 |
| 0.1 | 0.1 | ptsI | 1 | 2.5 | 0.248 |
| 0.1 | 0.1 | clcB | 1 | 2.5 | 0.246 |
| 0.1 | 0.1 | ycaM | 1 | 2.5 | 0.222 |
| 0.1 | 0.1 | yadI | 1 | 2.5 | 0.267 |
| 0.1 | 0.1 | AG1 | 1 | 2.5 | 0.246 |
| 0.1 | 0.1 | ptsI | 1 | 2.75 | 0.253 |
| 0.1 | 0.1 | clcB | 1 | 2.75 | 0.252 |
| 0.1 | 0.1 | ycaM | 1 | 2.75 | 0.228 |
| 0.1 | 0.1 | yadI | 1 | 2.75 | 0.273 |
| 0.1 | 0.1 | AG1 | 1 | 2.75 | 0.255 |
| 0.1 | 0.1 | ptsI | 1 | 3 | 0.264 |
| 0.1 | 0.1 | clcB | 1 | 3 | 0.255 |
| 0.1 | 0.1 | ycaM | 1 | 3 | 0.234 |
| 0.1 | 0.1 | yadI | 1 | 3 | 0.286 |
| 0.1 | 0.1 | AG1 | 1 | 3 | 0.258 |
| 0.1 | 0.1 | ptsI | 1 | 3.25 | 0.274 |
| 0.1 | 0.1 | clcB | 1 | 3.25 | 0.263 |
| 0.1 | 0.1 | ycaM | 1 | 3.25 | 0.242 |
| 0.1 | 0.1 | yadI | 1 | 3.25 | 0.293 |
| 0.1 | 0.1 | AG1 | 1 | 3.25 | 0.264 |
| 0.1 | 0.1 | ptsI | 1 | 3.5 | 0.281 |
| 0.1 | 0.1 | clcB | 1 | 3.5 | 0.268 |
| 0.1 | 0.1 | ycaM | 1 | 3.5 | 0.249 |
| 0.1 | 0.1 | yadI | 1 | 3.5 | 0.304 |
| 0.1 | 0.1 | AG1 | 1 | 3.5 | 0.274 |
| 0.1 | 0.1 | ptsI | 1 | 3.75 | 0.29 |
| 0.1 | 0.1 | clcB | 1 | 3.75 | 0.272 |
| 0.1 | 0.1 | ycaM | 1 | 3.75 | 0.252 |
| 0.1 | 0.1 | yadI | 1 | 3.75 | 0.318 |
| 0.1 | 0.1 | AG1 | 1 | 3.75 | 0.29 |
| 0.1 | 0.1 | ptsI | 1 | 4 | 0.307 |
| 0.1 | 0.1 | clcB | 1 | 4 | 0.279 |
| 0.1 | 0.1 | ycaM | 1 | 4 | 0.265 |
| 0.1 | 0.1 | yadI | 1 | 4 | 0.332 |
| 0.1 | 0.1 | AG1 | 1 | 4 | 0.3 |
| 0.1 | 0.1 | ptsI | 1 | 4.25 | 0.305 |
| 0.1 | 0.1 | clcB | 1 | 4.25 | 0.288 |
| 0.1 | 0.1 | ycaM | 1 | 4.25 | 0.269 |
| 0.1 | 0.1 | yadI | 1 | 4.25 | 0.339 |
| 0.1 | 0.1 | AG1 | 1 | 4.25 | 0.31 |
| 0.1 | 0.1 | ptsI | 1 | 4.5 | 0.316 |
| 0.1 | 0.1 | clcB | 1 | 4.5 | 0.295 |
| 0.1 | 0.1 | ycaM | 1 | 4.5 | 0.276 |
| 0.1 | 0.1 | yadI | 1 | 4.5 | 0.35 |
| 0.1 | 0.1 | AG1 | 1 | 4.5 | 0.319 |

| | | | | | |
|---|---|---|---|---|---|
| 0.1 | 0.1 | ptsI | 1 | 4.75 | 0.329 |
| 0.1 | 0.1 | clcB | 1 | 4.75 | 0.304 |
| 0.1 | 0.1 | ycaM | 1 | 4.75 | 0.28 |
| 0.1 | 0.1 | yadI | 1 | 4.75 | 0.362 |
| 0.1 | 0.1 | AG1 | 1 | 4.75 | 0.334 |
| 0.1 | 0.1 | ptsI | 1 | 5 | 0.343 |
| 0.1 | 0.1 | clcB | 1 | 5 | 0.31 |
| 0.1 | 0.1 | ycaM | 1 | 5 | 0.286 |
| 0.1 | 0.1 | yadI | 1 | 5 | 0.367 |
| 0.1 | 0.1 | AG1 | 1 | 5 | 0.34 |
| 0.1 | 0.1 | ptsI | 1 | 5.25 | 0.355 |
| 0.1 | 0.1 | clcB | 1 | 5.25 | 0.313 |
| 0.1 | 0.1 | ycaM | 1 | 5.25 | 0.291 |
| 0.1 | 0.1 | yadI | 1 | 5.25 | 0.38 |
| 0.1 | 0.1 | AG1 | 1 | 5.25 | 0.351 |
| 0.1 | 0.1 | ptsI | 1 | 5.5 | 0.365 |
| 0.1 | 0.1 | clcB | 1 | 5.5 | 0.321 |
| 0.1 | 0.1 | ycaM | 1 | 5.5 | 0.301 |
| 0.1 | 0.1 | yadI | 1 | 5.5 | 0.39 |
| 0.1 | 0.1 | AG1 | 1 | 5.5 | 0.362 |
| 0.1 | 0.1 | ptsI | 1 | 5.75 | 0.373 |
| 0.1 | 0.1 | clcB | 1 | 5.75 | 0.325 |
| 0.1 | 0.1 | ycaM | 1 | 5.75 | 0.306 |
| 0.1 | 0.1 | yadI | 1 | 5.75 | 0.398 |
| 0.1 | 0.1 | AG1 | 1 | 5.75 | 0.371 |
| 0.1 | 0.1 | ptsI | 1 | 6 | 0.375 |
| 0.1 | 0.1 | clcB | 1 | 6 | 0.334 |
| 0.1 | 0.1 | ycaM | 1 | 6 | 0.311 |
| 0.1 | 0.1 | yadI | 1 | 6 | 0.409 |
| 0.1 | 0.1 | AG1 | 1 | 6 | 0.387 |
| 0.1 | 0.1 | ptsI | 1 | 6.25 | 0.384 |
| 0.1 | 0.1 | clcB | 1 | 6.25 | 0.339 |
| 0.1 | 0.1 | ycaM | 1 | 6.25 | 0.318 |
| 0.1 | 0.1 | yadI | 1 | 6.25 | 0.42 |
| 0.1 | 0.1 | AG1 | 1 | 6.25 | 0.397 |
| 0.1 | 0.1 | ptsI | 1 | 6.5 | 0.394 |
| 0.1 | 0.1 | clcB | 1 | 6.5 | 0.349 |
| 0.1 | 0.1 | ycaM | 1 | 6.5 | 0.325 |
| 0.1 | 0.1 | yadI | 1 | 6.5 | 0.436 |
| 0.1 | 0.1 | AG1 | 1 | 6.5 | 0.415 |
| 0.1 | 0.1 | ptsI | 1 | 6.75 | 0.416 |
| 0.1 | 0.1 | clcB | 1 | 6.75 | 0.356 |
| 0.1 | 0.1 | ycaM | 1 | 6.75 | 0.333 |
| 0.1 | 0.1 | yadI | 1 | 6.75 | 0.444 |
| 0.1 | 0.1 | AG1 | 1 | 6.75 | 0.421 |
| 0.1 | 0.1 | ptsI | 1 | 7 | 0.438 |
| 0.1 | 0.1 | clcB | 1 | 7 | 0.362 |
| 0.1 | 0.1 | ycaM | 1 | 7 | 0.341 |

| | | | | | |
|---|---|---|---|---|---|
| 0.1 | 0.1 | yadI | 1 | 7 | 0.45 |
| 0.1 | 0.1 | AG1 | 1 | 7 | 0.427 |
| 0.1 | 0.1 | ptsI | 1 | 7.25 | 0.448 |
| 0.1 | 0.1 | clcB | 1 | 7.25 | 0.366 |
| 0.1 | 0.1 | ycaM | 1 | 7.25 | 0.349 |
| 0.1 | 0.1 | yadI | 1 | 7.25 | 0.461 |
| 0.1 | 0.1 | AG1 | 1 | 7.25 | 0.437 |
| 0.1 | 0.1 | ptsI | 1 | 7.5 | 0.465 |
| 0.1 | 0.1 | clcB | 1 | 7.5 | 0.374 |
| 0.1 | 0.1 | ycaM | 1 | 7.5 | 0.355 |
| 0.1 | 0.1 | yadI | 1 | 7.5 | 0.473 |
| 0.1 | 0.1 | AG1 | 1 | 7.5 | 0.444 |
| 0.1 | 0.1 | ptsI | 1 | 7.75 | 0.479 |
| 0.1 | 0.1 | clcB | 1 | 7.75 | 0.381 |
| 0.1 | 0.1 | ycaM | 1 | 7.75 | 0.362 |
| 0.1 | 0.1 | yadI | 1 | 7.75 | 0.484 |
| 0.1 | 0.1 | AG1 | 1 | 7.75 | 0.458 |
| 0.1 | 0.1 | ptsI | 1 | 8 | 0.496 |
| 0.1 | 0.1 | clcB | 1 | 8 | 0.391 |
| 0.1 | 0.1 | ycaM | 1 | 8 | 0.367 |
| 0.1 | 0.1 | yadI | 1 | 8 | 0.488 |
| 0.1 | 0.1 | AG1 | 1 | 8 | 0.466 |
| 0.1 | 0.1 | ptsI | 1 | 8.25 | 0.509 |
| 0.1 | 0.1 | clcB | 1 | 8.25 | 0.395 |
| 0.1 | 0.1 | ycaM | 1 | 8.25 | 0.371 |
| 0.1 | 0.1 | yadI | 1 | 8.25 | 0.5 |
| 0.1 | 0.1 | AG1 | 1 | 8.25 | 0.478 |
| 0.1 | 0.1 | ptsI | 1 | 8.5 | 0.525 |
| 0.1 | 0.1 | clcB | 1 | 8.5 | 0.405 |
| 0.1 | 0.1 | ycaM | 1 | 8.5 | 0.372 |
| 0.1 | 0.1 | yadI | 1 | 8.5 | 0.51 |
| 0.1 | 0.1 | AG1 | 1 | 8.5 | 0.492 |
| 0.1 | 0.1 | ptsI | 1 | 8.75 | 0.524 |
| 0.1 | 0.1 | clcB | 1 | 8.75 | 0.413 |
| 0.1 | 0.1 | ycaM | 1 | 8.75 | 0.374 |
| 0.1 | 0.1 | yadI | 1 | 8.75 | 0.518 |
| 0.1 | 0.1 | AG1 | 1 | 8.75 | 0.502 |
| 0.1 | 0.1 | ptsI | 1 | 9 | 0.537 |
| 0.1 | 0.1 | clcB | 1 | 9 | 0.418 |
| 0.1 | 0.1 | ycaM | 1 | 9 | 0.376 |
| 0.1 | 0.1 | yadI | 1 | 9 | 0.524 |
| 0.1 | 0.1 | AG1 | 1 | 9 | 0.513 |
| 0.1 | 0.1 | ptsI | 1 | 9.25 | 0.554 |
| 0.1 | 0.1 | clcB | 1 | 9.25 | 0.422 |
| 0.1 | 0.1 | ycaM | 1 | 9.25 | 0.384 |
| 0.1 | 0.1 | yadI | 1 | 9.25 | 0.524 |
| 0.1 | 0.1 | AG1 | 1 | 9.25 | 0.536 |
| 0.1 | 0.1 | ptsI | 1 | 9.5 | 0.626 |

| | | | | | |
|---|---|---|---|---|---|
| 0.1 | 0.1 | clcB | 1 | 9.5 | 0.437 |
| 0.1 | 0.1 | ycaM | 1 | 9.5 | 0.392 |
| 0.1 | 0.1 | yadI | 1 | 9.5 | 0.538 |
| 0.1 | 0.1 | AG1 | 1 | 9.5 | 0.58 |
| 0.1 | 0.1 | ptsI | 1 | 9.75 | 0.59 |
| 0.1 | 0.1 | clcB | 1 | 9.75 | 0.442 |
| 0.1 | 0.1 | ycaM | 1 | 9.75 | 0.398 |
| 0.1 | 0.1 | yadI | 1 | 9.75 | 0.546 |
| 0.1 | 0.1 | AG1 | 1 | 9.75 | 0.587 |
| 0.1 | 0.1 | ptsI | 1 | 10 | 0.564 |
| 0.1 | 0.1 | clcB | 1 | 10 | 0.443 |
| 0.1 | 0.1 | ycaM | 1 | 10 | 0.402 |
| 0.1 | 0.1 | yadI | 1 | 10 | 0.552 |
| 0.1 | 0.1 | AG1 | 1 | 10 | 0.583 |
| 0.1 | 0.1 | ptsI | 1 | 10.25 | 0.625 |
| 0.1 | 0.1 | clcB | 1 | 10.25 | 0.451 |
| 0.1 | 0.1 | ycaM | 1 | 10.25 | 0.407 |
| 0.1 | 0.1 | yadI | 1 | 10.25 | 0.562 |
| 0.1 | 0.1 | AG1 | 1 | 10.25 | 0.593 |
| 0.1 | 0.1 | ptsI | 1 | 10.5 | 0.627 |
| 0.1 | 0.1 | clcB | 1 | 10.5 | 0.459 |
| 0.1 | 0.1 | ycaM | 1 | 10.5 | 0.42 |
| 0.1 | 0.1 | yadI | 1 | 10.5 | 0.567 |
| 0.1 | 0.1 | AG1 | 1 | 10.5 | 0.603 |
| 0.1 | 0.1 | ptsI | 1 | 10.75 | 0.624 |
| 0.1 | 0.1 | clcB | 1 | 10.75 | 0.467 |
| 0.1 | 0.1 | ycaM | 1 | 10.75 | 0.423 |
| 0.1 | 0.1 | yadI | 1 | 10.75 | 0.576 |
| 0.1 | 0.1 | AG1 | 1 | 10.75 | 0.61 |
| 0.1 | 0.1 | ptsI | 1 | 11 | 0.678 |
| 0.1 | 0.1 | clcB | 1 | 11 | 0.485 |
| 0.1 | 0.1 | ycaM | 1 | 11 | 0.444 |
| 0.1 | 0.1 | yadI | 1 | 11 | 0.591 |
| 0.1 | 0.1 | AG1 | 1 | 11 | 0.697 |
| 0.1 | 0.1 | ptsI | 1 | 11.25 | 0.686 |
| 0.1 | 0.1 | clcB | 1 | 11.25 | 0.501 |
| 0.1 | 0.1 | ycaM | 1 | 11.25 | 0.444 |
| 0.1 | 0.1 | yadI | 1 | 11.25 | 0.597 |
| 0.1 | 0.1 | AG1 | 1 | 11.25 | 0.675 |
| 0.1 | 0.1 | ptsI | 1 | 11.5 | 0.658 |
| 0.1 | 0.1 | clcB | 1 | 11.5 | 0.508 |
| 0.1 | 0.1 | ycaM | 1 | 11.5 | 0.454 |
| 0.1 | 0.1 | yadI | 1 | 11.5 | 0.61 |
| 0.1 | 0.1 | AG1 | 1 | 11.5 | 0.738 |
| 0.1 | 0.1 | ptsI | 1 | 11.75 | 0.686 |
| 0.1 | 0.1 | clcB | 1 | 11.75 | 0.522 |
| 0.1 | 0.1 | ycaM | 1 | 11.75 | 0.459 |
| 0.1 | 0.1 | yadI | 1 | 11.75 | 0.634 |

| | | | | | |
|---|---|---|---|---|---|
| 0.1 | 0.1 | AG1 | 1 | 11.75 | 0.793 |
| 0.1 | 0.1 | ptsI | 1 | 12 | 0.699 |
| 0.1 | 0.1 | clcB | 1 | 12 | 0.549 |
| 0.1 | 0.1 | ycaM | 1 | 12 | 0.472 |
| 0.1 | 0.1 | yadI | 1 | 12 | 0.642 |
| 0.1 | 0.1 | AG1 | 1 | 12 | 0.784 |
| 0.1 | 0.1 | ptsI | 1 | 12.25 | 0.664 |
| 0.1 | 0.1 | clcB | 1 | 12.25 | 0.539 |
| 0.1 | 0.1 | ycaM | 1 | 12.25 | 0.462 |
| 0.1 | 0.1 | yadI | 1 | 12.25 | 0.635 |
| 0.1 | 0.1 | AG1 | 1 | 12.25 | 0.828 |
| 0.1 | 0.1 | ptsI | 1 | 12.5 | 0.683 |
| 0.1 | 0.1 | clcB | 1 | 12.5 | 0.566 |
| 0.1 | 0.1 | ycaM | 1 | 12.5 | 0.469 |
| 0.1 | 0.1 | yadI | 1 | 12.5 | 0.637 |
| 0.1 | 0.1 | AG1 | 1 | 12.5 | 0.814 |
| 0.1 | 0.1 | ptsI | 1 | 12.75 | 0.712 |
| 0.1 | 0.1 | clcB | 1 | 12.75 | 0.585 |
| 0.1 | 0.1 | ycaM | 1 | 12.75 | 0.473 |
| 0.1 | 0.1 | yadI | 1 | 12.75 | 0.64 |
| 0.1 | 0.1 | AG1 | 1 | 12.75 | 0.822 |
| 0.1 | 0.1 | ptsI | 1 | 13 | 0.712 |
| 0.1 | 0.1 | clcB | 1 | 13 | 0.6 |
| 0.1 | 0.1 | ycaM | 1 | 13 | 0.481 |
| 0.1 | 0.1 | yadI | 1 | 13 | 0.647 |
| 0.1 | 0.1 | AG1 | 1 | 13 | 0.834 |
| 0.1 | 0.1 | ptsI | 1 | 13.25 | 0.686 |
| 0.1 | 0.1 | clcB | 1 | 13.25 | 0.607 |
| 0.1 | 0.1 | ycaM | 1 | 13.25 | 0.482 |
| 0.1 | 0.1 | yadI | 1 | 13.25 | 0.648 |
| 0.1 | 0.1 | AG1 | 1 | 13.25 | 0.852 |
| 0.1 | 0.1 | ptsI | 1 | 13.5 | 0.733 |
| 0.1 | 0.1 | clcB | 1 | 13.5 | 0.615 |
| 0.1 | 0.1 | ycaM | 1 | 13.5 | 0.497 |
| 0.1 | 0.1 | yadI | 1 | 13.5 | 0.663 |
| 0.1 | 0.1 | AG1 | 1 | 13.5 | 0.854 |
| 0.1 | 0.1 | ptsI | 1 | 13.75 | 0.702 |
| 0.1 | 0.1 | clcB | 1 | 13.75 | 0.612 |
| 0.1 | 0.1 | ycaM | 1 | 13.75 | 0.501 |
| 0.1 | 0.1 | yadI | 1 | 13.75 | 0.661 |
| 0.1 | 0.1 | AG1 | 1 | 13.75 | 0.853 |
| 0.1 | 0.1 | ptsI | 1 | 14 | 0.742 |
| 0.1 | 0.1 | clcB | 1 | 14 | 0.615 |
| 0.1 | 0.1 | ycaM | 1 | 14 | 0.517 |
| 0.1 | 0.1 | yadI | 1 | 14 | 0.666 |
| 0.1 | 0.1 | AG1 | 1 | 14 | 0.875 |
| 0.1 | 0.1 | ptsI | 1 | 14.25 | 0.728 |
| 0.1 | 0.1 | clcB | 1 | 14.25 | 0.624 |

| | | | | | |
|---|---|---|---|---|---|
| 0.1 | 0.1 | ycaM | 1 | 14.25 | 0.534 |
| 0.1 | 0.1 | yadI | 1 | 14.25 | 0.673 |
| 0.1 | 0.1 | AG1 | 1 | 14.25 | 0.887 |
| 0.1 | 0.1 | ptsI | 1 | 14.5 | 0.726 |
| 0.1 | 0.1 | clcB | 1 | 14.5 | 0.651 |
| 0.1 | 0.1 | ycaM | 1 | 14.5 | 0.558 |
| 0.1 | 0.1 | yadI | 1 | 14.5 | 0.678 |
| 0.1 | 0.1 | AG1 | 1 | 14.5 | 0.882 |
| 0.1 | 0.1 | ptsI | 1 | 14.75 | 0.724 |
| 0.1 | 0.1 | clcB | 1 | 14.75 | 0.65 |
| 0.1 | 0.1 | ycaM | 1 | 14.75 | 0.561 |
| 0.1 | 0.1 | yadI | 1 | 14.75 | 0.675 |
| 0.1 | 0.1 | AG1 | 1 | 14.75 | 0.9 |
| 0.1 | 0.1 | ptsI | 1 | 15 | 0.758 |
| 0.1 | 0.1 | clcB | 1 | 15 | 0.7 |
| 0.1 | 0.1 | ycaM | 1 | 15 | 0.588 |
| 0.1 | 0.1 | yadI | 1 | 15 | 0.679 |
| 0.1 | 0.1 | AG1 | 1 | 15 | 0.901 |
| 0.1 | 0.1 | ptsI | 1 | 15.25 | 0.766 |
| 0.1 | 0.1 | clcB | 1 | 15.25 | 0.717 |
| 0.1 | 0.1 | ycaM | 1 | 15.25 | 0.612 |
| 0.1 | 0.1 | yadI | 1 | 15.25 | 0.68 |
| 0.1 | 0.1 | AG1 | 1 | 15.25 | 0.912 |
| 0.1 | 0.1 | ptsI | 1 | 15.5 | 0.74 |
| 0.1 | 0.1 | clcB | 1 | 15.5 | 0.714 |
| 0.1 | 0.1 | ycaM | 1 | 15.5 | 0.601 |
| 0.1 | 0.1 | yadI | 1 | 15.5 | 0.686 |
| 0.1 | 0.1 | AG1 | 1 | 15.5 | 0.899 |
| 0.1 | 0.1 | ptsI | 1 | 15.75 | 0.756 |
| 0.1 | 0.1 | clcB | 1 | 15.75 | 0.71 |
| 0.1 | 0.1 | ycaM | 1 | 15.75 | 0.615 |
| 0.1 | 0.1 | yadI | 1 | 15.75 | 0.69 |
| 0.1 | 0.1 | AG1 | 1 | 15.75 | 0.892 |
| 0.1 | 0.1 | ptsI | 1 | 16 | 0.75 |
| 0.1 | 0.1 | clcB | 1 | 16 | 0.702 |
| 0.1 | 0.1 | ycaM | 1 | 16 | 0.611 |
| 0.1 | 0.1 | yadI | 1 | 16 | 0.697 |
| 0.1 | 0.1 | AG1 | 1 | 16 | 0.908 |
| 0.1 | 0.1 | ptsI | 1 | 16.25 | 0.781 |
| 0.1 | 0.1 | clcB | 1 | 16.25 | 0.74 |
| 0.1 | 0.1 | ycaM | 1 | 16.25 | 0.619 |
| 0.1 | 0.1 | yadI | 1 | 16.25 | 0.704 |
| 0.1 | 0.1 | AG1 | 1 | 16.25 | 0.917 |
| 0.1 | 0.1 | ptsI | 1 | 16.5 | 0.78 |
| 0.1 | 0.1 | clcB | 1 | 16.5 | 0.705 |
| 0.1 | 0.1 | ycaM | 1 | 16.5 | 0.608 |
| 0.1 | 0.1 | yadI | 1 | 16.5 | 0.711 |
| 0.1 | 0.1 | AG1 | 1 | 16.5 | 0.909 |

| | | | | | |
|---|---|---|---|---|---|
| 0.1 | 0.1 | ptsI | 1 | 17.0667 | 0.67 |
| 0.1 | 0.1 | clcB | 1 | 17.0667 | 0.536 |
| 0.1 | 0.1 | ycaM | 1 | 17.0667 | 0.632 |
| 0.1 | 0.1 | yadI | 1 | 17.0667 | 0.842 |
| 0.1 | 0.1 | AG1 | 1 | 17.0667 | 0.781 |
| 0.1 | 0.1 | ptsI | 1 | 17.3167 | 0.661 |
| 0.1 | 0.1 | clcB | 1 | 17.3167 | 0.49 |
| 0.1 | 0.1 | ycaM | 1 | 17.3167 | 0.593 |
| 0.1 | 0.1 | yadI | 1 | 17.3167 | 0.821 |
| 0.1 | 0.1 | AG1 | 1 | 17.3167 | 0.747 |
| 0.1 | 0.1 | ptsI | 1 | 17.5667 | 0.661 |
| 0.1 | 0.1 | clcB | 1 | 17.5667 | 0.487 |
| 0.1 | 0.1 | ycaM | 1 | 17.5667 | 0.583 |
| 0.1 | 0.1 | yadI | 1 | 17.5667 | 0.826 |
| 0.1 | 0.1 | AG1 | 1 | 17.5667 | 0.754 |
| 0.1 | 0.1 | ptsI | 1 | 17.8167 | 0.731 |
| 0.1 | 0.1 | clcB | 1 | 17.8167 | 0.49 |
| 0.1 | 0.1 | ycaM | 1 | 17.8167 | 0.584 |
| 0.1 | 0.1 | yadI | 1 | 17.8167 | 0.838 |
| 0.1 | 0.1 | AG1 | 1 | 17.8167 | 0.807 |
| 0.1 | 0.1 | ptsI | 1 | 18.0667 | 0.714 |
| 0.1 | 0.1 | clcB | 1 | 18.0667 | 0.484 |
| 0.1 | 0.1 | ycaM | 1 | 18.0667 | 0.591 |
| 0.1 | 0.1 | yadI | 1 | 18.0667 | 0.853 |
| 0.1 | 0.1 | AG1 | 1 | 18.0667 | 0.788 |
| 0.1 | 0.1 | ptsI | 1 | 18.3167 | 0.737 |
| 0.1 | 0.1 | clcB | 1 | 18.3167 | 0.488 |
| 0.1 | 0.1 | ycaM | 1 | 18.3167 | 0.576 |
| 0.1 | 0.1 | yadI | 1 | 18.3167 | 0.85 |
| 0.1 | 0.1 | AG1 | 1 | 18.3167 | 0.802 |
| 0.1 | 0.1 | ptsI | 1 | 18.5667 | 0.78 |
| 0.1 | 0.1 | clcB | 1 | 18.5667 | 0.486 |
| 0.1 | 0.1 | ycaM | 1 | 18.5667 | 0.584 |
| 0.1 | 0.1 | yadI | 1 | 18.5667 | 0.86 |
| 0.1 | 0.1 | AG1 | 1 | 18.5667 | 0.817 |
| 0.1 | 0.1 | ptsI | 1 | 18.8167 | 0.787 |
| 0.1 | 0.1 | clcB | 1 | 18.8167 | 0.49 |
| 0.1 | 0.1 | ycaM | 1 | 18.8167 | 0.576 |
| 0.1 | 0.1 | yadI | 1 | 18.8167 | 0.864 |
| 0.1 | 0.1 | AG1 | 1 | 18.8167 | 0.823 |
| 0.1 | 0.1 | ptsI | 1 | 19.0667 | 0.812 |
| 0.1 | 0.1 | clcB | 1 | 19.0667 | 0.49 |
| 0.1 | 0.1 | ycaM | 1 | 19.0667 | 0.578 |
| 0.1 | 0.1 | yadI | 1 | 19.0667 | 0.875 |
| 0.1 | 0.1 | AG1 | 1 | 19.0667 | 0.847 |
| 0.1 | 0.1 | ptsI | 1 | 19.3167 | 0.793 |
| 0.1 | 0.1 | clcB | 1 | 19.3167 | 0.489 |
| 0.1 | 0.1 | ycaM | 1 | 19.3167 | 0.582 |

| | | | | | |
|---|---|---|---|---|---|
| 0.1 | 0.1 | yadI | 1 | 19.3167 | 0.874 |
| 0.1 | 0.1 | AG1 | 1 | 19.3167 | 0.834 |
| 0.1 | 0.1 | ptsI | 1 | 19.5667 | 0.813 |
| 0.1 | 0.1 | clcB | 1 | 19.5667 | 0.483 |
| 0.1 | 0.1 | ycaM | 1 | 19.5667 | 0.572 |
| 0.1 | 0.1 | yadI | 1 | 19.5667 | 0.881 |
| 0.1 | 0.1 | AG1 | 1 | 19.5667 | 0.849 |
| 0.1 | 0.1 | ptsI | 1 | 19.8167 | 0.816 |
| 0.1 | 0.1 | clcB | 1 | 19.8167 | 0.477 |
| 0.1 | 0.1 | ycaM | 1 | 19.8167 | 0.575 |
| 0.1 | 0.1 | yadI | 1 | 19.8167 | 0.884 |
| 0.1 | 0.1 | AG1 | 1 | 19.8167 | 0.84 |
| 0.1 | 0.1 | ptsI | 1 | 20.0667 | 0.818 |
| 0.1 | 0.1 | clcB | 1 | 20.0667 | 0.48 |
| 0.1 | 0.1 | ycaM | 1 | 20.0667 | 0.572 |
| 0.1 | 0.1 | yadI | 1 | 20.0667 | 0.888 |
| 0.1 | 0.1 | AG1 | 1 | 20.0667 | 0.842 |
| 0.1 | 0.1 | ptsI | 1 | 20.3167 | 0.82 |
| 0.1 | 0.1 | clcB | 1 | 20.3167 | 0.473 |
| 0.1 | 0.1 | ycaM | 1 | 20.3167 | 0.574 |
| 0.1 | 0.1 | yadI | 1 | 20.3167 | 0.889 |
| 0.1 | 0.1 | AG1 | 1 | 20.3167 | 0.841 |
| 0.1 | 0.1 | ptsI | 1 | 20.5667 | 0.807 |
| 0.1 | 0.1 | clcB | 1 | 20.5667 | 0.458 |
| 0.1 | 0.1 | ycaM | 1 | 20.5667 | 0.574 |
| 0.1 | 0.1 | yadI | 1 | 20.5667 | 0.896 |
| 0.1 | 0.1 | AG1 | 1 | 20.5667 | 0.838 |
| 0.1 | 0.1 | ptsI | 1 | 20.8167 | 0.81 |
| 0.1 | 0.1 | clcB | 1 | 20.8167 | 0.456 |
| 0.1 | 0.1 | ycaM | 1 | 20.8167 | 0.57 |
| 0.1 | 0.1 | yadI | 1 | 20.8167 | 0.899 |
| 0.1 | 0.1 | AG1 | 1 | 20.8167 | 0.845 |
| 0.1 | 0.1 | ptsI | 1 | 21.0667 | 0.828 |
| 0.1 | 0.1 | clcB | 1 | 21.0667 | 0.456 |
| 0.1 | 0.1 | ycaM | 1 | 21.0667 | 0.564 |
| 0.1 | 0.1 | yadI | 1 | 21.0667 | 0.898 |
| 0.1 | 0.1 | AG1 | 1 | 21.0667 | 0.858 |
| 0.1 | 0.1 | ptsI | 1 | 21.3167 | 0.825 |
| 0.1 | 0.1 | clcB | 1 | 21.3167 | 0.451 |
| 0.1 | 0.1 | ycaM | 1 | 21.3167 | 0.572 |
| 0.1 | 0.1 | yadI | 1 | 21.3167 | 0.903 |
| 0.1 | 0.1 | AG1 | 1 | 21.3167 | 0.856 |
| 0.1 | 0.1 | ptsI | 1 | 21.5667 | 0.827 |
| 0.1 | 0.1 | clcB | 1 | 21.5667 | 0.464 |
| 0.1 | 0.1 | ycaM | 1 | 21.5667 | 0.568 |
| 0.1 | 0.1 | yadI | 1 | 21.5667 | 0.912 |
| 0.1 | 0.1 | AG1 | 1 | 21.5667 | 0.859 |
| 0.1 | 0.1 | ptsI | 1 | 21.8167 | 0.83 |

| | | | | | |
|---|---|---|---|---|---|
| 0.1 | 0.1 | clcB | 1 | 21.8167 | 0.46 |
| 0.1 | 0.1 | ycaM | 1 | 21.8167 | 0.569 |
| 0.1 | 0.1 | yadI | 1 | 21.8167 | 0.913 |
| 0.1 | 0.1 | AG1 | 1 | 21.8167 | 0.869 |
| 0.1 | 0.1 | ptsI | 1 | 22.0667 | 0.838 |
| 0.1 | 0.1 | clcB | 1 | 22.0667 | 0.456 |
| 0.1 | 0.1 | ycaM | 1 | 22.0667 | 0.566 |
| 0.1 | 0.1 | yadI | 1 | 22.0667 | 0.914 |
| 0.1 | 0.1 | AG1 | 1 | 22.0667 | 0.874 |
| 0.1 | 0.1 | ptsI | 1 | 22.9333 | 0.827 |
| 0.1 | 0.1 | clcB | 1 | 22.9333 | 0.488 |
| 0.1 | 0.1 | ycaM | 1 | 22.9333 | 0.41 |
| 0.1 | 0.1 | yadI | 1 | 22.9333 | 0.953 |
| 0.1 | 0.1 | AG1 | 1 | 22.9333 | 0.865 |
| 0.1 | 0.1 | ptsI | 1 | 23.1833 | 0.798 |
| 0.1 | 0.1 | clcB | 1 | 23.1833 | 0.476 |
| 0.1 | 0.1 | ycaM | 1 | 23.1833 | 0.404 |
| 0.1 | 0.1 | yadI | 1 | 23.1833 | 0.933 |
| 0.1 | 0.1 | AG1 | 1 | 23.1833 | 0.841 |
| 0.1 | 0.1 | ptsI | 1 | 23.4333 | 0.792 |
| 0.1 | 0.1 | clcB | 1 | 23.4333 | 0.473 |
| 0.1 | 0.1 | ycaM | 1 | 23.4333 | 0.406 |
| 0.1 | 0.1 | yadI | 1 | 23.4333 | 0.941 |
| 0.1 | 0.1 | AG1 | 1 | 23.4333 | 0.847 |
| 0.1 | 0.1 | ptsI | 1 | 23.6833 | 0.807 |
| 0.1 | 0.1 | clcB | 1 | 23.6833 | 0.466 |
| 0.1 | 0.1 | ycaM | 1 | 23.6833 | 0.407 |
| 0.1 | 0.1 | yadI | 1 | 23.6833 | 0.953 |
| 0.1 | 0.1 | AG1 | 1 | 23.6833 | 0.865 |
| 0.1 | 0.1 | ptsI | 1 | 23.9333 | 0.807 |
| 0.1 | 0.1 | clcB | 1 | 23.9333 | 0.472 |
| 0.1 | 0.1 | ycaM | 1 | 23.9333 | 0.41 |
| 0.1 | 0.1 | yadI | 1 | 23.9333 | 0.955 |
| 0.1 | 0.1 | AG1 | 1 | 23.9333 | 0.867 |
| 0.1 | 0.1 | ptsI | 1 | 24.1833 | 0.808 |
| 0.1 | 0.1 | clcB | 1 | 24.1833 | 0.461 |
| 0.1 | 0.1 | ycaM | 1 | 24.1833 | 0.407 |
| 0.1 | 0.1 | yadI | 1 | 24.1833 | 0.964 |
| 0.1 | 0.1 | AG1 | 1 | 24.1833 | 0.868 |
| 0.1 | 0.1 | ptsI | 1 | 24.4333 | 0.833 |
| 0.1 | 0.1 | clcB | 1 | 24.4333 | 0.461 |
| 0.1 | 0.1 | ycaM | 1 | 24.4333 | 0.408 |
| 0.1 | 0.1 | yadI | 1 | 24.4333 | 0.968 |
| 0.1 | 0.1 | AG1 | 1 | 24.4333 | 0.868 |
| 0.1 | 0.1 | ptsI | 1 | 24.6833 | 0.831 |
| 0.1 | 0.1 | clcB | 1 | 24.6833 | 0.448 |
| 0.1 | 0.1 | ycaM | 1 | 24.6833 | 0.406 |
| 0.1 | 0.1 | yadI | 1 | 24.6833 | 0.974 |

| | | | | | |
|---|---|---|---|---|---|
| 0.1 | 0.1 | AG1 | 1 | 24.6833 | 0.868 |
| 0.1 | 0.1 | ptsI | 1 | 24.9333 | 0.839 |
| 0.1 | 0.1 | clcB | 1 | 24.9333 | 0.502 |
| 0.1 | 0.1 | ycaM | 1 | 24.9333 | 0.408 |
| 0.1 | 0.1 | yadI | 1 | 24.9333 | 0.976 |
| 0.1 | 0.1 | AG1 | 1 | 24.9333 | 0.876 |
| 0.1 | 0.1 | ptsI | 1 | 25.1833 | 0.837 |
| 0.1 | 0.1 | clcB | 1 | 25.1833 | 0.445 |
| 0.1 | 0.1 | ycaM | 1 | 25.1833 | 0.407 |
| 0.1 | 0.1 | yadI | 1 | 25.1833 | 0.976 |
| 0.1 | 0.1 | AG1 | 1 | 25.1833 | 0.87 |
| 0.1 | 0.1 | ptsI | 1 | 25.4333 | 0.844 |
| 0.1 | 0.1 | clcB | 1 | 25.4333 | 0.452 |
| 0.1 | 0.1 | ycaM | 1 | 25.4333 | 0.409 |
| 0.1 | 0.1 | yadI | 1 | 25.4333 | 0.984 |
| 0.1 | 0.1 | AG1 | 1 | 25.4333 | 0.876 |
| 0.1 | 0.1 | ptsI | 1 | 25.6833 | 0.84 |
| 0.1 | 0.1 | clcB | 1 | 25.6833 | 0.45 |
| 0.1 | 0.1 | ycaM | 1 | 25.6833 | 0.408 |
| 0.1 | 0.1 | yadI | 1 | 25.6833 | 0.988 |
| 0.1 | 0.1 | AG1 | 1 | 25.6833 | 0.873 |
| 0.1 | 0.1 | ptsI | 1 | 25.9333 | 0.854 |
| 0.1 | 0.1 | clcB | 1 | 25.9333 | 0.449 |
| 0.1 | 0.1 | ycaM | 1 | 25.9333 | 0.41 |
| 0.1 | 0.1 | yadI | 1 | 25.9333 | 0.982 |
| 0.1 | 0.1 | AG1 | 1 | 25.9333 | 0.881 |
| 0.1 | 0.1 | ptsI | 1 | 26.1833 | 0.855 |
| 0.1 | 0.1 | clcB | 1 | 26.1833 | 0.45 |
| 0.1 | 0.1 | ycaM | 1 | 26.1833 | 0.414 |
| 0.1 | 0.1 | yadI | 1 | 26.1833 | 0.998 |
| 0.1 | 0.1 | AG1 | 1 | 26.1833 | 0.884 |
| 0.1 | 0.1 | ptsI | 1 | 26.4333 | 0.853 |
| 0.1 | 0.1 | clcB | 1 | 26.4333 | 0.447 |
| 0.1 | 0.1 | ycaM | 1 | 26.4333 | 0.414 |
| 0.1 | 0.1 | yadI | 1 | 26.4333 | 0.994 |
| 0.1 | 0.1 | AG1 | 1 | 26.4333 | 0.877 |
| 0.1 | 0.1 | ptsI | 1 | 26.6833 | 0.856 |
| 0.1 | 0.1 | clcB | 1 | 26.6833 | 0.45 |
| 0.1 | 0.1 | ycaM | 1 | 26.6833 | 0.412 |
| 0.1 | 0.1 | yadI | 1 | 26.6833 | 1.005 |
| 0.1 | 0.1 | AG1 | 1 | 26.6833 | 0.884 |
| 0.1 | 0.1 | ptsI | 1 | 26.9333 | 0.862 |
| 0.1 | 0.1 | clcB | 1 | 26.9333 | 0.449 |
| 0.1 | 0.1 | ycaM | 1 | 26.9333 | 0.414 |
| 0.1 | 0.1 | yadI | 1 | 26.9333 | 1 |
| 0.1 | 0.1 | AG1 | 1 | 26.9333 | 0.883 |
| 0.1 | 0.1 | ptsI | 1 | 27.1833 | 0.866 |
| 0.1 | 0.1 | clcB | 1 | 27.1833 | 0.448 |

| | | | | | |
|---|---|---|---|---|---|
| 0.1 | 0.1 | ycaM | 1 | 27.1833 | 0.413 |
| 0.1 | 0.1 | yadI | 1 | 27.1833 | 1.005 |
| 0.1 | 0.1 | AG1 | 1 | 27.1833 | 0.89 |
| 0.1 | 0.1 | ptsI | 1 | 27.4333 | 0.859 |
| 0.1 | 0.1 | clcB | 1 | 27.4333 | 0.448 |
| 0.1 | 0.1 | ycaM | 1 | 27.4333 | 0.412 |
| 0.1 | 0.1 | yadI | 1 | 27.4333 | 1.01 |
| 0.1 | 0.1 | AG1 | 1 | 27.4333 | 0.887 |
| 0.1 | 0.1 | ptsI | 1 | 27.6833 | 0.867 |
| 0.1 | 0.1 | clcB | 1 | 27.6833 | 0.451 |
| 0.1 | 0.1 | ycaM | 1 | 27.6833 | 0.413 |
| 0.1 | 0.1 | yadI | 1 | 27.6833 | 1.011 |
| 0.1 | 0.1 | AG1 | 1 | 27.6833 | 0.894 |
| 0.1 | 0.1 | ptsI | 1 | 27.9333 | 0.86 |
| 0.1 | 0.1 | clcB | 1 | 27.9333 | 0.444 |
| 0.1 | 0.1 | ycaM | 1 | 27.9333 | 0.403 |
| 0.1 | 0.1 | yadI | 1 | 27.9333 | 0.992 |
| 0.1 | 0.1 | AG1 | 1 | 27.9333 | 0.892 |
| 0.1 | 0.1 | ptsI | 1 | 28.1833 | 0.88 |
| 0.1 | 0.1 | clcB | 1 | 28.1833 | 0.447 |
| 0.1 | 0.1 | ycaM | 1 | 28.1833 | 0.415 |
| 0.1 | 0.1 | yadI | 1 | 28.1833 | 1.013 |
| 0.1 | 0.1 | AG1 | 1 | 28.1833 | 0.906 |
| 0.1 | 0.1 | ptsI | 1 | 28.4333 | 0.879 |
| 0.1 | 0.1 | clcB | 1 | 28.4333 | 0.449 |
| 0.1 | 0.1 | ycaM | 1 | 28.4333 | 0.416 |
| 0.1 | 0.1 | yadI | 1 | 28.4333 | 1.013 |
| 0.1 | 0.1 | AG1 | 1 | 28.4333 | 0.905 |
| 0.1 | 0.1 | ptsI | 1 | 28.6833 | 0.873 |
| 0.1 | 0.1 | clcB | 1 | 28.6833 | 0.448 |
| 0.1 | 0.1 | ycaM | 1 | 28.6833 | 0.416 |
| 0.1 | 0.1 | yadI | 1 | 28.6833 | 1.018 |
| 0.1 | 0.1 | AG1 | 1 | 28.6833 | 0.912 |
| 0.1 | 0.1 | ptsI | 1 | 28.9333 | 0.881 |
| 0.1 | 0.1 | clcB | 1 | 28.9333 | 0.449 |
| 0.1 | 0.1 | ycaM | 1 | 28.9333 | 0.417 |
| 0.1 | 0.1 | yadI | 1 | 28.9333 | 1.02 |
| 0.1 | 0.1 | AG1 | 1 | 28.9333 | 0.918 |
| 0.1 | 0.1 | ptsI | 1 | 29.1833 | 0.87 |
| 0.1 | 0.1 | clcB | 1 | 29.1833 | 0.45 |
| 0.1 | 0.1 | ycaM | 1 | 29.1833 | 0.416 |
| 0.1 | 0.1 | yadI | 1 | 29.1833 | 1.025 |
| 0.1 | 0.1 | AG1 | 1 | 29.1833 | 0.914 |
| 0.1 | 0.1 | ptsI | 1 | 29.4333 | 0.878 |
| 0.1 | 0.1 | clcB | 1 | 29.4333 | 0.451 |
| 0.1 | 0.1 | ycaM | 1 | 29.4333 | 0.418 |
| 0.1 | 0.1 | yadI | 1 | 29.4333 | 1.026 |
| 0.1 | 0.1 | AG1 | 1 | 29.4333 | 0.919 |

| | | | | | |
|---|---|---|---|---|---|
| 0.1 | 0.1 | ptsI | 1 | 29.6833 | 0.876 |
| 0.1 | 0.1 | clcB | 1 | 29.6833 | 0.452 |
| 0.1 | 0.1 | ycaM | 1 | 29.6833 | 0.42 |
| 0.1 | 0.1 | yadI | 1 | 29.6833 | 1.029 |
| 0.1 | 0.1 | AG1 | 1 | 29.6833 | 0.924 |
| 0.1 | 0.1 | ptsI | 1 | 29.9333 | 0.88 |
| 0.1 | 0.1 | clcB | 1 | 29.9333 | 0.453 |
| 0.1 | 0.1 | ycaM | 1 | 29.9333 | 0.418 |
| 0.1 | 0.1 | yadI | 1 | 29.9333 | 1.028 |
| 0.1 | 0.1 | AG1 | 1 | 29.9333 | 0.931 |
| 0.1 | 0.1 | ptsI | 1 | 30.1833 | 0.884 |
| 0.1 | 0.1 | clcB | 1 | 30.1833 | 0.454 |
| 0.1 | 0.1 | ycaM | 1 | 30.1833 | 0.418 |
| 0.1 | 0.1 | yadI | 1 | 30.1833 | 1.03 |
| 0.1 | 0.1 | AG1 | 1 | 30.1833 | 0.938 |
| 0.1 | 0.1 | ptsI | 1 | 30.4333 | 0.887 |
| 0.1 | 0.1 | clcB | 1 | 30.4333 | 0.451 |
| 0.1 | 0.1 | ycaM | 1 | 30.4333 | 0.419 |
| 0.1 | 0.1 | yadI | 1 | 30.4333 | 1.031 |
| 0.1 | 0.1 | AG1 | 1 | 30.4333 | 0.944 |
| 0.1 | 0.1 | ptsI | 1 | 30.6833 | 0.877 |
| 0.1 | 0.1 | clcB | 1 | 30.6833 | 0.456 |
| 0.1 | 0.1 | ycaM | 1 | 30.6833 | 0.422 |
| 0.1 | 0.1 | yadI | 1 | 30.6833 | 1.041 |
| 0.1 | 0.1 | AG1 | 1 | 30.6833 | 0.943 |
| 0.1 | 0.1 | ptsI | 1 | 30.9333 | 0.884 |
| 0.1 | 0.1 | clcB | 1 | 30.9333 | 0.458 |
| 0.1 | 0.1 | ycaM | 1 | 30.9333 | 0.422 |
| 0.1 | 0.1 | yadI | 1 | 30.9333 | 1.042 |
| 0.1 | 0.1 | AG1 | 1 | 30.9333 | 0.951 |
| 0.1 | 0.1 | ptsI | 1 | 31.1833 | 0.88 |
| 0.1 | 0.1 | clcB | 1 | 31.1833 | 0.454 |
| 0.1 | 0.1 | ycaM | 1 | 31.1833 | 0.423 |
| 0.1 | 0.1 | yadI | 1 | 31.1833 | 1.042 |
| 0.1 | 0.1 | AG1 | 1 | 31.1833 | 0.952 |
| 0.1 | 0.1 | ptsI | 1 | 31.4333 | 0.888 |
| 0.1 | 0.1 | clcB | 1 | 31.4333 | 0.454 |
| 0.1 | 0.1 | ycaM | 1 | 31.4333 | 0.426 |
| 0.1 | 0.1 | yadI | 1 | 31.4333 | 1.044 |
| 0.1 | 0.1 | AG1 | 1 | 31.4333 | 0.962 |
| 0.1 | 0.1 | ptsI | 1 | 31.6833 | 0.886 |
| 0.1 | 0.1 | clcB | 1 | 31.6833 | 0.456 |
| 0.1 | 0.1 | ycaM | 1 | 31.6833 | 0.427 |
| 0.1 | 0.1 | yadI | 1 | 31.6833 | 1.046 |
| 0.1 | 0.1 | AG1 | 1 | 31.6833 | 0.967 |
| 0.1 | 0.1 | ptsI | 1 | 31.9333 | 0.888 |
| 0.1 | 0.1 | clcB | 1 | 31.9333 | 0.459 |
| 0.1 | 0.1 | ycaM | 1 | 31.9333 | 0.423 |

| | | | | | |
|---|---|---|---|---|---|
| 0.1 | 0.1 | yadI | 1 | 31.9333 | 1.049 |
| 0.1 | 0.1 | AG1 | 1 | 31.9333 | 0.972 |
| 0.1 | 0.1 | ptsI | 1 | 32.1833 | 0.891 |
| 0.1 | 0.1 | clcB | 1 | 32.1833 | 0.456 |
| 0.1 | 0.1 | ycaM | 1 | 32.1833 | 0.428 |
| 0.1 | 0.1 | yadI | 1 | 32.1833 | 1.048 |
| 0.1 | 0.1 | AG1 | 1 | 32.1833 | 0.976 |
| 0.1 | 0.1 | ptsI | 1 | 32.4333 | 0.89 |
| 0.1 | 0.1 | clcB | 1 | 32.4333 | 0.451 |
| 0.1 | 0.1 | ycaM | 1 | 32.4333 | 0.417 |
| 0.1 | 0.1 | yadI | 1 | 32.4333 | 1.032 |
| 0.1 | 0.1 | AG1 | 1 | 32.4333 | 0.982 |
| 0.1 | 0.1 | ptsI | 1 | 32.6833 | 0.89 |
| 0.1 | 0.1 | clcB | 1 | 32.6833 | 0.459 |
| 0.1 | 0.1 | ycaM | 1 | 32.6833 | 0.432 |
| 0.1 | 0.1 | yadI | 1 | 32.6833 | 1.055 |
| 0.1 | 0.1 | AG1 | 1 | 32.6833 | 0.987 |
| 0.1 | 0.1 | ptsI | 1 | 32.9333 | 0.886 |
| 0.1 | 0.1 | clcB | 1 | 32.9333 | 0.455 |
| 0.1 | 0.1 | ycaM | 1 | 32.9333 | 0.427 |
| 0.1 | 0.1 | yadI | 1 | 32.9333 | 1.057 |
| 0.1 | 0.1 | AG1 | 1 | 32.9333 | 0.995 |
| 0.1 | 0.1 | ptsI | 1 | 33.1833 | 0.886 |
| 0.1 | 0.1 | clcB | 1 | 33.1833 | 0.455 |
| 0.1 | 0.1 | ycaM | 1 | 33.1833 | 0.422 |
| 0.1 | 0.1 | yadI | 1 | 33.1833 | 1.049 |
| 0.1 | 0.1 | AG1 | 1 | 33.1833 | 1.001 |
| 0.1 | 0.1 | ptsI | 1 | 33.4333 | 0.891 |
| 0.1 | 0.1 | clcB | 1 | 33.4333 | 0.46 |
| 0.1 | 0.1 | ycaM | 1 | 33.4333 | 0.43 |
| 0.1 | 0.1 | yadI | 1 | 33.4333 | 1.06 |
| 0.1 | 0.1 | AG1 | 1 | 33.4333 | 1.013 |
| 0.1 | 0.1 | ptsI | 1 | 33.6833 | 0.886 |
| 0.1 | 0.1 | clcB | 1 | 33.6833 | 0.454 |
| 0.1 | 0.1 | ycaM | 1 | 33.6833 | 0.426 |
| 0.1 | 0.1 | yadI | 1 | 33.6833 | 1.046 |
| 0.1 | 0.1 | AG1 | 1 | 33.6833 | 1.016 |
| 0.1 | 0.1 | ptsI | 1 | 33.9333 | 0.888 |
| 0.1 | 0.1 | clcB | 1 | 33.9333 | 0.461 |
| 0.1 | 0.1 | ycaM | 1 | 33.9333 | 0.432 |
| 0.1 | 0.1 | yadI | 1 | 33.9333 | 1.063 |
| 0.1 | 0.1 | AG1 | 1 | 33.9333 | 1.026 |
| 0.1 | 0.1 | ptsI | 1 | 34.1833 | 0.881 |
| 0.1 | 0.1 | clcB | 1 | 34.1833 | 0.467 |
| 0.1 | 0.1 | ycaM | 1 | 34.1833 | 0.432 |
| 0.1 | 0.1 | yadI | 1 | 34.1833 | 1.069 |
| 0.1 | 0.1 | AG1 | 1 | 34.1833 | 1.028 |
| 0.1 | 0.1 | ptsI | 1 | 34.4333 | 0.891 |

| | | | | | |
|---|---|---|---|---|---|
| 0.1 | 0.1 | clcB | 1 | 34.4333 | 0.465 |
| 0.1 | 0.1 | ycaM | 1 | 34.4333 | 0.433 |
| 0.1 | 0.1 | yadI | 1 | 34.4333 | 1.07 |
| 0.1 | 0.1 | AG1 | 1 | 34.4333 | 1.042 |
| 0.1 | 0.1 | ptsI | 1 | 34.6833 | 0.892 |
| 0.1 | 0.1 | clcB | 1 | 34.6833 | 0.466 |
| 0.1 | 0.1 | ycaM | 1 | 34.6833 | 0.434 |
| 0.1 | 0.1 | yadI | 1 | 34.6833 | 1.066 |
| 0.1 | 0.1 | AG1 | 1 | 34.6833 | 1.054 |
| 0.1 | 0.1 | ptsI | 1 | 34.9333 | 0.888 |
| 0.1 | 0.1 | clcB | 1 | 34.9333 | 0.47 |
| 0.1 | 0.1 | ycaM | 1 | 34.9333 | 0.433 |
| 0.1 | 0.1 | yadI | 1 | 34.9333 | 1.065 |
| 0.1 | 0.1 | AG1 | 1 | 34.9333 | 1.056 |
| 0.1 | 0.1 | ptsI | 1 | 35.1833 | 0.886 |
| 0.1 | 0.1 | clcB | 1 | 35.1833 | 0.467 |
| 0.1 | 0.1 | ycaM | 1 | 35.1833 | 0.435 |
| 0.1 | 0.1 | yadI | 1 | 35.1833 | 1.067 |
| 0.1 | 0.1 | AG1 | 1 | 35.1833 | 1.062 |
| 0.1 | 0.1 | ptsI | 1 | 35.4333 | 0.888 |
| 0.1 | 0.1 | clcB | 1 | 35.4333 | 0.466 |
| 0.1 | 0.1 | ycaM | 1 | 35.4333 | 0.435 |
| 0.1 | 0.1 | yadI | 1 | 35.4333 | 1.068 |
| 0.1 | 0.1 | AG1 | 1 | 35.4333 | 1.072 |
| 0.1 | 0.1 | ptsI | 1 | 35.6833 | 0.888 |
| 0.1 | 0.1 | clcB | 1 | 35.6833 | 0.472 |
| 0.1 | 0.1 | ycaM | 1 | 35.6833 | 0.437 |
| 0.1 | 0.1 | yadI | 1 | 35.6833 | 1.069 |
| 0.1 | 0.1 | AG1 | 1 | 35.6833 | 1.076 |
| 0.1 | 0.1 | ptsI | 1 | 35.9333 | 0.883 |
| 0.1 | 0.1 | clcB | 1 | 35.9333 | 0.467 |
| 0.1 | 0.1 | ycaM | 1 | 35.9333 | 0.43 |
| 0.1 | 0.1 | yadI | 1 | 35.9333 | 1.061 |
| 0.1 | 0.1 | AG1 | 1 | 35.9333 | 1.083 |
| 0.1 | 0.1 | ptsI | 1 | 36.1833 | 0.889 |
| 0.1 | 0.1 | clcB | 1 | 36.1833 | 0.472 |
| 0.1 | 0.1 | ycaM | 1 | 36.1833 | 0.442 |
| 0.1 | 0.1 | yadI | 1 | 36.1833 | 1.073 |
| 0.1 | 0.1 | AG1 | 1 | 36.1833 | 1.092 |
| 0.1 | 0.1 | ptsI | 1 | 36.4333 | 0.888 |
| 0.1 | 0.1 | clcB | 1 | 36.4333 | 0.471 |
| 0.1 | 0.1 | ycaM | 1 | 36.4333 | 0.44 |
| 0.1 | 0.1 | yadI | 1 | 36.4333 | 1.073 |
| 0.1 | 0.1 | AG1 | 1 | 36.4333 | 1.099 |
| 0.1 | 0.1 | ptsI | 1 | 36.6833 | 0.885 |
| 0.1 | 0.1 | clcB | 1 | 36.6833 | 0.472 |
| 0.1 | 0.1 | ycaM | 1 | 36.6833 | 0.439 |
| 0.1 | 0.1 | yadI | 1 | 36.6833 | 1.076 |

| | | | | | |
|---|---|---|---|---|---|
| 0.1 | 0.1 | AG1 | 1 | 36.6833 | 1.103 |
| 0.2 | 0.1 | ptsI | 1 | 0 | 0.234 |
| 0.2 | 0.1 | clcB | 1 | 0 | 0.25 |
| 0.2 | 0.1 | ycaM | 1 | 0 | 0.232 |
| 0.2 | 0.1 | yadI | 1 | 0 | 0.28 |
| 0.2 | 0.1 | AG1 | 1 | 0 | 0.259 |
| 0.2 | 0.1 | ptsI | 1 | 0.25 | 0.224 |
| 0.2 | 0.1 | clcB | 1 | 0.25 | 0.238 |
| 0.2 | 0.1 | ycaM | 1 | 0.25 | 0.216 |
| 0.2 | 0.1 | yadI | 1 | 0.25 | 0.265 |
| 0.2 | 0.1 | AG1 | 1 | 0.25 | 0.245 |
| 0.2 | 0.1 | ptsI | 1 | 0.5 | 0.224 |
| 0.2 | 0.1 | clcB | 1 | 0.5 | 0.232 |
| 0.2 | 0.1 | ycaM | 1 | 0.5 | 0.213 |
| 0.2 | 0.1 | yadI | 1 | 0.5 | 0.262 |
| 0.2 | 0.1 | AG1 | 1 | 0.5 | 0.238 |
| 0.2 | 0.1 | ptsI | 1 | 0.75 | 0.223 |
| 0.2 | 0.1 | clcB | 1 | 0.75 | 0.231 |
| 0.2 | 0.1 | ycaM | 1 | 0.75 | 0.211 |
| 0.2 | 0.1 | yadI | 1 | 0.75 | 0.261 |
| 0.2 | 0.1 | AG1 | 1 | 0.75 | 0.235 |
| 0.2 | 0.1 | ptsI | 1 | 1 | 0.223 |
| 0.2 | 0.1 | clcB | 1 | 1 | 0.231 |
| 0.2 | 0.1 | ycaM | 1 | 1 | 0.213 |
| 0.2 | 0.1 | yadI | 1 | 1 | 0.261 |
| 0.2 | 0.1 | AG1 | 1 | 1 | 0.238 |
| 0.2 | 0.1 | ptsI | 1 | 1.25 | 0.225 |
| 0.2 | 0.1 | clcB | 1 | 1.25 | 0.233 |
| 0.2 | 0.1 | ycaM | 1 | 1.25 | 0.21 |
| 0.2 | 0.1 | yadI | 1 | 1.25 | 0.258 |
| 0.2 | 0.1 | AG1 | 1 | 1.25 | 0.237 |
| 0.2 | 0.1 | ptsI | 1 | 1.5 | 0.229 |
| 0.2 | 0.1 | clcB | 1 | 1.5 | 0.234 |
| 0.2 | 0.1 | ycaM | 1 | 1.5 | 0.21 |
| 0.2 | 0.1 | yadI | 1 | 1.5 | 0.26 |
| 0.2 | 0.1 | AG1 | 1 | 1.5 | 0.239 |
| 0.2 | 0.1 | ptsI | 1 | 1.75 | 0.233 |
| 0.2 | 0.1 | clcB | 1 | 1.75 | 0.235 |
| 0.2 | 0.1 | ycaM | 1 | 1.75 | 0.214 |
| 0.2 | 0.1 | yadI | 1 | 1.75 | 0.262 |
| 0.2 | 0.1 | AG1 | 1 | 1.75 | 0.245 |
| 0.2 | 0.1 | ptsI | 1 | 2 | 0.242 |
| 0.2 | 0.1 | clcB | 1 | 2 | 0.237 |
| 0.2 | 0.1 | ycaM | 1 | 2 | 0.217 |
| 0.2 | 0.1 | yadI | 1 | 2 | 0.268 |
| 0.2 | 0.1 | AG1 | 1 | 2 | 0.249 |
| 0.2 | 0.1 | ptsI | 1 | 2.25 | 0.247 |
| 0.2 | 0.1 | clcB | 1 | 2.25 | 0.24 |

| | | | | | |
|---|---|---|---|---|---|
| 0.2 | 0.1 | ycaM | 1 | 2.25 | 0.221 |
| 0.2 | 0.1 | yadI | 1 | 2.25 | 0.272 |
| 0.2 | 0.1 | AG1 | 1 | 2.25 | 0.255 |
| 0.2 | 0.1 | ptsI | 1 | 2.5 | 0.258 |
| 0.2 | 0.1 | clcB | 1 | 2.5 | 0.248 |
| 0.2 | 0.1 | ycaM | 1 | 2.5 | 0.226 |
| 0.2 | 0.1 | yadI | 1 | 2.5 | 0.28 |
| 0.2 | 0.1 | AG1 | 1 | 2.5 | 0.258 |
| 0.2 | 0.1 | ptsI | 1 | 2.75 | 0.263 |
| 0.2 | 0.1 | clcB | 1 | 2.75 | 0.255 |
| 0.2 | 0.1 | ycaM | 1 | 2.75 | 0.233 |
| 0.2 | 0.1 | yadI | 1 | 2.75 | 0.289 |
| 0.2 | 0.1 | AG1 | 1 | 2.75 | 0.263 |
| 0.2 | 0.1 | ptsI | 1 | 3 | 0.278 |
| 0.2 | 0.1 | clcB | 1 | 3 | 0.259 |
| 0.2 | 0.1 | ycaM | 1 | 3 | 0.236 |
| 0.2 | 0.1 | yadI | 1 | 3 | 0.301 |
| 0.2 | 0.1 | AG1 | 1 | 3 | 0.267 |
| 0.2 | 0.1 | ptsI | 1 | 3.25 | 0.291 |
| 0.2 | 0.1 | clcB | 1 | 3.25 | 0.265 |
| 0.2 | 0.1 | ycaM | 1 | 3.25 | 0.243 |
| 0.2 | 0.1 | yadI | 1 | 3.25 | 0.31 |
| 0.2 | 0.1 | AG1 | 1 | 3.25 | 0.275 |
| 0.2 | 0.1 | ptsI | 1 | 3.5 | 0.308 |
| 0.2 | 0.1 | clcB | 1 | 3.5 | 0.27 |
| 0.2 | 0.1 | ycaM | 1 | 3.5 | 0.248 |
| 0.2 | 0.1 | yadI | 1 | 3.5 | 0.321 |
| 0.2 | 0.1 | AG1 | 1 | 3.5 | 0.28 |
| 0.2 | 0.1 | ptsI | 1 | 3.75 | 0.327 |
| 0.2 | 0.1 | clcB | 1 | 3.75 | 0.278 |
| 0.2 | 0.1 | ycaM | 1 | 3.75 | 0.257 |
| 0.2 | 0.1 | yadI | 1 | 3.75 | 0.33 |
| 0.2 | 0.1 | AG1 | 1 | 3.75 | 0.288 |
| 0.2 | 0.1 | ptsI | 1 | 4 | 0.354 |
| 0.2 | 0.1 | clcB | 1 | 4 | 0.288 |
| 0.2 | 0.1 | ycaM | 1 | 4 | 0.272 |
| 0.2 | 0.1 | yadI | 1 | 4 | 0.357 |
| 0.2 | 0.1 | AG1 | 1 | 4 | 0.299 |
| 0.2 | 0.1 | ptsI | 1 | 4.25 | 0.363 |
| 0.2 | 0.1 | clcB | 1 | 4.25 | 0.292 |
| 0.2 | 0.1 | ycaM | 1 | 4.25 | 0.268 |
| 0.2 | 0.1 | yadI | 1 | 4.25 | 0.352 |
| 0.2 | 0.1 | AG1 | 1 | 4.25 | 0.305 |
| 0.2 | 0.1 | ptsI | 1 | 4.5 | 0.375 |
| 0.2 | 0.1 | clcB | 1 | 4.5 | 0.3 |
| 0.2 | 0.1 | ycaM | 1 | 4.5 | 0.275 |
| 0.2 | 0.1 | yadI | 1 | 4.5 | 0.364 |
| 0.2 | 0.1 | AG1 | 1 | 4.5 | 0.31 |

| | | | | | |
|---|---|---|---|---|---|
| 0.2 | 0.1 | ptsI | 1 | 4.75 | 0.38 |
| 0.2 | 0.1 | clcB | 1 | 4.75 | 0.31 |
| 0.2 | 0.1 | ycaM | 1 | 4.75 | 0.28 |
| 0.2 | 0.1 | yadI | 1 | 4.75 | 0.375 |
| 0.2 | 0.1 | AG1 | 1 | 4.75 | 0.32 |
| 0.2 | 0.1 | ptsI | 1 | 5 | 0.402 |
| 0.2 | 0.1 | clcB | 1 | 5 | 0.317 |
| 0.2 | 0.1 | ycaM | 1 | 5 | 0.289 |
| 0.2 | 0.1 | yadI | 1 | 5 | 0.385 |
| 0.2 | 0.1 | AG1 | 1 | 5 | 0.328 |
| 0.2 | 0.1 | ptsI | 1 | 5.25 | 0.411 |
| 0.2 | 0.1 | clcB | 1 | 5.25 | 0.326 |
| 0.2 | 0.1 | ycaM | 1 | 5.25 | 0.292 |
| 0.2 | 0.1 | yadI | 1 | 5.25 | 0.396 |
| 0.2 | 0.1 | AG1 | 1 | 5.25 | 0.333 |
| 0.2 | 0.1 | ptsI | 1 | 5.5 | 0.42 |
| 0.2 | 0.1 | clcB | 1 | 5.5 | 0.328 |
| 0.2 | 0.1 | ycaM | 1 | 5.5 | 0.298 |
| 0.2 | 0.1 | yadI | 1 | 5.5 | 0.407 |
| 0.2 | 0.1 | AG1 | 1 | 5.5 | 0.341 |
| 0.2 | 0.1 | ptsI | 1 | 5.75 | 0.43 |
| 0.2 | 0.1 | clcB | 1 | 5.75 | 0.335 |
| 0.2 | 0.1 | ycaM | 1 | 5.75 | 0.309 |
| 0.2 | 0.1 | yadI | 1 | 5.75 | 0.416 |
| 0.2 | 0.1 | AG1 | 1 | 5.75 | 0.35 |
| 0.2 | 0.1 | ptsI | 1 | 6 | 0.418 |
| 0.2 | 0.1 | clcB | 1 | 6 | 0.346 |
| 0.2 | 0.1 | ycaM | 1 | 6 | 0.31 |
| 0.2 | 0.1 | yadI | 1 | 6 | 0.427 |
| 0.2 | 0.1 | AG1 | 1 | 6 | 0.352 |
| 0.2 | 0.1 | ptsI | 1 | 6.25 | 0.425 |
| 0.2 | 0.1 | clcB | 1 | 6.25 | 0.352 |
| 0.2 | 0.1 | ycaM | 1 | 6.25 | 0.317 |
| 0.2 | 0.1 | yadI | 1 | 6.25 | 0.438 |
| 0.2 | 0.1 | AG1 | 1 | 6.25 | 0.353 |
| 0.2 | 0.1 | ptsI | 1 | 6.5 | 0.435 |
| 0.2 | 0.1 | clcB | 1 | 6.5 | 0.365 |
| 0.2 | 0.1 | ycaM | 1 | 6.5 | 0.32 |
| 0.2 | 0.1 | yadI | 1 | 6.5 | 0.45 |
| 0.2 | 0.1 | AG1 | 1 | 6.5 | 0.367 |
| 0.2 | 0.1 | ptsI | 1 | 6.75 | 0.456 |
| 0.2 | 0.1 | clcB | 1 | 6.75 | 0.369 |
| 0.2 | 0.1 | ycaM | 1 | 6.75 | 0.328 |
| 0.2 | 0.1 | yadI | 1 | 6.75 | 0.459 |
| 0.2 | 0.1 | AG1 | 1 | 6.75 | 0.378 |
| 0.2 | 0.1 | ptsI | 1 | 7 | 0.474 |
| 0.2 | 0.1 | clcB | 1 | 7 | 0.375 |
| 0.2 | 0.1 | ycaM | 1 | 7 | 0.334 |

| | | | | | |
|---|---|---|---|---|---|
| 0.2 | 0.1 | yadI | 1 | 7 | 0.468 |
| 0.2 | 0.1 | AG1 | 1 | 7 | 0.388 |
| 0.2 | 0.1 | ptsI | 1 | 7.25 | 0.491 |
| 0.2 | 0.1 | clcB | 1 | 7.25 | 0.384 |
| 0.2 | 0.1 | ycaM | 1 | 7.25 | 0.341 |
| 0.2 | 0.1 | yadI | 1 | 7.25 | 0.474 |
| 0.2 | 0.1 | AG1 | 1 | 7.25 | 0.398 |
| 0.2 | 0.1 | ptsI | 1 | 7.5 | 0.507 |
| 0.2 | 0.1 | clcB | 1 | 7.5 | 0.389 |
| 0.2 | 0.1 | ycaM | 1 | 7.5 | 0.347 |
| 0.2 | 0.1 | yadI | 1 | 7.5 | 0.487 |
| 0.2 | 0.1 | AG1 | 1 | 7.5 | 0.408 |
| 0.2 | 0.1 | ptsI | 1 | 7.75 | 0.51 |
| 0.2 | 0.1 | clcB | 1 | 7.75 | 0.403 |
| 0.2 | 0.1 | ycaM | 1 | 7.75 | 0.352 |
| 0.2 | 0.1 | yadI | 1 | 7.75 | 0.499 |
| 0.2 | 0.1 | AG1 | 1 | 7.75 | 0.422 |
| 0.2 | 0.1 | ptsI | 1 | 8 | 0.524 |
| 0.2 | 0.1 | clcB | 1 | 8 | 0.405 |
| 0.2 | 0.1 | ycaM | 1 | 8 | 0.355 |
| 0.2 | 0.1 | yadI | 1 | 8 | 0.506 |
| 0.2 | 0.1 | AG1 | 1 | 8 | 0.428 |
| 0.2 | 0.1 | ptsI | 1 | 8.25 | 0.534 |
| 0.2 | 0.1 | clcB | 1 | 8.25 | 0.411 |
| 0.2 | 0.1 | ycaM | 1 | 8.25 | 0.359 |
| 0.2 | 0.1 | yadI | 1 | 8.25 | 0.511 |
| 0.2 | 0.1 | AG1 | 1 | 8.25 | 0.439 |
| 0.2 | 0.1 | ptsI | 1 | 8.5 | 0.538 |
| 0.2 | 0.1 | clcB | 1 | 8.5 | 0.419 |
| 0.2 | 0.1 | ycaM | 1 | 8.5 | 0.359 |
| 0.2 | 0.1 | yadI | 1 | 8.5 | 0.518 |
| 0.2 | 0.1 | AG1 | 1 | 8.5 | 0.45 |
| 0.2 | 0.1 | ptsI | 1 | 8.75 | 0.536 |
| 0.2 | 0.1 | clcB | 1 | 8.75 | 0.424 |
| 0.2 | 0.1 | ycaM | 1 | 8.75 | 0.361 |
| 0.2 | 0.1 | yadI | 1 | 8.75 | 0.526 |
| 0.2 | 0.1 | AG1 | 1 | 8.75 | 0.456 |
| 0.2 | 0.1 | ptsI | 1 | 9 | 0.539 |
| 0.2 | 0.1 | clcB | 1 | 9 | 0.428 |
| 0.2 | 0.1 | ycaM | 1 | 9 | 0.364 |
| 0.2 | 0.1 | yadI | 1 | 9 | 0.536 |
| 0.2 | 0.1 | AG1 | 1 | 9 | 0.47 |
| 0.2 | 0.1 | ptsI | 1 | 9.25 | 0.524 |
| 0.2 | 0.1 | clcB | 1 | 9.25 | 0.438 |
| 0.2 | 0.1 | ycaM | 1 | 9.25 | 0.371 |
| 0.2 | 0.1 | yadI | 1 | 9.25 | 0.537 |
| 0.2 | 0.1 | AG1 | 1 | 9.25 | 0.496 |
| 0.2 | 0.1 | ptsI | 1 | 9.5 | 0.566 |

| | | | | | |
|---|---|---|---|---|---|
| 0.2 | 0.1 | clcB | 1 | 9.5 | 0.449 |
| 0.2 | 0.1 | ycaM | 1 | 9.5 | 0.382 |
| 0.2 | 0.1 | yadI | 1 | 9.5 | 0.565 |
| 0.2 | 0.1 | AG1 | 1 | 9.5 | 0.579 |
| 0.2 | 0.1 | ptsI | 1 | 9.75 | 0.579 |
| 0.2 | 0.1 | clcB | 1 | 9.75 | 0.45 |
| 0.2 | 0.1 | ycaM | 1 | 9.75 | 0.388 |
| 0.2 | 0.1 | yadI | 1 | 9.75 | 0.57 |
| 0.2 | 0.1 | AG1 | 1 | 9.75 | 0.596 |
| 0.2 | 0.1 | ptsI | 1 | 10 | 0.54 |
| 0.2 | 0.1 | clcB | 1 | 10 | 0.454 |
| 0.2 | 0.1 | ycaM | 1 | 10 | 0.391 |
| 0.2 | 0.1 | yadI | 1 | 10 | 0.566 |
| 0.2 | 0.1 | AG1 | 1 | 10 | 0.607 |
| 0.2 | 0.1 | ptsI | 1 | 10.25 | 0.591 |
| 0.2 | 0.1 | clcB | 1 | 10.25 | 0.462 |
| 0.2 | 0.1 | ycaM | 1 | 10.25 | 0.389 |
| 0.2 | 0.1 | yadI | 1 | 10.25 | 0.57 |
| 0.2 | 0.1 | AG1 | 1 | 10.25 | 0.625 |
| 0.2 | 0.1 | ptsI | 1 | 10.5 | 0.631 |
| 0.2 | 0.1 | clcB | 1 | 10.5 | 0.477 |
| 0.2 | 0.1 | ycaM | 1 | 10.5 | 0.398 |
| 0.2 | 0.1 | yadI | 1 | 10.5 | 0.58 |
| 0.2 | 0.1 | AG1 | 1 | 10.5 | 0.641 |
| 0.2 | 0.1 | ptsI | 1 | 10.75 | 0.597 |
| 0.2 | 0.1 | clcB | 1 | 10.75 | 0.482 |
| 0.2 | 0.1 | ycaM | 1 | 10.75 | 0.402 |
| 0.2 | 0.1 | yadI | 1 | 10.75 | 0.572 |
| 0.2 | 0.1 | AG1 | 1 | 10.75 | 0.572 |
| 0.2 | 0.1 | ptsI | 1 | 11 | 0.617 |
| 0.2 | 0.1 | clcB | 1 | 11 | 0.509 |
| 0.2 | 0.1 | ycaM | 1 | 11 | 0.415 |
| 0.2 | 0.1 | yadI | 1 | 11 | 0.6 |
| 0.2 | 0.1 | AG1 | 1 | 11 | 0.661 |
| 0.2 | 0.1 | ptsI | 1 | 11.25 | 0.663 |
| 0.2 | 0.1 | clcB | 1 | 11.25 | 0.514 |
| 0.2 | 0.1 | ycaM | 1 | 11.25 | 0.421 |
| 0.2 | 0.1 | yadI | 1 | 11.25 | 0.607 |
| 0.2 | 0.1 | AG1 | 1 | 11.25 | 0.69 |
| 0.2 | 0.1 | ptsI | 1 | 11.5 | 0.632 |
| 0.2 | 0.1 | clcB | 1 | 11.5 | 0.516 |
| 0.2 | 0.1 | ycaM | 1 | 11.5 | 0.426 |
| 0.2 | 0.1 | yadI | 1 | 11.5 | 0.616 |
| 0.2 | 0.1 | AG1 | 1 | 11.5 | 0.734 |
| 0.2 | 0.1 | ptsI | 1 | 11.75 | 0.638 |
| 0.2 | 0.1 | clcB | 1 | 11.75 | 0.533 |
| 0.2 | 0.1 | ycaM | 1 | 11.75 | 0.437 |
| 0.2 | 0.1 | yadI | 1 | 11.75 | 0.616 |

| | | | | | |
|---|---|---|---|---|---|
| 0.2 | 0.1 | AG1 | 1 | 11.75 | 0.713 |
| 0.2 | 0.1 | ptsI | 1 | 12 | 0.678 |
| 0.2 | 0.1 | clcB | 1 | 12 | 0.553 |
| 0.2 | 0.1 | ycaM | 1 | 12 | 0.445 |
| 0.2 | 0.1 | yadI | 1 | 12 | 0.622 |
| 0.2 | 0.1 | AG1 | 1 | 12 | 0.734 |
| 0.2 | 0.1 | ptsI | 1 | 12.25 | 0.641 |
| 0.2 | 0.1 | clcB | 1 | 12.25 | 0.551 |
| 0.2 | 0.1 | ycaM | 1 | 12.25 | 0.447 |
| 0.2 | 0.1 | yadI | 1 | 12.25 | 0.589 |
| 0.2 | 0.1 | AG1 | 1 | 12.25 | 0.697 |
| 0.2 | 0.1 | ptsI | 1 | 12.5 | 0.68 |
| 0.2 | 0.1 | clcB | 1 | 12.5 | 0.568 |
| 0.2 | 0.1 | ycaM | 1 | 12.5 | 0.448 |
| 0.2 | 0.1 | yadI | 1 | 12.5 | 0.585 |
| 0.2 | 0.1 | AG1 | 1 | 12.5 | 0.745 |
| 0.2 | 0.1 | ptsI | 1 | 12.75 | 0.694 |
| 0.2 | 0.1 | clcB | 1 | 12.75 | 0.596 |
| 0.2 | 0.1 | ycaM | 1 | 12.75 | 0.454 |
| 0.2 | 0.1 | yadI | 1 | 12.75 | 0.583 |
| 0.2 | 0.1 | AG1 | 1 | 12.75 | 0.767 |
| 0.2 | 0.1 | ptsI | 1 | 13 | 0.7 |
| 0.2 | 0.1 | clcB | 1 | 13 | 0.62 |
| 0.2 | 0.1 | ycaM | 1 | 13 | 0.462 |
| 0.2 | 0.1 | yadI | 1 | 13 | 0.588 |
| 0.2 | 0.1 | AG1 | 1 | 13 | 0.792 |
| 0.2 | 0.1 | ptsI | 1 | 13.25 | 0.687 |
| 0.2 | 0.1 | clcB | 1 | 13.25 | 0.628 |
| 0.2 | 0.1 | ycaM | 1 | 13.25 | 0.47 |
| 0.2 | 0.1 | yadI | 1 | 13.25 | 0.583 |
| 0.2 | 0.1 | AG1 | 1 | 13.25 | 0.798 |
| 0.2 | 0.1 | ptsI | 1 | 13.5 | 0.718 |
| 0.2 | 0.1 | clcB | 1 | 13.5 | 0.631 |
| 0.2 | 0.1 | ycaM | 1 | 13.5 | 0.487 |
| 0.2 | 0.1 | yadI | 1 | 13.5 | 0.595 |
| 0.2 | 0.1 | AG1 | 1 | 13.5 | 0.823 |
| 0.2 | 0.1 | ptsI | 1 | 13.75 | 0.681 |
| 0.2 | 0.1 | clcB | 1 | 13.75 | 0.616 |
| 0.2 | 0.1 | ycaM | 1 | 13.75 | 0.501 |
| 0.2 | 0.1 | yadI | 1 | 13.75 | 0.598 |
| 0.2 | 0.1 | AG1 | 1 | 13.75 | 0.816 |
| 0.2 | 0.1 | ptsI | 1 | 14 | 0.736 |
| 0.2 | 0.1 | clcB | 1 | 14 | 0.627 |
| 0.2 | 0.1 | ycaM | 1 | 14 | 0.53 |
| 0.2 | 0.1 | yadI | 1 | 14 | 0.617 |
| 0.2 | 0.1 | AG1 | 1 | 14 | 0.864 |
| 0.2 | 0.1 | ptsI | 1 | 14.25 | 0.703 |
| 0.2 | 0.1 | clcB | 1 | 14.25 | 0.669 |

| | | | | | |
|---|---|---|---|---|---|
| 0.2 | 0.1 | ycaM | 1 | 14.25 | 0.556 |
| 0.2 | 0.1 | yadI | 1 | 14.25 | 0.615 |
| 0.2 | 0.1 | AG1 | 1 | 14.25 | 0.868 |
| 0.2 | 0.1 | ptsI | 1 | 14.5 | 0.679 |
| 0.2 | 0.1 | clcB | 1 | 14.5 | 0.673 |
| 0.2 | 0.1 | ycaM | 1 | 14.5 | 0.58 |
| 0.2 | 0.1 | yadI | 1 | 14.5 | 0.619 |
| 0.2 | 0.1 | AG1 | 1 | 14.5 | 0.872 |
| 0.2 | 0.1 | ptsI | 1 | 14.75 | 0.698 |
| 0.2 | 0.1 | clcB | 1 | 14.75 | 0.68 |
| 0.2 | 0.1 | ycaM | 1 | 14.75 | 0.645 |
| 0.2 | 0.1 | yadI | 1 | 14.75 | 0.613 |
| 0.2 | 0.1 | AG1 | 1 | 14.75 | 0.902 |
| 0.2 | 0.1 | ptsI | 1 | 15 | 0.754 |
| 0.2 | 0.1 | clcB | 1 | 15 | 0.697 |
| 0.2 | 0.1 | ycaM | 1 | 15 | 0.586 |
| 0.2 | 0.1 | yadI | 1 | 15 | 0.616 |
| 0.2 | 0.1 | AG1 | 1 | 15 | 0.885 |
| 0.2 | 0.1 | ptsI | 1 | 15.25 | 0.763 |
| 0.2 | 0.1 | clcB | 1 | 15.25 | 0.712 |
| 0.2 | 0.1 | ycaM | 1 | 15.25 | 0.598 |
| 0.2 | 0.1 | yadI | 1 | 15.25 | 0.627 |
| 0.2 | 0.1 | AG1 | 1 | 15.25 | 0.904 |
| 0.2 | 0.1 | ptsI | 1 | 15.5 | 0.758 |
| 0.2 | 0.1 | clcB | 1 | 15.5 | 0.7 |
| 0.2 | 0.1 | ycaM | 1 | 15.5 | 0.569 |
| 0.2 | 0.1 | yadI | 1 | 15.5 | 0.629 |
| 0.2 | 0.1 | AG1 | 1 | 15.5 | 0.896 |
| 0.2 | 0.1 | ptsI | 1 | 15.75 | 0.741 |
| 0.2 | 0.1 | clcB | 1 | 15.75 | 0.694 |
| 0.2 | 0.1 | ycaM | 1 | 15.75 | 0.578 |
| 0.2 | 0.1 | yadI | 1 | 15.75 | 0.626 |
| 0.2 | 0.1 | AG1 | 1 | 15.75 | 0.922 |
| 0.2 | 0.1 | ptsI | 1 | 16 | 0.74 |
| 0.2 | 0.1 | clcB | 1 | 16 | 0.693 |
| 0.2 | 0.1 | ycaM | 1 | 16 | 0.574 |
| 0.2 | 0.1 | yadI | 1 | 16 | 0.607 |
| 0.2 | 0.1 | AG1 | 1 | 16 | 0.922 |
| 0.2 | 0.1 | ptsI | 1 | 16.25 | 0.798 |
| 0.2 | 0.1 | clcB | 1 | 16.25 | 0.714 |
| 0.2 | 0.1 | ycaM | 1 | 16.25 | 0.556 |
| 0.2 | 0.1 | yadI | 1 | 16.25 | 0.618 |
| 0.2 | 0.1 | AG1 | 1 | 16.25 | 0.923 |
| 0.2 | 0.1 | ptsI | 1 | 16.5 | 0.794 |
| 0.2 | 0.1 | clcB | 1 | 16.5 | 0.693 |
| 0.2 | 0.1 | ycaM | 1 | 16.5 | 0.559 |
| 0.2 | 0.1 | yadI | 1 | 16.5 | 0.63 |
| 0.2 | 0.1 | AG1 | 1 | 16.5 | 0.923 |

| | | | | | |
|---|---|---|---|---|---|
| 0.2 | 0.1 | ptsI | 1 | 17.0667 | 0.588 |
| 0.2 | 0.1 | clcB | 1 | 17.0667 | 0.52 |
| 0.2 | 0.1 | ycaM | 1 | 17.0667 | 0.498 |
| 0.2 | 0.1 | yadI | 1 | 17.0667 | 0.749 |
| 0.2 | 0.1 | AG1 | 1 | 17.0667 | 0.787 |
| 0.2 | 0.1 | ptsI | 1 | 17.3167 | 0.642 |
| 0.2 | 0.1 | clcB | 1 | 17.3167 | 0.48 |
| 0.2 | 0.1 | ycaM | 1 | 17.3167 | 0.47 |
| 0.2 | 0.1 | yadI | 1 | 17.3167 | 0.729 |
| 0.2 | 0.1 | AG1 | 1 | 17.3167 | 0.746 |
| 0.2 | 0.1 | ptsI | 1 | 17.5667 | 0.653 |
| 0.2 | 0.1 | clcB | 1 | 17.5667 | 0.474 |
| 0.2 | 0.1 | ycaM | 1 | 17.5667 | 0.469 |
| 0.2 | 0.1 | yadI | 1 | 17.5667 | 0.722 |
| 0.2 | 0.1 | AG1 | 1 | 17.5667 | 0.757 |
| 0.2 | 0.1 | ptsI | 1 | 17.8167 | 0.726 |
| 0.2 | 0.1 | clcB | 1 | 17.8167 | 0.467 |
| 0.2 | 0.1 | ycaM | 1 | 17.8167 | 0.477 |
| 0.2 | 0.1 | yadI | 1 | 17.8167 | 0.732 |
| 0.2 | 0.1 | AG1 | 1 | 17.8167 | 0.814 |
| 0.2 | 0.1 | ptsI | 1 | 18.0667 | 0.66 |
| 0.2 | 0.1 | clcB | 1 | 18.0667 | 0.471 |
| 0.2 | 0.1 | ycaM | 1 | 18.0667 | 0.474 |
| 0.2 | 0.1 | yadI | 1 | 18.0667 | 0.754 |
| 0.2 | 0.1 | AG1 | 1 | 18.0667 | 0.785 |
| 0.2 | 0.1 | ptsI | 1 | 18.3167 | 0.685 |
| 0.2 | 0.1 | clcB | 1 | 18.3167 | 0.475 |
| 0.2 | 0.1 | ycaM | 1 | 18.3167 | 0.47 |
| 0.2 | 0.1 | yadI | 1 | 18.3167 | 0.745 |
| 0.2 | 0.1 | AG1 | 1 | 18.3167 | 0.783 |
| 0.2 | 0.1 | ptsI | 1 | 18.5667 | 0.748 |
| 0.2 | 0.1 | clcB | 1 | 18.5667 | 0.481 |
| 0.2 | 0.1 | ycaM | 1 | 18.5667 | 0.471 |
| 0.2 | 0.1 | yadI | 1 | 18.5667 | 0.759 |
| 0.2 | 0.1 | AG1 | 1 | 18.5667 | 0.821 |
| 0.2 | 0.1 | ptsI | 1 | 18.8167 | 0.767 |
| 0.2 | 0.1 | clcB | 1 | 18.8167 | 0.479 |
| 0.2 | 0.1 | ycaM | 1 | 18.8167 | 0.471 |
| 0.2 | 0.1 | yadI | 1 | 18.8167 | 0.772 |
| 0.2 | 0.1 | AG1 | 1 | 18.8167 | 0.826 |
| 0.2 | 0.1 | ptsI | 1 | 19.0667 | 0.787 |
| 0.2 | 0.1 | clcB | 1 | 19.0667 | 0.48 |
| 0.2 | 0.1 | ycaM | 1 | 19.0667 | 0.478 |
| 0.2 | 0.1 | yadI | 1 | 19.0667 | 0.782 |
| 0.2 | 0.1 | AG1 | 1 | 19.0667 | 0.851 |
| 0.2 | 0.1 | ptsI | 1 | 19.3167 | 0.77 |
| 0.2 | 0.1 | clcB | 1 | 19.3167 | 0.47 |
| 0.2 | 0.1 | ycaM | 1 | 19.3167 | 0.474 |

| | | | | | |
|---|---|---|---|---|---|
| 0.2 | 0.1 | yadI | 1 | 19.3167 | 0.781 |
| 0.2 | 0.1 | AG1 | 1 | 19.3167 | 0.835 |
| 0.2 | 0.1 | ptsI | 1 | 19.5667 | 0.803 |
| 0.2 | 0.1 | clcB | 1 | 19.5667 | 0.486 |
| 0.2 | 0.1 | ycaM | 1 | 19.5667 | 0.473 |
| 0.2 | 0.1 | yadI | 1 | 19.5667 | 0.787 |
| 0.2 | 0.1 | AG1 | 1 | 19.5667 | 0.858 |
| 0.2 | 0.1 | ptsI | 1 | 19.8167 | 0.802 |
| 0.2 | 0.1 | clcB | 1 | 19.8167 | 0.466 |
| 0.2 | 0.1 | ycaM | 1 | 19.8167 | 0.476 |
| 0.2 | 0.1 | yadI | 1 | 19.8167 | 0.79 |
| 0.2 | 0.1 | AG1 | 1 | 19.8167 | 0.866 |
| 0.2 | 0.1 | ptsI | 1 | 20.0667 | 0.8 |
| 0.2 | 0.1 | clcB | 1 | 20.0667 | 0.463 |
| 0.2 | 0.1 | ycaM | 1 | 20.0667 | 0.474 |
| 0.2 | 0.1 | yadI | 1 | 20.0667 | 0.796 |
| 0.2 | 0.1 | AG1 | 1 | 20.0667 | 0.871 |
| 0.2 | 0.1 | ptsI | 1 | 20.3167 | 0.807 |
| 0.2 | 0.1 | clcB | 1 | 20.3167 | 0.459 |
| 0.2 | 0.1 | ycaM | 1 | 20.3167 | 0.476 |
| 0.2 | 0.1 | yadI | 1 | 20.3167 | 0.799 |
| 0.2 | 0.1 | AG1 | 1 | 20.3167 | 0.876 |
| 0.2 | 0.1 | ptsI | 1 | 20.5667 | 0.788 |
| 0.2 | 0.1 | clcB | 1 | 20.5667 | 0.446 |
| 0.2 | 0.1 | ycaM | 1 | 20.5667 | 0.469 |
| 0.2 | 0.1 | yadI | 1 | 20.5667 | 0.806 |
| 0.2 | 0.1 | AG1 | 1 | 20.5667 | 0.865 |
| 0.2 | 0.1 | ptsI | 1 | 20.8167 | 0.797 |
| 0.2 | 0.1 | clcB | 1 | 20.8167 | 0.439 |
| 0.2 | 0.1 | ycaM | 1 | 20.8167 | 0.472 |
| 0.2 | 0.1 | yadI | 1 | 20.8167 | 0.809 |
| 0.2 | 0.1 | AG1 | 1 | 20.8167 | 0.877 |
| 0.2 | 0.1 | ptsI | 1 | 21.0667 | 0.818 |
| 0.2 | 0.1 | clcB | 1 | 21.0667 | 0.448 |
| 0.2 | 0.1 | ycaM | 1 | 21.0667 | 0.466 |
| 0.2 | 0.1 | yadI | 1 | 21.0667 | 0.813 |
| 0.2 | 0.1 | AG1 | 1 | 21.0667 | 0.886 |
| 0.2 | 0.1 | ptsI | 1 | 21.3167 | 0.817 |
| 0.2 | 0.1 | clcB | 1 | 21.3167 | 0.439 |
| 0.2 | 0.1 | ycaM | 1 | 21.3167 | 0.47 |
| 0.2 | 0.1 | yadI | 1 | 21.3167 | 0.815 |
| 0.2 | 0.1 | AG1 | 1 | 21.3167 | 0.889 |
| 0.2 | 0.1 | ptsI | 1 | 21.5667 | 0.814 |
| 0.2 | 0.1 | clcB | 1 | 21.5667 | 0.44 |
| 0.2 | 0.1 | ycaM | 1 | 21.5667 | 0.468 |
| 0.2 | 0.1 | yadI | 1 | 21.5667 | 0.821 |
| 0.2 | 0.1 | AG1 | 1 | 21.5667 | 0.889 |
| 0.2 | 0.1 | ptsI | 1 | 21.8167 | 0.819 |

| | | | | | |
|---|---|---|---|---|---|
| 0.2 | 0.1 | clcB | 1 | 21.8167 | 0.436 |
| 0.2 | 0.1 | ycaM | 1 | 21.8167 | 0.474 |
| 0.2 | 0.1 | yadI | 1 | 21.8167 | 0.822 |
| 0.2 | 0.1 | AG1 | 1 | 21.8167 | 0.899 |
| 0.2 | 0.1 | ptsI | 1 | 22.0667 | 0.834 |
| 0.2 | 0.1 | clcB | 1 | 22.0667 | 0.442 |
| 0.2 | 0.1 | ycaM | 1 | 22.0667 | 0.47 |
| 0.2 | 0.1 | yadI | 1 | 22.0667 | 0.822 |
| 0.2 | 0.1 | AG1 | 1 | 22.0667 | 0.902 |
| 0.2 | 0.1 | ptsI | 1 | 22.9333 | 0.827 |
| 0.2 | 0.1 | clcB | 1 | 22.9333 | 0.452 |
| 0.2 | 0.1 | ycaM | 1 | 22.9333 | 0.456 |
| 0.2 | 0.1 | yadI | 1 | 22.9333 | 0.83 |
| 0.2 | 0.1 | AG1 | 1 | 22.9333 | 0.817 |
| 0.2 | 0.1 | ptsI | 1 | 23.1833 | 0.799 |
| 0.2 | 0.1 | clcB | 1 | 23.1833 | 0.447 |
| 0.2 | 0.1 | ycaM | 1 | 23.1833 | 0.449 |
| 0.2 | 0.1 | yadI | 1 | 23.1833 | 0.825 |
| 0.2 | 0.1 | AG1 | 1 | 23.1833 | 0.778 |
| 0.2 | 0.1 | ptsI | 1 | 23.4333 | 0.791 |
| 0.2 | 0.1 | clcB | 1 | 23.4333 | 0.44 |
| 0.2 | 0.1 | ycaM | 1 | 23.4333 | 0.447 |
| 0.2 | 0.1 | yadI | 1 | 23.4333 | 0.83 |
| 0.2 | 0.1 | AG1 | 1 | 23.4333 | 0.785 |
| 0.2 | 0.1 | ptsI | 1 | 23.6833 | 0.794 |
| 0.2 | 0.1 | clcB | 1 | 23.6833 | 0.44 |
| 0.2 | 0.1 | ycaM | 1 | 23.6833 | 0.449 |
| 0.2 | 0.1 | yadI | 1 | 23.6833 | 0.84 |
| 0.2 | 0.1 | AG1 | 1 | 23.6833 | 0.826 |
| 0.2 | 0.1 | ptsI | 1 | 23.9333 | 0.819 |
| 0.2 | 0.1 | clcB | 1 | 23.9333 | 0.44 |
| 0.2 | 0.1 | ycaM | 1 | 23.9333 | 0.444 |
| 0.2 | 0.1 | yadI | 1 | 23.9333 | 0.844 |
| 0.2 | 0.1 | AG1 | 1 | 23.9333 | 0.827 |
| 0.2 | 0.1 | ptsI | 1 | 24.1833 | 0.837 |
| 0.2 | 0.1 | clcB | 1 | 24.1833 | 0.438 |
| 0.2 | 0.1 | ycaM | 1 | 24.1833 | 0.443 |
| 0.2 | 0.1 | yadI | 1 | 24.1833 | 0.855 |
| 0.2 | 0.1 | AG1 | 1 | 24.1833 | 0.842 |
| 0.2 | 0.1 | ptsI | 1 | 24.4333 | 0.838 |
| 0.2 | 0.1 | clcB | 1 | 24.4333 | 0.435 |
| 0.2 | 0.1 | ycaM | 1 | 24.4333 | 0.442 |
| 0.2 | 0.1 | yadI | 1 | 24.4333 | 0.864 |
| 0.2 | 0.1 | AG1 | 1 | 24.4333 | 0.841 |
| 0.2 | 0.1 | ptsI | 1 | 24.6833 | 0.844 |
| 0.2 | 0.1 | clcB | 1 | 24.6833 | 0.435 |
| 0.2 | 0.1 | ycaM | 1 | 24.6833 | 0.443 |
| 0.2 | 0.1 | yadI | 1 | 24.6833 | 0.863 |

| | | | | | |
|---|---|---|---|---|---|
| 0.2 | 0.1 | AG1 | 1 | 24.6833 | 0.842 |
| 0.2 | 0.1 | ptsI | 1 | 24.9333 | 0.828 |
| 0.2 | 0.1 | clcB | 1 | 24.9333 | 0.433 |
| 0.2 | 0.1 | ycaM | 1 | 24.9333 | 0.441 |
| 0.2 | 0.1 | yadI | 1 | 24.9333 | 0.873 |
| 0.2 | 0.1 | AG1 | 1 | 24.9333 | 0.848 |
| 0.2 | 0.1 | ptsI | 1 | 25.1833 | 0.842 |
| 0.2 | 0.1 | clcB | 1 | 25.1833 | 0.428 |
| 0.2 | 0.1 | ycaM | 1 | 25.1833 | 0.44 |
| 0.2 | 0.1 | yadI | 1 | 25.1833 | 0.872 |
| 0.2 | 0.1 | AG1 | 1 | 25.1833 | 0.842 |
| 0.2 | 0.1 | ptsI | 1 | 25.4333 | 0.852 |
| 0.2 | 0.1 | clcB | 1 | 25.4333 | 0.437 |
| 0.2 | 0.1 | ycaM | 1 | 25.4333 | 0.441 |
| 0.2 | 0.1 | yadI | 1 | 25.4333 | 0.879 |
| 0.2 | 0.1 | AG1 | 1 | 25.4333 | 0.854 |
| 0.2 | 0.1 | ptsI | 1 | 25.6833 | 0.852 |
| 0.2 | 0.1 | clcB | 1 | 25.6833 | 0.432 |
| 0.2 | 0.1 | ycaM | 1 | 25.6833 | 0.444 |
| 0.2 | 0.1 | yadI | 1 | 25.6833 | 0.884 |
| 0.2 | 0.1 | AG1 | 1 | 25.6833 | 0.852 |
| 0.2 | 0.1 | ptsI | 1 | 25.9333 | 0.858 |
| 0.2 | 0.1 | clcB | 1 | 25.9333 | 0.425 |
| 0.2 | 0.1 | ycaM | 1 | 25.9333 | 0.441 |
| 0.2 | 0.1 | yadI | 1 | 25.9333 | 0.868 |
| 0.2 | 0.1 | AG1 | 1 | 25.9333 | 0.861 |
| 0.2 | 0.1 | ptsI | 1 | 26.1833 | 0.866 |
| 0.2 | 0.1 | clcB | 1 | 26.1833 | 0.43 |
| 0.2 | 0.1 | ycaM | 1 | 26.1833 | 0.443 |
| 0.2 | 0.1 | yadI | 1 | 26.1833 | 0.889 |
| 0.2 | 0.1 | AG1 | 1 | 26.1833 | 0.86 |
| 0.2 | 0.1 | ptsI | 1 | 26.4333 | 0.859 |
| 0.2 | 0.1 | clcB | 1 | 26.4333 | 0.433 |
| 0.2 | 0.1 | ycaM | 1 | 26.4333 | 0.445 |
| 0.2 | 0.1 | yadI | 1 | 26.4333 | 0.89 |
| 0.2 | 0.1 | AG1 | 1 | 26.4333 | 0.855 |
| 0.2 | 0.1 | ptsI | 1 | 26.6833 | 0.865 |
| 0.2 | 0.1 | clcB | 1 | 26.6833 | 0.433 |
| 0.2 | 0.1 | ycaM | 1 | 26.6833 | 0.446 |
| 0.2 | 0.1 | yadI | 1 | 26.6833 | 0.899 |
| 0.2 | 0.1 | AG1 | 1 | 26.6833 | 0.862 |
| 0.2 | 0.1 | ptsI | 1 | 26.9333 | 0.867 |
| 0.2 | 0.1 | clcB | 1 | 26.9333 | 0.437 |
| 0.2 | 0.1 | ycaM | 1 | 26.9333 | 0.444 |
| 0.2 | 0.1 | yadI | 1 | 26.9333 | 0.899 |
| 0.2 | 0.1 | AG1 | 1 | 26.9333 | 0.862 |
| 0.2 | 0.1 | ptsI | 1 | 27.1833 | 0.873 |
| 0.2 | 0.1 | clcB | 1 | 27.1833 | 0.436 |

| | | | | | |
|---|---|---|---|---|---|
| 0.2 | 0.1 | ycaM | 1 | 27.1833 | 0.445 |
| 0.2 | 0.1 | yadI | 1 | 27.1833 | 0.904 |
| 0.2 | 0.1 | AG1 | 1 | 27.1833 | 0.87 |
| 0.2 | 0.1 | ptsI | 1 | 27.4333 | 0.868 |
| 0.2 | 0.1 | clcB | 1 | 27.4333 | 0.433 |
| 0.2 | 0.1 | ycaM | 1 | 27.4333 | 0.446 |
| 0.2 | 0.1 | yadI | 1 | 27.4333 | 0.907 |
| 0.2 | 0.1 | AG1 | 1 | 27.4333 | 0.865 |
| 0.2 | 0.1 | ptsI | 1 | 27.6833 | 0.875 |
| 0.2 | 0.1 | clcB | 1 | 27.6833 | 0.435 |
| 0.2 | 0.1 | ycaM | 1 | 27.6833 | 0.444 |
| 0.2 | 0.1 | yadI | 1 | 27.6833 | 0.908 |
| 0.2 | 0.1 | AG1 | 1 | 27.6833 | 0.868 |
| 0.2 | 0.1 | ptsI | 1 | 27.9333 | 0.877 |
| 0.2 | 0.1 | clcB | 1 | 27.9333 | 0.43 |
| 0.2 | 0.1 | ycaM | 1 | 27.9333 | 0.443 |
| 0.2 | 0.1 | yadI | 1 | 27.9333 | 0.883 |
| 0.2 | 0.1 | AG1 | 1 | 27.9333 | 0.865 |
| 0.2 | 0.1 | ptsI | 1 | 28.1833 | 0.878 |
| 0.2 | 0.1 | clcB | 1 | 28.1833 | 0.437 |
| 0.2 | 0.1 | ycaM | 1 | 28.1833 | 0.445 |
| 0.2 | 0.1 | yadI | 1 | 28.1833 | 0.911 |
| 0.2 | 0.1 | AG1 | 1 | 28.1833 | 0.884 |
| 0.2 | 0.1 | ptsI | 1 | 28.4333 | 0.875 |
| 0.2 | 0.1 | clcB | 1 | 28.4333 | 0.434 |
| 0.2 | 0.1 | ycaM | 1 | 28.4333 | 0.446 |
| 0.2 | 0.1 | yadI | 1 | 28.4333 | 0.913 |
| 0.2 | 0.1 | AG1 | 1 | 28.4333 | 0.882 |
| 0.2 | 0.1 | ptsI | 1 | 28.6833 | 0.877 |
| 0.2 | 0.1 | clcB | 1 | 28.6833 | 0.438 |
| 0.2 | 0.1 | ycaM | 1 | 28.6833 | 0.447 |
| 0.2 | 0.1 | yadI | 1 | 28.6833 | 0.914 |
| 0.2 | 0.1 | AG1 | 1 | 28.6833 | 0.888 |
| 0.2 | 0.1 | ptsI | 1 | 28.9333 | 0.88 |
| 0.2 | 0.1 | clcB | 1 | 28.9333 | 0.437 |
| 0.2 | 0.1 | ycaM | 1 | 28.9333 | 0.447 |
| 0.2 | 0.1 | yadI | 1 | 28.9333 | 0.917 |
| 0.2 | 0.1 | AG1 | 1 | 28.9333 | 0.892 |
| 0.2 | 0.1 | ptsI | 1 | 29.1833 | 0.875 |
| 0.2 | 0.1 | clcB | 1 | 29.1833 | 0.438 |
| 0.2 | 0.1 | ycaM | 1 | 29.1833 | 0.447 |
| 0.2 | 0.1 | yadI | 1 | 29.1833 | 0.923 |
| 0.2 | 0.1 | AG1 | 1 | 29.1833 | 0.891 |
| 0.2 | 0.1 | ptsI | 1 | 29.4333 | 0.883 |
| 0.2 | 0.1 | clcB | 1 | 29.4333 | 0.44 |
| 0.2 | 0.1 | ycaM | 1 | 29.4333 | 0.447 |
| 0.2 | 0.1 | yadI | 1 | 29.4333 | 0.923 |
| 0.2 | 0.1 | AG1 | 1 | 29.4333 | 0.899 |

| | | | | | |
|---|---|---|---|---|---|
| 0.2 | 0.1 | ptsI | 1 | 29.6833 | 0.882 |
| 0.2 | 0.1 | clcB | 1 | 29.6833 | 0.444 |
| 0.2 | 0.1 | ycaM | 1 | 29.6833 | 0.448 |
| 0.2 | 0.1 | yadI | 1 | 29.6833 | 0.929 |
| 0.2 | 0.1 | AG1 | 1 | 29.6833 | 0.901 |
| 0.2 | 0.1 | ptsI | 1 | 29.9333 | 0.886 |
| 0.2 | 0.1 | clcB | 1 | 29.9333 | 0.444 |
| 0.2 | 0.1 | ycaM | 1 | 29.9333 | 0.449 |
| 0.2 | 0.1 | yadI | 1 | 29.9333 | 0.928 |
| 0.2 | 0.1 | AG1 | 1 | 29.9333 | 0.909 |
| 0.2 | 0.1 | ptsI | 1 | 30.1833 | 0.886 |
| 0.2 | 0.1 | clcB | 1 | 30.1833 | 0.445 |
| 0.2 | 0.1 | ycaM | 1 | 30.1833 | 0.448 |
| 0.2 | 0.1 | yadI | 1 | 30.1833 | 0.928 |
| 0.2 | 0.1 | AG1 | 1 | 30.1833 | 0.916 |
| 0.2 | 0.1 | ptsI | 1 | 30.4333 | 0.886 |
| 0.2 | 0.1 | clcB | 1 | 30.4333 | 0.444 |
| 0.2 | 0.1 | ycaM | 1 | 30.4333 | 0.45 |
| 0.2 | 0.1 | yadI | 1 | 30.4333 | 0.93 |
| 0.2 | 0.1 | AG1 | 1 | 30.4333 | 0.918 |
| 0.2 | 0.1 | ptsI | 1 | 30.6833 | 0.886 |
| 0.2 | 0.1 | clcB | 1 | 30.6833 | 0.445 |
| 0.2 | 0.1 | ycaM | 1 | 30.6833 | 0.451 |
| 0.2 | 0.1 | yadI | 1 | 30.6833 | 0.938 |
| 0.2 | 0.1 | AG1 | 1 | 30.6833 | 0.92 |
| 0.2 | 0.1 | ptsI | 1 | 30.9333 | 0.889 |
| 0.2 | 0.1 | clcB | 1 | 30.9333 | 0.446 |
| 0.2 | 0.1 | ycaM | 1 | 30.9333 | 0.452 |
| 0.2 | 0.1 | yadI | 1 | 30.9333 | 0.936 |
| 0.2 | 0.1 | AG1 | 1 | 30.9333 | 0.927 |
| 0.2 | 0.1 | ptsI | 1 | 31.1833 | 0.883 |
| 0.2 | 0.1 | clcB | 1 | 31.1833 | 0.442 |
| 0.2 | 0.1 | ycaM | 1 | 31.1833 | 0.452 |
| 0.2 | 0.1 | yadI | 1 | 31.1833 | 0.943 |
| 0.2 | 0.1 | AG1 | 1 | 31.1833 | 0.927 |
| 0.2 | 0.1 | ptsI | 1 | 31.4333 | 0.894 |
| 0.2 | 0.1 | clcB | 1 | 31.4333 | 0.447 |
| 0.2 | 0.1 | ycaM | 1 | 31.4333 | 0.455 |
| 0.2 | 0.1 | yadI | 1 | 31.4333 | 0.948 |
| 0.2 | 0.1 | AG1 | 1 | 31.4333 | 0.936 |
| 0.2 | 0.1 | ptsI | 1 | 31.6833 | 0.893 |
| 0.2 | 0.1 | clcB | 1 | 31.6833 | 0.446 |
| 0.2 | 0.1 | ycaM | 1 | 31.6833 | 0.457 |
| 0.2 | 0.1 | yadI | 1 | 31.6833 | 0.951 |
| 0.2 | 0.1 | AG1 | 1 | 31.6833 | 0.945 |
| 0.2 | 0.1 | ptsI | 1 | 31.9333 | 0.894 |
| 0.2 | 0.1 | clcB | 1 | 31.9333 | 0.451 |
| 0.2 | 0.1 | ycaM | 1 | 31.9333 | 0.453 |

| | | | | | |
|---|---|---|---|---|---|
| 0.2 | 0.1 | yadI | 1 | 31.9333 | 0.95 |
| 0.2 | 0.1 | AG1 | 1 | 31.9333 | 0.951 |
| 0.2 | 0.1 | ptsI | 1 | 32.1833 | 0.893 |
| 0.2 | 0.1 | clcB | 1 | 32.1833 | 0.451 |
| 0.2 | 0.1 | ycaM | 1 | 32.1833 | 0.46 |
| 0.2 | 0.1 | yadI | 1 | 32.1833 | 0.951 |
| 0.2 | 0.1 | AG1 | 1 | 32.1833 | 0.959 |
| 0.2 | 0.1 | ptsI | 1 | 32.4333 | 0.89 |
| 0.2 | 0.1 | clcB | 1 | 32.4333 | 0.435 |
| 0.2 | 0.1 | ycaM | 1 | 32.4333 | 0.449 |
| 0.2 | 0.1 | yadI | 1 | 32.4333 | 0.924 |
| 0.2 | 0.1 | AG1 | 1 | 32.4333 | 0.954 |
| 0.2 | 0.1 | ptsI | 1 | 32.6833 | 0.891 |
| 0.2 | 0.1 | clcB | 1 | 32.6833 | 0.45 |
| 0.2 | 0.1 | ycaM | 1 | 32.6833 | 0.46 |
| 0.2 | 0.1 | yadI | 1 | 32.6833 | 0.956 |
| 0.2 | 0.1 | AG1 | 1 | 32.6833 | 0.969 |
| 0.2 | 0.1 | ptsI | 1 | 32.9333 | 0.891 |
| 0.2 | 0.1 | clcB | 1 | 32.9333 | 0.447 |
| 0.2 | 0.1 | ycaM | 1 | 32.9333 | 0.456 |
| 0.2 | 0.1 | yadI | 1 | 32.9333 | 0.96 |
| 0.2 | 0.1 | AG1 | 1 | 32.9333 | 0.977 |
| 0.2 | 0.1 | ptsI | 1 | 33.1833 | 0.89 |
| 0.2 | 0.1 | clcB | 1 | 33.1833 | 0.442 |
| 0.2 | 0.1 | ycaM | 1 | 33.1833 | 0.456 |
| 0.2 | 0.1 | yadI | 1 | 33.1833 | 0.943 |
| 0.2 | 0.1 | AG1 | 1 | 33.1833 | 0.978 |
| 0.2 | 0.1 | ptsI | 1 | 33.4333 | 0.892 |
| 0.2 | 0.1 | clcB | 1 | 33.4333 | 0.453 |
| 0.2 | 0.1 | ycaM | 1 | 33.4333 | 0.459 |
| 0.2 | 0.1 | yadI | 1 | 33.4333 | 0.964 |
| 0.2 | 0.1 | AG1 | 1 | 33.4333 | 0.992 |
| 0.2 | 0.1 | ptsI | 1 | 33.6833 | 0.89 |
| 0.2 | 0.1 | clcB | 1 | 33.6833 | 0.449 |
| 0.2 | 0.1 | ycaM | 1 | 33.6833 | 0.46 |
| 0.2 | 0.1 | yadI | 1 | 33.6833 | 0.942 |
| 0.2 | 0.1 | AG1 | 1 | 33.6833 | 0.992 |
| 0.2 | 0.1 | ptsI | 1 | 33.9333 | 0.889 |
| 0.2 | 0.1 | clcB | 1 | 33.9333 | 0.452 |
| 0.2 | 0.1 | ycaM | 1 | 33.9333 | 0.46 |
| 0.2 | 0.1 | yadI | 1 | 33.9333 | 0.963 |
| 0.2 | 0.1 | AG1 | 1 | 33.9333 | 1.007 |
| 0.2 | 0.1 | ptsI | 1 | 34.1833 | 0.889 |
| 0.2 | 0.1 | clcB | 1 | 34.1833 | 0.461 |
| 0.2 | 0.1 | ycaM | 1 | 34.1833 | 0.458 |
| 0.2 | 0.1 | yadI | 1 | 34.1833 | 0.974 |
| 0.2 | 0.1 | AG1 | 1 | 34.1833 | 1.006 |
| 0.2 | 0.1 | ptsI | 1 | 34.4333 | 0.891 |

| | | | | | |
|---|---|---|---|---|---|
| 0.2 | 0.1 | clcB | 1 | 34.4333 | 0.456 |
| 0.2 | 0.1 | ycaM | 1 | 34.4333 | 0.461 |
| 0.2 | 0.1 | yadI | 1 | 34.4333 | 0.965 |
| 0.2 | 0.1 | AG1 | 1 | 34.4333 | 1.022 |
| 0.2 | 0.1 | ptsI | 1 | 34.6833 | 0.891 |
| 0.2 | 0.1 | clcB | 1 | 34.6833 | 0.46 |
| 0.2 | 0.1 | ycaM | 1 | 34.6833 | 0.464 |
| 0.2 | 0.1 | yadI | 1 | 34.6833 | 0.964 |
| 0.2 | 0.1 | AG1 | 1 | 34.6833 | 1.03 |
| 0.2 | 0.1 | ptsI | 1 | 34.9333 | 0.891 |
| 0.2 | 0.1 | clcB | 1 | 34.9333 | 0.457 |
| 0.2 | 0.1 | ycaM | 1 | 34.9333 | 0.461 |
| 0.2 | 0.1 | yadI | 1 | 34.9333 | 0.966 |
| 0.2 | 0.1 | AG1 | 1 | 34.9333 | 1.032 |
| 0.2 | 0.1 | ptsI | 1 | 35.1833 | 0.888 |
| 0.2 | 0.1 | clcB | 1 | 35.1833 | 0.458 |
| 0.2 | 0.1 | ycaM | 1 | 35.1833 | 0.462 |
| 0.2 | 0.1 | yadI | 1 | 35.1833 | 0.967 |
| 0.2 | 0.1 | AG1 | 1 | 35.1833 | 1.042 |
| 0.2 | 0.1 | ptsI | 1 | 35.4333 | 0.893 |
| 0.2 | 0.1 | clcB | 1 | 35.4333 | 0.458 |
| 0.2 | 0.1 | ycaM | 1 | 35.4333 | 0.46 |
| 0.2 | 0.1 | yadI | 1 | 35.4333 | 0.969 |
| 0.2 | 0.1 | AG1 | 1 | 35.4333 | 1.052 |
| 0.2 | 0.1 | ptsI | 1 | 35.6833 | 0.893 |
| 0.2 | 0.1 | clcB | 1 | 35.6833 | 0.462 |
| 0.2 | 0.1 | ycaM | 1 | 35.6833 | 0.465 |
| 0.2 | 0.1 | yadI | 1 | 35.6833 | 0.969 |
| 0.2 | 0.1 | AG1 | 1 | 35.6833 | 1.061 |
| 0.2 | 0.1 | ptsI | 1 | 35.9333 | 0.893 |
| 0.2 | 0.1 | clcB | 1 | 35.9333 | 0.455 |
| 0.2 | 0.1 | ycaM | 1 | 35.9333 | 0.463 |
| 0.2 | 0.1 | yadI | 1 | 35.9333 | 0.952 |
| 0.2 | 0.1 | AG1 | 1 | 35.9333 | 1.062 |
| 0.2 | 0.1 | ptsI | 1 | 36.1833 | 0.892 |
| 0.2 | 0.1 | clcB | 1 | 36.1833 | 0.463 |
| 0.2 | 0.1 | ycaM | 1 | 36.1833 | 0.47 |
| 0.2 | 0.1 | yadI | 1 | 36.1833 | 0.971 |
| 0.2 | 0.1 | AG1 | 1 | 36.1833 | 1.078 |
| 0.2 | 0.1 | ptsI | 1 | 36.4333 | 0.89 |
| 0.2 | 0.1 | clcB | 1 | 36.4333 | 0.461 |
| 0.2 | 0.1 | ycaM | 1 | 36.4333 | 0.466 |
| 0.2 | 0.1 | yadI | 1 | 36.4333 | 0.97 |
| 0.2 | 0.1 | AG1 | 1 | 36.4333 | 1.078 |
| 0.2 | 0.1 | ptsI | 1 | 36.6833 | 0.891 |
| 0.2 | 0.1 | clcB | 1 | 36.6833 | 0.46 |
| 0.2 | 0.1 | ycaM | 1 | 36.6833 | 0.466 |
| 0.2 | 0.1 | yadI | 1 | 36.6833 | 0.974 |

| | | | | | |
|---|---|---|---|---|---|
| 0.2 | 0.1 | AG1 | 1 | 36.6833 | 1.083 |
| 0.4 | 0.1 | ptsI | 1 | 0 | 0.232 |
| 0.4 | 0.1 | clcB | 1 | 0 | 0.253 |
| 0.4 | 0.1 | ycaM | 1 | 0 | 0.24 |
| 0.4 | 0.1 | yadI | 1 | 0 | 0.263 |
| 0.4 | 0.1 | AG1 | 1 | 0 | 0.254 |
| 0.4 | 0.1 | ptsI | 1 | 0.25 | 0.229 |
| 0.4 | 0.1 | clcB | 1 | 0.25 | 0.239 |
| 0.4 | 0.1 | ycaM | 1 | 0.25 | 0.23 |
| 0.4 | 0.1 | yadI | 1 | 0.25 | 0.252 |
| 0.4 | 0.1 | AG1 | 1 | 0.25 | 0.241 |
| 0.4 | 0.1 | ptsI | 1 | 0.5 | 0.228 |
| 0.4 | 0.1 | clcB | 1 | 0.5 | 0.233 |
| 0.4 | 0.1 | ycaM | 1 | 0.5 | 0.225 |
| 0.4 | 0.1 | yadI | 1 | 0.5 | 0.243 |
| 0.4 | 0.1 | AG1 | 1 | 0.5 | 0.237 |
| 0.4 | 0.1 | ptsI | 1 | 0.75 | 0.229 |
| 0.4 | 0.1 | clcB | 1 | 0.75 | 0.231 |
| 0.4 | 0.1 | ycaM | 1 | 0.75 | 0.223 |
| 0.4 | 0.1 | yadI | 1 | 0.75 | 0.243 |
| 0.4 | 0.1 | AG1 | 1 | 0.75 | 0.236 |
| 0.4 | 0.1 | ptsI | 1 | 1 | 0.228 |
| 0.4 | 0.1 | clcB | 1 | 1 | 0.233 |
| 0.4 | 0.1 | ycaM | 1 | 1 | 0.225 |
| 0.4 | 0.1 | yadI | 1 | 1 | 0.244 |
| 0.4 | 0.1 | AG1 | 1 | 1 | 0.235 |
| 0.4 | 0.1 | ptsI | 1 | 1.25 | 0.228 |
| 0.4 | 0.1 | clcB | 1 | 1.25 | 0.232 |
| 0.4 | 0.1 | ycaM | 1 | 1.25 | 0.225 |
| 0.4 | 0.1 | yadI | 1 | 1.25 | 0.246 |
| 0.4 | 0.1 | AG1 | 1 | 1.25 | 0.236 |
| 0.4 | 0.1 | ptsI | 1 | 1.5 | 0.235 |
| 0.4 | 0.1 | clcB | 1 | 1.5 | 0.233 |
| 0.4 | 0.1 | ycaM | 1 | 1.5 | 0.227 |
| 0.4 | 0.1 | yadI | 1 | 1.5 | 0.251 |
| 0.4 | 0.1 | AG1 | 1 | 1.5 | 0.239 |
| 0.4 | 0.1 | ptsI | 1 | 1.75 | 0.235 |
| 0.4 | 0.1 | clcB | 1 | 1.75 | 0.237 |
| 0.4 | 0.1 | ycaM | 1 | 1.75 | 0.231 |
| 0.4 | 0.1 | yadI | 1 | 1.75 | 0.252 |
| 0.4 | 0.1 | AG1 | 1 | 1.75 | 0.241 |
| 0.4 | 0.1 | ptsI | 1 | 2 | 0.239 |
| 0.4 | 0.1 | clcB | 1 | 2 | 0.239 |
| 0.4 | 0.1 | ycaM | 1 | 2 | 0.234 |
| 0.4 | 0.1 | yadI | 1 | 2 | 0.257 |
| 0.4 | 0.1 | AG1 | 1 | 2 | 0.246 |
| 0.4 | 0.1 | ptsI | 1 | 2.25 | 0.245 |
| 0.4 | 0.1 | clcB | 1 | 2.25 | 0.242 |

| | | | | | |
|---|---|---|---|---|---|
| 0.4 | 0.1 | ycaM | 1 | 2.25 | 0.24 |
| 0.4 | 0.1 | yadI | 1 | 2.25 | 0.264 |
| 0.4 | 0.1 | AG1 | 1 | 2.25 | 0.249 |
| 0.4 | 0.1 | ptsI | 1 | 2.5 | 0.251 |
| 0.4 | 0.1 | clcB | 1 | 2.5 | 0.246 |
| 0.4 | 0.1 | ycaM | 1 | 2.5 | 0.246 |
| 0.4 | 0.1 | yadI | 1 | 2.5 | 0.268 |
| 0.4 | 0.1 | AG1 | 1 | 2.5 | 0.254 |
| 0.4 | 0.1 | ptsI | 1 | 2.75 | 0.256 |
| 0.4 | 0.1 | clcB | 1 | 2.75 | 0.252 |
| 0.4 | 0.1 | ycaM | 1 | 2.75 | 0.253 |
| 0.4 | 0.1 | yadI | 1 | 2.75 | 0.276 |
| 0.4 | 0.1 | AG1 | 1 | 2.75 | 0.26 |
| 0.4 | 0.1 | ptsI | 1 | 3 | 0.262 |
| 0.4 | 0.1 | clcB | 1 | 3 | 0.254 |
| 0.4 | 0.1 | ycaM | 1 | 3 | 0.257 |
| 0.4 | 0.1 | yadI | 1 | 3 | 0.283 |
| 0.4 | 0.1 | AG1 | 1 | 3 | 0.268 |
| 0.4 | 0.1 | ptsI | 1 | 3.25 | 0.273 |
| 0.4 | 0.1 | clcB | 1 | 3.25 | 0.26 |
| 0.4 | 0.1 | ycaM | 1 | 3.25 | 0.266 |
| 0.4 | 0.1 | yadI | 1 | 3.25 | 0.295 |
| 0.4 | 0.1 | AG1 | 1 | 3.25 | 0.276 |
| 0.4 | 0.1 | ptsI | 1 | 3.5 | 0.278 |
| 0.4 | 0.1 | clcB | 1 | 3.5 | 0.269 |
| 0.4 | 0.1 | ycaM | 1 | 3.5 | 0.272 |
| 0.4 | 0.1 | yadI | 1 | 3.5 | 0.302 |
| 0.4 | 0.1 | AG1 | 1 | 3.5 | 0.278 |
| 0.4 | 0.1 | ptsI | 1 | 3.75 | 0.292 |
| 0.4 | 0.1 | clcB | 1 | 3.75 | 0.274 |
| 0.4 | 0.1 | ycaM | 1 | 3.75 | 0.276 |
| 0.4 | 0.1 | yadI | 1 | 3.75 | 0.31 |
| 0.4 | 0.1 | AG1 | 1 | 3.75 | 0.283 |
| 0.4 | 0.1 | ptsI | 1 | 4 | 0.31 |
| 0.4 | 0.1 | clcB | 1 | 4 | 0.285 |
| 0.4 | 0.1 | ycaM | 1 | 4 | 0.287 |
| 0.4 | 0.1 | yadI | 1 | 4 | 0.325 |
| 0.4 | 0.1 | AG1 | 1 | 4 | 0.292 |
| 0.4 | 0.1 | ptsI | 1 | 4.25 | 0.314 |
| 0.4 | 0.1 | clcB | 1 | 4.25 | 0.286 |
| 0.4 | 0.1 | ycaM | 1 | 4.25 | 0.29 |
| 0.4 | 0.1 | yadI | 1 | 4.25 | 0.327 |
| 0.4 | 0.1 | AG1 | 1 | 4.25 | 0.299 |
| 0.4 | 0.1 | ptsI | 1 | 4.5 | 0.328 |
| 0.4 | 0.1 | clcB | 1 | 4.5 | 0.292 |
| 0.4 | 0.1 | ycaM | 1 | 4.5 | 0.296 |
| 0.4 | 0.1 | yadI | 1 | 4.5 | 0.338 |
| 0.4 | 0.1 | AG1 | 1 | 4.5 | 0.305 |

| | | | | | |
|---|---|---|---|---|---|
| 0.4 | 0.1 | ptsI | 1 | 4.75 | 0.337 |
| 0.4 | 0.1 | clcB | 1 | 4.75 | 0.299 |
| 0.4 | 0.1 | ycaM | 1 | 4.75 | 0.301 |
| 0.4 | 0.1 | yadI | 1 | 4.75 | 0.342 |
| 0.4 | 0.1 | AG1 | 1 | 4.75 | 0.309 |
| 0.4 | 0.1 | ptsI | 1 | 5 | 0.354 |
| 0.4 | 0.1 | clcB | 1 | 5 | 0.302 |
| 0.4 | 0.1 | ycaM | 1 | 5 | 0.304 |
| 0.4 | 0.1 | yadI | 1 | 5 | 0.356 |
| 0.4 | 0.1 | AG1 | 1 | 5 | 0.317 |
| 0.4 | 0.1 | ptsI | 1 | 5.25 | 0.355 |
| 0.4 | 0.1 | clcB | 1 | 5.25 | 0.308 |
| 0.4 | 0.1 | ycaM | 1 | 5.25 | 0.309 |
| 0.4 | 0.1 | yadI | 1 | 5.25 | 0.363 |
| 0.4 | 0.1 | AG1 | 1 | 5.25 | 0.324 |
| 0.4 | 0.1 | ptsI | 1 | 5.5 | 0.365 |
| 0.4 | 0.1 | clcB | 1 | 5.5 | 0.315 |
| 0.4 | 0.1 | ycaM | 1 | 5.5 | 0.315 |
| 0.4 | 0.1 | yadI | 1 | 5.5 | 0.375 |
| 0.4 | 0.1 | AG1 | 1 | 5.5 | 0.332 |
| 0.4 | 0.1 | ptsI | 1 | 5.75 | 0.375 |
| 0.4 | 0.1 | clcB | 1 | 5.75 | 0.323 |
| 0.4 | 0.1 | ycaM | 1 | 5.75 | 0.322 |
| 0.4 | 0.1 | yadI | 1 | 5.75 | 0.385 |
| 0.4 | 0.1 | AG1 | 1 | 5.75 | 0.342 |
| 0.4 | 0.1 | ptsI | 1 | 6 | 0.368 |
| 0.4 | 0.1 | clcB | 1 | 6 | 0.328 |
| 0.4 | 0.1 | ycaM | 1 | 6 | 0.329 |
| 0.4 | 0.1 | yadI | 1 | 6 | 0.388 |
| 0.4 | 0.1 | AG1 | 1 | 6 | 0.341 |
| 0.4 | 0.1 | ptsI | 1 | 6.25 | 0.377 |
| 0.4 | 0.1 | clcB | 1 | 6.25 | 0.338 |
| 0.4 | 0.1 | ycaM | 1 | 6.25 | 0.328 |
| 0.4 | 0.1 | yadI | 1 | 6.25 | 0.382 |
| 0.4 | 0.1 | AG1 | 1 | 6.25 | 0.346 |
| 0.4 | 0.1 | ptsI | 1 | 6.5 | 0.388 |
| 0.4 | 0.1 | clcB | 1 | 6.5 | 0.34 |
| 0.4 | 0.1 | ycaM | 1 | 6.5 | 0.336 |
| 0.4 | 0.1 | yadI | 1 | 6.5 | 0.383 |
| 0.4 | 0.1 | AG1 | 1 | 6.5 | 0.353 |
| 0.4 | 0.1 | ptsI | 1 | 6.75 | 0.404 |
| 0.4 | 0.1 | clcB | 1 | 6.75 | 0.346 |
| 0.4 | 0.1 | ycaM | 1 | 6.75 | 0.342 |
| 0.4 | 0.1 | yadI | 1 | 6.75 | 0.393 |
| 0.4 | 0.1 | AG1 | 1 | 6.75 | 0.363 |
| 0.4 | 0.1 | ptsI | 1 | 7 | 0.42 |
| 0.4 | 0.1 | clcB | 1 | 7 | 0.356 |
| 0.4 | 0.1 | ycaM | 1 | 7 | 0.349 |

| | | | | | |
|---|---|---|---|---|---|
| 0.4 | 0.1 | yadI | 1 | 7 | 0.404 |
| 0.4 | 0.1 | AG1 | 1 | 7 | 0.374 |
| 0.4 | 0.1 | ptsI | 1 | 7.25 | 0.449 |
| 0.4 | 0.1 | clcB | 1 | 7.25 | 0.362 |
| 0.4 | 0.1 | ycaM | 1 | 7.25 | 0.354 |
| 0.4 | 0.1 | yadI | 1 | 7.25 | 0.409 |
| 0.4 | 0.1 | AG1 | 1 | 7.25 | 0.382 |
| 0.4 | 0.1 | ptsI | 1 | 7.5 | 0.46 |
| 0.4 | 0.1 | clcB | 1 | 7.5 | 0.369 |
| 0.4 | 0.1 | ycaM | 1 | 7.5 | 0.359 |
| 0.4 | 0.1 | yadI | 1 | 7.5 | 0.418 |
| 0.4 | 0.1 | AG1 | 1 | 7.5 | 0.392 |
| 0.4 | 0.1 | ptsI | 1 | 7.75 | 0.472 |
| 0.4 | 0.1 | clcB | 1 | 7.75 | 0.38 |
| 0.4 | 0.1 | ycaM | 1 | 7.75 | 0.368 |
| 0.4 | 0.1 | yadI | 1 | 7.75 | 0.426 |
| 0.4 | 0.1 | AG1 | 1 | 7.75 | 0.403 |
| 0.4 | 0.1 | ptsI | 1 | 8 | 0.498 |
| 0.4 | 0.1 | clcB | 1 | 8 | 0.386 |
| 0.4 | 0.1 | ycaM | 1 | 8 | 0.367 |
| 0.4 | 0.1 | yadI | 1 | 8 | 0.432 |
| 0.4 | 0.1 | AG1 | 1 | 8 | 0.409 |
| 0.4 | 0.1 | ptsI | 1 | 8.25 | 0.502 |
| 0.4 | 0.1 | clcB | 1 | 8.25 | 0.396 |
| 0.4 | 0.1 | ycaM | 1 | 8.25 | 0.368 |
| 0.4 | 0.1 | yadI | 1 | 8.25 | 0.437 |
| 0.4 | 0.1 | AG1 | 1 | 8.25 | 0.42 |
| 0.4 | 0.1 | ptsI | 1 | 8.5 | 0.517 |
| 0.4 | 0.1 | clcB | 1 | 8.5 | 0.402 |
| 0.4 | 0.1 | ycaM | 1 | 8.5 | 0.37 |
| 0.4 | 0.1 | yadI | 1 | 8.5 | 0.443 |
| 0.4 | 0.1 | AG1 | 1 | 8.5 | 0.426 |
| 0.4 | 0.1 | ptsI | 1 | 8.75 | 0.527 |
| 0.4 | 0.1 | clcB | 1 | 8.75 | 0.413 |
| 0.4 | 0.1 | ycaM | 1 | 8.75 | 0.373 |
| 0.4 | 0.1 | yadI | 1 | 8.75 | 0.448 |
| 0.4 | 0.1 | AG1 | 1 | 8.75 | 0.435 |
| 0.4 | 0.1 | ptsI | 1 | 9 | 0.537 |
| 0.4 | 0.1 | clcB | 1 | 9 | 0.417 |
| 0.4 | 0.1 | ycaM | 1 | 9 | 0.373 |
| 0.4 | 0.1 | yadI | 1 | 9 | 0.458 |
| 0.4 | 0.1 | AG1 | 1 | 9 | 0.443 |
| 0.4 | 0.1 | ptsI | 1 | 9.25 | 0.544 |
| 0.4 | 0.1 | clcB | 1 | 9.25 | 0.421 |
| 0.4 | 0.1 | ycaM | 1 | 9.25 | 0.376 |
| 0.4 | 0.1 | yadI | 1 | 9.25 | 0.446 |
| 0.4 | 0.1 | AG1 | 1 | 9.25 | 0.458 |
| 0.4 | 0.1 | ptsI | 1 | 9.5 | 0.564 |

| | | | | | |
|---|---|---|---|---|---|
| 0.4 | 0.1 | clcB | 1 | 9.5 | 0.435 |
| 0.4 | 0.1 | ycaM | 1 | 9.5 | 0.383 |
| 0.4 | 0.1 | yadI | 1 | 9.5 | 0.458 |
| 0.4 | 0.1 | AG1 | 1 | 9.5 | 0.494 |
| 0.4 | 0.1 | ptsI | 1 | 9.75 | 0.592 |
| 0.4 | 0.1 | clcB | 1 | 9.75 | 0.454 |
| 0.4 | 0.1 | ycaM | 1 | 9.75 | 0.39 |
| 0.4 | 0.1 | yadI | 1 | 9.75 | 0.465 |
| 0.4 | 0.1 | AG1 | 1 | 9.75 | 0.512 |
| 0.4 | 0.1 | ptsI | 1 | 10 | 0.565 |
| 0.4 | 0.1 | clcB | 1 | 10 | 0.459 |
| 0.4 | 0.1 | ycaM | 1 | 10 | 0.392 |
| 0.4 | 0.1 | yadI | 1 | 10 | 0.454 |
| 0.4 | 0.1 | AG1 | 1 | 10 | 0.522 |
| 0.4 | 0.1 | ptsI | 1 | 10.25 | 0.623 |
| 0.4 | 0.1 | clcB | 1 | 10.25 | 0.471 |
| 0.4 | 0.1 | ycaM | 1 | 10.25 | 0.399 |
| 0.4 | 0.1 | yadI | 1 | 10.25 | 0.464 |
| 0.4 | 0.1 | AG1 | 1 | 10.25 | 0.535 |
| 0.4 | 0.1 | ptsI | 1 | 10.5 | 0.641 |
| 0.4 | 0.1 | clcB | 1 | 10.5 | 0.495 |
| 0.4 | 0.1 | ycaM | 1 | 10.5 | 0.401 |
| 0.4 | 0.1 | yadI | 1 | 10.5 | 0.475 |
| 0.4 | 0.1 | AG1 | 1 | 10.5 | 0.549 |
| 0.4 | 0.1 | ptsI | 1 | 10.75 | 0.607 |
| 0.4 | 0.1 | clcB | 1 | 10.75 | 0.492 |
| 0.4 | 0.1 | ycaM | 1 | 10.75 | 0.404 |
| 0.4 | 0.1 | yadI | 1 | 10.75 | 0.456 |
| 0.4 | 0.1 | AG1 | 1 | 10.75 | 0.546 |
| 0.4 | 0.1 | ptsI | 1 | 11 | 0.613 |
| 0.4 | 0.1 | clcB | 1 | 11 | 0.51 |
| 0.4 | 0.1 | ycaM | 1 | 11 | 0.412 |
| 0.4 | 0.1 | yadI | 1 | 11 | 0.47 |
| 0.4 | 0.1 | AG1 | 1 | 11 | 0.582 |
| 0.4 | 0.1 | ptsI | 1 | 11.25 | 0.679 |
| 0.4 | 0.1 | clcB | 1 | 11.25 | 0.548 |
| 0.4 | 0.1 | ycaM | 1 | 11.25 | 0.411 |
| 0.4 | 0.1 | yadI | 1 | 11.25 | 0.473 |
| 0.4 | 0.1 | AG1 | 1 | 11.25 | 0.595 |
| 0.4 | 0.1 | ptsI | 1 | 11.5 | 0.657 |
| 0.4 | 0.1 | clcB | 1 | 11.5 | 0.538 |
| 0.4 | 0.1 | ycaM | 1 | 11.5 | 0.419 |
| 0.4 | 0.1 | yadI | 1 | 11.5 | 0.476 |
| 0.4 | 0.1 | AG1 | 1 | 11.5 | 0.624 |
| 0.4 | 0.1 | ptsI | 1 | 11.75 | 0.667 |
| 0.4 | 0.1 | clcB | 1 | 11.75 | 0.553 |
| 0.4 | 0.1 | ycaM | 1 | 11.75 | 0.419 |
| 0.4 | 0.1 | yadI | 1 | 11.75 | 0.47 |

| | | | | | |
|---|---|---|---|---|---|
| 0.4 | 0.1 | AG1 | 1 | 11.75 | 0.63 |
| 0.4 | 0.1 | ptsI | 1 | 12 | 0.703 |
| 0.4 | 0.1 | clcB | 1 | 12 | 0.599 |
| 0.4 | 0.1 | ycaM | 1 | 12 | 0.422 |
| 0.4 | 0.1 | yadI | 1 | 12 | 0.476 |
| 0.4 | 0.1 | AG1 | 1 | 12 | 0.66 |
| 0.4 | 0.1 | ptsI | 1 | 12.25 | 0.678 |
| 0.4 | 0.1 | clcB | 1 | 12.25 | 0.575 |
| 0.4 | 0.1 | ycaM | 1 | 12.25 | 0.422 |
| 0.4 | 0.1 | yadI | 1 | 12.25 | 0.469 |
| 0.4 | 0.1 | AG1 | 1 | 12.25 | 0.618 |
| 0.4 | 0.1 | ptsI | 1 | 12.5 | 0.713 |
| 0.4 | 0.1 | clcB | 1 | 12.5 | 0.588 |
| 0.4 | 0.1 | ycaM | 1 | 12.5 | 0.423 |
| 0.4 | 0.1 | yadI | 1 | 12.5 | 0.461 |
| 0.4 | 0.1 | AG1 | 1 | 12.5 | 0.63 |
| 0.4 | 0.1 | ptsI | 1 | 12.75 | 0.721 |
| 0.4 | 0.1 | clcB | 1 | 12.75 | 0.604 |
| 0.4 | 0.1 | ycaM | 1 | 12.75 | 0.429 |
| 0.4 | 0.1 | yadI | 1 | 12.75 | 0.47 |
| 0.4 | 0.1 | AG1 | 1 | 12.75 | 0.643 |
| 0.4 | 0.1 | ptsI | 1 | 13 | 0.728 |
| 0.4 | 0.1 | clcB | 1 | 13 | 0.608 |
| 0.4 | 0.1 | ycaM | 1 | 13 | 0.443 |
| 0.4 | 0.1 | yadI | 1 | 13 | 0.485 |
| 0.4 | 0.1 | AG1 | 1 | 13 | 0.69 |
| 0.4 | 0.1 | ptsI | 1 | 13.25 | 0.716 |
| 0.4 | 0.1 | clcB | 1 | 13.25 | 0.608 |
| 0.4 | 0.1 | ycaM | 1 | 13.25 | 0.439 |
| 0.4 | 0.1 | yadI | 1 | 13.25 | 0.48 |
| 0.4 | 0.1 | AG1 | 1 | 13.25 | 0.698 |
| 0.4 | 0.1 | ptsI | 1 | 13.5 | 0.748 |
| 0.4 | 0.1 | clcB | 1 | 13.5 | 0.637 |
| 0.4 | 0.1 | ycaM | 1 | 13.5 | 0.452 |
| 0.4 | 0.1 | yadI | 1 | 13.5 | 0.481 |
| 0.4 | 0.1 | AG1 | 1 | 13.5 | 0.732 |
| 0.4 | 0.1 | ptsI | 1 | 13.75 | 0.73 |
| 0.4 | 0.1 | clcB | 1 | 13.75 | 0.632 |
| 0.4 | 0.1 | ycaM | 1 | 13.75 | 0.455 |
| 0.4 | 0.1 | yadI | 1 | 13.75 | 0.477 |
| 0.4 | 0.1 | AG1 | 1 | 13.75 | 0.744 |
| 0.4 | 0.1 | ptsI | 1 | 14 | 0.768 |
| 0.4 | 0.1 | clcB | 1 | 14 | 0.669 |
| 0.4 | 0.1 | ycaM | 1 | 14 | 0.467 |
| 0.4 | 0.1 | yadI | 1 | 14 | 0.482 |
| 0.4 | 0.1 | AG1 | 1 | 14 | 0.787 |
| 0.4 | 0.1 | ptsI | 1 | 14.25 | 0.772 |
| 0.4 | 0.1 | clcB | 1 | 14.25 | 0.675 |

| | | | | | |
|---|---|---|---|---|---|
| 0.4 | 0.1 | ycaM | 1 | 14.25 | 0.471 |
| 0.4 | 0.1 | yadI | 1 | 14.25 | 0.49 |
| 0.4 | 0.1 | AG1 | 1 | 14.25 | 0.802 |
| 0.4 | 0.1 | ptsI | 1 | 14.5 | 0.767 |
| 0.4 | 0.1 | clcB | 1 | 14.5 | 0.683 |
| 0.4 | 0.1 | ycaM | 1 | 14.5 | 0.486 |
| 0.4 | 0.1 | yadI | 1 | 14.5 | 0.5 |
| 0.4 | 0.1 | AG1 | 1 | 14.5 | 0.812 |
| 0.4 | 0.1 | ptsI | 1 | 14.75 | 0.739 |
| 0.4 | 0.1 | clcB | 1 | 14.75 | 0.668 |
| 0.4 | 0.1 | ycaM | 1 | 14.75 | 0.489 |
| 0.4 | 0.1 | yadI | 1 | 14.75 | 0.504 |
| 0.4 | 0.1 | AG1 | 1 | 14.75 | 0.808 |
| 0.4 | 0.1 | ptsI | 1 | 15 | 0.792 |
| 0.4 | 0.1 | clcB | 1 | 15 | 0.707 |
| 0.4 | 0.1 | ycaM | 1 | 15 | 0.504 |
| 0.4 | 0.1 | yadI | 1 | 15 | 0.518 |
| 0.4 | 0.1 | AG1 | 1 | 15 | 0.818 |
| 0.4 | 0.1 | ptsI | 1 | 15.25 | 0.794 |
| 0.4 | 0.1 | clcB | 1 | 15.25 | 0.71 |
| 0.4 | 0.1 | ycaM | 1 | 15.25 | 0.543 |
| 0.4 | 0.1 | yadI | 1 | 15.25 | 0.523 |
| 0.4 | 0.1 | AG1 | 1 | 15.25 | 0.845 |
| 0.4 | 0.1 | ptsI | 1 | 15.5 | 0.78 |
| 0.4 | 0.1 | clcB | 1 | 15.5 | 0.701 |
| 0.4 | 0.1 | ycaM | 1 | 15.5 | 0.542 |
| 0.4 | 0.1 | yadI | 1 | 15.5 | 0.526 |
| 0.4 | 0.1 | AG1 | 1 | 15.5 | 0.832 |
| 0.4 | 0.1 | ptsI | 1 | 15.75 | 0.773 |
| 0.4 | 0.1 | clcB | 1 | 15.75 | 0.701 |
| 0.4 | 0.1 | ycaM | 1 | 15.75 | 0.536 |
| 0.4 | 0.1 | yadI | 1 | 15.75 | 0.513 |
| 0.4 | 0.1 | AG1 | 1 | 15.75 | 0.839 |
| 0.4 | 0.1 | ptsI | 1 | 16 | 0.77 |
| 0.4 | 0.1 | clcB | 1 | 16 | 0.696 |
| 0.4 | 0.1 | ycaM | 1 | 16 | 0.529 |
| 0.4 | 0.1 | yadI | 1 | 16 | 0.514 |
| 0.4 | 0.1 | AG1 | 1 | 16 | 0.833 |
| 0.4 | 0.1 | ptsI | 1 | 16.25 | 0.822 |
| 0.4 | 0.1 | clcB | 1 | 16.25 | 0.723 |
| 0.4 | 0.1 | ycaM | 1 | 16.25 | 0.534 |
| 0.4 | 0.1 | yadI | 1 | 16.25 | 0.527 |
| 0.4 | 0.1 | AG1 | 1 | 16.25 | 0.85 |
| 0.4 | 0.1 | ptsI | 1 | 16.5 | 0.815 |
| 0.4 | 0.1 | clcB | 1 | 16.5 | 0.73 |
| 0.4 | 0.1 | ycaM | 1 | 16.5 | 0.569 |
| 0.4 | 0.1 | yadI | 1 | 16.5 | 0.537 |
| 0.4 | 0.1 | AG1 | 1 | 16.5 | 0.864 |

| | | | | | |
|---|---|---|---|---|---|
| 0.4 | 0.1 | ptsI | 1 | 17.0667 | 0.629 |
| 0.4 | 0.1 | clcB | 1 | 17.0667 | 0.519 |
| 0.4 | 0.1 | ycaM | 1 | 17.0667 | 0.509 |
| 0.4 | 0.1 | yadI | 1 | 17.0667 | 0.645 |
| 0.4 | 0.1 | AG1 | 1 | 17.0667 | 0.759 |
| 0.4 | 0.1 | ptsI | 1 | 17.3167 | 0.659 |
| 0.4 | 0.1 | clcB | 1 | 17.3167 | 0.481 |
| 0.4 | 0.1 | ycaM | 1 | 17.3167 | 0.498 |
| 0.4 | 0.1 | yadI | 1 | 17.3167 | 0.622 |
| 0.4 | 0.1 | AG1 | 1 | 17.3167 | 0.756 |
| 0.4 | 0.1 | ptsI | 1 | 17.5667 | 0.654 |
| 0.4 | 0.1 | clcB | 1 | 17.5667 | 0.479 |
| 0.4 | 0.1 | ycaM | 1 | 17.5667 | 0.492 |
| 0.4 | 0.1 | yadI | 1 | 17.5667 | 0.624 |
| 0.4 | 0.1 | AG1 | 1 | 17.5667 | 0.746 |
| 0.4 | 0.1 | ptsI | 1 | 17.8167 | 0.742 |
| 0.4 | 0.1 | clcB | 1 | 17.8167 | 0.483 |
| 0.4 | 0.1 | ycaM | 1 | 17.8167 | 0.49 |
| 0.4 | 0.1 | yadI | 1 | 17.8167 | 0.641 |
| 0.4 | 0.1 | AG1 | 1 | 17.8167 | 0.773 |
| 0.4 | 0.1 | ptsI | 1 | 18.0667 | 0.69 |
| 0.4 | 0.1 | clcB | 1 | 18.0667 | 0.483 |
| 0.4 | 0.1 | ycaM | 1 | 18.0667 | 0.486 |
| 0.4 | 0.1 | yadI | 1 | 18.0667 | 0.652 |
| 0.4 | 0.1 | AG1 | 1 | 18.0667 | 0.775 |
| 0.4 | 0.1 | ptsI | 1 | 18.3167 | 0.7 |
| 0.4 | 0.1 | clcB | 1 | 18.3167 | 0.479 |
| 0.4 | 0.1 | ycaM | 1 | 18.3167 | 0.479 |
| 0.4 | 0.1 | yadI | 1 | 18.3167 | 0.644 |
| 0.4 | 0.1 | AG1 | 1 | 18.3167 | 0.794 |
| 0.4 | 0.1 | ptsI | 1 | 18.5667 | 0.764 |
| 0.4 | 0.1 | clcB | 1 | 18.5667 | 0.482 |
| 0.4 | 0.1 | ycaM | 1 | 18.5667 | 0.481 |
| 0.4 | 0.1 | yadI | 1 | 18.5667 | 0.659 |
| 0.4 | 0.1 | AG1 | 1 | 18.5667 | 0.807 |
| 0.4 | 0.1 | ptsI | 1 | 18.8167 | 0.766 |
| 0.4 | 0.1 | clcB | 1 | 18.8167 | 0.486 |
| 0.4 | 0.1 | ycaM | 1 | 18.8167 | 0.48 |
| 0.4 | 0.1 | yadI | 1 | 18.8167 | 0.67 |
| 0.4 | 0.1 | AG1 | 1 | 18.8167 | 0.818 |
| 0.4 | 0.1 | ptsI | 1 | 19.0667 | 0.783 |
| 0.4 | 0.1 | clcB | 1 | 19.0667 | 0.493 |
| 0.4 | 0.1 | ycaM | 1 | 19.0667 | 0.487 |
| 0.4 | 0.1 | yadI | 1 | 19.0667 | 0.686 |
| 0.4 | 0.1 | AG1 | 1 | 19.0667 | 0.839 |
| 0.4 | 0.1 | ptsI | 1 | 19.3167 | 0.764 |
| 0.4 | 0.1 | clcB | 1 | 19.3167 | 0.486 |
| 0.4 | 0.1 | ycaM | 1 | 19.3167 | 0.482 |

| | | | | | |
|---|---|---|---|---|---|
| 0.4 | 0.1 | yadI | 1 | 19.3167 | 0.685 |
| 0.4 | 0.1 | AG1 | 1 | 19.3167 | 0.833 |
| 0.4 | 0.1 | ptsI | 1 | 19.5667 | 0.784 |
| 0.4 | 0.1 | clcB | 1 | 19.5667 | 0.497 |
| 0.4 | 0.1 | ycaM | 1 | 19.5667 | 0.483 |
| 0.4 | 0.1 | yadI | 1 | 19.5667 | 0.693 |
| 0.4 | 0.1 | AG1 | 1 | 19.5667 | 0.836 |
| 0.4 | 0.1 | ptsI | 1 | 19.8167 | 0.786 |
| 0.4 | 0.1 | clcB | 1 | 19.8167 | 0.491 |
| 0.4 | 0.1 | ycaM | 1 | 19.8167 | 0.489 |
| 0.4 | 0.1 | yadI | 1 | 19.8167 | 0.697 |
| 0.4 | 0.1 | AG1 | 1 | 19.8167 | 0.838 |
| 0.4 | 0.1 | ptsI | 1 | 20.0667 | 0.786 |
| 0.4 | 0.1 | clcB | 1 | 20.0667 | 0.484 |
| 0.4 | 0.1 | ycaM | 1 | 20.0667 | 0.487 |
| 0.4 | 0.1 | yadI | 1 | 20.0667 | 0.705 |
| 0.4 | 0.1 | AG1 | 1 | 20.0667 | 0.847 |
| 0.4 | 0.1 | ptsI | 1 | 20.3167 | 0.789 |
| 0.4 | 0.1 | clcB | 1 | 20.3167 | 0.487 |
| 0.4 | 0.1 | ycaM | 1 | 20.3167 | 0.484 |
| 0.4 | 0.1 | yadI | 1 | 20.3167 | 0.706 |
| 0.4 | 0.1 | AG1 | 1 | 20.3167 | 0.85 |
| 0.4 | 0.1 | ptsI | 1 | 20.5667 | 0.778 |
| 0.4 | 0.1 | clcB | 1 | 20.5667 | 0.468 |
| 0.4 | 0.1 | ycaM | 1 | 20.5667 | 0.482 |
| 0.4 | 0.1 | yadI | 1 | 20.5667 | 0.705 |
| 0.4 | 0.1 | AG1 | 1 | 20.5667 | 0.852 |
| 0.4 | 0.1 | ptsI | 1 | 20.8167 | 0.786 |
| 0.4 | 0.1 | clcB | 1 | 20.8167 | 0.472 |
| 0.4 | 0.1 | ycaM | 1 | 20.8167 | 0.48 |
| 0.4 | 0.1 | yadI | 1 | 20.8167 | 0.708 |
| 0.4 | 0.1 | AG1 | 1 | 20.8167 | 0.85 |
| 0.4 | 0.1 | ptsI | 1 | 21.0667 | 0.806 |
| 0.4 | 0.1 | clcB | 1 | 21.0667 | 0.472 |
| 0.4 | 0.1 | ycaM | 1 | 21.0667 | 0.48 |
| 0.4 | 0.1 | yadI | 1 | 21.0667 | 0.715 |
| 0.4 | 0.1 | AG1 | 1 | 21.0667 | 0.861 |
| 0.4 | 0.1 | ptsI | 1 | 21.3167 | 0.803 |
| 0.4 | 0.1 | clcB | 1 | 21.3167 | 0.484 |
| 0.4 | 0.1 | ycaM | 1 | 21.3167 | 0.482 |
| 0.4 | 0.1 | yadI | 1 | 21.3167 | 0.716 |
| 0.4 | 0.1 | AG1 | 1 | 21.3167 | 0.857 |
| 0.4 | 0.1 | ptsI | 1 | 21.5667 | 0.801 |
| 0.4 | 0.1 | clcB | 1 | 21.5667 | 0.477 |
| 0.4 | 0.1 | ycaM | 1 | 21.5667 | 0.48 |
| 0.4 | 0.1 | yadI | 1 | 21.5667 | 0.716 |
| 0.4 | 0.1 | AG1 | 1 | 21.5667 | 0.855 |
| 0.4 | 0.1 | ptsI | 1 | 21.8167 | 0.805 |

| | | | | | |
|---|---|---|---|---|---|
| 0.4 | 0.1 | clcB | 1 | 21.8167 | 0.462 |
| 0.4 | 0.1 | ycaM | 1 | 21.8167 | 0.483 |
| 0.4 | 0.1 | yadI | 1 | 21.8167 | 0.724 |
| 0.4 | 0.1 | AG1 | 1 | 21.8167 | 0.865 |
| 0.4 | 0.1 | ptsI | 1 | 22.0667 | 0.823 |
| 0.4 | 0.1 | clcB | 1 | 22.0667 | 0.467 |
| 0.4 | 0.1 | ycaM | 1 | 22.0667 | 0.481 |
| 0.4 | 0.1 | yadI | 1 | 22.0667 | 0.726 |
| 0.4 | 0.1 | AG1 | 1 | 22.0667 | 0.874 |
| 0.4 | 0.1 | ptsI | 1 | 22.9333 | 0.654 |
| 0.4 | 0.1 | clcB | 1 | 22.9333 | 0.384 |
| 0.4 | 0.1 | ycaM | 1 | 22.9333 | 0.374 |
| 0.4 | 0.1 | yadI | 1 | 22.9333 | 0.792 |
| 0.4 | 0.1 | AG1 | 1 | 22.9333 | 0.687 |
| 0.4 | 0.1 | ptsI | 1 | 23.1833 | 0.722 |
| 0.4 | 0.1 | clcB | 1 | 23.1833 | 0.366 |
| 0.4 | 0.1 | ycaM | 1 | 23.1833 | 0.37 |
| 0.4 | 0.1 | yadI | 1 | 23.1833 | 0.77 |
| 0.4 | 0.1 | AG1 | 1 | 23.1833 | 0.703 |
| 0.4 | 0.1 | ptsI | 1 | 23.4333 | 0.727 |
| 0.4 | 0.1 | clcB | 1 | 23.4333 | 0.365 |
| 0.4 | 0.1 | ycaM | 1 | 23.4333 | 0.367 |
| 0.4 | 0.1 | yadI | 1 | 23.4333 | 0.777 |
| 0.4 | 0.1 | AG1 | 1 | 23.4333 | 0.718 |
| 0.4 | 0.1 | ptsI | 1 | 23.6833 | 0.769 |
| 0.4 | 0.1 | clcB | 1 | 23.6833 | 0.364 |
| 0.4 | 0.1 | ycaM | 1 | 23.6833 | 0.365 |
| 0.4 | 0.1 | yadI | 1 | 23.6833 | 0.779 |
| 0.4 | 0.1 | AG1 | 1 | 23.6833 | 0.742 |
| 0.4 | 0.1 | ptsI | 1 | 23.9333 | 0.76 |
| 0.4 | 0.1 | clcB | 1 | 23.9333 | 0.368 |
| 0.4 | 0.1 | ycaM | 1 | 23.9333 | 0.365 |
| 0.4 | 0.1 | yadI | 1 | 23.9333 | 0.783 |
| 0.4 | 0.1 | AG1 | 1 | 23.9333 | 0.763 |
| 0.4 | 0.1 | ptsI | 1 | 24.1833 | 0.798 |
| 0.4 | 0.1 | clcB | 1 | 24.1833 | 0.364 |
| 0.4 | 0.1 | ycaM | 1 | 24.1833 | 0.366 |
| 0.4 | 0.1 | yadI | 1 | 24.1833 | 0.788 |
| 0.4 | 0.1 | AG1 | 1 | 24.1833 | 0.777 |
| 0.4 | 0.1 | ptsI | 1 | 24.4333 | 0.805 |
| 0.4 | 0.1 | clcB | 1 | 24.4333 | 0.367 |
| 0.4 | 0.1 | ycaM | 1 | 24.4333 | 0.368 |
| 0.4 | 0.1 | yadI | 1 | 24.4333 | 0.797 |
| 0.4 | 0.1 | AG1 | 1 | 24.4333 | 0.778 |
| 0.4 | 0.1 | ptsI | 1 | 24.6833 | 0.804 |
| 0.4 | 0.1 | clcB | 1 | 24.6833 | 0.364 |
| 0.4 | 0.1 | ycaM | 1 | 24.6833 | 0.364 |
| 0.4 | 0.1 | yadI | 1 | 24.6833 | 0.8 |

| | | | | | |
|---|---|---|---|---|---|
| 0.4 | 0.1 | AG1 | 1 | 24.6833 | 0.782 |
| 0.4 | 0.1 | ptsI | 1 | 24.9333 | 0.797 |
| 0.4 | 0.1 | clcB | 1 | 24.9333 | 0.364 |
| 0.4 | 0.1 | ycaM | 1 | 24.9333 | 0.366 |
| 0.4 | 0.1 | yadI | 1 | 24.9333 | 0.807 |
| 0.4 | 0.1 | AG1 | 1 | 24.9333 | 0.797 |
| 0.4 | 0.1 | ptsI | 1 | 25.1833 | 0.799 |
| 0.4 | 0.1 | clcB | 1 | 25.1833 | 0.365 |
| 0.4 | 0.1 | ycaM | 1 | 25.1833 | 0.367 |
| 0.4 | 0.1 | yadI | 1 | 25.1833 | 0.812 |
| 0.4 | 0.1 | AG1 | 1 | 25.1833 | 0.794 |
| 0.4 | 0.1 | ptsI | 1 | 25.4333 | 0.814 |
| 0.4 | 0.1 | clcB | 1 | 25.4333 | 0.366 |
| 0.4 | 0.1 | ycaM | 1 | 25.4333 | 0.366 |
| 0.4 | 0.1 | yadI | 1 | 25.4333 | 0.821 |
| 0.4 | 0.1 | AG1 | 1 | 25.4333 | 0.803 |
| 0.4 | 0.1 | ptsI | 1 | 25.6833 | 0.812 |
| 0.4 | 0.1 | clcB | 1 | 25.6833 | 0.367 |
| 0.4 | 0.1 | ycaM | 1 | 25.6833 | 0.367 |
| 0.4 | 0.1 | yadI | 1 | 25.6833 | 0.823 |
| 0.4 | 0.1 | AG1 | 1 | 25.6833 | 0.803 |
| 0.4 | 0.1 | ptsI | 1 | 25.9333 | 0.824 |
| 0.4 | 0.1 | clcB | 1 | 25.9333 | 0.369 |
| 0.4 | 0.1 | ycaM | 1 | 25.9333 | 0.363 |
| 0.4 | 0.1 | yadI | 1 | 25.9333 | 0.836 |
| 0.4 | 0.1 | AG1 | 1 | 25.9333 | 0.81 |
| 0.4 | 0.1 | ptsI | 1 | 26.1833 | 0.826 |
| 0.4 | 0.1 | clcB | 1 | 26.1833 | 0.369 |
| 0.4 | 0.1 | ycaM | 1 | 26.1833 | 0.367 |
| 0.4 | 0.1 | yadI | 1 | 26.1833 | 0.835 |
| 0.4 | 0.1 | AG1 | 1 | 26.1833 | 0.812 |
| 0.4 | 0.1 | ptsI | 1 | 26.4333 | 0.819 |
| 0.4 | 0.1 | clcB | 1 | 26.4333 | 0.37 |
| 0.4 | 0.1 | ycaM | 1 | 26.4333 | 0.367 |
| 0.4 | 0.1 | yadI | 1 | 26.4333 | 0.834 |
| 0.4 | 0.1 | AG1 | 1 | 26.4333 | 0.81 |
| 0.4 | 0.1 | ptsI | 1 | 26.6833 | 0.823 |
| 0.4 | 0.1 | clcB | 1 | 26.6833 | 0.372 |
| 0.4 | 0.1 | ycaM | 1 | 26.6833 | 0.37 |
| 0.4 | 0.1 | yadI | 1 | 26.6833 | 0.841 |
| 0.4 | 0.1 | AG1 | 1 | 26.6833 | 0.818 |
| 0.4 | 0.1 | ptsI | 1 | 26.9333 | 0.829 |
| 0.4 | 0.1 | clcB | 1 | 26.9333 | 0.374 |
| 0.4 | 0.1 | ycaM | 1 | 26.9333 | 0.371 |
| 0.4 | 0.1 | yadI | 1 | 26.9333 | 0.837 |
| 0.4 | 0.1 | AG1 | 1 | 26.9333 | 0.817 |
| 0.4 | 0.1 | ptsI | 1 | 27.1833 | 0.836 |
| 0.4 | 0.1 | clcB | 1 | 27.1833 | 0.373 |

| | | | | | |
|---|---|---|---|---|---|
| 0.4 | 0.1 | ycaM | 1 | 27.1833 | 0.372 |
| 0.4 | 0.1 | yadI | 1 | 27.1833 | 0.84 |
| 0.4 | 0.1 | AG1 | 1 | 27.1833 | 0.824 |
| 0.4 | 0.1 | ptsI | 1 | 27.4333 | 0.823 |
| 0.4 | 0.1 | clcB | 1 | 27.4333 | 0.372 |
| 0.4 | 0.1 | ycaM | 1 | 27.4333 | 0.372 |
| 0.4 | 0.1 | yadI | 1 | 27.4333 | 0.844 |
| 0.4 | 0.1 | AG1 | 1 | 27.4333 | 0.817 |
| 0.4 | 0.1 | ptsI | 1 | 27.6833 | 0.833 |
| 0.4 | 0.1 | clcB | 1 | 27.6833 | 0.376 |
| 0.4 | 0.1 | ycaM | 1 | 27.6833 | 0.372 |
| 0.4 | 0.1 | yadI | 1 | 27.6833 | 0.846 |
| 0.4 | 0.1 | AG1 | 1 | 27.6833 | 0.832 |
| 0.4 | 0.1 | ptsI | 1 | 27.9333 | 0.824 |
| 0.4 | 0.1 | clcB | 1 | 27.9333 | 0.375 |
| 0.4 | 0.1 | ycaM | 1 | 27.9333 | 0.377 |
| 0.4 | 0.1 | yadI | 1 | 27.9333 | 0.811 |
| 0.4 | 0.1 | AG1 | 1 | 27.9333 | 0.828 |
| 0.4 | 0.1 | ptsI | 1 | 28.1833 | 0.841 |
| 0.4 | 0.1 | clcB | 1 | 28.1833 | 0.377 |
| 0.4 | 0.1 | ycaM | 1 | 28.1833 | 0.375 |
| 0.4 | 0.1 | yadI | 1 | 28.1833 | 0.852 |
| 0.4 | 0.1 | AG1 | 1 | 28.1833 | 0.838 |
| 0.4 | 0.1 | ptsI | 1 | 28.4333 | 0.838 |
| 0.4 | 0.1 | clcB | 1 | 28.4333 | 0.378 |
| 0.4 | 0.1 | ycaM | 1 | 28.4333 | 0.376 |
| 0.4 | 0.1 | yadI | 1 | 28.4333 | 0.85 |
| 0.4 | 0.1 | AG1 | 1 | 28.4333 | 0.838 |
| 0.4 | 0.1 | ptsI | 1 | 28.6833 | 0.842 |
| 0.4 | 0.1 | clcB | 1 | 28.6833 | 0.38 |
| 0.4 | 0.1 | ycaM | 1 | 28.6833 | 0.376 |
| 0.4 | 0.1 | yadI | 1 | 28.6833 | 0.854 |
| 0.4 | 0.1 | AG1 | 1 | 28.6833 | 0.843 |
| 0.4 | 0.1 | ptsI | 1 | 28.9333 | 0.847 |
| 0.4 | 0.1 | clcB | 1 | 28.9333 | 0.38 |
| 0.4 | 0.1 | ycaM | 1 | 28.9333 | 0.376 |
| 0.4 | 0.1 | yadI | 1 | 28.9333 | 0.858 |
| 0.4 | 0.1 | AG1 | 1 | 28.9333 | 0.849 |
| 0.4 | 0.1 | ptsI | 1 | 29.1833 | 0.838 |
| 0.4 | 0.1 | clcB | 1 | 29.1833 | 0.379 |
| 0.4 | 0.1 | ycaM | 1 | 29.1833 | 0.375 |
| 0.4 | 0.1 | yadI | 1 | 29.1833 | 0.857 |
| 0.4 | 0.1 | AG1 | 1 | 29.1833 | 0.843 |
| 0.4 | 0.1 | ptsI | 1 | 29.4333 | 0.847 |
| 0.4 | 0.1 | clcB | 1 | 29.4333 | 0.383 |
| 0.4 | 0.1 | ycaM | 1 | 29.4333 | 0.376 |
| 0.4 | 0.1 | yadI | 1 | 29.4333 | 0.86 |
| 0.4 | 0.1 | AG1 | 1 | 29.4333 | 0.857 |

| | | | | | |
|---|---|---|---|---|---|
| 0.4 | 0.1 | ptsI | 1 | 29.6833 | 0.845 |
| 0.4 | 0.1 | clcB | 1 | 29.6833 | 0.383 |
| 0.4 | 0.1 | ycaM | 1 | 29.6833 | 0.378 |
| 0.4 | 0.1 | yadI | 1 | 29.6833 | 0.862 |
| 0.4 | 0.1 | AG1 | 1 | 29.6833 | 0.858 |
| 0.4 | 0.1 | ptsI | 1 | 29.9333 | 0.851 |
| 0.4 | 0.1 | clcB | 1 | 29.9333 | 0.386 |
| 0.4 | 0.1 | ycaM | 1 | 29.9333 | 0.377 |
| 0.4 | 0.1 | yadI | 1 | 29.9333 | 0.864 |
| 0.4 | 0.1 | AG1 | 1 | 29.9333 | 0.868 |
| 0.4 | 0.1 | ptsI | 1 | 30.1833 | 0.857 |
| 0.4 | 0.1 | clcB | 1 | 30.1833 | 0.385 |
| 0.4 | 0.1 | ycaM | 1 | 30.1833 | 0.378 |
| 0.4 | 0.1 | yadI | 1 | 30.1833 | 0.867 |
| 0.4 | 0.1 | AG1 | 1 | 30.1833 | 0.875 |
| 0.4 | 0.1 | ptsI | 1 | 30.4333 | 0.853 |
| 0.4 | 0.1 | clcB | 1 | 30.4333 | 0.387 |
| 0.4 | 0.1 | ycaM | 1 | 30.4333 | 0.38 |
| 0.4 | 0.1 | yadI | 1 | 30.4333 | 0.864 |
| 0.4 | 0.1 | AG1 | 1 | 30.4333 | 0.878 |
| 0.4 | 0.1 | ptsI | 1 | 30.6833 | 0.848 |
| 0.4 | 0.1 | clcB | 1 | 30.6833 | 0.387 |
| 0.4 | 0.1 | ycaM | 1 | 30.6833 | 0.381 |
| 0.4 | 0.1 | yadI | 1 | 30.6833 | 0.876 |
| 0.4 | 0.1 | AG1 | 1 | 30.6833 | 0.874 |
| 0.4 | 0.1 | ptsI | 1 | 30.9333 | 0.86 |
| 0.4 | 0.1 | clcB | 1 | 30.9333 | 0.387 |
| 0.4 | 0.1 | ycaM | 1 | 30.9333 | 0.384 |
| 0.4 | 0.1 | yadI | 1 | 30.9333 | 0.879 |
| 0.4 | 0.1 | AG1 | 1 | 30.9333 | 0.884 |
| 0.4 | 0.1 | ptsI | 1 | 31.1833 | 0.853 |
| 0.4 | 0.1 | clcB | 1 | 31.1833 | 0.386 |
| 0.4 | 0.1 | ycaM | 1 | 31.1833 | 0.382 |
| 0.4 | 0.1 | yadI | 1 | 31.1833 | 0.88 |
| 0.4 | 0.1 | AG1 | 1 | 31.1833 | 0.884 |
| 0.4 | 0.1 | ptsI | 1 | 31.4333 | 0.862 |
| 0.4 | 0.1 | clcB | 1 | 31.4333 | 0.393 |
| 0.4 | 0.1 | ycaM | 1 | 31.4333 | 0.384 |
| 0.4 | 0.1 | yadI | 1 | 31.4333 | 0.884 |
| 0.4 | 0.1 | AG1 | 1 | 31.4333 | 0.9 |
| 0.4 | 0.1 | ptsI | 1 | 31.6833 | 0.864 |
| 0.4 | 0.1 | clcB | 1 | 31.6833 | 0.39 |
| 0.4 | 0.1 | ycaM | 1 | 31.6833 | 0.387 |
| 0.4 | 0.1 | yadI | 1 | 31.6833 | 0.888 |
| 0.4 | 0.1 | AG1 | 1 | 31.6833 | 0.903 |
| 0.4 | 0.1 | ptsI | 1 | 31.9333 | 0.865 |
| 0.4 | 0.1 | clcB | 1 | 31.9333 | 0.396 |
| 0.4 | 0.1 | ycaM | 1 | 31.9333 | 0.386 |

| | | | | | |
|---|---|---|---|---|---|
| 0.4 | 0.1 | yadI | 1 | 31.9333 | 0.881 |
| 0.4 | 0.1 | AG1 | 1 | 31.9333 | 0.909 |
| 0.4 | 0.1 | ptsI | 1 | 32.1833 | 0.864 |
| 0.4 | 0.1 | clcB | 1 | 32.1833 | 0.398 |
| 0.4 | 0.1 | ycaM | 1 | 32.1833 | 0.391 |
| 0.4 | 0.1 | yadI | 1 | 32.1833 | 0.885 |
| 0.4 | 0.1 | AG1 | 1 | 32.1833 | 0.917 |
| 0.4 | 0.1 | ptsI | 1 | 32.4333 | 0.862 |
| 0.4 | 0.1 | clcB | 1 | 32.4333 | 0.393 |
| 0.4 | 0.1 | ycaM | 1 | 32.4333 | 0.387 |
| 0.4 | 0.1 | yadI | 1 | 32.4333 | 0.873 |
| 0.4 | 0.1 | AG1 | 1 | 32.4333 | 0.913 |
| 0.4 | 0.1 | ptsI | 1 | 32.6833 | 0.864 |
| 0.4 | 0.1 | clcB | 1 | 32.6833 | 0.398 |
| 0.4 | 0.1 | ycaM | 1 | 32.6833 | 0.394 |
| 0.4 | 0.1 | yadI | 1 | 32.6833 | 0.894 |
| 0.4 | 0.1 | AG1 | 1 | 32.6833 | 0.927 |
| 0.4 | 0.1 | ptsI | 1 | 32.9333 | 0.865 |
| 0.4 | 0.1 | clcB | 1 | 32.9333 | 0.396 |
| 0.4 | 0.1 | ycaM | 1 | 32.9333 | 0.393 |
| 0.4 | 0.1 | yadI | 1 | 32.9333 | 0.891 |
| 0.4 | 0.1 | AG1 | 1 | 32.9333 | 0.937 |
| 0.4 | 0.1 | ptsI | 1 | 33.1833 | 0.862 |
| 0.4 | 0.1 | clcB | 1 | 33.1833 | 0.397 |
| 0.4 | 0.1 | ycaM | 1 | 33.1833 | 0.4 |
| 0.4 | 0.1 | yadI | 1 | 33.1833 | 0.879 |
| 0.4 | 0.1 | AG1 | 1 | 33.1833 | 0.942 |
| 0.4 | 0.1 | ptsI | 1 | 33.4333 | 0.875 |
| 0.4 | 0.1 | clcB | 1 | 33.4333 | 0.403 |
| 0.4 | 0.1 | ycaM | 1 | 33.4333 | 0.398 |
| 0.4 | 0.1 | yadI | 1 | 33.4333 | 0.902 |
| 0.4 | 0.1 | AG1 | 1 | 33.4333 | 0.955 |
| 0.4 | 0.1 | ptsI | 1 | 33.6833 | 0.866 |
| 0.4 | 0.1 | clcB | 1 | 33.6833 | 0.404 |
| 0.4 | 0.1 | ycaM | 1 | 33.6833 | 0.404 |
| 0.4 | 0.1 | yadI | 1 | 33.6833 | 0.881 |
| 0.4 | 0.1 | AG1 | 1 | 33.6833 | 0.962 |
| 0.4 | 0.1 | ptsI | 1 | 33.9333 | 0.872 |
| 0.4 | 0.1 | clcB | 1 | 33.9333 | 0.404 |
| 0.4 | 0.1 | ycaM | 1 | 33.9333 | 0.4 |
| 0.4 | 0.1 | yadI | 1 | 33.9333 | 0.906 |
| 0.4 | 0.1 | AG1 | 1 | 33.9333 | 0.974 |
| 0.4 | 0.1 | ptsI | 1 | 34.1833 | 0.864 |
| 0.4 | 0.1 | clcB | 1 | 34.1833 | 0.409 |
| 0.4 | 0.1 | ycaM | 1 | 34.1833 | 0.41 |
| 0.4 | 0.1 | yadI | 1 | 34.1833 | 0.901 |
| 0.4 | 0.1 | AG1 | 1 | 34.1833 | 0.981 |
| 0.4 | 0.1 | ptsI | 1 | 34.4333 | 0.876 |

| | | | | | |
|---|---|---|---|---|---|
| 0.4 | 0.1 | clcB | 1 | 34.4333 | 0.406 |
| 0.4 | 0.1 | ycaM | 1 | 34.4333 | 0.405 |
| 0.4 | 0.1 | yadI | 1 | 34.4333 | 0.911 |
| 0.4 | 0.1 | AG1 | 1 | 34.4333 | 0.992 |
| 0.4 | 0.1 | ptsI | 1 | 34.6833 | 0.879 |
| 0.4 | 0.1 | clcB | 1 | 34.6833 | 0.41 |
| 0.4 | 0.1 | ycaM | 1 | 34.6833 | 0.41 |
| 0.4 | 0.1 | yadI | 1 | 34.6833 | 0.91 |
| 0.4 | 0.1 | AG1 | 1 | 34.6833 | 1.006 |
| 0.4 | 0.1 | ptsI | 1 | 34.9333 | 0.876 |
| 0.4 | 0.1 | clcB | 1 | 34.9333 | 0.407 |
| 0.4 | 0.1 | ycaM | 1 | 34.9333 | 0.408 |
| 0.4 | 0.1 | yadI | 1 | 34.9333 | 0.909 |
| 0.4 | 0.1 | AG1 | 1 | 34.9333 | 1.014 |
| 0.4 | 0.1 | ptsI | 1 | 35.1833 | 0.878 |
| 0.4 | 0.1 | clcB | 1 | 35.1833 | 0.409 |
| 0.4 | 0.1 | ycaM | 1 | 35.1833 | 0.408 |
| 0.4 | 0.1 | yadI | 1 | 35.1833 | 0.911 |
| 0.4 | 0.1 | AG1 | 1 | 35.1833 | 1.025 |
| 0.4 | 0.1 | ptsI | 1 | 35.4333 | 0.878 |
| 0.4 | 0.1 | clcB | 1 | 35.4333 | 0.408 |
| 0.4 | 0.1 | ycaM | 1 | 35.4333 | 0.409 |
| 0.4 | 0.1 | yadI | 1 | 35.4333 | 0.914 |
| 0.4 | 0.1 | AG1 | 1 | 35.4333 | 1.032 |
| 0.4 | 0.1 | ptsI | 1 | 35.6833 | 0.876 |
| 0.4 | 0.1 | clcB | 1 | 35.6833 | 0.41 |
| 0.4 | 0.1 | ycaM | 1 | 35.6833 | 0.411 |
| 0.4 | 0.1 | yadI | 1 | 35.6833 | 0.916 |
| 0.4 | 0.1 | AG1 | 1 | 35.6833 | 1.041 |
| 0.4 | 0.1 | ptsI | 1 | 35.9333 | 0.866 |
| 0.4 | 0.1 | clcB | 1 | 35.9333 | 0.41 |
| 0.4 | 0.1 | ycaM | 1 | 35.9333 | 0.414 |
| 0.4 | 0.1 | yadI | 1 | 35.9333 | 0.9 |
| 0.4 | 0.1 | AG1 | 1 | 35.9333 | 1.041 |
| 0.4 | 0.1 | ptsI | 1 | 36.1833 | 0.882 |
| 0.4 | 0.1 | clcB | 1 | 36.1833 | 0.414 |
| 0.4 | 0.1 | ycaM | 1 | 36.1833 | 0.417 |
| 0.4 | 0.1 | yadI | 1 | 36.1833 | 0.918 |
| 0.4 | 0.1 | AG1 | 1 | 36.1833 | 1.052 |
| 0.4 | 0.1 | ptsI | 1 | 36.4333 | 0.878 |
| 0.4 | 0.1 | clcB | 1 | 36.4333 | 0.413 |
| 0.4 | 0.1 | ycaM | 1 | 36.4333 | 0.418 |
| 0.4 | 0.1 | yadI | 1 | 36.4333 | 0.926 |
| 0.4 | 0.1 | AG1 | 1 | 36.4333 | 1.066 |
| 0.4 | 0.1 | ptsI | 1 | 36.6833 | 0.88 |
| 0.4 | 0.1 | clcB | 1 | 36.6833 | 0.414 |
| 0.4 | 0.1 | ycaM | 1 | 36.6833 | 0.416 |
| 0.4 | 0.1 | yadI | 1 | 36.6833 | 0.924 |

| | | | | | |
|---|---|---|---|---|---|
| 0.4 | 0.1 | AG1 | 1 | 36.6833 | 1.074 |
| 0.8 | 0.1 | ptsI | 1 | 0 | 0.223 |
| 0.8 | 0.1 | clcB | 1 | 0 | 0.257 |
| 0.8 | 0.1 | ycaM | 1 | 0 | 0.243 |
| 0.8 | 0.1 | yadI | 1 | 0 | 0.253 |
| 0.8 | 0.1 | AG1 | 1 | 0 | 0.254 |
| 0.8 | 0.1 | ptsI | 1 | 0.25 | 0.222 |
| 0.8 | 0.1 | clcB | 1 | 0.25 | 0.242 |
| 0.8 | 0.1 | ycaM | 1 | 0.25 | 0.229 |
| 0.8 | 0.1 | yadI | 1 | 0.25 | 0.244 |
| 0.8 | 0.1 | AG1 | 1 | 0.25 | 0.239 |
| 0.8 | 0.1 | ptsI | 1 | 0.5 | 0.219 |
| 0.8 | 0.1 | clcB | 1 | 0.5 | 0.239 |
| 0.8 | 0.1 | ycaM | 1 | 0.5 | 0.228 |
| 0.8 | 0.1 | yadI | 1 | 0.5 | 0.241 |
| 0.8 | 0.1 | AG1 | 1 | 0.5 | 0.234 |
| 0.8 | 0.1 | ptsI | 1 | 0.75 | 0.219 |
| 0.8 | 0.1 | clcB | 1 | 0.75 | 0.237 |
| 0.8 | 0.1 | ycaM | 1 | 0.75 | 0.224 |
| 0.8 | 0.1 | yadI | 1 | 0.75 | 0.238 |
| 0.8 | 0.1 | AG1 | 1 | 0.75 | 0.232 |
| 0.8 | 0.1 | ptsI | 1 | 1 | 0.22 |
| 0.8 | 0.1 | clcB | 1 | 1 | 0.238 |
| 0.8 | 0.1 | ycaM | 1 | 1 | 0.224 |
| 0.8 | 0.1 | yadI | 1 | 1 | 0.239 |
| 0.8 | 0.1 | AG1 | 1 | 1 | 0.232 |
| 0.8 | 0.1 | ptsI | 1 | 1.25 | 0.22 |
| 0.8 | 0.1 | clcB | 1 | 1.25 | 0.24 |
| 0.8 | 0.1 | ycaM | 1 | 1.25 | 0.223 |
| 0.8 | 0.1 | yadI | 1 | 1.25 | 0.242 |
| 0.8 | 0.1 | AG1 | 1 | 1.25 | 0.232 |
| 0.8 | 0.1 | ptsI | 1 | 1.5 | 0.222 |
| 0.8 | 0.1 | clcB | 1 | 1.5 | 0.238 |
| 0.8 | 0.1 | ycaM | 1 | 1.5 | 0.226 |
| 0.8 | 0.1 | yadI | 1 | 1.5 | 0.242 |
| 0.8 | 0.1 | AG1 | 1 | 1.5 | 0.234 |
| 0.8 | 0.1 | ptsI | 1 | 1.75 | 0.226 |
| 0.8 | 0.1 | clcB | 1 | 1.75 | 0.24 |
| 0.8 | 0.1 | ycaM | 1 | 1.75 | 0.229 |
| 0.8 | 0.1 | yadI | 1 | 1.75 | 0.248 |
| 0.8 | 0.1 | AG1 | 1 | 1.75 | 0.234 |
| 0.8 | 0.1 | ptsI | 1 | 2 | 0.231 |
| 0.8 | 0.1 | clcB | 1 | 2 | 0.243 |
| 0.8 | 0.1 | ycaM | 1 | 2 | 0.233 |
| 0.8 | 0.1 | yadI | 1 | 2 | 0.252 |
| 0.8 | 0.1 | AG1 | 1 | 2 | 0.238 |
| 0.8 | 0.1 | ptsI | 1 | 2.25 | 0.236 |
| 0.8 | 0.1 | clcB | 1 | 2.25 | 0.248 |

| | | | | | |
|---|---|---|---|---|---|
| 0.8 | 0.1 | ycaM | 1 | 2.25 | 0.239 |
| 0.8 | 0.1 | yadI | 1 | 2.25 | 0.257 |
| 0.8 | 0.1 | AG1 | 1 | 2.25 | 0.242 |
| 0.8 | 0.1 | ptsI | 1 | 2.5 | 0.241 |
| 0.8 | 0.1 | clcB | 1 | 2.5 | 0.252 |
| 0.8 | 0.1 | ycaM | 1 | 2.5 | 0.244 |
| 0.8 | 0.1 | yadI | 1 | 2.5 | 0.262 |
| 0.8 | 0.1 | AG1 | 1 | 2.5 | 0.244 |
| 0.8 | 0.1 | ptsI | 1 | 2.75 | 0.248 |
| 0.8 | 0.1 | clcB | 1 | 2.75 | 0.256 |
| 0.8 | 0.1 | ycaM | 1 | 2.75 | 0.246 |
| 0.8 | 0.1 | yadI | 1 | 2.75 | 0.271 |
| 0.8 | 0.1 | AG1 | 1 | 2.75 | 0.248 |
| 0.8 | 0.1 | ptsI | 1 | 3 | 0.255 |
| 0.8 | 0.1 | clcB | 1 | 3 | 0.261 |
| 0.8 | 0.1 | ycaM | 1 | 3 | 0.255 |
| 0.8 | 0.1 | yadI | 1 | 3 | 0.28 |
| 0.8 | 0.1 | AG1 | 1 | 3 | 0.253 |
| 0.8 | 0.1 | ptsI | 1 | 3.25 | 0.263 |
| 0.8 | 0.1 | clcB | 1 | 3.25 | 0.265 |
| 0.8 | 0.1 | ycaM | 1 | 3.25 | 0.265 |
| 0.8 | 0.1 | yadI | 1 | 3.25 | 0.291 |
| 0.8 | 0.1 | AG1 | 1 | 3.25 | 0.261 |
| 0.8 | 0.1 | ptsI | 1 | 3.5 | 0.273 |
| 0.8 | 0.1 | clcB | 1 | 3.5 | 0.27 |
| 0.8 | 0.1 | ycaM | 1 | 3.5 | 0.266 |
| 0.8 | 0.1 | yadI | 1 | 3.5 | 0.299 |
| 0.8 | 0.1 | AG1 | 1 | 3.5 | 0.263 |
| 0.8 | 0.1 | ptsI | 1 | 3.75 | 0.279 |
| 0.8 | 0.1 | clcB | 1 | 3.75 | 0.272 |
| 0.8 | 0.1 | ycaM | 1 | 3.75 | 0.27 |
| 0.8 | 0.1 | yadI | 1 | 3.75 | 0.303 |
| 0.8 | 0.1 | AG1 | 1 | 3.75 | 0.266 |
| 0.8 | 0.1 | ptsI | 1 | 4 | 0.296 |
| 0.8 | 0.1 | clcB | 1 | 4 | 0.283 |
| 0.8 | 0.1 | ycaM | 1 | 4 | 0.282 |
| 0.8 | 0.1 | yadI | 1 | 4 | 0.316 |
| 0.8 | 0.1 | AG1 | 1 | 4 | 0.271 |
| 0.8 | 0.1 | ptsI | 1 | 4.25 | 0.289 |
| 0.8 | 0.1 | clcB | 1 | 4.25 | 0.286 |
| 0.8 | 0.1 | ycaM | 1 | 4.25 | 0.279 |
| 0.8 | 0.1 | yadI | 1 | 4.25 | 0.316 |
| 0.8 | 0.1 | AG1 | 1 | 4.25 | 0.28 |
| 0.8 | 0.1 | ptsI | 1 | 4.5 | 0.297 |
| 0.8 | 0.1 | clcB | 1 | 4.5 | 0.292 |
| 0.8 | 0.1 | ycaM | 1 | 4.5 | 0.286 |
| 0.8 | 0.1 | yadI | 1 | 4.5 | 0.326 |
| 0.8 | 0.1 | AG1 | 1 | 4.5 | 0.288 |

| | | | | | |
|---|---|---|---|---|---|
| 0.8 | 0.1 | ptsI | 1 | 4.75 | 0.303 |
| 0.8 | 0.1 | clcB | 1 | 4.75 | 0.298 |
| 0.8 | 0.1 | ycaM | 1 | 4.75 | 0.287 |
| 0.8 | 0.1 | yadI | 1 | 4.75 | 0.327 |
| 0.8 | 0.1 | AG1 | 1 | 4.75 | 0.29 |
| 0.8 | 0.1 | ptsI | 1 | 5 | 0.309 |
| 0.8 | 0.1 | clcB | 1 | 5 | 0.303 |
| 0.8 | 0.1 | ycaM | 1 | 5 | 0.295 |
| 0.8 | 0.1 | yadI | 1 | 5 | 0.336 |
| 0.8 | 0.1 | AG1 | 1 | 5 | 0.298 |
| 0.8 | 0.1 | ptsI | 1 | 5.25 | 0.315 |
| 0.8 | 0.1 | clcB | 1 | 5.25 | 0.31 |
| 0.8 | 0.1 | ycaM | 1 | 5.25 | 0.299 |
| 0.8 | 0.1 | yadI | 1 | 5.25 | 0.342 |
| 0.8 | 0.1 | AG1 | 1 | 5.25 | 0.301 |
| 0.8 | 0.1 | ptsI | 1 | 5.5 | 0.325 |
| 0.8 | 0.1 | clcB | 1 | 5.5 | 0.314 |
| 0.8 | 0.1 | ycaM | 1 | 5.5 | 0.305 |
| 0.8 | 0.1 | yadI | 1 | 5.5 | 0.352 |
| 0.8 | 0.1 | AG1 | 1 | 5.5 | 0.309 |
| 0.8 | 0.1 | ptsI | 1 | 5.75 | 0.334 |
| 0.8 | 0.1 | clcB | 1 | 5.75 | 0.322 |
| 0.8 | 0.1 | ycaM | 1 | 5.75 | 0.311 |
| 0.8 | 0.1 | yadI | 1 | 5.75 | 0.363 |
| 0.8 | 0.1 | AG1 | 1 | 5.75 | 0.317 |
| 0.8 | 0.1 | ptsI | 1 | 6 | 0.339 |
| 0.8 | 0.1 | clcB | 1 | 6 | 0.325 |
| 0.8 | 0.1 | ycaM | 1 | 6 | 0.311 |
| 0.8 | 0.1 | yadI | 1 | 6 | 0.36 |
| 0.8 | 0.1 | AG1 | 1 | 6 | 0.315 |
| 0.8 | 0.1 | ptsI | 1 | 6.25 | 0.344 |
| 0.8 | 0.1 | clcB | 1 | 6.25 | 0.332 |
| 0.8 | 0.1 | ycaM | 1 | 6.25 | 0.316 |
| 0.8 | 0.1 | yadI | 1 | 6.25 | 0.358 |
| 0.8 | 0.1 | AG1 | 1 | 6.25 | 0.321 |
| 0.8 | 0.1 | ptsI | 1 | 6.5 | 0.35 |
| 0.8 | 0.1 | clcB | 1 | 6.5 | 0.337 |
| 0.8 | 0.1 | ycaM | 1 | 6.5 | 0.322 |
| 0.8 | 0.1 | yadI | 1 | 6.5 | 0.363 |
| 0.8 | 0.1 | AG1 | 1 | 6.5 | 0.324 |
| 0.8 | 0.1 | ptsI | 1 | 6.75 | 0.358 |
| 0.8 | 0.1 | clcB | 1 | 6.75 | 0.344 |
| 0.8 | 0.1 | ycaM | 1 | 6.75 | 0.33 |
| 0.8 | 0.1 | yadI | 1 | 6.75 | 0.37 |
| 0.8 | 0.1 | AG1 | 1 | 6.75 | 0.334 |
| 0.8 | 0.1 | ptsI | 1 | 7 | 0.369 |
| 0.8 | 0.1 | clcB | 1 | 7 | 0.351 |
| 0.8 | 0.1 | ycaM | 1 | 7 | 0.337 |

| | | | | | |
|---|---|---|---|---|---|
| 0.8 | 0.1 | yadI | 1 | 7 | 0.377 |
| 0.8 | 0.1 | AG1 | 1 | 7 | 0.34 |
| 0.8 | 0.1 | ptsI | 1 | 7.25 | 0.377 |
| 0.8 | 0.1 | clcB | 1 | 7.25 | 0.353 |
| 0.8 | 0.1 | ycaM | 1 | 7.25 | 0.34 |
| 0.8 | 0.1 | yadI | 1 | 7.25 | 0.381 |
| 0.8 | 0.1 | AG1 | 1 | 7.25 | 0.348 |
| 0.8 | 0.1 | ptsI | 1 | 7.5 | 0.388 |
| 0.8 | 0.1 | clcB | 1 | 7.5 | 0.36 |
| 0.8 | 0.1 | ycaM | 1 | 7.5 | 0.348 |
| 0.8 | 0.1 | yadI | 1 | 7.5 | 0.392 |
| 0.8 | 0.1 | AG1 | 1 | 7.5 | 0.357 |
| 0.8 | 0.1 | ptsI | 1 | 7.75 | 0.402 |
| 0.8 | 0.1 | clcB | 1 | 7.75 | 0.373 |
| 0.8 | 0.1 | ycaM | 1 | 7.75 | 0.349 |
| 0.8 | 0.1 | yadI | 1 | 7.75 | 0.401 |
| 0.8 | 0.1 | AG1 | 1 | 7.75 | 0.369 |
| 0.8 | 0.1 | ptsI | 1 | 8 | 0.415 |
| 0.8 | 0.1 | clcB | 1 | 8 | 0.379 |
| 0.8 | 0.1 | ycaM | 1 | 8 | 0.347 |
| 0.8 | 0.1 | yadI | 1 | 8 | 0.402 |
| 0.8 | 0.1 | AG1 | 1 | 8 | 0.375 |
| 0.8 | 0.1 | ptsI | 1 | 8.25 | 0.436 |
| 0.8 | 0.1 | clcB | 1 | 8.25 | 0.382 |
| 0.8 | 0.1 | ycaM | 1 | 8.25 | 0.346 |
| 0.8 | 0.1 | yadI | 1 | 8.25 | 0.41 |
| 0.8 | 0.1 | AG1 | 1 | 8.25 | 0.383 |
| 0.8 | 0.1 | ptsI | 1 | 8.5 | 0.449 |
| 0.8 | 0.1 | clcB | 1 | 8.5 | 0.394 |
| 0.8 | 0.1 | ycaM | 1 | 8.5 | 0.35 |
| 0.8 | 0.1 | yadI | 1 | 8.5 | 0.415 |
| 0.8 | 0.1 | AG1 | 1 | 8.5 | 0.389 |
| 0.8 | 0.1 | ptsI | 1 | 8.75 | 0.462 |
| 0.8 | 0.1 | clcB | 1 | 8.75 | 0.401 |
| 0.8 | 0.1 | ycaM | 1 | 8.75 | 0.352 |
| 0.8 | 0.1 | yadI | 1 | 8.75 | 0.42 |
| 0.8 | 0.1 | AG1 | 1 | 8.75 | 0.399 |
| 0.8 | 0.1 | ptsI | 1 | 9 | 0.475 |
| 0.8 | 0.1 | clcB | 1 | 9 | 0.406 |
| 0.8 | 0.1 | ycaM | 1 | 9 | 0.354 |
| 0.8 | 0.1 | yadI | 1 | 9 | 0.433 |
| 0.8 | 0.1 | AG1 | 1 | 9 | 0.407 |
| 0.8 | 0.1 | ptsI | 1 | 9.25 | 0.489 |
| 0.8 | 0.1 | clcB | 1 | 9.25 | 0.41 |
| 0.8 | 0.1 | ycaM | 1 | 9.25 | 0.358 |
| 0.8 | 0.1 | yadI | 1 | 9.25 | 0.434 |
| 0.8 | 0.1 | AG1 | 1 | 9.25 | 0.422 |
| 0.8 | 0.1 | ptsI | 1 | 9.5 | 0.502 |

| | | | | | |
|---|---|---|---|---|---|
| 0.8 | 0.1 | clcB | 1 | 9.5 | 0.42 |
| 0.8 | 0.1 | ycaM | 1 | 9.5 | 0.367 |
| 0.8 | 0.1 | yadI | 1 | 9.5 | 0.434 |
| 0.8 | 0.1 | AG1 | 1 | 9.5 | 0.444 |
| 0.8 | 0.1 | ptsI | 1 | 9.75 | 0.538 |
| 0.8 | 0.1 | clcB | 1 | 9.75 | 0.432 |
| 0.8 | 0.1 | ycaM | 1 | 9.75 | 0.373 |
| 0.8 | 0.1 | yadI | 1 | 9.75 | 0.441 |
| 0.8 | 0.1 | AG1 | 1 | 9.75 | 0.459 |
| 0.8 | 0.1 | ptsI | 1 | 10 | 0.485 |
| 0.8 | 0.1 | clcB | 1 | 10 | 0.432 |
| 0.8 | 0.1 | ycaM | 1 | 10 | 0.37 |
| 0.8 | 0.1 | yadI | 1 | 10 | 0.421 |
| 0.8 | 0.1 | AG1 | 1 | 10 | 0.43 |
| 0.8 | 0.1 | ptsI | 1 | 10.25 | 0.517 |
| 0.8 | 0.1 | clcB | 1 | 10.25 | 0.437 |
| 0.8 | 0.1 | ycaM | 1 | 10.25 | 0.387 |
| 0.8 | 0.1 | yadI | 1 | 10.25 | 0.428 |
| 0.8 | 0.1 | AG1 | 1 | 10.25 | 0.448 |
| 0.8 | 0.1 | ptsI | 1 | 10.5 | 0.542 |
| 0.8 | 0.1 | clcB | 1 | 10.5 | 0.441 |
| 0.8 | 0.1 | ycaM | 1 | 10.5 | 0.385 |
| 0.8 | 0.1 | yadI | 1 | 10.5 | 0.435 |
| 0.8 | 0.1 | AG1 | 1 | 10.5 | 0.463 |
| 0.8 | 0.1 | ptsI | 1 | 10.75 | 0.522 |
| 0.8 | 0.1 | clcB | 1 | 10.75 | 0.448 |
| 0.8 | 0.1 | ycaM | 1 | 10.75 | 0.386 |
| 0.8 | 0.1 | yadI | 1 | 10.75 | 0.423 |
| 0.8 | 0.1 | AG1 | 1 | 10.75 | 0.431 |
| 0.8 | 0.1 | ptsI | 1 | 11 | 0.521 |
| 0.8 | 0.1 | clcB | 1 | 11 | 0.467 |
| 0.8 | 0.1 | ycaM | 1 | 11 | 0.39 |
| 0.8 | 0.1 | yadI | 1 | 11 | 0.435 |
| 0.8 | 0.1 | AG1 | 1 | 11 | 0.452 |
| 0.8 | 0.1 | ptsI | 1 | 11.25 | 0.595 |
| 0.8 | 0.1 | clcB | 1 | 11.25 | 0.493 |
| 0.8 | 0.1 | ycaM | 1 | 11.25 | 0.404 |
| 0.8 | 0.1 | yadI | 1 | 11.25 | 0.439 |
| 0.8 | 0.1 | AG1 | 1 | 11.25 | 0.49 |
| 0.8 | 0.1 | ptsI | 1 | 11.5 | 0.577 |
| 0.8 | 0.1 | clcB | 1 | 11.5 | 0.491 |
| 0.8 | 0.1 | ycaM | 1 | 11.5 | 0.4 |
| 0.8 | 0.1 | yadI | 1 | 11.5 | 0.44 |
| 0.8 | 0.1 | AG1 | 1 | 11.5 | 0.482 |
| 0.8 | 0.1 | ptsI | 1 | 11.75 | 0.579 |
| 0.8 | 0.1 | clcB | 1 | 11.75 | 0.499 |
| 0.8 | 0.1 | ycaM | 1 | 11.75 | 0.404 |
| 0.8 | 0.1 | yadI | 1 | 11.75 | 0.426 |

| | | | | | |
|---|---|---|---|---|---|
| 0.8 | 0.1 | AG1 | 1 | 11.75 | 0.475 |
| 0.8 | 0.1 | ptsI | 1 | 12 | 0.646 |
| 0.8 | 0.1 | clcB | 1 | 12 | 0.546 |
| 0.8 | 0.1 | ycaM | 1 | 12 | 0.423 |
| 0.8 | 0.1 | yadI | 1 | 12 | 0.43 |
| 0.8 | 0.1 | AG1 | 1 | 12 | 0.526 |
| 0.8 | 0.1 | ptsI | 1 | 12.25 | 0.6 |
| 0.8 | 0.1 | clcB | 1 | 12.25 | 0.525 |
| 0.8 | 0.1 | ycaM | 1 | 12.25 | 0.414 |
| 0.8 | 0.1 | yadI | 1 | 12.25 | 0.43 |
| 0.8 | 0.1 | AG1 | 1 | 12.25 | 0.507 |
| 0.8 | 0.1 | ptsI | 1 | 12.5 | 0.641 |
| 0.8 | 0.1 | clcB | 1 | 12.5 | 0.553 |
| 0.8 | 0.1 | ycaM | 1 | 12.5 | 0.423 |
| 0.8 | 0.1 | yadI | 1 | 12.5 | 0.429 |
| 0.8 | 0.1 | AG1 | 1 | 12.5 | 0.544 |
| 0.8 | 0.1 | ptsI | 1 | 12.75 | 0.658 |
| 0.8 | 0.1 | clcB | 1 | 12.75 | 0.583 |
| 0.8 | 0.1 | ycaM | 1 | 12.75 | 0.436 |
| 0.8 | 0.1 | yadI | 1 | 12.75 | 0.447 |
| 0.8 | 0.1 | AG1 | 1 | 12.75 | 0.569 |
| 0.8 | 0.1 | ptsI | 1 | 13 | 0.668 |
| 0.8 | 0.1 | clcB | 1 | 13 | 0.598 |
| 0.8 | 0.1 | ycaM | 1 | 13 | 0.465 |
| 0.8 | 0.1 | yadI | 1 | 13 | 0.447 |
| 0.8 | 0.1 | AG1 | 1 | 13 | 0.627 |
| 0.8 | 0.1 | ptsI | 1 | 13.25 | 0.657 |
| 0.8 | 0.1 | clcB | 1 | 13.25 | 0.582 |
| 0.8 | 0.1 | ycaM | 1 | 13.25 | 0.451 |
| 0.8 | 0.1 | yadI | 1 | 13.25 | 0.44 |
| 0.8 | 0.1 | AG1 | 1 | 13.25 | 0.613 |
| 0.8 | 0.1 | ptsI | 1 | 13.5 | 0.698 |
| 0.8 | 0.1 | clcB | 1 | 13.5 | 0.615 |
| 0.8 | 0.1 | ycaM | 1 | 13.5 | 0.471 |
| 0.8 | 0.1 | yadI | 1 | 13.5 | 0.444 |
| 0.8 | 0.1 | AG1 | 1 | 13.5 | 0.668 |
| 0.8 | 0.1 | ptsI | 1 | 13.75 | 0.669 |
| 0.8 | 0.1 | clcB | 1 | 13.75 | 0.602 |
| 0.8 | 0.1 | ycaM | 1 | 13.75 | 0.471 |
| 0.8 | 0.1 | yadI | 1 | 13.75 | 0.454 |
| 0.8 | 0.1 | AG1 | 1 | 13.75 | 0.688 |
| 0.8 | 0.1 | ptsI | 1 | 14 | 0.709 |
| 0.8 | 0.1 | clcB | 1 | 14 | 0.626 |
| 0.8 | 0.1 | ycaM | 1 | 14 | 0.485 |
| 0.8 | 0.1 | yadI | 1 | 14 | 0.447 |
| 0.8 | 0.1 | AG1 | 1 | 14 | 0.751 |
| 0.8 | 0.1 | ptsI | 1 | 14.25 | 0.716 |
| 0.8 | 0.1 | clcB | 1 | 14.25 | 0.655 |

| | | | | | |
|---|---|---|---|---|---|
| 0.8 | 0.1 | ycaM | 1 | 14.25 | 0.498 |
| 0.8 | 0.1 | yadI | 1 | 14.25 | 0.459 |
| 0.8 | 0.1 | AG1 | 1 | 14.25 | 0.777 |
| 0.8 | 0.1 | ptsI | 1 | 14.5 | 0.713 |
| 0.8 | 0.1 | clcB | 1 | 14.5 | 0.666 |
| 0.8 | 0.1 | ycaM | 1 | 14.5 | 0.503 |
| 0.8 | 0.1 | yadI | 1 | 14.5 | 0.454 |
| 0.8 | 0.1 | AG1 | 1 | 14.5 | 0.785 |
| 0.8 | 0.1 | ptsI | 1 | 14.75 | 0.703 |
| 0.8 | 0.1 | clcB | 1 | 14.75 | 0.646 |
| 0.8 | 0.1 | ycaM | 1 | 14.75 | 0.497 |
| 0.8 | 0.1 | yadI | 1 | 14.75 | 0.451 |
| 0.8 | 0.1 | AG1 | 1 | 14.75 | 0.76 |
| 0.8 | 0.1 | ptsI | 1 | 15 | 0.738 |
| 0.8 | 0.1 | clcB | 1 | 15 | 0.681 |
| 0.8 | 0.1 | ycaM | 1 | 15 | 0.549 |
| 0.8 | 0.1 | yadI | 1 | 15 | 0.456 |
| 0.8 | 0.1 | AG1 | 1 | 15 | 0.8 |
| 0.8 | 0.1 | ptsI | 1 | 15.25 | 0.746 |
| 0.8 | 0.1 | clcB | 1 | 15.25 | 0.701 |
| 0.8 | 0.1 | ycaM | 1 | 15.25 | 0.589 |
| 0.8 | 0.1 | yadI | 1 | 15.25 | 0.47 |
| 0.8 | 0.1 | AG1 | 1 | 15.25 | 0.838 |
| 0.8 | 0.1 | ptsI | 1 | 15.5 | 0.727 |
| 0.8 | 0.1 | clcB | 1 | 15.5 | 0.683 |
| 0.8 | 0.1 | ycaM | 1 | 15.5 | 0.575 |
| 0.8 | 0.1 | yadI | 1 | 15.5 | 0.466 |
| 0.8 | 0.1 | AG1 | 1 | 15.5 | 0.819 |
| 0.8 | 0.1 | ptsI | 1 | 15.75 | 0.728 |
| 0.8 | 0.1 | clcB | 1 | 15.75 | 0.694 |
| 0.8 | 0.1 | ycaM | 1 | 15.75 | 0.564 |
| 0.8 | 0.1 | yadI | 1 | 15.75 | 0.462 |
| 0.8 | 0.1 | AG1 | 1 | 15.75 | 0.799 |
| 0.8 | 0.1 | ptsI | 1 | 16 | 0.708 |
| 0.8 | 0.1 | clcB | 1 | 16 | 0.675 |
| 0.8 | 0.1 | ycaM | 1 | 16 | 0.548 |
| 0.8 | 0.1 | yadI | 1 | 16 | 0.458 |
| 0.8 | 0.1 | AG1 | 1 | 16 | 0.764 |
| 0.8 | 0.1 | ptsI | 1 | 16.25 | 0.759 |
| 0.8 | 0.1 | clcB | 1 | 16.25 | 0.725 |
| 0.8 | 0.1 | ycaM | 1 | 16.25 | 0.614 |
| 0.8 | 0.1 | yadI | 1 | 16.25 | 0.474 |
| 0.8 | 0.1 | AG1 | 1 | 16.25 | 0.82 |
| 0.8 | 0.1 | ptsI | 1 | 16.5 | 0.766 |
| 0.8 | 0.1 | clcB | 1 | 16.5 | 0.725 |
| 0.8 | 0.1 | ycaM | 1 | 16.5 | 0.64 |
| 0.8 | 0.1 | yadI | 1 | 16.5 | 0.476 |
| 0.8 | 0.1 | AG1 | 1 | 16.5 | 0.822 |

| | | | | | |
|---|---|---|---|---|---|
| 0.8 | 0.1 | ptsI | 1 | 17.0667 | 0.596 |
| 0.8 | 0.1 | clcB | 1 | 17.0667 | 0.493 |
| 0.8 | 0.1 | ycaM | 1 | 17.0667 | 0.517 |
| 0.8 | 0.1 | yadI | 1 | 17.0667 | 0.638 |
| 0.8 | 0.1 | AG1 | 1 | 17.0667 | 0.855 |
| 0.8 | 0.1 | ptsI | 1 | 17.3167 | 0.639 |
| 0.8 | 0.1 | clcB | 1 | 17.3167 | 0.467 |
| 0.8 | 0.1 | ycaM | 1 | 17.3167 | 0.484 |
| 0.8 | 0.1 | yadI | 1 | 17.3167 | 0.63 |
| 0.8 | 0.1 | AG1 | 1 | 17.3167 | 0.812 |
| 0.8 | 0.1 | ptsI | 1 | 17.5667 | 0.651 |
| 0.8 | 0.1 | clcB | 1 | 17.5667 | 0.46 |
| 0.8 | 0.1 | ycaM | 1 | 17.5667 | 0.476 |
| 0.8 | 0.1 | yadI | 1 | 17.5667 | 0.633 |
| 0.8 | 0.1 | AG1 | 1 | 17.5667 | 0.821 |
| 0.8 | 0.1 | ptsI | 1 | 17.8167 | 0.64 |
| 0.8 | 0.1 | clcB | 1 | 17.8167 | 0.465 |
| 0.8 | 0.1 | ycaM | 1 | 17.8167 | 0.477 |
| 0.8 | 0.1 | yadI | 1 | 17.8167 | 0.644 |
| 0.8 | 0.1 | AG1 | 1 | 17.8167 | 0.809 |
| 0.8 | 0.1 | ptsI | 1 | 18.0667 | 0.651 |
| 0.8 | 0.1 | clcB | 1 | 18.0667 | 0.464 |
| 0.8 | 0.1 | ycaM | 1 | 18.0667 | 0.471 |
| 0.8 | 0.1 | yadI | 1 | 18.0667 | 0.657 |
| 0.8 | 0.1 | AG1 | 1 | 18.0667 | 0.814 |
| 0.8 | 0.1 | ptsI | 1 | 18.3167 | 0.675 |
| 0.8 | 0.1 | clcB | 1 | 18.3167 | 0.45 |
| 0.8 | 0.1 | ycaM | 1 | 18.3167 | 0.468 |
| 0.8 | 0.1 | yadI | 1 | 18.3167 | 0.649 |
| 0.8 | 0.1 | AG1 | 1 | 18.3167 | 0.814 |
| 0.8 | 0.1 | ptsI | 1 | 18.5667 | 0.756 |
| 0.8 | 0.1 | clcB | 1 | 18.5667 | 0.453 |
| 0.8 | 0.1 | ycaM | 1 | 18.5667 | 0.469 |
| 0.8 | 0.1 | yadI | 1 | 18.5667 | 0.656 |
| 0.8 | 0.1 | AG1 | 1 | 18.5667 | 0.843 |
| 0.8 | 0.1 | ptsI | 1 | 18.8167 | 0.761 |
| 0.8 | 0.1 | clcB | 1 | 18.8167 | 0.454 |
| 0.8 | 0.1 | ycaM | 1 | 18.8167 | 0.465 |
| 0.8 | 0.1 | yadI | 1 | 18.8167 | 0.667 |
| 0.8 | 0.1 | AG1 | 1 | 18.8167 | 0.849 |
| 0.8 | 0.1 | ptsI | 1 | 19.0667 | 0.791 |
| 0.8 | 0.1 | clcB | 1 | 19.0667 | 0.46 |
| 0.8 | 0.1 | ycaM | 1 | 19.0667 | 0.466 |
| 0.8 | 0.1 | yadI | 1 | 19.0667 | 0.689 |
| 0.8 | 0.1 | AG1 | 1 | 19.0667 | 0.87 |
| 0.8 | 0.1 | ptsI | 1 | 19.3167 | 0.768 |
| 0.8 | 0.1 | clcB | 1 | 19.3167 | 0.454 |
| 0.8 | 0.1 | ycaM | 1 | 19.3167 | 0.462 |

| | | | | | |
|---|---|---|---|---|---|
| 0.8 | 0.1 | yadI | 1 | 19.3167 | 0.683 |
| 0.8 | 0.1 | AG1 | 1 | 19.3167 | 0.864 |
| 0.8 | 0.1 | ptsI | 1 | 19.5667 | 0.793 |
| 0.8 | 0.1 | clcB | 1 | 19.5667 | 0.465 |
| 0.8 | 0.1 | ycaM | 1 | 19.5667 | 0.467 |
| 0.8 | 0.1 | yadI | 1 | 19.5667 | 0.695 |
| 0.8 | 0.1 | AG1 | 1 | 19.5667 | 0.88 |
| 0.8 | 0.1 | ptsI | 1 | 19.8167 | 0.798 |
| 0.8 | 0.1 | clcB | 1 | 19.8167 | 0.452 |
| 0.8 | 0.1 | ycaM | 1 | 19.8167 | 0.47 |
| 0.8 | 0.1 | yadI | 1 | 19.8167 | 0.697 |
| 0.8 | 0.1 | AG1 | 1 | 19.8167 | 0.879 |
| 0.8 | 0.1 | ptsI | 1 | 20.0667 | 0.799 |
| 0.8 | 0.1 | clcB | 1 | 20.0667 | 0.452 |
| 0.8 | 0.1 | ycaM | 1 | 20.0667 | 0.473 |
| 0.8 | 0.1 | yadI | 1 | 20.0667 | 0.71 |
| 0.8 | 0.1 | AG1 | 1 | 20.0667 | 0.882 |
| 0.8 | 0.1 | ptsI | 1 | 20.3167 | 0.804 |
| 0.8 | 0.1 | clcB | 1 | 20.3167 | 0.453 |
| 0.8 | 0.1 | ycaM | 1 | 20.3167 | 0.471 |
| 0.8 | 0.1 | yadI | 1 | 20.3167 | 0.714 |
| 0.8 | 0.1 | AG1 | 1 | 20.3167 | 0.888 |
| 0.8 | 0.1 | ptsI | 1 | 20.5667 | 0.796 |
| 0.8 | 0.1 | clcB | 1 | 20.5667 | 0.431 |
| 0.8 | 0.1 | ycaM | 1 | 20.5667 | 0.469 |
| 0.8 | 0.1 | yadI | 1 | 20.5667 | 0.711 |
| 0.8 | 0.1 | AG1 | 1 | 20.5667 | 0.885 |
| 0.8 | 0.1 | ptsI | 1 | 20.8167 | 0.798 |
| 0.8 | 0.1 | clcB | 1 | 20.8167 | 0.435 |
| 0.8 | 0.1 | ycaM | 1 | 20.8167 | 0.468 |
| 0.8 | 0.1 | yadI | 1 | 20.8167 | 0.713 |
| 0.8 | 0.1 | AG1 | 1 | 20.8167 | 0.891 |
| 0.8 | 0.1 | ptsI | 1 | 21.0667 | 0.807 |
| 0.8 | 0.1 | clcB | 1 | 21.0667 | 0.428 |
| 0.8 | 0.1 | ycaM | 1 | 21.0667 | 0.464 |
| 0.8 | 0.1 | yadI | 1 | 21.0667 | 0.723 |
| 0.8 | 0.1 | AG1 | 1 | 21.0667 | 0.899 |
| 0.8 | 0.1 | ptsI | 1 | 21.3167 | 0.806 |
| 0.8 | 0.1 | clcB | 1 | 21.3167 | 0.426 |
| 0.8 | 0.1 | ycaM | 1 | 21.3167 | 0.467 |
| 0.8 | 0.1 | yadI | 1 | 21.3167 | 0.729 |
| 0.8 | 0.1 | AG1 | 1 | 21.3167 | 0.898 |
| 0.8 | 0.1 | ptsI | 1 | 21.5667 | 0.805 |
| 0.8 | 0.1 | clcB | 1 | 21.5667 | 0.423 |
| 0.8 | 0.1 | ycaM | 1 | 21.5667 | 0.469 |
| 0.8 | 0.1 | yadI | 1 | 21.5667 | 0.725 |
| 0.8 | 0.1 | AG1 | 1 | 21.5667 | 0.902 |
| 0.8 | 0.1 | ptsI | 1 | 21.8167 | 0.811 |

| | | | | | |
|---|---|---|---|---|---|
| 0.8 | 0.1 | clcB | 1 | 21.8167 | 0.41 |
| 0.8 | 0.1 | ycaM | 1 | 21.8167 | 0.47 |
| 0.8 | 0.1 | yadI | 1 | 21.8167 | 0.73 |
| 0.8 | 0.1 | AG1 | 1 | 21.8167 | 0.905 |
| 0.8 | 0.1 | ptsI | 1 | 22.0667 | 0.829 |
| 0.8 | 0.1 | clcB | 1 | 22.0667 | 0.418 |
| 0.8 | 0.1 | ycaM | 1 | 22.0667 | 0.466 |
| 0.8 | 0.1 | yadI | 1 | 22.0667 | 0.743 |
| 0.8 | 0.1 | AG1 | 1 | 22.0667 | 0.911 |
| 0.8 | 0.1 | ptsI | 1 | 22.9333 | 0.82 |
| 0.8 | 0.1 | clcB | 1 | 22.9333 | 0.394 |
| 0.8 | 0.1 | ycaM | 1 | 22.9333 | 0.431 |
| 0.8 | 0.1 | yadI | 1 | 22.9333 | 0.74 |
| 0.8 | 0.1 | AG1 | 1 | 22.9333 | 0.738 |
| 0.8 | 0.1 | ptsI | 1 | 23.1833 | 0.796 |
| 0.8 | 0.1 | clcB | 1 | 23.1833 | 0.378 |
| 0.8 | 0.1 | ycaM | 1 | 23.1833 | 0.42 |
| 0.8 | 0.1 | yadI | 1 | 23.1833 | 0.724 |
| 0.8 | 0.1 | AG1 | 1 | 23.1833 | 0.728 |
| 0.8 | 0.1 | ptsI | 1 | 23.4333 | 0.771 |
| 0.8 | 0.1 | clcB | 1 | 23.4333 | 0.375 |
| 0.8 | 0.1 | ycaM | 1 | 23.4333 | 0.414 |
| 0.8 | 0.1 | yadI | 1 | 23.4333 | 0.728 |
| 0.8 | 0.1 | AG1 | 1 | 23.4333 | 0.739 |
| 0.8 | 0.1 | ptsI | 1 | 23.6833 | 0.824 |
| 0.8 | 0.1 | clcB | 1 | 23.6833 | 0.372 |
| 0.8 | 0.1 | ycaM | 1 | 23.6833 | 0.415 |
| 0.8 | 0.1 | yadI | 1 | 23.6833 | 0.738 |
| 0.8 | 0.1 | AG1 | 1 | 23.6833 | 0.745 |
| 0.8 | 0.1 | ptsI | 1 | 23.9333 | 0.832 |
| 0.8 | 0.1 | clcB | 1 | 23.9333 | 0.375 |
| 0.8 | 0.1 | ycaM | 1 | 23.9333 | 0.416 |
| 0.8 | 0.1 | yadI | 1 | 23.9333 | 0.738 |
| 0.8 | 0.1 | AG1 | 1 | 23.9333 | 0.762 |
| 0.8 | 0.1 | ptsI | 1 | 24.1833 | 0.855 |
| 0.8 | 0.1 | clcB | 1 | 24.1833 | 0.377 |
| 0.8 | 0.1 | ycaM | 1 | 24.1833 | 0.416 |
| 0.8 | 0.1 | yadI | 1 | 24.1833 | 0.75 |
| 0.8 | 0.1 | AG1 | 1 | 24.1833 | 0.788 |
| 0.8 | 0.1 | ptsI | 1 | 24.4333 | 0.853 |
| 0.8 | 0.1 | clcB | 1 | 24.4333 | 0.378 |
| 0.8 | 0.1 | ycaM | 1 | 24.4333 | 0.416 |
| 0.8 | 0.1 | yadI | 1 | 24.4333 | 0.758 |
| 0.8 | 0.1 | AG1 | 1 | 24.4333 | 0.793 |
| 0.8 | 0.1 | ptsI | 1 | 24.6833 | 0.855 |
| 0.8 | 0.1 | clcB | 1 | 24.6833 | 0.382 |
| 0.8 | 0.1 | ycaM | 1 | 24.6833 | 0.412 |
| 0.8 | 0.1 | yadI | 1 | 24.6833 | 0.761 |

| | | | | | |
|---|---|---|---|---|---|
| 0.8 | 0.1 | AG1 | 1 | 24.6833 | 0.794 |
| 0.8 | 0.1 | ptsI | 1 | 24.9333 | 0.839 |
| 0.8 | 0.1 | clcB | 1 | 24.9333 | 0.427 |
| 0.8 | 0.1 | ycaM | 1 | 24.9333 | 0.412 |
| 0.8 | 0.1 | yadI | 1 | 24.9333 | 0.765 |
| 0.8 | 0.1 | AG1 | 1 | 24.9333 | 0.8 |
| 0.8 | 0.1 | ptsI | 1 | 25.1833 | 0.843 |
| 0.8 | 0.1 | clcB | 1 | 25.1833 | 0.443 |
| 0.8 | 0.1 | ycaM | 1 | 25.1833 | 0.413 |
| 0.8 | 0.1 | yadI | 1 | 25.1833 | 0.77 |
| 0.8 | 0.1 | AG1 | 1 | 25.1833 | 0.795 |
| 0.8 | 0.1 | ptsI | 1 | 25.4333 | 0.861 |
| 0.8 | 0.1 | clcB | 1 | 25.4333 | 0.385 |
| 0.8 | 0.1 | ycaM | 1 | 25.4333 | 0.415 |
| 0.8 | 0.1 | yadI | 1 | 25.4333 | 0.777 |
| 0.8 | 0.1 | AG1 | 1 | 25.4333 | 0.81 |
| 0.8 | 0.1 | ptsI | 1 | 25.6833 | 0.854 |
| 0.8 | 0.1 | clcB | 1 | 25.6833 | 0.414 |
| 0.8 | 0.1 | ycaM | 1 | 25.6833 | 0.412 |
| 0.8 | 0.1 | yadI | 1 | 25.6833 | 0.782 |
| 0.8 | 0.1 | AG1 | 1 | 25.6833 | 0.797 |
| 0.8 | 0.1 | ptsI | 1 | 25.9333 | 0.856 |
| 0.8 | 0.1 | clcB | 1 | 25.9333 | 0.388 |
| 0.8 | 0.1 | ycaM | 1 | 25.9333 | 0.412 |
| 0.8 | 0.1 | yadI | 1 | 25.9333 | 0.788 |
| 0.8 | 0.1 | AG1 | 1 | 25.9333 | 0.81 |
| 0.8 | 0.1 | ptsI | 1 | 26.1833 | 0.867 |
| 0.8 | 0.1 | clcB | 1 | 26.1833 | 0.393 |
| 0.8 | 0.1 | ycaM | 1 | 26.1833 | 0.416 |
| 0.8 | 0.1 | yadI | 1 | 26.1833 | 0.792 |
| 0.8 | 0.1 | AG1 | 1 | 26.1833 | 0.812 |
| 0.8 | 0.1 | ptsI | 1 | 26.4333 | 0.867 |
| 0.8 | 0.1 | clcB | 1 | 26.4333 | 0.386 |
| 0.8 | 0.1 | ycaM | 1 | 26.4333 | 0.414 |
| 0.8 | 0.1 | yadI | 1 | 26.4333 | 0.792 |
| 0.8 | 0.1 | AG1 | 1 | 26.4333 | 0.809 |
| 0.8 | 0.1 | ptsI | 1 | 26.6833 | 0.866 |
| 0.8 | 0.1 | clcB | 1 | 26.6833 | 0.383 |
| 0.8 | 0.1 | ycaM | 1 | 26.6833 | 0.415 |
| 0.8 | 0.1 | yadI | 1 | 26.6833 | 0.796 |
| 0.8 | 0.1 | AG1 | 1 | 26.6833 | 0.813 |
| 0.8 | 0.1 | ptsI | 1 | 26.9333 | 0.871 |
| 0.8 | 0.1 | clcB | 1 | 26.9333 | 0.386 |
| 0.8 | 0.1 | ycaM | 1 | 26.9333 | 0.414 |
| 0.8 | 0.1 | yadI | 1 | 26.9333 | 0.799 |
| 0.8 | 0.1 | AG1 | 1 | 26.9333 | 0.816 |
| 0.8 | 0.1 | ptsI | 1 | 27.1833 | 0.872 |
| 0.8 | 0.1 | clcB | 1 | 27.1833 | 0.382 |

| | | | | | |
|---|---|---|---|---|---|
| 0.8 | 0.1 | ycaM | 1 | 27.1833 | 0.417 |
| 0.8 | 0.1 | yadI | 1 | 27.1833 | 0.803 |
| 0.8 | 0.1 | AG1 | 1 | 27.1833 | 0.822 |
| 0.8 | 0.1 | ptsI | 1 | 27.4333 | 0.859 |
| 0.8 | 0.1 | clcB | 1 | 27.4333 | 0.373 |
| 0.8 | 0.1 | ycaM | 1 | 27.4333 | 0.417 |
| 0.8 | 0.1 | yadI | 1 | 27.4333 | 0.802 |
| 0.8 | 0.1 | AG1 | 1 | 27.4333 | 0.811 |
| 0.8 | 0.1 | ptsI | 1 | 27.6833 | 0.873 |
| 0.8 | 0.1 | clcB | 1 | 27.6833 | 0.376 |
| 0.8 | 0.1 | ycaM | 1 | 27.6833 | 0.417 |
| 0.8 | 0.1 | yadI | 1 | 27.6833 | 0.805 |
| 0.8 | 0.1 | AG1 | 1 | 27.6833 | 0.816 |
| 0.8 | 0.1 | ptsI | 1 | 27.9333 | 0.871 |
| 0.8 | 0.1 | clcB | 1 | 27.9333 | 0.366 |
| 0.8 | 0.1 | ycaM | 1 | 27.9333 | 0.412 |
| 0.8 | 0.1 | yadI | 1 | 27.9333 | 0.79 |
| 0.8 | 0.1 | AG1 | 1 | 27.9333 | 0.812 |
| 0.8 | 0.1 | ptsI | 1 | 28.1833 | 0.886 |
| 0.8 | 0.1 | clcB | 1 | 28.1833 | 0.385 |
| 0.8 | 0.1 | ycaM | 1 | 28.1833 | 0.417 |
| 0.8 | 0.1 | yadI | 1 | 28.1833 | 0.806 |
| 0.8 | 0.1 | AG1 | 1 | 28.1833 | 0.825 |
| 0.8 | 0.1 | ptsI | 1 | 28.4333 | 0.878 |
| 0.8 | 0.1 | clcB | 1 | 28.4333 | 0.385 |
| 0.8 | 0.1 | ycaM | 1 | 28.4333 | 0.418 |
| 0.8 | 0.1 | yadI | 1 | 28.4333 | 0.812 |
| 0.8 | 0.1 | AG1 | 1 | 28.4333 | 0.822 |
| 0.8 | 0.1 | ptsI | 1 | 28.6833 | 0.882 |
| 0.8 | 0.1 | clcB | 1 | 28.6833 | 0.386 |
| 0.8 | 0.1 | ycaM | 1 | 28.6833 | 0.418 |
| 0.8 | 0.1 | yadI | 1 | 28.6833 | 0.818 |
| 0.8 | 0.1 | AG1 | 1 | 28.6833 | 0.824 |
| 0.8 | 0.1 | ptsI | 1 | 28.9333 | 0.888 |
| 0.8 | 0.1 | clcB | 1 | 28.9333 | 0.389 |
| 0.8 | 0.1 | ycaM | 1 | 28.9333 | 0.419 |
| 0.8 | 0.1 | yadI | 1 | 28.9333 | 0.815 |
| 0.8 | 0.1 | AG1 | 1 | 28.9333 | 0.825 |
| 0.8 | 0.1 | ptsI | 1 | 29.1833 | 0.872 |
| 0.8 | 0.1 | clcB | 1 | 29.1833 | 0.387 |
| 0.8 | 0.1 | ycaM | 1 | 29.1833 | 0.418 |
| 0.8 | 0.1 | yadI | 1 | 29.1833 | 0.814 |
| 0.8 | 0.1 | AG1 | 1 | 29.1833 | 0.818 |
| 0.8 | 0.1 | ptsI | 1 | 29.4333 | 0.885 |
| 0.8 | 0.1 | clcB | 1 | 29.4333 | 0.386 |
| 0.8 | 0.1 | ycaM | 1 | 29.4333 | 0.42 |
| 0.8 | 0.1 | yadI | 1 | 29.4333 | 0.821 |
| 0.8 | 0.1 | AG1 | 1 | 29.4333 | 0.828 |

| | | | | | |
|---|---|---|---|---|---|
| 0.8 | 0.1 | ptsI | 1 | 29.6833 | 0.878 |
| 0.8 | 0.1 | clcB | 1 | 29.6833 | 0.388 |
| 0.8 | 0.1 | ycaM | 1 | 29.6833 | 0.42 |
| 0.8 | 0.1 | yadI | 1 | 29.6833 | 0.831 |
| 0.8 | 0.1 | AG1 | 1 | 29.6833 | 0.827 |
| 0.8 | 0.1 | ptsI | 1 | 29.9333 | 0.887 |
| 0.8 | 0.1 | clcB | 1 | 29.9333 | 0.389 |
| 0.8 | 0.1 | ycaM | 1 | 29.9333 | 0.419 |
| 0.8 | 0.1 | yadI | 1 | 29.9333 | 0.826 |
| 0.8 | 0.1 | AG1 | 1 | 29.9333 | 0.833 |
| 0.8 | 0.1 | ptsI | 1 | 30.1833 | 0.886 |
| 0.8 | 0.1 | clcB | 1 | 30.1833 | 0.39 |
| 0.8 | 0.1 | ycaM | 1 | 30.1833 | 0.42 |
| 0.8 | 0.1 | yadI | 1 | 30.1833 | 0.825 |
| 0.8 | 0.1 | AG1 | 1 | 30.1833 | 0.835 |
| 0.8 | 0.1 | ptsI | 1 | 30.4333 | 0.885 |
| 0.8 | 0.1 | clcB | 1 | 30.4333 | 0.392 |
| 0.8 | 0.1 | ycaM | 1 | 30.4333 | 0.423 |
| 0.8 | 0.1 | yadI | 1 | 30.4333 | 0.831 |
| 0.8 | 0.1 | AG1 | 1 | 30.4333 | 0.838 |
| 0.8 | 0.1 | ptsI | 1 | 30.6833 | 0.878 |
| 0.8 | 0.1 | clcB | 1 | 30.6833 | 0.393 |
| 0.8 | 0.1 | ycaM | 1 | 30.6833 | 0.422 |
| 0.8 | 0.1 | yadI | 1 | 30.6833 | 0.841 |
| 0.8 | 0.1 | AG1 | 1 | 30.6833 | 0.836 |
| 0.8 | 0.1 | ptsI | 1 | 30.9333 | 0.888 |
| 0.8 | 0.1 | clcB | 1 | 30.9333 | 0.394 |
| 0.8 | 0.1 | ycaM | 1 | 30.9333 | 0.424 |
| 0.8 | 0.1 | yadI | 1 | 30.9333 | 0.843 |
| 0.8 | 0.1 | AG1 | 1 | 30.9333 | 0.841 |
| 0.8 | 0.1 | ptsI | 1 | 31.1833 | 0.884 |
| 0.8 | 0.1 | clcB | 1 | 31.1833 | 0.394 |
| 0.8 | 0.1 | ycaM | 1 | 31.1833 | 0.423 |
| 0.8 | 0.1 | yadI | 1 | 31.1833 | 0.845 |
| 0.8 | 0.1 | AG1 | 1 | 31.1833 | 0.839 |
| 0.8 | 0.1 | ptsI | 1 | 31.4333 | 0.892 |
| 0.8 | 0.1 | clcB | 1 | 31.4333 | 0.397 |
| 0.8 | 0.1 | ycaM | 1 | 31.4333 | 0.425 |
| 0.8 | 0.1 | yadI | 1 | 31.4333 | 0.851 |
| 0.8 | 0.1 | AG1 | 1 | 31.4333 | 0.847 |
| 0.8 | 0.1 | ptsI | 1 | 31.6833 | 0.89 |
| 0.8 | 0.1 | clcB | 1 | 31.6833 | 0.398 |
| 0.8 | 0.1 | ycaM | 1 | 31.6833 | 0.424 |
| 0.8 | 0.1 | yadI | 1 | 31.6833 | 0.854 |
| 0.8 | 0.1 | AG1 | 1 | 31.6833 | 0.847 |
| 0.8 | 0.1 | ptsI | 1 | 31.9333 | 0.894 |
| 0.8 | 0.1 | clcB | 1 | 31.9333 | 0.399 |
| 0.8 | 0.1 | ycaM | 1 | 31.9333 | 0.426 |

| | | | | | |
|---|---|---|---|---|---|
| 0.8 | 0.1 | yadI | 1 | 31.9333 | 0.848 |
| 0.8 | 0.1 | AG1 | 1 | 31.9333 | 0.853 |
| 0.8 | 0.1 | ptsI | 1 | 32.1833 | 0.896 |
| 0.8 | 0.1 | clcB | 1 | 32.1833 | 0.4 |
| 0.8 | 0.1 | ycaM | 1 | 32.1833 | 0.427 |
| 0.8 | 0.1 | yadI | 1 | 32.1833 | 0.852 |
| 0.8 | 0.1 | AG1 | 1 | 32.1833 | 0.857 |
| 0.8 | 0.1 | ptsI | 1 | 32.4333 | 0.891 |
| 0.8 | 0.1 | clcB | 1 | 32.4333 | 0.385 |
| 0.8 | 0.1 | ycaM | 1 | 32.4333 | 0.42 |
| 0.8 | 0.1 | yadI | 1 | 32.4333 | 0.838 |
| 0.8 | 0.1 | AG1 | 1 | 32.4333 | 0.856 |
| 0.8 | 0.1 | ptsI | 1 | 32.6833 | 0.893 |
| 0.8 | 0.1 | clcB | 1 | 32.6833 | 0.406 |
| 0.8 | 0.1 | ycaM | 1 | 32.6833 | 0.428 |
| 0.8 | 0.1 | yadI | 1 | 32.6833 | 0.861 |
| 0.8 | 0.1 | AG1 | 1 | 32.6833 | 0.861 |
| 0.8 | 0.1 | ptsI | 1 | 32.9333 | 0.892 |
| 0.8 | 0.1 | clcB | 1 | 32.9333 | 0.404 |
| 0.8 | 0.1 | ycaM | 1 | 32.9333 | 0.428 |
| 0.8 | 0.1 | yadI | 1 | 32.9333 | 0.863 |
| 0.8 | 0.1 | AG1 | 1 | 32.9333 | 0.864 |
| 0.8 | 0.1 | ptsI | 1 | 33.1833 | 0.892 |
| 0.8 | 0.1 | clcB | 1 | 33.1833 | 0.397 |
| 0.8 | 0.1 | ycaM | 1 | 33.1833 | 0.425 |
| 0.8 | 0.1 | yadI | 1 | 33.1833 | 0.858 |
| 0.8 | 0.1 | AG1 | 1 | 33.1833 | 0.863 |
| 0.8 | 0.1 | ptsI | 1 | 33.4333 | 0.897 |
| 0.8 | 0.1 | clcB | 1 | 33.4333 | 0.408 |
| 0.8 | 0.1 | ycaM | 1 | 33.4333 | 0.429 |
| 0.8 | 0.1 | yadI | 1 | 33.4333 | 0.874 |
| 0.8 | 0.1 | AG1 | 1 | 33.4333 | 0.868 |
| 0.8 | 0.1 | ptsI | 1 | 33.6833 | 0.891 |
| 0.8 | 0.1 | clcB | 1 | 33.6833 | 0.402 |
| 0.8 | 0.1 | ycaM | 1 | 33.6833 | 0.427 |
| 0.8 | 0.1 | yadI | 1 | 33.6833 | 0.864 |
| 0.8 | 0.1 | AG1 | 1 | 33.6833 | 0.864 |
| 0.8 | 0.1 | ptsI | 1 | 33.9333 | 0.892 |
| 0.8 | 0.1 | clcB | 1 | 33.9333 | 0.408 |
| 0.8 | 0.1 | ycaM | 1 | 33.9333 | 0.433 |
| 0.8 | 0.1 | yadI | 1 | 33.9333 | 0.876 |
| 0.8 | 0.1 | AG1 | 1 | 33.9333 | 0.872 |
| 0.8 | 0.1 | ptsI | 1 | 34.1833 | 0.9 |
| 0.8 | 0.1 | clcB | 1 | 34.1833 | 0.416 |
| 0.8 | 0.1 | ycaM | 1 | 34.1833 | 0.434 |
| 0.8 | 0.1 | yadI | 1 | 34.1833 | 0.88 |
| 0.8 | 0.1 | AG1 | 1 | 34.1833 | 0.875 |
| 0.8 | 0.1 | ptsI | 1 | 34.4333 | 0.895 |

| | | | | | |
|---|---|---|---|---|---|
| 0.8 | 0.1 | clcB | 1 | 34.4333 | 0.41 |
| 0.8 | 0.1 | ycaM | 1 | 34.4333 | 0.434 |
| 0.8 | 0.1 | yadI | 1 | 34.4333 | 0.881 |
| 0.8 | 0.1 | AG1 | 1 | 34.4333 | 0.877 |
| 0.8 | 0.1 | ptsI | 1 | 34.6833 | 0.897 |
| 0.8 | 0.1 | clcB | 1 | 34.6833 | 0.413 |
| 0.8 | 0.1 | ycaM | 1 | 34.6833 | 0.434 |
| 0.8 | 0.1 | yadI | 1 | 34.6833 | 0.886 |
| 0.8 | 0.1 | AG1 | 1 | 34.6833 | 0.878 |
| 0.8 | 0.1 | ptsI | 1 | 34.9333 | 0.898 |
| 0.8 | 0.1 | clcB | 1 | 34.9333 | 0.412 |
| 0.8 | 0.1 | ycaM | 1 | 34.9333 | 0.434 |
| 0.8 | 0.1 | yadI | 1 | 34.9333 | 0.884 |
| 0.8 | 0.1 | AG1 | 1 | 34.9333 | 0.878 |
| 0.8 | 0.1 | ptsI | 1 | 35.1833 | 0.898 |
| 0.8 | 0.1 | clcB | 1 | 35.1833 | 0.411 |
| 0.8 | 0.1 | ycaM | 1 | 35.1833 | 0.434 |
| 0.8 | 0.1 | yadI | 1 | 35.1833 | 0.889 |
| 0.8 | 0.1 | AG1 | 1 | 35.1833 | 0.878 |
| 0.8 | 0.1 | ptsI | 1 | 35.4333 | 0.898 |
| 0.8 | 0.1 | clcB | 1 | 35.4333 | 0.413 |
| 0.8 | 0.1 | ycaM | 1 | 35.4333 | 0.436 |
| 0.8 | 0.1 | yadI | 1 | 35.4333 | 0.895 |
| 0.8 | 0.1 | AG1 | 1 | 35.4333 | 0.881 |
| 0.8 | 0.1 | ptsI | 1 | 35.6833 | 0.898 |
| 0.8 | 0.1 | clcB | 1 | 35.6833 | 0.416 |
| 0.8 | 0.1 | ycaM | 1 | 35.6833 | 0.437 |
| 0.8 | 0.1 | yadI | 1 | 35.6833 | 0.9 |
| 0.8 | 0.1 | AG1 | 1 | 35.6833 | 0.882 |
| 0.8 | 0.1 | ptsI | 1 | 35.9333 | 0.893 |
| 0.8 | 0.1 | clcB | 1 | 35.9333 | 0.409 |
| 0.8 | 0.1 | ycaM | 1 | 35.9333 | 0.431 |
| 0.8 | 0.1 | yadI | 1 | 35.9333 | 0.889 |
| 0.8 | 0.1 | AG1 | 1 | 35.9333 | 0.878 |
| 0.8 | 0.1 | ptsI | 1 | 36.1833 | 0.9 |
| 0.8 | 0.1 | clcB | 1 | 36.1833 | 0.421 |
| 0.8 | 0.1 | ycaM | 1 | 36.1833 | 0.44 |
| 0.8 | 0.1 | yadI | 1 | 36.1833 | 0.9 |
| 0.8 | 0.1 | AG1 | 1 | 36.1833 | 0.881 |
| 0.8 | 0.1 | ptsI | 1 | 36.4333 | 0.896 |
| 0.8 | 0.1 | clcB | 1 | 36.4333 | 0.416 |
| 0.8 | 0.1 | ycaM | 1 | 36.4333 | 0.439 |
| 0.8 | 0.1 | yadI | 1 | 36.4333 | 0.909 |
| 0.8 | 0.1 | AG1 | 1 | 36.4333 | 0.882 |
| 0.8 | 0.1 | ptsI | 1 | 36.6833 | 0.895 |
| 0.8 | 0.1 | clcB | 1 | 36.6833 | 0.419 |
| 0.8 | 0.1 | ycaM | 1 | 36.6833 | 0.44 |
| 0.8 | 0.1 | yadI | 1 | 36.6833 | 0.909 |

| | | | | | |
|---|---|---|---|---|---|
| 0.8 | 0.1 | AG1 | 1 | 36.6833 | 0.883 |
| 1.6 | 0.1 | ptsI | 1 | 0 | 0.228 |
| 1.6 | 0.1 | clcB | 1 | 0 | 0.285 |
| 1.6 | 0.1 | ycaM | 1 | 0 | 0.258 |
| 1.6 | 0.1 | yadI | 1 | 0 | 0.289 |
| 1.6 | 0.1 | AG1 | 1 | 0 | 0.236 |
| 1.6 | 0.1 | ptsI | 1 | 0.25 | 0.226 |
| 1.6 | 0.1 | clcB | 1 | 0.25 | 0.27 |
| 1.6 | 0.1 | ycaM | 1 | 0.25 | 0.243 |
| 1.6 | 0.1 | yadI | 1 | 0.25 | 0.271 |
| 1.6 | 0.1 | AG1 | 1 | 0.25 | 0.228 |
| 1.6 | 0.1 | ptsI | 1 | 0.5 | 0.227 |
| 1.6 | 0.1 | clcB | 1 | 0.5 | 0.268 |
| 1.6 | 0.1 | ycaM | 1 | 0.5 | 0.236 |
| 1.6 | 0.1 | yadI | 1 | 0.5 | 0.271 |
| 1.6 | 0.1 | AG1 | 1 | 0.5 | 0.227 |
| 1.6 | 0.1 | ptsI | 1 | 0.75 | 0.224 |
| 1.6 | 0.1 | clcB | 1 | 0.75 | 0.262 |
| 1.6 | 0.1 | ycaM | 1 | 0.75 | 0.233 |
| 1.6 | 0.1 | yadI | 1 | 0.75 | 0.27 |
| 1.6 | 0.1 | AG1 | 1 | 0.75 | 0.228 |
| 1.6 | 0.1 | ptsI | 1 | 1 | 0.227 |
| 1.6 | 0.1 | clcB | 1 | 1 | 0.266 |
| 1.6 | 0.1 | ycaM | 1 | 1 | 0.232 |
| 1.6 | 0.1 | yadI | 1 | 1 | 0.268 |
| 1.6 | 0.1 | AG1 | 1 | 1 | 0.23 |
| 1.6 | 0.1 | ptsI | 1 | 1.25 | 0.231 |
| 1.6 | 0.1 | clcB | 1 | 1.25 | 0.257 |
| 1.6 | 0.1 | ycaM | 1 | 1.25 | 0.228 |
| 1.6 | 0.1 | yadI | 1 | 1.25 | 0.267 |
| 1.6 | 0.1 | AG1 | 1 | 1.25 | 0.231 |
| 1.6 | 0.1 | ptsI | 1 | 1.5 | 0.232 |
| 1.6 | 0.1 | clcB | 1 | 1.5 | 0.262 |
| 1.6 | 0.1 | ycaM | 1 | 1.5 | 0.231 |
| 1.6 | 0.1 | yadI | 1 | 1.5 | 0.271 |
| 1.6 | 0.1 | AG1 | 1 | 1.5 | 0.228 |
| 1.6 | 0.1 | ptsI | 1 | 1.75 | 0.24 |
| 1.6 | 0.1 | clcB | 1 | 1.75 | 0.266 |
| 1.6 | 0.1 | ycaM | 1 | 1.75 | 0.232 |
| 1.6 | 0.1 | yadI | 1 | 1.75 | 0.273 |
| 1.6 | 0.1 | AG1 | 1 | 1.75 | 0.233 |
| 1.6 | 0.1 | ptsI | 1 | 2 | 0.245 |
| 1.6 | 0.1 | clcB | 1 | 2 | 0.272 |
| 1.6 | 0.1 | ycaM | 1 | 2 | 0.238 |
| 1.6 | 0.1 | yadI | 1 | 2 | 0.278 |
| 1.6 | 0.1 | AG1 | 1 | 2 | 0.234 |
| 1.6 | 0.1 | ptsI | 1 | 2.25 | 0.255 |
| 1.6 | 0.1 | clcB | 1 | 2.25 | 0.278 |

| | | | | | |
|---|---|---|---|---|---|
| 1.6 | 0.1 | ycaM | 1 | 2.25 | 0.238 |
| 1.6 | 0.1 | yadI | 1 | 2.25 | 0.28 |
| 1.6 | 0.1 | AG1 | 1 | 2.25 | 0.241 |
| 1.6 | 0.1 | ptsI | 1 | 2.5 | 0.261 |
| 1.6 | 0.1 | clcB | 1 | 2.5 | 0.284 |
| 1.6 | 0.1 | ycaM | 1 | 2.5 | 0.248 |
| 1.6 | 0.1 | yadI | 1 | 2.5 | 0.291 |
| 1.6 | 0.1 | AG1 | 1 | 2.5 | 0.241 |
| 1.6 | 0.1 | ptsI | 1 | 2.75 | 0.272 |
| 1.6 | 0.1 | clcB | 1 | 2.75 | 0.293 |
| 1.6 | 0.1 | ycaM | 1 | 2.75 | 0.247 |
| 1.6 | 0.1 | yadI | 1 | 2.75 | 0.293 |
| 1.6 | 0.1 | AG1 | 1 | 2.75 | 0.248 |
| 1.6 | 0.1 | ptsI | 1 | 3 | 0.282 |
| 1.6 | 0.1 | clcB | 1 | 3 | 0.299 |
| 1.6 | 0.1 | ycaM | 1 | 3 | 0.258 |
| 1.6 | 0.1 | yadI | 1 | 3 | 0.304 |
| 1.6 | 0.1 | AG1 | 1 | 3 | 0.251 |
| 1.6 | 0.1 | ptsI | 1 | 3.25 | 0.294 |
| 1.6 | 0.1 | clcB | 1 | 3.25 | 0.308 |
| 1.6 | 0.1 | ycaM | 1 | 3.25 | 0.266 |
| 1.6 | 0.1 | yadI | 1 | 3.25 | 0.316 |
| 1.6 | 0.1 | AG1 | 1 | 3.25 | 0.257 |
| 1.6 | 0.1 | ptsI | 1 | 3.5 | 0.306 |
| 1.6 | 0.1 | clcB | 1 | 3.5 | 0.318 |
| 1.6 | 0.1 | ycaM | 1 | 3.5 | 0.268 |
| 1.6 | 0.1 | yadI | 1 | 3.5 | 0.32 |
| 1.6 | 0.1 | AG1 | 1 | 3.5 | 0.264 |
| 1.6 | 0.1 | ptsI | 1 | 3.75 | 0.314 |
| 1.6 | 0.1 | clcB | 1 | 3.75 | 0.321 |
| 1.6 | 0.1 | ycaM | 1 | 3.75 | 0.265 |
| 1.6 | 0.1 | yadI | 1 | 3.75 | 0.328 |
| 1.6 | 0.1 | AG1 | 1 | 3.75 | 0.264 |
| 1.6 | 0.1 | ptsI | 1 | 4 | 0.33 |
| 1.6 | 0.1 | clcB | 1 | 4 | 0.339 |
| 1.6 | 0.1 | ycaM | 1 | 4 | 0.28 |
| 1.6 | 0.1 | yadI | 1 | 4 | 0.343 |
| 1.6 | 0.1 | AG1 | 1 | 4 | 0.273 |
| 1.6 | 0.1 | ptsI | 1 | 4.25 | 0.336 |
| 1.6 | 0.1 | clcB | 1 | 4.25 | 0.342 |
| 1.6 | 0.1 | ycaM | 1 | 4.25 | 0.278 |
| 1.6 | 0.1 | yadI | 1 | 4.25 | 0.343 |
| 1.6 | 0.1 | AG1 | 1 | 4.25 | 0.274 |
| 1.6 | 0.1 | ptsI | 1 | 4.5 | 0.348 |
| 1.6 | 0.1 | clcB | 1 | 4.5 | 0.356 |
| 1.6 | 0.1 | ycaM | 1 | 4.5 | 0.286 |
| 1.6 | 0.1 | yadI | 1 | 4.5 | 0.354 |
| 1.6 | 0.1 | AG1 | 1 | 4.5 | 0.279 |

| | | | | | |
|---|---|---|---|---|---|
| 1.6 | 0.1 | ptsI | 1 | 4.75 | 0.344 |
| 1.6 | 0.1 | clcB | 1 | 4.75 | 0.353 |
| 1.6 | 0.1 | ycaM | 1 | 4.75 | 0.282 |
| 1.6 | 0.1 | yadI | 1 | 4.75 | 0.356 |
| 1.6 | 0.1 | AG1 | 1 | 4.75 | 0.281 |
| 1.6 | 0.1 | ptsI | 1 | 5 | 0.36 |
| 1.6 | 0.1 | clcB | 1 | 5 | 0.37 |
| 1.6 | 0.1 | ycaM | 1 | 5 | 0.294 |
| 1.6 | 0.1 | yadI | 1 | 5 | 0.368 |
| 1.6 | 0.1 | AG1 | 1 | 5 | 0.287 |
| 1.6 | 0.1 | ptsI | 1 | 5.25 | 0.365 |
| 1.6 | 0.1 | clcB | 1 | 5.25 | 0.378 |
| 1.6 | 0.1 | ycaM | 1 | 5.25 | 0.293 |
| 1.6 | 0.1 | yadI | 1 | 5.25 | 0.372 |
| 1.6 | 0.1 | AG1 | 1 | 5.25 | 0.288 |
| 1.6 | 0.1 | ptsI | 1 | 5.5 | 0.376 |
| 1.6 | 0.1 | clcB | 1 | 5.5 | 0.399 |
| 1.6 | 0.1 | ycaM | 1 | 5.5 | 0.306 |
| 1.6 | 0.1 | yadI | 1 | 5.5 | 0.387 |
| 1.6 | 0.1 | AG1 | 1 | 5.5 | 0.295 |
| 1.6 | 0.1 | ptsI | 1 | 5.75 | 0.385 |
| 1.6 | 0.1 | clcB | 1 | 5.75 | 0.408 |
| 1.6 | 0.1 | ycaM | 1 | 5.75 | 0.308 |
| 1.6 | 0.1 | yadI | 1 | 5.75 | 0.394 |
| 1.6 | 0.1 | AG1 | 1 | 5.75 | 0.304 |
| 1.6 | 0.1 | ptsI | 1 | 6 | 0.373 |
| 1.6 | 0.1 | clcB | 1 | 6 | 0.401 |
| 1.6 | 0.1 | ycaM | 1 | 6 | 0.304 |
| 1.6 | 0.1 | yadI | 1 | 6 | 0.383 |
| 1.6 | 0.1 | AG1 | 1 | 6 | 0.309 |
| 1.6 | 0.1 | ptsI | 1 | 6.25 | 0.378 |
| 1.6 | 0.1 | clcB | 1 | 6.25 | 0.401 |
| 1.6 | 0.1 | ycaM | 1 | 6.25 | 0.307 |
| 1.6 | 0.1 | yadI | 1 | 6.25 | 0.386 |
| 1.6 | 0.1 | AG1 | 1 | 6.25 | 0.304 |
| 1.6 | 0.1 | ptsI | 1 | 6.5 | 0.388 |
| 1.6 | 0.1 | clcB | 1 | 6.5 | 0.4 |
| 1.6 | 0.1 | ycaM | 1 | 6.5 | 0.308 |
| 1.6 | 0.1 | yadI | 1 | 6.5 | 0.388 |
| 1.6 | 0.1 | AG1 | 1 | 6.5 | 0.311 |
| 1.6 | 0.1 | ptsI | 1 | 6.75 | 0.411 |
| 1.6 | 0.1 | clcB | 1 | 6.75 | 0.429 |
| 1.6 | 0.1 | ycaM | 1 | 6.75 | 0.322 |
| 1.6 | 0.1 | yadI | 1 | 6.75 | 0.4 |
| 1.6 | 0.1 | AG1 | 1 | 6.75 | 0.316 |
| 1.6 | 0.1 | ptsI | 1 | 7 | 0.426 |
| 1.6 | 0.1 | clcB | 1 | 7 | 0.446 |
| 1.6 | 0.1 | ycaM | 1 | 7 | 0.328 |

| | | | | | |
|---|---|---|---|---|---|
| 1.6 | 0.1 | yadI | 1 | 7 | 0.406 |
| 1.6 | 0.1 | AG1 | 1 | 7 | 0.327 |
| 1.6 | 0.1 | ptsI | 1 | 7.25 | 0.44 |
| 1.6 | 0.1 | clcB | 1 | 7.25 | 0.459 |
| 1.6 | 0.1 | ycaM | 1 | 7.25 | 0.339 |
| 1.6 | 0.1 | yadI | 1 | 7.25 | 0.416 |
| 1.6 | 0.1 | AG1 | 1 | 7.25 | 0.33 |
| 1.6 | 0.1 | ptsI | 1 | 7.5 | 0.456 |
| 1.6 | 0.1 | clcB | 1 | 7.5 | 0.468 |
| 1.6 | 0.1 | ycaM | 1 | 7.5 | 0.344 |
| 1.6 | 0.1 | yadI | 1 | 7.5 | 0.421 |
| 1.6 | 0.1 | AG1 | 1 | 7.5 | 0.345 |
| 1.6 | 0.1 | ptsI | 1 | 7.75 | 0.468 |
| 1.6 | 0.1 | clcB | 1 | 7.75 | 0.465 |
| 1.6 | 0.1 | ycaM | 1 | 7.75 | 0.35 |
| 1.6 | 0.1 | yadI | 1 | 7.75 | 0.428 |
| 1.6 | 0.1 | AG1 | 1 | 7.75 | 0.363 |
| 1.6 | 0.1 | ptsI | 1 | 8 | 0.48 |
| 1.6 | 0.1 | clcB | 1 | 8 | 0.489 |
| 1.6 | 0.1 | ycaM | 1 | 8 | 0.355 |
| 1.6 | 0.1 | yadI | 1 | 8 | 0.438 |
| 1.6 | 0.1 | AG1 | 1 | 8 | 0.359 |
| 1.6 | 0.1 | ptsI | 1 | 8.25 | 0.488 |
| 1.6 | 0.1 | clcB | 1 | 8.25 | 0.493 |
| 1.6 | 0.1 | ycaM | 1 | 8.25 | 0.352 |
| 1.6 | 0.1 | yadI | 1 | 8.25 | 0.442 |
| 1.6 | 0.1 | AG1 | 1 | 8.25 | 0.378 |
| 1.6 | 0.1 | ptsI | 1 | 8.5 | 0.498 |
| 1.6 | 0.1 | clcB | 1 | 8.5 | 0.502 |
| 1.6 | 0.1 | ycaM | 1 | 8.5 | 0.357 |
| 1.6 | 0.1 | yadI | 1 | 8.5 | 0.448 |
| 1.6 | 0.1 | AG1 | 1 | 8.5 | 0.381 |
| 1.6 | 0.1 | ptsI | 1 | 8.75 | 0.508 |
| 1.6 | 0.1 | clcB | 1 | 8.75 | 0.503 |
| 1.6 | 0.1 | ycaM | 1 | 8.75 | 0.36 |
| 1.6 | 0.1 | yadI | 1 | 8.75 | 0.456 |
| 1.6 | 0.1 | AG1 | 1 | 8.75 | 0.393 |
| 1.6 | 0.1 | ptsI | 1 | 9 | 0.511 |
| 1.6 | 0.1 | clcB | 1 | 9 | 0.513 |
| 1.6 | 0.1 | ycaM | 1 | 9 | 0.369 |
| 1.6 | 0.1 | yadI | 1 | 9 | 0.46 |
| 1.6 | 0.1 | AG1 | 1 | 9 | 0.389 |
| 1.6 | 0.1 | ptsI | 1 | 9.25 | 0.522 |
| 1.6 | 0.1 | clcB | 1 | 9.25 | 0.513 |
| 1.6 | 0.1 | ycaM | 1 | 9.25 | 0.371 |
| 1.6 | 0.1 | yadI | 1 | 9.25 | 0.457 |
| 1.6 | 0.1 | AG1 | 1 | 9.25 | 0.409 |
| 1.6 | 0.1 | ptsI | 1 | 9.5 | 0.544 |

| | | | | | |
|---|---|---|---|---|---|
| 1.6 | 0.1 | clcB | 1 | 9.5 | 0.515 |
| 1.6 | 0.1 | ycaM | 1 | 9.5 | 0.373 |
| 1.6 | 0.1 | yadI | 1 | 9.5 | 0.456 |
| 1.6 | 0.1 | AG1 | 1 | 9.5 | 0.418 |
| 1.6 | 0.1 | ptsI | 1 | 9.75 | 0.552 |
| 1.6 | 0.1 | clcB | 1 | 9.75 | 0.555 |
| 1.6 | 0.1 | ycaM | 1 | 9.75 | 0.388 |
| 1.6 | 0.1 | yadI | 1 | 9.75 | 0.474 |
| 1.6 | 0.1 | AG1 | 1 | 9.75 | 0.436 |
| 1.6 | 0.1 | ptsI | 1 | 10 | 0.503 |
| 1.6 | 0.1 | clcB | 1 | 10 | 0.514 |
| 1.6 | 0.1 | ycaM | 1 | 10 | 0.382 |
| 1.6 | 0.1 | yadI | 1 | 10 | 0.452 |
| 1.6 | 0.1 | AG1 | 1 | 10 | 0.408 |
| 1.6 | 0.1 | ptsI | 1 | 10.25 | 0.539 |
| 1.6 | 0.1 | clcB | 1 | 10.25 | 0.546 |
| 1.6 | 0.1 | ycaM | 1 | 10.25 | 0.388 |
| 1.6 | 0.1 | yadI | 1 | 10.25 | 0.461 |
| 1.6 | 0.1 | AG1 | 1 | 10.25 | 0.418 |
| 1.6 | 0.1 | ptsI | 1 | 10.5 | 0.581 |
| 1.6 | 0.1 | clcB | 1 | 10.5 | 0.551 |
| 1.6 | 0.1 | ycaM | 1 | 10.5 | 0.402 |
| 1.6 | 0.1 | yadI | 1 | 10.5 | 0.474 |
| 1.6 | 0.1 | AG1 | 1 | 10.5 | 0.44 |
| 1.6 | 0.1 | ptsI | 1 | 10.75 | 0.529 |
| 1.6 | 0.1 | clcB | 1 | 10.75 | 0.516 |
| 1.6 | 0.1 | ycaM | 1 | 10.75 | 0.397 |
| 1.6 | 0.1 | yadI | 1 | 10.75 | 0.453 |
| 1.6 | 0.1 | AG1 | 1 | 10.75 | 0.426 |
| 1.6 | 0.1 | ptsI | 1 | 11 | 0.526 |
| 1.6 | 0.1 | clcB | 1 | 11 | 0.492 |
| 1.6 | 0.1 | ycaM | 1 | 11 | 0.398 |
| 1.6 | 0.1 | yadI | 1 | 11 | 0.454 |
| 1.6 | 0.1 | AG1 | 1 | 11 | 0.442 |
| 1.6 | 0.1 | ptsI | 1 | 11.25 | 0.577 |
| 1.6 | 0.1 | clcB | 1 | 11.25 | 0.543 |
| 1.6 | 0.1 | ycaM | 1 | 11.25 | 0.423 |
| 1.6 | 0.1 | yadI | 1 | 11.25 | 0.465 |
| 1.6 | 0.1 | AG1 | 1 | 11.25 | 0.48 |
| 1.6 | 0.1 | ptsI | 1 | 11.5 | 0.572 |
| 1.6 | 0.1 | clcB | 1 | 11.5 | 0.511 |
| 1.6 | 0.1 | ycaM | 1 | 11.5 | 0.42 |
| 1.6 | 0.1 | yadI | 1 | 11.5 | 0.465 |
| 1.6 | 0.1 | AG1 | 1 | 11.5 | 0.47 |
| 1.6 | 0.1 | ptsI | 1 | 11.75 | 0.565 |
| 1.6 | 0.1 | clcB | 1 | 11.75 | 0.501 |
| 1.6 | 0.1 | ycaM | 1 | 11.75 | 0.414 |
| 1.6 | 0.1 | yadI | 1 | 11.75 | 0.451 |

| | | | | | |
|---|---|---|---|---|---|
| 1.6 | 0.1 | AG1 | 1 | 11.75 | 0.465 |
| 1.6 | 0.1 | ptsI | 1 | 12 | 0.601 |
| 1.6 | 0.1 | clcB | 1 | 12 | 0.546 |
| 1.6 | 0.1 | ycaM | 1 | 12 | 0.443 |
| 1.6 | 0.1 | yadI | 1 | 12 | 0.46 |
| 1.6 | 0.1 | AG1 | 1 | 12 | 0.516 |
| 1.6 | 0.1 | ptsI | 1 | 12.25 | 0.579 |
| 1.6 | 0.1 | clcB | 1 | 12.25 | 0.503 |
| 1.6 | 0.1 | ycaM | 1 | 12.25 | 0.426 |
| 1.6 | 0.1 | yadI | 1 | 12.25 | 0.454 |
| 1.6 | 0.1 | AG1 | 1 | 12.25 | 0.492 |
| 1.6 | 0.1 | ptsI | 1 | 12.5 | 0.612 |
| 1.6 | 0.1 | clcB | 1 | 12.5 | 0.535 |
| 1.6 | 0.1 | ycaM | 1 | 12.5 | 0.438 |
| 1.6 | 0.1 | yadI | 1 | 12.5 | 0.453 |
| 1.6 | 0.1 | AG1 | 1 | 12.5 | 0.541 |
| 1.6 | 0.1 | ptsI | 1 | 12.75 | 0.616 |
| 1.6 | 0.1 | clcB | 1 | 12.75 | 0.556 |
| 1.6 | 0.1 | ycaM | 1 | 12.75 | 0.474 |
| 1.6 | 0.1 | yadI | 1 | 12.75 | 0.463 |
| 1.6 | 0.1 | AG1 | 1 | 12.75 | 0.553 |
| 1.6 | 0.1 | ptsI | 1 | 13 | 0.644 |
| 1.6 | 0.1 | clcB | 1 | 13 | 0.564 |
| 1.6 | 0.1 | ycaM | 1 | 13 | 0.506 |
| 1.6 | 0.1 | yadI | 1 | 13 | 0.474 |
| 1.6 | 0.1 | AG1 | 1 | 13 | 0.614 |
| 1.6 | 0.1 | ptsI | 1 | 13.25 | 0.625 |
| 1.6 | 0.1 | clcB | 1 | 13.25 | 0.563 |
| 1.6 | 0.1 | ycaM | 1 | 13.25 | 0.478 |
| 1.6 | 0.1 | yadI | 1 | 13.25 | 0.466 |
| 1.6 | 0.1 | AG1 | 1 | 13.25 | 0.594 |
| 1.6 | 0.1 | ptsI | 1 | 13.5 | 0.663 |
| 1.6 | 0.1 | clcB | 1 | 13.5 | 0.589 |
| 1.6 | 0.1 | ycaM | 1 | 13.5 | 0.507 |
| 1.6 | 0.1 | yadI | 1 | 13.5 | 0.483 |
| 1.6 | 0.1 | AG1 | 1 | 13.5 | 0.685 |
| 1.6 | 0.1 | ptsI | 1 | 13.75 | 0.64 |
| 1.6 | 0.1 | clcB | 1 | 13.75 | 0.608 |
| 1.6 | 0.1 | ycaM | 1 | 13.75 | 0.487 |
| 1.6 | 0.1 | yadI | 1 | 13.75 | 0.484 |
| 1.6 | 0.1 | AG1 | 1 | 13.75 | 0.666 |
| 1.6 | 0.1 | ptsI | 1 | 14 | 0.671 |
| 1.6 | 0.1 | clcB | 1 | 14 | 0.638 |
| 1.6 | 0.1 | ycaM | 1 | 14 | 0.522 |
| 1.6 | 0.1 | yadI | 1 | 14 | 0.486 |
| 1.6 | 0.1 | AG1 | 1 | 14 | 0.744 |
| 1.6 | 0.1 | ptsI | 1 | 14.25 | 0.668 |
| 1.6 | 0.1 | clcB | 1 | 14.25 | 0.645 |

| | | | | | |
|---|---|---|---|---|---|
| 1.6 | 0.1 | ycaM | 1 | 14.25 | 0.545 |
| 1.6 | 0.1 | yadI | 1 | 14.25 | 0.494 |
| 1.6 | 0.1 | AG1 | 1 | 14.25 | 0.771 |
| 1.6 | 0.1 | ptsI | 1 | 14.5 | 0.683 |
| 1.6 | 0.1 | clcB | 1 | 14.5 | 0.652 |
| 1.6 | 0.1 | ycaM | 1 | 14.5 | 0.561 |
| 1.6 | 0.1 | yadI | 1 | 14.5 | 0.504 |
| 1.6 | 0.1 | AG1 | 1 | 14.5 | 0.79 |
| 1.6 | 0.1 | ptsI | 1 | 14.75 | 0.657 |
| 1.6 | 0.1 | clcB | 1 | 14.75 | 0.624 |
| 1.6 | 0.1 | ycaM | 1 | 14.75 | 0.537 |
| 1.6 | 0.1 | yadI | 1 | 14.75 | 0.495 |
| 1.6 | 0.1 | AG1 | 1 | 14.75 | 0.758 |
| 1.6 | 0.1 | ptsI | 1 | 15 | 0.701 |
| 1.6 | 0.1 | clcB | 1 | 15 | 0.662 |
| 1.6 | 0.1 | ycaM | 1 | 15 | 0.567 |
| 1.6 | 0.1 | yadI | 1 | 15 | 0.514 |
| 1.6 | 0.1 | AG1 | 1 | 15 | 0.823 |
| 1.6 | 0.1 | ptsI | 1 | 15.25 | 0.716 |
| 1.6 | 0.1 | clcB | 1 | 15.25 | 0.663 |
| 1.6 | 0.1 | ycaM | 1 | 15.25 | 0.583 |
| 1.6 | 0.1 | yadI | 1 | 15.25 | 0.522 |
| 1.6 | 0.1 | AG1 | 1 | 15.25 | 0.831 |
| 1.6 | 0.1 | ptsI | 1 | 15.5 | 0.698 |
| 1.6 | 0.1 | clcB | 1 | 15.5 | 0.674 |
| 1.6 | 0.1 | ycaM | 1 | 15.5 | 0.581 |
| 1.6 | 0.1 | yadI | 1 | 15.5 | 0.52 |
| 1.6 | 0.1 | AG1 | 1 | 15.5 | 0.822 |
| 1.6 | 0.1 | ptsI | 1 | 15.75 | 0.683 |
| 1.6 | 0.1 | clcB | 1 | 15.75 | 0.661 |
| 1.6 | 0.1 | ycaM | 1 | 15.75 | 0.551 |
| 1.6 | 0.1 | yadI | 1 | 15.75 | 0.506 |
| 1.6 | 0.1 | AG1 | 1 | 15.75 | 0.799 |
| 1.6 | 0.1 | ptsI | 1 | 16 | 0.672 |
| 1.6 | 0.1 | clcB | 1 | 16 | 0.666 |
| 1.6 | 0.1 | ycaM | 1 | 16 | 0.538 |
| 1.6 | 0.1 | yadI | 1 | 16 | 0.498 |
| 1.6 | 0.1 | AG1 | 1 | 16 | 0.755 |
| 1.6 | 0.1 | ptsI | 1 | 16.25 | 0.732 |
| 1.6 | 0.1 | clcB | 1 | 16.25 | 0.7 |
| 1.6 | 0.1 | ycaM | 1 | 16.25 | 0.581 |
| 1.6 | 0.1 | yadI | 1 | 16.25 | 0.512 |
| 1.6 | 0.1 | AG1 | 1 | 16.25 | 0.819 |
| 1.6 | 0.1 | ptsI | 1 | 16.5 | 0.725 |
| 1.6 | 0.1 | clcB | 1 | 16.5 | 0.712 |
| 1.6 | 0.1 | ycaM | 1 | 16.5 | 0.598 |
| 1.6 | 0.1 | yadI | 1 | 16.5 | 0.52 |
| 1.6 | 0.1 | AG1 | 1 | 16.5 | 0.834 |

| | | | | | |
|---|---|---|---|---|---|
| 1.6 | 0.1 | ptsI | 1 | 17.0667 | 0.746 |
| 1.6 | 0.1 | clcB | 1 | 17.0667 | 0.52 |
| 1.6 | 0.1 | ycaM | 1 | 17.0667 | 0.507 |
| 1.6 | 0.1 | yadI | 1 | 17.0667 | 0.673 |
| 1.6 | 0.1 | AG1 | 1 | 17.0667 | 0.851 |
| 1.6 | 0.1 | ptsI | 1 | 17.3167 | 0.68 |
| 1.6 | 0.1 | clcB | 1 | 17.3167 | 0.493 |
| 1.6 | 0.1 | ycaM | 1 | 17.3167 | 0.468 |
| 1.6 | 0.1 | yadI | 1 | 17.3167 | 0.651 |
| 1.6 | 0.1 | AG1 | 1 | 17.3167 | 0.817 |
| 1.6 | 0.1 | ptsI | 1 | 17.5667 | 0.669 |
| 1.6 | 0.1 | clcB | 1 | 17.5667 | 0.482 |
| 1.6 | 0.1 | ycaM | 1 | 17.5667 | 0.458 |
| 1.6 | 0.1 | yadI | 1 | 17.5667 | 0.652 |
| 1.6 | 0.1 | AG1 | 1 | 17.5667 | 0.831 |
| 1.6 | 0.1 | ptsI | 1 | 17.8167 | 0.745 |
| 1.6 | 0.1 | clcB | 1 | 17.8167 | 0.484 |
| 1.6 | 0.1 | ycaM | 1 | 17.8167 | 0.46 |
| 1.6 | 0.1 | yadI | 1 | 17.8167 | 0.658 |
| 1.6 | 0.1 | AG1 | 1 | 17.8167 | 0.849 |
| 1.6 | 0.1 | ptsI | 1 | 18.0667 | 0.67 |
| 1.6 | 0.1 | clcB | 1 | 18.0667 | 0.481 |
| 1.6 | 0.1 | ycaM | 1 | 18.0667 | 0.453 |
| 1.6 | 0.1 | yadI | 1 | 18.0667 | 0.678 |
| 1.6 | 0.1 | AG1 | 1 | 18.0667 | 0.824 |
| 1.6 | 0.1 | ptsI | 1 | 18.3167 | 0.682 |
| 1.6 | 0.1 | clcB | 1 | 18.3167 | 0.48 |
| 1.6 | 0.1 | ycaM | 1 | 18.3167 | 0.444 |
| 1.6 | 0.1 | yadI | 1 | 18.3167 | 0.663 |
| 1.6 | 0.1 | AG1 | 1 | 18.3167 | 0.798 |
| 1.6 | 0.1 | ptsI | 1 | 18.5667 | 0.757 |
| 1.6 | 0.1 | clcB | 1 | 18.5667 | 0.477 |
| 1.6 | 0.1 | ycaM | 1 | 18.5667 | 0.446 |
| 1.6 | 0.1 | yadI | 1 | 18.5667 | 0.677 |
| 1.6 | 0.1 | AG1 | 1 | 18.5667 | 0.827 |
| 1.6 | 0.1 | ptsI | 1 | 18.8167 | 0.745 |
| 1.6 | 0.1 | clcB | 1 | 18.8167 | 0.472 |
| 1.6 | 0.1 | ycaM | 1 | 18.8167 | 0.448 |
| 1.6 | 0.1 | yadI | 1 | 18.8167 | 0.69 |
| 1.6 | 0.1 | AG1 | 1 | 18.8167 | 0.832 |
| 1.6 | 0.1 | ptsI | 1 | 19.0667 | 0.751 |
| 1.6 | 0.1 | clcB | 1 | 19.0667 | 0.472 |
| 1.6 | 0.1 | ycaM | 1 | 19.0667 | 0.452 |
| 1.6 | 0.1 | yadI | 1 | 19.0667 | 0.703 |
| 1.6 | 0.1 | AG1 | 1 | 19.0667 | 0.851 |
| 1.6 | 0.1 | ptsI | 1 | 19.3167 | 0.74 |
| 1.6 | 0.1 | clcB | 1 | 19.3167 | 0.469 |
| 1.6 | 0.1 | ycaM | 1 | 19.3167 | 0.447 |

| | | | | | |
|---|---|---|---|---|---|
| 1.6 | 0.1 | yadI | 1 | 19.3167 | 0.706 |
| 1.6 | 0.1 | AG1 | 1 | 19.3167 | 0.844 |
| 1.6 | 0.1 | ptsI | 1 | 19.5667 | 0.757 |
| 1.6 | 0.1 | clcB | 1 | 19.5667 | 0.467 |
| 1.6 | 0.1 | ycaM | 1 | 19.5667 | 0.451 |
| 1.6 | 0.1 | yadI | 1 | 19.5667 | 0.714 |
| 1.6 | 0.1 | AG1 | 1 | 19.5667 | 0.859 |
| 1.6 | 0.1 | ptsI | 1 | 19.8167 | 0.765 |
| 1.6 | 0.1 | clcB | 1 | 19.8167 | 0.466 |
| 1.6 | 0.1 | ycaM | 1 | 19.8167 | 0.451 |
| 1.6 | 0.1 | yadI | 1 | 19.8167 | 0.72 |
| 1.6 | 0.1 | AG1 | 1 | 19.8167 | 0.861 |
| 1.6 | 0.1 | ptsI | 1 | 20.0667 | 0.761 |
| 1.6 | 0.1 | clcB | 1 | 20.0667 | 0.465 |
| 1.6 | 0.1 | ycaM | 1 | 20.0667 | 0.451 |
| 1.6 | 0.1 | yadI | 1 | 20.0667 | 0.728 |
| 1.6 | 0.1 | AG1 | 1 | 20.0667 | 0.865 |
| 1.6 | 0.1 | ptsI | 1 | 20.3167 | 0.768 |
| 1.6 | 0.1 | clcB | 1 | 20.3167 | 0.463 |
| 1.6 | 0.1 | ycaM | 1 | 20.3167 | 0.452 |
| 1.6 | 0.1 | yadI | 1 | 20.3167 | 0.734 |
| 1.6 | 0.1 | AG1 | 1 | 20.3167 | 0.872 |
| 1.6 | 0.1 | ptsI | 1 | 20.5667 | 0.761 |
| 1.6 | 0.1 | clcB | 1 | 20.5667 | 0.461 |
| 1.6 | 0.1 | ycaM | 1 | 20.5667 | 0.445 |
| 1.6 | 0.1 | yadI | 1 | 20.5667 | 0.732 |
| 1.6 | 0.1 | AG1 | 1 | 20.5667 | 0.867 |
| 1.6 | 0.1 | ptsI | 1 | 20.8167 | 0.767 |
| 1.6 | 0.1 | clcB | 1 | 20.8167 | 0.458 |
| 1.6 | 0.1 | ycaM | 1 | 20.8167 | 0.449 |
| 1.6 | 0.1 | yadI | 1 | 20.8167 | 0.737 |
| 1.6 | 0.1 | AG1 | 1 | 20.8167 | 0.868 |
| 1.6 | 0.1 | ptsI | 1 | 21.0667 | 0.774 |
| 1.6 | 0.1 | clcB | 1 | 21.0667 | 0.456 |
| 1.6 | 0.1 | ycaM | 1 | 21.0667 | 0.447 |
| 1.6 | 0.1 | yadI | 1 | 21.0667 | 0.743 |
| 1.6 | 0.1 | AG1 | 1 | 21.0667 | 0.879 |
| 1.6 | 0.1 | ptsI | 1 | 21.3167 | 0.775 |
| 1.6 | 0.1 | clcB | 1 | 21.3167 | 0.463 |
| 1.6 | 0.1 | ycaM | 1 | 21.3167 | 0.451 |
| 1.6 | 0.1 | yadI | 1 | 21.3167 | 0.744 |
| 1.6 | 0.1 | AG1 | 1 | 21.3167 | 0.882 |
| 1.6 | 0.1 | ptsI | 1 | 21.5667 | 0.772 |
| 1.6 | 0.1 | clcB | 1 | 21.5667 | 0.454 |
| 1.6 | 0.1 | ycaM | 1 | 21.5667 | 0.444 |
| 1.6 | 0.1 | yadI | 1 | 21.5667 | 0.738 |
| 1.6 | 0.1 | AG1 | 1 | 21.5667 | 0.886 |
| 1.6 | 0.1 | ptsI | 1 | 21.8167 | 0.776 |

| | | | | | |
|---|---|---|---|---|---|
| 1.6 | 0.1 | clcB | 1 | 21.8167 | 0.452 |
| 1.6 | 0.1 | ycaM | 1 | 21.8167 | 0.448 |
| 1.6 | 0.1 | yadI | 1 | 21.8167 | 0.749 |
| 1.6 | 0.1 | AG1 | 1 | 21.8167 | 0.886 |
| 1.6 | 0.1 | ptsI | 1 | 22.0667 | 0.785 |
| 1.6 | 0.1 | clcB | 1 | 22.0667 | 0.454 |
| 1.6 | 0.1 | ycaM | 1 | 22.0667 | 0.445 |
| 1.6 | 0.1 | yadI | 1 | 22.0667 | 0.749 |
| 1.6 | 0.1 | AG1 | 1 | 22.0667 | 0.893 |
| 1.6 | 0.1 | ptsI | 1 | 22.9333 | 0.811 |
| 1.6 | 0.1 | clcB | 1 | 22.9333 | 0.47 |
| 1.6 | 0.1 | ycaM | 1 | 22.9333 | 0.398 |
| 1.6 | 0.1 | yadI | 1 | 22.9333 | 0.776 |
| 1.6 | 0.1 | AG1 | 1 | 22.9333 | 0.872 |
| 1.6 | 0.1 | ptsI | 1 | 23.1833 | 0.772 |
| 1.6 | 0.1 | clcB | 1 | 23.1833 | 0.456 |
| 1.6 | 0.1 | ycaM | 1 | 23.1833 | 0.384 |
| 1.6 | 0.1 | yadI | 1 | 23.1833 | 0.759 |
| 1.6 | 0.1 | AG1 | 1 | 23.1833 | 0.844 |
| 1.6 | 0.1 | ptsI | 1 | 23.4333 | 0.748 |
| 1.6 | 0.1 | clcB | 1 | 23.4333 | 0.449 |
| 1.6 | 0.1 | ycaM | 1 | 23.4333 | 0.378 |
| 1.6 | 0.1 | yadI | 1 | 23.4333 | 0.768 |
| 1.6 | 0.1 | AG1 | 1 | 23.4333 | 0.833 |
| 1.6 | 0.1 | ptsI | 1 | 23.6833 | 0.786 |
| 1.6 | 0.1 | clcB | 1 | 23.6833 | 0.449 |
| 1.6 | 0.1 | ycaM | 1 | 23.6833 | 0.384 |
| 1.6 | 0.1 | yadI | 1 | 23.6833 | 0.779 |
| 1.6 | 0.1 | AG1 | 1 | 23.6833 | 0.84 |
| 1.6 | 0.1 | ptsI | 1 | 23.9333 | 0.789 |
| 1.6 | 0.1 | clcB | 1 | 23.9333 | 0.449 |
| 1.6 | 0.1 | ycaM | 1 | 23.9333 | 0.381 |
| 1.6 | 0.1 | yadI | 1 | 23.9333 | 0.785 |
| 1.6 | 0.1 | AG1 | 1 | 23.9333 | 0.843 |
| 1.6 | 0.1 | ptsI | 1 | 24.1833 | 0.808 |
| 1.6 | 0.1 | clcB | 1 | 24.1833 | 0.444 |
| 1.6 | 0.1 | ycaM | 1 | 24.1833 | 0.383 |
| 1.6 | 0.1 | yadI | 1 | 24.1833 | 0.79 |
| 1.6 | 0.1 | AG1 | 1 | 24.1833 | 0.847 |
| 1.6 | 0.1 | ptsI | 1 | 24.4333 | 0.818 |
| 1.6 | 0.1 | clcB | 1 | 24.4333 | 0.44 |
| 1.6 | 0.1 | ycaM | 1 | 24.4333 | 0.384 |
| 1.6 | 0.1 | yadI | 1 | 24.4333 | 0.795 |
| 1.6 | 0.1 | AG1 | 1 | 24.4333 | 0.853 |
| 1.6 | 0.1 | ptsI | 1 | 24.6833 | 0.816 |
| 1.6 | 0.1 | clcB | 1 | 24.6833 | 0.44 |
| 1.6 | 0.1 | ycaM | 1 | 24.6833 | 0.384 |
| 1.6 | 0.1 | yadI | 1 | 24.6833 | 0.798 |

| | | | | | |
|---|---|---|---|---|---|
| 1.6 | 0.1 | AG1 | 1 | 24.6833 | 0.849 |
| 1.6 | 0.1 | ptsI | 1 | 24.9333 | 0.815 |
| 1.6 | 0.1 | clcB | 1 | 24.9333 | 0.435 |
| 1.6 | 0.1 | ycaM | 1 | 24.9333 | 0.383 |
| 1.6 | 0.1 | yadI | 1 | 24.9333 | 0.798 |
| 1.6 | 0.1 | AG1 | 1 | 24.9333 | 0.846 |
| 1.6 | 0.1 | ptsI | 1 | 25.1833 | 0.815 |
| 1.6 | 0.1 | clcB | 1 | 25.1833 | 0.437 |
| 1.6 | 0.1 | ycaM | 1 | 25.1833 | 0.383 |
| 1.6 | 0.1 | yadI | 1 | 25.1833 | 0.805 |
| 1.6 | 0.1 | AG1 | 1 | 25.1833 | 0.85 |
| 1.6 | 0.1 | ptsI | 1 | 25.4333 | 0.819 |
| 1.6 | 0.1 | clcB | 1 | 25.4333 | 0.441 |
| 1.6 | 0.1 | ycaM | 1 | 25.4333 | 0.386 |
| 1.6 | 0.1 | yadI | 1 | 25.4333 | 0.813 |
| 1.6 | 0.1 | AG1 | 1 | 25.4333 | 0.859 |
| 1.6 | 0.1 | ptsI | 1 | 25.6833 | 0.822 |
| 1.6 | 0.1 | clcB | 1 | 25.6833 | 0.441 |
| 1.6 | 0.1 | ycaM | 1 | 25.6833 | 0.387 |
| 1.6 | 0.1 | yadI | 1 | 25.6833 | 0.812 |
| 1.6 | 0.1 | AG1 | 1 | 25.6833 | 0.858 |
| 1.6 | 0.1 | ptsI | 1 | 25.9333 | 0.822 |
| 1.6 | 0.1 | clcB | 1 | 25.9333 | 0.441 |
| 1.6 | 0.1 | ycaM | 1 | 25.9333 | 0.386 |
| 1.6 | 0.1 | yadI | 1 | 25.9333 | 0.808 |
| 1.6 | 0.1 | AG1 | 1 | 25.9333 | 0.863 |
| 1.6 | 0.1 | ptsI | 1 | 26.1833 | 0.839 |
| 1.6 | 0.1 | clcB | 1 | 26.1833 | 0.445 |
| 1.6 | 0.1 | ycaM | 1 | 26.1833 | 0.39 |
| 1.6 | 0.1 | yadI | 1 | 26.1833 | 0.816 |
| 1.6 | 0.1 | AG1 | 1 | 26.1833 | 0.868 |
| 1.6 | 0.1 | ptsI | 1 | 26.4333 | 0.831 |
| 1.6 | 0.1 | clcB | 1 | 26.4333 | 0.446 |
| 1.6 | 0.1 | ycaM | 1 | 26.4333 | 0.39 |
| 1.6 | 0.1 | yadI | 1 | 26.4333 | 0.816 |
| 1.6 | 0.1 | AG1 | 1 | 26.4333 | 0.863 |
| 1.6 | 0.1 | ptsI | 1 | 26.6833 | 0.836 |
| 1.6 | 0.1 | clcB | 1 | 26.6833 | 0.448 |
| 1.6 | 0.1 | ycaM | 1 | 26.6833 | 0.39 |
| 1.6 | 0.1 | yadI | 1 | 26.6833 | 0.818 |
| 1.6 | 0.1 | AG1 | 1 | 26.6833 | 0.869 |
| 1.6 | 0.1 | ptsI | 1 | 26.9333 | 0.836 |
| 1.6 | 0.1 | clcB | 1 | 26.9333 | 0.448 |
| 1.6 | 0.1 | ycaM | 1 | 26.9333 | 0.391 |
| 1.6 | 0.1 | yadI | 1 | 26.9333 | 0.822 |
| 1.6 | 0.1 | AG1 | 1 | 26.9333 | 0.87 |
| 1.6 | 0.1 | ptsI | 1 | 27.1833 | 0.844 |
| 1.6 | 0.1 | clcB | 1 | 27.1833 | 0.45 |

| | | | | | |
|---|---|---|---|---|---|
| 1.6 | 0.1 | ycaM | 1 | 27.1833 | 0.394 |
| 1.6 | 0.1 | yadI | 1 | 27.1833 | 0.823 |
| 1.6 | 0.1 | AG1 | 1 | 27.1833 | 0.872 |
| 1.6 | 0.1 | ptsI | 1 | 27.4333 | 0.835 |
| 1.6 | 0.1 | clcB | 1 | 27.4333 | 0.451 |
| 1.6 | 0.1 | ycaM | 1 | 27.4333 | 0.392 |
| 1.6 | 0.1 | yadI | 1 | 27.4333 | 0.818 |
| 1.6 | 0.1 | AG1 | 1 | 27.4333 | 0.87 |
| 1.6 | 0.1 | ptsI | 1 | 27.6833 | 0.842 |
| 1.6 | 0.1 | clcB | 1 | 27.6833 | 0.45 |
| 1.6 | 0.1 | ycaM | 1 | 27.6833 | 0.394 |
| 1.6 | 0.1 | yadI | 1 | 27.6833 | 0.82 |
| 1.6 | 0.1 | AG1 | 1 | 27.6833 | 0.878 |
| 1.6 | 0.1 | ptsI | 1 | 27.9333 | 0.84 |
| 1.6 | 0.1 | clcB | 1 | 27.9333 | 0.448 |
| 1.6 | 0.1 | ycaM | 1 | 27.9333 | 0.389 |
| 1.6 | 0.1 | yadI | 1 | 27.9333 | 0.814 |
| 1.6 | 0.1 | AG1 | 1 | 27.9333 | 0.868 |
| 1.6 | 0.1 | ptsI | 1 | 28.1833 | 0.846 |
| 1.6 | 0.1 | clcB | 1 | 28.1833 | 0.45 |
| 1.6 | 0.1 | ycaM | 1 | 28.1833 | 0.394 |
| 1.6 | 0.1 | yadI | 1 | 28.1833 | 0.824 |
| 1.6 | 0.1 | AG1 | 1 | 28.1833 | 0.888 |
| 1.6 | 0.1 | ptsI | 1 | 28.4333 | 0.846 |
| 1.6 | 0.1 | clcB | 1 | 28.4333 | 0.454 |
| 1.6 | 0.1 | ycaM | 1 | 28.4333 | 0.396 |
| 1.6 | 0.1 | yadI | 1 | 28.4333 | 0.826 |
| 1.6 | 0.1 | AG1 | 1 | 28.4333 | 0.887 |
| 1.6 | 0.1 | ptsI | 1 | 28.6833 | 0.846 |
| 1.6 | 0.1 | clcB | 1 | 28.6833 | 0.454 |
| 1.6 | 0.1 | ycaM | 1 | 28.6833 | 0.395 |
| 1.6 | 0.1 | yadI | 1 | 28.6833 | 0.822 |
| 1.6 | 0.1 | AG1 | 1 | 28.6833 | 0.892 |
| 1.6 | 0.1 | ptsI | 1 | 28.9333 | 0.849 |
| 1.6 | 0.1 | clcB | 1 | 28.9333 | 0.457 |
| 1.6 | 0.1 | ycaM | 1 | 28.9333 | 0.393 |
| 1.6 | 0.1 | yadI | 1 | 28.9333 | 0.828 |
| 1.6 | 0.1 | AG1 | 1 | 28.9333 | 0.893 |
| 1.6 | 0.1 | ptsI | 1 | 29.1833 | 0.847 |
| 1.6 | 0.1 | clcB | 1 | 29.1833 | 0.454 |
| 1.6 | 0.1 | ycaM | 1 | 29.1833 | 0.396 |
| 1.6 | 0.1 | yadI | 1 | 29.1833 | 0.827 |
| 1.6 | 0.1 | AG1 | 1 | 29.1833 | 0.892 |
| 1.6 | 0.1 | ptsI | 1 | 29.4333 | 0.857 |
| 1.6 | 0.1 | clcB | 1 | 29.4333 | 0.458 |
| 1.6 | 0.1 | ycaM | 1 | 29.4333 | 0.399 |
| 1.6 | 0.1 | yadI | 1 | 29.4333 | 0.831 |
| 1.6 | 0.1 | AG1 | 1 | 29.4333 | 0.903 |

| | | | | | |
|---|---|---|---|---|---|
| 1.6 | 0.1 | ptsI | 1 | 29.6833 | 0.853 |
| 1.6 | 0.1 | clcB | 1 | 29.6833 | 0.458 |
| 1.6 | 0.1 | ycaM | 1 | 29.6833 | 0.396 |
| 1.6 | 0.1 | yadI | 1 | 29.6833 | 0.833 |
| 1.6 | 0.1 | AG1 | 1 | 29.6833 | 0.902 |
| 1.6 | 0.1 | ptsI | 1 | 29.9333 | 0.855 |
| 1.6 | 0.1 | clcB | 1 | 29.9333 | 0.459 |
| 1.6 | 0.1 | ycaM | 1 | 29.9333 | 0.397 |
| 1.6 | 0.1 | yadI | 1 | 29.9333 | 0.832 |
| 1.6 | 0.1 | AG1 | 1 | 29.9333 | 0.907 |
| 1.6 | 0.1 | ptsI | 1 | 30.1833 | 0.863 |
| 1.6 | 0.1 | clcB | 1 | 30.1833 | 0.457 |
| 1.6 | 0.1 | ycaM | 1 | 30.1833 | 0.4 |
| 1.6 | 0.1 | yadI | 1 | 30.1833 | 0.83 |
| 1.6 | 0.1 | AG1 | 1 | 30.1833 | 0.913 |
| 1.6 | 0.1 | ptsI | 1 | 30.4333 | 0.86 |
| 1.6 | 0.1 | clcB | 1 | 30.4333 | 0.461 |
| 1.6 | 0.1 | ycaM | 1 | 30.4333 | 0.401 |
| 1.6 | 0.1 | yadI | 1 | 30.4333 | 0.831 |
| 1.6 | 0.1 | AG1 | 1 | 30.4333 | 0.914 |
| 1.6 | 0.1 | ptsI | 1 | 30.6833 | 0.863 |
| 1.6 | 0.1 | clcB | 1 | 30.6833 | 0.46 |
| 1.6 | 0.1 | ycaM | 1 | 30.6833 | 0.403 |
| 1.6 | 0.1 | yadI | 1 | 30.6833 | 0.838 |
| 1.6 | 0.1 | AG1 | 1 | 30.6833 | 0.916 |
| 1.6 | 0.1 | ptsI | 1 | 30.9333 | 0.868 |
| 1.6 | 0.1 | clcB | 1 | 30.9333 | 0.462 |
| 1.6 | 0.1 | ycaM | 1 | 30.9333 | 0.402 |
| 1.6 | 0.1 | yadI | 1 | 30.9333 | 0.836 |
| 1.6 | 0.1 | AG1 | 1 | 30.9333 | 0.928 |
| 1.6 | 0.1 | ptsI | 1 | 31.1833 | 0.871 |
| 1.6 | 0.1 | clcB | 1 | 31.1833 | 0.461 |
| 1.6 | 0.1 | ycaM | 1 | 31.1833 | 0.402 |
| 1.6 | 0.1 | yadI | 1 | 31.1833 | 0.842 |
| 1.6 | 0.1 | AG1 | 1 | 31.1833 | 0.926 |
| 1.6 | 0.1 | ptsI | 1 | 31.4333 | 0.872 |
| 1.6 | 0.1 | clcB | 1 | 31.4333 | 0.462 |
| 1.6 | 0.1 | ycaM | 1 | 31.4333 | 0.403 |
| 1.6 | 0.1 | yadI | 1 | 31.4333 | 0.842 |
| 1.6 | 0.1 | AG1 | 1 | 31.4333 | 0.934 |
| 1.6 | 0.1 | ptsI | 1 | 31.6833 | 0.874 |
| 1.6 | 0.1 | clcB | 1 | 31.6833 | 0.461 |
| 1.6 | 0.1 | ycaM | 1 | 31.6833 | 0.404 |
| 1.6 | 0.1 | yadI | 1 | 31.6833 | 0.844 |
| 1.6 | 0.1 | AG1 | 1 | 31.6833 | 0.943 |
| 1.6 | 0.1 | ptsI | 1 | 31.9333 | 0.877 |
| 1.6 | 0.1 | clcB | 1 | 31.9333 | 0.465 |
| 1.6 | 0.1 | ycaM | 1 | 31.9333 | 0.405 |

| | | | | | |
|---|---|---|---|---|---|
| 1.6 | 0.1 | yadI | 1 | 31.9333 | 0.848 |
| 1.6 | 0.1 | AG1 | 1 | 31.9333 | 0.944 |
| 1.6 | 0.1 | ptsI | 1 | 32.1833 | 0.879 |
| 1.6 | 0.1 | clcB | 1 | 32.1833 | 0.465 |
| 1.6 | 0.1 | ycaM | 1 | 32.1833 | 0.404 |
| 1.6 | 0.1 | yadI | 1 | 32.1833 | 0.84 |
| 1.6 | 0.1 | AG1 | 1 | 32.1833 | 0.95 |
| 1.6 | 0.1 | ptsI | 1 | 32.4333 | 0.878 |
| 1.6 | 0.1 | clcB | 1 | 32.4333 | 0.461 |
| 1.6 | 0.1 | ycaM | 1 | 32.4333 | 0.403 |
| 1.6 | 0.1 | yadI | 1 | 32.4333 | 0.831 |
| 1.6 | 0.1 | AG1 | 1 | 32.4333 | 0.946 |
| 1.6 | 0.1 | ptsI | 1 | 32.6833 | 0.883 |
| 1.6 | 0.1 | clcB | 1 | 32.6833 | 0.467 |
| 1.6 | 0.1 | ycaM | 1 | 32.6833 | 0.41 |
| 1.6 | 0.1 | yadI | 1 | 32.6833 | 0.845 |
| 1.6 | 0.1 | AG1 | 1 | 32.6833 | 0.961 |
| 1.6 | 0.1 | ptsI | 1 | 32.9333 | 0.881 |
| 1.6 | 0.1 | clcB | 1 | 32.9333 | 0.466 |
| 1.6 | 0.1 | ycaM | 1 | 32.9333 | 0.408 |
| 1.6 | 0.1 | yadI | 1 | 32.9333 | 0.845 |
| 1.6 | 0.1 | AG1 | 1 | 32.9333 | 0.969 |
| 1.6 | 0.1 | ptsI | 1 | 33.1833 | 0.888 |
| 1.6 | 0.1 | clcB | 1 | 33.1833 | 0.464 |
| 1.6 | 0.1 | ycaM | 1 | 33.1833 | 0.41 |
| 1.6 | 0.1 | yadI | 1 | 33.1833 | 0.847 |
| 1.6 | 0.1 | AG1 | 1 | 33.1833 | 0.963 |
| 1.6 | 0.1 | ptsI | 1 | 33.4333 | 0.885 |
| 1.6 | 0.1 | clcB | 1 | 33.4333 | 0.466 |
| 1.6 | 0.1 | ycaM | 1 | 33.4333 | 0.413 |
| 1.6 | 0.1 | yadI | 1 | 33.4333 | 0.847 |
| 1.6 | 0.1 | AG1 | 1 | 33.4333 | 0.981 |
| 1.6 | 0.1 | ptsI | 1 | 33.6833 | 0.887 |
| 1.6 | 0.1 | clcB | 1 | 33.6833 | 0.467 |
| 1.6 | 0.1 | ycaM | 1 | 33.6833 | 0.414 |
| 1.6 | 0.1 | yadI | 1 | 33.6833 | 0.847 |
| 1.6 | 0.1 | AG1 | 1 | 33.6833 | 0.981 |
| 1.6 | 0.1 | ptsI | 1 | 33.9333 | 0.883 |
| 1.6 | 0.1 | clcB | 1 | 33.9333 | 0.469 |
| 1.6 | 0.1 | ycaM | 1 | 33.9333 | 0.414 |
| 1.6 | 0.1 | yadI | 1 | 33.9333 | 0.849 |
| 1.6 | 0.1 | AG1 | 1 | 33.9333 | 0.992 |
| 1.6 | 0.1 | ptsI | 1 | 34.1833 | 0.886 |
| 1.6 | 0.1 | clcB | 1 | 34.1833 | 0.473 |
| 1.6 | 0.1 | ycaM | 1 | 34.1833 | 0.419 |
| 1.6 | 0.1 | yadI | 1 | 34.1833 | 0.855 |
| 1.6 | 0.1 | AG1 | 1 | 34.1833 | 0.994 |
| 1.6 | 0.1 | ptsI | 1 | 34.4333 | 0.884 |

| | | | | | |
|---|---|---|---|---|---|
| 1.6 | 0.1 | clcB | 1 | 34.4333 | 0.47 |
| 1.6 | 0.1 | ycaM | 1 | 34.4333 | 0.42 |
| 1.6 | 0.1 | yadI | 1 | 34.4333 | 0.852 |
| 1.6 | 0.1 | AG1 | 1 | 34.4333 | 1.002 |
| 1.6 | 0.1 | ptsI | 1 | 34.6833 | 0.887 |
| 1.6 | 0.1 | clcB | 1 | 34.6833 | 0.474 |
| 1.6 | 0.1 | ycaM | 1 | 34.6833 | 0.417 |
| 1.6 | 0.1 | yadI | 1 | 34.6833 | 0.851 |
| 1.6 | 0.1 | AG1 | 1 | 34.6833 | 1.003 |
| 1.6 | 0.1 | ptsI | 1 | 34.9333 | 0.883 |
| 1.6 | 0.1 | clcB | 1 | 34.9333 | 0.474 |
| 1.6 | 0.1 | ycaM | 1 | 34.9333 | 0.419 |
| 1.6 | 0.1 | yadI | 1 | 34.9333 | 0.854 |
| 1.6 | 0.1 | AG1 | 1 | 34.9333 | 1.008 |
| 1.6 | 0.1 | ptsI | 1 | 35.1833 | 0.885 |
| 1.6 | 0.1 | clcB | 1 | 35.1833 | 0.474 |
| 1.6 | 0.1 | ycaM | 1 | 35.1833 | 0.417 |
| 1.6 | 0.1 | yadI | 1 | 35.1833 | 0.858 |
| 1.6 | 0.1 | AG1 | 1 | 35.1833 | 1.015 |
| 1.6 | 0.1 | ptsI | 1 | 35.4333 | 0.885 |
| 1.6 | 0.1 | clcB | 1 | 35.4333 | 0.474 |
| 1.6 | 0.1 | ycaM | 1 | 35.4333 | 0.42 |
| 1.6 | 0.1 | yadI | 1 | 35.4333 | 0.858 |
| 1.6 | 0.1 | AG1 | 1 | 35.4333 | 1.019 |
| 1.6 | 0.1 | ptsI | 1 | 35.6833 | 0.886 |
| 1.6 | 0.1 | clcB | 1 | 35.6833 | 0.479 |
| 1.6 | 0.1 | ycaM | 1 | 35.6833 | 0.42 |
| 1.6 | 0.1 | yadI | 1 | 35.6833 | 0.854 |
| 1.6 | 0.1 | AG1 | 1 | 35.6833 | 1.022 |
| 1.6 | 0.1 | ptsI | 1 | 35.9333 | 0.885 |
| 1.6 | 0.1 | clcB | 1 | 35.9333 | 0.474 |
| 1.6 | 0.1 | ycaM | 1 | 35.9333 | 0.418 |
| 1.6 | 0.1 | yadI | 1 | 35.9333 | 0.856 |
| 1.6 | 0.1 | AG1 | 1 | 35.9333 | 1.018 |
| 1.6 | 0.1 | ptsI | 1 | 36.1833 | 0.884 |
| 1.6 | 0.1 | clcB | 1 | 36.1833 | 0.482 |
| 1.6 | 0.1 | ycaM | 1 | 36.1833 | 0.422 |
| 1.6 | 0.1 | yadI | 1 | 36.1833 | 0.858 |
| 1.6 | 0.1 | AG1 | 1 | 36.1833 | 1.028 |
| 1.6 | 0.1 | ptsI | 1 | 36.4333 | 0.882 |
| 1.6 | 0.1 | clcB | 1 | 36.4333 | 0.479 |
| 1.6 | 0.1 | ycaM | 1 | 36.4333 | 0.424 |
| 1.6 | 0.1 | yadI | 1 | 36.4333 | 0.862 |
| 1.6 | 0.1 | AG1 | 1 | 36.4333 | 1.034 |
| 1.6 | 0.1 | ptsI | 1 | 36.6833 | 0.883 |
| 1.6 | 0.1 | clcB | 1 | 36.6833 | 0.481 |
| 1.6 | 0.1 | ycaM | 1 | 36.6833 | 0.426 |
| 1.6 | 0.1 | yadI | 1 | 36.6833 | 0.863 |

| | | | | | |
|---|---|---|---|---|---|
| 1.6 | 0.1 | AG1 | 1 | 36.6833 | 1.039 |
| 3.2 | 0.1 | ptsI | 1 | 0 | 0.252 |
| 3.2 | 0.1 | clcB | 1 | 0 | 0.55 |
| 3.2 | 0.1 | ycaM | 1 | 0 | 0.228 |
| 3.2 | 0.1 | yadI | 1 | 0 | 0.275 |
| 3.2 | 0.1 | AG1 | 1 | 0 | 0.244 |
| 3.2 | 0.1 | ptsI | 1 | 0.25 | 0.24 |
| 3.2 | 0.1 | clcB | 1 | 0.25 | 0.508 |
| 3.2 | 0.1 | ycaM | 1 | 0.25 | 0.218 |
| 3.2 | 0.1 | yadI | 1 | 0.25 | 0.257 |
| 3.2 | 0.1 | AG1 | 1 | 0.25 | 0.234 |
| 3.2 | 0.1 | ptsI | 1 | 0.5 | 0.238 |
| 3.2 | 0.1 | clcB | 1 | 0.5 | 0.494 |
| 3.2 | 0.1 | ycaM | 1 | 0.5 | 0.215 |
| 3.2 | 0.1 | yadI | 1 | 0.5 | 0.254 |
| 3.2 | 0.1 | AG1 | 1 | 0.5 | 0.231 |
| 3.2 | 0.1 | ptsI | 1 | 0.75 | 0.236 |
| 3.2 | 0.1 | clcB | 1 | 0.75 | 0.492 |
| 3.2 | 0.1 | ycaM | 1 | 0.75 | 0.213 |
| 3.2 | 0.1 | yadI | 1 | 0.75 | 0.252 |
| 3.2 | 0.1 | AG1 | 1 | 0.75 | 0.228 |
| 3.2 | 0.1 | ptsI | 1 | 1 | 0.239 |
| 3.2 | 0.1 | clcB | 1 | 1 | 0.497 |
| 3.2 | 0.1 | ycaM | 1 | 1 | 0.212 |
| 3.2 | 0.1 | yadI | 1 | 1 | 0.255 |
| 3.2 | 0.1 | AG1 | 1 | 1 | 0.232 |
| 3.2 | 0.1 | ptsI | 1 | 1.25 | 0.244 |
| 3.2 | 0.1 | clcB | 1 | 1.25 | 0.499 |
| 3.2 | 0.1 | ycaM | 1 | 1.25 | 0.214 |
| 3.2 | 0.1 | yadI | 1 | 1.25 | 0.254 |
| 3.2 | 0.1 | AG1 | 1 | 1.25 | 0.227 |
| 3.2 | 0.1 | ptsI | 1 | 1.5 | 0.245 |
| 3.2 | 0.1 | clcB | 1 | 1.5 | 0.502 |
| 3.2 | 0.1 | ycaM | 1 | 1.5 | 0.215 |
| 3.2 | 0.1 | yadI | 1 | 1.5 | 0.258 |
| 3.2 | 0.1 | AG1 | 1 | 1.5 | 0.232 |
| 3.2 | 0.1 | ptsI | 1 | 1.75 | 0.252 |
| 3.2 | 0.1 | clcB | 1 | 1.75 | 0.51 |
| 3.2 | 0.1 | ycaM | 1 | 1.75 | 0.218 |
| 3.2 | 0.1 | yadI | 1 | 1.75 | 0.261 |
| 3.2 | 0.1 | AG1 | 1 | 1.75 | 0.232 |
| 3.2 | 0.1 | ptsI | 1 | 2 | 0.257 |
| 3.2 | 0.1 | clcB | 1 | 2 | 0.522 |
| 3.2 | 0.1 | ycaM | 1 | 2 | 0.223 |
| 3.2 | 0.1 | yadI | 1 | 2 | 0.265 |
| 3.2 | 0.1 | AG1 | 1 | 2 | 0.238 |
| 3.2 | 0.1 | ptsI | 1 | 2.25 | 0.267 |
| 3.2 | 0.1 | clcB | 1 | 2.25 | 0.528 |

| | | | | | |
|---|---|---|---|---|---|
| 3.2 | 0.1 | ycaM | 1 | 2.25 | 0.229 |
| 3.2 | 0.1 | yadI | 1 | 2.25 | 0.271 |
| 3.2 | 0.1 | AG1 | 1 | 2.25 | 0.242 |
| 3.2 | 0.1 | ptsI | 1 | 2.5 | 0.272 |
| 3.2 | 0.1 | clcB | 1 | 2.5 | 0.517 |
| 3.2 | 0.1 | ycaM | 1 | 2.5 | 0.231 |
| 3.2 | 0.1 | yadI | 1 | 2.5 | 0.275 |
| 3.2 | 0.1 | AG1 | 1 | 2.5 | 0.245 |
| 3.2 | 0.1 | ptsI | 1 | 2.75 | 0.278 |
| 3.2 | 0.1 | clcB | 1 | 2.75 | 0.526 |
| 3.2 | 0.1 | ycaM | 1 | 2.75 | 0.238 |
| 3.2 | 0.1 | yadI | 1 | 2.75 | 0.278 |
| 3.2 | 0.1 | AG1 | 1 | 2.75 | 0.248 |
| 3.2 | 0.1 | ptsI | 1 | 3 | 0.286 |
| 3.2 | 0.1 | clcB | 1 | 3 | 0.531 |
| 3.2 | 0.1 | ycaM | 1 | 3 | 0.244 |
| 3.2 | 0.1 | yadI | 1 | 3 | 0.285 |
| 3.2 | 0.1 | AG1 | 1 | 3 | 0.253 |
| 3.2 | 0.1 | ptsI | 1 | 3.25 | 0.296 |
| 3.2 | 0.1 | clcB | 1 | 3.25 | 0.546 |
| 3.2 | 0.1 | ycaM | 1 | 3.25 | 0.255 |
| 3.2 | 0.1 | yadI | 1 | 3.25 | 0.296 |
| 3.2 | 0.1 | AG1 | 1 | 3.25 | 0.26 |
| 3.2 | 0.1 | ptsI | 1 | 3.5 | 0.304 |
| 3.2 | 0.1 | clcB | 1 | 3.5 | 0.561 |
| 3.2 | 0.1 | ycaM | 1 | 3.5 | 0.257 |
| 3.2 | 0.1 | yadI | 1 | 3.5 | 0.306 |
| 3.2 | 0.1 | AG1 | 1 | 3.5 | 0.265 |
| 3.2 | 0.1 | ptsI | 1 | 3.75 | 0.309 |
| 3.2 | 0.1 | clcB | 1 | 3.75 | 0.57 |
| 3.2 | 0.1 | ycaM | 1 | 3.75 | 0.256 |
| 3.2 | 0.1 | yadI | 1 | 3.75 | 0.316 |
| 3.2 | 0.1 | AG1 | 1 | 3.75 | 0.262 |
| 3.2 | 0.1 | ptsI | 1 | 4 | 0.328 |
| 3.2 | 0.1 | clcB | 1 | 4 | 0.58 |
| 3.2 | 0.1 | ycaM | 1 | 4 | 0.266 |
| 3.2 | 0.1 | yadI | 1 | 4 | 0.33 |
| 3.2 | 0.1 | AG1 | 1 | 4 | 0.275 |
| 3.2 | 0.1 | ptsI | 1 | 4.25 | 0.326 |
| 3.2 | 0.1 | clcB | 1 | 4.25 | 0.624 |
| 3.2 | 0.1 | ycaM | 1 | 4.25 | 0.269 |
| 3.2 | 0.1 | yadI | 1 | 4.25 | 0.333 |
| 3.2 | 0.1 | AG1 | 1 | 4.25 | 0.276 |
| 3.2 | 0.1 | ptsI | 1 | 4.5 | 0.341 |
| 3.2 | 0.1 | clcB | 1 | 4.5 | 0.64 |
| 3.2 | 0.1 | ycaM | 1 | 4.5 | 0.278 |
| 3.2 | 0.1 | yadI | 1 | 4.5 | 0.345 |
| 3.2 | 0.1 | AG1 | 1 | 4.5 | 0.285 |

| | | | | | |
|---|---|---|---|---|---|
| 3.2 | 0.1 | ptsI | 1 | 4.75 | 0.338 |
| 3.2 | 0.1 | clcB | 1 | 4.75 | 0.655 |
| 3.2 | 0.1 | ycaM | 1 | 4.75 | 0.278 |
| 3.2 | 0.1 | yadI | 1 | 4.75 | 0.349 |
| 3.2 | 0.1 | AG1 | 1 | 4.75 | 0.283 |
| 3.2 | 0.1 | ptsI | 1 | 5 | 0.351 |
| 3.2 | 0.1 | clcB | 1 | 5 | 0.686 |
| 3.2 | 0.1 | ycaM | 1 | 5 | 0.281 |
| 3.2 | 0.1 | yadI | 1 | 5 | 0.358 |
| 3.2 | 0.1 | AG1 | 1 | 5 | 0.294 |
| 3.2 | 0.1 | ptsI | 1 | 5.25 | 0.356 |
| 3.2 | 0.1 | clcB | 1 | 5.25 | 0.715 |
| 3.2 | 0.1 | ycaM | 1 | 5.25 | 0.292 |
| 3.2 | 0.1 | yadI | 1 | 5.25 | 0.369 |
| 3.2 | 0.1 | AG1 | 1 | 5.25 | 0.294 |
| 3.2 | 0.1 | ptsI | 1 | 5.5 | 0.37 |
| 3.2 | 0.1 | clcB | 1 | 5.5 | 0.738 |
| 3.2 | 0.1 | ycaM | 1 | 5.5 | 0.299 |
| 3.2 | 0.1 | yadI | 1 | 5.5 | 0.382 |
| 3.2 | 0.1 | AG1 | 1 | 5.5 | 0.304 |
| 3.2 | 0.1 | ptsI | 1 | 5.75 | 0.386 |
| 3.2 | 0.1 | clcB | 1 | 5.75 | 0.774 |
| 3.2 | 0.1 | ycaM | 1 | 5.75 | 0.314 |
| 3.2 | 0.1 | yadI | 1 | 5.75 | 0.394 |
| 3.2 | 0.1 | AG1 | 1 | 5.75 | 0.31 |
| 3.2 | 0.1 | ptsI | 1 | 6 | 0.375 |
| 3.2 | 0.1 | clcB | 1 | 6 | 0.807 |
| 3.2 | 0.1 | ycaM | 1 | 6 | 0.31 |
| 3.2 | 0.1 | yadI | 1 | 6 | 0.396 |
| 3.2 | 0.1 | AG1 | 1 | 6 | 0.306 |
| 3.2 | 0.1 | ptsI | 1 | 6.25 | 0.379 |
| 3.2 | 0.1 | clcB | 1 | 6.25 | 0.844 |
| 3.2 | 0.1 | ycaM | 1 | 6.25 | 0.306 |
| 3.2 | 0.1 | yadI | 1 | 6.25 | 0.394 |
| 3.2 | 0.1 | AG1 | 1 | 6.25 | 0.306 |
| 3.2 | 0.1 | ptsI | 1 | 6.5 | 0.382 |
| 3.2 | 0.1 | clcB | 1 | 6.5 | 0.868 |
| 3.2 | 0.1 | ycaM | 1 | 6.5 | 0.312 |
| 3.2 | 0.1 | yadI | 1 | 6.5 | 0.397 |
| 3.2 | 0.1 | AG1 | 1 | 6.5 | 0.312 |
| 3.2 | 0.1 | ptsI | 1 | 6.75 | 0.406 |
| 3.2 | 0.1 | clcB | 1 | 6.75 | 0.861 |
| 3.2 | 0.1 | ycaM | 1 | 6.75 | 0.323 |
| 3.2 | 0.1 | yadI | 1 | 6.75 | 0.409 |
| 3.2 | 0.1 | AG1 | 1 | 6.75 | 0.322 |
| 3.2 | 0.1 | ptsI | 1 | 7 | 0.428 |
| 3.2 | 0.1 | clcB | 1 | 7 | 0.811 |
| 3.2 | 0.1 | ycaM | 1 | 7 | 0.337 |

| | | | | | |
|---|---|---|---|---|---|
| 3.2 | 0.1 | yadI | 1 | 7 | 0.419 |
| 3.2 | 0.1 | AG1 | 1 | 7 | 0.327 |
| 3.2 | 0.1 | ptsI | 1 | 7.25 | 0.441 |
| 3.2 | 0.1 | clcB | 1 | 7.25 | 0.734 |
| 3.2 | 0.1 | ycaM | 1 | 7.25 | 0.348 |
| 3.2 | 0.1 | yadI | 1 | 7.25 | 0.431 |
| 3.2 | 0.1 | AG1 | 1 | 7.25 | 0.334 |
| 3.2 | 0.1 | ptsI | 1 | 7.5 | 0.46 |
| 3.2 | 0.1 | clcB | 1 | 7.5 | 0.617 |
| 3.2 | 0.1 | ycaM | 1 | 7.5 | 0.366 |
| 3.2 | 0.1 | yadI | 1 | 7.5 | 0.449 |
| 3.2 | 0.1 | AG1 | 1 | 7.5 | 0.344 |
| 3.2 | 0.1 | ptsI | 1 | 7.75 | 0.473 |
| 3.2 | 0.1 | clcB | 1 | 7.75 | 0.498 |
| 3.2 | 0.1 | ycaM | 1 | 7.75 | 0.374 |
| 3.2 | 0.1 | yadI | 1 | 7.75 | 0.458 |
| 3.2 | 0.1 | AG1 | 1 | 7.75 | 0.352 |
| 3.2 | 0.1 | ptsI | 1 | 8 | 0.492 |
| 3.2 | 0.1 | clcB | 1 | 8 | 0.388 |
| 3.2 | 0.1 | ycaM | 1 | 8 | 0.377 |
| 3.2 | 0.1 | yadI | 1 | 8 | 0.464 |
| 3.2 | 0.1 | AG1 | 1 | 8 | 0.36 |
| 3.2 | 0.1 | ptsI | 1 | 8.25 | 0.507 |
| 3.2 | 0.1 | clcB | 1 | 8.25 | 0.388 |
| 3.2 | 0.1 | ycaM | 1 | 8.25 | 0.388 |
| 3.2 | 0.1 | yadI | 1 | 8.25 | 0.479 |
| 3.2 | 0.1 | AG1 | 1 | 8.25 | 0.368 |
| 3.2 | 0.1 | ptsI | 1 | 8.5 | 0.523 |
| 3.2 | 0.1 | clcB | 1 | 8.5 | 0.405 |
| 3.2 | 0.1 | ycaM | 1 | 8.5 | 0.398 |
| 3.2 | 0.1 | yadI | 1 | 8.5 | 0.497 |
| 3.2 | 0.1 | AG1 | 1 | 8.5 | 0.374 |
| 3.2 | 0.1 | ptsI | 1 | 8.75 | 0.534 |
| 3.2 | 0.1 | clcB | 1 | 8.75 | 0.421 |
| 3.2 | 0.1 | ycaM | 1 | 8.75 | 0.396 |
| 3.2 | 0.1 | yadI | 1 | 8.75 | 0.504 |
| 3.2 | 0.1 | AG1 | 1 | 8.75 | 0.38 |
| 3.2 | 0.1 | ptsI | 1 | 9 | 0.541 |
| 3.2 | 0.1 | clcB | 1 | 9 | 0.435 |
| 3.2 | 0.1 | ycaM | 1 | 9 | 0.407 |
| 3.2 | 0.1 | yadI | 1 | 9 | 0.513 |
| 3.2 | 0.1 | AG1 | 1 | 9 | 0.388 |
| 3.2 | 0.1 | ptsI | 1 | 9.25 | 0.535 |
| 3.2 | 0.1 | clcB | 1 | 9.25 | 0.454 |
| 3.2 | 0.1 | ycaM | 1 | 9.25 | 0.401 |
| 3.2 | 0.1 | yadI | 1 | 9.25 | 0.519 |
| 3.2 | 0.1 | AG1 | 1 | 9.25 | 0.4 |
| 3.2 | 0.1 | ptsI | 1 | 9.5 | 0.535 |

| | | | | | |
|---|---|---|---|---|---|
| 3.2 | 0.1 | clcB | 1 | 9.5 | 0.446 |
| 3.2 | 0.1 | ycaM | 1 | 9.5 | 0.407 |
| 3.2 | 0.1 | yadI | 1 | 9.5 | 0.51 |
| 3.2 | 0.1 | AG1 | 1 | 9.5 | 0.403 |
| 3.2 | 0.1 | ptsI | 1 | 9.75 | 0.586 |
| 3.2 | 0.1 | clcB | 1 | 9.75 | 0.499 |
| 3.2 | 0.1 | ycaM | 1 | 9.75 | 0.434 |
| 3.2 | 0.1 | yadI | 1 | 9.75 | 0.542 |
| 3.2 | 0.1 | AG1 | 1 | 9.75 | 0.426 |
| 3.2 | 0.1 | ptsI | 1 | 10 | 0.495 |
| 3.2 | 0.1 | clcB | 1 | 10 | 0.456 |
| 3.2 | 0.1 | ycaM | 1 | 10 | 0.422 |
| 3.2 | 0.1 | yadI | 1 | 10 | 0.507 |
| 3.2 | 0.1 | AG1 | 1 | 10 | 0.423 |
| 3.2 | 0.1 | ptsI | 1 | 10.25 | 0.554 |
| 3.2 | 0.1 | clcB | 1 | 10.25 | 0.497 |
| 3.2 | 0.1 | ycaM | 1 | 10.25 | 0.437 |
| 3.2 | 0.1 | yadI | 1 | 10.25 | 0.522 |
| 3.2 | 0.1 | AG1 | 1 | 10.25 | 0.44 |
| 3.2 | 0.1 | ptsI | 1 | 10.5 | 0.581 |
| 3.2 | 0.1 | clcB | 1 | 10.5 | 0.52 |
| 3.2 | 0.1 | ycaM | 1 | 10.5 | 0.458 |
| 3.2 | 0.1 | yadI | 1 | 10.5 | 0.538 |
| 3.2 | 0.1 | AG1 | 1 | 10.5 | 0.457 |
| 3.2 | 0.1 | ptsI | 1 | 10.75 | 0.507 |
| 3.2 | 0.1 | clcB | 1 | 10.75 | 0.475 |
| 3.2 | 0.1 | ycaM | 1 | 10.75 | 0.447 |
| 3.2 | 0.1 | yadI | 1 | 10.75 | 0.516 |
| 3.2 | 0.1 | AG1 | 1 | 10.75 | 0.448 |
| 3.2 | 0.1 | ptsI | 1 | 11 | 0.528 |
| 3.2 | 0.1 | clcB | 1 | 11 | 0.454 |
| 3.2 | 0.1 | ycaM | 1 | 11 | 0.452 |
| 3.2 | 0.1 | yadI | 1 | 11 | 0.523 |
| 3.2 | 0.1 | AG1 | 1 | 11 | 0.448 |
| 3.2 | 0.1 | ptsI | 1 | 11.25 | 0.61 |
| 3.2 | 0.1 | clcB | 1 | 11.25 | 0.528 |
| 3.2 | 0.1 | ycaM | 1 | 11.25 | 0.488 |
| 3.2 | 0.1 | yadI | 1 | 11.25 | 0.555 |
| 3.2 | 0.1 | AG1 | 1 | 11.25 | 0.505 |
| 3.2 | 0.1 | ptsI | 1 | 11.5 | 0.586 |
| 3.2 | 0.1 | clcB | 1 | 11.5 | 0.497 |
| 3.2 | 0.1 | ycaM | 1 | 11.5 | 0.494 |
| 3.2 | 0.1 | yadI | 1 | 11.5 | 0.528 |
| 3.2 | 0.1 | AG1 | 1 | 11.5 | 0.489 |
| 3.2 | 0.1 | ptsI | 1 | 11.75 | 0.572 |
| 3.2 | 0.1 | clcB | 1 | 11.75 | 0.498 |
| 3.2 | 0.1 | ycaM | 1 | 11.75 | 0.491 |
| 3.2 | 0.1 | yadI | 1 | 11.75 | 0.528 |

| | | | | | |
|---|---|---|---|---|---|
| 3.2 | 0.1 | AG1 | 1 | 11.75 | 0.488 |
| 3.2 | 0.1 | ptsI | 1 | 12 | 0.674 |
| 3.2 | 0.1 | clcB | 1 | 12 | 0.58 |
| 3.2 | 0.1 | ycaM | 1 | 12 | 0.533 |
| 3.2 | 0.1 | yadI | 1 | 12 | 0.539 |
| 3.2 | 0.1 | AG1 | 1 | 12 | 0.558 |
| 3.2 | 0.1 | ptsI | 1 | 12.25 | 0.606 |
| 3.2 | 0.1 | clcB | 1 | 12.25 | 0.519 |
| 3.2 | 0.1 | ycaM | 1 | 12.25 | 0.512 |
| 3.2 | 0.1 | yadI | 1 | 12.25 | 0.527 |
| 3.2 | 0.1 | AG1 | 1 | 12.25 | 0.516 |
| 3.2 | 0.1 | ptsI | 1 | 12.5 | 0.661 |
| 3.2 | 0.1 | clcB | 1 | 12.5 | 0.555 |
| 3.2 | 0.1 | ycaM | 1 | 12.5 | 0.537 |
| 3.2 | 0.1 | yadI | 1 | 12.5 | 0.53 |
| 3.2 | 0.1 | AG1 | 1 | 12.5 | 0.551 |
| 3.2 | 0.1 | ptsI | 1 | 12.75 | 0.697 |
| 3.2 | 0.1 | clcB | 1 | 12.75 | 0.623 |
| 3.2 | 0.1 | ycaM | 1 | 12.75 | 0.574 |
| 3.2 | 0.1 | yadI | 1 | 12.75 | 0.558 |
| 3.2 | 0.1 | AG1 | 1 | 12.75 | 0.596 |
| 3.2 | 0.1 | ptsI | 1 | 13 | 0.715 |
| 3.2 | 0.1 | clcB | 1 | 13 | 0.64 |
| 3.2 | 0.1 | ycaM | 1 | 13 | 0.595 |
| 3.2 | 0.1 | yadI | 1 | 13 | 0.586 |
| 3.2 | 0.1 | AG1 | 1 | 13 | 0.658 |
| 3.2 | 0.1 | ptsI | 1 | 13.25 | 0.664 |
| 3.2 | 0.1 | clcB | 1 | 13.25 | 0.625 |
| 3.2 | 0.1 | ycaM | 1 | 13.25 | 0.575 |
| 3.2 | 0.1 | yadI | 1 | 13.25 | 0.557 |
| 3.2 | 0.1 | AG1 | 1 | 13.25 | 0.644 |
| 3.2 | 0.1 | ptsI | 1 | 13.5 | 0.737 |
| 3.2 | 0.1 | clcB | 1 | 13.5 | 0.677 |
| 3.2 | 0.1 | ycaM | 1 | 13.5 | 0.624 |
| 3.2 | 0.1 | yadI | 1 | 13.5 | 0.591 |
| 3.2 | 0.1 | AG1 | 1 | 13.5 | 0.702 |
| 3.2 | 0.1 | ptsI | 1 | 13.75 | 0.687 |
| 3.2 | 0.1 | clcB | 1 | 13.75 | 0.653 |
| 3.2 | 0.1 | ycaM | 1 | 13.75 | 0.603 |
| 3.2 | 0.1 | yadI | 1 | 13.75 | 0.577 |
| 3.2 | 0.1 | AG1 | 1 | 13.75 | 0.688 |
| 3.2 | 0.1 | ptsI | 1 | 14 | 0.772 |
| 3.2 | 0.1 | clcB | 1 | 14 | 0.691 |
| 3.2 | 0.1 | ycaM | 1 | 14 | 0.635 |
| 3.2 | 0.1 | yadI | 1 | 14 | 0.602 |
| 3.2 | 0.1 | AG1 | 1 | 14 | 0.747 |
| 3.2 | 0.1 | ptsI | 1 | 14.25 | 0.782 |
| 3.2 | 0.1 | clcB | 1 | 14.25 | 0.705 |

| | | | | | |
|---|---|---|---|---|---|
| 3.2 | 0.1 | ycaM | 1 | 14.25 | 0.666 |
| 3.2 | 0.1 | yadI | 1 | 14.25 | 0.612 |
| 3.2 | 0.1 | AG1 | 1 | 14.25 | 0.764 |
| 3.2 | 0.1 | ptsI | 1 | 14.5 | 0.776 |
| 3.2 | 0.1 | clcB | 1 | 14.5 | 0.705 |
| 3.2 | 0.1 | ycaM | 1 | 14.5 | 0.68 |
| 3.2 | 0.1 | yadI | 1 | 14.5 | 0.622 |
| 3.2 | 0.1 | AG1 | 1 | 14.5 | 0.763 |
| 3.2 | 0.1 | ptsI | 1 | 14.75 | 0.732 |
| 3.2 | 0.1 | clcB | 1 | 14.75 | 0.658 |
| 3.2 | 0.1 | ycaM | 1 | 14.75 | 0.654 |
| 3.2 | 0.1 | yadI | 1 | 14.75 | 0.602 |
| 3.2 | 0.1 | AG1 | 1 | 14.75 | 0.757 |
| 3.2 | 0.1 | ptsI | 1 | 15 | 0.785 |
| 3.2 | 0.1 | clcB | 1 | 15 | 0.703 |
| 3.2 | 0.1 | ycaM | 1 | 15 | 0.683 |
| 3.2 | 0.1 | yadI | 1 | 15 | 0.631 |
| 3.2 | 0.1 | AG1 | 1 | 15 | 0.793 |
| 3.2 | 0.1 | ptsI | 1 | 15.25 | 0.795 |
| 3.2 | 0.1 | clcB | 1 | 15.25 | 0.718 |
| 3.2 | 0.1 | ycaM | 1 | 15.25 | 0.702 |
| 3.2 | 0.1 | yadI | 1 | 15.25 | 0.646 |
| 3.2 | 0.1 | AG1 | 1 | 15.25 | 0.806 |
| 3.2 | 0.1 | ptsI | 1 | 15.5 | 0.786 |
| 3.2 | 0.1 | clcB | 1 | 15.5 | 0.711 |
| 3.2 | 0.1 | ycaM | 1 | 15.5 | 0.686 |
| 3.2 | 0.1 | yadI | 1 | 15.5 | 0.634 |
| 3.2 | 0.1 | AG1 | 1 | 15.5 | 0.783 |
| 3.2 | 0.1 | ptsI | 1 | 15.75 | 0.761 |
| 3.2 | 0.1 | clcB | 1 | 15.75 | 0.677 |
| 3.2 | 0.1 | ycaM | 1 | 15.75 | 0.677 |
| 3.2 | 0.1 | yadI | 1 | 15.75 | 0.621 |
| 3.2 | 0.1 | AG1 | 1 | 15.75 | 0.799 |
| 3.2 | 0.1 | ptsI | 1 | 16 | 0.729 |
| 3.2 | 0.1 | clcB | 1 | 16 | 0.664 |
| 3.2 | 0.1 | ycaM | 1 | 16 | 0.66 |
| 3.2 | 0.1 | yadI | 1 | 16 | 0.616 |
| 3.2 | 0.1 | AG1 | 1 | 16 | 0.78 |
| 3.2 | 0.1 | ptsI | 1 | 16.25 | 0.779 |
| 3.2 | 0.1 | clcB | 1 | 16.25 | 0.699 |
| 3.2 | 0.1 | ycaM | 1 | 16.25 | 0.692 |
| 3.2 | 0.1 | yadI | 1 | 16.25 | 0.642 |
| 3.2 | 0.1 | AG1 | 1 | 16.25 | 0.814 |
| 3.2 | 0.1 | ptsI | 1 | 16.5 | 0.793 |
| 3.2 | 0.1 | clcB | 1 | 16.5 | 0.708 |
| 3.2 | 0.1 | ycaM | 1 | 16.5 | 0.702 |
| 3.2 | 0.1 | yadI | 1 | 16.5 | 0.662 |
| 3.2 | 0.1 | AG1 | 1 | 16.5 | 0.827 |

| | | | | | |
|---|---|---|---|---|---|
| 3.2 | 0.1 | ptsI | 1 | 17.0667 | 0.646 |
| 3.2 | 0.1 | clcB | 1 | 17.0667 | 0.596 |
| 3.2 | 0.1 | ycaM | 1 | 17.0667 | 0.6 |
| 3.2 | 0.1 | yadI | 1 | 17.0667 | 0.663 |
| 3.2 | 0.1 | AG1 | 1 | 17.0667 | 0.837 |
| 3.2 | 0.1 | ptsI | 1 | 17.3167 | 0.641 |
| 3.2 | 0.1 | clcB | 1 | 17.3167 | 0.576 |
| 3.2 | 0.1 | ycaM | 1 | 17.3167 | 0.551 |
| 3.2 | 0.1 | yadI | 1 | 17.3167 | 0.645 |
| 3.2 | 0.1 | AG1 | 1 | 17.3167 | 0.79 |
| 3.2 | 0.1 | ptsI | 1 | 17.5667 | 0.642 |
| 3.2 | 0.1 | clcB | 1 | 17.5667 | 0.564 |
| 3.2 | 0.1 | ycaM | 1 | 17.5667 | 0.539 |
| 3.2 | 0.1 | yadI | 1 | 17.5667 | 0.656 |
| 3.2 | 0.1 | AG1 | 1 | 17.5667 | 0.779 |
| 3.2 | 0.1 | ptsI | 1 | 17.8167 | 0.649 |
| 3.2 | 0.1 | clcB | 1 | 17.8167 | 0.552 |
| 3.2 | 0.1 | ycaM | 1 | 17.8167 | 0.537 |
| 3.2 | 0.1 | yadI | 1 | 17.8167 | 0.676 |
| 3.2 | 0.1 | AG1 | 1 | 17.8167 | 0.797 |
| 3.2 | 0.1 | ptsI | 1 | 18.0667 | 0.654 |
| 3.2 | 0.1 | clcB | 1 | 18.0667 | 0.557 |
| 3.2 | 0.1 | ycaM | 1 | 18.0667 | 0.529 |
| 3.2 | 0.1 | yadI | 1 | 18.0667 | 0.7 |
| 3.2 | 0.1 | AG1 | 1 | 18.0667 | 0.772 |
| 3.2 | 0.1 | ptsI | 1 | 18.3167 | 0.65 |
| 3.2 | 0.1 | clcB | 1 | 18.3167 | 0.537 |
| 3.2 | 0.1 | ycaM | 1 | 18.3167 | 0.521 |
| 3.2 | 0.1 | yadI | 1 | 18.3167 | 0.672 |
| 3.2 | 0.1 | AG1 | 1 | 18.3167 | 0.772 |
| 3.2 | 0.1 | ptsI | 1 | 18.5667 | 0.671 |
| 3.2 | 0.1 | clcB | 1 | 18.5667 | 0.546 |
| 3.2 | 0.1 | ycaM | 1 | 18.5667 | 0.519 |
| 3.2 | 0.1 | yadI | 1 | 18.5667 | 0.703 |
| 3.2 | 0.1 | AG1 | 1 | 18.5667 | 0.776 |
| 3.2 | 0.1 | ptsI | 1 | 18.8167 | 0.678 |
| 3.2 | 0.1 | clcB | 1 | 18.8167 | 0.539 |
| 3.2 | 0.1 | ycaM | 1 | 18.8167 | 0.519 |
| 3.2 | 0.1 | yadI | 1 | 18.8167 | 0.725 |
| 3.2 | 0.1 | AG1 | 1 | 18.8167 | 0.788 |
| 3.2 | 0.1 | ptsI | 1 | 19.0667 | 0.702 |
| 3.2 | 0.1 | clcB | 1 | 19.0667 | 0.536 |
| 3.2 | 0.1 | ycaM | 1 | 19.0667 | 0.519 |
| 3.2 | 0.1 | yadI | 1 | 19.0667 | 0.742 |
| 3.2 | 0.1 | AG1 | 1 | 19.0667 | 0.809 |
| 3.2 | 0.1 | ptsI | 1 | 19.3167 | 0.689 |
| 3.2 | 0.1 | clcB | 1 | 19.3167 | 0.537 |
| 3.2 | 0.1 | ycaM | 1 | 19.3167 | 0.511 |

| | | | | | |
|---|---|---|---|---|---|
| 3.2 | 0.1 | yadI | 1 | 19.3167 | 0.747 |
| 3.2 | 0.1 | AG1 | 1 | 19.3167 | 0.797 |
| 3.2 | 0.1 | ptsI | 1 | 19.5667 | 0.72 |
| 3.2 | 0.1 | clcB | 1 | 19.5667 | 0.539 |
| 3.2 | 0.1 | ycaM | 1 | 19.5667 | 0.511 |
| 3.2 | 0.1 | yadI | 1 | 19.5667 | 0.758 |
| 3.2 | 0.1 | AG1 | 1 | 19.5667 | 0.81 |
| 3.2 | 0.1 | ptsI | 1 | 19.8167 | 0.73 |
| 3.2 | 0.1 | clcB | 1 | 19.8167 | 0.533 |
| 3.2 | 0.1 | ycaM | 1 | 19.8167 | 0.509 |
| 3.2 | 0.1 | yadI | 1 | 19.8167 | 0.767 |
| 3.2 | 0.1 | AG1 | 1 | 19.8167 | 0.813 |
| 3.2 | 0.1 | ptsI | 1 | 20.0667 | 0.743 |
| 3.2 | 0.1 | clcB | 1 | 20.0667 | 0.534 |
| 3.2 | 0.1 | ycaM | 1 | 20.0667 | 0.512 |
| 3.2 | 0.1 | yadI | 1 | 20.0667 | 0.773 |
| 3.2 | 0.1 | AG1 | 1 | 20.0667 | 0.821 |
| 3.2 | 0.1 | ptsI | 1 | 20.3167 | 0.756 |
| 3.2 | 0.1 | clcB | 1 | 20.3167 | 0.532 |
| 3.2 | 0.1 | ycaM | 1 | 20.3167 | 0.512 |
| 3.2 | 0.1 | yadI | 1 | 20.3167 | 0.785 |
| 3.2 | 0.1 | AG1 | 1 | 20.3167 | 0.834 |
| 3.2 | 0.1 | ptsI | 1 | 20.5667 | 0.738 |
| 3.2 | 0.1 | clcB | 1 | 20.5667 | 0.531 |
| 3.2 | 0.1 | ycaM | 1 | 20.5667 | 0.515 |
| 3.2 | 0.1 | yadI | 1 | 20.5667 | 0.77 |
| 3.2 | 0.1 | AG1 | 1 | 20.5667 | 0.823 |
| 3.2 | 0.1 | ptsI | 1 | 20.8167 | 0.755 |
| 3.2 | 0.1 | clcB | 1 | 20.8167 | 0.534 |
| 3.2 | 0.1 | ycaM | 1 | 20.8167 | 0.512 |
| 3.2 | 0.1 | yadI | 1 | 20.8167 | 0.785 |
| 3.2 | 0.1 | AG1 | 1 | 20.8167 | 0.825 |
| 3.2 | 0.1 | ptsI | 1 | 21.0667 | 0.772 |
| 3.2 | 0.1 | clcB | 1 | 21.0667 | 0.533 |
| 3.2 | 0.1 | ycaM | 1 | 21.0667 | 0.513 |
| 3.2 | 0.1 | yadI | 1 | 21.0667 | 0.797 |
| 3.2 | 0.1 | AG1 | 1 | 21.0667 | 0.842 |
| 3.2 | 0.1 | ptsI | 1 | 21.3167 | 0.783 |
| 3.2 | 0.1 | clcB | 1 | 21.3167 | 0.536 |
| 3.2 | 0.1 | ycaM | 1 | 21.3167 | 0.514 |
| 3.2 | 0.1 | yadI | 1 | 21.3167 | 0.799 |
| 3.2 | 0.1 | AG1 | 1 | 21.3167 | 0.843 |
| 3.2 | 0.1 | ptsI | 1 | 21.5667 | 0.776 |
| 3.2 | 0.1 | clcB | 1 | 21.5667 | 0.535 |
| 3.2 | 0.1 | ycaM | 1 | 21.5667 | 0.514 |
| 3.2 | 0.1 | yadI | 1 | 21.5667 | 0.792 |
| 3.2 | 0.1 | AG1 | 1 | 21.5667 | 0.845 |
| 3.2 | 0.1 | ptsI | 1 | 21.8167 | 0.792 |

| | | | | | |
|---|---|---|---|---|---|
| 3.2 | 0.1 | clcB | 1 | 21.8167 | 0.536 |
| 3.2 | 0.1 | ycaM | 1 | 21.8167 | 0.516 |
| 3.2 | 0.1 | yadI | 1 | 21.8167 | 0.814 |
| 3.2 | 0.1 | AG1 | 1 | 21.8167 | 0.849 |
| 3.2 | 0.1 | ptsI | 1 | 22.0667 | 0.797 |
| 3.2 | 0.1 | clcB | 1 | 22.0667 | 0.536 |
| 3.2 | 0.1 | ycaM | 1 | 22.0667 | 0.514 |
| 3.2 | 0.1 | yadI | 1 | 22.0667 | 0.82 |
| 3.2 | 0.1 | AG1 | 1 | 22.0667 | 0.863 |
| 3.2 | 0.1 | ptsI | 1 | 22.9333 | 0.898 |
| 3.2 | 0.1 | clcB | 1 | 22.9333 | 0.472 |
| 3.2 | 0.1 | ycaM | 1 | 22.9333 | 0.523 |
| 3.2 | 0.1 | yadI | 1 | 22.9333 | 0.83 |
| 3.2 | 0.1 | AG1 | 1 | 22.9333 | 0.985 |
| 3.2 | 0.1 | ptsI | 1 | 23.1833 | 0.848 |
| 3.2 | 0.1 | clcB | 1 | 23.1833 | 0.465 |
| 3.2 | 0.1 | ycaM | 1 | 23.1833 | 0.513 |
| 3.2 | 0.1 | yadI | 1 | 23.1833 | 0.816 |
| 3.2 | 0.1 | AG1 | 1 | 23.1833 | 0.946 |
| 3.2 | 0.1 | ptsI | 1 | 23.4333 | 0.81 |
| 3.2 | 0.1 | clcB | 1 | 23.4333 | 0.462 |
| 3.2 | 0.1 | ycaM | 1 | 23.4333 | 0.518 |
| 3.2 | 0.1 | yadI | 1 | 23.4333 | 0.826 |
| 3.2 | 0.1 | AG1 | 1 | 23.4333 | 0.932 |
| 3.2 | 0.1 | ptsI | 1 | 23.6833 | 0.822 |
| 3.2 | 0.1 | clcB | 1 | 23.6833 | 0.44 |
| 3.2 | 0.1 | ycaM | 1 | 23.6833 | 0.517 |
| 3.2 | 0.1 | yadI | 1 | 23.6833 | 0.842 |
| 3.2 | 0.1 | AG1 | 1 | 23.6833 | 0.94 |
| 3.2 | 0.1 | ptsI | 1 | 23.9333 | 0.824 |
| 3.2 | 0.1 | clcB | 1 | 23.9333 | 0.446 |
| 3.2 | 0.1 | ycaM | 1 | 23.9333 | 0.516 |
| 3.2 | 0.1 | yadI | 1 | 23.9333 | 0.854 |
| 3.2 | 0.1 | AG1 | 1 | 23.9333 | 0.943 |
| 3.2 | 0.1 | ptsI | 1 | 24.1833 | 0.821 |
| 3.2 | 0.1 | clcB | 1 | 24.1833 | 0.446 |
| 3.2 | 0.1 | ycaM | 1 | 24.1833 | 0.516 |
| 3.2 | 0.1 | yadI | 1 | 24.1833 | 0.862 |
| 3.2 | 0.1 | AG1 | 1 | 24.1833 | 0.946 |
| 3.2 | 0.1 | ptsI | 1 | 24.4333 | 0.825 |
| 3.2 | 0.1 | clcB | 1 | 24.4333 | 0.449 |
| 3.2 | 0.1 | ycaM | 1 | 24.4333 | 0.514 |
| 3.2 | 0.1 | yadI | 1 | 24.4333 | 0.859 |
| 3.2 | 0.1 | AG1 | 1 | 24.4333 | 0.946 |
| 3.2 | 0.1 | ptsI | 1 | 24.6833 | 0.826 |
| 3.2 | 0.1 | clcB | 1 | 24.6833 | 0.448 |
| 3.2 | 0.1 | ycaM | 1 | 24.6833 | 0.514 |
| 3.2 | 0.1 | yadI | 1 | 24.6833 | 0.858 |

| | | | | | |
|---|---|---|---|---|---|
| 3.2 | 0.1 | AG1 | 1 | 24.6833 | 0.946 |
| 3.2 | 0.1 | ptsI | 1 | 24.9333 | 0.828 |
| 3.2 | 0.1 | clcB | 1 | 24.9333 | 0.442 |
| 3.2 | 0.1 | ycaM | 1 | 24.9333 | 0.515 |
| 3.2 | 0.1 | yadI | 1 | 24.9333 | 0.861 |
| 3.2 | 0.1 | AG1 | 1 | 24.9333 | 0.945 |
| 3.2 | 0.1 | ptsI | 1 | 25.1833 | 0.838 |
| 3.2 | 0.1 | clcB | 1 | 25.1833 | 0.443 |
| 3.2 | 0.1 | ycaM | 1 | 25.1833 | 0.516 |
| 3.2 | 0.1 | yadI | 1 | 25.1833 | 0.865 |
| 3.2 | 0.1 | AG1 | 1 | 25.1833 | 0.952 |
| 3.2 | 0.1 | ptsI | 1 | 25.4333 | 0.849 |
| 3.2 | 0.1 | clcB | 1 | 25.4333 | 0.446 |
| 3.2 | 0.1 | ycaM | 1 | 25.4333 | 0.514 |
| 3.2 | 0.1 | yadI | 1 | 25.4333 | 0.862 |
| 3.2 | 0.1 | AG1 | 1 | 25.4333 | 0.954 |
| 3.2 | 0.1 | ptsI | 1 | 25.6833 | 0.847 |
| 3.2 | 0.1 | clcB | 1 | 25.6833 | 0.447 |
| 3.2 | 0.1 | ycaM | 1 | 25.6833 | 0.513 |
| 3.2 | 0.1 | yadI | 1 | 25.6833 | 0.864 |
| 3.2 | 0.1 | AG1 | 1 | 25.6833 | 0.958 |
| 3.2 | 0.1 | ptsI | 1 | 25.9333 | 0.85 |
| 3.2 | 0.1 | clcB | 1 | 25.9333 | 0.449 |
| 3.2 | 0.1 | ycaM | 1 | 25.9333 | 0.505 |
| 3.2 | 0.1 | yadI | 1 | 25.9333 | 0.854 |
| 3.2 | 0.1 | AG1 | 1 | 25.9333 | 0.956 |
| 3.2 | 0.1 | ptsI | 1 | 26.1833 | 0.855 |
| 3.2 | 0.1 | clcB | 1 | 26.1833 | 0.449 |
| 3.2 | 0.1 | ycaM | 1 | 26.1833 | 0.512 |
| 3.2 | 0.1 | yadI | 1 | 26.1833 | 0.864 |
| 3.2 | 0.1 | AG1 | 1 | 26.1833 | 0.967 |
| 3.2 | 0.1 | ptsI | 1 | 26.4333 | 0.856 |
| 3.2 | 0.1 | clcB | 1 | 26.4333 | 0.45 |
| 3.2 | 0.1 | ycaM | 1 | 26.4333 | 0.51 |
| 3.2 | 0.1 | yadI | 1 | 26.4333 | 0.861 |
| 3.2 | 0.1 | AG1 | 1 | 26.4333 | 0.962 |
| 3.2 | 0.1 | ptsI | 1 | 26.6833 | 0.858 |
| 3.2 | 0.1 | clcB | 1 | 26.6833 | 0.45 |
| 3.2 | 0.1 | ycaM | 1 | 26.6833 | 0.509 |
| 3.2 | 0.1 | yadI | 1 | 26.6833 | 0.862 |
| 3.2 | 0.1 | AG1 | 1 | 26.6833 | 0.96 |
| 3.2 | 0.1 | ptsI | 1 | 26.9333 | 0.853 |
| 3.2 | 0.1 | clcB | 1 | 26.9333 | 0.448 |
| 3.2 | 0.1 | ycaM | 1 | 26.9333 | 0.509 |
| 3.2 | 0.1 | yadI | 1 | 26.9333 | 0.86 |
| 3.2 | 0.1 | AG1 | 1 | 26.9333 | 0.961 |
| 3.2 | 0.1 | ptsI | 1 | 27.1833 | 0.862 |
| 3.2 | 0.1 | clcB | 1 | 27.1833 | 0.453 |

| | | | | | |
|---|---|---|---|---|---|
| 3.2 | 0.1 | ycaM | 1 | 27.1833 | 0.508 |
| 3.2 | 0.1 | yadI | 1 | 27.1833 | 0.863 |
| 3.2 | 0.1 | AG1 | 1 | 27.1833 | 0.956 |
| 3.2 | 0.1 | ptsI | 1 | 27.4333 | 0.868 |
| 3.2 | 0.1 | clcB | 1 | 27.4333 | 0.451 |
| 3.2 | 0.1 | ycaM | 1 | 27.4333 | 0.507 |
| 3.2 | 0.1 | yadI | 1 | 27.4333 | 0.867 |
| 3.2 | 0.1 | AG1 | 1 | 27.4333 | 0.954 |
| 3.2 | 0.1 | ptsI | 1 | 27.6833 | 0.868 |
| 3.2 | 0.1 | clcB | 1 | 27.6833 | 0.452 |
| 3.2 | 0.1 | ycaM | 1 | 27.6833 | 0.504 |
| 3.2 | 0.1 | yadI | 1 | 27.6833 | 0.877 |
| 3.2 | 0.1 | AG1 | 1 | 27.6833 | 0.958 |
| 3.2 | 0.1 | ptsI | 1 | 27.9333 | 0.864 |
| 3.2 | 0.1 | clcB | 1 | 27.9333 | 0.448 |
| 3.2 | 0.1 | ycaM | 1 | 27.9333 | 0.506 |
| 3.2 | 0.1 | yadI | 1 | 27.9333 | 0.88 |
| 3.2 | 0.1 | AG1 | 1 | 27.9333 | 0.959 |
| 3.2 | 0.1 | ptsI | 1 | 28.1833 | 0.875 |
| 3.2 | 0.1 | clcB | 1 | 28.1833 | 0.451 |
| 3.2 | 0.1 | ycaM | 1 | 28.1833 | 0.507 |
| 3.2 | 0.1 | yadI | 1 | 28.1833 | 0.881 |
| 3.2 | 0.1 | AG1 | 1 | 28.1833 | 0.963 |
| 3.2 | 0.1 | ptsI | 1 | 28.4333 | 0.88 |
| 3.2 | 0.1 | clcB | 1 | 28.4333 | 0.454 |
| 3.2 | 0.1 | ycaM | 1 | 28.4333 | 0.507 |
| 3.2 | 0.1 | yadI | 1 | 28.4333 | 0.886 |
| 3.2 | 0.1 | AG1 | 1 | 28.4333 | 0.964 |
| 3.2 | 0.1 | ptsI | 1 | 28.6833 | 0.885 |
| 3.2 | 0.1 | clcB | 1 | 28.6833 | 0.453 |
| 3.2 | 0.1 | ycaM | 1 | 28.6833 | 0.506 |
| 3.2 | 0.1 | yadI | 1 | 28.6833 | 0.893 |
| 3.2 | 0.1 | AG1 | 1 | 28.6833 | 0.968 |
| 3.2 | 0.1 | ptsI | 1 | 28.9333 | 0.884 |
| 3.2 | 0.1 | clcB | 1 | 28.9333 | 0.45 |
| 3.2 | 0.1 | ycaM | 1 | 28.9333 | 0.51 |
| 3.2 | 0.1 | yadI | 1 | 28.9333 | 0.891 |
| 3.2 | 0.1 | AG1 | 1 | 28.9333 | 0.965 |
| 3.2 | 0.1 | ptsI | 1 | 29.1833 | 0.883 |
| 3.2 | 0.1 | clcB | 1 | 29.1833 | 0.45 |
| 3.2 | 0.1 | ycaM | 1 | 29.1833 | 0.508 |
| 3.2 | 0.1 | yadI | 1 | 29.1833 | 0.898 |
| 3.2 | 0.1 | AG1 | 1 | 29.1833 | 0.969 |
| 3.2 | 0.1 | ptsI | 1 | 29.4333 | 0.891 |
| 3.2 | 0.1 | clcB | 1 | 29.4333 | 0.452 |
| 3.2 | 0.1 | ycaM | 1 | 29.4333 | 0.509 |
| 3.2 | 0.1 | yadI | 1 | 29.4333 | 0.904 |
| 3.2 | 0.1 | AG1 | 1 | 29.4333 | 0.974 |

| | | | | | |
|---|---|---|---|---|---|
| 3.2 | 0.1 | ptsI | 1 | 29.6833 | 0.892 |
| 3.2 | 0.1 | clcB | 1 | 29.6833 | 0.45 |
| 3.2 | 0.1 | ycaM | 1 | 29.6833 | 0.505 |
| 3.2 | 0.1 | yadI | 1 | 29.6833 | 0.907 |
| 3.2 | 0.1 | AG1 | 1 | 29.6833 | 0.976 |
| 3.2 | 0.1 | ptsI | 1 | 29.9333 | 0.891 |
| 3.2 | 0.1 | clcB | 1 | 29.9333 | 0.449 |
| 3.2 | 0.1 | ycaM | 1 | 29.9333 | 0.511 |
| 3.2 | 0.1 | yadI | 1 | 29.9333 | 0.906 |
| 3.2 | 0.1 | AG1 | 1 | 29.9333 | 0.977 |
| 3.2 | 0.1 | ptsI | 1 | 30.1833 | 0.897 |
| 3.2 | 0.1 | clcB | 1 | 30.1833 | 0.448 |
| 3.2 | 0.1 | ycaM | 1 | 30.1833 | 0.508 |
| 3.2 | 0.1 | yadI | 1 | 30.1833 | 0.909 |
| 3.2 | 0.1 | AG1 | 1 | 30.1833 | 0.981 |
| 3.2 | 0.1 | ptsI | 1 | 30.4333 | 0.897 |
| 3.2 | 0.1 | clcB | 1 | 30.4333 | 0.446 |
| 3.2 | 0.1 | ycaM | 1 | 30.4333 | 0.506 |
| 3.2 | 0.1 | yadI | 1 | 30.4333 | 0.914 |
| 3.2 | 0.1 | AG1 | 1 | 30.4333 | 0.982 |
| 3.2 | 0.1 | ptsI | 1 | 30.6833 | 0.902 |
| 3.2 | 0.1 | clcB | 1 | 30.6833 | 0.443 |
| 3.2 | 0.1 | ycaM | 1 | 30.6833 | 0.508 |
| 3.2 | 0.1 | yadI | 1 | 30.6833 | 0.909 |
| 3.2 | 0.1 | AG1 | 1 | 30.6833 | 0.99 |
| 3.2 | 0.1 | ptsI | 1 | 30.9333 | 0.896 |
| 3.2 | 0.1 | clcB | 1 | 30.9333 | 0.442 |
| 3.2 | 0.1 | ycaM | 1 | 30.9333 | 0.506 |
| 3.2 | 0.1 | yadI | 1 | 30.9333 | 0.915 |
| 3.2 | 0.1 | AG1 | 1 | 30.9333 | 0.997 |
| 3.2 | 0.1 | ptsI | 1 | 31.1833 | 0.896 |
| 3.2 | 0.1 | clcB | 1 | 31.1833 | 0.44 |
| 3.2 | 0.1 | ycaM | 1 | 31.1833 | 0.505 |
| 3.2 | 0.1 | yadI | 1 | 31.1833 | 0.912 |
| 3.2 | 0.1 | AG1 | 1 | 31.1833 | 1.002 |
| 3.2 | 0.1 | ptsI | 1 | 31.4333 | 0.89 |
| 3.2 | 0.1 | clcB | 1 | 31.4333 | 0.439 |
| 3.2 | 0.1 | ycaM | 1 | 31.4333 | 0.506 |
| 3.2 | 0.1 | yadI | 1 | 31.4333 | 0.921 |
| 3.2 | 0.1 | AG1 | 1 | 31.4333 | 1.004 |
| 3.2 | 0.1 | ptsI | 1 | 31.6833 | 0.891 |
| 3.2 | 0.1 | clcB | 1 | 31.6833 | 0.441 |
| 3.2 | 0.1 | ycaM | 1 | 31.6833 | 0.507 |
| 3.2 | 0.1 | yadI | 1 | 31.6833 | 0.925 |
| 3.2 | 0.1 | AG1 | 1 | 31.6833 | 1.012 |
| 3.2 | 0.1 | ptsI | 1 | 31.9333 | 0.888 |
| 3.2 | 0.1 | clcB | 1 | 31.9333 | 0.438 |
| 3.2 | 0.1 | ycaM | 1 | 31.9333 | 0.506 |

| | | | | | |
|---|---|---|---|---|---|
| 3.2 | 0.1 | yadI | 1 | 31.9333 | 0.929 |
| 3.2 | 0.1 | AG1 | 1 | 31.9333 | 1.02 |
| 3.2 | 0.1 | ptsI | 1 | 32.1833 | 0.891 |
| 3.2 | 0.1 | clcB | 1 | 32.1833 | 0.436 |
| 3.2 | 0.1 | ycaM | 1 | 32.1833 | 0.507 |
| 3.2 | 0.1 | yadI | 1 | 32.1833 | 0.936 |
| 3.2 | 0.1 | AG1 | 1 | 32.1833 | 1.023 |
| 3.2 | 0.1 | ptsI | 1 | 32.4333 | 0.892 |
| 3.2 | 0.1 | clcB | 1 | 32.4333 | 0.431 |
| 3.2 | 0.1 | ycaM | 1 | 32.4333 | 0.507 |
| 3.2 | 0.1 | yadI | 1 | 32.4333 | 0.931 |
| 3.2 | 0.1 | AG1 | 1 | 32.4333 | 1.034 |
| 3.2 | 0.1 | ptsI | 1 | 32.6833 | 0.89 |
| 3.2 | 0.1 | clcB | 1 | 32.6833 | 0.439 |
| 3.2 | 0.1 | ycaM | 1 | 32.6833 | 0.507 |
| 3.2 | 0.1 | yadI | 1 | 32.6833 | 0.948 |
| 3.2 | 0.1 | AG1 | 1 | 32.6833 | 1.032 |
| 3.2 | 0.1 | ptsI | 1 | 32.9333 | 0.895 |
| 3.2 | 0.1 | clcB | 1 | 32.9333 | 0.435 |
| 3.2 | 0.1 | ycaM | 1 | 32.9333 | 0.509 |
| 3.2 | 0.1 | yadI | 1 | 32.9333 | 0.947 |
| 3.2 | 0.1 | AG1 | 1 | 32.9333 | 1.035 |
| 3.2 | 0.1 | ptsI | 1 | 33.1833 | 0.892 |
| 3.2 | 0.1 | clcB | 1 | 33.1833 | 0.431 |
| 3.2 | 0.1 | ycaM | 1 | 33.1833 | 0.508 |
| 3.2 | 0.1 | yadI | 1 | 33.1833 | 0.953 |
| 3.2 | 0.1 | AG1 | 1 | 33.1833 | 1.04 |
| 3.2 | 0.1 | ptsI | 1 | 33.4333 | 0.89 |
| 3.2 | 0.1 | clcB | 1 | 33.4333 | 0.434 |
| 3.2 | 0.1 | ycaM | 1 | 33.4333 | 0.508 |
| 3.2 | 0.1 | yadI | 1 | 33.4333 | 0.964 |
| 3.2 | 0.1 | AG1 | 1 | 33.4333 | 1.048 |
| 3.2 | 0.1 | ptsI | 1 | 33.6833 | 0.89 |
| 3.2 | 0.1 | clcB | 1 | 33.6833 | 0.432 |
| 3.2 | 0.1 | ycaM | 1 | 33.6833 | 0.506 |
| 3.2 | 0.1 | yadI | 1 | 33.6833 | 0.978 |
| 3.2 | 0.1 | AG1 | 1 | 33.6833 | 1.055 |
| 3.2 | 0.1 | ptsI | 1 | 33.9333 | 0.89 |
| 3.2 | 0.1 | clcB | 1 | 33.9333 | 0.44 |
| 3.2 | 0.1 | ycaM | 1 | 33.9333 | 0.507 |
| 3.2 | 0.1 | yadI | 1 | 33.9333 | 0.99 |
| 3.2 | 0.1 | AG1 | 1 | 33.9333 | 1.06 |
| 3.2 | 0.1 | ptsI | 1 | 34.1833 | 0.889 |
| 3.2 | 0.1 | clcB | 1 | 34.1833 | 0.437 |
| 3.2 | 0.1 | ycaM | 1 | 34.1833 | 0.512 |
| 3.2 | 0.1 | yadI | 1 | 34.1833 | 1.008 |
| 3.2 | 0.1 | AG1 | 1 | 34.1833 | 1.072 |
| 3.2 | 0.1 | ptsI | 1 | 34.4333 | 0.887 |

| | | | | | |
|---|---|---|---|---|---|
| 3.2 | 0.1 | clcB | 1 | 34.4333 | 0.436 |
| 3.2 | 0.1 | ycaM | 1 | 34.4333 | 0.509 |
| 3.2 | 0.1 | yadI | 1 | 34.4333 | 1.015 |
| 3.2 | 0.1 | AG1 | 1 | 34.4333 | 1.079 |
| 3.2 | 0.1 | ptsI | 1 | 34.6833 | 0.886 |
| 3.2 | 0.1 | clcB | 1 | 34.6833 | 0.434 |
| 3.2 | 0.1 | ycaM | 1 | 34.6833 | 0.512 |
| 3.2 | 0.1 | yadI | 1 | 34.6833 | 1.023 |
| 3.2 | 0.1 | AG1 | 1 | 34.6833 | 1.092 |
| 3.2 | 0.1 | ptsI | 1 | 34.9333 | 0.887 |
| 3.2 | 0.1 | clcB | 1 | 34.9333 | 0.436 |
| 3.2 | 0.1 | ycaM | 1 | 34.9333 | 0.506 |
| 3.2 | 0.1 | yadI | 1 | 34.9333 | 1.039 |
| 3.2 | 0.1 | AG1 | 1 | 34.9333 | 1.099 |
| 3.2 | 0.1 | ptsI | 1 | 35.1833 | 0.891 |
| 3.2 | 0.1 | clcB | 1 | 35.1833 | 0.438 |
| 3.2 | 0.1 | ycaM | 1 | 35.1833 | 0.508 |
| 3.2 | 0.1 | yadI | 1 | 35.1833 | 1.048 |
| 3.2 | 0.1 | AG1 | 1 | 35.1833 | 1.109 |
| 3.2 | 0.1 | ptsI | 1 | 35.4333 | 0.894 |
| 3.2 | 0.1 | clcB | 1 | 35.4333 | 0.438 |
| 3.2 | 0.1 | ycaM | 1 | 35.4333 | 0.509 |
| 3.2 | 0.1 | yadI | 1 | 35.4333 | 1.058 |
| 3.2 | 0.1 | AG1 | 1 | 35.4333 | 1.118 |
| 3.2 | 0.1 | ptsI | 1 | 35.6833 | 0.889 |
| 3.2 | 0.1 | clcB | 1 | 35.6833 | 0.438 |
| 3.2 | 0.1 | ycaM | 1 | 35.6833 | 0.511 |
| 3.2 | 0.1 | yadI | 1 | 35.6833 | 1.069 |
| 3.2 | 0.1 | AG1 | 1 | 35.6833 | 1.127 |
| 3.2 | 0.1 | ptsI | 1 | 35.9333 | 0.892 |
| 3.2 | 0.1 | clcB | 1 | 35.9333 | 0.438 |
| 3.2 | 0.1 | ycaM | 1 | 35.9333 | 0.511 |
| 3.2 | 0.1 | yadI | 1 | 35.9333 | 1.078 |
| 3.2 | 0.1 | AG1 | 1 | 35.9333 | 1.134 |
| 3.2 | 0.1 | ptsI | 1 | 36.1833 | 0.888 |
| 3.2 | 0.1 | clcB | 1 | 36.1833 | 0.441 |
| 3.2 | 0.1 | ycaM | 1 | 36.1833 | 0.519 |
| 3.2 | 0.1 | yadI | 1 | 36.1833 | 1.083 |
| 3.2 | 0.1 | AG1 | 1 | 36.1833 | 1.145 |
| 3.2 | 0.1 | ptsI | 1 | 36.4333 | 0.887 |
| 3.2 | 0.1 | clcB | 1 | 36.4333 | 0.442 |
| 3.2 | 0.1 | ycaM | 1 | 36.4333 | 0.517 |
| 3.2 | 0.1 | yadI | 1 | 36.4333 | 1.095 |
| 3.2 | 0.1 | AG1 | 1 | 36.4333 | 1.153 |
| 3.2 | 0.1 | ptsI | 1 | 36.6833 | 0.89 |
| 3.2 | 0.1 | clcB | 1 | 36.6833 | 0.446 |
| 3.2 | 0.1 | ycaM | 1 | 36.6833 | 0.512 |
| 3.2 | 0.1 | yadI | 1 | 36.6833 | 1.101 |

| | | | | | |
|---|---|---|---|---|---|
| 3.2 | 0.1 | AG1 | 1 | 36.6833 | 1.159 |
