## Supplementary material for "Membrane proteins ClcB, PtsI and YcaM mediate the bactericidal effects of colistin in *Escherichia coli*": Supp_table_1

| Strain | Growth_LB | Growth_LB | Growth_LB | Growth_LB | Growth_LB_colistin_5ug-ml |
|---|---|---|---|---|---|
| aroP | 1.678 | 0.861 | 1.695 | 0.358 | 1.905 |
| clcB | 1.485 | 1.259 | 1.768 | 1.716 | 1.757 |
| yadI | 1.959 | 0.975 | 1.718 | 1.808 | 1.625 |
| ycaM | 1.459 | 0.88 | 0.971 | 0.941 | 1.218 |
| rhtC | 2.126 | 1.151 | 1.439 | 0.695 | 1.144 |
| focB | 1.426 | 0.984 | 1.134 | 1.05 | 1.074 |
| ptsI | 1.96 | 1.13 | 1.69 | 1.527 | 0.897 |
| ycaD | 1.912 | 0.955 | 0.62 | 0.82 | 0.782 |
| yraQ | 1.658 | 0.243 | 0.347 | 0.504 | 0.727 |
| crr | 2.022 | 0.971 | 1.544 | 0.296 | 0.428 |
| phnK | 1.574 | 0.948 | 0.203 | 0.35 | 0.424 |
| sapC | 1.555 | 0.936 | 0.242 | 0.338 | 0.421 |
| oppB | 1.607 | 0.655 | 0.302 | 0.368 | 0.402 |
| ybhG | 1.44 | 0.797 | 1.593 | 0.36 | 0.396 |
| ulaB | 1.842 | 1 | 1.58 | 0.3 | 0.383 |
| artI | 1.94 | 0.76 | 0.934 | 0.297 | 0.359 |
| yidE | 1.767 | 0.814 | 0.256 | 0.278 | 0.356 |
| btuF | 2.081 | 0.557 | 0.263 | 0.282 | 0.336 |
| ydcT | 1.839 | 0.958 | 1.764 | 0.288 | 0.326 |
| gadC | 1.782 | 0.821 | 1.49 | 0.729 | 0.324 |
| pitA | 2.222 | 0.358 | 0.22 | 0.296 | 0.322 |
| ybhS | 2.036 | 0.334 | 0.255 | 0.274 | 0.32 |
| oppD | 1.834 | 0.918 | 1.466 | 0.279 | 0.311 |
| yicE | 0.761 | 0.948 | 0.291 | 0.27 | 0.311 |
| mgtA | 1.928 | 0.999 | 0.273 | 0.289 | 0.308 |
| putP | 1.764 | 0.878 | 1.258 | 0.301 | 0.306 |
| marA | 0.989 | 0.964 | 0.249 | 0.302 | 0.301 |
| uhpC | 2.016 | 0.968 | 0.261 | 0.288 | 0.297 |
| sapB | 1.793 | 0.895 | 0.253 | 0.296 | 0.294 |
| atpF | 1.514 | 0.994 | 0.275 | 0.291 | 0.288 |
| betT | 1.932 | 0.892 | 0.28 | 0.282 | 0.288 |
| yieG | 2.353 | 0.46 | 0.239 | 0.329 | 0.287 |
| ynjB | 1.75 | 0.838 | 0.223 | 0.276 | 0.287 |
| chaA | 1.68 | 0.812 | 1.457 | 0.312 | 0.285 |
| glvC | 2.29 | 1.11 | 0.24 | 0.258 | 0.283 |
| mtlA | 2.088 | 0.86 | 0.228 | 0.288 | 0.279 |
| trkD | 2.201 | 0.861 | 1.355 | 0.276 | 0.276 |
| narK | 1.921 | 1.069 | 0.224 | 0.253 | 0.274 |
| frwB | 1.64 | 0.974 | 0.223 | 0.382 | 0.258 |
| proP | 2.046 | 0.664 | 1.584 | 0.247 | 0.257 |
| frvB | 2.396 | 0.501 | 0.224 | 0.282 | 0.256 |
| fieF | 1.912 | 1.04 | 1.542 | 0.479 | 0.249 |
| yigM | 2.302 | 0.932 | 1.615 | 0.306 | 0.239 |
| hsrA | 2.055 | 0.749 | 0 | 0 | 0 |
| WT | 0.044 | 0.517 | 0 | 0 | 0 |
| garP | 2.126 | 0.435 | 0 | 0 | 0 |
| rbsB | 0.1 | 0.261 | 0 | 0 | 0 |

| | | | | | |
|---|---|---|---|---|---|
| yfhD | 1.432 | 0.232 | 0 | 0 | 0 |
| tolQ | 1.37 | 0.198 | 0 | 0 | 0 |
| yijE | 1.696 | 0.195 | 0 | 0 | 0 |
| wzxE | 1.826 | 0.194 | 0 | 0 | 0 |
| caiT | 1.806 | 0.192 | 0 | 0 | 0 |
| ybhF | 2.245 | 0.18 | 0 | 0 | 0 |
| ugpC | 2.133 | 0.178 | 0 | 0 | 0 |
| atoS | 0.076 | 0.175 | 0 | 0 | 0 |
| guaB | 1.327 | 0.172 | 0 | 0 | 0 |
| sbp | 1.975 | 0.168 | 0 | 0 | 0 |
| dppC | 1.856 | 0.162 | 0 | 0 | 0 |
| ydhK | 1.97 | 0.154 | 0 | 0 | 0 |
| malX | 1.994 | 0.142 | 0 | 0 | 0 |
| agaV | 1.903 | 0.141 | 0 | 0 | 0 |
| ybgH | 1.942 | 0.135 | 0 | 0 | 0 |
| yicL | 2.03 | 0.134 | 0 | 0 | 0 |
| ycjP | 2.213 | 0.13 | 0 | 0 | 0 |
| barA | 2.233 | 0.126 | 0 | 0 | 0 |
| yicG | 1.75 | 0.126 | 0 | 0 | 0 |
| dppD | 1.614 | 0.124 | 0 | 0 | 0 |
| yjiO | 2.124 | 0.123 | 0 | 0 | 0 |
| gntT | 0.13 | 0.123 | 0 | 0 | 0 |
| WT | 0.08 | 0.122 | 0 | 0 | 0 |
| oppA | 2.11 | 0.12 | 0 | 0 | 0 |
| ynfA | 1.857 | 0.119 | 0 | 0 | 0 |
| alsC | 1.735 | 0.118 | 0 | 0 | 0 |
| gntP | 0.102 | 0.118 | 0 | 0 | 0 |
| ulaA | 2.216 | 0.117 | 0 | 0 | 0 |
| rarD | 1.804 | 0.115 | 0 | 0 | 0 |
| rbbA | 1.775 | 0.115 | 0 | 0 | 0 |
| mhpT | 1.645 | 0.115 | 0 | 0 | 0 |
| yeiJ | 2.119 | 0.113 | 0 | 0 | 0 |
| emrD | 1.991 | 0.113 | 0 | 0 | 0 |
| emrB | 1.56 | 0.113 | 0 | 0 | 0 |
| bglF | 1.847 | 0.111 | 0 | 0 | 0 |
| cmtB | 1.624 | 0.111 | 0 | 0 | 0 |
| potH | 0.052 | 0.111 | 0 | 0 | 0 |
| ybaL | 2 | 0.11 | 0 | 0 | 0 |
| feoB | 2.226 | 0.109 | 0 | 0 | 0 |
| ydjK | 2.055 | 0.108 | 0 | 0 | 0 |
| mdtI | 1.97 | 0.108 | 0 | 0 | 0 |
| yejE | 1.95 | 0.108 | 0 | 0 | 0 |
| pheP | 1.726 | 0.108 | 0 | 0 | 0 |
| hisP | 1.698 | 0.108 | 0 | 0 | 0 |
| yedA | 1.648 | 0.108 | 0 | 0 | 0 |
| manX | 1.611 | 0.108 | 0 | 0 | 0 |
| yehZ | 1.979 | 0.107 | 0 | 0 | 0 |
| yqcE | 1.789 | 0.107 | 0 | 0 | 0 |

| | | | | | |
|---|---|---|---|---|---|
| ydeE | 1.738 | 0.107 | 0 | 0 | 0 |
| yddA | 1.692 | 0.107 | 0 | 0 | 0 |
| ybbM | 1.427 | 0.107 | 0 | 0 | 0 |
| yhdW | 1.9 | 0.106 | 0 | 0 | 0 |
| yhaO | 1.785 | 0.106 | 0 | 0 | 0 |
| ydhP | 1.527 | 0.106 | 0 | 0 | 0 |
| yfiK | 1.918 | 0.105 | 0 | 0 | 0 |
| cusA | 1.789 | 0.105 | 0 | 0 | 0 |
| yqeG | 1.677 | 0.105 | 0 | 0 | 0 |
| fsr | 2.232 | 0.104 | 0 | 0 | 0 |
| yphF | 1.994 | 0.104 | 0 | 0 | 0 |
| gntU | 1.945 | 0.104 | 0 | 0 | 0 |
| atoE | 1.764 | 0.103 | 0 | 0 | 0 |
| tsgA | 1.646 | 0.103 | 0 | 0 | 0 |
| ytfQ | 1.609 | 0.103 | 0 | 0 | 0 |
| ynjD | 1.591 | 0.103 | 0 | 0 | 0 |
| yebQ | 1.54 | 0.103 | 0 | 0 | 0 |
| ygcS | 2.077 | 0.102 | 0 | 0 | 0 |
| rhtB | 2.023 | 0.102 | 0 | 0 | 0 |
| pstA | 1.901 | 0.102 | 0 | 0 | 0 |
| uhpT | 1.886 | 0.102 | 0 | 0 | 0 |
| oppC | 1.8 | 0.102 | 0 | 0 | 0 |
| agaC | 1.778 | 0.102 | 0 | 0 | 0 |
| ycjO | 1.583 | 0.102 | 0 | 0 | 0 |
| livK | 1.488 | 0.102 | 0 | 0 | 0 |
| ygiS | 0.111 | 0.102 | 0 | 0 | 0 |
| yjcQ | 1.919 | 0.101 | 0 | 0 | 0 |
| dppF | 1.772 | 0.101 | 0 | 0 | 0 |
| manZ | 1.509 | 0.101 | 0 | 0 | 0 |
| hisM | 1.416 | 0.101 | 0 | 0 | 0 |
| ybbP | 1.977 | 0.1 | 0 | 0 | 0 |
| sufC | 1.905 | 0.1 | 0 | 0 | 0 |
| araE | 1.899 | 0.1 | 0 | 0 | 0 |
| acrE | 1.788 | 0.1 | 0 | 0 | 0 |
| alx | 1.756 | 0.1 | 0 | 0 | 0 |
| sufD | 1.656 | 0.1 | 0 | 0 | 0 |
| yrbE | 1.552 | 0.1 | 0 | 0 | 0 |
| ydgG | 1.489 | 0.1 | 0 | 0 | 0 |
| yhdX | 1.453 | 0.1 | 0 | 0 | 0 |
| yjjK | 1.416 | 0.1 | 0 | 0 | 0 |
| mdtD | 1.988 | 0.099 | 0 | 0 | 0 |
| ymjB | 1.897 | 0.099 | 0 | 0 | 0 |
| uraA | 1.868 | 0.099 | 0 | 0 | 0 |
| actP | 1.531 | 0.099 | 0 | 0 | 0 |
| yfdC | 1.41 | 0.099 | 0 | 0 | 0 |
| ygdQ | 0.042 | 0.099 | 0 | 0 | 0 |
| mdtA | 2.015 | 0.098 | 0 | 0 | 0 |
| yhjE | 1.895 | 0.098 | 0 | 0 | 0 |

| | | | | | |
|---|---|---|---|---|---|
| ydjE | 1.847 | 0.098 | 0 | 0 | 0 |
| yihN | 1.809 | 0.098 | 0 | 0 | 0 |
| chbA | 1.793 | 0.098 | 0 | 0 | 0 |
| frwD | 1.782 | 0.098 | 0 | 0 | 0 |
| nikA | 1.745 | 0.098 | 0 | 0 | 0 |
| agaB | 1.639 | 0.098 | 0 | 0 | 0 |
| yaaJ | 1.621 | 0.098 | 0 | 0 | 0 |
| ytfF | 1.586 | 0.098 | 0 | 0 | 0 |
| ybbY | 1.32 | 0.098 | 0 | 0 | 0 |
| argT | 2.102 | 0.097 | 0 | 0 | 0 |
| kefC | 2.024 | 0.097 | 0 | 0 | 0 |
| yhiP | 1.794 | 0.097 | 0 | 0 | 0 |
| sugE | 1.765 | 0.097 | 0 | 0 | 0 |
| dsdX | 1.762 | 0.097 | 0 | 0 | 0 |
| malG | 1.719 | 0.097 | 0 | 0 | 0 |
| metI | 1.594 | 0.097 | 0 | 0 | 0 |
| manY | 1.584 | 0.097 | 0 | 0 | 0 |
| tdcC | 2.105 | 0.096 | 0 | 0 | 0 |
| panF | 2.033 | 0.096 | 0 | 0 | 0 |
| yeeO | 1.95 | 0.096 | 0 | 0 | 0 |
| ptsA | 1.81 | 0.096 | 0 | 0 | 0 |
| fepB | 1.79 | 0.096 | 0 | 0 | 0 |
| yebA | 1.786 | 0.096 | 0 | 0 | 0 |
| kefB | 1.761 | 0.096 | 0 | 0 | 0 |
| livF | 1.734 | 0.096 | 0 | 0 | 0 |
| treB | 1.69 | 0.096 | 0 | 0 | 0 |
| yfeH | 1.683 | 0.096 | 0 | 0 | 0 |
| rbsC | 1.659 | 0.096 | 0 | 0 | 0 |
| yiaO | 1.561 | 0.096 | 0 | 0 | 0 |
| yhdZ | 1.539 | 0.096 | 0 | 0 | 0 |
| ygfO | 1.522 | 0.096 | 0 | 0 | 0 |
| gudP | 1.289 | 0.096 | 0 | 0 | 0 |
| ygjI | 1.278 | 0.096 | 0 | 0 | 0 |
| yicO | 2.054 | 0.095 | 0 | 0 | 0 |
| yehX | 2.029 | 0.095 | 0 | 0 | 0 |
| yeeF | 1.943 | 0.095 | 0 | 0 | 0 |
| yajR | 1.908 | 0.095 | 0 | 0 | 0 |
| malE | 1.869 | 0.095 | 0 | 0 | 0 |
| xylH | 1.847 | 0.095 | 0 | 0 | 0 |
| sapA | 1.844 | 0.095 | 0 | 0 | 0 |
| yjeH | 1.697 | 0.095 | 0 | 0 | 0 |
| tehA | 1.558 | 0.095 | 0 | 0 | 0 |
| lysP | 1.455 | 0.095 | 0 | 0 | 0 |
| gltS | 1.373 | 0.095 | 0 | 0 | 0 |
| ybbA | 1.311 | 0.095 | 0 | 0 | 0 |
| eamA | 2.33 | 0.094 | 0 | 0 | 0 |
| ybjL | 2.088 | 0.094 | 0 | 0 | 0 |
| hisQ | 1.97 | 0.094 | 0 | 0 | 0 |

| | | | | | |
|---|---|---|---|---|---|
| ygbN | 1.968 | 0.094 | 0 | 0 | 0 |
| ytfT | 1.954 | 0.094 | 0 | 0 | 0 |
| narU | 1.897 | 0.094 | 0 | 0 | 0 |
| mdlA | 1.89 | 0.094 | 0 | 0 | 0 |
| cycA | 1.756 | 0.094 | 0 | 0 | 0 |
| yfgO | 1.731 | 0.094 | 0 | 0 | 0 |
| cmtA | 1.726 | 0.094 | 0 | 0 | 0 |
| pstC | 1.724 | 0.094 | 0 | 0 | 0 |
| srlA | 1.709 | 0.094 | 0 | 0 | 0 |
| modA | 1.704 | 0.094 | 0 | 0 | 0 |
| ydgR | 1.649 | 0.094 | 0 | 0 | 0 |
| gltL | 1.641 | 0.094 | 0 | 0 | 0 |
| ddpB | 1.637 | 0.094 | 0 | 0 | 0 |
| thiP | 1.6 | 0.094 | 0 | 0 | 0 |
| setC | 1.553 | 0.094 | 0 | 0 | 0 |
| mdtL | 1.551 | 0.094 | 0 | 0 | 0 |
| ddpF | 1.519 | 0.094 | 0 | 0 | 0 |
| ynjC | 1.486 | 0.094 | 0 | 0 | 0 |
| cadB | 1.431 | 0.094 | 0 | 0 | 0 |
| gltJ | 1.342 | 0.094 | 0 | 0 | 0 |
| atpD | 0.059 | 0.094 | 0 | 0 | 0 |
| potI | 0.042 | 0.094 | 0 | 0 | 0 |
| ygfU | 2.192 | 0.093 | 0 | 0 | 0 |
| ydiK | 2.161 | 0.093 | 0 | 0 | 0 |
| yhdY | 2.116 | 0.093 | 0 | 0 | 0 |
| ydcS | 1.971 | 0.093 | 0 | 0 | 0 |
| yhhS | 1.962 | 0.093 | 0 | 0 | 0 |
| yfbJ | 1.891 | 0.093 | 0 | 0 | 0 |
| ddpA | 1.871 | 0.093 | 0 | 0 | 0 |
| yhhT | 1.865 | 0.093 | 0 | 0 | 0 |
| trkH | 1.797 | 0.093 | 0 | 0 | 0 |
| ynfM | 1.787 | 0.093 | 0 | 0 | 0 |
| mscS | 1.772 | 0.093 | 0 | 0 | 0 |
| dcuA | 1.759 | 0.093 | 0 | 0 | 0 |
| glpF | 1.581 | 0.093 | 0 | 0 | 0 |
| ydcO | 1.569 | 0.093 | 0 | 0 | 0 |
| yeaN | 1.561 | 0.093 | 0 | 0 | 0 |
| dctA | 1.546 | 0.093 | 0 | 0 | 0 |
| ypdG | 1.49 | 0.093 | 0 | 0 | 0 |
| acrA | 1.293 | 0.093 | 0 | 0 | 0 |
| kdpD | 1.208 | 0.093 | 0 | 0 | 0 |
| yjbB | 2.097 | 0.092 | 0 | 0 | 0 |
| kdgT | 2.072 | 0.092 | 0 | 0 | 0 |
| focA | 1.973 | 0.092 | 0 | 0 | 0 |
| ydeA | 1.961 | 0.092 | 0 | 0 | 0 |
| potA | 1.961 | 0.092 | 0 | 0 | 0 |
| yiaN | 1.944 | 0.092 | 0 | 0 | 0 |
| artM | 1.94 | 0.092 | 0 | 0 | 0 |

| | | | | | |
|---|---|---|---|---|---|
| yjfF | 1.921 | 0.092 | 0 | 0 | 0 |
| mngA | 1.892 | 0.092 | 0 | 0 | 0 |
| tbpA | 1.89 | 0.092 | 0 | 0 | 0 |
| setA | 1.885 | 0.092 | 0 | 0 | 0 |
| ycjN | 1.864 | 0.092 | 0 | 0 | 0 |
| ydiN | 1.831 | 0.092 | 0 | 0 | 0 |
| exbD | 1.814 | 0.092 | 0 | 0 | 0 |
| macB | 1.782 | 0.092 | 0 | 0 | 0 |
| ychM | 1.774 | 0.092 | 0 | 0 | 0 |
| kch | 1.752 | 0.092 | 0 | 0 | 0 |
| sdaC | 1.728 | 0.092 | 0 | 0 | 0 |
| yehY | 1.718 | 0.092 | 0 | 0 | 0 |
| codB | 1.681 | 0.092 | 0 | 0 | 0 |
| ybhN | 1.662 | 0.092 | 0 | 0 | 0 |
| shiA | 1.656 | 0.092 | 0 | 0 | 0 |
| murP | 1.628 | 0.092 | 0 | 0 | 0 |
| dcuC | 1.618 | 0.092 | 0 | 0 | 0 |
| yliC | 1.597 | 0.092 | 0 | 0 | 0 |
| araF | 1.472 | 0.092 | 0 | 0 | 0 |
| glnP | 1.472 | 0.092 | 0 | 0 | 0 |
| cysU | 1.426 | 0.092 | 0 | 0 | 0 |
| yjeP | 1.369 | 0.092 | 0 | 0 | 0 |
| mdlB | 2.218 | 0.091 | 0 | 0 | 0 |
| ytfL | 2.157 | 0.091 | 0 | 0 | 0 |
| yjcD | 2.076 | 0.091 | 0 | 0 | 0 |
| ydgI | 2.05 | 0.091 | 0 | 0 | 0 |
| nupC | 1.973 | 0.091 | 0 | 0 | 0 |
| nlpA | 1.927 | 0.091 | 0 | 0 | 0 |
| ybiR | 1.881 | 0.091 | 0 | 0 | 0 |
| yidK | 1.861 | 0.091 | 0 | 0 | 0 |
| livM | 1.789 | 0.091 | 0 | 0 | 0 |
| ydcV | 1.786 | 0.091 | 0 | 0 | 0 |
| ydcZ | 1.75 | 0.091 | 0 | 0 | 0 |
| lsrB | 1.746 | 0.091 | 0 | 0 | 0 |
| potD | 1.696 | 0.091 | 0 | 0 | 0 |
| xylF | 1.684 | 0.091 | 0 | 0 | 0 |
| ego | 1.661 | 0.091 | 0 | 0 | 0 |
| ygiE | 1.636 | 0.091 | 0 | 0 | 0 |
| nirC | 1.627 | 0.091 | 0 | 0 | 0 |
| ddpC | 1.626 | 0.091 | 0 | 0 | 0 |
| yoaE | 1.592 | 0.091 | 0 | 0 | 0 |
| artJ | 1.591 | 0.091 | 0 | 0 | 0 |
| ydhJ | 1.587 | 0.091 | 0 | 0 | 0 |
| yhjX | 1.585 | 0.091 | 0 | 0 | 0 |
| gltP | 1.55 | 0.091 | 0 | 0 | 0 |
| chbB | 1.513 | 0.091 | 0 | 0 | 0 |
| citT | 1.508 | 0.091 | 0 | 0 | 0 |
| modC | 1.409 | 0.091 | 0 | 0 | 0 |

| | | | | | |
|---|---|---|---|---|---|
| atpI | 2.25 | 0.09 | 0 | 0 | 0 |
| exbB | 2.214 | 0.09 | 0 | 0 | 0 |
| gltK | 2.093 | 0.09 | 0 | 0 | 0 |
| glnH | 2.092 | 0.09 | 0 | 0 | 0 |
| yheS | 2.074 | 0.09 | 0 | 0 | 0 |
| metN | 2.067 | 0.09 | 0 | 0 | 0 |
| lldP | 2.046 | 0.09 | 0 | 0 | 0 |
| ssuC | 2.022 | 0.09 | 0 | 0 | 0 |
| galP | 2.018 | 0.09 | 0 | 0 | 0 |
| fruB | 2.006 | 0.09 | 0 | 0 | 0 |
| ybhR | 1.93 | 0.09 | 0 | 0 | 0 |
| yjdL | 1.922 | 0.09 | 0 | 0 | 0 |
| yhfK | 1.916 | 0.09 | 0 | 0 | 0 |
| tatB | 1.892 | 0.09 | 0 | 0 | 0 |
| dcuD | 1.888 | 0.09 | 0 | 0 | 0 |
| yaaU | 1.887 | 0.09 | 0 | 0 | 0 |
| chbC | 1.885 | 0.09 | 0 | 0 | 0 |
| yeaV | 1.852 | 0.09 | 0 | 0 | 0 |
| ybhI | 1.85 | 0.09 | 0 | 0 | 0 |
| kdpC | 1.848 | 0.09 | 0 | 0 | 0 |
| mglC | 1.844 | 0.09 | 0 | 0 | 0 |
| yjcE | 1.829 | 0.09 | 0 | 0 | 0 |
| yojI | 1.819 | 0.09 | 0 | 0 | 0 |
| yegT | 1.768 | 0.09 | 0 | 0 | 0 |
| dppB | 1.722 | 0.09 | 0 | 0 | 0 |
| araH | 1.719 | 0.09 | 0 | 0 | 0 |
| dppA | 1.705 | 0.09 | 0 | 0 | 0 |
| yadS | 1.694 | 0.09 | 0 | 0 | 0 |
| yliA | 1.687 | 0.09 | 0 | 0 | 0 |
| potE | 1.665 | 0.09 | 0 | 0 | 0 |
| potF | 1.649 | 0.09 | 0 | 0 | 0 |
| mtr | 1.633 | 0.09 | 0 | 0 | 0 |
| ydcU | 1.616 | 0.09 | 0 | 0 | 0 |
| aqpZ | 1.612 | 0.09 | 0 | 0 | 0 |
| yegH | 1.588 | 0.09 | 0 | 0 | 0 |
| yfcJ | 1.575 | 0.09 | 0 | 0 | 0 |
| ptsP | 1.573 | 0.09 | 0 | 0 | 0 |
| ugpA | 1.569 | 0.09 | 0 | 0 | 0 |
| cysW | 1.553 | 0.09 | 0 | 0 | 0 |
| xapB | 1.522 | 0.09 | 0 | 0 | 0 |
| puuP | 1.486 | 0.09 | 0 | 0 | 0 |
| ptsN | 1.447 | 0.09 | 0 | 0 | 0 |
| hisJ | 1.435 | 0.09 | 0 | 0 | 0 |
| rbsA | 1.338 | 0.09 | 0 | 0 | 0 |
| artQ | 1.334 | 0.09 | 0 | 0 | 0 |
| yohN | 1.161 | 0.09 | 0 | 0 | 0 |
| yhbE | 0.049 | 0.09 | 0 | 0 | 0 |
| yhhJ | 2.115 | 0.089 | 0 | 0 | 0 |

| | | | | | |
|---|---|---|---|---|---|
| ccmA | 1.954 | 0.089 | 0 | 0 | 0 |
| phnC | 1.92 | 0.089 | 0 | 0 | 0 |
| yejB | 1.906 | 0.089 | 0 | 0 | 0 |
| mdtK | 1.892 | 0.089 | 0 | 0 | 0 |
| aaeB | 1.889 | 0.089 | 0 | 0 | 0 |
| btuC | 1.821 | 0.089 | 0 | 0 | 0 |
| ybiO | 1.797 | 0.089 | 0 | 0 | 0 |
| srlB | 1.762 | 0.089 | 0 | 0 | 0 |
| yphD | 1.725 | 0.089 | 0 | 0 | 0 |
| yneE | 1.72 | 0.089 | 0 | 0 | 0 |
| yrbF | 1.713 | 0.089 | 0 | 0 | 0 |
| livG | 1.683 | 0.089 | 0 | 0 | 0 |
| fruA | 1.674 | 0.089 | 0 | 0 | 0 |
| nupG | 1.668 | 0.089 | 0 | 0 | 0 |
| agaD | 1.655 | 0.089 | 0 | 0 | 0 |
| zitB | 1.64 | 0.089 | 0 | 0 | 0 |
| ssuA | 1.625 | 0.089 | 0 | 0 | 0 |
| yohM | 1.594 | 0.089 | 0 | 0 | 0 |
| yhjV | 1.583 | 0.089 | 0 | 0 | 0 |
| emrE | 1.574 | 0.089 | 0 | 0 | 0 |
| yliD | 1.515 | 0.089 | 0 | 0 | 0 |
| pstB | 1.508 | 0.089 | 0 | 0 | 0 |
| mdtE | 1.47 | 0.089 | 0 | 0 | 0 |
| potG | 1.461 | 0.089 | 0 | 0 | 0 |
| nagE | 1.456 | 0.089 | 0 | 0 | 0 |
| btuD | 1.447 | 0.089 | 0 | 0 | 0 |
| atpB | 1.447 | 0.089 | 0 | 0 | 0 |
| tnaB | 1.431 | 0.089 | 0 | 0 | 0 |
| lacY | 1.417 | 0.089 | 0 | 0 | 0 |
| cvrA | 1.417 | 0.089 | 0 | 0 | 0 |
| nikB | 1.402 | 0.089 | 0 | 0 | 0 |
| frlA | 1.38 | 0.089 | 0 | 0 | 0 |
| phnL | 1.338 | 0.089 | 0 | 0 | 0 |
| idnT | 1.335 | 0.089 | 0 | 0 | 0 |
| pstS | 1.322 | 0.089 | 0 | 0 | 0 |
| kgtP | 1.288 | 0.089 | 0 | 0 | 0 |
| hofC | 2.037 | 0.088 | 0 | 0 | 0 |
| ugpE | 1.971 | 0.088 | 0 | 0 | 0 |
| ydiM | 1.953 | 0.088 | 0 | 0 | 0 |
| yiaM | 1.95 | 0.088 | 0 | 0 | 0 |
| yicM | 1.93 | 0.088 | 0 | 0 | 0 |
| malK | 1.925 | 0.088 | 0 | 0 | 0 |
| cydD | 1.904 | 0.088 | 0 | 0 | 0 |
| brnQ | 1.868 | 0.088 | 0 | 0 | 0 |
| yjeM | 1.816 | 0.088 | 0 | 0 | 0 |
| sstT | 1.812 | 0.088 | 0 | 0 | 0 |
| ypdD | 1.803 | 0.088 | 0 | 0 | 0 |
| yadH | 1.774 | 0.088 | 0 | 0 | 0 |

| | | | | | |
|---|---|---|---|---|---|
| proY | 1.762 | 0.088 | 0 | 0 | 0 |
| yejA | 1.751 | 0.088 | 0 | 0 | 0 |
| mdtF | 1.75 | 0.088 | 0 | 0 | 0 |
| cysP | 1.749 | 0.088 | 0 | 0 | 0 |
| ampG | 1.745 | 0.088 | 0 | 0 | 0 |
| ygfQ | 1.709 | 0.088 | 0 | 0 | 0 |
| yeiM | 1.688 | 0.088 | 0 | 0 | 0 |
| kdpA | 1.686 | 0.088 | 0 | 0 | 0 |
| mdtG | 1.655 | 0.088 | 0 | 0 | 0 |
| proW | 1.647 | 0.088 | 0 | 0 | 0 |
| livH | 1.644 | 0.088 | 0 | 0 | 0 |
| bcr | 1.627 | 0.088 | 0 | 0 | 0 |
| malF | 1.619 | 0.088 | 0 | 0 | 0 |
| fepD | 1.588 | 0.088 | 0 | 0 | 0 |
| gltI | 1.578 | 0.088 | 0 | 0 | 0 |
| yrbD | 1.53 | 0.088 | 0 | 0 | 0 |
| acrD | 1.51 | 0.088 | 0 | 0 | 0 |
| nikD | 1.505 | 0.088 | 0 | 0 | 0 |
| nikC | 1.497 | 0.088 | 0 | 0 | 0 |
| lsrC | 1.477 | 0.088 | 0 | 0 | 0 |
| nanT | 1.438 | 0.088 | 0 | 0 | 0 |
| uup | 1.393 | 0.088 | 0 | 0 | 0 |
| clcA | 1.392 | 0.088 | 0 | 0 | 0 |
| nikE | 1.382 | 0.088 | 0 | 0 | 0 |
| ybaT | 1.356 | 0.088 | 0 | 0 | 0 |
| yfbS | 1.316 | 0.088 | 0 | 0 | 0 |
| dhaH | 1.31 | 0.088 | 0 | 0 | 0 |
| corA | 1.307 | 0.088 | 0 | 0 | 0 |
| torT | 1.3 | 0.088 | 0 | 0 | 0 |
| ycdG | 1.276 | 0.088 | 0 | 0 | 0 |
| yeaS | 1.248 | 0.088 | 0 | 0 | 0 |
| srlE | 1.141 | 0.088 | 0 | 0 | 0 |
| tatC | 0.996 | 0.088 | 0 | 0 | 0 |
| modB | 0.913 | 0.088 | 0 | 0 | 0 |
| dgoT | 0.174 | 0.088 | 0 | 0 | 0 |
| ccmC | 1.906 | 0.087 | 0 | 0 | 0 |
| yfjD | 1.894 | 0.087 | 0 | 0 | 0 |
| hcaT | 1.846 | 0.087 | 0 | 0 | 0 |
| gabP | 1.813 | 0.087 | 0 | 0 | 0 |
| yihP | 1.778 | 0.087 | 0 | 0 | 0 |
| mscL | 1.763 | 0.087 | 0 | 0 | 0 |
| alsA | 1.759 | 0.087 | 0 | 0 | 0 |
| kdpB | 1.751 | 0.087 | 0 | 0 | 0 |
| fepG | 1.75 | 0.087 | 0 | 0 | 0 |
| phnD | 1.698 | 0.087 | 0 | 0 | 0 |
| metQ | 1.694 | 0.087 | 0 | 0 | 0 |
| thiQ | 1.682 | 0.087 | 0 | 0 | 0 |
| ybiT | 1.65 | 0.087 | 0 | 0 | 0 |

| | | | | | |
|---|---|---|---|---|---|
| exuT | 1.643 | 0.087 | 0 | 0 | 0 |
| ansP | 1.609 | 0.087 | 0 | 0 | 0 |
| acrF | 1.597 | 0.087 | 0 | 0 | 0 |
| frvA | 1.578 | 0.087 | 0 | 0 | 0 |
| ybbW | 1.57 | 0.087 | 0 | 0 | 0 |
| yrbG | 1.566 | 0.087 | 0 | 0 | 0 |
| mglA | 1.552 | 0.087 | 0 | 0 | 0 |
| ulaC | 1.547 | 0.087 | 0 | 0 | 0 |
| yphE | 1.513 | 0.087 | 0 | 0 | 0 |
| glnQ | 1.509 | 0.087 | 0 | 0 | 0 |
| yehW | 1.491 | 0.087 | 0 | 0 | 0 |
| fepC | 1.463 | 0.087 | 0 | 0 | 0 |
| glpT | 1.458 | 0.087 | 0 | 0 | 0 |
| xylE | 1.419 | 0.087 | 0 | 0 | 0 |
| yahN | 1.394 | 0.087 | 0 | 0 | 0 |
| mglB | 1.308 | 0.087 | 0 | 0 | 0 |
| proV | 1.245 | 0.087 | 0 | 0 | 0 |
| yccS | 0.834 | 0.087 | 0 | 0 | 0 |
| phoR | 0.066 | 0.087 | 0 | 0 | 0 |
| amtB | 0.048 | 0.087 | 0 | 0 | 0 |
| ygeD | 2.234 | 0.086 | 0 | 0 | 0 |
| mdtH | 2.041 | 0.086 | 0 | 0 | 0 |
| mdtC | 2.027 | 0.086 | 0 | 0 | 0 |
| fucP | 1.909 | 0.086 | 0 | 0 | 0 |
| livJ | 1.829 | 0.086 | 0 | 0 | 0 |
| emrY | 1.814 | 0.086 | 0 | 0 | 0 |
| uidB | 1.791 | 0.086 | 0 | 0 | 0 |
| ygjE | 1.749 | 0.086 | 0 | 0 | 0 |
| yejF | 1.744 | 0.086 | 0 | 0 | 0 |
| yggT | 1.743 | 0.086 | 0 | 0 | 0 |
| yliB | 1.695 | 0.086 | 0 | 0 | 0 |
| yfaV | 1.692 | 0.086 | 0 | 0 | 0 |
| ygaZ | 1.69 | 0.086 | 0 | 0 | 0 |
| ascF | 1.679 | 0.086 | 0 | 0 | 0 |
| ypdH | 1.635 | 0.086 | 0 | 0 | 0 |
| proX | 1.629 | 0.086 | 0 | 0 | 0 |
| xylG | 1.612 | 0.086 | 0 | 0 | 0 |
| dcuB | 1.608 | 0.086 | 0 | 0 | 0 |
| ybbL | 1.581 | 0.086 | 0 | 0 | 0 |
| setB | 1.553 | 0.086 | 0 | 0 | 0 |
| aaeA | 1.455 | 0.086 | 0 | 0 | 0 |
| cysA | 1.451 | 0.086 | 0 | 0 | 0 |
| frwC | 1.413 | 0.086 | 0 | 0 | 0 |
| artP | 1.319 | 0.086 | 0 | 0 | 0 |
| sapF | 1.309 | 0.086 | 0 | 0 | 0 |
| yjjP | 1.02 | 0.086 | 0 | 0 | 0 |
| WT | 0.06 | 0.086 | 0 | 0 | 0 |
| melB | 0.054 | 0.086 | 0 | 0 | 0 |

| | | | | | |
|---|---|---|---|---|---|
| cmr | 0.052 | 0.086 | 0 | 0 | 0 |
| ssuB | 0.049 | 0.086 | 0 | 0 | 0 |
| lsrD | 1.908 | 0.085 | 0 | 0 | 0 |
| araJ | 1.824 | 0.085 | 0 | 0 | 0 |
| alsB | 1.8 | 0.085 | 0 | 0 | 0 |
| acrB | 1.738 | 0.085 | 0 | 0 | 0 |
| mdtB | 1.702 | 0.085 | 0 | 0 | 0 |
| argO | 1.558 | 0.085 | 0 | 0 | 0 |
| glvB | 1.489 | 0.085 | 0 | 0 | 0 |
| rhaT | 1.478 | 0.085 | 0 | 0 | 0 |
| tyrP | 1.422 | 0.085 | 0 | 0 | 0 |
| kefA | 1.418 | 0.085 | 0 | 0 | 0 |
| oppF | 1.375 | 0.085 | 0 | 0 | 0 |
| potB | 1.313 | 0.085 | 0 | 0 | 0 |
| nhaB | 1.288 | 0.085 | 0 | 0 | 0 |
| arsB | 1.137 | 0.085 | 0 | 0 | 0 |
| tolC | 0.89 | 0.085 | 0 | 0 | 0 |
| ybjJ | 0.748 | 0.085 | 0 | 0 | 0 |
| atpH | 0.668 | 0.085 | 0 | 0 | 0 |
| yddG | 0.068 | 0.085 | 0 | 0 | 0 |
| atpA | 0.047 | 0.085 | 0 | 0 | 0 |
| yfdV | 1.862 | 0.084 | 0 | 0 | 0 |
| zntA | 1.843 | 0.084 | 0 | 0 | 0 |
| yjcR | 1.824 | 0.084 | 0 | 0 | 0 |
| modF | 1.769 | 0.084 | 0 | 0 | 0 |
| adiC | 1.639 | 0.084 | 0 | 0 | 0 |
| ccmB | 1.611 | 0.084 | 0 | 0 | 0 |
| cynX | 1.54 | 0.084 | 0 | 0 | 0 |
| copA | 1.415 | 0.084 | 0 | 0 | 0 |
| sapD | 1.365 | 0.084 | 0 | 0 | 0 |
| yihO | 1.184 | 0.084 | 0 | 0 | 0 |
| atpC | 0.512 | 0.084 | 0 | 0 | 0 |
| ugpB | 0.085 | 0.084 | 0 | 0 | 0 |
| rhtA | 0.07 | 0.084 | 0 | 0 | 0 |
| ddpD | 0.061 | 0.084 | 0 | 0 | 0 |
| ompF | 0.048 | 0.084 | 0 | 0 | 0 |
| yfbW | 0.047 | 0.084 | 0 | 0 | 0 |
| tsr | 0.046 | 0.084 | 0 | 0 | 0 |
| yadG | 1.559 | 0.083 | 0 | 0 | 0 |
| znuB | 1.526 | 0.083 | 0 | 0 | 0 |
| ptsG | 1.342 | 0.083 | 0 | 0 | 0 |
| yrbC | 1.163 | 0.083 | 0 | 0 | 0 |
| sufB | 0.849 | 0.083 | 0 | 0 | 0 |
| atpG | 0.712 | 0.083 | 0 | 0 | 0 |
| tolR | 0.565 | 0.083 | 0 | 0 | 0 |
| WT | 0.278 | 0.083 | 0 | 0 | 0 |
| ydjN | 0.07 | 0.083 | 0 | 0 | 0 |
| RtolC | 0.045 | 0.083 | 0 | 0 | 0 |

| | | | | | |
|---|---|---|---|---|---|
| yhbG | 0.996 | 0.082 | 0 | 0 | 0 |
| ygaH | 0.12 | 0.082 | 0 | 0 | 0 |
| mdtJ | 0.086 | 0.082 | 0 | 0 | 0 |
| WT | 0.082 | 0.082 | 0 | 0 | 0 |
| ytfR | 0.07 | 0.082 | 0 | 0 | 0 |
| yeeA | 0.052 | 0.082 | 0 | 0 | 0 |
| yifK | 0.052 | 0.082 | 0 | 0 | 0 |
| yicJ | 0.05 | 0.082 | 0 | 0 | 0 |
| WT | 0.046 | 0.082 | 0 | 0 | 0 |
| WT | 0.046 | 0.082 | 0 | 0 | 0 |
| ompC | 0.044 | 0.082 | 0 | 0 | 0 |
| potC | 0.59 | 0.081 | 0 | 0 | 0 |
| WT | 0.044 | 0.081 | 0 | 0 | 0 |
| dinF | 0.043 | 0.081 | 0 | 0 | 0 |
| ompA | 0.041 | 0.081 | 0 | 0 | 0 |
| WT | 0.044 | 0.08 | 0 | 0 | 0 |
| WT | 0.04 | 0.08 | 0 | 0 | 0 |
